# Rapid accumulation of fluorophores and fast kill identify drugs with bactericidal effects against Gram-negative bacteria

**DOI:** 10.1101/2025.07.10.664096

**Authors:** J. Enrique Salcedo-Sora, Douglas B. Kell

## Abstract

Antimicrobial resistance is a massive threat, but it can take decades to develop a new antibiotic. That time could be drastically reduced if we were able to anticipate desirable properties of a chemical, such as its potential to target specific bacterial compartments. This would provide the opportunity to prioritise the development of molecules that target, for instance, the cell membrane, as this does not rely on transporters and usually results in a fast-acting bactericidal effect. We used flow cytometry and a set of fluorophores together with a group of antibiotics to discriminate between antimicrobials acting on cell membrane versus intracellularly against two Gram-negative bacteria *E. coli* and *A. baylyi*. We then chose Rhodamine 123 to screen a commercial library of chemical compounds. Using flow cytometry, several drugs present in the Prestwick library were observed to have cytotoxic effects towards *E. coli*. This was confirmed with growth inhibitory assays in both *E. coli* and *A. baylyi* for Pantoprazole, Theophylline and Zoledronic acid. This represents an approach to the large-scale screening of small molecules with the potential to deliver fast-acting molecules that target cell membranes in Gram-negative bacteria.

**Impact statement:** The discovery of novel antimicrobials is essential to build resilience against infectious diseases. Understanding the mechanism of action of known as well as novel compounds is equally crucial. Achieving this understanding is a long and complex process that is usually addressed on an individual basis for an antibiotic, or a class of antibiotics. Using high-throughput pipelines such as those possible with flow cytometry can help to shorten this process. We present a flow cytometry-based approach capable of determining if a given molecule has rapid antimicrobial effects that involve increasing the permeability of the cell membrane. The latter is a property that usually makes a chemical a very effective antimicrobial.

**Data summary:** The authors confirm all supporting data, code and protocols have been provided within the article or through supplementary data files.

## 5. Introduction

The membrane transport of natural substrates as well as xenobiotics in live cells is mediated by proteins embedded in the otherwise impermeable cell membrane (1, 2). Accordingly, antibiotics targeting the bacterial cell membrane, which make for a formidable barrier, are very effective bactericides (3). Molecules that preferentially target the cell membrane are much sought after, with the majority of antibiotics in preclinical and clinical development seeking to target cell wall and membrane synthesis (4-6). However, it still commonly takes a further 10 years for the first of any of the successful drugs to reach the market (4, 6).

Flow cytometry can interrogate a cell’s physiology rapidly and on an individual-cell basis. This technology has prospered throughout microbiological research ever since the technology was acknowledged to be applicable to the much smaller microbial cells (7-12). More recently, flow cytometry has been utilised for the study of membrane transport (13). With the wealth of fluorogenic dyes under continuous expansion, flow cytometry has allowed us to interrogate and compare, for instance, influx versus efflux membrane transport systems (14). The use of fluorogenic compounds as a proxy for transporter substrates has benefited from the structural similarities that often exist between available fluorophores and transporter substrates (15). This has facilitated the comparative measurement of transport of xenobiotics by specific membrane proteins (16, 17).

We leveraged these developments to assemble a screening method that detects lethal cell wall and cell membrane interacting compounds. Conventional antibiotic sensitisation screens often capture large numbers of hit molecules with unrelated activities that will require further characterisation. Given the multiparametric nature of flow cytometry, while detecting cell accumulation of fluorogenic dyes (18), we here exploited here the capacity of a palette of dyes to reveal cell damage in response to a short exposure to a range of known antibiotics in *E. coli* and *A. baylyi*. These antibiotics represented examples of antibacterials considered to affect either the cell membranes or intracellular targets. The most discriminatory parameters derived from these assays were then applied to interrogate the Prestwick chemical library of marketed drugs for their antimicrobial activity against *E. coli*. A number of these drugs increased the accumulation of the fluorophore and reduced the number of detectable cells beyond the threshold established by the control (polymyxin E or colistin). The bactericidal effects found in Pantoprazole, Theophylline and Zoledronic acid were then confirmed against both Gram-negative bacteria *E. coli* and *A. baylyi* by standard growth inhibitory assays.

## 6. Methods

### Bacterial strains and chemicals

Strains: *E. coli* strain BW25113 [Keio collection reference strain: *Δ(araD-araB)567, ΔlacZ4787(::rrnB-3), λ* −, *rph-1, Δ(rhaD-rhaB)568, hsdR514] and Acinetobacter. baylyi* strain ADP1 (ATCC 33305). Antibiotics were purchased from Merck KGaA (Darmsadt, Germany) and prepared in PBS (50 mg/mL stocks), except for nitrofurantoin, pentamidine and rifampicin which needed DMSO to dissolve, and chloramphenicol and tetracycline that were dissolved in ethanol.

### Growth and permeability effects measured by flow cytometry

Fluorophores were purchased from Sigma Aldrich (Merck KGaA, Darmsadt, Germany), ThermoFisher, TCI America, or ATTO-TEC GmbH. Other reagents were purchased from Sigma Aldrich. Fluorophore uptake assays started with the spread of a small flake from a frozen glycerol stock of bacteria onto a fresh plate of complex solid media (Merck LB 110283) and incubated overnight at 37⍰ ° C. A single colony from a fresh solid media plate was then incubated in 5 mL of complex media (Merck LB 110285) overnight at 37 °C with shaking at 250 rpm. in the absence of antibiotics. The overnight cultures were diluted 1:⍰1000 in fresh liquid complex media and grown for 2⍰h at 37⍰°C with shaking at 250 rpm. The cell density was then adjusted to approximately 2000⍰cells per microlitre, as inferred by turbidity at OD_600_. For antibiotic exposure, cells were first incubated in complex media for 30 min, 37°C, in 384-well plates, at a final volume of 50 µL. Antibiotics were present at final concentrations equivalent to twice their IC_50_ values for either *E. coli* or *A. baylyi*. Fluorophores were then added at 1 µM final concentration, and cells incubated for further 10 min at 37°C (similarly to our previous report (14)) before interrogation in the flow cytometry analyser as detailed below.

### Flow cytometry

We used a high-throughput flow cytometer, the Intellicyt iQue Screener Plus (Sartorius, Göttingen, Germany), with the following protocol: buffer equilibration (QSol, Sartorius) and plate shaking 2000 r.p.m. for 50⍰s, sampling for 2⍰s with 1⍰s upload time; 5 s wash in Qsol buffer every three wells, and further probe wash for 10 s every 12 wells. The instrument has three LED lasers (405⍰nm, 488⍰nm, 640⍰nm) and collects data for 2 light scattering channels and 13 fluorescence channels. Once the light from any of three lasers has reached the samples, these channels collect the fluorescent (emission) signals back from the samples in the following spectral ranges (channel name, emission range in nm): VL1 (445±45), VL2 (530±30), VL3 (572±28), VL4 (615±24), VL5 (675±30), VL6 (780±60), BL1 (530±30), BL2 (572±28), BL3 (615±24), BL4 (675±30), BL5 (780±60), RL1(675±30) and RL2(780±60). This instrument is highly resistant to the detection of extracellular fluorescence, a potential source of noise in fluorescent assays (19). As with any other transport or cell accumulation assays (e.g., filtration or flow dialysis (20)), flow cytometry would not discriminate directly between molecules that bind outward facing cell structures from those molecules that have actually reached intracellular compartments.

### Screening of chemical library

The Prestwick Chemical Library® of mostly FDA-approved small molecules were used to test for antibacterial activity of non-antibiotic drugs. A set of 1,280 compounds at 10 mM in DMSO (full list in Supplementary table 1) were aliquoted out in 384-multi-well plates for testing at final 1 µM concentration in PBS (0.01% DMSO final concentration) using in-house robotics for liquid handling (GeneMill Research Facility, Liverpool Shared Research Facilities (LIV-SRF), University of Liverpool). Bacteria cultures growing in logarithmic phase in complex media were added to 50 µL final volume per well. After 30 min incubation at 37°C, Rhodamine 123 (a dye normally excluded by Gram-negative bacteria)(21) was added to 1 µM final concentration for a further incubation of 15 min at 37°C.

**Table 1.** Inhibitory concentrations (IC_50_) for *E. coli* and *A. baylyi*. The average of at least three assays (Mean) and the standard deviation (SD) are listed. The first eight antibiotics (shaded) are categorized as cell membrane acting. The others are categorized as intracellularly acting antibiotics.

|  | *E. coli* |  | *A. baylyi* |  |
| --- | --- | --- | --- | --- |
|  | Mean (mg/L) | SD | Mean (mg/L) | SD |
| Ampicillin | 1.48 | 0.32 | 23.4 | 4.57 |
| Amoxicillin | 1.05 | 0.95 | 0.43 | 0.40 |
| Bacitracin | >1000 | NA | 48.4 | 2.56 |
| Colistin | 0.18 | 0.10 | 0.15 | 0.02 |
| Fosfomycin | 0.25 | 0.24 | 15.3 | 14.4 |
| Pentamidine | 71.1 | 2.79 | 46.4 | 2.13 |
| Polymyxin B | 85.3 | 5.3 | >500 | NA |
| Vancomycin | 68.9 | 55.4 | 46.0 | 42.6 |
| Azithromycin | 4.95 | 4.67 | 0.11 | 0.11 |
| Cefotaxime | 0.01 | <0.01 | 1.85 | 1.5 |
| Chloramphenicol | 0.70 | 0.21 | 0.82 | 0.12 |
| Ciprofloxacin | < 0.01 | NA | 0.040 | NA |
| Gentamicin | 1.29 | 0.06 | 1.11 | 0.12 |
| Nalidixic acid | 2.57 | 1.98 | 1.34 | 1.08 |
| Nitrofurantoin | 2.08 | 1.98 | 68.3 | 44.3 |
| Ofloxacin | 0.05 | 0.01 | 0.08 | 0.01 |
| Rifampicin | 0.55 | 0.14 | 0.08 | 0.01 |
| Tetracycline | 0.46 | 0.03 | 0.75 | 0.03 |
| Trimethoprim | 0.08 | 0.07 | 7.05 | 6.15 |

### Data analysis

End-point growth inhibitory concentrations were calculated from microtitration assays. The inhibitory concentrations of antibiotic that kill half of the bacteria population (IC_50_) were calculated with the four-parameter logistic model as implemented in py50 (22). Flow cytometry data were analysed using a combination of the instrument’s Forecyt software and routines written by the first author in R and Python. The principal component analyses were carried out using the R’s packages FactoMineR (23).

### Inhibitory concentrations assays

Bacterial cultures were carried out in complex media (lysogeny broth, Merck LB 110285). Overnight cultures in LB were diluted 1 :1000 in fresh LB and incubated for two hours at 37°C, shaking 250 rpm. One hundred microlitres of two-fold dilutions of drugs were prepared in flat bottom 96-well plates to which 100 µL of fresh bacterial culture were added. Endpoint read-outs of the media turbidity at 600 nm (OD_600_) were taken after an overnight incubation (approximately 20 h) at 37°C, using a BMG LabTech CLARIOstar Plus plate reader.

## 7. Results

### Testing the sensitivity of *E. coli* and *A. baylyi* strains to a panel of antibiotics

We tested 19 antibiotics in current clinical use (Table 1). The collated group had eight antibiotics considered to have cell membrane acting activity as their primary mechanism of action: ampicillin, amoxicillin, bacitracin, colistin (polymyxin E), fosfomycin, pentamidine, polymyxin B and vancomycin. The rest of the antibiotics in Table 1 were considered to have primary targets that are intracellular. The sensitivity of both *E. coli* BW25113 and *A. baylyi* ADP1 to this set of antibiotics was measured as growth inhibitory effect (IC_50_) in microtiter plates (Table 1). These values informed the later use of this set of antibiotics in the present work.

### Testing the permeability of *E. coli* and *A. baylyi* to fluorophores

An extensive variety of fluorogenic compounds have been previously shown to accumulate with different levels of permeability in *E. coli* (14). This provides the opportunity to have fluorescent-based assays to interrogate the natural chemical space membrane transporters have evolved under in Gram-negative bacteria (14). It was then logical to apply those resources to address issues of significance in antibiotic research. The inclusion of *A. baylyi* - as a model for pathogenic *Acinetobacter* such as *A. baumannii* - expanded the relevance of this work to other Gram-negative of current clinical importance (24). *E. coli* and *A. baylyi* were exposed to 67 fluorophores chosen from the previous palette tested in *E. coli* (14). The range of cell accumulation as measured by flow cytometry (Fig. 1a) corroborated the permeability profile previously reported for *E. coli* (14).

**Fig. 1.**
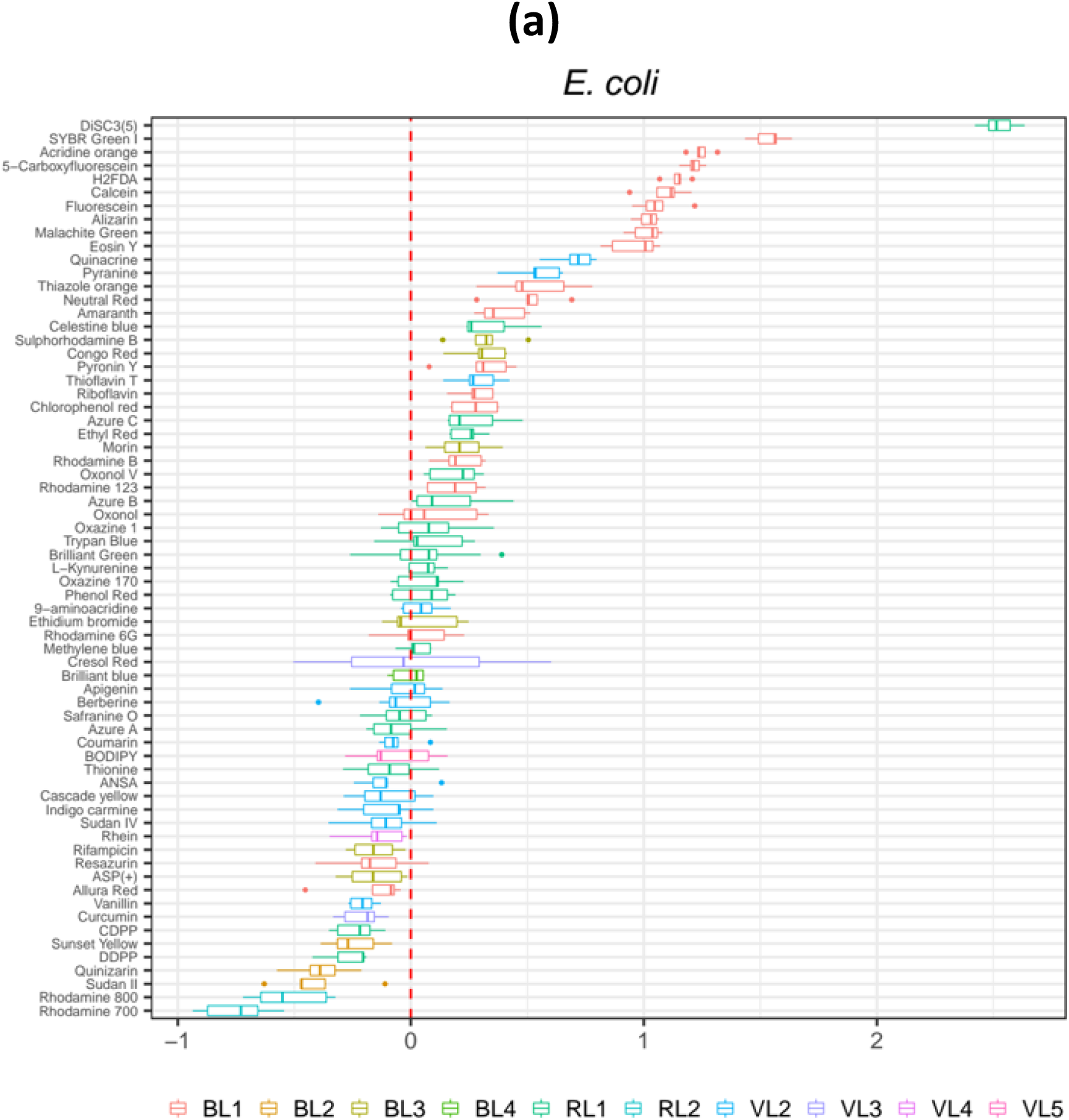
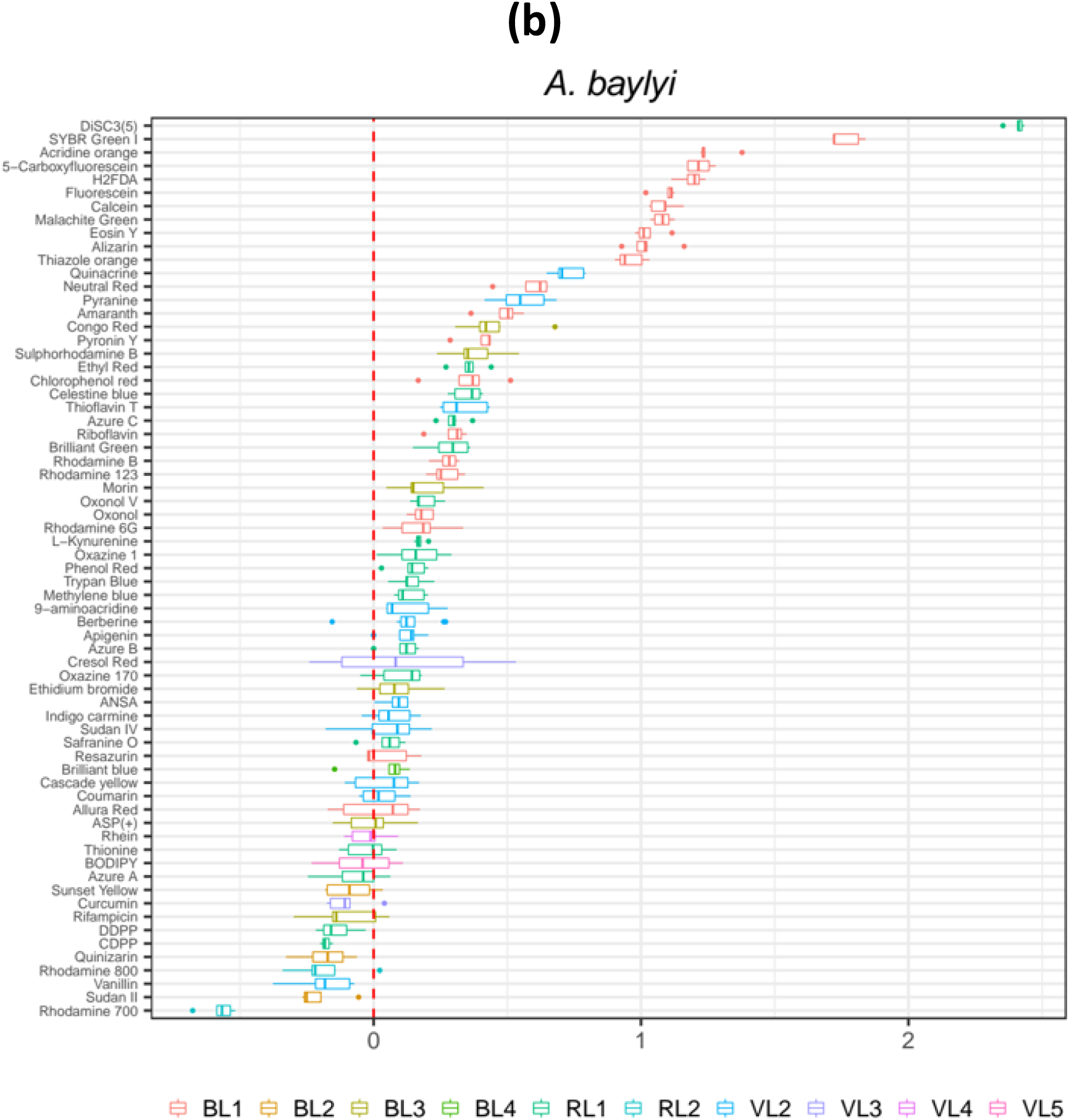
Cell accumulation of a panel of 67 fluorophores. **(a)** *E. coli* ; **(b)** *A. baylyi*. Fluorophores are listed on the ordinate. The bar plots represent the distribution of the data from cellular accumulation of those fluorophores in four or more assays. The colour code of the bar plots represents the flow cytometer channels (listed below each plot) appropriate for each fluorophore in the Intellicyt iQue Screener Plus cytometer (Methods). The abscissa represents the ratio, as log_10_ values, of the fluorescence signal generated by each fluorophore against the background signal (cells in media without fluorophores).

For *A. baylyi* this is a first description of its permeability profile to a set of fluorophores. *A. baylyi* is naturally transformable (25, 26), and generally had higher levels of permeability in comparison to *E. coli* (Fig. 1b). *A. baylyi* showed 52 compounds accumulating at levels greater than the background (Fig. 1b, ratios as log_10_ values equal or above 0 in the abscissa), while the corresponding value in *E. coli* was 37 compounds. This could be interpreted as some indication of the high levels of membrane permeability expected for bacteria known to be naturally competent (i.e., uninduced uptake of DNA) such as *A. baylyi* (27). The same ten compounds were accumulated 10-fold or higher in both species (Fig. 1, ratios as log_10_ values equal or greater than 1 in the abscissa) with the cyanine DiSC3(5) exhibiting the highest accumulation: two orders of magnitude above background.

### Interactivity of fluorophore accumulation and antibiotic exposure

Twenty-three fluorophores representing the range interrogated above were selected, and their accumulation studied in cells exposed to each of the 19 antibiotics chosen (Table 1). This was carried out against both *E. coli* and *A. baylyi*. Very apparent effects were observed on cell accumulation of fluorophores after 30 min exposure to antibiotics alone followed by 15 min of incubation with fluorophores. In *E. coli*, cell membrane-acting antibiotics such as colistin caused an increase of ten-fold and above in the cellular accumulation of almost all 23 fluorophores tested, except for Chlorophenol red and Congo red (Supplementary figure 1). Data represented here are the ratios of the fluorescence signals from antibiotic-exposed cells against the fluorescence signals from non-antibiotic controls. As usually acknowledged in cell membrane transport research, the cell accumulation of a given compound can be the result of net influx as well as the possibility of inhibited efflux.

Exposure to Pentamidine - another cell membrane-acting drug (28) - caused a ten-fold increase in the accumulation of SYBR green while vancomycin caused the accumulation of Allura red, Amaranth, and Calcein (Supplementary figure 1). In *A. baylyi* exposure to most antibiotics increased the cell accumulation of DiSC3(5). Cell membrane inhibitors such as Colistin or Pentamidine caused a ten-fold or higher accumulation of Alizarin, Rhodamine 123, and Thioflavin T (Supplementary figure 1). The accumulation of some of the fluorophores such as DiSC3(5), SYBR green and Thioflavin T illustrated the species-specific differences (Fig. 2). For instance, although most antibiotics induced higher permeability to DiSC3(5) in *A. baylyi* than in *E. coli*, Colistin and Pentamidine tended to standout in causing the specific accumulation of SYBR and Thioflavin T in both species (Fig. 2).

**Fig. 2.**
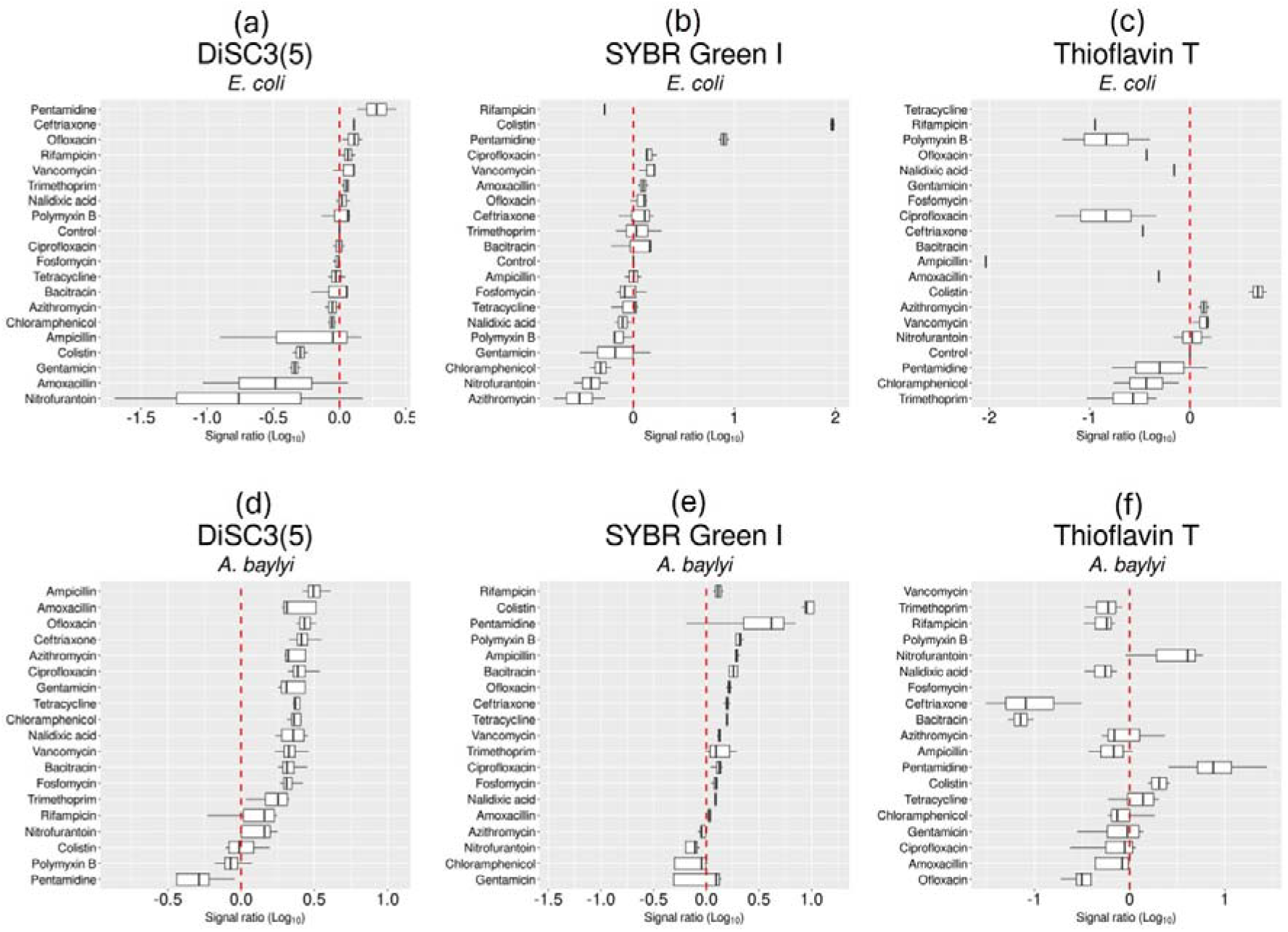
Examples of specific fluorophore uptake in *E. coli* and *A. baylyi* exposed to antibiotics. **(a-c)** *E. coli* ; **(d-f)** *A. baylyi*. Fluorescence signals from three fluorophores selected from the set of 67 molecules (Supplementary Figure 1). The signals (Signal ratio) for all fluorophores (Log10 values, abscissa) are shown against each antibiotic as ratios against non-antibiotic controls. The cellular accumulation of the fluorophore upon exposure to each antibiotic is represented by at least four biological replicates.

### Dimensional reduction of the multiple parameters detected in flow cytometry separates data according to antibiotic target as well as species

The data generated with the 23 fluorophores and 19 antibiotics were used to test for the parameters that differentiates cell membrane acting from intracellular acting antibiotics. The parameters used were species (*E. coli* and *A. baylyi*), antibiotic, fluor, events (median of cell count by cytometry), fluorescence signal (median of the emission) and fluorescence signal ratio (Supplementary table 2). When scoring the target (denoted as Intracellular or Wall) as the dependent variable, the PCA – despite being an unsupervised method with no knowledge of this – showed the data approximately segregating into two regions in one of the dimensions (Fig. 3a). Clear orthogonality was shown by the fluorophore signal (either as net fluorescence or as a ratio of exposed versus non-exposed to antibiotics), antibiotic, and the number of events detected by flow cytometry (Fig. 3a). Interestingly, even more apparent clusters were seen for the species, *E. coli* versus *A. baylyi* (Fig. 3b). The platform of the set of fluorophores and multiple known antibiotics, as implemented in this work, captured the differences in both cellular permeability and antibiotic sensitivity in these two Gram-negative bacteria.

**Fig. 3.**
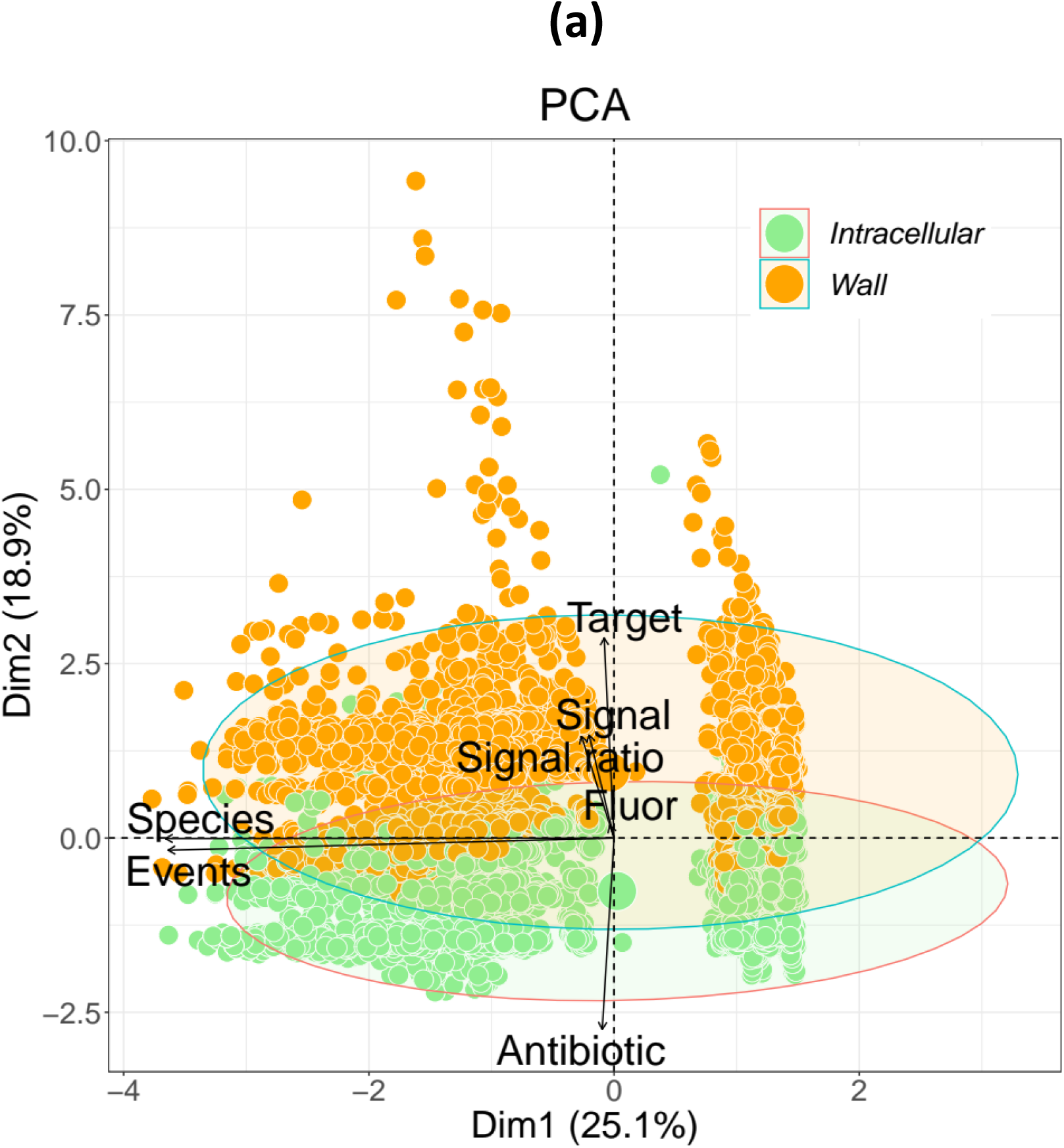
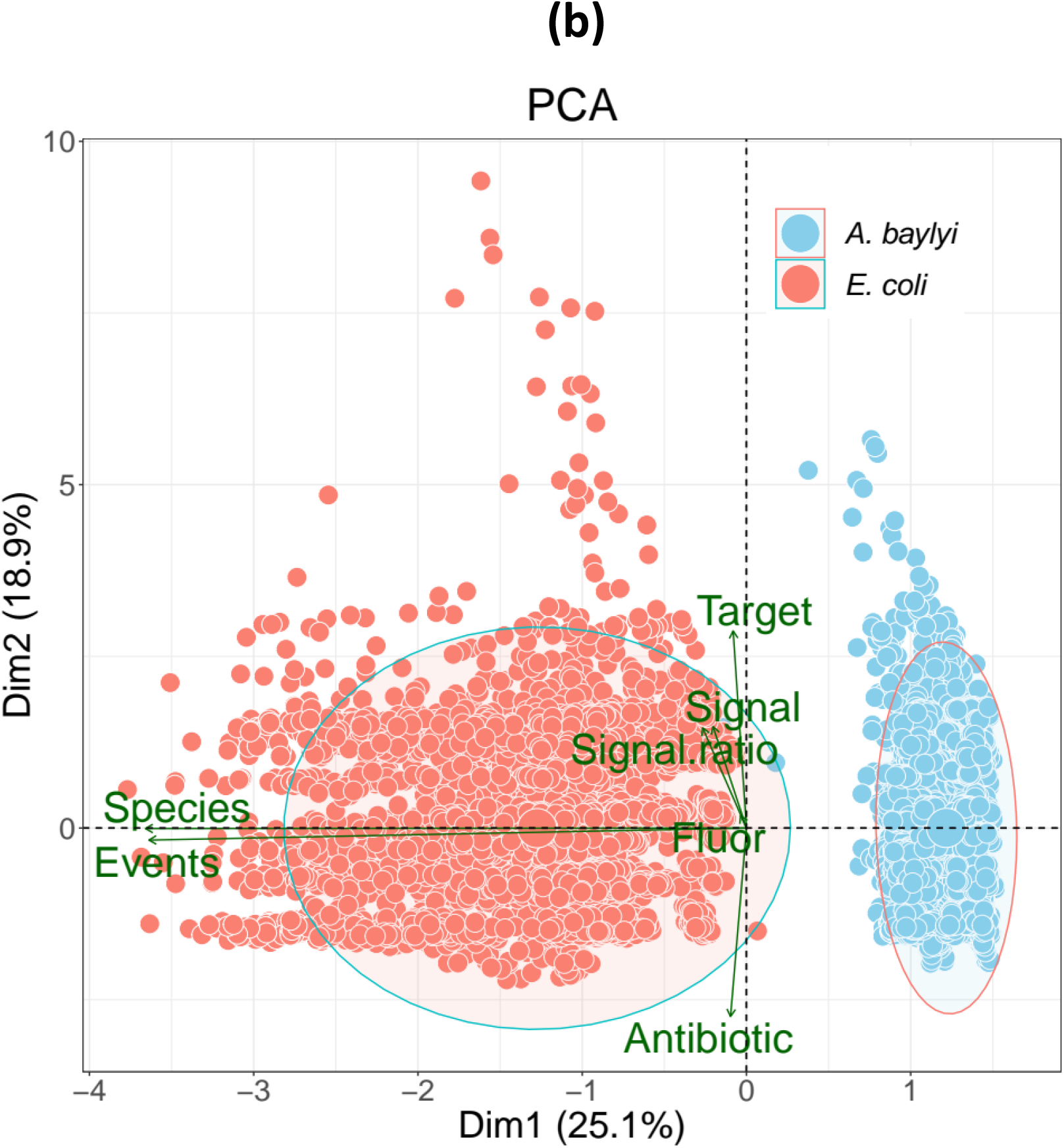
Principal components analysis (PCA) of fluorophore uptake in *E. coli* and *A. baylyi* exposed to antibiotics. **(a)** Intracellular versus cell membrane acting antibiotics; **(b)** *A. baylyi* versus *E. coli*. Fluorescence signals from 23 fluorophores (Supplementary Table 2) in cells exposed to 19 different antibiotics. Data generated from four or more biological replicates. Components consisted of species (*E. coli* and *A. baylyi*), antibiotics, fluorophore, events (median), signal (median) and signal ratio (fluorescence emission from samples exposed to antibiotic versus samples without exposure to antibiotics), and target type for each antibiotic (Supplementary tables 1 and 2).

### Screening of the Prestwick small molecule chemical library in *E. coli*

Five fluorophores were observed to accumulate in *E. coli* and *A. baylyi* at higher levels post-exposure to cell membrane-acting antibiotics such as Colistin and Pentamidine versus intracellularly acting antibiotics such as tetracycline (Fig. 4). From those fluorophores, Rhodamine 123 – to which viable cells are broadly and functionally impermeable – was the dye of choice for a proof of concept of flow cytometry screening for bactericidal compounds. We tested 1280 compounds from the Prestwick library of approved drugs (Supplementary Table 3) against *E. coli*. The cytotoxicity of those drugs against *E. coli* was estimated via the accumulation of Rhodamine 123 after 30 minutes post-exposure to each of the Prestwick chemical compounds. We ran two biological replicates for this screen (Supplementary Table 4). The trends were similar in both runs and the data for one replicate are illustrated in Figure 5.

**Fig. 4.**
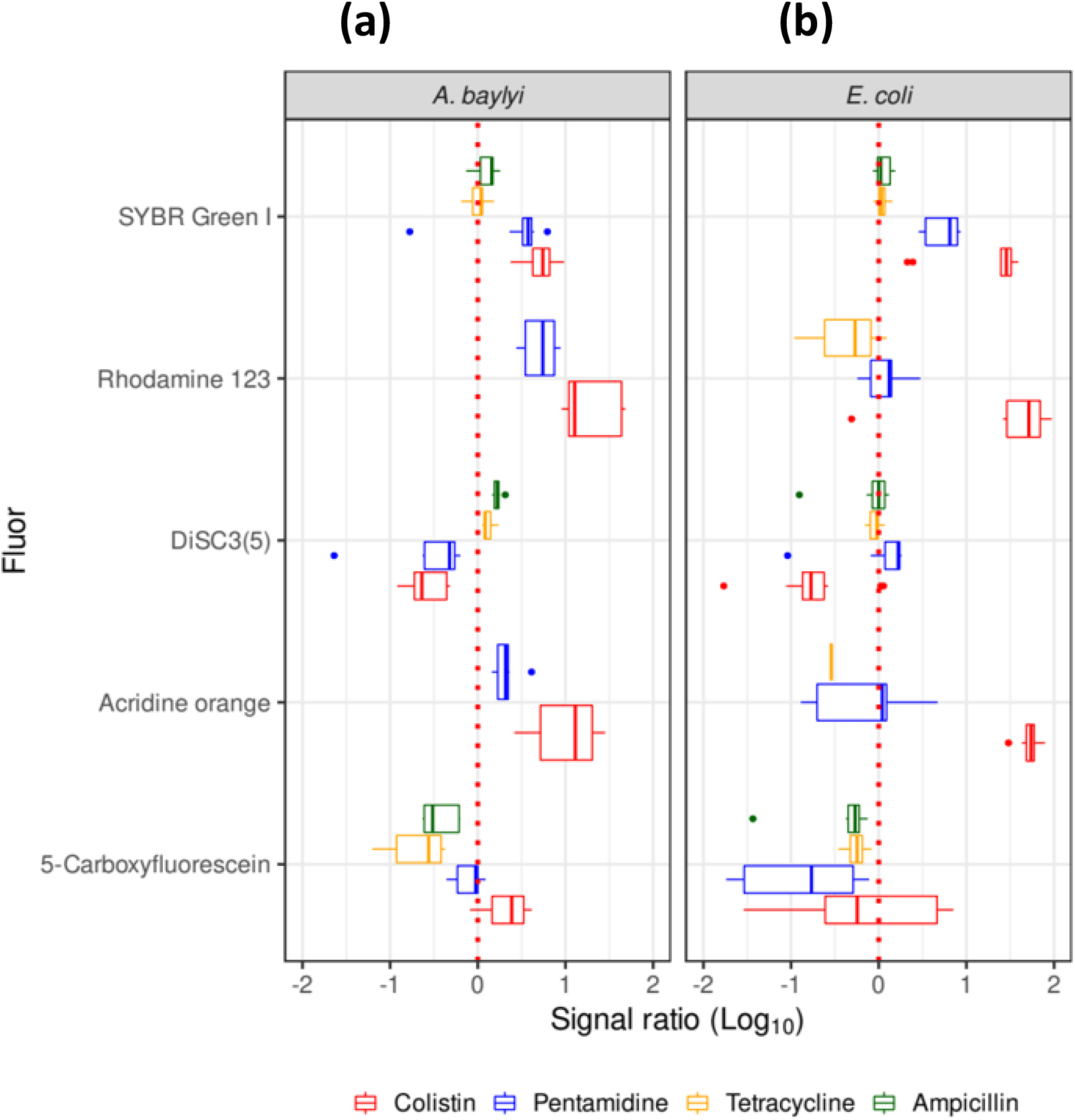
Antibiotics with the strongest effect on rapid fluorophore cell accumulation. **(a)** *A. baylyi*; **(b)** *E. coli*. Fluorophore uptake in *E. coli* and *A. baylyi* exposed to antibiotics is represented as signal ratio of fluorescent from exposed over non-exposed to antibiotics (Log10 values, abscissa). The four antibiotics and their effect of the accumulation of each of these five fluorophores are represented as box plots: the higher the permeability caused by the exposure to a given antibiotic the higher the signal ratio. Colour code: red Colistin, blue Pentamidine, yellow Tetracycline, green Ampicillin. The ratios for Rhodamine 123 and Acridine orange post-exposure to Tetracycline and Ampicillin in *A. baylyi* were lower than the cut off value of -2 for this graph. For the same reason the data on Rhodamine 123 and Acridine orange post-exposure to Ampicillin in *E. coli* were not visible in this graph.

**Fig. 5.**
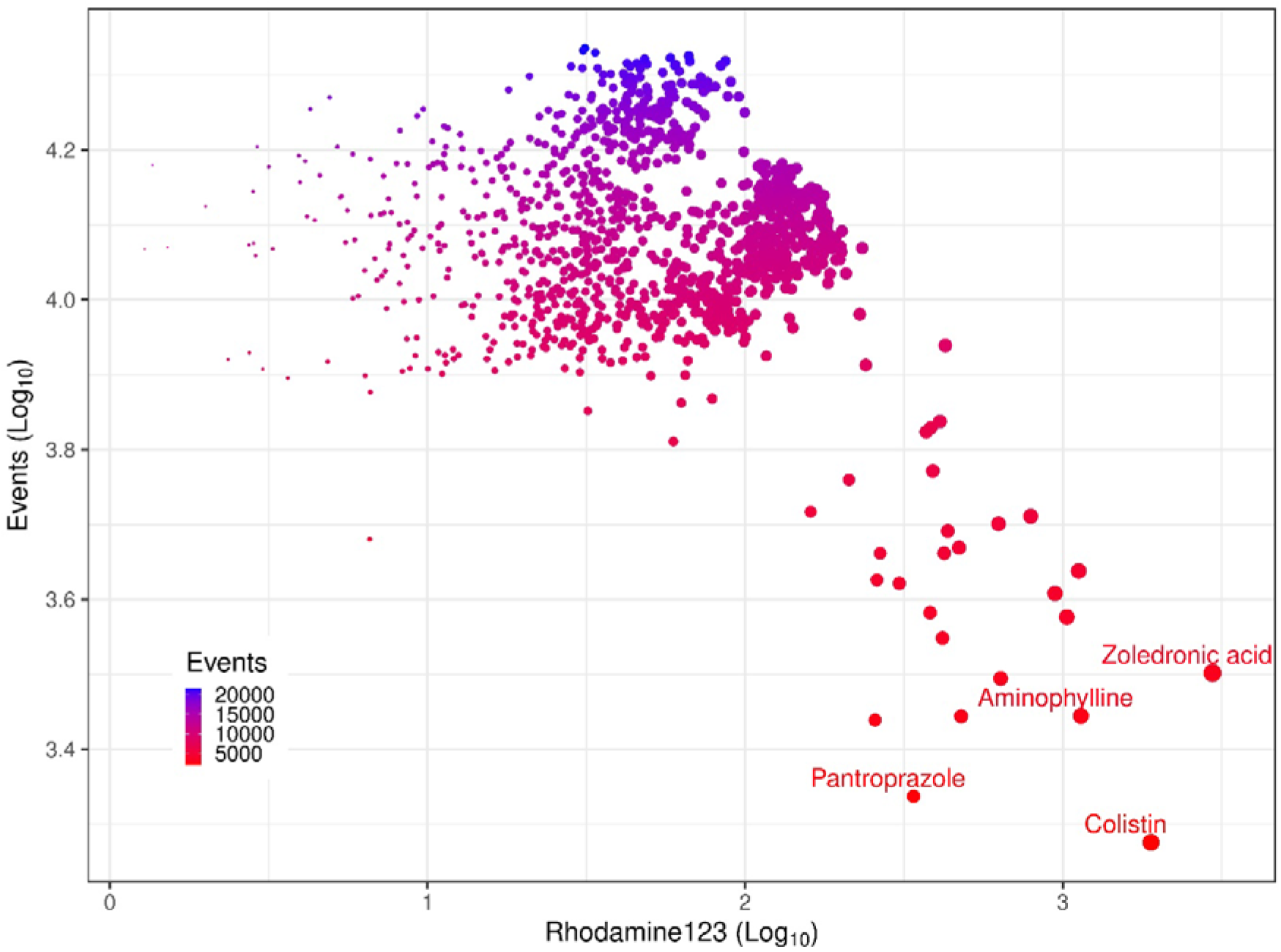
Distribution of the cytotoxic effect of a set of the Prestwick library. Flow cytometric detection of fluorophore uptake cell count (Events, ordinate) and dye uptake (Rhodamine 123, abscissa) in *E. coli* exposed to 1280 drugs from the Prestwick library. Compounds with minimal toxic effect (higher number of Events and low permeability to Rhodamine 123) were in the top and left areas of the plot while high cytotoxicity, similar to that of colistin, was represented towards the lower and right section of the plot. The size of the dots increases towards the higher signals of Rhodamine 123 and the colour gradient from blue to red follows the decreasing numbers of events (cell count).

Most drugs did not reduce the cell count (number of events by flow cytometry) below 7000 events (the negative control of 1% DMSO had counts of over 20000 events) (Fig. 5). The cellular accumulation of Rhodamine 123 on the other hand spread across two orders of magnitude (Fig. 5). Three compounds delimited the boundary of these data: Aminophylline, Pantoprazole sodium and Zoledronic acid hydrate (Fig. 5). Pantoprazole reduced the cell count to the lowest level of approximately 2000 events (Fig. 5).

### Pantoprazole, Theophylline and Zoledronic acid showed measurable growth inhibitory concentrations for Gram-negative bacteria *E. coli* and *A. baylyi*

The growth inhibitory concentrations were tested for the three drugs selected from the screening of the Prestwick library: Pantoprazole, Theophylline and Zoledronic acid (Fig. 6). Colistin was included as a known bactericidal (29). These assays were carried out for both *E. coli* and *A. baylyi* (Fig. 7). In *A. baylyi* Pantoprazole, Theophylline and Zoledronic acid displayed IC_50_ values in a narrow range of 0.57 to 0.59 mg/L (Fig. 7a). A similarly close range was seen for all three compounds in *E. coli* (0.68 – 0.72 mg/L) (Fig. 7b). All IC_50_ values observed were within the same order of magnitude of that of Colistin.

**Fig. 6.**
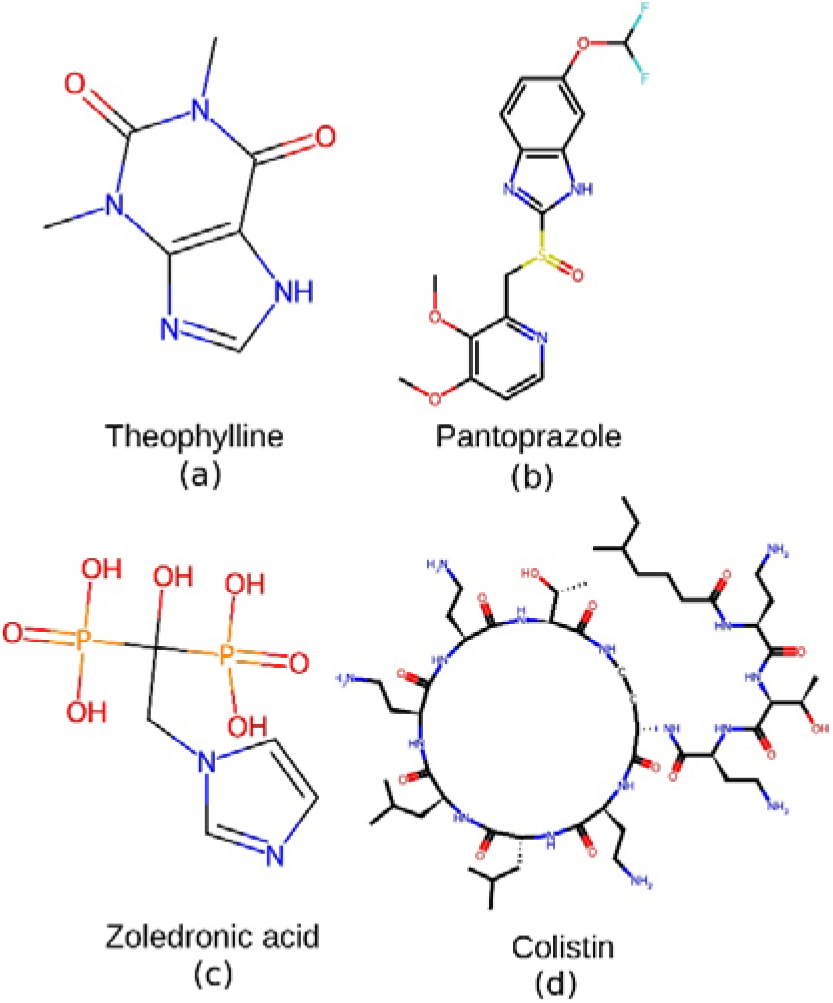
Compounds with apparent antimicrobial activity from the screen of the Prestwick library by flow cytometry. **(a)** Theophylline (MW 180), **(b)** Pantoprazole (MW 383), **(c)** Zoledronic acid (MW 272) and (d) Colistin (MW 1,155). Colistin was used as the control for a known bactericidal in the growth inhibitory assays (Fig. 7).

**Fig. 7.**
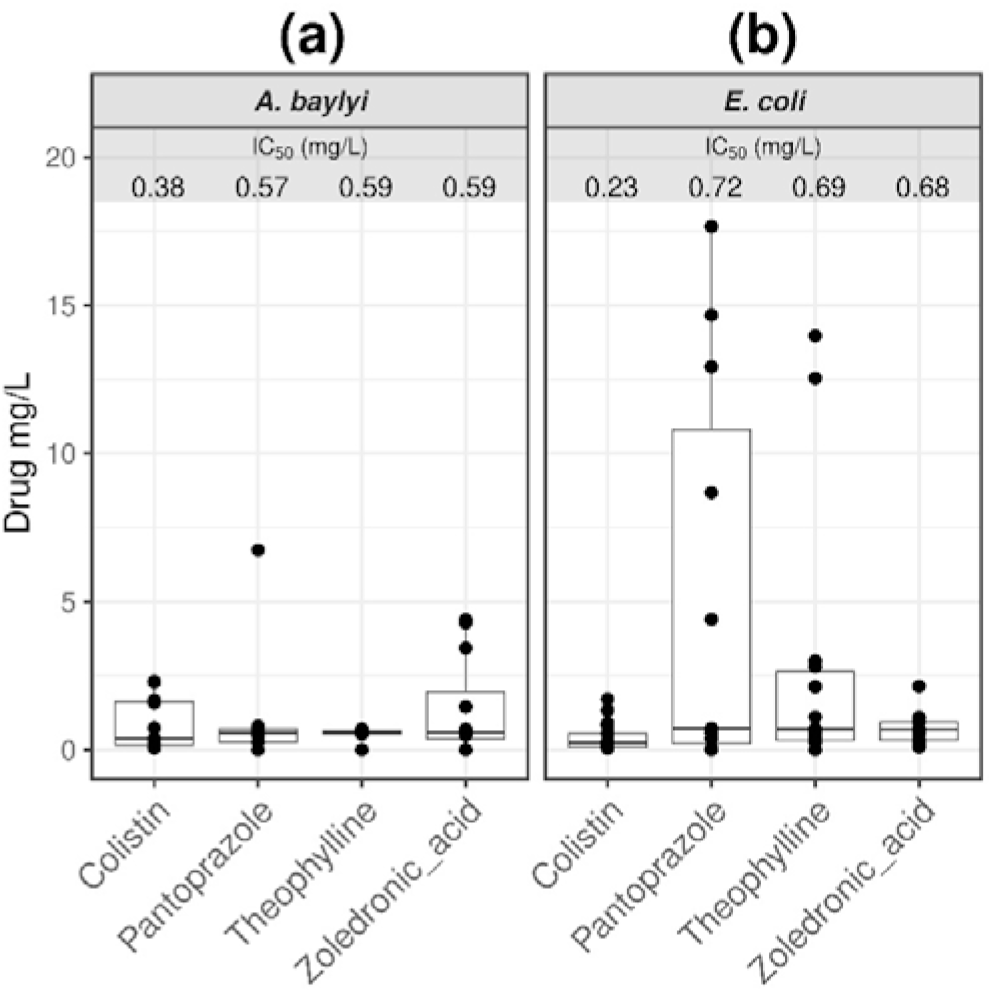
Growth inhibitory assays in *E. coli* and *A. baylyi*. **(a)** *A. baylyi*; **(b)** *E. coli*. The median values for the IC_50_ values (mg/L) of Colistin, Pantoprazole, Theophylline and Zoledronic acid are presented for *A. baylyi* and *E. coli*. Data represented at least five biological replicates with four technical replicates each (black dots).

## 8. Discussion

The discovery of new antibiotics is still a sporadic event measured in decades (30-32). At the same time infections caused by Gram-negative bacteria maintain their critical status. Carbapenem-resistant *Acinetobacter baumannii*, carbapenem-resistant Enterobacterales and third-generation cephalosporin-resistant Enterobacterales top the most recent of the WHO’s Bacterial Priority Pathogens List (33). The situation with *A. baumannii* is particularly challenging because, among other factors, of the limited antibiotic treatment options (33). The importance of targeted high-throughput platforms for the screening of antimicrobials is self-evident, particularly for the screening of highly effective bactericides. We consider cell membrane-acting antibiotics (direct interaction with prominent components of outer and inner cell membranes) to be a key trait of bactericidal antibiotics (28, 34-38). However, the list of antibiotics of current clinical relevance with such a property is limited (Table 1).

Antibiotic discovery can be done both phenotypically and/or by choosing particular targets (39). The former has the advantage that it can anticipate issues with cell efflux pumps (40-42). Along with flow cytometry-based methodology for bacteria phenotyping, bacterial cytological workflows have also been developed for antibiotics research. This methodology uses fluorescent microscopy of stained bacteria (including a membrane-impermeable reporter) (43). This methodology has been used to create classification models of antibiotic effects. (43). With the advantage of the high-throughput scales, single cell interrogation, and quantitative multiparametric outputs, flow cytometry continues to be a highly productive approach to investigate the mechanisms of action of antibiotics (44).

The work presented here focused on an approach to facilitate the discovery of rapid killing and cell-permeating agents. We built on our experience on cell phenotyping using extensive numbers of fluorogenic chemicals as reported before for *E. coli* (14) and complemented here for *A. baylyi*. Together with a representation of antibiotics of clinical relevance, we provided a flow cytometry platform to specifically identify cell membrane-binding molecules that seem to be in themselves rapid acting antimicrobials. We have illustrated its application in the screening of drugs which are considered primarily for their effects as antibiotics (45). Molecules with membrane-acting properties against bacteria can serve different purposes. For instance, polymyxins have been used to increase the in vitro accumulation of antibiotics with poor cellular penetration (46). Such a permeabilization strategy would be relevant in naturally resistance Gram-negative bacteria as in the case of P. aeruginosa. These bacteria have only one-eighth of the net permeability of a typical *E. coli* cell and three times the number of resistance-nodulation-division pumps (47).

The flow cytometry-based screening of a sample of the Prestwick library of mostly FDA-approved drugs here delivered three candidates for compounds with anti-Gram-negative activity. Aminophylline is a mix of the active compound Theophylline and ethylenediamine in a 2:1 ratio, used mainly for its bronchodilator effects via the interaction with several adenosine receptors, phosphodiesterases and histone deacetylases (48). Pantoprazole is a H+, K(+)-ATPase inhibitors (49), which as part of its effects in the treatment of gastric and duodenal ulcers seems to have a direct antimicrobial action on *Helicobacter pylori* (50), as well as other bacteria (51). Zoledronic acid is a synthetic imidazole bisphosphonate analogue of pyrophosphate with anti-bone-resorption activity. It binds to hydroxyapatite crystals in the bone matrix and also inhibits farnesyl pyrophosphate synthase, an enzyme involved in terpenoid biosynthesis (52). Zoledronic acid also inhibits the potassium ATP-sensitive channel (53). All three examples seemed to have the effects of being bactericidal as judged by flow cytometry parameters, as they were fast-acting and cell permeating. Equally relevant, all three examples represent chemical scaffolds that are different from known bactericidal antibiotics such as Colistin.

## 9. Author statements

### 9.1 Author contributions

Conceptualization, JES-S; methodology, JES-S; formal analysis, JES-S; resources, DBK.; writing—original draft preparation, JES-S; writing—review and editing, DBK; funding acquisition, DBK. Both authors have read and agreed to the published version of the manuscript.

### 9.4 Ethical approval

Non applicable

### 9.5 Consent for publication

Non applicable

## 9.6 Acknowledgements

We thank Miss Maria Lopez Rios and Mr Rhys Donafee for their assistance with the microtitration growth inhibitory assays.

## References

1. Kell DB. The Transporter-Mediated Cellular Uptake and Efflux of Pharmaceutical Drugs and Biotechnology Products: How and Why Phospholipid Bilayer Transport Is Negligible in Real Biomembranes. Molecules. 2021;26(18).

2. Kell DB. What would be the observable consequences if phospholipid bilayer diffusion of drugs into cells is negligible? Trends Pharmacol Sci. 2015;36(1):15–21.

3. Bush K. Antimicrobial agents targeting bacterial cell walls and cell membranes. Rev Sci Tech. 2012;31(1):43–56.

4. WHO. 2021 Antibacterial agents in clinical and preclinical development: an overview and analysis. Geneva: World Health Organization; 2021.

5. WHO/EMP/IAU/. Antibacterial Agents in Preclinical Development. Geneva: World Health Organization; 2019.

6. Theuretzbacher U, Outterson K, Engel A, Karlen A. The global preclinical antibacterial pipeline. Nat Rev Microbiol. 2020;18(5):275–85.

7. Steen HB. Flow cytometry of bacteria: glimpses from the past with a view to the future. J Microbiol Methods. 2000;42(1):65–74.

8. Davey HM, Kell DB. Flow cytometry and cell sorting of heterogeneous microbial populations: the importance of single-cell analyses. Microbiol Rev. 1996;60(4):641–96.

9. Lopez-Galvez J, Schiessl K, Besmer MD, Bruckmann C, Harms H, Muller S. Development of an Automated Online Flow Cytometry Method to Quantify Cell Density and Fingerprint Bacterial Communities. Cells. 2023;12(12).

10. Mao G, Wang Y, Hammes F. Automated flow cytometry as a flexible tool for comparing disinfection characteristics of indigenous bacterial communities and pure cultures. Ecotoxicol Environ Saf. 2021;225:112799.

11. Muller S, Davey H. Recent advances in the analysis of individual microbial cells. Cytometry A. 2009;75(2):83–5.

12. Van Nevel S, Koetzsch S, Proctor CR, Besmer MD, Prest EI, Vrouwenvelder JS, et al. Flow cytometric bacterial cell counts challenge conventional heterotrophic plate counts for routine microbiological drinking water monitoring. Water Res. 2017;113:191–206.

13. Jindal S, Yang L, Day PJ, Kell DB. Involvement of multiple influx and efflux transporters in the accumulation of cationic fluorescent dyes by Escherichia coli. BMC Microbiol. 2019;19(1):195.

14. Salcedo-Sora JE, Jindal S, O’Hagan S, Kell DB. A palette of fluorophores that are differentially accumulated by wild-type and mutant strains of Escherichia coli: surrogate ligands for profiling bacterial membrane transporters. Microbiology (Reading). 2021;167(2).

15. O’Hagan S, Kell DB. Structural Similarities between Some Common Fluorophores Used in Biology, Marketed Drugs, Endogenous Metabolites, and Natural Products. Mar Drugs. 2020;18(11).

16. Radi MS, Munro LJ, Salcedo-Sora JE, Kim SH, Feist AM, Kell DB. Understanding Functional Redundancy and Promiscuity of Multidrug Transporters in E. coli under Lipophilic Cation Stress. Membranes (Basel). 2022;12(12).

17. Radi MS, SalcedoSora JE, Kim SH, Sudarsan S, Sastry AV, Kell DB, et al. Membrane transporter identification and modulation via adaptive laboratory evolution. Metab Eng. 2022;72:376–90.

18. Davey HM, Jones A, Shaw AD, Kell DB. Variable selection and multivariate methods for the identification of microorganisms by flow cytometry. Cytometry. 1999;35(2):162–8.

19. Wurm M, Ilhan S, Jandt U, Zeng AP. Direct and highly sensitive measurement of fluorescent molecules in bulk solutions using flow cytometry. Anal Biochem. 2019;570:32–42.

20. Kell DB, Ferguson SJ, John P. Measurement by a flow dialysis technique of the steady-state proton-motive force in chromatophores from Rhodospirillum rubrum. Comparison with phosphorylation potential. Biochim Biophys Acta. 1978;502(1):111–26.

21. Diaper JP, Tither K, Edwards C. Rapid assessment of bacterial viability by flow cytometry. Appl Microbiol Biotechnol. 1992;38(2):268–72.

22. Lin TE. py50: Generate Dose-Response Curves (v1.0.4). Zenodo. 2024.

23. Lê S, Josse J, Husson F. FactoMineR: An R Package for Multivariate Analysis. Journal of Statistical Software. 2008;25(1):1–18.

24. WHO. Prioritization of pathogens to guide discovery, research and development of new antibiotics for drug resistant bacteria infections, including tuberculosis. Geneva: World Health Organization; 2017.

25. Biggs BW, Bedore SR, Arvay E, Huang S, Subramanian H, McIntyre EA, et al. Development of a genetic toolset for the highly engineerable and metabolically versatile Acinetobacter baylyi ADP1. Nucleic Acids Res. 2020;48(9):5169–82.

26. Jiang X, Palazzotto E, Wybraniec E, Munro LJ, Zhang H, Kell DB, et al. Automating Cloning by Natural Transformation. ACS Synth Biol. 2020;9(12):3228–35.

27. Young DM, Parke D, Ornston LN. Opportunities for genetic investigation afforded by Acinetobacter baylyi, a nutritionally versatile bacterial species that is highly competent for natural transformation. Annu Rev Microbiol. 2005;59:519–51.

28. Stokes JM, MacNair CR, Ilyas B, French S, Cote JP, Bouwman C, et al. Pentamidine sensitizes Gram-negative pathogens to antibiotics and overcomes acquired colistin resistance. Nat Microbiol. 2017;2:17028.

29. Jerala R. Synthetic lipopeptides: a novel class of anti-infectives. Expert Opin Investig Drugs. 2007;16(8):1159–69.

30. Lewis K. The Science of Antibiotic Discovery. Cell. 2020;181(1):29–45.

31. Jangra M, Travin DY, Aleksandrova EV, Kaur M, Darwish L, Koteva K, et al. A broad-spectrum lasso peptide antibiotic targeting the bacterial ribosome. Nature. 2025;640(8060):1022–30.

32. Stokes JM, Yang K, Swanson K, Jin W, Cubillos-Ruiz A, Donghia NM, et al. A Deep Learning Approach to Antibiotic Discovery. Cell. 2020;180(4):688–702 e13.

33. Sati H, Carrara E, Savoldi A, Hansen P, Garlasco J, Campagnaro E, et al. The WHO Bacterial Priority Pathogens List 2024: a prioritisation study to guide research, development, and public health strategies against antimicrobial resistance. Lancet Infect Dis. 2025.

34. Berglund NA, Piggot TJ, Jefferies D, Sessions RB, Bond PJ, Khalid S. Interaction of the antimicrobial peptide polymyxin B1 with both membranes of E. coli: a molecular dynamics study. PLoS Comput Biol. 2015;11(4):e1004180.

35. Deris ZZ, Swarbrick JD, Roberts KD, Azad MA, Akter J, Horne AS, et al. Probing the penetration of antimicrobial polymyxin lipopeptides into gram-negative bacteria. Bioconjug Chem. 2014;25(4):750–60.

36. Sabnis A, Hagart KL, Klockner A, Becce M, Evans LE, Furniss RCD, et al. Colistin kills bacteria by targeting lipopolysaccharide in the cytoplasmic membrane. Elife. 2021;10.

37. Buijs NP, Matheson EJ, Cochrane SA, Martin NI. Targeting membrane-bound bacterial cell wall precursors: a tried and true antibiotic strategy in nature and the clinic. Chem Commun (Camb). 2023;59(50):7685–703.

38. Bialvaei AZ, Samadi Kafil H. Colistin, mechanisms and prevalence of resistance. Curr Med Res Opin. 2015;31(4):707–21.

39. White TA, Kell DB. Comparative genomic assessment of novel broad-spectrum targets for antibacterial drugs. Comp Funct Genomics. 2004;5(4):304–27.

40. Piddock LJV. The 2019 Garrod Lecture: MDR efflux in Gram-negative bacteria-how understanding resistance led to a new tool for drug discovery. J Antimicrob Chemother. 2019;74(11):3128–34.

41. Kell DB. Control of metabolite efflux in microbial cell factories: current advances and future prospects. In: El-Mansi EMT, Nielsen, J., Mousdale, D., Allman, T.and Carlson, R., editor. Fermentation microbiology and biotechnology. Boca Raton: CRC Press; 2019. p. 117–38.

42. Mendes P, Girardi, E., Superti-Furga, G. and Kell, D. B. Why most transporter mutations that cause antibiotic resistance are to efflux pumps rather than to import transporters. bioRxiv. 2020;2020.2001.2016.909507v909501.

43. Salgado J, Rayner J, Ojkic N. Advancing antibiotic discovery with bacterial cytological profiling: a high-throughput solution to antimicrobial resistance. Front Microbiol. 2025;16:1536131.

44. Mermans F, De Baets H, Garcia-Timermans C, Teughels W, Boon N. Unlocking the mechanism of action: a cost-effective flow cytometry approach for accelerating antimicrobial drug development. Microbiol Spectr. 2024;12(4):e0393123.

45. Foletto VS, da Rosa TF, Serafin MB, Bottega A, Horner R. Repositioning of non-antibiotic drugs as an alternative to microbial resistance: a systematic review. Int J Antimicrob Agents. 2021;58(3):106380.

46. Huang Y, Zhu Y, Yue HY, Liu YY, Deng LM, Lv L, et al. Flavomycin restores colistin susceptibility in multidrug-resistant Gram-negative bacteria. mSystems. 2024;9(6):e0010924.

47. Zgurskaya HI, Lopez CA, Gnanakaran S. Permeability Barrier of Gram-Negative Cell Envelopes and Approaches To Bypass It. ACS Infect Dis. 2015;1(11):512–22.

48. Zafar Gondal A, Zulfiqar H. Aminophylline. StatPearls. Treasure Island (FL) 2024.

49. Parsons ME. Pantoprazole, a new proton-pump inhibitor, has a precise and predictable profile of activity. Eur J Gastroenterol Hepatol. 1996;8 Suppl 1:S15–20.

50. Nakao M, Malfertheiner P. Growth inhibitory and bactericidal activities of lansoprazole compared with those of omeprazole and pantoprazole against Helicobacter pylori. Helicobacter. 1998;3(1):21–7.

51. Ni WT, Cai XJ, Liang BB, Cai Y, Cui JC, Wang R. Effect of Proton Pump Inhibitors on Activity of Tigecycline against Several Common Clinical Pathogens. Plos One. 2014;9(1).

52. Green JR. Bisphosphonates: preclinical review. Oncologist. 2004;9 Suppl 4:3–13.

53. Scala R, Maqoud F, McClenaghan C, Harter TM, Perrone MG, Scilimati A, et al. Zoledronic Acid Blocks Overactive Kir6.1/SUR2-Dependent K(ATP) Channels in Skeletal Muscle and Osteoblasts in a Murine Model of Cantu Syndrome. Cells. 2023;12(6).

