## Supplementary table 1 for "Rapid accumulation of fluorophores and fast kill identify drugs with bactericidal effects against Gram-negative bacteria"

| Pre_incuba | Washes | Species | Antibiotic | Target | Code | Fluor | Events | Channel | BL1 |
|---|---|---|---|---|---|---|---|---|---|
| 40 | Yes | E. coli | None | None | NA | Autofluores | 18391 | NA | 764 |
| 40 | Yes | E. coli | None | None | F194 | Oxonol | 22395 | RL1 | 1309 |
| 40 | Yes | E. coli | None | None | F11 | Allura Red | 22329 | BL3 | 1749 |
| 40 | Yes | E. coli | None | None | F54 | DiSC3(5) | 26549 | RL1 | 1645 |
| 40 | Yes | E. coli | None | None | F124 | Thioflavin T | 26079 | VL2 | 1610 |
| 40 | Yes | E. coli | None | None | F101 | Quinacrine | 28108 | VL2 | 1725 |
| 40 | Yes | E. coli | None | None | F138 | Apigenin | 25169 | VL2 | 1814 |
| 40 | Yes | E. coli | None | None | F65 | H2FDA | 23974 | BL1 | 2814.5 |
| 40 | Yes | E. coli | None | None | F142 | Chlorophe | 23223 | BL3 | 3719 |
| 40 | Yes | E. coli | None | None | F39 | Congo Red | 20661 | BL3 | 3968 |
| 40 | Yes | E. coli | None | None | F116 | Sunset Yel | 24346 | BL2 | 4434 |
| 40 | Yes | E. coli | None | None | F10 | Alizarin | 22385 | BL1 | 3437 |
| 40 | Yes | E. coli | None | None | F84 | Neutral Re | 24029 | BL1 | 3055 |
| 40 | Yes | E. coli | None | None | F99 | Pyronin Y | 20228 | BL1 | 2513 |
| 40 | Yes | E. coli | None | None | F108 | Rhodamine | 26959 | BL1 | 2011 |
| 40 | Yes | E. coli | None | None | F109 | Riboflavin | 22236 | BL1 | 1865 |
| 40 | Yes | E. coli | None | None | F117 | SYBR Gree | 21591 | BL1 | 89582 |
| 40 | Yes | E. coli | None | None | F106 | Rhodamine | 26239 | BL1 | 3645 |
| 40 | Yes | E. coli | None | None | F73 | Malachite G | 22126 | BL1 | 3372 |
| 40 | Yes | E. coli | None | None | F7 | Acridine or | 14738 | BL1 | 4336 |
| 40 | Yes | E. coli | None | None | F192 | 5-Carboxyf | 21848 | BL1 | 407211.5 |
| 40 | Yes | E. coli | None | None | F57 | Eosin Y | 11306 | BL1 | 744235.5 |
| 40 | Yes | E. coli | None | None | F27 | Calcein | 24186 | BL1 | 30738.5 |
| 40 | Yes | E. coli | None | None | F12 | Amaranth | 18846 | BL1 | 29931.5 |
| 40 | Yes | E. coli | None | None | NA | Autofluores | 19461 | NA | 6163 |
| 40 | Yes | E. coli | Amoxacilli | Wall | F194 | Oxonol | 24115 | RL1 | 3680 |
| 40 | Yes | E. coli | Amoxacilli | Wall | F11 | Allura Red | 23627 | BL3 | 3500 |
| 40 | Yes | E. coli | Amoxacilli | Wall | F54 | DiSC3(5) | 14299 | RL1 | 4685 |
| 40 | Yes | E. coli | Amoxacilli | Wall | F124 | Thioflavin T | 23247 | VL2 | 3038 |
| 40 | Yes | E. coli | Amoxacilli | Wall | F101 | Quinacrine | 18528 | VL2 | 2516 |
| 40 | Yes | E. coli | Amoxacilli | Wall | F138 | Apigenin | 26866 | VL2 | 2644.5 |
| 40 | Yes | E. coli | Amoxacilli | Wall | F65 | H2FDA | 26458 | BL1 | 2886 |
| 40 | Yes | E. coli | Amoxacilli | Wall | F142 | Chlorophe | 27103 | BL3 | 4062 |
| 40 | Yes | E. coli | Amoxacilli | Wall | F39 | Congo Red | 25510 | BL3 | 5209 |
| 40 | Yes | E. coli | Amoxacilli | Wall | F116 | Sunset Yel | 21197 | BL2 | 3750 |
| 40 | Yes | E. coli | Amoxacilli | Wall | F10 | Alizarin | 26505 | BL1 | 3425 |
| 40 | Yes | E. coli | Amoxacilli | Wall | F84 | Neutral Re | 19651 | BL1 | 3409 |
| 40 | Yes | E. coli | Amoxacilli | Wall | F99 | Pyronin Y | 23483 | BL1 | 3051 |
| 40 | Yes | E. coli | Amoxacilli | Wall | F108 | Rhodamine | 23502 | BL1 | 2744 |
| 40 | Yes | E. coli | Amoxacilli | Wall | F109 | Riboflavin | 20388 | BL1 | 2967.5 |
| 40 | Yes | E. coli | Amoxacilli | Wall | F117 | SYBR Gree | 20182 | BL1 | 103846.5 |
| 40 | Yes | E. coli | Amoxacilli | Wall | F106 | Rhodamine | 23025 | BL1 | 7832 |
| 40 | Yes | E. coli | Amoxacilli | Wall | F73 | Malachite G | 19148 | BL1 | 5440 |
| 40 | Yes | E. coli | Amoxacilli | Wall | F7 | Acridine or | 16392 | BL1 | 6102 |
| 40 | Yes | E. coli | Amoxacilli | Wall | F192 | 5-Carboxyf | 16759 | BL1 | 15970 |
| 40 | Yes | E. coli | Amoxacilli | Wall | F57 | Eosin Y | 22973 | BL1 | 34432 |
| 40 | Yes | E. coli | Amoxacilli | Wall | F27 | Calcein | 23074 | BL1 | 21698.5 |

| | | | | | | | | | |
|---|---|---|---|---|---|---|---|---|---|
| 40 | Yes | E. coli | Amoxacilli | Wall | F12 | Amaranth | 22945 | BL1 | 63164 |
| 40 | Yes | E. coli | None | None | NA | Autofluore: | 21296 | NA | 5446 |
| 40 | Yes | E. coli | Azithromyc | Intracellul | F194 | Oxonol | 14696 | RL1 | 4654.5 |
| 40 | Yes | E. coli | Azithromyc | Intracellul | F11 | Allura Red | 12616 | BL3 | 4471 |
| 40 | Yes | E. coli | Azithromyc | Intracellul | F54 | DiSC3(5) | 11963 | RL1 | 3965 |
| 40 | Yes | E. coli | Azithromyc | Intracellul | F124 | Thioflavin T | 12538 | VL2 | 3653.5 |
| 40 | Yes | E. coli | Azithromyc | Intracellul | F101 | Quinacrine | 12884 | VL2 | 3634.5 |
| 40 | Yes | E. coli | Azithromyc | Intracellul | F138 | Apigenin | 11968 | VL2 | 3821.5 |
| 40 | Yes | E. coli | Azithromyc | Intracellul | F65 | H2FDA | 12962 | BL1 | 4813 |
| 40 | Yes | E. coli | Azithromyc | Intracellul | F142 | Chlorophe | 12558 | BL3 | 5831 |
| 40 | Yes | E. coli | Azithromyc | Intracellul | F39 | Congo Red | 12077 | BL3 | 6857 |
| 40 | Yes | E. coli | Azithromyc | Intracellul | F116 | Sunset Yel | 12571 | BL2 | 6069 |
| 40 | Yes | E. coli | Azithromyc | Intracellul | F10 | Alizarin | 11737 | BL1 | 5223 |
| 40 | Yes | E. coli | Azithromyc | Intracellul | F84 | Neutral Re | 11581 | BL1 | 4515 |
| 40 | Yes | E. coli | Azithromyc | Intracellul | F99 | Pyronin Y | 11632 | BL1 | 3806 |
| 40 | Yes | E. coli | Azithromyc | Intracellul | F108 | Rhodamine | 11735 | BL1 | 3671 |
| 40 | Yes | E. coli | Azithromyc | Intracellul | F109 | Riboflavin | 11664 | BL1 | 3965 |
| 40 | Yes | E. coli | Azithromyc | Intracellul | F117 | SYBR Gree | 11794 | BL1 | 81278 |
| 40 | Yes | E. coli | Azithromyc | Intracellul | F106 | Rhodamine | 11724 | BL1 | 5015.5 |
| 40 | Yes | E. coli | Azithromyc | Intracellul | F73 | Malachite ( | 12195 | BL1 | 4793 |
| 40 | Yes | E. coli | Azithromyc | Intracellul | F7 | Acridine or | 10975 | BL1 | 4454 |
| 40 | Yes | E. coli | Azithromyc | Intracellul | F192 | 5-Carboxyf | 11060 | BL1 | 16340.5 |
| 40 | Yes | E. coli | Azithromyc | Intracellul | F57 | Eosin Y | 9746 | BL1 | 22268.5 |
| 40 | Yes | E. coli | Azithromyc | Intracellul | F27 | Calcein | 13175 | BL1 | 31402 |
| 40 | Yes | E. coli | Azithromyc | Intracellul | F12 | Amaranth | 10733 | BL1 | 28894 |
| 40 | Yes | E. coli | None | None | NA | Autofluore: | 19832 | NA | 6743 |
| 40 | Yes | E. coli | Trimethopr | Intracellul | F194 | Oxonol | 18224 | RL1 | 4539 |
| 40 | Yes | E. coli | Trimethopr | Intracellul | F11 | Allura Red | 18011 | BL3 | 4081 |
| 40 | Yes | E. coli | Trimethopr | Intracellul | F54 | DiSC3(5) | 12656 | RL1 | 4407.5 |
| 40 | Yes | E. coli | Trimethopr | Intracellul | F124 | Thioflavin T | 17543 | VL2 | 3519 |
| 40 | Yes | E. coli | Trimethopr | Intracellul | F101 | Quinacrine | 15879 | VL2 | 3395 |
| 40 | Yes | E. coli | Trimethopr | Intracellul | F138 | Apigenin | 18063 | VL2 | 3263 |
| 40 | Yes | E. coli | Trimethopr | Intracellul | F65 | H2FDA | 15753 | BL1 | 4323 |
| 40 | Yes | E. coli | Trimethopr | Intracellul | F142 | Chlorophe | 16826 | BL3 | 5428 |
| 40 | Yes | E. coli | Trimethopr | Intracellul | F39 | Congo Red | 17208 | BL3 | 5759 |
| 40 | Yes | E. coli | Trimethopr | Intracellul | F116 | Sunset Yel | 17768 | BL2 | 5516 |
| 40 | Yes | E. coli | Trimethopr | Intracellul | F10 | Alizarin | 15887 | BL1 | 5252 |
| 40 | Yes | E. coli | Trimethopr | Intracellul | F84 | Neutral Re | 14141 | BL1 | 4678 |
| 40 | Yes | E. coli | Trimethopr | Intracellul | F99 | Pyronin Y | 17958 | BL1 | 3609.5 |
| 40 | Yes | E. coli | Trimethopr | Intracellul | F108 | Rhodamine | 16979 | BL1 | 3307 |
| 40 | Yes | E. coli | Trimethopr | Intracellul | F109 | Riboflavin | 15951 | BL1 | 4064 |
| 40 | Yes | E. coli | Trimethopr | Intracellul | F117 | SYBR Gree | 15587 | BL1 | 83947 |
| 40 | Yes | E. coli | Trimethopr | Intracellul | F106 | Rhodamine | 16875 | BL1 | 5343 |
| 40 | Yes | E. coli | Trimethopr | Intracellul | F73 | Malachite ( | 16883 | BL1 | 4104 |
| 40 | Yes | E. coli | Trimethopr | Intracellul | F7 | Acridine or | 15579 | BL1 | 4684 |
| 40 | Yes | E. coli | Trimethopr | Intracellul | F192 | 5-Carboxyf | 13941 | BL1 | 21718 |
| 40 | Yes | E. coli | Trimethopr | Intracellul | F57 | Eosin Y | 15043 | BL1 | 26183 |
| 40 | Yes | E. coli | Trimethopr | Intracellul | F27 | Calcein | 17985 | BL1 | 18529 |

| | | | | | | | | | |
|---|---|---|---|---|---|---|---|---|---|
| 40 | Yes | E. coli | Trimethopr | Intracellula | F12 | Amaranth | 16711 | BL1 | 19925 |
| 40 | Yes | E. coli | None | None | NA | Autofluores | 23825 | NA | 5000 |
| 40 | Yes | E. coli | Chloramph | Intracellula | F194 | Oxonol | 15897 | RL1 | 4367 |
| 40 | Yes | E. coli | Chloramph | Intracellula | F11 | Allura Red | 14503 | BL3 | 4143 |
| 40 | Yes | E. coli | Chloramph | Intracellula | F54 | DiSC3(5) | 12200 | RL1 | 3889 |
| 40 | Yes | E. coli | Chloramph | Intracellula | F124 | Thioflavin T | 13944 | VL2 | 3759.5 |
| 40 | Yes | E. coli | Chloramph | Intracellula | F101 | Quinacrine | 12967 | VL2 | 3751 |
| 40 | Yes | E. coli | Chloramph | Intracellula | F138 | Apigenin | 12892 | VL2 | 3901 |
| 40 | Yes | E. coli | Chloramph | Intracellula | F65 | H2FDA | 13514 | BL1 | 4537 |
| 40 | Yes | E. coli | Chloramph | Intracellula | F142 | Chlorophe | 13372 | BL3 | 6487.5 |
| 40 | Yes | E. coli | Chloramph | Intracellula | F39 | Congo Red | 11940 | BL3 | 6885.5 |
| 40 | Yes | E. coli | Chloramph | Intracellula | F116 | Sunset Yel | 13027 | BL2 | 6728 |
| 40 | Yes | E. coli | Chloramph | Intracellula | F10 | Alizarin | 12427 | BL1 | 4943 |
| 40 | Yes | E. coli | Chloramph | Intracellula | F84 | Neutral Re | 14931 | BL1 | 4050 |
| 40 | Yes | E. coli | Chloramph | Intracellula | F99 | Pyronin Y | 12954 | BL1 | 4426.5 |
| 40 | Yes | E. coli | Chloramph | Intracellula | F108 | Rhodamine | 11195 | BL1 | 4187 |
| 40 | Yes | E. coli | Chloramph | Intracellula | F109 | Riboflavin | 12447 | BL1 | 3616 |
| 40 | Yes | E. coli | Chloramph | Intracellula | F117 | SYBR Gree | 12777 | BL1 | 59956 |
| 40 | Yes | E. coli | Chloramph | Intracellula | F106 | Rhodamine | 13267 | BL1 | 4843 |
| 40 | Yes | E. coli | Chloramph | Intracellula | F73 | Malachite ( | 13745 | BL1 | 4000 |
| 40 | Yes | E. coli | Chloramph | Intracellula | F7 | Acridine or | 14172 | BL1 | 3847 |
| 40 | Yes | E. coli | Chloramph | Intracellula | F192 | 5-Carboxyf | 13373 | BL1 | 14514 |
| 40 | Yes | E. coli | Chloramph | Intracellula | F57 | Eosin Y | 11500 | BL1 | 19587.5 |
| 40 | Yes | E. coli | Chloramph | Intracellula | F27 | Calcein | 12971 | BL1 | 23987 |
| 40 | Yes | E. coli | Chloramph | Intracellula | F12 | Amaranth | 11846 | BL1 | 22995 |
| 40 | Yes | E. coli | None | None | NA | Autofluores | 21821 | NA | 5116 |
| 40 | Yes | E. coli | Colistin | Wall | F194 | Oxonol | 10040 | RL1 | 58318 |
| 40 | Yes | E. coli | Colistin | Wall | F11 | Allura Red | 7170 | BL3 | 65542 |
| 40 | Yes | E. coli | Colistin | Wall | F54 | DiSC3(5) | 5572 | RL1 | 57295 |
| 40 | Yes | E. coli | Colistin | Wall | F124 | Thioflavin T | 5529 | VL2 | 48354 |
| 40 | Yes | E. coli | Colistin | Wall | F101 | Quinacrine | 5069 | VL2 | 59710 |
| 40 | Yes | E. coli | Colistin | Wall | F138 | Apigenin | 4846 | VL2 | 47947.5 |
| 40 | Yes | E. coli | Colistin | Wall | F65 | H2FDA | 4608 | BL1 | 46939.5 |
| 40 | Yes | E. coli | Colistin | Wall | F142 | Chlorophe | 4945 | BL3 | 55459 |
| 40 | Yes | E. coli | Colistin | Wall | F39 | Congo Red | 4595 | BL3 | 36160 |
| 40 | Yes | E. coli | Colistin | Wall | F116 | Sunset Yel | 4487 | BL2 | 48655 |
| 40 | Yes | E. coli | Colistin | Wall | F10 | Alizarin | 4998 | BL1 | 42417 |
| 40 | Yes | E. coli | Colistin | Wall | F84 | Neutral Re | 4917 | BL1 | 40574 |
| 40 | Yes | E. coli | Colistin | Wall | F99 | Pyronin Y | 4475 | BL1 | 38679 |
| 40 | Yes | E. coli | Colistin | Wall | F108 | Rhodamine | 4472 | BL1 | 40059 |
| 40 | Yes | E. coli | Colistin | Wall | F109 | Riboflavin | 4632 | BL1 | 36714.5 |
| 40 | Yes | E. coli | Colistin | Wall | F117 | SYBR Gree | 5140 | BL1 | 3151302 |
| 40 | Yes | E. coli | Colistin | Wall | F106 | Rhodamine | 4143 | BL1 | 138937 |
| 40 | Yes | E. coli | Colistin | Wall | F73 | Malachite ( | 4547 | BL1 | 112095 |
| 40 | Yes | E. coli | Colistin | Wall | F7 | Acridine or | 4313 | BL1 | 193381 |
| 40 | Yes | E. coli | Colistin | Wall | F192 | 5-Carboxyf | 3952 | BL1 | 128968 |
| 40 | Yes | E. coli | Colistin | Wall | F57 | Eosin Y | 4369 | BL1 | 94103 |
| 40 | Yes | E. coli | Colistin | Wall | F27 | Calcein | 4220 | BL1 | 83219.5 |

| | | | | | | | | | |
|---|---|---|---|---|---|---|---|---|---|
| 40 | Yes | E. coli | Colistin | Wall | F12 | Amaranth | 4157 | BL1 | 74572 |
| 40 | Yes | E. coli | None | None | NA | Autofluores | 24763 | NA | 4860 |
| 40 | Yes | E. coli | Gentamici | Intracellula | F194 | Oxonol | 25581 | RL1 | 4133 |
| 40 | Yes | E. coli | Gentamici | Intracellula | F11 | Allura Red | 23359 | BL3 | 3924 |
| 40 | Yes | E. coli | Gentamici | Intracellula | F54 | DiSC3(5) | 26221 | RL1 | 3659 |
| 40 | Yes | E. coli | Gentamici | Intracellula | F124 | Thioflavin T | 22691 | VL2 | 3897 |
| 40 | Yes | E. coli | Gentamici | Intracellula | F101 | Quinacrine | 22865 | VL2 | 3888 |
| 40 | Yes | E. coli | Gentamici | Intracellula | F138 | Apigenin | 20459 | VL2 | 4797 |
| 40 | Yes | E. coli | Gentamici | Intracellula | F65 | H2FDA | 22961 | BL1 | 4432 |
| 40 | Yes | E. coli | Gentamici | Intracellula | F142 | Chlorophe | 20506 | BL3 | 6629.5 |
| 40 | Yes | E. coli | Gentamici | Intracellula | F39 | Congo Red | 22260 | BL3 | 5417 |
| 40 | Yes | E. coli | Gentamici | Intracellula | F116 | Sunset Yel | 23667 | BL2 | 5176 |
| 40 | Yes | E. coli | Gentamici | Intracellula | F10 | Alizarin | 20004 | BL1 | 5534 |
| 40 | Yes | E. coli | Gentamici | Intracellula | F84 | Neutral Re | 19715 | BL1 | 6962 |
| 40 | Yes | E. coli | Gentamici | Intracellula | F99 | Pyronin Y | 20243 | BL1 | 4155 |
| 40 | Yes | E. coli | Gentamici | Intracellula | F108 | Rhodamine | 21804 | BL1 | 3808 |
| 40 | Yes | E. coli | Gentamici | Intracellula | F109 | Riboflavin | 17721 | BL1 | 4936 |
| 40 | Yes | E. coli | Gentamici | Intracellula | F117 | SYBR Gree | 22211 | BL1 | 169250 |
| 40 | Yes | E. coli | Gentamici | Intracellula | F106 | Rhodamine | 16099 | BL1 | 6263 |
| 40 | Yes | E. coli | Gentamici | Intracellula | F73 | Malachite ( | 20637 | BL1 | 4301 |
| 40 | Yes | E. coli | Gentamici | Intracellula | F7 | Acridine or | 12067 | BL1 | 6048 |
| 40 | Yes | E. coli | Gentamici | Intracellula | F192 | 5-Carboxyf | 18482 | BL1 | 18494.5 |
| 40 | Yes | E. coli | Gentamici | Intracellula | F57 | Eosin Y | 17625 | BL1 | 19430 |
| 40 | Yes | E. coli | Gentamici | Intracellula | F27 | Calcein | 20053 | BL1 | 30615 |
| 40 | Yes | E. coli | Gentamici | Intracellula | F12 | Amaranth | 20517 | BL1 | 33702 |
| 40 | Yes | E. coli | None | None | NA | Autofluores | 22392 | NA | 5206 |
| 40 | Yes | E. coli | Nitrofurant | Intracellula | F194 | Oxonol | 14417 | RL1 | 4780 |
| 40 | Yes | E. coli | Nitrofurant | Intracellula | F11 | Allura Red | 13265 | BL3 | 4442 |
| 40 | Yes | E. coli | Nitrofurant | Intracellula | F54 | DiSC3(5) | 11673 | RL1 | 4618 |
| 40 | Yes | E. coli | Nitrofurant | Intracellula | F124 | Thioflavin T | 10609 | VL2 | 4702 |
| 40 | Yes | E. coli | Nitrofurant | Intracellula | F101 | Quinacrine | 10794 | VL2 | 4913 |
| 40 | Yes | E. coli | Nitrofurant | Intracellula | F138 | Apigenin | 10470 | VL2 | 4580 |
| 40 | Yes | E. coli | Nitrofurant | Intracellula | F65 | H2FDA | 10792 | BL1 | 5672 |
| 40 | Yes | E. coli | Nitrofurant | Intracellula | F142 | Chlorophe | 10849 | BL3 | 6083 |
| 40 | Yes | E. coli | Nitrofurant | Intracellula | F39 | Congo Red | 11456 | BL3 | 5895 |
| 40 | Yes | E. coli | Nitrofurant | Intracellula | F116 | Sunset Yel | 11291 | BL2 | 5732 |
| 40 | Yes | E. coli | Nitrofurant | Intracellula | F10 | Alizarin | 10931 | BL1 | 4801 |
| 40 | Yes | E. coli | Nitrofurant | Intracellula | F84 | Neutral Re | 10796 | BL1 | 4458 |
| 40 | Yes | E. coli | Nitrofurant | Intracellula | F99 | Pyronin Y | 12477 | BL1 | 4184 |
| 40 | Yes | E. coli | Nitrofurant | Intracellula | F108 | Rhodamine | 11066 | BL1 | 3831 |
| 40 | Yes | E. coli | Nitrofurant | Intracellula | F109 | Riboflavin | 9907 | BL1 | 4379 |
| 40 | Yes | E. coli | Nitrofurant | Intracellula | F117 | SYBR Gree | 10933 | BL1 | 101067 |
| 40 | Yes | E. coli | Nitrofurant | Intracellula | F106 | Rhodamine | 11061 | BL1 | 5480 |
| 40 | Yes | E. coli | Nitrofurant | Intracellula | F73 | Malachite ( | 10833 | BL1 | 4670 |
| 40 | Yes | E. coli | Nitrofurant | Intracellula | F7 | Acridine or | 10850 | BL1 | 4348 |
| 40 | Yes | E. coli | Nitrofurant | Intracellula | F192 | 5-Carboxyf | 11084 | BL1 | 17800.5 |
| 40 | Yes | E. coli | Nitrofurant | Intracellula | F57 | Eosin Y | 9265 | BL1 | 21241 |
| 40 | Yes | E. coli | Nitrofurant | Intracellula | F27 | Calcein | 7992 | BL1 | 29890 |

| | | | | | | | | | |
|---|---|---|---|---|---|---|---|---|---|
| 40 | Yes | E. coli | Nitrofurant | Intracellula | F12 | Amaranth | 9762 | BL1 | 34603 |
| 40 | Yes | E. coli | None | None | NA | Autofluore: | 23026 | NA | 5236 |
| 40 | Yes | E. coli | Pentamidi | Wall | F194 | Oxonol | 15700 | RL1 | 5636 |
| 40 | Yes | E. coli | Pentamidi | Wall | F11 | Allura Red | 14680 | BL3 | 5007 |
| 40 | Yes | E. coli | Pentamidi | Wall | F54 | DiSC3(5) | 12072 | RL1 | 5451 |
| 40 | Yes | E. coli | Pentamidi | Wall | F124 | Thioflavin T | 12103 | VL2 | 6139 |
| 40 | Yes | E. coli | Pentamidi | Wall | F101 | Quinacrine | 11367 | VL2 | 6848 |
| 40 | Yes | E. coli | Pentamidi | Wall | F138 | Apigenin | 11372 | VL2 | 5710 |
| 40 | Yes | E. coli | Pentamidi | Wall | F65 | H2FDA | 12371 | BL1 | 5489 |
| 40 | Yes | E. coli | Pentamidi | Wall | F142 | Chlorophe | 13558 | BL3 | 6783.5 |
| 40 | Yes | E. coli | Pentamidi | Wall | F39 | Congo Red | 12238 | BL3 | 7289 |
| 40 | Yes | E. coli | Pentamidi | Wall | F116 | Sunset Yel | 13349 | BL2 | 6621 |
| 40 | Yes | E. coli | Pentamidi | Wall | F10 | Alizarin | 12258 | BL1 | 6360 |
| 40 | Yes | E. coli | Pentamidi | Wall | F84 | Neutral Re | 12422 | BL1 | 5525 |
| 40 | Yes | E. coli | Pentamidi | Wall | F99 | Pyronin Y | 13955 | BL1 | 4979 |
| 40 | Yes | E. coli | Pentamidi | Wall | F108 | Rhodamine | 12439 | BL1 | 5490 |
| 40 | Yes | E. coli | Pentamidi | Wall | F109 | Riboflavin | 12137 | BL1 | 6019 |
| 40 | Yes | E. coli | Pentamidi | Wall | F117 | SYBR Gree | 12161 | BL1 | 744993 |
| 40 | Yes | E. coli | Pentamidi | Wall | F106 | Rhodamine | 12271 | BL1 | 7573 |
| 40 | Yes | E. coli | Pentamidi | Wall | F73 | Malachite ( | 12500 | BL1 | 7658 |
| 40 | Yes | E. coli | Pentamidi | Wall | F7 | Acridine or | 11178 | BL1 | 9115.5 |
| 40 | Yes | E. coli | Pentamidi | Wall | F192 | 5-Carboxyf | 11441 | BL1 | 16749 |
| 40 | Yes | E. coli | Pentamidi | Wall | F57 | Eosin Y | 10845 | BL1 | 22584 |
| 40 | Yes | E. coli | Pentamidi | Wall | F27 | Calcein | 11622 | BL1 | 21113.5 |
| 40 | Yes | E. coli | Pentamidi | Wall | F12 | Amaranth | 11611 | BL1 | 19532 |
| 40 | Yes | E. coli | None | None | NA | Autofluore: | 23390 | NA | 7530.5 |
| 40 | Yes | E. coli | Rifampicin | Intracellula | F194 | Oxonol | 27575 | RL1 | 5232 |
| 40 | Yes | E. coli | Rifampicin | Intracellula | F11 | Allura Red | 26549 | BL3 | 4703 |
| 40 | Yes | E. coli | Rifampicin | Intracellula | F54 | DiSC3(5) | 24914 | RL1 | 4360 |
| 40 | Yes | E. coli | Rifampicin | Intracellula | F124 | Thioflavin T | 22962 | VL2 | 4811.5 |
| 40 | Yes | E. coli | Rifampicin | Intracellula | F101 | Quinacrine | 20519 | VL2 | 5035 |
| 40 | Yes | E. coli | Rifampicin | Intracellula | F138 | Apigenin | 22972 | VL2 | 4276 |
| 40 | Yes | E. coli | Rifampicin | Intracellula | F65 | H2FDA | 22980 | BL1 | 4667 |
| 40 | Yes | E. coli | Rifampicin | Intracellula | F142 | Chlorophe | 23767 | BL3 | 5253 |
| 40 | Yes | E. coli | Rifampicin | Intracellula | F39 | Congo Red | 25362 | BL3 | 5135.5 |
| 40 | Yes | E. coli | Rifampicin | Intracellula | F116 | Sunset Yel | 24242 | BL2 | 4401 |
| 40 | Yes | E. coli | Rifampicin | Intracellula | F10 | Alizarin | 23636 | BL1 | 4688.5 |
| 40 | Yes | E. coli | Rifampicin | Intracellula | F84 | Neutral Re | 22192 | BL1 | 4138.5 |
| 40 | Yes | E. coli | Rifampicin | Intracellula | F99 | Pyronin Y | 27071 | BL1 | 3254 |
| 40 | Yes | E. coli | Rifampicin | Intracellula | F108 | Rhodamine | 22925 | BL1 | 3650 |
| 40 | Yes | E. coli | Rifampicin | Intracellula | F109 | Riboflavin | 25950 | BL1 | 3464.5 |
| 40 | Yes | E. coli | Rifampicin | Intracellula | F117 | SYBR Gree | 20222 | BL1 | 114866.5 |
| 40 | Yes | E. coli | Rifampicin | Intracellula | F106 | Rhodamine | 21213 | BL1 | 6901 |
| 40 | Yes | E. coli | Rifampicin | Intracellula | F73 | Malachite ( | 20891 | BL1 | 4821 |
| 40 | Yes | E. coli | Rifampicin | Intracellula | F7 | Acridine or | 20396 | BL1 | 4588 |
| 40 | Yes | E. coli | Rifampicin | Intracellula | F192 | 5-Carboxyf | 23840 | BL1 | 15202 |
| 40 | Yes | E. coli | Rifampicin | Intracellula | F57 | Eosin Y | 19112 | BL1 | 23605.5 |
| 40 | Yes | E. coli | Rifampicin | Intracellula | F27 | Calcein | 23955 | BL1 | 281440 |

| | | | | | | | | | |
|---|---|---|---|---|---|---|---|---|---|
| 40 | Yes | E. coli | Rifampicin | Intracellula | F12 | Amaranth | 19240 | BL1 | 468853.5 |
| 40 | Yes | E. coli | None | None | NA | Autofluore: | 23571 | NA | 6044 |
| 40 | Yes | E. coli | Trimethopr | Intracellula | F194 | Oxonol | 19168 | RL1 | 4807 |
| 40 | Yes | E. coli | Trimethopr | Intracellula | F11 | Allura Red | 20434 | BL3 | 4576 |
| 40 | Yes | E. coli | Trimethopr | Intracellula | F54 | DiSC3(5) | 17846 | RL1 | 4903 |
| 40 | Yes | E. coli | Trimethopr | Intracellula | F124 | Thioflavin T | 18043 | VL2 | 4420 |
| 40 | Yes | E. coli | Trimethopr | Intracellula | F101 | Quinacrine | 18314 | VL2 | 4022.5 |
| 40 | Yes | E. coli | Trimethopr | Intracellula | F138 | Apigenin | 18160 | VL2 | 3695.5 |
| 40 | Yes | E. coli | Trimethopr | Intracellula | F65 | H2FDA | 18270 | BL1 | 4897 |
| 40 | Yes | E. coli | Trimethopr | Intracellula | F142 | Chlorophe | 18311 | BL3 | 5625 |
| 40 | Yes | E. coli | Trimethopr | Intracellula | F39 | Congo Red | 18826 | BL3 | 5213 |
| 40 | Yes | E. coli | Trimethopr | Intracellula | F116 | Sunset Yel | 14198 | BL2 | 6387 |
| 40 | Yes | E. coli | Trimethopr | Intracellula | F10 | Alizarin | 15651 | BL1 | 4479 |
| 40 | Yes | E. coli | Trimethopr | Intracellula | F84 | Neutral Re | 17517 | BL1 | 4408 |
| 40 | Yes | E. coli | Trimethopr | Intracellula | F99 | Pyronin Y | 18018 | BL1 | 3952 |
| 40 | Yes | E. coli | Trimethopr | Intracellula | F108 | Rhodamine | 18189 | BL1 | 4455 |
| 40 | Yes | E. coli | Trimethopr | Intracellula | F109 | Riboflavin | 16218 | BL1 | 3934 |
| 40 | Yes | E. coli | Trimethopr | Intracellula | F117 | SYBR Gree | 17427 | BL1 | 78685 |
| 40 | Yes | E. coli | Trimethopr | Intracellula | F106 | Rhodamine | 17134 | BL1 | 6794 |
| 40 | Yes | E. coli | Trimethopr | Intracellula | F73 | Malachite ( | 19184 | BL1 | 4302 |
| 40 | Yes | E. coli | Trimethopr | Intracellula | F7 | Acridine or | 16178 | BL1 | 3921.5 |
| 40 | Yes | E. coli | Trimethopr | Intracellula | F192 | 5-Carboxyf | 20177 | BL1 | 13032 |
| 40 | Yes | E. coli | Trimethopr | Intracellula | F57 | Eosin Y | 15805 | BL1 | 491390 |
| 40 | Yes | E. coli | Trimethopr | Intracellula | F27 | Calcein | 18860 | BL1 | 20724.5 |
| 40 | Yes | E. coli | Trimethopr | Intracellula | F12 | Amaranth | 14492 | BL1 | 21572.5 |
| 40 | Yes | E. coli | None | None | NA | Autofluore: | 21899 | NA | 775 |
| 40 | Yes | E. coli | None | None | F194 | Oxonol | 21150 | RL1 | 1123 |
| 40 | Yes | E. coli | None | None | F11 | Allura Red | 25313 | BL3 | 1453 |
| 40 | Yes | E. coli | None | None | F54 | DiSC3(5) | 25313 | RL1 | 1800 |
| 40 | Yes | E. coli | None | None | F124 | Thioflavin T | 25369 | VL2 | 1895 |
| 40 | Yes | E. coli | None | None | F101 | Quinacrine | 24184 | VL2 | 2098 |
| 40 | Yes | E. coli | None | None | F138 | Apigenin | 24035 | VL2 | 2147 |
| 40 | Yes | E. coli | None | None | F65 | H2FDA | 23086 | BL1 | 3018 |
| 40 | Yes | E. coli | None | None | F142 | Chlorophe | 26034 | BL3 | 3688 |
| 40 | Yes | E. coli | None | None | F39 | Congo Red | 24953 | BL3 | 3817 |
| 40 | Yes | E. coli | None | None | F116 | Sunset Yel | 27947 | BL2 | 3549 |
| 40 | Yes | E. coli | None | None | F10 | Alizarin | 21925 | BL1 | 3403 |
| 40 | Yes | E. coli | None | None | F84 | Neutral Re | 23330 | BL1 | 2993 |
| 40 | Yes | E. coli | None | None | F99 | Pyronin Y | 24598 | BL1 | 2457 |
| 40 | Yes | E. coli | None | None | F108 | Rhodamine | 23958 | BL1 | 2401 |
| 40 | Yes | E. coli | None | None | F109 | Riboflavin | 23764 | BL1 | 2558 |
| 40 | Yes | E. coli | None | None | F117 | SYBR Gree | 24998 | BL1 | 89964.5 |
| 40 | Yes | E. coli | None | None | F106 | Rhodamine | 23658 | BL1 | 3528.5 |
| 40 | Yes | E. coli | None | None | F73 | Malachite ( | 23892 | BL1 | 2930 |
| 40 | Yes | E. coli | None | None | F7 | Acridine or | 22884 | BL1 | 2765.5 |
| 40 | Yes | E. coli | None | None | F192 | 5-Carboxyf | 23625 | BL1 | 17185 |
| 40 | Yes | E. coli | None | None | F57 | Eosin Y | 18630 | BL1 | 941330 |
| 40 | Yes | E. coli | None | None | F27 | Calcein | 19910 | BL1 | 1067810 |

| | | | | | | | | | |
|---|---|---|---|---|---|---|---|---|---|
| 40 | Yes | E. coli | None | None | F12 | Amaranth | 22873 | BL1 | 58414 |
| 40 | Yes | E. coli | None | None | NA | Autofluore: | 21200 | NA | 5170.5 |
| 40 | Yes | E. coli | Amoxacilli | Wall | F194 | Oxonol | 26217 | RL1 | 3855 |
| 40 | Yes | E. coli | Amoxacilli | Wall | F11 | Allura Red | 26966 | BL3 | 3191 |
| 40 | Yes | E. coli | Amoxacilli | Wall | F54 | DiSC3(5) | 24231 | RL1 | 3014 |
| 40 | Yes | E. coli | Amoxacilli | Wall | F124 | Thioflavin T | 25168 | VL2 | 2583 |
| 40 | Yes | E. coli | Amoxacilli | Wall | F101 | Quinacrine | 22912 | VL2 | 2578 |
| 40 | Yes | E. coli | Amoxacilli | Wall | F138 | Apigenin | 25091 | VL2 | 2635 |
| 40 | Yes | E. coli | Amoxacilli | Wall | F65 | H2FDA | 20415 | BL1 | 3267 |
| 40 | Yes | E. coli | Amoxacilli | Wall | F142 | Chlorophe | 24603 | BL3 | 4323 |
| 40 | Yes | E. coli | Amoxacilli | Wall | F39 | Congo Red | 22267 | BL3 | 5142 |
| 40 | Yes | E. coli | Amoxacilli | Wall | F116 | Sunset Yel | 24300 | BL2 | 4110 |
| 40 | Yes | E. coli | Amoxacilli | Wall | F10 | Alizarin | 22259 | BL1 | 3774 |
| 40 | Yes | E. coli | Amoxacilli | Wall | F84 | Neutral Re | 20193 | BL1 | 3149 |
| 40 | Yes | E. coli | Amoxacilli | Wall | F99 | Pyronin Y | 23348 | BL1 | 2906 |
| 40 | Yes | E. coli | Amoxacilli | Wall | F108 | Rhodamine | 22350 | BL1 | 2818 |
| 40 | Yes | E. coli | Amoxacilli | Wall | F109 | Riboflavin | 20732 | BL1 | 2847 |
| 40 | Yes | E. coli | Amoxacilli | Wall | F117 | SYBR Gree | 22939 | BL1 | 101942 |
| 40 | Yes | E. coli | Amoxacilli | Wall | F106 | Rhodamine | 25322 | BL1 | 3422 |
| 40 | Yes | E. coli | Amoxacilli | Wall | F73 | Malachite ( | 20664 | BL1 | 3479 |
| 40 | Yes | E. coli | Amoxacilli | Wall | F7 | Acridine or | 22170 | BL1 | 2923 |
| 40 | Yes | E. coli | Amoxacilli | Wall | F192 | 5-Carboxyf | 24225 | BL1 | 13254 |
| 40 | Yes | E. coli | Amoxacilli | Wall | F57 | Eosin Y | 21693 | BL1 | 445753 |
| 40 | Yes | E. coli | Amoxacilli | Wall | F27 | Calcein | 23708 | BL1 | 26937.5 |
| 40 | Yes | E. coli | Amoxacilli | Wall | F12 | Amaranth | 21526 | BL1 | 757545.5 |
| 40 | Yes | E. coli | None | None | NA | Autofluore: | 21429 | NA | 5668 |
| 40 | Yes | E. coli | Azithromyc | Intracellul: | F194 | Oxonol | 13337 | RL1 | 4568 |
| 40 | Yes | E. coli | Azithromyc | Intracellul: | F11 | Allura Red | 11886 | BL3 | 4062 |
| 40 | Yes | E. coli | Azithromyc | Intracellul: | F54 | DiSC3(5) | 11361 | RL1 | 3983 |
| 40 | Yes | E. coli | Azithromyc | Intracellul: | F124 | Thioflavin T | 10840 | VL2 | 3381 |
| 40 | Yes | E. coli | Azithromyc | Intracellul: | F101 | Quinacrine | 10404 | VL2 | 3369 |
| 40 | Yes | E. coli | Azithromyc | Intracellul: | F138 | Apigenin | 9674 | VL2 | 3402.5 |
| 40 | Yes | E. coli | Azithromyc | Intracellul: | F65 | H2FDA | 11067 | BL1 | 4454 |
| 40 | Yes | E. coli | Azithromyc | Intracellul: | F142 | Chlorophe | 10108 | BL3 | 5397 |
| 40 | Yes | E. coli | Azithromyc | Intracellul: | F39 | Congo Red | 10425 | BL3 | 5784 |
| 40 | Yes | E. coli | Azithromyc | Intracellul: | F116 | Sunset Yel | 10399 | BL2 | 5027 |
| 40 | Yes | E. coli | Azithromyc | Intracellul: | F10 | Alizarin | 10594 | BL1 | 4394.5 |
| 40 | Yes | E. coli | Azithromyc | Intracellul: | F84 | Neutral Re | 10753 | BL1 | 3828 |
| 40 | Yes | E. coli | Azithromyc | Intracellul: | F99 | Pyronin Y | 10606 | BL1 | 3583 |
| 40 | Yes | E. coli | Azithromyc | Intracellul: | F108 | Rhodamine | 10223 | BL1 | 3441 |
| 40 | Yes | E. coli | Azithromyc | Intracellul: | F109 | Riboflavin | 10093 | BL1 | 3621 |
| 40 | Yes | E. coli | Azithromyc | Intracellul: | F117 | SYBR Gree | 10152 | BL1 | 79396.5 |
| 40 | Yes | E. coli | Azithromyc | Intracellul: | F106 | Rhodamine | 10800 | BL1 | 4165 |
| 40 | Yes | E. coli | Azithromyc | Intracellul: | F73 | Malachite ( | 10603 | BL1 | 3801 |
| 40 | Yes | E. coli | Azithromyc | Intracellul: | F7 | Acridine or | 11096 | BL1 | 3535 |
| 40 | Yes | E. coli | Azithromyc | Intracellul: | F192 | 5-Carboxyf | 10354 | BL1 | 16162.5 |
| 40 | Yes | E. coli | Azithromyc | Intracellul: | F57 | Eosin Y | 9498 | BL1 | 25868.5 |
| 40 | Yes | E. coli | Azithromyc | Intracellul: | F27 | Calcein | 9608 | BL1 | 27665.5 |

| | | | | | | | | | |
|---|---|---|---|---|---|---|---|---|---|
| 40 | Yes | E. coli | Azithromyc | Intracellul | F12 | Amaranth | 10044 | BL1 | 28418 |
| 40 | Yes | E. coli | None | None | NA | Autofluore | 21878 | NA | 5517 |
| 40 | Yes | E. coli | Trimethopr | Intracellul | F194 | Oxonol | 19039 | RL1 | 4265 |
| 40 | Yes | E. coli | Trimethopr | Intracellul | F11 | Allura Red | 18006 | BL3 | 3727 |
| 40 | Yes | E. coli | Trimethopr | Intracellul | F54 | DiSC3(5) | 16816 | RL1 | 3575 |
| 40 | Yes | E. coli | Trimethopr | Intracellul | F124 | Thioflavin T | 14793 | VL2 | 3644 |
| 40 | Yes | E. coli | Trimethopr | Intracellul | F101 | Quinacrine | 17151 | VL2 | 3152 |
| 40 | Yes | E. coli | Trimethopr | Intracellul | F138 | Apigenin | 17623 | VL2 | 2916 |
| 40 | Yes | E. coli | Trimethopr | Intracellul | F65 | H2FDA | 16500 | BL1 | 3892 |
| 40 | Yes | E. coli | Trimethopr | Intracellul | F142 | Chlorophe | 16598 | BL3 | 4817 |
| 40 | Yes | E. coli | Trimethopr | Intracellul | F39 | Congo Red | 17153 | BL3 | 5251 |
| 40 | Yes | E. coli | Trimethopr | Intracellul | F116 | Sunset Yel | 16846 | BL2 | 4933 |
| 40 | Yes | E. coli | Trimethopr | Intracellul | F10 | Alizarin | 15885 | BL1 | 4186 |
| 40 | Yes | E. coli | Trimethopr | Intracellul | F84 | Neutral Re | 16087 | BL1 | 4045 |
| 40 | Yes | E. coli | Trimethopr | Intracellul | F99 | Pyronin Y | 17475 | BL1 | 3363 |
| 40 | Yes | E. coli | Trimethopr | Intracellul | F108 | Rhodamine | 16095 | BL1 | 3606 |
| 40 | Yes | E. coli | Trimethopr | Intracellul | F109 | Riboflavin | 15261 | BL1 | 4748 |
| 40 | Yes | E. coli | Trimethopr | Intracellul | F117 | SYBR Gree | 15615 | BL1 | 73706 |
| 40 | Yes | E. coli | Trimethopr | Intracellul | F106 | Rhodamine | 16059 | BL1 | 4198 |
| 40 | Yes | E. coli | Trimethopr | Intracellul | F73 | Malachite ( | 16444 | BL1 | 3936 |
| 40 | Yes | E. coli | Trimethopr | Intracellul | F7 | Acridine or | 14655 | BL1 | 3047 |
| 40 | Yes | E. coli | Trimethopr | Intracellul | F192 | 5-Carboxyf | 16764 | BL1 | 13146 |
| 40 | Yes | E. coli | Trimethopr | Intracellul | F57 | Eosin Y | 15699 | BL1 | 23232 |
| 40 | Yes | E. coli | Trimethopr | Intracellul | F27 | Calcein | 16424 | BL1 | 35674.5 |
| 40 | Yes | E. coli | Trimethopr | Intracellul | F12 | Amaranth | 19071 | BL1 | 21467 |
| 40 | Yes | E. coli | None | None | NA | Autofluore | 22744 | NA | 5010 |
| 40 | Yes | E. coli | Chloramph | Intracellul | F194 | Oxonol | 14079 | RL1 | 4456 |
| 40 | Yes | E. coli | Chloramph | Intracellul | F11 | Allura Red | 13055 | BL3 | 3890 |
| 40 | Yes | E. coli | Chloramph | Intracellul | F54 | DiSC3(5) | 12333 | RL1 | 3679 |
| 40 | Yes | E. coli | Chloramph | Intracellul | F124 | Thioflavin T | 12934 | VL2 | 3516 |
| 40 | Yes | E. coli | Chloramph | Intracellul | F101 | Quinacrine | 10841 | VL2 | 3788 |
| 40 | Yes | E. coli | Chloramph | Intracellul | F138 | Apigenin | 12087 | VL2 | 3824 |
| 40 | Yes | E. coli | Chloramph | Intracellul | F65 | H2FDA | 12526 | BL1 | 4439 |
| 40 | Yes | E. coli | Chloramph | Intracellul | F142 | Chlorophe | 12633 | BL3 | 5077 |
| 40 | Yes | E. coli | Chloramph | Intracellul | F39 | Congo Red | 13185 | BL3 | 5076 |
| 40 | Yes | E. coli | Chloramph | Intracellul | F116 | Sunset Yel | 12503 | BL2 | 5026 |
| 40 | Yes | E. coli | Chloramph | Intracellul | F10 | Alizarin | 12345 | BL1 | 4368 |
| 40 | Yes | E. coli | Chloramph | Intracellul | F84 | Neutral Re | 12361 | BL1 | 4105 |
| 40 | Yes | E. coli | Chloramph | Intracellul | F99 | Pyronin Y | 13373 | BL1 | 3312 |
| 40 | Yes | E. coli | Chloramph | Intracellul | F108 | Rhodamine | 11972 | BL1 | 3210.5 |
| 40 | Yes | E. coli | Chloramph | Intracellul | F109 | Riboflavin | 12159 | BL1 | 3955 |
| 40 | Yes | E. coli | Chloramph | Intracellul | F117 | SYBR Gree | 12522 | BL1 | 67186.5 |
| 40 | Yes | E. coli | Chloramph | Intracellul | F106 | Rhodamine | 12300 | BL1 | 4341 |
| 40 | Yes | E. coli | Chloramph | Intracellul | F73 | Malachite ( | 12318 | BL1 | 3766 |
| 40 | Yes | E. coli | Chloramph | Intracellul | F7 | Acridine or | 12235 | BL1 | 3817 |
| 40 | Yes | E. coli | Chloramph | Intracellul | F192 | 5-Carboxyf | 11572 | BL1 | 14609 |
| 40 | Yes | E. coli | Chloramph | Intracellul | F57 | Eosin Y | 12766 | BL1 | 20011.5 |
| 40 | Yes | E. coli | Chloramph | Intracellul | F27 | Calcein | 12157 | BL1 | 20466 |

| | | | | | | | | | |
|---|---|---|---|---|---|---|---|---|---|
| 40 | Yes | E. coli | Chloramph | Intracellul | F12 | Amaranth | 11887 | BL1 | 21454 |
| 40 | Yes | E. coli | None | None | NA | Autofluore | 23181 | NA | 5493 |
| 40 | Yes | E. coli | Colistin | Wall | F194 | Oxonol | 7337 | RL1 | 75360 |
| 40 | Yes | E. coli | Colistin | Wall | F11 | Allura Red | 5357 | BL3 | 53498 |
| 40 | Yes | E. coli | Colistin | Wall | F54 | DiSC3(5) | 4714 | RL1 | 52990.5 |
| 40 | Yes | E. coli | Colistin | Wall | F124 | Thioflavin T | 4815 | VL2 | 44867 |
| 40 | Yes | E. coli | Colistin | Wall | F101 | Quinacrine | 3829 | VL2 | 48363 |
| 40 | Yes | E. coli | Colistin | Wall | F138 | Apigenin | 4645 | VL2 | 44472 |
| 40 | Yes | E. coli | Colistin | Wall | F65 | H2FDA | 4409 | BL1 | 42009 |
| 40 | Yes | E. coli | Colistin | Wall | F142 | Chlorophe | 4304 | BL3 | 40940 |
| 40 | Yes | E. coli | Colistin | Wall | F39 | Congo Red | 4693 | BL3 | 25380 |
| 40 | Yes | E. coli | Colistin | Wall | F116 | Sunset Yel | 4518 | BL2 | 39359.5 |
| 40 | Yes | E. coli | Colistin | Wall | F10 | Alizarin | 4159 | BL1 | 33885 |
| 40 | Yes | E. coli | Colistin | Wall | F84 | Neutral Re | 4867 | BL1 | 34328 |
| 40 | Yes | E. coli | Colistin | Wall | F99 | Pyronin Y | 4649 | BL1 | 31731 |
| 40 | Yes | E. coli | Colistin | Wall | F108 | Rhodamine | 4741 | BL1 | 33697 |
| 40 | Yes | E. coli | Colistin | Wall | F109 | Riboflavin | 4221 | BL1 | 32499 |
| 40 | Yes | E. coli | Colistin | Wall | F117 | SYBR Gree | 4682 | BL1 | 3498376 |
| 40 | Yes | E. coli | Colistin | Wall | F106 | Rhodamine | 4607 | BL1 | 163115 |
| 40 | Yes | E. coli | Colistin | Wall | F73 | Malachite ( | 4677 | BL1 | 118563 |
| 40 | Yes | E. coli | Colistin | Wall | F7 | Acridine or | 4032 | BL1 | 162537 |
| 40 | Yes | E. coli | Colistin | Wall | F192 | 5-Carboxyf | 4763 | BL1 | 86861 |
| 40 | Yes | E. coli | Colistin | Wall | F57 | Eosin Y | 4631 | BL1 | 74049 |
| 40 | Yes | E. coli | Colistin | Wall | F27 | Calcein | 4400 | BL1 | 50407 |
| 40 | Yes | E. coli | Colistin | Wall | F12 | Amaranth | 4397 | BL1 | 60377 |
| 40 | Yes | E. coli | None | None | NA | Autofluore | 17141 | NA | 7874 |
| 40 | Yes | E. coli | Gentamici | Intracellul | F194 | Oxonol | 25688 | RL1 | 5013 |
| 40 | Yes | E. coli | Gentamici | Intracellul | F11 | Allura Red | 25234 | BL3 | 4657.5 |
| 40 | Yes | E. coli | Gentamici | Intracellul | F54 | DiSC3(5) | 25274 | RL1 | 3433 |
| 40 | Yes | E. coli | Gentamici | Intracellul | F124 | Thioflavin T | 26218 | VL2 | 4171 |
| 40 | Yes | E. coli | Gentamici | Intracellul | F101 | Quinacrine | 25536 | VL2 | 3783 |
| 40 | Yes | E. coli | Gentamici | Intracellul | F138 | Apigenin | 28396 | VL2 | 3198.5 |
| 40 | Yes | E. coli | Gentamici | Intracellul | F65 | H2FDA | 26091 | BL1 | 4378 |
| 40 | Yes | E. coli | Gentamici | Intracellul | F142 | Chlorophe | 30275 | BL3 | 4195 |
| 40 | Yes | E. coli | Gentamici | Intracellul | F39 | Congo Red | 28330 | BL3 | 4804 |
| 40 | Yes | E. coli | Gentamici | Intracellul | F116 | Sunset Yel | 22661 | BL2 | 6086 |
| 40 | Yes | E. coli | Gentamici | Intracellul | F10 | Alizarin | 25495 | BL1 | 5100 |
| 40 | Yes | E. coli | Gentamici | Intracellul | F84 | Neutral Re | 23656 | BL1 | 5034 |
| 40 | Yes | E. coli | Gentamici | Intracellul | F99 | Pyronin Y | 26622 | BL1 | 3305 |
| 40 | Yes | E. coli | Gentamici | Intracellul | F108 | Rhodamine | 23033 | BL1 | 3890 |
| 40 | Yes | E. coli | Gentamici | Intracellul | F109 | Riboflavin | 25518 | BL1 | 3181 |
| 40 | Yes | E. coli | Gentamici | Intracellul | F117 | SYBR Gree | 25839 | BL1 | 162058 |
| 40 | Yes | E. coli | Gentamici | Intracellul | F106 | Rhodamine | 25487 | BL1 | 4451 |
| 40 | Yes | E. coli | Gentamici | Intracellul | F73 | Malachite ( | 25036 | BL1 | 3675 |
| 40 | Yes | E. coli | Gentamici | Intracellul | F7 | Acridine or | 24263 | BL1 | 3907 |
| 40 | Yes | E. coli | Gentamici | Intracellul | F192 | 5-Carboxyf | 23877 | BL1 | 13139 |
| 40 | Yes | E. coli | Gentamici | Intracellul | F57 | Eosin Y | 21567 | BL1 | 109769 |
| 40 | Yes | E. coli | Gentamici | Intracellul | F27 | Calcein | 23787 | BL1 | 30633 |

| | | | | | | | | | |
|---|---|---|---|---|---|---|---|---|---|
| 40 | Yes | E. coli | Gentamici | Intracellula | F12 | Amaranth | 20842 | BL1 | 220603.5 |
| 40 | Yes | E. coli | None | None | NA | Autofluore: | 22672 | NA | 5529 |
| 40 | Yes | E. coli | Nitrofurant | Intracellula | F194 | Oxonol | 12599 | RL1 | 5620 |
| 40 | Yes | E. coli | Nitrofurant | Intracellula | F11 | Allura Red | 10186 | BL3 | 4633.5 |
| 40 | Yes | E. coli | Nitrofurant | Intracellula | F54 | DiSC3(5) | 9018 | RL1 | 4687 |
| 40 | Yes | E. coli | Nitrofurant | Intracellula | F124 | Thioflavin T | 11315 | VL2 | 4099 |
| 40 | Yes | E. coli | Nitrofurant | Intracellula | F101 | Quinacrine | 10187 | VL2 | 4015 |
| 40 | Yes | E. coli | Nitrofurant | Intracellula | F138 | Apigenin | 9897 | VL2 | 4691 |
| 40 | Yes | E. coli | Nitrofurant | Intracellula | F65 | H2FDA | 9449 | BL1 | 4757 |
| 40 | Yes | E. coli | Nitrofurant | Intracellula | F142 | Chlorophe | 10619 | BL3 | 6654 |
| 40 | Yes | E. coli | Nitrofurant | Intracellula | F39 | Congo Red | 8567 | BL3 | 6332 |
| 40 | Yes | E. coli | Nitrofurant | Intracellula | F116 | Sunset Yel | 9189 | BL2 | 5621 |
| 40 | Yes | E. coli | Nitrofurant | Intracellula | F10 | Alizarin | 9456 | BL1 | 5468.5 |
| 40 | Yes | E. coli | Nitrofurant | Intracellula | F84 | Neutral Re | 7991 | BL1 | 4764 |
| 40 | Yes | E. coli | Nitrofurant | Intracellula | F99 | Pyronin Y | 11215 | BL1 | 4313 |
| 40 | Yes | E. coli | Nitrofurant | Intracellula | F108 | Rhodamine | 10804 | BL1 | 3770.5 |
| 40 | Yes | E. coli | Nitrofurant | Intracellula | F109 | Riboflavin | 10960 | BL1 | 4076 |
| 40 | Yes | E. coli | Nitrofurant | Intracellula | F117 | SYBR Gree | 9358 | BL1 | 78195 |
| 40 | Yes | E. coli | Nitrofurant | Intracellula | F106 | Rhodamine | 9814 | BL1 | 5189 |
| 40 | Yes | E. coli | Nitrofurant | Intracellula | F73 | Malachite ( | 9880 | BL1 | 4699.5 |
| 40 | Yes | E. coli | Nitrofurant | Intracellula | F7 | Acridine or | 9183 | BL1 | 4410 |
| 40 | Yes | E. coli | Nitrofurant | Intracellula | F192 | 5-Carboxyf | 9976 | BL1 | 16162.5 |
| 40 | Yes | E. coli | Nitrofurant | Intracellula | F57 | Eosin Y | 8978 | BL1 | 23507 |
| 40 | Yes | E. coli | Nitrofurant | Intracellula | F27 | Calcein | 9364 | BL1 | 23942 |
| 40 | Yes | E. coli | Nitrofurant | Intracellula | F12 | Amaranth | 9122 | BL1 | 25662.5 |
| 40 | Yes | E. coli | None | None | NA | Autofluore: | 20491 | NA | 7267 |
| 40 | Yes | E. coli | Pentamidi | Wall | F194 | Oxonol | 15081 | RL1 | 5461 |
| 40 | Yes | E. coli | Pentamidi | Wall | F11 | Allura Red | 10968 | BL3 | 5236.5 |
| 40 | Yes | E. coli | Pentamidi | Wall | F54 | DiSC3(5) | 11431 | RL1 | 4414 |
| 40 | Yes | E. coli | Pentamidi | Wall | F124 | Thioflavin T | 11762 | VL2 | 4563.5 |
| 40 | Yes | E. coli | Pentamidi | Wall | F101 | Quinacrine | 12015 | VL2 | 5332 |
| 40 | Yes | E. coli | Pentamidi | Wall | F138 | Apigenin | 11359 | VL2 | 5206 |
| 40 | Yes | E. coli | Pentamidi | Wall | F65 | H2FDA | 11707 | BL1 | 6841 |
| 40 | Yes | E. coli | Pentamidi | Wall | F142 | Chlorophe | 11804 | BL3 | 6532 |
| 40 | Yes | E. coli | Pentamidi | Wall | F39 | Congo Red | 12305 | BL3 | 6632 |
| 40 | Yes | E. coli | Pentamidi | Wall | F116 | Sunset Yel | 11700 | BL2 | 6328.5 |
| 40 | Yes | E. coli | Pentamidi | Wall | F10 | Alizarin | 12598 | BL1 | 5388.5 |
| 40 | Yes | E. coli | Pentamidi | Wall | F84 | Neutral Re | 12290 | BL1 | 5123 |
| 40 | Yes | E. coli | Pentamidi | Wall | F99 | Pyronin Y | 12682 | BL1 | 4375.5 |
| 40 | Yes | E. coli | Pentamidi | Wall | F108 | Rhodamine | 12630 | BL1 | 4348.5 |
| 40 | Yes | E. coli | Pentamidi | Wall | F109 | Riboflavin | 11389 | BL1 | 4967 |
| 40 | Yes | E. coli | Pentamidi | Wall | F117 | SYBR Gree | 10890 | BL1 | 773955.5 |
| 40 | Yes | E. coli | Pentamidi | Wall | F106 | Rhodamine | 11588 | BL1 | 6841 |
| 40 | Yes | E. coli | Pentamidi | Wall | F73 | Malachite ( | 11724 | BL1 | 6554.5 |
| 40 | Yes | E. coli | Pentamidi | Wall | F7 | Acridine or | 10548 | BL1 | 5877 |
| 40 | Yes | E. coli | Pentamidi | Wall | F192 | 5-Carboxyf | 11802 | BL1 | 15081 |
| 40 | Yes | E. coli | Pentamidi | Wall | F57 | Eosin Y | 10949 | BL1 | 19821 |
| 40 | Yes | E. coli | Pentamidi | Wall | F27 | Calcein | 11938 | BL1 | 22357 |

| | | | | | | | | | |
|---|---|---|---|---|---|---|---|---|---|
| 40 | Yes | E. coli | Pentamidi | Wall | F12 | Amaranth | 10959 | BL1 | 20890 |
| 40 | Yes | E. coli | None | None | NA | Autofluore | 21666 | NA | 8586 |
| 40 | Yes | E. coli | Rifampicin | Intracellul | F194 | Oxonol | 23676 | RL1 | 6337 |
| 40 | Yes | E. coli | Rifampicin | Intracellul | F11 | Allura Red | 23119 | BL3 | 5401 |
| 40 | Yes | E. coli | Rifampicin | Intracellul | F54 | DiSC3(5) | 24508 | RL1 | 4226 |
| 40 | Yes | E. coli | Rifampicin | Intracellul | F124 | Thioflavin T | 31333 | VL2 | 3448 |
| 40 | Yes | E. coli | Rifampicin | Intracellul | F101 | Quinacrine | 25175 | VL2 | 3965 |
| 40 | Yes | E. coli | Rifampicin | Intracellul | F138 | Apigenin | 27985 | VL2 | 3480 |
| 40 | Yes | E. coli | Rifampicin | Intracellul | F65 | H2FDA | 23783 | BL1 | 5257 |
| 40 | Yes | E. coli | Rifampicin | Intracellul | F142 | Chlorophe | 25222 | BL3 | 6643 |
| 40 | Yes | E. coli | Rifampicin | Intracellul | F39 | Congo Red | 24837 | BL3 | 6628 |
| 40 | Yes | E. coli | Rifampicin | Intracellul | F116 | Sunset Yel | 21104 | BL2 | 7052 |
| 40 | Yes | E. coli | Rifampicin | Intracellul | F10 | Alizarin | 26648 | BL1 | 4321.5 |
| 40 | Yes | E. coli | Rifampicin | Intracellul | F84 | Neutral Re | 20586 | BL1 | 5481 |
| 40 | Yes | E. coli | Rifampicin | Intracellul | F99 | Pyronin Y | 26935 | BL1 | 3344 |
| 40 | Yes | E. coli | Rifampicin | Intracellul | F108 | Rhodamine | 24352 | BL1 | 3212 |
| 40 | Yes | E. coli | Rifampicin | Intracellul | F109 | Riboflavin | 25522 | BL1 | 3633.5 |
| 40 | Yes | E. coli | Rifampicin | Intracellul | F117 | SYBR Gree | 22747 | BL1 | 84280 |
| 40 | Yes | E. coli | Rifampicin | Intracellul | F106 | Rhodamine | 26301 | BL1 | 4383 |
| 40 | Yes | E. coli | Rifampicin | Intracellul | F73 | Malachite ( | 27045 | BL1 | 3713 |
| 40 | Yes | E. coli | Rifampicin | Intracellul | F7 | Acridine or | 22220 | BL1 | 4258 |
| 40 | Yes | E. coli | Rifampicin | Intracellul | F192 | 5-Carboxyf | 27398 | BL1 | 13857.5 |
| 40 | Yes | E. coli | Rifampicin | Intracellul | F57 | Eosin Y | 24061 | BL1 | 28690 |
| 40 | Yes | E. coli | Rifampicin | Intracellul | F27 | Calcein | 28314 | BL1 | 21601 |
| 40 | Yes | E. coli | Rifampicin | Intracellul | F12 | Amaranth | 27073 | BL1 | 19692 |
| 40 | Yes | E. coli | None | None | NA | Autofluore | 19424 | NA | 8198 |
| 40 | Yes | E. coli | Trimethopr | Intracellul | F194 | Oxonol | 23425 | RL1 | 5697 |
| 40 | Yes | E. coli | Trimethopr | Intracellul | F11 | Allura Red | 22542 | BL3 | 4153 |
| 40 | Yes | E. coli | Trimethopr | Intracellul | F54 | DiSC3(5) | 21251 | RL1 | 3853 |
| 40 | Yes | E. coli | Trimethopr | Intracellul | F124 | Thioflavin T | 23685 | VL2 | 3908 |
| 40 | Yes | E. coli | Trimethopr | Intracellul | F101 | Quinacrine | 21292 | VL2 | 3886.5 |
| 40 | Yes | E. coli | Trimethopr | Intracellul | F138 | Apigenin | 18879 | VL2 | 4351 |
| 40 | Yes | E. coli | Trimethopr | Intracellul | F65 | H2FDA | 20780 | BL1 | 4825 |
| 40 | Yes | E. coli | Trimethopr | Intracellul | F142 | Chlorophe | 20154 | BL3 | 5226 |
| 40 | Yes | E. coli | Trimethopr | Intracellul | F39 | Congo Red | 22011 | BL3 | 5144 |
| 40 | Yes | E. coli | Trimethopr | Intracellul | F116 | Sunset Yel | 18797 | BL2 | 5378 |
| 40 | Yes | E. coli | Trimethopr | Intracellul | F10 | Alizarin | 18757 | BL1 | 4726 |
| 40 | Yes | E. coli | Trimethopr | Intracellul | F84 | Neutral Re | 16858 | BL1 | 4399 |
| 40 | Yes | E. coli | Trimethopr | Intracellul | F99 | Pyronin Y | 19140 | BL1 | 3522 |
| 40 | Yes | E. coli | Trimethopr | Intracellul | F108 | Rhodamine | 17074 | BL1 | 3659 |
| 40 | Yes | E. coli | Trimethopr | Intracellul | F109 | Riboflavin | 16830 | BL1 | 4248 |
| 40 | Yes | E. coli | Trimethopr | Intracellul | F117 | SYBR Gree | 16590 | BL1 | 73983.5 |
| 40 | Yes | E. coli | Trimethopr | Intracellul | F106 | Rhodamine | 17331 | BL1 | 4352 |
| 40 | Yes | E. coli | Trimethopr | Intracellul | F73 | Malachite ( | 18219 | BL1 | 3880 |
| 40 | Yes | E. coli | Trimethopr | Intracellul | F7 | Acridine or | 16964 | BL1 | 3858 |
| 40 | Yes | E. coli | Trimethopr | Intracellul | F192 | 5-Carboxyf | 18920 | BL1 | 13154.5 |
| 40 | Yes | E. coli | Trimethopr | Intracellul | F57 | Eosin Y | 15773 | BL1 | 18085 |
| 40 | Yes | E. coli | Trimethopr | Intracellul | F27 | Calcein | 18791 | BL1 | 18157 |

| | | | | | | | | | |
|---|---|---|---|---|---|---|---|---|---|
| 40 | Yes | E. coli | Trimethopr | Intracellula | F12 | Amaranth | 16700 | BL1 | 20682.5 |
| 40 | Yes | E. coli | None | None | NA | Autofluores | 17824 | NA | 765 |
| 40 | Yes | E. coli | None | None | F194 | Oxonol | 25189 | RL1 | 1285 |
| 40 | Yes | E. coli | None | None | F11 | Allura Red | 21369 | BL3 | 1594 |
| 40 | Yes | E. coli | None | None | F54 | DiSC3(5) | 22684 | RL1 | 2036 |
| 40 | Yes | E. coli | None | None | F124 | Thioflavin T | 25215 | VL2 | 2045 |
| 40 | Yes | E. coli | None | None | F101 | Quinacrine | 20905 | VL2 | 2218 |
| 40 | Yes | E. coli | None | None | F138 | Apigenin | 21026 | VL2 | 2354 |
| 40 | Yes | E. coli | None | None | F65 | H2FDA | 22985 | BL1 | 3031 |
| 40 | Yes | E. coli | None | None | F142 | Chlorophe | 22213 | BL3 | 3839 |
| 40 | Yes | E. coli | None | None | F39 | Congo Red | 20853 | BL3 | 4037 |
| 40 | Yes | E. coli | None | None | F116 | Sunset Yel | 23501 | BL2 | 3918 |
| 40 | Yes | E. coli | None | None | F10 | Alizarin | 23744 | BL1 | 3520 |
| 40 | Yes | E. coli | None | None | F84 | Neutral Re | 19903 | BL1 | 3036 |
| 40 | Yes | E. coli | None | None | F99 | Pyronin Y | 21194 | BL1 | 2751.5 |
| 40 | Yes | E. coli | None | None | F108 | Rhodamine | 20268 | BL1 | 2600 |
| 40 | Yes | E. coli | None | None | F109 | Riboflavin | 22510 | BL1 | 2635 |
| 40 | Yes | E. coli | None | None | F117 | SYBR Gree | 20608 | BL1 | 92729 |
| 40 | Yes | E. coli | None | None | F106 | Rhodamine | 21914 | BL1 | 4119 |
| 40 | Yes | E. coli | None | None | F73 | Malachite ( | 20513 | BL1 | 3105 |
| 40 | Yes | E. coli | None | None | F7 | Acridine or | 20340 | BL1 | 3131 |
| 40 | Yes | E. coli | None | None | F192 | 5-Carboxyf | 21587 | BL1 | 15070 |
| 40 | Yes | E. coli | None | None | F57 | Eosin Y | 19698 | BL1 | 38477.5 |
| 40 | Yes | E. coli | None | None | F27 | Calcein | 17880 | BL1 | 909545.5 |
| 40 | Yes | E. coli | None | None | F12 | Amaranth | 17684 | BL1 | 406202 |
| 40 | Yes | E. coli | None | None | NA | Autofluores | 22226 | NA | 4728.5 |
| 40 | Yes | E. coli | Amoxacilli | Wall | F194 | Oxonol | 25328 | RL1 | 3733 |
| 40 | Yes | E. coli | Amoxacilli | Wall | F11 | Allura Red | 25883 | BL3 | 3040 |
| 40 | Yes | E. coli | Amoxacilli | Wall | F54 | DiSC3(5) | 23325 | RL1 | 2948 |
| 40 | Yes | E. coli | Amoxacilli | Wall | F124 | Thioflavin T | 21991 | VL2 | 2601 |
| 40 | Yes | E. coli | Amoxacilli | Wall | F101 | Quinacrine | 21019 | VL2 | 2698 |
| 40 | Yes | E. coli | Amoxacilli | Wall | F138 | Apigenin | 20858 | VL2 | 2914 |
| 40 | Yes | E. coli | Amoxacilli | Wall | F65 | H2FDA | 21575 | BL1 | 3371 |
| 40 | Yes | E. coli | Amoxacilli | Wall | F142 | Chlorophe | 21802 | BL3 | 3957 |
| 40 | Yes | E. coli | Amoxacilli | Wall | F39 | Congo Red | 23005 | BL3 | 4273 |
| 40 | Yes | E. coli | Amoxacilli | Wall | F116 | Sunset Yel | 22571 | BL2 | 4025 |
| 40 | Yes | E. coli | Amoxacilli | Wall | F10 | Alizarin | 18652 | BL1 | 4008 |
| 40 | Yes | E. coli | Amoxacilli | Wall | F84 | Neutral Re | 20050 | BL1 | 3427 |
| 40 | Yes | E. coli | Amoxacilli | Wall | F99 | Pyronin Y | 21445 | BL1 | 3056 |
| 40 | Yes | E. coli | Amoxacilli | Wall | F108 | Rhodamine | 19380 | BL1 | 3181 |
| 40 | Yes | E. coli | Amoxacilli | Wall | F109 | Riboflavin | 21913 | BL1 | 2888 |
| 40 | Yes | E. coli | Amoxacilli | Wall | F117 | SYBR Gree | 21321 | BL1 | 106359 |
| 40 | Yes | E. coli | Amoxacilli | Wall | F106 | Rhodamine | 23251 | BL1 | 3668 |
| 40 | Yes | E. coli | Amoxacilli | Wall | F73 | Malachite ( | 20433 | BL1 | 3764 |
| 40 | Yes | E. coli | Amoxacilli | Wall | F7 | Acridine or | 18398 | BL1 | 3393.5 |
| 40 | Yes | E. coli | Amoxacilli | Wall | F192 | 5-Carboxyf | 20477 | BL1 | 17452 |
| 40 | Yes | E. coli | Amoxacilli | Wall | F57 | Eosin Y | 18425 | BL1 | 953337 |
| 40 | Yes | E. coli | Amoxacilli | Wall | F27 | Calcein | 18231 | BL1 | 41033 |

| | | | | | | | | | |
|---|---|---|---|---|---|---|---|---|---|
| 40 | Yes | E. coli | Amoxacilli | Wall | F12 | Amaranth | 18722 | BL1 | 614137.5 |
| 40 | Yes | E. coli | None | None | NA | Autofluore | 22463 | NA | 5448 |
| 40 | Yes | E. coli | Azithromyc | Intracellul | F194 | Oxonol | 13116 | RL1 | 4840 |
| 40 | Yes | E. coli | Azithromyc | Intracellul | F11 | Allura Red | 12258 | BL3 | 3746 |
| 40 | Yes | E. coli | Azithromyc | Intracellul | F54 | DiSC3(5) | 11715 | RL1 | 3640 |
| 40 | Yes | E. coli | Azithromyc | Intracellul | F124 | Thioflavin T | 11153 | VL2 | 3319 |
| 40 | Yes | E. coli | Azithromyc | Intracellul | F101 | Quinacrine | 10594 | VL2 | 3441 |
| 40 | Yes | E. coli | Azithromyc | Intracellul | F138 | Apigenin | 10257 | VL2 | 3472 |
| 40 | Yes | E. coli | Azithromyc | Intracellul | F65 | H2FDA | 10157 | BL1 | 4031 |
| 40 | Yes | E. coli | Azithromyc | Intracellul | F142 | Chlorophe | 10785 | BL3 | 5070 |
| 40 | Yes | E. coli | Azithromyc | Intracellul | F39 | Congo Red | 11064 | BL3 | 5133 |
| 40 | Yes | E. coli | Azithromyc | Intracellul | F116 | Sunset Yel | 11271 | BL2 | 4690 |
| 40 | Yes | E. coli | Azithromyc | Intracellul | F10 | Alizarin | 9562 | BL1 | 4499.5 |
| 40 | Yes | E. coli | Azithromyc | Intracellul | F84 | Neutral Re | 10338 | BL1 | 4065 |
| 40 | Yes | E. coli | Azithromyc | Intracellul | F99 | Pyronin Y | 10555 | BL1 | 3680 |
| 40 | Yes | E. coli | Azithromyc | Intracellul | F108 | Rhodamine | 10111 | BL1 | 3465 |
| 40 | Yes | E. coli | Azithromyc | Intracellul | F109 | Riboflavin | 9955 | BL1 | 3488 |
| 40 | Yes | E. coli | Azithromyc | Intracellul | F117 | SYBR Gree | 10505 | BL1 | 69817 |
| 40 | Yes | E. coli | Azithromyc | Intracellul | F106 | Rhodamine | 10941 | BL1 | 4830 |
| 40 | Yes | E. coli | Azithromyc | Intracellul | F73 | Malachite ( | 10023 | BL1 | 4047 |
| 40 | Yes | E. coli | Azithromyc | Intracellul | F7 | Acridine or | 10572 | BL1 | 3808.5 |
| 40 | Yes | E. coli | Azithromyc | Intracellul | F192 | 5-Carboxyf | 9929 | BL1 | 16250 |
| 40 | Yes | E. coli | Azithromyc | Intracellul | F57 | Eosin Y | 8879 | BL1 | 20364 |
| 40 | Yes | E. coli | Azithromyc | Intracellul | F27 | Calcein | 10988 | BL1 | 23709.5 |
| 40 | Yes | E. coli | Azithromyc | Intracellul | F12 | Amaranth | 9895 | BL1 | 25382 |
| 40 | Yes | E. coli | None | None | NA | Autofluore | 21792 | NA | 5279 |
| 40 | Yes | E. coli | Trimethopr | Intracellul | F194 | Oxonol | 19349 | RL1 | 4262 |
| 40 | Yes | E. coli | Trimethopr | Intracellul | F11 | Allura Red | 17593 | BL3 | 3551 |
| 40 | Yes | E. coli | Trimethopr | Intracellul | F54 | DiSC3(5) | 16514 | RL1 | 3399 |
| 40 | Yes | E. coli | Trimethopr | Intracellul | F124 | Thioflavin T | 16576 | VL2 | 3035 |
| 40 | Yes | E. coli | Trimethopr | Intracellul | F101 | Quinacrine | 15515 | VL2 | 3179 |
| 40 | Yes | E. coli | Trimethopr | Intracellul | F138 | Apigenin | 14277 | VL2 | 3325 |
| 40 | Yes | E. coli | Trimethopr | Intracellul | F65 | H2FDA | 15342 | BL1 | 3760.5 |
| 40 | Yes | E. coli | Trimethopr | Intracellul | F142 | Chlorophe | 17668 | BL3 | 4830.5 |
| 40 | Yes | E. coli | Trimethopr | Intracellul | F39 | Congo Red | 15932 | BL3 | 5058 |
| 40 | Yes | E. coli | Trimethopr | Intracellul | F116 | Sunset Yel | 16672 | BL2 | 4172.5 |
| 40 | Yes | E. coli | Trimethopr | Intracellul | F10 | Alizarin | 16162 | BL1 | 4153.5 |
| 40 | Yes | E. coli | Trimethopr | Intracellul | F84 | Neutral Re | 14969 | BL1 | 3522 |
| 40 | Yes | E. coli | Trimethopr | Intracellul | F99 | Pyronin Y | 15792 | BL1 | 3329 |
| 40 | Yes | E. coli | Trimethopr | Intracellul | F108 | Rhodamine | 15958 | BL1 | 3282 |
| 40 | Yes | E. coli | Trimethopr | Intracellul | F109 | Riboflavin | 15922 | BL1 | 3345 |
| 40 | Yes | E. coli | Trimethopr | Intracellul | F117 | SYBR Gree | 15889 | BL1 | 78612 |
| 40 | Yes | E. coli | Trimethopr | Intracellul | F106 | Rhodamine | 16158 | BL1 | 4109.5 |
| 40 | Yes | E. coli | Trimethopr | Intracellul | F73 | Malachite ( | 14937 | BL1 | 3821 |
| 40 | Yes | E. coli | Trimethopr | Intracellul | F7 | Acridine or | 15821 | BL1 | 3726 |
| 40 | Yes | E. coli | Trimethopr | Intracellul | F192 | 5-Carboxyf | 16491 | BL1 | 11904 |
| 40 | Yes | E. coli | Trimethopr | Intracellul | F57 | Eosin Y | 16155 | BL1 | 16353 |
| 40 | Yes | E. coli | Trimethopr | Intracellul | F27 | Calcein | 20455 | BL1 | 28029 |

| | | | | | | | | | |
|---|---|---|---|---|---|---|---|---|---|
| 40 | Yes | E. coli | Trimethopr | Intracellula | F12 | Amaranth | 18143 | BL1 | 883067 |
| 40 | Yes | E. coli | None | None | NA | Autofluore: | 22240 | NA | 5355.5 |
| 40 | Yes | E. coli | Chloramph | Intracellula | F194 | Oxonol | 15727 | RL1 | 4374 |
| 40 | Yes | E. coli | Chloramph | Intracellula | F11 | Allura Red | 13958 | BL3 | 3653 |
| 40 | Yes | E. coli | Chloramph | Intracellula | F54 | DiSC3(5) | 13096 | RL1 | 3620.5 |
| 40 | Yes | E. coli | Chloramph | Intracellula | F124 | Thioflavin T | 10577 | VL2 | 3045 |
| 40 | Yes | E. coli | Chloramph | Intracellula | F101 | Quinacrine | 13516 | VL2 | 3258 |
| 40 | Yes | E. coli | Chloramph | Intracellula | F138 | Apigenin | 11912 | VL2 | 3458 |
| 40 | Yes | E. coli | Chloramph | Intracellula | F65 | H2FDA | 11072 | BL1 | 3799.5 |
| 40 | Yes | E. coli | Chloramph | Intracellula | F142 | Chlorophe | 12439 | BL3 | 5083 |
| 40 | Yes | E. coli | Chloramph | Intracellula | F39 | Congo Red | 12132 | BL3 | 5292 |
| 40 | Yes | E. coli | Chloramph | Intracellula | F116 | Sunset Yel | 11960 | BL2 | 4509 |
| 40 | Yes | E. coli | Chloramph | Intracellula | F10 | Alizarin | 12321 | BL1 | 4146 |
| 40 | Yes | E. coli | Chloramph | Intracellula | F84 | Neutral Re | 12354 | BL1 | 3547.5 |
| 40 | Yes | E. coli | Chloramph | Intracellula | F99 | Pyronin Y | 11590 | BL1 | 3277 |
| 40 | Yes | E. coli | Chloramph | Intracellula | F108 | Rhodamine | 12646 | BL1 | 3082.5 |
| 40 | Yes | E. coli | Chloramph | Intracellula | F109 | Riboflavin | 11590 | BL1 | 3151 |
| 40 | Yes | E. coli | Chloramph | Intracellula | F117 | SYBR Gree | 12495 | BL1 | 57482 |
| 40 | Yes | E. coli | Chloramph | Intracellula | F106 | Rhodamine | 12342 | BL1 | 3880 |
| 40 | Yes | E. coli | Chloramph | Intracellula | F73 | Malachite ( | 11458 | BL1 | 3543.5 |
| 40 | Yes | E. coli | Chloramph | Intracellula | F7 | Acridine or | 12460 | BL1 | 3342.5 |
| 40 | Yes | E. coli | Chloramph | Intracellula | F192 | 5-Carboxyf | 11472 | BL1 | 14240 |
| 40 | Yes | E. coli | Chloramph | Intracellula | F57 | Eosin Y | 11287 | BL1 | 18101 |
| 40 | Yes | E. coli | Chloramph | Intracellula | F27 | Calcein | 11038 | BL1 | 21098.5 |
| 40 | Yes | E. coli | Chloramph | Intracellula | F12 | Amaranth | 11325 | BL1 | 18060 |
| 40 | Yes | E. coli | None | None | NA | Autofluore: | 20692 | NA | 5670 |
| 40 | Yes | E. coli | Colistin | Wall | F194 | Oxonol | 6701 | RL1 | 64325 |
| 40 | Yes | E. coli | Colistin | Wall | F11 | Allura Red | 5161 | BL3 | 45842 |
| 40 | Yes | E. coli | Colistin | Wall | F54 | DiSC3(5) | 4435 | RL1 | 40498 |
| 40 | Yes | E. coli | Colistin | Wall | F124 | Thioflavin T | 4972 | VL2 | 40488 |
| 40 | Yes | E. coli | Colistin | Wall | F101 | Quinacrine | 4077 | VL2 | 35469 |
| 40 | Yes | E. coli | Colistin | Wall | F138 | Apigenin | 4329 | VL2 | 35249 |
| 40 | Yes | E. coli | Colistin | Wall | F65 | H2FDA | 4628 | BL1 | 32154.5 |
| 40 | Yes | E. coli | Colistin | Wall | F142 | Chlorophe | 4345 | BL3 | 30801 |
| 40 | Yes | E. coli | Colistin | Wall | F39 | Congo Red | 5131 | BL3 | 16146 |
| 40 | Yes | E. coli | Colistin | Wall | F116 | Sunset Yel | 4757 | BL2 | 29842 |
| 40 | Yes | E. coli | Colistin | Wall | F10 | Alizarin | 4770 | BL1 | 23718 |
| 40 | Yes | E. coli | Colistin | Wall | F84 | Neutral Re | 4195 | BL1 | 27771 |
| 40 | Yes | E. coli | Colistin | Wall | F99 | Pyronin Y | 4567 | BL1 | 24671 |
| 40 | Yes | E. coli | Colistin | Wall | F108 | Rhodamine | 4204 | BL1 | 24513 |
| 40 | Yes | E. coli | Colistin | Wall | F109 | Riboflavin | 11515 | BL1 | 4431 |
| 40 | Yes | E. coli | Colistin | Wall | F117 | SYBR Gree | 5264 | BL1 | 3084067 |
| 40 | Yes | E. coli | Colistin | Wall | F106 | Rhodamine | 4450 | BL1 | 120432.5 |
| 40 | Yes | E. coli | Colistin | Wall | F73 | Malachite ( | 4609 | BL1 | 97575 |
| 40 | Yes | E. coli | Colistin | Wall | F7 | Acridine or | 3896 | BL1 | 141534.5 |
| 40 | Yes | E. coli | Colistin | Wall | F192 | 5-Carboxyf | 3826 | BL1 | 75554 |
| 40 | Yes | E. coli | Colistin | Wall | F57 | Eosin Y | 3476 | BL1 | 51214.5 |
| 40 | Yes | E. coli | Colistin | Wall | F27 | Calcein | 15774 | BL1 | 20684.5 |

| | | | | | | | | | |
|---|---|---|---|---|---|---|---|---|---|
| 40 | Yes | E. coli | Colistin | Wall | F12 | Amaranth | 5697 | BL1 | 44743 |
| 40 | Yes | E. coli | None | None | NA | Autofluore: | 23307 | NA | 5536 |
| 40 | Yes | E. coli | Gentamici | Intracellul | F194 | Oxonol | 23534 | RL1 | 4571 |
| 40 | Yes | E. coli | Gentamici | Intracellul | F11 | Allura Red | 25001 | BL3 | 3606 |
| 40 | Yes | E. coli | Gentamici | Intracellul | F54 | DiSC3(5) | 23559 | RL1 | 3414 |
| 40 | Yes | E. coli | Gentamici | Intracellul | F124 | Thioflavin T | 21242 | VL2 | 3538 |
| 40 | Yes | E. coli | Gentamici | Intracellul | F101 | Quinacrine | 21540 | VL2 | 3598.5 |
| 40 | Yes | E. coli | Gentamici | Intracellul | F138 | Apigenin | 24856 | VL2 | 3665 |
| 40 | Yes | E. coli | Gentamici | Intracellul | F65 | H2FDA | 23459 | BL1 | 4305 |
| 40 | Yes | E. coli | Gentamici | Intracellul | F142 | Chlorophe | 24475 | BL3 | 4349 |
| 40 | Yes | E. coli | Gentamici | Intracellul | F39 | Congo Red | 27029 | BL3 | 4312 |
| 40 | Yes | E. coli | Gentamici | Intracellul | F116 | Sunset Yel | 24573 | BL2 | 4875 |
| 40 | Yes | E. coli | Gentamici | Intracellul | F10 | Alizarin | 24180 | BL1 | 4019.5 |
| 40 | Yes | E. coli | Gentamici | Intracellul | F84 | Neutral Re | 22765 | BL1 | 3749 |
| 40 | Yes | E. coli | Gentamici | Intracellul | F99 | Pyronin Y | 25228 | BL1 | 3078 |
| 40 | Yes | E. coli | Gentamici | Intracellul | F108 | Rhodamine | 23225 | BL1 | 3177 |
| 40 | Yes | E. coli | Gentamici | Intracellul | F109 | Riboflavin | 23358 | BL1 | 3788.5 |
| 40 | Yes | E. coli | Gentamici | Intracellul | F117 | SYBR Gree | 22936 | BL1 | 133836 |
| 40 | Yes | E. coli | Gentamici | Intracellul | F106 | Rhodamine | 23559 | BL1 | 4418 |
| 40 | Yes | E. coli | Gentamici | Intracellul | F73 | Malachite ( | 24074 | BL1 | 3967.5 |
| 40 | Yes | E. coli | Gentamici | Intracellul | F7 | Acridine or | 20399 | BL1 | 3636 |
| 40 | Yes | E. coli | Gentamici | Intracellul | F192 | 5-Carboxyf | 26153 | BL1 | 12664 |
| 40 | Yes | E. coli | Gentamici | Intracellul | F57 | Eosin Y | 19021 | BL1 | 44452 |
| 40 | Yes | E. coli | Gentamici | Intracellul | F27 | Calcein | 25159 | BL1 | 22330 |
| 40 | Yes | E. coli | Gentamici | Intracellul | F12 | Amaranth | 20519 | BL1 | 710259 |
| 40 | Yes | E. coli | None | None | NA | Autofluore: | 20315 | NA | 6044 |
| 40 | Yes | E. coli | Nitrofurant | Intracellul | F194 | Oxonol | 13608 | RL1 | 5116.5 |
| 40 | Yes | E. coli | Nitrofurant | Intracellul | F11 | Allura Red | 10163 | BL3 | 4167 |
| 40 | Yes | E. coli | Nitrofurant | Intracellul | F54 | DiSC3(5) | 9407 | RL1 | 4105 |
| 40 | Yes | E. coli | Nitrofurant | Intracellul | F124 | Thioflavin T | 11736 | VL2 | 4025.5 |
| 40 | Yes | E. coli | Nitrofurant | Intracellul | F101 | Quinacrine | 9938 | VL2 | 4034 |
| 40 | Yes | E. coli | Nitrofurant | Intracellul | F138 | Apigenin | 10022 | VL2 | 4489 |
| 40 | Yes | E. coli | Nitrofurant | Intracellul | F65 | H2FDA | 10880 | BL1 | 4791.5 |
| 40 | Yes | E. coli | Nitrofurant | Intracellul | F142 | Chlorophe | 9877 | BL3 | 5314 |
| 40 | Yes | E. coli | Nitrofurant | Intracellul | F39 | Congo Red | 10691 | BL3 | 5201 |
| 40 | Yes | E. coli | Nitrofurant | Intracellul | F116 | Sunset Yel | 9855 | BL2 | 5490 |
| 40 | Yes | E. coli | Nitrofurant | Intracellul | F10 | Alizarin | 10601 | BL1 | 5231 |
| 40 | Yes | E. coli | Nitrofurant | Intracellul | F84 | Neutral Re | 9585 | BL1 | 4451 |
| 40 | Yes | E. coli | Nitrofurant | Intracellul | F99 | Pyronin Y | 11077 | BL1 | 3815 |
| 40 | Yes | E. coli | Nitrofurant | Intracellul | F108 | Rhodamine | 9489 | BL1 | 3654 |
| 40 | Yes | E. coli | Nitrofurant | Intracellul | F109 | Riboflavin | 9088 | BL1 | 4734 |
| 40 | Yes | E. coli | Nitrofurant | Intracellul | F117 | SYBR Gree | 10436 | BL1 | 68996.5 |
| 40 | Yes | E. coli | Nitrofurant | Intracellul | F106 | Rhodamine | 9636 | BL1 | 5214 |
| 40 | Yes | E. coli | Nitrofurant | Intracellul | F73 | Malachite ( | 8831 | BL1 | 4195 |
| 40 | Yes | E. coli | Nitrofurant | Intracellul | F7 | Acridine or | 8911 | BL1 | 4158 |
| 40 | Yes | E. coli | Nitrofurant | Intracellul | F192 | 5-Carboxyf | 10612 | BL1 | 14340 |
| 40 | Yes | E. coli | Nitrofurant | Intracellul | F57 | Eosin Y | 9413 | BL1 | 19729 |
| 40 | Yes | E. coli | Nitrofurant | Intracellul | F27 | Calcein | 9135 | BL1 | 24391 |

| | | | | | | | | | |
|---|---|---|---|---|---|---|---|---|---|
| 40 | Yes | E. coli | Nitrofurant | Intracellula | F12 | Amaranth | 8361 | BL1 | 18900 |
| 40 | Yes | E. coli | None | None | NA | Autofluores | 21759 | NA | 6023 |
| 40 | Yes | E. coli | Pentamidi | Wall | F194 | Oxonol | 12475 | RL1 | 5629 |
| 40 | Yes | E. coli | Pentamidi | Wall | F11 | Allura Red | 12128 | BL3 | 4831 |
| 40 | Yes | E. coli | Pentamidi | Wall | F54 | DiSC3(5) | 11192 | RL1 | 4388 |
| 40 | Yes | E. coli | Pentamidi | Wall | F124 | Thioflavin T | 10405 | VL2 | 4497 |
| 40 | Yes | E. coli | Pentamidi | Wall | F101 | Quinacrine | 10517 | VL2 | 4624 |
| 40 | Yes | E. coli | Pentamidi | Wall | F138 | Apigenin | 10736 | VL2 | 4486 |
| 40 | Yes | E. coli | Pentamidi | Wall | F65 | H2FDA | 10563 | BL1 | 5090 |
| 40 | Yes | E. coli | Pentamidi | Wall | F142 | Chlorophe | 11356 | BL3 | 5585.5 |
| 40 | Yes | E. coli | Pentamidi | Wall | F39 | Congo Red | 11579 | BL3 | 5736 |
| 40 | Yes | E. coli | Pentamidi | Wall | F116 | Sunset Yel | 10718 | BL2 | 6729.5 |
| 40 | Yes | E. coli | Pentamidi | Wall | F10 | Alizarin | 11127 | BL1 | 5761 |
| 40 | Yes | E. coli | Pentamidi | Wall | F84 | Neutral Re | 11209 | BL1 | 5095 |
| 40 | Yes | E. coli | Pentamidi | Wall | F99 | Pyronin Y | 11796 | BL1 | 4524.5 |
| 40 | Yes | E. coli | Pentamidi | Wall | F108 | Rhodamine | 11697 | BL1 | 4549 |
| 40 | Yes | E. coli | Pentamidi | Wall | F109 | Riboflavin | 10275 | BL1 | 4676 |
| 40 | Yes | E. coli | Pentamidi | Wall | F117 | SYBR Gree | 10031 | BL1 | 656198 |
| 40 | Yes | E. coli | Pentamidi | Wall | F106 | Rhodamine | 11059 | BL1 | 7934 |
| 40 | Yes | E. coli | Pentamidi | Wall | F73 | Malachite ( | 9597 | BL1 | 6578 |
| 40 | Yes | E. coli | Pentamidi | Wall | F7 | Acridine or | 10480 | BL1 | 6405.5 |
| 40 | Yes | E. coli | Pentamidi | Wall | F192 | 5-Carboxyf | 10200 | BL1 | 15212.5 |
| 40 | Yes | E. coli | Pentamidi | Wall | F57 | Eosin Y | 9970 | BL1 | 20199.5 |
| 40 | Yes | E. coli | Pentamidi | Wall | F27 | Calcein | 10944 | BL1 | 20592 |
| 40 | Yes | E. coli | Pentamidi | Wall | F12 | Amaranth | 9698 | BL1 | 19868 |
| 40 | Yes | E. coli | None | None | NA | Autofluores | 22055 | NA | 7840 |
| 40 | Yes | E. coli | Rifampicin | Intracellula | F194 | Oxonol | 23634 | RL1 | 6134 |
| 40 | Yes | E. coli | Rifampicin | Intracellula | F11 | Allura Red | 21055 | BL3 | 4878 |
| 40 | Yes | E. coli | Rifampicin | Intracellula | F54 | DiSC3(5) | 23214 | RL1 | 4083 |
| 40 | Yes | E. coli | Rifampicin | Intracellula | F124 | Thioflavin T | 23199 | VL2 | 3710 |
| 40 | Yes | E. coli | Rifampicin | Intracellula | F101 | Quinacrine | 22367 | VL2 | 4392 |
| 40 | Yes | E. coli | Rifampicin | Intracellula | F138 | Apigenin | 19399 | VL2 | 4513 |
| 40 | Yes | E. coli | Rifampicin | Intracellula | F65 | H2FDA | 20677 | BL1 | 5010 |
| 40 | Yes | E. coli | Rifampicin | Intracellula | F142 | Chlorophe | 24907 | BL3 | 4618 |
| 40 | Yes | E. coli | Rifampicin | Intracellula | F39 | Congo Red | 20792 | BL3 | 5146.5 |
| 40 | Yes | E. coli | Rifampicin | Intracellula | F116 | Sunset Yel | 22653 | BL2 | 5345 |
| 40 | Yes | E. coli | Rifampicin | Intracellula | F10 | Alizarin | 22426 | BL1 | 4802 |
| 40 | Yes | E. coli | Rifampicin | Intracellula | F84 | Neutral Re | 21427 | BL1 | 4912 |
| 40 | Yes | E. coli | Rifampicin | Intracellula | F99 | Pyronin Y | 24301 | BL1 | 3349 |
| 40 | Yes | E. coli | Rifampicin | Intracellula | F108 | Rhodamine | 21603 | BL1 | 4020 |
| 40 | Yes | E. coli | Rifampicin | Intracellula | F109 | Riboflavin | 19314 | BL1 | 4374 |
| 40 | Yes | E. coli | Rifampicin | Intracellula | F117 | SYBR Gree | 21100 | BL1 | 93036.5 |
| 40 | Yes | E. coli | Rifampicin | Intracellula | F106 | Rhodamine | 22484 | BL1 | 5306 |
| 40 | Yes | E. coli | Rifampicin | Intracellula | F73 | Malachite ( | 22002 | BL1 | 4529.5 |
| 40 | Yes | E. coli | Rifampicin | Intracellula | F7 | Acridine or | 21997 | BL1 | 4214 |
| 40 | Yes | E. coli | Rifampicin | Intracellula | F192 | 5-Carboxyf | 21808 | BL1 | 20713.5 |
| 40 | Yes | E. coli | Rifampicin | Intracellula | F57 | Eosin Y | 20862 | BL1 | 74878 |
| 40 | Yes | E. coli | Rifampicin | Intracellula | F27 | Calcein | 21407 | BL1 | 303749 |

| | | | | | | | | | |
|---|---|---|---|---|---|---|---|---|---|
| 40 | Yes | E. coli | Rifampicin | Intracellula | F12 | Amaranth | 21988 | BL1 | 80441.5 |
| 40 | Yes | E. coli | None | None | NA | Autofluore: | 22539 | NA | 5909 |
| 40 | Yes | E. coli | Trimethopr | Intracellula | F194 | Oxonol | 26188 | RL1 | 4543 |
| 40 | Yes | E. coli | Trimethopr | Intracellula | F11 | Allura Red | 25176 | BL3 | 4134.5 |
| 40 | Yes | E. coli | Trimethopr | Intracellula | F54 | DiSC3(5) | 24733 | RL1 | 3510 |
| 40 | Yes | E. coli | Trimethopr | Intracellula | F124 | Thioflavin T | 24542 | VL2 | 3619.5 |
| 40 | Yes | E. coli | Trimethopr | Intracellula | F101 | Quinacrine | 23379 | VL2 | 4066 |
| 40 | Yes | E. coli | Trimethopr | Intracellula | F138 | Apigenin | 24799 | VL2 | 3637 |
| 40 | Yes | E. coli | Trimethopr | Intracellula | F65 | H2FDA | 22748 | BL1 | 4562 |
| 40 | Yes | E. coli | Trimethopr | Intracellula | F142 | Chlorophe | 27301 | BL3 | 5007 |
| 40 | Yes | E. coli | Trimethopr | Intracellula | F39 | Congo Red | 25884 | BL3 | 5090 |
| 40 | Yes | E. coli | Trimethopr | Intracellula | F116 | Sunset Yel | 24376 | BL2 | 5131 |
| 40 | Yes | E. coli | Trimethopr | Intracellula | F10 | Alizarin | 21088 | BL1 | 5316 |
| 40 | Yes | E. coli | Trimethopr | Intracellula | F84 | Neutral Re | 22410 | BL1 | 4571.5 |
| 40 | Yes | E. coli | Trimethopr | Intracellula | F99 | Pyronin Y | 21252 | BL1 | 3422.5 |
| 40 | Yes | E. coli | Trimethopr | Intracellula | F108 | Rhodamine | 19993 | BL1 | 3413 |
| 40 | Yes | E. coli | Trimethopr | Intracellula | F109 | Riboflavin | 18012 | BL1 | 4015.5 |
| 40 | Yes | E. coli | Trimethopr | Intracellula | F117 | SYBR Gree | 18145 | BL1 | 132123 |
| 40 | Yes | E. coli | Trimethopr | Intracellula | F106 | Rhodamine | 21223 | BL1 | 4846 |
| 40 | Yes | E. coli | Trimethopr | Intracellula | F73 | Malachite ( | 20968 | BL1 | 3668 |
| 40 | Yes | E. coli | Trimethopr | Intracellula | F7 | Acridine or | 19771 | BL1 | 3924 |
| 40 | Yes | E. coli | Trimethopr | Intracellula | F192 | 5-Carboxyf | 19133 | BL1 | 11701 |
| 40 | Yes | E. coli | Trimethopr | Intracellula | F57 | Eosin Y | 19164 | BL1 | 17087 |
| 40 | Yes | E. coli | Trimethopr | Intracellula | F27 | Calcein | 22580 | BL1 | 18745.5 |
| 40 | Yes | E. coli | Trimethopr | Intracellula | F12 | Amaranth | 19694 | BL1 | 21960.5 |
| 40 | Yes | E. coli | None | None | NA | Autofluore: | 20752 | NA | 856 |
| 40 | Yes | E. coli | None | None | F194 | Oxonol | 28134 | RL1 | 1495 |
| 40 | Yes | E. coli | None | None | F11 | Allura Red | 24863 | BL3 | 1889 |
| 40 | Yes | E. coli | None | None | F54 | DiSC3(5) | 21882 | RL1 | 2465.5 |
| 40 | Yes | E. coli | None | None | F124 | Thioflavin T | 25495 | VL2 | 2520 |
| 40 | Yes | E. coli | None | None | F101 | Quinacrine | 23924 | VL2 | 2913 |
| 40 | Yes | E. coli | None | None | F138 | Apigenin | 21287 | VL2 | 3084 |
| 40 | Yes | E. coli | None | None | F65 | H2FDA | 22651 | BL1 | 4072 |
| 40 | Yes | E. coli | None | None | F142 | Chlorophe | 21198 | BL3 | 4388 |
| 40 | Yes | E. coli | None | None | F39 | Congo Red | 27130 | BL3 | 4662 |
| 40 | Yes | E. coli | None | None | F116 | Sunset Yel | 24493 | BL2 | 4542 |
| 40 | Yes | E. coli | None | None | F10 | Alizarin | 22793 | BL1 | 4937 |
| 40 | Yes | E. coli | None | None | F84 | Neutral Re | 21867 | BL1 | 4301 |
| 40 | Yes | E. coli | None | None | F99 | Pyronin Y | 21854 | BL1 | 3494.5 |
| 40 | Yes | E. coli | None | None | F108 | Rhodamine | 21674 | BL1 | 3543 |
| 40 | Yes | E. coli | None | None | F109 | Riboflavin | 23430 | BL1 | 3527 |
| 40 | Yes | E. coli | None | None | F117 | SYBR Gree | 23840 | BL1 | 98775 |
| 40 | Yes | E. coli | None | None | F106 | Rhodamine | 23330 | BL1 | 4432.5 |
| 40 | Yes | E. coli | None | None | F73 | Malachite ( | 23601 | BL1 | 3679 |
| 40 | Yes | E. coli | None | None | F7 | Acridine or | 20585 | BL1 | 3936 |
| 40 | Yes | E. coli | None | None | F192 | 5-Carboxyf | 22274 | BL1 | 267081 |
| 40 | Yes | E. coli | None | None | F57 | Eosin Y | 19119 | BL1 | 773739 |
| 40 | Yes | E. coli | None | None | F27 | Calcein | 22387 | BL1 | 1012990 |

| | | | | | | | | | |
|---|---|---|---|---|---|---|---|---|---|
| 40 | Yes | E. coli | None | None | F12 | Amaranth | 23181 | BL1 | 536024 |
| 40 | Yes | E. coli | None | None | NA | Autofluores | 23346 | NA | 5498.5 |
| 40 | Yes | E. coli | Ampicillin | Wall | F194 | Oxonol | 25880 | RL1 | 4184 |
| 40 | Yes | E. coli | Ampicillin | Wall | F11 | Allura Red | 26529 | BL3 | 3572 |
| 40 | Yes | E. coli | Ampicillin | Wall | F54 | DiSC3(5) | 23622 | RL1 | 3405 |
| 40 | Yes | E. coli | Ampicillin | Wall | F124 | Thioflavin T | 21988 | VL2 | 3358 |
| 40 | Yes | E. coli | Ampicillin | Wall | F101 | Quinacrine | 19589 | VL2 | 3488 |
| 40 | Yes | E. coli | Ampicillin | Wall | F138 | Apigenin | 21095 | VL2 | 3516 |
| 40 | Yes | E. coli | Ampicillin | Wall | F65 | H2FDA | 20898 | BL1 | 4309 |
| 40 | Yes | E. coli | Ampicillin | Wall | F142 | Chlorophe | 23961 | BL3 | 4425 |
| 40 | Yes | E. coli | Ampicillin | Wall | F39 | Congo Red | 23090 | BL3 | 4703.5 |
| 40 | Yes | E. coli | Ampicillin | Wall | F116 | Sunset Yel | 22505 | BL2 | 4503 |
| 40 | Yes | E. coli | Ampicillin | Wall | F10 | Alizarin | 23711 | BL1 | 4546 |
| 40 | Yes | E. coli | Ampicillin | Wall | F84 | Neutral Re | 20331 | BL1 | 3884 |
| 40 | Yes | E. coli | Ampicillin | Wall | F99 | Pyronin Y | 21123 | BL1 | 3525 |
| 40 | Yes | E. coli | Ampicillin | Wall | F108 | Rhodamine | 20268 | BL1 | 3743 |
| 40 | Yes | E. coli | Ampicillin | Wall | F109 | Riboflavin | 22303 | BL1 | 3496 |
| 40 | Yes | E. coli | Ampicillin | Wall | F117 | SYBR Gree | 23630 | BL1 | 122596.5 |
| 40 | Yes | E. coli | Ampicillin | Wall | F106 | Rhodamine | 22277 | BL1 | 4449 |
| 40 | Yes | E. coli | Ampicillin | Wall | F73 | Malachite | 21700 | BL1 | 3914 |
| 40 | Yes | E. coli | Ampicillin | Wall | F7 | Acridine or | 19764 | BL1 | 4186.5 |
| 40 | Yes | E. coli | Ampicillin | Wall | F192 | 5-Carboxyf | 21950 | BL1 | 15257 |
| 40 | Yes | E. coli | Ampicillin | Wall | F57 | Eosin Y | 20650 | BL1 | 348769 |
| 40 | Yes | E. coli | Ampicillin | Wall | F27 | Calcein | 22158 | BL1 | 110735.5 |
| 40 | Yes | E. coli | Ampicillin | Wall | F12 | Amaranth | 23087 | BL1 | 106496 |
| 40 | Yes | E. coli | None | None | NA | Autofluores | 23587 | NA | 6048 |
| 40 | Yes | E. coli | Bacitracin | Wall | F194 | Oxonol | 27268 | RL1 | 4852 |
| 40 | Yes | E. coli | Bacitracin | Wall | F11 | Allura Red | 19076 | BL3 | 5588 |
| 40 | Yes | E. coli | Bacitracin | Wall | F54 | DiSC3(5) | 24846 | RL1 | 3652 |
| 40 | Yes | E. coli | Bacitracin | Wall | F124 | Thioflavin T | 25139 | VL2 | 3358 |
| 40 | Yes | E. coli | Bacitracin | Wall | F101 | Quinacrine | 22682 | VL2 | 3533 |
| 40 | Yes | E. coli | Bacitracin | Wall | F138 | Apigenin | 21520 | VL2 | 3852 |
| 40 | Yes | E. coli | Bacitracin | Wall | F65 | H2FDA | 19393 | BL1 | 4894 |
| 40 | Yes | E. coli | Bacitracin | Wall | F142 | Chlorophe | 26358 | BL3 | 4861 |
| 40 | Yes | E. coli | Bacitracin | Wall | F39 | Congo Red | 25916 | BL3 | 5109.5 |
| 40 | Yes | E. coli | Bacitracin | Wall | F116 | Sunset Yel | 24736 | BL2 | 4761 |
| 40 | Yes | E. coli | Bacitracin | Wall | F10 | Alizarin | 22297 | BL1 | 5345 |
| 40 | Yes | E. coli | Bacitracin | Wall | F84 | Neutral Re | 21008 | BL1 | 5073 |
| 40 | Yes | E. coli | Bacitracin | Wall | F99 | Pyronin Y | 21095 | BL1 | 4594 |
| 40 | Yes | E. coli | Bacitracin | Wall | F108 | Rhodamine | 20850 | BL1 | 4688 |
| 40 | Yes | E. coli | Bacitracin | Wall | F109 | Riboflavin | 22105 | BL1 | 4387 |
| 40 | Yes | E. coli | Bacitracin | Wall | F117 | SYBR Gree | 24899 | BL1 | 129772 |
| 40 | Yes | E. coli | Bacitracin | Wall | F106 | Rhodamine | 21483 | BL1 | 6952 |
| 40 | Yes | E. coli | Bacitracin | Wall | F73 | Malachite | 23128 | BL1 | 4288 |
| 40 | Yes | E. coli | Bacitracin | Wall | F7 | Acridine or | 22661 | BL1 | 4122 |
| 40 | Yes | E. coli | Bacitracin | Wall | F192 | 5-Carboxyf | 23156 | BL1 | 40659.5 |
| 40 | Yes | E. coli | Bacitracin | Wall | F57 | Eosin Y | 23269 | BL1 | 44457 |
| 40 | Yes | E. coli | Bacitracin | Wall | F27 | Calcein | 24539 | BL1 | 596872 |

| | | | | | | | | | |
|---|---|---|---|---|---|---|---|---|---|
| 40 | Yes | E. coli | Bacitracin | Wall | F12 | Amaranth | 21745 | BL1 | 79083 |
| 40 | Yes | E. coli | None | None | NA | Autofluore: | 24273 | NA | 6197 |
| 40 | Yes | E. coli | Cefotaxim | Wall | F194 | Oxonol | 27320 | RL1 | 4780 |
| 40 | Yes | E. coli | Cefotaxim | Wall | F11 | Allura Red | 24452 | BL3 | 4080.5 |
| 40 | Yes | E. coli | Cefotaxim | Wall | F54 | DiSC3(5) | 20294 | RL1 | 4185 |
| 40 | Yes | E. coli | Cefotaxim | Wall | F124 | Thioflavin T | 21498 | VL2 | 3729 |
| 40 | Yes | E. coli | Cefotaxim | Wall | F101 | Quinacrine | 19491 | VL2 | 4048 |
| 40 | Yes | E. coli | Cefotaxim | Wall | F138 | Apigenin | 18954 | VL2 | 3974.5 |
| 40 | Yes | E. coli | Cefotaxim | Wall | F65 | H2FDA | 19707 | BL1 | 4342 |
| 40 | Yes | E. coli | Cefotaxim | Wall | F142 | Chlorophe | 17740 | BL3 | 5158 |
| 40 | Yes | E. coli | Cefotaxim | Wall | F39 | Congo Red | 18879 | BL3 | 5424 |
| 40 | Yes | E. coli | Cefotaxim | Wall | F116 | Sunset Yel | 18207 | BL2 | 5498 |
| 40 | Yes | E. coli | Cefotaxim | Wall | F10 | Alizarin | 22696 | BL1 | 4484.5 |
| 40 | Yes | E. coli | Cefotaxim | Wall | F84 | Neutral Re | 20345 | BL1 | 4064 |
| 40 | Yes | E. coli | Cefotaxim | Wall | F99 | Pyronin Y | 22473 | BL1 | 3657 |
| 40 | Yes | E. coli | Cefotaxim | Wall | F108 | Rhodamine | 19376 | BL1 | 3570 |
| 40 | Yes | E. coli | Cefotaxim | Wall | F109 | Riboflavin | 19334 | BL1 | 4021.5 |
| 40 | Yes | E. coli | Cefotaxim | Wall | F117 | SYBR Gree | 20434 | BL1 | 152192.5 |
| 40 | Yes | E. coli | Cefotaxim | Wall | F106 | Rhodamine | 23091 | BL1 | 4454 |
| 40 | Yes | E. coli | Cefotaxim | Wall | F73 | Malachite ( | 16814 | BL1 | 5004 |
| 40 | Yes | E. coli | Cefotaxim | Wall | F7 | Acridine or | 20321 | BL1 | 4285 |
| 40 | Yes | E. coli | Cefotaxim | Wall | F192 | 5-Carboxyf | 22997 | BL1 | 13436 |
| 40 | Yes | E. coli | Cefotaxim | Wall | F57 | Eosin Y | 19643 | BL1 | 34415 |
| 40 | Yes | E. coli | Cefotaxim | Wall | F27 | Calcein | 21108 | BL1 | 34416 |
| 40 | Yes | E. coli | Cefotaxim | Wall | F12 | Amaranth | 20876 | BL1 | 990237.5 |
| 40 | Yes | E. coli | None | None | NA | Autofluore: | 23685 | NA | 6571 |
| 40 | Yes | E. coli | Ciprofloxa | Intracellula | F194 | Oxonol | 15374 | RL1 | 6465 |
| 40 | Yes | E. coli | Ciprofloxa | Intracellula | F11 | Allura Red | 18355 | BL3 | 4999 |
| 40 | Yes | E. coli | Ciprofloxa | Intracellula | F54 | DiSC3(5) | 16656 | RL1 | 4472 |
| 40 | Yes | E. coli | Ciprofloxa | Intracellula | F124 | Thioflavin T | 14491 | VL2 | 5067 |
| 40 | Yes | E. coli | Ciprofloxa | Intracellula | F101 | Quinacrine | 13497 | VL2 | 5042 |
| 40 | Yes | E. coli | Ciprofloxa | Intracellula | F138 | Apigenin | 14476 | VL2 | 4263 |
| 40 | Yes | E. coli | Ciprofloxa | Intracellula | F65 | H2FDA | 14036 | BL1 | 5667 |
| 40 | Yes | E. coli | Ciprofloxa | Intracellula | F142 | Chlorophe | 16993 | BL3 | 6235 |
| 40 | Yes | E. coli | Ciprofloxa | Intracellula | F39 | Congo Red | 16934 | BL3 | 6735 |
| 40 | Yes | E. coli | Ciprofloxa | Intracellula | F116 | Sunset Yel | 15488 | BL2 | 6527.5 |
| 40 | Yes | E. coli | Ciprofloxa | Intracellula | F10 | Alizarin | 15266 | BL1 | 4707.5 |
| 40 | Yes | E. coli | Ciprofloxa | Intracellula | F84 | Neutral Re | 16616 | BL1 | 4745 |
| 40 | Yes | E. coli | Ciprofloxa | Intracellula | F99 | Pyronin Y | 13863 | BL1 | 5094 |
| 40 | Yes | E. coli | Ciprofloxa | Intracellula | F108 | Rhodamine | 16991 | BL1 | 3784 |
| 40 | Yes | E. coli | Ciprofloxa | Intracellula | F109 | Riboflavin | 17497 | BL1 | 4645 |
| 40 | Yes | E. coli | Ciprofloxa | Intracellula | F117 | SYBR Gree | 16749 | BL1 | 170936 |
| 40 | Yes | E. coli | Ciprofloxa | Intracellula | F106 | Rhodamine | 17756 | BL1 | 5477 |
| 40 | Yes | E. coli | Ciprofloxa | Intracellula | F73 | Malachite ( | 14761 | BL1 | 4769 |
| 40 | Yes | E. coli | Ciprofloxa | Intracellula | F7 | Acridine or | 13736 | BL1 | 4745 |
| 40 | Yes | E. coli | Ciprofloxa | Intracellula | F192 | 5-Carboxyf | 14128 | BL1 | 17567 |
| 40 | Yes | E. coli | Ciprofloxa | Intracellula | F57 | Eosin Y | 14154 | BL1 | 26002.5 |
| 40 | Yes | E. coli | Ciprofloxa | Intracellula | F27 | Calcein | 14496 | BL1 | 29566 |

| | | | | | | | | | |
|---|---|---|---|---|---|---|---|---|---|
| 40 | Yes | E. coli | Ciprofloxa | Intracellula | F12 | Amaranth | 17024 | BL1 | 40972.5 |
| 40 | Yes | E. coli | None | None | NA | Autofluore: | 25300 | NA | 5620 |
| 40 | Yes | E. coli | Fosfomyci | Wall | F194 | Oxonol | 29917 | RL1 | 4475 |
| 40 | Yes | E. coli | Fosfomyci | Wall | F11 | Allura Red | 32545 | BL3 | 3670 |
| 40 | Yes | E. coli | Fosfomyci | Wall | F54 | DiSC3(5) | 28732 | RL1 | 3816 |
| 40 | Yes | E. coli | Fosfomyci | Wall | F124 | Thioflavin T | 28150 | VL2 | 3451 |
| 40 | Yes | E. coli | Fosfomyci | Wall | F101 | Quinacrine | 26553 | VL2 | 3749 |
| 40 | Yes | E. coli | Fosfomyci | Wall | F138 | Apigenin | 25140 | VL2 | 4160 |
| 40 | Yes | E. coli | Fosfomyci | Wall | F65 | H2FDA | 23554 | BL1 | 5288 |
| 40 | Yes | E. coli | Fosfomyci | Wall | F142 | Chlorophe | 23679 | BL3 | 5062 |
| 40 | Yes | E. coli | Fosfomyci | Wall | F39 | Congo Red | 30029 | BL3 | 4990 |
| 40 | Yes | E. coli | Fosfomyci | Wall | F116 | Sunset Yel | 26631 | BL2 | 5134 |
| 40 | Yes | E. coli | Fosfomyci | Wall | F10 | Alizarin | 24572 | BL1 | 5713.5 |
| 40 | Yes | E. coli | Fosfomyci | Wall | F84 | Neutral Re | 24003 | BL1 | 5062 |
| 40 | Yes | E. coli | Fosfomyci | Wall | F99 | Pyronin Y | 23691 | BL1 | 4385 |
| 40 | Yes | E. coli | Fosfomyci | Wall | F108 | Rhodamine | 23066 | BL1 | 5410.5 |
| 40 | Yes | E. coli | Fosfomyci | Wall | F109 | Riboflavin | 23557 | BL1 | 4511 |
| 40 | Yes | E. coli | Fosfomyci | Wall | F117 | SYBR Gree | 24123 | BL1 | 136652 |
| 40 | Yes | E. coli | Fosfomyci | Wall | F106 | Rhodamine | 23445 | BL1 | 5529 |
| 40 | Yes | E. coli | Fosfomyci | Wall | F73 | Malachite | 23683 | BL1 | 5150 |
| 40 | Yes | E. coli | Fosfomyci | Wall | F7 | Acridine or | 23259 | BL1 | 4842 |
| 40 | Yes | E. coli | Fosfomyci | Wall | F192 | 5-Carboxyf | 21577 | BL1 | 419029 |
| 40 | Yes | E. coli | Fosfomyci | Wall | F57 | Eosin Y | 22580 | BL1 | 821114 |
| 40 | Yes | E. coli | Fosfomyci | Wall | F27 | Calcein | 22174 | BL1 | 958479.5 |
| 40 | Yes | E. coli | Fosfomyci | Wall | F12 | Amaranth | 22288 | BL1 | 1098936 |
| 40 | Yes | E. coli | None | None | NA | Autofluore: | 26348 | NA | 6026 |
| 40 | Yes | E. coli | Nalidixic a | Intracellula | F194 | Oxonol | 21947 | RL1 | 4899 |
| 40 | Yes | E. coli | Nalidixic a | Intracellula | F11 | Allura Red | 26385 | BL3 | 4136 |
| 40 | Yes | E. coli | Nalidixic a | Intracellula | F54 | DiSC3(5) | 22764 | RL1 | 3960.5 |
| 40 | Yes | E. coli | Nalidixic a | Intracellula | F124 | Thioflavin T | 22511 | VL2 | 3922 |
| 40 | Yes | E. coli | Nalidixic a | Intracellula | F101 | Quinacrine | 18456 | VL2 | 4434.5 |
| 40 | Yes | E. coli | Nalidixic a | Intracellula | F138 | Apigenin | 21282 | VL2 | 4187 |
| 40 | Yes | E. coli | Nalidixic a | Intracellula | F65 | H2FDA | 20351 | BL1 | 4818 |
| 40 | Yes | E. coli | Nalidixic a | Intracellula | F142 | Chlorophe | 22664 | BL3 | 4992.5 |
| 40 | Yes | E. coli | Nalidixic a | Intracellula | F39 | Congo Red | 22621 | BL3 | 5145 |
| 40 | Yes | E. coli | Nalidixic a | Intracellula | F116 | Sunset Yel | 24165 | BL2 | 4937 |
| 40 | Yes | E. coli | Nalidixic a | Intracellula | F10 | Alizarin | 19128 | BL1 | 5358.5 |
| 40 | Yes | E. coli | Nalidixic a | Intracellula | F84 | Neutral Re | 18221 | BL1 | 4614 |
| 40 | Yes | E. coli | Nalidixic a | Intracellula | F99 | Pyronin Y | 16689 | BL1 | 4436 |
| 40 | Yes | E. coli | Nalidixic a | Intracellula | F108 | Rhodamine | 17792 | BL1 | 4411.5 |
| 40 | Yes | E. coli | Nalidixic a | Intracellula | F109 | Riboflavin | 21833 | BL1 | 4192 |
| 40 | Yes | E. coli | Nalidixic a | Intracellula | F117 | SYBR Gree | 21057 | BL1 | 129966 |
| 40 | Yes | E. coli | Nalidixic a | Intracellula | F106 | Rhodamine | 22358 | BL1 | 5006 |
| 40 | Yes | E. coli | Nalidixic a | Intracellula | F73 | Malachite | 21193 | BL1 | 4577 |
| 40 | Yes | E. coli | Nalidixic a | Intracellula | F7 | Acridine or | 21342 | BL1 | 4785 |
| 40 | Yes | E. coli | Nalidixic a | Intracellula | F192 | 5-Carboxyf | 18772 | BL1 | 18075 |
| 40 | Yes | E. coli | Nalidixic a | Intracellula | F57 | Eosin Y | 19046 | BL1 | 445558.5 |
| 40 | Yes | E. coli | Nalidixic a | Intracellula | F27 | Calcein | 19448 | BL1 | 42455.5 |

| | | | | | | | | | |
|---|---|---|---|---|---|---|---|---|---|
| 40 | Yes | E. coli | Nalidixic a | Intracellul | F12 | Amaranth | 18277 | BL1 | 292699 |
| 40 | Yes | E. coli | None | None | NA | Autofluore | 27246 | NA | 5726.5 |
| 40 | Yes | E. coli | Ofloxacin | Intracellul | F194 | Oxonol | 20023 | RL1 | 5580 |
| 40 | Yes | E. coli | Ofloxacin | Intracellul | F11 | Allura Red | 17337 | BL3 | 5117 |
| 40 | Yes | E. coli | Ofloxacin | Intracellul | F54 | DiSC3(5) | 16935 | RL1 | 4868 |
| 40 | Yes | E. coli | Ofloxacin | Intracellul | F124 | Thioflavin T | 16238 | VL2 | 4768.5 |
| 40 | Yes | E. coli | Ofloxacin | Intracellul | F101 | Quinacrine | 14222 | VL2 | 5009 |
| 40 | Yes | E. coli | Ofloxacin | Intracellul | F138 | Apigenin | 15751 | VL2 | 5034 |
| 40 | Yes | E. coli | Ofloxacin | Intracellul | F65 | H2FDA | 15905 | BL1 | 5442 |
| 40 | Yes | E. coli | Ofloxacin | Intracellul | F142 | Chlorophe | 15079 | BL3 | 6237 |
| 40 | Yes | E. coli | Ofloxacin | Intracellul | F39 | Congo Red | 16684 | BL3 | 6499.5 |
| 40 | Yes | E. coli | Ofloxacin | Intracellul | F116 | Sunset Yel | 15280 | BL2 | 6348.5 |
| 40 | Yes | E. coli | Ofloxacin | Intracellul | F10 | Alizarin | 14123 | BL1 | 6130 |
| 40 | Yes | E. coli | Ofloxacin | Intracellul | F84 | Neutral Re | 15080 | BL1 | 5626 |
| 40 | Yes | E. coli | Ofloxacin | Intracellul | F99 | Pyronin Y | 16564 | BL1 | 4817 |
| 40 | Yes | E. coli | Ofloxacin | Intracellul | F108 | Rhodamine | 13584 | BL1 | 5365.5 |
| 40 | Yes | E. coli | Ofloxacin | Intracellul | F109 | Riboflavin | 14195 | BL1 | 5044 |
| 40 | Yes | E. coli | Ofloxacin | Intracellul | F117 | SYBR Gree | 14219 | BL1 | 206243 |
| 40 | Yes | E. coli | Ofloxacin | Intracellul | F106 | Rhodamine | 15659 | BL1 | 6158 |
| 40 | Yes | E. coli | Ofloxacin | Intracellul | F73 | Malachite ( | 14652 | BL1 | 5389.5 |
| 40 | Yes | E. coli | Ofloxacin | Intracellul | F7 | Acridine or | 14106 | BL1 | 5898 |
| 40 | Yes | E. coli | Ofloxacin | Intracellul | F192 | 5-Carboxyf | 15161 | BL1 | 17648 |
| 40 | Yes | E. coli | Ofloxacin | Intracellul | F57 | Eosin Y | 12855 | BL1 | 23767 |
| 40 | Yes | E. coli | Ofloxacin | Intracellul | F27 | Calcein | 13194 | BL1 | 27174.5 |
| 40 | Yes | E. coli | Ofloxacin | Intracellul | F12 | Amaranth | 14947 | BL1 | 35770 |
| 40 | Yes | E. coli | None | None | NA | Autofluore | 27747 | NA | 5762 |
| 40 | Yes | E. coli | Polymyxin | Wall | F194 | Oxonol | 25908 | RL1 | 5851 |
| 40 | Yes | E. coli | Polymyxin | Wall | F11 | Allura Red | 24670 | BL3 | 4752.5 |
| 40 | Yes | E. coli | Polymyxin | Wall | F54 | DiSC3(5) | 24155 | RL1 | 4541 |
| 40 | Yes | E. coli | Polymyxin | Wall | F124 | Thioflavin T | 22594 | VL2 | 4284 |
| 40 | Yes | E. coli | Polymyxin | Wall | F101 | Quinacrine | 22362 | VL2 | 4508 |
| 40 | Yes | E. coli | Polymyxin | Wall | F138 | Apigenin | 20622 | VL2 | 4482.5 |
| 40 | Yes | E. coli | Polymyxin | Wall | F65 | H2FDA | 20365 | BL1 | 5498 |
| 40 | Yes | E. coli | Polymyxin | Wall | F142 | Chlorophe | 22017 | BL3 | 5756 |
| 40 | Yes | E. coli | Polymyxin | Wall | F39 | Congo Red | 23295 | BL3 | 5543 |
| 40 | Yes | E. coli | Polymyxin | Wall | F116 | Sunset Yel | 22414 | BL2 | 5615 |
| 40 | Yes | E. coli | Polymyxin | Wall | F10 | Alizarin | 21166 | BL1 | 5690 |
| 40 | Yes | E. coli | Polymyxin | Wall | F84 | Neutral Re | 20169 | BL1 | 4892 |
| 40 | Yes | E. coli | Polymyxin | Wall | F99 | Pyronin Y | 20631 | BL1 | 4601 |
| 40 | Yes | E. coli | Polymyxin | Wall | F108 | Rhodamine | 18988 | BL1 | 5120.5 |
| 40 | Yes | E. coli | Polymyxin | Wall | F109 | Riboflavin | 20258 | BL1 | 4964.5 |
| 40 | Yes | E. coli | Polymyxin | Wall | F117 | SYBR Gree | 21607 | BL1 | 201381 |
| 40 | Yes | E. coli | Polymyxin | Wall | F106 | Rhodamine | 20476 | BL1 | 5777 |
| 40 | Yes | E. coli | Polymyxin | Wall | F73 | Malachite ( | 21496 | BL1 | 5233 |
| 40 | Yes | E. coli | Polymyxin | Wall | F7 | Acridine or | 20308 | BL1 | 5404 |
| 40 | Yes | E. coli | Polymyxin | Wall | F192 | 5-Carboxyf | 18781 | BL1 | 28318 |
| 40 | Yes | E. coli | Polymyxin | Wall | F57 | Eosin Y | 19224 | BL1 | 81984 |
| 40 | Yes | E. coli | Polymyxin | Wall | F27 | Calcein | 20351 | BL1 | 270344 |

| | | | | | | | | | |
|---|---|---|---|---|---|---|---|---|---|
| 40 | Yes | E. coli | Polymyxin | Wall | F12 | Amaranth | 18770 | BL1 | 626132 |
| 40 | Yes | E. coli | None | None | NA | Autofluores | 26465 | NA | 5774 |
| 40 | Yes | E. coli | Tetracyclin | Intracellula | F194 | Oxonol | 29234 | RL1 | 4341.5 |
| 40 | Yes | E. coli | Tetracyclin | Intracellula | F11 | Allura Red | 28709 | BL3 | 3942 |
| 40 | Yes | E. coli | Tetracyclin | Intracellula | F54 | DiSC3(5) | 26454 | RL1 | 4044 |
| 40 | Yes | E. coli | Tetracyclin | Intracellula | F124 | Thioflavin T | 25775 | VL2 | 3846 |
| 40 | Yes | E. coli | Tetracyclin | Intracellula | F101 | Quinacrine | 23425 | VL2 | 4097 |
| 40 | Yes | E. coli | Tetracyclin | Intracellula | F138 | Apigenin | 22267 | VL2 | 4772 |
| 40 | Yes | E. coli | Tetracyclin | Intracellula | F65 | H2FDA | 22668 | BL1 | 5218.5 |
| 40 | Yes | E. coli | Tetracyclin | Intracellula | F142 | Chlorophe | 21624 | BL3 | 5934 |
| 40 | Yes | E. coli | Tetracyclin | Intracellula | F39 | Congo Red | 26085 | BL3 | 5151 |
| 40 | Yes | E. coli | Tetracyclin | Intracellula | F116 | Sunset Yel | 25285 | BL2 | 5022 |
| 40 | Yes | E. coli | Tetracyclin | Intracellula | F10 | Alizarin | 23988 | BL1 | 5085 |
| 40 | Yes | E. coli | Tetracyclin | Intracellula | F84 | Neutral Re | 21815 | BL1 | 4937 |
| 40 | Yes | E. coli | Tetracyclin | Intracellula | F99 | Pyronin Y | 21279 | BL1 | 4308 |
| 40 | Yes | E. coli | Tetracyclin | Intracellula | F108 | Rhodamine | 20956 | BL1 | 4568.5 |
| 40 | Yes | E. coli | Tetracyclin | Intracellula | F109 | Riboflavin | 22750 | BL1 | 4176 |
| 40 | Yes | E. coli | Tetracyclin | Intracellula | F117 | SYBR Gree | 22399 | BL1 | 110657 |
| 40 | Yes | E. coli | Tetracyclin | Intracellula | F106 | Rhodamine | 23141 | BL1 | 5393 |
| 40 | Yes | E. coli | Tetracyclin | Intracellula | F73 | Malachite ( | 24006 | BL1 | 5270 |
| 40 | Yes | E. coli | Tetracyclin | Intracellula | F7 | Acridine or | 22366 | BL1 | 5109 |
| 40 | Yes | E. coli | Tetracyclin | Intracellula | F192 | 5-Carboxyf | 21333 | BL1 | 184689 |
| 40 | Yes | E. coli | Tetracyclin | Intracellula | F57 | Eosin Y | 20082 | BL1 | 885532.5 |
| 40 | Yes | E. coli | Tetracyclin | Intracellula | F27 | Calcein | 20008 | BL1 | 1006448 |
| 40 | Yes | E. coli | Tetracyclin | Intracellula | F12 | Amaranth | 20593 | BL1 | 1126370 |
| 40 | Yes | E. coli | None | None | NA | Autofluores | 27709 | NA | 5921 |
| 40 | Yes | E. coli | Vancomyc | Wall | F194 | Oxonol | 31408 | RL1 | 4487 |
| 40 | Yes | E. coli | Vancomyc | Wall | F11 | Allura Red | 30403 | BL3 | 4067 |
| 40 | Yes | E. coli | Vancomyc | Wall | F54 | DiSC3(5) | 27115 | RL1 | 5157 |
| 40 | Yes | E. coli | Vancomyc | Wall | F124 | Thioflavin T | 23394 | VL2 | 5267 |
| 40 | Yes | E. coli | Vancomyc | Wall | F101 | Quinacrine | 25882 | VL2 | 4485 |
| 40 | Yes | E. coli | Vancomyc | Wall | F138 | Apigenin | 26412 | VL2 | 4241 |
| 40 | Yes | E. coli | Vancomyc | Wall | F65 | H2FDA | 23898 | BL1 | 5841 |
| 40 | Yes | E. coli | Vancomyc | Wall | F142 | Chlorophe | 23202 | BL3 | 10973 |
| 40 | Yes | E. coli | Vancomyc | Wall | F39 | Congo Red | 23781 | BL3 | 9688 |
| 40 | Yes | E. coli | Vancomyc | Wall | F116 | Sunset Yel | 27639 | BL2 | 5528 |
| 40 | Yes | E. coli | Vancomyc | Wall | F10 | Alizarin | 27885 | BL1 | 5066 |
| 40 | Yes | E. coli | Vancomyc | Wall | F84 | Neutral Re | 19754 | BL1 | 10005.5 |
| 40 | Yes | E. coli | Vancomyc | Wall | F99 | Pyronin Y | 21392 | BL1 | 7635.5 |
| 40 | Yes | E. coli | Vancomyc | Wall | F108 | Rhodamine | 21286 | BL1 | 7584.5 |
| 40 | Yes | E. coli | Vancomyc | Wall | F109 | Riboflavin | 19324 | BL1 | 7095 |
| 40 | Yes | E. coli | Vancomyc | Wall | F117 | SYBR Gree | 21809 | BL1 | 161644 |
| 40 | Yes | E. coli | Vancomyc | Wall | F106 | Rhodamine | 21373 | BL1 | 9993 |
| 40 | Yes | E. coli | Vancomyc | Wall | F73 | Malachite ( | 20698 | BL1 | 8250.5 |
| 40 | Yes | E. coli | Vancomyc | Wall | F7 | Acridine or | 18009 | BL1 | 8350 |
| 40 | Yes | E. coli | Vancomyc | Wall | F192 | 5-Carboxyf | 18868 | BL1 | 65761 |
| 40 | Yes | E. coli | Vancomyc | Wall | F57 | Eosin Y | 19134 | BL1 | 102932 |
| 40 | Yes | E. coli | Vancomyc | Wall | F27 | Calcein | 22207 | BL1 | 114136 |

| | | | | | | | | | |
|---|---|---|---|---|---|---|---|---|---|
| 40 | Yes | E. coli | Vancomyc | Wall | F12 | Amaranth | 22674 | BL1 | 73410 |
| 40 | Yes | E. coli | None | None | NA | Autofluore: | 13672 | NA | 863 |
| 40 | Yes | E. coli | None | None | F194 | Oxonol | 18573 | RL1 | 1480 |
| 40 | Yes | E. coli | None | None | F11 | Allura Red | 20562 | BL3 | 2271.5 |
| 40 | Yes | E. coli | None | None | F54 | DiSC3(5) | 18847 | RL1 | 2558 |
| 40 | Yes | E. coli | None | None | F124 | Thioflavin T | 20424 | VL2 | 2576 |
| 40 | Yes | E. coli | None | None | F101 | Quinacrine | 18997 | VL2 | 2875 |
| 40 | Yes | E. coli | None | None | F138 | Apigenin | 18328 | VL2 | 2776 |
| 40 | Yes | E. coli | None | None | F65 | H2FDA | 19206 | BL1 | 3425 |
| 40 | Yes | E. coli | None | None | F142 | Chlorophe | 19230 | BL3 | 4119 |
| 40 | Yes | E. coli | None | None | F39 | Congo Red | 18613 | BL3 | 4292 |
| 40 | Yes | E. coli | None | None | F116 | Sunset Yel | 19534 | BL2 | 4102 |
| 40 | Yes | E. coli | None | None | F10 | Alizarin | 17549 | BL1 | 4216 |
| 40 | Yes | E. coli | None | None | F84 | Neutral Re | 17459 | BL1 | 3557 |
| 40 | Yes | E. coli | None | None | F99 | Pyronin Y | 18610 | BL1 | 3381.5 |
| 40 | Yes | E. coli | None | None | F108 | Rhodamine | 18408 | BL1 | 2979 |
| 40 | Yes | E. coli | None | None | F109 | Riboflavin | 17417 | BL1 | 3398 |
| 40 | Yes | E. coli | None | None | F117 | SYBR Gree | 17987 | BL1 | 133379 |
| 40 | Yes | E. coli | None | None | F106 | Rhodamine | 15497 | BL1 | 4378 |
| 40 | Yes | E. coli | None | None | F73 | Malachite ( | 18242 | BL1 | 4077 |
| 40 | Yes | E. coli | None | None | F7 | Acridine or | 15772 | BL1 | 3938 |
| 40 | Yes | E. coli | None | None | F192 | 5-Carboxyf | 17217 | BL1 | 15110 |
| 40 | Yes | E. coli | None | None | F57 | Eosin Y | 17176 | BL1 | 21867 |
| 40 | Yes | E. coli | None | None | F27 | Calcein | 16768 | BL1 | 17594.5 |
| 40 | Yes | E. coli | None | None | F12 | Amaranth | 14779 | BL1 | 36848 |
| 40 | Yes | E. coli | None | None | NA | Autofluore: | 16288 | NA | 5790 |
| 40 | Yes | E. coli | Ampicillin | Wall | F194 | Oxonol | 18034 | RL1 | 4530.5 |
| 40 | Yes | E. coli | Ampicillin | Wall | F11 | Allura Red | 18485 | BL3 | 3551 |
| 40 | Yes | E. coli | Ampicillin | Wall | F54 | DiSC3(5) | 14623 | RL1 | 3492 |
| 40 | Yes | E. coli | Ampicillin | Wall | F124 | Thioflavin T | 16577 | VL2 | 3354 |
| 40 | Yes | E. coli | Ampicillin | Wall | F101 | Quinacrine | 14140 | VL2 | 3430 |
| 40 | Yes | E. coli | Ampicillin | Wall | F138 | Apigenin | 15235 | VL2 | 3375 |
| 40 | Yes | E. coli | Ampicillin | Wall | F65 | H2FDA | 14653 | BL1 | 4056 |
| 40 | Yes | E. coli | Ampicillin | Wall | F142 | Chlorophe | 15532 | BL3 | 4796 |
| 40 | Yes | E. coli | Ampicillin | Wall | F39 | Congo Red | 15579 | BL3 | 4969 |
| 40 | Yes | E. coli | Ampicillin | Wall | F116 | Sunset Yel | 16343 | BL2 | 4849 |
| 40 | Yes | E. coli | Ampicillin | Wall | F10 | Alizarin | 15667 | BL1 | 4462 |
| 40 | Yes | E. coli | Ampicillin | Wall | F84 | Neutral Re | 14767 | BL1 | 4245 |
| 40 | Yes | E. coli | Ampicillin | Wall | F99 | Pyronin Y | 15472 | BL1 | 3658 |
| 40 | Yes | E. coli | Ampicillin | Wall | F108 | Rhodamine | 14916 | BL1 | 3770.5 |
| 40 | Yes | E. coli | Ampicillin | Wall | F109 | Riboflavin | 15926 | BL1 | 3639 |
| 40 | Yes | E. coli | Ampicillin | Wall | F117 | SYBR Gree | 15613 | BL1 | 152767 |
| 40 | Yes | E. coli | Ampicillin | Wall | F106 | Rhodamine | 14553 | BL1 | 4685 |
| 40 | Yes | E. coli | Ampicillin | Wall | F73 | Malachite ( | 14883 | BL1 | 3868 |
| 40 | Yes | E. coli | Ampicillin | Wall | F7 | Acridine or | 14802 | BL1 | 4256.5 |
| 40 | Yes | E. coli | Ampicillin | Wall | F192 | 5-Carboxyf | 15394 | BL1 | 13418 |
| 40 | Yes | E. coli | Ampicillin | Wall | F57 | Eosin Y | 15294 | BL1 | 38581 |
| 40 | Yes | E. coli | Ampicillin | Wall | F27 | Calcein | 17145 | BL1 | 30101 |

| | | | | | | | | | |
|---|---|---|---|---|---|---|---|---|---|
| 40 | Yes | E. coli | Ampicillin | Wall | F12 | Amaranth | 16484 | BL1 | 23063.5 |
| 40 | Yes | E. coli | None | None | NA | Autofluore: | 18482 | NA | 5845.5 |
| 40 | Yes | E. coli | Bacitracin | Wall | F194 | Oxonol | 21026 | RL1 | 4475 |
| 40 | Yes | E. coli | Bacitracin | Wall | F11 | Allura Red | 19849 | BL3 | 3732 |
| 40 | Yes | E. coli | Bacitracin | Wall | F54 | DiSC3(5) | 18451 | RL1 | 3780 |
| 40 | Yes | E. coli | Bacitracin | Wall | F124 | Thioflavin T | 17023 | VL2 | 3174 |
| 40 | Yes | E. coli | Bacitracin | Wall | F101 | Quinacrine | 16740 | VL2 | 3382.5 |
| 40 | Yes | E. coli | Bacitracin | Wall | F138 | Apigenin | 16509 | VL2 | 3533 |
| 40 | Yes | E. coli | Bacitracin | Wall | F65 | H2FDA | 16951 | BL1 | 3821 |
| 40 | Yes | E. coli | Bacitracin | Wall | F142 | Chlorophe | 18021 | BL3 | 4668 |
| 40 | Yes | E. coli | Bacitracin | Wall | F39 | Congo Red | 16787 | BL3 | 4851 |
| 40 | Yes | E. coli | Bacitracin | Wall | F116 | Sunset Yel | 18040 | BL2 | 4428 |
| 40 | Yes | E. coli | Bacitracin | Wall | F10 | Alizarin | 18506 | BL1 | 4344 |
| 40 | Yes | E. coli | Bacitracin | Wall | F84 | Neutral Re | 16763 | BL1 | 4020 |
| 40 | Yes | E. coli | Bacitracin | Wall | F99 | Pyronin Y | 17732 | BL1 | 3619 |
| 40 | Yes | E. coli | Bacitracin | Wall | F108 | Rhodamine | 16533 | BL1 | 3528 |
| 40 | Yes | E. coli | Bacitracin | Wall | F109 | Riboflavin | 16847 | BL1 | 3704 |
| 40 | Yes | E. coli | Bacitracin | Wall | F117 | SYBR Gree | 17056 | BL1 | 152957.5 |
| 40 | Yes | E. coli | Bacitracin | Wall | F106 | Rhodamine | 17619 | BL1 | 5673 |
| 40 | Yes | E. coli | Bacitracin | Wall | F73 | Malachite ( | 17158 | BL1 | 4720.5 |
| 40 | Yes | E. coli | Bacitracin | Wall | F7 | Acridine or | 14527 | BL1 | 4363 |
| 40 | Yes | E. coli | Bacitracin | Wall | F192 | 5-Carboxyf | 15947 | BL1 | 23130 |
| 40 | Yes | E. coli | Bacitracin | Wall | F57 | Eosin Y | 16960 | BL1 | 137069 |
| 40 | Yes | E. coli | Bacitracin | Wall | F27 | Calcein | 17951 | BL1 | 21265 |
| 40 | Yes | E. coli | Bacitracin | Wall | F12 | Amaranth | 15037 | BL1 | 53575 |
| 40 | Yes | E. coli | None | None | NA | Autofluore: | 16730 | NA | 6356 |
| 40 | Yes | E. coli | Cefotaxim | Wall | F194 | Oxonol | 14508 | RL1 | 5436 |
| 40 | Yes | E. coli | Cefotaxim | Wall | F11 | Allura Red | 13145 | BL3 | 4703 |
| 40 | Yes | E. coli | Cefotaxim | Wall | F54 | DiSC3(5) | 12720 | RL1 | 4384 |
| 40 | Yes | E. coli | Cefotaxim | Wall | F124 | Thioflavin T | 12016 | VL2 | 4073 |
| 40 | Yes | E. coli | Cefotaxim | Wall | F101 | Quinacrine | 11495 | VL2 | 4407 |
| 40 | Yes | E. coli | Cefotaxim | Wall | F138 | Apigenin | 11737 | VL2 | 4156 |
| 40 | Yes | E. coli | Cefotaxim | Wall | F65 | H2FDA | 10837 | BL1 | 4590 |
| 40 | Yes | E. coli | Cefotaxim | Wall | F142 | Chlorophe | 11846 | BL3 | 5643.5 |
| 40 | Yes | E. coli | Cefotaxim | Wall | F39 | Congo Red | 10929 | BL3 | 6242 |
| 40 | Yes | E. coli | Cefotaxim | Wall | F116 | Sunset Yel | 12460 | BL2 | 5903 |
| 40 | Yes | E. coli | Cefotaxim | Wall | F10 | Alizarin | 11288 | BL1 | 5741.5 |
| 40 | Yes | E. coli | Cefotaxim | Wall | F84 | Neutral Re | 11525 | BL1 | 4999 |
| 40 | Yes | E. coli | Cefotaxim | Wall | F99 | Pyronin Y | 10749 | BL1 | 4803 |
| 40 | Yes | E. coli | Cefotaxim | Wall | F108 | Rhodamine | 12143 | BL1 | 4190 |
| 40 | Yes | E. coli | Cefotaxim | Wall | F109 | Riboflavin | 10885 | BL1 | 4486 |
| 40 | Yes | E. coli | Cefotaxim | Wall | F117 | SYBR Gree | 11376 | BL1 | 212905 |
| 40 | Yes | E. coli | Cefotaxim | Wall | F106 | Rhodamine | 12526 | BL1 | 5123.5 |
| 40 | Yes | E. coli | Cefotaxim | Wall | F73 | Malachite ( | 11778 | BL1 | 5168 |
| 40 | Yes | E. coli | Cefotaxim | Wall | F7 | Acridine or | 12183 | BL1 | 4683 |
| 40 | Yes | E. coli | Cefotaxim | Wall | F192 | 5-Carboxyf | 11822 | BL1 | 16409 |
| 40 | Yes | E. coli | Cefotaxim | Wall | F57 | Eosin Y | 13675 | BL1 | 17596 |
| 40 | Yes | E. coli | Cefotaxim | Wall | F27 | Calcein | 13227 | BL1 | 23246 |

| | | | | | | | | | |
|---|---|---|---|---|---|---|---|---|---|
| 40 | Yes | E. coli | Cefotaxim | Wall | F12 | Amaranth | 14553 | BL1 | 19910 |
| 40 | Yes | E. coli | None | None | NA | Autofluore | 18470 | NA | 5519 |
| 40 | Yes | E. coli | Ciprofloxa | Intracellul | F194 | Oxonol | 15575 | RL1 | 5039 |
| 40 | Yes | E. coli | Ciprofloxa | Intracellul | F11 | Allura Red | 13087 | BL3 | 4568 |
| 40 | Yes | E. coli | Ciprofloxa | Intracellul | F54 | DiSC3(5) | 11339 | RL1 | 4627 |
| 40 | Yes | E. coli | Ciprofloxa | Intracellul | F124 | Thioflavin T | 10220 | VL2 | 4082 |
| 40 | Yes | E. coli | Ciprofloxa | Intracellul | F101 | Quinacrine | 9543 | VL2 | 4409 |
| 40 | Yes | E. coli | Ciprofloxa | Intracellul | F138 | Apigenin | 11269 | VL2 | 4558 |
| 40 | Yes | E. coli | Ciprofloxa | Intracellul | F65 | H2FDA | 11158 | BL1 | 4952.5 |
| 40 | Yes | E. coli | Ciprofloxa | Intracellul | F142 | Chlorophe | 10238 | BL3 | 6388 |
| 40 | Yes | E. coli | Ciprofloxa | Intracellul | F39 | Congo Red | 10233 | BL3 | 6878 |
| 40 | Yes | E. coli | Ciprofloxa | Intracellul | F116 | Sunset Yel | 9782 | BL2 | 6108.5 |
| 40 | Yes | E. coli | Ciprofloxa | Intracellul | F10 | Alizarin | 9152 | BL1 | 6320 |
| 40 | Yes | E. coli | Ciprofloxa | Intracellul | F84 | Neutral Re | 8403 | BL1 | 5604 |
| 40 | Yes | E. coli | Ciprofloxa | Intracellul | F99 | Pyronin Y | 9225 | BL1 | 5126 |
| 40 | Yes | E. coli | Ciprofloxa | Intracellul | F108 | Rhodamine | 8600 | BL1 | 5151 |
| 40 | Yes | E. coli | Ciprofloxa | Intracellul | F109 | Riboflavin | 8247 | BL1 | 4793 |
| 40 | Yes | E. coli | Ciprofloxa | Intracellul | F117 | SYBR Gree | 9311 | BL1 | 249708 |
| 40 | Yes | E. coli | Ciprofloxa | Intracellul | F106 | Rhodamine | 8658 | BL1 | 6053 |
| 40 | Yes | E. coli | Ciprofloxa | Intracellul | F73 | Malachite ( | 8905 | BL1 | 5552 |
| 40 | Yes | E. coli | Ciprofloxa | Intracellul | F7 | Acridine or | 10866 | BL1 | 4937 |
| 40 | Yes | E. coli | Ciprofloxa | Intracellul | F192 | 5-Carboxyf | 9006 | BL1 | 19569.5 |
| 40 | Yes | E. coli | Ciprofloxa | Intracellul | F57 | Eosin Y | 8054 | BL1 | 28360 |
| 40 | Yes | E. coli | Ciprofloxa | Intracellul | F27 | Calcein | 8557 | BL1 | 29529 |
| 40 | Yes | E. coli | Ciprofloxa | Intracellul | F12 | Amaranth | 7923 | BL1 | 25194 |
| 40 | Yes | E. coli | None | None | NA | Autofluore | 16130 | NA | 6035 |
| 40 | Yes | E. coli | Fosfomyci | Wall | F194 | Oxonol | 22940 | RL1 | 4309.5 |
| 40 | Yes | E. coli | Fosfomyci | Wall | F11 | Allura Red | 21973 | BL3 | 3702 |
| 40 | Yes | E. coli | Fosfomyci | Wall | F54 | DiSC3(5) | 18545 | RL1 | 3713 |
| 40 | Yes | E. coli | Fosfomyci | Wall | F124 | Thioflavin T | 20198 | VL2 | 3423 |
| 40 | Yes | E. coli | Fosfomyci | Wall | F101 | Quinacrine | 16955 | VL2 | 3427 |
| 40 | Yes | E. coli | Fosfomyci | Wall | F138 | Apigenin | 19136 | VL2 | 3727 |
| 40 | Yes | E. coli | Fosfomyci | Wall | F65 | H2FDA | 16601 | BL1 | 4574 |
| 40 | Yes | E. coli | Fosfomyci | Wall | F142 | Chlorophe | 19792 | BL3 | 5085 |
| 40 | Yes | E. coli | Fosfomyci | Wall | F39 | Congo Red | 19218 | BL3 | 4664 |
| 40 | Yes | E. coli | Fosfomyci | Wall | F116 | Sunset Yel | 20800 | BL2 | 4750 |
| 40 | Yes | E. coli | Fosfomyci | Wall | F10 | Alizarin | 19054 | BL1 | 4423.5 |
| 40 | Yes | E. coli | Fosfomyci | Wall | F84 | Neutral Re | 18163 | BL1 | 4399 |
| 40 | Yes | E. coli | Fosfomyci | Wall | F99 | Pyronin Y | 14551 | BL1 | 3721 |
| 40 | Yes | E. coli | Fosfomyci | Wall | F108 | Rhodamine | 18678 | BL1 | 3854.5 |
| 40 | Yes | E. coli | Fosfomyci | Wall | F109 | Riboflavin | 18563 | BL1 | 4193 |
| 40 | Yes | E. coli | Fosfomyci | Wall | F117 | SYBR Gree | 18014 | BL1 | 145455.5 |
| 40 | Yes | E. coli | Fosfomyci | Wall | F106 | Rhodamine | 15179 | BL1 | 6552 |
| 40 | Yes | E. coli | Fosfomyci | Wall | F73 | Malachite ( | 20166 | BL1 | 4284 |
| 40 | Yes | E. coli | Fosfomyci | Wall | F7 | Acridine or | 17133 | BL1 | 4588 |
| 40 | Yes | E. coli | Fosfomyci | Wall | F192 | 5-Carboxyf | 18550 | BL1 | 13486.5 |
| 40 | Yes | E. coli | Fosfomyci | Wall | F57 | Eosin Y | 16093 | BL1 | 16651 |
| 40 | Yes | E. coli | Fosfomyci | Wall | F27 | Calcein | 14937 | BL1 | 42433 |

| | | | | | | | | | |
|---|---|---|---|---|---|---|---|---|---|
| 40 | Yes | E. coli | Fosfomycin | Wall | F12 | Amaranth | 16988 | BL1 | 34181 |
| 40 | Yes | E. coli | None | None | NA | Autofluores | 18273 | NA | 5881 |
| 40 | Yes | E. coli | Nalidixic ac | Intracellula | F194 | Oxonol | 18845 | RL1 | 4761 |
| 40 | Yes | E. coli | Nalidixic ac | Intracellula | F11 | Allura Red | 15745 | BL3 | 4001 |
| 40 | Yes | E. coli | Nalidixic ac | Intracellula | F54 | DiSC3(5) | 16229 | RL1 | 3977 |
| 40 | Yes | E. coli | Nalidixic ac | Intracellula | F124 | Thioflavin T | 14089 | VL2 | 3708 |
| 40 | Yes | E. coli | Nalidixic ac | Intracellula | F101 | Quinacrine | 14702 | VL2 | 3847 |
| 40 | Yes | E. coli | Nalidixic ac | Intracellula | F138 | Apigenin | 13246 | VL2 | 4002 |
| 40 | Yes | E. coli | Nalidixic ac | Intracellula | F65 | H2FDA | 13948 | BL1 | 4903 |
| 40 | Yes | E. coli | Nalidixic ac | Intracellula | F142 | Chlorophe | 13177 | BL3 | 5382 |
| 40 | Yes | E. coli | Nalidixic ac | Intracellula | F39 | Congo Red | 12062 | BL3 | 5919.5 |
| 40 | Yes | E. coli | Nalidixic ac | Intracellula | F116 | Sunset Yel | 13375 | BL2 | 5802 |
| 40 | Yes | E. coli | Nalidixic ac | Intracellula | F10 | Alizarin | 12686 | BL1 | 5249.5 |
| 40 | Yes | E. coli | Nalidixic ac | Intracellula | F84 | Neutral Re | 12773 | BL1 | 4872 |
| 40 | Yes | E. coli | Nalidixic ac | Intracellula | F99 | Pyronin Y | 12032 | BL1 | 4600 |
| 40 | Yes | E. coli | Nalidixic ac | Intracellula | F108 | Rhodamine | 11446 | BL1 | 4707 |
| 40 | Yes | E. coli | Nalidixic ac | Intracellula | F109 | Riboflavin | 13129 | BL1 | 4345 |
| 40 | Yes | E. coli | Nalidixic ac | Intracellula | F117 | SYBR Gree | 9857 | BL1 | 124217 |
| 40 | Yes | E. coli | Nalidixic ac | Intracellula | F106 | Rhodamine | 12769 | BL1 | 5302 |
| 40 | Yes | E. coli | Nalidixic ac | Intracellula | F73 | Malachite ( | 13085 | BL1 | 4846 |
| 40 | Yes | E. coli | Nalidixic ac | Intracellula | F7 | Acridine or | 9978 | BL1 | 5092 |
| 40 | Yes | E. coli | Nalidixic ac | Intracellula | F192 | 5-Carboxyf | 13123 | BL1 | 13795 |
| 40 | Yes | E. coli | Nalidixic ac | Intracellula | F57 | Eosin Y | 9464 | BL1 | 24801.5 |
| 40 | Yes | E. coli | Nalidixic ac | Intracellula | F27 | Calcein | 9520 | BL1 | 23560.5 |
| 40 | Yes | E. coli | Nalidixic ac | Intracellula | F12 | Amaranth | 14632 | BL1 | 21050 |
| 40 | Yes | E. coli | None | None | NA | Autofluores | 19344 | NA | 5517.5 |
| 40 | Yes | E. coli | Ofloxacin | Intracellula | F194 | Oxonol | 11457 | RL1 | 6011 |
| 40 | Yes | E. coli | Ofloxacin | Intracellula | F11 | Allura Red | 10649 | BL3 | 5562 |
| 40 | Yes | E. coli | Ofloxacin | Intracellula | F54 | DiSC3(5) | 9982 | RL1 | 5324.5 |
| 40 | Yes | E. coli | Ofloxacin | Intracellula | F124 | Thioflavin T | 9373 | VL2 | 4965 |
| 40 | Yes | E. coli | Ofloxacin | Intracellula | F101 | Quinacrine | 9332 | VL2 | 4817.5 |
| 40 | Yes | E. coli | Ofloxacin | Intracellula | F138 | Apigenin | 8861 | VL2 | 5116 |
| 40 | Yes | E. coli | Ofloxacin | Intracellula | F65 | H2FDA | 9204 | BL1 | 5954 |
| 40 | Yes | E. coli | Ofloxacin | Intracellula | F142 | Chlorophe | 8011 | BL3 | 7582 |
| 40 | Yes | E. coli | Ofloxacin | Intracellula | F39 | Congo Red | 9635 | BL3 | 7251 |
| 40 | Yes | E. coli | Ofloxacin | Intracellula | F116 | Sunset Yel | 9040 | BL2 | 7290 |
| 40 | Yes | E. coli | Ofloxacin | Intracellula | F10 | Alizarin | 8891 | BL1 | 6351 |
| 40 | Yes | E. coli | Ofloxacin | Intracellula | F84 | Neutral Re | 8873 | BL1 | 5685 |
| 40 | Yes | E. coli | Ofloxacin | Intracellula | F99 | Pyronin Y | 9990 | BL1 | 5281.5 |
| 40 | Yes | E. coli | Ofloxacin | Intracellula | F108 | Rhodamine | 8464 | BL1 | 5002.5 |
| 40 | Yes | E. coli | Ofloxacin | Intracellula | F109 | Riboflavin | 9925 | BL1 | 4920 |
| 40 | Yes | E. coli | Ofloxacin | Intracellula | F117 | SYBR Gree | 8150 | BL1 | 281485.5 |
| 40 | Yes | E. coli | Ofloxacin | Intracellula | F106 | Rhodamine | 9093 | BL1 | 6978 |
| 40 | Yes | E. coli | Ofloxacin | Intracellula | F73 | Malachite ( | 8509 | BL1 | 5646 |
| 40 | Yes | E. coli | Ofloxacin | Intracellula | F7 | Acridine or | 7894 | BL1 | 5964 |
| 40 | Yes | E. coli | Ofloxacin | Intracellula | F192 | 5-Carboxyf | 7539 | BL1 | 21643 |
| 40 | Yes | E. coli | Ofloxacin | Intracellula | F57 | Eosin Y | 8485 | BL1 | 29860 |
| 40 | Yes | E. coli | Ofloxacin | Intracellula | F27 | Calcein | 8222 | BL1 | 29479.5 |

| | | | | | | | | | |
|---|---|---|---|---|---|---|---|---|---|
| 40 | Yes | E. coli | Ofloxacin | Intracellula | F12 | Amaranth | 9778 | BL1 | 23243.5 |
| 40 | Yes | E. coli | None | None | NA | Autofluores | 18210 | NA | 5628.5 |
| 40 | Yes | E. coli | Polymyxin | Wall | F194 | Oxonol | 17442 | RL1 | 5493.5 |
| 40 | Yes | E. coli | Polymyxin | Wall | F11 | Allura Red | 16348 | BL3 | 4728 |
| 40 | Yes | E. coli | Polymyxin | Wall | F54 | DiSC3(5) | 15775 | RL1 | 4283 |
| 40 | Yes | E. coli | Polymyxin | Wall | F124 | Thioflavin T | 14527 | VL2 | 4322 |
| 40 | Yes | E. coli | Polymyxin | Wall | F101 | Quinacrine | 13917 | VL2 | 4544 |
| 40 | Yes | E. coli | Polymyxin | Wall | F138 | Apigenin | 13084 | VL2 | 4605.5 |
| 40 | Yes | E. coli | Polymyxin | Wall | F65 | H2FDA | 14562 | BL1 | 5460 |
| 40 | Yes | E. coli | Polymyxin | Wall | F142 | Chlorophe | 14669 | BL3 | 6033 |
| 40 | Yes | E. coli | Polymyxin | Wall | F39 | Congo Red | 13884 | BL3 | 6065.5 |
| 40 | Yes | E. coli | Polymyxin | Wall | F116 | Sunset Yel | 14190 | BL2 | 5934 |
| 40 | Yes | E. coli | Polymyxin | Wall | F10 | Alizarin | 12877 | BL1 | 5313 |
| 40 | Yes | E. coli | Polymyxin | Wall | F84 | Neutral Re | 14573 | BL1 | 4951 |
| 40 | Yes | E. coli | Polymyxin | Wall | F99 | Pyronin Y | 13428 | BL1 | 4380.5 |
| 40 | Yes | E. coli | Polymyxin | Wall | F108 | Rhodamine | 12317 | BL1 | 4867 |
| 40 | Yes | E. coli | Polymyxin | Wall | F109 | Riboflavin | 14920 | BL1 | 4495.5 |
| 40 | Yes | E. coli | Polymyxin | Wall | F117 | SYBR Gree | 13182 | BL1 | 259158.5 |
| 40 | Yes | E. coli | Polymyxin | Wall | F106 | Rhodamine | 12109 | BL1 | 5979 |
| 40 | Yes | E. coli | Polymyxin | Wall | F73 | Malachite ( | 15180 | BL1 | 5123 |
| 40 | Yes | E. coli | Polymyxin | Wall | F7 | Acridine or | 13497 | BL1 | 5921 |
| 40 | Yes | E. coli | Polymyxin | Wall | F192 | 5-Carboxyf | 12163 | BL1 | 16903 |
| 40 | Yes | E. coli | Polymyxin | Wall | F57 | Eosin Y | 11755 | BL1 | 49120 |
| 40 | Yes | E. coli | Polymyxin | Wall | F27 | Calcein | 13488 | BL1 | 22613 |
| 40 | Yes | E. coli | Polymyxin | Wall | F12 | Amaranth | 14358 | BL1 | 25283 |
| 40 | Yes | E. coli | None | None | NA | Autofluores | 18513 | NA | 5192 |
| 40 | Yes | E. coli | Tetracyclin | Intracellula | F194 | Oxonol | 18926 | RL1 | 4400.5 |
| 40 | Yes | E. coli | Tetracyclin | Intracellula | F11 | Allura Red | 18446 | BL3 | 3864 |
| 40 | Yes | E. coli | Tetracyclin | Intracellula | F54 | DiSC3(5) | 17006 | RL1 | 3938 |
| 40 | Yes | E. coli | Tetracyclin | Intracellula | F124 | Thioflavin T | 15911 | VL2 | 3670 |
| 40 | Yes | E. coli | Tetracyclin | Intracellula | F101 | Quinacrine | 16900 | VL2 | 3613 |
| 40 | Yes | E. coli | Tetracyclin | Intracellula | F138 | Apigenin | 16457 | VL2 | 3803 |
| 40 | Yes | E. coli | Tetracyclin | Intracellula | F65 | H2FDA | 16176 | BL1 | 4930 |
| 40 | Yes | E. coli | Tetracyclin | Intracellula | F142 | Chlorophe | 17431 | BL3 | 4937 |
| 40 | Yes | E. coli | Tetracyclin | Intracellula | F39 | Congo Red | 17121 | BL3 | 5285 |
| 40 | Yes | E. coli | Tetracyclin | Intracellula | F116 | Sunset Yel | 17352 | BL2 | 4858 |
| 40 | Yes | E. coli | Tetracyclin | Intracellula | F10 | Alizarin | 15221 | BL1 | 4747 |
| 40 | Yes | E. coli | Tetracyclin | Intracellula | F84 | Neutral Re | 17362 | BL1 | 3853.5 |
| 40 | Yes | E. coli | Tetracyclin | Intracellula | F99 | Pyronin Y | 15083 | BL1 | 4137 |
| 40 | Yes | E. coli | Tetracyclin | Intracellula | F108 | Rhodamine | 16150 | BL1 | 3803 |
| 40 | Yes | E. coli | Tetracyclin | Intracellula | F109 | Riboflavin | 16324 | BL1 | 3769 |
| 40 | Yes | E. coli | Tetracyclin | Intracellula | F117 | SYBR Gree | 14930 | BL1 | 144592 |
| 40 | Yes | E. coli | Tetracyclin | Intracellula | F106 | Rhodamine | 15215 | BL1 | 5009 |
| 40 | Yes | E. coli | Tetracyclin | Intracellula | F73 | Malachite ( | 17270 | BL1 | 4753 |
| 40 | Yes | E. coli | Tetracyclin | Intracellula | F7 | Acridine or | 14810 | BL1 | 4223 |
| 40 | Yes | E. coli | Tetracyclin | Intracellula | F192 | 5-Carboxyf | 15131 | BL1 | 13008 |
| 40 | Yes | E. coli | Tetracyclin | Intracellula | F57 | Eosin Y | 15203 | BL1 | 16168 |
| 40 | Yes | E. coli | Tetracyclin | Intracellula | F27 | Calcein | 15559 | BL1 | 14669 |

| | | | | | | | | | |
|---|---|---|---|---|---|---|---|---|---|
| 40 | Yes | E. coli | Tetracyclin | Intracellula | F12 | Amaranth | 14492 | BL1 | 30345 |
| 40 | Yes | E. coli | None | None | NA | Autofluore: | 19643 | NA | 5294 |
| 40 | Yes | E. coli | Vancomyc | Wall | F194 | Oxonol | 23555 | RL1 | 4325 |
| 40 | Yes | E. coli | Vancomyc | Wall | F11 | Allura Red | 19965 | BL3 | 3830 |
| 40 | Yes | E. coli | Vancomyc | Wall | F54 | DiSC3(5) | 22655 | RL1 | 3958 |
| 40 | Yes | E. coli | Vancomyc | Wall | F124 | Thioflavin T | 21449 | VL2 | 3613 |
| 40 | Yes | E. coli | Vancomyc | Wall | F101 | Quinacrine | 18780 | VL2 | 4113.5 |
| 40 | Yes | E. coli | Vancomyc | Wall | F138 | Apigenin | 15885 | VL2 | 4456 |
| 40 | Yes | E. coli | Vancomyc | Wall | F65 | H2FDA | 16388 | BL1 | 6044 |
| 40 | Yes | E. coli | Vancomyc | Wall | F142 | Chlorophe | 16214 | BL3 | 7683.5 |
| 40 | Yes | E. coli | Vancomyc | Wall | F39 | Congo Red | 15934 | BL3 | 9493.5 |
| 40 | Yes | E. coli | Vancomyc | Wall | F116 | Sunset Yel | 16324 | BL2 | 8433 |
| 40 | Yes | E. coli | Vancomyc | Wall | F10 | Alizarin | 14546 | BL1 | 6838.5 |
| 40 | Yes | E. coli | Vancomyc | Wall | F84 | Neutral Re | 15816 | BL1 | 5825 |
| 40 | Yes | E. coli | Vancomyc | Wall | F99 | Pyronin Y | 14582 | BL1 | 6345 |
| 40 | Yes | E. coli | Vancomyc | Wall | F108 | Rhodamine | 14933 | BL1 | 5564 |
| 40 | Yes | E. coli | Vancomyc | Wall | F109 | Riboflavin | 14933 | BL1 | 5402 |
| 40 | Yes | E. coli | Vancomyc | Wall | F117 | SYBR Gree | 15968 | BL1 | 179757.5 |
| 40 | Yes | E. coli | Vancomyc | Wall | F106 | Rhodamine | 13058 | BL1 | 8182.5 |
| 40 | Yes | E. coli | Vancomyc | Wall | F73 | Malachite ( | 15733 | BL1 | 6748 |
| 40 | Yes | E. coli | Vancomyc | Wall | F7 | Acridine or | 13315 | BL1 | 6047 |
| 40 | Yes | E. coli | Vancomyc | Wall | F192 | 5-Carboxyf | 14531 | BL1 | 40594 |
| 40 | Yes | E. coli | Vancomyc | Wall | F57 | Eosin Y | 15115 | BL1 | 67003 |
| 40 | Yes | E. coli | Vancomyc | Wall | F27 | Calcein | 16866 | BL1 | 67175.5 |
| 40 | Yes | E. coli | Vancomyc | Wall | F12 | Amaranth | 13663 | BL1 | 63823 |
| 40 | Yes | E. coli | None | None | NA | Autofluore: | 17516 | NA | 859 |
| 40 | Yes | E. coli | None | None | F194 | Oxonol | 21709 | RL1 | 1539 |
| 40 | Yes | E. coli | None | None | F11 | Allura Red | 20786 | BL3 | 2343 |
| 40 | Yes | E. coli | None | None | F54 | DiSC3(5) | 22897 | RL1 | 2499 |
| 40 | Yes | E. coli | None | None | F124 | Thioflavin T | 20437 | VL2 | 2618 |
| 40 | Yes | E. coli | None | None | F101 | Quinacrine | 18494 | VL2 | 2746 |
| 40 | Yes | E. coli | None | None | F138 | Apigenin | 22136 | VL2 | 2600 |
| 40 | Yes | E. coli | None | None | F65 | H2FDA | 21182 | BL1 | 3066.5 |
| 40 | Yes | E. coli | None | None | F142 | Chlorophe | 20030 | BL3 | 4793.5 |
| 40 | Yes | E. coli | None | None | F39 | Congo Red | 21270 | BL3 | 4625 |
| 40 | Yes | E. coli | None | None | F116 | Sunset Yel | 20285 | BL2 | 4294 |
| 40 | Yes | E. coli | None | None | F10 | Alizarin | 20514 | BL1 | 3647.5 |
| 40 | Yes | E. coli | None | None | F84 | Neutral Re | 19484 | BL1 | 3558 |
| 40 | Yes | E. coli | None | None | F99 | Pyronin Y | 19891 | BL1 | 3166 |
| 40 | Yes | E. coli | None | None | F108 | Rhodamine | 20315 | BL1 | 3110 |
| 40 | Yes | E. coli | None | None | F109 | Riboflavin | 19764 | BL1 | 3004 |
| 40 | Yes | E. coli | None | None | F117 | SYBR Gree | 22044 | BL1 | 121869.5 |
| 40 | Yes | E. coli | None | None | F106 | Rhodamine | 20751 | BL1 | 4102 |
| 40 | Yes | E. coli | None | None | F73 | Malachite ( | 17703 | BL1 | 3438 |
| 40 | Yes | E. coli | None | None | F7 | Acridine or | 19847 | BL1 | 3183 |
| 40 | Yes | E. coli | None | None | F192 | 5-Carboxyf | 20254 | BL1 | 11710 |
| 40 | Yes | E. coli | None | None | F57 | Eosin Y | 18748 | BL1 | 19247.5 |
| 40 | Yes | E. coli | None | None | F27 | Calcein | 17360 | BL1 | 26452.5 |

| | | | | | | | | | |
|---|---|---|---|---|---|---|---|---|---|
| 40 | Yes | E. coli | None | None | F12 | Amaranth | 18245 | BL1 | 21901 |
| 40 | Yes | E. coli | None | None | NA | Autofluore: | 15978 | NA | 6204.5 |
| 40 | Yes | E. coli | Ampicillin | Wall | F194 | Oxonol | 20609 | RL1 | 4230 |
| 40 | Yes | E. coli | Ampicillin | Wall | F11 | Allura Red | 20150 | BL3 | 3856.5 |
| 40 | Yes | E. coli | Ampicillin | Wall | F54 | DiSC3(5) | 19461 | RL1 | 3334 |
| 40 | Yes | E. coli | Ampicillin | Wall | F124 | Thioflavin T | 20952 | VL2 | 3490 |
| 40 | Yes | E. coli | Ampicillin | Wall | F101 | Quinacrine | 21236 | VL2 | 3467 |
| 40 | Yes | E. coli | Ampicillin | Wall | F138 | Apigenin | 22087 | VL2 | 3250 |
| 40 | Yes | E. coli | Ampicillin | Wall | F65 | H2FDA | 19055 | BL1 | 3409 |
| 40 | Yes | E. coli | Ampicillin | Wall | F142 | Chlorophe | 18282 | BL3 | 4953 |
| 40 | Yes | E. coli | Ampicillin | Wall | F39 | Congo Red | 20600 | BL3 | 4532 |
| 40 | Yes | E. coli | Ampicillin | Wall | F116 | Sunset Yel | 20016 | BL2 | 4397 |
| 40 | Yes | E. coli | Ampicillin | Wall | F10 | Alizarin | 20369 | BL1 | 3915 |
| 40 | Yes | E. coli | Ampicillin | Wall | F84 | Neutral Re | 22099 | BL1 | 3318 |
| 40 | Yes | E. coli | Ampicillin | Wall | F99 | Pyronin Y | 19039 | BL1 | 3470 |
| 40 | Yes | E. coli | Ampicillin | Wall | F108 | Rhodamine | 22287 | BL1 | 3196 |
| 40 | Yes | E. coli | Ampicillin | Wall | F109 | Riboflavin | 20608 | BL1 | 3326 |
| 40 | Yes | E. coli | Ampicillin | Wall | F117 | SYBR Gree | 23610 | BL1 | 122950 |
| 40 | Yes | E. coli | Ampicillin | Wall | F106 | Rhodamine | 21950 | BL1 | 4102 |
| 40 | Yes | E. coli | Ampicillin | Wall | F73 | Malachite ( | 22122 | BL1 | 3611 |
| 40 | Yes | E. coli | Ampicillin | Wall | F7 | Acridine or | 19913 | BL1 | 3298 |
| 40 | Yes | E. coli | Ampicillin | Wall | F192 | 5-Carboxyf | 18758 | BL1 | 12061 |
| 40 | Yes | E. coli | Ampicillin | Wall | F57 | Eosin Y | 20048 | BL1 | 15214 |
| 40 | Yes | E. coli | Ampicillin | Wall | F27 | Calcein | 20574 | BL1 | 17472.5 |
| 40 | Yes | E. coli | Ampicillin | Wall | F12 | Amaranth | 17919 | BL1 | 20678 |
| 40 | Yes | E. coli | None | None | NA | Autofluore: | 18082 | NA | 5930.5 |
| 40 | Yes | E. coli | Bacitracin | Wall | F194 | Oxonol | 23163 | RL1 | 4475 |
| 40 | Yes | E. coli | Bacitracin | Wall | F11 | Allura Red | 19490 | BL3 | 4135 |
| 40 | Yes | E. coli | Bacitracin | Wall | F54 | DiSC3(5) | 21436 | RL1 | 3561 |
| 40 | Yes | E. coli | Bacitracin | Wall | F124 | Thioflavin T | 18933 | VL2 | 3316 |
| 40 | Yes | E. coli | Bacitracin | Wall | F101 | Quinacrine | 18075 | VL2 | 3531 |
| 40 | Yes | E. coli | Bacitracin | Wall | F138 | Apigenin | 16601 | VL2 | 3637 |
| 40 | Yes | E. coli | Bacitracin | Wall | F65 | H2FDA | 18404 | BL1 | 4463.5 |
| 40 | Yes | E. coli | Bacitracin | Wall | F142 | Chlorophe | 20067 | BL3 | 4678 |
| 40 | Yes | E. coli | Bacitracin | Wall | F39 | Congo Red | 20780 | BL3 | 4791 |
| 40 | Yes | E. coli | Bacitracin | Wall | F116 | Sunset Yel | 19714 | BL2 | 4776.5 |
| 40 | Yes | E. coli | Bacitracin | Wall | F10 | Alizarin | 18905 | BL1 | 4642 |
| 40 | Yes | E. coli | Bacitracin | Wall | F84 | Neutral Re | 16874 | BL1 | 4567 |
| 40 | Yes | E. coli | Bacitracin | Wall | F99 | Pyronin Y | 16968 | BL1 | 4159 |
| 40 | Yes | E. coli | Bacitracin | Wall | F108 | Rhodamine | 17252 | BL1 | 3894.5 |
| 40 | Yes | E. coli | Bacitracin | Wall | F109 | Riboflavin | 18986 | BL1 | 3874 |
| 40 | Yes | E. coli | Bacitracin | Wall | F117 | SYBR Gree | 19677 | BL1 | 146867 |
| 40 | Yes | E. coli | Bacitracin | Wall | F106 | Rhodamine | 18990 | BL1 | 4379.5 |
| 40 | Yes | E. coli | Bacitracin | Wall | F73 | Malachite ( | 18318 | BL1 | 4052 |
| 40 | Yes | E. coli | Bacitracin | Wall | F7 | Acridine or | 19457 | BL1 | 3901 |
| 40 | Yes | E. coli | Bacitracin | Wall | F192 | 5-Carboxyf | 19351 | BL1 | 15150 |
| 40 | Yes | E. coli | Bacitracin | Wall | F57 | Eosin Y | 17861 | BL1 | 18287 |
| 40 | Yes | E. coli | Bacitracin | Wall | F27 | Calcein | 18941 | BL1 | 19224 |

| | | | | | | | | | |
|---|---|---|---|---|---|---|---|---|---|
| 40 | Yes | E. coli | Bacitracin | Wall | F12 | Amaranth | 18400 | BL1 | 59713 |
| 40 | Yes | E. coli | None | None | NA | Autofluore: | 17653 | NA | 6608 |
| 40 | Yes | E. coli | Cefotaxim | Wall | F194 | Oxonol | 19531 | RL1 | 5269 |
| 40 | Yes | E. coli | Cefotaxim | Wall | F11 | Allura Red | 19233 | BL3 | 4330 |
| 40 | Yes | E. coli | Cefotaxim | Wall | F54 | DiSC3(5) | 16025 | RL1 | 4265 |
| 40 | Yes | E. coli | Cefotaxim | Wall | F124 | Thioflavin T | 14256 | VL2 | 3875 |
| 40 | Yes | E. coli | Cefotaxim | Wall | F101 | Quinacrine | 15968 | VL2 | 3966 |
| 40 | Yes | E. coli | Cefotaxim | Wall | F138 | Apigenin | 16387 | VL2 | 3732 |
| 40 | Yes | E. coli | Cefotaxim | Wall | F65 | H2FDA | 13603 | BL1 | 4368 |
| 40 | Yes | E. coli | Cefotaxim | Wall | F142 | Chlorophe | 14648 | BL3 | 5123 |
| 40 | Yes | E. coli | Cefotaxim | Wall | F39 | Congo Red | 14091 | BL3 | 5746 |
| 40 | Yes | E. coli | Cefotaxim | Wall | F116 | Sunset Yel | 15225 | BL2 | 5468 |
| 40 | Yes | E. coli | Cefotaxim | Wall | F10 | Alizarin | 14942 | BL1 | 5173 |
| 40 | Yes | E. coli | Cefotaxim | Wall | F84 | Neutral Re | 14403 | BL1 | 4830 |
| 40 | Yes | E. coli | Cefotaxim | Wall | F99 | Pyronin Y | 13592 | BL1 | 3919 |
| 40 | Yes | E. coli | Cefotaxim | Wall | F108 | Rhodamine | 16569 | BL1 | 3598 |
| 40 | Yes | E. coli | Cefotaxim | Wall | F109 | Riboflavin | 17507 | BL1 | 3781 |
| 40 | Yes | E. coli | Cefotaxim | Wall | F117 | SYBR Gree | 16390 | BL1 | 161720 |
| 40 | Yes | E. coli | Cefotaxim | Wall | F106 | Rhodamine | 17286 | BL1 | 4822.5 |
| 40 | Yes | E. coli | Cefotaxim | Wall | F73 | Malachite ( | 16353 | BL1 | 4336 |
| 40 | Yes | E. coli | Cefotaxim | Wall | F7 | Acridine or | 16612 | BL1 | 4107.5 |
| 40 | Yes | E. coli | Cefotaxim | Wall | F192 | 5-Carboxyf | 16002 | BL1 | 14534 |
| 40 | Yes | E. coli | Cefotaxim | Wall | F57 | Eosin Y | 18708 | BL1 | 21329 |
| 40 | Yes | E. coli | Cefotaxim | Wall | F27 | Calcein | 18528 | BL1 | 19772 |
| 40 | Yes | E. coli | Cefotaxim | Wall | F12 | Amaranth | 19354 | BL1 | 34878.5 |
| 40 | Yes | E. coli | None | None | NA | Autofluore: | 17737 | NA | 5707 |
| 40 | Yes | E. coli | Ciprofloxa | Intracellula | F194 | Oxonol | 19299 | RL1 | 4938 |
| 40 | Yes | E. coli | Ciprofloxa | Intracellula | F11 | Allura Red | 15428 | BL3 | 4426.5 |
| 40 | Yes | E. coli | Ciprofloxa | Intracellula | F54 | DiSC3(5) | 13478 | RL1 | 4528 |
| 40 | Yes | E. coli | Ciprofloxa | Intracellula | F124 | Thioflavin T | 11790 | VL2 | 4284 |
| 40 | Yes | E. coli | Ciprofloxa | Intracellula | F101 | Quinacrine | 13527 | VL2 | 4549 |
| 40 | Yes | E. coli | Ciprofloxa | Intracellula | F138 | Apigenin | 12505 | VL2 | 4238 |
| 40 | Yes | E. coli | Ciprofloxa | Intracellula | F65 | H2FDA | 12061 | BL1 | 5012 |
| 40 | Yes | E. coli | Ciprofloxa | Intracellula | F142 | Chlorophe | 13350 | BL3 | 5687 |
| 40 | Yes | E. coli | Ciprofloxa | Intracellula | F39 | Congo Red | 13112 | BL3 | 5941.5 |
| 40 | Yes | E. coli | Ciprofloxa | Intracellula | F116 | Sunset Yel | 12217 | BL2 | 6493 |
| 40 | Yes | E. coli | Ciprofloxa | Intracellula | F10 | Alizarin | 11769 | BL1 | 5396 |
| 40 | Yes | E. coli | Ciprofloxa | Intracellula | F84 | Neutral Re | 12865 | BL1 | 5175 |
| 40 | Yes | E. coli | Ciprofloxa | Intracellula | F99 | Pyronin Y | 13002 | BL1 | 4433.5 |
| 40 | Yes | E. coli | Ciprofloxa | Intracellula | F108 | Rhodamine | 12007 | BL1 | 4452 |
| 40 | Yes | E. coli | Ciprofloxa | Intracellula | F109 | Riboflavin | 11790 | BL1 | 4388.5 |
| 40 | Yes | E. coli | Ciprofloxa | Intracellula | F117 | SYBR Gree | 11774 | BL1 | 217668.5 |
| 40 | Yes | E. coli | Ciprofloxa | Intracellula | F106 | Rhodamine | 11146 | BL1 | 5327 |
| 40 | Yes | E. coli | Ciprofloxa | Intracellula | F73 | Malachite ( | 11972 | BL1 | 4838.5 |
| 40 | Yes | E. coli | Ciprofloxa | Intracellula | F7 | Acridine or | 11592 | BL1 | 4687 |
| 40 | Yes | E. coli | Ciprofloxa | Intracellula | F192 | 5-Carboxyf | 11408 | BL1 | 18040 |
| 40 | Yes | E. coli | Ciprofloxa | Intracellula | F57 | Eosin Y | 11380 | BL1 | 21983.5 |
| 40 | Yes | E. coli | Ciprofloxa | Intracellula | F27 | Calcein | 13430 | BL1 | 23497 |

| | | | | | | | | | |
|---|---|---|---|---|---|---|---|---|---|
| 40 | Yes | E. coli | Ciprofloxa | Intracellula | F12 | Amaranth | 18270 | BL1 | 22882.5 |
| 40 | Yes | E. coli | None | None | NA | Autofluore: | 19401 | NA | 6178 |
| 40 | Yes | E. coli | Fosfomyci | Wall | F194 | Oxonol | 22219 | RL1 | 4323 |
| 40 | Yes | E. coli | Fosfomyci | Wall | F11 | Allura Red | 23924 | BL3 | 3819 |
| 40 | Yes | E. coli | Fosfomyci | Wall | F54 | DiSC3(5) | 21729 | RL1 | 3682 |
| 40 | Yes | E. coli | Fosfomyci | Wall | F124 | Thioflavin T | 22762 | VL2 | 3454.5 |
| 40 | Yes | E. coli | Fosfomyci | Wall | F101 | Quinacrine | 20288 | VL2 | 3820.5 |
| 40 | Yes | E. coli | Fosfomyci | Wall | F138 | Apigenin | 21792 | VL2 | 3859 |
| 40 | Yes | E. coli | Fosfomyci | Wall | F65 | H2FDA | 19728 | BL1 | 4198 |
| 40 | Yes | E. coli | Fosfomyci | Wall | F142 | Chlorophe | 21126 | BL3 | 4551 |
| 40 | Yes | E. coli | Fosfomyci | Wall | F39 | Congo Red | 20995 | BL3 | 4999 |
| 40 | Yes | E. coli | Fosfomyci | Wall | F116 | Sunset Yel | 19665 | BL2 | 4755 |
| 40 | Yes | E. coli | Fosfomyci | Wall | F10 | Alizarin | 21300 | BL1 | 4767.5 |
| 40 | Yes | E. coli | Fosfomyci | Wall | F84 | Neutral Re | 20561 | BL1 | 4308 |
| 40 | Yes | E. coli | Fosfomyci | Wall | F99 | Pyronin Y | 19169 | BL1 | 4135 |
| 40 | Yes | E. coli | Fosfomyci | Wall | F108 | Rhodamine | 18893 | BL1 | 4089 |
| 40 | Yes | E. coli | Fosfomyci | Wall | F109 | Riboflavin | 19667 | BL1 | 3952 |
| 40 | Yes | E. coli | Fosfomyci | Wall | F117 | SYBR Gree | 18420 | BL1 | 148922.5 |
| 40 | Yes | E. coli | Fosfomyci | Wall | F106 | Rhodamine | 20406 | BL1 | 4619.5 |
| 40 | Yes | E. coli | Fosfomyci | Wall | F73 | Malachite | 19422 | BL1 | 4449 |
| 40 | Yes | E. coli | Fosfomyci | Wall | F7 | Acridine or | 20630 | BL1 | 4130 |
| 40 | Yes | E. coli | Fosfomyci | Wall | F192 | 5-Carboxyf | 19039 | BL1 | 17761 |
| 40 | Yes | E. coli | Fosfomyci | Wall | F57 | Eosin Y | 20214 | BL1 | 30448 |
| 40 | Yes | E. coli | Fosfomyci | Wall | F27 | Calcein | 20457 | BL1 | 26162 |
| 40 | Yes | E. coli | Fosfomyci | Wall | F12 | Amaranth | 19661 | BL1 | 38304 |
| 40 | Yes | E. coli | None | None | NA | Autofluore: | 19792 | NA | 5773.5 |
| 40 | Yes | E. coli | Nalidixic a | Intracellula | F194 | Oxonol | 15358 | RL1 | 5148.5 |
| 40 | Yes | E. coli | Nalidixic a | Intracellula | F11 | Allura Red | 19450 | BL3 | 4094 |
| 40 | Yes | E. coli | Nalidixic a | Intracellula | F54 | DiSC3(5) | 17472 | RL1 | 4003.5 |
| 40 | Yes | E. coli | Nalidixic a | Intracellula | F124 | Thioflavin T | 17032 | VL2 | 3774 |
| 40 | Yes | E. coli | Nalidixic a | Intracellula | F101 | Quinacrine | 13138 | VL2 | 4480 |
| 40 | Yes | E. coli | Nalidixic a | Intracellula | F138 | Apigenin | 12466 | VL2 | 4879 |
| 40 | Yes | E. coli | Nalidixic a | Intracellula | F65 | H2FDA | 14748 | BL1 | 4611 |
| 40 | Yes | E. coli | Nalidixic a | Intracellula | F142 | Chlorophe | 15779 | BL3 | 5178 |
| 40 | Yes | E. coli | Nalidixic a | Intracellula | F39 | Congo Red | 13360 | BL3 | 6109 |
| 40 | Yes | E. coli | Nalidixic a | Intracellula | F116 | Sunset Yel | 14308 | BL2 | 5734.5 |
| 40 | Yes | E. coli | Nalidixic a | Intracellula | F10 | Alizarin | 12154 | BL1 | 6014.5 |
| 40 | Yes | E. coli | Nalidixic a | Intracellula | F84 | Neutral Re | 13431 | BL1 | 5286 |
| 40 | Yes | E. coli | Nalidixic a | Intracellula | F99 | Pyronin Y | 12735 | BL1 | 4519 |
| 40 | Yes | E. coli | Nalidixic a | Intracellula | F108 | Rhodamine | 11776 | BL1 | 4589.5 |
| 40 | Yes | E. coli | Nalidixic a | Intracellula | F109 | Riboflavin | 15361 | BL1 | 4298 |
| 40 | Yes | E. coli | Nalidixic a | Intracellula | F117 | SYBR Gree | 15324 | BL1 | 142353 |
| 40 | Yes | E. coli | Nalidixic a | Intracellula | F106 | Rhodamine | 12183 | BL1 | 5593 |
| 40 | Yes | E. coli | Nalidixic a | Intracellula | F73 | Malachite | 14011 | BL1 | 4579 |
| 40 | Yes | E. coli | Nalidixic a | Intracellula | F7 | Acridine or | 14559 | BL1 | 4610 |
| 40 | Yes | E. coli | Nalidixic a | Intracellula | F192 | 5-Carboxyf | 12913 | BL1 | 15907 |
| 40 | Yes | E. coli | Nalidixic a | Intracellula | F57 | Eosin Y | 12496 | BL1 | 18785 |
| 40 | Yes | E. coli | Nalidixic a | Intracellula | F27 | Calcein | 13763 | BL1 | 22786 |

| | | | | | | | | | |
|---|---|---|---|---|---|---|---|---|---|
| 40 | Yes | E. coli | Nalidixic ac | Intracellula | F12 | Amaranth | 14689 | BL1 | 20833 |
| 40 | Yes | E. coli | None | None | NA | Autofluores | 21525 | NA | 5548 |
| 40 | Yes | E. coli | Ofloxacin | Intracellula | F194 | Oxonol | 15153 | RL1 | 5831 |
| 40 | Yes | E. coli | Ofloxacin | Intracellula | F11 | Allura Red | 12462 | BL3 | 5300 |
| 40 | Yes | E. coli | Ofloxacin | Intracellula | F54 | DiSC3(5) | 11445 | RL1 | 4914 |
| 40 | Yes | E. coli | Ofloxacin | Intracellula | F124 | Thioflavin T | 11468 | VL2 | 4837.5 |
| 40 | Yes | E. coli | Ofloxacin | Intracellula | F101 | Quinacrine | 10924 | VL2 | 4950.5 |
| 40 | Yes | E. coli | Ofloxacin | Intracellula | F138 | Apigenin | 10529 | VL2 | 5142 |
| 40 | Yes | E. coli | Ofloxacin | Intracellula | F65 | H2FDA | 10825 | BL1 | 5961 |
| 40 | Yes | E. coli | Ofloxacin | Intracellula | F142 | Chlorophe | 13073 | BL3 | 5852 |
| 40 | Yes | E. coli | Ofloxacin | Intracellula | F39 | Congo Red | 14036 | BL3 | 6114 |
| 40 | Yes | E. coli | Ofloxacin | Intracellula | F116 | Sunset Yell | 12083 | BL2 | 6664 |
| 40 | Yes | E. coli | Ofloxacin | Intracellula | F10 | Alizarin | 13566 | BL1 | 5423 |
| 40 | Yes | E. coli | Ofloxacin | Intracellula | F84 | Neutral Re | 13574 | BL1 | 5354 |
| 40 | Yes | E. coli | Ofloxacin | Intracellula | F99 | Pyronin Y | 13577 | BL1 | 4986 |
| 40 | Yes | E. coli | Ofloxacin | Intracellula | F108 | Rhodamine | 10520 | BL1 | 5014.5 |
| 40 | Yes | E. coli | Ofloxacin | Intracellula | F109 | Riboflavin | 13626 | BL1 | 4809 |
| 40 | Yes | E. coli | Ofloxacin | Intracellula | F117 | SYBR Gree | 16706 | BL1 | 186336 |
| 40 | Yes | E. coli | Ofloxacin | Intracellula | F106 | Rhodamine | 13999 | BL1 | 5801 |
| 40 | Yes | E. coli | Ofloxacin | Intracellula | F73 | Malachite G | 12561 | BL1 | 5552 |
| 40 | Yes | E. coli | Ofloxacin | Intracellula | F7 | Acridine or | 12523 | BL1 | 5498 |
| 40 | Yes | E. coli | Ofloxacin | Intracellula | F192 | 5-Carboxyf | 12500 | BL1 | 17759 |
| 40 | Yes | E. coli | Ofloxacin | Intracellula | F57 | Eosin Y | 13678 | BL1 | 23847 |
| 40 | Yes | E. coli | Ofloxacin | Intracellula | F27 | Calcein | 18700 | BL1 | 26786 |
| 40 | Yes | E. coli | Ofloxacin | Intracellula | F12 | Amaranth | 19020 | BL1 | 333443 |
| 40 | Yes | E. coli | None | None | NA | Autofluores | 20468 | NA | 5688 |
| 40 | Yes | E. coli | Polymyxin | Wall | F194 | Oxonol | 18695 | RL1 | 5764 |
| 40 | Yes | E. coli | Polymyxin | Wall | F11 | Allura Red | 18952 | BL3 | 4652 |
| 40 | Yes | E. coli | Polymyxin | Wall | F54 | DiSC3(5) | 17598 | RL1 | 4306 |
| 40 | Yes | E. coli | Polymyxin | Wall | F124 | Thioflavin T | 17347 | VL2 | 4312 |
| 40 | Yes | E. coli | Polymyxin | Wall | F101 | Quinacrine | 16016 | VL2 | 4727.5 |
| 40 | Yes | E. coli | Polymyxin | Wall | F138 | Apigenin | 15438 | VL2 | 4955 |
| 40 | Yes | E. coli | Polymyxin | Wall | F65 | H2FDA | 16026 | BL1 | 4918 |
| 40 | Yes | E. coli | Polymyxin | Wall | F142 | Chlorophe | 16051 | BL3 | 5981 |
| 40 | Yes | E. coli | Polymyxin | Wall | F39 | Congo Red | 16460 | BL3 | 6074 |
| 40 | Yes | E. coli | Polymyxin | Wall | F116 | Sunset Yell | 17660 | BL2 | 5804.5 |
| 40 | Yes | E. coli | Polymyxin | Wall | F10 | Alizarin | 15633 | BL1 | 6002 |
| 40 | Yes | E. coli | Polymyxin | Wall | F84 | Neutral Re | 15906 | BL1 | 5126 |
| 40 | Yes | E. coli | Polymyxin | Wall | F99 | Pyronin Y | 15518 | BL1 | 5064 |
| 40 | Yes | E. coli | Polymyxin | Wall | F108 | Rhodamine | 15018 | BL1 | 5158 |
| 40 | Yes | E. coli | Polymyxin | Wall | F109 | Riboflavin | 15784 | BL1 | 4609 |
| 40 | Yes | E. coli | Polymyxin | Wall | F117 | SYBR Gree | 16351 | BL1 | 245267 |
| 40 | Yes | E. coli | Polymyxin | Wall | F106 | Rhodamine | 15654 | BL1 | 5766 |
| 40 | Yes | E. coli | Polymyxin | Wall | F73 | Malachite G | 16112 | BL1 | 5126 |
| 40 | Yes | E. coli | Polymyxin | Wall | F7 | Acridine or | 15472 | BL1 | 5018 |
| 40 | Yes | E. coli | Polymyxin | Wall | F192 | 5-Carboxyf | 15079 | BL1 | 16994 |
| 40 | Yes | E. coli | Polymyxin | Wall | F57 | Eosin Y | 15253 | BL1 | 23125 |
| 40 | Yes | E. coli | Polymyxin | Wall | F27 | Calcein | 16084 | BL1 | 25402.5 |

| | | | | | | | | | |
|---|---|---|---|---|---|---|---|---|---|
| 40 | Yes | E. coli | Polymyxin | Wall | F12 | Amaranth | 15228 | BL1 | 29273 |
| 40 | Yes | E. coli | None | None | NA | Autofluore: | 20673 | NA | 6077 |
| 40 | Yes | E. coli | Tetracyclin | Intracellul: | F194 | Oxonol | 21829 | RL1 | 4748 |
| 40 | Yes | E. coli | Tetracyclin | Intracellul: | F11 | Allura Red | 21073 | BL3 | 4073 |
| 40 | Yes | E. coli | Tetracyclin | Intracellul: | F54 | DiSC3(5) | 19573 | RL1 | 4136 |
| 40 | Yes | E. coli | Tetracyclin | Intracellul: | F124 | Thioflavin T | 19011 | VL2 | 3812 |
| 40 | Yes | E. coli | Tetracyclin | Intracellul: | F101 | Quinacrine | 17136 | VL2 | 4097 |
| 40 | Yes | E. coli | Tetracyclin | Intracellul: | F138 | Apigenin | 17591 | VL2 | 4035 |
| 40 | Yes | E. coli | Tetracyclin | Intracellul: | F65 | H2FDA | 19267 | BL1 | 4315 |
| 40 | Yes | E. coli | Tetracyclin | Intracellul: | F142 | Chlorophe | 19514 | BL3 | 4852 |
| 40 | Yes | E. coli | Tetracyclin | Intracellul: | F39 | Congo Red | 19016 | BL3 | 5346.5 |
| 40 | Yes | E. coli | Tetracyclin | Intracellul: | F116 | Sunset Yel | 17864 | BL2 | 5299 |
| 40 | Yes | E. coli | Tetracyclin | Intracellul: | F10 | Alizarin | 17793 | BL1 | 5520 |
| 40 | Yes | E. coli | Tetracyclin | Intracellul: | F84 | Neutral Re | 18770 | BL1 | 4780 |
| 40 | Yes | E. coli | Tetracyclin | Intracellul: | F99 | Pyronin Y | 17401 | BL1 | 4586 |
| 40 | Yes | E. coli | Tetracyclin | Intracellul: | F108 | Rhodamine | 17268 | BL1 | 4289 |
| 40 | Yes | E. coli | Tetracyclin | Intracellul: | F109 | Riboflavin | 18836 | BL1 | 4122.5 |
| 40 | Yes | E. coli | Tetracyclin | Intracellul: | F117 | SYBR Gree | 18724 | BL1 | 137565 |
| 40 | Yes | E. coli | Tetracyclin | Intracellul: | F106 | Rhodamine | 18889 | BL1 | 4999 |
| 40 | Yes | E. coli | Tetracyclin | Intracellul: | F73 | Malachite ( | 17976 | BL1 | 4485.5 |
| 40 | Yes | E. coli | Tetracyclin | Intracellul: | F7 | Acridine or | 17484 | BL1 | 4530 |
| 40 | Yes | E. coli | Tetracyclin | Intracellul: | F192 | 5-Carboxyf | 16899 | BL1 | 15066 |
| 40 | Yes | E. coli | Tetracyclin | Intracellul: | F57 | Eosin Y | 17753 | BL1 | 24154 |
| 40 | Yes | E. coli | Tetracyclin | Intracellul: | F27 | Calcein | 17743 | BL1 | 17828 |
| 40 | Yes | E. coli | Tetracyclin | Intracellul: | F12 | Amaranth | 16837 | BL1 | 19961 |
| 40 | Yes | E. coli | None | None | NA | Autofluore: | 21785 | NA | 5591 |
| 40 | Yes | E. coli | Vancomyc | Wall | F194 | Oxonol | 24857 | RL1 | 4386 |
| 40 | Yes | E. coli | Vancomyc | Wall | F11 | Allura Red | 24851 | BL3 | 3836 |
| 40 | Yes | E. coli | Vancomyc | Wall | F54 | DiSC3(5) | 24376 | RL1 | 3907 |
| 40 | Yes | E. coli | Vancomyc | Wall | F124 | Thioflavin T | 21956 | VL2 | 3909 |
| 40 | Yes | E. coli | Vancomyc | Wall | F101 | Quinacrine | 20576 | VL2 | 4290.5 |
| 40 | Yes | E. coli | Vancomyc | Wall | F138 | Apigenin | 22065 | VL2 | 3693 |
| 40 | Yes | E. coli | Vancomyc | Wall | F65 | H2FDA | 20260 | BL1 | 4198.5 |
| 40 | Yes | E. coli | Vancomyc | Wall | F142 | Chlorophe | 20592 | BL3 | 5179.5 |
| 40 | Yes | E. coli | Vancomyc | Wall | F39 | Congo Red | 21496 | BL3 | 5894 |
| 40 | Yes | E. coli | Vancomyc | Wall | F116 | Sunset Yel | 22520 | BL2 | 5784 |
| 40 | Yes | E. coli | Vancomyc | Wall | F10 | Alizarin | 23557 | BL1 | 4625 |
| 40 | Yes | E. coli | Vancomyc | Wall | F84 | Neutral Re | 23562 | BL1 | 4186 |
| 40 | Yes | E. coli | Vancomyc | Wall | F99 | Pyronin Y | 22055 | BL1 | 3944 |
| 40 | Yes | E. coli | Vancomyc | Wall | F108 | Rhodamine | 20785 | BL1 | 4264 |
| 40 | Yes | E. coli | Vancomyc | Wall | F109 | Riboflavin | 22388 | BL1 | 3920 |
| 40 | Yes | E. coli | Vancomyc | Wall | F117 | SYBR Gree | 21275 | BL1 | 149637 |
| 40 | Yes | E. coli | Vancomyc | Wall | F106 | Rhodamine | 22275 | BL1 | 4882 |
| 40 | Yes | E. coli | Vancomyc | Wall | F73 | Malachite ( | 20291 | BL1 | 4932 |
| 40 | Yes | E. coli | Vancomyc | Wall | F7 | Acridine or | 16804 | BL1 | 5607.5 |
| 40 | Yes | E. coli | Vancomyc | Wall | F192 | 5-Carboxyf | 16607 | BL1 | 33862 |
| 40 | Yes | E. coli | Vancomyc | Wall | F57 | Eosin Y | 16075 | BL1 | 60930 |
| 40 | Yes | E. coli | Vancomyc | Wall | F27 | Calcein | 16872 | BL1 | 59534.5 |

| | | | | | | | | | |
|---|---|---|---|---|---|---|---|---|---|
| 40 | Yes | E. coli | Vancomyc | Wall | F12 | Amaranth | 14840 | BL1 | 65318.5 |
| 40 | Yes | A. baylyi | None | None | NA | Autofluore: | 2451 | NA | 854 |
| 40 | Yes | A. baylyi | None | None | F194 | Oxonol | 2552 | RL1 | 1823 |
| 40 | Yes | A. baylyi | None | None | F11 | Allura Red | 2447 | BL3 | 1986 |
| 40 | Yes | A. baylyi | None | None | F54 | DiSC3(5) | 2574 | RL1 | 1904.5 |
| 40 | Yes | A. baylyi | None | None | F124 | Thioflavin T | 2368 | VL2 | 2491.5 |
| 40 | Yes | A. baylyi | None | None | F101 | Quinacrine | 2216 | VL2 | 2538.5 |
| 40 | Yes | A. baylyi | None | None | F138 | Apigenin | 2190 | VL2 | 2591.5 |
| 40 | Yes | A. baylyi | None | None | F65 | H2FDA | 2132 | BL1 | 4161 |
| 40 | Yes | A. baylyi | None | None | F142 | Chlorophe | 2134 | BL3 | 5113.5 |
| 40 | Yes | A. baylyi | None | None | F39 | Congo Red | 2190 | BL3 | 5118.5 |
| 40 | Yes | A. baylyi | None | None | F116 | Sunset Yel | 2268 | BL2 | 5411 |
| 40 | Yes | A. baylyi | None | None | F10 | Alizarin | 1893 | BL1 | 5563 |
| 40 | Yes | A. baylyi | None | None | F84 | Neutral Re | 2125 | BL1 | 3999 |
| 40 | Yes | A. baylyi | None | None | F99 | Pyronin Y | 2181 | BL1 | 3889 |
| 40 | Yes | A. baylyi | None | None | F108 | Rhodamine | 2148 | BL1 | 3263 |
| 40 | Yes | A. baylyi | None | None | F109 | Riboflavin | 2039 | BL1 | 2932 |
| 40 | Yes | A. baylyi | None | None | F117 | SYBR Gree | 1796 | BL1 | 237630 |
| 40 | Yes | A. baylyi | None | None | F106 | Rhodamine | 2067 | BL1 | 4694 |
| 40 | Yes | A. baylyi | None | None | F73 | Malachite ( | 2072 | BL1 | 4542 |
| 40 | Yes | A. baylyi | None | None | F7 | Acridine or | 2073 | BL1 | 3473 |
| 40 | Yes | A. baylyi | None | None | F192 | 5-Carboxyf | 2074 | BL1 | 12400.5 |
| 40 | Yes | A. baylyi | None | None | F57 | Eosin Y | 1929 | BL1 | 12595 |
| 40 | Yes | A. baylyi | None | None | F27 | Calcein | 1949 | BL1 | 13871 |
| 40 | Yes | A. baylyi | None | None | F12 | Amaranth | 2044 | BL1 | 12730 |
| 40 | Yes | A. baylyi | None | None | NA | Autofluore: | 2231 | NA | 5793 |
| 40 | Yes | A. baylyi | Amoxacilli | Wall | F194 | Oxonol | 2245 | RL1 | 4166 |
| 40 | Yes | A. baylyi | Amoxacilli | Wall | F11 | Allura Red | 2112 | BL3 | 4058 |
| 40 | Yes | A. baylyi | Amoxacilli | Wall | F54 | DiSC3(5) | 1927 | RL1 | 3335 |
| 40 | Yes | A. baylyi | Amoxacilli | Wall | F124 | Thioflavin T | 2139 | VL2 | 3653 |
| 40 | Yes | A. baylyi | Amoxacilli | Wall | F101 | Quinacrine | 2064 | VL2 | 3698 |
| 40 | Yes | A. baylyi | Amoxacilli | Wall | F138 | Apigenin | 2083 | VL2 | 3900 |
| 40 | Yes | A. baylyi | Amoxacilli | Wall | F65 | H2FDA | 2002 | BL1 | 4903.5 |
| 40 | Yes | A. baylyi | Amoxacilli | Wall | F142 | Chlorophe | 1958 | BL3 | 5705 |
| 40 | Yes | A. baylyi | Amoxacilli | Wall | F39 | Congo Red | 1767 | BL3 | 8742 |
| 40 | Yes | A. baylyi | Amoxacilli | Wall | F116 | Sunset Yel | 1780 | BL2 | 6935 |
| 40 | Yes | A. baylyi | Amoxacilli | Wall | F10 | Alizarin | 1746 | BL1 | 6605.5 |
| 40 | Yes | A. baylyi | Amoxacilli | Wall | F84 | Neutral Re | 1665 | BL1 | 5306 |
| 40 | Yes | A. baylyi | Amoxacilli | Wall | F99 | Pyronin Y | 2153 | BL1 | 4646 |
| 40 | Yes | A. baylyi | Amoxacilli | Wall | F108 | Rhodamine | 1908 | BL1 | 4319.5 |
| 40 | Yes | A. baylyi | Amoxacilli | Wall | F109 | Riboflavin | 2083 | BL1 | 3933 |
| 40 | Yes | A. baylyi | Amoxacilli | Wall | F117 | SYBR Gree | 1861 | BL1 | 281956 |
| 40 | Yes | A. baylyi | Amoxacilli | Wall | F106 | Rhodamine | 1931 | BL1 | 4577 |
| 40 | Yes | A. baylyi | Amoxacilli | Wall | F73 | Malachite ( | 2071 | BL1 | 4385 |
| 40 | Yes | A. baylyi | Amoxacilli | Wall | F7 | Acridine or | 1978 | BL1 | 4167 |
| 40 | Yes | A. baylyi | Amoxacilli | Wall | F192 | 5-Carboxyf | 1708 | BL1 | 12405 |
| 40 | Yes | A. baylyi | Amoxacilli | Wall | F57 | Eosin Y | 1758 | BL1 | 13799 |
| 40 | Yes | A. baylyi | Amoxacilli | Wall | F27 | Calcein | 1935 | BL1 | 14515 |

| | | | | | | | | | |
|---|---|---|---|---|---|---|---|---|---|
| 40 | Yes | A. baylyi | Amoxacillin | Wall | F12 | Amaranth | 1950 | BL1 | 15408.5 |
| 40 | Yes | A. baylyi | None | None | NA | Autofluores | 2228 | NA | 5629 |
| 40 | Yes | A. baylyi | Azithromyc | Intracellula | F194 | Oxonol | 2348 | RL1 | 4440.5 |
| 40 | Yes | A. baylyi | Azithromyc | Intracellula | F11 | Allura Red | 2258 | BL3 | 4306.5 |
| 40 | Yes | A. baylyi | Azithromyc | Intracellula | F54 | DiSC3(5) | 2222 | RL1 | 3718.5 |
| 40 | Yes | A. baylyi | Azithromyc | Intracellula | F124 | Thioflavin T | 2004 | VL2 | 3992 |
| 40 | Yes | A. baylyi | Azithromyc | Intracellula | F101 | Quinacrine | 2141 | VL2 | 4389 |
| 40 | Yes | A. baylyi | Azithromyc | Intracellula | F138 | Apigenin | 1880 | VL2 | 4542.5 |
| 40 | Yes | A. baylyi | Azithromyc | Intracellula | F65 | H2FDA | 1939 | BL1 | 5915 |
| 40 | Yes | A. baylyi | Azithromyc | Intracellula | F142 | Chlorophe | 2214 | BL3 | 7571 |
| 40 | Yes | A. baylyi | Azithromyc | Intracellula | F39 | Congo Red | 2108 | BL3 | 7078.5 |
| 40 | Yes | A. baylyi | Azithromyc | Intracellula | F116 | Sunset Yel | 2158 | BL2 | 5545 |
| 40 | Yes | A. baylyi | Azithromyc | Intracellula | F10 | Alizarin | 1840 | BL1 | 5617 |
| 40 | Yes | A. baylyi | Azithromyc | Intracellula | F84 | Neutral Re | 2148 | BL1 | 4579 |
| 40 | Yes | A. baylyi | Azithromyc | Intracellula | F99 | Pyronin Y | 1978 | BL1 | 5715.5 |
| 40 | Yes | A. baylyi | Azithromyc | Intracellula | F108 | Rhodamine | 2021 | BL1 | 3984 |
| 40 | Yes | A. baylyi | Azithromyc | Intracellula | F109 | Riboflavin | 1816 | BL1 | 3874.5 |
| 40 | Yes | A. baylyi | Azithromyc | Intracellula | F117 | SYBR Gree | 2102 | BL1 | 203606.5 |
| 40 | Yes | A. baylyi | Azithromyc | Intracellula | F106 | Rhodamine | 2093 | BL1 | 5190 |
| 40 | Yes | A. baylyi | Azithromyc | Intracellula | F73 | Malachite ( | 1939 | BL1 | 5271 |
| 40 | Yes | A. baylyi | Azithromyc | Intracellula | F7 | Acridine or | 1701 | BL1 | 4438 |
| 40 | Yes | A. baylyi | Azithromyc | Intracellula | F192 | 5-Carboxyf | 1980 | BL1 | 12201 |
| 40 | Yes | A. baylyi | Azithromyc | Intracellula | F57 | Eosin Y | 2022 | BL1 | 14486.5 |
| 40 | Yes | A. baylyi | Azithromyc | Intracellula | F27 | Calcein | 1923 | BL1 | 13287 |
| 40 | Yes | A. baylyi | Azithromyc | Intracellula | F12 | Amaranth | 2181 | BL1 | 16304 |
| 40 | Yes | A. baylyi | None | None | NA | Autofluores | 2172 | NA | 5167 |
| 40 | Yes | A. baylyi | Trimethopr | Intracellula | F194 | Oxonol | 2166 | RL1 | 4085 |
| 40 | Yes | A. baylyi | Trimethopr | Intracellula | F11 | Allura Red | 2540 | BL3 | 3876 |
| 40 | Yes | A. baylyi | Trimethopr | Intracellula | F54 | DiSC3(5) | 2329 | RL1 | 3905 |
| 40 | Yes | A. baylyi | Trimethopr | Intracellula | F124 | Thioflavin T | 2078 | VL2 | 4241.5 |
| 40 | Yes | A. baylyi | Trimethopr | Intracellula | F101 | Quinacrine | 1898 | VL2 | 4545.5 |
| 40 | Yes | A. baylyi | Trimethopr | Intracellula | F138 | Apigenin | 2051 | VL2 | 4554 |
| 40 | Yes | A. baylyi | Trimethopr | Intracellula | F65 | H2FDA | 2143 | BL1 | 5366 |
| 40 | Yes | A. baylyi | Trimethopr | Intracellula | F142 | Chlorophe | 2187 | BL3 | 4996 |
| 40 | Yes | A. baylyi | Trimethopr | Intracellula | F39 | Congo Red | 1987 | BL3 | 6861 |
| 40 | Yes | A. baylyi | Trimethopr | Intracellula | F116 | Sunset Yel | 2160 | BL2 | 8266 |
| 40 | Yes | A. baylyi | Trimethopr | Intracellula | F10 | Alizarin | 2181 | BL1 | 5368 |
| 40 | Yes | A. baylyi | Trimethopr | Intracellula | F84 | Neutral Re | 2155 | BL1 | 5839 |
| 40 | Yes | A. baylyi | Trimethopr | Intracellula | F99 | Pyronin Y | 1969 | BL1 | 4226 |
| 40 | Yes | A. baylyi | Trimethopr | Intracellula | F108 | Rhodamine | 2122 | BL1 | 3484.5 |
| 40 | Yes | A. baylyi | Trimethopr | Intracellula | F109 | Riboflavin | 1938 | BL1 | 3820 |
| 40 | Yes | A. baylyi | Trimethopr | Intracellula | F117 | SYBR Gree | 2120 | BL1 | 279880.5 |
| 40 | Yes | A. baylyi | Trimethopr | Intracellula | F106 | Rhodamine | 2180 | BL1 | 5020.5 |
| 40 | Yes | A. baylyi | Trimethopr | Intracellula | F73 | Malachite ( | 2057 | BL1 | 4715 |
| 40 | Yes | A. baylyi | Trimethopr | Intracellula | F7 | Acridine or | 2101 | BL1 | 4911 |
| 40 | Yes | A. baylyi | Trimethopr | Intracellula | F192 | 5-Carboxyf | 2157 | BL1 | 11982 |
| 40 | Yes | A. baylyi | Trimethopr | Intracellula | F57 | Eosin Y | 2179 | BL1 | 10658 |
| 40 | Yes | A. baylyi | Trimethopr | Intracellula | F27 | Calcein | 2025 | BL1 | 13867 |

| | | | | | | | | | |
|---|---|---|---|---|---|---|---|---|---|
| 40 | Yes | A. baylyi | Trimethopr | Intracellul | F12 | Amaranth | 1964 | BL1 | 13567 |
| 40 | Yes | A. baylyi | None | None | NA | Autofluore | 2371 | NA | 5909 |
| 40 | Yes | A. baylyi | Chloramph | Intracellul | F194 | Oxonol | 1637 | RL1 | 4892 |
| 40 | Yes | A. baylyi | Chloramph | Intracellul | F11 | Allura Red | 1827 | BL3 | 5285 |
| 40 | Yes | A. baylyi | Chloramph | Intracellul | F54 | DiSC3(5) | 1507 | RL1 | 5052 |
| 40 | Yes | A. baylyi | Chloramph | Intracellul | F124 | Thioflavin T | 1608 | VL2 | 5393.5 |
| 40 | Yes | A. baylyi | Chloramph | Intracellul | F101 | Quinacrine | 1545 | VL2 | 4500 |
| 40 | Yes | A. baylyi | Chloramph | Intracellul | F138 | Apigenin | 1525 | VL2 | 4084 |
| 40 | Yes | A. baylyi | Chloramph | Intracellul | F65 | H2FDA | 1629 | BL1 | 5025 |
| 40 | Yes | A. baylyi | Chloramph | Intracellul | F142 | Chlorophe | 1610 | BL3 | 5686.5 |
| 40 | Yes | A. baylyi | Chloramph | Intracellul | F39 | Congo Red | 1623 | BL3 | 6090 |
| 40 | Yes | A. baylyi | Chloramph | Intracellul | F116 | Sunset Yel | 1610 | BL2 | 5803 |
| 40 | Yes | A. baylyi | Chloramph | Intracellul | F10 | Alizarin | 1381 | BL1 | 7573 |
| 40 | Yes | A. baylyi | Chloramph | Intracellul | F84 | Neutral Re | 1436 | BL1 | 6912.5 |
| 40 | Yes | A. baylyi | Chloramph | Intracellul | F99 | Pyronin Y | 1069 | BL1 | 5467 |
| 40 | Yes | A. baylyi | Chloramph | Intracellul | F108 | Rhodamine | 1388 | BL1 | 6498 |
| 40 | Yes | A. baylyi | Chloramph | Intracellul | F109 | Riboflavin | 1411 | BL1 | 3998 |
| 40 | Yes | A. baylyi | Chloramph | Intracellul | F117 | SYBR Gree | 1292 | BL1 | 202917 |
| 40 | Yes | A. baylyi | Chloramph | Intracellul | F106 | Rhodamine | 1368 | BL1 | 5240 |
| 40 | Yes | A. baylyi | Chloramph | Intracellul | F73 | Malachite ( | 1391 | BL1 | 5013 |
| 40 | Yes | A. baylyi | Chloramph | Intracellul | F7 | Acridine or | 1364 | BL1 | 4731 |
| 40 | Yes | A. baylyi | Chloramph | Intracellul | F192 | 5-Carboxyf | 1364 | BL1 | 12627 |
| 40 | Yes | A. baylyi | Chloramph | Intracellul | F57 | Eosin Y | 1643 | BL1 | 17133 |
| 40 | Yes | A. baylyi | Chloramph | Intracellul | F27 | Calcein | 1539 | BL1 | 13121 |
| 40 | Yes | A. baylyi | Chloramph | Intracellul | F12 | Amaranth | 1140 | BL1 | 14536 |
| 40 | Yes | A. baylyi | None | None | NA | Autofluore | 2566 | NA | 6534.5 |
| 40 | Yes | A. baylyi | Colistin | Wall | F194 | Oxonol | 1205 | RL1 | 39134 |
| 40 | Yes | A. baylyi | Colistin | Wall | F11 | Allura Red | 1184 | BL3 | 24452 |
| 40 | Yes | A. baylyi | Colistin | Wall | F54 | DiSC3(5) | 1098 | RL1 | 30284 |
| 40 | Yes | A. baylyi | Colistin | Wall | F124 | Thioflavin T | 1025 | VL2 | 15334 |
| 40 | Yes | A. baylyi | Colistin | Wall | F101 | Quinacrine | 869 | VL2 | 19443 |
| 40 | Yes | A. baylyi | Colistin | Wall | F138 | Apigenin | 899 | VL2 | 17260 |
| 40 | Yes | A. baylyi | Colistin | Wall | F65 | H2FDA | 1040 | BL1 | 18129.5 |
| 40 | Yes | A. baylyi | Colistin | Wall | F142 | Chlorophe | 1079 | BL3 | 16297 |
| 40 | Yes | A. baylyi | Colistin | Wall | F39 | Congo Red | 1016 | BL3 | 13500.5 |
| 40 | Yes | A. baylyi | Colistin | Wall | F116 | Sunset Yel | 923 | BL2 | 15345 |
| 40 | Yes | A. baylyi | Colistin | Wall | F10 | Alizarin | 1019 | BL1 | 15779 |
| 40 | Yes | A. baylyi | Colistin | Wall | F84 | Neutral Re | 973 | BL1 | 16420 |
| 40 | Yes | A. baylyi | Colistin | Wall | F99 | Pyronin Y | 959 | BL1 | 10689 |
| 40 | Yes | A. baylyi | Colistin | Wall | F108 | Rhodamine | 973 | BL1 | 12742 |
| 40 | Yes | A. baylyi | Colistin | Wall | F109 | Riboflavin | 981 | BL1 | 13424 |
| 40 | Yes | A. baylyi | Colistin | Wall | F117 | SYBR Gree | 903 | BL1 | 2291640 |
| 40 | Yes | A. baylyi | Colistin | Wall | F106 | Rhodamine | 980 | BL1 | 55453.5 |
| 40 | Yes | A. baylyi | Colistin | Wall | F73 | Malachite ( | 969 | BL1 | 31126 |
| 40 | Yes | A. baylyi | Colistin | Wall | F7 | Acridine or | 912 | BL1 | 40431.5 |
| 40 | Yes | A. baylyi | Colistin | Wall | F192 | 5-Carboxyf | 899 | BL1 | 46788 |
| 40 | Yes | A. baylyi | Colistin | Wall | F57 | Eosin Y | 866 | BL1 | 36696 |
| 40 | Yes | A. baylyi | Colistin | Wall | F27 | Calcein | 891 | BL1 | 43135 |

| | | | | | | | | | |
|---|---|---|---|---|---|---|---|---|---|
| 40 | Yes | A. baylyi | Colistin | Wall | F12 | Amaranth | 907 | BL1 | 39837 |
| 40 | Yes | A. baylyi | None | None | NA | Autofluore: | 2285 | NA | 6526 |
| 40 | Yes | A. baylyi | Gentamici | Intracellul | F194 | Oxonol | 2159 | RL1 | 5766 |
| 40 | Yes | A. baylyi | Gentamici | Intracellul | F11 | Allura Red | 2034 | BL3 | 5372 |
| 40 | Yes | A. baylyi | Gentamici | Intracellul | F54 | DiSC3(5) | 1749 | RL1 | 5209 |
| 40 | Yes | A. baylyi | Gentamici | Intracellul | F124 | Thioflavin T | 1702 | VL2 | 5452.5 |
| 40 | Yes | A. baylyi | Gentamici | Intracellul | F101 | Quinacrine | 1543 | VL2 | 6307 |
| 40 | Yes | A. baylyi | Gentamici | Intracellul | F138 | Apigenin | 1656 | VL2 | 5460.5 |
| 40 | Yes | A. baylyi | Gentamici | Intracellul | F65 | H2FDA | 1789 | BL1 | 6166 |
| 40 | Yes | A. baylyi | Gentamici | Intracellul | F142 | Chlorophe | 1855 | BL3 | 6404 |
| 40 | Yes | A. baylyi | Gentamici | Intracellul | F39 | Congo Red | 1674 | BL3 | 6009.5 |
| 40 | Yes | A. baylyi | Gentamici | Intracellul | F116 | Sunset Yel | 2029 | BL2 | 5932 |
| 40 | Yes | A. baylyi | Gentamici | Intracellul | F10 | Alizarin | 1653 | BL1 | 5508 |
| 40 | Yes | A. baylyi | Gentamici | Intracellul | F84 | Neutral Re | 1807 | BL1 | 4958 |
| 40 | Yes | A. baylyi | Gentamici | Intracellul | F99 | Pyronin Y | 1789 | BL1 | 4870 |
| 40 | Yes | A. baylyi | Gentamici | Intracellul | F108 | Rhodamine | 1903 | BL1 | 4210 |
| 40 | Yes | A. baylyi | Gentamici | Intracellul | F109 | Riboflavin | 1641 | BL1 | 4996 |
| 40 | Yes | A. baylyi | Gentamici | Intracellul | F117 | SYBR Gree | 1685 | BL1 | 337365 |
| 40 | Yes | A. baylyi | Gentamici | Intracellul | F106 | Rhodamine | 1689 | BL1 | 5830 |
| 40 | Yes | A. baylyi | Gentamici | Intracellul | F73 | Malachite ( | 1812 | BL1 | 5783 |
| 40 | Yes | A. baylyi | Gentamici | Intracellul | F7 | Acridine or | 1829 | BL1 | 5971 |
| 40 | Yes | A. baylyi | Gentamici | Intracellul | F192 | 5-Carboxyf | 1692 | BL1 | 13460 |
| 40 | Yes | A. baylyi | Gentamici | Intracellul | F57 | Eosin Y | 1619 | BL1 | 11328 |
| 40 | Yes | A. baylyi | Gentamici | Intracellul | F27 | Calcein | 1648 | BL1 | 20948.5 |
| 40 | Yes | A. baylyi | Gentamici | Intracellul | F12 | Amaranth | 1766 | BL1 | 14585 |
| 40 | Yes | A. baylyi | None | None | NA | Autofluore: | 2290 | NA | 6917.5 |
| 40 | Yes | A. baylyi | Nitrofurant | Intracellul | F194 | Oxonol | 2098 | RL1 | 5730 |
| 40 | Yes | A. baylyi | Nitrofurant | Intracellul | F11 | Allura Red | 1984 | BL3 | 4900.5 |
| 40 | Yes | A. baylyi | Nitrofurant | Intracellul | F54 | DiSC3(5) | 1845 | RL1 | 5059 |
| 40 | Yes | A. baylyi | Nitrofurant | Intracellul | F124 | Thioflavin T | 1763 | VL2 | 5177 |
| 40 | Yes | A. baylyi | Nitrofurant | Intracellul | F101 | Quinacrine | 1812 | VL2 | 5705 |
| 40 | Yes | A. baylyi | Nitrofurant | Intracellul | F138 | Apigenin | 1706 | VL2 | 5278 |
| 40 | Yes | A. baylyi | Nitrofurant | Intracellul | F65 | H2FDA | 1898 | BL1 | 5858.5 |
| 40 | Yes | A. baylyi | Nitrofurant | Intracellul | F142 | Chlorophe | 1966 | BL3 | 6317 |
| 40 | Yes | A. baylyi | Nitrofurant | Intracellul | F39 | Congo Red | 1915 | BL3 | 5899 |
| 40 | Yes | A. baylyi | Nitrofurant | Intracellul | F116 | Sunset Yel | 1851 | BL2 | 5614 |
| 40 | Yes | A. baylyi | Nitrofurant | Intracellul | F10 | Alizarin | 1634 | BL1 | 5031 |
| 40 | Yes | A. baylyi | Nitrofurant | Intracellul | F84 | Neutral Re | 1656 | BL1 | 5040 |
| 40 | Yes | A. baylyi | Nitrofurant | Intracellul | F99 | Pyronin Y | 1917 | BL1 | 4239 |
| 40 | Yes | A. baylyi | Nitrofurant | Intracellul | F108 | Rhodamine | 1717 | BL1 | 3724 |
| 40 | Yes | A. baylyi | Nitrofurant | Intracellul | F109 | Riboflavin | 1908 | BL1 | 4492 |
| 40 | Yes | A. baylyi | Nitrofurant | Intracellul | F117 | SYBR Gree | 1545 | BL1 | 157362 |
| 40 | Yes | A. baylyi | Nitrofurant | Intracellul | F106 | Rhodamine | 1677 | BL1 | 4818 |
| 40 | Yes | A. baylyi | Nitrofurant | Intracellul | F73 | Malachite ( | 1594 | BL1 | 4899 |
| 40 | Yes | A. baylyi | Nitrofurant | Intracellul | F7 | Acridine or | 1854 | BL1 | 4860.5 |
| 40 | Yes | A. baylyi | Nitrofurant | Intracellul | F192 | 5-Carboxyf | 1540 | BL1 | 7290 |
| 40 | Yes | A. baylyi | Nitrofurant | Intracellul | F57 | Eosin Y | 1687 | BL1 | 9056 |
| 40 | Yes | A. baylyi | Nitrofurant | Intracellul | F27 | Calcein | 1361 | BL1 | 11904 |

| | | | | | | | | | |
|---|---|---|---|---|---|---|---|---|---|
| 40 | Yes | A. baylyi | Nitrofurant | Intracellula | F12 | Amaranth | 1547 | BL1 | 13417 |
| 40 | Yes | A. baylyi | None | None | NA | Autofluore: | 2243 | NA | 6540 |
| 40 | Yes | A. baylyi | Pentamidi | Wall | F194 | Oxonol | 1352 | RL1 | 12108 |
| 40 | Yes | A. baylyi | Pentamidi | Wall | F11 | Allura Red | 1218 | BL3 | 8040.5 |
| 40 | Yes | A. baylyi | Pentamidi | Wall | F54 | DiSC3(5) | 1137 | RL1 | 8196 |
| 40 | Yes | A. baylyi | Pentamidi | Wall | F124 | Thioflavin T | 1039 | VL2 | 11199 |
| 40 | Yes | A. baylyi | Pentamidi | Wall | F101 | Quinacrine | 1053 | VL2 | 8249 |
| 40 | Yes | A. baylyi | Pentamidi | Wall | F138 | Apigenin | 1024 | VL2 | 9628 |
| 40 | Yes | A. baylyi | Pentamidi | Wall | F65 | H2FDA | 1091 | BL1 | 8516 |
| 40 | Yes | A. baylyi | Pentamidi | Wall | F142 | Chlorophe | 1047 | BL3 | 7852 |
| 40 | Yes | A. baylyi | Pentamidi | Wall | F39 | Congo Red | 1360 | BL3 | 9158.5 |
| 40 | Yes | A. baylyi | Pentamidi | Wall | F116 | Sunset Yel | 1126 | BL2 | 8561.5 |
| 40 | Yes | A. baylyi | Pentamidi | Wall | F10 | Alizarin | 1068 | BL1 | 7177.5 |
| 40 | Yes | A. baylyi | Pentamidi | Wall | F84 | Neutral Re | 1062 | BL1 | 8205.5 |
| 40 | Yes | A. baylyi | Pentamidi | Wall | F99 | Pyronin Y | 961 | BL1 | 6404 |
| 40 | Yes | A. baylyi | Pentamidi | Wall | F108 | Rhodamine | 1059 | BL1 | 6583 |
| 40 | Yes | A. baylyi | Pentamidi | Wall | F109 | Riboflavin | 1024 | BL1 | 7375.5 |
| 40 | Yes | A. baylyi | Pentamidi | Wall | F117 | SYBR Gree | 946 | BL1 | 1479346 |
| 40 | Yes | A. baylyi | Pentamidi | Wall | F106 | Rhodamine | 1067 | BL1 | 27712 |
| 40 | Yes | A. baylyi | Pentamidi | Wall | F73 | Malachite ( | 1081 | BL1 | 10239 |
| 40 | Yes | A. baylyi | Pentamidi | Wall | F7 | Acridine or | 1030 | BL1 | 12320 |
| 40 | Yes | A. baylyi | Pentamidi | Wall | F192 | 5-Carboxyf | 956 | BL1 | 12937 |
| 40 | Yes | A. baylyi | Pentamidi | Wall | F57 | Eosin Y | 984 | BL1 | 15378.5 |
| 40 | Yes | A. baylyi | Pentamidi | Wall | F27 | Calcein | 1052 | BL1 | 13719 |
| 40 | Yes | A. baylyi | Pentamidi | Wall | F12 | Amaranth | 898 | BL1 | 15202 |
| 40 | Yes | A. baylyi | None | None | NA | Autofluore: | 2235 | NA | 9149 |
| 40 | Yes | A. baylyi | Rifampicin | Intracellula | F194 | Oxonol | 2510 | RL1 | 7221.5 |
| 40 | Yes | A. baylyi | Rifampicin | Intracellula | F11 | Allura Red | 2420 | BL3 | 6532.5 |
| 40 | Yes | A. baylyi | Rifampicin | Intracellula | F54 | DiSC3(5) | 2245 | RL1 | 5164 |
| 40 | Yes | A. baylyi | Rifampicin | Intracellula | F124 | Thioflavin T | 2315 | VL2 | 5974 |
| 40 | Yes | A. baylyi | Rifampicin | Intracellula | F101 | Quinacrine | 2007 | VL2 | 5721 |
| 40 | Yes | A. baylyi | Rifampicin | Intracellula | F138 | Apigenin | 2053 | VL2 | 5009 |
| 40 | Yes | A. baylyi | Rifampicin | Intracellula | F65 | H2FDA | 2030 | BL1 | 5537.5 |
| 40 | Yes | A. baylyi | Rifampicin | Intracellula | F142 | Chlorophe | 2291 | BL3 | 5752 |
| 40 | Yes | A. baylyi | Rifampicin | Intracellula | F39 | Congo Red | 2311 | BL3 | 6319 |
| 40 | Yes | A. baylyi | Rifampicin | Intracellula | F116 | Sunset Yel | 2153 | BL2 | 5671 |
| 40 | Yes | A. baylyi | Rifampicin | Intracellula | F10 | Alizarin | 2068 | BL1 | 4682 |
| 40 | Yes | A. baylyi | Rifampicin | Intracellula | F84 | Neutral Re | 2135 | BL1 | 5133 |
| 40 | Yes | A. baylyi | Rifampicin | Intracellula | F99 | Pyronin Y | 2028 | BL1 | 4569 |
| 40 | Yes | A. baylyi | Rifampicin | Intracellula | F108 | Rhodamine | 2018 | BL1 | 3999.5 |
| 40 | Yes | A. baylyi | Rifampicin | Intracellula | F109 | Riboflavin | 1957 | BL1 | 4490 |
| 40 | Yes | A. baylyi | Rifampicin | Intracellula | F117 | SYBR Gree | 1931 | BL1 | 443114 |
| 40 | Yes | A. baylyi | Rifampicin | Intracellula | F106 | Rhodamine | 2121 | BL1 | 5679 |
| 40 | Yes | A. baylyi | Rifampicin | Intracellula | F73 | Malachite ( | 1805 | BL1 | 5132 |
| 40 | Yes | A. baylyi | Rifampicin | Intracellula | F7 | Acridine or | 2092 | BL1 | 5283 |
| 40 | Yes | A. baylyi | Rifampicin | Intracellula | F192 | 5-Carboxyf | 1737 | BL1 | 6008 |
| 40 | Yes | A. baylyi | Rifampicin | Intracellula | F57 | Eosin Y | 1794 | BL1 | 6229.5 |
| 40 | Yes | A. baylyi | Rifampicin | Intracellula | F27 | Calcein | 1936 | BL1 | 13039 |

| | | | | | | | | | |
|---|---|---|---|---|---|---|---|---|---|
| 40 | Yes | A. baylyi | Rifampicin | Intracellul | F12 | Amaranth | 2081 | BL1 | 13252 |
| 40 | Yes | A. baylyi | None | None | NA | Autofluore | 2303 | NA | 7013 |
| 40 | Yes | A. baylyi | Trimethopr | Intracellul | F194 | Oxonol | 2766 | RL1 | 6090 |
| 40 | Yes | A. baylyi | Trimethopr | Intracellul | F11 | Allura Red | 2578 | BL3 | 5289 |
| 40 | Yes | A. baylyi | Trimethopr | Intracellul | F54 | DiSC3(5) | 2171 | RL1 | 5539 |
| 40 | Yes | A. baylyi | Trimethopr | Intracellul | F124 | Thioflavin T | 2265 | VL2 | 5658 |
| 40 | Yes | A. baylyi | Trimethopr | Intracellul | F101 | Quinacrine | 2415 | VL2 | 5423 |
| 40 | Yes | A. baylyi | Trimethopr | Intracellul | F138 | Apigenin | 2502 | VL2 | 4939 |
| 40 | Yes | A. baylyi | Trimethopr | Intracellul | F65 | H2FDA | 2359 | BL1 | 5839 |
| 40 | Yes | A. baylyi | Trimethopr | Intracellul | F142 | Chlorophe | 2518 | BL3 | 6771 |
| 40 | Yes | A. baylyi | Trimethopr | Intracellul | F39 | Congo Red | 2491 | BL3 | 6547 |
| 40 | Yes | A. baylyi | Trimethopr | Intracellul | F116 | Sunset Yel | 2625 | BL2 | 6218 |
| 40 | Yes | A. baylyi | Trimethopr | Intracellul | F10 | Alizarin | 2510 | BL1 | 5633 |
| 40 | Yes | A. baylyi | Trimethopr | Intracellul | F84 | Neutral Re | 2396 | BL1 | 5264.5 |
| 40 | Yes | A. baylyi | Trimethopr | Intracellul | F99 | Pyronin Y | 2440 | BL1 | 4713.5 |
| 40 | Yes | A. baylyi | Trimethopr | Intracellul | F108 | Rhodamine | 2481 | BL1 | 4569 |
| 40 | Yes | A. baylyi | Trimethopr | Intracellul | F109 | Riboflavin | 2585 | BL1 | 4577 |
| 40 | Yes | A. baylyi | Trimethopr | Intracellul | F117 | SYBR Gree | 2271 | BL1 | 322434 |
| 40 | Yes | A. baylyi | Trimethopr | Intracellul | F106 | Rhodamine | 2118 | BL1 | 5524 |
| 40 | Yes | A. baylyi | Trimethopr | Intracellul | F73 | Malachite ( | 2249 | BL1 | 5190 |
| 40 | Yes | A. baylyi | Trimethopr | Intracellul | F7 | Acridine or | 2367 | BL1 | 6005 |
| 40 | Yes | A. baylyi | Trimethopr | Intracellul | F192 | 5-Carboxyf | 2205 | BL1 | 5968 |
| 40 | Yes | A. baylyi | Trimethopr | Intracellul | F57 | Eosin Y | 2288 | BL1 | 7335.5 |
| 40 | Yes | A. baylyi | Trimethopr | Intracellul | F27 | Calcein | 2338 | BL1 | 11268 |
| 40 | Yes | A. baylyi | Trimethopr | Intracellul | F12 | Amaranth | 2092 | BL1 | 15565.5 |
| 40 | Yes | A. baylyi | None | None | NA | Autofluore | 2617 | NA | 784 |
| 40 | Yes | A. baylyi | None | None | F194 | Oxonol | 3141 | RL1 | 1628 |
| 40 | Yes | A. baylyi | None | None | F11 | Allura Red | 2637 | BL3 | 1810 |
| 40 | Yes | A. baylyi | None | None | F54 | DiSC3(5) | 2591 | RL1 | 2287 |
| 40 | Yes | A. baylyi | None | None | F124 | Thioflavin T | 2647 | VL2 | 2377 |
| 40 | Yes | A. baylyi | None | None | F101 | Quinacrine | 2436 | VL2 | 2610.5 |
| 40 | Yes | A. baylyi | None | None | F138 | Apigenin | 2546 | VL2 | 2692 |
| 40 | Yes | A. baylyi | None | None | F65 | H2FDA | 2137 | BL1 | 3740 |
| 40 | Yes | A. baylyi | None | None | F142 | Chlorophe | 2345 | BL3 | 4784 |
| 40 | Yes | A. baylyi | None | None | F39 | Congo Red | 2470 | BL3 | 4729 |
| 40 | Yes | A. baylyi | None | None | F116 | Sunset Yel | 2401 | BL2 | 4171 |
| 40 | Yes | A. baylyi | None | None | F10 | Alizarin | 2421 | BL1 | 3814 |
| 40 | Yes | A. baylyi | None | None | F84 | Neutral Re | 2196 | BL1 | 3488 |
| 40 | Yes | A. baylyi | None | None | F99 | Pyronin Y | 2254 | BL1 | 3295.5 |
| 40 | Yes | A. baylyi | None | None | F108 | Rhodamine | 2389 | BL1 | 3124 |
| 40 | Yes | A. baylyi | None | None | F109 | Riboflavin | 2130 | BL1 | 3176 |
| 40 | Yes | A. baylyi | None | None | F117 | SYBR Gree | 2085 | BL1 | 361526 |
| 40 | Yes | A. baylyi | None | None | F106 | Rhodamine | 2157 | BL1 | 3555 |
| 40 | Yes | A. baylyi | None | None | F73 | Malachite ( | 2286 | BL1 | 3420 |
| 40 | Yes | A. baylyi | None | None | F7 | Acridine or | 2141 | BL1 | 3449 |
| 40 | Yes | A. baylyi | None | None | F192 | 5-Carboxyf | 2037 | BL1 | 12011 |
| 40 | Yes | A. baylyi | None | None | F57 | Eosin Y | 1874 | BL1 | 13197.5 |
| 40 | Yes | A. baylyi | None | None | F27 | Calcein | 1851 | BL1 | 15463 |

| | | | | | | | | | |
|---|---|---|---|---|---|---|---|---|---|
| 40 | Yes | A. baylyi | None | None | F12 | Amaranth | 2144 | BL1 | 14735 |
| 40 | Yes | A. baylyi | None | None | NA | Autofluores | 2568 | NA | 6219.5 |
| 40 | Yes | A. baylyi | Amoxacilli | Wall | F194 | Oxonol | 2491 | RL1 | 4390 |
| 40 | Yes | A. baylyi | Amoxacilli | Wall | F11 | Allura Red | 2331 | BL3 | 3814 |
| 40 | Yes | A. baylyi | Amoxacilli | Wall | F54 | DiSC3(5) | 2278 | RL1 | 3672 |
| 40 | Yes | A. baylyi | Amoxacilli | Wall | F124 | Thioflavin T | 2189 | VL2 | 3327 |
| 40 | Yes | A. baylyi | Amoxacilli | Wall | F101 | Quinacrine | 1876 | VL2 | 3258.5 |
| 40 | Yes | A. baylyi | Amoxacilli | Wall | F138 | Apigenin | 2000 | VL2 | 3366 |
| 40 | Yes | A. baylyi | Amoxacilli | Wall | F65 | H2FDA | 2003 | BL1 | 4558 |
| 40 | Yes | A. baylyi | Amoxacilli | Wall | F142 | Chlorophe | 2047 | BL3 | 5041 |
| 40 | Yes | A. baylyi | Amoxacilli | Wall | F39 | Congo Red | 2145 | BL3 | 5481 |
| 40 | Yes | A. baylyi | Amoxacilli | Wall | F116 | Sunset Yel | 2165 | BL2 | 4747 |
| 40 | Yes | A. baylyi | Amoxacilli | Wall | F10 | Alizarin | 2028 | BL1 | 4542 |
| 40 | Yes | A. baylyi | Amoxacilli | Wall | F84 | Neutral Re | 1936 | BL1 | 3792.5 |
| 40 | Yes | A. baylyi | Amoxacilli | Wall | F99 | Pyronin Y | 1992 | BL1 | 3686.5 |
| 40 | Yes | A. baylyi | Amoxacilli | Wall | F108 | Rhodamine | 1780 | BL1 | 3573 |
| 40 | Yes | A. baylyi | Amoxacilli | Wall | F109 | Riboflavin | 1681 | BL1 | 3593 |
| 40 | Yes | A. baylyi | Amoxacilli | Wall | F117 | SYBR Gree | 1809 | BL1 | 359876 |
| 40 | Yes | A. baylyi | Amoxacilli | Wall | F106 | Rhodamine | 2113 | BL1 | 4322 |
| 40 | Yes | A. baylyi | Amoxacilli | Wall | F73 | Malachite ( | 1778 | BL1 | 4237.5 |
| 40 | Yes | A. baylyi | Amoxacilli | Wall | F7 | Acridine or | 1549 | BL1 | 4246 |
| 40 | Yes | A. baylyi | Amoxacilli | Wall | F192 | 5-Carboxyf | 1745 | BL1 | 12878 |
| 40 | Yes | A. baylyi | Amoxacilli | Wall | F57 | Eosin Y | 1646 | BL1 | 14385.5 |
| 40 | Yes | A. baylyi | Amoxacilli | Wall | F27 | Calcein | 1676 | BL1 | 17398 |
| 40 | Yes | A. baylyi | Amoxacilli | Wall | F12 | Amaranth | 1917 | BL1 | 16257 |
| 40 | Yes | A. baylyi | None | None | NA | Autofluores | 2538 | NA | 6346 |
| 40 | Yes | A. baylyi | Azithromyc | Intracellula | F194 | Oxonol | 2024 | RL1 | 5000 |
| 40 | Yes | A. baylyi | Azithromyc | Intracellula | F11 | Allura Red | 2377 | BL3 | 4160 |
| 40 | Yes | A. baylyi | Azithromyc | Intracellula | F54 | DiSC3(5) | 2087 | RL1 | 4119 |
| 40 | Yes | A. baylyi | Azithromyc | Intracellula | F124 | Thioflavin T | 2013 | VL2 | 3583 |
| 40 | Yes | A. baylyi | Azithromyc | Intracellula | F101 | Quinacrine | 1888 | VL2 | 3559.5 |
| 40 | Yes | A. baylyi | Azithromyc | Intracellula | F138 | Apigenin | 1835 | VL2 | 3500 |
| 40 | Yes | A. baylyi | Azithromyc | Intracellula | F65 | H2FDA | 1826 | BL1 | 4274 |
| 40 | Yes | A. baylyi | Azithromyc | Intracellula | F142 | Chlorophe | 1912 | BL3 | 5771.5 |
| 40 | Yes | A. baylyi | Azithromyc | Intracellula | F39 | Congo Red | 1984 | BL3 | 6078.5 |
| 40 | Yes | A. baylyi | Azithromyc | Intracellula | F116 | Sunset Yel | 1978 | BL2 | 4981.5 |
| 40 | Yes | A. baylyi | Azithromyc | Intracellula | F10 | Alizarin | 2167 | BL1 | 4557 |
| 40 | Yes | A. baylyi | Azithromyc | Intracellula | F84 | Neutral Re | 1822 | BL1 | 4313 |
| 40 | Yes | A. baylyi | Azithromyc | Intracellula | F99 | Pyronin Y | 1716 | BL1 | 4104.5 |
| 40 | Yes | A. baylyi | Azithromyc | Intracellula | F108 | Rhodamine | 1768 | BL1 | 3660 |
| 40 | Yes | A. baylyi | Azithromyc | Intracellula | F109 | Riboflavin | 1671 | BL1 | 3778 |
| 40 | Yes | A. baylyi | Azithromyc | Intracellula | F117 | SYBR Gree | 1672 | BL1 | 196875 |
| 40 | Yes | A. baylyi | Azithromyc | Intracellula | F106 | Rhodamine | 2074 | BL1 | 4805.5 |
| 40 | Yes | A. baylyi | Azithromyc | Intracellula | F73 | Malachite ( | 1759 | BL1 | 4452 |
| 40 | Yes | A. baylyi | Azithromyc | Intracellula | F7 | Acridine or | 1766 | BL1 | 4811.5 |
| 40 | Yes | A. baylyi | Azithromyc | Intracellula | F192 | 5-Carboxyf | 1687 | BL1 | 13599 |
| 40 | Yes | A. baylyi | Azithromyc | Intracellula | F57 | Eosin Y | 1696 | BL1 | 13311.5 |
| 40 | Yes | A. baylyi | Azithromyc | Intracellula | F27 | Calcein | 1781 | BL1 | 15040 |

| | | | | | | | | | |
|---|---|---|---|---|---|---|---|---|---|
| 40 | Yes | A. baylyi | Azithromyc | Intracellul | F12 | Amaranth | 1874 | BL1 | 15740 |
| 40 | Yes | A. baylyi | None | None | NA | Autofluore | 2763 | NA | 6390 |
| 40 | Yes | A. baylyi | Trimethopr | Intracellul | F194 | Oxonol | 2291 | RL1 | 4752 |
| 40 | Yes | A. baylyi | Trimethopr | Intracellul | F11 | Allura Red | 2365 | BL3 | 4183 |
| 40 | Yes | A. baylyi | Trimethopr | Intracellul | F54 | DiSC3(5) | 2155 | RL1 | 4100 |
| 40 | Yes | A. baylyi | Trimethopr | Intracellul | F124 | Thioflavin T | 2317 | VL2 | 3783 |
| 40 | Yes | A. baylyi | Trimethopr | Intracellul | F101 | Quinacrine | 2039 | VL2 | 3787 |
| 40 | Yes | A. baylyi | Trimethopr | Intracellul | F138 | Apigenin | 1991 | VL2 | 4068 |
| 40 | Yes | A. baylyi | Trimethopr | Intracellul | F65 | H2FDA | 1810 | BL1 | 4666 |
| 40 | Yes | A. baylyi | Trimethopr | Intracellul | F142 | Chlorophe | 1848 | BL3 | 6073.5 |
| 40 | Yes | A. baylyi | Trimethopr | Intracellul | F39 | Congo Red | 2069 | BL3 | 5783 |
| 40 | Yes | A. baylyi | Trimethopr | Intracellul | F116 | Sunset Yel | 2101 | BL2 | 5002 |
| 40 | Yes | A. baylyi | Trimethopr | Intracellul | F10 | Alizarin | 2357 | BL1 | 4609 |
| 40 | Yes | A. baylyi | Trimethopr | Intracellul | F84 | Neutral Re | 2052 | BL1 | 4297 |
| 40 | Yes | A. baylyi | Trimethopr | Intracellul | F99 | Pyronin Y | 2080 | BL1 | 4327 |
| 40 | Yes | A. baylyi | Trimethopr | Intracellul | F108 | Rhodamine | 1911 | BL1 | 4038 |
| 40 | Yes | A. baylyi | Trimethopr | Intracellul | F109 | Riboflavin | 1835 | BL1 | 4055 |
| 40 | Yes | A. baylyi | Trimethopr | Intracellul | F117 | SYBR Gree | 1827 | BL1 | 344026 |
| 40 | Yes | A. baylyi | Trimethopr | Intracellul | F106 | Rhodamine | 2117 | BL1 | 4622 |
| 40 | Yes | A. baylyi | Trimethopr | Intracellul | F73 | Malachite ( | 2040 | BL1 | 4695.5 |
| 40 | Yes | A. baylyi | Trimethopr | Intracellul | F7 | Acridine or | 1653 | BL1 | 4837 |
| 40 | Yes | A. baylyi | Trimethopr | Intracellul | F192 | 5-Carboxyf | 1787 | BL1 | 12152 |
| 40 | Yes | A. baylyi | Trimethopr | Intracellul | F57 | Eosin Y | 1781 | BL1 | 15422 |
| 40 | Yes | A. baylyi | Trimethopr | Intracellul | F27 | Calcein | 1853 | BL1 | 14436 |
| 40 | Yes | A. baylyi | Trimethopr | Intracellul | F12 | Amaranth | 1901 | BL1 | 15291 |
| 40 | Yes | A. baylyi | None | None | NA | Autofluore | 2554 | NA | 6990.5 |
| 40 | Yes | A. baylyi | Chloramph | Intracellul | F194 | Oxonol | 1620 | RL1 | 5595.5 |
| 40 | Yes | A. baylyi | Chloramph | Intracellul | F11 | Allura Red | 1564 | BL3 | 4729.5 |
| 40 | Yes | A. baylyi | Chloramph | Intracellul | F54 | DiSC3(5) | 1331 | RL1 | 4377 |
| 40 | Yes | A. baylyi | Chloramph | Intracellul | F124 | Thioflavin T | 1428 | VL2 | 4175 |
| 40 | Yes | A. baylyi | Chloramph | Intracellul | F101 | Quinacrine | 1481 | VL2 | 3804 |
| 40 | Yes | A. baylyi | Chloramph | Intracellul | F138 | Apigenin | 1376 | VL2 | 4354 |
| 40 | Yes | A. baylyi | Chloramph | Intracellul | F65 | H2FDA | 1179 | BL1 | 5382 |
| 40 | Yes | A. baylyi | Chloramph | Intracellul | F142 | Chlorophe | 1334 | BL3 | 5672.5 |
| 40 | Yes | A. baylyi | Chloramph | Intracellul | F39 | Congo Red | 1428 | BL3 | 6147.5 |
| 40 | Yes | A. baylyi | Chloramph | Intracellul | F116 | Sunset Yel | 1561 | BL2 | 5215 |
| 40 | Yes | A. baylyi | Chloramph | Intracellul | F10 | Alizarin | 1724 | BL1 | 4825 |
| 40 | Yes | A. baylyi | Chloramph | Intracellul | F84 | Neutral Re | 1579 | BL1 | 4584 |
| 40 | Yes | A. baylyi | Chloramph | Intracellul | F99 | Pyronin Y | 1377 | BL1 | 4616 |
| 40 | Yes | A. baylyi | Chloramph | Intracellul | F108 | Rhodamine | 1240 | BL1 | 4215.5 |
| 40 | Yes | A. baylyi | Chloramph | Intracellul | F109 | Riboflavin | 1169 | BL1 | 4280 |
| 40 | Yes | A. baylyi | Chloramph | Intracellul | F117 | SYBR Gree | 1205 | BL1 | 210970 |
| 40 | Yes | A. baylyi | Chloramph | Intracellul | F106 | Rhodamine | 1278 | BL1 | 4745 |
| 40 | Yes | A. baylyi | Chloramph | Intracellul | F73 | Malachite ( | 1247 | BL1 | 4608 |
| 40 | Yes | A. baylyi | Chloramph | Intracellul | F7 | Acridine or | 1188 | BL1 | 4969 |
| 40 | Yes | A. baylyi | Chloramph | Intracellul | F192 | 5-Carboxyf | 1328 | BL1 | 12339.5 |
| 40 | Yes | A. baylyi | Chloramph | Intracellul | F57 | Eosin Y | 1085 | BL1 | 14358 |
| 40 | Yes | A. baylyi | Chloramph | Intracellul | F27 | Calcein | 1211 | BL1 | 14931 |

| | | | | | | | | | |
|---|---|---|---|---|---|---|---|---|---|
| 40 | Yes | A. baylyi | Chloramph | Intracellula | F12 | Amaranth | 1210 | BL1 | 15610.5 |
| 40 | Yes | A. baylyi | None | None | NA | Autofluores | 2762 | NA | 7423 |
| 40 | Yes | A. baylyi | Colistin | Wall | F194 | Oxonol | 1440 | RL1 | 41115.5 |
| 40 | Yes | A. baylyi | Colistin | Wall | F11 | Allura Red | 1129 | BL3 | 25990 |
| 40 | Yes | A. baylyi | Colistin | Wall | F54 | DiSC3(5) | 1003 | RL1 | 24576 |
| 40 | Yes | A. baylyi | Colistin | Wall | F124 | Thioflavin T | 1098 | VL2 | 20819 |
| 40 | Yes | A. baylyi | Colistin | Wall | F101 | Quinacrine | 1007 | VL2 | 21013 |
| 40 | Yes | A. baylyi | Colistin | Wall | F138 | Apigenin | 1033 | VL2 | 17893 |
| 40 | Yes | A. baylyi | Colistin | Wall | F65 | H2FDA | 1004 | BL1 | 18771.5 |
| 40 | Yes | A. baylyi | Colistin | Wall | F142 | Chlorophe | 1140 | BL3 | 19111 |
| 40 | Yes | A. baylyi | Colistin | Wall | F39 | Congo Red | 1055 | BL3 | 12766 |
| 40 | Yes | A. baylyi | Colistin | Wall | F116 | Sunset Yel | 1134 | BL2 | 16296.5 |
| 40 | Yes | A. baylyi | Colistin | Wall | F10 | Alizarin | 1213 | BL1 | 15646 |
| 40 | Yes | A. baylyi | Colistin | Wall | F84 | Neutral Re | 1164 | BL1 | 16471.5 |
| 40 | Yes | A. baylyi | Colistin | Wall | F99 | Pyronin Y | 1073 | BL1 | 11990 |
| 40 | Yes | A. baylyi | Colistin | Wall | F108 | Rhodamine | 1066 | BL1 | 14357 |
| 40 | Yes | A. baylyi | Colistin | Wall | F109 | Riboflavin | 1061 | BL1 | 14694 |
| 40 | Yes | A. baylyi | Colistin | Wall | F117 | SYBR Gree | 1083 | BL1 | 2523923 |
| 40 | Yes | A. baylyi | Colistin | Wall | F106 | Rhodamine | 1069 | BL1 | 45258 |
| 40 | Yes | A. baylyi | Colistin | Wall | F73 | Malachite ( | 1025 | BL1 | 31068 |
| 40 | Yes | A. baylyi | Colistin | Wall | F7 | Acridine or | 937 | BL1 | 61247 |
| 40 | Yes | A. baylyi | Colistin | Wall | F192 | 5-Carboxyf | 920 | BL1 | 37779.5 |
| 40 | Yes | A. baylyi | Colistin | Wall | F57 | Eosin Y | 1004 | BL1 | 37251.5 |
| 40 | Yes | A. baylyi | Colistin | Wall | F27 | Calcein | 931 | BL1 | 28146 |
| 40 | Yes | A. baylyi | Colistin | Wall | F12 | Amaranth | 1102 | BL1 | 27688 |
| 40 | Yes | A. baylyi | None | None | NA | Autofluores | 2614 | NA | 7094.5 |
| 40 | Yes | A. baylyi | Gentamici | Intracellula | F194 | Oxonol | 2645 | RL1 | 5335 |
| 40 | Yes | A. baylyi | Gentamici | Intracellula | F11 | Allura Red | 2446 | BL3 | 4904.5 |
| 40 | Yes | A. baylyi | Gentamici | Intracellula | F54 | DiSC3(5) | 2381 | RL1 | 4539 |
| 40 | Yes | A. baylyi | Gentamici | Intracellula | F124 | Thioflavin T | 2261 | VL2 | 4052 |
| 40 | Yes | A. baylyi | Gentamici | Intracellula | F101 | Quinacrine | 2147 | VL2 | 4247 |
| 40 | Yes | A. baylyi | Gentamici | Intracellula | F138 | Apigenin | 2277 | VL2 | 4268 |
| 40 | Yes | A. baylyi | Gentamici | Intracellula | F65 | H2FDA | 2309 | BL1 | 5004 |
| 40 | Yes | A. baylyi | Gentamici | Intracellula | F142 | Chlorophe | 2375 | BL3 | 6435 |
| 40 | Yes | A. baylyi | Gentamici | Intracellula | F39 | Congo Red | 2435 | BL3 | 6225 |
| 40 | Yes | A. baylyi | Gentamici | Intracellula | F116 | Sunset Yel | 2391 | BL2 | 5591 |
| 40 | Yes | A. baylyi | Gentamici | Intracellula | F10 | Alizarin | 2549 | BL1 | 5283 |
| 40 | Yes | A. baylyi | Gentamici | Intracellula | F84 | Neutral Re | 1913 | BL1 | 5063 |
| 40 | Yes | A. baylyi | Gentamici | Intracellula | F99 | Pyronin Y | 2636 | BL1 | 4884 |
| 40 | Yes | A. baylyi | Gentamici | Intracellula | F108 | Rhodamine | 2387 | BL1 | 4493 |
| 40 | Yes | A. baylyi | Gentamici | Intracellula | F109 | Riboflavin | 2209 | BL1 | 4500 |
| 40 | Yes | A. baylyi | Gentamici | Intracellula | F117 | SYBR Gree | 2099 | BL1 | 370165 |
| 40 | Yes | A. baylyi | Gentamici | Intracellula | F106 | Rhodamine | 2385 | BL1 | 5468 |
| 40 | Yes | A. baylyi | Gentamici | Intracellula | F73 | Malachite ( | 2136 | BL1 | 5024.5 |
| 40 | Yes | A. baylyi | Gentamici | Intracellula | F7 | Acridine or | 1815 | BL1 | 5331 |
| 40 | Yes | A. baylyi | Gentamici | Intracellula | F192 | 5-Carboxyf | 1910 | BL1 | 14415 |
| 40 | Yes | A. baylyi | Gentamici | Intracellula | F57 | Eosin Y | 1908 | BL1 | 17157.5 |
| 40 | Yes | A. baylyi | Gentamici | Intracellula | F27 | Calcein | 1890 | BL1 | 17563 |

| | | | | | | | | | |
|---|---|---|---|---|---|---|---|---|---|
| 40 | Yes | A. baylyi | Gentamici | Intracellul | F12 | Amaranth | 1949 | BL1 | 16219 |
| 40 | Yes | A. baylyi | None | None | NA | Autofluore | 2597 | NA | 6347 |
| 40 | Yes | A. baylyi | Nitrofurant | Intracellul | F194 | Oxonol | 2170 | RL1 | 5441.5 |
| 40 | Yes | A. baylyi | Nitrofurant | Intracellul | F11 | Allura Red | 2078 | BL3 | 4845 |
| 40 | Yes | A. baylyi | Nitrofurant | Intracellul | F54 | DiSC3(5) | 1704 | RL1 | 4788.5 |
| 40 | Yes | A. baylyi | Nitrofurant | Intracellul | F124 | Thioflavin T | 1785 | VL2 | 4255 |
| 40 | Yes | A. baylyi | Nitrofurant | Intracellul | F101 | Quinacrine | 1863 | VL2 | 4114 |
| 40 | Yes | A. baylyi | Nitrofurant | Intracellul | F138 | Apigenin | 2035 | VL2 | 4390 |
| 40 | Yes | A. baylyi | Nitrofurant | Intracellul | F65 | H2FDA | 1841 | BL1 | 5208 |
| 40 | Yes | A. baylyi | Nitrofurant | Intracellul | F142 | Chlorophe | 1946 | BL3 | 6350.5 |
| 40 | Yes | A. baylyi | Nitrofurant | Intracellul | F39 | Congo Red | 2154 | BL3 | 6192.5 |
| 40 | Yes | A. baylyi | Nitrofurant | Intracellul | F116 | Sunset Yel | 2175 | BL2 | 5271 |
| 40 | Yes | A. baylyi | Nitrofurant | Intracellul | F10 | Alizarin | 1898 | BL1 | 5102 |
| 40 | Yes | A. baylyi | Nitrofurant | Intracellul | F84 | Neutral Re | 2058 | BL1 | 4647.5 |
| 40 | Yes | A. baylyi | Nitrofurant | Intracellul | F99 | Pyronin Y | 2060 | BL1 | 4400.5 |
| 40 | Yes | A. baylyi | Nitrofurant | Intracellul | F108 | Rhodamine | 1932 | BL1 | 4368 |
| 40 | Yes | A. baylyi | Nitrofurant | Intracellul | F109 | Riboflavin | 2035 | BL1 | 4590 |
| 40 | Yes | A. baylyi | Nitrofurant | Intracellul | F117 | SYBR Gree | 1691 | BL1 | 136949 |
| 40 | Yes | A. baylyi | Nitrofurant | Intracellul | F106 | Rhodamine | 2002 | BL1 | 5313 |
| 40 | Yes | A. baylyi | Nitrofurant | Intracellul | F73 | Malachite ( | 1752 | BL1 | 5054.5 |
| 40 | Yes | A. baylyi | Nitrofurant | Intracellul | F7 | Acridine or | 1713 | BL1 | 5565 |
| 40 | Yes | A. baylyi | Nitrofurant | Intracellul | F192 | 5-Carboxyf | 1649 | BL1 | 12062 |
| 40 | Yes | A. baylyi | Nitrofurant | Intracellul | F57 | Eosin Y | 1672 | BL1 | 13629 |
| 40 | Yes | A. baylyi | Nitrofurant | Intracellul | F27 | Calcein | 1626 | BL1 | 12036 |
| 40 | Yes | A. baylyi | Nitrofurant | Intracellul | F12 | Amaranth | 1731 | BL1 | 14005 |
| 40 | Yes | A. baylyi | None | None | NA | Autofluore | 2731 | NA | 6925 |
| 40 | Yes | A. baylyi | Pentamidi | Wall | F194 | Oxonol | 1461 | RL1 | 10239 |
| 40 | Yes | A. baylyi | Pentamidi | Wall | F11 | Allura Red | 1298 | BL3 | 6903 |
| 40 | Yes | A. baylyi | Pentamidi | Wall | F54 | DiSC3(5) | 1161 | RL1 | 6462 |
| 40 | Yes | A. baylyi | Pentamidi | Wall | F124 | Thioflavin T | 1166 | VL2 | 5815 |
| 40 | Yes | A. baylyi | Pentamidi | Wall | F101 | Quinacrine | 1187 | VL2 | 5948 |
| 40 | Yes | A. baylyi | Pentamidi | Wall | F138 | Apigenin | 1210 | VL2 | 11924.5 |
| 40 | Yes | A. baylyi | Pentamidi | Wall | F65 | H2FDA | 1154 | BL1 | 8752 |
| 40 | Yes | A. baylyi | Pentamidi | Wall | F142 | Chlorophe | 1168 | BL3 | 10152.5 |
| 40 | Yes | A. baylyi | Pentamidi | Wall | F39 | Congo Red | 1356 | BL3 | 11804.5 |
| 40 | Yes | A. baylyi | Pentamidi | Wall | F116 | Sunset Yel | 1147 | BL2 | 8580 |
| 40 | Yes | A. baylyi | Pentamidi | Wall | F10 | Alizarin | 1110 | BL1 | 8313.5 |
| 40 | Yes | A. baylyi | Pentamidi | Wall | F84 | Neutral Re | 1154 | BL1 | 8967 |
| 40 | Yes | A. baylyi | Pentamidi | Wall | F99 | Pyronin Y | 1191 | BL1 | 6817 |
| 40 | Yes | A. baylyi | Pentamidi | Wall | F108 | Rhodamine | 1115 | BL1 | 7152 |
| 40 | Yes | A. baylyi | Pentamidi | Wall | F109 | Riboflavin | 1174 | BL1 | 7422.5 |
| 40 | Yes | A. baylyi | Pentamidi | Wall | F117 | SYBR Gree | 1184 | BL1 | 1485568 |
| 40 | Yes | A. baylyi | Pentamidi | Wall | F106 | Rhodamine | 1130 | BL1 | 27267 |
| 40 | Yes | A. baylyi | Pentamidi | Wall | F73 | Malachite ( | 1133 | BL1 | 7713 |
| 40 | Yes | A. baylyi | Pentamidi | Wall | F7 | Acridine or | 1051 | BL1 | 11433 |
| 40 | Yes | A. baylyi | Pentamidi | Wall | F192 | 5-Carboxyf | 1036 | BL1 | 13860.5 |
| 40 | Yes | A. baylyi | Pentamidi | Wall | F57 | Eosin Y | 1073 | BL1 | 14480 |
| 40 | Yes | A. baylyi | Pentamidi | Wall | F27 | Calcein | 1247 | BL1 | 17556 |

| | | | | | | | | | |
|---|---|---|---|---|---|---|---|---|---|
| 40 | Yes | A. baylyi | Pentamidi | Wall | F12 | Amaranth | 1176 | BL1 | 13820 |
| 40 | Yes | A. baylyi | None | None | NA | Autofluore: | 2447 | NA | 8494 |
| 40 | Yes | A. baylyi | Rifampicin | Intracellul | F194 | Oxonol | 2390 | RL1 | 7728 |
| 40 | Yes | A. baylyi | Rifampicin | Intracellul | F11 | Allura Red | 2320 | BL3 | 6485 |
| 40 | Yes | A. baylyi | Rifampicin | Intracellul | F54 | DiSC3(5) | 2239 | RL1 | 5755 |
| 40 | Yes | A. baylyi | Rifampicin | Intracellul | F124 | Thioflavin T | 2227 | VL2 | 4964 |
| 40 | Yes | A. baylyi | Rifampicin | Intracellul | F101 | Quinacrine | 2085 | VL2 | 5017 |
| 40 | Yes | A. baylyi | Rifampicin | Intracellul | F138 | Apigenin | 2144 | VL2 | 5067 |
| 40 | Yes | A. baylyi | Rifampicin | Intracellul | F65 | H2FDA | 2015 | BL1 | 5987 |
| 40 | Yes | A. baylyi | Rifampicin | Intracellul | F142 | Chlorophe | 2212 | BL3 | 7118 |
| 40 | Yes | A. baylyi | Rifampicin | Intracellul | F39 | Congo Red | 2683 | BL3 | 6623 |
| 40 | Yes | A. baylyi | Rifampicin | Intracellul | F116 | Sunset Yel | 2507 | BL2 | 5963 |
| 40 | Yes | A. baylyi | Rifampicin | Intracellul | F10 | Alizarin | 2642 | BL1 | 5416 |
| 40 | Yes | A. baylyi | Rifampicin | Intracellul | F84 | Neutral Re | 2445 | BL1 | 5019 |
| 40 | Yes | A. baylyi | Rifampicin | Intracellul | F99 | Pyronin Y | 2288 | BL1 | 4902.5 |
| 40 | Yes | A. baylyi | Rifampicin | Intracellul | F108 | Rhodamine | 2255 | BL1 | 4560 |
| 40 | Yes | A. baylyi | Rifampicin | Intracellul | F109 | Riboflavin | 2062 | BL1 | 4696.5 |
| 40 | Yes | A. baylyi | Rifampicin | Intracellul | F117 | SYBR Gree | 1880 | BL1 | 416943.5 |
| 40 | Yes | A. baylyi | Rifampicin | Intracellul | F106 | Rhodamine | 2134 | BL1 | 5385 |
| 40 | Yes | A. baylyi | Rifampicin | Intracellul | F73 | Malachite ( | 2146 | BL1 | 5392.5 |
| 40 | Yes | A. baylyi | Rifampicin | Intracellul | F7 | Acridine or | 1927 | BL1 | 5016 |
| 40 | Yes | A. baylyi | Rifampicin | Intracellul | F192 | 5-Carboxyf | 1991 | BL1 | 13103 |
| 40 | Yes | A. baylyi | Rifampicin | Intracellul | F57 | Eosin Y | 1848 | BL1 | 14920.5 |
| 40 | Yes | A. baylyi | Rifampicin | Intracellul | F27 | Calcein | 1909 | BL1 | 14131 |
| 40 | Yes | A. baylyi | Rifampicin | Intracellul | F12 | Amaranth | 1961 | BL1 | 15446 |
| 40 | Yes | A. baylyi | None | None | NA | Autofluore: | 2832 | NA | 7114.5 |
| 40 | Yes | A. baylyi | Trimethopr | Intracellul | F194 | Oxonol | 2827 | RL1 | 5519 |
| 40 | Yes | A. baylyi | Trimethopr | Intracellul | F11 | Allura Red | 2466 | BL3 | 4794 |
| 40 | Yes | A. baylyi | Trimethopr | Intracellul | F54 | DiSC3(5) | 2279 | RL1 | 4894 |
| 40 | Yes | A. baylyi | Trimethopr | Intracellul | F124 | Thioflavin T | 2472 | VL2 | 4118.5 |
| 40 | Yes | A. baylyi | Trimethopr | Intracellul | F101 | Quinacrine | 2127 | VL2 | 4566 |
| 40 | Yes | A. baylyi | Trimethopr | Intracellul | F138 | Apigenin | 2272 | VL2 | 4733.5 |
| 40 | Yes | A. baylyi | Trimethopr | Intracellul | F65 | H2FDA | 2359 | BL1 | 5395 |
| 40 | Yes | A. baylyi | Trimethopr | Intracellul | F142 | Chlorophe | 2567 | BL3 | 6169 |
| 40 | Yes | A. baylyi | Trimethopr | Intracellul | F39 | Congo Red | 2702 | BL3 | 6073 |
| 40 | Yes | A. baylyi | Trimethopr | Intracellul | F116 | Sunset Yel | 2860 | BL2 | 6095 |
| 40 | Yes | A. baylyi | Trimethopr | Intracellul | F10 | Alizarin | 2744 | BL1 | 6131.5 |
| 40 | Yes | A. baylyi | Trimethopr | Intracellul | F84 | Neutral Re | 2461 | BL1 | 4935 |
| 40 | Yes | A. baylyi | Trimethopr | Intracellul | F99 | Pyronin Y | 2344 | BL1 | 4553.5 |
| 40 | Yes | A. baylyi | Trimethopr | Intracellul | F108 | Rhodamine | 2370 | BL1 | 4588.5 |
| 40 | Yes | A. baylyi | Trimethopr | Intracellul | F109 | Riboflavin | 2190 | BL1 | 5048.5 |
| 40 | Yes | A. baylyi | Trimethopr | Intracellul | F117 | SYBR Gree | 1949 | BL1 | 349406 |
| 40 | Yes | A. baylyi | Trimethopr | Intracellul | F106 | Rhodamine | 2360 | BL1 | 5128.5 |
| 40 | Yes | A. baylyi | Trimethopr | Intracellul | F73 | Malachite ( | 2357 | BL1 | 4955 |
| 40 | Yes | A. baylyi | Trimethopr | Intracellul | F7 | Acridine or | 2139 | BL1 | 5352 |
| 40 | Yes | A. baylyi | Trimethopr | Intracellul | F192 | 5-Carboxyf | 2117 | BL1 | 12259 |
| 40 | Yes | A. baylyi | Trimethopr | Intracellul | F57 | Eosin Y | 1840 | BL1 | 15357.5 |
| 40 | Yes | A. baylyi | Trimethopr | Intracellul | F27 | Calcein | 2091 | BL1 | 16513 |

| | | | | | | | | | | |
|---|---|---|---|---|---|---|---|---|---|---|
| 40 | Yes | A. baylyi | Trimethopr | Intracellula | F12 | Amaranth | 2180 | BL1 | | 17413.5 |
| 40 | Yes | A. baylyi | None | None | NA | Autofluore: | 2363 | NA | | 826 |
| 40 | Yes | A. baylyi | None | None | F194 | Oxonol | 3059 | RL1 | | 1665 |
| 40 | Yes | A. baylyi | None | None | F11 | Allura Red | 2814 | BL3 | | 2024.5 |
| 40 | Yes | A. baylyi | None | None | F54 | DiSC3(5) | 2554 | RL1 | | 2545.5 |
| 40 | Yes | A. baylyi | None | None | F124 | Thioflavin T | 2621 | VL2 | | 2764 |
| 40 | Yes | A. baylyi | None | None | F101 | Quinacrine | 2082 | VL2 | | 2708 |
| 40 | Yes | A. baylyi | None | None | F138 | Apigenin | 2651 | VL2 | | 2449 |
| 40 | Yes | A. baylyi | None | None | F65 | H2FDA | 2161 | BL1 | | 3386 |
| 40 | Yes | A. baylyi | None | None | F142 | Chlorophe | 2187 | BL3 | | 4539 |
| 40 | Yes | A. baylyi | None | None | F39 | Congo Red | 2085 | BL3 | | 5456 |
| 40 | Yes | A. baylyi | None | None | F116 | Sunset Yel | 2021 | BL2 | | 5662 |
| 40 | Yes | A. baylyi | None | None | F10 | Alizarin | 2174 | BL1 | | 4115 |
| 40 | Yes | A. baylyi | None | None | F84 | Neutral Re | 1917 | BL1 | | 3768 |
| 40 | Yes | A. baylyi | None | None | F99 | Pyronin Y | 2153 | BL1 | | 3822 |
| 40 | Yes | A. baylyi | None | None | F108 | Rhodamine | 2050 | BL1 | | 3249 |
| 40 | Yes | A. baylyi | None | None | F109 | Riboflavin | 1982 | BL1 | | 3142.5 |
| 40 | Yes | A. baylyi | None | None | F117 | SYBR Gree | 2013 | BL1 | | 366461 |
| 40 | Yes | A. baylyi | None | None | F106 | Rhodamine | 1956 | BL1 | | 3922 |
| 40 | Yes | A. baylyi | None | None | F73 | Malachite ( | 2027 | BL1 | | 4163 |
| 40 | Yes | A. baylyi | None | None | F7 | Acridine or | 1832 | BL1 | | 4136 |
| 40 | Yes | A. baylyi | None | None | F192 | 5-Carboxyf | 1863 | BL1 | | 12459 |
| 40 | Yes | A. baylyi | None | None | F57 | Eosin Y | 1912 | BL1 | | 14937.5 |
| 40 | Yes | A. baylyi | None | None | F27 | Calcein | 1785 | BL1 | | 15898 |
| 40 | Yes | A. baylyi | None | None | F12 | Amaranth | 1812 | BL1 | | 11722.5 |
| 40 | Yes | A. baylyi | None | None | NA | Autofluore: | 2272 | NA | | 6377 |
| 40 | Yes | A. baylyi | Amoxacilli | Wall | F194 | Oxonol | 2229 | RL1 | | 4540 |
| 40 | Yes | A. baylyi | Amoxacilli | Wall | F11 | Allura Red | 2272 | BL3 | | 3803 |
| 40 | Yes | A. baylyi | Amoxacilli | Wall | F54 | DiSC3(5) | 1940 | RL1 | | 3244.5 |
| 40 | Yes | A. baylyi | Amoxacilli | Wall | F124 | Thioflavin T | 1869 | VL2 | | 3214 |
| 40 | Yes | A. baylyi | Amoxacilli | Wall | F101 | Quinacrine | 1639 | VL2 | | 3490 |
| 40 | Yes | A. baylyi | Amoxacilli | Wall | F138 | Apigenin | 2021 | VL2 | | 3262 |
| 40 | Yes | A. baylyi | Amoxacilli | Wall | F65 | H2FDA | 1751 | BL1 | | 3700 |
| 40 | Yes | A. baylyi | Amoxacilli | Wall | F142 | Chlorophe | 1606 | BL3 | | 4482 |
| 40 | Yes | A. baylyi | Amoxacilli | Wall | F39 | Congo Red | 1778 | BL3 | | 6164.5 |
| 40 | Yes | A. baylyi | Amoxacilli | Wall | F116 | Sunset Yel | 1854 | BL2 | | 5397.5 |
| 40 | Yes | A. baylyi | Amoxacilli | Wall | F10 | Alizarin | 1934 | BL1 | | 4692 |
| 40 | Yes | A. baylyi | Amoxacilli | Wall | F84 | Neutral Re | 1712 | BL1 | | 4164 |
| 40 | Yes | A. baylyi | Amoxacilli | Wall | F99 | Pyronin Y | 1813 | BL1 | | 4177 |
| 40 | Yes | A. baylyi | Amoxacilli | Wall | F108 | Rhodamine | 1581 | BL1 | | 3560 |
| 40 | Yes | A. baylyi | Amoxacilli | Wall | F109 | Riboflavin | 1678 | BL1 | | 3684 |
| 40 | Yes | A. baylyi | Amoxacilli | Wall | F117 | SYBR Gree | 1537 | BL1 | | 358714 |
| 40 | Yes | A. baylyi | Amoxacilli | Wall | F106 | Rhodamine | 1963 | BL1 | | 4635 |
| 40 | Yes | A. baylyi | Amoxacilli | Wall | F73 | Malachite ( | 1695 | BL1 | | 4199 |
| 40 | Yes | A. baylyi | Amoxacilli | Wall | F7 | Acridine or | 1366 | BL1 | | 4180.5 |
| 40 | Yes | A. baylyi | Amoxacilli | Wall | F192 | 5-Carboxyf | 1881 | BL1 | | 12514 |
| 40 | Yes | A. baylyi | Amoxacilli | Wall | F57 | Eosin Y | 1638 | BL1 | | 15450.5 |
| 40 | Yes | A. baylyi | Amoxacilli | Wall | F27 | Calcein | 1649 | BL1 | | 14968 |

| | | | | | | | | | |
|---|---|---|---|---|---|---|---|---|---|
| 40 | Yes | A. baylyi | Amoxacilli | Wall | F12 | Amaranth | 1781 | BL1 | 15072 |
| 40 | Yes | A. baylyi | None | None | NA | Autofluore | 2378 | NA | 6362 |
| 40 | Yes | A. baylyi | Azithromyc | Intracellula | F194 | Oxonol | 2141 | RL1 | 4968 |
| 40 | Yes | A. baylyi | Azithromyc | Intracellula | F11 | Allura Red | 1976 | BL3 | 4000.5 |
| 40 | Yes | A. baylyi | Azithromyc | Intracellula | F54 | DiSC3(5) | 1695 | RL1 | 3891 |
| 40 | Yes | A. baylyi | Azithromyc | Intracellula | F124 | Thioflavin T | 1894 | VL2 | 3662.5 |
| 40 | Yes | A. baylyi | Azithromyc | Intracellula | F101 | Quinacrine | 1724 | VL2 | 3938.5 |
| 40 | Yes | A. baylyi | Azithromyc | Intracellula | F138 | Apigenin | 1812 | VL2 | 3477 |
| 40 | Yes | A. baylyi | Azithromyc | Intracellula | F65 | H2FDA | 1771 | BL1 | 3998 |
| 40 | Yes | A. baylyi | Azithromyc | Intracellula | F142 | Chlorophe | 1740 | BL3 | 4837.5 |
| 40 | Yes | A. baylyi | Azithromyc | Intracellula | F39 | Congo Red | 1803 | BL3 | 5660 |
| 40 | Yes | A. baylyi | Azithromyc | Intracellula | F116 | Sunset Yel | 1875 | BL2 | 6055 |
| 40 | Yes | A. baylyi | Azithromyc | Intracellula | F10 | Alizarin | 1922 | BL1 | 4794.5 |
| 40 | Yes | A. baylyi | Azithromyc | Intracellula | F84 | Neutral Re | 1642 | BL1 | 4109 |
| 40 | Yes | A. baylyi | Azithromyc | Intracellula | F99 | Pyronin Y | 1708 | BL1 | 4120.5 |
| 40 | Yes | A. baylyi | Azithromyc | Intracellula | F108 | Rhodamine | 1622 | BL1 | 3593 |
| 40 | Yes | A. baylyi | Azithromyc | Intracellula | F109 | Riboflavin | 1592 | BL1 | 3733.5 |
| 40 | Yes | A. baylyi | Azithromyc | Intracellula | F117 | SYBR Gree | 1612 | BL1 | 241597.5 |
| 40 | Yes | A. baylyi | Azithromyc | Intracellula | F106 | Rhodamine | 1773 | BL1 | 4549 |
| 40 | Yes | A. baylyi | Azithromyc | Intracellula | F73 | Malachite ( | 1709 | BL1 | 4407 |
| 40 | Yes | A. baylyi | Azithromyc | Intracellula | F7 | Acridine or | 1585 | BL1 | 4928 |
| 40 | Yes | A. baylyi | Azithromyc | Intracellula | F192 | 5-Carboxyf | 1546 | BL1 | 12805.5 |
| 40 | Yes | A. baylyi | Azithromyc | Intracellula | F57 | Eosin Y | 1523 | BL1 | 15718 |
| 40 | Yes | A. baylyi | Azithromyc | Intracellula | F27 | Calcein | 1596 | BL1 | 14641 |
| 40 | Yes | A. baylyi | Azithromyc | Intracellula | F12 | Amaranth | 1663 | BL1 | 14937 |
| 40 | Yes | A. baylyi | None | None | NA | Autofluore | 2258 | NA | 6169 |
| 40 | Yes | A. baylyi | Trimethopr | Intracellula | F194 | Oxonol | 2228 | RL1 | 4423.5 |
| 40 | Yes | A. baylyi | Trimethopr | Intracellula | F11 | Allura Red | 1988 | BL3 | 3674.5 |
| 40 | Yes | A. baylyi | Trimethopr | Intracellula | F54 | DiSC3(5) | 1952 | RL1 | 3873 |
| 40 | Yes | A. baylyi | Trimethopr | Intracellula | F124 | Thioflavin T | 1712 | VL2 | 3806.5 |
| 40 | Yes | A. baylyi | Trimethopr | Intracellula | F101 | Quinacrine | 2003 | VL2 | 3968 |
| 40 | Yes | A. baylyi | Trimethopr | Intracellula | F138 | Apigenin | 1727 | VL2 | 3688 |
| 40 | Yes | A. baylyi | Trimethopr | Intracellula | F65 | H2FDA | 2065 | BL1 | 4223 |
| 40 | Yes | A. baylyi | Trimethopr | Intracellula | F142 | Chlorophe | 1892 | BL3 | 4929.5 |
| 40 | Yes | A. baylyi | Trimethopr | Intracellula | F39 | Congo Red | 1925 | BL3 | 5685 |
| 40 | Yes | A. baylyi | Trimethopr | Intracellula | F116 | Sunset Yel | 1661 | BL2 | 5450 |
| 40 | Yes | A. baylyi | Trimethopr | Intracellula | F10 | Alizarin | 2049 | BL1 | 5555 |
| 40 | Yes | A. baylyi | Trimethopr | Intracellula | F84 | Neutral Re | 1797 | BL1 | 4657 |
| 40 | Yes | A. baylyi | Trimethopr | Intracellula | F99 | Pyronin Y | 1833 | BL1 | 4586 |
| 40 | Yes | A. baylyi | Trimethopr | Intracellula | F108 | Rhodamine | 1648 | BL1 | 4245 |
| 40 | Yes | A. baylyi | Trimethopr | Intracellula | F109 | Riboflavin | 1673 | BL1 | 4069 |
| 40 | Yes | A. baylyi | Trimethopr | Intracellula | F117 | SYBR Gree | 1514 | BL1 | 345718 |
| 40 | Yes | A. baylyi | Trimethopr | Intracellula | F106 | Rhodamine | 1652 | BL1 | 4957.5 |
| 40 | Yes | A. baylyi | Trimethopr | Intracellula | F73 | Malachite ( | 1926 | BL1 | 4712.5 |
| 40 | Yes | A. baylyi | Trimethopr | Intracellula | F7 | Acridine or | 1811 | BL1 | 4948 |
| 40 | Yes | A. baylyi | Trimethopr | Intracellula | F192 | 5-Carboxyf | 1732 | BL1 | 12105 |
| 40 | Yes | A. baylyi | Trimethopr | Intracellula | F57 | Eosin Y | 1720 | BL1 | 14656.5 |
| 40 | Yes | A. baylyi | Trimethopr | Intracellula | F27 | Calcein | 1836 | BL1 | 16395.5 |

| | | | | | | | | | |
|---|---|---|---|---|---|---|---|---|---|
| 40 | Yes | A. baylyi | Trimethopr | Intracellula | F12 | Amaranth | 1632 | BL1 | 15007.5 |
| 40 | Yes | A. baylyi | None | None | NA | Autofluores | 2312 | NA | 7743.5 |
| 40 | Yes | A. baylyi | Chloramph | Intracellula | F194 | Oxonol | 1551 | RL1 | 5500 |
| 40 | Yes | A. baylyi | Chloramph | Intracellula | F11 | Allura Red | 1400 | BL3 | 4581 |
| 40 | Yes | A. baylyi | Chloramph | Intracellula | F54 | DiSC3(5) | 1397 | RL1 | 4138 |
| 40 | Yes | A. baylyi | Chloramph | Intracellula | F124 | Thioflavin T | 1477 | VL2 | 3915 |
| 40 | Yes | A. baylyi | Chloramph | Intracellula | F101 | Quinacrine | 1386 | VL2 | 4297.5 |
| 40 | Yes | A. baylyi | Chloramph | Intracellula | F138 | Apigenin | 1316 | VL2 | 3998 |
| 40 | Yes | A. baylyi | Chloramph | Intracellula | F65 | H2FDA | 1152 | BL1 | 4494 |
| 40 | Yes | A. baylyi | Chloramph | Intracellula | F142 | Chlorophe | 1139 | BL3 | 5326 |
| 40 | Yes | A. baylyi | Chloramph | Intracellula | F39 | Congo Red | 1254 | BL3 | 5987.5 |
| 40 | Yes | A. baylyi | Chloramph | Intracellula | F116 | Sunset Yel | 1542 | BL2 | 6018 |
| 40 | Yes | A. baylyi | Chloramph | Intracellula | F10 | Alizarin | 1622 | BL1 | 5386 |
| 40 | Yes | A. baylyi | Chloramph | Intracellula | F84 | Neutral Re | 1427 | BL1 | 5048 |
| 40 | Yes | A. baylyi | Chloramph | Intracellula | F99 | Pyronin Y | 1259 | BL1 | 4890 |
| 40 | Yes | A. baylyi | Chloramph | Intracellula | F108 | Rhodamine | 1178 | BL1 | 4508.5 |
| 40 | Yes | A. baylyi | Chloramph | Intracellula | F109 | Riboflavin | 1141 | BL1 | 4566 |
| 40 | Yes | A. baylyi | Chloramph | Intracellula | F117 | SYBR Gree | 1257 | BL1 | 256956 |
| 40 | Yes | A. baylyi | Chloramph | Intracellula | F106 | Rhodamine | 1278 | BL1 | 5361 |
| 40 | Yes | A. baylyi | Chloramph | Intracellula | F73 | Malachite ( | 1079 | BL1 | 4981 |
| 40 | Yes | A. baylyi | Chloramph | Intracellula | F7 | Acridine or | 1121 | BL1 | 5001 |
| 40 | Yes | A. baylyi | Chloramph | Intracellula | F192 | 5-Carboxyf | 1193 | BL1 | 12525 |
| 40 | Yes | A. baylyi | Chloramph | Intracellula | F57 | Eosin Y | 1125 | BL1 | 14521 |
| 40 | Yes | A. baylyi | Chloramph | Intracellula | F27 | Calcein | 1095 | BL1 | 15149 |
| 40 | Yes | A. baylyi | Chloramph | Intracellula | F12 | Amaranth | 1071 | BL1 | 14553 |
| 40 | Yes | A. baylyi | None | None | NA | Autofluores | 2377 | NA | 8009 |
| 40 | Yes | A. baylyi | Colistin | Wall | F194 | Oxonol | 1308 | RL1 | 33117 |
| 40 | Yes | A. baylyi | Colistin | Wall | F11 | Allura Red | 1039 | BL3 | 22400 |
| 40 | Yes | A. baylyi | Colistin | Wall | F54 | DiSC3(5) | 823 | RL1 | 20746 |
| 40 | Yes | A. baylyi | Colistin | Wall | F124 | Thioflavin T | 1021 | VL2 | 15986 |
| 40 | Yes | A. baylyi | Colistin | Wall | F101 | Quinacrine | 938 | VL2 | 17324 |
| 40 | Yes | A. baylyi | Colistin | Wall | F138 | Apigenin | 1054 | VL2 | 13946.5 |
| 40 | Yes | A. baylyi | Colistin | Wall | F65 | H2FDA | 913 | BL1 | 15243 |
| 40 | Yes | A. baylyi | Colistin | Wall | F142 | Chlorophe | 945 | BL3 | 15441 |
| 40 | Yes | A. baylyi | Colistin | Wall | F39 | Congo Red | 1025 | BL3 | 12684 |
| 40 | Yes | A. baylyi | Colistin | Wall | F116 | Sunset Yel | 1101 | BL2 | 14091 |
| 40 | Yes | A. baylyi | Colistin | Wall | F10 | Alizarin | 1009 | BL1 | 14390 |
| 40 | Yes | A. baylyi | Colistin | Wall | F84 | Neutral Re | 1033 | BL1 | 13860 |
| 40 | Yes | A. baylyi | Colistin | Wall | F99 | Pyronin Y | 914 | BL1 | 10352.5 |
| 40 | Yes | A. baylyi | Colistin | Wall | F108 | Rhodamine | 957 | BL1 | 11240 |
| 40 | Yes | A. baylyi | Colistin | Wall | F109 | Riboflavin | 980 | BL1 | 12706 |
| 40 | Yes | A. baylyi | Colistin | Wall | F117 | SYBR Gree | 1002 | BL1 | 2470338 |
| 40 | Yes | A. baylyi | Colistin | Wall | F106 | Rhodamine | 873 | BL1 | 39849 |
| 40 | Yes | A. baylyi | Colistin | Wall | F73 | Malachite ( | 946 | BL1 | 27866 |
| 40 | Yes | A. baylyi | Colistin | Wall | F7 | Acridine or | 838 | BL1 | 46068 |
| 40 | Yes | A. baylyi | Colistin | Wall | F192 | 5-Carboxyf | 890 | BL1 | 32085 |
| 40 | Yes | A. baylyi | Colistin | Wall | F57 | Eosin Y | 928 | BL1 | 33398 |
| 40 | Yes | A. baylyi | Colistin | Wall | F27 | Calcein | 868 | BL1 | 28216 |

| | | | | | | | | | |
|---|---|---|---|---|---|---|---|---|---|
| 40 | Yes | A. baylyi | Colistin | Wall | F12 | Amaranth | 955 | BL1 | 24324 |
| 40 | Yes | A. baylyi | None | None | NA | Autofluore: | 2430 | NA | 7604.5 |
| 40 | Yes | A. baylyi | Gentamici | Intracellul | F194 | Oxonol | 2453 | RL1 | 5465 |
| 40 | Yes | A. baylyi | Gentamici | Intracellul | F11 | Allura Red | 2078 | BL3 | 4579.5 |
| 40 | Yes | A. baylyi | Gentamici | Intracellul | F54 | DiSC3(5) | 2005 | RL1 | 4442 |
| 40 | Yes | A. baylyi | Gentamici | Intracellul | F124 | Thioflavin T | 2187 | VL2 | 4406 |
| 40 | Yes | A. baylyi | Gentamici | Intracellul | F101 | Quinacrine | 1945 | VL2 | 4512 |
| 40 | Yes | A. baylyi | Gentamici | Intracellul | F138 | Apigenin | 2046 | VL2 | 4216.5 |
| 40 | Yes | A. baylyi | Gentamici | Intracellul | F65 | H2FDA | 2173 | BL1 | 4726 |
| 40 | Yes | A. baylyi | Gentamici | Intracellul | F142 | Chlorophe | 2125 | BL3 | 5576 |
| 40 | Yes | A. baylyi | Gentamici | Intracellul | F39 | Congo Red | 2196 | BL3 | 6091.5 |
| 40 | Yes | A. baylyi | Gentamici | Intracellul | F116 | Sunset Yel | 2484 | BL2 | 5973 |
| 40 | Yes | A. baylyi | Gentamici | Intracellul | F10 | Alizarin | 2517 | BL1 | 6017 |
| 40 | Yes | A. baylyi | Gentamici | Intracellul | F84 | Neutral Re | 2413 | BL1 | 5145 |
| 40 | Yes | A. baylyi | Gentamici | Intracellul | F99 | Pyronin Y | 2233 | BL1 | 4693 |
| 40 | Yes | A. baylyi | Gentamici | Intracellul | F108 | Rhodamine | 2096 | BL1 | 5162 |
| 40 | Yes | A. baylyi | Gentamici | Intracellul | F109 | Riboflavin | 1962 | BL1 | 4519.5 |
| 40 | Yes | A. baylyi | Gentamici | Intracellul | F117 | SYBR Gree | 2176 | BL1 | 440873.5 |
| 40 | Yes | A. baylyi | Gentamici | Intracellul | F106 | Rhodamine | 2170 | BL1 | 5544 |
| 40 | Yes | A. baylyi | Gentamici | Intracellul | F73 | Malachite | 1921 | BL1 | 6060 |
| 40 | Yes | A. baylyi | Gentamici | Intracellul | F7 | Acridine or | 1871 | BL1 | 5612 |
| 40 | Yes | A. baylyi | Gentamici | Intracellul | F192 | 5-Carboxyf | 1646 | BL1 | 12344 |
| 40 | Yes | A. baylyi | Gentamici | Intracellul | F57 | Eosin Y | 1661 | BL1 | 13240 |
| 40 | Yes | A. baylyi | Gentamici | Intracellul | F27 | Calcein | 1924 | BL1 | 15592 |
| 40 | Yes | A. baylyi | Gentamici | Intracellul | F12 | Amaranth | 1847 | BL1 | 14039 |
| 40 | Yes | A. baylyi | None | None | NA | Autofluore: | 2376 | NA | 7286.5 |
| 40 | Yes | A. baylyi | Nitrofurant | Intracellul | F194 | Oxonol | 2001 | RL1 | 5380 |
| 40 | Yes | A. baylyi | Nitrofurant | Intracellul | F11 | Allura Red | 1931 | BL3 | 4702 |
| 40 | Yes | A. baylyi | Nitrofurant | Intracellul | F54 | DiSC3(5) | 1727 | RL1 | 4362 |
| 40 | Yes | A. baylyi | Nitrofurant | Intracellul | F124 | Thioflavin T | 1775 | VL2 | 4370 |
| 40 | Yes | A. baylyi | Nitrofurant | Intracellul | F101 | Quinacrine | 1613 | VL2 | 4870 |
| 40 | Yes | A. baylyi | Nitrofurant | Intracellul | F138 | Apigenin | 1700 | VL2 | 4458 |
| 40 | Yes | A. baylyi | Nitrofurant | Intracellul | F65 | H2FDA | 1835 | BL1 | 4709 |
| 40 | Yes | A. baylyi | Nitrofurant | Intracellul | F142 | Chlorophe | 1628 | BL3 | 5355.5 |
| 40 | Yes | A. baylyi | Nitrofurant | Intracellul | F39 | Congo Red | 1852 | BL3 | 5616.5 |
| 40 | Yes | A. baylyi | Nitrofurant | Intracellul | F116 | Sunset Yel | 1719 | BL2 | 6184 |
| 40 | Yes | A. baylyi | Nitrofurant | Intracellul | F10 | Alizarin | 2103 | BL1 | 5805 |
| 40 | Yes | A. baylyi | Nitrofurant | Intracellul | F84 | Neutral Re | 1972 | BL1 | 5334.5 |
| 40 | Yes | A. baylyi | Nitrofurant | Intracellul | F99 | Pyronin Y | 1836 | BL1 | 5087 |
| 40 | Yes | A. baylyi | Nitrofurant | Intracellul | F108 | Rhodamine | 1859 | BL1 | 4977 |
| 40 | Yes | A. baylyi | Nitrofurant | Intracellul | F109 | Riboflavin | 1854 | BL1 | 4679.5 |
| 40 | Yes | A. baylyi | Nitrofurant | Intracellul | F117 | SYBR Gree | 1730 | BL1 | 184520 |
| 40 | Yes | A. baylyi | Nitrofurant | Intracellul | F106 | Rhodamine | 1793 | BL1 | 5945 |
| 40 | Yes | A. baylyi | Nitrofurant | Intracellul | F73 | Malachite | 1556 | BL1 | 5507 |
| 40 | Yes | A. baylyi | Nitrofurant | Intracellul | F7 | Acridine or | 1410 | BL1 | 5761.5 |
| 40 | Yes | A. baylyi | Nitrofurant | Intracellul | F192 | 5-Carboxyf | 1524 | BL1 | 11701 |
| 40 | Yes | A. baylyi | Nitrofurant | Intracellul | F57 | Eosin Y | 1667 | BL1 | 13756 |
| 40 | Yes | A. baylyi | Nitrofurant | Intracellul | F27 | Calcein | 1560 | BL1 | 13214 |

| | | | | | | | | | |
|---|---|---|---|---|---|---|---|---|---|
| 40 | Yes | A. baylyi | Nitrofurant | Intracellula | F12 | Amaranth | 1804 | BL1 | 15184 |
| 40 | Yes | A. baylyi | None | None | NA | Autofluores | 2408 | NA | 8312 |
| 40 | Yes | A. baylyi | Pentamidi | Wall | F194 | Oxonol | 1141 | RL1 | 9931 |
| 40 | Yes | A. baylyi | Pentamidi | Wall | F11 | Allura Red | 1131 | BL3 | 6328 |
| 40 | Yes | A. baylyi | Pentamidi | Wall | F54 | DiSC3(5) | 1031 | RL1 | 6666 |
| 40 | Yes | A. baylyi | Pentamidi | Wall | F124 | Thioflavin T | 946 | VL2 | 5741.5 |
| 40 | Yes | A. baylyi | Pentamidi | Wall | F101 | Quinacrine | 953 | VL2 | 5768 |
| 40 | Yes | A. baylyi | Pentamidi | Wall | F138 | Apigenin | 1113 | VL2 | 8530 |
| 40 | Yes | A. baylyi | Pentamidi | Wall | F65 | H2FDA | 1050 | BL1 | 7003.5 |
| 40 | Yes | A. baylyi | Pentamidi | Wall | F142 | Chlorophe | 1156 | BL3 | 7071 |
| 40 | Yes | A. baylyi | Pentamidi | Wall | F39 | Congo Red | 1307 | BL3 | 9642 |
| 40 | Yes | A. baylyi | Pentamidi | Wall | F116 | Sunset Yel | 1104 | BL2 | 9557.5 |
| 40 | Yes | A. baylyi | Pentamidi | Wall | F10 | Alizarin | 1111 | BL1 | 10120 |
| 40 | Yes | A. baylyi | Pentamidi | Wall | F84 | Neutral Re | 1156 | BL1 | 11590 |
| 40 | Yes | A. baylyi | Pentamidi | Wall | F99 | Pyronin Y | 1089 | BL1 | 9346 |
| 40 | Yes | A. baylyi | Pentamidi | Wall | F108 | Rhodamine | 1112 | BL1 | 9227.5 |
| 40 | Yes | A. baylyi | Pentamidi | Wall | F109 | Riboflavin | 1104 | BL1 | 9182 |
| 40 | Yes | A. baylyi | Pentamidi | Wall | F117 | SYBR Gree | 1018 | BL1 | 1609992 |
| 40 | Yes | A. baylyi | Pentamidi | Wall | F106 | Rhodamine | 1045 | BL1 | 21082 |
| 40 | Yes | A. baylyi | Pentamidi | Wall | F73 | Malachite ( | 1039 | BL1 | 9065 |
| 40 | Yes | A. baylyi | Pentamidi | Wall | F7 | Acridine or | 909 | BL1 | 13901 |
| 40 | Yes | A. baylyi | Pentamidi | Wall | F192 | 5-Carboxyf | 929 | BL1 | 13419 |
| 40 | Yes | A. baylyi | Pentamidi | Wall | F57 | Eosin Y | 1003 | BL1 | 15524 |
| 40 | Yes | A. baylyi | Pentamidi | Wall | F27 | Calcein | 945 | BL1 | 14685 |
| 40 | Yes | A. baylyi | Pentamidi | Wall | F12 | Amaranth | 998 | BL1 | 14433.5 |
| 40 | Yes | A. baylyi | None | None | NA | Autofluores | 2281 | NA | 10304 |
| 40 | Yes | A. baylyi | Rifampicin | Intracellula | F194 | Oxonol | 2171 | RL1 | 7681 |
| 40 | Yes | A. baylyi | Rifampicin | Intracellula | F11 | Allura Red | 1967 | BL3 | 6833 |
| 40 | Yes | A. baylyi | Rifampicin | Intracellula | F54 | DiSC3(5) | 1906 | RL1 | 5869.5 |
| 40 | Yes | A. baylyi | Rifampicin | Intracellula | F124 | Thioflavin T | 1986 | VL2 | 5782.5 |
| 40 | Yes | A. baylyi | Rifampicin | Intracellula | F101 | Quinacrine | 1954 | VL2 | 5797.5 |
| 40 | Yes | A. baylyi | Rifampicin | Intracellula | F138 | Apigenin | 1918 | VL2 | 4938.5 |
| 40 | Yes | A. baylyi | Rifampicin | Intracellula | F65 | H2FDA | 1888 | BL1 | 5299 |
| 40 | Yes | A. baylyi | Rifampicin | Intracellula | F142 | Chlorophe | 2372 | BL3 | 6074 |
| 40 | Yes | A. baylyi | Rifampicin | Intracellula | F39 | Congo Red | 2591 | BL3 | 6534 |
| 40 | Yes | A. baylyi | Rifampicin | Intracellula | F116 | Sunset Yel | 2396 | BL2 | 6509 |
| 40 | Yes | A. baylyi | Rifampicin | Intracellula | F10 | Alizarin | 2535 | BL1 | 6470 |
| 40 | Yes | A. baylyi | Rifampicin | Intracellula | F84 | Neutral Re | 2356 | BL1 | 5421 |
| 40 | Yes | A. baylyi | Rifampicin | Intracellula | F99 | Pyronin Y | 2195 | BL1 | 5116 |
| 40 | Yes | A. baylyi | Rifampicin | Intracellula | F108 | Rhodamine | 2547 | BL1 | 4858 |
| 40 | Yes | A. baylyi | Rifampicin | Intracellula | F109 | Riboflavin | 2095 | BL1 | 5013 |
| 40 | Yes | A. baylyi | Rifampicin | Intracellula | F117 | SYBR Gree | 1851 | BL1 | 525030 |
| 40 | Yes | A. baylyi | Rifampicin | Intracellula | F106 | Rhodamine | 1917 | BL1 | 5819 |
| 40 | Yes | A. baylyi | Rifampicin | Intracellula | F73 | Malachite ( | 1891 | BL1 | 6162 |
| 40 | Yes | A. baylyi | Rifampicin | Intracellula | F7 | Acridine or | 1820 | BL1 | 5890 |
| 40 | Yes | A. baylyi | Rifampicin | Intracellula | F192 | 5-Carboxyf | 1931 | BL1 | 11389 |
| 40 | Yes | A. baylyi | Rifampicin | Intracellula | F57 | Eosin Y | 1816 | BL1 | 14971 |
| 40 | Yes | A. baylyi | Rifampicin | Intracellula | F27 | Calcein | 1841 | BL1 | 14941 |

| | | | | | | | | | |
|---|---|---|---|---|---|---|---|---|---|
| 40 | Yes | A. baylyi | Rifampicin | Intracellula | F12 | Amaranth | 1987 | BL1 | 11456 |
| 40 | Yes | A. baylyi | None | None | NA | Autofluore: | 2376 | NA | 7989.5 |
| 40 | Yes | A. baylyi | Trimethopr | Intracellula | F194 | Oxonol | 2329 | RL1 | 5948 |
| 40 | Yes | A. baylyi | Trimethopr | Intracellula | F11 | Allura Red | 2178 | BL3 | 5326.5 |
| 40 | Yes | A. baylyi | Trimethopr | Intracellula | F54 | DiSC3(5) | 1952 | RL1 | 4433.5 |
| 40 | Yes | A. baylyi | Trimethopr | Intracellula | F124 | Thioflavin T | 1968 | VL2 | 4527 |
| 40 | Yes | A. baylyi | Trimethopr | Intracellula | F101 | Quinacrine | 2206 | VL2 | 4783.5 |
| 40 | Yes | A. baylyi | Trimethopr | Intracellula | F138 | Apigenin | 2053 | VL2 | 4575 |
| 40 | Yes | A. baylyi | Trimethopr | Intracellula | F65 | H2FDA | 2192 | BL1 | 4985 |
| 40 | Yes | A. baylyi | Trimethopr | Intracellula | F142 | Chlorophe | 2423 | BL3 | 5945 |
| 40 | Yes | A. baylyi | Trimethopr | Intracellula | F39 | Congo Red | 2385 | BL3 | 6355 |
| 40 | Yes | A. baylyi | Trimethopr | Intracellula | F116 | Sunset Yel | 2484 | BL2 | 6260 |
| 40 | Yes | A. baylyi | Trimethopr | Intracellula | F10 | Alizarin | 2287 | BL1 | 5481 |
| 40 | Yes | A. baylyi | Trimethopr | Intracellula | F84 | Neutral Re | 2220 | BL1 | 4707.5 |
| 40 | Yes | A. baylyi | Trimethopr | Intracellula | F99 | Pyronin Y | 2190 | BL1 | 4509.5 |
| 40 | Yes | A. baylyi | Trimethopr | Intracellula | F108 | Rhodamine | 2131 | BL1 | 4592 |
| 40 | Yes | A. baylyi | Trimethopr | Intracellula | F109 | Riboflavin | 2170 | BL1 | 4451.5 |
| 40 | Yes | A. baylyi | Trimethopr | Intracellula | F117 | SYBR Gree | 1879 | BL1 | 379985 |
| 40 | Yes | A. baylyi | Trimethopr | Intracellula | F106 | Rhodamine | 2138 | BL1 | 4909.5 |
| 40 | Yes | A. baylyi | Trimethopr | Intracellula | F73 | Malachite ( | 1956 | BL1 | 5051.5 |
| 40 | Yes | A. baylyi | Trimethopr | Intracellula | F7 | Acridine or | 1914 | BL1 | 4874 |
| 40 | Yes | A. baylyi | Trimethopr | Intracellula | F192 | 5-Carboxyf | 1792 | BL1 | 11578 |
| 40 | Yes | A. baylyi | Trimethopr | Intracellula | F57 | Eosin Y | 1790 | BL1 | 15723.5 |
| 40 | Yes | A. baylyi | Trimethopr | Intracellula | F27 | Calcein | 1778 | BL1 | 15936 |
| 40 | Yes | A. baylyi | Trimethopr | Intracellula | F12 | Amaranth | 1980 | BL1 | 16688 |
| 40 | Yes | A. baylyi | None | None | NA | Autofluore: | 3188 | NA | 810 |
| 40 | Yes | A. baylyi | None | None | F194 | Oxonol | 2878 | RL1 | 1679.5 |
| 40 | Yes | A. baylyi | None | None | F11 | Allura Red | 2513 | BL3 | 1963 |
| 40 | Yes | A. baylyi | None | None | F54 | DiSC3(5) | 2391 | RL1 | 2595 |
| 40 | Yes | A. baylyi | None | None | F124 | Thioflavin T | 2429 | VL2 | 2568 |
| 40 | Yes | A. baylyi | None | None | F101 | Quinacrine | 2234 | VL2 | 2760 |
| 40 | Yes | A. baylyi | None | None | F138 | Apigenin | 1851 | VL2 | 2578 |
| 40 | Yes | A. baylyi | None | None | F65 | H2FDA | 2025 | BL1 | 3461 |
| 40 | Yes | A. baylyi | None | None | F142 | Chlorophe | 1477 | BL3 | 4434 |
| 40 | Yes | A. baylyi | None | None | F39 | Congo Red | 1825 | BL3 | 5514 |
| 40 | Yes | A. baylyi | None | None | F116 | Sunset Yel | 1778 | BL2 | 5978 |
| 40 | Yes | A. baylyi | None | None | F10 | Alizarin | 2241 | BL1 | 4715 |
| 40 | Yes | A. baylyi | None | None | F84 | Neutral Re | 1933 | BL1 | 3978 |
| 40 | Yes | A. baylyi | None | None | F99 | Pyronin Y | 1953 | BL1 | 3862 |
| 40 | Yes | A. baylyi | None | None | F108 | Rhodamine | 1870 | BL1 | 3465 |
| 40 | Yes | A. baylyi | None | None | F109 | Riboflavin | 1740 | BL1 | 3274 |
| 40 | Yes | A. baylyi | None | None | F117 | SYBR Gree | 1777 | BL1 | 386787 |
| 40 | Yes | A. baylyi | None | None | F106 | Rhodamine | 1759 | BL1 | 3871 |
| 40 | Yes | A. baylyi | None | None | F73 | Malachite ( | 1864 | BL1 | 3741.5 |
| 40 | Yes | A. baylyi | None | None | F7 | Acridine or | 1754 | BL1 | 3877 |
| 40 | Yes | A. baylyi | None | None | F192 | 5-Carboxyf | 1560 | BL1 | 11577.5 |
| 40 | Yes | A. baylyi | None | None | F57 | Eosin Y | 1665 | BL1 | 14629 |
| 40 | Yes | A. baylyi | None | None | F27 | Calcein | 1568 | BL1 | 14032.5 |

| | | | | | | | | | |
|---|---|---|---|---|---|---|---|---|---|
| 40 | Yes | A. baylyi | None | None | F12 | Amaranth | 1783 | BL1 | 15304 |
| 40 | Yes | A. baylyi | None | None | NA | Autofluores | 1986 | NA | 7294 |
| 40 | Yes | A. baylyi | Ampicillin | Wall | F194 | Oxonol | 1646 | RL1 | 4796.5 |
| 40 | Yes | A. baylyi | Ampicillin | Wall | F11 | Allura Red | 1562 | BL3 | 4242.5 |
| 40 | Yes | A. baylyi | Ampicillin | Wall | F54 | DiSC3(5) | 1438 | RL1 | 3490 |
| 40 | Yes | A. baylyi | Ampicillin | Wall | F124 | Thioflavin T | 1447 | VL2 | 3547 |
| 40 | Yes | A. baylyi | Ampicillin | Wall | F101 | Quinacrine | 1256 | VL2 | 4308 |
| 40 | Yes | A. baylyi | Ampicillin | Wall | F138 | Apigenin | 1297 | VL2 | 3791 |
| 40 | Yes | A. baylyi | Ampicillin | Wall | F65 | H2FDA | 1097 | BL1 | 4290 |
| 40 | Yes | A. baylyi | Ampicillin | Wall | F142 | Chlorophe | 1150 | BL3 | 5364 |
| 40 | Yes | A. baylyi | Ampicillin | Wall | F39 | Congo Red | 1414 | BL3 | 5696 |
| 40 | Yes | A. baylyi | Ampicillin | Wall | F116 | Sunset Yel | 1350 | BL2 | 5701 |
| 40 | Yes | A. baylyi | Ampicillin | Wall | F10 | Alizarin | 1396 | BL1 | 5655 |
| 40 | Yes | A. baylyi | Ampicillin | Wall | F84 | Neutral Re | 1230 | BL1 | 4787.5 |
| 40 | Yes | A. baylyi | Ampicillin | Wall | F99 | Pyronin Y | 1257 | BL1 | 4814 |
| 40 | Yes | A. baylyi | Ampicillin | Wall | F108 | Rhodamine | 1142 | BL1 | 4429 |
| 40 | Yes | A. baylyi | Ampicillin | Wall | F109 | Riboflavin | 1037 | BL1 | 3755 |
| 40 | Yes | A. baylyi | Ampicillin | Wall | F117 | SYBR Gree | 1135 | BL1 | 597528 |
| 40 | Yes | A. baylyi | Ampicillin | Wall | F106 | Rhodamine | 1289 | BL1 | 4693 |
| 40 | Yes | A. baylyi | Ampicillin | Wall | F73 | Malachite ( | 1229 | BL1 | 4757 |
| 40 | Yes | A. baylyi | Ampicillin | Wall | F7 | Acridine or | 1133 | BL1 | 4987 |
| 40 | Yes | A. baylyi | Ampicillin | Wall | F192 | 5-Carboxyf | 1196 | BL1 | 13858 |
| 40 | Yes | A. baylyi | Ampicillin | Wall | F57 | Eosin Y | 1046 | BL1 | 17994.5 |
| 40 | Yes | A. baylyi | Ampicillin | Wall | F27 | Calcein | 1040 | BL1 | 16012 |
| 40 | Yes | A. baylyi | Ampicillin | Wall | F12 | Amaranth | 1163 | BL1 | 19067 |
| 40 | Yes | A. baylyi | None | None | NA | Autofluores | 2120 | NA | 6216 |
| 40 | Yes | A. baylyi | Bacitracin | Wall | F194 | Oxonol | 2319 | RL1 | 4903 |
| 40 | Yes | A. baylyi | Bacitracin | Wall | F11 | Allura Red | 2082 | BL3 | 4247.5 |
| 40 | Yes | A. baylyi | Bacitracin | Wall | F54 | DiSC3(5) | 1894 | RL1 | 3749 |
| 40 | Yes | A. baylyi | Bacitracin | Wall | F124 | Thioflavin T | 2163 | VL2 | 3456 |
| 40 | Yes | A. baylyi | Bacitracin | Wall | F101 | Quinacrine | 2080 | VL2 | 3982 |
| 40 | Yes | A. baylyi | Bacitracin | Wall | F138 | Apigenin | 1975 | VL2 | 4032 |
| 40 | Yes | A. baylyi | Bacitracin | Wall | F65 | H2FDA | 2103 | BL1 | 4654 |
| 40 | Yes | A. baylyi | Bacitracin | Wall | F142 | Chlorophe | 2111 | BL3 | 5197 |
| 40 | Yes | A. baylyi | Bacitracin | Wall | F39 | Congo Red | 2153 | BL3 | 5263 |
| 40 | Yes | A. baylyi | Bacitracin | Wall | F116 | Sunset Yel | 2157 | BL2 | 5198 |
| 40 | Yes | A. baylyi | Bacitracin | Wall | F10 | Alizarin | 2076 | BL1 | 5142 |
| 40 | Yes | A. baylyi | Bacitracin | Wall | F84 | Neutral Re | 1675 | BL1 | 4642 |
| 40 | Yes | A. baylyi | Bacitracin | Wall | F99 | Pyronin Y | 2022 | BL1 | 5015 |
| 40 | Yes | A. baylyi | Bacitracin | Wall | F108 | Rhodamine | 1993 | BL1 | 4436 |
| 40 | Yes | A. baylyi | Bacitracin | Wall | F109 | Riboflavin | 2038 | BL1 | 4171 |
| 40 | Yes | A. baylyi | Bacitracin | Wall | F117 | SYBR Gree | 1995 | BL1 | 632023 |
| 40 | Yes | A. baylyi | Bacitracin | Wall | F106 | Rhodamine | 2043 | BL1 | 5361 |
| 40 | Yes | A. baylyi | Bacitracin | Wall | F73 | Malachite ( | 1971 | BL1 | 4741 |
| 40 | Yes | A. baylyi | Bacitracin | Wall | F7 | Acridine or | 2019 | BL1 | 5228 |
| 40 | Yes | A. baylyi | Bacitracin | Wall | F192 | 5-Carboxyf | 2159 | BL1 | 13067 |
| 40 | Yes | A. baylyi | Bacitracin | Wall | F57 | Eosin Y | 2043 | BL1 | 15223 |
| 40 | Yes | A. baylyi | Bacitracin | Wall | F27 | Calcein | 2024 | BL1 | 15009.5 |

| | | | | | | | | | |
|---|---|---|---|---|---|---|---|---|---|
| 40 | Yes | A. baylyi | Bacitracin | Wall | F12 | Amaranth | 2073 | BL1 | 17024 |
| 40 | Yes | A. baylyi | None | None | NA | Autofluore: | 2015 | NA | 7789 |
| 40 | Yes | A. baylyi | Cefotaxim | Wall | F194 | Oxonol | 1613 | RL1 | 5162 |
| 40 | Yes | A. baylyi | Cefotaxim | Wall | F11 | Allura Red | 1641 | BL3 | 4712 |
| 40 | Yes | A. baylyi | Cefotaxim | Wall | F54 | DiSC3(5) | 1547 | RL1 | 3975 |
| 40 | Yes | A. baylyi | Cefotaxim | Wall | F124 | Thioflavin T | 1328 | VL2 | 3977.5 |
| 40 | Yes | A. baylyi | Cefotaxim | Wall | F101 | Quinacrine | 1433 | VL2 | 4525 |
| 40 | Yes | A. baylyi | Cefotaxim | Wall | F138 | Apigenin | 1245 | VL2 | 4029 |
| 40 | Yes | A. baylyi | Cefotaxim | Wall | F65 | H2FDA | 1354 | BL1 | 5051.5 |
| 40 | Yes | A. baylyi | Cefotaxim | Wall | F142 | Chlorophe | 1229 | BL3 | 5865 |
| 40 | Yes | A. baylyi | Cefotaxim | Wall | F39 | Congo Red | 1462 | BL3 | 6408.5 |
| 40 | Yes | A. baylyi | Cefotaxim | Wall | F116 | Sunset Yel | 1382 | BL2 | 6099 |
| 40 | Yes | A. baylyi | Cefotaxim | Wall | F10 | Alizarin | 1571 | BL1 | 6037 |
| 40 | Yes | A. baylyi | Cefotaxim | Wall | F84 | Neutral Re | 1311 | BL1 | 5475 |
| 40 | Yes | A. baylyi | Cefotaxim | Wall | F99 | Pyronin Y | 1408 | BL1 | 5417.5 |
| 40 | Yes | A. baylyi | Cefotaxim | Wall | F108 | Rhodamine | 1270 | BL1 | 4969 |
| 40 | Yes | A. baylyi | Cefotaxim | Wall | F109 | Riboflavin | 1224 | BL1 | 4826.5 |
| 40 | Yes | A. baylyi | Cefotaxim | Wall | F117 | SYBR Gree | 1286 | BL1 | 641357 |
| 40 | Yes | A. baylyi | Cefotaxim | Wall | F106 | Rhodamine | 1354 | BL1 | 5792 |
| 40 | Yes | A. baylyi | Cefotaxim | Wall | F73 | Malachite ( | 1339 | BL1 | 5352 |
| 40 | Yes | A. baylyi | Cefotaxim | Wall | F7 | Acridine or | 1285 | BL1 | 5748 |
| 40 | Yes | A. baylyi | Cefotaxim | Wall | F192 | 5-Carboxyf | 1089 | BL1 | 13570 |
| 40 | Yes | A. baylyi | Cefotaxim | Wall | F57 | Eosin Y | 1147 | BL1 | 16225 |
| 40 | Yes | A. baylyi | Cefotaxim | Wall | F27 | Calcein | 1149 | BL1 | 14833 |
| 40 | Yes | A. baylyi | Cefotaxim | Wall | F12 | Amaranth | 1356 | BL1 | 16122 |
| 40 | Yes | A. baylyi | None | None | NA | Autofluore: | 2153 | NA | 7236 |
| 40 | Yes | A. baylyi | Ciprofloxa | Intracellul | F194 | Oxonol | 1437 | RL1 | 5888 |
| 40 | Yes | A. baylyi | Ciprofloxa | Intracellul | F11 | Allura Red | 1424 | BL3 | 5001.5 |
| 40 | Yes | A. baylyi | Ciprofloxa | Intracellul | F54 | DiSC3(5) | 1187 | RL1 | 4771 |
| 40 | Yes | A. baylyi | Ciprofloxa | Intracellul | F124 | Thioflavin T | 1275 | VL2 | 4390 |
| 40 | Yes | A. baylyi | Ciprofloxa | Intracellul | F101 | Quinacrine | 1255 | VL2 | 4826 |
| 40 | Yes | A. baylyi | Ciprofloxa | Intracellul | F138 | Apigenin | 1234 | VL2 | 5143 |
| 40 | Yes | A. baylyi | Ciprofloxa | Intracellul | F65 | H2FDA | 1042 | BL1 | 5489.5 |
| 40 | Yes | A. baylyi | Ciprofloxa | Intracellul | F142 | Chlorophe | 1051 | BL3 | 6176 |
| 40 | Yes | A. baylyi | Ciprofloxa | Intracellul | F39 | Congo Red | 1209 | BL3 | 6547 |
| 40 | Yes | A. baylyi | Ciprofloxa | Intracellul | F116 | Sunset Yel | 1351 | BL2 | 6481 |
| 40 | Yes | A. baylyi | Ciprofloxa | Intracellul | F10 | Alizarin | 1400 | BL1 | 5988.5 |
| 40 | Yes | A. baylyi | Ciprofloxa | Intracellul | F84 | Neutral Re | 1416 | BL1 | 6131.5 |
| 40 | Yes | A. baylyi | Ciprofloxa | Intracellul | F99 | Pyronin Y | 1147 | BL1 | 5368 |
| 40 | Yes | A. baylyi | Ciprofloxa | Intracellul | F108 | Rhodamine | 1187 | BL1 | 5523 |
| 40 | Yes | A. baylyi | Ciprofloxa | Intracellul | F109 | Riboflavin | 1099 | BL1 | 5095 |
| 40 | Yes | A. baylyi | Ciprofloxa | Intracellul | F117 | SYBR Gree | 1050 | BL1 | 382501.5 |
| 40 | Yes | A. baylyi | Ciprofloxa | Intracellul | F106 | Rhodamine | 1340 | BL1 | 6547 |
| 40 | Yes | A. baylyi | Ciprofloxa | Intracellul | F73 | Malachite ( | 1166 | BL1 | 5980.5 |
| 40 | Yes | A. baylyi | Ciprofloxa | Intracellul | F7 | Acridine or | 1063 | BL1 | 6016 |
| 40 | Yes | A. baylyi | Ciprofloxa | Intracellul | F192 | 5-Carboxyf | 1154 | BL1 | 13611 |
| 40 | Yes | A. baylyi | Ciprofloxa | Intracellul | F57 | Eosin Y | 1066 | BL1 | 17449 |
| 40 | Yes | A. baylyi | Ciprofloxa | Intracellul | F27 | Calcein | 1050 | BL1 | 16876 |

| | | | | | | | | | |
|---|---|---|---|---|---|---|---|---|---|
| 40 | Yes | A. baylyi | Ciprofloxa | Intracellula | F12 | Amaranth | 1211 | BL1 | 17768 |
| 40 | Yes | A. baylyi | None | None | NA | Autofluore | 1874 | NA | 7275.5 |
| 40 | Yes | A. baylyi | Fosfomyci | Wall | F194 | Oxonol | 2293 | RL1 | 5426 |
| 40 | Yes | A. baylyi | Fosfomyci | Wall | F11 | Allura Red | 2089 | BL3 | 4871 |
| 40 | Yes | A. baylyi | Fosfomyci | Wall | F54 | DiSC3(5) | 1536 | RL1 | 4202 |
| 40 | Yes | A. baylyi | Fosfomyci | Wall | F124 | Thioflavin T | 2087 | VL2 | 4150 |
| 40 | Yes | A. baylyi | Fosfomyci | Wall | F101 | Quinacrine | 1670 | VL2 | 4727.5 |
| 40 | Yes | A. baylyi | Fosfomyci | Wall | F138 | Apigenin | 1836 | VL2 | 4781 |
| 40 | Yes | A. baylyi | Fosfomyci | Wall | F65 | H2FDA | 1892 | BL1 | 5269.5 |
| 40 | Yes | A. baylyi | Fosfomyci | Wall | F142 | Chlorophe | 1931 | BL3 | 5825 |
| 40 | Yes | A. baylyi | Fosfomyci | Wall | F39 | Congo Red | 2042 | BL3 | 6157.5 |
| 40 | Yes | A. baylyi | Fosfomyci | Wall | F116 | Sunset Yel | 2069 | BL2 | 6002 |
| 40 | Yes | A. baylyi | Fosfomyci | Wall | F10 | Alizarin | 2221 | BL1 | 5792 |
| 40 | Yes | A. baylyi | Fosfomyci | Wall | F84 | Neutral Re | 2146 | BL1 | 5480 |
| 40 | Yes | A. baylyi | Fosfomyci | Wall | F99 | Pyronin Y | 2051 | BL1 | 5161 |
| 40 | Yes | A. baylyi | Fosfomyci | Wall | F108 | Rhodamine | 1767 | BL1 | 4689 |
| 40 | Yes | A. baylyi | Fosfomyci | Wall | F109 | Riboflavin | 1806 | BL1 | 5074 |
| 40 | Yes | A. baylyi | Fosfomyci | Wall | F117 | SYBR Gree | 1728 | BL1 | 503529 |
| 40 | Yes | A. baylyi | Fosfomyci | Wall | F106 | Rhodamine | 1772 | BL1 | 5547.5 |
| 40 | Yes | A. baylyi | Fosfomyci | Wall | F73 | Malachite ( | 1695 | BL1 | 5314 |
| 40 | Yes | A. baylyi | Fosfomyci | Wall | F7 | Acridine or | 1636 | BL1 | 5838.5 |
| 40 | Yes | A. baylyi | Fosfomyci | Wall | F192 | 5-Carboxyf | 1542 | BL1 | 12504.5 |
| 40 | Yes | A. baylyi | Fosfomyci | Wall | F57 | Eosin Y | 1576 | BL1 | 13887 |
| 40 | Yes | A. baylyi | Fosfomyci | Wall | F27 | Calcein | 1650 | BL1 | 12824 |
| 40 | Yes | A. baylyi | Fosfomyci | Wall | F12 | Amaranth | 1839 | BL1 | 13769 |
| 40 | Yes | A. baylyi | None | None | NA | Autofluore | 2177 | NA | 8032 |
| 40 | Yes | A. baylyi | Nalidixic a | Intracellula | F194 | Oxonol | 1646 | RL1 | 6144 |
| 40 | Yes | A. baylyi | Nalidixic a | Intracellula | F11 | Allura Red | 1562 | BL3 | 5180 |
| 40 | Yes | A. baylyi | Nalidixic a | Intracellula | F54 | DiSC3(5) | 1366 | RL1 | 4897 |
| 40 | Yes | A. baylyi | Nalidixic a | Intracellula | F124 | Thioflavin T | 1437 | VL2 | 4683 |
| 40 | Yes | A. baylyi | Nalidixic a | Intracellula | F101 | Quinacrine | 1510 | VL2 | 5298.5 |
| 40 | Yes | A. baylyi | Nalidixic a | Intracellula | F138 | Apigenin | 1567 | VL2 | 5098 |
| 40 | Yes | A. baylyi | Nalidixic a | Intracellula | F65 | H2FDA | 1559 | BL1 | 4840 |
| 40 | Yes | A. baylyi | Nalidixic a | Intracellula | F142 | Chlorophe | 1548 | BL3 | 6848 |
| 40 | Yes | A. baylyi | Nalidixic a | Intracellula | F39 | Congo Red | 1706 | BL3 | 7941 |
| 40 | Yes | A. baylyi | Nalidixic a | Intracellula | F116 | Sunset Yel | 1809 | BL2 | 6584 |
| 40 | Yes | A. baylyi | Nalidixic a | Intracellula | F10 | Alizarin | 2030 | BL1 | 5684.5 |
| 40 | Yes | A. baylyi | Nalidixic a | Intracellula | F84 | Neutral Re | 2029 | BL1 | 5481 |
| 40 | Yes | A. baylyi | Nalidixic a | Intracellula | F99 | Pyronin Y | 1674 | BL1 | 5498 |
| 40 | Yes | A. baylyi | Nalidixic a | Intracellula | F108 | Rhodamine | 1548 | BL1 | 4981 |
| 40 | Yes | A. baylyi | Nalidixic a | Intracellula | F109 | Riboflavin | 1578 | BL1 | 4946.5 |
| 40 | Yes | A. baylyi | Nalidixic a | Intracellula | F117 | SYBR Gree | 1599 | BL1 | 415001 |
| 40 | Yes | A. baylyi | Nalidixic a | Intracellula | F106 | Rhodamine | 1584 | BL1 | 5642.5 |
| 40 | Yes | A. baylyi | Nalidixic a | Intracellula | F73 | Malachite ( | 1413 | BL1 | 5226 |
| 40 | Yes | A. baylyi | Nalidixic a | Intracellula | F7 | Acridine or | 1395 | BL1 | 5343 |
| 40 | Yes | A. baylyi | Nalidixic a | Intracellula | F192 | 5-Carboxyf | 1374 | BL1 | 13278.5 |
| 40 | Yes | A. baylyi | Nalidixic a | Intracellula | F57 | Eosin Y | 1387 | BL1 | 16416 |
| 40 | Yes | A. baylyi | Nalidixic a | Intracellula | F27 | Calcein | 1253 | BL1 | 16828 |

| | | | | | | | | | |
|---|---|---|---|---|---|---|---|---|---|
| 40 | Yes | A. baylyi | Nalidixic a | Intracellula | F12 | Amaranth | 1463 | BL1 | 18561 |
| 40 | Yes | A. baylyi | None | None | NA | Autofluore: | 2067 | NA | 7393 |
| 40 | Yes | A. baylyi | Ofloxacin | Intracellula | F194 | Oxonol | 1429 | RL1 | 6205 |
| 40 | Yes | A. baylyi | Ofloxacin | Intracellula | F11 | Allura Red | 1375 | BL3 | 6135 |
| 40 | Yes | A. baylyi | Ofloxacin | Intracellula | F54 | DiSC3(5) | 1278 | RL1 | 5721.5 |
| 40 | Yes | A. baylyi | Ofloxacin | Intracellula | F124 | Thioflavin T | 1297 | VL2 | 5035 |
| 40 | Yes | A. baylyi | Ofloxacin | Intracellula | F101 | Quinacrine | 1314 | VL2 | 5905.5 |
| 40 | Yes | A. baylyi | Ofloxacin | Intracellula | F138 | Apigenin | 1326 | VL2 | 7331 |
| 40 | Yes | A. baylyi | Ofloxacin | Intracellula | F65 | H2FDA | 1425 | BL1 | 6604 |
| 40 | Yes | A. baylyi | Ofloxacin | Intracellula | F142 | Chlorophe | 1582 | BL3 | 6460.5 |
| 40 | Yes | A. baylyi | Ofloxacin | Intracellula | F39 | Congo Red | 1444 | BL3 | 8442.5 |
| 40 | Yes | A. baylyi | Ofloxacin | Intracellula | F116 | Sunset Yel | 1567 | BL2 | 7224 |
| 40 | Yes | A. baylyi | Ofloxacin | Intracellula | F10 | Alizarin | 1462 | BL1 | 8243.5 |
| 40 | Yes | A. baylyi | Ofloxacin | Intracellula | F84 | Neutral Re | 1766 | BL1 | 6057.5 |
| 40 | Yes | A. baylyi | Ofloxacin | Intracellula | F99 | Pyronin Y | 1559 | BL1 | 6042 |
| 40 | Yes | A. baylyi | Ofloxacin | Intracellula | F108 | Rhodamine | 1418 | BL1 | 4359.5 |
| 40 | Yes | A. baylyi | Ofloxacin | Intracellula | F109 | Riboflavin | 1276 | BL1 | 5466 |
| 40 | Yes | A. baylyi | Ofloxacin | Intracellula | F117 | SYBR Gree | 1379 | BL1 | 542109 |
| 40 | Yes | A. baylyi | Ofloxacin | Intracellula | F106 | Rhodamine | 1306 | BL1 | 6208 |
| 40 | Yes | A. baylyi | Ofloxacin | Intracellula | F73 | Malachite ( | 1440 | BL1 | 6012 |
| 40 | Yes | A. baylyi | Ofloxacin | Intracellula | F7 | Acridine or | 1244 | BL1 | 5833.5 |
| 40 | Yes | A. baylyi | Ofloxacin | Intracellula | F192 | 5-Carboxyf | 1108 | BL1 | 14293.5 |
| 40 | Yes | A. baylyi | Ofloxacin | Intracellula | F57 | Eosin Y | 1138 | BL1 | 19301 |
| 40 | Yes | A. baylyi | Ofloxacin | Intracellula | F27 | Calcein | 1175 | BL1 | 20954 |
| 40 | Yes | A. baylyi | Ofloxacin | Intracellula | F12 | Amaranth | 1102 | BL1 | 16652.5 |
| 40 | Yes | A. baylyi | None | None | NA | Autofluore: | 2324 | NA | 6809 |
| 40 | Yes | A. baylyi | Polymyxin | Wall | F194 | Oxonol | 2366 | RL1 | 5990 |
| 40 | Yes | A. baylyi | Polymyxin | Wall | F11 | Allura Red | 2668 | BL3 | 4976.5 |
| 40 | Yes | A. baylyi | Polymyxin | Wall | F54 | DiSC3(5) | 2073 | RL1 | 7335 |
| 40 | Yes | A. baylyi | Polymyxin | Wall | F124 | Thioflavin T | 2663 | VL2 | 4597 |
| 40 | Yes | A. baylyi | Polymyxin | Wall | F101 | Quinacrine | 2252 | VL2 | 4608.5 |
| 40 | Yes | A. baylyi | Polymyxin | Wall | F138 | Apigenin | 2606 | VL2 | 4799.5 |
| 40 | Yes | A. baylyi | Polymyxin | Wall | F65 | H2FDA | 2568 | BL1 | 6046 |
| 40 | Yes | A. baylyi | Polymyxin | Wall | F142 | Chlorophe | 2615 | BL3 | 6465 |
| 40 | Yes | A. baylyi | Polymyxin | Wall | F39 | Congo Red | 2936 | BL3 | 5778.5 |
| 40 | Yes | A. baylyi | Polymyxin | Wall | F116 | Sunset Yel | 3068 | BL2 | 5935 |
| 40 | Yes | A. baylyi | Polymyxin | Wall | F10 | Alizarin | 2844 | BL1 | 5339 |
| 40 | Yes | A. baylyi | Polymyxin | Wall | F84 | Neutral Re | 2935 | BL1 | 5600 |
| 40 | Yes | A. baylyi | Polymyxin | Wall | F99 | Pyronin Y | 2509 | BL1 | 5045 |
| 40 | Yes | A. baylyi | Polymyxin | Wall | F108 | Rhodamine | 2661 | BL1 | 5006 |
| 40 | Yes | A. baylyi | Polymyxin | Wall | F109 | Riboflavin | 2562 | BL1 | 4923 |
| 40 | Yes | A. baylyi | Polymyxin | Wall | F117 | SYBR Gree | 2406 | BL1 | 885627 |
| 40 | Yes | A. baylyi | Polymyxin | Wall | F106 | Rhodamine | 2455 | BL1 | 5698 |
| 40 | Yes | A. baylyi | Polymyxin | Wall | F73 | Malachite ( | 2042 | BL1 | 5288.5 |
| 40 | Yes | A. baylyi | Polymyxin | Wall | F7 | Acridine or | 2121 | BL1 | 5667 |
| 40 | Yes | A. baylyi | Polymyxin | Wall | F192 | 5-Carboxyf | 2312 | BL1 | 11633 |
| 40 | Yes | A. baylyi | Polymyxin | Wall | F57 | Eosin Y | 2103 | BL1 | 15288 |
| 40 | Yes | A. baylyi | Polymyxin | Wall | F27 | Calcein | 2206 | BL1 | 14464.5 |

| | | | | | | | | | |
|---|---|---|---|---|---|---|---|---|---|
| 40 | Yes | A. baylyi | Polymyxin | Wall | F12 | Amaranth | 2245 | BL1 | 15416 |
| 40 | Yes | A. baylyi | None | None | NA | Autofluore: | 2240 | NA | 6837 |
| 40 | Yes | A. baylyi | Tetracyclin | Intracellula | F194 | Oxonol | 1208 | RL1 | 5911.5 |
| 40 | Yes | A. baylyi | Tetracyclin | Intracellula | F11 | Allura Red | 1143 | BL3 | 4950 |
| 40 | Yes | A. baylyi | Tetracyclin | Intracellula | F54 | DiSC3(5) | 1133 | RL1 | 4955 |
| 40 | Yes | A. baylyi | Tetracyclin | Intracellula | F124 | Thioflavin T | 1178 | VL2 | 4411 |
| 40 | Yes | A. baylyi | Tetracyclin | Intracellula | F101 | Quinacrine | 1219 | VL2 | 4918 |
| 40 | Yes | A. baylyi | Tetracyclin | Intracellula | F138 | Apigenin | 1114 | VL2 | 4884.5 |
| 40 | Yes | A. baylyi | Tetracyclin | Intracellula | F65 | H2FDA | 1234 | BL1 | 6329.5 |
| 40 | Yes | A. baylyi | Tetracyclin | Intracellula | F142 | Chlorophe | 1296 | BL3 | 6466.5 |
| 40 | Yes | A. baylyi | Tetracyclin | Intracellula | F39 | Congo Red | 1379 | BL3 | 6126 |
| 40 | Yes | A. baylyi | Tetracyclin | Intracellula | F116 | Sunset Yel | 1456 | BL2 | 6450.5 |
| 40 | Yes | A. baylyi | Tetracyclin | Intracellula | F10 | Alizarin | 1362 | BL1 | 6188.5 |
| 40 | Yes | A. baylyi | Tetracyclin | Intracellula | F84 | Neutral Re | 1441 | BL1 | 5298 |
| 40 | Yes | A. baylyi | Tetracyclin | Intracellula | F99 | Pyronin Y | 1317 | BL1 | 5823 |
| 40 | Yes | A. baylyi | Tetracyclin | Intracellula | F108 | Rhodamine | 1347 | BL1 | 5340 |
| 40 | Yes | A. baylyi | Tetracyclin | Intracellula | F109 | Riboflavin | 1091 | BL1 | 4966 |
| 40 | Yes | A. baylyi | Tetracyclin | Intracellula | F117 | SYBR Gree | 1139 | BL1 | 402808 |
| 40 | Yes | A. baylyi | Tetracyclin | Intracellula | F106 | Rhodamine | 1252 | BL1 | 5794.5 |
| 40 | Yes | A. baylyi | Tetracyclin | Intracellula | F73 | Malachite ( | 1187 | BL1 | 5382 |
| 40 | Yes | A. baylyi | Tetracyclin | Intracellula | F7 | Acridine or | 1060 | BL1 | 5077 |
| 40 | Yes | A. baylyi | Tetracyclin | Intracellula | F192 | 5-Carboxyf | 1047 | BL1 | 11400 |
| 40 | Yes | A. baylyi | Tetracyclin | Intracellula | F57 | Eosin Y | 958 | BL1 | 15217.5 |
| 40 | Yes | A. baylyi | Tetracyclin | Intracellula | F27 | Calcein | 1076 | BL1 | 13793.5 |
| 40 | Yes | A. baylyi | Tetracyclin | Intracellula | F12 | Amaranth | 994 | BL1 | 14118 |
| 40 | Yes | A. baylyi | None | None | NA | Autofluore: | 2348 | NA | 7769.5 |
| 40 | Yes | A. baylyi | Vancomyc | Wall | F194 | Oxonol | 1954 | RL1 | 6191.5 |
| 40 | Yes | A. baylyi | Vancomyc | Wall | F11 | Allura Red | 1909 | BL3 | 4811 |
| 40 | Yes | A. baylyi | Vancomyc | Wall | F54 | DiSC3(5) | 1851 | RL1 | 4634 |
| 40 | Yes | A. baylyi | Vancomyc | Wall | F124 | Thioflavin T | 2034 | VL2 | 4482.5 |
| 40 | Yes | A. baylyi | Vancomyc | Wall | F101 | Quinacrine | 1745 | VL2 | 5006 |
| 40 | Yes | A. baylyi | Vancomyc | Wall | F138 | Apigenin | 1693 | VL2 | 4507 |
| 40 | Yes | A. baylyi | Vancomyc | Wall | F65 | H2FDA | 1884 | BL1 | 6061 |
| 40 | Yes | A. baylyi | Vancomyc | Wall | F142 | Chlorophe | 1871 | BL3 | 7587 |
| 40 | Yes | A. baylyi | Vancomyc | Wall | F39 | Congo Red | 1991 | BL3 | 6702 |
| 40 | Yes | A. baylyi | Vancomyc | Wall | F116 | Sunset Yel | 1974 | BL2 | 6229.5 |
| 40 | Yes | A. baylyi | Vancomyc | Wall | F10 | Alizarin | 2213 | BL1 | 5980 |
| 40 | Yes | A. baylyi | Vancomyc | Wall | F84 | Neutral Re | 2078 | BL1 | 5651 |
| 40 | Yes | A. baylyi | Vancomyc | Wall | F99 | Pyronin Y | 1839 | BL1 | 5391 |
| 40 | Yes | A. baylyi | Vancomyc | Wall | F108 | Rhodamine | 1734 | BL1 | 5035.5 |
| 40 | Yes | A. baylyi | Vancomyc | Wall | F109 | Riboflavin | 1643 | BL1 | 5168 |
| 40 | Yes | A. baylyi | Vancomyc | Wall | F117 | SYBR Gree | 1623 | BL1 | 441193 |
| 40 | Yes | A. baylyi | Vancomyc | Wall | F106 | Rhodamine | 1878 | BL1 | 5975 |
| 40 | Yes | A. baylyi | Vancomyc | Wall | F73 | Malachite ( | 1634 | BL1 | 5045 |
| 40 | Yes | A. baylyi | Vancomyc | Wall | F7 | Acridine or | 1789 | BL1 | 5038 |
| 40 | Yes | A. baylyi | Vancomyc | Wall | F192 | 5-Carboxyf | 1601 | BL1 | 12871 |
| 40 | Yes | A. baylyi | Vancomyc | Wall | F57 | Eosin Y | 1460 | BL1 | 16788 |
| 40 | Yes | A. baylyi | Vancomyc | Wall | F27 | Calcein | 1636 | BL1 | 17496 |

| | | | | | | | | | |
|---|---|---|---|---|---|---|---|---|---|
| 40 | Yes | A. baylyi | Vancomyc | Wall | F12 | Amaranth | 1635 | BL1 | 18417 |
| 40 | Yes | A. baylyi | None | None | NA | Autofluore: | 2696 | NA | 889.5 |
| 40 | Yes | A. baylyi | None | None | F194 | Oxonol | 3381 | RL1 | 1534 |
| 40 | Yes | A. baylyi | None | None | F11 | Allura Red | 3054 | BL3 | 2387 |
| 40 | Yes | A. baylyi | None | None | F54 | DiSC3(5) | 2832 | RL1 | 2527 |
| 40 | Yes | A. baylyi | None | None | F124 | Thioflavin T | 2690 | VL2 | 2660.5 |
| 40 | Yes | A. baylyi | None | None | F101 | Quinacrine | 2541 | VL2 | 2735 |
| 40 | Yes | A. baylyi | None | None | F138 | Apigenin | 2259 | VL2 | 2580 |
| 40 | Yes | A. baylyi | None | None | F65 | H2FDA | 2441 | BL1 | 3299 |
| 40 | Yes | A. baylyi | None | None | F142 | Chlorophe | 2247 | BL3 | 4269 |
| 40 | Yes | A. baylyi | None | None | F39 | Congo Red | 2435 | BL3 | 4987 |
| 40 | Yes | A. baylyi | None | None | F116 | Sunset Yel | 2433 | BL2 | 4889 |
| 40 | Yes | A. baylyi | None | None | F10 | Alizarin | 2234 | BL1 | 4515.5 |
| 40 | Yes | A. baylyi | None | None | F84 | Neutral Re | 2302 | BL1 | 3914 |
| 40 | Yes | A. baylyi | None | None | F99 | Pyronin Y | 1828 | BL1 | 4107 |
| 40 | Yes | A. baylyi | None | None | F108 | Rhodamine | 2573 | BL1 | 3321 |
| 40 | Yes | A. baylyi | None | None | F109 | Riboflavin | 2183 | BL1 | 3221 |
| 40 | Yes | A. baylyi | None | None | F117 | SYBR Gree | 2126 | BL1 | 403874 |
| 40 | Yes | A. baylyi | None | None | F106 | Rhodamine | 2129 | BL1 | 3958 |
| 40 | Yes | A. baylyi | None | None | F73 | Malachite ( | 1971 | BL1 | 3790 |
| 40 | Yes | A. baylyi | None | None | F7 | Acridine or | 2150 | BL1 | 3723 |
| 40 | Yes | A. baylyi | None | None | F192 | 5-Carboxyf | 1968 | BL1 | 11492.5 |
| 40 | Yes | A. baylyi | None | None | F57 | Eosin Y | 1874 | BL1 | 15112 |
| 40 | Yes | A. baylyi | None | None | F27 | Calcein | 1995 | BL1 | 15746 |
| 40 | Yes | A. baylyi | None | None | F12 | Amaranth | 1846 | BL1 | 15971 |
| 40 | Yes | A. baylyi | None | None | NA | Autofluore: | 2485 | NA | 7221 |
| 40 | Yes | A. baylyi | Ampicillin | Wall | F194 | Oxonol | 1702 | RL1 | 5171.5 |
| 40 | Yes | A. baylyi | Ampicillin | Wall | F11 | Allura Red | 1614 | BL3 | 4259 |
| 40 | Yes | A. baylyi | Ampicillin | Wall | F54 | DiSC3(5) | 1504 | RL1 | 3699 |
| 40 | Yes | A. baylyi | Ampicillin | Wall | F124 | Thioflavin T | 1499 | VL2 | 3777 |
| 40 | Yes | A. baylyi | Ampicillin | Wall | F101 | Quinacrine | 1290 | VL2 | 4114 |
| 40 | Yes | A. baylyi | Ampicillin | Wall | F138 | Apigenin | 1307 | VL2 | 3524 |
| 40 | Yes | A. baylyi | Ampicillin | Wall | F65 | H2FDA | 1388 | BL1 | 4188 |
| 40 | Yes | A. baylyi | Ampicillin | Wall | F142 | Chlorophe | 1340 | BL3 | 5317.5 |
| 40 | Yes | A. baylyi | Ampicillin | Wall | F39 | Congo Red | 1518 | BL3 | 5867.5 |
| 40 | Yes | A. baylyi | Ampicillin | Wall | F116 | Sunset Yel | 1424 | BL2 | 5946.5 |
| 40 | Yes | A. baylyi | Ampicillin | Wall | F10 | Alizarin | 1457 | BL1 | 5268 |
| 40 | Yes | A. baylyi | Ampicillin | Wall | F84 | Neutral Re | 1324 | BL1 | 4647.5 |
| 40 | Yes | A. baylyi | Ampicillin | Wall | F99 | Pyronin Y | 1328 | BL1 | 4602.5 |
| 40 | Yes | A. baylyi | Ampicillin | Wall | F108 | Rhodamine | 996 | BL1 | 3855 |
| 40 | Yes | A. baylyi | Ampicillin | Wall | F109 | Riboflavin | 1221 | BL1 | 3987 |
| 40 | Yes | A. baylyi | Ampicillin | Wall | F117 | SYBR Gree | 1239 | BL1 | 732310 |
| 40 | Yes | A. baylyi | Ampicillin | Wall | F106 | Rhodamine | 1314 | BL1 | 4883 |
| 40 | Yes | A. baylyi | Ampicillin | Wall | F73 | Malachite ( | 1204 | BL1 | 5262.5 |
| 40 | Yes | A. baylyi | Ampicillin | Wall | F7 | Acridine or | 1189 | BL1 | 5073 |
| 40 | Yes | A. baylyi | Ampicillin | Wall | F192 | 5-Carboxyf | 1252 | BL1 | 13729.5 |
| 40 | Yes | A. baylyi | Ampicillin | Wall | F57 | Eosin Y | 1239 | BL1 | 18275 |
| 40 | Yes | A. baylyi | Ampicillin | Wall | F27 | Calcein | 1024 | BL1 | 19563.5 |

| | | | | | | | | | |
|---|---|---|---|---|---|---|---|---|---|
| 40 | Yes | A. baylyi | Ampicillin | Wall | F12 | Amaranth | 1195 | BL1 | 18571 |
| 40 | Yes | A. baylyi | None | None | NA | Autofluore: | 2542 | NA | 6097.5 |
| 40 | Yes | A. baylyi | Bacitracin | Wall | F194 | Oxonol | 2346 | RL1 | 4522 |
| 40 | Yes | A. baylyi | Bacitracin | Wall | F11 | Allura Red | 2633 | BL3 | 4209 |
| 40 | Yes | A. baylyi | Bacitracin | Wall | F54 | DiSC3(5) | 2176 | RL1 | 3754.5 |
| 40 | Yes | A. baylyi | Bacitracin | Wall | F124 | Thioflavin T | 2557 | VL2 | 3597 |
| 40 | Yes | A. baylyi | Bacitracin | Wall | F101 | Quinacrine | 2227 | VL2 | 3711 |
| 40 | Yes | A. baylyi | Bacitracin | Wall | F138 | Apigenin | 2245 | VL2 | 4157 |
| 40 | Yes | A. baylyi | Bacitracin | Wall | F65 | H2FDA | 2426 | BL1 | 4312.5 |
| 40 | Yes | A. baylyi | Bacitracin | Wall | F142 | Chlorophe | 2574 | BL3 | 4739 |
| 40 | Yes | A. baylyi | Bacitracin | Wall | F39 | Congo Red | 2235 | BL3 | 5326 |
| 40 | Yes | A. baylyi | Bacitracin | Wall | F116 | Sunset Yel | 2485 | BL2 | 5304 |
| 40 | Yes | A. baylyi | Bacitracin | Wall | F10 | Alizarin | 2526 | BL1 | 5329.5 |
| 40 | Yes | A. baylyi | Bacitracin | Wall | F84 | Neutral Re | 2337 | BL1 | 4785 |
| 40 | Yes | A. baylyi | Bacitracin | Wall | F99 | Pyronin Y | 2238 | BL1 | 4610.5 |
| 40 | Yes | A. baylyi | Bacitracin | Wall | F108 | Rhodamine | 2253 | BL1 | 4266 |
| 40 | Yes | A. baylyi | Bacitracin | Wall | F109 | Riboflavin | 2341 | BL1 | 4093 |
| 40 | Yes | A. baylyi | Bacitracin | Wall | F117 | SYBR Gree | 2345 | BL1 | 565184 |
| 40 | Yes | A. baylyi | Bacitracin | Wall | F106 | Rhodamine | 2515 | BL1 | 4674 |
| 40 | Yes | A. baylyi | Bacitracin | Wall | F73 | Malachite ( | 2314 | BL1 | 4443.5 |
| 40 | Yes | A. baylyi | Bacitracin | Wall | F7 | Acridine or | 2348 | BL1 | 4756.5 |
| 40 | Yes | A. baylyi | Bacitracin | Wall | F192 | 5-Carboxyf | 2322 | BL1 | 12582 |
| 40 | Yes | A. baylyi | Bacitracin | Wall | F57 | Eosin Y | 2437 | BL1 | 16939 |
| 40 | Yes | A. baylyi | Bacitracin | Wall | F27 | Calcein | 2325 | BL1 | 19562 |
| 40 | Yes | A. baylyi | Bacitracin | Wall | F12 | Amaranth | 2352 | BL1 | 14103 |
| 40 | Yes | A. baylyi | None | None | NA | Autofluore: | 2521 | NA | 7310 |
| 40 | Yes | A. baylyi | Cefotaxim | Wall | F194 | Oxonol | 1811 | RL1 | 5704 |
| 40 | Yes | A. baylyi | Cefotaxim | Wall | F11 | Allura Red | 1688 | BL3 | 4609 |
| 40 | Yes | A. baylyi | Cefotaxim | Wall | F54 | DiSC3(5) | 1852 | RL1 | 4058.5 |
| 40 | Yes | A. baylyi | Cefotaxim | Wall | F124 | Thioflavin T | 1642 | VL2 | 4201 |
| 40 | Yes | A. baylyi | Cefotaxim | Wall | F101 | Quinacrine | 1559 | VL2 | 4312 |
| 40 | Yes | A. baylyi | Cefotaxim | Wall | F138 | Apigenin | 1508 | VL2 | 4269.5 |
| 40 | Yes | A. baylyi | Cefotaxim | Wall | F65 | H2FDA | 1512 | BL1 | 4642 |
| 40 | Yes | A. baylyi | Cefotaxim | Wall | F142 | Chlorophe | 1368 | BL3 | 5553.5 |
| 40 | Yes | A. baylyi | Cefotaxim | Wall | F39 | Congo Red | 1575 | BL3 | 5904 |
| 40 | Yes | A. baylyi | Cefotaxim | Wall | F116 | Sunset Yel | 1603 | BL2 | 5996 |
| 40 | Yes | A. baylyi | Cefotaxim | Wall | F10 | Alizarin | 1550 | BL1 | 5630.5 |
| 40 | Yes | A. baylyi | Cefotaxim | Wall | F84 | Neutral Re | 1565 | BL1 | 5015 |
| 40 | Yes | A. baylyi | Cefotaxim | Wall | F99 | Pyronin Y | 1378 | BL1 | 4832.5 |
| 40 | Yes | A. baylyi | Cefotaxim | Wall | F108 | Rhodamine | 1404 | BL1 | 4599.5 |
| 40 | Yes | A. baylyi | Cefotaxim | Wall | F109 | Riboflavin | 1337 | BL1 | 4463 |
| 40 | Yes | A. baylyi | Cefotaxim | Wall | F117 | SYBR Gree | 1369 | BL1 | 603144 |
| 40 | Yes | A. baylyi | Cefotaxim | Wall | F106 | Rhodamine | 1567 | BL1 | 5242 |
| 40 | Yes | A. baylyi | Cefotaxim | Wall | F73 | Malachite ( | 1376 | BL1 | 5392 |
| 40 | Yes | A. baylyi | Cefotaxim | Wall | F7 | Acridine or | 1188 | BL1 | 5380 |
| 40 | Yes | A. baylyi | Cefotaxim | Wall | F192 | 5-Carboxyf | 1313 | BL1 | 14182 |
| 40 | Yes | A. baylyi | Cefotaxim | Wall | F57 | Eosin Y | 1325 | BL1 | 17865 |
| 40 | Yes | A. baylyi | Cefotaxim | Wall | F27 | Calcein | 1198 | BL1 | 17079.5 |

| | | | | | | | | | |
|---|---|---|---|---|---|---|---|---|---|
| 40 | Yes | A. baylyi | Cefotaxim | Wall | F12 | Amaranth | 1267 | BL1 | 16518 |
| 40 | Yes | A. baylyi | None | None | NA | Autofluore | 2455 | NA | 5854 |
| 40 | Yes | A. baylyi | Ciprofloxa | Intracellul | F194 | Oxonol | 1665 | RL1 | 5097 |
| 40 | Yes | A. baylyi | Ciprofloxa | Intracellul | F11 | Allura Red | 1507 | BL3 | 4673 |
| 40 | Yes | A. baylyi | Ciprofloxa | Intracellul | F54 | DiSC3(5) | 1209 | RL1 | 4110 |
| 40 | Yes | A. baylyi | Ciprofloxa | Intracellul | F124 | Thioflavin T | 1504 | VL2 | 4341.5 |
| 40 | Yes | A. baylyi | Ciprofloxa | Intracellul | F101 | Quinacrine | 1333 | VL2 | 4751 |
| 40 | Yes | A. baylyi | Ciprofloxa | Intracellul | F138 | Apigenin | 1386 | VL2 | 4583 |
| 40 | Yes | A. baylyi | Ciprofloxa | Intracellul | F65 | H2FDA | 1211 | BL1 | 5097 |
| 40 | Yes | A. baylyi | Ciprofloxa | Intracellul | F142 | Chlorophe | 1423 | BL3 | 5907 |
| 40 | Yes | A. baylyi | Ciprofloxa | Intracellul | F39 | Congo Red | 1379 | BL3 | 6464 |
| 40 | Yes | A. baylyi | Ciprofloxa | Intracellul | F116 | Sunset Yel | 1407 | BL2 | 6529 |
| 40 | Yes | A. baylyi | Ciprofloxa | Intracellul | F10 | Alizarin | 1610 | BL1 | 6290 |
| 40 | Yes | A. baylyi | Ciprofloxa | Intracellul | F84 | Neutral Re | 1524 | BL1 | 5593 |
| 40 | Yes | A. baylyi | Ciprofloxa | Intracellul | F99 | Pyronin Y | 1398 | BL1 | 5531 |
| 40 | Yes | A. baylyi | Ciprofloxa | Intracellul | F108 | Rhodamine | 1333 | BL1 | 5198 |
| 40 | Yes | A. baylyi | Ciprofloxa | Intracellul | F109 | Riboflavin | 1182 | BL1 | 4776.5 |
| 40 | Yes | A. baylyi | Ciprofloxa | Intracellul | F117 | SYBR Gree | 1156 | BL1 | 342272.5 |
| 40 | Yes | A. baylyi | Ciprofloxa | Intracellul | F106 | Rhodamine | 1299 | BL1 | 6168 |
| 40 | Yes | A. baylyi | Ciprofloxa | Intracellul | F73 | Malachite | 1278 | BL1 | 5600 |
| 40 | Yes | A. baylyi | Ciprofloxa | Intracellul | F7 | Acridine or | 1269 | BL1 | 6244 |
| 40 | Yes | A. baylyi | Ciprofloxa | Intracellul | F192 | 5-Carboxyf | 1202 | BL1 | 14684.5 |
| 40 | Yes | A. baylyi | Ciprofloxa | Intracellul | F57 | Eosin Y | 1149 | BL1 | 18863 |
| 40 | Yes | A. baylyi | Ciprofloxa | Intracellul | F27 | Calcein | 1169 | BL1 | 17914 |
| 40 | Yes | A. baylyi | Ciprofloxa | Intracellul | F12 | Amaranth | 1334 | BL1 | 17883.5 |
| 40 | Yes | A. baylyi | None | None | NA | Autofluore | 2296 | NA | 6756 |
| 40 | Yes | A. baylyi | Fosfomyci | Wall | F194 | Oxonol | 2351 | RL1 | 5019 |
| 40 | Yes | A. baylyi | Fosfomyci | Wall | F11 | Allura Red | 2251 | BL3 | 4309 |
| 40 | Yes | A. baylyi | Fosfomyci | Wall | F54 | DiSC3(5) | 1846 | RL1 | 3995.5 |
| 40 | Yes | A. baylyi | Fosfomyci | Wall | F124 | Thioflavin T | 2128 | VL2 | 4033.5 |
| 40 | Yes | A. baylyi | Fosfomyci | Wall | F101 | Quinacrine | 2234 | VL2 | 4859.5 |
| 40 | Yes | A. baylyi | Fosfomyci | Wall | F138 | Apigenin | 2003 | VL2 | 4173 |
| 40 | Yes | A. baylyi | Fosfomyci | Wall | F65 | H2FDA | 1987 | BL1 | 4657 |
| 40 | Yes | A. baylyi | Fosfomyci | Wall | F142 | Chlorophe | 2150 | BL3 | 5518 |
| 40 | Yes | A. baylyi | Fosfomyci | Wall | F39 | Congo Red | 2333 | BL3 | 5872 |
| 40 | Yes | A. baylyi | Fosfomyci | Wall | F116 | Sunset Yel | 2469 | BL2 | 6065 |
| 40 | Yes | A. baylyi | Fosfomyci | Wall | F10 | Alizarin | 2454 | BL1 | 5784 |
| 40 | Yes | A. baylyi | Fosfomyci | Wall | F84 | Neutral Re | 2427 | BL1 | 5223 |
| 40 | Yes | A. baylyi | Fosfomyci | Wall | F99 | Pyronin Y | 2273 | BL1 | 4984 |
| 40 | Yes | A. baylyi | Fosfomyci | Wall | F108 | Rhodamine | 2088 | BL1 | 4593.5 |
| 40 | Yes | A. baylyi | Fosfomyci | Wall | F109 | Riboflavin | 2037 | BL1 | 4697 |
| 40 | Yes | A. baylyi | Fosfomyci | Wall | F117 | SYBR Gree | 1946 | BL1 | 454051 |
| 40 | Yes | A. baylyi | Fosfomyci | Wall | F106 | Rhodamine | 2272 | BL1 | 5422 |
| 40 | Yes | A. baylyi | Fosfomyci | Wall | F73 | Malachite | 1838 | BL1 | 5453.5 |
| 40 | Yes | A. baylyi | Fosfomyci | Wall | F7 | Acridine or | 1816 | BL1 | 5646 |
| 40 | Yes | A. baylyi | Fosfomyci | Wall | F192 | 5-Carboxyf | 1729 | BL1 | 12507 |
| 40 | Yes | A. baylyi | Fosfomyci | Wall | F57 | Eosin Y | 1782 | BL1 | 14391.5 |
| 40 | Yes | A. baylyi | Fosfomyci | Wall | F27 | Calcein | 1805 | BL1 | 16672 |

| | | | | | | | | | |
|---|---|---|---|---|---|---|---|---|---|
| 40 | Yes | A. baylyi | Fosfomyci | Wall | F12 | Amaranth | 2213 | BL1 | 14388 |
| 40 | Yes | A. baylyi | None | None | NA | Autofluore | 2621 | NA | 7315 |
| 40 | Yes | A. baylyi | Nalidixic a | Intracellul | F194 | Oxonol | 2154 | RL1 | 5476 |
| 40 | Yes | A. baylyi | Nalidixic a | Intracellul | F11 | Allura Red | 2192 | BL3 | 4442 |
| 40 | Yes | A. baylyi | Nalidixic a | Intracellul | F54 | DiSC3(5) | 1773 | RL1 | 4330 |
| 40 | Yes | A. baylyi | Nalidixic a | Intracellul | F124 | Thioflavin T | 1758 | VL2 | 4580.5 |
| 40 | Yes | A. baylyi | Nalidixic a | Intracellul | F101 | Quinacrine | 1723 | VL2 | 4785 |
| 40 | Yes | A. baylyi | Nalidixic a | Intracellul | F138 | Apigenin | 1695 | VL2 | 4710 |
| 40 | Yes | A. baylyi | Nalidixic a | Intracellul | F65 | H2FDA | 1751 | BL1 | 5246 |
| 40 | Yes | A. baylyi | Nalidixic a | Intracellul | F142 | Chlorophe | 1810 | BL3 | 6051.5 |
| 40 | Yes | A. baylyi | Nalidixic a | Intracellul | F39 | Congo Red | 1890 | BL3 | 6453 |
| 40 | Yes | A. baylyi | Nalidixic a | Intracellul | F116 | Sunset Yel | 2033 | BL2 | 5980 |
| 40 | Yes | A. baylyi | Nalidixic a | Intracellul | F10 | Alizarin | 2127 | BL1 | 6359 |
| 40 | Yes | A. baylyi | Nalidixic a | Intracellul | F84 | Neutral Re | 2048 | BL1 | 5246.5 |
| 40 | Yes | A. baylyi | Nalidixic a | Intracellul | F99 | Pyronin Y | 1951 | BL1 | 5382 |
| 40 | Yes | A. baylyi | Nalidixic a | Intracellul | F108 | Rhodamine | 1833 | BL1 | 5111 |
| 40 | Yes | A. baylyi | Nalidixic a | Intracellul | F109 | Riboflavin | 1773 | BL1 | 5042 |
| 40 | Yes | A. baylyi | Nalidixic a | Intracellul | F117 | SYBR Gree | 1635 | BL1 | 462199 |
| 40 | Yes | A. baylyi | Nalidixic a | Intracellul | F106 | Rhodamine | 1761 | BL1 | 5567 |
| 40 | Yes | A. baylyi | Nalidixic a | Intracellul | F73 | Malachite ( | 1715 | BL1 | 5596 |
| 40 | Yes | A. baylyi | Nalidixic a | Intracellul | F7 | Acridine or | 1539 | BL1 | 5223 |
| 40 | Yes | A. baylyi | Nalidixic a | Intracellul | F192 | 5-Carboxyf | 1535 | BL1 | 12810 |
| 40 | Yes | A. baylyi | Nalidixic a | Intracellul | F57 | Eosin Y | 1377 | BL1 | 15659 |
| 40 | Yes | A. baylyi | Nalidixic a | Intracellul | F27 | Calcein | 1325 | BL1 | 15884 |
| 40 | Yes | A. baylyi | Nalidixic a | Intracellul | F12 | Amaranth | 1461 | BL1 | 15638 |
| 40 | Yes | A. baylyi | None | None | NA | Autofluore | 2538 | NA | 6766 |
| 40 | Yes | A. baylyi | Ofloxacin | Intracellul | F194 | Oxonol | 1723 | RL1 | 5803 |
| 40 | Yes | A. baylyi | Ofloxacin | Intracellul | F11 | Allura Red | 1597 | BL3 | 5027 |
| 40 | Yes | A. baylyi | Ofloxacin | Intracellul | F54 | DiSC3(5) | 1419 | RL1 | 4575 |
| 40 | Yes | A. baylyi | Ofloxacin | Intracellul | F124 | Thioflavin T | 1367 | VL2 | 4926 |
| 40 | Yes | A. baylyi | Ofloxacin | Intracellul | F101 | Quinacrine | 1396 | VL2 | 5475 |
| 40 | Yes | A. baylyi | Ofloxacin | Intracellul | F138 | Apigenin | 1414 | VL2 | 5220 |
| 40 | Yes | A. baylyi | Ofloxacin | Intracellul | F65 | H2FDA | 1466 | BL1 | 5454 |
| 40 | Yes | A. baylyi | Ofloxacin | Intracellul | F142 | Chlorophe | 1685 | BL3 | 6604 |
| 40 | Yes | A. baylyi | Ofloxacin | Intracellul | F39 | Congo Red | 1739 | BL3 | 6909 |
| 40 | Yes | A. baylyi | Ofloxacin | Intracellul | F116 | Sunset Yel | 1826 | BL2 | 6532 |
| 40 | Yes | A. baylyi | Ofloxacin | Intracellul | F10 | Alizarin | 1723 | BL1 | 6441 |
| 40 | Yes | A. baylyi | Ofloxacin | Intracellul | F84 | Neutral Re | 1820 | BL1 | 5720.5 |
| 40 | Yes | A. baylyi | Ofloxacin | Intracellul | F99 | Pyronin Y | 1740 | BL1 | 5624.5 |
| 40 | Yes | A. baylyi | Ofloxacin | Intracellul | F108 | Rhodamine | 1542 | BL1 | 5115.5 |
| 40 | Yes | A. baylyi | Ofloxacin | Intracellul | F109 | Riboflavin | 1519 | BL1 | 5461 |
| 40 | Yes | A. baylyi | Ofloxacin | Intracellul | F117 | SYBR Gree | 1514 | BL1 | 564022.5 |
| 40 | Yes | A. baylyi | Ofloxacin | Intracellul | F106 | Rhodamine | 1402 | BL1 | 6481 |
| 40 | Yes | A. baylyi | Ofloxacin | Intracellul | F73 | Malachite ( | 1384 | BL1 | 5502 |
| 40 | Yes | A. baylyi | Ofloxacin | Intracellul | F7 | Acridine or | 1094 | BL1 | 6916.5 |
| 40 | Yes | A. baylyi | Ofloxacin | Intracellul | F192 | 5-Carboxyf | 1243 | BL1 | 13710 |
| 40 | Yes | A. baylyi | Ofloxacin | Intracellul | F57 | Eosin Y | 1055 | BL1 | 20174 |
| 40 | Yes | A. baylyi | Ofloxacin | Intracellul | F27 | Calcein | 1696 | BL1 | 16666.5 |

| | | | | | | | | | |
|---|---|---|---|---|---|---|---|---|---|
| 40 | Yes | A. baylyi | Ofloxacin | Intracellula | F12 | Amaranth | 1919 | BL1 | 14857 |
| 40 | Yes | A. baylyi | None | None | NA | Autofluore: | 2478 | NA | 6842.5 |
| 40 | Yes | A. baylyi | Polymyxin | Wall | F194 | Oxonol | 2721 | RL1 | 5554 |
| 40 | Yes | A. baylyi | Polymyxin | Wall | F11 | Allura Red | 3150 | BL3 | 4789 |
| 40 | Yes | A. baylyi | Polymyxin | Wall | F54 | DiSC3(5) | 2466 | RL1 | 6530 |
| 40 | Yes | A. baylyi | Polymyxin | Wall | F124 | Thioflavin T | 2872 | VL2 | 4478.5 |
| 40 | Yes | A. baylyi | Polymyxin | Wall | F101 | Quinacrine | 2713 | VL2 | 4780 |
| 40 | Yes | A. baylyi | Polymyxin | Wall | F138 | Apigenin | 3106 | VL2 | 4659 |
| 40 | Yes | A. baylyi | Polymyxin | Wall | F65 | H2FDA | 3217 | BL1 | 5243 |
| 40 | Yes | A. baylyi | Polymyxin | Wall | F142 | Chlorophe | 3014 | BL3 | 5705 |
| 40 | Yes | A. baylyi | Polymyxin | Wall | F39 | Congo Red | 3110 | BL3 | 5940 |
| 40 | Yes | A. baylyi | Polymyxin | Wall | F116 | Sunset Yel | 3633 | BL2 | 5785 |
| 40 | Yes | A. baylyi | Polymyxin | Wall | F10 | Alizarin | 3412 | BL1 | 6155 |
| 40 | Yes | A. baylyi | Polymyxin | Wall | F84 | Neutral Re | 3430 | BL1 | 5184 |
| 40 | Yes | A. baylyi | Polymyxin | Wall | F99 | Pyronin Y | 2835 | BL1 | 4939 |
| 40 | Yes | A. baylyi | Polymyxin | Wall | F108 | Rhodamine | 3062 | BL1 | 4606 |
| 40 | Yes | A. baylyi | Polymyxin | Wall | F109 | Riboflavin | 2820 | BL1 | 4763.5 |
| 40 | Yes | A. baylyi | Polymyxin | Wall | F117 | SYBR Gree | 2747 | BL1 | 879279 |
| 40 | Yes | A. baylyi | Polymyxin | Wall | F106 | Rhodamine | 2967 | BL1 | 5598 |
| 40 | Yes | A. baylyi | Polymyxin | Wall | F73 | Malachite ( | 2146 | BL1 | 5186 |
| 40 | Yes | A. baylyi | Polymyxin | Wall | F7 | Acridine or | 2229 | BL1 | 5621 |
| 40 | Yes | A. baylyi | Polymyxin | Wall | F192 | 5-Carboxyf | 2680 | BL1 | 11648 |
| 40 | Yes | A. baylyi | Polymyxin | Wall | F57 | Eosin Y | 2290 | BL1 | 13808 |
| 40 | Yes | A. baylyi | Polymyxin | Wall | F27 | Calcein | 2675 | BL1 | 14399 |
| 40 | Yes | A. baylyi | Polymyxin | Wall | F12 | Amaranth | 2727 | BL1 | 14785 |
| 40 | Yes | A. baylyi | None | None | NA | Autofluore: | 2513 | NA | 7726 |
| 40 | Yes | A. baylyi | Tetracyclin | Intracellula | F194 | Oxonol | 1411 | RL1 | 5733 |
| 40 | Yes | A. baylyi | Tetracyclin | Intracellula | F11 | Allura Red | 1314 | BL3 | 4915.5 |
| 40 | Yes | A. baylyi | Tetracyclin | Intracellula | F54 | DiSC3(5) | 1233 | RL1 | 4695 |
| 40 | Yes | A. baylyi | Tetracyclin | Intracellula | F124 | Thioflavin T | 1321 | VL2 | 4416 |
| 40 | Yes | A. baylyi | Tetracyclin | Intracellula | F101 | Quinacrine | 1192 | VL2 | 5045 |
| 40 | Yes | A. baylyi | Tetracyclin | Intracellula | F138 | Apigenin | 1272 | VL2 | 4850.5 |
| 40 | Yes | A. baylyi | Tetracyclin | Intracellula | F65 | H2FDA | 1197 | BL1 | 5563 |
| 40 | Yes | A. baylyi | Tetracyclin | Intracellula | F142 | Chlorophe | 1354 | BL3 | 6144 |
| 40 | Yes | A. baylyi | Tetracyclin | Intracellula | F39 | Congo Red | 1401 | BL3 | 6355 |
| 40 | Yes | A. baylyi | Tetracyclin | Intracellula | F116 | Sunset Yel | 1416 | BL2 | 5917.5 |
| 40 | Yes | A. baylyi | Tetracyclin | Intracellula | F10 | Alizarin | 1577 | BL1 | 5963 |
| 40 | Yes | A. baylyi | Tetracyclin | Intracellula | F84 | Neutral Re | 1337 | BL1 | 4807 |
| 40 | Yes | A. baylyi | Tetracyclin | Intracellula | F99 | Pyronin Y | 1472 | BL1 | 5156 |
| 40 | Yes | A. baylyi | Tetracyclin | Intracellula | F108 | Rhodamine | 1386 | BL1 | 4935.5 |
| 40 | Yes | A. baylyi | Tetracyclin | Intracellula | F109 | Riboflavin | 1348 | BL1 | 4933 |
| 40 | Yes | A. baylyi | Tetracyclin | Intracellula | F117 | SYBR Gree | 1072 | BL1 | 446979 |
| 40 | Yes | A. baylyi | Tetracyclin | Intracellula | F106 | Rhodamine | 1069 | BL1 | 6180 |
| 40 | Yes | A. baylyi | Tetracyclin | Intracellula | F73 | Malachite ( | 1263 | BL1 | 5346 |
| 40 | Yes | A. baylyi | Tetracyclin | Intracellula | F7 | Acridine or | 1109 | BL1 | 5732 |
| 40 | Yes | A. baylyi | Tetracyclin | Intracellula | F192 | 5-Carboxyf | 1084 | BL1 | 11545 |
| 40 | Yes | A. baylyi | Tetracyclin | Intracellula | F57 | Eosin Y | 1080 | BL1 | 15704.5 |
| 40 | Yes | A. baylyi | Tetracyclin | Intracellula | F27 | Calcein | 1097 | BL1 | 15022 |

| | | | | | | | | | |
|---|---|---|---|---|---|---|---|---|---|
| 40 | Yes | A. baylyi | Tetracyclin | Intracellula | F12 | Amaranth | 1088 | BL1 | 13617.5 |
| 40 | Yes | A. baylyi | None | None | NA | Autofluore: | 2677 | NA | 7648 |
| 40 | Yes | A. baylyi | Vancomyc | Wall | F194 | Oxonol | 2513 | RL1 | 5370 |
| 40 | Yes | A. baylyi | Vancomyc | Wall | F11 | Allura Red | 2573 | BL3 | 4503 |
| 40 | Yes | A. baylyi | Vancomyc | Wall | F54 | DiSC3(5) | 1974 | RL1 | 4210 |
| 40 | Yes | A. baylyi | Vancomyc | Wall | F124 | Thioflavin T | 2269 | VL2 | 4470 |
| 40 | Yes | A. baylyi | Vancomyc | Wall | F101 | Quinacrine | 2232 | VL2 | 4403.5 |
| 40 | Yes | A. baylyi | Vancomyc | Wall | F138 | Apigenin | 2121 | VL2 | 4496 |
| 40 | Yes | A. baylyi | Vancomyc | Wall | F65 | H2FDA | 1902 | BL1 | 5291.5 |
| 40 | Yes | A. baylyi | Vancomyc | Wall | F142 | Chlorophe | 2281 | BL3 | 6294 |
| 40 | Yes | A. baylyi | Vancomyc | Wall | F39 | Congo Red | 2256 | BL3 | 7524 |
| 40 | Yes | A. baylyi | Vancomyc | Wall | F116 | Sunset Yel | 2317 | BL2 | 6211 |
| 40 | Yes | A. baylyi | Vancomyc | Wall | F10 | Alizarin | 2534 | BL1 | 5563 |
| 40 | Yes | A. baylyi | Vancomyc | Wall | F84 | Neutral Re | 2075 | BL1 | 5118 |
| 40 | Yes | A. baylyi | Vancomyc | Wall | F99 | Pyronin Y | 2204 | BL1 | 5259 |
| 40 | Yes | A. baylyi | Vancomyc | Wall | F108 | Rhodamine | 1576 | BL1 | 4824 |
| 40 | Yes | A. baylyi | Vancomyc | Wall | F109 | Riboflavin | 2099 | BL1 | 4434 |
| 40 | Yes | A. baylyi | Vancomyc | Wall | F117 | SYBR Gree | 1812 | BL1 | 454216.5 |
| 40 | Yes | A. baylyi | Vancomyc | Wall | F106 | Rhodamine | 2041 | BL1 | 5520 |
| 40 | Yes | A. baylyi | Vancomyc | Wall | F73 | Malachite ( | 1932 | BL1 | 5349.5 |
| 40 | Yes | A. baylyi | Vancomyc | Wall | F7 | Acridine or | 1582 | BL1 | 4840 |
| 40 | Yes | A. baylyi | Vancomyc | Wall | F192 | 5-Carboxyf | 2063 | BL1 | 12480 |
| 40 | Yes | A. baylyi | Vancomyc | Wall | F57 | Eosin Y | 1697 | BL1 | 16986 |
| 40 | Yes | A. baylyi | Vancomyc | Wall | F27 | Calcein | 1741 | BL1 | 18002 |
| 40 | Yes | A. baylyi | Vancomyc | Wall | F12 | Amaranth | 2045 | BL1 | 16502 |
| 40 | Yes | A. baylyi | None | None | NA | Autofluore: | 2517 | NA | 898 |
| 40 | Yes | A. baylyi | None | None | F194 | Oxonol | 2840 | RL1 | 1581 |
| 40 | Yes | A. baylyi | None | None | F11 | Allura Red | 2844 | BL3 | 2449 |
| 40 | Yes | A. baylyi | None | None | F54 | DiSC3(5) | 2486 | RL1 | 2558.5 |
| 40 | Yes | A. baylyi | None | None | F124 | Thioflavin T | 2472 | VL2 | 2605 |
| 40 | Yes | A. baylyi | None | None | F101 | Quinacrine | 2452 | VL2 | 2840 |
| 40 | Yes | A. baylyi | None | None | F138 | Apigenin | 1991 | VL2 | 2728 |
| 40 | Yes | A. baylyi | None | None | F65 | H2FDA | 2036 | BL1 | 3000 |
| 40 | Yes | A. baylyi | None | None | F142 | Chlorophe | 2230 | BL3 | 4535.5 |
| 40 | Yes | A. baylyi | None | None | F39 | Congo Red | 2152 | BL3 | 5391.5 |
| 40 | Yes | A. baylyi | None | None | F116 | Sunset Yel | 2027 | BL2 | 5600 |
| 40 | Yes | A. baylyi | None | None | F10 | Alizarin | 2113 | BL1 | 4623 |
| 40 | Yes | A. baylyi | None | None | F84 | Neutral Re | 2149 | BL1 | 4126 |
| 40 | Yes | A. baylyi | None | None | F99 | Pyronin Y | 2017 | BL1 | 3928 |
| 40 | Yes | A. baylyi | None | None | F108 | Rhodamine | 2000 | BL1 | 3381 |
| 40 | Yes | A. baylyi | None | None | F109 | Riboflavin | 1903 | BL1 | 3159 |
| 40 | Yes | A. baylyi | None | None | F117 | SYBR Gree | 1852 | BL1 | 328927.5 |
| 40 | Yes | A. baylyi | None | None | F106 | Rhodamine | 1918 | BL1 | 4006 |
| 40 | Yes | A. baylyi | None | None | F73 | Malachite ( | 1871 | BL1 | 3727 |
| 40 | Yes | A. baylyi | None | None | F7 | Acridine or | 1903 | BL1 | 3975 |
| 40 | Yes | A. baylyi | None | None | F192 | 5-Carboxyf | 1728 | BL1 | 11952 |
| 40 | Yes | A. baylyi | None | None | F57 | Eosin Y | 1707 | BL1 | 15775 |
| 40 | Yes | A. baylyi | None | None | F27 | Calcein | 1588 | BL1 | 15340 |

| | | | | | | | | | |
|---|---|---|---|---|---|---|---|---|---|
| 40 | Yes | A. baylyi | None | None | F12 | Amaranth | 1724 | BL1 | 15323.5 |
| 40 | Yes | A. baylyi | None | None | NA | Autofluore: | 2148 | NA | 7170.5 |
| 40 | Yes | A. baylyi | Ampicillin | Wall | F194 | Oxonol | 1586 | RL1 | 5475.5 |
| 40 | Yes | A. baylyi | Ampicillin | Wall | F11 | Allura Red | 1376 | BL3 | 4714 |
| 40 | Yes | A. baylyi | Ampicillin | Wall | F54 | DiSC3(5) | 1222 | RL1 | 3863 |
| 40 | Yes | A. baylyi | Ampicillin | Wall | F124 | Thioflavin T | 1219 | VL2 | 3757 |
| 40 | Yes | A. baylyi | Ampicillin | Wall | F101 | Quinacrine | 1130 | VL2 | 4247 |
| 40 | Yes | A. baylyi | Ampicillin | Wall | F138 | Apigenin | 1196 | VL2 | 3752.5 |
| 40 | Yes | A. baylyi | Ampicillin | Wall | F65 | H2FDA | 1122 | BL1 | 4378.5 |
| 40 | Yes | A. baylyi | Ampicillin | Wall | F142 | Chlorophe | 1138 | BL3 | 5880.5 |
| 40 | Yes | A. baylyi | Ampicillin | Wall | F39 | Congo Red | 1328 | BL3 | 6814.5 |
| 40 | Yes | A. baylyi | Ampicillin | Wall | F116 | Sunset Yel | 1312 | BL2 | 6615 |
| 40 | Yes | A. baylyi | Ampicillin | Wall | F10 | Alizarin | 1323 | BL1 | 5469 |
| 40 | Yes | A. baylyi | Ampicillin | Wall | F84 | Neutral Re | 1190 | BL1 | 4916 |
| 40 | Yes | A. baylyi | Ampicillin | Wall | F99 | Pyronin Y | 1316 | BL1 | 4728.5 |
| 40 | Yes | A. baylyi | Ampicillin | Wall | F108 | Rhodamine | 1203 | BL1 | 4011 |
| 40 | Yes | A. baylyi | Ampicillin | Wall | F109 | Riboflavin | 1059 | BL1 | 3870 |
| 40 | Yes | A. baylyi | Ampicillin | Wall | F117 | SYBR Gree | 1172 | BL1 | 489814.5 |
| 40 | Yes | A. baylyi | Ampicillin | Wall | F106 | Rhodamine | 1428 | BL1 | 4498.5 |
| 40 | Yes | A. baylyi | Ampicillin | Wall | F73 | Malachite ( | 1269 | BL1 | 4336 |
| 40 | Yes | A. baylyi | Ampicillin | Wall | F7 | Acridine or | 1262 | BL1 | 4407 |
| 40 | Yes | A. baylyi | Ampicillin | Wall | F192 | 5-Carboxyf | 1239 | BL1 | 14085 |
| 40 | Yes | A. baylyi | Ampicillin | Wall | F57 | Eosin Y | 1233 | BL1 | 17108 |
| 40 | Yes | A. baylyi | Ampicillin | Wall | F27 | Calcein | 1096 | BL1 | 18030.5 |
| 40 | Yes | A. baylyi | Ampicillin | Wall | F12 | Amaranth | 1218 | BL1 | 19730.5 |
| 40 | Yes | A. baylyi | None | None | NA | Autofluore: | 2042 | NA | 5856.5 |
| 40 | Yes | A. baylyi | Bacitracin | Wall | F194 | Oxonol | 2369 | RL1 | 4418 |
| 40 | Yes | A. baylyi | Bacitracin | Wall | F11 | Allura Red | 2073 | BL3 | 3725 |
| 40 | Yes | A. baylyi | Bacitracin | Wall | F54 | DiSC3(5) | 2065 | RL1 | 3555 |
| 40 | Yes | A. baylyi | Bacitracin | Wall | F124 | Thioflavin T | 1875 | VL2 | 3310 |
| 40 | Yes | A. baylyi | Bacitracin | Wall | F101 | Quinacrine | 1715 | VL2 | 3359 |
| 40 | Yes | A. baylyi | Bacitracin | Wall | F138 | Apigenin | 1976 | VL2 | 3656.5 |
| 40 | Yes | A. baylyi | Bacitracin | Wall | F65 | H2FDA | 1821 | BL1 | 4368 |
| 40 | Yes | A. baylyi | Bacitracin | Wall | F142 | Chlorophe | 1804 | BL3 | 4759 |
| 40 | Yes | A. baylyi | Bacitracin | Wall | F39 | Congo Red | 1596 | BL3 | 5479 |
| 40 | Yes | A. baylyi | Bacitracin | Wall | F116 | Sunset Yel | 2036 | BL2 | 5258 |
| 40 | Yes | A. baylyi | Bacitracin | Wall | F10 | Alizarin | 2096 | BL1 | 5953.5 |
| 40 | Yes | A. baylyi | Bacitracin | Wall | F84 | Neutral Re | 1689 | BL1 | 4493 |
| 40 | Yes | A. baylyi | Bacitracin | Wall | F99 | Pyronin Y | 1744 | BL1 | 4650.5 |
| 40 | Yes | A. baylyi | Bacitracin | Wall | F108 | Rhodamine | 1742 | BL1 | 4227 |
| 40 | Yes | A. baylyi | Bacitracin | Wall | F109 | Riboflavin | 1563 | BL1 | 4312 |
| 40 | Yes | A. baylyi | Bacitracin | Wall | F117 | SYBR Gree | 1704 | BL1 | 358092 |
| 40 | Yes | A. baylyi | Bacitracin | Wall | F106 | Rhodamine | 2044 | BL1 | 4603.5 |
| 40 | Yes | A. baylyi | Bacitracin | Wall | F73 | Malachite ( | 2129 | BL1 | 4540 |
| 40 | Yes | A. baylyi | Bacitracin | Wall | F7 | Acridine or | 2036 | BL1 | 4523.5 |
| 40 | Yes | A. baylyi | Bacitracin | Wall | F192 | 5-Carboxyf | 2124 | BL1 | 12509.5 |
| 40 | Yes | A. baylyi | Bacitracin | Wall | F57 | Eosin Y | 2187 | BL1 | 15716 |
| 40 | Yes | A. baylyi | Bacitracin | Wall | F27 | Calcein | 2066 | BL1 | 16792.5 |

| | | | | | | | | | |
|---|---|---|---|---|---|---|---|---|---|
| 40 | Yes | A. baylyi | Bacitracin | Wall | F12 | Amaranth | 2290 | BL1 | 17381 |
| 40 | Yes | A. baylyi | None | None | NA | Autofluore: | 2172 | NA | 7807 |
| 40 | Yes | A. baylyi | Cefotaxim | Wall | F194 | Oxonol | 1636 | RL1 | 6039.5 |
| 40 | Yes | A. baylyi | Cefotaxim | Wall | F11 | Allura Red | 1515 | BL3 | 4455 |
| 40 | Yes | A. baylyi | Cefotaxim | Wall | F54 | DiSC3(5) | 1605 | RL1 | 4353 |
| 40 | Yes | A. baylyi | Cefotaxim | Wall | F124 | Thioflavin T | 1323 | VL2 | 4369 |
| 40 | Yes | A. baylyi | Cefotaxim | Wall | F101 | Quinacrine | 1416 | VL2 | 4685.5 |
| 40 | Yes | A. baylyi | Cefotaxim | Wall | F138 | Apigenin | 1429 | VL2 | 3801 |
| 40 | Yes | A. baylyi | Cefotaxim | Wall | F65 | H2FDA | 1533 | BL1 | 4973 |
| 40 | Yes | A. baylyi | Cefotaxim | Wall | F142 | Chlorophe | 1203 | BL3 | 6536 |
| 40 | Yes | A. baylyi | Cefotaxim | Wall | F39 | Congo Red | 1568 | BL3 | 5739.5 |
| 40 | Yes | A. baylyi | Cefotaxim | Wall | F116 | Sunset Yel | 1476 | BL2 | 5933.5 |
| 40 | Yes | A. baylyi | Cefotaxim | Wall | F10 | Alizarin | 1587 | BL1 | 5471 |
| 40 | Yes | A. baylyi | Cefotaxim | Wall | F84 | Neutral Re | 1555 | BL1 | 4538 |
| 40 | Yes | A. baylyi | Cefotaxim | Wall | F99 | Pyronin Y | 1394 | BL1 | 4497 |
| 40 | Yes | A. baylyi | Cefotaxim | Wall | F108 | Rhodamine | 1317 | BL1 | 4043 |
| 40 | Yes | A. baylyi | Cefotaxim | Wall | F109 | Riboflavin | 1345 | BL1 | 3833 |
| 40 | Yes | A. baylyi | Cefotaxim | Wall | F117 | SYBR Gree | 1402 | BL1 | 587536.5 |
| 40 | Yes | A. baylyi | Cefotaxim | Wall | F106 | Rhodamine | 1219 | BL1 | 5391 |
| 40 | Yes | A. baylyi | Cefotaxim | Wall | F73 | Malachite ( | 1480 | BL1 | 4748 |
| 40 | Yes | A. baylyi | Cefotaxim | Wall | F7 | Acridine or | 1375 | BL1 | 4927 |
| 40 | Yes | A. baylyi | Cefotaxim | Wall | F192 | 5-Carboxyf | 1232 | BL1 | 12768.5 |
| 40 | Yes | A. baylyi | Cefotaxim | Wall | F57 | Eosin Y | 1627 | BL1 | 15817 |
| 40 | Yes | A. baylyi | Cefotaxim | Wall | F27 | Calcein | 1591 | BL1 | 14370 |
| 40 | Yes | A. baylyi | Cefotaxim | Wall | F12 | Amaranth | 1778 | BL1 | 15071 |
| 40 | Yes | A. baylyi | None | None | NA | Autofluore: | 1848 | NA | 8193.5 |
| 40 | Yes | A. baylyi | Ciprofloxa | Intracellula | F194 | Oxonol | 1664 | RL1 | 5941 |
| 40 | Yes | A. baylyi | Ciprofloxa | Intracellula | F11 | Allura Red | 1080 | BL3 | 4913.5 |
| 40 | Yes | A. baylyi | Ciprofloxa | Intracellula | F54 | DiSC3(5) | 1396 | RL1 | 5494.5 |
| 40 | Yes | A. baylyi | Ciprofloxa | Intracellula | F124 | Thioflavin T | 1397 | VL2 | 5206 |
| 40 | Yes | A. baylyi | Ciprofloxa | Intracellula | F101 | Quinacrine | 1319 | VL2 | 4331 |
| 40 | Yes | A. baylyi | Ciprofloxa | Intracellula | F138 | Apigenin | 1152 | VL2 | 4740.5 |
| 40 | Yes | A. baylyi | Ciprofloxa | Intracellula | F65 | H2FDA | 1108 | BL1 | 6339.5 |
| 40 | Yes | A. baylyi | Ciprofloxa | Intracellula | F142 | Chlorophe | 1199 | BL3 | 6410 |
| 40 | Yes | A. baylyi | Ciprofloxa | Intracellula | F39 | Congo Red | 1228 | BL3 | 6055 |
| 40 | Yes | A. baylyi | Ciprofloxa | Intracellula | F116 | Sunset Yel | 1396 | BL2 | 6564 |
| 40 | Yes | A. baylyi | Ciprofloxa | Intracellula | F10 | Alizarin | 1363 | BL1 | 5686 |
| 40 | Yes | A. baylyi | Ciprofloxa | Intracellula | F84 | Neutral Re | 1314 | BL1 | 5743 |
| 40 | Yes | A. baylyi | Ciprofloxa | Intracellula | F99 | Pyronin Y | 1069 | BL1 | 4901 |
| 40 | Yes | A. baylyi | Ciprofloxa | Intracellula | F108 | Rhodamine | 1215 | BL1 | 4368 |
| 40 | Yes | A. baylyi | Ciprofloxa | Intracellula | F109 | Riboflavin | 1033 | BL1 | 5401 |
| 40 | Yes | A. baylyi | Ciprofloxa | Intracellula | F117 | SYBR Gree | 1202 | BL1 | 325325.5 |
| 40 | Yes | A. baylyi | Ciprofloxa | Intracellula | F106 | Rhodamine | 1360 | BL1 | 5464 |
| 40 | Yes | A. baylyi | Ciprofloxa | Intracellula | F73 | Malachite ( | 1206 | BL1 | 5344.5 |
| 40 | Yes | A. baylyi | Ciprofloxa | Intracellula | F7 | Acridine or | 897 | BL1 | 5741 |
| 40 | Yes | A. baylyi | Ciprofloxa | Intracellula | F192 | 5-Carboxyf | 1051 | BL1 | 13669 |
| 40 | Yes | A. baylyi | Ciprofloxa | Intracellula | F57 | Eosin Y | 1012 | BL1 | 18650.5 |
| 40 | Yes | A. baylyi | Ciprofloxa | Intracellula | F27 | Calcein | 871 | BL1 | 16635 |

| | | | | | | | | | |
|---|---|---|---|---|---|---|---|---|---|
| 40 | Yes | A. baylyi | Ciprofloxa | Intracellula | F12 | Amaranth | 1132 | BL1 | 14681.5 |
| 40 | Yes | A. baylyi | None | None | NA | Autofluore | 2216 | NA | 7551 |
| 40 | Yes | A. baylyi | Fosfomyci | Wall | F194 | Oxonol | 2215 | RL1 | 5806 |
| 40 | Yes | A. baylyi | Fosfomyci | Wall | F11 | Allura Red | 2043 | BL3 | 4771 |
| 40 | Yes | A. baylyi | Fosfomyci | Wall | F54 | DiSC3(5) | 1520 | RL1 | 4540.5 |
| 40 | Yes | A. baylyi | Fosfomyci | Wall | F124 | Thioflavin T | 1992 | VL2 | 4556.5 |
| 40 | Yes | A. baylyi | Fosfomyci | Wall | F101 | Quinacrine | 1916 | VL2 | 4358.5 |
| 40 | Yes | A. baylyi | Fosfomyci | Wall | F138 | Apigenin | 1913 | VL2 | 4066 |
| 40 | Yes | A. baylyi | Fosfomyci | Wall | F65 | H2FDA | 1854 | BL1 | 4272 |
| 40 | Yes | A. baylyi | Fosfomyci | Wall | F142 | Chlorophe | 2018 | BL3 | 6221 |
| 40 | Yes | A. baylyi | Fosfomyci | Wall | F39 | Congo Red | 2213 | BL3 | 6897 |
| 40 | Yes | A. baylyi | Fosfomyci | Wall | F116 | Sunset Yel | 2279 | BL2 | 6824 |
| 40 | Yes | A. baylyi | Fosfomyci | Wall | F10 | Alizarin | 2290 | BL1 | 6498.5 |
| 40 | Yes | A. baylyi | Fosfomyci | Wall | F84 | Neutral Re | 2244 | BL1 | 5478 |
| 40 | Yes | A. baylyi | Fosfomyci | Wall | F99 | Pyronin Y | 2083 | BL1 | 5041 |
| 40 | Yes | A. baylyi | Fosfomyci | Wall | F108 | Rhodamine | 1957 | BL1 | 4550 |
| 40 | Yes | A. baylyi | Fosfomyci | Wall | F109 | Riboflavin | 1911 | BL1 | 4317 |
| 40 | Yes | A. baylyi | Fosfomyci | Wall | F117 | SYBR Gree | 1891 | BL1 | 434854 |
| 40 | Yes | A. baylyi | Fosfomyci | Wall | F106 | Rhodamine | 1744 | BL1 | 4977 |
| 40 | Yes | A. baylyi | Fosfomyci | Wall | F73 | Malachite ( | 1848 | BL1 | 4674 |
| 40 | Yes | A. baylyi | Fosfomyci | Wall | F7 | Acridine or | 1511 | BL1 | 5005 |
| 40 | Yes | A. baylyi | Fosfomyci | Wall | F192 | 5-Carboxyf | 1488 | BL1 | 12258 |
| 40 | Yes | A. baylyi | Fosfomyci | Wall | F57 | Eosin Y | 1475 | BL1 | 12875 |
| 40 | Yes | A. baylyi | Fosfomyci | Wall | F27 | Calcein | 1770 | BL1 | 15212.5 |
| 40 | Yes | A. baylyi | Fosfomyci | Wall | F12 | Amaranth | 1858 | BL1 | 15947.5 |
| 40 | Yes | A. baylyi | None | None | NA | Autofluore | 2278 | NA | 7269.5 |
| 40 | Yes | A. baylyi | Nalidixic a | Intracellula | F194 | Oxonol | 2103 | RL1 | 5733 |
| 40 | Yes | A. baylyi | Nalidixic a | Intracellula | F11 | Allura Red | 1958 | BL3 | 4428.5 |
| 40 | Yes | A. baylyi | Nalidixic a | Intracellula | F54 | DiSC3(5) | 1919 | RL1 | 4421 |
| 40 | Yes | A. baylyi | Nalidixic a | Intracellula | F124 | Thioflavin T | 1925 | VL2 | 4801 |
| 40 | Yes | A. baylyi | Nalidixic a | Intracellula | F101 | Quinacrine | 1982 | VL2 | 4564.5 |
| 40 | Yes | A. baylyi | Nalidixic a | Intracellula | F138 | Apigenin | 1908 | VL2 | 4814 |
| 40 | Yes | A. baylyi | Nalidixic a | Intracellula | F65 | H2FDA | 2075 | BL1 | 4915 |
| 40 | Yes | A. baylyi | Nalidixic a | Intracellula | F142 | Chlorophe | 2008 | BL3 | 5442.5 |
| 40 | Yes | A. baylyi | Nalidixic a | Intracellula | F39 | Congo Red | 2228 | BL3 | 6074 |
| 40 | Yes | A. baylyi | Nalidixic a | Intracellula | F116 | Sunset Yel | 2349 | BL2 | 6150 |
| 40 | Yes | A. baylyi | Nalidixic a | Intracellula | F10 | Alizarin | 2570 | BL1 | 6993.5 |
| 40 | Yes | A. baylyi | Nalidixic a | Intracellula | F84 | Neutral Re | 2266 | BL1 | 5579.5 |
| 40 | Yes | A. baylyi | Nalidixic a | Intracellula | F99 | Pyronin Y | 2059 | BL1 | 5440 |
| 40 | Yes | A. baylyi | Nalidixic a | Intracellula | F108 | Rhodamine | 1753 | BL1 | 5779 |
| 40 | Yes | A. baylyi | Nalidixic a | Intracellula | F109 | Riboflavin | 1733 | BL1 | 5268 |
| 40 | Yes | A. baylyi | Nalidixic a | Intracellula | F117 | SYBR Gree | 1422 | BL1 | 436445.5 |
| 40 | Yes | A. baylyi | Nalidixic a | Intracellula | F106 | Rhodamine | 1557 | BL1 | 6811 |
| 40 | Yes | A. baylyi | Nalidixic a | Intracellula | F73 | Malachite ( | 1505 | BL1 | 6142 |
| 40 | Yes | A. baylyi | Nalidixic a | Intracellula | F7 | Acridine or | 1332 | BL1 | 6219.5 |
| 40 | Yes | A. baylyi | Nalidixic a | Intracellula | F192 | 5-Carboxyf | 1226 | BL1 | 12646 |
| 40 | Yes | A. baylyi | Nalidixic a | Intracellula | F57 | Eosin Y | 1381 | BL1 | 16295 |
| 40 | Yes | A. baylyi | Nalidixic a | Intracellula | F27 | Calcein | 1292 | BL1 | 16257 |

| | | | | | | | | | |
|---|---|---|---|---|---|---|---|---|---|
| 40 | Yes | A. baylyi | Nalidixic a | Intracellula | F12 | Amaranth | 1219 | BL1 | 18518 |
| 40 | Yes | A. baylyi | None | None | NA | Autofluore | 2262 | NA | 7359 |
| 40 | Yes | A. baylyi | Ofloxacin | Intracellula | F194 | Oxonol | 1671 | RL1 | 6157 |
| 40 | Yes | A. baylyi | Ofloxacin | Intracellula | F11 | Allura Red | 1443 | BL3 | 5391 |
| 40 | Yes | A. baylyi | Ofloxacin | Intracellula | F54 | DiSC3(5) | 1444 | RL1 | 4961 |
| 40 | Yes | A. baylyi | Ofloxacin | Intracellula | F124 | Thioflavin T | 1552 | VL2 | 4870.5 |
| 40 | Yes | A. baylyi | Ofloxacin | Intracellula | F101 | Quinacrine | 1429 | VL2 | 5825 |
| 40 | Yes | A. baylyi | Ofloxacin | Intracellula | F138 | Apigenin | 1602 | VL2 | 4955.5 |
| 40 | Yes | A. baylyi | Ofloxacin | Intracellula | F65 | H2FDA | 1737 | BL1 | 5250 |
| 40 | Yes | A. baylyi | Ofloxacin | Intracellula | F142 | Chlorophe | 1846 | BL3 | 5896 |
| 40 | Yes | A. baylyi | Ofloxacin | Intracellula | F39 | Congo Red | 2077 | BL3 | 7044 |
| 40 | Yes | A. baylyi | Ofloxacin | Intracellula | F116 | Sunset Yel | 2176 | BL2 | 7531 |
| 40 | Yes | A. baylyi | Ofloxacin | Intracellula | F10 | Alizarin | 2227 | BL1 | 6226 |
| 40 | Yes | A. baylyi | Ofloxacin | Intracellula | F84 | Neutral Re | 1660 | BL1 | 6001 |
| 40 | Yes | A. baylyi | Ofloxacin | Intracellula | F99 | Pyronin Y | 2083 | BL1 | 6118 |
| 40 | Yes | A. baylyi | Ofloxacin | Intracellula | F108 | Rhodamine | 1899 | BL1 | 5393 |
| 40 | Yes | A. baylyi | Ofloxacin | Intracellula | F109 | Riboflavin | 1813 | BL1 | 5616 |
| 40 | Yes | A. baylyi | Ofloxacin | Intracellula | F117 | SYBR Gree | 1872 | BL1 | 492180.5 |
| 40 | Yes | A. baylyi | Ofloxacin | Intracellula | F106 | Rhodamine | 1916 | BL1 | 5725 |
| 40 | Yes | A. baylyi | Ofloxacin | Intracellula | F73 | Malachite ( | 1756 | BL1 | 5926 |
| 40 | Yes | A. baylyi | Ofloxacin | Intracellula | F7 | Acridine or | 1693 | BL1 | 5551 |
| 40 | Yes | A. baylyi | Ofloxacin | Intracellula | F192 | 5-Carboxyf | 1692 | BL1 | 13042.5 |
| 40 | Yes | A. baylyi | Ofloxacin | Intracellula | F57 | Eosin Y | 1762 | BL1 | 14862.5 |
| 40 | Yes | A. baylyi | Ofloxacin | Intracellula | F27 | Calcein | 1802 | BL1 | 15513.5 |
| 40 | Yes | A. baylyi | Ofloxacin | Intracellula | F12 | Amaranth | 1762 | BL1 | 11486.5 |
| 40 | Yes | A. baylyi | None | None | NA | Autofluore | 2429 | NA | 6839 |
| 40 | Yes | A. baylyi | Polymyxin | Wall | F194 | Oxonol | 2586 | RL1 | 6320 |
| 40 | Yes | A. baylyi | Polymyxin | Wall | F11 | Allura Red | 2284 | BL3 | 5185.5 |
| 40 | Yes | A. baylyi | Polymyxin | Wall | F54 | DiSC3(5) | 2276 | RL1 | 4868.5 |
| 40 | Yes | A. baylyi | Polymyxin | Wall | F124 | Thioflavin T | 2363 | VL2 | 4648 |
| 40 | Yes | A. baylyi | Polymyxin | Wall | F101 | Quinacrine | 2051 | VL2 | 5431 |
| 40 | Yes | A. baylyi | Polymyxin | Wall | F138 | Apigenin | 2381 | VL2 | 4816 |
| 40 | Yes | A. baylyi | Polymyxin | Wall | F65 | H2FDA | 2740 | BL1 | 5072.5 |
| 40 | Yes | A. baylyi | Polymyxin | Wall | F142 | Chlorophe | 2968 | BL3 | 5739.5 |
| 40 | Yes | A. baylyi | Polymyxin | Wall | F39 | Congo Red | 2937 | BL3 | 6915 |
| 40 | Yes | A. baylyi | Polymyxin | Wall | F116 | Sunset Yel | 3080 | BL2 | 7631 |
| 40 | Yes | A. baylyi | Polymyxin | Wall | F10 | Alizarin | 2625 | BL1 | 6294 |
| 40 | Yes | A. baylyi | Polymyxin | Wall | F84 | Neutral Re | 2937 | BL1 | 6090 |
| 40 | Yes | A. baylyi | Polymyxin | Wall | F99 | Pyronin Y | 2740 | BL1 | 5455.5 |
| 40 | Yes | A. baylyi | Polymyxin | Wall | F108 | Rhodamine | 2641 | BL1 | 5560 |
| 40 | Yes | A. baylyi | Polymyxin | Wall | F109 | Riboflavin | 2641 | BL1 | 5123 |
| 40 | Yes | A. baylyi | Polymyxin | Wall | F117 | SYBR Gree | 2031 | BL1 | 947392 |
| 40 | Yes | A. baylyi | Polymyxin | Wall | F106 | Rhodamine | 2537 | BL1 | 6853 |
| 40 | Yes | A. baylyi | Polymyxin | Wall | F73 | Malachite ( | 2120 | BL1 | 5881.5 |
| 40 | Yes | A. baylyi | Polymyxin | Wall | F7 | Acridine or | 2318 | BL1 | 6046 |
| 40 | Yes | A. baylyi | Polymyxin | Wall | F192 | 5-Carboxyf | 2101 | BL1 | 10911 |
| 40 | Yes | A. baylyi | Polymyxin | Wall | F57 | Eosin Y | 2057 | BL1 | 13671 |
| 40 | Yes | A. baylyi | Polymyxin | Wall | F27 | Calcein | 2233 | BL1 | 14107 |

| | | | | | | | | | |
|---|---|---|---|---|---|---|---|---|---|
| 40 | Yes | A. baylyi | Polymyxin | Wall | F12 | Amaranth | 2352 | BL1 | 12160.5 |
| 40 | Yes | A. baylyi | None | None | NA | Autofluore: | 2365 | NA | 7208 |
| 40 | Yes | A. baylyi | Tetracyclin | Intracellula | F194 | Oxonol | 1344 | RL1 | 5942 |
| 40 | Yes | A. baylyi | Tetracyclin | Intracellula | F11 | Allura Red | 1191 | BL3 | 4957 |
| 40 | Yes | A. baylyi | Tetracyclin | Intracellula | F54 | DiSC3(5) | 1047 | RL1 | 4681 |
| 40 | Yes | A. baylyi | Tetracyclin | Intracellula | F124 | Thioflavin T | 1024 | VL2 | 4618.5 |
| 40 | Yes | A. baylyi | Tetracyclin | Intracellula | F101 | Quinacrine | 1092 | VL2 | 5265 |
| 40 | Yes | A. baylyi | Tetracyclin | Intracellula | F138 | Apigenin | 1037 | VL2 | 4867 |
| 40 | Yes | A. baylyi | Tetracyclin | Intracellula | F65 | H2FDA | 976 | BL1 | 5213 |
| 40 | Yes | A. baylyi | Tetracyclin | Intracellula | F142 | Chlorophe | 1401 | BL3 | 6117 |
| 40 | Yes | A. baylyi | Tetracyclin | Intracellula | F39 | Congo Red | 1292 | BL3 | 5904.5 |
| 40 | Yes | A. baylyi | Tetracyclin | Intracellula | F116 | Sunset Yel | 1070 | BL2 | 6845.5 |
| 40 | Yes | A. baylyi | Tetracyclin | Intracellula | F10 | Alizarin | 1278 | BL1 | 7668.5 |
| 40 | Yes | A. baylyi | Tetracyclin | Intracellula | F84 | Neutral Re | 1293 | BL1 | 5857 |
| 40 | Yes | A. baylyi | Tetracyclin | Intracellula | F99 | Pyronin Y | 1226 | BL1 | 6642.5 |
| 40 | Yes | A. baylyi | Tetracyclin | Intracellula | F108 | Rhodamine | 1230 | BL1 | 5722.5 |
| 40 | Yes | A. baylyi | Tetracyclin | Intracellula | F109 | Riboflavin | 1111 | BL1 | 5519 |
| 40 | Yes | A. baylyi | Tetracyclin | Intracellula | F117 | SYBR Gree | 1050 | BL1 | 508451 |
| 40 | Yes | A. baylyi | Tetracyclin | Intracellula | F106 | Rhodamine | 1037 | BL1 | 6163 |
| 40 | Yes | A. baylyi | Tetracyclin | Intracellula | F73 | Malachite ( | 1220 | BL1 | 6444.5 |
| 40 | Yes | A. baylyi | Tetracyclin | Intracellula | F7 | Acridine or | 1079 | BL1 | 6234 |
| 40 | Yes | A. baylyi | Tetracyclin | Intracellula | F192 | 5-Carboxyf | 818 | BL1 | 11671 |
| 40 | Yes | A. baylyi | Tetracyclin | Intracellula | F57 | Eosin Y | 1050 | BL1 | 14317.5 |
| 40 | Yes | A. baylyi | Tetracyclin | Intracellula | F27 | Calcein | 1104 | BL1 | 14099 |
| 40 | Yes | A. baylyi | Tetracyclin | Intracellula | F12 | Amaranth | 1079 | BL1 | 15258 |
| 40 | Yes | A. baylyi | None | None | NA | Autofluore: | 2386 | NA | 7527 |
| 40 | Yes | A. baylyi | Vancomyc | Wall | F194 | Oxonol | 2124 | RL1 | 6080.5 |
| 40 | Yes | A. baylyi | Vancomyc | Wall | F11 | Allura Red | 1965 | BL3 | 5360 |
| 40 | Yes | A. baylyi | Vancomyc | Wall | F54 | DiSC3(5) | 1598 | RL1 | 4773 |
| 40 | Yes | A. baylyi | Vancomyc | Wall | F124 | Thioflavin T | 1906 | VL2 | 4988 |
| 40 | Yes | A. baylyi | Vancomyc | Wall | F101 | Quinacrine | 1913 | VL2 | 5650 |
| 40 | Yes | A. baylyi | Vancomyc | Wall | F138 | Apigenin | 1837 | VL2 | 5365 |
| 40 | Yes | A. baylyi | Vancomyc | Wall | F65 | H2FDA | 1522 | BL1 | 5368.5 |
| 40 | Yes | A. baylyi | Vancomyc | Wall | F142 | Chlorophe | 1890 | BL3 | 6886.5 |
| 40 | Yes | A. baylyi | Vancomyc | Wall | F39 | Congo Red | 1938 | BL3 | 8080 |
| 40 | Yes | A. baylyi | Vancomyc | Wall | F116 | Sunset Yel | 1699 | BL2 | 7425 |
| 40 | Yes | A. baylyi | Vancomyc | Wall | F10 | Alizarin | 2088 | BL1 | 5951.5 |
| 40 | Yes | A. baylyi | Vancomyc | Wall | F84 | Neutral Re | 2128 | BL1 | 5483.5 |
| 40 | Yes | A. baylyi | Vancomyc | Wall | F99 | Pyronin Y | 1799 | BL1 | 5528 |
| 40 | Yes | A. baylyi | Vancomyc | Wall | F108 | Rhodamine | 1824 | BL1 | 5469 |
| 40 | Yes | A. baylyi | Vancomyc | Wall | F109 | Riboflavin | 1700 | BL1 | 4976 |
| 40 | Yes | A. baylyi | Vancomyc | Wall | F117 | SYBR Gree | 1690 | BL1 | 480627.5 |
| 40 | Yes | A. baylyi | Vancomyc | Wall | F106 | Rhodamine | 1796 | BL1 | 6055 |
| 40 | Yes | A. baylyi | Vancomyc | Wall | F73 | Malachite ( | 1674 | BL1 | 5612.5 |
| 40 | Yes | A. baylyi | Vancomyc | Wall | F7 | Acridine or | 1598 | BL1 | 6048 |
| 40 | Yes | A. baylyi | Vancomyc | Wall | F192 | 5-Carboxyf | 1557 | BL1 | 12010 |
| 40 | Yes | A. baylyi | Vancomyc | Wall | F57 | Eosin Y | 1728 | BL1 | 17139 |
| 40 | Yes | A. baylyi | Vancomyc | Wall | F27 | Calcein | 1487 | BL1 | 16963 |

| | | | | | | | | | |
|---|---|---|---|---|---|---|---|---|---|
| 40 | Yes | A. baylyi | Vancomyc | Wall | F12 | Amaranth | 1744 | BL1 | 16742.5 |
| 40 | No | E. coli | None | None | NA | Autofluore: | 8779 | NA | 769 |
| 40 | No | E. coli | None | None | F194 | Oxonol | 9577 | RL1 | 23112 |
| 40 | No | E. coli | None | None | F11 | Allura Red | 6660 | BL3 | 18920 |
| 40 | No | E. coli | None | None | F54 | DiSC3(5) | 10007 | RL1 | 15664 |
| 40 | No | E. coli | None | None | F124 | Thioflavin T | 10679 | VL2 | 13757 |
| 40 | No | E. coli | None | None | F101 | Quinacrine | 10352 | VL2 | 10386 |
| 40 | No | E. coli | None | None | F138 | Apigenin | 10034 | VL2 | 9542 |
| 40 | No | E. coli | None | None | F65 | H2FDA | 9871 | BL1 | 6387 |
| 40 | No | E. coli | None | None | F142 | Chlorophe | 8705 | BL3 | 5600 |
| 40 | No | E. coli | None | None | F39 | Congo Red | 9440 | BL3 | 4682 |
| 40 | No | E. coli | None | None | F116 | Sunset Yel | 9652 | BL2 | 4918.5 |
| 40 | No | E. coli | None | None | F10 | Alizarin | 8735 | BL1 | 4887 |
| 40 | No | E. coli | None | None | F84 | Neutral Re | 8189 | BL1 | 5320 |
| 40 | No | E. coli | None | None | F99 | Pyronin Y | 8877 | BL1 | 4462 |
| 40 | No | E. coli | None | None | F108 | Rhodamine | 10633 | BL1 | 4320 |
| 40 | No | E. coli | None | None | F109 | Riboflavin | 7955 | BL1 | 3495 |
| 40 | No | E. coli | None | None | F117 | SYBR Gree | 8751 | BL1 | 67837 |
| 40 | No | E. coli | None | None | F106 | Rhodamine | 9225 | BL1 | 5350 |
| 40 | No | E. coli | None | None | F73 | Malachite ( | 8634 | BL1 | 4405 |
| 40 | No | E. coli | None | None | F7 | Acridine or | 10432 | BL1 | 4003.5 |
| 40 | No | E. coli | None | None | F192 | 5-Carboxyf | 8975 | BL1 | 12714 |
| 40 | No | E. coli | None | None | F57 | Eosin Y | 8426 | BL1 | 17470 |
| 40 | No | E. coli | None | None | F27 | Calcein | 10827 | BL1 | 19775 |
| 40 | No | E. coli | None | None | F12 | Amaranth | 9609 | BL1 | 19787 |
| 40 | No | E. coli | None | None | NA | Autofluore: | 8460 | NA | 19026 |
| 40 | No | E. coli | Amoxacilli | Wall | F194 | Oxonol | 9723 | RL1 | 19460 |
| 40 | No | E. coli | Amoxacilli | Wall | F11 | Allura Red | 8316 | BL3 | 21458.5 |
| 40 | No | E. coli | Amoxacilli | Wall | F54 | DiSC3(5) | 7139 | RL1 | 17586 |
| 40 | No | E. coli | Amoxacilli | Wall | F124 | Thioflavin T | 9549 | VL2 | 12551 |
| 40 | No | E. coli | Amoxacilli | Wall | F101 | Quinacrine | 8737 | VL2 | 12401 |
| 40 | No | E. coli | Amoxacilli | Wall | F138 | Apigenin | 8994 | VL2 | 10655.5 |
| 40 | No | E. coli | Amoxacilli | Wall | F65 | H2FDA | 8118 | BL1 | 5992.5 |
| 40 | No | E. coli | Amoxacilli | Wall | F142 | Chlorophe | 8252 | BL3 | 5492.5 |
| 40 | No | E. coli | Amoxacilli | Wall | F39 | Congo Red | 7743 | BL3 | 5040 |
| 40 | No | E. coli | Amoxacilli | Wall | F116 | Sunset Yel | 8650 | BL2 | 6125 |
| 40 | No | E. coli | Amoxacilli | Wall | F10 | Alizarin | 7712 | BL1 | 6010 |
| 40 | No | E. coli | Amoxacilli | Wall | F84 | Neutral Re | 7058 | BL1 | 8818.5 |
| 40 | No | E. coli | Amoxacilli | Wall | F99 | Pyronin Y | 7508 | BL1 | 6120 |
| 40 | No | E. coli | Amoxacilli | Wall | F108 | Rhodamine | 8492 | BL1 | 5034.5 |
| 40 | No | E. coli | Amoxacilli | Wall | F109 | Riboflavin | 8313 | BL1 | 6105 |
| 40 | No | E. coli | Amoxacilli | Wall | F117 | SYBR Gree | 8327 | BL1 | 108032 |
| 40 | No | E. coli | Amoxacilli | Wall | F106 | Rhodamine | 6419 | BL1 | 8709 |
| 40 | No | E. coli | Amoxacilli | Wall | F73 | Malachite ( | 9082 | BL1 | 5319 |
| 40 | No | E. coli | Amoxacilli | Wall | F7 | Acridine or | 8218 | BL1 | 6503 |
| 40 | No | E. coli | Amoxacilli | Wall | F192 | 5-Carboxyf | 7392 | BL1 | 11233.5 |
| 40 | No | E. coli | Amoxacilli | Wall | F57 | Eosin Y | 8403 | BL1 | 14595 |
| 40 | No | E. coli | Amoxacilli | Wall | F27 | Calcein | 9495 | BL1 | 21624 |

| | | | | | | | | | |
|---|---|---|---|---|---|---|---|---|---|
| 40 | No | E. coli | Amoxacilli | Wall | F12 | Amaranth | 8210 | BL1 | 21683.5 |
| 40 | No | E. coli | None | None | NA | Autofluore: | 10204 | NA | 20904.5 |
| 40 | No | E. coli | Azithromyc | Intracellul | F194 | Oxonol | 6822 | RL1 | 20932.5 |
| 40 | No | E. coli | Azithromyc | Intracellul | F11 | Allura Red | 5689 | BL3 | 17880 |
| 40 | No | E. coli | Azithromyc | Intracellul | F54 | DiSC3(5) | 6301 | RL1 | 17661 |
| 40 | No | E. coli | Azithromyc | Intracellul | F124 | Thioflavin T | 6055 | VL2 | 15406 |
| 40 | No | E. coli | Azithromyc | Intracellul | F101 | Quinacrine | 6715 | VL2 | 13239 |
| 40 | No | E. coli | Azithromyc | Intracellul | F138 | Apigenin | 6957 | VL2 | 8932 |
| 40 | No | E. coli | Azithromyc | Intracellul | F65 | H2FDA | 5741 | BL1 | 6622 |
| 40 | No | E. coli | Azithromyc | Intracellul | F142 | Chlorophe | 5423 | BL3 | 4832 |
| 40 | No | E. coli | Azithromyc | Intracellul | F39 | Congo Red | 6456 | BL3 | 6696 |
| 40 | No | E. coli | Azithromyc | Intracellul | F116 | Sunset Yel | 5757 | BL2 | 5639 |
| 40 | No | E. coli | Azithromyc | Intracellul | F10 | Alizarin | 4826 | BL1 | 7137 |
| 40 | No | E. coli | Azithromyc | Intracellul | F84 | Neutral Re | 6072 | BL1 | 6728 |
| 40 | No | E. coli | Azithromyc | Intracellul | F99 | Pyronin Y | 5998 | BL1 | 5241.5 |
| 40 | No | E. coli | Azithromyc | Intracellul | F108 | Rhodamine | 5518 | BL1 | 5344 |
| 40 | No | E. coli | Azithromyc | Intracellul | F109 | Riboflavin | 5268 | BL1 | 4771 |
| 40 | No | E. coli | Azithromyc | Intracellul | F117 | SYBR Gree | 5562 | BL1 | 18330.5 |
| 40 | No | E. coli | Azithromyc | Intracellul | F106 | Rhodamine | 6365 | BL1 | 7207 |
| 40 | No | E. coli | Azithromyc | Intracellul | F73 | Malachite ( | 6516 | BL1 | 5904.5 |
| 40 | No | E. coli | Azithromyc | Intracellul | F7 | Acridine or | 5320 | BL1 | 5454 |
| 40 | No | E. coli | Azithromyc | Intracellul | F192 | 5-Carboxyf | 5085 | BL1 | 17402 |
| 40 | No | E. coli | Azithromyc | Intracellul | F57 | Eosin Y | 4761 | BL1 | 12983 |
| 40 | No | E. coli | Azithromyc | Intracellul | F27 | Calcein | 5627 | BL1 | 18553 |
| 40 | No | E. coli | Azithromyc | Intracellul | F12 | Amaranth | 4947 | BL1 | 24123 |
| 40 | No | E. coli | None | None | NA | Autofluore: | 7710 | NA | 18805 |
| 40 | No | E. coli | Trimethopr | Intracellul | F194 | Oxonol | 6350 | RL1 | 18244.5 |
| 40 | No | E. coli | Trimethopr | Intracellul | F11 | Allura Red | 7651 | BL3 | 17833 |
| 40 | No | E. coli | Trimethopr | Intracellul | F54 | DiSC3(5) | 7121 | RL1 | 17796 |
| 40 | No | E. coli | Trimethopr | Intracellul | F124 | Thioflavin T | 7877 | VL2 | 13712 |
| 40 | No | E. coli | Trimethopr | Intracellul | F101 | Quinacrine | 6198 | VL2 | 12826 |
| 40 | No | E. coli | Trimethopr | Intracellul | F138 | Apigenin | 8474 | VL2 | 7901.5 |
| 40 | No | E. coli | Trimethopr | Intracellul | F65 | H2FDA | 5893 | BL1 | 6888 |
| 40 | No | E. coli | Trimethopr | Intracellul | F142 | Chlorophe | 8217 | BL3 | 4816 |
| 40 | No | E. coli | Trimethopr | Intracellul | F39 | Congo Red | 6337 | BL3 | 5880 |
| 40 | No | E. coli | Trimethopr | Intracellul | F116 | Sunset Yel | 7015 | BL2 | 5314 |
| 40 | No | E. coli | Trimethopr | Intracellul | F10 | Alizarin | 6623 | BL1 | 6601 |
| 40 | No | E. coli | Trimethopr | Intracellul | F84 | Neutral Re | 8610 | BL1 | 5633.5 |
| 40 | No | E. coli | Trimethopr | Intracellul | F99 | Pyronin Y | 6491 | BL1 | 6343 |
| 40 | No | E. coli | Trimethopr | Intracellul | F108 | Rhodamine | 8027 | BL1 | 5344 |
| 40 | No | E. coli | Trimethopr | Intracellul | F109 | Riboflavin | 6641 | BL1 | 5189 |
| 40 | No | E. coli | Trimethopr | Intracellul | F117 | SYBR Gree | 7182 | BL1 | 86369 |
| 40 | No | E. coli | Trimethopr | Intracellul | F106 | Rhodamine | 5394 | BL1 | 7515 |
| 40 | No | E. coli | Trimethopr | Intracellul | F73 | Malachite ( | 6560 | BL1 | 7532.5 |
| 40 | No | E. coli | Trimethopr | Intracellul | F7 | Acridine or | 8239 | BL1 | 5414 |
| 40 | No | E. coli | Trimethopr | Intracellul | F192 | 5-Carboxyf | 7859 | BL1 | 12262 |
| 40 | No | E. coli | Trimethopr | Intracellul | F57 | Eosin Y | 7445 | BL1 | 16036 |
| 40 | No | E. coli | Trimethopr | Intracellul | F27 | Calcein | 9131 | BL1 | 15967 |

| | | | | | | | | | |
|---|---|---|---|---|---|---|---|---|---|
| 40 | No | E. coli | Trimethopr | Intracellula | F12 | Amaranth | 7433 | BL1 | 22294 |
| 40 | No | E. coli | None | None | NA | Autofluores | 9406 | NA | 23036.5 |
| 40 | No | E. coli | Chloramph | Intracellula | F194 | Oxonol | 8078 | RL1 | 22215 |
| 40 | No | E. coli | Chloramph | Intracellula | F11 | Allura Red | 7342 | BL3 | 14041 |
| 40 | No | E. coli | Chloramph | Intracellula | F54 | DiSC3(5) | 7836 | RL1 | 16336.5 |
| 40 | No | E. coli | Chloramph | Intracellula | F124 | Thioflavin T | 8741 | VL2 | 14229 |
| 40 | No | E. coli | Chloramph | Intracellula | F101 | Quinacrine | 8037 | VL2 | 12557 |
| 40 | No | E. coli | Chloramph | Intracellula | F138 | Apigenin | 9005 | VL2 | 6802 |
| 40 | No | E. coli | Chloramph | Intracellula | F65 | H2FDA | 8460 | BL1 | 4829 |
| 40 | No | E. coli | Chloramph | Intracellula | F142 | Chlorophe | 9512 | BL3 | 5197.5 |
| 40 | No | E. coli | Chloramph | Intracellula | F39 | Congo Red | 6296 | BL3 | 4648.5 |
| 40 | No | E. coli | Chloramph | Intracellula | F116 | Sunset Yel | 8508 | BL2 | 6091 |
| 40 | No | E. coli | Chloramph | Intracellula | F10 | Alizarin | 7829 | BL1 | 6510 |
| 40 | No | E. coli | Chloramph | Intracellula | F84 | Neutral Re | 8502 | BL1 | 5942.5 |
| 40 | No | E. coli | Chloramph | Intracellula | F99 | Pyronin Y | 9056 | BL1 | 5079.5 |
| 40 | No | E. coli | Chloramph | Intracellula | F108 | Rhodamine | 8041 | BL1 | 5092 |
| 40 | No | E. coli | Chloramph | Intracellula | F109 | Riboflavin | 6436 | BL1 | 5400 |
| 40 | No | E. coli | Chloramph | Intracellula | F117 | SYBR Gree | 8424 | BL1 | 44650.5 |
| 40 | No | E. coli | Chloramph | Intracellula | F106 | Rhodamine | 8762 | BL1 | 6451 |
| 40 | No | E. coli | Chloramph | Intracellula | F73 | Malachite ( | 8307 | BL1 | 5475 |
| 40 | No | E. coli | Chloramph | Intracellula | F7 | Acridine or | 8644 | BL1 | 4717.5 |
| 40 | No | E. coli | Chloramph | Intracellula | F192 | 5-Carboxyf | 8847 | BL1 | 13175 |
| 40 | No | E. coli | Chloramph | Intracellula | F57 | Eosin Y | 8296 | BL1 | 17744.5 |
| 40 | No | E. coli | Chloramph | Intracellula | F27 | Calcein | 8010 | BL1 | 17089 |
| 40 | No | E. coli | Chloramph | Intracellula | F12 | Amaranth | 8783 | BL1 | 20655 |
| 40 | No | E. coli | None | None | NA | Autofluores | 8299 | NA | 23715 |
| 40 | No | E. coli | Colistin | Wall | F194 | Oxonol | 5868 | RL1 | 46248.5 |
| 40 | No | E. coli | Colistin | Wall | F11 | Allura Red | 3570 | BL3 | 44992.5 |
| 40 | No | E. coli | Colistin | Wall | F54 | DiSC3(5) | 2805 | RL1 | 55063 |
| 40 | No | E. coli | Colistin | Wall | F124 | Thioflavin T | 2465 | VL2 | 26043 |
| 40 | No | E. coli | Colistin | Wall | F101 | Quinacrine | 2889 | VL2 | 30232 |
| 40 | No | E. coli | Colistin | Wall | F138 | Apigenin | 2900 | VL2 | 19733 |
| 40 | No | E. coli | Colistin | Wall | F65 | H2FDA | 2934 | BL1 | 17685.5 |
| 40 | No | E. coli | Colistin | Wall | F142 | Chlorophe | 3254 | BL3 | 17824.5 |
| 40 | No | E. coli | Colistin | Wall | F39 | Congo Red | 2598 | BL3 | 14942 |
| 40 | No | E. coli | Colistin | Wall | F116 | Sunset Yel | 2236 | BL2 | 18235.5 |
| 40 | No | E. coli | Colistin | Wall | F10 | Alizarin | 3431 | BL1 | 15202 |
| 40 | No | E. coli | Colistin | Wall | F84 | Neutral Re | 4036 | BL1 | 14646 |
| 40 | No | E. coli | Colistin | Wall | F99 | Pyronin Y | 2956 | BL1 | 15925.5 |
| 40 | No | E. coli | Colistin | Wall | F108 | Rhodamine | 3371 | BL1 | 13709 |
| 40 | No | E. coli | Colistin | Wall | F109 | Riboflavin | 2585 | BL1 | 15528 |
| 40 | No | E. coli | Colistin | Wall | F117 | SYBR Gree | 3337 | BL1 | 2257512 |
| 40 | No | E. coli | Colistin | Wall | F106 | Rhodamine | 2510 | BL1 | 256579 |
| 40 | No | E. coli | Colistin | Wall | F73 | Malachite ( | 3625 | BL1 | 98446 |
| 40 | No | E. coli | Colistin | Wall | F7 | Acridine or | 3250 | BL1 | 112852.5 |
| 40 | No | E. coli | Colistin | Wall | F192 | 5-Carboxyf | 3051 | BL1 | 59961 |
| 40 | No | E. coli | Colistin | Wall | F57 | Eosin Y | 2403 | BL1 | 38706 |
| 40 | No | E. coli | Colistin | Wall | F27 | Calcein | 2447 | BL1 | 31394 |

| | | | | | | | | | |
|---|---|---|---|---|---|---|---|---|---|
| 40 | No | E. coli | Colistin | Wall | F12 | Amaranth | 2406 | BL1 | 29279.5 |
| 40 | No | E. coli | None | None | NA | Autofluore: | 4589 | NA | 25077 |
| 40 | No | E. coli | Gentamici | Intracellula | F194 | Oxonol | 5459 | RL1 | 22968 |
| 40 | No | E. coli | Gentamici | Intracellula | F11 | Allura Red | 7632 | BL3 | 23068.5 |
| 40 | No | E. coli | Gentamici | Intracellula | F54 | DiSC3(5) | 9802 | RL1 | 16982 |
| 40 | No | E. coli | Gentamici | Intracellula | F124 | Thioflavin T | 10430 | VL2 | 13487.5 |
| 40 | No | E. coli | Gentamici | Intracellula | F101 | Quinacrine | 9169 | VL2 | 10656 |
| 40 | No | E. coli | Gentamici | Intracellula | F138 | Apigenin | 9972 | VL2 | 8461.5 |
| 40 | No | E. coli | Gentamici | Intracellula | F65 | H2FDA | 8766 | BL1 | 5453.5 |
| 40 | No | E. coli | Gentamici | Intracellula | F142 | Chlorophe | 8863 | BL3 | 4402 |
| 40 | No | E. coli | Gentamici | Intracellula | F39 | Congo Red | 9932 | BL3 | 4979.5 |
| 40 | No | E. coli | Gentamici | Intracellula | F116 | Sunset Yel | 8995 | BL2 | 6913 |
| 40 | No | E. coli | Gentamici | Intracellula | F10 | Alizarin | 9071 | BL1 | 6948 |
| 40 | No | E. coli | Gentamici | Intracellula | F84 | Neutral Re | 9870 | BL1 | 7010 |
| 40 | No | E. coli | Gentamici | Intracellula | F99 | Pyronin Y | 8516 | BL1 | 7407.5 |
| 40 | No | E. coli | Gentamici | Intracellula | F108 | Rhodamine | 8439 | BL1 | 5940 |
| 40 | No | E. coli | Gentamici | Intracellula | F109 | Riboflavin | 10340 | BL1 | 4474 |
| 40 | No | E. coli | Gentamici | Intracellula | F117 | SYBR Gree | 10113 | BL1 | 78204 |
| 40 | No | E. coli | Gentamici | Intracellula | F106 | Rhodamine | 8412 | BL1 | 6689 |
| 40 | No | E. coli | Gentamici | Intracellula | F73 | Malachite ( | 8650 | BL1 | 4975 |
| 40 | No | E. coli | Gentamici | Intracellula | F7 | Acridine or | 7579 | BL1 | 5302 |
| 40 | No | E. coli | Gentamici | Intracellula | F192 | 5-Carboxyf | 9248 | BL1 | 6115 |
| 40 | No | E. coli | Gentamici | Intracellula | F57 | Eosin Y | 7235 | BL1 | 13758 |
| 40 | No | E. coli | Gentamici | Intracellula | F27 | Calcein | 9848 | BL1 | 19189 |
| 40 | No | E. coli | Gentamici | Intracellula | F12 | Amaranth | 8858 | BL1 | 14725 |
| 40 | No | E. coli | None | None | NA | Autofluore: | 8731 | NA | 17650 |
| 40 | No | E. coli | Nitrofurant | Intracellula | F194 | Oxonol | 8544 | RL1 | 20110 |
| 40 | No | E. coli | Nitrofurant | Intracellula | F11 | Allura Red | 9017 | BL3 | 16678 |
| 40 | No | E. coli | Nitrofurant | Intracellula | F54 | DiSC3(5) | 6072 | RL1 | 19759.5 |
| 40 | No | E. coli | Nitrofurant | Intracellula | F124 | Thioflavin T | 8195 | VL2 | 18058 |
| 40 | No | E. coli | Nitrofurant | Intracellula | F101 | Quinacrine | 6021 | VL2 | 15036 |
| 40 | No | E. coli | Nitrofurant | Intracellula | F138 | Apigenin | 9511 | VL2 | 6511 |
| 40 | No | E. coli | Nitrofurant | Intracellula | F65 | H2FDA | 7743 | BL1 | 5536 |
| 40 | No | E. coli | Nitrofurant | Intracellula | F142 | Chlorophe | 8000 | BL3 | 5207 |
| 40 | No | E. coli | Nitrofurant | Intracellula | F39 | Congo Red | 6962 | BL3 | 4776.5 |
| 40 | No | E. coli | Nitrofurant | Intracellula | F116 | Sunset Yel | 7557 | BL2 | 6638 |
| 40 | No | E. coli | Nitrofurant | Intracellula | F10 | Alizarin | 6244 | BL1 | 5933.5 |
| 40 | No | E. coli | Nitrofurant | Intracellula | F84 | Neutral Re | 6738 | BL1 | 6824 |
| 40 | No | E. coli | Nitrofurant | Intracellula | F99 | Pyronin Y | 7136 | BL1 | 7172 |
| 40 | No | E. coli | Nitrofurant | Intracellula | F108 | Rhodamine | 8097 | BL1 | 5571 |
| 40 | No | E. coli | Nitrofurant | Intracellula | F109 | Riboflavin | 7935 | BL1 | 4490 |
| 40 | No | E. coli | Nitrofurant | Intracellula | F117 | SYBR Gree | 6912 | BL1 | 27647.5 |
| 40 | No | E. coli | Nitrofurant | Intracellula | F106 | Rhodamine | 6163 | BL1 | 5702 |
| 40 | No | E. coli | Nitrofurant | Intracellula | F73 | Malachite ( | 6704 | BL1 | 5352.5 |
| 40 | No | E. coli | Nitrofurant | Intracellula | F7 | Acridine or | 5511 | BL1 | 5020 |
| 40 | No | E. coli | Nitrofurant | Intracellula | F192 | 5-Carboxyf | 6823 | BL1 | 10606 |
| 40 | No | E. coli | Nitrofurant | Intracellula | F57 | Eosin Y | 7324 | BL1 | 17742.5 |
| 40 | No | E. coli | Nitrofurant | Intracellula | F27 | Calcein | 7840 | BL1 | 22020.5 |

| | | | | | | | | | |
|---|---|---|---|---|---|---|---|---|---|
| 40 | No | E. coli | Nitrofurant | Intracellula | F12 | Amaranth | 5180 | BL1 | 20630.5 |
| 40 | No | E. coli | None | None | NA | Autofluore: | 5786 | NA | 21118.5 |
| 40 | No | E. coli | Pentamidi | Wall | F194 | Oxonol | 9334 | RL1 | 21795.5 |
| 40 | No | E. coli | Pentamidi | Wall | F11 | Allura Red | 8854 | BL3 | 18017.5 |
| 40 | No | E. coli | Pentamidi | Wall | F54 | DiSC3(5) | 6892 | RL1 | 17223 |
| 40 | No | E. coli | Pentamidi | Wall | F124 | Thioflavin T | 5977 | VL2 | 12575 |
| 40 | No | E. coli | Pentamidi | Wall | F101 | Quinacrine | 8808 | VL2 | 10601 |
| 40 | No | E. coli | Pentamidi | Wall | F138 | Apigenin | 8187 | VL2 | 13187 |
| 40 | No | E. coli | Pentamidi | Wall | F65 | H2FDA | 6482 | BL1 | 11242 |
| 40 | No | E. coli | Pentamidi | Wall | F142 | Chlorophe | 7045 | BL3 | 11609 |
| 40 | No | E. coli | Pentamidi | Wall | F39 | Congo Red | 7757 | BL3 | 12151 |
| 40 | No | E. coli | Pentamidi | Wall | F116 | Sunset Yel | 7317 | BL2 | 11218 |
| 40 | No | E. coli | Pentamidi | Wall | F10 | Alizarin | 6048 | BL1 | 10835 |
| 40 | No | E. coli | Pentamidi | Wall | F84 | Neutral Re | 6515 | BL1 | 12730 |
| 40 | No | E. coli | Pentamidi | Wall | F99 | Pyronin Y | 7331 | BL1 | 9175 |
| 40 | No | E. coli | Pentamidi | Wall | F108 | Rhodamine | 5612 | BL1 | 10666.5 |
| 40 | No | E. coli | Pentamidi | Wall | F109 | Riboflavin | 5023 | BL1 | 8009 |
| 40 | No | E. coli | Pentamidi | Wall | F117 | SYBR Gree | 4868 | BL1 | 276859 |
| 40 | No | E. coli | Pentamidi | Wall | F106 | Rhodamine | 8277 | BL1 | 9023 |
| 40 | No | E. coli | Pentamidi | Wall | F73 | Malachite ( | 7194 | BL1 | 6448 |
| 40 | No | E. coli | Pentamidi | Wall | F7 | Acridine or | 5999 | BL1 | 5994 |
| 40 | No | E. coli | Pentamidi | Wall | F192 | 5-Carboxyf | 7769 | BL1 | 12639 |
| 40 | No | E. coli | Pentamidi | Wall | F57 | Eosin Y | 6754 | BL1 | 13277 |
| 40 | No | E. coli | Pentamidi | Wall | F27 | Calcein | 4286 | BL1 | 28355 |
| 40 | No | E. coli | Pentamidi | Wall | F12 | Amaranth | 7143 | BL1 | 14322 |
| 40 | No | E. coli | None | None | NA | Autofluore: | 8349 | NA | 18745 |
| 40 | No | E. coli | Rifampicin | Intracellula | F194 | Oxonol | 6914 | RL1 | 19403.5 |
| 40 | No | E. coli | Rifampicin | Intracellula | F11 | Allura Red | 9799 | BL3 | 21015 |
| 40 | No | E. coli | Rifampicin | Intracellula | F54 | DiSC3(5) | 6508 | RL1 | 17371 |
| 40 | No | E. coli | Rifampicin | Intracellula | F124 | Thioflavin T | 9799 | VL2 | 14395 |
| 40 | No | E. coli | Rifampicin | Intracellula | F101 | Quinacrine | 8029 | VL2 | 11783 |
| 40 | No | E. coli | Rifampicin | Intracellula | F138 | Apigenin | 9247 | VL2 | 10175 |
| 40 | No | E. coli | Rifampicin | Intracellula | F65 | H2FDA | 7536 | BL1 | 10632.5 |
| 40 | No | E. coli | Rifampicin | Intracellula | F142 | Chlorophe | 8469 | BL3 | 5871 |
| 40 | No | E. coli | Rifampicin | Intracellula | F39 | Congo Red | 8486 | BL3 | 7072.5 |
| 40 | No | E. coli | Rifampicin | Intracellula | F116 | Sunset Yel | 8022 | BL2 | 8083.5 |
| 40 | No | E. coli | Rifampicin | Intracellula | F10 | Alizarin | 7327 | BL1 | 8584 |
| 40 | No | E. coli | Rifampicin | Intracellula | F84 | Neutral Re | 8213 | BL1 | 8453 |
| 40 | No | E. coli | Rifampicin | Intracellula | F99 | Pyronin Y | 7447 | BL1 | 8448 |
| 40 | No | E. coli | Rifampicin | Intracellula | F108 | Rhodamine | 7136 | BL1 | 6491 |
| 40 | No | E. coli | Rifampicin | Intracellula | F109 | Riboflavin | 8065 | BL1 | 4596 |
| 40 | No | E. coli | Rifampicin | Intracellula | F117 | SYBR Gree | 6990 | BL1 | 11371.5 |
| 40 | No | E. coli | Rifampicin | Intracellula | F106 | Rhodamine | 7648 | BL1 | 10670 |
| 40 | No | E. coli | Rifampicin | Intracellula | F73 | Malachite ( | 7108 | BL1 | 7656 |
| 40 | No | E. coli | Rifampicin | Intracellula | F7 | Acridine or | 7980 | BL1 | 6759.5 |
| 40 | No | E. coli | Rifampicin | Intracellula | F192 | 5-Carboxyf | 5730 | BL1 | 14273 |
| 40 | No | E. coli | Rifampicin | Intracellula | F57 | Eosin Y | 8364 | BL1 | 17528.5 |
| 40 | No | E. coli | Rifampicin | Intracellula | F27 | Calcein | 9362 | BL1 | 22371 |

| | | | | | | | | | |
|---|---|---|---|---|---|---|---|---|---|
| 40 | No | E. coli | Rifampicin | Intracellula | F12 | Amaranth | 6765 | BL1 | 22229 |
| 40 | No | E. coli | None | None | NA | Autofluore: | 9125 | NA | 20191 |
| 40 | No | E. coli | Trimethopr | Intracellula | F194 | Oxonol | 7879 | RL1 | 18231 |
| 40 | No | E. coli | Trimethopr | Intracellula | F11 | Allura Red | 8237 | BL3 | 21460 |
| 40 | No | E. coli | Trimethopr | Intracellula | F54 | DiSC3(5) | 7384 | RL1 | 35997 |
| 40 | No | E. coli | Trimethopr | Intracellula | F124 | Thioflavin T | 8959 | VL2 | 13160 |
| 40 | No | E. coli | Trimethopr | Intracellula | F101 | Quinacrine | 7083 | VL2 | 11860 |
| 40 | No | E. coli | Trimethopr | Intracellula | F138 | Apigenin | 8183 | VL2 | 8364 |
| 40 | No | E. coli | Trimethopr | Intracellula | F65 | H2FDA | 7986 | BL1 | 7835 |
| 40 | No | E. coli | Trimethopr | Intracellula | F142 | Chlorophe | 5734 | BL3 | 5821 |
| 40 | No | E. coli | Trimethopr | Intracellula | F39 | Congo Red | 7317 | BL3 | 7164 |
| 40 | No | E. coli | Trimethopr | Intracellula | F116 | Sunset Yel | 8917 | BL2 | 6880 |
| 40 | No | E. coli | Trimethopr | Intracellula | F10 | Alizarin | 7350 | BL1 | 9961 |
| 40 | No | E. coli | Trimethopr | Intracellula | F84 | Neutral Re | 7565 | BL1 | 7397 |
| 40 | No | E. coli | Trimethopr | Intracellula | F99 | Pyronin Y | 7066 | BL1 | 7460.5 |
| 40 | No | E. coli | Trimethopr | Intracellula | F108 | Rhodamine | 6647 | BL1 | 5402 |
| 40 | No | E. coli | Trimethopr | Intracellula | F109 | Riboflavin | 7465 | BL1 | 6949 |
| 40 | No | E. coli | Trimethopr | Intracellula | F117 | SYBR Gree | 6914 | BL1 | 116072.5 |
| 40 | No | E. coli | Trimethopr | Intracellula | F106 | Rhodamine | 7678 | BL1 | 7247 |
| 40 | No | E. coli | Trimethopr | Intracellula | F73 | Malachite ( | 6607 | BL1 | 7002 |
| 40 | No | E. coli | Trimethopr | Intracellula | F7 | Acridine or | 8303 | BL1 | 5558 |
| 40 | No | E. coli | Trimethopr | Intracellula | F192 | 5-Carboxyf | 8203 | BL1 | 11587 |
| 40 | No | E. coli | Trimethopr | Intracellula | F57 | Eosin Y | 6760 | BL1 | 16670 |
| 40 | No | E. coli | Trimethopr | Intracellula | F27 | Calcein | 8273 | BL1 | 20517 |
| 40 | No | E. coli | Trimethopr | Intracellula | F12 | Amaranth | 0 | BL1 | 0 |
| 40 | No | E. coli | None | None | NA | Autofluore: | 5888 | NA | 21447 |
| 40 | No | E. coli | None | None | F194 | Oxonol | 7610 | RL1 | 24421.5 |
| 40 | No | E. coli | None | None | F11 | Allura Red | 7422 | BL3 | 23842 |
| 40 | No | E. coli | None | None | F54 | DiSC3(5) | 7960 | RL1 | 15645 |
| 40 | No | E. coli | None | None | F124 | Thioflavin T | 8970 | VL2 | 15030.5 |
| 40 | No | E. coli | None | None | F101 | Quinacrine | 8813 | VL2 | 13788 |
| 40 | No | E. coli | None | None | F138 | Apigenin | 7054 | VL2 | 13466.5 |
| 40 | No | E. coli | None | None | F65 | H2FDA | 7543 | BL1 | 12924 |
| 40 | No | E. coli | None | None | F142 | Chlorophe | 8477 | BL3 | 13309 |
| 40 | No | E. coli | None | None | F39 | Congo Red | 8534 | BL3 | 11785.5 |
| 40 | No | E. coli | None | None | F116 | Sunset Yel | 6354 | BL2 | 12446.5 |
| 40 | No | E. coli | None | None | F10 | Alizarin | 8365 | BL1 | 11472 |
| 40 | No | E. coli | None | None | F84 | Neutral Re | 7002 | BL1 | 11453 |
| 40 | No | E. coli | None | None | F99 | Pyronin Y | 8832 | BL1 | 9130.5 |
| 40 | No | E. coli | None | None | F108 | Rhodamine | 7978 | BL1 | 9829 |
| 40 | No | E. coli | None | None | F109 | Riboflavin | 8257 | BL1 | 8326 |
| 40 | No | E. coli | None | None | F117 | SYBR Gree | 7771 | BL1 | 152780 |
| 40 | No | E. coli | None | None | F106 | Rhodamine | 8911 | BL1 | 10184 |
| 40 | No | E. coli | None | None | F73 | Malachite ( | 6521 | BL1 | 7915 |
| 40 | No | E. coli | None | None | F7 | Acridine or | 9104 | BL1 | 7231 |
| 40 | No | E. coli | None | None | F192 | 5-Carboxyf | 8151 | BL1 | 13553 |
| 40 | No | E. coli | None | None | F57 | Eosin Y | 7631 | BL1 | 15823 |
| 40 | No | E. coli | None | None | F27 | Calcein | 7100 | BL1 | 19246 |

| | | | | | | | | | |
|---|---|---|---|---|---|---|---|---|---|
| 40 | No | E. coli | None | None | F12 | Amaranth | 8987 | BL1 | 19465 |
| 40 | No | E. coli | None | None | NA | Autofluore: | 8431 | NA | 19740 |
| 40 | No | E. coli | Amoxacilli | Wall | F194 | Oxonol | 6914 | RL1 | 18180.5 |
| 40 | No | E. coli | Amoxacilli | Wall | F11 | Allura Red | 6555 | BL3 | 16216 |
| 40 | No | E. coli | Amoxacilli | Wall | F54 | DiSC3(5) | 5892 | RL1 | 15713.5 |
| 40 | No | E. coli | Amoxacilli | Wall | F124 | Thioflavin T | 8814 | VL2 | 14185.5 |
| 40 | No | E. coli | Amoxacilli | Wall | F101 | Quinacrine | 8500 | VL2 | 12785.5 |
| 40 | No | E. coli | Amoxacilli | Wall | F138 | Apigenin | 6762 | VL2 | 11712 |
| 40 | No | E. coli | Amoxacilli | Wall | F65 | H2FDA | 8188 | BL1 | 7056 |
| 40 | No | E. coli | Amoxacilli | Wall | F142 | Chlorophe | 8560 | BL3 | 6105 |
| 40 | No | E. coli | Amoxacilli | Wall | F39 | Congo Red | 6765 | BL3 | 5527 |
| 40 | No | E. coli | Amoxacilli | Wall | F116 | Sunset Yel | 8294 | BL2 | 6038 |
| 40 | No | E. coli | Amoxacilli | Wall | F10 | Alizarin | 6371 | BL1 | 6089 |
| 40 | No | E. coli | Amoxacilli | Wall | F84 | Neutral Re | 8760 | BL1 | 5676.5 |
| 40 | No | E. coli | Amoxacilli | Wall | F99 | Pyronin Y | 7407 | BL1 | 5364 |
| 40 | No | E. coli | Amoxacilli | Wall | F108 | Rhodamine | 7386 | BL1 | 5470.5 |
| 40 | No | E. coli | Amoxacilli | Wall | F109 | Riboflavin | 6749 | BL1 | 4635 |
| 40 | No | E. coli | Amoxacilli | Wall | F117 | SYBR Gree | 7906 | BL1 | 151986.5 |
| 40 | No | E. coli | Amoxacilli | Wall | F106 | Rhodamine | 7949 | BL1 | 6230 |
| 40 | No | E. coli | Amoxacilli | Wall | F73 | Malachite ( | 8072 | BL1 | 5142.5 |
| 40 | No | E. coli | Amoxacilli | Wall | F7 | Acridine or | 8590 | BL1 | 5405 |
| 40 | No | E. coli | Amoxacilli | Wall | F192 | 5-Carboxyf | 7819 | BL1 | 13314 |
| 40 | No | E. coli | Amoxacilli | Wall | F57 | Eosin Y | 6170 | BL1 | 15819.5 |
| 40 | No | E. coli | Amoxacilli | Wall | F27 | Calcein | 8717 | BL1 | 20359 |
| 40 | No | E. coli | Amoxacilli | Wall | F12 | Amaranth | 7986 | BL1 | 21201.5 |
| 40 | No | E. coli | None | None | NA | Autofluore: | 6917 | NA | 18001 |
| 40 | No | E. coli | Azithromyc | Intracellula | F194 | Oxonol | 5953 | RL1 | 18436 |
| 40 | No | E. coli | Azithromyc | Intracellula | F11 | Allura Red | 5322 | BL3 | 20246 |
| 40 | No | E. coli | Azithromyc | Intracellula | F54 | DiSC3(5) | 5408 | RL1 | 17774 |
| 40 | No | E. coli | Azithromyc | Intracellula | F124 | Thioflavin T | 6896 | VL2 | 14690 |
| 40 | No | E. coli | Azithromyc | Intracellula | F101 | Quinacrine | 5244 | VL2 | 13269.5 |
| 40 | No | E. coli | Azithromyc | Intracellula | F138 | Apigenin | 5404 | VL2 | 8654 |
| 40 | No | E. coli | Azithromyc | Intracellula | F65 | H2FDA | 6303 | BL1 | 5654 |
| 40 | No | E. coli | Azithromyc | Intracellula | F142 | Chlorophe | 5118 | BL3 | 5462.5 |
| 40 | No | E. coli | Azithromyc | Intracellula | F39 | Congo Red | 4825 | BL3 | 5522 |
| 40 | No | E. coli | Azithromyc | Intracellula | F116 | Sunset Yel | 4695 | BL2 | 6880 |
| 40 | No | E. coli | Azithromyc | Intracellula | F10 | Alizarin | 5469 | BL1 | 6254 |
| 40 | No | E. coli | Azithromyc | Intracellula | F84 | Neutral Re | 4985 | BL1 | 6330 |
| 40 | No | E. coli | Azithromyc | Intracellula | F99 | Pyronin Y | 5176 | BL1 | 5905 |
| 40 | No | E. coli | Azithromyc | Intracellula | F108 | Rhodamine | 5641 | BL1 | 4834 |
| 40 | No | E. coli | Azithromyc | Intracellula | F109 | Riboflavin | 5386 | BL1 | 4854.5 |
| 40 | No | E. coli | Azithromyc | Intracellula | F117 | SYBR Gree | 5499 | BL1 | 29185 |
| 40 | No | E. coli | Azithromyc | Intracellula | F106 | Rhodamine | 4937 | BL1 | 6369 |
| 40 | No | E. coli | Azithromyc | Intracellula | F73 | Malachite ( | 4468 | BL1 | 5639.5 |
| 40 | No | E. coli | Azithromyc | Intracellula | F7 | Acridine or | 6169 | BL1 | 5269 |
| 40 | No | E. coli | Azithromyc | Intracellula | F192 | 5-Carboxyf | 4441 | BL1 | 10048 |
| 40 | No | E. coli | Azithromyc | Intracellula | F57 | Eosin Y | 5015 | BL1 | 16907 |
| 40 | No | E. coli | Azithromyc | Intracellula | F27 | Calcein | 5531 | BL1 | 17032 |

| | | | | | | | | | |
|---|---|---|---|---|---|---|---|---|---|
| 40 | No | E. coli | Azithromyc | Intracellula | F12 | Amaranth | 4807 | BL1 | 19455 |
| 40 | No | E. coli | None | None | NA | Autofluores | 6716 | NA | 17761 |
| 40 | No | E. coli | Trimethopr | Intracellula | F194 | Oxonol | 5180 | RL1 | 16958 |
| 40 | No | E. coli | Trimethopr | Intracellula | F11 | Allura Red | 5518 | BL3 | 18679 |
| 40 | No | E. coli | Trimethopr | Intracellula | F54 | DiSC3(5) | 6533 | RL1 | 16428 |
| 40 | No | E. coli | Trimethopr | Intracellula | F124 | Thioflavin T | 7102 | VL2 | 13442.5 |
| 40 | No | E. coli | Trimethopr | Intracellula | F101 | Quinacrine | 7039 | VL2 | 12390 |
| 40 | No | E. coli | Trimethopr | Intracellula | F138 | Apigenin | 6587 | VL2 | 10014 |
| 40 | No | E. coli | Trimethopr | Intracellula | F65 | H2FDA | 7067 | BL1 | 7455 |
| 40 | No | E. coli | Trimethopr | Intracellula | F142 | Chlorophe | 7565 | BL3 | 5438 |
| 40 | No | E. coli | Trimethopr | Intracellula | F39 | Congo Red | 7465 | BL3 | 7014 |
| 40 | No | E. coli | Trimethopr | Intracellula | F116 | Sunset Yel | 6143 | BL2 | 6902 |
| 40 | No | E. coli | Trimethopr | Intracellula | F10 | Alizarin | 7841 | BL1 | 6798 |
| 40 | No | E. coli | Trimethopr | Intracellula | F84 | Neutral Re | 7268 | BL1 | 7343 |
| 40 | No | E. coli | Trimethopr | Intracellula | F99 | Pyronin Y | 7632 | BL1 | 6435.5 |
| 40 | No | E. coli | Trimethopr | Intracellula | F108 | Rhodamine | 7399 | BL1 | 5738 |
| 40 | No | E. coli | Trimethopr | Intracellula | F109 | Riboflavin | 6555 | BL1 | 5755 |
| 40 | No | E. coli | Trimethopr | Intracellula | F117 | SYBR Gree | 8456 | BL1 | 115451.5 |
| 40 | No | E. coli | Trimethopr | Intracellula | F106 | Rhodamine | 6084 | BL1 | 6924.5 |
| 40 | No | E. coli | Trimethopr | Intracellula | F73 | Malachite ( | 8519 | BL1 | 5849 |
| 40 | No | E. coli | Trimethopr | Intracellula | F7 | Acridine or | 7822 | BL1 | 5552.5 |
| 40 | No | E. coli | Trimethopr | Intracellula | F192 | 5-Carboxyf | 7430 | BL1 | 13277 |
| 40 | No | E. coli | Trimethopr | Intracellula | F57 | Eosin Y | 7794 | BL1 | 18400 |
| 40 | No | E. coli | Trimethopr | Intracellula | F27 | Calcein | 6357 | BL1 | 22217 |
| 40 | No | E. coli | Trimethopr | Intracellula | F12 | Amaranth | 5741 | BL1 | 21948 |
| 40 | No | E. coli | None | None | NA | Autofluores | 7350 | NA | 17229 |
| 40 | No | E. coli | Chloramph | Intracellula | F194 | Oxonol | 7046 | RL1 | 18936.5 |
| 40 | No | E. coli | Chloramph | Intracellula | F11 | Allura Red | 5136 | BL3 | 19602 |
| 40 | No | E. coli | Chloramph | Intracellula | F54 | DiSC3(5) | 6536 | RL1 | 18464 |
| 40 | No | E. coli | Chloramph | Intracellula | F124 | Thioflavin T | 6146 | VL2 | 13877.5 |
| 40 | No | E. coli | Chloramph | Intracellula | F101 | Quinacrine | 7992 | VL2 | 12656.5 |
| 40 | No | E. coli | Chloramph | Intracellula | F138 | Apigenin | 7583 | VL2 | 10234 |
| 40 | No | E. coli | Chloramph | Intracellula | F65 | H2FDA | 7360 | BL1 | 6675 |
| 40 | No | E. coli | Chloramph | Intracellula | F142 | Chlorophe | 7756 | BL3 | 6052 |
| 40 | No | E. coli | Chloramph | Intracellula | F39 | Congo Red | 5911 | BL3 | 5793 |
| 40 | No | E. coli | Chloramph | Intracellula | F116 | Sunset Yel | 7724 | BL2 | 7415.5 |
| 40 | No | E. coli | Chloramph | Intracellula | F10 | Alizarin | 6221 | BL1 | 7654 |
| 40 | No | E. coli | Chloramph | Intracellula | F84 | Neutral Re | 7332 | BL1 | 6479 |
| 40 | No | E. coli | Chloramph | Intracellula | F99 | Pyronin Y | 6292 | BL1 | 6141 |
| 40 | No | E. coli | Chloramph | Intracellula | F108 | Rhodamine | 7044 | BL1 | 5533 |
| 40 | No | E. coli | Chloramph | Intracellula | F109 | Riboflavin | 6219 | BL1 | 5470 |
| 40 | No | E. coli | Chloramph | Intracellula | F117 | SYBR Gree | 7143 | BL1 | 24121 |
| 40 | No | E. coli | Chloramph | Intracellula | F106 | Rhodamine | 7053 | BL1 | 10926 |
| 40 | No | E. coli | Chloramph | Intracellula | F73 | Malachite ( | 7288 | BL1 | 5540.5 |
| 40 | No | E. coli | Chloramph | Intracellula | F7 | Acridine or | 7102 | BL1 | 5347 |
| 40 | No | E. coli | Chloramph | Intracellula | F192 | 5-Carboxyf | 7951 | BL1 | 11700 |
| 40 | No | E. coli | Chloramph | Intracellula | F57 | Eosin Y | 7574 | BL1 | 14704 |
| 40 | No | E. coli | Chloramph | Intracellula | F27 | Calcein | 6714 | BL1 | 21974 |

| | | | | | | | | | |
|---|---|---|---|---|---|---|---|---|---|
| 40 | No | E. coli | Chloramph | Intracellula | F12 | Amaranth | 7177 | BL1 | 24542 |
| 40 | No | E. coli | None | None | NA | Autofluores | 6649 | NA | 19193 |
| 40 | No | E. coli | Colistin | Wall | F194 | Oxonol | 5039 | RL1 | 32050 |
| 40 | No | E. coli | Colistin | Wall | F11 | Allura Red | 3308 | BL3 | 48242 |
| 40 | No | E. coli | Colistin | Wall | F54 | DiSC3(5) | 2101 | RL1 | 32934 |
| 40 | No | E. coli | Colistin | Wall | F124 | Thioflavin T | 2478 | VL2 | 50551 |
| 40 | No | E. coli | Colistin | Wall | F101 | Quinacrine | 2483 | VL2 | 64894 |
| 40 | No | E. coli | Colistin | Wall | F138 | Apigenin | 2348 | VL2 | 55901.5 |
| 40 | No | E. coli | Colistin | Wall | F65 | H2FDA | 2835 | BL1 | 63999 |
| 40 | No | E. coli | Colistin | Wall | F142 | Chlorophe | 2768 | BL3 | 52018 |
| 40 | No | E. coli | Colistin | Wall | F39 | Congo Red | 3075 | BL3 | 44099 |
| 40 | No | E. coli | Colistin | Wall | F116 | Sunset Yell | 1688 | BL2 | 54567 |
| 40 | No | E. coli | Colistin | Wall | F10 | Alizarin | 3202 | BL1 | 41843.5 |
| 40 | No | E. coli | Colistin | Wall | F84 | Neutral Re | 2131 | BL1 | 47961 |
| 40 | No | E. coli | Colistin | Wall | F99 | Pyronin Y | 2394 | BL1 | 53197.5 |
| 40 | No | E. coli | Colistin | Wall | F108 | Rhodamine | 3178 | BL1 | 46801.5 |
| 40 | No | E. coli | Colistin | Wall | F109 | Riboflavin | 3332 | BL1 | 53682.5 |
| 40 | No | E. coli | Colistin | Wall | F117 | SYBR Gree | 2508 | BL1 | 2212247 |
| 40 | No | E. coli | Colistin | Wall | F106 | Rhodamine | 2185 | BL1 | 264715 |
| 40 | No | E. coli | Colistin | Wall | F73 | Malachite ( | 3646 | BL1 | 124763 |
| 40 | No | E. coli | Colistin | Wall | F7 | Acridine or | 3310 | BL1 | 131205.5 |
| 40 | No | E. coli | Colistin | Wall | F192 | 5-Carboxyf | 2839 | BL1 | 77489 |
| 40 | No | E. coli | Colistin | Wall | F57 | Eosin Y | 1649 | BL1 | 57291 |
| 40 | No | E. coli | Colistin | Wall | F27 | Calcein | 2193 | BL1 | 38609 |
| 40 | No | E. coli | Colistin | Wall | F12 | Amaranth | 2188 | BL1 | 33389.5 |
| 40 | No | E. coli | None | None | NA | Autofluores | 5013 | NA | 20329 |
| 40 | No | E. coli | Gentamici | Intracellula | F194 | Oxonol | 6139 | RL1 | 18047 |
| 40 | No | E. coli | Gentamici | Intracellula | F11 | Allura Red | 6903 | BL3 | 16138 |
| 40 | No | E. coli | Gentamici | Intracellula | F54 | DiSC3(5) | 7384 | RL1 | 16010.5 |
| 40 | No | E. coli | Gentamici | Intracellula | F124 | Thioflavin T | 7312 | VL2 | 13460.5 |
| 40 | No | E. coli | Gentamici | Intracellula | F101 | Quinacrine | 7012 | VL2 | 14326 |
| 40 | No | E. coli | Gentamici | Intracellula | F138 | Apigenin | 8472 | VL2 | 11379.5 |
| 40 | No | E. coli | Gentamici | Intracellula | F65 | H2FDA | 6896 | BL1 | 7623.5 |
| 40 | No | E. coli | Gentamici | Intracellula | F142 | Chlorophe | 8369 | BL3 | 5888 |
| 40 | No | E. coli | Gentamici | Intracellula | F39 | Congo Red | 7080 | BL3 | 5721 |
| 40 | No | E. coli | Gentamici | Intracellula | F116 | Sunset Yell | 9821 | BL2 | 5832 |
| 40 | No | E. coli | Gentamici | Intracellula | F10 | Alizarin | 6511 | BL1 | 7340 |
| 40 | No | E. coli | Gentamici | Intracellula | F84 | Neutral Re | 8863 | BL1 | 6877 |
| 40 | No | E. coli | Gentamici | Intracellula | F99 | Pyronin Y | 7891 | BL1 | 5889 |
| 40 | No | E. coli | Gentamici | Intracellula | F108 | Rhodamine | 6935 | BL1 | 5689 |
| 40 | No | E. coli | Gentamici | Intracellula | F109 | Riboflavin | 7224 | BL1 | 4815 |
| 40 | No | E. coli | Gentamici | Intracellula | F117 | SYBR Gree | 8612 | BL1 | 87406.5 |
| 40 | No | E. coli | Gentamici | Intracellula | F106 | Rhodamine | 7195 | BL1 | 6796 |
| 40 | No | E. coli | Gentamici | Intracellula | F73 | Malachite ( | 8619 | BL1 | 5602 |
| 40 | No | E. coli | Gentamici | Intracellula | F7 | Acridine or | 8072 | BL1 | 6311 |
| 40 | No | E. coli | Gentamici | Intracellula | F192 | 5-Carboxyf | 7247 | BL1 | 11601 |
| 40 | No | E. coli | Gentamici | Intracellula | F57 | Eosin Y | 9316 | BL1 | 18074 |
| 40 | No | E. coli | Gentamici | Intracellula | F27 | Calcein | 9800 | BL1 | 18639 |

| | | | | | | | | | |
|---|---|---|---|---|---|---|---|---|---|
| 40 | No | E. coli | Gentamici | Intracellula | F12 | Amaranth | 7951 | BL1 | 16808 |
| 40 | No | E. coli | None | None | NA | Autofluores | 7466 | NA | 21536.5 |
| 40 | No | E. coli | Nitrofurant | Intracellula | F194 | Oxonol | 5835 | RL1 | 19995 |
| 40 | No | E. coli | Nitrofurant | Intracellula | F11 | Allura Red | 6383 | BL3 | 20629 |
| 40 | No | E. coli | Nitrofurant | Intracellula | F54 | DiSC3(5) | 4033 | RL1 | 15562 |
| 40 | No | E. coli | Nitrofurant | Intracellula | F124 | Thioflavin T | 7901 | VL2 | 14277 |
| 40 | No | E. coli | Nitrofurant | Intracellula | F101 | Quinacrine | 6907 | VL2 | 10890 |
| 40 | No | E. coli | Nitrofurant | Intracellula | F138 | Apigenin | 5811 | VL2 | 11159 |
| 40 | No | E. coli | Nitrofurant | Intracellula | F65 | H2FDA | 7382 | BL1 | 6711 |
| 40 | No | E. coli | Nitrofurant | Intracellula | F142 | Chlorophe | 6041 | BL3 | 5764 |
| 40 | No | E. coli | Nitrofurant | Intracellula | F39 | Congo Red | 6292 | BL3 | 5709 |
| 40 | No | E. coli | Nitrofurant | Intracellula | F116 | Sunset Yel | 6906 | BL2 | 7186.5 |
| 40 | No | E. coli | Nitrofurant | Intracellula | F10 | Alizarin | 5634 | BL1 | 6893.5 |
| 40 | No | E. coli | Nitrofurant | Intracellula | F84 | Neutral Re | 6838 | BL1 | 6395 |
| 40 | No | E. coli | Nitrofurant | Intracellula | F99 | Pyronin Y | 5905 | BL1 | 6390 |
| 40 | No | E. coli | Nitrofurant | Intracellula | F108 | Rhodamine | 6150 | BL1 | 5529 |
| 40 | No | E. coli | Nitrofurant | Intracellula | F109 | Riboflavin | 4763 | BL1 | 5391 |
| 40 | No | E. coli | Nitrofurant | Intracellula | F117 | SYBR Gree | 5909 | BL1 | 31209 |
| 40 | No | E. coli | Nitrofurant | Intracellula | F106 | Rhodamine | 5808 | BL1 | 8981.5 |
| 40 | No | E. coli | Nitrofurant | Intracellula | F73 | Malachite ( | 5161 | BL1 | 6487 |
| 40 | No | E. coli | Nitrofurant | Intracellula | F7 | Acridine or | 4121 | BL1 | 5772 |
| 40 | No | E. coli | Nitrofurant | Intracellula | F192 | 5-Carboxyf | 5019 | BL1 | 7766 |
| 40 | No | E. coli | Nitrofurant | Intracellula | F57 | Eosin Y | 5804 | BL1 | 16028 |
| 40 | No | E. coli | Nitrofurant | Intracellula | F27 | Calcein | 5437 | BL1 | 18162 |
| 40 | No | E. coli | Nitrofurant | Intracellula | F12 | Amaranth | 6182 | BL1 | 21686.5 |
| 40 | No | E. coli | None | None | NA | Autofluores | 8557 | NA | 21325 |
| 40 | No | E. coli | Pentamidi | Wall | F194 | Oxonol | 6101 | RL1 | 15605 |
| 40 | No | E. coli | Pentamidi | Wall | F11 | Allura Red | 6639 | BL3 | 19951 |
| 40 | No | E. coli | Pentamidi | Wall | F54 | DiSC3(5) | 5197 | RL1 | 16721 |
| 40 | No | E. coli | Pentamidi | Wall | F124 | Thioflavin T | 6014 | VL2 | 15222.5 |
| 40 | No | E. coli | Pentamidi | Wall | F101 | Quinacrine | 4962 | VL2 | 16653.5 |
| 40 | No | E. coli | Pentamidi | Wall | F138 | Apigenin | 5019 | VL2 | 17663 |
| 40 | No | E. coli | Pentamidi | Wall | F65 | H2FDA | 6297 | BL1 | 10761 |
| 40 | No | E. coli | Pentamidi | Wall | F142 | Chlorophe | 7313 | BL3 | 11369 |
| 40 | No | E. coli | Pentamidi | Wall | F39 | Congo Red | 5895 | BL3 | 10601 |
| 40 | No | E. coli | Pentamidi | Wall | F116 | Sunset Yel | 6119 | BL2 | 8914 |
| 40 | No | E. coli | Pentamidi | Wall | F10 | Alizarin | 6541 | BL1 | 9744 |
| 40 | No | E. coli | Pentamidi | Wall | F84 | Neutral Re | 6779 | BL1 | 7572 |
| 40 | No | E. coli | Pentamidi | Wall | F99 | Pyronin Y | 7221 | BL1 | 6357 |
| 40 | No | E. coli | Pentamidi | Wall | F108 | Rhodamine | 5549 | BL1 | 5856 |
| 40 | No | E. coli | Pentamidi | Wall | F109 | Riboflavin | 7145 | BL1 | 4464 |
| 40 | No | E. coli | Pentamidi | Wall | F117 | SYBR Gree | 6604 | BL1 | 321297 |
| 40 | No | E. coli | Pentamidi | Wall | F106 | Rhodamine | 4839 | BL1 | 6740 |
| 40 | No | E. coli | Pentamidi | Wall | F73 | Malachite ( | 6931 | BL1 | 5447 |
| 40 | No | E. coli | Pentamidi | Wall | F7 | Acridine or | 6318 | BL1 | 4858 |
| 40 | No | E. coli | Pentamidi | Wall | F192 | 5-Carboxyf | 6213 | BL1 | 10074 |
| 40 | No | E. coli | Pentamidi | Wall | F57 | Eosin Y | 5712 | BL1 | 17531 |
| 40 | No | E. coli | Pentamidi | Wall | F27 | Calcein | 7496 | BL1 | 22647 |

| | | | | | | | | |
|---|---|---|---|---|---|---|---|---|
| 40 | No | E. coli | Pentamidi | Wall | F12 | Amaranth | 5253 | BL1 | 17404 |
| 40 | No | E. coli | None | None | NA | Autofluore | 6981 | NA | 20890 |
| 40 | No | E. coli | Rifampicin | Intracellul | F194 | Oxonol | 6764 | RL1 | 17564.5 |
| 40 | No | E. coli | Rifampicin | Intracellul | F11 | Allura Red | 7694 | BL3 | 18058 |
| 40 | No | E. coli | Rifampicin | Intracellul | F54 | DiSC3(5) | 8091 | RL1 | 16858 |
| 40 | No | E. coli | Rifampicin | Intracellul | F124 | Thioflavin T | 8944 | VL2 | 13589 |
| 40 | No | E. coli | Rifampicin | Intracellul | F101 | Quinacrine | 9673 | VL2 | 11933 |
| 40 | No | E. coli | Rifampicin | Intracellul | F138 | Apigenin | 9135 | VL2 | 8588 |
| 40 | No | E. coli | Rifampicin | Intracellul | F65 | H2FDA | 10029 | BL1 | 5551 |
| 40 | No | E. coli | Rifampicin | Intracellul | F142 | Chlorophe | 7576 | BL3 | 4621.5 |
| 40 | No | E. coli | Rifampicin | Intracellul | F39 | Congo Red | 9582 | BL3 | 5887.5 |
| 40 | No | E. coli | Rifampicin | Intracellul | F116 | Sunset Yel | 8030 | BL2 | 5173 |
| 40 | No | E. coli | Rifampicin | Intracellul | F10 | Alizarin | 9912 | BL1 | 5749.5 |
| 40 | No | E. coli | Rifampicin | Intracellul | F84 | Neutral Re | 7498 | BL1 | 5440 |
| 40 | No | E. coli | Rifampicin | Intracellul | F99 | Pyronin Y | 9555 | BL1 | 4554 |
| 40 | No | E. coli | Rifampicin | Intracellul | F108 | Rhodamine | 9104 | BL1 | 5093.5 |
| 40 | No | E. coli | Rifampicin | Intracellul | F109 | Riboflavin | 8576 | BL1 | 4716.5 |
| 40 | No | E. coli | Rifampicin | Intracellul | F117 | SYBR Gree | 8615 | BL1 | 27904 |
| 40 | No | E. coli | Rifampicin | Intracellul | F106 | Rhodamine | 8319 | BL1 | 6416 |
| 40 | No | E. coli | Rifampicin | Intracellul | F73 | Malachite | 7804 | BL1 | 5845.5 |
| 40 | No | E. coli | Rifampicin | Intracellul | F7 | Acridine or | 9612 | BL1 | 4312 |
| 40 | No | E. coli | Rifampicin | Intracellul | F192 | 5-Carboxyf | 9672 | BL1 | 10812.5 |
| 40 | No | E. coli | Rifampicin | Intracellul | F57 | Eosin Y | 9657 | BL1 | 15896 |
| 40 | No | E. coli | Rifampicin | Intracellul | F27 | Calcein | 7047 | BL1 | 16225 |
| 40 | No | E. coli | Rifampicin | Intracellul | F12 | Amaranth | 7547 | BL1 | 15753 |
| 40 | No | E. coli | None | None | NA | Autofluore | 6280 | NA | 24140 |
| 40 | No | E. coli | Trimethopr | Intracellul | F194 | Oxonol | 6395 | RL1 | 18440 |
| 40 | No | E. coli | Trimethopr | Intracellul | F11 | Allura Red | 6293 | BL3 | 18813 |
| 40 | No | E. coli | Trimethopr | Intracellul | F54 | DiSC3(5) | 7407 | RL1 | 15275 |
| 40 | No | E. coli | Trimethopr | Intracellul | F124 | Thioflavin T | 8570 | VL2 | 13935 |
| 40 | No | E. coli | Trimethopr | Intracellul | F101 | Quinacrine | 7861 | VL2 | 11388 |
| 40 | No | E. coli | Trimethopr | Intracellul | F138 | Apigenin | 6963 | VL2 | 8733 |
| 40 | No | E. coli | Trimethopr | Intracellul | F65 | H2FDA | 8485 | BL1 | 5520 |
| 40 | No | E. coli | Trimethopr | Intracellul | F142 | Chlorophe | 7205 | BL3 | 5501 |
| 40 | No | E. coli | Trimethopr | Intracellul | F39 | Congo Red | 6806 | BL3 | 5358 |
| 40 | No | E. coli | Trimethopr | Intracellul | F116 | Sunset Yel | 6793 | BL2 | 5593 |
| 40 | No | E. coli | Trimethopr | Intracellul | F10 | Alizarin | 6851 | BL1 | 5466 |
| 40 | No | E. coli | Trimethopr | Intracellul | F84 | Neutral Re | 5399 | BL1 | 6930 |
| 40 | No | E. coli | Trimethopr | Intracellul | F99 | Pyronin Y | 8359 | BL1 | 4791 |
| 40 | No | E. coli | Trimethopr | Intracellul | F108 | Rhodamine | 7270 | BL1 | 4970.5 |
| 40 | No | E. coli | Trimethopr | Intracellul | F109 | Riboflavin | 6298 | BL1 | 4916 |
| 40 | No | E. coli | Trimethopr | Intracellul | F117 | SYBR Gree | 8443 | BL1 | 11849 |
| 40 | No | E. coli | Trimethopr | Intracellul | F106 | Rhodamine | 8240 | BL1 | 7111.5 |
| 40 | No | E. coli | Trimethopr | Intracellul | F73 | Malachite | 8301 | BL1 | 5790 |
| 40 | No | E. coli | Trimethopr | Intracellul | F7 | Acridine or | 7167 | BL1 | 6938 |
| 40 | No | E. coli | Trimethopr | Intracellul | F192 | 5-Carboxyf | 7039 | BL1 | 6362 |
| 40 | No | E. coli | Trimethopr | Intracellul | F57 | Eosin Y | 7836 | BL1 | 12736 |
| 40 | No | E. coli | Trimethopr | Intracellul | F27 | Calcein | 8649 | BL1 | 16677 |

| | | | | | | | | | |
|---|---|---|---|---|---|---|---|---|---|
| 40 | No | E. coli | Trimethopr | Intracellula | F12 | Amaranth | 0 | BL1 | 0 |
| 40 | No | E. coli | None | None | NA | Autofluore: | 6191 | NA | 22531 |
| 40 | No | E. coli | None | None | F194 | Oxonol | 9511 | RL1 | 22471 |
| 40 | No | E. coli | None | None | F11 | Allura Red | 8456 | BL3 | 21182 |
| 40 | No | E. coli | None | None | F54 | DiSC3(5) | 8996 | RL1 | 16871 |
| 40 | No | E. coli | None | None | F124 | Thioflavin T | 9413 | VL2 | 16757 |
| 40 | No | E. coli | None | None | F101 | Quinacrine | 9299 | VL2 | 14842 |
| 40 | No | E. coli | None | None | F138 | Apigenin | 8575 | VL2 | 12689 |
| 40 | No | E. coli | None | None | F65 | H2FDA | 9246 | BL1 | 13999.5 |
| 40 | No | E. coli | None | None | F142 | Chlorophe | 8234 | BL3 | 13315.5 |
| 40 | No | E. coli | None | None | F39 | Congo Red | 8140 | BL3 | 11394 |
| 40 | No | E. coli | None | None | F116 | Sunset Yel | 8295 | BL2 | 11976 |
| 40 | No | E. coli | None | None | F10 | Alizarin | 7738 | BL1 | 11233 |
| 40 | No | E. coli | None | None | F84 | Neutral Re | 9160 | BL1 | 10300 |
| 40 | No | E. coli | None | None | F99 | Pyronin Y | 7225 | BL1 | 8855 |
| 40 | No | E. coli | None | None | F108 | Rhodamine | 9680 | BL1 | 8158 |
| 40 | No | E. coli | None | None | F109 | Riboflavin | 9408 | BL1 | 7830 |
| 40 | No | E. coli | None | None | F117 | SYBR Gree | 8923 | BL1 | 122758 |
| 40 | No | E. coli | None | None | F106 | Rhodamine | 9587 | BL1 | 7869 |
| 40 | No | E. coli | None | None | F73 | Malachite ( | 7958 | BL1 | 6988 |
| 40 | No | E. coli | None | None | F7 | Acridine or | 7908 | BL1 | 6482 |
| 40 | No | E. coli | None | None | F192 | 5-Carboxyf | 8769 | BL1 | 13572 |
| 40 | No | E. coli | None | None | F57 | Eosin Y | 8152 | BL1 | 17310.5 |
| 40 | No | E. coli | None | None | F27 | Calcein | 9142 | BL1 | 18277 |
| 40 | No | E. coli | None | None | F12 | Amaranth | 8435 | BL1 | 20025 |
| 40 | No | E. coli | None | None | NA | Autofluore: | 9236 | NA | 22341.5 |
| 40 | No | E. coli | Amoxacilli | Wall | F194 | Oxonol | 8579 | RL1 | 21081 |
| 40 | No | E. coli | Amoxacilli | Wall | F11 | Allura Red | 7078 | BL3 | 16848.5 |
| 40 | No | E. coli | Amoxacilli | Wall | F54 | DiSC3(5) | 6522 | RL1 | 17747 |
| 40 | No | E. coli | Amoxacilli | Wall | F124 | Thioflavin T | 11035 | VL2 | 13589 |
| 40 | No | E. coli | Amoxacilli | Wall | F101 | Quinacrine | 8227 | VL2 | 13025 |
| 40 | No | E. coli | Amoxacilli | Wall | F138 | Apigenin | 9872 | VL2 | 9246 |
| 40 | No | E. coli | Amoxacilli | Wall | F65 | H2FDA | 7919 | BL1 | 6291 |
| 40 | No | E. coli | Amoxacilli | Wall | F142 | Chlorophe | 9752 | BL3 | 5263 |
| 40 | No | E. coli | Amoxacilli | Wall | F39 | Congo Red | 7420 | BL3 | 4575 |
| 40 | No | E. coli | Amoxacilli | Wall | F116 | Sunset Yel | 7425 | BL2 | 5657 |
| 40 | No | E. coli | Amoxacilli | Wall | F10 | Alizarin | 7789 | BL1 | 6209 |
| 40 | No | E. coli | Amoxacilli | Wall | F84 | Neutral Re | 9588 | BL1 | 5888 |
| 40 | No | E. coli | Amoxacilli | Wall | F99 | Pyronin Y | 5944 | BL1 | 5713.5 |
| 40 | No | E. coli | Amoxacilli | Wall | F108 | Rhodamine | 8325 | BL1 | 4905 |
| 40 | No | E. coli | Amoxacilli | Wall | F109 | Riboflavin | 7892 | BL1 | 4919 |
| 40 | No | E. coli | Amoxacilli | Wall | F117 | SYBR Gree | 9284 | BL1 | 99569 |
| 40 | No | E. coli | Amoxacilli | Wall | F106 | Rhodamine | 8967 | BL1 | 5762 |
| 40 | No | E. coli | Amoxacilli | Wall | F73 | Malachite ( | 8701 | BL1 | 4779 |
| 40 | No | E. coli | Amoxacilli | Wall | F7 | Acridine or | 6522 | BL1 | 5439 |
| 40 | No | E. coli | Amoxacilli | Wall | F192 | 5-Carboxyf | 7930 | BL1 | 11105 |
| 40 | No | E. coli | Amoxacilli | Wall | F57 | Eosin Y | 7573 | BL1 | 17639 |
| 40 | No | E. coli | Amoxacilli | Wall | F27 | Calcein | 9733 | BL1 | 21290 |

| | | | | | | | | | |
|---|---|---|---|---|---|---|---|---|---|
| 40 | No | E. coli | Amoxacilli | Wall | F12 | Amaranth | 8042 | BL1 | 17657.5 |
| 40 | No | E. coli | None | None | NA | Autofluores | 7763 | NA | 16798 |
| 40 | No | E. coli | Azithromyc | Intracellula | F194 | Oxonol | 6554 | RL1 | 21002.5 |
| 40 | No | E. coli | Azithromyc | Intracellula | F11 | Allura Red | 5789 | BL3 | 18018 |
| 40 | No | E. coli | Azithromyc | Intracellula | F54 | DiSC3(5) | 5259 | RL1 | 17124 |
| 40 | No | E. coli | Azithromyc | Intracellula | F124 | Thioflavin T | 5329 | VL2 | 14018 |
| 40 | No | E. coli | Azithromyc | Intracellula | F101 | Quinacrine | 5754 | VL2 | 12367.5 |
| 40 | No | E. coli | Azithromyc | Intracellula | F138 | Apigenin | 5940 | VL2 | 9752 |
| 40 | No | E. coli | Azithromyc | Intracellula | F65 | H2FDA | 5489 | BL1 | 5453 |
| 40 | No | E. coli | Azithromyc | Intracellula | F142 | Chlorophe | 5318 | BL3 | 5797 |
| 40 | No | E. coli | Azithromyc | Intracellula | F39 | Congo Red | 5428 | BL3 | 5180.5 |
| 40 | No | E. coli | Azithromyc | Intracellula | F116 | Sunset Yel | 5562 | BL2 | 6068 |
| 40 | No | E. coli | Azithromyc | Intracellula | F10 | Alizarin | 4948 | BL1 | 5985.5 |
| 40 | No | E. coli | Azithromyc | Intracellula | F84 | Neutral Re | 5694 | BL1 | 5746 |
| 40 | No | E. coli | Azithromyc | Intracellula | F99 | Pyronin Y | 4607 | BL1 | 6007 |
| 40 | No | E. coli | Azithromyc | Intracellula | F108 | Rhodamine | 5916 | BL1 | 4846 |
| 40 | No | E. coli | Azithromyc | Intracellula | F109 | Riboflavin | 4310 | BL1 | 4829.5 |
| 40 | No | E. coli | Azithromyc | Intracellula | F117 | SYBR Gree | 5767 | BL1 | 29096 |
| 40 | No | E. coli | Azithromyc | Intracellula | F106 | Rhodamine | 5187 | BL1 | 5892 |
| 40 | No | E. coli | Azithromyc | Intracellula | F73 | Malachite ( | 5707 | BL1 | 4966 |
| 40 | No | E. coli | Azithromyc | Intracellula | F7 | Acridine or | 5049 | BL1 | 4921 |
| 40 | No | E. coli | Azithromyc | Intracellula | F192 | 5-Carboxyf | 5257 | BL1 | 13593 |
| 40 | No | E. coli | Azithromyc | Intracellula | F57 | Eosin Y | 4346 | BL1 | 17886 |
| 40 | No | E. coli | Azithromyc | Intracellula | F27 | Calcein | 4519 | BL1 | 18433 |
| 40 | No | E. coli | Azithromyc | Intracellula | F12 | Amaranth | 5315 | BL1 | 18781 |
| 40 | No | E. coli | None | None | NA | Autofluores | 6872 | NA | 16997 |
| 40 | No | E. coli | Trimethopr | Intracellula | F194 | Oxonol | 6221 | RL1 | 22240 |
| 40 | No | E. coli | Trimethopr | Intracellula | F11 | Allura Red | 7122 | BL3 | 18538.5 |
| 40 | No | E. coli | Trimethopr | Intracellula | F54 | DiSC3(5) | 5851 | RL1 | 16681 |
| 40 | No | E. coli | Trimethopr | Intracellula | F124 | Thioflavin T | 8343 | VL2 | 13362 |
| 40 | No | E. coli | Trimethopr | Intracellula | F101 | Quinacrine | 5893 | VL2 | 12786 |
| 40 | No | E. coli | Trimethopr | Intracellula | F138 | Apigenin | 9322 | VL2 | 7523.5 |
| 40 | No | E. coli | Trimethopr | Intracellula | F65 | H2FDA | 8063 | BL1 | 6213 |
| 40 | No | E. coli | Trimethopr | Intracellula | F142 | Chlorophe | 6558 | BL3 | 5118.5 |
| 40 | No | E. coli | Trimethopr | Intracellula | F39 | Congo Red | 7325 | BL3 | 5298 |
| 40 | No | E. coli | Trimethopr | Intracellula | F116 | Sunset Yel | 7932 | BL2 | 5605.5 |
| 40 | No | E. coli | Trimethopr | Intracellula | F10 | Alizarin | 5626 | BL1 | 5437.5 |
| 40 | No | E. coli | Trimethopr | Intracellula | F84 | Neutral Re | 8795 | BL1 | 6044 |
| 40 | No | E. coli | Trimethopr | Intracellula | F99 | Pyronin Y | 5893 | BL1 | 5660 |
| 40 | No | E. coli | Trimethopr | Intracellula | F108 | Rhodamine | 8885 | BL1 | 4987 |
| 40 | No | E. coli | Trimethopr | Intracellula | F109 | Riboflavin | 7695 | BL1 | 5068 |
| 40 | No | E. coli | Trimethopr | Intracellula | F117 | SYBR Gree | 6847 | BL1 | 81217 |
| 40 | No | E. coli | Trimethopr | Intracellula | F106 | Rhodamine | 6028 | BL1 | 11276 |
| 40 | No | E. coli | Trimethopr | Intracellula | F73 | Malachite ( | 9348 | BL1 | 5403.5 |
| 40 | No | E. coli | Trimethopr | Intracellula | F7 | Acridine or | 7856 | BL1 | 5635 |
| 40 | No | E. coli | Trimethopr | Intracellula | F192 | 5-Carboxyf | 6661 | BL1 | 11386 |
| 40 | No | E. coli | Trimethopr | Intracellula | F57 | Eosin Y | 7476 | BL1 | 16643 |
| 40 | No | E. coli | Trimethopr | Intracellula | F27 | Calcein | 7162 | BL1 | 21405 |

| | | | | | | | | | |
|---|---|---|---|---|---|---|---|---|---|
| 40 | No | E. coli | Trimethopr | Intracellula | F12 | Amaranth | 8810 | BL1 | 19016.5 |
| 40 | No | E. coli | None | None | NA | Autofluore: | 6660 | NA | 15806.5 |
| 40 | No | E. coli | Chloramph | Intracellula | F194 | Oxonol | 6141 | RL1 | 20840 |
| 40 | No | E. coli | Chloramph | Intracellula | F11 | Allura Red | 7166 | BL3 | 18658 |
| 40 | No | E. coli | Chloramph | Intracellula | F54 | DiSC3(5) | 5971 | RL1 | 18665 |
| 40 | No | E. coli | Chloramph | Intracellula | F124 | Thioflavin T | 8924 | VL2 | 14168.5 |
| 40 | No | E. coli | Chloramph | Intracellula | F101 | Quinacrine | 7482 | VL2 | 13226.5 |
| 40 | No | E. coli | Chloramph | Intracellula | F138 | Apigenin | 9025 | VL2 | 7616 |
| 40 | No | E. coli | Chloramph | Intracellula | F65 | H2FDA | 8122 | BL1 | 5932.5 |
| 40 | No | E. coli | Chloramph | Intracellula | F142 | Chlorophe | 6937 | BL3 | 5217 |
| 40 | No | E. coli | Chloramph | Intracellula | F39 | Congo Red | 8262 | BL3 | 5344 |
| 40 | No | E. coli | Chloramph | Intracellula | F116 | Sunset Yel | 6662 | BL2 | 5923 |
| 40 | No | E. coli | Chloramph | Intracellula | F10 | Alizarin | 7110 | BL1 | 5755.5 |
| 40 | No | E. coli | Chloramph | Intracellula | F84 | Neutral Re | 8234 | BL1 | 5471.5 |
| 40 | No | E. coli | Chloramph | Intracellula | F99 | Pyronin Y | 5727 | BL1 | 6214 |
| 40 | No | E. coli | Chloramph | Intracellula | F108 | Rhodamine | 7685 | BL1 | 4923 |
| 40 | No | E. coli | Chloramph | Intracellula | F109 | Riboflavin | 7261 | BL1 | 4416 |
| 40 | No | E. coli | Chloramph | Intracellula | F117 | SYBR Gree | 8514 | BL1 | 43913 |
| 40 | No | E. coli | Chloramph | Intracellula | F106 | Rhodamine | 7591 | BL1 | 6515 |
| 40 | No | E. coli | Chloramph | Intracellula | F73 | Malachite ( | 6337 | BL1 | 4894 |
| 40 | No | E. coli | Chloramph | Intracellula | F7 | Acridine or | 8902 | BL1 | 5521.5 |
| 40 | No | E. coli | Chloramph | Intracellula | F192 | 5-Carboxyf | 6512 | BL1 | 9865.5 |
| 40 | No | E. coli | Chloramph | Intracellula | F57 | Eosin Y | 8552 | BL1 | 16141 |
| 40 | No | E. coli | Chloramph | Intracellula | F27 | Calcein | 6679 | BL1 | 15180 |
| 40 | No | E. coli | Chloramph | Intracellula | F12 | Amaranth | 8068 | BL1 | 17954.5 |
| 40 | No | E. coli | None | None | NA | Autofluore: | 8125 | NA | 17506 |
| 40 | No | E. coli | Colistin | Wall | F194 | Oxonol | 7366 | RL1 | 24740.5 |
| 40 | No | E. coli | Colistin | Wall | F11 | Allura Red | 3951 | BL3 | 33551 |
| 40 | No | E. coli | Colistin | Wall | F54 | DiSC3(5) | 2238 | RL1 | 42373.5 |
| 40 | No | E. coli | Colistin | Wall | F124 | Thioflavin T | 4261 | VL2 | 52329 |
| 40 | No | E. coli | Colistin | Wall | F101 | Quinacrine | 2960 | VL2 | 72821 |
| 40 | No | E. coli | Colistin | Wall | F138 | Apigenin | 3976 | VL2 | 77323.5 |
| 40 | No | E. coli | Colistin | Wall | F65 | H2FDA | 3380 | BL1 | 72528.5 |
| 40 | No | E. coli | Colistin | Wall | F142 | Chlorophe | 3454 | BL3 | 55193 |
| 40 | No | E. coli | Colistin | Wall | F39 | Congo Red | 2371 | BL3 | 56921 |
| 40 | No | E. coli | Colistin | Wall | F116 | Sunset Yel | 1898 | BL2 | 59189 |
| 40 | No | E. coli | Colistin | Wall | F10 | Alizarin | 1947 | BL1 | 52800 |
| 40 | No | E. coli | Colistin | Wall | F84 | Neutral Re | 2936 | BL1 | 56842 |
| 40 | No | E. coli | Colistin | Wall | F99 | Pyronin Y | 2673 | BL1 | 60773 |
| 40 | No | E. coli | Colistin | Wall | F108 | Rhodamine | 2858 | BL1 | 56369.5 |
| 40 | No | E. coli | Colistin | Wall | F109 | Riboflavin | 2136 | BL1 | 64294.5 |
| 40 | No | E. coli | Colistin | Wall | F117 | SYBR Gree | 2841 | BL1 | 2289023 |
| 40 | No | E. coli | Colistin | Wall | F106 | Rhodamine | 2959 | BL1 | 240656 |
| 40 | No | E. coli | Colistin | Wall | F73 | Malachite ( | 2405 | BL1 | 141245 |
| 40 | No | E. coli | Colistin | Wall | F7 | Acridine or | 2590 | BL1 | 134706.5 |
| 40 | No | E. coli | Colistin | Wall | F192 | 5-Carboxyf | 2860 | BL1 | 107355.5 |
| 40 | No | E. coli | Colistin | Wall | F57 | Eosin Y | 2837 | BL1 | 55962 |
| 40 | No | E. coli | Colistin | Wall | F27 | Calcein | 2337 | BL1 | 50796 |

| | | | | | | | | | |
|---|---|---|---|---|---|---|---|---|---|
| 40 | No | E. coli | Colistin | Wall | F12 | Amaranth | 2031 | BL1 | 30136 |
| 40 | No | E. coli | None | None | NA | Autofluore: | 3575 | NA | 20226 |
| 40 | No | E. coli | Gentamici | Intracellula | F194 | Oxonol | 8066 | RL1 | 19380 |
| 40 | No | E. coli | Gentamici | Intracellula | F11 | Allura Red | 7685 | BL3 | 19939 |
| 40 | No | E. coli | Gentamici | Intracellula | F54 | DiSC3(5) | 7150 | RL1 | 15547.5 |
| 40 | No | E. coli | Gentamici | Intracellula | F124 | Thioflavin T | 9044 | VL2 | 13074 |
| 40 | No | E. coli | Gentamici | Intracellula | F101 | Quinacrine | 8396 | VL2 | 12634.5 |
| 40 | No | E. coli | Gentamici | Intracellula | F138 | Apigenin | 9034 | VL2 | 8179 |
| 40 | No | E. coli | Gentamici | Intracellula | F65 | H2FDA | 8192 | BL1 | 6787 |
| 40 | No | E. coli | Gentamici | Intracellula | F142 | Chlorophe | 9901 | BL3 | 5427 |
| 40 | No | E. coli | Gentamici | Intracellula | F39 | Congo Red | 9786 | BL3 | 5638 |
| 40 | No | E. coli | Gentamici | Intracellula | F116 | Sunset Yel | 6548 | BL2 | 6715.5 |
| 40 | No | E. coli | Gentamici | Intracellula | F10 | Alizarin | 8333 | BL1 | 6696 |
| 40 | No | E. coli | Gentamici | Intracellula | F84 | Neutral Re | 8529 | BL1 | 6147 |
| 40 | No | E. coli | Gentamici | Intracellula | F99 | Pyronin Y | 8962 | BL1 | 5290 |
| 40 | No | E. coli | Gentamici | Intracellula | F108 | Rhodamine | 9051 | BL1 | 5563 |
| 40 | No | E. coli | Gentamici | Intracellula | F109 | Riboflavin | 7582 | BL1 | 4732 |
| 40 | No | E. coli | Gentamici | Intracellula | F117 | SYBR Gree | 7430 | BL1 | 118064 |
| 40 | No | E. coli | Gentamici | Intracellula | F106 | Rhodamine | 8234 | BL1 | 8704 |
| 40 | No | E. coli | Gentamici | Intracellula | F73 | Malachite ( | 9379 | BL1 | 5515 |
| 40 | No | E. coli | Gentamici | Intracellula | F7 | Acridine or | 8661 | BL1 | 6142 |
| 40 | No | E. coli | Gentamici | Intracellula | F192 | 5-Carboxyf | 8488 | BL1 | 11760.5 |
| 40 | No | E. coli | Gentamici | Intracellula | F57 | Eosin Y | 7683 | BL1 | 27899 |
| 40 | No | E. coli | Gentamici | Intracellula | F27 | Calcein | 8167 | BL1 | 24511 |
| 40 | No | E. coli | Gentamici | Intracellula | F12 | Amaranth | 8595 | BL1 | 19339 |
| 40 | No | E. coli | None | None | NA | Autofluore: | 5696 | NA | 20863 |
| 40 | No | E. coli | Nitrofurant | Intracellula | F194 | Oxonol | 6473 | RL1 | 16873 |
| 40 | No | E. coli | Nitrofurant | Intracellula | F11 | Allura Red | 6326 | BL3 | 19588 |
| 40 | No | E. coli | Nitrofurant | Intracellula | F54 | DiSC3(5) | 5476 | RL1 | 20196 |
| 40 | No | E. coli | Nitrofurant | Intracellula | F124 | Thioflavin T | 8426 | VL2 | 13408.5 |
| 40 | No | E. coli | Nitrofurant | Intracellula | F101 | Quinacrine | 6212 | VL2 | 13581 |
| 40 | No | E. coli | Nitrofurant | Intracellula | F138 | Apigenin | 6915 | VL2 | 10350 |
| 40 | No | E. coli | Nitrofurant | Intracellula | F65 | H2FDA | 7238 | BL1 | 6538 |
| 40 | No | E. coli | Nitrofurant | Intracellula | F142 | Chlorophe | 7592 | BL3 | 6116.5 |
| 40 | No | E. coli | Nitrofurant | Intracellula | F39 | Congo Red | 5042 | BL3 | 6712 |
| 40 | No | E. coli | Nitrofurant | Intracellula | F116 | Sunset Yel | 6332 | BL2 | 6021.5 |
| 40 | No | E. coli | Nitrofurant | Intracellula | F10 | Alizarin | 5885 | BL1 | 6554 |
| 40 | No | E. coli | Nitrofurant | Intracellula | F84 | Neutral Re | 5708 | BL1 | 6939 |
| 40 | No | E. coli | Nitrofurant | Intracellula | F99 | Pyronin Y | 6850 | BL1 | 5594.5 |
| 40 | No | E. coli | Nitrofurant | Intracellula | F108 | Rhodamine | 6401 | BL1 | 5744 |
| 40 | No | E. coli | Nitrofurant | Intracellula | F109 | Riboflavin | 5766 | BL1 | 5344 |
| 40 | No | E. coli | Nitrofurant | Intracellula | F117 | SYBR Gree | 5931 | BL1 | 51897 |
| 40 | No | E. coli | Nitrofurant | Intracellula | F106 | Rhodamine | 6028 | BL1 | 6741 |
| 40 | No | E. coli | Nitrofurant | Intracellula | F73 | Malachite ( | 5611 | BL1 | 5803 |
| 40 | No | E. coli | Nitrofurant | Intracellula | F7 | Acridine or | 5872 | BL1 | 5950.5 |
| 40 | No | E. coli | Nitrofurant | Intracellula | F192 | 5-Carboxyf | 7555 | BL1 | 10226 |
| 40 | No | E. coli | Nitrofurant | Intracellula | F57 | Eosin Y | 6129 | BL1 | 19383 |
| 40 | No | E. coli | Nitrofurant | Intracellula | F27 | Calcein | 5310 | BL1 | 19918 |

| | | | | | | | | | |
|---|---|---|---|---|---|---|---|---|---|
| 40 | No | E. coli | Nitrofurant | Intracellula | F12 | Amaranth | 7304 | BL1 | 20863 |
| 40 | No | E. coli | None | None | NA | Autofluores | 7399 | NA | 22042 |
| 40 | No | E. coli | Pentamidi | Wall | F194 | Oxonol | 6689 | RL1 | 18563 |
| 40 | No | E. coli | Pentamidi | Wall | F11 | Allura Red | 5656 | BL3 | 18544.5 |
| 40 | No | E. coli | Pentamidi | Wall | F54 | DiSC3(5) | 4421 | RL1 | 17818 |
| 40 | No | E. coli | Pentamidi | Wall | F124 | Thioflavin T | 6372 | VL2 | 18568.5 |
| 40 | No | E. coli | Pentamidi | Wall | F101 | Quinacrine | 7062 | VL2 | 19912.5 |
| 40 | No | E. coli | Pentamidi | Wall | F138 | Apigenin | 6855 | VL2 | 14522 |
| 40 | No | E. coli | Pentamidi | Wall | F65 | H2FDA | 6199 | BL1 | 11552 |
| 40 | No | E. coli | Pentamidi | Wall | F142 | Chlorophe | 6240 | BL3 | 10989.5 |
| 40 | No | E. coli | Pentamidi | Wall | F39 | Congo Red | 7168 | BL3 | 11142 |
| 40 | No | E. coli | Pentamidi | Wall | F116 | Sunset Yel | 6053 | BL2 | 10321 |
| 40 | No | E. coli | Pentamidi | Wall | F10 | Alizarin | 5221 | BL1 | 10673 |
| 40 | No | E. coli | Pentamidi | Wall | F84 | Neutral Re | 5167 | BL1 | 8035 |
| 40 | No | E. coli | Pentamidi | Wall | F99 | Pyronin Y | 5886 | BL1 | 6690.5 |
| 40 | No | E. coli | Pentamidi | Wall | F108 | Rhodamine | 5545 | BL1 | 6283 |
| 40 | No | E. coli | Pentamidi | Wall | F109 | Riboflavin | 6371 | BL1 | 5744 |
| 40 | No | E. coli | Pentamidi | Wall | F117 | SYBR Gree | 6194 | BL1 | 311169.5 |
| 40 | No | E. coli | Pentamidi | Wall | F106 | Rhodamine | 7448 | BL1 | 8824.5 |
| 40 | No | E. coli | Pentamidi | Wall | F73 | Malachite ( | 5554 | BL1 | 6134.5 |
| 40 | No | E. coli | Pentamidi | Wall | F7 | Acridine or | 6170 | BL1 | 5215.5 |
| 40 | No | E. coli | Pentamidi | Wall | F192 | 5-Carboxyf | 5288 | BL1 | 10515 |
| 40 | No | E. coli | Pentamidi | Wall | F57 | Eosin Y | 6461 | BL1 | 15281 |
| 40 | No | E. coli | Pentamidi | Wall | F27 | Calcein | 6965 | BL1 | 18135 |
| 40 | No | E. coli | Pentamidi | Wall | F12 | Amaranth | 8495 | BL1 | 17984 |
| 40 | No | E. coli | None | None | NA | Autofluores | 5671 | NA | 15942 |
| 40 | No | E. coli | Rifampicin | Intracellula | F194 | Oxonol | 6213 | RL1 | 14592 |
| 40 | No | E. coli | Rifampicin | Intracellula | F11 | Allura Red | 7839 | BL3 | 22179 |
| 40 | No | E. coli | Rifampicin | Intracellula | F54 | DiSC3(5) | 7025 | RL1 | 17228 |
| 40 | No | E. coli | Rifampicin | Intracellula | F124 | Thioflavin T | 8714 | VL2 | 13095.5 |
| 40 | No | E. coli | Rifampicin | Intracellula | F101 | Quinacrine | 11214 | VL2 | 13277.5 |
| 40 | No | E. coli | Rifampicin | Intracellula | F138 | Apigenin | 9519 | VL2 | 8653 |
| 40 | No | E. coli | Rifampicin | Intracellula | F65 | H2FDA | 9657 | BL1 | 5893 |
| 40 | No | E. coli | Rifampicin | Intracellula | F142 | Chlorophe | 9542 | BL3 | 4710 |
| 40 | No | E. coli | Rifampicin | Intracellula | F39 | Congo Red | 8659 | BL3 | 5304 |
| 40 | No | E. coli | Rifampicin | Intracellula | F116 | Sunset Yel | 8469 | BL2 | 4906 |
| 40 | No | E. coli | Rifampicin | Intracellula | F10 | Alizarin | 8153 | BL1 | 5855 |
| 40 | No | E. coli | Rifampicin | Intracellula | F84 | Neutral Re | 5854 | BL1 | 6043.5 |
| 40 | No | E. coli | Rifampicin | Intracellula | F99 | Pyronin Y | 7305 | BL1 | 5544 |
| 40 | No | E. coli | Rifampicin | Intracellula | F108 | Rhodamine | 8603 | BL1 | 5027 |
| 40 | No | E. coli | Rifampicin | Intracellula | F109 | Riboflavin | 8519 | BL1 | 5836 |
| 40 | No | E. coli | Rifampicin | Intracellula | F117 | SYBR Gree | 9324 | BL1 | 62337 |
| 40 | No | E. coli | Rifampicin | Intracellula | F106 | Rhodamine | 7365 | BL1 | 6496 |
| 40 | No | E. coli | Rifampicin | Intracellula | F73 | Malachite ( | 7584 | BL1 | 5297 |
| 40 | No | E. coli | Rifampicin | Intracellula | F7 | Acridine or | 8000 | BL1 | 5316.5 |
| 40 | No | E. coli | Rifampicin | Intracellula | F192 | 5-Carboxyf | 8735 | BL1 | 12674 |
| 40 | No | E. coli | Rifampicin | Intracellula | F57 | Eosin Y | 7319 | BL1 | 17793 |
| 40 | No | E. coli | Rifampicin | Intracellula | F27 | Calcein | 8663 | BL1 | 21607 |

| | | | | | | | | | |
|---|---|---|---|---|---|---|---|---|---|
| 40 | No | E. coli | Rifampicin | Intracellul | F12 | Amaranth | 8558 | BL1 | 17035 |
| 40 | No | E. coli | None | None | NA | Autofluore: | 7798 | NA | 17089.5 |
| 40 | No | E. coli | Trimethopr | Intracellul | F194 | Oxonol | 5209 | RL1 | 13701 |
| 40 | No | E. coli | Trimethopr | Intracellul | F11 | Allura Red | 7101 | BL3 | 18076 |
| 40 | No | E. coli | Trimethopr | Intracellul | F54 | DiSC3(5) | 8675 | RL1 | 20033 |
| 40 | No | E. coli | Trimethopr | Intracellul | F124 | Thioflavin T | 7810 | VL2 | 13143.5 |
| 40 | No | E. coli | Trimethopr | Intracellul | F101 | Quinacrine | 7140 | VL2 | 10947.5 |
| 40 | No | E. coli | Trimethopr | Intracellul | F138 | Apigenin | 7968 | VL2 | 7309.5 |
| 40 | No | E. coli | Trimethopr | Intracellul | F65 | H2FDA | 8654 | BL1 | 6655 |
| 40 | No | E. coli | Trimethopr | Intracellul | F142 | Chlorophe | 7669 | BL3 | 6385 |
| 40 | No | E. coli | Trimethopr | Intracellul | F39 | Congo Red | 8185 | BL3 | 5527 |
| 40 | No | E. coli | Trimethopr | Intracellul | F116 | Sunset Yel | 6655 | BL2 | 5983 |
| 40 | No | E. coli | Trimethopr | Intracellul | F10 | Alizarin | 8907 | BL1 | 6786 |
| 40 | No | E. coli | Trimethopr | Intracellul | F84 | Neutral Re | 7578 | BL1 | 6857 |
| 40 | No | E. coli | Trimethopr | Intracellul | F99 | Pyronin Y | 5726 | BL1 | 6251.5 |
| 40 | No | E. coli | Trimethopr | Intracellul | F108 | Rhodamine | 8377 | BL1 | 5247 |
| 40 | No | E. coli | Trimethopr | Intracellul | F109 | Riboflavin | 6221 | BL1 | 4790 |
| 40 | No | E. coli | Trimethopr | Intracellul | F117 | SYBR Gree | 8057 | BL1 | 71960 |
| 40 | No | E. coli | Trimethopr | Intracellul | F106 | Rhodamine | 7116 | BL1 | 7807 |
| 40 | No | E. coli | Trimethopr | Intracellul | F73 | Malachite ( | 7767 | BL1 | 5873 |
| 40 | No | E. coli | Trimethopr | Intracellul | F7 | Acridine or | 8568 | BL1 | 5419.5 |
| 40 | No | E. coli | Trimethopr | Intracellul | F192 | 5-Carboxyf | 7202 | BL1 | 10296.5 |
| 40 | No | E. coli | Trimethopr | Intracellul | F57 | Eosin Y | 7061 | BL1 | 13341 |
| 40 | No | E. coli | Trimethopr | Intracellul | F27 | Calcein | 9145 | BL1 | 20442 |
| 40 | No | E. coli | Trimethopr | Intracellul | F12 | Amaranth | 7912 | BL1 | 15229.5 |
| 40 | No | E. coli | None | None | NA | Autofluore: | 7699 | NA | 959 |
| 40 | No | E. coli | None | None | F194 | Oxonol | 6417 | RL1 | 18681 |
| 40 | No | E. coli | None | None | F11 | Allura Red | 10527 | BL3 | 17231 |
| 40 | No | E. coli | None | None | F54 | DiSC3(5) | 8816 | RL1 | 9793 |
| 40 | No | E. coli | None | None | F124 | Thioflavin T | 9548 | VL2 | 8263 |
| 40 | No | E. coli | None | None | F101 | Quinacrine | 7438 | VL2 | 6609.5 |
| 40 | No | E. coli | None | None | F138 | Apigenin | 10417 | VL2 | 5295 |
| 40 | No | E. coli | None | None | F65 | H2FDA | 9051 | BL1 | 4725 |
| 40 | No | E. coli | None | None | F142 | Chlorophe | 8875 | BL3 | 4316 |
| 40 | No | E. coli | None | None | F39 | Congo Red | 8660 | BL3 | 4303 |
| 40 | No | E. coli | None | None | F116 | Sunset Yel | 8813 | BL2 | 4732 |
| 40 | No | E. coli | None | None | F10 | Alizarin | 9578 | BL1 | 4645 |
| 40 | No | E. coli | None | None | F84 | Neutral Re | 7961 | BL1 | 4940 |
| 40 | No | E. coli | None | None | F99 | Pyronin Y | 8145 | BL1 | 4588 |
| 40 | No | E. coli | None | None | F108 | Rhodamine | 10424 | BL1 | 4146 |
| 40 | No | E. coli | None | None | F109 | Riboflavin | 7116 | BL1 | 3840 |
| 40 | No | E. coli | None | None | F117 | SYBR Gree | 8177 | BL1 | 107541 |
| 40 | No | E. coli | None | None | F106 | Rhodamine | 9062 | BL1 | 5626 |
| 40 | No | E. coli | None | None | F73 | Malachite ( | 9873 | BL1 | 4424 |
| 40 | No | E. coli | None | None | F7 | Acridine or | 9755 | BL1 | 4193 |
| 40 | No | E. coli | None | None | F192 | 5-Carboxyf | 9212 | BL1 | 12061 |
| 40 | No | E. coli | None | None | F57 | Eosin Y | 7118 | BL1 | 16365.5 |
| 40 | No | E. coli | None | None | F27 | Calcein | 9145 | BL1 | 15889 |

| | | | | | | | | | |
|---|---|---|---|---|---|---|---|---|---|
| 40 | No | E. coli | None | None | F12 | Amaranth | 8557 | BL1 | 19182 |
| 40 | No | E. coli | None | None | NA | Autofluore: | 7829 | NA | 15515 |
| 40 | No | E. coli | Amoxacilli | Wall | F194 | Oxonol | 8900 | RL1 | 15685.5 |
| 40 | No | E. coli | Amoxacilli | Wall | F11 | Allura Red | 9033 | BL3 | 19484 |
| 40 | No | E. coli | Amoxacilli | Wall | F54 | DiSC3(5) | 6328 | RL1 | 15372 |
| 40 | No | E. coli | Amoxacilli | Wall | F124 | Thioflavin T | 8602 | VL2 | 13541 |
| 40 | No | E. coli | Amoxacilli | Wall | F101 | Quinacrine | 10347 | VL2 | 12505 |
| 40 | No | E. coli | Amoxacilli | Wall | F138 | Apigenin | 8162 | VL2 | 9474 |
| 40 | No | E. coli | Amoxacilli | Wall | F65 | H2FDA | 9487 | BL1 | 6312 |
| 40 | No | E. coli | Amoxacilli | Wall | F142 | Chlorophe | 9609 | BL3 | 5626 |
| 40 | No | E. coli | Amoxacilli | Wall | F39 | Congo Red | 8154 | BL3 | 5130 |
| 40 | No | E. coli | Amoxacilli | Wall | F116 | Sunset Yel | 10155 | BL2 | 5640 |
| 40 | No | E. coli | Amoxacilli | Wall | F10 | Alizarin | 8118 | BL1 | 5654 |
| 40 | No | E. coli | Amoxacilli | Wall | F84 | Neutral Re | 9370 | BL1 | 5403.5 |
| 40 | No | E. coli | Amoxacilli | Wall | F99 | Pyronin Y | 8301 | BL1 | 5099 |
| 40 | No | E. coli | Amoxacilli | Wall | F108 | Rhodamine | 9645 | BL1 | 4722 |
| 40 | No | E. coli | Amoxacilli | Wall | F109 | Riboflavin | 9086 | BL1 | 4981 |
| 40 | No | E. coli | Amoxacilli | Wall | F117 | SYBR Gree | 8906 | BL1 | 146437.5 |
| 40 | No | E. coli | Amoxacilli | Wall | F106 | Rhodamine | 8784 | BL1 | 5344 |
| 40 | No | E. coli | Amoxacilli | Wall | F73 | Malachite ( | 7564 | BL1 | 5142.5 |
| 40 | No | E. coli | Amoxacilli | Wall | F7 | Acridine or | 9029 | BL1 | 4492 |
| 40 | No | E. coli | Amoxacilli | Wall | F192 | 5-Carboxyf | 10434 | BL1 | 11855.5 |
| 40 | No | E. coli | Amoxacilli | Wall | F57 | Eosin Y | 6642 | BL1 | 16117.5 |
| 40 | No | E. coli | Amoxacilli | Wall | F27 | Calcein | 8042 | BL1 | 16656 |
| 40 | No | E. coli | Amoxacilli | Wall | F12 | Amaranth | 9679 | BL1 | 20693 |
| 40 | No | E. coli | None | None | NA | Autofluore: | 9262 | NA | 17231 |
| 40 | No | E. coli | Azithromyc | Intracellula | F194 | Oxonol | 5200 | RL1 | 19377.5 |
| 40 | No | E. coli | Azithromyc | Intracellula | F11 | Allura Red | 6003 | BL3 | 21030 |
| 40 | No | E. coli | Azithromyc | Intracellula | F54 | DiSC3(5) | 4334 | RL1 | 17941.5 |
| 40 | No | E. coli | Azithromyc | Intracellula | F124 | Thioflavin T | 5552 | VL2 | 14108 |
| 40 | No | E. coli | Azithromyc | Intracellula | F101 | Quinacrine | 5210 | VL2 | 12734 |
| 40 | No | E. coli | Azithromyc | Intracellula | F138 | Apigenin | 5755 | VL2 | 9611 |
| 40 | No | E. coli | Azithromyc | Intracellula | F65 | H2FDA | 5795 | BL1 | 7046 |
| 40 | No | E. coli | Azithromyc | Intracellula | F142 | Chlorophe | 5062 | BL3 | 5905 |
| 40 | No | E. coli | Azithromyc | Intracellula | F39 | Congo Red | 5700 | BL3 | 6617.5 |
| 40 | No | E. coli | Azithromyc | Intracellula | F116 | Sunset Yel | 4012 | BL2 | 7791.5 |
| 40 | No | E. coli | Azithromyc | Intracellula | F10 | Alizarin | 4955 | BL1 | 6874 |
| 40 | No | E. coli | Azithromyc | Intracellula | F84 | Neutral Re | 4753 | BL1 | 6867 |
| 40 | No | E. coli | Azithromyc | Intracellula | F99 | Pyronin Y | 5127 | BL1 | 6263 |
| 40 | No | E. coli | Azithromyc | Intracellula | F108 | Rhodamine | 4811 | BL1 | 5426 |
| 40 | No | E. coli | Azithromyc | Intracellula | F109 | Riboflavin | 4270 | BL1 | 5379 |
| 40 | No | E. coli | Azithromyc | Intracellula | F117 | SYBR Gree | 4436 | BL1 | 38729.5 |
| 40 | No | E. coli | Azithromyc | Intracellula | F106 | Rhodamine | 5075 | BL1 | 7482 |
| 40 | No | E. coli | Azithromyc | Intracellula | F73 | Malachite ( | 4928 | BL1 | 6052 |
| 40 | No | E. coli | Azithromyc | Intracellula | F7 | Acridine or | 4593 | BL1 | 5514 |
| 40 | No | E. coli | Azithromyc | Intracellula | F192 | 5-Carboxyf | 4664 | BL1 | 9933.5 |
| 40 | No | E. coli | Azithromyc | Intracellula | F57 | Eosin Y | 5289 | BL1 | 18143 |
| 40 | No | E. coli | Azithromyc | Intracellula | F27 | Calcein | 4456 | BL1 | 18663 |

| | | | | | | | | | |
|---|---|---|---|---|---|---|---|---|---|
| 40 | No | E. coli | Azithromyc | Intracellula | F12 | Amaranth | 5033 | BL1 | 19058 |
| 40 | No | E. coli | None | None | NA | Autofluore: | 6225 | NA | 20196 |
| 40 | No | E. coli | Trimethopr | Intracellula | F194 | Oxonol | 5999 | RL1 | 17161 |
| 40 | No | E. coli | Trimethopr | Intracellula | F11 | Allura Red | 7236 | BL3 | 17710 |
| 40 | No | E. coli | Trimethopr | Intracellula | F54 | DiSC3(5) | 6731 | RL1 | 17175 |
| 40 | No | E. coli | Trimethopr | Intracellula | F124 | Thioflavin T | 7351 | VL2 | 13897 |
| 40 | No | E. coli | Trimethopr | Intracellula | F101 | Quinacrine | 6607 | VL2 | 11885 |
| 40 | No | E. coli | Trimethopr | Intracellula | F138 | Apigenin | 7039 | VL2 | 10354 |
| 40 | No | E. coli | Trimethopr | Intracellula | F65 | H2FDA | 7573 | BL1 | 6689 |
| 40 | No | E. coli | Trimethopr | Intracellula | F142 | Chlorophe | 8352 | BL3 | 5781 |
| 40 | No | E. coli | Trimethopr | Intracellula | F39 | Congo Red | 6536 | BL3 | 5688 |
| 40 | No | E. coli | Trimethopr | Intracellula | F116 | Sunset Yel | 7131 | BL2 | 6431 |
| 40 | No | E. coli | Trimethopr | Intracellula | F10 | Alizarin | 7266 | BL1 | 6142 |
| 40 | No | E. coli | Trimethopr | Intracellula | F84 | Neutral Re | 6664 | BL1 | 6292 |
| 40 | No | E. coli | Trimethopr | Intracellula | F99 | Pyronin Y | 7734 | BL1 | 5759.5 |
| 40 | No | E. coli | Trimethopr | Intracellula | F108 | Rhodamine | 8200 | BL1 | 5240.5 |
| 40 | No | E. coli | Trimethopr | Intracellula | F109 | Riboflavin | 5800 | BL1 | 5503 |
| 40 | No | E. coli | Trimethopr | Intracellula | F117 | SYBR Gree | 8741 | BL1 | 105011 |
| 40 | No | E. coli | Trimethopr | Intracellula | F106 | Rhodamine | 5620 | BL1 | 6126 |
| 40 | No | E. coli | Trimethopr | Intracellula | F73 | Malachite ( | 8678 | BL1 | 5873.5 |
| 40 | No | E. coli | Trimethopr | Intracellula | F7 | Acridine or | 8078 | BL1 | 5515.5 |
| 40 | No | E. coli | Trimethopr | Intracellula | F192 | 5-Carboxyf | 7536 | BL1 | 11979.5 |
| 40 | No | E. coli | Trimethopr | Intracellula | F57 | Eosin Y | 5879 | BL1 | 14388 |
| 40 | No | E. coli | Trimethopr | Intracellula | F27 | Calcein | 6871 | BL1 | 15895 |
| 40 | No | E. coli | Trimethopr | Intracellula | F12 | Amaranth | 6845 | BL1 | 14812 |
| 40 | No | E. coli | None | None | NA | Autofluore: | 7159 | NA | 17667 |
| 40 | No | E. coli | Chloramph | Intracellula | F194 | Oxonol | 6381 | RL1 | 21476 |
| 40 | No | E. coli | Chloramph | Intracellula | F11 | Allura Red | 6562 | BL3 | 15362 |
| 40 | No | E. coli | Chloramph | Intracellula | F54 | DiSC3(5) | 6574 | RL1 | 17675 |
| 40 | No | E. coli | Chloramph | Intracellula | F124 | Thioflavin T | 6491 | VL2 | 13735 |
| 40 | No | E. coli | Chloramph | Intracellula | F101 | Quinacrine | 7464 | VL2 | 11003 |
| 40 | No | E. coli | Chloramph | Intracellula | F138 | Apigenin | 8051 | VL2 | 9767 |
| 40 | No | E. coli | Chloramph | Intracellula | F65 | H2FDA | 6230 | BL1 | 7330 |
| 40 | No | E. coli | Chloramph | Intracellula | F142 | Chlorophe | 6708 | BL3 | 5747 |
| 40 | No | E. coli | Chloramph | Intracellula | F39 | Congo Red | 6913 | BL3 | 5813 |
| 40 | No | E. coli | Chloramph | Intracellula | F116 | Sunset Yel | 5790 | BL2 | 6768.5 |
| 40 | No | E. coli | Chloramph | Intracellula | F10 | Alizarin | 7613 | BL1 | 6619 |
| 40 | No | E. coli | Chloramph | Intracellula | F84 | Neutral Re | 8087 | BL1 | 6603 |
| 40 | No | E. coli | Chloramph | Intracellula | F99 | Pyronin Y | 6115 | BL1 | 6066 |
| 40 | No | E. coli | Chloramph | Intracellula | F108 | Rhodamine | 7306 | BL1 | 5542 |
| 40 | No | E. coli | Chloramph | Intracellula | F109 | Riboflavin | 7018 | BL1 | 5140 |
| 40 | No | E. coli | Chloramph | Intracellula | F117 | SYBR Gree | 7347 | BL1 | 75943 |
| 40 | No | E. coli | Chloramph | Intracellula | F106 | Rhodamine | 5809 | BL1 | 6341 |
| 40 | No | E. coli | Chloramph | Intracellula | F73 | Malachite ( | 6744 | BL1 | 5530 |
| 40 | No | E. coli | Chloramph | Intracellula | F7 | Acridine or | 6522 | BL1 | 5350.5 |
| 40 | No | E. coli | Chloramph | Intracellula | F192 | 5-Carboxyf | 6606 | BL1 | 10932.5 |
| 40 | No | E. coli | Chloramph | Intracellula | F57 | Eosin Y | 6637 | BL1 | 15628 |
| 40 | No | E. coli | Chloramph | Intracellula | F27 | Calcein | 6871 | BL1 | 17819 |

| | | | | | | | | | |
|---|---|---|---|---|---|---|---|---|---|
| 40 | No | E. coli | Chloramph | Intracellula | F12 | Amaranth | 7297 | BL1 | 20439 |
| 40 | No | E. coli | None | None | NA | Autofluores | 8195 | NA | 14853 |
| 40 | No | E. coli | Colistin | Wall | F194 | Oxonol | 3909 | RL1 | 46029 |
| 40 | No | E. coli | Colistin | Wall | F11 | Allura Red | 2276 | BL3 | 55806.5 |
| 40 | No | E. coli | Colistin | Wall | F54 | DiSC3(5) | 1869 | RL1 | 52254 |
| 40 | No | E. coli | Colistin | Wall | F124 | Thioflavin T | 2290 | VL2 | 35486.5 |
| 40 | No | E. coli | Colistin | Wall | F101 | Quinacrine | 2578 | VL2 | 45841 |
| 40 | No | E. coli | Colistin | Wall | F138 | Apigenin | 3842 | VL2 | 33733 |
| 40 | No | E. coli | Colistin | Wall | F65 | H2FDA | 2527 | BL1 | 32882 |
| 40 | No | E. coli | Colistin | Wall | F142 | Chlorophe | 3669 | BL3 | 28321 |
| 40 | No | E. coli | Colistin | Wall | F39 | Congo Red | 2618 | BL3 | 22968.5 |
| 40 | No | E. coli | Colistin | Wall | F116 | Sunset Yel | 2828 | BL2 | 26064 |
| 40 | No | E. coli | Colistin | Wall | F10 | Alizarin | 2113 | BL1 | 22575 |
| 40 | No | E. coli | Colistin | Wall | F84 | Neutral Re | 3219 | BL1 | 23334 |
| 40 | No | E. coli | Colistin | Wall | F99 | Pyronin Y | 1519 | BL1 | 22993 |
| 40 | No | E. coli | Colistin | Wall | F108 | Rhodamine | 2068 | BL1 | 24354 |
| 40 | No | E. coli | Colistin | Wall | F109 | Riboflavin | 1667 | BL1 | 26583 |
| 40 | No | E. coli | Colistin | Wall | F117 | SYBR Gree | 2172 | BL1 | 2804762 |
| 40 | No | E. coli | Colistin | Wall | F106 | Rhodamine | 1726 | BL1 | 246430 |
| 40 | No | E. coli | Colistin | Wall | F73 | Malachite ( | 2279 | BL1 | 148426 |
| 40 | No | E. coli | Colistin | Wall | F7 | Acridine or | 1531 | BL1 | 137876 |
| 40 | No | E. coli | Colistin | Wall | F192 | 5-Carboxyf | 2140 | BL1 | 69874.5 |
| 40 | No | E. coli | Colistin | Wall | F57 | Eosin Y | 1970 | BL1 | 37328 |
| 40 | No | E. coli | Colistin | Wall | F27 | Calcein | 1794 | BL1 | 24486 |
| 40 | No | E. coli | Colistin | Wall | F12 | Amaranth | 1572 | BL1 | 21047.5 |
| 40 | No | E. coli | None | None | NA | Autofluores | 4306 | NA | 18403.5 |
| 40 | No | E. coli | Gentamici | Intracellula | F194 | Oxonol | 7602 | RL1 | 18638.5 |
| 40 | No | E. coli | Gentamici | Intracellula | F11 | Allura Red | 6020 | BL3 | 18115 |
| 40 | No | E. coli | Gentamici | Intracellula | F54 | DiSC3(5) | 6666 | RL1 | 15170.5 |
| 40 | No | E. coli | Gentamici | Intracellula | F124 | Thioflavin T | 9868 | VL2 | 13588.5 |
| 40 | No | E. coli | Gentamici | Intracellula | F101 | Quinacrine | 7836 | VL2 | 11751.5 |
| 40 | No | E. coli | Gentamici | Intracellula | F138 | Apigenin | 7580 | VL2 | 10614.5 |
| 40 | No | E. coli | Gentamici | Intracellula | F65 | H2FDA | 8536 | BL1 | 6474.5 |
| 40 | No | E. coli | Gentamici | Intracellula | F142 | Chlorophe | 7085 | BL3 | 5704 |
| 40 | No | E. coli | Gentamici | Intracellula | F39 | Congo Red | 9041 | BL3 | 5521 |
| 40 | No | E. coli | Gentamici | Intracellula | F116 | Sunset Yel | 8568 | BL2 | 6088 |
| 40 | No | E. coli | Gentamici | Intracellula | F10 | Alizarin | 7033 | BL1 | 5848 |
| 40 | No | E. coli | Gentamici | Intracellula | F84 | Neutral Re | 9251 | BL1 | 6114 |
| 40 | No | E. coli | Gentamici | Intracellula | F99 | Pyronin Y | 6975 | BL1 | 5718 |
| 40 | No | E. coli | Gentamici | Intracellula | F108 | Rhodamine | 9198 | BL1 | 5116 |
| 40 | No | E. coli | Gentamici | Intracellula | F109 | Riboflavin | 8112 | BL1 | 5102 |
| 40 | No | E. coli | Gentamici | Intracellula | F117 | SYBR Gree | 6348 | BL1 | 149608 |
| 40 | No | E. coli | Gentamici | Intracellula | F106 | Rhodamine | 8453 | BL1 | 6896 |
| 40 | No | E. coli | Gentamici | Intracellula | F73 | Malachite ( | 8226 | BL1 | 5720.5 |
| 40 | No | E. coli | Gentamici | Intracellula | F7 | Acridine or | 9194 | BL1 | 5359.5 |
| 40 | No | E. coli | Gentamici | Intracellula | F192 | 5-Carboxyf | 7613 | BL1 | 10591 |
| 40 | No | E. coli | Gentamici | Intracellula | F57 | Eosin Y | 9225 | BL1 | 17482 |
| 40 | No | E. coli | Gentamici | Intracellula | F27 | Calcein | 7547 | BL1 | 15542 |

| | | | | | | | | | |
|---|---|---|---|---|---|---|---|---|---|
| 40 | No | E. coli | Gentamici | Intracellul | F12 | Amaranth | 7975 | BL1 | 17008 |
| 40 | No | E. coli | None | None | NA | Autofluore: | 6746 | NA | 14635 |
| 40 | No | E. coli | Nitrofurant | Intracellul | F194 | Oxonol | 7212 | RL1 | 18587.5 |
| 40 | No | E. coli | Nitrofurant | Intracellul | F11 | Allura Red | 7052 | BL3 | 16549.5 |
| 40 | No | E. coli | Nitrofurant | Intracellul | F54 | DiSC3(5) | 5649 | RL1 | 17993 |
| 40 | No | E. coli | Nitrofurant | Intracellul | F124 | Thioflavin T | 5121 | VL2 | 15875 |
| 40 | No | E. coli | Nitrofurant | Intracellul | F101 | Quinacrine | 6162 | VL2 | 12513.5 |
| 40 | No | E. coli | Nitrofurant | Intracellul | F138 | Apigenin | 7607 | VL2 | 11387 |
| 40 | No | E. coli | Nitrofurant | Intracellul | F65 | H2FDA | 4633 | BL1 | 8853 |
| 40 | No | E. coli | Nitrofurant | Intracellul | F142 | Chlorophe | 5849 | BL3 | 6579 |
| 40 | No | E. coli | Nitrofurant | Intracellul | F39 | Congo Red | 6255 | BL3 | 6991 |
| 40 | No | E. coli | Nitrofurant | Intracellul | F116 | Sunset Yel | 6605 | BL2 | 6226 |
| 40 | No | E. coli | Nitrofurant | Intracellul | F10 | Alizarin | 4770 | BL1 | 6650 |
| 40 | No | E. coli | Nitrofurant | Intracellul | F84 | Neutral Re | 6931 | BL1 | 6282 |
| 40 | No | E. coli | Nitrofurant | Intracellul | F99 | Pyronin Y | 3945 | BL1 | 6200 |
| 40 | No | E. coli | Nitrofurant | Intracellul | F108 | Rhodamine | 7252 | BL1 | 5531 |
| 40 | No | E. coli | Nitrofurant | Intracellul | F109 | Riboflavin | 4317 | BL1 | 5001 |
| 40 | No | E. coli | Nitrofurant | Intracellul | F117 | SYBR Gree | 6588 | BL1 | 62795 |
| 40 | No | E. coli | Nitrofurant | Intracellul | F106 | Rhodamine | 5803 | BL1 | 6466 |
| 40 | No | E. coli | Nitrofurant | Intracellul | F73 | Malachite ( | 5021 | BL1 | 5865 |
| 40 | No | E. coli | Nitrofurant | Intracellul | F7 | Acridine or | 5077 | BL1 | 5484 |
| 40 | No | E. coli | Nitrofurant | Intracellul | F192 | 5-Carboxyf | 4971 | BL1 | 8835 |
| 40 | No | E. coli | Nitrofurant | Intracellul | F57 | Eosin Y | 4996 | BL1 | 16204.5 |
| 40 | No | E. coli | Nitrofurant | Intracellul | F27 | Calcein | 5881 | BL1 | 17342 |
| 40 | No | E. coli | Nitrofurant | Intracellul | F12 | Amaranth | 5309 | BL1 | 17445 |
| 40 | No | E. coli | None | None | NA | Autofluore: | 6552 | NA | 15449.5 |
| 40 | No | E. coli | Pentamidi | Wall | F194 | Oxonol | 7587 | RL1 | 15718 |
| 40 | No | E. coli | Pentamidi | Wall | F11 | Allura Red | 6713 | BL3 | 16715 |
| 40 | No | E. coli | Pentamidi | Wall | F54 | DiSC3(5) | 4328 | RL1 | 15470 |
| 40 | No | E. coli | Pentamidi | Wall | F124 | Thioflavin T | 6040 | VL2 | 15791 |
| 40 | No | E. coli | Pentamidi | Wall | F101 | Quinacrine | 5134 | VL2 | 14388 |
| 40 | No | E. coli | Pentamidi | Wall | F138 | Apigenin | 5609 | VL2 | 12572 |
| 40 | No | E. coli | Pentamidi | Wall | F65 | H2FDA | 6258 | BL1 | 9973 |
| 40 | No | E. coli | Pentamidi | Wall | F142 | Chlorophe | 5511 | BL3 | 9133 |
| 40 | No | E. coli | Pentamidi | Wall | F39 | Congo Red | 6411 | BL3 | 8372 |
| 40 | No | E. coli | Pentamidi | Wall | F116 | Sunset Yel | 5638 | BL2 | 8691 |
| 40 | No | E. coli | Pentamidi | Wall | F10 | Alizarin | 5860 | BL1 | 7285 |
| 40 | No | E. coli | Pentamidi | Wall | F84 | Neutral Re | 5313 | BL1 | 7756 |
| 40 | No | E. coli | Pentamidi | Wall | F99 | Pyronin Y | 6362 | BL1 | 7093 |
| 40 | No | E. coli | Pentamidi | Wall | F108 | Rhodamine | 6313 | BL1 | 7047 |
| 40 | No | E. coli | Pentamidi | Wall | F109 | Riboflavin | 5044 | BL1 | 6370.5 |
| 40 | No | E. coli | Pentamidi | Wall | F117 | SYBR Gree | 5398 | BL1 | 638948.5 |
| 40 | No | E. coli | Pentamidi | Wall | F106 | Rhodamine | 5447 | BL1 | 8597 |
| 40 | No | E. coli | Pentamidi | Wall | F73 | Malachite ( | 5914 | BL1 | 6922 |
| 40 | No | E. coli | Pentamidi | Wall | F7 | Acridine or | 7450 | BL1 | 6160 |
| 40 | No | E. coli | Pentamidi | Wall | F192 | 5-Carboxyf | 6111 | BL1 | 11686 |
| 40 | No | E. coli | Pentamidi | Wall | F57 | Eosin Y | 6129 | BL1 | 16549 |
| 40 | No | E. coli | Pentamidi | Wall | F27 | Calcein | 8218 | BL1 | 20962.5 |

| | | | | | | | | | |
|---|---|---|---|---|---|---|---|---|---|
| 40 | No | E. coli | Pentamidi | Wall | F12 | Amaranth | 6945 | BL1 | 18815 |
| 40 | No | E. coli | None | None | NA | Autofluore | 6799 | NA | 20933 |
| 40 | No | E. coli | Rifampicin | Intracellul | F194 | Oxonol | 6090 | RL1 | 16826 |
| 40 | No | E. coli | Rifampicin | Intracellul | F11 | Allura Red | 6845 | BL3 | 15014 |
| 40 | No | E. coli | Rifampicin | Intracellul | F54 | DiSC3(5) | 6580 | RL1 | 17676.5 |
| 40 | No | E. coli | Rifampicin | Intracellul | F124 | Thioflavin T | 10544 | VL2 | 13045.5 |
| 40 | No | E. coli | Rifampicin | Intracellul | F101 | Quinacrine | 9508 | VL2 | 12577 |
| 40 | No | E. coli | Rifampicin | Intracellul | F138 | Apigenin | 7423 | VL2 | 10730 |
| 40 | No | E. coli | Rifampicin | Intracellul | F65 | H2FDA | 10642 | BL1 | 6628.5 |
| 40 | No | E. coli | Rifampicin | Intracellul | F142 | Chlorophe | 9333 | BL3 | 6171 |
| 40 | No | E. coli | Rifampicin | Intracellul | F39 | Congo Red | 6361 | BL3 | 5612 |
| 40 | No | E. coli | Rifampicin | Intracellul | F116 | Sunset Yel | 8994 | BL2 | 6103.5 |
| 40 | No | E. coli | Rifampicin | Intracellul | F10 | Alizarin | 9684 | BL1 | 6223.5 |
| 40 | No | E. coli | Rifampicin | Intracellul | F84 | Neutral Re | 7172 | BL1 | 5980 |
| 40 | No | E. coli | Rifampicin | Intracellul | F99 | Pyronin Y | 8131 | BL1 | 5610 |
| 40 | No | E. coli | Rifampicin | Intracellul | F108 | Rhodamine | 9145 | BL1 | 5252 |
| 40 | No | E. coli | Rifampicin | Intracellul | F109 | Riboflavin | 7529 | BL1 | 4762 |
| 40 | No | E. coli | Rifampicin | Intracellul | F117 | SYBR Gree | 10016 | BL1 | 98650 |
| 40 | No | E. coli | Rifampicin | Intracellul | F106 | Rhodamine | 7951 | BL1 | 5831 |
| 40 | No | E. coli | Rifampicin | Intracellul | F73 | Malachite ( | 9925 | BL1 | 5572 |
| 40 | No | E. coli | Rifampicin | Intracellul | F7 | Acridine or | 8673 | BL1 | 5191 |
| 40 | No | E. coli | Rifampicin | Intracellul | F192 | 5-Carboxyf | 7848 | BL1 | 13098 |
| 40 | No | E. coli | Rifampicin | Intracellul | F57 | Eosin Y | 7744 | BL1 | 15279 |
| 40 | No | E. coli | Rifampicin | Intracellul | F27 | Calcein | 8758 | BL1 | 17733.5 |
| 40 | No | E. coli | Rifampicin | Intracellul | F12 | Amaranth | 9828 | BL1 | 18531 |
| 40 | No | E. coli | None | None | NA | Autofluore | 7161 | NA | 14734 |
| 40 | No | E. coli | Trimethopr | Intracellul | F194 | Oxonol | 8058 | RL1 | 14745 |
| 40 | No | E. coli | Trimethopr | Intracellul | F11 | Allura Red | 7295 | BL3 | 18845 |
| 40 | No | E. coli | Trimethopr | Intracellul | F54 | DiSC3(5) | 7276 | RL1 | 16023 |
| 40 | No | E. coli | Trimethopr | Intracellul | F124 | Thioflavin T | 7992 | VL2 | 12805 |
| 40 | No | E. coli | Trimethopr | Intracellul | F101 | Quinacrine | 7113 | VL2 | 12673 |
| 40 | No | E. coli | Trimethopr | Intracellul | F138 | Apigenin | 7790 | VL2 | 11297 |
| 40 | No | E. coli | Trimethopr | Intracellul | F65 | H2FDA | 8387 | BL1 | 7501 |
| 40 | No | E. coli | Trimethopr | Intracellul | F142 | Chlorophe | 7863 | BL3 | 5727 |
| 40 | No | E. coli | Trimethopr | Intracellul | F39 | Congo Red | 8628 | BL3 | 6106.5 |
| 40 | No | E. coli | Trimethopr | Intracellul | F116 | Sunset Yel | 8214 | BL2 | 6921.5 |
| 40 | No | E. coli | Trimethopr | Intracellul | F10 | Alizarin | 8074 | BL1 | 6180 |
| 40 | No | E. coli | Trimethopr | Intracellul | F84 | Neutral Re | 8099 | BL1 | 6286 |
| 40 | No | E. coli | Trimethopr | Intracellul | F99 | Pyronin Y | 6280 | BL1 | 6614.5 |
| 40 | No | E. coli | Trimethopr | Intracellul | F108 | Rhodamine | 7755 | BL1 | 5843 |
| 40 | No | E. coli | Trimethopr | Intracellul | F109 | Riboflavin | 8732 | BL1 | 5388 |
| 40 | No | E. coli | Trimethopr | Intracellul | F117 | SYBR Gree | 7555 | BL1 | 106015 |
| 40 | No | E. coli | Trimethopr | Intracellul | F106 | Rhodamine | 9245 | BL1 | 6966 |
| 40 | No | E. coli | Trimethopr | Intracellul | F73 | Malachite ( | 6863 | BL1 | 5752 |
| 40 | No | E. coli | Trimethopr | Intracellul | F7 | Acridine or | 8798 | BL1 | 5322.5 |
| 40 | No | E. coli | Trimethopr | Intracellul | F192 | 5-Carboxyf | 7891 | BL1 | 12097 |
| 40 | No | E. coli | Trimethopr | Intracellul | F57 | Eosin Y | 6993 | BL1 | 15718 |
| 40 | No | E. coli | Trimethopr | Intracellul | F27 | Calcein | 8444 | BL1 | 21299.5 |

| | | | | | | | | | |
|---|---|---|---|---|---|---|---|---|---|
| 40 | No | E. coli | Trimethopr | Intracellula | F12 | Amaranth | 8155 | BL1 | 18687 |
| 40 | No | E. coli | None | None | NA | Autofluore: | 7660 | NA | 899 |
| 40 | No | E. coli | None | None | F194 | Oxonol | 14820 | RL1 | 17230 |
| 40 | No | E. coli | None | None | F11 | Allura Red | 15993 | BL3 | 16995 |
| 40 | No | E. coli | None | None | F54 | DiSC3(5) | 12883 | RL1 | 11544 |
| 40 | No | E. coli | None | None | F124 | Thioflavin T | 17484 | VL2 | 8610 |
| 40 | No | E. coli | None | None | F101 | Quinacrine | 15973 | VL2 | 7290 |
| 40 | No | E. coli | None | None | F138 | Apigenin | 15542 | VL2 | 4958 |
| 40 | No | E. coli | None | None | F65 | H2FDA | 15993 | BL1 | 4622 |
| 40 | No | E. coli | None | None | F142 | Chlorophe | 15653 | BL3 | 4316 |
| 40 | No | E. coli | None | None | F39 | Congo Red | 15021 | BL3 | 4318 |
| 40 | No | E. coli | None | None | F116 | Sunset Yel | 15480 | BL2 | 4232.5 |
| 40 | No | E. coli | None | None | F10 | Alizarin | 14283 | BL1 | 4561 |
| 40 | No | E. coli | None | None | F84 | Neutral Re | 14392 | BL1 | 3678 |
| 40 | No | E. coli | None | None | F99 | Pyronin Y | 16314 | BL1 | 3364 |
| 40 | No | E. coli | None | None | F108 | Rhodamine | 14815 | BL1 | 2901 |
| 40 | No | E. coli | None | None | F109 | Riboflavin | 16280 | BL1 | 2793.5 |
| 40 | No | E. coli | None | None | F117 | SYBR Gree | 11256 | BL1 | 183836.5 |
| 40 | No | E. coli | None | None | F106 | Rhodamine | 18973 | BL1 | 3767 |
| 40 | No | E. coli | None | None | F73 | Malachite ( | 14816 | BL1 | 3747 |
| 40 | No | E. coli | None | None | F7 | Acridine or | 12393 | BL1 | 3288 |
| 40 | No | E. coli | None | None | F192 | 5-Carboxyf | 15747 | BL1 | 7675 |
| 40 | No | E. coli | None | None | F57 | Eosin Y | 17780 | BL1 | 17085 |
| 40 | No | E. coli | None | None | F27 | Calcein | 16476 | BL1 | 18721 |
| 40 | No | E. coli | None | None | F12 | Amaranth | 12066 | BL1 | 24515.5 |
| 40 | No | E. coli | None | None | NA | Autofluore: | 12835 | NA | 19114 |
| 40 | No | E. coli | Ampicillin | Wall | F194 | Oxonol | 14273 | RL1 | 19885 |
| 40 | No | E. coli | Ampicillin | Wall | F11 | Allura Red | 10972 | BL3 | 21646 |
| 40 | No | E. coli | Ampicillin | Wall | F54 | DiSC3(5) | 13581 | RL1 | 16325 |
| 40 | No | E. coli | Ampicillin | Wall | F124 | Thioflavin T | 17328 | VL2 | 12312.5 |
| 40 | No | E. coli | Ampicillin | Wall | F101 | Quinacrine | 13966 | VL2 | 12437 |
| 40 | No | E. coli | Ampicillin | Wall | F138 | Apigenin | 12180 | VL2 | 6653.5 |
| 40 | No | E. coli | Ampicillin | Wall | F65 | H2FDA | 12507 | BL1 | 5233 |
| 40 | No | E. coli | Ampicillin | Wall | F142 | Chlorophe | 14747 | BL3 | 4521 |
| 40 | No | E. coli | Ampicillin | Wall | F39 | Congo Red | 15122 | BL3 | 5501.5 |
| 40 | No | E. coli | Ampicillin | Wall | F116 | Sunset Yel | 12754 | BL2 | 5375 |
| 40 | No | E. coli | Ampicillin | Wall | F10 | Alizarin | 14675 | BL1 | 4986 |
| 40 | No | E. coli | Ampicillin | Wall | F84 | Neutral Re | 11962 | BL1 | 4501 |
| 40 | No | E. coli | Ampicillin | Wall | F99 | Pyronin Y | 15419 | BL1 | 4116 |
| 40 | No | E. coli | Ampicillin | Wall | F108 | Rhodamine | 14684 | BL1 | 3591 |
| 40 | No | E. coli | Ampicillin | Wall | F109 | Riboflavin | 14269 | BL1 | 3470 |
| 40 | No | E. coli | Ampicillin | Wall | F117 | SYBR Gree | 10806 | BL1 | 193016.5 |
| 40 | No | E. coli | Ampicillin | Wall | F106 | Rhodamine | 15674 | BL1 | 5283 |
| 40 | No | E. coli | Ampicillin | Wall | F73 | Malachite ( | 11495 | BL1 | 4408 |
| 40 | No | E. coli | Ampicillin | Wall | F7 | Acridine or | 15351 | BL1 | 3935 |
| 40 | No | E. coli | Ampicillin | Wall | F192 | 5-Carboxyf | 14432 | BL1 | 12944 |
| 40 | No | E. coli | Ampicillin | Wall | F57 | Eosin Y | 10173 | BL1 | 16711 |
| 40 | No | E. coli | Ampicillin | Wall | F27 | Calcein | 15305 | BL1 | 18925 |

| | | | | | | | | | |
|---|---|---|---|---|---|---|---|---|---|
| 40 | No | E. coli | Ampicillin | Wall | F12 | Amaranth | 13547 | BL1 | 17743 |
| 40 | No | E. coli | None | None | NA | Autofluore: | 10357 | NA | 26729 |
| 40 | No | E. coli | Bacitracin | Wall | F194 | Oxonol | 14894 | RL1 | 17490 |
| 40 | No | E. coli | Bacitracin | Wall | F11 | Allura Red | 10099 | BL3 | 17409 |
| 40 | No | E. coli | Bacitracin | Wall | F54 | DiSC3(5) | 13284 | RL1 | 19035 |
| 40 | No | E. coli | Bacitracin | Wall | F124 | Thioflavin T | 14327 | VL2 | 13162 |
| 40 | No | E. coli | Bacitracin | Wall | F101 | Quinacrine | 16934 | VL2 | 11299.5 |
| 40 | No | E. coli | Bacitracin | Wall | F138 | Apigenin | 12389 | VL2 | 8014 |
| 40 | No | E. coli | Bacitracin | Wall | F65 | H2FDA | 14119 | BL1 | 5953 |
| 40 | No | E. coli | Bacitracin | Wall | F142 | Chlorophe | 17326 | BL3 | 5229 |
| 40 | No | E. coli | Bacitracin | Wall | F39 | Congo Red | 14972 | BL3 | 5114 |
| 40 | No | E. coli | Bacitracin | Wall | F116 | Sunset Yel | 15086 | BL2 | 5204 |
| 40 | No | E. coli | Bacitracin | Wall | F10 | Alizarin | 13170 | BL1 | 4875.5 |
| 40 | No | E. coli | Bacitracin | Wall | F84 | Neutral Re | 15451 | BL1 | 4771 |
| 40 | No | E. coli | Bacitracin | Wall | F99 | Pyronin Y | 13116 | BL1 | 4068 |
| 40 | No | E. coli | Bacitracin | Wall | F108 | Rhodamine | 16195 | BL1 | 3919 |
| 40 | No | E. coli | Bacitracin | Wall | F109 | Riboflavin | 13996 | BL1 | 3905 |
| 40 | No | E. coli | Bacitracin | Wall | F117 | SYBR Gree | 14622 | BL1 | 179694 |
| 40 | No | E. coli | Bacitracin | Wall | F106 | Rhodamine | 14327 | BL1 | 4627 |
| 40 | No | E. coli | Bacitracin | Wall | F73 | Malachite ( | 13843 | BL1 | 4372 |
| 40 | No | E. coli | Bacitracin | Wall | F7 | Acridine or | 16296 | BL1 | 4291 |
| 40 | No | E. coli | Bacitracin | Wall | F192 | 5-Carboxyf | 14451 | BL1 | 11913 |
| 40 | No | E. coli | Bacitracin | Wall | F57 | Eosin Y | 16710 | BL1 | 17463 |
| 40 | No | E. coli | Bacitracin | Wall | F27 | Calcein | 14170 | BL1 | 20777.5 |
| 40 | No | E. coli | Bacitracin | Wall | F12 | Amaranth | 13153 | BL1 | 22167 |
| 40 | No | E. coli | None | None | NA | Autofluore: | 15323 | NA | 23910 |
| 40 | No | E. coli | Cefotaxim | Wall | F194 | Oxonol | 13842 | RL1 | 28902.5 |
| 40 | No | E. coli | Cefotaxim | Wall | F11 | Allura Red | 10639 | BL3 | 45067 |
| 40 | No | E. coli | Cefotaxim | Wall | F54 | DiSC3(5) | 11443 | RL1 | 19181 |
| 40 | No | E. coli | Cefotaxim | Wall | F124 | Thioflavin T | 13287 | VL2 | 13646 |
| 40 | No | E. coli | Cefotaxim | Wall | F101 | Quinacrine | 13429 | VL2 | 12794 |
| 40 | No | E. coli | Cefotaxim | Wall | F138 | Apigenin | 13637 | VL2 | 9822 |
| 40 | No | E. coli | Cefotaxim | Wall | F65 | H2FDA | 12029 | BL1 | 7170 |
| 40 | No | E. coli | Cefotaxim | Wall | F142 | Chlorophe | 11003 | BL3 | 5678 |
| 40 | No | E. coli | Cefotaxim | Wall | F39 | Congo Red | 13088 | BL3 | 6394.5 |
| 40 | No | E. coli | Cefotaxim | Wall | F116 | Sunset Yel | 12660 | BL2 | 6385 |
| 40 | No | E. coli | Cefotaxim | Wall | F10 | Alizarin | 9900 | BL1 | 6386 |
| 40 | No | E. coli | Cefotaxim | Wall | F84 | Neutral Re | 10254 | BL1 | 5547 |
| 40 | No | E. coli | Cefotaxim | Wall | F99 | Pyronin Y | 10173 | BL1 | 5443 |
| 40 | No | E. coli | Cefotaxim | Wall | F108 | Rhodamine | 11300 | BL1 | 4624.5 |
| 40 | No | E. coli | Cefotaxim | Wall | F109 | Riboflavin | 11342 | BL1 | 4605.5 |
| 40 | No | E. coli | Cefotaxim | Wall | F117 | SYBR Gree | 9893 | BL1 | 130008 |
| 40 | No | E. coli | Cefotaxim | Wall | F106 | Rhodamine | 12064 | BL1 | 8406.5 |
| 40 | No | E. coli | Cefotaxim | Wall | F73 | Malachite ( | 13007 | BL1 | 5065 |
| 40 | No | E. coli | Cefotaxim | Wall | F7 | Acridine or | 10473 | BL1 | 4843 |
| 40 | No | E. coli | Cefotaxim | Wall | F192 | 5-Carboxyf | 10160 | BL1 | 7833 |
| 40 | No | E. coli | Cefotaxim | Wall | F57 | Eosin Y | 11966 | BL1 | 18719.5 |
| 40 | No | E. coli | Cefotaxim | Wall | F27 | Calcein | 11006 | BL1 | 20187 |

| | | | | | | | | | |
|---|---|---|---|---|---|---|---|---|---|
| 40 | No | E. coli | Cefotaxim | Wall | F12 | Amaranth | 12409 | BL1 | 24099 |
| 40 | No | E. coli | None | None | NA | Autofluore | 10362 | NA | 20132.5 |
| 40 | No | E. coli | Ciprofloxa | Intracellul | F194 | Oxonol | 12663 | RL1 | 18101 |
| 40 | No | E. coli | Ciprofloxa | Intracellul | F11 | Allura Red | 10748 | BL3 | 20955.5 |
| 40 | No | E. coli | Ciprofloxa | Intracellul | F54 | DiSC3(5) | 11080 | RL1 | 14919.5 |
| 40 | No | E. coli | Ciprofloxa | Intracellul | F124 | Thioflavin T | 11327 | VL2 | 14371 |
| 40 | No | E. coli | Ciprofloxa | Intracellul | F101 | Quinacrine | 13867 | VL2 | 12114 |
| 40 | No | E. coli | Ciprofloxa | Intracellul | F138 | Apigenin | 10242 | VL2 | 9761 |
| 40 | No | E. coli | Ciprofloxa | Intracellul | F65 | H2FDA | 10264 | BL1 | 6039 |
| 40 | No | E. coli | Ciprofloxa | Intracellul | F142 | Chlorophe | 10667 | BL3 | 5537 |
| 40 | No | E. coli | Ciprofloxa | Intracellul | F39 | Congo Red | 7837 | BL3 | 6303 |
| 40 | No | E. coli | Ciprofloxa | Intracellul | F116 | Sunset Yel | 12152 | BL2 | 6308 |
| 40 | No | E. coli | Ciprofloxa | Intracellul | F10 | Alizarin | 7547 | BL1 | 6149 |
| 40 | No | E. coli | Ciprofloxa | Intracellul | F84 | Neutral Re | 12801 | BL1 | 5439 |
| 40 | No | E. coli | Ciprofloxa | Intracellul | F99 | Pyronin Y | 8343 | BL1 | 4697 |
| 40 | No | E. coli | Ciprofloxa | Intracellul | F108 | Rhodamine | 12662 | BL1 | 4939 |
| 40 | No | E. coli | Ciprofloxa | Intracellul | F109 | Riboflavin | 13350 | BL1 | 4849 |
| 40 | No | E. coli | Ciprofloxa | Intracellul | F117 | SYBR Gree | 8266 | BL1 | 244342 |
| 40 | No | E. coli | Ciprofloxa | Intracellul | F106 | Rhodamine | 11114 | BL1 | 6862.5 |
| 40 | No | E. coli | Ciprofloxa | Intracellul | F73 | Malachite ( | 9987 | BL1 | 5612 |
| 40 | No | E. coli | Ciprofloxa | Intracellul | F7 | Acridine or | 9979 | BL1 | 4701 |
| 40 | No | E. coli | Ciprofloxa | Intracellul | F192 | 5-Carboxyf | 9634 | BL1 | 6754.5 |
| 40 | No | E. coli | Ciprofloxa | Intracellul | F57 | Eosin Y | 12121 | BL1 | 21376 |
| 40 | No | E. coli | Ciprofloxa | Intracellul | F27 | Calcein | 10320 | BL1 | 22311.5 |
| 40 | No | E. coli | Ciprofloxa | Intracellul | F12 | Amaranth | 11128 | BL1 | 22689 |
| 40 | No | E. coli | None | None | NA | Autofluore | 10569 | NA | 18491 |
| 40 | No | E. coli | Fosfomyci | Wall | F194 | Oxonol | 12848 | RL1 | 17558 |
| 40 | No | E. coli | Fosfomyci | Wall | F11 | Allura Red | 15225 | BL3 | 15950 |
| 40 | No | E. coli | Fosfomyci | Wall | F54 | DiSC3(5) | 11661 | RL1 | 16847 |
| 40 | No | E. coli | Fosfomyci | Wall | F124 | Thioflavin T | 14440 | VL2 | 16176.5 |
| 40 | No | E. coli | Fosfomyci | Wall | F101 | Quinacrine | 10142 | VL2 | 16595.5 |
| 40 | No | E. coli | Fosfomyci | Wall | F138 | Apigenin | 19559 | VL2 | 7653 |
| 40 | No | E. coli | Fosfomyci | Wall | F65 | H2FDA | 11212 | BL1 | 6006 |
| 40 | No | E. coli | Fosfomyci | Wall | F142 | Chlorophe | 15212 | BL3 | 5522 |
| 40 | No | E. coli | Fosfomyci | Wall | F39 | Congo Red | 15521 | BL3 | 6429 |
| 40 | No | E. coli | Fosfomyci | Wall | F116 | Sunset Yel | 10303 | BL2 | 6188 |
| 40 | No | E. coli | Fosfomyci | Wall | F10 | Alizarin | 13252 | BL1 | 5530 |
| 40 | No | E. coli | Fosfomyci | Wall | F84 | Neutral Re | 11026 | BL1 | 5745.5 |
| 40 | No | E. coli | Fosfomyci | Wall | F99 | Pyronin Y | 12650 | BL1 | 5021 |
| 40 | No | E. coli | Fosfomyci | Wall | F108 | Rhodamine | 15181 | BL1 | 4247 |
| 40 | No | E. coli | Fosfomyci | Wall | F109 | Riboflavin | 13802 | BL1 | 4320 |
| 40 | No | E. coli | Fosfomyci | Wall | F117 | SYBR Gree | 13141 | BL1 | 167210 |
| 40 | No | E. coli | Fosfomyci | Wall | F106 | Rhodamine | 11028 | BL1 | 6404 |
| 40 | No | E. coli | Fosfomyci | Wall | F73 | Malachite ( | 12051 | BL1 | 5465 |
| 40 | No | E. coli | Fosfomyci | Wall | F7 | Acridine or | 14991 | BL1 | 4804 |
| 40 | No | E. coli | Fosfomyci | Wall | F192 | 5-Carboxyf | 18385 | BL1 | 11703 |
| 40 | No | E. coli | Fosfomyci | Wall | F57 | Eosin Y | 9615 | BL1 | 19259 |
| 40 | No | E. coli | Fosfomyci | Wall | F27 | Calcein | 17615 | BL1 | 24747 |

| | | | | | | | | | | |
|---|---|---|---|---|---|---|---|---|---|---|
| 40 | No | E. coli | Fosfomyci | Wall | F12 | Amaranth | 13163 | BL1 | | 18622 |
| 40 | No | E. coli | None | None | NA | Autofluore: | 16662 | NA | | 30537.5 |
| 40 | No | E. coli | Nalidixic ac | Intracellula | F194 | Oxonol | 11034 | RL1 | | 21240.5 |
| 40 | No | E. coli | Nalidixic ac | Intracellula | F11 | Allura Red | 12686 | BL3 | | 14956 |
| 40 | No | E. coli | Nalidixic ac | Intracellula | F54 | DiSC3(5) | 10039 | RL1 | | 16091 |
| 40 | No | E. coli | Nalidixic ac | Intracellula | F124 | Thioflavin T | 13124 | VL2 | | 13683 |
| 40 | No | E. coli | Nalidixic ac | Intracellula | F101 | Quinacrine | 10818 | VL2 | | 12078.5 |
| 40 | No | E. coli | Nalidixic ac | Intracellula | F138 | Apigenin | 13632 | VL2 | | 8644 |
| 40 | No | E. coli | Nalidixic ac | Intracellula | F65 | H2FDA | 13598 | BL1 | | 6681.5 |
| 40 | No | E. coli | Nalidixic ac | Intracellula | F142 | Chlorophe | 12174 | BL3 | | 5983.5 |
| 40 | No | E. coli | Nalidixic ac | Intracellula | F39 | Congo Red | 13313 | BL3 | | 6414 |
| 40 | No | E. coli | Nalidixic ac | Intracellula | F116 | Sunset Yel | 9713 | BL2 | | 6350 |
| 40 | No | E. coli | Nalidixic ac | Intracellula | F10 | Alizarin | 9786 | BL1 | | 6215 |
| 40 | No | E. coli | Nalidixic ac | Intracellula | F84 | Neutral Re | 12437 | BL1 | | 5573 |
| 40 | No | E. coli | Nalidixic ac | Intracellula | F99 | Pyronin Y | 9849 | BL1 | | 5227 |
| 40 | No | E. coli | Nalidixic ac | Intracellula | F108 | Rhodamine | 8445 | BL1 | | 4615 |
| 40 | No | E. coli | Nalidixic ac | Intracellula | F109 | Riboflavin | 13231 | BL1 | | 4781 |
| 40 | No | E. coli | Nalidixic ac | Intracellula | F117 | SYBR Gree | 9263 | BL1 | | 134711 |
| 40 | No | E. coli | Nalidixic ac | Intracellula | F106 | Rhodamine | 13421 | BL1 | | 6098 |
| 40 | No | E. coli | Nalidixic ac | Intracellula | F73 | Malachite ( | 11612 | BL1 | | 5822.5 |
| 40 | No | E. coli | Nalidixic ac | Intracellula | F7 | Acridine or | 8696 | BL1 | | 5149 |
| 40 | No | E. coli | Nalidixic ac | Intracellula | F192 | 5-Carboxyf | 12024 | BL1 | | 10584 |
| 40 | No | E. coli | Nalidixic ac | Intracellula | F57 | Eosin Y | 11161 | BL1 | | 16670 |
| 40 | No | E. coli | Nalidixic ac | Intracellula | F27 | Calcein | 11320 | BL1 | | 19912.5 |
| 40 | No | E. coli | Nalidixic ac | Intracellula | F12 | Amaranth | 11248 | BL1 | | 17103.5 |
| 40 | No | E. coli | None | None | NA | Autofluore: | 15419 | NA | | 19120 |
| 40 | No | E. coli | Ofloxacin | Intracellula | F194 | Oxonol | 9737 | RL1 | | 39709 |
| 40 | No | E. coli | Ofloxacin | Intracellula | F11 | Allura Red | 11443 | BL3 | | 19979 |
| 40 | No | E. coli | Ofloxacin | Intracellula | F54 | DiSC3(5) | 8257 | RL1 | | 20506 |
| 40 | No | E. coli | Ofloxacin | Intracellula | F124 | Thioflavin T | 13604 | VL2 | | 13245 |
| 40 | No | E. coli | Ofloxacin | Intracellula | F101 | Quinacrine | 10752 | VL2 | | 13692 |
| 40 | No | E. coli | Ofloxacin | Intracellula | F138 | Apigenin | 10259 | VL2 | | 9670 |
| 40 | No | E. coli | Ofloxacin | Intracellula | F65 | H2FDA | 8730 | BL1 | | 6598 |
| 40 | No | E. coli | Ofloxacin | Intracellula | F142 | Chlorophe | 13811 | BL3 | | 6525 |
| 40 | No | E. coli | Ofloxacin | Intracellula | F39 | Congo Red | 8035 | BL3 | | 6935 |
| 40 | No | E. coli | Ofloxacin | Intracellula | F116 | Sunset Yel | 10809 | BL2 | | 6364 |
| 40 | No | E. coli | Ofloxacin | Intracellula | F10 | Alizarin | 9840 | BL1 | | 6859.5 |
| 40 | No | E. coli | Ofloxacin | Intracellula | F84 | Neutral Re | 9261 | BL1 | | 6167 |
| 40 | No | E. coli | Ofloxacin | Intracellula | F99 | Pyronin Y | 8765 | BL1 | | 5296 |
| 40 | No | E. coli | Ofloxacin | Intracellula | F108 | Rhodamine | 12228 | BL1 | | 5052 |
| 40 | No | E. coli | Ofloxacin | Intracellula | F109 | Riboflavin | 6848 | BL1 | | 4862 |
| 40 | No | E. coli | Ofloxacin | Intracellula | F117 | SYBR Gree | 12796 | BL1 | | 209546.5 |
| 40 | No | E. coli | Ofloxacin | Intracellula | F106 | Rhodamine | 8880 | BL1 | | 5934.5 |
| 40 | No | E. coli | Ofloxacin | Intracellula | F73 | Malachite ( | 9876 | BL1 | | 5265 |
| 40 | No | E. coli | Ofloxacin | Intracellula | F7 | Acridine or | 10057 | BL1 | | 5198 |
| 40 | No | E. coli | Ofloxacin | Intracellula | F192 | 5-Carboxyf | 10944 | BL1 | | 12360.5 |
| 40 | No | E. coli | Ofloxacin | Intracellula | F57 | Eosin Y | 8655 | BL1 | | 20980 |
| 40 | No | E. coli | Ofloxacin | Intracellula | F27 | Calcein | 10466 | BL1 | | 21622.5 |

| | | | | | | | | | |
|---|---|---|---|---|---|---|---|---|---|
| 40 | No | E. coli | Ofloxacin | Intracellula | F12 | Amaranth | 9096 | BL1 | 21531 |
| 40 | No | E. coli | None | None | NA | Autofluore: | 13060 | NA | 16861 |
| 40 | No | E. coli | Polymyxin | Wall | F194 | Oxonol | 15605 | RL1 | 38390 |
| 40 | No | E. coli | Polymyxin | Wall | F11 | Allura Red | 10273 | BL3 | 21260 |
| 40 | No | E. coli | Polymyxin | Wall | F54 | DiSC3(5) | 17155 | RL1 | 16704 |
| 40 | No | E. coli | Polymyxin | Wall | F124 | Thioflavin T | 13998 | VL2 | 13278 |
| 40 | No | E. coli | Polymyxin | Wall | F101 | Quinacrine | 16508 | VL2 | 12477 |
| 40 | No | E. coli | Polymyxin | Wall | F138 | Apigenin | 11302 | VL2 | 10270 |
| 40 | No | E. coli | Polymyxin | Wall | F65 | H2FDA | 13487 | BL1 | 6559 |
| 40 | No | E. coli | Polymyxin | Wall | F142 | Chlorophe | 16087 | BL3 | 6246 |
| 40 | No | E. coli | Polymyxin | Wall | F39 | Congo Red | 11081 | BL3 | 5602 |
| 40 | No | E. coli | Polymyxin | Wall | F116 | Sunset Yel | 14297 | BL2 | 5948 |
| 40 | No | E. coli | Polymyxin | Wall | F10 | Alizarin | 15239 | BL1 | 5298 |
| 40 | No | E. coli | Polymyxin | Wall | F84 | Neutral Re | 10962 | BL1 | 5372.5 |
| 40 | No | E. coli | Polymyxin | Wall | F99 | Pyronin Y | 12546 | BL1 | 4994.5 |
| 40 | No | E. coli | Polymyxin | Wall | F108 | Rhodamine | 13193 | BL1 | 4690 |
| 40 | No | E. coli | Polymyxin | Wall | F109 | Riboflavin | 11562 | BL1 | 4474 |
| 40 | No | E. coli | Polymyxin | Wall | F117 | SYBR Gree | 13451 | BL1 | 159757 |
| 40 | No | E. coli | Polymyxin | Wall | F106 | Rhodamine | 14822 | BL1 | 5706 |
| 40 | No | E. coli | Polymyxin | Wall | F73 | Malachite ( | 14474 | BL1 | 5676 |
| 40 | No | E. coli | Polymyxin | Wall | F7 | Acridine or | 13432 | BL1 | 4944.5 |
| 40 | No | E. coli | Polymyxin | Wall | F192 | 5-Carboxyf | 10870 | BL1 | 6881 |
| 40 | No | E. coli | Polymyxin | Wall | F57 | Eosin Y | 13125 | BL1 | 36403 |
| 40 | No | E. coli | Polymyxin | Wall | F27 | Calcein | 12376 | BL1 | 33330 |
| 40 | No | E. coli | Polymyxin | Wall | F12 | Amaranth | 17485 | BL1 | 25474 |
| 40 | No | E. coli | None | None | NA | Autofluore: | 13559 | NA | 30947 |
| 40 | No | E. coli | Tetracyclin | Intracellula | F194 | Oxonol | 11747 | RL1 | 16257 |
| 40 | No | E. coli | Tetracyclin | Intracellula | F11 | Allura Red | 13706 | BL3 | 13593 |
| 40 | No | E. coli | Tetracyclin | Intracellula | F54 | DiSC3(5) | 12128 | RL1 | 15811 |
| 40 | No | E. coli | Tetracyclin | Intracellula | F124 | Thioflavin T | 16759 | VL2 | 11330 |
| 40 | No | E. coli | Tetracyclin | Intracellula | F101 | Quinacrine | 11392 | VL2 | 12501 |
| 40 | No | E. coli | Tetracyclin | Intracellula | F138 | Apigenin | 19026 | VL2 | 7978.5 |
| 40 | No | E. coli | Tetracyclin | Intracellula | F65 | H2FDA | 11860 | BL1 | 5724 |
| 40 | No | E. coli | Tetracyclin | Intracellula | F142 | Chlorophe | 15156 | BL3 | 4843 |
| 40 | No | E. coli | Tetracyclin | Intracellula | F39 | Congo Red | 10410 | BL3 | 5939 |
| 40 | No | E. coli | Tetracyclin | Intracellula | F116 | Sunset Yel | 15561 | BL2 | 6225 |
| 40 | No | E. coli | Tetracyclin | Intracellula | F10 | Alizarin | 9955 | BL1 | 5794 |
| 40 | No | E. coli | Tetracyclin | Intracellula | F84 | Neutral Re | 15128 | BL1 | 5250.5 |
| 40 | No | E. coli | Tetracyclin | Intracellula | F99 | Pyronin Y | 10863 | BL1 | 4904 |
| 40 | No | E. coli | Tetracyclin | Intracellula | F108 | Rhodamine | 13211 | BL1 | 4484 |
| 40 | No | E. coli | Tetracyclin | Intracellula | F109 | Riboflavin | 12273 | BL1 | 3994 |
| 40 | No | E. coli | Tetracyclin | Intracellula | F117 | SYBR Gree | 12526 | BL1 | 151423.5 |
| 40 | No | E. coli | Tetracyclin | Intracellula | F106 | Rhodamine | 11670 | BL1 | 10076.5 |
| 40 | No | E. coli | Tetracyclin | Intracellula | F73 | Malachite ( | 16389 | BL1 | 5097 |
| 40 | No | E. coli | Tetracyclin | Intracellula | F7 | Acridine or | 13388 | BL1 | 4711.5 |
| 40 | No | E. coli | Tetracyclin | Intracellula | F192 | 5-Carboxyf | 13819 | BL1 | 9832 |
| 40 | No | E. coli | Tetracyclin | Intracellula | F57 | Eosin Y | 10729 | BL1 | 16054 |
| 40 | No | E. coli | Tetracyclin | Intracellula | F27 | Calcein | 14288 | BL1 | 26281.5 |

| | | | | | | | | | |
|---|---|---|---|---|---|---|---|---|---|
| 40 | No | E. coli | Tetracyclin | Intracellula | F12 | Amaranth | 10724 | BL1 | 30245 |
| 40 | No | E. coli | None | None | NA | Autofluore: | 15419 | NA | 21564 |
| 40 | No | E. coli | Vancomyc | Wall | F194 | Oxonol | 15566 | RL1 | 58451.5 |
| 40 | No | E. coli | Vancomyc | Wall | F11 | Allura Red | 10313 | BL3 | 50846 |
| 40 | No | E. coli | Vancomyc | Wall | F54 | DiSC3(5) | 14070 | RL1 | 38012 |
| 40 | No | E. coli | Vancomyc | Wall | F124 | Thioflavin T | 11481 | VL2 | 24723 |
| 40 | No | E. coli | Vancomyc | Wall | F101 | Quinacrine | 13779 | VL2 | 22647 |
| 40 | No | E. coli | Vancomyc | Wall | F138 | Apigenin | 13502 | VL2 | 13756.5 |
| 40 | No | E. coli | Vancomyc | Wall | F65 | H2FDA | 11549 | BL1 | 8752 |
| 40 | No | E. coli | Vancomyc | Wall | F142 | Chlorophe | 10363 | BL3 | 7552 |
| 40 | No | E. coli | Vancomyc | Wall | F39 | Congo Red | 11309 | BL3 | 7729 |
| 40 | No | E. coli | Vancomyc | Wall | F116 | Sunset Yel | 13317 | BL2 | 8271 |
| 40 | No | E. coli | Vancomyc | Wall | F10 | Alizarin | 9682 | BL1 | 8269.5 |
| 40 | No | E. coli | Vancomyc | Wall | F84 | Neutral Re | 10066 | BL1 | 8158 |
| 40 | No | E. coli | Vancomyc | Wall | F99 | Pyronin Y | 12245 | BL1 | 7123 |
| 40 | No | E. coli | Vancomyc | Wall | F108 | Rhodamine | 10924 | BL1 | 5575 |
| 40 | No | E. coli | Vancomyc | Wall | F109 | Riboflavin | 13807 | BL1 | 4816 |
| 40 | No | E. coli | Vancomyc | Wall | F117 | SYBR Gree | 9143 | BL1 | 181981 |
| 40 | No | E. coli | Vancomyc | Wall | F106 | Rhodamine | 11639 | BL1 | 13118 |
| 40 | No | E. coli | Vancomyc | Wall | F73 | Malachite ( | 10877 | BL1 | 6715 |
| 40 | No | E. coli | Vancomyc | Wall | F7 | Acridine or | 14732 | BL1 | 5225 |
| 40 | No | E. coli | Vancomyc | Wall | F192 | 5-Carboxyf | 12580 | BL1 | 14752 |
| 40 | No | E. coli | Vancomyc | Wall | F57 | Eosin Y | 11269 | BL1 | 38736 |
| 40 | No | E. coli | Vancomyc | Wall | F27 | Calcein | 11212 | BL1 | 57185 |
| 40 | No | E. coli | Vancomyc | Wall | F12 | Amaranth | 14278 | BL1 | 93077.5 |
| 40 | No | E. coli | None | None | NA | Autofluore: | 10273 | NA | 793 |
| 40 | No | E. coli | None | None | F194 | Oxonol | 9633 | RL1 | 19153 |
| 40 | No | E. coli | None | None | F11 | Allura Red | 15149 | BL3 | 14829 |
| 40 | No | E. coli | None | None | F54 | DiSC3(5) | 13530 | RL1 | 7942.5 |
| 40 | No | E. coli | None | None | F124 | Thioflavin T | 12987 | VL2 | 5772 |
| 40 | No | E. coli | None | None | F101 | Quinacrine | 15281 | VL2 | 4113 |
| 40 | No | E. coli | None | None | F138 | Apigenin | 13004 | VL2 | 3553 |
| 40 | No | E. coli | None | None | F65 | H2FDA | 14949 | BL1 | 3295 |
| 40 | No | E. coli | None | None | F142 | Chlorophe | 13479 | BL3 | 3655 |
| 40 | No | E. coli | None | None | F39 | Congo Red | 10841 | BL3 | 3876 |
| 40 | No | E. coli | None | None | F116 | Sunset Yel | 15619 | BL2 | 4000 |
| 40 | No | E. coli | None | None | F10 | Alizarin | 11592 | BL1 | 4189 |
| 40 | No | E. coli | None | None | F84 | Neutral Re | 15771 | BL1 | 3725 |
| 40 | No | E. coli | None | None | F99 | Pyronin Y | 12264 | BL1 | 3455 |
| 40 | No | E. coli | None | None | F108 | Rhodamine | 16566 | BL1 | 2933 |
| 40 | No | E. coli | None | None | F109 | Riboflavin | 14191 | BL1 | 2865 |
| 40 | No | E. coli | None | None | F117 | SYBR Gree | 14939 | BL1 | 176018 |
| 40 | No | E. coli | None | None | F106 | Rhodamine | 13743 | BL1 | 3420 |
| 40 | No | E. coli | None | None | F73 | Malachite ( | 14678 | BL1 | 3384 |
| 40 | No | E. coli | None | None | F7 | Acridine or | 13095 | BL1 | 3194 |
| 40 | No | E. coli | None | None | F192 | 5-Carboxyf | 13840 | BL1 | 12477 |
| 40 | No | E. coli | None | None | F57 | Eosin Y | 14001 | BL1 | 16820 |
| 40 | No | E. coli | None | None | F27 | Calcein | 14631 | BL1 | 18822 |

| | | | | | | | | | |
|---|---|---|---|---|---|---|---|---|---|
| 40 | No | E. coli | None | None | F12 | Amaranth | 13439 | BL1 | 19449 |
| 40 | No | E. coli | None | None | NA | Autofluores | 9398 | NA | 16290.5 |
| 40 | No | E. coli | Ampicillin | Wall | F194 | Oxonol | 12308 | RL1 | 16493.5 |
| 40 | No | E. coli | Ampicillin | Wall | F11 | Allura Red | 11456 | BL3 | 15668 |
| 40 | No | E. coli | Ampicillin | Wall | F54 | DiSC3(5) | 11015 | RL1 | 16680 |
| 40 | No | E. coli | Ampicillin | Wall | F124 | Thioflavin T | 16187 | VL2 | 11814 |
| 40 | No | E. coli | Ampicillin | Wall | F101 | Quinacrine | 14076 | VL2 | 12554 |
| 40 | No | E. coli | Ampicillin | Wall | F138 | Apigenin | 15244 | VL2 | 8656 |
| 40 | No | E. coli | Ampicillin | Wall | F65 | H2FDA | 9925 | BL1 | 5487 |
| 40 | No | E. coli | Ampicillin | Wall | F142 | Chlorophe | 17385 | BL3 | 5351 |
| 40 | No | E. coli | Ampicillin | Wall | F39 | Congo Red | 12103 | BL3 | 5964 |
| 40 | No | E. coli | Ampicillin | Wall | F116 | Sunset Yell | 13256 | BL2 | 5576 |
| 40 | No | E. coli | Ampicillin | Wall | F10 | Alizarin | 11244 | BL1 | 5776 |
| 40 | No | E. coli | Ampicillin | Wall | F84 | Neutral Re | 13143 | BL1 | 4817 |
| 40 | No | E. coli | Ampicillin | Wall | F99 | Pyronin Y | 12182 | BL1 | 4644 |
| 40 | No | E. coli | Ampicillin | Wall | F108 | Rhodamine | 12516 | BL1 | 4013 |
| 40 | No | E. coli | Ampicillin | Wall | F109 | Riboflavin | 10164 | BL1 | 3713.5 |
| 40 | No | E. coli | Ampicillin | Wall | F117 | SYBR Gree | 10568 | BL1 | 166497.5 |
| 40 | No | E. coli | Ampicillin | Wall | F106 | Rhodamine | 12856 | BL1 | 5549 |
| 40 | No | E. coli | Ampicillin | Wall | F73 | Malachite ( | 14410 | BL1 | 4568 |
| 40 | No | E. coli | Ampicillin | Wall | F7 | Acridine or | 11862 | BL1 | 4615.5 |
| 40 | No | E. coli | Ampicillin | Wall | F192 | 5-Carboxyf | 13629 | BL1 | 13343 |
| 40 | No | E. coli | Ampicillin | Wall | F57 | Eosin Y | 9848 | BL1 | 17914 |
| 40 | No | E. coli | Ampicillin | Wall | F27 | Calcein | 12899 | BL1 | 19550 |
| 40 | No | E. coli | Ampicillin | Wall | F12 | Amaranth | 12700 | BL1 | 22776 |
| 40 | No | E. coli | None | None | NA | Autofluores | 12413 | NA | 19119 |
| 40 | No | E. coli | Bacitracin | Wall | F194 | Oxonol | 11350 | RL1 | 19706.5 |
| 40 | No | E. coli | Bacitracin | Wall | F11 | Allura Red | 9435 | BL3 | 15599 |
| 40 | No | E. coli | Bacitracin | Wall | F54 | DiSC3(5) | 11059 | RL1 | 16450 |
| 40 | No | E. coli | Bacitracin | Wall | F124 | Thioflavin T | 14615 | VL2 | 12808 |
| 40 | No | E. coli | Bacitracin | Wall | F101 | Quinacrine | 13476 | VL2 | 12323.5 |
| 40 | No | E. coli | Bacitracin | Wall | F138 | Apigenin | 11299 | VL2 | 9238 |
| 40 | No | E. coli | Bacitracin | Wall | F65 | H2FDA | 13246 | BL1 | 6135.5 |
| 40 | No | E. coli | Bacitracin | Wall | F142 | Chlorophe | 14123 | BL3 | 5693 |
| 40 | No | E. coli | Bacitracin | Wall | F39 | Congo Red | 13430 | BL3 | 5381 |
| 40 | No | E. coli | Bacitracin | Wall | F116 | Sunset Yell | 12716 | BL2 | 5749.5 |
| 40 | No | E. coli | Bacitracin | Wall | F10 | Alizarin | 12792 | BL1 | 4983 |
| 40 | No | E. coli | Bacitracin | Wall | F84 | Neutral Re | 12568 | BL1 | 5149 |
| 40 | No | E. coli | Bacitracin | Wall | F99 | Pyronin Y | 12671 | BL1 | 4744 |
| 40 | No | E. coli | Bacitracin | Wall | F108 | Rhodamine | 13798 | BL1 | 4118 |
| 40 | No | E. coli | Bacitracin | Wall | F109 | Riboflavin | 12657 | BL1 | 4054 |
| 40 | No | E. coli | Bacitracin | Wall | F117 | SYBR Gree | 11027 | BL1 | 213297 |
| 40 | No | E. coli | Bacitracin | Wall | F106 | Rhodamine | 12963 | BL1 | 5224 |
| 40 | No | E. coli | Bacitracin | Wall | F73 | Malachite ( | 13797 | BL1 | 4707 |
| 40 | No | E. coli | Bacitracin | Wall | F7 | Acridine or | 11899 | BL1 | 4303 |
| 40 | No | E. coli | Bacitracin | Wall | F192 | 5-Carboxyf | 13033 | BL1 | 12228 |
| 40 | No | E. coli | Bacitracin | Wall | F57 | Eosin Y | 10632 | BL1 | 17782 |
| 40 | No | E. coli | Bacitracin | Wall | F27 | Calcein | 13017 | BL1 | 20562 |

| | | | | | | | | | |
|---|---|---|---|---|---|---|---|---|---|
| 40 | No | E. coli | Bacitracin | Wall | F12 | Amaranth | 11582 | BL1 | 19320 |
| 40 | No | E. coli | None | None | NA | Autofluores | 11699 | NA | 19097 |
| 40 | No | E. coli | Cefotaxime | Wall | F194 | Oxonol | 10440 | RL1 | 19110 |
| 40 | No | E. coli | Cefotaxime | Wall | F11 | Allura Red | 13411 | BL3 | 19644 |
| 40 | No | E. coli | Cefotaxime | Wall | F54 | DiSC3(5) | 10580 | RL1 | 15885.5 |
| 40 | No | E. coli | Cefotaxime | Wall | F124 | Thioflavin T | 13158 | VL2 | 12802 |
| 40 | No | E. coli | Cefotaxime | Wall | F101 | Quinacrine | 11888 | VL2 | 13300 |
| 40 | No | E. coli | Cefotaxime | Wall | F138 | Apigenin | 13532 | VL2 | 7855 |
| 40 | No | E. coli | Cefotaxime | Wall | F65 | H2FDA | 10511 | BL1 | 6271 |
| 40 | No | E. coli | Cefotaxime | Wall | F142 | Chlorophe | 10495 | BL3 | 5908 |
| 40 | No | E. coli | Cefotaxime | Wall | F39 | Congo Red | 9773 | BL3 | 6511 |
| 40 | No | E. coli | Cefotaxime | Wall | F116 | Sunset Yel | 12480 | BL2 | 6222.5 |
| 40 | No | E. coli | Cefotaxime | Wall | F10 | Alizarin | 10526 | BL1 | 6384 |
| 40 | No | E. coli | Cefotaxime | Wall | F84 | Neutral Re | 11869 | BL1 | 5112 |
| 40 | No | E. coli | Cefotaxime | Wall | F99 | Pyronin Y | 10074 | BL1 | 5046 |
| 40 | No | E. coli | Cefotaxime | Wall | F108 | Rhodamine | 10841 | BL1 | 4480 |
| 40 | No | E. coli | Cefotaxime | Wall | F109 | Riboflavin | 11171 | BL1 | 4346 |
| 40 | No | E. coli | Cefotaxime | Wall | F117 | SYBR Gree | 10894 | BL1 | 221301.5 |
| 40 | No | E. coli | Cefotaxime | Wall | F106 | Rhodamine | 9258 | BL1 | 6292 |
| 40 | No | E. coli | Cefotaxime | Wall | F73 | Malachite ( | 10804 | BL1 | 5113 |
| 40 | No | E. coli | Cefotaxime | Wall | F7 | Acridine or | 10035 | BL1 | 4778 |
| 40 | No | E. coli | Cefotaxime | Wall | F192 | 5-Carboxyf | 11582 | BL1 | 10315.5 |
| 40 | No | E. coli | Cefotaxime | Wall | F57 | Eosin Y | 9617 | BL1 | 19005 |
| 40 | No | E. coli | Cefotaxime | Wall | F27 | Calcein | 11759 | BL1 | 23223 |
| 40 | No | E. coli | Cefotaxime | Wall | F12 | Amaranth | 9864 | BL1 | 27482 |
| 40 | No | E. coli | None | None | NA | Autofluores | 9939 | NA | 17658 |
| 40 | No | E. coli | Ciprofloxac | Intracellula | F194 | Oxonol | 12053 | RL1 | 17816 |
| 40 | No | E. coli | Ciprofloxac | Intracellula | F11 | Allura Red | 10750 | BL3 | 18757.5 |
| 40 | No | E. coli | Ciprofloxac | Intracellula | F54 | DiSC3(5) | 12877 | RL1 | 15986 |
| 40 | No | E. coli | Ciprofloxac | Intracellula | F124 | Thioflavin T | 11534 | VL2 | 13102 |
| 40 | No | E. coli | Ciprofloxac | Intracellula | F101 | Quinacrine | 11276 | VL2 | 13501 |
| 40 | No | E. coli | Ciprofloxac | Intracellula | F138 | Apigenin | 11117 | VL2 | 8253 |
| 40 | No | E. coli | Ciprofloxac | Intracellula | F65 | H2FDA | 9757 | BL1 | 5472 |
| 40 | No | E. coli | Ciprofloxac | Intracellula | F142 | Chlorophe | 11595 | BL3 | 5113 |
| 40 | No | E. coli | Ciprofloxac | Intracellula | F39 | Congo Red | 10672 | BL3 | 5670.5 |
| 40 | No | E. coli | Ciprofloxac | Intracellula | F116 | Sunset Yel | 13020 | BL2 | 5390 |
| 40 | No | E. coli | Ciprofloxac | Intracellula | F10 | Alizarin | 10041 | BL1 | 5077 |
| 40 | No | E. coli | Ciprofloxac | Intracellula | F84 | Neutral Re | 11513 | BL1 | 5179 |
| 40 | No | E. coli | Ciprofloxac | Intracellula | F99 | Pyronin Y | 8835 | BL1 | 4826 |
| 40 | No | E. coli | Ciprofloxac | Intracellula | F108 | Rhodamine | 8960 | BL1 | 4559 |
| 40 | No | E. coli | Ciprofloxac | Intracellula | F109 | Riboflavin | 10615 | BL1 | 4012 |
| 40 | No | E. coli | Ciprofloxac | Intracellula | F117 | SYBR Gree | 11959 | BL1 | 206001 |
| 40 | No | E. coli | Ciprofloxac | Intracellula | F106 | Rhodamine | 8366 | BL1 | 4972 |
| 40 | No | E. coli | Ciprofloxac | Intracellula | F73 | Malachite ( | 9763 | BL1 | 4557 |
| 40 | No | E. coli | Ciprofloxac | Intracellula | F7 | Acridine or | 10431 | BL1 | 4416 |
| 40 | No | E. coli | Ciprofloxac | Intracellula | F192 | 5-Carboxyf | 12524 | BL1 | 14622.5 |
| 40 | No | E. coli | Ciprofloxac | Intracellula | F57 | Eosin Y | 9069 | BL1 | 18224 |
| 40 | No | E. coli | Ciprofloxac | Intracellula | F27 | Calcein | 9819 | BL1 | 24103 |

| | | | | | | | | | |
|---|---|---|---|---|---|---|---|---|---|
| 40 | No | E. coli | Ciprofloxa | Intracellul | F12 | Amaranth | 10604 | BL1 | 19343.5 |
| 40 | No | E. coli | None | None | NA | Autofluore | 10554 | NA | 15919.5 |
| 40 | No | E. coli | Fosfomyci | Wall | F194 | Oxonol | 9890 | RL1 | 22949.5 |
| 40 | No | E. coli | Fosfomyci | Wall | F11 | Allura Red | 12210 | BL3 | 17050.5 |
| 40 | No | E. coli | Fosfomyci | Wall | F54 | DiSC3(5) | 13035 | RL1 | 17086 |
| 40 | No | E. coli | Fosfomyci | Wall | F124 | Thioflavin T | 14534 | VL2 | 13346 |
| 40 | No | E. coli | Fosfomyci | Wall | F101 | Quinacrine | 12997 | VL2 | 12366 |
| 40 | No | E. coli | Fosfomyci | Wall | F138 | Apigenin | 14562 | VL2 | 7619.5 |
| 40 | No | E. coli | Fosfomyci | Wall | F65 | H2FDA | 11288 | BL1 | 5966 |
| 40 | No | E. coli | Fosfomyci | Wall | F142 | Chlorophe | 13428 | BL3 | 5143.5 |
| 40 | No | E. coli | Fosfomyci | Wall | F39 | Congo Red | 11972 | BL3 | 5472.5 |
| 40 | No | E. coli | Fosfomyci | Wall | F116 | Sunset Yel | 15640 | BL2 | 5347.5 |
| 40 | No | E. coli | Fosfomyci | Wall | F10 | Alizarin | 10565 | BL1 | 5462 |
| 40 | No | E. coli | Fosfomyci | Wall | F84 | Neutral Re | 14838 | BL1 | 4643 |
| 40 | No | E. coli | Fosfomyci | Wall | F99 | Pyronin Y | 11575 | BL1 | 4192 |
| 40 | No | E. coli | Fosfomyci | Wall | F108 | Rhodamine | 15131 | BL1 | 4244 |
| 40 | No | E. coli | Fosfomyci | Wall | F109 | Riboflavin | 14444 | BL1 | 4095 |
| 40 | No | E. coli | Fosfomyci | Wall | F117 | SYBR Gree | 11576 | BL1 | 201814 |
| 40 | No | E. coli | Fosfomyci | Wall | F106 | Rhodamine | 13443 | BL1 | 5314 |
| 40 | No | E. coli | Fosfomyci | Wall | F73 | Malachite ( | 11721 | BL1 | 4807 |
| 40 | No | E. coli | Fosfomyci | Wall | F7 | Acridine or | 13494 | BL1 | 4344.5 |
| 40 | No | E. coli | Fosfomyci | Wall | F192 | 5-Carboxyf | 11907 | BL1 | 7897 |
| 40 | No | E. coli | Fosfomyci | Wall | F57 | Eosin Y | 14184 | BL1 | 17319 |
| 40 | No | E. coli | Fosfomyci | Wall | F27 | Calcein | 10988 | BL1 | 21227 |
| 40 | No | E. coli | Fosfomyci | Wall | F12 | Amaranth | 11417 | BL1 | 18130 |
| 40 | No | E. coli | None | None | NA | Autofluore | 14772 | NA | 20058 |
| 40 | No | E. coli | Nalidixic a | Intracellul | F194 | Oxonol | 10760 | RL1 | 18602 |
| 40 | No | E. coli | Nalidixic a | Intracellul | F11 | Allura Red | 13508 | BL3 | 17988.5 |
| 40 | No | E. coli | Nalidixic a | Intracellul | F54 | DiSC3(5) | 11100 | RL1 | 16599.5 |
| 40 | No | E. coli | Nalidixic a | Intracellul | F124 | Thioflavin T | 16421 | VL2 | 12093 |
| 40 | No | E. coli | Nalidixic a | Intracellul | F101 | Quinacrine | 12069 | VL2 | 12040 |
| 40 | No | E. coli | Nalidixic a | Intracellul | F138 | Apigenin | 15802 | VL2 | 7085.5 |
| 40 | No | E. coli | Nalidixic a | Intracellul | F65 | H2FDA | 9883 | BL1 | 5941 |
| 40 | No | E. coli | Nalidixic a | Intracellul | F142 | Chlorophe | 11176 | BL3 | 5627 |
| 40 | No | E. coli | Nalidixic a | Intracellul | F39 | Congo Red | 9598 | BL3 | 5660.5 |
| 40 | No | E. coli | Nalidixic a | Intracellul | F116 | Sunset Yel | 10426 | BL2 | 5647 |
| 40 | No | E. coli | Nalidixic a | Intracellul | F10 | Alizarin | 11242 | BL1 | 5521 |
| 40 | No | E. coli | Nalidixic a | Intracellul | F84 | Neutral Re | 10630 | BL1 | 5031 |
| 40 | No | E. coli | Nalidixic a | Intracellul | F99 | Pyronin Y | 12107 | BL1 | 4824 |
| 40 | No | E. coli | Nalidixic a | Intracellul | F108 | Rhodamine | 8700 | BL1 | 4624.5 |
| 40 | No | E. coli | Nalidixic a | Intracellul | F109 | Riboflavin | 10281 | BL1 | 4159 |
| 40 | No | E. coli | Nalidixic a | Intracellul | F117 | SYBR Gree | 10020 | BL1 | 126263 |
| 40 | No | E. coli | Nalidixic a | Intracellul | F106 | Rhodamine | 10582 | BL1 | 5766.5 |
| 40 | No | E. coli | Nalidixic a | Intracellul | F73 | Malachite ( | 11060 | BL1 | 4767.5 |
| 40 | No | E. coli | Nalidixic a | Intracellul | F7 | Acridine or | 8894 | BL1 | 4616 |
| 40 | No | E. coli | Nalidixic a | Intracellul | F192 | 5-Carboxyf | 10722 | BL1 | 7323.5 |
| 40 | No | E. coli | Nalidixic a | Intracellul | F57 | Eosin Y | 10197 | BL1 | 18590 |
| 40 | No | E. coli | Nalidixic a | Intracellul | F27 | Calcein | 10135 | BL1 | 22103 |

| | | | | | | | | | |
|---|---|---|---|---|---|---|---|---|---|
| 40 | No | E. coli | Nalidixic a | Intracellula | F12 | Amaranth | 10271 | BL1 | 21432 |
| 40 | No | E. coli | None | None | NA | Autofluore: | 11572 | NA | 18337.5 |
| 40 | No | E. coli | Ofloxacin | Intracellula | F194 | Oxonol | 12485 | RL1 | 19019 |
| 40 | No | E. coli | Ofloxacin | Intracellula | F11 | Allura Red | 11870 | BL3 | 18689.5 |
| 40 | No | E. coli | Ofloxacin | Intracellula | F54 | DiSC3(5) | 10846 | RL1 | 17355.5 |
| 40 | No | E. coli | Ofloxacin | Intracellula | F124 | Thioflavin T | 17107 | VL2 | 13558 |
| 40 | No | E. coli | Ofloxacin | Intracellula | F101 | Quinacrine | 12608 | VL2 | 11844 |
| 40 | No | E. coli | Ofloxacin | Intracellula | F138 | Apigenin | 15093 | VL2 | 8314 |
| 40 | No | E. coli | Ofloxacin | Intracellula | F65 | H2FDA | 11585 | BL1 | 6481 |
| 40 | No | E. coli | Ofloxacin | Intracellula | F142 | Chlorophe | 11666 | BL3 | 5938.5 |
| 40 | No | E. coli | Ofloxacin | Intracellula | F39 | Congo Red | 10032 | BL3 | 6235.5 |
| 40 | No | E. coli | Ofloxacin | Intracellula | F116 | Sunset Yel | 9786 | BL2 | 6036.5 |
| 40 | No | E. coli | Ofloxacin | Intracellula | F10 | Alizarin | 9407 | BL1 | 6321 |
| 40 | No | E. coli | Ofloxacin | Intracellula | F84 | Neutral Re | 9691 | BL1 | 5625 |
| 40 | No | E. coli | Ofloxacin | Intracellula | F99 | Pyronin Y | 10517 | BL1 | 5316 |
| 40 | No | E. coli | Ofloxacin | Intracellula | F108 | Rhodamine | 10053 | BL1 | 4374 |
| 40 | No | E. coli | Ofloxacin | Intracellula | F109 | Riboflavin | 6770 | BL1 | 4797.5 |
| 40 | No | E. coli | Ofloxacin | Intracellula | F117 | SYBR Gree | 11896 | BL1 | 209941 |
| 40 | No | E. coli | Ofloxacin | Intracellula | F106 | Rhodamine | 9710 | BL1 | 5996 |
| 40 | No | E. coli | Ofloxacin | Intracellula | F73 | Malachite ( | 9300 | BL1 | 5196 |
| 40 | No | E. coli | Ofloxacin | Intracellula | F7 | Acridine or | 10674 | BL1 | 5002.5 |
| 40 | No | E. coli | Ofloxacin | Intracellula | F192 | 5-Carboxyf | 8664 | BL1 | 12296 |
| 40 | No | E. coli | Ofloxacin | Intracellula | F57 | Eosin Y | 10037 | BL1 | 21640 |
| 40 | No | E. coli | Ofloxacin | Intracellula | F27 | Calcein | 11129 | BL1 | 23533 |
| 40 | No | E. coli | Ofloxacin | Intracellula | F12 | Amaranth | 7121 | BL1 | 28145 |
| 40 | No | E. coli | None | None | NA | Autofluore: | 9382 | NA | 21560.5 |
| 40 | No | E. coli | Polymyxin | Wall | F194 | Oxonol | 14803 | RL1 | 22804 |
| 40 | No | E. coli | Polymyxin | Wall | F11 | Allura Red | 10175 | BL3 | 18341 |
| 40 | No | E. coli | Polymyxin | Wall | F54 | DiSC3(5) | 10833 | RL1 | 18728 |
| 40 | No | E. coli | Polymyxin | Wall | F124 | Thioflavin T | 14584 | VL2 | 13798.5 |
| 40 | No | E. coli | Polymyxin | Wall | F101 | Quinacrine | 11221 | VL2 | 13460 |
| 40 | No | E. coli | Polymyxin | Wall | F138 | Apigenin | 13152 | VL2 | 8001 |
| 40 | No | E. coli | Polymyxin | Wall | F65 | H2FDA | 12596 | BL1 | 5653.5 |
| 40 | No | E. coli | Polymyxin | Wall | F142 | Chlorophe | 14219 | BL3 | 5265 |
| 40 | No | E. coli | Polymyxin | Wall | F39 | Congo Red | 10733 | BL3 | 5478 |
| 40 | No | E. coli | Polymyxin | Wall | F116 | Sunset Yel | 11059 | BL2 | 5586 |
| 40 | No | E. coli | Polymyxin | Wall | F10 | Alizarin | 12739 | BL1 | 5514 |
| 40 | No | E. coli | Polymyxin | Wall | F84 | Neutral Re | 14303 | BL1 | 4736 |
| 40 | No | E. coli | Polymyxin | Wall | F99 | Pyronin Y | 11014 | BL1 | 4624.5 |
| 40 | No | E. coli | Polymyxin | Wall | F108 | Rhodamine | 13944 | BL1 | 4560 |
| 40 | No | E. coli | Polymyxin | Wall | F109 | Riboflavin | 10599 | BL1 | 4309 |
| 40 | No | E. coli | Polymyxin | Wall | F117 | SYBR Gree | 10305 | BL1 | 140960 |
| 40 | No | E. coli | Polymyxin | Wall | F106 | Rhodamine | 12113 | BL1 | 6129 |
| 40 | No | E. coli | Polymyxin | Wall | F73 | Malachite ( | 12493 | BL1 | 4534 |
| 40 | No | E. coli | Polymyxin | Wall | F7 | Acridine or | 12617 | BL1 | 4532 |
| 40 | No | E. coli | Polymyxin | Wall | F192 | 5-Carboxyf | 11333 | BL1 | 9751 |
| 40 | No | E. coli | Polymyxin | Wall | F57 | Eosin Y | 13773 | BL1 | 20740 |
| 40 | No | E. coli | Polymyxin | Wall | F27 | Calcein | 11414 | BL1 | 21290 |

| | | | | | | | | | |
|---|---|---|---|---|---|---|---|---|---|
| 40 | No | E. coli | Polymyxin | Wall | F12 | Amaranth | 12865 | BL1 | 24467 |
| 40 | No | E. coli | None | None | NA | Autofluore: | 11862 | NA | 20624 |
| 40 | No | E. coli | Tetracyclin | Intracellul | F194 | Oxonol | 11264 | RL1 | 18614.5 |
| 40 | No | E. coli | Tetracyclin | Intracellul | F11 | Allura Red | 12210 | BL3 | 16597.5 |
| 40 | No | E. coli | Tetracyclin | Intracellul | F54 | DiSC3(5) | 13701 | RL1 | 16325 |
| 40 | No | E. coli | Tetracyclin | Intracellul | F124 | Thioflavin T | 15267 | VL2 | 12673 |
| 40 | No | E. coli | Tetracyclin | Intracellul | F101 | Quinacrine | 10445 | VL2 | 11654 |
| 40 | No | E. coli | Tetracyclin | Intracellul | F138 | Apigenin | 13968 | VL2 | 8470 |
| 40 | No | E. coli | Tetracyclin | Intracellul | F65 | H2FDA | 12387 | BL1 | 6260 |
| 40 | No | E. coli | Tetracyclin | Intracellul | F142 | Chlorophe | 13214 | BL3 | 5384 |
| 40 | No | E. coli | Tetracyclin | Intracellul | F39 | Congo Red | 10604 | BL3 | 5598 |
| 40 | No | E. coli | Tetracyclin | Intracellul | F116 | Sunset Yel | 13540 | BL2 | 5756 |
| 40 | No | E. coli | Tetracyclin | Intracellul | F10 | Alizarin | 11279 | BL1 | 5424 |
| 40 | No | E. coli | Tetracyclin | Intracellul | F84 | Neutral Re | 11496 | BL1 | 4905.5 |
| 40 | No | E. coli | Tetracyclin | Intracellul | F99 | Pyronin Y | 11672 | BL1 | 4366 |
| 40 | No | E. coli | Tetracyclin | Intracellul | F108 | Rhodamine | 11745 | BL1 | 4111 |
| 40 | No | E. coli | Tetracyclin | Intracellul | F109 | Riboflavin | 14661 | BL1 | 3842 |
| 40 | No | E. coli | Tetracyclin | Intracellul | F117 | SYBR Gree | 12934 | BL1 | 177914 |
| 40 | No | E. coli | Tetracyclin | Intracellul | F106 | Rhodamine | 11957 | BL1 | 6133 |
| 40 | No | E. coli | Tetracyclin | Intracellul | F73 | Malachite ( | 14741 | BL1 | 4835 |
| 40 | No | E. coli | Tetracyclin | Intracellul | F7 | Acridine or | 10180 | BL1 | 4535.5 |
| 40 | No | E. coli | Tetracyclin | Intracellul | F192 | 5-Carboxyf | 14607 | BL1 | 12470 |
| 40 | No | E. coli | Tetracyclin | Intracellul | F57 | Eosin Y | 12504 | BL1 | 15682.5 |
| 40 | No | E. coli | Tetracyclin | Intracellul | F27 | Calcein | 14486 | BL1 | 19550.5 |
| 40 | No | E. coli | Tetracyclin | Intracellul | F12 | Amaranth | 10515 | BL1 | 18665 |
| 40 | No | E. coli | None | None | NA | Autofluore: | 14584 | NA | 19130 |
| 40 | No | E. coli | Vancomyc | Wall | F194 | Oxonol | 11518 | RL1 | 57174 |
| 40 | No | E. coli | Vancomyc | Wall | F11 | Allura Red | 10889 | BL3 | 61415 |
| 40 | No | E. coli | Vancomyc | Wall | F54 | DiSC3(5) | 10391 | RL1 | 43848 |
| 40 | No | E. coli | Vancomyc | Wall | F124 | Thioflavin T | 10564 | VL2 | 35611 |
| 40 | No | E. coli | Vancomyc | Wall | F101 | Quinacrine | 10945 | VL2 | 31912 |
| 40 | No | E. coli | Vancomyc | Wall | F138 | Apigenin | 14489 | VL2 | 13231 |
| 40 | No | E. coli | Vancomyc | Wall | F65 | H2FDA | 10767 | BL1 | 9059 |
| 40 | No | E. coli | Vancomyc | Wall | F142 | Chlorophe | 12214 | BL3 | 7874 |
| 40 | No | E. coli | Vancomyc | Wall | F39 | Congo Red | 11527 | BL3 | 7483 |
| 40 | No | E. coli | Vancomyc | Wall | F116 | Sunset Yel | 13673 | BL2 | 7455 |
| 40 | No | E. coli | Vancomyc | Wall | F10 | Alizarin | 11204 | BL1 | 6909.5 |
| 40 | No | E. coli | Vancomyc | Wall | F84 | Neutral Re | 10139 | BL1 | 7491 |
| 40 | No | E. coli | Vancomyc | Wall | F99 | Pyronin Y | 12856 | BL1 | 6152.5 |
| 40 | No | E. coli | Vancomyc | Wall | F108 | Rhodamine | 10413 | BL1 | 5499 |
| 40 | No | E. coli | Vancomyc | Wall | F109 | Riboflavin | 12758 | BL1 | 5262 |
| 40 | No | E. coli | Vancomyc | Wall | F117 | SYBR Gree | 11236 | BL1 | 230726 |
| 40 | No | E. coli | Vancomyc | Wall | F106 | Rhodamine | 9818 | BL1 | 8973.5 |
| 40 | No | E. coli | Vancomyc | Wall | F73 | Malachite ( | 12712 | BL1 | 6114 |
| 40 | No | E. coli | Vancomyc | Wall | F7 | Acridine or | 9133 | BL1 | 5826 |
| 40 | No | E. coli | Vancomyc | Wall | F192 | 5-Carboxyf | 9423 | BL1 | 9607 |
| 40 | No | E. coli | Vancomyc | Wall | F57 | Eosin Y | 12334 | BL1 | 36086 |
| 40 | No | E. coli | Vancomyc | Wall | F27 | Calcein | 10315 | BL1 | 55520 |

| | | | | | | | | | |
|---|---|---|---|---|---|---|---|---|---|
| 40 | No | E. coli | Vancomyc | Wall | F12 | Amaranth | 10911 | BL1 | 48322 |
| 40 | No | E. coli | None | None | NA | Autofluore: | 10110 | NA | 776 |
| 40 | No | E. coli | None | None | F194 | Oxonol | 13643 | RL1 | 13790 |
| 40 | No | E. coli | None | None | F11 | Allura Red | 17024 | BL3 | 11053 |
| 40 | No | E. coli | None | None | F54 | DiSC3(5) | 16885 | RL1 | 7139 |
| 40 | No | E. coli | None | None | F124 | Thioflavin T | 12984 | VL2 | 5189.5 |
| 40 | No | E. coli | None | None | F101 | Quinacrine | 17202 | VL2 | 3959 |
| 40 | No | E. coli | None | None | F138 | Apigenin | 14242 | VL2 | 3256.5 |
| 40 | No | E. coli | None | None | F65 | H2FDA | 18256 | BL1 | 3224.5 |
| 40 | No | E. coli | None | None | F142 | Chlorophe | 15238 | BL3 | 3389 |
| 40 | No | E. coli | None | None | F39 | Congo Red | 14998 | BL3 | 4198 |
| 40 | No | E. coli | None | None | F116 | Sunset Yel | 18181 | BL2 | 4345 |
| 40 | No | E. coli | None | None | F10 | Alizarin | 17743 | BL1 | 4269 |
| 40 | No | E. coli | None | None | F84 | Neutral Re | 12676 | BL1 | 3972 |
| 40 | No | E. coli | None | None | F99 | Pyronin Y | 16806 | BL1 | 3514 |
| 40 | No | E. coli | None | None | F108 | Rhodamine | 13000 | BL1 | 3162.5 |
| 40 | No | E. coli | None | None | F109 | Riboflavin | 17049 | BL1 | 2803 |
| 40 | No | E. coli | None | None | F117 | SYBR Gree | 16875 | BL1 | 139899 |
| 40 | No | E. coli | None | None | F106 | Rhodamine | 16445 | BL1 | 3874 |
| 40 | No | E. coli | None | None | F73 | Malachite ( | 13570 | BL1 | 3299 |
| 40 | No | E. coli | None | None | F7 | Acridine or | 17813 | BL1 | 3260 |
| 40 | No | E. coli | None | None | F192 | 5-Carboxyf | 13841 | BL1 | 9473 |
| 40 | No | E. coli | None | None | F57 | Eosin Y | 17756 | BL1 | 17265.5 |
| 40 | No | E. coli | None | None | F27 | Calcein | 13798 | BL1 | 20139 |
| 40 | No | E. coli | None | None | F12 | Amaranth | 15652 | BL1 | 19748.5 |
| 40 | No | E. coli | None | None | NA | Autofluore: | 15084 | NA | 18934.5 |
| 40 | No | E. coli | Ampicillin | Wall | F194 | Oxonol | 13296 | RL1 | 19171 |
| 40 | No | E. coli | Ampicillin | Wall | F11 | Allura Red | 15253 | BL3 | 18280 |
| 40 | No | E. coli | Ampicillin | Wall | F54 | DiSC3(5) | 10764 | RL1 | 16367.5 |
| 40 | No | E. coli | Ampicillin | Wall | F124 | Thioflavin T | 17542 | VL2 | 12252.5 |
| 40 | No | E. coli | Ampicillin | Wall | F101 | Quinacrine | 16054 | VL2 | 11271 |
| 40 | No | E. coli | Ampicillin | Wall | F138 | Apigenin | 15147 | VL2 | 6972 |
| 40 | No | E. coli | Ampicillin | Wall | F65 | H2FDA | 15232 | BL1 | 4756 |
| 40 | No | E. coli | Ampicillin | Wall | F142 | Chlorophe | 16171 | BL3 | 4313 |
| 40 | No | E. coli | Ampicillin | Wall | F39 | Congo Red | 16394 | BL3 | 4722 |
| 40 | No | E. coli | Ampicillin | Wall | F116 | Sunset Yel | 13720 | BL2 | 4885 |
| 40 | No | E. coli | Ampicillin | Wall | F10 | Alizarin | 14297 | BL1 | 4798 |
| 40 | No | E. coli | Ampicillin | Wall | F84 | Neutral Re | 14374 | BL1 | 4664 |
| 40 | No | E. coli | Ampicillin | Wall | F99 | Pyronin Y | 16508 | BL1 | 3844 |
| 40 | No | E. coli | Ampicillin | Wall | F108 | Rhodamine | 14013 | BL1 | 3491 |
| 40 | No | E. coli | Ampicillin | Wall | F109 | Riboflavin | 10436 | BL1 | 3627 |
| 40 | No | E. coli | Ampicillin | Wall | F117 | SYBR Gree | 17585 | BL1 | 140313 |
| 40 | No | E. coli | Ampicillin | Wall | F106 | Rhodamine | 15502 | BL1 | 4517 |
| 40 | No | E. coli | Ampicillin | Wall | F73 | Malachite ( | 14148 | BL1 | 4253 |
| 40 | No | E. coli | Ampicillin | Wall | F7 | Acridine or | 13768 | BL1 | 4000.5 |
| 40 | No | E. coli | Ampicillin | Wall | F192 | 5-Carboxyf | 12487 | BL1 | 8293 |
| 40 | No | E. coli | Ampicillin | Wall | F57 | Eosin Y | 15106 | BL1 | 17246.5 |
| 40 | No | E. coli | Ampicillin | Wall | F27 | Calcein | 14749 | BL1 | 21597 |

| | | | | | | | | | |
|---|---|---|---|---|---|---|---|---|---|
| 40 | No | E. coli | Ampicillin | Wall | F12 | Amaranth | 13680 | BL1 | 20800.5 |
| 40 | No | E. coli | None | None | NA | Autofluore: | 16297 | NA | 23673 |
| 40 | No | E. coli | Bacitracin | Wall | F194 | Oxonol | 13757 | RL1 | 18402 |
| 40 | No | E. coli | Bacitracin | Wall | F11 | Allura Red | 13583 | BL3 | 16418 |
| 40 | No | E. coli | Bacitracin | Wall | F54 | DiSC3(5) | 14486 | RL1 | 18020 |
| 40 | No | E. coli | Bacitracin | Wall | F124 | Thioflavin T | 15992 | VL2 | 12628 |
| 40 | No | E. coli | Bacitracin | Wall | F101 | Quinacrine | 15707 | VL2 | 12809 |
| 40 | No | E. coli | Bacitracin | Wall | F138 | Apigenin | 14608 | VL2 | 8161 |
| 40 | No | E. coli | Bacitracin | Wall | F65 | H2FDA | 14987 | BL1 | 6246 |
| 40 | No | E. coli | Bacitracin | Wall | F142 | Chlorophe | 14653 | BL3 | 4994 |
| 40 | No | E. coli | Bacitracin | Wall | F39 | Congo Red | 13312 | BL3 | 4957 |
| 40 | No | E. coli | Bacitracin | Wall | F116 | Sunset Yel | 14232 | BL2 | 4992.5 |
| 40 | No | E. coli | Bacitracin | Wall | F10 | Alizarin | 15050 | BL1 | 4858 |
| 40 | No | E. coli | Bacitracin | Wall | F84 | Neutral Re | 15531 | BL1 | 4467 |
| 40 | No | E. coli | Bacitracin | Wall | F99 | Pyronin Y | 15481 | BL1 | 4008 |
| 40 | No | E. coli | Bacitracin | Wall | F108 | Rhodamine | 14272 | BL1 | 4050.5 |
| 40 | No | E. coli | Bacitracin | Wall | F109 | Riboflavin | 15016 | BL1 | 3710 |
| 40 | No | E. coli | Bacitracin | Wall | F117 | SYBR Gree | 11632 | BL1 | 155726 |
| 40 | No | E. coli | Bacitracin | Wall | F106 | Rhodamine | 16838 | BL1 | 5294 |
| 40 | No | E. coli | Bacitracin | Wall | F73 | Malachite ( | 14919 | BL1 | 4186 |
| 40 | No | E. coli | Bacitracin | Wall | F7 | Acridine or | 14410 | BL1 | 3855 |
| 40 | No | E. coli | Bacitracin | Wall | F192 | 5-Carboxyf | 14935 | BL1 | 12178 |
| 40 | No | E. coli | Bacitracin | Wall | F57 | Eosin Y | 17028 | BL1 | 17336 |
| 40 | No | E. coli | Bacitracin | Wall | F27 | Calcein | 15481 | BL1 | 21667 |
| 40 | No | E. coli | Bacitracin | Wall | F12 | Amaranth | 14083 | BL1 | 37810 |
| 40 | No | E. coli | None | None | NA | Autofluore: | 15367 | NA | 21406 |
| 40 | No | E. coli | Cefotaxim( | Wall | F194 | Oxonol | 13852 | RL1 | 19314.5 |
| 40 | No | E. coli | Cefotaxim( | Wall | F11 | Allura Red | 14097 | BL3 | 22970 |
| 40 | No | E. coli | Cefotaxim( | Wall | F54 | DiSC3(5) | 12766 | RL1 | 17862 |
| 40 | No | E. coli | Cefotaxim( | Wall | F124 | Thioflavin T | 17670 | VL2 | 12472 |
| 40 | No | E. coli | Cefotaxim( | Wall | F101 | Quinacrine | 13050 | VL2 | 11768.5 |
| 40 | No | E. coli | Cefotaxim( | Wall | F138 | Apigenin | 17800 | VL2 | 8390.5 |
| 40 | No | E. coli | Cefotaxim( | Wall | F65 | H2FDA | 12078 | BL1 | 5464.5 |
| 40 | No | E. coli | Cefotaxim( | Wall | F142 | Chlorophe | 18133 | BL3 | 5256 |
| 40 | No | E. coli | Cefotaxim( | Wall | F39 | Congo Red | 12841 | BL3 | 5408 |
| 40 | No | E. coli | Cefotaxim( | Wall | F116 | Sunset Yel | 17891 | BL2 | 5295 |
| 40 | No | E. coli | Cefotaxim( | Wall | F10 | Alizarin | 11286 | BL1 | 5555.5 |
| 40 | No | E. coli | Cefotaxim( | Wall | F84 | Neutral Re | 13319 | BL1 | 5232 |
| 40 | No | E. coli | Cefotaxim( | Wall | F99 | Pyronin Y | 14373 | BL1 | 4381 |
| 40 | No | E. coli | Cefotaxim( | Wall | F108 | Rhodamine | 12870 | BL1 | 3962.5 |
| 40 | No | E. coli | Cefotaxim( | Wall | F109 | Riboflavin | 12479 | BL1 | 3864 |
| 40 | No | E. coli | Cefotaxim( | Wall | F117 | SYBR Gree | 13561 | BL1 | 163319 |
| 40 | No | E. coli | Cefotaxim( | Wall | F106 | Rhodamine | 9560 | BL1 | 6350.5 |
| 40 | No | E. coli | Cefotaxim( | Wall | F73 | Malachite ( | 16164 | BL1 | 4715 |
| 40 | No | E. coli | Cefotaxim( | Wall | F7 | Acridine or | 13671 | BL1 | 4523 |
| 40 | No | E. coli | Cefotaxim( | Wall | F192 | 5-Carboxyf | 10648 | BL1 | 9257.5 |
| 40 | No | E. coli | Cefotaxim( | Wall | F57 | Eosin Y | 11889 | BL1 | 17310 |
| 40 | No | E. coli | Cefotaxim( | Wall | F27 | Calcein | 14107 | BL1 | 23293 |

| | | | | | | | | | |
|---|---|---|---|---|---|---|---|---|---|
| 40 | No | E. coli | Cefotaxim | Wall | F12 | Amaranth | 11289 | BL1 | 25475 |
| 40 | No | E. coli | None | None | NA | Autofluore: | 11864 | NA | 23218 |
| 40 | No | E. coli | Ciprofloxa | Intracellul | F194 | Oxonol | 16165 | RL1 | 19661 |
| 40 | No | E. coli | Ciprofloxa | Intracellul | F11 | Allura Red | 12989 | BL3 | 18140 |
| 40 | No | E. coli | Ciprofloxa | Intracellul | F54 | DiSC3(5) | 15052 | RL1 | 18215.5 |
| 40 | No | E. coli | Ciprofloxa | Intracellul | F124 | Thioflavin T | 13863 | VL2 | 13721 |
| 40 | No | E. coli | Ciprofloxa | Intracellul | F101 | Quinacrine | 11748 | VL2 | 12488.5 |
| 40 | No | E. coli | Ciprofloxa | Intracellul | F138 | Apigenin | 12975 | VL2 | 8007 |
| 40 | No | E. coli | Ciprofloxa | Intracellul | F65 | H2FDA | 12393 | BL1 | 5833 |
| 40 | No | E. coli | Ciprofloxa | Intracellul | F142 | Chlorophe | 10340 | BL3 | 5384 |
| 40 | No | E. coli | Ciprofloxa | Intracellul | F39 | Congo Red | 13402 | BL3 | 5605.5 |
| 40 | No | E. coli | Ciprofloxa | Intracellul | F116 | Sunset Yel | 9308 | BL2 | 5629.5 |
| 40 | No | E. coli | Ciprofloxa | Intracellul | F10 | Alizarin | 10491 | BL1 | 5583 |
| 40 | No | E. coli | Ciprofloxa | Intracellul | F84 | Neutral Re | 11741 | BL1 | 5113 |
| 40 | No | E. coli | Ciprofloxa | Intracellul | F99 | Pyronin Y | 10564 | BL1 | 4740.5 |
| 40 | No | E. coli | Ciprofloxa | Intracellul | F108 | Rhodamine | 12633 | BL1 | 4259 |
| 40 | No | E. coli | Ciprofloxa | Intracellul | F109 | Riboflavin | 9515 | BL1 | 4340 |
| 40 | No | E. coli | Ciprofloxa | Intracellul | F117 | SYBR Gree | 12443 | BL1 | 163863 |
| 40 | No | E. coli | Ciprofloxa | Intracellul | F106 | Rhodamine | 12588 | BL1 | 5204 |
| 40 | No | E. coli | Ciprofloxa | Intracellul | F73 | Malachite ( | 11765 | BL1 | 4557 |
| 40 | No | E. coli | Ciprofloxa | Intracellul | F7 | Acridine or | 11798 | BL1 | 4475 |
| 40 | No | E. coli | Ciprofloxa | Intracellul | F192 | 5-Carboxyf | 12612 | BL1 | 14907.5 |
| 40 | No | E. coli | Ciprofloxa | Intracellul | F57 | Eosin Y | 9849 | BL1 | 18606 |
| 40 | No | E. coli | Ciprofloxa | Intracellul | F27 | Calcein | 12340 | BL1 | 22211 |
| 40 | No | E. coli | Ciprofloxa | Intracellul | F12 | Amaranth | 11176 | BL1 | 21281.5 |
| 40 | No | E. coli | None | None | NA | Autofluore: | 12392 | NA | 18041 |
| 40 | No | E. coli | Fosfomyci | Wall | F194 | Oxonol | 11134 | RL1 | 23188.5 |
| 40 | No | E. coli | Fosfomyci | Wall | F11 | Allura Red | 13416 | BL3 | 20539 |
| 40 | No | E. coli | Fosfomyci | Wall | F54 | DiSC3(5) | 14222 | RL1 | 15459 |
| 40 | No | E. coli | Fosfomyci | Wall | F124 | Thioflavin T | 18871 | VL2 | 12678 |
| 40 | No | E. coli | Fosfomyci | Wall | F101 | Quinacrine | 13290 | VL2 | 12350 |
| 40 | No | E. coli | Fosfomyci | Wall | F138 | Apigenin | 16951 | VL2 | 7167 |
| 40 | No | E. coli | Fosfomyci | Wall | F65 | H2FDA | 15263 | BL1 | 5662 |
| 40 | No | E. coli | Fosfomyci | Wall | F142 | Chlorophe | 11206 | BL3 | 5106.5 |
| 40 | No | E. coli | Fosfomyci | Wall | F39 | Congo Red | 11832 | BL3 | 5529.5 |
| 40 | No | E. coli | Fosfomyci | Wall | F116 | Sunset Yel | 16719 | BL2 | 5275 |
| 40 | No | E. coli | Fosfomyci | Wall | F10 | Alizarin | 12308 | BL1 | 5337.5 |
| 40 | No | E. coli | Fosfomyci | Wall | F84 | Neutral Re | 14662 | BL1 | 5061 |
| 40 | No | E. coli | Fosfomyci | Wall | F99 | Pyronin Y | 13715 | BL1 | 4354 |
| 40 | No | E. coli | Fosfomyci | Wall | F108 | Rhodamine | 16605 | BL1 | 4040 |
| 40 | No | E. coli | Fosfomyci | Wall | F109 | Riboflavin | 12903 | BL1 | 3782 |
| 40 | No | E. coli | Fosfomyci | Wall | F117 | SYBR Gree | 16016 | BL1 | 131229 |
| 40 | No | E. coli | Fosfomyci | Wall | F106 | Rhodamine | 15314 | BL1 | 5035 |
| 40 | No | E. coli | Fosfomyci | Wall | F73 | Malachite ( | 18969 | BL1 | 4477 |
| 40 | No | E. coli | Fosfomyci | Wall | F7 | Acridine or | 14888 | BL1 | 4249 |
| 40 | No | E. coli | Fosfomyci | Wall | F192 | 5-Carboxyf | 19177 | BL1 | 13054 |
| 40 | No | E. coli | Fosfomyci | Wall | F57 | Eosin Y | 11886 | BL1 | 18102.5 |
| 40 | No | E. coli | Fosfomyci | Wall | F27 | Calcein | 16714 | BL1 | 27464.5 |

| | | | | | | | | | |
|---|---|---|---|---|---|---|---|---|---|
| 40 | No | E. coli | Fosfomyci | Wall | F12 | Amaranth | 16801 | BL1 | 27397 |
| 40 | No | E. coli | None | None | NA | Autofluore | 15749 | NA | 29337 |
| 40 | No | E. coli | Nalidixic a | Intracellul | F194 | Oxonol | 16018 | RL1 | 19184.5 |
| 40 | No | E. coli | Nalidixic a | Intracellul | F11 | Allura Red | 14699 | BL3 | 17866 |
| 40 | No | E. coli | Nalidixic a | Intracellul | F54 | DiSC3(5) | 12550 | RL1 | 15582.5 |
| 40 | No | E. coli | Nalidixic a | Intracellul | F124 | Thioflavin T | 18535 | VL2 | 13479 |
| 40 | No | E. coli | Nalidixic a | Intracellul | F101 | Quinacrine | 15793 | VL2 | 14845 |
| 40 | No | E. coli | Nalidixic a | Intracellul | F138 | Apigenin | 14071 | VL2 | 8235 |
| 40 | No | E. coli | Nalidixic a | Intracellul | F65 | H2FDA | 16175 | BL1 | 6821 |
| 40 | No | E. coli | Nalidixic a | Intracellul | F142 | Chlorophe | 14633 | BL3 | 5328 |
| 40 | No | E. coli | Nalidixic a | Intracellul | F39 | Congo Red | 12884 | BL3 | 5553.5 |
| 40 | No | E. coli | Nalidixic a | Intracellul | F116 | Sunset Yel | 12242 | BL2 | 6027 |
| 40 | No | E. coli | Nalidixic a | Intracellul | F10 | Alizarin | 9101 | BL1 | 5241 |
| 40 | No | E. coli | Nalidixic a | Intracellul | F84 | Neutral Re | 10091 | BL1 | 5442 |
| 40 | No | E. coli | Nalidixic a | Intracellul | F99 | Pyronin Y | 10580 | BL1 | 4805 |
| 40 | No | E. coli | Nalidixic a | Intracellul | F108 | Rhodamine | 11402 | BL1 | 4552 |
| 40 | No | E. coli | Nalidixic a | Intracellul | F109 | Riboflavin | 10229 | BL1 | 4225 |
| 40 | No | E. coli | Nalidixic a | Intracellul | F117 | SYBR Gree | 12232 | BL1 | 87988.5 |
| 40 | No | E. coli | Nalidixic a | Intracellul | F106 | Rhodamine | 14037 | BL1 | 5468 |
| 40 | No | E. coli | Nalidixic a | Intracellul | F73 | Malachite ( | 11313 | BL1 | 4790 |
| 40 | No | E. coli | Nalidixic a | Intracellul | F7 | Acridine or | 11329 | BL1 | 4459 |
| 40 | No | E. coli | Nalidixic a | Intracellul | F192 | 5-Carboxyf | 12525 | BL1 | 11133 |
| 40 | No | E. coli | Nalidixic a | Intracellul | F57 | Eosin Y | 10457 | BL1 | 17715 |
| 40 | No | E. coli | Nalidixic a | Intracellul | F27 | Calcein | 12621 | BL1 | 19660 |
| 40 | No | E. coli | Nalidixic a | Intracellul | F12 | Amaranth | 13642 | BL1 | 23860 |
| 40 | No | E. coli | None | None | NA | Autofluore | 10237 | NA | 16537 |
| 40 | No | E. coli | Ofloxacin | Intracellul | F194 | Oxonol | 16262 | RL1 | 20991.5 |
| 40 | No | E. coli | Ofloxacin | Intracellul | F11 | Allura Red | 11855 | BL3 | 17285 |
| 40 | No | E. coli | Ofloxacin | Intracellul | F54 | DiSC3(5) | 12986 | RL1 | 17792.5 |
| 40 | No | E. coli | Ofloxacin | Intracellul | F124 | Thioflavin T | 16656 | VL2 | 13393 |
| 40 | No | E. coli | Ofloxacin | Intracellul | F101 | Quinacrine | 17416 | VL2 | 11947.5 |
| 40 | No | E. coli | Ofloxacin | Intracellul | F138 | Apigenin | 14952 | VL2 | 7621 |
| 40 | No | E. coli | Ofloxacin | Intracellul | F65 | H2FDA | 9000 | BL1 | 5926 |
| 40 | No | E. coli | Ofloxacin | Intracellul | F142 | Chlorophe | 12728 | BL3 | 5609.5 |
| 40 | No | E. coli | Ofloxacin | Intracellul | F39 | Congo Red | 10756 | BL3 | 6175 |
| 40 | No | E. coli | Ofloxacin | Intracellul | F116 | Sunset Yel | 12797 | BL2 | 5626 |
| 40 | No | E. coli | Ofloxacin | Intracellul | F10 | Alizarin | 8799 | BL1 | 6378 |
| 40 | No | E. coli | Ofloxacin | Intracellul | F84 | Neutral Re | 9506 | BL1 | 5705.5 |
| 40 | No | E. coli | Ofloxacin | Intracellul | F99 | Pyronin Y | 10614 | BL1 | 5335.5 |
| 40 | No | E. coli | Ofloxacin | Intracellul | F108 | Rhodamine | 12878 | BL1 | 4524 |
| 40 | No | E. coli | Ofloxacin | Intracellul | F109 | Riboflavin | 8801 | BL1 | 4258 |
| 40 | No | E. coli | Ofloxacin | Intracellul | F117 | SYBR Gree | 13587 | BL1 | 164913 |
| 40 | No | E. coli | Ofloxacin | Intracellul | F106 | Rhodamine | 8587 | BL1 | 6139 |
| 40 | No | E. coli | Ofloxacin | Intracellul | F73 | Malachite ( | 13565 | BL1 | 4739 |
| 40 | No | E. coli | Ofloxacin | Intracellul | F7 | Acridine or | 9503 | BL1 | 4843 |
| 40 | No | E. coli | Ofloxacin | Intracellul | F192 | 5-Carboxyf | 11531 | BL1 | 8985 |
| 40 | No | E. coli | Ofloxacin | Intracellul | F57 | Eosin Y | 12195 | BL1 | 22322 |
| 40 | No | E. coli | Ofloxacin | Intracellul | F27 | Calcein | 9969 | BL1 | 22297 |

| | | | | | | | | | |
|---|---|---|---|---|---|---|---|---|---|
| 40 | No | E. coli | Ofloxacin | Intracellula | F12 | Amaranth | 9129 | BL1 | 27409 |
| 40 | No | E. coli | None | None | NA | Autofluore: | 13964 | NA | 24619 |
| 40 | No | E. coli | Polymyxin | Wall | F194 | Oxonol | 12928 | RL1 | 22881.5 |
| 40 | No | E. coli | Polymyxin | Wall | F11 | Allura Red | 13987 | BL3 | 20380 |
| 40 | No | E. coli | Polymyxin | Wall | F54 | DiSC3(5) | 14805 | RL1 | 18289 |
| 40 | No | E. coli | Polymyxin | Wall | F124 | Thioflavin T | 14316 | VL2 | 15822.5 |
| 40 | No | E. coli | Polymyxin | Wall | F101 | Quinacrine | 17644 | VL2 | 12199 |
| 40 | No | E. coli | Polymyxin | Wall | F138 | Apigenin | 13573 | VL2 | 7274 |
| 40 | No | E. coli | Polymyxin | Wall | F65 | H2FDA | 12494 | BL1 | 5698 |
| 40 | No | E. coli | Polymyxin | Wall | F142 | Chlorophe | 15374 | BL3 | 4749.5 |
| 40 | No | E. coli | Polymyxin | Wall | F39 | Congo Red | 11881 | BL3 | 5879 |
| 40 | No | E. coli | Polymyxin | Wall | F116 | Sunset Yel | 13057 | BL2 | 5264 |
| 40 | No | E. coli | Polymyxin | Wall | F10 | Alizarin | 15740 | BL1 | 5090.5 |
| 40 | No | E. coli | Polymyxin | Wall | F84 | Neutral Re | 16617 | BL1 | 4942 |
| 40 | No | E. coli | Polymyxin | Wall | F99 | Pyronin Y | 14455 | BL1 | 4254 |
| 40 | No | E. coli | Polymyxin | Wall | F108 | Rhodamine | 16036 | BL1 | 4177 |
| 40 | No | E. coli | Polymyxin | Wall | F109 | Riboflavin | 11790 | BL1 | 3789 |
| 40 | No | E. coli | Polymyxin | Wall | F117 | SYBR Gree | 16566 | BL1 | 115874 |
| 40 | No | E. coli | Polymyxin | Wall | F106 | Rhodamine | 15961 | BL1 | 5000 |
| 40 | No | E. coli | Polymyxin | Wall | F73 | Malachite ( | 15887 | BL1 | 4307 |
| 40 | No | E. coli | Polymyxin | Wall | F7 | Acridine or | 14814 | BL1 | 4285 |
| 40 | No | E. coli | Polymyxin | Wall | F192 | 5-Carboxyf | 14333 | BL1 | 13596 |
| 40 | No | E. coli | Polymyxin | Wall | F57 | Eosin Y | 14714 | BL1 | 19940 |
| 40 | No | E. coli | Polymyxin | Wall | F27 | Calcein | 14184 | BL1 | 22674.5 |
| 40 | No | E. coli | Polymyxin | Wall | F12 | Amaranth | 13436 | BL1 | 72681.5 |
| 40 | No | E. coli | None | None | NA | Autofluore: | 14569 | NA | 23603 |
| 40 | No | E. coli | Tetracyclin | Intracellula | F194 | Oxonol | 12356 | RL1 | 22173 |
| 40 | No | E. coli | Tetracyclin | Intracellula | F11 | Allura Red | 13954 | BL3 | 21803.5 |
| 40 | No | E. coli | Tetracyclin | Intracellula | F54 | DiSC3(5) | 11805 | RL1 | 17174 |
| 40 | No | E. coli | Tetracyclin | Intracellula | F124 | Thioflavin T | 19222 | VL2 | 13071 |
| 40 | No | E. coli | Tetracyclin | Intracellula | F101 | Quinacrine | 12084 | VL2 | 12419 |
| 40 | No | E. coli | Tetracyclin | Intracellula | F138 | Apigenin | 17181 | VL2 | 7858 |
| 40 | No | E. coli | Tetracyclin | Intracellula | F65 | H2FDA | 11836 | BL1 | 5922 |
| 40 | No | E. coli | Tetracyclin | Intracellula | F142 | Chlorophe | 13962 | BL3 | 5602 |
| 40 | No | E. coli | Tetracyclin | Intracellula | F39 | Congo Red | 12544 | BL3 | 5564 |
| 40 | No | E. coli | Tetracyclin | Intracellula | F116 | Sunset Yel | 13514 | BL2 | 5265.5 |
| 40 | No | E. coli | Tetracyclin | Intracellula | F10 | Alizarin | 14348 | BL1 | 6037 |
| 40 | No | E. coli | Tetracyclin | Intracellula | F84 | Neutral Re | 15550 | BL1 | 4546 |
| 40 | No | E. coli | Tetracyclin | Intracellula | F99 | Pyronin Y | 11693 | BL1 | 4501 |
| 40 | No | E. coli | Tetracyclin | Intracellula | F108 | Rhodamine | 16669 | BL1 | 4200 |
| 40 | No | E. coli | Tetracyclin | Intracellula | F109 | Riboflavin | 14014 | BL1 | 3927 |
| 40 | No | E. coli | Tetracyclin | Intracellula | F117 | SYBR Gree | 13417 | BL1 | 144730 |
| 40 | No | E. coli | Tetracyclin | Intracellula | F106 | Rhodamine | 12760 | BL1 | 5607.5 |
| 40 | No | E. coli | Tetracyclin | Intracellula | F73 | Malachite ( | 16125 | BL1 | 4429 |
| 40 | No | E. coli | Tetracyclin | Intracellula | F7 | Acridine or | 14531 | BL1 | 4308 |
| 40 | No | E. coli | Tetracyclin | Intracellula | F192 | 5-Carboxyf | 17759 | BL1 | 10209 |
| 40 | No | E. coli | Tetracyclin | Intracellula | F57 | Eosin Y | 14216 | BL1 | 16424.5 |
| 40 | No | E. coli | Tetracyclin | Intracellula | F27 | Calcein | 13683 | BL1 | 23803 |

| | | | | | | | | | |
|---|---|---|---|---|---|---|---|---|---|
| 40 | No | E. coli | Tetracyclin | Intracellula | F12 | Amaranth | 13071 | BL1 | 20151 |
| 40 | No | E. coli | None | None | NA | Autofluore: | 17537 | NA | 18249 |
| 40 | No | E. coli | Vancomyc | Wall | F194 | Oxonol | 11216 | RL1 | 52851 |
| 40 | No | E. coli | Vancomyc | Wall | F11 | Allura Red | 13086 | BL3 | 54618 |
| 40 | No | E. coli | Vancomyc | Wall | F54 | DiSC3(5) | 11672 | RL1 | 61522 |
| 40 | No | E. coli | Vancomyc | Wall | F124 | Thioflavin T | 13691 | VL2 | 30042 |
| 40 | No | E. coli | Vancomyc | Wall | F101 | Quinacrine | 15798 | VL2 | 20624 |
| 40 | No | E. coli | Vancomyc | Wall | F138 | Apigenin | 13021 | VL2 | 13773 |
| 40 | No | E. coli | Vancomyc | Wall | F65 | H2FDA | 13614 | BL1 | 7961 |
| 40 | No | E. coli | Vancomyc | Wall | F142 | Chlorophe | 14072 | BL3 | 6984.5 |
| 40 | No | E. coli | Vancomyc | Wall | F39 | Congo Red | 10888 | BL3 | 6338 |
| 40 | No | E. coli | Vancomyc | Wall | F116 | Sunset Yel | 15204 | BL2 | 6455.5 |
| 40 | No | E. coli | Vancomyc | Wall | F10 | Alizarin | 15183 | BL1 | 6313 |
| 40 | No | E. coli | Vancomyc | Wall | F84 | Neutral Re | 12359 | BL1 | 6920 |
| 40 | No | E. coli | Vancomyc | Wall | F99 | Pyronin Y | 13170 | BL1 | 6271 |
| 40 | No | E. coli | Vancomyc | Wall | F108 | Rhodamine | 13417 | BL1 | 4921 |
| 40 | No | E. coli | Vancomyc | Wall | F109 | Riboflavin | 14791 | BL1 | 4894 |
| 40 | No | E. coli | Vancomyc | Wall | F117 | SYBR Gree | 11685 | BL1 | 167190 |
| 40 | No | E. coli | Vancomyc | Wall | F106 | Rhodamine | 14779 | BL1 | 7780 |
| 40 | No | E. coli | Vancomyc | Wall | F73 | Malachite ( | 12854 | BL1 | 6098.5 |
| 40 | No | E. coli | Vancomyc | Wall | F7 | Acridine or | 15600 | BL1 | 5298 |
| 40 | No | E. coli | Vancomyc | Wall | F192 | 5-Carboxyf | 13735 | BL1 | 20679 |
| 40 | No | E. coli | Vancomyc | Wall | F57 | Eosin Y | 15112 | BL1 | 37804.5 |
| 40 | No | E. coli | Vancomyc | Wall | F27 | Calcein | 12145 | BL1 | 53739 |
| 40 | No | E. coli | Vancomyc | Wall | F12 | Amaranth | 14618 | BL1 | 55316 |
| 40 | No | A. baylyi | None | None | NA | Autofluore: | 793 | NA | 819 |
| 40 | No | A. baylyi | None | None | F194 | Oxonol | 1298 | RL1 | 393453 |
| 40 | No | A. baylyi | None | None | F11 | Allura Red | 936 | BL3 | 393224.5 |
| 40 | No | A. baylyi | None | None | F54 | DiSC3(5) | 654 | RL1 | 313382.5 |
| 40 | No | A. baylyi | None | None | F124 | Thioflavin T | 992 | VL2 | 401740.5 |
| 40 | No | A. baylyi | None | None | F101 | Quinacrine | 647 | VL2 | 54751 |
| 40 | No | A. baylyi | None | None | F138 | Apigenin | 576 | VL2 | 26389 |
| 40 | No | A. baylyi | None | None | F65 | H2FDA | 485 | BL1 | 12371 |
| 40 | No | A. baylyi | None | None | F142 | Chlorophe | 503 | BL3 | 9327 |
| 40 | No | A. baylyi | None | None | F39 | Congo Red | 503 | BL3 | 7824 |
| 40 | No | A. baylyi | None | None | F116 | Sunset Yel | 415 | BL2 | 7617 |
| 40 | No | A. baylyi | None | None | F10 | Alizarin | 405 | BL1 | 8367 |
| 40 | No | A. baylyi | None | None | F84 | Neutral Re | 534 | BL1 | 7311 |
| 40 | No | A. baylyi | None | None | F99 | Pyronin Y | 389 | BL1 | 7628 |
| 40 | No | A. baylyi | None | None | F108 | Rhodamine | 330 | BL1 | 5412 |
| 40 | No | A. baylyi | None | None | F109 | Riboflavin | 324 | BL1 | 4739 |
| 40 | No | A. baylyi | None | None | F117 | SYBR Gree | 361 | BL1 | 111770 |
| 40 | No | A. baylyi | None | None | F106 | Rhodamine | 420 | BL1 | 6573 |
| 40 | No | A. baylyi | None | None | F73 | Malachite ( | 392 | BL1 | 5174.5 |
| 40 | No | A. baylyi | None | None | F7 | Acridine or | 299 | BL1 | 5235 |
| 40 | No | A. baylyi | None | None | F192 | 5-Carboxyf | 313 | BL1 | 12064 |
| 40 | No | A. baylyi | None | None | F57 | Eosin Y | 388 | BL1 | 20859.5 |
| 40 | No | A. baylyi | None | None | F27 | Calcein | 376 | BL1 | 17049 |

| | | | | | | | | | |
|---|---|---|---|---|---|---|---|---|---|
| 40 | No | A. baylyi | None | None | F12 | Amaranth | 302 | BL1 | 17387.5 |
| 40 | No | A. baylyi | None | None | NA | Autofluore: | 330 | NA | 23045.5 |
| 40 | No | A. baylyi | Amoxacilli | Wall | F194 | Oxonol | 943 | RL1 | 11984 |
| 40 | No | A. baylyi | Amoxacilli | Wall | F11 | Allura Red | 381 | BL3 | 21243 |
| 40 | No | A. baylyi | Amoxacilli | Wall | F54 | DiSC3(5) | 142 | RL1 | 16150 |
| 40 | No | A. baylyi | Amoxacilli | Wall | F124 | Thioflavin T | 389 | VL2 | 16396 |
| 40 | No | A. baylyi | Amoxacilli | Wall | F101 | Quinacrine | 221 | VL2 | 13446 |
| 40 | No | A. baylyi | Amoxacilli | Wall | F138 | Apigenin | 219 | VL2 | 10993 |
| 40 | No | A. baylyi | Amoxacilli | Wall | F65 | H2FDA | 225 | BL1 | 6554 |
| 40 | No | A. baylyi | Amoxacilli | Wall | F142 | Chlorophe | 229 | BL3 | 6124 |
| 40 | No | A. baylyi | Amoxacilli | Wall | F39 | Congo Red | 302 | BL3 | 6508 |
| 40 | No | A. baylyi | Amoxacilli | Wall | F116 | Sunset Yel | 237 | BL2 | 6637 |
| 40 | No | A. baylyi | Amoxacilli | Wall | F10 | Alizarin | 236 | BL1 | 6897.5 |
| 40 | No | A. baylyi | Amoxacilli | Wall | F84 | Neutral Re | 331 | BL1 | 6282 |
| 40 | No | A. baylyi | Amoxacilli | Wall | F99 | Pyronin Y | 209 | BL1 | 6119 |
| 40 | No | A. baylyi | Amoxacilli | Wall | F108 | Rhodamin | 371 | BL1 | 4973 |
| 40 | No | A. baylyi | Amoxacilli | Wall | F109 | Riboflavin | 165 | BL1 | 4442 |
| 40 | No | A. baylyi | Amoxacilli | Wall | F117 | SYBR Gree | 371 | BL1 | 145247 |
| 40 | No | A. baylyi | Amoxacilli | Wall | F106 | Rhodamin | 292 | BL1 | 6263.5 |
| 40 | No | A. baylyi | Amoxacilli | Wall | F73 | Malachite ( | 307 | BL1 | 6128 |
| 40 | No | A. baylyi | Amoxacilli | Wall | F7 | Acridine or | 262 | BL1 | 5349.5 |
| 40 | No | A. baylyi | Amoxacilli | Wall | F192 | 5-Carboxyf | 262 | BL1 | 7211 |
| 40 | No | A. baylyi | Amoxacilli | Wall | F57 | Eosin Y | 335 | BL1 | 16163 |
| 40 | No | A. baylyi | Amoxacilli | Wall | F27 | Calcein | 397 | BL1 | 18179 |
| 40 | No | A. baylyi | Amoxacilli | Wall | F12 | Amaranth | 140 | BL1 | 18661.5 |
| 40 | No | A. baylyi | None | None | NA | Autofluore: | 395 | NA | 18983 |
| 40 | No | A. baylyi | Azithromyc | Intracellul: | F194 | Oxonol | 534 | RL1 | 17454.5 |
| 40 | No | A. baylyi | Azithromyc | Intracellul: | F11 | Allura Red | 361 | BL3 | 14175 |
| 40 | No | A. baylyi | Azithromyc | Intracellul: | F54 | DiSC3(5) | 306 | RL1 | 15494 |
| 40 | No | A. baylyi | Azithromyc | Intracellul: | F124 | Thioflavin T | 238 | VL2 | 12108.5 |
| 40 | No | A. baylyi | Azithromyc | Intracellul: | F101 | Quinacrine | 219 | VL2 | 15703 |
| 40 | No | A. baylyi | Azithromyc | Intracellul: | F138 | Apigenin | 341 | VL2 | 7780 |
| 40 | No | A. baylyi | Azithromyc | Intracellul: | F65 | H2FDA | 410 | BL1 | 6954 |
| 40 | No | A. baylyi | Azithromyc | Intracellul: | F142 | Chlorophe | 273 | BL3 | 6693 |
| 40 | No | A. baylyi | Azithromyc | Intracellul: | F39 | Congo Red | 392 | BL3 | 6465 |
| 40 | No | A. baylyi | Azithromyc | Intracellul: | F116 | Sunset Yel | 275 | BL2 | 5954 |
| 40 | No | A. baylyi | Azithromyc | Intracellul: | F10 | Alizarin | 376 | BL1 | 5699 |
| 40 | No | A. baylyi | Azithromyc | Intracellul: | F84 | Neutral Re | 351 | BL1 | 6285 |
| 40 | No | A. baylyi | Azithromyc | Intracellul: | F99 | Pyronin Y | 204 | BL1 | 5671.5 |
| 40 | No | A. baylyi | Azithromyc | Intracellul: | F108 | Rhodamin | 399 | BL1 | 4910 |
| 40 | No | A. baylyi | Azithromyc | Intracellul: | F109 | Riboflavin | 293 | BL1 | 4677 |
| 40 | No | A. baylyi | Azithromyc | Intracellul: | F117 | SYBR Gree | 252 | BL1 | 109039 |
| 40 | No | A. baylyi | Azithromyc | Intracellul: | F106 | Rhodamin | 345 | BL1 | 9563 |
| 40 | No | A. baylyi | Azithromyc | Intracellul: | F73 | Malachite ( | 306 | BL1 | 6159 |
| 40 | No | A. baylyi | Azithromyc | Intracellul: | F7 | Acridine or | 279 | BL1 | 5750 |
| 40 | No | A. baylyi | Azithromyc | Intracellul: | F192 | 5-Carboxyf | 402 | BL1 | 12811.5 |
| 40 | No | A. baylyi | Azithromyc | Intracellul: | F57 | Eosin Y | 250 | BL1 | 17075.5 |
| 40 | No | A. baylyi | Azithromyc | Intracellul: | F27 | Calcein | 406 | BL1 | 20089 |

| | | | | | | | | | |
|---|---|---|---|---|---|---|---|---|---|
| 40 | No | A. baylyi | Azithromyc | Intracellul | F12 | Amaranth | 250 | BL1 | 14348 |
| 40 | No | A. baylyi | None | None | NA | Autofluore | 245 | NA | 18147 |
| 40 | No | A. baylyi | Trimethopr | Intracellul | F194 | Oxonol | 1029 | RL1 | 13795 |
| 40 | No | A. baylyi | Trimethopr | Intracellul | F11 | Allura Red | 358 | BL3 | 17230.5 |
| 40 | No | A. baylyi | Trimethopr | Intracellul | F54 | DiSC3(5) | 241 | RL1 | 14251 |
| 40 | No | A. baylyi | Trimethopr | Intracellul | F124 | Thioflavin T | 344 | VL2 | 14941.5 |
| 40 | No | A. baylyi | Trimethopr | Intracellul | F101 | Quinacrine | 204 | VL2 | 16575.5 |
| 40 | No | A. baylyi | Trimethopr | Intracellul | F138 | Apigenin | 364 | VL2 | 13486.5 |
| 40 | No | A. baylyi | Trimethopr | Intracellul | F65 | H2FDA | 192 | BL1 | 7235.5 |
| 40 | No | A. baylyi | Trimethopr | Intracellul | F142 | Chlorophe | 328 | BL3 | 6407 |
| 40 | No | A. baylyi | Trimethopr | Intracellul | F39 | Congo Red | 287 | BL3 | 6326 |
| 40 | No | A. baylyi | Trimethopr | Intracellul | F116 | Sunset Yel | 365 | BL2 | 7510 |
| 40 | No | A. baylyi | Trimethopr | Intracellul | F10 | Alizarin | 280 | BL1 | 7397 |
| 40 | No | A. baylyi | Trimethopr | Intracellul | F84 | Neutral Re | 423 | BL1 | 7987 |
| 40 | No | A. baylyi | Trimethopr | Intracellul | F99 | Pyronin Y | 206 | BL1 | 6572 |
| 40 | No | A. baylyi | Trimethopr | Intracellul | F108 | Rhodamine | 225 | BL1 | 5673 |
| 40 | No | A. baylyi | Trimethopr | Intracellul | F109 | Riboflavin | 332 | BL1 | 5360.5 |
| 40 | No | A. baylyi | Trimethopr | Intracellul | F117 | SYBR Gree | 263 | BL1 | 131628 |
| 40 | No | A. baylyi | Trimethopr | Intracellul | F106 | Rhodamine | 318 | BL1 | 9854.5 |
| 40 | No | A. baylyi | Trimethopr | Intracellul | F73 | Malachite ( | 308 | BL1 | 6098 |
| 40 | No | A. baylyi | Trimethopr | Intracellul | F7 | Acridine or | 240 | BL1 | 7198.5 |
| 40 | No | A. baylyi | Trimethopr | Intracellul | F192 | 5-Carboxyf | 393 | BL1 | 11986 |
| 40 | No | A. baylyi | Trimethopr | Intracellul | F57 | Eosin Y | 300 | BL1 | 17005 |
| 40 | No | A. baylyi | Trimethopr | Intracellul | F27 | Calcein | 327 | BL1 | 19672 |
| 40 | No | A. baylyi | Trimethopr | Intracellul | F12 | Amaranth | 242 | BL1 | 14478 |
| 40 | No | A. baylyi | None | None | NA | Autofluore | 325 | NA | 14564 |
| 40 | No | A. baylyi | Chloramph | Intracellul | F194 | Oxonol | 565 | RL1 | 15938 |
| 40 | No | A. baylyi | Chloramph | Intracellul | F11 | Allura Red | 257 | BL3 | 15473 |
| 40 | No | A. baylyi | Chloramph | Intracellul | F54 | DiSC3(5) | 424 | RL1 | 16850.5 |
| 40 | No | A. baylyi | Chloramph | Intracellul | F124 | Thioflavin T | 318 | VL2 | 14592.5 |
| 40 | No | A. baylyi | Chloramph | Intracellul | F101 | Quinacrine | 300 | VL2 | 16203.5 |
| 40 | No | A. baylyi | Chloramph | Intracellul | F138 | Apigenin | 200 | VL2 | 14722.5 |
| 40 | No | A. baylyi | Chloramph | Intracellul | F65 | H2FDA | 362 | BL1 | 7849 |
| 40 | No | A. baylyi | Chloramph | Intracellul | F142 | Chlorophe | 296 | BL3 | 5970.5 |
| 40 | No | A. baylyi | Chloramph | Intracellul | F39 | Congo Red | 284 | BL3 | 5958 |
| 40 | No | A. baylyi | Chloramph | Intracellul | F116 | Sunset Yel | 300 | BL2 | 6224 |
| 40 | No | A. baylyi | Chloramph | Intracellul | F10 | Alizarin | 273 | BL1 | 6935 |
| 40 | No | A. baylyi | Chloramph | Intracellul | F84 | Neutral Re | 277 | BL1 | 6246 |
| 40 | No | A. baylyi | Chloramph | Intracellul | F99 | Pyronin Y | 272 | BL1 | 5764.5 |
| 40 | No | A. baylyi | Chloramph | Intracellul | F108 | Rhodamine | 336 | BL1 | 5213 |
| 40 | No | A. baylyi | Chloramph | Intracellul | F109 | Riboflavin | 277 | BL1 | 5477 |
| 40 | No | A. baylyi | Chloramph | Intracellul | F117 | SYBR Gree | 378 | BL1 | 29021.5 |
| 40 | No | A. baylyi | Chloramph | Intracellul | F106 | Rhodamine | 360 | BL1 | 9453 |
| 40 | No | A. baylyi | Chloramph | Intracellul | F73 | Malachite ( | 251 | BL1 | 6737 |
| 40 | No | A. baylyi | Chloramph | Intracellul | F7 | Acridine or | 239 | BL1 | 6286 |
| 40 | No | A. baylyi | Chloramph | Intracellul | F192 | 5-Carboxyf | 331 | BL1 | 8867 |
| 40 | No | A. baylyi | Chloramph | Intracellul | F57 | Eosin Y | 299 | BL1 | 18038 |
| 40 | No | A. baylyi | Chloramph | Intracellul | F27 | Calcein | 314 | BL1 | 16780 |

| | | | | | | | | | |
|---|---|---|---|---|---|---|---|---|---|
| 40 | No | A. baylyi | Chloramph | Intracellula | F12 | Amaranth | 346 | BL1 | 19521.5 |
| 40 | No | A. baylyi | None | None | NA | Autofluores | 333 | NA | 17417 |
| 40 | No | A. baylyi | Colistin | Wall | F194 | Oxonol | 435 | RL1 | 16092 |
| 40 | No | A. baylyi | Colistin | Wall | F11 | Allura Red | 348 | BL3 | 36244.5 |
| 40 | No | A. baylyi | Colistin | Wall | F54 | DiSC3(5) | 197 | RL1 | 43371 |
| 40 | No | A. baylyi | Colistin | Wall | F124 | Thioflavin T | 249 | VL2 | 34481 |
| 40 | No | A. baylyi | Colistin | Wall | F101 | Quinacrine | 153 | VL2 | 46180 |
| 40 | No | A. baylyi | Colistin | Wall | F138 | Apigenin | 233 | VL2 | 35454 |
| 40 | No | A. baylyi | Colistin | Wall | F65 | H2FDA | 196 | BL1 | 32802.5 |
| 40 | No | A. baylyi | Colistin | Wall | F142 | Chlorophe | 193 | BL3 | 24460 |
| 40 | No | A. baylyi | Colistin | Wall | F39 | Congo Red | 155 | BL3 | 15763 |
| 40 | No | A. baylyi | Colistin | Wall | F116 | Sunset Yel | 214 | BL2 | 22298 |
| 40 | No | A. baylyi | Colistin | Wall | F10 | Alizarin | 173 | BL1 | 25418 |
| 40 | No | A. baylyi | Colistin | Wall | F84 | Neutral Re | 230 | BL1 | 25742 |
| 40 | No | A. baylyi | Colistin | Wall | F99 | Pyronin Y | 183 | BL1 | 29774 |
| 40 | No | A. baylyi | Colistin | Wall | F108 | Rhodamine | 215 | BL1 | 27683 |
| 40 | No | A. baylyi | Colistin | Wall | F109 | Riboflavin | 200 | BL1 | 23872.5 |
| 40 | No | A. baylyi | Colistin | Wall | F117 | SYBR Gree | 168 | BL1 | 874005 |
| 40 | No | A. baylyi | Colistin | Wall | F106 | Rhodamine | 174 | BL1 | 157326 |
| 40 | No | A. baylyi | Colistin | Wall | F73 | Malachite ( | 198 | BL1 | 68650.5 |
| 40 | No | A. baylyi | Colistin | Wall | F7 | Acridine or | 218 | BL1 | 60508.5 |
| 40 | No | A. baylyi | Colistin | Wall | F192 | 5-Carboxyf | 255 | BL1 | 56025 |
| 40 | No | A. baylyi | Colistin | Wall | F57 | Eosin Y | 154 | BL1 | 36863.5 |
| 40 | No | A. baylyi | Colistin | Wall | F27 | Calcein | 178 | BL1 | 40424 |
| 40 | No | A. baylyi | Colistin | Wall | F12 | Amaranth | 208 | BL1 | 27426.5 |
| 40 | No | A. baylyi | None | None | NA | Autofluores | 357 | NA | 18362 |
| 40 | No | A. baylyi | Gentamici | Intracellula | F194 | Oxonol | 550 | RL1 | 13154.5 |
| 40 | No | A. baylyi | Gentamici | Intracellula | F11 | Allura Red | 351 | BL3 | 17426 |
| 40 | No | A. baylyi | Gentamici | Intracellula | F54 | DiSC3(5) | 363 | RL1 | 17245 |
| 40 | No | A. baylyi | Gentamici | Intracellula | F124 | Thioflavin T | 290 | VL2 | 17571 |
| 40 | No | A. baylyi | Gentamici | Intracellula | F101 | Quinacrine | 393 | VL2 | 17339 |
| 40 | No | A. baylyi | Gentamici | Intracellula | F138 | Apigenin | 338 | VL2 | 10964.5 |
| 40 | No | A. baylyi | Gentamici | Intracellula | F65 | H2FDA | 339 | BL1 | 8628 |
| 40 | No | A. baylyi | Gentamici | Intracellula | F142 | Chlorophe | 286 | BL3 | 6070.5 |
| 40 | No | A. baylyi | Gentamici | Intracellula | F39 | Congo Red | 334 | BL3 | 5981.5 |
| 40 | No | A. baylyi | Gentamici | Intracellula | F116 | Sunset Yel | 274 | BL2 | 6462 |
| 40 | No | A. baylyi | Gentamici | Intracellula | F10 | Alizarin | 248 | BL1 | 5980 |
| 40 | No | A. baylyi | Gentamici | Intracellula | F84 | Neutral Re | 400 | BL1 | 6190.5 |
| 40 | No | A. baylyi | Gentamici | Intracellula | F99 | Pyronin Y | 331 | BL1 | 5804 |
| 40 | No | A. baylyi | Gentamici | Intracellula | F108 | Rhodamine | 245 | BL1 | 5743 |
| 40 | No | A. baylyi | Gentamici | Intracellula | F109 | Riboflavin | 278 | BL1 | 5036.5 |
| 40 | No | A. baylyi | Gentamici | Intracellula | F117 | SYBR Gree | 233 | BL1 | 14539 |
| 40 | No | A. baylyi | Gentamici | Intracellula | F106 | Rhodamine | 374 | BL1 | 11425.5 |
| 40 | No | A. baylyi | Gentamici | Intracellula | F73 | Malachite ( | 299 | BL1 | 6872 |
| 40 | No | A. baylyi | Gentamici | Intracellula | F7 | Acridine or | 342 | BL1 | 6452 |
| 40 | No | A. baylyi | Gentamici | Intracellula | F192 | 5-Carboxyf | 328 | BL1 | 13710.5 |
| 40 | No | A. baylyi | Gentamici | Intracellula | F57 | Eosin Y | 336 | BL1 | 16672.5 |
| 40 | No | A. baylyi | Gentamici | Intracellula | F27 | Calcein | 212 | BL1 | 16017.5 |

| | | | | | | | | | |
|---|---|---|---|---|---|---|---|---|---|
| 40 | No | A. baylyi | Gentamici | Intracellula | F12 | Amaranth | 361 | BL1 | 16419 |
| 40 | No | A. baylyi | None | None | NA | Autofluore: | 388 | NA | 13781.5 |
| 40 | No | A. baylyi | Nitrofurant | Intracellula | F194 | Oxonol | 741 | RL1 | 12462 |
| 40 | No | A. baylyi | Nitrofurant | Intracellula | F11 | Allura Red | 487 | BL3 | 12922 |
| 40 | No | A. baylyi | Nitrofurant | Intracellula | F54 | DiSC3(5) | 154 | RL1 | 15486.5 |
| 40 | No | A. baylyi | Nitrofurant | Intracellula | F124 | Thioflavin T | 387 | VL2 | 14422 |
| 40 | No | A. baylyi | Nitrofurant | Intracellula | F101 | Quinacrine | 255 | VL2 | 12230 |
| 40 | No | A. baylyi | Nitrofurant | Intracellula | F138 | Apigenin | 264 | VL2 | 12315 |
| 40 | No | A. baylyi | Nitrofurant | Intracellula | F65 | H2FDA | 300 | BL1 | 7139 |
| 40 | No | A. baylyi | Nitrofurant | Intracellula | F142 | Chlorophe | 279 | BL3 | 6529 |
| 40 | No | A. baylyi | Nitrofurant | Intracellula | F39 | Congo Red | 343 | BL3 | 7116 |
| 40 | No | A. baylyi | Nitrofurant | Intracellula | F116 | Sunset Yel | 224 | BL2 | 6871.5 |
| 40 | No | A. baylyi | Nitrofurant | Intracellula | F10 | Alizarin | 266 | BL1 | 6779 |
| 40 | No | A. baylyi | Nitrofurant | Intracellula | F84 | Neutral Re | 270 | BL1 | 6507 |
| 40 | No | A. baylyi | Nitrofurant | Intracellula | F99 | Pyronin Y | 194 | BL1 | 6409 |
| 40 | No | A. baylyi | Nitrofurant | Intracellula | F108 | Rhodamine | 384 | BL1 | 5661.5 |
| 40 | No | A. baylyi | Nitrofurant | Intracellula | F109 | Riboflavin | 216 | BL1 | 4976 |
| 40 | No | A. baylyi | Nitrofurant | Intracellula | F117 | SYBR Gree | 393 | BL1 | 106687 |
| 40 | No | A. baylyi | Nitrofurant | Intracellula | F106 | Rhodamine | 232 | BL1 | 9020 |
| 40 | No | A. baylyi | Nitrofurant | Intracellula | F73 | Malachite ( | 296 | BL1 | 7582 |
| 40 | No | A. baylyi | Nitrofurant | Intracellula | F7 | Acridine or | 267 | BL1 | 6620 |
| 40 | No | A. baylyi | Nitrofurant | Intracellula | F192 | 5-Carboxyf | 270 | BL1 | 11449 |
| 40 | No | A. baylyi | Nitrofurant | Intracellula | F57 | Eosin Y | 318 | BL1 | 16055.5 |
| 40 | No | A. baylyi | Nitrofurant | Intracellula | F27 | Calcein | 353 | BL1 | 19783 |
| 40 | No | A. baylyi | Nitrofurant | Intracellula | F12 | Amaranth | 236 | BL1 | 19066 |
| 40 | No | A. baylyi | None | None | NA | Autofluore: | 385 | NA | 16201 |
| 40 | No | A. baylyi | Pentamidi | Wall | F194 | Oxonol | 1224 | RL1 | 9349.5 |
| 40 | No | A. baylyi | Pentamidi | Wall | F11 | Allura Red | 486 | BL3 | 13513.5 |
| 40 | No | A. baylyi | Pentamidi | Wall | F54 | DiSC3(5) | 271 | RL1 | 16418 |
| 40 | No | A. baylyi | Pentamidi | Wall | F124 | Thioflavin T | 287 | VL2 | 22094 |
| 40 | No | A. baylyi | Pentamidi | Wall | F101 | Quinacrine | 350 | VL2 | 23628.5 |
| 40 | No | A. baylyi | Pentamidi | Wall | F138 | Apigenin | 287 | VL2 | 15689 |
| 40 | No | A. baylyi | Pentamidi | Wall | F65 | H2FDA | 269 | BL1 | 10310 |
| 40 | No | A. baylyi | Pentamidi | Wall | F142 | Chlorophe | 220 | BL3 | 14361.5 |
| 40 | No | A. baylyi | Pentamidi | Wall | F39 | Congo Red | 295 | BL3 | 8883 |
| 40 | No | A. baylyi | Pentamidi | Wall | F116 | Sunset Yel | 238 | BL2 | 8421 |
| 40 | No | A. baylyi | Pentamidi | Wall | F10 | Alizarin | 294 | BL1 | 7954.5 |
| 40 | No | A. baylyi | Pentamidi | Wall | F84 | Neutral Re | 323 | BL1 | 8155 |
| 40 | No | A. baylyi | Pentamidi | Wall | F99 | Pyronin Y | 223 | BL1 | 8666 |
| 40 | No | A. baylyi | Pentamidi | Wall | F108 | Rhodamine | 386 | BL1 | 9359.5 |
| 40 | No | A. baylyi | Pentamidi | Wall | F109 | Riboflavin | 449 | BL1 | 10646 |
| 40 | No | A. baylyi | Pentamidi | Wall | F117 | SYBR Gree | 288 | BL1 | 69493 |
| 40 | No | A. baylyi | Pentamidi | Wall | F106 | Rhodamine | 236 | BL1 | 31793.5 |
| 40 | No | A. baylyi | Pentamidi | Wall | F73 | Malachite ( | 288 | BL1 | 15315 |
| 40 | No | A. baylyi | Pentamidi | Wall | F7 | Acridine or | 213 | BL1 | 13273 |
| 40 | No | A. baylyi | Pentamidi | Wall | F192 | 5-Carboxyf | 386 | BL1 | 14743.5 |
| 40 | No | A. baylyi | Pentamidi | Wall | F57 | Eosin Y | 333 | BL1 | 33020 |
| 40 | No | A. baylyi | Pentamidi | Wall | F27 | Calcein | 604 | BL1 | 15295 |

| | | | | | | | | | |
|---|---|---|---|---|---|---|---|---|---|
| 40 | No | A. baylyi | Pentamidi | Wall | F12 | Amaranth | 265 | BL1 | 17033 |
| 40 | No | A. baylyi | None | None | NA | Autofluore: | 371 | NA | 21601 |
| 40 | No | A. baylyi | Rifampicin | Intracellul | F194 | Oxonol | 2192 | RL1 | 8766 |
| 40 | No | A. baylyi | Rifampicin | Intracellul | F11 | Allura Red | 772 | BL3 | 13919.5 |
| 40 | No | A. baylyi | Rifampicin | Intracellul | F54 | DiSC3(5) | 284 | RL1 | 18364 |
| 40 | No | A. baylyi | Rifampicin | Intracellul | F124 | Thioflavin T | 398 | VL2 | 19098.5 |
| 40 | No | A. baylyi | Rifampicin | Intracellul | F101 | Quinacrine | 313 | VL2 | 16127 |
| 40 | No | A. baylyi | Rifampicin | Intracellul | F138 | Apigenin | 323 | VL2 | 15597 |
| 40 | No | A. baylyi | Rifampicin | Intracellul | F65 | H2FDA | 202 | BL1 | 11140 |
| 40 | No | A. baylyi | Rifampicin | Intracellul | F142 | Chlorophe | 296 | BL3 | 8860 |
| 40 | No | A. baylyi | Rifampicin | Intracellul | F39 | Congo Red | 210 | BL3 | 7735 |
| 40 | No | A. baylyi | Rifampicin | Intracellul | F116 | Sunset Yel | 374 | BL2 | 9235 |
| 40 | No | A. baylyi | Rifampicin | Intracellul | F10 | Alizarin | 229 | BL1 | 7236 |
| 40 | No | A. baylyi | Rifampicin | Intracellul | F84 | Neutral Re | 251 | BL1 | 6824 |
| 40 | No | A. baylyi | Rifampicin | Intracellul | F99 | Pyronin Y | 297 | BL1 | 8002 |
| 40 | No | A. baylyi | Rifampicin | Intracellul | F108 | Rhodamine | 353 | BL1 | 5463 |
| 40 | No | A. baylyi | Rifampicin | Intracellul | F109 | Riboflavin | 315 | BL1 | 5409 |
| 40 | No | A. baylyi | Rifampicin | Intracellul | F117 | SYBR Gree | 208 | BL1 | 8023.5 |
| 40 | No | A. baylyi | Rifampicin | Intracellul | F106 | Rhodamine | 262 | BL1 | 33489.5 |
| 40 | No | A. baylyi | Rifampicin | Intracellul | F73 | Malachite ( | 342 | BL1 | 6584 |
| 40 | No | A. baylyi | Rifampicin | Intracellul | F7 | Acridine or | 220 | BL1 | 6319.5 |
| 40 | No | A. baylyi | Rifampicin | Intracellul | F192 | 5-Carboxyf | 414 | BL1 | 14298.5 |
| 40 | No | A. baylyi | Rifampicin | Intracellul | F57 | Eosin Y | 263 | BL1 | 18038 |
| 40 | No | A. baylyi | Rifampicin | Intracellul | F27 | Calcein | 276 | BL1 | 25385.5 |
| 40 | No | A. baylyi | Rifampicin | Intracellul | F12 | Amaranth | 275 | BL1 | 17237 |
| 40 | No | A. baylyi | None | None | NA | Autofluore: | 361 | NA | 16412 |
| 40 | No | A. baylyi | Trimethopr | Intracellul | F194 | Oxonol | 1461 | RL1 | 13645 |
| 40 | No | A. baylyi | Trimethopr | Intracellul | F11 | Allura Red | 517 | BL3 | 14677 |
| 40 | No | A. baylyi | Trimethopr | Intracellul | F54 | DiSC3(5) | 356 | RL1 | 12628.5 |
| 40 | No | A. baylyi | Trimethopr | Intracellul | F124 | Thioflavin T | 353 | VL2 | 14260 |
| 40 | No | A. baylyi | Trimethopr | Intracellul | F101 | Quinacrine | 323 | VL2 | 14369 |
| 40 | No | A. baylyi | Trimethopr | Intracellul | F138 | Apigenin | 375 | VL2 | 10447 |
| 40 | No | A. baylyi | Trimethopr | Intracellul | F65 | H2FDA | 294 | BL1 | 9058 |
| 40 | No | A. baylyi | Trimethopr | Intracellul | F142 | Chlorophe | 236 | BL3 | 7064.5 |
| 40 | No | A. baylyi | Trimethopr | Intracellul | F39 | Congo Red | 242 | BL3 | 6051 |
| 40 | No | A. baylyi | Trimethopr | Intracellul | F116 | Sunset Yel | 429 | BL2 | 6682 |
| 40 | No | A. baylyi | Trimethopr | Intracellul | F10 | Alizarin | 331 | BL1 | 7064 |
| 40 | No | A. baylyi | Trimethopr | Intracellul | F84 | Neutral Re | 362 | BL1 | 6758 |
| 40 | No | A. baylyi | Trimethopr | Intracellul | F99 | Pyronin Y | 329 | BL1 | 6997 |
| 40 | No | A. baylyi | Trimethopr | Intracellul | F108 | Rhodamine | 292 | BL1 | 5757 |
| 40 | No | A. baylyi | Trimethopr | Intracellul | F109 | Riboflavin | 378 | BL1 | 5426 |
| 40 | No | A. baylyi | Trimethopr | Intracellul | F117 | SYBR Gree | 262 | BL1 | 99010 |
| 40 | No | A. baylyi | Trimethopr | Intracellul | F106 | Rhodamine | 312 | BL1 | 8273.5 |
| 40 | No | A. baylyi | Trimethopr | Intracellul | F73 | Malachite ( | 251 | BL1 | 7675 |
| 40 | No | A. baylyi | Trimethopr | Intracellul | F7 | Acridine or | 397 | BL1 | 7029 |
| 40 | No | A. baylyi | Trimethopr | Intracellul | F192 | 5-Carboxyf | 240 | BL1 | 7813 |
| 40 | No | A. baylyi | Trimethopr | Intracellul | F57 | Eosin Y | 277 | BL1 | 16686 |
| 40 | No | A. baylyi | Trimethopr | Intracellul | F27 | Calcein | 247 | BL1 | 18289 |

| | | | | | | | | | |
|---|---|---|---|---|---|---|---|---|---|
| 40 | No | A. baylyi | Trimethopr | Intracellula | F12 | Amaranth | 298 | BL1 | 15386 |
| 40 | No | A. baylyi | None | None | NA | Autofluore: | 2007 | NA | 1401 |
| 40 | No | A. baylyi | None | None | F194 | Oxonol | 2541 | RL1 | 59696 |
| 40 | No | A. baylyi | None | None | F11 | Allura Red | 1494 | BL3 | 53235.5 |
| 40 | No | A. baylyi | None | None | F54 | DiSC3(5) | 1326 | RL1 | 9659 |
| 40 | No | A. baylyi | None | None | F124 | Thioflavin T | 1153 | VL2 | 7085 |
| 40 | No | A. baylyi | None | None | F101 | Quinacrine | 1083 | VL2 | 5420 |
| 40 | No | A. baylyi | None | None | F138 | Apigenin | 1244 | VL2 | 4891.5 |
| 40 | No | A. baylyi | None | None | F65 | H2FDA | 1023 | BL1 | 6813 |
| 40 | No | A. baylyi | None | None | F142 | Chlorophe | 1036 | BL3 | 8790.5 |
| 40 | No | A. baylyi | None | None | F39 | Congo Red | 1040 | BL3 | 7922 |
| 40 | No | A. baylyi | None | None | F116 | Sunset Yel | 932 | BL2 | 6599.5 |
| 40 | No | A. baylyi | None | None | F10 | Alizarin | 1022 | BL1 | 5749 |
| 40 | No | A. baylyi | None | None | F84 | Neutral Re | 999 | BL1 | 4586 |
| 40 | No | A. baylyi | None | None | F99 | Pyronin Y | 877 | BL1 | 4438 |
| 40 | No | A. baylyi | None | None | F108 | Rhodamine | 940 | BL1 | 3835.5 |
| 40 | No | A. baylyi | None | None | F109 | Riboflavin | 923 | BL1 | 4218 |
| 40 | No | A. baylyi | None | None | F117 | SYBR Gree | 876 | BL1 | 253573 |
| 40 | No | A. baylyi | None | None | F106 | Rhodamine | 1070 | BL1 | 6177.5 |
| 40 | No | A. baylyi | None | None | F73 | Malachite ( | 914 | BL1 | 4439 |
| 40 | No | A. baylyi | None | None | F7 | Acridine or | 897 | BL1 | 5476 |
| 40 | No | A. baylyi | None | None | F192 | 5-Carboxyf | 1018 | BL1 | 18394 |
| 40 | No | A. baylyi | None | None | F57 | Eosin Y | 942 | BL1 | 17785 |
| 40 | No | A. baylyi | None | None | F27 | Calcein | 943 | BL1 | 16549 |
| 40 | No | A. baylyi | None | None | F12 | Amaranth | 939 | BL1 | 18026 |
| 40 | No | A. baylyi | None | None | NA | Autofluore: | 1054 | NA | 18344.5 |
| 40 | No | A. baylyi | Amoxacilli | Wall | F194 | Oxonol | 2701 | RL1 | 14644 |
| 40 | No | A. baylyi | Amoxacilli | Wall | F11 | Allura Red | 923 | BL3 | 14042 |
| 40 | No | A. baylyi | Amoxacilli | Wall | F54 | DiSC3(5) | 904 | RL1 | 12763 |
| 40 | No | A. baylyi | Amoxacilli | Wall | F124 | Thioflavin T | 917 | VL2 | 11426 |
| 40 | No | A. baylyi | Amoxacilli | Wall | F101 | Quinacrine | 884 | VL2 | 7942.5 |
| 40 | No | A. baylyi | Amoxacilli | Wall | F138 | Apigenin | 921 | VL2 | 6711 |
| 40 | No | A. baylyi | Amoxacilli | Wall | F65 | H2FDA | 921 | BL1 | 8686 |
| 40 | No | A. baylyi | Amoxacilli | Wall | F142 | Chlorophe | 942 | BL3 | 8937 |
| 40 | No | A. baylyi | Amoxacilli | Wall | F39 | Congo Red | 928 | BL3 | 9607 |
| 40 | No | A. baylyi | Amoxacilli | Wall | F116 | Sunset Yel | 956 | BL2 | 7078 |
| 40 | No | A. baylyi | Amoxacilli | Wall | F10 | Alizarin | 910 | BL1 | 5841 |
| 40 | No | A. baylyi | Amoxacilli | Wall | F84 | Neutral Re | 999 | BL1 | 6218 |
| 40 | No | A. baylyi | Amoxacilli | Wall | F99 | Pyronin Y | 909 | BL1 | 5919 |
| 40 | No | A. baylyi | Amoxacilli | Wall | F108 | Rhodamine | 881 | BL1 | 4329 |
| 40 | No | A. baylyi | Amoxacilli | Wall | F109 | Riboflavin | 901 | BL1 | 5310 |
| 40 | No | A. baylyi | Amoxacilli | Wall | F117 | SYBR Gree | 920 | BL1 | 289610.5 |
| 40 | No | A. baylyi | Amoxacilli | Wall | F106 | Rhodamine | 907 | BL1 | 7824 |
| 40 | No | A. baylyi | Amoxacilli | Wall | F73 | Malachite ( | 902 | BL1 | 5694.5 |
| 40 | No | A. baylyi | Amoxacilli | Wall | F7 | Acridine or | 816 | BL1 | 5866.5 |
| 40 | No | A. baylyi | Amoxacilli | Wall | F192 | 5-Carboxyf | 920 | BL1 | 17286.5 |
| 40 | No | A. baylyi | Amoxacilli | Wall | F57 | Eosin Y | 865 | BL1 | 19314 |
| 40 | No | A. baylyi | Amoxacilli | Wall | F27 | Calcein | 865 | BL1 | 17986 |

| | | | | | | | | | |
|---|---|---|---|---|---|---|---|---|---|
| 40 | No | A. baylyi | Amoxacilli | Wall | F12 | Amaranth | 760 | BL1 | 18059 |
| 40 | No | A. baylyi | None | None | NA | Autofluore: | 916 | NA | 17435 |
| 40 | No | A. baylyi | Azithromyc | Intracellul | F194 | Oxonol | 2195 | RL1 | 14969 |
| 40 | No | A. baylyi | Azithromyc | Intracellul | F11 | Allura Red | 927 | BL3 | 14096 |
| 40 | No | A. baylyi | Azithromyc | Intracellul | F54 | DiSC3(5) | 805 | RL1 | 13056 |
| 40 | No | A. baylyi | Azithromyc | Intracellul | F124 | Thioflavin T | 850 | VL2 | 11865 |
| 40 | No | A. baylyi | Azithromyc | Intracellul | F101 | Quinacrine | 948 | VL2 | 7988 |
| 40 | No | A. baylyi | Azithromyc | Intracellul | F138 | Apigenin | 1060 | VL2 | 6569.5 |
| 40 | No | A. baylyi | Azithromyc | Intracellul | F65 | H2FDA | 994 | BL1 | 8134.5 |
| 40 | No | A. baylyi | Azithromyc | Intracellul | F142 | Chlorophe | 988 | BL3 | 8021 |
| 40 | No | A. baylyi | Azithromyc | Intracellul | F39 | Congo Red | 1011 | BL3 | 7348 |
| 40 | No | A. baylyi | Azithromyc | Intracellul | F116 | Sunset Yel | 907 | BL2 | 6293 |
| 40 | No | A. baylyi | Azithromyc | Intracellul | F10 | Alizarin | 949 | BL1 | 5684 |
| 40 | No | A. baylyi | Azithromyc | Intracellul | F84 | Neutral Re | 893 | BL1 | 5250 |
| 40 | No | A. baylyi | Azithromyc | Intracellul | F99 | Pyronin Y | 837 | BL1 | 5352 |
| 40 | No | A. baylyi | Azithromyc | Intracellul | F108 | Rhodamine | 849 | BL1 | 5015 |
| 40 | No | A. baylyi | Azithromyc | Intracellul | F109 | Riboflavin | 912 | BL1 | 5094 |
| 40 | No | A. baylyi | Azithromyc | Intracellul | F117 | SYBR Gree | 880 | BL1 | 247398 |
| 40 | No | A. baylyi | Azithromyc | Intracellul | F106 | Rhodamine | 928 | BL1 | 7182 |
| 40 | No | A. baylyi | Azithromyc | Intracellul | F73 | Malachite ( | 880 | BL1 | 5321.5 |
| 40 | No | A. baylyi | Azithromyc | Intracellul | F7 | Acridine or | 1055 | BL1 | 6533 |
| 40 | No | A. baylyi | Azithromyc | Intracellul | F192 | 5-Carboxyf | 941 | BL1 | 18623 |
| 40 | No | A. baylyi | Azithromyc | Intracellul | F57 | Eosin Y | 879 | BL1 | 15790 |
| 40 | No | A. baylyi | Azithromyc | Intracellul | F27 | Calcein | 903 | BL1 | 17234 |
| 40 | No | A. baylyi | Azithromyc | Intracellul | F12 | Amaranth | 1002 | BL1 | 18306 |
| 40 | No | A. baylyi | None | None | NA | Autofluore: | 927 | NA | 17831 |
| 40 | No | A. baylyi | Trimethopr | Intracellul | F194 | Oxonol | 4020 | RL1 | 12852.5 |
| 40 | No | A. baylyi | Trimethopr | Intracellul | F11 | Allura Red | 962 | BL3 | 13513 |
| 40 | No | A. baylyi | Trimethopr | Intracellul | F54 | DiSC3(5) | 879 | RL1 | 11579 |
| 40 | No | A. baylyi | Trimethopr | Intracellul | F124 | Thioflavin T | 914 | VL2 | 10410 |
| 40 | No | A. baylyi | Trimethopr | Intracellul | F101 | Quinacrine | 897 | VL2 | 7631 |
| 40 | No | A. baylyi | Trimethopr | Intracellul | F138 | Apigenin | 977 | VL2 | 6726 |
| 40 | No | A. baylyi | Trimethopr | Intracellul | F65 | H2FDA | 903 | BL1 | 7935 |
| 40 | No | A. baylyi | Trimethopr | Intracellul | F142 | Chlorophe | 915 | BL3 | 9322 |
| 40 | No | A. baylyi | Trimethopr | Intracellul | F39 | Congo Red | 852 | BL3 | 8342 |
| 40 | No | A. baylyi | Trimethopr | Intracellul | F116 | Sunset Yel | 923 | BL2 | 7414 |
| 40 | No | A. baylyi | Trimethopr | Intracellul | F10 | Alizarin | 906 | BL1 | 6421 |
| 40 | No | A. baylyi | Trimethopr | Intracellul | F84 | Neutral Re | 927 | BL1 | 5625 |
| 40 | No | A. baylyi | Trimethopr | Intracellul | F99 | Pyronin Y | 821 | BL1 | 5578 |
| 40 | No | A. baylyi | Trimethopr | Intracellul | F108 | Rhodamine | 911 | BL1 | 4724 |
| 40 | No | A. baylyi | Trimethopr | Intracellul | F109 | Riboflavin | 818 | BL1 | 5159.5 |
| 40 | No | A. baylyi | Trimethopr | Intracellul | F117 | SYBR Gree | 827 | BL1 | 275826 |
| 40 | No | A. baylyi | Trimethopr | Intracellul | F106 | Rhodamine | 870 | BL1 | 8001.5 |
| 40 | No | A. baylyi | Trimethopr | Intracellul | F73 | Malachite ( | 1001 | BL1 | 6443 |
| 40 | No | A. baylyi | Trimethopr | Intracellul | F7 | Acridine or | 792 | BL1 | 6105 |
| 40 | No | A. baylyi | Trimethopr | Intracellul | F192 | 5-Carboxyf | 848 | BL1 | 18061 |
| 40 | No | A. baylyi | Trimethopr | Intracellul | F57 | Eosin Y | 887 | BL1 | 15450 |
| 40 | No | A. baylyi | Trimethopr | Intracellul | F27 | Calcein | 926 | BL1 | 15488.5 |

| | | | | | | | | | |
|---|---|---|---|---|---|---|---|---|---|
| 40 | No | A. baylyi | Trimethopr | Intracellula | F12 | Amaranth | 911 | BL1 | 17329 |
| 40 | No | A. baylyi | None | None | NA | Autofluore: | 938 | NA | 18595.5 |
| 40 | No | A. baylyi | Chloramph | Intracellula | F194 | Oxonol | 2431 | RL1 | 15442 |
| 40 | No | A. baylyi | Chloramph | Intracellula | F11 | Allura Red | 853 | BL3 | 15089 |
| 40 | No | A. baylyi | Chloramph | Intracellula | F54 | DiSC3(5) | 753 | RL1 | 12825 |
| 40 | No | A. baylyi | Chloramph | Intracellula | F124 | Thioflavin T | 883 | VL2 | 11799 |
| 40 | No | A. baylyi | Chloramph | Intracellula | F101 | Quinacrine | 857 | VL2 | 8344 |
| 40 | No | A. baylyi | Chloramph | Intracellula | F138 | Apigenin | 748 | VL2 | 6242 |
| 40 | No | A. baylyi | Chloramph | Intracellula | F65 | H2FDA | 871 | BL1 | 8806 |
| 40 | No | A. baylyi | Chloramph | Intracellula | F142 | Chlorophe | 837 | BL3 | 8794 |
| 40 | No | A. baylyi | Chloramph | Intracellula | F39 | Congo Red | 826 | BL3 | 8230.5 |
| 40 | No | A. baylyi | Chloramph | Intracellula | F116 | Sunset Yel | 796 | BL2 | 7874.5 |
| 40 | No | A. baylyi | Chloramph | Intracellula | F10 | Alizarin | 773 | BL1 | 6174 |
| 40 | No | A. baylyi | Chloramph | Intracellula | F84 | Neutral Re | 787 | BL1 | 5607 |
| 40 | No | A. baylyi | Chloramph | Intracellula | F99 | Pyronin Y | 899 | BL1 | 6424 |
| 40 | No | A. baylyi | Chloramph | Intracellula | F108 | Rhodamine | 838 | BL1 | 4903 |
| 40 | No | A. baylyi | Chloramph | Intracellula | F109 | Riboflavin | 811 | BL1 | 5643 |
| 40 | No | A. baylyi | Chloramph | Intracellula | F117 | SYBR Gree | 843 | BL1 | 221812 |
| 40 | No | A. baylyi | Chloramph | Intracellula | F106 | Rhodamine | 863 | BL1 | 10286 |
| 40 | No | A. baylyi | Chloramph | Intracellula | F73 | Malachite ( | 829 | BL1 | 5937 |
| 40 | No | A. baylyi | Chloramph | Intracellula | F7 | Acridine or | 902 | BL1 | 7877.5 |
| 40 | No | A. baylyi | Chloramph | Intracellula | F192 | 5-Carboxyf | 806 | BL1 | 19386 |
| 40 | No | A. baylyi | Chloramph | Intracellula | F57 | Eosin Y | 790 | BL1 | 17644.5 |
| 40 | No | A. baylyi | Chloramph | Intracellula | F27 | Calcein | 865 | BL1 | 17258 |
| 40 | No | A. baylyi | Chloramph | Intracellula | F12 | Amaranth | 765 | BL1 | 19931 |
| 40 | No | A. baylyi | None | None | NA | Autofluore: | 900 | NA | 18539.5 |
| 40 | No | A. baylyi | Colistin | Wall | F194 | Oxonol | 4008 | RL1 | 12366.5 |
| 40 | No | A. baylyi | Colistin | Wall | F11 | Allura Red | 676 | BL3 | 72837 |
| 40 | No | A. baylyi | Colistin | Wall | F54 | DiSC3(5) | 532 | RL1 | 82078.5 |
| 40 | No | A. baylyi | Colistin | Wall | F124 | Thioflavin T | 615 | VL2 | 98525 |
| 40 | No | A. baylyi | Colistin | Wall | F101 | Quinacrine | 687 | VL2 | 73589 |
| 40 | No | A. baylyi | Colistin | Wall | F138 | Apigenin | 646 | VL2 | 62718 |
| 40 | No | A. baylyi | Colistin | Wall | F65 | H2FDA | 635 | BL1 | 76391 |
| 40 | No | A. baylyi | Colistin | Wall | F142 | Chlorophe | 540 | BL3 | 53412.5 |
| 40 | No | A. baylyi | Colistin | Wall | F39 | Congo Red | 652 | BL3 | 42489.5 |
| 40 | No | A. baylyi | Colistin | Wall | F116 | Sunset Yel | 687 | BL2 | 55203 |
| 40 | No | A. baylyi | Colistin | Wall | F10 | Alizarin | 624 | BL1 | 48375.5 |
| 40 | No | A. baylyi | Colistin | Wall | F84 | Neutral Re | 665 | BL1 | 47997 |
| 40 | No | A. baylyi | Colistin | Wall | F99 | Pyronin Y | 557 | BL1 | 60792 |
| 40 | No | A. baylyi | Colistin | Wall | F108 | Rhodamine | 593 | BL1 | 49112 |
| 40 | No | A. baylyi | Colistin | Wall | F109 | Riboflavin | 554 | BL1 | 54717.5 |
| 40 | No | A. baylyi | Colistin | Wall | F117 | SYBR Gree | 608 | BL1 | 1600107 |
| 40 | No | A. baylyi | Colistin | Wall | F106 | Rhodamine | 564 | BL1 | 150708 |
| 40 | No | A. baylyi | Colistin | Wall | F73 | Malachite ( | 682 | BL1 | 97254 |
| 40 | No | A. baylyi | Colistin | Wall | F7 | Acridine or | 620 | BL1 | 94279.5 |
| 40 | No | A. baylyi | Colistin | Wall | F192 | 5-Carboxyf | 597 | BL1 | 33525 |
| 40 | No | A. baylyi | Colistin | Wall | F57 | Eosin Y | 594 | BL1 | 28615.5 |
| 40 | No | A. baylyi | Colistin | Wall | F27 | Calcein | 615 | BL1 | 29795 |

| | | | | | | | | | |
|---|---|---|---|---|---|---|---|---|---|
| 40 | No | A. baylyi | Colistin | Wall | F12 | Amaranth | 644 | BL1 | 29999.5 |
| 40 | No | A. baylyi | None | None | NA | Autofluores | 883 | NA | 17875 |
| 40 | No | A. baylyi | Gentamici | Intracellula | F194 | Oxonol | 3322 | RL1 | 14252 |
| 40 | No | A. baylyi | Gentamici | Intracellula | F11 | Allura Red | 949 | BL3 | 14755 |
| 40 | No | A. baylyi | Gentamici | Intracellula | F54 | DiSC3(5) | 943 | RL1 | 12136 |
| 40 | No | A. baylyi | Gentamici | Intracellula | F124 | Thioflavin T | 967 | VL2 | 12536 |
| 40 | No | A. baylyi | Gentamici | Intracellula | F101 | Quinacrine | 906 | VL2 | 8775.5 |
| 40 | No | A. baylyi | Gentamici | Intracellula | F138 | Apigenin | 991 | VL2 | 7449 |
| 40 | No | A. baylyi | Gentamici | Intracellula | F65 | H2FDA | 915 | BL1 | 7996 |
| 40 | No | A. baylyi | Gentamici | Intracellula | F142 | Chlorophe | 1031 | BL3 | 10258 |
| 40 | No | A. baylyi | Gentamici | Intracellula | F39 | Congo Red | 977 | BL3 | 8598 |
| 40 | No | A. baylyi | Gentamici | Intracellula | F116 | Sunset Yel | 896 | BL2 | 7743.5 |
| 40 | No | A. baylyi | Gentamici | Intracellula | F10 | Alizarin | 929 | BL1 | 6774 |
| 40 | No | A. baylyi | Gentamici | Intracellula | F84 | Neutral Re | 880 | BL1 | 6205 |
| 40 | No | A. baylyi | Gentamici | Intracellula | F99 | Pyronin Y | 769 | BL1 | 6025 |
| 40 | No | A. baylyi | Gentamici | Intracellula | F108 | Rhodamine | 866 | BL1 | 5080 |
| 40 | No | A. baylyi | Gentamici | Intracellula | F109 | Riboflavin | 981 | BL1 | 5612 |
| 40 | No | A. baylyi | Gentamici | Intracellula | F117 | SYBR Gree | 856 | BL1 | 339929.5 |
| 40 | No | A. baylyi | Gentamici | Intracellula | F106 | Rhodamine | 877 | BL1 | 8395 |
| 40 | No | A. baylyi | Gentamici | Intracellula | F73 | Malachite ( | 968 | BL1 | 6031 |
| 40 | No | A. baylyi | Gentamici | Intracellula | F7 | Acridine or | 840 | BL1 | 6291 |
| 40 | No | A. baylyi | Gentamici | Intracellula | F192 | 5-Carboxyf | 926 | BL1 | 19372.5 |
| 40 | No | A. baylyi | Gentamici | Intracellula | F57 | Eosin Y | 928 | BL1 | 17127.5 |
| 40 | No | A. baylyi | Gentamici | Intracellula | F27 | Calcein | 904 | BL1 | 15919.5 |
| 40 | No | A. baylyi | Gentamici | Intracellula | F12 | Amaranth | 927 | BL1 | 19231 |
| 40 | No | A. baylyi | None | None | NA | Autofluores | 905 | NA | 20530 |
| 40 | No | A. baylyi | Nitrofurant | Intracellula | F194 | Oxonol | 3819 | RL1 | 12579 |
| 40 | No | A. baylyi | Nitrofurant | Intracellula | F11 | Allura Red | 909 | BL3 | 13630 |
| 40 | No | A. baylyi | Nitrofurant | Intracellula | F54 | DiSC3(5) | 782 | RL1 | 11880.5 |
| 40 | No | A. baylyi | Nitrofurant | Intracellula | F124 | Thioflavin T | 839 | VL2 | 12814 |
| 40 | No | A. baylyi | Nitrofurant | Intracellula | F101 | Quinacrine | 824 | VL2 | 8899 |
| 40 | No | A. baylyi | Nitrofurant | Intracellula | F138 | Apigenin | 721 | VL2 | 6896 |
| 40 | No | A. baylyi | Nitrofurant | Intracellula | F65 | H2FDA | 793 | BL1 | 8719 |
| 40 | No | A. baylyi | Nitrofurant | Intracellula | F142 | Chlorophe | 844 | BL3 | 10466.5 |
| 40 | No | A. baylyi | Nitrofurant | Intracellula | F39 | Congo Red | 749 | BL3 | 8455 |
| 40 | No | A. baylyi | Nitrofurant | Intracellula | F116 | Sunset Yel | 806 | BL2 | 7852 |
| 40 | No | A. baylyi | Nitrofurant | Intracellula | F10 | Alizarin | 778 | BL1 | 6700.5 |
| 40 | No | A. baylyi | Nitrofurant | Intracellula | F84 | Neutral Re | 911 | BL1 | 5737 |
| 40 | No | A. baylyi | Nitrofurant | Intracellula | F99 | Pyronin Y | 744 | BL1 | 6475 |
| 40 | No | A. baylyi | Nitrofurant | Intracellula | F108 | Rhodamine | 777 | BL1 | 5265 |
| 40 | No | A. baylyi | Nitrofurant | Intracellula | F109 | Riboflavin | 759 | BL1 | 5765 |
| 40 | No | A. baylyi | Nitrofurant | Intracellula | F117 | SYBR Gree | 850 | BL1 | 109921 |
| 40 | No | A. baylyi | Nitrofurant | Intracellula | F106 | Rhodamine | 677 | BL1 | 15039 |
| 40 | No | A. baylyi | Nitrofurant | Intracellula | F73 | Malachite ( | 829 | BL1 | 6042 |
| 40 | No | A. baylyi | Nitrofurant | Intracellula | F7 | Acridine or | 781 | BL1 | 7905 |
| 40 | No | A. baylyi | Nitrofurant | Intracellula | F192 | 5-Carboxyf | 843 | BL1 | 17425 |
| 40 | No | A. baylyi | Nitrofurant | Intracellula | F57 | Eosin Y | 806 | BL1 | 13468 |
| 40 | No | A. baylyi | Nitrofurant | Intracellula | F27 | Calcein | 941 | BL1 | 14875 |

| | | | | | | | | | |
|---|---|---|---|---|---|---|---|---|---|
| 40 | No | A. baylyi | Nitrofurant | Intracellula | F12 | Amaranth | 771 | BL1 | 16208 |
| 40 | No | A. baylyi | None | None | NA | Autofluore: | 986 | NA | 18483 |
| 40 | No | A. baylyi | Pentamidi | Wall | F194 | Oxonol | 9856 | RL1 | 17634.5 |
| 40 | No | A. baylyi | Pentamidi | Wall | F11 | Allura Red | 1568 | BL3 | 12552.5 |
| 40 | No | A. baylyi | Pentamidi | Wall | F54 | DiSC3(5) | 855 | RL1 | 14752 |
| 40 | No | A. baylyi | Pentamidi | Wall | F124 | Thioflavin T | 730 | VL2 | 36644 |
| 40 | No | A. baylyi | Pentamidi | Wall | F101 | Quinacrine | 765 | VL2 | 12058 |
| 40 | No | A. baylyi | Pentamidi | Wall | F138 | Apigenin | 711 | VL2 | 9061 |
| 40 | No | A. baylyi | Pentamidi | Wall | F65 | H2FDA | 787 | BL1 | 11043 |
| 40 | No | A. baylyi | Pentamidi | Wall | F142 | Chlorophe | 781 | BL3 | 10737 |
| 40 | No | A. baylyi | Pentamidi | Wall | F39 | Congo Red | 1085 | BL3 | 11242 |
| 40 | No | A. baylyi | Pentamidi | Wall | F116 | Sunset Yel | 761 | BL2 | 9170 |
| 40 | No | A. baylyi | Pentamidi | Wall | F10 | Alizarin | 806 | BL1 | 7446 |
| 40 | No | A. baylyi | Pentamidi | Wall | F84 | Neutral Re | 795 | BL1 | 7093 |
| 40 | No | A. baylyi | Pentamidi | Wall | F99 | Pyronin Y | 806 | BL1 | 6983.5 |
| 40 | No | A. baylyi | Pentamidi | Wall | F108 | Rhodamine | 705 | BL1 | 6139 |
| 40 | No | A. baylyi | Pentamidi | Wall | F109 | Riboflavin | 778 | BL1 | 6897 |
| 40 | No | A. baylyi | Pentamidi | Wall | F117 | SYBR Gree | 825 | BL1 | 1209562 |
| 40 | No | A. baylyi | Pentamidi | Wall | F106 | Rhodamine | 715 | BL1 | 22945 |
| 40 | No | A. baylyi | Pentamidi | Wall | F73 | Malachite ( | 804 | BL1 | 8912.5 |
| 40 | No | A. baylyi | Pentamidi | Wall | F7 | Acridine or | 669 | BL1 | 12450 |
| 40 | No | A. baylyi | Pentamidi | Wall | F192 | 5-Carboxyf | 655 | BL1 | 17532 |
| 40 | No | A. baylyi | Pentamidi | Wall | F57 | Eosin Y | 743 | BL1 | 14892 |
| 40 | No | A. baylyi | Pentamidi | Wall | F27 | Calcein | 694 | BL1 | 14803 |
| 40 | No | A. baylyi | Pentamidi | Wall | F12 | Amaranth | 750 | BL1 | 15363.5 |
| 40 | No | A. baylyi | None | None | NA | Autofluore: | 888 | NA | 18073 |
| 40 | No | A. baylyi | Rifampicin | Intracellula | F194 | Oxonol | 5801 | RL1 | 11922 |
| 40 | No | A. baylyi | Rifampicin | Intracellula | F11 | Allura Red | 1247 | BL3 | 18324 |
| 40 | No | A. baylyi | Rifampicin | Intracellula | F54 | DiSC3(5) | 973 | RL1 | 12791 |
| 40 | No | A. baylyi | Rifampicin | Intracellula | F124 | Thioflavin T | 944 | VL2 | 11960 |
| 40 | No | A. baylyi | Rifampicin | Intracellula | F101 | Quinacrine | 1063 | VL2 | 8226 |
| 40 | No | A. baylyi | Rifampicin | Intracellula | F138 | Apigenin | 1049 | VL2 | 6924 |
| 40 | No | A. baylyi | Rifampicin | Intracellula | F65 | H2FDA | 979 | BL1 | 8493 |
| 40 | No | A. baylyi | Rifampicin | Intracellula | F142 | Chlorophe | 1042 | BL3 | 9480 |
| 40 | No | A. baylyi | Rifampicin | Intracellula | F39 | Congo Red | 999 | BL3 | 9363 |
| 40 | No | A. baylyi | Rifampicin | Intracellula | F116 | Sunset Yel | 1006 | BL2 | 7651 |
| 40 | No | A. baylyi | Rifampicin | Intracellula | F10 | Alizarin | 908 | BL1 | 7426 |
| 40 | No | A. baylyi | Rifampicin | Intracellula | F84 | Neutral Re | 1018 | BL1 | 6587 |
| 40 | No | A. baylyi | Rifampicin | Intracellula | F99 | Pyronin Y | 926 | BL1 | 6813.5 |
| 40 | No | A. baylyi | Rifampicin | Intracellula | F108 | Rhodamine | 1025 | BL1 | 5907 |
| 40 | No | A. baylyi | Rifampicin | Intracellula | F109 | Riboflavin | 918 | BL1 | 7065 |
| 40 | No | A. baylyi | Rifampicin | Intracellula | F117 | SYBR Gree | 995 | BL1 | 352194 |
| 40 | No | A. baylyi | Rifampicin | Intracellula | F106 | Rhodamine | 948 | BL1 | 9335.5 |
| 40 | No | A. baylyi | Rifampicin | Intracellula | F73 | Malachite ( | 974 | BL1 | 7191 |
| 40 | No | A. baylyi | Rifampicin | Intracellula | F7 | Acridine or | 860 | BL1 | 7197 |
| 40 | No | A. baylyi | Rifampicin | Intracellula | F192 | 5-Carboxyf | 1018 | BL1 | 18511 |
| 40 | No | A. baylyi | Rifampicin | Intracellula | F57 | Eosin Y | 904 | BL1 | 17752 |
| 40 | No | A. baylyi | Rifampicin | Intracellula | F27 | Calcein | 1006 | BL1 | 16855.5 |

| | | | | | | | | | |
|---|---|---|---|---|---|---|---|---|---|
| 40 | No | A. baylyi | Rifampicin | Intracellul | F12 | Amaranth | 904 | BL1 | 18943 |
| 40 | No | A. baylyi | None | None | NA | Autofluore | 1035 | NA | 18433 |
| 40 | No | A. baylyi | Trimethopr | Intracellul | F194 | Oxonol | 3856 | RL1 | 13735.5 |
| 40 | No | A. baylyi | Trimethopr | Intracellul | F11 | Allura Red | 1131 | BL3 | 14281 |
| 40 | No | A. baylyi | Trimethopr | Intracellul | F54 | DiSC3(5) | 965 | RL1 | 11703 |
| 40 | No | A. baylyi | Trimethopr | Intracellul | F124 | Thioflavin T | 976 | VL2 | 12146 |
| 40 | No | A. baylyi | Trimethopr | Intracellul | F101 | Quinacrine | 1110 | VL2 | 9395 |
| 40 | No | A. baylyi | Trimethopr | Intracellul | F138 | Apigenin | 1144 | VL2 | 7651.5 |
| 40 | No | A. baylyi | Trimethopr | Intracellul | F65 | H2FDA | 1015 | BL1 | 9426 |
| 40 | No | A. baylyi | Trimethopr | Intracellul | F142 | Chlorophe | 1038 | BL3 | 9749 |
| 40 | No | A. baylyi | Trimethopr | Intracellul | F39 | Congo Red | 1054 | BL3 | 10634 |
| 40 | No | A. baylyi | Trimethopr | Intracellul | F116 | Sunset Yel | 973 | BL2 | 9032 |
| 40 | No | A. baylyi | Trimethopr | Intracellul | F10 | Alizarin | 948 | BL1 | 7339 |
| 40 | No | A. baylyi | Trimethopr | Intracellul | F84 | Neutral Re | 1064 | BL1 | 6317.5 |
| 40 | No | A. baylyi | Trimethopr | Intracellul | F99 | Pyronin Y | 993 | BL1 | 6512 |
| 40 | No | A. baylyi | Trimethopr | Intracellul | F108 | Rhodamine | 1098 | BL1 | 5987 |
| 40 | No | A. baylyi | Trimethopr | Intracellul | F109 | Riboflavin | 993 | BL1 | 6560 |
| 40 | No | A. baylyi | Trimethopr | Intracellul | F117 | SYBR Gree | 954 | BL1 | 554733.5 |
| 40 | No | A. baylyi | Trimethopr | Intracellul | F106 | Rhodamine | 985 | BL1 | 9954 |
| 40 | No | A. baylyi | Trimethopr | Intracellul | F73 | Malachite ( | 1061 | BL1 | 5776 |
| 40 | No | A. baylyi | Trimethopr | Intracellul | F7 | Acridine or | 997 | BL1 | 7210 |
| 40 | No | A. baylyi | Trimethopr | Intracellul | F192 | 5-Carboxyf | 1038 | BL1 | 18759 |
| 40 | No | A. baylyi | Trimethopr | Intracellul | F57 | Eosin Y | 972 | BL1 | 19846 |
| 40 | No | A. baylyi | Trimethopr | Intracellul | F27 | Calcein | 988 | BL1 | 18343.5 |
| 40 | No | A. baylyi | Trimethopr | Intracellul | F12 | Amaranth | 864 | BL1 | 20566.5 |
| 40 | No | A. baylyi | None | None | NA | Autofluore | 1817 | NA | 1403 |
| 40 | No | A. baylyi | None | None | F194 | Oxonol | 2002 | RL1 | 103281.5 |
| 40 | No | A. baylyi | None | None | F11 | Allura Red | 1446 | BL3 | 19104 |
| 40 | No | A. baylyi | None | None | F54 | DiSC3(5) | 1431 | RL1 | 10064 |
| 40 | No | A. baylyi | None | None | F124 | Thioflavin T | 1392 | VL2 | 5435.5 |
| 40 | No | A. baylyi | None | None | F101 | Quinacrine | 1171 | VL2 | 4559 |
| 40 | No | A. baylyi | None | None | F138 | Apigenin | 1143 | VL2 | 4115 |
| 40 | No | A. baylyi | None | None | F65 | H2FDA | 1147 | BL1 | 6335 |
| 40 | No | A. baylyi | None | None | F142 | Chlorophe | 1148 | BL3 | 7582.5 |
| 40 | No | A. baylyi | None | None | F39 | Congo Red | 1143 | BL3 | 6734 |
| 40 | No | A. baylyi | None | None | F116 | Sunset Yel | 1008 | BL2 | 5969 |
| 40 | No | A. baylyi | None | None | F10 | Alizarin | 1047 | BL1 | 4680 |
| 40 | No | A. baylyi | None | None | F84 | Neutral Re | 1247 | BL1 | 4342 |
| 40 | No | A. baylyi | None | None | F99 | Pyronin Y | 1049 | BL1 | 4266 |
| 40 | No | A. baylyi | None | None | F108 | Rhodamine | 1110 | BL1 | 3768 |
| 40 | No | A. baylyi | None | None | F109 | Riboflavin | 1083 | BL1 | 4117 |
| 40 | No | A. baylyi | None | None | F117 | SYBR Gree | 1049 | BL1 | 247834 |
| 40 | No | A. baylyi | None | None | F106 | Rhodamine | 1057 | BL1 | 5407 |
| 40 | No | A. baylyi | None | None | F73 | Malachite ( | 1075 | BL1 | 4418 |
| 40 | No | A. baylyi | None | None | F7 | Acridine or | 1039 | BL1 | 4957 |
| 40 | No | A. baylyi | None | None | F192 | 5-Carboxyf | 1049 | BL1 | 19168 |
| 40 | No | A. baylyi | None | None | F57 | Eosin Y | 1046 | BL1 | 18022.5 |
| 40 | No | A. baylyi | None | None | F27 | Calcein | 995 | BL1 | 15734 |

| | | | | | | | | | |
|---|---|---|---|---|---|---|---|---|---|
| 40 | No | A. baylyi | None | None | F12 | Amaranth | 1105 | BL1 | 18554 |
| 40 | No | A. baylyi | None | None | NA | Autofluore: | 1111 | NA | 18163 |
| 40 | No | A. baylyi | Amoxacilli | Wall | F194 | Oxonol | 3509 | RL1 | 13747 |
| 40 | No | A. baylyi | Amoxacilli | Wall | F11 | Allura Red | 952 | BL3 | 13707 |
| 40 | No | A. baylyi | Amoxacilli | Wall | F54 | DiSC3(5) | 1009 | RL1 | 13767 |
| 40 | No | A. baylyi | Amoxacilli | Wall | F124 | Thioflavin T | 942 | VL2 | 11075.5 |
| 40 | No | A. baylyi | Amoxacilli | Wall | F101 | Quinacrine | 943 | VL2 | 7421 |
| 40 | No | A. baylyi | Amoxacilli | Wall | F138 | Apigenin | 1072 | VL2 | 6222.5 |
| 40 | No | A. baylyi | Amoxacilli | Wall | F65 | H2FDA | 978 | BL1 | 7789 |
| 40 | No | A. baylyi | Amoxacilli | Wall | F142 | Chlorophe | 1055 | BL3 | 8151 |
| 40 | No | A. baylyi | Amoxacilli | Wall | F39 | Congo Red | 1054 | BL3 | 7336 |
| 40 | No | A. baylyi | Amoxacilli | Wall | F116 | Sunset Yel | 1059 | BL2 | 7079 |
| 40 | No | A. baylyi | Amoxacilli | Wall | F10 | Alizarin | 890 | BL1 | 5882 |
| 40 | No | A. baylyi | Amoxacilli | Wall | F84 | Neutral Re | 965 | BL1 | 5069 |
| 40 | No | A. baylyi | Amoxacilli | Wall | F99 | Pyronin Y | 910 | BL1 | 5296.5 |
| 40 | No | A. baylyi | Amoxacilli | Wall | F108 | Rhodamine | 1053 | BL1 | 4618 |
| 40 | No | A. baylyi | Amoxacilli | Wall | F109 | Riboflavin | 897 | BL1 | 4944 |
| 40 | No | A. baylyi | Amoxacilli | Wall | F117 | SYBR Gree | 944 | BL1 | 259491 |
| 40 | No | A. baylyi | Amoxacilli | Wall | F106 | Rhodamine | 948 | BL1 | 6701 |
| 40 | No | A. baylyi | Amoxacilli | Wall | F73 | Malachite ( | 949 | BL1 | 4938 |
| 40 | No | A. baylyi | Amoxacilli | Wall | F7 | Acridine or | 930 | BL1 | 5639 |
| 40 | No | A. baylyi | Amoxacilli | Wall | F192 | 5-Carboxyf | 963 | BL1 | 19101 |
| 40 | No | A. baylyi | Amoxacilli | Wall | F57 | Eosin Y | 996 | BL1 | 17901 |
| 40 | No | A. baylyi | Amoxacilli | Wall | F27 | Calcein | 955 | BL1 | 19950 |
| 40 | No | A. baylyi | Amoxacilli | Wall | F12 | Amaranth | 995 | BL1 | 19037 |
| 40 | No | A. baylyi | None | None | NA | Autofluore: | 1068 | NA | 18193 |
| 40 | No | A. baylyi | Azithromyc | Intracellul: | F194 | Oxonol | 1327 | RL1 | 19108 |
| 40 | No | A. baylyi | Azithromyc | Intracellul: | F11 | Allura Red | 963 | BL3 | 13068 |
| 40 | No | A. baylyi | Azithromyc | Intracellul: | F54 | DiSC3(5) | 987 | RL1 | 13354 |
| 40 | No | A. baylyi | Azithromyc | Intracellul: | F124 | Thioflavin T | 1017 | VL2 | 9733 |
| 40 | No | A. baylyi | Azithromyc | Intracellul: | F101 | Quinacrine | 1046 | VL2 | 7698.5 |
| 40 | No | A. baylyi | Azithromyc | Intracellul: | F138 | Apigenin | 1076 | VL2 | 6512 |
| 40 | No | A. baylyi | Azithromyc | Intracellul: | F65 | H2FDA | 965 | BL1 | 6764 |
| 40 | No | A. baylyi | Azithromyc | Intracellul: | F142 | Chlorophe | 1044 | BL3 | 7577.5 |
| 40 | No | A. baylyi | Azithromyc | Intracellul: | F39 | Congo Red | 951 | BL3 | 6410 |
| 40 | No | A. baylyi | Azithromyc | Intracellul: | F116 | Sunset Yel | 967 | BL2 | 6388 |
| 40 | No | A. baylyi | Azithromyc | Intracellul: | F10 | Alizarin | 1016 | BL1 | 5285 |
| 40 | No | A. baylyi | Azithromyc | Intracellul: | F84 | Neutral Re | 1008 | BL1 | 5027 |
| 40 | No | A. baylyi | Azithromyc | Intracellul: | F99 | Pyronin Y | 1010 | BL1 | 5041 |
| 40 | No | A. baylyi | Azithromyc | Intracellul: | F108 | Rhodamine | 948 | BL1 | 4562.5 |
| 40 | No | A. baylyi | Azithromyc | Intracellul: | F109 | Riboflavin | 960 | BL1 | 4750 |
| 40 | No | A. baylyi | Azithromyc | Intracellul: | F117 | SYBR Gree | 1000 | BL1 | 229557.5 |
| 40 | No | A. baylyi | Azithromyc | Intracellul: | F106 | Rhodamine | 996 | BL1 | 6555.5 |
| 40 | No | A. baylyi | Azithromyc | Intracellul: | F73 | Malachite ( | 1031 | BL1 | 5093 |
| 40 | No | A. baylyi | Azithromyc | Intracellul: | F7 | Acridine or | 1006 | BL1 | 5852.5 |
| 40 | No | A. baylyi | Azithromyc | Intracellul: | F192 | 5-Carboxyf | 954 | BL1 | 20089 |
| 40 | No | A. baylyi | Azithromyc | Intracellul: | F57 | Eosin Y | 978 | BL1 | 19267 |
| 40 | No | A. baylyi | Azithromyc | Intracellul: | F27 | Calcein | 985 | BL1 | 20287 |

| | | | | | | | | | |
|---|---|---|---|---|---|---|---|---|---|
| 40 | No | A. baylyi | Azithromyc | Intracellula | F12 | Amaranth | 994 | BL1 | 18768.5 |
| 40 | No | A. baylyi | None | None | NA | Autofluore: | 1049 | NA | 18499 |
| 40 | No | A. baylyi | Trimethopr | Intracellula | F194 | Oxonol | 3620 | RL1 | 12523 |
| 40 | No | A. baylyi | Trimethopr | Intracellula | F11 | Allura Red | 985 | BL3 | 13838 |
| 40 | No | A. baylyi | Trimethopr | Intracellula | F54 | DiSC3(5) | 852 | RL1 | 12450 |
| 40 | No | A. baylyi | Trimethopr | Intracellula | F124 | Thioflavin T | 895 | VL2 | 9600 |
| 40 | No | A. baylyi | Trimethopr | Intracellula | F101 | Quinacrine | 862 | VL2 | 6942 |
| 40 | No | A. baylyi | Trimethopr | Intracellula | F138 | Apigenin | 973 | VL2 | 5994 |
| 40 | No | A. baylyi | Trimethopr | Intracellula | F65 | H2FDA | 844 | BL1 | 7156.5 |
| 40 | No | A. baylyi | Trimethopr | Intracellula | F142 | Chlorophe | 1039 | BL3 | 7846 |
| 40 | No | A. baylyi | Trimethopr | Intracellula | F39 | Congo Red | 962 | BL3 | 7456.5 |
| 40 | No | A. baylyi | Trimethopr | Intracellula | F116 | Sunset Yel | 955 | BL2 | 6444 |
| 40 | No | A. baylyi | Trimethopr | Intracellula | F10 | Alizarin | 882 | BL1 | 5542 |
| 40 | No | A. baylyi | Trimethopr | Intracellula | F84 | Neutral Re | 956 | BL1 | 5299.5 |
| 40 | No | A. baylyi | Trimethopr | Intracellula | F99 | Pyronin Y | 886 | BL1 | 5085 |
| 40 | No | A. baylyi | Trimethopr | Intracellula | F108 | Rhodamine | 896 | BL1 | 4706.5 |
| 40 | No | A. baylyi | Trimethopr | Intracellula | F109 | Riboflavin | 915 | BL1 | 5264 |
| 40 | No | A. baylyi | Trimethopr | Intracellula | F117 | SYBR Gree | 954 | BL1 | 265335.5 |
| 40 | No | A. baylyi | Trimethopr | Intracellula | F106 | Rhodamine | 903 | BL1 | 7225 |
| 40 | No | A. baylyi | Trimethopr | Intracellula | F73 | Malachite ( | 1001 | BL1 | 6073 |
| 40 | No | A. baylyi | Trimethopr | Intracellula | F7 | Acridine or | 962 | BL1 | 6233 |
| 40 | No | A. baylyi | Trimethopr | Intracellula | F192 | 5-Carboxyf | 902 | BL1 | 18503.5 |
| 40 | No | A. baylyi | Trimethopr | Intracellula | F57 | Eosin Y | 964 | BL1 | 18408 |
| 40 | No | A. baylyi | Trimethopr | Intracellula | F27 | Calcein | 982 | BL1 | 19190.5 |
| 40 | No | A. baylyi | Trimethopr | Intracellula | F12 | Amaranth | 913 | BL1 | 16432 |
| 40 | No | A. baylyi | None | None | NA | Autofluore: | 966 | NA | 18060 |
| 40 | No | A. baylyi | Chloramph | Intracellula | F194 | Oxonol | 1264 | RL1 | 16380.5 |
| 40 | No | A. baylyi | Chloramph | Intracellula | F11 | Allura Red | 869 | BL3 | 14149 |
| 40 | No | A. baylyi | Chloramph | Intracellula | F54 | DiSC3(5) | 842 | RL1 | 12406 |
| 40 | No | A. baylyi | Chloramph | Intracellula | F124 | Thioflavin T | 866 | VL2 | 9554 |
| 40 | No | A. baylyi | Chloramph | Intracellula | F101 | Quinacrine | 858 | VL2 | 7259 |
| 40 | No | A. baylyi | Chloramph | Intracellula | F138 | Apigenin | 915 | VL2 | 6843 |
| 40 | No | A. baylyi | Chloramph | Intracellula | F65 | H2FDA | 962 | BL1 | 7726.5 |
| 40 | No | A. baylyi | Chloramph | Intracellula | F142 | Chlorophe | 879 | BL3 | 8698 |
| 40 | No | A. baylyi | Chloramph | Intracellula | F39 | Congo Red | 703 | BL3 | 7265 |
| 40 | No | A. baylyi | Chloramph | Intracellula | F116 | Sunset Yel | 904 | BL2 | 6493 |
| 40 | No | A. baylyi | Chloramph | Intracellula | F10 | Alizarin | 869 | BL1 | 6102 |
| 40 | No | A. baylyi | Chloramph | Intracellula | F84 | Neutral Re | 852 | BL1 | 5284 |
| 40 | No | A. baylyi | Chloramph | Intracellula | F99 | Pyronin Y | 861 | BL1 | 5974 |
| 40 | No | A. baylyi | Chloramph | Intracellula | F108 | Rhodamine | 690 | BL1 | 4909 |
| 40 | No | A. baylyi | Chloramph | Intracellula | F109 | Riboflavin | 1001 | BL1 | 5357 |
| 40 | No | A. baylyi | Chloramph | Intracellula | F117 | SYBR Gree | 893 | BL1 | 220593 |
| 40 | No | A. baylyi | Chloramph | Intracellula | F106 | Rhodamine | 880 | BL1 | 8704.5 |
| 40 | No | A. baylyi | Chloramph | Intracellula | F73 | Malachite ( | 849 | BL1 | 5914 |
| 40 | No | A. baylyi | Chloramph | Intracellula | F7 | Acridine or | 798 | BL1 | 6266.5 |
| 40 | No | A. baylyi | Chloramph | Intracellula | F192 | 5-Carboxyf | 888 | BL1 | 19680 |
| 40 | No | A. baylyi | Chloramph | Intracellula | F57 | Eosin Y | 807 | BL1 | 19290 |
| 40 | No | A. baylyi | Chloramph | Intracellula | F27 | Calcein | 854 | BL1 | 19033 |

| | | | | | | | | | |
|---|---|---|---|---|---|---|---|---|---|
| 40 | No | A. baylyi | Chloramph | Intracellula | F12 | Amaranth | 860 | BL1 | 20472 |
| 40 | No | A. baylyi | None | None | NA | Autofluore: | 986 | NA | 16396 |
| 40 | No | A. baylyi | Colistin | Wall | F194 | Oxonol | 3099 | RL1 | 11535 |
| 40 | No | A. baylyi | Colistin | Wall | F11 | Allura Red | 752 | BL3 | 76717 |
| 40 | No | A. baylyi | Colistin | Wall | F54 | DiSC3(5) | 503 | RL1 | 91313 |
| 40 | No | A. baylyi | Colistin | Wall | F124 | Thioflavin T | 708 | VL2 | 99136.5 |
| 40 | No | A. baylyi | Colistin | Wall | F101 | Quinacrine | 632 | VL2 | 66481 |
| 40 | No | A. baylyi | Colistin | Wall | F138 | Apigenin | 673 | VL2 | 54026 |
| 40 | No | A. baylyi | Colistin | Wall | F65 | H2FDA | 680 | BL1 | 48043 |
| 40 | No | A. baylyi | Colistin | Wall | F142 | Chlorophe | 641 | BL3 | 48867 |
| 40 | No | A. baylyi | Colistin | Wall | F39 | Congo Red | 669 | BL3 | 26448 |
| 40 | No | A. baylyi | Colistin | Wall | F116 | Sunset Yel | 705 | BL2 | 32572 |
| 40 | No | A. baylyi | Colistin | Wall | F10 | Alizarin | 623 | BL1 | 41942 |
| 40 | No | A. baylyi | Colistin | Wall | F84 | Neutral Re | 727 | BL1 | 49434 |
| 40 | No | A. baylyi | Colistin | Wall | F99 | Pyronin Y | 654 | BL1 | 52209 |
| 40 | No | A. baylyi | Colistin | Wall | F108 | Rhodamine | 738 | BL1 | 46415.5 |
| 40 | No | A. baylyi | Colistin | Wall | F109 | Riboflavin | 635 | BL1 | 37381 |
| 40 | No | A. baylyi | Colistin | Wall | F117 | SYBR Gree | 656 | BL1 | 1379246 |
| 40 | No | A. baylyi | Colistin | Wall | F106 | Rhodamine | 602 | BL1 | 145673 |
| 40 | No | A. baylyi | Colistin | Wall | F73 | Malachite ( | 580 | BL1 | 97698.5 |
| 40 | No | A. baylyi | Colistin | Wall | F7 | Acridine or | 652 | BL1 | 64017 |
| 40 | No | A. baylyi | Colistin | Wall | F192 | 5-Carboxyf | 734 | BL1 | 40759 |
| 40 | No | A. baylyi | Colistin | Wall | F57 | Eosin Y | 620 | BL1 | 36247.5 |
| 40 | No | A. baylyi | Colistin | Wall | F27 | Calcein | 611 | BL1 | 41387 |
| 40 | No | A. baylyi | Colistin | Wall | F12 | Amaranth | 600 | BL1 | 35565 |
| 40 | No | A. baylyi | None | None | NA | Autofluore: | 1083 | NA | 18451 |
| 40 | No | A. baylyi | Gentamici | Intracellula | F194 | Oxonol | 1639 | RL1 | 16228 |
| 40 | No | A. baylyi | Gentamici | Intracellula | F11 | Allura Red | 926 | BL3 | 14398 |
| 40 | No | A. baylyi | Gentamici | Intracellula | F54 | DiSC3(5) | 949 | RL1 | 13087 |
| 40 | No | A. baylyi | Gentamici | Intracellula | F124 | Thioflavin T | 910 | VL2 | 10484 |
| 40 | No | A. baylyi | Gentamici | Intracellula | F101 | Quinacrine | 1028 | VL2 | 7338.5 |
| 40 | No | A. baylyi | Gentamici | Intracellula | F138 | Apigenin | 1083 | VL2 | 7252 |
| 40 | No | A. baylyi | Gentamici | Intracellula | F65 | H2FDA | 921 | BL1 | 7821 |
| 40 | No | A. baylyi | Gentamici | Intracellula | F142 | Chlorophe | 873 | BL3 | 7988 |
| 40 | No | A. baylyi | Gentamici | Intracellula | F39 | Congo Red | 1061 | BL3 | 7816 |
| 40 | No | A. baylyi | Gentamici | Intracellula | F116 | Sunset Yel | 856 | BL2 | 7220.5 |
| 40 | No | A. baylyi | Gentamici | Intracellula | F10 | Alizarin | 982 | BL1 | 5646.5 |
| 40 | No | A. baylyi | Gentamici | Intracellula | F84 | Neutral Re | 994 | BL1 | 5444.5 |
| 40 | No | A. baylyi | Gentamici | Intracellula | F99 | Pyronin Y | 1018 | BL1 | 5459.5 |
| 40 | No | A. baylyi | Gentamici | Intracellula | F108 | Rhodamine | 975 | BL1 | 4972 |
| 40 | No | A. baylyi | Gentamici | Intracellula | F109 | Riboflavin | 888 | BL1 | 5190.5 |
| 40 | No | A. baylyi | Gentamici | Intracellula | F117 | SYBR Gree | 1032 | BL1 | 303533.5 |
| 40 | No | A. baylyi | Gentamici | Intracellula | F106 | Rhodamine | 828 | BL1 | 8124.5 |
| 40 | No | A. baylyi | Gentamici | Intracellula | F73 | Malachite ( | 1037 | BL1 | 5632 |
| 40 | No | A. baylyi | Gentamici | Intracellula | F7 | Acridine or | 967 | BL1 | 6132 |
| 40 | No | A. baylyi | Gentamici | Intracellula | F192 | 5-Carboxyf | 910 | BL1 | 18809 |
| 40 | No | A. baylyi | Gentamici | Intracellula | F57 | Eosin Y | 1014 | BL1 | 18257.5 |
| 40 | No | A. baylyi | Gentamici | Intracellula | F27 | Calcein | 921 | BL1 | 17288 |

| | | | | | | | | | |
|---|---|---|---|---|---|---|---|---|---|
| 40 | No | A. baylyi | Gentamici | Intracellul | F12 | Amaranth | 871 | BL1 | 19161 |
| 40 | No | A. baylyi | None | None | NA | Autofluore | 1135 | NA | 19029 |
| 40 | No | A. baylyi | Nitrofurant | Intracellul | F194 | Oxonol | 3285 | RL1 | 11342 |
| 40 | No | A. baylyi | Nitrofurant | Intracellul | F11 | Allura Red | 1006 | BL3 | 13840 |
| 40 | No | A. baylyi | Nitrofurant | Intracellul | F54 | DiSC3(5) | 703 | RL1 | 11000 |
| 40 | No | A. baylyi | Nitrofurant | Intracellul | F124 | Thioflavin T | 984 | VL2 | 12369 |
| 40 | No | A. baylyi | Nitrofurant | Intracellul | F101 | Quinacrine | 815 | VL2 | 7926 |
| 40 | No | A. baylyi | Nitrofurant | Intracellul | F138 | Apigenin | 895 | VL2 | 6839 |
| 40 | No | A. baylyi | Nitrofurant | Intracellul | F65 | H2FDA | 937 | BL1 | 8815 |
| 40 | No | A. baylyi | Nitrofurant | Intracellul | F142 | Chlorophe | 816 | BL3 | 8015.5 |
| 40 | No | A. baylyi | Nitrofurant | Intracellul | F39 | Congo Red | 777 | BL3 | 7706 |
| 40 | No | A. baylyi | Nitrofurant | Intracellul | F116 | Sunset Yel | 987 | BL2 | 7317 |
| 40 | No | A. baylyi | Nitrofurant | Intracellul | F10 | Alizarin | 711 | BL1 | 5593 |
| 40 | No | A. baylyi | Nitrofurant | Intracellul | F84 | Neutral Re | 973 | BL1 | 5185 |
| 40 | No | A. baylyi | Nitrofurant | Intracellul | F99 | Pyronin Y | 888 | BL1 | 5295 |
| 40 | No | A. baylyi | Nitrofurant | Intracellul | F108 | Rhodamine | 958 | BL1 | 4654 |
| 40 | No | A. baylyi | Nitrofurant | Intracellul | F109 | Riboflavin | 896 | BL1 | 6079 |
| 40 | No | A. baylyi | Nitrofurant | Intracellul | F117 | SYBR Gree | 960 | BL1 | 196414 |
| 40 | No | A. baylyi | Nitrofurant | Intracellul | F106 | Rhodamine | 809 | BL1 | 7772 |
| 40 | No | A. baylyi | Nitrofurant | Intracellul | F73 | Malachite ( | 979 | BL1 | 5988 |
| 40 | No | A. baylyi | Nitrofurant | Intracellul | F7 | Acridine or | 833 | BL1 | 6884 |
| 40 | No | A. baylyi | Nitrofurant | Intracellul | F192 | 5-Carboxyf | 1034 | BL1 | 18100 |
| 40 | No | A. baylyi | Nitrofurant | Intracellul | F57 | Eosin Y | 783 | BL1 | 14780 |
| 40 | No | A. baylyi | Nitrofurant | Intracellul | F27 | Calcein | 840 | BL1 | 18109 |
| 40 | No | A. baylyi | Nitrofurant | Intracellul | F12 | Amaranth | 1032 | BL1 | 16388.5 |
| 40 | No | A. baylyi | None | None | NA | Autofluore | 1015 | NA | 18489 |
| 40 | No | A. baylyi | Pentamidi | Wall | F194 | Oxonol | 11909 | RL1 | 848593 |
| 40 | No | A. baylyi | Pentamidi | Wall | F11 | Allura Red | 1332 | BL3 | 17296.5 |
| 40 | No | A. baylyi | Pentamidi | Wall | F54 | DiSC3(5) | 696 | RL1 | 20601 |
| 40 | No | A. baylyi | Pentamidi | Wall | F124 | Thioflavin T | 809 | VL2 | 24724 |
| 40 | No | A. baylyi | Pentamidi | Wall | F101 | Quinacrine | 717 | VL2 | 12557 |
| 40 | No | A. baylyi | Pentamidi | Wall | F138 | Apigenin | 741 | VL2 | 7996 |
| 40 | No | A. baylyi | Pentamidi | Wall | F65 | H2FDA | 778 | BL1 | 10443 |
| 40 | No | A. baylyi | Pentamidi | Wall | F142 | Chlorophe | 720 | BL3 | 11445.5 |
| 40 | No | A. baylyi | Pentamidi | Wall | F39 | Congo Red | 828 | BL3 | 12590 |
| 40 | No | A. baylyi | Pentamidi | Wall | F116 | Sunset Yel | 681 | BL2 | 10509 |
| 40 | No | A. baylyi | Pentamidi | Wall | F10 | Alizarin | 730 | BL1 | 9204.5 |
| 40 | No | A. baylyi | Pentamidi | Wall | F84 | Neutral Re | 813 | BL1 | 9489 |
| 40 | No | A. baylyi | Pentamidi | Wall | F99 | Pyronin Y | 744 | BL1 | 8046 |
| 40 | No | A. baylyi | Pentamidi | Wall | F108 | Rhodamine | 798 | BL1 | 8569 |
| 40 | No | A. baylyi | Pentamidi | Wall | F109 | Riboflavin | 711 | BL1 | 8270 |
| 40 | No | A. baylyi | Pentamidi | Wall | F117 | SYBR Gree | 1129 | BL1 | 935470 |
| 40 | No | A. baylyi | Pentamidi | Wall | F106 | Rhodamine | 790 | BL1 | 32174.5 |
| 40 | No | A. baylyi | Pentamidi | Wall | F73 | Malachite ( | 809 | BL1 | 14897 |
| 40 | No | A. baylyi | Pentamidi | Wall | F7 | Acridine or | 722 | BL1 | 12766.5 |
| 40 | No | A. baylyi | Pentamidi | Wall | F192 | 5-Carboxyf | 756 | BL1 | 19870.5 |
| 40 | No | A. baylyi | Pentamidi | Wall | F57 | Eosin Y | 886 | BL1 | 14242 |
| 40 | No | A. baylyi | Pentamidi | Wall | F27 | Calcein | 914 | BL1 | 17973 |

| | | | | | | | | | |
|---|---|---|---|---|---|---|---|---|---|
| 40 | No | A. baylyi | Pentamidi | Wall | F12 | Amaranth | 829 | BL1 | 16153 |
| 40 | No | A. baylyi | None | None | NA | Autofluore: | 1055 | NA | 16097 |
| 40 | No | A. baylyi | Rifampicin | Intracellul | F194 | Oxonol | 5305 | RL1 | 11992 |
| 40 | No | A. baylyi | Rifampicin | Intracellul | F11 | Allura Red | 1445 | BL3 | 13145 |
| 40 | No | A. baylyi | Rifampicin | Intracellul | F54 | DiSC3(5) | 1087 | RL1 | 13996 |
| 40 | No | A. baylyi | Rifampicin | Intracellul | F124 | Thioflavin T | 1026 | VL2 | 9174.5 |
| 40 | No | A. baylyi | Rifampicin | Intracellul | F101 | Quinacrine | 1080 | VL2 | 6763.5 |
| 40 | No | A. baylyi | Rifampicin | Intracellul | F138 | Apigenin | 1103 | VL2 | 6101 |
| 40 | No | A. baylyi | Rifampicin | Intracellul | F65 | H2FDA | 1121 | BL1 | 7275 |
| 40 | No | A. baylyi | Rifampicin | Intracellul | F142 | Chlorophe | 995 | BL3 | 8221 |
| 40 | No | A. baylyi | Rifampicin | Intracellul | F39 | Congo Red | 954 | BL3 | 8037.5 |
| 40 | No | A. baylyi | Rifampicin | Intracellul | F116 | Sunset Yel | 1034 | BL2 | 7578.5 |
| 40 | No | A. baylyi | Rifampicin | Intracellul | F10 | Alizarin | 953 | BL1 | 6413 |
| 40 | No | A. baylyi | Rifampicin | Intracellul | F84 | Neutral Re | 984 | BL1 | 6126.5 |
| 40 | No | A. baylyi | Rifampicin | Intracellul | F99 | Pyronin Y | 881 | BL1 | 6361 |
| 40 | No | A. baylyi | Rifampicin | Intracellul | F108 | Rhodamine | 1033 | BL1 | 5572 |
| 40 | No | A. baylyi | Rifampicin | Intracellul | F109 | Riboflavin | 1119 | BL1 | 5950 |
| 40 | No | A. baylyi | Rifampicin | Intracellul | F117 | SYBR Gree | 1064 | BL1 | 342452 |
| 40 | No | A. baylyi | Rifampicin | Intracellul | F106 | Rhodamine | 1025 | BL1 | 7828 |
| 40 | No | A. baylyi | Rifampicin | Intracellul | F73 | Malachite | 1206 | BL1 | 6827 |
| 40 | No | A. baylyi | Rifampicin | Intracellul | F7 | Acridine or | 991 | BL1 | 7147 |
| 40 | No | A. baylyi | Rifampicin | Intracellul | F192 | 5-Carboxyf | 1078 | BL1 | 19975.5 |
| 40 | No | A. baylyi | Rifampicin | Intracellul | F57 | Eosin Y | 1089 | BL1 | 19212 |
| 40 | No | A. baylyi | Rifampicin | Intracellul | F27 | Calcein | 1037 | BL1 | 19142 |
| 40 | No | A. baylyi | Rifampicin | Intracellul | F12 | Amaranth | 1066 | BL1 | 19560 |
| 40 | No | A. baylyi | None | None | NA | Autofluore: | 1135 | NA | 18400 |
| 40 | No | A. baylyi | Trimethopr | Intracellul | F194 | Oxonol | 2443 | RL1 | 15806 |
| 40 | No | A. baylyi | Trimethopr | Intracellul | F11 | Allura Red | 1030 | BL3 | 13701.5 |
| 40 | No | A. baylyi | Trimethopr | Intracellul | F54 | DiSC3(5) | 884 | RL1 | 11831 |
| 40 | No | A. baylyi | Trimethopr | Intracellul | F124 | Thioflavin T | 912 | VL2 | 8661.5 |
| 40 | No | A. baylyi | Trimethopr | Intracellul | F101 | Quinacrine | 954 | VL2 | 7121.5 |
| 40 | No | A. baylyi | Trimethopr | Intracellul | F138 | Apigenin | 1123 | VL2 | 6074 |
| 40 | No | A. baylyi | Trimethopr | Intracellul | F65 | H2FDA | 1040 | BL1 | 6980.5 |
| 40 | No | A. baylyi | Trimethopr | Intracellul | F142 | Chlorophe | 992 | BL3 | 8395 |
| 40 | No | A. baylyi | Trimethopr | Intracellul | F39 | Congo Red | 1254 | BL3 | 8024 |
| 40 | No | A. baylyi | Trimethopr | Intracellul | F116 | Sunset Yel | 944 | BL2 | 8198 |
| 40 | No | A. baylyi | Trimethopr | Intracellul | F10 | Alizarin | 958 | BL1 | 6753 |
| 40 | No | A. baylyi | Trimethopr | Intracellul | F84 | Neutral Re | 1031 | BL1 | 6283 |
| 40 | No | A. baylyi | Trimethopr | Intracellul | F99 | Pyronin Y | 982 | BL1 | 6295 |
| 40 | No | A. baylyi | Trimethopr | Intracellul | F108 | Rhodamine | 1118 | BL1 | 5625 |
| 40 | No | A. baylyi | Trimethopr | Intracellul | F109 | Riboflavin | 1040 | BL1 | 5500.5 |
| 40 | No | A. baylyi | Trimethopr | Intracellul | F117 | SYBR Gree | 1170 | BL1 | 470547.5 |
| 40 | No | A. baylyi | Trimethopr | Intracellul | F106 | Rhodamine | 1024 | BL1 | 7319 |
| 40 | No | A. baylyi | Trimethopr | Intracellul | F73 | Malachite | 1148 | BL1 | 6134 |
| 40 | No | A. baylyi | Trimethopr | Intracellul | F7 | Acridine or | 1166 | BL1 | 6027 |
| 40 | No | A. baylyi | Trimethopr | Intracellul | F192 | 5-Carboxyf | 1161 | BL1 | 18241 |
| 40 | No | A. baylyi | Trimethopr | Intracellul | F57 | Eosin Y | 1180 | BL1 | 18859 |
| 40 | No | A. baylyi | Trimethopr | Intracellul | F27 | Calcein | 1184 | BL1 | 17520.5 |

| | | | | | | | | | |
|---|---|---|---|---|---|---|---|---|---|
| 40 | No | A. baylyi | Trimethopr | Intracellul | F12 | Amaranth | 1084 | BL1 | 19153 |
| 40 | No | A. baylyi | None | None | NA | Autofluore | 1877 | NA | 1440 |
| 40 | No | A. baylyi | None | None | F194 | Oxonol | 2200 | RL1 | 92730 |
| 40 | No | A. baylyi | None | None | F11 | Allura Red | 1458 | BL3 | 14320.5 |
| 40 | No | A. baylyi | None | None | F54 | DiSC3(5) | 1358 | RL1 | 7500 |
| 40 | No | A. baylyi | None | None | F124 | Thioflavin T | 1466 | VL2 | 7553.5 |
| 40 | No | A. baylyi | None | None | F101 | Quinacrine | 1226 | VL2 | 4447 |
| 40 | No | A. baylyi | None | None | F138 | Apigenin | 1110 | VL2 | 4072.5 |
| 40 | No | A. baylyi | None | None | F65 | H2FDA | 1114 | BL1 | 5876 |
| 40 | No | A. baylyi | None | None | F142 | Chlorophe | 1082 | BL3 | 6924 |
| 40 | No | A. baylyi | None | None | F39 | Congo Red | 1145 | BL3 | 6507 |
| 40 | No | A. baylyi | None | None | F116 | Sunset Yel | 1103 | BL2 | 5122 |
| 40 | No | A. baylyi | None | None | F10 | Alizarin | 1155 | BL1 | 4648 |
| 40 | No | A. baylyi | None | None | F84 | Neutral Re | 1042 | BL1 | 3633.5 |
| 40 | No | A. baylyi | None | None | F99 | Pyronin Y | 973 | BL1 | 4064 |
| 40 | No | A. baylyi | None | None | F108 | Rhodamine | 1057 | BL1 | 3448 |
| 40 | No | A. baylyi | None | None | F109 | Riboflavin | 1058 | BL1 | 3882 |
| 40 | No | A. baylyi | None | None | F117 | SYBR Gree | 1039 | BL1 | 271939 |
| 40 | No | A. baylyi | None | None | F106 | Rhodamine | 963 | BL1 | 4904 |
| 40 | No | A. baylyi | None | None | F73 | Malachite ( | 1040 | BL1 | 4227 |
| 40 | No | A. baylyi | None | None | F7 | Acridine or | 1023 | BL1 | 4788 |
| 40 | No | A. baylyi | None | None | F192 | 5-Carboxyf | 1003 | BL1 | 18995 |
| 40 | No | A. baylyi | None | None | F57 | Eosin Y | 993 | BL1 | 19680 |
| 40 | No | A. baylyi | None | None | F27 | Calcein | 939 | BL1 | 18043 |
| 40 | No | A. baylyi | None | None | F12 | Amaranth | 1071 | BL1 | 17682 |
| 40 | No | A. baylyi | None | None | NA | Autofluore | 1026 | NA | 17278 |
| 40 | No | A. baylyi | Amoxacilli | Wall | F194 | Oxonol | 2298 | RL1 | 15199.5 |
| 40 | No | A. baylyi | Amoxacilli | Wall | F11 | Allura Red | 961 | BL3 | 13739 |
| 40 | No | A. baylyi | Amoxacilli | Wall | F54 | DiSC3(5) | 918 | RL1 | 13278.5 |
| 40 | No | A. baylyi | Amoxacilli | Wall | F124 | Thioflavin T | 926 | VL2 | 7904.5 |
| 40 | No | A. baylyi | Amoxacilli | Wall | F101 | Quinacrine | 960 | VL2 | 5839 |
| 40 | No | A. baylyi | Amoxacilli | Wall | F138 | Apigenin | 1056 | VL2 | 5558 |
| 40 | No | A. baylyi | Amoxacilli | Wall | F65 | H2FDA | 979 | BL1 | 6392 |
| 40 | No | A. baylyi | Amoxacilli | Wall | F142 | Chlorophe | 1013 | BL3 | 6988 |
| 40 | No | A. baylyi | Amoxacilli | Wall | F39 | Congo Red | 982 | BL3 | 6294 |
| 40 | No | A. baylyi | Amoxacilli | Wall | F116 | Sunset Yel | 1109 | BL2 | 5915 |
| 40 | No | A. baylyi | Amoxacilli | Wall | F10 | Alizarin | 962 | BL1 | 4701 |
| 40 | No | A. baylyi | Amoxacilli | Wall | F84 | Neutral Re | 977 | BL1 | 4583 |
| 40 | No | A. baylyi | Amoxacilli | Wall | F99 | Pyronin Y | 951 | BL1 | 4880 |
| 40 | No | A. baylyi | Amoxacilli | Wall | F108 | Rhodamine | 943 | BL1 | 3967 |
| 40 | No | A. baylyi | Amoxacilli | Wall | F109 | Riboflavin | 911 | BL1 | 4585 |
| 40 | No | A. baylyi | Amoxacilli | Wall | F117 | SYBR Gree | 951 | BL1 | 292078 |
| 40 | No | A. baylyi | Amoxacilli | Wall | F106 | Rhodamine | 1029 | BL1 | 6297 |
| 40 | No | A. baylyi | Amoxacilli | Wall | F73 | Malachite ( | 1030 | BL1 | 4470 |
| 40 | No | A. baylyi | Amoxacilli | Wall | F7 | Acridine or | 977 | BL1 | 5326 |
| 40 | No | A. baylyi | Amoxacilli | Wall | F192 | 5-Carboxyf | 1005 | BL1 | 19823 |
| 40 | No | A. baylyi | Amoxacilli | Wall | F57 | Eosin Y | 988 | BL1 | 18577.5 |
| 40 | No | A. baylyi | Amoxacilli | Wall | F27 | Calcein | 1026 | BL1 | 20058.5 |

| | | | | | | | | | |
|---|---|---|---|---|---|---|---|---|---|
| 40 | No | A. baylyi | Amoxacilli | Wall | F12 | Amaranth | 1047 | BL1 | 19575 |
| 40 | No | A. baylyi | None | None | NA | Autofluore: | 1121 | NA | 18448 |
| 40 | No | A. baylyi | Azithromyc | Intracellula | F194 | Oxonol | 1922 | RL1 | 15489.5 |
| 40 | No | A. baylyi | Azithromyc | Intracellula | F11 | Allura Red | 998 | BL3 | 13984.5 |
| 40 | No | A. baylyi | Azithromyc | Intracellula | F54 | DiSC3(5) | 912 | RL1 | 12742 |
| 40 | No | A. baylyi | Azithromyc | Intracellula | F124 | Thioflavin T | 1005 | VL2 | 8535 |
| 40 | No | A. baylyi | Azithromyc | Intracellula | F101 | Quinacrine | 1042 | VL2 | 7005.5 |
| 40 | No | A. baylyi | Azithromyc | Intracellula | F138 | Apigenin | 1034 | VL2 | 5842.5 |
| 40 | No | A. baylyi | Azithromyc | Intracellula | F65 | H2FDA | 979 | BL1 | 6475 |
| 40 | No | A. baylyi | Azithromyc | Intracellula | F142 | Chlorophe | 1033 | BL3 | 7076 |
| 40 | No | A. baylyi | Azithromyc | Intracellula | F39 | Congo Red | 977 | BL3 | 6695 |
| 40 | No | A. baylyi | Azithromyc | Intracellula | F116 | Sunset Yel | 1131 | BL2 | 6195 |
| 40 | No | A. baylyi | Azithromyc | Intracellula | F10 | Alizarin | 1043 | BL1 | 5297 |
| 40 | No | A. baylyi | Azithromyc | Intracellula | F84 | Neutral Re | 953 | BL1 | 4374 |
| 40 | No | A. baylyi | Azithromyc | Intracellula | F99 | Pyronin Y | 976 | BL1 | 4800 |
| 40 | No | A. baylyi | Azithromyc | Intracellula | F108 | Rhodamine | 949 | BL1 | 4036 |
| 40 | No | A. baylyi | Azithromyc | Intracellula | F109 | Riboflavin | 1029 | BL1 | 4984 |
| 40 | No | A. baylyi | Azithromyc | Intracellula | F117 | SYBR Gree | 972 | BL1 | 226369.5 |
| 40 | No | A. baylyi | Azithromyc | Intracellula | F106 | Rhodamine | 1064 | BL1 | 6537.5 |
| 40 | No | A. baylyi | Azithromyc | Intracellula | F73 | Malachite ( | 1060 | BL1 | 4827 |
| 40 | No | A. baylyi | Azithromyc | Intracellula | F7 | Acridine or | 957 | BL1 | 5534 |
| 40 | No | A. baylyi | Azithromyc | Intracellula | F192 | 5-Carboxyf | 1007 | BL1 | 19422 |
| 40 | No | A. baylyi | Azithromyc | Intracellula | F57 | Eosin Y | 979 | BL1 | 19583 |
| 40 | No | A. baylyi | Azithromyc | Intracellula | F27 | Calcein | 1018 | BL1 | 18559 |
| 40 | No | A. baylyi | Azithromyc | Intracellula | F12 | Amaranth | 899 | BL1 | 18692 |
| 40 | No | A. baylyi | None | None | NA | Autofluore: | 1037 | NA | 18904 |
| 40 | No | A. baylyi | Trimethopr | Intracellula | F194 | Oxonol | 3305 | RL1 | 13382 |
| 40 | No | A. baylyi | Trimethopr | Intracellula | F11 | Allura Red | 1030 | BL3 | 13669 |
| 40 | No | A. baylyi | Trimethopr | Intracellula | F54 | DiSC3(5) | 867 | RL1 | 12674 |
| 40 | No | A. baylyi | Trimethopr | Intracellula | F124 | Thioflavin T | 938 | VL2 | 8105 |
| 40 | No | A. baylyi | Trimethopr | Intracellula | F101 | Quinacrine | 951 | VL2 | 6669 |
| 40 | No | A. baylyi | Trimethopr | Intracellula | F138 | Apigenin | 1015 | VL2 | 6059 |
| 40 | No | A. baylyi | Trimethopr | Intracellula | F65 | H2FDA | 955 | BL1 | 6427 |
| 40 | No | A. baylyi | Trimethopr | Intracellula | F142 | Chlorophe | 1040 | BL3 | 7244.5 |
| 40 | No | A. baylyi | Trimethopr | Intracellula | F39 | Congo Red | 915 | BL3 | 7068 |
| 40 | No | A. baylyi | Trimethopr | Intracellula | F116 | Sunset Yel | 1024 | BL2 | 6393.5 |
| 40 | No | A. baylyi | Trimethopr | Intracellula | F10 | Alizarin | 920 | BL1 | 5167 |
| 40 | No | A. baylyi | Trimethopr | Intracellula | F84 | Neutral Re | 950 | BL1 | 4859 |
| 40 | No | A. baylyi | Trimethopr | Intracellula | F99 | Pyronin Y | 904 | BL1 | 5296 |
| 40 | No | A. baylyi | Trimethopr | Intracellula | F108 | Rhodamine | 912 | BL1 | 4309 |
| 40 | No | A. baylyi | Trimethopr | Intracellula | F109 | Riboflavin | 824 | BL1 | 4374.5 |
| 40 | No | A. baylyi | Trimethopr | Intracellula | F117 | SYBR Gree | 1042 | BL1 | 285831.5 |
| 40 | No | A. baylyi | Trimethopr | Intracellula | F106 | Rhodamine | 842 | BL1 | 6110.5 |
| 40 | No | A. baylyi | Trimethopr | Intracellula | F73 | Malachite ( | 1023 | BL1 | 5247 |
| 40 | No | A. baylyi | Trimethopr | Intracellula | F7 | Acridine or | 806 | BL1 | 5685 |
| 40 | No | A. baylyi | Trimethopr | Intracellula | F192 | 5-Carboxyf | 939 | BL1 | 18943 |
| 40 | No | A. baylyi | Trimethopr | Intracellula | F57 | Eosin Y | 878 | BL1 | 20589 |
| 40 | No | A. baylyi | Trimethopr | Intracellula | F27 | Calcein | 940 | BL1 | 17383 |

| | | | | | | | | | |
|---|---|---|---|---|---|---|---|---|---|
| 40 | No | A. baylyi | Trimethopr | Intracellula | F12 | Amaranth | 879 | BL1 | 18143 |
| 40 | No | A. baylyi | None | None | NA | Autofluores | 1012 | NA | 15180 |
| 40 | No | A. baylyi | Chloramph | Intracellula | F194 | Oxonol | 1847 | RL1 | 13954 |
| 40 | No | A. baylyi | Chloramph | Intracellula | F11 | Allura Red | 795 | BL3 | 14429 |
| 40 | No | A. baylyi | Chloramph | Intracellula | F54 | DiSC3(5) | 832 | RL1 | 13136 |
| 40 | No | A. baylyi | Chloramph | Intracellula | F124 | Thioflavin T | 875 | VL2 | 8169 |
| 40 | No | A. baylyi | Chloramph | Intracellula | F101 | Quinacrine | 912 | VL2 | 7667.5 |
| 40 | No | A. baylyi | Chloramph | Intracellula | F138 | Apigenin | 902 | VL2 | 6393.5 |
| 40 | No | A. baylyi | Chloramph | Intracellula | F65 | H2FDA | 848 | BL1 | 7082 |
| 40 | No | A. baylyi | Chloramph | Intracellula | F142 | Chlorophe | 843 | BL3 | 7636 |
| 40 | No | A. baylyi | Chloramph | Intracellula | F39 | Congo Red | 812 | BL3 | 7220 |
| 40 | No | A. baylyi | Chloramph | Intracellula | F116 | Sunset Yel | 879 | BL2 | 6812 |
| 40 | No | A. baylyi | Chloramph | Intracellula | F10 | Alizarin | 797 | BL1 | 5292 |
| 40 | No | A. baylyi | Chloramph | Intracellula | F84 | Neutral Re | 771 | BL1 | 4674 |
| 40 | No | A. baylyi | Chloramph | Intracellula | F99 | Pyronin Y | 724 | BL1 | 5425 |
| 40 | No | A. baylyi | Chloramph | Intracellula | F108 | Rhodamine | 748 | BL1 | 4603.5 |
| 40 | No | A. baylyi | Chloramph | Intracellula | F109 | Riboflavin | 862 | BL1 | 4896 |
| 40 | No | A. baylyi | Chloramph | Intracellula | F117 | SYBR Gree | 907 | BL1 | 258801 |
| 40 | No | A. baylyi | Chloramph | Intracellula | F106 | Rhodamine | 754 | BL1 | 6256 |
| 40 | No | A. baylyi | Chloramph | Intracellula | F73 | Malachite ( | 855 | BL1 | 5404 |
| 40 | No | A. baylyi | Chloramph | Intracellula | F7 | Acridine or | 820 | BL1 | 5671 |
| 40 | No | A. baylyi | Chloramph | Intracellula | F192 | 5-Carboxyf | 811 | BL1 | 19864 |
| 40 | No | A. baylyi | Chloramph | Intracellula | F57 | Eosin Y | 784 | BL1 | 20010.5 |
| 40 | No | A. baylyi | Chloramph | Intracellula | F27 | Calcein | 802 | BL1 | 18385.5 |
| 40 | No | A. baylyi | Chloramph | Intracellula | F12 | Amaranth | 833 | BL1 | 20902 |
| 40 | No | A. baylyi | None | None | NA | Autofluores | 1058 | NA | 19953 |
| 40 | No | A. baylyi | Colistin | Wall | F194 | Oxonol | 2796 | RL1 | 14611 |
| 40 | No | A. baylyi | Colistin | Wall | F11 | Allura Red | 672 | BL3 | 49428.5 |
| 40 | No | A. baylyi | Colistin | Wall | F54 | DiSC3(5) | 511 | RL1 | 116472 |
| 40 | No | A. baylyi | Colistin | Wall | F124 | Thioflavin T | 752 | VL2 | 96769 |
| 40 | No | A. baylyi | Colistin | Wall | F101 | Quinacrine | 616 | VL2 | 76996 |
| 40 | No | A. baylyi | Colistin | Wall | F138 | Apigenin | 818 | VL2 | 66849.5 |
| 40 | No | A. baylyi | Colistin | Wall | F65 | H2FDA | 612 | BL1 | 68086.5 |
| 40 | No | A. baylyi | Colistin | Wall | F142 | Chlorophe | 732 | BL3 | 59574.5 |
| 40 | No | A. baylyi | Colistin | Wall | F39 | Congo Red | 630 | BL3 | 34468 |
| 40 | No | A. baylyi | Colistin | Wall | F116 | Sunset Yel | 662 | BL2 | 54570.5 |
| 40 | No | A. baylyi | Colistin | Wall | F10 | Alizarin | 697 | BL1 | 42635 |
| 40 | No | A. baylyi | Colistin | Wall | F84 | Neutral Re | 649 | BL1 | 54876 |
| 40 | No | A. baylyi | Colistin | Wall | F99 | Pyronin Y | 637 | BL1 | 61251 |
| 40 | No | A. baylyi | Colistin | Wall | F108 | Rhodamine | 599 | BL1 | 53620 |
| 40 | No | A. baylyi | Colistin | Wall | F109 | Riboflavin | 619 | BL1 | 40219 |
| 40 | No | A. baylyi | Colistin | Wall | F117 | SYBR Gree | 683 | BL1 | 1432589 |
| 40 | No | A. baylyi | Colistin | Wall | F106 | Rhodamine | 586 | BL1 | 138306 |
| 40 | No | A. baylyi | Colistin | Wall | F73 | Malachite ( | 693 | BL1 | 100251 |
| 40 | No | A. baylyi | Colistin | Wall | F7 | Acridine or | 674 | BL1 | 76041.5 |
| 40 | No | A. baylyi | Colistin | Wall | F192 | 5-Carboxyf | 583 | BL1 | 47614 |
| 40 | No | A. baylyi | Colistin | Wall | F57 | Eosin Y | 641 | BL1 | 39052 |
| 40 | No | A. baylyi | Colistin | Wall | F27 | Calcein | 648 | BL1 | 37779.5 |

| | | | | | | | | | |
|---|---|---|---|---|---|---|---|---|---|
| 40 | No | A. baylyi | Colistin | Wall | F12 | Amaranth | 623 | BL1 | 40618 |
| 40 | No | A. baylyi | None | None | NA | Autofluore: | 941 | NA | 18251 |
| 40 | No | A. baylyi | Gentamici | Intracellul | F194 | Oxonol | 2130 | RL1 | 15133 |
| 40 | No | A. baylyi | Gentamici | Intracellul | F11 | Allura Red | 1002 | BL3 | 13792.5 |
| 40 | No | A. baylyi | Gentamici | Intracellul | F54 | DiSC3(5) | 910 | RL1 | 12948 |
| 40 | No | A. baylyi | Gentamici | Intracellul | F124 | Thioflavin T | 964 | VL2 | 8403 |
| 40 | No | A. baylyi | Gentamici | Intracellul | F101 | Quinacrine | 1028 | VL2 | 6877 |
| 40 | No | A. baylyi | Gentamici | Intracellul | F138 | Apigenin | 1017 | VL2 | 6124 |
| 40 | No | A. baylyi | Gentamici | Intracellul | F65 | H2FDA | 1039 | BL1 | 7595 |
| 40 | No | A. baylyi | Gentamici | Intracellul | F142 | Chlorophe | 984 | BL3 | 7630.5 |
| 40 | No | A. baylyi | Gentamici | Intracellul | F39 | Congo Red | 935 | BL3 | 7133 |
| 40 | No | A. baylyi | Gentamici | Intracellul | F116 | Sunset Yel | 1034 | BL2 | 6720.5 |
| 40 | No | A. baylyi | Gentamici | Intracellul | F10 | Alizarin | 919 | BL1 | 5847 |
| 40 | No | A. baylyi | Gentamici | Intracellul | F84 | Neutral Re | 1000 | BL1 | 5156.5 |
| 40 | No | A. baylyi | Gentamici | Intracellul | F99 | Pyronin Y | 978 | BL1 | 5650.5 |
| 40 | No | A. baylyi | Gentamici | Intracellul | F108 | Rhodamine | 914 | BL1 | 4516 |
| 40 | No | A. baylyi | Gentamici | Intracellul | F109 | Riboflavin | 944 | BL1 | 5038.5 |
| 40 | No | A. baylyi | Gentamici | Intracellul | F117 | SYBR Gree | 1031 | BL1 | 397132 |
| 40 | No | A. baylyi | Gentamici | Intracellul | F106 | Rhodamine | 1058 | BL1 | 8708 |
| 40 | No | A. baylyi | Gentamici | Intracellul | F73 | Malachite ( | 979 | BL1 | 5582 |
| 40 | No | A. baylyi | Gentamici | Intracellul | F7 | Acridine or | 910 | BL1 | 5547 |
| 40 | No | A. baylyi | Gentamici | Intracellul | F192 | 5-Carboxyf | 1009 | BL1 | 20301 |
| 40 | No | A. baylyi | Gentamici | Intracellul | F57 | Eosin Y | 967 | BL1 | 18347 |
| 40 | No | A. baylyi | Gentamici | Intracellul | F27 | Calcein | 909 | BL1 | 18181 |
| 40 | No | A. baylyi | Gentamici | Intracellul | F12 | Amaranth | 991 | BL1 | 20631 |
| 40 | No | A. baylyi | None | None | NA | Autofluore: | 1052 | NA | 18490.5 |
| 40 | No | A. baylyi | Nitrofurant | Intracellul | F194 | Oxonol | 2290 | RL1 | 12631.5 |
| 40 | No | A. baylyi | Nitrofurant | Intracellul | F11 | Allura Red | 1074 | BL3 | 14439.5 |
| 40 | No | A. baylyi | Nitrofurant | Intracellul | F54 | DiSC3(5) | 1026 | RL1 | 13149 |
| 40 | No | A. baylyi | Nitrofurant | Intracellul | F124 | Thioflavin T | 998 | VL2 | 8338 |
| 40 | No | A. baylyi | Nitrofurant | Intracellul | F101 | Quinacrine | 971 | VL2 | 6595 |
| 40 | No | A. baylyi | Nitrofurant | Intracellul | F138 | Apigenin | 1083 | VL2 | 5940 |
| 40 | No | A. baylyi | Nitrofurant | Intracellul | F65 | H2FDA | 1022 | BL1 | 7191 |
| 40 | No | A. baylyi | Nitrofurant | Intracellul | F142 | Chlorophe | 1123 | BL3 | 8297 |
| 40 | No | A. baylyi | Nitrofurant | Intracellul | F39 | Congo Red | 990 | BL3 | 7141.5 |
| 40 | No | A. baylyi | Nitrofurant | Intracellul | F116 | Sunset Yel | 1161 | BL2 | 7329 |
| 40 | No | A. baylyi | Nitrofurant | Intracellul | F10 | Alizarin | 874 | BL1 | 6045 |
| 40 | No | A. baylyi | Nitrofurant | Intracellul | F84 | Neutral Re | 1059 | BL1 | 4961 |
| 40 | No | A. baylyi | Nitrofurant | Intracellul | F99 | Pyronin Y | 1000 | BL1 | 5601 |
| 40 | No | A. baylyi | Nitrofurant | Intracellul | F108 | Rhodamine | 935 | BL1 | 4541 |
| 40 | No | A. baylyi | Nitrofurant | Intracellul | F109 | Riboflavin | 948 | BL1 | 5053 |
| 40 | No | A. baylyi | Nitrofurant | Intracellul | F117 | SYBR Gree | 1051 | BL1 | 256776 |
| 40 | No | A. baylyi | Nitrofurant | Intracellul | F106 | Rhodamine | 1040 | BL1 | 7720.5 |
| 40 | No | A. baylyi | Nitrofurant | Intracellul | F73 | Malachite ( | 1095 | BL1 | 5798 |
| 40 | No | A. baylyi | Nitrofurant | Intracellul | F7 | Acridine or | 849 | BL1 | 6208 |
| 40 | No | A. baylyi | Nitrofurant | Intracellul | F192 | 5-Carboxyf | 923 | BL1 | 18712 |
| 40 | No | A. baylyi | Nitrofurant | Intracellul | F57 | Eosin Y | 836 | BL1 | 17253 |
| 40 | No | A. baylyi | Nitrofurant | Intracellul | F27 | Calcein | 944 | BL1 | 16746.5 |

| | | | | | | | | |
|---|---|---|---|---|---|---|---|---|
| 40 | No | A. baylyi | Nitrofurant | Intracellula | F12 | Amaranth | 936 | BL1 | 17630.5 |
| 40 | No | A. baylyi | None | None | NA | Autofluores | 1077 | NA | 18380 |
| 40 | No | A. baylyi | Pentamidi | Wall | F194 | Oxonol | 17047 | RL1 | 312420 |
| 40 | No | A. baylyi | Pentamidi | Wall | F11 | Allura Red | 1214 | BL3 | 14136 |
| 40 | No | A. baylyi | Pentamidi | Wall | F54 | DiSC3(5) | 866 | RL1 | 21224 |
| 40 | No | A. baylyi | Pentamidi | Wall | F124 | Thioflavin T | 723 | VL2 | 32955 |
| 40 | No | A. baylyi | Pentamidi | Wall | F101 | Quinacrine | 941 | VL2 | 13047 |
| 40 | No | A. baylyi | Pentamidi | Wall | F138 | Apigenin | 813 | VL2 | 9530 |
| 40 | No | A. baylyi | Pentamidi | Wall | F65 | H2FDA | 859 | BL1 | 9800 |
| 40 | No | A. baylyi | Pentamidi | Wall | F142 | Chlorophe | 739 | BL3 | 13679 |
| 40 | No | A. baylyi | Pentamidi | Wall | F39 | Congo Red | 924 | BL3 | 13446 |
| 40 | No | A. baylyi | Pentamidi | Wall | F116 | Sunset Yel | 788 | BL2 | 11622.5 |
| 40 | No | A. baylyi | Pentamidi | Wall | F10 | Alizarin | 802 | BL1 | 11692.5 |
| 40 | No | A. baylyi | Pentamidi | Wall | F84 | Neutral Re | 726 | BL1 | 14404 |
| 40 | No | A. baylyi | Pentamidi | Wall | F99 | Pyronin Y | 680 | BL1 | 11538.5 |
| 40 | No | A. baylyi | Pentamidi | Wall | F108 | Rhodamine | 823 | BL1 | 12428 |
| 40 | No | A. baylyi | Pentamidi | Wall | F109 | Riboflavin | 817 | BL1 | 12675 |
| 40 | No | A. baylyi | Pentamidi | Wall | F117 | SYBR Gree | 682 | BL1 | 1263176 |
| 40 | No | A. baylyi | Pentamidi | Wall | F106 | Rhodamine | 771 | BL1 | 34186 |
| 40 | No | A. baylyi | Pentamidi | Wall | F73 | Malachite ( | 782 | BL1 | 13693 |
| 40 | No | A. baylyi | Pentamidi | Wall | F7 | Acridine or | 833 | BL1 | 11289 |
| 40 | No | A. baylyi | Pentamidi | Wall | F192 | 5-Carboxyf | 771 | BL1 | 18882 |
| 40 | No | A. baylyi | Pentamidi | Wall | F57 | Eosin Y | 734 | BL1 | 18009 |
| 40 | No | A. baylyi | Pentamidi | Wall | F27 | Calcein | 775 | BL1 | 16373 |
| 40 | No | A. baylyi | Pentamidi | Wall | F12 | Amaranth | 753 | BL1 | 17022 |
| 40 | No | A. baylyi | None | None | NA | Autofluores | 1099 | NA | 19895 |
| 40 | No | A. baylyi | Rifampicin | Intracellula | F194 | Oxonol | 5050 | RL1 | 12362.5 |
| 40 | No | A. baylyi | Rifampicin | Intracellula | F11 | Allura Red | 1353 | BL3 | 18866 |
| 40 | No | A. baylyi | Rifampicin | Intracellula | F54 | DiSC3(5) | 1238 | RL1 | 15117 |
| 40 | No | A. baylyi | Rifampicin | Intracellula | F124 | Thioflavin T | 1102 | VL2 | 9623.5 |
| 40 | No | A. baylyi | Rifampicin | Intracellula | F101 | Quinacrine | 1176 | VL2 | 7491 |
| 40 | No | A. baylyi | Rifampicin | Intracellula | F138 | Apigenin | 1049 | VL2 | 5693 |
| 40 | No | A. baylyi | Rifampicin | Intracellula | F65 | H2FDA | 1117 | BL1 | 7776 |
| 40 | No | A. baylyi | Rifampicin | Intracellula | F142 | Chlorophe | 1103 | BL3 | 8328 |
| 40 | No | A. baylyi | Rifampicin | Intracellula | F39 | Congo Red | 990 | BL3 | 8573 |
| 40 | No | A. baylyi | Rifampicin | Intracellula | F116 | Sunset Yel | 1163 | BL2 | 7876 |
| 40 | No | A. baylyi | Rifampicin | Intracellula | F10 | Alizarin | 992 | BL1 | 6530.5 |
| 40 | No | A. baylyi | Rifampicin | Intracellula | F84 | Neutral Re | 1262 | BL1 | 6467 |
| 40 | No | A. baylyi | Rifampicin | Intracellula | F99 | Pyronin Y | 1070 | BL1 | 6801.5 |
| 40 | No | A. baylyi | Rifampicin | Intracellula | F108 | Rhodamine | 1107 | BL1 | 6246 |
| 40 | No | A. baylyi | Rifampicin | Intracellula | F109 | Riboflavin | 980 | BL1 | 5836.5 |
| 40 | No | A. baylyi | Rifampicin | Intracellula | F117 | SYBR Gree | 1001 | BL1 | 401670 |
| 40 | No | A. baylyi | Rifampicin | Intracellula | F106 | Rhodamine | 1031 | BL1 | 9351 |
| 40 | No | A. baylyi | Rifampicin | Intracellula | F73 | Malachite ( | 1093 | BL1 | 6091 |
| 40 | No | A. baylyi | Rifampicin | Intracellula | F7 | Acridine or | 1088 | BL1 | 7647.5 |
| 40 | No | A. baylyi | Rifampicin | Intracellula | F192 | 5-Carboxyf | 1238 | BL1 | 17610 |
| 40 | No | A. baylyi | Rifampicin | Intracellula | F57 | Eosin Y | 1097 | BL1 | 17271 |
| 40 | No | A. baylyi | Rifampicin | Intracellula | F27 | Calcein | 1058 | BL1 | 18556 |

| | | | | | | | | | |
|---|---|---|---|---|---|---|---|---|---|
| 40 | No | A. baylyi | Rifampicin | Intracellula | F12 | Amaranth | 1051 | BL1 | 18094 |
| 40 | No | A. baylyi | None | None | NA | Autofluore: | 1176 | NA | 18211.5 |
| 40 | No | A. baylyi | Trimethopr | Intracellula | F194 | Oxonol | 2867 | RL1 | 13679 |
| 40 | No | A. baylyi | Trimethopr | Intracellula | F11 | Allura Red | 1455 | BL3 | 12521 |
| 40 | No | A. baylyi | Trimethopr | Intracellula | F54 | DiSC3(5) | 1069 | RL1 | 12757 |
| 40 | No | A. baylyi | Trimethopr | Intracellula | F124 | Thioflavin T | 1223 | VL2 | 8309 |
| 40 | No | A. baylyi | Trimethopr | Intracellula | F101 | Quinacrine | 1189 | VL2 | 6533 |
| 40 | No | A. baylyi | Trimethopr | Intracellula | F138 | Apigenin | 1249 | VL2 | 6657 |
| 40 | No | A. baylyi | Trimethopr | Intracellula | F65 | H2FDA | 1232 | BL1 | 7670 |
| 40 | No | A. baylyi | Trimethopr | Intracellula | F142 | Chlorophe | 1180 | BL3 | 8372 |
| 40 | No | A. baylyi | Trimethopr | Intracellula | F39 | Congo Red | 1243 | BL3 | 9483 |
| 40 | No | A. baylyi | Trimethopr | Intracellula | F116 | Sunset Yel | 1154 | BL2 | 7617 |
| 40 | No | A. baylyi | Trimethopr | Intracellula | F10 | Alizarin | 1170 | BL1 | 6860 |
| 40 | No | A. baylyi | Trimethopr | Intracellula | F84 | Neutral Re | 1016 | BL1 | 5915 |
| 40 | No | A. baylyi | Trimethopr | Intracellula | F99 | Pyronin Y | 1066 | BL1 | 6859 |
| 40 | No | A. baylyi | Trimethopr | Intracellula | F108 | Rhodamine | 1247 | BL1 | 5826 |
| 40 | No | A. baylyi | Trimethopr | Intracellula | F109 | Riboflavin | 965 | BL1 | 5405 |
| 40 | No | A. baylyi | Trimethopr | Intracellula | F117 | SYBR Gree | 982 | BL1 | 458793.5 |
| 40 | No | A. baylyi | Trimethopr | Intracellula | F106 | Rhodamine | 1096 | BL1 | 7493.5 |
| 40 | No | A. baylyi | Trimethopr | Intracellula | F73 | Malachite ( | 1171 | BL1 | 7121 |
| 40 | No | A. baylyi | Trimethopr | Intracellula | F7 | Acridine or | 1035 | BL1 | 6735 |
| 40 | No | A. baylyi | Trimethopr | Intracellula | F192 | 5-Carboxyf | 1115 | BL1 | 19021 |
| 40 | No | A. baylyi | Trimethopr | Intracellula | F57 | Eosin Y | 988 | BL1 | 13951.5 |
| 40 | No | A. baylyi | Trimethopr | Intracellula | F27 | Calcein | 1175 | BL1 | 17557 |
| 40 | No | A. baylyi | Trimethopr | Intracellula | F12 | Amaranth | 1103 | BL1 | 18694 |
| 40 | No | A. baylyi | None | None | NA | Autofluore: | 2082 | NA | 1365.5 |
| 40 | No | A. baylyi | None | None | F194 | Oxonol | 3386 | RL1 | 658925 |
| 40 | No | A. baylyi | None | None | F11 | Allura Red | 1781 | BL3 | 45682 |
| 40 | No | A. baylyi | None | None | F54 | DiSC3(5) | 1483 | RL1 | 7058 |
| 40 | No | A. baylyi | None | None | F124 | Thioflavin T | 1438 | VL2 | 5277 |
| 40 | No | A. baylyi | None | None | F101 | Quinacrine | 1338 | VL2 | 4219.5 |
| 40 | No | A. baylyi | None | None | F138 | Apigenin | 1194 | VL2 | 3668 |
| 40 | No | A. baylyi | None | None | F65 | H2FDA | 1279 | BL1 | 5450 |
| 40 | No | A. baylyi | None | None | F142 | Chlorophe | 1276 | BL3 | 6234 |
| 40 | No | A. baylyi | None | None | F39 | Congo Red | 1107 | BL3 | 5638 |
| 40 | No | A. baylyi | None | None | F116 | Sunset Yel | 1182 | BL2 | 5385.5 |
| 40 | No | A. baylyi | None | None | F10 | Alizarin | 1184 | BL1 | 4277 |
| 40 | No | A. baylyi | None | None | F84 | Neutral Re | 1176 | BL1 | 3699.5 |
| 40 | No | A. baylyi | None | None | F99 | Pyronin Y | 1160 | BL1 | 4202 |
| 40 | No | A. baylyi | None | None | F108 | Rhodamine | 1193 | BL1 | 3404 |
| 40 | No | A. baylyi | None | None | F109 | Riboflavin | 1150 | BL1 | 3715 |
| 40 | No | A. baylyi | None | None | F117 | SYBR Gree | 1106 | BL1 | 413749.5 |
| 40 | No | A. baylyi | None | None | F106 | Rhodamine | 1128 | BL1 | 5096 |
| 40 | No | A. baylyi | None | None | F73 | Malachite ( | 1060 | BL1 | 3922.5 |
| 40 | No | A. baylyi | None | None | F7 | Acridine or | 1305 | BL1 | 4491 |
| 40 | No | A. baylyi | None | None | F192 | 5-Carboxyf | 1090 | BL1 | 18249 |
| 40 | No | A. baylyi | None | None | F57 | Eosin Y | 1096 | BL1 | 19002 |
| 40 | No | A. baylyi | None | None | F27 | Calcein | 1184 | BL1 | 18017.5 |

| | | | | | | | | | |
|---|---|---|---|---|---|---|---|---|---|
| 40 | No | A. baylyi | None | None | F12 | Amaranth | 1102 | BL1 | 16472 |
| 40 | No | A. baylyi | None | None | NA | Autofluore: | 1183 | NA | 17709 |
| 40 | No | A. baylyi | Ampicillin | Wall | F194 | Oxonol | 2036 | RL1 | 13874 |
| 40 | No | A. baylyi | Ampicillin | Wall | F11 | Allura Red | 899 | BL3 | 15302 |
| 40 | No | A. baylyi | Ampicillin | Wall | F54 | DiSC3(5) | 810 | RL1 | 13053 |
| 40 | No | A. baylyi | Ampicillin | Wall | F124 | Thioflavin T | 812 | VL2 | 9075 |
| 40 | No | A. baylyi | Ampicillin | Wall | F101 | Quinacrine | 766 | VL2 | 6770.5 |
| 40 | No | A. baylyi | Ampicillin | Wall | F138 | Apigenin | 892 | VL2 | 5918.5 |
| 40 | No | A. baylyi | Ampicillin | Wall | F65 | H2FDA | 792 | BL1 | 7009 |
| 40 | No | A. baylyi | Ampicillin | Wall | F142 | Chlorophe | 883 | BL3 | 8182 |
| 40 | No | A. baylyi | Ampicillin | Wall | F39 | Congo Red | 767 | BL3 | 6791 |
| 40 | No | A. baylyi | Ampicillin | Wall | F116 | Sunset Yell | 836 | BL2 | 5758 |
| 40 | No | A. baylyi | Ampicillin | Wall | F10 | Alizarin | 854 | BL1 | 5559 |
| 40 | No | A. baylyi | Ampicillin | Wall | F84 | Neutral Re | 922 | BL1 | 4983.5 |
| 40 | No | A. baylyi | Ampicillin | Wall | F99 | Pyronin Y | 718 | BL1 | 5082 |
| 40 | No | A. baylyi | Ampicillin | Wall | F108 | Rhodamine | 878 | BL1 | 4603 |
| 40 | No | A. baylyi | Ampicillin | Wall | F109 | Riboflavin | 803 | BL1 | 4810 |
| 40 | No | A. baylyi | Ampicillin | Wall | F117 | SYBR Gree | 839 | BL1 | 622781 |
| 40 | No | A. baylyi | Ampicillin | Wall | F106 | Rhodamine | 854 | BL1 | 7316 |
| 40 | No | A. baylyi | Ampicillin | Wall | F73 | Malachite ( | 862 | BL1 | 5469.5 |
| 40 | No | A. baylyi | Ampicillin | Wall | F7 | Acridine or | 809 | BL1 | 5908 |
| 40 | No | A. baylyi | Ampicillin | Wall | F192 | 5-Carboxyf | 852 | BL1 | 21996 |
| 40 | No | A. baylyi | Ampicillin | Wall | F57 | Eosin Y | 809 | BL1 | 23516 |
| 40 | No | A. baylyi | Ampicillin | Wall | F27 | Calcein | 792 | BL1 | 21783.5 |
| 40 | No | A. baylyi | Ampicillin | Wall | F12 | Amaranth | 802 | BL1 | 22525 |
| 40 | No | A. baylyi | None | None | NA | Autofluore: | 1130 | NA | 17425.5 |
| 40 | No | A. baylyi | Bacitracin | Wall | F194 | Oxonol | 1659 | RL1 | 19280 |
| 40 | No | A. baylyi | Bacitracin | Wall | F11 | Allura Red | 1107 | BL3 | 12247 |
| 40 | No | A. baylyi | Bacitracin | Wall | F54 | DiSC3(5) | 1026 | RL1 | 13017.5 |
| 40 | No | A. baylyi | Bacitracin | Wall | F124 | Thioflavin T | 1060 | VL2 | 8531 |
| 40 | No | A. baylyi | Bacitracin | Wall | F101 | Quinacrine | 1078 | VL2 | 7014 |
| 40 | No | A. baylyi | Bacitracin | Wall | F138 | Apigenin | 1203 | VL2 | 6793 |
| 40 | No | A. baylyi | Bacitracin | Wall | F65 | H2FDA | 1102 | BL1 | 7069 |
| 40 | No | A. baylyi | Bacitracin | Wall | F142 | Chlorophe | 1110 | BL3 | 6921 |
| 40 | No | A. baylyi | Bacitracin | Wall | F39 | Congo Red | 1082 | BL3 | 6869.5 |
| 40 | No | A. baylyi | Bacitracin | Wall | F116 | Sunset Yell | 1075 | BL2 | 5653 |
| 40 | No | A. baylyi | Bacitracin | Wall | F10 | Alizarin | 1186 | BL1 | 5777.5 |
| 40 | No | A. baylyi | Bacitracin | Wall | F84 | Neutral Re | 1153 | BL1 | 5028 |
| 40 | No | A. baylyi | Bacitracin | Wall | F99 | Pyronin Y | 1024 | BL1 | 4744.5 |
| 40 | No | A. baylyi | Bacitracin | Wall | F108 | Rhodamine | 1157 | BL1 | 4838 |
| 40 | No | A. baylyi | Bacitracin | Wall | F109 | Riboflavin | 1119 | BL1 | 4785 |
| 40 | No | A. baylyi | Bacitracin | Wall | F117 | SYBR Gree | 988 | BL1 | 614720.5 |
| 40 | No | A. baylyi | Bacitracin | Wall | F106 | Rhodamine | 1028 | BL1 | 6845 |
| 40 | No | A. baylyi | Bacitracin | Wall | F73 | Malachite ( | 1084 | BL1 | 5297.5 |
| 40 | No | A. baylyi | Bacitracin | Wall | F7 | Acridine or | 1109 | BL1 | 5507 |
| 40 | No | A. baylyi | Bacitracin | Wall | F192 | 5-Carboxyf | 1101 | BL1 | 18545 |
| 40 | No | A. baylyi | Bacitracin | Wall | F57 | Eosin Y | 1142 | BL1 | 19670.5 |
| 40 | No | A. baylyi | Bacitracin | Wall | F27 | Calcein | 1110 | BL1 | 19284.5 |

| | | | | | | | | | |
|---|---|---|---|---|---|---|---|---|---|
| 40 | No | A. baylyi | Bacitracin | Wall | F12 | Amaranth | 1078 | BL1 | 19698.5 |
| 40 | No | A. baylyi | None | None | NA | Autofluore: | 1088 | NA | 17229.5 |
| 40 | No | A. baylyi | Cefotaxim | Wall | F194 | Oxonol | 2960 | RL1 | 14583 |
| 40 | No | A. baylyi | Cefotaxim | Wall | F11 | Allura Red | 1098 | BL3 | 14051 |
| 40 | No | A. baylyi | Cefotaxim | Wall | F54 | DiSC3(5) | 884 | RL1 | 13167.5 |
| 40 | No | A. baylyi | Cefotaxim | Wall | F124 | Thioflavin T | 961 | VL2 | 9503 |
| 40 | No | A. baylyi | Cefotaxim | Wall | F101 | Quinacrine | 933 | VL2 | 7041 |
| 40 | No | A. baylyi | Cefotaxim | Wall | F138 | Apigenin | 954 | VL2 | 5950.5 |
| 40 | No | A. baylyi | Cefotaxim | Wall | F65 | H2FDA | 826 | BL1 | 7253.5 |
| 40 | No | A. baylyi | Cefotaxim | Wall | F142 | Chlorophe | 1062 | BL3 | 7854.5 |
| 40 | No | A. baylyi | Cefotaxim | Wall | F39 | Congo Red | 873 | BL3 | 8145 |
| 40 | No | A. baylyi | Cefotaxim | Wall | F116 | Sunset Yel | 1025 | BL2 | 7278 |
| 40 | No | A. baylyi | Cefotaxim | Wall | F10 | Alizarin | 914 | BL1 | 5552 |
| 40 | No | A. baylyi | Cefotaxim | Wall | F84 | Neutral Re | 966 | BL1 | 5222.5 |
| 40 | No | A. baylyi | Cefotaxim | Wall | F99 | Pyronin Y | 825 | BL1 | 5349 |
| 40 | No | A. baylyi | Cefotaxim | Wall | F108 | Rhodamine | 902 | BL1 | 4949.5 |
| 40 | No | A. baylyi | Cefotaxim | Wall | F109 | Riboflavin | 950 | BL1 | 5243.5 |
| 40 | No | A. baylyi | Cefotaxim | Wall | F117 | SYBR Gree | 914 | BL1 | 558460.5 |
| 40 | No | A. baylyi | Cefotaxim | Wall | F106 | Rhodamine | 847 | BL1 | 7961 |
| 40 | No | A. baylyi | Cefotaxim | Wall | F73 | Malachite ( | 989 | BL1 | 5319 |
| 40 | No | A. baylyi | Cefotaxim | Wall | F7 | Acridine or | 893 | BL1 | 6591 |
| 40 | No | A. baylyi | Cefotaxim | Wall | F192 | 5-Carboxyf | 819 | BL1 | 21293 |
| 40 | No | A. baylyi | Cefotaxim | Wall | F57 | Eosin Y | 969 | BL1 | 19267 |
| 40 | No | A. baylyi | Cefotaxim | Wall | F27 | Calcein | 908 | BL1 | 18515.5 |
| 40 | No | A. baylyi | Cefotaxim | Wall | F12 | Amaranth | 853 | BL1 | 20728 |
| 40 | No | A. baylyi | None | None | NA | Autofluore: | 1179 | NA | 17435 |
| 40 | No | A. baylyi | Ciprofloxa | Intracellul | F194 | Oxonol | 1322 | RL1 | 17887.5 |
| 40 | No | A. baylyi | Ciprofloxa | Intracellul | F11 | Allura Red | 808 | BL3 | 14213.5 |
| 40 | No | A. baylyi | Ciprofloxa | Intracellul | F54 | DiSC3(5) | 803 | RL1 | 15011 |
| 40 | No | A. baylyi | Ciprofloxa | Intracellul | F124 | Thioflavin T | 832 | VL2 | 9262 |
| 40 | No | A. baylyi | Ciprofloxa | Intracellul | F101 | Quinacrine | 826 | VL2 | 7060.5 |
| 40 | No | A. baylyi | Ciprofloxa | Intracellul | F138 | Apigenin | 918 | VL2 | 6375.5 |
| 40 | No | A. baylyi | Ciprofloxa | Intracellul | F65 | H2FDA | 887 | BL1 | 7471 |
| 40 | No | A. baylyi | Ciprofloxa | Intracellul | F142 | Chlorophe | 849 | BL3 | 7972 |
| 40 | No | A. baylyi | Ciprofloxa | Intracellul | F39 | Congo Red | 850 | BL3 | 7629.5 |
| 40 | No | A. baylyi | Ciprofloxa | Intracellul | F116 | Sunset Yel | 847 | BL2 | 6980 |
| 40 | No | A. baylyi | Ciprofloxa | Intracellul | F10 | Alizarin | 831 | BL1 | 5687 |
| 40 | No | A. baylyi | Ciprofloxa | Intracellul | F84 | Neutral Re | 844 | BL1 | 5393 |
| 40 | No | A. baylyi | Ciprofloxa | Intracellul | F99 | Pyronin Y | 844 | BL1 | 5676 |
| 40 | No | A. baylyi | Ciprofloxa | Intracellul | F108 | Rhodamine | 900 | BL1 | 4815 |
| 40 | No | A. baylyi | Ciprofloxa | Intracellul | F109 | Riboflavin | 878 | BL1 | 5731.5 |
| 40 | No | A. baylyi | Ciprofloxa | Intracellul | F117 | SYBR Gree | 841 | BL1 | 356385 |
| 40 | No | A. baylyi | Ciprofloxa | Intracellul | F106 | Rhodamine | 852 | BL1 | 6647 |
| 40 | No | A. baylyi | Ciprofloxa | Intracellul | F73 | Malachite ( | 888 | BL1 | 6311 |
| 40 | No | A. baylyi | Ciprofloxa | Intracellul | F7 | Acridine or | 848 | BL1 | 6731 |
| 40 | No | A. baylyi | Ciprofloxa | Intracellul | F192 | 5-Carboxyf | 785 | BL1 | 21601 |
| 40 | No | A. baylyi | Ciprofloxa | Intracellul | F57 | Eosin Y | 840 | BL1 | 21682.5 |
| 40 | No | A. baylyi | Ciprofloxa | Intracellul | F27 | Calcein | 811 | BL1 | 21114 |

| | | | | | | | | | |
|---|---|---|---|---|---|---|---|---|---|
| 40 | No | A. baylyi | Ciprofloxa | Intracellula | F12 | Amaranth | 886 | BL1 | 19825 |
| 40 | No | A. baylyi | None | None | NA | Autofluore | 1087 | NA | 16252 |
| 40 | No | A. baylyi | Fosfomyci | Wall | F194 | Oxonol | 1546 | RL1 | 18304 |
| 40 | No | A. baylyi | Fosfomyci | Wall | F11 | Allura Red | 1153 | BL3 | 12165 |
| 40 | No | A. baylyi | Fosfomyci | Wall | F54 | DiSC3(5) | 1003 | RL1 | 11760 |
| 40 | No | A. baylyi | Fosfomyci | Wall | F124 | Thioflavin T | 1072 | VL2 | 7288 |
| 40 | No | A. baylyi | Fosfomyci | Wall | F101 | Quinacrine | 1040 | VL2 | 6286.5 |
| 40 | No | A. baylyi | Fosfomyci | Wall | F138 | Apigenin | 1104 | VL2 | 5520.5 |
| 40 | No | A. baylyi | Fosfomyci | Wall | F65 | H2FDA | 1183 | BL1 | 6627 |
| 40 | No | A. baylyi | Fosfomyci | Wall | F142 | Chlorophe | 1166 | BL3 | 7262 |
| 40 | No | A. baylyi | Fosfomyci | Wall | F39 | Congo Red | 1181 | BL3 | 7059 |
| 40 | No | A. baylyi | Fosfomyci | Wall | F116 | Sunset Yel | 1050 | BL2 | 6552.5 |
| 40 | No | A. baylyi | Fosfomyci | Wall | F10 | Alizarin | 1188 | BL1 | 5222 |
| 40 | No | A. baylyi | Fosfomyci | Wall | F84 | Neutral Re | 1174 | BL1 | 4744.5 |
| 40 | No | A. baylyi | Fosfomyci | Wall | F99 | Pyronin Y | 1092 | BL1 | 5232 |
| 40 | No | A. baylyi | Fosfomyci | Wall | F108 | Rhodamine | 1052 | BL1 | 4753 |
| 40 | No | A. baylyi | Fosfomyci | Wall | F109 | Riboflavin | 1082 | BL1 | 4651 |
| 40 | No | A. baylyi | Fosfomyci | Wall | F117 | SYBR Gree | 928 | BL1 | 454276 |
| 40 | No | A. baylyi | Fosfomyci | Wall | F106 | Rhodamine | 1084 | BL1 | 6769 |
| 40 | No | A. baylyi | Fosfomyci | Wall | F73 | Malachite | 1092 | BL1 | 5024.5 |
| 40 | No | A. baylyi | Fosfomyci | Wall | F7 | Acridine or | 1167 | BL1 | 5842 |
| 40 | No | A. baylyi | Fosfomyci | Wall | F192 | 5-Carboxyf | 1163 | BL1 | 17641 |
| 40 | No | A. baylyi | Fosfomyci | Wall | F57 | Eosin Y | 1094 | BL1 | 15105 |
| 40 | No | A. baylyi | Fosfomyci | Wall | F27 | Calcein | 1224 | BL1 | 16932.5 |
| 40 | No | A. baylyi | Fosfomyci | Wall | F12 | Amaranth | 1117 | BL1 | 17881 |
| 40 | No | A. baylyi | None | None | NA | Autofluore | 1186 | NA | 16654.5 |
| 40 | No | A. baylyi | Nalidixic a | Intracellula | F194 | Oxonol | 1549 | RL1 | 15696 |
| 40 | No | A. baylyi | Nalidixic a | Intracellula | F11 | Allura Red | 1036 | BL3 | 13291.5 |
| 40 | No | A. baylyi | Nalidixic a | Intracellula | F54 | DiSC3(5) | 795 | RL1 | 11922 |
| 40 | No | A. baylyi | Nalidixic a | Intracellula | F124 | Thioflavin T | 888 | VL2 | 7607.5 |
| 40 | No | A. baylyi | Nalidixic a | Intracellula | F101 | Quinacrine | 966 | VL2 | 7074.5 |
| 40 | No | A. baylyi | Nalidixic a | Intracellula | F138 | Apigenin | 950 | VL2 | 6355.5 |
| 40 | No | A. baylyi | Nalidixic a | Intracellula | F65 | H2FDA | 908 | BL1 | 6872.5 |
| 40 | No | A. baylyi | Nalidixic a | Intracellula | F142 | Chlorophe | 1114 | BL3 | 8840.5 |
| 40 | No | A. baylyi | Nalidixic a | Intracellula | F39 | Congo Red | 916 | BL3 | 7694 |
| 40 | No | A. baylyi | Nalidixic a | Intracellula | F116 | Sunset Yel | 910 | BL2 | 7101.5 |
| 40 | No | A. baylyi | Nalidixic a | Intracellula | F10 | Alizarin | 930 | BL1 | 6140 |
| 40 | No | A. baylyi | Nalidixic a | Intracellula | F84 | Neutral Re | 972 | BL1 | 5158.5 |
| 40 | No | A. baylyi | Nalidixic a | Intracellula | F99 | Pyronin Y | 935 | BL1 | 5680 |
| 40 | No | A. baylyi | Nalidixic a | Intracellula | F108 | Rhodamine | 1028 | BL1 | 5018 |
| 40 | No | A. baylyi | Nalidixic a | Intracellula | F109 | Riboflavin | 941 | BL1 | 5241 |
| 40 | No | A. baylyi | Nalidixic a | Intracellula | F117 | SYBR Gree | 924 | BL1 | 404692 |
| 40 | No | A. baylyi | Nalidixic a | Intracellula | F106 | Rhodamine | 975 | BL1 | 7231 |
| 40 | No | A. baylyi | Nalidixic a | Intracellula | F73 | Malachite | 923 | BL1 | 6006 |
| 40 | No | A. baylyi | Nalidixic a | Intracellula | F7 | Acridine or | 967 | BL1 | 6370 |
| 40 | No | A. baylyi | Nalidixic a | Intracellula | F192 | 5-Carboxyf | 908 | BL1 | 19179 |
| 40 | No | A. baylyi | Nalidixic a | Intracellula | F57 | Eosin Y | 986 | BL1 | 18209 |
| 40 | No | A. baylyi | Nalidixic a | Intracellula | F27 | Calcein | 955 | BL1 | 17716 |

| | | | | | | | | | |
|---|---|---|---|---|---|---|---|---|---|
| 40 | No | A. baylyi | Nalidixic a | Intracellula | F12 | Amaranth | 853 | BL1 | 18880 |
| 40 | No | A. baylyi | None | None | NA | Autofluore: | 1195 | NA | 18283 |
| 40 | No | A. baylyi | Ofloxacin | Intracellula | F194 | Oxonol | 1847 | RL1 | 12704 |
| 40 | No | A. baylyi | Ofloxacin | Intracellula | F11 | Allura Red | 1010 | BL3 | 15060.5 |
| 40 | No | A. baylyi | Ofloxacin | Intracellula | F54 | DiSC3(5) | 788 | RL1 | 12511 |
| 40 | No | A. baylyi | Ofloxacin | Intracellula | F124 | Thioflavin T | 907 | VL2 | 8928 |
| 40 | No | A. baylyi | Ofloxacin | Intracellula | F101 | Quinacrine | 872 | VL2 | 7470.5 |
| 40 | No | A. baylyi | Ofloxacin | Intracellula | F138 | Apigenin | 833 | VL2 | 6278 |
| 40 | No | A. baylyi | Ofloxacin | Intracellula | F65 | H2FDA | 856 | BL1 | 7548 |
| 40 | No | A. baylyi | Ofloxacin | Intracellula | F142 | Chlorophe | 941 | BL3 | 8767 |
| 40 | No | A. baylyi | Ofloxacin | Intracellula | F39 | Congo Red | 871 | BL3 | 8273 |
| 40 | No | A. baylyi | Ofloxacin | Intracellula | F116 | Sunset Yel | 951 | BL2 | 6913 |
| 40 | No | A. baylyi | Ofloxacin | Intracellula | F10 | Alizarin | 859 | BL1 | 6171 |
| 40 | No | A. baylyi | Ofloxacin | Intracellula | F84 | Neutral Re | 911 | BL1 | 5637 |
| 40 | No | A. baylyi | Ofloxacin | Intracellula | F99 | Pyronin Y | 822 | BL1 | 5629 |
| 40 | No | A. baylyi | Ofloxacin | Intracellula | F108 | Rhodamine | 915 | BL1 | 5217 |
| 40 | No | A. baylyi | Ofloxacin | Intracellula | F109 | Riboflavin | 810 | BL1 | 4913.5 |
| 40 | No | A. baylyi | Ofloxacin | Intracellula | F117 | SYBR Gree | 722 | BL1 | 461077.5 |
| 40 | No | A. baylyi | Ofloxacin | Intracellula | F106 | Rhodamine | 954 | BL1 | 8763.5 |
| 40 | No | A. baylyi | Ofloxacin | Intracellula | F73 | Malachite ( | 970 | BL1 | 5889 |
| 40 | No | A. baylyi | Ofloxacin | Intracellula | F7 | Acridine or | 765 | BL1 | 6242 |
| 40 | No | A. baylyi | Ofloxacin | Intracellula | F192 | 5-Carboxyf | 915 | BL1 | 22321 |
| 40 | No | A. baylyi | Ofloxacin | Intracellula | F57 | Eosin Y | 888 | BL1 | 20344.5 |
| 40 | No | A. baylyi | Ofloxacin | Intracellula | F27 | Calcein | 841 | BL1 | 20680 |
| 40 | No | A. baylyi | Ofloxacin | Intracellula | F12 | Amaranth | 883 | BL1 | 21029 |
| 40 | No | A. baylyi | None | None | NA | Autofluore: | 1125 | NA | 17088 |
| 40 | No | A. baylyi | Polymyxin | Wall | F194 | Oxonol | 1736 | RL1 | 17918.5 |
| 40 | No | A. baylyi | Polymyxin | Wall | F11 | Allura Red | 1283 | BL3 | 13433 |
| 40 | No | A. baylyi | Polymyxin | Wall | F54 | DiSC3(5) | 1139 | RL1 | 10947 |
| 40 | No | A. baylyi | Polymyxin | Wall | F124 | Thioflavin T | 1174 | VL2 | 7553 |
| 40 | No | A. baylyi | Polymyxin | Wall | F101 | Quinacrine | 1206 | VL2 | 6720.5 |
| 40 | No | A. baylyi | Polymyxin | Wall | F138 | Apigenin | 1358 | VL2 | 5980.5 |
| 40 | No | A. baylyi | Polymyxin | Wall | F65 | H2FDA | 1227 | BL1 | 7073 |
| 40 | No | A. baylyi | Polymyxin | Wall | F142 | Chlorophe | 1300 | BL3 | 7524 |
| 40 | No | A. baylyi | Polymyxin | Wall | F39 | Congo Red | 1175 | BL3 | 7693 |
| 40 | No | A. baylyi | Polymyxin | Wall | F116 | Sunset Yel | 1193 | BL2 | 6184 |
| 40 | No | A. baylyi | Polymyxin | Wall | F10 | Alizarin | 1258 | BL1 | 5382 |
| 40 | No | A. baylyi | Polymyxin | Wall | F84 | Neutral Re | 1198 | BL1 | 5419 |
| 40 | No | A. baylyi | Polymyxin | Wall | F99 | Pyronin Y | 1076 | BL1 | 5129 |
| 40 | No | A. baylyi | Polymyxin | Wall | F108 | Rhodamine | 1194 | BL1 | 4588 |
| 40 | No | A. baylyi | Polymyxin | Wall | F109 | Riboflavin | 1097 | BL1 | 4985 |
| 40 | No | A. baylyi | Polymyxin | Wall | F117 | SYBR Gree | 1104 | BL1 | 631274.5 |
| 40 | No | A. baylyi | Polymyxin | Wall | F106 | Rhodamine | 1225 | BL1 | 6518 |
| 40 | No | A. baylyi | Polymyxin | Wall | F73 | Malachite ( | 1149 | BL1 | 5354 |
| 40 | No | A. baylyi | Polymyxin | Wall | F7 | Acridine or | 1053 | BL1 | 5497 |
| 40 | No | A. baylyi | Polymyxin | Wall | F192 | 5-Carboxyf | 1280 | BL1 | 20056.5 |
| 40 | No | A. baylyi | Polymyxin | Wall | F57 | Eosin Y | 1253 | BL1 | 19044 |
| 40 | No | A. baylyi | Polymyxin | Wall | F27 | Calcein | 1214 | BL1 | 19363 |

| | | | | | | | | | |
|---|---|---|---|---|---|---|---|---|---|
| 40 | No | A. baylyi | Polymyxin | Wall | F12 | Amaranth | 1205 | BL1 | 17888 |
| 40 | No | A. baylyi | None | None | NA | Autofluore: | 1160 | NA | 15151 |
| 40 | No | A. baylyi | Tetracyclin | Intracellula | F194 | Oxonol | 1972 | RL1 | 16101.5 |
| 40 | No | A. baylyi | Tetracyclin | Intracellula | F11 | Allura Red | 909 | BL3 | 14751 |
| 40 | No | A. baylyi | Tetracyclin | Intracellula | F54 | DiSC3(5) | 754 | RL1 | 10918 |
| 40 | No | A. baylyi | Tetracyclin | Intracellula | F124 | Thioflavin T | 893 | VL2 | 8389 |
| 40 | No | A. baylyi | Tetracyclin | Intracellula | F101 | Quinacrine | 820 | VL2 | 7220 |
| 40 | No | A. baylyi | Tetracyclin | Intracellula | F138 | Apigenin | 909 | VL2 | 6253 |
| 40 | No | A. baylyi | Tetracyclin | Intracellula | F65 | H2FDA | 780 | BL1 | 7169.5 |
| 40 | No | A. baylyi | Tetracyclin | Intracellula | F142 | Chlorophe | 846 | BL3 | 8236.5 |
| 40 | No | A. baylyi | Tetracyclin | Intracellula | F39 | Congo Red | 812 | BL3 | 7372.5 |
| 40 | No | A. baylyi | Tetracyclin | Intracellula | F116 | Sunset Yel | 896 | BL2 | 6927 |
| 40 | No | A. baylyi | Tetracyclin | Intracellula | F10 | Alizarin | 884 | BL1 | 6340.5 |
| 40 | No | A. baylyi | Tetracyclin | Intracellula | F84 | Neutral Re | 792 | BL1 | 5094 |
| 40 | No | A. baylyi | Tetracyclin | Intracellula | F99 | Pyronin Y | 730 | BL1 | 6214 |
| 40 | No | A. baylyi | Tetracyclin | Intracellula | F108 | Rhodamine | 927 | BL1 | 4879 |
| 40 | No | A. baylyi | Tetracyclin | Intracellula | F109 | Riboflavin | 694 | BL1 | 5148 |
| 40 | No | A. baylyi | Tetracyclin | Intracellula | F117 | SYBR Gree | 870 | BL1 | 523643 |
| 40 | No | A. baylyi | Tetracyclin | Intracellula | F106 | Rhodamine | 806 | BL1 | 7469.5 |
| 40 | No | A. baylyi | Tetracyclin | Intracellula | F73 | Malachite ( | 795 | BL1 | 6141 |
| 40 | No | A. baylyi | Tetracyclin | Intracellula | F7 | Acridine or | 804 | BL1 | 6620.5 |
| 40 | No | A. baylyi | Tetracyclin | Intracellula | F192 | 5-Carboxyf | 796 | BL1 | 19789 |
| 40 | No | A. baylyi | Tetracyclin | Intracellula | F57 | Eosin Y | 795 | BL1 | 19423 |
| 40 | No | A. baylyi | Tetracyclin | Intracellula | F27 | Calcein | 709 | BL1 | 18885 |
| 40 | No | A. baylyi | Tetracyclin | Intracellula | F12 | Amaranth | 920 | BL1 | 18376.5 |
| 40 | No | A. baylyi | None | None | NA | Autofluore: | 1179 | NA | 17319 |
| 40 | No | A. baylyi | Vancomyc | Wall | F194 | Oxonol | 1411 | RL1 | 19151 |
| 40 | No | A. baylyi | Vancomyc | Wall | F11 | Allura Red | 994 | BL3 | 13698 |
| 40 | No | A. baylyi | Vancomyc | Wall | F54 | DiSC3(5) | 961 | RL1 | 11733 |
| 40 | No | A. baylyi | Vancomyc | Wall | F124 | Thioflavin T | 1137 | VL2 | 7531 |
| 40 | No | A. baylyi | Vancomyc | Wall | F101 | Quinacrine | 1176 | VL2 | 6790 |
| 40 | No | A. baylyi | Vancomyc | Wall | F138 | Apigenin | 1086 | VL2 | 5988 |
| 40 | No | A. baylyi | Vancomyc | Wall | F65 | H2FDA | 1131 | BL1 | 6565 |
| 40 | No | A. baylyi | Vancomyc | Wall | F142 | Chlorophe | 1139 | BL3 | 7982 |
| 40 | No | A. baylyi | Vancomyc | Wall | F39 | Congo Red | 1092 | BL3 | 7886 |
| 40 | No | A. baylyi | Vancomyc | Wall | F116 | Sunset Yel | 1095 | BL2 | 6472 |
| 40 | No | A. baylyi | Vancomyc | Wall | F10 | Alizarin | 1139 | BL1 | 6629 |
| 40 | No | A. baylyi | Vancomyc | Wall | F84 | Neutral Re | 1148 | BL1 | 5389 |
| 40 | No | A. baylyi | Vancomyc | Wall | F99 | Pyronin Y | 979 | BL1 | 5439 |
| 40 | No | A. baylyi | Vancomyc | Wall | F108 | Rhodamine | 1135 | BL1 | 4781 |
| 40 | No | A. baylyi | Vancomyc | Wall | F109 | Riboflavin | 1185 | BL1 | 5397 |
| 40 | No | A. baylyi | Vancomyc | Wall | F117 | SYBR Gree | 1047 | BL1 | 506969 |
| 40 | No | A. baylyi | Vancomyc | Wall | F106 | Rhodamine | 1087 | BL1 | 8204 |
| 40 | No | A. baylyi | Vancomyc | Wall | F73 | Malachite ( | 1082 | BL1 | 5144.5 |
| 40 | No | A. baylyi | Vancomyc | Wall | F7 | Acridine or | 1116 | BL1 | 6209.5 |
| 40 | No | A. baylyi | Vancomyc | Wall | F192 | 5-Carboxyf | 832 | BL1 | 16424 |
| 40 | No | A. baylyi | Vancomyc | Wall | F57 | Eosin Y | 1188 | BL1 | 16978.5 |
| 40 | No | A. baylyi | Vancomyc | Wall | F27 | Calcein | 1075 | BL1 | 18616 |

| | | | | | | | | | |
|---|---|---|---|---|---|---|---|---|---|
| 40 | No | A. baylyi | Vancomyc | Wall | F12 | Amaranth | 1004 | BL1 | 20468 |
| 40 | No | A. baylyi | None | None | NA | Autofluores | 2788 | NA | 1287 |
| 40 | No | A. baylyi | None | None | F194 | Oxonol | 2594 | RL1 | 354223 |
| 40 | No | A. baylyi | None | None | F11 | Allura Red | 1743 | BL3 | 27307 |
| 40 | No | A. baylyi | None | None | F54 | DiSC3(5) | 1492 | RL1 | 6440.5 |
| 40 | No | A. baylyi | None | None | F124 | Thioflavin T | 1380 | VL2 | 4980.5 |
| 40 | No | A. baylyi | None | None | F101 | Quinacrine | 1386 | VL2 | 4333 |
| 40 | No | A. baylyi | None | None | F138 | Apigenin | 1154 | VL2 | 3463 |
| 40 | No | A. baylyi | None | None | F65 | H2FDA | 1281 | BL1 | 5623 |
| 40 | No | A. baylyi | None | None | F142 | Chlorophe | 1219 | BL3 | 6421 |
| 40 | No | A. baylyi | None | None | F39 | Congo Red | 1394 | BL3 | 7160.5 |
| 40 | No | A. baylyi | None | None | F116 | Sunset Yel | 1296 | BL2 | 5696.5 |
| 40 | No | A. baylyi | None | None | F10 | Alizarin | 1129 | BL1 | 4220 |
| 40 | No | A. baylyi | None | None | F84 | Neutral Re | 1205 | BL1 | 4106 |
| 40 | No | A. baylyi | None | None | F99 | Pyronin Y | 1109 | BL1 | 3948 |
| 40 | No | A. baylyi | None | None | F108 | Rhodamine | 1245 | BL1 | 3617 |
| 40 | No | A. baylyi | None | None | F109 | Riboflavin | 1126 | BL1 | 3549 |
| 40 | No | A. baylyi | None | None | F117 | SYBR Gree | 1172 | BL1 | 284082 |
| 40 | No | A. baylyi | None | None | F106 | Rhodamine | 1142 | BL1 | 4627 |
| 40 | No | A. baylyi | None | None | F73 | Malachite ( | 1160 | BL1 | 4295 |
| 40 | No | A. baylyi | None | None | F7 | Acridine or | 1086 | BL1 | 4480 |
| 40 | No | A. baylyi | None | None | F192 | 5-Carboxyf | 1197 | BL1 | 17273 |
| 40 | No | A. baylyi | None | None | F57 | Eosin Y | 1149 | BL1 | 17008 |
| 40 | No | A. baylyi | None | None | F27 | Calcein | 1118 | BL1 | 18239.5 |
| 40 | No | A. baylyi | None | None | F12 | Amaranth | 1158 | BL1 | 17220.5 |
| 40 | No | A. baylyi | None | None | NA | Autofluores | 1158 | NA | 17435 |
| 40 | No | A. baylyi | Ampicillin | Wall | F194 | Oxonol | 3192 | RL1 | 14464 |
| 40 | No | A. baylyi | Ampicillin | Wall | F11 | Allura Red | 1039 | BL3 | 13396 |
| 40 | No | A. baylyi | Ampicillin | Wall | F54 | DiSC3(5) | 871 | RL1 | 14150 |
| 40 | No | A. baylyi | Ampicillin | Wall | F124 | Thioflavin T | 924 | VL2 | 9102.5 |
| 40 | No | A. baylyi | Ampicillin | Wall | F101 | Quinacrine | 890 | VL2 | 6419.5 |
| 40 | No | A. baylyi | Ampicillin | Wall | F138 | Apigenin | 883 | VL2 | 5025 |
| 40 | No | A. baylyi | Ampicillin | Wall | F65 | H2FDA | 827 | BL1 | 6871 |
| 40 | No | A. baylyi | Ampicillin | Wall | F142 | Chlorophe | 919 | BL3 | 6963 |
| 40 | No | A. baylyi | Ampicillin | Wall | F39 | Congo Red | 956 | BL3 | 7076.5 |
| 40 | No | A. baylyi | Ampicillin | Wall | F116 | Sunset Yel | 988 | BL2 | 6356 |
| 40 | No | A. baylyi | Ampicillin | Wall | F10 | Alizarin | 892 | BL1 | 5601.5 |
| 40 | No | A. baylyi | Ampicillin | Wall | F84 | Neutral Re | 942 | BL1 | 4840.5 |
| 40 | No | A. baylyi | Ampicillin | Wall | F99 | Pyronin Y | 807 | BL1 | 5243 |
| 40 | No | A. baylyi | Ampicillin | Wall | F108 | Rhodamine | 979 | BL1 | 4310 |
| 40 | No | A. baylyi | Ampicillin | Wall | F109 | Riboflavin | 793 | BL1 | 4662 |
| 40 | No | A. baylyi | Ampicillin | Wall | F117 | SYBR Gree | 938 | BL1 | 442678 |
| 40 | No | A. baylyi | Ampicillin | Wall | F106 | Rhodamine | 850 | BL1 | 6157.5 |
| 40 | No | A. baylyi | Ampicillin | Wall | F73 | Malachite ( | 929 | BL1 | 5038 |
| 40 | No | A. baylyi | Ampicillin | Wall | F7 | Acridine or | 828 | BL1 | 5357.5 |
| 40 | No | A. baylyi | Ampicillin | Wall | F192 | 5-Carboxyf | 944 | BL1 | 21200.5 |
| 40 | No | A. baylyi | Ampicillin | Wall | F57 | Eosin Y | 851 | BL1 | 22007 |
| 40 | No | A. baylyi | Ampicillin | Wall | F27 | Calcein | 829 | BL1 | 21491 |

| | | | | | | | | | |
|---|---|---|---|---|---|---|---|---|---|
| 40 | No | A. baylyi | Ampicillin | Wall | F12 | Amaranth | 836 | BL1 | 21535.5 |
| 40 | No | A. baylyi | None | None | NA | Autofluore: | 1144 | NA | 17203.5 |
| 40 | No | A. baylyi | Bacitracin | Wall | F194 | Oxonol | 1487 | RL1 | 18442 |
| 40 | No | A. baylyi | Bacitracin | Wall | F11 | Allura Red | 1110 | BL3 | 15048.5 |
| 40 | No | A. baylyi | Bacitracin | Wall | F54 | DiSC3(5) | 1014 | RL1 | 14190.5 |
| 40 | No | A. baylyi | Bacitracin | Wall | F124 | Thioflavin T | 1015 | VL2 | 8295 |
| 40 | No | A. baylyi | Bacitracin | Wall | F101 | Quinacrine | 1163 | VL2 | 7073 |
| 40 | No | A. baylyi | Bacitracin | Wall | F138 | Apigenin | 1156 | VL2 | 6102 |
| 40 | No | A. baylyi | Bacitracin | Wall | F65 | H2FDA | 1081 | BL1 | 6327 |
| 40 | No | A. baylyi | Bacitracin | Wall | F142 | Chlorophe | 1116 | BL3 | 7286 |
| 40 | No | A. baylyi | Bacitracin | Wall | F39 | Congo Red | 1007 | BL3 | 6398 |
| 40 | No | A. baylyi | Bacitracin | Wall | F116 | Sunset Yel | 1184 | BL2 | 6266.5 |
| 40 | No | A. baylyi | Bacitracin | Wall | F10 | Alizarin | 1171 | BL1 | 5237 |
| 40 | No | A. baylyi | Bacitracin | Wall | F84 | Neutral Re | 1160 | BL1 | 4552 |
| 40 | No | A. baylyi | Bacitracin | Wall | F99 | Pyronin Y | 1085 | BL1 | 4971 |
| 40 | No | A. baylyi | Bacitracin | Wall | F108 | Rhodamine | 1073 | BL1 | 4251 |
| 40 | No | A. baylyi | Bacitracin | Wall | F109 | Riboflavin | 1056 | BL1 | 4670.5 |
| 40 | No | A. baylyi | Bacitracin | Wall | F117 | SYBR Gree | 1093 | BL1 | 456312 |
| 40 | No | A. baylyi | Bacitracin | Wall | F106 | Rhodamine | 1102 | BL1 | 6186 |
| 40 | No | A. baylyi | Bacitracin | Wall | F73 | Malachite ( | 1165 | BL1 | 5239 |
| 40 | No | A. baylyi | Bacitracin | Wall | F7 | Acridine or | 1012 | BL1 | 5106.5 |
| 40 | No | A. baylyi | Bacitracin | Wall | F192 | 5-Carboxyf | 1123 | BL1 | 19917 |
| 40 | No | A. baylyi | Bacitracin | Wall | F57 | Eosin Y | 1161 | BL1 | 20449 |
| 40 | No | A. baylyi | Bacitracin | Wall | F27 | Calcein | 980 | BL1 | 20009.5 |
| 40 | No | A. baylyi | Bacitracin | Wall | F12 | Amaranth | 1080 | BL1 | 19911.5 |
| 40 | No | A. baylyi | None | None | NA | Autofluore: | 1155 | NA | 17978 |
| 40 | No | A. baylyi | Cefotaxim | Wall | F194 | Oxonol | 5116 | RL1 | 12072 |
| 40 | No | A. baylyi | Cefotaxim | Wall | F11 | Allura Red | 1342 | BL3 | 14835.5 |
| 40 | No | A. baylyi | Cefotaxim | Wall | F54 | DiSC3(5) | 949 | RL1 | 11743 |
| 40 | No | A. baylyi | Cefotaxim | Wall | F124 | Thioflavin T | 1000 | VL2 | 8236.5 |
| 40 | No | A. baylyi | Cefotaxim | Wall | F101 | Quinacrine | 986 | VL2 | 6876 |
| 40 | No | A. baylyi | Cefotaxim | Wall | F138 | Apigenin | 1065 | VL2 | 5872 |
| 40 | No | A. baylyi | Cefotaxim | Wall | F65 | H2FDA | 994 | BL1 | 6995.5 |
| 40 | No | A. baylyi | Cefotaxim | Wall | F142 | Chlorophe | 1076 | BL3 | 8315.5 |
| 40 | No | A. baylyi | Cefotaxim | Wall | F39 | Congo Red | 965 | BL3 | 7588 |
| 40 | No | A. baylyi | Cefotaxim | Wall | F116 | Sunset Yel | 1073 | BL2 | 7012 |
| 40 | No | A. baylyi | Cefotaxim | Wall | F10 | Alizarin | 952 | BL1 | 5277.5 |
| 40 | No | A. baylyi | Cefotaxim | Wall | F84 | Neutral Re | 1088 | BL1 | 5217.5 |
| 40 | No | A. baylyi | Cefotaxim | Wall | F99 | Pyronin Y | 874 | BL1 | 5204.5 |
| 40 | No | A. baylyi | Cefotaxim | Wall | F108 | Rhodamine | 1130 | BL1 | 4916 |
| 40 | No | A. baylyi | Cefotaxim | Wall | F109 | Riboflavin | 985 | BL1 | 5150 |
| 40 | No | A. baylyi | Cefotaxim | Wall | F117 | SYBR Gree | 1005 | BL1 | 367591 |
| 40 | No | A. baylyi | Cefotaxim | Wall | F106 | Rhodamine | 1009 | BL1 | 6778 |
| 40 | No | A. baylyi | Cefotaxim | Wall | F73 | Malachite ( | 966 | BL1 | 5533 |
| 40 | No | A. baylyi | Cefotaxim | Wall | F7 | Acridine or | 933 | BL1 | 5773 |
| 40 | No | A. baylyi | Cefotaxim | Wall | F192 | 5-Carboxyf | 1092 | BL1 | 18982.5 |
| 40 | No | A. baylyi | Cefotaxim | Wall | F57 | Eosin Y | 936 | BL1 | 20471 |
| 40 | No | A. baylyi | Cefotaxim | Wall | F27 | Calcein | 1034 | BL1 | 19525.5 |

| | | | | | | | | | |
|---|---|---|---|---|---|---|---|---|---|
| 40 | No | A. baylyi | Cefotaxim | Wall | F12 | Amaranth | 981 | BL1 | 20233 |
| 40 | No | A. baylyi | None | None | NA | Autofluore: | 1037 | NA | 17024 |
| 40 | No | A. baylyi | Ciprofloxa | Intracellul | F194 | Oxonol | 2228 | RL1 | 15003.5 |
| 40 | No | A. baylyi | Ciprofloxa | Intracellul | F11 | Allura Red | 990 | BL3 | 13916 |
| 40 | No | A. baylyi | Ciprofloxa | Intracellul | F54 | DiSC3(5) | 883 | RL1 | 13856 |
| 40 | No | A. baylyi | Ciprofloxa | Intracellul | F124 | Thioflavin T | 944 | VL2 | 8941.5 |
| 40 | No | A. baylyi | Ciprofloxa | Intracellul | F101 | Quinacrine | 923 | VL2 | 6268 |
| 40 | No | A. baylyi | Ciprofloxa | Intracellul | F138 | Apigenin | 958 | VL2 | 5764.5 |
| 40 | No | A. baylyi | Ciprofloxa | Intracellul | F65 | H2FDA | 967 | BL1 | 6891 |
| 40 | No | A. baylyi | Ciprofloxa | Intracellul | F142 | Chlorophe | 892 | BL3 | 7314 |
| 40 | No | A. baylyi | Ciprofloxa | Intracellul | F39 | Congo Red | 871 | BL3 | 7321 |
| 40 | No | A. baylyi | Ciprofloxa | Intracellul | F116 | Sunset Yel | 894 | BL2 | 5937.5 |
| 40 | No | A. baylyi | Ciprofloxa | Intracellul | F10 | Alizarin | 971 | BL1 | 5849 |
| 40 | No | A. baylyi | Ciprofloxa | Intracellul | F84 | Neutral Re | 943 | BL1 | 4965 |
| 40 | No | A. baylyi | Ciprofloxa | Intracellul | F99 | Pyronin Y | 897 | BL1 | 5274 |
| 40 | No | A. baylyi | Ciprofloxa | Intracellul | F108 | Rhodamine | 948 | BL1 | 4679.5 |
| 40 | No | A. baylyi | Ciprofloxa | Intracellul | F109 | Riboflavin | 888 | BL1 | 5038.5 |
| 40 | No | A. baylyi | Ciprofloxa | Intracellul | F117 | SYBR Gree | 895 | BL1 | 298099 |
| 40 | No | A. baylyi | Ciprofloxa | Intracellul | F106 | Rhodamine | 821 | BL1 | 6464 |
| 40 | No | A. baylyi | Ciprofloxa | Intracellul | F73 | Malachite ( | 955 | BL1 | 5672 |
| 40 | No | A. baylyi | Ciprofloxa | Intracellul | F7 | Acridine or | 855 | BL1 | 5819 |
| 40 | No | A. baylyi | Ciprofloxa | Intracellul | F192 | 5-Carboxyf | 905 | BL1 | 20330 |
| 40 | No | A. baylyi | Ciprofloxa | Intracellul | F57 | Eosin Y | 899 | BL1 | 19661 |
| 40 | No | A. baylyi | Ciprofloxa | Intracellul | F27 | Calcein | 929 | BL1 | 20448 |
| 40 | No | A. baylyi | Ciprofloxa | Intracellul | F12 | Amaranth | 872 | BL1 | 21761 |
| 40 | No | A. baylyi | None | None | NA | Autofluore: | 1211 | NA | 18891 |
| 40 | No | A. baylyi | Fosfomyci | Wall | F194 | Oxonol | 3904 | RL1 | 12650.5 |
| 40 | No | A. baylyi | Fosfomyci | Wall | F11 | Allura Red | 1166 | BL3 | 11780 |
| 40 | No | A. baylyi | Fosfomyci | Wall | F54 | DiSC3(5) | 945 | RL1 | 12196 |
| 40 | No | A. baylyi | Fosfomyci | Wall | F124 | Thioflavin T | 1080 | VL2 | 7098 |
| 40 | No | A. baylyi | Fosfomyci | Wall | F101 | Quinacrine | 1115 | VL2 | 6080 |
| 40 | No | A. baylyi | Fosfomyci | Wall | F138 | Apigenin | 1180 | VL2 | 5975.5 |
| 40 | No | A. baylyi | Fosfomyci | Wall | F65 | H2FDA | 1061 | BL1 | 6307 |
| 40 | No | A. baylyi | Fosfomyci | Wall | F142 | Chlorophe | 1276 | BL3 | 8250 |
| 40 | No | A. baylyi | Fosfomyci | Wall | F39 | Congo Red | 1130 | BL3 | 7875 |
| 40 | No | A. baylyi | Fosfomyci | Wall | F116 | Sunset Yel | 1178 | BL2 | 6033.5 |
| 40 | No | A. baylyi | Fosfomyci | Wall | F10 | Alizarin | 1052 | BL1 | 5323 |
| 40 | No | A. baylyi | Fosfomyci | Wall | F84 | Neutral Re | 1128 | BL1 | 4710.5 |
| 40 | No | A. baylyi | Fosfomyci | Wall | F99 | Pyronin Y | 1004 | BL1 | 4927 |
| 40 | No | A. baylyi | Fosfomyci | Wall | F108 | Rhodamine | 1049 | BL1 | 4581 |
| 40 | No | A. baylyi | Fosfomyci | Wall | F109 | Riboflavin | 1182 | BL1 | 4679.5 |
| 40 | No | A. baylyi | Fosfomyci | Wall | F117 | SYBR Gree | 1085 | BL1 | 348826 |
| 40 | No | A. baylyi | Fosfomyci | Wall | F106 | Rhodamine | 1114 | BL1 | 6386 |
| 40 | No | A. baylyi | Fosfomyci | Wall | F73 | Malachite ( | 1106 | BL1 | 5218.5 |
| 40 | No | A. baylyi | Fosfomyci | Wall | F7 | Acridine or | 1102 | BL1 | 5319.5 |
| 40 | No | A. baylyi | Fosfomyci | Wall | F192 | 5-Carboxyf | 1209 | BL1 | 18183 |
| 40 | No | A. baylyi | Fosfomyci | Wall | F57 | Eosin Y | 1105 | BL1 | 17900 |
| 40 | No | A. baylyi | Fosfomyci | Wall | F27 | Calcein | 1129 | BL1 | 16305 |

| | | | | | | | | | |
|---|---|---|---|---|---|---|---|---|---|
| 40 | No | A. baylyi | Fosfomyci | Wall | F12 | Amaranth | 1084 | BL1 | 18367.5 |
| 40 | No | A. baylyi | None | None | NA | Autofluore: | 1264 | NA | 16125.5 |
| 40 | No | A. baylyi | Nalidixic a | Intracellula | F194 | Oxonol | 2234 | RL1 | 13655.5 |
| 40 | No | A. baylyi | Nalidixic a | Intracellula | F11 | Allura Red | 1105 | BL3 | 12998 |
| 40 | No | A. baylyi | Nalidixic a | Intracellula | F54 | DiSC3(5) | 1042 | RL1 | 9453 |
| 40 | No | A. baylyi | Nalidixic a | Intracellula | F124 | Thioflavin T | 1005 | VL2 | 7704 |
| 40 | No | A. baylyi | Nalidixic a | Intracellula | F101 | Quinacrine | 1109 | VL2 | 6522 |
| 40 | No | A. baylyi | Nalidixic a | Intracellula | F138 | Apigenin | 1148 | VL2 | 6025.5 |
| 40 | No | A. baylyi | Nalidixic a | Intracellula | F65 | H2FDA | 1031 | BL1 | 7152 |
| 40 | No | A. baylyi | Nalidixic a | Intracellula | F142 | Chlorophe | 1014 | BL3 | 7149.5 |
| 40 | No | A. baylyi | Nalidixic a | Intracellula | F39 | Congo Red | 1155 | BL3 | 7541 |
| 40 | No | A. baylyi | Nalidixic a | Intracellula | F116 | Sunset Yel | 1055 | BL2 | 6127 |
| 40 | No | A. baylyi | Nalidixic a | Intracellula | F10 | Alizarin | 1102 | BL1 | 5166 |
| 40 | No | A. baylyi | Nalidixic a | Intracellula | F84 | Neutral Re | 1065 | BL1 | 4866 |
| 40 | No | A. baylyi | Nalidixic a | Intracellula | F99 | Pyronin Y | 970 | BL1 | 4797 |
| 40 | No | A. baylyi | Nalidixic a | Intracellula | F108 | Rhodamine | 1128 | BL1 | 4500.5 |
| 40 | No | A. baylyi | Nalidixic a | Intracellula | F109 | Riboflavin | 1089 | BL1 | 4863 |
| 40 | No | A. baylyi | Nalidixic a | Intracellula | F117 | SYBR Gree | 1004 | BL1 | 316544.5 |
| 40 | No | A. baylyi | Nalidixic a | Intracellula | F106 | Rhodamine | 1119 | BL1 | 6687 |
| 40 | No | A. baylyi | Nalidixic a | Intracellula | F73 | Malachite ( | 1047 | BL1 | 5436 |
| 40 | No | A. baylyi | Nalidixic a | Intracellula | F7 | Acridine or | 1043 | BL1 | 5642 |
| 40 | No | A. baylyi | Nalidixic a | Intracellula | F192 | 5-Carboxyf | 1118 | BL1 | 19218 |
| 40 | No | A. baylyi | Nalidixic a | Intracellula | F57 | Eosin Y | 1067 | BL1 | 21158 |
| 40 | No | A. baylyi | Nalidixic a | Intracellula | F27 | Calcein | 1002 | BL1 | 19610 |
| 40 | No | A. baylyi | Nalidixic a | Intracellula | F12 | Amaranth | 1051 | BL1 | 20339 |
| 40 | No | A. baylyi | None | None | NA | Autofluore: | 1242 | NA | 18842 |
| 40 | No | A. baylyi | Ofloxacin | Intracellula | F194 | Oxonol | 3259 | RL1 | 12078 |
| 40 | No | A. baylyi | Ofloxacin | Intracellula | F11 | Allura Red | 996 | BL3 | 14657 |
| 40 | No | A. baylyi | Ofloxacin | Intracellula | F54 | DiSC3(5) | 828 | RL1 | 10873 |
| 40 | No | A. baylyi | Ofloxacin | Intracellula | F124 | Thioflavin T | 1015 | VL2 | 7302 |
| 40 | No | A. baylyi | Ofloxacin | Intracellula | F101 | Quinacrine | 946 | VL2 | 6899 |
| 40 | No | A. baylyi | Ofloxacin | Intracellula | F138 | Apigenin | 1025 | VL2 | 6454 |
| 40 | No | A. baylyi | Ofloxacin | Intracellula | F65 | H2FDA | 925 | BL1 | 6808 |
| 40 | No | A. baylyi | Ofloxacin | Intracellula | F142 | Chlorophe | 1066 | BL3 | 8397 |
| 40 | No | A. baylyi | Ofloxacin | Intracellula | F39 | Congo Red | 960 | BL3 | 8913 |
| 40 | No | A. baylyi | Ofloxacin | Intracellula | F116 | Sunset Yel | 941 | BL2 | 6707 |
| 40 | No | A. baylyi | Ofloxacin | Intracellula | F10 | Alizarin | 995 | BL1 | 6194 |
| 40 | No | A. baylyi | Ofloxacin | Intracellula | F84 | Neutral Re | 880 | BL1 | 4864 |
| 40 | No | A. baylyi | Ofloxacin | Intracellula | F99 | Pyronin Y | 913 | BL1 | 6334 |
| 40 | No | A. baylyi | Ofloxacin | Intracellula | F108 | Rhodamine | 1000 | BL1 | 4843.5 |
| 40 | No | A. baylyi | Ofloxacin | Intracellula | F109 | Riboflavin | 1011 | BL1 | 5886 |
| 40 | No | A. baylyi | Ofloxacin | Intracellula | F117 | SYBR Gree | 926 | BL1 | 406537.5 |
| 40 | No | A. baylyi | Ofloxacin | Intracellula | F106 | Rhodamine | 951 | BL1 | 7966 |
| 40 | No | A. baylyi | Ofloxacin | Intracellula | F73 | Malachite ( | 979 | BL1 | 6137 |
| 40 | No | A. baylyi | Ofloxacin | Intracellula | F7 | Acridine or | 884 | BL1 | 6521 |
| 40 | No | A. baylyi | Ofloxacin | Intracellula | F192 | 5-Carboxyf | 979 | BL1 | 21267 |
| 40 | No | A. baylyi | Ofloxacin | Intracellula | F57 | Eosin Y | 887 | BL1 | 22017 |
| 40 | No | A. baylyi | Ofloxacin | Intracellula | F27 | Calcein | 975 | BL1 | 22090 |

| | | | | | | | | | |
|---|---|---|---|---|---|---|---|---|---|
| 40 | No | A. baylyi | Ofloxacin | Intracellul | F12 | Amaranth | 950 | BL1 | 21174.5 |
| 40 | No | A. baylyi | None | None | NA | Autofluore | 1212 | NA | 19764 |
| 40 | No | A. baylyi | Polymyxin | Wall | F194 | Oxonol | 2664 | RL1 | 14120 |
| 40 | No | A. baylyi | Polymyxin | Wall | F11 | Allura Red | 1366 | BL3 | 12312 |
| 40 | No | A. baylyi | Polymyxin | Wall | F54 | DiSC3(5) | 1085 | RL1 | 14085 |
| 40 | No | A. baylyi | Polymyxin | Wall | F124 | Thioflavin T | 1162 | VL2 | 8128.5 |
| 40 | No | A. baylyi | Polymyxin | Wall | F101 | Quinacrine | 1199 | VL2 | 7191 |
| 40 | No | A. baylyi | Polymyxin | Wall | F138 | Apigenin | 1268 | VL2 | 6130 |
| 40 | No | A. baylyi | Polymyxin | Wall | F65 | H2FDA | 1169 | BL1 | 6790 |
| 40 | No | A. baylyi | Polymyxin | Wall | F142 | Chlorophe | 1244 | BL3 | 8206 |
| 40 | No | A. baylyi | Polymyxin | Wall | F39 | Congo Red | 1173 | BL3 | 7375 |
| 40 | No | A. baylyi | Polymyxin | Wall | F116 | Sunset Yel | 1301 | BL2 | 7405 |
| 40 | No | A. baylyi | Polymyxin | Wall | F10 | Alizarin | 1106 | BL1 | 5637.5 |
| 40 | No | A. baylyi | Polymyxin | Wall | F84 | Neutral Re | 1132 | BL1 | 5012.5 |
| 40 | No | A. baylyi | Polymyxin | Wall | F99 | Pyronin Y | 1161 | BL1 | 5425 |
| 40 | No | A. baylyi | Polymyxin | Wall | F108 | Rhodamine | 1062 | BL1 | 4666.5 |
| 40 | No | A. baylyi | Polymyxin | Wall | F109 | Riboflavin | 1209 | BL1 | 5300 |
| 40 | No | A. baylyi | Polymyxin | Wall | F117 | SYBR Gree | 1057 | BL1 | 548094 |
| 40 | No | A. baylyi | Polymyxin | Wall | F106 | Rhodamine | 1139 | BL1 | 6257 |
| 40 | No | A. baylyi | Polymyxin | Wall | F73 | Malachite | 1186 | BL1 | 5257.5 |
| 40 | No | A. baylyi | Polymyxin | Wall | F7 | Acridine or | 1145 | BL1 | 5581 |
| 40 | No | A. baylyi | Polymyxin | Wall | F192 | 5-Carboxyf | 1075 | BL1 | 18414 |
| 40 | No | A. baylyi | Polymyxin | Wall | F57 | Eosin Y | 1072 | BL1 | 18717.5 |
| 40 | No | A. baylyi | Polymyxin | Wall | F27 | Calcein | 1233 | BL1 | 19310 |
| 40 | No | A. baylyi | Polymyxin | Wall | F12 | Amaranth | 1141 | BL1 | 16626 |
| 40 | No | A. baylyi | None | None | NA | Autofluore | 1129 | NA | 18755 |
| 40 | No | A. baylyi | Tetracyclin | Intracellul | F194 | Oxonol | 2602 | RL1 | 13126 |
| 40 | No | A. baylyi | Tetracyclin | Intracellul | F11 | Allura Red | 950 | BL3 | 13648 |
| 40 | No | A. baylyi | Tetracyclin | Intracellul | F54 | DiSC3(5) | 777 | RL1 | 11085 |
| 40 | No | A. baylyi | Tetracyclin | Intracellul | F124 | Thioflavin T | 919 | VL2 | 7390 |
| 40 | No | A. baylyi | Tetracyclin | Intracellul | F101 | Quinacrine | 886 | VL2 | 6876.5 |
| 40 | No | A. baylyi | Tetracyclin | Intracellul | F138 | Apigenin | 934 | VL2 | 6062 |
| 40 | No | A. baylyi | Tetracyclin | Intracellul | F65 | H2FDA | 949 | BL1 | 7399 |
| 40 | No | A. baylyi | Tetracyclin | Intracellul | F142 | Chlorophe | 877 | BL3 | 7670 |
| 40 | No | A. baylyi | Tetracyclin | Intracellul | F39 | Congo Red | 858 | BL3 | 8093.5 |
| 40 | No | A. baylyi | Tetracyclin | Intracellul | F116 | Sunset Yel | 916 | BL2 | 7405.5 |
| 40 | No | A. baylyi | Tetracyclin | Intracellul | F10 | Alizarin | 885 | BL1 | 5919 |
| 40 | No | A. baylyi | Tetracyclin | Intracellul | F84 | Neutral Re | 978 | BL1 | 5579 |
| 40 | No | A. baylyi | Tetracyclin | Intracellul | F99 | Pyronin Y | 774 | BL1 | 5629.5 |
| 40 | No | A. baylyi | Tetracyclin | Intracellul | F108 | Rhodamine | 922 | BL1 | 5292.5 |
| 40 | No | A. baylyi | Tetracyclin | Intracellul | F109 | Riboflavin | 829 | BL1 | 5149 |
| 40 | No | A. baylyi | Tetracyclin | Intracellul | F117 | SYBR Gree | 895 | BL1 | 336586 |
| 40 | No | A. baylyi | Tetracyclin | Intracellul | F106 | Rhodamine | 849 | BL1 | 7174 |
| 40 | No | A. baylyi | Tetracyclin | Intracellul | F73 | Malachite | 863 | BL1 | 5377 |
| 40 | No | A. baylyi | Tetracyclin | Intracellul | F7 | Acridine or | 793 | BL1 | 6111 |
| 40 | No | A. baylyi | Tetracyclin | Intracellul | F192 | 5-Carboxyf | 958 | BL1 | 18169 |
| 40 | No | A. baylyi | Tetracyclin | Intracellul | F57 | Eosin Y | 807 | BL1 | 17046 |
| 40 | No | A. baylyi | Tetracyclin | Intracellul | F27 | Calcein | 991 | BL1 | 17381 |

| | | | | | | | | | |
|---|---|---|---|---|---|---|---|---|---|
| 40 | No | A. baylyi | Tetracyclin | Intracellula | F12 | Amaranth | 749 | BL1 | 16803 |
| 40 | No | A. baylyi | None | None | NA | Autofluore: | 1167 | NA | 19605 |
| 40 | No | A. baylyi | Vancomyc | Wall | F194 | Oxonol | 1384 | RL1 | 17609.5 |
| 40 | No | A. baylyi | Vancomyc | Wall | F11 | Allura Red | 1140 | BL3 | 14220 |
| 40 | No | A. baylyi | Vancomyc | Wall | F54 | DiSC3(5) | 1043 | RL1 | 11711 |
| 40 | No | A. baylyi | Vancomyc | Wall | F124 | Thioflavin T | 1035 | VL2 | 8077 |
| 40 | No | A. baylyi | Vancomyc | Wall | F101 | Quinacrine | 1120 | VL2 | 6877.5 |
| 40 | No | A. baylyi | Vancomyc | Wall | F138 | Apigenin | 1166 | VL2 | 5800.5 |
| 40 | No | A. baylyi | Vancomyc | Wall | F65 | H2FDA | 1141 | BL1 | 7056 |
| 40 | No | A. baylyi | Vancomyc | Wall | F142 | Chlorophe | 1125 | BL3 | 8006 |
| 40 | No | A. baylyi | Vancomyc | Wall | F39 | Congo Red | 1205 | BL3 | 8719 |
| 40 | No | A. baylyi | Vancomyc | Wall | F116 | Sunset Yel | 1023 | BL2 | 6347 |
| 40 | No | A. baylyi | Vancomyc | Wall | F10 | Alizarin | 1073 | BL1 | 5845 |
| 40 | No | A. baylyi | Vancomyc | Wall | F84 | Neutral Re | 1086 | BL1 | 5667.5 |
| 40 | No | A. baylyi | Vancomyc | Wall | F99 | Pyronin Y | 1034 | BL1 | 5396 |
| 40 | No | A. baylyi | Vancomyc | Wall | F108 | Rhodamine | 1148 | BL1 | 4978 |
| 40 | No | A. baylyi | Vancomyc | Wall | F109 | Riboflavin | 1046 | BL1 | 5431 |
| 40 | No | A. baylyi | Vancomyc | Wall | F117 | SYBR Gree | 1055 | BL1 | 355169 |
| 40 | No | A. baylyi | Vancomyc | Wall | F106 | Rhodamine | 1052 | BL1 | 6728 |
| 40 | No | A. baylyi | Vancomyc | Wall | F73 | Malachite ( | 1097 | BL1 | 6007 |
| 40 | No | A. baylyi | Vancomyc | Wall | F7 | Acridine or | 1162 | BL1 | 6568 |
| 40 | No | A. baylyi | Vancomyc | Wall | F192 | 5-Carboxyf | 1099 | BL1 | 19833 |
| 40 | No | A. baylyi | Vancomyc | Wall | F57 | Eosin Y | 1011 | BL1 | 16342 |
| 40 | No | A. baylyi | Vancomyc | Wall | F27 | Calcein | 1096 | BL1 | 16826.5 |
| 40 | No | A. baylyi | Vancomyc | Wall | F12 | Amaranth | 1084 | BL1 | 19980.5 |
| 40 | No | A. baylyi | None | None | NA | Autofluore: | 2153 | NA | 1451 |
| 40 | No | A. baylyi | None | None | F194 | Oxonol | 2964 | RL1 | 232257.5 |
| 40 | No | A. baylyi | None | None | F11 | Allura Red | 1697 | BL3 | 48069 |
| 40 | No | A. baylyi | None | None | F54 | DiSC3(5) | 1593 | RL1 | 9333 |
| 40 | No | A. baylyi | None | None | F124 | Thioflavin T | 1450 | VL2 | 5244 |
| 40 | No | A. baylyi | None | None | F101 | Quinacrine | 1377 | VL2 | 4546 |
| 40 | No | A. baylyi | None | None | F138 | Apigenin | 1448 | VL2 | 4273 |
| 40 | No | A. baylyi | None | None | F65 | H2FDA | 1253 | BL1 | 4930 |
| 40 | No | A. baylyi | None | None | F142 | Chlorophe | 1286 | BL3 | 5713.5 |
| 40 | No | A. baylyi | None | None | F39 | Congo Red | 1249 | BL3 | 5567 |
| 40 | No | A. baylyi | None | None | F116 | Sunset Yel | 1244 | BL2 | 4796 |
| 40 | No | A. baylyi | None | None | F10 | Alizarin | 1153 | BL1 | 4114 |
| 40 | No | A. baylyi | None | None | F84 | Neutral Re | 1173 | BL1 | 3817 |
| 40 | No | A. baylyi | None | None | F99 | Pyronin Y | 1128 | BL1 | 3889 |
| 40 | No | A. baylyi | None | None | F108 | Rhodamine | 1188 | BL1 | 3289.5 |
| 40 | No | A. baylyi | None | None | F109 | Riboflavin | 1150 | BL1 | 3763 |
| 40 | No | A. baylyi | None | None | F117 | SYBR Gree | 1215 | BL1 | 378314 |
| 40 | No | A. baylyi | None | None | F106 | Rhodamine | 1138 | BL1 | 4591 |
| 40 | No | A. baylyi | None | None | F73 | Malachite ( | 1140 | BL1 | 4070 |
| 40 | No | A. baylyi | None | None | F7 | Acridine or | 1188 | BL1 | 4664 |
| 40 | No | A. baylyi | None | None | F192 | 5-Carboxyf | 1070 | BL1 | 17765 |
| 40 | No | A. baylyi | None | None | F57 | Eosin Y | 1140 | BL1 | 19635.5 |
| 40 | No | A. baylyi | None | None | F27 | Calcein | 1135 | BL1 | 18732 |

| | | | | | | | | | |
|---|---|---|---|---|---|---|---|---|---|
| 40 | No | A. baylyi | None | None | F12 | Amaranth | 1099 | BL1 | 19814 |
| 40 | No | A. baylyi | None | None | NA | Autofluore: | 1065 | NA | 18402 |
| 40 | No | A. baylyi | Ampicillin | Wall | F194 | Oxonol | 3912 | RL1 | 13372 |
| 40 | No | A. baylyi | Ampicillin | Wall | F11 | Allura Red | 963 | BL3 | 14614 |
| 40 | No | A. baylyi | Ampicillin | Wall | F54 | DiSC3(5) | 802 | RL1 | 12273 |
| 40 | No | A. baylyi | Ampicillin | Wall | F124 | Thioflavin T | 886 | VL2 | 7173.5 |
| 40 | No | A. baylyi | Ampicillin | Wall | F101 | Quinacrine | 800 | VL2 | 6163 |
| 40 | No | A. baylyi | Ampicillin | Wall | F138 | Apigenin | 816 | VL2 | 6018 |
| 40 | No | A. baylyi | Ampicillin | Wall | F65 | H2FDA | 767 | BL1 | 6519 |
| 40 | No | A. baylyi | Ampicillin | Wall | F142 | Chlorophe | 838 | BL3 | 7214 |
| 40 | No | A. baylyi | Ampicillin | Wall | F39 | Congo Red | 885 | BL3 | 7300 |
| 40 | No | A. baylyi | Ampicillin | Wall | F116 | Sunset Yel | 825 | BL2 | 5824 |
| 40 | No | A. baylyi | Ampicillin | Wall | F10 | Alizarin | 823 | BL1 | 5636 |
| 40 | No | A. baylyi | Ampicillin | Wall | F84 | Neutral Re | 890 | BL1 | 4874.5 |
| 40 | No | A. baylyi | Ampicillin | Wall | F99 | Pyronin Y | 740 | BL1 | 5198.5 |
| 40 | No | A. baylyi | Ampicillin | Wall | F108 | Rhodamine | 925 | BL1 | 4586 |
| 40 | No | A. baylyi | Ampicillin | Wall | F109 | Riboflavin | 812 | BL1 | 4781 |
| 40 | No | A. baylyi | Ampicillin | Wall | F117 | SYBR Gree | 848 | BL1 | 561938 |
| 40 | No | A. baylyi | Ampicillin | Wall | F106 | Rhodamine | 773 | BL1 | 6966 |
| 40 | No | A. baylyi | Ampicillin | Wall | F73 | Malachite ( | 832 | BL1 | 5212 |
| 40 | No | A. baylyi | Ampicillin | Wall | F7 | Acridine or | 781 | BL1 | 5868 |
| 40 | No | A. baylyi | Ampicillin | Wall | F192 | 5-Carboxyf | 799 | BL1 | 23391 |
| 40 | No | A. baylyi | Ampicillin | Wall | F57 | Eosin Y | 778 | BL1 | 23232 |
| 40 | No | A. baylyi | Ampicillin | Wall | F27 | Calcein | 853 | BL1 | 21745 |
| 40 | No | A. baylyi | Ampicillin | Wall | F12 | Amaranth | 761 | BL1 | 22440 |
| 40 | No | A. baylyi | None | None | NA | Autofluore: | 1024 | NA | 14842 |
| 40 | No | A. baylyi | Bacitracin | Wall | F194 | Oxonol | 2347 | RL1 | 20681 |
| 40 | No | A. baylyi | Bacitracin | Wall | F11 | Allura Red | 964 | BL3 | 12563 |
| 40 | No | A. baylyi | Bacitracin | Wall | F54 | DiSC3(5) | 920 | RL1 | 11995.5 |
| 40 | No | A. baylyi | Bacitracin | Wall | F124 | Thioflavin T | 1100 | VL2 | 7859.5 |
| 40 | No | A. baylyi | Bacitracin | Wall | F101 | Quinacrine | 1182 | VL2 | 8091 |
| 40 | No | A. baylyi | Bacitracin | Wall | F138 | Apigenin | 1178 | VL2 | 6227.5 |
| 40 | No | A. baylyi | Bacitracin | Wall | F65 | H2FDA | 980 | BL1 | 7005.5 |
| 40 | No | A. baylyi | Bacitracin | Wall | F142 | Chlorophe | 1118 | BL3 | 7631.5 |
| 40 | No | A. baylyi | Bacitracin | Wall | F39 | Congo Red | 1021 | BL3 | 6698 |
| 40 | No | A. baylyi | Bacitracin | Wall | F116 | Sunset Yel | 1135 | BL2 | 5990 |
| 40 | No | A. baylyi | Bacitracin | Wall | F10 | Alizarin | 1030 | BL1 | 5595 |
| 40 | No | A. baylyi | Bacitracin | Wall | F84 | Neutral Re | 1098 | BL1 | 4937 |
| 40 | No | A. baylyi | Bacitracin | Wall | F99 | Pyronin Y | 960 | BL1 | 4946.5 |
| 40 | No | A. baylyi | Bacitracin | Wall | F108 | Rhodamine | 1212 | BL1 | 4936 |
| 40 | No | A. baylyi | Bacitracin | Wall | F109 | Riboflavin | 1149 | BL1 | 4973 |
| 40 | No | A. baylyi | Bacitracin | Wall | F117 | SYBR Gree | 969 | BL1 | 614347 |
| 40 | No | A. baylyi | Bacitracin | Wall | F106 | Rhodamine | 1090 | BL1 | 6544.5 |
| 40 | No | A. baylyi | Bacitracin | Wall | F73 | Malachite ( | 1089 | BL1 | 5131 |
| 40 | No | A. baylyi | Bacitracin | Wall | F7 | Acridine or | 1056 | BL1 | 5077 |
| 40 | No | A. baylyi | Bacitracin | Wall | F192 | 5-Carboxyf | 1089 | BL1 | 21502 |
| 40 | No | A. baylyi | Bacitracin | Wall | F57 | Eosin Y | 1040 | BL1 | 19277 |
| 40 | No | A. baylyi | Bacitracin | Wall | F27 | Calcein | 1007 | BL1 | 21117 |

| | | | | | | | | | |
|---|---|---|---|---|---|---|---|---|---|
| 40 | No | A. baylyi | Bacitracin | Wall | F12 | Amaranth | 974 | BL1 | 22021.5 |
| 40 | No | A. baylyi | None | None | NA | Autofluores | 1068 | NA | 16952.5 |
| 40 | No | A. baylyi | Cefotaxime | Wall | F194 | Oxonol | 4974 | RL1 | 12745.5 |
| 40 | No | A. baylyi | Cefotaxime | Wall | F11 | Allura Red | 1189 | BL3 | 14260 |
| 40 | No | A. baylyi | Cefotaxime | Wall | F54 | DiSC3(5) | 832 | RL1 | 12190.5 |
| 40 | No | A. baylyi | Cefotaxime | Wall | F124 | Thioflavin T | 1001 | VL2 | 7771 |
| 40 | No | A. baylyi | Cefotaxime | Wall | F101 | Quinacrine | 943 | VL2 | 7114 |
| 40 | No | A. baylyi | Cefotaxime | Wall | F138 | Apigenin | 1033 | VL2 | 6356 |
| 40 | No | A. baylyi | Cefotaxime | Wall | F65 | H2FDA | 913 | BL1 | 7371 |
| 40 | No | A. baylyi | Cefotaxime | Wall | F142 | Chlorophe | 974 | BL3 | 7947 |
| 40 | No | A. baylyi | Cefotaxime | Wall | F39 | Congo Red | 965 | BL3 | 8400 |
| 40 | No | A. baylyi | Cefotaxime | Wall | F116 | Sunset Yell | 941 | BL2 | 6800 |
| 40 | No | A. baylyi | Cefotaxime | Wall | F10 | Alizarin | 885 | BL1 | 5441 |
| 40 | No | A. baylyi | Cefotaxime | Wall | F84 | Neutral Re | 1038 | BL1 | 5164 |
| 40 | No | A. baylyi | Cefotaxime | Wall | F99 | Pyronin Y | 784 | BL1 | 5446.5 |
| 40 | No | A. baylyi | Cefotaxime | Wall | F108 | Rhodamine | 977 | BL1 | 4769 |
| 40 | No | A. baylyi | Cefotaxime | Wall | F109 | Riboflavin | 944 | BL1 | 5488 |
| 40 | No | A. baylyi | Cefotaxime | Wall | F117 | SYBR Gree | 968 | BL1 | 489559.5 |
| 40 | No | A. baylyi | Cefotaxime | Wall | F106 | Rhodamine | 932 | BL1 | 7015.5 |
| 40 | No | A. baylyi | Cefotaxime | Wall | F73 | Malachite ( | 1015 | BL1 | 5650 |
| 40 | No | A. baylyi | Cefotaxime | Wall | F7 | Acridine or | 871 | BL1 | 5882 |
| 40 | No | A. baylyi | Cefotaxime | Wall | F192 | 5-Carboxyf | 899 | BL1 | 23114 |
| 40 | No | A. baylyi | Cefotaxime | Wall | F57 | Eosin Y | 868 | BL1 | 20880.5 |
| 40 | No | A. baylyi | Cefotaxime | Wall | F27 | Calcein | 950 | BL1 | 20603.5 |
| 40 | No | A. baylyi | Cefotaxime | Wall | F12 | Amaranth | 894 | BL1 | 21457.5 |
| 40 | No | A. baylyi | None | None | NA | Autofluores | 1103 | NA | 15886 |
| 40 | No | A. baylyi | Ciprofloxac | Intracellula | F194 | Oxonol | 2377 | RL1 | 13013 |
| 40 | No | A. baylyi | Ciprofloxac | Intracellula | F11 | Allura Red | 886 | BL3 | 15677 |
| 40 | No | A. baylyi | Ciprofloxac | Intracellula | F54 | DiSC3(5) | 800 | RL1 | 13771.5 |
| 40 | No | A. baylyi | Ciprofloxac | Intracellula | F124 | Thioflavin T | 828 | VL2 | 9839.5 |
| 40 | No | A. baylyi | Ciprofloxac | Intracellula | F101 | Quinacrine | 842 | VL2 | 7399 |
| 40 | No | A. baylyi | Ciprofloxac | Intracellula | F138 | Apigenin | 891 | VL2 | 6160 |
| 40 | No | A. baylyi | Ciprofloxac | Intracellula | F65 | H2FDA | 911 | BL1 | 7743 |
| 40 | No | A. baylyi | Ciprofloxac | Intracellula | F142 | Chlorophe | 911 | BL3 | 7937 |
| 40 | No | A. baylyi | Ciprofloxac | Intracellula | F39 | Congo Red | 844 | BL3 | 6721.5 |
| 40 | No | A. baylyi | Ciprofloxac | Intracellula | F116 | Sunset Yell | 894 | BL2 | 6919 |
| 40 | No | A. baylyi | Ciprofloxac | Intracellula | F10 | Alizarin | 856 | BL1 | 6302 |
| 40 | No | A. baylyi | Ciprofloxac | Intracellula | F84 | Neutral Re | 838 | BL1 | 5297.5 |
| 40 | No | A. baylyi | Ciprofloxac | Intracellula | F99 | Pyronin Y | 903 | BL1 | 6017 |
| 40 | No | A. baylyi | Ciprofloxac | Intracellula | F108 | Rhodamine | 882 | BL1 | 5151 |
| 40 | No | A. baylyi | Ciprofloxac | Intracellula | F109 | Riboflavin | 795 | BL1 | 5347 |
| 40 | No | A. baylyi | Ciprofloxac | Intracellula | F117 | SYBR Gree | 841 | BL1 | 367231 |
| 40 | No | A. baylyi | Ciprofloxac | Intracellula | F106 | Rhodamine | 877 | BL1 | 7343 |
| 40 | No | A. baylyi | Ciprofloxac | Intracellula | F73 | Malachite ( | 813 | BL1 | 5494 |
| 40 | No | A. baylyi | Ciprofloxac | Intracellula | F7 | Acridine or | 802 | BL1 | 5974 |
| 40 | No | A. baylyi | Ciprofloxac | Intracellula | F192 | 5-Carboxyf | 859 | BL1 | 23240 |
| 40 | No | A. baylyi | Ciprofloxac | Intracellula | F57 | Eosin Y | 777 | BL1 | 22275 |
| 40 | No | A. baylyi | Ciprofloxac | Intracellula | F27 | Calcein | 845 | BL1 | 22590 |

| | | | | | | | | | |
|---|---|---|---|---|---|---|---|---|---|
| 40 | No | A. baylyi | Ciprofloxa | Intracellula | F12 | Amaranth | 886 | BL1 | 23750.5 |
| 40 | No | A. baylyi | None | None | NA | Autofluore: | 1151 | NA | 19847 |
| 40 | No | A. baylyi | Fosfomyci | Wall | F194 | Oxonol | 3182 | RL1 | 12606.5 |
| 40 | No | A. baylyi | Fosfomyci | Wall | F11 | Allura Red | 1265 | BL3 | 13227 |
| 40 | No | A. baylyi | Fosfomyci | Wall | F54 | DiSC3(5) | 955 | RL1 | 13091 |
| 40 | No | A. baylyi | Fosfomyci | Wall | F124 | Thioflavin T | 1208 | VL2 | 8913.5 |
| 40 | No | A. baylyi | Fosfomyci | Wall | F101 | Quinacrine | 1109 | VL2 | 6164 |
| 40 | No | A. baylyi | Fosfomyci | Wall | F138 | Apigenin | 1128 | VL2 | 5628.5 |
| 40 | No | A. baylyi | Fosfomyci | Wall | F65 | H2FDA | 1060 | BL1 | 6864 |
| 40 | No | A. baylyi | Fosfomyci | Wall | F142 | Chlorophe | 1111 | BL3 | 7369 |
| 40 | No | A. baylyi | Fosfomyci | Wall | F39 | Congo Red | 1153 | BL3 | 7769 |
| 40 | No | A. baylyi | Fosfomyci | Wall | F116 | Sunset Yel | 1157 | BL2 | 6663 |
| 40 | No | A. baylyi | Fosfomyci | Wall | F10 | Alizarin | 1109 | BL1 | 5107 |
| 40 | No | A. baylyi | Fosfomyci | Wall | F84 | Neutral Re | 1134 | BL1 | 4876.5 |
| 40 | No | A. baylyi | Fosfomyci | Wall | F99 | Pyronin Y | 991 | BL1 | 5241 |
| 40 | No | A. baylyi | Fosfomyci | Wall | F108 | Rhodamine | 1221 | BL1 | 4803 |
| 40 | No | A. baylyi | Fosfomyci | Wall | F109 | Riboflavin | 1128 | BL1 | 4993.5 |
| 40 | No | A. baylyi | Fosfomyci | Wall | F117 | SYBR Gree | 1185 | BL1 | 427157 |
| 40 | No | A. baylyi | Fosfomyci | Wall | F106 | Rhodamine | 1090 | BL1 | 5769.5 |
| 40 | No | A. baylyi | Fosfomyci | Wall | F73 | Malachite ( | 1122 | BL1 | 5338 |
| 40 | No | A. baylyi | Fosfomyci | Wall | F7 | Acridine or | 1162 | BL1 | 5912.5 |
| 40 | No | A. baylyi | Fosfomyci | Wall | F192 | 5-Carboxyf | 1174 | BL1 | 18966 |
| 40 | No | A. baylyi | Fosfomyci | Wall | F57 | Eosin Y | 1046 | BL1 | 17451.5 |
| 40 | No | A. baylyi | Fosfomyci | Wall | F27 | Calcein | 1188 | BL1 | 21224.5 |
| 40 | No | A. baylyi | Fosfomyci | Wall | F12 | Amaranth | 1105 | BL1 | 19358 |
| 40 | No | A. baylyi | None | None | NA | Autofluore: | 1234 | NA | 19011.5 |
| 40 | No | A. baylyi | Nalidixic a | Intracellula | F194 | Oxonol | 2877 | RL1 | 14751 |
| 40 | No | A. baylyi | Nalidixic a | Intracellula | F11 | Allura Red | 1034 | BL3 | 13701 |
| 40 | No | A. baylyi | Nalidixic a | Intracellula | F54 | DiSC3(5) | 918 | RL1 | 13528.5 |
| 40 | No | A. baylyi | Nalidixic a | Intracellula | F124 | Thioflavin T | 1045 | VL2 | 9107 |
| 40 | No | A. baylyi | Nalidixic a | Intracellula | F101 | Quinacrine | 1095 | VL2 | 7360 |
| 40 | No | A. baylyi | Nalidixic a | Intracellula | F138 | Apigenin | 1007 | VL2 | 6173 |
| 40 | No | A. baylyi | Nalidixic a | Intracellula | F65 | H2FDA | 1078 | BL1 | 7387 |
| 40 | No | A. baylyi | Nalidixic a | Intracellula | F142 | Chlorophe | 1031 | BL3 | 7380 |
| 40 | No | A. baylyi | Nalidixic a | Intracellula | F39 | Congo Red | 955 | BL3 | 6993 |
| 40 | No | A. baylyi | Nalidixic a | Intracellula | F116 | Sunset Yel | 1063 | BL2 | 6440 |
| 40 | No | A. baylyi | Nalidixic a | Intracellula | F10 | Alizarin | 986 | BL1 | 5439 |
| 40 | No | A. baylyi | Nalidixic a | Intracellula | F84 | Neutral Re | 1034 | BL1 | 5168 |
| 40 | No | A. baylyi | Nalidixic a | Intracellula | F99 | Pyronin Y | 926 | BL1 | 5354.5 |
| 40 | No | A. baylyi | Nalidixic a | Intracellula | F108 | Rhodamine | 1065 | BL1 | 5258 |
| 40 | No | A. baylyi | Nalidixic a | Intracellula | F109 | Riboflavin | 910 | BL1 | 4685 |
| 40 | No | A. baylyi | Nalidixic a | Intracellula | F117 | SYBR Gree | 992 | BL1 | 393006.5 |
| 40 | No | A. baylyi | Nalidixic a | Intracellula | F106 | Rhodamine | 1019 | BL1 | 6961 |
| 40 | No | A. baylyi | Nalidixic a | Intracellula | F73 | Malachite ( | 932 | BL1 | 5011.5 |
| 40 | No | A. baylyi | Nalidixic a | Intracellula | F7 | Acridine or | 1071 | BL1 | 6301 |
| 40 | No | A. baylyi | Nalidixic a | Intracellula | F192 | 5-Carboxyf | 1020 | BL1 | 20797.5 |
| 40 | No | A. baylyi | Nalidixic a | Intracellula | F57 | Eosin Y | 953 | BL1 | 21239 |
| 40 | No | A. baylyi | Nalidixic a | Intracellula | F27 | Calcein | 1101 | BL1 | 20180 |

| | | | | | | | | | |
|---|---|---|---|---|---|---|---|---|---|
| 40 | No | A. baylyi | Nalidixic a | Intracellula | F12 | Amaranth | 987 | BL1 | 21507 |
| 40 | No | A. baylyi | None | None | NA | Autofluore: | 1151 | NA | 18935 |
| 40 | No | A. baylyi | Ofloxacin | Intracellula | F194 | Oxonol | 3337 | RL1 | 12829 |
| 40 | No | A. baylyi | Ofloxacin | Intracellula | F11 | Allura Red | 1026 | BL3 | 16137 |
| 40 | No | A. baylyi | Ofloxacin | Intracellula | F54 | DiSC3(5) | 838 | RL1 | 14192.5 |
| 40 | No | A. baylyi | Ofloxacin | Intracellula | F124 | Thioflavin T | 928 | VL2 | 8679 |
| 40 | No | A. baylyi | Ofloxacin | Intracellula | F101 | Quinacrine | 898 | VL2 | 7017 |
| 40 | No | A. baylyi | Ofloxacin | Intracellula | F138 | Apigenin | 953 | VL2 | 6559 |
| 40 | No | A. baylyi | Ofloxacin | Intracellula | F65 | H2FDA | 882 | BL1 | 8005 |
| 40 | No | A. baylyi | Ofloxacin | Intracellula | F142 | Chlorophe | 840 | BL3 | 7828 |
| 40 | No | A. baylyi | Ofloxacin | Intracellula | F39 | Congo Red | 869 | BL3 | 8697 |
| 40 | No | A. baylyi | Ofloxacin | Intracellula | F116 | Sunset Yel | 980 | BL2 | 7340.5 |
| 40 | No | A. baylyi | Ofloxacin | Intracellula | F10 | Alizarin | 851 | BL1 | 6138 |
| 40 | No | A. baylyi | Ofloxacin | Intracellula | F84 | Neutral Re | 850 | BL1 | 5654 |
| 40 | No | A. baylyi | Ofloxacin | Intracellula | F99 | Pyronin Y | 790 | BL1 | 6242 |
| 40 | No | A. baylyi | Ofloxacin | Intracellula | F108 | Rhodamine | 927 | BL1 | 4780 |
| 40 | No | A. baylyi | Ofloxacin | Intracellula | F109 | Riboflavin | 789 | BL1 | 5365 |
| 40 | No | A. baylyi | Ofloxacin | Intracellula | F117 | SYBR Gree | 957 | BL1 | 474373 |
| 40 | No | A. baylyi | Ofloxacin | Intracellula | F106 | Rhodamine | 872 | BL1 | 7237 |
| 40 | No | A. baylyi | Ofloxacin | Intracellula | F73 | Malachite ( | 853 | BL1 | 5696 |
| 40 | No | A. baylyi | Ofloxacin | Intracellula | F7 | Acridine or | 830 | BL1 | 7033 |
| 40 | No | A. baylyi | Ofloxacin | Intracellula | F192 | 5-Carboxyf | 796 | BL1 | 24067 |
| 40 | No | A. baylyi | Ofloxacin | Intracellula | F57 | Eosin Y | 826 | BL1 | 22642 |
| 40 | No | A. baylyi | Ofloxacin | Intracellula | F27 | Calcein | 928 | BL1 | 23231.5 |
| 40 | No | A. baylyi | Ofloxacin | Intracellula | F12 | Amaranth | 839 | BL1 | 22018 |
| 40 | No | A. baylyi | None | None | NA | Autofluore: | 1267 | NA | 19121 |
| 40 | No | A. baylyi | Polymyxin | Wall | F194 | Oxonol | 2981 | RL1 | 13305 |
| 40 | No | A. baylyi | Polymyxin | Wall | F11 | Allura Red | 1306 | BL3 | 12536.5 |
| 40 | No | A. baylyi | Polymyxin | Wall | F54 | DiSC3(5) | 1034 | RL1 | 13358.5 |
| 40 | No | A. baylyi | Polymyxin | Wall | F124 | Thioflavin T | 1206 | VL2 | 8436.5 |
| 40 | No | A. baylyi | Polymyxin | Wall | F101 | Quinacrine | 1197 | VL2 | 6569 |
| 40 | No | A. baylyi | Polymyxin | Wall | F138 | Apigenin | 1249 | VL2 | 6118 |
| 40 | No | A. baylyi | Polymyxin | Wall | F65 | H2FDA | 1269 | BL1 | 6721 |
| 40 | No | A. baylyi | Polymyxin | Wall | F142 | Chlorophe | 1253 | BL3 | 7790 |
| 40 | No | A. baylyi | Polymyxin | Wall | F39 | Congo Red | 1266 | BL3 | 7438.5 |
| 40 | No | A. baylyi | Polymyxin | Wall | F116 | Sunset Yel | 1154 | BL2 | 6334.5 |
| 40 | No | A. baylyi | Polymyxin | Wall | F10 | Alizarin | 1171 | BL1 | 6067 |
| 40 | No | A. baylyi | Polymyxin | Wall | F84 | Neutral Re | 1264 | BL1 | 5475 |
| 40 | No | A. baylyi | Polymyxin | Wall | F99 | Pyronin Y | 1134 | BL1 | 4912 |
| 40 | No | A. baylyi | Polymyxin | Wall | F108 | Rhodamine | 1215 | BL1 | 4954 |
| 40 | No | A. baylyi | Polymyxin | Wall | F109 | Riboflavin | 1232 | BL1 | 5379 |
| 40 | No | A. baylyi | Polymyxin | Wall | F117 | SYBR Gree | 1019 | BL1 | 735454 |
| 40 | No | A. baylyi | Polymyxin | Wall | F106 | Rhodamine | 1266 | BL1 | 6919.5 |
| 40 | No | A. baylyi | Polymyxin | Wall | F73 | Malachite ( | 1139 | BL1 | 5695 |
| 40 | No | A. baylyi | Polymyxin | Wall | F7 | Acridine or | 1111 | BL1 | 5502 |
| 40 | No | A. baylyi | Polymyxin | Wall | F192 | 5-Carboxyf | 1167 | BL1 | 19318 |
| 40 | No | A. baylyi | Polymyxin | Wall | F57 | Eosin Y | 1220 | BL1 | 20084 |
| 40 | No | A. baylyi | Polymyxin | Wall | F27 | Calcein | 1153 | BL1 | 17460 |

| | | | | | | | | | |
|---|---|---|---|---|---|---|---|---|---|
| 40 | No | A. baylyi | Polymyxin | Wall | F12 | Amaranth | 1110 | BL1 | 17712.5 |
| 40 | No | A. baylyi | None | None | NA | Autofluore: | 1163 | NA | 17224 |
| 40 | No | A. baylyi | Tetracyclin | Intracellul | F194 | Oxonol | 3470 | RL1 | 12411 |
| 40 | No | A. baylyi | Tetracyclin | Intracellul | F11 | Allura Red | 1074 | BL3 | 13854.5 |
| 40 | No | A. baylyi | Tetracyclin | Intracellul | F54 | DiSC3(5) | 734 | RL1 | 11596 |
| 40 | No | A. baylyi | Tetracyclin | Intracellul | F124 | Thioflavin T | 818 | VL2 | 8568.5 |
| 40 | No | A. baylyi | Tetracyclin | Intracellul | F101 | Quinacrine | 878 | VL2 | 7434.5 |
| 40 | No | A. baylyi | Tetracyclin | Intracellul | F138 | Apigenin | 815 | VL2 | 6207 |
| 40 | No | A. baylyi | Tetracyclin | Intracellul | F65 | H2FDA | 750 | BL1 | 6977.5 |
| 40 | No | A. baylyi | Tetracyclin | Intracellul | F142 | Chlorophe | 946 | BL3 | 8427.5 |
| 40 | No | A. baylyi | Tetracyclin | Intracellul | F39 | Congo Red | 832 | BL3 | 8226.5 |
| 40 | No | A. baylyi | Tetracyclin | Intracellul | F116 | Sunset Yel | 740 | BL2 | 6611 |
| 40 | No | A. baylyi | Tetracyclin | Intracellul | F10 | Alizarin | 816 | BL1 | 6273.5 |
| 40 | No | A. baylyi | Tetracyclin | Intracellul | F84 | Neutral Re | 931 | BL1 | 5769 |
| 40 | No | A. baylyi | Tetracyclin | Intracellul | F99 | Pyronin Y | 666 | BL1 | 5945.5 |
| 40 | No | A. baylyi | Tetracyclin | Intracellul | F108 | Rhodamine | 798 | BL1 | 5048 |
| 40 | No | A. baylyi | Tetracyclin | Intracellul | F109 | Riboflavin | 808 | BL1 | 5846.5 |
| 40 | No | A. baylyi | Tetracyclin | Intracellul | F117 | SYBR Gree | 870 | BL1 | 442955.5 |
| 40 | No | A. baylyi | Tetracyclin | Intracellul | F106 | Rhodamine | 833 | BL1 | 8757 |
| 40 | No | A. baylyi | Tetracyclin | Intracellul | F73 | Malachite ( | 872 | BL1 | 5805 |
| 40 | No | A. baylyi | Tetracyclin | Intracellul | F7 | Acridine or | 761 | BL1 | 6565 |
| 40 | No | A. baylyi | Tetracyclin | Intracellul | F192 | 5-Carboxyf | 870 | BL1 | 18557.5 |
| 40 | No | A. baylyi | Tetracyclin | Intracellul | F57 | Eosin Y | 772 | BL1 | 18269 |
| 40 | No | A. baylyi | Tetracyclin | Intracellul | F27 | Calcein | 836 | BL1 | 17933 |
| 40 | No | A. baylyi | Tetracyclin | Intracellul | F12 | Amaranth | 725 | BL1 | 19222 |
| 40 | No | A. baylyi | None | None | NA | Autofluore: | 1180 | NA | 20919 |
| 40 | No | A. baylyi | Vancomyc | Wall | F194 | Oxonol | 1511 | RL1 | 19599 |
| 40 | No | A. baylyi | Vancomyc | Wall | F11 | Allura Red | 1132 | BL3 | 13311.5 |
| 40 | No | A. baylyi | Vancomyc | Wall | F54 | DiSC3(5) | 894 | RL1 | 11532.5 |
| 40 | No | A. baylyi | Vancomyc | Wall | F124 | Thioflavin T | 1181 | VL2 | 7871 |
| 40 | No | A. baylyi | Vancomyc | Wall | F101 | Quinacrine | 1129 | VL2 | 6681 |
| 40 | No | A. baylyi | Vancomyc | Wall | F138 | Apigenin | 1112 | VL2 | 5908 |
| 40 | No | A. baylyi | Vancomyc | Wall | F65 | H2FDA | 1053 | BL1 | 7443 |
| 40 | No | A. baylyi | Vancomyc | Wall | F142 | Chlorophe | 1166 | BL3 | 7947.5 |
| 40 | No | A. baylyi | Vancomyc | Wall | F39 | Congo Red | 1023 | BL3 | 7990 |
| 40 | No | A. baylyi | Vancomyc | Wall | F116 | Sunset Yel | 1048 | BL2 | 6894 |
| 40 | No | A. baylyi | Vancomyc | Wall | F10 | Alizarin | 1168 | BL1 | 6380 |
| 40 | No | A. baylyi | Vancomyc | Wall | F84 | Neutral Re | 1049 | BL1 | 5678 |
| 40 | No | A. baylyi | Vancomyc | Wall | F99 | Pyronin Y | 964 | BL1 | 5530.5 |
| 40 | No | A. baylyi | Vancomyc | Wall | F108 | Rhodamine | 1109 | BL1 | 5072 |
| 40 | No | A. baylyi | Vancomyc | Wall | F109 | Riboflavin | 1102 | BL1 | 5324 |
| 40 | No | A. baylyi | Vancomyc | Wall | F117 | SYBR Gree | 984 | BL1 | 466189.5 |
| 40 | No | A. baylyi | Vancomyc | Wall | F106 | Rhodamine | 1021 | BL1 | 7496 |
| 40 | No | A. baylyi | Vancomyc | Wall | F73 | Malachite ( | 1003 | BL1 | 5806 |
| 40 | No | A. baylyi | Vancomyc | Wall | F7 | Acridine or | 1084 | BL1 | 6296 |
| 40 | No | A. baylyi | Vancomyc | Wall | F192 | 5-Carboxyf | 1082 | BL1 | 20722.5 |
| 40 | No | A. baylyi | Vancomyc | Wall | F57 | Eosin Y | 1051 | BL1 | 15949 |
| 40 | No | A. baylyi | Vancomyc | Wall | F27 | Calcein | 1067 | BL1 | 20417 |

| | | | | | | | | | |
|---|---|---|---|---|---|---|---|---|---|
| 40 | No | A. baylyi | Vancomyc | Wall | F12 | Amaranth | 1090 | BL1 | 21083 |
| 40 | No | A. baylyi | None | None | NA | Autofluore: | 2632 | NA | 1481.5 |
| 40 | No | A. baylyi | None | None | F194 | Oxonol | 2264 | RL1 | 362466.5 |
| 40 | No | A. baylyi | None | None | F11 | Allura Red | 1701 | BL3 | 51598 |
| 40 | No | A. baylyi | None | None | F54 | DiSC3(5) | 1359 | RL1 | 6954 |
| 40 | No | A. baylyi | None | None | F124 | Thioflavin T | 1309 | VL2 | 5833 |
| 40 | No | A. baylyi | None | None | F101 | Quinacrine | 1458 | VL2 | 5690.5 |
| 40 | No | A. baylyi | None | None | F138 | Apigenin | 1203 | VL2 | 3957 |
| 40 | No | A. baylyi | None | None | F65 | H2FDA | 1244 | BL1 | 5487.5 |
| 40 | No | A. baylyi | None | None | F142 | Chlorophe | 1190 | BL3 | 6232.5 |
| 40 | No | A. baylyi | None | None | F39 | Congo Red | 1035 | BL3 | 5743 |
| 40 | No | A. baylyi | None | None | F116 | Sunset Yel | 1009 | BL2 | 4750 |
| 40 | No | A. baylyi | None | None | F10 | Alizarin | 1094 | BL1 | 4051 |
| 40 | No | A. baylyi | None | None | F84 | Neutral Re | 1042 | BL1 | 3610 |
| 40 | No | A. baylyi | None | None | F99 | Pyronin Y | 999 | BL1 | 3867 |
| 40 | No | A. baylyi | None | None | F108 | Rhodamine | 1036 | BL1 | 3419 |
| 40 | No | A. baylyi | None | None | F109 | Riboflavin | 1141 | BL1 | 3880 |
| 40 | No | A. baylyi | None | None | F117 | SYBR Gree | 1113 | BL1 | 329149 |
| 40 | No | A. baylyi | None | None | F106 | Rhodamine | 1033 | BL1 | 4778 |
| 40 | No | A. baylyi | None | None | F73 | Malachite ( | 1042 | BL1 | 4014.5 |
| 40 | No | A. baylyi | None | None | F7 | Acridine or | 1052 | BL1 | 4751.5 |
| 40 | No | A. baylyi | None | None | F192 | 5-Carboxyf | 1037 | BL1 | 18442 |
| 40 | No | A. baylyi | None | None | F57 | Eosin Y | 979 | BL1 | 18229 |
| 40 | No | A. baylyi | None | None | F27 | Calcein | 984 | BL1 | 18294.5 |
| 40 | No | A. baylyi | None | None | F12 | Amaranth | 976 | BL1 | 17985.5 |
| 40 | No | A. baylyi | None | None | NA | Autofluore: | 1014 | NA | 17862 |
| 40 | No | A. baylyi | Ampicillin | Wall | F194 | Oxonol | 3286 | RL1 | 13638.5 |
| 40 | No | A. baylyi | Ampicillin | Wall | F11 | Allura Red | 877 | BL3 | 15597 |
| 40 | No | A. baylyi | Ampicillin | Wall | F54 | DiSC3(5) | 771 | RL1 | 15000 |
| 40 | No | A. baylyi | Ampicillin | Wall | F124 | Thioflavin T | 749 | VL2 | 8333 |
| 40 | No | A. baylyi | Ampicillin | Wall | F101 | Quinacrine | 730 | VL2 | 6252.5 |
| 40 | No | A. baylyi | Ampicillin | Wall | F138 | Apigenin | 894 | VL2 | 5859 |
| 40 | No | A. baylyi | Ampicillin | Wall | F65 | H2FDA | 784 | BL1 | 7027.5 |
| 40 | No | A. baylyi | Ampicillin | Wall | F142 | Chlorophe | 740 | BL3 | 7375.5 |
| 40 | No | A. baylyi | Ampicillin | Wall | F39 | Congo Red | 872 | BL3 | 7596 |
| 40 | No | A. baylyi | Ampicillin | Wall | F116 | Sunset Yel | 836 | BL2 | 6519 |
| 40 | No | A. baylyi | Ampicillin | Wall | F10 | Alizarin | 787 | BL1 | 5309 |
| 40 | No | A. baylyi | Ampicillin | Wall | F84 | Neutral Re | 808 | BL1 | 4772 |
| 40 | No | A. baylyi | Ampicillin | Wall | F99 | Pyronin Y | 776 | BL1 | 5479.5 |
| 40 | No | A. baylyi | Ampicillin | Wall | F108 | Rhodamine | 832 | BL1 | 4436 |
| 40 | No | A. baylyi | Ampicillin | Wall | F109 | Riboflavin | 738 | BL1 | 4971 |
| 40 | No | A. baylyi | Ampicillin | Wall | F117 | SYBR Gree | 831 | BL1 | 482882 |
| 40 | No | A. baylyi | Ampicillin | Wall | F106 | Rhodamine | 717 | BL1 | 6289 |
| 40 | No | A. baylyi | Ampicillin | Wall | F73 | Malachite ( | 816 | BL1 | 5357 |
| 40 | No | A. baylyi | Ampicillin | Wall | F7 | Acridine or | 816 | BL1 | 6138 |
| 40 | No | A. baylyi | Ampicillin | Wall | F192 | 5-Carboxyf | 729 | BL1 | 21872 |
| 40 | No | A. baylyi | Ampicillin | Wall | F57 | Eosin Y | 736 | BL1 | 17199.5 |
| 40 | No | A. baylyi | Ampicillin | Wall | F27 | Calcein | 823 | BL1 | 20179 |

| | | | | | | | | | |
|---|---|---|---|---|---|---|---|---|---|
| 40 | No | A. baylyi | Ampicillin | Wall | F12 | Amaranth | 660 | BL1 | 21669 |
| 40 | No | A. baylyi | None | None | NA | Autofluore: | 994 | NA | 16697 |
| 40 | No | A. baylyi | Bacitracin | Wall | F194 | Oxonol | 1456 | RL1 | 19440 |
| 40 | No | A. baylyi | Bacitracin | Wall | F11 | Allura Red | 948 | BL3 | 12593.5 |
| 40 | No | A. baylyi | Bacitracin | Wall | F54 | DiSC3(5) | 826 | RL1 | 14100 |
| 40 | No | A. baylyi | Bacitracin | Wall | F124 | Thioflavin T | 1099 | VL2 | 11311 |
| 40 | No | A. baylyi | Bacitracin | Wall | F101 | Quinacrine | 1078 | VL2 | 7542.5 |
| 40 | No | A. baylyi | Bacitracin | Wall | F138 | Apigenin | 942 | VL2 | 6009.5 |
| 40 | No | A. baylyi | Bacitracin | Wall | F65 | H2FDA | 1039 | BL1 | 7298 |
| 40 | No | A. baylyi | Bacitracin | Wall | F142 | Chlorophe | 1031 | BL3 | 7519 |
| 40 | No | A. baylyi | Bacitracin | Wall | F39 | Congo Red | 960 | BL3 | 7011.5 |
| 40 | No | A. baylyi | Bacitracin | Wall | F116 | Sunset Yel | 1024 | BL2 | 6588 |
| 40 | No | A. baylyi | Bacitracin | Wall | F10 | Alizarin | 1025 | BL1 | 5628 |
| 40 | No | A. baylyi | Bacitracin | Wall | F84 | Neutral Re | 951 | BL1 | 5115 |
| 40 | No | A. baylyi | Bacitracin | Wall | F99 | Pyronin Y | 891 | BL1 | 5280 |
| 40 | No | A. baylyi | Bacitracin | Wall | F108 | Rhodamine | 1000 | BL1 | 4517.5 |
| 40 | No | A. baylyi | Bacitracin | Wall | F109 | Riboflavin | 985 | BL1 | 4712 |
| 40 | No | A. baylyi | Bacitracin | Wall | F117 | SYBR Gree | 901 | BL1 | 539503 |
| 40 | No | A. baylyi | Bacitracin | Wall | F106 | Rhodamine | 1032 | BL1 | 6938 |
| 40 | No | A. baylyi | Bacitracin | Wall | F73 | Malachite ( | 909 | BL1 | 5438 |
| 40 | No | A. baylyi | Bacitracin | Wall | F7 | Acridine or | 880 | BL1 | 5752 |
| 40 | No | A. baylyi | Bacitracin | Wall | F192 | 5-Carboxyf | 900 | BL1 | 20650 |
| 40 | No | A. baylyi | Bacitracin | Wall | F57 | Eosin Y | 1028 | BL1 | 18143.5 |
| 40 | No | A. baylyi | Bacitracin | Wall | F27 | Calcein | 984 | BL1 | 19324.5 |
| 40 | No | A. baylyi | Bacitracin | Wall | F12 | Amaranth | 858 | BL1 | 19814.5 |
| 40 | No | A. baylyi | None | None | NA | Autofluore: | 984 | NA | 19610 |
| 40 | No | A. baylyi | Cefotaxim | Wall | F194 | Oxonol | 3655 | RL1 | 12370 |
| 40 | No | A. baylyi | Cefotaxim | Wall | F11 | Allura Red | 1055 | BL3 | 14654 |
| 40 | No | A. baylyi | Cefotaxim | Wall | F54 | DiSC3(5) | 786 | RL1 | 14776.5 |
| 40 | No | A. baylyi | Cefotaxim | Wall | F124 | Thioflavin T | 883 | VL2 | 8344 |
| 40 | No | A. baylyi | Cefotaxim | Wall | F101 | Quinacrine | 887 | VL2 | 7161 |
| 40 | No | A. baylyi | Cefotaxim | Wall | F138 | Apigenin | 963 | VL2 | 6356 |
| 40 | No | A. baylyi | Cefotaxim | Wall | F65 | H2FDA | 990 | BL1 | 7496 |
| 40 | No | A. baylyi | Cefotaxim | Wall | F142 | Chlorophe | 913 | BL3 | 7451 |
| 40 | No | A. baylyi | Cefotaxim | Wall | F39 | Congo Red | 835 | BL3 | 7932 |
| 40 | No | A. baylyi | Cefotaxim | Wall | F116 | Sunset Yel | 915 | BL2 | 7147 |
| 40 | No | A. baylyi | Cefotaxim | Wall | F10 | Alizarin | 867 | BL1 | 5636 |
| 40 | No | A. baylyi | Cefotaxim | Wall | F84 | Neutral Re | 1041 | BL1 | 6012 |
| 40 | No | A. baylyi | Cefotaxim | Wall | F99 | Pyronin Y | 778 | BL1 | 5441 |
| 40 | No | A. baylyi | Cefotaxim | Wall | F108 | Rhodamine | 831 | BL1 | 4533 |
| 40 | No | A. baylyi | Cefotaxim | Wall | F109 | Riboflavin | 989 | BL1 | 5931 |
| 40 | No | A. baylyi | Cefotaxim | Wall | F117 | SYBR Gree | 931 | BL1 | 481502 |
| 40 | No | A. baylyi | Cefotaxim | Wall | F106 | Rhodamine | 890 | BL1 | 6839.5 |
| 40 | No | A. baylyi | Cefotaxim | Wall | F73 | Malachite ( | 897 | BL1 | 5745 |
| 40 | No | A. baylyi | Cefotaxim | Wall | F7 | Acridine or | 880 | BL1 | 6240 |
| 40 | No | A. baylyi | Cefotaxim | Wall | F192 | 5-Carboxyf | 926 | BL1 | 21326 |
| 40 | No | A. baylyi | Cefotaxim | Wall | F57 | Eosin Y | 868 | BL1 | 21694 |
| 40 | No | A. baylyi | Cefotaxim | Wall | F27 | Calcein | 930 | BL1 | 21385 |

| | | | | | | | | | |
|---|---|---|---|---|---|---|---|---|---|
| 40 | No | A. baylyi | Cefotaxim | Wall | F12 | Amaranth | 836 | BL1 | 21631.5 |
| 40 | No | A. baylyi | None | None | NA | Autofluore: | 1081 | NA | 18587 |
| 40 | No | A. baylyi | Ciprofloxa | Intracellul | F194 | Oxonol | 1365 | RL1 | 17397 |
| 40 | No | A. baylyi | Ciprofloxa | Intracellul | F11 | Allura Red | 881 | BL3 | 16089 |
| 40 | No | A. baylyi | Ciprofloxa | Intracellul | F54 | DiSC3(5) | 821 | RL1 | 14588 |
| 40 | No | A. baylyi | Ciprofloxa | Intracellul | F124 | Thioflavin T | 795 | VL2 | 8857 |
| 40 | No | A. baylyi | Ciprofloxa | Intracellul | F101 | Quinacrine | 792 | VL2 | 7106.5 |
| 40 | No | A. baylyi | Ciprofloxa | Intracellul | F138 | Apigenin | 894 | VL2 | 6942 |
| 40 | No | A. baylyi | Ciprofloxa | Intracellul | F65 | H2FDA | 836 | BL1 | 7494.5 |
| 40 | No | A. baylyi | Ciprofloxa | Intracellul | F142 | Chlorophe | 820 | BL3 | 7938.5 |
| 40 | No | A. baylyi | Ciprofloxa | Intracellul | F39 | Congo Red | 797 | BL3 | 7928 |
| 40 | No | A. baylyi | Ciprofloxa | Intracellul | F116 | Sunset Yel | 855 | BL2 | 6961 |
| 40 | No | A. baylyi | Ciprofloxa | Intracellul | F10 | Alizarin | 797 | BL1 | 5561 |
| 40 | No | A. baylyi | Ciprofloxa | Intracellul | F84 | Neutral Re | 801 | BL1 | 5481 |
| 40 | No | A. baylyi | Ciprofloxa | Intracellul | F99 | Pyronin Y | 819 | BL1 | 5683 |
| 40 | No | A. baylyi | Ciprofloxa | Intracellul | F108 | Rhodamine | 856 | BL1 | 4994 |
| 40 | No | A. baylyi | Ciprofloxa | Intracellul | F109 | Riboflavin | 793 | BL1 | 5065 |
| 40 | No | A. baylyi | Ciprofloxa | Intracellul | F117 | SYBR Gree | 794 | BL1 | 347757.5 |
| 40 | No | A. baylyi | Ciprofloxa | Intracellul | F106 | Rhodamine | 850 | BL1 | 7927.5 |
| 40 | No | A. baylyi | Ciprofloxa | Intracellul | F73 | Malachite ( | 794 | BL1 | 5189 |
| 40 | No | A. baylyi | Ciprofloxa | Intracellul | F7 | Acridine or | 811 | BL1 | 6912 |
| 40 | No | A. baylyi | Ciprofloxa | Intracellul | F192 | 5-Carboxyf | 838 | BL1 | 21535 |
| 40 | No | A. baylyi | Ciprofloxa | Intracellul | F57 | Eosin Y | 770 | BL1 | 22717.5 |
| 40 | No | A. baylyi | Ciprofloxa | Intracellul | F27 | Calcein | 873 | BL1 | 21487 |
| 40 | No | A. baylyi | Ciprofloxa | Intracellul | F12 | Amaranth | 822 | BL1 | 22625 |
| 40 | No | A. baylyi | None | None | NA | Autofluore: | 1006 | NA | 18234.5 |
| 40 | No | A. baylyi | Fosfomyci | Wall | F194 | Oxonol | 2586 | RL1 | 14970.5 |
| 40 | No | A. baylyi | Fosfomyci | Wall | F11 | Allura Red | 1036 | BL3 | 14015 |
| 40 | No | A. baylyi | Fosfomyci | Wall | F54 | DiSC3(5) | 885 | RL1 | 13177 |
| 40 | No | A. baylyi | Fosfomyci | Wall | F124 | Thioflavin T | 1047 | VL2 | 8763 |
| 40 | No | A. baylyi | Fosfomyci | Wall | F101 | Quinacrine | 1009 | VL2 | 6111 |
| 40 | No | A. baylyi | Fosfomyci | Wall | F138 | Apigenin | 1114 | VL2 | 6257 |
| 40 | No | A. baylyi | Fosfomyci | Wall | F65 | H2FDA | 1034 | BL1 | 7123.5 |
| 40 | No | A. baylyi | Fosfomyci | Wall | F142 | Chlorophe | 986 | BL3 | 6991.5 |
| 40 | No | A. baylyi | Fosfomyci | Wall | F39 | Congo Red | 1022 | BL3 | 7819.5 |
| 40 | No | A. baylyi | Fosfomyci | Wall | F116 | Sunset Yel | 1038 | BL2 | 6619 |
| 40 | No | A. baylyi | Fosfomyci | Wall | F10 | Alizarin | 977 | BL1 | 5081 |
| 40 | No | A. baylyi | Fosfomyci | Wall | F84 | Neutral Re | 1067 | BL1 | 4910 |
| 40 | No | A. baylyi | Fosfomyci | Wall | F99 | Pyronin Y | 906 | BL1 | 5293 |
| 40 | No | A. baylyi | Fosfomyci | Wall | F108 | Rhodamine | 1083 | BL1 | 4651 |
| 40 | No | A. baylyi | Fosfomyci | Wall | F109 | Riboflavin | 930 | BL1 | 4963.5 |
| 40 | No | A. baylyi | Fosfomyci | Wall | F117 | SYBR Gree | 1044 | BL1 | 393851 |
| 40 | No | A. baylyi | Fosfomyci | Wall | F106 | Rhodamine | 1029 | BL1 | 6577 |
| 40 | No | A. baylyi | Fosfomyci | Wall | F73 | Malachite ( | 966 | BL1 | 5073.5 |
| 40 | No | A. baylyi | Fosfomyci | Wall | F7 | Acridine or | 1082 | BL1 | 6034.5 |
| 40 | No | A. baylyi | Fosfomyci | Wall | F192 | 5-Carboxyf | 964 | BL1 | 19869.5 |
| 40 | No | A. baylyi | Fosfomyci | Wall | F57 | Eosin Y | 947 | BL1 | 21205 |
| 40 | No | A. baylyi | Fosfomyci | Wall | F27 | Calcein | 1085 | BL1 | 18650 |

| | | | | | | | | | |
|---|---|---|---|---|---|---|---|---|---|
| 40 | No | A. baylyi | Fosfomyci | Wall | F12 | Amaranth | 1015 | BL1 | 19876 |
| 40 | No | A. baylyi | None | None | NA | Autofluore: | 1071 | NA | 18077 |
| 40 | No | A. baylyi | Nalidixic a | Intracellula | F194 | Oxonol | 1708 | RL1 | 15875 |
| 40 | No | A. baylyi | Nalidixic a | Intracellula | F11 | Allura Red | 998 | BL3 | 15037 |
| 40 | No | A. baylyi | Nalidixic a | Intracellula | F54 | DiSC3(5) | 903 | RL1 | 12584 |
| 40 | No | A. baylyi | Nalidixic a | Intracellula | F124 | Thioflavin T | 1009 | VL2 | 8241 |
| 40 | No | A. baylyi | Nalidixic a | Intracellula | F101 | Quinacrine | 937 | VL2 | 6595 |
| 40 | No | A. baylyi | Nalidixic a | Intracellula | F138 | Apigenin | 927 | VL2 | 6422 |
| 40 | No | A. baylyi | Nalidixic a | Intracellula | F65 | H2FDA | 988 | BL1 | 7260 |
| 40 | No | A. baylyi | Nalidixic a | Intracellula | F142 | Chlorophe | 959 | BL3 | 7672 |
| 40 | No | A. baylyi | Nalidixic a | Intracellula | F39 | Congo Red | 884 | BL3 | 7542 |
| 40 | No | A. baylyi | Nalidixic a | Intracellula | F116 | Sunset Yel | 938 | BL2 | 6763.5 |
| 40 | No | A. baylyi | Nalidixic a | Intracellula | F10 | Alizarin | 929 | BL1 | 5814 |
| 40 | No | A. baylyi | Nalidixic a | Intracellula | F84 | Neutral Re | 972 | BL1 | 5145.5 |
| 40 | No | A. baylyi | Nalidixic a | Intracellula | F99 | Pyronin Y | 838 | BL1 | 5518 |
| 40 | No | A. baylyi | Nalidixic a | Intracellula | F108 | Rhodamine | 947 | BL1 | 5111 |
| 40 | No | A. baylyi | Nalidixic a | Intracellula | F109 | Riboflavin | 893 | BL1 | 4939 |
| 40 | No | A. baylyi | Nalidixic a | Intracellula | F117 | SYBR Gree | 947 | BL1 | 362559 |
| 40 | No | A. baylyi | Nalidixic a | Intracellula | F106 | Rhodamine | 896 | BL1 | 7170.5 |
| 40 | No | A. baylyi | Nalidixic a | Intracellula | F73 | Malachite ( | 889 | BL1 | 5221 |
| 40 | No | A. baylyi | Nalidixic a | Intracellula | F7 | Acridine or | 991 | BL1 | 6870 |
| 40 | No | A. baylyi | Nalidixic a | Intracellula | F192 | 5-Carboxyf | 939 | BL1 | 22545 |
| 40 | No | A. baylyi | Nalidixic a | Intracellula | F57 | Eosin Y | 830 | BL1 | 20472 |
| 40 | No | A. baylyi | Nalidixic a | Intracellula | F27 | Calcein | 966 | BL1 | 20335.5 |
| 40 | No | A. baylyi | Nalidixic a | Intracellula | F12 | Amaranth | 924 | BL1 | 22846 |
| 40 | No | A. baylyi | None | None | NA | Autofluore: | 1120 | NA | 17991 |
| 40 | No | A. baylyi | Ofloxacin | Intracellula | F194 | Oxonol | 2907 | RL1 | 13005 |
| 40 | No | A. baylyi | Ofloxacin | Intracellula | F11 | Allura Red | 983 | BL3 | 15507 |
| 40 | No | A. baylyi | Ofloxacin | Intracellula | F54 | DiSC3(5) | 771 | RL1 | 13282 |
| 40 | No | A. baylyi | Ofloxacin | Intracellula | F124 | Thioflavin T | 926 | VL2 | 9604.5 |
| 40 | No | A. baylyi | Ofloxacin | Intracellula | F101 | Quinacrine | 788 | VL2 | 6754 |
| 40 | No | A. baylyi | Ofloxacin | Intracellula | F138 | Apigenin | 888 | VL2 | 6100.5 |
| 40 | No | A. baylyi | Ofloxacin | Intracellula | F65 | H2FDA | 830 | BL1 | 8200.5 |
| 40 | No | A. baylyi | Ofloxacin | Intracellula | F142 | Chlorophe | 905 | BL3 | 8819 |
| 40 | No | A. baylyi | Ofloxacin | Intracellula | F39 | Congo Red | 891 | BL3 | 8676 |
| 40 | No | A. baylyi | Ofloxacin | Intracellula | F116 | Sunset Yel | 925 | BL2 | 7706 |
| 40 | No | A. baylyi | Ofloxacin | Intracellula | F10 | Alizarin | 815 | BL1 | 5774 |
| 40 | No | A. baylyi | Ofloxacin | Intracellula | F84 | Neutral Re | 878 | BL1 | 5556.5 |
| 40 | No | A. baylyi | Ofloxacin | Intracellula | F99 | Pyronin Y | 730 | BL1 | 6105.5 |
| 40 | No | A. baylyi | Ofloxacin | Intracellula | F108 | Rhodamine | 896 | BL1 | 5166.5 |
| 40 | No | A. baylyi | Ofloxacin | Intracellula | F109 | Riboflavin | 745 | BL1 | 5272 |
| 40 | No | A. baylyi | Ofloxacin | Intracellula | F117 | SYBR Gree | 844 | BL1 | 420978 |
| 40 | No | A. baylyi | Ofloxacin | Intracellula | F106 | Rhodamine | 784 | BL1 | 7085.5 |
| 40 | No | A. baylyi | Ofloxacin | Intracellula | F73 | Malachite ( | 851 | BL1 | 5588 |
| 40 | No | A. baylyi | Ofloxacin | Intracellula | F7 | Acridine or | 805 | BL1 | 7362 |
| 40 | No | A. baylyi | Ofloxacin | Intracellula | F192 | 5-Carboxyf | 818 | BL1 | 24274.5 |
| 40 | No | A. baylyi | Ofloxacin | Intracellula | F57 | Eosin Y | 752 | BL1 | 23046 |
| 40 | No | A. baylyi | Ofloxacin | Intracellula | F27 | Calcein | 795 | BL1 | 24621 |

| | | | | | | | | | |
|---|---|---|---|---|---|---|---|---|---|
| 40 | No | A. baylyi | Ofloxacin | Intracellul | F12 | Amaranth | 739 | BL1 | 20790 |
| 40 | No | A. baylyi | None | None | NA | Autofluore | 1126 | NA | 19005.5 |
| 40 | No | A. baylyi | Polymyxin | Wall | F194 | Oxonol | 1581 | RL1 | 17788 |
| 40 | No | A. baylyi | Polymyxin | Wall | F11 | Allura Red | 1064 | BL3 | 13771 |
| 40 | No | A. baylyi | Polymyxin | Wall | F54 | DiSC3(5) | 1012 | RL1 | 14669.5 |
| 40 | No | A. baylyi | Polymyxin | Wall | F124 | Thioflavin T | 1063 | VL2 | 9219 |
| 40 | No | A. baylyi | Polymyxin | Wall | F101 | Quinacrine | 1033 | VL2 | 6524 |
| 40 | No | A. baylyi | Polymyxin | Wall | F138 | Apigenin | 1133 | VL2 | 6280 |
| 40 | No | A. baylyi | Polymyxin | Wall | F65 | H2FDA | 1106 | BL1 | 7521.5 |
| 40 | No | A. baylyi | Polymyxin | Wall | F142 | Chlorophe | 1100 | BL3 | 7729 |
| 40 | No | A. baylyi | Polymyxin | Wall | F39 | Congo Red | 1025 | BL3 | 7234 |
| 40 | No | A. baylyi | Polymyxin | Wall | F116 | Sunset Yel | 1153 | BL2 | 6381 |
| 40 | No | A. baylyi | Polymyxin | Wall | F10 | Alizarin | 1053 | BL1 | 6165 |
| 40 | No | A. baylyi | Polymyxin | Wall | F84 | Neutral Re | 992 | BL1 | 5147 |
| 40 | No | A. baylyi | Polymyxin | Wall | F99 | Pyronin Y | 1084 | BL1 | 5460 |
| 40 | No | A. baylyi | Polymyxin | Wall | F108 | Rhodamine | 1068 | BL1 | 4962 |
| 40 | No | A. baylyi | Polymyxin | Wall | F109 | Riboflavin | 976 | BL1 | 5094 |
| 40 | No | A. baylyi | Polymyxin | Wall | F117 | SYBR Gree | 1019 | BL1 | 644645 |
| 40 | No | A. baylyi | Polymyxin | Wall | F106 | Rhodamine | 1102 | BL1 | 7045.5 |
| 40 | No | A. baylyi | Polymyxin | Wall | F73 | Malachite ( | 989 | BL1 | 5277 |
| 40 | No | A. baylyi | Polymyxin | Wall | F7 | Acridine or | 1101 | BL1 | 5973 |
| 40 | No | A. baylyi | Polymyxin | Wall | F192 | 5-Carboxyf | 1097 | BL1 | 18718 |
| 40 | No | A. baylyi | Polymyxin | Wall | F57 | Eosin Y | 996 | BL1 | 18124.5 |
| 40 | No | A. baylyi | Polymyxin | Wall | F27 | Calcein | 1033 | BL1 | 17496 |
| 40 | No | A. baylyi | Polymyxin | Wall | F12 | Amaranth | 1013 | BL1 | 20206 |
| 40 | No | A. baylyi | None | None | NA | Autofluore | 1155 | NA | 19602 |
| 40 | No | A. baylyi | Tetracyclin | Intracellul | F194 | Oxonol | 2512 | RL1 | 12206.5 |
| 40 | No | A. baylyi | Tetracyclin | Intracellul | F11 | Allura Red | 894 | BL3 | 13679 |
| 40 | No | A. baylyi | Tetracyclin | Intracellul | F54 | DiSC3(5) | 682 | RL1 | 12845 |
| 40 | No | A. baylyi | Tetracyclin | Intracellul | F124 | Thioflavin T | 792 | VL2 | 9164 |
| 40 | No | A. baylyi | Tetracyclin | Intracellul | F101 | Quinacrine | 765 | VL2 | 6619 |
| 40 | No | A. baylyi | Tetracyclin | Intracellul | F138 | Apigenin | 842 | VL2 | 6388.5 |
| 40 | No | A. baylyi | Tetracyclin | Intracellul | F65 | H2FDA | 796 | BL1 | 7427 |
| 40 | No | A. baylyi | Tetracyclin | Intracellul | F142 | Chlorophe | 844 | BL3 | 8633 |
| 40 | No | A. baylyi | Tetracyclin | Intracellul | F39 | Congo Red | 711 | BL3 | 8167 |
| 40 | No | A. baylyi | Tetracyclin | Intracellul | F116 | Sunset Yel | 873 | BL2 | 8168 |
| 40 | No | A. baylyi | Tetracyclin | Intracellul | F10 | Alizarin | 754 | BL1 | 6053.5 |
| 40 | No | A. baylyi | Tetracyclin | Intracellul | F84 | Neutral Re | 741 | BL1 | 5057 |
| 40 | No | A. baylyi | Tetracyclin | Intracellul | F99 | Pyronin Y | 874 | BL1 | 6624.5 |
| 40 | No | A. baylyi | Tetracyclin | Intracellul | F108 | Rhodamine | 802 | BL1 | 5316.5 |
| 40 | No | A. baylyi | Tetracyclin | Intracellul | F109 | Riboflavin | 721 | BL1 | 5370 |
| 40 | No | A. baylyi | Tetracyclin | Intracellul | F117 | SYBR Gree | 798 | BL1 | 386498.5 |
| 40 | No | A. baylyi | Tetracyclin | Intracellul | F106 | Rhodamine | 772 | BL1 | 8696.5 |
| 40 | No | A. baylyi | Tetracyclin | Intracellul | F73 | Malachite ( | 721 | BL1 | 5338 |
| 40 | No | A. baylyi | Tetracyclin | Intracellul | F7 | Acridine or | 777 | BL1 | 6610 |
| 40 | No | A. baylyi | Tetracyclin | Intracellul | F192 | 5-Carboxyf | 888 | BL1 | 20659.5 |
| 40 | No | A. baylyi | Tetracyclin | Intracellul | F57 | Eosin Y | 760 | BL1 | 20026.5 |
| 40 | No | A. baylyi | Tetracyclin | Intracellul | F27 | Calcein | 777 | BL1 | 17625 |

| | | | | | | | | | |
|---|---|---|---|---|---|---|---|---|---|
| 40 | No | A. baylyi | Tetracyclin | Intracellula | F12 | Amaranth | 732 | BL1 | 19365.5 |
| 40 | No | A. baylyi | None | None | NA | Autofluore: | 1101 | NA | 19455 |
| 40 | No | A. baylyi | Vancomyc | Wall | F194 | Oxonol | 1070 | RL1 | 17944.5 |
| 40 | No | A. baylyi | Vancomyc | Wall | F11 | Allura Red | 981 | BL3 | 14879 |
| 40 | No | A. baylyi | Vancomyc | Wall | F54 | DiSC3(5) | 853 | RL1 | 12946 |
| 40 | No | A. baylyi | Vancomyc | Wall | F124 | Thioflavin T | 1010 | VL2 | 8807 |
| 40 | No | A. baylyi | Vancomyc | Wall | F101 | Quinacrine | 917 | VL2 | 6249 |
| 40 | No | A. baylyi | Vancomyc | Wall | F138 | Apigenin | 1001 | VL2 | 6113 |
| 40 | No | A. baylyi | Vancomyc | Wall | F65 | H2FDA | 999 | BL1 | 7439 |
| 40 | No | A. baylyi | Vancomyc | Wall | F142 | Chlorophe | 998 | BL3 | 8156 |
| 40 | No | A. baylyi | Vancomyc | Wall | F39 | Congo Red | 987 | BL3 | 7807 |
| 40 | No | A. baylyi | Vancomyc | Wall | F116 | Sunset Yel | 1062 | BL2 | 7472.5 |
| 40 | No | A. baylyi | Vancomyc | Wall | F10 | Alizarin | 1034 | BL1 | 6095 |
| 40 | No | A. baylyi | Vancomyc | Wall | F84 | Neutral Re | 937 | BL1 | 5633 |
| 40 | No | A. baylyi | Vancomyc | Wall | F99 | Pyronin Y | 955 | BL1 | 5710 |
| 40 | No | A. baylyi | Vancomyc | Wall | F108 | Rhodamine | 1087 | BL1 | 5357 |
| 40 | No | A. baylyi | Vancomyc | Wall | F109 | Riboflavin | 930 | BL1 | 5339.5 |
| 40 | No | A. baylyi | Vancomyc | Wall | F117 | SYBR Gree | 954 | BL1 | 391584.5 |
| 40 | No | A. baylyi | Vancomyc | Wall | F106 | Rhodamine | 1029 | BL1 | 8014 |
| 40 | No | A. baylyi | Vancomyc | Wall | F73 | Malachite ( | 961 | BL1 | 5939 |
| 40 | No | A. baylyi | Vancomyc | Wall | F7 | Acridine or | 991 | BL1 | 5978 |
| 40 | No | A. baylyi | Vancomyc | Wall | F192 | 5-Carboxyf | 933 | BL1 | 21444 |
| 40 | No | A. baylyi | Vancomyc | Wall | F57 | Eosin Y | 897 | BL1 | 16491 |
| 40 | No | A. baylyi | Vancomyc | Wall | F27 | Calcein | 1043 | BL1 | 20304 |
| 40 | No | A. baylyi | Vancomyc | Wall | F12 | Amaranth | 899 | BL1 | 22145 |
| 20 | Yes | E. coli | None | None | NA | Autofluore: | 3634 | NA | 891 |
| 20 | Yes | E. coli | None | None | F194 | Oxonol | 4930 | RL1 | 2418 |
| 20 | Yes | E. coli | None | None | F11 | Allura Red | 3803 | BL3 | 2087 |
| 20 | Yes | E. coli | None | None | F54 | DiSC3(5) | 4347 | RL1 | 2770 |
| 20 | Yes | E. coli | None | None | F124 | Thioflavin T | 4064 | VL2 | 2717 |
| 20 | Yes | E. coli | None | None | F101 | Quinacrine | 4060 | VL2 | 2590 |
| 20 | Yes | E. coli | None | None | F138 | Apigenin | 4055 | VL2 | 3342 |
| 20 | Yes | E. coli | None | None | F65 | H2FDA | 3814 | BL1 | 4477 |
| 20 | Yes | E. coli | None | None | F142 | Chlorophe | 3493 | BL3 | 6221 |
| 20 | Yes | E. coli | None | None | F39 | Congo Red | 4765 | BL3 | 5972 |
| 20 | Yes | E. coli | None | None | F116 | Sunset Yel | 1938 | BL2 | 6633.5 |
| 20 | Yes | E. coli | None | None | F10 | Alizarin | 4078 | BL1 | 4824.5 |
| 20 | Yes | E. coli | None | None | F84 | Neutral Re | 2800 | BL1 | 4720 |
| 20 | Yes | E. coli | None | None | F99 | Pyronin Y | 3280 | BL1 | 3568 |
| 20 | Yes | E. coli | None | None | F108 | Rhodamine | 5138 | BL1 | 4081 |
| 20 | Yes | E. coli | None | None | F109 | Riboflavin | 4632 | BL1 | 120661.5 |
| 20 | Yes | E. coli | None | None | F117 | SYBR Gree | 2096 | BL1 | 3441.5 |
| 20 | Yes | E. coli | None | None | F106 | Rhodamine | 3296 | BL1 | 5058.5 |
| 20 | Yes | E. coli | None | None | F73 | Malachite ( | 3434 | BL1 | 4660 |
| 20 | Yes | E. coli | None | None | F7 | Acridine or | 3313 | BL1 | 11719 |
| 20 | Yes | E. coli | None | None | F192 | 5-Carboxyf | 2625 | BL1 | 12779 |
| 20 | Yes | E. coli | None | None | F57 | Eosin Y | 1780 | BL1 | 18193 |
| 20 | Yes | E. coli | None | None | F27 | Calcein | 2005 | BL1 | 16869 |

| | | | | | | | | | |
|---|---|---|---|---|---|---|---|---|---|
| 20 | Yes | E. coli | None | None | F12 | Amaranth | 3217 | BL1 | 18980 |
| 20 | Yes | E. coli | None | None | NA | Autofluores | 3198 | NA | 7259 |
| 20 | Yes | E. coli | Amoxacilli | Wall | F194 | Oxonol | 3760 | RL1 | 5995 |
| 20 | Yes | E. coli | Amoxacilli | Wall | F11 | Allura Red | 2800 | BL3 | 4207 |
| 20 | Yes | E. coli | Amoxacilli | Wall | F54 | DiSC3(5) | 3267 | RL1 | 4495 |
| 20 | Yes | E. coli | Amoxacilli | Wall | F124 | Thioflavin T | 4345 | VL2 | 4092 |
| 20 | Yes | E. coli | Amoxacilli | Wall | F101 | Quinacrine | 3713 | VL2 | 4625 |
| 20 | Yes | E. coli | Amoxacilli | Wall | F138 | Apigenin | 3214 | VL2 | 4757 |
| 20 | Yes | E. coli | Amoxacilli | Wall | F65 | H2FDA | 3074 | BL1 | 6574 |
| 20 | Yes | E. coli | Amoxacilli | Wall | F142 | Chlorophe | 3495 | BL3 | 7842 |
| 20 | Yes | E. coli | Amoxacilli | Wall | F39 | Congo Red | 2961 | BL3 | 8960 |
| 20 | Yes | E. coli | Amoxacilli | Wall | F116 | Sunset Yel | 3477 | BL2 | 7691 |
| 20 | Yes | E. coli | Amoxacilli | Wall | F10 | Alizarin | 2885 | BL1 | 6827 |
| 20 | Yes | E. coli | Amoxacilli | Wall | F84 | Neutral Re | 3196 | BL1 | 6173 |
| 20 | Yes | E. coli | Amoxacilli | Wall | F99 | Pyronin Y | 3330 | BL1 | 6137.5 |
| 20 | Yes | E. coli | Amoxacilli | Wall | F108 | Rhodamine | 3347 | BL1 | 4813 |
| 20 | Yes | E. coli | Amoxacilli | Wall | F109 | Riboflavin | 2444 | BL1 | 5228.5 |
| 20 | Yes | E. coli | Amoxacilli | Wall | F117 | SYBR Gree | 2773 | BL1 | 179759 |
| 20 | Yes | E. coli | Amoxacilli | Wall | F106 | Rhodamine | 3333 | BL1 | 6890 |
| 20 | Yes | E. coli | Amoxacilli | Wall | F73 | Malachite ( | 3043 | BL1 | 5433 |
| 20 | Yes | E. coli | Amoxacilli | Wall | F7 | Acridine or | 2860 | BL1 | 5376.5 |
| 20 | Yes | E. coli | Amoxacilli | Wall | F192 | 5-Carboxyf | 2962 | BL1 | 12420 |
| 20 | Yes | E. coli | Amoxacilli | Wall | F57 | Eosin Y | 3511 | BL1 | 13519 |
| 20 | Yes | E. coli | Amoxacilli | Wall | F27 | Calcein | 3299 | BL1 | 18623 |
| 20 | Yes | E. coli | Amoxacilli | Wall | F12 | Amaranth | 3433 | BL1 | 18983 |
| 20 | Yes | E. coli | None | None | NA | Autofluores | 3190 | NA | 7759.5 |
| 20 | Yes | E. coli | Azithromyc | Intracellula | F194 | Oxonol | 2692 | RL1 | 6048.5 |
| 20 | Yes | E. coli | Azithromyc | Intracellula | F11 | Allura Red | 2560 | BL3 | 5730 |
| 20 | Yes | E. coli | Azithromyc | Intracellula | F54 | DiSC3(5) | 2526 | RL1 | 4589.5 |
| 20 | Yes | E. coli | Azithromyc | Intracellula | F124 | Thioflavin T | 2588 | VL2 | 5623.5 |
| 20 | Yes | E. coli | Azithromyc | Intracellula | F101 | Quinacrine | 2590 | VL2 | 4151.5 |
| 20 | Yes | E. coli | Azithromyc | Intracellula | F138 | Apigenin | 2484 | VL2 | 5240.5 |
| 20 | Yes | E. coli | Azithromyc | Intracellula | F65 | H2FDA | 1970 | BL1 | 5686 |
| 20 | Yes | E. coli | Azithromyc | Intracellula | F142 | Chlorophe | 2495 | BL3 | 6730 |
| 20 | Yes | E. coli | Azithromyc | Intracellula | F39 | Congo Red | 2309 | BL3 | 7238 |
| 20 | Yes | E. coli | Azithromyc | Intracellula | F116 | Sunset Yel | 2482 | BL2 | 7846 |
| 20 | Yes | E. coli | Azithromyc | Intracellula | F10 | Alizarin | 2439 | BL1 | 7147 |
| 20 | Yes | E. coli | Azithromyc | Intracellula | F84 | Neutral Re | 2062 | BL1 | 5877 |
| 20 | Yes | E. coli | Azithromyc | Intracellula | F99 | Pyronin Y | 2252 | BL1 | 5924 |
| 20 | Yes | E. coli | Azithromyc | Intracellula | F108 | Rhodamine | 2489 | BL1 | 7216 |
| 20 | Yes | E. coli | Azithromyc | Intracellula | F109 | Riboflavin | 2242 | BL1 | 7914.5 |
| 20 | Yes | E. coli | Azithromyc | Intracellula | F117 | SYBR Gree | 1972 | BL1 | 67135.5 |
| 20 | Yes | E. coli | Azithromyc | Intracellula | F106 | Rhodamine | 2346 | BL1 | 7029.5 |
| 20 | Yes | E. coli | Azithromyc | Intracellula | F73 | Malachite ( | 1978 | BL1 | 7891.5 |
| 20 | Yes | E. coli | Azithromyc | Intracellula | F7 | Acridine or | 2012 | BL1 | 5951.5 |
| 20 | Yes | E. coli | Azithromyc | Intracellula | F192 | 5-Carboxyf | 2128 | BL1 | 12797 |
| 20 | Yes | E. coli | Azithromyc | Intracellula | F57 | Eosin Y | 2169 | BL1 | 15789 |
| 20 | Yes | E. coli | Azithromyc | Intracellula | F27 | Calcein | 2616 | BL1 | 20577 |

| | | | | | | | | | |
|---|---|---|---|---|---|---|---|---|---|
| 20 | Yes | E. coli | Azithromyc | Intracellul | F12 | Amaranth | 2623 | BL1 | 18778 |
| 20 | Yes | E. coli | None | None | NA | Autofluore | 3059 | NA | 7413 |
| 20 | Yes | E. coli | Trimethopr | Intracellul | F194 | Oxonol | 3527 | RL1 | 6421 |
| 20 | Yes | E. coli | Trimethopr | Intracellul | F11 | Allura Red | 3126 | BL3 | 4866 |
| 20 | Yes | E. coli | Trimethopr | Intracellul | F54 | DiSC3(5) | 3337 | RL1 | 4587 |
| 20 | Yes | E. coli | Trimethopr | Intracellul | F124 | Thioflavin T | 3246 | VL2 | 4629.5 |
| 20 | Yes | E. coli | Trimethopr | Intracellul | F101 | Quinacrine | 3356 | VL2 | 5014.5 |
| 20 | Yes | E. coli | Trimethopr | Intracellul | F138 | Apigenin | 2964 | VL2 | 5784 |
| 20 | Yes | E. coli | Trimethopr | Intracellul | F65 | H2FDA | 3309 | BL1 | 4933 |
| 20 | Yes | E. coli | Trimethopr | Intracellul | F142 | Chlorophe | 2974 | BL3 | 6346 |
| 20 | Yes | E. coli | Trimethopr | Intracellul | F39 | Congo Red | 3174 | BL3 | 6207 |
| 20 | Yes | E. coli | Trimethopr | Intracellul | F116 | Sunset Yel | 2665 | BL2 | 6695 |
| 20 | Yes | E. coli | Trimethopr | Intracellul | F10 | Alizarin | 2499 | BL1 | 5444 |
| 20 | Yes | E. coli | Trimethopr | Intracellul | F84 | Neutral Re | 2766 | BL1 | 4703 |
| 20 | Yes | E. coli | Trimethopr | Intracellul | F99 | Pyronin Y | 2956 | BL1 | 4365.5 |
| 20 | Yes | E. coli | Trimethopr | Intracellul | F108 | Rhodamine | 2933 | BL1 | 4765 |
| 20 | Yes | E. coli | Trimethopr | Intracellul | F109 | Riboflavin | 2920 | BL1 | 4002.5 |
| 20 | Yes | E. coli | Trimethopr | Intracellul | F117 | SYBR Gree | 2770 | BL1 | 117532.5 |
| 20 | Yes | E. coli | Trimethopr | Intracellul | F106 | Rhodamine | 2967 | BL1 | 5313 |
| 20 | Yes | E. coli | Trimethopr | Intracellul | F73 | Malachite | 2575 | BL1 | 7247 |
| 20 | Yes | E. coli | Trimethopr | Intracellul | F7 | Acridine or | 2807 | BL1 | 5480 |
| 20 | Yes | E. coli | Trimethopr | Intracellul | F192 | 5-Carboxyf | 2695 | BL1 | 9189 |
| 20 | Yes | E. coli | Trimethopr | Intracellul | F57 | Eosin Y | 2880 | BL1 | 14687 |
| 20 | Yes | E. coli | Trimethopr | Intracellul | F27 | Calcein | 2672 | BL1 | 19488 |
| 20 | Yes | E. coli | Trimethopr | Intracellul | F12 | Amaranth | 2868 | BL1 | 15001.5 |
| 20 | Yes | E. coli | None | None | NA | Autofluore | 3205 | NA | 7580 |
| 20 | Yes | E. coli | Chloramph | Intracellul | F194 | Oxonol | 2731 | RL1 | 6754 |
| 20 | Yes | E. coli | Chloramph | Intracellul | F11 | Allura Red | 2704 | BL3 | 4936 |
| 20 | Yes | E. coli | Chloramph | Intracellul | F54 | DiSC3(5) | 2666 | RL1 | 5395.5 |
| 20 | Yes | E. coli | Chloramph | Intracellul | F124 | Thioflavin T | 2859 | VL2 | 6169 |
| 20 | Yes | E. coli | Chloramph | Intracellul | F101 | Quinacrine | 3058 | VL2 | 4789.5 |
| 20 | Yes | E. coli | Chloramph | Intracellul | F138 | Apigenin | 2768 | VL2 | 5043 |
| 20 | Yes | E. coli | Chloramph | Intracellul | F65 | H2FDA | 3092 | BL1 | 7669.5 |
| 20 | Yes | E. coli | Chloramph | Intracellul | F142 | Chlorophe | 2814 | BL3 | 6024 |
| 20 | Yes | E. coli | Chloramph | Intracellul | F39 | Congo Red | 2520 | BL3 | 8395.5 |
| 20 | Yes | E. coli | Chloramph | Intracellul | F116 | Sunset Yel | 2624 | BL2 | 6586 |
| 20 | Yes | E. coli | Chloramph | Intracellul | F10 | Alizarin | 2071 | BL1 | 7490 |
| 20 | Yes | E. coli | Chloramph | Intracellul | F84 | Neutral Re | 2630 | BL1 | 5744 |
| 20 | Yes | E. coli | Chloramph | Intracellul | F99 | Pyronin Y | 2558 | BL1 | 5011.5 |
| 20 | Yes | E. coli | Chloramph | Intracellul | F108 | Rhodamine | 2575 | BL1 | 5523 |
| 20 | Yes | E. coli | Chloramph | Intracellul | F109 | Riboflavin | 2743 | BL1 | 5919 |
| 20 | Yes | E. coli | Chloramph | Intracellul | F117 | SYBR Gree | 2457 | BL1 | 108756 |
| 20 | Yes | E. coli | Chloramph | Intracellul | F106 | Rhodamine | 2645 | BL1 | 7185 |
| 20 | Yes | E. coli | Chloramph | Intracellul | F73 | Malachite | 2411 | BL1 | 5479 |
| 20 | Yes | E. coli | Chloramph | Intracellul | F7 | Acridine or | 2238 | BL1 | 7073.5 |
| 20 | Yes | E. coli | Chloramph | Intracellul | F192 | 5-Carboxyf | 2627 | BL1 | 13807 |
| 20 | Yes | E. coli | Chloramph | Intracellul | F57 | Eosin Y | 2123 | BL1 | 11371 |
| 20 | Yes | E. coli | Chloramph | Intracellul | F27 | Calcein | 2499 | BL1 | 15688 |

| | | | | | | | | | |
|---|---|---|---|---|---|---|---|---|---|
| 20 | Yes | E. coli | Chloramph | Intracellula | F12 | Amaranth | 2350 | BL1 | 16145.5 |
| 20 | Yes | E. coli | None | None | NA | Autofluore: | 3122 | NA | 7300.5 |
| 20 | Yes | E. coli | Colistin | Wall | F194 | Oxonol | 1404 | RL1 | 95817.5 |
| 20 | Yes | E. coli | Colistin | Wall | F11 | Allura Red | 1564 | BL3 | 61714.5 |
| 20 | Yes | E. coli | Colistin | Wall | F54 | DiSC3(5) | 1429 | RL1 | 57958 |
| 20 | Yes | E. coli | Colistin | Wall | F124 | Thioflavin T | 1632 | VL2 | 54593 |
| 20 | Yes | E. coli | Colistin | Wall | F101 | Quinacrine | 1345 | VL2 | 63472 |
| 20 | Yes | E. coli | Colistin | Wall | F138 | Apigenin | 1098 | VL2 | 61149.5 |
| 20 | Yes | E. coli | Colistin | Wall | F65 | H2FDA | 1238 | BL1 | 60417 |
| 20 | Yes | E. coli | Colistin | Wall | F142 | Chlorophe | 1112 | BL3 | 52852 |
| 20 | Yes | E. coli | Colistin | Wall | F39 | Congo Red | 1317 | BL3 | 47135 |
| 20 | Yes | E. coli | Colistin | Wall | F116 | Sunset Yel | 962 | BL2 | 51150.5 |
| 20 | Yes | E. coli | Colistin | Wall | F10 | Alizarin | 1135 | BL1 | 41843 |
| 20 | Yes | E. coli | Colistin | Wall | F84 | Neutral Re | 943 | BL1 | 40530 |
| 20 | Yes | E. coli | Colistin | Wall | F99 | Pyronin Y | 1239 | BL1 | 32045 |
| 20 | Yes | E. coli | Colistin | Wall | F108 | Rhodamine | 916 | BL1 | 34033 |
| 20 | Yes | E. coli | Colistin | Wall | F109 | Riboflavin | 1007 | BL1 | 34223 |
| 20 | Yes | E. coli | Colistin | Wall | F117 | SYBR Gree | 940 | BL1 | 2699515 |
| 20 | Yes | E. coli | Colistin | Wall | F106 | Rhodamine | 1123 | BL1 | 84015 |
| 20 | Yes | E. coli | Colistin | Wall | F73 | Malachite ( | 1064 | BL1 | 81362.5 |
| 20 | Yes | E. coli | Colistin | Wall | F7 | Acridine or | 1069 | BL1 | 177692 |
| 20 | Yes | E. coli | Colistin | Wall | F192 | 5-Carboxyf | 952 | BL1 | 106220 |
| 20 | Yes | E. coli | Colistin | Wall | F57 | Eosin Y | 1030 | BL1 | 49651 |
| 20 | Yes | E. coli | Colistin | Wall | F27 | Calcein | 1088 | BL1 | 84257.5 |
| 20 | Yes | E. coli | Colistin | Wall | F12 | Amaranth | 937 | BL1 | 67448 |
| 20 | Yes | E. coli | None | None | NA | Autofluore: | 3279 | NA | 7727 |
| 20 | Yes | E. coli | Gentamicii | Intracellula | F194 | Oxonol | 3546 | RL1 | 6385.5 |
| 20 | Yes | E. coli | Gentamicii | Intracellula | F11 | Allura Red | 3454 | BL3 | 5808.5 |
| 20 | Yes | E. coli | Gentamicii | Intracellula | F54 | DiSC3(5) | 3528 | RL1 | 5155 |
| 20 | Yes | E. coli | Gentamicii | Intracellula | F124 | Thioflavin T | 3544 | VL2 | 5360.5 |
| 20 | Yes | E. coli | Gentamicii | Intracellula | F101 | Quinacrine | 3696 | VL2 | 5458 |
| 20 | Yes | E. coli | Gentamicii | Intracellula | F138 | Apigenin | 3595 | VL2 | 5108 |
| 20 | Yes | E. coli | Gentamicii | Intracellula | F65 | H2FDA | 3262 | BL1 | 5968 |
| 20 | Yes | E. coli | Gentamicii | Intracellula | F142 | Chlorophe | 3651 | BL3 | 6558 |
| 20 | Yes | E. coli | Gentamicii | Intracellula | F39 | Congo Red | 3497 | BL3 | 7485 |
| 20 | Yes | E. coli | Gentamicii | Intracellula | F116 | Sunset Yel | 3587 | BL2 | 6066 |
| 20 | Yes | E. coli | Gentamicii | Intracellula | F10 | Alizarin | 3286 | BL1 | 8669.5 |
| 20 | Yes | E. coli | Gentamicii | Intracellula | F84 | Neutral Re | 3409 | BL1 | 5270 |
| 20 | Yes | E. coli | Gentamicii | Intracellula | F99 | Pyronin Y | 3391 | BL1 | 7177 |
| 20 | Yes | E. coli | Gentamicii | Intracellula | F108 | Rhodamine | 3520 | BL1 | 5905.5 |
| 20 | Yes | E. coli | Gentamicii | Intracellula | F109 | Riboflavin | 3128 | BL1 | 5916.5 |
| 20 | Yes | E. coli | Gentamicii | Intracellula | F117 | SYBR Gree | 3376 | BL1 | 194743 |
| 20 | Yes | E. coli | Gentamicii | Intracellula | F106 | Rhodamine | 3635 | BL1 | 8684 |
| 20 | Yes | E. coli | Gentamicii | Intracellula | F73 | Malachite ( | 2894 | BL1 | 4864.5 |
| 20 | Yes | E. coli | Gentamicii | Intracellula | F7 | Acridine or | 2488 | BL1 | 8952.5 |
| 20 | Yes | E. coli | Gentamicii | Intracellula | F192 | 5-Carboxyf | 3672 | BL1 | 12316 |
| 20 | Yes | E. coli | Gentamicii | Intracellula | F57 | Eosin Y | 2749 | BL1 | 12447 |
| 20 | Yes | E. coli | Gentamicii | Intracellula | F27 | Calcein | 3640 | BL1 | 14617.5 |

| | | | | | | | | | |
|---|---|---|---|---|---|---|---|---|---|
| 20 | Yes | E. coli | Gentamici | Intracellul | F12 | Amaranth | 3260 | BL1 | 16500 |
| 20 | Yes | E. coli | None | None | NA | Autofluore | 2765 | NA | 7440 |
| 20 | Yes | E. coli | Nitrofurant | Intracellul | F194 | Oxonol | 2817 | RL1 | 6866 |
| 20 | Yes | E. coli | Nitrofurant | Intracellul | F11 | Allura Red | 2932 | BL3 | 5498 |
| 20 | Yes | E. coli | Nitrofurant | Intracellul | F54 | DiSC3(5) | 2669 | RL1 | 5265 |
| 20 | Yes | E. coli | Nitrofurant | Intracellul | F124 | Thioflavin T | 2758 | VL2 | 5283.5 |
| 20 | Yes | E. coli | Nitrofurant | Intracellul | F101 | Quinacrine | 2373 | VL2 | 4904 |
| 20 | Yes | E. coli | Nitrofurant | Intracellul | F138 | Apigenin | 2746 | VL2 | 4992.5 |
| 20 | Yes | E. coli | Nitrofurant | Intracellul | F65 | H2FDA | 2856 | BL1 | 5721.5 |
| 20 | Yes | E. coli | Nitrofurant | Intracellul | F142 | Chlorophe | 2337 | BL3 | 6665 |
| 20 | Yes | E. coli | Nitrofurant | Intracellul | F39 | Congo Red | 2448 | BL3 | 6362.5 |
| 20 | Yes | E. coli | Nitrofurant | Intracellul | F116 | Sunset Yel | 2205 | BL2 | 6952 |
| 20 | Yes | E. coli | Nitrofurant | Intracellul | F10 | Alizarin | 2309 | BL1 | 4853 |
| 20 | Yes | E. coli | Nitrofurant | Intracellul | F84 | Neutral Re | 2579 | BL1 | 5354 |
| 20 | Yes | E. coli | Nitrofurant | Intracellul | F99 | Pyronin Y | 2521 | BL1 | 6134 |
| 20 | Yes | E. coli | Nitrofurant | Intracellul | F108 | Rhodamine | 2159 | BL1 | 4975 |
| 20 | Yes | E. coli | Nitrofurant | Intracellul | F109 | Riboflavin | 2374 | BL1 | 4809 |
| 20 | Yes | E. coli | Nitrofurant | Intracellul | F117 | SYBR Gree | 2329 | BL1 | 63334 |
| 20 | Yes | E. coli | Nitrofurant | Intracellul | F106 | Rhodamine | 2110 | BL1 | 6777.5 |
| 20 | Yes | E. coli | Nitrofurant | Intracellul | F73 | Malachite ( | 2549 | BL1 | 7048 |
| 20 | Yes | E. coli | Nitrofurant | Intracellul | F7 | Acridine or | 2483 | BL1 | 7743 |
| 20 | Yes | E. coli | Nitrofurant | Intracellul | F192 | 5-Carboxyf | 2326 | BL1 | 12950 |
| 20 | Yes | E. coli | Nitrofurant | Intracellul | F57 | Eosin Y | 2372 | BL1 | 14971 |
| 20 | Yes | E. coli | Nitrofurant | Intracellul | F27 | Calcein | 1731 | BL1 | 16384 |
| 20 | Yes | E. coli | Nitrofurant | Intracellul | F12 | Amaranth | 2426 | BL1 | 16073.5 |
| 20 | Yes | E. coli | None | None | NA | Autofluore | 3025 | NA | 7403 |
| 20 | Yes | E. coli | Pentamidi | Wall | F194 | Oxonol | 2601 | RL1 | 11789 |
| 20 | Yes | E. coli | Pentamidi | Wall | F11 | Allura Red | 2318 | BL3 | 7073 |
| 20 | Yes | E. coli | Pentamidi | Wall | F54 | DiSC3(5) | 2205 | RL1 | 6740 |
| 20 | Yes | E. coli | Pentamidi | Wall | F124 | Thioflavin T | 2352 | VL2 | 7226.5 |
| 20 | Yes | E. coli | Pentamidi | Wall | F101 | Quinacrine | 2191 | VL2 | 8286 |
| 20 | Yes | E. coli | Pentamidi | Wall | F138 | Apigenin | 2251 | VL2 | 7651 |
| 20 | Yes | E. coli | Pentamidi | Wall | F65 | H2FDA | 2208 | BL1 | 8536.5 |
| 20 | Yes | E. coli | Pentamidi | Wall | F142 | Chlorophe | 2206 | BL3 | 8583.5 |
| 20 | Yes | E. coli | Pentamidi | Wall | F39 | Congo Red | 2011 | BL3 | 9734 |
| 20 | Yes | E. coli | Pentamidi | Wall | F116 | Sunset Yel | 2172 | BL2 | 8409 |
| 20 | Yes | E. coli | Pentamidi | Wall | F10 | Alizarin | 1997 | BL1 | 7917 |
| 20 | Yes | E. coli | Pentamidi | Wall | F84 | Neutral Re | 1901 | BL1 | 6315 |
| 20 | Yes | E. coli | Pentamidi | Wall | F99 | Pyronin Y | 2071 | BL1 | 5685 |
| 20 | Yes | E. coli | Pentamidi | Wall | F108 | Rhodamine | 2238 | BL1 | 5949 |
| 20 | Yes | E. coli | Pentamidi | Wall | F109 | Riboflavin | 2198 | BL1 | 6144 |
| 20 | Yes | E. coli | Pentamidi | Wall | F117 | SYBR Gree | 2085 | BL1 | 691912 |
| 20 | Yes | E. coli | Pentamidi | Wall | F106 | Rhodamine | 2152 | BL1 | 9261 |
| 20 | Yes | E. coli | Pentamidi | Wall | F73 | Malachite ( | 2107 | BL1 | 10474 |
| 20 | Yes | E. coli | Pentamidi | Wall | F7 | Acridine or | 2088 | BL1 | 9929 |
| 20 | Yes | E. coli | Pentamidi | Wall | F192 | 5-Carboxyf | 2279 | BL1 | 17438 |
| 20 | Yes | E. coli | Pentamidi | Wall | F57 | Eosin Y | 2301 | BL1 | 16396 |
| 20 | Yes | E. coli | Pentamidi | Wall | F27 | Calcein | 2155 | BL1 | 16547 |

| | | | | | | | | | |
|---|---|---|---|---|---|---|---|---|---|
| 20 | Yes | E. coli | Pentamidi | Wall | F12 | Amaranth | 2061 | BL1 | 14945 |
| 20 | Yes | E. coli | None | None | NA | Autofluore | 3074 | NA | 8940 |
| 20 | Yes | E. coli | Rifampicin | Intracellul | F194 | Oxonol | 3592 | RL1 | 8868 |
| 20 | Yes | E. coli | Rifampicin | Intracellul | F11 | Allura Red | 3406 | BL3 | 8179 |
| 20 | Yes | E. coli | Rifampicin | Intracellul | F54 | DiSC3(5) | 3473 | RL1 | 6377 |
| 20 | Yes | E. coli | Rifampicin | Intracellul | F124 | Thioflavin T | 3669 | VL2 | 5842 |
| 20 | Yes | E. coli | Rifampicin | Intracellul | F101 | Quinacrine | 3447 | VL2 | 5375 |
| 20 | Yes | E. coli | Rifampicin | Intracellul | F138 | Apigenin | 3297 | VL2 | 6641 |
| 20 | Yes | E. coli | Rifampicin | Intracellul | F65 | H2FDA | 3161 | BL1 | 7031 |
| 20 | Yes | E. coli | Rifampicin | Intracellul | F142 | Chlorophe | 3390 | BL3 | 7724.5 |
| 20 | Yes | E. coli | Rifampicin | Intracellul | F39 | Congo Red | 3332 | BL3 | 8724 |
| 20 | Yes | E. coli | Rifampicin | Intracellul | F116 | Sunset Yel | 3545 | BL2 | 6531 |
| 20 | Yes | E. coli | Rifampicin | Intracellul | F10 | Alizarin | 2632 | BL1 | 10731.5 |
| 20 | Yes | E. coli | Rifampicin | Intracellul | F84 | Neutral Re | 2946 | BL1 | 6373.5 |
| 20 | Yes | E. coli | Rifampicin | Intracellul | F99 | Pyronin Y | 2677 | BL1 | 8428 |
| 20 | Yes | E. coli | Rifampicin | Intracellul | F108 | Rhodamine | 2588 | BL1 | 6863.5 |
| 20 | Yes | E. coli | Rifampicin | Intracellul | F109 | Riboflavin | 2858 | BL1 | 6118.5 |
| 20 | Yes | E. coli | Rifampicin | Intracellul | F117 | SYBR Gree | 2825 | BL1 | 156948 |
| 20 | Yes | E. coli | Rifampicin | Intracellul | F106 | Rhodamine | 3415 | BL1 | 8200 |
| 20 | Yes | E. coli | Rifampicin | Intracellul | F73 | Malachite ( | 3079 | BL1 | 5451 |
| 20 | Yes | E. coli | Rifampicin | Intracellul | F7 | Acridine or | 2848 | BL1 | 6818 |
| 20 | Yes | E. coli | Rifampicin | Intracellul | F192 | 5-Carboxyf | 3250 | BL1 | 13230 |
| 20 | Yes | E. coli | Rifampicin | Intracellul | F57 | Eosin Y | 3016 | BL1 | 12616.5 |
| 20 | Yes | E. coli | Rifampicin | Intracellul | F27 | Calcein | 3402 | BL1 | 17131 |
| 20 | Yes | E. coli | Rifampicin | Intracellul | F12 | Amaranth | 3332 | BL1 | 17610 |
| 20 | Yes | E. coli | None | None | NA | Autofluore | 3069 | NA | 10181 |
| 20 | Yes | E. coli | Trimethopr | Intracellul | F194 | Oxonol | 2867 | RL1 | 6917 |
| 20 | Yes | E. coli | Trimethopr | Intracellul | F11 | Allura Red | 2979 | BL3 | 6807 |
| 20 | Yes | E. coli | Trimethopr | Intracellul | F54 | DiSC3(5) | 3183 | RL1 | 6426 |
| 20 | Yes | E. coli | Trimethopr | Intracellul | F124 | Thioflavin T | 3265 | VL2 | 5662 |
| 20 | Yes | E. coli | Trimethopr | Intracellul | F101 | Quinacrine | 3235 | VL2 | 5866 |
| 20 | Yes | E. coli | Trimethopr | Intracellul | F138 | Apigenin | 3082 | VL2 | 5788 |
| 20 | Yes | E. coli | Trimethopr | Intracellul | F65 | H2FDA | 2682 | BL1 | 7058.5 |
| 20 | Yes | E. coli | Trimethopr | Intracellul | F142 | Chlorophe | 3036 | BL3 | 8931.5 |
| 20 | Yes | E. coli | Trimethopr | Intracellul | F39 | Congo Red | 3070 | BL3 | 7643.5 |
| 20 | Yes | E. coli | Trimethopr | Intracellul | F116 | Sunset Yel | 3087 | BL2 | 8559 |
| 20 | Yes | E. coli | Trimethopr | Intracellul | F10 | Alizarin | 2769 | BL1 | 6582 |
| 20 | Yes | E. coli | Trimethopr | Intracellul | F84 | Neutral Re | 2518 | BL1 | 7222.5 |
| 20 | Yes | E. coli | Trimethopr | Intracellul | F99 | Pyronin Y | 2925 | BL1 | 8006 |
| 20 | Yes | E. coli | Trimethopr | Intracellul | F108 | Rhodamine | 2466 | BL1 | 6619.5 |
| 20 | Yes | E. coli | Trimethopr | Intracellul | F109 | Riboflavin | 2728 | BL1 | 8111.5 |
| 20 | Yes | E. coli | Trimethopr | Intracellul | F117 | SYBR Gree | 2631 | BL1 | 145296 |
| 20 | Yes | E. coli | Trimethopr | Intracellul | F106 | Rhodamine | 2778 | BL1 | 7231 |
| 20 | Yes | E. coli | Trimethopr | Intracellul | F73 | Malachite ( | 2411 | BL1 | 8560 |
| 20 | Yes | E. coli | Trimethopr | Intracellul | F7 | Acridine or | 2678 | BL1 | 5916 |
| 20 | Yes | E. coli | Trimethopr | Intracellul | F192 | 5-Carboxyf | 2388 | BL1 | 11386.5 |
| 20 | Yes | E. coli | Trimethopr | Intracellul | F57 | Eosin Y | 2873 | BL1 | 13072 |
| 20 | Yes | E. coli | Trimethopr | Intracellul | F27 | Calcein | 2987 | BL1 | 16631 |

| | | | | | | | | | |
|---|---|---|---|---|---|---|---|---|---|
| 20 | Yes | E. coli | Trimethopr | Intracellula | F12 | Amaranth | 3116 | BL1 | 16723.5 |
| 20 | Yes | E. coli | None | None | NA | Autofluore: | 14835 | NA | 1071 |
| 20 | Yes | E. coli | None | None | F194 | Oxonol | 17198 | RL1 | 1841 |
| 20 | Yes | E. coli | None | None | F11 | Allura Red | 16539 | BL3 | 2824 |
| 20 | Yes | E. coli | None | None | F54 | DiSC3(5) | 17122 | RL1 | 2361 |
| 20 | Yes | E. coli | None | None | F124 | Thioflavin T | 19740 | VL2 | 2754 |
| 20 | Yes | E. coli | None | None | F101 | Quinacrine | 18979 | VL2 | 2593 |
| 20 | Yes | E. coli | None | None | F138 | Apigenin | 16254 | VL2 | 2986.5 |
| 20 | Yes | E. coli | None | None | F65 | H2FDA | 17460 | BL1 | 4131 |
| 20 | Yes | E. coli | None | None | F142 | Chlorophe | 17157 | BL3 | 5813 |
| 20 | Yes | E. coli | None | None | F39 | Congo Red | 16213 | BL3 | 7286 |
| 20 | Yes | E. coli | None | None | F116 | Sunset Yel | 18015 | BL2 | 5788 |
| 20 | Yes | E. coli | None | None | F10 | Alizarin | 17307 | BL1 | 4999 |
| 20 | Yes | E. coli | None | None | F84 | Neutral Re | 17073 | BL1 | 3801 |
| 20 | Yes | E. coli | None | None | F99 | Pyronin Y | 13080 | BL1 | 3883.5 |
| 20 | Yes | E. coli | None | None | F108 | Rhodamine | 19184 | BL1 | 2963.5 |
| 20 | Yes | E. coli | None | None | F109 | Riboflavin | 17348 | BL1 | 3503.5 |
| 20 | Yes | E. coli | None | None | F117 | SYBR Gree | 15829 | BL1 | 196674 |
| 20 | Yes | E. coli | None | None | F106 | Rhodamine | 18457 | BL1 | 4185 |
| 20 | Yes | E. coli | None | None | F73 | Malachite ( | 17039 | BL1 | 4466 |
| 20 | Yes | E. coli | None | None | F7 | Acridine or | 17656 | BL1 | 3675 |
| 20 | Yes | E. coli | None | None | F192 | 5-Carboxyf | 17076 | BL1 | 13305.5 |
| 20 | Yes | E. coli | None | None | F57 | Eosin Y | 18294 | BL1 | 16779.5 |
| 20 | Yes | E. coli | None | None | F27 | Calcein | 18163 | BL1 | 19981 |
| 20 | Yes | E. coli | None | None | F12 | Amaranth | 19560 | BL1 | 17836 |
| 20 | Yes | E. coli | None | None | NA | Autofluore: | 14016 | NA | 7431 |
| 20 | Yes | E. coli | Amoxacilli | Wall | F194 | Oxonol | 17626 | RL1 | 5676 |
| 20 | Yes | E. coli | Amoxacilli | Wall | F11 | Allura Red | 13869 | BL3 | 4482 |
| 20 | Yes | E. coli | Amoxacilli | Wall | F54 | DiSC3(5) | 15143 | RL1 | 3970 |
| 20 | Yes | E. coli | Amoxacilli | Wall | F124 | Thioflavin T | 15915 | VL2 | 3531 |
| 20 | Yes | E. coli | Amoxacilli | Wall | F101 | Quinacrine | 17825 | VL2 | 3453 |
| 20 | Yes | E. coli | Amoxacilli | Wall | F138 | Apigenin | 18315 | VL2 | 3436 |
| 20 | Yes | E. coli | Amoxacilli | Wall | F65 | H2FDA | 17429 | BL1 | 4665 |
| 20 | Yes | E. coli | Amoxacilli | Wall | F142 | Chlorophe | 16698 | BL3 | 6219 |
| 20 | Yes | E. coli | Amoxacilli | Wall | F39 | Congo Red | 17531 | BL3 | 7336 |
| 20 | Yes | E. coli | Amoxacilli | Wall | F116 | Sunset Yel | 16577 | BL2 | 5350 |
| 20 | Yes | E. coli | Amoxacilli | Wall | F10 | Alizarin | 15357 | BL1 | 5167 |
| 20 | Yes | E. coli | Amoxacilli | Wall | F84 | Neutral Re | 15382 | BL1 | 4481 |
| 20 | Yes | E. coli | Amoxacilli | Wall | F99 | Pyronin Y | 17769 | BL1 | 3872 |
| 20 | Yes | E. coli | Amoxacilli | Wall | F108 | Rhodamine | 13970 | BL1 | 3579 |
| 20 | Yes | E. coli | Amoxacilli | Wall | F109 | Riboflavin | 15274 | BL1 | 3592 |
| 20 | Yes | E. coli | Amoxacilli | Wall | F117 | SYBR Gree | 17001 | BL1 | 171259 |
| 20 | Yes | E. coli | Amoxacilli | Wall | F106 | Rhodamine | 18600 | BL1 | 6052 |
| 20 | Yes | E. coli | Amoxacilli | Wall | F73 | Malachite ( | 16772 | BL1 | 4514 |
| 20 | Yes | E. coli | Amoxacilli | Wall | F7 | Acridine or | 15973 | BL1 | 4042 |
| 20 | Yes | E. coli | Amoxacilli | Wall | F192 | 5-Carboxyf | 17371 | BL1 | 13644 |
| 20 | Yes | E. coli | Amoxacilli | Wall | F57 | Eosin Y | 16309 | BL1 | 15981 |
| 20 | Yes | E. coli | Amoxacilli | Wall | F27 | Calcein | 18390 | BL1 | 19694.5 |

| | | | | | | | | | |
|---|---|---|---|---|---|---|---|---|---|
| 20 | Yes | E. coli | Amoxacilli | Wall | F12 | Amaranth | 18129 | BL1 | 18397 |
| 20 | Yes | E. coli | None | None | NA | Autofluore: | 14961 | NA | 6648 |
| 20 | Yes | E. coli | Azithromyc | Intracellul: | F194 | Oxonol | 15453 | RL1 | 5848 |
| 20 | Yes | E. coli | Azithromyc | Intracellul: | F11 | Allura Red | 13902 | BL3 | 6318.5 |
| 20 | Yes | E. coli | Azithromyc | Intracellul: | F54 | DiSC3(5) | 14131 | RL1 | 4327 |
| 20 | Yes | E. coli | Azithromyc | Intracellul: | F124 | Thioflavin T | 15941 | VL2 | 3881 |
| 20 | Yes | E. coli | Azithromyc | Intracellul: | F101 | Quinacrine | 13702 | VL2 | 3815 |
| 20 | Yes | E. coli | Azithromyc | Intracellul: | F138 | Apigenin | 13825 | VL2 | 3971 |
| 20 | Yes | E. coli | Azithromyc | Intracellul: | F65 | H2FDA | 17226 | BL1 | 4734.5 |
| 20 | Yes | E. coli | Azithromyc | Intracellul: | F142 | Chlorophe | 14649 | BL3 | 5999 |
| 20 | Yes | E. coli | Azithromyc | Intracellul: | F39 | Congo Red | 13894 | BL3 | 6240 |
| 20 | Yes | E. coli | Azithromyc | Intracellul: | F116 | Sunset Yel | 14732 | BL2 | 5838.5 |
| 20 | Yes | E. coli | Azithromyc | Intracellul: | F10 | Alizarin | 15471 | BL1 | 5090 |
| 20 | Yes | E. coli | Azithromyc | Intracellul: | F84 | Neutral Re | 13266 | BL1 | 4695.5 |
| 20 | Yes | E. coli | Azithromyc | Intracellul: | F99 | Pyronin Y | 15932 | BL1 | 4161 |
| 20 | Yes | E. coli | Azithromyc | Intracellul: | F108 | Rhodamine | 15683 | BL1 | 3634 |
| 20 | Yes | E. coli | Azithromyc | Intracellul: | F109 | Riboflavin | 15387 | BL1 | 4117 |
| 20 | Yes | E. coli | Azithromyc | Intracellul: | F117 | SYBR Gree | 15187 | BL1 | 75786 |
| 20 | Yes | E. coli | Azithromyc | Intracellul: | F106 | Rhodamine | 14355 | BL1 | 5699 |
| 20 | Yes | E. coli | Azithromyc | Intracellul: | F73 | Malachite ( | 15309 | BL1 | 4238 |
| 20 | Yes | E. coli | Azithromyc | Intracellul: | F7 | Acridine or | 14285 | BL1 | 5404 |
| 20 | Yes | E. coli | Azithromyc | Intracellul: | F192 | 5-Carboxyf | 13578 | BL1 | 12848.5 |
| 20 | Yes | E. coli | Azithromyc | Intracellul: | F57 | Eosin Y | 15222 | BL1 | 16153 |
| 20 | Yes | E. coli | Azithromyc | Intracellul: | F27 | Calcein | 14715 | BL1 | 17417 |
| 20 | Yes | E. coli | Azithromyc | Intracellul: | F12 | Amaranth | 16596 | BL1 | 16573 |
| 20 | Yes | E. coli | None | None | NA | Autofluore: | 13883 | NA | 7606 |
| 20 | Yes | E. coli | Trimethopr | Intracellul: | F194 | Oxonol | 15209 | RL1 | 6450 |
| 20 | Yes | E. coli | Trimethopr | Intracellul: | F11 | Allura Red | 15237 | BL3 | 5979 |
| 20 | Yes | E. coli | Trimethopr | Intracellul: | F54 | DiSC3(5) | 14356 | RL1 | 4457 |
| 20 | Yes | E. coli | Trimethopr | Intracellul: | F124 | Thioflavin T | 16807 | VL2 | 4184 |
| 20 | Yes | E. coli | Trimethopr | Intracellul: | F101 | Quinacrine | 16384 | VL2 | 4126.5 |
| 20 | Yes | E. coli | Trimethopr | Intracellul: | F138 | Apigenin | 15138 | VL2 | 3814 |
| 20 | Yes | E. coli | Trimethopr | Intracellul: | F65 | H2FDA | 14564 | BL1 | 4579.5 |
| 20 | Yes | E. coli | Trimethopr | Intracellul: | F142 | Chlorophe | 16939 | BL3 | 5904 |
| 20 | Yes | E. coli | Trimethopr | Intracellul: | F39 | Congo Red | 16925 | BL3 | 6161 |
| 20 | Yes | E. coli | Trimethopr | Intracellul: | F116 | Sunset Yel | 15730 | BL2 | 6687 |
| 20 | Yes | E. coli | Trimethopr | Intracellul: | F10 | Alizarin | 15559 | BL1 | 5291 |
| 20 | Yes | E. coli | Trimethopr | Intracellul: | F84 | Neutral Re | 16247 | BL1 | 5469 |
| 20 | Yes | E. coli | Trimethopr | Intracellul: | F99 | Pyronin Y | 14197 | BL1 | 4988 |
| 20 | Yes | E. coli | Trimethopr | Intracellul: | F108 | Rhodamine | 13539 | BL1 | 4611 |
| 20 | Yes | E. coli | Trimethopr | Intracellul: | F109 | Riboflavin | 16350 | BL1 | 4485.5 |
| 20 | Yes | E. coli | Trimethopr | Intracellul: | F117 | SYBR Gree | 16033 | BL1 | 170959 |
| 20 | Yes | E. coli | Trimethopr | Intracellul: | F106 | Rhodamine | 15908 | BL1 | 6351.5 |
| 20 | Yes | E. coli | Trimethopr | Intracellul: | F73 | Malachite ( | 15689 | BL1 | 4983 |
| 20 | Yes | E. coli | Trimethopr | Intracellul: | F7 | Acridine or | 13621 | BL1 | 5623 |
| 20 | Yes | E. coli | Trimethopr | Intracellul: | F192 | 5-Carboxyf | 16736 | BL1 | 12532.5 |
| 20 | Yes | E. coli | Trimethopr | Intracellul: | F57 | Eosin Y | 15984 | BL1 | 15661.5 |
| 20 | Yes | E. coli | Trimethopr | Intracellul: | F27 | Calcein | 15280 | BL1 | 16822 |

| | | | | | | | | | |
|---|---|---|---|---|---|---|---|---|---|
| 20 | Yes | E. coli | Trimethopr | Intracellula | F12 | Amaranth | 17790 | BL1 | 16716.5 |
| 20 | Yes | E. coli | None | None | NA | Autofluores | 14237 | NA | 7935 |
| 20 | Yes | E. coli | Chloramph | Intracellula | F194 | Oxonol | 17094 | RL1 | 5524 |
| 20 | Yes | E. coli | Chloramph | Intracellula | F11 | Allura Red | 14015 | BL3 | 6123 |
| 20 | Yes | E. coli | Chloramph | Intracellula | F54 | DiSC3(5) | 14488 | RL1 | 5429.5 |
| 20 | Yes | E. coli | Chloramph | Intracellula | F124 | Thioflavin T | 17901 | VL2 | 4665 |
| 20 | Yes | E. coli | Chloramph | Intracellula | F101 | Quinacrine | 18284 | VL2 | 4526 |
| 20 | Yes | E. coli | Chloramph | Intracellula | F138 | Apigenin | 17579 | VL2 | 4389 |
| 20 | Yes | E. coli | Chloramph | Intracellula | F65 | H2FDA | 18144 | BL1 | 4875 |
| 20 | Yes | E. coli | Chloramph | Intracellula | F142 | Chlorophe | 16998 | BL3 | 6106 |
| 20 | Yes | E. coli | Chloramph | Intracellula | F39 | Congo Red | 14684 | BL3 | 7799.5 |
| 20 | Yes | E. coli | Chloramph | Intracellula | F116 | Sunset Yel | 17093 | BL2 | 6609 |
| 20 | Yes | E. coli | Chloramph | Intracellula | F10 | Alizarin | 14873 | BL1 | 6452 |
| 20 | Yes | E. coli | Chloramph | Intracellula | F84 | Neutral Re | 15514 | BL1 | 5121.5 |
| 20 | Yes | E. coli | Chloramph | Intracellula | F99 | Pyronin Y | 14393 | BL1 | 6597 |
| 20 | Yes | E. coli | Chloramph | Intracellula | F108 | Rhodamine | 13886 | BL1 | 4293 |
| 20 | Yes | E. coli | Chloramph | Intracellula | F109 | Riboflavin | 15128 | BL1 | 5753.5 |
| 20 | Yes | E. coli | Chloramph | Intracellula | F117 | SYBR Gree | 15582 | BL1 | 135790.5 |
| 20 | Yes | E. coli | Chloramph | Intracellula | F106 | Rhodamine | 16107 | BL1 | 5591 |
| 20 | Yes | E. coli | Chloramph | Intracellula | F73 | Malachite ( | 14862 | BL1 | 6508 |
| 20 | Yes | E. coli | Chloramph | Intracellula | F7 | Acridine or | 15965 | BL1 | 5023 |
| 20 | Yes | E. coli | Chloramph | Intracellula | F192 | 5-Carboxyf | 15214 | BL1 | 14852.5 |
| 20 | Yes | E. coli | Chloramph | Intracellula | F57 | Eosin Y | 16542 | BL1 | 16327.5 |
| 20 | Yes | E. coli | Chloramph | Intracellula | F27 | Calcein | 15560 | BL1 | 17122.5 |
| 20 | Yes | E. coli | Chloramph | Intracellula | F12 | Amaranth | 17932 | BL1 | 15716.5 |
| 20 | Yes | E. coli | None | None | NA | Autofluores | 14224 | NA | 9491 |
| 20 | Yes | E. coli | Colistin | Wall | F194 | Oxonol | 9238 | RL1 | 82335.5 |
| 20 | Yes | E. coli | Colistin | Wall | F11 | Allura Red | 7845 | BL3 | 59232 |
| 20 | Yes | E. coli | Colistin | Wall | F54 | DiSC3(5) | 8102 | RL1 | 70454 |
| 20 | Yes | E. coli | Colistin | Wall | F124 | Thioflavin T | 7984 | VL2 | 55636.5 |
| 20 | Yes | E. coli | Colistin | Wall | F101 | Quinacrine | 8416 | VL2 | 53000.5 |
| 20 | Yes | E. coli | Colistin | Wall | F138 | Apigenin | 7483 | VL2 | 47905 |
| 20 | Yes | E. coli | Colistin | Wall | F65 | H2FDA | 7097 | BL1 | 47482 |
| 20 | Yes | E. coli | Colistin | Wall | F142 | Chlorophe | 7910 | BL3 | 48214.5 |
| 20 | Yes | E. coli | Colistin | Wall | F39 | Congo Red | 7802 | BL3 | 48771.5 |
| 20 | Yes | E. coli | Colistin | Wall | F116 | Sunset Yel | 7457 | BL2 | 48963 |
| 20 | Yes | E. coli | Colistin | Wall | F10 | Alizarin | 7615 | BL1 | 40464 |
| 20 | Yes | E. coli | Colistin | Wall | F84 | Neutral Re | 7568 | BL1 | 49347.5 |
| 20 | Yes | E. coli | Colistin | Wall | F99 | Pyronin Y | 7876 | BL1 | 36534 |
| 20 | Yes | E. coli | Colistin | Wall | F108 | Rhodamine | 7627 | BL1 | 33699 |
| 20 | Yes | E. coli | Colistin | Wall | F109 | Riboflavin | 7821 | BL1 | 31373 |
| 20 | Yes | E. coli | Colistin | Wall | F117 | SYBR Gree | 7919 | BL1 | 2800471 |
| 20 | Yes | E. coli | Colistin | Wall | F106 | Rhodamine | 7972 | BL1 | 77489 |
| 20 | Yes | E. coli | Colistin | Wall | F73 | Malachite ( | 6509 | BL1 | 66937 |
| 20 | Yes | E. coli | Colistin | Wall | F7 | Acridine or | 7294 | BL1 | 123319 |
| 20 | Yes | E. coli | Colistin | Wall | F192 | 5-Carboxyf | 7462 | BL1 | 67414 |
| 20 | Yes | E. coli | Colistin | Wall | F57 | Eosin Y | 7522 | BL1 | 56031 |
| 20 | Yes | E. coli | Colistin | Wall | F27 | Calcein | 7128 | BL1 | 60286 |

| | | | | | | | | | |
|---|---|---|---|---|---|---|---|---|---|
| 20 | Yes | E. coli | Colistin | Wall | F12 | Amaranth | 7526 | BL1 | 58171 |
| 20 | Yes | E. coli | None | None | NA | Autofluore: | 15475 | NA | 9482 |
| 20 | Yes | E. coli | Gentamici | Intracellula | F194 | Oxonol | 17390 | RL1 | 7359 |
| 20 | Yes | E. coli | Gentamici | Intracellula | F11 | Allura Red | 16491 | BL3 | 6681 |
| 20 | Yes | E. coli | Gentamici | Intracellula | F54 | DiSC3(5) | 13469 | RL1 | 6213 |
| 20 | Yes | E. coli | Gentamici | Intracellula | F124 | Thioflavin T | 18646 | VL2 | 5042.5 |
| 20 | Yes | E. coli | Gentamici | Intracellula | F101 | Quinacrine | 18416 | VL2 | 5046.5 |
| 20 | Yes | E. coli | Gentamici | Intracellula | F138 | Apigenin | 17774 | VL2 | 4393 |
| 20 | Yes | E. coli | Gentamici | Intracellula | F65 | H2FDA | 16968 | BL1 | 5073 |
| 20 | Yes | E. coli | Gentamici | Intracellula | F142 | Chlorophe | 18160 | BL3 | 6230.5 |
| 20 | Yes | E. coli | Gentamici | Intracellula | F39 | Congo Red | 17974 | BL3 | 8555 |
| 20 | Yes | E. coli | Gentamici | Intracellula | F116 | Sunset Yel | 17631 | BL2 | 6503 |
| 20 | Yes | E. coli | Gentamici | Intracellula | F10 | Alizarin | 15680 | BL1 | 6828.5 |
| 20 | Yes | E. coli | Gentamici | Intracellula | F84 | Neutral Re | 16831 | BL1 | 6095 |
| 20 | Yes | E. coli | Gentamici | Intracellula | F99 | Pyronin Y | 16121 | BL1 | 6364 |
| 20 | Yes | E. coli | Gentamici | Intracellula | F108 | Rhodamine | 14791 | BL1 | 4947 |
| 20 | Yes | E. coli | Gentamici | Intracellula | F109 | Riboflavin | 18074 | BL1 | 5225 |
| 20 | Yes | E. coli | Gentamici | Intracellula | F117 | SYBR Gree | 16003 | BL1 | 214692 |
| 20 | Yes | E. coli | Gentamici | Intracellula | F106 | Rhodamine | 16737 | BL1 | 7506 |
| 20 | Yes | E. coli | Gentamici | Intracellula | F73 | Malachite ( | 16721 | BL1 | 5916 |
| 20 | Yes | E. coli | Gentamici | Intracellula | F7 | Acridine or | 18181 | BL1 | 6091 |
| 20 | Yes | E. coli | Gentamici | Intracellula | F192 | 5-Carboxyf | 17269 | BL1 | 14275 |
| 20 | Yes | E. coli | Gentamici | Intracellula | F57 | Eosin Y | 16420 | BL1 | 15352 |
| 20 | Yes | E. coli | Gentamici | Intracellula | F27 | Calcein | 19237 | BL1 | 18021 |
| 20 | Yes | E. coli | Gentamici | Intracellula | F12 | Amaranth | 18245 | BL1 | 18550 |
| 20 | Yes | E. coli | None | None | NA | Autofluore: | 15424 | NA | 8722.5 |
| 20 | Yes | E. coli | Nitrofurant | Intracellula | F194 | Oxonol | 14383 | RL1 | 6932 |
| 20 | Yes | E. coli | Nitrofurant | Intracellula | F11 | Allura Red | 12404 | BL3 | 6691.5 |
| 20 | Yes | E. coli | Nitrofurant | Intracellula | F54 | DiSC3(5) | 13881 | RL1 | 6240 |
| 20 | Yes | E. coli | Nitrofurant | Intracellula | F124 | Thioflavin T | 13717 | VL2 | 4908 |
| 20 | Yes | E. coli | Nitrofurant | Intracellula | F101 | Quinacrine | 14441 | VL2 | 4928 |
| 20 | Yes | E. coli | Nitrofurant | Intracellula | F138 | Apigenin | 16429 | VL2 | 4428 |
| 20 | Yes | E. coli | Nitrofurant | Intracellula | F65 | H2FDA | 13785 | BL1 | 5398 |
| 20 | Yes | E. coli | Nitrofurant | Intracellula | F142 | Chlorophe | 14945 | BL3 | 6173 |
| 20 | Yes | E. coli | Nitrofurant | Intracellula | F39 | Congo Red | 14764 | BL3 | 8952.5 |
| 20 | Yes | E. coli | Nitrofurant | Intracellula | F116 | Sunset Yel | 12737 | BL2 | 7904 |
| 20 | Yes | E. coli | Nitrofurant | Intracellula | F10 | Alizarin | 15391 | BL1 | 5506 |
| 20 | Yes | E. coli | Nitrofurant | Intracellula | F84 | Neutral Re | 14517 | BL1 | 5856 |
| 20 | Yes | E. coli | Nitrofurant | Intracellula | F99 | Pyronin Y | 13580 | BL1 | 6431 |
| 20 | Yes | E. coli | Nitrofurant | Intracellula | F108 | Rhodamine | 13811 | BL1 | 5132 |
| 20 | Yes | E. coli | Nitrofurant | Intracellula | F109 | Riboflavin | 13840 | BL1 | 6721.5 |
| 20 | Yes | E. coli | Nitrofurant | Intracellula | F117 | SYBR Gree | 13256 | BL1 | 110338 |
| 20 | Yes | E. coli | Nitrofurant | Intracellula | F106 | Rhodamine | 13998 | BL1 | 7425.5 |
| 20 | Yes | E. coli | Nitrofurant | Intracellula | F73 | Malachite ( | 12728 | BL1 | 6952 |
| 20 | Yes | E. coli | Nitrofurant | Intracellula | F7 | Acridine or | 14303 | BL1 | 6203 |
| 20 | Yes | E. coli | Nitrofurant | Intracellula | F192 | 5-Carboxyf | 14203 | BL1 | 13534 |
| 20 | Yes | E. coli | Nitrofurant | Intracellula | F57 | Eosin Y | 12763 | BL1 | 14891 |
| 20 | Yes | E. coli | Nitrofurant | Intracellula | F27 | Calcein | 13766 | BL1 | 16295 |

| | | | | | | | | | |
|---|---|---|---|---|---|---|---|---|---|
| 20 | Yes | E. coli | Nitrofurant | Intracellula | F12 | Amaranth | 15035 | BL1 | 16923 |
| 20 | Yes | E. coli | None | None | NA | Autofluore: | 13994 | NA | 9994.5 |
| 20 | Yes | E. coli | Pentamidi | Wall | F194 | Oxonol | 12922 | RL1 | 10182 |
| 20 | Yes | E. coli | Pentamidi | Wall | F11 | Allura Red | 13500 | BL3 | 6982 |
| 20 | Yes | E. coli | Pentamidi | Wall | F54 | DiSC3(5) | 12604 | RL1 | 7468 |
| 20 | Yes | E. coli | Pentamidi | Wall | F124 | Thioflavin T | 13527 | VL2 | 8121 |
| 20 | Yes | E. coli | Pentamidi | Wall | F101 | Quinacrine | 13226 | VL2 | 6097 |
| 20 | Yes | E. coli | Pentamidi | Wall | F138 | Apigenin | 13278 | VL2 | 6719.5 |
| 20 | Yes | E. coli | Pentamidi | Wall | F65 | H2FDA | 13661 | BL1 | 7839 |
| 20 | Yes | E. coli | Pentamidi | Wall | F142 | Chlorophe | 15278 | BL3 | 10588 |
| 20 | Yes | E. coli | Pentamidi | Wall | F39 | Congo Red | 13948 | BL3 | 10929.5 |
| 20 | Yes | E. coli | Pentamidi | Wall | F116 | Sunset Yel | 13494 | BL2 | 11412 |
| 20 | Yes | E. coli | Pentamidi | Wall | F10 | Alizarin | 14574 | BL1 | 10640 |
| 20 | Yes | E. coli | Pentamidi | Wall | F84 | Neutral Re | 12649 | BL1 | 9480 |
| 20 | Yes | E. coli | Pentamidi | Wall | F99 | Pyronin Y | 14428 | BL1 | 10119 |
| 20 | Yes | E. coli | Pentamidi | Wall | F108 | Rhodamine | 15822 | BL1 | 9637 |
| 20 | Yes | E. coli | Pentamidi | Wall | F109 | Riboflavin | 14578 | BL1 | 8117.5 |
| 20 | Yes | E. coli | Pentamidi | Wall | F117 | SYBR Gree | 15427 | BL1 | 728159 |
| 20 | Yes | E. coli | Pentamidi | Wall | F106 | Rhodamine | 14105 | BL1 | 15470 |
| 20 | Yes | E. coli | Pentamidi | Wall | F73 | Malachite ( | 15065 | BL1 | 10357 |
| 20 | Yes | E. coli | Pentamidi | Wall | F7 | Acridine or | 15495 | BL1 | 16621 |
| 20 | Yes | E. coli | Pentamidi | Wall | F192 | 5-Carboxyf | 15149 | BL1 | 20023 |
| 20 | Yes | E. coli | Pentamidi | Wall | F57 | Eosin Y | 15558 | BL1 | 19845 |
| 20 | Yes | E. coli | Pentamidi | Wall | F27 | Calcein | 16217 | BL1 | 22634 |
| 20 | Yes | E. coli | Pentamidi | Wall | F12 | Amaranth | 13846 | BL1 | 18690.5 |
| 20 | Yes | E. coli | None | None | NA | Autofluore: | 14144 | NA | 12684 |
| 20 | Yes | E. coli | Rifampicin | Intracellula | F194 | Oxonol | 15486 | RL1 | 8691 |
| 20 | Yes | E. coli | Rifampicin | Intracellula | F11 | Allura Red | 16197 | BL3 | 6814 |
| 20 | Yes | E. coli | Rifampicin | Intracellula | F54 | DiSC3(5) | 16843 | RL1 | 6919 |
| 20 | Yes | E. coli | Rifampicin | Intracellula | F124 | Thioflavin T | 17171 | VL2 | 6570 |
| 20 | Yes | E. coli | Rifampicin | Intracellula | F101 | Quinacrine | 17034 | VL2 | 5505.5 |
| 20 | Yes | E. coli | Rifampicin | Intracellula | F138 | Apigenin | 18514 | VL2 | 5018 |
| 20 | Yes | E. coli | Rifampicin | Intracellula | F65 | H2FDA | 16533 | BL1 | 5773 |
| 20 | Yes | E. coli | Rifampicin | Intracellula | F142 | Chlorophe | 20353 | BL3 | 6287 |
| 20 | Yes | E. coli | Rifampicin | Intracellula | F39 | Congo Red | 17109 | BL3 | 6845 |
| 20 | Yes | E. coli | Rifampicin | Intracellula | F116 | Sunset Yel | 18562 | BL2 | 5796.5 |
| 20 | Yes | E. coli | Rifampicin | Intracellula | F10 | Alizarin | 16875 | BL1 | 5931 |
| 20 | Yes | E. coli | Rifampicin | Intracellula | F84 | Neutral Re | 17695 | BL1 | 5107 |
| 20 | Yes | E. coli | Rifampicin | Intracellula | F99 | Pyronin Y | 14906 | BL1 | 5301 |
| 20 | Yes | E. coli | Rifampicin | Intracellula | F108 | Rhodamine | 16652 | BL1 | 5120.5 |
| 20 | Yes | E. coli | Rifampicin | Intracellula | F109 | Riboflavin | 15873 | BL1 | 5311 |
| 20 | Yes | E. coli | Rifampicin | Intracellula | F117 | SYBR Gree | 16620 | BL1 | 194508.5 |
| 20 | Yes | E. coli | Rifampicin | Intracellula | F106 | Rhodamine | 17809 | BL1 | 7174 |
| 20 | Yes | E. coli | Rifampicin | Intracellula | F73 | Malachite ( | 16161 | BL1 | 6591 |
| 20 | Yes | E. coli | Rifampicin | Intracellula | F7 | Acridine or | 15128 | BL1 | 7081.5 |
| 20 | Yes | E. coli | Rifampicin | Intracellula | F192 | 5-Carboxyf | 17292 | BL1 | 13471 |
| 20 | Yes | E. coli | Rifampicin | Intracellula | F57 | Eosin Y | 16906 | BL1 | 18219 |
| 20 | Yes | E. coli | Rifampicin | Intracellula | F27 | Calcein | 15453 | BL1 | 16129 |

| | | | | | | | | | |
|---|---|---|---|---|---|---|---|---|---|
| 20 | Yes | E. coli | Rifampicin | Intracellula | F12 | Amaranth | 17220 | BL1 | 16384.5 |
| 20 | Yes | E. coli | None | None | NA | Autofluore: | 13860 | NA | 8807 |
| 20 | Yes | E. coli | Trimethopr | Intracellula | F194 | Oxonol | 17490 | RL1 | 6622.5 |
| 20 | Yes | E. coli | Trimethopr | Intracellula | F11 | Allura Red | 14501 | BL3 | 5508 |
| 20 | Yes | E. coli | Trimethopr | Intracellula | F54 | DiSC3(5) | 15758 | RL1 | 6386.5 |
| 20 | Yes | E. coli | Trimethopr | Intracellula | F124 | Thioflavin T | 18609 | VL2 | 5136 |
| 20 | Yes | E. coli | Trimethopr | Intracellula | F101 | Quinacrine | 17865 | VL2 | 5324 |
| 20 | Yes | E. coli | Trimethopr | Intracellula | F138 | Apigenin | 17856 | VL2 | 4631 |
| 20 | Yes | E. coli | Trimethopr | Intracellula | F65 | H2FDA | 16451 | BL1 | 7200 |
| 20 | Yes | E. coli | Trimethopr | Intracellula | F142 | Chlorophe | 16165 | BL3 | 6247 |
| 20 | Yes | E. coli | Trimethopr | Intracellula | F39 | Congo Red | 15539 | BL3 | 8609 |
| 20 | Yes | E. coli | Trimethopr | Intracellula | F116 | Sunset Yel | 15576 | BL2 | 7002.5 |
| 20 | Yes | E. coli | Trimethopr | Intracellula | F10 | Alizarin | 18062 | BL1 | 5819.5 |
| 20 | Yes | E. coli | Trimethopr | Intracellula | F84 | Neutral Re | 14584 | BL1 | 4940.5 |
| 20 | Yes | E. coli | Trimethopr | Intracellula | F99 | Pyronin Y | 17196 | BL1 | 5623.5 |
| 20 | Yes | E. coli | Trimethopr | Intracellula | F108 | Rhodamine | 13441 | BL1 | 4896 |
| 20 | Yes | E. coli | Trimethopr | Intracellula | F109 | Riboflavin | 15966 | BL1 | 5534 |
| 20 | Yes | E. coli | Trimethopr | Intracellula | F117 | SYBR Gree | 14377 | BL1 | 176584 |
| 20 | Yes | E. coli | Trimethopr | Intracellula | F106 | Rhodamine | 17669 | BL1 | 6089 |
| 20 | Yes | E. coli | Trimethopr | Intracellula | F73 | Malachite ( | 15012 | BL1 | 5598.5 |
| 20 | Yes | E. coli | Trimethopr | Intracellula | F7 | Acridine or | 16215 | BL1 | 5170 |
| 20 | Yes | E. coli | Trimethopr | Intracellula | F192 | 5-Carboxyf | 15817 | BL1 | 12791 |
| 20 | Yes | E. coli | Trimethopr | Intracellula | F57 | Eosin Y | 14239 | BL1 | 15548 |
| 20 | Yes | E. coli | Trimethopr | Intracellula | F27 | Calcein | 14965 | BL1 | 15438 |
| 20 | Yes | E. coli | Trimethopr | Intracellula | F12 | Amaranth | 17638 | BL1 | 18142.5 |
| 20 | Yes | E. coli | None | None | NA | Autofluore: | 7518 | NA | 965 |
| 20 | Yes | E. coli | None | None | F194 | Oxonol | 10498 | RL1 | 1889 |
| 20 | Yes | E. coli | None | None | F11 | Allura Red | 8973 | BL3 | 2442 |
| 20 | Yes | E. coli | None | None | F54 | DiSC3(5) | 8357 | RL1 | 2589 |
| 20 | Yes | E. coli | None | None | F124 | Thioflavin T | 8870 | VL2 | 3019 |
| 20 | Yes | E. coli | None | None | F101 | Quinacrine | 8832 | VL2 | 3082.5 |
| 20 | Yes | E. coli | None | None | F138 | Apigenin | 8004 | VL2 | 3245 |
| 20 | Yes | E. coli | None | None | F65 | H2FDA | 8320 | BL1 | 4512 |
| 20 | Yes | E. coli | None | None | F142 | Chlorophe | 7571 | BL3 | 6284 |
| 20 | Yes | E. coli | None | None | F39 | Congo Red | 8915 | BL3 | 5498 |
| 20 | Yes | E. coli | None | None | F116 | Sunset Yel | 8160 | BL2 | 5262 |
| 20 | Yes | E. coli | None | None | F10 | Alizarin | 7381 | BL1 | 5089 |
| 20 | Yes | E. coli | None | None | F84 | Neutral Re | 8405 | BL1 | 3721 |
| 20 | Yes | E. coli | None | None | F99 | Pyronin Y | 7948 | BL1 | 3531 |
| 20 | Yes | E. coli | None | None | F108 | Rhodamine | 8714 | BL1 | 3048 |
| 20 | Yes | E. coli | None | None | F109 | Riboflavin | 8346 | BL1 | 3433 |
| 20 | Yes | E. coli | None | None | F117 | SYBR Gree | 7513 | BL1 | 163907 |
| 20 | Yes | E. coli | None | None | F106 | Rhodamine | 8657 | BL1 | 4444 |
| 20 | Yes | E. coli | None | None | F73 | Malachite ( | 8606 | BL1 | 3693 |
| 20 | Yes | E. coli | None | None | F7 | Acridine or | 8232 | BL1 | 3646 |
| 20 | Yes | E. coli | None | None | F192 | 5-Carboxyf | 8347 | BL1 | 12775 |
| 20 | Yes | E. coli | None | None | F57 | Eosin Y | 8283 | BL1 | 16884 |
| 20 | Yes | E. coli | None | None | F27 | Calcein | 8416 | BL1 | 18370.5 |

| | | | | | | | | | |
|---|---|---|---|---|---|---|---|---|---|
| 20 | Yes | E. coli | None | None | F12 | Amaranth | 8215 | BL1 | 17718 |
| 20 | Yes | E. coli | None | None | NA | Autofluore: | 7668 | NA | 7551 |
| 20 | Yes | E. coli | Amoxacillii | Wall | F194 | Oxonol | 8463 | RL1 | 5803 |
| 20 | Yes | E. coli | Amoxacillii | Wall | F11 | Allura Red | 7836 | BL3 | 4617.5 |
| 20 | Yes | E. coli | Amoxacillii | Wall | F54 | DiSC3(5) | 7925 | RL1 | 3884 |
| 20 | Yes | E. coli | Amoxacillii | Wall | F124 | Thioflavin T | 8560 | VL2 | 3702.5 |
| 20 | Yes | E. coli | Amoxacillii | Wall | F101 | Quinacrine | 7883 | VL2 | 3626 |
| 20 | Yes | E. coli | Amoxacillii | Wall | F138 | Apigenin | 7297 | VL2 | 3769 |
| 20 | Yes | E. coli | Amoxacillii | Wall | F65 | H2FDA | 8117 | BL1 | 5612 |
| 20 | Yes | E. coli | Amoxacillii | Wall | F142 | Chlorophe | 7970 | BL3 | 6545 |
| 20 | Yes | E. coli | Amoxacillii | Wall | F39 | Congo Red | 7613 | BL3 | 7097 |
| 20 | Yes | E. coli | Amoxacillii | Wall | F116 | Sunset Yel | 8182 | BL2 | 5852 |
| 20 | Yes | E. coli | Amoxacillii | Wall | F10 | Alizarin | 7740 | BL1 | 5748.5 |
| 20 | Yes | E. coli | Amoxacillii | Wall | F84 | Neutral Re | 7379 | BL1 | 4360 |
| 20 | Yes | E. coli | Amoxacillii | Wall | F99 | Pyronin Y | 7872 | BL1 | 3575.5 |
| 20 | Yes | E. coli | Amoxacillii | Wall | F108 | Rhodamine | 7223 | BL1 | 3494 |
| 20 | Yes | E. coli | Amoxacillii | Wall | F109 | Riboflavin | 8426 | BL1 | 3676 |
| 20 | Yes | E. coli | Amoxacillii | Wall | F117 | SYBR Gree | 7944 | BL1 | 181563.5 |
| 20 | Yes | E. coli | Amoxacillii | Wall | F106 | Rhodamine | 7027 | BL1 | 4113 |
| 20 | Yes | E. coli | Amoxacillii | Wall | F73 | Malachite ( | 8086 | BL1 | 4417.5 |
| 20 | Yes | E. coli | Amoxacillii | Wall | F7 | Acridine or | 8030 | BL1 | 4059.5 |
| 20 | Yes | E. coli | Amoxacillii | Wall | F192 | 5-Carboxyf | 7814 | BL1 | 13087 |
| 20 | Yes | E. coli | Amoxacillii | Wall | F57 | Eosin Y | 7620 | BL1 | 14648.5 |
| 20 | Yes | E. coli | Amoxacillii | Wall | F27 | Calcein | 7998 | BL1 | 16543.5 |
| 20 | Yes | E. coli | Amoxacillii | Wall | F12 | Amaranth | 8470 | BL1 | 17976 |
| 20 | Yes | E. coli | None | None | NA | Autofluore: | 7431 | NA | 7256 |
| 20 | Yes | E. coli | Azithromyc | Intracellula | F194 | Oxonol | 6909 | RL1 | 5394 |
| 20 | Yes | E. coli | Azithromyc | Intracellula | F11 | Allura Red | 6172 | BL3 | 4958 |
| 20 | Yes | E. coli | Azithromyc | Intracellula | F54 | DiSC3(5) | 6196 | RL1 | 4293 |
| 20 | Yes | E. coli | Azithromyc | Intracellula | F124 | Thioflavin T | 6709 | VL2 | 4188 |
| 20 | Yes | E. coli | Azithromyc | Intracellula | F101 | Quinacrine | 6264 | VL2 | 4241 |
| 20 | Yes | E. coli | Azithromyc | Intracellula | F138 | Apigenin | 6135 | VL2 | 4614 |
| 20 | Yes | E. coli | Azithromyc | Intracellula | F65 | H2FDA | 6410 | BL1 | 6054 |
| 20 | Yes | E. coli | Azithromyc | Intracellula | F142 | Chlorophe | 6029 | BL3 | 7624 |
| 20 | Yes | E. coli | Azithromyc | Intracellula | F39 | Congo Red | 6013 | BL3 | 6176 |
| 20 | Yes | E. coli | Azithromyc | Intracellula | F116 | Sunset Yel | 6387 | BL2 | 5920 |
| 20 | Yes | E. coli | Azithromyc | Intracellula | F10 | Alizarin | 6204 | BL1 | 5388 |
| 20 | Yes | E. coli | Azithromyc | Intracellula | F84 | Neutral Re | 6316 | BL1 | 4519.5 |
| 20 | Yes | E. coli | Azithromyc | Intracellula | F99 | Pyronin Y | 5428 | BL1 | 3938 |
| 20 | Yes | E. coli | Azithromyc | Intracellula | F108 | Rhodamine | 6612 | BL1 | 3852.5 |
| 20 | Yes | E. coli | Azithromyc | Intracellula | F109 | Riboflavin | 6296 | BL1 | 4070 |
| 20 | Yes | E. coli | Azithromyc | Intracellula | F117 | SYBR Gree | 5644 | BL1 | 54559.5 |
| 20 | Yes | E. coli | Azithromyc | Intracellula | F106 | Rhodamine | 6474 | BL1 | 4879 |
| 20 | Yes | E. coli | Azithromyc | Intracellula | F73 | Malachite ( | 5881 | BL1 | 5011 |
| 20 | Yes | E. coli | Azithromyc | Intracellula | F7 | Acridine or | 6247 | BL1 | 4224 |
| 20 | Yes | E. coli | Azithromyc | Intracellula | F192 | 5-Carboxyf | 6027 | BL1 | 12551 |
| 20 | Yes | E. coli | Azithromyc | Intracellula | F57 | Eosin Y | 6101 | BL1 | 17651 |
| 20 | Yes | E. coli | Azithromyc | Intracellula | F27 | Calcein | 5992 | BL1 | 19104.5 |

| | | | | | | | | | |
|---|---|---|---|---|---|---|---|---|---|
| 20 | Yes | E. coli | Azithromyc | Intracellula | F12 | Amaranth | 7927 | BL1 | 17568 |
| 20 | Yes | E. coli | None | None | NA | Autofluore: | 7620 | NA | 7887.5 |
| 20 | Yes | E. coli | Trimethopr | Intracellula | F194 | Oxonol | 8196 | RL1 | 6336.5 |
| 20 | Yes | E. coli | Trimethopr | Intracellula | F11 | Allura Red | 7630 | BL3 | 5025 |
| 20 | Yes | E. coli | Trimethopr | Intracellula | F54 | DiSC3(5) | 7405 | RL1 | 4282 |
| 20 | Yes | E. coli | Trimethopr | Intracellula | F124 | Thioflavin T | 8286 | VL2 | 4205 |
| 20 | Yes | E. coli | Trimethopr | Intracellula | F101 | Quinacrine | 8291 | VL2 | 4195 |
| 20 | Yes | E. coli | Trimethopr | Intracellula | F138 | Apigenin | 7366 | VL2 | 4689 |
| 20 | Yes | E. coli | Trimethopr | Intracellula | F65 | H2FDA | 8100 | BL1 | 5747 |
| 20 | Yes | E. coli | Trimethopr | Intracellula | F142 | Chlorophe | 7869 | BL3 | 8113 |
| 20 | Yes | E. coli | Trimethopr | Intracellula | F39 | Congo Red | 7928 | BL3 | 6777 |
| 20 | Yes | E. coli | Trimethopr | Intracellula | F116 | Sunset Yel | 8007 | BL2 | 6844 |
| 20 | Yes | E. coli | Trimethopr | Intracellula | F10 | Alizarin | 8163 | BL1 | 5129 |
| 20 | Yes | E. coli | Trimethopr | Intracellula | F84 | Neutral Re | 7642 | BL1 | 4878.5 |
| 20 | Yes | E. coli | Trimethopr | Intracellula | F99 | Pyronin Y | 8615 | BL1 | 4031 |
| 20 | Yes | E. coli | Trimethopr | Intracellula | F108 | Rhodamine | 8069 | BL1 | 3594 |
| 20 | Yes | E. coli | Trimethopr | Intracellula | F109 | Riboflavin | 7021 | BL1 | 4698 |
| 20 | Yes | E. coli | Trimethopr | Intracellula | F117 | SYBR Gree | 7921 | BL1 | 171211 |
| 20 | Yes | E. coli | Trimethopr | Intracellula | F106 | Rhodamine | 7930 | BL1 | 5684.5 |
| 20 | Yes | E. coli | Trimethopr | Intracellula | F73 | Malachite ( | 8178 | BL1 | 4554.5 |
| 20 | Yes | E. coli | Trimethopr | Intracellula | F7 | Acridine or | 8160 | BL1 | 4609.5 |
| 20 | Yes | E. coli | Trimethopr | Intracellula | F192 | 5-Carboxyf | 8056 | BL1 | 11704 |
| 20 | Yes | E. coli | Trimethopr | Intracellula | F57 | Eosin Y | 8027 | BL1 | 16436 |
| 20 | Yes | E. coli | Trimethopr | Intracellula | F27 | Calcein | 7764 | BL1 | 16407.5 |
| 20 | Yes | E. coli | Trimethopr | Intracellula | F12 | Amaranth | 8252 | BL1 | 18337.5 |
| 20 | Yes | E. coli | None | None | NA | Autofluore: | 7222 | NA | 7167 |
| 20 | Yes | E. coli | Chloramph | Intracellula | F194 | Oxonol | 8692 | RL1 | 6271 |
| 20 | Yes | E. coli | Chloramph | Intracellula | F11 | Allura Red | 8017 | BL3 | 5935 |
| 20 | Yes | E. coli | Chloramph | Intracellula | F54 | DiSC3(5) | 7737 | RL1 | 4831 |
| 20 | Yes | E. coli | Chloramph | Intracellula | F124 | Thioflavin T | 8814 | VL2 | 4258 |
| 20 | Yes | E. coli | Chloramph | Intracellula | F101 | Quinacrine | 8028 | VL2 | 4579.5 |
| 20 | Yes | E. coli | Chloramph | Intracellula | F138 | Apigenin | 8430 | VL2 | 4589 |
| 20 | Yes | E. coli | Chloramph | Intracellula | F65 | H2FDA | 7775 | BL1 | 6632 |
| 20 | Yes | E. coli | Chloramph | Intracellula | F142 | Chlorophe | 8247 | BL3 | 7530 |
| 20 | Yes | E. coli | Chloramph | Intracellula | F39 | Congo Red | 7834 | BL3 | 8343.5 |
| 20 | Yes | E. coli | Chloramph | Intracellula | F116 | Sunset Yel | 8208 | BL2 | 6549 |
| 20 | Yes | E. coli | Chloramph | Intracellula | F10 | Alizarin | 8340 | BL1 | 6135 |
| 20 | Yes | E. coli | Chloramph | Intracellula | F84 | Neutral Re | 7366 | BL1 | 4695 |
| 20 | Yes | E. coli | Chloramph | Intracellula | F99 | Pyronin Y | 8601 | BL1 | 4167 |
| 20 | Yes | E. coli | Chloramph | Intracellula | F108 | Rhodamine | 6379 | BL1 | 3879 |
| 20 | Yes | E. coli | Chloramph | Intracellula | F109 | Riboflavin | 7017 | BL1 | 4995 |
| 20 | Yes | E. coli | Chloramph | Intracellula | F117 | SYBR Gree | 6219 | BL1 | 105541 |
| 20 | Yes | E. coli | Chloramph | Intracellula | F106 | Rhodamine | 7630 | BL1 | 5838.5 |
| 20 | Yes | E. coli | Chloramph | Intracellula | F73 | Malachite ( | 8111 | BL1 | 4800 |
| 20 | Yes | E. coli | Chloramph | Intracellula | F7 | Acridine or | 7827 | BL1 | 5230 |
| 20 | Yes | E. coli | Chloramph | Intracellula | F192 | 5-Carboxyf | 8152 | BL1 | 12376 |
| 20 | Yes | E. coli | Chloramph | Intracellula | F57 | Eosin Y | 7674 | BL1 | 15789.5 |
| 20 | Yes | E. coli | Chloramph | Intracellula | F27 | Calcein | 7667 | BL1 | 13972 |

| | | | | | | | | | |
|---|---|---|---|---|---|---|---|---|---|
| 20 | Yes | E. coli | Chloramph | Intracellula | F12 | Amaranth | 8603 | BL1 | 15434 |
| 20 | Yes | E. coli | None | None | NA | Autofluore: | 7725 | NA | 7955 |
| 20 | Yes | E. coli | Colistin | Wall | F194 | Oxonol | 8558 | RL1 | 6076 |
| 20 | Yes | E. coli | Colistin | Wall | F11 | Allura Red | 8024 | BL3 | 5698.5 |
| 20 | Yes | E. coli | Colistin | Wall | F54 | DiSC3(5) | 7395 | RL1 | 4859 |
| 20 | Yes | E. coli | Colistin | Wall | F124 | Thioflavin T | 8365 | VL2 | 4607 |
| 20 | Yes | E. coli | Colistin | Wall | F101 | Quinacrine | 7921 | VL2 | 4917 |
| 20 | Yes | E. coli | Colistin | Wall | F138 | Apigenin | 8129 | VL2 | 5276 |
| 20 | Yes | E. coli | Colistin | Wall | F65 | H2FDA | 8254 | BL1 | 6261.5 |
| 20 | Yes | E. coli | Colistin | Wall | F142 | Chlorophe | 7287 | BL3 | 7161 |
| 20 | Yes | E. coli | Colistin | Wall | F39 | Congo Red | 7951 | BL3 | 6788 |
| 20 | Yes | E. coli | Colistin | Wall | F116 | Sunset Yel | 8342 | BL2 | 6247 |
| 20 | Yes | E. coli | Colistin | Wall | F10 | Alizarin | 7992 | BL1 | 5918.5 |
| 20 | Yes | E. coli | Colistin | Wall | F84 | Neutral Re | 7701 | BL1 | 4689 |
| 20 | Yes | E. coli | Colistin | Wall | F99 | Pyronin Y | 7343 | BL1 | 4584 |
| 20 | Yes | E. coli | Colistin | Wall | F108 | Rhodamine | 8165 | BL1 | 3917 |
| 20 | Yes | E. coli | Colistin | Wall | F109 | Riboflavin | 8209 | BL1 | 4529 |
| 20 | Yes | E. coli | Colistin | Wall | F117 | SYBR Gree | 8113 | BL1 | 200756 |
| 20 | Yes | E. coli | Colistin | Wall | F106 | Rhodamine | 8396 | BL1 | 5185.5 |
| 20 | Yes | E. coli | Colistin | Wall | F73 | Malachite ( | 8322 | BL1 | 4786 |
| 20 | Yes | E. coli | Colistin | Wall | F7 | Acridine or | 8187 | BL1 | 5059 |
| 20 | Yes | E. coli | Colistin | Wall | F192 | 5-Carboxyf | 7767 | BL1 | 13815 |
| 20 | Yes | E. coli | Colistin | Wall | F57 | Eosin Y | 8466 | BL1 | 16539.5 |
| 20 | Yes | E. coli | Colistin | Wall | F27 | Calcein | 7307 | BL1 | 17377 |
| 20 | Yes | E. coli | Colistin | Wall | F12 | Amaranth | 9468 | BL1 | 17880 |
| 20 | Yes | E. coli | None | None | NA | Autofluore: | 7402 | NA | 7706 |
| 20 | Yes | E. coli | Gentamici | Intracellula | F194 | Oxonol | 8421 | RL1 | 6154 |
| 20 | Yes | E. coli | Gentamici | Intracellula | F11 | Allura Red | 7821 | BL3 | 5685 |
| 20 | Yes | E. coli | Gentamici | Intracellula | F54 | DiSC3(5) | 7306 | RL1 | 5102 |
| 20 | Yes | E. coli | Gentamici | Intracellula | F124 | Thioflavin T | 8596 | VL2 | 5046 |
| 20 | Yes | E. coli | Gentamici | Intracellula | F101 | Quinacrine | 8687 | VL2 | 4326 |
| 20 | Yes | E. coli | Gentamici | Intracellula | F138 | Apigenin | 8106 | VL2 | 4688 |
| 20 | Yes | E. coli | Gentamici | Intracellula | F65 | H2FDA | 8677 | BL1 | 6135 |
| 20 | Yes | E. coli | Gentamici | Intracellula | F142 | Chlorophe | 6992 | BL3 | 8261.5 |
| 20 | Yes | E. coli | Gentamici | Intracellula | F39 | Congo Red | 8650 | BL3 | 8945.5 |
| 20 | Yes | E. coli | Gentamici | Intracellula | F116 | Sunset Yel | 8227 | BL2 | 8767 |
| 20 | Yes | E. coli | Gentamici | Intracellula | F10 | Alizarin | 7668 | BL1 | 6022 |
| 20 | Yes | E. coli | Gentamici | Intracellula | F84 | Neutral Re | 8193 | BL1 | 6162 |
| 20 | Yes | E. coli | Gentamici | Intracellula | F99 | Pyronin Y | 7579 | BL1 | 5162 |
| 20 | Yes | E. coli | Gentamici | Intracellula | F108 | Rhodamine | 8611 | BL1 | 4228 |
| 20 | Yes | E. coli | Gentamici | Intracellula | F109 | Riboflavin | 8149 | BL1 | 4882 |
| 20 | Yes | E. coli | Gentamici | Intracellula | F117 | SYBR Gree | 8043 | BL1 | 211304 |
| 20 | Yes | E. coli | Gentamici | Intracellula | F106 | Rhodamine | 8113 | BL1 | 5510 |
| 20 | Yes | E. coli | Gentamici | Intracellula | F73 | Malachite ( | 8313 | BL1 | 5213 |
| 20 | Yes | E. coli | Gentamici | Intracellula | F7 | Acridine or | 7946 | BL1 | 4721.5 |
| 20 | Yes | E. coli | Gentamici | Intracellula | F192 | 5-Carboxyf | 8349 | BL1 | 13380 |
| 20 | Yes | E. coli | Gentamici | Intracellula | F57 | Eosin Y | 8142 | BL1 | 16319 |
| 20 | Yes | E. coli | Gentamici | Intracellula | F27 | Calcein | 8358 | BL1 | 18484 |

| | | | | | | | | | |
|---|---|---|---|---|---|---|---|---|---|
| 20 | Yes | E. coli | Gentamici | Intracellul | F12 | Amaranth | 8891 | BL1 | 17762 |
| 20 | Yes | E. coli | None | None | NA | Autofluore | 7446 | NA | 8045 |
| 20 | Yes | E. coli | Nitrofurant | Intracellul | F194 | Oxonol | 5821 | RL1 | 6563 |
| 20 | Yes | E. coli | Nitrofurant | Intracellul | F11 | Allura Red | 5541 | BL3 | 7666 |
| 20 | Yes | E. coli | Nitrofurant | Intracellul | F54 | DiSC3(5) | 5869 | RL1 | 5694 |
| 20 | Yes | E. coli | Nitrofurant | Intracellul | F124 | Thioflavin T | 5893 | VL2 | 4816 |
| 20 | Yes | E. coli | Nitrofurant | Intracellul | F101 | Quinacrine | 6020 | VL2 | 4703.5 |
| 20 | Yes | E. coli | Nitrofurant | Intracellul | F138 | Apigenin | 6196 | VL2 | 4629.5 |
| 20 | Yes | E. coli | Nitrofurant | Intracellul | F65 | H2FDA | 6123 | BL1 | 6035 |
| 20 | Yes | E. coli | Nitrofurant | Intracellul | F142 | Chlorophe | 5722 | BL3 | 8637.5 |
| 20 | Yes | E. coli | Nitrofurant | Intracellul | F39 | Congo Red | 5967 | BL3 | 9598 |
| 20 | Yes | E. coli | Nitrofurant | Intracellul | F116 | Sunset Yel | 6033 | BL2 | 7690 |
| 20 | Yes | E. coli | Nitrofurant | Intracellul | F10 | Alizarin | 5300 | BL1 | 8551.5 |
| 20 | Yes | E. coli | Nitrofurant | Intracellul | F84 | Neutral Re | 6187 | BL1 | 6648 |
| 20 | Yes | E. coli | Nitrofurant | Intracellul | F99 | Pyronin Y | 5827 | BL1 | 5740 |
| 20 | Yes | E. coli | Nitrofurant | Intracellul | F108 | Rhodamine | 5187 | BL1 | 5005 |
| 20 | Yes | E. coli | Nitrofurant | Intracellul | F109 | Riboflavin | 6103 | BL1 | 5499 |
| 20 | Yes | E. coli | Nitrofurant | Intracellul | F117 | SYBR Gree | 7542 | BL1 | 224490.5 |
| 20 | Yes | E. coli | Nitrofurant | Intracellul | F106 | Rhodamine | 7992 | BL1 | 7760 |
| 20 | Yes | E. coli | Nitrofurant | Intracellul | F73 | Malachite | 7880 | BL1 | 5062.5 |
| 20 | Yes | E. coli | Nitrofurant | Intracellul | F7 | Acridine or | 7040 | BL1 | 5217 |
| 20 | Yes | E. coli | Nitrofurant | Intracellul | F192 | 5-Carboxyf | 7932 | BL1 | 10385.5 |
| 20 | Yes | E. coli | Nitrofurant | Intracellul | F57 | Eosin Y | 8461 | BL1 | 17176 |
| 20 | Yes | E. coli | Nitrofurant | Intracellul | F27 | Calcein | 7549 | BL1 | 16706 |
| 20 | Yes | E. coli | Nitrofurant | Intracellul | F12 | Amaranth | 8260 | BL1 | 13963 |
| 20 | Yes | E. coli | None | None | NA | Autofluore | 6880 | NA | 10883.5 |
| 20 | Yes | E. coli | Pentamidi | Wall | F194 | Oxonol | 4840 | RL1 | 10495 |
| 20 | Yes | E. coli | Pentamidi | Wall | F11 | Allura Red | 4867 | BL3 | 8037 |
| 20 | Yes | E. coli | Pentamidi | Wall | F54 | DiSC3(5) | 5396 | RL1 | 6008 |
| 20 | Yes | E. coli | Pentamidi | Wall | F124 | Thioflavin T | 5866 | VL2 | 5973 |
| 20 | Yes | E. coli | Pentamidi | Wall | F101 | Quinacrine | 5573 | VL2 | 5283 |
| 20 | Yes | E. coli | Pentamidi | Wall | F138 | Apigenin | 5407 | VL2 | 6506 |
| 20 | Yes | E. coli | Pentamidi | Wall | F65 | H2FDA | 5738 | BL1 | 7296.5 |
| 20 | Yes | E. coli | Pentamidi | Wall | F142 | Chlorophe | 5418 | BL3 | 9371.5 |
| 20 | Yes | E. coli | Pentamidi | Wall | F39 | Congo Red | 5205 | BL3 | 11065 |
| 20 | Yes | E. coli | Pentamidi | Wall | F116 | Sunset Yel | 5444 | BL2 | 10926 |
| 20 | Yes | E. coli | Pentamidi | Wall | F10 | Alizarin | 5351 | BL1 | 11217 |
| 20 | Yes | E. coli | Pentamidi | Wall | F84 | Neutral Re | 5229 | BL1 | 10884 |
| 20 | Yes | E. coli | Pentamidi | Wall | F99 | Pyronin Y | 6419 | BL1 | 10989 |
| 20 | Yes | E. coli | Pentamidi | Wall | F108 | Rhodamine | 6823 | BL1 | 10801 |
| 20 | Yes | E. coli | Pentamidi | Wall | F109 | Riboflavin | 6741 | BL1 | 10162 |
| 20 | Yes | E. coli | Pentamidi | Wall | F117 | SYBR Gree | 7999 | BL1 | 558735 |
| 20 | Yes | E. coli | Pentamidi | Wall | F106 | Rhodamine | 6601 | BL1 | 10883 |
| 20 | Yes | E. coli | Pentamidi | Wall | F73 | Malachite | 8404 | BL1 | 8545.5 |
| 20 | Yes | E. coli | Pentamidi | Wall | F7 | Acridine or | 8272 | BL1 | 8855.5 |
| 20 | Yes | E. coli | Pentamidi | Wall | F192 | 5-Carboxyf | 7986 | BL1 | 13862.5 |
| 20 | Yes | E. coli | Pentamidi | Wall | F57 | Eosin Y | 8292 | BL1 | 16969.5 |
| 20 | Yes | E. coli | Pentamidi | Wall | F27 | Calcein | 8190 | BL1 | 17882 |

| | | | | | | | | | |
|---|---|---|---|---|---|---|---|---|---|
| 20 | Yes | E. coli | Pentamidi | Wall | F12 | Amaranth | 8787 | BL1 | 17858 |
| 20 | Yes | E. coli | None | None | NA | Autofluore: | 7039 | NA | 12444 |
| 20 | Yes | E. coli | Rifampicin | Intracellula | F194 | Oxonol | 7627 | RL1 | 9479 |
| 20 | Yes | E. coli | Rifampicin | Intracellula | F11 | Allura Red | 6993 | BL3 | 7755 |
| 20 | Yes | E. coli | Rifampicin | Intracellula | F54 | DiSC3(5) | 6158 | RL1 | 8274 |
| 20 | Yes | E. coli | Rifampicin | Intracellula | F124 | Thioflavin T | 8873 | VL2 | 5386 |
| 20 | Yes | E. coli | Rifampicin | Intracellula | F101 | Quinacrine | 7906 | VL2 | 5724.5 |
| 20 | Yes | E. coli | Rifampicin | Intracellula | F138 | Apigenin | 7877 | VL2 | 5122 |
| 20 | Yes | E. coli | Rifampicin | Intracellula | F65 | H2FDA | 8292 | BL1 | 5876.5 |
| 20 | Yes | E. coli | Rifampicin | Intracellula | F142 | Chlorophe | 8179 | BL3 | 6251 |
| 20 | Yes | E. coli | Rifampicin | Intracellula | F39 | Congo Red | 7611 | BL3 | 8017 |
| 20 | Yes | E. coli | Rifampicin | Intracellula | F116 | Sunset Yel | 7929 | BL2 | 7268 |
| 20 | Yes | E. coli | Rifampicin | Intracellula | F10 | Alizarin | 7198 | BL1 | 6235.5 |
| 20 | Yes | E. coli | Rifampicin | Intracellula | F84 | Neutral Re | 7755 | BL1 | 5895 |
| 20 | Yes | E. coli | Rifampicin | Intracellula | F99 | Pyronin Y | 7216 | BL1 | 6984 |
| 20 | Yes | E. coli | Rifampicin | Intracellula | F108 | Rhodamine | 7879 | BL1 | 5090 |
| 20 | Yes | E. coli | Rifampicin | Intracellula | F109 | Riboflavin | 7780 | BL1 | 6717 |
| 20 | Yes | E. coli | Rifampicin | Intracellula | F117 | SYBR Gree | 7032 | BL1 | 220497 |
| 20 | Yes | E. coli | Rifampicin | Intracellula | F106 | Rhodamine | 8178 | BL1 | 7258 |
| 20 | Yes | E. coli | Rifampicin | Intracellula | F73 | Malachite ( | 6743 | BL1 | 6117 |
| 20 | Yes | E. coli | Rifampicin | Intracellula | F7 | Acridine or | 8456 | BL1 | 5111 |
| 20 | Yes | E. coli | Rifampicin | Intracellula | F192 | 5-Carboxyf | 8066 | BL1 | 13980 |
| 20 | Yes | E. coli | Rifampicin | Intracellula | F57 | Eosin Y | 8008 | BL1 | 16606 |
| 20 | Yes | E. coli | Rifampicin | Intracellula | F27 | Calcein | 7461 | BL1 | 16218 |
| 20 | Yes | E. coli | Rifampicin | Intracellula | F12 | Amaranth | 8953 | BL1 | 18541 |
| 20 | Yes | E. coli | None | None | NA | Autofluore: | 7195 | NA | 10848 |
| 20 | Yes | E. coli | Trimethopr | Intracellula | F194 | Oxonol | 7813 | RL1 | 8330 |
| 20 | Yes | E. coli | Trimethopr | Intracellula | F11 | Allura Red | 7424 | BL3 | 6821 |
| 20 | Yes | E. coli | Trimethopr | Intracellula | F54 | DiSC3(5) | 7683 | RL1 | 6250 |
| 20 | Yes | E. coli | Trimethopr | Intracellula | F124 | Thioflavin T | 8541 | VL2 | 5374 |
| 20 | Yes | E. coli | Trimethopr | Intracellula | F101 | Quinacrine | 8556 | VL2 | 5070 |
| 20 | Yes | E. coli | Trimethopr | Intracellula | F138 | Apigenin | 7902 | VL2 | 5434.5 |
| 20 | Yes | E. coli | Trimethopr | Intracellula | F65 | H2FDA | 8438 | BL1 | 5536 |
| 20 | Yes | E. coli | Trimethopr | Intracellula | F142 | Chlorophe | 8140 | BL3 | 7312.5 |
| 20 | Yes | E. coli | Trimethopr | Intracellula | F39 | Congo Red | 8094 | BL3 | 9475 |
| 20 | Yes | E. coli | Trimethopr | Intracellula | F116 | Sunset Yel | 8545 | BL2 | 9001 |
| 20 | Yes | E. coli | Trimethopr | Intracellula | F10 | Alizarin | 8205 | BL1 | 6845 |
| 20 | Yes | E. coli | Trimethopr | Intracellula | F84 | Neutral Re | 8179 | BL1 | 5088 |
| 20 | Yes | E. coli | Trimethopr | Intracellula | F99 | Pyronin Y | 8907 | BL1 | 5057 |
| 20 | Yes | E. coli | Trimethopr | Intracellula | F108 | Rhodamine | 8483 | BL1 | 4551 |
| 20 | Yes | E. coli | Trimethopr | Intracellula | F109 | Riboflavin | 7394 | BL1 | 5348 |
| 20 | Yes | E. coli | Trimethopr | Intracellula | F117 | SYBR Gree | 7532 | BL1 | 231387 |
| 20 | Yes | E. coli | Trimethopr | Intracellula | F106 | Rhodamine | 7728 | BL1 | 6731 |
| 20 | Yes | E. coli | Trimethopr | Intracellula | F73 | Malachite ( | 7312 | BL1 | 5541.5 |
| 20 | Yes | E. coli | Trimethopr | Intracellula | F7 | Acridine or | 7088 | BL1 | 5224 |
| 20 | Yes | E. coli | Trimethopr | Intracellula | F192 | 5-Carboxyf | 7816 | BL1 | 12971 |
| 20 | Yes | E. coli | Trimethopr | Intracellula | F57 | Eosin Y | 7659 | BL1 | 17313 |
| 20 | Yes | E. coli | Trimethopr | Intracellula | F27 | Calcein | 8280 | BL1 | 17157 |

| | | | | | | | | | |
|---|---|---|---|---|---|---|---|---|---|
| 20 | Yes | E. coli | Trimethopr | Intracellula | F12 | Amaranth | 8076 | BL1 | 13718 |
| 20 | Yes | E. coli | None | None | NA | Autofluore: | 10115 | NA | 1053 |
| 20 | Yes | E. coli | None | None | F194 | Oxonol | 12235 | RL1 | 1878 |
| 20 | Yes | E. coli | None | None | F11 | Allura Red | 10536 | BL3 | 2574 |
| 20 | Yes | E. coli | None | None | F54 | DiSC3(5) | 10137 | RL1 | 2909 |
| 20 | Yes | E. coli | None | None | F124 | Thioflavin T | 11011 | VL2 | 2892 |
| 20 | Yes | E. coli | None | None | F101 | Quinacrine | 9955 | VL2 | 2888 |
| 20 | Yes | E. coli | None | None | F138 | Apigenin | 11191 | VL2 | 2740 |
| 20 | Yes | E. coli | None | None | F65 | H2FDA | 10075 | BL1 | 3969 |
| 20 | Yes | E. coli | None | None | F142 | Chlorophe | 9934 | BL3 | 4995 |
| 20 | Yes | E. coli | None | None | F39 | Congo Red | 9906 | BL3 | 6234 |
| 20 | Yes | E. coli | None | None | F116 | Sunset Yel | 10696 | BL2 | 5891.5 |
| 20 | Yes | E. coli | None | None | F10 | Alizarin | 9533 | BL1 | 4887 |
| 20 | Yes | E. coli | None | None | F84 | Neutral Re | 9291 | BL1 | 4068 |
| 20 | Yes | E. coli | None | None | F99 | Pyronin Y | 10298 | BL1 | 3597 |
| 20 | Yes | E. coli | None | None | F108 | Rhodamine | 9646 | BL1 | 3644.5 |
| 20 | Yes | E. coli | None | None | F109 | Riboflavin | 10022 | BL1 | 3976.5 |
| 20 | Yes | E. coli | None | None | F117 | SYBR Gree | 8905 | BL1 | 172481 |
| 20 | Yes | E. coli | None | None | F106 | Rhodamine | 9524 | BL1 | 4177 |
| 20 | Yes | E. coli | None | None | F73 | Malachite ( | 9282 | BL1 | 4540.5 |
| 20 | Yes | E. coli | None | None | F7 | Acridine or | 10511 | BL1 | 3965 |
| 20 | Yes | E. coli | None | None | F192 | 5-Carboxyf | 9159 | BL1 | 11992 |
| 20 | Yes | E. coli | None | None | F57 | Eosin Y | 11064 | BL1 | 16212 |
| 20 | Yes | E. coli | None | None | F27 | Calcein | 10325 | BL1 | 17989 |
| 20 | Yes | E. coli | None | None | F12 | Amaranth | 9037 | BL1 | 14113 |
| 20 | Yes | E. coli | None | None | NA | Autofluore: | 8969 | NA | 6830 |
| 20 | Yes | E. coli | Amoxacilli | Wall | F194 | Oxonol | 9851 | RL1 | 5197 |
| 20 | Yes | E. coli | Amoxacilli | Wall | F11 | Allura Red | 9400 | BL3 | 4021 |
| 20 | Yes | E. coli | Amoxacilli | Wall | F54 | DiSC3(5) | 7837 | RL1 | 3918 |
| 20 | Yes | E. coli | Amoxacilli | Wall | F124 | Thioflavin T | 10773 | VL2 | 3665 |
| 20 | Yes | E. coli | Amoxacilli | Wall | F101 | Quinacrine | 9466 | VL2 | 3577 |
| 20 | Yes | E. coli | Amoxacilli | Wall | F138 | Apigenin | 8921 | VL2 | 3460 |
| 20 | Yes | E. coli | Amoxacilli | Wall | F65 | H2FDA | 10521 | BL1 | 4145 |
| 20 | Yes | E. coli | Amoxacilli | Wall | F142 | Chlorophe | 9917 | BL3 | 5975 |
| 20 | Yes | E. coli | Amoxacilli | Wall | F39 | Congo Red | 9538 | BL3 | 5562.5 |
| 20 | Yes | E. coli | Amoxacilli | Wall | F116 | Sunset Yel | 8762 | BL2 | 6688 |
| 20 | Yes | E. coli | Amoxacilli | Wall | F10 | Alizarin | 8308 | BL1 | 4974 |
| 20 | Yes | E. coli | Amoxacilli | Wall | F84 | Neutral Re | 10049 | BL1 | 4425 |
| 20 | Yes | E. coli | Amoxacilli | Wall | F99 | Pyronin Y | 9512 | BL1 | 3790 |
| 20 | Yes | E. coli | Amoxacilli | Wall | F108 | Rhodamine | 8911 | BL1 | 3784 |
| 20 | Yes | E. coli | Amoxacilli | Wall | F109 | Riboflavin | 9315 | BL1 | 4599 |
| 20 | Yes | E. coli | Amoxacilli | Wall | F117 | SYBR Gree | 9117 | BL1 | 202184 |
| 20 | Yes | E. coli | Amoxacilli | Wall | F106 | Rhodamine | 8258 | BL1 | 4734 |
| 20 | Yes | E. coli | Amoxacilli | Wall | F73 | Malachite ( | 9427 | BL1 | 4424 |
| 20 | Yes | E. coli | Amoxacilli | Wall | F7 | Acridine or | 9802 | BL1 | 5033.5 |
| 20 | Yes | E. coli | Amoxacilli | Wall | F192 | 5-Carboxyf | 8704 | BL1 | 10466 |
| 20 | Yes | E. coli | Amoxacilli | Wall | F57 | Eosin Y | 9924 | BL1 | 15899.5 |
| 20 | Yes | E. coli | Amoxacilli | Wall | F27 | Calcein | 8787 | BL1 | 14936 |

| | | | | | | | | | |
|---|---|---|---|---|---|---|---|---|---|
| 20 | Yes | E. coli | Amoxacilli | Wall | F12 | Amaranth | 10752 | BL1 | 17234.5 |
| 20 | Yes | E. coli | None | None | NA | Autofluore: | 9658 | NA | 7109 |
| 20 | Yes | E. coli | Azithromyc | Intracellul | F194 | Oxonol | 9344 | RL1 | 5408 |
| 20 | Yes | E. coli | Azithromyc | Intracellul | F11 | Allura Red | 9907 | BL3 | 4973 |
| 20 | Yes | E. coli | Azithromyc | Intracellul | F54 | DiSC3(5) | 8869 | RL1 | 4448 |
| 20 | Yes | E. coli | Azithromyc | Intracellul | F124 | Thioflavin T | 9956 | VL2 | 4192 |
| 20 | Yes | E. coli | Azithromyc | Intracellul | F101 | Quinacrine | 9288 | VL2 | 3867 |
| 20 | Yes | E. coli | Azithromyc | Intracellul | F138 | Apigenin | 9822 | VL2 | 3715 |
| 20 | Yes | E. coli | Azithromyc | Intracellul | F65 | H2FDA | 9276 | BL1 | 4438 |
| 20 | Yes | E. coli | Azithromyc | Intracellul | F142 | Chlorophe | 9514 | BL3 | 5634 |
| 20 | Yes | E. coli | Azithromyc | Intracellul | F39 | Congo Red | 9263 | BL3 | 6523 |
| 20 | Yes | E. coli | Azithromyc | Intracellul | F116 | Sunset Yel | 9641 | BL2 | 7409 |
| 20 | Yes | E. coli | Azithromyc | Intracellul | F10 | Alizarin | 9640 | BL1 | 5419 |
| 20 | Yes | E. coli | Azithromyc | Intracellul | F84 | Neutral Re | 8208 | BL1 | 4787 |
| 20 | Yes | E. coli | Azithromyc | Intracellul | F99 | Pyronin Y | 9284 | BL1 | 3903.5 |
| 20 | Yes | E. coli | Azithromyc | Intracellul | F108 | Rhodamine | 9177 | BL1 | 3774 |
| 20 | Yes | E. coli | Azithromyc | Intracellul | F109 | Riboflavin | 9442 | BL1 | 4991.5 |
| 20 | Yes | E. coli | Azithromyc | Intracellul | F117 | SYBR Gree | 9428 | BL1 | 203448 |
| 20 | Yes | E. coli | Azithromyc | Intracellul | F106 | Rhodamine | 9377 | BL1 | 6206 |
| 20 | Yes | E. coli | Azithromyc | Intracellul | F73 | Malachite ( | 8607 | BL1 | 4781 |
| 20 | Yes | E. coli | Azithromyc | Intracellul | F7 | Acridine or | 10163 | BL1 | 5063 |
| 20 | Yes | E. coli | Azithromyc | Intracellul | F192 | 5-Carboxyf | 8224 | BL1 | 10737 |
| 20 | Yes | E. coli | Azithromyc | Intracellul | F57 | Eosin Y | 10826 | BL1 | 15572.5 |
| 20 | Yes | E. coli | Azithromyc | Intracellul | F27 | Calcein | 7755 | BL1 | 18485 |
| 20 | Yes | E. coli | Azithromyc | Intracellul | F12 | Amaranth | 10087 | BL1 | 17363 |
| 20 | Yes | E. coli | None | None | NA | Autofluore: | 9882 | NA | 7464 |
| 20 | Yes | E. coli | Trimethopr | Intracellul | F194 | Oxonol | 10354 | RL1 | 5688 |
| 20 | Yes | E. coli | Trimethopr | Intracellul | F11 | Allura Red | 9018 | BL3 | 5255 |
| 20 | Yes | E. coli | Trimethopr | Intracellul | F54 | DiSC3(5) | 9398 | RL1 | 4619.5 |
| 20 | Yes | E. coli | Trimethopr | Intracellul | F124 | Thioflavin T | 9737 | VL2 | 4735 |
| 20 | Yes | E. coli | Trimethopr | Intracellul | F101 | Quinacrine | 10184 | VL2 | 4287.5 |
| 20 | Yes | E. coli | Trimethopr | Intracellul | F138 | Apigenin | 10137 | VL2 | 3965 |
| 20 | Yes | E. coli | Trimethopr | Intracellul | F65 | H2FDA | 9981 | BL1 | 4870 |
| 20 | Yes | E. coli | Trimethopr | Intracellul | F142 | Chlorophe | 10117 | BL3 | 6362 |
| 20 | Yes | E. coli | Trimethopr | Intracellul | F39 | Congo Red | 9802 | BL3 | 6364.5 |
| 20 | Yes | E. coli | Trimethopr | Intracellul | F116 | Sunset Yel | 8269 | BL2 | 6627 |
| 20 | Yes | E. coli | Trimethopr | Intracellul | F10 | Alizarin | 9733 | BL1 | 5385 |
| 20 | Yes | E. coli | Trimethopr | Intracellul | F84 | Neutral Re | 8811 | BL1 | 4771 |
| 20 | Yes | E. coli | Trimethopr | Intracellul | F99 | Pyronin Y | 9966 | BL1 | 4387 |
| 20 | Yes | E. coli | Trimethopr | Intracellul | F108 | Rhodamine | 9192 | BL1 | 4275.5 |
| 20 | Yes | E. coli | Trimethopr | Intracellul | F109 | Riboflavin | 8833 | BL1 | 4989 |
| 20 | Yes | E. coli | Trimethopr | Intracellul | F117 | SYBR Gree | 9254 | BL1 | 208297.5 |
| 20 | Yes | E. coli | Trimethopr | Intracellul | F106 | Rhodamine | 8933 | BL1 | 7149 |
| 20 | Yes | E. coli | Trimethopr | Intracellul | F73 | Malachite ( | 8161 | BL1 | 4847 |
| 20 | Yes | E. coli | Trimethopr | Intracellul | F7 | Acridine or | 10309 | BL1 | 4832 |
| 20 | Yes | E. coli | Trimethopr | Intracellul | F192 | 5-Carboxyf | 9495 | BL1 | 11563 |
| 20 | Yes | E. coli | Trimethopr | Intracellul | F57 | Eosin Y | 10720 | BL1 | 16184 |
| 20 | Yes | E. coli | Trimethopr | Intracellul | F27 | Calcein | 9561 | BL1 | 17033 |

| | | | | | | | | | |
|---|---|---|---|---|---|---|---|---|---|
| 20 | Yes | E. coli | Trimethopr | Intracellul | F12 | Amaranth | 11031 | BL1 | 17697 |
| 20 | Yes | E. coli | None | None | NA | Autofluore | 8843 | NA | 7191 |
| 20 | Yes | E. coli | Chloramph | Intracellul | F194 | Oxonol | 10135 | RL1 | 5991 |
| 20 | Yes | E. coli | Chloramph | Intracellul | F11 | Allura Red | 9435 | BL3 | 4895 |
| 20 | Yes | E. coli | Chloramph | Intracellul | F54 | DiSC3(5) | 8962 | RL1 | 5025.5 |
| 20 | Yes | E. coli | Chloramph | Intracellul | F124 | Thioflavin T | 8767 | VL2 | 4557 |
| 20 | Yes | E. coli | Chloramph | Intracellul | F101 | Quinacrine | 9263 | VL2 | 4476 |
| 20 | Yes | E. coli | Chloramph | Intracellul | F138 | Apigenin | 8574 | VL2 | 4413 |
| 20 | Yes | E. coli | Chloramph | Intracellul | F65 | H2FDA | 10291 | BL1 | 5128 |
| 20 | Yes | E. coli | Chloramph | Intracellul | F142 | Chlorophe | 9978 | BL3 | 6119 |
| 20 | Yes | E. coli | Chloramph | Intracellul | F39 | Congo Red | 9902 | BL3 | 6313 |
| 20 | Yes | E. coli | Chloramph | Intracellul | F116 | Sunset Yel | 9535 | BL2 | 6524 |
| 20 | Yes | E. coli | Chloramph | Intracellul | F10 | Alizarin | 9386 | BL1 | 5482 |
| 20 | Yes | E. coli | Chloramph | Intracellul | F84 | Neutral Re | 9552 | BL1 | 5199 |
| 20 | Yes | E. coli | Chloramph | Intracellul | F99 | Pyronin Y | 9506 | BL1 | 4390.5 |
| 20 | Yes | E. coli | Chloramph | Intracellul | F108 | Rhodamine | 9623 | BL1 | 4567 |
| 20 | Yes | E. coli | Chloramph | Intracellul | F109 | Riboflavin | 8844 | BL1 | 4442.5 |
| 20 | Yes | E. coli | Chloramph | Intracellul | F117 | SYBR Gree | 9354 | BL1 | 235990 |
| 20 | Yes | E. coli | Chloramph | Intracellul | F106 | Rhodamine | 8126 | BL1 | 5948.5 |
| 20 | Yes | E. coli | Chloramph | Intracellul | F73 | Malachite G | 8115 | BL1 | 6275 |
| 20 | Yes | E. coli | Chloramph | Intracellul | F7 | Acridine or | 8938 | BL1 | 4862.5 |
| 20 | Yes | E. coli | Chloramph | Intracellul | F192 | 5-Carboxyf | 9990 | BL1 | 13914.5 |
| 20 | Yes | E. coli | Chloramph | Intracellul | F57 | Eosin Y | 8932 | BL1 | 14597.5 |
| 20 | Yes | E. coli | Chloramph | Intracellul | F27 | Calcein | 8994 | BL1 | 17923 |
| 20 | Yes | E. coli | Chloramph | Intracellul | F12 | Amaranth | 10404 | BL1 | 16123 |
| 20 | Yes | E. coli | None | None | NA | Autofluore | 9343 | NA | 7546 |
| 20 | Yes | E. coli | Colistin | Wall | F194 | Oxonol | 10320 | RL1 | 6255.5 |
| 20 | Yes | E. coli | Colistin | Wall | F11 | Allura Red | 9863 | BL3 | 5410 |
| 20 | Yes | E. coli | Colistin | Wall | F54 | DiSC3(5) | 8952 | RL1 | 4843 |
| 20 | Yes | E. coli | Colistin | Wall | F124 | Thioflavin T | 9571 | VL2 | 5261 |
| 20 | Yes | E. coli | Colistin | Wall | F101 | Quinacrine | 10279 | VL2 | 4830 |
| 20 | Yes | E. coli | Colistin | Wall | F138 | Apigenin | 10065 | VL2 | 4322 |
| 20 | Yes | E. coli | Colistin | Wall | F65 | H2FDA | 9311 | BL1 | 5412 |
| 20 | Yes | E. coli | Colistin | Wall | F142 | Chlorophe | 9871 | BL3 | 6426 |
| 20 | Yes | E. coli | Colistin | Wall | F39 | Congo Red | 8140 | BL3 | 9088 |
| 20 | Yes | E. coli | Colistin | Wall | F116 | Sunset Yel | 9441 | BL2 | 6192 |
| 20 | Yes | E. coli | Colistin | Wall | F10 | Alizarin | 8674 | BL1 | 6645.5 |
| 20 | Yes | E. coli | Colistin | Wall | F84 | Neutral Re | 9520 | BL1 | 5778 |
| 20 | Yes | E. coli | Colistin | Wall | F99 | Pyronin Y | 4060 | BL1 | 25341.5 |
| 20 | Yes | E. coli | Colistin | Wall | F108 | Rhodamine | 3264 | BL1 | 30765.5 |
| 20 | Yes | E. coli | Colistin | Wall | F109 | Riboflavin | 3457 | BL1 | 34642 |
| 20 | Yes | E. coli | Colistin | Wall | F117 | SYBR Gree | 8484 | BL1 | 245257 |
| 20 | Yes | E. coli | Colistin | Wall | F106 | Rhodamine | 9103 | BL1 | 7370 |
| 20 | Yes | E. coli | Colistin | Wall | F73 | Malachite G | 9087 | BL1 | 5515 |
| 20 | Yes | E. coli | Colistin | Wall | F7 | Acridine or | 10057 | BL1 | 5230 |
| 20 | Yes | E. coli | Colistin | Wall | F192 | 5-Carboxyf | 9628 | BL1 | 13284.5 |
| 20 | Yes | E. coli | Colistin | Wall | F57 | Eosin Y | 10204 | BL1 | 16239.5 |
| 20 | Yes | E. coli | Colistin | Wall | F27 | Calcein | 10765 | BL1 | 16896 |

| | | | | | | | | | |
|---|---|---|---|---|---|---|---|---|---|
| 20 | Yes | E. coli | Colistin | Wall | F12 | Amaranth | 10391 | BL1 | 17077 |
| 20 | Yes | E. coli | None | None | NA | Autofluores | 9187 | NA | 7552 |
| 20 | Yes | E. coli | Gentamicin | Intracellula | F194 | Oxonol | 10547 | RL1 | 6490 |
| 20 | Yes | E. coli | Gentamicin | Intracellula | F11 | Allura Red | 9654 | BL3 | 5435 |
| 20 | Yes | E. coli | Gentamicin | Intracellula | F54 | DiSC3(5) | 9127 | RL1 | 5371 |
| 20 | Yes | E. coli | Gentamicin | Intracellula | F124 | Thioflavin T | 10098 | VL2 | 4786 |
| 20 | Yes | E. coli | Gentamicin | Intracellula | F101 | Quinacrine | 10159 | VL2 | 4848 |
| 20 | Yes | E. coli | Gentamicin | Intracellula | F138 | Apigenin | 9924 | VL2 | 4240.5 |
| 20 | Yes | E. coli | Gentamicin | Intracellula | F65 | H2FDA | 9841 | BL1 | 5213 |
| 20 | Yes | E. coli | Gentamicin | Intracellula | F142 | Chlorophe | 10459 | BL3 | 6241 |
| 20 | Yes | E. coli | Gentamicin | Intracellula | F39 | Congo Red | 10122 | BL3 | 6719 |
| 20 | Yes | E. coli | Gentamicin | Intracellula | F116 | Sunset Yell | 10335 | BL2 | 6547 |
| 20 | Yes | E. coli | Gentamicin | Intracellula | F10 | Alizarin | 9761 | BL1 | 6815 |
| 20 | Yes | E. coli | Gentamicin | Intracellula | F84 | Neutral Re | 10157 | BL1 | 4956 |
| 20 | Yes | E. coli | Gentamicin | Intracellula | F99 | Pyronin Y | 9460 | BL1 | 4581 |
| 20 | Yes | E. coli | Gentamicin | Intracellula | F108 | Rhodamine | 9717 | BL1 | 4463 |
| 20 | Yes | E. coli | Gentamicin | Intracellula | F109 | Riboflavin | 10376 | BL1 | 4806.5 |
| 20 | Yes | E. coli | Gentamicin | Intracellula | F117 | SYBR Gree | 8938 | BL1 | 239855.5 |
| 20 | Yes | E. coli | Gentamicin | Intracellula | F106 | Rhodamine | 9916 | BL1 | 6014.5 |
| 20 | Yes | E. coli | Gentamicin | Intracellula | F73 | Malachite G | 9615 | BL1 | 6099 |
| 20 | Yes | E. coli | Gentamicin | Intracellula | F7 | Acridine or | 10267 | BL1 | 4906 |
| 20 | Yes | E. coli | Gentamicin | Intracellula | F192 | 5-Carboxyf | 10033 | BL1 | 13071 |
| 20 | Yes | E. coli | Gentamicin | Intracellula | F57 | Eosin Y | 9031 | BL1 | 13849 |
| 20 | Yes | E. coli | Gentamicin | Intracellula | F27 | Calcein | 10672 | BL1 | 15956.5 |
| 20 | Yes | E. coli | Gentamicin | Intracellula | F12 | Amaranth | 10021 | BL1 | 14205 |
| 20 | Yes | E. coli | None | None | NA | Autofluores | 9007 | NA | 7069 |
| 20 | Yes | E. coli | Nitrofurant | Intracellula | F194 | Oxonol | 8039 | RL1 | 6104 |
| 20 | Yes | E. coli | Nitrofurant | Intracellula | F11 | Allura Red | 7411 | BL3 | 5498 |
| 20 | Yes | E. coli | Nitrofurant | Intracellula | F54 | DiSC3(5) | 6719 | RL1 | 5231 |
| 20 | Yes | E. coli | Nitrofurant | Intracellula | F124 | Thioflavin T | 7441 | VL2 | 5063 |
| 20 | Yes | E. coli | Nitrofurant | Intracellula | F101 | Quinacrine | 7446 | VL2 | 5445.5 |
| 20 | Yes | E. coli | Nitrofurant | Intracellula | F138 | Apigenin | 6775 | VL2 | 5252 |
| 20 | Yes | E. coli | Nitrofurant | Intracellula | F65 | H2FDA | 7670 | BL1 | 6152 |
| 20 | Yes | E. coli | Nitrofurant | Intracellula | F142 | Chlorophe | 7313 | BL3 | 6644 |
| 20 | Yes | E. coli | Nitrofurant | Intracellula | F39 | Congo Red | 6718 | BL3 | 7375 |
| 20 | Yes | E. coli | Nitrofurant | Intracellula | F116 | Sunset Yell | 7163 | BL2 | 7503 |
| 20 | Yes | E. coli | Nitrofurant | Intracellula | F10 | Alizarin | 6872 | BL1 | 6066.5 |
| 20 | Yes | E. coli | Nitrofurant | Intracellula | F84 | Neutral Re | 7515 | BL1 | 6521 |
| 20 | Yes | E. coli | Nitrofurant | Intracellula | F99 | Pyronin Y | 7167 | BL1 | 4991 |
| 20 | Yes | E. coli | Nitrofurant | Intracellula | F108 | Rhodamine | 6283 | BL1 | 4812 |
| 20 | Yes | E. coli | Nitrofurant | Intracellula | F109 | Riboflavin | 7279 | BL1 | 6037 |
| 20 | Yes | E. coli | Nitrofurant | Intracellula | F117 | SYBR Gree | 9741 | BL1 | 253908 |
| 20 | Yes | E. coli | Nitrofurant | Intracellula | F106 | Rhodamine | 9621 | BL1 | 7186 |
| 20 | Yes | E. coli | Nitrofurant | Intracellula | F73 | Malachite G | 8218 | BL1 | 6059.5 |
| 20 | Yes | E. coli | Nitrofurant | Intracellula | F7 | Acridine or | 10258 | BL1 | 6254 |
| 20 | Yes | E. coli | Nitrofurant | Intracellula | F192 | 5-Carboxyf | 9566 | BL1 | 12837.5 |
| 20 | Yes | E. coli | Nitrofurant | Intracellula | F57 | Eosin Y | 10499 | BL1 | 14650 |
| 20 | Yes | E. coli | Nitrofurant | Intracellula | F27 | Calcein | 10294 | BL1 | 17770.5 |

| | | | | | | | | | |
|---|---|---|---|---|---|---|---|---|---|
| 20 | Yes | E. coli | Nitrofurant | Intracellula | F12 | Amaranth | 10399 | BL1 | 17215 |
| 20 | Yes | E. coli | None | None | NA | Autofluores | 8795 | NA | 8104 |
| 20 | Yes | E. coli | Pentamidi | Wall | F194 | Oxonol | 7099 | RL1 | 9744 |
| 20 | Yes | E. coli | Pentamidi | Wall | F11 | Allura Red | 6845 | BL3 | 7137 |
| 20 | Yes | E. coli | Pentamidi | Wall | F54 | DiSC3(5) | 6096 | RL1 | 6184.5 |
| 20 | Yes | E. coli | Pentamidi | Wall | F124 | Thioflavin T | 6913 | VL2 | 6629 |
| 20 | Yes | E. coli | Pentamidi | Wall | F101 | Quinacrine | 6223 | VL2 | 6119 |
| 20 | Yes | E. coli | Pentamidi | Wall | F138 | Apigenin | 6697 | VL2 | 5818 |
| 20 | Yes | E. coli | Pentamidi | Wall | F65 | H2FDA | 6290 | BL1 | 6480 |
| 20 | Yes | E. coli | Pentamidi | Wall | F142 | Chlorophe | 7428 | BL3 | 8444 |
| 20 | Yes | E. coli | Pentamidi | Wall | F39 | Congo Red | 7313 | BL3 | 11177 |
| 20 | Yes | E. coli | Pentamidi | Wall | F116 | Sunset Yel | 6951 | BL2 | 11839 |
| 20 | Yes | E. coli | Pentamidi | Wall | F10 | Alizarin | 6878 | BL1 | 11355.5 |
| 20 | Yes | E. coli | Pentamidi | Wall | F84 | Neutral Re | 8080 | BL1 | 12296.5 |
| 20 | Yes | E. coli | Pentamidi | Wall | F99 | Pyronin Y | 8419 | BL1 | 10342 |
| 20 | Yes | E. coli | Pentamidi | Wall | F108 | Rhodamine | 8304 | BL1 | 9385.5 |
| 20 | Yes | E. coli | Pentamidi | Wall | F109 | Riboflavin | 7258 | BL1 | 9877 |
| 20 | Yes | E. coli | Pentamidi | Wall | F117 | SYBR Gree | 8828 | BL1 | 285460 |
| 20 | Yes | E. coli | Pentamidi | Wall | F106 | Rhodamine | 8715 | BL1 | 10503 |
| 20 | Yes | E. coli | Pentamidi | Wall | F73 | Malachite ( | 9627 | BL1 | 9953 |
| 20 | Yes | E. coli | Pentamidi | Wall | F7 | Acridine or | 9187 | BL1 | 9176 |
| 20 | Yes | E. coli | Pentamidi | Wall | F192 | 5-Carboxyf | 7777 | BL1 | 15988 |
| 20 | Yes | E. coli | Pentamidi | Wall | F57 | Eosin Y | 7069 | BL1 | 16117 |
| 20 | Yes | E. coli | Pentamidi | Wall | F27 | Calcein | 7224 | BL1 | 16330 |
| 20 | Yes | E. coli | Pentamidi | Wall | F12 | Amaranth | 10223 | BL1 | 16150 |
| 20 | Yes | E. coli | None | None | NA | Autofluores | 9260 | NA | 9735.5 |
| 20 | Yes | E. coli | Rifampicin | Intracellula | F194 | Oxonol | 10143 | RL1 | 7470 |
| 20 | Yes | E. coli | Rifampicin | Intracellula | F11 | Allura Red | 9895 | BL3 | 6599 |
| 20 | Yes | E. coli | Rifampicin | Intracellula | F54 | DiSC3(5) | 9197 | RL1 | 6411 |
| 20 | Yes | E. coli | Rifampicin | Intracellula | F124 | Thioflavin T | 9857 | VL2 | 6452 |
| 20 | Yes | E. coli | Rifampicin | Intracellula | F101 | Quinacrine | 9539 | VL2 | 5999 |
| 20 | Yes | E. coli | Rifampicin | Intracellula | F138 | Apigenin | 9373 | VL2 | 6042 |
| 20 | Yes | E. coli | Rifampicin | Intracellula | F65 | H2FDA | 9300 | BL1 | 6161.5 |
| 20 | Yes | E. coli | Rifampicin | Intracellula | F142 | Chlorophe | 9742 | BL3 | 6004.5 |
| 20 | Yes | E. coli | Rifampicin | Intracellula | F39 | Congo Red | 9663 | BL3 | 6318 |
| 20 | Yes | E. coli | Rifampicin | Intracellula | F116 | Sunset Yel | 9895 | BL2 | 6375 |
| 20 | Yes | E. coli | Rifampicin | Intracellula | F10 | Alizarin | 10621 | BL1 | 6077 |
| 20 | Yes | E. coli | Rifampicin | Intracellula | F84 | Neutral Re | 9790 | BL1 | 5305 |
| 20 | Yes | E. coli | Rifampicin | Intracellula | F99 | Pyronin Y | 10136 | BL1 | 4712 |
| 20 | Yes | E. coli | Rifampicin | Intracellula | F108 | Rhodamine | 9658 | BL1 | 5071 |
| 20 | Yes | E. coli | Rifampicin | Intracellula | F109 | Riboflavin | 9969 | BL1 | 5075 |
| 20 | Yes | E. coli | Rifampicin | Intracellula | F117 | SYBR Gree | 10132 | BL1 | 275067 |
| 20 | Yes | E. coli | Rifampicin | Intracellula | F106 | Rhodamine | 9616 | BL1 | 6006 |
| 20 | Yes | E. coli | Rifampicin | Intracellula | F73 | Malachite ( | 10199 | BL1 | 6544 |
| 20 | Yes | E. coli | Rifampicin | Intracellula | F7 | Acridine or | 9117 | BL1 | 5305 |
| 20 | Yes | E. coli | Rifampicin | Intracellula | F192 | 5-Carboxyf | 10595 | BL1 | 12812 |
| 20 | Yes | E. coli | Rifampicin | Intracellula | F57 | Eosin Y | 10501 | BL1 | 14351 |
| 20 | Yes | E. coli | Rifampicin | Intracellula | F27 | Calcein | 11012 | BL1 | 15351 |

| | | | | | | | | | |
|---|---|---|---|---|---|---|---|---|---|
| 20 | Yes | E. coli | Rifampicin | Intracellula | F12 | Amaranth | 9978 | BL1 | 14232.5 |
| 20 | Yes | E. coli | None | None | NA | Autofluore: | 9959 | NA | 8309 |
| 20 | Yes | E. coli | Trimethopr | Intracellula | F194 | Oxonol | 11033 | RL1 | 6266 |
| 20 | Yes | E. coli | Trimethopr | Intracellula | F11 | Allura Red | 9897 | BL3 | 5532 |
| 20 | Yes | E. coli | Trimethopr | Intracellula | F54 | DiSC3(5) | 9717 | RL1 | 5682 |
| 20 | Yes | E. coli | Trimethopr | Intracellula | F124 | Thioflavin T | 9992 | VL2 | 5361.5 |
| 20 | Yes | E. coli | Trimethopr | Intracellula | F101 | Quinacrine | 9770 | VL2 | 4567 |
| 20 | Yes | E. coli | Trimethopr | Intracellula | F138 | Apigenin | 11123 | VL2 | 4680 |
| 20 | Yes | E. coli | Trimethopr | Intracellula | F65 | H2FDA | 10132 | BL1 | 6023.5 |
| 20 | Yes | E. coli | Trimethopr | Intracellula | F142 | Chlorophe | 10190 | BL3 | 6905.5 |
| 20 | Yes | E. coli | Trimethopr | Intracellula | F39 | Congo Red | 9110 | BL3 | 8904.5 |
| 20 | Yes | E. coli | Trimethopr | Intracellula | F116 | Sunset Yel | 9260 | BL2 | 8084.5 |
| 20 | Yes | E. coli | Trimethopr | Intracellula | F10 | Alizarin | 11270 | BL1 | 5968 |
| 20 | Yes | E. coli | Trimethopr | Intracellula | F84 | Neutral Re | 9962 | BL1 | 5248 |
| 20 | Yes | E. coli | Trimethopr | Intracellula | F99 | Pyronin Y | 9679 | BL1 | 4631 |
| 20 | Yes | E. coli | Trimethopr | Intracellula | F108 | Rhodamine | 8510 | BL1 | 4636.5 |
| 20 | Yes | E. coli | Trimethopr | Intracellula | F109 | Riboflavin | 8883 | BL1 | 4909 |
| 20 | Yes | E. coli | Trimethopr | Intracellula | F117 | SYBR Gree | 9554 | BL1 | 242210.5 |
| 20 | Yes | E. coli | Trimethopr | Intracellula | F106 | Rhodamine | 9067 | BL1 | 5737 |
| 20 | Yes | E. coli | Trimethopr | Intracellula | F73 | Malachite ( | 10667 | BL1 | 5238 |
| 20 | Yes | E. coli | Trimethopr | Intracellula | F7 | Acridine or | 10154 | BL1 | 4822 |
| 20 | Yes | E. coli | Trimethopr | Intracellula | F192 | 5-Carboxyf | 9821 | BL1 | 12073 |
| 20 | Yes | E. coli | Trimethopr | Intracellula | F57 | Eosin Y | 11070 | BL1 | 15458 |
| 20 | Yes | E. coli | Trimethopr | Intracellula | F27 | Calcein | 10419 | BL1 | 15197 |
| 20 | Yes | E. coli | Trimethopr | Intracellula | F12 | Amaranth | 10384 | BL1 | 16696.5 |
| 20 | Yes | E. coli | None | None | NA | Autofluore: | 12303 | NA | 965 |
| 20 | Yes | E. coli | None | None | F194 | Oxonol | 13218 | RL1 | 1890 |
| 20 | Yes | E. coli | None | None | F11 | Allura Red | 11579 | BL3 | 2643 |
| 20 | Yes | E. coli | None | None | F54 | DiSC3(5) | 10809 | RL1 | 2999 |
| 20 | Yes | E. coli | None | None | F124 | Thioflavin T | 11975 | VL2 | 3115 |
| 20 | Yes | E. coli | None | None | F101 | Quinacrine | 11512 | VL2 | 2826 |
| 20 | Yes | E. coli | None | None | F138 | Apigenin | 11300 | VL2 | 2918 |
| 20 | Yes | E. coli | None | None | F65 | H2FDA | 11030 | BL1 | 3804.5 |
| 20 | Yes | E. coli | None | None | F142 | Chlorophe | 11737 | BL3 | 5123 |
| 20 | Yes | E. coli | None | None | F39 | Congo Red | 11198 | BL3 | 6112 |
| 20 | Yes | E. coli | None | None | F116 | Sunset Yel | 9625 | BL2 | 6106 |
| 20 | Yes | E. coli | None | None | F10 | Alizarin | 11220 | BL1 | 4650.5 |
| 20 | Yes | E. coli | None | None | F84 | Neutral Re | 10036 | BL1 | 3977 |
| 20 | Yes | E. coli | None | None | F99 | Pyronin Y | 11206 | BL1 | 3664.5 |
| 20 | Yes | E. coli | None | None | F108 | Rhodamine | 10871 | BL1 | 3733 |
| 20 | Yes | E. coli | None | None | F109 | Riboflavin | 9128 | BL1 | 3895.5 |
| 20 | Yes | E. coli | None | None | F117 | SYBR Gree | 11699 | BL1 | 169472 |
| 20 | Yes | E. coli | None | None | F106 | Rhodamine | 10200 | BL1 | 4433 |
| 20 | Yes | E. coli | None | None | F73 | Malachite ( | 11355 | BL1 | 4649 |
| 20 | Yes | E. coli | None | None | F7 | Acridine or | 9635 | BL1 | 3945 |
| 20 | Yes | E. coli | None | None | F192 | 5-Carboxyf | 11664 | BL1 | 13150 |
| 20 | Yes | E. coli | None | None | F57 | Eosin Y | 10917 | BL1 | 14748 |
| 20 | Yes | E. coli | None | None | F27 | Calcein | 12175 | BL1 | 16449 |

| | | | | | | | | | |
|---|---|---|---|---|---|---|---|---|---|
| 20 | Yes | E. coli | None | None | F12 | Amaranth | 11150 | BL1 | 15274.5 |
| 20 | Yes | E. coli | None | None | NA | Autofluore: | 10307 | NA | 6994 |
| 20 | Yes | E. coli | Ampicillin | Wall | F194 | Oxonol | 11361 | RL1 | 5513 |
| 20 | Yes | E. coli | Ampicillin | Wall | F11 | Allura Red | 11219 | BL3 | 4544 |
| 20 | Yes | E. coli | Ampicillin | Wall | F54 | DiSC3(5) | 10035 | RL1 | 3978 |
| 20 | Yes | E. coli | Ampicillin | Wall | F124 | Thioflavin T | 10890 | VL2 | 3928 |
| 20 | Yes | E. coli | Ampicillin | Wall | F101 | Quinacrine | 10644 | VL2 | 3587.5 |
| 20 | Yes | E. coli | Ampicillin | Wall | F138 | Apigenin | 10888 | VL2 | 3423 |
| 20 | Yes | E. coli | Ampicillin | Wall | F65 | H2FDA | 11220 | BL1 | 4298.5 |
| 20 | Yes | E. coli | Ampicillin | Wall | F142 | Chlorophe | 10894 | BL3 | 5356 |
| 20 | Yes | E. coli | Ampicillin | Wall | F39 | Congo Red | 10947 | BL3 | 6457 |
| 20 | Yes | E. coli | Ampicillin | Wall | F116 | Sunset Yel | 10726 | BL2 | 6016.5 |
| 20 | Yes | E. coli | Ampicillin | Wall | F10 | Alizarin | 10550 | BL1 | 5464 |
| 20 | Yes | E. coli | Ampicillin | Wall | F84 | Neutral Re | 9463 | BL1 | 4582 |
| 20 | Yes | E. coli | Ampicillin | Wall | F99 | Pyronin Y | 11277 | BL1 | 3690 |
| 20 | Yes | E. coli | Ampicillin | Wall | F108 | Rhodamine | 8973 | BL1 | 4425 |
| 20 | Yes | E. coli | Ampicillin | Wall | F109 | Riboflavin | 11600 | BL1 | 4097 |
| 20 | Yes | E. coli | Ampicillin | Wall | F117 | SYBR Gree | 10232 | BL1 | 209758.5 |
| 20 | Yes | E. coli | Ampicillin | Wall | F106 | Rhodamine | 10596 | BL1 | 5390.5 |
| 20 | Yes | E. coli | Ampicillin | Wall | F73 | Malachite ( | 9945 | BL1 | 4467 |
| 20 | Yes | E. coli | Ampicillin | Wall | F7 | Acridine or | 10116 | BL1 | 4724 |
| 20 | Yes | E. coli | Ampicillin | Wall | F192 | 5-Carboxyf | 9912 | BL1 | 11806.5 |
| 20 | Yes | E. coli | Ampicillin | Wall | F57 | Eosin Y | 10183 | BL1 | 14116 |
| 20 | Yes | E. coli | Ampicillin | Wall | F27 | Calcein | 9721 | BL1 | 15115 |
| 20 | Yes | E. coli | Ampicillin | Wall | F12 | Amaranth | 12713 | BL1 | 16593 |
| 20 | Yes | E. coli | None | None | NA | Autofluore: | 10042 | NA | 6857 |
| 20 | Yes | E. coli | Bacitracin | Wall | F194 | Oxonol | 11623 | RL1 | 5522 |
| 20 | Yes | E. coli | Bacitracin | Wall | F11 | Allura Red | 11314 | BL3 | 4761.5 |
| 20 | Yes | E. coli | Bacitracin | Wall | F54 | DiSC3(5) | 9944 | RL1 | 3879 |
| 20 | Yes | E. coli | Bacitracin | Wall | F124 | Thioflavin T | 10540 | VL2 | 3698 |
| 20 | Yes | E. coli | Bacitracin | Wall | F101 | Quinacrine | 10640 | VL2 | 4216.5 |
| 20 | Yes | E. coli | Bacitracin | Wall | F138 | Apigenin | 10546 | VL2 | 3982.5 |
| 20 | Yes | E. coli | Bacitracin | Wall | F65 | H2FDA | 11020 | BL1 | 5755.5 |
| 20 | Yes | E. coli | Bacitracin | Wall | F142 | Chlorophe | 10522 | BL3 | 6851 |
| 20 | Yes | E. coli | Bacitracin | Wall | F39 | Congo Red | 9992 | BL3 | 5818 |
| 20 | Yes | E. coli | Bacitracin | Wall | F116 | Sunset Yel | 10382 | BL2 | 6296 |
| 20 | Yes | E. coli | Bacitracin | Wall | F10 | Alizarin | 9964 | BL1 | 5440 |
| 20 | Yes | E. coli | Bacitracin | Wall | F84 | Neutral Re | 9263 | BL1 | 4706 |
| 20 | Yes | E. coli | Bacitracin | Wall | F99 | Pyronin Y | 10001 | BL1 | 4001 |
| 20 | Yes | E. coli | Bacitracin | Wall | F108 | Rhodamine | 9459 | BL1 | 3749 |
| 20 | Yes | E. coli | Bacitracin | Wall | F109 | Riboflavin | 10046 | BL1 | 4337.5 |
| 20 | Yes | E. coli | Bacitracin | Wall | F117 | SYBR Gree | 9316 | BL1 | 238355.5 |
| 20 | Yes | E. coli | Bacitracin | Wall | F106 | Rhodamine | 10116 | BL1 | 5885 |
| 20 | Yes | E. coli | Bacitracin | Wall | F73 | Malachite ( | 8777 | BL1 | 5146 |
| 20 | Yes | E. coli | Bacitracin | Wall | F7 | Acridine or | 9975 | BL1 | 4941 |
| 20 | Yes | E. coli | Bacitracin | Wall | F192 | 5-Carboxyf | 10535 | BL1 | 13324 |
| 20 | Yes | E. coli | Bacitracin | Wall | F57 | Eosin Y | 9727 | BL1 | 16988 |
| 20 | Yes | E. coli | Bacitracin | Wall | F27 | Calcein | 10496 | BL1 | 18584.5 |

| | | | | | | | | | |
|---|---|---|---|---|---|---|---|---|---|
| 20 | Yes | E. coli | Bacitracin | Wall | F12 | Amaranth | 10566 | BL1 | 18006 |
| 20 | Yes | E. coli | None | None | NA | Autofluore: | 10607 | NA | 7140 |
| 20 | Yes | E. coli | Cefotaxim | Wall | F194 | Oxonol | 8913 | RL1 | 5570 |
| 20 | Yes | E. coli | Cefotaxim | Wall | F11 | Allura Red | 8288 | BL3 | 4932 |
| 20 | Yes | E. coli | Cefotaxim | Wall | F54 | DiSC3(5) | 7704 | RL1 | 5160.5 |
| 20 | Yes | E. coli | Cefotaxim | Wall | F124 | Thioflavin T | 8209 | VL2 | 4637 |
| 20 | Yes | E. coli | Cefotaxim | Wall | F101 | Quinacrine | 8188 | VL2 | 5287 |
| 20 | Yes | E. coli | Cefotaxim | Wall | F138 | Apigenin | 8229 | VL2 | 4736 |
| 20 | Yes | E. coli | Cefotaxim | Wall | F65 | H2FDA | 8196 | BL1 | 5563 |
| 20 | Yes | E. coli | Cefotaxim | Wall | F142 | Chlorophe | 7931 | BL3 | 6586 |
| 20 | Yes | E. coli | Cefotaxim | Wall | F39 | Congo Red | 8335 | BL3 | 6987 |
| 20 | Yes | E. coli | Cefotaxim | Wall | F116 | Sunset Yel | 8109 | BL2 | 7681 |
| 20 | Yes | E. coli | Cefotaxim | Wall | F10 | Alizarin | 8080 | BL1 | 6120 |
| 20 | Yes | E. coli | Cefotaxim | Wall | F84 | Neutral Re | 7274 | BL1 | 5679 |
| 20 | Yes | E. coli | Cefotaxim | Wall | F99 | Pyronin Y | 8645 | BL1 | 4674 |
| 20 | Yes | E. coli | Cefotaxim | Wall | F108 | Rhodamine | 7731 | BL1 | 4562 |
| 20 | Yes | E. coli | Cefotaxim | Wall | F109 | Riboflavin | 7659 | BL1 | 5118 |
| 20 | Yes | E. coli | Cefotaxim | Wall | F117 | SYBR Gree | 7989 | BL1 | 272340 |
| 20 | Yes | E. coli | Cefotaxim | Wall | F106 | Rhodamine | 7206 | BL1 | 6071 |
| 20 | Yes | E. coli | Cefotaxim | Wall | F73 | Malachite ( | 8162 | BL1 | 5572 |
| 20 | Yes | E. coli | Cefotaxim | Wall | F7 | Acridine or | 7950 | BL1 | 4876.5 |
| 20 | Yes | E. coli | Cefotaxim | Wall | F192 | 5-Carboxyf | 8311 | BL1 | 15354 |
| 20 | Yes | E. coli | Cefotaxim | Wall | F57 | Eosin Y | 7531 | BL1 | 18623 |
| 20 | Yes | E. coli | Cefotaxim | Wall | F27 | Calcein | 8388 | BL1 | 22260 |
| 20 | Yes | E. coli | Cefotaxim | Wall | F12 | Amaranth | 7718 | BL1 | 21228 |
| 20 | Yes | E. coli | None | None | NA | Autofluore: | 10028 | NA | 7424 |
| 20 | Yes | E. coli | Ciprofloxa | Intracellul | F194 | Oxonol | 8840 | RL1 | 5943 |
| 20 | Yes | E. coli | Ciprofloxa | Intracellul | F11 | Allura Red | 7985 | BL3 | 5199 |
| 20 | Yes | E. coli | Ciprofloxa | Intracellul | F54 | DiSC3(5) | 7581 | RL1 | 4397 |
| 20 | Yes | E. coli | Ciprofloxa | Intracellul | F124 | Thioflavin T | 8586 | VL2 | 4626.5 |
| 20 | Yes | E. coli | Ciprofloxa | Intracellul | F101 | Quinacrine | 8217 | VL2 | 4573 |
| 20 | Yes | E. coli | Ciprofloxa | Intracellul | F138 | Apigenin | 8974 | VL2 | 4849 |
| 20 | Yes | E. coli | Ciprofloxa | Intracellul | F65 | H2FDA | 8542 | BL1 | 5731 |
| 20 | Yes | E. coli | Ciprofloxa | Intracellul | F142 | Chlorophe | 8403 | BL3 | 8905 |
| 20 | Yes | E. coli | Ciprofloxa | Intracellul | F39 | Congo Red | 7167 | BL3 | 7194 |
| 20 | Yes | E. coli | Ciprofloxa | Intracellul | F116 | Sunset Yel | 8557 | BL2 | 6894 |
| 20 | Yes | E. coli | Ciprofloxa | Intracellul | F10 | Alizarin | 7494 | BL1 | 6154.5 |
| 20 | Yes | E. coli | Ciprofloxa | Intracellul | F84 | Neutral Re | 8365 | BL1 | 5223 |
| 20 | Yes | E. coli | Ciprofloxa | Intracellul | F99 | Pyronin Y | 8322 | BL1 | 5434.5 |
| 20 | Yes | E. coli | Ciprofloxa | Intracellul | F108 | Rhodamine | 7703 | BL1 | 4107 |
| 20 | Yes | E. coli | Ciprofloxa | Intracellul | F109 | Riboflavin | 7757 | BL1 | 4702 |
| 20 | Yes | E. coli | Ciprofloxa | Intracellul | F117 | SYBR Gree | 7680 | BL1 | 176905.5 |
| 20 | Yes | E. coli | Ciprofloxa | Intracellul | F106 | Rhodamine | 8026 | BL1 | 5310.5 |
| 20 | Yes | E. coli | Ciprofloxa | Intracellul | F73 | Malachite ( | 8842 | BL1 | 4942.5 |
| 20 | Yes | E. coli | Ciprofloxa | Intracellul | F7 | Acridine or | 7838 | BL1 | 5652 |
| 20 | Yes | E. coli | Ciprofloxa | Intracellul | F192 | 5-Carboxyf | 7389 | BL1 | 14328 |
| 20 | Yes | E. coli | Ciprofloxa | Intracellul | F57 | Eosin Y | 8659 | BL1 | 17457 |
| 20 | Yes | E. coli | Ciprofloxa | Intracellul | F27 | Calcein | 8199 | BL1 | 19486 |

| | | | | | | | | | |
|---|---|---|---|---|---|---|---|---|---|
| 20 | Yes | E. coli | Ciprofloxa | Intracellula | F12 | Amaranth | 8503 | BL1 | 21174 |
| 20 | Yes | E. coli | None | None | NA | Autofluore | 9405 | NA | 7563 |
| 20 | Yes | E. coli | Fosfomyci | Wall | F194 | Oxonol | 11237 | RL1 | 6373 |
| 20 | Yes | E. coli | Fosfomyci | Wall | F11 | Allura Red | 9721 | BL3 | 5383 |
| 20 | Yes | E. coli | Fosfomyci | Wall | F54 | DiSC3(5) | 10722 | RL1 | 4702 |
| 20 | Yes | E. coli | Fosfomyci | Wall | F124 | Thioflavin T | 11240 | VL2 | 4253 |
| 20 | Yes | E. coli | Fosfomyci | Wall | F101 | Quinacrine | 10501 | VL2 | 4505 |
| 20 | Yes | E. coli | Fosfomyci | Wall | F138 | Apigenin | 11632 | VL2 | 4681.5 |
| 20 | Yes | E. coli | Fosfomyci | Wall | F65 | H2FDA | 11493 | BL1 | 5523 |
| 20 | Yes | E. coli | Fosfomyci | Wall | F142 | Chlorophe | 10327 | BL3 | 8723 |
| 20 | Yes | E. coli | Fosfomyci | Wall | F39 | Congo Red | 11015 | BL3 | 7218 |
| 20 | Yes | E. coli | Fosfomyci | Wall | F116 | Sunset Yel | 9932 | BL2 | 9080 |
| 20 | Yes | E. coli | Fosfomyci | Wall | F10 | Alizarin | 10581 | BL1 | 5729 |
| 20 | Yes | E. coli | Fosfomyci | Wall | F84 | Neutral Re | 10539 | BL1 | 6251 |
| 20 | Yes | E. coli | Fosfomyci | Wall | F99 | Pyronin Y | 10769 | BL1 | 4418 |
| 20 | Yes | E. coli | Fosfomyci | Wall | F108 | Rhodamine | 10192 | BL1 | 4834 |
| 20 | Yes | E. coli | Fosfomyci | Wall | F109 | Riboflavin | 10250 | BL1 | 4308 |
| 20 | Yes | E. coli | Fosfomyci | Wall | F117 | SYBR Gree | 9549 | BL1 | 217515 |
| 20 | Yes | E. coli | Fosfomyci | Wall | F106 | Rhodamine | 11363 | BL1 | 5263 |
| 20 | Yes | E. coli | Fosfomyci | Wall | F73 | Malachite | 11273 | BL1 | 5446 |
| 20 | Yes | E. coli | Fosfomyci | Wall | F7 | Acridine or | 10305 | BL1 | 4363 |
| 20 | Yes | E. coli | Fosfomyci | Wall | F192 | 5-Carboxyf | 10937 | BL1 | 12385 |
| 20 | Yes | E. coli | Fosfomyci | Wall | F57 | Eosin Y | 10674 | BL1 | 14196 |
| 20 | Yes | E. coli | Fosfomyci | Wall | F27 | Calcein | 11545 | BL1 | 16632 |
| 20 | Yes | E. coli | Fosfomyci | Wall | F12 | Amaranth | 11749 | BL1 | 17845 |
| 20 | Yes | E. coli | None | None | NA | Autofluore | 10556 | NA | 7613.5 |
| 20 | Yes | E. coli | Nalidixic a | Intracellula | F194 | Oxonol | 10284 | RL1 | 6485.5 |
| 20 | Yes | E. coli | Nalidixic a | Intracellula | F11 | Allura Red | 9467 | BL3 | 5609 |
| 20 | Yes | E. coli | Nalidixic a | Intracellula | F54 | DiSC3(5) | 9353 | RL1 | 4658 |
| 20 | Yes | E. coli | Nalidixic a | Intracellula | F124 | Thioflavin T | 11089 | VL2 | 4877 |
| 20 | Yes | E. coli | Nalidixic a | Intracellula | F101 | Quinacrine | 9193 | VL2 | 4723 |
| 20 | Yes | E. coli | Nalidixic a | Intracellula | F138 | Apigenin | 10871 | VL2 | 4831 |
| 20 | Yes | E. coli | Nalidixic a | Intracellula | F65 | H2FDA | 9694 | BL1 | 6076.5 |
| 20 | Yes | E. coli | Nalidixic a | Intracellula | F142 | Chlorophe | 9864 | BL3 | 8580 |
| 20 | Yes | E. coli | Nalidixic a | Intracellula | F39 | Congo Red | 9871 | BL3 | 7601 |
| 20 | Yes | E. coli | Nalidixic a | Intracellula | F116 | Sunset Yel | 9747 | BL2 | 6824 |
| 20 | Yes | E. coli | Nalidixic a | Intracellula | F10 | Alizarin | 9733 | BL1 | 6646 |
| 20 | Yes | E. coli | Nalidixic a | Intracellula | F84 | Neutral Re | 9358 | BL1 | 5063 |
| 20 | Yes | E. coli | Nalidixic a | Intracellula | F99 | Pyronin Y | 10212 | BL1 | 4817 |
| 20 | Yes | E. coli | Nalidixic a | Intracellula | F108 | Rhodamine | 9508 | BL1 | 3933.5 |
| 20 | Yes | E. coli | Nalidixic a | Intracellula | F109 | Riboflavin | 9414 | BL1 | 4572 |
| 20 | Yes | E. coli | Nalidixic a | Intracellula | F117 | SYBR Gree | 10116 | BL1 | 180895 |
| 20 | Yes | E. coli | Nalidixic a | Intracellula | F106 | Rhodamine | 10380 | BL1 | 5138 |
| 20 | Yes | E. coli | Nalidixic a | Intracellula | F73 | Malachite | 9461 | BL1 | 5543 |
| 20 | Yes | E. coli | Nalidixic a | Intracellula | F7 | Acridine or | 8184 | BL1 | 4532 |
| 20 | Yes | E. coli | Nalidixic a | Intracellula | F192 | 5-Carboxyf | 10531 | BL1 | 12340 |
| 20 | Yes | E. coli | Nalidixic a | Intracellula | F57 | Eosin Y | 9004 | BL1 | 15178 |
| 20 | Yes | E. coli | Nalidixic a | Intracellula | F27 | Calcein | 10095 | BL1 | 18853 |

| | | | | | | | | | |
|---|---|---|---|---|---|---|---|---|---|
| 20 | Yes | E. coli | Nalidixic a | Intracellula | F12 | Amaranth | 9534 | BL1 | 19172 |
| 20 | Yes | E. coli | None | None | NA | Autofluore: | 10222 | NA | 6941.5 |
| 20 | Yes | E. coli | Ofloxacin | Intracellula | F194 | Oxonol | 8678 | RL1 | 6750 |
| 20 | Yes | E. coli | Ofloxacin | Intracellula | F11 | Allura Red | 8329 | BL3 | 6290 |
| 20 | Yes | E. coli | Ofloxacin | Intracellula | F54 | DiSC3(5) | 8297 | RL1 | 5075 |
| 20 | Yes | E. coli | Ofloxacin | Intracellula | F124 | Thioflavin T | 8613 | VL2 | 5149 |
| 20 | Yes | E. coli | Ofloxacin | Intracellula | F101 | Quinacrine | 8463 | VL2 | 4715 |
| 20 | Yes | E. coli | Ofloxacin | Intracellula | F138 | Apigenin | 9351 | VL2 | 5323 |
| 20 | Yes | E. coli | Ofloxacin | Intracellula | F65 | H2FDA | 9231 | BL1 | 5973 |
| 20 | Yes | E. coli | Ofloxacin | Intracellula | F142 | Chlorophe | 8769 | BL3 | 9118 |
| 20 | Yes | E. coli | Ofloxacin | Intracellula | F39 | Congo Red | 8348 | BL3 | 9579.5 |
| 20 | Yes | E. coli | Ofloxacin | Intracellula | F116 | Sunset Yel | 8597 | BL2 | 9345 |
| 20 | Yes | E. coli | Ofloxacin | Intracellula | F10 | Alizarin | 8130 | BL1 | 6561 |
| 20 | Yes | E. coli | Ofloxacin | Intracellula | F84 | Neutral Re | 8524 | BL1 | 6626 |
| 20 | Yes | E. coli | Ofloxacin | Intracellula | F99 | Pyronin Y | 8667 | BL1 | 5627 |
| 20 | Yes | E. coli | Ofloxacin | Intracellula | F108 | Rhodamine | 8328 | BL1 | 4606.5 |
| 20 | Yes | E. coli | Ofloxacin | Intracellula | F109 | Riboflavin | 8424 | BL1 | 4954 |
| 20 | Yes | E. coli | Ofloxacin | Intracellula | F117 | SYBR Gree | 8562 | BL1 | 224175.5 |
| 20 | Yes | E. coli | Ofloxacin | Intracellula | F106 | Rhodamine | 8357 | BL1 | 5199 |
| 20 | Yes | E. coli | Ofloxacin | Intracellula | F73 | Malachite ( | 9100 | BL1 | 5437 |
| 20 | Yes | E. coli | Ofloxacin | Intracellula | F7 | Acridine or | 8375 | BL1 | 5051 |
| 20 | Yes | E. coli | Ofloxacin | Intracellula | F192 | 5-Carboxyf | 8847 | BL1 | 13465 |
| 20 | Yes | E. coli | Ofloxacin | Intracellula | F57 | Eosin Y | 7368 | BL1 | 17538.5 |
| 20 | Yes | E. coli | Ofloxacin | Intracellula | F27 | Calcein | 9158 | BL1 | 20993.5 |
| 20 | Yes | E. coli | Ofloxacin | Intracellula | F12 | Amaranth | 8322 | BL1 | 21369 |
| 20 | Yes | E. coli | None | None | NA | Autofluore: | 10152 | NA | 8173.5 |
| 20 | Yes | E. coli | Polymyxin | Wall | F194 | Oxonol | 10553 | RL1 | 6493 |
| 20 | Yes | E. coli | Polymyxin | Wall | F11 | Allura Red | 8864 | BL3 | 5782.5 |
| 20 | Yes | E. coli | Polymyxin | Wall | F54 | DiSC3(5) | 8864 | RL1 | 5174 |
| 20 | Yes | E. coli | Polymyxin | Wall | F124 | Thioflavin T | 8757 | VL2 | 5258 |
| 20 | Yes | E. coli | Polymyxin | Wall | F101 | Quinacrine | 8905 | VL2 | 5454 |
| 20 | Yes | E. coli | Polymyxin | Wall | F138 | Apigenin | 8734 | VL2 | 5617 |
| 20 | Yes | E. coli | Polymyxin | Wall | F65 | H2FDA | 8614 | BL1 | 6666 |
| 20 | Yes | E. coli | Polymyxin | Wall | F142 | Chlorophe | 9048 | BL3 | 9432.5 |
| 20 | Yes | E. coli | Polymyxin | Wall | F39 | Congo Red | 9130 | BL3 | 7831 |
| 20 | Yes | E. coli | Polymyxin | Wall | F116 | Sunset Yel | 8935 | BL2 | 9425 |
| 20 | Yes | E. coli | Polymyxin | Wall | F10 | Alizarin | 9381 | BL1 | 6798 |
| 20 | Yes | E. coli | Polymyxin | Wall | F84 | Neutral Re | 8626 | BL1 | 6088.5 |
| 20 | Yes | E. coli | Polymyxin | Wall | F99 | Pyronin Y | 9326 | BL1 | 5553.5 |
| 20 | Yes | E. coli | Polymyxin | Wall | F108 | Rhodamine | 9045 | BL1 | 4892 |
| 20 | Yes | E. coli | Polymyxin | Wall | F109 | Riboflavin | 7944 | BL1 | 5000.5 |
| 20 | Yes | E. coli | Polymyxin | Wall | F117 | SYBR Gree | 9302 | BL1 | 280057.5 |
| 20 | Yes | E. coli | Polymyxin | Wall | F106 | Rhodamine | 8618 | BL1 | 6472.5 |
| 20 | Yes | E. coli | Polymyxin | Wall | F73 | Malachite ( | 8527 | BL1 | 5447 |
| 20 | Yes | E. coli | Polymyxin | Wall | F7 | Acridine or | 9151 | BL1 | 5777 |
| 20 | Yes | E. coli | Polymyxin | Wall | F192 | 5-Carboxyf | 9068 | BL1 | 14455 |
| 20 | Yes | E. coli | Polymyxin | Wall | F57 | Eosin Y | 9093 | BL1 | 17635 |
| 20 | Yes | E. coli | Polymyxin | Wall | F27 | Calcein | 9435 | BL1 | 18529 |

| | | | | | | | | | |
|---|---|---|---|---|---|---|---|---|---|
| 20 | Yes | E. coli | Polymyxin | Wall | F12 | Amaranth | 9708 | BL1 | 17555 |
| 20 | Yes | E. coli | None | None | NA | Autofluore: | 9997 | NA | 7687 |
| 20 | Yes | E. coli | Tetracyclin | Intracellula | F194 | Oxonol | 9869 | RL1 | 6051 |
| 20 | Yes | E. coli | Tetracyclin | Intracellula | F11 | Allura Red | 10162 | BL3 | 5612.5 |
| 20 | Yes | E. coli | Tetracyclin | Intracellula | F54 | DiSC3(5) | 9258 | RL1 | 5001 |
| 20 | Yes | E. coli | Tetracyclin | Intracellula | F124 | Thioflavin T | 9539 | VL2 | 5408 |
| 20 | Yes | E. coli | Tetracyclin | Intracellula | F101 | Quinacrine | 10818 | VL2 | 5652.5 |
| 20 | Yes | E. coli | Tetracyclin | Intracellula | F138 | Apigenin | 10681 | VL2 | 5310 |
| 20 | Yes | E. coli | Tetracyclin | Intracellula | F65 | H2FDA | 9365 | BL1 | 6828 |
| 20 | Yes | E. coli | Tetracyclin | Intracellula | F142 | Chlorophe | 10731 | BL3 | 7670 |
| 20 | Yes | E. coli | Tetracyclin | Intracellula | F39 | Congo Red | 10279 | BL3 | 7017 |
| 20 | Yes | E. coli | Tetracyclin | Intracellula | F116 | Sunset Yel | 10785 | BL2 | 7590 |
| 20 | Yes | E. coli | Tetracyclin | Intracellula | F10 | Alizarin | 10016 | BL1 | 6581 |
| 20 | Yes | E. coli | Tetracyclin | Intracellula | F84 | Neutral Re | 10102 | BL1 | 5210 |
| 20 | Yes | E. coli | Tetracyclin | Intracellula | F99 | Pyronin Y | 10421 | BL1 | 4972 |
| 20 | Yes | E. coli | Tetracyclin | Intracellula | F108 | Rhodamine | 10018 | BL1 | 4583 |
| 20 | Yes | E. coli | Tetracyclin | Intracellula | F109 | Riboflavin | 10726 | BL1 | 5331 |
| 20 | Yes | E. coli | Tetracyclin | Intracellula | F117 | SYBR Gree | 9818 | BL1 | 195171.5 |
| 20 | Yes | E. coli | Tetracyclin | Intracellula | F106 | Rhodamine | 10689 | BL1 | 7080 |
| 20 | Yes | E. coli | Tetracyclin | Intracellula | F73 | Malachite ( | 9660 | BL1 | 5120 |
| 20 | Yes | E. coli | Tetracyclin | Intracellula | F7 | Acridine or | 10678 | BL1 | 6076 |
| 20 | Yes | E. coli | Tetracyclin | Intracellula | F192 | 5-Carboxyf | 10778 | BL1 | 11612.5 |
| 20 | Yes | E. coli | Tetracyclin | Intracellula | F57 | Eosin Y | 10058 | BL1 | 16039.5 |
| 20 | Yes | E. coli | Tetracyclin | Intracellula | F27 | Calcein | 10663 | BL1 | 16894 |
| 20 | Yes | E. coli | Tetracyclin | Intracellula | F12 | Amaranth | 10227 | BL1 | 13146 |
| 20 | Yes | E. coli | None | None | NA | Autofluore: | 9855 | NA | 7791 |
| 20 | Yes | E. coli | Vancomyc | Wall | F194 | Oxonol | 10246 | RL1 | 5919 |
| 20 | Yes | E. coli | Vancomyc | Wall | F11 | Allura Red | 11836 | BL3 | 5379 |
| 20 | Yes | E. coli | Vancomyc | Wall | F54 | DiSC3(5) | 11108 | RL1 | 5618 |
| 20 | Yes | E. coli | Vancomyc | Wall | F124 | Thioflavin T | 11981 | VL2 | 5675 |
| 20 | Yes | E. coli | Vancomyc | Wall | F101 | Quinacrine | 11320 | VL2 | 4947 |
| 20 | Yes | E. coli | Vancomyc | Wall | F138 | Apigenin | 10890 | VL2 | 5107.5 |
| 20 | Yes | E. coli | Vancomyc | Wall | F65 | H2FDA | 10818 | BL1 | 7039 |
| 20 | Yes | E. coli | Vancomyc | Wall | F142 | Chlorophe | 12012 | BL3 | 7915.5 |
| 20 | Yes | E. coli | Vancomyc | Wall | F39 | Congo Red | 9953 | BL3 | 9359 |
| 20 | Yes | E. coli | Vancomyc | Wall | F116 | Sunset Yel | 11314 | BL2 | 7518 |
| 20 | Yes | E. coli | Vancomyc | Wall | F10 | Alizarin | 11434 | BL1 | 6267.5 |
| 20 | Yes | E. coli | Vancomyc | Wall | F84 | Neutral Re | 10004 | BL1 | 6265 |
| 20 | Yes | E. coli | Vancomyc | Wall | F99 | Pyronin Y | 12161 | BL1 | 5486 |
| 20 | Yes | E. coli | Vancomyc | Wall | F108 | Rhodamine | 10495 | BL1 | 5326 |
| 20 | Yes | E. coli | Vancomyc | Wall | F109 | Riboflavin | 9999 | BL1 | 4808 |
| 20 | Yes | E. coli | Vancomyc | Wall | F117 | SYBR Gree | 10947 | BL1 | 231096 |
| 20 | Yes | E. coli | Vancomyc | Wall | F106 | Rhodamine | 9322 | BL1 | 10320.5 |
| 20 | Yes | E. coli | Vancomyc | Wall | F73 | Malachite ( | 8790 | BL1 | 8855.5 |
| 20 | Yes | E. coli | Vancomyc | Wall | F7 | Acridine or | 9146 | BL1 | 8667.5 |
| 20 | Yes | E. coli | Vancomyc | Wall | F192 | 5-Carboxyf | 9179 | BL1 | 29963 |
| 20 | Yes | E. coli | Vancomyc | Wall | F57 | Eosin Y | 8441 | BL1 | 38315 |
| 20 | Yes | E. coli | Vancomyc | Wall | F27 | Calcein | 9796 | BL1 | 49983.5 |

| | | | | | | | | | |
|---|---|---|---|---|---|---|---|---|---|
| 20 | Yes | E. coli | Vancomyc | Wall | F12 | Amaranth | 9881 | BL1 | 45544 |
| 20 | Yes | E. coli | None | None | NA | Autofluore: | 8106 | NA | 1049 |
| 20 | Yes | E. coli | None | None | F194 | Oxonol | 11032 | RL1 | 1756 |
| 20 | Yes | E. coli | None | None | F11 | Allura Red | 9804 | BL3 | 2375 |
| 20 | Yes | E. coli | None | None | F54 | DiSC3(5) | 8592 | RL1 | 2732.5 |
| 20 | Yes | E. coli | None | None | F124 | Thioflavin T | 10555 | VL2 | 3150 |
| 20 | Yes | E. coli | None | None | F101 | Quinacrine | 8507 | VL2 | 3112 |
| 20 | Yes | E. coli | None | None | F138 | Apigenin | 9193 | VL2 | 3400 |
| 20 | Yes | E. coli | None | None | F65 | H2FDA | 7758 | BL1 | 4294 |
| 20 | Yes | E. coli | None | None | F142 | Chlorophe | 10090 | BL3 | 6315 |
| 20 | Yes | E. coli | None | None | F39 | Congo Red | 8317 | BL3 | 6654 |
| 20 | Yes | E. coli | None | None | F116 | Sunset Yel | 9583 | BL2 | 7306 |
| 20 | Yes | E. coli | None | None | F10 | Alizarin | 8582 | BL1 | 5551 |
| 20 | Yes | E. coli | None | None | F84 | Neutral Re | 8965 | BL1 | 4389 |
| 20 | Yes | E. coli | None | None | F99 | Pyronin Y | 8781 | BL1 | 3845 |
| 20 | Yes | E. coli | None | None | F108 | Rhodamine | 8582 | BL1 | 3017.5 |
| 20 | Yes | E. coli | None | None | F109 | Riboflavin | 8645 | BL1 | 4220 |
| 20 | Yes | E. coli | None | None | F117 | SYBR Gree | 8616 | BL1 | 144864.5 |
| 20 | Yes | E. coli | None | None | F106 | Rhodamine | 8816 | BL1 | 5229 |
| 20 | Yes | E. coli | None | None | F73 | Malachite ( | 8768 | BL1 | 4474 |
| 20 | Yes | E. coli | None | None | F7 | Acridine or | 8857 | BL1 | 4262 |
| 20 | Yes | E. coli | None | None | F192 | 5-Carboxyf | 9132 | BL1 | 13304 |
| 20 | Yes | E. coli | None | None | F57 | Eosin Y | 8763 | BL1 | 16596 |
| 20 | Yes | E. coli | None | None | F27 | Calcein | 9271 | BL1 | 16236 |
| 20 | Yes | E. coli | None | None | F12 | Amaranth | 8957 | BL1 | 14012 |
| 20 | Yes | E. coli | None | None | NA | Autofluore: | 8331 | NA | 6346 |
| 20 | Yes | E. coli | Ampicillin | Wall | F194 | Oxonol | 8521 | RL1 | 5497 |
| 20 | Yes | E. coli | Ampicillin | Wall | F11 | Allura Red | 8365 | BL3 | 4463 |
| 20 | Yes | E. coli | Ampicillin | Wall | F54 | DiSC3(5) | 7473 | RL1 | 4172 |
| 20 | Yes | E. coli | Ampicillin | Wall | F124 | Thioflavin T | 7812 | VL2 | 4292 |
| 20 | Yes | E. coli | Ampicillin | Wall | F101 | Quinacrine | 7507 | VL2 | 3932 |
| 20 | Yes | E. coli | Ampicillin | Wall | F138 | Apigenin | 7056 | VL2 | 4312.5 |
| 20 | Yes | E. coli | Ampicillin | Wall | F65 | H2FDA | 8674 | BL1 | 5347 |
| 20 | Yes | E. coli | Ampicillin | Wall | F142 | Chlorophe | 7993 | BL3 | 6932 |
| 20 | Yes | E. coli | Ampicillin | Wall | F39 | Congo Red | 7534 | BL3 | 7965 |
| 20 | Yes | E. coli | Ampicillin | Wall | F116 | Sunset Yel | 8215 | BL2 | 6781 |
| 20 | Yes | E. coli | Ampicillin | Wall | F10 | Alizarin | 7279 | BL1 | 6140 |
| 20 | Yes | E. coli | Ampicillin | Wall | F84 | Neutral Re | 8011 | BL1 | 4686 |
| 20 | Yes | E. coli | Ampicillin | Wall | F99 | Pyronin Y | 7127 | BL1 | 3922 |
| 20 | Yes | E. coli | Ampicillin | Wall | F108 | Rhodamine | 7313 | BL1 | 3757 |
| 20 | Yes | E. coli | Ampicillin | Wall | F109 | Riboflavin | 7677 | BL1 | 3791 |
| 20 | Yes | E. coli | Ampicillin | Wall | F117 | SYBR Gree | 7576 | BL1 | 176376 |
| 20 | Yes | E. coli | Ampicillin | Wall | F106 | Rhodamine | 6804 | BL1 | 4193 |
| 20 | Yes | E. coli | Ampicillin | Wall | F73 | Malachite ( | 7766 | BL1 | 5206 |
| 20 | Yes | E. coli | Ampicillin | Wall | F7 | Acridine or | 7094 | BL1 | 3849 |
| 20 | Yes | E. coli | Ampicillin | Wall | F192 | 5-Carboxyf | 7999 | BL1 | 14322 |
| 20 | Yes | E. coli | Ampicillin | Wall | F57 | Eosin Y | 7616 | BL1 | 16847 |
| 20 | Yes | E. coli | Ampicillin | Wall | F27 | Calcein | 8655 | BL1 | 18791 |

| | | | | | | | | | |
|---|---|---|---|---|---|---|---|---|---|
| 20 | Yes | E. coli | Ampicillin | Wall | F12 | Amaranth | 8372 | BL1 | 16883 |
| 20 | Yes | E. coli | None | None | NA | Autofluore: | 7698 | NA | 6921.5 |
| 20 | Yes | E. coli | Bacitracin | Wall | F194 | Oxonol | 9693 | RL1 | 5281 |
| 20 | Yes | E. coli | Bacitracin | Wall | F11 | Allura Red | 8443 | BL3 | 4794 |
| 20 | Yes | E. coli | Bacitracin | Wall | F54 | DiSC3(5) | 8148 | RL1 | 3879 |
| 20 | Yes | E. coli | Bacitracin | Wall | F124 | Thioflavin T | 8008 | VL2 | 3825.5 |
| 20 | Yes | E. coli | Bacitracin | Wall | F101 | Quinacrine | 9300 | VL2 | 4259 |
| 20 | Yes | E. coli | Bacitracin | Wall | F138 | Apigenin | 9068 | VL2 | 4107 |
| 20 | Yes | E. coli | Bacitracin | Wall | F65 | H2FDA | 7862 | BL1 | 5775.5 |
| 20 | Yes | E. coli | Bacitracin | Wall | F142 | Chlorophe | 8152 | BL3 | 8038.5 |
| 20 | Yes | E. coli | Bacitracin | Wall | F39 | Congo Red | 8343 | BL3 | 8460 |
| 20 | Yes | E. coli | Bacitracin | Wall | F116 | Sunset Yel | 8197 | BL2 | 7479 |
| 20 | Yes | E. coli | Bacitracin | Wall | F10 | Alizarin | 7782 | BL1 | 5768 |
| 20 | Yes | E. coli | Bacitracin | Wall | F84 | Neutral Re | 8118 | BL1 | 4801.5 |
| 20 | Yes | E. coli | Bacitracin | Wall | F99 | Pyronin Y | 8056 | BL1 | 4084.5 |
| 20 | Yes | E. coli | Bacitracin | Wall | F108 | Rhodamine | 8732 | BL1 | 3943.5 |
| 20 | Yes | E. coli | Bacitracin | Wall | F109 | Riboflavin | 7377 | BL1 | 3817 |
| 20 | Yes | E. coli | Bacitracin | Wall | F117 | SYBR Gree | 8516 | BL1 | 177547.5 |
| 20 | Yes | E. coli | Bacitracin | Wall | F106 | Rhodamine | 7753 | BL1 | 4717 |
| 20 | Yes | E. coli | Bacitracin | Wall | F73 | Malachite ( | 8127 | BL1 | 5050 |
| 20 | Yes | E. coli | Bacitracin | Wall | F7 | Acridine or | 8296 | BL1 | 5236.5 |
| 20 | Yes | E. coli | Bacitracin | Wall | F192 | 5-Carboxyf | 7335 | BL1 | 12331 |
| 20 | Yes | E. coli | Bacitracin | Wall | F57 | Eosin Y | 8214 | BL1 | 17000 |
| 20 | Yes | E. coli | Bacitracin | Wall | F27 | Calcein | 8145 | BL1 | 16763 |
| 20 | Yes | E. coli | Bacitracin | Wall | F12 | Amaranth | 8499 | BL1 | 16625 |
| 20 | Yes | E. coli | None | None | NA | Autofluore: | 8372 | NA | 6449.5 |
| 20 | Yes | E. coli | Cefotaxim | Wall | F194 | Oxonol | 7274 | RL1 | 5235 |
| 20 | Yes | E. coli | Cefotaxim | Wall | F11 | Allura Red | 6860 | BL3 | 4681.5 |
| 20 | Yes | E. coli | Cefotaxim | Wall | F54 | DiSC3(5) | 6370 | RL1 | 4611.5 |
| 20 | Yes | E. coli | Cefotaxim | Wall | F124 | Thioflavin T | 7463 | VL2 | 5007 |
| 20 | Yes | E. coli | Cefotaxim | Wall | F101 | Quinacrine | 6609 | VL2 | 5016 |
| 20 | Yes | E. coli | Cefotaxim | Wall | F138 | Apigenin | 7315 | VL2 | 4669 |
| 20 | Yes | E. coli | Cefotaxim | Wall | F65 | H2FDA | 7002 | BL1 | 6571 |
| 20 | Yes | E. coli | Cefotaxim | Wall | F142 | Chlorophe | 7214 | BL3 | 7710.5 |
| 20 | Yes | E. coli | Cefotaxim | Wall | F39 | Congo Red | 6810 | BL3 | 7857 |
| 20 | Yes | E. coli | Cefotaxim | Wall | F116 | Sunset Yel | 7484 | BL2 | 8688.5 |
| 20 | Yes | E. coli | Cefotaxim | Wall | F10 | Alizarin | 7218 | BL1 | 6909 |
| 20 | Yes | E. coli | Cefotaxim | Wall | F84 | Neutral Re | 6471 | BL1 | 5631 |
| 20 | Yes | E. coli | Cefotaxim | Wall | F99 | Pyronin Y | 7687 | BL1 | 4551 |
| 20 | Yes | E. coli | Cefotaxim | Wall | F108 | Rhodamine | 6588 | BL1 | 4827 |
| 20 | Yes | E. coli | Cefotaxim | Wall | F109 | Riboflavin | 7622 | BL1 | 5268 |
| 20 | Yes | E. coli | Cefotaxim | Wall | F117 | SYBR Gree | 6987 | BL1 | 213913 |
| 20 | Yes | E. coli | Cefotaxim | Wall | F106 | Rhodamine | 6242 | BL1 | 6537.5 |
| 20 | Yes | E. coli | Cefotaxim | Wall | F73 | Malachite ( | 6843 | BL1 | 5720 |
| 20 | Yes | E. coli | Cefotaxim | Wall | F7 | Acridine or | 6892 | BL1 | 6364 |
| 20 | Yes | E. coli | Cefotaxim | Wall | F192 | 5-Carboxyf | 6322 | BL1 | 12671 |
| 20 | Yes | E. coli | Cefotaxim | Wall | F57 | Eosin Y | 7546 | BL1 | 18445 |
| 20 | Yes | E. coli | Cefotaxim | Wall | F27 | Calcein | 8568 | BL1 | 14125.5 |

| | | | | | | | | | |
|---|---|---|---|---|---|---|---|---|---|
| 20 | Yes | E. coli | Cefotaxim | Wall | F12 | Amaranth | 9054 | BL1 | 15646 |
| 20 | Yes | E. coli | None | None | NA | Autofluore | 8456 | NA | 7629.5 |
| 20 | Yes | E. coli | Ciprofloxa | Intracellul | F194 | Oxonol | 9299 | RL1 | 5702 |
| 20 | Yes | E. coli | Ciprofloxa | Intracellul | F11 | Allura Red | 8831 | BL3 | 5091 |
| 20 | Yes | E. coli | Ciprofloxa | Intracellul | F54 | DiSC3(5) | 7925 | RL1 | 4421 |
| 20 | Yes | E. coli | Ciprofloxa | Intracellul | F124 | Thioflavin T | 7627 | VL2 | 5287 |
| 20 | Yes | E. coli | Ciprofloxa | Intracellul | F101 | Quinacrine | 8587 | VL2 | 5105 |
| 20 | Yes | E. coli | Ciprofloxa | Intracellul | F138 | Apigenin | 8167 | VL2 | 5511 |
| 20 | Yes | E. coli | Ciprofloxa | Intracellul | F65 | H2FDA | 7343 | BL1 | 6134 |
| 20 | Yes | E. coli | Ciprofloxa | Intracellul | F142 | Chlorophe | 8439 | BL3 | 7703 |
| 20 | Yes | E. coli | Ciprofloxa | Intracellul | F39 | Congo Red | 7160 | BL3 | 7633.5 |
| 20 | Yes | E. coli | Ciprofloxa | Intracellul | F116 | Sunset Yel | 7955 | BL2 | 8481 |
| 20 | Yes | E. coli | Ciprofloxa | Intracellul | F10 | Alizarin | 7626 | BL1 | 6330 |
| 20 | Yes | E. coli | Ciprofloxa | Intracellul | F84 | Neutral Re | 7540 | BL1 | 5480 |
| 20 | Yes | E. coli | Ciprofloxa | Intracellul | F99 | Pyronin Y | 7872 | BL1 | 4865 |
| 20 | Yes | E. coli | Ciprofloxa | Intracellul | F108 | Rhodamine | 7660 | BL1 | 4918.5 |
| 20 | Yes | E. coli | Ciprofloxa | Intracellul | F109 | Riboflavin | 7573 | BL1 | 6164 |
| 20 | Yes | E. coli | Ciprofloxa | Intracellul | F117 | SYBR Gree | 7099 | BL1 | 152957 |
| 20 | Yes | E. coli | Ciprofloxa | Intracellul | F106 | Rhodamine | 7532 | BL1 | 7793 |
| 20 | Yes | E. coli | Ciprofloxa | Intracellul | F73 | Malachite ( | 7027 | BL1 | 5687 |
| 20 | Yes | E. coli | Ciprofloxa | Intracellul | F7 | Acridine or | 7920 | BL1 | 5913 |
| 20 | Yes | E. coli | Ciprofloxa | Intracellul | F192 | 5-Carboxyf | 7964 | BL1 | 13725 |
| 20 | Yes | E. coli | Ciprofloxa | Intracellul | F57 | Eosin Y | 7463 | BL1 | 16354 |
| 20 | Yes | E. coli | Ciprofloxa | Intracellul | F27 | Calcein | 7906 | BL1 | 17534 |
| 20 | Yes | E. coli | Ciprofloxa | Intracellul | F12 | Amaranth | 7836 | BL1 | 17606.5 |
| 20 | Yes | E. coli | None | None | NA | Autofluore | 7710 | NA | 7579.5 |
| 20 | Yes | E. coli | Fosfomyci | Wall | F194 | Oxonol | 9310 | RL1 | 6020 |
| 20 | Yes | E. coli | Fosfomyci | Wall | F11 | Allura Red | 8970 | BL3 | 5111 |
| 20 | Yes | E. coli | Fosfomyci | Wall | F54 | DiSC3(5) | 8377 | RL1 | 5092 |
| 20 | Yes | E. coli | Fosfomyci | Wall | F124 | Thioflavin T | 8931 | VL2 | 4851 |
| 20 | Yes | E. coli | Fosfomyci | Wall | F101 | Quinacrine | 9188 | VL2 | 5358.5 |
| 20 | Yes | E. coli | Fosfomyci | Wall | F138 | Apigenin | 9012 | VL2 | 5171 |
| 20 | Yes | E. coli | Fosfomyci | Wall | F65 | H2FDA | 8136 | BL1 | 6383.5 |
| 20 | Yes | E. coli | Fosfomyci | Wall | F142 | Chlorophe | 8697 | BL3 | 7625 |
| 20 | Yes | E. coli | Fosfomyci | Wall | F39 | Congo Red | 8993 | BL3 | 8868 |
| 20 | Yes | E. coli | Fosfomyci | Wall | F116 | Sunset Yel | 8295 | BL2 | 7616 |
| 20 | Yes | E. coli | Fosfomyci | Wall | F10 | Alizarin | 8845 | BL1 | 6584 |
| 20 | Yes | E. coli | Fosfomyci | Wall | F84 | Neutral Re | 8398 | BL1 | 5158 |
| 20 | Yes | E. coli | Fosfomyci | Wall | F99 | Pyronin Y | 8563 | BL1 | 4598 |
| 20 | Yes | E. coli | Fosfomyci | Wall | F108 | Rhodamine | 9313 | BL1 | 4895 |
| 20 | Yes | E. coli | Fosfomyci | Wall | F109 | Riboflavin | 8602 | BL1 | 4465.5 |
| 20 | Yes | E. coli | Fosfomyci | Wall | F117 | SYBR Gree | 8844 | BL1 | 224818 |
| 20 | Yes | E. coli | Fosfomyci | Wall | F106 | Rhodamine | 8077 | BL1 | 5418 |
| 20 | Yes | E. coli | Fosfomyci | Wall | F73 | Malachite ( | 8646 | BL1 | 6756.5 |
| 20 | Yes | E. coli | Fosfomyci | Wall | F7 | Acridine or | 8484 | BL1 | 4935 |
| 20 | Yes | E. coli | Fosfomyci | Wall | F192 | 5-Carboxyf | 8141 | BL1 | 12135 |
| 20 | Yes | E. coli | Fosfomyci | Wall | F57 | Eosin Y | 9311 | BL1 | 14990 |
| 20 | Yes | E. coli | Fosfomyci | Wall | F27 | Calcein | 9247 | BL1 | 16867 |

| | | | | | | | | | |
|---|---|---|---|---|---|---|---|---|---|
| 20 | Yes | E. coli | Fosfomycii | Wall | F12 | Amaranth | 8417 | BL1 | 14854 |
| 20 | Yes | E. coli | None | None | NA | Autofluore: | 8145 | NA | 7601 |
| 20 | Yes | E. coli | Nalidixic ac | Intracellul: | F194 | Oxonol | 9746 | RL1 | 5775 |
| 20 | Yes | E. coli | Nalidixic ac | Intracellul: | F11 | Allura Red | 8574 | BL3 | 5111 |
| 20 | Yes | E. coli | Nalidixic ac | Intracellul: | F54 | DiSC3(5) | 8396 | RL1 | 5191.5 |
| 20 | Yes | E. coli | Nalidixic ac | Intracellul: | F124 | Thioflavin T | 9327 | VL2 | 5422 |
| 20 | Yes | E. coli | Nalidixic ac | Intracellul: | F101 | Quinacrine | 8746 | VL2 | 5344 |
| 20 | Yes | E. coli | Nalidixic ac | Intracellul: | F138 | Apigenin | 8755 | VL2 | 5005 |
| 20 | Yes | E. coli | Nalidixic ac | Intracellul: | F65 | H2FDA | 9040 | BL1 | 6494 |
| 20 | Yes | E. coli | Nalidixic ac | Intracellul: | F142 | Chlorophe | 8363 | BL3 | 7144 |
| 20 | Yes | E. coli | Nalidixic ac | Intracellul: | F39 | Congo Red | 9347 | BL3 | 9576 |
| 20 | Yes | E. coli | Nalidixic ac | Intracellul: | F116 | Sunset Yel | 8346 | BL2 | 7395 |
| 20 | Yes | E. coli | Nalidixic ac | Intracellul: | F10 | Alizarin | 8632 | BL1 | 7667.5 |
| 20 | Yes | E. coli | Nalidixic ac | Intracellul: | F84 | Neutral Re | 8037 | BL1 | 5684 |
| 20 | Yes | E. coli | Nalidixic ac | Intracellul: | F99 | Pyronin Y | 8746 | BL1 | 5338 |
| 20 | Yes | E. coli | Nalidixic ac | Intracellul: | F108 | Rhodamine | 8014 | BL1 | 5165 |
| 20 | Yes | E. coli | Nalidixic ac | Intracellul: | F109 | Riboflavin | 8784 | BL1 | 5420.5 |
| 20 | Yes | E. coli | Nalidixic ac | Intracellul: | F117 | SYBR Gree | 8118 | BL1 | 172358 |
| 20 | Yes | E. coli | Nalidixic ac | Intracellul: | F106 | Rhodamine | 8196 | BL1 | 6270 |
| 20 | Yes | E. coli | Nalidixic ac | Intracellul: | F73 | Malachite ( | 8538 | BL1 | 6390.5 |
| 20 | Yes | E. coli | Nalidixic ac | Intracellul: | F7 | Acridine or | 8771 | BL1 | 6009 |
| 20 | Yes | E. coli | Nalidixic ac | Intracellul: | F192 | 5-Carboxyf | 8271 | BL1 | 12203 |
| 20 | Yes | E. coli | Nalidixic ac | Intracellul: | F57 | Eosin Y | 8756 | BL1 | 15367 |
| 20 | Yes | E. coli | Nalidixic ac | Intracellul: | F27 | Calcein | 8403 | BL1 | 14765 |
| 20 | Yes | E. coli | Nalidixic ac | Intracellul: | F12 | Amaranth | 8872 | BL1 | 17810.5 |
| 20 | Yes | E. coli | None | None | NA | Autofluore: | 9193 | NA | 7380 |
| 20 | Yes | E. coli | Ofloxacin | Intracellul: | F194 | Oxonol | 8138 | RL1 | 5870.5 |
| 20 | Yes | E. coli | Ofloxacin | Intracellul: | F11 | Allura Red | 8220 | BL3 | 5556 |
| 20 | Yes | E. coli | Ofloxacin | Intracellul: | F54 | DiSC3(5) | 7069 | RL1 | 5002 |
| 20 | Yes | E. coli | Ofloxacin | Intracellul: | F124 | Thioflavin T | 7747 | VL2 | 6232 |
| 20 | Yes | E. coli | Ofloxacin | Intracellul: | F101 | Quinacrine | 8414 | VL2 | 5540 |
| 20 | Yes | E. coli | Ofloxacin | Intracellul: | F138 | Apigenin | 7922 | VL2 | 5469 |
| 20 | Yes | E. coli | Ofloxacin | Intracellul: | F65 | H2FDA | 7175 | BL1 | 7324 |
| 20 | Yes | E. coli | Ofloxacin | Intracellul: | F142 | Chlorophe | 8102 | BL3 | 9778 |
| 20 | Yes | E. coli | Ofloxacin | Intracellul: | F39 | Congo Red | 7062 | BL3 | 9208.5 |
| 20 | Yes | E. coli | Ofloxacin | Intracellul: | F116 | Sunset Yel | 7746 | BL2 | 8195.5 |
| 20 | Yes | E. coli | Ofloxacin | Intracellul: | F10 | Alizarin | 7822 | BL1 | 7101.5 |
| 20 | Yes | E. coli | Ofloxacin | Intracellul: | F84 | Neutral Re | 8026 | BL1 | 5983 |
| 20 | Yes | E. coli | Ofloxacin | Intracellul: | F99 | Pyronin Y | 7911 | BL1 | 5285 |
| 20 | Yes | E. coli | Ofloxacin | Intracellul: | F108 | Rhodamine | 8096 | BL1 | 5669 |
| 20 | Yes | E. coli | Ofloxacin | Intracellul: | F109 | Riboflavin | 7842 | BL1 | 6203 |
| 20 | Yes | E. coli | Ofloxacin | Intracellul: | F117 | SYBR Gree | 7198 | BL1 | 182253 |
| 20 | Yes | E. coli | Ofloxacin | Intracellul: | F106 | Rhodamine | 8070 | BL1 | 8019 |
| 20 | Yes | E. coli | Ofloxacin | Intracellul: | F73 | Malachite ( | 6992 | BL1 | 5616.5 |
| 20 | Yes | E. coli | Ofloxacin | Intracellul: | F7 | Acridine or | 8298 | BL1 | 6311 |
| 20 | Yes | E. coli | Ofloxacin | Intracellul: | F192 | 5-Carboxyf | 8395 | BL1 | 12821 |
| 20 | Yes | E. coli | Ofloxacin | Intracellul: | F57 | Eosin Y | 8767 | BL1 | 15255 |
| 20 | Yes | E. coli | Ofloxacin | Intracellul: | F27 | Calcein | 8773 | BL1 | 14209 |

| | | | | | | | | | |
|---|---|---|---|---|---|---|---|---|---|
| 20 | Yes | E. coli | Ofloxacin | Intracellula | F12 | Amaranth | 9746 | BL1 | 16021 |
| 20 | Yes | E. coli | None | None | NA | Autofluore: | 8139 | NA | 8592 |
| 20 | Yes | E. coli | Polymyxin | Wall | F194 | Oxonol | 9475 | RL1 | 6100 |
| 20 | Yes | E. coli | Polymyxin | Wall | F11 | Allura Red | 8307 | BL3 | 5490 |
| 20 | Yes | E. coli | Polymyxin | Wall | F54 | DiSC3(5) | 8558 | RL1 | 5730 |
| 20 | Yes | E. coli | Polymyxin | Wall | F124 | Thioflavin T | 8954 | VL2 | 5447.5 |
| 20 | Yes | E. coli | Polymyxin | Wall | F101 | Quinacrine | 8716 | VL2 | 5516.5 |
| 20 | Yes | E. coli | Polymyxin | Wall | F138 | Apigenin | 9213 | VL2 | 5107 |
| 20 | Yes | E. coli | Polymyxin | Wall | F65 | H2FDA | 8991 | BL1 | 6338 |
| 20 | Yes | E. coli | Polymyxin | Wall | F142 | Chlorophe | 9917 | BL3 | 7652 |
| 20 | Yes | E. coli | Polymyxin | Wall | F39 | Congo Red | 8843 | BL3 | 8215 |
| 20 | Yes | E. coli | Polymyxin | Wall | F116 | Sunset Yel | 8983 | BL2 | 9166 |
| 20 | Yes | E. coli | Polymyxin | Wall | F10 | Alizarin | 9479 | BL1 | 6946 |
| 20 | Yes | E. coli | Polymyxin | Wall | F84 | Neutral Re | 7189 | BL1 | 6187 |
| 20 | Yes | E. coli | Polymyxin | Wall | F99 | Pyronin Y | 9229 | BL1 | 4949 |
| 20 | Yes | E. coli | Polymyxin | Wall | F108 | Rhodamine | 8632 | BL1 | 4945 |
| 20 | Yes | E. coli | Polymyxin | Wall | F109 | Riboflavin | 8803 | BL1 | 4866 |
| 20 | Yes | E. coli | Polymyxin | Wall | F117 | SYBR Gree | 8395 | BL1 | 240490 |
| 20 | Yes | E. coli | Polymyxin | Wall | F106 | Rhodamine | 7013 | BL1 | 6562 |
| 20 | Yes | E. coli | Polymyxin | Wall | F73 | Malachite ( | 8717 | BL1 | 6575 |
| 20 | Yes | E. coli | Polymyxin | Wall | F7 | Acridine or | 8673 | BL1 | 5919 |
| 20 | Yes | E. coli | Polymyxin | Wall | F192 | 5-Carboxyf | 8143 | BL1 | 13678 |
| 20 | Yes | E. coli | Polymyxin | Wall | F57 | Eosin Y | 8005 | BL1 | 16660 |
| 20 | Yes | E. coli | Polymyxin | Wall | F27 | Calcein | 8146 | BL1 | 18229.5 |
| 20 | Yes | E. coli | Polymyxin | Wall | F12 | Amaranth | 8179 | BL1 | 16282 |
| 20 | Yes | E. coli | None | None | NA | Autofluore: | 8166 | NA | 7733.5 |
| 20 | Yes | E. coli | Tetracyclin | Intracellula | F194 | Oxonol | 8578 | RL1 | 6674.5 |
| 20 | Yes | E. coli | Tetracyclin | Intracellula | F11 | Allura Red | 8900 | BL3 | 5645 |
| 20 | Yes | E. coli | Tetracyclin | Intracellula | F54 | DiSC3(5) | 8997 | RL1 | 5397 |
| 20 | Yes | E. coli | Tetracyclin | Intracellula | F124 | Thioflavin T | 9150 | VL2 | 5589 |
| 20 | Yes | E. coli | Tetracyclin | Intracellula | F101 | Quinacrine | 9072 | VL2 | 5466.5 |
| 20 | Yes | E. coli | Tetracyclin | Intracellula | F138 | Apigenin | 7834 | VL2 | 5450 |
| 20 | Yes | E. coli | Tetracyclin | Intracellula | F65 | H2FDA | 8560 | BL1 | 6235.5 |
| 20 | Yes | E. coli | Tetracyclin | Intracellula | F142 | Chlorophe | 9153 | BL3 | 8803 |
| 20 | Yes | E. coli | Tetracyclin | Intracellula | F39 | Congo Red | 8227 | BL3 | 8783 |
| 20 | Yes | E. coli | Tetracyclin | Intracellula | F116 | Sunset Yel | 8992 | BL2 | 7723.5 |
| 20 | Yes | E. coli | Tetracyclin | Intracellula | F10 | Alizarin | 8842 | BL1 | 6722 |
| 20 | Yes | E. coli | Tetracyclin | Intracellula | F84 | Neutral Re | 9359 | BL1 | 6017 |
| 20 | Yes | E. coli | Tetracyclin | Intracellula | F99 | Pyronin Y | 8602 | BL1 | 4907 |
| 20 | Yes | E. coli | Tetracyclin | Intracellula | F108 | Rhodamine | 8457 | BL1 | 4869 |
| 20 | Yes | E. coli | Tetracyclin | Intracellula | F109 | Riboflavin | 8866 | BL1 | 4783.5 |
| 20 | Yes | E. coli | Tetracyclin | Intracellula | F117 | SYBR Gree | 8044 | BL1 | 178148.5 |
| 20 | Yes | E. coli | Tetracyclin | Intracellula | F106 | Rhodamine | 8927 | BL1 | 6429 |
| 20 | Yes | E. coli | Tetracyclin | Intracellula | F73 | Malachite ( | 7405 | BL1 | 6228 |
| 20 | Yes | E. coli | Tetracyclin | Intracellula | F7 | Acridine or | 9148 | BL1 | 5804.5 |
| 20 | Yes | E. coli | Tetracyclin | Intracellula | F192 | 5-Carboxyf | 8962 | BL1 | 12453.5 |
| 20 | Yes | E. coli | Tetracyclin | Intracellula | F57 | Eosin Y | 8937 | BL1 | 17260 |
| 20 | Yes | E. coli | Tetracyclin | Intracellula | F27 | Calcein | 9559 | BL1 | 17871 |

| | | | | | | | | | | |
|---|---|---|---|---|---|---|---|---|---|---|
| 20 | Yes | E. coli | Tetracyclin | Intracellula | F12 | Amaranth | 9452 | BL1 | | 17157.5 |
| 20 | Yes | E. coli | None | None | NA | Autofluore: | 7711 | NA | | 7243 |
| 20 | Yes | E. coli | Vancomyc | Wall | F194 | Oxonol | 9186 | RL1 | | 6246 |
| 20 | Yes | E. coli | Vancomyc | Wall | F11 | Allura Red | 9081 | BL3 | | 5568 |
| 20 | Yes | E. coli | Vancomyc | Wall | F54 | DiSC3(5) | 7935 | RL1 | | 5446 |
| 20 | Yes | E. coli | Vancomyc | Wall | F124 | Thioflavin T | 9398 | VL2 | | 5572 |
| 20 | Yes | E. coli | Vancomyc | Wall | F101 | Quinacrine | 9563 | VL2 | | 5342 |
| 20 | Yes | E. coli | Vancomyc | Wall | F138 | Apigenin | 8410 | VL2 | | 5657.5 |
| 20 | Yes | E. coli | Vancomyc | Wall | F65 | H2FDA | 9095 | BL1 | | 8063 |
| 20 | Yes | E. coli | Vancomyc | Wall | F142 | Chlorophe | 9586 | BL3 | | 9803 |
| 20 | Yes | E. coli | Vancomyc | Wall | F39 | Congo Red | 9443 | BL3 | | 10429 |
| 20 | Yes | E. coli | Vancomyc | Wall | F116 | Sunset Yel | 8460 | BL2 | | 9731.5 |
| 20 | Yes | E. coli | Vancomyc | Wall | F10 | Alizarin | 9896 | BL1 | | 6856 |
| 20 | Yes | E. coli | Vancomyc | Wall | F84 | Neutral Re | 8563 | BL1 | | 6399 |
| 20 | Yes | E. coli | Vancomyc | Wall | F99 | Pyronin Y | 9397 | BL1 | | 6380 |
| 20 | Yes | E. coli | Vancomyc | Wall | F108 | Rhodamine | 8873 | BL1 | | 5397 |
| 20 | Yes | E. coli | Vancomyc | Wall | F109 | Riboflavin | 8445 | BL1 | | 5197 |
| 20 | Yes | E. coli | Vancomyc | Wall | F117 | SYBR Gree | 8090 | BL1 | | 241507.5 |
| 20 | Yes | E. coli | Vancomyc | Wall | F106 | Rhodamine | 6664 | BL1 | | 11019.5 |
| 20 | Yes | E. coli | Vancomyc | Wall | F73 | Malachite ( | 6643 | BL1 | | 11086 |
| 20 | Yes | E. coli | Vancomyc | Wall | F7 | Acridine or | 6334 | BL1 | | 8935.5 |
| 20 | Yes | E. coli | Vancomyc | Wall | F192 | 5-Carboxyf | 7393 | BL1 | | 25787 |
| 20 | Yes | E. coli | Vancomyc | Wall | F57 | Eosin Y | 7217 | BL1 | | 36282 |
| 20 | Yes | E. coli | Vancomyc | Wall | F27 | Calcein | 7319 | BL1 | | 48932 |
| 20 | Yes | E. coli | Vancomyc | Wall | F12 | Amaranth | 7912 | BL1 | | 47147.5 |
| 20 | Yes | E. coli | None | None | NA | Autofluore: | 12262 | NA | | 1050 |
| 20 | Yes | E. coli | None | None | F194 | Oxonol | 11410 | RL1 | | 1843 |
| 20 | Yes | E. coli | None | None | F11 | Allura Red | 10518 | BL3 | | 2536 |
| 20 | Yes | E. coli | None | None | F54 | DiSC3(5) | 11481 | RL1 | | 2843 |
| 20 | Yes | E. coli | None | None | F124 | Thioflavin T | 11853 | VL2 | | 2955 |
| 20 | Yes | E. coli | None | None | F101 | Quinacrine | 11584 | VL2 | | 2831 |
| 20 | Yes | E. coli | None | None | F138 | Apigenin | 11268 | VL2 | | 2819 |
| 20 | Yes | E. coli | None | None | F65 | H2FDA | 11132 | BL1 | | 3897 |
| 20 | Yes | E. coli | None | None | F142 | Chlorophe | 10825 | BL3 | | 5093 |
| 20 | Yes | E. coli | None | None | F39 | Congo Red | 11422 | BL3 | | 5769.5 |
| 20 | Yes | E. coli | None | None | F116 | Sunset Yel | 11144 | BL2 | | 5899 |
| 20 | Yes | E. coli | None | None | F10 | Alizarin | 10198 | BL1 | | 5461 |
| 20 | Yes | E. coli | None | None | F84 | Neutral Re | 10820 | BL1 | | 4064 |
| 20 | Yes | E. coli | None | None | F99 | Pyronin Y | 10549 | BL1 | | 3659 |
| 20 | Yes | E. coli | None | None | F108 | Rhodamine | 9290 | BL1 | | 3338.5 |
| 20 | Yes | E. coli | None | None | F109 | Riboflavin | 10152 | BL1 | | 3747.5 |
| 20 | Yes | E. coli | None | None | F117 | SYBR Gree | 10930 | BL1 | | 170085.5 |
| 20 | Yes | E. coli | None | None | F106 | Rhodamine | 10550 | BL1 | | 4687 |
| 20 | Yes | E. coli | None | None | F73 | Malachite ( | 8787 | BL1 | | 4042 |
| 20 | Yes | E. coli | None | None | F7 | Acridine or | 11629 | BL1 | | 3867 |
| 20 | Yes | E. coli | None | None | F192 | 5-Carboxyf | 9857 | BL1 | | 12694 |
| 20 | Yes | E. coli | None | None | F57 | Eosin Y | 11015 | BL1 | | 16934 |
| 20 | Yes | E. coli | None | None | F27 | Calcein | 11785 | BL1 | | 17801 |

| | | | | | | | | | |
|---|---|---|---|---|---|---|---|---|---|
| 20 | Yes | E. coli | None | None | F12 | Amaranth | 10943 | BL1 | 16437 |
| 20 | Yes | E. coli | None | None | NA | Autofluore: | 10912 | NA | 6883.5 |
| 20 | Yes | E. coli | Ampicillin | Wall | F194 | Oxonol | 10209 | RL1 | 5113 |
| 20 | Yes | E. coli | Ampicillin | Wall | F11 | Allura Red | 9261 | BL3 | 4069 |
| 20 | Yes | E. coli | Ampicillin | Wall | F54 | DiSC3(5) | 8581 | RL1 | 3988 |
| 20 | Yes | E. coli | Ampicillin | Wall | F124 | Thioflavin T | 10461 | VL2 | 3927 |
| 20 | Yes | E. coli | Ampicillin | Wall | F101 | Quinacrine | 9449 | VL2 | 3611 |
| 20 | Yes | E. coli | Ampicillin | Wall | F138 | Apigenin | 8813 | VL2 | 3675 |
| 20 | Yes | E. coli | Ampicillin | Wall | F65 | H2FDA | 10779 | BL1 | 4761 |
| 20 | Yes | E. coli | Ampicillin | Wall | F142 | Chlorophe | 9430 | BL3 | 5929 |
| 20 | Yes | E. coli | Ampicillin | Wall | F39 | Congo Red | 8019 | BL3 | 6190 |
| 20 | Yes | E. coli | Ampicillin | Wall | F116 | Sunset Yel | 9831 | BL2 | 6859 |
| 20 | Yes | E. coli | Ampicillin | Wall | F10 | Alizarin | 7775 | BL1 | 5690 |
| 20 | Yes | E. coli | Ampicillin | Wall | F84 | Neutral Re | 9966 | BL1 | 4610 |
| 20 | Yes | E. coli | Ampicillin | Wall | F99 | Pyronin Y | 9129 | BL1 | 4220 |
| 20 | Yes | E. coli | Ampicillin | Wall | F108 | Rhodamine | 9182 | BL1 | 3996.5 |
| 20 | Yes | E. coli | Ampicillin | Wall | F109 | Riboflavin | 8798 | BL1 | 4555 |
| 20 | Yes | E. coli | Ampicillin | Wall | F117 | SYBR Gree | 8655 | BL1 | 188941 |
| 20 | Yes | E. coli | Ampicillin | Wall | F106 | Rhodamine | 9391 | BL1 | 5718 |
| 20 | Yes | E. coli | Ampicillin | Wall | F73 | Malachite ( | 7527 | BL1 | 4183 |
| 20 | Yes | E. coli | Ampicillin | Wall | F7 | Acridine or | 10875 | BL1 | 4568 |
| 20 | Yes | E. coli | Ampicillin | Wall | F192 | 5-Carboxyf | 9650 | BL1 | 12103.5 |
| 20 | Yes | E. coli | Ampicillin | Wall | F57 | Eosin Y | 11360 | BL1 | 15522 |
| 20 | Yes | E. coli | Ampicillin | Wall | F27 | Calcein | 11225 | BL1 | 16386 |
| 20 | Yes | E. coli | Ampicillin | Wall | F12 | Amaranth | 9793 | BL1 | 14295 |
| 20 | Yes | E. coli | None | None | NA | Autofluore: | 10403 | NA | 6511 |
| 20 | Yes | E. coli | Bacitracin | Wall | F194 | Oxonol | 10989 | RL1 | 5031 |
| 20 | Yes | E. coli | Bacitracin | Wall | F11 | Allura Red | 11096 | BL3 | 4442.5 |
| 20 | Yes | E. coli | Bacitracin | Wall | F54 | DiSC3(5) | 10259 | RL1 | 3688 |
| 20 | Yes | E. coli | Bacitracin | Wall | F124 | Thioflavin T | 10505 | VL2 | 3718 |
| 20 | Yes | E. coli | Bacitracin | Wall | F101 | Quinacrine | 10507 | VL2 | 4248 |
| 20 | Yes | E. coli | Bacitracin | Wall | F138 | Apigenin | 10573 | VL2 | 3982 |
| 20 | Yes | E. coli | Bacitracin | Wall | F65 | H2FDA | 10168 | BL1 | 5612 |
| 20 | Yes | E. coli | Bacitracin | Wall | F142 | Chlorophe | 10293 | BL3 | 6214 |
| 20 | Yes | E. coli | Bacitracin | Wall | F39 | Congo Red | 9914 | BL3 | 6093 |
| 20 | Yes | E. coli | Bacitracin | Wall | F116 | Sunset Yel | 9369 | BL2 | 5723 |
| 20 | Yes | E. coli | Bacitracin | Wall | F10 | Alizarin | 9928 | BL1 | 5030.5 |
| 20 | Yes | E. coli | Bacitracin | Wall | F84 | Neutral Re | 9689 | BL1 | 4455 |
| 20 | Yes | E. coli | Bacitracin | Wall | F99 | Pyronin Y | 10287 | BL1 | 4012 |
| 20 | Yes | E. coli | Bacitracin | Wall | F108 | Rhodamine | 9556 | BL1 | 3837 |
| 20 | Yes | E. coli | Bacitracin | Wall | F109 | Riboflavin | 10024 | BL1 | 3957.5 |
| 20 | Yes | E. coli | Bacitracin | Wall | F117 | SYBR Gree | 9778 | BL1 | 219185.5 |
| 20 | Yes | E. coli | Bacitracin | Wall | F106 | Rhodamine | 8604 | BL1 | 4866 |
| 20 | Yes | E. coli | Bacitracin | Wall | F73 | Malachite ( | 10108 | BL1 | 4872.5 |
| 20 | Yes | E. coli | Bacitracin | Wall | F7 | Acridine or | 8893 | BL1 | 4761 |
| 20 | Yes | E. coli | Bacitracin | Wall | F192 | 5-Carboxyf | 9116 | BL1 | 12384 |
| 20 | Yes | E. coli | Bacitracin | Wall | F57 | Eosin Y | 9745 | BL1 | 16992 |
| 20 | Yes | E. coli | Bacitracin | Wall | F27 | Calcein | 10221 | BL1 | 16103 |

| | | | | | | | | | |
|---|---|---|---|---|---|---|---|---|---|
| 20 | Yes | E. coli | Bacitracin | Wall | F12 | Amaranth | 9859 | BL1 | 17862 |
| 20 | Yes | E. coli | None | None | NA | Autofluore: | 9533 | NA | 6672 |
| 20 | Yes | E. coli | Cefotaxim | Wall | F194 | Oxonol | 9608 | RL1 | 5434 |
| 20 | Yes | E. coli | Cefotaxim | Wall | F11 | Allura Red | 8678 | BL3 | 4785 |
| 20 | Yes | E. coli | Cefotaxim | Wall | F54 | DiSC3(5) | 8111 | RL1 | 4447 |
| 20 | Yes | E. coli | Cefotaxim | Wall | F124 | Thioflavin T | 8471 | VL2 | 4380 |
| 20 | Yes | E. coli | Cefotaxim | Wall | F101 | Quinacrine | 8054 | VL2 | 4354 |
| 20 | Yes | E. coli | Cefotaxim | Wall | F138 | Apigenin | 8500 | VL2 | 4442.5 |
| 20 | Yes | E. coli | Cefotaxim | Wall | F65 | H2FDA | 8923 | BL1 | 5904 |
| 20 | Yes | E. coli | Cefotaxim | Wall | F142 | Chlorophe | 8587 | BL3 | 6912 |
| 20 | Yes | E. coli | Cefotaxim | Wall | F39 | Congo Red | 8327 | BL3 | 6660 |
| 20 | Yes | E. coli | Cefotaxim | Wall | F116 | Sunset Yel | 7939 | BL2 | 6656 |
| 20 | Yes | E. coli | Cefotaxim | Wall | F10 | Alizarin | 8736 | BL1 | 5981 |
| 20 | Yes | E. coli | Cefotaxim | Wall | F84 | Neutral Re | 7763 | BL1 | 4838 |
| 20 | Yes | E. coli | Cefotaxim | Wall | F99 | Pyronin Y | 8489 | BL1 | 4341 |
| 20 | Yes | E. coli | Cefotaxim | Wall | F108 | Rhodamine | 8827 | BL1 | 4020 |
| 20 | Yes | E. coli | Cefotaxim | Wall | F109 | Riboflavin | 9009 | BL1 | 4056 |
| 20 | Yes | E. coli | Cefotaxim | Wall | F117 | SYBR Gree | 9679 | BL1 | 210497 |
| 20 | Yes | E. coli | Cefotaxim | Wall | F106 | Rhodamine | 9256 | BL1 | 4723 |
| 20 | Yes | E. coli | Cefotaxim | Wall | F73 | Malachite ( | 10419 | BL1 | 4624 |
| 20 | Yes | E. coli | Cefotaxim | Wall | F7 | Acridine or | 9713 | BL1 | 3989 |
| 20 | Yes | E. coli | Cefotaxim | Wall | F192 | 5-Carboxyf | 10178 | BL1 | 12100.5 |
| 20 | Yes | E. coli | Cefotaxim | Wall | F57 | Eosin Y | 9836 | BL1 | 15926.5 |
| 20 | Yes | E. coli | Cefotaxim | Wall | F27 | Calcein | 11081 | BL1 | 16196 |
| 20 | Yes | E. coli | Cefotaxim | Wall | F12 | Amaranth | 10030 | BL1 | 15320.5 |
| 20 | Yes | E. coli | None | None | NA | Autofluore: | 8244 | NA | 7459.5 |
| 20 | Yes | E. coli | Ciprofloxa | Intracellul | F194 | Oxonol | 12320 | RL1 | 5609.5 |
| 20 | Yes | E. coli | Ciprofloxa | Intracellul | F11 | Allura Red | 10462 | BL3 | 4923 |
| 20 | Yes | E. coli | Ciprofloxa | Intracellul | F54 | DiSC3(5) | 10244 | RL1 | 4402.5 |
| 20 | Yes | E. coli | Ciprofloxa | Intracellul | F124 | Thioflavin T | 11373 | VL2 | 4124 |
| 20 | Yes | E. coli | Ciprofloxa | Intracellul | F101 | Quinacrine | 10189 | VL2 | 4397 |
| 20 | Yes | E. coli | Ciprofloxa | Intracellul | F138 | Apigenin | 10805 | VL2 | 4325 |
| 20 | Yes | E. coli | Ciprofloxa | Intracellul | F65 | H2FDA | 10500 | BL1 | 6638.5 |
| 20 | Yes | E. coli | Ciprofloxa | Intracellul | F142 | Chlorophe | 9603 | BL3 | 6247 |
| 20 | Yes | E. coli | Ciprofloxa | Intracellul | F39 | Congo Red | 10783 | BL3 | 7730 |
| 20 | Yes | E. coli | Ciprofloxa | Intracellul | F116 | Sunset Yel | 9249 | BL2 | 6582 |
| 20 | Yes | E. coli | Ciprofloxa | Intracellul | F10 | Alizarin | 9812 | BL1 | 5055 |
| 20 | Yes | E. coli | Ciprofloxa | Intracellul | F84 | Neutral Re | 11324 | BL1 | 4847 |
| 20 | Yes | E. coli | Ciprofloxa | Intracellul | F99 | Pyronin Y | 9832 | BL1 | 4173 |
| 20 | Yes | E. coli | Ciprofloxa | Intracellul | F108 | Rhodamine | 9939 | BL1 | 3943 |
| 20 | Yes | E. coli | Ciprofloxa | Intracellul | F109 | Riboflavin | 9699 | BL1 | 4504 |
| 20 | Yes | E. coli | Ciprofloxa | Intracellul | F117 | SYBR Gree | 8808 | BL1 | 195076 |
| 20 | Yes | E. coli | Ciprofloxa | Intracellul | F106 | Rhodamine | 8721 | BL1 | 5717 |
| 20 | Yes | E. coli | Ciprofloxa | Intracellul | F73 | Malachite ( | 9946 | BL1 | 5060 |
| 20 | Yes | E. coli | Ciprofloxa | Intracellul | F7 | Acridine or | 9161 | BL1 | 6126 |
| 20 | Yes | E. coli | Ciprofloxa | Intracellul | F192 | 5-Carboxyf | 7640 | BL1 | 11711.5 |
| 20 | Yes | E. coli | Ciprofloxa | Intracellul | F57 | Eosin Y | 9860 | BL1 | 15917 |
| 20 | Yes | E. coli | Ciprofloxa | Intracellul | F27 | Calcein | 10191 | BL1 | 15758 |

| | | | | | | | | | |
|---|---|---|---|---|---|---|---|---|---|
| 20 | Yes | E. coli | Ciprofloxa | Intracellula | F12 | Amaranth | 9579 | BL1 | 13594 |
| 20 | Yes | E. coli | None | None | NA | Autofluore | 9884 | NA | 8069.5 |
| 20 | Yes | E. coli | Fosfomyci | Wall | F194 | Oxonol | 10045 | RL1 | 5970 |
| 20 | Yes | E. coli | Fosfomyci | Wall | F11 | Allura Red | 10565 | BL3 | 5343 |
| 20 | Yes | E. coli | Fosfomyci | Wall | F54 | DiSC3(5) | 9923 | RL1 | 4983 |
| 20 | Yes | E. coli | Fosfomyci | Wall | F124 | Thioflavin T | 9207 | VL2 | 4953 |
| 20 | Yes | E. coli | Fosfomyci | Wall | F101 | Quinacrine | 10916 | VL2 | 4942 |
| 20 | Yes | E. coli | Fosfomyci | Wall | F138 | Apigenin | 11096 | VL2 | 4265 |
| 20 | Yes | E. coli | Fosfomyci | Wall | F65 | H2FDA | 11203 | BL1 | 5372 |
| 20 | Yes | E. coli | Fosfomyci | Wall | F142 | Chlorophe | 9236 | BL3 | 6196 |
| 20 | Yes | E. coli | Fosfomyci | Wall | F39 | Congo Red | 11935 | BL3 | 7099 |
| 20 | Yes | E. coli | Fosfomyci | Wall | F116 | Sunset Yel | 9357 | BL2 | 7309 |
| 20 | Yes | E. coli | Fosfomyci | Wall | F10 | Alizarin | 11742 | BL1 | 6397.5 |
| 20 | Yes | E. coli | Fosfomyci | Wall | F84 | Neutral Re | 10282 | BL1 | 4978 |
| 20 | Yes | E. coli | Fosfomyci | Wall | F99 | Pyronin Y | 10632 | BL1 | 4878.5 |
| 20 | Yes | E. coli | Fosfomyci | Wall | F108 | Rhodamine | 9128 | BL1 | 4323 |
| 20 | Yes | E. coli | Fosfomyci | Wall | F109 | Riboflavin | 11595 | BL1 | 4820 |
| 20 | Yes | E. coli | Fosfomyci | Wall | F117 | SYBR Gree | 9697 | BL1 | 226104 |
| 20 | Yes | E. coli | Fosfomyci | Wall | F106 | Rhodamine | 11139 | BL1 | 5557 |
| 20 | Yes | E. coli | Fosfomyci | Wall | F73 | Malachite ( | 9588 | BL1 | 5986 |
| 20 | Yes | E. coli | Fosfomyci | Wall | F7 | Acridine or | 10033 | BL1 | 4543 |
| 20 | Yes | E. coli | Fosfomyci | Wall | F192 | 5-Carboxyf | 11260 | BL1 | 12879 |
| 20 | Yes | E. coli | Fosfomyci | Wall | F57 | Eosin Y | 10580 | BL1 | 16805.5 |
| 20 | Yes | E. coli | Fosfomyci | Wall | F27 | Calcein | 11714 | BL1 | 17088.5 |
| 20 | Yes | E. coli | Fosfomyci | Wall | F12 | Amaranth | 10679 | BL1 | 16325 |
| 20 | Yes | E. coli | None | None | NA | Autofluore | 10088 | NA | 8339 |
| 20 | Yes | E. coli | Nalidixic a | Intracellula | F194 | Oxonol | 11067 | RL1 | 6551 |
| 20 | Yes | E. coli | Nalidixic a | Intracellula | F11 | Allura Red | 9862 | BL3 | 5399 |
| 20 | Yes | E. coli | Nalidixic a | Intracellula | F54 | DiSC3(5) | 10483 | RL1 | 4849 |
| 20 | Yes | E. coli | Nalidixic a | Intracellula | F124 | Thioflavin T | 10872 | VL2 | 4721 |
| 20 | Yes | E. coli | Nalidixic a | Intracellula | F101 | Quinacrine | 9789 | VL2 | 4421 |
| 20 | Yes | E. coli | Nalidixic a | Intracellula | F138 | Apigenin | 11660 | VL2 | 4253.5 |
| 20 | Yes | E. coli | Nalidixic a | Intracellula | F65 | H2FDA | 10581 | BL1 | 5103 |
| 20 | Yes | E. coli | Nalidixic a | Intracellula | F142 | Chlorophe | 10728 | BL3 | 6120 |
| 20 | Yes | E. coli | Nalidixic a | Intracellula | F39 | Congo Red | 11445 | BL3 | 6498 |
| 20 | Yes | E. coli | Nalidixic a | Intracellula | F116 | Sunset Yel | 10442 | BL2 | 6924 |
| 20 | Yes | E. coli | Nalidixic a | Intracellula | F10 | Alizarin | 10519 | BL1 | 5424 |
| 20 | Yes | E. coli | Nalidixic a | Intracellula | F84 | Neutral Re | 9726 | BL1 | 5067.5 |
| 20 | Yes | E. coli | Nalidixic a | Intracellula | F99 | Pyronin Y | 10854 | BL1 | 4310.5 |
| 20 | Yes | E. coli | Nalidixic a | Intracellula | F108 | Rhodamine | 11488 | BL1 | 4612 |
| 20 | Yes | E. coli | Nalidixic a | Intracellula | F109 | Riboflavin | 9494 | BL1 | 4513.5 |
| 20 | Yes | E. coli | Nalidixic a | Intracellula | F117 | SYBR Gree | 10461 | BL1 | 204411 |
| 20 | Yes | E. coli | Nalidixic a | Intracellula | F106 | Rhodamine | 10864 | BL1 | 6230 |
| 20 | Yes | E. coli | Nalidixic a | Intracellula | F73 | Malachite ( | 9774 | BL1 | 4858 |
| 20 | Yes | E. coli | Nalidixic a | Intracellula | F7 | Acridine or | 10029 | BL1 | 5477 |
| 20 | Yes | E. coli | Nalidixic a | Intracellula | F192 | 5-Carboxyf | 9613 | BL1 | 12396 |
| 20 | Yes | E. coli | Nalidixic a | Intracellula | F57 | Eosin Y | 9751 | BL1 | 12868 |
| 20 | Yes | E. coli | Nalidixic a | Intracellula | F27 | Calcein | 9920 | BL1 | 13234 |

| | | | | | | | | | |
|---|---|---|---|---|---|---|---|---|---|
| 20 | Yes | E. coli | Nalidixic a | Intracellul | F12 | Amaranth | 9944 | BL1 | 13335.5 |
| 20 | Yes | E. coli | None | None | NA | Autofluore | 10445 | NA | 8152 |
| 20 | Yes | E. coli | Ofloxacin | Intracellul | F194 | Oxonol | 11361 | RL1 | 6534 |
| 20 | Yes | E. coli | Ofloxacin | Intracellul | F11 | Allura Red | 10781 | BL3 | 5821 |
| 20 | Yes | E. coli | Ofloxacin | Intracellul | F54 | DiSC3(5) | 9704 | RL1 | 5524.5 |
| 20 | Yes | E. coli | Ofloxacin | Intracellul | F124 | Thioflavin T | 10861 | VL2 | 5086 |
| 20 | Yes | E. coli | Ofloxacin | Intracellul | F101 | Quinacrine | 10250 | VL2 | 4667.5 |
| 20 | Yes | E. coli | Ofloxacin | Intracellul | F138 | Apigenin | 10828 | VL2 | 4495 |
| 20 | Yes | E. coli | Ofloxacin | Intracellul | F65 | H2FDA | 11982 | BL1 | 5298 |
| 20 | Yes | E. coli | Ofloxacin | Intracellul | F142 | Chlorophe | 10928 | BL3 | 6036.5 |
| 20 | Yes | E. coli | Ofloxacin | Intracellul | F39 | Congo Red | 10880 | BL3 | 7264 |
| 20 | Yes | E. coli | Ofloxacin | Intracellul | F116 | Sunset Yel | 8970 | BL2 | 7194 |
| 20 | Yes | E. coli | Ofloxacin | Intracellul | F10 | Alizarin | 11642 | BL1 | 6211.5 |
| 20 | Yes | E. coli | Ofloxacin | Intracellul | F84 | Neutral Re | 10317 | BL1 | 5471 |
| 20 | Yes | E. coli | Ofloxacin | Intracellul | F99 | Pyronin Y | 10965 | BL1 | 4485 |
| 20 | Yes | E. coli | Ofloxacin | Intracellul | F108 | Rhodamine | 10931 | BL1 | 4740 |
| 20 | Yes | E. coli | Ofloxacin | Intracellul | F109 | Riboflavin | 10752 | BL1 | 4911.5 |
| 20 | Yes | E. coli | Ofloxacin | Intracellul | F117 | SYBR Gree | 10242 | BL1 | 234847.5 |
| 20 | Yes | E. coli | Ofloxacin | Intracellul | F106 | Rhodamine | 9518 | BL1 | 5742.5 |
| 20 | Yes | E. coli | Ofloxacin | Intracellul | F73 | Malachite ( | 11167 | BL1 | 5818 |
| 20 | Yes | E. coli | Ofloxacin | Intracellul | F7 | Acridine or | 10995 | BL1 | 5572 |
| 20 | Yes | E. coli | Ofloxacin | Intracellul | F192 | 5-Carboxyf | 10417 | BL1 | 11951 |
| 20 | Yes | E. coli | Ofloxacin | Intracellul | F57 | Eosin Y | 11531 | BL1 | 15092 |
| 20 | Yes | E. coli | Ofloxacin | Intracellul | F27 | Calcein | 10271 | BL1 | 12877 |
| 20 | Yes | E. coli | Ofloxacin | Intracellul | F12 | Amaranth | 11149 | BL1 | 14117 |
| 20 | Yes | E. coli | None | None | NA | Autofluore | 9874 | NA | 7420 |
| 20 | Yes | E. coli | Polymyxin | Wall | F194 | Oxonol | 11493 | RL1 | 6247 |
| 20 | Yes | E. coli | Polymyxin | Wall | F11 | Allura Red | 11476 | BL3 | 5518.5 |
| 20 | Yes | E. coli | Polymyxin | Wall | F54 | DiSC3(5) | 10236 | RL1 | 5166 |
| 20 | Yes | E. coli | Polymyxin | Wall | F124 | Thioflavin T | 10219 | VL2 | 5126 |
| 20 | Yes | E. coli | Polymyxin | Wall | F101 | Quinacrine | 10311 | VL2 | 4547 |
| 20 | Yes | E. coli | Polymyxin | Wall | F138 | Apigenin | 11049 | VL2 | 4742 |
| 20 | Yes | E. coli | Polymyxin | Wall | F65 | H2FDA | 10463 | BL1 | 5282 |
| 20 | Yes | E. coli | Polymyxin | Wall | F142 | Chlorophe | 10727 | BL3 | 6438 |
| 20 | Yes | E. coli | Polymyxin | Wall | F39 | Congo Red | 10820 | BL3 | 6623.5 |
| 20 | Yes | E. coli | Polymyxin | Wall | F116 | Sunset Yel | 11159 | BL2 | 6871 |
| 20 | Yes | E. coli | Polymyxin | Wall | F10 | Alizarin | 10061 | BL1 | 5847 |
| 20 | Yes | E. coli | Polymyxin | Wall | F84 | Neutral Re | 11384 | BL1 | 5018.5 |
| 20 | Yes | E. coli | Polymyxin | Wall | F99 | Pyronin Y | 9737 | BL1 | 4557 |
| 20 | Yes | E. coli | Polymyxin | Wall | F108 | Rhodamine | 11270 | BL1 | 4228 |
| 20 | Yes | E. coli | Polymyxin | Wall | F109 | Riboflavin | 11056 | BL1 | 5139.5 |
| 20 | Yes | E. coli | Polymyxin | Wall | F117 | SYBR Gree | 10696 | BL1 | 218590.5 |
| 20 | Yes | E. coli | Polymyxin | Wall | F106 | Rhodamine | 10376 | BL1 | 6087 |
| 20 | Yes | E. coli | Polymyxin | Wall | F73 | Malachite ( | 10912 | BL1 | 5536.5 |
| 20 | Yes | E. coli | Polymyxin | Wall | F7 | Acridine or | 10259 | BL1 | 4954 |
| 20 | Yes | E. coli | Polymyxin | Wall | F192 | 5-Carboxyf | 10355 | BL1 | 12012 |
| 20 | Yes | E. coli | Polymyxin | Wall | F57 | Eosin Y | 10486 | BL1 | 13592.5 |
| 20 | Yes | E. coli | Polymyxin | Wall | F27 | Calcein | 11382 | BL1 | 14705 |

| | | | | | | | | | |
|---|---|---|---|---|---|---|---|---|---|
| 20 | Yes | E. coli | Polymyxin | Wall | F12 | Amaranth | 11480 | BL1 | 16440.5 |
| 20 | Yes | E. coli | None | None | NA | Autofluore: | 10514 | NA | 7405 |
| 20 | Yes | E. coli | Tetracyclin | Intracellula | F194 | Oxonol | 11175 | RL1 | 6068 |
| 20 | Yes | E. coli | Tetracyclin | Intracellula | F11 | Allura Red | 10216 | BL3 | 5182 |
| 20 | Yes | E. coli | Tetracyclin | Intracellula | F54 | DiSC3(5) | 10366 | RL1 | 4883.5 |
| 20 | Yes | E. coli | Tetracyclin | Intracellula | F124 | Thioflavin T | 10460 | VL2 | 4524.5 |
| 20 | Yes | E. coli | Tetracyclin | Intracellula | F101 | Quinacrine | 10501 | VL2 | 4571 |
| 20 | Yes | E. coli | Tetracyclin | Intracellula | F138 | Apigenin | 11041 | VL2 | 4908 |
| 20 | Yes | E. coli | Tetracyclin | Intracellula | F65 | H2FDA | 10240 | BL1 | 5965 |
| 20 | Yes | E. coli | Tetracyclin | Intracellula | F142 | Chlorophe | 10329 | BL3 | 7675 |
| 20 | Yes | E. coli | Tetracyclin | Intracellula | F39 | Congo Red | 10776 | BL3 | 8790 |
| 20 | Yes | E. coli | Tetracyclin | Intracellula | F116 | Sunset Yel | 10079 | BL2 | 8792 |
| 20 | Yes | E. coli | Tetracyclin | Intracellula | F10 | Alizarin | 10243 | BL1 | 6737 |
| 20 | Yes | E. coli | Tetracyclin | Intracellula | F84 | Neutral Re | 10390 | BL1 | 5907.5 |
| 20 | Yes | E. coli | Tetracyclin | Intracellula | F99 | Pyronin Y | 10488 | BL1 | 5415 |
| 20 | Yes | E. coli | Tetracyclin | Intracellula | F108 | Rhodamine | 10319 | BL1 | 4361 |
| 20 | Yes | E. coli | Tetracyclin | Intracellula | F109 | Riboflavin | 11083 | BL1 | 5002 |
| 20 | Yes | E. coli | Tetracyclin | Intracellula | F117 | SYBR Gree | 10436 | BL1 | 194245 |
| 20 | Yes | E. coli | Tetracyclin | Intracellula | F106 | Rhodamine | 10132 | BL1 | 5565.5 |
| 20 | Yes | E. coli | Tetracyclin | Intracellula | F73 | Malachite ( | 10172 | BL1 | 4866.5 |
| 20 | Yes | E. coli | Tetracyclin | Intracellula | F7 | Acridine or | 9617 | BL1 | 4954 |
| 20 | Yes | E. coli | Tetracyclin | Intracellula | F192 | 5-Carboxyf | 10768 | BL1 | 11111.5 |
| 20 | Yes | E. coli | Tetracyclin | Intracellula | F57 | Eosin Y | 9045 | BL1 | 15217 |
| 20 | Yes | E. coli | Tetracyclin | Intracellula | F27 | Calcein | 9294 | BL1 | 16432.5 |
| 20 | Yes | E. coli | Tetracyclin | Intracellula | F12 | Amaranth | 11671 | BL1 | 17981 |
| 20 | Yes | E. coli | None | None | NA | Autofluore: | 9676 | NA | 7839 |
| 20 | Yes | E. coli | Vancomyc | Wall | F194 | Oxonol | 11645 | RL1 | 5628 |
| 20 | Yes | E. coli | Vancomyc | Wall | F11 | Allura Red | 11069 | BL3 | 4804 |
| 20 | Yes | E. coli | Vancomyc | Wall | F54 | DiSC3(5) | 11003 | RL1 | 5180 |
| 20 | Yes | E. coli | Vancomyc | Wall | F124 | Thioflavin T | 12160 | VL2 | 5184.5 |
| 20 | Yes | E. coli | Vancomyc | Wall | F101 | Quinacrine | 10255 | VL2 | 4809 |
| 20 | Yes | E. coli | Vancomyc | Wall | F138 | Apigenin | 11483 | VL2 | 4830 |
| 20 | Yes | E. coli | Vancomyc | Wall | F65 | H2FDA | 11346 | BL1 | 5641.5 |
| 20 | Yes | E. coli | Vancomyc | Wall | F142 | Chlorophe | 11322 | BL3 | 7581.5 |
| 20 | Yes | E. coli | Vancomyc | Wall | F39 | Congo Red | 10740 | BL3 | 8199 |
| 20 | Yes | E. coli | Vancomyc | Wall | F116 | Sunset Yel | 10352 | BL2 | 6813 |
| 20 | Yes | E. coli | Vancomyc | Wall | F10 | Alizarin | 12222 | BL1 | 5914.5 |
| 20 | Yes | E. coli | Vancomyc | Wall | F84 | Neutral Re | 10372 | BL1 | 5518 |
| 20 | Yes | E. coli | Vancomyc | Wall | F99 | Pyronin Y | 9671 | BL1 | 4820 |
| 20 | Yes | E. coli | Vancomyc | Wall | F108 | Rhodamine | 11670 | BL1 | 4818.5 |
| 20 | Yes | E. coli | Vancomyc | Wall | F109 | Riboflavin | 10709 | BL1 | 5140 |
| 20 | Yes | E. coli | Vancomyc | Wall | F117 | SYBR Gree | 9231 | BL1 | 250473 |
| 20 | Yes | E. coli | Vancomyc | Wall | F106 | Rhodamine | 8717 | BL1 | 7223 |
| 20 | Yes | E. coli | Vancomyc | Wall | F73 | Malachite ( | 8948 | BL1 | 7010.5 |
| 20 | Yes | E. coli | Vancomyc | Wall | F7 | Acridine or | 8809 | BL1 | 8014 |
| 20 | Yes | E. coli | Vancomyc | Wall | F192 | 5-Carboxyf | 8904 | BL1 | 26394.5 |
| 20 | Yes | E. coli | Vancomyc | Wall | F57 | Eosin Y | 8850 | BL1 | 45504 |
| 20 | Yes | E. coli | Vancomyc | Wall | F27 | Calcein | 8430 | BL1 | 64544 |

| | | | | | | | | | |
|---|---|---|---|---|---|---|---|---|---|
| 20 | Yes | E. coli | Vancomyc | Wall | F12 | Amaranth | 8643 | BL1 | 49200 |
| 20 | Yes | E. coli | None | None | NA | Autofluore: | 9595 | NA | 1015 |
| 20 | Yes | E. coli | None | None | F194 | Oxonol | 13313 | RL1 | 1877 |
| 20 | Yes | E. coli | None | None | F11 | Allura Red | 11270 | BL3 | 2634 |
| 20 | Yes | E. coli | None | None | F54 | DiSC3(5) | 9009 | RL1 | 2862 |
| 20 | Yes | E. coli | None | None | F124 | Thioflavin 1 | 12848 | VL2 | 2706.5 |
| 20 | Yes | E. coli | None | None | F101 | Quinacrine | 10907 | VL2 | 2822 |
| 20 | Yes | E. coli | None | None | F138 | Apigenin | 10966 | VL2 | 2782 |
| 20 | Yes | E. coli | None | None | F65 | H2FDA | 8947 | BL1 | 3695 |
| 20 | Yes | E. coli | None | None | F142 | Chlorophe | 11341 | BL3 | 5410 |
| 20 | Yes | E. coli | None | None | F39 | Congo Red | 10823 | BL3 | 5769 |
| 20 | Yes | E. coli | None | None | F116 | Sunset Yel | 10207 | BL2 | 5842 |
| 20 | Yes | E. coli | None | None | F10 | Alizarin | 10662 | BL1 | 4421.5 |
| 20 | Yes | E. coli | None | None | F84 | Neutral Re | 10348 | BL1 | 4034.5 |
| 20 | Yes | E. coli | None | None | F99 | Pyronin Y | 10366 | BL1 | 3304 |
| 20 | Yes | E. coli | None | None | F108 | Rhodamin | 10423 | BL1 | 3476 |
| 20 | Yes | E. coli | None | None | F109 | Riboflavin | 9699 | BL1 | 3605 |
| 20 | Yes | E. coli | None | None | F117 | SYBR Gree | 10783 | BL1 | 142424 |
| 20 | Yes | E. coli | None | None | F106 | Rhodamin | 10185 | BL1 | 4102 |
| 20 | Yes | E. coli | None | None | F73 | Malachite ( | 10973 | BL1 | 3678 |
| 20 | Yes | E. coli | None | None | F7 | Acridine or | 9484 | BL1 | 3477 |
| 20 | Yes | E. coli | None | None | F192 | 5-Carboxyf | 10858 | BL1 | 12013.5 |
| 20 | Yes | E. coli | None | None | F57 | Eosin Y | 10768 | BL1 | 17273 |
| 20 | Yes | E. coli | None | None | F27 | Calcein | 11122 | BL1 | 17168 |
| 20 | Yes | E. coli | None | None | F12 | Amaranth | 11332 | BL1 | 17584 |
| 20 | Yes | E. coli | None | None | NA | Autofluore: | 9652 | NA | 6758.5 |
| 20 | Yes | E. coli | Ampicillin | Wall | F194 | Oxonol | 10265 | RL1 | 5704 |
| 20 | Yes | E. coli | Ampicillin | Wall | F11 | Allura Red | 9751 | BL3 | 4211 |
| 20 | Yes | E. coli | Ampicillin | Wall | F54 | DiSC3(5) | 9146 | RL1 | 4234 |
| 20 | Yes | E. coli | Ampicillin | Wall | F124 | Thioflavin 1 | 9364 | VL2 | 3504 |
| 20 | Yes | E. coli | Ampicillin | Wall | F101 | Quinacrine | 9184 | VL2 | 3256 |
| 20 | Yes | E. coli | Ampicillin | Wall | F138 | Apigenin | 9398 | VL2 | 3543 |
| 20 | Yes | E. coli | Ampicillin | Wall | F65 | H2FDA | 10153 | BL1 | 4182 |
| 20 | Yes | E. coli | Ampicillin | Wall | F142 | Chlorophe | 9268 | BL3 | 5782 |
| 20 | Yes | E. coli | Ampicillin | Wall | F39 | Congo Red | 10419 | BL3 | 6489 |
| 20 | Yes | E. coli | Ampicillin | Wall | F116 | Sunset Yel | 10089 | BL2 | 5817 |
| 20 | Yes | E. coli | Ampicillin | Wall | F10 | Alizarin | 10419 | BL1 | 5252 |
| 20 | Yes | E. coli | Ampicillin | Wall | F84 | Neutral Re | 10058 | BL1 | 4098 |
| 20 | Yes | E. coli | Ampicillin | Wall | F99 | Pyronin Y | 9196 | BL1 | 3717 |
| 20 | Yes | E. coli | Ampicillin | Wall | F108 | Rhodamin | 10715 | BL1 | 3774 |
| 20 | Yes | E. coli | Ampicillin | Wall | F109 | Riboflavin | 9804 | BL1 | 4286.5 |
| 20 | Yes | E. coli | Ampicillin | Wall | F117 | SYBR Gree | 9113 | BL1 | 158125 |
| 20 | Yes | E. coli | Ampicillin | Wall | F106 | Rhodamin | 11180 | BL1 | 4635 |
| 20 | Yes | E. coli | Ampicillin | Wall | F73 | Malachite ( | 8764 | BL1 | 4087 |
| 20 | Yes | E. coli | Ampicillin | Wall | F7 | Acridine or | 10532 | BL1 | 3683 |
| 20 | Yes | E. coli | Ampicillin | Wall | F192 | 5-Carboxyf | 10330 | BL1 | 12508 |
| 20 | Yes | E. coli | Ampicillin | Wall | F57 | Eosin Y | 10957 | BL1 | 16949 |
| 20 | Yes | E. coli | Ampicillin | Wall | F27 | Calcein | 10311 | BL1 | 17970 |

| | | | | | | | | | |
|---|---|---|---|---|---|---|---|---|---|
| 20 | Yes | E. coli | Ampicillin | Wall | F12 | Amaranth | 10780 | BL1 | 17753 |
| 20 | Yes | E. coli | None | None | NA | Autofluore: | 9516 | NA | 7118 |
| 20 | Yes | E. coli | Bacitracin | Wall | F194 | Oxonol | 10691 | RL1 | 5781 |
| 20 | Yes | E. coli | Bacitracin | Wall | F11 | Allura Red | 10044 | BL3 | 4368.5 |
| 20 | Yes | E. coli | Bacitracin | Wall | F54 | DiSC3(5) | 10476 | RL1 | 4139.5 |
| 20 | Yes | E. coli | Bacitracin | Wall | F124 | Thioflavin T | 10759 | VL2 | 4074 |
| 20 | Yes | E. coli | Bacitracin | Wall | F101 | Quinacrine | 9291 | VL2 | 4631 |
| 20 | Yes | E. coli | Bacitracin | Wall | F138 | Apigenin | 9383 | VL2 | 4047 |
| 20 | Yes | E. coli | Bacitracin | Wall | F65 | H2FDA | 11330 | BL1 | 4600 |
| 20 | Yes | E. coli | Bacitracin | Wall | F142 | Chlorophe | 9363 | BL3 | 5467 |
| 20 | Yes | E. coli | Bacitracin | Wall | F39 | Congo Red | 10574 | BL3 | 5870.5 |
| 20 | Yes | E. coli | Bacitracin | Wall | F116 | Sunset Yel | 9512 | BL2 | 6227 |
| 20 | Yes | E. coli | Bacitracin | Wall | F10 | Alizarin | 8964 | BL1 | 5620 |
| 20 | Yes | E. coli | Bacitracin | Wall | F84 | Neutral Re | 9668 | BL1 | 5159 |
| 20 | Yes | E. coli | Bacitracin | Wall | F99 | Pyronin Y | 8529 | BL1 | 4199 |
| 20 | Yes | E. coli | Bacitracin | Wall | F108 | Rhodamine | 10507 | BL1 | 4020 |
| 20 | Yes | E. coli | Bacitracin | Wall | F109 | Riboflavin | 8804 | BL1 | 4342 |
| 20 | Yes | E. coli | Bacitracin | Wall | F117 | SYBR Gree | 10343 | BL1 | 193380 |
| 20 | Yes | E. coli | Bacitracin | Wall | F106 | Rhodamine | 9033 | BL1 | 4586 |
| 20 | Yes | E. coli | Bacitracin | Wall | F73 | Malachite ( | 9874 | BL1 | 4427.5 |
| 20 | Yes | E. coli | Bacitracin | Wall | F7 | Acridine or | 10189 | BL1 | 4105 |
| 20 | Yes | E. coli | Bacitracin | Wall | F192 | 5-Carboxyf | 9289 | BL1 | 12872 |
| 20 | Yes | E. coli | Bacitracin | Wall | F57 | Eosin Y | 9723 | BL1 | 18483 |
| 20 | Yes | E. coli | Bacitracin | Wall | F27 | Calcein | 9481 | BL1 | 16875 |
| 20 | Yes | E. coli | Bacitracin | Wall | F12 | Amaranth | 10907 | BL1 | 18395 |
| 20 | Yes | E. coli | None | None | NA | Autofluore: | 9489 | NA | 7160 |
| 20 | Yes | E. coli | Cefotaxime | Wall | F194 | Oxonol | 9267 | RL1 | 6032 |
| 20 | Yes | E. coli | Cefotaxime | Wall | F11 | Allura Red | 9030 | BL3 | 4893.5 |
| 20 | Yes | E. coli | Cefotaxime | Wall | F54 | DiSC3(5) | 8998 | RL1 | 5219 |
| 20 | Yes | E. coli | Cefotaxime | Wall | F124 | Thioflavin T | 9078 | VL2 | 4153.5 |
| 20 | Yes | E. coli | Cefotaxime | Wall | F101 | Quinacrine | 8814 | VL2 | 4078 |
| 20 | Yes | E. coli | Cefotaxime | Wall | F138 | Apigenin | 9785 | VL2 | 3691 |
| 20 | Yes | E. coli | Cefotaxime | Wall | F65 | H2FDA | 7971 | BL1 | 4748 |
| 20 | Yes | E. coli | Cefotaxime | Wall | F142 | Chlorophe | 11053 | BL3 | 6020 |
| 20 | Yes | E. coli | Cefotaxime | Wall | F39 | Congo Red | 9894 | BL3 | 6675 |
| 20 | Yes | E. coli | Cefotaxime | Wall | F116 | Sunset Yel | 9823 | BL2 | 6549 |
| 20 | Yes | E. coli | Cefotaxime | Wall | F10 | Alizarin | 9732 | BL1 | 5240 |
| 20 | Yes | E. coli | Cefotaxime | Wall | F84 | Neutral Re | 9768 | BL1 | 4667 |
| 20 | Yes | E. coli | Cefotaxime | Wall | F99 | Pyronin Y | 9569 | BL1 | 3968 |
| 20 | Yes | E. coli | Cefotaxime | Wall | F108 | Rhodamine | 10216 | BL1 | 4292.5 |
| 20 | Yes | E. coli | Cefotaxime | Wall | F109 | Riboflavin | 8857 | BL1 | 4203 |
| 20 | Yes | E. coli | Cefotaxime | Wall | F117 | SYBR Gree | 10849 | BL1 | 182109 |
| 20 | Yes | E. coli | Cefotaxime | Wall | F106 | Rhodamine | 9823 | BL1 | 4724 |
| 20 | Yes | E. coli | Cefotaxime | Wall | F73 | Malachite ( | 10846 | BL1 | 4452 |
| 20 | Yes | E. coli | Cefotaxime | Wall | F7 | Acridine or | 9636 | BL1 | 4025.5 |
| 20 | Yes | E. coli | Cefotaxime | Wall | F192 | 5-Carboxyf | 10545 | BL1 | 12900 |
| 20 | Yes | E. coli | Cefotaxime | Wall | F57 | Eosin Y | 10386 | BL1 | 17130.5 |
| 20 | Yes | E. coli | Cefotaxime | Wall | F27 | Calcein | 11000 | BL1 | 18310 |

| | | | | | | | | | |
|---|---|---|---|---|---|---|---|---|---|
| 20 | Yes | E. coli | Cefotaxim | Wall | F12 | Amaranth | 11246 | BL1 | 17785.5 |
| 20 | Yes | E. coli | None | None | NA | Autofluore | 9404 | NA | 7475.5 |
| 20 | Yes | E. coli | Ciprofloxa | Intracellul | F194 | Oxonol | 10272 | RL1 | 6200 |
| 20 | Yes | E. coli | Ciprofloxa | Intracellul | F11 | Allura Red | 10960 | BL3 | 4951 |
| 20 | Yes | E. coli | Ciprofloxa | Intracellul | F54 | DiSC3(5) | 9816 | RL1 | 5077 |
| 20 | Yes | E. coli | Ciprofloxa | Intracellul | F124 | Thioflavin T | 10933 | VL2 | 4059 |
| 20 | Yes | E. coli | Ciprofloxa | Intracellul | F101 | Quinacrine | 10174 | VL2 | 3808 |
| 20 | Yes | E. coli | Ciprofloxa | Intracellul | F138 | Apigenin | 11106 | VL2 | 3676 |
| 20 | Yes | E. coli | Ciprofloxa | Intracellul | F65 | H2FDA | 10936 | BL1 | 4666.5 |
| 20 | Yes | E. coli | Ciprofloxa | Intracellul | F142 | Chlorophe | 10154 | BL3 | 5511 |
| 20 | Yes | E. coli | Ciprofloxa | Intracellul | F39 | Congo Red | 10137 | BL3 | 6485 |
| 20 | Yes | E. coli | Ciprofloxa | Intracellul | F116 | Sunset Yel | 10141 | BL2 | 6545 |
| 20 | Yes | E. coli | Ciprofloxa | Intracellul | F10 | Alizarin | 10097 | BL1 | 5936 |
| 20 | Yes | E. coli | Ciprofloxa | Intracellul | F84 | Neutral Re | 10250 | BL1 | 5044 |
| 20 | Yes | E. coli | Ciprofloxa | Intracellul | F99 | Pyronin Y | 10482 | BL1 | 4570 |
| 20 | Yes | E. coli | Ciprofloxa | Intracellul | F108 | Rhodamine | 10224 | BL1 | 4781.5 |
| 20 | Yes | E. coli | Ciprofloxa | Intracellul | F109 | Riboflavin | 9763 | BL1 | 4203 |
| 20 | Yes | E. coli | Ciprofloxa | Intracellul | F117 | SYBR Gree | 10538 | BL1 | 201596.5 |
| 20 | Yes | E. coli | Ciprofloxa | Intracellul | F106 | Rhodamine | 8540 | BL1 | 4872 |
| 20 | Yes | E. coli | Ciprofloxa | Intracellul | F73 | Malachite ( | 10312 | BL1 | 4934 |
| 20 | Yes | E. coli | Ciprofloxa | Intracellul | F7 | Acridine or | 8969 | BL1 | 4258 |
| 20 | Yes | E. coli | Ciprofloxa | Intracellul | F192 | 5-Carboxyf | 10021 | BL1 | 12821 |
| 20 | Yes | E. coli | Ciprofloxa | Intracellul | F57 | Eosin Y | 10450 | BL1 | 16458.5 |
| 20 | Yes | E. coli | Ciprofloxa | Intracellul | F27 | Calcein | 10541 | BL1 | 16848 |
| 20 | Yes | E. coli | Ciprofloxa | Intracellul | F12 | Amaranth | 11232 | BL1 | 15124.5 |
| 20 | Yes | E. coli | None | None | NA | Autofluore | 9415 | NA | 7219 |
| 20 | Yes | E. coli | Fosfomyci | Wall | F194 | Oxonol | 10705 | RL1 | 5978 |
| 20 | Yes | E. coli | Fosfomyci | Wall | F11 | Allura Red | 10786 | BL3 | 5052.5 |
| 20 | Yes | E. coli | Fosfomyci | Wall | F54 | DiSC3(5) | 9538 | RL1 | 5391.5 |
| 20 | Yes | E. coli | Fosfomyci | Wall | F124 | Thioflavin T | 10713 | VL2 | 4222 |
| 20 | Yes | E. coli | Fosfomyci | Wall | F101 | Quinacrine | 10545 | VL2 | 4145 |
| 20 | Yes | E. coli | Fosfomyci | Wall | F138 | Apigenin | 10697 | VL2 | 3811 |
| 20 | Yes | E. coli | Fosfomyci | Wall | F65 | H2FDA | 10862 | BL1 | 4802.5 |
| 20 | Yes | E. coli | Fosfomyci | Wall | F142 | Chlorophe | 9578 | BL3 | 6151.5 |
| 20 | Yes | E. coli | Fosfomyci | Wall | F39 | Congo Red | 11862 | BL3 | 7121 |
| 20 | Yes | E. coli | Fosfomyci | Wall | F116 | Sunset Yel | 10224 | BL2 | 6486.5 |
| 20 | Yes | E. coli | Fosfomyci | Wall | F10 | Alizarin | 10546 | BL1 | 5893 |
| 20 | Yes | E. coli | Fosfomyci | Wall | F84 | Neutral Re | 10474 | BL1 | 4938.5 |
| 20 | Yes | E. coli | Fosfomyci | Wall | F99 | Pyronin Y | 9430 | BL1 | 4385 |
| 20 | Yes | E. coli | Fosfomyci | Wall | F108 | Rhodamine | 11052 | BL1 | 4440 |
| 20 | Yes | E. coli | Fosfomyci | Wall | F109 | Riboflavin | 9804 | BL1 | 4627.5 |
| 20 | Yes | E. coli | Fosfomyci | Wall | F117 | SYBR Gree | 10065 | BL1 | 204754 |
| 20 | Yes | E. coli | Fosfomyci | Wall | F106 | Rhodamine | 9962 | BL1 | 5128.5 |
| 20 | Yes | E. coli | Fosfomyci | Wall | F73 | Malachite ( | 10885 | BL1 | 5181 |
| 20 | Yes | E. coli | Fosfomyci | Wall | F7 | Acridine or | 10425 | BL1 | 4528 |
| 20 | Yes | E. coli | Fosfomyci | Wall | F192 | 5-Carboxyf | 10328 | BL1 | 13558.5 |
| 20 | Yes | E. coli | Fosfomyci | Wall | F57 | Eosin Y | 10851 | BL1 | 16433 |
| 20 | Yes | E. coli | Fosfomyci | Wall | F27 | Calcein | 11023 | BL1 | 17451 |

| | | | | | | | | | |
|---|---|---|---|---|---|---|---|---|---|
| 20 | Yes | E. coli | Fosfomyci | Wall | F12 | Amaranth | 11068 | BL1 | 17062 |
| 20 | Yes | E. coli | None | None | NA | Autofluore | 9733 | NA | 8355 |
| 20 | Yes | E. coli | Nalidixic a | Intracellul | F194 | Oxonol | 11078 | RL1 | 6117 |
| 20 | Yes | E. coli | Nalidixic a | Intracellul | F11 | Allura Red | 10741 | BL3 | 5774 |
| 20 | Yes | E. coli | Nalidixic a | Intracellul | F54 | DiSC3(5) | 9599 | RL1 | 4781 |
| 20 | Yes | E. coli | Nalidixic a | Intracellul | F124 | Thioflavin T | 10912 | VL2 | 4444.5 |
| 20 | Yes | E. coli | Nalidixic a | Intracellul | F101 | Quinacrine | 10962 | VL2 | 4203 |
| 20 | Yes | E. coli | Nalidixic a | Intracellul | F138 | Apigenin | 10471 | VL2 | 3991 |
| 20 | Yes | E. coli | Nalidixic a | Intracellul | F65 | H2FDA | 11126 | BL1 | 5006 |
| 20 | Yes | E. coli | Nalidixic a | Intracellul | F142 | Chlorophe | 11675 | BL3 | 6380 |
| 20 | Yes | E. coli | Nalidixic a | Intracellul | F39 | Congo Red | 10845 | BL3 | 6643 |
| 20 | Yes | E. coli | Nalidixic a | Intracellul | F116 | Sunset Yel | 10505 | BL2 | 7013 |
| 20 | Yes | E. coli | Nalidixic a | Intracellul | F10 | Alizarin | 10896 | BL1 | 5766.5 |
| 20 | Yes | E. coli | Nalidixic a | Intracellul | F84 | Neutral Re | 10613 | BL1 | 5217 |
| 20 | Yes | E. coli | Nalidixic a | Intracellul | F99 | Pyronin Y | 10949 | BL1 | 4630 |
| 20 | Yes | E. coli | Nalidixic a | Intracellul | F108 | Rhodamine | 10331 | BL1 | 5022 |
| 20 | Yes | E. coli | Nalidixic a | Intracellul | F109 | Riboflavin | 10199 | BL1 | 4820 |
| 20 | Yes | E. coli | Nalidixic a | Intracellul | F117 | SYBR Gree | 9976 | BL1 | 163716.5 |
| 20 | Yes | E. coli | Nalidixic a | Intracellul | F106 | Rhodamine | 9146 | BL1 | 5701.5 |
| 20 | Yes | E. coli | Nalidixic a | Intracellul | F73 | Malachite ( | 10254 | BL1 | 4750.5 |
| 20 | Yes | E. coli | Nalidixic a | Intracellul | F7 | Acridine or | 10372 | BL1 | 4797 |
| 20 | Yes | E. coli | Nalidixic a | Intracellul | F192 | 5-Carboxyf | 9761 | BL1 | 12480 |
| 20 | Yes | E. coli | Nalidixic a | Intracellul | F57 | Eosin Y | 10430 | BL1 | 16114.5 |
| 20 | Yes | E. coli | Nalidixic a | Intracellul | F27 | Calcein | 9399 | BL1 | 14986 |
| 20 | Yes | E. coli | Nalidixic a | Intracellul | F12 | Amaranth | 11330 | BL1 | 16508 |
| 20 | Yes | E. coli | None | None | NA | Autofluore | 9678 | NA | 7617 |
| 20 | Yes | E. coli | Ofloxacin | Intracellul | F194 | Oxonol | 10848 | RL1 | 7055 |
| 20 | Yes | E. coli | Ofloxacin | Intracellul | F11 | Allura Red | 10190 | BL3 | 5722 |
| 20 | Yes | E. coli | Ofloxacin | Intracellul | F54 | DiSC3(5) | 10046 | RL1 | 5350 |
| 20 | Yes | E. coli | Ofloxacin | Intracellul | F124 | Thioflavin T | 10728 | VL2 | 4859 |
| 20 | Yes | E. coli | Ofloxacin | Intracellul | F101 | Quinacrine | 10076 | VL2 | 4332.5 |
| 20 | Yes | E. coli | Ofloxacin | Intracellul | F138 | Apigenin | 10826 | VL2 | 4148.5 |
| 20 | Yes | E. coli | Ofloxacin | Intracellul | F65 | H2FDA | 10945 | BL1 | 4843 |
| 20 | Yes | E. coli | Ofloxacin | Intracellul | F142 | Chlorophe | 11059 | BL3 | 6157 |
| 20 | Yes | E. coli | Ofloxacin | Intracellul | F39 | Congo Red | 11112 | BL3 | 6630 |
| 20 | Yes | E. coli | Ofloxacin | Intracellul | F116 | Sunset Yel | 10230 | BL2 | 7779 |
| 20 | Yes | E. coli | Ofloxacin | Intracellul | F10 | Alizarin | 10674 | BL1 | 6029.5 |
| 20 | Yes | E. coli | Ofloxacin | Intracellul | F84 | Neutral Re | 10738 | BL1 | 5701 |
| 20 | Yes | E. coli | Ofloxacin | Intracellul | F99 | Pyronin Y | 10784 | BL1 | 4450.5 |
| 20 | Yes | E. coli | Ofloxacin | Intracellul | F108 | Rhodamine | 10278 | BL1 | 5192 |
| 20 | Yes | E. coli | Ofloxacin | Intracellul | F109 | Riboflavin | 10701 | BL1 | 5067 |
| 20 | Yes | E. coli | Ofloxacin | Intracellul | F117 | SYBR Gree | 10431 | BL1 | 227154 |
| 20 | Yes | E. coli | Ofloxacin | Intracellul | F106 | Rhodamine | 10996 | BL1 | 6373.5 |
| 20 | Yes | E. coli | Ofloxacin | Intracellul | F73 | Malachite ( | 10100 | BL1 | 4948 |
| 20 | Yes | E. coli | Ofloxacin | Intracellul | F7 | Acridine or | 11222 | BL1 | 4886 |
| 20 | Yes | E. coli | Ofloxacin | Intracellul | F192 | 5-Carboxyf | 9725 | BL1 | 11327 |
| 20 | Yes | E. coli | Ofloxacin | Intracellul | F57 | Eosin Y | 11216 | BL1 | 16463 |
| 20 | Yes | E. coli | Ofloxacin | Intracellul | F27 | Calcein | 11043 | BL1 | 16930 |

| | | | | | | | | | |
|---|---|---|---|---|---|---|---|---|---|
| 20 | Yes | E. coli | Ofloxacin | Intracellula | F12 | Amaranth | 10366 | BL1 | 13652 |
| 20 | Yes | E. coli | None | None | NA | Autofluores | 9120 | NA | 7075 |
| 20 | Yes | E. coli | Polymyxin | Wall | F194 | Oxonol | 10796 | RL1 | 6724.5 |
| 20 | Yes | E. coli | Polymyxin | Wall | F11 | Allura Red | 10636 | BL3 | 5311.5 |
| 20 | Yes | E. coli | Polymyxin | Wall | F54 | DiSC3(5) | 10015 | RL1 | 5690 |
| 20 | Yes | E. coli | Polymyxin | Wall | F124 | Thioflavin T | 11132 | VL2 | 4745 |
| 20 | Yes | E. coli | Polymyxin | Wall | F101 | Quinacrine | 11190 | VL2 | 4321.5 |
| 20 | Yes | E. coli | Polymyxin | Wall | F138 | Apigenin | 10480 | VL2 | 4349 |
| 20 | Yes | E. coli | Polymyxin | Wall | F65 | H2FDA | 11285 | BL1 | 5023 |
| 20 | Yes | E. coli | Polymyxin | Wall | F142 | Chlorophe | 11033 | BL3 | 6449 |
| 20 | Yes | E. coli | Polymyxin | Wall | F39 | Congo Red | 10370 | BL3 | 6864 |
| 20 | Yes | E. coli | Polymyxin | Wall | F116 | Sunset Yel | 10897 | BL2 | 7210 |
| 20 | Yes | E. coli | Polymyxin | Wall | F10 | Alizarin | 10132 | BL1 | 6008.5 |
| 20 | Yes | E. coli | Polymyxin | Wall | F84 | Neutral Re | 11598 | BL1 | 5189 |
| 20 | Yes | E. coli | Polymyxin | Wall | F99 | Pyronin Y | 9505 | BL1 | 4620 |
| 20 | Yes | E. coli | Polymyxin | Wall | F108 | Rhodamine | 11463 | BL1 | 4820 |
| 20 | Yes | E. coli | Polymyxin | Wall | F109 | Riboflavin | 9436 | BL1 | 4902 |
| 20 | Yes | E. coli | Polymyxin | Wall | F117 | SYBR Gree | 10147 | BL1 | 248664 |
| 20 | Yes | E. coli | Polymyxin | Wall | F106 | Rhodamine | 9695 | BL1 | 5724 |
| 20 | Yes | E. coli | Polymyxin | Wall | F73 | Malachite ( | 10655 | BL1 | 5417 |
| 20 | Yes | E. coli | Polymyxin | Wall | F7 | Acridine or | 10059 | BL1 | 4865 |
| 20 | Yes | E. coli | Polymyxin | Wall | F192 | 5-Carboxyf | 11320 | BL1 | 13426 |
| 20 | Yes | E. coli | Polymyxin | Wall | F57 | Eosin Y | 10285 | BL1 | 15798 |
| 20 | Yes | E. coli | Polymyxin | Wall | F27 | Calcein | 11345 | BL1 | 16654 |
| 20 | Yes | E. coli | Polymyxin | Wall | F12 | Amaranth | 10693 | BL1 | 15363 |
| 20 | Yes | E. coli | None | None | NA | Autofluores | 9781 | NA | 8641 |
| 20 | Yes | E. coli | Tetracyclin | Intracellula | F194 | Oxonol | 10302 | RL1 | 6373 |
| 20 | Yes | E. coli | Tetracyclin | Intracellula | F11 | Allura Red | 10046 | BL3 | 6163.5 |
| 20 | Yes | E. coli | Tetracyclin | Intracellula | F54 | DiSC3(5) | 9038 | RL1 | 5121.5 |
| 20 | Yes | E. coli | Tetracyclin | Intracellula | F124 | Thioflavin T | 11278 | VL2 | 4895.5 |
| 20 | Yes | E. coli | Tetracyclin | Intracellula | F101 | Quinacrine | 11118 | VL2 | 4668 |
| 20 | Yes | E. coli | Tetracyclin | Intracellula | F138 | Apigenin | 10856 | VL2 | 4163.5 |
| 20 | Yes | E. coli | Tetracyclin | Intracellula | F65 | H2FDA | 10911 | BL1 | 5160 |
| 20 | Yes | E. coli | Tetracyclin | Intracellula | F142 | Chlorophe | 11184 | BL3 | 6518.5 |
| 20 | Yes | E. coli | Tetracyclin | Intracellula | F39 | Congo Red | 10628 | BL3 | 7203.5 |
| 20 | Yes | E. coli | Tetracyclin | Intracellula | F116 | Sunset Yel | 9631 | BL2 | 7615 |
| 20 | Yes | E. coli | Tetracyclin | Intracellula | F10 | Alizarin | 11293 | BL1 | 5957 |
| 20 | Yes | E. coli | Tetracyclin | Intracellula | F84 | Neutral Re | 10626 | BL1 | 6286 |
| 20 | Yes | E. coli | Tetracyclin | Intracellula | F99 | Pyronin Y | 10681 | BL1 | 4622 |
| 20 | Yes | E. coli | Tetracyclin | Intracellula | F108 | Rhodamine | 9957 | BL1 | 5292 |
| 20 | Yes | E. coli | Tetracyclin | Intracellula | F109 | Riboflavin | 9316 | BL1 | 5193.5 |
| 20 | Yes | E. coli | Tetracyclin | Intracellula | F117 | SYBR Gree | 10554 | BL1 | 173781.5 |
| 20 | Yes | E. coli | Tetracyclin | Intracellula | F106 | Rhodamine | 10289 | BL1 | 6924 |
| 20 | Yes | E. coli | Tetracyclin | Intracellula | F73 | Malachite ( | 9654 | BL1 | 5121 |
| 20 | Yes | E. coli | Tetracyclin | Intracellula | F7 | Acridine or | 10696 | BL1 | 5652 |
| 20 | Yes | E. coli | Tetracyclin | Intracellula | F192 | 5-Carboxyf | 9334 | BL1 | 11871 |
| 20 | Yes | E. coli | Tetracyclin | Intracellula | F57 | Eosin Y | 10736 | BL1 | 16484 |
| 20 | Yes | E. coli | Tetracyclin | Intracellula | F27 | Calcein | 10154 | BL1 | 15376 |

| | | | | | | | | | |
|---|---|---|---|---|---|---|---|---|---|
| 20 | Yes | E. coli | Tetracyclin | Intracellula | F12 | Amaranth | 10816 | BL1 | 14809 |
| 20 | Yes | E. coli | None | None | NA | Autofluore: | 9601 | NA | 7679 |
| 20 | Yes | E. coli | Vancomyc | Wall | F194 | Oxonol | 10805 | RL1 | 6789 |
| 20 | Yes | E. coli | Vancomyc | Wall | F11 | Allura Red | 10832 | BL3 | 5525.5 |
| 20 | Yes | E. coli | Vancomyc | Wall | F54 | DiSC3(5) | 9935 | RL1 | 5488 |
| 20 | Yes | E. coli | Vancomyc | Wall | F124 | Thioflavin T | 11874 | VL2 | 5012 |
| 20 | Yes | E. coli | Vancomyc | Wall | F101 | Quinacrine | 10433 | VL2 | 4601 |
| 20 | Yes | E. coli | Vancomyc | Wall | F138 | Apigenin | 10256 | VL2 | 4394 |
| 20 | Yes | E. coli | Vancomyc | Wall | F65 | H2FDA | 11206 | BL1 | 5346.5 |
| 20 | Yes | E. coli | Vancomyc | Wall | F142 | Chlorophe | 11766 | BL3 | 6577.5 |
| 20 | Yes | E. coli | Vancomyc | Wall | F39 | Congo Red | 10953 | BL3 | 7479 |
| 20 | Yes | E. coli | Vancomyc | Wall | F116 | Sunset Yel | 10696 | BL2 | 7726 |
| 20 | Yes | E. coli | Vancomyc | Wall | F10 | Alizarin | 9967 | BL1 | 5924 |
| 20 | Yes | E. coli | Vancomyc | Wall | F84 | Neutral Re | 11357 | BL1 | 5278 |
| 20 | Yes | E. coli | Vancomyc | Wall | F99 | Pyronin Y | 10863 | BL1 | 4251 |
| 20 | Yes | E. coli | Vancomyc | Wall | F108 | Rhodamine | 10348 | BL1 | 4546 |
| 20 | Yes | E. coli | Vancomyc | Wall | F109 | Riboflavin | 10605 | BL1 | 5323 |
| 20 | Yes | E. coli | Vancomyc | Wall | F117 | SYBR Gree | 10830 | BL1 | 226150 |
| 20 | Yes | E. coli | Vancomyc | Wall | F106 | Rhodamine | 7617 | BL1 | 8407 |
| 20 | Yes | E. coli | Vancomyc | Wall | F73 | Malachite ( | 9331 | BL1 | 7122 |
| 20 | Yes | E. coli | Vancomyc | Wall | F7 | Acridine or | 8238 | BL1 | 6168.5 |
| 20 | Yes | E. coli | Vancomyc | Wall | F192 | 5-Carboxyf | 9548 | BL1 | 26955.5 |
| 20 | Yes | E. coli | Vancomyc | Wall | F57 | Eosin Y | 8530 | BL1 | 48812 |
| 20 | Yes | E. coli | Vancomyc | Wall | F27 | Calcein | 9088 | BL1 | 64051 |
| 20 | Yes | E. coli | Vancomyc | Wall | F12 | Amaranth | 9414 | BL1 | 58893 |
| 20 | Yes | A. baylyi | None | None | NA | Autofluore: | 5081 | NA | 1300 |
| 20 | Yes | A. baylyi | None | None | F194 | Oxonol | 5215 | RL1 | 2637 |
| 20 | Yes | A. baylyi | None | None | F11 | Allura Red | 4986 | BL3 | 2968.5 |
| 20 | Yes | A. baylyi | None | None | F54 | DiSC3(5) | 5073 | RL1 | 2523 |
| 20 | Yes | A. baylyi | None | None | F124 | Thioflavin T | 5111 | VL2 | 2693 |
| 20 | Yes | A. baylyi | None | None | F101 | Quinacrine | 4991 | VL2 | 2701 |
| 20 | Yes | A. baylyi | None | None | F138 | Apigenin | 4922 | VL2 | 2780.5 |
| 20 | Yes | A. baylyi | None | None | F65 | H2FDA | 4705 | BL1 | 3733 |
| 20 | Yes | A. baylyi | None | None | F142 | Chlorophe | 4644 | BL3 | 6368 |
| 20 | Yes | A. baylyi | None | None | F39 | Congo Red | 4071 | BL3 | 6357 |
| 20 | Yes | A. baylyi | None | None | F116 | Sunset Yel | 5169 | BL2 | 5555 |
| 20 | Yes | A. baylyi | None | None | F10 | Alizarin | 4631 | BL1 | 4539 |
| 20 | Yes | A. baylyi | None | None | F84 | Neutral Re | 4049 | BL1 | 4703 |
| 20 | Yes | A. baylyi | None | None | F99 | Pyronin Y | 4465 | BL1 | 3849 |
| 20 | Yes | A. baylyi | None | None | F108 | Rhodamine | 4145 | BL1 | 3074 |
| 20 | Yes | A. baylyi | None | None | F109 | Riboflavin | 4207 | BL1 | 3088 |
| 20 | Yes | A. baylyi | None | None | F117 | SYBR Gree | 4296 | BL1 | 250854.5 |
| 20 | Yes | A. baylyi | None | None | F106 | Rhodamine | 4340 | BL1 | 4330.5 |
| 20 | Yes | A. baylyi | None | None | F73 | Malachite ( | 4722 | BL1 | 4110 |
| 20 | Yes | A. baylyi | None | None | F7 | Acridine or | 3906 | BL1 | 4804 |
| 20 | Yes | A. baylyi | None | None | F192 | 5-Carboxyf | 3937 | BL1 | 13912 |
| 20 | Yes | A. baylyi | None | None | F57 | Eosin Y | 4063 | BL1 | 13902 |
| 20 | Yes | A. baylyi | None | None | F27 | Calcein | 3837 | BL1 | 13942 |

| | | | | | | | | | | |
|---|---|---|---|---|---|---|---|---|---|---|
| 20 | Yes | A. baylyi | None | None | F12 | Amaranth | 4110 | BL1 | | 11756.5 |
| 20 | Yes | A. baylyi | None | None | NA | Autofluore: | 4655 | NA | | 5488 |
| 20 | Yes | A. baylyi | Amoxacilli | Wall | F194 | Oxonol | 4701 | RL1 | | 5154 |
| 20 | Yes | A. baylyi | Amoxacilli | Wall | F11 | Allura Red | 4267 | BL3 | | 5170 |
| 20 | Yes | A. baylyi | Amoxacilli | Wall | F54 | DiSC3(5) | 4643 | RL1 | | 5578 |
| 20 | Yes | A. baylyi | Amoxacilli | Wall | F124 | Thioflavin T | 4260 | VL2 | | 4146.5 |
| 20 | Yes | A. baylyi | Amoxacilli | Wall | F101 | Quinacrine | 4542 | VL2 | | 4340.5 |
| 20 | Yes | A. baylyi | Amoxacilli | Wall | F138 | Apigenin | 4451 | VL2 | | 4171 |
| 20 | Yes | A. baylyi | Amoxacilli | Wall | F65 | H2FDA | 4335 | BL1 | | 5134 |
| 20 | Yes | A. baylyi | Amoxacilli | Wall | F142 | Chlorophe | 4558 | BL3 | | 6571.5 |
| 20 | Yes | A. baylyi | Amoxacilli | Wall | F39 | Congo Red | 3390 | BL3 | | 8037.5 |
| 20 | Yes | A. baylyi | Amoxacilli | Wall | F116 | Sunset Yel | 4312 | BL2 | | 5256.5 |
| 20 | Yes | A. baylyi | Amoxacilli | Wall | F10 | Alizarin | 4184 | BL1 | | 5957.5 |
| 20 | Yes | A. baylyi | Amoxacilli | Wall | F84 | Neutral Re | 4261 | BL1 | | 5616 |
| 20 | Yes | A. baylyi | Amoxacilli | Wall | F99 | Pyronin Y | 3581 | BL1 | | 3963 |
| 20 | Yes | A. baylyi | Amoxacilli | Wall | F108 | Rhodamine | 3739 | BL1 | | 3534 |
| 20 | Yes | A. baylyi | Amoxacilli | Wall | F109 | Riboflavin | 3193 | BL1 | | 3733 |
| 20 | Yes | A. baylyi | Amoxacilli | Wall | F117 | SYBR Gree | 3474 | BL1 | | 284958.5 |
| 20 | Yes | A. baylyi | Amoxacilli | Wall | F106 | Rhodamine | 3845 | BL1 | | 8032 |
| 20 | Yes | A. baylyi | Amoxacilli | Wall | F73 | Malachite ( | 3632 | BL1 | | 5440.5 |
| 20 | Yes | A. baylyi | Amoxacilli | Wall | F7 | Acridine or | 3571 | BL1 | | 5755 |
| 20 | Yes | A. baylyi | Amoxacilli | Wall | F192 | 5-Carboxyf | 3756 | BL1 | | 14374 |
| 20 | Yes | A. baylyi | Amoxacilli | Wall | F57 | Eosin Y | 3708 | BL1 | | 13947.5 |
| 20 | Yes | A. baylyi | Amoxacilli | Wall | F27 | Calcein | 3724 | BL1 | | 14152.5 |
| 20 | Yes | A. baylyi | Amoxacilli | Wall | F12 | Amaranth | 3901 | BL1 | | 11031 |
| 20 | Yes | A. baylyi | None | None | NA | Autofluore: | 4970 | NA | | 5948 |
| 20 | Yes | A. baylyi | Azithromyc | Intracellul | F194 | Oxonol | 4446 | RL1 | | 5462.5 |
| 20 | Yes | A. baylyi | Azithromyc | Intracellul | F11 | Allura Red | 4067 | BL3 | | 5320 |
| 20 | Yes | A. baylyi | Azithromyc | Intracellul | F54 | DiSC3(5) | 3902 | RL1 | | 5858 |
| 20 | Yes | A. baylyi | Azithromyc | Intracellul | F124 | Thioflavin T | 4389 | VL2 | | 4644 |
| 20 | Yes | A. baylyi | Azithromyc | Intracellul | F101 | Quinacrine | 4184 | VL2 | | 4432 |
| 20 | Yes | A. baylyi | Azithromyc | Intracellul | F138 | Apigenin | 4044 | VL2 | | 4604 |
| 20 | Yes | A. baylyi | Azithromyc | Intracellul | F65 | H2FDA | 4307 | BL1 | | 5650 |
| 20 | Yes | A. baylyi | Azithromyc | Intracellul | F142 | Chlorophe | 3875 | BL3 | | 6542 |
| 20 | Yes | A. baylyi | Azithromyc | Intracellul | F39 | Congo Red | 4010 | BL3 | | 6801 |
| 20 | Yes | A. baylyi | Azithromyc | Intracellul | F116 | Sunset Yel | 4148 | BL2 | | 7686.5 |
| 20 | Yes | A. baylyi | Azithromyc | Intracellul | F10 | Alizarin | 3864 | BL1 | | 6251 |
| 20 | Yes | A. baylyi | Azithromyc | Intracellul | F84 | Neutral Re | 4192 | BL1 | | 4361.5 |
| 20 | Yes | A. baylyi | Azithromyc | Intracellul | F99 | Pyronin Y | 3650 | BL1 | | 4212 |
| 20 | Yes | A. baylyi | Azithromyc | Intracellul | F108 | Rhodamine | 3263 | BL1 | | 3621 |
| 20 | Yes | A. baylyi | Azithromyc | Intracellul | F109 | Riboflavin | 3800 | BL1 | | 4201.5 |
| 20 | Yes | A. baylyi | Azithromyc | Intracellul | F117 | SYBR Gree | 3709 | BL1 | | 257184 |
| 20 | Yes | A. baylyi | Azithromyc | Intracellul | F106 | Rhodamine | 3939 | BL1 | | 5450 |
| 20 | Yes | A. baylyi | Azithromyc | Intracellul | F73 | Malachite ( | 3894 | BL1 | | 6407.5 |
| 20 | Yes | A. baylyi | Azithromyc | Intracellul | F7 | Acridine or | 3435 | BL1 | | 6084 |
| 20 | Yes | A. baylyi | Azithromyc | Intracellul | F192 | 5-Carboxyf | 3750 | BL1 | | 14422.5 |
| 20 | Yes | A. baylyi | Azithromyc | Intracellul | F57 | Eosin Y | 3799 | BL1 | | 14473 |
| 20 | Yes | A. baylyi | Azithromyc | Intracellul | F27 | Calcein | 3498 | BL1 | | 14516.5 |

| | | | | | | | | | |
|---|---|---|---|---|---|---|---|---|---|
| 20 | Yes | A. baylyi | Azithromyc | Intracellula | F12 | Amaranth | 3895 | BL1 | 10830 |
| 20 | Yes | A. baylyi | None | None | NA | Autofluore: | 4606 | NA | 5785.5 |
| 20 | Yes | A. baylyi | Trimethopr | Intracellula | F194 | Oxonol | 4159 | RL1 | 6197 |
| 20 | Yes | A. baylyi | Trimethopr | Intracellula | F11 | Allura Red | 4043 | BL3 | 6939 |
| 20 | Yes | A. baylyi | Trimethopr | Intracellula | F54 | DiSC3(5) | 4498 | RL1 | 5570.5 |
| 20 | Yes | A. baylyi | Trimethopr | Intracellula | F124 | Thioflavin T | 4194 | VL2 | 5713.5 |
| 20 | Yes | A. baylyi | Trimethopr | Intracellula | F101 | Quinacrine | 4646 | VL2 | 6771 |
| 20 | Yes | A. baylyi | Trimethopr | Intracellula | F138 | Apigenin | 4808 | VL2 | 5127 |
| 20 | Yes | A. baylyi | Trimethopr | Intracellula | F65 | H2FDA | 4179 | BL1 | 6860 |
| 20 | Yes | A. baylyi | Trimethopr | Intracellula | F142 | Chlorophe | 3801 | BL3 | 9419 |
| 20 | Yes | A. baylyi | Trimethopr | Intracellula | F39 | Congo Red | 4266 | BL3 | 8283 |
| 20 | Yes | A. baylyi | Trimethopr | Intracellula | F116 | Sunset Yel | 4400 | BL2 | 5816 |
| 20 | Yes | A. baylyi | Trimethopr | Intracellula | F10 | Alizarin | 4943 | BL1 | 4386 |
| 20 | Yes | A. baylyi | Trimethopr | Intracellula | F84 | Neutral Re | 4615 | BL1 | 3937 |
| 20 | Yes | A. baylyi | Trimethopr | Intracellula | F99 | Pyronin Y | 4015 | BL1 | 4106 |
| 20 | Yes | A. baylyi | Trimethopr | Intracellula | F108 | Rhodamine | 4084 | BL1 | 4526 |
| 20 | Yes | A. baylyi | Trimethopr | Intracellula | F109 | Riboflavin | 3340 | BL1 | 5648.5 |
| 20 | Yes | A. baylyi | Trimethopr | Intracellula | F117 | SYBR Gree | 3315 | BL1 | 296804 |
| 20 | Yes | A. baylyi | Trimethopr | Intracellula | F106 | Rhodamine | 4437 | BL1 | 6156 |
| 20 | Yes | A. baylyi | Trimethopr | Intracellula | F73 | Malachite ( | 4106 | BL1 | 6075.5 |
| 20 | Yes | A. baylyi | Trimethopr | Intracellula | F7 | Acridine or | 4086 | BL1 | 6667 |
| 20 | Yes | A. baylyi | Trimethopr | Intracellula | F192 | 5-Carboxyf | 3821 | BL1 | 14931 |
| 20 | Yes | A. baylyi | Trimethopr | Intracellula | F57 | Eosin Y | 3814 | BL1 | 11597.5 |
| 20 | Yes | A. baylyi | Trimethopr | Intracellula | F27 | Calcein | 3735 | BL1 | 14817 |
| 20 | Yes | A. baylyi | Trimethopr | Intracellula | F12 | Amaranth | 3703 | BL1 | 10458 |
| 20 | Yes | A. baylyi | None | None | NA | Autofluore: | 4925 | NA | 6347 |
| 20 | Yes | A. baylyi | Chloramph | Intracellula | F194 | Oxonol | 3782 | RL1 | 6297 |
| 20 | Yes | A. baylyi | Chloramph | Intracellula | F11 | Allura Red | 3353 | BL3 | 6591 |
| 20 | Yes | A. baylyi | Chloramph | Intracellula | F54 | DiSC3(5) | 3480 | RL1 | 6360 |
| 20 | Yes | A. baylyi | Chloramph | Intracellula | F124 | Thioflavin T | 3367 | VL2 | 5452 |
| 20 | Yes | A. baylyi | Chloramph | Intracellula | F101 | Quinacrine | 3813 | VL2 | 5141 |
| 20 | Yes | A. baylyi | Chloramph | Intracellula | F138 | Apigenin | 3818 | VL2 | 4917.5 |
| 20 | Yes | A. baylyi | Chloramph | Intracellula | F65 | H2FDA | 3583 | BL1 | 6832 |
| 20 | Yes | A. baylyi | Chloramph | Intracellula | F142 | Chlorophe | 3293 | BL3 | 9027 |
| 20 | Yes | A. baylyi | Chloramph | Intracellula | F39 | Congo Red | 3717 | BL3 | 8635 |
| 20 | Yes | A. baylyi | Chloramph | Intracellula | F116 | Sunset Yel | 3799 | BL2 | 8603 |
| 20 | Yes | A. baylyi | Chloramph | Intracellula | F10 | Alizarin | 4052 | BL1 | 5295 |
| 20 | Yes | A. baylyi | Chloramph | Intracellula | F84 | Neutral Re | 4341 | BL1 | 4104 |
| 20 | Yes | A. baylyi | Chloramph | Intracellula | F99 | Pyronin Y | 3816 | BL1 | 4447.5 |
| 20 | Yes | A. baylyi | Chloramph | Intracellula | F108 | Rhodamine | 3474 | BL1 | 4734.5 |
| 20 | Yes | A. baylyi | Chloramph | Intracellula | F109 | Riboflavin | 3477 | BL1 | 4704 |
| 20 | Yes | A. baylyi | Chloramph | Intracellula | F117 | SYBR Gree | 3099 | BL1 | 216249 |
| 20 | Yes | A. baylyi | Chloramph | Intracellula | F106 | Rhodamine | 3029 | BL1 | 8018 |
| 20 | Yes | A. baylyi | Chloramph | Intracellula | F73 | Malachite ( | 3198 | BL1 | 6583 |
| 20 | Yes | A. baylyi | Chloramph | Intracellula | F7 | Acridine or | 3399 | BL1 | 9265 |
| 20 | Yes | A. baylyi | Chloramph | Intracellula | F192 | 5-Carboxyf | 3146 | BL1 | 15650 |
| 20 | Yes | A. baylyi | Chloramph | Intracellula | F57 | Eosin Y | 3160 | BL1 | 13507 |
| 20 | Yes | A. baylyi | Chloramph | Intracellula | F27 | Calcein | 3155 | BL1 | 15006 |

| | | | | | | | | | |
|---|---|---|---|---|---|---|---|---|---|
| 20 | Yes | A. baylyi | Chloramph | Intracellula | F12 | Amaranth | 3481 | BL1 | 13344 |
| 20 | Yes | A. baylyi | None | None | NA | Autofluore: | 4868 | NA | 6371.5 |
| 20 | Yes | A. baylyi | Colistin | Wall | F194 | Oxonol | 3413 | RL1 | 13115 |
| 20 | Yes | A. baylyi | Colistin | Wall | F11 | Allura Red | 3115 | BL3 | 10878 |
| 20 | Yes | A. baylyi | Colistin | Wall | F54 | DiSC3(5) | 3060 | RL1 | 18138 |
| 20 | Yes | A. baylyi | Colistin | Wall | F124 | Thioflavin T | 3155 | VL2 | 9551 |
| 20 | Yes | A. baylyi | Colistin | Wall | F101 | Quinacrine | 3543 | VL2 | 8374 |
| 20 | Yes | A. baylyi | Colistin | Wall | F138 | Apigenin | 3303 | VL2 | 9150 |
| 20 | Yes | A. baylyi | Colistin | Wall | F65 | H2FDA | 3277 | BL1 | 10137 |
| 20 | Yes | A. baylyi | Colistin | Wall | F142 | Chlorophe | 3289 | BL3 | 13322 |
| 20 | Yes | A. baylyi | Colistin | Wall | F39 | Congo Red | 3675 | BL3 | 11239 |
| 20 | Yes | A. baylyi | Colistin | Wall | F116 | Sunset Yel | 3376 | BL2 | 11272 |
| 20 | Yes | A. baylyi | Colistin | Wall | F10 | Alizarin | 3638 | BL1 | 10556 |
| 20 | Yes | A. baylyi | Colistin | Wall | F84 | Neutral Re | 3411 | BL1 | 11469 |
| 20 | Yes | A. baylyi | Colistin | Wall | F99 | Pyronin Y | 3358 | BL1 | 7166 |
| 20 | Yes | A. baylyi | Colistin | Wall | F108 | Rhodamine | 3101 | BL1 | 7928 |
| 20 | Yes | A. baylyi | Colistin | Wall | F109 | Riboflavin | 2969 | BL1 | 9240 |
| 20 | Yes | A. baylyi | Colistin | Wall | F117 | SYBR Gree | 3237 | BL1 | 1922431 |
| 20 | Yes | A. baylyi | Colistin | Wall | F106 | Rhodamine | 2819 | BL1 | 35953 |
| 20 | Yes | A. baylyi | Colistin | Wall | F73 | Malachite ( | 2885 | BL1 | 9191 |
| 20 | Yes | A. baylyi | Colistin | Wall | F7 | Acridine or | 3237 | BL1 | 16676 |
| 20 | Yes | A. baylyi | Colistin | Wall | F192 | 5-Carboxyf | 3081 | BL1 | 17481 |
| 20 | Yes | A. baylyi | Colistin | Wall | F57 | Eosin Y | 2950 | BL1 | 13643 |
| 20 | Yes | A. baylyi | Colistin | Wall | F27 | Calcein | 3336 | BL1 | 21340.5 |
| 20 | Yes | A. baylyi | Colistin | Wall | F12 | Amaranth | 3293 | BL1 | 14331 |
| 20 | Yes | A. baylyi | None | None | NA | Autofluore: | 4760 | NA | 7522.5 |
| 20 | Yes | A. baylyi | Gentamici | Intracellula | F194 | Oxonol | 4399 | RL1 | 7593 |
| 20 | Yes | A. baylyi | Gentamici | Intracellula | F11 | Allura Red | 3918 | BL3 | 6909 |
| 20 | Yes | A. baylyi | Gentamici | Intracellula | F54 | DiSC3(5) | 4011 | RL1 | 7062 |
| 20 | Yes | A. baylyi | Gentamici | Intracellula | F124 | Thioflavin T | 4035 | VL2 | 6974 |
| 20 | Yes | A. baylyi | Gentamici | Intracellula | F101 | Quinacrine | 4664 | VL2 | 6004.5 |
| 20 | Yes | A. baylyi | Gentamici | Intracellula | F138 | Apigenin | 4731 | VL2 | 6094 |
| 20 | Yes | A. baylyi | Gentamici | Intracellula | F65 | H2FDA | 4722 | BL1 | 6753.5 |
| 20 | Yes | A. baylyi | Gentamici | Intracellula | F142 | Chlorophe | 4915 | BL3 | 9456 |
| 20 | Yes | A. baylyi | Gentamici | Intracellula | F39 | Congo Red | 5128 | BL3 | 9297 |
| 20 | Yes | A. baylyi | Gentamici | Intracellula | F116 | Sunset Yel | 5497 | BL2 | 7887 |
| 20 | Yes | A. baylyi | Gentamici | Intracellula | F10 | Alizarin | 5449 | BL1 | 6142 |
| 20 | Yes | A. baylyi | Gentamici | Intracellula | F84 | Neutral Re | 5419 | BL1 | 5799 |
| 20 | Yes | A. baylyi | Gentamici | Intracellula | F99 | Pyronin Y | 4775 | BL1 | 4467 |
| 20 | Yes | A. baylyi | Gentamici | Intracellula | F108 | Rhodamine | 4947 | BL1 | 4610 |
| 20 | Yes | A. baylyi | Gentamici | Intracellula | F109 | Riboflavin | 4791 | BL1 | 4844 |
| 20 | Yes | A. baylyi | Gentamici | Intracellula | F117 | SYBR Gree | 4028 | BL1 | 245447 |
| 20 | Yes | A. baylyi | Gentamici | Intracellula | F106 | Rhodamine | 4778 | BL1 | 6618 |
| 20 | Yes | A. baylyi | Gentamici | Intracellula | F73 | Malachite ( | 4284 | BL1 | 8201.5 |
| 20 | Yes | A. baylyi | Gentamici | Intracellula | F7 | Acridine or | 4070 | BL1 | 7014.5 |
| 20 | Yes | A. baylyi | Gentamici | Intracellula | F192 | 5-Carboxyf | 4153 | BL1 | 13730 |
| 20 | Yes | A. baylyi | Gentamici | Intracellula | F57 | Eosin Y | 3904 | BL1 | 13098.5 |
| 20 | Yes | A. baylyi | Gentamici | Intracellula | F27 | Calcein | 4364 | BL1 | 15446.5 |

| | | | | | | | | | |
|---|---|---|---|---|---|---|---|---|---|
| 20 | Yes | A. baylyi | Gentamici | Intracellula | F12 | Amaranth | 4068 | BL1 | 11473 |
| 20 | Yes | A. baylyi | None | None | NA | Autofluore: | 4683 | NA | 7438 |
| 20 | Yes | A. baylyi | Nitrofurant | Intracellula | F194 | Oxonol | 3926 | RL1 | 6656 |
| 20 | Yes | A. baylyi | Nitrofurant | Intracellula | F11 | Allura Red | 4146 | BL3 | 6875 |
| 20 | Yes | A. baylyi | Nitrofurant | Intracellula | F54 | DiSC3(5) | 4168 | RL1 | 7224 |
| 20 | Yes | A. baylyi | Nitrofurant | Intracellula | F124 | Thioflavin T | 3959 | VL2 | 6245 |
| 20 | Yes | A. baylyi | Nitrofurant | Intracellula | F101 | Quinacrine | 4404 | VL2 | 6574.5 |
| 20 | Yes | A. baylyi | Nitrofurant | Intracellula | F138 | Apigenin | 4523 | VL2 | 6163 |
| 20 | Yes | A. baylyi | Nitrofurant | Intracellula | F65 | H2FDA | 5160 | BL1 | 6832 |
| 20 | Yes | A. baylyi | Nitrofurant | Intracellula | F142 | Chlorophe | 4921 | BL3 | 8789 |
| 20 | Yes | A. baylyi | Nitrofurant | Intracellula | F39 | Congo Red | 4896 | BL3 | 9708.5 |
| 20 | Yes | A. baylyi | Nitrofurant | Intracellula | F116 | Sunset Yel | 5665 | BL2 | 6553 |
| 20 | Yes | A. baylyi | Nitrofurant | Intracellula | F10 | Alizarin | 5454 | BL1 | 5189 |
| 20 | Yes | A. baylyi | Nitrofurant | Intracellula | F84 | Neutral Re | 5403 | BL1 | 4692 |
| 20 | Yes | A. baylyi | Nitrofurant | Intracellula | F99 | Pyronin Y | 4776 | BL1 | 4505.5 |
| 20 | Yes | A. baylyi | Nitrofurant | Intracellula | F108 | Rhodamine | 5396 | BL1 | 5728 |
| 20 | Yes | A. baylyi | Nitrofurant | Intracellula | F109 | Riboflavin | 4548 | BL1 | 4960 |
| 20 | Yes | A. baylyi | Nitrofurant | Intracellula | F117 | SYBR Gree | 4540 | BL1 | 244564.5 |
| 20 | Yes | A. baylyi | Nitrofurant | Intracellula | F106 | Rhodamine | 4741 | BL1 | 7919 |
| 20 | Yes | A. baylyi | Nitrofurant | Intracellula | F73 | Malachite ( | 3969 | BL1 | 8413 |
| 20 | Yes | A. baylyi | Nitrofurant | Intracellula | F7 | Acridine or | 4051 | BL1 | 6875 |
| 20 | Yes | A. baylyi | Nitrofurant | Intracellula | F192 | 5-Carboxyf | 3884 | BL1 | 13549 |
| 20 | Yes | A. baylyi | Nitrofurant | Intracellula | F57 | Eosin Y | 3523 | BL1 | 14720 |
| 20 | Yes | A. baylyi | Nitrofurant | Intracellula | F27 | Calcein | 4187 | BL1 | 13752 |
| 20 | Yes | A. baylyi | Nitrofurant | Intracellula | F12 | Amaranth | 4077 | BL1 | 13042 |
| 20 | Yes | A. baylyi | None | None | NA | Autofluore: | 5161 | NA | 6818 |
| 20 | Yes | A. baylyi | Pentamidi | Wall | F194 | Oxonol | 3098 | RL1 | 11343.5 |
| 20 | Yes | A. baylyi | Pentamidi | Wall | F11 | Allura Red | 2989 | BL3 | 9278 |
| 20 | Yes | A. baylyi | Pentamidi | Wall | F54 | DiSC3(5) | 3089 | RL1 | 12065 |
| 20 | Yes | A. baylyi | Pentamidi | Wall | F124 | Thioflavin T | 3168 | VL2 | 11029.5 |
| 20 | Yes | A. baylyi | Pentamidi | Wall | F101 | Quinacrine | 3366 | VL2 | 8186 |
| 20 | Yes | A. baylyi | Pentamidi | Wall | F138 | Apigenin | 3723 | VL2 | 10725 |
| 20 | Yes | A. baylyi | Pentamidi | Wall | F65 | H2FDA | 3631 | BL1 | 9903 |
| 20 | Yes | A. baylyi | Pentamidi | Wall | F142 | Chlorophe | 4281 | BL3 | 10830 |
| 20 | Yes | A. baylyi | Pentamidi | Wall | F39 | Congo Red | 6048 | BL3 | 10924 |
| 20 | Yes | A. baylyi | Pentamidi | Wall | F116 | Sunset Yel | 5391 | BL2 | 9985 |
| 20 | Yes | A. baylyi | Pentamidi | Wall | F10 | Alizarin | 4038 | BL1 | 7861 |
| 20 | Yes | A. baylyi | Pentamidi | Wall | F84 | Neutral Re | 4674 | BL1 | 7336 |
| 20 | Yes | A. baylyi | Pentamidi | Wall | F99 | Pyronin Y | 5016 | BL1 | 6107 |
| 20 | Yes | A. baylyi | Pentamidi | Wall | F108 | Rhodamine | 4202 | BL1 | 6053 |
| 20 | Yes | A. baylyi | Pentamidi | Wall | F109 | Riboflavin | 4496 | BL1 | 7349.5 |
| 20 | Yes | A. baylyi | Pentamidi | Wall | F117 | SYBR Gree | 4120 | BL1 | 1252297 |
| 20 | Yes | A. baylyi | Pentamidi | Wall | F106 | Rhodamine | 4436 | BL1 | 29317 |
| 20 | Yes | A. baylyi | Pentamidi | Wall | F73 | Malachite ( | 4833 | BL1 | 14230 |
| 20 | Yes | A. baylyi | Pentamidi | Wall | F7 | Acridine or | 4195 | BL1 | 21892 |
| 20 | Yes | A. baylyi | Pentamidi | Wall | F192 | 5-Carboxyf | 4241 | BL1 | 19315 |
| 20 | Yes | A. baylyi | Pentamidi | Wall | F57 | Eosin Y | 3914 | BL1 | 14528.5 |
| 20 | Yes | A. baylyi | Pentamidi | Wall | F27 | Calcein | 4207 | BL1 | 17614 |

| | | | | | | | | | |
|---|---|---|---|---|---|---|---|---|---|
| 20 | Yes | A. baylyi | Pentamidi | Wall | F12 | Amaranth | 3180 | BL1 | 13131 |
| 20 | Yes | A. baylyi | None | None | NA | Autofluore | 4980 | NA | 7493.5 |
| 20 | Yes | A. baylyi | Rifampicin | Intracellul | F194 | Oxonol | 4589 | RL1 | 6978 |
| 20 | Yes | A. baylyi | Rifampicin | Intracellul | F11 | Allura Red | 4154 | BL3 | 7313.5 |
| 20 | Yes | A. baylyi | Rifampicin | Intracellul | F54 | DiSC3(5) | 4345 | RL1 | 5317 |
| 20 | Yes | A. baylyi | Rifampicin | Intracellul | F124 | Thioflavin T | 4439 | VL2 | 5912 |
| 20 | Yes | A. baylyi | Rifampicin | Intracellul | F101 | Quinacrine | 4718 | VL2 | 5652 |
| 20 | Yes | A. baylyi | Rifampicin | Intracellul | F138 | Apigenin | 4649 | VL2 | 4994 |
| 20 | Yes | A. baylyi | Rifampicin | Intracellul | F65 | H2FDA | 4507 | BL1 | 5859 |
| 20 | Yes | A. baylyi | Rifampicin | Intracellul | F142 | Chlorophe | 4458 | BL3 | 8672 |
| 20 | Yes | A. baylyi | Rifampicin | Intracellul | F39 | Congo Red | 5522 | BL3 | 9367 |
| 20 | Yes | A. baylyi | Rifampicin | Intracellul | F116 | Sunset Yel | 5791 | BL2 | 6539 |
| 20 | Yes | A. baylyi | Rifampicin | Intracellul | F10 | Alizarin | 5381 | BL1 | 6629 |
| 20 | Yes | A. baylyi | Rifampicin | Intracellul | F84 | Neutral Re | 4981 | BL1 | 5513 |
| 20 | Yes | A. baylyi | Rifampicin | Intracellul | F99 | Pyronin Y | 5250 | BL1 | 4442 |
| 20 | Yes | A. baylyi | Rifampicin | Intracellul | F108 | Rhodamine | 5297 | BL1 | 4102 |
| 20 | Yes | A. baylyi | Rifampicin | Intracellul | F109 | Riboflavin | 4639 | BL1 | 5209 |
| 20 | Yes | A. baylyi | Rifampicin | Intracellul | F117 | SYBR Gree | 4064 | BL1 | 331536 |
| 20 | Yes | A. baylyi | Rifampicin | Intracellul | F106 | Rhodamine | 5061 | BL1 | 7704 |
| 20 | Yes | A. baylyi | Rifampicin | Intracellul | F73 | Malachite | 4253 | BL1 | 8483 |
| 20 | Yes | A. baylyi | Rifampicin | Intracellul | F7 | Acridine or | 4358 | BL1 | 6042 |
| 20 | Yes | A. baylyi | Rifampicin | Intracellul | F192 | 5-Carboxyf | 4335 | BL1 | 12184 |
| 20 | Yes | A. baylyi | Rifampicin | Intracellul | F57 | Eosin Y | 3525 | BL1 | 13221 |
| 20 | Yes | A. baylyi | Rifampicin | Intracellul | F27 | Calcein | 4158 | BL1 | 12392.5 |
| 20 | Yes | A. baylyi | Rifampicin | Intracellul | F12 | Amaranth | 4363 | BL1 | 12637 |
| 20 | Yes | A. baylyi | None | None | NA | Autofluore | 5348 | NA | 6899 |
| 20 | Yes | A. baylyi | Trimethopr | Intracellul | F194 | Oxonol | 4756 | RL1 | 8140 |
| 20 | Yes | A. baylyi | Trimethopr | Intracellul | F11 | Allura Red | 4268 | BL3 | 8624 |
| 20 | Yes | A. baylyi | Trimethopr | Intracellul | F54 | DiSC3(5) | 4433 | RL1 | 6339 |
| 20 | Yes | A. baylyi | Trimethopr | Intracellul | F124 | Thioflavin T | 4418 | VL2 | 6906 |
| 20 | Yes | A. baylyi | Trimethopr | Intracellul | F101 | Quinacrine | 4693 | VL2 | 7148 |
| 20 | Yes | A. baylyi | Trimethopr | Intracellul | F138 | Apigenin | 4466 | VL2 | 5965.5 |
| 20 | Yes | A. baylyi | Trimethopr | Intracellul | F65 | H2FDA | 4984 | BL1 | 7937.5 |
| 20 | Yes | A. baylyi | Trimethopr | Intracellul | F142 | Chlorophe | 4788 | BL3 | 9839.5 |
| 20 | Yes | A. baylyi | Trimethopr | Intracellul | F39 | Congo Red | 5334 | BL3 | 9741 |
| 20 | Yes | A. baylyi | Trimethopr | Intracellul | F116 | Sunset Yel | 5428 | BL2 | 7920 |
| 20 | Yes | A. baylyi | Trimethopr | Intracellul | F10 | Alizarin | 5600 | BL1 | 5484 |
| 20 | Yes | A. baylyi | Trimethopr | Intracellul | F84 | Neutral Re | 5102 | BL1 | 5104 |
| 20 | Yes | A. baylyi | Trimethopr | Intracellul | F99 | Pyronin Y | 4937 | BL1 | 5149 |
| 20 | Yes | A. baylyi | Trimethopr | Intracellul | F108 | Rhodamine | 4446 | BL1 | 6311.5 |
| 20 | Yes | A. baylyi | Trimethopr | Intracellul | F109 | Riboflavin | 4739 | BL1 | 5163 |
| 20 | Yes | A. baylyi | Trimethopr | Intracellul | F117 | SYBR Gree | 4396 | BL1 | 299585 |
| 20 | Yes | A. baylyi | Trimethopr | Intracellul | F106 | Rhodamine | 4279 | BL1 | 6928 |
| 20 | Yes | A. baylyi | Trimethopr | Intracellul | F73 | Malachite | 4573 | BL1 | 6125 |
| 20 | Yes | A. baylyi | Trimethopr | Intracellul | F7 | Acridine or | 4119 | BL1 | 6636 |
| 20 | Yes | A. baylyi | Trimethopr | Intracellul | F192 | 5-Carboxyf | 3910 | BL1 | 13105 |
| 20 | Yes | A. baylyi | Trimethopr | Intracellul | F57 | Eosin Y | 4379 | BL1 | 19017 |
| 20 | Yes | A. baylyi | Trimethopr | Intracellul | F27 | Calcein | 4423 | BL1 | 16258 |

| | | | | | | | | | |
|---|---|---|---|---|---|---|---|---|---|
| 20 | Yes | A. baylyi | Trimethopr | Intracellula | F12 | Amaranth | 4233 | BL1 | 14814 |
| 20 | Yes | A. baylyi | None | None | NA | Autofluore: | 4864 | NA | 1280 |
| 20 | Yes | A. baylyi | None | None | F194 | Oxonol | 4761 | RL1 | 2514 |
| 20 | Yes | A. baylyi | None | None | F11 | Allura Red | 5007 | BL3 | 3115 |
| 20 | Yes | A. baylyi | None | None | F54 | DiSC3(5) | 4641 | RL1 | 2967 |
| 20 | Yes | A. baylyi | None | None | F124 | Thioflavin T | 4657 | VL2 | 2950 |
| 20 | Yes | A. baylyi | None | None | F101 | Quinacrine | 4825 | VL2 | 2920 |
| 20 | Yes | A. baylyi | None | None | F138 | Apigenin | 5104 | VL2 | 2860.5 |
| 20 | Yes | A. baylyi | None | None | F65 | H2FDA | 4534 | BL1 | 4468.5 |
| 20 | Yes | A. baylyi | None | None | F142 | Chlorophe | 4484 | BL3 | 6415 |
| 20 | Yes | A. baylyi | None | None | F39 | Congo Red | 4817 | BL3 | 6257 |
| 20 | Yes | A. baylyi | None | None | F116 | Sunset Yel | 4637 | BL2 | 5476 |
| 20 | Yes | A. baylyi | None | None | F10 | Alizarin | 4287 | BL1 | 4337 |
| 20 | Yes | A. baylyi | None | None | F84 | Neutral Re | 4529 | BL1 | 3592 |
| 20 | Yes | A. baylyi | None | None | F99 | Pyronin Y | 4141 | BL1 | 3161 |
| 20 | Yes | A. baylyi | None | None | F108 | Rhodamine | 3857 | BL1 | 3184 |
| 20 | Yes | A. baylyi | None | None | F109 | Riboflavin | 4119 | BL1 | 2999 |
| 20 | Yes | A. baylyi | None | None | F117 | SYBR Gree | 3808 | BL1 | 273371 |
| 20 | Yes | A. baylyi | None | None | F106 | Rhodamine | 3857 | BL1 | 7214 |
| 20 | Yes | A. baylyi | None | None | F73 | Malachite ( | 4477 | BL1 | 6268 |
| 20 | Yes | A. baylyi | None | None | F7 | Acridine or | 3906 | BL1 | 4845 |
| 20 | Yes | A. baylyi | None | None | F192 | 5-Carboxyf | 4138 | BL1 | 12732 |
| 20 | Yes | A. baylyi | None | None | F57 | Eosin Y | 4230 | BL1 | 12712 |
| 20 | Yes | A. baylyi | None | None | F27 | Calcein | 4230 | BL1 | 15408.5 |
| 20 | Yes | A. baylyi | None | None | F12 | Amaranth | 3456 | BL1 | 11688 |
| 20 | Yes | A. baylyi | None | None | NA | Autofluore: | 4438 | NA | 5172.5 |
| 20 | Yes | A. baylyi | Amoxacilli | Wall | F194 | Oxonol | 4150 | RL1 | 4483 |
| 20 | Yes | A. baylyi | Amoxacilli | Wall | F11 | Allura Red | 3612 | BL3 | 4348 |
| 20 | Yes | A. baylyi | Amoxacilli | Wall | F54 | DiSC3(5) | 3893 | RL1 | 4169 |
| 20 | Yes | A. baylyi | Amoxacilli | Wall | F124 | Thioflavin T | 4319 | VL2 | 4613 |
| 20 | Yes | A. baylyi | Amoxacilli | Wall | F101 | Quinacrine | 3991 | VL2 | 3933 |
| 20 | Yes | A. baylyi | Amoxacilli | Wall | F138 | Apigenin | 4079 | VL2 | 3632 |
| 20 | Yes | A. baylyi | Amoxacilli | Wall | F65 | H2FDA | 4293 | BL1 | 5065 |
| 20 | Yes | A. baylyi | Amoxacilli | Wall | F142 | Chlorophe | 3701 | BL3 | 7426 |
| 20 | Yes | A. baylyi | Amoxacilli | Wall | F39 | Congo Red | 4306 | BL3 | 7148.5 |
| 20 | Yes | A. baylyi | Amoxacilli | Wall | F116 | Sunset Yel | 4024 | BL2 | 7519.5 |
| 20 | Yes | A. baylyi | Amoxacilli | Wall | F10 | Alizarin | 3206 | BL1 | 5759.5 |
| 20 | Yes | A. baylyi | Amoxacilli | Wall | F84 | Neutral Re | 4199 | BL1 | 5163 |
| 20 | Yes | A. baylyi | Amoxacilli | Wall | F99 | Pyronin Y | 4006 | BL1 | 4676 |
| 20 | Yes | A. baylyi | Amoxacilli | Wall | F108 | Rhodamine | 3571 | BL1 | 4612 |
| 20 | Yes | A. baylyi | Amoxacilli | Wall | F109 | Riboflavin | 3424 | BL1 | 3844 |
| 20 | Yes | A. baylyi | Amoxacilli | Wall | F117 | SYBR Gree | 3222 | BL1 | 298316.5 |
| 20 | Yes | A. baylyi | Amoxacilli | Wall | F106 | Rhodamine | 3666 | BL1 | 7839 |
| 20 | Yes | A. baylyi | Amoxacilli | Wall | F73 | Malachite ( | 3352 | BL1 | 7483.5 |
| 20 | Yes | A. baylyi | Amoxacilli | Wall | F7 | Acridine or | 3709 | BL1 | 6768 |
| 20 | Yes | A. baylyi | Amoxacilli | Wall | F192 | 5-Carboxyf | 3967 | BL1 | 11870 |
| 20 | Yes | A. baylyi | Amoxacilli | Wall | F57 | Eosin Y | 3685 | BL1 | 13814 |
| 20 | Yes | A. baylyi | Amoxacilli | Wall | F27 | Calcein | 3386 | BL1 | 14728.5 |

| | | | | | | | | | |
|---|---|---|---|---|---|---|---|---|---|
| 20 | Yes | A. baylyi | Amoxacilli | Wall | F12 | Amaranth | 3085 | BL1 | 11177 |
| 20 | Yes | A. baylyi | None | None | NA | Autofluore | 4436 | NA | 5894 |
| 20 | Yes | A. baylyi | Azithromyc | Intracellul | F194 | Oxonol | 4167 | RL1 | 4936 |
| 20 | Yes | A. baylyi | Azithromyc | Intracellul | F11 | Allura Red | 3753 | BL3 | 4534 |
| 20 | Yes | A. baylyi | Azithromyc | Intracellul | F54 | DiSC3(5) | 3966 | RL1 | 5377 |
| 20 | Yes | A. baylyi | Azithromyc | Intracellul | F124 | Thioflavin T | 4004 | VL2 | 5273 |
| 20 | Yes | A. baylyi | Azithromyc | Intracellul | F101 | Quinacrine | 4073 | VL2 | 5056 |
| 20 | Yes | A. baylyi | Azithromyc | Intracellul | F138 | Apigenin | 3781 | VL2 | 4780 |
| 20 | Yes | A. baylyi | Azithromyc | Intracellul | F65 | H2FDA | 3939 | BL1 | 5872 |
| 20 | Yes | A. baylyi | Azithromyc | Intracellul | F142 | Chlorophe | 4081 | BL3 | 8191 |
| 20 | Yes | A. baylyi | Azithromyc | Intracellul | F39 | Congo Red | 3763 | BL3 | 8084 |
| 20 | Yes | A. baylyi | Azithromyc | Intracellul | F116 | Sunset Yel | 4203 | BL2 | 5903 |
| 20 | Yes | A. baylyi | Azithromyc | Intracellul | F10 | Alizarin | 4160 | BL1 | 5066 |
| 20 | Yes | A. baylyi | Azithromyc | Intracellul | F84 | Neutral Re | 4325 | BL1 | 4214 |
| 20 | Yes | A. baylyi | Azithromyc | Intracellul | F99 | Pyronin Y | 3459 | BL1 | 4079 |
| 20 | Yes | A. baylyi | Azithromyc | Intracellul | F108 | Rhodamine | 3889 | BL1 | 4475 |
| 20 | Yes | A. baylyi | Azithromyc | Intracellul | F109 | Riboflavin | 3135 | BL1 | 4568 |
| 20 | Yes | A. baylyi | Azithromyc | Intracellul | F117 | SYBR Gree | 2789 | BL1 | 281925 |
| 20 | Yes | A. baylyi | Azithromyc | Intracellul | F106 | Rhodamine | 3331 | BL1 | 8686 |
| 20 | Yes | A. baylyi | Azithromyc | Intracellul | F73 | Malachite | 3928 | BL1 | 7412 |
| 20 | Yes | A. baylyi | Azithromyc | Intracellul | F7 | Acridine or | 3224 | BL1 | 7517.5 |
| 20 | Yes | A. baylyi | Azithromyc | Intracellul | F192 | 5-Carboxyf | 3375 | BL1 | 12947 |
| 20 | Yes | A. baylyi | Azithromyc | Intracellul | F57 | Eosin Y | 3521 | BL1 | 13660 |
| 20 | Yes | A. baylyi | Azithromyc | Intracellul | F27 | Calcein | 3965 | BL1 | 16382 |
| 20 | Yes | A. baylyi | Azithromyc | Intracellul | F12 | Amaranth | 3575 | BL1 | 13384 |
| 20 | Yes | A. baylyi | None | None | NA | Autofluore | 4328 | NA | 5948 |
| 20 | Yes | A. baylyi | Trimethopr | Intracellul | F194 | Oxonol | 4053 | RL1 | 5748 |
| 20 | Yes | A. baylyi | Trimethopr | Intracellul | F11 | Allura Red | 3970 | BL3 | 5542 |
| 20 | Yes | A. baylyi | Trimethopr | Intracellul | F54 | DiSC3(5) | 4103 | RL1 | 4925 |
| 20 | Yes | A. baylyi | Trimethopr | Intracellul | F124 | Thioflavin T | 4035 | VL2 | 5316 |
| 20 | Yes | A. baylyi | Trimethopr | Intracellul | F101 | Quinacrine | 4226 | VL2 | 4854 |
| 20 | Yes | A. baylyi | Trimethopr | Intracellul | F138 | Apigenin | 4052 | VL2 | 4743 |
| 20 | Yes | A. baylyi | Trimethopr | Intracellul | F65 | H2FDA | 3703 | BL1 | 5536 |
| 20 | Yes | A. baylyi | Trimethopr | Intracellul | F142 | Chlorophe | 4029 | BL3 | 8164 |
| 20 | Yes | A. baylyi | Trimethopr | Intracellul | F39 | Congo Red | 3944 | BL3 | 8635.5 |
| 20 | Yes | A. baylyi | Trimethopr | Intracellul | F116 | Sunset Yel | 3936 | BL2 | 7195.5 |
| 20 | Yes | A. baylyi | Trimethopr | Intracellul | F10 | Alizarin | 4260 | BL1 | 5734 |
| 20 | Yes | A. baylyi | Trimethopr | Intracellul | F84 | Neutral Re | 3828 | BL1 | 5743.5 |
| 20 | Yes | A. baylyi | Trimethopr | Intracellul | F99 | Pyronin Y | 4120 | BL1 | 4173.5 |
| 20 | Yes | A. baylyi | Trimethopr | Intracellul | F108 | Rhodamine | 3882 | BL1 | 4119 |
| 20 | Yes | A. baylyi | Trimethopr | Intracellul | F109 | Riboflavin | 3676 | BL1 | 4297 |
| 20 | Yes | A. baylyi | Trimethopr | Intracellul | F117 | SYBR Gree | 3877 | BL1 | 306987 |
| 20 | Yes | A. baylyi | Trimethopr | Intracellul | F106 | Rhodamine | 3701 | BL1 | 7377 |
| 20 | Yes | A. baylyi | Trimethopr | Intracellul | F73 | Malachite | 3371 | BL1 | 8119 |
| 20 | Yes | A. baylyi | Trimethopr | Intracellul | F7 | Acridine or | 3905 | BL1 | 7884 |
| 20 | Yes | A. baylyi | Trimethopr | Intracellul | F192 | 5-Carboxyf | 3616 | BL1 | 12725.5 |
| 20 | Yes | A. baylyi | Trimethopr | Intracellul | F57 | Eosin Y | 3244 | BL1 | 16317.5 |
| 20 | Yes | A. baylyi | Trimethopr | Intracellul | F27 | Calcein | 3458 | BL1 | 15694 |

| | | | | | | | | | |
|---|---|---|---|---|---|---|---|---|---|
| 20 | Yes | A. baylyi | Trimethopr | Intracellula | F12 | Amaranth | 3817 | BL1 | 14298 |
| 20 | Yes | A. baylyi | None | None | NA | Autofluore: | 4342 | NA | 5994.5 |
| 20 | Yes | A. baylyi | Chloramph | Intracellula | F194 | Oxonol | 3458 | RL1 | 6114.5 |
| 20 | Yes | A. baylyi | Chloramph | Intracellula | F11 | Allura Red | 3228 | BL3 | 6150.5 |
| 20 | Yes | A. baylyi | Chloramph | Intracellula | F54 | DiSC3(5) | 3376 | RL1 | 5537 |
| 20 | Yes | A. baylyi | Chloramph | Intracellula | F124 | Thioflavin T | 3364 | VL2 | 5907.5 |
| 20 | Yes | A. baylyi | Chloramph | Intracellula | F101 | Quinacrine | 3711 | VL2 | 5825 |
| 20 | Yes | A. baylyi | Chloramph | Intracellula | F138 | Apigenin | 3804 | VL2 | 5139 |
| 20 | Yes | A. baylyi | Chloramph | Intracellula | F65 | H2FDA | 3210 | BL1 | 6427.5 |
| 20 | Yes | A. baylyi | Chloramph | Intracellula | F142 | Chlorophe | 3450 | BL3 | 9390 |
| 20 | Yes | A. baylyi | Chloramph | Intracellula | F39 | Congo Red | 3322 | BL3 | 7213.5 |
| 20 | Yes | A. baylyi | Chloramph | Intracellula | F116 | Sunset Yel | 3610 | BL2 | 6333.5 |
| 20 | Yes | A. baylyi | Chloramph | Intracellula | F10 | Alizarin | 3741 | BL1 | 5458 |
| 20 | Yes | A. baylyi | Chloramph | Intracellula | F84 | Neutral Re | 3894 | BL1 | 4263.5 |
| 20 | Yes | A. baylyi | Chloramph | Intracellula | F99 | Pyronin Y | 3601 | BL1 | 4665 |
| 20 | Yes | A. baylyi | Chloramph | Intracellula | F108 | Rhodamine | 3427 | BL1 | 4695 |
| 20 | Yes | A. baylyi | Chloramph | Intracellula | F109 | Riboflavin | 3095 | BL1 | 4646 |
| 20 | Yes | A. baylyi | Chloramph | Intracellula | F117 | SYBR Gree | 2441 | BL1 | 228324 |
| 20 | Yes | A. baylyi | Chloramph | Intracellula | F106 | Rhodamine | 3183 | BL1 | 8673 |
| 20 | Yes | A. baylyi | Chloramph | Intracellula | F73 | Malachite ( | 2793 | BL1 | 7622 |
| 20 | Yes | A. baylyi | Chloramph | Intracellula | F7 | Acridine or | 2874 | BL1 | 7758.5 |
| 20 | Yes | A. baylyi | Chloramph | Intracellula | F192 | 5-Carboxyf | 3311 | BL1 | 14244 |
| 20 | Yes | A. baylyi | Chloramph | Intracellula | F57 | Eosin Y | 3197 | BL1 | 15954 |
| 20 | Yes | A. baylyi | Chloramph | Intracellula | F27 | Calcein | 2895 | BL1 | 13420 |
| 20 | Yes | A. baylyi | Chloramph | Intracellula | F12 | Amaranth | 3449 | BL1 | 14891 |
| 20 | Yes | A. baylyi | None | None | NA | Autofluore: | 4405 | NA | 7038 |
| 20 | Yes | A. baylyi | Colistin | Wall | F194 | Oxonol | 2805 | RL1 | 13879 |
| 20 | Yes | A. baylyi | Colistin | Wall | F11 | Allura Red | 2661 | BL3 | 11955 |
| 20 | Yes | A. baylyi | Colistin | Wall | F54 | DiSC3(5) | 2672 | RL1 | 21620.5 |
| 20 | Yes | A. baylyi | Colistin | Wall | F124 | Thioflavin T | 2692 | VL2 | 10839.5 |
| 20 | Yes | A. baylyi | Colistin | Wall | F101 | Quinacrine | 2937 | VL2 | 9497 |
| 20 | Yes | A. baylyi | Colistin | Wall | F138 | Apigenin | 2674 | VL2 | 8873 |
| 20 | Yes | A. baylyi | Colistin | Wall | F65 | H2FDA | 2750 | BL1 | 12112.5 |
| 20 | Yes | A. baylyi | Colistin | Wall | F142 | Chlorophe | 2883 | BL3 | 13774 |
| 20 | Yes | A. baylyi | Colistin | Wall | F39 | Congo Red | 3097 | BL3 | 9634 |
| 20 | Yes | A. baylyi | Colistin | Wall | F116 | Sunset Yel | 3186 | BL2 | 9988 |
| 20 | Yes | A. baylyi | Colistin | Wall | F10 | Alizarin | 2913 | BL1 | 8557 |
| 20 | Yes | A. baylyi | Colistin | Wall | F84 | Neutral Re | 2997 | BL1 | 10127 |
| 20 | Yes | A. baylyi | Colistin | Wall | F99 | Pyronin Y | 3123 | BL1 | 6916 |
| 20 | Yes | A. baylyi | Colistin | Wall | F108 | Rhodamine | 2587 | BL1 | 7411 |
| 20 | Yes | A. baylyi | Colistin | Wall | F109 | Riboflavin | 2746 | BL1 | 9174 |
| 20 | Yes | A. baylyi | Colistin | Wall | F117 | SYBR Gree | 2450 | BL1 | 2215436 |
| 20 | Yes | A. baylyi | Colistin | Wall | F106 | Rhodamine | 2666 | BL1 | 41174.5 |
| 20 | Yes | A. baylyi | Colistin | Wall | F73 | Malachite ( | 2525 | BL1 | 10919 |
| 20 | Yes | A. baylyi | Colistin | Wall | F7 | Acridine or | 2188 | BL1 | 20792 |
| 20 | Yes | A. baylyi | Colistin | Wall | F192 | 5-Carboxyf | 2452 | BL1 | 22225 |
| 20 | Yes | A. baylyi | Colistin | Wall | F57 | Eosin Y | 2556 | BL1 | 12593.5 |
| 20 | Yes | A. baylyi | Colistin | Wall | F27 | Calcein | 2773 | BL1 | 22609 |

| | | | | | | | | | |
|---|---|---|---|---|---|---|---|---|---|
| 20 | Yes | A. baylyi | Colistin | Wall | F12 | Amaranth | 2598 | BL1 | 18635 |
| 20 | Yes | A. baylyi | None | None | NA | Autofluore: | 4524 | NA | 6539.5 |
| 20 | Yes | A. baylyi | Gentamici | Intracellul | F194 | Oxonol | 3992 | RL1 | 6227 |
| 20 | Yes | A. baylyi | Gentamici | Intracellul | F11 | Allura Red | 4013 | BL3 | 6463 |
| 20 | Yes | A. baylyi | Gentamici | Intracellul | F54 | DiSC3(5) | 3978 | RL1 | 5352 |
| 20 | Yes | A. baylyi | Gentamici | Intracellul | F124 | Thioflavin T | 4254 | VL2 | 5766.5 |
| 20 | Yes | A. baylyi | Gentamici | Intracellul | F101 | Quinacrine | 4292 | VL2 | 5416 |
| 20 | Yes | A. baylyi | Gentamici | Intracellul | F138 | Apigenin | 4193 | VL2 | 5533 |
| 20 | Yes | A. baylyi | Gentamici | Intracellul | F65 | H2FDA | 4284 | BL1 | 6074.5 |
| 20 | Yes | A. baylyi | Gentamici | Intracellul | F142 | Chlorophe | 4596 | BL3 | 7716.5 |
| 20 | Yes | A. baylyi | Gentamici | Intracellul | F39 | Congo Red | 4570 | BL3 | 8906.5 |
| 20 | Yes | A. baylyi | Gentamici | Intracellul | F116 | Sunset Yel | 4827 | BL2 | 7834 |
| 20 | Yes | A. baylyi | Gentamici | Intracellul | F10 | Alizarin | 4723 | BL1 | 5817 |
| 20 | Yes | A. baylyi | Gentamici | Intracellul | F84 | Neutral Re | 4902 | BL1 | 5259.5 |
| 20 | Yes | A. baylyi | Gentamici | Intracellul | F99 | Pyronin Y | 4549 | BL1 | 5275 |
| 20 | Yes | A. baylyi | Gentamici | Intracellul | F108 | Rhodamine | 4371 | BL1 | 4836 |
| 20 | Yes | A. baylyi | Gentamici | Intracellul | F109 | Riboflavin | 4406 | BL1 | 4561 |
| 20 | Yes | A. baylyi | Gentamici | Intracellul | F117 | SYBR Gree | 3625 | BL1 | 312924 |
| 20 | Yes | A. baylyi | Gentamici | Intracellul | F106 | Rhodamine | 2755 | BL1 | 7586 |
| 20 | Yes | A. baylyi | Gentamici | Intracellul | F73 | Malachite ( | 3332 | BL1 | 7914 |
| 20 | Yes | A. baylyi | Gentamici | Intracellul | F7 | Acridine or | 3783 | BL1 | 5531 |
| 20 | Yes | A. baylyi | Gentamici | Intracellul | F192 | 5-Carboxyf | 3277 | BL1 | 10771 |
| 20 | Yes | A. baylyi | Gentamici | Intracellul | F57 | Eosin Y | 3933 | BL1 | 12863 |
| 20 | Yes | A. baylyi | Gentamici | Intracellul | F27 | Calcein | 3645 | BL1 | 15776 |
| 20 | Yes | A. baylyi | Gentamici | Intracellul | F12 | Amaranth | 4130 | BL1 | 16146 |
| 20 | Yes | A. baylyi | None | None | NA | Autofluore: | 4532 | NA | 5905 |
| 20 | Yes | A. baylyi | Nitrofurant | Intracellul | F194 | Oxonol | 4056 | RL1 | 6439.5 |
| 20 | Yes | A. baylyi | Nitrofurant | Intracellul | F11 | Allura Red | 3772 | BL3 | 5253 |
| 20 | Yes | A. baylyi | Nitrofurant | Intracellul | F54 | DiSC3(5) | 4061 | RL1 | 5699 |
| 20 | Yes | A. baylyi | Nitrofurant | Intracellul | F124 | Thioflavin T | 4341 | VL2 | 5706 |
| 20 | Yes | A. baylyi | Nitrofurant | Intracellul | F101 | Quinacrine | 3850 | VL2 | 5280 |
| 20 | Yes | A. baylyi | Nitrofurant | Intracellul | F138 | Apigenin | 4173 | VL2 | 5235 |
| 20 | Yes | A. baylyi | Nitrofurant | Intracellul | F65 | H2FDA | 4350 | BL1 | 7114.5 |
| 20 | Yes | A. baylyi | Nitrofurant | Intracellul | F142 | Chlorophe | 4596 | BL3 | 8817 |
| 20 | Yes | A. baylyi | Nitrofurant | Intracellul | F39 | Congo Red | 4603 | BL3 | 6366 |
| 20 | Yes | A. baylyi | Nitrofurant | Intracellul | F116 | Sunset Yel | 4798 | BL2 | 5546 |
| 20 | Yes | A. baylyi | Nitrofurant | Intracellul | F10 | Alizarin | 4826 | BL1 | 4937.5 |
| 20 | Yes | A. baylyi | Nitrofurant | Intracellul | F84 | Neutral Re | 4635 | BL1 | 3904 |
| 20 | Yes | A. baylyi | Nitrofurant | Intracellul | F99 | Pyronin Y | 4714 | BL1 | 4318 |
| 20 | Yes | A. baylyi | Nitrofurant | Intracellul | F108 | Rhodamine | 3845 | BL1 | 4908 |
| 20 | Yes | A. baylyi | Nitrofurant | Intracellul | F109 | Riboflavin | 4179 | BL1 | 4828 |
| 20 | Yes | A. baylyi | Nitrofurant | Intracellul | F117 | SYBR Gree | 4119 | BL1 | 289771 |
| 20 | Yes | A. baylyi | Nitrofurant | Intracellul | F106 | Rhodamine | 3833 | BL1 | 7263 |
| 20 | Yes | A. baylyi | Nitrofurant | Intracellul | F73 | Malachite ( | 3939 | BL1 | 7255 |
| 20 | Yes | A. baylyi | Nitrofurant | Intracellul | F7 | Acridine or | 3558 | BL1 | 7249.5 |
| 20 | Yes | A. baylyi | Nitrofurant | Intracellul | F192 | 5-Carboxyf | 3813 | BL1 | 12950 |
| 20 | Yes | A. baylyi | Nitrofurant | Intracellul | F57 | Eosin Y | 3806 | BL1 | 12183.5 |
| 20 | Yes | A. baylyi | Nitrofurant | Intracellul | F27 | Calcein | 3570 | BL1 | 12991 |

| | | | | | | | | | |
|---|---|---|---|---|---|---|---|---|---|
| 20 | Yes | A. baylyi | Nitrofurant | Intracellula | F12 | Amaranth | 3656 | BL1 | 13088.5 |
| 20 | Yes | A. baylyi | None | None | NA | Autofluores | 4338 | NA | 6338.5 |
| 20 | Yes | A. baylyi | Pentamidi | Wall | F194 | Oxonol | 2626 | RL1 | 9532 |
| 20 | Yes | A. baylyi | Pentamidi | Wall | F11 | Allura Red | 2740 | BL3 | 8859 |
| 20 | Yes | A. baylyi | Pentamidi | Wall | F54 | DiSC3(5) | 2951 | RL1 | 10088 |
| 20 | Yes | A. baylyi | Pentamidi | Wall | F124 | Thioflavin T | 3001 | VL2 | 10421 |
| 20 | Yes | A. baylyi | Pentamidi | Wall | F101 | Quinacrine | 2634 | VL2 | 10485 |
| 20 | Yes | A. baylyi | Pentamidi | Wall | F138 | Apigenin | 3213 | VL2 | 10811 |
| 20 | Yes | A. baylyi | Pentamidi | Wall | F65 | H2FDA | 2986 | BL1 | 11199.5 |
| 20 | Yes | A. baylyi | Pentamidi | Wall | F142 | Chlorophe | 2873 | BL3 | 11742 |
| 20 | Yes | A. baylyi | Pentamidi | Wall | F39 | Congo Red | 4160 | BL3 | 12973 |
| 20 | Yes | A. baylyi | Pentamidi | Wall | F116 | Sunset Yel | 3216 | BL2 | 7874.5 |
| 20 | Yes | A. baylyi | Pentamidi | Wall | F10 | Alizarin | 2970 | BL1 | 6977.5 |
| 20 | Yes | A. baylyi | Pentamidi | Wall | F84 | Neutral Re | 3102 | BL1 | 7170 |
| 20 | Yes | A. baylyi | Pentamidi | Wall | F99 | Pyronin Y | 3110 | BL1 | 6301 |
| 20 | Yes | A. baylyi | Pentamidi | Wall | F108 | Rhodamine | 3937 | BL1 | 7652 |
| 20 | Yes | A. baylyi | Pentamidi | Wall | F109 | Riboflavin | 2974 | BL1 | 8121.5 |
| 20 | Yes | A. baylyi | Pentamidi | Wall | F117 | SYBR Gree | 2919 | BL1 | 1450828 |
| 20 | Yes | A. baylyi | Pentamidi | Wall | F106 | Rhodamine | 2862 | BL1 | 15807.5 |
| 20 | Yes | A. baylyi | Pentamidi | Wall | F73 | Malachite ( | 3277 | BL1 | 13408 |
| 20 | Yes | A. baylyi | Pentamidi | Wall | F7 | Acridine or | 2983 | BL1 | 13832 |
| 20 | Yes | A. baylyi | Pentamidi | Wall | F192 | 5-Carboxyf | 3292 | BL1 | 17565 |
| 20 | Yes | A. baylyi | Pentamidi | Wall | F57 | Eosin Y | 3089 | BL1 | 16963 |
| 20 | Yes | A. baylyi | Pentamidi | Wall | F27 | Calcein | 3133 | BL1 | 17448 |
| 20 | Yes | A. baylyi | Pentamidi | Wall | F12 | Amaranth | 2880 | BL1 | 17589 |
| 20 | Yes | A. baylyi | None | None | NA | Autofluores | 4542 | NA | 7049 |
| 20 | Yes | A. baylyi | Rifampicin | Intracellula | F194 | Oxonol | 3956 | RL1 | 6174 |
| 20 | Yes | A. baylyi | Rifampicin | Intracellula | F11 | Allura Red | 3862 | BL3 | 5981 |
| 20 | Yes | A. baylyi | Rifampicin | Intracellula | F54 | DiSC3(5) | 3977 | RL1 | 5093 |
| 20 | Yes | A. baylyi | Rifampicin | Intracellula | F124 | Thioflavin T | 4261 | VL2 | 5426 |
| 20 | Yes | A. baylyi | Rifampicin | Intracellula | F101 | Quinacrine | 4425 | VL2 | 5151 |
| 20 | Yes | A. baylyi | Rifampicin | Intracellula | F138 | Apigenin | 4351 | VL2 | 4875 |
| 20 | Yes | A. baylyi | Rifampicin | Intracellula | F65 | H2FDA | 3936 | BL1 | 6225 |
| 20 | Yes | A. baylyi | Rifampicin | Intracellula | F142 | Chlorophe | 4317 | BL3 | 7800 |
| 20 | Yes | A. baylyi | Rifampicin | Intracellula | F39 | Congo Red | 4610 | BL3 | 8421 |
| 20 | Yes | A. baylyi | Rifampicin | Intracellula | F116 | Sunset Yel | 4711 | BL2 | 8087 |
| 20 | Yes | A. baylyi | Rifampicin | Intracellula | F10 | Alizarin | 4246 | BL1 | 5450 |
| 20 | Yes | A. baylyi | Rifampicin | Intracellula | F84 | Neutral Re | 4556 | BL1 | 4408 |
| 20 | Yes | A. baylyi | Rifampicin | Intracellula | F99 | Pyronin Y | 4144 | BL1 | 4063.5 |
| 20 | Yes | A. baylyi | Rifampicin | Intracellula | F108 | Rhodamine | 4329 | BL1 | 4146 |
| 20 | Yes | A. baylyi | Rifampicin | Intracellula | F109 | Riboflavin | 3962 | BL1 | 4428 |
| 20 | Yes | A. baylyi | Rifampicin | Intracellula | F117 | SYBR Gree | 3701 | BL1 | 420720 |
| 20 | Yes | A. baylyi | Rifampicin | Intracellula | F106 | Rhodamine | 4557 | BL1 | 6030 |
| 20 | Yes | A. baylyi | Rifampicin | Intracellula | F73 | Malachite ( | 3879 | BL1 | 7098 |
| 20 | Yes | A. baylyi | Rifampicin | Intracellula | F7 | Acridine or | 2997 | BL1 | 7070 |
| 20 | Yes | A. baylyi | Rifampicin | Intracellula | F192 | 5-Carboxyf | 3901 | BL1 | 14366 |
| 20 | Yes | A. baylyi | Rifampicin | Intracellula | F57 | Eosin Y | 3639 | BL1 | 14170 |
| 20 | Yes | A. baylyi | Rifampicin | Intracellula | F27 | Calcein | 3324 | BL1 | 11654.5 |

| | | | | | | | | | |
|---|---|---|---|---|---|---|---|---|---|
| 20 | Yes | A. baylyi | Rifampicin | Intracellula | F12 | Amaranth | 3817 | BL1 | 12387 |
| 20 | Yes | A. baylyi | None | None | NA | Autofluores | 4269 | NA | 5324 |
| 20 | Yes | A. baylyi | Trimethopr | Intracellula | F194 | Oxonol | 4502 | RL1 | 5244.5 |
| 20 | Yes | A. baylyi | Trimethopr | Intracellula | F11 | Allura Red | 4355 | BL3 | 5308 |
| 20 | Yes | A. baylyi | Trimethopr | Intracellula | F54 | DiSC3(5) | 4188 | RL1 | 5088.5 |
| 20 | Yes | A. baylyi | Trimethopr | Intracellula | F124 | Thioflavin T | 4246 | VL2 | 4864 |
| 20 | Yes | A. baylyi | Trimethopr | Intracellula | F101 | Quinacrine | 4472 | VL2 | 5564.5 |
| 20 | Yes | A. baylyi | Trimethopr | Intracellula | F138 | Apigenin | 3949 | VL2 | 4960 |
| 20 | Yes | A. baylyi | Trimethopr | Intracellula | F65 | H2FDA | 4300 | BL1 | 6269 |
| 20 | Yes | A. baylyi | Trimethopr | Intracellula | F142 | Chlorophe | 4509 | BL3 | 9136 |
| 20 | Yes | A. baylyi | Trimethopr | Intracellula | F39 | Congo Red | 4695 | BL3 | 7059 |
| 20 | Yes | A. baylyi | Trimethopr | Intracellula | F116 | Sunset Yel | 4476 | BL2 | 6135 |
| 20 | Yes | A. baylyi | Trimethopr | Intracellula | F10 | Alizarin | 4460 | BL1 | 5102.5 |
| 20 | Yes | A. baylyi | Trimethopr | Intracellula | F84 | Neutral Re | 4555 | BL1 | 4944 |
| 20 | Yes | A. baylyi | Trimethopr | Intracellula | F99 | Pyronin Y | 3847 | BL1 | 5544 |
| 20 | Yes | A. baylyi | Trimethopr | Intracellula | F108 | Rhodamine | 4123 | BL1 | 5332 |
| 20 | Yes | A. baylyi | Trimethopr | Intracellula | F109 | Riboflavin | 3868 | BL1 | 4632.5 |
| 20 | Yes | A. baylyi | Trimethopr | Intracellula | F117 | SYBR Gree | 3746 | BL1 | 380135.5 |
| 20 | Yes | A. baylyi | Trimethopr | Intracellula | F106 | Rhodamine | 4402 | BL1 | 5882.5 |
| 20 | Yes | A. baylyi | Trimethopr | Intracellula | F73 | Malachite ( | 4084 | BL1 | 5117.5 |
| 20 | Yes | A. baylyi | Trimethopr | Intracellula | F7 | Acridine or | 3792 | BL1 | 5995.5 |
| 20 | Yes | A. baylyi | Trimethopr | Intracellula | F192 | 5-Carboxyf | 3800 | BL1 | 13595 |
| 20 | Yes | A. baylyi | Trimethopr | Intracellula | F57 | Eosin Y | 3690 | BL1 | 16064.5 |
| 20 | Yes | A. baylyi | Trimethopr | Intracellula | F27 | Calcein | 3968 | BL1 | 17834.5 |
| 20 | Yes | A. baylyi | Trimethopr | Intracellula | F12 | Amaranth | 4245 | BL1 | 16415 |
| 20 | Yes | A. baylyi | None | None | NA | Autofluores | 4559 | NA | 1114 |
| 20 | Yes | A. baylyi | None | None | F194 | Oxonol | 5706 | RL1 | 2368 |
| 20 | Yes | A. baylyi | None | None | F11 | Allura Red | 5586 | BL3 | 2763 |
| 20 | Yes | A. baylyi | None | None | F54 | DiSC3(5) | 5342 | RL1 | 3008.5 |
| 20 | Yes | A. baylyi | None | None | F124 | Thioflavin T | 5270 | VL2 | 2807 |
| 20 | Yes | A. baylyi | None | None | F101 | Quinacrine | 5332 | VL2 | 2840 |
| 20 | Yes | A. baylyi | None | None | F138 | Apigenin | 5030 | VL2 | 2911 |
| 20 | Yes | A. baylyi | None | None | F65 | H2FDA | 4871 | BL1 | 4138 |
| 20 | Yes | A. baylyi | None | None | F142 | Chlorophe | 4726 | BL3 | 5896.5 |
| 20 | Yes | A. baylyi | None | None | F39 | Congo Red | 4792 | BL3 | 5798.5 |
| 20 | Yes | A. baylyi | None | None | F116 | Sunset Yel | 4495 | BL2 | 5376 |
| 20 | Yes | A. baylyi | None | None | F10 | Alizarin | 4572 | BL1 | 4831 |
| 20 | Yes | A. baylyi | None | None | F84 | Neutral Re | 4269 | BL1 | 4214 |
| 20 | Yes | A. baylyi | None | None | F99 | Pyronin Y | 4229 | BL1 | 3124 |
| 20 | Yes | A. baylyi | None | None | F108 | Rhodamine | 4654 | BL1 | 2909 |
| 20 | Yes | A. baylyi | None | None | F109 | Riboflavin | 4672 | BL1 | 3284 |
| 20 | Yes | A. baylyi | None | None | F117 | SYBR Gree | 4283 | BL1 | 287453 |
| 20 | Yes | A. baylyi | None | None | F106 | Rhodamine | 4346 | BL1 | 5532 |
| 20 | Yes | A. baylyi | None | None | F73 | Malachite ( | 3792 | BL1 | 4998 |
| 20 | Yes | A. baylyi | None | None | F7 | Acridine or | 4138 | BL1 | 5425.5 |
| 20 | Yes | A. baylyi | None | None | F192 | 5-Carboxyf | 3623 | BL1 | 10753 |
| 20 | Yes | A. baylyi | None | None | F57 | Eosin Y | 4640 | BL1 | 16311 |
| 20 | Yes | A. baylyi | None | None | F27 | Calcein | 4195 | BL1 | 13664 |

| | | | | | | | | | |
|---|---|---|---|---|---|---|---|---|---|
| 20 | Yes | A. baylyi | None | None | F12 | Amaranth | 4771 | BL1 | 13022 |
| 20 | Yes | A. baylyi | None | None | NA | Autofluore: | 4759 | NA | 4361 |
| 20 | Yes | A. baylyi | Amoxacilli | Wall | F194 | Oxonol | 4682 | RL1 | 4226 |
| 20 | Yes | A. baylyi | Amoxacilli | Wall | F11 | Allura Red | 4541 | BL3 | 3945 |
| 20 | Yes | A. baylyi | Amoxacilli | Wall | F54 | DiSC3(5) | 4486 | RL1 | 3685 |
| 20 | Yes | A. baylyi | Amoxacilli | Wall | F124 | Thioflavin T | 4747 | VL2 | 3864 |
| 20 | Yes | A. baylyi | Amoxacilli | Wall | F101 | Quinacrine | 4397 | VL2 | 3663 |
| 20 | Yes | A. baylyi | Amoxacilli | Wall | F138 | Apigenin | 4365 | VL2 | 3711 |
| 20 | Yes | A. baylyi | Amoxacilli | Wall | F65 | H2FDA | 4614 | BL1 | 4367 |
| 20 | Yes | A. baylyi | Amoxacilli | Wall | F142 | Chlorophe | 4481 | BL3 | 6339 |
| 20 | Yes | A. baylyi | Amoxacilli | Wall | F39 | Congo Red | 4274 | BL3 | 6720 |
| 20 | Yes | A. baylyi | Amoxacilli | Wall | F116 | Sunset Yel | 4330 | BL2 | 5355.5 |
| 20 | Yes | A. baylyi | Amoxacilli | Wall | F10 | Alizarin | 4594 | BL1 | 4969 |
| 20 | Yes | A. baylyi | Amoxacilli | Wall | F84 | Neutral Re | 4373 | BL1 | 4902 |
| 20 | Yes | A. baylyi | Amoxacilli | Wall | F99 | Pyronin Y | 4314 | BL1 | 3865 |
| 20 | Yes | A. baylyi | Amoxacilli | Wall | F108 | Rhodamine | 4530 | BL1 | 3317 |
| 20 | Yes | A. baylyi | Amoxacilli | Wall | F109 | Riboflavin | 4000 | BL1 | 3750 |
| 20 | Yes | A. baylyi | Amoxacilli | Wall | F117 | SYBR Gree | 4246 | BL1 | 289040.5 |
| 20 | Yes | A. baylyi | Amoxacilli | Wall | F106 | Rhodamine | 3868 | BL1 | 6404.5 |
| 20 | Yes | A. baylyi | Amoxacilli | Wall | F73 | Malachite ( | 3920 | BL1 | 7180.5 |
| 20 | Yes | A. baylyi | Amoxacilli | Wall | F7 | Acridine or | 3048 | BL1 | 5412.5 |
| 20 | Yes | A. baylyi | Amoxacilli | Wall | F192 | 5-Carboxyf | 4019 | BL1 | 12634 |
| 20 | Yes | A. baylyi | Amoxacilli | Wall | F57 | Eosin Y | 3538 | BL1 | 12943.5 |
| 20 | Yes | A. baylyi | Amoxacilli | Wall | F27 | Calcein | 3883 | BL1 | 15590 |
| 20 | Yes | A. baylyi | Amoxacilli | Wall | F12 | Amaranth | 4190 | BL1 | 13681 |
| 20 | Yes | A. baylyi | None | None | NA | Autofluore: | 4766 | NA | 5809.5 |
| 20 | Yes | A. baylyi | Azithromyc | Intracellula | F194 | Oxonol | 4595 | RL1 | 4763 |
| 20 | Yes | A. baylyi | Azithromyc | Intracellula | F11 | Allura Red | 4583 | BL3 | 4680 |
| 20 | Yes | A. baylyi | Azithromyc | Intracellula | F54 | DiSC3(5) | 4491 | RL1 | 4702 |
| 20 | Yes | A. baylyi | Azithromyc | Intracellula | F124 | Thioflavin T | 4618 | VL2 | 4595 |
| 20 | Yes | A. baylyi | Azithromyc | Intracellula | F101 | Quinacrine | 4500 | VL2 | 4372.5 |
| 20 | Yes | A. baylyi | Azithromyc | Intracellula | F138 | Apigenin | 4292 | VL2 | 4477 |
| 20 | Yes | A. baylyi | Azithromyc | Intracellula | F65 | H2FDA | 4242 | BL1 | 5258 |
| 20 | Yes | A. baylyi | Azithromyc | Intracellula | F142 | Chlorophe | 4051 | BL3 | 6636 |
| 20 | Yes | A. baylyi | Azithromyc | Intracellula | F39 | Congo Red | 4513 | BL3 | 7260 |
| 20 | Yes | A. baylyi | Azithromyc | Intracellula | F116 | Sunset Yel | 4565 | BL2 | 7276 |
| 20 | Yes | A. baylyi | Azithromyc | Intracellula | F10 | Alizarin | 4203 | BL1 | 5092 |
| 20 | Yes | A. baylyi | Azithromyc | Intracellula | F84 | Neutral Re | 4421 | BL1 | 4852 |
| 20 | Yes | A. baylyi | Azithromyc | Intracellula | F99 | Pyronin Y | 3348 | BL1 | 4782 |
| 20 | Yes | A. baylyi | Azithromyc | Intracellula | F108 | Rhodamine | 3768 | BL1 | 4175 |
| 20 | Yes | A. baylyi | Azithromyc | Intracellula | F109 | Riboflavin | 4479 | BL1 | 4157 |
| 20 | Yes | A. baylyi | Azithromyc | Intracellula | F117 | SYBR Gree | 3668 | BL1 | 255017.5 |
| 20 | Yes | A. baylyi | Azithromyc | Intracellula | F106 | Rhodamine | 4008 | BL1 | 7021.5 |
| 20 | Yes | A. baylyi | Azithromyc | Intracellula | F73 | Malachite ( | 3500 | BL1 | 7229.5 |
| 20 | Yes | A. baylyi | Azithromyc | Intracellula | F7 | Acridine or | 4212 | BL1 | 7422 |
| 20 | Yes | A. baylyi | Azithromyc | Intracellula | F192 | 5-Carboxyf | 3442 | BL1 | 12558 |
| 20 | Yes | A. baylyi | Azithromyc | Intracellula | F57 | Eosin Y | 3858 | BL1 | 11634.5 |
| 20 | Yes | A. baylyi | Azithromyc | Intracellula | F27 | Calcein | 4302 | BL1 | 16702 |

| | | | | | | | | | |
|---|---|---|---|---|---|---|---|---|---|
| 20 | Yes | A. baylyi | Azithromyc | Intracellula | F12 | Amaranth | 4529 | BL1 | 13276 |
| 20 | Yes | A. baylyi | None | None | NA | Autofluores | 4366 | NA | 6441 |
| 20 | Yes | A. baylyi | Trimethopr | Intracellula | F194 | Oxonol | 4447 | RL1 | 5785 |
| 20 | Yes | A. baylyi | Trimethopr | Intracellula | F11 | Allura Red | 4270 | BL3 | 5196.5 |
| 20 | Yes | A. baylyi | Trimethopr | Intracellula | F54 | DiSC3(5) | 4452 | RL1 | 5396 |
| 20 | Yes | A. baylyi | Trimethopr | Intracellula | F124 | Thioflavin T | 4251 | VL2 | 5019 |
| 20 | Yes | A. baylyi | Trimethopr | Intracellula | F101 | Quinacrine | 5118 | VL2 | 4936 |
| 20 | Yes | A. baylyi | Trimethopr | Intracellula | F138 | Apigenin | 4553 | VL2 | 4678 |
| 20 | Yes | A. baylyi | Trimethopr | Intracellula | F65 | H2FDA | 4651 | BL1 | 6373 |
| 20 | Yes | A. baylyi | Trimethopr | Intracellula | F142 | Chlorophe | 4189 | BL3 | 6842 |
| 20 | Yes | A. baylyi | Trimethopr | Intracellula | F39 | Congo Red | 4672 | BL3 | 8679.5 |
| 20 | Yes | A. baylyi | Trimethopr | Intracellula | F116 | Sunset Yel | 4527 | BL2 | 6470 |
| 20 | Yes | A. baylyi | Trimethopr | Intracellula | F10 | Alizarin | 4659 | BL1 | 5744 |
| 20 | Yes | A. baylyi | Trimethopr | Intracellula | F84 | Neutral Re | 4661 | BL1 | 4939 |
| 20 | Yes | A. baylyi | Trimethopr | Intracellula | F99 | Pyronin Y | 4358 | BL1 | 4464.5 |
| 20 | Yes | A. baylyi | Trimethopr | Intracellula | F108 | Rhodamine | 4168 | BL1 | 4118 |
| 20 | Yes | A. baylyi | Trimethopr | Intracellula | F109 | Riboflavin | 3975 | BL1 | 4144 |
| 20 | Yes | A. baylyi | Trimethopr | Intracellula | F117 | SYBR Gree | 3953 | BL1 | 309982 |
| 20 | Yes | A. baylyi | Trimethopr | Intracellula | F106 | Rhodamine | 4475 | BL1 | 5974 |
| 20 | Yes | A. baylyi | Trimethopr | Intracellula | F73 | Malachite ( | 4482 | BL1 | 5684.5 |
| 20 | Yes | A. baylyi | Trimethopr | Intracellula | F7 | Acridine or | 4193 | BL1 | 5975 |
| 20 | Yes | A. baylyi | Trimethopr | Intracellula | F192 | 5-Carboxyf | 3811 | BL1 | 14855 |
| 20 | Yes | A. baylyi | Trimethopr | Intracellula | F57 | Eosin Y | 4044 | BL1 | 12852 |
| 20 | Yes | A. baylyi | Trimethopr | Intracellula | F27 | Calcein | 4258 | BL1 | 13814.5 |
| 20 | Yes | A. baylyi | Trimethopr | Intracellula | F12 | Amaranth | 3836 | BL1 | 10576 |
| 20 | Yes | A. baylyi | None | None | NA | Autofluores | 4913 | NA | 6559 |
| 20 | Yes | A. baylyi | Chloramph | Intracellula | F194 | Oxonol | 3302 | RL1 | 5246 |
| 20 | Yes | A. baylyi | Chloramph | Intracellula | F11 | Allura Red | 3658 | BL3 | 5262.5 |
| 20 | Yes | A. baylyi | Chloramph | Intracellula | F54 | DiSC3(5) | 3710 | RL1 | 4890 |
| 20 | Yes | A. baylyi | Chloramph | Intracellula | F124 | Thioflavin T | 3968 | VL2 | 5056.5 |
| 20 | Yes | A. baylyi | Chloramph | Intracellula | F101 | Quinacrine | 4111 | VL2 | 4857 |
| 20 | Yes | A. baylyi | Chloramph | Intracellula | F138 | Apigenin | 3721 | VL2 | 4744 |
| 20 | Yes | A. baylyi | Chloramph | Intracellula | F65 | H2FDA | 3453 | BL1 | 5661 |
| 20 | Yes | A. baylyi | Chloramph | Intracellula | F142 | Chlorophe | 3933 | BL3 | 8130 |
| 20 | Yes | A. baylyi | Chloramph | Intracellula | F39 | Congo Red | 3174 | BL3 | 8230.5 |
| 20 | Yes | A. baylyi | Chloramph | Intracellula | F116 | Sunset Yel | 4163 | BL2 | 6161 |
| 20 | Yes | A. baylyi | Chloramph | Intracellula | F10 | Alizarin | 3849 | BL1 | 5009 |
| 20 | Yes | A. baylyi | Chloramph | Intracellula | F84 | Neutral Re | 4067 | BL1 | 4417 |
| 20 | Yes | A. baylyi | Chloramph | Intracellula | F99 | Pyronin Y | 3692 | BL1 | 4122 |
| 20 | Yes | A. baylyi | Chloramph | Intracellula | F108 | Rhodamine | 3548 | BL1 | 4306.5 |
| 20 | Yes | A. baylyi | Chloramph | Intracellula | F109 | Riboflavin | 3441 | BL1 | 4256 |
| 20 | Yes | A. baylyi | Chloramph | Intracellula | F117 | SYBR Gree | 3416 | BL1 | 243231 |
| 20 | Yes | A. baylyi | Chloramph | Intracellula | F106 | Rhodamine | 3559 | BL1 | 7462 |
| 20 | Yes | A. baylyi | Chloramph | Intracellula | F73 | Malachite ( | 3311 | BL1 | 6727 |
| 20 | Yes | A. baylyi | Chloramph | Intracellula | F7 | Acridine or | 3030 | BL1 | 6542.5 |
| 20 | Yes | A. baylyi | Chloramph | Intracellula | F192 | 5-Carboxyf | 3262 | BL1 | 14932 |
| 20 | Yes | A. baylyi | Chloramph | Intracellula | F57 | Eosin Y | 3445 | BL1 | 12522 |
| 20 | Yes | A. baylyi | Chloramph | Intracellula | F27 | Calcein | 3142 | BL1 | 17984.5 |

| | | | | | | | | | |
|---|---|---|---|---|---|---|---|---|---|
| 20 | Yes | A. baylyi | Chloramph | Intracellula | F12 | Amaranth | 3828 | BL1 | 18243 |
| 20 | Yes | A. baylyi | None | None | NA | Autofluore: | 4490 | NA | 6212 |
| 20 | Yes | A. baylyi | Colistin | Wall | F194 | Oxonol | 2834 | RL1 | 16061.5 |
| 20 | Yes | A. baylyi | Colistin | Wall | F11 | Allura Red | 2743 | BL3 | 12182 |
| 20 | Yes | A. baylyi | Colistin | Wall | F54 | DiSC3(5) | 3012 | RL1 | 24268 |
| 20 | Yes | A. baylyi | Colistin | Wall | F124 | Thioflavin T | 3056 | VL2 | 10653.5 |
| 20 | Yes | A. baylyi | Colistin | Wall | F101 | Quinacrine | 2939 | VL2 | 9608 |
| 20 | Yes | A. baylyi | Colistin | Wall | F138 | Apigenin | 2890 | VL2 | 10279.5 |
| 20 | Yes | A. baylyi | Colistin | Wall | F65 | H2FDA | 2883 | BL1 | 10972 |
| 20 | Yes | A. baylyi | Colistin | Wall | F142 | Chlorophe | 2852 | BL3 | 13986.5 |
| 20 | Yes | A. baylyi | Colistin | Wall | F39 | Congo Red | 2658 | BL3 | 12976 |
| 20 | Yes | A. baylyi | Colistin | Wall | F116 | Sunset Yel | 2974 | BL2 | 12480.5 |
| 20 | Yes | A. baylyi | Colistin | Wall | F10 | Alizarin | 3054 | BL1 | 9727 |
| 20 | Yes | A. baylyi | Colistin | Wall | F84 | Neutral Re | 2933 | BL1 | 11773 |
| 20 | Yes | A. baylyi | Colistin | Wall | F99 | Pyronin Y | 2900 | BL1 | 7242.5 |
| 20 | Yes | A. baylyi | Colistin | Wall | F108 | Rhodamine | 1944 | BL1 | 7762 |
| 20 | Yes | A. baylyi | Colistin | Wall | F109 | Riboflavin | 2966 | BL1 | 9002.5 |
| 20 | Yes | A. baylyi | Colistin | Wall | F117 | SYBR Gree | 2466 | BL1 | 2229571 |
| 20 | Yes | A. baylyi | Colistin | Wall | F106 | Rhodamine | 2511 | BL1 | 44001 |
| 20 | Yes | A. baylyi | Colistin | Wall | F73 | Malachite ( | 2705 | BL1 | 10570 |
| 20 | Yes | A. baylyi | Colistin | Wall | F7 | Acridine or | 2632 | BL1 | 23348 |
| 20 | Yes | A. baylyi | Colistin | Wall | F192 | 5-Carboxyf | 2443 | BL1 | 22856 |
| 20 | Yes | A. baylyi | Colistin | Wall | F57 | Eosin Y | 2356 | BL1 | 18231 |
| 20 | Yes | A. baylyi | Colistin | Wall | F27 | Calcein | 3412 | BL1 | 23190 |
| 20 | Yes | A. baylyi | Colistin | Wall | F12 | Amaranth | 3571 | BL1 | 17476 |
| 20 | Yes | A. baylyi | None | None | NA | Autofluore: | 4684 | NA | 6678.5 |
| 20 | Yes | A. baylyi | Gentamici | Intracellula | F194 | Oxonol | 3582 | RL1 | 5313.5 |
| 20 | Yes | A. baylyi | Gentamici | Intracellula | F11 | Allura Red | 4771 | BL3 | 5188 |
| 20 | Yes | A. baylyi | Gentamici | Intracellula | F54 | DiSC3(5) | 4408 | RL1 | 5060.5 |
| 20 | Yes | A. baylyi | Gentamici | Intracellula | F124 | Thioflavin T | 4693 | VL2 | 5075 |
| 20 | Yes | A. baylyi | Gentamici | Intracellula | F101 | Quinacrine | 4807 | VL2 | 5082 |
| 20 | Yes | A. baylyi | Gentamici | Intracellula | F138 | Apigenin | 4815 | VL2 | 5258 |
| 20 | Yes | A. baylyi | Gentamici | Intracellula | F65 | H2FDA | 5013 | BL1 | 6448 |
| 20 | Yes | A. baylyi | Gentamici | Intracellula | F142 | Chlorophe | 4818 | BL3 | 7994 |
| 20 | Yes | A. baylyi | Gentamici | Intracellula | F39 | Congo Red | 4948 | BL3 | 8538 |
| 20 | Yes | A. baylyi | Gentamici | Intracellula | F116 | Sunset Yel | 5299 | BL2 | 6295 |
| 20 | Yes | A. baylyi | Gentamici | Intracellula | F10 | Alizarin | 4455 | BL1 | 4815 |
| 20 | Yes | A. baylyi | Gentamici | Intracellula | F84 | Neutral Re | 5637 | BL1 | 4254 |
| 20 | Yes | A. baylyi | Gentamici | Intracellula | F99 | Pyronin Y | 4725 | BL1 | 4519 |
| 20 | Yes | A. baylyi | Gentamici | Intracellula | F108 | Rhodamine | 5191 | BL1 | 4933 |
| 20 | Yes | A. baylyi | Gentamici | Intracellula | F109 | Riboflavin | 4806 | BL1 | 5622 |
| 20 | Yes | A. baylyi | Gentamici | Intracellula | F117 | SYBR Gree | 4409 | BL1 | 335992 |
| 20 | Yes | A. baylyi | Gentamici | Intracellula | F106 | Rhodamine | 4878 | BL1 | 7711 |
| 20 | Yes | A. baylyi | Gentamici | Intracellula | F73 | Malachite ( | 4118 | BL1 | 6854.5 |
| 20 | Yes | A. baylyi | Gentamici | Intracellula | F7 | Acridine or | 4161 | BL1 | 7397 |
| 20 | Yes | A. baylyi | Gentamici | Intracellula | F192 | 5-Carboxyf | 4293 | BL1 | 14901 |
| 20 | Yes | A. baylyi | Gentamici | Intracellula | F57 | Eosin Y | 4581 | BL1 | 12454 |
| 20 | Yes | A. baylyi | Gentamici | Intracellula | F27 | Calcein | 4178 | BL1 | 14971.5 |

| | | | | | | | | | |
|---|---|---|---|---|---|---|---|---|---|
| 20 | Yes | A. baylyi | Gentamicii | Intracellul | F12 | Amaranth | 4713 | BL1 | 13736 |
| 20 | Yes | A. baylyi | None | None | NA | Autofluore | 4671 | NA | 6051 |
| 20 | Yes | A. baylyi | Nitrofurant | Intracellul | F194 | Oxonol | 4310 | RL1 | 5519 |
| 20 | Yes | A. baylyi | Nitrofurant | Intracellul | F11 | Allura Red | 4186 | BL3 | 5287 |
| 20 | Yes | A. baylyi | Nitrofurant | Intracellul | F54 | DiSC3(5) | 3990 | RL1 | 5198 |
| 20 | Yes | A. baylyi | Nitrofurant | Intracellul | F124 | Thioflavin T | 4156 | VL2 | 5235 |
| 20 | Yes | A. baylyi | Nitrofurant | Intracellul | F101 | Quinacrine | 4488 | VL2 | 5071 |
| 20 | Yes | A. baylyi | Nitrofurant | Intracellul | F138 | Apigenin | 4452 | VL2 | 4972.5 |
| 20 | Yes | A. baylyi | Nitrofurant | Intracellul | F65 | H2FDA | 4049 | BL1 | 6514 |
| 20 | Yes | A. baylyi | Nitrofurant | Intracellul | F142 | Chlorophe | 4642 | BL3 | 8911 |
| 20 | Yes | A. baylyi | Nitrofurant | Intracellul | F39 | Congo Red | 4653 | BL3 | 7812 |
| 20 | Yes | A. baylyi | Nitrofurant | Intracellul | F116 | Sunset Yel | 5063 | BL2 | 7214 |
| 20 | Yes | A. baylyi | Nitrofurant | Intracellul | F10 | Alizarin | 5027 | BL1 | 5255 |
| 20 | Yes | A. baylyi | Nitrofurant | Intracellul | F84 | Neutral Re | 4764 | BL1 | 4229 |
| 20 | Yes | A. baylyi | Nitrofurant | Intracellul | F99 | Pyronin Y | 4146 | BL1 | 4018.5 |
| 20 | Yes | A. baylyi | Nitrofurant | Intracellul | F108 | Rhodamine | 4473 | BL1 | 3935 |
| 20 | Yes | A. baylyi | Nitrofurant | Intracellul | F109 | Riboflavin | 4270 | BL1 | 4448.5 |
| 20 | Yes | A. baylyi | Nitrofurant | Intracellul | F117 | SYBR Gree | 3658 | BL1 | 188719 |
| 20 | Yes | A. baylyi | Nitrofurant | Intracellul | F106 | Rhodamine | 4300 | BL1 | 6673.5 |
| 20 | Yes | A. baylyi | Nitrofurant | Intracellul | F73 | Malachite | 4416 | BL1 | 6750.5 |
| 20 | Yes | A. baylyi | Nitrofurant | Intracellul | F7 | Acridine or | 3615 | BL1 | 6026 |
| 20 | Yes | A. baylyi | Nitrofurant | Intracellul | F192 | 5-Carboxyf | 4320 | BL1 | 13192.5 |
| 20 | Yes | A. baylyi | Nitrofurant | Intracellul | F57 | Eosin Y | 4324 | BL1 | 12795.5 |
| 20 | Yes | A. baylyi | Nitrofurant | Intracellul | F27 | Calcein | 3726 | BL1 | 15754 |
| 20 | Yes | A. baylyi | Nitrofurant | Intracellul | F12 | Amaranth | 4145 | BL1 | 12718 |
| 20 | Yes | A. baylyi | None | None | NA | Autofluore | 4782 | NA | 6059.5 |
| 20 | Yes | A. baylyi | Pentamidi | Wall | F194 | Oxonol | 2923 | RL1 | 10546 |
| 20 | Yes | A. baylyi | Pentamidi | Wall | F11 | Allura Red | 2499 | BL3 | 8530 |
| 20 | Yes | A. baylyi | Pentamidi | Wall | F54 | DiSC3(5) | 2704 | RL1 | 8908 |
| 20 | Yes | A. baylyi | Pentamidi | Wall | F124 | Thioflavin T | 2893 | VL2 | 9416 |
| 20 | Yes | A. baylyi | Pentamidi | Wall | F101 | Quinacrine | 2964 | VL2 | 9064 |
| 20 | Yes | A. baylyi | Pentamidi | Wall | F138 | Apigenin | 3118 | VL2 | 10537.5 |
| 20 | Yes | A. baylyi | Pentamidi | Wall | F65 | H2FDA | 3045 | BL1 | 13189 |
| 20 | Yes | A. baylyi | Pentamidi | Wall | F142 | Chlorophe | 2928 | BL3 | 12638 |
| 20 | Yes | A. baylyi | Pentamidi | Wall | F39 | Congo Red | 4050 | BL3 | 12771 |
| 20 | Yes | A. baylyi | Pentamidi | Wall | F116 | Sunset Yel | 2907 | BL2 | 9529 |
| 20 | Yes | A. baylyi | Pentamidi | Wall | F10 | Alizarin | 2995 | BL1 | 7744 |
| 20 | Yes | A. baylyi | Pentamidi | Wall | F84 | Neutral Re | 3180 | BL1 | 7978 |
| 20 | Yes | A. baylyi | Pentamidi | Wall | F99 | Pyronin Y | 2960 | BL1 | 6366.5 |
| 20 | Yes | A. baylyi | Pentamidi | Wall | F108 | Rhodamine | 3030 | BL1 | 6755 |
| 20 | Yes | A. baylyi | Pentamidi | Wall | F109 | Riboflavin | 2995 | BL1 | 7368 |
| 20 | Yes | A. baylyi | Pentamidi | Wall | F117 | SYBR Gree | 3072 | BL1 | 1420348 |
| 20 | Yes | A. baylyi | Pentamidi | Wall | F106 | Rhodamine | 2771 | BL1 | 15269 |
| 20 | Yes | A. baylyi | Pentamidi | Wall | F73 | Malachite | 2686 | BL1 | 10429.5 |
| 20 | Yes | A. baylyi | Pentamidi | Wall | F7 | Acridine or | 2937 | BL1 | 12729 |
| 20 | Yes | A. baylyi | Pentamidi | Wall | F192 | 5-Carboxyf | 2672 | BL1 | 17312 |
| 20 | Yes | A. baylyi | Pentamidi | Wall | F57 | Eosin Y | 2646 | BL1 | 15393.5 |
| 20 | Yes | A. baylyi | Pentamidi | Wall | F27 | Calcein | 2828 | BL1 | 17272.5 |

| | | | | | | | | | |
|---|---|---|---|---|---|---|---|---|---|
| 20 | Yes | A. baylyi | Pentamidi | Wall | F12 | Amaranth | 2782 | BL1 | 14601.5 |
| 20 | Yes | A. baylyi | None | None | NA | Autofluore: | 4543 | NA | 7379 |
| 20 | Yes | A. baylyi | Rifampicin | Intracellul | F194 | Oxonol | 4284 | RL1 | 6324 |
| 20 | Yes | A. baylyi | Rifampicin | Intracellul | F11 | Allura Red | 4059 | BL3 | 5972 |
| 20 | Yes | A. baylyi | Rifampicin | Intracellul | F54 | DiSC3(5) | 4561 | RL1 | 5819 |
| 20 | Yes | A. baylyi | Rifampicin | Intracellul | F124 | Thioflavin T | 4584 | VL2 | 5377.5 |
| 20 | Yes | A. baylyi | Rifampicin | Intracellul | F101 | Quinacrine | 4333 | VL2 | 5186 |
| 20 | Yes | A. baylyi | Rifampicin | Intracellul | F138 | Apigenin | 4477 | VL2 | 4886 |
| 20 | Yes | A. baylyi | Rifampicin | Intracellul | F65 | H2FDA | 4806 | BL1 | 5888 |
| 20 | Yes | A. baylyi | Rifampicin | Intracellul | F142 | Chlorophe | 4753 | BL3 | 8584 |
| 20 | Yes | A. baylyi | Rifampicin | Intracellul | F39 | Congo Red | 5038 | BL3 | 9148 |
| 20 | Yes | A. baylyi | Rifampicin | Intracellul | F116 | Sunset Yel | 4843 | BL2 | 7376 |
| 20 | Yes | A. baylyi | Rifampicin | Intracellul | F10 | Alizarin | 5025 | BL1 | 5289 |
| 20 | Yes | A. baylyi | Rifampicin | Intracellul | F84 | Neutral Re | 4932 | BL1 | 4744.5 |
| 20 | Yes | A. baylyi | Rifampicin | Intracellul | F99 | Pyronin Y | 4524 | BL1 | 4956.5 |
| 20 | Yes | A. baylyi | Rifampicin | Intracellul | F108 | Rhodamine | 4504 | BL1 | 4216.5 |
| 20 | Yes | A. baylyi | Rifampicin | Intracellul | F109 | Riboflavin | 4703 | BL1 | 4292 |
| 20 | Yes | A. baylyi | Rifampicin | Intracellul | F117 | SYBR Gree | 3694 | BL1 | 442724.5 |
| 20 | Yes | A. baylyi | Rifampicin | Intracellul | F106 | Rhodamine | 4826 | BL1 | 5275 |
| 20 | Yes | A. baylyi | Rifampicin | Intracellul | F73 | Malachite ( | 4332 | BL1 | 7153.5 |
| 20 | Yes | A. baylyi | Rifampicin | Intracellul | F7 | Acridine or | 3493 | BL1 | 6710 |
| 20 | Yes | A. baylyi | Rifampicin | Intracellul | F192 | 5-Carboxyf | 4256 | BL1 | 13538 |
| 20 | Yes | A. baylyi | Rifampicin | Intracellul | F57 | Eosin Y | 4375 | BL1 | 14432 |
| 20 | Yes | A. baylyi | Rifampicin | Intracellul | F27 | Calcein | 4191 | BL1 | 12746 |
| 20 | Yes | A. baylyi | Rifampicin | Intracellul | F12 | Amaranth | 3938 | BL1 | 14562 |
| 20 | Yes | A. baylyi | None | None | NA | Autofluore: | 4743 | NA | 6826 |
| 20 | Yes | A. baylyi | Trimethopr | Intracellul | F194 | Oxonol | 4421 | RL1 | 5923 |
| 20 | Yes | A. baylyi | Trimethopr | Intracellul | F11 | Allura Red | 4905 | BL3 | 5511 |
| 20 | Yes | A. baylyi | Trimethopr | Intracellul | F54 | DiSC3(5) | 4778 | RL1 | 5735.5 |
| 20 | Yes | A. baylyi | Trimethopr | Intracellul | F124 | Thioflavin T | 4981 | VL2 | 5539 |
| 20 | Yes | A. baylyi | Trimethopr | Intracellul | F101 | Quinacrine | 4804 | VL2 | 4951 |
| 20 | Yes | A. baylyi | Trimethopr | Intracellul | F138 | Apigenin | 4846 | VL2 | 4862.5 |
| 20 | Yes | A. baylyi | Trimethopr | Intracellul | F65 | H2FDA | 5192 | BL1 | 6287 |
| 20 | Yes | A. baylyi | Trimethopr | Intracellul | F142 | Chlorophe | 5074 | BL3 | 9647.5 |
| 20 | Yes | A. baylyi | Trimethopr | Intracellul | F39 | Congo Red | 5042 | BL3 | 7688.5 |
| 20 | Yes | A. baylyi | Trimethopr | Intracellul | F116 | Sunset Yel | 5199 | BL2 | 6344 |
| 20 | Yes | A. baylyi | Trimethopr | Intracellul | F10 | Alizarin | 5148 | BL1 | 5361.5 |
| 20 | Yes | A. baylyi | Trimethopr | Intracellul | F84 | Neutral Re | 5133 | BL1 | 5086 |
| 20 | Yes | A. baylyi | Trimethopr | Intracellul | F99 | Pyronin Y | 4846 | BL1 | 5158 |
| 20 | Yes | A. baylyi | Trimethopr | Intracellul | F108 | Rhodamine | 4845 | BL1 | 5131 |
| 20 | Yes | A. baylyi | Trimethopr | Intracellul | F109 | Riboflavin | 4692 | BL1 | 4945 |
| 20 | Yes | A. baylyi | Trimethopr | Intracellul | F117 | SYBR Gree | 4053 | BL1 | 350237 |
| 20 | Yes | A. baylyi | Trimethopr | Intracellul | F106 | Rhodamine | 5058 | BL1 | 5448 |
| 20 | Yes | A. baylyi | Trimethopr | Intracellul | F73 | Malachite ( | 4276 | BL1 | 5231.5 |
| 20 | Yes | A. baylyi | Trimethopr | Intracellul | F7 | Acridine or | 4362 | BL1 | 6007 |
| 20 | Yes | A. baylyi | Trimethopr | Intracellul | F192 | 5-Carboxyf | 4747 | BL1 | 15935 |
| 20 | Yes | A. baylyi | Trimethopr | Intracellul | F57 | Eosin Y | 4367 | BL1 | 13688 |
| 20 | Yes | A. baylyi | Trimethopr | Intracellul | F27 | Calcein | 4309 | BL1 | 15371 |

| | | | | | | | | |
|---|---|---|---|---|---|---|---|---|
| 20 | Yes | A. baylyi | Trimethopr | Intracellula | F12 | Amaranth | 4812 | BL1 | 17250 |
| 20 | Yes | A. baylyi | None | None | NA | Autofluores | 4969 | NA | 1137 |
| 20 | Yes | A. baylyi | None | None | F194 | Oxonol | 5101 | RL1 | 2587 |
| 20 | Yes | A. baylyi | None | None | F11 | Allura Red | 5480 | BL3 | 2796.5 |
| 20 | Yes | A. baylyi | None | None | F54 | DiSC3(5) | 4879 | RL1 | 2998 |
| 20 | Yes | A. baylyi | None | None | F124 | Thioflavin T | 4701 | VL2 | 2964 |
| 20 | Yes | A. baylyi | None | None | F101 | Quinacrine | 4520 | VL2 | 3258 |
| 20 | Yes | A. baylyi | None | None | F138 | Apigenin | 4319 | VL2 | 3117 |
| 20 | Yes | A. baylyi | None | None | F65 | H2FDA | 4442 | BL1 | 4482.5 |
| 20 | Yes | A. baylyi | None | None | F142 | Chlorophe | 4006 | BL3 | 6320 |
| 20 | Yes | A. baylyi | None | None | F39 | Congo Red | 4369 | BL3 | 6499 |
| 20 | Yes | A. baylyi | None | None | F116 | Sunset Yel | 4410 | BL2 | 6103.5 |
| 20 | Yes | A. baylyi | None | None | F10 | Alizarin | 4146 | BL1 | 4203 |
| 20 | Yes | A. baylyi | None | None | F84 | Neutral Re | 4245 | BL1 | 3154 |
| 20 | Yes | A. baylyi | None | None | F99 | Pyronin Y | 3701 | BL1 | 3045 |
| 20 | Yes | A. baylyi | None | None | F108 | Rhodamine | 4180 | BL1 | 2916 |
| 20 | Yes | A. baylyi | None | None | F109 | Riboflavin | 4262 | BL1 | 3163 |
| 20 | Yes | A. baylyi | None | None | F117 | SYBR Gree | 4150 | BL1 | 264581.5 |
| 20 | Yes | A. baylyi | None | None | F106 | Rhodamine | 4021 | BL1 | 5890 |
| 20 | Yes | A. baylyi | None | None | F73 | Malachite ( | 3929 | BL1 | 5010 |
| 20 | Yes | A. baylyi | None | None | F7 | Acridine or | 3722 | BL1 | 5886 |
| 20 | Yes | A. baylyi | None | None | F192 | 5-Carboxyf | 4004 | BL1 | 12510 |
| 20 | Yes | A. baylyi | None | None | F57 | Eosin Y | 3988 | BL1 | 11480.5 |
| 20 | Yes | A. baylyi | None | None | F27 | Calcein | 3649 | BL1 | 13256 |
| 20 | Yes | A. baylyi | None | None | F12 | Amaranth | 3926 | BL1 | 14384.5 |
| 20 | Yes | A. baylyi | None | None | NA | Autofluores | 4082 | NA | 4795.5 |
| 20 | Yes | A. baylyi | Amoxacilli | Wall | F194 | Oxonol | 4054 | RL1 | 4401 |
| 20 | Yes | A. baylyi | Amoxacilli | Wall | F11 | Allura Red | 4175 | BL3 | 4298 |
| 20 | Yes | A. baylyi | Amoxacilli | Wall | F54 | DiSC3(5) | 4342 | RL1 | 3763.5 |
| 20 | Yes | A. baylyi | Amoxacilli | Wall | F124 | Thioflavin T | 4395 | VL2 | 3792 |
| 20 | Yes | A. baylyi | Amoxacilli | Wall | F101 | Quinacrine | 4444 | VL2 | 3800.5 |
| 20 | Yes | A. baylyi | Amoxacilli | Wall | F138 | Apigenin | 3991 | VL2 | 3814 |
| 20 | Yes | A. baylyi | Amoxacilli | Wall | F65 | H2FDA | 4051 | BL1 | 4588 |
| 20 | Yes | A. baylyi | Amoxacilli | Wall | F142 | Chlorophe | 4102 | BL3 | 6918 |
| 20 | Yes | A. baylyi | Amoxacilli | Wall | F39 | Congo Red | 4077 | BL3 | 6342 |
| 20 | Yes | A. baylyi | Amoxacilli | Wall | F116 | Sunset Yel | 3922 | BL2 | 5460.5 |
| 20 | Yes | A. baylyi | Amoxacilli | Wall | F10 | Alizarin | 3876 | BL1 | 5215.5 |
| 20 | Yes | A. baylyi | Amoxacilli | Wall | F84 | Neutral Re | 3463 | BL1 | 6379 |
| 20 | Yes | A. baylyi | Amoxacilli | Wall | F99 | Pyronin Y | 3976 | BL1 | 4372.5 |
| 20 | Yes | A. baylyi | Amoxacilli | Wall | F108 | Rhodamine | 3390 | BL1 | 3539 |
| 20 | Yes | A. baylyi | Amoxacilli | Wall | F109 | Riboflavin | 3681 | BL1 | 3937 |
| 20 | Yes | A. baylyi | Amoxacilli | Wall | F117 | SYBR Gree | 3669 | BL1 | 243967 |
| 20 | Yes | A. baylyi | Amoxacilli | Wall | F106 | Rhodamine | 3727 | BL1 | 6072 |
| 20 | Yes | A. baylyi | Amoxacilli | Wall | F73 | Malachite ( | 3825 | BL1 | 6841 |
| 20 | Yes | A. baylyi | Amoxacilli | Wall | F7 | Acridine or | 3318 | BL1 | 6258.5 |
| 20 | Yes | A. baylyi | Amoxacilli | Wall | F192 | 5-Carboxyf | 3466 | BL1 | 13069 |
| 20 | Yes | A. baylyi | Amoxacilli | Wall | F57 | Eosin Y | 3835 | BL1 | 13860 |
| 20 | Yes | A. baylyi | Amoxacilli | Wall | F27 | Calcein | 3685 | BL1 | 14028 |

| | | | | | | | | | |
|---|---|---|---|---|---|---|---|---|---|
| 20 | Yes | A. baylyi | Amoxacilli | Wall | F12 | Amaranth | 3398 | BL1 | 12001.5 |
| 20 | Yes | A. baylyi | None | None | NA | Autofluore | 4212 | NA | 5409.5 |
| 20 | Yes | A. baylyi | Azithromyc | Intracellula | F194 | Oxonol | 4028 | RL1 | 4697 |
| 20 | Yes | A. baylyi | Azithromyc | Intracellula | F11 | Allura Red | 3889 | BL3 | 4595 |
| 20 | Yes | A. baylyi | Azithromyc | Intracellula | F54 | DiSC3(5) | 3953 | RL1 | 4384 |
| 20 | Yes | A. baylyi | Azithromyc | Intracellula | F124 | Thioflavin T | 3996 | VL2 | 4506.5 |
| 20 | Yes | A. baylyi | Azithromyc | Intracellula | F101 | Quinacrine | 4138 | VL2 | 4578.5 |
| 20 | Yes | A. baylyi | Azithromyc | Intracellula | F138 | Apigenin | 3992 | VL2 | 4280.5 |
| 20 | Yes | A. baylyi | Azithromyc | Intracellula | F65 | H2FDA | 4189 | BL1 | 5561 |
| 20 | Yes | A. baylyi | Azithromyc | Intracellula | F142 | Chlorophe | 3779 | BL3 | 7611 |
| 20 | Yes | A. baylyi | Azithromyc | Intracellula | F39 | Congo Red | 4217 | BL3 | 8457 |
| 20 | Yes | A. baylyi | Azithromyc | Intracellula | F116 | Sunset Yel | 4088 | BL2 | 6932 |
| 20 | Yes | A. baylyi | Azithromyc | Intracellula | F10 | Alizarin | 3972 | BL1 | 5201.5 |
| 20 | Yes | A. baylyi | Azithromyc | Intracellula | F84 | Neutral Re | 3840 | BL1 | 4690 |
| 20 | Yes | A. baylyi | Azithromyc | Intracellula | F99 | Pyronin Y | 4062 | BL1 | 4072 |
| 20 | Yes | A. baylyi | Azithromyc | Intracellula | F108 | Rhodamine | 3786 | BL1 | 4148.5 |
| 20 | Yes | A. baylyi | Azithromyc | Intracellula | F109 | Riboflavin | 3658 | BL1 | 4170 |
| 20 | Yes | A. baylyi | Azithromyc | Intracellula | F117 | SYBR Gree | 3701 | BL1 | 216155 |
| 20 | Yes | A. baylyi | Azithromyc | Intracellula | F106 | Rhodamine | 3671 | BL1 | 7202 |
| 20 | Yes | A. baylyi | Azithromyc | Intracellula | F73 | Malachite ( | 3891 | BL1 | 5375 |
| 20 | Yes | A. baylyi | Azithromyc | Intracellula | F7 | Acridine or | 3674 | BL1 | 6718.5 |
| 20 | Yes | A. baylyi | Azithromyc | Intracellula | F192 | 5-Carboxyf | 3850 | BL1 | 14253 |
| 20 | Yes | A. baylyi | Azithromyc | Intracellula | F57 | Eosin Y | 3413 | BL1 | 10172 |
| 20 | Yes | A. baylyi | Azithromyc | Intracellula | F27 | Calcein | 3416 | BL1 | 13862 |
| 20 | Yes | A. baylyi | Azithromyc | Intracellula | F12 | Amaranth | 3732 | BL1 | 11394 |
| 20 | Yes | A. baylyi | None | None | NA | Autofluore | 4225 | NA | 5523 |
| 20 | Yes | A. baylyi | Trimethopr | Intracellula | F194 | Oxonol | 4079 | RL1 | 4797 |
| 20 | Yes | A. baylyi | Trimethopr | Intracellula | F11 | Allura Red | 3806 | BL3 | 4817.5 |
| 20 | Yes | A. baylyi | Trimethopr | Intracellula | F54 | DiSC3(5) | 4355 | RL1 | 4339 |
| 20 | Yes | A. baylyi | Trimethopr | Intracellula | F124 | Thioflavin T | 4083 | VL2 | 4788 |
| 20 | Yes | A. baylyi | Trimethopr | Intracellula | F101 | Quinacrine | 4288 | VL2 | 4842.5 |
| 20 | Yes | A. baylyi | Trimethopr | Intracellula | F138 | Apigenin | 4156 | VL2 | 4669 |
| 20 | Yes | A. baylyi | Trimethopr | Intracellula | F65 | H2FDA | 4145 | BL1 | 5612 |
| 20 | Yes | A. baylyi | Trimethopr | Intracellula | F142 | Chlorophe | 4163 | BL3 | 7349 |
| 20 | Yes | A. baylyi | Trimethopr | Intracellula | F39 | Congo Red | 4168 | BL3 | 6689 |
| 20 | Yes | A. baylyi | Trimethopr | Intracellula | F116 | Sunset Yel | 4085 | BL2 | 7915 |
| 20 | Yes | A. baylyi | Trimethopr | Intracellula | F10 | Alizarin | 4188 | BL1 | 7281.5 |
| 20 | Yes | A. baylyi | Trimethopr | Intracellula | F84 | Neutral Re | 3754 | BL1 | 5072.5 |
| 20 | Yes | A. baylyi | Trimethopr | Intracellula | F99 | Pyronin Y | 3805 | BL1 | 4693 |
| 20 | Yes | A. baylyi | Trimethopr | Intracellula | F108 | Rhodamine | 3887 | BL1 | 4228 |
| 20 | Yes | A. baylyi | Trimethopr | Intracellula | F109 | Riboflavin | 3876 | BL1 | 3982 |
| 20 | Yes | A. baylyi | Trimethopr | Intracellula | F117 | SYBR Gree | 3432 | BL1 | 271118.5 |
| 20 | Yes | A. baylyi | Trimethopr | Intracellula | F106 | Rhodamine | 3990 | BL1 | 5430 |
| 20 | Yes | A. baylyi | Trimethopr | Intracellula | F73 | Malachite ( | 3625 | BL1 | 5349 |
| 20 | Yes | A. baylyi | Trimethopr | Intracellula | F7 | Acridine or | 4243 | BL1 | 6271 |
| 20 | Yes | A. baylyi | Trimethopr | Intracellula | F192 | 5-Carboxyf | 3359 | BL1 | 12600 |
| 20 | Yes | A. baylyi | Trimethopr | Intracellula | F57 | Eosin Y | 3605 | BL1 | 9275 |
| 20 | Yes | A. baylyi | Trimethopr | Intracellula | F27 | Calcein | 3448 | BL1 | 14903 |

| | | | | | | | | | |
|---|---|---|---|---|---|---|---|---|---|
| 20 | Yes | A. baylyi | Trimethopr | Intracellula | F12 | Amaranth | 3516 | BL1 | 13445 |
| 20 | Yes | A. baylyi | None | None | NA | Autofluore: | 4190 | NA | 6089.5 |
| 20 | Yes | A. baylyi | Chloramph | Intracellula | F194 | Oxonol | 3198 | RL1 | 5438.5 |
| 20 | Yes | A. baylyi | Chloramph | Intracellula | F11 | Allura Red | 3062 | BL3 | 5115.5 |
| 20 | Yes | A. baylyi | Chloramph | Intracellula | F54 | DiSC3(5) | 2989 | RL1 | 5051 |
| 20 | Yes | A. baylyi | Chloramph | Intracellula | F124 | Thioflavin T | 3814 | VL2 | 5162 |
| 20 | Yes | A. baylyi | Chloramph | Intracellula | F101 | Quinacrine | 3563 | VL2 | 5104 |
| 20 | Yes | A. baylyi | Chloramph | Intracellula | F138 | Apigenin | 3403 | VL2 | 4588 |
| 20 | Yes | A. baylyi | Chloramph | Intracellula | F65 | H2FDA | 3467 | BL1 | 6177 |
| 20 | Yes | A. baylyi | Chloramph | Intracellula | F142 | Chlorophe | 3472 | BL3 | 8951 |
| 20 | Yes | A. baylyi | Chloramph | Intracellula | F39 | Congo Red | 3500 | BL3 | 8873.5 |
| 20 | Yes | A. baylyi | Chloramph | Intracellula | F116 | Sunset Yel | 3569 | BL2 | 7863 |
| 20 | Yes | A. baylyi | Chloramph | Intracellula | F10 | Alizarin | 3594 | BL1 | 5922 |
| 20 | Yes | A. baylyi | Chloramph | Intracellula | F84 | Neutral Re | 3158 | BL1 | 5149 |
| 20 | Yes | A. baylyi | Chloramph | Intracellula | F99 | Pyronin Y | 3587 | BL1 | 4297 |
| 20 | Yes | A. baylyi | Chloramph | Intracellula | F108 | Rhodamine | 3383 | BL1 | 4898 |
| 20 | Yes | A. baylyi | Chloramph | Intracellula | F109 | Riboflavin | 3247 | BL1 | 4802 |
| 20 | Yes | A. baylyi | Chloramph | Intracellula | F117 | SYBR Gree | 3247 | BL1 | 215806 |
| 20 | Yes | A. baylyi | Chloramph | Intracellula | F106 | Rhodamine | 3027 | BL1 | 7781 |
| 20 | Yes | A. baylyi | Chloramph | Intracellula | F73 | Malachite ( | 3022 | BL1 | 7709 |
| 20 | Yes | A. baylyi | Chloramph | Intracellula | F7 | Acridine or | 3252 | BL1 | 7572.5 |
| 20 | Yes | A. baylyi | Chloramph | Intracellula | F192 | 5-Carboxyf | 3171 | BL1 | 14137 |
| 20 | Yes | A. baylyi | Chloramph | Intracellula | F57 | Eosin Y | 3045 | BL1 | 10105 |
| 20 | Yes | A. baylyi | Chloramph | Intracellula | F27 | Calcein | 2867 | BL1 | 18432 |
| 20 | Yes | A. baylyi | Chloramph | Intracellula | F12 | Amaranth | 3317 | BL1 | 13271 |
| 20 | Yes | A. baylyi | None | None | NA | Autofluore: | 4226 | NA | 6308 |
| 20 | Yes | A. baylyi | Colistin | Wall | F194 | Oxonol | 2416 | RL1 | 15971.5 |
| 20 | Yes | A. baylyi | Colistin | Wall | F11 | Allura Red | 2324 | BL3 | 12848 |
| 20 | Yes | A. baylyi | Colistin | Wall | F54 | DiSC3(5) | 2624 | RL1 | 21886.5 |
| 20 | Yes | A. baylyi | Colistin | Wall | F124 | Thioflavin T | 2552 | VL2 | 10587 |
| 20 | Yes | A. baylyi | Colistin | Wall | F101 | Quinacrine | 2525 | VL2 | 10529 |
| 20 | Yes | A. baylyi | Colistin | Wall | F138 | Apigenin | 2585 | VL2 | 9753 |
| 20 | Yes | A. baylyi | Colistin | Wall | F65 | H2FDA | 2184 | BL1 | 11554 |
| 20 | Yes | A. baylyi | Colistin | Wall | F142 | Chlorophe | 2520 | BL3 | 15063 |
| 20 | Yes | A. baylyi | Colistin | Wall | F39 | Congo Red | 2358 | BL3 | 13338.5 |
| 20 | Yes | A. baylyi | Colistin | Wall | F116 | Sunset Yel | 2527 | BL2 | 10115 |
| 20 | Yes | A. baylyi | Colistin | Wall | F10 | Alizarin | 2541 | BL1 | 9657 |
| 20 | Yes | A. baylyi | Colistin | Wall | F84 | Neutral Re | 2545 | BL1 | 10383 |
| 20 | Yes | A. baylyi | Colistin | Wall | F99 | Pyronin Y | 2594 | BL1 | 7125 |
| 20 | Yes | A. baylyi | Colistin | Wall | F108 | Rhodamine | 2453 | BL1 | 8091 |
| 20 | Yes | A. baylyi | Colistin | Wall | F109 | Riboflavin | 2574 | BL1 | 9366 |
| 20 | Yes | A. baylyi | Colistin | Wall | F117 | SYBR Gree | 2223 | BL1 | 2120890 |
| 20 | Yes | A. baylyi | Colistin | Wall | F106 | Rhodamine | 2509 | BL1 | 40940 |
| 20 | Yes | A. baylyi | Colistin | Wall | F73 | Malachite ( | 2448 | BL1 | 11295.5 |
| 20 | Yes | A. baylyi | Colistin | Wall | F7 | Acridine or | 2405 | BL1 | 21715 |
| 20 | Yes | A. baylyi | Colistin | Wall | F192 | 5-Carboxyf | 2219 | BL1 | 20345 |
| 20 | Yes | A. baylyi | Colistin | Wall | F57 | Eosin Y | 2596 | BL1 | 19260.5 |
| 20 | Yes | A. baylyi | Colistin | Wall | F27 | Calcein | 2558 | BL1 | 20751 |

| | | | | | | | | | |
|---|---|---|---|---|---|---|---|---|---|
| 20 | Yes | A. baylyi | Colistin | Wall | F12 | Amaranth | 2789 | BL1 | 20119 |
| 20 | Yes | A. baylyi | None | None | NA | Autofluore: | 4130 | NA | 6174.5 |
| 20 | Yes | A. baylyi | Gentamici | Intracellula | F194 | Oxonol | 4080 | RL1 | 5739 |
| 20 | Yes | A. baylyi | Gentamici | Intracellula | F11 | Allura Red | 4248 | BL3 | 5611.5 |
| 20 | Yes | A. baylyi | Gentamici | Intracellula | F54 | DiSC3(5) | 4251 | RL1 | 5037 |
| 20 | Yes | A. baylyi | Gentamici | Intracellula | F124 | Thioflavin T | 4406 | VL2 | 5475 |
| 20 | Yes | A. baylyi | Gentamici | Intracellula | F101 | Quinacrine | 4442 | VL2 | 5459.5 |
| 20 | Yes | A. baylyi | Gentamici | Intracellula | F138 | Apigenin | 4279 | VL2 | 5180 |
| 20 | Yes | A. baylyi | Gentamici | Intracellula | F65 | H2FDA | 4351 | BL1 | 6445 |
| 20 | Yes | A. baylyi | Gentamici | Intracellula | F142 | Chlorophe | 4459 | BL3 | 9338 |
| 20 | Yes | A. baylyi | Gentamici | Intracellula | F39 | Congo Red | 3958 | BL3 | 8997.5 |
| 20 | Yes | A. baylyi | Gentamici | Intracellula | F116 | Sunset Yel | 4716 | BL2 | 6084.5 |
| 20 | Yes | A. baylyi | Gentamici | Intracellula | F10 | Alizarin | 4389 | BL1 | 5137 |
| 20 | Yes | A. baylyi | Gentamici | Intracellula | F84 | Neutral Re | 4591 | BL1 | 4405 |
| 20 | Yes | A. baylyi | Gentamici | Intracellula | F99 | Pyronin Y | 4434 | BL1 | 4396.5 |
| 20 | Yes | A. baylyi | Gentamici | Intracellula | F108 | Rhodamine | 4075 | BL1 | 4388 |
| 20 | Yes | A. baylyi | Gentamici | Intracellula | F109 | Riboflavin | 4200 | BL1 | 4748 |
| 20 | Yes | A. baylyi | Gentamici | Intracellula | F117 | SYBR Gree | 3735 | BL1 | 297611 |
| 20 | Yes | A. baylyi | Gentamici | Intracellula | F106 | Rhodamine | 4529 | BL1 | 6595 |
| 20 | Yes | A. baylyi | Gentamici | Intracellula | F73 | Malachite ( | 4157 | BL1 | 6562 |
| 20 | Yes | A. baylyi | Gentamici | Intracellula | F7 | Acridine or | 4016 | BL1 | 7036.5 |
| 20 | Yes | A. baylyi | Gentamici | Intracellula | F192 | 5-Carboxyf | 3161 | BL1 | 12806 |
| 20 | Yes | A. baylyi | Gentamici | Intracellula | F57 | Eosin Y | 4268 | BL1 | 13882 |
| 20 | Yes | A. baylyi | Gentamici | Intracellula | F27 | Calcein | 3937 | BL1 | 13846 |
| 20 | Yes | A. baylyi | Gentamici | Intracellula | F12 | Amaranth | 3860 | BL1 | 14058.5 |
| 20 | Yes | A. baylyi | None | None | NA | Autofluore: | 4092 | NA | 6757 |
| 20 | Yes | A. baylyi | Nitrofurant | Intracellula | F194 | Oxonol | 4191 | RL1 | 6020 |
| 20 | Yes | A. baylyi | Nitrofurant | Intracellula | F11 | Allura Red | 3733 | BL3 | 5719 |
| 20 | Yes | A. baylyi | Nitrofurant | Intracellula | F54 | DiSC3(5) | 4079 | RL1 | 5079 |
| 20 | Yes | A. baylyi | Nitrofurant | Intracellula | F124 | Thioflavin T | 3814 | VL2 | 5485 |
| 20 | Yes | A. baylyi | Nitrofurant | Intracellula | F101 | Quinacrine | 3885 | VL2 | 5246 |
| 20 | Yes | A. baylyi | Nitrofurant | Intracellula | F138 | Apigenin | 4226 | VL2 | 5091.5 |
| 20 | Yes | A. baylyi | Nitrofurant | Intracellula | F65 | H2FDA | 4060 | BL1 | 6915 |
| 20 | Yes | A. baylyi | Nitrofurant | Intracellula | F142 | Chlorophe | 3848 | BL3 | 9062.5 |
| 20 | Yes | A. baylyi | Nitrofurant | Intracellula | F39 | Congo Red | 4298 | BL3 | 7727.5 |
| 20 | Yes | A. baylyi | Nitrofurant | Intracellula | F116 | Sunset Yel | 3986 | BL2 | 5867 |
| 20 | Yes | A. baylyi | Nitrofurant | Intracellula | F10 | Alizarin | 4324 | BL1 | 5109.5 |
| 20 | Yes | A. baylyi | Nitrofurant | Intracellula | F84 | Neutral Re | 4233 | BL1 | 4277 |
| 20 | Yes | A. baylyi | Nitrofurant | Intracellula | F99 | Pyronin Y | 3828 | BL1 | 4210 |
| 20 | Yes | A. baylyi | Nitrofurant | Intracellula | F108 | Rhodamine | 3951 | BL1 | 4598 |
| 20 | Yes | A. baylyi | Nitrofurant | Intracellula | F109 | Riboflavin | 4114 | BL1 | 5005 |
| 20 | Yes | A. baylyi | Nitrofurant | Intracellula | F117 | SYBR Gree | 3947 | BL1 | 202526 |
| 20 | Yes | A. baylyi | Nitrofurant | Intracellula | F106 | Rhodamine | 4202 | BL1 | 7224 |
| 20 | Yes | A. baylyi | Nitrofurant | Intracellula | F73 | Malachite ( | 3823 | BL1 | 6914 |
| 20 | Yes | A. baylyi | Nitrofurant | Intracellula | F7 | Acridine or | 3228 | BL1 | 6958 |
| 20 | Yes | A. baylyi | Nitrofurant | Intracellula | F192 | 5-Carboxyf | 3672 | BL1 | 12696 |
| 20 | Yes | A. baylyi | Nitrofurant | Intracellula | F57 | Eosin Y | 3734 | BL1 | 12800 |
| 20 | Yes | A. baylyi | Nitrofurant | Intracellula | F27 | Calcein | 4204 | BL1 | 14249 |

| | | | | | | | | | |
|---|---|---|---|---|---|---|---|---|---|
| 20 | Yes | A. baylyi | Nitrofurant | Intracellula | F12 | Amaranth | 4027 | BL1 | 12037 |
| 20 | Yes | A. baylyi | None | None | NA | Autofluores | 4392 | NA | 6216.5 |
| 20 | Yes | A. baylyi | Pentamidi | Wall | F194 | Oxonol | 2709 | RL1 | 10562 |
| 20 | Yes | A. baylyi | Pentamidi | Wall | F11 | Allura Red | 2488 | BL3 | 7728 |
| 20 | Yes | A. baylyi | Pentamidi | Wall | F54 | DiSC3(5) | 2904 | RL1 | 8060.5 |
| 20 | Yes | A. baylyi | Pentamidi | Wall | F124 | Thioflavin T | 2714 | VL2 | 8676.5 |
| 20 | Yes | A. baylyi | Pentamidi | Wall | F101 | Quinacrine | 2715 | VL2 | 8836 |
| 20 | Yes | A. baylyi | Pentamidi | Wall | F138 | Apigenin | 2738 | VL2 | 11976 |
| 20 | Yes | A. baylyi | Pentamidi | Wall | F65 | H2FDA | 2619 | BL1 | 13297 |
| 20 | Yes | A. baylyi | Pentamidi | Wall | F142 | Chlorophe | 2673 | BL3 | 11422 |
| 20 | Yes | A. baylyi | Pentamidi | Wall | F39 | Congo Red | 3668 | BL3 | 13769.5 |
| 20 | Yes | A. baylyi | Pentamidi | Wall | F116 | Sunset Yel | 2798 | BL2 | 9095 |
| 20 | Yes | A. baylyi | Pentamidi | Wall | F10 | Alizarin | 2688 | BL1 | 8281.5 |
| 20 | Yes | A. baylyi | Pentamidi | Wall | F84 | Neutral Re | 2643 | BL1 | 7928 |
| 20 | Yes | A. baylyi | Pentamidi | Wall | F99 | Pyronin Y | 2787 | BL1 | 6545 |
| 20 | Yes | A. baylyi | Pentamidi | Wall | F108 | Rhodamine | 2727 | BL1 | 7395 |
| 20 | Yes | A. baylyi | Pentamidi | Wall | F109 | Riboflavin | 2702 | BL1 | 7681 |
| 20 | Yes | A. baylyi | Pentamidi | Wall | F117 | SYBR Gree | 2608 | BL1 | 1359427 |
| 20 | Yes | A. baylyi | Pentamidi | Wall | F106 | Rhodamine | 2541 | BL1 | 15601 |
| 20 | Yes | A. baylyi | Pentamidi | Wall | F73 | Malachite ( | 2412 | BL1 | 11520.5 |
| 20 | Yes | A. baylyi | Pentamidi | Wall | F7 | Acridine or | 2400 | BL1 | 13696.5 |
| 20 | Yes | A. baylyi | Pentamidi | Wall | F192 | 5-Carboxyf | 2363 | BL1 | 14433 |
| 20 | Yes | A. baylyi | Pentamidi | Wall | F57 | Eosin Y | 2367 | BL1 | 12721 |
| 20 | Yes | A. baylyi | Pentamidi | Wall | F27 | Calcein | 2575 | BL1 | 14837 |
| 20 | Yes | A. baylyi | Pentamidi | Wall | F12 | Amaranth | 2558 | BL1 | 16923.5 |
| 20 | Yes | A. baylyi | None | None | NA | Autofluores | 4334 | NA | 8014 |
| 20 | Yes | A. baylyi | Rifampicin | Intracellula | F194 | Oxonol | 3843 | RL1 | 7292 |
| 20 | Yes | A. baylyi | Rifampicin | Intracellula | F11 | Allura Red | 3946 | BL3 | 6627 |
| 20 | Yes | A. baylyi | Rifampicin | Intracellula | F54 | DiSC3(5) | 4188 | RL1 | 5850.5 |
| 20 | Yes | A. baylyi | Rifampicin | Intracellula | F124 | Thioflavin T | 4295 | VL2 | 5774 |
| 20 | Yes | A. baylyi | Rifampicin | Intracellula | F101 | Quinacrine | 4258 | VL2 | 5164.5 |
| 20 | Yes | A. baylyi | Rifampicin | Intracellula | F138 | Apigenin | 4260 | VL2 | 5347.5 |
| 20 | Yes | A. baylyi | Rifampicin | Intracellula | F65 | H2FDA | 3983 | BL1 | 5984 |
| 20 | Yes | A. baylyi | Rifampicin | Intracellula | F142 | Chlorophe | 4246 | BL3 | 8898 |
| 20 | Yes | A. baylyi | Rifampicin | Intracellula | F39 | Congo Red | 4505 | BL3 | 8273 |
| 20 | Yes | A. baylyi | Rifampicin | Intracellula | F116 | Sunset Yel | 4312 | BL2 | 8629 |
| 20 | Yes | A. baylyi | Rifampicin | Intracellula | F10 | Alizarin | 4385 | BL1 | 5880 |
| 20 | Yes | A. baylyi | Rifampicin | Intracellula | F84 | Neutral Re | 4580 | BL1 | 4596 |
| 20 | Yes | A. baylyi | Rifampicin | Intracellula | F99 | Pyronin Y | 4348 | BL1 | 4282 |
| 20 | Yes | A. baylyi | Rifampicin | Intracellula | F108 | Rhodamine | 4399 | BL1 | 4258 |
| 20 | Yes | A. baylyi | Rifampicin | Intracellula | F109 | Riboflavin | 4519 | BL1 | 4493 |
| 20 | Yes | A. baylyi | Rifampicin | Intracellula | F117 | SYBR Gree | 3701 | BL1 | 369116 |
| 20 | Yes | A. baylyi | Rifampicin | Intracellula | F106 | Rhodamine | 4034 | BL1 | 7246.5 |
| 20 | Yes | A. baylyi | Rifampicin | Intracellula | F73 | Malachite ( | 3912 | BL1 | 6878 |
| 20 | Yes | A. baylyi | Rifampicin | Intracellula | F7 | Acridine or | 4198 | BL1 | 6824.5 |
| 20 | Yes | A. baylyi | Rifampicin | Intracellula | F192 | 5-Carboxyf | 3988 | BL1 | 14925 |
| 20 | Yes | A. baylyi | Rifampicin | Intracellula | F57 | Eosin Y | 3576 | BL1 | 11091 |
| 20 | Yes | A. baylyi | Rifampicin | Intracellula | F27 | Calcein | 3722 | BL1 | 15467.5 |

| | | | | | | | | | |
|---|---|---|---|---|---|---|---|---|---|
| 20 | Yes | A. baylyi | Rifampicin | Intracellula | F12 | Amaranth | 3881 | BL1 | 14311 |
| 20 | Yes | A. baylyi | None | None | NA | Autofluore: | 4061 | NA | 5753 |
| 20 | Yes | A. baylyi | Trimethopr | Intracellula | F194 | Oxonol | 3749 | RL1 | 5427 |
| 20 | Yes | A. baylyi | Trimethopr | Intracellula | F11 | Allura Red | 4352 | BL3 | 5845.5 |
| 20 | Yes | A. baylyi | Trimethopr | Intracellula | F54 | DiSC3(5) | 4289 | RL1 | 4885 |
| 20 | Yes | A. baylyi | Trimethopr | Intracellula | F124 | Thioflavin T | 4477 | VL2 | 5161 |
| 20 | Yes | A. baylyi | Trimethopr | Intracellula | F101 | Quinacrine | 4557 | VL2 | 5204 |
| 20 | Yes | A. baylyi | Trimethopr | Intracellula | F138 | Apigenin | 4310 | VL2 | 4820.5 |
| 20 | Yes | A. baylyi | Trimethopr | Intracellula | F65 | H2FDA | 4363 | BL1 | 6375 |
| 20 | Yes | A. baylyi | Trimethopr | Intracellula | F142 | Chlorophe | 4330 | BL3 | 9345 |
| 20 | Yes | A. baylyi | Trimethopr | Intracellula | F39 | Congo Red | 4365 | BL3 | 7232 |
| 20 | Yes | A. baylyi | Trimethopr | Intracellula | F116 | Sunset Yel | 4464 | BL2 | 6377.5 |
| 20 | Yes | A. baylyi | Trimethopr | Intracellula | F10 | Alizarin | 4627 | BL1 | 5761 |
| 20 | Yes | A. baylyi | Trimethopr | Intracellula | F84 | Neutral Re | 4144 | BL1 | 5379.5 |
| 20 | Yes | A. baylyi | Trimethopr | Intracellula | F99 | Pyronin Y | 3855 | BL1 | 4626 |
| 20 | Yes | A. baylyi | Trimethopr | Intracellula | F108 | Rhodamine | 4460 | BL1 | 4314.5 |
| 20 | Yes | A. baylyi | Trimethopr | Intracellula | F109 | Riboflavin | 4333 | BL1 | 4598 |
| 20 | Yes | A. baylyi | Trimethopr | Intracellula | F117 | SYBR Gree | 3873 | BL1 | 302737 |
| 20 | Yes | A. baylyi | Trimethopr | Intracellula | F106 | Rhodamine | 4461 | BL1 | 6048 |
| 20 | Yes | A. baylyi | Trimethopr | Intracellula | F73 | Malachite ( | 4420 | BL1 | 6074 |
| 20 | Yes | A. baylyi | Trimethopr | Intracellula | F7 | Acridine or | 3625 | BL1 | 6473 |
| 20 | Yes | A. baylyi | Trimethopr | Intracellula | F192 | 5-Carboxyf | 4022 | BL1 | 13748 |
| 20 | Yes | A. baylyi | Trimethopr | Intracellula | F57 | Eosin Y | 4238 | BL1 | 14666 |
| 20 | Yes | A. baylyi | Trimethopr | Intracellula | F27 | Calcein | 4380 | BL1 | 15641 |
| 20 | Yes | A. baylyi | Trimethopr | Intracellula | F12 | Amaranth | 4469 | BL1 | 14637 |
| 20 | Yes | A. baylyi | None | None | NA | Autofluore: | 4984 | NA | 1077.5 |
| 20 | Yes | A. baylyi | None | None | F194 | Oxonol | 5792 | RL1 | 2456 |
| 20 | Yes | A. baylyi | None | None | F11 | Allura Red | 4919 | BL3 | 2533 |
| 20 | Yes | A. baylyi | None | None | F54 | DiSC3(5) | 5657 | RL1 | 2761 |
| 20 | Yes | A. baylyi | None | None | F124 | Thioflavin T | 5119 | VL2 | 2605 |
| 20 | Yes | A. baylyi | None | None | F101 | Quinacrine | 5261 | VL2 | 2824 |
| 20 | Yes | A. baylyi | None | None | F138 | Apigenin | 4970 | VL2 | 2915.5 |
| 20 | Yes | A. baylyi | None | None | F65 | H2FDA | 5076 | BL1 | 4095 |
| 20 | Yes | A. baylyi | None | None | F142 | Chlorophe | 4872 | BL3 | 5500 |
| 20 | Yes | A. baylyi | None | None | F39 | Congo Red | 4871 | BL3 | 5511 |
| 20 | Yes | A. baylyi | None | None | F116 | Sunset Yel | 5067 | BL2 | 4383 |
| 20 | Yes | A. baylyi | None | None | F10 | Alizarin | 4795 | BL1 | 4270 |
| 20 | Yes | A. baylyi | None | None | F84 | Neutral Re | 4616 | BL1 | 4000.5 |
| 20 | Yes | A. baylyi | None | None | F99 | Pyronin Y | 4371 | BL1 | 3823 |
| 20 | Yes | A. baylyi | None | None | F108 | Rhodamine | 4420 | BL1 | 3955.5 |
| 20 | Yes | A. baylyi | None | None | F109 | Riboflavin | 4429 | BL1 | 4623 |
| 20 | Yes | A. baylyi | None | None | F117 | SYBR Gree | 4367 | BL1 | 310396 |
| 20 | Yes | A. baylyi | None | None | F106 | Rhodamine | 3681 | BL1 | 4075 |
| 20 | Yes | A. baylyi | None | None | F73 | Malachite ( | 4641 | BL1 | 4018 |
| 20 | Yes | A. baylyi | None | None | F7 | Acridine or | 4032 | BL1 | 4345.5 |
| 20 | Yes | A. baylyi | None | None | F192 | 5-Carboxyf | 3338 | BL1 | 14744 |
| 20 | Yes | A. baylyi | None | None | F57 | Eosin Y | 4377 | BL1 | 13689 |
| 20 | Yes | A. baylyi | None | None | F27 | Calcein | 4301 | BL1 | 18974 |

| | | | | | | | | | |
|---|---|---|---|---|---|---|---|---|---|
| 20 | Yes | A. baylyi | None | None | F12 | Amaranth | 4247 | BL1 | 13334 |
| 20 | Yes | A. baylyi | None | None | NA | Autofluore: | 4301 | NA | 5259 |
| 20 | Yes | A. baylyi | Ampicillin | Wall | F194 | Oxonol | 3800 | RL1 | 4451.5 |
| 20 | Yes | A. baylyi | Ampicillin | Wall | F11 | Allura Red | 3701 | BL3 | 4222 |
| 20 | Yes | A. baylyi | Ampicillin | Wall | F54 | DiSC3(5) | 3864 | RL1 | 4031.5 |
| 20 | Yes | A. baylyi | Ampicillin | Wall | F124 | Thioflavin T | 3915 | VL2 | 4165 |
| 20 | Yes | A. baylyi | Ampicillin | Wall | F101 | Quinacrine | 3573 | VL2 | 4380 |
| 20 | Yes | A. baylyi | Ampicillin | Wall | F138 | Apigenin | 3494 | VL2 | 4065 |
| 20 | Yes | A. baylyi | Ampicillin | Wall | F65 | H2FDA | 3595 | BL1 | 5010 |
| 20 | Yes | A. baylyi | Ampicillin | Wall | F142 | Chlorophe | 3770 | BL3 | 6417 |
| 20 | Yes | A. baylyi | Ampicillin | Wall | F39 | Congo Red | 3722 | BL3 | 6611 |
| 20 | Yes | A. baylyi | Ampicillin | Wall | F116 | Sunset Yel | 3766 | BL2 | 5740 |
| 20 | Yes | A. baylyi | Ampicillin | Wall | F10 | Alizarin | 3751 | BL1 | 4782 |
| 20 | Yes | A. baylyi | Ampicillin | Wall | F84 | Neutral Re | 3528 | BL1 | 4542 |
| 20 | Yes | A. baylyi | Ampicillin | Wall | F99 | Pyronin Y | 3604 | BL1 | 4632.5 |
| 20 | Yes | A. baylyi | Ampicillin | Wall | F108 | Rhodamine | 3517 | BL1 | 4127 |
| 20 | Yes | A. baylyi | Ampicillin | Wall | F109 | Riboflavin | 3506 | BL1 | 3908 |
| 20 | Yes | A. baylyi | Ampicillin | Wall | F117 | SYBR Gree | 3566 | BL1 | 321335 |
| 20 | Yes | A. baylyi | Ampicillin | Wall | F106 | Rhodamine | 3547 | BL1 | 4902 |
| 20 | Yes | A. baylyi | Ampicillin | Wall | F73 | Malachite ( | 3725 | BL1 | 4500 |
| 20 | Yes | A. baylyi | Ampicillin | Wall | F7 | Acridine or | 3340 | BL1 | 5000.5 |
| 20 | Yes | A. baylyi | Ampicillin | Wall | F192 | 5-Carboxyf | 3269 | BL1 | 15128 |
| 20 | Yes | A. baylyi | Ampicillin | Wall | F57 | Eosin Y | 3611 | BL1 | 16204 |
| 20 | Yes | A. baylyi | Ampicillin | Wall | F27 | Calcein | 3355 | BL1 | 19161 |
| 20 | Yes | A. baylyi | Ampicillin | Wall | F12 | Amaranth | 3574 | BL1 | 17333.5 |
| 20 | Yes | A. baylyi | None | None | NA | Autofluore: | 4192 | NA | 6104.5 |
| 20 | Yes | A. baylyi | Bacitracin | Wall | F194 | Oxonol | 4297 | RL1 | 6210 |
| 20 | Yes | A. baylyi | Bacitracin | Wall | F11 | Allura Red | 3862 | BL3 | 5732.5 |
| 20 | Yes | A. baylyi | Bacitracin | Wall | F54 | DiSC3(5) | 4321 | RL1 | 5474 |
| 20 | Yes | A. baylyi | Bacitracin | Wall | F124 | Thioflavin T | 4415 | VL2 | 5774 |
| 20 | Yes | A. baylyi | Bacitracin | Wall | F101 | Quinacrine | 4052 | VL2 | 6717.5 |
| 20 | Yes | A. baylyi | Bacitracin | Wall | F138 | Apigenin | 4446 | VL2 | 5587 |
| 20 | Yes | A. baylyi | Bacitracin | Wall | F65 | H2FDA | 4229 | BL1 | 6358 |
| 20 | Yes | A. baylyi | Bacitracin | Wall | F142 | Chlorophe | 4139 | BL3 | 6645 |
| 20 | Yes | A. baylyi | Bacitracin | Wall | F39 | Congo Red | 4609 | BL3 | 6883 |
| 20 | Yes | A. baylyi | Bacitracin | Wall | F116 | Sunset Yel | 4338 | BL2 | 5789.5 |
| 20 | Yes | A. baylyi | Bacitracin | Wall | F10 | Alizarin | 4151 | BL1 | 5759 |
| 20 | Yes | A. baylyi | Bacitracin | Wall | F84 | Neutral Re | 4407 | BL1 | 6293 |
| 20 | Yes | A. baylyi | Bacitracin | Wall | F99 | Pyronin Y | 3675 | BL1 | 4618 |
| 20 | Yes | A. baylyi | Bacitracin | Wall | F108 | Rhodamine | 4703 | BL1 | 5460 |
| 20 | Yes | A. baylyi | Bacitracin | Wall | F109 | Riboflavin | 4300 | BL1 | 4577.5 |
| 20 | Yes | A. baylyi | Bacitracin | Wall | F117 | SYBR Gree | 3991 | BL1 | 361411 |
| 20 | Yes | A. baylyi | Bacitracin | Wall | F106 | Rhodamine | 4416 | BL1 | 5511 |
| 20 | Yes | A. baylyi | Bacitracin | Wall | F73 | Malachite ( | 3647 | BL1 | 4893 |
| 20 | Yes | A. baylyi | Bacitracin | Wall | F7 | Acridine or | 4139 | BL1 | 5370 |
| 20 | Yes | A. baylyi | Bacitracin | Wall | F192 | 5-Carboxyf | 4377 | BL1 | 16134 |
| 20 | Yes | A. baylyi | Bacitracin | Wall | F57 | Eosin Y | 3985 | BL1 | 17100 |
| 20 | Yes | A. baylyi | Bacitracin | Wall | F27 | Calcein | 4393 | BL1 | 17054 |

| | | | | | | | | | |
|---|---|---|---|---|---|---|---|---|---|
| 20 | Yes | A. baylyi | Bacitracin | Wall | F12 | Amaranth | 4243 | BL1 | 16673 |
| 20 | Yes | A. baylyi | None | None | NA | Autofluore: | 4432 | NA | 4976.5 |
| 20 | Yes | A. baylyi | Cefotaxim | Wall | F194 | Oxonol | 4064 | RL1 | 4810 |
| 20 | Yes | A. baylyi | Cefotaxim | Wall | F11 | Allura Red | 3557 | BL3 | 5014 |
| 20 | Yes | A. baylyi | Cefotaxim | Wall | F54 | DiSC3(5) | 3789 | RL1 | 4752 |
| 20 | Yes | A. baylyi | Cefotaxim | Wall | F124 | Thioflavin T | 3910 | VL2 | 5285.5 |
| 20 | Yes | A. baylyi | Cefotaxim | Wall | F101 | Quinacrine | 3882 | VL2 | 5199.5 |
| 20 | Yes | A. baylyi | Cefotaxim | Wall | F138 | Apigenin | 3417 | VL2 | 6157 |
| 20 | Yes | A. baylyi | Cefotaxim | Wall | F65 | H2FDA | 3602 | BL1 | 5862.5 |
| 20 | Yes | A. baylyi | Cefotaxim | Wall | F142 | Chlorophe | 3900 | BL3 | 6526.5 |
| 20 | Yes | A. baylyi | Cefotaxim | Wall | F39 | Congo Red | 3673 | BL3 | 6186 |
| 20 | Yes | A. baylyi | Cefotaxim | Wall | F116 | Sunset Yel | 3882 | BL2 | 5914.5 |
| 20 | Yes | A. baylyi | Cefotaxim | Wall | F10 | Alizarin | 3555 | BL1 | 5246 |
| 20 | Yes | A. baylyi | Cefotaxim | Wall | F84 | Neutral Re | 3130 | BL1 | 5666.5 |
| 20 | Yes | A. baylyi | Cefotaxim | Wall | F99 | Pyronin Y | 3442 | BL1 | 4816 |
| 20 | Yes | A. baylyi | Cefotaxim | Wall | F108 | Rhodamin | 3885 | BL1 | 4376 |
| 20 | Yes | A. baylyi | Cefotaxim | Wall | F109 | Riboflavin | 3494 | BL1 | 4431.5 |
| 20 | Yes | A. baylyi | Cefotaxim | Wall | F117 | SYBR Gree | 3782 | BL1 | 334439.5 |
| 20 | Yes | A. baylyi | Cefotaxim | Wall | F106 | Rhodamin | 3923 | BL1 | 5367 |
| 20 | Yes | A. baylyi | Cefotaxim | Wall | F73 | Malachite | 3604 | BL1 | 4881.5 |
| 20 | Yes | A. baylyi | Cefotaxim | Wall | F7 | Acridine or | 3463 | BL1 | 4644 |
| 20 | Yes | A. baylyi | Cefotaxim | Wall | F192 | 5-Carboxyf | 3691 | BL1 | 16666 |
| 20 | Yes | A. baylyi | Cefotaxim | Wall | F57 | Eosin Y | 3474 | BL1 | 17904 |
| 20 | Yes | A. baylyi | Cefotaxim | Wall | F27 | Calcein | 3478 | BL1 | 16191 |
| 20 | Yes | A. baylyi | Cefotaxim | Wall | F12 | Amaranth | 3396 | BL1 | 19271.5 |
| 20 | Yes | A. baylyi | None | None | NA | Autofluore: | 4758 | NA | 5402 |
| 20 | Yes | A. baylyi | Ciprofloxa | Intracellul | F194 | Oxonol | 3605 | RL1 | 5427 |
| 20 | Yes | A. baylyi | Ciprofloxa | Intracellul | F11 | Allura Red | 3352 | BL3 | 5072 |
| 20 | Yes | A. baylyi | Ciprofloxa | Intracellul | F54 | DiSC3(5) | 3523 | RL1 | 4962 |
| 20 | Yes | A. baylyi | Ciprofloxa | Intracellul | F124 | Thioflavin T | 3387 | VL2 | 5295 |
| 20 | Yes | A. baylyi | Ciprofloxa | Intracellul | F101 | Quinacrine | 3515 | VL2 | 5130 |
| 20 | Yes | A. baylyi | Ciprofloxa | Intracellul | F138 | Apigenin | 3481 | VL2 | 4846 |
| 20 | Yes | A. baylyi | Ciprofloxa | Intracellul | F65 | H2FDA | 3392 | BL1 | 6097 |
| 20 | Yes | A. baylyi | Ciprofloxa | Intracellul | F142 | Chlorophe | 3632 | BL3 | 7715.5 |
| 20 | Yes | A. baylyi | Ciprofloxa | Intracellul | F39 | Congo Red | 3372 | BL3 | 7346 |
| 20 | Yes | A. baylyi | Ciprofloxa | Intracellul | F116 | Sunset Yel | 3344 | BL2 | 7064.5 |
| 20 | Yes | A. baylyi | Ciprofloxa | Intracellul | F10 | Alizarin | 3482 | BL1 | 6334 |
| 20 | Yes | A. baylyi | Ciprofloxa | Intracellul | F84 | Neutral Re | 3435 | BL1 | 4929 |
| 20 | Yes | A. baylyi | Ciprofloxa | Intracellul | F99 | Pyronin Y | 3368 | BL1 | 4637.5 |
| 20 | Yes | A. baylyi | Ciprofloxa | Intracellul | F108 | Rhodamin | 3192 | BL1 | 4489.5 |
| 20 | Yes | A. baylyi | Ciprofloxa | Intracellul | F109 | Riboflavin | 3391 | BL1 | 4747 |
| 20 | Yes | A. baylyi | Ciprofloxa | Intracellul | F117 | SYBR Gree | 3127 | BL1 | 265266 |
| 20 | Yes | A. baylyi | Ciprofloxa | Intracellul | F106 | Rhodamin | 3081 | BL1 | 5304 |
| 20 | Yes | A. baylyi | Ciprofloxa | Intracellul | F73 | Malachite | 3362 | BL1 | 4939.5 |
| 20 | Yes | A. baylyi | Ciprofloxa | Intracellul | F7 | Acridine or | 3133 | BL1 | 6016 |
| 20 | Yes | A. baylyi | Ciprofloxa | Intracellul | F192 | 5-Carboxyf | 2761 | BL1 | 16281 |
| 20 | Yes | A. baylyi | Ciprofloxa | Intracellul | F57 | Eosin Y | 2884 | BL1 | 18508.5 |
| 20 | Yes | A. baylyi | Ciprofloxa | Intracellul | F27 | Calcein | 3060 | BL1 | 16637.5 |

| | | | | | | | | | |
|---|---|---|---|---|---|---|---|---|---|
| 20 | Yes | A. baylyi | Ciprofloxa | Intracellula | F12 | Amaranth | 3269 | BL1 | 19972 |
| 20 | Yes | A. baylyi | None | None | NA | Autofluore: | 4454 | NA | 5231 |
| 20 | Yes | A. baylyi | Fosfomyci | Wall | F194 | Oxonol | 4231 | RL1 | 4861 |
| 20 | Yes | A. baylyi | Fosfomyci | Wall | F11 | Allura Red | 4340 | BL3 | 4989 |
| 20 | Yes | A. baylyi | Fosfomyci | Wall | F54 | DiSC3(5) | 4342 | RL1 | 4697 |
| 20 | Yes | A. baylyi | Fosfomyci | Wall | F124 | Thioflavin T | 4546 | VL2 | 4825.5 |
| 20 | Yes | A. baylyi | Fosfomyci | Wall | F101 | Quinacrine | 4468 | VL2 | 4763.5 |
| 20 | Yes | A. baylyi | Fosfomyci | Wall | F138 | Apigenin | 3855 | VL2 | 4765 |
| 20 | Yes | A. baylyi | Fosfomyci | Wall | F65 | H2FDA | 4418 | BL1 | 5740.5 |
| 20 | Yes | A. baylyi | Fosfomyci | Wall | F142 | Chlorophe | 4193 | BL3 | 7373 |
| 20 | Yes | A. baylyi | Fosfomyci | Wall | F39 | Congo Red | 4490 | BL3 | 7634.5 |
| 20 | Yes | A. baylyi | Fosfomyci | Wall | F116 | Sunset Yel | 4736 | BL2 | 6613 |
| 20 | Yes | A. baylyi | Fosfomyci | Wall | F10 | Alizarin | 4541 | BL1 | 5388 |
| 20 | Yes | A. baylyi | Fosfomyci | Wall | F84 | Neutral Re | 4326 | BL1 | 4839 |
| 20 | Yes | A. baylyi | Fosfomyci | Wall | F99 | Pyronin Y | 4399 | BL1 | 4860 |
| 20 | Yes | A. baylyi | Fosfomyci | Wall | F108 | Rhodamine | 4108 | BL1 | 4280.5 |
| 20 | Yes | A. baylyi | Fosfomyci | Wall | F109 | Riboflavin | 4740 | BL1 | 4567.5 |
| 20 | Yes | A. baylyi | Fosfomyci | Wall | F117 | SYBR Gree | 4131 | BL1 | 285199 |
| 20 | Yes | A. baylyi | Fosfomyci | Wall | F106 | Rhodamine | 4397 | BL1 | 5480 |
| 20 | Yes | A. baylyi | Fosfomyci | Wall | F73 | Malachite ( | 4322 | BL1 | 5002.5 |
| 20 | Yes | A. baylyi | Fosfomyci | Wall | F7 | Acridine or | 4035 | BL1 | 5113 |
| 20 | Yes | A. baylyi | Fosfomyci | Wall | F192 | 5-Carboxyf | 3290 | BL1 | 14659 |
| 20 | Yes | A. baylyi | Fosfomyci | Wall | F57 | Eosin Y | 4387 | BL1 | 12871 |
| 20 | Yes | A. baylyi | Fosfomyci | Wall | F27 | Calcein | 4068 | BL1 | 15741.5 |
| 20 | Yes | A. baylyi | Fosfomyci | Wall | F12 | Amaranth | 4433 | BL1 | 15562 |
| 20 | Yes | A. baylyi | None | None | NA | Autofluore: | 4395 | NA | 5986 |
| 20 | Yes | A. baylyi | Nalidixic a | Intracellula | F194 | Oxonol | 3814 | RL1 | 5136.5 |
| 20 | Yes | A. baylyi | Nalidixic a | Intracellula | F11 | Allura Red | 3864 | BL3 | 4994.5 |
| 20 | Yes | A. baylyi | Nalidixic a | Intracellula | F54 | DiSC3(5) | 3964 | RL1 | 4784 |
| 20 | Yes | A. baylyi | Nalidixic a | Intracellula | F124 | Thioflavin T | 4162 | VL2 | 5390.5 |
| 20 | Yes | A. baylyi | Nalidixic a | Intracellula | F101 | Quinacrine | 4248 | VL2 | 5085.5 |
| 20 | Yes | A. baylyi | Nalidixic a | Intracellula | F138 | Apigenin | 3971 | VL2 | 5213 |
| 20 | Yes | A. baylyi | Nalidixic a | Intracellula | F65 | H2FDA | 4005 | BL1 | 6006 |
| 20 | Yes | A. baylyi | Nalidixic a | Intracellula | F142 | Chlorophe | 4276 | BL3 | 8143.5 |
| 20 | Yes | A. baylyi | Nalidixic a | Intracellula | F39 | Congo Red | 4312 | BL3 | 8716.5 |
| 20 | Yes | A. baylyi | Nalidixic a | Intracellula | F116 | Sunset Yel | 4102 | BL2 | 7205 |
| 20 | Yes | A. baylyi | Nalidixic a | Intracellula | F10 | Alizarin | 4379 | BL1 | 5707 |
| 20 | Yes | A. baylyi | Nalidixic a | Intracellula | F84 | Neutral Re | 4236 | BL1 | 4998 |
| 20 | Yes | A. baylyi | Nalidixic a | Intracellula | F99 | Pyronin Y | 4115 | BL1 | 4759 |
| 20 | Yes | A. baylyi | Nalidixic a | Intracellula | F108 | Rhodamine | 4364 | BL1 | 4579 |
| 20 | Yes | A. baylyi | Nalidixic a | Intracellula | F109 | Riboflavin | 3608 | BL1 | 4639 |
| 20 | Yes | A. baylyi | Nalidixic a | Intracellula | F117 | SYBR Gree | 4147 | BL1 | 283907 |
| 20 | Yes | A. baylyi | Nalidixic a | Intracellula | F106 | Rhodamine | 4145 | BL1 | 5309 |
| 20 | Yes | A. baylyi | Nalidixic a | Intracellula | F73 | Malachite ( | 3670 | BL1 | 5017 |
| 20 | Yes | A. baylyi | Nalidixic a | Intracellula | F7 | Acridine or | 3769 | BL1 | 5554 |
| 20 | Yes | A. baylyi | Nalidixic a | Intracellula | F192 | 5-Carboxyf | 4109 | BL1 | 16285 |
| 20 | Yes | A. baylyi | Nalidixic a | Intracellula | F57 | Eosin Y | 3869 | BL1 | 17531 |
| 20 | Yes | A. baylyi | Nalidixic a | Intracellula | F27 | Calcein | 3989 | BL1 | 16946 |

| | | | | | | | | | |
|---|---|---|---|---|---|---|---|---|---|
| 20 | Yes | A. baylyi | Nalidixic a | Intracellul | F12 | Amaranth | 3592 | BL1 | 17894 |
| 20 | Yes | A. baylyi | None | None | NA | Autofluore | 4524 | NA | 5512 |
| 20 | Yes | A. baylyi | Ofloxacin | Intracellul | F194 | Oxonol | 3267 | RL1 | 5666 |
| 20 | Yes | A. baylyi | Ofloxacin | Intracellul | F11 | Allura Red | 3488 | BL3 | 5653 |
| 20 | Yes | A. baylyi | Ofloxacin | Intracellul | F54 | DiSC3(5) | 3516 | RL1 | 5410 |
| 20 | Yes | A. baylyi | Ofloxacin | Intracellul | F124 | Thioflavin T | 3498 | VL2 | 5475.5 |
| 20 | Yes | A. baylyi | Ofloxacin | Intracellul | F101 | Quinacrine | 3708 | VL2 | 5485.5 |
| 20 | Yes | A. baylyi | Ofloxacin | Intracellul | F138 | Apigenin | 3507 | VL2 | 5494 |
| 20 | Yes | A. baylyi | Ofloxacin | Intracellul | F65 | H2FDA | 3841 | BL1 | 6528 |
| 20 | Yes | A. baylyi | Ofloxacin | Intracellul | F142 | Chlorophe | 3780 | BL3 | 8312 |
| 20 | Yes | A. baylyi | Ofloxacin | Intracellul | F39 | Congo Red | 3804 | BL3 | 9390 |
| 20 | Yes | A. baylyi | Ofloxacin | Intracellul | F116 | Sunset Yel | 3682 | BL2 | 7832.5 |
| 20 | Yes | A. baylyi | Ofloxacin | Intracellul | F10 | Alizarin | 3674 | BL1 | 6191.5 |
| 20 | Yes | A. baylyi | Ofloxacin | Intracellul | F84 | Neutral Re | 4056 | BL1 | 5887 |
| 20 | Yes | A. baylyi | Ofloxacin | Intracellul | F99 | Pyronin Y | 3739 | BL1 | 5172 |
| 20 | Yes | A. baylyi | Ofloxacin | Intracellul | F108 | Rhodamine | 3553 | BL1 | 4852 |
| 20 | Yes | A. baylyi | Ofloxacin | Intracellul | F109 | Riboflavin | 3472 | BL1 | 4982.5 |
| 20 | Yes | A. baylyi | Ofloxacin | Intracellul | F117 | SYBR Gree | 3485 | BL1 | 324962 |
| 20 | Yes | A. baylyi | Ofloxacin | Intracellul | F106 | Rhodamine | 3582 | BL1 | 5870 |
| 20 | Yes | A. baylyi | Ofloxacin | Intracellul | F73 | Malachite ( | 3191 | BL1 | 5349 |
| 20 | Yes | A. baylyi | Ofloxacin | Intracellul | F7 | Acridine or | 3615 | BL1 | 5643 |
| 20 | Yes | A. baylyi | Ofloxacin | Intracellul | F192 | 5-Carboxyf | 3348 | BL1 | 19119.5 |
| 20 | Yes | A. baylyi | Ofloxacin | Intracellul | F57 | Eosin Y | 2902 | BL1 | 18325 |
| 20 | Yes | A. baylyi | Ofloxacin | Intracellul | F27 | Calcein | 3273 | BL1 | 22400 |
| 20 | Yes | A. baylyi | Ofloxacin | Intracellul | F12 | Amaranth | 3355 | BL1 | 19814 |
| 20 | Yes | A. baylyi | None | None | NA | Autofluore | 4258 | NA | 6276.5 |
| 20 | Yes | A. baylyi | Polymyxin | Wall | F194 | Oxonol | 4045 | RL1 | 5891 |
| 20 | Yes | A. baylyi | Polymyxin | Wall | F11 | Allura Red | 4557 | BL3 | 5525 |
| 20 | Yes | A. baylyi | Polymyxin | Wall | F54 | DiSC3(5) | 4414 | RL1 | 6861.5 |
| 20 | Yes | A. baylyi | Polymyxin | Wall | F124 | Thioflavin T | 4691 | VL2 | 5836 |
| 20 | Yes | A. baylyi | Polymyxin | Wall | F101 | Quinacrine | 4890 | VL2 | 5486 |
| 20 | Yes | A. baylyi | Polymyxin | Wall | F138 | Apigenin | 4630 | VL2 | 5553 |
| 20 | Yes | A. baylyi | Polymyxin | Wall | F65 | H2FDA | 4487 | BL1 | 6641 |
| 20 | Yes | A. baylyi | Polymyxin | Wall | F142 | Chlorophe | 4620 | BL3 | 7499.5 |
| 20 | Yes | A. baylyi | Polymyxin | Wall | F39 | Congo Red | 4738 | BL3 | 9365 |
| 20 | Yes | A. baylyi | Polymyxin | Wall | F116 | Sunset Yel | 4605 | BL2 | 8502 |
| 20 | Yes | A. baylyi | Polymyxin | Wall | F10 | Alizarin | 4716 | BL1 | 5761.5 |
| 20 | Yes | A. baylyi | Polymyxin | Wall | F84 | Neutral Re | 4576 | BL1 | 5162.5 |
| 20 | Yes | A. baylyi | Polymyxin | Wall | F99 | Pyronin Y | 4563 | BL1 | 4672 |
| 20 | Yes | A. baylyi | Polymyxin | Wall | F108 | Rhodamine | 4792 | BL1 | 4657 |
| 20 | Yes | A. baylyi | Polymyxin | Wall | F109 | Riboflavin | 4821 | BL1 | 4838 |
| 20 | Yes | A. baylyi | Polymyxin | Wall | F117 | SYBR Gree | 4388 | BL1 | 662152.5 |
| 20 | Yes | A. baylyi | Polymyxin | Wall | F106 | Rhodamine | 4402 | BL1 | 6836 |
| 20 | Yes | A. baylyi | Polymyxin | Wall | F73 | Malachite ( | 3915 | BL1 | 5695 |
| 20 | Yes | A. baylyi | Polymyxin | Wall | F7 | Acridine or | 4132 | BL1 | 6601 |
| 20 | Yes | A. baylyi | Polymyxin | Wall | F192 | 5-Carboxyf | 4250 | BL1 | 16241.5 |
| 20 | Yes | A. baylyi | Polymyxin | Wall | F57 | Eosin Y | 4427 | BL1 | 15562 |
| 20 | Yes | A. baylyi | Polymyxin | Wall | F27 | Calcein | 4105 | BL1 | 15287 |

| | | | | | | | | | |
|---|---|---|---|---|---|---|---|---|---|
| 20 | Yes | A. baylyi | Polymyxin | Wall | F12 | Amaranth | 4172 | BL1 | 15265.5 |
| 20 | Yes | A. baylyi | None | None | NA | Autofluore: | 4386 | NA | 5768 |
| 20 | Yes | A. baylyi | Tetracyclin | Intracellul: | F194 | Oxonol | 3212 | RL1 | 5778 |
| 20 | Yes | A. baylyi | Tetracyclin | Intracellul: | F11 | Allura Red | 3100 | BL3 | 5700.5 |
| 20 | Yes | A. baylyi | Tetracyclin | Intracellul: | F54 | DiSC3(5) | 2870 | RL1 | 5773 |
| 20 | Yes | A. baylyi | Tetracyclin | Intracellul: | F124 | Thioflavin T | 3542 | VL2 | 5673 |
| 20 | Yes | A. baylyi | Tetracyclin | Intracellul: | F101 | Quinacrine | 3406 | VL2 | 5760 |
| 20 | Yes | A. baylyi | Tetracyclin | Intracellul: | F138 | Apigenin | 3419 | VL2 | 5803 |
| 20 | Yes | A. baylyi | Tetracyclin | Intracellul: | F65 | H2FDA | 3260 | BL1 | 6825.5 |
| 20 | Yes | A. baylyi | Tetracyclin | Intracellul: | F142 | Chlorophe | 3369 | BL3 | 8322 |
| 20 | Yes | A. baylyi | Tetracyclin | Intracellul: | F39 | Congo Red | 3529 | BL3 | 8874 |
| 20 | Yes | A. baylyi | Tetracyclin | Intracellul: | F116 | Sunset Yel | 3332 | BL2 | 7374.5 |
| 20 | Yes | A. baylyi | Tetracyclin | Intracellul: | F10 | Alizarin | 3300 | BL1 | 7042 |
| 20 | Yes | A. baylyi | Tetracyclin | Intracellul: | F84 | Neutral Re | 3126 | BL1 | 7365.5 |
| 20 | Yes | A. baylyi | Tetracyclin | Intracellul: | F99 | Pyronin Y | 3161 | BL1 | 5284 |
| 20 | Yes | A. baylyi | Tetracyclin | Intracellul: | F108 | Rhodamine | 3315 | BL1 | 5074 |
| 20 | Yes | A. baylyi | Tetracyclin | Intracellul: | F109 | Riboflavin | 3491 | BL1 | 5262 |
| 20 | Yes | A. baylyi | Tetracyclin | Intracellul: | F117 | SYBR Gree | 3199 | BL1 | 281027 |
| 20 | Yes | A. baylyi | Tetracyclin | Intracellul: | F106 | Rhodamine | 3333 | BL1 | 6395 |
| 20 | Yes | A. baylyi | Tetracyclin | Intracellul: | F73 | Malachite ( | 3215 | BL1 | 5481 |
| 20 | Yes | A. baylyi | Tetracyclin | Intracellul: | F7 | Acridine or | 3189 | BL1 | 6184 |
| 20 | Yes | A. baylyi | Tetracyclin | Intracellul: | F192 | 5-Carboxyf | 3312 | BL1 | 18039 |
| 20 | Yes | A. baylyi | Tetracyclin | Intracellul: | F57 | Eosin Y | 2928 | BL1 | 17019 |
| 20 | Yes | A. baylyi | Tetracyclin | Intracellul: | F27 | Calcein | 3369 | BL1 | 20633 |
| 20 | Yes | A. baylyi | Tetracyclin | Intracellul: | F12 | Amaranth | 2815 | BL1 | 21731 |
| 20 | Yes | A. baylyi | None | None | NA | Autofluore: | 4501 | NA | 6048 |
| 20 | Yes | A. baylyi | Vancomyc | Wall | F194 | Oxonol | 3606 | RL1 | 5742 |
| 20 | Yes | A. baylyi | Vancomyc | Wall | F11 | Allura Red | 3537 | BL3 | 5757 |
| 20 | Yes | A. baylyi | Vancomyc | Wall | F54 | DiSC3(5) | 4509 | RL1 | 5530 |
| 20 | Yes | A. baylyi | Vancomyc | Wall | F124 | Thioflavin T | 4511 | VL2 | 5649 |
| 20 | Yes | A. baylyi | Vancomyc | Wall | F101 | Quinacrine | 4133 | VL2 | 5732 |
| 20 | Yes | A. baylyi | Vancomyc | Wall | F138 | Apigenin | 4491 | VL2 | 5471 |
| 20 | Yes | A. baylyi | Vancomyc | Wall | F65 | H2FDA | 4095 | BL1 | 6461 |
| 20 | Yes | A. baylyi | Vancomyc | Wall | F142 | Chlorophe | 4327 | BL3 | 8405 |
| 20 | Yes | A. baylyi | Vancomyc | Wall | F39 | Congo Red | 3899 | BL3 | 8814 |
| 20 | Yes | A. baylyi | Vancomyc | Wall | F116 | Sunset Yel | 4696 | BL2 | 7291 |
| 20 | Yes | A. baylyi | Vancomyc | Wall | F10 | Alizarin | 4511 | BL1 | 6363 |
| 20 | Yes | A. baylyi | Vancomyc | Wall | F84 | Neutral Re | 4074 | BL1 | 5767 |
| 20 | Yes | A. baylyi | Vancomyc | Wall | F99 | Pyronin Y | 3731 | BL1 | 5270 |
| 20 | Yes | A. baylyi | Vancomyc | Wall | F108 | Rhodamine | 4991 | BL1 | 4538 |
| 20 | Yes | A. baylyi | Vancomyc | Wall | F109 | Riboflavin | 4299 | BL1 | 5082 |
| 20 | Yes | A. baylyi | Vancomyc | Wall | F117 | SYBR Gree | 4006 | BL1 | 352989.5 |
| 20 | Yes | A. baylyi | Vancomyc | Wall | F106 | Rhodamine | 4435 | BL1 | 6671 |
| 20 | Yes | A. baylyi | Vancomyc | Wall | F73 | Malachite ( | 4104 | BL1 | 5726.5 |
| 20 | Yes | A. baylyi | Vancomyc | Wall | F7 | Acridine or | 3619 | BL1 | 5772 |
| 20 | Yes | A. baylyi | Vancomyc | Wall | F192 | 5-Carboxyf | 4634 | BL1 | 16650 |
| 20 | Yes | A. baylyi | Vancomyc | Wall | F57 | Eosin Y | 3772 | BL1 | 19250 |
| 20 | Yes | A. baylyi | Vancomyc | Wall | F27 | Calcein | 3905 | BL1 | 17532 |

| | | | | | | | | | |
|---|---|---|---|---|---|---|---|---|---|
| 20 | Yes | A. baylyi | Vancomyc | Wall | F12 | Amaranth | 4012 | BL1 | 15026.5 |
| 20 | Yes | A. baylyi | None | None | NA | Autofluore: | 4966 | NA | 974 |
| 20 | Yes | A. baylyi | None | None | F194 | Oxonol | 5387 | RL1 | 1847 |
| 20 | Yes | A. baylyi | None | None | F11 | Allura Red | 4944 | BL3 | 2494.5 |
| 20 | Yes | A. baylyi | None | None | F54 | DiSC3(5) | 5014 | RL1 | 2638 |
| 20 | Yes | A. baylyi | None | None | F124 | Thioflavin T | 4755 | VL2 | 2755 |
| 20 | Yes | A. baylyi | None | None | F101 | Quinacrine | 4707 | VL2 | 2898 |
| 20 | Yes | A. baylyi | None | None | F138 | Apigenin | 5042 | VL2 | 2944 |
| 20 | Yes | A. baylyi | None | None | F65 | H2FDA | 4777 | BL1 | 4047 |
| 20 | Yes | A. baylyi | None | None | F142 | Chlorophe | 5039 | BL3 | 5074 |
| 20 | Yes | A. baylyi | None | None | F39 | Congo Red | 4844 | BL3 | 5192.5 |
| 20 | Yes | A. baylyi | None | None | F116 | Sunset Yel | 4900 | BL2 | 4553 |
| 20 | Yes | A. baylyi | None | None | F10 | Alizarin | 4382 | BL1 | 3826 |
| 20 | Yes | A. baylyi | None | None | F84 | Neutral Re | 4288 | BL1 | 3841 |
| 20 | Yes | A. baylyi | None | None | F99 | Pyronin Y | 4461 | BL1 | 5092 |
| 20 | Yes | A. baylyi | None | None | F108 | Rhodamine | 4344 | BL1 | 3297.5 |
| 20 | Yes | A. baylyi | None | None | F109 | Riboflavin | 4322 | BL1 | 3617.5 |
| 20 | Yes | A. baylyi | None | None | F117 | SYBR Gree | 3343 | BL1 | 264119 |
| 20 | Yes | A. baylyi | None | None | F106 | Rhodamine | 4692 | BL1 | 4070.5 |
| 20 | Yes | A. baylyi | None | None | F73 | Malachite ( | 4642 | BL1 | 4218.5 |
| 20 | Yes | A. baylyi | None | None | F7 | Acridine or | 4608 | BL1 | 4230.5 |
| 20 | Yes | A. baylyi | None | None | F192 | 5-Carboxyf | 4007 | BL1 | 12791 |
| 20 | Yes | A. baylyi | None | None | F57 | Eosin Y | 4288 | BL1 | 14178.5 |
| 20 | Yes | A. baylyi | None | None | F27 | Calcein | 4133 | BL1 | 15260 |
| 20 | Yes | A. baylyi | None | None | F12 | Amaranth | 3971 | BL1 | 14428 |
| 20 | Yes | A. baylyi | None | None | NA | Autofluore: | 4464 | NA | 5729 |
| 20 | Yes | A. baylyi | Ampicillin | Wall | F194 | Oxonol | 3288 | RL1 | 5320 |
| 20 | Yes | A. baylyi | Ampicillin | Wall | F11 | Allura Red | 3543 | BL3 | 4380 |
| 20 | Yes | A. baylyi | Ampicillin | Wall | F54 | DiSC3(5) | 3471 | RL1 | 4485 |
| 20 | Yes | A. baylyi | Ampicillin | Wall | F124 | Thioflavin T | 3495 | VL2 | 4568 |
| 20 | Yes | A. baylyi | Ampicillin | Wall | F101 | Quinacrine | 3449 | VL2 | 4278 |
| 20 | Yes | A. baylyi | Ampicillin | Wall | F138 | Apigenin | 3382 | VL2 | 4382.5 |
| 20 | Yes | A. baylyi | Ampicillin | Wall | F65 | H2FDA | 3392 | BL1 | 5279.5 |
| 20 | Yes | A. baylyi | Ampicillin | Wall | F142 | Chlorophe | 3546 | BL3 | 6564 |
| 20 | Yes | A. baylyi | Ampicillin | Wall | F39 | Congo Red | 3363 | BL3 | 6436 |
| 20 | Yes | A. baylyi | Ampicillin | Wall | F116 | Sunset Yel | 3432 | BL2 | 6111.5 |
| 20 | Yes | A. baylyi | Ampicillin | Wall | F10 | Alizarin | 3405 | BL1 | 4685 |
| 20 | Yes | A. baylyi | Ampicillin | Wall | F84 | Neutral Re | 3422 | BL1 | 4484 |
| 20 | Yes | A. baylyi | Ampicillin | Wall | F99 | Pyronin Y | 3379 | BL1 | 5233 |
| 20 | Yes | A. baylyi | Ampicillin | Wall | F108 | Rhodamine | 3168 | BL1 | 4131.5 |
| 20 | Yes | A. baylyi | Ampicillin | Wall | F109 | Riboflavin | 3231 | BL1 | 4635 |
| 20 | Yes | A. baylyi | Ampicillin | Wall | F117 | SYBR Gree | 3419 | BL1 | 342361 |
| 20 | Yes | A. baylyi | Ampicillin | Wall | F106 | Rhodamine | 3203 | BL1 | 4971 |
| 20 | Yes | A. baylyi | Ampicillin | Wall | F73 | Malachite ( | 3496 | BL1 | 4697.5 |
| 20 | Yes | A. baylyi | Ampicillin | Wall | F7 | Acridine or | 3257 | BL1 | 4884 |
| 20 | Yes | A. baylyi | Ampicillin | Wall | F192 | 5-Carboxyf | 3494 | BL1 | 16860 |
| 20 | Yes | A. baylyi | Ampicillin | Wall | F57 | Eosin Y | 2954 | BL1 | 13857.5 |
| 20 | Yes | A. baylyi | Ampicillin | Wall | F27 | Calcein | 3502 | BL1 | 18571.5 |

| | | | | | | | | | |
|---|---|---|---|---|---|---|---|---|---|
| 20 | Yes | A. baylyi | Ampicillin | Wall | F12 | Amaranth | 3301 | BL1 | 21117 |
| 20 | Yes | A. baylyi | None | None | NA | Autofluore: | 4398 | NA | 5730 |
| 20 | Yes | A. baylyi | Bacitracin | Wall | F194 | Oxonol | 3909 | RL1 | 5735 |
| 20 | Yes | A. baylyi | Bacitracin | Wall | F11 | Allura Red | 4167 | BL3 | 5182 |
| 20 | Yes | A. baylyi | Bacitracin | Wall | F54 | DiSC3(5) | 4437 | RL1 | 5313 |
| 20 | Yes | A. baylyi | Bacitracin | Wall | F124 | Thioflavin T | 4239 | VL2 | 6082 |
| 20 | Yes | A. baylyi | Bacitracin | Wall | F101 | Quinacrine | 3917 | VL2 | 6129 |
| 20 | Yes | A. baylyi | Bacitracin | Wall | F138 | Apigenin | 4079 | VL2 | 4874 |
| 20 | Yes | A. baylyi | Bacitracin | Wall | F65 | H2FDA | 3973 | BL1 | 7994 |
| 20 | Yes | A. baylyi | Bacitracin | Wall | F142 | Chlorophe | 3942 | BL3 | 7092 |
| 20 | Yes | A. baylyi | Bacitracin | Wall | F39 | Congo Red | 4465 | BL3 | 6535 |
| 20 | Yes | A. baylyi | Bacitracin | Wall | F116 | Sunset Yel | 3967 | BL2 | 5798 |
| 20 | Yes | A. baylyi | Bacitracin | Wall | F10 | Alizarin | 4333 | BL1 | 5639 |
| 20 | Yes | A. baylyi | Bacitracin | Wall | F84 | Neutral Re | 4032 | BL1 | 6239.5 |
| 20 | Yes | A. baylyi | Bacitracin | Wall | F99 | Pyronin Y | 3870 | BL1 | 5426.5 |
| 20 | Yes | A. baylyi | Bacitracin | Wall | F108 | Rhodamine | 4287 | BL1 | 4869 |
| 20 | Yes | A. baylyi | Bacitracin | Wall | F109 | Riboflavin | 4209 | BL1 | 4639 |
| 20 | Yes | A. baylyi | Bacitracin | Wall | F117 | SYBR Gree | 4063 | BL1 | 391810 |
| 20 | Yes | A. baylyi | Bacitracin | Wall | F106 | Rhodamine | 4096 | BL1 | 5229 |
| 20 | Yes | A. baylyi | Bacitracin | Wall | F73 | Malachite ( | 3889 | BL1 | 5472 |
| 20 | Yes | A. baylyi | Bacitracin | Wall | F7 | Acridine or | 4134 | BL1 | 4940.5 |
| 20 | Yes | A. baylyi | Bacitracin | Wall | F192 | 5-Carboxyf | 4317 | BL1 | 15288 |
| 20 | Yes | A. baylyi | Bacitracin | Wall | F57 | Eosin Y | 4202 | BL1 | 16848 |
| 20 | Yes | A. baylyi | Bacitracin | Wall | F27 | Calcein | 3998 | BL1 | 17007 |
| 20 | Yes | A. baylyi | Bacitracin | Wall | F12 | Amaranth | 3786 | BL1 | 16510 |
| 20 | Yes | A. baylyi | None | None | NA | Autofluore: | 4409 | NA | 5358 |
| 20 | Yes | A. baylyi | Cefotaxim | Wall | F194 | Oxonol | 3345 | RL1 | 5495 |
| 20 | Yes | A. baylyi | Cefotaxim | Wall | F11 | Allura Red | 3476 | BL3 | 5518 |
| 20 | Yes | A. baylyi | Cefotaxim | Wall | F54 | DiSC3(5) | 3622 | RL1 | 5436 |
| 20 | Yes | A. baylyi | Cefotaxim | Wall | F124 | Thioflavin T | 3740 | VL2 | 5308.5 |
| 20 | Yes | A. baylyi | Cefotaxim | Wall | F101 | Quinacrine | 3802 | VL2 | 5185 |
| 20 | Yes | A. baylyi | Cefotaxim | Wall | F138 | Apigenin | 3374 | VL2 | 5377.5 |
| 20 | Yes | A. baylyi | Cefotaxim | Wall | F65 | H2FDA | 3556 | BL1 | 6224.5 |
| 20 | Yes | A. baylyi | Cefotaxim | Wall | F142 | Chlorophe | 3649 | BL3 | 7092 |
| 20 | Yes | A. baylyi | Cefotaxim | Wall | F39 | Congo Red | 3552 | BL3 | 6687.5 |
| 20 | Yes | A. baylyi | Cefotaxim | Wall | F116 | Sunset Yel | 3158 | BL2 | 5964 |
| 20 | Yes | A. baylyi | Cefotaxim | Wall | F10 | Alizarin | 3741 | BL1 | 4994 |
| 20 | Yes | A. baylyi | Cefotaxim | Wall | F84 | Neutral Re | 3389 | BL1 | 5210 |
| 20 | Yes | A. baylyi | Cefotaxim | Wall | F99 | Pyronin Y | 3181 | BL1 | 5736 |
| 20 | Yes | A. baylyi | Cefotaxim | Wall | F108 | Rhodamine | 3420 | BL1 | 5709.5 |
| 20 | Yes | A. baylyi | Cefotaxim | Wall | F109 | Riboflavin | 3303 | BL1 | 5808 |
| 20 | Yes | A. baylyi | Cefotaxim | Wall | F117 | SYBR Gree | 3320 | BL1 | 383786 |
| 20 | Yes | A. baylyi | Cefotaxim | Wall | F106 | Rhodamine | 3888 | BL1 | 5328 |
| 20 | Yes | A. baylyi | Cefotaxim | Wall | F73 | Malachite ( | 3380 | BL1 | 5226 |
| 20 | Yes | A. baylyi | Cefotaxim | Wall | F7 | Acridine or | 3372 | BL1 | 5232 |
| 20 | Yes | A. baylyi | Cefotaxim | Wall | F192 | 5-Carboxyf | 2925 | BL1 | 12711 |
| 20 | Yes | A. baylyi | Cefotaxim | Wall | F57 | Eosin Y | 3122 | BL1 | 14398 |
| 20 | Yes | A. baylyi | Cefotaxim | Wall | F27 | Calcein | 3653 | BL1 | 19082 |

| | | | | | | | | | |
|---|---|---|---|---|---|---|---|---|---|
| 20 | Yes | A. baylyi | Cefotaxim | Wall | F12 | Amaranth | 3180 | BL1 | 18700 |
| 20 | Yes | A. baylyi | None | None | NA | Autofluore | 4627 | NA | 6033 |
| 20 | Yes | A. baylyi | Ciprofloxa | Intracellul | F194 | Oxonol | 3064 | RL1 | 5521.5 |
| 20 | Yes | A. baylyi | Ciprofloxa | Intracellul | F11 | Allura Red | 3108 | BL3 | 5616 |
| 20 | Yes | A. baylyi | Ciprofloxa | Intracellul | F54 | DiSC3(5) | 3161 | RL1 | 5144 |
| 20 | Yes | A. baylyi | Ciprofloxa | Intracellul | F124 | Thioflavin T | 3256 | VL2 | 5641.5 |
| 20 | Yes | A. baylyi | Ciprofloxa | Intracellul | F101 | Quinacrine | 3153 | VL2 | 5329 |
| 20 | Yes | A. baylyi | Ciprofloxa | Intracellul | F138 | Apigenin | 3051 | VL2 | 5578 |
| 20 | Yes | A. baylyi | Ciprofloxa | Intracellul | F65 | H2FDA | 3191 | BL1 | 6314 |
| 20 | Yes | A. baylyi | Ciprofloxa | Intracellul | F142 | Chlorophe | 3359 | BL3 | 7388 |
| 20 | Yes | A. baylyi | Ciprofloxa | Intracellul | F39 | Congo Red | 3220 | BL3 | 7510 |
| 20 | Yes | A. baylyi | Ciprofloxa | Intracellul | F116 | Sunset Yel | 3267 | BL2 | 7681 |
| 20 | Yes | A. baylyi | Ciprofloxa | Intracellul | F10 | Alizarin | 3256 | BL1 | 7022 |
| 20 | Yes | A. baylyi | Ciprofloxa | Intracellul | F84 | Neutral Re | 3546 | BL1 | 5600 |
| 20 | Yes | A. baylyi | Ciprofloxa | Intracellul | F99 | Pyronin Y | 3198 | BL1 | 5110 |
| 20 | Yes | A. baylyi | Ciprofloxa | Intracellul | F108 | Rhodamine | 3000 | BL1 | 4892.5 |
| 20 | Yes | A. baylyi | Ciprofloxa | Intracellul | F109 | Riboflavin | 2939 | BL1 | 5172 |
| 20 | Yes | A. baylyi | Ciprofloxa | Intracellul | F117 | SYBR Gree | 3008 | BL1 | 279714.5 |
| 20 | Yes | A. baylyi | Ciprofloxa | Intracellul | F106 | Rhodamine | 3253 | BL1 | 5789 |
| 20 | Yes | A. baylyi | Ciprofloxa | Intracellul | F73 | Malachite | 3498 | BL1 | 4866 |
| 20 | Yes | A. baylyi | Ciprofloxa | Intracellul | F7 | Acridine or | 2985 | BL1 | 5582 |
| 20 | Yes | A. baylyi | Ciprofloxa | Intracellul | F192 | 5-Carboxyf | 2955 | BL1 | 17376 |
| 20 | Yes | A. baylyi | Ciprofloxa | Intracellul | F57 | Eosin Y | 2797 | BL1 | 11047 |
| 20 | Yes | A. baylyi | Ciprofloxa | Intracellul | F27 | Calcein | 2889 | BL1 | 24317 |
| 20 | Yes | A. baylyi | Ciprofloxa | Intracellul | F12 | Amaranth | 2994 | BL1 | 23002 |
| 20 | Yes | A. baylyi | None | None | NA | Autofluore | 4260 | NA | 5938.5 |
| 20 | Yes | A. baylyi | Fosfomyci | Wall | F194 | Oxonol | 4384 | RL1 | 5271 |
| 20 | Yes | A. baylyi | Fosfomyci | Wall | F11 | Allura Red | 4282 | BL3 | 5063 |
| 20 | Yes | A. baylyi | Fosfomyci | Wall | F54 | DiSC3(5) | 4460 | RL1 | 4908 |
| 20 | Yes | A. baylyi | Fosfomyci | Wall | F124 | Thioflavin T | 4232 | VL2 | 5111.5 |
| 20 | Yes | A. baylyi | Fosfomyci | Wall | F101 | Quinacrine | 4644 | VL2 | 5032 |
| 20 | Yes | A. baylyi | Fosfomyci | Wall | F138 | Apigenin | 4329 | VL2 | 5101 |
| 20 | Yes | A. baylyi | Fosfomyci | Wall | F65 | H2FDA | 4378 | BL1 | 5654 |
| 20 | Yes | A. baylyi | Fosfomyci | Wall | F142 | Chlorophe | 4460 | BL3 | 7019 |
| 20 | Yes | A. baylyi | Fosfomyci | Wall | F39 | Congo Red | 4566 | BL3 | 7515.5 |
| 20 | Yes | A. baylyi | Fosfomyci | Wall | F116 | Sunset Yel | 4430 | BL2 | 6568 |
| 20 | Yes | A. baylyi | Fosfomyci | Wall | F10 | Alizarin | 4580 | BL1 | 5485.5 |
| 20 | Yes | A. baylyi | Fosfomyci | Wall | F84 | Neutral Re | 4592 | BL1 | 4917 |
| 20 | Yes | A. baylyi | Fosfomyci | Wall | F99 | Pyronin Y | 4420 | BL1 | 4525 |
| 20 | Yes | A. baylyi | Fosfomyci | Wall | F108 | Rhodamine | 4204 | BL1 | 4692 |
| 20 | Yes | A. baylyi | Fosfomyci | Wall | F109 | Riboflavin | 3848 | BL1 | 4452 |
| 20 | Yes | A. baylyi | Fosfomyci | Wall | F117 | SYBR Gree | 4434 | BL1 | 299747.5 |
| 20 | Yes | A. baylyi | Fosfomyci | Wall | F106 | Rhodamine | 4648 | BL1 | 5511.5 |
| 20 | Yes | A. baylyi | Fosfomyci | Wall | F73 | Malachite | 3911 | BL1 | 4846 |
| 20 | Yes | A. baylyi | Fosfomyci | Wall | F7 | Acridine or | 4379 | BL1 | 5283 |
| 20 | Yes | A. baylyi | Fosfomyci | Wall | F192 | 5-Carboxyf | 4082 | BL1 | 15366.5 |
| 20 | Yes | A. baylyi | Fosfomyci | Wall | F57 | Eosin Y | 4181 | BL1 | 13223 |
| 20 | Yes | A. baylyi | Fosfomyci | Wall | F27 | Calcein | 4113 | BL1 | 16030 |

| | | | | | | | | | |
|---|---|---|---|---|---|---|---|---|---|
| 20 | Yes | A. baylyi | Fosfomyci | Wall | F12 | Amaranth | 4336 | BL1 | 16745 |
| 20 | Yes | A. baylyi | None | None | NA | Autofluore: | 4428 | NA | 5941 |
| 20 | Yes | A. baylyi | Nalidixic a | Intracellul | F194 | Oxonol | 3521 | RL1 | 5207 |
| 20 | Yes | A. baylyi | Nalidixic a | Intracellul | F11 | Allura Red | 3746 | BL3 | 5482.5 |
| 20 | Yes | A. baylyi | Nalidixic a | Intracellul | F54 | DiSC3(5) | 3945 | RL1 | 5185 |
| 20 | Yes | A. baylyi | Nalidixic a | Intracellul | F124 | Thioflavin T | 3715 | VL2 | 5358 |
| 20 | Yes | A. baylyi | Nalidixic a | Intracellul | F101 | Quinacrine | 3908 | VL2 | 5396 |
| 20 | Yes | A. baylyi | Nalidixic a | Intracellul | F138 | Apigenin | 3896 | VL2 | 5396.5 |
| 20 | Yes | A. baylyi | Nalidixic a | Intracellul | F65 | H2FDA | 3499 | BL1 | 5896 |
| 20 | Yes | A. baylyi | Nalidixic a | Intracellul | F142 | Chlorophe | 3723 | BL3 | 8385 |
| 20 | Yes | A. baylyi | Nalidixic a | Intracellul | F39 | Congo Red | 3892 | BL3 | 7504.5 |
| 20 | Yes | A. baylyi | Nalidixic a | Intracellul | F116 | Sunset Yel | 4084 | BL2 | 6891.5 |
| 20 | Yes | A. baylyi | Nalidixic a | Intracellul | F10 | Alizarin | 3892 | BL1 | 6479 |
| 20 | Yes | A. baylyi | Nalidixic a | Intracellul | F84 | Neutral Re | 3832 | BL1 | 5698.5 |
| 20 | Yes | A. baylyi | Nalidixic a | Intracellul | F99 | Pyronin Y | 3732 | BL1 | 5065.5 |
| 20 | Yes | A. baylyi | Nalidixic a | Intracellul | F108 | Rhodamine | 3742 | BL1 | 4588.5 |
| 20 | Yes | A. baylyi | Nalidixic a | Intracellul | F109 | Riboflavin | 3893 | BL1 | 5231 |
| 20 | Yes | A. baylyi | Nalidixic a | Intracellul | F117 | SYBR Gree | 3683 | BL1 | 294161 |
| 20 | Yes | A. baylyi | Nalidixic a | Intracellul | F106 | Rhodamine | 3658 | BL1 | 5485 |
| 20 | Yes | A. baylyi | Nalidixic a | Intracellul | F73 | Malachite ( | 3499 | BL1 | 5047 |
| 20 | Yes | A. baylyi | Nalidixic a | Intracellul | F7 | Acridine or | 3575 | BL1 | 5545 |
| 20 | Yes | A. baylyi | Nalidixic a | Intracellul | F192 | 5-Carboxyf | 3357 | BL1 | 16373 |
| 20 | Yes | A. baylyi | Nalidixic a | Intracellul | F57 | Eosin Y | 3340 | BL1 | 17859 |
| 20 | Yes | A. baylyi | Nalidixic a | Intracellul | F27 | Calcein | 3612 | BL1 | 17757.5 |
| 20 | Yes | A. baylyi | Nalidixic a | Intracellul | F12 | Amaranth | 3414 | BL1 | 18457 |
| 20 | Yes | A. baylyi | None | None | NA | Autofluore: | 4276 | NA | 5967 |
| 20 | Yes | A. baylyi | Ofloxacin | Intracellul | F194 | Oxonol | 3104 | RL1 | 5718.5 |
| 20 | Yes | A. baylyi | Ofloxacin | Intracellul | F11 | Allura Red | 3630 | BL3 | 5691 |
| 20 | Yes | A. baylyi | Ofloxacin | Intracellul | F54 | DiSC3(5) | 3664 | RL1 | 5391 |
| 20 | Yes | A. baylyi | Ofloxacin | Intracellul | F124 | Thioflavin T | 3529 | VL2 | 5735 |
| 20 | Yes | A. baylyi | Ofloxacin | Intracellul | F101 | Quinacrine | 3617 | VL2 | 5709 |
| 20 | Yes | A. baylyi | Ofloxacin | Intracellul | F138 | Apigenin | 3578 | VL2 | 5740.5 |
| 20 | Yes | A. baylyi | Ofloxacin | Intracellul | F65 | H2FDA | 3769 | BL1 | 6367 |
| 20 | Yes | A. baylyi | Ofloxacin | Intracellul | F142 | Chlorophe | 3634 | BL3 | 7982 |
| 20 | Yes | A. baylyi | Ofloxacin | Intracellul | F39 | Congo Red | 3692 | BL3 | 8434.5 |
| 20 | Yes | A. baylyi | Ofloxacin | Intracellul | F116 | Sunset Yel | 3578 | BL2 | 8039.5 |
| 20 | Yes | A. baylyi | Ofloxacin | Intracellul | F10 | Alizarin | 3582 | BL1 | 6672.5 |
| 20 | Yes | A. baylyi | Ofloxacin | Intracellul | F84 | Neutral Re | 3746 | BL1 | 5526.5 |
| 20 | Yes | A. baylyi | Ofloxacin | Intracellul | F99 | Pyronin Y | 3664 | BL1 | 4972 |
| 20 | Yes | A. baylyi | Ofloxacin | Intracellul | F108 | Rhodamine | 3583 | BL1 | 5006 |
| 20 | Yes | A. baylyi | Ofloxacin | Intracellul | F109 | Riboflavin | 3627 | BL1 | 5276 |
| 20 | Yes | A. baylyi | Ofloxacin | Intracellul | F117 | SYBR Gree | 3686 | BL1 | 342622.5 |
| 20 | Yes | A. baylyi | Ofloxacin | Intracellul | F106 | Rhodamine | 3612 | BL1 | 5880 |
| 20 | Yes | A. baylyi | Ofloxacin | Intracellul | F73 | Malachite ( | 3754 | BL1 | 5598.5 |
| 20 | Yes | A. baylyi | Ofloxacin | Intracellul | F7 | Acridine or | 3456 | BL1 | 5663 |
| 20 | Yes | A. baylyi | Ofloxacin | Intracellul | F192 | 5-Carboxyf | 3312 | BL1 | 15876 |
| 20 | Yes | A. baylyi | Ofloxacin | Intracellul | F57 | Eosin Y | 3632 | BL1 | 16346.5 |
| 20 | Yes | A. baylyi | Ofloxacin | Intracellul | F27 | Calcein | 3880 | BL1 | 17200 |

| | | | | | | | | | |
|---|---|---|---|---|---|---|---|---|---|
| 20 | Yes | A. baylyi | Ofloxacin | Intracellula | F12 | Amaranth | 4295 | BL1 | 18042 |
| 20 | Yes | A. baylyi | None | None | NA | Autofluore: | 4435 | NA | 6509 |
| 20 | Yes | A. baylyi | Polymyxin | Wall | F194 | Oxonol | 4484 | RL1 | 6299.5 |
| 20 | Yes | A. baylyi | Polymyxin | Wall | F11 | Allura Red | 4691 | BL3 | 5847 |
| 20 | Yes | A. baylyi | Polymyxin | Wall | F54 | DiSC3(5) | 4337 | RL1 | 6869 |
| 20 | Yes | A. baylyi | Polymyxin | Wall | F124 | Thioflavin T | 4916 | VL2 | 5833.5 |
| 20 | Yes | A. baylyi | Polymyxin | Wall | F101 | Quinacrine | 4566 | VL2 | 5798.5 |
| 20 | Yes | A. baylyi | Polymyxin | Wall | F138 | Apigenin | 4367 | VL2 | 5767 |
| 20 | Yes | A. baylyi | Polymyxin | Wall | F65 | H2FDA | 4911 | BL1 | 6238 |
| 20 | Yes | A. baylyi | Polymyxin | Wall | F142 | Chlorophe | 4997 | BL3 | 7591 |
| 20 | Yes | A. baylyi | Polymyxin | Wall | F39 | Congo Red | 5023 | BL3 | 8375 |
| 20 | Yes | A. baylyi | Polymyxin | Wall | F116 | Sunset Yel | 4744 | BL2 | 8448.5 |
| 20 | Yes | A. baylyi | Polymyxin | Wall | F10 | Alizarin | 4389 | BL1 | 6172 |
| 20 | Yes | A. baylyi | Polymyxin | Wall | F84 | Neutral Re | 4787 | BL1 | 5109 |
| 20 | Yes | A. baylyi | Polymyxin | Wall | F99 | Pyronin Y | 4516 | BL1 | 4926 |
| 20 | Yes | A. baylyi | Polymyxin | Wall | F108 | Rhodamine | 4945 | BL1 | 4708 |
| 20 | Yes | A. baylyi | Polymyxin | Wall | F109 | Riboflavin | 4672 | BL1 | 4711.5 |
| 20 | Yes | A. baylyi | Polymyxin | Wall | F117 | SYBR Gree | 4351 | BL1 | 694653 |
| 20 | Yes | A. baylyi | Polymyxin | Wall | F106 | Rhodamine | 4593 | BL1 | 6419 |
| 20 | Yes | A. baylyi | Polymyxin | Wall | F73 | Malachite ( | 4255 | BL1 | 5748 |
| 20 | Yes | A. baylyi | Polymyxin | Wall | F7 | Acridine or | 4415 | BL1 | 6480 |
| 20 | Yes | A. baylyi | Polymyxin | Wall | F192 | 5-Carboxyf | 4512 | BL1 | 15121 |
| 20 | Yes | A. baylyi | Polymyxin | Wall | F57 | Eosin Y | 4666 | BL1 | 13198.5 |
| 20 | Yes | A. baylyi | Polymyxin | Wall | F27 | Calcein | 4207 | BL1 | 15981 |
| 20 | Yes | A. baylyi | Polymyxin | Wall | F12 | Amaranth | 4670 | BL1 | 16818.5 |
| 20 | Yes | A. baylyi | None | None | NA | Autofluore: | 4505 | NA | 6154 |
| 20 | Yes | A. baylyi | Tetracyclin | Intracellula | F194 | Oxonol | 3000 | RL1 | 6099.5 |
| 20 | Yes | A. baylyi | Tetracyclin | Intracellula | F11 | Allura Red | 2956 | BL3 | 6103 |
| 20 | Yes | A. baylyi | Tetracyclin | Intracellula | F54 | DiSC3(5) | 2913 | RL1 | 5963 |
| 20 | Yes | A. baylyi | Tetracyclin | Intracellula | F124 | Thioflavin T | 3121 | VL2 | 6178 |
| 20 | Yes | A. baylyi | Tetracyclin | Intracellula | F101 | Quinacrine | 3078 | VL2 | 6141 |
| 20 | Yes | A. baylyi | Tetracyclin | Intracellula | F138 | Apigenin | 2846 | VL2 | 5907.5 |
| 20 | Yes | A. baylyi | Tetracyclin | Intracellula | F65 | H2FDA | 3037 | BL1 | 6931 |
| 20 | Yes | A. baylyi | Tetracyclin | Intracellula | F142 | Chlorophe | 3043 | BL3 | 8515 |
| 20 | Yes | A. baylyi | Tetracyclin | Intracellula | F39 | Congo Red | 3018 | BL3 | 8700.5 |
| 20 | Yes | A. baylyi | Tetracyclin | Intracellula | F116 | Sunset Yel | 2929 | BL2 | 8314 |
| 20 | Yes | A. baylyi | Tetracyclin | Intracellula | F10 | Alizarin | 3032 | BL1 | 7330 |
| 20 | Yes | A. baylyi | Tetracyclin | Intracellula | F84 | Neutral Re | 3071 | BL1 | 5633 |
| 20 | Yes | A. baylyi | Tetracyclin | Intracellula | F99 | Pyronin Y | 3057 | BL1 | 5257 |
| 20 | Yes | A. baylyi | Tetracyclin | Intracellula | F108 | Rhodamine | 2905 | BL1 | 4702 |
| 20 | Yes | A. baylyi | Tetracyclin | Intracellula | F109 | Riboflavin | 3075 | BL1 | 5287 |
| 20 | Yes | A. baylyi | Tetracyclin | Intracellula | F117 | SYBR Gree | 3048 | BL1 | 282491.5 |
| 20 | Yes | A. baylyi | Tetracyclin | Intracellula | F106 | Rhodamine | 3016 | BL1 | 6417 |
| 20 | Yes | A. baylyi | Tetracyclin | Intracellula | F73 | Malachite ( | 2891 | BL1 | 6168 |
| 20 | Yes | A. baylyi | Tetracyclin | Intracellula | F7 | Acridine or | 2984 | BL1 | 6471.5 |
| 20 | Yes | A. baylyi | Tetracyclin | Intracellula | F192 | 5-Carboxyf | 3048 | BL1 | 15994.5 |
| 20 | Yes | A. baylyi | Tetracyclin | Intracellula | F57 | Eosin Y | 2931 | BL1 | 17611 |
| 20 | Yes | A. baylyi | Tetracyclin | Intracellula | F27 | Calcein | 2958 | BL1 | 21436.5 |

| | | | | | | | | | |
|---|---|---|---|---|---|---|---|---|---|
| 20 | Yes | A. baylyi | Tetracyclin | Intracellula | F12 | Amaranth | 3038 | BL1 | 21032.5 |
| 20 | Yes | A. baylyi | None | None | NA | Autofluores | 4499 | NA | 6229 |
| 20 | Yes | A. baylyi | Vancomyc | Wall | F194 | Oxonol | 4120 | RL1 | 5640.5 |
| 20 | Yes | A. baylyi | Vancomyc | Wall | F11 | Allura Red | 4778 | BL3 | 5630.5 |
| 20 | Yes | A. baylyi | Vancomyc | Wall | F54 | DiSC3(5) | 4692 | RL1 | 5455.5 |
| 20 | Yes | A. baylyi | Vancomyc | Wall | F124 | Thioflavin T | 4359 | VL2 | 5835 |
| 20 | Yes | A. baylyi | Vancomyc | Wall | F101 | Quinacrine | 4181 | VL2 | 5709 |
| 20 | Yes | A. baylyi | Vancomyc | Wall | F138 | Apigenin | 4166 | VL2 | 5763 |
| 20 | Yes | A. baylyi | Vancomyc | Wall | F65 | H2FDA | 3748 | BL1 | 6599.5 |
| 20 | Yes | A. baylyi | Vancomyc | Wall | F142 | Chlorophe | 3681 | BL3 | 8352 |
| 20 | Yes | A. baylyi | Vancomyc | Wall | F39 | Congo Red | 4156 | BL3 | 8395 |
| 20 | Yes | A. baylyi | Vancomyc | Wall | F116 | Sunset Yel | 4457 | BL2 | 7413 |
| 20 | Yes | A. baylyi | Vancomyc | Wall | F10 | Alizarin | 4173 | BL1 | 6603 |
| 20 | Yes | A. baylyi | Vancomyc | Wall | F84 | Neutral Re | 3962 | BL1 | 5710.5 |
| 20 | Yes | A. baylyi | Vancomyc | Wall | F99 | Pyronin Y | 4036 | BL1 | 5725 |
| 20 | Yes | A. baylyi | Vancomyc | Wall | F108 | Rhodamine | 3863 | BL1 | 5112 |
| 20 | Yes | A. baylyi | Vancomyc | Wall | F109 | Riboflavin | 3673 | BL1 | 5427 |
| 20 | Yes | A. baylyi | Vancomyc | Wall | F117 | SYBR Gree | 3689 | BL1 | 340699 |
| 20 | Yes | A. baylyi | Vancomyc | Wall | F106 | Rhodamine | 4013 | BL1 | 7148 |
| 20 | Yes | A. baylyi | Vancomyc | Wall | F73 | Malachite ( | 4119 | BL1 | 5814 |
| 20 | Yes | A. baylyi | Vancomyc | Wall | F7 | Acridine or | 3726 | BL1 | 6523 |
| 20 | Yes | A. baylyi | Vancomyc | Wall | F192 | 5-Carboxyf | 3869 | BL1 | 17322 |
| 20 | Yes | A. baylyi | Vancomyc | Wall | F57 | Eosin Y | 3826 | BL1 | 19059 |
| 20 | Yes | A. baylyi | Vancomyc | Wall | F27 | Calcein | 3712 | BL1 | 18999.5 |
| 20 | Yes | A. baylyi | Vancomyc | Wall | F12 | Amaranth | 3897 | BL1 | 15620 |
| 20 | Yes | A. baylyi | None | None | NA | Autofluores | 4891 | NA | 1092 |
| 20 | Yes | A. baylyi | None | None | F194 | Oxonol | 5049 | RL1 | 1865 |
| 20 | Yes | A. baylyi | None | None | F11 | Allura Red | 4403 | BL3 | 2407 |
| 20 | Yes | A. baylyi | None | None | F54 | DiSC3(5) | 4920 | RL1 | 2785 |
| 20 | Yes | A. baylyi | None | None | F124 | Thioflavin T | 4594 | VL2 | 2790.5 |
| 20 | Yes | A. baylyi | None | None | F101 | Quinacrine | 4661 | VL2 | 2899 |
| 20 | Yes | A. baylyi | None | None | F138 | Apigenin | 4517 | VL2 | 3024 |
| 20 | Yes | A. baylyi | None | None | F65 | H2FDA | 4279 | BL1 | 4245 |
| 20 | Yes | A. baylyi | None | None | F142 | Chlorophe | 4395 | BL3 | 5300 |
| 20 | Yes | A. baylyi | None | None | F39 | Congo Red | 4495 | BL3 | 5764 |
| 20 | Yes | A. baylyi | None | None | F116 | Sunset Yel | 4349 | BL2 | 5142 |
| 20 | Yes | A. baylyi | None | None | F10 | Alizarin | 4401 | BL1 | 4832 |
| 20 | Yes | A. baylyi | None | None | F84 | Neutral Re | 3952 | BL1 | 3655 |
| 20 | Yes | A. baylyi | None | None | F99 | Pyronin Y | 4156 | BL1 | 3550 |
| 20 | Yes | A. baylyi | None | None | F108 | Rhodamine | 4178 | BL1 | 2945.5 |
| 20 | Yes | A. baylyi | None | None | F109 | Riboflavin | 4122 | BL1 | 3273 |
| 20 | Yes | A. baylyi | None | None | F117 | SYBR Gree | 4213 | BL1 | 269566 |
| 20 | Yes | A. baylyi | None | None | F106 | Rhodamine | 4317 | BL1 | 4272 |
| 20 | Yes | A. baylyi | None | None | F73 | Malachite ( | 4201 | BL1 | 3865 |
| 20 | Yes | A. baylyi | None | None | F7 | Acridine or | 4016 | BL1 | 4006 |
| 20 | Yes | A. baylyi | None | None | F192 | 5-Carboxyf | 4257 | BL1 | 14203 |
| 20 | Yes | A. baylyi | None | None | F57 | Eosin Y | 3929 | BL1 | 16035 |
| 20 | Yes | A. baylyi | None | None | F27 | Calcein | 3945 | BL1 | 16432 |

| | | | | | | | | | |
|---|---|---|---|---|---|---|---|---|---|
| 20 | Yes | A. baylyi | None | None | F12 | Amaranth | 3927 | BL1 | 16126 |
| 20 | Yes | A. baylyi | None | None | NA | Autofluores | 4095 | NA | 5150 |
| 20 | Yes | A. baylyi | Ampicillin | Wall | F194 | Oxonol | 3585 | RL1 | 4720 |
| 20 | Yes | A. baylyi | Ampicillin | Wall | F11 | Allura Red | 3326 | BL3 | 4557.5 |
| 20 | Yes | A. baylyi | Ampicillin | Wall | F54 | DiSC3(5) | 3601 | RL1 | 4159 |
| 20 | Yes | A. baylyi | Ampicillin | Wall | F124 | Thioflavin T | 3581 | VL2 | 4291 |
| 20 | Yes | A. baylyi | Ampicillin | Wall | F101 | Quinacrine | 3489 | VL2 | 4826 |
| 20 | Yes | A. baylyi | Ampicillin | Wall | F138 | Apigenin | 3509 | VL2 | 4558 |
| 20 | Yes | A. baylyi | Ampicillin | Wall | F65 | H2FDA | 3678 | BL1 | 5316.5 |
| 20 | Yes | A. baylyi | Ampicillin | Wall | F142 | Chlorophe | 3466 | BL3 | 6246 |
| 20 | Yes | A. baylyi | Ampicillin | Wall | F39 | Congo Red | 3451 | BL3 | 6216 |
| 20 | Yes | A. baylyi | Ampicillin | Wall | F116 | Sunset Yel | 3489 | BL2 | 5374 |
| 20 | Yes | A. baylyi | Ampicillin | Wall | F10 | Alizarin | 3399 | BL1 | 4840 |
| 20 | Yes | A. baylyi | Ampicillin | Wall | F84 | Neutral Re | 3486 | BL1 | 4778.5 |
| 20 | Yes | A. baylyi | Ampicillin | Wall | F99 | Pyronin Y | 3277 | BL1 | 4811 |
| 20 | Yes | A. baylyi | Ampicillin | Wall | F108 | Rhodamine | 3318 | BL1 | 4667.5 |
| 20 | Yes | A. baylyi | Ampicillin | Wall | F109 | Riboflavin | 3432 | BL1 | 4477 |
| 20 | Yes | A. baylyi | Ampicillin | Wall | F117 | SYBR Gree | 3367 | BL1 | 338447 |
| 20 | Yes | A. baylyi | Ampicillin | Wall | F106 | Rhodamine | 2880 | BL1 | 4980.5 |
| 20 | Yes | A. baylyi | Ampicillin | Wall | F73 | Malachite ( | 3304 | BL1 | 5039 |
| 20 | Yes | A. baylyi | Ampicillin | Wall | F7 | Acridine or | 3122 | BL1 | 5242 |
| 20 | Yes | A. baylyi | Ampicillin | Wall | F192 | 5-Carboxyf | 3036 | BL1 | 11786 |
| 20 | Yes | A. baylyi | Ampicillin | Wall | F57 | Eosin Y | 3071 | BL1 | 12993 |
| 20 | Yes | A. baylyi | Ampicillin | Wall | F27 | Calcein | 3616 | BL1 | 16608.5 |
| 20 | Yes | A. baylyi | Ampicillin | Wall | F12 | Amaranth | 3681 | BL1 | 16887 |
| 20 | Yes | A. baylyi | None | None | NA | Autofluores | 4066 | NA | 6398.5 |
| 20 | Yes | A. baylyi | Bacitracin | Wall | F194 | Oxonol | 3872 | RL1 | 5144 |
| 20 | Yes | A. baylyi | Bacitracin | Wall | F11 | Allura Red | 3662 | BL3 | 4929 |
| 20 | Yes | A. baylyi | Bacitracin | Wall | F54 | DiSC3(5) | 3901 | RL1 | 5177 |
| 20 | Yes | A. baylyi | Bacitracin | Wall | F124 | Thioflavin T | 3980 | VL2 | 5779 |
| 20 | Yes | A. baylyi | Bacitracin | Wall | F101 | Quinacrine | 3882 | VL2 | 5612.5 |
| 20 | Yes | A. baylyi | Bacitracin | Wall | F138 | Apigenin | 3822 | VL2 | 5180 |
| 20 | Yes | A. baylyi | Bacitracin | Wall | F65 | H2FDA | 3888 | BL1 | 7204.5 |
| 20 | Yes | A. baylyi | Bacitracin | Wall | F142 | Chlorophe | 3763 | BL3 | 6585 |
| 20 | Yes | A. baylyi | Bacitracin | Wall | F39 | Congo Red | 3996 | BL3 | 6099 |
| 20 | Yes | A. baylyi | Bacitracin | Wall | F116 | Sunset Yel | 3942 | BL2 | 6038 |
| 20 | Yes | A. baylyi | Bacitracin | Wall | F10 | Alizarin | 3932 | BL1 | 5651 |
| 20 | Yes | A. baylyi | Bacitracin | Wall | F84 | Neutral Re | 3770 | BL1 | 5517.5 |
| 20 | Yes | A. baylyi | Bacitracin | Wall | F99 | Pyronin Y | 3854 | BL1 | 5425 |
| 20 | Yes | A. baylyi | Bacitracin | Wall | F108 | Rhodamine | 3695 | BL1 | 4818 |
| 20 | Yes | A. baylyi | Bacitracin | Wall | F109 | Riboflavin | 3999 | BL1 | 5311 |
| 20 | Yes | A. baylyi | Bacitracin | Wall | F117 | SYBR Gree | 3924 | BL1 | 369998 |
| 20 | Yes | A. baylyi | Bacitracin | Wall | F106 | Rhodamine | 4002 | BL1 | 5091 |
| 20 | Yes | A. baylyi | Bacitracin | Wall | F73 | Malachite ( | 3657 | BL1 | 5783 |
| 20 | Yes | A. baylyi | Bacitracin | Wall | F7 | Acridine or | 3895 | BL1 | 5622 |
| 20 | Yes | A. baylyi | Bacitracin | Wall | F192 | 5-Carboxyf | 3763 | BL1 | 14779 |
| 20 | Yes | A. baylyi | Bacitracin | Wall | F57 | Eosin Y | 3892 | BL1 | 15725 |
| 20 | Yes | A. baylyi | Bacitracin | Wall | F27 | Calcein | 4117 | BL1 | 18245 |

| | | | | | | | | | |
|---|---|---|---|---|---|---|---|---|---|
| 20 | Yes | A. baylyi | Bacitracin | Wall | F12 | Amaranth | 3725 | BL1 | 17270 |
| 20 | Yes | A. baylyi | None | None | NA | Autofluore: | 4311 | NA | 5930 |
| 20 | Yes | A. baylyi | Cefotaxim | Wall | F194 | Oxonol | 3372 | RL1 | 5410 |
| 20 | Yes | A. baylyi | Cefotaxim | Wall | F11 | Allura Red | 3385 | BL3 | 5492 |
| 20 | Yes | A. baylyi | Cefotaxim | Wall | F54 | DiSC3(5) | 3689 | RL1 | 4956 |
| 20 | Yes | A. baylyi | Cefotaxim | Wall | F124 | Thioflavin T | 3741 | VL2 | 5358 |
| 20 | Yes | A. baylyi | Cefotaxim | Wall | F101 | Quinacrine | 3484 | VL2 | 5609.5 |
| 20 | Yes | A. baylyi | Cefotaxim | Wall | F138 | Apigenin | 3322 | VL2 | 5497.5 |
| 20 | Yes | A. baylyi | Cefotaxim | Wall | F65 | H2FDA | 3323 | BL1 | 6120 |
| 20 | Yes | A. baylyi | Cefotaxim | Wall | F142 | Chlorophe | 3655 | BL3 | 6794 |
| 20 | Yes | A. baylyi | Cefotaxim | Wall | F39 | Congo Red | 3185 | BL3 | 6831 |
| 20 | Yes | A. baylyi | Cefotaxim | Wall | F116 | Sunset Yel | 3721 | BL2 | 6097 |
| 20 | Yes | A. baylyi | Cefotaxim | Wall | F10 | Alizarin | 3620 | BL1 | 5884.5 |
| 20 | Yes | A. baylyi | Cefotaxim | Wall | F84 | Neutral Re | 3570 | BL1 | 6257 |
| 20 | Yes | A. baylyi | Cefotaxim | Wall | F99 | Pyronin Y | 3593 | BL1 | 6961 |
| 20 | Yes | A. baylyi | Cefotaxim | Wall | F108 | Rhodamine | 3352 | BL1 | 5390.5 |
| 20 | Yes | A. baylyi | Cefotaxim | Wall | F109 | Riboflavin | 3390 | BL1 | 6014 |
| 20 | Yes | A. baylyi | Cefotaxim | Wall | F117 | SYBR Gree | 3010 | BL1 | 347505 |
| 20 | Yes | A. baylyi | Cefotaxim | Wall | F106 | Rhodamine | 2990 | BL1 | 6295.5 |
| 20 | Yes | A. baylyi | Cefotaxim | Wall | F73 | Malachite ( | 2917 | BL1 | 5647 |
| 20 | Yes | A. baylyi | Cefotaxim | Wall | F7 | Acridine or | 3070 | BL1 | 5569 |
| 20 | Yes | A. baylyi | Cefotaxim | Wall | F192 | 5-Carboxyf | 3178 | BL1 | 12458 |
| 20 | Yes | A. baylyi | Cefotaxim | Wall | F57 | Eosin Y | 3386 | BL1 | 13479.5 |
| 20 | Yes | A. baylyi | Cefotaxim | Wall | F27 | Calcein | 3637 | BL1 | 17746 |
| 20 | Yes | A. baylyi | Cefotaxim | Wall | F12 | Amaranth | 3713 | BL1 | 17333 |
| 20 | Yes | A. baylyi | None | None | NA | Autofluore: | 3873 | NA | 6545 |
| 20 | Yes | A. baylyi | Ciprofloxa | Intracellul | F194 | Oxonol | 2948 | RL1 | 5923 |
| 20 | Yes | A. baylyi | Ciprofloxa | Intracellul | F11 | Allura Red | 3063 | BL3 | 5839 |
| 20 | Yes | A. baylyi | Ciprofloxa | Intracellul | F54 | DiSC3(5) | 3452 | RL1 | 5631 |
| 20 | Yes | A. baylyi | Ciprofloxa | Intracellul | F124 | Thioflavin T | 3355 | VL2 | 6086 |
| 20 | Yes | A. baylyi | Ciprofloxa | Intracellul | F101 | Quinacrine | 3312 | VL2 | 6852 |
| 20 | Yes | A. baylyi | Ciprofloxa | Intracellul | F138 | Apigenin | 2967 | VL2 | 6027 |
| 20 | Yes | A. baylyi | Ciprofloxa | Intracellul | F65 | H2FDA | 3425 | BL1 | 6754 |
| 20 | Yes | A. baylyi | Ciprofloxa | Intracellul | F142 | Chlorophe | 3283 | BL3 | 7563 |
| 20 | Yes | A. baylyi | Ciprofloxa | Intracellul | F39 | Congo Red | 3050 | BL3 | 7833.5 |
| 20 | Yes | A. baylyi | Ciprofloxa | Intracellul | F116 | Sunset Yel | 3091 | BL2 | 7286 |
| 20 | Yes | A. baylyi | Ciprofloxa | Intracellul | F10 | Alizarin | 3107 | BL1 | 6731 |
| 20 | Yes | A. baylyi | Ciprofloxa | Intracellul | F84 | Neutral Re | 3073 | BL1 | 6025 |
| 20 | Yes | A. baylyi | Ciprofloxa | Intracellul | F99 | Pyronin Y | 3084 | BL1 | 5561 |
| 20 | Yes | A. baylyi | Ciprofloxa | Intracellul | F108 | Rhodamine | 2792 | BL1 | 5363 |
| 20 | Yes | A. baylyi | Ciprofloxa | Intracellul | F109 | Riboflavin | 3408 | BL1 | 5235.5 |
| 20 | Yes | A. baylyi | Ciprofloxa | Intracellul | F117 | SYBR Gree | 2555 | BL1 | 277288 |
| 20 | Yes | A. baylyi | Ciprofloxa | Intracellul | F106 | Rhodamine | 3307 | BL1 | 5871 |
| 20 | Yes | A. baylyi | Ciprofloxa | Intracellul | F73 | Malachite ( | 3236 | BL1 | 5318 |
| 20 | Yes | A. baylyi | Ciprofloxa | Intracellul | F7 | Acridine or | 2961 | BL1 | 5879 |
| 20 | Yes | A. baylyi | Ciprofloxa | Intracellul | F192 | 5-Carboxyf | 2963 | BL1 | 15653 |
| 20 | Yes | A. baylyi | Ciprofloxa | Intracellul | F57 | Eosin Y | 2408 | BL1 | 18379 |
| 20 | Yes | A. baylyi | Ciprofloxa | Intracellul | F27 | Calcein | 3278 | BL1 | 20203.5 |

| | | | | | | | | | |
|---|---|---|---|---|---|---|---|---|---|
| 20 | Yes | A. baylyi | Ciprofloxa | Intracellula | F12 | Amaranth | 2963 | BL1 | 18900 |
| 20 | Yes | A. baylyi | None | None | NA | Autofluore | 4089 | NA | 6186 |
| 20 | Yes | A. baylyi | Fosfomyci | Wall | F194 | Oxonol | 3861 | RL1 | 5477 |
| 20 | Yes | A. baylyi | Fosfomyci | Wall | F11 | Allura Red | 4041 | BL3 | 5275 |
| 20 | Yes | A. baylyi | Fosfomyci | Wall | F54 | DiSC3(5) | 4190 | RL1 | 4963.5 |
| 20 | Yes | A. baylyi | Fosfomyci | Wall | F124 | Thioflavin T | 4214 | VL2 | 5429 |
| 20 | Yes | A. baylyi | Fosfomyci | Wall | F101 | Quinacrine | 4489 | VL2 | 5088 |
| 20 | Yes | A. baylyi | Fosfomyci | Wall | F138 | Apigenin | 4318 | VL2 | 4926.5 |
| 20 | Yes | A. baylyi | Fosfomyci | Wall | F65 | H2FDA | 4288 | BL1 | 5889 |
| 20 | Yes | A. baylyi | Fosfomyci | Wall | F142 | Chlorophe | 4335 | BL3 | 7602 |
| 20 | Yes | A. baylyi | Fosfomyci | Wall | F39 | Congo Red | 4263 | BL3 | 7093 |
| 20 | Yes | A. baylyi | Fosfomyci | Wall | F116 | Sunset Yel | 4322 | BL2 | 6777.5 |
| 20 | Yes | A. baylyi | Fosfomyci | Wall | F10 | Alizarin | 4058 | BL1 | 6193.5 |
| 20 | Yes | A. baylyi | Fosfomyci | Wall | F84 | Neutral Re | 4117 | BL1 | 5239 |
| 20 | Yes | A. baylyi | Fosfomyci | Wall | F99 | Pyronin Y | 4045 | BL1 | 4709 |
| 20 | Yes | A. baylyi | Fosfomyci | Wall | F108 | Rhodamine | 4031 | BL1 | 4432 |
| 20 | Yes | A. baylyi | Fosfomyci | Wall | F109 | Riboflavin | 4545 | BL1 | 4638 |
| 20 | Yes | A. baylyi | Fosfomyci | Wall | F117 | SYBR Gree | 3937 | BL1 | 291544 |
| 20 | Yes | A. baylyi | Fosfomyci | Wall | F106 | Rhodamine | 4266 | BL1 | 5362.5 |
| 20 | Yes | A. baylyi | Fosfomyci | Wall | F73 | Malachite | 3830 | BL1 | 5256 |
| 20 | Yes | A. baylyi | Fosfomyci | Wall | F7 | Acridine or | 3924 | BL1 | 5359 |
| 20 | Yes | A. baylyi | Fosfomyci | Wall | F192 | 5-Carboxyf | 4109 | BL1 | 14115 |
| 20 | Yes | A. baylyi | Fosfomyci | Wall | F57 | Eosin Y | 4136 | BL1 | 16242.5 |
| 20 | Yes | A. baylyi | Fosfomyci | Wall | F27 | Calcein | 4126 | BL1 | 16903 |
| 20 | Yes | A. baylyi | Fosfomyci | Wall | F12 | Amaranth | 4289 | BL1 | 16083 |
| 20 | Yes | A. baylyi | None | None | NA | Autofluore | 4248 | NA | 6239.5 |
| 20 | Yes | A. baylyi | Nalidixic a | Intracellula | F194 | Oxonol | 3772 | RL1 | 5396 |
| 20 | Yes | A. baylyi | Nalidixic a | Intracellula | F11 | Allura Red | 4310 | BL3 | 5059.5 |
| 20 | Yes | A. baylyi | Nalidixic a | Intracellula | F54 | DiSC3(5) | 4262 | RL1 | 5247.5 |
| 20 | Yes | A. baylyi | Nalidixic a | Intracellula | F124 | Thioflavin T | 4074 | VL2 | 5798.5 |
| 20 | Yes | A. baylyi | Nalidixic a | Intracellula | F101 | Quinacrine | 3866 | VL2 | 5786.5 |
| 20 | Yes | A. baylyi | Nalidixic a | Intracellula | F138 | Apigenin | 3539 | VL2 | 5422 |
| 20 | Yes | A. baylyi | Nalidixic a | Intracellula | F65 | H2FDA | 4000 | BL1 | 6229 |
| 20 | Yes | A. baylyi | Nalidixic a | Intracellula | F142 | Chlorophe | 3817 | BL3 | 7552 |
| 20 | Yes | A. baylyi | Nalidixic a | Intracellula | F39 | Congo Red | 3998 | BL3 | 8087.5 |
| 20 | Yes | A. baylyi | Nalidixic a | Intracellula | F116 | Sunset Yel | 3906 | BL2 | 7280.5 |
| 20 | Yes | A. baylyi | Nalidixic a | Intracellula | F10 | Alizarin | 3984 | BL1 | 6815.5 |
| 20 | Yes | A. baylyi | Nalidixic a | Intracellula | F84 | Neutral Re | 3953 | BL1 | 6004 |
| 20 | Yes | A. baylyi | Nalidixic a | Intracellula | F99 | Pyronin Y | 3819 | BL1 | 5088 |
| 20 | Yes | A. baylyi | Nalidixic a | Intracellula | F108 | Rhodamine | 3973 | BL1 | 4531 |
| 20 | Yes | A. baylyi | Nalidixic a | Intracellula | F109 | Riboflavin | 3902 | BL1 | 4799 |
| 20 | Yes | A. baylyi | Nalidixic a | Intracellula | F117 | SYBR Gree | 3660 | BL1 | 286220.5 |
| 20 | Yes | A. baylyi | Nalidixic a | Intracellula | F106 | Rhodamine | 3876 | BL1 | 5387 |
| 20 | Yes | A. baylyi | Nalidixic a | Intracellula | F73 | Malachite | 3890 | BL1 | 5221 |
| 20 | Yes | A. baylyi | Nalidixic a | Intracellula | F7 | Acridine or | 3792 | BL1 | 5393 |
| 20 | Yes | A. baylyi | Nalidixic a | Intracellula | F192 | 5-Carboxyf | 3789 | BL1 | 15916 |
| 20 | Yes | A. baylyi | Nalidixic a | Intracellula | F57 | Eosin Y | 3457 | BL1 | 15534 |
| 20 | Yes | A. baylyi | Nalidixic a | Intracellula | F27 | Calcein | 3572 | BL1 | 16527 |

| | | | | | | | | | |
|---|---|---|---|---|---|---|---|---|---|
| 20 | Yes | A. baylyi | Nalidixic a | Intracellul | F12 | Amaranth | 3850 | BL1 | 18060.5 |
| 20 | Yes | A. baylyi | None | None | NA | Autofluore | 3890 | NA | 6394 |
| 20 | Yes | A. baylyi | Ofloxacin | Intracellul | F194 | Oxonol | 3436 | RL1 | 6058 |
| 20 | Yes | A. baylyi | Ofloxacin | Intracellul | F11 | Allura Red | 3451 | BL3 | 5760 |
| 20 | Yes | A. baylyi | Ofloxacin | Intracellul | F54 | DiSC3(5) | 3619 | RL1 | 5397 |
| 20 | Yes | A. baylyi | Ofloxacin | Intracellul | F124 | Thioflavin T | 3740 | VL2 | 5679 |
| 20 | Yes | A. baylyi | Ofloxacin | Intracellul | F101 | Quinacrine | 3474 | VL2 | 5933 |
| 20 | Yes | A. baylyi | Ofloxacin | Intracellul | F138 | Apigenin | 3560 | VL2 | 5791 |
| 20 | Yes | A. baylyi | Ofloxacin | Intracellul | F65 | H2FDA | 3807 | BL1 | 6378 |
| 20 | Yes | A. baylyi | Ofloxacin | Intracellul | F142 | Chlorophe | 3696 | BL3 | 7596 |
| 20 | Yes | A. baylyi | Ofloxacin | Intracellul | F39 | Congo Red | 3987 | BL3 | 7655 |
| 20 | Yes | A. baylyi | Ofloxacin | Intracellul | F116 | Sunset Yel | 3268 | BL2 | 7886.5 |
| 20 | Yes | A. baylyi | Ofloxacin | Intracellul | F10 | Alizarin | 3877 | BL1 | 7099 |
| 20 | Yes | A. baylyi | Ofloxacin | Intracellul | F84 | Neutral Re | 3318 | BL1 | 5698.5 |
| 20 | Yes | A. baylyi | Ofloxacin | Intracellul | F99 | Pyronin Y | 3986 | BL1 | 4943 |
| 20 | Yes | A. baylyi | Ofloxacin | Intracellul | F108 | Rhodamine | 3912 | BL1 | 4516 |
| 20 | Yes | A. baylyi | Ofloxacin | Intracellul | F109 | Riboflavin | 4018 | BL1 | 4779 |
| 20 | Yes | A. baylyi | Ofloxacin | Intracellul | F117 | SYBR Gree | 3940 | BL1 | 320176.5 |
| 20 | Yes | A. baylyi | Ofloxacin | Intracellul | F106 | Rhodamine | 3860 | BL1 | 5574 |
| 20 | Yes | A. baylyi | Ofloxacin | Intracellul | F73 | Malachite ( | 3852 | BL1 | 5366 |
| 20 | Yes | A. baylyi | Ofloxacin | Intracellul | F7 | Acridine or | 3654 | BL1 | 5321.5 |
| 20 | Yes | A. baylyi | Ofloxacin | Intracellul | F192 | 5-Carboxyf | 3777 | BL1 | 16089 |
| 20 | Yes | A. baylyi | Ofloxacin | Intracellul | F57 | Eosin Y | 3798 | BL1 | 16224 |
| 20 | Yes | A. baylyi | Ofloxacin | Intracellul | F27 | Calcein | 3819 | BL1 | 15926 |
| 20 | Yes | A. baylyi | Ofloxacin | Intracellul | F12 | Amaranth | 4165 | BL1 | 16143 |
| 20 | Yes | A. baylyi | None | None | NA | Autofluore | 4136 | NA | 6570 |
| 20 | Yes | A. baylyi | Polymyxin | Wall | F194 | Oxonol | 3810 | RL1 | 6113 |
| 20 | Yes | A. baylyi | Polymyxin | Wall | F11 | Allura Red | 4652 | BL3 | 5552.5 |
| 20 | Yes | A. baylyi | Polymyxin | Wall | F54 | DiSC3(5) | 4351 | RL1 | 6004 |
| 20 | Yes | A. baylyi | Polymyxin | Wall | F124 | Thioflavin T | 4124 | VL2 | 5785 |
| 20 | Yes | A. baylyi | Polymyxin | Wall | F101 | Quinacrine | 4231 | VL2 | 5698 |
| 20 | Yes | A. baylyi | Polymyxin | Wall | F138 | Apigenin | 4335 | VL2 | 5539 |
| 20 | Yes | A. baylyi | Polymyxin | Wall | F65 | H2FDA | 4083 | BL1 | 6261 |
| 20 | Yes | A. baylyi | Polymyxin | Wall | F142 | Chlorophe | 4728 | BL3 | 6972.5 |
| 20 | Yes | A. baylyi | Polymyxin | Wall | F39 | Congo Red | 4504 | BL3 | 8202.5 |
| 20 | Yes | A. baylyi | Polymyxin | Wall | F116 | Sunset Yel | 4666 | BL2 | 7438 |
| 20 | Yes | A. baylyi | Polymyxin | Wall | F10 | Alizarin | 3742 | BL1 | 6982.5 |
| 20 | Yes | A. baylyi | Polymyxin | Wall | F84 | Neutral Re | 4488 | BL1 | 5910.5 |
| 20 | Yes | A. baylyi | Polymyxin | Wall | F99 | Pyronin Y | 4147 | BL1 | 5234 |
| 20 | Yes | A. baylyi | Polymyxin | Wall | F108 | Rhodamine | 4419 | BL1 | 5216 |
| 20 | Yes | A. baylyi | Polymyxin | Wall | F109 | Riboflavin | 4394 | BL1 | 5129 |
| 20 | Yes | A. baylyi | Polymyxin | Wall | F117 | SYBR Gree | 3910 | BL1 | 606786 |
| 20 | Yes | A. baylyi | Polymyxin | Wall | F106 | Rhodamine | 3877 | BL1 | 6323 |
| 20 | Yes | A. baylyi | Polymyxin | Wall | F73 | Malachite ( | 3913 | BL1 | 5157 |
| 20 | Yes | A. baylyi | Polymyxin | Wall | F7 | Acridine or | 4265 | BL1 | 5936 |
| 20 | Yes | A. baylyi | Polymyxin | Wall | F192 | 5-Carboxyf | 4248 | BL1 | 15508.5 |
| 20 | Yes | A. baylyi | Polymyxin | Wall | F57 | Eosin Y | 3588 | BL1 | 16627 |
| 20 | Yes | A. baylyi | Polymyxin | Wall | F27 | Calcein | 4220 | BL1 | 16639.5 |

| | | | | | | | | | |
|---|---|---|---|---|---|---|---|---|---|
| 20 | Yes | A. baylyi | Polymyxin | Wall | F12 | Amaranth | 4072 | BL1 | 16031 |
| 20 | Yes | A. baylyi | None | None | NA | Autofluore: | 3671 | NA | 6421 |
| 20 | Yes | A. baylyi | Tetracyclin | Intracellula | F194 | Oxonol | 3231 | RL1 | 6014 |
| 20 | Yes | A. baylyi | Tetracyclin | Intracellula | F11 | Allura Red | 2885 | BL3 | 5846 |
| 20 | Yes | A. baylyi | Tetracyclin | Intracellula | F54 | DiSC3(5) | 3119 | RL1 | 6092 |
| 20 | Yes | A. baylyi | Tetracyclin | Intracellula | F124 | Thioflavin T | 3152 | VL2 | 6118.5 |
| 20 | Yes | A. baylyi | Tetracyclin | Intracellula | F101 | Quinacrine | 2922 | VL2 | 6980 |
| 20 | Yes | A. baylyi | Tetracyclin | Intracellula | F138 | Apigenin | 2962 | VL2 | 6451 |
| 20 | Yes | A. baylyi | Tetracyclin | Intracellula | F65 | H2FDA | 3253 | BL1 | 6919 |
| 20 | Yes | A. baylyi | Tetracyclin | Intracellula | F142 | Chlorophe | 3078 | BL3 | 7749.5 |
| 20 | Yes | A. baylyi | Tetracyclin | Intracellula | F39 | Congo Red | 2978 | BL3 | 8198 |
| 20 | Yes | A. baylyi | Tetracyclin | Intracellula | F116 | Sunset Yel | 3102 | BL2 | 7573.5 |
| 20 | Yes | A. baylyi | Tetracyclin | Intracellula | F10 | Alizarin | 3200 | BL1 | 7439 |
| 20 | Yes | A. baylyi | Tetracyclin | Intracellula | F84 | Neutral Re | 2931 | BL1 | 6666 |
| 20 | Yes | A. baylyi | Tetracyclin | Intracellula | F99 | Pyronin Y | 3185 | BL1 | 5698 |
| 20 | Yes | A. baylyi | Tetracyclin | Intracellula | F108 | Rhodamine | 3063 | BL1 | 5169 |
| 20 | Yes | A. baylyi | Tetracyclin | Intracellula | F109 | Riboflavin | 3106 | BL1 | 5243 |
| 20 | Yes | A. baylyi | Tetracyclin | Intracellula | F117 | SYBR Gree | 3155 | BL1 | 279120 |
| 20 | Yes | A. baylyi | Tetracyclin | Intracellula | F106 | Rhodamine | 2871 | BL1 | 5997 |
| 20 | Yes | A. baylyi | Tetracyclin | Intracellula | F73 | Malachite ( | 2844 | BL1 | 5025 |
| 20 | Yes | A. baylyi | Tetracyclin | Intracellula | F7 | Acridine or | 3042 | BL1 | 5812 |
| 20 | Yes | A. baylyi | Tetracyclin | Intracellula | F192 | 5-Carboxyf | 2953 | BL1 | 16019 |
| 20 | Yes | A. baylyi | Tetracyclin | Intracellula | F57 | Eosin Y | 2700 | BL1 | 15357 |
| 20 | Yes | A. baylyi | Tetracyclin | Intracellula | F27 | Calcein | 2903 | BL1 | 17862 |
| 20 | Yes | A. baylyi | Tetracyclin | Intracellula | F12 | Amaranth | 2942 | BL1 | 19149.5 |
| 20 | Yes | A. baylyi | None | None | NA | Autofluore: | 4189 | NA | 6360 |
| 20 | Yes | A. baylyi | Vancomyc | Wall | F194 | Oxonol | 3931 | RL1 | 5709 |
| 20 | Yes | A. baylyi | Vancomyc | Wall | F11 | Allura Red | 4499 | BL3 | 6087 |
| 20 | Yes | A. baylyi | Vancomyc | Wall | F54 | DiSC3(5) | 4465 | RL1 | 5580 |
| 20 | Yes | A. baylyi | Vancomyc | Wall | F124 | Thioflavin T | 4250 | VL2 | 5714.5 |
| 20 | Yes | A. baylyi | Vancomyc | Wall | F101 | Quinacrine | 4016 | VL2 | 5776 |
| 20 | Yes | A. baylyi | Vancomyc | Wall | F138 | Apigenin | 4141 | VL2 | 5894 |
| 20 | Yes | A. baylyi | Vancomyc | Wall | F65 | H2FDA | 4542 | BL1 | 6076.5 |
| 20 | Yes | A. baylyi | Vancomyc | Wall | F142 | Chlorophe | 4222 | BL3 | 8143 |
| 20 | Yes | A. baylyi | Vancomyc | Wall | F39 | Congo Red | 4364 | BL3 | 10071 |
| 20 | Yes | A. baylyi | Vancomyc | Wall | F116 | Sunset Yel | 4421 | BL2 | 8478 |
| 20 | Yes | A. baylyi | Vancomyc | Wall | F10 | Alizarin | 4221 | BL1 | 7103 |
| 20 | Yes | A. baylyi | Vancomyc | Wall | F84 | Neutral Re | 4619 | BL1 | 5742 |
| 20 | Yes | A. baylyi | Vancomyc | Wall | F99 | Pyronin Y | 4124 | BL1 | 5154.5 |
| 20 | Yes | A. baylyi | Vancomyc | Wall | F108 | Rhodamine | 3608 | BL1 | 4838.5 |
| 20 | Yes | A. baylyi | Vancomyc | Wall | F109 | Riboflavin | 3556 | BL1 | 5198 |
| 20 | Yes | A. baylyi | Vancomyc | Wall | F117 | SYBR Gree | 3503 | BL1 | 319624 |
| 20 | Yes | A. baylyi | Vancomyc | Wall | F106 | Rhodamine | 3892 | BL1 | 6410 |
| 20 | Yes | A. baylyi | Vancomyc | Wall | F73 | Malachite ( | 3804 | BL1 | 5465.5 |
| 20 | Yes | A. baylyi | Vancomyc | Wall | F7 | Acridine or | 3780 | BL1 | 5556 |
| 20 | Yes | A. baylyi | Vancomyc | Wall | F192 | 5-Carboxyf | 3987 | BL1 | 15682 |
| 20 | Yes | A. baylyi | Vancomyc | Wall | F57 | Eosin Y | 4047 | BL1 | 15804 |
| 20 | Yes | A. baylyi | Vancomyc | Wall | F27 | Calcein | 4226 | BL1 | 13960 |

| | | | | | | | | | |
|---|---|---|---|---|---|---|---|---|---|
| 20 | Yes | A. baylyi | Vancomyc | Wall | F12 | Amaranth | 3763 | BL1 | 11903 |
| 20 | Yes | A. baylyi | None | None | NA | Autofluore: | 6527 | NA | 1051 |
| 20 | Yes | A. baylyi | None | None | F194 | Oxonol | 5928 | RL1 | 1942 |
| 20 | Yes | A. baylyi | None | None | F11 | Allura Red | 5292 | BL3 | 2491 |
| 20 | Yes | A. baylyi | None | None | F54 | DiSC3(5) | 5161 | RL1 | 2661 |
| 20 | Yes | A. baylyi | None | None | F124 | Thioflavin T | 5385 | VL2 | 2758 |
| 20 | Yes | A. baylyi | None | None | F101 | Quinacrine | 4775 | VL2 | 2901 |
| 20 | Yes | A. baylyi | None | None | F138 | Apigenin | 5212 | VL2 | 2937 |
| 20 | Yes | A. baylyi | None | None | F65 | H2FDA | 4788 | BL1 | 4076.5 |
| 20 | Yes | A. baylyi | None | None | F142 | Chlorophe | 5144 | BL3 | 5428 |
| 20 | Yes | A. baylyi | None | None | F39 | Congo Red | 4962 | BL3 | 5285 |
| 20 | Yes | A. baylyi | None | None | F116 | Sunset Yel | 5131 | BL2 | 4562 |
| 20 | Yes | A. baylyi | None | None | F10 | Alizarin | 4777 | BL1 | 4130 |
| 20 | Yes | A. baylyi | None | None | F84 | Neutral Re | 4877 | BL1 | 3830 |
| 20 | Yes | A. baylyi | None | None | F99 | Pyronin Y | 4593 | BL1 | 3461 |
| 20 | Yes | A. baylyi | None | None | F108 | Rhodamine | 4554 | BL1 | 3122 |
| 20 | Yes | A. baylyi | None | None | F109 | Riboflavin | 4896 | BL1 | 3606.5 |
| 20 | Yes | A. baylyi | None | None | F117 | SYBR Gree | 4324 | BL1 | 287343.5 |
| 20 | Yes | A. baylyi | None | None | F106 | Rhodamine | 4895 | BL1 | 3877 |
| 20 | Yes | A. baylyi | None | None | F73 | Malachite ( | 4668 | BL1 | 3684.5 |
| 20 | Yes | A. baylyi | None | None | F7 | Acridine or | 4447 | BL1 | 3958 |
| 20 | Yes | A. baylyi | None | None | F192 | 5-Carboxyf | 4481 | BL1 | 14879 |
| 20 | Yes | A. baylyi | None | None | F57 | Eosin Y | 4716 | BL1 | 14409.5 |
| 20 | Yes | A. baylyi | None | None | F27 | Calcein | 4814 | BL1 | 17688.5 |
| 20 | Yes | A. baylyi | None | None | F12 | Amaranth | 4888 | BL1 | 16855.5 |
| 20 | Yes | A. baylyi | None | None | NA | Autofluore: | 4725 | NA | 5651 |
| 20 | Yes | A. baylyi | Ampicillin | Wall | F194 | Oxonol | 3776 | RL1 | 4755 |
| 20 | Yes | A. baylyi | Ampicillin | Wall | F11 | Allura Red | 3843 | BL3 | 4273 |
| 20 | Yes | A. baylyi | Ampicillin | Wall | F54 | DiSC3(5) | 4414 | RL1 | 4169 |
| 20 | Yes | A. baylyi | Ampicillin | Wall | F124 | Thioflavin T | 4217 | VL2 | 4458 |
| 20 | Yes | A. baylyi | Ampicillin | Wall | F101 | Quinacrine | 4049 | VL2 | 4314 |
| 20 | Yes | A. baylyi | Ampicillin | Wall | F138 | Apigenin | 3991 | VL2 | 4247 |
| 20 | Yes | A. baylyi | Ampicillin | Wall | F65 | H2FDA | 4092 | BL1 | 5250 |
| 20 | Yes | A. baylyi | Ampicillin | Wall | F142 | Chlorophe | 3990 | BL3 | 6214 |
| 20 | Yes | A. baylyi | Ampicillin | Wall | F39 | Congo Red | 4064 | BL3 | 5929 |
| 20 | Yes | A. baylyi | Ampicillin | Wall | F116 | Sunset Yel | 3831 | BL2 | 5311 |
| 20 | Yes | A. baylyi | Ampicillin | Wall | F10 | Alizarin | 4091 | BL1 | 4954 |
| 20 | Yes | A. baylyi | Ampicillin | Wall | F84 | Neutral Re | 3837 | BL1 | 4912 |
| 20 | Yes | A. baylyi | Ampicillin | Wall | F99 | Pyronin Y | 3866 | BL1 | 4359.5 |
| 20 | Yes | A. baylyi | Ampicillin | Wall | F108 | Rhodamine | 3820 | BL1 | 4042 |
| 20 | Yes | A. baylyi | Ampicillin | Wall | F109 | Riboflavin | 3694 | BL1 | 4344.5 |
| 20 | Yes | A. baylyi | Ampicillin | Wall | F117 | SYBR Gree | 3905 | BL1 | 345702 |
| 20 | Yes | A. baylyi | Ampicillin | Wall | F106 | Rhodamine | 4073 | BL1 | 4313 |
| 20 | Yes | A. baylyi | Ampicillin | Wall | F73 | Malachite ( | 4027 | BL1 | 4270 |
| 20 | Yes | A. baylyi | Ampicillin | Wall | F7 | Acridine or | 4198 | BL1 | 4006.5 |
| 20 | Yes | A. baylyi | Ampicillin | Wall | F192 | 5-Carboxyf | 4494 | BL1 | 13951.5 |
| 20 | Yes | A. baylyi | Ampicillin | Wall | F57 | Eosin Y | 4290 | BL1 | 15801 |
| 20 | Yes | A. baylyi | Ampicillin | Wall | F27 | Calcein | 4961 | BL1 | 17789 |

| | | | | | | | | | |
|---|---|---|---|---|---|---|---|---|---|
| 20 | Yes | A. baylyi | Ampicillin | Wall | F12 | Amaranth | 4648 | BL1 | 17001.5 |
| 20 | Yes | A. baylyi | None | None | NA | Autofluore: | 4667 | NA | 6391 |
| 20 | Yes | A. baylyi | Bacitracin | Wall | F194 | Oxonol | 4476 | RL1 | 4584 |
| 20 | Yes | A. baylyi | Bacitracin | Wall | F11 | Allura Red | 4412 | BL3 | 4505.5 |
| 20 | Yes | A. baylyi | Bacitracin | Wall | F54 | DiSC3(5) | 4891 | RL1 | 4185 |
| 20 | Yes | A. baylyi | Bacitracin | Wall | F124 | Thioflavin T | 4636 | VL2 | 4700 |
| 20 | Yes | A. baylyi | Bacitracin | Wall | F101 | Quinacrine | 4789 | VL2 | 4459 |
| 20 | Yes | A. baylyi | Bacitracin | Wall | F138 | Apigenin | 4686 | VL2 | 4958.5 |
| 20 | Yes | A. baylyi | Bacitracin | Wall | F65 | H2FDA | 4705 | BL1 | 6831 |
| 20 | Yes | A. baylyi | Bacitracin | Wall | F142 | Chlorophe | 4411 | BL3 | 7371 |
| 20 | Yes | A. baylyi | Bacitracin | Wall | F39 | Congo Red | 4746 | BL3 | 6477 |
| 20 | Yes | A. baylyi | Bacitracin | Wall | F116 | Sunset Yel | 4833 | BL2 | 5760 |
| 20 | Yes | A. baylyi | Bacitracin | Wall | F10 | Alizarin | 4316 | BL1 | 4841.5 |
| 20 | Yes | A. baylyi | Bacitracin | Wall | F84 | Neutral Re | 4203 | BL1 | 5833 |
| 20 | Yes | A. baylyi | Bacitracin | Wall | F99 | Pyronin Y | 4188 | BL1 | 5234 |
| 20 | Yes | A. baylyi | Bacitracin | Wall | F108 | Rhodamine | 4704 | BL1 | 4499.5 |
| 20 | Yes | A. baylyi | Bacitracin | Wall | F109 | Riboflavin | 4324 | BL1 | 4704 |
| 20 | Yes | A. baylyi | Bacitracin | Wall | F117 | SYBR Gree | 4388 | BL1 | 391701.5 |
| 20 | Yes | A. baylyi | Bacitracin | Wall | F106 | Rhodamine | 4603 | BL1 | 4812 |
| 20 | Yes | A. baylyi | Bacitracin | Wall | F73 | Malachite ( | 4466 | BL1 | 5017.5 |
| 20 | Yes | A. baylyi | Bacitracin | Wall | F7 | Acridine or | 4497 | BL1 | 4578 |
| 20 | Yes | A. baylyi | Bacitracin | Wall | F192 | 5-Carboxyf | 4552 | BL1 | 15077 |
| 20 | Yes | A. baylyi | Bacitracin | Wall | F57 | Eosin Y | 4386 | BL1 | 18350.5 |
| 20 | Yes | A. baylyi | Bacitracin | Wall | F27 | Calcein | 4680 | BL1 | 16736.5 |
| 20 | Yes | A. baylyi | Bacitracin | Wall | F12 | Amaranth | 4628 | BL1 | 15592.5 |
| 20 | Yes | A. baylyi | None | None | NA | Autofluore: | 4957 | NA | 5811 |
| 20 | Yes | A. baylyi | Cefotaxim | Wall | F194 | Oxonol | 3751 | RL1 | 5486 |
| 20 | Yes | A. baylyi | Cefotaxim | Wall | F11 | Allura Red | 4105 | BL3 | 5236 |
| 20 | Yes | A. baylyi | Cefotaxim | Wall | F54 | DiSC3(5) | 4046 | RL1 | 5080.5 |
| 20 | Yes | A. baylyi | Cefotaxim | Wall | F124 | Thioflavin T | 4231 | VL2 | 5439 |
| 20 | Yes | A. baylyi | Cefotaxim | Wall | F101 | Quinacrine | 4169 | VL2 | 5254 |
| 20 | Yes | A. baylyi | Cefotaxim | Wall | F138 | Apigenin | 4080 | VL2 | 5495 |
| 20 | Yes | A. baylyi | Cefotaxim | Wall | F65 | H2FDA | 4164 | BL1 | 5930.5 |
| 20 | Yes | A. baylyi | Cefotaxim | Wall | F142 | Chlorophe | 4034 | BL3 | 7024 |
| 20 | Yes | A. baylyi | Cefotaxim | Wall | F39 | Congo Red | 4155 | BL3 | 6685 |
| 20 | Yes | A. baylyi | Cefotaxim | Wall | F116 | Sunset Yel | 4087 | BL2 | 5636 |
| 20 | Yes | A. baylyi | Cefotaxim | Wall | F10 | Alizarin | 4346 | BL1 | 5226.5 |
| 20 | Yes | A. baylyi | Cefotaxim | Wall | F84 | Neutral Re | 4032 | BL1 | 5694.5 |
| 20 | Yes | A. baylyi | Cefotaxim | Wall | F99 | Pyronin Y | 4281 | BL1 | 5455 |
| 20 | Yes | A. baylyi | Cefotaxim | Wall | F108 | Rhodamine | 3977 | BL1 | 4520 |
| 20 | Yes | A. baylyi | Cefotaxim | Wall | F109 | Riboflavin | 4329 | BL1 | 5392 |
| 20 | Yes | A. baylyi | Cefotaxim | Wall | F117 | SYBR Gree | 4367 | BL1 | 310880 |
| 20 | Yes | A. baylyi | Cefotaxim | Wall | F106 | Rhodamine | 4513 | BL1 | 5088 |
| 20 | Yes | A. baylyi | Cefotaxim | Wall | F73 | Malachite ( | 4644 | BL1 | 4543 |
| 20 | Yes | A. baylyi | Cefotaxim | Wall | F7 | Acridine or | 3946 | BL1 | 4861.5 |
| 20 | Yes | A. baylyi | Cefotaxim | Wall | F192 | 5-Carboxyf | 4372 | BL1 | 15422.5 |
| 20 | Yes | A. baylyi | Cefotaxim | Wall | F57 | Eosin Y | 3855 | BL1 | 12256 |
| 20 | Yes | A. baylyi | Cefotaxim | Wall | F27 | Calcein | 4005 | BL1 | 15911 |

| | | | | | | | | | |
|---|---|---|---|---|---|---|---|---|---|
| 20 | Yes | A. baylyi | Cefotaxim | Wall | F12 | Amaranth | 4137 | BL1 | 19906 |
| 20 | Yes | A. baylyi | None | None | NA | Autofluore | 4592 | NA | 6077 |
| 20 | Yes | A. baylyi | Ciprofloxa | Intracellul | F194 | Oxonol | 3536 | RL1 | 5750.5 |
| 20 | Yes | A. baylyi | Ciprofloxa | Intracellul | F11 | Allura Red | 3455 | BL3 | 5499 |
| 20 | Yes | A. baylyi | Ciprofloxa | Intracellul | F54 | DiSC3(5) | 3817 | RL1 | 5435 |
| 20 | Yes | A. baylyi | Ciprofloxa | Intracellul | F124 | Thioflavin T | 3876 | VL2 | 5570 |
| 20 | Yes | A. baylyi | Ciprofloxa | Intracellul | F101 | Quinacrine | 3460 | VL2 | 5423 |
| 20 | Yes | A. baylyi | Ciprofloxa | Intracellul | F138 | Apigenin | 3688 | VL2 | 5322.5 |
| 20 | Yes | A. baylyi | Ciprofloxa | Intracellul | F65 | H2FDA | 3864 | BL1 | 5974.5 |
| 20 | Yes | A. baylyi | Ciprofloxa | Intracellul | F142 | Chlorophe | 3905 | BL3 | 7291 |
| 20 | Yes | A. baylyi | Ciprofloxa | Intracellul | F39 | Congo Red | 3690 | BL3 | 7794 |
| 20 | Yes | A. baylyi | Ciprofloxa | Intracellul | F116 | Sunset Yel | 3569 | BL2 | 6976 |
| 20 | Yes | A. baylyi | Ciprofloxa | Intracellul | F10 | Alizarin | 3600 | BL1 | 5920.5 |
| 20 | Yes | A. baylyi | Ciprofloxa | Intracellul | F84 | Neutral Re | 3891 | BL1 | 4991 |
| 20 | Yes | A. baylyi | Ciprofloxa | Intracellul | F99 | Pyronin Y | 3763 | BL1 | 4733 |
| 20 | Yes | A. baylyi | Ciprofloxa | Intracellul | F108 | Rhodamine | 3650 | BL1 | 4443.5 |
| 20 | Yes | A. baylyi | Ciprofloxa | Intracellul | F109 | Riboflavin | 3740 | BL1 | 4796 |
| 20 | Yes | A. baylyi | Ciprofloxa | Intracellul | F117 | SYBR Gree | 3385 | BL1 | 283945 |
| 20 | Yes | A. baylyi | Ciprofloxa | Intracellul | F106 | Rhodamine | 3890 | BL1 | 5185.5 |
| 20 | Yes | A. baylyi | Ciprofloxa | Intracellul | F73 | Malachite ( | 3734 | BL1 | 4854.5 |
| 20 | Yes | A. baylyi | Ciprofloxa | Intracellul | F7 | Acridine or | 3632 | BL1 | 5181 |
| 20 | Yes | A. baylyi | Ciprofloxa | Intracellul | F192 | 5-Carboxyf | 3430 | BL1 | 16724.5 |
| 20 | Yes | A. baylyi | Ciprofloxa | Intracellul | F57 | Eosin Y | 3655 | BL1 | 17280 |
| 20 | Yes | A. baylyi | Ciprofloxa | Intracellul | F27 | Calcein | 3518 | BL1 | 19671 |
| 20 | Yes | A. baylyi | Ciprofloxa | Intracellul | F12 | Amaranth | 3679 | BL1 | 18840 |
| 20 | Yes | A. baylyi | None | None | NA | Autofluore | 4510 | NA | 6115.5 |
| 20 | Yes | A. baylyi | Fosfomyci | Wall | F194 | Oxonol | 4582 | RL1 | 5267 |
| 20 | Yes | A. baylyi | Fosfomyci | Wall | F11 | Allura Red | 4696 | BL3 | 5154 |
| 20 | Yes | A. baylyi | Fosfomyci | Wall | F54 | DiSC3(5) | 4706 | RL1 | 4948 |
| 20 | Yes | A. baylyi | Fosfomyci | Wall | F124 | Thioflavin T | 4716 | VL2 | 5170.5 |
| 20 | Yes | A. baylyi | Fosfomyci | Wall | F101 | Quinacrine | 4984 | VL2 | 5259.5 |
| 20 | Yes | A. baylyi | Fosfomyci | Wall | F138 | Apigenin | 4795 | VL2 | 5314 |
| 20 | Yes | A. baylyi | Fosfomyci | Wall | F65 | H2FDA | 4796 | BL1 | 5598.5 |
| 20 | Yes | A. baylyi | Fosfomyci | Wall | F142 | Chlorophe | 4777 | BL3 | 7478 |
| 20 | Yes | A. baylyi | Fosfomyci | Wall | F39 | Congo Red | 4526 | BL3 | 6945.5 |
| 20 | Yes | A. baylyi | Fosfomyci | Wall | F116 | Sunset Yel | 4924 | BL2 | 7676 |
| 20 | Yes | A. baylyi | Fosfomyci | Wall | F10 | Alizarin | 4896 | BL1 | 6435.5 |
| 20 | Yes | A. baylyi | Fosfomyci | Wall | F84 | Neutral Re | 4953 | BL1 | 5214 |
| 20 | Yes | A. baylyi | Fosfomyci | Wall | F99 | Pyronin Y | 4695 | BL1 | 4400 |
| 20 | Yes | A. baylyi | Fosfomyci | Wall | F108 | Rhodamine | 4882 | BL1 | 4361.5 |
| 20 | Yes | A. baylyi | Fosfomyci | Wall | F109 | Riboflavin | 4495 | BL1 | 4442 |
| 20 | Yes | A. baylyi | Fosfomyci | Wall | F117 | SYBR Gree | 4571 | BL1 | 317579 |
| 20 | Yes | A. baylyi | Fosfomyci | Wall | F106 | Rhodamine | 4908 | BL1 | 5041 |
| 20 | Yes | A. baylyi | Fosfomyci | Wall | F73 | Malachite ( | 4672 | BL1 | 4532 |
| 20 | Yes | A. baylyi | Fosfomyci | Wall | F7 | Acridine or | 4464 | BL1 | 4937 |
| 20 | Yes | A. baylyi | Fosfomyci | Wall | F192 | 5-Carboxyf | 4575 | BL1 | 15027 |
| 20 | Yes | A. baylyi | Fosfomyci | Wall | F57 | Eosin Y | 4521 | BL1 | 14303 |
| 20 | Yes | A. baylyi | Fosfomyci | Wall | F27 | Calcein | 4684 | BL1 | 19008 |

| | | | | | | | | | |
|---|---|---|---|---|---|---|---|---|---|
| 20 | Yes | A. baylyi | Fosfomyci | Wall | F12 | Amaranth | 4848 | BL1 | 16861.5 |
| 20 | Yes | A. baylyi | None | None | NA | Autofluore: | 4679 | NA | 6478 |
| 20 | Yes | A. baylyi | Nalidixic a | Intracellul | F194 | Oxonol | 4744 | RL1 | 5491.5 |
| 20 | Yes | A. baylyi | Nalidixic a | Intracellul | F11 | Allura Red | 4929 | BL3 | 5147 |
| 20 | Yes | A. baylyi | Nalidixic a | Intracellul | F54 | DiSC3(5) | 4762 | RL1 | 5142 |
| 20 | Yes | A. baylyi | Nalidixic a | Intracellul | F124 | Thioflavin T | 4968 | VL2 | 5417 |
| 20 | Yes | A. baylyi | Nalidixic a | Intracellul | F101 | Quinacrine | 4691 | VL2 | 5479 |
| 20 | Yes | A. baylyi | Nalidixic a | Intracellul | F138 | Apigenin | 4641 | VL2 | 5164 |
| 20 | Yes | A. baylyi | Nalidixic a | Intracellul | F65 | H2FDA | 4927 | BL1 | 6158 |
| 20 | Yes | A. baylyi | Nalidixic a | Intracellul | F142 | Chlorophe | 4420 | BL3 | 7225 |
| 20 | Yes | A. baylyi | Nalidixic a | Intracellul | F39 | Congo Red | 4741 | BL3 | 7414 |
| 20 | Yes | A. baylyi | Nalidixic a | Intracellul | F116 | Sunset Yel | 4598 | BL2 | 6667 |
| 20 | Yes | A. baylyi | Nalidixic a | Intracellul | F10 | Alizarin | 4473 | BL1 | 6599 |
| 20 | Yes | A. baylyi | Nalidixic a | Intracellul | F84 | Neutral Re | 4264 | BL1 | 5521.5 |
| 20 | Yes | A. baylyi | Nalidixic a | Intracellul | F99 | Pyronin Y | 4142 | BL1 | 4867 |
| 20 | Yes | A. baylyi | Nalidixic a | Intracellul | F108 | Rhodamine | 4483 | BL1 | 5455 |
| 20 | Yes | A. baylyi | Nalidixic a | Intracellul | F109 | Riboflavin | 4377 | BL1 | 4911 |
| 20 | Yes | A. baylyi | Nalidixic a | Intracellul | F117 | SYBR Gree | 4093 | BL1 | 296677 |
| 20 | Yes | A. baylyi | Nalidixic a | Intracellul | F106 | Rhodamine | 4568 | BL1 | 5570.5 |
| 20 | Yes | A. baylyi | Nalidixic a | Intracellul | F73 | Malachite ( | 3888 | BL1 | 4707.5 |
| 20 | Yes | A. baylyi | Nalidixic a | Intracellul | F7 | Acridine or | 4211 | BL1 | 5492 |
| 20 | Yes | A. baylyi | Nalidixic a | Intracellul | F192 | 5-Carboxyf | 3986 | BL1 | 16174.5 |
| 20 | Yes | A. baylyi | Nalidixic a | Intracellul | F57 | Eosin Y | 4194 | BL1 | 16091 |
| 20 | Yes | A. baylyi | Nalidixic a | Intracellul | F27 | Calcein | 4063 | BL1 | 18542 |
| 20 | Yes | A. baylyi | Nalidixic a | Intracellul | F12 | Amaranth | 4398 | BL1 | 16792.5 |
| 20 | Yes | A. baylyi | None | None | NA | Autofluore: | 4501 | NA | 6060 |
| 20 | Yes | A. baylyi | Ofloxacin | Intracellul | F194 | Oxonol | 3975 | RL1 | 5836 |
| 20 | Yes | A. baylyi | Ofloxacin | Intracellul | F11 | Allura Red | 4161 | BL3 | 6009 |
| 20 | Yes | A. baylyi | Ofloxacin | Intracellul | F54 | DiSC3(5) | 3993 | RL1 | 5705 |
| 20 | Yes | A. baylyi | Ofloxacin | Intracellul | F124 | Thioflavin T | 4092 | VL2 | 5848 |
| 20 | Yes | A. baylyi | Ofloxacin | Intracellul | F101 | Quinacrine | 4035 | VL2 | 5646 |
| 20 | Yes | A. baylyi | Ofloxacin | Intracellul | F138 | Apigenin | 3919 | VL2 | 5560 |
| 20 | Yes | A. baylyi | Ofloxacin | Intracellul | F65 | H2FDA | 4409 | BL1 | 6332 |
| 20 | Yes | A. baylyi | Ofloxacin | Intracellul | F142 | Chlorophe | 4734 | BL3 | 7072.5 |
| 20 | Yes | A. baylyi | Ofloxacin | Intracellul | F39 | Congo Red | 4623 | BL3 | 7819 |
| 20 | Yes | A. baylyi | Ofloxacin | Intracellul | F116 | Sunset Yel | 4794 | BL2 | 7427 |
| 20 | Yes | A. baylyi | Ofloxacin | Intracellul | F10 | Alizarin | 4572 | BL1 | 6643.5 |
| 20 | Yes | A. baylyi | Ofloxacin | Intracellul | F84 | Neutral Re | 4604 | BL1 | 5799.5 |
| 20 | Yes | A. baylyi | Ofloxacin | Intracellul | F99 | Pyronin Y | 4251 | BL1 | 4966 |
| 20 | Yes | A. baylyi | Ofloxacin | Intracellul | F108 | Rhodamine | 4481 | BL1 | 4544 |
| 20 | Yes | A. baylyi | Ofloxacin | Intracellul | F109 | Riboflavin | 4609 | BL1 | 4735 |
| 20 | Yes | A. baylyi | Ofloxacin | Intracellul | F117 | SYBR Gree | 4041 | BL1 | 330922 |
| 20 | Yes | A. baylyi | Ofloxacin | Intracellul | F106 | Rhodamine | 4778 | BL1 | 5685 |
| 20 | Yes | A. baylyi | Ofloxacin | Intracellul | F73 | Malachite ( | 4487 | BL1 | 5037 |
| 20 | Yes | A. baylyi | Ofloxacin | Intracellul | F7 | Acridine or | 4410 | BL1 | 5571 |
| 20 | Yes | A. baylyi | Ofloxacin | Intracellul | F192 | 5-Carboxyf | 4614 | BL1 | 15441.5 |
| 20 | Yes | A. baylyi | Ofloxacin | Intracellul | F57 | Eosin Y | 4562 | BL1 | 16834 |
| 20 | Yes | A. baylyi | Ofloxacin | Intracellul | F27 | Calcein | 4622 | BL1 | 18324.5 |

| | | | | | | | | | |
|---|---|---|---|---|---|---|---|---|---|
| 20 | Yes | A. baylyi | Ofloxacin | Intracellul | F12 | Amaranth | 4786 | BL1 | 17618.5 |
| 20 | Yes | A. baylyi | None | None | NA | Autofluore | 4582 | NA | 6367 |
| 20 | Yes | A. baylyi | Polymyxin | Wall | F194 | Oxonol | 4662 | RL1 | 6155 |
| 20 | Yes | A. baylyi | Polymyxin | Wall | F11 | Allura Red | 4732 | BL3 | 5990 |
| 20 | Yes | A. baylyi | Polymyxin | Wall | F54 | DiSC3(5) | 4664 | RL1 | 6350 |
| 20 | Yes | A. baylyi | Polymyxin | Wall | F124 | Thioflavin T | 5263 | VL2 | 6098 |
| 20 | Yes | A. baylyi | Polymyxin | Wall | F101 | Quinacrine | 5015 | VL2 | 6200 |
| 20 | Yes | A. baylyi | Polymyxin | Wall | F138 | Apigenin | 4809 | VL2 | 6852 |
| 20 | Yes | A. baylyi | Polymyxin | Wall | F65 | H2FDA | 4939 | BL1 | 6420 |
| 20 | Yes | A. baylyi | Polymyxin | Wall | F142 | Chlorophe | 5195 | BL3 | 7619 |
| 20 | Yes | A. baylyi | Polymyxin | Wall | F39 | Congo Red | 5183 | BL3 | 7539 |
| 20 | Yes | A. baylyi | Polymyxin | Wall | F116 | Sunset Yel | 5043 | BL2 | 7037 |
| 20 | Yes | A. baylyi | Polymyxin | Wall | F10 | Alizarin | 4912 | BL1 | 6320 |
| 20 | Yes | A. baylyi | Polymyxin | Wall | F84 | Neutral Re | 4994 | BL1 | 6218 |
| 20 | Yes | A. baylyi | Polymyxin | Wall | F99 | Pyronin Y | 4793 | BL1 | 5892 |
| 20 | Yes | A. baylyi | Polymyxin | Wall | F108 | Rhodamine | 4666 | BL1 | 4911.5 |
| 20 | Yes | A. baylyi | Polymyxin | Wall | F109 | Riboflavin | 4982 | BL1 | 5178 |
| 20 | Yes | A. baylyi | Polymyxin | Wall | F117 | SYBR Gree | 4472 | BL1 | 659595.5 |
| 20 | Yes | A. baylyi | Polymyxin | Wall | F106 | Rhodamine | 5016 | BL1 | 5529 |
| 20 | Yes | A. baylyi | Polymyxin | Wall | F73 | Malachite | 4591 | BL1 | 5230 |
| 20 | Yes | A. baylyi | Polymyxin | Wall | F7 | Acridine or | 4675 | BL1 | 5621 |
| 20 | Yes | A. baylyi | Polymyxin | Wall | F192 | 5-Carboxyf | 4674 | BL1 | 15975 |
| 20 | Yes | A. baylyi | Polymyxin | Wall | F57 | Eosin Y | 4652 | BL1 | 14677.5 |
| 20 | Yes | A. baylyi | Polymyxin | Wall | F27 | Calcein | 4790 | BL1 | 18176 |
| 20 | Yes | A. baylyi | Polymyxin | Wall | F12 | Amaranth | 4837 | BL1 | 17289 |
| 20 | Yes | A. baylyi | None | None | NA | Autofluore | 4618 | NA | 6453 |
| 20 | Yes | A. baylyi | Tetracyclin | Intracellul | F194 | Oxonol | 3352 | RL1 | 5868 |
| 20 | Yes | A. baylyi | Tetracyclin | Intracellul | F11 | Allura Red | 3557 | BL3 | 5721 |
| 20 | Yes | A. baylyi | Tetracyclin | Intracellul | F54 | DiSC3(5) | 3884 | RL1 | 5667 |
| 20 | Yes | A. baylyi | Tetracyclin | Intracellul | F124 | Thioflavin T | 3551 | VL2 | 6208 |
| 20 | Yes | A. baylyi | Tetracyclin | Intracellul | F101 | Quinacrine | 3562 | VL2 | 6096 |
| 20 | Yes | A. baylyi | Tetracyclin | Intracellul | F138 | Apigenin | 3543 | VL2 | 7110 |
| 20 | Yes | A. baylyi | Tetracyclin | Intracellul | F65 | H2FDA | 3553 | BL1 | 7048 |
| 20 | Yes | A. baylyi | Tetracyclin | Intracellul | F142 | Chlorophe | 3839 | BL3 | 8249 |
| 20 | Yes | A. baylyi | Tetracyclin | Intracellul | F39 | Congo Red | 3723 | BL3 | 8785 |
| 20 | Yes | A. baylyi | Tetracyclin | Intracellul | F116 | Sunset Yel | 3261 | BL2 | 8100 |
| 20 | Yes | A. baylyi | Tetracyclin | Intracellul | F10 | Alizarin | 3708 | BL1 | 6891.5 |
| 20 | Yes | A. baylyi | Tetracyclin | Intracellul | F84 | Neutral Re | 3686 | BL1 | 5927 |
| 20 | Yes | A. baylyi | Tetracyclin | Intracellul | F99 | Pyronin Y | 3663 | BL1 | 5200 |
| 20 | Yes | A. baylyi | Tetracyclin | Intracellul | F108 | Rhodamine | 3604 | BL1 | 4748.5 |
| 20 | Yes | A. baylyi | Tetracyclin | Intracellul | F109 | Riboflavin | 3305 | BL1 | 5033 |
| 20 | Yes | A. baylyi | Tetracyclin | Intracellul | F117 | SYBR Gree | 3546 | BL1 | 288981.5 |
| 20 | Yes | A. baylyi | Tetracyclin | Intracellul | F106 | Rhodamine | 3666 | BL1 | 5843.5 |
| 20 | Yes | A. baylyi | Tetracyclin | Intracellul | F73 | Malachite | 3628 | BL1 | 5301.5 |
| 20 | Yes | A. baylyi | Tetracyclin | Intracellul | F7 | Acridine or | 3399 | BL1 | 5774 |
| 20 | Yes | A. baylyi | Tetracyclin | Intracellul | F192 | 5-Carboxyf | 3282 | BL1 | 17050 |
| 20 | Yes | A. baylyi | Tetracyclin | Intracellul | F57 | Eosin Y | 3263 | BL1 | 15568 |
| 20 | Yes | A. baylyi | Tetracyclin | Intracellul | F27 | Calcein | 3495 | BL1 | 21341 |

| | | | | | | | | | |
|---|---|---|---|---|---|---|---|---|---|
| 20 | Yes | A. baylyi | Tetracyclin | Intracellula | F12 | Amaranth | 3250 | BL1 | 19182.5 |
| 20 | Yes | A. baylyi | None | None | NA | Autofluores | 4560 | NA | 6283.5 |
| 20 | Yes | A. baylyi | Vancomyc | Wall | F194 | Oxonol | 4922 | RL1 | 5668.5 |
| 20 | Yes | A. baylyi | Vancomyc | Wall | F11 | Allura Red | 5218 | BL3 | 5540 |
| 20 | Yes | A. baylyi | Vancomyc | Wall | F54 | DiSC3(5) | 4895 | RL1 | 5485 |
| 20 | Yes | A. baylyi | Vancomyc | Wall | F124 | Thioflavin T | 4901 | VL2 | 6055 |
| 20 | Yes | A. baylyi | Vancomyc | Wall | F101 | Quinacrine | 5071 | VL2 | 5550 |
| 20 | Yes | A. baylyi | Vancomyc | Wall | F138 | Apigenin | 4941 | VL2 | 6087 |
| 20 | Yes | A. baylyi | Vancomyc | Wall | F65 | H2FDA | 5071 | BL1 | 6709 |
| 20 | Yes | A. baylyi | Vancomyc | Wall | F142 | Chlorophe | 4597 | BL3 | 8384 |
| 20 | Yes | A. baylyi | Vancomyc | Wall | F39 | Congo Red | 4951 | BL3 | 10201 |
| 20 | Yes | A. baylyi | Vancomyc | Wall | F116 | Sunset Yel | 5018 | BL2 | 9056 |
| 20 | Yes | A. baylyi | Vancomyc | Wall | F10 | Alizarin | 5256 | BL1 | 7059 |
| 20 | Yes | A. baylyi | Vancomyc | Wall | F84 | Neutral Re | 5187 | BL1 | 5643 |
| 20 | Yes | A. baylyi | Vancomyc | Wall | F99 | Pyronin Y | 4303 | BL1 | 4882 |
| 20 | Yes | A. baylyi | Vancomyc | Wall | F108 | Rhodamine | 5187 | BL1 | 4601 |
| 20 | Yes | A. baylyi | Vancomyc | Wall | F109 | Riboflavin | 5066 | BL1 | 4955 |
| 20 | Yes | A. baylyi | Vancomyc | Wall | F117 | SYBR Gree | 4229 | BL1 | 315222 |
| 20 | Yes | A. baylyi | Vancomyc | Wall | F106 | Rhodamine | 4890 | BL1 | 7193 |
| 20 | Yes | A. baylyi | Vancomyc | Wall | F73 | Malachite ( | 4855 | BL1 | 5573 |
| 20 | Yes | A. baylyi | Vancomyc | Wall | F7 | Acridine or | 4926 | BL1 | 4853.5 |
| 20 | Yes | A. baylyi | Vancomyc | Wall | F192 | 5-Carboxyf | 4719 | BL1 | 15326 |
| 20 | Yes | A. baylyi | Vancomyc | Wall | F57 | Eosin Y | 4257 | BL1 | 15047 |
| 20 | Yes | A. baylyi | Vancomyc | Wall | F27 | Calcein | 4338 | BL1 | 16752 |
| 20 | Yes | A. baylyi | Vancomyc | Wall | F12 | Amaranth | 4664 | BL1 | 15564.5 |

| BL2 | BL3 | BL4 | BL5 | RL1 | RL2 | VL1 | VL2 | VL3 | VL4 |
|---|---|---|---|---|---|---|---|---|---|
| 690 | 896 | 734 | 572 | 1319 | 1114 | 257 | 2272 | 1339 | 1773 |
| 874 | 1418 | 1261 | 678 | 2002 | 1278 | 419 | 3843 | 1953 | 2878 |
| 1051 | 1850 | 1703 | 783 | 2561 | 1444 | 548 | 5704 | 2516 | 3878 |
| 1069 | 1901 | 4699 | 1543 | 812677 | 79086 | 572 | 5485 | 2591 | 3935 |
| 1070 | 1869 | 2361 | 929 | 5122 | 2009 | 564 | 5425 | 2562 | 3904 |
| 1093 | 1964 | 2012 | 876 | 3504 | 1628 | 584 | 5727 | 2613 | 4003 |
| 1102 | 1989 | 1946 | 857 | 3232 | 1575 | 587 | 6174 | 2639 | 4064 |
| 1289 | 2212 | 2087 | 887 | 3291.5 | 1610.5 | 618 | 7199 | 2927 | 4372.5 |
| 1450 | 2322 | 2003 | 880 | 3079 | 1564 | 598 | 7773 | 2984 | 4367 |
| 1505 | 2411 | 2018 | 881 | 2966 | 1565 | 600 | 7667 | 2981 | 4340 |
| 1635 | 2761 | 2340 | 953 | 3451.5 | 1681 | 691 | 9765 | 3613 | 5335 |
| 1424 | 2430 | 2102 | 912 | 3080 | 1604 | 632 | 8321 | 3157 | 4718 |
| 1383 | 2552 | 2294 | 960 | 3488 | 1688 | 702 | 10023 | 3530 | 5368 |
| 1329 | 2308 | 2109 | 913 | 3054 | 1588 | 647 | 7584.5 | 3038 | 4652 |
| 1267 | 2260 | 2048 | 884 | 2995 | 1560 | 602 | 6349 | 2823 | 4311 |
| 1194 | 2031 | 1778.5 | 803 | 2606 | 1414 | 516 | 5298 | 2512 | 3773 |
| 23041 | 13694 | 5005 | 1156 | 3867 | 1769 | 695 | 13059 | 4831 | 6616 |
| 1964 | 3563 | 2842 | 1007 | 3716 | 1685 | 636 | 8409 | 3699 | 5596 |
| 1863 | 3402 | 2780 | 1037 | 3926 | 1784 | 715 | 11223 | 4339 | 6500.5 |
| 2168 | 3863 | 3089 | 1096.5 | 3811 | 1879.5 | 858 | 16922.5 | 5240.5 | 7884 |
| 9882 | 19152 | 5018 | 1279 | 5481 | 2040 | 781 | 24635 | 7725.5 | 19315 |
| 25526.5 | 23714 | 7862.5 | 1649.5 | 6831 | 2390 | 910 | 20212.5 | 10983 | 20846.5 |
| 8977 | 20495.5 | 6151.5 | 1428 | 6613 | 2131 | 638 | 16161.5 | 7995.5 | 21121.5 |
| 12200.5 | 22669 | 6664 | 1479 | 6803 | 2176.5 | 693 | 16154 | 9985.5 | 22992 |
| 2512 | 3477 | 1867 | 704 | 2531 | 1289 | 353 | 7845 | 3393 | 4331 |
| 1723 | 2639 | 1703 | 734 | 2430 | 1322 | 435 | 6081 | 2742 | 3779 |
| 1649 | 2733 | 2038 | 826 | 2905 | 1476 | 550 | 7354 | 3059 | 4464 |
| 2124 | 3341 | 6450 | 1606 | 865094 | 79134 | 759 | 11791 | 4214 | 6090 |
| 1584 | 2697 | 2975 | 1025 | 6184 | 2270 | 631 | 7642 | 3192 | 4803 |
| 1451 | 2393 | 2243 | 909 | 3972.5 | 1734 | 587 | 6053.5 | 2886 | 4305 |
| 1409 | 2464 | 2185 | 898 | 3668.5 | 1646.5 | 599 | 6761 | 2918.5 | 4461.5 |
| 1425 | 2404 | 2040 | 869 | 3267 | 1559 | 583 | 6686 | 2870 | 4299 |
| 1611 | 2574 | 2021 | 851 | 3150 | 1533 | 576 | 7507 | 3008 | 4372 |
| 1836.5 | 2904 | 2330 | 936.5 | 3399.5 | 1616 | 629 | 9749.5 | 3528 | 5058 |
| 1531 | 2381 | 1968 | 863 | 2855 | 1515 | 561 | 7378 | 2870 | 4180 |
| 1484 | 2530 | 2132 | 887 | 3172 | 1566 | 599 | 7532 | 3059 | 4539 |
| 1556 | 2709 | 2307 | 946 | 3349 | 1664 | 658 | 8345 | 3287 | 5000 |
| 1680 | 2709 | 2282 | 936 | 3188 | 1641 | 688 | 7773 | 3405 | 5153 |
| 1550 | 2713 | 2304 | 944 | 3380 | 1635 | 651 | 7556.5 | 3259 | 4992 |
| 1594 | 2863 | 2372.5 | 944 | 3467 | 1641 | 645 | 8010 | 3366.5 | 5145.5 |
| 25415.5 | 16240 | 5384 | 1229 | 4324 | 1883 | 763 | 23986.5 | 5908 | 8867 |
| 3279 | 5321 | 3632 | 1117 | 4308 | 1844 | 743 | 23262 | 5783 | 8117 |
| 2528 | 4432.5 | 3172 | 1094 | 4158 | 1841 | 790 | 19154 | 5650 | 8189 |
| 2580 | 4288 | 3211.5 | 1087 | 3928 | 1869 | 889.5 | 139120.5 | 6200.5 | 8847.5 |
| 4761 | 7466 | 3791 | 1136 | 4506 | 1861 | 708 | 16911 | 6319 | 9836 |
| 9874 | 22077 | 6612 | 1517 | 6788 | 2231 | 701 | 17235 | 8603 | 22754 |
| 7935.5 | 19250.5 | 6000.5 | 1459 | 6735.5 | 2142 | 617 | 14032 | 7467 | 20274 |

| | | | | | | | | | |
|---|---|---|---|---|---|---|---|---|---|
| 11147 | 22440 | 6191 | 1459 | 6889 | 2182 | 656 | 13744 | 9261 | 22394 |
| 2239 | 3109 | 1683 | 677 | 2343.5 | 1263 | 301 | 6976 | 3019 | 3847 |
| 2175.5 | 3157 | 2122.5 | 850 | 2922.5 | 1485 | 518 | 7424.5 | 3279 | 4536.5 |
| 2163 | 3255.5 | 2465.5 | 986 | 3339.5 | 1657 | 664 | 8537.5 | 3743 | 5236.5 |
| 1897 | 2890 | 6360 | 1731 | 1184403 | 127436 | 688 | 8480 | 3461 | 4891 |
| 1801 | 2875 | 2813 | 1039 | 4832.5 | 1913 | 698 | 8232.5 | 3428.5 | 4970.5 |
| 1800 | 2927.5 | 2612 | 1019 | 4285 | 1798 | 717 | 8469 | 3506.5 | 5159.5 |
| 1911.5 | 3147 | 2812 | 1091 | 4348 | 1862 | 783.5 | 9050 | 3811 | 5596 |
| 2011 | 3191 | 2768 | 1067 | 4075.5 | 1791 | 778 | 9599 | 3874 | 5602 |
| 2327 | 3460 | 2862.5 | 1104 | 4180.5 | 1822.5 | 782 | 10494.5 | 4307 | 5834 |
| 2649 | 4087 | 3288 | 1237 | 4708 | 2006 | 860 | 12056 | 4956 | 6863 |
| 2370 | 3791 | 3006 | 1178 | 4290 | 1915 | 825 | 11463 | 4594 | 6469 |
| 2196 | 3510 | 3076 | 1181 | 4303 | 1924 | 847 | 10894 | 4406 | 6378 |
| 2169 | 3602 | 3078 | 1215 | 4331 | 1958 | 858 | 10381 | 4534 | 6485 |
| 2014 | 3175 | 2756 | 1079.5 | 3838 | 1807 | 793 | 9331 | 3946.5 | 5758 |
| 2030 | 3351 | 2833 | 1121 | 3968 | 1830 | 799 | 9434 | 4108 | 6004 |
| 2264.5 | 3677.5 | 3138.5 | 1188 | 4336.5 | 1897 | 805.5 | 9670.5 | 4463.5 | 6354.5 |
| 20215 | 12658.5 | 5531.5 | 1381 | 4834 | 2023 | 830 | 12584.5 | 5477 | 7897.5 |
| 2713 | 4355 | 3444 | 1204.5 | 4468 | 1922 | 789 | 10369 | 4814 | 6901 |
| 2530 | 4276 | 3471 | 1278 | 4776 | 2024 | 858 | 11023 | 4991 | 7282 |
| 2261 | 3836 | 3096 | 1186 | 4312 | 1907 | 833 | 10461 | 4513 | 6596 |
| 5698 | 11172 | 5006.5 | 1494 | 6145 | 2231 | 844 | 15944 | 6882.5 | 14021 |
| 7439.5 | 18786 | 6431.5 | 1679 | 7327.5 | 2371 | 780.5 | 14441.5 | 6849.5 | 20418.5 |
| 10857 | 21687 | 6502 | 1601 | 7007 | 2238 | 736 | 15919 | 8620 | 21503 |
| 12689 | 23596 | 7202 | 1728 | 7812 | 2394 | 825 | 12971 | 9605 | 23288 |
| 2541 | 3491 | 1902 | 709 | 2637 | 1310 | 333 | 8109 | 3368 | 4333.5 |
| 2072 | 3190 | 2099.5 | 824 | 2970 | 1441 | 486 | 7461 | 3272 | 4499 |
| 1972 | 3168 | 2391 | 929 | 3343 | 1607 | 615 | 7970 | 3478 | 5053 |
| 2167.5 | 3412.5 | 5819 | 1646 | 805043 | 85290 | 703 | 9291 | 3902.5 | 5682.5 |
| 1743 | 2887 | 2850 | 1044 | 5087 | 2013 | 638 | 7654 | 3291 | 4868 |
| 1696 | 2804 | 2508 | 978 | 4076 | 1758 | 638 | 7582 | 3267 | 4855 |
| 1631 | 2724 | 2371 | 949 | 3652 | 1649 | 632 | 7473 | 3151 | 4709 |
| 1956 | 3279 | 2763 | 1061 | 4262 | 1822 | 726 | 9436 | 3863 | 5751 |
| 2137.5 | 3342 | 2690 | 1038 | 4025 | 1778.5 | 692 | 9945 | 4008.5 | 5583 |
| 2293.5 | 3549 | 2792 | 1065.5 | 3962.5 | 1783 | 707 | 10338 | 4195 | 5875 |
| 2202 | 3519 | 2899 | 1093 | 4161 | 1848 | 738 | 10625 | 4293.5 | 6033 |
| 2175 | 3587 | 2883 | 1134 | 4144 | 1865 | 773 | 10878 | 4431 | 6428 |
| 2062 | 3535 | 2891 | 1098 | 4091 | 1853 | 778 | 10831 | 4216 | 6482 |
| 1870 | 3181 | 2693.5 | 1045 | 3850 | 1795 | 721 | 8859 | 3799 | 5671 |
| 1720 | 2982 | 2502 | 1009 | 3625 | 1711 | 671 | 7905 | 3442 | 5180 |
| 2222 | 3911 | 3092 | 1146 | 4293 | 1892 | 758 | 10106 | 4530 | 6935 |
| 20424 | 11860 | 4930 | 1248 | 4419 | 1888 | 718 | 11555 | 4953 | 6929 |
| 2761 | 4626 | 3522 | 1181 | 4609 | 1929 | 742 | 10719 | 4803 | 7168 |
| 2228 | 3695 | 3019 | 1099 | 4172 | 1813 | 698 | 8922 | 4209 | 6147 |
| 2392 | 4090 | 3303 | 1196 | 4609 | 1964 | 808 | 12016 | 4958 | 7264 |
| 6681 | 11795 | 4445 | 1320 | 5429 | 2068 | 754 | 15026 | 7445 | 13698 |
| 8398 | 19170 | 6635 | 1640 | 7225 | 2329 | 739 | 16186 | 8089 | 20602 |
| 6256 | 18027 | 5927 | 1489 | 6838 | 2140 | 634 | 14179 | 6360 | 19946 |

| | | | | | | | | | |
|---|---|---|---|---|---|---|---|---|---|
| 8736 | 18139 | 5665 | 1410 | 6593 | 2096 | 629 | 13960 | 8489 | 19672 |
| 2128 | 3080 | 1703 | 697 | 2437 | 1269 | 303 | 6608 | 2936 | 3882 |
| 2073 | 3060 | 2045 | 823 | 2837 | 1452 | 488 | 6877 | 3164 | 4337 |
| 2052 | 3190 | 2418 | 949 | 3331 | 1616 | 609 | 7837 | 3510 | 5013 |
| 1889 | 2946 | 5526 | 1588.5 | 865782.5 | 97816.5 | 634 | 7852 | 3345 | 4828 |
| 1852.5 | 3031 | 2882.5 | 1064 | 4682 | 1939.5 | 674 | 7959.5 | 3484 | 5084.5 |
| 1842 | 3060 | 2720 | 1020 | 4255 | 1817 | 685 | 8230 | 3542 | 5221 |
| 1892 | 3158.5 | 2722 | 1063 | 4107 | 1797 | 715.5 | 8671 | 3654 | 5466 |
| 2009 | 3266 | 2741.5 | 1062 | 4086 | 1821 | 724 | 8983.5 | 3804 | 5613 |
| 2486 | 3762 | 3070 | 1141 | 4413 | 1889 | 766 | 11703.5 | 4580.5 | 6272 |
| 2686 | 4146 | 3160.5 | 1182 | 4444 | 1959 | 800 | 12259 | 5068.5 | 7028 |
| 2569 | 4103 | 3211 | 1220 | 4611 | 1987 | 808 | 12782 | 4864 | 6943 |
| 2080 | 3470 | 2876 | 1117 | 4036 | 1842 | 758 | 10169 | 4156 | 5994 |
| 1880 | 3180 | 2735 | 1070 | 3884 | 1798 | 735 | 8951 | 3779 | 5601 |
| 2391 | 3764.5 | 3176 | 1191 | 4434 | 1971 | 818 | 10505.5 | 4589 | 6717 |
| 2286 | 3885 | 3193 | 1184 | 4367 | 1959 | 828 | 10608 | 4528 | 6822 |
| 2007 | 3352 | 2788 | 1067 | 3870 | 1772 | 704 | 8435 | 3841 | 5641 |
| 15361 | 9523 | 4639 | 1257 | 4375 | 1895 | 729 | 10700 | 4778 | 6805 |
| 2596 | 4230 | 3263 | 1132 | 4090 | 1802 | 689 | 9269 | 4194 | 6144 |
| 2089 | 3527 | 2923 | 1086 | 3981 | 1799 | 693 | 8749 | 3929 | 5981 |
| 1981 | 3292 | 2707.5 | 1038 | 3797 | 1742 | 693 | 8506 | 3809 | 5573 |
| 5527 | 13913 | 5066 | 1428 | 6022 | 2132 | 756 | 14729 | 6658 | 16833 |
| 7680.5 | 17204 | 5693.5 | 1490 | 6522 | 2164 | 694 | 15434 | 7909 | 19016 |
| 10517 | 22158 | 6835 | 1687 | 7597 | 2359 | 723 | 14980 | 9596 | 23265 |
| 9980 | 19971 | 6272.5 | 1559 | 7029.5 | 2215 | 706 | 13997.5 | 9159 | 21422 |
| 2099 | 3035 | 1671 | 682 | 2422 | 1271 | 290 | 6663 | 2905 | 3795 |
| 12480 | 7617.5 | 3304.5 | 900 | 3096 | 1513 | 514 | 8594.5 | 3710 | 4690 |
| 17856.5 | 10304 | 3955.5 | 1006.5 | 3471 | 1635 | 621 | 9787 | 4184.5 | 5302 |
| 15446 | 9397 | 6304.5 | 1555 | 176211.5 | 19320.5 | 657.5 | 9377.5 | 4060 | 5460.5 |
| 12961 | 8022 | 4040 | 1111 | 15716 | 2774 | 710 | 9657 | 4039 | 5223 |
| 16288 | 10169 | 4192 | 1091 | 6758 | 2025 | 747 | 10474 | 4368 | 5657 |
| 13047.5 | 8234.5 | 4134 | 1156 | 5961.5 | 2064 | 803 | 11397 | 4593 | 6073 |
| 13305.5 | 8305.5 | 3903.5 | 1093 | 4947.5 | 1910 | 757 | 11163.5 | 4511 | 5880.5 |
| 14520 | 9554 | 4676 | 1310 | 5487 | 2142 | 881 | 16273 | 6048 | 7823 |
| 10142 | 9912 | 12259 | 5574 | 5584 | 2192 | 754 | 14055 | 5156 | 7412 |
| 14249 | 9864 | 4742 | 1414 | 5068 | 2121 | 886 | 14459 | 5992 | 7962 |
| 12407 | 7993.5 | 4035 | 1187.5 | 4453 | 1941 | 812.5 | 12115 | 5051.5 | 6523 |
| 12241 | 7974 | 4011 | 1206 | 4359 | 1950 | 838 | 11782 | 4996 | 6564 |
| 15915 | 10344 | 4242 | 1093 | 3777 | 1732 | 752 | 10158 | 5433 | 6502 |
| 15859.5 | 10519.5 | 4168 | 1038.5 | 3660.5 | 1672 | 736.5 | 9906.5 | 5215.5 | 6156.5 |
| 15810.5 | 10361 | 4306.5 | 1145.5 | 4103.5 | 1799 | 742 | 9934 | 5627 | 6776 |
| 1350611 | 1054850 | 306546 | 22343 | 5417 | 2236 | 879 | 128967.5 | 40452 | 31179.5 |
| 41056 | 23840 | 7860 | 1449 | 4649 | 1927 | 780 | 13280 | 6450 | 7873 |
| 32351 | 18664 | 6421 | 1281 | 4268 | 1850 | 753 | 11655 | 5595 | 6870 |
| 52652 | 29938 | 8883 | 1341 | 4100 | 1819 | 761 | 15875 | 6580 | 7218 |
| 38515 | 33717.5 | 8206.5 | 1479 | 5331.5 | 1938 | 744 | 17913 | 7580.5 | 13301 |
| 37938 | 45484 | 9932 | 1810 | 7836 | 2459 | 804 | 18257 | 9213 | 22752 |
| 37700.5 | 42696 | 9756.5 | 1789 | 7598.5 | 2356.5 | 737 | 16413.5 | 10180 | 23468 |

| | | | | | | | | | |
|---|---|---|---|---|---|---|---|---|---|
| 34959 | 42412 | 9333 | 1706 | 7267 | 2230 | 738 | 15675 | 9480 | 22332 |
| 2062 | 3019 | 1719 | 690 | 2496 | 1279 | 310 | 6595 | 2896 | 3831 |
| 1883 | 2904 | 1902 | 765 | 2717 | 1380 | 446 | 6595 | 2919 | 4049 |
| 1844 | 2938 | 2111 | 839 | 3021 | 1481 | 526 | 7245 | 3115 | 4483 |
| 1746 | 2792 | 4984 | 1425 | 781011 | 73411 | 576 | 7256 | 3144 | 4559 |
| 1894 | 3163 | 2865 | 1013 | 5287 | 1964 | 647 | 8562 | 3617 | 5260 |
| 1868 | 3147 | 2686 | 997 | 4592 | 1815 | 660 | 8610 | 3591 | 5317 |
| 2193 | 3565 | 2923 | 1048 | 4723 | 1890 | 745 | 13459 | 4696 | 6498 |
| 1938 | 3144 | 2554 | 961 | 4085 | 1738 | 649 | 8976 | 3671 | 5301 |
| 2490 | 3815 | 2862 | 1021 | 4291 | 1797.5 | 743 | 11857 | 4538.5 | 6393.5 |
| 2041 | 3150 | 2408 | 948 | 3525 | 1631 | 625 | 9510 | 3731 | 5218 |
| 2079 | 3336 | 2587 | 990 | 3790 | 1723 | 675 | 9546 | 3940 | 5643 |
| 2158 | 3494 | 2788.5 | 1037.5 | 3989.5 | 1781 | 733 | 11954.5 | 4366 | 6319 |
| 2372 | 3768 | 2942 | 1056 | 4138 | 1840 | 817 | 80518 | 5595 | 7608 |
| 2077 | 3270 | 2571 | 981 | 3659 | 1718 | 701 | 9580 | 3968 | 5732 |
| 2006 | 3183 | 2472 | 957 | 3587 | 1681 | 672 | 10020.5 | 3979.5 | 5582 |
| 2429 | 4106 | 3029 | 1041 | 4117 | 1814 | 773 | 11902 | 4785 | 7033 |
| 41604 | 23945 | 7014 | 1298 | 4224 | 1821 | 686 | 13853 | 5493 | 7310 |
| 2964 | 4859 | 3600 | 1143 | 4272 | 1862 | 751 | 13816 | 5305 | 7701 |
| 2157 | 3744 | 2771 | 995 | 3758 | 1715 | 653 | 9423 | 4070 | 6208 |
| 2971 | 4850 | 3400 | 1113 | 3840 | 1878 | 854 | 13717 | 5247 | 8001 |
| 4217 | 7333 | 4037.5 | 1171 | 4856.5 | 1922 | 672 | 16741.5 | 5396.5 | 9418.5 |
| 7870 | 17933 | 5990 | 1491 | 6828 | 2207 | 673 | 13854 | 7981 | 19992 |
| 10862 | 24083 | 7086 | 1570 | 7627 | 2317 | 679 | 14982 | 9393 | 24569 |
| 10409 | 23454 | 6475 | 1540 | 7320 | 2270 | 675 | 12775 | 8910 | 23271 |
| 2120 | 3018.5 | 1664 | 679 | 2464 | 1261 | 286 | 6793 | 2866 | 3786 |
| 2203 | 3235 | 2241 | 861 | 2983 | 1470 | 553 | 7867 | 3346 | 4625 |
| 2153 | 3246 | 2516 | 966 | 3285 | 1600 | 676 | 8507 | 3621 | 5058 |
| 2211 | 3360 | 6371 | 1791 | 950204 | 104793 | 749 | 9421 | 3935 | 5532 |
| 2221 | 3514 | 3207 | 1162 | 5458 | 2060 | 780 | 9829 | 4034 | 5745 |
| 2344 | 3792 | 3343 | 1207.5 | 5254.5 | 2018 | 847 | 10677 | 4472 | 6352 |
| 2241 | 3579 | 3116 | 1164 | 4753.5 | 1938.5 | 832 | 10174 | 4383 | 6259 |
| 2559.5 | 3952 | 3312.5 | 1192 | 4871.5 | 1963 | 861 | 11276.5 | 4784.5 | 6660.5 |
| 2507 | 3758 | 3011 | 1141 | 4343 | 1880 | 822 | 10973 | 4489 | 6203 |
| 2438.5 | 3583 | 2897.5 | 1116 | 3926 | 1792 | 786 | 10443.5 | 4295.5 | 5842 |
| 2380 | 3645 | 2979 | 1121 | 4116 | 1852 | 803 | 10628 | 4370 | 6123 |
| 2089 | 3241 | 2696 | 1069 | 3702 | 1725 | 765 | 9818 | 3919 | 5464 |
| 2165 | 3462.5 | 2939 | 1130 | 4096 | 1847 | 808 | 9835 | 4248 | 5939.5 |
| 2060 | 3297 | 2770 | 1083 | 3843 | 1787 | 788 | 9698 | 3990 | 5682 |
| 2010 | 3133 | 2664 | 1041 | 3591.5 | 1718.5 | 752 | 8920 | 3772.5 | 5351 |
| 2341 | 3717 | 3096 | 1150 | 4100 | 1839 | 793 | 10142 | 4480 | 6275 |
| 24463 | 14251 | 5634 | 1278 | 4164 | 1812 | 773 | 12207 | 5040 | 6672 |
| 2843 | 4386 | 3454 | 1148 | 4037 | 1772 | 763 | 10485 | 4594 | 6380 |
| 2377 | 3746 | 3169 | 1125 | 3997 | 1792 | 784 | 9927 | 4352 | 6204 |
| 2208 | 3471 | 2861 | 1076 | 3758.5 | 1737 | 757 | 9453 | 4002.5 | 5804.5 |
| 5187.5 | 12559 | 4785 | 1343.5 | 5374.5 | 2004.5 | 778 | 14812 | 5689 | 14970.5 |
| 9094 | 18847 | 6464 | 1633 | 7008 | 2273 | 796 | 15077 | 8703 | 20716 |
| 10791 | 25006.5 | 8464.5 | 1931 | 9011 | 2615 | 857 | 11981.5 | 8351.5 | 24692 |

| | | | | | | | | | |
|---|---|---|---|---|---|---|---|---|---|
| 12471.5 | 21953 | 6853.5 | 1636 | 7244.5 | 2271.5 | 815 | 15427 | 9517 | 22269 |
| 2152 | 3112 | 1793 | 694 | 2585 | 1293 | 303 | 6929 | 3005 | 3967 |
| 2247.5 | 3186 | 2156 | 826 | 3220.5 | 1496 | 467 | 7342.5 | 3162 | 4334.5 |
| 2172 | 3255 | 2351 | 912 | 3745.5 | 1614 | 569 | 7825.5 | 3310.5 | 4753 |
| 2438 | 3609 | 7121 | 1947 | 1388141 | 117681.5 | 650 | 8951.5 | 3794.5 | 5436.5 |
| 2730 | 4049 | 3764 | 1198 | 17085 | 3090 | 682 | 9874 | 4379 | 6034 |
| 2933 | 4573 | 3678 | 1218 | 13482 | 2749 | 754 | 11314 | 4784 | 6717 |
| 2589.5 | 4014 | 3214 | 1130 | 10267 | 2395 | 707 | 9939 | 4323 | 6187.5 |
| 2372 | 3559 | 2820 | 1041 | 9342 | 2195 | 666 | 9066 | 3890 | 5463 |
| 2726 | 3897 | 3044 | 1074 | 8135.5 | 2137 | 699 | 10527 | 4437 | 5910 |
| 2902.5 | 4621 | 3991 | 1378.5 | 8125.5 | 2362 | 723 | 11257 | 4673 | 6577.5 |
| 2733 | 4077 | 3175 | 1169 | 7208 | 2146 | 728 | 10811 | 4561 | 6271 |
| 2637 | 3989.5 | 3263 | 1149.5 | 6505.5 | 2112 | 744 | 10864.5 | 4542 | 6247.5 |
| 2455 | 3797.5 | 3155 | 1129 | 6193 | 2067 | 751 | 9903 | 4325 | 6082.5 |
| 4884 | 5165 | 3461 | 1115 | 5653 | 1975 | 711 | 9213 | 5059 | 6496 |
| 3076 | 4689 | 3494 | 1188 | 5905 | 2065 | 761 | 10482 | 4711 | 6939 |
| 2960 | 5604 | 3764 | 1261 | 6181 | 2156 | 783 | 11351 | 4802 | 8430 |
| 140738 | 122474 | 24077 | 2283 | 6347 | 2186 | 778 | 29114 | 7321 | 13286 |
| 3393 | 6187 | 4081 | 1338 | 6134 | 2120 | 750 | 11484 | 5090 | 8886 |
| 3172 | 5064 | 3675 | 1225.5 | 5568 | 2011 | 732 | 11382.5 | 4749 | 7378.5 |
| 2991 | 5870.5 | 3780 | 1246.5 | 5760.5 | 2084 | 761.5 | 13480 | 4581 | 8374.5 |
| 6996 | 16550 | 5869 | 1586 | 7599 | 2372 | 772 | 15366 | 7831 | 19199 |
| 9791 | 24834 | 8377 | 1949 | 10300 | 2808 | 809 | 14226 | 9455 | 27126 |
| 9954.5 | 22960.5 | 8212.5 | 1860 | 9703 | 2635 | 745 | 14479.5 | 9369.5 | 24466 |
| 9325 | 22977 | 8033 | 1845 | 9775 | 2645 | 769 | 13895 | 9269 | 25083 |
| 2300 | 3196 | 1847 | 719 | 2671 | 1328 | 338 | 8545 | 3134 | 4045.5 |
| 1946 | 2934 | 1988 | 774 | 2830 | 1403 | 445 | 7587 | 2995 | 4147 |
| 1925 | 3121 | 2264 | 853 | 3270 | 1545 | 546 | 8216 | 3323 | 4775 |
| 1917 | 3097 | 5493.5 | 1526 | 784712.5 | 73408 | 606 | 8230 | 3429 | 4962 |
| 2156 | 3442 | 3008 | 1060.5 | 5205 | 2003 | 670 | 10098 | 4068 | 5732.5 |
| 2177 | 3536 | 2866 | 1019 | 4569 | 1868 | 702 | 11374 | 4222 | 6023 |
| 1946 | 3169 | 2548 | 952 | 4010.5 | 1727 | 656 | 9055.5 | 3748.5 | 5420 |
| 1989 | 3179 | 2501 | 946 | 3838 | 1690 | 642 | 9166.5 | 3694 | 5291.5 |
| 2123 | 3229 | 2460 | 931 | 3649 | 1662 | 633 | 9181 | 3726 | 5291 |
| 2054 | 3177.5 | 2538 | 1011 | 3610 | 1743 | 611 | 8902.5 | 3641 | 5112 |
| 1810 | 2907.5 | 2293.5 | 903 | 3364 | 1590 | 581 | 8050 | 3281 | 4741 |
| 1959 | 3135.5 | 2420.5 | 937 | 3454.5 | 1632 | 642 | 9017 | 3607 | 5170 |
| 1901 | 3059.5 | 2452 | 948 | 3481 | 1662 | 656 | 8775 | 3642 | 5233 |
| 1642 | 2710 | 2187 | 890 | 3181 | 1547 | 581 | 7209 | 3085 | 4563 |
| 1887 | 3133 | 2452 | 936 | 3404 | 1634 | 652 | 7924 | 3517 | 5149 |
| 1823 | 3120 | 2460 | 923 | 3484 | 1626 | 602 | 7611.5 | 3468.5 | 5136 |
| 26654 | 15226 | 5242 | 1170.5 | 4178 | 1816 | 697 | 14414.5 | 5206 | 7363.5 |
| 3173 | 5026 | 3392 | 1069 | 4166 | 1804 | 706 | 13280 | 5059 | 7419 |
| 2427 | 3991 | 3009 | 1048 | 4004 | 1772 | 696 | 10280 | 4440 | 6455 |
| 2209 | 3542 | 2737 | 993.5 | 3493 | 1699 | 668 | 9171 | 3843 | 5727 |
| 4468 | 9195.5 | 3854.5 | 1117 | 4821.5 | 1822.5 | 616.5 | 14549.5 | 5616 | 11148 |
| 9190.5 | 20337.5 | 6296 | 1526 | 6994 | 2248 | 698 | 15510 | 8506.5 | 21142.5 |
| 13707 | 27990 | 7141 | 1599 | 7692 | 2326 | 659 | 13497 | 9360 | 24437 |

| | | | | | | | | | |
|---|---|---|---|---|---|---|---|---|---|
| 16046 | 34373 | 6820.5 | 1559 | 7219.5 | 2251 | 672 | 13848.5 | 8726.5 | 27727.5 |
| 2374 | 3428 | 1922 | 725 | 2752 | 1337 | 323 | 7725 | 3241 | 4286 |
| 2205 | 3384 | 2249 | 833 | 3099 | 1471 | 477 | 7433 | 3334 | 4636 |
| 2141 | 3360 | 2499 | 927 | 3447 | 1601 | 582 | 8201 | 3548.5 | 5077 |
| 2291 | 3582 | 6170 | 1710 | 844912.5 | 85052.5 | 683 | 9224 | 3996 | 5664 |
| 2150 | 3394 | 2990 | 1058 | 4977 | 1959 | 670 | 8839 | 3832 | 5461 |
| 1930.5 | 3166.5 | 2666 | 1004 | 4160 | 1774 | 637 | 7968 | 3470 | 5152 |
| 1789 | 2922 | 2419 | 942 | 3686.5 | 1646.5 | 604 | 7438 | 3273 | 4738 |
| 2130 | 3364 | 2684.5 | 1034 | 4013 | 1753 | 669 | 9058.5 | 3922 | 5559 |
| 2272 | 3447 | 2767 | 1038 | 4010 | 1749 | 671 | 9716 | 3994 | 5590 |
| 2065.5 | 3123 | 2435.5 | 981 | 3447.5 | 1653 | 616 | 8726.5 | 3557.5 | 4965 |
| 2541 | 3924 | 2968.5 | 1100 | 4108.5 | 1871 | 740 | 11412 | 4527 | 6580.5 |
| 1938 | 3075 | 2519 | 975 | 3495 | 1675 | 632 | 8403 | 3469 | 5052 |
| 1945 | 3227 | 2615 | 1017 | 3715 | 1714 | 676 | 9091 | 3720 | 5388 |
| 1890 | 2941.5 | 2347 | 964 | 3348 | 1644 | 627 | 8466.5 | 3421.5 | 4884 |
| 2239 | 3782 | 2950 | 1112 | 4188 | 1846 | 745 | 10198 | 4275 | 6349 |
| 2044 | 3383.5 | 2730 | 1021.5 | 3831 | 1716 | 653 | 8341 | 3763 | 5576 |
| 18918 | 10563 | 4602 | 1149 | 4069 | 1772 | 646 | 10659 | 4662 | 6174 |
| 3388 | 5663 | 3657.5 | 1186 | 4554 | 1929 | 726 | 12511 | 5251.5 | 7990 |
| 2246 | 3651 | 2917.5 | 1052 | 3872.5 | 1746 | 649 | 8418 | 3962 | 5697 |
| 1950 | 3258 | 2620 | 974 | 3575.5 | 1662 | 619 | 7800 | 3430 | 5162.5 |
| 4353 | 11909 | 4440 | 1252 | 5408 | 1929 | 646 | 13558 | 5345 | 14455 |
| 12451 | 28853 | 8292 | 1802 | 8888 | 2605 | 783 | 17690 | 9624 | 27696 |
| 9056.5 | 20945 | 6628 | 1612.5 | 7594 | 2326 | 676 | 14166 | 9000.5 | 22614.5 |
| 9293 | 20624 | 7992 | 1705 | 8969.5 | 2441.5 | 743.5 | 14035.5 | 9049 | 21897 |
| 701 | 870 | 742 | 576 | 1317 | 1114 | 247 | 2154 | 1333 | 1743 |
| 838 | 1297 | 1176 | 668 | 1884.5 | 1250 | 382 | 3368 | 1810 | 2608 |
| 1006 | 1723 | 1634 | 760 | 2464 | 1388 | 514 | 4610 | 2337 | 3465 |
| 1111 | 1964 | 4794 | 1522 | 754647 | 76310 | 592 | 6111 | 2658 | 4065 |
| 1120 | 2010 | 2237 | 922 | 3792 | 1813 | 593 | 6307 | 2655 | 4061 |
| 1210 | 2257 | 2292 | 944 | 3605 | 1712 | 651 | 7024.5 | 2962 | 4575 |
| 1231 | 2318 | 2282 | 951 | 3535 | 1692 | 681 | 7953 | 3140 | 4778 |
| 1388 | 2451 | 2288 | 936 | 3430 | 1668.5 | 668 | 7900.5 | 3175 | 4821.5 |
| 1489 | 2500 | 2208.5 | 908 | 3251 | 1614 | 630 | 7842 | 3113 | 4603 |
| 1490 | 2448 | 2136 | 914 | 3140 | 1597 | 616 | 7751 | 3050 | 4440 |
| 1455 | 2434 | 2138 | 915 | 3118 | 1573 | 616 | 7563 | 3024 | 4452 |
| 1440 | 2500 | 2263 | 950 | 3317 | 1633 | 652 | 8111 | 3162 | 4793 |
| 1371 | 2467 | 2269 | 950 | 3384 | 1648 | 673 | 8251.5 | 3181.5 | 4880 |
| 1319 | 2315 | 2174 | 916 | 3148.5 | 1605 | 637 | 7037.5 | 2971 | 4490 |
| 1338 | 2461 | 2298 | 934 | 3404 | 1651 | 679 | 7799 | 3198 | 4874.5 |
| 1426 | 2640 | 2388 | 952 | 3491 | 1666 | 661 | 7866 | 3327 | 5108 |
| 21027 | 12452.5 | 4669.5 | 1086 | 3510 | 1635 | 619 | 9417 | 3769 | 5348.5 |
| 1831 | 3266.5 | 2790 | 1022 | 3765.5 | 1735 | 678 | 9598.5 | 3835 | 5724.5 |
| 1561.5 | 2893 | 2535 | 973 | 3615 | 1683 | 658 | 8189 | 3461 | 5329.5 |
| 1524 | 2816 | 2524 | 984 | 3692 | 1691 | 685 | 8422 | 3496 | 5377 |
| 4551 | 11746 | 4151 | 1200 | 5149 | 1917 | 695 | 14010 | 5579 | 14465 |
| 180779.5 | 131441 | 7449 | 1578 | 7149.5 | 2306 | 797 | 25934 | 11646.5 | 29217 |
| 239740.5 | 217561 | 8723.5 | 1637.5 | 7753 | 2420 | 780 | 20158.5 | 11120 | 34824.5 |

| | | | | | | | | | |
|---|---|---|---|---|---|---|---|---|---|
| 11919 | 22959 | 6017 | 1421 | 6639 | 2114 | 683 | 16059 | 9446 | 22566 |
| 1991 | 2855 | 1561 | 664 | 2186 | 1227 | 297 | 6743 | 2824.5 | 3609.5 |
| 1687 | 2584 | 1733 | 736 | 2472 | 1336 | 436 | 6621 | 2746 | 3830 |
| 1582.5 | 2619.5 | 2062 | 842 | 2960 | 1498 | 569 | 7203 | 3052 | 4438 |
| 1541 | 2573 | 5281 | 1561 | 817405 | 79863 | 622 | 7551 | 3157 | 4620 |
| 1367 | 2288 | 2311 | 914 | 3987.5 | 1743 | 586 | 6447 | 2796 | 4150 |
| 1373 | 2344 | 2225 | 907.5 | 3578 | 1646 | 606 | 6640 | 2892 | 4315 |
| 1416 | 2472 | 2236 | 925 | 3534 | 1625 | 631 | 7057 | 3058 | 4572 |
| 1529 | 2596 | 2316 | 945 | 3461 | 1665 | 651 | 8273 | 3237 | 4807 |
| 1694 | 2812 | 2394 | 948 | 3551 | 1646 | 659 | 8612 | 3375 | 4982 |
| 1880 | 3085 | 2530 | 990 | 3699 | 1711 | 690 | 10547 | 3813 | 5489 |
| 1664.5 | 2721 | 2300 | 938.5 | 3348 | 1634 | 648 | 8477 | 3310 | 4866 |
| 1588 | 2685 | 2292 | 937 | 3356 | 1629 | 657 | 8516 | 3360 | 4918 |
| 1478 | 2572 | 2266 | 950 | 3403 | 1662 | 651 | 7778 | 3214 | 4805 |
| 1504.5 | 2574 | 2281 | 939 | 3326 | 1623 | 674 | 8446 | 3338.5 | 4961.5 |
| 1496 | 2634.5 | 2359 | 970 | 3428 | 1665 | 690 | 8409 | 3436 | 5093 |
| 1555 | 2685 | 2322 | 939 | 3380.5 | 1639 | 652 | 8043 | 3421 | 5106 |
| 24995 | 14432 | 5090 | 1134 | 3746 | 1695 | 658 | 11161 | 4415 | 6085 |
| 1784 | 3074 | 2566.5 | 965 | 3456 | 1622.5 | 619 | 8174 | 3505 | 5172.5 |
| 1791.5 | 3074 | 2547 | 977.5 | 3530 | 1681 | 674 | 8920.5 | 3805.5 | 5613 |
| 1592.5 | 2681.5 | 2305 | 921 | 3213.5 | 1602 | 647 | 7679.5 | 3346 | 4954.5 |
| 3820 | 6288 | 3487 | 1080 | 4390 | 1766 | 625 | 14581 | 5062 | 8299 |
| 17348 | 30482 | 6635 | 1538 | 6977 | 2284 | 714 | 17827 | 9755 | 25572 |
| 9976.5 | 20663.5 | 6030 | 1459 | 6885 | 2170 | 646 | 14763.5 | 8874 | 21165.5 |
| 24911 | 32169.5 | 6687 | 1505 | 7046.5 | 2238 | 716 | 16061.5 | 9878 | 25059 |
| 2199 | 3105 | 1714 | 698 | 2409 | 1261 | 323 | 7375 | 3028 | 3904 |
| 2028 | 2858 | 1996 | 827 | 2694 | 1433 | 505 | 7163 | 3062 | 4179 |
| 1923 | 2969.5 | 2367 | 959 | 3201.5 | 1588 | 649 | 7940 | 3377 | 4871.5 |
| 1971 | 3073 | 6750 | 1864 | 1163427 | 124607 | 767 | 8964 | 3767 | 5382 |
| 1734 | 2686 | 2602.5 | 1024 | 4259.5 | 1816 | 695 | 8108 | 3342 | 4781 |
| 1716 | 2718 | 2488 | 985 | 3785 | 1715 | 729 | 8266 | 3395 | 4921 |
| 1730 | 2827.5 | 2601.5 | 1051.5 | 3813 | 1741 | 760 | 8666.5 | 3554 | 5110.5 |
| 1923 | 3115 | 2745 | 1072 | 3949 | 1788 | 787 | 9491 | 3850 | 5535 |
| 2094 | 3219 | 2757 | 1075 | 3885 | 1765 | 783 | 10215 | 3937.5 | 5594.5 |
| 2188 | 3336 | 2841 | 1113 | 3944 | 1809 | 784 | 10301 | 4013 | 5803 |
| 2060 | 3119 | 2663 | 1058 | 3688 | 1754 | 764 | 9529 | 3840 | 5405 |
| 1887 | 2954.5 | 2604 | 1065 | 3580 | 1744 | 763 | 9151 | 3723.5 | 5351 |
| 1797 | 2905 | 2606 | 1054 | 3590 | 1740 | 774 | 8869 | 3645 | 5314 |
| 1920 | 2947 | 2648 | 1056.5 | 3624.5 | 1760 | 775 | 8791 | 3709 | 5349.5 |
| 1831 | 3028 | 2754 | 1103 | 3773 | 1772 | 791 | 9157 | 3856 | 5639 |
| 1969 | 3268 | 2839 | 1111 | 3934 | 1823 | 787 | 9303 | 4087 | 5917 |
| 19016.5 | 11451 | 4940 | 1225 | 3903.5 | 1772.5 | 743.5 | 11140 | 4518.5 | 6142 |
| 2171 | 3528 | 3054.5 | 1119.5 | 3888 | 1795 | 760.5 | 9528.5 | 4132.5 | 5993.5 |
| 2015 | 3408 | 2965 | 1123 | 3973 | 1814 | 783 | 9408 | 4084 | 5947 |
| 1878 | 3108 | 2768.5 | 1093 | 3780.5 | 1766.5 | 784 | 9043 | 3865.5 | 5594.5 |
| 4720 | 10629 | 4124 | 1198.5 | 4746 | 1859.5 | 721 | 14275.5 | 5186 | 12920 |
| 10202.5 | 20468 | 6764 | 1705 | 7360.5 | 2391 | 824 | 16279 | 9228 | 21674 |
| 12026 | 23034.5 | 7211 | 1762.5 | 7612 | 2399 | 816 | 14392.5 | 9651 | 23211.5 |

| | | | | | | | | | |
|---|---|---|---|---|---|---|---|---|---|
| 11892 | 21655 | 6796.5 | 1673 | 7335.5 | 2321 | 815 | 15450 | 9543 | 22378.5 |
| 2153 | 3160 | 1782 | 689 | 2533.5 | 1278 | 331 | 7359 | 3079 | 4019 |
| 1946 | 2930 | 2041 | 814 | 2817 | 1432 | 491 | 7008 | 3078 | 4280 |
| 1773 | 2857.5 | 2231 | 886 | 3137.5 | 1531.5 | 575 | 7334 | 3169 | 4624.5 |
| 1783 | 2886.5 | 5547 | 1639 | 781681.5 | 83727.5 | 651.5 | 7872.5 | 3404 | 4976 |
| 1796 | 2965 | 3079 | 1108 | 5276 | 2111 | 693 | 8703 | 3648 | 5286 |
| 1604 | 2621 | 2383 | 973 | 3799 | 1680 | 632 | 7325 | 3132 | 4633 |
| 1529 | 2487 | 2227 | 920 | 3441 | 1606 | 617 | 6918 | 3018 | 4406 |
| 1736 | 2883 | 2505 | 995 | 3803.5 | 1706.5 | 672 | 8219.5 | 3423 | 5009 |
| 1934.5 | 3160 | 2600.5 | 1014 | 3812 | 1737 | 691 | 9053.5 | 3587 | 5310.5 |
| 1998 | 3139 | 2631 | 1033 | 3708 | 1716 | 684 | 9497 | 3659 | 5367 |
| 2036 | 3234 | 2722.5 | 1052 | 3953 | 1786 | 712 | 9432.5 | 3933.5 | 5547 |
| 1813 | 2986 | 2605 | 1028 | 3661 | 1713 | 692 | 8690 | 3575 | 5250 |
| 1797 | 3021 | 2665 | 1064 | 3827 | 1776 | 730 | 9177 | 3735 | 5489 |
| 1677 | 2821 | 2490 | 1012 | 3557 | 1696 | 679 | 7980 | 3421 | 5113 |
| 1846 | 3068 | 2692 | 1060 | 3898 | 1785 | 730 | 9051 | 3830 | 5562 |
| 2296 | 3955 | 3351 | 1245 | 4741 | 2020 | 844 | 12964 | 4973 | 7306 |
| 17485 | 10512 | 4557 | 1164 | 3926 | 1763 | 674 | 9956 | 4129 | 5905 |
| 2187 | 3578 | 2970 | 1063 | 3801 | 1752 | 662 | 8767 | 3919 | 5665 |
| 2087 | 3354.5 | 2857.5 | 1077 | 3946 | 1783 | 696 | 9337.5 | 4208 | 5763 |
| 1616 | 2681 | 2330 | 937 | 3262 | 1595 | 621 | 7401 | 3180 | 4662 |
| 4581 | 11784.5 | 4224.5 | 1229 | 5109 | 1920 | 674 | 12923.5 | 5541 | 14401 |
| 7921 | 18493 | 6038 | 1564 | 6943 | 2272 | 716 | 15210 | 7850 | 20005 |
| 10878.5 | 22018.5 | 8637.5 | 1790 | 9184 | 2495 | 783 | 15337 | 9315 | 22806.5 |
| 9571 | 20730 | 6122 | 1473 | 6949 | 2167 | 682 | 13327 | 8864 | 21903 |
| 2015 | 2867 | 1630 | 682 | 2349 | 1247 | 295 | 6735.5 | 2814 | 3655 |
| 1999 | 2941 | 2014 | 822 | 2816 | 1417 | 479 | 6942 | 3051 | 4177 |
| 1867 | 2869 | 2227 | 903 | 3058 | 1536 | 576 | 7335 | 3181 | 4544 |
| 1820 | 2793 | 5247 | 1593 | 766726 | 91043 | 633 | 7652 | 3262 | 4661 |
| 1772 | 2846 | 2713 | 1023 | 4255.5 | 1808 | 672 | 7873 | 3378 | 4907.5 |
| 1890 | 3256 | 2978 | 1109 | 4399 | 1874 | 747 | 8930 | 3793 | 5633 |
| 1892 | 3200 | 2827 | 1086 | 4234 | 1851 | 752 | 9043 | 3797 | 5653 |
| 1996 | 3245 | 2765 | 1067 | 4045 | 1793 | 729.5 | 9105.5 | 3906 | 5573 |
| 2061 | 3274 | 2647 | 1056 | 3904 | 1758 | 711 | 9347 | 3872 | 5453 |
| 1988 | 3073 | 2581 | 1019 | 3619 | 1696 | 687 | 9011 | 3602 | 5221 |
| 2019 | 3197 | 2696 | 1037 | 3856 | 1757 | 726 | 9428 | 3811 | 5510 |
| 1839 | 2967 | 2617 | 1024 | 3692 | 1728 | 705 | 8985 | 3552 | 5296 |
| 1873 | 3144 | 2778 | 1092 | 3924 | 1805 | 753 | 9169 | 3855 | 5623 |
| 1865 | 2849 | 2464 | 990 | 3434 | 1661 | 675 | 7973 | 3428 | 4983 |
| 1706 | 2877 | 2532 | 1021 | 3546 | 1699.5 | 692 | 7994 | 3441 | 5098.5 |
| 2169 | 3454 | 3020 | 1131 | 4178 | 1851 | 760 | 9800 | 4459 | 6139 |
| 16676.5 | 9902 | 4678.5 | 1219 | 4206 | 1840.5 | 729 | 10765.5 | 4629 | 6478.5 |
| 2316.5 | 3673 | 3077 | 1118 | 4170.5 | 1821 | 717 | 9628.5 | 4395 | 6075.5 |
| 1952.5 | 3209.5 | 2751 | 1043 | 3800 | 1728 | 695 | 8807 | 3828 | 5665 |
| 1876 | 3211 | 2715 | 1052 | 3839 | 1732 | 718 | 8922 | 3806 | 5584 |
| 4370 | 9423.5 | 4308.5 | 1306 | 5266.5 | 1979 | 729 | 14849.5 | 5501 | 11820.5 |
| 9456 | 20227 | 6731.5 | 1713 | 7709 | 2390 | 763 | 15438.5 | 9720 | 22607.5 |
| 9315 | 20343 | 7134 | 1719 | 7888 | 2391 | 738 | 14234 | 8946 | 22128 |

| | | | | | | | | | |
|---|---|---|---|---|---|---|---|---|---|
| 9717 | 21180 | 6499 | 1655 | 7344 | 2314 | 750 | 14642 | 9271 | 22688 |
| 2113 | 3036 | 1733 | 705 | 2493 | 1280 | 310 | 7219 | 2968 | 3844 |
| 17154 | 9251 | 3366 | 886 | 2946 | 1438 | 489 | 8399 | 3556 | 4423 |
| 14526 | 8838 | 3616 | 982 | 3349 | 1640 | 630 | 9254 | 3991 | 5111 |
| 14252 | 8739.5 | 5644 | 1499 | 129477.5 | 14421.5 | 654.5 | 9125.5 | 3830 | 5010.5 |
| 12575 | 7958 | 3974 | 1129 | 11392 | 2437 | 720 | 9572 | 4053 | 5432 |
| 13220 | 8443 | 3877 | 1112 | 5564 | 1977 | 750 | 10064 | 4245 | 5533 |
| 12651 | 8028 | 4187 | 1221 | 5169 | 2000 | 830 | 11146 | 4743 | 6432 |
| 11967 | 7599 | 3876 | 1140 | 4658 | 1891 | 806 | 11323 | 4715 | 6133 |
| 11906 | 7489 | 3730 | 1085 | 4149 | 1788 | 776.5 | 10970.5 | 4534.5 | 5892 |
| 7930 | 8004 | 10233 | 4837 | 4563 | 1947 | 674 | 9811 | 3980 | 5893 |
| 10691 | 7101 | 3606.5 | 1086.5 | 3995.5 | 1734.5 | 753.5 | 11660.5 | 4442 | 5853 |
| 9399 | 6349 | 3535 | 1112 | 4109 | 1818 | 790 | 11505 | 4544 | 5889 |
| 9748 | 6575 | 3387 | 1059 | 3713 | 1712 | 766 | 10588 | 4228 | 5578 |
| 13096 | 8779 | 3853 | 1012 | 3505 | 1661 | 737 | 10081 | 5158 | 6090 |
| 12259 | 8218 | 3641 | 1072 | 3606 | 1692 | 745 | 10287 | 4849 | 5858 |
| 11812 | 7771 | 3659 | 1080 | 3697 | 1706 | 737 | 10601 | 4847 | 6110 |
| 1387742 | 1005932 | 315156.5 | 23486.5 | 4564.5 | 1900 | 832 | 148328 | 40643 | 28871.5 |
| 43275 | 25206 | 7937 | 1255 | 3546 | 1665 | 706 | 12752 | 5490 | 6318 |
| 32736 | 18516 | 6318 | 1240 | 4039 | 1799 | 760 | 12480 | 5587 | 6534 |
| 43009.5 | 24899.5 | 7723 | 1243.5 | 3773.5 | 1732.5 | 756 | 15141.5 | 5858.5 | 6532 |
| 25714 | 30799 | 7112 | 1323 | 5196 | 1912 | 716 | 15154 | 6366 | 14808 |
| 33659 | 37409 | 8746 | 1768 | 7470 | 2405 | 792 | 16038 | 10864 | 22420 |
| 20664 | 26778 | 7551.5 | 1659 | 8053 | 2505 | 797 | 15737 | 9660 | 21499.5 |
| 29351 | 36417 | 8796 | 1814 | 7795 | 2418 | 784 | 14620 | 10077 | 24024 |
| 2540 | 3408 | 1873 | 712 | 2606 | 1326 | 319 | 8751 | 3252 | 4204 |
| 2042.5 | 3053 | 2005 | 766 | 2834.5 | 1400 | 462 | 7992 | 3185.5 | 4332 |
| 1990.5 | 3148 | 2317 | 873 | 3274 | 1542.5 | 583 | 9358 | 3655 | 5086.5 |
| 1659.5 | 2666 | 5373.5 | 1526 | 835866.5 | 79033.5 | 565 | 7050 | 3068 | 4414.5 |
| 1955 | 3205 | 2999 | 1048 | 5300.5 | 1997 | 687 | 10098.5 | 3954 | 5616.5 |
| 1835.5 | 3122.5 | 2643 | 992 | 4455 | 1808 | 678.5 | 9832.5 | 3830 | 5511.5 |
| 1609 | 2743 | 2330 | 910.5 | 3820 | 1648.5 | 615 | 7461 | 3196 | 4740 |
| 1884 | 3141 | 2573 | 985 | 4098 | 1744 | 673 | 9629 | 3733 | 5393 |
| 1709 | 2784 | 2223 | 881 | 3436 | 1581 | 589 | 7775 | 3168 | 4599 |
| 1848 | 3042 | 2440 | 946.5 | 3606 | 1615 | 629 | 8801.5 | 3446 | 5015.5 |
| 2235 | 3288 | 2603 | 992 | 3867 | 1721 | 696 | 12715 | 4414 | 5803 |
| 1966 | 3182 | 2575 | 988 | 3836 | 1700 | 709 | 12713 | 4212 | 5855 |
| 2066 | 3355.5 | 2727 | 1021 | 3923 | 1771 | 758 | 14562 | 4584 | 6523 |
| 1697 | 2832.5 | 2394 | 953 | 3474 | 1639 | 647 | 7962 | 3366 | 5046 |
| 1889 | 3178 | 2659 | 996 | 3844 | 1749 | 723 | 10977 | 4091 | 5976 |
| 1687 | 2919 | 2410 | 931.5 | 3513 | 1629 | 617 | 7592 | 3366 | 5000.5 |
| 40952 | 21932 | 6919 | 1229 | 4063 | 1737 | 658 | 12237 | 5031 | 6350 |
| 2382 | 3727 | 3044 | 1032 | 3984 | 1721 | 649 | 9375 | 4374 | 5836 |
| 1894 | 3189 | 2676 | 981 | 3705 | 1673 | 632 | 8054.5 | 3584 | 5388.5 |
| 1957 | 3273 | 2708 | 1014 | 3830 | 1727 | 686 | 9541 | 4064 | 5684 |
| 4319 | 11786 | 4237 | 1212 | 5284 | 1916 | 659 | 13434 | 5493 | 14510 |
| 12079 | 24816 | 6783 | 1536 | 7119 | 2278 | 721 | 19023 | 10169 | 25193 |
| 9859 | 20790 | 6221 | 1497 | 7080 | 2207 | 660 | 15362 | 8850 | 21598 |

| | | | | | | | | | |
|---|---|---|---|---|---|---|---|---|---|
| 11972 | 25828.5 | 7136 | 1554 | 7771 | 2294 | 732 | 16200.5 | 9950 | 25457 |
| 2090 | 3032 | 1715 | 710 | 2476 | 1270 | 301 | 7129.5 | 2932 | 3748 |
| 2487 | 3585 | 2509 | 933 | 3361 | 1560 | 591 | 9272 | 3899 | 5147 |
| 2213 | 3311 | 2606 | 1007 | 3404.5 | 1633.5 | 697 | 9041 | 3770 | 5275.5 |
| 2315 | 3538 | 6974 | 1925 | 1011228 | 108649 | 801 | 9897.5 | 4264 | 5994.5 |
| 2031 | 3148 | 3134 | 1115 | 5310 | 2053 | 766 | 9191 | 3832 | 5411 |
| 1983 | 3026 | 2730 | 1063 | 4243 | 1809 | 766 | 9133 | 3723 | 5281 |
| 2295 | 3544 | 3171 | 1173 | 4719 | 1958 | 863 | 10907 | 4487 | 6179 |
| 2150 | 3392 | 2961 | 1118 | 4221 | 1843 | 821 | 10035 | 4094 | 5824 |
| 2648 | 3810 | 3220 | 1172 | 4467 | 1937 | 855 | 12418 | 4899 | 6403 |
| 2506 | 3721 | 3139 | 1164 | 4160 | 1841 | 830 | 11479 | 4546 | 6172 |
| 2286 | 3445 | 2896 | 1100 | 3929 | 1781 | 823 | 10604 | 4160 | 5850 |
| 2416 | 3716.5 | 3187 | 1205 | 4269.5 | 1893 | 872 | 11136 | 4614.5 | 6443 |
| 2113 | 3323 | 2916 | 1124 | 3976 | 1803 | 802 | 10414 | 4111 | 5928 |
| 2122 | 3382 | 2948 | 1121 | 4050 | 1824 | 844 | 10196 | 4239 | 6063 |
| 1935.5 | 3147 | 2741 | 1082 | 3708 | 1747 | 782.5 | 9428.5 | 3864.5 | 5603.5 |
| 2125 | 3419 | 2968 | 1099.5 | 3912 | 1780 | 787 | 9789 | 4188 | 5936.5 |
| 19030.5 | 11427 | 5087 | 1315 | 4295 | 1863 | 827.5 | 12556.5 | 5173 | 6800.5 |
| 2746 | 4045.5 | 3409 | 1201 | 4357 | 1903 | 837 | 11594 | 5173 | 6717.5 |
| 2479 | 3756.5 | 3262 | 1173 | 4182 | 1857.5 | 847 | 11089.5 | 4915 | 6438.5 |
| 2268 | 3576 | 3070 | 1155 | 3992 | 1815 | 828 | 10453 | 4365 | 6168 |
| 5165 | 12546.5 | 4912 | 1391 | 5542 | 2024.5 | 811 | 14959 | 5900 | 15073.5 |
| 9660.5 | 18302.5 | 6322.5 | 1606.5 | 6916.5 | 2273.5 | 825 | 17066.5 | 9228.5 | 20413 |
| 10505.5 | 21134.5 | 7611.5 | 1765 | 8201 | 2457 | 861 | 14862 | 9364.5 | 22606.5 |
| 11483 | 22498.5 | 6986 | 1733 | 7699 | 2404 | 865 | 15632.5 | 10284.5 | 23909 |
| 2431 | 3310 | 1845 | 728 | 2656 | 1324 | 329 | 8269 | 3227 | 4096 |
| 2140 | 3076 | 2090 | 823 | 3146 | 1461 | 464 | 7367 | 3076 | 4226 |
| 2328.5 | 3450 | 2559 | 984 | 4047 | 1696 | 591 | 8375.5 | 3775.5 | 5206 |
| 2032 | 3098 | 7028 | 1829 | 1374782 | 116349 | 619 | 7916 | 3415 | 4870 |
| 2134 | 3342 | 3036.5 | 1075 | 12899 | 2598 | 661 | 8503 | 3636 | 5272 |
| 2488 | 3713 | 3102 | 1100 | 10696 | 2407 | 710 | 9688 | 4361 | 5851 |
| 2502 | 3755 | 3062 | 1112 | 9340 | 2289 | 706 | 9208 | 4280 | 5853 |
| 2783 | 4293 | 3411 | 1173 | 9443 | 2387 | 760 | 11593 | 4714 | 6633 |
| 2718 | 3997 | 3082.5 | 1102 | 8605.5 | 2230 | 704 | 10523 | 4462 | 6097 |
| 2731 | 4442 | 3925 | 1376 | 8029 | 2346 | 712 | 10490 | 4377 | 6396 |
| 2568.5 | 3707 | 2946.5 | 1082 | 7079 | 2064.5 | 695.5 | 10133 | 4242.5 | 5740 |
| 2310 | 3508 | 2855.5 | 1052 | 6119.5 | 1951 | 680 | 9123 | 3968 | 5508 |
| 2207 | 3465 | 2825 | 1070 | 5897 | 1952 | 702.5 | 9391 | 3845 | 5621.5 |
| 4651 | 4769.5 | 3078.5 | 1015 | 5243 | 1830.5 | 652 | 8169.5 | 4324 | 5753.5 |
| 2619.5 | 3714.5 | 2829.5 | 1031 | 5166 | 1852 | 663 | 8487.5 | 4014 | 5558.5 |
| 2869 | 4224 | 3264 | 1151 | 5468 | 1977 | 713 | 9719 | 4726 | 6440 |
| 173163 | 97421 | 26950.5 | 2138 | 5331 | 1918 | 669 | 25761 | 7453.5 | 7996 |
| 3184 | 4465 | 3460 | 1191 | 5267 | 1931 | 682 | 10412 | 4819 | 6525.5 |
| 3036 | 4163 | 3245.5 | 1121 | 5189.5 | 1931 | 703 | 10048 | 4645 | 6343.5 |
| 2721.5 | 3905 | 2995 | 1066 | 4785 | 1845.5 | 661 | 9396.5 | 4308 | 5902.5 |
| 5318.5 | 13179.5 | 4837 | 1341 | 6390 | 2112 | 696 | 13835.5 | 5990.5 | 15414.5 |
| 8813 | 17979 | 6304 | 1616 | 7899 | 2375 | 725 | 14908 | 8747 | 19812 |
| 10095 | 19935 | 6485 | 1595 | 7855 | 2344.5 | 691 | 14840 | 9177 | 21067 |

| | | | | | | | | | |
|---|---|---|---|---|---|---|---|---|---|
| 9520 | 22477 | 7196 | 1728 | 8688 | 2513 | 754 | 14335 | 9433 | 24681 |
| 2472 | 3267 | 1881.5 | 726 | 2702.5 | 1330 | 355 | 9457.5 | 3256.5 | 4077 |
| 2150 | 3081 | 2027 | 793 | 2872 | 1414 | 469.5 | 8576 | 3236 | 4372 |
| 1958 | 3070 | 2229 | 859 | 3181 | 1523 | 549 | 9319 | 3388 | 4834 |
| 1839 | 2945.5 | 4997 | 1467.5 | 679189 | 70190.5 | 594 | 8058 | 3303 | 4795 |
| 1651 | 2736 | 2582 | 930 | 4437 | 1826 | 573 | 7067 | 3068 | 4485 |
| 1842 | 3119 | 2701 | 993 | 4312 | 1785 | 660 | 8740 | 3599 | 5264 |
| 1672 | 2837 | 2374 | 915 | 3745 | 1632 | 609 | 7575 | 3221 | 4739 |
| 2189 | 3380 | 2785 | 1017 | 4175 | 1781 | 700 | 11788 | 4332 | 5885 |
| 2410 | 3657 | 2868 | 1036 | 4225 | 1794 | 715 | 12891.5 | 4655.5 | 6241.5 |
| 2334 | 3555 | 2840 | 1133 | 4072 | 1882 | 695 | 12673 | 4470 | 6091 |
| 2470 | 3765.5 | 2891 | 1062 | 4160 | 1809 | 746 | 15075 | 4860.5 | 6582 |
| 1821 | 3068.5 | 2477 | 959.5 | 3619 | 1651 | 643 | 8579 | 3527 | 5158 |
| 2180 | 3472.5 | 2728 | 1027 | 3962 | 1773 | 750 | 15191.5 | 4718.5 | 6523 |
| 1671 | 2743 | 2258 | 901 | 3284 | 1587 | 601 | 7514 | 3201 | 4666 |
| 1639 | 2809 | 2304 | 918 | 3331 | 1597 | 608 | 7524.5 | 3209 | 4741.5 |
| 1858 | 3190 | 2597.5 | 990 | 3737 | 1682 | 645 | 8532 | 3680 | 5450 |
| 20331 | 11594 | 4634 | 1122 | 3961 | 1729 | 653 | 11565 | 4598 | 6217 |
| 2219 | 3599 | 2738 | 981 | 3636 | 1655 | 611 | 8864 | 3859 | 5534 |
| 1868 | 3204 | 2580 | 948 | 3541 | 1640 | 604 | 8044 | 3522 | 5233 |
| 2016 | 3348 | 2697.5 | 983 | 3722 | 1694 | 670 | 10668 | 4047 | 5928.5 |
| 4044.5 | 10567 | 3870 | 1134 | 4815 | 1815 | 616 | 13526 | 4886.5 | 12907 |
| 8996 | 19162 | 6025 | 1492 | 6786 | 2181 | 680 | 16563 | 9054 | 21012 |
| 8644 | 18080.5 | 5519 | 1388 | 6464 | 2073.5 | 609 | 14507.5 | 8458.5 | 19497 |
| 7798 | 16854 | 5267 | 1334 | 6150 | 2017 | 607 | 13672 | 7905 | 18246 |
| 2605.5 | 3374.5 | 1819.5 | 730 | 2570.5 | 1288 | 311 | 8620 | 3313 | 4131 |
| 2249 | 3252 | 2127 | 799 | 2961 | 1424 | 470 | 8613 | 3441 | 4583 |
| 1979 | 3133 | 2308 | 887 | 3245 | 1542 | 551 | 7541 | 3356 | 4765 |
| 1843 | 2832 | 5414 | 1508 | 835332 | 83778 | 576 | 7405 | 3205 | 4536 |
| 1897 | 3141 | 2938 | 1039 | 5142 | 1986 | 643 | 8103 | 3492 | 5141 |
| 1962 | 3171 | 2760 | 1027 | 4458 | 1824 | 662 | 8308 | 3643 | 5291.5 |
| 2155 | 3450 | 2955 | 1080 | 4596 | 1885 | 723 | 9431 | 4153 | 5819 |
| 2113 | 3356 | 2763 | 1034.5 | 4157.5 | 1771 | 682 | 9421.5 | 4004 | 5580 |
| 2252 | 3415 | 2751 | 1027 | 4089 | 1760 | 684 | 9493.5 | 4076 | 5557 |
| 2045 | 3256 | 2587 | 994 | 3712 | 1670 | 646 | 9051 | 3656 | 5204 |
| 2244 | 3497 | 2813 | 1059 | 4023 | 1781 | 700 | 9724 | 4163 | 5732 |
| 2080 | 3310 | 2736 | 1046 | 3919 | 1769 | 696 | 9128 | 3943 | 5541 |
| 1991 | 3235.5 | 2730 | 1025 | 3857 | 1738 | 695 | 9110 | 3801 | 5528 |
| 1782.5 | 2873.5 | 2414.5 | 951 | 3369 | 1611 | 625 | 7547 | 3284 | 4823.5 |
| 1887 | 3101.5 | 2665 | 1012 | 3654 | 1705 | 674 | 8147 | 3587 | 5258 |
| 2241 | 3646.5 | 2990 | 1105 | 4249 | 1824 | 714 | 9396 | 4380.5 | 6182.5 |
| 18210.5 | 10005.5 | 4304.5 | 1106 | 3847 | 1701 | 632 | 9596 | 4318 | 5660.5 |
| 2247 | 3615 | 2927 | 1042 | 3903 | 1719 | 644 | 8682 | 3949 | 5730 |
| 2019 | 3281 | 2706 | 1014 | 3773 | 1690 | 633 | 8203 | 3711 | 5465 |
| 1909 | 3205 | 2649 | 1021.5 | 3681 | 1702 | 645 | 8175 | 3600 | 5316.5 |
| 4166 | 11057 | 4154.5 | 1209 | 5062.5 | 1879 | 631 | 13699 | 5085.5 | 13542 |
| 6612 | 17356 | 5815 | 1498 | 6705 | 2189 | 686 | 14847 | 7278 | 19381 |
| 8342 | 17777 | 5859 | 1489 | 6804 | 2173 | 643 | 14701 | 8553 | 19701 |

| | | | | | | | | | |
|---|---|---|---|---|---|---|---|---|---|
| 9101.5 | 19225.5 | 6038 | 1481.5 | 6911 | 2207.5 | 679 | 14374.5 | 9039 | 21201 |
| 706 | 832 | 653 | 540 | 1165 | 1100 | 230 | 2045 | 1282 | 1552 |
| 896 | 1428 | 1293 | 668 | 2028 | 1267 | 413 | 3714 | 1933 | 2820 |
| 1052 | 1766 | 1652 | 766 | 2513 | 1395 | 518 | 5134 | 2438 | 3582 |
| 1180 | 2088 | 5185 | 1557.5 | 804151 | 83725.5 | 610 | 6497.5 | 2787 | 4193 |
| 1175 | 2063 | 2302 | 935 | 3718 | 1824 | 595 | 6421 | 2724 | 4154 |
| 1245 | 2260 | 2257 | 936 | 3492 | 1702 | 648 | 7592 | 3051 | 4590 |
| 1327 | 2455 | 2392.5 | 990 | 3579 | 1733.5 | 703 | 8301.5 | 3318 | 5071.5 |
| 1387 | 2443 | 2260 | 928 | 3411 | 1635 | 654 | 7699 | 3121 | 4730 |
| 1522 | 2554 | 2243 | 926 | 3322 | 1627 | 637 | 8112 | 3217 | 4716 |
| 1527 | 2502 | 2152 | 909 | 3127 | 1599 | 613 | 8109 | 3119 | 4524 |
| 1507 | 2521 | 2202 | 912 | 3171 | 1591 | 630 | 8372 | 3140 | 4669 |
| 1476 | 2576 | 2316.5 | 949 | 3392 | 1644 | 672 | 8483 | 3274 | 4929 |
| 1388 | 2450 | 2292 | 952 | 3336 | 1655 | 663 | 8326 | 3165 | 4823 |
| 1427 | 2553 | 2384.5 | 968 | 3447 | 1672 | 700 | 8651.5 | 3356 | 5093.5 |
| 1403 | 2574 | 2415 | 977 | 3560 | 1707 | 701 | 8544 | 3362 | 5170 |
| 1441.5 | 2635 | 2438 | 977 | 3537 | 1676 | 666 | 8210 | 3363 | 5140 |
| 21686.5 | 12758 | 4875.5 | 1129 | 3680 | 1691 | 645 | 10128 | 4006 | 5633.5 |
| 2156 | 3752.5 | 3080.5 | 1057 | 3952 | 1758 | 694 | 10474 | 4263 | 6114.5 |
| 1640 | 2975 | 2692 | 1022 | 3785 | 1735 | 677 | 8648 | 3638 | 5584 |
| 1602 | 3001.5 | 2725 | 1059 | 3941 | 1791.5 | 733 | 11292 | 4001 | 6016 |
| 4305 | 11065 | 4145 | 1219 | 5215 | 1928 | 698 | 14412 | 5517 | 13977 |
| 9908.5 | 21265.5 | 6223 | 1524 | 6960.5 | 2258 | 730.5 | 16575 | 9503 | 22657 |
| 92488.5 | 59889 | 8377.5 | 1706 | 7860 | 2450 | 783 | 16971 | 10412.5 | 28993 |
| 13617.5 | 29837.5 | 7284 | 1589 | 7486.5 | 2314.5 | 747 | 12450.5 | 8862.5 | 27016 |
| 1951 | 2702 | 1464.5 | 641 | 2067 | 1194 | 277 | 6317 | 2702 | 3416 |
| 1676 | 2508 | 1684 | 712 | 2376 | 1312 | 414 | 6174 | 2672 | 3642 |
| 1500 | 2418 | 1918 | 811 | 2758 | 1431 | 530 | 6465 | 2801 | 4022 |
| 1510 | 2508 | 5461 | 1557 | 835436 | 82870 | 608 | 7299 | 3046 | 4514 |
| 1405 | 2333 | 2459 | 948 | 4277 | 1828 | 597 | 6746 | 2894 | 4322 |
| 1457 | 2566 | 2379 | 951 | 3755 | 1702 | 652 | 7453 | 3175 | 4786 |
| 1503 | 2719.5 | 2501.5 | 994 | 3885 | 1750.5 | 696 | 8239 | 3361.5 | 5155.5 |
| 1550 | 2660 | 2374 | 955 | 3563 | 1675 | 668 | 8579 | 3326 | 4983 |
| 1635 | 2682 | 2275 | 914 | 3325 | 1604.5 | 626 | 8053 | 3214 | 4708 |
| 1643 | 2646 | 2271 | 916 | 3279 | 1604 | 624 | 8029 | 3191 | 4666 |
| 1616 | 2636 | 2276 | 911 | 3290 | 1598 | 621 | 8000 | 3175 | 4675 |
| 1622 | 2711 | 2371 | 958.5 | 3453.5 | 1666 | 671 | 9110.5 | 3396 | 5047 |
| 1544 | 2729 | 2453 | 975 | 3542 | 1696 | 694 | 8680.5 | 3441.5 | 5162 |
| 1519 | 2692 | 2442 | 986 | 3595 | 1697 | 686 | 8306 | 3323 | 5108 |
| 1585.5 | 2907 | 2644 | 1032 | 3889.5 | 1788 | 745 | 11723.5 | 3829 | 5819.5 |
| 1522 | 2754 | 2459 | 975 | 3637 | 1693 | 663 | 8018 | 3412 | 5181 |
| 25113 | 14266 | 5341 | 1180 | 4019 | 1780 | 686 | 11937 | 4533 | 6170 |
| 1858 | 3289 | 2839 | 1007 | 3834 | 1698 | 664 | 8704 | 3764 | 5617 |
| 1962 | 3376 | 2914 | 1077 | 4173 | 1820 | 723 | 10414 | 4426 | 6194 |
| 1737 | 3176.5 | 2775 | 1046.5 | 3957 | 1821 | 743 | 10467 | 4063 | 6044 |
| 4816 | 11683 | 4488 | 1288 | 5534 | 2020 | 728 | 16066 | 6059 | 14581 |
| 179688 | 133132 | 7266 | 1642 | 7482 | 2396 | 798 | 21593 | 11138 | 30208 |
| 11892 | 25993 | 7010 | 1577 | 7321 | 2281 | 719 | 14414 | 9398 | 25562 |

| | | | | | | | | | |
|---|---|---|---|---|---|---|---|---|---|
| 14226.5 | 27848 | 7722 | 1599 | 7792 | 2337 | 755 | 16881 | 9959.5 | 26348 |
| 2132 | 3006 | 1678 | 682 | 2324 | 1243 | 301 | 7121 | 2978 | 3806 |
| 2089 | 3010 | 2079.5 | 814.5 | 2770 | 1429 | 503 | 7402 | 3131.5 | 4291.5 |
| 1800 | 2772 | 2198.5 | 894 | 2999 | 1511.5 | 594 | 7459.5 | 3200 | 4459 |
| 1830 | 2838 | 6249 | 1760 | 1029350 | 117075 | 722 | 8560 | 3557 | 4976 |
| 1691 | 2725 | 2660 | 1017 | 4244 | 1834 | 714 | 8241 | 3433 | 4910 |
| 1742 | 2878 | 2689.5 | 1073 | 4069 | 1811 | 768 | 8793 | 3652 | 5369.5 |
| 1759 | 2963 | 2758 | 1070 | 4044 | 1809 | 778 | 9113 | 3733 | 5458 |
| 1857 | 3113 | 2789 | 1101 | 4106 | 1840 | 789 | 9654 | 3843 | 5662 |
| 1984 | 3039 | 2616 | 1031 | 3745 | 1727 | 746 | 9505 | 3784 | 5366 |
| 1963 | 2890.5 | 2492 | 1010 | 3433.5 | 1657 | 714 | 9238.5 | 3629 | 5077 |
| 1864 | 2832 | 2433 | 986 | 3371 | 1642 | 709 | 9051 | 3473 | 4960 |
| 1875 | 2975 | 2623 | 1064 | 3716 | 1754 | 775 | 9347.5 | 3735.5 | 5472 |
| 1825.5 | 3078.5 | 2834 | 1109.5 | 3894 | 1828 | 804 | 9763 | 3889 | 5749 |
| 1926 | 3051 | 2818 | 1085 | 3842 | 1808 | 800 | 9350 | 3817 | 5637 |
| 1832 | 3149 | 2919 | 1145 | 4061 | 1869 | 828 | 9439 | 3922 | 5835 |
| 1834 | 3083 | 2820 | 1092 | 3857 | 1806 | 778 | 9251 | 3862 | 5653 |
| 16612 | 10074 | 4675 | 1239 | 4084 | 1828 | 759 | 11287 | 4525 | 6266 |
| 2544 | 4029 | 3448 | 1200 | 4360 | 1911 | 807 | 10972 | 4853 | 6718 |
| 2090 | 3491 | 3121 | 1149 | 4209 | 1894 | 801 | 10126 | 4484 | 6368 |
| 2007 | 3384 | 3065.5 | 1176 | 4212.5 | 1907 | 835 | 10220.5 | 4351.5 | 6358 |
| 4927 | 12429 | 4954 | 1430 | 5794 | 2105 | 810 | 15032 | 5757 | 15323 |
| 8686 | 19778 | 6647 | 1733 | 7731 | 2427 | 832 | 13680 | 8446 | 21790 |
| 11099 | 24866.5 | 7607.5 | 1855 | 8316 | 2541.5 | 836 | 14584.5 | 9923 | 25939 |
| 10623 | 24231 | 7463 | 1783 | 8284 | 2458 | 844 | 15321 | 9397 | 25523 |
| 2107 | 3013 | 1682 | 692 | 2348.5 | 1246 | 297 | 6985.5 | 2940 | 3761 |
| 1950 | 2865 | 1973 | 780 | 2715 | 1398 | 474 | 6898 | 3024 | 4128 |
| 1747 | 2766 | 2182 | 877 | 3076 | 1525 | 562 | 7142 | 3116 | 4480 |
| 1701 | 2746.5 | 5231.5 | 1584 | 748493 | 84811 | 636 | 7615.5 | 3266.5 | 4730 |
| 1560 | 2526 | 2596 | 970 | 4470 | 1856.5 | 616 | 7013 | 3047 | 4439 |
| 1628 | 2751 | 2583 | 1013 | 4089 | 1790 | 666 | 7789 | 3354 | 5045 |
| 1671 | 2888 | 2636 | 1065 | 4068 | 1788 | 703 | 8412 | 3508 | 5270 |
| 1703 | 2830 | 2567.5 | 1004 | 3829 | 1725 | 682 | 8246 | 3432 | 5085.5 |
| 1966 | 3176.5 | 2687 | 1034 | 3953 | 1742 | 715 | 9360 | 3840 | 5474 |
| 1964 | 3106.5 | 2601 | 1009 | 3763 | 1744 | 686.5 | 9399.5 | 3678.5 | 5251.5 |
| 1699 | 2695 | 2311 | 929 | 3280 | 1592 | 629 | 8000 | 3213 | 4682.5 |
| 1737 | 2879 | 2514 | 997 | 3608 | 1709.5 | 685 | 8692.5 | 3460 | 5123 |
| 1629 | 2740 | 2416 | 985 | 3473 | 1675 | 666 | 8094 | 3266 | 4934 |
| 1665.5 | 2829 | 2562 | 1026 | 3692 | 1744 | 701 | 8413 | 3437 | 5215 |
| 1704.5 | 2944.5 | 2690.5 | 1062 | 3862 | 1797 | 735 | 8675.5 | 3624 | 5425.5 |
| 1715.5 | 3007 | 2715.5 | 1040 | 3819 | 1763.5 | 701 | 8406 | 3658.5 | 5440 |
| 18358 | 10855 | 4514 | 1127 | 3814 | 1725 | 655 | 9598 | 3954 | 5679 |
| 2171 | 3510.5 | 3012 | 1091 | 4035.5 | 1785.5 | 704 | 9313.5 | 4273 | 5991 |
| 2109 | 3420 | 2976 | 1112 | 4189 | 1847 | 735 | 9233 | 4387 | 6077 |
| 1896 | 3283 | 2935 | 1119 | 4100 | 1844 | 742 | 9601 | 4113 | 5960 |
| 4127 | 10568 | 4409 | 1294 | 5445 | 2011 | 724 | 13777 | 5482 | 13407 |
| 7885 | 18137 | 6117 | 1585 | 7185 | 2314 | 728 | 14644 | 8920 | 20966 |
| 9641 | 21010 | 6719 | 1546 | 7560 | 2274 | 687 | 14922 | 9188 | 22302 |

| | | | | | | | | | |
|---|---|---|---|---|---|---|---|---|---|
| 22796 | 34397 | 7913 | 1633 | 7982 | 2367 | 762 | 16789 | 10062 | 26771 |
| 2098 | 2984.5 | 1681 | 670 | 2360 | 1258 | 298 | 7008 | 2951 | 3767 |
| 1955 | 2892 | 1951 | 783 | 2682 | 1389 | 459 | 6875 | 2979 | 4072 |
| 1765 | 2775 | 2149 | 874 | 2955.5 | 1523 | 562 | 7125.5 | 3079.5 | 4366.5 |
| 1780 | 2845 | 5376.5 | 1575 | 748123 | 89035 | 660 | 7907.5 | 3384 | 4836.5 |
| 1579 | 2493 | 2491 | 981 | 3998 | 1765 | 604 | 6958 | 2984 | 4332 |
| 1660 | 2752.5 | 2570 | 999 | 3814.5 | 1730 | 677 | 7853 | 3330 | 4927.5 |
| 1725 | 2905 | 2649.5 | 1030 | 3922 | 1777 | 720 | 8451 | 3541.5 | 5285.5 |
| 1758 | 2948.5 | 2565 | 1015 | 3800 | 1747 | 706 | 8542 | 3513 | 5144.5 |
| 2057 | 3251 | 2707 | 1055 | 3873 | 1761 | 719 | 9497 | 3827 | 5492 |
| 2073 | 3198 | 2695 | 1051 | 3751 | 1766 | 710 | 9548 | 3832 | 5436 |
| 1819 | 2814 | 2420 | 958 | 3340 | 1641 | 666 | 8395.5 | 3369 | 4826 |
| 1737 | 2767 | 2443 | 983 | 3424 | 1662 | 673 | 8249 | 3359 | 4906 |
| 1619 | 2645 | 2353 | 972 | 3303 | 1646 | 664 | 7889 | 3263 | 4789 |
| 1844 | 2826 | 2511 | 1006 | 3512.5 | 1678 | 686 | 7956 | 3397.5 | 4983 |
| 1646 | 2719 | 2464.5 | 993.5 | 3483 | 1679 | 688 | 7859.5 | 3371 | 4996.5 |
| 1666 | 2904.5 | 2563 | 1020 | 3615.5 | 1695 | 690 | 8044 | 3479 | 5194.5 |
| 13890 | 8610 | 4140 | 1120 | 3762 | 1711 | 670 | 9395 | 3942 | 5742 |
| 1952.5 | 3295.5 | 2825 | 1038 | 3700.5 | 1697.5 | 677 | 8646 | 3721 | 5523 |
| 1854 | 3077 | 2686 | 1044 | 3759 | 1742 | 684 | 8486 | 3705 | 5521 |
| 1708.5 | 2947 | 2621.5 | 1029 | 3681 | 1702 | 695 | 8290 | 3548.5 | 5336 |
| 4293 | 10761.5 | 4219.5 | 1255 | 5090 | 1916 | 687 | 13543 | 5123 | 13304.5 |
| 6543 | 16234 | 5695 | 1494 | 6432 | 2138 | 706 | 14336 | 6857 | 18240 |
| 9646.5 | 19386.5 | 6315 | 1593 | 7078.5 | 2274 | 713 | 14946.5 | 9136.5 | 21208 |
| 8878 | 21991 | 7159 | 1772 | 8136 | 2494 | 803 | 14339 | 9262 | 24177 |
| 2182 | 3120 | 1760.5 | 694 | 2484 | 1279 | 304 | 7365.5 | 3070 | 3908 |
| 14400 | 7835 | 3064 | 828 | 2752 | 1411 | 477 | 8476 | 3468 | 4276 |
| 12093 | 7418 | 3170 | 907 | 3004 | 1492 | 576 | 8928 | 3664 | 4651 |
| 10918 | 6931 | 5035 | 1404 | 119156 | 13330 | 651 | 8872 | 3766 | 4993 |
| 11242 | 7129 | 3761.5 | 1118 | 13443.5 | 2616.5 | 729 | 9928 | 4151 | 5654 |
| 9800 | 6382 | 3660 | 1106 | 6277 | 2085 | 752 | 9955 | 4156 | 5639 |
| 9725 | 6340 | 3502 | 1079 | 5357 | 1930 | 756 | 9926 | 4085 | 5582 |
| 9203.5 | 6076 | 3432 | 1066 | 4840 | 1897 | 771 | 10093.5 | 4144 | 5784.5 |
| 9032 | 6081 | 3418 | 1065 | 4268 | 1825 | 771 | 10807 | 4412 | 5772 |
| 4860 | 5217 | 5711 | 2780 | 6135 | 2476 | 775 | 11101 | 4129 | 5724 |
| 8504 | 5817 | 3417 | 1076 | 3938 | 1766 | 757 | 10448 | 4166 | 5566 |
| 6942 | 5070.5 | 3198 | 1075.5 | 4088.5 | 1864.5 | 776 | 10058.5 | 4082 | 5725.5 |
| 7914 | 5648 | 3278 | 1029 | 3829 | 1740 | 766 | 10034 | 4110 | 5556 |
| 10305 | 7084 | 3631 | 1076 | 3614 | 1715 | 752 | 9578 | 4744 | 6058 |
| 9023 | 6165.5 | 3166.5 | 1026 | 3564 | 1689.5 | 725.5 | 9138 | 4220 | 5490.5 |
| 2318 | 3590 | 2968 | 1085 | 14599 | 2786 | 760 | 9724 | 4309 | 5929 |
| 1172550 | 916397.5 | 260086 | 17720.5 | 5712 | 2291 | 880 | 128773.5 | 35653 | 27730 |
| 32970 | 18453 | 6385.5 | 1319 | 4362.5 | 1886.5 | 779 | 13103.5 | 5653 | 6947 |
| 26401 | 15231 | 5475 | 1211 | 4114 | 1788 | 762 | 12131 | 5142 | 6361 |
| 36425.5 | 20853.5 | 6702.5 | 1181 | 3895.5 | 1767 | 746 | 14774 | 5537.5 | 6415 |
| 22024 | 23596 | 6107.5 | 1325.5 | 5101 | 1908 | 732 | 15356.5 | 6288.5 | 13634.5 |
| 26218 | 30747 | 8350 | 1702.5 | 7647.5 | 2405.5 | 774 | 14797 | 9745.5 | 21823 |
| 9058 | 20616.5 | 6679.5 | 1580 | 11843 | 2720 | 696 | 11598 | 8358 | 21677 |

| | | | | | | | | | |
|---|---|---|---|---|---|---|---|---|---|
| 20416 | 27569 | 7988 | 1689 | 7884 | 2353 | 763 | 13987 | 9505 | 22063 |
| 2164 | 3102 | 1747 | 702 | 2513 | 1277 | 304 | 7217 | 3038 | 3907 |
| 1921.5 | 2941.5 | 1899 | 750 | 2718 | 1366 | 436 | 7147 | 3041 | 4084 |
| 1723 | 2755 | 2072 | 818 | 2943 | 1447 | 521 | 6777 | 3003 | 4311 |
| 1675 | 2712 | 5419 | 1526 | 834665 | 80965 | 574 | 7199 | 3093 | 4504 |
| 1711 | 2889 | 2785.5 | 1003 | 5001.5 | 1957 | 645 | 8781 | 3466 | 5049 |
| 1733 | 2955 | 2645 | 972 | 4381 | 1783 | 670 | 9329.5 | 3618 | 5267.5 |
| 1809 | 3035 | 2651 | 991 | 4236 | 1762 | 702 | 10067 | 3872 | 5563 |
| 1854 | 3034 | 2548 | 980 | 3981 | 1721 | 676 | 10194 | 3914 | 5441 |
| 1811 | 2937 | 2424 | 939 | 3715 | 1666 | 634 | 8422 | 3428 | 5013 |
| 1717 | 2792 | 2292 | 903 | 3345 | 1579 | 597 | 8044 | 3249 | 4709 |
| 1907 | 3067 | 2532 | 959 | 3731 | 1677 | 662 | 9456 | 3742 | 5286 |
| 1715 | 2874 | 2408.5 | 945 | 3536 | 1661 | 648 | 8423 | 3425 | 4984.5 |
| 1691 | 2902 | 2460 | 968 | 3654 | 1657 | 667 | 8804 | 3508 | 5175 |
| 1598 | 2683 | 2307 | 911 | 3356.5 | 1604 | 626 | 7444.5 | 3169.5 | 4733 |
| 1618 | 2785 | 2372 | 955 | 3528 | 1653 | 666 | 8631 | 3410 | 5037 |
| 1929.5 | 3270 | 2789 | 1033.5 | 4004 | 1771 | 712 | 11112 | 4253.5 | 5939.5 |
| 31930.5 | 17245 | 5978 | 1228 | 4255 | 1809.5 | 700 | 13760 | 5093 | 6625 |
| 2270 | 3661 | 2985 | 1042 | 4095 | 1763 | 690 | 11068 | 4526 | 6285 |
| 1986 | 3341 | 2805 | 1018 | 4029 | 1736.5 | 682 | 9956 | 4151 | 5902 |
| 1761 | 3069 | 2605 | 980 | 3705 | 1703 | 663 | 9769 | 3763 | 5576 |
| 4116 | 10998 | 4067 | 1162 | 5082 | 1861 | 646 | 13074 | 5213 | 13739 |
| 10091 | 20855 | 6024 | 1525 | 6777 | 2239 | 706 | 15278 | 8976 | 21549 |
| 9006 | 19078 | 5792 | 1424 | 6669 | 2114 | 637 | 13930 | 8725 | 20305 |
| 17071 | 28678 | 6981 | 1540 | 7359 | 2286 | 718 | 16584 | 10005 | 25052 |
| 2304 | 3219 | 1816 | 708 | 2592 | 1302 | 307 | 7819 | 3171 | 4012 |
| 2193 | 3208 | 2175 | 823 | 2906 | 1426.5 | 518 | 8024.5 | 3325 | 4433.5 |
| 1987 | 3024 | 2291 | 897 | 3056 | 1526 | 619 | 7901 | 3388 | 4654 |
| 2013 | 3065 | 6148 | 1694 | 876308 | 101686 | 713 | 8885 | 3700 | 5130 |
| 1997 | 3238.5 | 3350.5 | 1186 | 5437 | 2111 | 776 | 9310 | 3895 | 5562.5 |
| 2045 | 3343 | 3022.5 | 1124 | 4656 | 1900 | 811 | 9608 | 4089 | 5846.5 |
| 2284 | 3483 | 3185 | 1182 | 4717 | 1967 | 856 | 11054.5 | 4684.5 | 6297.5 |
| 2257 | 3490 | 3082.5 | 1160.5 | 4467 | 1896.5 | 826.5 | 10448 | 4443 | 6108 |
| 2279 | 3489 | 2914 | 1102 | 4145 | 1845 | 803 | 10433 | 4297 | 5886 |
| 2075 | 3113 | 2643 | 1019 | 3574 | 1675 | 736 | 9619 | 3806 | 5285 |
| 2302 | 3440 | 2953 | 1111 | 4004 | 1831 | 816 | 10455 | 4325 | 5880 |
| 2267 | 3525 | 3043 | 1143 | 4215 | 1856 | 849 | 11192 | 4506 | 6263 |
| 2150 | 3468 | 3082 | 1180 | 4271 | 1921 | 847 | 10324 | 4447 | 6148 |
| 1913 | 3121 | 2758 | 1074 | 3810 | 1769 | 787 | 9359 | 3874 | 5602 |
| 1882 | 3144 | 2811 | 1104 | 3830 | 1783 | 795 | 9284 | 3889 | 5713 |
| 2398.5 | 3866 | 3382 | 1267.5 | 4784.5 | 2030.5 | 882 | 11994.5 | 5054.5 | 7113.5 |
| 16624 | 10223.5 | 5049 | 1357 | 4788 | 2028 | 872 | 13525 | 5470.5 | 7541 |
| 2672 | 4237 | 3596 | 1267 | 4707.5 | 2002.5 | 870 | 11927.5 | 5204.5 | 7256 |
| 2310 | 3572 | 3144 | 1141 | 4236 | 1858 | 807 | 10570 | 4826 | 6324 |
| 2247 | 3596 | 3199 | 1190 | 4343 | 1919 | 859 | 10193 | 4565 | 6368 |
| 5071 | 11945.5 | 4890.5 | 1398 | 5817.5 | 2052 | 809.5 | 14776 | 6243 | 14814 |
| 9539 | 21488 | 7226 | 1891 | 8125 | 2572 | 898 | 15612 | 9978 | 24394 |
| 10481 | 21354 | 6968 | 1732 | 7587 | 2393 | 829 | 15074 | 9624 | 22877 |

| | | | | | | | | | |
|---|---|---|---|---|---|---|---|---|---|
| 9192 | 21346 | 8594 | 1940 | 9751 | 2587 | 897 | 13085 | 8942 | 23369 |
| 2263 | 3192 | 1816 | 698 | 2588 | 1282 | 302 | 7615 | 3148 | 3991 |
| 2216 | 3162 | 2092 | 825 | 3122 | 1467 | 453 | 7694 | 3196 | 4268 |
| 2115.5 | 3242 | 2414.5 | 927.5 | 3786 | 1661 | 597 | 8104 | 3511 | 4965 |
| 2046.5 | 3217 | 7374 | 1905 | 1415340 | 124156 | 655 | 8382.5 | 3569 | 5150 |
| 2154 | 3318 | 3129 | 1118 | 11373 | 2500 | 689 | 8841 | 3873 | 5489 |
| 2275 | 3581 | 3040 | 1118 | 9501 | 2363 | 736 | 9198 | 4192 | 5744 |
| 2228.5 | 3466.5 | 2911 | 1074.5 | 7893 | 2139 | 709 | 8854.5 | 4022 | 5637.5 |
| 2295 | 3497 | 2872 | 1072 | 7557 | 2087 | 697 | 9268 | 4065 | 5591 |
| 2378 | 3449 | 2763.5 | 1028 | 6930 | 2007 | 668 | 9297.5 | 3981 | 5441.5 |
| 2430 | 3926 | 3501 | 1250 | 6786 | 2174 | 652 | 9191 | 3902 | 5654 |
| 2672 | 3965 | 3212 | 1129 | 6837 | 2094.5 | 719 | 11219.5 | 4624.5 | 6195 |
| 2515 | 3879 | 3094 | 1118 | 6320 | 2050 | 721 | 9966 | 4378 | 6112 |
| 2286 | 3461 | 2877 | 1062 | 5802 | 1939 | 705 | 9448 | 4004 | 5606 |
| 5025.5 | 4983 | 3273 | 1076.5 | 5400.5 | 1890 | 678 | 8610 | 4783 | 6054.5 |
| 2773 | 3868 | 3049 | 1081 | 5477 | 1925 | 715 | 9093 | 4358 | 5906 |
| 2738 | 4152 | 3290 | 1156 | 5698 | 2021 | 736 | 9380 | 4648 | 6498 |
| 163598 | 102829 | 24496 | 2208 | 5943 | 2095 | 728 | 24413 | 7938 | 9489 |
| 3231 | 5539 | 4159 | 1327 | 5870 | 2094 | 756 | 12954 | 5038 | 7884 |
| 3066 | 4918 | 3658 | 1238 | 6051 | 2112 | 772 | 10699 | 4895 | 7554 |
| 2916 | 4414.5 | 3549 | 1221 | 5571.5 | 2057 | 774 | 11118.5 | 4810 | 6995.5 |
| 6163 | 14578 | 5259.5 | 1462 | 7051 | 2300.5 | 768 | 14750 | 7228 | 17444 |
| 9243 | 20784.5 | 6714.5 | 1733 | 8704.5 | 2580 | 768 | 14713 | 9161 | 22796.5 |
| 9964.5 | 22400 | 6995 | 1686 | 8590.5 | 2480 | 726 | 14253.5 | 9344 | 23810.5 |
| 9570.5 | 21812 | 7361.5 | 1753 | 8822.5 | 2523.5 | 774 | 12992 | 9006 | 23150.5 |
| 2363 | 3117 | 1765 | 695 | 2558 | 1298 | 332 | 8472 | 3097 | 3870 |
| 2039 | 2877 | 1888 | 756 | 2692 | 1362.5 | 443 | 7940 | 2977 | 4056 |
| 1892 | 2915 | 2124 | 832 | 3022 | 1488 | 526 | 8040 | 3189 | 4477 |
| 1830 | 2932 | 5095 | 1438 | 708351 | 71731.5 | 600 | 7879.5 | 3324 | 4780.5 |
| 1764 | 2931 | 2723 | 976 | 4660 | 1874 | 614 | 8253 | 3334 | 4828 |
| 2049 | 3352 | 2877 | 1041 | 4645 | 1861 | 705 | 10558 | 4157 | 5828 |
| 2100 | 3359 | 2800 | 1035 | 4377 | 1841 | 735 | 11139 | 4444 | 6132 |
| 2115 | 3389 | 2772 | 1025 | 4286 | 1802 | 712 | 12414 | 4342 | 5987 |
| 1904 | 3032 | 2418 | 939 | 3689 | 1654 | 623 | 8656 | 3530 | 5016 |
| 2043 | 3180 | 2620 | 1075 | 3720.5 | 1798.5 | 634 | 9192 | 3795 | 5215 |
| 2109 | 3264 | 2634 | 991 | 3803 | 1695 | 664 | 10031 | 4002 | 5427 |
| 2022.5 | 3220 | 2594 | 1007 | 3815.5 | 1723 | 672 | 10106.5 | 4063 | 5533 |
| 2087 | 3325 | 2781 | 1027 | 3918 | 1761 | 725 | 12182 | 4428 | 5994 |
| 1664 | 2864 | 2392 | 935 | 3533 | 1643 | 630 | 7668 | 3303 | 4858 |
| 1991 | 3293 | 2774 | 1041 | 4027 | 1777 | 725 | 10660 | 4194 | 5916 |
| 2170 | 3627 | 2983.5 | 1105.5 | 4232.5 | 1859 | 735 | 11556.5 | 4673 | 6416.5 |
| 21880.5 | 12890.5 | 5095 | 1212 | 4451 | 1881 | 733 | 14714.5 | 5299.5 | 7478 |
| 2560 | 4248.5 | 3257 | 1109 | 4344.5 | 1828 | 707 | 11535 | 4668.5 | 6870 |
| 2301 | 3722 | 2955 | 1067 | 4137 | 1814 | 718 | 11460 | 4668 | 6474.5 |
| 2095 | 3458 | 2840 | 1047 | 4072 | 1792 | 702 | 10412 | 4271 | 6048 |
| 5500.5 | 13243 | 4495.5 | 1273 | 5602 | 1998 | 698 | 15063 | 6415.5 | 15753 |
| 11699.5 | 23046 | 6167 | 1550 | 7055 | 2287.5 | 721 | 16975 | 9523.5 | 23140.5 |
| 14455 | 27757 | 6757 | 1602 | 7522 | 2317 | 706 | 15073 | 9472 | 24461 |

| | | | | | | | | | |
|---|---|---|---|---|---|---|---|---|---|
| 11231 | 23192.5 | 6445.5 | 1510 | 7107.5 | 2219 | 692 | 15266.5 | 9437.5 | 23446 |
| 2291 | 3236 | 1822 | 707 | 2575 | 1282 | 297 | 7571 | 3169 | 4015 |
| 1960 | 2948 | 1872 | 751 | 2693.5 | 1358 | 415 | 6838 | 2961 | 4015 |
| 1862 | 2959 | 2119 | 821 | 3019.5 | 1460 | 518 | 7489 | 3198.5 | 4478 |
| 1716 | 2694 | 5108 | 1454 | 780451 | 76549 | 551 | 6929 | 3031 | 4308 |
| 1764 | 2898 | 2593 | 961 | 4593 | 1833 | 604 | 8024.5 | 3335 | 4874 |
| 1967 | 3186 | 2707 | 992 | 4500 | 1822 | 676 | 9793 | 4035 | 5665 |
| 1795 | 3047 | 2581 | 972 | 4149 | 1735 | 663 | 8797 | 3635 | 5240 |
| 2025 | 3225.5 | 2657 | 1002 | 4133 | 1743 | 680 | 9846 | 3979 | 5477 |
| 1990 | 3175 | 2539 | 954 | 3894 | 1690 | 644 | 9453 | 3733 | 5250 |
| 2050 | 3167 | 2513 | 972.5 | 3698.5 | 1673 | 638 | 9339.5 | 3815 | 5253 |
| 2043.5 | 3249.5 | 2599 | 992 | 3855 | 1710.5 | 666 | 10061 | 3914 | 5428 |
| 2158 | 3337 | 2716 | 1033 | 3966 | 1760 | 702 | 11039.5 | 4274 | 5740 |
| 2047 | 3230 | 2667 | 1013 | 3889.5 | 1751 | 703.5 | 10410 | 4226 | 5703.5 |
| 1716 | 2786 | 2337 | 926 | 3411 | 1608 | 633 | 7723.5 | 3281.5 | 4792.5 |
| 1717 | 2910 | 2489 | 975 | 3619 | 1679 | 658 | 8323 | 3463 | 5123 |
| 2048 | 3450 | 2813.5 | 1054 | 4056 | 1792 | 711 | 11242 | 4380 | 6301 |
| 32312 | 17269 | 5999 | 1240 | 4341 | 1837 | 696 | 12411 | 4982 | 6509 |
| 2509 | 3986 | 3209 | 1077 | 4122 | 1776 | 677 | 9813 | 4439 | 6076 |
| 1866 | 3146.5 | 2601 | 975.5 | 3649 | 1679 | 615 | 7852 | 3539 | 5283.5 |
| 2014 | 3394 | 2816 | 1057 | 3992 | 1786 | 697 | 9181 | 4098 | 5931 |
| 4028 | 10532 | 4215 | 1228 | 5266 | 1913 | 668 | 13165 | 5318 | 13197 |
| 7186 | 16107.5 | 5473 | 1426 | 6448 | 2119 | 657 | 15170 | 8080 | 18434 |
| 7595 | 17013 | 5544 | 1413 | 6616 | 2094 | 623 | 14066.5 | 7908 | 18748 |
| 9115 | 20555.5 | 6140 | 1535 | 7304 | 2275.5 | 689 | 14695 | 9265.5 | 22387 |
| 718 | 905 | 717 | 555 | 1259 | 1087 | 238 | 2191.5 | 1351 | 1674 |
| 985 | 1569 | 1372.5 | 670 | 2068 | 1260 | 399 | 3823 | 2017 | 2937 |
| 1190 | 1998 | 1751 | 754 | 2572 | 1382 | 502 | 5444 | 2557 | 3762 |
| 1340 | 2333.5 | 5074 | 1546 | 800308.5 | 81825.5 | 594 | 8440.5 | 3113 | 4599.5 |
| 1385 | 2455 | 2792 | 1025 | 4882 | 2177 | 614 | 7331 | 3091 | 4668 |
| 1500 | 2774 | 2602 | 971 | 3912.5 | 1779 | 699 | 19627.5 | 3869 | 5678.5 |
| 1593 | 2914 | 2612 | 982 | 3823 | 1758 | 745 | 14504 | 4378 | 6344 |
| 1715 | 2956 | 2548 | 971 | 3682 | 1707 | 721 | 19432 | 4251 | 6073 |
| 1689 | 2785 | 2378 | 916 | 3374 | 1627 | 646 | 10304 | 3582 | 5246 |
| 1737.5 | 2862.5 | 2391 | 943 | 3401 | 1624 | 637 | 9377.5 | 3490 | 5040 |
| 1728 | 2916 | 2431 | 956 | 3496 | 1641 | 647 | 9619 | 3578 | 5230 |
| 1825 | 3119 | 2662 | 1004 | 3789 | 1720 | 714 | 14976 | 4238 | 6050 |
| 1736 | 3056 | 2666 | 992 | 3771 | 1728 | 745 | 20282 | 4332 | 6216 |
| 1676 | 2942 | 2588 | 988 | 3682.5 | 1684 | 704 | 11133 | 3933 | 5776.5 |
| 1722 | 3087 | 2681 | 1005 | 3816 | 1751 | 753 | 20810.5 | 4487 | 6425 |
| 1746 | 3147 | 2720.5 | 1006 | 3835 | 1726 | 706 | 11408.5 | 4294 | 6220 |
| 23921.5 | 13730.5 | 5155.5 | 1143 | 3941 | 1724 | 663 | 12606.5 | 4723.5 | 6369 |
| 2246 | 3859 | 3169 | 1069 | 4070.5 | 1763 | 709 | 12812.5 | 4770 | 6740.5 |
| 1888 | 3332 | 2833 | 1024 | 3969 | 1749 | 690 | 11410 | 4363 | 6324 |
| 1957 | 3434 | 2929 | 1043 | 4039 | 1786 | 765 | 75638 | 5244 | 7169 |
| 6415.5 | 13556.5 | 4661 | 1244 | 5389 | 1989 | 743 | 22331.5 | 6863 | 16258 |
| 168409 | 146060 | 7901 | 1495 | 6618 | 2239 | 821 | 26616 | 11772 | 32452 |
| 130656 | 95514 | 8866 | 1629 | 7965 | 2355 | 775 | 21188 | 11808 | 31294 |

| | | | | | | | | | |
|---|---|---|---|---|---|---|---|---|---|
| 16501 | 25926 | 6621 | 1424 | 6662 | 2122 | 694 | 18948 | 9939 | 23492 |
| 2088 | 2975 | 1672.5 | 670 | 2315 | 1226 | 308 | 7384 | 2959 | 3783 |
| 1813 | 2724 | 1844.5 | 737 | 2574 | 1339 | 443 | 6747 | 2908 | 3960 |
| 1701 | 2778 | 2163 | 846 | 3020 | 1476 | 556 | 7278 | 3145 | 4517 |
| 1706 | 2835.5 | 5658 | 1644 | 879960.5 | 89152 | 623 | 8435.5 | 3431 | 4966.5 |
| 1697 | 2963 | 2965 | 1050 | 4974 | 2005 | 676 | 9101 | 3629 | 5287.5 |
| 1743 | 3045 | 2803 | 1016 | 4380 | 1829 | 696 | 9580 | 3777 | 5485 |
| 1757 | 3040 | 2653 | 1000 | 3934 | 1727 | 704 | 10095 | 3947 | 5659 |
| 1918 | 3233 | 2740.5 | 1019 | 3980.5 | 1742.5 | 729 | 11074 | 4236 | 5923 |
| 1827 | 2987 | 2470 | 949 | 3546 | 1608 | 650 | 8780 | 3546 | 5146 |
| 1872.5 | 3029 | 2490 | 962.5 | 3505 | 1641 | 651 | 8736 | 3549 | 5120 |
| 1829 | 3016 | 2523 | 958 | 3498 | 1646 | 649 | 8794 | 3548 | 5167 |
| 1867 | 3179 | 2689 | 1008 | 3808 | 1725 | 699 | 9998 | 3852 | 5644 |
| 1725 | 3000 | 2609 | 977 | 3686 | 1682 | 693 | 9217 | 3650 | 5404 |
| 1757 | 3013 | 2602 | 998 | 3639 | 1678 | 686 | 8721 | 3595 | 5342 |
| 1820 | 3190 | 2763 | 1030 | 3886 | 1755 | 747 | 11248 | 4169.5 | 6034 |
| 1780 | 3128 | 2720 | 1004 | 3826 | 1707 | 694 | 9022 | 3781 | 5646 |
| 29699 | 16159 | 5836 | 1188 | 4048 | 1754 | 676 | 12297.5 | 4931 | 6325 |
| 2375 | 3855 | 3163 | 1070 | 4105 | 1771 | 697 | 10554 | 4674 | 6355 |
| 2039 | 3379.5 | 2892 | 1040 | 4000.5 | 1740 | 684 | 9684.5 | 4252.5 | 5954 |
| 2124 | 3527 | 3006 | 1099 | 4203 | 1830 | 763 | 12125.5 | 4860 | 6610 |
| 4496 | 11490 | 4322.5 | 1233 | 5345.5 | 1959 | 704 | 14426.5 | 5616 | 14450 |
| 12465 | 25069.5 | 6317.5 | 1507 | 7038 | 2246 | 727 | 18697.5 | 9219 | 23725 |
| 13330 | 26391.5 | 7022 | 1604 | 7572 | 2318 | 735 | 15937 | 9823.5 | 25287 |
| 12718 | 23370 | 6318 | 1489 | 6927 | 2203 | 698 | 15986 | 9505 | 22905 |
| 2220 | 3130 | 1762 | 680 | 2437 | 1250 | 317 | 7689 | 3105 | 4027 |
| 1966 | 2939 | 1954 | 765 | 2731 | 1354 | 473 | 7990.5 | 3161 | 4321 |
| 2162.5 | 3217.5 | 2303 | 850 | 3123.5 | 1513 | 608 | 16810 | 4288 | 5777 |
| 1756 | 2933.5 | 5438 | 1690 | 902237.5 | 80652.5 | 627 | 9199 | 3593 | 5091 |
| 1693 | 2822 | 2959 | 1013 | 5878 | 2087 | 632 | 9029 | 3468 | 4959 |
| 1754 | 3039 | 2748 | 986 | 4594 | 1816 | 687 | 10136.5 | 3768 | 5451.5 |
| 1822.5 | 3155 | 2751 | 1013 | 4431 | 1808 | 716 | 12497.5 | 4149 | 5853 |
| 2034 | 3267 | 2716 | 979 | 4098 | 1743 | 761 | 16155 | 4796 | 6392 |
| 1882 | 3038 | 2458.5 | 923 | 3708 | 1630.5 | 647 | 10476.5 | 3781 | 5277 |
| 1946 | 3149 | 2595.5 | 1020 | 3770 | 1736 | 660 | 10126.5 | 3738.5 | 5362.5 |
| 1839.5 | 3043 | 2494 | 961 | 3688 | 1663 | 659 | 9873.5 | 3644 | 5229.5 |
| 2030 | 3265 | 2704 | 993 | 3914 | 1748 | 719 | 12981 | 4399 | 5932 |
| 2001.5 | 3340 | 2778 | 1017 | 4011.5 | 1770 | 759 | 18359.5 | 4713 | 6327 |
| 2031 | 3372 | 2771 | 1012 | 4004 | 1767 | 774 | 20114 | 4832 | 6514 |
| 2084 | 3434 | 2815 | 1038 | 4082 | 1792 | 812 | 88234 | 5790 | 7184 |
| 2118 | 3537 | 2897 | 1045 | 4097 | 1763 | 782 | 17612 | 5164 | 7003 |
| 31371 | 17385 | 5856 | 1154 | 3916 | 1698 | 652 | 13092 | 4781 | 6181 |
| 2912 | 4533 | 3333 | 1076 | 4224 | 1808 | 781 | 72737 | 7266 | 8455 |
| 2122 | 3560 | 2887 | 1026 | 4025.5 | 1742 | 721 | 14131 | 4798 | 6544.5 |
| 2026 | 3469 | 2837 | 1023 | 3966 | 1753 | 728 | 13125 | 4726 | 6492 |
| 5351 | 9232.5 | 4195 | 1150 | 4970 | 1872 | 707 | 20961 | 5613 | 11140 |
| 8729 | 18138 | 5362 | 1351 | 6124 | 2062 | 666 | 17192 | 7575 | 18642 |
| 26376 | 31490 | 6711 | 1486 | 6945 | 2183 | 717 | 19008 | 10093 | 24430 |

| | | | | | | | | | |
|---|---|---|---|---|---|---|---|---|---|
| 13419 | 22155 | 5869 | 1371 | 6461 | 2087 | 678 | 16637 | 8999 | 21131 |
| 2314 | 3277 | 1856 | 686 | 2540 | 1270 | 320 | 7876 | 3211 | 4116 |
| 1982 | 2977 | 2022 | 768 | 2785 | 1372 | 467 | 7397.5 | 3118 | 4252.5 |
| 1881 | 2965.5 | 2286 | 882 | 3179.5 | 1513 | 579 | 7891 | 3347 | 4722 |
| 1980 | 3112 | 6548 | 1791 | 1101846 | 109978 | 693 | 8939 | 3719 | 5304.5 |
| 1828 | 3024 | 2944 | 1037 | 5032 | 1923 | 677 | 8484.5 | 3582 | 5207.5 |
| 1904 | 3266 | 2909 | 1061 | 4590 | 1830 | 722 | 9391 | 3884 | 5699 |
| 1974 | 3323 | 2933 | 1066 | 4461 | 1843 | 766 | 9482 | 4086.5 | 5827 |
| 1964 | 3122 | 2683 | 992 | 3932 | 1714 | 722 | 9115 | 3810 | 5429 |
| 2114 | 3377.5 | 2749 | 1017 | 3980.5 | 1739 | 721 | 9766.5 | 3998 | 5609 |
| 2098 | 3229 | 2682 | 1009 | 3691 | 1684 | 706 | 9744 | 3849 | 5461 |
| 2262 | 3481 | 2849 | 1049 | 4006 | 1760 | 754 | 10489 | 4335 | 5952 |
| 1902 | 3105 | 2596 | 978 | 3690 | 1664 | 681 | 9166.5 | 3699 | 5349 |
| 1858 | 3042 | 2626 | 995 | 3655 | 1675 | 704 | 8855 | 3687 | 5306 |
| 1832 | 3049 | 2563 | 972 | 3683 | 1655 | 671 | 8590 | 3618 | 5333 |
| 1817 | 3004 | 2543 | 991 | 3546 | 1661 | 715 | 8659.5 | 3680.5 | 5266 |
| 2026 | 3377 | 2876.5 | 1052 | 3977 | 1761 | 746 | 9645 | 4109.5 | 5968 |
| 37458.5 | 19627 | 6655 | 1260 | 4050 | 1747 | 689 | 12580 | 5233.5 | 6382.5 |
| 2182 | 3614 | 3016 | 1030 | 3872 | 1683 | 680 | 9580 | 4095 | 5873 |
| 2456 | 3851 | 3232.5 | 1144 | 4335 | 1834 | 783 | 11390.5 | 4883 | 6642 |
| 2077 | 3600 | 2937 | 1039 | 4019 | 1775 | 740 | 11022 | 4431 | 6467 |
| 4457 | 11212 | 4262 | 1191 | 5127 | 1877 | 682 | 13525 | 5381 | 13778 |
| 9603 | 19851 | 6284 | 1576 | 7021 | 2262 | 761 | 16465 | 8515 | 20871 |
| 11599 | 23486.5 | 6567 | 1563 | 7252 | 2259.5 | 704 | 15311 | 9483 | 23457 |
| 123460 | 76454 | 8714.5 | 1611 | 7688 | 2331 | 779 | 21512.5 | 12201.5 | 32912 |
| 2321 | 3288 | 1887 | 716 | 2609 | 1295 | 335 | 8233 | 3246 | 4163 |
| 2566 | 3712.5 | 2559 | 897 | 3359.5 | 1540 | 562 | 9638.5 | 3885 | 5167 |
| 2261 | 3433 | 2691 | 989 | 3537 | 1643 | 678 | 9166 | 3866 | 5435 |
| 2181.5 | 3345 | 7309 | 2000.5 | 1300318 | 129633 | 745 | 9481.5 | 4043.5 | 5684.5 |
| 2501 | 3771 | 3870 | 1270 | 6647 | 2321 | 837 | 11647 | 4969 | 6580 |
| 2494 | 3835 | 3461 | 1211 | 5472 | 2056 | 868 | 11508 | 4925 | 6600 |
| 2115.5 | 3349 | 2916 | 1068 | 4444 | 1810 | 792 | 9825 | 4139.5 | 5832 |
| 2588 | 3800.5 | 3220 | 1166 | 4720 | 1910.5 | 860 | 12040.5 | 5045 | 6596.5 |
| 2394 | 3520 | 2864 | 1056 | 3997 | 1759 | 775 | 10751 | 4222 | 5842 |
| 2517 | 3595.5 | 2979.5 | 1099 | 3948 | 1777 | 782 | 11379.5 | 4476 | 5936 |
| 2696 | 3873 | 3243 | 1176 | 4339 | 1866 | 859 | 11832.5 | 5024 | 6509 |
| 2022 | 3028 | 2565 | 988 | 3457 | 1633 | 733 | 9406.5 | 3766 | 5235 |
| 2100 | 3339 | 2849 | 1068 | 3926.5 | 1768 | 796 | 10355 | 4176 | 5889 |
| 2748 | 4171 | 3552 | 1269 | 4778 | 2044 | 927 | 12104 | 5343 | 7200 |
| 1937 | 3040 | 2618 | 1024 | 3522 | 1669 | 761 | 9043 | 3814 | 5373 |
| 2380 | 3771 | 3196 | 1153 | 4259 | 1848 | 830 | 11001 | 4812 | 6605 |
| 43505 | 22749 | 7482 | 1354 | 4328 | 1826 | 786 | 14316 | 5876 | 7081 |
| 2774.5 | 4103 | 3396.5 | 1154 | 4302 | 1820 | 800 | 11653 | 5186 | 6731 |
| 2429 | 3784 | 3223 | 1132 | 4222 | 1805 | 804 | 10878 | 4687 | 6482 |
| 2334 | 3698 | 3185 | 1155 | 4167 | 1837 | 835 | 11026 | 4637 | 6452.5 |
| 5384.5 | 12244.5 | 4679.5 | 1333 | 5402 | 1984 | 808 | 14733.5 | 5776.5 | 14707.5 |
| 9386 | 18031 | 6116 | 1517 | 6600.5 | 2159.5 | 795 | 16640 | 8387 | 19184 |
| 12164.5 | 24071.5 | 7754.5 | 1840 | 8216.5 | 2520.5 | 872 | 15419 | 9787.5 | 25095 |

| | | | | | | | | | |
|---|---|---|---|---|---|---|---|---|---|
| 12926.5 | 23554.5 | 7161 | 1718.5 | 7769 | 2372 | 860 | 15907 | 9669 | 23633.5 |
| 2116 | 3025 | 1754 | 688 | 2471 | 1250 | 310 | 7445 | 2992 | 3855.5 |
| 1873 | 2871 | 1921 | 749 | 2710 | 1357 | 436 | 7079 | 2986 | 4118 |
| 1728 | 2836 | 2130 | 820 | 3034 | 1462 | 525 | 7276 | 3098 | 4465 |
| 1797 | 3038 | 5700 | 1631 | 850748.5 | 79575.5 | 600 | 8527 | 3493 | 5073 |
| 1691 | 2913 | 2770 | 961 | 4938 | 1918 | 597 | 7821 | 3346.5 | 4855 |
| 1783 | 3066 | 2652 | 960 | 4310 | 1761 | 632 | 9499 | 3625 | 5340 |
| 1940.5 | 3255.5 | 2749 | 982 | 4218.5 | 1748 | 686 | 12978.5 | 4244.5 | 5903 |
| 2129 | 3470 | 2825 | 998 | 4187 | 1768 | 704 | 14930.5 | 4630 | 6378 |
| 1976 | 3147 | 2515 | 927 | 3718 | 1627 | 639 | 10557 | 3827 | 5391 |
| 1907 | 3094 | 2474 | 926 | 3591 | 1613 | 610 | 9295 | 3576 | 5146 |
| 1929 | 3187 | 2550 | 940 | 3704 | 1645 | 637 | 10017 | 3773 | 5400 |
| 2188 | 3419 | 2721 | 972 | 3865 | 1710 | 692 | 14050 | 4558.5 | 6116.5 |
| 2065 | 3339 | 2688 | 991 | 3911 | 1723 | 710 | 16345 | 4597 | 6200 |
| 2026 | 3311 | 2727 | 989 | 3830 | 1718 | 702 | 12905 | 4371 | 6051 |
| 2251.5 | 3623 | 2872.5 | 1016 | 4045 | 1768 | 786 | 110221 | 5921 | 7593 |
| 2179 | 3607 | 2900 | 1034 | 4086 | 1763 | 715 | 27490 | 4979 | 6795 |
| 32395 | 18024 | 6131 | 1194 | 4245 | 1789 | 698 | 18535 | 5753 | 7388 |
| 2547 | 4165 | 3158 | 1040 | 4182 | 1750 | 685 | 15653 | 5168 | 7121 |
| 2384 | 3884 | 3053 | 1053 | 4102 | 1773 | 713 | 16647 | 5178 | 7072 |
| 2273 | 3689 | 2907 | 1033 | 4049 | 1746 | 721 | 17959 | 5170 | 6949 |
| 13567 | 22243 | 5188 | 1299 | 5662 | 2011 | 756 | 35112 | 7570 | 21535 |
| 152387.5 | 118650 | 7135 | 1511 | 6943 | 2251.5 | 744 | 25404 | 11179.5 | 28904 |
| 198090 | 151550.5 | 8245.5 | 1575 | 7574.5 | 2300.5 | 735 | 22108 | 11811.5 | 32636.5 |
| 199792.5 | 160367.5 | 7910.5 | 1504 | 7158.5 | 2213 | 741 | 23116.5 | 11729 | 31360 |
| 2128 | 3014 | 1758 | 688 | 2486 | 1267 | 303 | 7624 | 3018 | 3878 |
| 2046 | 2993 | 2021 | 785 | 2777 | 1377 | 451 | 7324 | 3088 | 4184 |
| 1900 | 3059 | 2333 | 876 | 3260 | 1520 | 564 | 7907 | 3383 | 4832 |
| 1935 | 3083 | 5971.5 | 1736 | 903043 | 94572.5 | 625 | 8259 | 3520 | 5099 |
| 1938 | 3213 | 3135 | 1083 | 5502 | 2022 | 658 | 8467 | 3735 | 5372 |
| 2174.5 | 3461.5 | 3060 | 1091.5 | 4874 | 1892.5 | 741 | 9846 | 4203 | 5963.5 |
| 2074 | 3400 | 2861 | 1038 | 4446.5 | 1793 | 708 | 9956.5 | 4213 | 5819.5 |
| 2206 | 3553 | 2948 | 1071 | 4442 | 1822 | 739 | 9870 | 4276 | 5986 |
| 2087 | 3310 | 2684.5 | 971 | 3972 | 1708 | 657 | 9328 | 3891 | 5495.5 |
| 2048 | 3221 | 2629 | 976 | 3643 | 1660 | 653 | 9158 | 3711 | 5243 |
| 1982 | 3180 | 2556 | 951 | 3677 | 1648 | 641 | 9136 | 3686 | 5276 |
| 2301 | 3473 | 2914 | 1060 | 4056 | 1791 | 742 | 10542.5 | 4338 | 5898 |
| 2082 | 3346 | 2797 | 1042 | 3929 | 1742 | 728 | 9752 | 4036 | 5780 |
| 2216 | 3463 | 2884 | 1058 | 4046 | 1754 | 742 | 9782 | 4207 | 5893 |
| 2264.5 | 3663.5 | 3128 | 1142 | 4346 | 1865 | 799 | 10912 | 4554 | 6405 |
| 2158 | 3528 | 2963 | 1066 | 4143 | 1774 | 710 | 9940 | 4337 | 6183 |
| 29672 | 16691 | 6029 | 1291 | 4533 | 1841 | 723 | 13638 | 5245 | 7058 |
| 2512.5 | 4068 | 3251 | 1087 | 4292.5 | 1805 | 700 | 10935 | 4782 | 6544.5 |
| 2374 | 3769 | 3080 | 1079 | 4288 | 1797 | 707 | 10966 | 4695 | 6473 |
| 2312 | 3707 | 3016 | 1080 | 4197 | 1800 | 724 | 12220 | 4824 | 6616.5 |
| 5755 | 13965.5 | 4960.5 | 1374 | 6029.5 | 2092 | 772 | 15918 | 6779 | 17012.5 |
| 13224.5 | 26741.5 | 6797.5 | 1641 | 7495 | 2389 | 777.5 | 17642 | 10204.5 | 26085 |
| 11622 | 26020 | 7342 | 1657.5 | 7896 | 2406 | 747 | 15438.5 | 9591 | 25923 |

| | | | | | | | | | |
|---|---|---|---|---|---|---|---|---|---|
| 14595 | 29016 | 7421 | 1664 | 8043 | 2436 | 783 | 15572 | 9931 | 26442 |
| 2084 | 2976 | 1740 | 688 | 2483 | 1261 | 302 | 7367 | 2945 | 3809.5 |
| 2334 | 3265 | 2273 | 854 | 3030 | 1443 | 524 | 8039 | 3417 | 4611 |
| 2306 | 3355 | 2527 | 930 | 3357 | 1576 | 644 | 8929 | 3748 | 5126 |
| 2286 | 3457 | 7368 | 2030 | 1305280 | 127080 | 732 | 9608 | 4102 | 5745 |
| 2303 | 3497 | 3347 | 1144 | 5824 | 2086 | 780 | 9894.5 | 4159.5 | 5836 |
| 2355 | 3643.5 | 3202.5 | 1126 | 5175 | 1956 | 808 | 10717 | 4403.5 | 6091 |
| 2503 | 3800 | 3292 | 1160 | 5082 | 1973 | 831 | 10955 | 4764 | 6410 |
| 2512 | 3793 | 3168 | 1146 | 4845 | 1917 | 813 | 10825 | 4692 | 6381 |
| 2573 | 3684 | 2978 | 1072 | 4394 | 1806 | 814 | 10924 | 4502 | 6044 |
| 2500 | 3652 | 2950.5 | 1076 | 3968 | 1746 | 779 | 11019.5 | 4360.5 | 5892 |
| 2487 | 3601 | 2882 | 1057 | 3984.5 | 1752.5 | 792 | 11067 | 4362 | 5867 |
| 2611 | 3703 | 3087 | 1109 | 4211 | 1803 | 820 | 11583 | 4770 | 6277 |
| 2478.5 | 3724.5 | 3127.5 | 1131 | 4310.5 | 1857 | 837 | 11447 | 4695.5 | 6354 |
| 2410 | 3602.5 | 3003 | 1096 | 4028 | 1799 | 797 | 10498.5 | 4368 | 6052 |
| 2661 | 4011 | 3392 | 1195 | 4608 | 1937 | 895 | 12146.5 | 5166 | 6889 |
| 2632 | 3963 | 3306 | 1165 | 4454 | 1881 | 840 | 11478 | 5015 | 6747 |
| 51726 | 26970 | 8529 | 1407 | 4404 | 1850 | 806 | 15328 | 6069 | 7212 |
| 3019 | 4613 | 3752 | 1226 | 4697 | 1942 | 839 | 12676 | 5327 | 7400 |
| 2734 | 4017 | 3277 | 1148.5 | 4381 | 1859 | 807 | 11440.5 | 5144.5 | 6760 |
| 2735.5 | 4340.5 | 3556.5 | 1235 | 4775.5 | 1974 | 874 | 13298.5 | 5340 | 7425.5 |
| 6151 | 13616 | 5100 | 1412 | 5916 | 2080 | 826 | 14958 | 6709 | 16195 |
| 11128 | 22148 | 7545 | 1873 | 8227 | 2563 | 902 | 15644 | 10172 | 23796 |
| 11668 | 24104.5 | 9127 | 2001 | 9811.5 | 2697 | 927 | 15036.5 | 9695.5 | 24889.5 |
| 13906 | 24818 | 7883 | 1831 | 8582 | 2496 | 895 | 15406 | 10352 | 24704 |
| 2122 | 2996 | 1756 | 694 | 2521 | 1269 | 308 | 7456 | 2962 | 3850 |
| 2282.5 | 3267 | 2225 | 814 | 3217 | 1452.5 | 492 | 8074.5 | 3328 | 4501.5 |
| 2108 | 3128.5 | 2328 | 881 | 3391.5 | 1535 | 572 | 8471.5 | 3513.5 | 4838 |
| 2133 | 3360 | 6888 | 1812 | 1278935 | 104978 | 662 | 9598 | 3914 | 5498 |
| 2046 | 3244 | 2991 | 1028 | 7553 | 2136 | 669.5 | 8739.5 | 3757 | 5315 |
| 2118 | 3419 | 2897 | 1037 | 6181 | 1958 | 719 | 9714.5 | 4008.5 | 5685.5 |
| 2146 | 3449 | 2849 | 1036 | 5490 | 1889 | 721 | 9675 | 4089 | 5790.5 |
| 2488 | 3725 | 3069 | 1072 | 5434 | 1931 | 751 | 11185 | 4749 | 6261 |
| 2374 | 3606 | 2866 | 1029 | 4999 | 1823 | 714 | 10511 | 4314 | 5909 |
| 2234 | 3407 | 2745 | 1003 | 4057 | 1725 | 677 | 9684 | 4001 | 5480 |
| 2208 | 3470 | 2754 | 1001 | 4283 | 1741 | 696 | 10042 | 4019 | 5665.5 |
| 2401 | 3616 | 2910 | 1036 | 4372 | 1789 | 737 | 11662 | 4606 | 6099.5 |
| 2172 | 3461 | 2797 | 1017 | 4200 | 1739 | 722 | 10509 | 4203 | 5884 |
| 2317 | 3560 | 2920 | 1056 | 4325 | 1789 | 739 | 10238 | 4370 | 6060 |
| 2466 | 3816 | 3073 | 1094 | 4578.5 | 1862 | 798 | 13912 | 5071 | 6668 |
| 2502 | 4003 | 3225.5 | 1140 | 4739 | 1882.5 | 791 | 12538 | 5157.5 | 6883 |
| 45353 | 25503 | 8106 | 1366 | 4729 | 1867 | 738 | 16223 | 5607 | 7335 |
| 2769 | 4392 | 3513 | 1148 | 4718 | 1860 | 750 | 11720 | 4947 | 6898 |
| 2561 | 4011 | 3193 | 1110 | 4606 | 1869 | 757 | 11821.5 | 5010.5 | 6751 |
| 2614 | 4104.5 | 3336.5 | 1140 | 4732 | 1911 | 798 | 13416 | 5201.5 | 7036 |
| 7105 | 15299 | 5255 | 1437 | 6359 | 2155 | 810 | 16826 | 7593 | 17868 |
| 12812 | 24911.5 | 6874 | 1683.5 | 7876 | 2427 | 811 | 16958.5 | 10030.5 | 24942.5 |
| 16864 | 29627 | 7384 | 1708 | 8066 | 2427 | 790 | 15909 | 9857 | 25944 |

| | | | | | | | | | |
|---|---|---|---|---|---|---|---|---|---|
| 25947.5 | 36563.5 | 7815 | 1666 | 8263 | 2394.5 | 819 | 17041 | 10418.5 | 27961 |
| 2057 | 2908 | 1675 | 680 | 2443 | 1263 | 290 | 7166 | 2864 | 3671 |
| 1868 | 2909 | 1976 | 759 | 2794 | 1359 | 435 | 7035.5 | 3005 | 4198 |
| 1835 | 2986 | 2218 | 836 | 3144 | 1470 | 522 | 7544 | 3237 | 4663 |
| 1908 | 3133 | 5652.5 | 1655 | 789817.5 | 78722.5 | 596 | 8263.5 | 3534 | 5074.5 |
| 1853 | 3138 | 2841 | 993 | 4844 | 1881 | 602 | 8682 | 3590 | 5243 |
| 2053 | 3332 | 2853 | 1004 | 4571 | 1818 | 647 | 9258 | 4041 | 5598 |
| 2268 | 3653 | 2974 | 1028 | 4480 | 1786 | 712 | 14342 | 4855 | 6547 |
| 2245 | 3558 | 2867 | 998 | 4301 | 1764 | 682 | 12681.5 | 4557 | 6228.5 |
| 2243 | 3486 | 2736 | 964 | 3959 | 1705 | 655 | 11585 | 4303 | 5941 |
| 2016 | 3234 | 2548 | 945 | 3651 | 1624 | 603 | 9270 | 3711 | 5245 |
| 1996 | 3240 | 2590 | 947 | 3753 | 1641 | 613 | 9297 | 3755 | 5321 |
| 2229 | 3430 | 2746.5 | 988 | 3935 | 1697 | 653 | 10520.5 | 4295 | 5738 |
| 2182 | 3423 | 2753 | 995 | 3989 | 1732 | 672 | 11478 | 4434 | 5919 |
| 2165 | 3370 | 2731 | 1006 | 3927 | 1713 | 672 | 11013 | 4306 | 5911 |
| 2236 | 3594 | 2882 | 1030 | 4062.5 | 1740 | 697 | 12870 | 4689.5 | 6264.5 |
| 2213 | 3598 | 2916 | 1027 | 4084 | 1748 | 671 | 10255.5 | 4475 | 6183.5 |
| 25397 | 14198 | 5378 | 1184 | 4288 | 1792 | 660 | 12882 | 4960 | 6690 |
| 2557 | 4183 | 3205 | 1069 | 4322 | 1781 | 667 | 11670 | 4779 | 6681 |
| 2420 | 4046 | 3159 | 1070 | 4386 | 1808.5 | 697 | 14352.5 | 5049 | 7031 |
| 2387 | 3845 | 3066 | 1056 | 4245 | 1790 | 725 | 14081.5 | 5062 | 6880.5 |
| 7584 | 16374 | 4848 | 1274 | 5668 | 2025 | 732 | 19870 | 7366 | 18968 |
| 195587.5 | 150273.5 | 7643 | 1568 | 7440.5 | 2355 | 759 | 24877 | 11840.5 | 32276 |
| 208273 | 166925 | 9093.5 | 1601 | 7447.5 | 2319.5 | 743 | 21828.5 | 12072.5 | 34756 |
| 108975 | 68607 | 8094 | 1523 | 7527 | 2256 | 717 | 21832 | 11390 | 30433 |
| 2110 | 3025 | 1756 | 688 | 2549 | 1272 | 298 | 7419 | 2971 | 3819 |
| 1899 | 2884 | 1937 | 748 | 2756 | 1346.5 | 422 | 6892 | 2964 | 4033 |
| 1882 | 2973 | 2206 | 827 | 3124 | 1471 | 518 | 7401 | 3194 | 4533 |
| 2256 | 3324 | 6057 | 1696 | 975068 | 87074 | 757 | 11083 | 4147 | 5428 |
| 2281 | 3403 | 2931.5 | 989 | 5242 | 1923 | 838 | 13490.5 | 4516 | 5670 |
| 2005 | 3326 | 2763 | 974 | 4538 | 1776.5 | 667 | 11445 | 4038.5 | 5670 |
| 1930 | 3178 | 2572 | 935.5 | 4106.5 | 1687 | 656 | 11207 | 3888 | 5435 |
| 2301 | 3591 | 2813 | 990.5 | 4301 | 1761.5 | 692 | 14722.5 | 4643 | 6332.5 |
| 3370 | 4373 | 3126 | 1049.5 | 4564 | 1827 | 1130 | 22858 | 6612 | 7501.5 |
| 2976 | 3956 | 2948 | 1024 | 4071 | 1740 | 940 | 18149 | 5722 | 6559 |
| 2105 | 3347 | 2625 | 948 | 3798 | 1654 | 645 | 10415 | 3937 | 5455 |
| 2052 | 3333 | 2609 | 950 | 3792 | 1649 | 645 | 10295 | 3943 | 5516 |
| 3226 | 4145.5 | 2998 | 1038 | 4079.5 | 1758 | 1304 | 27561 | 7473.5 | 7996.5 |
| 3387 | 4383.5 | 3158 | 1051 | 4130 | 1754 | 1429 | 21184 | 7468 | 7870 |
| 3378 | 4410 | 3194 | 1062 | 4221 | 1788 | 1512 | 21567 | 7292.5 | 7835 |
| 3202 | 4294.5 | 3072 | 1019 | 3997 | 1719 | 1341.5 | 19628.5 | 6764.5 | 7492 |
| 40213 | 21835 | 7165 | 1303 | 4311 | 1799 | 1341 | 25558 | 8791 | 8970 |
| 4533 | 5449 | 3634 | 1110 | 4402 | 1808 | 1404 | 22143 | 8078 | 8599 |
| 3884 | 4682.5 | 3285 | 1065 | 4178 | 1746 | 1333.5 | 21388.5 | 7889.5 | 7984 |
| 3804 | 4654 | 3285 | 1071 | 3970 | 1732 | 1402 | 22301 | 7881 | 7994 |
| 11191 | 16507.5 | 5133 | 1299 | 5487 | 2000 | 1406 | 25261 | 7530.5 | 15456 |
| 26746.5 | 35570.5 | 7612 | 1634 | 7403 | 2345 | 1399 | 21856.5 | 11643 | 25378.5 |
| 32026 | 38015 | 7664 | 1574 | 7031 | 2207 | 1207 | 19858 | 11464 | 24335 |

| | | | | | | | | | |
|---|---|---|---|---|---|---|---|---|---|
| 21249.5 | 28188.5 | 6489 | 1468 | 6771 | 2131 | 1023 | 17716.5 | 10266 | 21581.5 |
| 733 | 842.5 | 677 | 554 | 1210 | 1083.5 | 224 | 2101 | 1306 | 1604.5 |
| 1000 | 1566 | 1348 | 680 | 2074 | 1253 | 391 | 3732 | 1999 | 2894 |
| 1258.5 | 2181.5 | 1910 | 812 | 2791 | 1444 | 524 | 6077 | 2645 | 3978 |
| 1391 | 2449 | 5576 | 1652 | 910109 | 94445 | 598 | 7271 | 3044 | 4517 |
| 1447 | 2551 | 2766 | 1024 | 4525 | 2053 | 615 | 7161.5 | 3115 | 4693.5 |
| 1533 | 2802 | 2673 | 1007 | 3995 | 1809 | 666 | 7965 | 3407 | 5182 |
| 1516 | 2722 | 2545 | 978 | 3713 | 1708 | 661 | 7719.5 | 3355.5 | 5036 |
| 1603 | 2797.5 | 2494 | 964 | 3617 | 1680 | 650 | 7959 | 3415 | 5061.5 |
| 1703 | 2853 | 2443 | 944 | 3473.5 | 1627 | 634 | 8233.5 | 3372 | 4940.5 |
| 1715 | 2785 | 2385 | 943 | 3302 | 1615 | 606 | 8141 | 3286 | 4768 |
| 1681 | 2781 | 2391 | 930 | 3369 | 1598 | 617 | 8127 | 3312 | 4809.5 |
| 1731 | 2947 | 2545 | 1000 | 3637 | 1690 | 669 | 8847 | 3573 | 5273 |
| 1642 | 2847 | 2575 | 1005 | 3624 | 1700 | 672 | 8456 | 3453 | 5193 |
| 1665 | 2946 | 2593 | 1008 | 3718 | 1700 | 683 | 8846.5 | 3623 | 5402 |
| 1564 | 2763 | 2497 | 985 | 3541 | 1670 | 659 | 7743 | 3342 | 5120 |
| 1745 | 3140 | 2776 | 1040 | 3910 | 1754 | 692 | 9022 | 3817 | 5797 |
| 32536 | 17211 | 6071 | 1224 | 4049 | 1770 | 672 | 11587 | 4768 | 6225 |
| 2194 | 3798 | 3208 | 1093 | 4105 | 1793 | 691 | 10760 | 4415 | 6271 |
| 2096 | 3589 | 3085 | 1130.5 | 4362 | 1876 | 735 | 10697 | 4602 | 6511 |
| 2040 | 3576 | 3119.5 | 1146 | 4434 | 1911 | 774 | 12316 | 4676 | 6777 |
| 4777 | 12116 | 4698 | 1330 | 5737 | 2047 | 716 | 14964 | 6200 | 15530 |
| 7123 | 17458 | 6065.5 | 1522 | 7087.5 | 2276 | 714 | 16378 | 7670 | 19834.5 |
| 8229 | 21254 | 6714 | 1671 | 7862.5 | 2401 | 710 | 13119.5 | 8846 | 23563.5 |
| 10021 | 24278 | 7448 | 1708 | 8668 | 2475 | 752 | 13533 | 9240 | 25107 |
| 2231 | 3094 | 1736 | 689 | 2380 | 1228.5 | 313 | 7671.5 | 3105 | 3917.5 |
| 1932 | 2828 | 1952 | 761.5 | 2651 | 1365 | 461 | 7019 | 3022 | 4107 |
| 1724 | 2670 | 2100 | 833 | 2901 | 1449 | 541 | 6948 | 3058 | 4355 |
| 1752 | 2805 | 6235 | 1609 | 1089651 | 101619 | 628 | 7827 | 3371 | 4881 |
| 1702 | 2872 | 3062 | 1094 | 5153 | 2075 | 657 | 7981 | 3451 | 5021 |
| 1711.5 | 2902 | 2713 | 1015 | 4065.5 | 1779 | 688 | 8319 | 3536.5 | 5180 |
| 1742 | 2971 | 2707 | 1021 | 3977 | 1749 | 695 | 8299 | 3577 | 5316 |
| 1843 | 3058 | 2674 | 1008 | 3820 | 1723 | 692 | 8791 | 3672 | 5373 |
| 1961 | 3143.5 | 2626 | 1018 | 3724 | 1686 | 681 | 9132.5 | 3758 | 5387.5 |
| 1964 | 3081 | 2544 | 983 | 3555 | 1663 | 664 | 9161 | 3639 | 5187 |
| 1900 | 3030 | 2553 | 978 | 3518 | 1649 | 658 | 9046 | 3576 | 5128 |
| 1899 | 3113 | 2705 | 1026 | 3706 | 1720 | 693 | 9243 | 3782 | 5448 |
| 1845 | 3109 | 2734 | 1041 | 3804 | 1733 | 721 | 9487 | 3829 | 5581 |
| 1842 | 3109 | 2703.5 | 1041 | 3878.5 | 1744 | 716.5 | 8832.5 | 3820 | 5577.5 |
| 1860 | 3276 | 2857 | 1081 | 4022 | 1802 | 738 | 9489.5 | 3962.5 | 5903.5 |
| 1900 | 3289.5 | 2890 | 1049 | 4002 | 1770 | 708 | 8971.5 | 3980 | 5865 |
| 36677 | 20042 | 6863 | 1286 | 4076 | 1759 | 694 | 12410 | 5000 | 6422 |
| 2420 | 3925 | 3321 | 1134 | 4340 | 1858 | 729 | 10378 | 4694 | 6508 |
| 1945 | 3406 | 2935 | 1078 | 4024 | 1766 | 695 | 9134 | 4040 | 5904 |
| 2142 | 3587 | 3096 | 1116 | 4328 | 1856 | 755.5 | 10699.5 | 4567.5 | 6377.5 |
| 5169.5 | 12980 | 4954 | 1427 | 6086 | 2143 | 761 | 14691.5 | 6722 | 16422.5 |
| 10763.5 | 23297.5 | 6633.5 | 1555 | 7430 | 2321 | 732 | 16502 | 9439 | 24071.5 |
| 9514 | 22419 | 7136 | 1666 | 8247 | 2371 | 710 | 13805 | 9049 | 23479 |

| | | | | | | | | | |
|---|---|---|---|---|---|---|---|---|---|
| 9445 | 23315 | 7212 | 1640 | 8086.5 | 2398 | 739 | 14634 | 9402.5 | 24827 |
| 2244 | 3167 | 1799 | 689 | 2456 | 1248 | 311 | 7666 | 3151 | 4013.5 |
| 1945 | 2931 | 1989 | 771 | 2755 | 1371 | 455 | 7120.5 | 3081.5 | 4214.5 |
| 1756 | 2822 | 2160 | 843 | 3023 | 1487 | 540 | 7321 | 3176 | 4485 |
| 1870 | 3061 | 5939 | 1799 | 1045813 | 92246 | 644 | 8644 | 3662 | 5217 |
| 1626 | 2703 | 2754 | 986 | 5128 | 1914 | 594 | 7378 | 3196 | 4656 |
| 1719 | 2985 | 2738.5 | 1006 | 4510.5 | 1820 | 654.5 | 8303 | 3567.5 | 5206 |
| 1774 | 3062 | 2715 | 996 | 4268 | 1800 | 687 | 8718 | 3691 | 5433 |
| 1785 | 3039 | 2638 | 981 | 3973 | 1708 | 662 | 8565 | 3610 | 5279 |
| 1964 | 3157 | 2626 | 984 | 3874 | 1715 | 657 | 9190 | 3757 | 5382 |
| 1927 | 3087 | 2570 | 1062 | 3717 | 1729 | 630 | 9028 | 3652 | 5173 |
| 1779 | 2880.5 | 2406 | 954 | 3515 | 1630 | 612 | 8542.5 | 3380 | 4858.5 |
| 1828 | 3041 | 2574 | 993 | 3747 | 1692 | 659 | 8999 | 3639.5 | 5300 |
| 1785 | 3063 | 2629 | 998 | 3780 | 1701 | 682 | 8931 | 3645 | 5398 |
| 1748 | 2987 | 2602 | 980 | 3726 | 1682 | 670 | 8655.5 | 3555 | 5245 |
| 1762 | 3093 | 2687 | 1016 | 3859 | 1731 | 694 | 9021 | 3718 | 5506 |
| 1881 | 3265 | 2840 | 1059 | 4028 | 1762 | 710 | 9388 | 4001 | 5890 |
| 35506 | 20048.5 | 6421 | 1190.5 | 3833 | 1722 | 643 | 11811.5 | 4443 | 5992.5 |
| 2607 | 4404 | 3619 | 1161 | 4546 | 1905 | 745 | 12315 | 4834 | 6887 |
| 2364 | 3965.5 | 3331 | 1169 | 4596 | 1928 | 763 | 11434 | 4919.5 | 6809 |
| 2208 | 3834 | 3169 | 1136 | 4478 | 1899 | 776 | 14186 | 5108 | 7120 |
| 6040 | 10214 | 4218 | 1247 | 5303 | 1957 | 729 | 17280 | 7420 | 12936 |
| 11658 | 25147 | 6718 | 1577 | 7477 | 2315 | 755 | 18473 | 8353.5 | 24593.5 |
| 8627 | 18931 | 6158 | 1535 | 7185 | 2249 | 676 | 14636 | 8806 | 20938 |
| 10715 | 23913 | 6593 | 1557 | 7403 | 2269 | 704 | 14090 | 8278 | 23222 |
| 2506 | 3498 | 1981 | 719 | 2677 | 1309 | 326 | 8454 | 3490 | 4436 |
| 2341 | 3274 | 2283.5 | 850 | 2959 | 1453 | 520 | 8068.5 | 3502 | 4659 |
| 2179 | 3212 | 2541 | 958 | 3342 | 1600 | 658 | 8872 | 3733 | 5200 |
| 2110 | 3216 | 7234 | 1849 | 1309347 | 123484 | 726 | 9541 | 3933.5 | 5425 |
| 2013 | 3143 | 3027 | 1086 | 5123 | 1952 | 744 | 9193 | 3840 | 5414 |
| 2118 | 3407 | 3126 | 1112 | 4726 | 1896 | 803 | 10413 | 4237 | 5970 |
| 2081 | 3373 | 2999 | 1110 | 4400 | 1828 | 808 | 9958 | 4219 | 5990 |
| 2117 | 3377 | 2913 | 1086 | 4222 | 1824 | 804 | 10035 | 4125 | 5816 |
| 2220 | 3277 | 2742 | 1030 | 3815 | 1697 | 757 | 10246.5 | 4019.5 | 5533.5 |
| 2293 | 3395 | 2821 | 1082 | 3806 | 1752 | 759 | 10828 | 4133 | 5641 |
| 2248 | 3385.5 | 2810 | 1056 | 3850 | 1739 | 770 | 10591.5 | 4094 | 5742 |
| 2442.5 | 3595.5 | 3085.5 | 1134.5 | 4201.5 | 1838 | 826 | 11349.5 | 4642 | 6255.5 |
| 2181 | 3446 | 3015 | 1110 | 4099 | 1810 | 821 | 10882 | 4329 | 6049 |
| 2350 | 3676 | 3171 | 1174 | 4286 | 1878 | 853 | 11076 | 4643 | 6425 |
| 2108 | 3487 | 3044 | 1131 | 4154 | 1837 | 833 | 10173 | 4327 | 6184 |
| 2283 | 3687 | 3198 | 1167 | 4288 | 1862 | 840 | 10918 | 4701 | 6628 |
| 51868.5 | 26997.5 | 8723.5 | 1430 | 4414 | 1863 | 798 | 15480 | 5962 | 7146 |
| 2665.5 | 4091 | 3489 | 1191 | 4430 | 1883 | 795 | 11275 | 5138.5 | 6782 |
| 2563 | 4098 | 3475.5 | 1217 | 4745 | 1957 | 832 | 11874 | 5174 | 7115 |
| 2375 | 3823 | 3384 | 1206 | 4593 | 1952 | 827 | 11100 | 4867 | 6826 |
| 5559.5 | 12231.5 | 5044 | 1449 | 6101 | 2140 | 830 | 16012 | 6614 | 15045 |
| 8748 | 20652 | 6820 | 1737 | 7913 | 2472 | 791 | 15072 | 9389 | 23456 |
| 9473 | 23387 | 8903 | 1903 | 10087 | 2659 | 825 | 12135 | 8548 | 24686 |

| | | | | | | | | | |
|---|---|---|---|---|---|---|---|---|---|
| 9146 | 22972 | 7229 | 1738 | 8241 | 2463 | 771 | 14662 | 9377 | 25356 |
| 2136 | 2983 | 1709 | 694 | 2385 | 1260 | 295 | 7274 | 2966 | 3793 |
| 2158 | 3123 | 2151 | 820 | 2900 | 1422 | 489 | 7591 | 3268 | 4452 |
| 2113 | 3128 | 2465 | 926 | 3241 | 1578 | 629 | 8432 | 3589 | 4871 |
| 2220 | 3278 | 7509 | 1991 | 1426946 | 138138 | 751 | 9670 | 4081 | 5622 |
| 2014 | 3098.5 | 2938 | 1065 | 5114.5 | 1917 | 735 | 9141.5 | 3770.5 | 5271 |
| 2203 | 3410 | 3025 | 1112 | 4658 | 1891 | 824 | 10294 | 4206 | 5864 |
| 2206 | 3546 | 3206 | 1145 | 4697 | 1903 | 827 | 10647 | 4389 | 6249 |
| 2344 | 3627.5 | 3116.5 | 1150 | 4506.5 | 1876.5 | 820.5 | 10418 | 4470 | 6208 |
| 2594 | 3776.5 | 3118 | 1139 | 4382.5 | 1857 | 840 | 11459.5 | 4723.5 | 6271.5 |
| 2511 | 3595 | 3053 | 1111 | 3955 | 1805 | 813 | 11585 | 4479 | 5982 |
| 2402.5 | 3485.5 | 2945 | 1091 | 3946 | 1769 | 811 | 10889.5 | 4292 | 5876.5 |
| 2621.5 | 3814 | 3295 | 1191 | 4337 | 1893 | 899 | 12525.5 | 4923.5 | 6516.5 |
| 2415 | 3699 | 3284 | 1196 | 4314 | 1897 | 905 | 11877 | 4590 | 6508 |
| 2579 | 3771 | 3265 | 1221 | 4364 | 1906 | 923 | 11991 | 4916 | 6618 |
| 2587 | 3921.5 | 3503 | 1279.5 | 4578.5 | 1979 | 946 | 12472.5 | 5239.5 | 6926.5 |
| 2451 | 3829 | 3409 | 1229 | 4466 | 1933 | 906 | 11648 | 4774 | 6692 |
| 60304 | 31469 | 9897 | 1553 | 4643 | 1927 | 882 | 16922 | 6513 | 7716 |
| 3026 | 4479.5 | 3862 | 1321 | 4852.5 | 2011 | 915 | 13360.5 | 5854.5 | 7643 |
| 2865 | 4214 | 3619 | 1287 | 4767 | 1996 | 919 | 12647 | 5599 | 7328 |
| 2459.5 | 3959.5 | 3417 | 1231 | 4605.5 | 1962 | 839 | 11593.5 | 5030.5 | 6818 |
| 6868 | 14538 | 5570 | 1535 | 6303.5 | 2218 | 921 | 16387 | 7356 | 17532.5 |
| 12047.5 | 23003.5 | 7477.5 | 1911.5 | 7979 | 2549.5 | 952 | 17219 | 10407.5 | 24408.5 |
| 12996 | 25663 | 8978 | 2071 | 9209 | 2735 | 975 | 15122 | 10104 | 26375 |
| 12164 | 27773 | 8557 | 1957 | 9192 | 2666 | 965 | 12348 | 9513 | 28029 |
| 2267.5 | 3157 | 1830.5 | 704 | 2507 | 1279 | 306 | 7820.5 | 3189.5 | 4013.5 |
| 1894 | 2860 | 1963 | 756 | 2721 | 1360 | 433 | 6881 | 3009 | 4090 |
| 1767 | 2856 | 2183 | 840 | 3051 | 1468 | 525 | 7049 | 3108 | 4433 |
| 1833 | 2982 | 5627 | 1594 | 897800 | 82531 | 600 | 8062 | 3439 | 4919 |
| 1720 | 2877 | 2934 | 1017 | 5118 | 1968.5 | 601 | 7524.5 | 3347 | 4875 |
| 1736 | 2975 | 2663 | 988 | 4308 | 1752 | 626 | 7720 | 3389 | 5060 |
| 1861 | 3163 | 2754.5 | 1007 | 4249 | 1753 | 670 | 8696 | 3698 | 5489 |
| 2096 | 3466 | 2921 | 1056 | 4381 | 1816 | 709 | 10645 | 4306 | 6094 |
| 2059 | 3320.5 | 2693 | 977 | 3958.5 | 1700 | 642 | 9547.5 | 3880 | 5484 |
| 1875 | 2995 | 2441 | 939 | 3478 | 1612.5 | 602 | 8348 | 3410 | 4847 |
| 1903 | 3103 | 2543 | 957 | 3676 | 1648 | 625 | 8725.5 | 3571.5 | 5144.5 |
| 1870.5 | 3095 | 2591 | 970 | 3705 | 1653 | 634 | 8811.5 | 3578.5 | 5205 |
| 1972 | 3286 | 2788 | 1025 | 3997 | 1748 | 686 | 9630 | 4033 | 5723 |
| 1794 | 3065 | 2573 | 982 | 3747 | 1681 | 641 | 8440 | 3616 | 5262 |
| 1907 | 3311 | 2802 | 1046 | 4027 | 1768 | 703 | 9175 | 3914 | 5754 |
| 2159 | 3588 | 3074 | 1093 | 4292 | 1824 | 710 | 10113 | 4501 | 6289 |
| 36166 | 19503.5 | 6463.5 | 1233 | 4140.5 | 1762 | 648 | 11694 | 4855.5 | 6340 |
| 3049 | 5045 | 3964 | 1222 | 4745 | 1957 | 758 | 12926 | 5143 | 7413 |
| 2212 | 3743 | 3090 | 1104.5 | 4270.5 | 1808 | 682 | 9682 | 4412.5 | 6236 |
| 2216 | 3741 | 3110 | 1110 | 4433 | 1853 | 725 | 11291 | 4745 | 6638 |
| 4664 | 12244 | 4659 | 1341 | 5862.5 | 2040.5 | 700 | 14239 | 6204 | 15511 |
| 8091 | 19627 | 6496 | 1607 | 7545 | 2336 | 725 | 15129 | 9261 | 22905 |
| 9237 | 23032 | 7743 | 1767 | 8940 | 2509 | 742 | 13263 | 8636 | 23608 |

| | | | | | | | | | |
|---|---|---|---|---|---|---|---|---|---|
| 10889 | 25369 | 7312.5 | 1631 | 8053.5 | 2367 | 725 | 15870.5 | 9563 | 26455.5 |
| 2267 | 3171 | 1847 | 712 | 2563 | 1291 | 311 | 7701 | 3121 | 3996 |
| 2024 | 3021 | 2081 | 795 | 2838 | 1407 | 462 | 7313 | 3114 | 4277 |
| 1917 | 2991 | 2304 | 900 | 3211 | 1523 | 553 | 7419 | 3305 | 4698 |
| 1938 | 3097 | 6316 | 1650 | 978617 | 90849 | 631 | 8223 | 3565 | 5122 |
| 1881 | 2998 | 2909 | 1021 | 4820 | 1900 | 643 | 7922 | 3509 | 4975 |
| 1908.5 | 3123 | 2794 | 1019 | 4273 | 1777 | 673 | 8429.5 | 3680 | 5275 |
| 1977.5 | 3296 | 2848 | 1039 | 4314.5 | 1792 | 706 | 8758.5 | 3801 | 5588 |
| 2184 | 3556.5 | 3011 | 1074.5 | 4411.5 | 1832 | 718 | 9660 | 4168 | 5942.5 |
| 2238 | 3426 | 2821 | 1030 | 4019 | 1734 | 710 | 9724 | 4057 | 5590 |
| 2299 | 3423 | 2835 | 1047 | 3885.5 | 1742.5 | 704 | 10195.5 | 4035 | 5592 |
| 2283 | 3512 | 2910 | 1071 | 4018 | 1776 | 720 | 10348 | 4151 | 5840 |
| 2240 | 3447.5 | 2904 | 1066 | 4057 | 1759 | 727 | 10058 | 4129.5 | 5821.5 |
| 2226 | 3392 | 2956 | 1093 | 4106 | 1774 | 754 | 10254 | 4293 | 5919 |
| 2233 | 3559.5 | 3021.5 | 1125 | 4189 | 1833 | 755 | 9998.5 | 4292 | 6052 |
| 2244 | 3706.5 | 3171 | 1159.5 | 4341.5 | 1878 | 808 | 10782 | 4478 | 6437 |
| 2236 | 3652 | 3092 | 1119 | 4256 | 1844 | 746 | 10052 | 4388 | 6194 |
| 30459 | 15790 | 5979 | 1325 | 4328 | 1868 | 789 | 13099 | 5541 | 6829 |
| 2570 | 4129 | 3438 | 1183 | 4583 | 1895 | 739 | 11215 | 4835 | 6768 |
| 2503 | 3825 | 3202 | 1154 | 4435 | 1846 | 755 | 10550 | 4828 | 6517 |
| 2621 | 4051 | 3469.5 | 1247.5 | 4715 | 1963 | 851 | 11393.5 | 5228 | 6927.5 |
| 5537 | 13146 | 5022 | 1433 | 6135 | 2131 | 770 | 14700 | 7132 | 16545 |
| 10202 | 19290 | 6558.5 | 1702.5 | 7568.5 | 2404 | 807 | 15152.5 | 9436 | 21465.5 |
| 10952.5 | 24504.5 | 8123.5 | 1950.5 | 8875 | 2626 | 863 | 13572.5 | 9428.5 | 25299 |
| 8933 | 22867 | 7277.5 | 1758.5 | 8570 | 2494 | 763 | 13461.5 | 9190 | 25210.5 |
| 2129 | 3001 | 1743 | 698 | 2460 | 1259 | 295.5 | 7207 | 2955 | 3802 |
| 2540 | 3367 | 2355 | 901 | 3025 | 1465 | 552 | 8391 | 3595 | 4723 |
| 2528 | 3500 | 2687 | 993 | 3484 | 1639 | 714 | 9650 | 4009 | 5394 |
| 2526.5 | 3584 | 8191 | 2125 | 1548526 | 149855 | 805 | 10472.5 | 4419.5 | 5984.5 |
| 2344 | 3426 | 3290 | 1157 | 5776 | 2037 | 817 | 10335 | 4277 | 5725 |
| 2332 | 3523.5 | 3148 | 1130 | 4935 | 1904.5 | 832 | 10494.5 | 4316 | 5990.5 |
| 2404 | 3669 | 3274 | 1147 | 4901 | 1922 | 899 | 11439 | 4617 | 6345 |
| 2614 | 3840 | 3266.5 | 1184 | 4723 | 1916 | 874 | 11790 | 4861.5 | 6517.5 |
| 3052 | 4195 | 3469 | 1226 | 4821 | 1985 | 941 | 13038 | 5319 | 6930 |
| 2727 | 3768 | 3124 | 1146 | 4029 | 1805 | 860 | 11851 | 4612 | 6144 |
| 2764 | 3877.5 | 3148 | 1124.5 | 4213 | 1837 | 881 | 12302 | 4763 | 6371.5 |
| 2601 | 3730 | 3122 | 1164 | 4235 | 1851 | 852 | 11807 | 4756 | 6269 |
| 2608 | 3775 | 3304 | 1200 | 4435 | 1902 | 891 | 11605 | 4995 | 6569 |
| 2619 | 3831 | 3302 | 1182 | 4324.5 | 1889 | 893 | 11569.5 | 4835 | 6552 |
| 2470.5 | 3735 | 3305.5 | 1213 | 4378 | 1912 | 873 | 11443.5 | 4780 | 6500.5 |
| 2522 | 3843 | 3307 | 1186 | 4447 | 1901 | 822 | 11170 | 4869 | 6593 |
| 66716 | 34746.5 | 10613 | 1559.5 | 4592 | 1909.5 | 873 | 17727 | 6607 | 7638 |
| 3149 | 4894 | 4103 | 1369 | 5057 | 2056 | 933 | 14735 | 5820 | 8078 |
| 2983 | 4299 | 3639 | 1266 | 4769 | 1980 | 905 | 12281 | 5685 | 7234 |
| 2972.5 | 4418.5 | 3793 | 1303 | 4938 | 2084.5 | 956 | 13272 | 5753.5 | 7587.5 |
| 6291 | 13394 | 5429 | 1542 | 6293 | 2240 | 942 | 16903 | 6664 | 16164 |
| 12803 | 24880 | 8078 | 1972 | 8485 | 2658 | 971 | 16893 | 10678 | 25885 |
| 12746 | 27060 | 8364 | 1969.5 | 8905.5 | 2646 | 917.5 | 14082.5 | 9694 | 27636.5 |

| | | | | | | | | | |
|---|---|---|---|---|---|---|---|---|---|
| 10676 | 26249 | 8655.5 | 2013 | 9584 | 2697.5 | 930 | 12965.5 | 9455.5 | 27864.5 |
| 2202 | 3106 | 1820 | 721 | 2569 | 1289 | 302 | 7422.5 | 3039 | 3932 |
| 2257 | 3148 | 2155 | 817 | 3093 | 1442 | 477 | 7580.5 | 3219.5 | 4349 |
| 2108.5 | 3112 | 2366 | 890 | 3428 | 1548 | 573 | 8042.5 | 3443 | 4776.5 |
| 2070 | 3226 | 6998 | 1805 | 1307509 | 110420 | 631 | 8472 | 3718 | 5241 |
| 2097 | 3200 | 3015 | 1052 | 8762 | 2233 | 681 | 8726 | 3699 | 5234 |
| 2186 | 3399 | 2919 | 1066 | 6593 | 2027 | 731 | 9429 | 4025 | 5742 |
| 2229 | 3484 | 2988 | 1081.5 | 5887 | 1955 | 751 | 9844 | 4181 | 5867 |
| 2459 | 3775.5 | 3135 | 1117 | 5760 | 1970 | 775 | 10599 | 4586.5 | 6238.5 |
| 2505 | 3704 | 3000 | 1074 | 5227 | 1883 | 749 | 10625 | 4429 | 6002 |
| 2320 | 3491.5 | 2817 | 1057.5 | 4188.5 | 1735.5 | 699 | 10114.5 | 4055 | 5503 |
| 2324 | 3434.5 | 2835 | 1025 | 4380 | 1753 | 713.5 | 10326.5 | 4025 | 5602 |
| 2260 | 3459 | 2892 | 1074 | 4375 | 1782 | 727 | 10009 | 4149 | 5794 |
| 2249 | 3481 | 2979 | 1085 | 4432 | 1826 | 758 | 10079 | 4282 | 5918 |
| 2220 | 3430.5 | 2837 | 1040 | 4195 | 1750 | 734 | 9457.5 | 3981 | 5709 |
| 2482 | 3747 | 3164 | 1160 | 4688 | 1914 | 807 | 10954 | 4795 | 6382 |
| 2321 | 3673.5 | 3116 | 1121 | 4514 | 1841 | 749 | 10090 | 4502 | 6241.5 |
| 62643 | 32550 | 9939 | 1501 | 4657 | 1886 | 730 | 16000.5 | 6017.5 | 7091 |
| 2885 | 4355 | 3550 | 1188 | 4798 | 1907 | 760 | 11544 | 5085 | 6827 |
| 2563 | 3937 | 3254.5 | 1136.5 | 4655 | 1887 | 748 | 10831.5 | 4898 | 6560.5 |
| 2624 | 4234 | 3469 | 1208 | 5029 | 2008 | 825 | 12734 | 5066 | 7142 |
| 5805 | 13276 | 5063 | 1430 | 6254 | 2185 | 801 | 15145 | 6820 | 16371 |
| 12983 | 26307 | 8201 | 1873 | 9223 | 2684 | 880 | 14085 | 9612 | 26408 |
| 10356.5 | 23187 | 7698 | 1851.5 | 8691 | 2551 | 817 | 14282 | 9416.5 | 24659.5 |
| 10647.5 | 22495 | 7692 | 1795 | 8924 | 2567 | 824 | 14265 | 9766.5 | 23652.5 |
| 2069 | 2946 | 1732 | 686 | 2495 | 1259 | 290 | 6922 | 2909 | 3725 |
| 1972 | 3023 | 2120 | 794 | 2923 | 1404 | 441 | 7090.5 | 3122 | 4358 |
| 1863 | 3003 | 2279 | 856 | 3206 | 1510 | 527 | 7252 | 3254.5 | 4660 |
| 1931.5 | 3095 | 5440.5 | 1567 | 794010 | 79209.5 | 599 | 7972 | 3525.5 | 5018 |
| 1813 | 3005 | 2867 | 1008 | 4786 | 1891 | 588 | 7650 | 3378 | 4939 |
| 1810 | 3074 | 2707 | 992 | 4324.5 | 1749.5 | 613 | 7788.5 | 3499 | 5114 |
| 1926 | 3278 | 2800 | 1009 | 4331 | 1782 | 651 | 8355 | 3749 | 5490 |
| 2277.5 | 3671 | 3028.5 | 1075 | 4549 | 1859 | 697 | 9979.5 | 4397 | 6072 |
| 2132 | 3377 | 2719 | 995 | 3990 | 1706 | 632 | 9043 | 3935 | 5459 |
| 2174 | 3438 | 2760 | 1002 | 3931 | 1711 | 638 | 9448 | 3937 | 5519 |
| 2007 | 3275.5 | 2636 | 970 | 3754 | 1675 | 619 | 8906 | 3706 | 5335 |
| 2160 | 3381 | 2787 | 1036 | 3957 | 1740 | 660 | 9145 | 4179 | 5635 |
| 1778 | 3015 | 2519 | 947.5 | 3584 | 1621 | 609 | 8015 | 3455 | 5087 |
| 2151 | 3371 | 2801 | 1027 | 3964 | 1730 | 671 | 9033 | 4073 | 5691 |
| 1936 | 3307 | 2766 | 1030 | 3988.5 | 1729 | 665 | 8738 | 3819.5 | 5573 |
| 2058 | 3450 | 2888 | 1056 | 4142.5 | 1759 | 660 | 8600 | 4084 | 5834.5 |
| 36021 | 18359.5 | 6281 | 1252 | 4254 | 1780 | 648 | 11705.5 | 4992.5 | 6374.5 |
| 2637 | 4210 | 3351 | 1114 | 4334 | 1804 | 669 | 10024 | 4607 | 6337 |
| 2336.5 | 3911 | 3225.5 | 1104 | 4465.5 | 1840 | 692 | 10135.5 | 4507 | 6461 |
| 2293.5 | 3742 | 3082 | 1118 | 4352.5 | 1837.5 | 699 | 9267.5 | 4567 | 6340.5 |
| 5431 | 13452 | 4898 | 1391 | 6159 | 2114 | 725 | 14265 | 7179 | 16546 |
| 7650 | 19458 | 6521 | 1666 | 7683 | 2435 | 728 | 14171 | 9083 | 22898 |
| 7771 | 20604 | 7179 | 1735 | 8386 | 2472 | 710 | 12531 | 8494 | 23502 |

| | | | | | | | | | |
|---|---|---|---|---|---|---|---|---|---|
| 9528 | 24718.5 | 7275 | 1714 | 8541 | 2456 | 730 | 12752 | 8957.5 | 25522 |
| 2108 | 2993 | 1738 | 695 | 2499 | 1273 | 284 | 6975 | 2949 | 3751 |
| 1935 | 2950 | 2040 | 778 | 2843 | 1390 | 434 | 6882 | 3017 | 4198 |
| 1846 | 2931 | 2211 | 849 | 3097 | 1475 | 508 | 7098 | 3140 | 4468 |
| 1920 | 3068 | 5727 | 1523 | 888725 | 75998 | 587 | 7766 | 3435 | 4928 |
| 1801 | 2981 | 2733 | 965 | 4712 | 1831 | 576 | 7416 | 3296 | 4801 |
| 2102 | 3348 | 2884 | 1018 | 4590.5 | 1813 | 656 | 8804 | 3954 | 5527 |
| 2076 | 3212 | 2616 | 967 | 4001 | 1686 | 803 | 10988 | 4132 | 5392 |
| 2503 | 3744 | 2960 | 1062 | 4469.5 | 1806 | 921 | 13043 | 4898 | 6248.5 |
| 2848.5 | 3910.5 | 3048.5 | 1073 | 4415 | 1809 | 971 | 14346.5 | 5469.5 | 6523 |
| 3307.5 | 4266 | 3255 | 1141 | 4369 | 1833 | 1084.5 | 15832 | 6136 | 7006 |
| 2883 | 3904 | 2944 | 1036 | 4088.5 | 1739 | 1024.5 | 15341.5 | 5329 | 6508 |
| 2485 | 3629.5 | 2796 | 1015 | 3910.5 | 1690 | 966 | 13975 | 4924 | 6182.5 |
| 2457.5 | 3583 | 2901 | 1027 | 3963 | 1702 | 982 | 13694 | 4984.5 | 6185 |
| 2974 | 4109.5 | 3233 | 1104 | 4253 | 1831 | 1173 | 16144.5 | 6197.5 | 7104 |
| 2543 | 3747 | 2975 | 1064 | 4047 | 1753 | 1066 | 14983 | 5487 | 6638 |
| 2558 | 3757 | 2922 | 1039 | 3958 | 1726 | 1057 | 14506 | 5441 | 6651 |
| 44439 | 24055 | 7704 | 1325 | 4338 | 1817 | 1067 | 19165 | 7074 | 7935 |
| 3774 | 4995.5 | 3678.5 | 1179 | 4509 | 1882 | 1090 | 18611 | 6746 | 7825.5 |
| 3327 | 4361 | 3307 | 1111 | 4287 | 1820 | 1137 | 17199 | 6679 | 7405 |
| 2840 | 4019 | 3118 | 1081 | 4026 | 1739 | 1114 | 16820 | 5976 | 7036 |
| 7988 | 10645 | 4778 | 1290 | 5413 | 1965 | 1094 | 19735 | 6482 | 11524 |
| 19845 | 25780 | 7016 | 1620 | 7347 | 2322 | 1157 | 19095 | 10519 | 21576 |
| 21922.5 | 29128 | 6985 | 1554 | 7082.5 | 2224 | 1032 | 18693 | 10588.5 | 22189 |
| 20767 | 28341 | 6793 | 1567 | 7050 | 2206 | 1033 | 16220 | 10111 | 21604 |
| 747 | 867 | 682 | 545 | 1218 | 1073 | 227 | 2124 | 1320 | 1629.5 |
| 1030 | 1616 | 1410 | 677 | 2135 | 1265 | 412 | 3958 | 2087 | 2999 |
| 1262 | 2160 | 1920 | 794 | 2799 | 1433 | 536 | 6509 | 2695 | 4001 |
| 1375 | 2401 | 5549 | 1621 | 897611 | 93229 | 591 | 6884 | 3016 | 4478 |
| 1438 | 2553 | 2634 | 981 | 4168 | 1893 | 622 | 7159 | 3142 | 4689 |
| 1495 | 2680 | 2539 | 972 | 3804 | 1746 | 654 | 7795 | 3343 | 5027 |
| 1427 | 2555.5 | 2339 | 924 | 3371 | 1614 | 623 | 7100.5 | 3096 | 4684 |
| 1444 | 2413 | 2163 | 881 | 3100.5 | 1524.5 | 584 | 6926 | 2960 | 4326.5 |
| 1849 | 3117 | 2726 | 1001 | 3799 | 1747 | 690 | 9716 | 3827.5 | 5527 |
| 1854 | 3049 | 2575.5 | 1012 | 3687 | 1702 | 650 | 8963 | 3662 | 5274.5 |
| 1743 | 2873 | 2470 | 963 | 3472 | 1637 | 641 | 8374 | 3450 | 5038 |
| 1600 | 2628 | 2285.5 | 905 | 3226 | 1566 | 611 | 7583 | 3185 | 4664.5 |
| 1599 | 2785 | 2493 | 954 | 3533.5 | 1651.5 | 646 | 8251.5 | 3404 | 5043.5 |
| 1640 | 2818 | 2482 | 965 | 3557 | 1658 | 662 | 7791 | 3392 | 5128 |
| 1581 | 2823 | 2566 | 989 | 3643 | 1692 | 671 | 8095 | 3432 | 5216 |
| 1579.5 | 2778 | 2467 | 957 | 3492.5 | 1632 | 630 | 7674 | 3361 | 5064 |
| 28825 | 16211.5 | 5614 | 1137 | 3683.5 | 1651 | 616 | 10244 | 4133 | 5655 |
| 2169 | 3679 | 3173 | 1081 | 4103 | 1782 | 679 | 9493 | 4331 | 6131 |
| 1852 | 3305 | 2870 | 1022 | 4076 | 1755 | 677 | 9129 | 4006 | 5910 |
| 1656 | 2970 | 2632 | 989 | 3723 | 1671 | 658 | 8240 | 3561 | 5341 |
| 3756 | 9903.5 | 3950 | 1139 | 4840.5 | 1810.5 | 623 | 13092 | 4879 | 12694.5 |
| 8318.5 | 19037 | 6110 | 1578 | 7176 | 2286.5 | 724 | 14495 | 9051 | 21394.5 |
| 9929 | 22801 | 6657 | 1606 | 7659.5 | 2349 | 699.5 | 14278.5 | 9315 | 24230.5 |

| | | | | | | | | | |
|---|---|---|---|---|---|---|---|---|---|
| 8894 | 20462 | 6128 | 1552 | 7131 | 2247 | 681 | 13550 | 9034 | 22372 |
| 2384 | 3303.5 | 1892 | 696 | 2527.5 | 1262 | 326 | 8165.5 | 3365 | 4216 |
| 1849 | 2764 | 1887 | 746 | 2612 | 1339 | 436 | 6852 | 2952 | 3998 |
| 1849 | 2972 | 2337.5 | 898 | 3256.5 | 1535 | 581 | 8020.5 | 3382 | 4869 |
| 1672 | 2737 | 5752 | 1600 | 947303 | 94336 | 614 | 7714 | 3279 | 4738 |
| 1763 | 3052 | 3053 | 1070 | 4894 | 1976.5 | 669 | 8553 | 3663.5 | 5427 |
| 1717 | 3029.5 | 2756 | 1019 | 4144 | 1783 | 674 | 8563 | 3636 | 5407 |
| 1649 | 2911 | 2588 | 978 | 3891 | 1701 | 662 | 8051 | 3499 | 5233 |
| 1623 | 2693 | 2314 | 914 | 3340 | 1572 | 596 | 7506 | 3194 | 4700 |
| 2036 | 3168 | 2678 | 985.5 | 3741 | 1697 | 672 | 9650.5 | 3899 | 5394 |
| 1825 | 2902.5 | 2429 | 938 | 3374.5 | 1603 | 611 | 8444.5 | 3412 | 4903 |
| 1801 | 2902 | 2447 | 935.5 | 3400 | 1616 | 627.5 | 8423.5 | 3445 | 4923 |
| 1704 | 2840 | 2410 | 954 | 3450 | 1622 | 630 | 8188 | 3409 | 5019 |
| 1555 | 2642 | 2277 | 904 | 3235 | 1551 | 599 | 7360 | 3144 | 4675 |
| 1733 | 2952 | 2540 | 992 | 3664 | 1696 | 671 | 8342 | 3564 | 5245 |
| 1638 | 2895 | 2539 | 981 | 3680 | 1667 | 664 | 8097 | 3509 | 5175 |
| 1735 | 3051 | 2668 | 990 | 3662 | 1673 | 642 | 8112.5 | 3676 | 5365 |
| 28831.5 | 16275.5 | 5669.5 | 1157 | 3711 | 1654 | 616 | 10763 | 4289.5 | 5770 |
| 2246 | 3539 | 3010 | 1038 | 3909.5 | 1720 | 647 | 9041.5 | 4290 | 5808 |
| 1825.5 | 3168 | 2746.5 | 993 | 3820 | 1699 | 651 | 8594 | 3837 | 5585 |
| 1724 | 2989 | 2638 | 988 | 3732 | 1663 | 652 | 8091 | 3616 | 5396 |
| 3931 | 10504.5 | 4120 | 1190.5 | 5141 | 1884 | 655 | 13627.5 | 5167.5 | 13515 |
| 5913 | 14596.5 | 5171 | 1338 | 6191 | 2029 | 626 | 14486.5 | 6821.5 | 17156 |
| 6978 | 15892.5 | 5186.5 | 1324 | 6127 | 2000 | 597 | 13784.5 | 7396 | 17917.5 |
| 7772 | 16973 | 5519 | 1342 | 6340 | 2037 | 628 | 14478 | 7831 | 18843 |
| 2270 | 3162 | 1801 | 700 | 2463 | 1273 | 315 | 7739 | 3192 | 4037.5 |
| 1943 | 2918 | 1981 | 765 | 2738 | 1364 | 451 | 7156 | 3091 | 4193 |
| 1852 | 2949 | 2198.5 | 836 | 3079 | 1495 | 557 | 8894 | 3440 | 4816 |
| 1788 | 2915 | 5639 | 1737 | 976468.5 | 88291 | 624 | 7987 | 3481 | 5027 |
| 1686 | 2868 | 3016 | 1037 | 5758 | 2092 | 615 | 7796 | 3371 | 4927 |
| 1796 | 3049 | 2887 | 1023 | 4777 | 1872 | 680 | 8681 | 3678 | 5407 |
| 1811 | 3111 | 2734 | 1010 | 4227 | 1770 | 709 | 10286 | 3912 | 5640 |
| 2021 | 3297 | 2785.5 | 1021 | 4253.5 | 1792 | 708 | 10592 | 4154 | 5818 |
| 1969 | 3182 | 2630 | 989 | 3914 | 1713 | 668 | 9260 | 3845 | 5421 |
| 1921 | 3095.5 | 2590.5 | 1041 | 3717 | 1750.5 | 634.5 | 8950 | 3638.5 | 5173 |
| 1897 | 3127.5 | 2573.5 | 986 | 3799.5 | 1690 | 653 | 9279.5 | 3713 | 5313 |
| 1908 | 3142 | 2667 | 1005 | 3846 | 1722 | 679 | 9686 | 3801 | 5476 |
| 2038 | 3311 | 2840.5 | 1049 | 4049 | 1802.5 | 740.5 | 11495 | 4305.5 | 6014.5 |
| 1984 | 3355 | 2820.5 | 1040 | 4081.5 | 1797 | 733 | 10939.5 | 4251.5 | 6044 |
| 1969 | 3366 | 2811 | 1065 | 4141 | 1814 | 763 | 13390 | 4621.5 | 6354 |
| 1987 | 3425 | 2922 | 1076 | 4133.5 | 1798.5 | 723 | 10561.5 | 4377 | 6210 |
| 36292 | 19870 | 6430 | 1212 | 3912 | 1720 | 651 | 11864 | 4717 | 6082 |
| 2292 | 3787.5 | 3176.5 | 1071 | 4081.5 | 1771 | 674 | 9575 | 4302 | 6075 |
| 2070 | 3543.5 | 2979.5 | 1060 | 4123 | 1790 | 691 | 9648 | 4252.5 | 6090 |
| 1984 | 3377 | 2866 | 1053 | 4110 | 1784 | 701 | 10254 | 4275 | 6003 |
| 4134 | 10191 | 4276 | 1240 | 5315 | 1929 | 686 | 14998 | 5272 | 12975 |
| 7091 | 16246 | 5447 | 1412 | 6510 | 2144 | 683 | 14550 | 7963 | 18650 |
| 8364 | 18165 | 5891 | 1492 | 6908 | 2193 | 671 | 14481 | 8758 | 20336 |

| | | | | | | | | | |
|---|---|---|---|---|---|---|---|---|---|
| 10359 | 20689 | 6170 | 1533 | 7234 | 2238 | 705.5 | 14589.5 | 8942 | 21568.5 |
| 2468 | 3414 | 1989 | 715 | 2667 | 1295 | 331 | 8788 | 3446 | 4301 |
| 2140 | 3142 | 2129 | 801 | 2882 | 1400 | 478 | 8001 | 3280 | 4409 |
| 1984 | 3075 | 2388 | 909 | 3260 | 1553 | 594 | 8163 | 3485 | 4882 |
| 2036 | 3153 | 6691 | 1831 | 1139523 | 108837 | 696 | 9096 | 3817 | 5424 |
| 1926.5 | 3150 | 3046 | 1061 | 5076 | 1958 | 697 | 8791.5 | 3772 | 5386.5 |
| 1971 | 3202.5 | 2905 | 1050 | 4494 | 1827 | 731 | 9093.5 | 3919.5 | 5609.5 |
| 1907 | 3105 | 2791 | 1031 | 4119 | 1748 | 718 | 8763 | 3842 | 5480 |
| 1979 | 3045 | 2616 | 990 | 3790 | 1688 | 718 | 9161 | 3783 | 5296 |
| 2060 | 3221 | 2655.5 | 1009 | 3791 | 1670 | 694 | 9471.5 | 3842 | 5378 |
| 2228 | 3393 | 2766 | 1069 | 3840 | 1739 | 718 | 10320 | 4011 | 5634 |
| 2136 | 3316 | 2791 | 1024 | 3900 | 1723 | 718 | 10198 | 3980 | 5590 |
| 2223 | 3440 | 2948 | 1094 | 4089.5 | 1818 | 764 | 10304 | 4334 | 5974 |
| 2181 | 3492 | 3053 | 1125 | 4227 | 1841 | 791 | 10472 | 4437 | 6199 |
| 1957 | 3221 | 2762 | 1025 | 3864.5 | 1729 | 730 | 9045 | 3922 | 5573 |
| 1874 | 3161 | 2766 | 1044 | 3896 | 1750 | 704 | 8680 | 3865 | 5559 |
| 2036 | 3371 | 2896 | 1060 | 4058 | 1766 | 713 | 9162 | 4168 | 5927 |
| 40201.5 | 21708.5 | 7139 | 1304 | 4214 | 1792 | 693.5 | 12874 | 5221 | 6598 |
| 2472 | 3972 | 3320.5 | 1113 | 4184 | 1810 | 713 | 10046.5 | 4576 | 6340 |
| 2255 | 3651 | 3081 | 1125 | 4247 | 1821 | 728 | 9941 | 4465 | 6274 |
| 2198 | 3662 | 3156 | 1122 | 4390.5 | 1872 | 747 | 10078.5 | 4536.5 | 6420.5 |
| 4640.5 | 11992 | 4822.5 | 1373 | 5823 | 2064 | 743.5 | 14851.5 | 5808.5 | 14966 |
| 7736 | 17120.5 | 5657.5 | 1472 | 6738 | 2167.5 | 670.5 | 15098 | 8316 | 19274 |
| 8648.5 | 19815.5 | 6138 | 1530 | 7286.5 | 2252.5 | 667 | 14044 | 9002 | 21982.5 |
| 9171 | 20968.5 | 6212 | 1524 | 7216 | 2240.5 | 682 | 14395.5 | 8945.5 | 22141 |
| 2208 | 3082 | 1761 | 685 | 2414 | 1260 | 306 | 7474 | 3061 | 3867 |
| 2112 | 3069 | 2140 | 813 | 2878 | 1414 | 482 | 7621 | 3268 | 4382 |
| 2075 | 3073.5 | 2414 | 915 | 3171 | 1536 | 609 | 8150.5 | 3490 | 4820.5 |
| 2135 | 3225.5 | 7244 | 1882 | 1318451 | 126615 | 713 | 9375 | 3933.5 | 5450.5 |
| 2127.5 | 3249.5 | 3068.5 | 1099 | 5150 | 1923 | 753 | 9461.5 | 3984 | 5543 |
| 2224 | 3558 | 3164 | 1126 | 4891 | 1931 | 786 | 10384 | 4398 | 6162 |
| 2142 | 3393 | 2985 | 1094 | 4397 | 1852 | 780 | 9764 | 4201 | 5893 |
| 2407 | 3659 | 3130 | 1142 | 4585 | 1918 | 833 | 10784 | 4700 | 6239 |
| 2234 | 3326 | 2768 | 1021 | 3846 | 1704 | 725 | 10130 | 4032 | 5558 |
| 2273 | 3353 | 2758 | 1054 | 3680 | 1699 | 720 | 10182 | 3997 | 5512.5 |
| 2608 | 3688 | 3106 | 1135 | 4187 | 1803 | 807 | 11585 | 4724 | 6205 |
| 2389 | 3585 | 3035 | 1105 | 4165 | 1829 | 804 | 10708 | 4546 | 6108 |
| 2347 | 3681 | 3198 | 1159 | 4389 | 1874 | 820 | 11202 | 4703 | 6408 |
| 2283 | 3511 | 3056.5 | 1131 | 4167.5 | 1842 | 792 | 10166 | 4404.5 | 6141 |
| 2278 | 3658 | 3148 | 1176 | 4288 | 1883 | 850 | 10556 | 4636 | 6478 |
| 2292.5 | 3622 | 3121 | 1129 | 4135 | 1853 | 815 | 10414 | 4607 | 6288.5 |
| 54210 | 28586 | 8992 | 1426 | 4246.5 | 1845 | 785 | 14613.5 | 5785 | 7062 |
| 2726.5 | 4106 | 3533 | 1206 | 4421 | 1865 | 815 | 11419.5 | 5104.5 | 6826 |
| 2460.5 | 3787 | 3202.5 | 1148 | 4345 | 1842 | 795 | 10741.5 | 4808.5 | 6572 |
| 2481 | 3832 | 3310 | 1183 | 4440 | 1883.5 | 849 | 10855.5 | 4898.5 | 6665 |
| 5421.5 | 12542 | 5295 | 1491.5 | 6144.5 | 2189.5 | 884 | 16877.5 | 6309 | 16008.5 |
| 9787.5 | 20083 | 6832.5 | 1733 | 7661.5 | 2407.5 | 830 | 15999.5 | 9499 | 22058.5 |
| 11005 | 22800.5 | 7399 | 1779 | 8229 | 2470 | 814 | 14097 | 9404 | 23672 |

| | | | | | | | | | |
|---|---|---|---|---|---|---|---|---|---|
| 9545 | 22095 | 6582.5 | 1575 | 7533.5 | 2292.5 | 700 | 14402.5 | 9264 | 23701 |
| 2285 | 3188 | 1885 | 706 | 2567 | 1293 | 322 | 8112 | 3180 | 4071 |
| 1910 | 2889 | 1980 | 775 | 2792 | 1360 | 439 | 7041 | 3064 | 4139 |
| 1826.5 | 2982 | 2284 | 864 | 3184 | 1520 | 549 | 7393 | 3284 | 4701 |
| 1806 | 2943 | 5605 | 1569 | 876956 | 80159 | 588 | 7729 | 3373 | 4873 |
| 1746.5 | 2941 | 3006.5 | 1022 | 5275 | 2043 | 603 | 7586 | 3365 | 4900 |
| 1889 | 3249 | 2882 | 1017 | 4571 | 1839 | 658 | 8520 | 3754 | 5486.5 |
| 1929 | 3240 | 2779 | 1013 | 4326 | 1788.5 | 677 | 9118 | 3955 | 5614 |
| 1956 | 3247 | 2715 | 995 | 4098 | 1736 | 657 | 9116.5 | 3854.5 | 5545 |
| 1905 | 3083 | 2506 | 926 | 3703 | 1641 | 609 | 8576 | 3531.5 | 5101 |
| 2006 | 3219 | 2642 | 985 | 3727 | 1679 | 623 | 9145 | 3685 | 5261 |
| 1942 | 3147 | 2561 | 946 | 3658 | 1660 | 626 | 8888 | 3640 | 5188 |
| 2072 | 3316 | 2756 | 999 | 3911 | 1704 | 673 | 9734.5 | 4081 | 5603.5 |
| 2011 | 3273 | 2761 | 1019 | 3980 | 1733 | 682 | 9533 | 4102 | 5647 |
| 2068 | 3341 | 2811 | 1042 | 4027 | 1758 | 699 | 9782 | 4166 | 5809 |
| 2095 | 3476 | 2917 | 1061 | 4200 | 1799 | 730 | 10301 | 4443 | 6210 |
| 2074 | 3471 | 2920 | 1057 | 4199 | 1789 | 698 | 10073 | 4393 | 6167 |
| 37140.5 | 19751.5 | 6716 | 1253 | 4302.5 | 1814 | 674 | 12423 | 5169 | 6629 |
| 2414 | 3954.5 | 3207.5 | 1075 | 4216 | 1776 | 664 | 9656.5 | 4504 | 6219 |
| 2283 | 3754 | 3090 | 1077 | 4304 | 1811 | 692 | 10330.5 | 4666 | 6422 |
| 2176.5 | 3591 | 3019 | 1082 | 4295.5 | 1802 | 706 | 10152 | 4459 | 6298.5 |
| 4888 | 11876 | 4541 | 1268 | 5608 | 2002 | 687 | 15841 | 5922 | 14741 |
| 8896 | 19549.5 | 6030.5 | 1525 | 7095 | 2252 | 691 | 16122.5 | 8703 | 21187 |
| 9463 | 20962 | 6297 | 1534 | 7384 | 2255 | 671 | 14501 | 9127 | 22267 |
| 10003 | 22217 | 6390 | 1533 | 7274 | 2244 | 680 | 14671 | 9080 | 23218 |
| 2154 | 3058 | 1806 | 698.5 | 2509 | 1284.5 | 311 | 7468 | 3037.5 | 3922 |
| 2180 | 3072 | 2139 | 818 | 2843.5 | 1412 | 481 | 7644.5 | 3245 | 4301.5 |
| 1919 | 3008 | 2356 | 885 | 3233 | 1523 | 567 | 7642.5 | 3352.5 | 4711 |
| 1948 | 3079 | 6278 | 1739 | 959540 | 93334.5 | 627 | 8288 | 3567 | 5132 |
| 1900.5 | 3110 | 3025 | 1049 | 5103 | 1956 | 642 | 8112.5 | 3563.5 | 5128.5 |
| 2172.5 | 3479.5 | 3117.5 | 1097 | 4788 | 1919 | 747 | 9809 | 4233 | 5887 |
| 2452.5 | 3731 | 3279 | 1163 | 4870 | 1967 | 822 | 10912 | 4773 | 6441.5 |
| 2114 | 3369 | 2872 | 1045 | 4163.5 | 1776 | 708 | 9298 | 3997.5 | 5578.5 |
| 2153 | 3302 | 2703 | 996 | 3899 | 1696 | 677 | 9316 | 3836 | 5430 |
| 2372 | 3528.5 | 2925 | 1086 | 3936.5 | 1782 | 729 | 10542.5 | 4182 | 5831.5 |
| 2239.5 | 3438 | 2831 | 1041.5 | 3894.5 | 1718 | 708 | 10218.5 | 4056 | 5651 |
| 2545 | 3745 | 3112.5 | 1137 | 4281.5 | 1843 | 782 | 11567.5 | 4775 | 6295 |
| 2362 | 3631 | 3118 | 1139 | 4295 | 1851 | 791 | 10923 | 4639 | 6260 |
| 2341 | 3616 | 3101 | 1137 | 4255 | 1852 | 794 | 9983 | 4484 | 6198 |
| 2378 | 3720.5 | 3181 | 1164 | 4425.5 | 1888 | 824 | 10832 | 4734.5 | 6521.5 |
| 2260 | 3652 | 3176 | 1119 | 4240 | 1812 | 748 | 9723 | 4463 | 6263 |
| 35390.5 | 19022.5 | 6640.5 | 1312 | 4338 | 1834 | 719 | 12110 | 5258 | 6735.5 |
| 2966 | 4443 | 3686 | 1235 | 4615 | 1916 | 805 | 11512 | 5253 | 7144 |
| 2408 | 3856 | 3210 | 1138 | 4469 | 1854 | 728 | 10020 | 4717 | 6532 |
| 2404 | 3875 | 3271 | 1163 | 4474 | 1891 | 777 | 10348 | 4723 | 6616 |
| 5130 | 12663 | 5142 | 1456 | 6048 | 2124 | 795 | 15655 | 6068 | 15778 |
| 7898 | 18828 | 6563 | 1687 | 7569.5 | 2385.5 | 790 | 14033.5 | 8149.5 | 21049 |
| 10657 | 23260 | 7255 | 1767 | 8000 | 2405 | 774 | 15480 | 9856 | 24693 |

| | | | | | | | | | |
|---|---|---|---|---|---|---|---|---|---|
| 9633 | 23401 | 7320 | 1759 | 8386 | 2486 | 789 | 14379 | 9656 | 25167 |
| 2109 | 2946 | 1735 | 688 | 2451 | 1269 | 296 | 7304 | 2957 | 3739 |
| 2378 | 3301 | 2312 | 873 | 3033 | 1478 | 527 | 8304 | 3486 | 4670 |
| 2405.5 | 3412.5 | 2672.5 | 975 | 3466.5 | 1616 | 665 | 9327.5 | 3899 | 5306 |
| 2353 | 3489 | 7812 | 2029 | 1459500 | 138775 | 746 | 9646 | 4128 | 5689 |
| 2322 | 3496 | 3427.5 | 1182 | 5954 | 2142 | 799.5 | 10234 | 4322 | 5854.5 |
| 2407 | 3615 | 3250.5 | 1151 | 5104.5 | 1972 | 821 | 10694.5 | 4515 | 6181.5 |
| 2608 | 3846 | 3355 | 1177 | 5006 | 1951 | 878 | 11194 | 4854 | 6528 |
| 2736 | 3948 | 3333 | 1181 | 4877 | 1958 | 880 | 11740 | 5017 | 6673 |
| 2382 | 3518 | 2910 | 1063 | 4163 | 1767 | 733 | 10164 | 4211 | 5726 |
| 2357.5 | 3482.5 | 2860.5 | 1047.5 | 3855 | 1730.5 | 714 | 10248 | 4047 | 5635.5 |
| 2533 | 3703 | 2999 | 1089 | 4056 | 1787 | 803 | 11291 | 4448 | 6023 |
| 2474 | 3545.5 | 2930 | 1082 | 4123.5 | 1778 | 742 | 10371 | 4571.5 | 5934 |
| 2395 | 3671.5 | 3115.5 | 1120 | 4235.5 | 1835 | 814.5 | 11091 | 4599 | 6257 |
| 2530 | 3754 | 3125 | 1143 | 4293 | 1842 | 830 | 10969 | 4765 | 6379 |
| 2622 | 3936.5 | 3343 | 1212 | 4515 | 1926 | 868 | 11433.5 | 5113.5 | 6742 |
| 2519.5 | 3925.5 | 3333 | 1189 | 4545 | 1892.5 | 812 | 10957.5 | 5016 | 6686 |
| 45577 | 23834 | 7633 | 1319 | 4346.5 | 1803 | 688 | 13035 | 5336 | 6747.5 |
| 2981 | 4557 | 3725 | 1207 | 4528 | 1895 | 766 | 11450 | 5100 | 6922 |
| 2748 | 4209 | 3488 | 1205 | 4584 | 1924 | 822 | 11796 | 5178 | 6941 |
| 2631 | 4324 | 3505 | 1225 | 4709 | 1965 | 838 | 12128 | 5099 | 7403 |
| 5999.5 | 14105 | 5427 | 1502.5 | 6359 | 2208 | 837 | 15629 | 6762 | 17194.5 |
| 9126 | 19569 | 6599 | 1671 | 7591.5 | 2387 | 763 | 15012 | 8872 | 21374.5 |
| 9453 | 23540 | 7063 | 1668 | 8218 | 2409 | 711 | 13321.5 | 9117.5 | 24627 |
| 10858 | 25271.5 | 6723 | 1602 | 7813 | 2358 | 696 | 14387.5 | 9420.5 | 25405.5 |
| 2170.5 | 3061 | 1790 | 695 | 2510 | 1281 | 307 | 7524 | 3041 | 3864 |
| 2323 | 3224 | 2218 | 825 | 3100 | 1453 | 489 | 8055 | 3333 | 4500 |
| 2138 | 3162 | 2441.5 | 886 | 3431 | 1533 | 584 | 8114.5 | 3509 | 4873.5 |
| 2083 | 3187.5 | 7048 | 1796.5 | 1335913 | 110027 | 628 | 8524 | 3702 | 5217.5 |
| 2087 | 3217 | 3080 | 1061 | 7233 | 2156 | 668 | 8772 | 3732 | 5243 |
| 2265.5 | 3531 | 3047 | 1080 | 6169.5 | 2019.5 | 745 | 9972.5 | 4208 | 5916 |
| 2440 | 3800 | 3254.5 | 1141 | 6054.5 | 2047.5 | 785 | 10823.5 | 4757 | 6410 |
| 2272 | 3454.5 | 2860 | 1040 | 5175 | 1844 | 728 | 9506 | 4121 | 5702.5 |
| 2416 | 3655 | 2982 | 1049 | 4982 | 1845 | 733 | 10614 | 4329 | 5977 |
| 2369 | 3503 | 2857 | 1034 | 4166 | 1757.5 | 714 | 10259 | 4096 | 5648 |
| 2248 | 3342.5 | 2707.5 | 987 | 4196.5 | 1704 | 690 | 10018 | 3975 | 5487 |
| 2571 | 3698 | 3067 | 1101 | 4515 | 1844 | 758 | 11276 | 4733 | 6210 |
| 2308.5 | 3454 | 2864 | 1055 | 4281 | 1767 | 739 | 10479 | 4373 | 5851.5 |
| 2517 | 3768 | 3101 | 1097.5 | 4510 | 1854 | 779.5 | 11176.5 | 4724 | 6224 |
| 2549 | 3898.5 | 3292 | 1177 | 4832.5 | 1931 | 835.5 | 12077 | 5020 | 6667 |
| 2499.5 | 3873 | 3249 | 1163 | 4662 | 1889 | 792.5 | 10814 | 4896.5 | 6691.5 |
| 56345 | 30806 | 9514 | 1465 | 4759 | 1912 | 747 | 16528 | 5953 | 7437 |
| 2848 | 4364.5 | 3572.5 | 1180 | 4726 | 1899 | 760 | 11579 | 5098 | 6926 |
| 2722 | 4227 | 3507 | 1180 | 4926 | 1970.5 | 795 | 11484.5 | 5259 | 7115.5 |
| 2538 | 3913 | 3245.5 | 1145 | 4594 | 1884 | 790 | 10714 | 4879 | 6584 |
| 5792 | 13383 | 5186 | 1434 | 6374 | 2181 | 814 | 16030 | 6945 | 16561 |
| 9453 | 19612 | 6397 | 1637 | 7461 | 2342 | 785 | 16044 | 9157 | 21402 |
| 10412 | 21347 | 6950 | 1679.5 | 8000.5 | 2396 | 770 | 15173.5 | 9409 | 22798.5 |

| | | | | | | | | | |
|---|---|---|---|---|---|---|---|---|---|
| 10884 | 22783 | 6957.5 | 1671 | 7949 | 2364.5 | 769 | 14297 | 9489.5 | 23887.5 |
| 2252 | 3205 | 1912 | 711 | 2658 | 1310 | 322 | 7949 | 3186 | 4051 |
| 1996 | 3050 | 2114 | 785 | 2963 | 1410 | 453 | 7535 | 3203 | 4359 |
| 1883 | 3047 | 2261 | 860 | 3247 | 1499 | 527 | 7487 | 3339 | 4722 |
| 1983 | 3179 | 5858 | 1647 | 857576 | 78184 | 603 | 8267 | 3610 | 5140 |
| 1869 | 3139 | 2970 | 1030 | 4995 | 1937 | 599 | 7938 | 3564 | 5155 |
| 2069 | 3372 | 2925 | 1031 | 4606 | 1827.5 | 649 | 8842.5 | 4041 | 5604 |
| 2092 | 3478 | 2913 | 1051 | 4483 | 1841 | 679 | 8854 | 4154 | 5854 |
| 2039 | 3325 | 2749 | 1005 | 4155 | 1738 | 647 | 8726 | 3898 | 5562 |
| 2006 | 3229.5 | 2590 | 954 | 3827 | 1652 | 614 | 8849 | 3678 | 5188 |
| 2158 | 3443 | 2778.5 | 1011 | 3921.5 | 1728 | 635 | 9527.5 | 3951 | 5502.5 |
| 2216 | 3438.5 | 2778 | 994 | 3931 | 1690 | 642 | 9726 | 4138.5 | 5594.5 |
| 2331 | 3652 | 2967 | 1055 | 4243 | 1809 | 688 | 10816 | 4488 | 5987 |
| 2218.5 | 3528.5 | 2914 | 1042 | 4163.5 | 1790 | 692 | 10085 | 4437 | 5937 |
| 2265 | 3588 | 2953 | 1070 | 4187 | 1811 | 710 | 10234 | 4473 | 6113 |
| 2194 | 3618 | 3012 | 1088 | 4295.5 | 1827 | 709 | 10060 | 4486 | 6180.5 |
| 2183 | 3636.5 | 2990 | 1085 | 4277 | 1798 | 686 | 9479.5 | 4378 | 6219.5 |
| 33647 | 18044 | 6265 | 1251 | 4360 | 1809 | 669 | 11920.5 | 5013 | 6652 |
| 2554 | 4186 | 3340 | 1113 | 4460 | 1836 | 679 | 9998 | 4623 | 6496 |
| 2275 | 3834 | 3123 | 1088 | 4333 | 1803 | 675 | 9710 | 4456 | 6303.5 |
| 2327 | 3858.5 | 3173 | 1097 | 4443 | 1869 | 708 | 10215 | 4613.5 | 6531 |
| 4877 | 12692 | 4685 | 1339 | 5903 | 2063 | 703 | 15024 | 6315 | 15958 |
| 8390 | 20168 | 6145 | 1562 | 7398 | 2310 | 710 | 15186 | 9032 | 22369 |
| 8466 | 21517 | 6470 | 1601 | 7720 | 2319 | 669 | 13549 | 8883 | 23800 |
| 8696 | 22491 | 6766 | 1627 | 7916 | 2374 | 701 | 13846 | 8959 | 24820 |
| 2134 | 3032 | 1773 | 708 | 2579 | 1290 | 305 | 7398 | 3040 | 3850 |
| 1953 | 2932 | 2026 | 768 | 2843 | 1388 | 434 | 6976 | 3062 | 4164 |
| 1842 | 2927 | 2193 | 828 | 3100 | 1476 | 513 | 7176 | 3195 | 4485 |
| 1894 | 3073 | 5641 | 1530 | 853605.5 | 77215.5 | 585 | 7799.5 | 3437.5 | 4947 |
| 1892 | 3108 | 2862.5 | 996 | 4872 | 1878 | 606 | 8092.5 | 3507 | 5135 |
| 2192 | 3539 | 3027 | 1034 | 4748 | 1854 | 675.5 | 10087.5 | 4320 | 5916 |
| 1840 | 3117 | 2594 | 951 | 4097 | 1700 | 615 | 7965 | 3515 | 5157 |
| 1968 | 3200 | 2661 | 949 | 4005 | 1685 | 625 | 8456 | 3666.5 | 5287.5 |
| 2230 | 3459 | 2775 | 984 | 4135 | 1718 | 654.5 | 9597.5 | 4097 | 5632 |
| 2355 | 3630 | 2842 | 1021 | 4060 | 1735 | 659 | 10594.5 | 4426 | 5869 |
| 2413 | 3592.5 | 2845 | 1005 | 4089 | 1729 | 669 | 10451 | 4425.5 | 5823 |
| 1945 | 3193 | 2585 | 941 | 3715 | 1642 | 621 | 8725 | 3648 | 5242 |
| 1936 | 3199 | 2621.5 | 977 | 3786 | 1667 | 635 | 8613 | 3773 | 5368 |
| 2030 | 3283 | 2650 | 977 | 3835 | 1674 | 647 | 8532 | 3875 | 5503 |
| 2207 | 3545 | 2894 | 1030 | 4089 | 1793 | 709 | 11939 | 4517 | 6173 |
| 2052 | 3405 | 2751.5 | 979 | 3996 | 1715.5 | 642 | 8861.5 | 4044.5 | 5667 |
| 37545 | 19307 | 6406 | 1179 | 4093 | 1709 | 633 | 11569 | 4934 | 6264 |
| 2506 | 3999 | 3274 | 1065 | 4221 | 1773 | 662 | 9965 | 4539 | 6267 |
| 2525 | 3854 | 3104 | 1072 | 4266 | 1783 | 760 | 11801 | 4969 | 6401 |
| 2750 | 3869 | 3071 | 1064 | 4051 | 1736 | 963 | 14680 | 5717 | 6632 |
| 8234 | 14653 | 4690 | 1245 | 5217 | 1916 | 1066 | 17619 | 6526 | 15288 |
| 18202 | 25427 | 6583 | 1529 | 6695 | 2173 | 1122 | 19679 | 9772 | 20873 |
| 19768 | 27370.5 | 6866 | 1568 | 7114 | 2225 | 1070 | 18044 | 10395.5 | 22268 |

| | | | | | | | | | |
|---|---|---|---|---|---|---|---|---|---|
| 18762 | 26167.5 | 6603 | 1530.5 | 6961 | 2229 | 1088 | 18510 | 10209 | 22195 |
| 754 | 1027 | 889 | 642 | 1445 | 1184 | 284 | 2393 | 1486 | 1989 |
| 1006.5 | 1555 | 1383.5 | 766 | 2148 | 1306 | 410 | 3728.5 | 1968 | 2995 |
| 1238 | 2051 | 1917 | 850 | 2781 | 1478 | 525 | 5020 | 2591 | 3882 |
| 1210 | 2125.5 | 4931.5 | 1496 | 752453.5 | 63641.5 | 540.5 | 5243.5 | 2549.5 | 4085.5 |
| 1359.5 | 2607 | 3029.5 | 1064.5 | 5783 | 2238 | 647 | 6856 | 3102.5 | 4848.5 |
| 1428 | 2675 | 2802.5 | 1051.5 | 4579.5 | 1906.5 | 654 | 7036 | 3211.5 | 4980.5 |
| 1427.5 | 2713 | 2902 | 1069.5 | 4441 | 1857.5 | 688.5 | 7364 | 3284 | 5211.5 |
| 1880 | 3416 | 3085.5 | 1195.5 | 4818 | 1992.5 | 760 | 9359.5 | 4109.5 | 6239.5 |
| 1994 | 3322.5 | 2915.5 | 1134 | 4351.5 | 1894.5 | 700 | 10016 | 3988 | 5788.5 |
| 2102.5 | 3566 | 2860.5 | 1182 | 4217.5 | 1972.5 | 690 | 9420.5 | 3995 | 5821 |
| 2122 | 3577 | 3124.5 | 1234.5 | 4488.5 | 1985 | 753.5 | 10707.5 | 4488.5 | 6323.5 |
| 2272 | 4387 | 3525 | 1348 | 5220 | 2251 | 880 | 11491 | 4801 | 7963 |
| 1881 | 3386 | 3052 | 1233 | 4318 | 2014 | 747 | 9479 | 4038 | 6256 |
| 14684 | 11662 | 4978 | 1331 | 4341 | 1979 | 755 | 8878 | 7295 | 8815 |
| 2338 | 3880.5 | 3265 | 1247 | 4485.5 | 1977 | 789 | 8581.5 | 4473.5 | 6748 |
| 1899 | 3132 | 2706 | 1052 | 3766 | 1726 | 635 | 7581 | 3576 | 5370 |
| 65422 | 37816 | 10723.5 | 1861.5 | 5476.5 | 2304 | 819 | 15319.5 | 7031 | 9494.5 |
| 2717 | 5075 | 3966 | 1316 | 4897 | 2067 | 737 | 9505 | 4946 | 7571 |
| 2522.5 | 4541 | 3575.5 | 1326 | 5053.5 | 2072 | 762 | 9541 | 4866 | 7627 |
| 1754 | 3024 | 2546 | 990 | 3647 | 1629 | 610 | 7276 | 3417 | 5099 |
| 4125 | 10654 | 4219 | 1253.5 | 5256.5 | 1988 | 621 | 13882.5 | 5353.5 | 13019.5 |
| 7077 | 18573 | 6552 | 1699 | 7862 | 2441 | 685 | 12501 | 8331 | 21882 |
| 9112 | 20747 | 6830 | 1689 | 8002 | 2418 | 668 | 12135 | 9094 | 23267 |
| 7482.5 | 21042 | 6658 | 1628 | 7665 | 2276 | 658 | 11884 | 8713.5 | 23970 |
| 2380 | 3212 | 1881 | 767 | 2587 | 1320 | 330 | 7282 | 3226 | 4205 |
| 2015 | 2909 | 1938 | 831 | 2583 | 1377 | 425 | 6687 | 2959 | 3956 |
| 1943 | 3030.5 | 2278 | 910 | 3076 | 1513 | 550.5 | 7090.5 | 3310.5 | 4804.5 |
| 1602 | 2579 | 4599 | 1388 | 693742 | 64639 | 511 | 6589 | 2911 | 4290 |
| 1831 | 3029 | 2730 | 1032 | 5482 | 1981 | 613 | 7372 | 3367 | 4931 |
| 1869.5 | 3036.5 | 2721 | 1058.5 | 4540 | 1791.5 | 630.5 | 7507 | 3392.5 | 5079 |
| 1977 | 3421 | 2879 | 1138 | 4632 | 1893 | 686 | 8470 | 3839 | 5589 |
| 2113 | 3435.5 | 3003 | 1095 | 4434.5 | 1820 | 677 | 9462 | 4024.5 | 5694.5 |
| 2313 | 3658.5 | 2846.5 | 1117 | 4244.5 | 1894.5 | 681.5 | 9488 | 4232 | 5828 |
| 3042 | 5466 | 3885 | 1468 | 5576 | 2243 | 852 | 14660 | 5492 | 8884 |
| 2701.5 | 4427.5 | 3485 | 1275.5 | 4936 | 2096.5 | 800 | 12537.5 | 5020.5 | 7344.5 |
| 2800 | 5440.5 | 3732 | 1403 | 5141 | 2170 | 849.5 | 11923.5 | 5459.5 | 9443 |
| 2341 | 4273 | 3317 | 1288 | 4925 | 2022 | 807 | 10233 | 4691 | 7267 |
| 14712 | 10893 | 4781 | 1239 | 4186 | 1885 | 685 | 9556 | 7158 | 8081 |
| 2747 | 4505.5 | 3322.5 | 1212 | 4420 | 1923 | 727.5 | 9285 | 4706 | 6701 |
| 2451 | 3815 | 3141 | 1127 | 4187 | 1894 | 693 | 8129 | 4150 | 6128 |
| 75545 | 43730 | 11961 | 1853 | 5095 | 2098 | 763 | 15515 | 7230 | 9389 |
| 2498 | 3729 | 3111 | 1108 | 3917 | 1757 | 596 | 8496 | 4174 | 5955 |
| 2280 | 3590 | 3091 | 1082 | 4226 | 1813 | 637 | 8479 | 4145 | 5881 |
| 2190 | 3448.5 | 2789.5 | 1054 | 3795 | 1769.5 | 624.5 | 8279 | 3811 | 5578 |
| 3847 | 10321 | 4024.5 | 1262.5 | 5158.5 | 1953 | 654 | 14215 | 5261 | 12954.5 |
| 8063.5 | 19922.5 | 7121 | 1773 | 8242.5 | 2480.5 | 691 | 12034 | 8712.5 | 22392 |
| 8605 | 20368 | 7543 | 1724 | 8261 | 2409 | 668 | 12257 | 8721 | 23175 |

| | | | | | | | | | |
|---|---|---|---|---|---|---|---|---|---|
| 7569 | 21596 | 7486.5 | 1793 | 8977.5 | 2593 | 695.5 | 11023 | 8111 | 23979 |
| 2346.5 | 3383 | 1932 | 763 | 2613.5 | 1346.5 | 331 | 7235 | 3180 | 4170.5 |
| 2050 | 3131 | 2054.5 | 835 | 2809 | 1436.5 | 452.5 | 6899.5 | 3123 | 4342 |
| 2061 | 3189.5 | 2378.5 | 969.5 | 3199 | 1549 | 553.5 | 7421.5 | 3327 | 4924.5 |
| 1801.5 | 2837 | 4752.5 | 1466.5 | 694296 | 61768.5 | 537 | 6798 | 3094 | 4430 |
| 1892 | 3177.5 | 2910 | 1057 | 6099.5 | 2060.5 | 601.5 | 7686.5 | 3426.5 | 5157 |
| 2123 | 3614 | 3044 | 1110 | 5379 | 2041 | 693 | 8975 | 3916 | 5680 |
| 2086.5 | 3464 | 3031 | 1112.5 | 5003 | 1965 | 669 | 8570 | 3960 | 5675 |
| 2451 | 4188 | 3401 | 1256 | 5403 | 2172 | 772 | 10663 | 4930 | 6822 |
| 2767.5 | 4510 | 3477.5 | 1290 | 5095 | 2078 | 763 | 12672.5 | 4881 | 7300 |
| 2578 | 4307 | 3196 | 1177.5 | 4600 | 1985 | 718 | 11789.5 | 4689 | 6853 |
| 2407 | 3838.5 | 3028 | 1143.5 | 4385.5 | 1858.5 | 700 | 9936.5 | 4436.5 | 6185 |
| 2356 | 4023.5 | 3151 | 1267 | 4585 | 2015 | 726 | 10426 | 4666 | 6789 |
| 2113.5 | 3531 | 2864.5 | 1113.5 | 4233.5 | 1901.5 | 698.5 | 9111.5 | 4082.5 | 6060 |
| 14571.5 | 11234 | 5036.5 | 1349.5 | 4919.5 | 2066.5 | 784.5 | 11521 | 7540 | 9191 |
| 2447 | 3868 | 2886 | 1110 | 3976 | 1791 | 670 | 8398 | 4118 | 5735 |
| 2250.5 | 3660 | 2890 | 1076 | 4011.5 | 1698 | 620 | 8099.5 | 3889.5 | 5748 |
| 54231 | 32030 | 10036 | 1827 | 5924.5 | 2308.5 | 807 | 14817 | 6972.5 | 10516 |
| 2843 | 4982 | 3558 | 1199 | 4505 | 1901 | 694 | 9844 | 4904 | 7441 |
| 2909 | 5724 | 3953 | 1361 | 5392 | 2223 | 753 | 10385 | 5538 | 8962 |
| 2224 | 3497 | 2832 | 1053 | 3883 | 1714 | 627 | 8554 | 3798 | 5616 |
| 4801.5 | 10619.5 | 4563 | 1407 | 5880.5 | 2064 | 687.5 | 14820 | 6649 | 13380 |
| 7348 | 19347.5 | 6602 | 1691.5 | 7732 | 2388.5 | 696.5 | 14003.5 | 8345 | 22070 |
| 8489 | 21978 | 7122 | 1769 | 8344 | 2448 | 648 | 11866 | 8983 | 24772 |
| 9212 | 19943 | 6619 | 1633 | 7876 | 2315 | 663 | 13091 | 8363 | 21700 |
| 1979 | 2864 | 1732.5 | 745.5 | 2471.5 | 1302 | 328 | 6772 | 2817.5 | 3703 |
| 1824.5 | 2761 | 1962.5 | 818.5 | 2750.5 | 1432.5 | 442 | 6400 | 2816.5 | 3947 |
| 1817 | 2915 | 2248 | 898.5 | 3188 | 1524.5 | 527.5 | 6840.5 | 3078.5 | 4558.5 |
| 1862 | 3065 | 5253 | 1650 | 721630 | 60770 | 578 | 7603 | 3370 | 4853 |
| 2023.5 | 3370.5 | 3400 | 1130.5 | 6842.5 | 2273 | 654 | 8501 | 3635 | 5419.5 |
| 2220 | 3681 | 3270 | 1160.5 | 5616.5 | 2062 | 692 | 8707.5 | 4002.5 | 6024.5 |
| 2122 | 3669 | 3094 | 1162 | 5212 | 2011 | 674 | 9137 | 4078 | 5813 |
| 2362 | 3728 | 3152 | 1156 | 5035 | 1949 | 710 | 10024 | 4246 | 6134 |
| 2039 | 3158 | 2473 | 948 | 3864 | 1675 | 596 | 8305 | 3498 | 5050 |
| 2639 | 4306 | 3451 | 1323 | 4823 | 2061 | 707 | 11559 | 4845 | 7099 |
| 3049 | 5606 | 4203.5 | 1518.5 | 5971.5 | 2329 | 906 | 14675 | 5766 | 9253.5 |
| 2308 | 3855 | 3109 | 1181 | 4486 | 1920 | 701 | 9824 | 4435 | 6591 |
| 2436 | 4458 | 3258 | 1264 | 4904 | 2075 | 801 | 11417 | 4847 | 7525 |
| 14672 | 10957 | 4523 | 1135 | 3764 | 1718 | 632 | 7888 | 6823 | 7446 |
| 2253.5 | 3382.5 | 2692 | 1022 | 3555 | 1640 | 594.5 | 7305.5 | 3618 | 5452 |
| 2323 | 3719.5 | 2898 | 1048 | 4035.5 | 1766 | 640.5 | 8111.5 | 3968.5 | 5728.5 |
| 72616 | 38533.5 | 11024.5 | 1479.5 | 4315.5 | 1792.5 | 624 | 13435.5 | 5730 | 7054.5 |
| 2715 | 4184 | 3394.5 | 1171 | 4507.5 | 1871.5 | 632 | 9152 | 4559 | 6540.5 |
| 2467 | 3746 | 3137 | 1112 | 4231 | 1795 | 627 | 8754 | 4288 | 6062 |
| 2455 | 3988 | 3182 | 1163 | 4509 | 1872 | 696 | 9108 | 4264 | 6572 |
| 4508 | 12620 | 5017 | 1406 | 6050 | 2089 | 643 | 13518 | 5928 | 15681 |
| 6143 | 24243 | 8853 | 2087 | 11168 | 3000 | 807 | 11374 | 7435 | 27748 |
| 7699 | 18240 | 9329 | 1979 | 10917 | 2825 | 731 | 12740 | 8799 | 21070 |

| | | | | | | | | | |
|---|---|---|---|---|---|---|---|---|---|
| 8062.5 | 21862 | 7382.5 | 1803 | 8680 | 2453.5 | 680.5 | 11688 | 8724.5 | 24709.5 |
| 2306 | 3241 | 1977 | 775 | 2655 | 1344 | 329 | 7659 | 3116 | 3986 |
| 2130 | 3229 | 2175 | 872 | 2923 | 1436 | 453 | 7086 | 3132 | 4387 |
| 2534 | 3913 | 2925 | 1063 | 3833 | 1720 | 617 | 9094 | 4074 | 6000 |
| 2478 | 3908 | 6260 | 1908 | 868329 | 68969 | 681 | 9108 | 4414 | 6092 |
| 2687 | 4387.5 | 3965.5 | 1298 | 7363 | 2452 | 739 | 9952 | 4684 | 6931 |
| 2111 | 3480 | 3069 | 1114 | 5258 | 1922 | 653 | 8611 | 3795 | 5530 |
| 1927 | 3061 | 2605 | 992 | 4449 | 1721 | 602 | 7713 | 3463 | 5046 |
| 2170 | 3567 | 2867 | 1023 | 4459 | 1758 | 627 | 8624 | 3782 | 5599 |
| 2469.5 | 3604 | 2882.5 | 1040 | 4305 | 1815.5 | 652.5 | 9529 | 4200 | 5544 |
| 2367 | 3609 | 3046 | 1130 | 4127 | 1851 | 650 | 9993 | 4116 | 5802 |
| 2407.5 | 3753.5 | 2966 | 1128 | 4184.5 | 1838.5 | 674 | 9982.5 | 4330 | 6142 |
| 2699 | 4720 | 3617 | 1307 | 5189 | 2116 | 794 | 13050 | 5195 | 7846 |
| 2881.5 | 5157.5 | 3830.5 | 1350 | 5545.5 | 2172 | 857 | 13008 | 5608.5 | 8631.5 |
| 27588 | 19918 | 6611 | 1370 | 4447 | 1944 | 764 | 10364 | 11294 | 10674 |
| 4395.5 | 7578.5 | 5373.5 | 1726 | 6647 | 2646 | 1019.5 | 12579 | 7202.5 | 10792 |
| 2355 | 3669 | 2879 | 1059 | 3796 | 1694 | 627 | 8303 | 4052 | 5868 |
| 53165.5 | 27597 | 8341.5 | 1369.5 | 4115 | 1736 | 612 | 12516.5 | 5308.5 | 6556.5 |
| 3005.5 | 4386.5 | 3512 | 1119 | 4415 | 1816 | 641 | 9947 | 4749.5 | 6589.5 |
| 2805 | 4177 | 3311 | 1166 | 4467 | 1907 | 682 | 9529 | 4666 | 6338 |
| 2267.5 | 3664.5 | 2908.5 | 1040.5 | 3892.5 | 1759 | 621.5 | 8610 | 3978 | 5716.5 |
| 4708.5 | 9970.5 | 4604 | 1403 | 5915 | 2077 | 711 | 14322.5 | 6091 | 12531.5 |
| 7394 | 20943 | 6821 | 1742 | 7957 | 2470 | 674 | 14421 | 8326 | 23881 |
| 8071 | 23250 | 9043 | 2114 | 10606 | 2776 | 732 | 11260 | 8659 | 27134 |
| 8333 | 22848.5 | 8032.5 | 1915.5 | 9396 | 2714.5 | 748.5 | 8820 | 8007.5 | 25315 |
| 2685 | 3522.5 | 2146 | 810.5 | 2865 | 1391 | 335.5 | 8172 | 3403 | 4486.5 |
| 10764 | 6759 | 3397 | 983 | 4731 | 2005 | 493 | 9060 | 4008 | 4934 |
| 7215.5 | 5531 | 3119.5 | 1019 | 5979.5 | 2390 | 612 | 9114.5 | 4273.5 | 5398 |
| 7895 | 5487 | 6643.5 | 1852.5 | 236302 | 23805 | 631 | 9875 | 4035.5 | 5553.5 |
| 5712 | 5113 | 5974 | 1410 | 513052 | 55259 | 760 | 10377 | 4544 | 6102 |
| 6330 | 5313 | 4678 | 1309 | 29213 | 4946 | 793 | 10973 | 4997 | 6285 |
| 6137 | 5176 | 4742 | 1321 | 173284 | 19561 | 792 | 11426 | 5218 | 6929 |
| 5904.5 | 6052.5 | 4852.5 | 1435 | 212206.5 | 23749 | 864.5 | 13575 | 5396 | 7813.5 |
| 5634 | 5691 | 4413 | 1371 | 190246 | 21657 | 811 | 12362 | 5405 | 7212 |
| 4992 | 5461.5 | 4677.5 | 1699 | 72652.5 | 7291 | 749.5 | 12022 | 4721 | 6417.5 |
| 5140 | 5079 | 3681 | 1236 | 8446 | 3135 | 773 | 11970 | 5087 | 6811 |
| 5002 | 4795 | 3416 | 1166 | 6634 | 2472 | 770 | 10920 | 4740 | 6343 |
| 5539 | 5169 | 3583 | 1278 | 8222 | 3143 | 808 | 11458 | 5034 | 6934 |
| 43618 | 32231 | 11094 | 1913 | 5233 | 2306 | 763 | 10049 | 17049 | 14679 |
| 10998 | 8802 | 4550 | 1157 | 7089 | 2843 | 740 | 9965 | 6028 | 7025 |
| 12964 | 8765 | 4425 | 1188 | 59636 | 6149 | 730 | 10345 | 6113 | 7318 |
| 906294 | 665695 | 198719 | 15254 | 6502 | 2381 | 786 | 86314 | 29176 | 22323 |
| 21187.5 | 13129.5 | 5868 | 1355 | 59260.5 | 6339.5 | 762.5 | 12306 | 6295.5 | 7624.5 |
| 9389 | 6571 | 3813 | 1176 | 4645 | 1991 | 688 | 10233 | 4914 | 6569 |
| 14851 | 9985 | 4994 | 1243 | 52156.5 | 5374.5 | 722.5 | 11696 | 5505.5 | 6713 |
| 14817 | 20337 | 6077 | 1443 | 10141 | 3044 | 773 | 15317 | 6520 | 16232 |
| 20842.5 | 30074 | 8855.5 | 1837 | 17857 | 4089 | 804 | 13896.5 | 9929.5 | 25383 |
| 19777 | 30518 | 8519 | 1841 | 13696 | 3424 | 721 | 15007 | 9918 | 25470 |

| | | | | | | | | | |
|---|---|---|---|---|---|---|---|---|---|
| 20149 | 28052 | 7646 | 1784 | 12598 | 3261 | 777 | 14539 | 9720 | 24225 |
| 2631 | 3450 | 2155 | 782 | 2963 | 1415 | 326 | 7930 | 3370 | 4408 |
| 2408 | 3377 | 2316 | 852 | 3092 | 1453 | 466 | 7684 | 3288 | 4541 |
| 2468 | 3548 | 2523 | 947 | 3488.5 | 1618 | 544.5 | 8089.5 | 3712.5 | 5283.5 |
| 2293 | 3418 | 6212 | 1854 | 754130 | 67164 | 600 | 8596 | 3743 | 5379 |
| 2447 | 3736 | 3365 | 1121 | 7004.5 | 2261 | 637 | 9087.5 | 4136 | 5651 |
| 2794 | 4089 | 3544 | 1200 | 6361 | 2133 | 706 | 11192 | 4769 | 6467 |
| 2587.5 | 3913 | 3372.5 | 1179 | 5561 | 2059 | 732 | 9844 | 4479.5 | 6251 |
| 2676 | 4062 | 3311 | 1145 | 5194 | 2012 | 695 | 10067 | 4518 | 6353 |
| 2747 | 4068 | 3064 | 1088 | 4685 | 1867 | 705 | 10497 | 4579 | 6324 |
| 2501.5 | 3779 | 2899 | 1098 | 4021 | 1747.5 | 621 | 9927 | 4152.5 | 5835 |
| 2435 | 3678 | 2924 | 1033 | 4133 | 1788 | 641 | 9809 | 4138 | 5780 |
| 2392 | 3817 | 2977 | 1073 | 4309 | 1798 | 681 | 10023 | 4279 | 6093 |
| 2216 | 3516 | 2953 | 1120 | 4126 | 1814 | 683 | 8925 | 4019 | 5709 |
| 28887 | 21955 | 6822 | 1135 | 3607 | 1735 | 659 | 8327 | 11177 | 9895 |
| 2568 | 3659 | 2903 | 1034 | 3806 | 1700 | 637 | 8060 | 3963 | 5585 |
| 2837 | 4191 | 3322 | 1160 | 4576 | 1891 | 706 | 9618 | 4690 | 6657 |
| 85167 | 45753 | 13292 | 1571 | 4305 | 1812 | 657 | 15846 | 6276 | 7158 |
| 3007 | 4229 | 3358 | 1075 | 4282 | 1775 | 633 | 9152 | 4514 | 6260 |
| 2959.5 | 4599.5 | 3495.5 | 1199.5 | 5273.5 | 2004.5 | 699 | 9844.5 | 4882 | 7189.5 |
| 2840 | 4218 | 3187 | 1146 | 4331 | 1855 | 690 | 9496 | 4549 | 6362 |
| 5734.5 | 12458 | 5160.5 | 1373 | 6131 | 2125.5 | 684.5 | 14705.5 | 6742 | 14716.5 |
| 8707 | 21863 | 10121 | 2181 | 11818 | 3028 | 845 | 11394 | 8244 | 24632 |
| 9142.5 | 23165.5 | 7102 | 1792.5 | 8647.5 | 2604 | 683.5 | 11705 | 7775.5 | 21002.5 |
| 7285 | 20916 | 10318 | 2293.5 | 12068.5 | 3076.5 | 838 | 9785.5 | 8321 | 22568 |
| 2683.5 | 3644 | 2183.5 | 783 | 3016.5 | 1417 | 340.5 | 8439 | 3519.5 | 4501 |
| 2354.5 | 3550 | 2516.5 | 906 | 3331 | 1543 | 452.5 | 8004 | 3610 | 5017.5 |
| 2265 | 3501.5 | 2693.5 | 1038 | 3889.5 | 1802.5 | 540.5 | 8346 | 3734 | 5650.5 |
| 2539 | 3838 | 5739 | 1870 | 559542 | 52757 | 611 | 8909 | 4405 | 6452 |
| 2468 | 3992 | 3762 | 1323 | 6773 | 2457 | 630 | 9226 | 4698 | 6809 |
| 2460.5 | 4878 | 3636 | 1395 | 6555.5 | 2422.5 | 690 | 10781 | 4609.5 | 8718 |
| 2366.5 | 4507.5 | 3545.5 | 1358.5 | 6198.5 | 2380 | 711.5 | 10090.5 | 4682 | 8369.5 |
| 2598 | 4674 | 3616.5 | 1375 | 5873.5 | 2274.5 | 694.5 | 10697.5 | 4941.5 | 8306.5 |
| 2754.5 | 4797 | 3489 | 1342.5 | 5781.5 | 2203 | 676 | 10461.5 | 5009.5 | 8778.5 |
| 2470 | 4028 | 3347 | 1310 | 4956 | 2157 | 625 | 10167 | 4552 | 7106 |
| 2519 | 4077 | 3169 | 1301 | 5074 | 2134 | 649 | 10509 | 4633 | 7636 |
| 2160 | 3548.5 | 3079 | 1189 | 4714.5 | 2050.5 | 616.5 | 9093.5 | 4408 | 6753 |
| 2404.5 | 3979.5 | 3325 | 1374 | 5331 | 2205.5 | 681.5 | 10089.5 | 4795.5 | 7676.5 |
| 5955 | 5892 | 3853 | 1242 | 4561 | 2005 | 610 | 8333 | 5427 | 7304 |
| 2221 | 3313 | 2792 | 1137 | 4223 | 1900 | 558 | 7595 | 4012 | 6108 |
| 2723.5 | 4747 | 3543 | 1357.5 | 5312 | 2270.5 | 662 | 9878.5 | 5132 | 8389.5 |
| 38627 | 22200 | 7393 | 1492 | 4946 | 2079 | 615 | 12823 | 5481 | 7720 |
| 2540 | 4120 | 3502 | 1283 | 4947 | 1990 | 587 | 9195 | 4611 | 7243 |
| 2497 | 4793 | 3721.5 | 1339.5 | 5418.5 | 2232.5 | 666 | 9986 | 4822.5 | 8422 |
| 2428 | 3986 | 3333 | 1293.5 | 4870.5 | 2161 | 641 | 9187 | 4595.5 | 7102.5 |
| 2892 | 5938.5 | 4100.5 | 1494.5 | 5915.5 | 2351.5 | 709.5 | 11940.5 | 4828.5 | 10344.5 |
| 3736 | 11257 | 5321 | 1775 | 7622 | 2884 | 783 | 13020 | 6264 | 17585 |
| 4302 | 8961 | 4191 | 1481 | 5954 | 2268 | 677 | 14181 | 6062 | 12565 |

| | | | | | | | | | |
|---|---|---|---|---|---|---|---|---|---|
| 5014 | 13846 | 5202 | 1573 | 6833 | 2357 | 661 | 13700 | 6653 | 18597 |
| 2694 | 3586 | 2265 | 816 | 3070 | 1448 | 342 | 8410 | 3563 | 4657 |
| 3985.5 | 4778 | 3067.5 | 1043.5 | 8514 | 2109.5 | 509.5 | 9934.5 | 4280 | 5552 |
| 3152.5 | 4163.5 | 3013.5 | 1046.5 | 14063.5 | 2631.5 | 599.5 | 9492.5 | 4131 | 5685 |
| 3196 | 4004 | 7108 | 2217 | 456271 | 68629 | 595 | 9234 | 4383 | 5818 |
| 3298 | 6106 | 5516 | 1731 | 167493 | 24705 | 760 | 15196 | 4812 | 8442 |
| 3482 | 5564 | 4551 | 1396 | 135642 | 18927 | 697 | 11543 | 4998 | 7819 |
| 3907 | 5401 | 4711 | 1413 | 106327.5 | 14890 | 739 | 13908.5 | 7125 | 9401.5 |
| 3411 | 4709 | 3903 | 1255 | 102779 | 13888 | 661 | 10973 | 5117 | 6657 |
| 3282 | 4514 | 3692 | 1232 | 83226 | 11377 | 667 | 10894 | 4853 | 6625 |
| 3768 | 5824.5 | 5726 | 1781.5 | 70606.5 | 10476 | 764.5 | 12548 | 5633.5 | 7586 |
| 3180 | 4785.5 | 3880 | 1301.5 | 62972 | 8642 | 712.5 | 12930.5 | 4769.5 | 6927 |
| 3113 | 4613.5 | 4129.5 | 1309 | 38450.5 | 5449.5 | 711 | 10872.5 | 4728 | 6360 |
| 3216.5 | 4428.5 | 3753 | 1256 | 41107.5 | 5491.5 | 725 | 11062 | 4679 | 6641 |
| 57082 | 44987 | 46636 | 5927 | 33803 | 4814 | 654 | 9365 | 21993 | 18391 |
| 4742 | 7132 | 4698 | 1419 | 30938 | 4596 | 720 | 10885 | 5370 | 8969 |
| 3682.5 | 7046 | 4156.5 | 1295.5 | 28462.5 | 4268.5 | 715.5 | 10640.5 | 4665.5 | 10083 |
| 426767.5 | 437098.5 | 147038.5 | 14254.5 | 27151 | 4148 | 707 | 52283 | 14130 | 18515 |
| 7438 | 8871 | 5071 | 1403 | 24788 | 3829 | 761 | 13282 | 4987 | 10276 |
| 3333 | 8598 | 4673 | 1484 | 96501 | 19938 | 769 | 14573 | 5039 | 12473 |
| 4299.5 | 6825.5 | 4339 | 1363 | 23303 | 3721.5 | 698 | 12384.5 | 5128 | 9332 |
| 4170 | 8206 | 4436.5 | 1371.5 | 20226.5 | 3390.5 | 755.5 | 14184.5 | 5278.5 | 10658.5 |
| 4501 | 9593.5 | 4921.5 | 1471 | 17260 | 3267.5 | 780.5 | 15909.5 | 5290 | 11739 |
| 5048.5 | 11277.5 | 5013 | 1537 | 16198.5 | 3043 | 750.5 | 14426 | 5689 | 14373.5 |
| 5418.5 | 13740 | 5432 | 1539.5 | 15939 | 3058.5 | 739 | 14842 | 6452.5 | 16830 |
| 2956 | 3689 | 2254 | 818 | 3471 | 1464 | 348 | 10058 | 3642 | 4485 |
| 2634 | 3722.5 | 2495.5 | 901 | 3499.5 | 1579.5 | 477 | 8866 | 3581 | 4802.5 |
| 2652 | 3933.5 | 2852.5 | 1024 | 3974 | 1762 | 584 | 9143 | 4002.5 | 5416 |
| 2275 | 3605 | 5312 | 1651 | 668375 | 56493 | 601 | 8604 | 3737 | 5366 |
| 2657 | 3993 | 3771 | 1245 | 8537 | 2440 | 670 | 10183 | 4485 | 6128 |
| 2633 | 4316 | 3466 | 1206 | 6518 | 2157 | 702 | 9722 | 4660 | 6583 |
| 2316 | 3756 | 2956 | 1148 | 5323 | 2015 | 652 | 8633 | 4013 | 5841 |
| 2460.5 | 4036.5 | 3077.5 | 1118 | 5073.5 | 1969.5 | 688.5 | 9664.5 | 4291.5 | 6279 |
| 2358 | 3690 | 2858 | 1017 | 4464 | 1780 | 613 | 9180 | 4003 | 5539 |
| 2561 | 3942 | 3285 | 1311 | 4497 | 1963 | 635 | 10140 | 4366 | 5965 |
| 2372 | 3839 | 2919 | 1125 | 4253 | 1849 | 648 | 9693 | 4113 | 5900 |
| 2114.5 | 3312.5 | 2760.5 | 1004 | 3905 | 1710 | 615.5 | 8399.5 | 3672.5 | 5086.5 |
| 2293 | 3679 | 3048 | 1106 | 4477 | 1862 | 673 | 9461 | 4138 | 5968 |
| 22478 | 16124 | 5483 | 1128.5 | 3866.5 | 1710.5 | 618 | 7772 | 8488.5 | 8110.5 |
| 2546.5 | 3684 | 2766 | 991 | 3840 | 1662.5 | 603 | 7943 | 3874 | 5422 |
| 2845 | 4119 | 3141 | 1141 | 4209 | 1834 | 639 | 8710 | 4443 | 6212 |
| 117601 | 64451 | 17360 | 1923 | 5050 | 1991 | 701 | 18584 | 7416 | 8840 |
| 3017 | 4587 | 3648 | 1179 | 4732 | 1939 | 664 | 10099 | 4639 | 6730 |
| 2840 | 4552 | 3475 | 1193 | 4708 | 1956 | 663 | 9589 | 4765 | 6952 |
| 2727 | 4382.5 | 3338.5 | 1161.5 | 4480 | 1921.5 | 657 | 9481.5 | 4516.5 | 6489 |
| 2739 | 4240 | 3133 | 1194 | 4562 | 1891 | 690 | 10409 | 4627 | 6612 |
| 2823 | 4563.5 | 3468.5 | 1232.5 | 4687 | 1919 | 708 | 10582 | 4696 | 6906.5 |
| 3954 | 8468.5 | 4149 | 1270 | 5559 | 2012.5 | 717 | 14508.5 | 5508 | 10802.5 |

| | | | | | | | | | |
|---|---|---|---|---|---|---|---|---|---|
| 4768 | 12789 | 4613 | 1297 | 5931 | 2048 | 688 | 13365 | 5796 | 15547 |
| 2646 | 3652 | 2366 | 820 | 3103 | 1430 | 353 | 8316 | 3539 | 4669 |
| 2684.5 | 3837.5 | 2589.5 | 930.5 | 3562.5 | 1567.5 | 490 | 8452 | 3795.5 | 5207 |
| 2393.5 | 3582 | 2629 | 971 | 3570.5 | 1659.5 | 539.5 | 8198.5 | 3600.5 | 5160.5 |
| 2635 | 4029 | 6149 | 1891 | 757436 | 59559 | 618 | 9648 | 4406 | 5897 |
| 2684 | 4080 | 3472 | 1215 | 7175 | 2303 | 647 | 9419 | 4401 | 6101 |
| 2616 | 4031 | 3287 | 1135 | 5793 | 2041 | 665 | 9574 | 4462 | 6198 |
| 2294 | 3639.5 | 2986 | 1056 | 5068 | 1839 | 620 | 8623.5 | 3841.5 | 5527 |
| 2525 | 4051 | 3083 | 1096 | 5021 | 1918 | 664 | 9645 | 4450 | 6215 |
| 2704.5 | 4154.5 | 3188 | 1114 | 4949 | 1930 | 670 | 10597.5 | 4561.5 | 6232 |
| 2721 | 3923 | 3140 | 1138 | 4313 | 1867 | 628 | 10116 | 4548 | 5933 |
| 2623 | 4040 | 3075 | 1175 | 4528 | 1894 | 660 | 9856 | 4509 | 6066 |
| 2524 | 4051 | 3149 | 1143.5 | 4433.5 | 1856.5 | 690 | 9649 | 4459 | 6408 |
| 2413.5 | 3877 | 3099.5 | 1088 | 4345 | 1820 | 690 | 9434.5 | 4370 | 6153.5 |
| 13271 | 10080.5 | 4447.5 | 1102.5 | 3860.5 | 1721 | 610 | 7991.5 | 6467.5 | 7136.5 |
| 2857 | 4275 | 3169 | 1154 | 4352 | 1823 | 658 | 8790 | 4463 | 6190 |
| 2677 | 4009 | 3154 | 1112 | 4232 | 1795 | 630 | 8558 | 4213 | 6042 |
| 73465 | 44333 | 12513 | 1613 | 4880 | 2021 | 670 | 16358 | 5760 | 8047 |
| 2722.5 | 4282 | 3328.5 | 1139.5 | 4504.5 | 1781.5 | 612 | 9335 | 4378 | 6404.5 |
| 2722 | 4529 | 3263 | 1139 | 4581 | 1834 | 651 | 9401 | 4646 | 6952 |
| 2798 | 4299 | 3420 | 1193 | 4672 | 1894 | 693 | 10189 | 4720 | 6599 |
| 2755 | 4083 | 3013 | 1048 | 4386 | 1826 | 653 | 9686 | 4537 | 6237 |
| 3118 | 5005 | 3312 | 1149 | 4467.5 | 1843 | 665 | 10862 | 5087.5 | 6953 |
| 3880 | 9426.5 | 3700.5 | 1161.5 | 4815 | 1835 | 622 | 11470.5 | 4706 | 12042 |
| 4543.5 | 10592 | 4241.5 | 1159.5 | 5233.5 | 1863.5 | 623 | 15571.5 | 5505.5 | 13217 |
| 746 | 898 | 767 | 622 | 1308 | 1145 | 253 | 2213 | 1379 | 1722 |
| 967 | 1526 | 1367 | 702 | 2029 | 1312 | 401 | 3591 | 1972 | 2838 |
| 1202 | 2000 | 1876 | 814 | 2733 | 1413 | 498 | 4784 | 2485 | 3766 |
| 1274 | 2313 | 4283 | 1415 | 650618 | 58944 | 572 | 6301 | 2785 | 4289 |
| 1325 | 2356 | 2732 | 1047 | 5084 | 2190 | 598 | 6587 | 2876 | 4364 |
| 1476.5 | 2690.5 | 2740.5 | 1106 | 4250.5 | 1949 | 660 | 7413.5 | 3159 | 4989.5 |
| 1503.5 | 2785 | 2652 | 1075 | 4167.5 | 1881.5 | 679.5 | 7434 | 3200 | 4984 |
| 1674 | 2908 | 2737 | 1097 | 4075 | 1860 | 685 | 8313 | 3501 | 5494 |
| 1866 | 3137 | 2792 | 1098 | 4024 | 1866 | 697 | 9136 | 3738 | 5611 |
| 1877 | 3057 | 2686 | 1108 | 3743 | 1765.5 | 636 | 8916.5 | 3606 | 5192.5 |
| 1696 | 2871 | 2419 | 960 | 3390 | 1656 | 619 | 7900 | 3233 | 4894 |
| 1625 | 2745 | 2457 | 1018 | 3417 | 1650 | 628 | 7769 | 3149 | 4735 |
| 1530.5 | 2781.5 | 2517.5 | 996 | 3480 | 1694 | 637 | 7683 | 3191.5 | 4866 |
| 16089 | 11850 | 4544.5 | 1110.5 | 3536 | 1694 | 637 | 7396.5 | 6453 | 6942.5 |
| 1913 | 3216 | 2925 | 1104 | 3914 | 1833 | 695 | 8135 | 3575 | 5646 |
| 1820.5 | 3119 | 2908.5 | 1137 | 3974 | 1819 | 680 | 7952 | 3694 | 5564 |
| 88292 | 48043 | 13806 | 1581 | 4157 | 1862 | 651 | 15869 | 5255 | 6521 |
| 1811 | 3148 | 2832 | 1037 | 3698 | 1702 | 613 | 7645 | 3605 | 5371 |
| 1704 | 2964.5 | 2632.5 | 1023.5 | 4139 | 1759 | 600 | 7611.5 | 3375.5 | 5251 |
| 1713 | 2998 | 2563 | 1059 | 3800 | 1679 | 625 | 7900 | 3464 | 5239 |
| 3733 | 10033 | 4164 | 1218 | 5193 | 1851 | 627 | 13204 | 4880 | 12935 |
| 7500 | 19136.5 | 6676.5 | 1760.5 | 7795.5 | 2396.5 | 706.5 | 13255 | 8720.5 | 22208 |
| 8505 | 19440 | 6353 | 1640 | 7254 | 2282 | 643 | 12326 | 8606 | 21394 |

| | | | | | | | | | |
|---|---|---|---|---|---|---|---|---|---|
| 8010 | 20890 | 6871 | 1717.5 | 8204 | 2418.5 | 675 | 13185 | 9358 | 24140 |
| 2630.5 | 3657 | 2056 | 773.5 | 2667 | 1333 | 355 | 8331.5 | 3522.5 | 4515.5 |
| 2020 | 2991 | 2018 | 784 | 2676 | 1361 | 440 | 6879 | 3072 | 4247 |
| 1859 | 2942 | 2270 | 934 | 3207 | 1561 | 547 | 7120 | 3227 | 4703 |
| 1873.5 | 3140.5 | 5062 | 1489 | 698902 | 64868 | 617.5 | 8057 | 3523.5 | 5318.5 |
| 1729 | 2874 | 2668 | 1001 | 4588 | 1878 | 610 | 7307 | 3195 | 4734 |
| 1653.5 | 2811.5 | 2468 | 1006 | 4005 | 1774.5 | 618.5 | 7168 | 3180 | 4803 |
| 1700 | 2886.5 | 2672 | 1039 | 4029 | 1756 | 640.5 | 7487.5 | 3392 | 5068.5 |
| 1979 | 3364 | 2929 | 1127 | 4255 | 1867 | 685 | 9046 | 3770 | 5635 |
| 2055 | 3386 | 2801 | 1084 | 3986 | 1774 | 641 | 9244 | 3822 | 5485 |
| 2158 | 3520 | 2994 | 1132 | 4022 | 1852 | 669 | 9651 | 4038 | 5720 |
| 1882 | 2985 | 2572 | 994 | 3542 | 1696 | 626 | 8304 | 3498 | 5116 |
| 1900 | 3136 | 2751 | 1110 | 3855 | 1771 | 673 | 8844.5 | 3568.5 | 5371.5 |
| 1693.5 | 2934.5 | 2561.5 | 1011 | 3490.5 | 1665 | 635 | 7828 | 3397.5 | 5049.5 |
| 16044.5 | 11853.5 | 4475.5 | 1119.5 | 3711.5 | 1768 | 656 | 8013.5 | 6566 | 7264.5 |
| 2184.5 | 3449 | 2892 | 1170 | 4016 | 1832.5 | 681 | 8482 | 3895.5 | 5787 |
| 1992 | 3306 | 2895 | 1093 | 3955 | 1778 | 642 | 8114 | 3768 | 5621 |
| 87255 | 47487 | 13562 | 1580 | 4141 | 1839 | 668 | 15486 | 5617 | 6914 |
| 2278 | 3784 | 3262 | 1141 | 4313 | 1786 | 664 | 9252 | 4152 | 6133 |
| 2119.5 | 3449.5 | 2948.5 | 1094 | 4658 | 1898 | 664 | 9280.5 | 4007.5 | 5876 |
| 2174 | 3473 | 3119 | 1170 | 4202 | 1822 | 709 | 8859 | 4253 | 5904 |
| 5078 | 12408 | 5051 | 1354 | 6065 | 2047 | 679 | 14360 | 6452 | 15544 |
| 8135 | 20370.5 | 6773 | 1687 | 8127.5 | 2454 | 700 | 13595 | 9337 | 23549 |
| 8551.5 | 21062 | 7195.5 | 1774 | 8417 | 2475 | 661 | 13896 | 9191.5 | 24370 |
| 8441 | 22026 | 6748 | 1758 | 8221 | 2463 | 699 | 13225 | 9196 | 24809 |
| 2571.5 | 3505 | 2058 | 777.5 | 2776 | 1368 | 331 | 8199 | 3483.5 | 4526.5 |
| 2302 | 3362.5 | 2253 | 841 | 3048.5 | 1434 | 474 | 7532.5 | 3460.5 | 4819 |
| 2035 | 3244 | 2508 | 946 | 3486 | 1584 | 570 | 7572 | 3451 | 4969 |
| 1983 | 3345 | 5030 | 1512 | 663115 | 60780 | 638 | 8317 | 3699 | 5350 |
| 1723 | 2871 | 2672 | 964 | 4716 | 1862 | 595 | 7520 | 3245 | 4771 |
| 1829 | 3022 | 2700.5 | 1038.5 | 4364 | 1829.5 | 649.5 | 7946 | 3295.5 | 5004.5 |
| 1735 | 2843 | 2632 | 962 | 3932 | 1728 | 616 | 7763 | 3433 | 4899 |
| 2026 | 3279 | 2758 | 1078.5 | 4247 | 1883 | 690 | 8887 | 3657.5 | 5393 |
| 2298.5 | 3657.5 | 3016.5 | 1038 | 4202 | 1779 | 679 | 9962 | 4166.5 | 5846.5 |
| 2453.5 | 3814 | 3132 | 1173 | 4409.5 | 1887 | 703 | 9982.5 | 4276.5 | 6252.5 |
| 2058 | 3188 | 2755.5 | 1033.5 | 3677.5 | 1712 | 616 | 8831.5 | 3730 | 5227.5 |
| 1895 | 3170 | 2720 | 1064 | 3746 | 1758 | 640 | 8494 | 3667 | 5390 |
| 1944.5 | 3149.5 | 2727.5 | 1059 | 3961.5 | 1716.5 | 686.5 | 8492 | 3691.5 | 5477.5 |
| 13221.5 | 10101.5 | 4139.5 | 1068.5 | 3727.5 | 1718.5 | 635 | 7820.5 | 6100.5 | 6848 |
| 2146 | 3284.5 | 2847 | 1076.5 | 3740.5 | 1799 | 675.5 | 8060.5 | 3692.5 | 5536 |
| 2047 | 3429 | 2965 | 1080 | 4069 | 1763 | 653 | 8575 | 3961 | 5883 |
| 49630 | 26874.5 | 8437.5 | 1310 | 4043 | 1768 | 614.5 | 11875 | 4903.5 | 6324 |
| 2630.5 | 4120 | 3392 | 1225 | 4523.5 | 1921 | 676.5 | 10113 | 4731 | 6596.5 |
| 2193 | 3645 | 3106 | 1165 | 4642 | 1911 | 645 | 9208 | 4058 | 6018 |
| 2223.5 | 3516.5 | 2988 | 1186.5 | 4227 | 1836 | 666 | 9139 | 4013.5 | 6103 |
| 4709 | 12039 | 4893 | 1379 | 5893 | 2040 | 672 | 13703 | 6020 | 15261 |
| 7795.5 | 18887 | 6524 | 1745 | 7778.5 | 2413 | 685 | 13484.5 | 8997.5 | 22202.5 |
| 7393 | 20874 | 8944 | 2079 | 10942 | 2804 | 791 | 10717 | 7663 | 22649 |

| | | | | | | | | | |
|---|---|---|---|---|---|---|---|---|---|
| 8590.5 | 21062 | 6853.5 | 1713 | 8190 | 2517.5 | 716 | 13222.5 | 9519.5 | 23849 |
| 2578 | 3638 | 2113 | 801 | 2830 | 1369 | 339 | 8316 | 3655 | 4588 |
| 2153 | 3116 | 2163 | 802 | 2886 | 1422 | 424 | 7356 | 3117 | 4317 |
| 2017 | 3118 | 2358 | 915 | 3271 | 1568 | 534 | 7643 | 3395 | 4865 |
| 2027 | 3238 | 5019 | 1530 | 641506 | 58998 | 612 | 8079 | 3671 | 5393 |
| 1863 | 2988 | 2766 | 990 | 4952 | 1902 | 589 | 7430 | 3343 | 4921 |
| 1892 | 3054 | 2584 | 1009 | 4257 | 1789 | 602 | 7974 | 3434 | 5119 |
| 1931 | 3363 | 2919 | 1070 | 4458 | 1880 | 651 | 8918 | 3571 | 5449 |
| 2095 | 3400.5 | 2809.5 | 1058.5 | 4247 | 1801 | 668 | 9109.5 | 3905.5 | 5561 |
| 2481.5 | 3840 | 3074 | 1131 | 4290 | 1761 | 688 | 10591 | 4329.5 | 5916 |
| 2272 | 3502 | 2909 | 1075 | 3981 | 1798 | 648 | 9607 | 3960 | 5700 |
| 2008 | 3151 | 2538 | 978 | 3670 | 1664 | 591 | 8845 | 3496 | 5082 |
| 2020 | 3220 | 2687 | 1013 | 3821 | 1721 | 644 | 9164 | 3545 | 5351 |
| 1905 | 3202 | 2637.5 | 998.5 | 3792 | 1679.5 | 629.5 | 8386.5 | 3556 | 5294 |
| 14171.5 | 10838.5 | 4237.5 | 1152 | 3770 | 1703 | 649.5 | 8311 | 6319.5 | 6838 |
| 2262 | 3482 | 2883 | 1054 | 3991 | 1800 | 669 | 8377 | 3865 | 5783 |
| 2134 | 3508 | 2849 | 1032 | 3952 | 1743 | 651 | 8653 | 3935 | 5611 |
| 85428 | 45605 | 13027 | 1579 | 4243 | 1794 | 645 | 15084 | 5662 | 6846 |
| 2333 | 3846 | 3140 | 1163 | 4354 | 1839 | 655 | 9518 | 4202 | 6109 |
| 2484.5 | 3731 | 3075 | 1093 | 4655 | 1909.5 | 679.5 | 9560.5 | 4482 | 6024.5 |
| 2294 | 3683 | 2893 | 1128 | 4108 | 1875 | 686 | 9189 | 4194 | 6045 |
| 4636 | 12911 | 4994 | 1429 | 6227 | 2125 | 685 | 13985 | 6057 | 16309 |
| 8303 | 19512 | 6511 | 1642 | 7686 | 2419 | 693 | 13633 | 9251 | 21843 |
| 7967 | 20942 | 8531 | 1923 | 10174 | 2715 | 735 | 11608 | 8895 | 23590 |
| 8354 | 21657 | 7329 | 1832 | 8451 | 2551 | 725 | 11850 | 9229 | 24955 |
| 2823 | 3731.5 | 2175 | 805 | 2890 | 1370.5 | 350 | 8830.5 | 3706 | 4609 |
| 2251.5 | 3473.5 | 2261 | 914 | 3165 | 1449 | 476.5 | 7935.5 | 3498.5 | 4777.5 |
| 2087 | 3324 | 2604.5 | 947 | 3391.5 | 1596 | 563.5 | 8202.5 | 3596.5 | 5069 |
| 2028 | 3264 | 4951 | 1522 | 722240 | 63335 | 598 | 8322 | 3613 | 5235 |
| 2036 | 3339 | 3063.5 | 1133 | 5144 | 2005 | 624.5 | 7935 | 3633.5 | 5240.5 |
| 1883 | 2998 | 2635 | 997 | 4094 | 1716 | 592 | 7641 | 3401 | 4923 |
| 2051.5 | 3271 | 2931.5 | 1088.5 | 4488.5 | 1855 | 668 | 8571.5 | 3757 | 5660.5 |
| 2307 | 3514 | 3123 | 1084 | 4534 | 1852 | 690 | 9808 | 4176 | 5923 |
| 2448 | 3668.5 | 2874 | 1063 | 4079 | 1791.5 | 664 | 9792.5 | 4080 | 5697.5 |
| 2378 | 3665.5 | 3040.5 | 1132.5 | 4281 | 1844 | 649 | 10326.5 | 4230.5 | 5805.5 |
| 2119 | 3290 | 2643 | 1044 | 3847 | 1702 | 623 | 9140 | 3857 | 5370 |
| 2003.5 | 3197.5 | 2771.5 | 989.5 | 3673.5 | 1692.5 | 615 | 8694.5 | 3563 | 5179 |
| 1956 | 3235 | 2709 | 1077 | 3798 | 1737 | 647 | 8631 | 3546 | 5410 |
| 27473 | 20691 | 6617 | 1146 | 3599 | 1641 | 633 | 8287 | 10692 | 9368 |
| 2310 | 3697.5 | 3016.5 | 1072.5 | 3962.5 | 1750 | 691 | 8607.5 | 3934 | 5952 |
| 2268 | 3527 | 2967 | 1041 | 3987 | 1783 | 662 | 8976 | 4087 | 5860 |
| 53526 | 28558 | 8876 | 1475 | 4486 | 1859 | 686 | 12821 | 5397 | 6839 |
| 2403 | 3854 | 3115.5 | 1122.5 | 4106.5 | 1732.5 | 643.5 | 8954 | 4149 | 5993 |
| 2348 | 3898 | 3182 | 1079 | 4724 | 1948 | 641 | 9393 | 4268 | 6446 |
| 2497.5 | 3664.5 | 2963.5 | 1131 | 4191.5 | 1818 | 675 | 9462.5 | 4500 | 6065.5 |
| 4920 | 13067 | 4931 | 1411.5 | 6197 | 2145.5 | 691 | 13861 | 6455 | 16393.5 |
| 7721 | 20245 | 6438 | 1696 | 8008 | 2394 | 700 | 14449 | 9178 | 23734 |
| 7773 | 20369 | 7983 | 1807 | 10009 | 2721 | 747 | 12066 | 8459 | 22492 |

| | | | | | | | | | |
|---|---|---|---|---|---|---|---|---|---|
| 8388.5 | 21642.5 | 7550.5 | 1795.5 | 8508.5 | 2418.5 | 707 | 12605.5 | 9218 | 24295.5 |
| 2997.5 | 3828 | 2342 | 839 | 3097.5 | 1440 | 359 | 9495.5 | 3898 | 4823.5 |
| 10221 | 6429.5 | 3288.5 | 950.5 | 3736.5 | 1666 | 519 | 9461.5 | 3998 | 5182.5 |
| 7296 | 5426 | 3321 | 1064 | 4282 | 1867 | 644 | 9992 | 4399 | 5796 |
| 6516 | 5041 | 5354 | 1665 | 109588 | 12402 | 717 | 10392 | 4333 | 5765 |
| 6098 | 4846.5 | 3590 | 1157 | 20972.5 | 3280.5 | 692 | 9809 | 4223 | 5385.5 |
| 5891 | 4768 | 3414 | 1131 | 12105 | 2501 | 712 | 10137 | 4077 | 5516 |
| 5440 | 5031 | 3903 | 1176 | 10777 | 3191 | 792 | 11042 | 4545 | 6131 |
| 5795.5 | 4974.5 | 3705.5 | 1190 | 8344 | 2478.5 | 812.5 | 11803.5 | 4757.5 | 6458 |
| 5906 | 5139.5 | 3912.5 | 1231.5 | 7283 | 2624 | 807 | 13122 | 5314 | 6990 |
| 4999 | 5520 | 4929 | 1842 | 40435 | 4565 | 789 | 12126 | 5108 | 7096 |
| 4839 | 4213 | 3081.5 | 1077.5 | 4829.5 | 1948.5 | 682.5 | 10671.5 | 4227 | 5557 |
| 4619 | 4275 | 3112 | 1125 | 4537 | 1894 | 730 | 10721 | 4267 | 5898 |
| 4702 | 4323.5 | 3185 | 1106.5 | 4293.5 | 1916.5 | 735.5 | 10441.5 | 4192 | 5856 |
| 30029 | 21920 | 7015 | 1343 | 4295 | 1877 | 745 | 9921 | 12061 | 11018 |
| 6942.5 | 6024 | 3907.5 | 1257 | 4928.5 | 2069 | 822 | 11006 | 5345.5 | 7059.5 |
| 7446 | 5836 | 3912 | 1179 | 5644 | 2300 | 801 | 11630 | 5526 | 6955 |
| 1073692 | 642386 | 206700 | 16087 | 4619 | 1911 | 744 | 89948 | 31026 | 19791 |
| 14233 | 8770 | 4622 | 1333 | 5831 | 2395 | 815 | 13292 | 6120 | 7588 |
| 9376 | 6298 | 3831 | 1283 | 4860 | 1996 | 739 | 11335 | 5279 | 6858 |
| 17327 | 10155 | 4733 | 1297 | 5908 | 2436 | 776 | 14217 | 5948 | 7189 |
| 12981.5 | 18584.5 | 6313 | 1644 | 7248.5 | 2356.5 | 794 | 16548 | 7697.5 | 16785.5 |
| 19988 | 28174 | 8332 | 1963 | 10154 | 2890 | 821 | 15502 | 10391.5 | 26943.5 |
| 14643 | 26466 | 8588 | 1923 | 9722 | 2625 | 753 | 13983 | 9159 | 24713 |
| 14093.5 | 26480 | 8026 | 1886.5 | 9011 | 2622.5 | 793 | 13753.5 | 9849 | 26210 |
| 2736 | 3661 | 2181.5 | 799.5 | 3003 | 1395 | 339 | 8905.5 | 3525.5 | 4644 |
| 2397 | 3445 | 2375 | 865 | 3073 | 1502 | 464 | 7830 | 3424 | 4639 |
| 2215.5 | 3445 | 2593 | 939 | 3531.5 | 1604 | 551 | 8111.5 | 3636.5 | 5112 |
| 2118 | 3559 | 5250 | 1669 | 681393 | 62913 | 623 | 8415 | 3774 | 5639 |
| 1898 | 3011 | 2723 | 1007 | 5261 | 1887 | 559 | 7606 | 3467 | 4922 |
| 1979 | 3234 | 2850 | 1032 | 4755 | 1819 | 634 | 8002 | 3523 | 5225 |
| 2024 | 3370 | 2754 | 1026 | 4502 | 1793 | 627 | 8694 | 3602 | 5307 |
| 2260 | 3543 | 2968 | 1083 | 4553 | 1830 | 647 | 9308 | 3984 | 5712 |
| 2627 | 4002 | 3185 | 1164 | 4684 | 1869 | 695 | 11260 | 4731 | 6248 |
| 2550 | 3901 | 3147 | 1145 | 4379 | 1838 | 668 | 10558 | 4370 | 6017 |
| 2259 | 3271 | 2643 | 1007 | 3787 | 1702 | 607 | 9101 | 3737 | 5139 |
| 2213 | 3499 | 2880 | 1042 | 3909 | 1708 | 647 | 9208 | 3865 | 5491 |
| 2039 | 3197 | 2648 | 1001 | 3678 | 1636 | 616 | 8876 | 3701 | 5332 |
| 15648.5 | 11790 | 4485 | 1100 | 3925 | 1746.5 | 647.5 | 8809 | 6848 | 7423.5 |
| 2565 | 3788 | 2988 | 1073 | 4161 | 1762 | 678 | 9073 | 4216 | 5921 |
| 2374 | 3625 | 3027 | 1080 | 4094 | 1764 | 643 | 8962 | 4126 | 6031 |
| 90743 | 47907 | 13642 | 1579 | 4104 | 1684 | 620 | 15283 | 5646 | 6630 |
| 2785 | 4137 | 3357 | 1158 | 4456 | 1787 | 662 | 9796 | 4570 | 6528 |
| 2525 | 3773 | 3046.5 | 1086.5 | 4683.5 | 1824 | 655.5 | 9374.5 | 4304.5 | 5950 |
| 2489 | 3777 | 2947 | 1069 | 4050 | 1762 | 649 | 9705 | 4259 | 5889 |
| 5282.5 | 12433.5 | 4815 | 1342 | 5726 | 1944.5 | 663 | 14797 | 6193 | 15604 |
| 8395.5 | 18972.5 | 6584.5 | 1626 | 7651.5 | 2433.5 | 702 | 15313 | 9215 | 21778 |
| 9190 | 22927 | 7375 | 1811.5 | 8684 | 2478.5 | 710 | 12939.5 | 9017 | 24767.5 |

| | | | | | | | | | |
|---|---|---|---|---|---|---|---|---|---|
| 8922 | 22197 | 7051 | 1772 | 8379 | 2500 | 709 | 12955 | 9148 | 25267 |
| 2599 | 3502 | 2018 | 783 | 2823 | 1343 | 333 | 8364 | 3480 | 4377 |
| 2301.5 | 3366 | 2426 | 863.5 | 3223 | 1548 | 446 | 8143 | 3591 | 4808 |
| 2186 | 3483 | 2748.5 | 1023 | 3918 | 1803.5 | 551 | 8424 | 3789.5 | 5740.5 |
| 2282.5 | 3538 | 5670.5 | 1688 | 494195.5 | 50766.5 | 591.5 | 8583.5 | 4207.5 | 6213 |
| 1955 | 3326 | 3112 | 1179 | 5743 | 2177 | 582 | 8585 | 3868 | 6026 |
| 1986 | 3227 | 2951 | 1155 | 5096 | 2027 | 588 | 8316 | 3934 | 6095 |
| 2154 | 3336 | 3155 | 1276 | 5147 | 2131 | 611 | 8971 | 4361 | 6537 |
| 2301 | 3869 | 3492 | 1352 | 5411 | 2271 | 670 | 10624 | 4539 | 7127 |
| 2426.5 | 4066.5 | 3254.5 | 1285 | 5316 | 2157 | 655 | 10877 | 4425.5 | 7475 |
| 2417.5 | 4244 | 3578.5 | 1368.5 | 5270.5 | 2227.5 | 657 | 10880.5 | 4687.5 | 7572.5 |
| 2223 | 3403 | 3011 | 1186 | 4521 | 1994 | 604 | 9519 | 4162 | 6477 |
| 2233 | 3388 | 3177.5 | 1204 | 4705.5 | 2097.5 | 593 | 9518 | 4574.5 | 6868.5 |
| 2142.5 | 3303.5 | 3108 | 1234.5 | 4708.5 | 2045.5 | 624 | 9200.5 | 4340 | 6554.5 |
| 5119 | 4981.5 | 3741.5 | 1245.5 | 4629 | 2030 | 626 | 8716 | 5363 | 7069.5 |
| 2435 | 3901 | 3348 | 1296.5 | 5024.5 | 2174 | 659.5 | 9289 | 4716 | 7536 |
| 2302 | 3963 | 3494 | 1332 | 5062 | 2168 | 649 | 9797 | 4560 | 7747 |
| 32002 | 19805 | 6459 | 1456 | 5168 | 2100 | 626 | 12425 | 5164 | 8390 |
| 2407 | 5141 | 3657 | 1309.5 | 5184 | 2210.5 | 645.5 | 10496 | 4602.5 | 9184 |
| 2293 | 4295 | 3643.5 | 1312 | 5665 | 2250 | 652 | 10265.5 | 4496 | 7924.5 |
| 2282 | 5489 | 3774 | 1413 | 5303 | 2277 | 687 | 11003 | 4586 | 10247 |
| 5322 | 13734 | 5233 | 1641 | 6843 | 2429 | 671 | 15312 | 7444 | 17911 |
| 7396.5 | 19259 | 6936 | 1922 | 8883 | 2779.5 | 659 | 15035.5 | 9428 | 24002.5 |
| 6555 | 18035.5 | 8383.5 | 2078 | 10272.5 | 2876 | 703 | 11847.5 | 8065 | 22554 |
| 7444 | 21007 | 9362 | 2159 | 10856 | 2955 | 741 | 12021 | 9151 | 25483 |
| 2530 | 3599 | 2142 | 787 | 3013 | 1411 | 333 | 8591 | 3419 | 4539 |
| 3400 | 4201 | 2682 | 978 | 5608 | 1783 | 504 | 9518 | 3877 | 5134 |
| 2773.5 | 3854.5 | 2853.5 | 1010 | 8721.5 | 2158.5 | 600 | 9203 | 3967.5 | 5545 |
| 2494 | 3694 | 5956 | 2081 | 284929 | 57666 | 619 | 8664 | 3859 | 5577 |
| 2562.5 | 3762.5 | 3929 | 1220.5 | 106474 | 14394 | 648.5 | 9185.5 | 3961.5 | 5611.5 |
| 2565 | 3755 | 3444 | 1177 | 87104 | 11797 | 645 | 9612 | 3931 | 5586 |
| 4034.5 | 4356.5 | 4155 | 1173.5 | 79968 | 10723 | 702 | 12651.5 | 5692.5 | 7619 |
| 3387 | 4418 | 3748 | 1222 | 64603.5 | 8490.5 | 695.5 | 11274.5 | 4942 | 6598.5 |
| 3210.5 | 5204.5 | 3872 | 1240 | 55725 | 7552 | 722 | 14432.5 | 4768 | 7172.5 |
| 3581 | 6589 | 5637 | 1813.5 | 46455 | 6703 | 776.5 | 16539.5 | 5256.5 | 8781.5 |
| 3208 | 4501 | 3331 | 1136 | 44178 | 5917 | 672 | 11684 | 4835 | 6335 |
| 3321.5 | 5287 | 4497 | 1407.5 | 29303 | 4363.5 | 701 | 12151.5 | 5076 | 7329.5 |
| 3604 | 4522.5 | 3737 | 1187.5 | 31777 | 4578.5 | 705.5 | 11053.5 | 4956.5 | 6574.5 |
| 71047 | 53391 | 36386 | 3990 | 26910 | 3885 | 703 | 9888 | 29052 | 22443 |
| 4923 | 5726 | 4073 | 1240 | 26202 | 3741 | 695 | 10386 | 5645 | 7353 |
| 3801.5 | 5183.5 | 3622 | 1221.5 | 22105 | 3413 | 691.5 | 10517 | 5009.5 | 7241.5 |
| 464601 | 328745.5 | 139374 | 12841 | 20112 | 3361.5 | 673.5 | 50082 | 15909.5 | 12999.5 |
| 7858 | 6377.5 | 4302.5 | 1313 | 17904.5 | 3044 | 672 | 11999.5 | 5027.5 | 7453.5 |
| 3058 | 4827 | 3851 | 1307 | 124895 | 24529 | 701 | 11200 | 4715 | 7216 |
| 4061 | 5086 | 3789 | 1252 | 20222 | 3178 | 692 | 11250 | 5077 | 6941 |
| 6168 | 14128.5 | 5668 | 1613 | 17665.5 | 3252.5 | 733 | 13755 | 6977 | 17760 |
| 8103 | 20434 | 7605 | 1986 | 16372 | 3318 | 744 | 13430 | 9147 | 23769 |
| 8736 | 21922 | 7896 | 1853 | 14668 | 3136 | 745 | 13178 | 9084 | 25108 |

| | | | | | | | | | |
|---|---|---|---|---|---|---|---|---|---|
| 8455 | 23220.5 | 8084.5 | 2001 | 14680.5 | 3221.5 | 771 | 12284 | 9430 | 25425 |
| 2731 | 3462 | 2053 | 756 | 3159 | 1376 | 321 | 9231 | 3360 | 4271 |
| 2692 | 3616.5 | 2405 | 864 | 3374.5 | 1535.5 | 470 | 9181.5 | 3591.5 | 4829 |
| 2520.5 | 3777 | 2683 | 990.5 | 3771.5 | 1639 | 563.5 | 9102 | 3925.5 | 5453.5 |
| 2524 | 3748 | 5332 | 1569 | 626268 | 59195 | 598 | 9314 | 4058 | 5848 |
| 2246 | 3462 | 2988 | 1039 | 7963 | 2079 | 585 | 8071 | 3621 | 5196 |
| 2338 | 3579 | 2976 | 1069 | 5827 | 1953 | 633 | 9020 | 4027 | 5665 |
| 2294.5 | 3788.5 | 3011.5 | 1117.5 | 5087 | 1924.5 | 651 | 9421.5 | 4022.5 | 5625 |
| 2659 | 3815 | 3111 | 1141 | 5120 | 1977 | 673 | 10218 | 4464 | 6160 |
| 2689 | 4166.5 | 3316 | 1132.5 | 4920.5 | 1921 | 688 | 11330 | 4657 | 6334.5 |
| 2574 | 3974 | 3356 | 1299 | 4509 | 2014 | 643 | 10604 | 4316 | 6159 |
| 2435 | 3612 | 2870 | 1084 | 4133 | 1748 | 628 | 9865 | 4027 | 5723 |
| 2259.5 | 3437.5 | 2787.5 | 1065 | 4001 | 1737 | 630 | 9516.5 | 3878 | 5541.5 |
| 2189 | 3461 | 2870 | 1069 | 3953 | 1764 | 650 | 9431 | 3939 | 5707 |
| 16895 | 12347.5 | 4697 | 1127 | 3954 | 1793.5 | 659 | 8892.5 | 7030 | 7412.5 |
| 2510 | 3901 | 3154 | 1120 | 4067 | 1785 | 673 | 9171 | 4252 | 5956 |
| 2501 | 3858.5 | 3107 | 1095 | 4203.5 | 1744.5 | 638.5 | 8974.5 | 4221 | 6012.5 |
| 103379 | 53583.5 | 15143 | 1641.5 | 4380.5 | 1838 | 621.5 | 17176.5 | 6223.5 | 7074.5 |
| 2742.5 | 4205 | 3327 | 1163 | 4362.5 | 1817.5 | 631 | 10153 | 4620 | 6361.5 |
| 2722.5 | 4001.5 | 3226 | 1111 | 4674.5 | 1940 | 682 | 10136.5 | 4589.5 | 6440 |
| 2520 | 3784 | 3139 | 1045 | 4184 | 1822 | 656 | 9145 | 4168 | 6120 |
| 4771 | 12580 | 4918 | 1375 | 6101 | 2090 | 682 | 14332 | 6288 | 15624 |
| 8061.5 | 19810.5 | 6487.5 | 1730.5 | 7894.5 | 2425.5 | 696 | 14516.5 | 9394 | 23451 |
| 8493 | 21700 | 6975 | 1766 | 8042 | 2424 | 670 | 11825 | 8633 | 23936 |
| 8135 | 21199 | 6774 | 1692 | 7966 | 2396 | 682 | 13702 | 9052 | 24348 |
| 2783 | 3626 | 2255 | 818 | 3008.5 | 1385.5 | 350 | 9193.5 | 3637 | 4636 |
| 2332 | 3347 | 2262 | 845 | 3061 | 1442 | 450 | 7676 | 3461 | 4660 |
| 2230.5 | 3495 | 2527 | 933.5 | 3423 | 1598 | 538.5 | 7755 | 3567.5 | 5021 |
| 2339 | 3680 | 5274 | 1582 | 623519 | 57115 | 608 | 8501 | 4097 | 5720 |
| 2043.5 | 3212.5 | 2804 | 1001.5 | 5156.5 | 1863 | 588 | 7582 | 3459.5 | 4901 |
| 2211 | 3568 | 2970 | 1038 | 5005 | 1893 | 639 | 8930 | 3970 | 5570 |
| 2299.5 | 3732.5 | 3078 | 1124.5 | 4848.5 | 1912.5 | 682.5 | 8986 | 4125.5 | 5871 |
| 2444 | 3989 | 3050 | 1115 | 4797 | 1884 | 672 | 9862 | 4455 | 6085 |
| 2471 | 3717 | 3023 | 1057 | 4365 | 1824 | 642 | 10062 | 4221 | 5821 |
| 2387.5 | 3640.5 | 2904 | 1079.5 | 4063 | 1748.5 | 609.5 | 9724.5 | 4116 | 5611 |
| 2537.5 | 3797.5 | 3091.5 | 1123.5 | 4249 | 1854 | 674 | 10195.5 | 4337.5 | 5966 |
| 2631 | 3964 | 3235.5 | 1133 | 4461.5 | 1852 | 705 | 10881 | 4592.5 | 6342 |
| 2248 | 3620 | 2876 | 1095 | 4175 | 1774 | 658 | 9275 | 4070 | 5672 |
| 11749.5 | 8756.5 | 3979.5 | 1068 | 3732 | 1629.5 | 624 | 8418.5 | 5813.5 | 6662.5 |
| 2651.5 | 3971.5 | 3169 | 1107 | 4264.5 | 1861 | 679 | 8826 | 4151.5 | 6054.5 |
| 2708.5 | 4058 | 3274.5 | 1178 | 4563.5 | 1864 | 679 | 9650.5 | 4650.5 | 6660 |
| 86456 | 46678 | 13407 | 1660 | 4734 | 1882 | 684 | 16025 | 6015 | 7635 |
| 2631.5 | 4046 | 3312 | 1122 | 4319 | 1833.5 | 636 | 9515.5 | 4470 | 6390 |
| 2580 | 4282 | 3330 | 1175 | 4895 | 1991 | 667 | 9407 | 4492 | 6677 |
| 2692 | 4329 | 3524 | 1248 | 4739 | 1915 | 723 | 10221 | 4882 | 7116 |
| 4328 | 10454 | 4347 | 1387 | 5879 | 2023 | 659 | 14684 | 5760 | 12874 |
| 7719 | 18067.5 | 6138.5 | 1548.5 | 7314.5 | 2186.5 | 641 | 13680.5 | 8475 | 20305 |
| 8130 | 18260 | 6088 | 1531 | 7114 | 2176 | 592 | 13768 | 8871 | 20192 |

| | | | | | | | | | |
|---|---|---|---|---|---|---|---|---|---|
| 8504 | 20245.5 | 6700.5 | 1668.5 | 8014 | 2393.5 | 665.5 | 13998 | 9157.5 | 23131.5 |
| 750 | 923 | 769 | 574 | 1270 | 1112 | 261 | 2322 | 1407 | 1774 |
| 1037 | 1733 | 1505 | 738 | 2286 | 1306 | 453 | 4117 | 2142 | 3156 |
| 1224 | 2088.5 | 1907 | 827.5 | 2816 | 1455 | 529 | 5344 | 2541.5 | 3878 |
| 1389 | 2475.5 | 4693.5 | 1438 | 645398 | 59356.5 | 622 | 6976 | 3023 | 4735.5 |
| 1538 | 2840 | 2956 | 1137 | 5357 | 2282 | 679 | 7973 | 3454 | 5222 |
| 1492.5 | 2681 | 2689 | 1123 | 4120 | 1905 | 640 | 7195 | 3206.5 | 4952 |
| 1354 | 2447 | 2338 | 960 | 3454 | 1654 | 616 | 6897 | 3080 | 4636 |
| 1564 | 2650 | 2434 | 996 | 3495 | 1737 | 638 | 7586 | 3222 | 4858 |
| 1788 | 3086 | 2601 | 1050 | 3874 | 1779 | 673 | 8873 | 3599 | 5321 |
| 1976 | 3158 | 2849 | 1151 | 3887 | 1840 | 660 | 9279 | 3670 | 5572 |
| 1983 | 3307 | 2858 | 1139 | 4064 | 1894 | 696 | 10650 | 3968 | 5678 |
| 1748.5 | 2952.5 | 2607 | 1014 | 3655.5 | 1704.5 | 652 | 8336 | 3446 | 5103 |
| 1657 | 2850 | 2714 | 1031 | 3647 | 1693 | 664 | 8036 | 3389 | 5124 |
| 17239 | 12664 | 4907 | 1208 | 3984 | 1787 | 717 | 8469 | 6959 | 7481 |
| 2002.5 | 3320.5 | 2968 | 1176 | 4077 | 1862 | 712 | 8718.5 | 3840.5 | 5835 |
| 1839 | 3112.5 | 2832.5 | 1083.5 | 3884 | 1751 | 657 | 7880.5 | 3659.5 | 5479.5 |
| 92770 | 50145 | 14191 | 1617 | 4343 | 1800 | 680 | 16031 | 5690 | 6835 |
| 2084.5 | 3577 | 3049 | 1149 | 4352.5 | 1786.5 | 669 | 8896 | 4121.5 | 6103 |
| 2059 | 3646 | 3265 | 1214 | 5002 | 1991 | 705 | 9480 | 4507 | 6318 |
| 2157.5 | 3579 | 3213 | 1175 | 4463 | 1921 | 704.5 | 9809.5 | 4337 | 6389 |
| 4595 | 12186 | 4885 | 1443 | 6131 | 2155 | 698 | 13895 | 5686 | 15217 |
| 8028 | 18991 | 6550.5 | 1693.5 | 7841.5 | 2403 | 742 | 14385 | 9471 | 22935.5 |
| 8468 | 21435 | 7246 | 1743 | 8239 | 2452 | 691 | 13414 | 9254 | 25449 |
| 7321 | 23133.5 | 7734.5 | 1858 | 9068.5 | 2554 | 734.5 | 10652 | 8436 | 27222 |
| 2635 | 3431.5 | 1911.5 | 752 | 2520.5 | 1330 | 328 | 7913 | 3378 | 4322.5 |
| 2024 | 3063 | 2050 | 790 | 2809 | 1397 | 452 | 7004 | 3097 | 4208 |
| 1899 | 3001 | 2307.5 | 897.5 | 3200.5 | 1544.5 | 574 | 7133 | 3243.5 | 4778 |
| 1685.5 | 2882.5 | 4975.5 | 1431 | 710465 | 66655 | 608 | 7595.5 | 3296.5 | 4845.5 |
| 1670 | 2817 | 2721 | 1007 | 4585 | 1868 | 637 | 7534 | 3285 | 4927 |
| 1770 | 3056 | 2876 | 1141 | 4431 | 1919 | 705 | 8493 | 3676 | 5625 |
| 1737 | 2900 | 2728 | 1085 | 4075 | 1798 | 656 | 7833 | 3555 | 5125 |
| 1739 | 2909 | 2499 | 1026 | 3702 | 1691 | 640 | 7905 | 3317 | 5032 |
| 1932.5 | 2980 | 2532.5 | 978 | 3574.5 | 1710 | 625 | 8614.5 | 3569 | 4983 |
| 2170.5 | 3519 | 2972 | 1184.5 | 4072 | 1847.5 | 683.5 | 10705 | 3978.5 | 5866 |
| 2105.5 | 3283 | 2782 | 1091 | 3979 | 1750.5 | 667 | 9931 | 3830.5 | 5761.5 |
| 2014.5 | 3332 | 2900 | 1135.5 | 4038 | 1825 | 706 | 9637.5 | 3882 | 5802 |
| 1857 | 3176 | 2831.5 | 1079 | 3920 | 1827 | 693.5 | 8743 | 3813 | 5560 |
| 18135 | 13351 | 4883 | 1177 | 3863 | 1760 | 692 | 8486 | 7221 | 7667 |
| 2201 | 3505 | 3048 | 1179 | 4202 | 1811 | 733 | 8837 | 3958 | 5905 |
| 2093 | 3442.5 | 2865 | 1166 | 4172 | 1872.5 | 689 | 8720.5 | 3944.5 | 5903 |
| 87795 | 48360 | 14142 | 1653 | 4195 | 1839 | 651 | 15992 | 5593 | 6798 |
| 2348 | 3971 | 3456 | 1190 | 4328 | 1835 | 683 | 10271 | 4266 | 6171 |
| 2099 | 3464 | 3138 | 1159 | 4869 | 1979 | 660 | 9598 | 4197 | 6115 |
| 2144.5 | 3504.5 | 3111.5 | 1134.5 | 4248 | 1838 | 698 | 9193.5 | 4090 | 5940.5 |
| 4501 | 11481 | 4672 | 1354 | 5768 | 2062 | 692 | 13567 | 5364 | 14388 |
| 7520 | 17055.5 | 5962 | 1560 | 7096 | 2290.5 | 695 | 13887 | 8426 | 19830 |
| 8323 | 20436 | 6956 | 1741 | 8053 | 2504 | 715 | 13369 | 9060 | 24575 |

| | | | | | | | | | |
|---|---|---|---|---|---|---|---|---|---|
| 8203 | 21990 | 7165 | 1781 | 8311 | 2490 | 713 | 13033 | 9321 | 24566 |
| 2831.5 | 3638.5 | 2093.5 | 753 | 2744.5 | 1351.5 | 338 | 8394 | 3774 | 4596 |
| 2172 | 3299 | 2183 | 846 | 2905 | 1421 | 480 | 7493 | 3295 | 4646 |
| 1945 | 3073.5 | 2470 | 956 | 3357 | 1602 | 565 | 7728 | 3405.5 | 4784 |
| 1962 | 3095 | 5292 | 1555 | 673691 | 61185 | 642 | 8256 | 3632 | 5362 |
| 1839 | 3106 | 3175.5 | 1104 | 5652.5 | 2106.5 | 651 | 8270 | 3523.5 | 5163 |
| 1944.5 | 3285.5 | 3106 | 1177 | 4833 | 1977 | 717.5 | 9202.5 | 3765.5 | 5598.5 |
| 1801.5 | 3109.5 | 2703.5 | 1072.5 | 4155 | 1860 | 663 | 8414.5 | 3506.5 | 5318 |
| 1827 | 3014 | 2738 | 1048 | 3941 | 1785 | 641 | 8779 | 3475 | 5211 |
| 1904.5 | 3188 | 2636.5 | 1062.5 | 4022 | 1759 | 654.5 | 8924.5 | 3662 | 5256.5 |
| 2190 | 3449 | 2881 | 1109 | 4012 | 1856 | 657 | 10218 | 3995 | 5847 |
| 2271 | 3563 | 3006 | 1152 | 4206 | 1902 | 688 | 10604 | 4142 | 5813 |
| 2031.5 | 3417 | 2884.5 | 1140 | 4160 | 1803 | 700 | 9322.5 | 3956.5 | 5883 |
| 1960.5 | 3173 | 2817.5 | 1068.5 | 3975 | 1776.5 | 695.5 | 9003 | 3826 | 5581 |
| 14810 | 11106.5 | 4604 | 1134.5 | 3967 | 1849 | 690 | 8631.5 | 6549.5 | 7351.5 |
| 2150 | 3360 | 2924 | 1088.5 | 3875.5 | 1822.5 | 693 | 8481.5 | 3842 | 5499.5 |
| 2148 | 3477.5 | 2962 | 1113 | 4177.5 | 1892 | 685 | 8843 | 4054.5 | 5961.5 |
| 60943.5 | 32822 | 10122 | 1517.5 | 4528 | 1923 | 699 | 14200.5 | 5610 | 6867 |
| 2515 | 4107 | 3470 | 1242 | 4773 | 1932 | 705 | 10339 | 4598 | 6666 |
| 2296 | 3828 | 3283 | 1136 | 4966 | 2004 | 700 | 10053 | 4436 | 6334 |
| 2326 | 3824 | 3355 | 1227 | 4646 | 2056 | 742 | 10645 | 4477 | 6495 |
| 5032.5 | 13197.5 | 5024.5 | 1454.5 | 6302 | 2222 | 757.5 | 14367 | 6996 | 16434 |
| 8560 | 19608 | 6557 | 1735 | 7813 | 2464 | 749 | 14676 | 9384 | 23247 |
| 8248 | 23141.5 | 8168 | 1930 | 9619 | 2695 | 763 | 12390 | 8918.5 | 26246 |
| 8114 | 22695 | 7614 | 1802 | 9010 | 2552 | 755 | 12432 | 9138 | 25501 |
| 2546.5 | 3535 | 1996 | 766.5 | 2623.5 | 1321.5 | 314 | 8018 | 3440 | 4450 |
| 2114.5 | 3073 | 2073 | 832.5 | 2865.5 | 1445 | 451 | 7029.5 | 3287 | 4329.5 |
| 1842 | 2978.5 | 2210 | 885.5 | 3115.5 | 1488.5 | 514 | 7072 | 3110 | 4491 |
| 1828.5 | 3130.5 | 4938.5 | 1448 | 649722 | 59924.5 | 602 | 7869 | 3547 | 5027.5 |
| 1991.5 | 3223 | 3197 | 1144.5 | 5703 | 2135 | 663.5 | 8455 | 3637.5 | 5545 |
| 1915 | 3161 | 2764 | 1068 | 4620 | 1857 | 653 | 8366 | 3658 | 5192 |
| 1807 | 3073 | 2925 | 1098 | 4374 | 1869 | 662 | 8227 | 3563 | 5418 |
| 1916 | 3162 | 2852 | 1041 | 3941 | 1781 | 655 | 8711 | 3701 | 5365 |
| 1982 | 3238 | 2695.5 | 1022.5 | 3919 | 1697.5 | 636.5 | 8823 | 3725.5 | 5181.5 |
| 2158 | 3423 | 2904 | 1138 | 3997 | 1829 | 671 | 9800 | 3823 | 5565 |
| 2128 | 3322 | 2831 | 1022 | 3950 | 1749 | 665 | 9715 | 3972 | 5456 |
| 2224 | 3522 | 3117 | 1167 | 4318 | 1854 | 720 | 10875 | 4181 | 6037 |
| 2078 | 3377 | 2978 | 1100 | 4121 | 1876 | 691 | 9415 | 4002 | 5801 |
| 15848 | 11491 | 4785 | 1167 | 4072 | 1779 | 700 | 8905 | 6907 | 7557 |
| 2520.5 | 3856 | 3274 | 1191 | 4418 | 1922 | 742.5 | 9515 | 4449 | 6131.5 |
| 2343 | 3816 | 3223 | 1198 | 4348 | 1866 | 719 | 9425 | 4313 | 6306 |
| 88237 | 47000.5 | 13885 | 1685 | 4221 | 1778.5 | 669 | 15723.5 | 5950 | 6934 |
| 2616 | 4105.5 | 3417 | 1240.5 | 4418.5 | 1894 | 690 | 10480 | 4768 | 6793 |
| 2360 | 3726 | 3243.5 | 1210 | 4942.5 | 2018.5 | 693 | 10119.5 | 4425 | 6376 |
| 2417 | 3762 | 3258 | 1214 | 4528 | 1936 | 716 | 10639 | 4648 | 6644 |
| 4583.5 | 11728 | 4945 | 1403 | 6143 | 2149 | 704 | 14140.5 | 6198.5 | 15049.5 |
| 7628 | 18784 | 6670 | 1720 | 8107 | 2464 | 726 | 14311 | 8920 | 21871.5 |
| 8697 | 23102 | 7163.5 | 1822.5 | 8547.5 | 2534 | 695 | 12745.5 | 8970 | 26102.5 |

| | | | | | | | | | |
|---|---|---|---|---|---|---|---|---|---|
| 8019 | 21961.5 | 6991 | 1672 | 8196 | 2443.5 | 675 | 13184 | 9164.5 | 24949.5 |
| 3208.5 | 3895 | 2274 | 815 | 3027 | 1422.5 | 353 | 9679.5 | 4016.5 | 4899 |
| 2248 | 3502 | 2206 | 866 | 2990 | 1488 | 443 | 7901 | 3509 | 4461 |
| 2194.5 | 3301 | 2509.5 | 967.5 | 3466 | 1561 | 560 | 8027.5 | 3577 | 4913 |
| 1930 | 3330 | 5020 | 1584 | 723747 | 63490 | 618 | 8016 | 3675 | 5289 |
| 1973 | 3185 | 3291 | 1150 | 5554 | 2125 | 645 | 8140 | 3633 | 5342 |
| 2027.5 | 3299.5 | 3089 | 1132 | 4904.5 | 2007.5 | 692 | 9113 | 3874 | 5652 |
| 1953 | 3307 | 3036.5 | 1111.5 | 4483.5 | 1867.5 | 679 | 8611 | 3691 | 5439.5 |
| 1933 | 3167.5 | 2533.5 | 1015 | 3999 | 1766.5 | 627 | 8498 | 3507 | 5253.5 |
| 2151 | 3448 | 2778 | 1097 | 4162 | 1741 | 663 | 9858 | 3752 | 5572 |
| 2227 | 3549 | 2892.5 | 1115 | 3909 | 1752.5 | 646.5 | 10008 | 4018.5 | 5627 |
| 2459.5 | 3518 | 2957 | 1153.5 | 4185 | 1835.5 | 696 | 10539 | 4314 | 5728 |
| 2330.5 | 3636 | 3020.5 | 1188.5 | 4364.5 | 1890.5 | 707 | 10076 | 4281.5 | 5821.5 |
| 2177 | 3589 | 3062 | 1121 | 4370 | 1900 | 712 | 9795 | 4232 | 6057 |
| 32684 | 24081 | 7589 | 1289 | 4265 | 1864 | 697 | 9250 | 12656 | 10932 |
| 2366 | 3765 | 3166.5 | 1121.5 | 4520 | 1880 | 723 | 9343 | 4461 | 6290.5 |
| 2352 | 3716 | 3303 | 1238 | 4458 | 1916 | 744 | 9915 | 4479 | 6495 |
| 65129 | 34748 | 10813 | 1505 | 4471 | 1896 | 700 | 14360 | 5581 | 6907 |
| 2678.5 | 4321 | 3751.5 | 1241 | 4892.5 | 1966.5 | 711 | 10472.5 | 4743.5 | 6891.5 |
| 2486 | 3972 | 3373 | 1167 | 5057 | 2023 | 717 | 9500 | 4715 | 6580 |
| 2469 | 3933 | 3246 | 1231 | 4677 | 1960 | 742 | 10333 | 4590 | 6559 |
| 5394 | 13852 | 5326 | 1502 | 6733 | 2264 | 746 | 14590 | 6875 | 16972 |
| 8096 | 22209 | 6935 | 1878 | 8436 | 2587 | 776 | 13145 | 9217 | 24587 |
| 8257 | 22952 | 7409 | 1828 | 8959 | 2566 | 734 | 12915 | 9198 | 26365 |
| 7814 | 23146 | 8014 | 1800 | 9674 | 2611 | 708 | 12663 | 9275 | 25831 |
| 3272 | 3922 | 2262 | 848 | 3057 | 1429 | 350 | 9570 | 4092 | 4923 |
| 8662.5 | 6047 | 3163.5 | 923.5 | 3597.5 | 1606.5 | 492.5 | 9209.5 | 3874.5 | 4720.5 |
| 5976 | 4986 | 3240 | 1038 | 4435 | 1894 | 651 | 10160 | 4326 | 5749 |
| 5481 | 4455 | 4908 | 1609 | 84926 | 11114 | 644 | 8923 | 3846 | 5541 |
| 4824 | 4477 | 4114 | 1302 | 28045 | 4328 | 721 | 10423 | 4255 | 5985 |
| 5523 | 4804 | 4035.5 | 1232.5 | 12896 | 3188.5 | 852 | 11919 | 4981.5 | 6338 |
| 4738 | 4410.5 | 3524 | 1207 | 10032 | 2911 | 781.5 | 10517.5 | 4515 | 6017 |
| 4886 | 4289 | 3452 | 1084 | 9319 | 2749 | 751 | 10523 | 4261 | 5783 |
| 4725 | 4496 | 3325 | 1141 | 6739 | 2468 | 780 | 11500 | 4645 | 6112 |
| 4695 | 5131 | 4542 | 1697 | 26211 | 3730 | 819 | 12899 | 5250 | 6723 |
| 4521 | 4471 | 3325 | 1091 | 5827 | 2350 | 771 | 11412 | 4812 | 6146 |
| 4427 | 4541 | 3550 | 1225 | 5136 | 2110 | 803 | 12409 | 5045 | 6513 |
| 4456 | 4324 | 3309 | 1210 | 5043 | 2090 | 783 | 11025 | 4662 | 6253 |
| 34262.5 | 23707 | 7770.5 | 1472 | 4731.5 | 2014 | 799.5 | 11346.5 | 14743 | 12233 |
| 7030 | 6311 | 4163 | 1353 | 5697 | 2395 | 840 | 11973 | 6161 | 7798 |
| 6161.5 | 5751 | 3916.5 | 1271 | 5951.5 | 2387.5 | 849.5 | 11677.5 | 5447.5 | 7077 |
| 1019116 | 613733 | 203785 | 15945.5 | 5292 | 1934 | 753 | 88853.5 | 30081 | 19016 |
| 12744 | 8116 | 4480 | 1359 | 6953 | 2583 | 845 | 13191 | 5901 | 7462 |
| 8306.5 | 6074 | 3986.5 | 1276 | 5095 | 2132.5 | 772.5 | 12616.5 | 5191 | 6910.5 |
| 13999 | 8620.5 | 4730.5 | 1331 | 7637 | 2743.5 | 837 | 14754.5 | 5943.5 | 7595.5 |
| 10906 | 17950.5 | 6256 | 1677 | 8947 | 2843.5 | 862 | 15834 | 7415.5 | 18072 |
| 18544 | 27615.5 | 8262 | 1867 | 12367.5 | 3117.5 | 840.5 | 15473 | 10333.5 | 24968.5 |
| 14303 | 27066.5 | 7739.5 | 1932.5 | 8901.5 | 2788.5 | 794.5 | 13724.5 | 9545.5 | 25916 |

| | | | | | | | | | |
|---|---|---|---|---|---|---|---|---|---|
| 12742 | 27628 | 8074 | 1883 | 10912 | 2932 | 817 | 11528 | 9415 | 27139 |
| 3013 | 3882.5 | 2230 | 810 | 3044 | 1399.5 | 343 | 9418 | 3919 | 4737.5 |
| 2373 | 3489 | 2329 | 876 | 3123 | 1472 | 453 | 7927 | 3447 | 4774 |
| 2116 | 3089 | 2387.5 | 879 | 3181 | 1521 | 533 | 7695.5 | 3368 | 4727.5 |
| 2024 | 3354 | 5294 | 1588 | 685374 | 61719 | 614 | 8538 | 3643 | 5377 |
| 2055 | 3364 | 3039 | 1133 | 5517 | 2061 | 637 | 8727 | 3627 | 5530 |
| 2329 | 3573 | 3201 | 1138 | 4928 | 1927 | 686 | 8999 | 4379 | 5713 |
| 2040.5 | 3428 | 2956 | 1065.5 | 4526.5 | 1864 | 666 | 8618 | 3958 | 5666.5 |
| 2139 | 3415 | 2834 | 1040 | 4307 | 1824 | 659 | 8951 | 3858 | 5584 |
| 2252 | 3326 | 2657 | 996 | 4051 | 1724 | 644 | 9434 | 3853 | 5463 |
| 2368.5 | 3613.5 | 2821.5 | 1124 | 4130 | 1822 | 640 | 9580 | 4084 | 5626 |
| 2402 | 3616.5 | 2875.5 | 1097.5 | 4076.5 | 1840 | 664.5 | 10611.5 | 4233 | 5681 |
| 2432 | 3727 | 3122 | 1153 | 4395 | 1883 | 708 | 10841 | 4526 | 6127 |
| 2302 | 3677 | 3160 | 1152 | 4279 | 1872 | 727 | 10197 | 4331 | 5955 |
| 16345 | 11888 | 4948 | 1202 | 4047 | 1794 | 683 | 9043 | 7162 | 7631 |
| 2907 | 4153 | 3400 | 1226 | 4603.5 | 1966 | 761 | 10445.5 | 4849.5 | 6750 |
| 2516.5 | 3960 | 3263 | 1173.5 | 4477.5 | 1905.5 | 719 | 9954 | 4505.5 | 6429 |
| 110947.5 | 58285 | 16595 | 1815.5 | 4660 | 1961 | 685 | 18240 | 6678 | 7656.5 |
| 2717 | 4229 | 3584.5 | 1216.5 | 4632 | 1882.5 | 685 | 10702.5 | 4745.5 | 6637 |
| 2678 | 4399 | 3736 | 1279 | 5669 | 2125 | 754 | 12213 | 4902 | 7360 |
| 2643 | 4196 | 3393 | 1223 | 4664 | 1972 | 734 | 11253 | 4641 | 6700 |
| 5351.5 | 11192 | 4846.5 | 1429 | 6143 | 2167 | 706.5 | 14800.5 | 7411 | 14762 |
| 7667 | 19742 | 7022 | 1820 | 8363 | 2548 | 734 | 12572 | 8834 | 23367 |
| 8335 | 23041.5 | 8168.5 | 1982 | 9172 | 2672.5 | 729.5 | 12528 | 8683 | 26647 |
| 8270 | 24198 | 9023 | 2001 | 10200 | 2701 | 767 | 11982 | 9015 | 27703 |
| 2959.5 | 3872.5 | 2156 | 752 | 2967 | 1370.5 | 324.5 | 9351 | 4076 | 4720 |
| 2375 | 3389 | 2324 | 889 | 3226 | 1551 | 466 | 8023 | 3601 | 4871 |
| 2171 | 3379 | 2729 | 1032 | 3915 | 1729 | 553 | 8486 | 3720 | 5527 |
| 2112 | 3253 | 5192 | 1548 | 491937 | 50547 | 597 | 8856 | 4007 | 5844 |
| 2177 | 3289 | 3282 | 1249 | 5813 | 2254 | 614 | 9189 | 4327 | 6204 |
| 2234 | 3626 | 3290 | 1244 | 5579 | 2192 | 644 | 9985 | 4491 | 6964 |
| 2248.5 | 3714 | 3574 | 1375.5 | 5684.5 | 2244.5 | 684 | 9598 | 4606.5 | 7388.5 |
| 2169 | 3570 | 3222 | 1252 | 5109 | 2107 | 635 | 9455 | 4462 | 6820 |
| 2255 | 3408 | 2890 | 1195 | 4670 | 1983.5 | 615 | 9832 | 4480.5 | 6438 |
| 2393 | 3519 | 3216 | 1256.5 | 4768 | 2089.5 | 603.5 | 9411 | 4559.5 | 6629.5 |
| 2491 | 3766 | 3392 | 1322 | 5128 | 2209 | 676 | 11298 | 4623 | 7029 |
| 2244 | 3828 | 3528 | 1361 | 5219 | 2271 | 695 | 11636 | 4555 | 7662 |
| 2218.5 | 4238.5 | 3503.5 | 1373 | 5363.5 | 2334.5 | 695.5 | 10881.5 | 4471 | 8433 |
| 5495.5 | 5758.5 | 4132 | 1394.5 | 5313 | 2258.5 | 683 | 10542.5 | 5702 | 8109 |
| 2334 | 4471 | 3698 | 1445 | 5626 | 2330 | 736 | 11352 | 4726 | 8952 |
| 2273.5 | 4370 | 3646.5 | 1407 | 5223.5 | 2259 | 683 | 10112 | 4644.5 | 8402 |
| 42685 | 31104 | 8473.5 | 1609.5 | 5514.5 | 2230.5 | 663.5 | 13595.5 | 5363.5 | 10413 |
| 2418 | 6366 | 4133 | 1440 | 5804 | 2348 | 717 | 12169 | 4532 | 11506 |
| 2223.5 | 5180 | 3649 | 1387 | 5880.5 | 2390.5 | 678 | 11632.5 | 4564.5 | 10073.5 |
| 2323.5 | 5305.5 | 3941 | 1524 | 5752.5 | 2392 | 744.5 | 11725.5 | 4658 | 9631 |
| 5538.5 | 13125 | 5532.5 | 1682.5 | 7083.5 | 2650.5 | 712.5 | 13536.5 | 7306.5 | 18343 |
| 7561 | 20076 | 7083 | 1979 | 9132 | 2939 | 714 | 13713 | 9309 | 24778 |
| 7500.5 | 22541.5 | 7897.5 | 2128.5 | 9956 | 2894 | 717 | 12279 | 9083 | 26240.5 |

| | | | | | | | | | |
|---|---|---|---|---|---|---|---|---|---|
| 8275 | 23380.5 | 7916.5 | 1972.5 | 9622 | 2821.5 | 702.5 | 13258 | 9316 | 27279 |
| 3237.5 | 4027.5 | 2393.5 | 879.5 | 3207 | 1461 | 357.5 | 9970 | 4382.5 | 4965.5 |
| 3397 | 4171 | 2654 | 998 | 4788 | 1667 | 500 | 9881 | 4263 | 5208 |
| 2728 | 4064 | 3008 | 1032 | 6968 | 2046 | 637 | 9396 | 4142 | 5704 |
| 2797 | 3855 | 6056 | 1932 | 348849 | 65874 | 659 | 9225 | 4110 | 5664 |
| 2670.5 | 3640 | 3553.5 | 1196.5 | 76213 | 10171 | 644 | 9476.5 | 4396 | 5622.5 |
| 2624 | 3655 | 3331 | 1172 | 63207 | 8642 | 696 | 9889 | 4188 | 5965 |
| 3306 | 4189 | 3576 | 1237 | 56211 | 7742 | 730 | 13116 | 6304 | 8235 |
| 2834 | 3923.5 | 3247 | 1163 | 49163 | 6602 | 690 | 9690.5 | 4165 | 5893.5 |
| 2993.5 | 4035.5 | 3195.5 | 1121 | 41036.5 | 5606.5 | 688 | 10495.5 | 4477 | 5963 |
| 3675 | 5522 | 4825 | 1677 | 34834 | 5673 | 793 | 13866 | 5407 | 7722 |
| 3111 | 4606 | 3694 | 1259.5 | 33757 | 4771.5 | 747 | 13259.5 | 4786.5 | 6919.5 |
| 3174 | 4950 | 4455 | 1401 | 21438 | 3551 | 719 | 14639 | 4766 | 7277 |
| 3438.5 | 5231 | 4137.5 | 1282.5 | 25515 | 4070.5 | 771 | 15463.5 | 5016 | 7800.5 |
| 57878 | 47796 | 35633 | 4327 | 21397 | 3583 | 766 | 13284 | 22445 | 19743 |
| 4821.5 | 6312.5 | 4614 | 1383 | 20788 | 3486.5 | 775.5 | 14225 | 5738.5 | 8529.5 |
| 3698 | 5919 | 4178 | 1290.5 | 18693 | 3312 | 767.5 | 13120.5 | 4925 | 8448.5 |
| 480063 | 441999 | 169677.5 | 16766 | 18037.5 | 3262 | 711.5 | 59833.5 | 15997 | 17380.5 |
| 6086 | 7041 | 4613 | 1471 | 16097 | 3126 | 742 | 13649 | 5201 | 9111 |
| 3160 | 5229 | 4236 | 1337 | 99182 | 20655 | 741 | 12843 | 4689 | 8227 |
| 4221 | 6270 | 4395 | 1448 | 17976 | 3339 | 769 | 14101 | 5107 | 7993 |
| 6182 | 15454 | 5619 | 1650 | 16047 | 3154 | 774 | 14567 | 7639 | 18173 |
| 8827 | 22734 | 7936 | 2010 | 15613 | 3333 | 788 | 13592 | 9253 | 25977 |
| 8876 | 24753 | 8422 | 2148 | 15642 | 3347 | 805 | 12591 | 9239 | 26080 |
| 8581 | 24028 | 8489.5 | 1968 | 14159 | 3166 | 780 | 13641.5 | 9233.5 | 26573 |
| 3309 | 3818 | 2205 | 821 | 3244 | 1458 | 373 | 10914 | 3910 | 4732 |
| 2761 | 3711 | 2496 | 866 | 3502 | 1510 | 477 | 10128 | 3796 | 4870 |
| 2695 | 3762 | 2761 | 949 | 3808 | 1641 | 572 | 9245 | 3886 | 5428 |
| 2587 | 3795 | 5593.5 | 1489.5 | 654730.5 | 62285 | 616.5 | 9855 | 4258.5 | 5742.5 |
| 2573 | 3846.5 | 3285.5 | 1161 | 9725.5 | 2384 | 655 | 9610.5 | 4293.5 | 5871 |
| 2611.5 | 3821 | 3201 | 1149.5 | 6392.5 | 2013.5 | 685 | 9925.5 | 4536.5 | 6105 |
| 2482 | 3764.5 | 3249.5 | 1097.5 | 5225 | 1922 | 676 | 9308 | 4303.5 | 5908 |
| 2335 | 3616.5 | 3043.5 | 1096.5 | 4627.5 | 1800.5 | 645 | 9386.5 | 3896.5 | 5734.5 |
| 2495 | 3722 | 2962 | 1072 | 4418.5 | 1782.5 | 657 | 10051 | 4169 | 5730.5 |
| 2695 | 3927 | 3289 | 1264 | 4570 | 2047 | 645 | 10664 | 4395 | 5907 |
| 2702 | 3853.5 | 3158.5 | 1135.5 | 4302.5 | 1851.5 | 670.5 | 10576 | 4516.5 | 6008.5 |
| 2634 | 3962 | 3285 | 1191 | 4583 | 1903 | 716 | 11380 | 4532 | 6431 |
| 2461.5 | 3773 | 3161.5 | 1098 | 4369.5 | 1864 | 688 | 10523.5 | 4432 | 6195.5 |
| 16179 | 11184 | 4645 | 1193 | 4203 | 1772 | 699 | 9889 | 7406 | 7712 |
| 2802 | 4128 | 3188 | 1158 | 4454 | 1919 | 710 | 9813 | 4625 | 6381 |
| 2667 | 4038 | 3294 | 1194 | 4526 | 1886 | 695 | 9722 | 4745 | 6363 |
| 122907 | 67647 | 18823 | 1863 | 4771 | 1947 | 711 | 20576 | 6716 | 7915 |
| 2871 | 4295 | 3500 | 1225 | 4672 | 1920 | 678 | 11073 | 4879 | 6769 |
| 2596 | 4630 | 3660 | 1297 | 5597 | 2098 | 721 | 12350 | 4696 | 7186 |
| 2793 | 4319.5 | 3452 | 1274 | 4916 | 1999.5 | 733.5 | 11552.5 | 4889.5 | 6917 |
| 5081 | 11435 | 4882 | 1502 | 6225 | 2169 | 711 | 14465 | 6732 | 14551 |
| 8211.5 | 22702.5 | 7914 | 1943.5 | 9696.5 | 2691 | 768 | 12877 | 9194.5 | 24294 |
| 7929 | 22743 | 8876 | 2012 | 10551 | 2842 | 763 | 12079 | 8499 | 25145 |

| | | | | | | | | | |
|---|---|---|---|---|---|---|---|---|---|
| 7224 | 22118 | 8415 | 1887 | 10282 | 2658 | 716 | 10654 | 7996 | 24800 |
| 3338.5 | 4104.5 | 2401 | 815.5 | 3102 | 1425.5 | 346 | 9637.5 | 4183 | 4918.5 |
| 2582 | 3786 | 2473 | 893 | 3114 | 1435 | 449 | 8386 | 3632 | 4963 |
| 2373 | 3600.5 | 2723.5 | 1012.5 | 3502.5 | 1629.5 | 547 | 8609.5 | 3794 | 5257.5 |
| 2153 | 3411 | 4950.5 | 1472.5 | 608383 | 56375.5 | 618.5 | 8464 | 3686.5 | 5206.5 |
| 2285.5 | 3524 | 3222.5 | 1099.5 | 5444 | 2031 | 636.5 | 8880.5 | 3841.5 | 5409 |
| 2381 | 3749 | 3272 | 1155 | 5278.5 | 2040 | 691 | 9100.5 | 4101 | 6031.5 |
| 2351 | 3616 | 3116 | 1154 | 4710 | 1893 | 692 | 9173 | 4069 | 5764 |
| 2188 | 3479 | 2970.5 | 1082.5 | 4391.5 | 1838.5 | 658 | 9073 | 3864.5 | 5699.5 |
| 2527 | 3841 | 3005 | 1087 | 4445 | 1810 | 657 | 10158 | 4161 | 5969 |
| 2483 | 3850 | 3179 | 1222 | 4310 | 1939 | 667 | 10158 | 4282 | 6168 |
| 2475 | 3834 | 3011.5 | 1159 | 4444 | 1854 | 669 | 10729 | 4460.5 | 5887 |
| 2424 | 3537 | 2927 | 1071 | 4006 | 1830 | 666 | 10174 | 4220 | 5625 |
| 2138.5 | 3457 | 2896.5 | 1068 | 3961 | 1735 | 657 | 9072.5 | 3825.5 | 5596 |
| 13790 | 10148 | 4428 | 1135 | 3964 | 1798.5 | 660.5 | 8792 | 6352 | 7077.5 |
| 2611 | 3768 | 3192 | 1100 | 4293 | 1877 | 695 | 9454 | 4562 | 5970 |
| 2431 | 3661 | 3103.5 | 1126 | 4124 | 1804 | 678 | 9158.5 | 4387 | 5950.5 |
| 98836 | 51457 | 14895 | 1646 | 4158 | 1782 | 630 | 16286 | 6070 | 6895 |
| 2543.5 | 3998.5 | 3378.5 | 1156.5 | 4378 | 1870 | 662 | 9567.5 | 4501 | 6358.5 |
| 2623.5 | 4023 | 3345 | 1204 | 5169.5 | 1981.5 | 717 | 10630 | 4646.5 | 6651.5 |
| 2423.5 | 3910 | 3200 | 1228 | 4416.5 | 1873 | 683 | 10097.5 | 4610.5 | 6237 |
| 4369.5 | 11670 | 4509.5 | 1306.5 | 5582 | 2016.5 | 664 | 12626.5 | 5506 | 14370.5 |
| 6142.5 | 15709 | 6206.5 | 1541.5 | 6861 | 2229.5 | 656 | 14991.5 | 7110 | 18800 |
| 8482.5 | 22053.5 | 7327.5 | 1758.5 | 8593.5 | 2503.5 | 689 | 13845.5 | 9581.5 | 24728.5 |
| 8497.5 | 22240 | 7827 | 1828 | 9142.5 | 2522 | 729 | 12597 | 9219 | 25518.5 |
| 725.5 | 905 | 744 | 581.5 | 1304 | 1106 | 272 | 2363.5 | 1399.5 | 1763 |
| 1044 | 1719.5 | 1580 | 768 | 2305.5 | 1358 | 460 | 4032 | 2160 | 3208 |
| 1230 | 2085 | 1926 | 836 | 2855 | 1461 | 539 | 5327 | 2618 | 3915 |
| 1436 | 2562 | 4750 | 1412 | 642670 | 59326 | 623 | 6874 | 3106 | 4752 |
| 1447 | 2603 | 2629 | 1018 | 4673 | 1975 | 639 | 7362 | 3157 | 4835 |
| 1514.5 | 2822 | 2807 | 1082 | 4232.5 | 1868 | 676.5 | 7583.5 | 3414 | 5435.5 |
| 1408 | 2649 | 2442 | 967 | 3652 | 1748 | 629 | 7306 | 3245 | 4953 |
| 1627 | 2771 | 2730 | 1047 | 3918 | 1734 | 669 | 7840 | 3443 | 5203 |
| 1702 | 2941 | 2606 | 1017 | 3713 | 1767 | 658 | 8600 | 3418 | 4913 |
| 1994 | 3247 | 2784 | 1114 | 3901 | 1828 | 668 | 9719 | 3736 | 5480 |
| 2062.5 | 3412 | 3021.5 | 1135 | 4108.5 | 1912.5 | 717 | 10909.5 | 4313 | 5924.5 |
| 1980 | 3313 | 2916 | 1156 | 4185 | 1834 | 729 | 9747 | 3951 | 5716 |
| 1822 | 3112 | 3055 | 1154 | 4143 | 1872 | 733 | 9235 | 3741 | 5771 |
| 18327 | 13858 | 5101 | 1229 | 4204 | 1815 | 732 | 8550 | 7433 | 7900 |
| 1996.5 | 3345 | 3131.5 | 1142 | 4078 | 1836.5 | 704 | 8540.5 | 3819 | 5781.5 |
| 1766.5 | 3258.5 | 2924.5 | 1134 | 3948 | 1785 | 690 | 8638 | 3678 | 5690.5 |
| 97388 | 51424 | 14702 | 1687 | 4286 | 1894 | 661 | 16292 | 5769 | 6785 |
| 2165 | 3643 | 3347 | 1187 | 4209 | 1871 | 681 | 9369 | 4330 | 6116 |
| 1944.5 | 3343.5 | 2928.5 | 1123 | 4726.5 | 1924.5 | 678 | 8744 | 3850.5 | 5726 |
| 1972.5 | 3406 | 3051 | 1122 | 4315 | 1884 | 698.5 | 9172.5 | 4085.5 | 6069 |
| 3769.5 | 8104 | 4440.5 | 1308.5 | 5500.5 | 1977 | 684.5 | 14094 | 5629.5 | 10770 |
| 8006 | 18611 | 6441 | 1753 | 7817 | 2490 | 722 | 13984 | 9276 | 21448 |
| 8169.5 | 22345.5 | 7578.5 | 1855 | 8562 | 2468.5 | 730.5 | 13809 | 9394.5 | 25797 |

| | | | | | | | | | |
|---|---|---|---|---|---|---|---|---|---|
| 8141 | 22295 | 7523 | 1801 | 8481 | 2548 | 715 | 12872 | 9138 | 24854 |
| 2780 | 3708 | 2140.5 | 773 | 2793 | 1377 | 362.5 | 9443.5 | 3793 | 4727 |
| 2225 | 3140.5 | 2169.5 | 847.5 | 2845.5 | 1457.5 | 471.5 | 7257 | 3212 | 4342.5 |
| 2053 | 3162.5 | 2518 | 994 | 3384.5 | 1667 | 613 | 8131 | 3587 | 4929.5 |
| 1804 | 2850.5 | 5644 | 1557 | 942111.5 | 88478.5 | 619.5 | 7789.5 | 3322.5 | 4990 |
| 1802 | 2956 | 2995 | 1098 | 5196 | 2036 | 680 | 8264 | 3452 | 5041 |
| 2054 | 3700.5 | 3375 | 1235.5 | 5355 | 2160 | 787 | 9886.5 | 4139.5 | 6061.5 |
| 1930 | 3299 | 3082 | 1148 | 4604 | 1949 | 749 | 9115 | 3921 | 5870 |
| 1852 | 3091 | 2760 | 1000 | 4017 | 1820 | 685 | 9013 | 3731 | 5280 |
| 2111.5 | 3202 | 2898.5 | 1128 | 3920.5 | 1733.5 | 716.5 | 9925 | 3876.5 | 5492.5 |
| 2165.5 | 3490 | 2959.5 | 1148.5 | 4006 | 1766.5 | 726.5 | 10123 | 4053 | 5714 |
| 2169.5 | 3528.5 | 2935 | 1131.5 | 4194 | 1866.5 | 720 | 10611 | 4175 | 5876.5 |
| 2416.5 | 3767 | 3317 | 1215 | 4490 | 1998 | 774.5 | 12188.5 | 4596 | 6486.5 |
| 2317.5 | 3483 | 3120 | 1223.5 | 4542.5 | 1933 | 802.5 | 10545 | 4300.5 | 6346.5 |
| 22882 | 16329 | 5804 | 1285 | 4552 | 1974 | 781 | 10649 | 9131 | 8962 |
| 2570 | 3808 | 3332.5 | 1260 | 4537 | 2029 | 813 | 11068 | 4602.5 | 6428.5 |
| 2180 | 3442 | 3244 | 1178 | 4254 | 1890 | 736 | 9272 | 4232 | 6128 |
| 146986 | 75370 | 21403 | 1980 | 4747 | 2043 | 760 | 22396 | 7640 | 8054 |
| 2583 | 3972 | 3432 | 1258 | 4542 | 1907 | 736 | 10312 | 4946 | 6663 |
| 2532 | 3954 | 3448 | 1236 | 5631 | 2134 | 769 | 10836 | 4781 | 6801 |
| 2479 | 3829 | 3453 | 1250 | 4813 | 1908 | 768 | 11081 | 4813 | 6759 |
| 5641.5 | 13170.5 | 5304 | 1539.5 | 6531 | 2274.5 | 777 | 14936 | 7233 | 17245.5 |
| 9484 | 21216 | 6747.5 | 1797.5 | 8128 | 2567.5 | 768 | 13954 | 9181 | 22419.5 |
| 8534.5 | 23449.5 | 9193.5 | 2219.5 | 10498.5 | 2800 | 820 | 11386.5 | 8045.5 | 24885.5 |
| 9772 | 23563 | 8166 | 1888 | 9048 | 2631 | 801 | 12899 | 9626 | 26574 |
| 2629.5 | 3524.5 | 2174 | 769.5 | 2854.5 | 1323 | 349 | 8434 | 3648 | 4686.5 |
| 2256 | 3400 | 2299 | 838 | 2939 | 1455 | 496 | 7759 | 3465 | 4639 |
| 2033.5 | 3236.5 | 2464.5 | 949 | 3369 | 1573 | 570.5 | 7997.5 | 3473.5 | 4979 |
| 1869.5 | 3162.5 | 5717 | 1829 | 837858.5 | 69323 | 611 | 7750 | 3535.5 | 5295 |
| 1757 | 2895 | 3089 | 1086 | 6668 | 2200 | 614 | 7561 | 3372 | 4824 |
| 1949 | 3279.5 | 3128.5 | 1136 | 5455 | 2047.5 | 693.5 | 8841 | 3694 | 5435.5 |
| 1934 | 3253 | 3086 | 1152 | 4916 | 1962 | 693 | 9255 | 3776 | 5767 |
| 2148 | 3511 | 3087 | 1157 | 5025 | 1991 | 722 | 9815 | 4194 | 5901 |
| 2144 | 3359 | 2851 | 1109 | 4315 | 1818 | 678 | 9604 | 3849 | 5573 |
| 2092 | 3303 | 2848 | 1160 | 4005 | 1955 | 652 | 9534 | 3695 | 5384 |
| 2061 | 3273 | 2782 | 1098 | 3968 | 1776 | 653 | 9699 | 3774 | 5395 |
| 2296.5 | 3594 | 3165 | 1191.5 | 4338 | 1940.5 | 718 | 10522 | 4194 | 5845 |
| 2116 | 3256 | 2976 | 1152 | 4078 | 1820 | 704 | 9785 | 4171 | 5688 |
| 19137 | 13444.5 | 5202 | 1271.5 | 4533.5 | 1951 | 772.5 | 10155.5 | 8273 | 8200.5 |
| 2615 | 3868 | 3394 | 1217 | 4501 | 1948 | 763 | 10617 | 4726 | 6451 |
| 2432 | 3870 | 3440 | 1203.5 | 4674.5 | 1908 | 767 | 10488.5 | 4552 | 6282.5 |
| 158184 | 81825 | 22780 | 2110 | 4779 | 1986 | 730 | 24613 | 7698 | 8136 |
| 2671 | 4236 | 3590 | 1298 | 4885 | 1965 | 746 | 11134 | 5034 | 7207 |
| 2498 | 3903 | 3362 | 1188 | 5556 | 2125 | 739 | 10087 | 4675 | 6674 |
| 2519 | 3851 | 3534 | 1225 | 4778 | 1972 | 753 | 10815 | 4648 | 6664 |
| 4472 | 11805 | 4857 | 1434 | 6394 | 2202 | 761 | 14263 | 6198 | 15466 |
| 7372 | 18630 | 7027 | 1723 | 7973 | 2575 | 751 | 14074 | 8562 | 21567 |
| 8226 | 23575 | 7758.5 | 1854 | 9320 | 2594 | 740.5 | 12499.5 | 8994 | 25979.5 |

| | | | | | | | | | |
|---|---|---|---|---|---|---|---|---|---|
| 8533 | 22459 | 7617 | 1777 | 8739 | 2594 | 737 | 13322 | 9342 | 25655 |
| 2845 | 3671 | 2190 | 816 | 2840 | 1403 | 351 | 9692 | 3708 | 4643 |
| 2266 | 3211 | 2248 | 877 | 2879 | 1446 | 475 | 7769 | 3316 | 4492 |
| 2149 | 3321 | 2646 | 1010 | 3498 | 1650 | 601 | 8546 | 3615 | 5119 |
| 1978 | 3091 | 5678 | 1585 | 918164 | 84852 | 632 | 8265 | 3572 | 5138 |
| 1954.5 | 3035.5 | 3007 | 1079 | 5411 | 1949.5 | 657 | 8365.5 | 3688.5 | 5176.5 |
| 2105 | 3571 | 3081 | 1157 | 5187 | 1984 | 738 | 9553 | 3979 | 5871 |
| 2058 | 3303 | 3100 | 1148 | 4602 | 1897 | 739 | 8946 | 3934 | 5892 |
| 2248.5 | 3518.5 | 3089.5 | 1127 | 4588.5 | 1948.5 | 712.5 | 10091 | 4194 | 5906.5 |
| 2293 | 3417 | 2834 | 1139 | 4109 | 1785 | 701 | 10322 | 4031 | 5968 |
| 2463.5 | 3747 | 3126 | 1269 | 4059.5 | 1864.5 | 698.5 | 10938.5 | 4320 | 5804 |
| 2402 | 3624.5 | 3039 | 1170.5 | 4307 | 1809 | 709.5 | 11155.5 | 4350.5 | 5914 |
| 2479 | 3781 | 3327 | 1260 | 4708 | 1926 | 779 | 11967 | 4845 | 6303 |
| 2548 | 3685 | 3296 | 1238 | 4556 | 1923 | 781 | 11114 | 4636 | 6224 |
| 19780 | 14031.5 | 5457 | 1341 | 4565 | 1971 | 778 | 11263 | 8360.5 | 8453 |
| 2847.5 | 3994.5 | 3406.5 | 1279 | 4592 | 1976 | 782 | 11121 | 4809 | 6585.5 |
| 2662.5 | 3844.5 | 3357 | 1257.5 | 4567 | 1985.5 | 746 | 10251.5 | 4818.5 | 6648.5 |
| 156944.5 | 80398 | 22330.5 | 2072.5 | 4743.5 | 1926.5 | 734 | 23925.5 | 7680.5 | 7856 |
| 2937 | 4615 | 3803.5 | 1346.5 | 4919 | 2078 | 770.5 | 11929 | 5094.5 | 7472.5 |
| 2703 | 4080 | 3391 | 1238 | 5555 | 2136 | 734 | 10894 | 4987 | 6875 |
| 2634 | 4117 | 3530 | 1260 | 4742 | 2025 | 788 | 11508 | 4803 | 7050 |
| 5480 | 12959 | 4926 | 1422 | 6178 | 2176 | 727 | 13982 | 6544 | 15990 |
| 8668 | 22666 | 7914 | 1974 | 9144 | 2770 | 821 | 13139 | 9144 | 24658 |
| 8352 | 22086 | 9288 | 2073 | 10733 | 2872 | 812 | 12176 | 8444 | 23991 |
| 8646 | 23145.5 | 8483.5 | 1985 | 9755 | 2679 | 777 | 12917 | 9319 | 26669 |
| 2776 | 3679 | 2197 | 828 | 2922 | 1420 | 361 | 9227 | 3816 | 4595 |
| 2351 | 3498 | 2352 | 877 | 2999 | 1501 | 497 | 8405 | 3550 | 4684 |
| 2289 | 3345 | 2532.5 | 992 | 3378.5 | 1642.5 | 584 | 8346 | 3603 | 5006.5 |
| 2196 | 3354 | 5920 | 1680 | 801602 | 81257 | 664 | 9463 | 3965 | 5586 |
| 2106 | 3156 | 3114 | 1085 | 5344 | 1965 | 644 | 8524 | 3770 | 5222 |
| 2261 | 3350 | 3119 | 1177 | 5007 | 1916 | 724 | 9544 | 4079 | 5802 |
| 2524 | 3777 | 3403 | 1294 | 4904 | 2057 | 791 | 10589 | 4723.5 | 6275 |
| 2451.5 | 3812 | 3226.5 | 1198.5 | 4574 | 1964 | 742.5 | 10638.5 | 4434.5 | 6106.5 |
| 2423 | 3778 | 3107 | 1110 | 4036 | 1813 | 691 | 10512 | 4277 | 5677 |
| 2378 | 3600 | 3007 | 1124 | 3999 | 1793 | 666 | 10450 | 3939 | 5602 |
| 2614 | 3703 | 3128 | 1144 | 4352 | 1842 | 718 | 11044 | 4604 | 6025 |
| 2518 | 3623 | 3064.5 | 1149 | 4195.5 | 1856 | 730 | 11208 | 4452.5 | 6142.5 |
| 2612.5 | 4038.5 | 3303 | 1204.5 | 4661 | 1944 | 809 | 11676 | 4863 | 6457 |
| 24637 | 16678 | 5713 | 1233 | 4056 | 1835 | 694 | 9832 | 9759 | 8759 |
| 3170 | 4356 | 3703 | 1295 | 4658 | 2027 | 804 | 10997 | 5133 | 6908 |
| 2720 | 4099 | 3435 | 1209 | 4635 | 1937 | 795 | 11199 | 4882 | 6634 |
| 95989 | 50660.5 | 14190 | 1654.5 | 4707.5 | 1948.5 | 727 | 17209.5 | 6444 | 7243 |
| 2943.5 | 4871.5 | 4122 | 1419 | 5168 | 2073.5 | 810 | 12849 | 5158.5 | 8011.5 |
| 2855 | 4294.5 | 3707 | 1253 | 5420.5 | 2136 | 788.5 | 11550.5 | 4999 | 7149 |
| 2741 | 4231 | 3464 | 1272 | 4833 | 2027 | 786 | 11807 | 4912 | 6893 |
| 5845.5 | 12487.5 | 5414.5 | 1579 | 6454.5 | 2276 | 777 | 13992 | 7052 | 17089.5 |
| 9943 | 22807.5 | 7419 | 2023 | 9057 | 2728.5 | 842 | 13669.5 | 9520.5 | 24574 |
| 8928 | 23412.5 | 8790.5 | 2130 | 10212 | 2838 | 825.5 | 13287.5 | 8861 | 25164 |

| | | | | | | | | | |
|---|---|---|---|---|---|---|---|---|---|
| 8928 | 23094 | 8956 | 2048 | 10010 | 2739 | 788 | 12784 | 9349 | 26427 |
| 2830 | 3717 | 2127 | 814 | 2898 | 1364.5 | 369 | 9233.5 | 3745.5 | 4597 |
| 2334 | 3358 | 2346 | 847 | 3004 | 1446 | 457 | 7944 | 3395 | 4663 |
| 2243 | 3424 | 2706 | 951 | 3499 | 1630 | 564 | 8282 | 3649 | 5263 |
| 1908 | 2980 | 4977.5 | 1505 | 721983.5 | 65956 | 569 | 7651 | 3378.5 | 4949.5 |
| 2009 | 3260 | 3129 | 1052 | 5399 | 1995 | 628 | 8235 | 3571 | 5395 |
| 2245.5 | 3391.5 | 2923.5 | 1113.5 | 4905.5 | 1891 | 646 | 9229 | 4080.5 | 5614.5 |
| 2364.5 | 3478.5 | 3179.5 | 1189 | 4816.5 | 1956.5 | 676.5 | 9614.5 | 4183 | 5754 |
| 2470 | 3697.5 | 3218.5 | 1172 | 4710 | 1927.5 | 704 | 9687 | 4420 | 6196 |
| 2357 | 3603 | 3077 | 1025 | 4214 | 1837 | 661 | 9867 | 3989 | 5834 |
| 2480.5 | 3468 | 2895.5 | 1125 | 3984 | 1775 | 653 | 10164 | 4229.5 | 5740 |
| 2408 | 3624 | 2905 | 1072 | 4091 | 1795 | 654 | 10566 | 4207 | 5679 |
| 2496 | 3777 | 3138 | 1184 | 4307 | 1861 | 731 | 10910 | 4416 | 6039 |
| 2436.5 | 3615 | 3075.5 | 1143 | 4316 | 1943 | 698 | 10148.5 | 4448.5 | 6121 |
| 15946 | 11226 | 4761 | 1227 | 4245 | 1848 | 717 | 10748 | 7424 | 7687 |
| 2760 | 3902 | 3365 | 1184 | 4394 | 1843 | 732 | 9624 | 4742 | 6338 |
| 2692 | 4045.5 | 3299.5 | 1195 | 4609 | 1907.5 | 726 | 10446.5 | 4849.5 | 6521 |
| 119692 | 63547 | 17967 | 1726 | 4766.5 | 1965 | 704 | 20164 | 6493.5 | 7569 |
| 2606.5 | 4355.5 | 3652 | 1200 | 4859 | 1971.5 | 715.5 | 10870.5 | 4511 | 6951 |
| 2566 | 4145 | 3404 | 1215 | 5252 | 2030 | 694 | 9940 | 4775 | 6607 |
| 2617.5 | 4256 | 3519 | 1283 | 4815 | 2010.5 | 739.5 | 11447 | 4697 | 6725.5 |
| 5450.5 | 13687.5 | 5193.5 | 1497 | 6402 | 2220 | 757 | 13911 | 7385.5 | 17283 |
| 8278.5 | 21071 | 7419 | 1940.5 | 9328 | 2685 | 781.5 | 12994 | 9176 | 25063.5 |
| 7453.5 | 21931 | 8971.5 | 2083.5 | 10638 | 2834.5 | 751.5 | 11514 | 8595 | 24358.5 |
| 7925 | 22683 | 8405 | 1957 | 9900 | 2715 | 758 | 12243 | 8708 | 25266 |
| 3026 | 3745 | 2266 | 835 | 2993 | 1411 | 347 | 9746 | 3814 | 4761 |
| 2575.5 | 3663.5 | 2459 | 894 | 3115 | 1445.5 | 491 | 8659 | 3637 | 4952 |
| 2404.5 | 3459 | 2708.5 | 988.5 | 3439 | 1616 | 573.5 | 8329.5 | 3707 | 5087.5 |
| 2284 | 3519 | 5637 | 1598.5 | 783395.5 | 75893 | 619 | 8978 | 3814.5 | 5399.5 |
| 2134 | 3319 | 3142 | 1118 | 5338 | 2032 | 650 | 9213 | 3739 | 5420 |
| 2647 | 3806 | 3291 | 1184.5 | 5356 | 2047.5 | 742.5 | 10367 | 4604 | 6076.5 |
| 2440 | 3820 | 3376 | 1186 | 5058 | 2021 | 739 | 10324 | 4557 | 6099 |
| 2062 | 3324 | 2652 | 1010 | 4071 | 1764 | 639 | 8632 | 3771 | 5381 |
| 2801 | 3977.5 | 3217 | 1189.5 | 4589 | 1949.5 | 719 | 11226 | 4814.5 | 6205 |
| 2830.5 | 4260 | 3559 | 1280.5 | 4723.5 | 2033 | 763.5 | 13317.5 | 4848.5 | 6647 |
| 2643 | 3805 | 3136 | 1134 | 4263 | 1857 | 686 | 10980 | 4625 | 6209 |
| 2432 | 3784.5 | 3257.5 | 1140.5 | 4271.5 | 1836 | 694.5 | 9841.5 | 4414.5 | 6055.5 |
| 2290 | 3720 | 3211 | 1118 | 4246 | 1849 | 712 | 10269 | 4288 | 6233 |
| 19469 | 13868 | 5324.5 | 1266 | 4495 | 1899 | 750 | 10337.5 | 8141 | 8290 |
| 2879 | 3806.5 | 3223.5 | 1141 | 4305 | 1854.5 | 744 | 9536.5 | 4534 | 6349.5 |
| 2747 | 3993 | 3296 | 1193.5 | 4454.5 | 1900 | 734.5 | 9748 | 4704.5 | 6487 |
| 103018 | 53660 | 15208 | 1747 | 4604 | 1994 | 706 | 17835 | 6498 | 7573 |
| 2914 | 4403.5 | 3690.5 | 1233.5 | 4811 | 1936 | 713 | 10398.5 | 4865 | 6823.5 |
| 2768 | 4097 | 3356 | 1220 | 5175 | 1996 | 723 | 9759 | 4855 | 6611 |
| 2567 | 3903 | 3179 | 1153 | 4421 | 1932 | 703 | 10119 | 4551 | 6208 |
| 4713.5 | 11638 | 4626.5 | 1340 | 5713.5 | 1995 | 676.5 | 13459 | 5605 | 14407.5 |
| 8638 | 19095 | 6769 | 1727 | 8055 | 2441 | 756 | 15157 | 9532 | 22556 |
| 9119 | 22655 | 7715 | 1902 | 9117 | 2665 | 728 | 13736 | 9063 | 26306 |

| | | | | | | | | | |
|---|---|---|---|---|---|---|---|---|---|
| 9049 | 19034 | 6102 | 1634 | 7449 | 2217 | 663 | 13217 | 8675 | 21033 |
| 2934 | 3737 | 2210 | 827 | 2963 | 1372 | 350 | 9392 | 3843 | 4743 |
| 2586 | 3609 | 2424 | 903 | 3074 | 1504 | 485 | 8396 | 3639 | 4821 |
| 2868 | 4025 | 3125 | 1068 | 3953 | 1719 | 641 | 10271 | 4620 | 5886 |
| 2749.5 | 3869.5 | 6334.5 | 1817 | 840270.5 | 82835 | 668.5 | 11172.5 | 4609 | 6126 |
| 2367 | 3482 | 3350 | 1168 | 5432 | 2045 | 671 | 9608 | 4064 | 5597 |
| 2776 | 4095.5 | 3667.5 | 1260 | 5676 | 2114 | 773 | 11680.5 | 4885 | 6630.5 |
| 2916.5 | 4861.5 | 4020 | 1387 | 5697 | 2257 | 869 | 13324.5 | 5124 | 7644 |
| 2875 | 4112 | 3432 | 1256 | 4938 | 2010 | 796 | 11999 | 4941 | 6630 |
| 2529 | 3687 | 2973.5 | 1112.5 | 4218 | 1825.5 | 697 | 10304.5 | 4303 | 5739 |
| 3139 | 4294 | 3625.5 | 1331.5 | 4819.5 | 2070 | 765.5 | 13372 | 5047.5 | 6926.5 |
| 2942 | 4248 | 3395 | 1283 | 4729 | 2044 | 804 | 11785 | 4982 | 6823 |
| 3097.5 | 4297.5 | 3610 | 1339 | 5125.5 | 2155 | 813 | 14691.5 | 5269 | 7260 |
| 2684.5 | 3809 | 3271 | 1153.5 | 4383.5 | 1883.5 | 778 | 10908 | 4799.5 | 6476 |
| 26367 | 17391 | 6425 | 1382 | 4693 | 2049 | 776 | 11148 | 10649 | 9452 |
| 2698 | 3470.5 | 2741 | 1078.5 | 3594 | 1676.5 | 651.5 | 8584 | 3953.5 | 5465.5 |
| 2867 | 4025 | 3552.5 | 1249 | 4644 | 1893.5 | 737 | 10845.5 | 4903.5 | 6824 |
| 148691 | 75877 | 21054 | 1884 | 4644 | 2010 | 753 | 22034 | 8342 | 7980 |
| 3164.5 | 4589 | 3728.5 | 1336 | 4783.5 | 2002 | 786 | 11427.5 | 5299.5 | 7301.5 |
| 2858 | 4049 | 3393 | 1203.5 | 5285 | 2019 | 735 | 10974.5 | 5104 | 6531 |
| 3079 | 4007 | 3453 | 1254 | 4743 | 1974 | 755 | 11338 | 5247 | 6728.5 |
| 5123.5 | 12166.5 | 4847.5 | 1345 | 5835.5 | 2009 | 702.5 | 13452 | 5691.5 | 14558.5 |
| 8533 | 17263 | 6079.5 | 1573.5 | 6979.5 | 2232 | 691.5 | 14892.5 | 8553 | 19312 |
| 9837 | 24783 | 10286 | 2127 | 10901 | 2822 | 798 | 13612 | 8510 | 26924 |
| 8903 | 22761 | 9264.5 | 2098 | 10507 | 2856 | 827 | 12269 | 8743 | 24915 |
| 2584 | 3428.5 | 2023 | 797 | 2686.5 | 1379.5 | 326 | 8323.5 | 3432.5 | 4411.5 |
| 2568.5 | 3518.5 | 2352 | 880 | 3531.5 | 1511.5 | 470 | 8351.5 | 3624.5 | 4756.5 |
| 2244.5 | 3448.5 | 2531.5 | 899 | 3868.5 | 1599.5 | 527.5 | 8072 | 3552 | 4935 |
| 2761 | 3594 | 5202 | 1567 | 334070 | 37519 | 570 | 7968 | 3629 | 5180 |
| 2162 | 3391 | 3055 | 1068 | 29964 | 3811 | 623 | 8375 | 3627 | 5347 |
| 2163.5 | 3428 | 2892.5 | 1088 | 21412 | 3064 | 639.5 | 8592.5 | 3712.5 | 5405 |
| 2385 | 3543 | 2948.5 | 1078.5 | 16815.5 | 2716.5 | 663 | 9055 | 4239 | 5797 |
| 2538.5 | 3932.5 | 3132 | 1153 | 13904.5 | 2593.5 | 699 | 10607 | 4563 | 6430.5 |
| 2622 | 3848 | 3172 | 1091 | 11191 | 2289 | 672 | 10747 | 4409 | 6088 |
| 2370.5 | 3545 | 2846.5 | 1066 | 5205.5 | 1872 | 612 | 9273 | 3985.5 | 5412.5 |
| 2538.5 | 3567.5 | 2833.5 | 1050 | 7651.5 | 2034.5 | 631 | 9736.5 | 4304 | 5591 |
| 2333.5 | 3485.5 | 2791 | 1035.5 | 6578 | 1929 | 630 | 9182.5 | 4193.5 | 5542.5 |
| 2464 | 3748 | 3248 | 1158 | 6368 | 2044 | 695 | 10503 | 4447 | 5919 |
| 7547 | 6126 | 3578 | 1133 | 6339 | 1924 | 676 | 9391 | 5265 | 6437 |
| 2639 | 3753 | 3240 | 1166 | 6022 | 2003 | 694 | 10348 | 4451 | 6176 |
| 2603.5 | 3929 | 3315 | 1184.5 | 5997 | 2023.5 | 705.5 | 9763 | 4676 | 6251.5 |
| 225267.5 | 118647 | 32824.5 | 2451 | 6563.5 | 2031.5 | 678 | 30245.5 | 8975 | 8234 |
| 2585 | 4094 | 3324 | 1218 | 5700 | 1946 | 652 | 9545 | 4453 | 6570 |
| 2576 | 3880 | 3314.5 | 1180 | 14110 | 3090.5 | 675 | 10563.5 | 4465 | 6498.5 |
| 2617 | 3945 | 3187 | 1167 | 5794 | 2065 | 706 | 10376 | 4447 | 6150 |
| 5301 | 13200.5 | 5045.5 | 1445 | 7269 | 2268.5 | 697 | 12658 | 6752 | 16286.5 |
| 7971 | 18556 | 6368 | 1655 | 8599 | 2420 | 670 | 14036 | 9310 | 21217 |
| 7583.5 | 20998 | 8000 | 1884.5 | 10537.5 | 2646.5 | 689 | 11962.5 | 8987 | 23360 |

| | | | | | | | | | |
|---|---|---|---|---|---|---|---|---|---|
| 8333 | 20220 | 7395 | 1819 | 9740 | 2567 | 697 | 11985 | 8785 | 23271 |
| 2748 | 3693.5 | 2171 | 783 | 2839.5 | 1361 | 343 | 8577 | 3650 | 4628.5 |
| 2434.5 | 3715.5 | 2444 | 891 | 3132 | 1457.5 | 444 | 8325.5 | 3776.5 | 4951.5 |
| 2282 | 3425 | 2517 | 983 | 3413 | 1555 | 523 | 8305 | 3771 | 5092 |
| 2396 | 3558 | 5456 | 1603 | 764780 | 68368 | 622 | 9253 | 4374 | 5769 |
| 2093.5 | 3210.5 | 2967.5 | 1020.5 | 4914.5 | 1898 | 588 | 8004 | 3624 | 5184 |
| 2481 | 3830 | 3161 | 1146 | 5069 | 1991 | 673 | 10331 | 4517 | 6006 |
| 2522 | 3629 | 3127.5 | 1162 | 4467 | 1865 | 663 | 8740.5 | 4398.5 | 5903.5 |
| 2584.5 | 4078.5 | 3398.5 | 1137.5 | 4839.5 | 1993.5 | 709 | 10999 | 4770 | 6519 |
| 2616 | 3983 | 3179 | 1175.5 | 4586 | 1921.5 | 697 | 11145 | 4540.5 | 6441.5 |
| 2602 | 3760 | 3051 | 1117 | 4121 | 1818 | 624 | 10039 | 4390 | 5870 |
| 2573.5 | 3709 | 2972.5 | 1106.5 | 4328 | 1840.5 | 673 | 10504 | 4581.5 | 6020.5 |
| 2567.5 | 3789.5 | 3250 | 1142.5 | 4286 | 1875.5 | 679 | 10905.5 | 4572 | 6193 |
| 2482 | 3797 | 3069 | 1147 | 4283 | 1834 | 673 | 10087 | 4507 | 5974 |
| 57814 | 41095 | 11852 | 1525 | 4353 | 1964 | 713 | 10570 | 22752 | 17009 |
| 2791 | 4043 | 3352 | 1229 | 4406 | 1892 | 700 | 10131 | 4745 | 6524 |
| 2583 | 4000 | 3238 | 1167 | 4356 | 1798 | 718 | 9636 | 4721 | 6405 |
| 97105 | 52968 | 15033 | 1778 | 4755 | 1956 | 684 | 17890 | 6373 | 7626 |
| 2647 | 4302 | 3541.5 | 1198.5 | 4789.5 | 1878 | 692.5 | 10947.5 | 4771.5 | 6670.5 |
| 2653 | 4091 | 3362 | 1152 | 4794 | 1926 | 693 | 10367 | 4754 | 6614 |
| 2645.5 | 3965 | 3207.5 | 1134 | 4276.5 | 1901.5 | 684 | 10310 | 4730.5 | 6371.5 |
| 4318 | 11990 | 4698 | 1285 | 5699 | 2030 | 659 | 13059 | 5852 | 14592 |
| 8677.5 | 19037 | 6455.5 | 1713.5 | 7961 | 2530 | 702.5 | 13562 | 9248.5 | 22339 |
| 7488.5 | 21715 | 8421 | 2015 | 10486.5 | 2625.5 | 706.5 | 12131 | 8241 | 23603.5 |
| 8320.5 | 20368.5 | 6718 | 1643 | 7778 | 2300.5 | 668.5 | 11532 | 8925.5 | 23091 |
| 2822 | 3631.5 | 2148 | 830 | 2911 | 1372.5 | 335.5 | 9469 | 3595 | 4573.5 |
| 2470.5 | 3580.5 | 2297 | 839.5 | 3009 | 1415.5 | 441.5 | 8188.5 | 3581.5 | 4607.5 |
| 2209 | 3306 | 2344 | 925 | 3250 | 1528 | 512 | 7623 | 3435 | 4931 |
| 2190 | 3474 | 5284 | 1584 | 666343 | 60059 | 592 | 8352 | 3702 | 5180 |
| 2139 | 3321.5 | 2972.5 | 1080 | 5205 | 1956.5 | 597 | 8174.5 | 3589 | 5056 |
| 2418 | 3649 | 3165 | 1153 | 5135 | 1950 | 658 | 9555 | 4171 | 5745 |
| 2256 | 3483 | 2884 | 1097 | 4509 | 1807 | 648 | 8920 | 3921 | 5576 |
| 2691 | 3946 | 3232.5 | 1145 | 4790.5 | 1957 | 715.5 | 10391.5 | 4642 | 6233.5 |
| 2756 | 4378 | 3471 | 1260 | 4953 | 1956 | 742 | 12596 | 4773 | 6731 |
| 2789 | 4000 | 3205 | 1154 | 4264 | 1888 | 677 | 10867 | 4740 | 6020 |
| 2591.5 | 3687 | 2915.5 | 1066 | 4103 | 1774.5 | 649.5 | 9809.5 | 4443 | 5795 |
| 2593 | 3957 | 3144 | 1159 | 4444 | 1863 | 707 | 10509 | 4573 | 6090 |
| 2603 | 3924.5 | 3264 | 1200 | 4460.5 | 1960.5 | 727 | 10577.5 | 4633 | 6101.5 |
| 14033 | 9941 | 4452 | 1211 | 4120 | 1835 | 702 | 9730 | 6942 | 7416 |
| 2856 | 4042 | 3258 | 1183.5 | 4353.5 | 1885 | 726 | 9707 | 4668 | 6304 |
| 2843 | 4198 | 3438 | 1244 | 4536 | 1942 | 737 | 10202 | 4759 | 6659 |
| 113710 | 59530 | 16932 | 1845 | 4754 | 1914 | 681 | 18269 | 6738 | 7719 |
| 2886 | 4523 | 3802.5 | 1289.5 | 4838 | 1955.5 | 706 | 11390.5 | 4904.5 | 7392 |
| 2573.5 | 3669 | 3048.5 | 1086.5 | 4846.5 | 1899 | 654.5 | 9325 | 4693 | 6175 |
| 2562 | 3917 | 3256 | 1140 | 4393 | 1836 | 698 | 9556 | 4360 | 6180 |
| 4228 | 11025 | 4848 | 1358 | 5870 | 2042 | 697 | 14169 | 5510 | 14025 |
| 7978.5 | 18305 | 6408 | 1712.5 | 7748.5 | 2387.5 | 674 | 14389 | 8975 | 21002 |
| 9239.5 | 21852 | 7285.5 | 1814 | 8461 | 2463 | 693 | 13396.5 | 9198 | 24426 |

| | | | | | | | | | |
|---|---|---|---|---|---|---|---|---|---|
| 9417 | 21047 | 6752 | 1661 | 7800 | 2475 | 682 | 13587 | 9109 | 22680 |
| 790 | 965 | 828 | 582 | 1385.5 | 1126 | 278 | 2479.5 | 1470 | 1835.5 |
| 1010 | 1588 | 1411 | 702 | 2112 | 1284 | 405 | 3757 | 2024 | 2912 |
| 1294.5 | 2222 | 2037 | 894 | 2951 | 1483.5 | 561 | 6142 | 2749 | 4100 |
| 1384 | 2509.5 | 4592 | 1448 | 675380 | 61772 | 621 | 6859.5 | 2986 | 4557 |
| 1476 | 2686.5 | 2801.5 | 1075.5 | 4895 | 2179 | 668.5 | 7189 | 3198 | 4960.5 |
| 1516 | 2813 | 2759 | 1050 | 4200 | 1913 | 682 | 7645 | 3430 | 5156 |
| 1500 | 2508 | 2460 | 1028 | 3723 | 1748 | 643 | 7173 | 3098 | 4818 |
| 1609 | 2683 | 2421 | 989 | 3571 | 1695 | 637 | 7534 | 3156 | 4814 |
| 1743 | 2818 | 2536 | 1038 | 3578 | 1705 | 613 | 8438 | 3350 | 4896 |
| 1858 | 3105 | 2689 | 1063 | 3701 | 1800 | 655 | 9055 | 3589 | 5181 |
| 1879 | 3155 | 2753 | 1116 | 3921 | 1762 | 677 | 9303 | 3644 | 5276 |
| 1903 | 3163.5 | 2882 | 1091 | 4027.5 | 1845 | 713 | 9385.5 | 3846.5 | 5668 |
| 1814 | 3020 | 2821.5 | 1108.5 | 4000.5 | 1796 | 713 | 8614 | 3695 | 5468 |
| 20994.5 | 15220 | 5601.5 | 1287.5 | 4251 | 1850 | 719.5 | 8806 | 8258.5 | 8367 |
| 2199 | 3613 | 3205 | 1165 | 4083 | 1820 | 714 | 8213 | 4055 | 5937 |
| 1955 | 3267 | 2933 | 1130 | 4104 | 1889 | 685 | 8127 | 3854 | 5766 |
| 100705.5 | 54098 | 15138.5 | 1637 | 4063 | 1763.5 | 639 | 15667 | 5642.5 | 6621.5 |
| 2109 | 3650 | 3225 | 1119 | 4261 | 1850 | 678 | 8964 | 4162 | 5839 |
| 1956 | 3437 | 3031 | 1157 | 4829 | 1959 | 678 | 8808 | 4097 | 6068 |
| 1975 | 3288.5 | 3051.5 | 1126.5 | 4300.5 | 1852 | 675 | 9021.5 | 3806.5 | 5633 |
| 4090 | 11037.5 | 4753.5 | 1332 | 5753 | 2028 | 715.5 | 13536.5 | 5393 | 14446 |
| 7292.5 | 16707 | 6023 | 1634.5 | 7131.5 | 2346.5 | 686.5 | 14304.5 | 8686.5 | 20107 |
| 8113 | 20026 | 6708 | 1743 | 8053 | 2369 | 689 | 12798 | 8813 | 22188 |
| 8196 | 19832 | 6477 | 1703 | 7652 | 2424 | 705 | 12925 | 8728 | 22309 |
| 2724 | 3585 | 2129 | 763 | 2727 | 1330 | 350 | 9305 | 3671 | 4656 |
| 2252 | 3141.5 | 2094.5 | 847 | 2763.5 | 1408.5 | 474 | 7515 | 3265.5 | 4376 |
| 2040.5 | 3155 | 2443 | 973 | 3272 | 1612 | 606 | 7524.5 | 3541 | 4767 |
| 1968.5 | 2920.5 | 6141 | 1646 | 1067839 | 98341.5 | 649 | 8124.5 | 3525 | 5023 |
| 1930 | 3058 | 3294 | 1206 | 5924 | 2178 | 714 | 9111 | 3793 | 5388 |
| 2066 | 3345.5 | 3103 | 1198.5 | 4892 | 2083 | 767 | 9570.5 | 4067 | 5866.5 |
| 1826 | 3002 | 2836 | 1093 | 4319 | 1846 | 720 | 8776 | 3715 | 5389 |
| 1931.5 | 3034.5 | 2766 | 1068 | 3936 | 1780.5 | 696 | 9053.5 | 3639.5 | 5204.5 |
| 2146.5 | 3239.5 | 2827.5 | 1079 | 3917.5 | 1754 | 709 | 9596 | 3877.5 | 5527.5 |
| 2295.5 | 3583 | 3115.5 | 1203.5 | 4164 | 1888 | 729.5 | 10586.5 | 4151.5 | 5814.5 |
| 2258 | 3517 | 3131 | 1152.5 | 4083 | 1854.5 | 751.5 | 10703.5 | 4236.5 | 5903 |
| 2304 | 3591 | 3029 | 1160 | 4324 | 1922 | 767 | 10220 | 4232 | 6089 |
| 2073 | 3437 | 2966.5 | 1157.5 | 4294 | 1922 | 762.5 | 9983.5 | 4068.5 | 5939 |
| 22867 | 16459 | 5794.5 | 1272 | 4293 | 1903 | 762 | 9693.5 | 8960 | 8801.5 |
| 2475.5 | 3580 | 3279.5 | 1193 | 4214 | 1897.5 | 788 | 9615.5 | 4290.5 | 6120 |
| 2345 | 3619 | 3211 | 1234 | 4402 | 1945 | 755 | 9360 | 4389 | 6494 |
| 188626 | 95527 | 26635 | 2152 | 4357 | 1895 | 727 | 25289 | 7768 | 7767 |
| 2586 | 3957.5 | 3575.5 | 1239.5 | 4514.5 | 1936 | 761 | 10667.5 | 4866.5 | 6666.5 |
| 2547 | 3984 | 3545.5 | 1331.5 | 6068 | 2306 | 785.5 | 11789 | 5046.5 | 6987 |
| 2541 | 4021 | 3537 | 1334 | 4883 | 2018 | 825 | 11189 | 4862 | 6665 |
| 5766 | 13704.5 | 5639 | 1549 | 6747.5 | 2324 | 794.5 | 14384.5 | 7530 | 17091.5 |
| 9390 | 19928 | 6904 | 1882 | 8186 | 2522 | 790 | 13861 | 9550 | 22712 |
| 9639 | 24049.5 | 7806 | 2097.5 | 8807.5 | 2838 | 856 | 12744 | 9134.5 | 25865.5 |

| | | | | | | | | | |
|---|---|---|---|---|---|---|---|---|---|
| 9963 | 23647 | 7813 | 1955 | 8560 | 2638 | 788 | 12870 | 9785 | 25222 |
| 2661 | 3489.5 | 2067 | 773 | 2727.5 | 1322 | 342.5 | 8266.5 | 3602.5 | 4409 |
| 2071 | 2974.5 | 2048.5 | 796 | 2679.5 | 1368.5 | 431 | 7074.5 | 3036.5 | 4126 |
| 2085 | 3185 | 2485 | 964 | 3517 | 1603 | 599 | 8037 | 3627 | 5081 |
| 1895 | 3095 | 5968 | 1795.5 | 921513.5 | 74361 | 630 | 7918.5 | 3542 | 5348.5 |
| 1820 | 2956 | 3130 | 1108 | 6007 | 2124 | 660 | 7957 | 3466 | 5103 |
| 1814 | 3099 | 2968 | 1112 | 5112 | 2010 | 656 | 8721 | 3604 | 5361 |
| 2077 | 3375 | 3274 | 1185 | 5205 | 2011 | 738 | 10253 | 4279 | 6191 |
| 2001.5 | 3302.5 | 2964.5 | 1089 | 4421.5 | 1825.5 | 706 | 9065 | 3890.5 | 5586.5 |
| 1973 | 3153 | 2712.5 | 1056 | 3865 | 1701 | 651.5 | 8843 | 3657.5 | 5216 |
| 2067 | 3259 | 2834 | 1142 | 3900 | 1814 | 641 | 9578 | 3691 | 5459 |
| 2156 | 3313 | 2875 | 1112 | 4103 | 1876 | 684 | 9941 | 3923 | 5592 |
| 2237 | 3469 | 3023 | 1203 | 4326 | 1906.5 | 733.5 | 10059 | 4215 | 5922.5 |
| 2128 | 3527 | 3120 | 1183 | 4334 | 1892 | 738 | 10411 | 4165 | 6074 |
| 18868 | 13479 | 5341.5 | 1282 | 4067 | 1905.5 | 721.5 | 9687.5 | 7875.5 | 8008.5 |
| 2480 | 3873 | 3494 | 1240 | 4569 | 1985 | 764 | 9882 | 4628 | 6392 |
| 2279 | 3707 | 3194 | 1195 | 4491 | 1945 | 723 | 9803 | 4476 | 6226 |
| 146078 | 76820 | 20968 | 1925 | 4549 | 1932 | 701 | 20851 | 7184 | 7349 |
| 2491 | 3866 | 3289 | 1180 | 4389 | 1893 | 684 | 10125 | 4451 | 6470 |
| 2342 | 3663 | 3211.5 | 1159.5 | 5323.5 | 2022.5 | 699 | 9726.5 | 4455.5 | 6268.5 |
| 2334 | 3752 | 3301.5 | 1172 | 4462.5 | 1960 | 730 | 9773.5 | 4737.5 | 6412 |
| 4658 | 11568 | 5039.5 | 1473 | 6096.5 | 2161 | 747.5 | 14068.5 | 5853.5 | 15271 |
| 8019 | 18407 | 6370 | 1677 | 7487 | 2377 | 750 | 14508 | 9148 | 21028 |
| 9421 | 23294 | 7245 | 1858 | 8577 | 2530 | 725 | 13568 | 9330 | 24143 |
| 7998 | 21860.5 | 7324.5 | 1825.5 | 8849.5 | 2570 | 745 | 11162.5 | 8610.5 | 24918.5 |
| 3003 | 3746 | 2246 | 818 | 2876 | 1397 | 351 | 9463 | 3826 | 4785 |
| 2400 | 3370 | 2324 | 900 | 2963 | 1432 | 497 | 8057 | 3523 | 4558 |
| 2188 | 3349 | 2611.5 | 1003 | 3390.5 | 1637.5 | 582 | 8250 | 3663 | 5284 |
| 1922.5 | 3082 | 5837 | 1625.5 | 877243.5 | 81137 | 642.5 | 8052 | 3481 | 5134 |
| 2006.5 | 3281.5 | 3236 | 1121 | 5931 | 2109 | 699 | 8368 | 3780 | 5521.5 |
| 2123 | 3363 | 3335 | 1160 | 5293 | 2027 | 737 | 9586 | 4078 | 5734 |
| 2021 | 3389 | 3020 | 1164 | 4778 | 1946 | 724 | 9460.5 | 4154 | 5934.5 |
| 2079 | 3260 | 2948.5 | 1124 | 4227.5 | 1818 | 712.5 | 9258 | 3991 | 5666.5 |
| 2189.5 | 3374 | 2930 | 1093 | 4023.5 | 1767.5 | 707 | 9619.5 | 3914 | 5555.5 |
| 2365 | 3518 | 3053 | 1178 | 4179 | 1897 | 714 | 10549 | 4124 | 5834 |
| 2286 | 3492 | 3002 | 1109 | 4178 | 1837 | 735 | 10648 | 4119 | 5856 |
| 2350.5 | 3629.5 | 3114.5 | 1169.5 | 4287.5 | 1893 | 766 | 10865 | 4577.5 | 6117.5 |
| 2281 | 3551 | 3222 | 1181 | 4328 | 1960 | 759 | 10041 | 4182 | 6106 |
| 19981 | 14559 | 5473.5 | 1294 | 4370.5 | 1870 | 757 | 10112 | 8245 | 8620.5 |
| 2638.5 | 3975 | 3494 | 1216.5 | 4489.5 | 1982.5 | 798 | 10125.5 | 4671.5 | 6562.5 |
| 2535 | 3814 | 3488 | 1270 | 4705 | 2003 | 764 | 10267 | 4556 | 6619 |
| 156429 | 79478 | 22385 | 1973 | 4474 | 1894 | 721 | 21408 | 7556 | 7583 |
| 2719 | 4164 | 3578 | 1263 | 4775 | 1992 | 766 | 11079 | 5186 | 6929 |
| 2639 | 4049 | 3627 | 1307 | 5586 | 2065 | 754.5 | 10953 | 4909.5 | 6715.5 |
| 2523 | 3745.5 | 3320.5 | 1191.5 | 4384.5 | 1891 | 742.5 | 10798 | 4772.5 | 6503.5 |
| 5910 | 13482 | 5390 | 1480 | 6712 | 2228 | 783 | 14490 | 7660 | 17175 |
| 9470 | 20881 | 6926 | 1836 | 8591 | 2622 | 786 | 13049 | 9227 | 23591 |
| 9003.5 | 22343.5 | 7640 | 1916.5 | 8830 | 2485.5 | 774 | 13719 | 9672 | 25892.5 |

| | | | | | | | | | |
|---|---|---|---|---|---|---|---|---|---|
| 9125 | 24159 | 8586 | 2020 | 9826 | 2700 | 814 | 12490 | 9305 | 25726 |
| 2493 | 3279 | 1942 | 764 | 2640 | 1323 | 324 | 7722 | 3426 | 4251 |
| 2350 | 3218 | 2272 | 878 | 2913 | 1455 | 474 | 7782 | 3429 | 4520 |
| 2231 | 3258 | 2599 | 1022 | 3466 | 1623 | 617 | 8497 | 3767 | 5166 |
| 1993 | 3050 | 5665 | 1651 | 838921 | 85640 | 634 | 8257 | 3393 | 5234 |
| 2065 | 3306.5 | 3266 | 1175.5 | 5741 | 2118.5 | 676.5 | 8946 | 3861.5 | 5346 |
| 2353 | 3605 | 3362 | 1263 | 5609 | 2111 | 758 | 10607 | 4238 | 6203 |
| 2248 | 3612.5 | 3213 | 1231.5 | 5007 | 1988 | 764.5 | 9601 | 4358 | 6116 |
| 2332 | 3567 | 3191 | 1248 | 4555 | 1907 | 747 | 9990 | 4097 | 6024 |
| 2354 | 3613 | 3027 | 1162 | 4246 | 1810 | 712 | 10201 | 4073 | 5812 |
| 2440 | 3607 | 3042 | 1148 | 4031 | 1896 | 699 | 10696 | 4198 | 5864 |
| 2594 | 3630 | 3000 | 1123 | 4241 | 1803 | 718 | 11035 | 4509 | 5913 |
| 2650.5 | 3732 | 3293 | 1265 | 4628 | 1970 | 796 | 11315.5 | 4766.5 | 6499 |
| 2421 | 3702 | 3214.5 | 1220 | 4368.5 | 1918 | 775 | 11194.5 | 4591 | 6224.5 |
| 26431.5 | 18244.5 | 6407.5 | 1357 | 4350.5 | 1995.5 | 778 | 10444 | 10152 | 9285.5 |
| 3034 | 4312 | 3748 | 1321 | 4749 | 2073 | 827 | 11219 | 5167 | 6958 |
| 2689.5 | 3923.5 | 3387 | 1239.5 | 4541 | 1928.5 | 792 | 10019.5 | 4704.5 | 6417 |
| 91213 | 46824.5 | 13276 | 1587.5 | 4606.5 | 1869 | 746 | 16387 | 6514.5 | 7351.5 |
| 2992 | 4473 | 3904 | 1331 | 5087 | 2041 | 788 | 12814 | 5162 | 7437 |
| 2851.5 | 4269.5 | 3614.5 | 1291 | 5384.5 | 2099 | 779.5 | 11328 | 5121.5 | 7030.5 |
| 2931 | 4310 | 3749 | 1327 | 4893 | 2043 | 827 | 12175 | 5062 | 7040 |
| 6030 | 13736 | 5605.5 | 1574.5 | 6532 | 2207.5 | 774 | 14783.5 | 7304.5 | 16971 |
| 9558 | 21319 | 7408 | 2042 | 8416 | 2564 | 840 | 13938 | 9560 | 23984 |
| 9423 | 23851 | 8270 | 2061 | 9515 | 2794 | 815 | 12411 | 8971 | 25738 |
| 9529 | 23292 | 8079.5 | 2032.5 | 9159 | 2731.5 | 815.5 | 12763 | 9323 | 25881.5 |
| 2770 | 3657.5 | 2164.5 | 787 | 2750 | 1298.5 | 318 | 8795.5 | 3612.5 | 4573 |
| 2233 | 3223 | 2235 | 803 | 2956 | 1396 | 453 | 7501 | 3299 | 4450 |
| 2015 | 3163 | 2492 | 945 | 3363 | 1553 | 547 | 7737 | 3402 | 4944 |
| 2036 | 3149.5 | 5227 | 1515 | 734282 | 66854 | 596.5 | 8146 | 3453.5 | 5266 |
| 1940.5 | 3187 | 3045 | 1128 | 5245 | 1972.5 | 613 | 7990 | 3532 | 5054.5 |
| 2333 | 3694 | 3308.5 | 1195.5 | 5279 | 2023 | 721 | 9601 | 4433.5 | 5980.5 |
| 2231 | 3499 | 3049 | 1116 | 4686 | 1952 | 712 | 8879 | 4174 | 5800 |
| 2101 | 3466 | 2999 | 1069 | 4321 | 1851 | 660 | 8682 | 3920 | 5518 |
| 2198.5 | 3457.5 | 2937.5 | 1093.5 | 4096 | 1749 | 645 | 9412 | 3998.5 | 5499 |
| 2317 | 3614 | 3049 | 1156 | 4153 | 1847 | 653 | 10207 | 3982 | 5770 |
| 2554 | 3801 | 3086 | 1166 | 4413 | 1869 | 692 | 10698 | 4485 | 5951 |
| 2421 | 3752 | 3132 | 1171 | 4402 | 1860 | 713 | 10852 | 4453.5 | 6150.5 |
| 2310 | 3676 | 3117 | 1158 | 4397 | 1900 | 705 | 10281 | 4362 | 6140 |
| 15842 | 11452 | 4800 | 1229 | 4389 | 1874 | 719 | 9825 | 7090 | 7658 |
| 2697.5 | 3854 | 3264 | 1207 | 4350.5 | 1853 | 716 | 9895 | 4455 | 6239 |
| 2486 | 3967 | 3229 | 1209 | 4637 | 1952 | 744 | 10486 | 4661 | 6448 |
| 120135.5 | 62735 | 17269 | 1783.5 | 4652.5 | 1912 | 684 | 18076.5 | 6780 | 7500 |
| 2711 | 4260 | 3655 | 1265.5 | 4754 | 1940 | 723 | 10943 | 4800.5 | 6816 |
| 2657.5 | 4121 | 3499.5 | 1225 | 5407.5 | 2074.5 | 732.5 | 10642.5 | 4670 | 7008 |
| 2633 | 4228 | 3543.5 | 1275.5 | 4955 | 2046.5 | 762.5 | 11195 | 4855 | 6717 |
| 5219 | 12130 | 4975 | 1491 | 6125 | 2196 | 731 | 13831 | 6711 | 15754 |
| 8195.5 | 21126.5 | 6892.5 | 1764.5 | 8311.5 | 2535.5 | 755.5 | 13406.5 | 9248 | 24445 |
| 8685 | 23099 | 7471 | 1920 | 9082 | 2538 | 721 | 13311 | 9325 | 25380 |

| | | | | | | | | | |
|---|---|---|---|---|---|---|---|---|---|
| 8122 | 22969 | 7469 | 1807 | 8735 | 2534 | 713 | 12912 | 9331 | 26429 |
| 2859 | 3916 | 2209 | 819 | 2957 | 1380 | 351 | 9427 | 3871 | 4805 |
| 2283 | 3375 | 2241 | 880.5 | 2933.5 | 1453.5 | 458 | 7871.5 | 3536 | 4543 |
| 2063 | 3133 | 2463 | 909.5 | 3268.5 | 1523 | 536 | 7818 | 3453.5 | 4887.5 |
| 2061 | 3340 | 5411 | 1614 | 716414 | 67616 | 622 | 8540 | 3572 | 5177 |
| 2196 | 3447.5 | 3206.5 | 1137.5 | 5510.5 | 2128 | 670.5 | 9044.5 | 3871.5 | 5629.5 |
| 2358 | 3715 | 3494 | 1236 | 5153 | 2059 | 736 | 9701 | 4428 | 6118 |
| 2338 | 3636 | 3089 | 1155 | 4716 | 1971 | 720 | 9459 | 4234 | 6006 |
| 2440 | 3760 | 3173 | 1143 | 4638 | 1920 | 724 | 9668 | 4363 | 5900 |
| 2381 | 3516 | 2869 | 1077.5 | 4191.5 | 1800.5 | 669.5 | 9884 | 4074.5 | 5675 |
| 2511 | 3672.5 | 3132 | 1184.5 | 4207 | 1893.5 | 680 | 10520.5 | 4356 | 5918.5 |
| 2356 | 3487 | 2928 | 1067 | 3972 | 1795 | 672 | 9960 | 4078 | 5621 |
| 2732 | 3912 | 3365 | 1219 | 4611 | 1932 | 742 | 11358 | 4734 | 6555 |
| 2358.5 | 3645 | 3191.5 | 1137 | 4300 | 1892 | 733 | 10182.5 | 4254 | 5975.5 |
| 20411 | 14122 | 5439 | 1253 | 4277 | 1902 | 763 | 10086 | 8489 | 8362 |
| 2914 | 4005 | 3469 | 1234 | 4415 | 1886 | 775 | 10259 | 4896 | 6475 |
| 2764 | 4003 | 3374 | 1208 | 4587 | 1970 | 771 | 10091 | 4699 | 6328 |
| 109777 | 58040 | 16842 | 1863 | 4999 | 2038 | 753 | 20561 | 6726 | 7770 |
| 2830 | 4251 | 3723 | 1241 | 4684 | 1918 | 717 | 11221 | 4830 | 6860 |
| 2676 | 4160 | 3495 | 1281 | 5359 | 2058 | 739 | 11166 | 4765 | 6637 |
| 2602 | 3727 | 3139 | 1177 | 4321 | 1848 | 698 | 9775 | 4594 | 6166 |
| 5862 | 13259 | 5165 | 1524 | 6433 | 2231 | 744 | 13432 | 7165 | 16529 |
| 8945 | 20909 | 7121 | 1913 | 8321 | 2578 | 786 | 13651 | 9284 | 23401 |
| 8402 | 22118 | 8245 | 1902 | 9488 | 2544 | 763 | 13086 | 9076 | 25488 |
| 8584 | 22712 | 8371 | 1972 | 9996 | 2636 | 769 | 12319 | 9626 | 24847 |
| 2782 | 3769.5 | 2089.5 | 804 | 2861 | 1353.5 | 336.5 | 8734 | 3697.5 | 4622 |
| 2582 | 3588 | 2491 | 875 | 3151 | 1490 | 477 | 8235 | 3679 | 4759 |
| 2372 | 3368 | 2529 | 992 | 3432 | 1609 | 592 | 8437 | 3785 | 5089 |
| 2250 | 3374 | 5936 | 1693 | 903469 | 89850 | 630 | 8282 | 3779 | 5292 |
| 2308 | 3468 | 3381 | 1206 | 6003 | 2172 | 690 | 9381 | 3941 | 5657 |
| 2675.5 | 3895 | 3467 | 1306.5 | 5621 | 2063.5 | 745.5 | 10919 | 4724.5 | 6290 |
| 2643 | 3903.5 | 3398.5 | 1284 | 5223.5 | 2018 | 763.5 | 10538 | 4753 | 6385 |
| 2558 | 3589.5 | 3238 | 1145.5 | 4764 | 1850 | 733.5 | 10061.5 | 4474.5 | 6075.5 |
| 2646 | 3873 | 3247 | 1146 | 4550 | 1861 | 738 | 11003 | 4429 | 6168 |
| 2667 | 3747 | 3174 | 1180 | 4199 | 1879 | 707 | 10892 | 4389 | 5938 |
| 2672.5 | 3766 | 3090 | 1113.5 | 4156.5 | 1865.5 | 706 | 10758 | 4445 | 5998 |
| 2772 | 3858 | 3334 | 1224 | 4382 | 1924 | 756 | 10934 | 4730 | 6124 |
| 2531 | 3772.5 | 3339.5 | 1230.5 | 4355.5 | 1896 | 775 | 11159 | 4518 | 6280.5 |
| 24063.5 | 16984 | 6005 | 1272 | 4237.5 | 1857.5 | 727 | 10349.5 | 9340 | 8903 |
| 3045.5 | 4088.5 | 3425.5 | 1145.5 | 4256.5 | 1901.5 | 778 | 10199.5 | 4634 | 6426.5 |
| 2994 | 4281 | 3519 | 1271 | 4641 | 1983 | 798 | 11170 | 5050 | 6949 |
| 145670.5 | 73074 | 20727 | 1918 | 4554.5 | 1916.5 | 736 | 21799.5 | 7420.5 | 7669 |
| 2988 | 4456 | 3658 | 1261.5 | 4767 | 1905 | 753 | 12031 | 5057.5 | 7070.5 |
| 2712 | 3908.5 | 3360 | 1153.5 | 5081 | 2019.5 | 723 | 10215 | 4884.5 | 6513.5 |
| 2909 | 4506 | 3826.5 | 1305 | 4924 | 2029.5 | 784.5 | 12659.5 | 4970 | 7260 |
| 5798 | 13375 | 5170 | 1530 | 6435 | 2212 | 763 | 14823 | 6782 | 16586 |
| 10219 | 21162 | 7361 | 1891 | 8705 | 2603 | 808 | 13744 | 9302 | 24214 |
| 8488 | 21261 | 8016 | 1919 | 9398.5 | 2623 | 716 | 12367 | 8599.5 | 23534.5 |

| | | | | | | | | | |
|---|---|---|---|---|---|---|---|---|---|
| 8095 | 22064 | 7475 | 1889 | 8902 | 2565 | 723 | 12233 | 8544 | 24105 |
| 2683 | 3727.5 | 2112 | 772 | 2861.5 | 1346 | 307.5 | 8560.5 | 3681 | 4628 |
| 2397 | 3422 | 2278 | 840 | 3444 | 1487 | 420 | 7699 | 3416 | 4514 |
| 2191 | 3333 | 2475 | 919 | 3794.5 | 1564.5 | 525 | 7561.5 | 3470.5 | 4835.5 |
| 2602.5 | 3484 | 4996.5 | 1558.5 | 318928 | 35775.5 | 568.5 | 8040.5 | 3456 | 5137 |
| 2093.5 | 3303.5 | 2912.5 | 1036.5 | 27894 | 3589.5 | 597 | 8132 | 3715.5 | 5273 |
| 2311 | 3608 | 3045 | 1095 | 21721 | 3180 | 679 | 9434 | 4179 | 5926 |
| 2251.5 | 3533.5 | 2984.5 | 1074 | 16056 | 2682.5 | 666 | 9265.5 | 4150.5 | 5872.5 |
| 2430 | 3759 | 3000 | 1112 | 14437 | 2560 | 673 | 9585 | 4170 | 5872 |
| 2361.5 | 3659.5 | 2905.5 | 1104.5 | 10817 | 2242 | 644.5 | 9559.5 | 4057 | 5671 |
| 2398.5 | 3503.5 | 2848.5 | 1049 | 5207.5 | 1898 | 624 | 9790.5 | 4141.5 | 5483.5 |
| 2398 | 3589 | 2890 | 1077 | 8047 | 2081 | 645 | 9788 | 4189 | 5624 |
| 2477.5 | 3609 | 2961 | 1117 | 6984 | 2055.5 | 666.5 | 10184.5 | 4409 | 5820.5 |
| 2353 | 3539 | 2984 | 1084 | 6422 | 2005 | 678 | 9797 | 4271.5 | 5840 |
| 6932 | 5823 | 3623 | 1136 | 6519 | 2036 | 681 | 9761 | 5244 | 6496 |
| 2626.5 | 3860.5 | 3115.5 | 1160.5 | 6133.5 | 2030 | 703 | 9227 | 4566.5 | 6249.5 |
| 2523 | 3845 | 3203 | 1137 | 5914.5 | 2044.5 | 694 | 9668.5 | 4528.5 | 6400 |
| 221457 | 115861 | 32414 | 2416 | 6836 | 2076 | 679 | 29982 | 9007 | 8268 |
| 2580 | 4125 | 3376 | 1207 | 5927 | 2015 | 664 | 10326 | 4620 | 6423 |
| 2424.5 | 3640 | 3093.5 | 1117.5 | 13500.5 | 2997.5 | 644 | 9831 | 4468.5 | 6310.5 |
| 2531 | 3981 | 3259 | 1205 | 5878 | 2029 | 711 | 10498 | 4493 | 6425 |
| 5184 | 13021.5 | 4943.5 | 1414 | 7202 | 2300 | 691 | 13059 | 6716 | 16456 |
| 8404 | 19819.5 | 6634.5 | 1807.5 | 8983 | 2555.5 | 741.5 | 12295 | 8725 | 22482.5 |
| 7736 | 20718 | 7034 | 1834 | 9689 | 2602 | 709 | 12073 | 8493 | 23452 |
| 7847 | 20946 | 7023 | 1784 | 9334 | 2543 | 699 | 12166 | 8681 | 23875 |
| 2958 | 3848 | 2360 | 826 | 3043 | 1420 | 340 | 9514 | 3850 | 4934 |
| 2442 | 3511 | 2424 | 887 | 3120 | 1432 | 455 | 8138 | 3612 | 4718 |
| 2233 | 3441 | 2563 | 940 | 3436.5 | 1600 | 535.5 | 8201 | 3619 | 5155 |
| 2154 | 3503 | 5523 | 1624 | 794098 | 71031 | 607 | 8457 | 3705 | 5441 |
| 2176 | 3369 | 3168 | 1112 | 5567 | 1998 | 615 | 8439 | 3816 | 5406 |
| 2520 | 3696 | 3227.5 | 1155 | 5177.5 | 2021.5 | 697 | 9557.5 | 4468.5 | 5879 |
| 2411.5 | 3777.5 | 3245.5 | 1124.5 | 4933.5 | 1948 | 695.5 | 9641.5 | 4364.5 | 5952 |
| 2557 | 3743 | 3263 | 1182 | 4794 | 1967 | 685 | 9945 | 4651 | 6068 |
| 2593.5 | 3971.5 | 3305.5 | 1133.5 | 4732 | 1904.5 | 682 | 10735.5 | 4555.5 | 6281 |
| 2520 | 3637 | 3015 | 1080 | 4038 | 1723 | 633 | 9759 | 4212 | 5645 |
| 2562.5 | 3725 | 2966.5 | 1120 | 4109.5 | 1780.5 | 648.5 | 10075 | 4533.5 | 5939 |
| 2555 | 3790 | 3201 | 1095 | 4385 | 1878 | 706 | 10710 | 4572 | 6202 |
| 2118 | 3377 | 2863 | 1078 | 3896 | 1688 | 662 | 8592 | 3790 | 5493 |
| 53698 | 40601 | 11733.5 | 1370 | 4208.5 | 1838 | 666 | 9208 | 20705 | 16291 |
| 2799 | 4102 | 3502.5 | 1218 | 4522.5 | 1958.5 | 708.5 | 10251 | 4640.5 | 6555.5 |
| 2519 | 3897.5 | 3380.5 | 1183 | 4413 | 1906.5 | 706 | 9888.5 | 4672 | 6528 |
| 114134.5 | 60657 | 17174 | 1899.5 | 4686.5 | 1934 | 715.5 | 18914.5 | 6938 | 7501 |
| 2672 | 4750 | 3832 | 1354 | 4907 | 2095 | 728 | 11680 | 4787 | 7148 |
| 2684 | 4283 | 3505 | 1200 | 5009 | 1968 | 710 | 9989 | 4764 | 6840 |
| 2562 | 4145 | 3499 | 1319 | 4984 | 2004 | 729 | 11281 | 4852 | 6638 |
| 5659.5 | 13574.5 | 5183 | 1558.5 | 6407.5 | 2191 | 724.5 | 12858.5 | 7016 | 16609.5 |
| 8665.5 | 21672.5 | 6802 | 1932.5 | 8790 | 2638.5 | 771.5 | 12501 | 9120.5 | 23764 |
| 8091 | 21892 | 8689 | 2047 | 10145 | 2870 | 738 | 11711 | 9267 | 24060 |

| | | | | | | | | | |
|---|---|---|---|---|---|---|---|---|---|
| 7529.5 | 21035 | 7894.5 | 1863.5 | 9129 | 2595 | 723.5 | 12657.5 | 8921 | 24809 |
| 2965 | 3736 | 2230 | 810 | 2925 | 1397 | 346 | 9359 | 3746 | 4670 |
| 2320 | 3330 | 2149 | 839 | 2915 | 1413 | 407 | 7710 | 3361 | 4537 |
| 2148 | 3316 | 2458 | 906 | 3285 | 1520 | 530 | 7652 | 3414 | 4914 |
| 2068 | 3232 | 5154.5 | 1549 | 698299.5 | 63801.5 | 575 | 7830 | 3491 | 4869.5 |
| 2111 | 3322 | 3043 | 1089 | 5335 | 1999 | 605 | 8514 | 3697 | 5421 |
| 2209.5 | 3356 | 2897.5 | 1087.5 | 4682 | 1885.5 | 640 | 8515.5 | 3904 | 5420 |
| 2160 | 3534 | 3051 | 1136 | 4608 | 1825 | 663 | 8881 | 3925 | 5525 |
| 2489 | 3584.5 | 3089.5 | 1115 | 4528.5 | 1789.5 | 675 | 9696.5 | 4210 | 5903 |
| 2703 | 3859 | 3200 | 1161 | 4454 | 1873 | 686 | 10410 | 4605 | 6177 |
| 2758 | 4184 | 3405 | 1239 | 4480 | 1969 | 698 | 12135.5 | 4676.5 | 6467.5 |
| 2684 | 3479 | 2881 | 1084 | 4046 | 1737 | 647 | 10035 | 4488 | 5724 |
| 2385.5 | 3677.5 | 2971.5 | 1083 | 4206 | 1810 | 681.5 | 9726.5 | 4165 | 5837 |
| 2329 | 3619 | 3067 | 1127 | 4176 | 1795 | 681 | 9191 | 4129 | 5816 |
| 14012 | 10183 | 4533 | 1187.5 | 4331 | 1834.5 | 712.5 | 9631 | 6897 | 7528 |
| 2650.5 | 3750.5 | 3082.5 | 1149.5 | 4076 | 1797.5 | 690 | 9913 | 4464 | 5928.5 |
| 2459 | 3717 | 3105 | 1166 | 4091 | 1797 | 689 | 9009 | 4178 | 5957 |
| 117411.5 | 60899 | 17011.5 | 1760 | 4548 | 1876.5 | 680.5 | 18316 | 6646 | 7456.5 |
| 2759 | 4353 | 3697 | 1251 | 4662 | 1956 | 723 | 10533 | 4947 | 6760 |
| 2744 | 4162.5 | 3402.5 | 1162 | 5350 | 2058.5 | 713 | 10443 | 4813 | 6458 |
| 2576 | 3795 | 3133.5 | 1153.5 | 4372 | 1826 | 683 | 9893 | 4369 | 6054 |
| 4397 | 11314 | 4546 | 1301 | 5541 | 2025 | 664 | 12970 | 5369 | 13885 |
| 8100 | 18282 | 6525 | 1660 | 7384 | 2352 | 719 | 14829 | 9075 | 20816 |
| 9230 | 22218 | 7076 | 1815 | 8551 | 2515 | 735 | 13058 | 9089 | 24108 |
| 8778 | 21042 | 6953 | 1799 | 8080 | 2433 | 687 | 12794 | 9393 | 23296 |
| 783 | 920 | 826 | 585 | 1368 | 1132 | 275 | 2421 | 1474 | 1835 |
| 1039.5 | 1613.5 | 1424 | 701.5 | 2229 | 1300.5 | 416.5 | 3852.5 | 2080.5 | 3044.5 |
| 1355.5 | 2323.5 | 2152.5 | 896 | 3125 | 1522.5 | 573.5 | 6563.5 | 2779 | 4234 |
| 1376 | 2439 | 4580 | 1433 | 658168 | 60032 | 597 | 6771.5 | 3089 | 4672.5 |
| 1486 | 2638 | 2708.5 | 1073.5 | 4451 | 2042.5 | 653 | 7205 | 3249.5 | 4877.5 |
| 1567 | 2851 | 2842 | 1092 | 4333 | 1950 | 710.5 | 7911 | 3463 | 5370.5 |
| 1497 | 2799 | 2690 | 1052 | 4001 | 1823 | 668 | 7613 | 3341 | 5121 |
| 1489 | 2589 | 2389 | 965 | 3522.5 | 1657.5 | 622.5 | 7574.5 | 3250.5 | 4850 |
| 1758 | 2968.5 | 2646 | 1020 | 3782 | 1760.5 | 660 | 8958 | 3511 | 5162.5 |
| 2024 | 3207 | 2843 | 1162 | 4051 | 1850 | 675 | 9908 | 3841.5 | 5483 |
| 2165 | 3383 | 3019 | 1174 | 4187 | 1886 | 721 | 10718 | 4169 | 5797 |
| 1937 | 3302 | 2930 | 1141 | 4138 | 1817 | 703 | 9109 | 3912 | 5749 |
| 1798 | 3203 | 2957 | 1148 | 4186 | 1850 | 727 | 9263 | 3888 | 5725 |
| 15585 | 11278 | 4692 | 1159 | 4094 | 1793 | 720 | 8530 | 6772 | 7435 |
| 2025 | 3311.5 | 3026 | 1140 | 4093.5 | 1859 | 707.5 | 8906 | 3900 | 5725.5 |
| 1811 | 3144 | 2800 | 1032 | 3907 | 1776 | 682 | 8174 | 3660 | 5387 |
| 83801 | 44603.5 | 12836.5 | 1509.5 | 4048 | 1787.5 | 669 | 13794.5 | 5171.5 | 6404 |
| 2231 | 3735 | 3249 | 1154.5 | 4273.5 | 1849.5 | 671.5 | 8822.5 | 4354 | 6239 |
| 1898 | 3339 | 3066 | 1117 | 4808 | 1936 | 661 | 8532 | 4091 | 6073 |
| 1978 | 3439 | 3080 | 1156 | 4350 | 1909 | 709 | 9203 | 4147 | 5934 |
| 3873.5 | 9019 | 4484.5 | 1293.5 | 5323.5 | 2097.5 | 688 | 13658 | 5191 | 12752.5 |
| 6874 | 16082 | 5527 | 1520 | 6677 | 2227 | 667 | 14023 | 8009 | 18301 |
| 8244.5 | 20045.5 | 6966.5 | 1733 | 8098 | 2398 | 670 | 13498.5 | 8905 | 23591 |

| | | | | | | | | | |
|---|---|---|---|---|---|---|---|---|---|
| 7842 | 20351.5 | 6654 | 1734.5 | 8350.5 | 2415 | 706.5 | 13478 | 9108 | 23200 |
| 2807 | 3674 | 2161.5 | 777 | 2780.5 | 1337 | 373 | 9099 | 3610.5 | 4625 |
| 2415.5 | 3315.5 | 2294 | 888 | 2986.5 | 1442 | 505.5 | 8425.5 | 3546 | 4828.5 |
| 2240 | 3371.5 | 2714 | 1050 | 3533.5 | 1657 | 655 | 8728.5 | 3821 | 5256 |
| 2036 | 3219.5 | 6391 | 1726 | 1111152 | 102557.5 | 671 | 8524.5 | 3839.5 | 5578.5 |
| 1974 | 2998 | 3112 | 1128 | 5426 | 2126 | 714 | 8877 | 3773 | 5457 |
| 2150 | 3479.5 | 3392.5 | 1226 | 5345 | 2151.5 | 765.5 | 9790 | 4281 | 5991 |
| 1890 | 3174 | 2926.5 | 1105 | 4353 | 1904.5 | 739.5 | 8753.5 | 3689.5 | 5485 |
| 1955 | 3129 | 2780 | 1052.5 | 4023 | 1784 | 705 | 9175 | 3800.5 | 5512.5 |
| 2301 | 3287.5 | 2925.5 | 1136 | 4055.5 | 1832.5 | 708 | 10369.5 | 4128 | 5706.5 |
| 2637.5 | 3849 | 3284 | 1231.5 | 4432.5 | 1996 | 753.5 | 11862.5 | 4650 | 6264 |
| 2658 | 3890.5 | 3417.5 | 1248.5 | 4578.5 | 1976 | 791.5 | 12212.5 | 4884 | 6429 |
| 2316 | 3660 | 3187 | 1218 | 4334 | 1937 | 785 | 10801 | 4498 | 6212 |
| 2216 | 3575.5 | 3191 | 1204 | 4384.5 | 1935.5 | 786 | 10329 | 4338.5 | 6176.5 |
| 19933 | 13947 | 5229 | 1248 | 4254 | 1922.5 | 750 | 9584 | 8366 | 8332.5 |
| 2396 | 3575 | 3257 | 1205 | 4341 | 1947 | 749 | 9763 | 4525 | 6288 |
| 2175 | 3504 | 3150 | 1179 | 4204 | 1842 | 725 | 9547 | 4172 | 5940 |
| 124574.5 | 64345 | 18858 | 1803 | 4225 | 1945 | 709 | 19706.5 | 6668 | 7181.5 |
| 2462.5 | 3781 | 3553.5 | 1234 | 4558.5 | 1872 | 721 | 10018.5 | 4798.5 | 6443 |
| 2354 | 3543 | 3367 | 1211 | 5461 | 2071 | 723 | 9896 | 4432 | 6389 |
| 2281.5 | 3804 | 3299 | 1166 | 4591 | 1968 | 753.5 | 9885 | 4584.5 | 6352 |
| 4639 | 11580 | 5124 | 1452 | 6205 | 2169 | 720 | 15436 | 5827 | 14729 |
| 8368 | 18453 | 6395 | 1734 | 7658 | 2358 | 747 | 14922 | 9214 | 21318 |
| 9343.5 | 23538.5 | 7394.5 | 1872.5 | 8775.5 | 2523.5 | 762.5 | 14365 | 9519 | 25121.5 |
| 9321.5 | 22632 | 7475 | 1794.5 | 8574 | 2630.5 | 771 | 13597.5 | 9387.5 | 25697.5 |
| 2364.5 | 3333.5 | 1925 | 746.5 | 2583.5 | 1300 | 323 | 7695.5 | 3354.5 | 4257.5 |
| 2154 | 3055 | 2172 | 840 | 2884 | 1434 | 473 | 7172 | 3238 | 4372 |
| 1872 | 2920 | 2377 | 919 | 3200 | 1551 | 562 | 7271 | 3374 | 4720 |
| 1783 | 3046 | 5105 | 1471 | 706521 | 64080 | 612 | 7732 | 3488 | 5085 |
| 1702 | 2848 | 2762 | 1024 | 4598 | 1905 | 610 | 7593 | 3216 | 4849 |
| 1775 | 3024 | 2737 | 1028 | 4317 | 1770 | 654 | 7691 | 3494 | 5197 |
| 1882 | 3201 | 2950.5 | 1110 | 4434.5 | 1930.5 | 713.5 | 8573.5 | 3876.5 | 5535 |
| 2092 | 3297 | 2973 | 1100 | 4263 | 1822 | 696 | 8938 | 3962 | 5745 |
| 1998.5 | 3138 | 2686 | 1032 | 3872 | 1703 | 640 | 9059 | 3773 | 5388.5 |
| 2183.5 | 3510.5 | 2869 | 1153.5 | 4022.5 | 1799.5 | 677 | 10258.5 | 4026 | 5558 |
| 2186.5 | 3379 | 2789.5 | 1109.5 | 3954.5 | 1806 | 675 | 9978 | 3987 | 5652 |
| 2472 | 3790 | 3243 | 1172 | 4617 | 1962.5 | 772.5 | 12153 | 4681 | 6504.5 |
| 2094 | 3390 | 3089 | 1104 | 4213 | 1826 | 727 | 9269 | 4208 | 5825 |
| 13866.5 | 9767.5 | 4516.5 | 1286.5 | 4596 | 1959 | 744 | 10346 | 6858.5 | 7651 |
| 2440.5 | 3689.5 | 3412 | 1233 | 4531 | 1981 | 759.5 | 10010 | 4483 | 6313 |
| 2201 | 3719 | 3396 | 1229 | 4471 | 1946 | 754 | 10181 | 4528 | 6692 |
| 90518 | 46238 | 13734.5 | 1767 | 4735 | 1932 | 718.5 | 16945.5 | 6413 | 7448 |
| 2482.5 | 3960.5 | 3412.5 | 1238.5 | 4518.5 | 1932.5 | 699 | 10212.5 | 4751.5 | 6493.5 |
| 2400 | 3923 | 3448 | 1206 | 5544 | 2083 | 725 | 10194 | 4562 | 6636 |
| 2439 | 3680 | 3268.5 | 1209.5 | 4523 | 1929 | 733 | 10002.5 | 4700 | 6391 |
| 4210.5 | 9950 | 4487.5 | 1328 | 5782.5 | 2100 | 715.5 | 13982.5 | 5390 | 12695.5 |
| 7096 | 15679 | 5762 | 1540 | 6828 | 2217 | 693 | 14593 | 7884 | 18504 |
| 9021.5 | 18871 | 6713.5 | 1722 | 7890.5 | 2399.5 | 694 | 13826.5 | 8877.5 | 21916 |

| | | | | | | | | | |
|---|---|---|---|---|---|---|---|---|---|
| 8545.5 | 19805.5 | 6569 | 1717 | 7795 | 2375.5 | 701 | 13168 | 8801.5 | 21936 |
| 3057 | 3904 | 2322 | 790 | 3096.5 | 1420 | 378.5 | 9976.5 | 4121 | 4926 |
| 2565.5 | 3663 | 2535.5 | 919 | 3312.5 | 1555 | 522 | 8831.5 | 3831 | 5200.5 |
| 2152 | 3298 | 2512 | 947 | 3402 | 1615 | 599 | 8051 | 3688 | 5089 |
| 2072 | 3433 | 6109 | 1710 | 940421 | 85907 | 695 | 8677 | 3838 | 5557 |
| 2146 | 3429 | 3395 | 1202 | 6154 | 2200 | 711 | 9592 | 4148 | 5708 |
| 2338 | 3735.5 | 3424.5 | 1265 | 5519 | 2030 | 795 | 10818.5 | 4531 | 6285 |
| 1865 | 3060 | 2833 | 1098 | 4418 | 1816 | 690 | 8281 | 3685 | 5427 |
| 2209 | 3589 | 3060 | 1128 | 4469 | 1932 | 719 | 9697 | 4241 | 5804 |
| 2637 | 3991 | 3221 | 1178 | 4702 | 1960 | 803 | 12410 | 4690 | 6432 |
| 2190.5 | 3369.5 | 2996.5 | 1144 | 3977 | 1842 | 683.5 | 10287.5 | 4040.5 | 5632.5 |
| 2439.5 | 3672 | 3138.5 | 1203.5 | 4303.5 | 1894 | 727.5 | 10782 | 4556 | 6071 |
| 2334 | 3583 | 3032 | 1126 | 4136 | 1881 | 724 | 10319 | 4261 | 5944 |
| 1969 | 3140 | 2809 | 1055 | 3908 | 1767 | 698 | 9148 | 3726 | 5478 |
| 16721.5 | 11984.5 | 4904.5 | 1240.5 | 4099 | 1818.5 | 753 | 9215 | 7317 | 7571 |
| 2220 | 3515 | 2992 | 1055 | 3879 | 1771 | 718 | 8947 | 3947 | 5688 |
| 2166 | 3432 | 2942 | 1155 | 4144 | 1866 | 692 | 8940 | 3974 | 5785 |
| 144321 | 73204 | 20878.5 | 2079 | 4918.5 | 2058.5 | 743 | 22653 | 7322 | 7894.5 |
| 2725 | 4188 | 3709 | 1299 | 4783 | 2055 | 770 | 11757 | 4967 | 6931 |
| 2459 | 3846 | 3449.5 | 1185 | 5263.5 | 2061.5 | 724.5 | 10350.5 | 4574.5 | 6470 |
| 2317 | 3688 | 3204 | 1170 | 4408 | 1902 | 710 | 9695 | 4441 | 6147 |
| 4696.5 | 10918.5 | 4634.5 | 1349 | 5620.5 | 2075 | 712 | 12793 | 5524.5 | 14056.5 |
| 7626 | 16598 | 5784 | 1575 | 7205 | 2209 | 675 | 14476 | 8492 | 19380 |
| 8622 | 22374 | 7114 | 1824 | 8630 | 2604 | 746 | 12247 | 9051 | 25209 |
| 8702.5 | 20927.5 | 6900.5 | 1657 | 8046 | 2336 | 692 | 12874.5 | 9257.5 | 23446.5 |
| 3176 | 3850.5 | 2221 | 858.5 | 2955 | 1440 | 362 | 9661 | 4220 | 4846 |
| 2489.5 | 3419.5 | 2337.5 | 846.5 | 3096 | 1520.5 | 504 | 8364.5 | 3534.5 | 4587.5 |
| 2281 | 3268 | 2527 | 933.5 | 3293.5 | 1549.5 | 561 | 8665 | 3515.5 | 4747 |
| 2646.5 | 4013.5 | 6408.5 | 1899 | 828464 | 80639.5 | 777 | 10865 | 4904 | 6510.5 |
| 2483 | 3797 | 3858 | 1286 | 6642 | 2407 | 773 | 10585 | 4769 | 6526 |
| 2071 | 3249 | 2939 | 1151 | 4715 | 1943 | 683 | 8943 | 3891 | 5522 |
| 2395 | 3560.5 | 3262 | 1155.5 | 5020.5 | 1973.5 | 764.5 | 10353 | 4488 | 6276 |
| 2699.5 | 4008.5 | 3472 | 1279 | 5048.5 | 2056.5 | 796.5 | 12459.5 | 5172 | 6633 |
| 2767 | 3887 | 3368 | 1226 | 4730 | 1962 | 766 | 11695 | 4778 | 6250 |
| 2458 | 3678.5 | 3082.5 | 1225.5 | 4206 | 1819.5 | 708 | 10320 | 4279.5 | 5712 |
| 2475 | 3584.5 | 3118 | 1094.5 | 4277 | 1874 | 723 | 11086 | 4443.5 | 5835 |
| 2391 | 3761 | 3154 | 1125 | 4101 | 1887 | 762 | 10093 | 4386 | 5897 |
| 2571.5 | 3976 | 3494 | 1297.5 | 4744.5 | 2083 | 810 | 11451.5 | 5136 | 6875 |
| 22691 | 16342 | 5454 | 1155 | 3794 | 1744 | 680 | 9195 | 8706 | 8285 |
| 2577 | 3564 | 3026 | 1146 | 3874 | 1836 | 714 | 9022 | 4296 | 5607 |
| 2996 | 4543 | 3958 | 1405 | 5388 | 2164 | 834 | 11982 | 5502 | 7835 |
| 84428.5 | 44925.5 | 12866 | 1722 | 4858 | 1950.5 | 785.5 | 16127.5 | 6275 | 7719.5 |
| 2911.5 | 4385 | 3781.5 | 1297.5 | 4802.5 | 2012 | 746 | 10604 | 5236 | 7093 |
| 2679.5 | 4019.5 | 3637.5 | 1260.5 | 5103 | 2048.5 | 763.5 | 10564.5 | 4917.5 | 6708.5 |
| 2834 | 4124 | 3483 | 1303 | 4744 | 2073 | 807 | 11393 | 5033 | 6852 |
| 4779 | 11178 | 4923 | 1458 | 5911 | 2096 | 757 | 14162 | 5716 | 14252 |
| 8291 | 18107.5 | 6804.5 | 1741 | 7440 | 2448 | 745.5 | 15235 | 8524 | 21089.5 |
| 9414 | 24236 | 8493 | 2056 | 9325 | 2754 | 826 | 11597 | 9077 | 26894 |

| | | | | | | | | | |
|---|---|---|---|---|---|---|---|---|---|
| 8689.5 | 23447 | 7890.5 | 1868.5 | 8772 | 2591.5 | 772.5 | 11083.5 | 8919 | 26767 |
| 3125.5 | 3841.5 | 2319.5 | 839.5 | 2928 | 1433 | 363 | 9910 | 4047.5 | 4749 |
| 2511 | 3679 | 2504 | 875 | 3139 | 1484 | 484 | 8619 | 3708 | 4870 |
| 2156 | 3415 | 2558 | 970 | 3451 | 1622 | 567 | 8205 | 3742 | 5220 |
| 2160.5 | 3360.5 | 5486.5 | 1555 | 742649.5 | 66514 | 616 | 8644.5 | 3870.5 | 5327 |
| 2288 | 3484.5 | 3488.5 | 1180.5 | 5869.5 | 2150.5 | 680 | 9451 | 4127.5 | 5800 |
| 2273.5 | 3337 | 2970 | 1128.5 | 4941.5 | 1894.5 | 676.5 | 8887.5 | 4159.5 | 5745 |
| 2004 | 3220 | 2877 | 1092 | 4477 | 1826 | 680 | 8505 | 3815 | 5539 |
| 1990 | 3208 | 2908.5 | 1050.5 | 3923.5 | 1709 | 644 | 8505 | 3721 | 5254.5 |
| 2549.5 | 3802 | 3085.5 | 1142 | 4460.5 | 1852.5 | 682 | 10696 | 4451.5 | 6173 |
| 2605 | 3988 | 3259 | 1189 | 4436 | 1925 | 695 | 11706 | 4532 | 6051 |
| 2582 | 3802 | 3265 | 1190 | 4574 | 1925 | 724 | 12105 | 4510 | 6145 |
| 2603.5 | 4060.5 | 3321.5 | 1217 | 4808.5 | 1945.5 | 743 | 12435 | 4703.5 | 6542 |
| 2427 | 3813 | 3269 | 1179.5 | 4475 | 1938.5 | 716.5 | 10889 | 4524 | 6354.5 |
| 13463 | 9655 | 4524 | 1249 | 4500 | 1917 | 731 | 10645 | 6788 | 7447 |
| 2764 | 4024 | 3319 | 1228 | 4457 | 1922 | 759 | 10078 | 4550 | 6449 |
| 2553 | 3848 | 3259 | 1211 | 4477 | 1910 | 729 | 9619 | 4543 | 6356 |
| 113022 | 59664 | 16365 | 1747 | 4606 | 1859 | 679 | 17924 | 6501 | 7284 |
| 2628 | 3945 | 3455 | 1196 | 4479 | 1842.5 | 679 | 10006.5 | 4721 | 6453.5 |
| 2447.5 | 3818 | 3342 | 1203.5 | 5070 | 2029.5 | 684 | 9675.5 | 4442 | 6281.5 |
| 2398 | 3847 | 3229 | 1157 | 4445 | 1898 | 695 | 9944 | 4544 | 6329 |
| 4154.5 | 10302.5 | 4458 | 1378 | 5704.5 | 2099 | 697 | 14010 | 5438.5 | 13394 |
| 6091 | 17052 | 5936 | 1614 | 7117 | 2309 | 692 | 12058 | 7301 | 19260 |
| 8479 | 22153 | 7430 | 1798.5 | 8838.5 | 2532 | 691 | 13210 | 9174 | 25554.5 |
| 8089.5 | 22568 | 7882.5 | 1787 | 8647.5 | 2525 | 722 | 12718.5 | 9056.5 | 25368.5 |
| 3201.5 | 3817 | 2353.5 | 816 | 3023 | 1359 | 350.5 | 9641 | 4271 | 4715 |
| 2554 | 3654 | 2444 | 867 | 3186 | 1445 | 466 | 8617 | 3802 | 4980 |
| 2145 | 3334 | 2467.5 | 949 | 3409.5 | 1576 | 551 | 7886.5 | 3529 | 4910 |
| 2155 | 3440 | 5366 | 1605 | 693590 | 63043 | 629 | 8669 | 3853 | 5512 |
| 2385 | 3686 | 3519 | 1260 | 6008 | 2165 | 681 | 9655 | 4362 | 5859 |
| 2244.5 | 3513.5 | 3132.5 | 1177 | 4980 | 1954.5 | 677 | 8774.5 | 4316 | 5787 |
| 2316.5 | 3650 | 3284.5 | 1137 | 4898.5 | 2004 | 734.5 | 9608.5 | 4350 | 6038.5 |
| 2224 | 3513 | 3053 | 1075 | 4509 | 1889 | 683 | 9475 | 4145 | 5841 |
| 2247 | 3488 | 2854 | 1079 | 4062.5 | 1750 | 655.5 | 9520.5 | 4062.5 | 5720 |
| 2462 | 3501 | 2966 | 1096 | 4071 | 1832 | 629.5 | 9814 | 4312.5 | 5596 |
| 2525 | 3675 | 2940 | 1105 | 4228 | 1804 | 679 | 10535 | 4481 | 5894 |
| 2666 | 4022.5 | 3380 | 1215.5 | 4726.5 | 1970.5 | 750 | 12449.5 | 4749.5 | 6807 |
| 2500 | 3753.5 | 3213 | 1162 | 4516.5 | 1996.5 | 720.5 | 11336.5 | 4612 | 6161 |
| 15319 | 10688 | 4736 | 1192 | 4470 | 1899 | 723 | 10521 | 7369 | 7798 |
| 3003 | 4256 | 3397 | 1231 | 4667 | 1995 | 779 | 11769 | 5032 | 6661 |
| 2749 | 4192 | 3471 | 1246 | 4756 | 2032 | 772 | 11062 | 4702 | 6759 |
| 113884.5 | 58229.5 | 16316 | 1775.5 | 4394.5 | 1880 | 689.5 | 17429.5 | 6488 | 7513 |
| 3049 | 5167 | 4012 | 1369 | 5165 | 2071 | 788 | 13599 | 5086 | 8243 |
| 2807 | 4304 | 3772 | 1296 | 5456 | 2105 | 748 | 11717 | 4952 | 7014 |
| 2855.5 | 4430 | 3660.5 | 1303 | 4999 | 2070.5 | 780.5 | 12247 | 5019.5 | 7132 |
| 4002 | 8309.5 | 4508 | 1364.5 | 5659.5 | 2123 | 758.5 | 15296.5 | 5592 | 11440 |
| 8691 | 20787 | 6690 | 1778 | 7991 | 2510 | 755 | 14472 | 9322 | 23307 |
| 8583.5 | 23208.5 | 8077.5 | 2098.5 | 9678 | 2777.5 | 782.5 | 12547.5 | 8714 | 25657.5 |

| | | | | | | | | | |
|---|---|---|---|---|---|---|---|---|---|
| 9174 | 23428 | 8645 | 1911 | 9987 | 2735 | 780 | 13089 | 9438 | 26284 |
| 3225 | 3800.5 | 2283.5 | 774.5 | 3027 | 1376.5 | 354 | 9207.5 | 4258 | 4922.5 |
| 2898 | 3755 | 2577 | 893 | 3312 | 1522 | 491 | 8879 | 4081 | 5136 |
| 2468 | 3554 | 2643 | 924 | 3548 | 1598 | 580 | 8812 | 3824 | 5312 |
| 2466 | 3625 | 6026 | 1718 | 780994.5 | 75807 | 643.5 | 9060.5 | 4162 | 5887.5 |
| 2501.5 | 3535 | 3395 | 1185.5 | 5648.5 | 2061.5 | 682 | 9456 | 4402 | 5822 |
| 2753 | 4140 | 3658 | 1299 | 5875 | 2201 | 774 | 11688 | 4934 | 6404 |
| 2653.5 | 3733.5 | 3331 | 1180.5 | 5057 | 2045.5 | 740.5 | 9940.5 | 4620 | 6185.5 |
| 2472 | 3672 | 3085 | 1140 | 4764 | 1901 | 697 | 9805 | 4373 | 5911 |
| 2416.5 | 3551 | 2958.5 | 1091 | 4239.5 | 1810 | 661.5 | 9996 | 4202 | 5727 |
| 2692 | 3889 | 3185 | 1219 | 4269 | 1961 | 682 | 11483 | 4672 | 6145 |
| 2712 | 4094 | 3402.5 | 1188 | 4645 | 1940.5 | 713 | 12351 | 4699.5 | 6306 |
| 2528 | 3768 | 3142 | 1136 | 4348 | 1864 | 705 | 11244 | 4559 | 6055 |
| 2589.5 | 3976.5 | 3257 | 1231.5 | 4691.5 | 1988 | 754.5 | 11860.5 | 4743.5 | 6426.5 |
| 15239 | 11019 | 4927 | 1319 | 4744 | 1978 | 777 | 11485 | 7454 | 8125 |
| 3027 | 4365 | 3610 | 1287 | 4846 | 2000 | 766 | 11571 | 5091 | 6804 |
| 2688 | 4235 | 3628 | 1293 | 4788 | 2034 | 765 | 11354 | 4917 | 7040 |
| 118980 | 62114.5 | 17554.5 | 1846.5 | 4836 | 1965 | 707 | 20182.5 | 6766 | 7859.5 |
| 2807 | 4457.5 | 3628.5 | 1274 | 4774 | 1950 | 729 | 10901.5 | 4894 | 6935 |
| 2673 | 4512.5 | 3649.5 | 1338.5 | 5606.5 | 2091.5 | 751 | 11521.5 | 4803.5 | 7305 |
| 2635 | 4073 | 3486 | 1208 | 4621 | 1923 | 720 | 10566 | 4727 | 6586 |
| 5144.5 | 12853.5 | 5246 | 1461 | 6389.5 | 2253 | 757 | 14564 | 6670 | 16603.5 |
| 7842.5 | 19178 | 6823 | 1734.5 | 8037.5 | 2472 | 725 | 14143 | 9143.5 | 22174 |
| 8482 | 23319 | 7705 | 1922.5 | 9076 | 2647.5 | 736.5 | 12954.5 | 9439.5 | 26277 |
| 7389.5 | 23147.5 | 7464 | 1868 | 9041 | 2552 | 725.5 | 10532 | 8272 | 26498.5 |
| 3028 | 3630 | 2196 | 771 | 2962 | 1377 | 342 | 8699 | 4071 | 4659 |
| 2810.5 | 3750 | 2523.5 | 913.5 | 3584 | 1547.5 | 485 | 8835.5 | 3916.5 | 5059 |
| 2317.5 | 3538.5 | 2726 | 975.5 | 3754 | 1625.5 | 563 | 8346 | 3716.5 | 5303 |
| 2275.5 | 3531.5 | 5217 | 1566 | 469424 | 45889 | 603 | 9041.5 | 3733.5 | 5346 |
| 2314 | 3418 | 3119 | 1094 | 17039 | 2824 | 640 | 8474 | 4167 | 5438 |
| 2535 | 3883 | 3394 | 1236 | 10200 | 2412 | 710 | 10572 | 4437 | 6156 |
| 2421 | 3631 | 3055 | 1151 | 8496 | 2212 | 691 | 9037 | 4514 | 5861 |
| 2310 | 3602 | 3067 | 1083 | 9815.5 | 2178.5 | 658 | 9392 | 4108 | 5773.5 |
| 2428 | 3631.5 | 2923 | 1045 | 9591.5 | 2187.5 | 636.5 | 9763 | 4155.5 | 5684 |
| 2730 | 3835 | 3168 | 1211 | 5204 | 1954 | 673 | 11518 | 4517 | 6093 |
| 2615 | 4077 | 3217.5 | 1185 | 7403.5 | 2134.5 | 701.5 | 12063.5 | 4449.5 | 6267.5 |
| 2343 | 3840 | 3138 | 1142 | 6683 | 2035 | 683 | 11205 | 4303 | 6030 |
| 2556 | 4009 | 3313 | 1192 | 5831 | 2080 | 733 | 12053 | 4630 | 6489 |
| 6825 | 5971 | 3847.5 | 1242.5 | 6415 | 2115 | 736.5 | 10665.5 | 5552.5 | 6955 |
| 2698 | 4152 | 3441 | 1223 | 6153 | 2114 | 764 | 10971 | 4795 | 6621 |
| 2643 | 4035 | 3330 | 1209 | 5794 | 1991 | 720 | 10374 | 4763 | 6627 |
| 242053 | 129101 | 35552 | 2456 | 6116 | 2003 | 654 | 32099 | 9071 | 8145 |
| 2812 | 4806 | 3845 | 1319 | 5827 | 2068 | 723 | 12564 | 4915 | 7791 |
| 2561.5 | 4309 | 3591 | 1239.5 | 13824.5 | 3115 | 715.5 | 11251.5 | 4608.5 | 6852.5 |
| 2715 | 4100 | 3313 | 1246 | 5746.5 | 2099.5 | 720 | 11575 | 4687.5 | 6650 |
| 4635 | 11816 | 4619 | 1324 | 6539 | 2115 | 669 | 13506 | 6246 | 15391 |
| 7631 | 19496 | 6572 | 1842 | 8825 | 2555 | 735 | 13659 | 9732 | 21951 |
| 8081 | 22289 | 7439 | 1881 | 9878 | 2717 | 719 | 12329 | 8916 | 24233 |

| | | | | | | | | | |
|---|---|---|---|---|---|---|---|---|---|
| 7329.5 | 20724.5 | 7949 | 1814.5 | 10374.5 | 2618 | 715.5 | 10752 | 8265 | 23921.5 |
| 2852 | 3779 | 2246 | 782 | 2936 | 1351 | 329 | 9046 | 3814 | 4637 |
| 2528.5 | 3658.5 | 2552 | 893 | 3277.5 | 1481 | 448 | 8405 | 3729 | 4809.5 |
| 2393 | 3637 | 2648 | 976 | 3430 | 1637 | 539 | 8168 | 3679 | 5222 |
| 2268 | 3391 | 5469 | 1532 | 799755 | 70827 | 583 | 8465 | 3987 | 5240 |
| 2206 | 3566.5 | 3085 | 1119.5 | 5251 | 2012.5 | 659 | 8583 | 3820 | 5611 |
| 2532.5 | 3748.5 | 3352.5 | 1186.5 | 5298 | 1918.5 | 714.5 | 10695 | 4479 | 5933 |
| 2414 | 3655 | 3061 | 1164 | 4619 | 1912 | 672 | 9112 | 4450 | 6195 |
| 2408 | 3753.5 | 3100.5 | 1100 | 4569.5 | 1850 | 687 | 9369 | 4577 | 5964.5 |
| 2504 | 3726 | 2962 | 1081 | 4388 | 1860 | 660 | 9940 | 4452 | 5767 |
| 2624 | 3669 | 2931 | 1157.5 | 4091.5 | 1798.5 | 610 | 9993.5 | 4396 | 5776.5 |
| 2611.5 | 4012.5 | 3096.5 | 1159.5 | 4537.5 | 1881 | 692 | 11786 | 4517 | 6356.5 |
| 2682 | 4399.5 | 3556 | 1273 | 4834.5 | 2031 | 748 | 13570 | 4906.5 | 6855.5 |
| 2598 | 3854 | 3371 | 1142 | 4499 | 1916 | 707 | 11946 | 4629 | 6366 |
| 61914 | 45235 | 13480.5 | 1663 | 4852 | 2005 | 745.5 | 11772 | 24023 | 17900 |
| 3041.5 | 4667.5 | 3787.5 | 1346 | 4895.5 | 2004.5 | 808 | 12200.5 | 5417.5 | 7282 |
| 2656 | 4245 | 3428 | 1263 | 4759 | 2005 | 738 | 11186 | 4862 | 6845 |
| 122749 | 65990 | 18638.5 | 1976 | 4793.5 | 1960.5 | 733 | 21283 | 7000 | 7881.5 |
| 2871 | 4552 | 3692 | 1287 | 5005 | 2080 | 740 | 12702 | 4781 | 7446 |
| 2643.5 | 5258.5 | 3769.5 | 1388.5 | 5582.5 | 2238.5 | 746.5 | 13326.5 | 4822 | 8752 |
| 2703 | 4305 | 3618 | 1284 | 4910 | 2069 | 767 | 11479 | 4952 | 7134 |
| 4606.5 | 11473 | 4602.5 | 1392 | 5679 | 2256 | 742.5 | 12905.5 | 6083.5 | 15022.5 |
| 7960.5 | 19967.5 | 6972.5 | 1814.5 | 8354 | 2521.5 | 740 | 14656 | 9687.5 | 24286 |
| 7919.5 | 23368.5 | 8538.5 | 2029.5 | 9784 | 2736 | 748.5 | 12647.5 | 9413 | 26733 |
| 8081 | 22523 | 8788 | 1973 | 10204 | 2704 | 764 | 12853 | 8978 | 26196 |
| 3283 | 3886.5 | 2358 | 815.5 | 3046.5 | 1417 | 354 | 9415.5 | 4263.5 | 4928.5 |
| 2879 | 3796 | 2616.5 | 915.5 | 3277.5 | 1532.5 | 474.5 | 8964.5 | 4165.5 | 5064.5 |
| 2561 | 3654 | 2745 | 1006 | 3590 | 1660 | 586 | 8728 | 4019 | 5362 |
| 2193 | 3382 | 5396 | 1555 | 697996 | 64987.5 | 585 | 8346 | 3774.5 | 5326.5 |
| 2497.5 | 3681 | 3487.5 | 1211.5 | 6309 | 2257 | 669 | 9228 | 4355.5 | 5862.5 |
| 2661 | 4096 | 3611 | 1282 | 5563 | 2149 | 744 | 10750 | 4716 | 6359 |
| 2628 | 3920 | 3398 | 1215 | 5182 | 2041 | 705 | 10302 | 4502 | 6214 |
| 2456 | 3583 | 2988 | 1168 | 4614.5 | 1891 | 671.5 | 9646.5 | 4257 | 5940 |
| 2729 | 3797.5 | 3118 | 1110 | 4477 | 1845.5 | 661 | 10935 | 4623 | 6109 |
| 2813 | 4203 | 3474.5 | 1253.5 | 4718 | 1934.5 | 699 | 12876 | 4804 | 6534.5 |
| 2788 | 4148 | 3380 | 1249 | 4582 | 1927 | 715 | 11940 | 4819 | 6368 |
| 2743 | 3823 | 3166.5 | 1121 | 4294 | 1828 | 695 | 10461.5 | 4672 | 6271.5 |
| 2455 | 3635 | 3148 | 1131.5 | 4202.5 | 1794 | 700 | 10057.5 | 4235 | 5829 |
| 12071 | 8949 | 4483 | 1251 | 4610 | 1886 | 712 | 10162 | 6758 | 7663 |
| 3024.5 | 4128 | 3469.5 | 1241.5 | 4575 | 1910.5 | 739 | 10727.5 | 4924.5 | 6526.5 |
| 2649.5 | 3776.5 | 3223 | 1113.5 | 4437.5 | 1840.5 | 683.5 | 9910.5 | 4571.5 | 6132 |
| 120665.5 | 61134.5 | 17171.5 | 1750.5 | 4449 | 1972 | 683.5 | 19363 | 6852 | 7250 |
| 2880.5 | 4559 | 3761 | 1243 | 4852 | 1954 | 725 | 10942.5 | 5047.5 | 7283 |
| 2696.5 | 4211 | 3516.5 | 1256.5 | 5448 | 2061 | 705.5 | 10358.5 | 4750.5 | 6921.5 |
| 2761.5 | 4176 | 3559 | 1232 | 4787.5 | 1996 | 745 | 11200.5 | 4846.5 | 6807.5 |
| 4465 | 11152 | 4440 | 1357 | 5687 | 2005 | 695 | 12620 | 5605 | 14238 |
| 6491.5 | 16529 | 5694.5 | 1517 | 6766.5 | 2265.5 | 670 | 14430.5 | 7125 | 18179 |
| 8872 | 21937 | 7213 | 1738 | 8189 | 2401 | 679 | 14789 | 9481 | 25174 |

| | | | | | | | | | |
|---|---|---|---|---|---|---|---|---|---|
| 8923.5 | 22041 | 7266.5 | 1795.5 | 8352.5 | 2577.5 | 739 | 12848 | 9626 | 23759 |
| 693 | 826 | 630 | 558 | 1139 | 1105 | 184 | 1809 | 1216 | 1496 |
| 5476 | 7862 | 3597 | 1162 | 4119 | 1723 | 273 | 14031 | 5200 | 7970 |
| 3645 | 4862.5 | 2545.5 | 862 | 3121.5 | 1472 | 360 | 12773 | 4142.5 | 5568 |
| 3388 | 4168 | 10160 | 3491 | 697138 | 64417 | 515 | 14848 | 4416 | 5491 |
| 2874 | 3689 | 4222 | 1491 | 6437 | 2623 | 577 | 15676 | 4410 | 5633 |
| 2321 | 3371.5 | 3118.5 | 1121 | 4479 | 1992 | 612 | 13441.5 | 3861.5 | 5494 |
| 2346 | 3405 | 3019 | 1097 | 4329 | 1902.5 | 675 | 13798 | 4247.5 | 5785 |
| 1869 | 3093 | 2722 | 1021 | 3809 | 1774 | 648 | 10391 | 3621 | 5376 |
| 1735 | 3031 | 2610 | 1005 | 3779 | 1736 | 670 | 10391 | 3610 | 5469 |
| 1695 | 2949 | 2583.5 | 1014 | 3673 | 1740 | 668.5 | 9519.5 | 3507 | 5407 |
| 1747 | 3077.5 | 2633 | 1015 | 3724 | 1744 | 669.5 | 9795 | 3600.5 | 5516.5 |
| 1730 | 2929 | 2419 | 959 | 3488 | 1665 | 631 | 9496 | 3370 | 5100 |
| 1803 | 3159 | 2617 | 1028 | 3763 | 1751 | 675 | 10565 | 3697 | 5609 |
| 1801 | 2937 | 2472 | 974 | 3451 | 1688 | 637 | 9227 | 3388 | 5123 |
| 1761 | 3224 | 2762 | 1073 | 3891 | 1774 | 715 | 9968 | 3800 | 5855 |
| 1547 | 2746 | 2382 | 976 | 3418 | 1658 | 632 | 8343 | 3198 | 5032 |
| 13887 | 8026 | 3847 | 1047 | 3645 | 1684 | 628 | 9876 | 3562 | 5377 |
| 2546 | 4546 | 3444 | 1112 | 3957 | 1784 | 672 | 9991 | 3970 | 5985 |
| 2006 | 3630.5 | 3061.5 | 1074 | 3884 | 1787.5 | 668 | 9872 | 3851 | 6004.5 |
| 1928 | 3509 | 2918 | 1071 | 3955 | 1813 | 690 | 9612 | 4060.5 | 6115 |
| 3508 | 6720 | 3727 | 1192 | 4794 | 1939 | 728 | 16381 | 5276 | 9483 |
| 5785 | 16928 | 5538.5 | 1576 | 6753.5 | 2344 | 764 | 14897 | 6958 | 20200 |
| 7931 | 21189 | 6630 | 1647 | 7698 | 2362 | 660 | 12917 | 7970 | 23437 |
| 9460 | 27070 | 8343 | 1903 | 9501 | 2609 | 698 | 12804 | 9032 | 29138 |
| 9213 | 25761 | 8962.5 | 1980 | 9889 | 2733 | 755 | 12248.5 | 9067 | 28029 |
| 8756 | 20235 | 6630 | 1689 | 7485 | 2398 | 679 | 13808 | 8921 | 22937 |
| 6123.5 | 16977.5 | 5305.5 | 1416 | 6267 | 2131.5 | 685 | 14500.5 | 6944.5 | 19589.5 |
| 5870 | 13871 | 6560 | 1777 | 54332 | 7421 | 715 | 14622 | 6957 | 17218 |
| 3743 | 10231 | 4923 | 1496 | 8635 | 2996 | 743 | 13336 | 5194 | 13846 |
| 3038 | 5939 | 4024 | 1328 | 6027 | 2339 | 794 | 16443 | 4898 | 8937 |
| 2782.5 | 4448 | 3503 | 1236 | 5164.5 | 2088.5 | 760 | 16229 | 5071.5 | 7291 |
| 2192 | 3734.5 | 3095.5 | 1160.5 | 4471 | 1962 | 737 | 11222 | 4398.5 | 6530.5 |
| 2125.5 | 3699 | 3015 | 1116.5 | 4290 | 1914.5 | 759 | 12058.5 | 4350 | 6666 |
| 1924 | 3353 | 2816 | 1115 | 4089 | 1854 | 709 | 10846 | 3999 | 5990 |
| 2284.5 | 4050 | 3048.5 | 1153 | 4473 | 1945 | 754 | 11851 | 4583 | 7030 |
| 2234.5 | 3733 | 2908 | 1111.5 | 4147 | 1830 | 739 | 11267 | 4370.5 | 6394 |
| 2696.5 | 4735.5 | 3361.5 | 1250 | 4827 | 2083 | 840 | 16889.5 | 5684.5 | 8995 |
| 2335 | 3659.5 | 2936 | 1090 | 4168 | 1839.5 | 723 | 12079 | 4425 | 6375 |
| 2075.5 | 3528 | 2873.5 | 1092 | 4075.5 | 1830 | 728.5 | 10500 | 4120 | 6214.5 |
| 2441 | 4585 | 3380 | 1240 | 4772 | 2080 | 864 | 15127 | 5467 | 8744 |
| 22955 | 12772 | 4856 | 1202 | 4185 | 1872 | 736 | 12400 | 4575 | 6674 |
| 3679 | 6181 | 4464 | 1311 | 4878 | 2077 | 808 | 13350 | 5673 | 7656 |
| 2646.5 | 4714.5 | 3650 | 1202 | 4711 | 1984 | 752 | 11226 | 4908.5 | 7479.5 |
| 2694.5 | 5036 | 3687.5 | 1276.5 | 5052.5 | 2065.5 | 827 | 17492 | 5803 | 9237.5 |
| 3500 | 6595 | 3822.5 | 1242 | 4784 | 1987 | 754.5 | 15386.5 | 5733 | 9856 |
| 5681 | 13389 | 5505 | 1519 | 6654 | 2282 | 747 | 13091 | 6838 | 16506 |
| 8487 | 23642 | 7397 | 1728 | 8552 | 2474 | 692 | 14246 | 8725 | 26185 |

| | | | | | | | | | |
|---|---|---|---|---|---|---|---|---|---|
| 9754 | 28016.5 | 8306.5 | 1902.5 | 9346.5 | 2605 | 690 | 12045.5 | 8487.5 | 29052.5 |
| 8878 | 27873.5 | 9037.5 | 1983.5 | 10308 | 2793 | 744 | 11315 | 8542 | 29581 |
| 8464.5 | 22556 | 7009 | 1767 | 8062 | 2496 | 723 | 14401 | 8461.5 | 24594.5 |
| 7763 | 17973 | 5720 | 1519 | 6839 | 2212 | 690 | 14198 | 8528 | 20859 |
| 4584 | 12344 | 5408 | 1449 | 346251 | 41193 | 658 | 15712 | 5443 | 15202 |
| 3974 | 10364 | 4364 | 1332 | 6473 | 2430 | 707 | 16009 | 5352 | 13733 |
| 3081 | 4852 | 3429 | 1133 | 4660 | 1905 | 685 | 16435 | 4562 | 6945 |
| 2864 | 4477 | 3355 | 1195 | 4684 | 1962 | 741 | 13932 | 5011 | 7142 |
| 2568 | 4193 | 3228 | 1203 | 4658 | 1993 | 768 | 12179 | 4875 | 7185 |
| 1957 | 3313 | 2676 | 1013 | 3802 | 1757 | 653 | 9695 | 3704 | 5611 |
| 2431 | 4201.5 | 3177 | 1189.5 | 4481 | 1965 | 761 | 12552.5 | 4774.5 | 7115.5 |
| 2149 | 3454 | 2736 | 1063 | 3821 | 1767 | 678 | 10088 | 3912 | 5777 |
| 2524.5 | 4326 | 3251 | 1231 | 4696.5 | 2003.5 | 756 | 13137.5 | 4978 | 7376 |
| 2702.5 | 4332.5 | 3260 | 1205 | 4588.5 | 1954 | 783.5 | 12079.5 | 5078.5 | 7434.5 |
| 2683.5 | 3774.5 | 2859 | 1036 | 3900.5 | 1740.5 | 675 | 9940 | 4138 | 5960 |
| 2334 | 3712.5 | 2942 | 1077.5 | 3972.5 | 1827.5 | 702.5 | 10524 | 4144.5 | 6231 |
| 2193 | 3592.5 | 2872 | 1068.5 | 4005.5 | 1791 | 708.5 | 10038.5 | 4173 | 6150 |
| 4297.5 | 4366 | 3067.5 | 1068 | 3852 | 1773 | 663 | 9778.5 | 4020 | 5929.5 |
| 3209 | 4594 | 3250 | 1171 | 4303 | 1899 | 720 | 10851 | 4603 | 6907 |
| 2811 | 4698 | 3517.5 | 1248 | 4670 | 2016.5 | 770 | 11447 | 5059.5 | 7886.5 |
| 2512.5 | 4128 | 3258 | 1145 | 4329 | 1929.5 | 736 | 11091 | 4684 | 6883.5 |
| 5519 | 9933 | 4818 | 1489 | 6204 | 2329 | 850 | 20116 | 7894 | 13986 |
| 7237 | 19573 | 7028 | 1934 | 8657 | 2715 | 891 | 12838 | 8899 | 24067 |
| 9050 | 24636 | 7827 | 1842 | 9132 | 2598 | 736 | 15037 | 9598 | 27824 |
| 9306 | 22012 | 9210 | 1952 | 10521 | 2696 | 744 | 13081 | 8753 | 23102 |
| 9312.5 | 23241.5 | 7552 | 1782 | 8464.5 | 2504.5 | 689.5 | 11755.5 | 8520 | 24799.5 |
| 8002 | 19594.5 | 6556 | 1766 | 7420.5 | 2461 | 668 | 11339 | 7447.5 | 21617 |
| 7184 | 16806 | 5629 | 1522 | 6392 | 2217 | 711 | 14341 | 8089 | 19671 |
| 3912 | 10606 | 5458 | 1420 | 552964 | 53430 | 612 | 15758 | 4808 | 13121 |
| 3514 | 9920 | 4655 | 1373 | 7102 | 2558 | 737 | 15496 | 4950 | 13592 |
| 2851.5 | 4619.5 | 3398 | 1150 | 4843.5 | 2023 | 650 | 15007.5 | 4396 | 6876 |
| 2819 | 4281.5 | 3292 | 1170 | 4719 | 1987 | 720 | 12586.5 | 4935 | 6721.5 |
| 2537 | 3973 | 3238 | 1184 | 4585 | 1974 | 727 | 12505 | 4649 | 6462 |
| 2014 | 3323 | 2724 | 1033 | 3769 | 1743 | 670 | 9378 | 3766 | 5677 |
| 2375 | 3987 | 3228 | 1207 | 4436 | 1956 | 778 | 12255 | 4681 | 6678 |
| 2034 | 3213 | 2541 | 978 | 3457 | 1681 | 626 | 9913 | 3684 | 5366 |
| 2554 | 4233 | 3199 | 1154 | 4440 | 1927 | 745 | 12079 | 4889 | 6970 |
| 2239.5 | 3635 | 2863 | 1070 | 3957 | 1786 | 686 | 10469.5 | 4117 | 5974.5 |
| 2554 | 4304 | 3204 | 1193 | 4546 | 1959 | 772 | 12101 | 4750 | 7063 |
| 2344 | 3651 | 2945 | 1087 | 4063 | 1832 | 714 | 10860 | 4453 | 6234 |
| 2299 | 3761 | 3017 | 1107 | 4194 | 1869 | 717 | 10690 | 4399 | 6421 |
| 19152 | 10276.5 | 4333 | 1168 | 3951 | 1814.5 | 681 | 12147.5 | 4771.5 | 6282.5 |
| 3299.5 | 5400 | 3798.5 | 1208 | 4548 | 1959 | 718 | 11762 | 4805.5 | 7004.5 |
| 3025 | 5326.5 | 3808.5 | 1307 | 4928.5 | 2114 | 778 | 14489.5 | 5165 | 8184.5 |
| 2470 | 4055 | 3105 | 1117 | 4078 | 1858 | 682 | 11420 | 4743 | 6635 |
| 4489 | 10217 | 4518 | 1449 | 5976 | 2247 | 823 | 16947 | 6977 | 14976 |
| 5130 | 13732 | 4991 | 1362 | 5862 | 2067 | 687 | 14912 | 6095 | 16478 |
| 9009 | 23031 | 7915 | 1784 | 9236 | 2519 | 697 | 13536 | 9338 | 25737 |

| | | | | | | | | | |
|---|---|---|---|---|---|---|---|---|---|
| 9953 | 22458 | 6972 | 1682 | 7740 | 2323 | 641 | 14015 | 9696 | 24105 |
| 9332.5 | 19483.5 | 5908.5 | 1474 | 6783 | 2114 | 582 | 15035 | 8677 | 21167.5 |
| 9304.5 | 20776.5 | 6504 | 1657.5 | 7423 | 2396 | 671 | 15152.5 | 9614 | 23583 |
| 7056.5 | 17300.5 | 5796 | 1534.5 | 7037 | 2244 | 691 | 12239 | 8109 | 20640.5 |
| 4655 | 12146.5 | 5664 | 1491 | 525258 | 55622.5 | 641 | 14551 | 5580.5 | 14949 |
| 3370 | 6732 | 4108 | 1238 | 6043 | 2263 | 647 | 15881 | 4618 | 8985 |
| 2819 | 4577 | 3437 | 1148 | 4857 | 2028 | 653 | 15652 | 4377 | 6714 |
| 2511 | 3720 | 2954 | 1059 | 4146 | 1829 | 662 | 10994 | 4361 | 5975 |
| 1834.5 | 2999.5 | 2458.5 | 947 | 3473 | 1660 | 605 | 8700 | 3389 | 5045 |
| 2080.5 | 3401 | 2783 | 1039 | 3954.5 | 1811 | 680 | 10370 | 4093.5 | 5766 |
| 1782.5 | 2839.5 | 2294 | 909 | 3205.5 | 1588 | 584 | 8960 | 3332 | 4822 |
| 2424.5 | 3833 | 2977 | 1089 | 4206.5 | 1852 | 705 | 11223 | 4491 | 6373 |
| 2231 | 3446 | 2672 | 1010 | 3630 | 1685 | 634 | 11066 | 3994 | 5610 |
| 2155 | 3382 | 2600.5 | 983 | 3591.5 | 1672 | 639 | 10230 | 3911 | 5541 |
| 2207 | 3352 | 2604 | 970 | 3497.5 | 1648.5 | 631 | 9381 | 3732 | 5482.5 |
| 2192 | 3562 | 2790 | 1042 | 3987 | 1743 | 684 | 9962 | 4171 | 6008 |
| 2404.5 | 3872.5 | 3129.5 | 1158 | 4229 | 1933.5 | 731.5 | 11116 | 4597.5 | 6348.5 |
| 9557 | 6539 | 3558.5 | 1047.5 | 3750 | 1705 | 647 | 10071 | 4062 | 5784.5 |
| 3006 | 4749.5 | 3572 | 1187 | 4587 | 1940 | 739 | 11308.5 | 4890 | 7019 |
| 2534 | 3973 | 3128 | 1095 | 4129 | 1791 | 683 | 10805 | 4549 | 6394 |
| 2166 | 3581.5 | 2855 | 1050.5 | 3952 | 1765.5 | 659 | 9926.5 | 4197 | 5982.5 |
| 3974 | 8279 | 3989 | 1243 | 5183 | 2011 | 725 | 17146 | 5918 | 11462 |
| 5239 | 14653 | 5213 | 1424 | 6269.5 | 2121.5 | 684 | 16272 | 6183 | 17707.5 |
| 7851 | 23679.5 | 7426.5 | 1770 | 8549.5 | 2516.5 | 717 | 12847.5 | 8210 | 26447 |
| 9344 | 22517 | 6650 | 1646 | 7674 | 2303 | 626 | 12404 | 8324 | 23619 |
| 9279 | 26865 | 7883 | 1842 | 8907 | 2556 | 710 | 13427 | 8736 | 28044 |
| 16500 | 34528.5 | 13414.5 | 3412.5 | 8501.5 | 2650.5 | 693 | 14633.5 | 9663.5 | 24797.5 |
| 9611.5 | 19862.5 | 6320 | 1634.5 | 7204 | 2284.5 | 766.5 | 15297 | 7645 | 20848 |
| 8929 | 14388 | 15500 | 3332 | 115766 | 16288 | 694 | 14124 | 5861 | 15592 |
| 5031 | 6852 | 4964 | 1440 | 31630 | 4473 | 774 | 15925 | 5434 | 8748 |
| 4870 | 6074 | 4018 | 1307 | 18850 | 3286 | 784 | 16580 | 5157 | 8013 |
| 4316.5 | 4175 | 3198.5 | 1143 | 9495 | 2587.5 | 732 | 12195 | 4755.5 | 6230.5 |
| 3824.5 | 4064 | 3071 | 1129 | 7109.5 | 2263.5 | 766.5 | 11317.5 | 4432 | 6204 |
| 3839.5 | 3923.5 | 3015 | 1098.5 | 6570 | 2266.5 | 736 | 10891.5 | 4148 | 6027 |
| 3308.5 | 4191 | 3777 | 1374.5 | 5913 | 2194.5 | 727 | 11447 | 4336.5 | 6292.5 |
| 4230.5 | 4581 | 3517 | 1263 | 6688.5 | 2295.5 | 806.5 | 14498.5 | 5252.5 | 6957.5 |
| 3389 | 3825 | 2894 | 1043 | 4635 | 1883 | 677 | 11257 | 3995 | 5569 |
| 3618.5 | 3969 | 2857.5 | 1030 | 4913 | 1898 | 689 | 11081 | 4312.5 | 5884.5 |
| 7401 | 6389.5 | 3477.5 | 1057 | 4019.5 | 1753.5 | 669 | 10559 | 5499.5 | 6515.5 |
| 5695 | 5008 | 3232 | 1051 | 5599 | 2008 | 735 | 10807 | 4972 | 6406 |
| 4159 | 4262 | 3079 | 1063 | 4631 | 1936 | 741 | 12001 | 4506 | 5997 |
| 858945 | 515605 | 123300 | 7758 | 7632 | 2303 | 811 | 87753 | 25928 | 17576 |
| 62123 | 33818.5 | 9459.5 | 1560.5 | 5906 | 2218.5 | 822 | 16156.5 | 6756 | 8637 |
| 22251 | 12655 | 5161 | 1305 | 6152 | 2315 | 744 | 12511 | 5153 | 7257 |
| 23601.5 | 14351 | 5543.5 | 1428.5 | 5740.5 | 2221.5 | 849 | 16494 | 6009.5 | 8836 |
| 11489 | 12706 | 5231 | 1539 | 7010 | 2351 | 844 | 18481 | 6366 | 12594 |
| 15733 | 24864 | 7116 | 1742 | 7595 | 2473 | 828 | 16409 | 9725 | 23094 |
| 13420 | 25285 | 7421 | 1751 | 8184 | 2437 | 711 | 13583 | 9199 | 24102 |

| | | | | | | | | | |
|---|---|---|---|---|---|---|---|---|---|
| 13292.5 | 26385.5 | 8029.5 | 1869 | 8778 | 2500 | 703 | 13464 | 9275.5 | 25640 |
| 10467 | 19961 | 5950 | 1420 | 6640 | 2063 | 561 | 14159 | 8879 | 20842 |
| 10568 | 23871 | 7610 | 1790 | 8151 | 2489 | 689 | 11116 | 8640 | 23860 |
| 6421.5 | 16005.5 | 5232.5 | 1368 | 5921.5 | 2026.5 | 605 | 14386 | 6195 | 17652.5 |
| 4544 | 11651 | 5927.5 | 1501 | 297635 | 27551 | 626 | 15408.5 | 5399 | 14044 |
| 3321 | 8276 | 4187 | 1220 | 6675 | 2334 | 653 | 14219.5 | 4559 | 11103 |
| 3252 | 4480 | 3255 | 1091 | 4780 | 1948 | 646 | 13569 | 4854 | 6441 |
| 2842 | 4560 | 3354 | 1158 | 4953 | 1993 | 724 | 13720 | 4931.5 | 7202.5 |
| 2032.5 | 3212.5 | 2567.5 | 966 | 3708.5 | 1714 | 625 | 9535.5 | 3663 | 5323 |
| 1789 | 2905 | 2350 | 937 | 3400 | 1635 | 591 | 8387 | 3345 | 4946 |
| 1851 | 2999 | 2373.5 | 934 | 3392 | 1606 | 591 | 9180 | 3377.5 | 4970.5 |
| 2491 | 3884 | 2915 | 1073 | 4086 | 1810 | 691 | 11675 | 4399 | 6172 |
| 2519 | 3647 | 2741 | 992 | 3748 | 1708 | 645 | 11320 | 4334 | 5765 |
| 2616 | 4003 | 3062.5 | 1070 | 4195.5 | 1821 | 710.5 | 11860 | 4691 | 6458 |
| 2672.5 | 4114 | 3015 | 1089 | 4171.5 | 1834 | 712.5 | 12813 | 4637 | 6608 |
| 2394 | 4226 | 2899 | 1082 | 4171 | 1811 | 700 | 11357 | 4522 | 6961 |
| 1988 | 3237.5 | 2587 | 985 | 3612 | 1689.5 | 636 | 8947.5 | 3781 | 5371 |
| 16772 | 9682 | 3901 | 1041 | 3675 | 1672 | 619 | 10174 | 3890 | 5607 |
| 3161.5 | 5699 | 3673.5 | 1244.5 | 4773 | 2015 | 749 | 11788.5 | 5107.5 | 8068.5 |
| 2282 | 3814 | 3003 | 1039 | 3914 | 1753.5 | 634 | 9795 | 4023.5 | 6082 |
| 2286 | 3916 | 3078 | 1087 | 4272 | 1841 | 685 | 11308 | 4421 | 6414 |
| 2345.5 | 3920 | 2981.5 | 1064 | 4025.5 | 1767 | 643.5 | 10689.5 | 4195 | 6030 |
| 5720 | 11448 | 5221 | 1445 | 6210 | 2185 | 744 | 13007 | 7023 | 14016 |
| 7633 | 21089.5 | 6583 | 1667 | 7559.5 | 2340.5 | 660 | 14668 | 8125 | 23413.5 |
| 8650.5 | 24691.5 | 8042 | 1845 | 9356.5 | 2532.5 | 659 | 11433 | 8621 | 27028 |
| 8041 | 22269 | 7868 | 1746 | 8812 | 2516 | 671 | 10991 | 7589 | 24176 |
| 8554.5 | 21464 | 7167 | 1759 | 8292 | 2448 | 695 | 11667.5 | 7990.5 | 23232.5 |
| 6085 | 15662 | 5375 | 1447 | 6577 | 2183 | 684 | 14339 | 7254 | 18820 |
| 5382.5 | 11982 | 6150 | 1849 | 14563 | 4973.5 | 690.5 | 14886 | 6542 | 14935 |
| 4653 | 8059 | 5832 | 1706 | 16587 | 4694 | 764 | 17359 | 6298 | 10703 |
| 3623 | 5525 | 4352 | 1340 | 7121 | 2479 | 736 | 16402 | 5351 | 7719 |
| 2193 | 3107 | 2618 | 961 | 3872 | 1740 | 608 | 10097 | 3761 | 5077 |
| 2061 | 3266 | 2714 | 1054 | 3987 | 1771 | 662 | 10083 | 3823 | 5489 |
| 2107 | 3427 | 2861 | 1097 | 4178 | 1844 | 710 | 10402.5 | 4056.5 | 5983.5 |
| 1881 | 3075.5 | 2459 | 977.5 | 3569 | 1665 | 630.5 | 9507 | 3574.5 | 5127 |
| 2384 | 3783 | 2922 | 1072 | 4063 | 1827 | 712 | 11855 | 4359 | 6339 |
| 2089 | 3263.5 | 2635.5 | 986 | 3554 | 1631.5 | 618 | 10226.5 | 3732.5 | 5424.5 |
| 2571 | 4051.5 | 3043 | 1152.5 | 4344.5 | 1874.5 | 724 | 11842.5 | 4710 | 6620 |
| 2457.5 | 4100 | 2945 | 1108 | 4177 | 1889 | 727 | 13166.5 | 4551 | 6627.5 |
| 2393 | 3913 | 2983 | 1137 | 4305 | 1843 | 746 | 11085 | 4677 | 6582 |
| 1957 | 3215 | 2617 | 1011 | 3687 | 1718 | 662 | 9278 | 3747 | 5534 |
| 6153 | 4830.5 | 2918 | 959 | 3262.5 | 1580 | 622 | 8873 | 3527 | 5117.5 |
| 2712 | 4112 | 3014 | 1084 | 3956 | 1773 | 682 | 9872 | 4015 | 5854 |
| 2516.5 | 4026.5 | 3186.5 | 1119 | 4249 | 1843 | 694 | 10441.5 | 4579.5 | 6376.5 |
| 2429 | 3948 | 3182 | 1145 | 4263 | 1903 | 745 | 11180 | 4704 | 6713 |
| 3110 | 4913 | 3566 | 1214 | 4752 | 1957 | 750 | 16067 | 5281 | 7620 |
| 5346 | 14357.5 | 5421.5 | 1475.5 | 6423 | 2229 | 743 | 16800.5 | 6450 | 17488 |
| 7019.5 | 17495.5 | 6188 | 1541 | 7241.5 | 2249 | 642 | 14288 | 7089.5 | 19586.5 |

| | | | | | | | | | |
|---|---|---|---|---|---|---|---|---|---|
| 10891.5 | 26983 | 8132.5 | 1886 | 8718.5 | 2626 | 742.5 | 11476 | 9596.5 | 28453 |
| 10450.5 | 23152 | 7730 | 1775.5 | 8405 | 2465.5 | 693 | 11801.5 | 9412 | 24494.5 |
| 11427 | 38106 | 38406 | 10082.5 | 9433.5 | 3487.5 | 621 | 14091 | 9000.5 | 25463 |
| 5981 | 18169.5 | 11083.5 | 3305 | 6657 | 2619.5 | 628 | 13635 | 6128.5 | 17901 |
| 4489 | 12929 | 8750 | 2693.5 | 660322 | 55955.5 | 675 | 14330.5 | 5420 | 15171.5 |
| 3863 | 9908 | 6336 | 2294 | 13328 | 3804 | 700 | 13451 | 5263 | 12436 |
| 3223 | 4655.5 | 3878.5 | 1303 | 9698.5 | 2496 | 642 | 13016 | 4805 | 6700 |
| 3160 | 4896 | 4058 | 1492 | 9243 | 2630 | 784 | 16012 | 5200 | 7689 |
| 2711.5 | 3663.5 | 3169.5 | 1186 | 8797.5 | 2305 | 681.5 | 11041 | 4194.5 | 5822 |
| 2521 | 3511 | 2945 | 1091 | 7709 | 2171 | 663 | 10220 | 3841 | 5659 |
| 3053 | 4083 | 3468 | 1240 | 7187 | 2247 | 735 | 11724 | 4678 | 6415 |
| 2582 | 3492 | 2776 | 1036 | 5805 | 1909 | 646 | 10356 | 3799 | 5612 |
| 2448.5 | 3433 | 2719 | 1034.5 | 5108.5 | 1852 | 646 | 11311 | 3813.5 | 5483 |
| 2913 | 3933 | 3145 | 1182 | 5344 | 2000 | 728 | 13418 | 4808 | 6620 |
| 3825 | 3913 | 2652 | 948 | 4006 | 1639 | 588 | 9080 | 3799 | 5271 |
| 3067.5 | 4056 | 3330 | 1178 | 5127.5 | 2000 | 764 | 13545 | 5011 | 6792.5 |
| 2592 | 3668 | 3007 | 1121 | 4538 | 1896 | 717 | 11086 | 4521 | 6423 |
| 57252 | 30637 | 8264.5 | 1502 | 5429 | 2143.5 | 825 | 18113 | 6170.5 | 8158 |
| 3266 | 5601 | 4351 | 1271 | 5145 | 2038 | 750 | 12222 | 5004 | 7459 |
| 2751 | 4652 | 3738.5 | 1286 | 5317.5 | 2080 | 772 | 11336 | 5225.5 | 7692 |
| 2359 | 4111 | 3486 | 1204 | 4890 | 2028 | 750 | 11440 | 4685 | 7000 |
| 3513 | 7199 | 3724 | 1217 | 5245 | 1988 | 712 | 16690 | 5583 | 10302 |
| 7129 | 20655 | 8435.5 | 2016 | 10113.5 | 2886.5 | 926 | 12539 | 8482 | 24439 |
| 9093 | 26320.5 | 7831.5 | 2060 | 9651.5 | 2916 | 765 | 10388 | 7605.5 | 26074 |
| 7880 | 26183 | 10784 | 2283 | 12931 | 3157 | 830 | 10077 | 7794 | 28621 |
| 9436 | 21272 | 6469 | 1601 | 7644 | 2331 | 637 | 11862 | 8739 | 22614 |
| 10004.5 | 23983.5 | 15475 | 5204.5 | 10857.5 | 4391 | 699 | 13411.5 | 9666.5 | 24994 |
| 6239 | 16901 | 9238 | 3323 | 7370 | 3096 | 637 | 15308 | 6540 | 18568 |
| 6077 | 14604 | 8986 | 3547.5 | 599395 | 49609 | 689 | 13506 | 6797.5 | 16959 |
| 3938 | 9098 | 6916 | 2936 | 12020 | 4023 | 698 | 15326 | 5260 | 11758 |
| 3231 | 6234 | 5882 | 2367 | 7718 | 3030 | 704 | 13773 | 4864 | 8840 |
| 2902 | 3923 | 3892 | 1503 | 5338 | 2226 | 624 | 12980 | 4638 | 5824 |
| 2944.5 | 6097.5 | 4793.5 | 1750 | 6271.5 | 2543.5 | 856 | 17106.5 | 5512 | 10357 |
| 2160 | 3511 | 3236 | 1239 | 4598 | 1995 | 692 | 10695 | 4081 | 5906 |
| 2610 | 4300.5 | 3641.5 | 1369 | 5070 | 2163 | 773 | 12857.5 | 5068 | 7307.5 |
| 2618 | 4144 | 3323.5 | 1233 | 4620.5 | 1989.5 | 731 | 13179 | 4812.5 | 6774.5 |
| 2700 | 4141 | 3243 | 1198 | 4489 | 1924 | 703 | 13374 | 4842 | 6691 |
| 2728 | 4088 | 3106 | 1152 | 4325 | 1885 | 708 | 13758 | 4902 | 6593 |
| 2828 | 4576 | 3486 | 1246 | 4838 | 2031 | 779 | 15166 | 5153 | 7536 |
| 2373.5 | 4138 | 3012 | 1154.5 | 4292.5 | 1884.5 | 711 | 12822 | 4500.5 | 6896 |
| 1921 | 3259 | 2697 | 1023 | 3761 | 1734 | 663 | 9324 | 3784 | 5557 |
| 4428 | 6270.5 | 4027.5 | 1262.5 | 4634.5 | 1993 | 735 | 13702 | 4937 | 7029 |
| 4367 | 6254 | 4011 | 1232 | 4599.5 | 1986 | 714 | 12191 | 4804.5 | 6792 |
| 3101.5 | 5884 | 3845 | 1303 | 4850 | 2077 | 787 | 14688 | 5343 | 9092.5 |
| 2576 | 4766 | 3643.5 | 1266 | 4793.5 | 2027.5 | 744 | 14168.5 | 4800.5 | 7946.5 |
| 4108 | 8950.5 | 4382 | 1431 | 5654.5 | 2272.5 | 903 | 20118.5 | 6857.5 | 14345 |
| 7193.5 | 17426 | 5832.5 | 1645 | 7178 | 2425 | 799.5 | 14632.5 | 8290.5 | 21676.5 |
| 9141.5 | 21963 | 6627.5 | 1677 | 7736.5 | 2395 | 660.5 | 14216.5 | 9117 | 24156 |

| | | | | | | | | | |
|---|---|---|---|---|---|---|---|---|---|
| 10292 | 20874 | 7823 | 1618 | 8708 | 2355 | 625 | 14687 | 9512 | 22612 |
| 9880 | 22144 | 6722 | 1611 | 7695 | 2327 | 627 | 12730 | 9058 | 23581 |
| 8885 | 27308 | 14507 | 4455 | 13337 | 4838 | 769 | 11310 | 8372 | 28917 |
| 6202 | 15308 | 6147 | 1974 | 6297 | 2539 | 618 | 15131 | 6597 | 17432 |
| 7997.5 | 19293.5 | 10301 | 3002 | 612871.5 | 48359.5 | 797 | 15656 | 8419.5 | 21803.5 |
| 3816 | 10293 | 5348 | 2006 | 8021 | 3151 | 722 | 14861 | 5264 | 13614 |
| 3010 | 5983 | 4188 | 1520 | 5602 | 2339 | 675 | 13635 | 4454 | 8465 |
| 2847 | 4324 | 3403 | 1276 | 4790 | 2096 | 662 | 11920 | 4681 | 6531 |
| 2766 | 4299 | 3390 | 1282.5 | 4858 | 2064 | 738 | 13374 | 4931.5 | 6968 |
| 2412.5 | 3706 | 3074 | 1144 | 4274.5 | 1873.5 | 684 | 10738 | 4419.5 | 6042.5 |
| 2650 | 4143 | 3316 | 1193 | 4485 | 1958 | 730 | 13281 | 4924 | 6809 |
| 2456 | 3750 | 2895 | 1057 | 3944 | 1781 | 651 | 11243 | 4309 | 5925 |
| 3197.5 | 4960.5 | 3507 | 1245.5 | 4898.5 | 2036.5 | 773.5 | 16212 | 5669 | 7866 |
| 2816 | 4224 | 3190 | 1141 | 4306 | 1859 | 701 | 12162 | 4892 | 6727 |
| 3139.5 | 4773 | 3516.5 | 1269 | 4839.5 | 2036 | 782 | 13247.5 | 5640 | 7763 |
| 2109 | 3366 | 2653 | 996 | 3715 | 1694 | 634 | 9934 | 3851 | 5559 |
| 2765 | 4837 | 3474 | 1248 | 4966 | 2078 | 797 | 13157 | 5342 | 7908 |
| 24511.5 | 12891 | 5234 | 1350 | 4758 | 2035.5 | 759 | 14143.5 | 5384.5 | 7239 |
| 3291 | 5285 | 3733 | 1146 | 4198.5 | 1806 | 679 | 11140.5 | 4614.5 | 6471.5 |
| 3175 | 5117 | 3918 | 1307 | 4877 | 2027 | 750 | 12406 | 5178 | 7418 |
| 2403 | 4086 | 3072 | 1093 | 4233 | 1822 | 673 | 10942 | 4380 | 6528 |
| 3955 | 7217 | 3846 | 1261 | 5132 | 2032 | 744 | 16066 | 6370 | 10576 |
| 6762 | 16725.5 | 5764 | 1556 | 6720 | 2275 | 739 | 15828 | 8213 | 20175.5 |
| 8012 | 21180 | 6591 | 1648 | 7551 | 2385 | 673 | 14845 | 8112 | 23270 |
| 0 | 0 | 0 | 0 | 0 | 0 | 0 | 0 | 0 | 0 |
| 4857.5 | 5903.5 | 2924.5 | 931 | 3732.5 | 1498.5 | 190 | 13205 | 4872 | 6222.5 |
| 6049 | 13633.5 | 5775.5 | 2064 | 6624 | 2641.5 | 269 | 14659.5 | 5850 | 13579.5 |
| 5728 | 10658 | 4587 | 1425.5 | 5390.5 | 2061 | 427 | 14573 | 5965.5 | 11460 |
| 4162.5 | 6343.5 | 31161 | 11005.5 | 751794.5 | 75524.5 | 553 | 15017 | 5208 | 7934 |
| 3670.5 | 6300.5 | 10528 | 3508 | 15144 | 5176.5 | 607 | 14219.5 | 4795.5 | 8140 |
| 3707 | 6781 | 7424 | 2453 | 9819 | 3512 | 683 | 14097 | 5143 | 9145 |
| 3297 | 6663 | 5894 | 2042.5 | 7562.5 | 2917 | 728.5 | 14054 | 4801 | 9574.5 |
| 3115 | 5595 | 5204 | 1816 | 6906 | 2671 | 739 | 13996 | 4685 | 8159 |
| 2927 | 4509 | 4196 | 1454 | 5385 | 2206 | 692 | 15716 | 4499 | 6768 |
| 2778.5 | 4287 | 3885 | 1375.5 | 5156 | 2172 | 732 | 16351.5 | 4715 | 6826 |
| 2860.5 | 4481 | 3835 | 1381 | 5037.5 | 2114 | 731.5 | 16127 | 4666.5 | 6807 |
| 2717 | 3973 | 3282 | 1174 | 4361 | 1894 | 681 | 15705 | 4599 | 6353 |
| 2733 | 4169 | 3337.5 | 1234 | 4595.5 | 1951 | 716 | 15269 | 4612 | 6540 |
| 2629 | 3725 | 2961.5 | 1093 | 4126.5 | 1795 | 681.5 | 13618.5 | 4514 | 6052 |
| 2612 | 3862.5 | 3231.5 | 1163 | 4358 | 1911 | 724 | 14676.5 | 4727.5 | 6470 |
| 2417 | 3596 | 3001 | 1122 | 4142 | 1821 | 697 | 12941 | 4459 | 6082 |
| 34812 | 18218 | 5853 | 1268 | 4320 | 1857 | 707 | 13741 | 4943 | 6551 |
| 3009 | 4754 | 3750 | 1238 | 4659 | 1964 | 740 | 14578 | 4927 | 6983 |
| 2611 | 4170 | 3457 | 1222 | 4513 | 1973 | 721 | 12806 | 4884 | 6676 |
| 2383.5 | 3828 | 3190 | 1156 | 4290.5 | 1894 | 715 | 12586.5 | 4647 | 6524.5 |
| 4429 | 8973 | 4116 | 1290 | 5321 | 2070 | 737 | 16664 | 6503 | 12326 |
| 7682 | 18248 | 5822 | 1548 | 7134 | 2306 | 720 | 14200 | 8839 | 21682 |
| 8853 | 22450 | 6805.5 | 1728 | 7937.5 | 2438.5 | 689 | 12495.5 | 8809 | 24704.5 |

| | | | | | | | | | |
|---|---|---|---|---|---|---|---|---|---|
| 8953 | 21530 | 8546 | 1825 | 9677 | 2523 | 674 | 12930 | 8754 | 23492 |
| 9573 | 24361 | 7509 | 1821 | 8474 | 2558 | 684 | 12791 | 9221 | 26013 |
| 9613 | 22512.5 | 7344 | 2117 | 7980.5 | 2702 | 674.5 | 11603 | 9064.5 | 24600.5 |
| 8122 | 18323 | 5999 | 1582 | 6992 | 2296 | 718 | 13719 | 9072 | 21957 |
| 6044 | 13903 | 8789.5 | 2652.5 | 752073 | 61843.5 | 689.5 | 13243 | 6998 | 16969.5 |
| 3690.5 | 8309.5 | 5500.5 | 1791 | 8146 | 2938.5 | 679 | 15183 | 5096 | 11061.5 |
| 3104.5 | 5469 | 4260 | 1392 | 5938.5 | 2278 | 697.5 | 15893.5 | 4577 | 7937 |
| 2709.5 | 4110.5 | 3547 | 1270.5 | 4921.5 | 2060 | 689 | 15582 | 4475 | 6268.5 |
| 2344 | 3512 | 3167 | 1164 | 4395.5 | 1872 | 675 | 11786.5 | 4257 | 5818 |
| 2044 | 3411.5 | 2932.5 | 1105.5 | 4176 | 1836.5 | 680 | 10697.5 | 4008.5 | 5706.5 |
| 1989 | 3279 | 2928 | 1140 | 4073 | 1836 | 695 | 10229 | 3741 | 5607 |
| 2102.5 | 3395.5 | 2857 | 1091 | 4052 | 1787 | 685.5 | 11022.5 | 4040 | 5839 |
| 2178 | 3511 | 2981 | 1122 | 4152 | 1848 | 708 | 11163 | 4197 | 5883 |
| 2008 | 3309 | 2819.5 | 1048.5 | 3843.5 | 1747 | 687 | 10479 | 3918.5 | 5702 |
| 2044 | 3367 | 2847 | 1106 | 3913 | 1801 | 683 | 9925 | 3903 | 5743 |
| 2152 | 3381.5 | 2914 | 1127 | 4106 | 1839 | 720 | 10564 | 4197 | 6026.5 |
| 1878 | 3183 | 2742 | 1097 | 3933 | 1792 | 685 | 9795 | 3623 | 5531 |
| 33353 | 18221 | 5861.5 | 1235 | 4196.5 | 1870 | 702 | 12354.5 | 4508 | 6239 |
| 2567 | 4159 | 3585 | 1180 | 4208 | 1859 | 693 | 11117 | 4419 | 6272 |
| 2347 | 3819.5 | 3240 | 1163 | 4302 | 1859.5 | 707 | 10482.5 | 4543 | 6448 |
| 2251 | 3768.5 | 3244 | 1156.5 | 4327 | 1879 | 709 | 11460 | 4579.5 | 6388.5 |
| 3683 | 7778 | 3737 | 1199 | 4876 | 1889 | 686 | 16691 | 5416 | 10618 |
| 5996 | 15007 | 5420.5 | 1499 | 6391 | 2182 | 714.5 | 14332.5 | 6967 | 18216 |
| 9597 | 21438 | 6762 | 1668 | 7528 | 2331 | 662 | 14422 | 9372 | 23351 |
| 9927.5 | 24871 | 7567 | 1780 | 8417.5 | 2460 | 663 | 13485 | 9455 | 26596.5 |
| 9155 | 24530 | 7951 | 1791 | 8722 | 2560 | 695 | 10814 | 8455 | 25898 |
| 8682 | 19855 | 6152 | 1621 | 7369 | 2321 | 652 | 13035 | 8906 | 21925 |
| 7550.5 | 17978.5 | 6006.5 | 1632 | 7020.5 | 2324 | 722 | 13305.5 | 8379 | 20947.5 |
| 4645.5 | 12420.5 | 6388 | 1722 | 539365 | 58670 | 685.5 | 14100 | 5403 | 15412 |
| 3190 | 5470.5 | 4062.5 | 1230 | 5708.5 | 2246 | 661 | 15896.5 | 4398.5 | 7386 |
| 3265 | 6350 | 4009.5 | 1327 | 5567.5 | 2253 | 730 | 16599 | 5026.5 | 9289.5 |
| 2644.5 | 3756 | 3147 | 1158 | 4397 | 1909 | 680 | 13244.5 | 4732.5 | 5841 |
| 2140 | 3446 | 2908 | 1072 | 4011 | 1816 | 695 | 10362 | 4002 | 5785 |
| 2100.5 | 3438 | 2983.5 | 1127 | 4135.5 | 1812 | 699.5 | 10469.5 | 3961.5 | 5783 |
| 2059 | 3415 | 2994 | 1098 | 3979 | 1787 | 706 | 10542 | 3931 | 5832 |
| 2435 | 3753 | 3187 | 1168 | 4386 | 1922 | 728 | 12642 | 4599 | 6460 |
| 2226 | 3541 | 2862 | 1091 | 3911 | 1776 | 696 | 11136 | 4149 | 5922 |
| 2355 | 3715 | 3077 | 1153 | 4318 | 1875 | 750 | 12119 | 4512 | 6307 |
| 2637 | 3770.5 | 3016.5 | 1117.5 | 4125 | 1825 | 727 | 11390.5 | 4428.5 | 6146 |
| 2088 | 3389 | 2765 | 1054 | 3836 | 1762 | 692 | 9669 | 3776 | 5605 |
| 2076 | 3505.5 | 2956 | 1121.5 | 4116 | 1827 | 724 | 10245.5 | 4059 | 6065 |
| 6555 | 5137 | 3408 | 1148 | 4185 | 1829 | 706 | 10428 | 4323 | 6287 |
| 2928 | 4380 | 3548 | 1246 | 4716 | 2019 | 779 | 11147 | 4789 | 7011 |
| 2543.5 | 3958.5 | 3340.5 | 1184 | 4420.5 | 1902 | 714 | 11188.5 | 4682.5 | 6545 |
| 2389 | 3978 | 3262 | 1171 | 4455 | 1901 | 743 | 10866 | 4704 | 6658 |
| 3151 | 4826 | 3600 | 1229 | 4654 | 1926 | 712 | 14464 | 5027 | 7359 |
| 5959 | 15273 | 5492 | 1528 | 6509 | 2267 | 734 | 13907 | 6774 | 18436 |
| 9232 | 24342 | 7354 | 1824 | 8748 | 2527 | 699 | 12559 | 8920 | 26665 |

| | | | | | | | | | |
|---|---|---|---|---|---|---|---|---|---|
| 9432 | 27832 | 9135 | 2058 | 10079 | 2734 | 725 | 9561 | 8165 | 28956 |
| 8700.5 | 24710.5 | 8831.5 | 1935.5 | 10244.5 | 2660.5 | 727 | 12029 | 8604 | 26914 |
| 9126.5 | 24387 | 8486 | 2308.5 | 8900.5 | 2917 | 709 | 11605 | 8962.5 | 26644.5 |
| 5929 | 17232 | 5662 | 1509.5 | 6587 | 2194.5 | 688 | 12904 | 6443.5 | 19503 |
| 4932 | 12348 | 6394 | 1654 | 504895 | 49214 | 664 | 14915 | 5881 | 15222 |
| 3597 | 8126 | 4347 | 1316 | 6504 | 2424 | 684 | 14143.5 | 4835.5 | 10736.5 |
| 3365 | 6760 | 4052 | 1306 | 5704 | 2192 | 729 | 15927 | 5185 | 9786 |
| 2793 | 4349 | 3297 | 1180 | 4680 | 1969 | 699 | 14169 | 4725 | 6570 |
| 2780 | 4427 | 3419 | 1244 | 4850 | 2017 | 765 | 12799 | 5250 | 7287 |
| 2390 | 3692 | 3023 | 1125 | 4282 | 1882 | 704 | 10461 | 4536 | 6317 |
| 2627 | 4225 | 3422 | 1236 | 4775 | 2005 | 782 | 12863 | 4985 | 7213 |
| 2546 | 4016 | 3237 | 1202 | 4498 | 1941 | 738 | 12179 | 4808 | 6554 |
| 2611 | 4099 | 3198 | 1176 | 4472 | 1874 | 742 | 11981 | 4852 | 6752 |
| 2668.5 | 4139 | 3226 | 1173 | 4536 | 1934.5 | 750 | 13090.5 | 4916 | 6994.5 |
| 2607.5 | 4060.5 | 3111.5 | 1157.5 | 4476.5 | 1895 | 731 | 11572 | 4625 | 6600 |
| 2406 | 3787 | 3120 | 1177 | 4295 | 1888 | 723 | 10853 | 4560 | 6427 |
| 2439 | 3905 | 3120 | 1175 | 4469 | 1932 | 741 | 11325 | 4826 | 6609 |
| 24945 | 13167.5 | 5154.5 | 1269.5 | 4455.5 | 1913 | 745.5 | 13507 | 5014 | 6908 |
| 2936 | 4675 | 3717.5 | 1227 | 4532.5 | 1931 | 739.5 | 12216.5 | 4936 | 6938 |
| 2683 | 4841 | 3543 | 1222 | 4714 | 1965 | 747 | 11369 | 5009 | 7779 |
| 2507 | 3992.5 | 3222 | 1179.5 | 4474 | 1881.5 | 722 | 11012 | 4791 | 6688 |
| 4743 | 10841 | 4789.5 | 1460 | 6258 | 2244 | 803 | 16758.5 | 7210.5 | 15443 |
| 6531 | 16586.5 | 5562.5 | 1509 | 6623 | 2179 | 687 | 16299.5 | 7480.5 | 19532.5 |
| 10404 | 24981 | 7582 | 1843 | 8436 | 2515 | 707 | 14836 | 9963 | 26925 |
| 8752 | 26943 | 10866 | 2252 | 11702 | 3035 | 780 | 8756 | 7343 | 26395 |
| 8735 | 25798.5 | 8068.5 | 1874 | 9160.5 | 2601.5 | 688 | 11492 | 8490.5 | 28143 |
| 9679 | 24024.5 | 7617 | 1908.5 | 8570 | 2572.5 | 713 | 13089 | 9678.5 | 26874 |
| 8614 | 19444 | 6134 | 1621 | 7113 | 2314 | 723.5 | 14494.5 | 9247.5 | 22180.5 |
| 5205 | 13623.5 | 6714 | 1723 | 562268 | 62632.5 | 711 | 16171.5 | 6320.5 | 16513.5 |
| 3686 | 9864 | 4487 | 1343 | 6285 | 2352.5 | 711 | 13172 | 4875.5 | 13343 |
| 2975 | 5376 | 3771.5 | 1215 | 5012.5 | 2016 | 728 | 15868 | 4457 | 7904.5 |
| 2816 | 4549 | 3363 | 1162 | 4726 | 1967 | 723 | 15408 | 4797 | 7128 |
| 2627 | 4044 | 3181 | 1154 | 4482 | 1914 | 727.5 | 11568 | 4790 | 6604 |
| 2493.5 | 3790 | 3108 | 1134 | 4290.5 | 1878 | 720 | 11136 | 4661.5 | 6447 |
| 2446 | 3816 | 3126 | 1150 | 4291 | 1893 | 738 | 11011 | 4638 | 6483 |
| 2756 | 4522.5 | 3429 | 1242.5 | 4707.5 | 1961 | 792 | 13351.5 | 5212 | 7529.5 |
| 2662 | 4179 | 3263 | 1223 | 4573 | 1986 | 740 | 13367 | 5218 | 6872 |
| 2653 | 4448 | 3375 | 1233 | 4817.5 | 1969 | 779 | 11838.5 | 5162 | 7457.5 |
| 2453 | 3968 | 3107 | 1183 | 4344.5 | 1904 | 737 | 11083.5 | 4512 | 6485 |
| 2307.5 | 3521 | 2886 | 1111.5 | 4073 | 1855.5 | 684 | 11498 | 4174.5 | 5890.5 |
| 2413 | 3878 | 3180 | 1197 | 4411 | 1897 | 734 | 11239 | 4495 | 6389 |
| 5911 | 5888 | 3703 | 1158 | 4147 | 1844 | 722 | 11738 | 4673 | 6274 |
| 4135 | 7270 | 4300 | 1407 | 5397 | 2188 | 868 | 15994 | 5857 | 9888 |
| 2477.5 | 4040.5 | 3206 | 1109 | 4181.5 | 1843 | 708 | 10591.5 | 4437.5 | 6443 |
| 2519 | 4158.5 | 3336.5 | 1182 | 4557.5 | 1923 | 745 | 10886 | 4823 | 6950.5 |
| 3375 | 6485 | 3789 | 1227 | 4909 | 1963 | 703 | 16077 | 5144 | 9311 |
| 7922 | 20447.5 | 7603 | 1866 | 9267.5 | 2626 | 789 | 14591.5 | 9660.5 | 23976 |
| 10296.5 | 22085.5 | 7013.5 | 1710 | 7843.5 | 2380 | 685 | 14117.5 | 9573 | 23295.5 |

| | | | | | | | | | |
|---|---|---|---|---|---|---|---|---|---|
| 10261 | 24240 | 9167 | 1943 | 10165 | 2636 | 702 | 12779 | 9080 | 25273 |
| 9348 | 25717 | 8175 | 1914 | 9214 | 2624 | 725 | 12160 | 9254 | 28485 |
| 15316 | 75390 | 16212 | 3720 | 9643 | 2848 | 734 | 10905 | 8372 | 34343 |
| 10697.5 | 18962.5 | 5896 | 1532.5 | 6417.5 | 2122.5 | 686 | 16169 | 7433 | 19021.5 |
| 12270 | 21229 | 13423 | 3197 | 101012 | 10572 | 910 | 13262 | 10238 | 23508 |
| 7862.5 | 12926.5 | 5940.5 | 1652 | 19706 | 3515.5 | 785.5 | 15938 | 5572.5 | 14513 |
| 9799 | 8582 | 4807 | 1438 | 11590 | 2810 | 863 | 17652 | 6595 | 10250 |
| 7414 | 5622 | 3911 | 1269 | 8722.5 | 2538 | 749.5 | 15402 | 5020 | 6916.5 |
| 8391 | 5484 | 3875 | 1271 | 7812 | 2529 | 836 | 15676 | 5551 | 7119 |
| 7672.5 | 5150.5 | 3582.5 | 1227 | 8221 | 2602 | 809 | 13268.5 | 5203 | 6702 |
| 7207 | 5801 | 4922 | 1634 | 5386 | 2140 | 790 | 11914 | 4986 | 6994 |
| 7309.5 | 5465 | 3891 | 1402 | 6840 | 2394.5 | 809 | 15712.5 | 5176 | 7319.5 |
| 6357.5 | 4814.5 | 3372 | 1170 | 5211 | 1995 | 770 | 12387.5 | 4882 | 6397.5 |
| 7229 | 5375 | 3797 | 1274 | 6497 | 2279 | 826 | 13540 | 5361 | 7149 |
| 14961.5 | 9519.5 | 4751.5 | 1294 | 5974 | 2137.5 | 813 | 13950.5 | 7262 | 8489 |
| 9755.5 | 6418 | 3914.5 | 1257 | 6269.5 | 2240 | 831 | 12479.5 | 5703 | 7147.5 |
| 8652 | 5650.5 | 3837.5 | 1299.5 | 6474 | 2250 | 885.5 | 13600.5 | 5683.5 | 7531.5 |
| 862119 | 559491 | 138174 | 8529.5 | 8059.5 | 2506 | 794 | 97419 | 25975.5 | 18864.5 |
| 59109 | 31357 | 9273 | 1737 | 6771 | 2272 | 829 | 17200 | 6415 | 8351 |
| 25457.5 | 14308 | 5563 | 1441 | 6851.5 | 2538.5 | 833.5 | 13965 | 5905.5 | 8866 |
| 25551 | 14564.5 | 5351 | 1391.5 | 5501.5 | 2111.5 | 799 | 14979.5 | 5889.5 | 8360.5 |
| 14012 | 14665 | 5603 | 1587 | 7639 | 2490 | 906 | 17641 | 6598 | 14348 |
| 16003 | 22942 | 5930 | 1718 | 7263 | 2312 | 746 | 14088 | 8048 | 18650 |
| 15060 | 29878 | 8960 | 1925 | 9901 | 2641 | 786 | 14760 | 9873 | 28410 |
| 14118.5 | 29626 | 10389.5 | 2130 | 11000.5 | 2715.5 | 790.5 | 13060.5 | 9961.5 | 28302.5 |
| 10093 | 26554 | 8787 | 1928 | 9841 | 2640 | 701 | 12427 | 9180 | 28580 |
| 9526 | 23483 | 7537 | 1873 | 8456 | 2489 | 681 | 11613 | 9119 | 25748 |
| 8128 | 22088 | 7002 | 1761 | 8383 | 2526 | 780 | 13128 | 8959 | 25415 |
| 4850.5 | 13591.5 | 7190.5 | 2139.5 | 806981 | 62197.5 | 684 | 15145.5 | 6046 | 16574 |
| 4154.5 | 10859.5 | 4994 | 1436 | 8097 | 2640 | 703 | 13922 | 5500.5 | 13993 |
| 3659 | 6476 | 4255 | 1326 | 6533.5 | 2351.5 | 732 | 16496 | 5146 | 9070.5 |
| 2912.5 | 5033 | 3743 | 1255.5 | 5570 | 2135 | 759 | 15523.5 | 4937.5 | 7780 |
| 2579.5 | 3953 | 3254 | 1156.5 | 4694 | 1923 | 724 | 13095.5 | 4606.5 | 6371 |
| 2506 | 3857 | 3176 | 1164 | 4580 | 1910 | 740 | 10979 | 4634 | 6534 |
| 2272 | 3445 | 2889.5 | 1082 | 4062 | 1781 | 689 | 10837 | 4167 | 5871 |
| 2347 | 3568 | 2880 | 1063 | 4077 | 1781 | 676 | 10311 | 4340 | 5926 |
| 2599 | 3819 | 3011 | 1112 | 4262 | 1827 | 709 | 12359 | 4703 | 6263 |
| 2588 | 3889 | 3080 | 1118 | 4306 | 1854 | 715 | 12408 | 4781 | 6433 |
| 2485 | 3810 | 3032 | 1104 | 4223 | 1853 | 718 | 11028 | 4525 | 6259 |
| 2452 | 3818 | 3103 | 1146 | 4375 | 1916 | 728 | 10865 | 4645 | 6460 |
| 2069 | 3458.5 | 2823.5 | 1084 | 4034 | 1742 | 689 | 9819 | 3969.5 | 5903.5 |
| 19255.5 | 10388.5 | 4460.5 | 1186 | 4250 | 1831 | 697 | 11839.5 | 4774.5 | 6378 |
| 3001 | 4607 | 3492 | 1176 | 4399 | 1921 | 711 | 10869 | 4990 | 6558 |
| 2587 | 4089 | 3292 | 1141 | 4378 | 1863 | 699 | 10959 | 4793 | 6545 |
| 2635 | 4465 | 3429.5 | 1209 | 4758 | 1993.5 | 759.5 | 12782.5 | 5022 | 7541.5 |
| 3724 | 6115 | 3679 | 1229 | 4913 | 1918 | 706 | 16556 | 6044 | 9088 |
| 7661 | 18660 | 5950.5 | 1566 | 7118 | 2293 | 695 | 16356 | 8774.5 | 22154 |
| 9331 | 23416 | 7733 | 1826 | 8794.5 | 2517 | 698 | 14052.5 | 9289.5 | 25590.5 |

| | | | | | | | | | |
|---|---|---|---|---|---|---|---|---|---|
| 8102 | 23290 | 7995 | 1924 | 9923 | 2636 | 660 | 9696 | 7641 | 24521 |
| 9889.5 | 22322.5 | 6856 | 1667.5 | 7511 | 2312 | 633 | 13237 | 9109 | 23702.5 |
| 9775 | 22992 | 7623 | 1982 | 8174 | 2582 | 710 | 12095 | 8989 | 24541 |
| 5564 | 14187 | 4827 | 1321 | 5612 | 1951 | 626 | 14943 | 5874 | 16451 |
| 7896 | 17115 | 8334 | 2150 | 21470 | 6274 | 820 | 12157 | 8755 | 20592 |
| 3997 | 10328 | 4976 | 1482 | 9116 | 3065 | 706 | 15415 | 5316 | 13278 |
| 3431 | 7636 | 4492 | 1405 | 6831 | 2534 | 771 | 14525 | 5335 | 11261 |
| 2871 | 5367 | 4102 | 1378 | 5924 | 2320 | 808 | 15959 | 4956 | 8489 |
| 2621.5 | 4265.5 | 3362.5 | 1235.5 | 4841.5 | 1988.5 | 762 | 12163 | 4901 | 6975 |
| 2337 | 3765 | 3220 | 1211 | 4665 | 1973 | 734 | 11221 | 4350 | 6168 |
| 2296.5 | 3689.5 | 3112 | 1171.5 | 4266 | 1924 | 744 | 11244 | 4360 | 6223 |
| 2625.5 | 4337.5 | 3362.5 | 1235.5 | 4654.5 | 1983 | 797 | 13267 | 5038.5 | 7384 |
| 2634 | 4033.5 | 3262 | 1183.5 | 4496 | 1926.5 | 771 | 12631.5 | 4853 | 6922 |
| 2417.5 | 3773 | 3009 | 1153 | 4196.5 | 1821.5 | 732 | 12091.5 | 4543.5 | 6324.5 |
| 2750 | 4447 | 3563 | 1260 | 4850 | 2015 | 835 | 12509 | 5235 | 7426 |
| 2363 | 3835.5 | 3154.5 | 1179 | 4244.5 | 1871.5 | 763 | 11148 | 4497.5 | 6476 |
| 2490 | 4098 | 3481 | 1288 | 4728 | 2059 | 810 | 11414 | 4824 | 6619 |
| 6995 | 7461 | 4527 | 1363 | 4809 | 2052 | 858 | 15888 | 5791 | 8613 |
| 3709.5 | 7205.5 | 4558 | 1487 | 5741 | 2275 | 940 | 17349.5 | 6352.5 | 11205 |
| 3164 | 5473 | 3984 | 1327 | 4910 | 2032 | 838 | 13632 | 5843 | 8637 |
| 2643 | 4351 | 3491 | 1285 | 4650 | 1960 | 785 | 12723 | 5227 | 7184 |
| 3256 | 5935 | 4096 | 1399 | 5293 | 2179 | 818 | 13722 | 6296 | 9297 |
| 6228.5 | 17000.5 | 6053 | 1655 | 7162 | 2389.5 | 825 | 15669 | 7905.5 | 21478.5 |
| 8393 | 20537 | 6682 | 1697 | 7711 | 2372 | 702 | 13259 | 8463 | 23059 |
| 9523.5 | 27775.5 | 8644.5 | 2023.5 | 9855.5 | 2768 | 723.5 | 11477 | 8357 | 29624.5 |
| 9275 | 24318 | 7924 | 1762 | 9139 | 2485 | 687 | 14577 | 9226 | 26100 |
| 11129 | 61516 | 68933 | 17407 | 14638 | 5062 | 724 | 10342 | 8500 | 35173 |
| 7051 | 20012 | 8403 | 2433 | 7500 | 2613 | 694 | 15813 | 7641 | 21799 |
| 5310 | 15781 | 9813 | 2869 | 843612 | 72431 | 759 | 13915 | 6260 | 18397 |
| 4313 | 9681 | 5941 | 1803 | 20317 | 3849.5 | 714 | 15361 | 5520 | 12609 |
| 3900 | 8026 | 5677 | 1869 | 19156.5 | 3675.5 | 781 | 14900 | 5470 | 11018.5 |
| 3398 | 4829 | 4394 | 1540 | 14473 | 3133 | 755 | 15467 | 4988 | 6990 |
| 2835 | 3978 | 3495 | 1287 | 8744 | 2444 | 711 | 11395 | 4587 | 6450 |
| 2899 | 4311 | 3872 | 1422 | 7832 | 2441 | 800 | 13313 | 4920 | 7129 |
| 2738 | 3946 | 3699 | 1415 | 6824 | 2333 | 728 | 11510 | 4646 | 6480 |
| 2352 | 3315 | 2910 | 1161 | 5392 | 1984 | 656 | 9933 | 3872 | 5520 |
| 2573 | 3791 | 3326 | 1204 | 5418 | 2040 | 727 | 13521 | 4686 | 6403 |
| 2348 | 3580 | 3007 | 1141 | 4901 | 1928 | 706 | 11287 | 4200 | 6043 |
| 2839 | 3755 | 3126 | 1131 | 4700 | 1923 | 709 | 10980 | 4592 | 6158 |
| 2340 | 3629 | 3159 | 1183 | 4942 | 1966 | 734 | 10423 | 4243 | 6188 |
| 1954 | 3232 | 2795 | 1066 | 4644 | 1864 | 696 | 9400 | 3795 | 5678 |
| 75057 | 39413.5 | 10442.5 | 1383 | 4901.5 | 1882 | 688 | 14640.5 | 5188 | 6476.5 |
| 2800 | 4244 | 3817 | 1189 | 5115 | 1920 | 681 | 11420 | 4456 | 6086 |
| 2351 | 3790 | 3124 | 1116 | 5142 | 1916 | 660 | 10398 | 4213 | 6089 |
| 2152.5 | 3577 | 3004.5 | 1111 | 4768 | 1854 | 672 | 9837 | 4131 | 6068 |
| 3297 | 4890 | 3462 | 1166 | 5102 | 1946 | 709 | 14806 | 5454 | 7369 |
| 7037.5 | 17638.5 | 6391 | 1702 | 7787.5 | 2468 | 795 | 14149 | 7915 | 20724 |
| 8919 | 19829 | 6332.5 | 1549.5 | 7630.5 | 2273 | 636 | 15930 | 8765 | 21955 |

| | | | | | | | | | |
|---|---|---|---|---|---|---|---|---|---|
| 7858 | 31875 | 10378 | 2162 | 12533 | 3080 | 772 | 9516 | 7014 | 28978 |
| 9267 | 21923 | 6885 | 1625 | 7928 | 2372 | 636 | 14181 | 9035 | 24126 |
| 8211 | 22800.5 | 12796 | 4155 | 10155 | 3904 | 642 | 11702.5 | 7653 | 23835 |
| 7124 | 19560.5 | 9255 | 3193 | 8383 | 3373.5 | 688 | 12721.5 | 7676.5 | 21761.5 |
| 4242 | 11297 | 6811 | 2631 | 30865 | 7763 | 586 | 14000 | 4978 | 13617 |
| 3723.5 | 9694.5 | 6566.5 | 2553 | 10246.5 | 3644 | 703 | 13969.5 | 5024.5 | 12622.5 |
| 3349 | 5258 | 4690 | 1793 | 6944 | 2661 | 651 | 14072 | 5008 | 7541 |
| 2655 | 3950 | 3879 | 1471 | 5530 | 2252 | 644 | 12328 | 4428 | 6159 |
| 1927 | 3159 | 2991 | 1128 | 4218 | 1871 | 623 | 9700 | 3685 | 5221 |
| 1769.5 | 2934.5 | 2715 | 1058 | 3907.5 | 1754 | 617 | 9073.5 | 3439 | 5068.5 |
| 2198.5 | 3883 | 3230 | 1247.5 | 4723.5 | 2043 | 719.5 | 11771.5 | 4395.5 | 6635.5 |
| 2018 | 3289 | 2802 | 1096 | 3926.5 | 1812 | 646 | 9573.5 | 3810 | 5464 |
| 2104.5 | 3405 | 2767.5 | 1055.5 | 3943.5 | 1776 | 651 | 10393.5 | 3949 | 5706.5 |
| 2195.5 | 3444 | 2782.5 | 1073 | 4035 | 1764 | 660 | 10206 | 4088 | 5745.5 |
| 1803 | 2889 | 2400 | 949 | 3446 | 1613 | 590 | 8794 | 3348 | 4901 |
| 2076 | 3443.5 | 2900 | 1091 | 4125.5 | 1810 | 699 | 10510 | 4123 | 5913.5 |
| 1862.5 | 3182.5 | 2681.5 | 1030 | 3879.5 | 1737 | 640 | 10102.5 | 3689.5 | 5535.5 |
| 6365 | 5478 | 3531 | 1127 | 4163 | 1817 | 683 | 10556 | 4276 | 6185 |
| 2800 | 4370 | 3202 | 1083 | 4050 | 1800 | 652 | 10176 | 4057 | 5991 |
| 2569 | 4340.5 | 3334 | 1187.5 | 4501 | 1910 | 684 | 11346.5 | 4476 | 6607 |
| 2012 | 3436.5 | 2856.5 | 1037 | 3951 | 1727 | 630 | 9233 | 3823 | 5769.5 |
| 3342 | 5994 | 3500.5 | 1150 | 4734.5 | 1884 | 653 | 15329 | 5121 | 8523.5 |
| 7794 | 16902 | 6098 | 1602 | 7494 | 2348 | 714 | 15921 | 9459 | 20158 |
| 8408 | 25064 | 8518 | 2014 | 9519 | 2712 | 776 | 11236 | 8358 | 27286 |
| 7998 | 26705 | 9248 | 2056 | 10871 | 2823 | 735 | 10557 | 8027 | 29030 |
| 8890.5 | 21427.5 | 6936.5 | 1659 | 7758.5 | 2383 | 619 | 12721.5 | 7458 | 22662 |
| 7750 | 20814 | 7563 | 2149 | 8213 | 2820 | 633 | 12042 | 7409 | 22679 |
| 6043 | 16694 | 5834 | 1676 | 7009 | 2407 | 679 | 12313 | 6554 | 19233 |
| 4477 | 12245 | 6449 | 1832 | 522459 | 44913 | 687 | 13732 | 5769 | 15180 |
| 4235.5 | 6613.5 | 4700 | 1401.5 | 8114.5 | 2734 | 693 | 15531 | 5868 | 8768 |
| 3046 | 6751 | 4317 | 1426 | 6758 | 2471 | 770 | 15497 | 4906 | 10044 |
| 2764 | 4001 | 3450 | 1155 | 5060 | 2013 | 652 | 13642 | 4679 | 6185 |
| 2186 | 3472 | 2883 | 1055 | 4198 | 1805 | 646 | 10144 | 4002 | 5618 |
| 2169 | 3512 | 2957 | 1094 | 4189 | 1843 | 668 | 10460 | 4034 | 5878 |
| 2243.5 | 3663.5 | 2979 | 1172 | 4258.5 | 1899 | 699 | 10272.5 | 4294.5 | 6181.5 |
| 2220 | 3662 | 2984 | 1130 | 4231 | 1840 | 680 | 10569 | 4288 | 6110 |
| 2136 | 3414 | 2775 | 1022 | 3909 | 1741 | 658 | 9865 | 4030 | 5654 |
| 2672 | 4312 | 3320 | 1239 | 4693 | 2015 | 766 | 13407 | 4991 | 7048 |
| 1962 | 3112 | 2573 | 987 | 3599 | 1662 | 617 | 9313 | 3637 | 5262 |
| 2272.5 | 3683 | 3013.5 | 1129.5 | 4324 | 1887 | 701 | 10204 | 4417.5 | 6364 |
| 2136.5 | 3489.5 | 2880.5 | 1098 | 4089 | 1828 | 697 | 10121 | 4018 | 5917 |
| 4812 | 5852 | 3622 | 1107 | 3902 | 1751 | 647 | 10127 | 4070 | 5713 |
| 3026 | 4624 | 3344 | 1095 | 3978 | 1775.5 | 662 | 11056 | 4350 | 6081 |
| 2610 | 4128 | 3213 | 1138 | 4282 | 1863 | 675 | 10765 | 4434 | 6260 |
| 2920 | 4995 | 3810 | 1332 | 5272 | 2135 | 812 | 13608 | 5584 | 8348 |
| 2630 | 4049 | 3138 | 1125 | 4234 | 1831 | 669 | 11344 | 4562 | 6288 |
| 5343 | 11435 | 5055 | 1459 | 6465 | 2158.5 | 701 | 13788.5 | 6737.5 | 14403 |
| 8411 | 19675 | 6828 | 1710 | 8010 | 2409 | 654 | 13136 | 9006 | 22380 |

| | | | | | | | | | |
|---|---|---|---|---|---|---|---|---|---|
| 0 | 0 | 0 | 0 | 0 | 0 | 0 | 0 | 0 | 0 |
| 5268 | 6714 | 3181 | 1015 | 4116 | 1614 | 193 | 13191 | 5307 | 6967 |
| 7512 | 15826 | 6347 | 2188 | 7059 | 2733 | 295 | 14359 | 7136 | 15741 |
| 4640.5 | 7446 | 3998.5 | 1289.5 | 4801 | 1866 | 408 | 14989.5 | 4889 | 8187.5 |
| 3745 | 5426 | 25975 | 8420.5 | 825755 | 77052 | 505 | 15082.5 | 4567 | 6595 |
| 3844 | 6356 | 8210 | 2665 | 11433 | 3948 | 601 | 15104 | 4751 | 7926 |
| 3476 | 6128 | 5719 | 1929 | 7476 | 2823 | 654 | 15089 | 4739 | 8322 |
| 3260 | 6535 | 5076 | 1765 | 6756 | 2621 | 731 | 14194 | 4721 | 9370 |
| 3050.5 | 4908 | 3991.5 | 1418 | 5411 | 2150.5 | 689 | 15714.5 | 4523.5 | 7245.5 |
| 3074.5 | 5223.5 | 4050.5 | 1426 | 5407.5 | 2209.5 | 751 | 15629.5 | 4873.5 | 7786.5 |
| 2708 | 4000 | 3391.5 | 1270 | 4595 | 1953 | 690 | 15705.5 | 4652 | 6277.5 |
| 2730 | 4871 | 3543 | 1329 | 4934 | 2057 | 769 | 15899 | 4636 | 7749 |
| 2795 | 4105.5 | 3298 | 1256 | 4583 | 1934.5 | 726 | 14721 | 4725 | 6529 |
| 2602 | 3830.5 | 3050 | 1169 | 4232 | 1830 | 700 | 14750 | 4519.5 | 6195.5 |
| 2559 | 3622 | 2935 | 1116 | 4130 | 1802 | 679 | 12979 | 4488 | 5884 |
| 2383 | 3532 | 2855.5 | 1114.5 | 4034 | 1773 | 682 | 12291 | 4255.5 | 5842.5 |
| 2335 | 3602 | 2982 | 1140 | 4156.5 | 1815.5 | 696 | 12585.5 | 4420 | 6026.5 |
| 26308 | 13820 | 4972 | 1230 | 4057 | 1826 | 685 | 13099 | 4636 | 6206 |
| 2650 | 4004 | 3204 | 1163 | 4151 | 1823 | 687 | 11930 | 4473 | 6222 |
| 2485 | 3810 | 3121.5 | 1142 | 4122.5 | 1832.5 | 684 | 11136.5 | 4429 | 6205.5 |
| 2390 | 3614 | 3060.5 | 1151 | 4214.5 | 1838 | 692 | 11166.5 | 4403.5 | 6092.5 |
| 3146 | 5831 | 3662 | 1251 | 4796 | 1964 | 733 | 16696 | 4884 | 8569 |
| 5258 | 14924.5 | 5391.5 | 1491.5 | 6532.5 | 2167 | 681 | 15547.5 | 6245 | 18064 |
| 8958 | 23774 | 7302.5 | 1808.5 | 8517 | 2563 | 706 | 12679 | 9003 | 25750.5 |
| 9725 | 22672 | 6739 | 1652 | 7929 | 2314 | 631 | 13719 | 9282 | 24630 |
| 9771.5 | 22530 | 6813 | 1669 | 7739.5 | 2328 | 648 | 13178 | 9121.5 | 24070 |
| 8513 | 17587 | 6088 | 1765 | 6902 | 2315 | 605 | 15011 | 8708 | 19685 |
| 8013 | 18303 | 5977.5 | 1582.5 | 6955 | 2279 | 710 | 12400.5 | 8813 | 21405 |
| 5225.5 | 12683 | 7014 | 2071 | 517143 | 50424 | 667 | 14075.5 | 6154.5 | 15515.5 |
| 3637 | 7434 | 4296 | 1353 | 6107 | 2358 | 656 | 15047 | 4977 | 9928 |
| 2931 | 4441 | 3446 | 1209 | 4927 | 2021 | 657 | 15217 | 4354 | 6475 |
| 2654 | 3810 | 3096 | 1118 | 4355 | 1883 | 680 | 13815.5 | 4619.5 | 6130 |
| 2060 | 3278 | 2818 | 1115 | 3998 | 1759 | 661 | 10577 | 3852 | 5523 |
| 1936 | 3247.5 | 2795 | 1088 | 3914.5 | 1789 | 676 | 9984 | 3835.5 | 5644 |
| 1802 | 3063 | 2590.5 | 1054.5 | 3706 | 1704 | 648 | 9230.5 | 3546 | 5332 |
| 1942 | 3340 | 2742 | 1123 | 3863 | 1804 | 693 | 10354 | 3780 | 5639 |
| 2044 | 3409 | 2762 | 1095 | 3885 | 1785 | 689 | 10848 | 3986 | 5706 |
| 2051 | 3312 | 2747 | 1080 | 3923.5 | 1768 | 680.5 | 10747.5 | 4052 | 5784.5 |
| 2089 | 3508 | 2911 | 1139 | 3924 | 1810 | 704 | 10854 | 4005 | 5856.5 |
| 1880 | 3153 | 2690 | 1083 | 3737 | 1728 | 669 | 9508 | 3622 | 5420 |
| 1845.5 | 3228 | 2790.5 | 1142 | 4007.5 | 1830 | 700 | 10195.5 | 3845.5 | 5776 |
| 21215 | 11868.5 | 4495 | 1189.5 | 3917 | 1785 | 684 | 11402.5 | 4128.5 | 5938 |
| 2547 | 4132 | 3376 | 1152 | 4130 | 1817 | 696 | 10424 | 4437 | 6238 |
| 1994 | 3502 | 2853 | 1077 | 3869 | 1726 | 648 | 9800 | 3831 | 5728 |
| 2322 | 3775 | 3220 | 1193 | 4288.5 | 1905 | 722 | 11369.5 | 4607.5 | 6631.5 |
| 3023 | 4666 | 3376 | 1181 | 4451.5 | 1849 | 698 | 15867.5 | 4821.5 | 7108 |
| 5867 | 13935 | 5164 | 1442 | 6215 | 2158 | 704 | 15548 | 7230 | 16978 |
| 9456 | 21652 | 6784 | 1670 | 7698 | 2376 | 680 | 14959 | 9699 | 24036 |

| | | | | | | | | | |
|---|---|---|---|---|---|---|---|---|---|
| 9151.5 | 23443 | 7557 | 1771 | 8703.5 | 2456.5 | 668 | 12451.5 | 9124 | 26031.5 |
| 9104 | 23363 | 6966 | 1704 | 8104 | 2422 | 653 | 12697 | 9188 | 24774 |
| 9696 | 20874.5 | 6645.5 | 1736 | 7527.5 | 2370.5 | 690 | 11560 | 8590 | 22701 |
| 7594 | 16413 | 5501 | 1492 | 6226 | 2147 | 705 | 13213 | 8076 | 18990 |
| 5131 | 12881 | 6063 | 1685 | 334399 | 38077 | 740 | 15109 | 6126 | 16037 |
| 3487 | 8286 | 4124 | 1315 | 5936 | 2308 | 707 | 13616 | 4775 | 11015 |
| 2947.5 | 4429 | 3365 | 1179 | 4686.5 | 1971 | 696.5 | 15545.5 | 4781.5 | 6590.5 |
| 2716 | 3984 | 3170 | 1203 | 4466.5 | 1923 | 742 | 14701 | 4654.5 | 6498.5 |
| 2031 | 3337 | 2871 | 1125 | 3970 | 1821 | 687 | 10089 | 3785 | 5574 |
| 2308 | 3710.5 | 3144 | 1218.5 | 4507.5 | 1975.5 | 765 | 11175.5 | 4576 | 6692.5 |
| 1974 | 3301.5 | 2781 | 1111 | 3906 | 1778 | 702 | 10111.5 | 3833 | 5516.5 |
| 2309 | 3517 | 2937 | 1129 | 4166 | 1826 | 725 | 11338.5 | 4410 | 6101 |
| 2189.5 | 3524 | 2851 | 1167 | 4020 | 1833 | 729 | 10964.5 | 4066.5 | 5958 |
| 2288 | 3707 | 3087 | 1192 | 4328 | 1882.5 | 755 | 10968 | 4455.5 | 6358 |
| 2560 | 3808 | 3100 | 1181 | 4193 | 1892 | 745 | 11495 | 4388 | 6359 |
| 2142.5 | 3490.5 | 2842 | 1131 | 3971 | 1818 | 741 | 10102.5 | 4057.5 | 5958.5 |
| 2121.5 | 3536 | 2947 | 1191 | 4186 | 1929.5 | 749 | 10282.5 | 4339 | 6349.5 |
| 6445 | 4952 | 3318 | 1149 | 3964 | 1828 | 698 | 10298 | 4185 | 6031 |
| 2638 | 3964 | 3227 | 1202 | 4377 | 1863 | 730 | 10727 | 4524 | 6550 |
| 2203 | 3620 | 2930 | 1158 | 4170 | 1826 | 715 | 10517 | 4274 | 6170 |
| 2110 | 3555 | 3008 | 1141 | 4137 | 1796 | 696 | 10608 | 4236 | 6162 |
| 3994 | 7005 | 3851 | 1303 | 4968 | 1971 | 735 | 16761 | 6017 | 10141 |
| 6478 | 16224.5 | 5696.5 | 1586 | 7066 | 2312.5 | 750 | 14628.5 | 7497 | 19396 |
| 9001 | 21626 | 6590 | 1742 | 7976 | 2416 | 680 | 12056 | 8467 | 23634 |
| 9565 | 23056 | 8710 | 1886 | 9796 | 2557 | 726 | 11876 | 9090 | 24504 |
| 8742.5 | 26783 | 8985.5 | 1984 | 10221.5 | 2814 | 765 | 11082.5 | 8616.5 | 29323.5 |
| 6754 | 17034 | 6239 | 1897 | 6419 | 2361 | 592 | 14225 | 6457 | 18505 |
| 5913 | 14334.5 | 4927 | 1342 | 5699 | 1956.5 | 631 | 14984 | 6657.5 | 16970.5 |
| 5208 | 13030 | 6364 | 1700 | 575049 | 56204 | 677 | 14450 | 6274 | 15894 |
| 3360 | 7724 | 4159 | 1290 | 5904 | 2215 | 676 | 14872 | 4536 | 10196 |
| 2866 | 4618 | 3449 | 1190 | 4783 | 1961 | 648 | 15908 | 4417 | 6675 |
| 2557 | 3713.5 | 2875 | 1093 | 4021.5 | 1763 | 660 | 11334.5 | 4382 | 5850 |
| 2331 | 3606 | 2904 | 1130 | 4125 | 1802 | 703 | 10720 | 4331 | 5986 |
| 1952.5 | 3197 | 2613 | 1042 | 3696 | 1709.5 | 633.5 | 9577.5 | 3631 | 5366.5 |
| 2076 | 3357 | 2806 | 1088 | 3921 | 1796 | 677 | 9874 | 3942 | 5816 |
| 2194 | 3501 | 2793.5 | 1115 | 3895 | 1774.5 | 690 | 10370 | 4086.5 | 5921 |
| 2169.5 | 3370 | 2675.5 | 1085 | 3784 | 1723.5 | 661 | 9712.5 | 3939 | 5427.5 |
| 2451 | 3807 | 3034 | 1163 | 4285 | 1888 | 730 | 10981 | 4554 | 6340 |
| 2257 | 3642 | 2913 | 1172 | 4123 | 1875 | 707 | 10542 | 4160 | 6221 |
| 2109 | 3445 | 2837 | 1097 | 4002 | 1790 | 705 | 9928 | 4102 | 5863 |
| 2246 | 3605 | 2988 | 1126 | 4172 | 1868 | 722 | 10568 | 4435 | 6228 |
| 17711 | 9522 | 4473 | 1254 | 4364 | 1903 | 748 | 12772 | 4881 | 6703 |
| 4091.5 | 5807 | 3896 | 1240.5 | 4294 | 1897 | 725 | 12121 | 4660 | 6542 |
| 2457 | 4046.5 | 3193.5 | 1147.5 | 4263 | 1834 | 689 | 10509 | 4524 | 6509.5 |
| 2582 | 4107.5 | 3377 | 1236 | 4566 | 1970 | 743 | 11013 | 4887 | 6983 |
| 3827 | 6332 | 3909 | 1330 | 5175 | 2057 | 751 | 15527 | 6428 | 9216 |
| 5773 | 14438.5 | 5410.5 | 1518 | 6513.5 | 2180 | 717 | 15205 | 7156 | 17586 |
| 7646 | 21398 | 6624 | 1687.5 | 7734 | 2396 | 662 | 13197.5 | 7847 | 23360 |

| | | | | | | | | | |
|---|---|---|---|---|---|---|---|---|---|
| 9314.5 | 23901.5 | 7591 | 1806 | 8649 | 2464 | 669 | 14060.5 | 9611 | 26029.5 |
| 8850 | 25004.5 | 7785.5 | 1810 | 8585 | 2505 | 702 | 10715.5 | 8324 | 27009.5 |
| 7929 | 20152 | 6354 | 1665 | 7327 | 2317 | 650 | 11327 | 7654 | 22138 |
| 7213 | 16052.5 | 5360.5 | 1466.5 | 6218 | 2100 | 653 | 14329 | 7762.5 | 18200 |
| 4643 | 12295 | 5883 | 1637 | 386967 | 45088 | 683 | 14709 | 5453 | 15172 |
| 3773 | 6981.5 | 4288.5 | 1328 | 6411.5 | 2404 | 670 | 15261 | 5096 | 9312.5 |
| 2914.5 | 5714 | 3786 | 1294 | 5278 | 2117.5 | 744 | 16308 | 4554.5 | 8387.5 |
| 2542 | 3657 | 2820 | 1082 | 3952 | 1736 | 648 | 11613 | 4282 | 5763 |
| 2379 | 3679 | 2977 | 1136 | 4202 | 1817 | 702 | 10546.5 | 4378.5 | 6050 |
| 2079 | 3387 | 2781 | 1084 | 3919 | 1758 | 677 | 10060 | 3894 | 5595 |
| 2152 | 3453 | 2801 | 1080 | 3921.5 | 1782 | 694 | 10020.5 | 4039 | 5911 |
| 2291.5 | 3578.5 | 2845.5 | 1121 | 4072.5 | 1777 | 699 | 10830.5 | 4198.5 | 6046 |
| 2207.5 | 3544.5 | 2774.5 | 1099 | 3886 | 1769.5 | 685 | 10331 | 4073 | 5939 |
| 2158 | 3466 | 2753 | 1098 | 3837 | 1744 | 667 | 10015.5 | 4015.5 | 5758 |
| 2453 | 3757 | 2978 | 1169 | 4166 | 1862 | 722 | 11854 | 4455 | 6366 |
| 2045 | 3231 | 2650 | 1054 | 3705 | 1728 | 677 | 9677 | 3768 | 5555 |
| 1951 | 3225 | 2708 | 1075 | 3746 | 1741 | 670 | 9428 | 3765 | 5639 |
| 9260.5 | 6341 | 3649.5 | 1130 | 3832 | 1746.5 | 665 | 10563.5 | 4010 | 5972.5 |
| 2851 | 4312 | 3352 | 1175 | 4237 | 1849 | 721 | 11579 | 4579 | 6478 |
| 2198 | 3624 | 2856 | 1121 | 3871 | 1764 | 649 | 9462 | 3909 | 5752 |
| 2477 | 4370.5 | 3266.5 | 1214.5 | 4616 | 1958 | 764 | 11161.5 | 4908 | 7335.5 |
| 3381.5 | 5554.5 | 3747 | 1294.5 | 4892.5 | 2026 | 757 | 14463 | 5776 | 8439 |
| 7227.5 | 16911 | 6886.5 | 1774 | 8470 | 2561 | 806 | 15387 | 9191 | 20635.5 |
| 8235 | 23567 | 7650 | 1938 | 8991 | 2657 | 762 | 10958 | 8348 | 26623 |
| 9572.5 | 25538 | 8274 | 1921 | 9267 | 2625.5 | 713 | 12028 | 9255.5 | 27525.5 |
| 9180 | 25081 | 8525 | 1926 | 9709 | 2646 | 748 | 11846 | 9002 | 27678 |
| 10367 | 24091.5 | 7962 | 2177 | 7587 | 2410 | 666 | 15113 | 8971.5 | 24217.5 |
| 8619 | 21162 | 6740 | 1697 | 7199 | 2378 | 770 | 15450 | 7838 | 22529 |
| 8332.5 | 15608 | 8285 | 2312 | 16178.5 | 4552 | 755 | 15288.5 | 6661 | 16961.5 |
| 7963 | 11166 | 5622 | 1609 | 17168 | 3279 | 788 | 16129 | 5530 | 12496 |
| 8700 | 7902.5 | 4632 | 1420.5 | 9505 | 2602 | 826 | 18706 | 5375.5 | 9084.5 |
| 9312 | 6319.5 | 4101 | 1273 | 7270.5 | 2375 | 799.5 | 17469.5 | 5433 | 7389 |
| 9556 | 5697.5 | 3685.5 | 1253 | 5814.5 | 2142 | 795 | 15160.5 | 5241.5 | 6960.5 |
| 8664.5 | 5502.5 | 3750 | 1273 | 7153 | 2397.5 | 824.5 | 12543.5 | 5270.5 | 6854 |
| 8318 | 6266 | 5470 | 1848 | 5432 | 2167 | 831 | 13050 | 5176 | 7470 |
| 8783 | 5421 | 3537.5 | 1297.5 | 5942 | 2168.5 | 792 | 13149.5 | 4795 | 6495 |
| 7412 | 5017 | 3278 | 1212 | 6012 | 2205 | 781 | 12370 | 5003 | 6371 |
| 8424.5 | 5620 | 3656.5 | 1302 | 6477.5 | 2234 | 820 | 12792.5 | 5256.5 | 7187 |
| 14170 | 8966 | 4544 | 1204 | 5533 | 2081 | 794 | 12140 | 6690 | 7805 |
| 10056 | 6205.5 | 3674 | 1247.5 | 5995 | 2180.5 | 837 | 12497.5 | 5464 | 7108 |
| 9384 | 5443 | 3343.5 | 1185 | 4814 | 1969.5 | 759.5 | 12436 | 4681.5 | 6349.5 |
| 713465 | 404439 | 107409 | 6479 | 6938 | 2398 | 784 | 87034 | 23202 | 15324 |
| 52706 | 26492 | 7945 | 1522 | 4932 | 2005 | 785 | 14886 | 6002 | 7489 |
| 26042 | 14204 | 5203 | 1323 | 5547 | 2129 | 765 | 13972 | 5363 | 7910 |
| 26484.5 | 13367 | 5161.5 | 1359.5 | 5308.5 | 2084 | 765 | 14046.5 | 5921.5 | 7487 |
| 13663.5 | 10396 | 4831.5 | 1434 | 7884 | 2431 | 844.5 | 18802 | 5409 | 9579 |
| 20203 | 30607 | 9013 | 2090 | 10739 | 2896 | 933 | 14389 | 9789 | 26892 |
| 16994 | 26784 | 7945 | 1857 | 9473 | 2601 | 747 | 14576 | 9167 | 25005 |

| | | | | | | | | | |
|---|---|---|---|---|---|---|---|---|---|
| 12508 | 30462 | 10771 | 2329 | 11933 | 3026 | 847 | 10535 | 7925 | 28764 |
| 9788 | 23512 | 7211 | 1833 | 8496 | 2473 | 667 | 9388 | 7920 | 23680 |
| 9708 | 22372 | 7139 | 1859.5 | 8024 | 2510 | 720 | 12688 | 9147.5 | 24641 |
| 7203 | 16339 | 5404 | 1459 | 6256 | 2126 | 694 | 15419 | 7942 | 18719 |
| 5060.5 | 13636.5 | 6984 | 1814 | 586650.5 | 57044 | 737 | 12398 | 6152.5 | 16328.5 |
| 3505.5 | 8243.5 | 4511.5 | 1364.5 | 6861 | 2503 | 705 | 14222.5 | 4855 | 11127 |
| 2981.5 | 5432.5 | 3635 | 1267 | 5268 | 2074 | 710 | 14604 | 4814.5 | 7663.5 |
| 2920 | 4228 | 3170.5 | 1176 | 4581 | 1940 | 717 | 12507 | 5015 | 6562 |
| 2557.5 | 3946 | 3114.5 | 1161 | 4454.5 | 1909.5 | 735 | 11748.5 | 4787 | 6451 |
| 2175 | 3469 | 2781 | 1090 | 3988 | 1765 | 679 | 10289 | 4017 | 5779 |
| 2376.5 | 3773.5 | 2982 | 1161 | 4322 | 1893 | 729 | 10835 | 4541 | 6378.5 |
| 2340 | 3772 | 3004 | 1158.5 | 4171 | 1883 | 712.5 | 12166.5 | 4564 | 6117 |
| 2469 | 3813 | 3029 | 1165 | 4253 | 1872 | 728 | 12230 | 4788 | 6326 |
| 2394 | 3783 | 2975 | 1164 | 4216 | 1852 | 719 | 11041 | 4529 | 6119 |
| 2152.5 | 3492 | 2785.5 | 1077 | 3943.5 | 1774 | 694 | 9926.5 | 4065.5 | 5876 |
| 2371 | 3700 | 3043 | 1156 | 4238 | 1860 | 735 | 11208 | 4614 | 6389 |
| 2154 | 3504 | 2818 | 1111.5 | 4042 | 1811 | 697 | 10104 | 4295.5 | 5942.5 |
| 27836.5 | 14358.5 | 5030 | 1234 | 4158 | 1824 | 697 | 11504 | 4703.5 | 6443 |
| 3545 | 5601 | 3972 | 1270.5 | 4541 | 1939.5 | 745 | 12486.5 | 5004.5 | 7127.5 |
| 2591 | 4240 | 3339 | 1176 | 4416 | 1885 | 714 | 10890 | 4812 | 6741 |
| 2606 | 4355 | 3441 | 1247 | 4699 | 1975 | 777 | 12592 | 5083 | 7282 |
| 3261 | 5805 | 3719 | 1277 | 4974.5 | 1999 | 739 | 16521 | 5460.5 | 8759.5 |
| 8830 | 21699 | 7666 | 1793 | 8816 | 2586 | 829 | 15935 | 9764 | 25347 |
| 9846 | 23309 | 6882 | 1726 | 8206 | 2447 | 679 | 14379 | 9299 | 25394 |
| 8541 | 22609 | 9420 | 1964 | 10588 | 2746 | 739 | 12726 | 8940 | 24991 |
| 9693 | 22304 | 6956 | 1691 | 7903 | 2411 | 642.5 | 13743.5 | 9126.5 | 24739 |
| 8001 | 19242 | 6435 | 1769 | 7607 | 2438 | 663 | 10895 | 7926 | 20979 |
| 8155 | 17516 | 6043.5 | 1581 | 6964 | 2250 | 748.5 | 14268.5 | 8528.5 | 19904.5 |
| 5620.5 | 13772.5 | 6281.5 | 1674 | 13365 | 4259.5 | 733.5 | 15446.5 | 6485.5 | 17235.5 |
| 3459 | 8571.5 | 4543.5 | 1400.5 | 6951 | 2555.5 | 744.5 | 15011.5 | 4685.5 | 11520 |
| 3319.5 | 5397.5 | 3930.5 | 1282.5 | 5560 | 2205 | 760 | 16806.5 | 4993.5 | 7804.5 |
| 2812 | 4317 | 3338 | 1225 | 4775 | 2031 | 742 | 14710 | 4874 | 6893 |
| 2326 | 3589 | 2909.5 | 1106 | 4089 | 1795 | 699 | 11231 | 4168 | 5991.5 |
| 2610.5 | 4119 | 3279.5 | 1248 | 4732 | 1994 | 798 | 11901.5 | 5056.5 | 7005.5 |
| 2596 | 4170.5 | 3426 | 1285.5 | 4706 | 2035 | 808 | 13009 | 4865.5 | 6860 |
| 2255 | 3580 | 2890 | 1113 | 3993.5 | 1793.5 | 721 | 11435 | 4306 | 6006 |
| 2430 | 3729 | 2978 | 1162 | 4211 | 1836 | 746 | 12101 | 4568 | 6153 |
| 2534.5 | 4026 | 3392.5 | 1268 | 4678 | 1997.5 | 771 | 12413 | 4913.5 | 6634.5 |
| 2276.5 | 3549 | 2878 | 1121 | 3996 | 1786.5 | 736 | 10721 | 4312.5 | 6066 |
| 2481 | 3959 | 3139 | 1208 | 4492 | 1907 | 799 | 11862 | 4846 | 6784 |
| 2246 | 3670 | 3032 | 1188 | 4155.5 | 1866 | 772 | 11037.5 | 4379.5 | 6348 |
| 11617 | 7393 | 3789 | 1161 | 4050 | 1795 | 722 | 10836 | 4309 | 6109 |
| 3131.5 | 4733 | 3578 | 1220 | 4360 | 1885 | 740 | 11245.5 | 4472.5 | 6471 |
| 2573 | 4115 | 3340 | 1229 | 4378 | 1918 | 730 | 11466 | 4743 | 6763 |
| 2649 | 4423.5 | 3504.5 | 1275 | 4834 | 2028 | 783.5 | 12489.5 | 5344.5 | 7486 |
| 3160 | 4905 | 3516 | 1198 | 4568 | 1827 | 699 | 15035 | 5069 | 7343 |
| 6837 | 17586 | 6136 | 1646 | 7145 | 2340 | 822 | 17362 | 8209 | 21273 |
| 9388 | 20208 | 6580 | 1656 | 7461 | 2345.5 | 724.5 | 13369.5 | 8935.5 | 22453.5 |

| | | | | | | | | | |
|---|---|---|---|---|---|---|---|---|---|
| 9871.5 | 27221 | 8755.5 | 1983.5 | 9952 | 2685.5 | 769 | 13080 | 9099.5 | 28760 |
| 9473 | 27235 | 9387 | 2045 | 10549 | 2769 | 770 | 12228 | 9064 | 29495 |
| 9869 | 39464 | 36650 | 8193 | 11155 | 3981 | 666 | 12620 | 8017 | 28985 |
| 6065.5 | 18182 | 7858.5 | 2305 | 7332 | 2546.5 | 689 | 13952 | 6519.5 | 20278.5 |
| 4943 | 13514 | 8432 | 2343 | 75873 | 11462 | 671 | 14454 | 5928 | 15864 |
| 3861 | 7320 | 5392.5 | 1678.5 | 11040.5 | 3079 | 701 | 15857.5 | 5132.5 | 9800 |
| 3658.5 | 5320 | 4436.5 | 1539 | 8762 | 2545.5 | 741 | 17729 | 4994 | 7632 |
| 3382 | 4610 | 4006 | 1419 | 7300 | 2356 | 757 | 14937 | 5099 | 6948 |
| 2955 | 3954 | 3461 | 1300 | 6528 | 2190 | 719 | 11857 | 4447 | 6257 |
| 2997 | 4271.5 | 3615 | 1362 | 6783.5 | 2292 | 790 | 11837 | 5001.5 | 7087.5 |
| 2876 | 4216 | 3878 | 1426.5 | 6046.5 | 2209 | 768.5 | 11296.5 | 4739 | 6763.5 |
| 2747 | 3682 | 3141 | 1217 | 5385 | 2009 | 727 | 11136 | 4418 | 6280 |
| 2992 | 4719 | 3718 | 1381 | 5969 | 2265 | 823 | 13041 | 5224 | 8005 |
| 2563 | 3786 | 3111 | 1221 | 5065 | 1962 | 750 | 11321 | 4541 | 6339 |
| 4397.5 | 4946.5 | 3573.5 | 1223 | 5189.5 | 2015 | 743.5 | 10587 | 5295 | 6962.5 |
| 2915 | 4135 | 3355 | 1214 | 5125 | 2014 | 737 | 10634 | 4817 | 6522 |
| 2350 | 3653 | 3104 | 1158 | 4863 | 1916 | 749 | 10665 | 4276 | 6136 |
| 69442.5 | 36357 | 9526 | 1453 | 4943.5 | 1959.5 | 748.5 | 15654.5 | 5650.5 | 7112 |
| 3758 | 5156 | 4166.5 | 1274.5 | 5073.5 | 1957.5 | 746.5 | 12600 | 5178 | 6946.5 |
| 2605.5 | 4191 | 3529.5 | 1208 | 4824 | 1919 | 710 | 11752.5 | 4609.5 | 6532 |
| 2459 | 3999.5 | 3345.5 | 1222 | 4869.5 | 2016 | 763 | 10971.5 | 4987.5 | 6921 |
| 3194 | 5052 | 3726 | 1309 | 5093.5 | 2057 | 755 | 16908.5 | 5438 | 7764 |
| 6025 | 14473 | 5323 | 1539 | 6854 | 2246 | 766 | 13083 | 7170 | 17760 |
| 8570 | 20488 | 6704 | 1647 | 7951 | 2354 | 658 | 13393 | 8619 | 23178 |
| 8770 | 26576 | 9359 | 2095 | 10829 | 2860 | 746 | 12292 | 8976 | 29093 |
| 8535 | 27631 | 9424 | 2065 | 10546 | 2803 | 749 | 11352 | 8416 | 29988 |
| 8877 | 24073 | 11566 | 3788 | 10254 | 3724 | 679 | 9228 | 7986 | 24884 |
| 7594 | 19786 | 9558 | 3267 | 7968 | 3225 | 702 | 13788 | 7919 | 21985 |
| 5599 | 13115 | 10254 | 3945 | 507233 | 45451 | 651 | 14252 | 6409 | 15749 |
| 3964.5 | 11087.5 | 6979 | 2535 | 7298 | 3076.5 | 738 | 13305 | 5285 | 14512 |
| 2925 | 4739 | 4609 | 1612.5 | 5218.5 | 2286.5 | 661 | 16306.5 | 4346.5 | 6739.5 |
| 2551 | 3924 | 3548 | 1359 | 4527 | 2036 | 685 | 12785 | 4392 | 6168 |
| 2164 | 3427 | 3058 | 1179 | 4131 | 1872 | 668 | 10550 | 4039 | 5611 |
| 1830.5 | 3056.5 | 2679.5 | 1080.5 | 3782 | 1750 | 655 | 9220 | 3568 | 5310 |
| 1927 | 3235 | 2840 | 1145 | 3965 | 1828 | 679 | 10282 | 3748 | 5575 |
| 1918 | 3165 | 2700 | 1054 | 3827 | 1751 | 647 | 9477 | 3706 | 5472 |
| 2149 | 3446 | 2818 | 1083 | 4017 | 1788 | 673 | 10959 | 4125 | 5733 |
| 2172.5 | 3541.5 | 2884 | 1163 | 4160 | 1898.5 | 708 | 11315.5 | 4359 | 6032.5 |
| 2102 | 3431 | 2895 | 1138 | 4106 | 1808 | 683 | 10720 | 3996 | 5785 |
| 2017 | 3297 | 2769 | 1096 | 3938 | 1785 | 668 | 10117 | 3964 | 5654 |
| 2212 | 3844 | 3046 | 1171 | 4440 | 1918 | 751 | 12467 | 4543 | 6769 |
| 11730 | 7656.5 | 4158.5 | 1210 | 4162 | 1844 | 704.5 | 11257 | 4617.5 | 6310.5 |
| 3013 | 4525 | 3568 | 1217 | 4374 | 1912 | 708 | 11350 | 4886 | 6481 |
| 2490 | 4172 | 3420.5 | 1204 | 4510 | 1902 | 723.5 | 10904.5 | 4738 | 6842 |
| 2402 | 4591 | 3311 | 1212 | 4714 | 1918 | 731 | 11316.5 | 4818 | 7502 |
| 3673 | 6782 | 3688 | 1249 | 4993 | 1956 | 713 | 16718 | 5971 | 9844 |
| 7198 | 18000 | 6002 | 1626 | 7463 | 2373 | 758 | 15274 | 8725 | 22174 |
| 7753 | 19812 | 6654 | 1647 | 7789 | 2342 | 666 | 15373 | 8146 | 22834 |

| | | | | | | | | | |
|---|---|---|---|---|---|---|---|---|---|
| 9132 | 27298 | 8666 | 1983 | 9826 | 2693 | 730 | 11806.5 | 9030.5 | 29151 |
| 9597.5 | 24113 | 7535.5 | 1776.5 | 8959 | 2518 | 684.5 | 11693.5 | 8812 | 26241 |
| 7615 | 19250 | 7471 | 2168 | 8189 | 2687 | 639 | 8905 | 7261 | 21486 |
| 7656 | 18614 | 6439 | 1816 | 7148 | 2442 | 681 | 10710 | 7438 | 20433 |
| 4866 | 12263 | 6524 | 1876 | 14568 | 4665 | 682 | 15783 | 5702 | 14717 |
| 3722 | 8788.5 | 4746 | 1509 | 7241 | 2760 | 686 | 12484.5 | 5151 | 11583.5 |
| 2959.5 | 4782.5 | 3520 | 1267 | 5035 | 2121 | 644 | 13238.5 | 4483 | 7008 |
| 2522 | 3733 | 3064 | 1120 | 4306 | 1884 | 644 | 11327.5 | 4338 | 5917 |
| 2468.5 | 3824.5 | 3143.5 | 1183 | 4522.5 | 1936 | 718 | 11400.5 | 4520 | 6245.5 |
| 2625 | 4255 | 3426 | 1276 | 4938 | 2044 | 789 | 12289 | 4999 | 7233 |
| 2228 | 3560 | 2900 | 1122 | 4106 | 1824 | 685 | 10346 | 4237 | 5984 |
| 2426 | 3732 | 3145 | 1189 | 4350 | 1883 | 707 | 11063 | 4417 | 6273 |
| 2411 | 3750 | 2909 | 1123 | 4192 | 1859 | 709 | 12047 | 4495 | 6300 |
| 2502 | 3894.5 | 3171.5 | 1206 | 4452 | 1900 | 725 | 12130.5 | 4725 | 6452 |
| 2516 | 3899.5 | 3096 | 1188.5 | 4350.5 | 1899.5 | 734 | 11290.5 | 4677.5 | 6432.5 |
| 2198 | 3432 | 2795 | 1083 | 3894 | 1760 | 674 | 9948 | 4038 | 5743 |
| 1980 | 3251 | 2613 | 1066 | 3841 | 1740 | 651 | 9393 | 3807 | 5452 |
| 13782 | 8207 | 4245 | 1229 | 4439 | 1894 | 707 | 11943 | 4941 | 6449 |
| 3215 | 4809.5 | 3638 | 1178 | 4418.5 | 1895.5 | 694 | 12173.5 | 4783 | 6505.5 |
| 2738 | 4280 | 3353 | 1176 | 4427 | 1870 | 700 | 10865 | 4743 | 6597 |
| 2576 | 4185.5 | 3264.5 | 1194 | 4562 | 1923 | 724 | 10798 | 4722 | 6811 |
| 3443 | 5864 | 3691 | 1278 | 5035 | 1996 | 709 | 14549 | 5744 | 8627.5 |
| 4746 | 8552 | 4281 | 1303 | 5402 | 1990 | 661 | 13316 | 6067 | 10806 |
| 6850 | 17294 | 5775 | 1479 | 6801 | 2137 | 591 | 15091 | 6893 | 19361 |
| 8289 | 23069 | 8068.5 | 1899 | 9513.5 | 2611.5 | 685 | 11677.5 | 8632.5 | 25986 |
| 736 | 850 | 614 | 563 | 1142 | 1107 | 176 | 1901 | 1255 | 1466 |
| 3926 | 4254 | 2176 | 877 | 2678 | 1421 | 230 | 13842 | 4235 | 4491 |
| 3248 | 3794 | 2231 | 804 | 2792 | 1370 | 353 | 15492 | 3886 | 4536 |
| 2378.5 | 3103.5 | 7626.5 | 2672 | 753304 | 72036 | 468 | 10637 | 3351.5 | 4394 |
| 2175 | 3177 | 4001 | 1371 | 6586.5 | 2724 | 541 | 10637.5 | 3516.5 | 4860 |
| 1966.5 | 3082.5 | 2961 | 1057 | 4311.5 | 1944 | 588 | 9952 | 3482 | 5069.5 |
| 1817 | 3062 | 2724 | 1013 | 3924 | 1758 | 621 | 9521 | 3545 | 5221 |
| 1850 | 3154 | 2798 | 1049 | 4025 | 1773 | 662 | 9571 | 3691 | 5595 |
| 1731 | 2984 | 2626 | 989 | 3759 | 1685 | 622 | 8757 | 3491 | 5284 |
| 1817 | 3199 | 2782.5 | 1063 | 3961 | 1749 | 667 | 9242.5 | 3756 | 5755 |
| 1914 | 3285 | 2792 | 1043 | 3869 | 1752 | 666 | 9730 | 3857 | 5692 |
| 1854 | 3139.5 | 2627.5 | 999 | 3695.5 | 1667 | 634.5 | 9121.5 | 3669 | 5436 |
| 1948 | 3267 | 2743 | 1060 | 3930 | 1741 | 671 | 9764 | 3863 | 5693 |
| 1921 | 3272 | 2755 | 1051 | 3851 | 1714 | 662 | 9559 | 3810 | 5577 |
| 1823 | 3178 | 2716 | 1030 | 3793 | 1728 | 661 | 9235.5 | 3708 | 5701.5 |
| 1815 | 3162 | 2744 | 1038 | 3835.5 | 1744.5 | 660 | 8867 | 3702.5 | 5542 |
| 23509 | 13157 | 4855 | 1179 | 4066 | 1777 | 665 | 11270 | 4227 | 5997 |
| 2703.5 | 4713.5 | 3768.5 | 1157 | 4301 | 1820 | 693 | 10322.5 | 4354 | 6325.5 |
| 2218 | 3820 | 3193 | 1109 | 4264 | 1789 | 677 | 9866 | 4414 | 6454 |
| 2035 | 3554 | 3025 | 1068 | 4135 | 1799 | 678 | 9699 | 4189 | 6241 |
| 3881 | 8878.5 | 4148.5 | 1208 | 5279.5 | 1958.5 | 687 | 15214 | 5736 | 12125.5 |
| 5832.5 | 14895 | 5308 | 1436 | 6219.5 | 2121 | 669 | 15879 | 7056 | 17782 |
| 8126 | 21708 | 7026 | 1729 | 8108 | 2470 | 680 | 12290 | 8488 | 24458 |

| | | | | | | | | | |
|---|---|---|---|---|---|---|---|---|---|
| 9528 | 24962 | 7760 | 1800 | 8664 | 2492 | 656 | 11755 | 8737 | 26892 |
| 7874 | 23402 | 8376 | 1923 | 10017 | 2593 | 681 | 9784 | 7452 | 24617 |
| 8706.5 | 23047.5 | 7283.5 | 1877 | 8035.5 | 2550.5 | 697 | 10051.5 | 8364 | 25580.5 |
| 5678 | 16094 | 5336 | 1403 | 6322 | 2110 | 683 | 16050 | 6606 | 18736 |
| 4828 | 12487.5 | 6496.5 | 1654 | 634729.5 | 60890 | 669 | 14166 | 5965.5 | 15518.5 |
| 3714 | 8778 | 4921 | 1395 | 8222 | 2800.5 | 696 | 14395 | 5178 | 11554.5 |
| 2888 | 4900 | 3599 | 1184 | 5066 | 2014 | 695 | 16584 | 4672 | 7386 |
| 2675 | 4226.5 | 3334.5 | 1129 | 4652.5 | 1938 | 708 | 14352 | 4676.5 | 6642 |
| 2387 | 3748 | 3083 | 1093 | 4366 | 1859 | 707 | 11873 | 4501 | 6369 |
| 2289 | 3697 | 3096 | 1112 | 4382 | 1867 | 715 | 11294 | 4612 | 6383 |
| 2236.5 | 3670 | 3075.5 | 1129 | 4301 | 1882.5 | 720 | 10467 | 4577.5 | 6415.5 |
| 2213 | 3525 | 2940 | 1059 | 4171 | 1786 | 689 | 11024 | 4253 | 5986 |
| 2341 | 3733.5 | 3137 | 1115 | 4334 | 1868 | 720.5 | 10950 | 4650.5 | 6543.5 |
| 2131 | 3547.5 | 2925 | 1044.5 | 4065.5 | 1778 | 688 | 10509.5 | 4238 | 6046 |
| 2279 | 3641 | 3000 | 1098 | 4220 | 1826 | 717 | 10431 | 4483 | 6356 |
| 2103 | 3535 | 2911 | 1082 | 4084 | 1802 | 691 | 10025 | 4186 | 6068 |
| 2141 | 3652 | 3101.5 | 1124 | 4282 | 1844.5 | 715 | 10862 | 4443.5 | 6400.5 |
| 33226 | 17999 | 5745.5 | 1226 | 4065.5 | 1798 | 681 | 11844 | 4675 | 6429 |
| 2692 | 4337.5 | 3452 | 1109 | 4241 | 1840 | 689.5 | 10570 | 4526 | 6530.5 |
| 2508.5 | 4179 | 3471 | 1168 | 4509 | 1901 | 702 | 11041.5 | 4736.5 | 6716 |
| 2192 | 3639 | 3032 | 1093 | 4153 | 1796 | 664 | 9987 | 4387 | 6243 |
| 3762.5 | 9204.5 | 4181.5 | 1267 | 5358 | 1980 | 685 | 15166 | 5477.5 | 12276.5 |
| 5317 | 14609.5 | 5281.5 | 1438 | 6274 | 2167 | 674 | 14634 | 6254.5 | 17635 |
| 8226 | 23056 | 7155.5 | 1732 | 8527.5 | 2471 | 677 | 11965.5 | 8596.5 | 25643.5 |
| 9985 | 24871 | 8389 | 1861 | 9491 | 2577 | 673 | 14018 | 9578 | 26778 |
| 8839 | 25136 | 9075.5 | 1989 | 10492 | 2743 | 720 | 12324 | 9020 | 27692 |
| 9313.5 | 24295.5 | 8459.5 | 1939.5 | 9292 | 2663 | 750 | 9939 | 8228 | 25340.5 |
| 7093 | 18217 | 6073 | 1521 | 7082 | 2233 | 709 | 15572 | 7707 | 21326 |
| 6256 | 15301 | 6670.5 | 1674.5 | 377668.5 | 42656 | 705 | 13286 | 6923 | 17594.5 |
| 4202.5 | 10432 | 4749.5 | 1344 | 6842 | 2497 | 715 | 14904.5 | 5419.5 | 13911 |
| 3216 | 5243 | 3764 | 1207 | 4973 | 1984 | 712.5 | 16149.5 | 4702 | 7508 |
| 2968 | 4268 | 3280 | 1136 | 4368 | 1876 | 706 | 14383 | 4936 | 6688 |
| 2722 | 4110 | 3286 | 1151 | 4548 | 1878 | 731 | 12325 | 4847 | 6631 |
| 2382 | 3717.5 | 2992.5 | 1133 | 4218.5 | 1826.5 | 709 | 10942.5 | 4486.5 | 6185 |
| 2618 | 3999.5 | 3319 | 1168.5 | 4406 | 1899 | 750.5 | 12462.5 | 4847.5 | 6615 |
| 2820.5 | 4358.5 | 3541 | 1228 | 4657.5 | 1977 | 778 | 14181.5 | 5159 | 7042 |
| 2711 | 4057 | 3177 | 1159 | 4321 | 1864 | 733 | 11824 | 4955 | 6625 |
| 2684 | 4103 | 3190 | 1180 | 4339 | 1872 | 741 | 12023 | 4811 | 6583 |
| 3006 | 4198 | 3327 | 1174 | 4392 | 1866 | 730 | 11382 | 4896 | 6800 |
| 2431 | 3788 | 3006 | 1112 | 4099 | 1818 | 703 | 10770 | 4394 | 6374 |
| 2468.5 | 3915.5 | 3198 | 1188 | 4417 | 1870.5 | 726 | 10812.5 | 4613.5 | 6669.5 |
| 8732 | 5889 | 3707 | 1227 | 4441 | 1860 | 701 | 11680.5 | 4842.5 | 6845 |
| 3528 | 5033 | 3717 | 1227 | 4768 | 1947 | 742 | 11649 | 5192 | 7242 |
| 2891 | 4487 | 3532 | 1228 | 4666 | 1943.5 | 750.5 | 11789.5 | 5242.5 | 7325 |
| 2740 | 4162 | 3357 | 1186 | 4592 | 1953 | 736 | 11425 | 5022 | 6924 |
| 3388.5 | 5583.5 | 3785.5 | 1260.5 | 4950 | 1977 | 723 | 14919 | 5644.5 | 8097.5 |
| 7353 | 17598 | 6177 | 1598 | 7082 | 2341 | 735 | 14641 | 8027 | 21334 |
| 9597 | 26377.5 | 8089.5 | 1924 | 9043 | 2564 | 721 | 11687.5 | 9013 | 28257 |

| | | | | | | | | | |
|---|---|---|---|---|---|---|---|---|---|
| 9977 | 25155 | 9660 | 2106 | 10445 | 2755 | 740 | 12422 | 9560 | 26962 |
| 9629 | 23890 | 8851 | 1843 | 9665 | 2631 | 698 | 12175 | 8806 | 25493 |
| 8944 | 23329 | 7479 | 1944 | 8360 | 2581 | 686 | 10971 | 9080 | 25429 |
| 8065 | 18402 | 6206 | 1607 | 7045.5 | 2313 | 714 | 14632 | 9110.5 | 21478.5 |
| 6025 | 14110 | 6666 | 1686 | 603343 | 60172 | 701 | 16422 | 7253 | 17111 |
| 3852 | 10971 | 4766 | 1381 | 6905 | 2419 | 715 | 14973 | 5246 | 14682 |
| 3091 | 5551 | 3726 | 1206 | 5173 | 2024 | 692 | 15163 | 4628 | 7936 |
| 2757 | 4584 | 3428 | 1184 | 4788 | 1962 | 707 | 15005 | 4688 | 7025 |
| 2613 | 4054 | 3198 | 1144 | 4437 | 1856 | 705 | 11648 | 4739 | 6455 |
| 2349 | 3757.5 | 3040.5 | 1109 | 4289 | 1818 | 690 | 11040.5 | 4458.5 | 6335.5 |
| 2408 | 3768.5 | 3068 | 1109 | 4192 | 1841 | 677 | 10458 | 4475.5 | 6157 |
| 2605 | 4089 | 3293 | 1172 | 4506 | 1906 | 733 | 11738 | 4800 | 6799 |
| 2450 | 3765.5 | 3029.5 | 1113 | 4111.5 | 1785 | 679 | 11122 | 4454.5 | 6253 |
| 2520 | 3896.5 | 3125 | 1134.5 | 4331 | 1867 | 701.5 | 11329.5 | 4694.5 | 6373.5 |
| 2437 | 3805 | 2997 | 1098 | 4130 | 1785 | 689 | 11002 | 4557 | 6325 |
| 2356.5 | 3746.5 | 3006 | 1090 | 4149.5 | 1800.5 | 685 | 10652 | 4442 | 6188.5 |
| 2366 | 3785 | 3163 | 1124.5 | 4302 | 1841.5 | 702 | 11135.5 | 4616 | 6471.5 |
| 23925 | 12658 | 4891 | 1197 | 4260 | 1809 | 680 | 11911 | 4852 | 6647 |
| 2750 | 4557 | 3629 | 1175 | 4444 | 1833 | 695.5 | 11527 | 4823.5 | 6681 |
| 2610 | 4371 | 3510 | 1191 | 4601 | 1925 | 725 | 11912.5 | 4939 | 7197.5 |
| 2539 | 4247.5 | 3440 | 1177 | 4697 | 1925 | 714.5 | 11563 | 4978.5 | 7175 |
| 3619 | 9051 | 4205 | 1271 | 5284 | 1979.5 | 697.5 | 15318 | 5228.5 | 12304 |
| 5759 | 15130 | 5482 | 1489 | 6686 | 2243 | 709 | 14548 | 7020 | 18463 |
| 8226 | 23755 | 7558 | 1874 | 8748 | 2579 | 708 | 11469 | 8405 | 26616 |
| 7730 | 23540 | 9249 | 2058 | 10704 | 2739 | 714 | 10386 | 7873 | 25981 |
| 8912 | 21797 | 8545 | 1818 | 9644 | 2489 | 665 | 10797 | 8440 | 23473 |
| 9161 | 23142 | 7382 | 1992 | 8300 | 2599 | 698 | 11527 | 8740 | 25164 |
| 7892 | 18938 | 6051 | 1578 | 7032.5 | 2281 | 700.5 | 12429.5 | 8561.5 | 21741.5 |
| 5791 | 13836.5 | 6468 | 1638.5 | 464810 | 52461 | 702 | 16267 | 6919.5 | 16838 |
| 4540 | 11396 | 5135 | 1465 | 7559 | 2640 | 730 | 14157 | 6166 | 14969 |
| 2916 | 7629 | 3817 | 1207.5 | 5208 | 2039 | 708 | 14297.5 | 4584 | 10921.5 |
| 2907 | 4469 | 3388 | 1166 | 4665 | 1929 | 720 | 15105 | 4993 | 6983 |
| 2552 | 3994.5 | 3215.5 | 1144 | 4481.5 | 1870 | 714 | 12750.5 | 4774.5 | 6677 |
| 2485.5 | 3841 | 3154.5 | 1140.5 | 4259.5 | 1849.5 | 720 | 11411 | 4675.5 | 6328.5 |
| 2483 | 3813 | 3172 | 1152 | 4353 | 1844 | 712 | 11341 | 4612 | 6423 |
| 2638 | 3990 | 3250.5 | 1152.5 | 4252.5 | 1857 | 713.5 | 12289 | 4973.5 | 6634 |
| 2658 | 4097 | 3139 | 1157 | 4303 | 1873 | 716 | 11843 | 4794 | 6549 |
| 2524 | 3869 | 3118 | 1085 | 4188 | 1794 | 697 | 11811 | 4675 | 6362 |
| 2716 | 4049 | 3102 | 1129 | 4278 | 1828 | 711 | 11054 | 4701 | 6314 |
| 2536 | 3935.5 | 3180 | 1150 | 4403.5 | 1868 | 709 | 11102 | 4771 | 6624 |
| 2236 | 3734.5 | 3125 | 1126 | 4218 | 1799.5 | 708 | 10963 | 4475 | 6322 |
| 17328 | 9281 | 4471 | 1204 | 4265 | 1820 | 701 | 11981 | 4868 | 6583 |
| 3080 | 4744 | 3620 | 1212 | 4602 | 1906 | 707 | 11323 | 5012 | 7075 |
| 2630.5 | 4186.5 | 3385.5 | 1151 | 4496 | 1869.5 | 699 | 11300.5 | 4811.5 | 6827.5 |
| 2488 | 4024 | 3195 | 1146.5 | 4407.5 | 1817 | 690 | 10863 | 4826 | 6714 |
| 3257.5 | 5814.5 | 3726 | 1214 | 4800.5 | 1924 | 707 | 15577 | 5166 | 8524 |
| 6732 | 16812 | 5816 | 1600 | 7152 | 2323 | 706 | 14409 | 8007 | 20068 |
| 8656 | 22171 | 8868 | 1909 | 10225 | 2619 | 714 | 12638 | 8797 | 23828 |

| | | | | | | | | | |
|---|---|---|---|---|---|---|---|---|---|
| 9644 | 26412 | 9271 | 2012 | 10735 | 2770 | 711 | 13534 | 9444 | 28951 |
| 8613 | 23676 | 8413 | 1843 | 9729 | 2558 | 692 | 11879 | 8741 | 26310 |
| 23860 | 97503 | 49471 | 4540 | 9071 | 2734 | 708 | 11101 | 9860 | 31368 |
| 17626.5 | 25037 | 7862 | 1882.5 | 7526.5 | 2380 | 747 | 11866 | 9635 | 24542.5 |
| 13038 | 18272 | 9609 | 2399 | 72726 | 8981 | 772 | 12751 | 7970 | 18317 |
| 7018.5 | 11763 | 4745 | 1436 | 15172.5 | 3287.5 | 776.5 | 14975.5 | 5532 | 14140 |
| 6780.5 | 6929 | 4214 | 1323.5 | 8124 | 2341 | 794 | 19136.5 | 5415.5 | 8714.5 |
| 5326.5 | 5177.5 | 3602.5 | 1180 | 6277 | 2222 | 772.5 | 17192 | 4966 | 7044 |
| 5849 | 4967 | 3334 | 1151 | 5389 | 2068 | 799 | 14655 | 5294 | 6858 |
| 5648 | 4993 | 3627 | 1182 | 5830 | 2096 | 799 | 13295 | 5319 | 6891 |
| 4982.5 | 5458 | 4964.5 | 1610 | 5043.5 | 1964.5 | 789.5 | 12819 | 5342.5 | 7083.5 |
| 5288 | 4865.5 | 3728.5 | 1217 | 5420.5 | 2030 | 813 | 14026 | 5447 | 7060.5 |
| 5001 | 4741 | 3350 | 1158 | 4556 | 1869 | 798 | 13141 | 5358 | 6938 |
| 5115 | 4518 | 3285 | 1127 | 5022 | 1865 | 776 | 12176 | 5186 | 6592 |
| 8660 | 7037 | 4349 | 1233 | 4377 | 1833 | 742 | 11922 | 7230 | 8026 |
| 8239 | 5963.5 | 3669.5 | 1169 | 4940.5 | 1918.5 | 772.5 | 12786 | 5963 | 7097.5 |
| 6181 | 5193 | 3544 | 1203 | 4541 | 1895 | 772 | 13130 | 5437 | 6874 |
| 1273094 | 741392.5 | 213002.5 | 14460.5 | 4817.5 | 1939.5 | 747.5 | 112308 | 36135 | 21473.5 |
| 60574 | 31402.5 | 9578.5 | 1503.5 | 4555 | 1889 | 725 | 15471 | 6189.5 | 7633.5 |
| 29878 | 17194 | 6187 | 1414 | 5450 | 2120 | 817 | 16428 | 6014 | 8297 |
| 29667 | 16258 | 5737 | 1353 | 4751 | 1920 | 736 | 16158 | 5832 | 7525 |
| 15722.5 | 14857.5 | 5334.5 | 1392.5 | 5871 | 2084.5 | 784.5 | 19241.5 | 7088.5 | 12936 |
| 12942.5 | 21462.5 | 6963.5 | 1645.5 | 7476.5 | 2378 | 762 | 16907 | 8191 | 20715 |
| 12706 | 28129.5 | 8187.5 | 1815 | 8959 | 2544 | 762 | 11971 | 9074 | 26579 |
| 11555 | 26174.5 | 10154 | 2023 | 10802.5 | 2706 | 754 | 9783 | 7766 | 24939.5 |
| 8583 | 22756 | 8578 | 2005 | 10118 | 2722.5 | 697.5 | 10772.5 | 8115.5 | 24921.5 |
| 9534 | 23456 | 7854 | 1874.5 | 8615.5 | 2533 | 697.5 | 12692 | 9387.5 | 25184 |
| 7901.5 | 21111 | 6647.5 | 1651.5 | 7752.5 | 2367 | 726 | 12802 | 8379 | 23877.5 |
| 6477.5 | 14854.5 | 6736 | 1704.5 | 304677.5 | 33815.5 | 697 | 12221.5 | 7573.5 | 17462.5 |
| 3770.5 | 10428 | 4753 | 1329.5 | 7165.5 | 2496 | 707 | 14866.5 | 5218.5 | 13757 |
| 3378 | 7332 | 3880 | 1215 | 5513 | 2129.5 | 702.5 | 14629 | 4936.5 | 10199.5 |
| 2806.5 | 4504.5 | 3459.5 | 1202 | 4739.5 | 1983 | 708.5 | 14711 | 4725 | 6795.5 |
| 2564.5 | 3956 | 3106.5 | 1109 | 4461.5 | 1872 | 693 | 11796 | 4662 | 6611 |
| 2332 | 3736 | 3040 | 1084 | 4281 | 1798 | 691 | 10862 | 4537 | 6197 |
| 2343 | 3739 | 2973 | 1083 | 4219 | 1795 | 671 | 10682 | 4491 | 6192 |
| 2495 | 3953 | 3110.5 | 1107.5 | 4332.5 | 1826 | 710 | 11475 | 4685 | 6625.5 |
| 2374 | 3681 | 2912 | 1071 | 4037 | 1748 | 659 | 10578 | 4387 | 5980 |
| 2436 | 3823 | 3032 | 1087 | 4237 | 1812 | 690 | 11121 | 4522 | 6363 |
| 2489 | 3773 | 2941 | 1070 | 4029 | 1791 | 679 | 10563 | 4367 | 6208 |
| 2331 | 3682 | 2945.5 | 1063 | 4061 | 1769 | 675 | 10203 | 4380 | 6160.5 |
| 2308 | 3754.5 | 3052.5 | 1102 | 4188 | 1790 | 699 | 10565.5 | 4522 | 6302 |
| 35097.5 | 17990.5 | 5888.5 | 1258 | 4354 | 1852 | 700.5 | 12807 | 5158.5 | 6639 |
| 3275 | 5112 | 4052 | 1173 | 4344 | 1863 | 688 | 11475 | 4913 | 6708 |
| 2614.5 | 4262.5 | 3392 | 1153 | 4453 | 1863 | 680 | 11466.5 | 4820.5 | 6703.5 |
| 2506.5 | 4157 | 3364 | 1127 | 4447.5 | 1848 | 702 | 11001 | 4781 | 6850.5 |
| 3443 | 5937 | 3711 | 1187 | 4802 | 1927 | 684 | 15260 | 5610 | 8514 |
| 7583 | 18340 | 6108 | 1556 | 7242 | 2242 | 689 | 15223 | 8672 | 21495 |
| 8557 | 23660 | 7460 | 1765 | 8846 | 2527 | 676 | 11836 | 8627 | 26559 |

| | | | | | | | | | |
|---|---|---|---|---|---|---|---|---|---|
| 8682 | 24937 | 8527 | 1929 | 9744 | 2617 | 685 | 11115 | 8751 | 26862 |
| 7750.5 | 22222.5 | 8335.5 | 1860 | 9395.5 | 2526 | 663 | 10172.5 | 7806 | 24003.5 |
| 9539 | 22964 | 7983.5 | 1928 | 8758 | 2589 | 707.5 | 12324.5 | 9337.5 | 25673.5 |
| 8627.5 | 21182.5 | 6629.5 | 1684 | 7539 | 2381 | 740 | 14170.5 | 9541.5 | 24428 |
| 6898 | 15984 | 7585 | 2110 | 731799 | 62443 | 725 | 15438 | 8278 | 18518 |
| 4586 | 11677 | 5257 | 1479 | 8501 | 2771 | 718 | 14931 | 5890 | 14977 |
| 3345 | 8050.5 | 4312.5 | 1334 | 5970.5 | 2212.5 | 760 | 14992.5 | 5024.5 | 11510 |
| 2981 | 4707 | 3639 | 1194 | 5156 | 2005 | 758 | 17053 | 4931 | 7304 |
| 2856 | 4369 | 3583 | 1242 | 4829 | 1964 | 774 | 15017 | 5035 | 7050 |
| 2585 | 4037 | 3401 | 1193 | 4692 | 1933 | 774 | 12500 | 4963 | 6740 |
| 2606 | 4153 | 3413 | 1211 | 4605 | 1939 | 762 | 13599 | 4991 | 6996 |
| 2631 | 3977 | 3329 | 1199 | 4583 | 1892 | 739 | 11520 | 4948 | 6640 |
| 2650.5 | 4105 | 3356 | 1188.5 | 4563 | 1928.5 | 746 | 12050.5 | 5029 | 6729.5 |
| 2668 | 4039 | 3197 | 1155 | 4415 | 1844 | 754 | 11605 | 4901 | 6665 |
| 2544 | 3955 | 3198 | 1142 | 4396 | 1900 | 753 | 11998 | 4837 | 6656 |
| 2478.5 | 3889 | 3235 | 1130 | 4384 | 1862.5 | 762 | 11278 | 4731 | 6569.5 |
| 2288 | 3606 | 3132 | 1154 | 4286 | 1897 | 719 | 10735 | 4464 | 6239 |
| 14832 | 8498 | 4407.5 | 1232.5 | 4420 | 1897 | 756.5 | 12197.5 | 5165 | 6909.5 |
| 2989 | 4670 | 3677 | 1243 | 4518 | 1906 | 756 | 11877 | 5089 | 7158 |
| 2820 | 4479 | 3688 | 1267 | 4794 | 1975 | 768 | 12205 | 5199 | 7325 |
| 2668 | 4294 | 3574 | 1217 | 4692 | 1947 | 747 | 11589 | 5080 | 7087 |
| 3352 | 5333 | 3915 | 1285 | 4936 | 2047 | 776 | 13329 | 5511 | 7989 |
| 5397.5 | 13627 | 5214.5 | 1478 | 6077.5 | 2139 | 726.5 | 14883 | 6489.5 | 16338.5 |
| 9340 | 25069 | 7923 | 1922 | 9098 | 2639 | 776 | 12245 | 8950 | 28445 |
| 9526 | 25003 | 8863 | 1989 | 9933 | 2656 | 749 | 11172 | 8825 | 27351 |
| 8644.5 | 23521 | 9373.5 | 1993 | 10501 | 2715 | 717 | 9909 | 7945.5 | 25799.5 |
| 10219 | 32394 | 25896 | 7174 | 11308 | 3892 | 690 | 12693 | 9778 | 28765 |
| 8490 | 19779 | 7820 | 2161 | 7735 | 2601 | 696 | 12494 | 8863 | 22149 |
| 6148.5 | 15548.5 | 10139 | 2766 | 997683 | 104948 | 687 | 12746.5 | 7297 | 18020 |
| 4584.5 | 10331 | 6400 | 1909 | 14859.5 | 3611 | 758 | 16977 | 6059 | 13876.5 |
| 3655 | 7323.5 | 5091.5 | 1608 | 10259.5 | 2759 | 744 | 16384.5 | 5193 | 10288 |
| 2877 | 4606 | 3889 | 1318 | 8018 | 2291 | 711 | 16118 | 4546 | 7069 |
| 3016 | 4314 | 3589.5 | 1272.5 | 7736.5 | 2286 | 738 | 13878 | 5111 | 6991.5 |
| 2846 | 4091 | 3432 | 1206 | 7090 | 2164 | 731 | 12069 | 4899 | 6742 |
| 2836 | 4313 | 3848 | 1430 | 6641 | 2305 | 728 | 11518 | 4894 | 6802 |
| 2709.5 | 3929.5 | 3265 | 1182 | 6195 | 2082 | 706 | 12293.5 | 4661.5 | 6454.5 |
| 2609.5 | 3877.5 | 3191.5 | 1170 | 5712 | 2020.5 | 695 | 10956 | 4634.5 | 6347 |
| 2620 | 3895 | 3314 | 1164 | 5630 | 2026 | 717 | 11693 | 4840 | 6510 |
| 8350.5 | 6840 | 4081 | 1176.5 | 5683 | 1984 | 709 | 10799 | 6262.5 | 7438.5 |
| 2923 | 4246 | 3482 | 1161 | 5441 | 1974 | 729 | 11431 | 5040 | 6774 |
| 2532.5 | 3929 | 3291.5 | 1151 | 5413.5 | 2011 | 733 | 10918.5 | 4799.5 | 6791 |
| 163847.5 | 82118 | 21890 | 1864.5 | 5349.5 | 1990 | 697 | 22048 | 7483 | 7573 |
| 3442 | 5059 | 4414 | 1396 | 5454 | 2050 | 735 | 13223 | 5608 | 7290 |
| 2804 | 4376.5 | 3664 | 1272 | 5426 | 2038 | 730 | 11725.5 | 5098 | 7092 |
| 2474 | 3934.5 | 3380 | 1177 | 4855 | 1955.5 | 718 | 11326 | 4700 | 6722 |
| 3956 | 6858 | 4006 | 1266 | 5670 | 2064 | 734 | 15550 | 6388 | 10182 |
| 6141 | 16373 | 5941 | 1600 | 7373 | 2352 | 722 | 14166 | 6958 | 19122 |
| 9493.5 | 25778 | 8466.5 | 1866.5 | 9746 | 2615.5 | 698.5 | 15000 | 9601 | 27851.5 |

| | | | | | | | | | |
|---|---|---|---|---|---|---|---|---|---|
| 9723 | 25117 | 9327 | 1977 | 10568 | 2769 | 720 | 13085 | 9353 | 27041 |
| 9364 | 23420 | 8354 | 1851 | 9502 | 2593 | 691 | 11739 | 8731 | 25198 |
| 9087.5 | 25427.5 | 15404 | 4974.5 | 12426 | 4662 | 695.5 | 10042.5 | 8567 | 26682.5 |
| 8339 | 18898 | 7810 | 2542 | 7681 | 2889 | 679 | 10850 | 8406 | 21784 |
| 6063.5 | 14455 | 8170 | 2574 | 588241 | 51978.5 | 673 | 13146.5 | 6756 | 17068.5 |
| 3479.5 | 9870 | 5716 | 1907 | 9101 | 3121 | 698 | 14667.5 | 4916 | 13252.5 |
| 3222 | 6222 | 4661.5 | 1621 | 6644.5 | 2469 | 711 | 16417 | 4955 | 8782 |
| 2681 | 4509 | 3825 | 1347 | 5461 | 2118 | 714 | 15854 | 4483 | 6944 |
| 2508 | 4007.5 | 3466 | 1223 | 4971 | 1965 | 707 | 11692 | 4741.5 | 6651 |
| 2417 | 3841 | 3363 | 1203 | 4692 | 1969 | 717 | 11783 | 4711 | 6528 |
| 2310 | 3733 | 3279 | 1212 | 4434 | 1972 | 690 | 10768 | 4461 | 6246 |
| 2415 | 3736 | 3169.5 | 1153 | 4360 | 1874.5 | 687 | 11183.5 | 4565 | 6273 |
| 2401 | 3762 | 3164.5 | 1101.5 | 4383.5 | 1826 | 695 | 11357 | 4570 | 6314 |
| 2402 | 3800 | 3079.5 | 1136 | 4268.5 | 1856.5 | 682 | 11343.5 | 4452 | 6188.5 |
| 2239 | 3659 | 3004 | 1095 | 4185 | 1796 | 675 | 10906 | 4151 | 5962 |
| 2244 | 3705 | 3018 | 1097 | 4284 | 1827 | 694 | 10581 | 4341 | 6217 |
| 2072 | 3550 | 2934 | 1092 | 4085 | 1785 | 681 | 10055 | 4063 | 5886 |
| 22915.5 | 12116.5 | 4885 | 1187.5 | 4233.5 | 1844.5 | 667 | 11402.5 | 4810.5 | 6491 |
| 2765 | 4397 | 3539 | 1159 | 4386 | 1846 | 681 | 10577 | 4700 | 6573 |
| 2502 | 4363 | 3469 | 1150 | 4562 | 1895 | 693 | 11185 | 4803 | 7015 |
| 2450 | 4056 | 3218 | 1116 | 4407 | 1855 | 678 | 10689 | 4628 | 6757 |
| 4135.5 | 7453 | 3997 | 1237 | 5187 | 1961 | 676 | 16283 | 6286 | 10218 |
| 4895 | 13641 | 5124 | 1411 | 6092.5 | 2102.5 | 659 | 14591 | 5978 | 16447.5 |
| 8500.5 | 22322.5 | 7001 | 1751 | 8120 | 2466 | 681 | 12593.5 | 8664 | 24836 |
| 9661 | 24405.5 | 7881.5 | 1819 | 9000.5 | 2521.5 | 660 | 13397 | 9597.5 | 26833 |
| 8296 | 22357 | 8672 | 1879 | 10131 | 2629 | 690 | 9968 | 8121 | 24198 |
| 8774 | 24308.5 | 12552.5 | 4051 | 10968.5 | 4264 | 683.5 | 11296.5 | 9137.5 | 26224 |
| 7374 | 16763 | 6488 | 2072 | 6777 | 2540 | 637 | 16232 | 8269 | 19505 |
| 6268 | 14951.5 | 8327 | 2317.5 | 734791.5 | 62128.5 | 719 | 14443 | 7490.5 | 17943.5 |
| 4157 | 10746 | 5368.5 | 1802.5 | 7146.5 | 2757.5 | 687 | 14325 | 5622.5 | 13995.5 |
| 3212 | 5417 | 4169 | 1389 | 5459 | 2162 | 679 | 15722 | 4753 | 7686 |
| 2913 | 5287 | 4049.5 | 1404 | 5316 | 2132.5 | 738 | 15836 | 4782.5 | 7915.5 |
| 2728 | 4102 | 3425 | 1188 | 4620 | 1929 | 696 | 12991 | 4827 | 6723 |
| 2513 | 3764 | 3070 | 1104 | 4214 | 1828 | 662 | 10436 | 4651 | 6155 |
| 2592 | 4026 | 3283 | 1177 | 4473.5 | 1902 | 695 | 10930.5 | 4715 | 6593 |
| 2634 | 3998 | 3172 | 1122 | 4257.5 | 1832.5 | 675 | 12125 | 4670.5 | 6359.5 |
| 2546 | 3874 | 3052 | 1094 | 4128.5 | 1787.5 | 661 | 10538 | 4582 | 6199 |
| 2487 | 3794 | 2968 | 1048 | 4097 | 1743 | 661 | 11111 | 4470 | 6140 |
| 2741.5 | 4083 | 3225.5 | 1131 | 4333 | 1832 | 696 | 11503 | 4820 | 6520.5 |
| 2537 | 3749 | 3002 | 1077 | 4111 | 1795 | 676 | 10969 | 4579 | 6086 |
| 2476 | 3819 | 3063.5 | 1113 | 4230 | 1798.5 | 668 | 10377.5 | 4562.5 | 6260 |
| 24850 | 13035 | 5046 | 1208 | 4270 | 1850 | 672 | 11866 | 4791 | 6611 |
| 3222 | 4978 | 3794 | 1177 | 4480 | 1862 | 689 | 11471 | 4976 | 6921 |
| 2792 | 4279 | 3459 | 1138 | 4403 | 1842 | 666 | 10777 | 4658 | 6589 |
| 2438 | 3791 | 3019.5 | 1056 | 4046.5 | 1710 | 625 | 10245 | 4396 | 6052.5 |
| 3485 | 9044 | 4085 | 1269 | 5286 | 1969 | 682 | 15431 | 5101 | 12327 |
| 5618 | 15016 | 5457 | 1487 | 6557 | 2193 | 669 | 14876 | 6817 | 18423 |
| 9582 | 21024 | 6859.5 | 1679 | 7738 | 2355 | 648 | 15028.5 | 9495 | 23179.5 |

| | | | | | | | | | |
|---|---|---|---|---|---|---|---|---|---|
| 9329 | 24522 | 8336 | 1913 | 9555 | 2599 | 666 | 13678 | 9605 | 26511 |
| 698 | 752 | 533 | 537 | 1001 | 1086 | 169 | 1665.5 | 1139 | 1241.5 |
| 3996.5 | 4942 | 2485 | 817 | 2984.5 | 1398 | 266 | 13503.5 | 4436 | 5281.5 |
| 3246 | 3721 | 2065 | 761 | 2678 | 1353 | 421 | 15437 | 4043 | 4489 |
| 2348 | 3040 | 6871 | 2397 | 801454 | 76709 | 533 | 13288 | 3455 | 4411 |
| 2053 | 2960.5 | 3313 | 1172 | 5408 | 2407 | 634 | 11868 | 3488.5 | 4861 |
| 1860 | 2868 | 2817 | 1059 | 4204 | 1928 | 700 | 11397 | 3599 | 5225 |
| 1625 | 2734 | 2566 | 1018 | 3844.5 | 1787 | 732 | 9726 | 3508 | 5191 |
| 1606 | 2783 | 2635 | 1035 | 3839 | 1788 | 759 | 9950 | 3575 | 5409 |
| 1583 | 2741 | 2494 | 991 | 3723 | 1758 | 745 | 9524 | 3487 | 5268 |
| 1611 | 2781 | 2544 | 1011 | 3661 | 1748 | 734 | 9408 | 3479 | 5214 |
| 1571 | 2660.5 | 2347 | 958 | 3378 | 1659 | 702 | 8975.5 | 3296.5 | 4907 |
| 1634 | 2865 | 2555 | 1027 | 3737 | 1749 | 749 | 10089 | 3569 | 5456 |
| 1519 | 2633 | 2386 | 982 | 3503.5 | 1691 | 722 | 8673 | 3320 | 5091 |
| 1541.5 | 2618.5 | 2334 | 984 | 3447 | 1677 | 710 | 8529 | 3265 | 5015 |
| 1425 | 2519 | 2358 | 984 | 3406 | 1684 | 718 | 8185 | 3220 | 4945 |
| 1401 | 2536 | 2365.5 | 990 | 3459.5 | 1684 | 711 | 8206.5 | 3261 | 5020.5 |
| 40833 | 23206 | 6442 | 1163 | 3478 | 1689.5 | 705 | 11374.5 | 3898.5 | 5378.5 |
| 1914 | 3353 | 3039 | 999 | 3346 | 1633 | 666 | 8756 | 3431 | 5074 |
| 1786 | 3305 | 2967.5 | 1071 | 3948.5 | 1798 | 754 | 9885 | 3843 | 5831 |
| 1636 | 3029 | 2689 | 1043 | 3773 | 1755 | 733 | 9183 | 3603 | 5542 |
| 2631 | 4093 | 2960 | 1052 | 4016 | 1743 | 711 | 12707 | 4627 | 6322 |
| 6861 | 16259.5 | 5387.5 | 1417 | 6453 | 2125 | 724 | 15678 | 8192 | 18968.5 |
| 8973.5 | 21141.5 | 6393.5 | 1601 | 7375 | 2304 | 706 | 15145.5 | 9439.5 | 23542 |
| 10470 | 26200.5 | 7615 | 1775 | 8419.5 | 2495 | 745 | 11827 | 8662 | 26258.5 |
| 9686 | 23571 | 7031 | 1704 | 7974 | 2433 | 729 | 11809 | 8544 | 25286 |
| 9289 | 21453 | 6509 | 1650 | 7235 | 2329 | 744 | 14561 | 9558 | 23763 |
| 8167.5 | 18214 | 5538.5 | 1452 | 6267 | 2168 | 754 | 13323.5 | 8156 | 20257 |
| 5091 | 12324 | 5758 | 1514 | 113816 | 11854 | 766 | 15699 | 6414 | 15309 |
| 3169 | 5792 | 4053 | 1220 | 6597.5 | 2505 | 743 | 14382 | 4584 | 8088 |
| 2799 | 4168.5 | 3173.5 | 1101 | 4553.5 | 1963 | 770 | 16829.5 | 4617 | 6431 |
| 2259.5 | 3326 | 2692 | 1024 | 3866 | 1778.5 | 726 | 11199 | 4086 | 5501.5 |
| 1830 | 3005 | 2585 | 1043 | 3777 | 1754 | 754 | 10480 | 3639 | 5402 |
| 1682 | 2794 | 2387 | 970 | 3505 | 1675 | 705 | 9520 | 3413 | 5087 |
| 1887 | 3123 | 2628 | 1023 | 3696 | 1738 | 753 | 10667 | 3783 | 5579 |
| 1825 | 2923 | 2450.5 | 985.5 | 3443.5 | 1682 | 700 | 10161.5 | 3560 | 5180 |
| 1748 | 2946 | 2423 | 972 | 3507 | 1664 | 716 | 9838 | 3529 | 5232 |
| 1701.5 | 2789.5 | 2383.5 | 995 | 3400 | 1668 | 709 | 9233 | 3426.5 | 5096 |
| 1747 | 2863 | 2469 | 981 | 3512 | 1702 | 732 | 9536 | 3511 | 5255 |
| 1586.5 | 2764 | 2418 | 992 | 3509 | 1696.5 | 736 | 8996.5 | 3447 | 5155.5 |
| 1614 | 2829 | 2500 | 997 | 3580 | 1710 | 745 | 9090 | 3485 | 5327 |
| 42553.5 | 24201.5 | 6780 | 1193.5 | 3624.5 | 1709 | 721 | 12087.5 | 4116 | 5657 |
| 2447 | 4181.5 | 3536.5 | 1086 | 3752 | 1732 | 726 | 10334.5 | 3981.5 | 5775.5 |
| 2060 | 3540 | 3058 | 1093 | 3918 | 1769 | 751 | 10361 | 4015 | 5982 |
| 1891 | 3315 | 2796 | 1052 | 3884 | 1758 | 740 | 9627 | 4015 | 5959 |
| 3149.5 | 6068.5 | 3603.5 | 1151 | 4573.5 | 1851 | 743 | 16445.5 | 4573 | 8404 |
| 5157 | 13466 | 4925 | 1370 | 5925 | 2083 | 769 | 14790 | 6168 | 16325 |
| 9192 | 22422 | 6850 | 1674 | 7796 | 2371 | 745 | 13826 | 9162 | 24581 |

| | | | | | | | | | |
|---|---|---|---|---|---|---|---|---|---|
| 9645 | 25843 | 7999 | 1861 | 9139 | 2569 | 757 | 11786 | 8971 | 27416 |
| 10366 | 24462 | 7127 | 1709 | 7930 | 2421 | 719 | 11666 | 8328 | 24812 |
| 8819.5 | 20034 | 6068 | 1560 | 7029.5 | 2261 | 724 | 12226.5 | 8940 | 22017 |
| 6652 | 15676 | 5033 | 1346 | 5987 | 2065 | 728 | 12227 | 7285 | 17792 |
| 4983.5 | 12297 | 6042 | 1518.5 | 514432.5 | 46023.5 | 756 | 14232.5 | 5977.5 | 15186 |
| 3521 | 8147 | 4222 | 1251 | 7699 | 2762 | 737 | 14584 | 4931 | 11127 |
| 2722 | 4118 | 3071 | 1060 | 4494 | 1896 | 732 | 15906 | 4428 | 6241.5 |
| 2517 | 3554 | 2801 | 1031 | 4110 | 1808 | 737 | 12759 | 4474 | 5723 |
| 2131 | 3294 | 2697 | 1032 | 3901 | 1768 | 747 | 11255 | 4117 | 5673 |
| 1893 | 3115 | 2593 | 992 | 3702 | 1700 | 730 | 10432 | 3747 | 5462 |
| 1987.5 | 3159.5 | 2598 | 1021 | 3750 | 1720.5 | 745 | 10121.5 | 3900 | 5496.5 |
| 1888 | 3073 | 2553 | 986 | 3609 | 1696.5 | 717 | 10139 | 3661 | 5345.5 |
| 1856 | 3082 | 2518 | 991.5 | 3593 | 1702 | 712 | 9806 | 3670.5 | 5323 |
| 1824 | 3041 | 2504 | 985 | 3592 | 1701 | 728 | 9807 | 3664 | 5441 |
| 1742 | 2835.5 | 2372 | 941 | 3320 | 1633 | 682.5 | 8802 | 3440 | 5028.5 |
| 1703 | 2892 | 2416 | 974 | 3521 | 1678 | 710 | 9051 | 3484 | 5133 |
| 1717 | 2973.5 | 2525.5 | 1006 | 3604 | 1704 | 728 | 9351.5 | 3573.5 | 5364.5 |
| 38609 | 21847 | 6452.5 | 1176 | 3686.5 | 1684 | 712 | 12688 | 4301 | 5767 |
| 2115 | 3512 | 2925 | 1017 | 3551 | 1669 | 688 | 9411 | 3703 | 5387 |
| 1995 | 3376 | 2834 | 1038 | 3807 | 1726 | 708 | 9846 | 3903 | 5772 |
| 1918 | 3257.5 | 2716 | 1010 | 3768.5 | 1729 | 722 | 9750 | 3917 | 5744.5 |
| 3201 | 5446 | 3367 | 1103 | 4495 | 1829 | 717 | 16614 | 4860 | 7737 |
| 5075 | 12899.5 | 4686 | 1275 | 5679 | 1967 | 688 | 16343 | 5909.5 | 15369 |
| 8446 | 18315 | 5855 | 1480 | 6822.5 | 2182.5 | 692.5 | 15619 | 8820.5 | 20698.5 |
| 9437 | 21610 | 6574 | 1607 | 7578 | 2284 | 702 | 13378 | 8926 | 23304 |
| 8886 | 24309 | 7265 | 1705 | 8569 | 2460 | 754 | 14208 | 8936 | 26650 |
| 11096 | 28121 | 8746 | 2309 | 8752.5 | 2773 | 779 | 14077.5 | 10092.5 | 28376 |
| 9599 | 20354 | 6087 | 1606 | 7093 | 2345 | 853 | 12536 | 8602 | 22185 |
| 5880 | 14153 | 8156 | 1879 | 933178 | 76130 | 846 | 14292 | 6950 | 17020 |
| 3664 | 9432 | 4413 | 1303 | 6480 | 2432 | 807 | 14400 | 5049 | 12702 |
| 3156 | 5123 | 3518 | 1181 | 4978 | 2021 | 829 | 16825 | 4883 | 7669 |
| 2790 | 4099 | 3204 | 1155 | 4572 | 1911 | 828 | 15390 | 4935 | 6634 |
| 2514 | 3601 | 2912 | 1095 | 4176 | 1848 | 826 | 12736 | 4741 | 6206 |
| 2148 | 3370 | 2770 | 1081 | 3981 | 1795 | 775 | 11000 | 4135 | 5902 |
| 2278 | 3410 | 2785 | 1056 | 3841 | 1788 | 791 | 11996 | 4344 | 5892.5 |
| 2265 | 3452 | 2781 | 1058 | 3917.5 | 1801 | 789 | 11943 | 4370 | 5963 |
| 2457 | 3617.5 | 2917 | 1132 | 4138 | 1862 | 857 | 12100.5 | 4757.5 | 6342.5 |
| 2070 | 3331 | 2706.5 | 1043 | 3877.5 | 1788.5 | 793 | 11059 | 4054.5 | 5751 |
| 2062 | 3248 | 2669 | 1072 | 3779 | 1766 | 774 | 11426 | 4043 | 5742 |
| 2007 | 3262.5 | 2717 | 1068 | 3912 | 1798 | 804 | 11059 | 4122.5 | 5923.5 |
| 1980 | 3334.5 | 2847 | 1100 | 4054.5 | 1845 | 822.5 | 11078 | 4201 | 6107 |
| 29246 | 16371 | 5615 | 1238 | 4071 | 1838 | 797 | 12793 | 4625 | 6273 |
| 3807.5 | 5794.5 | 4324 | 1192 | 4253 | 1836.5 | 814 | 13003.5 | 4872 | 6660 |
| 2352 | 3843 | 3173 | 1124 | 4235 | 1852 | 773 | 11105 | 4610 | 6353 |
| 2194 | 3647 | 3027 | 1116 | 4194 | 1859 | 811 | 11488 | 4587 | 6359 |
| 2869 | 4289.5 | 3276 | 1172 | 4403.5 | 1916.5 | 793 | 13757 | 5104.5 | 6883.5 |
| 6499 | 15846 | 5697 | 1542 | 6721.5 | 2260 | 831 | 16146.5 | 7556.5 | 18767 |
| 9562 | 23905 | 7240 | 1797 | 8152 | 2475.5 | 815.5 | 12194 | 8610.5 | 25551.5 |

| | | | | | | | | | |
|---|---|---|---|---|---|---|---|---|---|
| 10211 | 26510 | 9260 | 2000 | 10597 | 2738 | 836 | 13606 | 8838 | 27063 |
| 9447 | 23617.5 | 7727 | 1771.5 | 8815.5 | 2501 | 753 | 11695 | 8732.5 | 25367.5 |
| 9507 | 21139 | 6767 | 1777 | 7612 | 2394 | 742 | 11909 | 8810 | 22802 |
| 8506 | 17219.5 | 5559.5 | 1499 | 6406.5 | 2202 | 830 | 12981.5 | 8403 | 19587 |
| 5005 | 11317.5 | 6494.5 | 1664.5 | 694130.5 | 69942 | 819 | 13793.5 | 6068.5 | 14043.5 |
| 3187 | 5092 | 3877 | 1220 | 5753 | 2279 | 753 | 15191 | 4273 | 7070 |
| 2937 | 4311 | 3202 | 1122 | 4554 | 1905 | 795 | 16677 | 4720 | 6499 |
| 2632 | 4087 | 3224.5 | 1178 | 4480 | 1943 | 873 | 15305.5 | 4689.5 | 6602 |
| 2336 | 3305 | 2704 | 1068 | 3869.5 | 1794 | 793 | 11235 | 4351.5 | 5713 |
| 2126 | 3378 | 2775 | 1064 | 3920 | 1807 | 834 | 11019 | 4177 | 5825 |
| 2300 | 3618 | 2908 | 1157 | 4109 | 1925 | 837 | 12299 | 4398 | 6111 |
| 2340.5 | 3509 | 2863 | 1086 | 4026 | 1816 | 824 | 12091.5 | 4484 | 6157 |
| 2078 | 3301 | 2720 | 1112 | 3749 | 1821 | 769 | 11358 | 3914 | 5577 |
| 2246 | 3435 | 2830 | 1097 | 4004 | 1810 | 844 | 10979 | 4404 | 6116 |
| 1997 | 3179 | 2672 | 1048 | 3718 | 1765 | 774 | 10187 | 3814 | 5572 |
| 2084.5 | 3340.5 | 2836 | 1101 | 4062.5 | 1815.5 | 829 | 11225.5 | 4297 | 5986.5 |
| 2060 | 3311 | 2779 | 1089 | 4045 | 1815 | 827 | 11288.5 | 4301 | 5999 |
| 56517.5 | 30655.5 | 8608.5 | 1372 | 4086.5 | 1838 | 823.5 | 14886 | 5251.5 | 6580 |
| 3212 | 4892.5 | 4039.5 | 1214.5 | 4183.5 | 1866 | 837 | 12605 | 4986 | 6592.5 |
| 2612 | 3997 | 3331 | 1181 | 4384 | 1927 | 854 | 12712 | 5076 | 6734 |
| 2184 | 3575 | 2984 | 1137 | 4082 | 1855 | 809 | 10851 | 4349 | 6277 |
| 2693 | 4153 | 3219 | 1167 | 4319.5 | 1871 | 797.5 | 12450.5 | 4786 | 6557 |
| 7254 | 17686 | 5712 | 1559 | 6710 | 2272 | 858 | 16226 | 7970 | 20653 |
| 10417 | 25163 | 7532 | 1853.5 | 8681.5 | 2571 | 838 | 12654.5 | 8829.5 | 26293.5 |
| 11023.5 | 28268 | 8773 | 1996 | 9883.5 | 2714 | 829 | 11675 | 8975.5 | 27649 |
| 9496 | 22911 | 6292 | 1602 | 7414 | 2276 | 688 | 11605 | 8480 | 23950 |
| 9297 | 23608.5 | 7588.5 | 1945 | 7795.5 | 2507 | 738 | 11001 | 8815.5 | 25510 |
| 7033 | 16075 | 5273 | 1400 | 6238 | 2128 | 750 | 13410 | 8006 | 18570 |
| 5092 | 12332 | 6438 | 1588 | 654631 | 59296 | 740 | 15524 | 6376 | 15239 |
| 3512.5 | 9209.5 | 4415 | 1270 | 6823.5 | 2571 | 788.5 | 15053.5 | 4902.5 | 12688 |
| 3354.5 | 4961 | 3584 | 1183.5 | 5163 | 2165 | 764 | 17724 | 5110 | 7408 |
| 2402 | 3603 | 2824 | 1049 | 4173 | 1818 | 755 | 13267 | 4444 | 5970 |
| 2149 | 3376 | 2742.5 | 1069 | 3991.5 | 1839 | 761 | 11497 | 4174 | 5813 |
| 2124 | 3277 | 2715 | 1041.5 | 3902.5 | 1755 | 763 | 11199.5 | 4253.5 | 5785 |
| 2209 | 3372 | 2713 | 1022 | 3870 | 1773 | 755 | 12628 | 4346 | 5959 |
| 2237 | 3483 | 2758 | 1050 | 3966 | 1833 | 785 | 11778 | 4285 | 6002 |
| 2153 | 3384 | 2688.5 | 1051 | 3947 | 1765 | 758 | 10665.5 | 4302 | 5892 |
| 2069.5 | 3372.5 | 2746.5 | 1045 | 3844 | 1791.5 | 757 | 11689.5 | 4182 | 5904 |
| 2014 | 3280.5 | 2718.5 | 1042.5 | 3829 | 1784 | 763 | 10747 | 3979 | 5829 |
| 1783 | 3020 | 2483 | 998 | 3631 | 1701 | 715 | 9595 | 3627 | 5392 |
| 1876 | 3160 | 2614 | 1025 | 3878.5 | 1756.5 | 750 | 10052 | 3871 | 5685 |
| 39568 | 21366 | 6189 | 1191 | 3862 | 1765 | 744 | 13304 | 4819 | 6265 |
| 2939 | 4612.5 | 3730.5 | 1152 | 4183 | 1858 | 747.5 | 12089 | 4684.5 | 6341.5 |
| 2458 | 4032 | 3287 | 1150 | 4308 | 1896 | 776 | 12922 | 4903 | 6847 |
| 2196 | 3658 | 2989 | 1099 | 4192 | 1811 | 752 | 11174 | 4513 | 6331 |
| 3649 | 8451 | 3733 | 1183 | 5070 | 1932 | 758 | 15983 | 5614 | 11807 |
| 5009 | 13571 | 4941 | 1371 | 6005 | 2122 | 763 | 14823 | 6158 | 16404 |
| 9328 | 25223 | 7838 | 1769 | 8967 | 2524 | 788 | 15036 | 9270 | 26865 |

| | | | | | | | | | |
|---|---|---|---|---|---|---|---|---|---|
| 8516 | 20781 | 7716 | 1725 | 9005 | 2476 | 722 | 11693 | 8155 | 22670 |
| 9633 | 26935 | 8471 | 1839 | 9889.5 | 2603 | 799 | 13337.5 | 9303 | 27899 |
| 10109.5 | 23402 | 7187.5 | 1904 | 7709 | 2512 | 738 | 10683 | 8901 | 24216 |
| 7798 | 18747 | 5972 | 1575 | 7007.5 | 2294.5 | 829 | 10931 | 8332.5 | 21686.5 |
| 5308 | 11994 | 6598 | 1685 | 592443 | 61389 | 798 | 13051 | 6416 | 14931 |
| 3578.5 | 8186 | 4424.5 | 1324 | 6898.5 | 2665.5 | 808 | 14725.5 | 4948.5 | 11294 |
| 2829.5 | 4419.5 | 3405 | 1173 | 4762 | 2010.5 | 816.5 | 15390 | 4496 | 6827.5 |
| 2701 | 3912 | 3107.5 | 1144 | 4490 | 1917 | 851 | 14761 | 4936.5 | 6602.5 |
| 2408 | 3674.5 | 3033 | 1140 | 4415 | 1906 | 840 | 12765 | 4670 | 6350.5 |
| 2278 | 3613 | 2945.5 | 1108 | 4272 | 1877.5 | 836 | 11753.5 | 4593.5 | 6209 |
| 2270 | 3418 | 2813 | 1065 | 3994 | 1785 | 793 | 12362 | 4351 | 5929 |
| 2271 | 3599 | 2882 | 1102 | 4002 | 1815 | 812 | 11746 | 4307 | 6091 |
| 2291 | 3540 | 2898.5 | 1089 | 3999.5 | 1811.5 | 802.5 | 11469.5 | 4280 | 6050 |
| 2192 | 3430 | 2819 | 1084 | 3941 | 1806 | 815 | 10805 | 4242 | 5929 |
| 2127 | 3420 | 2827 | 1102 | 3991 | 1815 | 804 | 10633 | 4143 | 5870 |
| 2000 | 3296 | 2802 | 1102 | 3987 | 1815 | 787 | 10589 | 3949 | 5758 |
| 2011 | 3270 | 2745 | 1063 | 3956 | 1793 | 790 | 10948 | 4177 | 5845 |
| 30851 | 16960 | 5552 | 1171 | 3898 | 1781 | 770 | 12375 | 4484 | 6063 |
| 2899 | 4578 | 3615 | 1156 | 4285 | 1892 | 809 | 11345 | 4799 | 6535 |
| 2447 | 3915 | 3258.5 | 1162 | 4390 | 1877 | 808 | 12368 | 4755.5 | 6540 |
| 2411.5 | 3975 | 3295.5 | 1218 | 4631.5 | 1984.5 | 828 | 11592.5 | 4712 | 6707.5 |
| 3342.5 | 5766 | 3483.5 | 1174 | 4605 | 1883 | 802 | 15987.5 | 5699 | 8752.5 |
| 5630 | 12708 | 5048 | 1424 | 6086 | 2153 | 818 | 15590 | 6805 | 15371 |
| 9490.5 | 20617.5 | 6924 | 1716 | 7982.5 | 2457.5 | 794 | 12432 | 8408.5 | 22257.5 |
| 9135.5 | 25369 | 9318 | 2016 | 10233.5 | 2729.5 | 834 | 10628.5 | 8149 | 26633 |
| 9109 | 22263 | 7740 | 1748 | 9085 | 2490 | 737 | 14209 | 9382 | 25017 |
| 12480 | 31125 | 8634 | 2105 | 8758 | 2725 | 818 | 10734 | 8793 | 28285 |
| 8406 | 16919 | 5505 | 1455 | 6170 | 2128 | 809 | 13132 | 8214 | 19239 |
| 5374 | 12560 | 7060 | 1810 | 766678 | 71426 | 838 | 14482 | 6172 | 15135 |
| 3444.5 | 7524.5 | 4381 | 1313 | 6551 | 2453 | 841 | 14858 | 4809.5 | 10361 |
| 3286 | 5173 | 3645.5 | 1234.5 | 5193 | 2081 | 863 | 17916 | 5175.5 | 7920.5 |
| 2664 | 3869 | 3094 | 1132 | 4319 | 1883 | 823 | 15050 | 4551 | 6357 |
| 2385.5 | 3597 | 2922 | 1148 | 4254.5 | 1902 | 837 | 12262 | 4611.5 | 6134.5 |
| 2396 | 3676 | 3082 | 1144 | 4389 | 1909 | 879 | 12987 | 4778 | 6486 |
| 2409 | 3575 | 2920 | 1140 | 4138 | 1861 | 841 | 12926 | 4588 | 6132 |
| 2344 | 3505 | 2826 | 1080 | 3951 | 1796 | 811 | 11397 | 4413 | 5938 |
| 2516 | 3754 | 3059.5 | 1134.5 | 4211.5 | 1902 | 897 | 12642.5 | 4835 | 6547 |
| 2382 | 3562 | 2962 | 1141 | 4168 | 1882 | 841 | 11960 | 4639 | 6224 |
| 2141 | 3237 | 2617 | 1043 | 3737 | 1779 | 805 | 10590 | 4085 | 5716 |
| 2071 | 3358.5 | 2855 | 1097 | 4016 | 1817 | 837 | 11144.5 | 4181 | 6067 |
| 2057 | 3339 | 2812.5 | 1145 | 3972.5 | 1863 | 827 | 11200 | 4114 | 5929.5 |
| 46762.5 | 24692.5 | 6911 | 1281 | 4069 | 1840.5 | 823 | 14182.5 | 5050 | 6483 |
| 2691.5 | 4165 | 3563 | 1197 | 4157 | 1858 | 839 | 12177.5 | 4614 | 6388 |
| 2459 | 3873 | 3290.5 | 1174.5 | 4388 | 1906 | 841 | 11708 | 4819 | 6696 |
| 2373 | 3869 | 3298 | 1184 | 4508 | 1943 | 873 | 12372 | 4904 | 6918 |
| 3766 | 7100.5 | 3722 | 1226.5 | 4982 | 1977 | 829 | 16614.5 | 5841 | 10166 |
| 5892 | 12123 | 5039 | 1460 | 6177 | 2173 | 852 | 15899 | 6467 | 14231 |
| 9924 | 24004 | 7358.5 | 1792 | 8318 | 2545.5 | 836 | 12843.5 | 8705 | 25334.5 |

| | | | | | | | | | |
|---|---|---|---|---|---|---|---|---|---|
| 10687 | 29862.5 | 9365 | 2031.5 | 10089.5 | 2756 | 873 | 9568 | 7959.5 | 28649 |
| 8666 | 23476.5 | 8383.5 | 1793 | 9769 | 2529.5 | 759 | 10795 | 8191.5 | 25064 |
| 11825 | 27390 | 8072 | 1882 | 8601 | 2549 | 805 | 14611 | 9968 | 27149 |
| 8067 | 18023 | 5575 | 1475 | 6515 | 2187 | 789 | 10844 | 7845 | 19720 |
| 5040 | 11333 | 7157 | 1651 | 791278 | 70321 | 787 | 15179 | 6349 | 14196 |
| 3398 | 9297 | 3999 | 1206 | 5857.5 | 2160.5 | 783 | 14156 | 4672 | 12570.5 |
| 2930 | 4439 | 3234 | 1082.5 | 4540 | 1867 | 756 | 16162 | 4539 | 6632 |
| 2613 | 4294 | 3304 | 1159.5 | 4630.5 | 1943 | 839 | 15255 | 4529 | 6947 |
| 2392 | 3520 | 2877 | 1086 | 4062 | 1838 | 822 | 12359 | 4668 | 6131 |
| 2223 | 3407 | 2798 | 1060 | 4109 | 1816 | 807 | 12455 | 4441 | 5963 |
| 2129 | 3336 | 2742 | 1064 | 3870 | 1797 | 789 | 10869 | 4045 | 5769 |
| 2191 | 3267 | 2655 | 1010 | 3784 | 1726 | 760 | 11260 | 4153 | 5712 |
| 2048 | 3187 | 2552 | 998 | 3682 | 1702 | 748 | 10070 | 3951 | 5578 |
| 2060.5 | 3329.5 | 2729 | 1063 | 3901.5 | 1786 | 797 | 10579.5 | 4085.5 | 5841 |
| 1968 | 3191 | 2638 | 1041 | 3691 | 1738 | 766 | 10567 | 3840.5 | 5616 |
| 1872 | 3105 | 2573 | 1018 | 3629 | 1718 | 766 | 10470 | 3771 | 5486 |
| 1912 | 3228 | 2653.5 | 1056 | 3835.5 | 1783 | 803 | 10809.5 | 4003 | 5815 |
| 36563 | 18988 | 5857 | 1200 | 4020 | 1760 | 768 | 12855 | 4748 | 6176 |
| 2649 | 4213 | 3431 | 1111.5 | 4075 | 1822 | 784 | 11070.5 | 4534 | 6240 |
| 2404 | 3891.5 | 3243 | 1129 | 4369 | 1872 | 806 | 12356.5 | 4732.5 | 6601 |
| 2279.5 | 3681.5 | 3027 | 1118 | 4313.5 | 1884 | 800 | 11221.5 | 4576.5 | 6290 |
| 2613 | 4083.5 | 3052 | 1092 | 4178 | 1805 | 756.5 | 12755.5 | 4622.5 | 6591.5 |
| 7485 | 14220 | 5208 | 1411 | 6163 | 2160 | 813 | 17605 | 7853 | 16583 |
| 9859 | 22306.5 | 6586 | 1633 | 7722 | 2381 | 747 | 13388 | 8309 | 22848 |
| 10223 | 24237 | 8552 | 1841 | 9888 | 2598 | 794 | 13727 | 9397 | 25218 |
| 9172 | 23792 | 8668 | 1847 | 9996 | 2609 | 794 | 11916 | 8513 | 24215 |
| 8965 | 22984 | 7864 | 1982 | 8264 | 2535 | 741 | 11830 | 9122 | 25178 |
| 7346 | 16106.5 | 5504.5 | 1506 | 6728 | 2188 | 758 | 11380 | 8230 | 18862.5 |
| 4813.5 | 12039.5 | 6342 | 1577 | 626181.5 | 57879 | 731 | 13526 | 5951.5 | 14899.5 |
| 2869 | 6672 | 3857 | 1207 | 6164 | 2359 | 746 | 13550 | 4282 | 9439 |
| 3086.5 | 4526 | 3427.5 | 1148 | 4993.5 | 2076 | 765 | 16518 | 5004.5 | 6908.5 |
| 2441 | 3676 | 2934 | 1069 | 4254.5 | 1811 | 757 | 13627.5 | 4512.5 | 6024.5 |
| 2133 | 3293 | 2668 | 1021 | 3926.5 | 1762 | 720 | 10884.5 | 4130.5 | 5516.5 |
| 1844.5 | 3013 | 2481.5 | 973 | 3555.5 | 1685 | 700 | 9492.5 | 3604 | 5219 |
| 2257 | 3359 | 2784 | 1078 | 3894.5 | 1807.5 | 744 | 11467.5 | 4341.5 | 5774 |
| 2300 | 3550 | 2824 | 1058 | 4045 | 1810 | 777 | 11767 | 4524 | 6031 |
| 2083 | 3314 | 2638 | 1015 | 3791 | 1765 | 730 | 10746 | 3980 | 5592 |
| 1921 | 3136 | 2585.5 | 993 | 3666 | 1701 | 724 | 10507.5 | 3806 | 5480 |
| 2012 | 3263 | 2627 | 1034 | 3775 | 1783 | 749 | 10142 | 3921 | 5734 |
| 1882 | 3127 | 2581 | 1013 | 3707 | 1732 | 746 | 9911 | 3799 | 5606 |
| 1742 | 2931 | 2474 | 978 | 3552 | 1694 | 713 | 9272 | 3569 | 5287 |
| 35134.5 | 18706 | 5792 | 1226.5 | 4145.5 | 1838.5 | 787 | 13877.5 | 4925 | 6539 |
| 4035.5 | 6272 | 4534 | 1203 | 4109 | 1849 | 776 | 12796 | 4770 | 6547.5 |
| 2326 | 3838 | 3200 | 1102 | 4290 | 1841 | 760 | 11351 | 4649 | 6489 |
| 2195 | 3527 | 2944.5 | 1076.5 | 4145.5 | 1811 | 735.5 | 10640 | 4455 | 6172.5 |
| 3199 | 5013 | 3461 | 1160 | 4620 | 1920 | 795 | 16448 | 5656 | 7873 |
| 5589 | 13479 | 4854 | 1357 | 5925 | 2099 | 778 | 15011 | 7026 | 16819 |
| 9968 | 25589 | 7306 | 1707.5 | 8215.5 | 2455.5 | 783 | 12571 | 8848.5 | 26554 |

| | | | | | | | | | |
|---|---|---|---|---|---|---|---|---|---|
| 11727 | 25126 | 7826 | 1804 | 8935 | 2564 | 727 | 10373.5 | 8056.5 | 25238.5 |
| 8723 | 25691 | 7324 | 1699 | 8216 | 2420 | 713 | 10925 | 7872 | 26732 |
| 17639 | 28216.5 | 7082.5 | 1739 | 7693.5 | 2395 | 1030 | 15622.5 | 9982.5 | 24246 |
| 15097 | 22661 | 5937 | 1449 | 6335 | 2142 | 1086 | 15681 | 9202 | 20489 |
| 6864.5 | 12186 | 7035 | 1603 | 790904.5 | 67550.5 | 1082 | 18514 | 5814.5 | 13639.5 |
| 5017 | 9894 | 4223 | 1221 | 6176 | 2273 | 1036 | 16393 | 5666 | 12536 |
| 3944 | 4712 | 3315 | 1078 | 4595 | 1895 | 1146 | 22008 | 5511 | 7067 |
| 3585 | 4283.5 | 3139 | 1096 | 4371 | 1870 | 1299 | 19015.5 | 6186 | 7076.5 |
| 2743 | 3613 | 2880 | 1066 | 4086 | 1816 | 1192 | 16778 | 5532 | 6487 |
| 2328 | 3399 | 2724 | 1040 | 3831 | 1759 | 1183 | 16462 | 5169 | 6399 |
| 2433 | 3506 | 2826 | 1081 | 3815 | 1787 | 1177 | 16978 | 5147 | 6502 |
| 2524 | 3598 | 2749 | 1038 | 3803 | 1742 | 1253 | 17093 | 5303 | 6591 |
| 2422 | 3513 | 2661 | 1034.5 | 3651 | 1724 | 1187.5 | 18106.5 | 5040 | 6370 |
| 2448 | 3628.5 | 2830.5 | 1108 | 3894 | 1847 | 1182 | 18304 | 5201.5 | 6582.5 |
| 2498 | 3710 | 2964 | 1111 | 3969 | 1806 | 1365 | 18735 | 5777 | 6970 |
| 2101 | 3268.5 | 2634.5 | 1037 | 3643 | 1739 | 1181 | 15379.5 | 4719.5 | 6082 |
| 1912 | 2962 | 2423 | 950 | 3430 | 1648 | 1060 | 13536 | 4356 | 5549 |
| 40610 | 22756 | 6494 | 1253 | 3772 | 1792 | 1036 | 17468 | 5183 | 6298 |
| 5001 | 6029 | 4206 | 1167 | 4075 | 1821 | 1258 | 20691 | 6247 | 7170 |
| 2671 | 3915 | 3054 | 1074 | 3856 | 1776 | 1151 | 17921 | 5716 | 6889 |
| 2234 | 3534 | 2801 | 1016.5 | 3813.5 | 1710 | 1049 | 14588 | 4977.5 | 6349.5 |
| 4025.5 | 5443.5 | 3516 | 1136 | 4517.5 | 1862 | 1236 | 21866 | 6911.5 | 8499 |
| 8578 | 13990 | 4741 | 1317 | 5710 | 2014 | 1113 | 19447 | 6854 | 15330 |
| 16467.5 | 24084.5 | 6582 | 1556.5 | 7135.5 | 2234 | 991 | 16999.5 | 8578 | 20828.5 |
| 26985 | 44598.5 | 9229.5 | 1888 | 9285 | 2568 | 1213 | 15689 | 10557.5 | 29427.5 |
| 693 | 749 | 561 | 550 | 1025 | 1079 | 175 | 1713 | 1172 | 1307 |
| 3613 | 3847 | 1895 | 760 | 2319 | 1295 | 222 | 14251 | 3770 | 3999 |
| 2683 | 3255 | 1939 | 751 | 2491 | 1331 | 414 | 13827 | 3334 | 4043 |
| 1963.5 | 2618 | 5548 | 1994 | 739401 | 79467.5 | 516 | 9659 | 3010.5 | 4018.5 |
| 1688 | 2477 | 2696 | 1044 | 4070 | 1982 | 602 | 8740 | 3090 | 4336 |
| 1473 | 2389 | 2298 | 942 | 3368 | 1682 | 636 | 8106 | 3022 | 4434 |
| 1446 | 2448 | 2334 | 958 | 3468.5 | 1684 | 699 | 8374.5 | 3208.5 | 4793 |
| 1428 | 2510 | 2368 | 974 | 3545 | 1705 | 721 | 8470 | 3266 | 4960 |
| 1512 | 2616 | 2441 | 995 | 3585 | 1716 | 739 | 8860 | 3369 | 5101 |
| 1590 | 2696 | 2496 | 1032 | 3591 | 1758 | 731 | 9312 | 3416 | 5164 |
| 1556 | 2633 | 2342 | 970 | 3410 | 1635 | 709 | 8839 | 3303 | 4932 |
| 1612 | 2736 | 2493 | 1014.5 | 3547.5 | 1732 | 742 | 9454 | 3414 | 5143.5 |
| 1524 | 2685 | 2475 | 996 | 3598 | 1717 | 733 | 9076 | 3414 | 5178 |
| 1612 | 2683.5 | 2490.5 | 1013 | 3518.5 | 1708 | 735 | 8835 | 3353.5 | 5067 |
| 1427 | 2500 | 2345 | 966 | 3456.5 | 1679 | 719 | 8278.5 | 3248 | 4976 |
| 1460 | 2606 | 2458 | 994 | 3611 | 1722 | 738 | 8397 | 3354 | 5152 |
| 39222 | 21914 | 6417 | 1178 | 3690 | 1718 | 738 | 11713 | 4135 | 5724 |
| 1758 | 3123 | 2917 | 1037 | 3591 | 1693 | 713 | 8847 | 3579 | 5389 |
| 1697 | 3072 | 2809 | 1052 | 3832 | 1769.5 | 745 | 9087 | 3726.5 | 5709.5 |
| 1639 | 3010 | 2710 | 1047 | 3826 | 1766 | 750 | 9061 | 3719 | 5667 |
| 3245 | 6977 | 3721 | 1180 | 4764.5 | 1909 | 750 | 15784 | 4782.5 | 9601 |
| 6538 | 16015 | 5520 | 1466 | 6519 | 2181 | 753 | 16036 | 7660 | 18931 |
| 8406 | 20182 | 6729 | 1685 | 8061 | 2430 | 750 | 15439 | 9071 | 22980 |

| | | | | | | | | | |
|---|---|---|---|---|---|---|---|---|---|
| 8734 | 24359 | 8373 | 1911 | 9501 | 2630 | 793 | 13256 | 9049 | 26428 |
| 8690 | 24294.5 | 7299 | 1755 | 8369.5 | 2495 | 751 | 10591.5 | 8382.5 | 26161 |
| 9005.5 | 22991 | 6914 | 1821 | 7379.5 | 2423 | 732 | 11469 | 8841 | 25075.5 |
| 6774.5 | 15894 | 5186.5 | 1382.5 | 6020 | 2067.5 | 730 | 12410.5 | 7368 | 18438.5 |
| 5405 | 12663 | 6186 | 1583 | 544903 | 52884 | 768 | 14552 | 6507 | 15590 |
| 3324 | 7529 | 4271 | 1284 | 6764 | 2522 | 763 | 14192 | 4767 | 10250 |
| 2891 | 4464 | 3408 | 1154 | 4805 | 2001 | 780.5 | 16834.5 | 4671 | 6802 |
| 2523 | 3671 | 2947 | 1092 | 4249.5 | 1864 | 782 | 14268 | 4607 | 6061 |
| 1890 | 3041 | 2570 | 1014 | 3746 | 1738 | 724 | 10436 | 3729 | 5339 |
| 1918 | 3146 | 2676 | 1033 | 3875 | 1748 | 774 | 10802 | 3870 | 5666 |
| 2076 | 3317 | 2749 | 1070 | 3918 | 1788 | 780 | 11558 | 4127 | 5777 |
| 2072 | 3195 | 2711 | 1042 | 3863 | 1778 | 770 | 10681 | 4081 | 5753 |
| 2025 | 3242 | 2742.5 | 1087 | 3791 | 1780 | 778 | 11331 | 3954 | 5750.5 |
| 1864 | 3095 | 2634 | 1028 | 3786 | 1752 | 770 | 9888 | 3768 | 5547 |
| 1897 | 3141 | 2658.5 | 1041.5 | 3704 | 1761 | 772 | 10361 | 3808 | 5656 |
| 1697 | 2902.5 | 2571 | 1020 | 3694 | 1746 | 770 | 9552.5 | 3610.5 | 5453.5 |
| 1700.5 | 2948 | 2605.5 | 1032 | 3807 | 1761 | 769 | 9150 | 3686 | 5462.5 |
| 37367 | 20759.5 | 6154 | 1224 | 3819.5 | 1770 | 757 | 12594.5 | 4399.5 | 5998.5 |
| 2749.5 | 4525 | 3769 | 1179.5 | 4229.5 | 1846 | 779 | 10989.5 | 4558 | 6281 |
| 2239 | 3697 | 3139 | 1107 | 4135 | 1788.5 | 757 | 10381.5 | 4431 | 6284 |
| 2195 | 3577 | 3039.5 | 1114 | 4137 | 1822.5 | 776 | 10782.5 | 4555.5 | 6333.5 |
| 3919 | 8678 | 3969 | 1231 | 5055 | 1946 | 771 | 16645 | 5691 | 11853 |
| 5369 | 15163 | 5490 | 1507.5 | 6599 | 2221 | 794 | 14613 | 6162 | 18046 |
| 9638 | 23454 | 7009 | 1762 | 7958 | 2464 | 768 | 12971 | 9103 | 25182 |
| 10119.5 | 26338.5 | 8695 | 1876 | 9542.5 | 2598.5 | 786 | 14416 | 9352.5 | 28183.5 |
| 9444 | 24402 | 7348 | 1783 | 8298 | 2477 | 767 | 13831 | 9572 | 26908 |
| 9209 | 22033 | 6586 | 1707 | 7603.5 | 2424 | 785 | 13792 | 9440.5 | 24420.5 |
| 7311 | 15955 | 5233 | 1437 | 6217 | 2096 | 731 | 12459 | 8003 | 18310 |
| 5463 | 13249 | 6946 | 1718 | 698337 | 60326 | 792 | 14013 | 6518 | 16135 |
| 3249 | 8672 | 4129 | 1249 | 6515 | 2393 | 753 | 14015 | 4684 | 11895 |
| 2804 | 4585.5 | 3494.5 | 1167 | 5041 | 2030.5 | 785 | 16358 | 4455 | 6944 |
| 2536 | 3761 | 2940 | 1063 | 4197 | 1851 | 762 | 14619 | 4464 | 6078 |
| 2271 | 3391.5 | 2755 | 1055 | 3995.5 | 1817 | 770 | 11613 | 4437 | 5901 |
| 2085 | 3293 | 2770 | 1046 | 3951 | 1785 | 788 | 11493 | 4234 | 5876 |
| 2181 | 3310 | 2795 | 1051.5 | 4032 | 1779 | 784 | 11053 | 4376 | 5874.5 |
| 2092 | 3277 | 2728 | 1053 | 3857 | 1773 | 778 | 11226 | 4181.5 | 5750 |
| 1977 | 3195.5 | 2660 | 1019 | 3754 | 1757 | 758 | 9930 | 3919.5 | 5628.5 |
| 2035 | 3262.5 | 2756 | 1076 | 3903 | 1781.5 | 770 | 10879 | 4120 | 5816 |
| 1985 | 3248 | 2688 | 1058 | 3844 | 1786 | 779 | 10482 | 4013 | 5692 |
| 1750 | 2968 | 2570 | 1009 | 3688 | 1706.5 | 741 | 9510.5 | 3603 | 5434 |
| 1803 | 3088 | 2612 | 1023 | 3699 | 1734 | 753 | 9854 | 3827 | 5617 |
| 47560 | 24804 | 7280 | 1323 | 4098 | 1836 | 791 | 14964 | 5222 | 6559 |
| 2452 | 3959 | 3359 | 1112 | 3819 | 1746 | 741 | 10535 | 4241 | 5782 |
| 2257 | 3578 | 3005 | 1073 | 4017 | 1786 | 747 | 10528 | 4433 | 6132 |
| 2119 | 3494 | 2937 | 1111 | 4084 | 1803 | 765 | 10028 | 4320 | 6146 |
| 3230 | 7886 | 3668 | 1172 | 4718 | 1882 | 756 | 15616 | 4689 | 10984 |
| 5251.5 | 14484 | 4978.5 | 1372 | 6103.5 | 2092 | 757 | 15964.5 | 6112 | 17049 |
| 9381 | 21439 | 6662 | 1643 | 7479 | 2349 | 746 | 14502 | 9348 | 23722 |

| | | | | | | | | | |
|---|---|---|---|---|---|---|---|---|---|
| 9822.5 | 24172 | 7950.5 | 1791.5 | 8975 | 2484 | 765 | 12829.5 | 9319 | 25737 |
| 9624 | 22236 | 6746 | 1612 | 7507 | 2316 | 719 | 13696 | 9536 | 24456 |
| 10002.5 | 26518.5 | 8683.5 | 2267 | 8394 | 2695.5 | 726 | 11815.5 | 9517.5 | 27843.5 |
| 7315 | 16282 | 5342 | 1450 | 6287 | 2152 | 769 | 14927 | 8098 | 18792 |
| 4703 | 12273 | 6935 | 1716 | 749147.5 | 64707 | 746 | 14306.5 | 5791 | 15152 |
| 3225 | 8726.5 | 3984.5 | 1228 | 5978 | 2292 | 737 | 14202 | 4528 | 11921 |
| 2875 | 4377 | 3280 | 1144.5 | 4721 | 2004.5 | 776 | 15764.5 | 4339 | 6588.5 |
| 2685 | 3713 | 3001 | 1092 | 4226 | 1868 | 792.5 | 12601 | 4785.5 | 6109.5 |
| 2358 | 3491 | 2909 | 1106 | 4067 | 1854 | 822 | 11649 | 4393 | 6023 |
| 2178 | 3376 | 2798 | 1081 | 3989 | 1809 | 803 | 11556 | 4225 | 5887 |
| 2312 | 3531 | 2867 | 1123 | 4066 | 1844 | 816 | 12319 | 4361 | 5937 |
| 2166 | 3327 | 2710 | 1029 | 3773.5 | 1745 | 771 | 11245.5 | 4007 | 5647 |
| 2199 | 3392 | 2764.5 | 1062 | 3770 | 1773 | 798 | 11511 | 4135.5 | 5774.5 |
| 1935 | 3058 | 2509 | 994 | 3518 | 1662 | 738 | 9823 | 3652 | 5344 |
| 2141 | 3307.5 | 2775 | 1090 | 3862 | 1807 | 807 | 10569.5 | 4055.5 | 5863.5 |
| 1924 | 3095 | 2598 | 1040 | 3667 | 1719 | 778 | 10102 | 3731 | 5457 |
| 1870 | 3108 | 2622 | 1031 | 3597 | 1727 | 787 | 9953 | 3822 | 5532 |
| 48966.5 | 27168 | 7689 | 1290 | 3733 | 1734.5 | 751.5 | 13318 | 4584 | 6093.5 |
| 2729 | 4286 | 3594.5 | 1123 | 3782.5 | 1748 | 771 | 11580 | 4205 | 5810 |
| 2322 | 3718 | 3152.5 | 1119 | 4050.5 | 1809 | 774 | 11197.5 | 4402.5 | 6110 |
| 2152 | 3488 | 2991 | 1076 | 3960 | 1792 | 793 | 10963 | 4273 | 6080 |
| 3491 | 5469 | 3498.5 | 1158 | 4543 | 1865 | 783 | 14892.5 | 5549.5 | 8051.5 |
| 5992 | 14654 | 5158 | 1396 | 6047 | 2109 | 801 | 15120 | 6404 | 17138 |
| 10612 | 23877 | 7123 | 1733 | 7893 | 2428 | 792 | 14802 | 9740 | 25246 |
| 12138.5 | 28383 | 8654 | 1899 | 9218 | 2608 | 832 | 15254.5 | 9989 | 28706 |
| 9211 | 21884 | 6701 | 1671 | 7822 | 2353 | 725 | 12073 | 8842 | 23934 |
| 8930 | 20544 | 6365 | 1636 | 7081 | 2307 | 730 | 12197 | 8855 | 22362 |
| 6427 | 15907.5 | 5154 | 1387.5 | 6103 | 2097 | 735 | 13386.5 | 6882 | 18285.5 |
| 4396 | 11448 | 5960 | 1499 | 636945 | 60111 | 725 | 14720 | 5293 | 14130 |
| 3432.5 | 8592.5 | 4179 | 1248 | 6170.5 | 2392 | 801 | 14277 | 4708 | 11868 |
| 3090 | 4423.5 | 3378 | 1138 | 4639.5 | 1997 | 809 | 16895.5 | 4799 | 6688 |
| 2695 | 3711 | 3008 | 1106 | 4191 | 1868 | 812 | 13408 | 4712 | 6093 |
| 2064 | 3234 | 2732 | 1046 | 3852 | 1801 | 779 | 10513 | 3913 | 5659 |
| 1948 | 3118 | 2618 | 1028 | 3643 | 1714 | 779 | 10241 | 3788 | 5470 |
| 2121.5 | 3309.5 | 2812.5 | 1070 | 3863.5 | 1767 | 808 | 11266.5 | 4033.5 | 5832 |
| 1937 | 3083.5 | 2529.5 | 996 | 3539.5 | 1671 | 746 | 10306 | 3756.5 | 5297 |
| 1949 | 3025 | 2548 | 999 | 3516 | 1662 | 751 | 9787 | 3665 | 5192 |
| 1969 | 3160 | 2645 | 1022 | 3705 | 1730 | 786 | 10659 | 3865 | 5597 |
| 2036 | 3188 | 2676 | 1064 | 3675 | 1764 | 799 | 10709 | 3882 | 5598 |
| 1969 | 3233 | 2725.5 | 1102 | 3830 | 1825.5 | 806 | 10480 | 3907 | 5749.5 |
| 1819 | 2994 | 2604 | 1015 | 3615 | 1697 | 767 | 9489 | 3660 | 5387 |
| 44462 | 24407 | 7042 | 1247 | 3797 | 1749 | 775 | 13921 | 4684 | 6090 |
| 2295 | 3663.5 | 3115 | 1106 | 3724 | 1731 | 763 | 10594 | 4021 | 5719.5 |
| 2046 | 3392 | 2869 | 1061 | 3861 | 1747 | 761 | 10393 | 4033 | 5826 |
| 2032 | 3375 | 2839 | 1070 | 3865 | 1794 | 777 | 10162 | 4035 | 5918 |
| 3772.5 | 7461.5 | 3802 | 1207.5 | 4868 | 1918.5 | 818 | 18359 | 5436 | 10540 |
| 5687 | 13647 | 5085 | 1403 | 6049 | 2125 | 786 | 15370 | 6380 | 16187 |
| 10815 | 22456 | 7103 | 1734 | 8120 | 2442 | 812 | 13467 | 9197 | 23628 |

| | | | | | | | | | |
|---|---|---|---|---|---|---|---|---|---|
| 10090.5 | 24549.5 | 7677 | 1883 | 8824 | 2536 | 786 | 11349.5 | 8586 | 25880.5 |
| 9313 | 23723.5 | 6989 | 1698 | 7879 | 2412 | 745 | 11010 | 8690 | 25645.5 |
| 10113 | 24188 | 7846 | 2054 | 7618 | 2590 | 718 | 11543 | 8593 | 24697 |
| 6888.5 | 15422 | 5085 | 1378 | 6013.5 | 2033.5 | 705 | 12721 | 7524.5 | 18095 |
| 4593 | 12260 | 6603 | 1555 | 711419 | 61104 | 745 | 15724 | 5564 | 15128 |
| 3249 | 8585 | 3922 | 1188 | 5818 | 2194 | 736 | 14349.5 | 4539 | 11608 |
| 2804 | 4489 | 3325 | 1101 | 4760 | 1923 | 749 | 16059 | 4393 | 6668 |
| 2581.5 | 3656.5 | 2884.5 | 1041 | 4079 | 1787.5 | 747.5 | 12294 | 4590 | 5891 |
| 2190 | 3269 | 2667 | 1017 | 3888 | 1746 | 734 | 11218 | 4213.5 | 5615 |
| 1970 | 3181 | 2638.5 | 1015 | 3810 | 1743 | 742 | 10267.5 | 3770 | 5553 |
| 1985.5 | 3179 | 2637 | 1020 | 3698.5 | 1719 | 727 | 10581 | 3807 | 5451.5 |
| 1982 | 3172 | 2589.5 | 985.5 | 3674 | 1690 | 722 | 10107.5 | 3753.5 | 5433 |
| 1990 | 3158 | 2579 | 1003 | 3634 | 1712 | 719 | 10690 | 3774 | 5409 |
| 1821 | 2998 | 2484.5 | 961 | 3554 | 1676 | 707 | 9296 | 3593 | 5229 |
| 1780 | 2891 | 2379 | 964 | 3448 | 1639 | 683 | 8955 | 3440 | 5026 |
| 1783 | 2977 | 2504 | 987 | 3602 | 1692 | 722 | 9368 | 3575 | 5325 |
| 1766 | 2964.5 | 2526.5 | 1004 | 3630.5 | 1676 | 707 | 9353.5 | 3558 | 5327 |
| 45747.5 | 25073.5 | 7051 | 1230 | 3795 | 1749 | 720.5 | 12656.5 | 4486 | 5902.5 |
| 2414 | 4005 | 3360 | 1064 | 3643 | 1692 | 710 | 10349 | 3993 | 5613 |
| 2166 | 3613 | 3061 | 1089 | 3964 | 1744 | 733 | 10327 | 4136 | 5969 |
| 2032 | 3424 | 2876.5 | 1056 | 3906.5 | 1774 | 741 | 9944.5 | 4190 | 5953 |
| 2884 | 4282 | 3114 | 1077 | 4163 | 1783 | 707 | 12845 | 4826 | 6596 |
| 5755 | 13447.5 | 4977 | 1338 | 5922 | 2031.5 | 708 | 16466.5 | 6823.5 | 15864.5 |
| 9151 | 20558.5 | 6364 | 1613 | 7361.5 | 2317 | 728.5 | 13252.5 | 8461.5 | 22423 |
| 9709 | 23672 | 7388 | 1816 | 8635 | 2504 | 746 | 12214 | 9013 | 25714 |
| 9686 | 23004.5 | 6908.5 | 1671 | 7879.5 | 2363 | 720 | 13880.5 | 9466 | 25272.5 |
| 9190.5 | 23117.5 | 6722 | 1748.5 | 7587.5 | 2410 | 756.5 | 11160.5 | 8663 | 24903 |
| 6586.5 | 15462.5 | 5194.5 | 1354 | 6160 | 2063 | 733 | 14513 | 7386 | 18006.5 |
| 4556.5 | 11838.5 | 6302 | 1587 | 637421.5 | 60929.5 | 746 | 14170.5 | 5483.5 | 14368.5 |
| 3065 | 5605 | 4055 | 1206 | 6348 | 2400 | 735 | 14082 | 4376 | 7722 |
| 2769 | 4487 | 3316 | 1111 | 4774 | 2000 | 754 | 15782 | 4311 | 6702 |
| 2566.5 | 3703 | 2907 | 1055 | 4249 | 1815 | 758 | 11845 | 4594 | 6061.5 |
| 2196 | 3377 | 2804 | 1070 | 3973 | 1838 | 769 | 11203 | 4098 | 5915 |
| 2139.5 | 3364 | 2882 | 1101 | 4075 | 1839 | 807.5 | 11321 | 4203 | 6024 |
| 2149 | 3425 | 2844 | 1112 | 3910 | 1829 | 807 | 10956 | 4081 | 5846 |
| 2072 | 3231 | 2642 | 1025.5 | 3669 | 1717 | 758.5 | 10605 | 3861 | 5527.5 |
| 2083.5 | 3244 | 2680.5 | 1032.5 | 3737.5 | 1763 | 754 | 10519 | 3836.5 | 5493 |
| 1955.5 | 3119.5 | 2611 | 1022 | 3652.5 | 1721 | 754 | 9918 | 3741.5 | 5484 |
| 1955 | 3148 | 2640 | 1016 | 3660 | 1706 | 767 | 10139 | 3786 | 5497 |
| 1967 | 3204 | 2718.5 | 1080 | 3823 | 1775.5 | 785 | 10350.5 | 3839 | 5679.5 |
| 1800 | 2949 | 2498 | 999 | 3531 | 1707 | 743 | 9445 | 3655 | 5297 |
| 27836.5 | 15522 | 5293 | 1206 | 3848 | 1776 | 778 | 12364.5 | 4348 | 5943 |
| 2539 | 4073.5 | 3441 | 1108 | 3867 | 1753 | 755 | 10740 | 4069 | 5798 |
| 2100 | 3531 | 2943.5 | 1082.5 | 3812 | 1756.5 | 751 | 10352.5 | 4022 | 5915 |
| 2105 | 3415.5 | 2840.5 | 1075 | 3888 | 1784.5 | 776 | 10745 | 4132.5 | 5943.5 |
| 2795.5 | 4193.5 | 3098.5 | 1109 | 4208 | 1838 | 768 | 12549 | 4866 | 6602 |
| 6013 | 14142 | 5081 | 1394 | 5959 | 2092 | 757 | 15743 | 6691 | 16847 |
| 9212 | 20501 | 6533 | 1605 | 7530 | 2305 | 742 | 14106 | 8510 | 21874 |

| | | | | | | | | | |
|---|---|---|---|---|---|---|---|---|---|
| 10524 | 25481 | 8235 | 1887 | 9125 | 2642 | 801 | 11538 | 8753 | 26301 |
| 9549.5 | 20910 | 6542 | 1637.5 | 7466 | 2319 | 700 | 12319.5 | 8954 | 22906 |
| 10054 | 24475 | 7658 | 1957 | 7678 | 2494 | 713 | 12021 | 9326 | 25614 |
| 6092.5 | 15195 | 4965 | 1336.5 | 5775.5 | 1972 | 691 | 14617.5 | 6660 | 17382 |
| 4678 | 12147.5 | 7261 | 1666 | 773261 | 63802.5 | 768 | 16187 | 5696.5 | 14967 |
| 3687 | 7826 | 4224 | 1235 | 6419 | 2336 | 738 | 14922 | 5066 | 10421 |
| 2847 | 4647.5 | 3414 | 1131 | 4879.5 | 1960 | 740 | 15247.5 | 4308 | 6746 |
| 2733 | 3861 | 2993 | 1064 | 4306 | 1848 | 763 | 12768 | 4701 | 6155 |
| 2338 | 3580 | 2929 | 1094 | 4234 | 1876 | 770 | 11462 | 4318 | 6006 |
| 2089 | 3356 | 2786 | 1052.5 | 3931 | 1791.5 | 796 | 11160.5 | 4090 | 5822.5 |
| 2237.5 | 3449.5 | 2827 | 1076 | 3883 | 1783 | 812 | 11522.5 | 4161.5 | 5873 |
| 2133.5 | 3284 | 2643.5 | 1054 | 3695.5 | 1730 | 775 | 10535 | 3943 | 5570.5 |
| 2200 | 3320 | 2704 | 1070 | 3736 | 1731 | 812 | 11273 | 4089 | 5756 |
| 2049 | 3218 | 2607 | 1034 | 3663 | 1729 | 784 | 10996 | 3876 | 5511 |
| 2127 | 3267 | 2738 | 1064 | 3758 | 1766 | 811 | 10934 | 4033 | 5704 |
| 1887 | 3056 | 2612 | 1031 | 3666 | 1739 | 788 | 9740 | 3782 | 5518 |
| 1975 | 3215.5 | 2745 | 1101 | 3888.5 | 1820.5 | 838.5 | 11108 | 4038.5 | 5777 |
| 47858.5 | 25999.5 | 7270 | 1280 | 3976 | 1794 | 799 | 13417 | 4796.5 | 6277 |
| 2655.5 | 4136.5 | 3522.5 | 1197.5 | 4308.5 | 1894 | 852 | 12307 | 4804.5 | 6495 |
| 2384.5 | 3662.5 | 3180 | 1125 | 4082 | 1815 | 792.5 | 11351.5 | 4532 | 6274 |
| 2153 | 3485.5 | 2996 | 1097 | 4119 | 1813 | 808.5 | 11404 | 4417.5 | 6262 |
| 3713.5 | 5270.5 | 3599.5 | 1157 | 4668.5 | 1871.5 | 811 | 16155.5 | 5778 | 7775.5 |
| 6392 | 13116 | 5300 | 1445 | 6134 | 2134 | 825 | 17507 | 7011 | 15320 |
| 10638 | 22600 | 6875 | 1708 | 7723 | 2390 | 795 | 16278 | 9933 | 24188 |
| 12043 | 26370 | 8053 | 1897 | 8560 | 2565 | 846 | 12601 | 9160 | 26591 |
| 9988.5 | 25097.5 | 7476 | 1710 | 8025.5 | 2411 | 743 | 10907.5 | 8382 | 26011.5 |
| 9462 | 20084 | 6488 | 1643 | 7221 | 2319 | 756 | 15623 | 9171 | 21902 |
| 8051 | 16468 | 5356 | 1425 | 6296 | 2091 | 762 | 12785 | 8071 | 18824 |
| 5225 | 12372 | 6673 | 1637 | 692558 | 64650 | 768 | 14152 | 5770 | 15055 |
| 3771 | 6880 | 4316 | 1276 | 7235 | 2599 | 787 | 14861.5 | 5005.5 | 9164.5 |
| 2985 | 4754 | 3497 | 1167 | 4998 | 2013 | 802 | 15935 | 4495 | 6941 |
| 2737 | 3713 | 2975 | 1095 | 4212 | 1829 | 786 | 12434.5 | 4751.5 | 6044 |
| 2055.5 | 3246 | 2678 | 1013 | 3861.5 | 1748 | 769 | 10648.5 | 3955.5 | 5607.5 |
| 1945 | 3060 | 2558 | 1011 | 3663 | 1694 | 754 | 10378 | 3765 | 5410 |
| 2052 | 3245 | 2683 | 1051 | 3833 | 1771 | 785 | 10675 | 3974 | 5615 |
| 1999 | 3167 | 2541 | 1006 | 3655 | 1703 | 750 | 10430 | 3764 | 5383 |
| 2030 | 3230 | 2624 | 1024 | 3698 | 1733 | 767 | 10438 | 3887 | 5531 |
| 1886 | 3037 | 2546 | 992 | 3538 | 1689 | 746 | 9612 | 3708 | 5284 |
| 1935 | 3102.5 | 2593 | 1007 | 3640.5 | 1734 | 753.5 | 9867 | 3721.5 | 5379 |
| 1883 | 3133 | 2627 | 1022 | 3710 | 1708 | 779 | 10122.5 | 3810 | 5569 |
| 1884 | 3075 | 2640 | 1057 | 3722 | 1744 | 768 | 9867 | 3718 | 5422 |
| 30991 | 17346 | 5323 | 1138 | 3598 | 1713 | 744 | 12033 | 4244 | 5647 |
| 2715 | 4260 | 3462 | 1089 | 3834 | 1749 | 761 | 11097 | 4159 | 5878 |
| 2136 | 3500 | 2911 | 1058 | 3865 | 1754 | 757 | 10103 | 4054 | 5883 |
| 2025 | 3367 | 2802 | 1055 | 3813 | 1744 | 750 | 10149 | 4048 | 5869 |
| 3208 | 4486 | 3214 | 1114 | 4227 | 1771 | 753 | 14181 | 5053 | 6682 |
| 7531 | 15460 | 5506 | 1440 | 6343 | 2118 | 786 | 17108 | 8391 | 18075 |
| 9537 | 20350 | 6340.5 | 1596 | 7330 | 2297 | 756 | 14392 | 9009 | 22133.5 |

| | | | | | | | | | |
|---|---|---|---|---|---|---|---|---|---|
| 10924 | 23369 | 7107 | 1682 | 7896 | 2408 | 754 | 15222 | 9764 | 24491 |
| 9812 | 23434 | 6869.5 | 1650.5 | 7914 | 2342 | 733 | 12657 | 9168.5 | 24373.5 |
| 9320.5 | 24580.5 | 7535 | 1987 | 7693.5 | 2493 | 710 | 11192.5 | 8897 | 25666.5 |
| 7259.5 | 15931.5 | 5413 | 1440 | 6267 | 2130 | 726 | 13352 | 8004 | 18798 |
| 4104 | 11194 | 6215 | 1536 | 710059 | 63705 | 704 | 15279 | 5057 | 13647 |
| 3601 | 7897 | 4145 | 1225 | 6082 | 2219 | 736 | 14300 | 4902 | 10589 |
| 2920 | 4727 | 3466 | 1130 | 4874 | 1941 | 734 | 14645 | 4394 | 6776 |
| 2616 | 3644 | 2791 | 1012 | 3968.5 | 1774 | 720 | 12905.5 | 4527.5 | 5776 |
| 2334 | 3462 | 2793 | 1065 | 4055 | 1813 | 755 | 11560 | 4399 | 5809 |
| 1993 | 3171 | 2622 | 1011 | 3815 | 1722 | 723 | 10397 | 3842 | 5417 |
| 1968.5 | 3183 | 2583 | 1029 | 3707 | 1726.5 | 730 | 10741.5 | 3855 | 5551.5 |
| 2006 | 3160 | 2572.5 | 998.5 | 3638 | 1701 | 711 | 10463.5 | 3792 | 5388 |
| 2016 | 3255 | 2611 | 1029 | 3685 | 1739 | 738 | 10308 | 3878 | 5569 |
| 1852 | 2985.5 | 2436 | 967 | 3458 | 1667 | 691 | 9422 | 3550.5 | 5107 |
| 1875 | 3020.5 | 2520 | 984.5 | 3571.5 | 1688 | 709.5 | 9328 | 3585.5 | 5298.5 |
| 1773 | 2938 | 2466 | 983 | 3521 | 1686 | 713 | 9224 | 3551 | 5199 |
| 1710 | 2864 | 2435 | 970 | 3442 | 1651 | 706 | 8780 | 3475 | 5163 |
| 38749 | 21393.5 | 6188.5 | 1192 | 3689 | 1735 | 714.5 | 12054.5 | 4218.5 | 5760 |
| 2713 | 4409 | 3684 | 1085 | 3697 | 1690 | 694 | 10613 | 4009 | 5655 |
| 2161 | 3563 | 3023 | 1059 | 3940 | 1744 | 724 | 10390 | 4110 | 5938 |
| 1999.5 | 3447.5 | 2870 | 1071 | 3860 | 1752 | 713 | 10185 | 3997 | 5872.5 |
| 2902 | 4905 | 3332 | 1089 | 4360 | 1786 | 709 | 16622 | 4402 | 7102 |
| 4749 | 11715.5 | 4781.5 | 1320 | 5773.5 | 2038.5 | 717 | 16018.5 | 5828.5 | 14436.5 |
| 8786.5 | 19200 | 6155 | 1548 | 7152.5 | 2215 | 699 | 15409.5 | 9180.5 | 21716 |
| 9631 | 22653 | 6862 | 1653 | 7808 | 2369 | 726 | 12847 | 9436 | 25128 |
| 9077 | 20145.5 | 6385 | 1548.5 | 7291 | 2216 | 687 | 14232.5 | 9152.5 | 22151.5 |
| 18594.5 | 27319.5 | 6873 | 1613 | 7100.5 | 2263.5 | 949 | 14142.5 | 9827 | 22848.5 |
| 15679 | 22908 | 6047 | 1515 | 6672 | 2213 | 1150 | 17250 | 9502 | 20834 |
| 9611 | 16184 | 7386 | 1716 | 733879 | 63541 | 1135 | 16111 | 7495 | 16992 |
| 6624 | 11554 | 4779 | 1335.5 | 7262 | 2603 | 1141 | 18195.5 | 6425 | 13903.5 |
| 4600 | 5792 | 3841 | 1214 | 5321 | 2048 | 1256 | 21734 | 5920 | 8036 |
| 3542 | 4417 | 3244 | 1125 | 4584 | 1908 | 1259 | 19078 | 6249 | 7234 |
| 2756 | 3790 | 3003 | 1116 | 4179 | 1832 | 1140 | 16622 | 5347 | 6555 |
| 2501 | 3591.5 | 2776.5 | 1058 | 3880 | 1761 | 1182 | 17130 | 5139.5 | 6449.5 |
| 2424 | 3512 | 2825 | 1070 | 3853 | 1774 | 1097 | 16509 | 5029 | 6328 |
| 2354 | 3450 | 2672 | 1003 | 3691 | 1718 | 1086 | 15003 | 4821 | 6094 |
| 2338 | 3381 | 2711 | 1019 | 3686 | 1725 | 1074 | 14407.5 | 4725 | 6047.5 |
| 2492 | 3662 | 2901 | 1087 | 3935 | 1811 | 1144 | 18049 | 5071 | 6446 |
| 2442 | 3522 | 2792 | 1077.5 | 3863 | 1766.5 | 1189 | 15871.5 | 5127 | 6487 |
| 2161 | 3360 | 2692 | 1022 | 3652 | 1717 | 1126 | 14538 | 4662 | 6020 |
| 2070 | 3220.5 | 2662 | 1008 | 3667.5 | 1693 | 1061 | 14294 | 4563.5 | 5964 |
| 51933 | 27746.5 | 7853 | 1298 | 3924 | 1781.5 | 1076.5 | 18467.5 | 5805.5 | 6714 |
| 3625.5 | 4767 | 3725 | 1137.5 | 3961.5 | 1788 | 1061 | 16034 | 5493 | 6683 |
| 2666 | 3868.5 | 3053.5 | 1086 | 3942 | 1745 | 1050 | 15091 | 5394 | 6690.5 |
| 2457 | 3762 | 3075 | 1116 | 4088 | 1802 | 973 | 15411 | 5063 | 6408 |
| 3329 | 4493 | 3278 | 1123 | 4297 | 1812 | 933 | 16839 | 5562 | 6887 |
| 7861 | 14537 | 4827 | 1306 | 5617 | 1984 | 952 | 18490.5 | 6364.5 | 15889 |
| 18321 | 30801 | 7245 | 1725 | 7636 | 2414 | 1013 | 14225 | 9266 | 25042 |

| | | | | | | | | | |
|---|---|---|---|---|---|---|---|---|---|
| 18524 | 30065 | 7561 | 1718 | 8142 | 2377 | 922 | 13439 | 9395 | 25119 |
| 687 | 730 | 522 | 529 | 995 | 1058 | 170 | 1632 | 1160 | 1263 |
| 2691 | 2970 | 1456 | 687 | 1990 | 1250 | 238 | 11179 | 3109 | 3327 |
| 2303 | 2853 | 1707 | 702 | 2327 | 1300 | 404 | 10833 | 2991 | 3706.5 |
| 1828 | 2508 | 5717 | 2064 | 737724 | 78870 | 539 | 9294 | 2990 | 4035 |
| 1622 | 2462 | 2593.5 | 1024 | 3981 | 1940 | 602 | 8630.5 | 3044 | 4362.5 |
| 1462 | 2444 | 2385 | 958 | 3604 | 1729 | 686 | 8441.5 | 3138 | 4691 |
| 1423 | 2475 | 2407 | 971 | 3582 | 1736 | 719 | 8669 | 3297 | 4962 |
| 1457 | 2586 | 2465 | 987 | 3654.5 | 1724 | 748 | 8980.5 | 3396 | 5176.5 |
| 1455 | 2557 | 2404 | 987 | 3573 | 1699 | 735 | 8899.5 | 3353 | 5049 |
| 1633 | 2792 | 2550 | 1020 | 3674 | 1737 | 759.5 | 9455 | 3545.5 | 5348.5 |
| 1647 | 2814 | 2488 | 987 | 3588 | 1706 | 746 | 9557 | 3534 | 5293 |
| 1615 | 2782 | 2484 | 1006 | 3608 | 1720 | 748 | 9611 | 3496 | 5238 |
| 1577.5 | 2817.5 | 2549.5 | 1021 | 3674 | 1714.5 | 774 | 9681 | 3492.5 | 5289 |
| 1560 | 2747 | 2526.5 | 1015 | 3689.5 | 1731 | 773 | 9437 | 3551.5 | 5364 |
| 1453 | 2600.5 | 2445.5 | 1005 | 3561 | 1726.5 | 754 | 8628.5 | 3327 | 5087 |
| 1431 | 2549 | 2439 | 981 | 3537 | 1704 | 740 | 8535 | 3376 | 5180 |
| 30414 | 17355 | 5342 | 1141 | 3652 | 1720 | 733 | 11194 | 4015 | 5568 |
| 1919 | 3469 | 3088 | 1079 | 3896 | 1784 | 758 | 9970 | 3864 | 5803 |
| 1667 | 3019.5 | 2725 | 1031 | 3721 | 1737 | 722.5 | 8993 | 3596 | 5504.5 |
| 1634 | 3057 | 2755 | 1036 | 3926 | 1768 | 764 | 9329 | 3800 | 5739 |
| 2897 | 4288 | 3107 | 1049 | 4114 | 1758 | 722 | 14633 | 4840 | 6537 |
| 6347.5 | 15011.5 | 5116 | 1351 | 6018 | 2069 | 724 | 15694 | 7320 | 17739 |
| 8814.5 | 20409.5 | 6309 | 1598 | 7283 | 2290 | 731 | 13674 | 8859 | 22404 |
| 9474 | 23405.5 | 7160 | 1712 | 8053.5 | 2419 | 738 | 11917 | 8858.5 | 24647.5 |
| 9314.5 | 21727 | 6379.5 | 1567 | 7408 | 2254.5 | 709 | 11795 | 8685.5 | 23459 |
| 9167.5 | 20068 | 6272.5 | 1643 | 7035 | 2313.5 | 728 | 11916.5 | 8807.5 | 21600 |
| 6488 | 14674 | 4914 | 1320 | 5878 | 2007 | 723 | 15380 | 7451 | 17271 |
| 5575.5 | 13297 | 6718.5 | 1666 | 690658 | 65099.5 | 772 | 13289 | 6504.5 | 16002 |
| 3057 | 7617.5 | 3915 | 1193 | 6020.5 | 2365 | 722 | 13798.5 | 4326.5 | 10504 |
| 2537 | 3802 | 2983.5 | 1035 | 4268 | 1856 | 723 | 15448 | 4205.5 | 5932 |
| 2323 | 3326 | 2767 | 1034 | 3972 | 1796 | 744 | 11738 | 4359 | 5649 |
| 1744 | 2877 | 2507 | 986 | 3616 | 1714 | 736 | 9692 | 3558.5 | 5129.5 |
| 1701 | 2839 | 2432 | 970 | 3567 | 1711 | 733 | 9239 | 3482 | 5188 |
| 1753 | 2924.5 | 2499 | 988 | 3592.5 | 1694 | 729 | 9696.5 | 3608 | 5297 |
| 1811 | 2954.5 | 2484 | 984 | 3609 | 1704 | 728 | 9673 | 3599.5 | 5263 |
| 1737 | 2844 | 2403 | 952 | 3395 | 1638 | 710 | 9649 | 3458 | 5031 |
| 1769 | 2975 | 2530.5 | 1009 | 3591 | 1737 | 757 | 10097.5 | 3689.5 | 5390.5 |
| 1662 | 2734 | 2396 | 948.5 | 3408 | 1644 | 720 | 8982 | 3437 | 5022.5 |
| 1575 | 2703 | 2391 | 964 | 3445 | 1663 | 724 | 8444 | 3339 | 4978 |
| 1650 | 2787 | 2519.5 | 1032 | 3622.5 | 1739 | 758 | 9611 | 3504 | 5197 |
| 30870 | 17306 | 5388 | 1166 | 3835 | 1751 | 762 | 11815 | 4337 | 5939 |
| 2114 | 3649 | 3126.5 | 1069 | 3909 | 1761 | 739 | 10221 | 4043 | 5845.5 |
| 2031 | 3502 | 2964 | 1076 | 4002 | 1788 | 754 | 10243.5 | 4171 | 6051 |
| 1941 | 3308 | 2862 | 1061 | 3962.5 | 1777 | 760 | 10018 | 4099 | 5981.5 |
| 3128 | 4535 | 3278 | 1115 | 4369 | 1847 | 788 | 13570 | 5327 | 7237 |
| 5535 | 13881 | 5027 | 1394.5 | 6015 | 2079 | 762 | 15717 | 6442 | 16785.5 |
| 9642 | 21708 | 6702 | 1677 | 7595 | 2382 | 756 | 13868 | 9235 | 23662 |

| | | | | | | | | | |
|---|---|---|---|---|---|---|---|---|---|
| 10383.5 | 24661 | 7235 | 1752 | 8201.5 | 2422.5 | 752 | 12711.5 | 9385.5 | 25840.5 |
| 9691 | 20475 | 6325 | 1539 | 7399 | 2241 | 716 | 15760 | 9360 | 22367 |
| 9114 | 20023 | 6205 | 1651 | 7076 | 2313 | 737 | 12534 | 8928 | 22384 |
| 6896 | 15752 | 5244 | 1379 | 6189 | 2079 | 743 | 11806 | 7408 | 18183 |
| 4597.5 | 11709 | 6533 | 1604 | 723916 | 62559.5 | 739 | 14304 | 5410 | 14555 |
| 3478.5 | 6895.5 | 4252.5 | 1235 | 6950 | 2499 | 759 | 14260.5 | 4890 | 9409 |
| 2770 | 4236 | 3193 | 1101 | 4686 | 1922 | 742 | 16392 | 4395 | 6375 |
| 2583.5 | 3552 | 2837.5 | 1042 | 4080 | 1783.5 | 759 | 13317 | 4608 | 5828 |
| 2196 | 3333 | 2751 | 1023 | 3981 | 1778 | 758 | 11720 | 4232 | 5736 |
| 2007 | 3160 | 2661 | 1012 | 3823 | 1761 | 771 | 10143 | 4036 | 5611 |
| 1919 | 3176 | 2594.5 | 1043 | 3779.5 | 1751 | 752 | 10361 | 3841.5 | 5582 |
| 1892.5 | 3039 | 2540.5 | 1007 | 3656 | 1704 | 741 | 10285 | 3815 | 5423.5 |
| 1850.5 | 2979.5 | 2498 | 980 | 3524 | 1682 | 720 | 9940.5 | 3610 | 5195.5 |
| 1790 | 2987 | 2495 | 989 | 3573 | 1670 | 736 | 9701 | 3578 | 5251 |
| 1771 | 2887 | 2450 | 966 | 3512 | 1666 | 730 | 9173 | 3603 | 5290 |
| 1748.5 | 2967.5 | 2497.5 | 994 | 3577 | 1706.5 | 734 | 9560 | 3618 | 5353.5 |
| 1668 | 2851 | 2444 | 956 | 3444 | 1669.5 | 716 | 8972 | 3502.5 | 5136.5 |
| 34922.5 | 19450 | 5983 | 1190 | 3715 | 1720 | 744 | 12953.5 | 4460 | 5785 |
| 2400 | 3905 | 3184 | 1016 | 3594 | 1664 | 709 | 10053.5 | 3829.5 | 5489 |
| 2003 | 3378 | 2887 | 1039 | 3865 | 1733 | 737 | 9689 | 3936 | 5747 |
| 1785 | 3057.5 | 2544 | 967.5 | 3569 | 1660 | 700 | 9380.5 | 3665 | 5412 |
| 3265 | 5310 | 3380 | 1096 | 4454 | 1815 | 742 | 16633 | 5124 | 7766 |
| 5593 | 13242 | 4854 | 1297 | 5797.5 | 2014 | 723 | 16231.5 | 6519.5 | 15634 |
| 9164 | 19275 | 6111 | 1531 | 7030 | 2236 | 740 | 16293 | 9253 | 21684 |
| 11622 | 24901 | 7100 | 1701 | 7925 | 2424 | 744 | 13447 | 9409 | 25406 |
| 9410 | 19990 | 6298 | 1533 | 7283 | 2234 | 703 | 14394 | 9116 | 21504 |
| 9507.5 | 20701 | 6893 | 1957 | 7413.5 | 2536.5 | 732 | 13798 | 9381.5 | 22642.5 |
| 6899 | 16125 | 5151 | 1364 | 6095 | 2085 | 743 | 14209 | 7284 | 18392 |
| 4508 | 12133 | 6671.5 | 1648 | 716826 | 63372 | 742 | 13894 | 5384 | 14758.5 |
| 3000 | 5786.5 | 3989 | 1196 | 6186 | 2314 | 740 | 14525.5 | 4333.5 | 7964.5 |
| 2611 | 4348 | 3240 | 1118 | 4640 | 1949 | 748 | 15257.5 | 4145.5 | 6513 |
| 2519 | 3650 | 2872 | 1046 | 4291 | 1822 | 760.5 | 13901.5 | 4555 | 6022 |
| 2029 | 3165 | 2624 | 1021 | 3853.5 | 1756 | 738 | 10392 | 3801 | 5443.5 |
| 1980 | 3150 | 2652 | 1003 | 3889 | 1729 | 747 | 10444 | 3942 | 5550 |
| 1963 | 3158 | 2601 | 1017 | 3745 | 1746 | 738 | 10442 | 3781 | 5436 |
| 1881 | 2983 | 2428 | 945 | 3455 | 1640 | 694 | 9903 | 3632 | 5173 |
| 1986.5 | 3167 | 2621.5 | 1016.5 | 3665.5 | 1742 | 739 | 10678.5 | 3786.5 | 5399 |
| 1904 | 3100 | 2550 | 999 | 3563 | 1710 | 750 | 10473 | 3783 | 5395 |
| 1811 | 2907 | 2468 | 970 | 3462 | 1661 | 725 | 9573 | 3548 | 5091 |
| 1730.5 | 2820.5 | 2424 | 968.5 | 3472.5 | 1654.5 | 731 | 9072.5 | 3465 | 5068 |
| 1748 | 2858 | 2487 | 995 | 3479 | 1699 | 758 | 9393 | 3582 | 5130 |
| 36935 | 20273 | 6094 | 1182 | 3779 | 1711 | 762 | 12464 | 4485 | 5911 |
| 2769.5 | 4495 | 3750.5 | 1177 | 3919 | 1796 | 793 | 11978.5 | 4386 | 6147.5 |
| 2158.5 | 3525 | 2960 | 1054 | 3950 | 1783 | 770 | 10589 | 4283 | 6109 |
| 2091 | 3330 | 2841 | 1045 | 3924 | 1733 | 766 | 10528 | 4292 | 5954 |
| 3250.5 | 4473.5 | 3217 | 1097 | 4219 | 1774.5 | 754 | 13712.5 | 5154 | 6683 |
| 5180 | 12450 | 4840 | 1321 | 5699 | 2027 | 776 | 15457 | 5859 | 15189 |
| 10015 | 20683 | 6488 | 1623 | 7323 | 2275 | 773 | 14685 | 9114 | 21881 |

| | | | | | | | | | |
|---|---|---|---|---|---|---|---|---|---|
| 11841 | 25006 | 7334 | 1781 | 7980 | 2409 | 779 | 13007 | 9370 | 24766 |
| 10061.5 | 20542.5 | 6292.5 | 1555 | 7109.5 | 2186 | 686 | 14042.5 | 8960.5 | 21861 |
| 9246 | 19735 | 6057 | 1548 | 6941 | 2251 | 734 | 12512 | 8832 | 21325 |
| 6306 | 14527 | 4788 | 1291 | 5718 | 1960 | 709 | 13811 | 7061 | 16935 |
| 4517 | 11156.5 | 6826 | 1615 | 728321 | 70316 | 787 | 15944 | 5338 | 13729.5 |
| 3754 | 7621 | 4105 | 1240 | 5718 | 2174 | 803 | 14839 | 5055 | 10210 |
| 2903 | 4682 | 3351 | 1127 | 4581.5 | 1909 | 816 | 15595.5 | 4367 | 6746.5 |
| 2680 | 3529 | 2777 | 1008 | 3860 | 1732 | 776 | 12044 | 4557 | 5798 |
| 2114 | 3270 | 2728 | 1040 | 3812 | 1754 | 800 | 10957 | 4014 | 5696 |
| 1992 | 3123.5 | 2637 | 1034 | 3711 | 1743 | 787 | 10589 | 3888.5 | 5520 |
| 2066 | 3241 | 2706.5 | 1050 | 3812 | 1757 | 792 | 10979 | 3995 | 5653 |
| 2040 | 3158.5 | 2621 | 1040 | 3654.5 | 1733 | 780.5 | 10899 | 4003 | 5521.5 |
| 2104 | 3234 | 2660 | 1034 | 3688 | 1736 | 800 | 10807 | 4008 | 5657 |
| 2028 | 3205 | 2673 | 1032 | 3763 | 1760 | 808 | 10578 | 3929 | 5674 |
| 1948 | 3091 | 2627.5 | 1041 | 3635 | 1725.5 | 785 | 10486.5 | 3793.5 | 5488.5 |
| 1832 | 2993 | 2562 | 1022 | 3536 | 1701 | 778 | 9736 | 3736 | 5377 |
| 1988 | 3254 | 2786 | 1105 | 3955 | 1833 | 823 | 10430 | 3990 | 5820 |
| 36484 | 20076 | 6047 | 1259 | 3988 | 1817 | 828 | 13497 | 4793 | 6286 |
| 2369 | 3850 | 3228 | 1081 | 3901 | 1739 | 786.5 | 10878 | 4231 | 5969 |
| 2157 | 3551 | 3030 | 1098 | 4033 | 1799 | 823 | 10520 | 4376 | 6147 |
| 2062 | 3455 | 2967 | 1094.5 | 4106 | 1845 | 824 | 10617 | 4346.5 | 6159 |
| 3528 | 6365 | 3756.5 | 1203.5 | 4808 | 1898.5 | 816 | 17511.5 | 4903 | 8809.5 |
| 5490 | 12083 | 4642 | 1318 | 5501 | 1973 | 788 | 16319 | 6113 | 14393 |
| 9903.5 | 21313 | 6725 | 1692 | 7511.5 | 2384 | 819 | 13965 | 8713 | 22820 |
| 11494 | 26414 | 7827 | 1863.5 | 8559.5 | 2521 | 836 | 12705.5 | 9505 | 26870.5 |
| 9335.5 | 22102 | 6722.5 | 1627.5 | 7497 | 2335 | 735 | 11006 | 8703.5 | 23200 |
| 9363.5 | 23197.5 | 6978 | 1760 | 7546 | 2432 | 756.5 | 11511.5 | 8505 | 24680.5 |
| 6532 | 15774.5 | 5177.5 | 1365 | 6170.5 | 2086 | 746.5 | 14301 | 7066 | 18171.5 |
| 4613.5 | 10563 | 5851.5 | 1483 | 630883 | 56438 | 711.5 | 14452 | 5665 | 13050.5 |
| 3229 | 5768 | 4060 | 1200 | 6616 | 2386 | 734 | 14607 | 4633 | 7895 |
| 2782.5 | 4241 | 3179 | 1071 | 4706 | 1948 | 732 | 16458 | 4422.5 | 6440.5 |
| 2478 | 3528 | 2817 | 1015 | 4118 | 1769 | 746 | 12043 | 4505 | 5796 |
| 1972 | 3097 | 2586 | 988 | 3767 | 1711 | 722 | 10583 | 3792 | 5404 |
| 1849 | 3026 | 2523.5 | 1003.5 | 3704 | 1728 | 712.5 | 10092 | 3641.5 | 5249.5 |
| 1929 | 3210 | 2624 | 1024 | 3694 | 1752 | 742 | 11158 | 3903 | 5695.5 |
| 1972 | 3144 | 2537 | 972 | 3633 | 1692 | 726 | 10304 | 3786 | 5377 |
| 1883.5 | 3046.5 | 2500 | 986 | 3536.5 | 1664 | 701.5 | 10144.5 | 3635 | 5189 |
| 1929 | 3112 | 2562.5 | 998 | 3704 | 1722.5 | 736 | 10191 | 3776.5 | 5407.5 |
| 1807 | 2925 | 2444 | 960 | 3467 | 1650 | 704 | 9597 | 3553 | 5190 |
| 1742 | 2926 | 2505 | 981 | 3570 | 1687 | 726 | 9044 | 3597 | 5237 |
| 1692 | 2868 | 2419 | 976 | 3483 | 1674 | 722 | 9229 | 3479 | 5201 |
| 30354.5 | 16702 | 5260 | 1120 | 3629 | 1681 | 717 | 11424 | 4164 | 5584.5 |
| 2266.5 | 3799 | 3195 | 1058 | 3785 | 1719 | 712 | 10158.5 | 3977.5 | 5727 |
| 2030 | 3460 | 2909 | 1044 | 3948 | 1748 | 728 | 10084 | 4088 | 5898 |
| 1916 | 3291 | 2785.5 | 1034 | 3872 | 1733 | 720 | 9757 | 3938 | 5802 |
| 3413 | 6723 | 3685 | 1180 | 4853 | 1891 | 747 | 16881 | 5088 | 9372 |
| 5663.5 | 11459.5 | 4568.5 | 1305 | 5631 | 1989.5 | 729 | 17259 | 6864.5 | 14097 |
| 8593 | 21831 | 6550 | 1604.5 | 7365 | 2321 | 751.5 | 15188.5 | 7757.5 | 22807 |

| | | | | | | | | | |
|---|---|---|---|---|---|---|---|---|---|
| 9623 | 23466 | 7197 | 1676 | 8269 | 2394 | 723 | 14469 | 9117 | 24532 |
| 10186 | 24883 | 7570 | 1755 | 8723 | 2497 | 771 | 13788 | 9440 | 26172 |
| 8865 | 20466.5 | 6489 | 1666 | 7578.5 | 2384.5 | 752 | 14901.5 | 9441 | 22747 |
| 6810 | 16842 | 5533 | 1468 | 6846 | 2207 | 791 | 15754 | 8154 | 19860 |
| 4223 | 12323 | 6713 | 1592 | 666396.5 | 62877.5 | 755 | 15351.5 | 5407.5 | 15395 |
| 3461 | 7493 | 4341 | 1259 | 7275 | 2626 | 752 | 14901 | 4773 | 10065 |
| 2924 | 4927 | 3484 | 1152 | 4989 | 2014 | 810 | 18599 | 4731 | 7519 |
| 2535 | 3569 | 2832 | 1039 | 4083 | 1795 | 756 | 13721 | 4672 | 6060 |
| 2254 | 3525 | 2873 | 1070 | 4258 | 1830 | 811 | 13627 | 4606 | 6209 |
| 2080 | 3271 | 2730 | 1037 | 3961 | 1772 | 785 | 11017 | 4199 | 5830 |
| 2102.5 | 3352 | 2791 | 1056 | 3912 | 1776 | 798.5 | 11035 | 4160 | 5829 |
| 2263.5 | 3499.5 | 2867.5 | 1095 | 4058.5 | 1842 | 835 | 11607 | 4433.5 | 6130.5 |
| 2073 | 3277 | 2720 | 1065 | 3844 | 1765 | 805 | 10766 | 4046 | 5772 |
| 2087 | 3298 | 2732 | 1066 | 3841 | 1783 | 806 | 11373 | 4067 | 5705 |
| 1996 | 3230 | 2747 | 1075 | 3879 | 1777 | 807 | 10486.5 | 3971 | 5717.5 |
| 1916 | 3143 | 2692 | 1058 | 3837 | 1792.5 | 799 | 10469.5 | 3879 | 5644 |
| 1887 | 3124 | 2678 | 1058 | 3801 | 1782 | 814 | 10296 | 3995 | 5687 |
| 19978 | 11143.5 | 4676 | 1189 | 4004.5 | 1813 | 812 | 11803 | 4461 | 6098.5 |
| 2614 | 4093 | 3354 | 1094 | 3953 | 1804 | 795 | 10634 | 4367 | 6069 |
| 2193 | 3580 | 2979 | 1081 | 4008 | 1778 | 782 | 10667 | 4238 | 6053 |
| 2052 | 3427 | 2846 | 1090 | 3992 | 1807 | 779 | 10224 | 4097 | 5987 |
| 3580 | 5877 | 3538 | 1189 | 4620 | 1899 | 814 | 14983 | 5672 | 8464 |
| 5402 | 11275 | 4705 | 1328 | 5646 | 2012 | 779 | 16056 | 6303 | 13936 |
| 9533 | 19032 | 6459 | 1616 | 7362 | 2299 | 774 | 14321 | 9230 | 21067 |
| 10478 | 24234.5 | 7726.5 | 1788 | 8492.5 | 2497 | 795 | 14895.5 | 9373.5 | 25789 |
| 8638 | 20605 | 6532 | 1580 | 7538 | 2293 | 697 | 11205 | 8180 | 22316 |
| 9464 | 22296.5 | 6910 | 1829 | 7481 | 2418.5 | 743 | 14377.5 | 9679.5 | 24120.5 |
| 7139 | 16494 | 5425 | 1418 | 6412 | 2135 | 742 | 12229 | 7683 | 19027 |
| 5280 | 12505 | 6245 | 1549 | 601505 | 54005.5 | 759 | 14395.5 | 6148.5 | 15337 |
| 3673 | 8465.5 | 4376 | 1294 | 7537.5 | 2704 | 779.5 | 14553 | 5044 | 11353 |
| 2881 | 4241 | 3235 | 1091.5 | 4733 | 1947 | 757 | 16450 | 4695 | 6467 |
| 2624 | 3684.5 | 2995 | 1090 | 4266.5 | 1872 | 819 | 12390 | 4737.5 | 6100.5 |
| 2241 | 3333 | 2804 | 1066 | 4017.5 | 1819 | 804 | 11210 | 4243.5 | 5680 |
| 2087.5 | 3220 | 2762 | 1053.5 | 3817.5 | 1764 | 815 | 11263 | 4015 | 5727 |
| 2187 | 3317.5 | 2772 | 1072 | 3811 | 1789 | 849 | 11903.5 | 4209 | 5818 |
| 2063 | 3154 | 2630 | 1010 | 3666 | 1697 | 773 | 10489 | 3896 | 5499 |
| 2223 | 3334 | 2790 | 1071 | 3773 | 1766 | 836 | 11628 | 4161 | 5792 |
| 2069 | 3197.5 | 2689.5 | 1061.5 | 3783 | 1735 | 801 | 11237.5 | 4013.5 | 5728 |
| 2056 | 3238.5 | 2685.5 | 1066 | 3749 | 1772 | 827 | 11060.5 | 3991.5 | 5636.5 |
| 1885 | 2992 | 2587 | 992 | 3607 | 1694 | 782 | 10063.5 | 3759.5 | 5481 |
| 1891 | 3022 | 2651 | 1058 | 3703 | 1720 | 835 | 10069 | 3910 | 5532 |
| 37641 | 20839 | 6352 | 1209 | 3864 | 1757 | 790 | 12670 | 4607 | 6047 |
| 2803 | 4475 | 3681 | 1166 | 4117 | 1828 | 868 | 12085 | 4666 | 6448 |
| 2183 | 3563 | 3058 | 1077 | 3976 | 1777 | 800 | 10648 | 4272 | 6034 |
| 2152 | 3544 | 3022 | 1100 | 4019 | 1820 | 827 | 11126 | 4290 | 6173 |
| 3150 | 4483 | 3271 | 1111 | 4282 | 1815 | 830 | 14167 | 5166 | 6925 |
| 6947 | 14106 | 5209 | 1396 | 6088 | 2111 | 832 | 18034 | 7297 | 16379 |
| 9808 | 22185 | 6739 | 1696 | 7734 | 2345 | 815 | 13569 | 8571 | 23183 |

| | | | | | | | | | |
|---|---|---|---|---|---|---|---|---|---|
| 12382 | 27007 | 7940 | 1840 | 8474 | 2525 | 869 | 12647 | 9324 | 26315 |
| 10702.5 | 23705 | 6822.5 | 1637 | 7703 | 2331 | 744 | 14583 | 9706 | 24943 |
| 9820 | 20815 | 6357.5 | 1639 | 7120.5 | 2315 | 747 | 12565 | 9174 | 22238.5 |
| 8265 | 17031 | 5473 | 1421 | 6478 | 2120 | 776 | 12673 | 8197 | 18696 |
| 4928 | 11797 | 6972 | 1619 | 790518 | 73496 | 772 | 14857 | 5560 | 14449 |
| 3585.5 | 9020 | 4149.5 | 1242 | 6159 | 2296 | 786 | 15337 | 4774.5 | 12206 |
| 2979 | 4071 | 3077 | 1064.5 | 4392 | 1838 | 767 | 16408.5 | 4640.5 | 6239 |
| 2478 | 3343 | 2703 | 985 | 3867 | 1718 | 749 | 11616 | 4401 | 5584 |
| 2026 | 3211.5 | 2656 | 1032 | 3810.5 | 1748.5 | 764 | 10946 | 3902.5 | 5598 |
| 1846 | 2938 | 2451 | 964 | 3533 | 1653.5 | 743.5 | 9594.5 | 3629 | 5183.5 |
| 2051 | 3300 | 2740 | 1060 | 3908 | 1765 | 792 | 11835 | 4100 | 5795 |
| 1911 | 2972 | 2483 | 984 | 3436 | 1662 | 725 | 10069 | 3583 | 5168 |
| 1917 | 3041 | 2511.5 | 985 | 3558.5 | 1665 | 742 | 10050.5 | 3701 | 5275.5 |
| 1869 | 3068 | 2567 | 997 | 3657 | 1718 | 756 | 10213 | 3810 | 5424 |
| 1843 | 2945 | 2479 | 979 | 3484 | 1687 | 751 | 9241 | 3617 | 5270 |
| 1766.5 | 2871 | 2462 | 976 | 3591.5 | 1676.5 | 741 | 9427 | 3601 | 5195 |
| 1671 | 2777 | 2395 | 956 | 3426 | 1636 | 733 | 8878 | 3469.5 | 5052.5 |
| 26342 | 14570.5 | 4976 | 1146 | 3812 | 1735 | 763 | 11952 | 4403 | 5902 |
| 2254 | 3644 | 3042 | 1033 | 3679 | 1682 | 730 | 9942 | 3892 | 5582 |
| 1951 | 3296 | 2774 | 1015 | 3782 | 1705 | 742 | 9648 | 3946 | 5673 |
| 2011.5 | 3334 | 2789 | 1042 | 3883 | 1772.5 | 774 | 9903.5 | 4152 | 5956 |
| 3168 | 5492 | 3527 | 1154 | 4571 | 1859 | 777 | 17243 | 4735 | 7882 |
| 6788.5 | 13141.5 | 4865 | 1329 | 5881 | 2025 | 771 | 17215 | 7723 | 15643 |
| 9682.5 | 20328.5 | 6406 | 1572 | 7188.5 | 2300 | 782 | 16285.5 | 9472 | 22087 |
| 11407.5 | 26104.5 | 7463 | 1741 | 8488 | 2458 | 796 | 14150.5 | 8869 | 25999 |
| 10086 | 23271 | 6876 | 1653 | 7698 | 2371 | 746 | 12635 | 9224 | 24495 |
| 9699.5 | 21141.5 | 6643 | 1772 | 7259 | 2390 | 706 | 12429 | 8877 | 22083 |
| 7255.5 | 16438 | 5318 | 1408.5 | 6353.5 | 2107 | 759 | 13512.5 | 7750 | 18738.5 |
| 4952 | 12300 | 6154 | 1536 | 578100 | 55367 | 735 | 14385 | 5954 | 15151 |
| 3364 | 6774 | 4277 | 1247 | 7298.5 | 2625 | 757 | 14857 | 4796.5 | 9237.5 |
| 2836.5 | 4501 | 3379 | 1126 | 4845 | 1998 | 729 | 15841.5 | 4453 | 6534.5 |
| 2497 | 3589 | 2812 | 1023 | 4064 | 1782 | 733 | 12876 | 4510 | 5863 |
| 2171 | 3338.5 | 2769 | 1041 | 4086.5 | 1809 | 742 | 11221.5 | 4187 | 5739 |
| 2040 | 3223.5 | 2750 | 1059 | 3960 | 1788 | 770 | 11436.5 | 4105 | 5749 |
| 2023 | 3311 | 2696.5 | 1045.5 | 3950.5 | 1777 | 775 | 11011 | 4049.5 | 5773.5 |
| 1924 | 3079 | 2531 | 996.5 | 3611 | 1695 | 716 | 10162.5 | 3756 | 5412 |
| 2019 | 3227 | 2629.5 | 1016 | 3745 | 1749 | 738 | 11820.5 | 4113 | 5707.5 |
| 1800 | 2974 | 2477 | 963 | 3532 | 1664 | 707 | 9357 | 3627 | 5206.5 |
| 1837 | 3011 | 2531 | 1002 | 3613 | 1707 | 721 | 9886 | 3708 | 5346 |
| 1776 | 2969 | 2540 | 1002 | 3674 | 1726 | 737 | 9650 | 3675 | 5349 |
| 1724 | 2964 | 2537.5 | 1009 | 3689 | 1722 | 739.5 | 9367.5 | 3668 | 5421 |
| 33403 | 17943 | 5812 | 1196 | 3960 | 1763 | 751 | 12699 | 4625 | 6139 |
| 2635.5 | 4312 | 3485.5 | 1087.5 | 3877.5 | 1772.5 | 730 | 10828 | 4199.5 | 5899 |
| 2023 | 3421 | 2873 | 1022 | 3811 | 1718 | 698 | 9959 | 3955 | 5720 |
| 1978 | 3377 | 2834 | 1044 | 3887 | 1761 | 730 | 10761 | 4148 | 5955 |
| 2959 | 5066 | 3329 | 1106 | 4335 | 1820 | 731 | 15088 | 4928 | 7568 |
| 4838.5 | 11663.5 | 4536 | 1265.5 | 5494.5 | 1982.5 | 722 | 15722 | 6054 | 13973 |
| 7016 | 18497 | 6237 | 1584 | 7369 | 2282 | 736 | 13669 | 7230 | 20120 |

| | | | | | | | | | |
|---|---|---|---|---|---|---|---|---|---|
| 9430 | 21358 | 6597 | 1609 | 7548 | 2287 | 682 | 12793 | 8813 | 23160 |
| 9110 | 21768 | 6419 | 1554 | 7413 | 2250 | 701 | 13202 | 9250 | 23892 |
| 17912.5 | 27637 | 7223.5 | 1753.5 | 7684 | 2404 | 935.5 | 12891.5 | 8982 | 24254 |
| 15118.5 | 21882 | 5778 | 1455.5 | 6374.5 | 2139 | 1064 | 15044 | 9210.5 | 20151 |
| 11252 | 17284.5 | 7376 | 1722.5 | 661382 | 62313.5 | 1175 | 16519 | 7923 | 18187 |
| 5002 | 6787 | 4573 | 1313 | 8038 | 2805 | 1030 | 17593 | 5467 | 8776 |
| 3823.5 | 4721 | 3320 | 1104 | 4725 | 1924 | 1072 | 20177 | 5509 | 7087 |
| 3095 | 4066 | 3054 | 1085 | 4256 | 1823 | 1114 | 19395 | 5407 | 6602 |
| 2561 | 3517.5 | 2774 | 1024 | 3948 | 1773 | 1155 | 15862.5 | 5394 | 6370.5 |
| 2288 | 3366 | 2696.5 | 1033 | 3800 | 1746 | 1088.5 | 15539 | 4900.5 | 6096 |
| 2186.5 | 3258.5 | 2623.5 | 1001 | 3731 | 1708 | 1075.5 | 14509.5 | 4536.5 | 5882 |
| 2095.5 | 3124 | 2509 | 980 | 3484.5 | 1662 | 1026 | 13914 | 4464 | 5691 |
| 2145 | 3236 | 2634 | 1013 | 3664 | 1701 | 1022 | 13957 | 4566 | 5877 |
| 2361 | 3438 | 2794 | 1047 | 3849 | 1774 | 1203 | 16364 | 5328 | 6494 |
| 2313.5 | 3439 | 2808.5 | 1052 | 3812 | 1784 | 1178.5 | 16437.5 | 5252.5 | 6469 |
| 1972 | 3107 | 2593 | 994 | 3580 | 1693 | 1055 | 13516 | 4541 | 5796 |
| 1979 | 3151 | 2679 | 1019 | 3823 | 1733 | 1087 | 14096 | 4676 | 6024 |
| 37443 | 20321 | 6237 | 1243 | 4084 | 1831 | 1128 | 19244 | 5773 | 6929 |
| 3272 | 4573 | 3485 | 1078 | 3815 | 1745 | 1093 | 15937 | 5328 | 6521 |
| 2482.5 | 3754 | 2975 | 1042 | 3775 | 1744 | 1078 | 16941.5 | 5107.5 | 6405.5 |
| 2339 | 3605 | 2921 | 1065 | 3894 | 1746 | 1095 | 15727.5 | 5466.5 | 6636.5 |
| 4571 | 5481 | 3491 | 1115 | 4444 | 1826 | 1132 | 21820 | 6417 | 7945 |
| 7804 | 12986.5 | 4737 | 1259 | 5379.5 | 1948 | 1020 | 19435.5 | 6420 | 14302.5 |
| 17208 | 26414 | 6692 | 1600 | 7408 | 2298 | 1082 | 16292 | 9303 | 23092 |
| 19461.5 | 30935.5 | 8078 | 1738 | 8331 | 2425 | 1016 | 15006.5 | 10034 | 25552 |
| 704 | 747 | 603 | 555 | 1030 | 1085 | 169 | 1506 | 1128 | 1273 |
| 78156 | 51617 | 12081.5 | 1640 | 4088 | 1618 | 233 | 16388.5 | 6481 | 8396.5 |
| 54774 | 21269.5 | 4868.5 | 1021 | 3312 | 1447 | 347 | 12674 | 4795 | 6100.5 |
| 33704.5 | 10998.5 | 60706 | 11268 | 157785 | 26678 | 421 | 14653.5 | 4130.5 | 4646.5 |
| 41794 | 12690.5 | 10612.5 | 2483 | 13441 | 3572.5 | 552.5 | 16237 | 4790 | 5714.5 |
| 5488 | 5621 | 4602 | 1378 | 5765 | 2133 | 543 | 13669 | 3916 | 5024 |
| 4076 | 4968 | 4085.5 | 1200 | 4859.5 | 1815.5 | 613.5 | 12925.5 | 3867.5 | 5383 |
| 2896 | 3741 | 3160 | 1021 | 3872 | 1756 | 596 | 11165 | 3459 | 4973 |
| 2450 | 3757 | 2965 | 1022 | 3875 | 1776 | 627 | 11089 | 3894 | 5139 |
| 2526 | 3759 | 2894 | 1138 | 3686 | 1635 | 647 | 9546 | 3429 | 5228 |
| 2220 | 3751 | 2812 | 1048 | 3438 | 1582 | 643 | 9723 | 3653 | 5256 |
| 2541 | 3555 | 2822 | 1043 | 3632 | 1612 | 639 | 10698 | 3708 | 5374 |
| 2373 | 3675.5 | 2830.5 | 1000.5 | 3485 | 1655 | 574.5 | 9341.5 | 3597.5 | 5326 |
| 20008 | 13586 | 4940 | 1146 | 3851 | 1735 | 678 | 10322 | 7167 | 7835 |
| 2361.5 | 3769 | 3135 | 1094 | 3720.5 | 1727.5 | 665.5 | 9201.5 | 3849 | 5924.5 |
| 2032.5 | 3480.5 | 2745.5 | 1070.5 | 3622 | 1659 | 675 | 8971.5 | 3593 | 5283 |
| 22207 | 12617 | 4841 | 1102 | 4010 | 1866 | 648 | 11746 | 4330 | 6078 |
| 2852.5 | 4653 | 3361.5 | 1131 | 4090.5 | 1732 | 728 | 10009 | 4031 | 6246.5 |
| 2444.5 | 3953 | 3017.5 | 1011 | 4509.5 | 1869.5 | 691.5 | 10140 | 4230.5 | 6241 |
| 2241 | 3615 | 3190 | 1033 | 3969 | 1637 | 688 | 9505 | 3894 | 5929 |
| 4138 | 5297 | 3547 | 1208 | 4502 | 1791 | 641 | 13953 | 5591 | 7054 |
| 6424.5 | 15070.5 | 5399.5 | 1484.5 | 6524.5 | 2174 | 693.5 | 15174 | 7008 | 18464.5 |
| 9988 | 22861 | 9296 | 1990.5 | 10687 | 2815 | 713 | 10837.5 | 8418 | 24108.5 |

| | | | | | | | | | |
|---|---|---|---|---|---|---|---|---|---|
| 8214 | 24302.5 | 9055 | 1974.5 | 10603 | 2730 | 640.5 | 9106 | 7092.5 | 23069.5 |
| 11179 | 25167.5 | 9579 | 1957.5 | 10831 | 2560 | 672 | 10170 | 8322 | 24628 |
| 13292 | 127250 | 177911 | 66961 | 9128 | 2796 | 620 | 11720 | 11449 | 65456 |
| 11208 | 23009 | 7178 | 1915 | 7036 | 2232 | 737 | 14625 | 10152 | 24785 |
| 5470 | 12885 | 6519 | 1632 | 394900 | 36465.5 | 569 | 13415 | 5137 | 15822 |
| 4129 | 10967 | 5584 | 1510 | 9385 | 2909 | 751 | 16083 | 5521 | 13759 |
| 3924 | 5067 | 3603 | 1189 | 5561 | 2058 | 636 | 16036 | 5485 | 7529 |
| 2693 | 4055 | 3269 | 1164 | 4141 | 1962 | 664 | 13896 | 4725 | 6382 |
| 2201 | 3446 | 2439 | 1120 | 3813 | 1844 | 582 | 10491 | 3928 | 5645 |
| 2221 | 3689 | 3051 | 1234 | 4447 | 1890 | 720 | 11210 | 4359 | 6539 |
| 2234.5 | 3620 | 2600.5 | 1151 | 3989.5 | 1845 | 713.5 | 11051.5 | 4049.5 | 6144.5 |
| 2200 | 4001 | 2775 | 1051 | 3825 | 1867 | 662 | 11406 | 3745 | 6009 |
| 2122.5 | 3627 | 2824.5 | 1091.5 | 3684 | 1699 | 732 | 10343.5 | 3792.5 | 6128.5 |
| 2377 | 3908 | 3026 | 1020 | 4058 | 1878 | 723 | 11214 | 4279 | 6072 |
| 13707 | 10608 | 5070 | 1214 | 3871 | 1720 | 717 | 10455 | 7915 | 8023 |
| 2487 | 3988 | 3040 | 1142 | 4309 | 1832 | 706 | 9410 | 4670 | 6380 |
| 1942 | 3452 | 2549 | 1098 | 3899 | 1604 | 638 | 8943 | 3788 | 5933 |
| 33830 | 18951 | 5626 | 1223 | 3940 | 1593 | 696 | 12014 | 4601 | 6156 |
| 3093 | 4340 | 3277.5 | 993 | 3862 | 1667 | 647.5 | 10589.5 | 4287.5 | 5862 |
| 2622 | 4324 | 3589 | 1203 | 4675 | 1858 | 685 | 10383 | 4906 | 6727 |
| 2527 | 3740 | 3070 | 1094.5 | 4262 | 1802.5 | 634 | 9807.5 | 4135 | 6092 |
| 2736.5 | 4115 | 3046 | 1134 | 4483 | 1820.5 | 727 | 11143.5 | 4815.5 | 6646 |
| 5664 | 11191 | 4792 | 1247 | 5478 | 1844 | 675 | 14870 | 5983 | 12908 |
| 8737 | 27227 | 9469 | 2162 | 10988 | 2811 | 754 | 13748 | 8979 | 30080 |
| 8865 | 27770.5 | 8048 | 2087.5 | 9629.5 | 3022 | 674 | 10234.5 | 8349.5 | 27489 |
| 10209 | 24315 | 11438 | 2177 | 11986 | 2984 | 722 | 10762 | 8634 | 23728 |
| 10605 | 32016.5 | 11371.5 | 2612 | 10096.5 | 2682.5 | 679.5 | 13700.5 | 10460.5 | 33187 |
| 10074 | 21888 | 7258 | 1717 | 8023 | 2379 | 722 | 11856 | 9428 | 25556 |
| 5880 | 13786.5 | 6492 | 1666 | 423743 | 40327 | 660 | 13243.5 | 6516 | 16745 |
| 3899.5 | 11115 | 4962 | 1426.5 | 7643.5 | 2337 | 781 | 14550 | 5577.5 | 15000 |
| 3860 | 4961 | 3206 | 1004 | 4450 | 1819 | 685 | 17565 | 5186 | 7243 |
| 2774 | 4193 | 2973 | 1047 | 4320 | 1735 | 670 | 11904 | 4685 | 6146 |
| 2783.5 | 4032.5 | 3128 | 1173 | 4242.5 | 1909.5 | 733.5 | 11327 | 4411 | 6271.5 |
| 2501 | 3794 | 3255 | 1120 | 4209 | 1687 | 660 | 11286 | 4523 | 6190 |
| 2326.5 | 3608 | 2999 | 1086 | 4099 | 1770.5 | 676.5 | 10405.5 | 4231 | 6087 |
| 2372 | 3680 | 3105 | 1133 | 3660 | 1685 | 669 | 10843 | 3814 | 5604 |
| 2314.5 | 3590 | 2774.5 | 1084.5 | 3689 | 1702 | 690 | 10308 | 4105 | 5890.5 |
| 2420 | 4023 | 2753 | 998 | 4174 | 1635 | 655 | 10260 | 4347 | 5764 |
| 9572 | 7228 | 3809 | 1168 | 3876 | 1674.5 | 642.5 | 9875 | 5546 | 7069.5 |
| 2860 | 4183 | 3189 | 1102 | 3649 | 1703 | 667 | 9243 | 4443 | 5771 |
| 2204 | 3430 | 2794 | 1013 | 3810 | 1661 | 592 | 8865 | 4012 | 5718 |
| 22696 | 11809.5 | 5113.5 | 1268.5 | 4151.5 | 1843.5 | 684.5 | 12468 | 4714.5 | 6446 |
| 4396 | 5468 | 3685 | 1166 | 4128 | 1824 | 691 | 11034 | 5100 | 6716 |
| 2766.5 | 4266.5 | 3236 | 1184 | 4670 | 1901.5 | 641 | 10398.5 | 4245 | 6253 |
| 2629 | 4114 | 2687 | 1083 | 4000 | 1873 | 632 | 10174 | 3758 | 5966 |
| 3607 | 5659 | 3179.5 | 1143 | 4379.5 | 1668 | 643 | 15377 | 5192.5 | 7418 |
| 4988.5 | 11505.5 | 4597.5 | 1220 | 5167 | 1752.5 | 623 | 15324.5 | 5774.5 | 12532 |
| 9349 | 23134 | 7447 | 1656 | 8187.5 | 2477.5 | 630 | 13786.5 | 8924.5 | 26005.5 |

| | | | | | | | | | |
|---|---|---|---|---|---|---|---|---|---|
| 9199 | 24227.5 | 9526.5 | 2157 | 11688 | 2891.5 | 665 | 9332.5 | 8304.5 | 25856 |
| 9284 | 24633 | 8602 | 1993 | 10710 | 2780 | 572 | 10409 | 8455 | 24824 |
| 11936 | 104571 | 171747 | 58351 | 11185 | 2832 | 640 | 12636 | 11439 | 57737 |
| 9386.5 | 23373 | 7790.5 | 1828 | 7928 | 2396.5 | 680 | 13448.5 | 9302.5 | 25961 |
| 6928 | 16474 | 6672 | 1592 | 409375 | 37788 | 619 | 12594 | 6495 | 18179 |
| 4831.5 | 10663 | 4856.5 | 1384 | 7582 | 2610.5 | 769.5 | 16288 | 6189 | 14526 |
| 4210 | 6565.5 | 4090.5 | 1222 | 6475.5 | 2064.5 | 721.5 | 17123.5 | 5885.5 | 9045 |
| 3590.5 | 4452 | 3211 | 1226 | 4498 | 1857 | 681.5 | 16339 | 5324.5 | 6850 |
| 2812.5 | 3920 | 2832.5 | 1020.5 | 4393 | 1860.5 | 656 | 11715 | 4647.5 | 5913 |
| 2685.5 | 4503 | 3222.5 | 1181.5 | 4648 | 1987.5 | 706 | 12461 | 4936 | 6532.5 |
| 2234 | 4036 | 2931 | 1107 | 3760 | 1809 | 685 | 10350 | 4402 | 6234 |
| 2723 | 3960 | 3099 | 1126 | 4135 | 1745 | 696 | 11111 | 4265 | 6250 |
| 2398 | 3605.5 | 2742 | 1098.5 | 4140 | 1768.5 | 638 | 9902.5 | 3942 | 5594.5 |
| 2660 | 3833 | 3254 | 1164 | 4264 | 1879 | 732 | 12521 | 4658 | 6508 |
| 16270.5 | 10213.5 | 4274.5 | 1286 | 4095 | 1796 | 685 | 10929 | 6903 | 8452 |
| 3164 | 4714 | 3529 | 1061 | 3730 | 1548 | 631 | 9877 | 4652 | 6280 |
| 2299.5 | 3888.5 | 3002 | 1054 | 3695.5 | 1667.5 | 680.5 | 11165.5 | 4148 | 5971.5 |
| 32631 | 16228 | 6012 | 1234 | 4159 | 1803 | 739 | 12524 | 4913 | 6761 |
| 4442 | 6340.5 | 4288 | 1166 | 4151.5 | 1731 | 669 | 12020.5 | 4711.5 | 6736 |
| 2886 | 4367.5 | 3369 | 1080 | 4634.5 | 1793 | 690.5 | 11275.5 | 4327 | 6727.5 |
| 2926.5 | 4570 | 3213.5 | 1037 | 4456 | 1920 | 619 | 11090 | 4390.5 | 6505.5 |
| 4077 | 6529 | 3685 | 1061 | 4688 | 1808 | 681 | 14440 | 5911 | 8049 |
| 6334 | 13099.5 | 5100 | 1374 | 6768 | 2205.5 | 651.5 | 15866 | 6663.5 | 14996 |
| 9812 | 22848 | 7100 | 1669 | 8583 | 2502 | 664 | 13875 | 9407 | 24842 |
| 7956 | 26676 | 9273 | 1878 | 9871 | 2710.5 | 635 | 10366.5 | 7189 | 29903.5 |
| 8425 | 24439 | 11814 | 2169 | 12019 | 2836 | 670 | 9002 | 7128 | 24523 |
| 12331 | 46963 | 15844 | 3987 | 10380 | 2776 | 696 | 13367 | 10885 | 39236 |
| 10675 | 24108 | 8008 | 1789 | 9096 | 2449 | 753 | 12527 | 9444 | 26519 |
| 6200 | 15049 | 6777.5 | 1609.5 | 355799.5 | 36630 | 658.5 | 14480.5 | 6987.5 | 17520.5 |
| 4464.5 | 13698.5 | 5076.5 | 1382 | 6667 | 2393.5 | 848 | 17287.5 | 6745.5 | 16593 |
| 3431.5 | 6404.5 | 4023.5 | 1185 | 5092 | 1968 | 689.5 | 17135.5 | 4717.5 | 8698.5 |
| 3097.5 | 4328.5 | 3112 | 1223 | 4551 | 1926 | 761.5 | 15630 | 5040 | 6786.5 |
| 2578.5 | 4099 | 3144 | 1134.5 | 3937 | 1881.5 | 713 | 11696.5 | 4669 | 6252 |
| 2375.5 | 3589.5 | 2845 | 1098 | 4000.5 | 1712.5 | 649 | 10196.5 | 4173 | 5978.5 |
| 2388.5 | 3409 | 2995 | 1170.5 | 3723.5 | 1731 | 661 | 9669.5 | 3823 | 5673.5 |
| 2499.5 | 3793.5 | 2974.5 | 1082 | 4506 | 1839 | 707 | 11040.5 | 4413 | 6302.5 |
| 2573 | 3878 | 2847 | 1155 | 4029 | 1788 | 699 | 10813 | 4380 | 6293 |
| 2400 | 3488 | 3026 | 1050 | 4203 | 1729 | 666 | 10674 | 4198 | 6085 |
| 11442 | 8450 | 3975 | 1225 | 3870.5 | 1829 | 671.5 | 9808.5 | 6024 | 7600.5 |
| 2708.5 | 4044 | 3204 | 1109 | 3840 | 1759 | 690 | 9778 | 4814 | 6334 |
| 2830 | 4140 | 3449 | 1190 | 4333 | 1950 | 753 | 11242 | 4750 | 6411 |
| 10042 | 8921 | 4293 | 1236 | 3951 | 1749 | 699 | 11156.5 | 4409.5 | 6448.5 |
| 3725.5 | 4759.5 | 3657.5 | 1035 | 3701 | 1806 | 603 | 9626.5 | 4006 | 5852.5 |
| 2689 | 4452 | 3085 | 1167 | 4423 | 1872 | 689 | 11075 | 4610 | 6159 |
| 2931 | 4296 | 3060 | 1116 | 4490 | 2015 | 671 | 10613 | 4588 | 6635 |
| 3305 | 4851 | 3610 | 1326 | 4819 | 1819 | 731 | 13334 | 5308 | 7215 |
| 6524 | 13413 | 4973 | 1347 | 6018 | 2005 | 641 | 16266 | 6875 | 15829 |
| 9477 | 23790.5 | 7271 | 1865 | 8646 | 2315.5 | 680.5 | 11660 | 8701 | 25576 |

| | | | | | | | | | |
|---|---|---|---|---|---|---|---|---|---|
| 10480.5 | 24128.5 | 10599.5 | 2058 | 10391.5 | 3083.5 | 703.5 | 12495 | 8655 | 23468 |
| 9993 | 25293 | 9927 | 2046 | 10971 | 2730 | 665 | 10841 | 8306 | 25179 |
| 17576 | 145848 | 215735 | 77172 | 8740 | 2313 | 525 | 10253 | 11686 | 74928 |
| 15867.5 | 38637.5 | 14199 | 3731.5 | 8147 | 2488 | 702 | 12695.5 | 10154.5 | 26217 |
| 13602 | 18536 | 17337 | 7407 | 80610 | 15920 | 756 | 13554 | 8402 | 19370 |
| 7227 | 13803 | 6952 | 2054 | 32009 | 5070 | 885 | 17653 | 8349 | 17008 |
| 5961 | 6103 | 4181 | 1222 | 11510 | 2500 | 797 | 18938 | 5849 | 8224 |
| 5529 | 4782 | 3582 | 1110 | 7630 | 2240 | 735 | 16047 | 5474 | 6801 |
| 5190.5 | 4618.5 | 3805.5 | 1216.5 | 5848 | 2127 | 878 | 14535 | 5282 | 6894.5 |
| 4705 | 4256 | 2914 | 1062 | 5131 | 1957 | 763 | 14347 | 4949 | 5933 |
| 3617 | 4127 | 3310 | 1198 | 6255 | 1893 | 799 | 11534 | 4571 | 6203 |
| 4125 | 4396.5 | 3382 | 1114 | 4836 | 1934 | 715 | 11674 | 4387 | 6711.5 |
| 4022 | 4156 | 3218 | 1178 | 4303 | 1741 | 749 | 11632 | 4635 | 6116 |
| 3877.5 | 4108.5 | 3493 | 1132.5 | 4713.5 | 1791 | 697.5 | 11507 | 4343 | 6140 |
| 8669 | 7636 | 4157 | 1382 | 4180 | 1924 | 738 | 11849 | 6221 | 7351 |
| 6146 | 5215 | 3654 | 1244 | 4540 | 1849 | 711 | 11793 | 5495 | 7155 |
| 5239 | 5005 | 3645 | 1227 | 5087 | 1935 | 723 | 11667 | 4736 | 6468 |
| 216556.5 | 123971 | 34085 | 3128 | 4444 | 1950.5 | 745.5 | 40828.5 | 10875.5 | 9365 |
| 36496 | 18527 | 6839 | 1855.5 | 5110 | 1944 | 764 | 15764.5 | 7388.5 | 8376.5 |
| 13808.5 | 7156 | 3912.5 | 1234 | 5315 | 2093 | 680.5 | 12276 | 5063.5 | 6947 |
| 11963 | 7697.5 | 4203 | 1241 | 4975 | 2089.5 | 740.5 | 12719.5 | 4776 | 6908.5 |
| 9498 | 7699 | 4395 | 1339 | 5821 | 2176 | 703 | 15983 | 6003 | 8385 |
| 11587.5 | 20543.5 | 6577 | 1514 | 7557.5 | 2383.5 | 781 | 15680 | 7262 | 18093 |
| 13143.5 | 22113.5 | 6960.5 | 1348.5 | 8344 | 2392.5 | 722.5 | 15542.5 | 9460 | 22012.5 |
| 14062.5 | 28623 | 8177 | 1691 | 10143.5 | 2597.5 | 668 | 13228 | 9548 | 27152.5 |
| 9826 | 23691 | 10852 | 2400 | 11992 | 3125 | 695 | 11053 | 8869 | 25671 |
| 9910.5 | 37317 | 19624.5 | 3808.5 | 11991.5 | 2920 | 684.5 | 8939.5 | 8133 | 33265 |
| 10784 | 24803 | 8441 | 1901 | 8178 | 2555 | 678 | 12327 | 8834 | 26661 |
| 5681 | 15958 | 7180 | 1790 | 482987 | 45648 | 677 | 15358 | 6724 | 18619 |
| 5598 | 11614.5 | 4858 | 1412.5 | 7932.5 | 2759 | 717.5 | 14835.5 | 6931 | 15868 |
| 3718 | 5826 | 3814 | 1169 | 5411 | 2024 | 683 | 18296 | 5054 | 7381 |
| 3197.5 | 4494 | 3284.5 | 1206 | 4611 | 1932.5 | 713 | 13446.5 | 5190.5 | 6982 |
| 3032 | 4506 | 3172 | 1098 | 4538 | 1829 | 711 | 14003 | 5110 | 6537 |
| 2533 | 3616 | 3242 | 1113 | 4173 | 1819.5 | 728 | 10425 | 4632 | 5801 |
| 2224 | 3477 | 3031 | 1055 | 3666.5 | 1765.5 | 646 | 10263 | 3645.5 | 5606.5 |
| 2380 | 3763 | 2740 | 1064.5 | 4156.5 | 1751.5 | 648 | 10242.5 | 4456.5 | 5827 |
| 2332 | 3598 | 2803.5 | 1073 | 3766.5 | 1705 | 649 | 9969 | 4064 | 5589 |
| 2336 | 3630 | 2925 | 1066 | 3786.5 | 1786.5 | 654.5 | 10095.5 | 4267.5 | 5671 |
| 11600 | 8616 | 4191 | 1096 | 4224 | 1860 | 696 | 10169 | 5937 | 7427 |
| 2940 | 4259 | 3236 | 1161 | 4291 | 1740 | 706 | 10588 | 4590 | 6417 |
| 2149 | 3417 | 2912.5 | 1091.5 | 3805.5 | 1668 | 652 | 9050 | 3713 | 5030 |
| 5255 | 6878 | 4162 | 1092 | 3768 | 1750 | 642 | 10653 | 4567 | 6021 |
| 4495.5 | 5557 | 3754 | 1214.5 | 4179 | 1795 | 695 | 11388 | 4364.5 | 6195 |
| 2779 | 4090 | 3141 | 1100 | 4804 | 1910 | 727 | 10837 | 5084 | 6522 |
| 2867.5 | 4209.5 | 3117.5 | 1037 | 4468 | 1784.5 | 667 | 9199 | 4418 | 6016.5 |
| 4082 | 6115.5 | 3474.5 | 1240 | 4878 | 1866.5 | 688.5 | 15897.5 | 5566.5 | 8447.5 |
| 6214 | 12753 | 5121.5 | 1420.5 | 6177 | 2143.5 | 681.5 | 16374 | 7313.5 | 15383.5 |
| 7634.5 | 21898.5 | 7293 | 1770 | 8802 | 2613 | 671.5 | 11893.5 | 7861 | 24655.5 |

| | | | | | | | | | |
|---|---|---|---|---|---|---|---|---|---|
| 9080 | 30782 | 10021 | 2074 | 11707 | 2804 | 683 | 10725 | 7990 | 32784 |
| 9785.5 | 23534 | 10424 | 2098.5 | 11285.5 | 2886.5 | 666 | 9918.5 | 8591.5 | 24935.5 |
| 12787 | 132046 | 201917 | 70484 | 10489 | 2763 | 616 | 10207 | 11958 | 76823 |
| 9174 | 26955 | 8441 | 2276 | 8874 | 2629 | 711 | 12726 | 10463 | 29396 |
| 8491.5 | 15309 | 7245.5 | 2008.5 | 18766.5 | 5901 | 563.5 | 9649.5 | 6990 | 18775 |
| 4766 | 12152 | 5251 | 1676 | 8604 | 2854 | 801 | 14942 | 6647 | 16729 |
| 3062 | 7697 | 4654 | 1510 | 6499 | 2599 | 703 | 16595 | 5325 | 11345 |
| 3351.5 | 6420 | 4360.5 | 1514 | 6800.5 | 2646.5 | 699.5 | 17045.5 | 5624.5 | 10761 |
| 2659 | 4173.5 | 3714 | 1407.5 | 5559 | 2169 | 658 | 12661 | 5267.5 | 8241 |
| 2572 | 3907 | 3197 | 1318 | 5382 | 2193 | 672 | 11371 | 4840 | 7151 |
| 2428 | 4045 | 3469 | 1506 | 5620 | 2333 | 648 | 12436 | 5152 | 7294 |
| 2425.5 | 4048 | 3406.5 | 1277 | 4701 | 2026.5 | 639 | 11431 | 5057 | 7785 |
| 2589 | 4092 | 3730.5 | 1281.5 | 5274.5 | 2212.5 | 664.5 | 11371 | 4510 | 7939 |
| 2562.5 | 3751 | 3454.5 | 1416.5 | 5097.5 | 2237 | 652.5 | 12071 | 4964 | 7785 |
| 5034 | 5935 | 4447.5 | 1429.5 | 4648.5 | 2211 | 601 | 10268.5 | 6509 | 8358 |
| 2850.5 | 4680.5 | 3838.5 | 1273 | 5301.5 | 2231.5 | 669.5 | 11139.5 | 5567.5 | 8612.5 |
| 2286.5 | 3720.5 | 3484 | 1231 | 4797 | 2096.5 | 589 | 9935 | 4435 | 6782.5 |
| 23897 | 12459 | 5562 | 1496 | 5464 | 2420 | 670 | 13200 | 5719 | 8378 |
| 3802 | 5349.5 | 3885 | 1392.5 | 5027.5 | 2267.5 | 639 | 11056.5 | 5661.5 | 8104 |
| 3205 | 6165.5 | 4315 | 1580.5 | 6090.5 | 2441.5 | 660 | 11301 | 5654.5 | 10873 |
| 2942 | 4206 | 4021 | 1354 | 5093 | 2209 | 619 | 11342 | 5382 | 7715 |
| 3448.5 | 5963.5 | 3981.5 | 1419 | 5620.5 | 2297 | 655 | 13929.5 | 5597 | 10498.5 |
| 5658 | 12923.5 | 5492 | 1517 | 6462.5 | 2443.5 | 691 | 17058.5 | 6513.5 | 17016 |
| 8852 | 21985 | 7276 | 2100 | 9075 | 2854 | 642 | 16413 | 8804 | 24034 |
| 9546 | 27673.5 | 9332.5 | 2058.5 | 10341 | 3091.5 | 609.5 | 12940.5 | 8918 | 27939 |
| 10488 | 27476 | 10613 | 2520 | 12922 | 3621 | 707 | 11244 | 8918 | 30810 |
| 9540 | 72728 | 101675 | 25204 | 13325.5 | 3824.5 | 575 | 9929 | 9139 | 50841 |
| 10621.5 | 40043 | 36228.5 | 9452.5 | 11526.5 | 3288 | 629 | 10208.5 | 9895 | 36524 |
| 6917 | 18412 | 9237 | 2863 | 17993 | 6060 | 654 | 14482 | 7443 | 21351 |
| 5707 | 12868 | 9926 | 2238 | 73252 | 10237 | 812 | 19046 | 7009 | 18611 |
| 4667.5 | 5888.5 | 5678 | 1658.5 | 60131.5 | 8202.5 | 733 | 18585.5 | 6034.5 | 8466 |
| 3853 | 4583 | 4164 | 1386 | 44971 | 5755 | 658 | 12718 | 4663 | 6435 |
| 3171 | 4073 | 3622 | 1253 | 42398 | 5873 | 694 | 11798 | 4801 | 6029 |
| 3434 | 3969.5 | 3533 | 1321 | 34119.5 | 4676 | 682 | 11762.5 | 4572.5 | 7077.5 |
| 2780 | 4265 | 3869 | 1400 | 25536 | 3953 | 658 | 10152 | 3978 | 5878 |
| 2752.5 | 4123.5 | 3245 | 1256.5 | 19512 | 2951 | 664 | 9651 | 3929.5 | 5650 |
| 2443 | 3930.5 | 2997.5 | 1108 | 13018 | 2516 | 659.5 | 9777.5 | 4059.5 | 5492.5 |
| 2617 | 3826 | 3247 | 1090 | 12481 | 2462 | 705 | 9819 | 4021 | 5885 |
| 11594 | 9257 | 5928 | 1537 | 14774 | 2713 | 667 | 9947 | 6048 | 7759 |
| 4750 | 5351 | 4517 | 1264.5 | 12887 | 2697.5 | 744 | 10132 | 5017.5 | 7600.5 |
| 3919 | 5002 | 3584 | 1161 | 10840 | 2280 | 850 | 12400 | 4708 | 6381 |
| 20431.5 | 14701 | 14475.5 | 2796.5 | 10554.5 | 2353 | 696 | 21360.5 | 6751.5 | 7464.5 |
| 6566 | 6258.5 | 4773.5 | 1580 | 9727 | 2152 | 685 | 13737.5 | 4799.5 | 6381.5 |
| 3907.5 | 4795 | 3708.5 | 1252.5 | 13000 | 2794 | 649.5 | 8955 | 4094.5 | 6491.5 |
| 3593 | 3871 | 3350 | 1168 | 9440 | 2361 | 615 | 9581 | 4074 | 5641 |
| 4431.5 | 5548.5 | 3850.5 | 1284 | 10799.5 | 2254.5 | 680 | 14577 | 5360 | 6958.5 |
| 15875 | 22917 | 7605 | 1712 | 9169 | 2472 | 706 | 15875 | 9020 | 18588 |
| 9329.5 | 29855 | 10414.5 | 2200.5 | 14132 | 3144.5 | 950.5 | 13729.5 | 10005 | 31159.5 |

| | | | | | | | | | |
|---|---|---|---|---|---|---|---|---|---|
| 9330 | 27096 | 10510 | 1947 | 12772 | 2911 | 867 | 11141 | 8837 | 27133 |
| 10402 | 26704 | 11095 | 2390 | 12739 | 2936 | 698 | 9917 | 8440 | 26196 |
| 15268 | 197689.5 | 342566 | 106828 | 14539.5 | 4877 | 615 | 10491 | 12246 | 98370 |
| 14159.5 | 112816.5 | 186456.5 | 59464.5 | 11780 | 3917 | 658.5 | 10722.5 | 10597.5 | 53590.5 |
| 9467 | 21830 | 20777.5 | 7164.5 | 49905 | 10540 | 716.5 | 12401.5 | 10147 | 24269.5 |
| 5469 | 14690.5 | 8900.5 | 3330 | 21817.5 | 4215.5 | 783 | 14705 | 6070 | 17163.5 |
| 3725 | 6425 | 4553 | 1673 | 9361 | 2544 | 683 | 15025 | 5280 | 7843 |
| 3638 | 5266 | 4769 | 1559 | 6846 | 2290 | 705 | 17480 | 5187 | 7306 |
| 2712.5 | 4034 | 3930.5 | 1551 | 6432.5 | 2066.5 | 638.5 | 12201.5 | 4527 | 5694 |
| 2493.5 | 3940 | 3790.5 | 1312.5 | 5209.5 | 1998.5 | 724 | 11986 | 4270 | 6336.5 |
| 2226 | 3517 | 3591 | 1148 | 4848 | 1850.5 | 639.5 | 10999 | 4014.5 | 6405 |
| 2574.5 | 4191.5 | 3457 | 1281 | 5252.5 | 1956 | 702.5 | 14334.5 | 4724.5 | 6926.5 |
| 2344 | 3850 | 3320 | 1160 | 4607 | 1919 | 670 | 10613 | 4093 | 5775 |
| 2049 | 3802 | 2912 | 1084 | 4123 | 1804 | 672 | 10370 | 3865 | 5878 |
| 4220 | 5883 | 4786 | 1379 | 4734 | 1908 | 741 | 11976 | 5676 | 7742 |
| 2853 | 4202 | 3381 | 1198 | 4271 | 1903 | 656 | 9784 | 4757 | 6478 |
| 2321 | 3919 | 3259 | 1203 | 4229 | 1810 | 678 | 10570 | 4346 | 6168 |
| 4288.5 | 6774.5 | 4560.5 | 1268 | 4084.5 | 1871.5 | 640.5 | 12018.5 | 4453 | 6017 |
| 6061 | 6996 | 4868 | 1273 | 4120.5 | 1714 | 663.5 | 11161.5 | 4598 | 6582 |
| 2715 | 4200.5 | 3435.5 | 1179 | 4824 | 1882.5 | 625 | 10709 | 4311 | 6341.5 |
| 2508.5 | 3903 | 3061.5 | 1081 | 4117.5 | 1680.5 | 622 | 9862.5 | 4294 | 5821.5 |
| 3996.5 | 6425 | 3723.5 | 1186 | 4795 | 1794 | 669.5 | 17630 | 5565.5 | 8243 |
| 7054 | 13990 | 4908 | 1389 | 6889 | 2040 | 676 | 17395 | 7665 | 16347 |
| 8664.5 | 18623 | 6038 | 1577.5 | 6866 | 2088.5 | 605.5 | 14675 | 7603 | 19750.5 |
| 8922 | 25867 | 8468 | 1960 | 9926 | 2800 | 650 | 9900 | 8289 | 28268 |
| 10716 | 27125 | 11345 | 2181 | 11820 | 2886 | 673 | 9957 | 8646 | 25104 |
| 17210 | 242513 | 378513 | 121499 | 13888 | 4178 | 641 | 12113 | 13088 | 108130 |
| 12642 | 37218 | 17878 | 5276 | 10449 | 3173 | 715 | 12844 | 10884 | 36545 |
| 7623 | 18416.5 | 9992 | 3203.5 | 103814 | 33356 | 702 | 10609 | 8625.5 | 20271 |
| 4380 | 10991 | 5574 | 1845 | 7056 | 2741 | 657 | 13608 | 5841 | 14299 |
| 3616 | 5372 | 3829 | 1232 | 5293 | 2141 | 697 | 17220 | 5009 | 7759 |
| 3642 | 4994 | 3810 | 1323 | 4478 | 1948 | 654 | 14225 | 5328 | 7039 |
| 2941 | 4575.5 | 3551 | 1237 | 4840.5 | 2025 | 701.5 | 13711 | 5109 | 6675.5 |
| 2462.5 | 3912 | 2955.5 | 1123.5 | 4329 | 1856 | 680.5 | 10932.5 | 4720.5 | 6193.5 |
| 2446.5 | 3579 | 3018.5 | 1123.5 | 3870.5 | 1705.5 | 609.5 | 9944 | 4075.5 | 5532.5 |
| 2763 | 3944 | 2887 | 1139 | 4143 | 1852 | 666 | 10761 | 4256 | 6149 |
| 2551 | 3808 | 2987 | 1166 | 4414 | 1864 | 670 | 11820 | 4293 | 6239 |
| 2499 | 3776.5 | 3235.5 | 1088 | 4040 | 1794 | 636 | 11122.5 | 4459 | 5970 |
| 7459 | 6601 | 4072 | 1166 | 4587 | 1799 | 635 | 10517 | 5283 | 6402 |
| 2532.5 | 3854 | 3060 | 995 | 3701 | 1777 | 657 | 9571.5 | 4138.5 | 5914.5 |
| 2575.5 | 3929.5 | 3311 | 1009.5 | 4263 | 1802 | 693 | 9461.5 | 4217 | 6119 |
| 26341 | 13641 | 6213.5 | 1191 | 4063 | 1675 | 605.5 | 12261 | 4835 | 5754.5 |
| 3652.5 | 4885 | 3923.5 | 1101.5 | 3776 | 1711 | 638.5 | 10458 | 4306.5 | 6588 |
| 3597 | 4492 | 3777 | 1218 | 4389 | 1865 | 662 | 11572 | 4752 | 6957 |
| 2898 | 3943 | 3339 | 1130 | 4549 | 1901 | 644 | 10993 | 4603 | 6351 |
| 3200 | 4234 | 3157.5 | 1042.5 | 4075.5 | 1723.5 | 615.5 | 11993.5 | 4876.5 | 5972 |
| 5957 | 10430 | 4674 | 1433 | 5855 | 1950 | 669 | 15278 | 6712 | 12374 |
| 11099 | 19442 | 7225 | 1581 | 7942 | 2125 | 581 | 12488 | 9101 | 20511 |

| | | | | | | | | | |
|---|---|---|---|---|---|---|---|---|---|
| 10435.5 | 26342 | 9022.5 | 1845 | 9789.5 | 2747 | 660 | 11483 | 9102 | 29397 |
| 940 | 1399 | 1189 | 677 | 1855 | 1223 | 326 | 3286 | 1796 | 2671 |
| 23773 | 48557 | 15217 | 2436 | 4538 | 1907 | 497 | 15098 | 5513 | 8589 |
| 6605.5 | 6589 | 3865 | 1084 | 3944 | 1692 | 627.5 | 15068.5 | 4617.5 | 5777.5 |
| 3174.5 | 4429.5 | 10064 | 3797.5 | 459879 | 47032.5 | 667.5 | 10291 | 3923 | 5601 |
| 2706 | 4155 | 3843 | 1409 | 5956 | 2297 | 836 | 11841 | 4860 | 6746 |
| 2503 | 3811 | 3349 | 1140 | 4944 | 2011 | 793 | 10854 | 4478 | 6209 |
| 2433.5 | 3885 | 3342.5 | 1177.5 | 4428 | 1910.5 | 730.5 | 9752 | 4130 | 5983.5 |
| 2631 | 4145 | 3210 | 1129 | 4461 | 1860 | 760 | 11249 | 4650 | 6422 |
| 2805 | 4372 | 3464.5 | 1171 | 4530.5 | 1875 | 744 | 12793 | 4672 | 6767 |
| 2601 | 4156 | 3414.5 | 1277 | 4581 | 1946 | 715.5 | 12177.5 | 4446.5 | 6743 |
| 2513 | 3982.5 | 3159 | 1207 | 4203 | 1906 | 763.5 | 10984 | 4629.5 | 6556 |
| 2339.5 | 3858.5 | 3120 | 1199.5 | 4336.5 | 1870.5 | 752.5 | 10819 | 4288.5 | 6455.5 |
| 2096 | 3400 | 3134 | 1200 | 4299 | 1853 | 758 | 9720 | 4179 | 6521 |
| 26368 | 18764 | 6050 | 1302 | 4370 | 1878 | 736 | 9784 | 9483 | 8996 |
| 2172.5 | 3644 | 3127.5 | 1105 | 4229.5 | 1863.5 | 726 | 9313 | 4172 | 6339 |
| 2223 | 3677 | 3270 | 1169 | 4460 | 1920 | 764 | 10047 | 4713 | 6842 |
| 60609.5 | 33373 | 9795.5 | 1489.5 | 4732 | 1946 | 770.5 | 14778 | 5725.5 | 7938.5 |
| 2950.5 | 4342 | 3530 | 1167 | 4560 | 1922.5 | 718 | 10298.5 | 4648 | 6854 |
| 2175 | 3773 | 3233.5 | 1204 | 5232 | 2047 | 759 | 10271 | 4499 | 6561 |
| 2719 | 4068 | 3471 | 1190 | 4493 | 1890 | 795 | 10882 | 4911 | 6587 |
| 9882 | 22196.5 | 9346.5 | 2223 | 10825 | 2868 | 721 | 13774 | 9952 | 24404 |
| 10109.5 | 23745 | 11897 | 2702.5 | 13481 | 3435.5 | 748.5 | 12231 | 8960 | 23862 |
| 9638 | 23876 | 12431 | 2708 | 13880 | 3484 | 762 | 11404 | 9081 | 24836 |
| 9662 | 21632 | 10749 | 2258 | 11655 | 2989 | 747 | 12973 | 9420 | 23400 |
| 9074.5 | 23256.5 | 8236.5 | 1797 | 9702 | 2593.5 | 739.5 | 13843.5 | 9695.5 | 26060 |
| 13347 | 114686 | 169774 | 66184 | 7580 | 2649 | 685 | 15742 | 11776 | 62650 |
| 4453 | 12023 | 4871 | 1452 | 5867 | 2118 | 757 | 14391 | 5963 | 16097 |
| 3124 | 7681 | 6651.5 | 1950 | 600283 | 56639 | 726 | 15844.5 | 4488 | 10829.5 |
| 2816 | 5201 | 4142 | 1387 | 5922 | 2257 | 894 | 19650 | 5203 | 8106 |
| 2805 | 4752.5 | 3856 | 1298 | 5133.5 | 1954 | 797.5 | 14603.5 | 4763 | 7532.5 |
| 2602 | 4363 | 3535 | 1246 | 4819 | 2026 | 778 | 12653 | 4670 | 7250 |
| 2715 | 4563 | 3540 | 1229 | 4623 | 1899 | 780 | 14233 | 4541 | 7133 |
| 2704 | 4655.5 | 3506 | 1196.5 | 4624 | 1996 | 769.5 | 13847 | 4525 | 7005.5 |
| 2606.5 | 5045 | 3857.5 | 1395.5 | 5087.5 | 2064 | 749.5 | 14342 | 4381.5 | 7426.5 |
| 2685 | 4374 | 3682.5 | 1256.5 | 4554.5 | 1854 | 769.5 | 12053 | 4419.5 | 7136.5 |
| 2539 | 3873.5 | 3290.5 | 1126.5 | 4592 | 1870.5 | 751.5 | 10787 | 4758.5 | 6541 |
| 2503 | 3947 | 3352 | 1218 | 4582 | 1937 | 787 | 11177 | 4484 | 6688 |
| 31886 | 22558 | 6734 | 1475 | 4565 | 1970 | 776 | 11167 | 11325 | 10265 |
| 2476 | 3996 | 3261 | 1181 | 4168 | 1837 | 754 | 9794 | 4513 | 6535 |
| 2534 | 4180 | 3499 | 1237 | 4729 | 1986 | 779 | 11199 | 4775 | 7122 |
| 59500.5 | 45009 | 10349 | 1644.5 | 5131 | 2133.5 | 765.5 | 16798.5 | 5586 | 10141.5 |
| 3047 | 5928 | 3951 | 1270 | 5091 | 2040 | 796 | 12512 | 4660 | 8444 |
| 2411 | 4223.5 | 3422.5 | 1272 | 5687.5 | 2154.5 | 771.5 | 11287 | 4666.5 | 7233 |
| 2583 | 4363 | 3568 | 1173 | 4625 | 1950.5 | 741.5 | 11254.5 | 4613 | 6768 |
| 9676 | 20008 | 10260.5 | 2334 | 11464.5 | 3044 | 767 | 13785.5 | 9773.5 | 22135 |
| 10524 | 25132 | 12892 | 2809 | 14475 | 3477 | 753 | 11579 | 8763 | 24822 |
| 10236 | 24524 | 12437 | 2691 | 13468 | 3318 | 740 | 12320 | 9177 | 24602 |

| | | | | | | | | | |
|---|---|---|---|---|---|---|---|---|---|
| 9702.5 | 21070.5 | 10507 | 2363 | 12205 | 3213 | 769 | 13068.5 | 9445.5 | 22713 |
| 8798.5 | 22136 | 8236.5 | 1753 | 9411 | 2446.5 | 767 | 14734.5 | 9427.5 | 26310.5 |
| 13769 | 110833 | 163652 | 60559 | 7636 | 2591 | 706 | 15633 | 11875 | 59135 |
| 4622 | 12756 | 4723 | 1452 | 5735 | 2032 | 764 | 15226 | 5842 | 15945 |
| 3174 | 8123 | 6513 | 1828 | 574385 | 54173 | 746 | 16181 | 4734 | 11261 |
| 2868.5 | 5145 | 4015.5 | 1337 | 5992.5 | 2282 | 918.5 | 18740.5 | 5198.5 | 8611 |
| 2769 | 4436.5 | 3773 | 1315 | 5292.5 | 2120.5 | 775 | 12631 | 4477.5 | 6955 |
| 2573 | 4421.5 | 3631 | 1263 | 4880 | 1936 | 758.5 | 12434.5 | 4501.5 | 7066.5 |
| 2554 | 4680.5 | 3525.5 | 1266.5 | 4912 | 2055.5 | 755 | 13409.5 | 4618 | 7309.5 |
| 2816 | 4575 | 3562 | 1259 | 4603 | 1917 | 746.5 | 11875.5 | 4530.5 | 6999 |
| 2586 | 4458 | 3440 | 1316 | 4587 | 2002 | 727 | 11565 | 4574 | 7169 |
| 2627 | 4229 | 3453 | 1205 | 4668 | 1910 | 757 | 11026 | 4636 | 6722 |
| 2424 | 3949 | 3184 | 1185 | 4443 | 1931 | 734 | 10630 | 4536 | 6623 |
| 2403 | 4001 | 3381 | 1181 | 4528 | 1855 | 785 | 10825 | 4570 | 6795 |
| 19964 | 13475 | 5635 | 1382 | 4370 | 1949 | 784 | 10479 | 8305 | 8931 |
| 2594 | 4115 | 3469 | 1262 | 4526 | 1926 | 819 | 10644 | 4903 | 6769 |
| 2330 | 4025 | 3472.5 | 1287 | 4575 | 1929.5 | 766.5 | 10543.5 | 4663 | 7135.5 |
| 54439 | 30992.5 | 9630 | 1584 | 4818.5 | 1993.5 | 783 | 15000 | 5328.5 | 7991 |
| 2999 | 4748 | 3667 | 1306.5 | 4840 | 2029.5 | 761.5 | 11991 | 4870 | 7179 |
| 2458 | 4069 | 3420.5 | 1198.5 | 5342 | 2073.5 | 753.5 | 10310 | 4624 | 6719 |
| 2604 | 3902 | 3374 | 1153 | 4470 | 1824 | 728 | 10587 | 4612 | 6454 |
| 10220 | 21258 | 10533 | 2392 | 11530 | 3028 | 754 | 13652 | 9268 | 23568 |
| 10013 | 25702 | 13318 | 2776 | 14520 | 3578 | 780 | 11356 | 8887 | 25618 |
| 10092 | 23272 | 12244 | 2812 | 14016 | 3444 | 765 | 11740 | 9098 | 24276 |
| 10048 | 21405 | 10575.5 | 2420 | 12183.5 | 2999.5 | 765.5 | 13252 | 9227.5 | 23032 |
| 8895 | 23288 | 8671 | 1874 | 9690 | 2583 | 762 | 13806 | 9239 | 25653 |
| 14998 | 135802.5 | 205183 | 76445.5 | 7912 | 2711.5 | 668 | 15309 | 12872 | 80833 |
| 5170 | 13069.5 | 5177.5 | 1519 | 5709 | 2023.5 | 745.5 | 14721 | 5990.5 | 16790 |
| 3357 | 7813 | 6388 | 1826 | 532430 | 49260 | 779 | 14835 | 4564 | 10936 |
| 2443.5 | 4704 | 4092.5 | 1330 | 5948 | 2233.5 | 837.5 | 18341.5 | 4506.5 | 7806.5 |
| 2784 | 4297 | 3850 | 1299 | 5439 | 2092 | 810 | 13213 | 4732 | 7158 |
| 2439 | 4316 | 3775 | 1272 | 5102 | 2004 | 788 | 11912 | 4474 | 6969 |
| 2524 | 4412 | 3601 | 1234 | 4729 | 1988 | 772 | 13082 | 4598 | 7009 |
| 2775 | 5160 | 3644 | 1319 | 4976 | 2033 | 758 | 14298 | 4564 | 7925 |
| 2654.5 | 4562 | 3567 | 1299 | 4870 | 2049 | 770 | 12917 | 4379 | 7072 |
| 2668 | 4492 | 3630 | 1291 | 4581 | 1977 | 744 | 11712 | 4458 | 6829 |
| 2597.5 | 4035 | 3279.5 | 1291 | 4626 | 1987.5 | 780 | 11056.5 | 4500 | 6653.5 |
| 2476 | 3958 | 3370 | 1235 | 4441 | 1833 | 786 | 10451 | 4636 | 6442 |
| 23538 | 16230 | 5833 | 1390 | 4667 | 1932 | 786 | 10935 | 9109 | 9241 |
| 2704 | 3848 | 3421 | 1197 | 4535 | 1832 | 765 | 10191 | 4656 | 6490 |
| 2464 | 4246.5 | 3490 | 1244 | 4891 | 1969 | 773.5 | 11027 | 4884.5 | 7101.5 |
| 56118 | 39565 | 10500 | 1537 | 5099 | 2064 | 776 | 16411 | 5336 | 9590 |
| 3281.5 | 6164.5 | 4165 | 1312.5 | 5047 | 2128 | 752 | 12626.5 | 4626 | 8442.5 |
| 2673 | 4528 | 3801 | 1319 | 5885 | 2209 | 762 | 11695 | 4574 | 7305 |
| 2718.5 | 4511.5 | 3608.5 | 1218 | 4597.5 | 1945 | 728 | 10681 | 4741 | 6923 |
| 9514.5 | 21226 | 9978.5 | 2256 | 11436.5 | 2961 | 774 | 13568 | 9021.5 | 23158 |
| 9428 | 23639 | 13152 | 2761 | 14607 | 3561 | 772 | 11447 | 8564 | 25843 |
| 9371 | 24040 | 12568 | 2646 | 14394.5 | 3480 | 752 | 11152 | 9036 | 26009.5 |

| | | | | | | | | | |
|---|---|---|---|---|---|---|---|---|---|
| 9659 | 20960 | 11234 | 2243 | 12033 | 3060 | 780 | 13318 | 9374 | 23052 |
| 9450 | 23283 | 8183.5 | 2056 | 9997.5 | 2691 | 731 | 13885.5 | 9256 | 25676 |
| 16635 | 141322 | 204508 | 75763 | 7902 | 2656 | 677 | 15573 | 12827 | 75001 |
| 4788 | 13153 | 5121 | 1489 | 5922 | 2111 | 767 | 15392 | 6059 | 16237 |
| 3467 | 8554 | 6424 | 1749 | 598885 | 55156 | 764 | 15448 | 4542 | 12180 |
| 2843 | 4932 | 4378 | 1409 | 6313 | 2365 | 868 | 19341 | 5304 | 7999 |
| 2707 | 4506 | 4041 | 1389 | 5542 | 2182 | 792 | 14313 | 4792 | 7219 |
| 2684.5 | 4297 | 3400 | 1169 | 4881 | 1960 | 785.5 | 11505 | 4416.5 | 6832.5 |
| 2827 | 4414 | 3835 | 1314 | 5049 | 1957 | 737 | 12837 | 4330 | 7068 |
| 2740 | 4707 | 3726 | 1289 | 5061 | 1965 | 776 | 13561 | 4564 | 7316 |
| 2698 | 4555 | 3838 | 1396.5 | 4832.5 | 1967 | 762 | 12763.5 | 4535 | 7303.5 |
| 2639 | 4411 | 3474.5 | 1265 | 4802.5 | 1941 | 769 | 11832.5 | 4477.5 | 6663 |
| 2640 | 4155 | 3276 | 1216 | 4677 | 1905 | 737 | 11409 | 4697 | 6659 |
| 2412 | 3969 | 3341 | 1216 | 4466 | 1892 | 741 | 10357 | 4718 | 6881 |
| 26907 | 18599 | 6620 | 1480 | 4791 | 2040 | 773 | 10932 | 9990 | 9496 |
| 2752.5 | 4148.5 | 3432.5 | 1178.5 | 4516.5 | 1927 | 743 | 10254.5 | 4654.5 | 6725 |
| 2447 | 4194 | 3554 | 1374 | 4990 | 1878 | 773 | 11438 | 4687 | 6856 |
| 44096 | 26848 | 8304 | 1571 | 5003 | 2056 | 781 | 15224 | 5054 | 7894 |
| 3749 | 5452 | 4155 | 1320 | 5196 | 2010 | 759 | 12207 | 4614 | 7266 |
| 2556 | 4262 | 3548 | 1289 | 5907 | 2139 | 737 | 11237 | 4631 | 7002 |
| 3137.5 | 4388 | 3679.5 | 1281 | 4902.5 | 1912.5 | 699.5 | 11120 | 4751 | 6560 |
| 10449 | 20124.5 | 10692.5 | 2192.5 | 11909 | 3003.5 | 782.5 | 13821 | 9506.5 | 21901 |
| 9905 | 25175.5 | 13307 | 2872 | 14825.5 | 3558.5 | 758 | 11851.5 | 8461 | 24150.5 |
| 9329 | 22966 | 12008 | 2921 | 14233 | 3610 | 762 | 11220 | 8876 | 24266 |
| 10078 | 21160 | 10738 | 2316 | 12436 | 2969 | 760 | 12894 | 9745 | 22915 |
| 9162 | 23252.5 | 8943 | 1906 | 10134.5 | 2623.5 | 755 | 14265.5 | 9030 | 26035.5 |
| 16869.5 | 147489 | 225236.5 | 83793.5 | 8036 | 2859 | 649.5 | 15338 | 12698 | 84739.5 |
| 13053.5 | 18382.5 | 6410 | 1836.5 | 6278.5 | 2064 | 791.5 | 16114 | 6535 | 17259.5 |
| 8044.5 | 11236 | 27336.5 | 9337.5 | 150353.5 | 23144 | 739 | 15031.5 | 4360.5 | 12979.5 |
| 9057 | 6592 | 7092 | 1973 | 40381 | 5129 | 956 | 19102 | 5431 | 8159 |
| 8194 | 5807 | 5159 | 1604 | 17885 | 3206 | 799 | 15042 | 5002 | 7071 |
| 7597.5 | 5408 | 4448.5 | 1460 | 9456 | 2731.5 | 803 | 13682 | 4973.5 | 7031 |
| 8077 | 5617 | 4083 | 1340 | 7073 | 2204 | 841 | 15370 | 4653 | 7234 |
| 5949.5 | 5774.5 | 4227.5 | 1323.5 | 6428 | 2225 | 797.5 | 15188.5 | 4661 | 7766 |
| 5431.5 | 5777 | 4802.5 | 1684.5 | 5629 | 2134 | 793.5 | 14714 | 4824 | 7663.5 |
| 6540 | 5518 | 4265 | 1303 | 5881 | 2181 | 815 | 13678 | 4705 | 7300 |
| 6287 | 4734 | 3693 | 1255.5 | 4999 | 1926 | 808.5 | 11685.5 | 4733.5 | 6771 |
| 6466 | 5103 | 4072 | 1229 | 5739 | 1995 | 850 | 13212 | 4940 | 6844 |
| 18468 | 10487 | 5556 | 1688 | 5120 | 2015 | 829 | 12081 | 6770 | 8421 |
| 9187 | 5444 | 3825 | 1232 | 5325 | 2022 | 802 | 11160 | 5365 | 7091 |
| 7310.5 | 5517.5 | 4191 | 1302 | 5378 | 2156 | 886.5 | 13498.5 | 5145 | 7544 |
| 425234.5 | 326546 | 83465 | 5774 | 5581.5 | 2092 | 818 | 52350 | 13108.5 | 14262 |
| 25687 | 16226 | 6049 | 1501 | 5819 | 2062.5 | 806.5 | 14909 | 5304.5 | 8964 |
| 14360.5 | 7777 | 4561 | 1430 | 6611.5 | 2578 | 832 | 13512 | 4887 | 7728 |
| 17091.5 | 8837 | 4785.5 | 1412 | 5323.5 | 2066.5 | 829.5 | 13466.5 | 5297 | 7414 |
| 14508 | 24798 | 12403 | 2512 | 13443 | 3367 | 876 | 14525 | 9936 | 23844 |
| 15082 | 28683.5 | 14928 | 2952.5 | 15478.5 | 3677.5 | 841 | 11881 | 9636.5 | 26000.5 |
| 13954 | 26514 | 14207 | 2729 | 15980 | 3442 | 810 | 13042 | 9500 | 26514 |

| | | | | | | | | | |
|---|---|---|---|---|---|---|---|---|---|
| 13172.5 | 24278 | 12163.5 | 2460.5 | 14127.5 | 3378.5 | 857.5 | 13393.5 | 9623.5 | 23531.5 |
| 8652 | 22180 | 8952 | 1940 | 10051 | 2477 | 784 | 14349 | 9460 | 24267 |
| 16563.5 | 145297.5 | 216917 | 79338.5 | 8038 | 2709 | 688 | 15524.5 | 13284.5 | 74714.5 |
| 5340 | 13346 | 5143 | 1580 | 6075 | 2117 | 751 | 16289 | 6245 | 16681 |
| 3185 | 8204 | 6628 | 1877 | 596268 | 56757 | 728 | 14956 | 4711 | 11702 |
| 2914 | 5947 | 4219 | 1399 | 6061 | 2374 | 925 | 21886 | 5579 | 9549 |
| 2619.5 | 4772 | 3919 | 1373 | 5223.5 | 2085.5 | 787.5 | 15558.5 | 4606 | 7247.5 |
| 2611 | 4503 | 3767 | 1298 | 5185 | 2117 | 763 | 12648 | 4370 | 6845 |
| 2729 | 4523 | 3695 | 1249 | 4736 | 1912 | 745 | 12619 | 4323 | 6990 |
| 2965 | 4859 | 3940 | 1333 | 4966 | 2062 | 764 | 14316 | 4411 | 7338 |
| 2779 | 4551 | 3924 | 1368 | 4826 | 2078 | 765 | 13007 | 4380 | 7064 |
| 2615 | 4496.5 | 3593 | 1307 | 4739 | 1974.5 | 761.5 | 12624 | 4497.5 | 6956.5 |
| 2669 | 4196 | 3534 | 1212 | 4736 | 1963 | 760 | 11190 | 4572 | 6745 |
| 2592.5 | 4080 | 3413 | 1258.5 | 4605 | 1945 | 774 | 11254 | 4652.5 | 6478 |
| 27187 | 18579 | 6199 | 1411 | 4778 | 1925 | 752 | 10449 | 10062 | 9433 |
| 2836 | 3959.5 | 3262.5 | 1231.5 | 4583 | 1938 | 778.5 | 10496.5 | 4841.5 | 6568 |
| 2487 | 4269 | 3719 | 1308 | 4761 | 1995 | 774 | 11298 | 4659 | 7354 |
| 70351.5 | 50394.5 | 12326.5 | 1724 | 5492.5 | 2078.5 | 753 | 16625.5 | 5704.5 | 9767 |
| 3366 | 5200 | 4131 | 1294 | 4872 | 1977 | 759 | 11972 | 4726 | 7308 |
| 2467 | 4365.5 | 3509 | 1327.5 | 5999.5 | 2142.5 | 758.5 | 11365 | 4805 | 6931 |
| 2867.5 | 4314 | 3666 | 1278.5 | 4814 | 1952 | 755.5 | 10524.5 | 4744.5 | 6754 |
| 10117.5 | 23298.5 | 10449.5 | 2297.5 | 11795.5 | 3004.5 | 766.5 | 14522 | 9634.5 | 24101.5 |
| 9803.5 | 23949.5 | 13149 | 2755 | 14486.5 | 3566.5 | 773 | 12124.5 | 8725.5 | 24423 |
| 9651.5 | 24433 | 12554 | 2832 | 14117 | 3466 | 758.5 | 11361.5 | 9002 | 24480.5 |
| 10335 | 21046 | 10496 | 2388 | 11677 | 3007 | 748 | 12719 | 9764 | 23244 |
| 10260 | 23377 | 7650 | 1751 | 8564 | 2602 | 733 | 14041 | 9164 | 26412 |
| 15477 | 139163 | 212313 | 78574 | 8548 | 3071 | 656 | 15093 | 13028 | 77378 |
| 4630 | 12863 | 5287 | 1668 | 6272 | 2394 | 715 | 13977 | 5480 | 16700 |
| 3181.5 | 8676 | 7424.5 | 2382 | 378677 | 37536.5 | 689 | 15387.5 | 4836 | 14556 |
| 2930 | 8063 | 5176 | 1869 | 8049 | 3140 | 1383 | 55629 | 8509 | 18374 |
| 2487.5 | 7401 | 4903 | 1825.5 | 7216.5 | 2895.5 | 753.5 | 16147 | 4849.5 | 14531.5 |
| 2263 | 7002 | 4388 | 1671 | 6664 | 2571 | 708 | 12759 | 4490 | 13478 |
| 2357 | 7158 | 4610 | 1791 | 6861 | 2726 | 769 | 14690 | 4321 | 14467 |
| 2612.5 | 7677.5 | 4889 | 1775 | 7064 | 2846.5 | 732.5 | 16522 | 4625 | 14797 |
| 2468 | 7123 | 4709 | 1677 | 6599 | 2677 | 693 | 13481 | 4722 | 14814 |
| 2271.5 | 7190.5 | 4590 | 1720.5 | 6679 | 2706.5 | 705 | 12773.5 | 4439.5 | 14825 |
| 2376.5 | 6802 | 4605 | 1718 | 6339 | 2689.5 | 762 | 12554.5 | 4265.5 | 14596 |
| 2133 | 6160 | 4507 | 1857 | 6591 | 2727 | 709 | 11045 | 4643 | 13844 |
| 28016.5 | 35193.5 | 8653.5 | 2128 | 6630.5 | 2677.5 | 723 | 11789.5 | 9669.5 | 18526.5 |
| 2154 | 6792 | 4393 | 1646 | 6389 | 2638 | 736 | 10823 | 4617 | 14494 |
| 2171 | 6836 | 4688 | 1696 | 6502 | 2676 | 693 | 12170 | 4671 | 14301 |
| 19991 | 21901.5 | 7242.5 | 1984 | 6783 | 2755 | 713.5 | 15693 | 5272.5 | 15775.5 |
| 4015 | 9542 | 5279 | 1863 | 7022 | 2903 | 745 | 14270 | 4813 | 15419 |
| 2100 | 6935 | 4734 | 1661 | 6972 | 2832 | 733 | 11502 | 4382 | 15306 |
| 2744 | 7007 | 4788 | 1835 | 6765 | 2799 | 700 | 12351 | 4542 | 13958 |
| 9102 | 20327 | 10688 | 2583 | 12311 | 3457 | 710 | 13722 | 9692 | 23193 |
| 8677 | 24659 | 13243.5 | 3228.5 | 14849.5 | 4025.5 | 711 | 12435 | 9186 | 26072 |
| 8720 | 22976 | 13145 | 3128 | 14962 | 3938 | 706 | 11614 | 9165 | 25726 |

| | | | | | | | | | |
|---|---|---|---|---|---|---|---|---|---|
| 8942 | 20430 | 11453 | 2799 | 13056 | 3517 | 734 | 13051 | 9521 | 23850 |
| 9225 | 21361.5 | 9144.5 | 2338.5 | 11150.5 | 3172.5 | 756 | 13834.5 | 9842.5 | 24560.5 |
| 14695.5 | 97107.5 | 108630.5 | 26585.5 | 12935 | 4903 | 624 | 13995 | 12976.5 | 71691.5 |
| 6697 | 42118 | 52831 | 13821.5 | 8777 | 2928 | 699.5 | 14372.5 | 7022.5 | 31270 |
| 4815 | 12017 | 27163 | 6643 | 158500 | 25698 | 743 | 14136 | 5265 | 15249 |
| 5490 | 6926.5 | 15343 | 3072.5 | 76984 | 10581.5 | 1520.5 | 65714 | 10539.5 | 15513.5 |
| 3615 | 5008 | 5310 | 1680 | 67980 | 8590 | 811 | 13651 | 4804 | 7187 |
| 2924 | 4656 | 4281 | 1430 | 31044 | 4517 | 773 | 13523 | 5020 | 7596 |
| 3250 | 4773 | 4410 | 1553 | 34699 | 4630 | 734 | 12918 | 4688 | 7205 |
| 3125 | 4641 | 4137 | 1344 | 21426 | 3471 | 754 | 12845 | 4627 | 6937 |
| 4698 | 10087 | 11787 | 3461 | 12601 | 3310 | 922 | 15518 | 6182 | 10895 |
| 2966 | 4547 | 3849 | 1451 | 10889 | 2545 | 747 | 12221 | 4657 | 6906 |
| 2872 | 4287.5 | 3818 | 1413 | 6925.5 | 2292.5 | 729 | 10829 | 4516 | 6598 |
| 2689 | 4182 | 3698 | 1330 | 7657 | 2282 | 755 | 10828 | 4592 | 6607 |
| 30012 | 21427.5 | 11819 | 2129 | 7334 | 2287.5 | 768 | 10239 | 10793.5 | 10398.5 |
| 3137 | 4464 | 3895 | 1363 | 6940 | 2167 | 735 | 10571 | 5161 | 6977 |
| 2943.5 | 4539 | 4063 | 1341 | 6908 | 2200.5 | 726.5 | 11042.5 | 4737 | 7133.5 |
| 290294 | 198658 | 56299 | 4131 | 6839 | 2263 | 752 | 38684 | 9757 | 11209 |
| 6126 | 6578 | 4637 | 1472 | 6832 | 2203 | 755 | 11646 | 4779 | 8418 |
| 3055 | 4742 | 4281.5 | 1396.5 | 19134 | 4224.5 | 758.5 | 11152 | 4384.5 | 6855 |
| 3775 | 4539 | 3674 | 1331 | 14454 | 2799 | 755 | 10717 | 4748 | 6814 |
| 9300 | 19910 | 10847 | 2476 | 18188 | 3778 | 794 | 13959 | 9745 | 22448 |
| 8770 | 24191 | 12588 | 2921 | 17164 | 3732 | 742 | 10885 | 8652 | 25365 |
| 7935.5 | 24217 | 13341 | 3037 | 16221.5 | 3750.5 | 745 | 10552 | 8164 | 25274.5 |
| 9136 | 21701.5 | 10367 | 2395.5 | 14772 | 3262.5 | 781 | 12748 | 9633.5 | 23389 |
| 9257.5 | 23810.5 | 8623 | 1918.5 | 9941.5 | 2692 | 764 | 13721.5 | 9511 | 26835 |
| 19996 | 180766 | 279477 | 91670 | 12670 | 5764 | 654 | 15444 | 14832 | 101176 |
| 6336 | 17017 | 10471 | 3546 | 6483 | 2608 | 734 | 15002 | 6475 | 18447 |
| 3475 | 9049 | 8238 | 2644 | 569897 | 55507 | 764 | 14598 | 5000 | 12622 |
| 2649.5 | 4779 | 4758.5 | 1655.5 | 7677.5 | 2519 | 864.5 | 19270.5 | 4742.5 | 8063 |
| 2906 | 4755 | 4439 | 1557 | 6054 | 2329 | 782 | 13064 | 4638 | 7299 |
| 2624 | 4446 | 3944 | 1354 | 5104 | 2057 | 762 | 11521 | 4604 | 6855 |
| 2717 | 4494 | 3886 | 1309 | 5263 | 1998 | 762 | 13077 | 4513 | 7280 |
| 2950 | 4887 | 4077.5 | 1394 | 5321 | 2020 | 789.5 | 13716.5 | 4508 | 7331.5 |
| 2812 | 4838 | 4248 | 1574 | 5511 | 2212 | 742 | 13404 | 4517 | 7797 |
| 2675.5 | 4638 | 3787 | 1344 | 5047 | 2079 | 754 | 12131 | 4521 | 7322 |
| 2721.5 | 4543 | 3763 | 1280.5 | 4838 | 1965 | 780.5 | 12211 | 4546.5 | 7311.5 |
| 2590 | 4456 | 3758 | 1324 | 5044.5 | 1886 | 786 | 11845 | 4534 | 7290 |
| 28222.5 | 21241 | 7174 | 1502.5 | 4826 | 1922 | 737.5 | 11923 | 9761 | 10280 |
| 2813 | 4466 | 3635 | 1255 | 4700 | 1944 | 777 | 11190 | 4687 | 7187 |
| 2717.5 | 5553.5 | 4141 | 1325 | 5286.5 | 2101 | 795 | 12140.5 | 4842.5 | 8933.5 |
| 67378 | 54257 | 13390 | 1819 | 5268 | 2032 | 741 | 17957 | 5526 | 10994 |
| 3455.5 | 7613.5 | 4459 | 1415.5 | 5459.5 | 2090 | 763 | 13507.5 | 4817 | 10367.5 |
| 2618.5 | 5084 | 4035 | 1323.5 | 5901.5 | 2177.5 | 760 | 12439.5 | 4605.5 | 7972.5 |
| 2903.5 | 4961.5 | 3824.5 | 1290.5 | 5232.5 | 1994.5 | 764.5 | 11650.5 | 4689.5 | 7318.5 |
| 9897.5 | 21588.5 | 10911.5 | 2420.5 | 12131 | 3151 | 812.5 | 13881 | 9336.5 | 23365.5 |
| 10706 | 26124 | 14314 | 2888.5 | 14674 | 3692.5 | 787.5 | 11973.5 | 9266.5 | 26849 |
| 9640.5 | 24539 | 12786.5 | 2734.5 | 14175.5 | 3427 | 752.5 | 11318 | 8757.5 | 24871 |

| | | | | | | | | | |
|---|---|---|---|---|---|---|---|---|---|
| 10628 | 22743 | 11342 | 2460 | 12472.5 | 3010.5 | 761.5 | 12672.5 | 9008 | 24563.5 |
| 9916 | 24682 | 9342 | 2089 | 10299 | 2645 | 744 | 14341 | 9629 | 25980 |
| 21560 | 188577 | 309853 | 100223.5 | 12128 | 4595.5 | 690 | 15812.5 | 15269.5 | 98211 |
| 5886 | 15727 | 8298 | 2578 | 6649 | 2555 | 740 | 15200 | 6693 | 18548 |
| 3234 | 10139 | 7585 | 2802 | 482357 | 49148 | 714 | 14683 | 4509 | 14146 |
| 3056.5 | 6026 | 5108 | 1712.5 | 6873.5 | 2674 | 859 | 18498 | 5339.5 | 9369 |
| 3036 | 5167 | 5140.5 | 1716.5 | 6486.5 | 2460 | 783 | 13263.5 | 4666 | 7550.5 |
| 3004.5 | 5027.5 | 4263.5 | 1578 | 5771 | 2279.5 | 748.5 | 12137 | 4668 | 7250 |
| 3141 | 5329 | 4413 | 1480 | 5718 | 2192 | 753 | 13782 | 4689 | 7955 |
| 3192.5 | 5365.5 | 4177.5 | 1516.5 | 5303 | 2082 | 704.5 | 12888.5 | 4585.5 | 7904.5 |
| 3093.5 | 5947 | 4642.5 | 1601.5 | 5453 | 2226 | 735.5 | 14395 | 4397 | 8623.5 |
| 2933 | 5093 | 4137 | 1465 | 5266 | 2121 | 734 | 13833 | 4281 | 7452 |
| 2952.5 | 4349 | 3604.5 | 1346 | 4778.5 | 2009 | 700.5 | 11652.5 | 4546.5 | 6607 |
| 2818.5 | 4382.5 | 3616 | 1274 | 4619 | 2036.5 | 707 | 10580 | 4676 | 6484 |
| 16956 | 12409 | 5795 | 1434 | 4797 | 2028 | 731 | 10499 | 7143 | 8377 |
| 3005.5 | 4535 | 3897.5 | 1351.5 | 4994 | 1991 | 762.5 | 10872.5 | 4856 | 6895 |
| 2670 | 4622 | 4016 | 1383 | 5156 | 2068 | 726 | 11528 | 4641 | 7176 |
| 115965.5 | 68106.5 | 21774.5 | 2103.5 | 5103 | 1988 | 728 | 21772.5 | 6289 | 8455.5 |
| 3737 | 5103 | 4133 | 1422 | 4888 | 2026 | 710 | 11630 | 4940 | 6781 |
| 2607 | 4121 | 3581 | 1247 | 5375 | 2035 | 738 | 10872 | 4634 | 6513 |
| 2906 | 4627 | 3717 | 1208 | 4995 | 1965 | 747 | 11007 | 4710 | 6748 |
| 10119.5 | 21175 | 10894.5 | 2492 | 12004 | 3082.5 | 732.5 | 13099.5 | 9409 | 22005.5 |
| 10010.5 | 23142.5 | 12769 | 2639.5 | 13905.5 | 3363.5 | 731 | 12073.5 | 8944.5 | 22419 |
| 10068.5 | 22423 | 11851.5 | 2480 | 13236.5 | 3140 | 698 | 12390 | 8805 | 21124 |
| 10996.5 | 23937.5 | 10404 | 2227.5 | 11352.5 | 2798 | 737.5 | 14284.5 | 9555.5 | 23796 |
| 922 | 1343 | 1253 | 667 | 1858 | 1226 | 351 | 3307 | 1842 | 2559 |
| 20953 | 22660 | 6823 | 1420 | 3835.5 | 1707 | 514 | 16945 | 4979 | 6269.5 |
| 4166.5 | 4946.5 | 3346 | 989 | 3521 | 1679.5 | 621.5 | 11058 | 3891.5 | 5229 |
| 3080 | 4316 | 8691 | 2694 | 374882 | 39466 | 685 | 9681 | 3725 | 5574 |
| 2375.5 | 3934 | 3502.5 | 1232 | 5185 | 2175 | 803 | 10722.5 | 4401 | 6255 |
| 2161 | 3527 | 3124 | 1144 | 4618 | 1865 | 753 | 10474 | 4371 | 6231 |
| 2054 | 3485 | 3020 | 1101 | 4346 | 1860 | 710 | 9475 | 4106 | 5944 |
| 2472 | 3892 | 3371 | 1158 | 4657 | 1925 | 747 | 10873 | 4604 | 6503 |
| 2610 | 3876.5 | 3067 | 1061 | 4190 | 1892 | 722 | 11303.5 | 4242.5 | 6258 |
| 2525 | 3864 | 3288 | 1230 | 4365 | 1876 | 716 | 10851 | 4510 | 6203 |
| 2421 | 3854.5 | 3017 | 1135.5 | 4301.5 | 1828.5 | 753 | 10632 | 4486.5 | 6145.5 |
| 2105 | 3413 | 2984 | 1112 | 4125 | 1819 | 705 | 9587 | 4117 | 5713 |
| 2075 | 3393 | 2997 | 1103 | 4092 | 1782 | 732 | 9153 | 4039 | 5880 |
| 21411 | 15110 | 5777 | 1274 | 4015 | 1825 | 724 | 9273 | 8571 | 8591 |
| 2256.5 | 3568 | 3065 | 1083 | 3898 | 1804 | 745 | 9146.5 | 4324 | 6182.5 |
| 2252 | 3755 | 3190 | 1112 | 4261 | 1872 | 760 | 9878 | 4585 | 6595 |
| 60360 | 32793 | 9597 | 1554 | 4717 | 1944 | 783 | 14343 | 5699 | 7534 |
| 2925 | 4432 | 3512 | 1217 | 4540 | 1876 | 694 | 9853 | 4635 | 6557 |
| 2273 | 3692 | 3246 | 1115 | 5082 | 1975 | 731 | 10063 | 4411 | 6335 |
| 2459 | 3816 | 3111 | 1165 | 4241 | 1806 | 732 | 10047 | 4487 | 6315 |
| 9766 | 22514 | 9720 | 2031 | 10736 | 2934 | 738 | 14402 | 9746 | 24255 |
| 10422.5 | 24852.5 | 11911 | 2704 | 13810 | 3477.5 | 742.5 | 11903 | 9024 | 24072 |
| 9704 | 22562 | 11661 | 2605 | 13519 | 3349 | 731 | 11852 | 9271 | 23536 |

| | | | | | | | | | |
|---|---|---|---|---|---|---|---|---|---|
| 10016 | 22362 | 9768 | 2174 | 11364 | 2822 | 743 | 13455 | 9822 | 23846 |
| 9112 | 24241 | 8057 | 1802 | 9444 | 2542 | 754 | 14495 | 9723 | 26486 |
| 12581 | 106328 | 164638 | 63875 | 7792 | 2723 | 684 | 15384 | 11748 | 59401 |
| 4165.5 | 12115.5 | 4586.5 | 1386 | 5448.5 | 2006 | 726 | 15049.5 | 5235 | 14997.5 |
| 3206 | 7166 | 7101 | 1986 | 570541 | 55148 | 743 | 16715 | 4583 | 10096 |
| 2717.5 | 5087.5 | 4330.5 | 1356 | 6172 | 2398.5 | 817 | 17834 | 4662.5 | 7981 |
| 2596 | 4562 | 3677 | 1230 | 5119 | 2014 | 799 | 13438 | 4794 | 7664 |
| 2600 | 4343.5 | 3393 | 1195.5 | 4686.5 | 1858 | 770.5 | 10926.5 | 4591.5 | 6610 |
| 2616 | 4390.5 | 3546 | 1207.5 | 4946.5 | 1915 | 759 | 11570.5 | 4655 | 7024 |
| 2631 | 4622 | 3418 | 1227 | 4602 | 1980 | 744 | 12820 | 4386 | 7258 |
| 2645.5 | 4522.5 | 3485.5 | 1369 | 4705 | 2044 | 751.5 | 11382 | 4545.5 | 6953.5 |
| 2653 | 4313 | 3541 | 1342 | 4773 | 1965 | 770 | 11495 | 4481 | 6859 |
| 2407.5 | 3778.5 | 3232.5 | 1166.5 | 4407 | 1945.5 | 752 | 10775 | 4572.5 | 6514.5 |
| 2356 | 3703 | 3169 | 1208 | 4433 | 1796 | 779 | 10087 | 4705 | 6412 |
| 28443.5 | 19532 | 6466.5 | 1369 | 4288.5 | 1890.5 | 739 | 9740.5 | 10394 | 9566.5 |
| 2465 | 3919 | 3300 | 1170 | 4188 | 1785 | 733 | 9687 | 4785 | 6574 |
| 2353 | 3936 | 3382 | 1209 | 4587 | 1878 | 770 | 10952 | 4576 | 6756 |
| 57675 | 34767.5 | 9879 | 1591 | 5126 | 1997.5 | 767 | 14750.5 | 5516 | 8359 |
| 2980.5 | 4985 | 3917.5 | 1209 | 4820 | 1956.5 | 771.5 | 11339 | 4797.5 | 7724 |
| 2352 | 3903 | 3287 | 1164 | 5093 | 1960 | 744 | 10642 | 4544 | 6713 |
| 2603.5 | 3942.5 | 3339 | 1160 | 4293 | 1866 | 717 | 10358.5 | 4551 | 6193 |
| 9855 | 23500 | 9155 | 2076 | 10070 | 2716 | 744 | 14651 | 10254 | 25108 |
| 9859 | 23828 | 12353 | 2660 | 14059.5 | 3541 | 764.5 | 12157 | 9124.5 | 24410.5 |
| 10443 | 23427 | 12142 | 2692 | 13305 | 3406 | 741 | 12630 | 9252 | 24270 |
| 10842 | 23594 | 10433 | 2223 | 11817 | 3038 | 752 | 13001 | 9714 | 24882 |
| 9721 | 25178.5 | 8173 | 2006 | 9770 | 2674.5 | 766.5 | 14387 | 10078.5 | 27415 |
| 10433 | 23633 | 8712 | 2567 | 7698 | 2486 | 751 | 15625 | 10206 | 25590 |
| 4052 | 11792 | 4777 | 1390 | 5714 | 2071 | 784 | 14357 | 5479 | 15483 |
| 3164 | 6573 | 6489 | 1783 | 567517 | 51889 | 725 | 16658 | 4678 | 9435 |
| 2864 | 4974 | 3970 | 1361 | 6000 | 2295 | 890 | 16755 | 5395 | 8172 |
| 2783 | 4716 | 3807.5 | 1282 | 5406.5 | 2135.5 | 801.5 | 13033 | 4863.5 | 7244.5 |
| 2715.5 | 4162.5 | 3536.5 | 1206 | 4801 | 1963.5 | 760.5 | 11183 | 4613.5 | 6652.5 |
| 2546 | 4451 | 3536 | 1177 | 4790 | 1950 | 766 | 11602 | 4518 | 6818 |
| 2825.5 | 4176 | 3363.5 | 1140.5 | 4479.5 | 1788 | 726 | 11832 | 4620 | 6546 |
| 2614 | 4005 | 3271 | 1197 | 4290 | 1787 | 709 | 10626 | 4339 | 6451 |
| 2567 | 3836 | 3138 | 1232 | 4433 | 1966 | 732 | 10481 | 4482 | 6484 |
| 2316.5 | 3604 | 3136.5 | 1105.5 | 4174.5 | 1880 | 702.5 | 10490.5 | 4422.5 | 6364 |
| 2269 | 3702 | 3141 | 1161 | 4278.5 | 1869 | 734 | 9867 | 4385.5 | 6100 |
| 17097.5 | 11986.5 | 5040.5 | 1242 | 4168 | 1760 | 746.5 | 9574 | 7746.5 | 8221 |
| 2321.5 | 3702 | 3124.5 | 1122 | 4222.5 | 1836.5 | 739 | 9626.5 | 4463.5 | 6224 |
| 2504.5 | 3816 | 3373.5 | 1162 | 4488 | 1871 | 734.5 | 10168 | 4768.5 | 6462 |
| 54607.5 | 28654 | 9059.5 | 1522.5 | 4540 | 1920 | 749.5 | 14944.5 | 5676.5 | 7543 |
| 3074 | 4688.5 | 3647 | 1286 | 4489 | 1979 | 761 | 10913.5 | 4769.5 | 7037.5 |
| 2404 | 3857 | 3223 | 1157 | 5339 | 1896 | 705 | 10177 | 4604 | 6587 |
| 2633 | 3978.5 | 3098 | 1180.5 | 4501 | 1897.5 | 759 | 10504.5 | 4416 | 6303 |
| 10487.5 | 22076.5 | 9626 | 2130 | 10603.5 | 2876.5 | 727 | 14330.5 | 10298.5 | 24716 |
| 10800.5 | 23954.5 | 12277 | 2739 | 13955.5 | 3312.5 | 761 | 12558.5 | 9334 | 23969 |
| 10982 | 23726 | 12431 | 2614 | 13412 | 3432 | 764 | 12084 | 9473 | 24231 |

| | | | | | | | | | |
|---|---|---|---|---|---|---|---|---|---|
| 10474.5 | 22544 | 9925 | 2224 | 11631 | 2865 | 749 | 13484.5 | 10085 | 23751 |
| 9239 | 23886 | 8110 | 1864 | 8925 | 2557 | 762 | 14386 | 9747 | 27279 |
| 12798 | 114636.5 | 175046 | 65007.5 | 7847.5 | 2781 | 679 | 15573.5 | 12347 | 69000 |
| 4889 | 13182 | 5197 | 1579 | 5809 | 2042 | 752 | 15065 | 5932 | 16629 |
| 2992.5 | 7068.5 | 6430.5 | 1898 | 482956 | 46333.5 | 749.5 | 15829 | 4515 | 10127 |
| 2695 | 4742 | 4109 | 1375 | 6266 | 2372 | 875 | 16690 | 4961 | 7915 |
| 2542 | 4310.5 | 3598 | 1311 | 5333.5 | 2116.5 | 761.5 | 12678.5 | 4772 | 6948.5 |
| 2474 | 4157 | 3668 | 1169 | 4985 | 1867 | 743 | 11306 | 4497 | 6597 |
| 2626.5 | 4187.5 | 3300.5 | 1174.5 | 4663.5 | 1892 | 735 | 11505.5 | 4420.5 | 6774.5 |
| 2754 | 4338 | 3371 | 1202 | 4638 | 1794 | 726 | 12246 | 4458 | 6668 |
| 2813.5 | 4351.5 | 3680 | 1324.5 | 4555 | 1980 | 706.5 | 11563.5 | 4517.5 | 7008.5 |
| 2581 | 3891 | 3315 | 1158 | 4529 | 1822 | 706 | 10826 | 4373 | 6285 |
| 2433.5 | 3705 | 3073 | 1140.5 | 4326 | 1869.5 | 731.5 | 10197 | 4386 | 6232 |
| 2393.5 | 3730.5 | 3003.5 | 1125.5 | 4498.5 | 1834 | 767.5 | 10251 | 4543 | 6317 |
| 22250 | 15408 | 5774.5 | 1245 | 4263 | 1885 | 722 | 9929.5 | 8463.5 | 8402 |
| 2448.5 | 3750.5 | 3163 | 1202 | 4345.5 | 1879 | 734 | 9601.5 | 4490.5 | 6265.5 |
| 2639 | 3926 | 3184 | 1183 | 4559 | 1891 | 726 | 10069 | 4600 | 6771 |
| 57534.5 | 38268.5 | 10527.5 | 1589.5 | 4926.5 | 2049 | 754.5 | 15241.5 | 5589.5 | 9054.5 |
| 3130 | 5188 | 3934 | 1285 | 4943 | 1970 | 733 | 11273 | 4482 | 7502 |
| 2533 | 4179 | 3392 | 1228 | 5264 | 2056 | 736 | 10351 | 4549 | 6820 |
| 2757.5 | 4198 | 3426 | 1158.5 | 4524.5 | 1863.5 | 735 | 10404 | 4613.5 | 6121 |
| 9852.5 | 22551.5 | 9364 | 2256.5 | 10779 | 2849.5 | 741.5 | 13954 | 9870.5 | 25274.5 |
| 9764.5 | 23649.5 | 12822.5 | 2726 | 14028 | 3456.5 | 741 | 11934 | 8885 | 24184.5 |
| 10060 | 22749.5 | 11770.5 | 2514.5 | 13496.5 | 3304 | 734 | 12051.5 | 8540.5 | 24028.5 |
| 9401 | 21529 | 10668 | 2221 | 12072 | 2997 | 711 | 12931 | 9389 | 23266 |
| 8999.5 | 23992 | 8127 | 1902 | 9533 | 2591 | 794 | 14201 | 9841.5 | 27869 |
| 10765.5 | 26557.5 | 11080 | 3207 | 7287.5 | 2601.5 | 719 | 14647 | 10008.5 | 27719 |
| 4065 | 10672 | 4393 | 1423 | 5656 | 2024 | 791 | 14902 | 5518 | 13029 |
| 2997.5 | 6794 | 6697.5 | 1981.5 | 521516 | 47779.5 | 735.5 | 15907 | 4131.5 | 9869.5 |
| 3049 | 4852 | 4135.5 | 1456 | 6498.5 | 2383.5 | 889 | 15739.5 | 5338 | 8140.5 |
| 2641.5 | 4666 | 4142.5 | 1328.5 | 5574 | 2133 | 773 | 12385.5 | 4502.5 | 7298 |
| 2726 | 4446 | 3755 | 1288 | 5254 | 1934 | 771 | 11378 | 4593 | 6812 |
| 2748.5 | 4378.5 | 3575 | 1303.5 | 4940 | 1945.5 | 783.5 | 12876.5 | 4678.5 | 7341 |
| 2994 | 4432 | 3678 | 1224 | 5039 | 1906 | 741 | 11934 | 4698 | 6735 |
| 2743 | 4074 | 3385 | 1214 | 4635 | 1855 | 729 | 11156 | 4693 | 6523 |
| 2514.5 | 3993 | 3204 | 1212.5 | 4402.5 | 1922.5 | 755 | 11437.5 | 4563 | 6663.5 |
| 2540 | 3828 | 3222 | 1121 | 4410 | 1843 | 738 | 10209 | 4367 | 6285 |
| 2474 | 3795 | 3264 | 1221.5 | 4452.5 | 1917.5 | 760 | 9964 | 4564 | 6231 |
| 29085 | 20080 | 6649 | 1363 | 4379 | 1869 | 753 | 9869 | 10486 | 9535 |
| 2625.5 | 3785 | 3364.5 | 1239 | 4464 | 1968.5 | 727 | 10054 | 4696 | 6386 |
| 2605 | 4030 | 3548 | 1291 | 4783 | 2012 | 746 | 10448 | 4558 | 7009 |
| 50986 | 27926 | 9207 | 1562 | 4644 | 1959 | 759 | 14781 | 5563 | 7629 |
| 3422.5 | 5116.5 | 4007 | 1231 | 4598.5 | 1953 | 734 | 11339.5 | 4787.5 | 6920 |
| 2671 | 4185 | 3480 | 1273 | 5757 | 2108 | 733 | 10097 | 4552 | 6734 |
| 2756.5 | 4063.5 | 3287 | 1195 | 4424 | 1809.5 | 736 | 10032 | 4719.5 | 6365 |
| 10321 | 23391.5 | 10090 | 2226.5 | 11514.5 | 2980 | 761 | 14157 | 9572.5 | 23964 |
| 10378 | 24253 | 12981 | 2694 | 13935 | 3498 | 773 | 11498 | 8525 | 24835 |
| 11378.5 | 24323.5 | 12301.5 | 2627 | 13597.5 | 3366 | 740.5 | 12328 | 9231.5 | 24057.5 |

| | | | | | | | | | |
|---|---|---|---|---|---|---|---|---|---|
| 10608 | 23544 | 10815 | 2306.5 | 11853.5 | 3034.5 | 744 | 13844 | 9501.5 | 23301 |
| 9028.5 | 23904 | 8468 | 2032.5 | 9859 | 2609.5 | 753 | 13315.5 | 9358 | 25610.5 |
| 17357 | 137580 | 207873 | 77604 | 7898 | 2896 | 678 | 13869 | 12160 | 74539 |
| 12788.5 | 15478.5 | 6387.5 | 1843.5 | 5847.5 | 2107 | 812.5 | 15257.5 | 6063 | 15149.5 |
| 8935 | 9561 | 30075 | 9890 | 152027 | 22506 | 771 | 17675 | 4954 | 11097 |
| 9213 | 6738.5 | 7051 | 2034.5 | 34486 | 4568 | 980 | 19467.5 | 5583.5 | 8457 |
| 8159.5 | 6117 | 5116 | 1571 | 17448.5 | 3351 | 942.5 | 17332.5 | 5407.5 | 7776 |
| 7713 | 5581 | 4611 | 1464 | 9580 | 2558 | 853 | 14408 | 5218 | 7353 |
| 6890.5 | 5585.5 | 4404.5 | 1332 | 7244.5 | 2364.5 | 867 | 14597 | 5254 | 7484 |
| 6778 | 6073 | 4434 | 1389 | 6025 | 2026 | 831 | 14472 | 5016 | 7721 |
| 4987 | 5435 | 4477 | 1559 | 5857 | 2150 | 832 | 13513 | 5068 | 7596 |
| 5845 | 5017 | 3785 | 1260 | 5488 | 1986 | 779 | 12565 | 4870 | 6907 |
| 5934 | 4801 | 3643 | 1326 | 4806 | 1957 | 822 | 11328 | 5026 | 6803 |
| 7056 | 5202 | 3756 | 1280 | 5060 | 1948 | 799 | 11852 | 4988 | 6681 |
| 17214.5 | 9750 | 5279.5 | 1489.5 | 4623.5 | 1955.5 | 807.5 | 11159 | 6730 | 7851.5 |
| 9492 | 6023.5 | 3913.5 | 1300.5 | 4977 | 1996.5 | 805.5 | 11699 | 5096 | 6995.5 |
| 7264 | 5399 | 3955 | 1262 | 5660 | 2000 | 857 | 12093 | 5400 | 7199 |
| 374142 | 246039.5 | 67465.5 | 4843.5 | 5126 | 1979.5 | 812 | 45390 | 12790 | 11691 |
| 27995.5 | 15828.5 | 6180.5 | 1447.5 | 5901 | 2140 | 821.5 | 14545.5 | 5768.5 | 8537 |
| 15693 | 7883.5 | 4333.5 | 1336 | 6540.5 | 2483.5 | 820 | 12504.5 | 5327 | 7211.5 |
| 13836 | 7712 | 4365 | 1202 | 5627 | 2078 | 830.5 | 12463 | 5467.5 | 7072 |
| 15119.5 | 28170.5 | 10126 | 2148.5 | 12732.5 | 3047 | 799 | 15521 | 9869 | 26264.5 |
| 16351 | 29268.5 | 14907.5 | 2948 | 16040 | 3539 | 823.5 | 13042.5 | 9437.5 | 24634.5 |
| 18300 | 31593 | 14797 | 2861 | 16647 | 3613 | 924 | 12935 | 9900 | 24859 |
| 14176.5 | 27826.5 | 12481.5 | 2536 | 15971 | 3396 | 865 | 13781.5 | 9585 | 24104 |
| 9394 | 25368 | 8446 | 1885 | 9567 | 2615 | 780 | 14148 | 9532 | 26261 |
| 10326 | 61661 | 66183 | 19883 | 7422 | 2605 | 732 | 16211 | 9493 | 35432 |
| 4301 | 11785.5 | 4827.5 | 1458 | 5397 | 2021.5 | 750.5 | 14620 | 5467.5 | 14493 |
| 3195 | 7615 | 6756 | 1913 | 549031 | 52768 | 757 | 16497 | 4533 | 10183 |
| 2618 | 5002.5 | 4362.5 | 1533.5 | 6437.5 | 2473 | 927 | 18371 | 5072.5 | 8625 |
| 2709 | 4463.5 | 3738 | 1382 | 5477 | 2059 | 766.5 | 12300 | 4795 | 6996 |
| 2801 | 4400 | 3879 | 1351 | 5297 | 2024 | 732 | 11279 | 4514 | 6495 |
| 2928 | 4402 | 3759 | 1253 | 4900 | 1934 | 748 | 12092 | 4686 | 7089 |
| 2979 | 4312 | 3407 | 1218 | 4634 | 1858 | 719 | 12006 | 4710 | 6575 |
| 2768 | 4194 | 3470 | 1228 | 4272 | 1833 | 699 | 10944 | 4517 | 6325 |
| 2691 | 3995 | 3320.5 | 1142 | 4420 | 1908 | 726 | 11323 | 4520 | 6166.5 |
| 2440 | 3734 | 3038.5 | 1158 | 4197.5 | 1777 | 718.5 | 10253.5 | 4466 | 6034 |
| 2509.5 | 3821.5 | 3299 | 1189 | 4096 | 1823.5 | 708 | 10335.5 | 4381.5 | 6143.5 |
| 24310.5 | 16633 | 6065.5 | 1360.5 | 4341.5 | 1772 | 730 | 9694.5 | 9111.5 | 8959.5 |
| 2659 | 4004 | 3340 | 1108 | 4258 | 1857 | 731 | 9536 | 4557 | 6411 |
| 2482.5 | 3644.5 | 3307.5 | 1185 | 4535.5 | 1843 | 711.5 | 9907 | 4449.5 | 6534 |
| 68282 | 37795.5 | 11774 | 1669 | 4913 | 1888.5 | 709.5 | 16217.5 | 5844.5 | 7779 |
| 3429.5 | 5043 | 3928 | 1291.5 | 4790.5 | 1950 | 704.5 | 11127 | 4799 | 7027 |
| 2547 | 4038 | 3324 | 1224 | 5484 | 2036 | 722 | 10201 | 4569 | 6617 |
| 2760 | 4029 | 3472 | 1173 | 4540 | 1893 | 751 | 10397 | 4563 | 6365 |
| 10146.5 | 25280 | 9364.5 | 2152 | 10328 | 2714 | 725 | 13333 | 9900 | 26625.5 |
| 10840.5 | 26375.5 | 13126 | 2834.5 | 14292.5 | 3603.5 | 761 | 11728 | 8786.5 | 24853.5 |
| 10070 | 23815 | 12815 | 2693 | 14042 | 3387 | 714 | 11927 | 8887 | 24806 |

| | | | | | | | | | |
|---|---|---|---|---|---|---|---|---|---|
| 9317 | 22253 | 10839 | 2305 | 12290 | 2994 | 722 | 12278 | 8664 | 23468 |
| 9401 | 24655 | 7909 | 1837 | 9004 | 2534 | 766 | 14208 | 9847 | 27172 |
| 15277 | 131512 | 206528 | 77012 | 8118 | 2972 | 652 | 13882 | 12139 | 72435 |
| 4815 | 12780.5 | 5620.5 | 1824.5 | 6416 | 2425 | 722.5 | 14888.5 | 5699 | 17541.5 |
| 2972 | 8438 | 6477 | 2171 | 331339 | 34144 | 718 | 15838 | 4756 | 14393 |
| 2766 | 7565 | 5487.5 | 1953 | 8857 | 3484.5 | 1083 | 35865.5 | 6782 | 16270.5 |
| 2431 | 6863 | 4893 | 1848 | 7525 | 2890 | 751 | 13372 | 4542 | 13387 |
| 2356 | 6162 | 4425 | 1822 | 6718 | 2580 | 739 | 11489 | 4462 | 12627 |
| 2527 | 6360 | 4583 | 1601 | 6397 | 2426 | 702 | 13214 | 4539 | 11902 |
| 2459.5 | 5672.5 | 3944 | 1502 | 5796.5 | 2438 | 702.5 | 12134.5 | 4460 | 11240 |
| 2367 | 5576 | 3972 | 1621 | 6103 | 2451 | 695 | 12257 | 4551 | 10932 |
| 2450 | 5881 | 4157 | 1627 | 6000 | 2536 | 707 | 12047 | 4792 | 11175 |
| 2327 | 4687 | 3613 | 1480 | 5667 | 2326 | 688 | 10786 | 4431 | 9552 |
| 2207 | 4365 | 3843 | 1511 | 5583 | 2348 | 723 | 10719 | 4751 | 8913 |
| 8580 | 7921.5 | 4785 | 1528.5 | 5307.5 | 2197 | 709 | 10782.5 | 6168 | 9192 |
| 2599 | 3869.5 | 3596.5 | 1329 | 4956.5 | 2134 | 739 | 10548 | 4972 | 7148.5 |
| 2703 | 4205.5 | 3995 | 1394.5 | 5046.5 | 2130 | 681 | 10693 | 4650 | 7370.5 |
| 40261 | 28316 | 8320 | 1669.5 | 5410 | 2232 | 733 | 14709.5 | 5348.5 | 10129.5 |
| 2979 | 6307 | 4384 | 1458 | 5458 | 2212 | 702 | 11713 | 4550 | 9582 |
| 2381 | 4531 | 3804 | 1389 | 6098 | 2326 | 739 | 11615 | 4492 | 8173 |
| 2667 | 4301 | 3749 | 1322 | 5161 | 2178 | 720 | 10446 | 4593 | 7253 |
| 9328 | 21808 | 9790 | 2287 | 11169 | 3005 | 738 | 14271.5 | 9512.5 | 24162.5 |
| 8773 | 24389 | 12031 | 2751 | 13696 | 3472 | 698 | 10736 | 8382 | 25240 |
| 10040 | 23349.5 | 12983 | 2953 | 14058 | 3393 | 724.5 | 11340.5 | 8315 | 25252 |
| 8973 | 21028.5 | 10062 | 2296.5 | 12098 | 3096 | 722.5 | 13593 | 9558 | 23036.5 |
| 9057 | 24492 | 7954 | 2049 | 9364 | 2752 | 714 | 14697 | 10051 | 27740 |
| 242018 | 498932 | 554023 | 141888 | 15914 | 6258 | 739 | 123464 | 59738 | 192366 |
| 18919 | 72236 | 89295 | 25873 | 8574.5 | 2969 | 695 | 14726.5 | 6959 | 31319.5 |
| 4432 | 10271 | 21466 | 6403.5 | 171254.5 | 27024.5 | 708.5 | 17089 | 5056 | 12320 |
| 5183 | 6057 | 11743 | 2672 | 66948 | 8857 | 1241 | 42776 | 9827 | 11098 |
| 3972 | 4885 | 5287 | 1707 | 50459 | 6587 | 777 | 12389 | 5059 | 6720 |
| 3087 | 4197 | 4022 | 1345 | 34246 | 4721 | 781 | 11964 | 5319 | 6944 |
| 3360.5 | 4557 | 3894 | 1413 | 25996.5 | 3822.5 | 793.5 | 12094 | 4754.5 | 7316 |
| 3377.5 | 4573 | 3839 | 1389.5 | 22208 | 3397.5 | 707 | 11787 | 4622.5 | 6731.5 |
| 4560.5 | 7186 | 7258 | 2338 | 16600 | 3400 | 819.5 | 12347 | 5524 | 8212.5 |
| 3246 | 4449 | 3557 | 1314 | 15579 | 2784 | 707 | 11266 | 4474 | 6655 |
| 2985.5 | 4306 | 3744 | 1427.5 | 10004.5 | 2447.5 | 732.5 | 10521 | 4441 | 6511 |
| 3160 | 4218 | 3715 | 1257 | 11417 | 2504 | 685 | 10195 | 4469 | 6165 |
| 25051 | 17969 | 10026 | 1820.5 | 8929 | 2290.5 | 730.5 | 9697.5 | 9595 | 9360.5 |
| 3699 | 4722 | 3963.5 | 1383.5 | 9605 | 2313 | 722 | 9674.5 | 5143.5 | 7015.5 |
| 3104 | 4323 | 3580 | 1288 | 8561 | 2323 | 724 | 10327 | 4608 | 6641 |
| 324485 | 232118 | 81906 | 8092 | 8861 | 2411 | 722 | 35399 | 12581 | 11517 |
| 8649 | 7005 | 5214 | 1634 | 7715.5 | 2317 | 707.5 | 12432.5 | 5240.5 | 7770.5 |
| 3990 | 5005 | 4635 | 1566 | 10529 | 3071 | 741 | 10834 | 4849 | 7305 |
| 3789 | 4787 | 3793 | 1335 | 8623 | 2313 | 752.5 | 10308 | 4606 | 6571.5 |
| 10587 | 24634.5 | 9104 | 2142.5 | 13269 | 3089 | 735.5 | 14418.5 | 9935.5 | 25888 |
| 9195 | 24628.5 | 13654 | 2969.5 | 15888.5 | 3640 | 734.5 | 10237.5 | 8838.5 | 26438.5 |
| 10235 | 23480.5 | 12838.5 | 2834 | 15029.5 | 3645.5 | 739 | 11761.5 | 8550 | 23511 |

| | | | | | | | | | |
|---|---|---|---|---|---|---|---|---|---|
| 9634 | 19668 | 10313 | 2433 | 12603 | 3069 | 725 | 12174 | 8844 | 21766 |
| 9272 | 23566 | 7244 | 1768 | 8554 | 2487 | 772 | 12711 | 9476 | 25952 |
| 14833 | 162490 | 266282 | 88274 | 12892 | 5896 | 669 | 15870 | 10783 | 90636 |
| 4991 | 14954 | 10278 | 3802 | 6262 | 2656 | 747 | 13566 | 5651 | 17546 |
| 3262 | 5461 | 8734 | 3161 | 509106 | 49203 | 710 | 16846 | 4805 | 7869 |
| 2905.5 | 4525.5 | 4596 | 1648 | 7983.5 | 2609 | 857.5 | 14341.5 | 5398.5 | 7638.5 |
| 2780.5 | 4292 | 3853.5 | 1450.5 | 5513.5 | 2080.5 | 746 | 11728 | 4782.5 | 6748 |
| 2651 | 4298 | 3882 | 1261 | 5194 | 2036 | 771 | 10518 | 4717 | 7178 |
| 2687 | 4434 | 3651 | 1272 | 5093 | 1977 | 738 | 11161 | 4639 | 6906 |
| 2867 | 4390 | 3668 | 1266 | 4802 | 1890 | 728 | 12373 | 4520 | 6734 |
| 2822.5 | 4857 | 4063.5 | 1424 | 4838.5 | 2028 | 724 | 12195 | 4445.5 | 7048.5 |
| 2874.5 | 4400 | 3688 | 1241.5 | 4592.5 | 1865 | 700.5 | 11280 | 4529.5 | 6727.5 |
| 2635 | 4280 | 3380 | 1177 | 4514 | 1829 | 708 | 10239 | 4427 | 6816 |
| 2868 | 4102.5 | 3491.5 | 1258 | 4419 | 1828.5 | 694 | 10057.5 | 4621.5 | 6418.5 |
| 25433 | 17454 | 6228 | 1437 | 4324 | 1780 | 695 | 10151 | 9112 | 8628 |
| 2997 | 4330 | 3610 | 1169 | 4424 | 1817 | 735 | 10339 | 4978 | 6845 |
| 2752 | 4364 | 3610 | 1196 | 4809 | 1833 | 707 | 10435 | 4717 | 7034 |
| 70834.5 | 41236.5 | 12784 | 1727.5 | 4801.5 | 1950 | 717 | 17248 | 5743.5 | 8078 |
| 3418 | 5577 | 4003 | 1368 | 4771 | 1891 | 704 | 10937 | 4799 | 7541 |
| 2786 | 4372 | 3747 | 1202.5 | 5334 | 2068 | 708.5 | 10949.5 | 4576 | 6692 |
| 3142 | 4486 | 3448 | 1197 | 4650 | 1925 | 719 | 11029 | 4702 | 6712 |
| 9777 | 24337.5 | 9673.5 | 2120 | 11075.5 | 2793.5 | 704.5 | 14468.5 | 9802 | 26092 |
| 10608 | 23036 | 11836 | 2615 | 13208 | 3244 | 730 | 12570 | 9006 | 21998 |
| 9981 | 23028 | 11749 | 2503 | 12611 | 3190 | 699 | 12013 | 8865 | 23505 |
| 10850.5 | 23390 | 9889 | 2238.5 | 11410 | 2868 | 701 | 13410.5 | 9622 | 23320.5 |
| 9963 | 22685 | 7652 | 1841 | 8567 | 2518 | 730 | 14731 | 10090 | 24323 |
| 12953 | 123948 | 201050 | 65767 | 10705 | 4537 | 699 | 15969 | 9386 | 61216 |
| 4379 | 13188 | 6504.5 | 2340 | 6052.5 | 2381 | 704 | 14073 | 5353 | 15735 |
| 2786 | 5032.5 | 6927.5 | 2425 | 479550.5 | 46249 | 696 | 15436 | 4243 | 7631.5 |
| 2827.5 | 4633.5 | 4826.5 | 1857.5 | 6451.5 | 2653 | 845.5 | 14042.5 | 5325 | 7881.5 |
| 3014.5 | 4586.5 | 4239.5 | 1484.5 | 5492 | 2249 | 741 | 11687.5 | 4852 | 7165.5 |
| 2754 | 4316 | 3712 | 1301 | 4853 | 1947 | 717 | 10800 | 4702 | 6734 |
| 2767 | 4427.5 | 3495 | 1307 | 4747 | 1954.5 | 712.5 | 10687.5 | 4592.5 | 6741 |
| 2863 | 4237 | 3237.5 | 1114 | 4407.5 | 1838 | 650.5 | 11698.5 | 4471.5 | 6403.5 |
| 2721 | 4524.5 | 3644 | 1383 | 4685 | 1961.5 | 694 | 11947 | 4408.5 | 6911 |
| 2824 | 4535 | 3850.5 | 1358 | 4977.5 | 1964.5 | 707 | 11203 | 4445 | 6769.5 |
| 2648 | 4125 | 3652 | 1223.5 | 4850 | 2007.5 | 728.5 | 11365.5 | 4370.5 | 6744.5 |
| 2737 | 4184 | 3610 | 1256 | 4772 | 1912 | 717 | 10637 | 4484 | 6478 |
| 14561.5 | 10557 | 5142 | 1350.5 | 4347 | 1889.5 | 721.5 | 10030.5 | 7051.5 | 7990.5 |
| 2883 | 4083.5 | 3523 | 1250 | 4450 | 1899.5 | 726.5 | 9819 | 4826.5 | 6405 |
| 2559 | 4055.5 | 3454 | 1262 | 4729.5 | 1865.5 | 728 | 10206.5 | 4775.5 | 6807.5 |
| 98208.5 | 72383.5 | 18266 | 2038 | 5357.5 | 2058 | 740 | 19779 | 6105 | 10511.5 |
| 3111 | 5333.5 | 3951.5 | 1322.5 | 4912.5 | 1919 | 750 | 11813 | 4618 | 7303 |
| 2640.5 | 4117 | 3489 | 1207 | 5552.5 | 2052.5 | 720 | 10727 | 4610.5 | 6404.5 |
| 2891 | 4018 | 3451.5 | 1161.5 | 4575.5 | 1802 | 721 | 9908.5 | 4510.5 | 6378.5 |
| 10041 | 22477 | 10017 | 2260 | 11038 | 2888 | 730 | 14217 | 9671 | 24543 |
| 10314.5 | 24212 | 13907.5 | 2822.5 | 14687 | 3495.5 | 737.5 | 11901 | 8896.5 | 24721 |
| 9399 | 23439.5 | 12828.5 | 2726.5 | 13854.5 | 3354 | 738.5 | 11787.5 | 8722 | 23234 |

| | | | | | | | | | |
|---|---|---|---|---|---|---|---|---|---|
| 9707 | 21914 | 10468.5 | 2264 | 11639.5 | 2812 | 682.5 | 13744.5 | 8936.5 | 22870 |
| 970 | 1311 | 1196 | 658 | 1812 | 1246 | 347 | 3242 | 1850 | 2595 |
| 27532 | 33720 | 8996 | 1544 | 3512 | 1638 | 475.5 | 17810 | 5056.5 | 6165.5 |
| 3551 | 4353 | 3175 | 1022.5 | 3356 | 1581.5 | 592.5 | 10001 | 3629.5 | 5015.5 |
| 2892.5 | 4109.5 | 7980 | 2391 | 449086.5 | 44383 | 647.5 | 8675.5 | 3656.5 | 5385.5 |
| 2698.5 | 3910 | 3743 | 1268 | 5542.5 | 2128 | 723 | 9829 | 3982 | 5621.5 |
| 2160.5 | 3526.5 | 3190 | 1126 | 4241.5 | 1871.5 | 726 | 9939.5 | 4104.5 | 5810.5 |
| 2050 | 3380.5 | 2986.5 | 1156.5 | 4285 | 1828.5 | 733 | 8844.5 | 3973.5 | 5776 |
| 2437.5 | 3625 | 3138.5 | 1132.5 | 4269.5 | 1812 | 748.5 | 9887 | 4499 | 6235 |
| 2608 | 3734.5 | 2947 | 1052 | 3958.5 | 1729.5 | 708 | 10739.5 | 4458.5 | 6074 |
| 2545 | 3755 | 3248 | 1264 | 4152 | 1844 | 678 | 10421 | 4310 | 5831 |
| 2147 | 3459 | 2895 | 1065 | 3862 | 1800 | 656 | 9444 | 3814 | 5807 |
| 2044 | 3304 | 2876 | 1085 | 3832 | 1817 | 673 | 9079 | 3750 | 5523 |
| 1806.5 | 3107.5 | 2801 | 1040 | 3916 | 1681 | 672.5 | 8399 | 3741 | 5654.5 |
| 17438 | 13258 | 4992 | 1262 | 3835 | 1755 | 722 | 8671 | 7508 | 7759 |
| 2168 | 3386 | 2934 | 1096 | 3996 | 1717 | 693 | 8705 | 3997 | 5808 |
| 2173 | 3391 | 3015.5 | 1171.5 | 4101 | 1848 | 728.5 | 9025 | 4197 | 6118 |
| 69897 | 35231 | 10484 | 1506 | 4401 | 1797 | 709 | 14421 | 5838 | 6940 |
| 2633 | 3997 | 3268 | 1152 | 4342 | 1850 | 715 | 9789 | 4572 | 6268 |
| 2148 | 3431 | 2981.5 | 1118 | 5106 | 1925 | 711 | 9582 | 4088.5 | 6066 |
| 2208 | 3598 | 3150 | 1107 | 4035 | 1769 | 704 | 9785 | 4078 | 5824 |
| 9681 | 22264 | 9370 | 2077 | 10696 | 2754 | 721 | 13866 | 9015 | 24258 |
| 10767 | 22512 | 12304 | 2568 | 13208 | 3220 | 705 | 12180 | 9154 | 22353 |
| 10108 | 21276 | 11261 | 2405 | 12726 | 3096 | 678 | 13053 | 9627 | 22432 |
| 9624 | 22067 | 10220 | 2055 | 11137 | 2749 | 686 | 13417 | 9690 | 22818 |
| 9449.5 | 22093.5 | 7262 | 1691.5 | 8521 | 2484 | 735 | 14705 | 10066 | 24260.5 |
| 9915 | 95563.5 | 149113.5 | 57256 | 7170.5 | 2617.5 | 664 | 15433.5 | 8788 | 52140 |
| 3853 | 9912 | 4310 | 1243 | 5023 | 1939 | 699 | 15302 | 4943 | 12022 |
| 2876.5 | 5470.5 | 6420.5 | 1861.5 | 607620.5 | 57649.5 | 719 | 16533.5 | 4331.5 | 7790.5 |
| 2605.5 | 4302.5 | 3736 | 1214.5 | 5780.5 | 2116 | 829.5 | 13704.5 | 4846.5 | 7203.5 |
| 2574.5 | 3910.5 | 3269.5 | 1144.5 | 4490 | 1828 | 732.5 | 11483 | 4874.5 | 6300.5 |
| 2474 | 3777.5 | 3225.5 | 1127 | 4379.5 | 1851 | 730.5 | 10640 | 4465 | 6175 |
| 2589 | 4082 | 3246 | 1122 | 4340 | 1781 | 727 | 10881 | 4532 | 6474 |
| 2698 | 3922 | 3043 | 1106 | 4233 | 1805 | 692 | 11470 | 4479 | 6169 |
| 2662.5 | 3742.5 | 3100.5 | 1154 | 3906.5 | 1839 | 704.5 | 10493 | 4564.5 | 6078 |
| 2503 | 3643 | 2933 | 1102 | 4065 | 1772 | 695 | 9954 | 4496 | 6000 |
| 1985.5 | 3310.5 | 2682.5 | 1098 | 3974.5 | 1725 | 674 | 8990 | 3940 | 5721 |
| 2008 | 3402 | 2966 | 1030 | 3909 | 1780 | 700 | 9326 | 4020 | 5711 |
| 26887 | 19378 | 6392 | 1298 | 4025 | 1765 | 720 | 9518 | 9708 | 8964 |
| 2076 | 3324 | 2838 | 1066 | 3818 | 1697 | 687 | 8656 | 4002 | 5863 |
| 2360 | 3909 | 3301 | 1161 | 4308 | 1842 | 721 | 10153 | 4543 | 6538 |
| 72856 | 38335 | 10819 | 1488 | 4550 | 1914 | 697 | 14690 | 5972 | 7244 |
| 2804 | 4447 | 3423 | 1227 | 4431 | 1884 | 695 | 10844 | 4689 | 6627 |
| 2307 | 3619 | 3104.5 | 1146 | 4976 | 1921 | 707.5 | 9683.5 | 4395.5 | 6126 |
| 2404 | 3772 | 3035 | 1083 | 4200 | 1807 | 702 | 9892 | 4170 | 5862 |
| 9587 | 21691 | 9401 | 2079 | 10464 | 2760 | 701 | 13718 | 9263 | 24046 |
| 10576 | 22894.5 | 11978.5 | 2497.5 | 12641.5 | 3153.5 | 730 | 13215 | 9362.5 | 24207.5 |
| 10358 | 22550.5 | 11944 | 2479.5 | 13039.5 | 3229 | 714.5 | 12305 | 8779 | 22174.5 |

| | | | | | | | | | |
|---|---|---|---|---|---|---|---|---|---|
| 9915 | 21281 | 9701 | 2150 | 10968 | 2809 | 712 | 13111 | 9558 | 22787 |
| 9142 | 21309 | 6977 | 1677 | 8067 | 2379 | 725 | 15579 | 10146 | 23698 |
| 9643 | 84574.5 | 128323.5 | 47051 | 7037 | 2412 | 656 | 15733 | 8346 | 42613 |
| 3995 | 9439 | 4373 | 1238.5 | 5178.5 | 1887.5 | 686 | 15580 | 5047.5 | 12158 |
| 2896 | 5567.5 | 6582 | 1825.5 | 601007.5 | 54676.5 | 721.5 | 16827 | 4411.5 | 7989 |
| 2905 | 4407 | 3932 | 1288 | 5642 | 2186 | 782 | 13017 | 4900 | 7223 |
| 2821 | 4068 | 3644.5 | 1192.5 | 5149.5 | 1930 | 760 | 11456 | 4867 | 6600.5 |
| 2612 | 3879.5 | 3312.5 | 1143 | 4502.5 | 1936.5 | 731 | 10757 | 4614.5 | 6281 |
| 2653 | 3932 | 3109 | 1119 | 4451 | 1791 | 723 | 10487 | 4594 | 6529 |
| 2811 | 4113 | 3185 | 1127 | 4171 | 1794 | 719 | 11501 | 4539 | 6100 |
| 2626 | 3896 | 3322 | 1153 | 4226 | 1857 | 671 | 10715 | 4575 | 6294 |
| 2537 | 3612 | 3151 | 1163 | 4185 | 1791 | 688 | 10099 | 4485 | 6043 |
| 2323 | 3526 | 3013 | 1128 | 4066 | 1777 | 729 | 9729 | 4376 | 5894 |
| 2083 | 3323 | 2996 | 1076 | 3882 | 1744 | 676 | 9525 | 3931 | 5729 |
| 15452 | 11092 | 4927.5 | 1198.5 | 4087 | 1735 | 694 | 9389.5 | 6983.5 | 7671 |
| 2195 | 3526 | 2979 | 1078 | 3971 | 1731 | 699 | 9000 | 4158 | 6023 |
| 2375 | 3504 | 3171 | 1132 | 4169 | 1776 | 668 | 9621 | 4513 | 6245 |
| 55476.5 | 28371 | 8672.5 | 1436 | 4316 | 1761.5 | 696 | 13661 | 5658 | 6885.5 |
| 3046.5 | 4495.5 | 3588.5 | 1183.5 | 4284.5 | 1866.5 | 712.5 | 10440.5 | 4792.5 | 6647.5 |
| 2454.5 | 3756 | 3196.5 | 1156 | 5163.5 | 1952.5 | 676 | 10175.5 | 4442 | 6163.5 |
| 2399 | 3708 | 3039 | 1067 | 4167 | 1768 | 736 | 9429 | 4215 | 5989 |
| 9665 | 21955 | 9722 | 2016 | 10871 | 2724 | 694 | 14098 | 9619 | 22793 |
| 10280 | 21984 | 11522 | 2581 | 13164 | 3281 | 688 | 12601 | 9190 | 22472 |
| 10071 | 21377 | 11587.5 | 2430 | 12349 | 3172 | 691 | 12444 | 8945 | 22043 |
| 10359 | 21660 | 9355 | 2095 | 10801 | 2627 | 702 | 13550 | 8818 | 21890 |
| 9359 | 19847 | 6753 | 1636 | 7362 | 2239 | 710 | 15752 | 9839 | 22008 |
| 10293 | 111842 | 174501 | 64810 | 6970 | 2512 | 633 | 15744 | 9347 | 66918 |
| 4280.5 | 9903 | 4660.5 | 1458 | 5236 | 1956 | 718.5 | 14598.5 | 5396 | 12399 |
| 2918 | 5089 | 6164 | 1838 | 536219 | 51178 | 693 | 16970 | 4399 | 7516 |
| 2822 | 4109 | 3712.5 | 1253.5 | 5834 | 2119.5 | 842.5 | 13887 | 5214.5 | 7218.5 |
| 2707 | 3980 | 3448 | 1275 | 5254 | 2035 | 705 | 11797 | 4738 | 6406 |
| 2677 | 3933 | 3489 | 1175 | 4547 | 1881 | 727 | 10729 | 4711 | 6298 |
| 2649 | 3756 | 3302 | 1112 | 4277 | 1767 | 707 | 10696 | 4452 | 6159 |
| 2790.5 | 3938.5 | 3252.5 | 1163 | 4171 | 1815 | 701 | 11377 | 4561.5 | 6063 |
| 2604 | 3989 | 3381 | 1302 | 4465 | 1898 | 689 | 11520 | 4452 | 6347 |
| 2632.5 | 3723 | 3245 | 1172.5 | 4210 | 1803 | 691.5 | 10932.5 | 4509 | 6156.5 |
| 2258.5 | 3396 | 2834 | 1050.5 | 3935.5 | 1766 | 671 | 9654.5 | 4048.5 | 5608.5 |
| 2190 | 3589 | 3056 | 1096 | 4016.5 | 1786.5 | 715 | 9705.5 | 4067.5 | 5822 |
| 23963.5 | 16267 | 5748.5 | 1312 | 4137.5 | 1805 | 695 | 9848.5 | 8488.5 | 8573 |
| 2316 | 3517.5 | 2883 | 1097.5 | 4024 | 1774.5 | 668 | 9028.5 | 4076 | 5866.5 |
| 2318.5 | 3621 | 3151.5 | 1115 | 4194.5 | 1808.5 | 681.5 | 9597 | 4690 | 6410.5 |
| 65699.5 | 35003.5 | 10518.5 | 1566.5 | 4600 | 1901 | 726 | 15046 | 5600.5 | 7345 |
| 2887 | 4762.5 | 3605.5 | 1116 | 4385 | 1813.5 | 671.5 | 10301 | 4667.5 | 6253.5 |
| 2378 | 3796 | 3226 | 1131 | 5002 | 1938 | 673 | 10119 | 4549 | 6246 |
| 2573.5 | 3760 | 3246 | 1080 | 4097 | 1710 | 694.5 | 9989.5 | 4299.5 | 6040 |
| 9997 | 21969 | 9455 | 2007 | 10684 | 2718 | 709 | 15006 | 9886 | 23050 |
| 10590 | 22862.5 | 12185 | 2717.5 | 13645.5 | 3458 | 721.5 | 12558.5 | 9158 | 23218 |
| 9699.5 | 22201 | 11585 | 2431.5 | 13110 | 3157.5 | 695 | 12878 | 9128.5 | 23115.5 |

| | | | | | | | | | |
|---|---|---|---|---|---|---|---|---|---|
| 9344 | 21125 | 10186 | 2133 | 10998 | 2798 | 681 | 13329 | 9147 | 23083 |
| 8831 | 23456.5 | 7389 | 1688 | 8699 | 2403 | 729.5 | 12470.5 | 9329 | 26330 |
| 10517 | 99429 | 150174 | 53263 | 7179 | 2494 | 699 | 15119 | 8725 | 51231 |
| 4340 | 11255 | 4754 | 1402 | 5234 | 1942 | 744 | 14746 | 4992 | 14075 |
| 2929 | 5231.5 | 6465.5 | 1757 | 570968 | 52027 | 713.5 | 16158 | 4268.5 | 7113.5 |
| 2899 | 4400 | 4193 | 1422 | 6167 | 2254 | 855 | 13679 | 5038 | 7475 |
| 3076.5 | 4320.5 | 4073.5 | 1408.5 | 5440.5 | 2150 | 760 | 12524 | 4953.5 | 6608 |
| 2725.5 | 4126.5 | 3580 | 1182.5 | 5055.5 | 1869.5 | 747 | 10401.5 | 4613 | 6166.5 |
| 2735 | 4075 | 3595.5 | 1257.5 | 4704 | 1881.5 | 742 | 11654.5 | 4435 | 6645 |
| 2908 | 4031 | 3299 | 1241 | 4340 | 1872 | 702 | 11236 | 4537 | 6433 |
| 2737.5 | 3805 | 3330 | 1233 | 4261.5 | 1824 | 673 | 10356 | 4383 | 6063 |
| 2684 | 3749 | 3238 | 1223 | 4316 | 1863 | 676 | 10557 | 4314 | 6125 |
| 2293 | 3608 | 2882 | 1117 | 4184 | 1769 | 693 | 9740 | 4188 | 5998 |
| 2063 | 3350 | 3010 | 1097 | 3979 | 1827 | 744 | 9500 | 3977 | 5644 |
| 29427 | 20489 | 6904 | 1294.5 | 4046.5 | 1709 | 730 | 9462 | 11067 | 9496 |
| 2588.5 | 3741 | 3273.5 | 1084 | 4019.5 | 1831 | 678.5 | 9177 | 4339.5 | 6145 |
| 2466 | 3705 | 3338 | 1098.5 | 4270.5 | 1863.5 | 694 | 9738.5 | 4358.5 | 6297.5 |
| 59968 | 31619 | 10141 | 1593 | 4507 | 1875 | 687 | 15133 | 5878 | 7384 |
| 3054.5 | 4413 | 3400.5 | 1230.5 | 4316 | 1841.5 | 703 | 10221.5 | 4778.5 | 6454 |
| 2657 | 3738 | 3361 | 1133 | 5075 | 1969 | 711 | 9660 | 4448 | 6124 |
| 2597.5 | 3824.5 | 3188.5 | 1137 | 4325.5 | 1760 | 737 | 9636.5 | 4315 | 6104 |
| 10385 | 20948 | 9756 | 1990 | 11002 | 2721 | 708 | 15200 | 9671 | 22825 |
| 10316.5 | 22275 | 12027.5 | 2503.5 | 12935 | 3443.5 | 702 | 12401 | 8764.5 | 23546 |
| 10304.5 | 23314.5 | 11492.5 | 2484 | 13191 | 3218.5 | 704 | 12492.5 | 9135 | 23129 |
| 10496 | 22376 | 9952 | 2060 | 10816 | 2728 | 713 | 13517 | 9737 | 23452 |
| 9958 | 22615.5 | 7332 | 1784.5 | 8796.5 | 2479 | 714.5 | 14541 | 9832.5 | 24787.5 |
| 14350 | 131696.5 | 201152 | 73980.5 | 7790.5 | 2719 | 663 | 15238.5 | 10339.5 | 72138.5 |
| 9283 | 14864 | 5229 | 1468.5 | 5434 | 1990 | 786.5 | 15723 | 5440 | 15286 |
| 10511 | 7417 | 30660 | 8337 | 146130 | 20697 | 756 | 17985 | 4610 | 8579 |
| 10967.5 | 6762.5 | 7790 | 2119 | 36338.5 | 4996 | 906 | 16146.5 | 5273 | 7994 |
| 10197.5 | 5821.5 | 4499.5 | 1402.5 | 13323 | 2716 | 808 | 13467 | 5238 | 6711.5 |
| 9366 | 5537 | 4789.5 | 1467 | 8165 | 2401.5 | 803 | 12497.5 | 5399.5 | 6554 |
| 9210 | 5511 | 3969.5 | 1288.5 | 5989 | 2074 | 775 | 12451.5 | 5017.5 | 6863.5 |
| 7992 | 5386.5 | 3928 | 1223.5 | 5444 | 1928.5 | 743 | 13326.5 | 4910 | 6683 |
| 5738.5 | 5679 | 4628 | 1557 | 4765 | 1954 | 754 | 11882 | 4772 | 6462.5 |
| 7894.5 | 5144.5 | 3694 | 1237.5 | 4699.5 | 1924 | 749 | 12019 | 4826 | 6593.5 |
| 5979 | 4459 | 3396 | 1188 | 4399 | 1851 | 804 | 11499 | 4468 | 6260 |
| 7518 | 5106 | 3603 | 1218 | 4704 | 1904 | 766 | 11219 | 4500 | 6235 |
| 16972 | 9206 | 4885 | 1419 | 4608 | 1940 | 759 | 10855 | 5905 | 7458 |
| 9770 | 5542 | 3638 | 1195 | 4646 | 1848 | 769 | 10630 | 4838 | 6763 |
| 7956 | 4923 | 3705 | 1218 | 4882 | 1904 | 755 | 11353 | 5263 | 6769 |
| 406382 | 250982 | 73612 | 4892 | 4896 | 1889 | 748 | 47846 | 13296 | 10826 |
| 28117 | 14040.5 | 5469.5 | 1349 | 4749.5 | 1918.5 | 798 | 13906.5 | 5742 | 7356.5 |
| 16435 | 7497 | 4224 | 1225 | 5095 | 2176 | 802 | 11983 | 5142 | 6626 |
| 16066 | 8227 | 4217 | 1290 | 5117 | 1947.5 | 777.5 | 11713.5 | 5206 | 6680.5 |
| 16744 | 25485 | 10511 | 2247 | 11787 | 2812 | 796 | 16359 | 10535 | 23141 |
| 17342 | 29959 | 13854 | 2802 | 13962 | 3419 | 809 | 13381 | 9780 | 23188 |
| 15292 | 26586.5 | 13444 | 2625.5 | 13759.5 | 3389.5 | 793 | 13570 | 10089 | 23816.5 |

| | | | | | | | | | |
|---|---|---|---|---|---|---|---|---|---|
| 15446 | 24673 | 10613 | 2191 | 11241 | 2827 | 811 | 13455 | 9747 | 22059 |
| 9611 | 23171 | 7492 | 1849 | 8784 | 2522 | 711 | 14751 | 10466 | 25141 |
| 10099.5 | 98279 | 148198 | 53788.5 | 6988 | 2396.5 | 655 | 16187 | 8638.5 | 51826.5 |
| 4528 | 11061.5 | 4587 | 1400 | 5461 | 1986.5 | 713 | 13378 | 5270 | 14024 |
| 2836 | 5141.5 | 6172 | 1777.5 | 582745.5 | 56141 | 703.5 | 16011.5 | 4134 | 7434 |
| 2787 | 4186 | 3825.5 | 1413 | 6182.5 | 2197.5 | 862 | 16470 | 5316 | 7312 |
| 2910.5 | 4183 | 3441.5 | 1251.5 | 5241.5 | 2044.5 | 740 | 12063.5 | 4949.5 | 6748 |
| 2778 | 3939 | 3509 | 1185 | 4478 | 1926 | 711 | 10319 | 4479 | 6126 |
| 2900 | 4021 | 3532 | 1208 | 4734 | 1902 | 715 | 11361 | 4620 | 6349 |
| 2773 | 3948 | 3357.5 | 1135 | 4381.5 | 1815.5 | 687.5 | 11788.5 | 4304.5 | 6202.5 |
| 2758 | 3936 | 3201 | 1229 | 4223 | 1823 | 678 | 10618 | 4354 | 6215 |
| 2710 | 3873.5 | 3266.5 | 1218.5 | 4330.5 | 1907.5 | 699 | 11355.5 | 4528.5 | 6254.5 |
| 2472 | 3699 | 3194 | 1078 | 4118 | 1737 | 692 | 9933 | 4479 | 5992 |
| 2285 | 3496 | 2964.5 | 1155 | 4015 | 1709.5 | 685.5 | 9350 | 4074.5 | 5867 |
| 28149 | 18669 | 6383 | 1343 | 4225.5 | 1828.5 | 721 | 9830 | 10615 | 8983 |
| 2430.5 | 3736.5 | 3192 | 1102.5 | 4069.5 | 1696 | 710 | 9475.5 | 4108 | 5884.5 |
| 2533.5 | 3801 | 3370.5 | 1165 | 4260.5 | 1831.5 | 684.5 | 9764 | 4458 | 6147 |
| 91113 | 49594 | 14305 | 1695 | 4499 | 1883 | 712 | 17819 | 5999 | 7563 |
| 3479 | 4766.5 | 3830.5 | 1245 | 4517.5 | 1892 | 718.5 | 10778.5 | 4831.5 | 6390.5 |
| 2561 | 3896 | 3307 | 1200 | 5377 | 2048 | 711 | 9881 | 4614 | 6271 |
| 2642.5 | 3870.5 | 3198.5 | 1084 | 4116 | 1726 | 718 | 9798.5 | 4512.5 | 6089 |
| 10377 | 21868 | 9755 | 2091 | 10658 | 2652 | 688 | 14089 | 9021 | 22937 |
| 10132 | 22150 | 11803 | 2539 | 13065 | 3303 | 708 | 12612 | 8917 | 22975 |
| 10774 | 21881 | 11184 | 2585 | 13017 | 3189 | 703 | 12727 | 9293 | 21775 |
| 9847 | 21717 | 9812 | 1916 | 10819 | 2730 | 707 | 13769 | 9094 | 22052 |
| 9909 | 21868.5 | 7122.5 | 1764 | 8250.5 | 2381 | 679.5 | 14766.5 | 10145 | 24479.5 |
| 10030.5 | 105159 | 163968 | 59944 | 7457.5 | 2756 | 655 | 15278.5 | 8633.5 | 59424.5 |
| 4405 | 11034.5 | 4938.5 | 1603.5 | 5637.5 | 2087.5 | 705 | 14667 | 5267 | 14577.5 |
| 2812.5 | 5580 | 6745.5 | 1981 | 522186 | 49078 | 698 | 17132.5 | 4427 | 8846.5 |
| 2491.5 | 4498 | 4124 | 1494 | 6482 | 2498.5 | 782 | 13823 | 4622.5 | 8055.5 |
| 2666 | 4024 | 3890 | 1463 | 5902 | 2276 | 749 | 12325 | 4819 | 7149 |
| 2347 | 3902 | 3585 | 1336 | 5269 | 2056 | 704 | 10537 | 4495 | 7085 |
| 2557 | 4141.5 | 3735 | 1384.5 | 5082.5 | 2045.5 | 730.5 | 12176 | 4685.5 | 7287.5 |
| 2730 | 4104 | 3550 | 1272 | 4812 | 1990 | 681 | 12064 | 4590 | 6878 |
| 2376.5 | 3953.5 | 3353 | 1297.5 | 4635.5 | 1942 | 665 | 10421.5 | 4093.5 | 6396.5 |
| 2623 | 3950 | 3565 | 1270 | 4910 | 2018 | 657 | 11183 | 4377 | 6509 |
| 2376 | 3772.5 | 3349 | 1265.5 | 4650.5 | 1997.5 | 709.5 | 11373.5 | 4550.5 | 6802.5 |
| 2436 | 3541 | 3374 | 1284 | 4524 | 1948 | 685 | 9973 | 4631 | 6596 |
| 7991.5 | 6596 | 4373 | 1439.5 | 4715 | 2047.5 | 711.5 | 10559 | 6289.5 | 7319 |
| 2494 | 3702 | 3341 | 1208 | 4486 | 1927 | 712 | 9765 | 4801 | 6679 |
| 2228.5 | 3792.5 | 3375 | 1304 | 4842.5 | 2027.5 | 667.5 | 10369 | 4323.5 | 6877 |
| 52461 | 47138 | 9809 | 1647 | 5155 | 2151 | 693 | 16454 | 5337 | 12339 |
| 3001.5 | 7107.5 | 4010.5 | 1323 | 5051.5 | 2097.5 | 699 | 11703.5 | 4387.5 | 10401 |
| 2332 | 3967 | 3478 | 1268 | 5631 | 2159 | 722 | 10999 | 4413 | 7051 |
| 2615 | 4139 | 3326 | 1227 | 4569 | 2016 | 704 | 10504 | 4418 | 6727 |
| 9598 | 19635 | 10140 | 2233 | 10882 | 2854 | 696 | 15138 | 9384 | 20973 |
| 9907.5 | 22191 | 13202.5 | 2958 | 14704 | 3571.5 | 707.5 | 12213.5 | 8931.5 | 23759 |
| 9528 | 21039.5 | 11702.5 | 2543.5 | 13282 | 3303.5 | 704.5 | 12865.5 | 9100.5 | 23735 |

| | | | | | | | | | |
|---|---|---|---|---|---|---|---|---|---|
| 9068.5 | 20102.5 | 10577.5 | 2464.5 | 11853 | 3067 | 682.5 | 13000.5 | 8996.5 | 20924.5 |
| 9519 | 22081 | 7227 | 1853 | 8313 | 2532 | 710 | 14539 | 9941 | 24888 |
| 50485 | 356406 | 470364 | 129552 | 13251 | 5181 | 614 | 19630 | 12765 | 90357 |
| 7832.5 | 34071 | 24985 | 6446 | 7477 | 2539.5 | 684.5 | 13032 | 6226 | 21201.5 |
| 4262.5 | 5996.5 | 17773.5 | 4923 | 157206 | 27770 | 664 | 15885 | 4432 | 7382.5 |
| 6920 | 6345 | 10458 | 2250 | 47760 | 6687 | 1828 | 110193 | 19252 | 17563 |
| 4065 | 4749 | 5426 | 1741 | 34027 | 4808 | 727 | 11584 | 4910 | 6759 |
| 3379 | 4326 | 3930 | 1423 | 25366 | 3687 | 689 | 11912 | 5642 | 7279 |
| 3255 | 4353 | 3731 | 1322 | 22249 | 3378 | 710 | 11058 | 4655 | 6155 |
| 3806 | 4715 | 4058 | 1462 | 16867 | 3042 | 722 | 12221 | 5000 | 6541 |
| 4243.5 | 6383 | 6397.5 | 2076 | 15853.5 | 3367.5 | 791 | 13388 | 5188.5 | 7810.5 |
| 3432 | 4518.5 | 3945.5 | 1363 | 13030.5 | 2610 | 703.5 | 11470.5 | 4527.5 | 6461.5 |
| 3296.5 | 4251 | 3781 | 1406.5 | 8843 | 2365 | 660 | 10298 | 4460 | 6303.5 |
| 3827.5 | 4324 | 3927 | 1423 | 11299 | 2559 | 717 | 10510.5 | 4727 | 6422.5 |
| 34020.5 | 23987.5 | 19853.5 | 2734 | 10606 | 2458 | 694 | 9071 | 12187.5 | 10647 |
| 4806 | 5297 | 4863 | 1625 | 9387 | 2443 | 707 | 9718 | 4910 | 6868 |
| 3689 | 4678 | 4155 | 1412 | 9778 | 2386 | 732 | 10587 | 4699 | 6655 |
| 366043.5 | 219592.5 | 81106 | 7825 | 8801.5 | 2357.5 | 696.5 | 42247 | 12392 | 10280.5 |
| 8880 | 6417 | 4760 | 1507 | 8568 | 2331 | 695 | 11248 | 4767 | 7002 |
| 3854.5 | 4774.5 | 4168.5 | 1457.5 | 21854.5 | 5439 | 720 | 10391 | 4624 | 6670.5 |
| 3763 | 4552 | 4032 | 1412 | 7124 | 2168 | 709 | 10099 | 4625 | 6100 |
| 10068 | 21137 | 10627 | 2286 | 13698 | 3291 | 699 | 13285 | 9137 | 21602 |
| 10934 | 23871.5 | 13265 | 2944.5 | 16299 | 3654.5 | 710 | 12321.5 | 9401 | 23469 |
| 9727 | 22345 | 13048 | 2962 | 15791 | 3456 | 703 | 12379 | 8752 | 21823 |
| 9539 | 21439 | 10476 | 2358 | 12529 | 3030 | 678 | 12898 | 8846 | 21581 |
| 10179 | 21173 | 7442 | 1820 | 8144 | 2447 | 720 | 14860 | 9815 | 22982 |
| 14258 | 151508 | 254960.5 | 84396.5 | 12789 | 5777 | 620 | 15480 | 10409.5 | 85973 |
| 5583 | 13398 | 9026 | 3066 | 5955 | 2459 | 687 | 14240 | 5656 | 15174 |
| 3412.5 | 5577.5 | 9020 | 2841 | 521575 | 52850 | 667 | 15968.5 | 4376 | 7464.5 |
| 3102 | 4518 | 4547.5 | 1624 | 8106.5 | 2526.5 | 805.5 | 14690.5 | 5485 | 7465 |
| 2983.5 | 4095.5 | 3941.5 | 1434 | 5530.5 | 2179 | 735 | 11598 | 4860.5 | 6680.5 |
| 2844 | 3890 | 3570 | 1257 | 4788 | 2001 | 747 | 9706 | 4862 | 6333 |
| 2982 | 4288 | 3634 | 1318 | 4781 | 1921 | 708 | 11490 | 4729 | 6610 |
| 2845 | 4520 | 3494 | 1241 | 4701 | 1916 | 723 | 11913 | 4533 | 6865 |
| 2645.5 | 4759 | 3918.5 | 1513 | 4867 | 2128 | 702.5 | 12846 | 4223.5 | 6779.5 |
| 2784 | 4559 | 3911 | 1343 | 4924 | 2050 | 737 | 12797 | 4313 | 6818 |
| 2588 | 3994 | 3384.5 | 1194 | 4316.5 | 1875 | 694 | 10565 | 4286 | 6342 |
| 2677 | 4099 | 3268 | 1267.5 | 4576 | 1875 | 707.5 | 10732.5 | 4321 | 6500 |
| 26001.5 | 19344.5 | 6956 | 1538 | 4633.5 | 1951 | 731.5 | 11388 | 9161.5 | 9655.5 |
| 2878 | 4479 | 3546 | 1295 | 4368 | 1947 | 725 | 10830 | 4558 | 6747 |
| 2456 | 4427.5 | 3577 | 1188.5 | 4626.5 | 1854 | 701 | 10904 | 4331.5 | 7103.5 |
| 79683 | 51305 | 14808 | 1759 | 4906 | 1885 | 708 | 18284 | 5882 | 8209 |
| 3401 | 7947 | 4436 | 1390 | 5144 | 2017 | 762 | 13158 | 4522 | 10631 |
| 2431 | 4232 | 3490 | 1242 | 5382 | 2034 | 694 | 11383 | 4238 | 6740 |
| 3055.5 | 4332 | 3672 | 1202.5 | 4571 | 1855 | 676 | 11065.5 | 4581 | 6640.5 |
| 10349.5 | 21311.5 | 10382 | 2455 | 11740 | 2980.5 | 717.5 | 13440 | 8810.5 | 21495 |
| 9936 | 23803 | 13592 | 2890 | 14505 | 3406 | 704 | 12010 | 8596 | 22569 |
| 9819.5 | 21741 | 11571 | 2546 | 12802.5 | 3160 | 721.5 | 12458 | 8756.5 | 21870 |

| | | | | | | | | | |
|---|---|---|---|---|---|---|---|---|---|
| 9391 | 21600 | 10523 | 2217 | 11659 | 2767 | 709 | 13083 | 8602 | 21853 |
| 9322.5 | 20912 | 6554 | 1682.5 | 7502 | 2321 | 696 | 15634.5 | 10017.5 | 23403 |
| 16001 | 146668 | 234628 | 83955 | 8990 | 3514 | 642 | 15549 | 9797 | 69838 |
| 4328 | 11882 | 5264 | 1735 | 5363 | 2039 | 664 | 13577 | 5154 | 15138 |
| 3176 | 5603 | 7160 | 2220 | 528760 | 52079 | 687 | 15991 | 4502 | 7686 |
| 2929 | 4458 | 4205 | 1481 | 5988 | 2338 | 750 | 13474 | 4904 | 7484 |
| 2876 | 4287 | 3736 | 1303 | 4901 | 1983 | 707 | 10896 | 4573 | 6401 |
| 2828 | 4336 | 3633 | 1358 | 5082 | 2011 | 734 | 10784 | 4740 | 6753 |
| 2870.5 | 4348 | 3540 | 1280 | 4692.5 | 1917 | 686.5 | 11236.5 | 4485.5 | 6466 |
| 2735 | 4442.5 | 3392 | 1200.5 | 4487 | 1944.5 | 669.5 | 11804.5 | 4317.5 | 6366 |
| 2885 | 5030 | 4026 | 1524 | 5145 | 2155 | 722 | 13865 | 4255 | 7374 |
| 2951 | 4375 | 3559.5 | 1306 | 4623.5 | 1939 | 697.5 | 11378.5 | 4634.5 | 6489 |
| 2735 | 4278 | 3721.5 | 1259 | 4727 | 1975.5 | 719.5 | 10792 | 4631 | 6816.5 |
| 2697.5 | 4241.5 | 3615 | 1293 | 4812.5 | 1927 | 746 | 9928.5 | 4416 | 6729.5 |
| 12283 | 9912 | 5463.5 | 1579.5 | 4886.5 | 2000.5 | 763 | 10406 | 6501.5 | 8435.5 |
| 2967 | 4414 | 3633 | 1305 | 4728 | 1981 | 747 | 10948 | 4737 | 6898 |
| 2738 | 4034 | 3543 | 1335 | 4538 | 1879 | 674 | 10019 | 4481 | 6429 |
| 110464.5 | 60586.5 | 17944 | 1932 | 4811 | 2011 | 697 | 18372.5 | 6549.5 | 7917.5 |
| 3333.5 | 5936 | 4380.5 | 1431 | 5339 | 2064.5 | 749.5 | 11489 | 4647 | 8012.5 |
| 2685 | 4607 | 3976 | 1484 | 6006 | 2198 | 759 | 11583 | 4359 | 7217 |
| 3034 | 4394 | 3626 | 1255 | 4556 | 1874 | 693 | 10784 | 4670 | 6352 |
| 10202 | 20547 | 10862 | 2351 | 11510 | 2876 | 711 | 13992 | 9441 | 20539 |
| 8864.5 | 26205 | 13643.5 | 2995 | 15139.5 | 3739.5 | 730.5 | 10104.5 | 8283.5 | 26912.5 |
| 9765 | 24185 | 13617 | 3035 | 14584 | 3530 | 718 | 11354 | 8405 | 24670 |
| 9663 | 20308 | 10172 | 2216 | 11148 | 2829 | 677 | 12470 | 8734 | 20913 |
| 942.5 | 1378 | 1150.5 | 663.5 | 1768 | 1231 | 339 | 3186.5 | 1833 | 2474 |
| 81209 | 44710.5 | 13062.5 | 1763 | 3573.5 | 1602 | 474 | 18576.5 | 5010 | 6055 |
| 5894 | 5426 | 3442 | 1011 | 3364 | 1573 | 577 | 10637 | 3690 | 5035 |
| 2783 | 3939 | 7984 | 2390 | 466092 | 45121 | 642 | 8955 | 3595 | 5094 |
| 2237.5 | 3652.5 | 3441 | 1170 | 5002.5 | 2030.5 | 727.5 | 9472.5 | 4042.5 | 5675 |
| 2042.5 | 3395.5 | 2947 | 1088.5 | 4270 | 1857 | 699 | 8831.5 | 3908.5 | 5490 |
| 1829.5 | 3129.5 | 2780.5 | 1024 | 3929.5 | 1743 | 677.5 | 8223 | 3637 | 5308.5 |
| 2353 | 3693 | 2906 | 1140 | 4153 | 1812 | 699 | 9879 | 4122 | 5951 |
| 2379 | 3605 | 3027.5 | 1093 | 3984 | 1794.5 | 650.5 | 10237 | 4127.5 | 5702.5 |
| 2448 | 3627 | 3169 | 1257 | 3992 | 1810 | 670 | 9827 | 4314 | 5880 |
| 2170 | 3353 | 2936.5 | 1171 | 4090.5 | 1834.5 | 704 | 9528.5 | 4035.5 | 5805.5 |
| 1856.5 | 3285.5 | 2933 | 1118 | 3906 | 1771 | 694.5 | 9373.5 | 3737 | 5668 |
| 1870 | 3028.5 | 2790.5 | 1037 | 3976.5 | 1736.5 | 690 | 8735 | 3694 | 5519 |
| 18918.5 | 13372.5 | 5122 | 1238 | 3961 | 1796.5 | 727.5 | 8819 | 7651 | 7813.5 |
| 2098 | 3401 | 3044 | 1089 | 3906 | 1766 | 721 | 8330 | 3904 | 5875 |
| 2002.5 | 3421.5 | 2859.5 | 1094.5 | 3970 | 1734.5 | 673.5 | 8600.5 | 4126.5 | 6042.5 |
| 102917 | 52490.5 | 15551.5 | 1652 | 4397.5 | 1898 | 720 | 17379 | 6246 | 7104 |
| 2677 | 4117.5 | 3376.5 | 1170.5 | 4202.5 | 1804 | 687 | 10111.5 | 4511.5 | 6155.5 |
| 2165.5 | 3439 | 2972 | 1106 | 5175 | 1933.5 | 668 | 9062.5 | 4426.5 | 5764.5 |
| 2245 | 3361 | 2931 | 1033 | 4161 | 1776 | 702 | 9332 | 4041 | 5647 |
| 9748.5 | 19034 | 9502 | 2070 | 10701.5 | 2821.5 | 681.5 | 14775.5 | 9456 | 20582 |
| 9986 | 22415 | 12514 | 2642.5 | 13850.5 | 3486 | 696.5 | 12274.5 | 8883 | 22584.5 |
| 9769.5 | 21168.5 | 11603 | 2532.5 | 13079.5 | 3160.5 | 654 | 12421.5 | 8848.5 | 22223 |

| | | | | | | | | | |
|---|---|---|---|---|---|---|---|---|---|
| 8692.5 | 19875 | 8984 | 1932.5 | 10208 | 2606.5 | 674 | 13162 | 9139.5 | 21913 |
| 8756 | 19272 | 6497 | 1627 | 7621 | 2312 | 700 | 14743 | 9569 | 22460 |
| 9528 | 97779.5 | 153879.5 | 60238 | 6549 | 2551.5 | 661 | 15179 | 8169.5 | 55395 |
| 4410 | 9204 | 4695 | 1426 | 5009 | 1969 | 742 | 14852 | 5303 | 11330 |
| 3171 | 4944 | 7557.5 | 2053 | 835369 | 78344 | 712.5 | 16483.5 | 4455 | 7029 |
| 2990.5 | 4395 | 3990 | 1402 | 6153 | 2331 | 856 | 14522 | 5250.5 | 7188.5 |
| 2868 | 3964.5 | 3498 | 1227 | 4803 | 1968.5 | 760.5 | 11754 | 5007.5 | 6512 |
| 2618 | 3874.5 | 3279.5 | 1177 | 4661 | 1959.5 | 752.5 | 10759 | 4556 | 6195.5 |
| 2820 | 3874.5 | 3208.5 | 1121 | 4304.5 | 1761 | 729.5 | 11056.5 | 4652 | 6159 |
| 2999 | 4099 | 3186 | 1235 | 4104 | 1806 | 713 | 11642 | 4815 | 6135 |
| 2628 | 4121 | 3131 | 1270 | 4326 | 1890 | 717 | 10581 | 4444 | 6234 |
| 2531 | 3659.5 | 3015 | 1173.5 | 4092.5 | 1873.5 | 729.5 | 10058.5 | 4694 | 5986 |
| 2212 | 3640.5 | 3013.5 | 1150.5 | 4075 | 1830.5 | 739 | 9881.5 | 4172.5 | 6074 |
| 2251.5 | 3504 | 3049 | 1109.5 | 3975 | 1852.5 | 738 | 9337 | 4040 | 5829.5 |
| 36390 | 25884.5 | 7994.5 | 1410 | 4010.5 | 1721 | 747.5 | 9668 | 12522.5 | 10562.5 |
| 2429.5 | 3616.5 | 3108.5 | 1193.5 | 4007.5 | 1813 | 726 | 9690.5 | 4379 | 6100.5 |
| 2500 | 3877 | 3245 | 1213 | 4415 | 1887 | 766 | 10698 | 4967 | 6558 |
| 155183 | 80179 | 22747 | 1923 | 4572 | 1864 | 735 | 22393 | 7485 | 7947 |
| 3424 | 4849 | 3626.5 | 1288.5 | 4356 | 1850.5 | 752.5 | 11756.5 | 4950 | 6660.5 |
| 2579.5 | 3746 | 3377.5 | 1218.5 | 6492.5 | 2177 | 749.5 | 10689.5 | 4857.5 | 6409 |
| 2624 | 3875 | 3168 | 1148 | 4139 | 1810 | 721 | 10133 | 4427 | 6132 |
| 10529 | 22321 | 9855 | 2133.5 | 10849 | 2809 | 756.5 | 14383.5 | 9776.5 | 23380.5 |
| 12217 | 24101 | 13349 | 2551 | 13828 | 3361 | 748 | 13001 | 9452 | 23606 |
| 11721 | 24167.5 | 11905 | 2499 | 13439.5 | 3281 | 726 | 12916.5 | 9463.5 | 22511.5 |
| 11702 | 22864.5 | 10475 | 2138 | 11085.5 | 2942 | 776.5 | 13982 | 9607 | 24623 |
| 9210 | 23010.5 | 7764.5 | 1791.5 | 8736.5 | 2522 | 740 | 14715 | 10059 | 26522 |
| 11298 | 34391 | 19647 | 6231 | 7608 | 2613 | 809 | 16212 | 9732 | 31154 |
| 3696 | 10461 | 4259 | 1318 | 5328 | 1966 | 743 | 14594 | 5040 | 13936 |
| 2979.5 | 5987.5 | 6600 | 2045.5 | 644280.5 | 60764 | 751 | 17224.5 | 4605 | 8320 |
| 2945.5 | 4705.5 | 4588.5 | 1585 | 7454 | 2582 | 857 | 14084.5 | 5074 | 7521.5 |
| 2837 | 4391 | 3940 | 1415 | 5662 | 2233 | 760.5 | 11691.5 | 4769.5 | 6961 |
| 2750 | 4168 | 3990 | 1402 | 5527 | 2083 | 745 | 11068 | 4604 | 6411 |
| 2797 | 4256.5 | 3505.5 | 1217 | 4906 | 1978 | 750 | 11777 | 4724 | 6887.5 |
| 2753 | 4083.5 | 3280.5 | 1160 | 4399.5 | 1825 | 711 | 11097 | 4716 | 6380.5 |
| 2758.5 | 4180 | 3517 | 1354 | 4513 | 2003.5 | 723.5 | 11169 | 4511.5 | 6405 |
| 2410 | 3818 | 3294 | 1199 | 4362 | 1927 | 709 | 9936 | 4507 | 6134 |
| 2387 | 3640.5 | 3161.5 | 1150.5 | 4315.5 | 1867.5 | 713 | 9987 | 4153 | 6125 |
| 2273 | 3546 | 3169 | 1150 | 4221 | 1873 | 718 | 9787 | 4079 | 5916 |
| 22113 | 15293 | 5604 | 1226 | 4121 | 1787 | 698.5 | 8961.5 | 8427 | 8452 |
| 2464 | 3743 | 3262 | 1143 | 4224 | 1863 | 705 | 9060 | 4283 | 5997 |
| 2461 | 3791 | 3434 | 1155 | 4312 | 1821 | 727 | 9875 | 4642 | 6343 |
| 160980.5 | 82886.5 | 23237 | 2024 | 4431.5 | 1853 | 705.5 | 21760 | 7427.5 | 7761 |
| 3105 | 4707.5 | 3663.5 | 1213 | 4445.5 | 1884 | 711.5 | 10946 | 4902.5 | 6594.5 |
| 2574 | 3590.5 | 3141.5 | 1115 | 5752 | 2057 | 709.5 | 9867 | 4758.5 | 6288 |
| 2452 | 3737 | 2959 | 1120 | 4309 | 1798 | 693 | 9529 | 4264 | 6127 |
| 9603 | 21815 | 9560 | 2060 | 11134 | 2722 | 702 | 14131 | 9151 | 24043 |
| 10480.5 | 22224 | 12261 | 2647 | 13410 | 3191 | 722 | 13040 | 9377.5 | 22775.5 |
| 10406 | 21751 | 11976 | 2498.5 | 12837 | 3306 | 717 | 12409.5 | 8965 | 23025.5 |

| | | | | | | | | | |
|---|---|---|---|---|---|---|---|---|---|
| 10357 | 21679 | 10174.5 | 2211 | 11532.5 | 2976.5 | 708.5 | 14278.5 | 9406.5 | 22496.5 |
| 8869.5 | 21209 | 6976.5 | 1719.5 | 7980 | 2368 | 688 | 14354.5 | 9827 | 23760 |
| 11004.5 | 112940.5 | 179242.5 | 66141 | 8665.5 | 3800.5 | 663 | 15198.5 | 9182 | 60492 |
| 4545.5 | 12270.5 | 5044 | 1585.5 | 5348.5 | 2101 | 740.5 | 14159 | 5120.5 | 15535.5 |
| 3014.5 | 5275 | 7042 | 2315.5 | 736184 | 68034.5 | 684.5 | 16631.5 | 4431 | 7275.5 |
| 3082 | 4728 | 4853 | 1842 | 7255 | 2856 | 859 | 14206 | 5044 | 7472 |
| 2976 | 4250 | 4113 | 1434 | 5840 | 2265 | 731 | 11590 | 4894 | 6781 |
| 2644 | 3928 | 3505.5 | 1255 | 4666.5 | 1933 | 723.5 | 10521 | 4728.5 | 6390 |
| 3046.5 | 3924 | 3594.5 | 1225.5 | 4605.5 | 1847.5 | 724 | 10633 | 4734.5 | 6304 |
| 3001 | 4030 | 3383.5 | 1148 | 4427 | 1837 | 705 | 11874.5 | 4577 | 6263.5 |
| 2961 | 4019 | 3587 | 1363 | 4485 | 1920 | 717 | 11640 | 4741 | 6153 |
| 2944 | 3917 | 3356 | 1161 | 4219 | 1841 | 737 | 11196 | 4630 | 6325 |
| 2322.5 | 3508.5 | 2965.5 | 1138.5 | 4026 | 1730.5 | 703.5 | 9924.5 | 4121 | 5917.5 |
| 2402 | 3487 | 3076.5 | 1155 | 4110 | 1835 | 692.5 | 9686 | 4151.5 | 5965.5 |
| 28747 | 19990 | 6474 | 1344 | 3974 | 1781 | 709 | 9278 | 10289 | 9061 |
| 2485 | 3685.5 | 3219.5 | 1141 | 3977.5 | 1782.5 | 707.5 | 9501.5 | 4226.5 | 6069.5 |
| 2412.5 | 3816.5 | 3469.5 | 1171 | 4320 | 1860 | 754.5 | 10237.5 | 4597.5 | 6417.5 |
| 141275.5 | 74288.5 | 21251 | 1943 | 4568 | 1864.5 | 696 | 20760 | 7229.5 | 7892.5 |
| 3519 | 5064 | 3849 | 1175 | 4776 | 1830 | 756 | 11523 | 5022 | 6729 |
| 2572 | 3989 | 3422 | 1227 | 5810 | 2106 | 732 | 10508 | 4865 | 6322 |
| 2699 | 3974 | 3328 | 1107 | 4193 | 1825 | 749 | 10847 | 4605 | 6175 |
| 10046 | 23978 | 7913 | 1928 | 9275 | 2578 | 726 | 14867 | 9628 | 26024 |
| 11347 | 24030 | 12924 | 2730 | 13993 | 3380 | 732 | 12187 | 9614 | 24252 |
| 11035 | 23735.5 | 12224.5 | 2630.5 | 13281.5 | 3329 | 732 | 12870.5 | 9470 | 24445.5 |
| 10931 | 22088 | 10323 | 2205 | 11245 | 2775 | 733 | 13726 | 9612 | 22730 |
| 8998 | 22213 | 7426 | 1821 | 8452 | 2523 | 728 | 14832 | 10007 | 25294 |
| 8643.5 | 25321 | 13115.5 | 4653 | 7172 | 2767.5 | 721.5 | 16692 | 7585 | 26736.5 |
| 4154.5 | 9855 | 4677 | 1416.5 | 5233 | 1903.5 | 733 | 15619.5 | 5284 | 11847 |
| 3448 | 6569 | 7580 | 2099 | 643980 | 64375 | 739 | 17424 | 4737 | 9206 |
| 3056 | 4769 | 4633 | 1601.5 | 6956 | 2401 | 872.5 | 14862 | 5166 | 7226 |
| 2951.5 | 4306.5 | 3818 | 1379.5 | 5531.5 | 2160.5 | 767.5 | 12069 | 4971 | 6872.5 |
| 2755.5 | 4099 | 3461 | 1208.5 | 4894 | 1894 | 768.5 | 11065.5 | 4816.5 | 6920.5 |
| 2978 | 4124 | 3430 | 1249 | 4714 | 1904 | 777 | 11713 | 4720 | 6600 |
| 3141 | 4222 | 3513 | 1228 | 4685 | 1826 | 736 | 11770 | 4784 | 6598 |
| 2845.5 | 4107 | 3451.5 | 1296.5 | 4347 | 1849.5 | 728.5 | 10734 | 4731 | 6376 |
| 2881 | 3970 | 3356 | 1225 | 4463 | 1817 | 716 | 11232 | 4778 | 6308 |
| 2491 | 3625 | 3117 | 1121 | 4080 | 1835 | 746 | 9945 | 4316 | 6116 |
| 2378.5 | 3583 | 3177 | 1190 | 4159.5 | 1829.5 | 724.5 | 10311 | 4419 | 6072.5 |
| 24686 | 16362.5 | 5897 | 1367.5 | 4264 | 1898.5 | 724.5 | 9479.5 | 9217 | 8546 |
| 2641.5 | 3644.5 | 3327.5 | 1198.5 | 4299.5 | 1793.5 | 745.5 | 9411.5 | 4523 | 6145 |
| 2941.5 | 3948 | 3550.5 | 1212.5 | 4518.5 | 1935 | 744 | 10302.5 | 4736.5 | 6478 |
| 89364 | 46784 | 13630 | 1668 | 4589 | 1920 | 726 | 16754 | 6171 | 7613 |
| 3288.5 | 4513.5 | 3798.5 | 1219 | 4551 | 1840 | 747.5 | 11286.5 | 5062.5 | 7062 |
| 2914.5 | 3832.5 | 3485.5 | 1158.5 | 5454.5 | 2080.5 | 731 | 10253.5 | 4682 | 6293.5 |
| 3026.5 | 4127 | 3531 | 1226.5 | 4422.5 | 1909 | 746 | 10856 | 4705 | 6496 |
| 10155 | 24214 | 9321 | 2062 | 10072 | 2713 | 738 | 15963 | 9721 | 25207 |
| 11499.5 | 25267 | 14620.5 | 2975.5 | 14988.5 | 3612 | 765.5 | 11853 | 8864 | 25599.5 |
| 11163 | 24451 | 12886 | 2777 | 14226 | 3494 | 756 | 12675 | 9333 | 24586 |

| | | | | | | | | | |
|---|---|---|---|---|---|---|---|---|---|
| 10995.5 | 22228 | 11155 | 2331.5 | 12376.5 | 3033 | 738 | 12995.5 | 9611.5 | 22739.5 |
| 8830 | 24170 | 7735 | 1788 | 9087 | 2599 | 730 | 15669 | 9663 | 27016 |
| 10545 | 25275.5 | 11401 | 3861 | 7561.5 | 2925 | 714.5 | 15859 | 10030.5 | 26357.5 |
| 4033 | 11001 | 4495 | 1419 | 5527 | 2135 | 744 | 13674 | 4980 | 14498 |
| 2665 | 5251 | 6500 | 1842 | 635364 | 57856 | 738 | 16173 | 4093 | 8106 |
| 2551 | 4319 | 4216.5 | 1422 | 6449.5 | 2497 | 794 | 12620 | 4707.5 | 7197.5 |
| 2687 | 4209 | 3815 | 1361 | 5531.5 | 2196.5 | 789.5 | 11209.5 | 4756.5 | 6974.5 |
| 2603 | 3806 | 3499 | 1249.5 | 4871 | 1971 | 742 | 10473.5 | 4679.5 | 6624.5 |
| 2560 | 3897 | 3443 | 1222 | 4576 | 1912 | 730 | 11443 | 4456 | 6649 |
| 2798 | 3979 | 3259.5 | 1158 | 4550.5 | 1855 | 703 | 11616.5 | 4480.5 | 6602.5 |
| 2706 | 4149 | 3486 | 1274 | 4467 | 1846 | 698 | 10916 | 4397 | 6425 |
| 2705 | 3918.5 | 3374.5 | 1240 | 4454.5 | 1856 | 700 | 10391 | 4591 | 6522 |
| 2496.5 | 3563.5 | 3002.5 | 1134.5 | 4259 | 1812.5 | 682.5 | 9639.5 | 4294 | 6077 |
| 2322 | 3576.5 | 3220.5 | 1150.5 | 4241.5 | 1819 | 724.5 | 9474.5 | 4343 | 6135 |
| 19157 | 13193 | 5215.5 | 1309 | 4135 | 1814.5 | 698.5 | 9380.5 | 7570 | 8230.5 |
| 2568 | 3748.5 | 3152 | 1141.5 | 4249 | 1816.5 | 713 | 10010.5 | 4424 | 6375.5 |
| 2343 | 3671.5 | 3282 | 1135 | 4287 | 1873.5 | 702.5 | 9399 | 4473.5 | 6315.5 |
| 110601 | 59639.5 | 16927 | 1771.5 | 4495 | 1913 | 728 | 17963.5 | 6316.5 | 7509.5 |
| 3106 | 4869.5 | 3866 | 1259 | 4599.5 | 1872 | 734 | 11532 | 4750.5 | 6808 |
| 2482.5 | 3853 | 3282 | 1138.5 | 5289.5 | 2059 | 703 | 10039 | 4537 | 6302 |
| 2568 | 3703 | 3220 | 1117 | 4363 | 1860 | 695 | 10187 | 4519 | 6243 |
| 9517 | 23294 | 8265 | 1963 | 9954 | 2655 | 735 | 15168 | 9535 | 25672 |
| 9455 | 25158 | 13188 | 2870 | 14556.5 | 3579.5 | 710 | 10502.5 | 8598.5 | 25560.5 |
| 9593 | 23509 | 13215 | 2774 | 14060.5 | 3379.5 | 726.5 | 11774 | 8683 | 23311 |
| 9549 | 20726 | 10252 | 2240 | 11430 | 2941 | 718 | 13824 | 9459 | 22817 |
| 9051.5 | 23321.5 | 8557.5 | 1825.5 | 9093 | 2557 | 745.5 | 13731.5 | 9585.5 | 27074 |
| 11685 | 59656 | 44440 | 15064 | 7613 | 2780 | 715 | 15257 | 10906 | 34685 |
| 4317.5 | 11767.5 | 4852.5 | 1539.5 | 5576.5 | 2019 | 766.5 | 14434.5 | 5287.5 | 14412.5 |
| 2893 | 5476 | 6869 | 2004 | 661028 | 62369 | 758 | 15793 | 4469 | 7912 |
| 2746 | 4597.5 | 4230.5 | 1516 | 6594 | 2445 | 858.5 | 13590.5 | 5224 | 7485 |
| 2676 | 4215.5 | 4070.5 | 1352.5 | 5739.5 | 2130.5 | 723 | 11966 | 4733 | 7052.5 |
| 2736.5 | 4155.5 | 3705 | 1268.5 | 5128.5 | 2007 | 736 | 11031 | 4702.5 | 6646 |
| 2733 | 4613.5 | 3605 | 1321.5 | 5054 | 1982.5 | 769 | 11359 | 4654 | 6810 |
| 3045 | 4427 | 3454 | 1221.5 | 4753.5 | 1873 | 717 | 11982.5 | 4527 | 6713 |
| 2891 | 4323.5 | 3570.5 | 1259.5 | 4569 | 1980 | 757 | 11468.5 | 4599.5 | 6666.5 |
| 2800.5 | 4061 | 3542.5 | 1259 | 4473.5 | 1939.5 | 705.5 | 10963.5 | 4578 | 6354.5 |
| 2617 | 3827 | 3323.5 | 1179.5 | 4392.5 | 1913.5 | 738 | 10280.5 | 4564.5 | 6234 |
| 2448.5 | 3637 | 3153 | 1147.5 | 4389 | 1952 | 771.5 | 9832 | 4519 | 6347 |
| 21295 | 14072 | 5605 | 1414 | 4383 | 1850 | 757 | 9950 | 8600 | 8607 |
| 2715.5 | 3939.5 | 3439 | 1235 | 4260.5 | 1846 | 737 | 10107.5 | 4622.5 | 6203.5 |
| 2558 | 3831 | 3338 | 1169 | 4669 | 1977 | 758 | 10364 | 4807 | 6820 |
| 96218 | 52144 | 15016.5 | 1770.5 | 4559.5 | 1906 | 735 | 17476.5 | 5963.5 | 7885.5 |
| 3174 | 5015 | 4067 | 1318 | 4803 | 1936 | 722 | 11148 | 4680 | 7109 |
| 2714 | 3878 | 3409 | 1208 | 5581 | 2159 | 759 | 10497 | 4969 | 6867 |
| 2894 | 4067 | 3462 | 1249 | 4516 | 1911 | 741 | 10542 | 4707 | 6221 |
| 9298.5 | 18604 | 9174.5 | 2031.5 | 10516 | 2649.5 | 767.5 | 15164 | 8739 | 19528.5 |
| 11096.5 | 26308.5 | 13956.5 | 2836.5 | 14547 | 3619 | 753 | 11591.5 | 9252 | 25799 |
| 9848 | 25466 | 13265 | 2869 | 14506 | 3583 | 737 | 10852 | 8253 | 25311 |

| | | | | | | | | | |
|---|---|---|---|---|---|---|---|---|---|
| 9540 | 21933 | 10399 | 2231 | 12040 | 2973 | 701 | 11629 | 8525 | 22157 |
| 9228 | 21215 | 7108 | 1824 | 8138 | 2507 | 739 | 14712 | 9914 | 24305 |
| 10701 | 98427 | 153931 | 55906 | 7773 | 3010 | 701 | 13831 | 9069 | 54125 |
| 4465 | 10043.5 | 4773.5 | 1573.5 | 5573 | 2069.5 | 743 | 15433.5 | 5238.5 | 12926.5 |
| 3129.5 | 5231 | 7191.5 | 2040.5 | 712580.5 | 69496.5 | 729.5 | 15952.5 | 4416.5 | 7383 |
| 3098 | 4698 | 4672 | 1687 | 6725 | 2700 | 861 | 14690 | 5357 | 7432 |
| 2924 | 4182.5 | 4044.5 | 1451 | 5615.5 | 2193.5 | 770.5 | 12640.5 | 4969.5 | 6780.5 |
| 2839 | 4332 | 3801 | 1369 | 5246 | 1994 | 814 | 11384 | 4784 | 6736 |
| 2673.5 | 3964 | 3519.5 | 1210 | 4561.5 | 1907 | 771 | 11717 | 4713 | 6550.5 |
| 3142 | 4259 | 3806 | 1251 | 4709 | 1908 | 740 | 12290 | 4781 | 6553 |
| 2924 | 4225 | 3565 | 1317 | 4457 | 1952 | 749 | 11712 | 4707 | 6489 |
| 2756 | 3722 | 3346 | 1217 | 4276 | 1838 | 731 | 11320 | 4628 | 6130 |
| 2673 | 3714 | 3241 | 1154 | 4307 | 1774 | 715 | 10451 | 4605 | 6253 |
| 2355 | 3692 | 3288 | 1178 | 4260 | 1785 | 743 | 9920 | 4347 | 6229 |
| 31981.5 | 21704.5 | 7109.5 | 1399.5 | 4143.5 | 1755 | 731 | 9957 | 10453.5 | 9864.5 |
| 2664 | 3921 | 3294 | 1169 | 4348 | 1847 | 743 | 9892 | 4651 | 6314 |
| 2762.5 | 3935.5 | 3339.5 | 1142.5 | 4263.5 | 1823 | 764 | 9996.5 | 4921.5 | 6671 |
| 110815.5 | 59419.5 | 17319 | 1789.5 | 4554.5 | 1942 | 726.5 | 18979 | 6691 | 8228.5 |
| 3698.5 | 5219 | 4269.5 | 1323.5 | 4652.5 | 1923 | 707 | 11801 | 5055.5 | 6828.5 |
| 2718.5 | 3934.5 | 3455.5 | 1188 | 5781.5 | 2121 | 783 | 10836 | 4794.5 | 6713 |
| 2859 | 4051 | 3362 | 1147 | 4379 | 1838 | 748 | 10833 | 4625 | 6112 |
| 10647 | 24836 | 9231 | 2044 | 10084 | 2700 | 745 | 15276 | 10235 | 27119 |
| 11827.5 | 26136 | 13533 | 2780.5 | 13930.5 | 3454.5 | 753 | 12407 | 9604 | 25600.5 |
| 11668 | 25620 | 13227 | 2817 | 14140 | 3494 | 714 | 12652 | 9358 | 24489 |
| 11713 | 22543 | 11596 | 2468 | 12196 | 3094 | 797 | 12486 | 8841 | 22249 |
| 9064 | 21123 | 7083 | 1660 | 7961 | 2407 | 693 | 14892 | 9425 | 22739 |
| 11032.5 | 86017.5 | 112894 | 35501.5 | 6935.5 | 2383.5 | 661.5 | 15209 | 7548 | 35110.5 |
| 4148 | 8542 | 4741 | 1478 | 5232 | 1976 | 695 | 14954 | 5035 | 10984 |
| 2709 | 4806 | 6383 | 2173 | 244238 | 27458 | 673 | 13357 | 4145 | 6935 |
| 2821.5 | 4305 | 3806.5 | 1332.5 | 12870 | 2747 | 731.5 | 12430 | 4981.5 | 6887 |
| 2732.5 | 4194 | 3970 | 1337.5 | 7170.5 | 2232.5 | 703.5 | 10639.5 | 4517 | 6624 |
| 2640 | 3780 | 3350.5 | 1195 | 5376.5 | 1971.5 | 703 | 10355 | 4504.5 | 6188 |
| 2762 | 4159 | 3357 | 1198 | 4836 | 1906 | 697 | 11034 | 4557 | 6389 |
| 2695 | 4159.5 | 3241 | 1147.5 | 4610 | 1842 | 681.5 | 11447.5 | 4378 | 6220.5 |
| 2664 | 4027 | 3448 | 1228 | 4452 | 1928 | 691 | 10975 | 4426 | 6361 |
| 2589 | 3755 | 3228 | 1112 | 4422 | 1809 | 679 | 9865 | 4455 | 6281 |
| 2352 | 3573.5 | 2941.5 | 1112 | 4271.5 | 1741.5 | 687 | 9595.5 | 4261.5 | 5757 |
| 2271.5 | 3601 | 3130 | 1112 | 4275 | 1856 | 698 | 9472.5 | 4163.5 | 5954.5 |
| 6976 | 5982 | 4007.5 | 1152 | 4406.5 | 1793.5 | 685 | 9309.5 | 5373.5 | 6805.5 |
| 2219 | 3500.5 | 3018.5 | 1097 | 4259.5 | 1766 | 677.5 | 8918.5 | 4128.5 | 5832.5 |
| 2512 | 3846 | 3251 | 1161 | 4524 | 1839 | 696 | 9867 | 4571 | 6147 |
| 156330.5 | 84788.5 | 23384.5 | 2086.5 | 4852 | 1880.5 | 718 | 23446 | 7382.5 | 8032.5 |
| 2968 | 4192 | 3509 | 1174 | 4700 | 1893 | 693 | 10503 | 4519 | 6584 |
| 2492 | 3676 | 3243 | 1100 | 13129 | 3024 | 703 | 9556 | 4607 | 5961 |
| 2465 | 3690 | 3085 | 1061 | 4448 | 1796 | 665 | 9488 | 4369 | 5991 |
| 10013.5 | 21728 | 9633.5 | 2077.5 | 11282.5 | 2859 | 716 | 13998.5 | 9381 | 22571 |
| 10335 | 22526 | 12211 | 2699 | 13462 | 3276 | 689 | 12489 | 9118 | 22745 |
| 10402.5 | 21780.5 | 11880 | 2559.5 | 13433 | 3345.5 | 693 | 12447 | 9148 | 22669.5 |

| | | | | | | | | | |
|---|---|---|---|---|---|---|---|---|---|
| 9812 | 21904 | 9707 | 2121 | 10871 | 2798 | 684 | 14336 | 9553 | 23222 |
| 8806.5 | 21084.5 | 7105.5 | 1729.5 | 8062.5 | 2468.5 | 718.5 | 13773 | 9883.5 | 23470 |
| 10753.5 | 100417 | 153391 | 55552 | 7721 | 2883.5 | 650 | 15419 | 8642 | 53438 |
| 4437 | 10788 | 4683 | 1497 | 5497 | 1997 | 698 | 14774 | 5271 | 13918 |
| 2644.5 | 4692.5 | 6123.5 | 1857 | 567051 | 55604.5 | 698.5 | 15670 | 4149.5 | 7247.5 |
| 3008 | 4363 | 4070 | 1500 | 6418 | 2416 | 945 | 19783 | 6359 | 7906 |
| 3004 | 4038.5 | 3765.5 | 1511.5 | 5805.5 | 2167.5 | 757.5 | 11958.5 | 5092.5 | 6743 |
| 2805 | 4092 | 3725 | 1289 | 5005 | 1968 | 735 | 10941 | 4724 | 6515 |
| 2782.5 | 4104.5 | 3596.5 | 1259.5 | 4489.5 | 1878 | 709 | 11463.5 | 4663.5 | 6528.5 |
| 3069 | 4203.5 | 3619 | 1211 | 4550.5 | 1824.5 | 686 | 11856 | 4738.5 | 6469.5 |
| 2902 | 4054 | 3348 | 1323.5 | 4450.5 | 1953.5 | 685.5 | 11263.5 | 4703 | 6643.5 |
| 2780.5 | 3880 | 3174 | 1225.5 | 4194 | 1843 | 709 | 10924 | 4666.5 | 6421 |
| 2602.5 | 3798 | 3270 | 1211.5 | 4434 | 1945.5 | 734.5 | 10169 | 4593.5 | 6376.5 |
| 2331 | 3650 | 3141.5 | 1193.5 | 4209 | 1834 | 698 | 9921.5 | 4148.5 | 6019 |
| 52020 | 38055 | 12288 | 1695 | 4369 | 1875.5 | 690.5 | 10029.5 | 19291.5 | 15603 |
| 2732 | 3963 | 3580 | 1191 | 4309 | 1813 | 682 | 9473 | 4634 | 6506 |
| 2585.5 | 3920 | 3600.5 | 1180 | 4336 | 1811.5 | 734 | 10443 | 4647.5 | 6579.5 |
| 123957 | 68220.5 | 19703.5 | 1899.5 | 4678 | 1949 | 715.5 | 19970.5 | 6552.5 | 8008.5 |
| 3361.5 | 5106 | 3971 | 1249 | 4709 | 1877 | 740.5 | 11513.5 | 4957 | 6786.5 |
| 2622 | 4131 | 3383 | 1264 | 5413 | 2113 | 711 | 10358 | 4605 | 6407 |
| 2917.5 | 4050.5 | 3329 | 1161.5 | 4473 | 1916.5 | 731.5 | 10600 | 4635 | 6226.5 |
| 10835.5 | 20761.5 | 9456.5 | 2100 | 10805.5 | 2775.5 | 701.5 | 15433 | 10184.5 | 21916 |
| 11028 | 23966 | 13226 | 2777 | 13494 | 3378 | 732 | 12567 | 9294 | 23409 |
| 9608 | 23930 | 12852 | 2775 | 14111 | 3417 | 697 | 12105 | 8934 | 23821 |
| 10082 | 20925 | 9816.5 | 2284 | 11640.5 | 2946.5 | 698.5 | 13425.5 | 9496 | 22663.5 |
| 9104 | 22103 | 7318 | 1804 | 8760 | 2447 | 728 | 14433 | 9639 | 24688 |
| 6854 | 17539 | 6891 | 1961 | 6904 | 2363 | 728 | 16117 | 6918 | 19702 |
| 3980 | 10655 | 4371 | 1338.5 | 5584 | 2000.5 | 729.5 | 14474 | 5028.5 | 13367.5 |
| 3036 | 5058 | 6447 | 1863 | 628427 | 57138 | 700 | 15518 | 4434 | 7152 |
| 2708 | 4396 | 4140 | 1385 | 6275 | 2478 | 819 | 12516 | 5100 | 7005 |
| 2871 | 4131 | 3830 | 1322.5 | 5378.5 | 2163.5 | 723.5 | 11523 | 4914.5 | 6589.5 |
| 2748.5 | 4018.5 | 3664.5 | 1238.5 | 4896 | 1965 | 740 | 10782.5 | 4710.5 | 6501.5 |
| 2719 | 4103 | 3368 | 1180 | 4670 | 1872 | 725 | 10689 | 4527 | 6194 |
| 3087 | 4008 | 3298 | 1130 | 4376 | 1772 | 680 | 11167 | 4645 | 5957 |
| 2643 | 4468.5 | 3526 | 1317.5 | 4598 | 1872.5 | 691 | 11345.5 | 4254.5 | 6565 |
| 2675 | 3989 | 3455 | 1287 | 4592 | 1915 | 717 | 10987 | 4479 | 6571 |
| 2584 | 4020 | 3646 | 1264 | 4805 | 2031 | 747 | 11314 | 4444 | 6689 |
| 2502.5 | 3779.5 | 3314 | 1204 | 4436.5 | 1873 | 714.5 | 10168 | 4549 | 6197 |
| 12572 | 9083 | 4620 | 1248 | 4300 | 1881 | 731 | 9721 | 7002 | 7719 |
| 2568 | 3611 | 3325 | 1193 | 4328 | 1825 | 717 | 9289 | 4432 | 6073 |
| 2665 | 3999 | 3472 | 1258 | 4622 | 1903 | 732 | 10468 | 4690 | 6721 |
| 106950 | 64988 | 17768 | 1990 | 4905 | 1888 | 741 | 19897 | 6396 | 8545 |
| 3158 | 5158 | 4150 | 1344 | 4926 | 2041 | 744 | 12199 | 4684 | 6979 |
| 2542.5 | 3653 | 3283.5 | 1194 | 5572.5 | 2015.5 | 721 | 10117 | 4700.5 | 6185 |
| 2821 | 4117.5 | 3380.5 | 1218.5 | 4389.5 | 1872 | 703 | 10335.5 | 4567 | 6267 |
| 8984 | 20689.5 | 9994.5 | 2181 | 11445 | 2992 | 778 | 13736.5 | 9667.5 | 21776.5 |
| 10044 | 26112 | 14484.5 | 3129.5 | 16101 | 3808.5 | 768 | 10377.5 | 8426 | 25400 |
| 10183 | 24258 | 13572 | 2909 | 14632 | 3531 | 731 | 11118 | 8283 | 24475 |

| | | | | | | | | | |
|---|---|---|---|---|---|---|---|---|---|
| 10190.5 | 21605.5 | 9935 | 2129.5 | 11080 | 2730 | 699.5 | 13801.5 | 8982.5 | 22250 |
| 896 | 1193 | 1033 | 630.5 | 1621 | 1150 | 320 | 2972.5 | 1669.5 | 2229.5 |
| 47667 | 36270.5 | 10322 | 1490 | 3417 | 1566 | 462.5 | 19004.5 | 5105.5 | 5904 |
| 4245 | 5253 | 3437 | 1027 | 3404 | 1610 | 598 | 10463 | 3694 | 5022 |
| 2458 | 3648.5 | 6894 | 2039 | 484483 | 44985 | 629 | 8473.5 | 3531 | 4925 |
| 2189.5 | 3441.5 | 3275.5 | 1160 | 4873 | 2041.5 | 739 | 9387 | 3847.5 | 5465 |
| 2060.5 | 3438 | 2949.5 | 1075 | 4270.5 | 1822.5 | 700.5 | 9081.5 | 3864.5 | 5560.5 |
| 1849 | 3051.5 | 2712 | 1040.5 | 3989 | 1727 | 677 | 8163.5 | 3656 | 5271 |
| 2253 | 3442 | 2931 | 1075 | 4093 | 1779 | 721 | 10055 | 4134 | 5665 |
| 2487 | 3576 | 2912 | 1043 | 3878 | 1721 | 663 | 10038 | 4373 | 5682 |
| 2572 | 4028 | 3557.5 | 1294 | 4302 | 1915 | 721 | 10941.5 | 4351.5 | 6162.5 |
| 2385.5 | 3762 | 3161.5 | 1218 | 4210.5 | 1828 | 691.5 | 10585.5 | 4429 | 6096 |
| 2037 | 3276 | 2901 | 1061 | 4013 | 1749 | 697 | 8873 | 4059 | 5475 |
| 1969 | 3413 | 2922 | 1150 | 4141 | 1844 | 693 | 9184 | 4121 | 5761 |
| 17836 | 12639 | 4872 | 1163 | 3937 | 1762 | 698 | 8631 | 7538 | 7725 |
| 2096 | 3463 | 2963 | 1096 | 4158 | 1792 | 725 | 8897 | 4056 | 5811 |
| 2216.5 | 3428 | 3047 | 1067 | 4145 | 1742 | 705.5 | 8781 | 4523 | 6239 |
| 66838.5 | 36013.5 | 10865 | 1569.5 | 4523.5 | 1899.5 | 719 | 15290 | 5562.5 | 7015.5 |
| 2591 | 4016 | 3329 | 1173 | 4359.5 | 1826.5 | 679.5 | 9862.5 | 4603.5 | 6283 |
| 2291.5 | 3684 | 3059 | 1079 | 5197.5 | 2044 | 735 | 9807.5 | 4680.5 | 6175.5 |
| 2423 | 3407 | 3139 | 1095.5 | 3968.5 | 1762.5 | 696.5 | 8932.5 | 4433.5 | 5889.5 |
| 8600 | 19515 | 8952 | 1974 | 10160 | 2728 | 674 | 14716 | 9164 | 21376 |
| 9543 | 23274 | 13317 | 2828 | 14753 | 3547 | 705 | 11501 | 8409 | 23682 |
| 10009 | 22019.5 | 11694.5 | 2574.5 | 13654.5 | 3206.5 | 706.5 | 11638.5 | 8855 | 22104 |
| 9433 | 19895.5 | 9765 | 2235 | 11513 | 2911.5 | 720 | 12786.5 | 8963 | 20621.5 |
| 9376 | 20893.5 | 6967 | 1730.5 | 8001 | 2333.5 | 696.5 | 14569.5 | 9667 | 23692 |
| 10559 | 104094.5 | 165778.5 | 64570.5 | 6856 | 2476.5 | 630.5 | 15759 | 9313.5 | 58874 |
| 4375 | 10918 | 4616 | 1294 | 4949 | 1892 | 691 | 14517 | 5303 | 13582 |
| 3064 | 4893 | 7078 | 1978 | 747547 | 68059 | 696 | 16682 | 4341 | 6862 |
| 3095.5 | 4414 | 4182.5 | 1418 | 5982.5 | 2361.5 | 748 | 13574 | 4942 | 6750 |
| 2783 | 3926.5 | 3450 | 1216.5 | 4838.5 | 1942.5 | 741.5 | 11199 | 4908.5 | 6406.5 |
| 2525 | 3586 | 2887 | 1138 | 4070 | 1787 | 685 | 9688 | 4642 | 5937 |
| 2773 | 3605 | 3086 | 1092 | 4161 | 1800 | 698 | 10652 | 4729 | 6097 |
| 2698 | 3747 | 3214 | 1065 | 4090 | 1808 | 698 | 10907 | 4601 | 6050 |
| 2608.5 | 3853 | 3318 | 1259.5 | 4161.5 | 1909.5 | 675.5 | 10096.5 | 4314.5 | 6127 |
| 2512 | 3923 | 3318.5 | 1200 | 4369 | 1926 | 724.5 | 11262 | 4476 | 6266 |
| 2418 | 3468.5 | 3116 | 1119.5 | 4067.5 | 1727.5 | 700.5 | 10046 | 4326 | 5898 |
| 2122.5 | 3439 | 2984 | 1088.5 | 4032 | 1784 | 742 | 9434.5 | 3910.5 | 5787 |
| 35458 | 25234 | 7783 | 1396 | 4070 | 1785 | 712 | 9523 | 12444 | 10316 |
| 2452 | 3614 | 3053 | 1126 | 4183 | 1793 | 749 | 9374 | 4123 | 5910 |
| 2356 | 3646 | 3243 | 1114 | 4272 | 1788 | 692 | 9518 | 4657 | 6291 |
| 105097.5 | 55529 | 16192.5 | 1763 | 4577.5 | 1874.5 | 713.5 | 19501.5 | 6381.5 | 7724 |
| 2837.5 | 4529.5 | 3556.5 | 1139.5 | 4317.5 | 1772 | 690 | 10673 | 4691 | 6540.5 |
| 2537 | 3901 | 3321 | 1148 | 5540 | 2054 | 723 | 10047 | 4682 | 6278 |
| 2451 | 3672.5 | 3234 | 1125.5 | 4058.5 | 1849.5 | 729.5 | 10075.5 | 4197 | 6210 |
| 10595 | 22087 | 9684.5 | 2169.5 | 10954 | 2896 | 728.5 | 13963.5 | 9284 | 22261 |
| 11775 | 23327 | 12777 | 2677 | 14082 | 3477 | 739 | 12531 | 9536 | 23636 |
| 11359 | 22085 | 11591 | 2395 | 12910 | 3111 | 694 | 12912 | 9125 | 22156 |

| | | | | | | | | | |
|---|---|---|---|---|---|---|---|---|---|
| 10696.5 | 22055.5 | 9998.5 | 2068.5 | 10856.5 | 2787 | 692.5 | 13731.5 | 9631 | 23380.5 |
| 8616.5 | 21657.5 | 7218.5 | 1745 | 8662 | 2475.5 | 707.5 | 14241 | 9643.5 | 24737.5 |
| 7562 | 21154 | 8172 | 2541 | 6980 | 2491 | 718 | 15733 | 7239 | 23728 |
| 4488 | 10464.5 | 4520 | 1412.5 | 5475 | 1995.5 | 718 | 14962.5 | 5445 | 12782 |
| 3060.5 | 5957 | 6848 | 2081.5 | 594070.5 | 56383 | 724.5 | 16897 | 4418 | 8274 |
| 2741 | 4456 | 4348 | 1422 | 6808 | 2432 | 770 | 13203 | 4678 | 7267 |
| 2935 | 4085 | 3797 | 1348 | 5648 | 2161 | 757 | 11850 | 4635 | 6577 |
| 2661 | 3854.5 | 3581 | 1228.5 | 4888.5 | 1986.5 | 722.5 | 10690 | 4644.5 | 6430.5 |
| 2739 | 4092 | 3304 | 1161 | 4591 | 1959 | 714 | 11200 | 4551 | 6594 |
| 2743 | 3929.5 | 3182 | 1141 | 4439 | 1810.5 | 702.5 | 10851 | 4671.5 | 6394.5 |
| 2775 | 3883 | 3360 | 1300 | 4302 | 1868 | 673 | 10147 | 4470 | 6178 |
| 2615 | 3784.5 | 3226 | 1202 | 4286.5 | 1874.5 | 673 | 9829 | 4364 | 5985.5 |
| 2351 | 3632 | 3064 | 1125 | 4114 | 1813 | 693 | 9546 | 4167 | 5890 |
| 2122 | 3354.5 | 2982.5 | 1124 | 3994 | 1758 | 687 | 9166.5 | 3997 | 5756 |
| 20446 | 14187 | 5286 | 1220 | 4201 | 1814 | 690 | 9033 | 7959 | 7885 |
| 2355 | 3599 | 3024 | 1091 | 3986 | 1719 | 692 | 8830 | 4154 | 5885 |
| 2477 | 3795 | 3220.5 | 1093.5 | 4326.5 | 1812.5 | 722 | 9772 | 4678.5 | 6264 |
| 119544 | 60495 | 17085 | 1812 | 4428 | 1818 | 692 | 17474 | 6547 | 7162 |
| 2942 | 4420 | 3807.5 | 1182.5 | 4431.5 | 1844 | 718 | 10503.5 | 4674 | 6377 |
| 2527 | 3806 | 3121 | 1117 | 5339 | 2023 | 723 | 10196 | 4694 | 6150 |
| 2264 | 3645.5 | 3101.5 | 1093 | 4126.5 | 1799.5 | 705.5 | 9258.5 | 4164.5 | 5920 |
| 10255 | 20796 | 9699 | 2057 | 10936 | 2752 | 689 | 13978 | 9386 | 22588 |
| 10989 | 22781 | 12424 | 2651 | 13492 | 3302 | 703 | 12633 | 9227 | 23244 |
| 10235.5 | 22070 | 11445 | 2382.5 | 13026.5 | 3173 | 689.5 | 12587.5 | 9112.5 | 22123 |
| 9979.5 | 21122.5 | 10056 | 2157 | 11104.5 | 2892.5 | 690.5 | 13599 | 8823 | 21975.5 |
| 9631 | 21557 | 7172 | 1687 | 8161 | 2439 | 681 | 15015 | 9412 | 23961 |
| 11604.5 | 132868 | 213728 | 80009 | 7763.5 | 3387 | 605 | 15742.5 | 9427.5 | 73233.5 |
| 5297.5 | 13107 | 5423 | 1755 | 5311.5 | 2092 | 674.5 | 13693.5 | 5407 | 14797.5 |
| 2776 | 4877 | 6652 | 2187 | 660122 | 60291 | 661 | 15555 | 4128 | 6936 |
| 2923.5 | 4119.5 | 4400 | 1583.5 | 6628 | 2652 | 808 | 13176 | 4904.5 | 6633.5 |
| 2870.5 | 3930.5 | 3887 | 1452.5 | 5652 | 2263 | 702 | 11408.5 | 4818.5 | 6356 |
| 2608 | 3671 | 3509 | 1244 | 4691 | 1931 | 676 | 10051 | 4509 | 6128 |
| 2903 | 4009 | 3468.5 | 1171.5 | 4711.5 | 1886 | 723 | 10788.5 | 4694 | 6248 |
| 2936.5 | 3980.5 | 3354.5 | 1115 | 4316 | 1800.5 | 681 | 11703.5 | 4576 | 6187.5 |
| 2736 | 3951 | 3344 | 1238 | 4146 | 1887 | 657 | 10719 | 4365 | 6055 |
| 2726 | 3893 | 3286 | 1253 | 4487 | 1930 | 684 | 11088 | 4452 | 6078 |
| 2382 | 3375.5 | 2890.5 | 1084 | 4163.5 | 1772 | 693 | 9688 | 4432 | 5774 |
| 2394 | 3482.5 | 3070 | 1109 | 4120.5 | 1832 | 695.5 | 9921.5 | 4449 | 5729 |
| 35148.5 | 24738.5 | 7703 | 1372 | 3921 | 1808 | 667 | 9106 | 12120.5 | 9998 |
| 2625.5 | 3656.5 | 3233 | 1148.5 | 3920.5 | 1782 | 697.5 | 9147 | 4173.5 | 5806.5 |
| 2554 | 3982 | 3261 | 1156 | 4460 | 1840 | 723 | 10496 | 4696 | 6381 |
| 85814 | 45698 | 13057 | 1638 | 4716 | 1913 | 678 | 17104 | 6072 | 7424 |
| 2964 | 4540 | 3659 | 1179 | 4581 | 1906 | 708 | 10754 | 4809 | 6754 |
| 2538 | 3739 | 3192.5 | 1131 | 5387 | 2024.5 | 693 | 10377.5 | 4322.5 | 6312 |
| 2658 | 3760 | 3195 | 1109 | 4284 | 1770 | 680 | 9427 | 4092 | 5899 |
| 9089.5 | 22251.5 | 8440.5 | 1910.5 | 9281 | 2606.5 | 694 | 14964 | 8661 | 22489.5 |
| 11423.5 | 23833.5 | 12728 | 2749.5 | 14212 | 3324.5 | 685 | 12449 | 9469 | 24388 |
| 9952.5 | 21473.5 | 12238.5 | 2536.5 | 13309.5 | 3202.5 | 672.5 | 12157 | 8788.5 | 22194 |

| | | | | | | | | | |
|---|---|---|---|---|---|---|---|---|---|
| 10449 | 21458 | 10078 | 2154 | 10697 | 2792 | 673 | 14063 | 9559 | 21966 |
| 9426 | 19649 | 6713 | 1496 | 7571 | 2252 | 691 | 14222 | 9780 | 22005 |
| 10385.5 | 97156.5 | 151400 | 55337.5 | 6874.5 | 2576 | 671.5 | 15398.5 | 8330.5 | 50643 |
| 3884 | 10115.5 | 4236.5 | 1341 | 4993 | 1908 | 697.5 | 14592.5 | 4662.5 | 12728.5 |
| 3048 | 5247 | 6891 | 2020 | 603892 | 59241 | 720 | 16664 | 4324 | 7499 |
| 2907 | 4428.5 | 4557.5 | 1546 | 6714.5 | 2569 | 813.5 | 14232.5 | 4898.5 | 6905 |
| 2890 | 3935 | 3528 | 1229 | 4889 | 2017 | 739 | 11197 | 4790 | 6433 |
| 2702.5 | 3900.5 | 3322 | 1235 | 4699.5 | 1817 | 715 | 10421.5 | 4689 | 6079.5 |
| 2930 | 4102 | 3367 | 1169 | 4527 | 1879 | 707 | 11263 | 4655 | 6271 |
| 2969 | 3901 | 3276 | 1159.5 | 4173.5 | 1817.5 | 685 | 11154.5 | 4598.5 | 6128.5 |
| 2829 | 4128 | 3364 | 1295 | 4368 | 1825 | 696 | 11217 | 4436 | 6181 |
| 2489 | 3631.5 | 3059.5 | 1135 | 4033.5 | 1825 | 708 | 10347 | 4480 | 5958 |
| 2643 | 3707 | 3121 | 1129 | 4132 | 1819 | 706 | 10226 | 4576 | 5947 |
| 2185 | 3357 | 2959 | 1061 | 3915 | 1782 | 716 | 9247 | 3961 | 5599 |
| 22183 | 15250 | 5763 | 1352 | 3944 | 1743 | 716 | 9634 | 8187 | 8204 |
| 2475.5 | 3592.5 | 3253 | 1080.5 | 4023.5 | 1776.5 | 696 | 9319.5 | 4225.5 | 6045 |
| 2506 | 3623.5 | 3238.5 | 1159.5 | 4195.5 | 1861.5 | 687.5 | 9815 | 4683 | 6087 |
| 74383 | 37846 | 11317 | 1519 | 4395 | 1860 | 714 | 15451 | 5912 | 7339 |
| 3190 | 4521 | 3637 | 1193 | 4418 | 1838 | 689 | 10827 | 4865 | 6484 |
| 2746 | 3897 | 3360 | 1117 | 5093 | 2003 | 708 | 9884 | 4656 | 6134 |
| 2736 | 3797 | 3299 | 1151 | 4186 | 1814 | 716 | 10054 | 4462 | 6039 |
| 10493 | 20733 | 9567 | 2080 | 10702 | 2695 | 758 | 15227 | 9819 | 22796 |
| 10942 | 23660 | 13049 | 2594 | 13350 | 3316 | 721 | 11979 | 9216 | 24334 |
| 11345 | 22340 | 12145 | 2606 | 13397 | 3492 | 692 | 12929 | 9701 | 22657 |
| 10914 | 21779.5 | 11005 | 2227 | 11246.5 | 2746.5 | 721.5 | 13597 | 9604 | 22863 |
| 9433 | 21932 | 7396 | 1771 | 8257 | 2477 | 695 | 14409 | 9398 | 24547 |
| 12504 | 124136 | 194374.5 | 71680 | 7494 | 2778.5 | 626.5 | 14629.5 | 10543.5 | 67698.5 |
| 3806 | 10705 | 4643 | 1370 | 5165 | 1947.5 | 706.5 | 13414.5 | 5010.5 | 13953.5 |
| 2881 | 4802 | 6308 | 1814 | 569364 | 52068 | 713 | 15482 | 4324 | 7021 |
| 2692.5 | 4130 | 4183.5 | 1516.5 | 6645.5 | 2537 | 751 | 12061.5 | 4623.5 | 6615 |
| 2651 | 3717 | 3340 | 1209 | 4829 | 2024 | 683 | 10703 | 4745 | 6025 |
| 2644.5 | 3691 | 3532 | 1222 | 4663.5 | 1891.5 | 718 | 10540 | 4617.5 | 6190 |
| 2857 | 3743 | 3149 | 1194 | 4311 | 1820 | 679 | 10316 | 4422 | 6027 |
| 2947.5 | 4192 | 3381 | 1204 | 4413.5 | 1817 | 680.5 | 11310.5 | 4359.5 | 6088 |
| 2693 | 4036 | 3266 | 1258 | 4419.5 | 1901 | 691 | 11549.5 | 4425 | 6306.5 |
| 2465.5 | 3701 | 3226 | 1147.5 | 4069.5 | 1834.5 | 640 | 10161 | 4411 | 5802 |
| 2482.5 | 3358 | 2970 | 1107.5 | 3881 | 1755.5 | 667 | 9255.5 | 4339 | 5494 |
| 2094 | 3355 | 2850 | 1080.5 | 3989.5 | 1769 | 677.5 | 9109.5 | 4005 | 5736.5 |
| 26987 | 18894.5 | 6263.5 | 1245.5 | 4043.5 | 1768.5 | 656 | 9212 | 9649 | 8841 |
| 2643 | 3745 | 3222 | 1133 | 4099 | 1872 | 700 | 9073 | 4449 | 5998 |
| 2376.5 | 3619.5 | 3194.5 | 1140.5 | 4262 | 1761.5 | 702 | 9564 | 4485 | 6023 |
| 77275 | 43738 | 12851 | 1630 | 4488 | 1830 | 673 | 16002 | 5485 | 7173 |
| 2973.5 | 4636 | 3938 | 1216.5 | 4530 | 1863.5 | 684 | 11055.5 | 4431 | 6657.5 |
| 2528 | 3668 | 3177.5 | 1176.5 | 5077.5 | 1939 | 694 | 9796.5 | 4362 | 6316.5 |
| 2605.5 | 3667.5 | 3009 | 1114 | 4268.5 | 1833 | 682.5 | 9640 | 4450.5 | 5842 |
| 9356 | 21578 | 9532 | 2054 | 10612 | 2721 | 697 | 14391 | 9560 | 22710 |
| 10295 | 22923 | 12164 | 2631 | 13865 | 3257 | 683 | 12080 | 9405 | 23642 |
| 9034 | 21448 | 11578 | 2547 | 13438 | 3178 | 655 | 11977 | 8695 | 22845 |

| | | | | | | | | | |
|---|---|---|---|---|---|---|---|---|---|
| 9563.5 | 20857.5 | 10241.5 | 2112.5 | 11278 | 2786.5 | 675 | 13689 | 9542.5 | 22631 |
| 8837 | 20633 | 6629 | 1673 | 7636.5 | 2352.5 | 722.5 | 13249.5 | 9713 | 23155.5 |
| 10505.5 | 101789.5 | 155849 | 56135.5 | 6912.5 | 2577 | 675 | 14298.5 | 8620.5 | 52701.5 |
| 3576 | 10265 | 4244 | 1330 | 5018 | 1888 | 689 | 14528 | 4843 | 13251 |
| 2741.5 | 4559 | 6131.5 | 1732 | 585347.5 | 55833 | 624 | 12669.5 | 4400 | 6753 |
| 2762 | 4234 | 4214 | 1369 | 5916 | 2273 | 757 | 12461 | 4630 | 6454 |
| 2683 | 4028 | 3665 | 1251 | 5107 | 2028 | 704 | 11262 | 4771 | 6446 |
| 2679.5 | 3811 | 3365.5 | 1186.5 | 4580.5 | 1804 | 701 | 10199.5 | 4470 | 6054.5 |
| 2878 | 4020 | 3345 | 1252 | 4649 | 1887 | 685 | 10900 | 4614 | 6342 |
| 2691.5 | 3901.5 | 3265.5 | 1094.5 | 4085 | 1687.5 | 657 | 10820.5 | 4548.5 | 5885.5 |
| 2750 | 4103 | 3456 | 1284 | 4411 | 1936 | 696 | 11080 | 4486 | 6594 |
| 2660 | 3874 | 3214 | 1157 | 4265 | 1807 | 677 | 10221 | 4489 | 5825 |
| 2402 | 3500 | 2943.5 | 1135 | 3956 | 1768.5 | 686.5 | 9863.5 | 4388.5 | 5848 |
| 2300 | 3399 | 3059 | 1157 | 3937 | 1723 | 710 | 8924 | 4015 | 5754 |
| 15645 | 10865 | 4630 | 1177 | 3929 | 1763.5 | 671 | 8743 | 6668.5 | 7245.5 |
| 2309 | 3601.5 | 3088 | 1080 | 3955 | 1741.5 | 694 | 9134 | 4118 | 5786 |
| 2483 | 3741 | 3273 | 1152 | 4375 | 1836 | 697 | 10549 | 4670 | 6302 |
| 75252.5 | 40915.5 | 11947 | 1614 | 4333 | 1855 | 704 | 15452.5 | 5663 | 7335.5 |
| 2880 | 4194 | 3711 | 1127 | 4152 | 1799 | 691 | 10492 | 4823 | 6153 |
| 2522 | 3887 | 3309 | 1184 | 5249 | 2005 | 723 | 10444 | 4603 | 6180 |
| 2648 | 3840 | 3141 | 1112 | 4073 | 1805 | 690 | 10131 | 4523 | 6031 |
| 9822.5 | 20192 | 9556 | 2145.5 | 11103.5 | 2948 | 722.5 | 13145 | 8773.5 | 21849 |
| 10614 | 23073 | 12623 | 2771 | 14315 | 3609 | 724 | 12278 | 8865 | 23041 |
| 10313.5 | 21968.5 | 11466 | 2568.5 | 13119 | 3336.5 | 722.5 | 12789 | 9212.5 | 22002.5 |
| 10475 | 20849 | 9612 | 2217 | 10921 | 2815 | 693 | 13506 | 9019 | 21238 |
| 9773.5 | 21486.5 | 7110 | 1770 | 7950 | 2426.5 | 676 | 14436.5 | 9903.5 | 23976 |
| 11982 | 119987 | 185393 | 68723 | 7016 | 2744 | 618 | 14492 | 9495 | 64775 |
| 4489 | 10127 | 4927 | 1527.5 | 5372 | 1937 | 708 | 14321.5 | 5489 | 13221 |
| 2711 | 4482 | 6286.5 | 1784 | 641516.5 | 59739.5 | 665.5 | 13585 | 4087 | 6652 |
| 2901 | 4045 | 3747 | 1329 | 5768 | 2197 | 756 | 12890 | 5083 | 6684 |
| 2858 | 3900 | 3814.5 | 1395 | 5324.5 | 2092.5 | 706 | 11603 | 4893 | 6233.5 |
| 2798 | 3971 | 3666 | 1325 | 4873 | 1947 | 735 | 10441 | 4523 | 6071 |
| 2809 | 4033 | 3365 | 1153 | 4346 | 1802 | 697 | 11151 | 4612 | 6244 |
| 2848.5 | 4209.5 | 3355 | 1228.5 | 4322.5 | 1838.5 | 710 | 12075.5 | 4440.5 | 6481.5 |
| 2873.5 | 4402.5 | 3952.5 | 1354.5 | 4765 | 2058.5 | 720 | 12650.5 | 4227.5 | 6579 |
| 2665 | 3962 | 3270 | 1245 | 4413 | 1832 | 708 | 11120 | 4467 | 6274 |
| 2725 | 3821 | 3388 | 1217 | 4458 | 1926 | 724 | 11005 | 4427 | 6086 |
| 2453 | 3524 | 2957.5 | 1158.5 | 4076 | 1847.5 | 696.5 | 9700 | 4562 | 5893.5 |
| 31134 | 19987 | 6995 | 1454 | 4284 | 1882 | 751 | 10328 | 10521 | 9361 |
| 2815 | 3823 | 3435.5 | 1127.5 | 4361 | 1814.5 | 720 | 9679.5 | 4785.5 | 6175.5 |
| 2642 | 4062 | 3660 | 1228 | 4608 | 1913 | 736 | 10871 | 4472 | 6624 |
| 87935 | 50773 | 14603.5 | 1655 | 4828 | 1868.5 | 718.5 | 17758.5 | 5915.5 | 7700.5 |
| 3158 | 5080 | 4076 | 1239 | 4805 | 1952 | 729 | 12146 | 4717 | 6945 |
| 2550 | 4044 | 3577 | 1219 | 5558 | 2129 | 739 | 10796 | 4461 | 6799 |
| 2972 | 4346 | 3640 | 1241.5 | 4600 | 1948.5 | 748.5 | 11002 | 4802.5 | 6426 |
| 10619 | 20586 | 10638 | 2280 | 11589 | 2988 | 746 | 13852 | 9523 | 21237 |
| 10933 | 24470 | 14691 | 2896 | 14720 | 3629 | 726 | 11829 | 8562 | 22892 |
| 11367 | 22920 | 12650 | 2654 | 13652 | 3368 | 734 | 12795 | 9339 | 23594 |

| | | | | | | | | | |
|---|---|---|---|---|---|---|---|---|---|
| 11116.5 | 20928.5 | 10893 | 2275 | 12058 | 2861.5 | 717.5 | 13713 | 9439.5 | 21851.5 |
| 9940 | 22135.5 | 7132.5 | 1739 | 7908 | 2348.5 | 688.5 | 14815.5 | 9573 | 24324 |
| 10689.5 | 105307 | 160589.5 | 57776.5 | 7021 | 2511 | 630.5 | 16036.5 | 9037 | 56545.5 |
| 4199 | 10567 | 4631 | 1373 | 5236 | 1910 | 685.5 | 12748 | 5270.5 | 13105.5 |
| 3103 | 5054 | 6238 | 2073 | 294586 | 31978 | 655 | 16119 | 4372 | 7245 |
| 2716.5 | 4072 | 3699.5 | 1257.5 | 9542.5 | 2353 | 730.5 | 12515.5 | 4450.5 | 6505 |
| 2769 | 4081 | 3596 | 1232 | 6265 | 2083 | 705 | 11839 | 4611 | 6323 |
| 2660 | 3911.5 | 3288 | 1163.5 | 4895 | 1880 | 687 | 10139 | 4555 | 6022 |
| 2784 | 3988 | 3333 | 1154 | 4594 | 1816 | 659 | 10391 | 4447 | 6127 |
| 2762.5 | 3993.5 | 3234 | 1145 | 4426 | 1794 | 662 | 11457.5 | 4395.5 | 6050 |
| 2510 | 4043 | 3146 | 1237 | 4227 | 1920 | 671 | 10925 | 4143 | 6259 |
| 2774 | 4124 | 3348 | 1208 | 4510 | 1887 | 671 | 10543 | 4401 | 6232 |
| 2493.5 | 3518.5 | 3025.5 | 1095 | 4393.5 | 1777 | 683 | 9671.5 | 4370 | 5900 |
| 2290 | 3436 | 3015 | 1084 | 4048 | 1705.5 | 660.5 | 8633 | 4268.5 | 5497.5 |
| 10694 | 7456 | 4162 | 1269 | 4382 | 1784 | 691 | 9186 | 6303 | 7124 |
| 2387 | 3612 | 3145.5 | 1150 | 4144.5 | 1795 | 696 | 9191 | 4172.5 | 5813.5 |
| 2561 | 3759 | 3301 | 1123 | 4364 | 1855 | 660 | 9484 | 4500 | 6152 |
| 136196 | 71736 | 20750 | 1903 | 4521 | 1869 | 660 | 20156 | 6895 | 7450 |
| 2719 | 4155 | 3392 | 1199 | 4545 | 1801 | 699 | 10443 | 4531 | 6452 |
| 2450 | 3630 | 3229 | 1136.5 | 11005.5 | 2701 | 679 | 9268.5 | 4475 | 6031 |
| 2600 | 3647 | 3416 | 1155 | 4357 | 1821 | 683 | 9973 | 4506 | 5992 |
| 9536 | 19509 | 9926 | 2210 | 11535 | 2874 | 721 | 13462 | 8956 | 21596 |
| 10461.5 | 23262.5 | 12772 | 2655 | 13548.5 | 3295 | 669 | 12108.5 | 9096.5 | 23575.5 |
| 9863 | 23359 | 12261 | 2737 | 13454 | 3398 | 716 | 11714 | 8258 | 22633 |
| 8776 | 19189 | 9425 | 2116 | 11063 | 2702 | 663 | 12873 | 8668 | 20666 |
| 9294 | 19133 | 6699 | 1630 | 7726 | 2274 | 675 | 15270 | 9924 | 22370 |
| 11414.5 | 118438.5 | 187572.5 | 68469.5 | 7204 | 2708 | 617 | 16025.5 | 9291 | 65666.5 |
| 4386 | 10962 | 4908.5 | 1473 | 5475.5 | 1968.5 | 715.5 | 13746 | 5193.5 | 14150.5 |
| 2563 | 6168 | 6200 | 1799 | 588394 | 53645 | 633 | 14561 | 4057 | 9007 |
| 2705 | 4157 | 3677 | 1238 | 5508 | 2143 | 828 | 15546 | 5164 | 6944 |
| 2819.5 | 3960 | 3773.5 | 1332.5 | 5221.5 | 2098.5 | 711 | 11756 | 4747 | 6338 |
| 2692 | 3897.5 | 3415.5 | 1232.5 | 4797.5 | 1954 | 688.5 | 10848 | 4474 | 6216.5 |
| 2900 | 4005 | 3501 | 1189 | 4578 | 1789 | 646 | 11382 | 4388 | 6315 |
| 2729 | 4075 | 3524 | 1177 | 4528 | 1845 | 693 | 11169 | 4372 | 6304 |
| 2748 | 4262 | 3575.5 | 1329 | 4729.5 | 2007 | 701 | 12171.5 | 4336 | 6723.5 |
| 2715.5 | 4142 | 3520 | 1284.5 | 4623.5 | 1915 | 696 | 11712.5 | 4423 | 6180.5 |
| 2492 | 3800 | 3174 | 1202 | 4390 | 1915 | 713 | 10118 | 4496 | 6129 |
| 2556.5 | 3792 | 3383.5 | 1188.5 | 4427.5 | 1864 | 705 | 10561 | 4424.5 | 6122 |
| 49639.5 | 34579.5 | 11060.5 | 1499 | 4158 | 1875.5 | 675 | 9447 | 18099.5 | 13589 |
| 2891 | 4020.5 | 3507.5 | 1228.5 | 4240.5 | 1854.5 | 712 | 9708 | 4729 | 6374.5 |
| 2554 | 3879 | 3451 | 1204 | 4407 | 1828 | 691 | 9882 | 4654 | 6290 |
| 71524 | 40867 | 12732 | 1758 | 4853 | 1942 | 697 | 16358 | 5680 | 7734 |
| 3186 | 4696 | 3805 | 1224 | 4670 | 1822 | 694 | 11402 | 4670 | 6680 |
| 2396 | 3758 | 3260 | 1138 | 4952 | 1898 | 696 | 10025 | 4401 | 6271 |
| 2969 | 3991 | 3240 | 1180 | 4078 | 1813 | 697 | 9451 | 4645 | 6220 |
| 9429.5 | 20987 | 10109 | 2234.5 | 11229.5 | 2930 | 686.5 | 12920.5 | 8878 | 21793.5 |
| 10013 | 22129 | 12062 | 2535 | 13655 | 3276 | 678 | 12602 | 9361 | 22999 |
| 9260 | 21357 | 11954 | 2572 | 13700 | 3243 | 682 | 12190 | 8670 | 22288 |

| | | | | | | | | | |
|---|---|---|---|---|---|---|---|---|---|
| 9247 | 19733 | 9514 | 2090 | 10988 | 2761 | 688 | 12810 | 9031 | 20624 |
| 9886 | 21040 | 6976 | 1749 | 7982 | 2377 | 689 | 14783 | 10117 | 22982 |
| 7598 | 19067.5 | 7724.5 | 2235.5 | 7101 | 2368 | 695.5 | 15500 | 7428 | 21834.5 |
| 4304.5 | 10530.5 | 4460 | 1356 | 5423.5 | 1986.5 | 696 | 14192.5 | 5287 | 13669.5 |
| 2637 | 4665 | 6026 | 1672 | 584481 | 54040 | 658 | 15607 | 3927 | 6631 |
| 2665 | 4397 | 3849 | 1291 | 5298 | 2153 | 791 | 12919 | 4946 | 6535 |
| 2759.5 | 4090.5 | 3425 | 1241.5 | 5038 | 1959 | 695 | 11507 | 4714 | 6150 |
| 2667.5 | 3805 | 3215.5 | 1180 | 4243.5 | 1838 | 704 | 10105.5 | 4495.5 | 5950.5 |
| 2773 | 3984 | 3267 | 1151 | 4520 | 1876 | 708 | 11117 | 4555 | 6174 |
| 2880 | 4019 | 3339 | 1163 | 4183 | 1761 | 674 | 11267 | 4448 | 6037 |
| 2713 | 4382 | 3814 | 1369 | 4827 | 1977 | 734 | 12691 | 4160 | 6815 |
| 2592 | 3978 | 3277 | 1242 | 4357 | 1870 | 709 | 10578 | 4544 | 6228 |
| 2649 | 3898 | 3312 | 1186 | 4314 | 1914 | 730 | 10010 | 4614 | 6039 |
| 2664.5 | 4155.5 | 3625 | 1278 | 4765.5 | 1891 | 744 | 10212.5 | 4637 | 6617.5 |
| 14287 | 10103.5 | 4855 | 1402 | 4492.5 | 1938 | 773.5 | 10287.5 | 6947.5 | 7969 |
| 2771 | 4009.5 | 3496 | 1222.5 | 4537.5 | 1910.5 | 706 | 9939.5 | 4708 | 6426.5 |
| 2583.5 | 4002 | 3317 | 1235.5 | 4594 | 1954 | 722 | 10420.5 | 4679.5 | 6507.5 |
| 82153 | 44547 | 13809 | 1755 | 4741 | 2006 | 718 | 16261 | 5971 | 7412 |
| 2896.5 | 5879 | 3987 | 1361 | 5045.5 | 2020 | 757 | 11620 | 4629 | 8744.5 |
| 2533 | 4148 | 3662 | 1284 | 5663 | 2111 | 747 | 11319 | 4501 | 6832 |
| 3000 | 3951.5 | 3368.5 | 1223 | 4491 | 1925 | 699 | 10251.5 | 4515.5 | 6267 |
| 10830 | 21861 | 9986 | 2128 | 10946 | 2844 | 702 | 15154 | 10022 | 22890 |
| 10332 | 26280 | 13984 | 2961 | 15410 | 3630 | 745 | 11125 | 8984 | 26682 |
| 9253 | 23569 | 13358.5 | 2793 | 14357.5 | 3421.5 | 740.5 | 11794 | 8641 | 23962 |
| 9990.5 | 21070 | 10813.5 | 2195.5 | 11267.5 | 2878 | 705 | 12929.5 | 9322 | 21972.5 |
| 941 | 1361 | 1236 | 692 | 1851 | 1240 | 356 | 3235 | 1797 | 2503 |
| 39010 | 47055.5 | 15380.5 | 2154.5 | 3841.5 | 1668.5 | 482 | 16032 | 5335 | 7191.5 |
| 6457 | 6159 | 3701 | 1014 | 3372 | 1544 | 596 | 11398 | 3911 | 5137 |
| 2953 | 4185 | 8213 | 2502 | 402804 | 41493 | 647 | 8964 | 3643 | 5261 |
| 2452 | 3610 | 3356.5 | 1167 | 5008 | 2054 | 666 | 8474.5 | 3555 | 5289 |
| 2195 | 3377 | 3101 | 1104 | 4167 | 1830 | 709 | 8792 | 3721 | 5512 |
| 2165 | 3437.5 | 3014.5 | 1085 | 4137 | 1794 | 682.5 | 8271.5 | 3781 | 5449.5 |
| 2087 | 3451 | 3050 | 1074 | 4089 | 1748 | 692 | 9145 | 4017 | 5763 |
| 2184.5 | 3322.5 | 2866 | 1035.5 | 3880.5 | 1715 | 672 | 9350.5 | 3902 | 5494 |
| 2408 | 3487 | 3189 | 1161 | 3898 | 1835 | 695 | 9347 | 4114 | 5666 |
| 2139.5 | 3308.5 | 3048 | 1111 | 4061.5 | 1738 | 689.5 | 9274.5 | 3814.5 | 5547.5 |
| 1848 | 3173 | 2844 | 1065 | 3908 | 1752 | 704 | 9080 | 3721 | 5530 |
| 1863 | 3104 | 2941 | 1087 | 3879 | 1690 | 712 | 8697 | 3716 | 5602 |
| 16505.5 | 11888.5 | 4819 | 1164.5 | 3753.5 | 1746.5 | 712 | 8416 | 7105 | 7471 |
| 2077.5 | 3324 | 2946 | 1040 | 3741.5 | 1744.5 | 689 | 8177 | 3796 | 5548 |
| 2174 | 3462.5 | 3065 | 1134.5 | 4178 | 1861.5 | 704 | 8968.5 | 4192 | 5999 |
| 100739 | 49246 | 14250 | 1667 | 4386 | 1829 | 710 | 16335 | 6155 | 6954 |
| 2535 | 3661.5 | 3193.5 | 1078.5 | 4116.5 | 1709 | 681 | 8974 | 4512.5 | 6092 |
| 2003 | 3311.5 | 2916.5 | 1119.5 | 4895 | 1913.5 | 682 | 8675 | 4065.5 | 5763 |
| 2259 | 3520.5 | 3003 | 1086.5 | 4001 | 1766 | 696 | 8772 | 3935 | 5648 |
| 9564.5 | 18804.5 | 9455 | 1988.5 | 10880 | 2745.5 | 685 | 14482 | 9417 | 21135.5 |
| 10425.5 | 23100.5 | 12878 | 2716 | 14310.5 | 3497.5 | 696 | 11617.5 | 8432.5 | 23135 |
| 9940 | 21832 | 12119 | 2700 | 13680 | 3284 | 715 | 12149 | 8910 | 21906 |

| | | | | | | | | | |
|---|---|---|---|---|---|---|---|---|---|
| 10070 | 21072 | 9640 | 2068 | 10620 | 2768 | 674 | 14314 | 9570 | 21647 |
| 9509 | 20344 | 7167 | 1731 | 7682 | 2448 | 710 | 14822 | 10141 | 23661 |
| 11415 | 118157 | 189152.5 | 73657.5 | 7130 | 2703 | 643.5 | 15216 | 9757 | 69235 |
| 4717 | 12157 | 4847 | 1440 | 5207 | 1940 | 721 | 14194 | 5187 | 14481 |
| 3096.5 | 4866.5 | 7402.5 | 2059.5 | 832160.5 | 74775.5 | 733.5 | 15831.5 | 4610.5 | 7142 |
| 2877 | 4247.5 | 4157.5 | 1447 | 6086 | 2328 | 732.5 | 11089 | 4689 | 6561 |
| 2854.5 | 3969 | 3430.5 | 1224.5 | 4884.5 | 2026 | 746 | 11360 | 4881 | 6209 |
| 2661 | 3839.5 | 3436.5 | 1234.5 | 4783 | 1860 | 765 | 10519.5 | 4579.5 | 6157 |
| 2766 | 3786 | 3226 | 1156 | 4293 | 1786 | 741 | 10864 | 4787 | 5968 |
| 2754 | 3784 | 3050.5 | 1035 | 4069 | 1738 | 724.5 | 11074.5 | 4716.5 | 6021.5 |
| 2815 | 4146 | 3387 | 1269 | 4240 | 1897 | 737 | 11670 | 4581 | 6310 |
| 2571 | 3601 | 3111 | 1095 | 4058 | 1758 | 698 | 10183 | 4686 | 5966 |
| 2300 | 3374 | 2817 | 1081 | 4105 | 1812 | 704 | 10079 | 4067 | 5710 |
| 2243 | 3553.5 | 3132 | 1148.5 | 4020.5 | 1792 | 719.5 | 9927.5 | 4098 | 5961 |
| 36738 | 26154 | 7921.5 | 1343 | 3973 | 1708 | 747 | 9507.5 | 13113 | 10958 |
| 2407 | 3486 | 3269 | 1108 | 4003 | 1724 | 733 | 9202 | 4140 | 5970 |
| 2595.5 | 3766 | 3278.5 | 1173.5 | 4361 | 1768.5 | 744 | 9884.5 | 4842 | 6346 |
| 147160.5 | 75361.5 | 21027 | 1933.5 | 4644 | 1889.5 | 730.5 | 20651 | 7406.5 | 7587.5 |
| 3246 | 4374 | 3501 | 1195 | 4367 | 1828 | 737 | 11440 | 5038 | 6689 |
| 2559 | 3782.5 | 3332.5 | 1230.5 | 6182.5 | 2157 | 724 | 10433.5 | 4797.5 | 6380.5 |
| 2569 | 3833 | 3137 | 1021 | 4040 | 1785 | 708 | 10092 | 4299 | 5897 |
| 10826 | 22358 | 10433 | 2183 | 11151 | 2873 | 728 | 13952 | 9830 | 22639 |
| 12919.5 | 25154 | 13138 | 2841.5 | 13817 | 3294.5 | 764.5 | 13068.5 | 9393.5 | 25254 |
| 11867 | 24131 | 12571 | 2651 | 13384 | 3462 | 762 | 12396 | 9057 | 23868 |
| 11496 | 21787 | 10741 | 2225 | 11293 | 2828 | 753 | 14292 | 10053 | 23282 |
| 8788 | 22458.5 | 7301.5 | 1757.5 | 8414.5 | 2591 | 718 | 13058.5 | 9461.5 | 26242 |
| 12363 | 112247 | 192562 | 75330 | 7998 | 2828 | 779 | 16731 | 9274 | 47469 |
| 3959 | 11773.5 | 4578.5 | 1344 | 5535 | 2038 | 750 | 14083 | 5188.5 | 15143.5 |
| 2936 | 5742.5 | 7037 | 2101 | 664148 | 62610.5 | 678 | 15921.5 | 4498.5 | 8485 |
| 2957 | 4558 | 4620.5 | 1538.5 | 7829 | 2647.5 | 826.5 | 13179 | 4988 | 7227 |
| 2996 | 4481 | 4610 | 1693.5 | 6948 | 2515 | 733 | 11695.5 | 4792.5 | 6684 |
| 2744.5 | 4158 | 3697.5 | 1303 | 5452 | 2117.5 | 745 | 10830.5 | 4651.5 | 6675.5 |
| 2715 | 4359 | 3789.5 | 1278.5 | 4933 | 1937 | 776 | 11355.5 | 4586 | 6593 |
| 2945 | 4205 | 3531.5 | 1246 | 4618.5 | 1896 | 738 | 11564 | 4773 | 6551 |
| 2766 | 3963 | 3597 | 1344 | 4510 | 2032 | 712 | 10561 | 4529 | 6474 |
| 2663 | 3995 | 3311 | 1231 | 4293 | 1805 | 726 | 10624 | 4360 | 6093 |
| 2405.5 | 3440 | 3018.5 | 1133.5 | 4123 | 1812.5 | 720.5 | 9826 | 4310 | 5814 |
| 2431 | 3443.5 | 3197.5 | 1176.5 | 4167 | 1884.5 | 707.5 | 9850 | 4375.5 | 5981 |
| 20786.5 | 13904 | 5310.5 | 1297 | 4123 | 1843.5 | 707 | 9382.5 | 7967.5 | 8089 |
| 2488 | 3798.5 | 3242 | 1155 | 4202 | 1783.5 | 686 | 9442.5 | 4235.5 | 6008 |
| 2516 | 3835 | 3364 | 1180 | 4613 | 1881 | 701 | 9976 | 4577 | 6210 |
| 155967 | 82420 | 22946 | 1994 | 4481 | 1828 | 703 | 21788 | 7525 | 7515 |
| 3147.5 | 4563 | 3830.5 | 1252 | 4542 | 1833 | 701.5 | 10785.5 | 4925 | 6665 |
| 2435 | 3841 | 3125 | 1169 | 5777 | 2033 | 707 | 10044 | 4514 | 6206 |
| 2493 | 3878.5 | 3073.5 | 1137.5 | 4210 | 1803 | 703.5 | 9638 | 4233 | 5895.5 |
| 10233 | 22494 | 10020 | 2145 | 11509 | 2928 | 726 | 14493 | 9372 | 23539 |
| 10414.5 | 23043.5 | 12498.5 | 2667 | 14101 | 3395 | 690 | 12616.5 | 9483.5 | 22767.5 |
| 11271 | 22742 | 12081 | 2427 | 13161 | 3205 | 698 | 13112 | 9418 | 23385 |

| | | | | | | | | | |
|---|---|---|---|---|---|---|---|---|---|
| 11338.5 | 21742 | 10329 | 2142 | 11815 | 2920.5 | 731.5 | 13454.5 | 9066.5 | 22586 |
| 8933.5 | 21178 | 7147.5 | 1754 | 8292 | 2390 | 724.5 | 14119.5 | 9753 | 24551.5 |
| 12380 | 139364 | 232868 | 82226.5 | 8835.5 | 3846.5 | 639 | 15431 | 10081.5 | 80332.5 |
| 5154 | 13340 | 5744 | 1846 | 5435 | 2119 | 718 | 14509 | 5498 | 16799 |
| 2821 | 4979 | 7165 | 2169.5 | 750304.5 | 69850 | 695 | 15347 | 4115 | 7191 |
| 2916 | 4319 | 4540 | 1782 | 7071 | 2810 | 797 | 12623 | 4939 | 6962 |
| 3080 | 4074 | 4357 | 1572 | 5981 | 2406 | 716 | 11083 | 4654 | 6140 |
| 2835 | 3931 | 3985 | 1353 | 5245 | 2058 | 728 | 10991 | 4570 | 6223 |
| 2979 | 3887 | 3679 | 1176 | 4662 | 1923 | 726 | 10948 | 4591 | 6267 |
| 3046 | 3887 | 3255.5 | 1106.5 | 4292 | 1732.5 | 715 | 11552 | 4438.5 | 5953.5 |
| 2862 | 3991 | 3482 | 1325 | 4455 | 1928 | 735 | 11825 | 4767 | 6369 |
| 2707 | 3825 | 3098 | 1211 | 4217 | 1811 | 693 | 10662 | 4631 | 6047 |
| 2278 | 3439 | 3061 | 1079 | 3942 | 1758 | 703 | 9983 | 4190 | 5856 |
| 2319.5 | 3538 | 3091.5 | 1146 | 4045 | 1806 | 689.5 | 9427 | 4087.5 | 5874 |
| 38697 | 27759 | 8694 | 1356 | 4133.5 | 1699.5 | 675.5 | 9090.5 | 13395.5 | 10790.5 |
| 2586 | 3615 | 3278 | 1127 | 4112 | 1845 | 717 | 9565 | 4423 | 6126 |
| 2724.5 | 3834 | 3394.5 | 1133 | 4547.5 | 1846 | 686 | 10120 | 4802 | 6530 |
| 118163 | 62594.5 | 17559 | 1870.5 | 4662 | 1896 | 723.5 | 20936.5 | 6613.5 | 7614 |
| 3240 | 4460 | 3836 | 1267.5 | 4624 | 1922 | 718.5 | 11303 | 4819.5 | 6766 |
| 2699 | 3739 | 3403 | 1150 | 5555 | 2011 | 680 | 10113 | 4631 | 6290 |
| 2587 | 3787 | 3013 | 1080 | 4104 | 1768 | 709 | 9640 | 4269 | 5966 |
| 10755 | 22294 | 10387 | 2174 | 11549 | 2935 | 754 | 13865 | 9268 | 22806 |
| 11878 | 24788.5 | 13121.5 | 2658 | 14016 | 3551 | 739.5 | 12399.5 | 9436 | 23894.5 |
| 10865 | 22633 | 12473.5 | 2592 | 13438.5 | 3249.5 | 698 | 12652.5 | 9404.5 | 23066 |
| 10286.5 | 21351 | 9847 | 2031.5 | 10933 | 2707.5 | 710.5 | 13657 | 9193.5 | 22217.5 |
| 9128 | 21288 | 7051 | 1746 | 8006 | 2363 | 725 | 13636 | 9550 | 24698 |
| 11354 | 114087 | 174384 | 64211 | 7379 | 2971 | 676 | 15138 | 8971 | 61632 |
| 4619 | 8862.5 | 4810 | 1434.5 | 5156.5 | 1970 | 731 | 15017 | 5332 | 10516 |
| 3197 | 5236 | 7002 | 1961.5 | 701140 | 68513.5 | 671 | 16551.5 | 4542.5 | 7457 |
| 3092.5 | 4737.5 | 4512 | 1553 | 6413.5 | 2475.5 | 834 | 14425 | 5222 | 7231 |
| 2847.5 | 4185 | 3939 | 1365 | 5307 | 2081.5 | 772 | 11781.5 | 4645 | 6533 |
| 2900 | 3990 | 3466 | 1157 | 4805 | 1916 | 769 | 10353 | 4868 | 6237 |
| 2969 | 4178 | 3658 | 1264 | 4777 | 1871 | 743 | 11362 | 4620 | 6383 |
| 3077 | 4111 | 3331 | 1145 | 4231 | 1833 | 734 | 11529 | 4641 | 6114 |
| 2996.5 | 3952.5 | 3303 | 1218 | 4182 | 1819 | 706.5 | 10749 | 4692 | 6133 |
| 2824 | 3967 | 3381 | 1126 | 4285.5 | 1793 | 693.5 | 10945.5 | 4754.5 | 6213.5 |
| 2579.5 | 3703 | 3046 | 1117.5 | 4007 | 1848.5 | 718.5 | 10368 | 4335.5 | 5962 |
| 2400 | 3476 | 3071.5 | 1124 | 4091.5 | 1690 | 714.5 | 9559.5 | 4295.5 | 5706 |
| 19630 | 13551 | 5466 | 1275 | 4117 | 1845 | 697 | 9420 | 7532 | 7841 |
| 2652 | 3858 | 3221 | 1182.5 | 4078.5 | 1808 | 753.5 | 9879 | 4319.5 | 6149.5 |
| 2616 | 3587 | 3350 | 1196 | 4209 | 1826 | 743 | 10136 | 4820 | 6452 |
| 91149 | 47183 | 13946 | 1658 | 4437 | 1849 | 718 | 16931 | 6346 | 7526 |
| 3173 | 4633 | 3687 | 1262 | 4541 | 1872 | 736 | 11420 | 4968 | 6867 |
| 2759 | 3921 | 3289 | 1202 | 5308 | 2028 | 719 | 10190 | 4966 | 6156 |
| 2699 | 3898.5 | 3311 | 1119 | 4407.5 | 1785.5 | 696 | 10035 | 4371.5 | 5960.5 |
| 11818 | 21803 | 10378 | 2258 | 11379 | 2899 | 752 | 14037 | 9503 | 23228 |
| 12098 | 24944 | 12677 | 2812 | 14150 | 3521 | 712 | 12989 | 9967 | 24796 |
| 11865 | 23437 | 13204 | 2663 | 13495 | 3332 | 726 | 12618 | 9124 | 23028 |

| | | | | | | | | | |
|---|---|---|---|---|---|---|---|---|---|
| 11309 | 22075 | 10325 | 2324.5 | 11812 | 2928.5 | 752.5 | 14185 | 9610 | 22775 |
| 9806 | 22043 | 7518 | 1797 | 8404 | 2383 | 719 | 14920 | 9443 | 25636 |
| 11326 | 118800.5 | 186235.5 | 68672.5 | 7443 | 2891 | 644 | 15317 | 9509.5 | 65136.5 |
| 4060 | 9176 | 4627 | 1394 | 5286 | 2040 | 661 | 14355 | 5056 | 11709 |
| 2931 | 5064 | 6354 | 1893 | 643248 | 57474 | 706 | 16431 | 4345 | 7372 |
| 2959.5 | 4443 | 4291 | 1529 | 6440.5 | 2480.5 | 789.5 | 14062.5 | 4889.5 | 7020 |
| 2708 | 3955 | 3737 | 1270 | 5162 | 2056 | 689 | 10553 | 4644 | 6478 |
| 2647 | 3903.5 | 3383 | 1259.5 | 4860 | 1931 | 715.5 | 9834.5 | 4598 | 6305 |
| 2755.5 | 4088 | 3408 | 1158 | 4506.5 | 1853.5 | 699 | 11231.5 | 4489.5 | 6169.5 |
| 2847 | 3937 | 3299 | 1081 | 4223 | 1836 | 689 | 10900 | 4621 | 6323 |
| 2657 | 3975 | 3441 | 1226 | 4374 | 1855 | 703 | 11890 | 4571 | 6176 |
| 2708 | 3844 | 3177 | 1104 | 4205 | 1806 | 725 | 10366 | 4580 | 5934 |
| 2343 | 3425 | 2931 | 1108 | 4041 | 1779 | 675 | 9260 | 4380 | 5712 |
| 2247.5 | 3487.5 | 2858 | 1046 | 3966 | 1749 | 669.5 | 9335 | 4100 | 5825.5 |
| 23758 | 16404 | 5810 | 1272 | 4048 | 1757 | 664 | 8918 | 8787 | 8274 |
| 2648 | 3701 | 3433 | 1163 | 4034 | 1811 | 703 | 9369 | 4251 | 6028 |
| 2491.5 | 3736.5 | 3266.5 | 1139 | 4214.5 | 1797.5 | 681 | 9590.5 | 4617.5 | 6391.5 |
| 97748 | 53166 | 15678 | 1831 | 4662 | 1896 | 712 | 18345 | 5920 | 7839 |
| 2597.5 | 4334.5 | 3384 | 1173 | 4331.5 | 1788 | 683.5 | 10484 | 4468.5 | 6564 |
| 2466.5 | 3650.5 | 3148.5 | 1128.5 | 5174 | 1938.5 | 669.5 | 9673 | 4670 | 6172.5 |
| 2847 | 3804.5 | 3365.5 | 1159 | 4376 | 1859 | 693 | 9980 | 4552 | 5955 |
| 10230 | 20861.5 | 10289.5 | 2275.5 | 11302 | 2865.5 | 680.5 | 13491 | 9344 | 22356.5 |
| 10482.5 | 23098 | 12805 | 2676.5 | 13808.5 | 3374 | 690 | 11816.5 | 8888 | 24074 |
| 10721 | 22198 | 12104 | 2637 | 13722 | 3369 | 687.5 | 11984 | 8681 | 22916.5 |
| 9896 | 20661 | 9605 | 2255 | 11235 | 2850 | 699 | 13636 | 9141 | 21193 |
| 9429.5 | 22239.5 | 7270.5 | 1732 | 8685 | 2411 | 711.5 | 13846.5 | 9752 | 24857.5 |
| 11596 | 110717 | 170735 | 62488 | 7425 | 2663 | 641 | 15513 | 9605 | 61901 |
| 4202 | 11014 | 4797 | 1442.5 | 5095.5 | 1952 | 704.5 | 14087.5 | 5164 | 14650.5 |
| 2993 | 5030 | 6317 | 1757.5 | 668612.5 | 62526.5 | 693.5 | 16416 | 4474.5 | 7334 |
| 3034 | 4405 | 4187 | 1459 | 6272 | 2417 | 833 | 14238 | 4853 | 7088 |
| 2902 | 4068 | 3814 | 1329 | 5434 | 2069 | 728 | 12178 | 4787 | 6640 |
| 2764 | 3935 | 3449 | 1201 | 4758 | 1947 | 718 | 10348 | 4739 | 6162 |
| 2971 | 4206.5 | 3595 | 1287 | 4658.5 | 1905.5 | 717 | 11154.5 | 4595 | 6280 |
| 2864 | 3921 | 3266 | 1135 | 4235 | 1751 | 706 | 11392 | 4543 | 6164 |
| 2653 | 4009 | 3463 | 1196 | 4297 | 1849 | 702 | 10870 | 4516 | 6364 |
| 2717 | 3684 | 3098 | 1156 | 4183 | 1800 | 689 | 10353 | 4550 | 5795 |
| 2736 | 3517.5 | 3015.5 | 1114 | 4098.5 | 1759 | 698.5 | 9562 | 4435.5 | 5878 |
| 2407 | 3457 | 3218.5 | 1101 | 4082 | 1784.5 | 723 | 9769 | 4369.5 | 5748.5 |
| 15404.5 | 10751.5 | 4857 | 1309.5 | 4144 | 1810.5 | 705.5 | 9497 | 6816.5 | 7931 |
| 2857 | 3889 | 3128 | 1220 | 4259 | 1823 | 730 | 10160 | 4970 | 6135 |
| 2453.5 | 3700 | 3354.5 | 1169 | 4187.5 | 1865.5 | 663 | 9355 | 4681 | 6188 |
| 92873 | 49162.5 | 14809 | 1750 | 4535.5 | 1827 | 734 | 17534 | 6078.5 | 7476.5 |
| 3213 | 4784 | 3717 | 1270 | 4507 | 1894 | 745 | 11560 | 4883 | 6788 |
| 2671.5 | 3599 | 3312.5 | 1146.5 | 5177 | 1929.5 | 673 | 9421.5 | 4795.5 | 6134.5 |
| 2971 | 3870 | 3271 | 1176 | 4248 | 1848 | 714 | 10336 | 4389 | 5926 |
| 10750.5 | 20997 | 10470.5 | 2251.5 | 11136.5 | 2947 | 731.5 | 13876.5 | 9567.5 | 22935.5 |
| 11698 | 23894 | 12899 | 2633 | 13874 | 3251 | 714 | 13097 | 9740 | 24115 |
| 10772 | 22639 | 12484 | 2612 | 13670 | 3458 | 704 | 12150 | 8621 | 21776 |

| | | | | | | | | | |
|---|---|---|---|---|---|---|---|---|---|
| 11074 | 21164 | 10132 | 2184 | 10993 | 2769 | 727 | 13660 | 9470 | 21933 |
| 9650 | 21565 | 7261 | 1712 | 8101 | 2431 | 699 | 14669 | 9925 | 22556 |
| 11586 | 127042 | 199596 | 73051 | 7610 | 2907 | 648 | 15882 | 9311 | 71970 |
| 4769 | 10846 | 5172.5 | 1594.5 | 5349 | 2062.5 | 750 | 14742.5 | 5468 | 12765 |
| 3330.5 | 5397 | 7406.5 | 2102.5 | 696144.5 | 66231.5 | 721 | 16384.5 | 4490 | 7379.5 |
| 2918.5 | 4514.5 | 4354 | 1513 | 6156 | 2385.5 | 809.5 | 13092.5 | 4962.5 | 6908 |
| 2870 | 4014.5 | 3932 | 1357.5 | 5297 | 2159 | 742.5 | 11814.5 | 4989.5 | 6426.5 |
| 2957 | 4119 | 4005 | 1330 | 5114 | 1962 | 738 | 11017 | 4646 | 6272 |
| 2939.5 | 4165.5 | 3799.5 | 1244 | 4725 | 1947 | 741 | 11888 | 4962 | 6430.5 |
| 3133 | 3998 | 3314.5 | 1212.5 | 4277.5 | 1824 | 692 | 11680.5 | 4905.5 | 6072.5 |
| 3024 | 4349 | 3757 | 1368 | 4617 | 1967 | 759 | 12075 | 4640 | 6616 |
| 2935 | 3957 | 3358.5 | 1187 | 4390.5 | 1927.5 | 728.5 | 11237.5 | 4646 | 6438 |
| 2648 | 3641 | 3111 | 1092 | 3995 | 1732 | 718 | 9688 | 4493 | 5745 |
| 2509.5 | 3603.5 | 3216 | 1210 | 4087.5 | 1818.5 | 745 | 10470 | 4360.5 | 6101.5 |
| 33947.5 | 23594 | 7704 | 1426 | 4070 | 1831.5 | 729 | 9821.5 | 11382.5 | 9841 |
| 2625 | 3830 | 3245 | 1201 | 4224 | 1815 | 716 | 9381 | 4209 | 6091 |
| 2738 | 3929 | 3563 | 1204 | 4483 | 1911 | 733 | 10519 | 4974 | 6673 |
| 107291 | 58895 | 17705 | 1856 | 4819 | 1939 | 710 | 19913 | 6165 | 7778 |
| 3264.5 | 4616.5 | 3838 | 1289.5 | 4464.5 | 1834.5 | 726.5 | 10882 | 4959.5 | 7052 |
| 2660 | 3990 | 3365 | 1216 | 5661 | 2039 | 734 | 11160 | 4726 | 6639 |
| 3096 | 4259.5 | 3516 | 1268.5 | 4292.5 | 1832.5 | 731.5 | 11012.5 | 5001.5 | 6434.5 |
| 11276 | 20611 | 10216.5 | 2151 | 10855 | 2832.5 | 735 | 15324 | 9856.5 | 20924.5 |
| 11868.5 | 23789.5 | 13033 | 2750 | 14116.5 | 3550.5 | 707.5 | 12101.5 | 9067.5 | 22979.5 |
| 12234 | 23826.5 | 12699.5 | 2883.5 | 13761 | 3530 | 745 | 12499.5 | 9082.5 | 23719.5 |
| 10898 | 22246 | 10921 | 2246 | 11546 | 2908 | 717 | 14224 | 9569 | 23443 |
| 9702 | 21970 | 7188 | 1854 | 8528 | 2440 | 683 | 14636 | 10101 | 24368 |
| 10504 | 107356 | 162083 | 58381 | 7327 | 2563 | 615 | 15875 | 8965 | 62072 |
| 4192.5 | 11548.5 | 4895 | 1547.5 | 5353.5 | 2008.5 | 664.5 | 13727.5 | 5289.5 | 14161 |
| 2912 | 4862 | 6483 | 2128 | 315930 | 34592.5 | 641 | 15233.5 | 3970.5 | 6770 |
| 2974 | 4452 | 4021.5 | 1444.5 | 9979.5 | 2611.5 | 761 | 13134 | 4877.5 | 7007 |
| 2832 | 4195 | 3618 | 1323 | 6325 | 2094 | 707 | 11124 | 4650 | 6334 |
| 2705 | 3919 | 3413 | 1272 | 4982 | 1896 | 666 | 10220 | 4637 | 6198 |
| 2748 | 4085 | 3394 | 1166 | 4876 | 1862 | 677 | 10376 | 4420 | 6085 |
| 2761 | 3951 | 3184 | 1119 | 4370 | 1880 | 676 | 11562 | 4553 | 6031 |
| 2669 | 4013 | 3396 | 1232 | 4346 | 1863.5 | 668 | 11440 | 4362.5 | 6187 |
| 2647.5 | 3820 | 3007 | 1109 | 4478 | 1777.5 | 669.5 | 10017.5 | 4414.5 | 5659.5 |
| 2633 | 3566 | 3165 | 1142 | 4174 | 1859 | 705 | 9835 | 4386 | 6035 |
| 2408 | 3526 | 2979 | 1139 | 4186.5 | 1821.5 | 668.5 | 9399 | 4012 | 5708 |
| 6602 | 5873.5 | 3845.5 | 1151.5 | 4104.5 | 1765.5 | 683 | 8809 | 5078 | 6405.5 |
| 2376 | 3661 | 3130 | 1154 | 4383 | 1823 | 688 | 9422 | 4113 | 6020 |
| 2445.5 | 3859 | 3249 | 1121.5 | 4529.5 | 1900 | 700 | 10199.5 | 4592 | 6371 |
| 186500 | 98303 | 27308 | 2170 | 4700 | 1876 | 660 | 24986 | 7972 | 7961 |
| 3104.5 | 4197 | 3527.5 | 1220 | 4515 | 1881 | 667.5 | 10423.5 | 4613.5 | 6383.5 |
| 2593 | 3633 | 3161 | 1200 | 12141 | 2831 | 688 | 9889 | 4544 | 6145 |
| 2456 | 3618 | 3179 | 1128 | 4181 | 1732 | 685 | 8972 | 4245 | 5870 |
| 9848 | 20401 | 9842 | 2080 | 11139 | 2894 | 678 | 14496 | 9921 | 22407 |
| 10885 | 23675 | 13425.5 | 2727 | 14467 | 3446 | 691 | 12109.5 | 8903.5 | 22808.5 |
| 9956 | 21531 | 11432 | 2588 | 13046 | 3213 | 699 | 13318 | 9178 | 22917 |

| | | | | | | | | | |
|---|---|---|---|---|---|---|---|---|---|
| 10048.5 | 21235 | 10053 | 2192 | 10992.5 | 2783.5 | 682.5 | 13903.5 | 9622 | 22022.5 |
| 9315 | 20157 | 7040 | 1681 | 7946 | 2315 | 703 | 14937 | 10031 | 22941 |
| 13132 | 133920 | 209983 | 77094.5 | 7841.5 | 2940 | 626 | 14491 | 9926 | 76398.5 |
| 4842 | 11232.5 | 5500.5 | 1717 | 5467 | 2008.5 | 689.5 | 14262.5 | 5542.5 | 14071.5 |
| 3013 | 4833.5 | 6570 | 1905.5 | 558511 | 52827.5 | 643 | 14321 | 4134.5 | 6787.5 |
| 2887.5 | 4462.5 | 4025 | 1417 | 5849.5 | 2346.5 | 858.5 | 16601.5 | 5361.5 | 7068.5 |
| 2848.5 | 4149 | 3847 | 1411 | 5404.5 | 2089 | 732.5 | 12402 | 4851 | 6685 |
| 2696 | 3932 | 3496 | 1357 | 5024 | 2053 | 705 | 10621 | 4631 | 6292 |
| 2706 | 3907 | 3413 | 1190.5 | 4497.5 | 1840.5 | 681.5 | 10899 | 4415.5 | 6302 |
| 3226.5 | 4044.5 | 3529.5 | 1251.5 | 4662.5 | 1873 | 688 | 11350.5 | 4558 | 6066.5 |
| 2846.5 | 4159 | 3338 | 1228 | 4341.5 | 1899 | 679 | 11539 | 4281.5 | 6386.5 |
| 2623 | 3811.5 | 3156.5 | 1215 | 4206.5 | 1810.5 | 724.5 | 11118 | 4446.5 | 6037 |
| 2785.5 | 3572 | 3272.5 | 1149.5 | 4210.5 | 1782 | 687 | 10118.5 | 4469 | 5997 |
| 2692 | 3636 | 3125 | 1122 | 4164 | 1869 | 691 | 10074 | 4596 | 5977 |
| 55096 | 41971.5 | 13670.5 | 1711.5 | 4131 | 1809 | 674.5 | 9146.5 | 19994.5 | 16171.5 |
| 2792 | 3906.5 | 3566.5 | 1257.5 | 4318 | 1796 | 707 | 9801 | 4658 | 6361 |
| 2601.5 | 3934 | 3392.5 | 1186.5 | 4510.5 | 1852.5 | 697.5 | 10193 | 4589.5 | 6451 |
| 106636 | 56555 | 16804 | 1746.5 | 4641.5 | 1898 | 699 | 18297.5 | 6313 | 7717.5 |
| 3535 | 5048 | 4022 | 1358 | 4691 | 1889 | 720 | 11085 | 4606 | 6522 |
| 2642 | 4048 | 3616 | 1232.5 | 5288.5 | 2018.5 | 720 | 10478 | 4661 | 6590 |
| 2778 | 3912 | 3284 | 1128 | 4342 | 1752 | 685 | 9994 | 4694 | 5989 |
| 9392.5 | 19285 | 9681 | 2261 | 11136.5 | 2905.5 | 711 | 13760 | 9431 | 22472 |
| 10921.5 | 23532 | 12700.5 | 2690.5 | 13219.5 | 3332 | 712 | 12785.5 | 9357 | 24149 |
| 10115.5 | 21585 | 12896.5 | 2557.5 | 13812.5 | 3408.5 | 691 | 12006 | 8790 | 22879.5 |
| 10496 | 21749 | 10273 | 2190 | 11348 | 2803 | 712 | 13307 | 9166 | 21271 |
| 9868 | 23064 | 7635 | 1741.5 | 8186.5 | 2416.5 | 667.5 | 15000 | 9930 | 24340.5 |
| 8587 | 21866 | 9193 | 2668 | 7363 | 2424 | 739 | 15854 | 8012 | 23361 |
| 3976 | 10036 | 4324.5 | 1285 | 5000 | 1843 | 674 | 14186 | 4935 | 12302.5 |
| 2606.5 | 4510 | 6324 | 1710 | 657391.5 | 60122.5 | 674.5 | 14838 | 3940.5 | 6787 |
| 2797 | 4267 | 3905 | 1391 | 5974 | 2294 | 733 | 12518 | 4641 | 6794 |
| 2807 | 4222 | 3730 | 1298 | 5189 | 2070 | 694 | 11464 | 4692 | 6379 |
| 2617 | 3856 | 3342.5 | 1212.5 | 4480 | 1863.5 | 702 | 10116.5 | 4588.5 | 6074 |
| 2829 | 4231 | 3529 | 1258 | 4674 | 1918 | 675 | 10976 | 4577 | 6060 |
| 2883.5 | 4186 | 3446 | 1175 | 4303.5 | 1793 | 680.5 | 11483.5 | 4482 | 6397 |
| 2578 | 4106 | 3596 | 1323 | 4331 | 1898 | 677 | 11436 | 4105 | 6200 |
| 2804 | 3960.5 | 3398.5 | 1183 | 4348 | 1791 | 713.5 | 11030 | 4694.5 | 6432.5 |
| 2694 | 3983.5 | 3475.5 | 1213 | 4497 | 1939.5 | 741.5 | 10363 | 4533.5 | 6109 |
| 2560 | 4017 | 3577 | 1263 | 4742 | 1941 | 772 | 10908 | 4538 | 6748 |
| 11982.5 | 8586.5 | 4823.5 | 1367.5 | 4403 | 1787.5 | 744 | 10052 | 6754.5 | 7722.5 |
| 2820 | 3806 | 3238 | 1142 | 4146 | 1838 | 690 | 9430 | 4629 | 5880 |
| 2751.5 | 3878.5 | 3450.5 | 1232.5 | 4588.5 | 1865 | 730.5 | 10124 | 4781.5 | 6666 |
| 107596 | 60341 | 18075 | 1931 | 4949 | 1935 | 734 | 18715 | 6380.5 | 7995 |
| 3053 | 6107 | 4299 | 1407 | 5181 | 2105 | 787 | 12317 | 4656 | 8827 |
| 2554 | 3894 | 3510 | 1199 | 5480 | 2112 | 731 | 10930 | 4677 | 6773 |
| 2801.5 | 4025 | 3474.5 | 1269 | 4260 | 1839.5 | 708 | 10271 | 4651 | 6118 |
| 10837 | 22108.5 | 10638 | 2258 | 11821 | 3052 | 729 | 14277.5 | 9681.5 | 24166 |
| 10364 | 26037 | 14257 | 3048 | 15300 | 3829 | 731 | 10853 | 8846 | 27388 |
| 10374 | 23194 | 13084 | 2802 | 13976 | 3470 | 723 | 12008 | 8911 | 23023 |

| | | | | | | | | | |
|---|---|---|---|---|---|---|---|---|---|
| 10236.5 | 21979 | 9765.5 | 2136.5 | 11188.5 | 2706 | 701 | 14283 | 9117.5 | 22302.5 |
| 960 | 1351 | 1185.5 | 651.5 | 1782.5 | 1163 | 353.5 | 3259.5 | 1810.5 | 2475.5 |
| 48437.5 | 30938.5 | 8163.5 | 1320.5 | 3499.5 | 1573 | 490 | 19539.5 | 5177 | 5775.5 |
| 8078 | 6168 | 3455 | 1030 | 3469 | 1611 | 609 | 10396 | 3837 | 5242 |
| 2743 | 4091 | 7868 | 2567 | 470805 | 46615 | 647 | 8815 | 3769 | 5395 |
| 2491 | 3769 | 3426 | 1219 | 5511 | 2105 | 733 | 9541 | 4056 | 5709 |
| 2656.5 | 3684.5 | 3223 | 1083 | 4510.5 | 1928 | 680.5 | 9121 | 3851.5 | 5568 |
| 2080 | 3296 | 2931 | 1086 | 4258 | 1819 | 710 | 8634 | 3957 | 5507 |
| 2423.5 | 3543 | 3180.5 | 1104.5 | 4190 | 1839.5 | 740.5 | 9599.5 | 4307 | 5902.5 |
| 2467 | 3482 | 2897.5 | 1062 | 3993.5 | 1769.5 | 686 | 10070 | 4155 | 5729.5 |
| 2232 | 3630 | 3207 | 1198 | 4027 | 1833 | 710 | 9435 | 4169 | 6008 |
| 2038 | 3319 | 3030 | 1076 | 3963 | 1752 | 687 | 9030 | 3887 | 5571 |
| 1806.5 | 3116 | 2807.5 | 1058 | 3847 | 1717 | 701.5 | 8120.5 | 3660 | 5261 |
| 1923.5 | 3053.5 | 2885 | 1071 | 3972.5 | 1769.5 | 690 | 8220 | 3803.5 | 5500.5 |
| 17462 | 12306 | 4754 | 1150 | 3626 | 1689 | 676 | 8386 | 7352 | 7672 |
| 2136.5 | 3151.5 | 2995 | 1063.5 | 3985.5 | 1753 | 709 | 8750 | 4071 | 5628 |
| 2142 | 3329 | 2942 | 1121 | 4119 | 1765 | 700 | 8505 | 3875 | 5714 |
| 85167 | 43099 | 12532 | 1595 | 4380 | 1805 | 721 | 14671 | 6027 | 6935 |
| 2627 | 3807 | 3348 | 1121 | 4345 | 1810 | 690 | 9132 | 4517 | 6145 |
| 2136 | 3383 | 3090 | 1084 | 5273.5 | 2043.5 | 694 | 9428 | 4271 | 5791 |
| 2278.5 | 3636 | 3110 | 1048 | 4140 | 1823 | 686 | 9249 | 4159.5 | 5867 |
| 10016 | 19541 | 9986 | 2041 | 10952 | 2919 | 687 | 14581 | 9881 | 21497 |
| 10559 | 23184 | 12018 | 2683 | 13950 | 3402 | 711 | 12187 | 9279 | 23545 |
| 10210 | 22046 | 11866.5 | 2600 | 13282 | 3198 | 700.5 | 12581 | 9030.5 | 22736 |
| 9614.5 | 19806.5 | 9003.5 | 2043.5 | 10768.5 | 2765.5 | 685 | 13055 | 9434 | 21728 |
| 9362 | 21464.5 | 7164.5 | 1763.5 | 8575 | 2374 | 714.5 | 14498.5 | 9828 | 23747.5 |
| 10350 | 109059.5 | 174259.5 | 69096.5 | 7118 | 2660 | 635 | 15635 | 9138 | 61817 |
| 4779 | 10664 | 4844 | 1440 | 5334 | 2002 | 713 | 14320 | 5508 | 13688 |
| 3374 | 5644 | 7702 | 2205 | 796683 | 74970 | 745 | 17221 | 4562 | 8177 |
| 2861 | 4492 | 4177 | 1398 | 6205 | 2345 | 842 | 13793 | 4821 | 7419 |
| 2727 | 3987.5 | 3672 | 1239 | 4716.5 | 1932 | 782 | 11078 | 5011 | 6233.5 |
| 2670 | 3900.5 | 3468.5 | 1141 | 4687.5 | 1845.5 | 774.5 | 10562.5 | 4755.5 | 6353.5 |
| 2742.5 | 4004 | 3292.5 | 1148 | 4473 | 1809.5 | 752.5 | 11175 | 4701 | 6317.5 |
| 2930 | 3980 | 3188 | 1105.5 | 4057 | 1756 | 711.5 | 11174 | 4659.5 | 6034.5 |
| 2860.5 | 3885 | 3215 | 1271.5 | 4274 | 1884 | 684.5 | 10764.5 | 4697.5 | 6327 |
| 2652 | 3908.5 | 3172.5 | 1168 | 4371.5 | 1813 | 751.5 | 10737 | 4605 | 6236.5 |
| 2242 | 3462 | 3014 | 1135 | 3897 | 1686 | 715 | 9732 | 4213 | 5856 |
| 2252.5 | 3497 | 3111 | 1114 | 4046 | 1781 | 721.5 | 9447 | 4120.5 | 5977.5 |
| 36340 | 25651.5 | 7941 | 1378 | 4197 | 1868 | 729.5 | 9443 | 12647.5 | 10461 |
| 2585.5 | 3783.5 | 3234 | 1074 | 3998 | 1757 | 777.5 | 9428 | 4457 | 6288 |
| 2629 | 3962.5 | 3455 | 1118 | 4207 | 1800 | 766 | 10189 | 4698 | 6713 |
| 124395 | 63287 | 17647 | 1791 | 4703 | 1943 | 755 | 18532 | 6852 | 7837 |
| 3205 | 4629 | 3800 | 1243 | 4748 | 1836 | 738 | 10926 | 4951 | 6774 |
| 2395.5 | 3782 | 3365.5 | 1185.5 | 5983.5 | 2078 | 728 | 10093 | 4585 | 6266.5 |
| 2874 | 3863.5 | 3362 | 1136 | 4354.5 | 1819.5 | 730 | 10471.5 | 4459.5 | 6221.5 |
| 11594 | 21626 | 10772 | 2385 | 12289 | 2949 | 780 | 14187 | 9774 | 21557 |
| 10122 | 25548.5 | 12860.5 | 2819.5 | 14668 | 3708.5 | 705.5 | 10787 | 8755 | 26146.5 |
| 11722 | 24451 | 13364 | 2667 | 13776 | 3389 | 763 | 12121 | 9325 | 25551 |

| | | | | | | | | | |
|---|---|---|---|---|---|---|---|---|---|
| 11213 | 22077.5 | 11204.5 | 2261 | 12236 | 3021 | 737 | 13697.5 | 9787 | 23475.5 |
| 8957 | 23763 | 8174.5 | 1877.5 | 9527.5 | 2599.5 | 729.5 | 14644.5 | 9728 | 27390 |
| 10859 | 27868 | 10981 | 3303.5 | 7702.5 | 2505 | 754.5 | 16664.5 | 9781.5 | 27375.5 |
| 4038 | 11895.5 | 4840.5 | 1328.5 | 5595 | 2030.5 | 752.5 | 14061 | 5250 | 15727 |
| 3299 | 6727.5 | 7072 | 2108 | 618178 | 59437.5 | 750 | 16384.5 | 4752 | 9314 |
| 3339 | 5019 | 5789 | 2065 | 9378 | 3357 | 794 | 13884 | 4829 | 7405 |
| 2948.5 | 4667.5 | 4073.5 | 1434 | 6082.5 | 2362 | 764.5 | 12079 | 4682.5 | 7106.5 |
| 2750 | 4267 | 3603 | 1263 | 5462 | 2053.5 | 769.5 | 10893 | 4607.5 | 6942.5 |
| 2943 | 4482 | 3863 | 1246 | 5092 | 1979 | 786 | 12347 | 4596 | 6820 |
| 2806 | 4236 | 3696 | 1212 | 4595 | 1877 | 737 | 10987 | 4791 | 6770 |
| 2791 | 4251.5 | 3547 | 1299 | 4496 | 1952.5 | 759 | 10635 | 4565 | 6540 |
| 2614 | 4093.5 | 3494 | 1175 | 4494.5 | 1862 | 704 | 9745 | 4436 | 6150 |
| 2455 | 3656 | 3204 | 1121 | 4346 | 1839 | 738 | 9572 | 4509 | 6150 |
| 2446 | 3648 | 3200 | 1227 | 4336 | 1845 | 765 | 9559 | 4457 | 6334 |
| 22474 | 14350 | 5816 | 1347 | 4186 | 1784 | 730 | 9545 | 8708 | 8650 |
| 2561 | 3777.5 | 3186 | 1152 | 4054.5 | 1837.5 | 733 | 9402 | 4497 | 6069 |
| 2499 | 3808 | 3191 | 1203 | 4486 | 1809 | 713 | 9544 | 4528 | 6591 |
| 139320 | 70073 | 20097 | 1936 | 4597 | 1839 | 697 | 20498 | 7230 | 7773 |
| 3046.5 | 4460 | 3698.5 | 1195.5 | 4541.5 | 1879 | 696.5 | 10323 | 4677.5 | 6356.5 |
| 2530 | 3908 | 3343 | 1125 | 5783 | 2048 | 710 | 9615 | 4638 | 6295 |
| 2621 | 3660.5 | 3165.5 | 1183 | 4154 | 1829.5 | 740 | 9606 | 4412 | 6242.5 |
| 10517.5 | 21667 | 10139 | 2099.5 | 11585 | 2904.5 | 698 | 14183.5 | 9466.5 | 23325.5 |
| 10102 | 23114 | 12601.5 | 2634 | 13840 | 3310 | 731 | 12064.5 | 9029 | 23443.5 |
| 10865 | 22061 | 12336 | 2430.5 | 13447.5 | 3214 | 707.5 | 12614.5 | 9388.5 | 22664.5 |
| 10079.5 | 21175.5 | 9971 | 2186.5 | 11230 | 2824.5 | 738.5 | 13567 | 9673 | 21779 |
| 10053 | 23126 | 7533.5 | 1833.5 | 8941.5 | 2467 | 745 | 14135 | 9966 | 25626 |
| 11025 | 122407 | 203139 | 73461 | 8719 | 3798 | 636 | 15909 | 9366 | 70003 |
| 5195 | 11003 | 5665 | 1729 | 5290 | 2097 | 726 | 14753 | 5683 | 13076 |
| 3304.5 | 5608.5 | 7651.5 | 2459 | 695230 | 63464.5 | 699 | 16431 | 4550.5 | 7801.5 |
| 2860 | 4519 | 4765 | 1678 | 7339 | 2802 | 815 | 13116 | 4807 | 6976 |
| 2883 | 4159 | 3894 | 1428 | 5641 | 2176 | 733 | 11658 | 4808 | 6375 |
| 2898 | 3888 | 3779 | 1361 | 5259 | 2015 | 704 | 10546 | 4651 | 6357 |
| 2965 | 4078.5 | 3547.5 | 1215 | 4667 | 1952.5 | 743 | 11527 | 4717.5 | 6338.5 |
| 2914 | 3956 | 3332 | 1130 | 4524 | 1833 | 713 | 11041 | 4587 | 5909 |
| 2939 | 4079 | 3494 | 1264 | 4342 | 1902 | 713 | 11333 | 4787 | 6275 |
| 2790 | 3925 | 3320 | 1178 | 4396 | 1834 | 747 | 10828 | 4718 | 6144 |
| 2442 | 3638 | 3023 | 1139 | 3930 | 1843 | 736 | 9766 | 4400 | 6000 |
| 2627 | 3598 | 3100 | 1181 | 4095 | 1728 | 707 | 9204 | 4006 | 5813 |
| 34073 | 23132.5 | 7542.5 | 1347.5 | 4038.5 | 1746.5 | 684.5 | 9081 | 11202 | 9775.5 |
| 2589 | 3680 | 3047 | 1096 | 3935 | 1750 | 709 | 9168 | 4320 | 5903 |
| 2997 | 3896 | 3493 | 1206 | 4511 | 1884 | 710 | 10328 | 4750 | 6486 |
| 110084 | 60748 | 17089 | 1784 | 4648 | 1919 | 706 | 19726 | 6357 | 7797 |
| 3103 | 4548 | 3694 | 1224 | 4554 | 1771.5 | 724.5 | 10908 | 4817.5 | 6711.5 |
| 2565 | 3702 | 3139 | 1103 | 5208 | 2004 | 695 | 9854 | 4658 | 6162 |
| 2703.5 | 4022.5 | 3302 | 1167.5 | 4244 | 1832 | 708.5 | 10132.5 | 4442.5 | 5968.5 |
| 11310 | 21770.5 | 10665 | 2242 | 11383 | 2860 | 715.5 | 13760.5 | 9468 | 23163.5 |
| 11680 | 24290 | 13200.5 | 2748.5 | 13723.5 | 3522.5 | 737 | 12252 | 8998 | 22176 |
| 11113 | 22938 | 12268 | 2675.5 | 13569 | 3432 | 721 | 12785.5 | 9040 | 22921.5 |

| | | | | | | | | | |
|---|---|---|---|---|---|---|---|---|---|
| 11090 | 21226 | 10380 | 2138.5 | 11078 | 2775 | 731 | 14022 | 9677 | 21880 |
| 9757 | 22710 | 7497 | 1806 | 8958 | 2506 | 711 | 14271 | 9926 | 24867 |
| 10230 | 88252 | 130008 | 46445 | 7217 | 2566 | 692 | 16499 | 8493 | 43167 |
| 4647 | 11142 | 4769 | 1507 | 5255 | 2028 | 724 | 14975 | 5277 | 13662 |
| 3279 | 5552 | 7052 | 2005 | 671459 | 66017 | 716 | 16654 | 4500 | 7826 |
| 2984 | 4550 | 4245 | 1508 | 6434 | 2368 | 858 | 14117 | 4857 | 7108 |
| 2970 | 4241.5 | 3830.5 | 1359.5 | 5225.5 | 2094 | 776 | 11586.5 | 5245.5 | 6427.5 |
| 3108.5 | 4198.5 | 3711 | 1257 | 4961 | 1988.5 | 754 | 10786 | 4905.5 | 6402 |
| 3092 | 4034 | 3435.5 | 1214 | 4482 | 1869 | 740 | 10965.5 | 4649 | 6480 |
| 2988 | 3947 | 3300 | 1125.5 | 4191 | 1857.5 | 733 | 11023 | 4678.5 | 6025 |
| 3038 | 4116 | 3635 | 1279 | 4502 | 1884 | 726 | 11434 | 4841 | 6315 |
| 3084 | 4004 | 3311 | 1170 | 4381 | 1885 | 738 | 11013 | 4914 | 6197 |
| 2475 | 3581 | 2926 | 1100 | 3990 | 1781 | 720 | 9229 | 4292 | 5778 |
| 2381 | 3671 | 3247 | 1139 | 4278 | 1859 | 737 | 10676 | 4328 | 5832 |
| 23669 | 15620 | 5830 | 1273 | 4107 | 1820 | 689 | 9713 | 8476 | 8091 |
| 2743.5 | 3955.5 | 3255.5 | 1168 | 4040.5 | 1832.5 | 773 | 9504 | 4514 | 6319.5 |
| 2638 | 4015 | 3453 | 1105 | 4219 | 1867 | 738 | 9995 | 4798 | 6264 |
| 85332.5 | 44887 | 13581.5 | 1616.5 | 4393 | 1861 | 732 | 16996.5 | 6285 | 7357.5 |
| 3536 | 4688.5 | 3852.5 | 1224 | 4523.5 | 1883.5 | 759 | 11664.5 | 4958.5 | 6665.5 |
| 2708 | 3659 | 3236 | 1126.5 | 5350.5 | 1968 | 731.5 | 10379.5 | 4727.5 | 6285 |
| 2870 | 3917 | 3141 | 1142 | 4236 | 1790 | 710 | 10183 | 4334 | 5978 |
| 11459 | 22044.5 | 11147.5 | 2266.5 | 11397.5 | 3089 | 740 | 14366 | 9668.5 | 21597.5 |
| 12234.5 | 25744.5 | 14113 | 2842.5 | 14233 | 3419 | 719.5 | 12863.5 | 9649.5 | 24831.5 |
| 10956 | 23425 | 12778 | 2747 | 13996 | 3495 | 730 | 12023 | 9167 | 23126 |
| 11041 | 21904.5 | 10629 | 2149.5 | 11614 | 2801.5 | 709 | 13920 | 9623 | 22989.5 |
| 9476.5 | 22880 | 7675 | 1756 | 8665 | 2530 | 694 | 14072 | 10168 | 24962 |
| 11216.5 | 108779.5 | 169876 | 62467 | 7625.5 | 2827.5 | 666.5 | 15169 | 9203.5 | 58736 |
| 4427 | 10603.5 | 4873.5 | 1460 | 5485 | 1972.5 | 706.5 | 14690.5 | 5335 | 13816.5 |
| 3131 | 5336 | 6553 | 1898 | 651371 | 58645 | 701 | 16510 | 4386 | 7671 |
| 2971 | 4432 | 4577 | 1576 | 6872 | 2491 | 758 | 13391 | 4923 | 6656 |
| 2730 | 3914 | 3598 | 1290 | 5034 | 1950 | 705 | 11480 | 4787 | 6149 |
| 2858 | 3897 | 3810.5 | 1296 | 5034.5 | 1982 | 693.5 | 9756.5 | 4592.5 | 6038.5 |
| 2729.5 | 4012.5 | 3379.5 | 1140 | 4662 | 1890 | 705.5 | 10818 | 4550.5 | 6240 |
| 2718.5 | 3971.5 | 3140.5 | 1141 | 4303 | 1803 | 683.5 | 10956 | 4596 | 6165 |
| 2740.5 | 3841 | 3528.5 | 1261 | 4571 | 1870.5 | 705 | 10903.5 | 4523 | 6109 |
| 2620.5 | 3827.5 | 3375.5 | 1245.5 | 4429.5 | 1922.5 | 699 | 10807 | 4778.5 | 6320.5 |
| 2416 | 3435 | 2931 | 1094 | 3988 | 1692 | 666 | 9193 | 4423 | 5636 |
| 2251 | 3596 | 3037 | 1157 | 4055 | 1826 | 711 | 9918 | 3976 | 5633 |
| 27643.5 | 18738 | 6370 | 1283.5 | 4068.5 | 1767.5 | 696 | 9505.5 | 9733 | 8815 |
| 2587 | 3776 | 3225 | 1152 | 4242 | 1767 | 714 | 9493 | 4282 | 6065 |
| 2580 | 3698.5 | 3358.5 | 1215 | 4486.5 | 1845 | 714.5 | 9933.5 | 4508 | 6437 |
| 88011 | 48029.5 | 14277.5 | 1788 | 4858.5 | 1929 | 717 | 17222 | 5801 | 7470 |
| 3001 | 4529 | 3687 | 1184 | 4557 | 1844 | 687 | 11203 | 4649 | 6611 |
| 2488.5 | 3875.5 | 3236.5 | 1122.5 | 5304.5 | 1945 | 690.5 | 9537 | 4531.5 | 6161 |
| 2856 | 3787.5 | 3283.5 | 1127.5 | 4177.5 | 1813 | 687.5 | 10184.5 | 4558 | 5967 |
| 10232 | 21748.5 | 10390.5 | 2254 | 11459 | 2972.5 | 709 | 13698 | 9042.5 | 22244.5 |
| 10908 | 24593 | 13345 | 2746 | 14561 | 3495 | 704 | 12536 | 8656 | 22791 |
| 10292 | 22158 | 11530 | 2597 | 13416 | 3323 | 703 | 12164 | 8761 | 22890 |

| | | | | | | | | | |
|---|---|---|---|---|---|---|---|---|---|
| 10163 | 20631 | 10134 | 2196 | 11390 | 2946 | 701 | 13036 | 8957 | 21585 |
| 9756 | 22861 | 7717 | 1787 | 8514 | 2463 | 693 | 14862 | 10139 | 25207 |
| 10334.5 | 75082 | 115648.5 | 38276 | 7220.5 | 2578.5 | 677 | 15181 | 8849 | 37866 |
| 4307.5 | 10811 | 4568 | 1380.5 | 5407.5 | 2004.5 | 737 | 14220.5 | 5208.5 | 13332 |
| 2849 | 5113 | 6643 | 1878 | 619322 | 56882 | 690 | 15746 | 4139 | 7187 |
| 3088 | 4338 | 4383 | 1474 | 6539 | 2407 | 795 | 12771 | 5042 | 6877 |
| 2840 | 4298 | 3789 | 1334 | 5373 | 2135 | 731 | 11347 | 4877 | 6626 |
| 2836 | 4032 | 3662 | 1267 | 4933 | 2035 | 714 | 10958 | 4649 | 6619 |
| 2839 | 4231.5 | 3459.5 | 1213.5 | 4565.5 | 1880 | 733.5 | 11352 | 4506 | 6436 |
| 3000 | 4030 | 3320 | 1203 | 4196 | 1728 | 682 | 10801 | 4661 | 5965 |
| 2846.5 | 4145 | 3437.5 | 1241.5 | 4374.5 | 1933.5 | 700 | 10962 | 4433.5 | 6279 |
| 2701.5 | 3863 | 3335.5 | 1214.5 | 4282.5 | 1848.5 | 701 | 10667 | 4575.5 | 5959.5 |
| 2691 | 3687 | 3249 | 1187 | 4113 | 1785 | 708 | 10018 | 4762 | 5764 |
| 2299.5 | 3530.5 | 3089 | 1134 | 4070.5 | 1739.5 | 703 | 9674.5 | 4202.5 | 5844.5 |
| 16595.5 | 11657.5 | 4903.5 | 1237 | 4138 | 1815 | 716.5 | 9260 | 6777 | 7588 |
| 2530 | 3859 | 3255 | 1140 | 4185 | 1773 | 708 | 9635 | 4427 | 6107 |
| 2537 | 3774 | 3148 | 1137 | 4307 | 1799 | 727 | 9684 | 4734 | 6186 |
| 86186 | 46312 | 13710 | 1690 | 4454 | 1838 | 730 | 17457 | 5849 | 7114 |
| 3371.5 | 4769 | 3824.5 | 1239.5 | 4547.5 | 1842.5 | 756 | 11119.5 | 4892 | 6877.5 |
| 2511 | 3812 | 3305 | 1140 | 5327 | 2035 | 706 | 10130 | 4742 | 6149 |
| 3048 | 3894 | 3281 | 1170 | 4145 | 1818 | 708 | 10223 | 4542 | 6019 |
| 10868 | 20994 | 10158 | 2158 | 11639 | 2845 | 704 | 14446 | 9384 | 21337 |
| 11579.5 | 23756 | 13207 | 2670.5 | 13730 | 3483 | 725.5 | 12803.5 | 9292 | 23862 |
| 10777 | 22485 | 12178 | 2592 | 14029.5 | 3400 | 708 | 12343.5 | 9148 | 23069 |
| 11499 | 22262 | 10897.5 | 2184 | 11524.5 | 2866.5 | 712 | 13074 | 9307 | 22619.5 |
| 9632 | 23640.5 | 7876.5 | 1788 | 8774.5 | 2433 | 707 | 14497.5 | 9682.5 | 25784.5 |
| 11910 | 122109 | 189303 | 69849 | 7627 | 2802 | 643 | 15239 | 9692 | 67223 |
| 4948 | 10231 | 5125 | 1597 | 5481 | 1984 | 707 | 14572 | 5498 | 12882 |
| 3250 | 5281 | 6858 | 2016 | 690167 | 66430 | 721 | 16886 | 4427 | 7377 |
| 3202 | 4752.5 | 4948.5 | 1606.5 | 6931 | 2541.5 | 808.5 | 13744 | 5143 | 7444 |
| 3024.5 | 4106 | 3788 | 1391.5 | 5364 | 2141.5 | 766 | 11373.5 | 4816.5 | 6487 |
| 2824.5 | 4099 | 3653 | 1254 | 5088.5 | 1921 | 742 | 11187 | 4665 | 6333.5 |
| 3093.5 | 4314 | 3718 | 1316.5 | 4851 | 1915.5 | 740.5 | 12002.5 | 4942 | 6583.5 |
| 3188 | 4222 | 3380 | 1266 | 4315 | 1863 | 696 | 11423 | 4686 | 6282 |
| 2834 | 4148 | 3497 | 1310 | 4487 | 1918 | 719 | 11909 | 4421 | 6283 |
| 2901 | 4079 | 3444 | 1230 | 4513 | 1898 | 726 | 11924 | 4772 | 6368 |
| 2599 | 3700 | 3157 | 1175 | 3975 | 1798 | 742 | 9834 | 4591 | 5919 |
| 2419.5 | 3727 | 3300.5 | 1192 | 4230.5 | 1834.5 | 736.5 | 9945.5 | 4230 | 6089.5 |
| 32877 | 22063.5 | 7377 | 1385.5 | 4268 | 1724.5 | 730 | 9808.5 | 11318 | 9523 |
| 2733 | 3943 | 3211 | 1251.5 | 4256 | 1891 | 762 | 10140.5 | 4518.5 | 6211 |
| 2759 | 4143 | 3303 | 1187 | 4450 | 1787 | 754 | 10424 | 4744 | 6498 |
| 95351 | 52028.5 | 16025.5 | 1852 | 4667 | 1954.5 | 731 | 19416 | 6307.5 | 7689.5 |
| 2965 | 4579 | 3493 | 1202 | 4639 | 1935.5 | 704 | 11003 | 4635.5 | 6735 |
| 2754 | 3869 | 3370 | 1144 | 5328 | 2056 | 734 | 10629 | 4829 | 6402 |
| 3057 | 4021 | 3402 | 1178 | 4380 | 1926 | 737 | 10883 | 4872 | 6091 |
| 11861.5 | 23184.5 | 11033.5 | 2361 | 11792 | 2887 | 747.5 | 13583 | 9528 | 22661 |
| 11816 | 25231.5 | 13081.5 | 2819.5 | 14794 | 3426 | 755 | 12243.5 | 9036.5 | 23247.5 |
| 11119 | 23500 | 13688 | 2852 | 14336 | 3524 | 741 | 11778 | 8643 | 22153 |

| | | | | | | | | | |
|---|---|---|---|---|---|---|---|---|---|
| 10451 | 21546 | 10207 | 2127 | 10661 | 2725 | 695 | 13890 | 9368 | 21847 |
| 9320.5 | 20717 | 7040 | 1641.5 | 7976 | 2363 | 683.5 | 14868.5 | 9971 | 23264 |
| 9810 | 38572 | 18224 | 4721 | 7136 | 2460 | 682 | 15642 | 8149 | 28167 |
| 4359.5 | 10931 | 4676 | 1421 | 5475.5 | 2014.5 | 687.5 | 13605 | 5315.5 | 13673.5 |
| 3066 | 5217.5 | 6824 | 2201 | 313755.5 | 33783.5 | 666 | 15921.5 | 4310 | 7222.5 |
| 2947 | 4492 | 4222 | 1450 | 10059 | 2673 | 763 | 13050 | 4670 | 7085 |
| 2564 | 4206 | 3498 | 1214 | 5867 | 2019 | 686 | 10518 | 4462 | 6559 |
| 2795 | 3863 | 3481 | 1183 | 4950 | 1980 | 678 | 10262 | 4729 | 6228 |
| 2883.5 | 4136 | 3455 | 1179 | 4745 | 1832 | 690.5 | 10744 | 4465.5 | 6215.5 |
| 2852 | 4096.5 | 3269 | 1160.5 | 4431 | 1803.5 | 680.5 | 11226 | 4474.5 | 6181.5 |
| 2629 | 3972 | 3555 | 1201 | 4289 | 1849 | 660 | 10473 | 4331 | 5925 |
| 2755 | 3851 | 3214 | 1140 | 4203 | 1835 | 678 | 10129 | 4373 | 6050 |
| 2720 | 3613 | 3213 | 1180 | 4429 | 1842 | 707 | 9766 | 4539 | 5927 |
| 2239 | 3591.5 | 2990.5 | 1054.5 | 4098 | 1724.5 | 672 | 9089.5 | 4158.5 | 5620 |
| 8137 | 6583.5 | 4144 | 1246.5 | 4193.5 | 1812.5 | 696.5 | 9518 | 5593 | 6800.5 |
| 2637 | 3810 | 3176.5 | 1141.5 | 4286 | 1799.5 | 692 | 9549.5 | 4181 | 5921.5 |
| 2607 | 3915.5 | 3326 | 1125.5 | 4561 | 1834 | 699.5 | 9413 | 4778.5 | 6401.5 |
| 153937 | 82224 | 24135 | 2014 | 4632 | 1827 | 701 | 22859 | 7273 | 7664 |
| 2941 | 4482.5 | 3799 | 1201.5 | 4680.5 | 1918.5 | 696.5 | 10984 | 4691.5 | 6734 |
| 2495 | 3720 | 3271 | 1147 | 10911 | 2656 | 679 | 9506 | 4651 | 5788 |
| 2684 | 3976 | 3319 | 1129 | 4481 | 1861 | 715 | 10086 | 4548 | 5844 |
| 9659 | 21002 | 10229 | 2377 | 11748 | 2978 | 709 | 13896 | 9135 | 22025 |
| 10662 | 23174.5 | 12775.5 | 2807.5 | 14033.5 | 3404.5 | 663.5 | 12032.5 | 9312.5 | 24025 |
| 10097 | 23044 | 13035 | 2606 | 13384 | 3311 | 697 | 12679 | 9386 | 23567 |
| 10332 | 21397 | 10629 | 2235 | 11591 | 2879 | 691 | 13944 | 9277 | 21525 |
| 9834 | 22418 | 8047 | 1849 | 8578 | 2462 | 713 | 14276 | 9788 | 24933 |
| 12411 | 122971.5 | 192736.5 | 71318.5 | 7623.5 | 2936 | 643 | 14847.5 | 9886.5 | 65640 |
| 4388 | 8084.5 | 4781.5 | 1596.5 | 5136 | 1911 | 671 | 14443.5 | 5097 | 10261.5 |
| 3299.5 | 4892 | 6589 | 1985 | 535517.5 | 52762.5 | 684 | 15633.5 | 4418 | 7076.5 |
| 3133.5 | 4450.5 | 4526 | 1503 | 6576.5 | 2562.5 | 802 | 15125.5 | 5336 | 7040 |
| 2787 | 4062 | 3725 | 1381 | 5117 | 1980 | 723 | 11534 | 5005 | 6472 |
| 2878 | 4093.5 | 3872.5 | 1320.5 | 5019 | 1958 | 734 | 11211.5 | 4813.5 | 6355.5 |
| 3045 | 4192 | 3716.5 | 1269 | 4960 | 1948 | 707.5 | 10725 | 4526 | 6085.5 |
| 2857 | 4106.5 | 3407 | 1163 | 4418 | 1849 | 723.5 | 11482.5 | 4561 | 6136 |
| 2920 | 4120 | 3397 | 1298 | 4242 | 1849 | 708 | 11690 | 4679 | 6154 |
| 3115 | 4053 | 3484 | 1242 | 4489 | 1902 | 677 | 11270 | 4447 | 6209 |
| 2717.5 | 3703 | 3296.5 | 1118 | 4185.5 | 1869 | 693.5 | 10537 | 4740 | 6164.5 |
| 2293 | 3556 | 3078 | 1098 | 4021 | 1816 | 691 | 9070 | 4222 | 5613 |
| 40836 | 29122.5 | 9730.5 | 1536 | 4352.5 | 1751.5 | 707.5 | 9386 | 14655 | 12109.5 |
| 2775 | 4027 | 3291.5 | 1224.5 | 4392.5 | 1793 | 755 | 10326 | 4743 | 6378.5 |
| 2616 | 3931 | 3468 | 1251 | 4422 | 1862 | 676 | 9872 | 4712 | 6426 |
| 85280.5 | 47413.5 | 15147 | 1855 | 4795 | 1961 | 688.5 | 18373 | 5972 | 8081.5 |
| 3674.5 | 5196.5 | 4215.5 | 1341 | 4737 | 1893 | 713 | 11971.5 | 5128.5 | 6746 |
| 2539 | 3986 | 3413 | 1135 | 5371 | 1946 | 709 | 10258 | 4723 | 6482 |
| 3079 | 4071 | 3381 | 1187 | 4327 | 1822 | 685 | 10093 | 4864 | 5982 |
| 10113 | 20464 | 10645 | 2269.5 | 11217.5 | 2940.5 | 723.5 | 14141.5 | 9014 | 21156 |
| 11044.5 | 23362 | 13485.5 | 2657.5 | 13984.5 | 3246 | 698.5 | 13212 | 9305.5 | 24615.5 |
| 10012 | 22055 | 12634 | 2631 | 13310 | 3355 | 712 | 11991 | 9001 | 23740 |

| | | | | | | | | | |
|---|---|---|---|---|---|---|---|---|---|
| 9803.5 | 21326 | 10642.5 | 2173.5 | 10658.5 | 2629 | 697 | 13601.5 | 9437.5 | 22642.5 |
| 9835 | 21737 | 7356 | 1854 | 8160 | 2472 | 692 | 14809 | 10402 | 23290 |
| 6668 | 17584 | 6876.5 | 2007.5 | 6641.5 | 2292.5 | 685.5 | 15530.5 | 6790.5 | 20520 |
| 4216 | 9678 | 4346 | 1281 | 5224 | 1966 | 704 | 14599 | 5169 | 11824 |
| 2870 | 4989 | 6308 | 1752 | 647086 | 58850 | 686 | 16306 | 4335 | 6915 |
| 2833 | 4361 | 3974 | 1416.5 | 6063 | 2257 | 760 | 13077 | 4772 | 6881 |
| 2730 | 3989 | 3628 | 1260 | 4802 | 1904 | 689 | 11076 | 4763 | 6117 |
| 2817 | 4055 | 3503 | 1207 | 4867 | 1996 | 717 | 10642 | 4832 | 6520 |
| 2900 | 4059 | 3641 | 1168 | 4617 | 1842 | 712 | 11277 | 4811 | 6169 |
| 2956 | 4162 | 3418 | 1165 | 4390.5 | 1802 | 683.5 | 11535 | 4556.5 | 6051 |
| 2727 | 4357 | 3669 | 1272 | 4677 | 1941 | 721 | 11898 | 4446 | 6487 |
| 3003.5 | 4095 | 3525.5 | 1214.5 | 4354 | 1873 | 700.5 | 10966 | 4722.5 | 6619 |
| 2699.5 | 3898 | 3342.5 | 1219 | 4571 | 1876.5 | 748.5 | 10620.5 | 4738.5 | 6334.5 |
| 2667 | 3945 | 3529 | 1203 | 4662 | 1882 | 764 | 10206 | 4642 | 6304 |
| 13818 | 10045 | 5014 | 1356 | 4452 | 1830 | 745 | 10395 | 7095 | 7803 |
| 2913 | 4033 | 3408 | 1199 | 4278 | 1883 | 743 | 10244 | 4986 | 6344 |
| 2705.5 | 4032 | 3611.5 | 1235.5 | 4461 | 1886.5 | 717 | 10192 | 4785 | 6373.5 |
| 90467.5 | 52996.5 | 15024.5 | 1872.5 | 5116.5 | 2000.5 | 764 | 18113 | 6041.5 | 8697 |
| 3298 | 5026 | 4099 | 1351 | 4951 | 2022 | 746 | 11910 | 4528 | 7182 |
| 2789 | 4094 | 3414 | 1215 | 5565 | 2111 | 714 | 11028 | 4683 | 6707 |
| 3012 | 4053 | 3522 | 1189 | 4566 | 1813 | 751 | 10342 | 4663 | 6596 |
| 10893 | 22962 | 11210 | 2494 | 12595 | 2965 | 721 | 13898 | 9386 | 23598 |
| 10509 | 26099 | 14615 | 3019 | 15539 | 3709 | 751 | 10593 | 8759 | 27051 |
| 10427 | 24043 | 12836 | 2775 | 14835 | 3506 | 724 | 11639 | 8580 | 23941 |
| 11453 | 22240 | 10689 | 2331 | 11884 | 2990 | 703 | 13680 | 9247 | 23693 |
| 765.5 | 1026 | 842 | 641 | 1391.5 | 1143.5 | 234 | 2227.5 | 1410 | 1875.5 |
| 1205.5 | 1901.5 | 1631 | 782 | 2313 | 1349 | 416 | 4313.5 | 2236 | 3324.5 |
| 1196 | 1938 | 1631 | 790 | 2350 | 1370 | 462 | 4420 | 2280 | 3460 |
| 1392 | 2411 | 4988 | 1490 | 773097 | 69639 | 555 | 6190 | 2783 | 4309 |
| 1414.5 | 2446 | 2666 | 1025 | 5033.5 | 2076.5 | 574 | 6655.5 | 2852 | 4327 |
| 1494.5 | 2645 | 2526 | 1032 | 4051.5 | 1848 | 617 | 6613.5 | 3139.5 | 4865 |
| 1805 | 3660 | 3039 | 1207 | 4708 | 2060 | 721 | 9190 | 3922 | 6826 |
| 1882.5 | 3276.5 | 2812.5 | 1139.5 | 4213 | 1839 | 672.5 | 9640 | 3975 | 5789.5 |
| 2407 | 4644 | 3308 | 1272 | 4823 | 2002 | 733 | 12006 | 4896 | 7890 |
| 2226 | 4925 | 3104 | 1250 | 4623 | 1993 | 718 | 10705 | 4350 | 8197 |
| 2734 | 5292.5 | 3643 | 1398 | 5412.5 | 2219.5 | 827.5 | 11821 | 5406.5 | 9487 |
| 2140.5 | 4378 | 3200.5 | 1289.5 | 4500.5 | 2000 | 748 | 9725 | 4450 | 7768 |
| 2206.5 | 4330.5 | 3509 | 1265 | 5075 | 2096 | 799 | 10976 | 4827.5 | 7701.5 |
| 1923 | 3413.5 | 3056 | 1185.5 | 4061.5 | 1877 | 717 | 8672.5 | 3944.5 | 6067.5 |
| 1991 | 5169 | 3459.5 | 1338 | 4891.5 | 2123.5 | 777 | 10171 | 4113 | 9276.5 |
| 27122.5 | 21333.5 | 7280.5 | 1690.5 | 6023.5 | 2421 | 830.5 | 13027.5 | 5993.5 | 11711 |
| 1896 | 3178 | 2669.5 | 971 | 3167.5 | 1592 | 541 | 7408 | 3378 | 5018.5 |
| 2442.5 | 5253 | 3737 | 1329 | 5046.5 | 2067 | 778 | 11460.5 | 4827.5 | 8732.5 |
| 2543 | 5446.5 | 3779.5 | 1370.5 | 5249 | 2180 | 810 | 10401 | 5263 | 9225 |
| 4713 | 11734 | 4699 | 1401 | 5784 | 2075 | 693 | 13081 | 6210 | 14829 |
| 7307 | 21766 | 11016 | 2427 | 12763 | 3267 | 924 | 11487 | 8244 | 26148 |
| 6638 | 17672.5 | 5873 | 1490.5 | 6701.5 | 2191 | 605.5 | 15383 | 6618.5 | 19583 |
| 7032 | 25441 | 7529 | 1934 | 9221 | 2701 | 696 | 11127 | 6999 | 27616 |

| | | | | | | | | | |
|---|---|---|---|---|---|---|---|---|---|
| 9141 | 20351 | 6625 | 1634 | 7595 | 2313 | 679 | 14762 | 9422 | 23558 |
| 3065.5 | 4222.5 | 2365.5 | 821.5 | 3014 | 1393 | 343 | 9008.5 | 4081.5 | 5189.5 |
| 2784.5 | 3922 | 2581 | 915.5 | 3349.5 | 1503.5 | 479.5 | 8162.5 | 3892.5 | 5222.5 |
| 2072.5 | 3082.5 | 2179 | 903 | 2939.5 | 1498.5 | 494.5 | 6830.5 | 3146 | 4500 |
| 2220 | 3618 | 5406 | 1649 | 757423 | 68398 | 629 | 8330 | 3858 | 5665 |
| 2084 | 3520 | 3417 | 1143 | 6954 | 2415 | 618 | 7986 | 3628 | 5506 |
| 2360 | 4066 | 3521 | 1219 | 5398 | 2113 | 730 | 9310 | 4402 | 6911 |
| 2468.5 | 4657.5 | 3593.5 | 1332.5 | 5496.5 | 2134 | 774.5 | 9644.5 | 4773.5 | 8010 |
| 2887 | 5755 | 3706.5 | 1364 | 5704.5 | 2200 | 780 | 11897 | 5441.5 | 9924 |
| 3136 | 6097 | 3896 | 1405 | 5635 | 2164 | 823 | 12928 | 5544 | 10007 |
| 3215 | 6628 | 3965 | 1429 | 5764 | 2232 | 833 | 14042 | 5908 | 10719 |
| 3749 | 8008 | 4935 | 1621 | 7119 | 2658 | 1005 | 13162 | 7029 | 13509 |
| 3125 | 6777 | 4282 | 1566 | 6255 | 2415 | 912 | 11693 | 5993 | 11379 |
| 2896 | 6049.5 | 3947 | 1414 | 5735.5 | 2280 | 881.5 | 12118 | 5918.5 | 10605.5 |
| 2996 | 6860.5 | 4216 | 1512.5 | 6030.5 | 2302.5 | 909 | 12725 | 5817 | 10842 |
| 2451 | 5207 | 3620 | 1350 | 5142 | 2187 | 809 | 8763 | 4631 | 8751 |
| 2800 | 6049 | 3711.5 | 1369.5 | 5436 | 2163.5 | 766 | 10230.5 | 5739.5 | 9500 |
| 44115 | 28717 | 8159 | 1590 | 5279 | 2164 | 757 | 13877 | 5823 | 9253 |
| 3760 | 8395 | 4890 | 1534 | 5521 | 2249 | 791 | 11906 | 5972 | 10397 |
| 2416 | 6567 | 3936 | 1352 | 5245 | 2083 | 773 | 10888 | 4436 | 10796 |
| 2637 | 5210.5 | 3769.5 | 1320.5 | 5174.5 | 2119 | 766 | 10353 | 5084.5 | 8688 |
| 6221 | 13780.5 | 5194.5 | 1546.5 | 6899.5 | 2278.5 | 751 | 14837 | 8352.5 | 17716 |
| 7383 | 20948 | 8948 | 2043 | 10214 | 2831 | 804 | 13272 | 8612 | 24662 |
| 6820 | 19265 | 6436 | 1577 | 7283 | 2247 | 622 | 15221 | 7104 | 21615 |
| 9700 | 20170 | 6418 | 1604 | 7370 | 2279 | 657 | 14390 | 9725 | 22477 |
| 3484 | 4490 | 2577 | 851 | 3244.5 | 1435 | 338 | 9476 | 4328 | 5513.5 |
| 2746.5 | 3970 | 2484 | 873 | 3290 | 1485 | 463.5 | 8395.5 | 3972.5 | 5210.5 |
| 2627.5 | 3992.5 | 2867.5 | 1043.5 | 3804 | 1679 | 584.5 | 9016.5 | 4183 | 5636 |
| 2187 | 3331 | 5433.5 | 1596.5 | 758483 | 80503.5 | 600 | 8157 | 3584.5 | 5243.5 |
| 2548 | 4266.5 | 3698.5 | 1255.5 | 6211.5 | 2304.5 | 716 | 10749 | 4620 | 6715 |
| 2119.5 | 3426.5 | 2963 | 1062.5 | 4359 | 1870 | 638 | 8099.5 | 3816.5 | 5565 |
| 2598.5 | 4530.5 | 3481 | 1219.5 | 5051 | 2050.5 | 738 | 10051 | 4656.5 | 7374 |
| 2413.5 | 4049.5 | 2982.5 | 1186 | 4359.5 | 1921.5 | 691 | 10755.5 | 4212.5 | 6540 |
| 2717 | 4566 | 3352 | 1211 | 4636 | 1948 | 713 | 10985 | 4752 | 7185 |
| 2794 | 4714 | 3376 | 1227 | 4594 | 1922 | 710 | 11599 | 4788 | 7233 |
| 3149 | 5288.5 | 3863 | 1371.5 | 5356.5 | 2146 | 817 | 12691.5 | 5601.5 | 8540.5 |
| 2805 | 5197 | 3698 | 1337 | 5117 | 2091 | 812 | 12746 | 5096 | 8570 |
| 2626 | 4526 | 3365.5 | 1306.5 | 4785.5 | 2062.5 | 761.5 | 11244.5 | 4833.5 | 7380.5 |
| 2693.5 | 5517.5 | 3415.5 | 1295 | 4719.5 | 2032 | 753 | 11333 | 4644.5 | 9081 |
| 4022 | 10043 | 5815 | 1974 | 8337 | 3009 | 1133 | 13311 | 7967 | 16013 |
| 4578 | 9727.5 | 6095.5 | 1959 | 9187.5 | 3090 | 1090 | 12876 | 8203 | 14814.5 |
| 13120.5 | 16390.5 | 4867.5 | 1319 | 4628.5 | 1969.5 | 686 | 11689.5 | 4110 | 11208.5 |
| 3827 | 7765.5 | 4436.5 | 1510 | 5827 | 2305 | 804.5 | 11090.5 | 5963 | 10911.5 |
| 4061.5 | 8488.5 | 5387 | 1825.5 | 7210.5 | 2676.5 | 980 | 13976 | 7415 | 14253 |
| 2976 | 5610 | 4023 | 1456 | 5465 | 2167 | 800.5 | 11188 | 5525.5 | 9348 |
| 5322.5 | 10461 | 4855 | 1418.5 | 6054.5 | 2205.5 | 726 | 14477.5 | 7020.5 | 13632.5 |
| 7681 | 21582 | 7577 | 1828 | 8494 | 2561 | 721 | 14260 | 8465 | 24617 |
| 7009 | 19757.5 | 6606 | 1594.5 | 7301 | 2241 | 658 | 15737.5 | 7190.5 | 21401 |

| | | | | | | | | | |
|---|---|---|---|---|---|---|---|---|---|
| 8946 | 18928 | 6670 | 1595 | 7653 | 2264 | 673 | 13963 | 8745 | 20976 |
| 3094 | 4062 | 2248 | 809 | 2936 | 1389 | 300 | 8367 | 3800 | 4910 |
| 2975 | 4051 | 2490 | 892 | 3320 | 1506 | 446 | 8085 | 3804 | 5090 |
| 2266 | 3364 | 2322 | 881 | 3100.5 | 1484.5 | 501.5 | 7302.5 | 3355 | 4694 |
| 2165 | 3223 | 5141 | 1514 | 682134 | 68645 | 541 | 7644 | 3313 | 4646 |
| 2273.5 | 3533 | 3213.5 | 1142.5 | 5721 | 2061 | 612.5 | 8155.5 | 3717 | 5445.5 |
| 2327.5 | 3891 | 3044.5 | 1147 | 5029.5 | 1988.5 | 654 | 9557.5 | 4137 | 6143 |
| 2550 | 4787 | 3539.5 | 1283.5 | 5405.5 | 2047 | 754 | 11315 | 4806 | 7812.5 |
| 2267 | 3642 | 2818 | 1091 | 4124 | 1786 | 619 | 8721 | 3909 | 5843 |
| 2659.5 | 4376.5 | 3199.5 | 1157 | 4518 | 1864.5 | 667 | 10502 | 4503.5 | 6620.5 |
| 2508 | 3966.5 | 2919 | 1133.5 | 4083.5 | 1838 | 642.5 | 9723 | 4079.5 | 6129.5 |
| 2783 | 5104 | 3464 | 1240 | 4819 | 2009 | 723 | 11123 | 4825 | 8255 |
| 2346 | 3917 | 2895 | 1133 | 4146 | 1810 | 669 | 9560 | 4287 | 5975 |
| 2285.5 | 3890 | 3069.5 | 1156.5 | 4117.5 | 1852.5 | 688 | 9049.5 | 4223.5 | 6388 |
| 2264.5 | 3761.5 | 3051.5 | 1149.5 | 4116 | 1786 | 663 | 8555 | 4017.5 | 6120.5 |
| 2482 | 4212 | 3244 | 1223 | 4446 | 1902 | 708 | 9534 | 4414 | 6644 |
| 2241 | 3543 | 2821.5 | 1043.5 | 3862.5 | 1699 | 598 | 7975.5 | 3965.5 | 5595.5 |
| 29856.5 | 19145.5 | 6482.5 | 1448.5 | 5086 | 2029 | 722 | 12189 | 5735 | 8892.5 |
| 2775 | 4552 | 3575 | 1156 | 4256 | 1825 | 628 | 9199 | 4452 | 6551 |
| 3579 | 8377 | 5011 | 1698 | 6631 | 2566 | 894 | 11909 | 6253 | 12454 |
| 2795 | 5349 | 3654 | 1285 | 4936 | 1997 | 727 | 10384 | 4952 | 8506 |
| 5292 | 16107 | 7441 | 1995 | 9635 | 2843 | 882 | 11627 | 7487 | 20476 |
| 8154.5 | 23232.5 | 8260.5 | 1980.5 | 9885.5 | 2678 | 758 | 14150.5 | 9232 | 26558 |
| 6864.5 | 19357.5 | 6497.5 | 1584 | 7325 | 2254 | 620 | 15608.5 | 7053 | 21634 |
| 7987 | 21802.5 | 6883.5 | 1715.5 | 7813.5 | 2370.5 | 676 | 12956.5 | 8568 | 24431 |
| 3356 | 4239 | 2432 | 855 | 3192 | 1429 | 326 | 9163 | 4242 | 5232 |
| 3100 | 4436 | 2800 | 977 | 3694 | 1547 | 486 | 9202 | 4309 | 5766 |
| 2274 | 3446.5 | 2400 | 924.5 | 3229 | 1493 | 508 | 7481 | 3441 | 4893 |
| 2434.5 | 3559.5 | 5652.5 | 1675 | 807801.5 | 83352.5 | 602 | 8999 | 3875 | 5416.5 |
| 2620 | 4987 | 3883 | 1310 | 6850 | 2358 | 706 | 11114 | 4429 | 7631 |
| 2244 | 3530.5 | 2904.5 | 1056.5 | 4464 | 1815 | 627 | 8427.5 | 3879.5 | 5440.5 |
| 2490 | 4032 | 3124 | 1165.5 | 4801 | 1931 | 683 | 8828.5 | 4333.5 | 6245.5 |
| 3185 | 6685 | 3967 | 1443 | 6000.5 | 2271 | 844.5 | 13620.5 | 5662.5 | 10500 |
| 2424.5 | 3782.5 | 2875 | 1060 | 4064 | 1761 | 630 | 9373.5 | 4182.5 | 5839 |
| 3567 | 8053.5 | 4441 | 1545 | 6163.5 | 2323.5 | 880.5 | 11700 | 6061.5 | 12300.5 |
| 2429.5 | 4771 | 2929.5 | 1089 | 4224 | 1806.5 | 649.5 | 10786 | 4295 | 7453.5 |
| 2747 | 6153 | 3740 | 1336 | 5390 | 2200 | 775 | 12519 | 5178 | 9842 |
| 2433 | 4834.5 | 3190 | 1171 | 4415.5 | 1946 | 712.5 | 11450 | 4372.5 | 7796 |
| 2657.5 | 4114.5 | 3218 | 1196.5 | 4443.5 | 1897 | 715 | 9807.5 | 4471.5 | 6562.5 |
| 2677 | 5304 | 3513 | 1281 | 4784 | 1981 | 752 | 10716 | 4795 | 8340 |
| 3189 | 6187 | 4070 | 1455 | 5753 | 2292 | 820 | 11165 | 5837 | 9808 |
| 27457 | 18964 | 6185 | 1450 | 5026 | 2047 | 712 | 12112 | 5626 | 9947 |
| 3287 | 7899 | 4352 | 1412 | 5375 | 2157 | 770 | 12360 | 5141 | 11252 |
| 2610 | 5252 | 3650 | 1271 | 4786 | 1947 | 709 | 10123 | 4783 | 7959 |
| 3195 | 7342.5 | 4352 | 1483 | 5889 | 2294.5 | 854.5 | 13992.5 | 5762 | 11585.5 |
| 5259 | 11739 | 4894 | 1391 | 5853 | 2041 | 688 | 15258 | 6584 | 15070 |
| 7149 | 20488 | 9281 | 2377 | 11664 | 3194 | 851 | 10450 | 7548 | 24578 |
| 9100 | 23981 | 8010 | 1913 | 9196 | 2650 | 707 | 12868 | 9373 | 26681 |

| | | | | | | | | | |
|---|---|---|---|---|---|---|---|---|---|
| 8963.5 | 20867.5 | 6916.5 | 1743 | 8114 | 2463.5 | 708 | 12929 | 9158.5 | 23988 |
| 2973.5 | 4026 | 2216.5 | 800 | 2950 | 1399 | 306 | 8743.5 | 3716.5 | 4843 |
| 25173.5 | 12581.5 | 4454 | 1014 | 3588.5 | 1520.5 | 480 | 11075.5 | 5084.5 | 5766.5 |
| 17846 | 10009.5 | 4276 | 1143 | 4436.5 | 1801.5 | 619.5 | 10811 | 5220.5 | 6671 |
| 16450 | 9368 | 6926 | 1625 | 144529 | 15004 | 625 | 10536 | 4790 | 6045 |
| 15576 | 9563.5 | 5039 | 1286.5 | 18902.5 | 3258.5 | 720 | 11476.5 | 5273.5 | 7064 |
| 16315 | 12469 | 5318 | 1424 | 11297 | 2771 | 792 | 14047 | 5666 | 10199 |
| 17640 | 14876.5 | 5270.5 | 1409 | 7763.5 | 2357.5 | 836.5 | 13243.5 | 6188 | 11728 |
| 16152.5 | 11717.5 | 4933.5 | 1350.5 | 6617 | 2327 | 814.5 | 15646 | 6307.5 | 9842.5 |
| 16987.5 | 13093 | 5844.5 | 1591.5 | 7344 | 2529 | 923 | 15718 | 7959 | 12327 |
| 13893 | 9786 | 4458 | 1225 | 5206 | 2010 | 716 | 14291 | 5796 | 8255 |
| 15079 | 11583 | 4951.5 | 1429 | 5920 | 2208.5 | 847.5 | 15811.5 | 7041 | 10709.5 |
| 12546 | 8213 | 4261 | 1206 | 4679 | 1921 | 715 | 12601 | 5454 | 7155 |
| 12009 | 7759 | 3863 | 1172 | 4685 | 1907 | 709 | 12266 | 5201 | 6934 |
| 23694 | 17866 | 5819 | 1365 | 5172 | 1992 | 756 | 12097 | 9274 | 11219 |
| 20603.5 | 18748 | 8040.5 | 2080 | 8899.5 | 2884 | 1209.5 | 17173 | 10383.5 | 17974.5 |
| 17340 | 11259 | 4508 | 1080 | 4060 | 1739 | 664 | 10423 | 6336 | 7137 |
| 1442626 | 1262620 | 347895 | 24188 | 10522.5 | 3237.5 | 1171.5 | 93645.5 | 46213.5 | 39560 |
| 28541 | 19371 | 7022 | 1725 | 5789 | 2315 | 816 | 12836 | 7706 | 10883 |
| 23962 | 18809 | 5943 | 1386 | 5126.5 | 2015.5 | 768 | 11983 | 5851 | 10040 |
| 50015 | 34100 | 9033 | 1871 | 7242 | 2599 | 1007 | 18239 | 9511 | 15557 |
| 38934 | 37112 | 8171 | 1534 | 5944 | 2120.5 | 686 | 17414.5 | 9052.5 | 17582.5 |
| 25984 | 39013.5 | 14087.5 | 2624.5 | 13200.5 | 3359 | 907 | 12830 | 9416.5 | 28664.5 |
| 28597.5 | 42834 | 9785 | 1774.5 | 8160.5 | 2454.5 | 662 | 16986 | 8254 | 24844 |
| 35382 | 38630 | 8785 | 1619 | 7382 | 2257 | 647 | 14814 | 9987 | 21744 |
| 3447 | 4533 | 2589 | 872 | 3489 | 1468 | 354 | 9244 | 4469 | 5537 |
| 2999.5 | 4134 | 2643 | 907 | 3525 | 1530 | 440 | 8504.5 | 4047.5 | 5333 |
| 2734 | 4033.5 | 2766 | 994.5 | 3794.5 | 1627.5 | 554 | 8585 | 4122 | 5689.5 |
| 2432 | 3563 | 5541 | 1623.5 | 856536 | 76372 | 556.5 | 8249.5 | 3732 | 5239 |
| 2538 | 4011 | 3377 | 1120 | 6201 | 2131 | 634 | 9136 | 4329 | 6039 |
| 2479 | 3988 | 3151 | 1131.5 | 5310 | 1998.5 | 631 | 9363.5 | 4027.5 | 6084 |
| 2442 | 4075 | 3175 | 1122 | 4945 | 1920 | 664 | 9062 | 4235 | 6380 |
| 2731 | 4457.5 | 3337.5 | 1216.5 | 5095.5 | 1979.5 | 693 | 9670.5 | 4720.5 | 6830.5 |
| 2620 | 4161 | 3110 | 1126 | 4473 | 1835 | 639 | 10137 | 4299 | 6340 |
| 3098 | 5244 | 3770 | 1325 | 5243 | 2055 | 731 | 11242 | 5068 | 8193 |
| 2502 | 4288 | 2955 | 1108 | 4190 | 1804 | 627 | 9876 | 4075 | 6553 |
| 3577.5 | 7333 | 4556.5 | 1592 | 6555.5 | 2440.5 | 884 | 13377 | 6428.5 | 11679.5 |
| 2313 | 3728 | 2915 | 1046 | 4053 | 1746 | 618 | 9512 | 4146 | 5908 |
| 3258 | 6292 | 4208 | 1481 | 5880 | 2294 | 852 | 13336 | 5850 | 10003 |
| 2884.5 | 5872.5 | 3781 | 1347 | 5212.5 | 2106 | 743 | 11247 | 5149.5 | 9186.5 |
| 3306 | 6939 | 4549 | 1492 | 6145.5 | 2284.5 | 855 | 10865.5 | 5810 | 10491.5 |
| 47751 | 29376.5 | 8115 | 1483.5 | 4809.5 | 1967 | 670.5 | 12544 | 5497 | 8291.5 |
| 5080 | 13073 | 7524 | 2035 | 9897 | 3160 | 1064 | 13274 | 7682 | 18847 |
| 2442.5 | 4151.5 | 2992 | 1081 | 4052 | 1760.5 | 607.5 | 8566 | 4130.5 | 6141 |
| 4076 | 7749.5 | 5317.5 | 1708.5 | 6786 | 2608.5 | 944 | 13628 | 6861 | 12180 |
| 3995 | 11052 | 4846.5 | 1377.5 | 5964.5 | 2065 | 660 | 14383.5 | 5390.5 | 13476 |
| 6713 | 21566 | 9071 | 2207 | 10727 | 2957 | 795 | 11025 | 7322 | 25270 |
| 6839 | 22659 | 8169 | 1935.5 | 9501.5 | 2717 | 726 | 13157 | 7536 | 26429 |

| | | | | | | | | | |
|---|---|---|---|---|---|---|---|---|---|
| 8615 | 19132 | 6340.5 | 1552 | 7198 | 2265 | 645 | 13377 | 8845 | 21543.5 |
| 3174 | 4129 | 2384 | 818 | 3239 | 1420 | 309 | 8872 | 4048 | 5120 |
| 3259 | 4446 | 2899 | 942 | 3827 | 1614 | 500 | 9254 | 4474 | 5786 |
| 2539.5 | 3645 | 2462.5 | 945 | 3393 | 1524.5 | 524.5 | 8010.5 | 3598.5 | 5025.5 |
| 2497 | 3708 | 5410 | 1576 | 598507 | 67425 | 604 | 8702 | 3888 | 5441 |
| 2464 | 3794.5 | 3236 | 1099.5 | 5521.5 | 2033.5 | 618 | 9008.5 | 4034.5 | 5668.5 |
| 2400 | 3588 | 3079 | 1058 | 4566 | 1818 | 631 | 8852 | 3907 | 5569 |
| 2418.5 | 3838 | 3131.5 | 1131.5 | 4649.5 | 1872 | 655.5 | 8906 | 4087.5 | 5953.5 |
| 2548.5 | 3857 | 3044 | 1080.5 | 4231 | 1765.5 | 633.5 | 9503 | 4122 | 5866.5 |
| 2732 | 4272 | 3126 | 1149 | 4414 | 1801 | 670 | 10502 | 4467 | 6599 |
| 2585 | 3914.5 | 3018 | 1054.5 | 4018.5 | 1777 | 621 | 10103.5 | 4277 | 5946.5 |
| 2734 | 4263 | 3327 | 1130 | 4495 | 1861 | 673 | 10997 | 4594 | 6524 |
| 2071 | 3303 | 2638 | 1004 | 3553 | 1636 | 597 | 8562 | 3610 | 5128 |
| 2429 | 3874 | 3156 | 1149 | 4269 | 1830 | 679 | 9942 | 4303 | 6283 |
| 2607 | 5382 | 3593 | 1290 | 4916 | 1998 | 744 | 12056 | 4656 | 8397 |
| 2731 | 4646 | 3363 | 1282 | 4547 | 1932 | 714 | 9976 | 4629 | 7269 |
| 2329 | 4233.5 | 3243.5 | 1131 | 4308.5 | 1832.5 | 656 | 9437.5 | 4131 | 6403 |
| 15233 | 9718 | 4768 | 1234 | 4416 | 1806 | 636 | 10645 | 4686 | 6763 |
| 3167 | 6529 | 4304 | 1466 | 5875.5 | 2218.5 | 786.5 | 11638 | 5297.5 | 10040 |
| 3563 | 7396 | 4590 | 1498 | 6152 | 2360 | 852 | 12111 | 6277 | 11651 |
| 3440 | 7986 | 4495 | 1534 | 6186 | 2372 | 872 | 14086 | 5989 | 12476 |
| 7155 | 16645.5 | 6615 | 1846 | 8181.5 | 2587.5 | 825.5 | 14206.5 | 9033.5 | 21047.5 |
| 7866.5 | 22293 | 8075.5 | 1928.5 | 9802.5 | 2609.5 | 752 | 14094 | 9110.5 | 26062 |
| 6962 | 21777 | 7796 | 1914 | 9591 | 2642 | 720 | 12710 | 7528 | 25777 |
| 8507.5 | 22536.5 | 8533.5 | 1870 | 9820 | 2586 | 744 | 14223.5 | 8968 | 26138.5 |
| 3389 | 4299 | 2471 | 841 | 3362 | 1427 | 314 | 9218 | 4053 | 5226 |
| 3800 | 5334 | 3288 | 994 | 4551 | 1708 | 481 | 11772 | 4491 | 6068 |
| 2805.5 | 3870 | 2571 | 960.5 | 3951 | 1633.5 | 503.5 | 8366 | 3699.5 | 5092 |
| 2980 | 4258 | 7889 | 2003 | 1343769 | 131721 | 593 | 9828 | 4408 | 6229 |
| 3066 | 4235.5 | 3652 | 1184.5 | 22650.5 | 3326 | 599 | 9747.5 | 4525.5 | 6003 |
| 3333 | 5589 | 3938 | 1313 | 17236 | 3008 | 661 | 11312 | 4905 | 7525 |
| 3185 | 4757 | 3515 | 1103 | 12063 | 2438 | 612 | 10197 | 4505 | 6489 |
| 3165.5 | 5202 | 3631.5 | 1182.5 | 11688.5 | 2410 | 646.5 | 11368.5 | 4615 | 7283 |
| 3201 | 4952 | 3494 | 1156 | 9843 | 2269.5 | 631.5 | 11412.5 | 4538 | 6937.5 |
| 3622 | 6897 | 4100 | 1351 | 10120 | 2415 | 697 | 13371 | 5418 | 9411 |
| 3275 | 5726.5 | 3805 | 1268.5 | 8999.5 | 2354.5 | 671 | 12149 | 4937 | 8064 |
| 3254 | 5944 | 3869 | 1274 | 7978 | 2275 | 705 | 11927 | 5190 | 8776 |
| 2840 | 4587 | 3344 | 1146 | 7035 | 2056 | 644 | 9771 | 4443 | 6653 |
| 6761 | 6209 | 3966 | 1159 | 6278 | 1936 | 618 | 8928 | 5624 | 6953 |
| 3460 | 5557 | 3837 | 1231.5 | 6716.5 | 2138.5 | 682 | 9640.5 | 5099 | 7622.5 |
| 3028.5 | 6707 | 3899 | 1265.5 | 6585.5 | 2123.5 | 684 | 10257 | 4579 | 9635 |
| 157392 | 110805 | 26759 | 2244 | 6538 | 2157 | 652 | 23344 | 7228 | 10292 |
| 3754.5 | 7826.5 | 4565.5 | 1342 | 6796 | 2176.5 | 660 | 11486.5 | 4863.5 | 9754.5 |
| 3314 | 8646 | 4722 | 1457 | 7186 | 2353 | 740 | 13830 | 4688 | 12044 |
| 3779 | 7153.5 | 4588 | 1540 | 6688 | 2169 | 682 | 12604.5 | 5009 | 9514 |
| 7951 | 20041 | 7195 | 1913 | 9138 | 2677 | 777 | 16740 | 8769 | 20862 |
| 9661 | 23537 | 9491 | 2146 | 11716 | 2913 | 745 | 12628 | 9525 | 24766 |
| 9104 | 25575 | 8728 | 2077 | 10552 | 2771 | 728 | 13285 | 9184 | 27472 |

| | | | | | | | | | |
|---|---|---|---|---|---|---|---|---|---|
| 8494 | 23740 | 8801 | 1923 | 10847 | 2724 | 719 | 12072 | 8843 | 26334 |
| 3289.5 | 4089 | 2361 | 821 | 3245.5 | 1443 | 311 | 9649 | 4071.5 | 4937 |
| 3371 | 4270 | 2670.5 | 933 | 3759 | 1571 | 452.5 | 10099 | 4278.5 | 5337.5 |
| 3260 | 4800 | 3349 | 1136.5 | 4651 | 1868 | 614 | 11406 | 4665.5 | 6707.5 |
| 2808 | 4109 | 5963 | 1715 | 836031 | 74279 | 637 | 9464 | 4431 | 6226 |
| 2657 | 4201 | 3659 | 1189 | 6258 | 2111 | 648 | 9987 | 4382 | 6271 |
| 2564 | 4107 | 3350 | 1157 | 5234 | 1952 | 662 | 9315 | 4408 | 6274 |
| 2830 | 5441 | 3927 | 1259 | 5721 | 2152 | 744 | 11823 | 4862 | 8420 |
| 2943 | 5200 | 3614 | 1255 | 5367 | 2076 | 732 | 11644 | 4842 | 8003 |
| 3055.5 | 5660.5 | 3798.5 | 1314.5 | 5306 | 2032.5 | 725.5 | 12073.5 | 5097.5 | 8490 |
| 2865.5 | 7254.5 | 3768.5 | 1299 | 5263.5 | 2048 | 748 | 14665 | 4829.5 | 10929.5 |
| 2705 | 4296 | 3353 | 1133 | 4668 | 1896 | 684 | 10699 | 4460 | 6569 |
| 5518.5 | 11484.5 | 7007.5 | 2120 | 9435 | 3256 | 1236.5 | 18527.5 | 9842.5 | 19896.5 |
| 2812 | 5000 | 3449.5 | 1281.5 | 5040.5 | 2023 | 738 | 11299 | 4996.5 | 7687 |
| 4333 | 9421 | 5741 | 1823 | 7944 | 2851 | 1075 | 14922 | 7837 | 14860 |
| 3708.5 | 8601 | 5065 | 1727 | 6929.5 | 2666 | 941 | 11732.5 | 6131.5 | 13814.5 |
| 3008 | 6196 | 3703.5 | 1297 | 5089.5 | 1985 | 717 | 11721 | 5344.5 | 9872.5 |
| 38925 | 25963 | 9433 | 1999 | 8483 | 2846 | 952 | 14921 | 8357 | 14948 |
| 4174 | 9017 | 5048 | 1544 | 5867 | 2293 | 764 | 11964 | 6097 | 10796 |
| 2648 | 4720 | 3431 | 1209 | 4752 | 1938 | 677 | 9868 | 4415 | 7196 |
| 3465.5 | 7788.5 | 4569.5 | 1564 | 6228.5 | 2395.5 | 819 | 11723 | 5937.5 | 12033.5 |
| 4036 | 8835 | 4176 | 1273 | 5156.5 | 1939.5 | 616 | 14568.5 | 5183 | 11531.5 |
| 7195.5 | 20616 | 10218.5 | 2272.5 | 11870.5 | 3075 | 831 | 11107 | 8177.5 | 23282.5 |
| 9233 | 23174 | 7845 | 1879 | 9250 | 2559 | 694 | 14261.5 | 9523 | 26042.5 |
| 8652.5 | 23269 | 7520.5 | 1819.5 | 8936.5 | 2463 | 694 | 14630 | 9297.5 | 26401 |
| 3607 | 5179 | 3020 | 935 | 4075 | 1566 | 376 | 12141 | 4545 | 6315 |
| 3278 | 4227 | 2701 | 931 | 3731 | 1534 | 448 | 9078 | 4396 | 5513 |
| 3081 | 4412 | 3117 | 1002 | 4159 | 1718 | 573 | 9743 | 4385 | 6030 |
| 3028 | 4049 | 5886 | 1615 | 794295 | 71678 | 629 | 9870 | 4520 | 6073 |
| 2727 | 4110 | 3536 | 1171 | 6385 | 2237 | 632 | 9017 | 4190 | 5948 |
| 2733 | 5132 | 3638 | 1227 | 5678 | 2047 | 675 | 9890 | 4543 | 7733 |
| 2599.5 | 4162 | 3377 | 1140 | 4958 | 1909 | 666.5 | 10227 | 4395 | 6345.5 |
| 3127 | 5573 | 3754 | 1300 | 5502.5 | 2098 | 730.5 | 11756.5 | 5343.5 | 8346 |
| 3435 | 6354 | 4012 | 1363.5 | 5665 | 2208.5 | 735 | 13180 | 5732 | 9638.5 |
| 2843 | 4324.5 | 3372.5 | 1120 | 4582 | 1829 | 646 | 11169.5 | 4398 | 6692 |
| 2691 | 4663 | 3489 | 1200 | 4944 | 1890 | 682 | 13469 | 4509 | 7131 |
| 2808 | 4474 | 3463 | 1170 | 4887 | 1925 | 698 | 11065 | 4847 | 7075 |
| 3008 | 6850.5 | 3777 | 1400.5 | 5652 | 2181.5 | 760.5 | 11459.5 | 5637.5 | 11216.5 |
| 4042 | 8910 | 5068 | 1715 | 7323 | 2654 | 993 | 14278 | 7051 | 13831 |
| 3070.5 | 6327 | 3642 | 1374.5 | 5281 | 2138 | 766.5 | 11432.5 | 5286 | 9685.5 |
| 4213 | 9631.5 | 5414 | 1753 | 7262.5 | 2625.5 | 929 | 14589.5 | 7508.5 | 14517 |
| 32775 | 24191 | 7897 | 1843 | 7340 | 2620 | 897 | 16094 | 7661 | 15380 |
| 3960.5 | 8152.5 | 4604 | 1431.5 | 5480 | 2079.5 | 711.5 | 11087.5 | 5527.5 | 10454.5 |
| 3946 | 9955 | 5032 | 1654 | 6666 | 2504 | 869 | 14371 | 6468 | 14987 |
| 2813.5 | 4888.5 | 3580 | 1255 | 4968.5 | 1955.5 | 699 | 10368.5 | 4693.5 | 7482.5 |
| 4453.5 | 8767.5 | 4391.5 | 1346 | 5504.5 | 2062.5 | 672.5 | 13601.5 | 6048.5 | 11201 |
| 8329 | 21127 | 9119 | 2073 | 10621 | 2809 | 746 | 12217 | 9238 | 24035 |
| 8940 | 19642 | 7108 | 1713 | 7845 | 2333 | 632 | 14469 | 9496 | 22290 |

| | | | | | | | | | |
|---|---|---|---|---|---|---|---|---|---|
| 7718 | 19454.5 | 6616.5 | 1706 | 7908 | 2332 | 671 | 15080.5 | 8600 | 22164 |
| 817 | 1192 | 975 | 614 | 1604 | 1159 | 247 | 2611 | 1569 | 2165 |
| 1119.5 | 1841.5 | 1548 | 725 | 2361 | 1325 | 403 | 4117 | 2194 | 3267 |
| 1412 | 2670 | 2313 | 903 | 3362 | 1567 | 566 | 6991 | 3071 | 4752 |
| 1335 | 2298 | 4820 | 1483 | 829442.5 | 77585 | 523 | 5866 | 2731 | 4145 |
| 1478 | 2719 | 2856 | 1041 | 4905 | 2089 | 599 | 7154.5 | 3182.5 | 4872.5 |
| 1421 | 2588 | 2431 | 935 | 3763 | 1698 | 576 | 6712 | 3041 | 4672 |
| 1622 | 2973 | 2744 | 1044 | 4130 | 1798 | 652 | 7801.5 | 3563 | 5415.5 |
| 1854 | 3118.5 | 2691 | 1019 | 3882 | 1735 | 652 | 8785.5 | 3864 | 5426.5 |
| 2164 | 3598 | 2997 | 1066 | 4232 | 1817 | 689 | 11105 | 4244 | 6072 |
| 2276 | 4441 | 3306 | 1230 | 4751 | 2013 | 741 | 13588 | 4396 | 7588 |
| 2138 | 3646 | 3046 | 1123 | 4396 | 1874 | 708 | 11130 | 4279 | 6296 |
| 2066 | 3582 | 3076 | 1139 | 4439 | 1872 | 719 | 10584 | 4315 | 6329 |
| 1896 | 3298 | 2844 | 1085 | 4134 | 1805 | 691 | 8567 | 4021 | 5931 |
| 2124 | 3658.5 | 3156.5 | 1175 | 4444 | 1929 | 759.5 | 10766 | 4558.5 | 6560 |
| 1609 | 2947 | 2597 | 1004 | 3713 | 1678 | 632 | 7521 | 3428 | 5283 |
| 1940.5 | 3533 | 3025 | 1101 | 4339 | 1821.5 | 684 | 8638.5 | 4155 | 6154.5 |
| 43266 | 26729 | 8000 | 1452 | 4744 | 1914 | 714 | 14169 | 5232 | 7954 |
| 2277 | 3857 | 3290 | 1109 | 4181 | 1758 | 637 | 9101 | 4321 | 6105 |
| 2085 | 4635 | 3331 | 1161 | 4692 | 1913 | 715 | 10394 | 4371 | 8104 |
| 1915 | 3389 | 2914.5 | 1051 | 4091.5 | 1743 | 654 | 8556 | 4040.5 | 5868.5 |
| 4182 | 11011 | 4476 | 1273 | 5563 | 1983 | 634.5 | 14911 | 5488 | 13751 |
| 8135 | 18350 | 6138 | 1593 | 7408.5 | 2311 | 671 | 15604 | 9459 | 21339.5 |
| 6396 | 17240 | 5785 | 1428 | 6796 | 2097 | 598 | 15716 | 6885 | 19515 |
| 7339.5 | 17541 | 5812.5 | 1438 | 6877.5 | 2151 | 621 | 15476.5 | 8233 | 20124 |
| 3194 | 4397 | 2435 | 780 | 3191 | 1374 | 344 | 9832.5 | 4395.5 | 5511.5 |
| 2651 | 3790.5 | 2490 | 846 | 3347 | 1468 | 473 | 8549.5 | 4006 | 5243 |
| 2265 | 3598 | 2649 | 952 | 3655 | 1602 | 564 | 8331 | 3960 | 5426 |
| 2098 | 3449 | 5704 | 1741 | 882678 | 74330 | 635 | 8522 | 3961 | 5687 |
| 1823 | 3145 | 3259 | 1107 | 5865 | 2177 | 618 | 7930 | 3536 | 5343 |
| 1785 | 3083 | 2755 | 1003 | 4336 | 1788 | 625 | 8015 | 3544 | 5265 |
| 1748 | 3081 | 2691 | 992 | 4055 | 1717 | 627 | 7887 | 3508 | 5363 |
| 2137 | 3462 | 2914 | 1060 | 4282 | 1803 | 670 | 9579 | 4196 | 5827 |
| 2341 | 3857.5 | 3103 | 1107 | 4455 | 1862 | 694 | 11443.5 | 4344 | 6315.5 |
| 2315 | 5143 | 3299 | 1204 | 4731 | 1957 | 732 | 13341 | 4451 | 8306 |
| 2253 | 3731 | 3020 | 1100 | 4279 | 1819 | 688 | 9895 | 4321 | 6155 |
| 2216 | 3869 | 3226 | 1157 | 4586 | 1921 | 718 | 10618 | 4528 | 6511 |
| 2115 | 3667 | 3100 | 1136 | 4430.5 | 1885 | 712 | 9904.5 | 4338.5 | 6336.5 |
| 2048 | 3444 | 2910 | 1087 | 4171 | 1787 | 674 | 9022 | 4105 | 5953 |
| 1963 | 3343 | 2857 | 1094 | 4061 | 1808 | 676 | 8553.5 | 3993.5 | 5819.5 |
| 1915 | 3412 | 2875 | 1062 | 4109 | 1769 | 654 | 8527 | 3905 | 5885 |
| 42025 | 22257 | 6942 | 1261 | 4115 | 1729 | 615 | 11575 | 4848 | 6378 |
| 2740 | 7060 | 4090 | 1268 | 5031.5 | 2012.5 | 731 | 11896 | 4768.5 | 10307 |
| 2249 | 3842 | 3219.5 | 1108 | 4449.5 | 1834 | 669 | 9656.5 | 4377.5 | 6457 |
| 2090 | 3543 | 2961 | 1074 | 4167 | 1793 | 665 | 8842 | 4201 | 6094 |
| 4588 | 12544 | 4813 | 1330 | 5953 | 2039 | 655 | 14476 | 5862 | 15582 |
| 7127 | 16863 | 5762 | 1500 | 6889 | 2187 | 637 | 14524 | 7976 | 19692 |
| 8118 | 17777 | 5937.5 | 1466 | 6945 | 2141 | 603 | 15999.5 | 8579 | 19926 |

| | | | | | | | | | |
|---|---|---|---|---|---|---|---|---|---|
| 8950 | 20145 | 6383 | 1556 | 7338 | 2267 | 660 | 14687 | 9672 | 22668 |
| 3024 | 4044 | 2244 | 754 | 2967 | 1337 | 300 | 8648 | 4156 | 4999 |
| 2790 | 3971 | 2514 | 842 | 3365 | 1480 | 457 | 8567 | 4100 | 5339 |
| 2801 | 4706.5 | 3087 | 1095 | 4338 | 1779 | 630 | 11207.5 | 4735 | 7115 |
| 2157 | 3440 | 5703 | 1739 | 763129 | 78471 | 597 | 8483 | 3881 | 5484 |
| 2050 | 3339 | 3290 | 1076 | 5568 | 2124 | 617 | 8212 | 3875 | 5509 |
| 2036 | 3333 | 2933 | 1069 | 4505 | 1840 | 632 | 8290.5 | 3862 | 5492.5 |
| 2064 | 3380 | 2952 | 1066 | 4294 | 1815 | 650 | 8625 | 3993 | 5732 |
| 2171 | 3494.5 | 2905 | 1045 | 4158.5 | 1781 | 647 | 9360.5 | 4122 | 5810 |
| 2412 | 4172 | 3187 | 1129 | 4503 | 1887 | 698 | 10533 | 4460 | 6773 |
| 2367.5 | 4265 | 3258 | 1159.5 | 4550 | 1907 | 685 | 10624.5 | 4319 | 6877 |
| 2365.5 | 3822.5 | 3082.5 | 1101 | 4344 | 1811.5 | 675 | 10225.5 | 4424 | 6237 |
| 2244 | 3901 | 3216 | 1136 | 4535 | 1882 | 701 | 9829 | 4334 | 6566 |
| 2186 | 3780.5 | 3089.5 | 1118 | 4424.5 | 1870.5 | 710 | 9733 | 4333.5 | 6355.5 |
| 2231 | 3657 | 3024 | 1099 | 4267 | 1799 | 685 | 9163.5 | 4182 | 6083 |
| 1884 | 3244 | 2681 | 1006 | 3774 | 1677 | 628 | 8013 | 3643 | 5470 |
| 2196 | 3862 | 3209 | 1122 | 4477 | 1850 | 684 | 9124 | 4294 | 6478 |
| 18853 | 10663 | 4669 | 1170 | 4174 | 1752 | 611 | 10272 | 4639 | 6270 |
| 2422 | 5548 | 3602 | 1208 | 4877 | 1946 | 697 | 11394 | 4461 | 8823 |
| 2224 | 3651 | 2963 | 1058 | 4112 | 1737 | 630 | 9064 | 4238 | 6080 |
| 2306 | 4852 | 3472 | 1222 | 4929 | 1969 | 735 | 11474 | 4587 | 8145 |
| 3814.5 | 9473.5 | 4142 | 1174 | 5119 | 1833 | 596 | 14548 | 5099 | 11794 |
| 8050.5 | 19121 | 6590.5 | 1633 | 7890.5 | 2350 | 670 | 15362.5 | 9221.5 | 22246 |
| 8767 | 18388 | 6107 | 1530 | 7217 | 2218 | 617 | 14437 | 9084 | 20627 |
| 8195 | 18849 | 6098.5 | 1521 | 7305.5 | 2231 | 639 | 14676 | 9200 | 21672.5 |
| 3193 | 4509 | 2479 | 784 | 3261 | 1388 | 320 | 9762 | 4332 | 5475 |
| 2814 | 4042 | 2560 | 860 | 3472 | 1503 | 468 | 9152 | 4172 | 5444 |
| 2638 | 4151 | 2955 | 1005 | 4079 | 1695 | 586 | 10406 | 4423 | 6224 |
| 2215 | 3497 | 5588 | 1717 | 721949 | 69381.5 | 599 | 8339.5 | 3904.5 | 5507 |
| 2067 | 3434 | 3313 | 1092 | 6041 | 2148 | 612 | 8199 | 3829 | 5495 |
| 2088 | 3471 | 3025.5 | 1057 | 4813 | 1880 | 636.5 | 8427.5 | 3900 | 5690.5 |
| 1862 | 3165 | 2711 | 995 | 4128 | 1708 | 604 | 8032.5 | 3530 | 5303.5 |
| 2139 | 3423.5 | 2838.5 | 1049 | 4207 | 1761 | 627 | 8951 | 4000.5 | 5602.5 |
| 2406 | 3681 | 2909 | 1038 | 4201 | 1749 | 636 | 10051 | 4206 | 5831 |
| 2389 | 3743 | 2961 | 1084 | 4100 | 1778 | 634 | 9976 | 4188 | 5825 |
| 2158 | 3964.5 | 3070 | 1113 | 4343.5 | 1842 | 665 | 11543 | 3890 | 6387.5 |
| 2237 | 3636 | 2977 | 1068 | 4106 | 1785 | 661 | 9982 | 4143 | 6006 |
| 2196 | 4554 | 3211 | 1159 | 4581 | 1923 | 712 | 11315 | 4336 | 7692 |
| 2289 | 4925 | 3356 | 1197 | 4778 | 1969 | 752 | 10653 | 4543 | 8249 |
| 2236 | 3924 | 3253 | 1166 | 4599 | 1890 | 721 | 10121 | 4448 | 6580 |
| 2216 | 4530.5 | 3285 | 1159 | 4589 | 1892 | 699 | 9704 | 4361.5 | 7626 |
| 34007 | 24837 | 7017 | 1373 | 4781 | 1904 | 677 | 13787 | 4789 | 8733 |
| 2557 | 6935 | 3841.5 | 1265 | 5198 | 2028 | 733 | 12360.5 | 4676 | 10779.5 |
| 2320 | 3990 | 3287 | 1127 | 4454 | 1829 | 667 | 10078 | 4404 | 6486 |
| 2242 | 5579 | 3595 | 1245 | 4966 | 1993 | 748 | 11795 | 4340 | 9425 |
| 4217.5 | 11892.5 | 4691 | 1342 | 5851.5 | 2037 | 665 | 14658.5 | 5622 | 15063.5 |
| 7334.5 | 18682 | 7224 | 1739.5 | 8922.5 | 2486 | 711 | 15256.5 | 8761 | 21945 |
| 8417.5 | 19367.5 | 6581 | 1635 | 7812.5 | 2348.5 | 641 | 14638 | 9505 | 22225 |

| | | | | | | | | | |
|---|---|---|---|---|---|---|---|---|---|
| 7884.5 | 18989 | 6208 | 1575 | 7489.5 | 2238 | 645 | 13891 | 8692 | 21365 |
| 3160 | 4423 | 2468 | 793 | 3288 | 1387 | 320 | 10463 | 4328 | 5518 |
| 2601.5 | 3633.5 | 2306 | 813 | 3184 | 1435 | 421 | 7981 | 3870 | 4899 |
| 2717 | 4040 | 2868 | 978 | 3905 | 1660 | 580 | 10064 | 4348 | 6014 |
| 2534 | 4148 | 6194 | 1877 | 866621 | 72171 | 641 | 10148 | 4474 | 6492 |
| 2301 | 3823 | 3868 | 1242 | 7421 | 2532 | 648 | 9272 | 4253 | 6158 |
| 2216 | 3759 | 3193.5 | 1119 | 5245 | 1966 | 664 | 9033.5 | 4258 | 6174 |
| 2215 | 3599 | 3014 | 1051 | 4660 | 1829 | 655 | 8957 | 4165 | 5909 |
| 2190 | 3435.5 | 2754 | 987 | 4149 | 1720 | 611 | 8901.5 | 3966 | 5512 |
| 2378.5 | 3636 | 2845 | 1009 | 4235 | 1714 | 618 | 10307 | 4144 | 5717.5 |
| 2284.5 | 5954 | 3227 | 1138 | 4613 | 1890.5 | 682 | 12773 | 3998 | 9521 |
| 2328 | 4001 | 3068 | 1101 | 4376 | 1802 | 666 | 11703 | 4172 | 6337 |
| 2196 | 4268 | 3327 | 1161 | 4697 | 1901 | 709 | 11975 | 4082 | 6833 |
| 2246 | 3679 | 2994 | 1082 | 4281.5 | 1800.5 | 671 | 10074 | 4228 | 6097.5 |
| 2715 | 6561 | 3759 | 1305 | 5324 | 2108 | 810 | 13910 | 5086 | 10891 |
| 2262 | 3634 | 3010 | 1085 | 4225 | 1789 | 664 | 9015 | 4293 | 6001 |
| 2298 | 5764 | 3515.5 | 1218 | 4937 | 1983 | 727 | 12368 | 4544 | 9545 |
| 28388.5 | 21255.5 | 6438 | 1369 | 4878 | 1944 | 694 | 13108 | 4805 | 8945.5 |
| 2514 | 4463 | 3475 | 1134 | 4518 | 1820 | 644 | 10361 | 4381 | 6764 |
| 2548 | 6724 | 3769 | 1305 | 5355 | 2066 | 749 | 13353.5 | 4840 | 10826 |
| 2347 | 4508 | 3290 | 1144 | 4607 | 1863 | 687 | 10168 | 4427 | 7338 |
| 5276 | 13260 | 5143.5 | 1417 | 6426.5 | 2140 | 684 | 15560 | 6854 | 16633.5 |
| 7397 | 17693 | 6373.5 | 1584 | 7626.5 | 2332 | 669 | 15329 | 8575.5 | 20689 |
| 7852 | 18975.5 | 6338.5 | 1577 | 7585.5 | 2289 | 633 | 14320.5 | 8714 | 21595 |
| 8284.5 | 18806 | 6190.5 | 1518 | 7429 | 2249 | 642 | 13287.5 | 9004 | 21314 |
| 3101 | 4678 | 2652 | 830 | 3510 | 1441 | 331 | 12287 | 4195 | 5816 |
| 20315.5 | 10731.5 | 4139 | 974 | 3989.5 | 1690 | 488 | 11732.5 | 4964 | 5833 |
| 16255 | 8885 | 3823 | 994 | 4317 | 1838 | 571 | 11162 | 4914 | 5791 |
| 16273 | 13507.5 | 8361 | 2202.5 | 201629 | 19890 | 796 | 15566 | 5693 | 10612.5 |
| 14206 | 8872.5 | 5238.5 | 1358 | 19468.5 | 3556 | 788 | 15274.5 | 5585 | 7444.5 |
| 13979.5 | 8210 | 4767 | 1249 | 9982.5 | 2750.5 | 765 | 13142 | 5322 | 6920 |
| 13329 | 9108 | 4660 | 1293 | 8158 | 2528 | 782 | 12195 | 5274 | 8001 |
| 13358 | 7632 | 4266 | 1178 | 7355 | 2446 | 746 | 12040 | 5240 | 6615 |
| 14676 | 8230 | 4161 | 1107 | 5325 | 2038 | 706 | 12151.5 | 5376 | 6443.5 |
| 10707.5 | 10432 | 5666 | 2203 | 7701 | 2847 | 816 | 17156 | 5358 | 10274.5 |
| 13253 | 8240 | 4143 | 1167 | 5016 | 2028 | 734 | 13272 | 5225 | 6726 |
| 11260 | 7173 | 3964 | 1161 | 5832 | 2314 | 750 | 12657 | 5231 | 6782 |
| 10961 | 9590.5 | 4342 | 1234 | 5788.5 | 2246.5 | 820 | 15248 | 5024.5 | 9222 |
| 36232 | 26754 | 8313 | 1378 | 5838.5 | 2308 | 821 | 14582.5 | 13687.5 | 13013 |
| 13188 | 11322 | 4960 | 1318 | 7365 | 2806 | 872 | 13939 | 6683 | 10433 |
| 10751 | 9212 | 4532 | 1295 | 7559 | 2917 | 803 | 13951 | 5945 | 9150 |
| 1016291 | 1077958 | 239732 | 16265 | 7403 | 2884 | 759 | 121422 | 33220 | 35133 |
| 18254.5 | 16146 | 5664.5 | 1419.5 | 6820 | 2568 | 813 | 16293 | 5902.5 | 11552 |
| 16042 | 13634 | 5154 | 1317 | 6443 | 2557 | 777 | 14662 | 5314 | 9711 |
| 30192 | 19629 | 6476 | 1374 | 6898 | 2661 | 808 | 17235 | 6399.5 | 8810.5 |
| 21918.5 | 25929.5 | 7392.5 | 1620.5 | 8164.5 | 2717.5 | 810 | 18664 | 8281.5 | 19118.5 |
| 29192 | 35282 | 9775 | 1959.5 | 10669.5 | 3002.5 | 806 | 17170.5 | 11342 | 25753.5 |
| 26490.5 | 32526.5 | 8537.5 | 1756 | 8330 | 2498.5 | 705.5 | 17334 | 10136 | 23077.5 |

| | | | | | | | | | |
|---|---|---|---|---|---|---|---|---|---|
| 23319.5 | 31356.5 | 8254.5 | 1688.5 | 8968.5 | 2669.5 | 731 | 17515 | 9515 | 23248.5 |
| 3232 | 4728 | 2636 | 805 | 3572 | 1426 | 332 | 12135 | 4362 | 5676 |
| 3005.5 | 4401.5 | 2782 | 892 | 3763 | 1547 | 482 | 10298 | 4377.5 | 5821 |
| 2748 | 4285 | 3009 | 981 | 4081 | 1696 | 586 | 10688 | 4372 | 6152 |
| 2799 | 4912 | 6634 | 2026 | 918162 | 75050 | 666 | 10863 | 4733 | 7415 |
| 2385 | 4044 | 3949 | 1232 | 8069 | 2606.5 | 637 | 9223.5 | 4205 | 6349.5 |
| 2385 | 3962 | 3362 | 1130 | 5527 | 2002.5 | 661 | 9735.5 | 4322 | 6279 |
| 2224 | 3518 | 2864 | 1007 | 4506.5 | 1764 | 617 | 8490 | 3986 | 5601 |
| 2340 | 3678 | 2993.5 | 1049 | 4557.5 | 1808 | 623 | 9127.5 | 4257.5 | 5799 |
| 2500 | 3820 | 2920 | 1022 | 4354 | 1756 | 620 | 10123 | 4266 | 5844.5 |
| 2589 | 6409 | 3452.5 | 1199.5 | 4985 | 1951.5 | 712 | 14066 | 4493.5 | 9866.5 |
| 2464 | 4055 | 3055 | 1060 | 4420 | 1801 | 652 | 10918 | 4297 | 6296 |
| 2317 | 5207 | 3273 | 1144 | 4761 | 1885 | 695 | 12293.5 | 4165 | 8275 |
| 2291 | 4621 | 3234 | 1122 | 4571 | 1867 | 706 | 11686 | 4246 | 7480 |
| 2564 | 6378 | 3536 | 1242 | 5145 | 2039 | 757 | 12921 | 4755 | 10294 |
| 2360 | 4160 | 3287 | 1160 | 4695 | 1895 | 705 | 9887 | 4390 | 6720 |
| 2320 | 4498 | 3304.5 | 1119 | 4687.5 | 1863 | 680 | 10615.5 | 4321.5 | 7337 |
| 48474 | 38279 | 8804 | 1437 | 4876 | 1920 | 670 | 13382 | 5125 | 9832 |
| 2985 | 7182 | 4127 | 1293 | 5162 | 2000 | 710 | 13096 | 4803 | 10325 |
| 2252 | 6361 | 3539 | 1181 | 4851 | 1933 | 691 | 11211 | 4007 | 9880 |
| 2383 | 5687 | 3480 | 1154 | 4862 | 1912 | 702 | 11691 | 4394 | 9091 |
| 5908 | 14357 | 5288 | 1467 | 6667 | 2212 | 707 | 15298 | 7666 | 17818 |
| 8065 | 20283.5 | 7350.5 | 1772 | 9019 | 2530 | 715 | 14734 | 9619.5 | 23547 |
| 6541 | 18545 | 6234 | 1556 | 7517 | 2283 | 627 | 15409 | 7175 | 21085 |
| 8356 | 20922 | 6814 | 1643 | 8197 | 2388 | 673 | 14797 | 9191 | 23765 |
| 3374 | 4909.5 | 2752 | 827.5 | 3764 | 1455.5 | 344 | 11004.5 | 4521.5 | 6066 |
| 2956 | 4339 | 2860 | 908 | 3746 | 1542 | 512 | 10417 | 4479 | 5903 |
| 2692 | 4381.5 | 3039 | 1016 | 4145 | 1707 | 603 | 11124 | 4343.5 | 6375.5 |
| 2738 | 4812 | 6716 | 2028 | 715511 | 67127 | 715 | 11598 | 4667 | 7280 |
| 2503 | 3986 | 3738 | 1191 | 6221 | 2264 | 668 | 9483 | 4435 | 6236 |
| 2429 | 3950 | 3293 | 1124 | 5033 | 1927 | 687 | 9664 | 4402 | 6338 |
| 2182 | 3488 | 2899 | 1019 | 4179 | 1707 | 631 | 8765 | 3982 | 5564 |
| 2482 | 3973 | 3209 | 1095 | 4549 | 1836 | 691 | 9942 | 4482 | 6389 |
| 2465 | 3917 | 3067 | 1064 | 4229 | 1747 | 657 | 10471 | 4244 | 6033 |
| 2792 | 6553 | 3666 | 1269 | 5028.5 | 2019 | 752 | 14847.5 | 4838.5 | 10069 |
| 2319 | 4729 | 3447 | 1175 | 4751 | 1925 | 725 | 13317 | 4079 | 7500 |
| 2383 | 3686 | 2985 | 1043 | 4025 | 1717 | 660 | 9919 | 4271 | 5913 |
| 2316 | 4925 | 3199 | 1116 | 4361 | 1813 | 713 | 11562 | 4287 | 8180 |
| 2426.5 | 6455 | 3580 | 1228 | 4915 | 1971 | 780 | 13104.5 | 4502.5 | 10448.5 |
| 2410 | 4050 | 3293 | 1148 | 4534 | 1864 | 718 | 10496 | 4517 | 6602 |
| 2560 | 7255 | 3860 | 1305 | 5307.5 | 2068 | 799 | 14193 | 4848 | 11894 |
| 24035 | 21224.5 | 6075 | 1335 | 4811 | 1917 | 707 | 12730 | 4904 | 10629.5 |
| 2900 | 8020.5 | 4101 | 1289 | 5226 | 2039 | 757.5 | 13588.5 | 4861 | 12018.5 |
| 2354 | 7819 | 3896 | 1300 | 5307.5 | 2043 | 773 | 13705.5 | 4325 | 12531 |
| 2632 | 6646 | 3827 | 1282 | 5314 | 2065 | 779 | 12278 | 4918 | 11025 |
| 6212 | 15061 | 5456 | 1549 | 6895 | 2278 | 753 | 16160 | 8366 | 18969 |
| 7840 | 21315 | 7360 | 1829 | 8922 | 2584 | 763 | 13348 | 9176 | 25204 |
| 8485.5 | 21762.5 | 7410 | 1745 | 8743.5 | 2447 | 701 | 14594.5 | 9465 | 24916.5 |

| | | | | | | | | | |
|---|---|---|---|---|---|---|---|---|---|
| 7794 | 20651 | 6707 | 1684 | 8084 | 2363 | 710 | 15133 | 8802 | 23792 |
| 3235 | 4882 | 2718 | 838.5 | 3703 | 1439.5 | 320.5 | 12562 | 4355 | 5918.5 |
| 3098.5 | 4541 | 2863.5 | 920 | 4047 | 1595 | 448 | 12218.5 | 4259 | 5856 |
| 2788 | 4078 | 2837 | 939 | 4245.5 | 1671 | 543 | 9947 | 4200.5 | 5680 |
| 2801.5 | 5048 | 8244.5 | 2367 | 1357953 | 129087.5 | 663 | 12007 | 4572.5 | 7558 |
| 2892 | 6124 | 4391 | 1370 | 12663 | 2936 | 766 | 13537 | 4964 | 9410 |
| 2746 | 4391 | 3421.5 | 1129.5 | 9827 | 2352.5 | 658 | 9925.5 | 4450 | 6592 |
| 2831.5 | 5830 | 3613 | 1177 | 8805 | 2295 | 674 | 10753.5 | 4571 | 8491.5 |
| 2871 | 5184 | 3709 | 1156 | 9251 | 2294 | 651 | 11319 | 4263 | 7245 |
| 3128 | 7634 | 3763 | 1157 | 9059 | 2264 | 632.5 | 14334.5 | 4486.5 | 10422 |
| 3207 | 8470.5 | 4445 | 1431 | 8770.5 | 2436 | 623 | 14527 | 4490 | 11063 |
| 3553.5 | 9612 | 4432 | 1376 | 9247 | 2489 | 703 | 15058.5 | 5099.5 | 12833 |
| 3149 | 8686.5 | 3968.5 | 1248 | 7914.5 | 2232 | 677 | 14840 | 4568.5 | 11945.5 |
| 2940 | 7293 | 3729 | 1200 | 7569 | 2197 | 659 | 13550 | 4308 | 10099 |
| 6237.5 | 12003 | 4980 | 1371 | 7943 | 2386 | 747 | 14813 | 6326 | 14346 |
| 3972.5 | 9552.5 | 4610.5 | 1350.5 | 7775.5 | 2388 | 737 | 14360 | 5551 | 12764 |
| 3328 | 8470 | 4088 | 1249 | 7022 | 2212.5 | 689 | 12540 | 4922.5 | 11661 |
| 152232 | 149202 | 26596 | 2575 | 8166 | 2576 | 719 | 28816 | 8484 | 17471 |
| 5453 | 13593 | 5980 | 1607 | 8143 | 2533 | 777 | 17049 | 6189 | 15435 |
| 4053 | 9465 | 4879 | 1380 | 7651 | 2455 | 730 | 12851 | 5269 | 12017 |
| 5534 | 13243 | 5744 | 1719 | 8348 | 2594 | 774 | 18495 | 6535 | 15152 |
| 6233 | 16879 | 6424 | 1617 | 8856 | 2569 | 748 | 16477 | 6728 | 16552 |
| 10623.5 | 23833.5 | 9611.5 | 2024 | 11755.5 | 2880 | 724 | 15843 | 9994.5 | 23111.5 |
| 11159 | 27013 | 9036 | 1925 | 11453 | 2854 | 739 | 14848 | 10076 | 26391 |
| 9631 | 24596 | 8189 | 1855 | 10026.5 | 2663 | 698 | 14147.5 | 9748 | 26013 |
| 3521 | 4771.5 | 2682.5 | 833 | 3681 | 1482 | 333 | 13383 | 4202 | 5622 |
| 3137 | 4332 | 2736 | 896 | 3741.5 | 1558 | 449 | 10620.5 | 4218 | 5639 |
| 2836 | 4193 | 2947 | 961 | 4015 | 1655 | 549 | 10036 | 4262 | 5919 |
| 2666 | 4940 | 6059 | 1763 | 837104 | 70666 | 632 | 11340 | 4341 | 7339 |
| 2595 | 5526 | 3979 | 1243 | 6813 | 2304 | 686 | 11605 | 4466 | 8390 |
| 2427.5 | 4183 | 3463 | 1129.5 | 5493.5 | 2011 | 666 | 10023 | 4198 | 6515 |
| 2409 | 3989 | 3224.5 | 1087.5 | 4945.5 | 1894 | 654 | 9391 | 4261 | 6273.5 |
| 2546 | 4222 | 3327 | 1138 | 4940 | 1908 | 675 | 10285 | 4420 | 6552 |
| 2507 | 3847 | 2922 | 1006 | 4322 | 1744 | 610 | 10317 | 4235 | 5890 |
| 2490 | 4137 | 3269 | 1143 | 4559 | 1880 | 639 | 11196 | 4247 | 6384 |
| 2435 | 3602 | 2777 | 997 | 3974 | 1679 | 593 | 9637 | 4212 | 5557 |
| 2444 | 4025 | 3124 | 1105 | 4463 | 1819 | 658 | 10505 | 4353 | 6346 |
| 2309 | 3542 | 2825 | 1018 | 3991 | 1714 | 613 | 9521 | 4138 | 5686 |
| 2354.5 | 4038 | 3232.5 | 1116 | 4593 | 1841 | 682 | 10309.5 | 4297.5 | 6466 |
| 2365 | 4008.5 | 3145 | 1093 | 4406 | 1817 | 668 | 10322 | 4333.5 | 6385 |
| 2372 | 5875 | 3431 | 1177 | 4916 | 1930 | 695 | 10845 | 4431 | 9514 |
| 37439 | 32500 | 7457 | 1391 | 4908 | 1924 | 671 | 14800 | 4924 | 10954.5 |
| 2912 | 7694 | 4100 | 1239 | 5169 | 1989 | 697 | 12540 | 4727 | 10802 |
| 2532 | 7178 | 3690 | 1227 | 5108 | 1989 | 713 | 12630 | 4530 | 11265 |
| 2514 | 7389.5 | 3777.5 | 1279 | 5319 | 2068 | 759.5 | 14173 | 4613.5 | 11834.5 |
| 5710 | 13443 | 5095 | 1410.5 | 6484.5 | 2163.5 | 688 | 15499.5 | 7497 | 16849 |
| 8741.5 | 19213 | 7866 | 1807 | 9472 | 2584 | 733.5 | 15336.5 | 9933.5 | 22032.5 |
| 8154 | 21687 | 6933 | 1677 | 8493 | 2415 | 653 | 13711 | 9105 | 24644 |

| | | | | | | | | | |
|---|---|---|---|---|---|---|---|---|---|
| 8227 | 21792 | 7374 | 1701 | 8809 | 2439 | 678 | 14292.5 | 9190 | 24802 |
| 3112 | 4352 | 2463.5 | 787.5 | 3429 | 1395 | 299 | 10767 | 4048 | 5335 |
| 2996.5 | 4040 | 2555 | 854 | 3569 | 1499 | 435 | 9065 | 4189 | 5363.5 |
| 2684 | 3856 | 2683 | 925 | 3757 | 1585 | 515 | 8500 | 4183 | 5453 |
| 2811 | 5958.5 | 6598.5 | 1870 | 918232 | 72175 | 674 | 11367.5 | 4796 | 8998 |
| 2413 | 3827 | 3492 | 1130 | 6173 | 2144 | 627 | 9417 | 4250 | 5990 |
| 2418 | 4117 | 3404 | 1127 | 5472 | 1991 | 664 | 10133 | 4354 | 6421 |
| 2374 | 3630.5 | 2982 | 1035 | 4637 | 1768.5 | 624 | 8969.5 | 4216 | 5742 |
| 2557 | 5970 | 3550 | 1198 | 5354 | 2019 | 732 | 13020 | 4515 | 9311 |
| 2440 | 3904 | 3003 | 1035 | 4448 | 1766 | 620 | 10147 | 4166 | 5970 |
| 2597 | 6875 | 3443 | 1186 | 4990 | 1954 | 695 | 13943 | 4401 | 10454 |
| 2459 | 4120 | 3041 | 1061 | 4338.5 | 1768 | 632 | 11461 | 4082 | 6344 |
| 2439 | 3992 | 3101 | 1072 | 4406.5 | 1783 | 639 | 10125.5 | 4311 | 6259.5 |
| 2351.5 | 3582 | 2804 | 1008 | 4016 | 1701 | 610 | 8866 | 4192 | 5618.5 |
| 2424.5 | 4317 | 3286 | 1127 | 4761 | 1862 | 689 | 11058 | 4431 | 6909.5 |
| 2407 | 3953 | 3110 | 1117 | 4436 | 1832 | 659 | 9681 | 4393 | 6275 |
| 2339.5 | 5028.5 | 3361 | 1161 | 4737 | 1906 | 686 | 10968 | 4287.5 | 7996.5 |
| 36917 | 22108 | 7035 | 1303 | 4584 | 1833 | 629 | 13039 | 4724 | 7120 |
| 2915 | 4922 | 3800 | 1154 | 4669 | 1833 | 635 | 10171 | 4488 | 6902 |
| 2526 | 4473 | 3446 | 1156 | 4770.5 | 1867 | 657 | 10408 | 4470 | 6901.5 |
| 2431 | 4032 | 3171 | 1084 | 4374 | 1769 | 634 | 9596 | 4288 | 6366 |
| 5543 | 12757 | 4948 | 1368 | 6259 | 2118 | 678 | 14719 | 7344 | 15862 |
| 8049 | 20515 | 7043 | 1715 | 8563 | 2473 | 695 | 13955 | 9284 | 23610 |
| 8373 | 19781 | 7403 | 1750 | 8933 | 2472 | 669 | 13628 | 9554 | 22472 |
| 8441 | 18238.5 | 6040.5 | 1497 | 7222 | 2190 | 619 | 15367 | 9358 | 20887.5 |
| 814 | 1096 | 877 | 604 | 1450.5 | 1159 | 232 | 2350 | 1460 | 1933 |
| 1118 | 1801 | 1545 | 701 | 2265 | 1289 | 392 | 3927.5 | 2145 | 3130 |
| 1346 | 2310 | 2059 | 828 | 2904 | 1455 | 508 | 5817 | 2705 | 4066 |
| 1413 | 2440 | 4711 | 1460 | 746482 | 77657 | 549 | 6406 | 2894 | 4344 |
| 1623 | 2937.5 | 3110 | 1125.5 | 5256 | 2235 | 638 | 7699 | 3470.5 | 5299 |
| 1656 | 3053 | 2872.5 | 1056 | 4297.5 | 1887.5 | 654 | 7885 | 3544 | 5425.5 |
| 1727 | 3059.5 | 2825 | 1070 | 4189 | 1843 | 673 | 8171 | 3675.5 | 5514.5 |
| 2033.5 | 3322.5 | 3016.5 | 1105 | 4229 | 1834.5 | 697 | 9323 | 4150 | 5888 |
| 2318 | 3812 | 3224 | 1164 | 4576 | 1903 | 726 | 11193 | 4430 | 6475 |
| 2075 | 3396 | 2820 | 1070 | 3881 | 1754 | 632 | 9479 | 3837 | 5510 |
| 2118 | 3366 | 2932 | 1048 | 3959 | 1764 | 654 | 9461 | 4084 | 5753 |
| 2176 | 3505 | 3084 | 1124 | 4238 | 1848 | 709 | 9912 | 4309 | 6108 |
| 1696 | 3056 | 2716 | 1013 | 3744 | 1683 | 638 | 8020 | 3547 | 5327 |
| 1937 | 3319 | 2942.5 | 1116.5 | 4086.5 | 1826.5 | 697.5 | 8405 | 3921.5 | 5959 |
| 1659 | 2913 | 2620.5 | 997 | 3667 | 1672 | 637.5 | 7478.5 | 3350.5 | 5214.5 |
| 1789.5 | 3150 | 2796 | 1024 | 3857.5 | 1689 | 621 | 8126 | 3690 | 5473.5 |
| 41011 | 21632 | 7048 | 1314 | 4204 | 1765 | 639 | 11526 | 4914 | 6476 |
| 2352 | 3968 | 3470 | 1156 | 4358 | 1842 | 665 | 9606 | 4478 | 6495 |
| 1867 | 3349 | 2938 | 1036 | 3963 | 1742 | 628 | 8323 | 3755 | 5779.5 |
| 1822.5 | 3248.5 | 2856.5 | 1053 | 3971 | 1718.5 | 641 | 8414.5 | 3718.5 | 5631 |
| 4718 | 12452 | 4795 | 1353 | 5857 | 2016 | 660 | 13986 | 6124 | 15456 |
| 8067 | 17241 | 6001 | 1547 | 7126 | 2213 | 642 | 16157 | 9123 | 19993 |
| 8900 | 18449 | 6212 | 1556 | 7289.5 | 2198.5 | 623 | 16034 | 9718 | 20993 |

| | | | | | | | | | |
|---|---|---|---|---|---|---|---|---|---|
| 9141 | 19159 | 6308 | 1571 | 7392 | 2260 | 655 | 14484 | 9596 | 22269 |
| 3203 | 4099.5 | 2295 | 758 | 2866 | 1316 | 309 | 8901.5 | 4027.5 | 5008.5 |
| 2685 | 3735 | 2412 | 863 | 3252 | 1449 | 461 | 8272 | 3902 | 5117 |
| 2287 | 3519.5 | 2630.5 | 948 | 3543.5 | 1595.5 | 553 | 8065 | 3752 | 5265 |
| 1931 | 3166 | 5477 | 1623 | 814679 | 75529 | 583 | 7816 | 3525 | 5183 |
| 1827 | 3114 | 3077.5 | 1047 | 5115 | 2003 | 616 | 8017 | 3538 | 5239 |
| 1829 | 3110 | 2834 | 1034 | 4294 | 1797 | 632 | 8210 | 3541 | 5308 |
| 2014 | 3344 | 2993 | 1121 | 4468 | 1861 | 680 | 8550 | 4060 | 5849 |
| 2291 | 3764 | 3229 | 1149 | 4713 | 1911 | 731 | 10878 | 4359 | 6365 |
| 2388.5 | 3737 | 3085.5 | 1093.5 | 4335.5 | 1828 | 685 | 11551 | 4367 | 6144.5 |
| 2469 | 4096 | 3295 | 1180 | 4600 | 1911 | 702 | 11957 | 4459 | 6596 |
| 2399 | 3691.5 | 3023.5 | 1111.5 | 4253 | 1818.5 | 678 | 10441.5 | 4346 | 6095.5 |
| 2289 | 3770.5 | 3196.5 | 1153.5 | 4516 | 1866 | 714 | 11126.5 | 4386.5 | 6334.5 |
| 2049 | 3432 | 3007 | 1096 | 4224 | 1798 | 683 | 9065 | 4134 | 6005 |
| 1826 | 3064 | 2692 | 996.5 | 3743 | 1666 | 638 | 8023.5 | 3521.5 | 5320.5 |
| 1825 | 3154 | 2808 | 1056 | 3890 | 1750 | 652 | 8070 | 3685 | 5606 |
| 1894 | 3320 | 2904.5 | 1069 | 3967.5 | 1726 | 630 | 8426.5 | 3768.5 | 5682 |
| 41042.5 | 22183.5 | 7235.5 | 1357.5 | 4498.5 | 1845 | 660 | 13019 | 5059 | 6848 |
| 2279 | 3586 | 2972 | 1049 | 3946 | 1715 | 606 | 8547 | 4232 | 5831 |
| 2256.5 | 3767.5 | 3206.5 | 1146 | 4473 | 1838 | 667 | 9721 | 4419 | 6431.5 |
| 2158 | 3455 | 2993.5 | 1099 | 4080 | 1773 | 646.5 | 8763.5 | 4203 | 5945.5 |
| 4646.5 | 11646 | 4595 | 1282 | 5636.5 | 1973 | 635 | 14542 | 5989.5 | 14551 |
| 8348.5 | 19769 | 6724.5 | 1715 | 7967 | 2421 | 693 | 14180.5 | 9601.5 | 23297 |
| 8981 | 19675.5 | 6530.5 | 1622 | 7579 | 2288 | 638 | 13820 | 9574 | 22349 |
| 8986 | 19412.5 | 6627.5 | 1636 | 7882 | 2316.5 | 664 | 15090.5 | 9815.5 | 22314 |
| 3158 | 4163 | 2255 | 745 | 2906 | 1329 | 299 | 8917 | 4034 | 5020 |
| 2465 | 3468 | 2269 | 809 | 2938 | 1409 | 429 | 7564 | 3436 | 4616 |
| 2453 | 3713 | 2803 | 957 | 3699.5 | 1657 | 580 | 8559 | 4014 | 5582.5 |
| 2089.5 | 3313.5 | 5671.5 | 1636 | 784571 | 78200.5 | 603 | 8205 | 3657.5 | 5423 |
| 2126 | 3318 | 3151 | 1065 | 5087 | 2016 | 626 | 8506 | 3888 | 5419 |
| 2193 | 3451.5 | 3050.5 | 1083 | 4674.5 | 1919.5 | 660 | 9084.5 | 4263 | 5713.5 |
| 2238 | 3716 | 3363 | 1173 | 4804 | 1950 | 714 | 9842 | 4325 | 6300 |
| 2310 | 3944 | 3343 | 1183.5 | 4753.5 | 1936 | 728 | 11565.5 | 4359 | 6574.5 |
| 2461 | 4703 | 3546 | 1206 | 4904 | 1967 | 736 | 13295 | 4471 | 7459 |
| 2524 | 3760 | 2982 | 1094 | 4186 | 1795 | 660 | 10168 | 4340 | 5997 |
| 2341 | 3555 | 2891 | 1050 | 3934 | 1738 | 650 | 10127 | 4214 | 5863 |
| 2326 | 3724.5 | 3135 | 1109 | 4375 | 1821 | 695 | 9895 | 4464.5 | 6150 |
| 2098.5 | 3348 | 2838 | 1077 | 3964 | 1752 | 669 | 9263 | 4026.5 | 5685.5 |
| 2124.5 | 3397 | 2840.5 | 1044 | 3869.5 | 1700 | 658 | 8419.5 | 3809.5 | 5601.5 |
| 1915 | 3320 | 2835.5 | 1050 | 3945.5 | 1767.5 | 674 | 8479.5 | 3778.5 | 5595.5 |
| 2119 | 3420 | 2907 | 1052 | 3975.5 | 1728 | 634 | 8881 | 4181.5 | 5680 |
| 13919 | 8440 | 4516.5 | 1186 | 4383 | 1863 | 670 | 10323 | 4686 | 6546 |
| 2469 | 4000.5 | 3213.5 | 1114 | 4226.5 | 1819.5 | 648 | 9399 | 4362 | 6229.5 |
| 2353 | 4268 | 3552 | 1230 | 4787 | 1959 | 719 | 10644 | 4588 | 7195 |
| 2179 | 3421 | 2947 | 1071 | 4001 | 1756 | 649 | 8878 | 4235 | 5884 |
| 4798 | 12130 | 4855 | 1374 | 5854 | 2035 | 679 | 14394 | 6285 | 15150 |
| 8701 | 18710 | 6626 | 1700 | 7870 | 2439 | 699 | 15260 | 9706 | 22056 |
| 9326 | 20597.5 | 6654.5 | 1650 | 7571 | 2329 | 651 | 15658 | 9608 | 22781 |

| | | | | | | | | | |
|---|---|---|---|---|---|---|---|---|---|
| 8579 | 20355 | 6712 | 1654 | 7851 | 2361 | 679 | 13403 | 9135 | 23193 |
| 3327 | 4313.5 | 2390.5 | 787.5 | 3135.5 | 1358 | 317 | 9597 | 4329 | 5287.5 |
| 2875 | 4045 | 2596.5 | 878 | 3488 | 1516 | 463 | 9180.5 | 4204 | 5442.5 |
| 2435.5 | 3632.5 | 2678 | 945.5 | 3569.5 | 1600 | 531.5 | 8306 | 3963.5 | 5422.5 |
| 2047 | 3238 | 5607 | 1559 | 852673 | 73821 | 564 | 7901 | 3507 | 5085 |
| 2068.5 | 3306.5 | 3106 | 1054.5 | 5259 | 1977 | 604 | 8257 | 3732 | 5300 |
| 2105 | 3311 | 2873 | 1048 | 4613 | 1841 | 627 | 8521 | 3953 | 5530 |
| 2180 | 3741.5 | 3170 | 1154.5 | 4836 | 1946.5 | 699 | 9895 | 4235 | 6283 |
| 2357 | 3879.5 | 3200.5 | 1149.5 | 4674 | 1884.5 | 684 | 10797 | 4323 | 6253 |
| 2454 | 5488 | 3509 | 1188 | 4956 | 1968 | 731 | 13752 | 4428 | 8807 |
| 2498 | 4089 | 3221 | 1155 | 4573.5 | 1875 | 678 | 11144.5 | 4320.5 | 6485.5 |
| 2386 | 3919 | 3198 | 1121 | 4489 | 1861 | 687 | 11628 | 4406 | 6298 |
| 2243 | 3473 | 2887 | 1032 | 4044 | 1720 | 637 | 9304 | 4129 | 5749 |
| 2246 | 3702 | 3104.5 | 1135 | 4443 | 1847.5 | 689 | 9515.5 | 4408.5 | 6213.5 |
| 2031 | 3416 | 2868 | 1035 | 3997 | 1719 | 646 | 8443 | 3836 | 5734 |
| 1800 | 3009 | 2594 | 979 | 3566 | 1638 | 602 | 7764 | 3396 | 5133 |
| 2321 | 3954 | 3281 | 1202 | 4690 | 1950 | 706 | 10101 | 4518 | 6725 |
| 42590 | 21572 | 6902 | 1237 | 4111 | 1739 | 609 | 11754 | 4954 | 6204 |
| 2482.5 | 4590.5 | 3703.5 | 1196.5 | 4792.5 | 1905 | 683 | 11015.5 | 4493 | 7170.5 |
| 2339 | 3693 | 3042 | 1070 | 4177 | 1774 | 625 | 9111 | 4370.5 | 6030.5 |
| 2284.5 | 3763 | 3203 | 1136.5 | 4418 | 1830 | 669 | 9453.5 | 4438 | 6343.5 |
| 4084.5 | 11142.5 | 4400 | 1264 | 5415 | 1927 | 625.5 | 12908.5 | 5276 | 14111 |
| 7991 | 18350 | 6318 | 1630 | 7413 | 2319 | 669 | 16125 | 9162 | 21245 |
| 9043 | 18218 | 6273.5 | 1587 | 7352 | 2258 | 626 | 13319.5 | 8881 | 21216 |
| 8821.5 | 19022 | 6269.5 | 1549 | 7388.5 | 2238.5 | 637 | 13610 | 9206.5 | 21613 |
| 3220 | 4114 | 2290 | 775 | 3048 | 1339.5 | 289 | 8752 | 4246 | 5049.5 |
| 2956 | 3978 | 2557 | 847 | 3402 | 1457 | 448.5 | 8604 | 4110 | 5318.5 |
| 2813 | 4078 | 2946 | 996 | 3981 | 1689 | 577 | 9646 | 4406 | 5961 |
| 2224 | 3530 | 5840 | 1624 | 835707 | 71877 | 595 | 8480 | 3779 | 5455 |
| 2013 | 3268.5 | 3056 | 1013.5 | 5152 | 1914.5 | 583 | 8185.5 | 3620 | 5170.5 |
| 2328 | 3659 | 3178.5 | 1120 | 5020 | 1923 | 665 | 9106.5 | 4334 | 5973 |
| 2293.5 | 3505.5 | 3030 | 1076.5 | 4666.5 | 1850 | 659 | 9116.5 | 4233 | 5800.5 |
| 2412 | 4883 | 3426 | 1195 | 5080 | 1976 | 722 | 11976 | 4405 | 7995 |
| 2450 | 4152 | 3305 | 1141 | 4719 | 1875 | 672 | 12403 | 4295 | 6561 |
| 2523 | 4229.5 | 3417 | 1199 | 4756 | 1904 | 690 | 13375 | 4368 | 6649 |
| 2507 | 3844 | 3088 | 1083 | 4355 | 1800.5 | 656 | 10864 | 4421.5 | 6165 |
| 2347 | 3955.5 | 3161 | 1137 | 4526 | 1844 | 689 | 11083 | 4401 | 6337.5 |
| 2226 | 3435.5 | 2821 | 1042 | 3970 | 1721 | 641.5 | 8902 | 4147.5 | 5609 |
| 2043 | 3356 | 2851 | 1047 | 3901 | 1718 | 631 | 8641 | 3734 | 5577 |
| 1924 | 3139 | 2671 | 977 | 3721 | 1680 | 607 | 8125 | 3566 | 5245 |
| 2321 | 4064 | 3350 | 1178 | 4571 | 1899 | 713 | 10299 | 4430 | 6667 |
| 26894 | 13794 | 5294 | 1197 | 4040 | 1735 | 613 | 10621 | 4762 | 6107 |
| 2518.5 | 4952.5 | 3632.5 | 1176 | 4748 | 1905 | 677 | 11164.5 | 4422 | 7827.5 |
| 2364 | 3774 | 3091 | 1093 | 4216 | 1780 | 636 | 9377 | 4326 | 6099 |
| 2300 | 3949 | 3301 | 1161 | 4557 | 1861 | 692 | 10449 | 4371 | 6608 |
| 4470.5 | 11907 | 4660 | 1300.5 | 5663.5 | 1998 | 631 | 13050 | 5807.5 | 14796 |
| 8285 | 19819 | 7265.5 | 1751 | 8829 | 2499.5 | 713 | 14701.5 | 9784 | 23555 |
| 8417 | 19287 | 6688 | 1672 | 7987 | 2366 | 658 | 12835 | 9173 | 21968 |

| | | | | | | | | | |
|---|---|---|---|---|---|---|---|---|---|
| 8362 | 20562 | 7082 | 1661 | 8376 | 2384 | 683 | 13764 | 9333 | 23467 |
| 3347 | 4269 | 2429 | 790 | 3133 | 1383 | 316 | 9563 | 4365 | 5176 |
| 2681.5 | 3775 | 2434.5 | 822 | 3216.5 | 1438 | 422 | 8389 | 3811.5 | 4953 |
| 2752 | 3995.5 | 2921 | 972.5 | 3975.5 | 1652 | 563 | 9138 | 4392 | 5787.5 |
| 2352 | 3576 | 5815 | 1649 | 848781 | 73167 | 583 | 8577 | 3951 | 5514 |
| 2299 | 3435 | 3172 | 1063 | 5378 | 2005 | 602 | 8611 | 4076 | 5484 |
| 2344 | 3706 | 3254 | 1070 | 5026 | 1923 | 655 | 9636 | 4249 | 6020 |
| 2292 | 4016 | 3415 | 1169 | 5136 | 1992 | 711 | 10577 | 4263 | 6638 |
| 2374 | 3991 | 3247 | 1144 | 4852.5 | 1901.5 | 679 | 11193 | 4294 | 6506 |
| 2594 | 4829 | 3464 | 1165 | 4954 | 1947 | 712 | 11621 | 4546 | 7566 |
| 2550 | 4150 | 3246 | 1154 | 4590 | 1844 | 664 | 10976 | 4337 | 6422 |
| 2504.5 | 3890.5 | 3148 | 1078.5 | 4400 | 1782 | 654 | 10222.5 | 4340 | 6103.5 |
| 2444.5 | 3846 | 3153 | 1100 | 4436 | 1796 | 667 | 10377.5 | 4429 | 6205 |
| 2289 | 3466 | 2821 | 1037 | 4076 | 1733 | 637 | 8848 | 4205 | 5640 |
| 2362 | 3798 | 3144 | 1120 | 4429 | 1833 | 687 | 9281 | 4405 | 6247 |
| 1917 | 3192 | 2669 | 986 | 3708 | 1664 | 603 | 8111 | 3571 | 5217 |
| 2303 | 3719 | 3066 | 1087 | 4250 | 1803 | 640 | 9446 | 4361 | 6289 |
| 46548 | 25339 | 7963 | 1362 | 4541 | 1839 | 638 | 12929 | 5057 | 6836 |
| 2580 | 4030 | 3352.5 | 1105 | 4307 | 1794 | 627 | 9815.5 | 4440 | 6281.5 |
| 2414 | 3955.5 | 3233 | 1117.5 | 4424 | 1852 | 648 | 9541 | 4453.5 | 6413.5 |
| 2383 | 3999 | 3296 | 1146 | 4464 | 1842 | 662 | 10190 | 4443 | 6467 |
| 4550 | 11135 | 4496 | 1228 | 5481 | 1898 | 626 | 14998 | 5797 | 13561 |
| 8572 | 19190.5 | 6647 | 1674 | 7785.5 | 2352 | 661.5 | 15994.5 | 9707.5 | 22415.5 |
| 8584 | 19246 | 6662 | 1646 | 7748 | 2347 | 636 | 13316 | 9122 | 22359 |
| 7904 | 18602.5 | 6047.5 | 1528 | 7245 | 2203 | 630 | 15391.5 | 8590.5 | 21178 |
| 3268 | 4425.5 | 2527.5 | 799 | 3334 | 1396.5 | 317 | 9589.5 | 4284 | 5428.5 |
| 2838 | 4021 | 2548 | 835 | 3451 | 1470 | 434 | 8599 | 4084 | 5304 |
| 2759 | 4161 | 2907 | 987 | 3983 | 1650 | 554 | 9151 | 4379 | 5912 |
| 2513.5 | 3748 | 5866 | 1633 | 851301.5 | 72063 | 600 | 8797.5 | 4180.5 | 5693 |
| 2461 | 3771 | 3388 | 1126 | 5738.5 | 2091 | 633 | 9301.5 | 4295 | 5894 |
| 2020 | 3268 | 2734 | 975 | 4495 | 1768 | 585 | 8042 | 3603 | 5203 |
| 2343 | 3663 | 3116 | 1094 | 4695 | 1852 | 653 | 9195 | 4310 | 5909 |
| 2448 | 3928 | 3153 | 1096 | 4678 | 1850 | 663 | 10792 | 4276 | 6258 |
| 2499 | 5408 | 3480 | 1190 | 5032 | 1956 | 693 | 13387 | 4371 | 8494.5 |
| 2429 | 4811 | 3480 | 1184.5 | 4807.5 | 1919 | 687 | 13874 | 4203 | 7411.5 |
| 2600 | 6567 | 3693 | 1256 | 5157 | 2010 | 733 | 14591 | 4515 | 10093 |
| 2462 | 4068.5 | 3266 | 1149.5 | 4642 | 1884 | 687 | 10663.5 | 4405 | 6650 |
| 2366 | 4489 | 3416 | 1180 | 4819 | 1929 | 714 | 11621 | 4409 | 7178 |
| 2584 | 4521 | 3478 | 1214 | 4888 | 1947 | 725 | 10172 | 4761 | 7352 |
| 2155 | 3499 | 2858 | 1025 | 3990 | 1732 | 628 | 8480 | 3914 | 5673 |
| 2413 | 4058 | 3300 | 1128 | 4634 | 1877 | 680 | 9893 | 4488 | 6586 |
| 47895 | 26572 | 8178 | 1384 | 4576 | 1857 | 647 | 13067 | 5125 | 7013 |
| 2621 | 4329 | 3562 | 1164 | 4550 | 1824 | 641 | 10093 | 4472 | 6624 |
| 2409 | 4265 | 3386 | 1165 | 4707 | 1900 | 661 | 10169 | 4415 | 6799 |
| 2433.5 | 3856.5 | 3108 | 1091 | 4215.5 | 1807.5 | 640 | 9147.5 | 4371.5 | 6218 |
| 5093 | 13073 | 5194 | 1414 | 6391 | 2109 | 673 | 15371 | 6494 | 16552 |
| 7425.5 | 16683 | 5897 | 1515 | 7130 | 2243 | 620 | 15379.5 | 8548.5 | 19263 |
| 8067.5 | 18395.5 | 6469.5 | 1576 | 7459.5 | 2262 | 623.5 | 15932.5 | 8747.5 | 21280 |

| | | | | | | | | | |
|---|---|---|---|---|---|---|---|---|---|
| 9166 | 19402 | 6411 | 1579 | 7567 | 2291 | 640 | 14173 | 9437 | 22030 |
| 3221 | 4410.5 | 2467.5 | 783 | 3323 | 1390 | 312 | 9954 | 4237 | 5326.5 |
| 3105 | 4250 | 2735 | 912 | 3545 | 1504 | 498 | 9499 | 4480 | 5623 |
| 3035 | 4934 | 3462 | 1130 | 4629 | 1834 | 658 | 12585 | 4815 | 7009 |
| 2773 | 4245 | 6276 | 1757 | 666232 | 66462 | 678 | 10272 | 4715 | 6430 |
| 2379 | 3607 | 3292 | 1098 | 5249 | 1965 | 640 | 9038 | 4174 | 5675 |
| 2372.5 | 3589 | 3104 | 1065 | 4501.5 | 1836.5 | 654 | 9297 | 4290.5 | 5780 |
| 2498.5 | 3799.5 | 3173.5 | 1095.5 | 4515.5 | 1836 | 693 | 9424.5 | 4459 | 6085.5 |
| 2536 | 4028 | 3290 | 1114 | 4581 | 1866 | 706 | 10989 | 4495 | 6267 |
| 2670 | 6402 | 3672 | 1217 | 5042.5 | 1951 | 749 | 14203.5 | 4606 | 9876 |
| 2982 | 7837 | 4309 | 1448 | 5797 | 2204 | 833 | 15728 | 5126 | 12212 |
| 2555 | 4238 | 3359 | 1173 | 4565 | 1854 | 717 | 13138 | 4414 | 6659 |
| 2924 | 7019.5 | 4000.5 | 1357 | 5371 | 2167.5 | 821 | 15362.5 | 5296 | 11134.5 |
| 2427 | 4899 | 3563 | 1238 | 4951 | 1974 | 766 | 12848 | 4511 | 7954 |
| 2505 | 5090 | 3713 | 1293 | 5090 | 2031 | 787 | 11705 | 4830 | 8241 |
| 2612 | 4241 | 3481 | 1222 | 4769 | 1943 | 766 | 10265 | 4985 | 7136 |
| 2445 | 4412 | 3594 | 1218 | 4917 | 1944 | 734 | 11296 | 4581 | 7238 |
| 48871 | 30811.5 | 8803 | 1430 | 4827 | 1897 | 657 | 13974 | 5148 | 7927 |
| 2966.5 | 8499 | 4482.5 | 1453 | 5997 | 2211.5 | 767 | 14519.5 | 5137 | 12710 |
| 2553 | 4168.5 | 3320.5 | 1140 | 4592 | 1834.5 | 649.5 | 9695 | 4593.5 | 6645 |
| 2506 | 4196.5 | 3419.5 | 1186 | 4737 | 1900.5 | 684.5 | 10072 | 4495 | 6760 |
| 5643.5 | 12638 | 5321.5 | 1514 | 6879 | 2285 | 723 | 13043.5 | 7789 | 15931.5 |
| 8001 | 18598 | 6399 | 1601 | 7605 | 2308 | 643 | 16258 | 9140 | 21564 |
| 8201 | 21611 | 7252 | 1747 | 8397 | 2484 | 669 | 14404 | 9069 | 24716 |
| 8279.5 | 21213 | 6986 | 1696 | 8219 | 2366.5 | 674 | 12177 | 9097 | 24088 |
| 3237 | 5267.5 | 2991 | 896 | 3954.5 | 1522 | 366 | 13977.5 | 4424 | 6372.5 |
| 3451 | 4845.5 | 3063 | 956 | 4157 | 1622.5 | 461 | 12778 | 4576 | 6190.5 |
| 3099 | 4818 | 3292 | 1064 | 4801 | 1829 | 601 | 11612 | 4587 | 6818 |
| 2808 | 4142.5 | 7718.5 | 1978 | 1340072 | 137515 | 626 | 9580 | 4437 | 6205.5 |
| 2791 | 3933 | 3377.5 | 1114 | 8669.5 | 2243 | 653.5 | 9711.5 | 4581.5 | 6163 |
| 2515 | 3678 | 2959 | 1036 | 7068 | 2004 | 588 | 8489 | 3968 | 5456 |
| 2739 | 4496 | 3463 | 1178 | 7063 | 2113 | 675 | 11043 | 4395 | 6682 |
| 2794 | 4408.5 | 3259 | 1096.5 | 6590 | 2015 | 634 | 10939 | 4234.5 | 6309 |
| 2962 | 6852 | 3591 | 1103 | 6774.5 | 2062 | 635.5 | 13085.5 | 4421 | 9759.5 |
| 3276 | 9482 | 4365 | 1398 | 7548 | 2324 | 712 | 14761 | 4798 | 13156 |
| 3319 | 9570.5 | 4242 | 1294.5 | 7509 | 2289 | 715 | 14544.5 | 4762 | 12841.5 |
| 2964 | 8479 | 4055 | 1231 | 6824 | 2158 | 673 | 14889 | 4298 | 11636 |
| 3360 | 9882 | 4448 | 1377 | 7349 | 2333 | 748 | 15169 | 4916 | 13287 |
| 8785 | 13791 | 5985 | 1495 | 7634 | 2368 | 854 | 15520 | 7209 | 15083 |
| 4559 | 10042 | 5081 | 1411 | 7635 | 2380 | 778 | 14681 | 6029 | 13057 |
| 4009 | 9704 | 4944 | 1398 | 7708 | 2432 | 780 | 13672 | 5477 | 12249 |
| 98414 | 88879 | 17024 | 1849 | 6922 | 2221 | 708 | 23213 | 6732 | 13761 |
| 4110 | 9201 | 4899 | 1317 | 6818 | 2179 | 695 | 12266 | 4968 | 10795 |
| 3235 | 7218 | 4169 | 1251 | 5687.5 | 2013 | 680 | 12194.5 | 4770.5 | 10180.5 |
| 2957 | 6834.5 | 4022.5 | 1291 | 5452 | 2029.5 | 708 | 13385.5 | 4569.5 | 9766.5 |
| 4921 | 12421.5 | 4928.5 | 1363.5 | 6229 | 2070 | 648 | 15146.5 | 6141 | 15104 |
| 8758 | 20468.5 | 7056 | 1730 | 8619 | 2504 | 664 | 16065.5 | 9900 | 23413 |
| 9071.5 | 21561 | 7123.5 | 1732.5 | 8749.5 | 2512 | 662 | 14845.5 | 9737 | 23980 |

| | | | | | | | | | |
|---|---|---|---|---|---|---|---|---|---|
| 7678 | 19707 | 6684 | 1636 | 7870 | 2348 | 635 | 15031 | 8223 | 22695 |
| 3538 | 5111 | 3008 | 877 | 3873 | 1522 | 370 | 14495 | 4562 | 6313 |
| 3149 | 4649 | 3000 | 952 | 3967 | 1589 | 479 | 11766 | 4241 | 5952 |
| 2859 | 4369 | 3070 | 1001 | 4132 | 1674 | 549 | 10912 | 4266 | 6071 |
| 2940 | 5561 | 6438.5 | 1846 | 833457 | 71086.5 | 658 | 12915 | 4671 | 8199.5 |
| 2588 | 3833 | 3593 | 1125 | 6260 | 2191 | 595 | 9037 | 4371 | 5760 |
| 2596 | 4253.5 | 3578 | 1170 | 5687 | 2055 | 671 | 9858 | 4446 | 6608 |
| 2506 | 3775 | 3136 | 1073 | 4736 | 1838 | 627 | 9120 | 4354 | 5922 |
| 2550 | 3900 | 3112 | 1054.5 | 4516 | 1805 | 628 | 9839.5 | 4243.5 | 5949 |
| 2628 | 3820 | 3054 | 1055 | 4299 | 1756 | 609 | 10064 | 4241 | 5824 |
| 2337 | 6367 | 3503 | 1225 | 4731 | 1942 | 662 | 12508 | 3867 | 9760 |
| 2479 | 4555 | 3190 | 1127 | 4628 | 1857 | 662 | 11848 | 4155 | 7063 |
| 2352.5 | 3933 | 3120 | 1101.5 | 4373.5 | 1804.5 | 648 | 10581 | 4096.5 | 6126 |
| 2411 | 4041 | 3186 | 1132 | 4418 | 1827 | 657 | 10776 | 4320 | 6379 |
| 2712 | 7025.5 | 3896 | 1348 | 5480.5 | 2142 | 774 | 13069 | 4821 | 11374.5 |
| 2465 | 3959 | 3295 | 1104 | 4524 | 1842 | 676 | 9745 | 4409 | 6420 |
| 2636.5 | 7068.5 | 3914.5 | 1300 | 5558 | 2096 | 760 | 13221.5 | 4774 | 11061.5 |
| 49281.5 | 29373 | 8798 | 1415 | 4822 | 1909.5 | 647 | 13365 | 5086 | 7445.5 |
| 2885 | 7282.5 | 4222 | 1326.5 | 5286 | 2041.5 | 710 | 12847 | 4759.5 | 10263.5 |
| 2717 | 6065 | 3951 | 1294 | 5318 | 2053 | 726 | 11394 | 4661 | 9393 |
| 2582 | 4300.5 | 3377.5 | 1142.5 | 4701.5 | 1884 | 674 | 9540 | 4626.5 | 6735.5 |
| 4759.5 | 12696 | 4958 | 1429 | 6278 | 2131.5 | 659 | 15762 | 6225 | 15863.5 |
| 8630 | 19379.5 | 6589 | 1722.5 | 7831.5 | 2395 | 677 | 16127 | 9958 | 22659.5 |
| 8156 | 20097 | 6898 | 1703 | 8100 | 2398 | 653 | 14207 | 8943 | 22703 |
| 7985 | 18626 | 6282 | 1546 | 7355 | 2227 | 636 | 15418 | 8676 | 20999 |
| 3443 | 5195 | 2960 | 882 | 3971 | 1512 | 358 | 13607 | 4560 | 6420 |
| 3104 | 4821 | 3070 | 957 | 4136 | 1607 | 499 | 11461 | 4453 | 6276 |
| 2775.5 | 4529.5 | 3115 | 1048 | 4353 | 1747.5 | 576 | 10675 | 4292 | 6410 |
| 2797 | 4564 | 6266 | 1773 | 811360 | 67422 | 665 | 10547 | 4692 | 6871 |
| 2575 | 4242 | 3748 | 1183 | 6386 | 2189 | 650 | 9476 | 4435 | 6446 |
| 2485.5 | 3928 | 3232 | 1104 | 5194 | 1932 | 640 | 9289 | 4272 | 6117 |
| 2375 | 4041 | 3349.5 | 1145 | 5133.5 | 1951 | 677 | 10155.5 | 4261 | 6305 |
| 2510 | 3689.5 | 2969 | 1041 | 4387 | 1735.5 | 607 | 9488.5 | 4248 | 5707 |
| 2625.5 | 4349.5 | 3341 | 1123.5 | 4849 | 1882.5 | 673 | 11582.5 | 4281.5 | 6620 |
| 2625 | 4621.5 | 3447 | 1194.5 | 4837 | 1921 | 677 | 14217 | 4274.5 | 6874 |
| 2738 | 6803 | 3692 | 1243 | 5158 | 2011 | 728 | 14438 | 4660 | 10451 |
| 2454 | 4124 | 3205 | 1101 | 4522 | 1815 | 647 | 11393 | 4167 | 6368 |
| 2357 | 3500 | 2799 | 995 | 3922 | 1676 | 601 | 8965 | 4123 | 5459 |
| 2524 | 3872 | 3155 | 1089 | 4362 | 1797 | 649 | 9487 | 4483 | 6145 |
| 2299 | 3644 | 2947 | 1052 | 4145 | 1719 | 634 | 8691 | 4144 | 5843 |
| 2400 | 4899 | 3524.5 | 1200 | 4934.5 | 1951 | 707 | 10404 | 4388 | 7845 |
| 48428.5 | 33269.5 | 8594.5 | 1411.5 | 4753 | 1880.5 | 647 | 14347 | 5055 | 8342.5 |
| 2839.5 | 7171 | 4025 | 1243 | 4991.5 | 1969.5 | 676 | 11653.5 | 4536 | 10270 |
| 2538 | 4308 | 3447 | 1163 | 4609.5 | 1857 | 655.5 | 10316 | 4333.5 | 6684 |
| 2423 | 4045.5 | 3248.5 | 1112.5 | 4457.5 | 1800 | 653 | 9944 | 4356 | 6330 |
| 4405 | 11412.5 | 4652 | 1278 | 5712 | 1987 | 633 | 14421 | 5640 | 14535.5 |
| 7111 | 17847 | 6247 | 1599 | 7503 | 2276 | 650 | 16497 | 8203 | 20937 |
| 8878 | 21560.5 | 7202 | 1797 | 8459.5 | 2442 | 671 | 14372.5 | 9958.5 | 24542 |

| | | | | | | | | | |
|---|---|---|---|---|---|---|---|---|---|
| 8044.5 | 20961.5 | 6924.5 | 1701.5 | 8369 | 2466 | 676.5 | 11350 | 8752.5 | 24253 |
| 842 | 1153 | 958 | 609 | 1542 | 1148 | 244 | 2556 | 1558 | 2107 |
| 1154 | 1900 | 1633 | 723 | 2388 | 1317 | 419 | 4224 | 2234 | 3326 |
| 1395.5 | 2427 | 2159.5 | 866 | 3082 | 1498 | 529 | 6094.5 | 2851 | 4372 |
| 1583 | 2834 | 5495 | 1609 | 840812 | 79186 | 621 | 7376 | 3345 | 5135 |
| 1584 | 2844 | 3033 | 1105 | 5069 | 2193 | 626 | 7462 | 3377 | 5130 |
| 1570 | 2771 | 2652 | 991 | 4014 | 1784 | 618 | 7373 | 3370 | 5038 |
| 1501 | 2660 | 2449 | 938 | 3650 | 1683 | 589 | 7025 | 3171 | 4771 |
| 1688 | 2895 | 2525 | 971 | 3535 | 1649 | 608 | 7966 | 3354 | 4960 |
| 1946.5 | 3130 | 2596 | 973 | 3643 | 1682 | 613 | 8761 | 3610.5 | 5267 |
| 2373 | 3605.5 | 2990.5 | 1096 | 4095.5 | 1802.5 | 668.5 | 10821.5 | 4380 | 5990 |
| 2312 | 3702.5 | 3127 | 1126 | 4312 | 1860 | 690 | 10650 | 4435.5 | 6167 |
| 2095 | 3386 | 2938 | 1060 | 4096 | 1764 | 678 | 9490 | 4105 | 5840 |
| 1918 | 3294 | 2910 | 1078 | 4035 | 1779 | 681 | 8822 | 3918 | 5723 |
| 1848 | 3184 | 2837 | 1058 | 3977.5 | 1767 | 670 | 8356 | 3702 | 5673.5 |
| 1906.5 | 3306.5 | 3002.5 | 1109 | 4206 | 1811 | 708 | 8963 | 4112 | 5920.5 |
| 2081.5 | 3616.5 | 3176 | 1169 | 4434.5 | 1856 | 693 | 9569 | 4364 | 6349 |
| 43310 | 24217 | 7736 | 1427 | 4813 | 1963 | 712 | 12505 | 5310 | 7554 |
| 2321 | 3918 | 3338 | 1133.5 | 4314.5 | 1823 | 660 | 9026 | 4500 | 6317.5 |
| 2241.5 | 5129 | 3652.5 | 1291 | 5086 | 2018 | 759 | 10232 | 4620.5 | 8941 |
| 2084 | 3579 | 3165 | 1137 | 4363 | 1854 | 701 | 9097 | 4290 | 6211 |
| 4026 | 10411 | 4676 | 1395 | 6021 | 2145 | 711 | 14595 | 5825 | 13765 |
| 8280.5 | 19008.5 | 6568 | 1678.5 | 7846 | 2374 | 680 | 15103 | 9649.5 | 22322 |
| 8828 | 20019 | 6596 | 1619 | 7667 | 2323 | 640 | 15677 | 9658 | 22557 |
| 8215 | 20254 | 6692 | 1683 | 7881 | 2371 | 676 | 11470 | 8877 | 22989 |
| 2876 | 3759 | 2124 | 729 | 2710 | 1294 | 299 | 8584 | 3822 | 4634 |
| 2370 | 3451 | 2321 | 821 | 3062 | 1422 | 451 | 7674 | 3502 | 4767 |
| 1997 | 3152.5 | 2426.5 | 921 | 3302 | 1533 | 542.5 | 7388.5 | 3442 | 4891.5 |
| 1923 | 3254 | 5405 | 1592 | 815978 | 79561 | 617 | 8122 | 3601 | 5277 |
| 1913 | 3217 | 3394 | 1159 | 5863 | 2244 | 642 | 8180 | 3790 | 5513 |
| 1861.5 | 3190 | 2910 | 1043 | 4482.5 | 1844 | 650 | 8134.5 | 3702 | 5440 |
| 1834 | 3164 | 2847 | 1026 | 4154 | 1767 | 648 | 8027 | 3641 | 5363 |
| 1843 | 3058 | 2628 | 1006 | 3754 | 1682 | 614 | 8079 | 3533 | 5098 |
| 2326 | 3487 | 2891 | 1045 | 4002 | 1752 | 643 | 10129 | 4148 | 5692 |
| 2103.5 | 3293 | 2661 | 976 | 3664.5 | 1641 | 600 | 9099 | 3684 | 5289 |
| 2384.5 | 3891 | 3251 | 1163.5 | 4499.5 | 1892 | 714 | 11719.5 | 4444.5 | 6610.5 |
| 2123.5 | 3362.5 | 2834 | 1063 | 3928 | 1723 | 649 | 9531.5 | 4133 | 5620.5 |
| 2083 | 3415 | 2986 | 1102 | 4196 | 1781 | 693 | 9521 | 4194 | 5867 |
| 1980 | 3258.5 | 2788 | 1036 | 3950 | 1720 | 668 | 8597.5 | 3855.5 | 5558.5 |
| 2097 | 3434 | 3034 | 1131 | 4254 | 1850 | 704 | 9045 | 4300 | 6104 |
| 2202 | 3934 | 3423 | 1215 | 4836 | 1949 | 738 | 10506 | 4523 | 6925 |
| 44147 | 24786 | 7992 | 1425 | 4654 | 1920 | 684 | 13843 | 5099 | 7174 |
| 2249 | 5596 | 3704 | 1276 | 4902 | 1991.5 | 706 | 10242.5 | 4343 | 8961 |
| 2293 | 3951 | 3396 | 1193 | 4640 | 1906 | 701 | 10105 | 4522 | 6720 |
| 2203.5 | 4361.5 | 3480 | 1225 | 4882 | 1990 | 741.5 | 11303 | 4556.5 | 7578.5 |
| 4296.5 | 8748 | 4428.5 | 1330 | 5674 | 2021.5 | 682 | 13230 | 6056 | 11093.5 |
| 8429 | 19384 | 7591.5 | 1797 | 9196 | 2551 | 703.5 | 15342.5 | 9774 | 22547 |
| 8292 | 20162 | 6892 | 1695 | 8025 | 2360 | 643 | 12353 | 9027 | 23141 |

| | | | | | | | | | |
|---|---|---|---|---|---|---|---|---|---|
| 7889.5 | 21056 | 6768 | 1634 | 8029.5 | 2339 | 666 | 14791 | 8744.5 | 23956.5 |
| 3092 | 4038.5 | 2283.5 | 762 | 3015 | 1308.5 | 321 | 9161.5 | 4228 | 4956 |
| 2413 | 3527 | 2301 | 817 | 3100 | 1415 | 443 | 8088 | 3601 | 4856.5 |
| 2289 | 3505 | 2612 | 947 | 3532 | 1591 | 548 | 8383 | 3768 | 5221 |
| 2163 | 3468 | 5882 | 1693 | 887072 | 78150 | 635 | 8672 | 3897 | 5677 |
| 2175 | 3509 | 3389 | 1137 | 5822.5 | 2144.5 | 658 | 8730 | 4129 | 5790.5 |
| 1967 | 3316.5 | 3009 | 1056 | 4711 | 1851 | 644 | 8269.5 | 3828 | 5588.5 |
| 1884 | 3183 | 2792 | 1024 | 4304.5 | 1781 | 639 | 8126 | 3694.5 | 5458 |
| 1927 | 3210.5 | 2705 | 981 | 3931 | 1666 | 608.5 | 8500.5 | 3663.5 | 5292 |
| 2284 | 3435 | 2818 | 1005 | 3945 | 1683 | 625 | 9553.5 | 4005.5 | 5581.5 |
| 2498 | 3728 | 2922 | 1085 | 4178 | 1795 | 646 | 10494 | 4384 | 5954 |
| 2399 | 4041 | 3276 | 1160 | 4600 | 1890 | 709 | 12797 | 4401 | 6529 |
| 2249 | 3572 | 3034 | 1097 | 4188 | 1780 | 668 | 10050.5 | 4295 | 5922 |
| 2300 | 3641 | 3119.5 | 1147 | 4351.5 | 1858 | 709 | 9807 | 4501 | 6173.5 |
| 2048 | 3328 | 2819 | 1051 | 3985.5 | 1758.5 | 649 | 8403.5 | 3879.5 | 5662.5 |
| 2118 | 3436 | 2968 | 1091 | 4152 | 1810 | 678 | 8642 | 4208 | 5922 |
| 2240 | 4807 | 3568 | 1279 | 5003 | 2010 | 761 | 11254.5 | 4526.5 | 8159.5 |
| 44808 | 25098 | 8072.5 | 1409 | 4560 | 1860 | 681 | 13421 | 5030 | 7056 |
| 2550 | 6582 | 3972 | 1324 | 5295 | 2087 | 750 | 12935 | 4764 | 10587 |
| 2253 | 4162 | 3492 | 1218 | 4867 | 1973 | 709 | 10218 | 4294 | 6966 |
| 2226 | 4146 | 3384 | 1203 | 4792 | 1951 | 723 | 10786 | 4461 | 7121 |
| 4578 | 9365.5 | 4422.5 | 1382 | 5797 | 2041.5 | 687 | 13144 | 6487 | 12395.5 |
| 8482.5 | 19985.5 | 6942 | 1748 | 8296.5 | 2464.5 | 683 | 15365 | 9802 | 23165.5 |
| 8320 | 20659 | 6697 | 1693 | 7883 | 2341 | 654 | 15655 | 8936 | 23081 |
| 8856 | 20882 | 6676 | 1643 | 7839 | 2358 | 666 | 15070 | 9837 | 23897 |
| 3087.5 | 4069 | 2289.5 | 759 | 3008 | 1347.5 | 305 | 9526 | 4149 | 5026.5 |
| 2466 | 3506 | 2292 | 805 | 3035.5 | 1412 | 424 | 8025.5 | 3545.5 | 4780 |
| 2570.5 | 3664 | 2698 | 944 | 3639 | 1586 | 559 | 9071 | 4216.5 | 5526 |
| 2217 | 3459 | 5845.5 | 1645 | 890918 | 75940 | 607 | 8959 | 3920 | 5552.5 |
| 2299 | 3751 | 3590 | 1174 | 5922 | 2143 | 678 | 9504 | 4315 | 6193 |
| 2227 | 3650 | 3259.5 | 1130 | 5147.5 | 1993 | 678 | 8828 | 4233 | 6130 |
| 2054 | 3427 | 3006 | 1059 | 4547 | 1808 | 653 | 8500 | 3994 | 5718 |
| 2167 | 3472 | 2948 | 1046 | 4284 | 1779 | 638 | 8960 | 4063 | 5688 |
| 2469 | 3833 | 3125 | 1104 | 4464 | 1795 | 661 | 10642 | 4447 | 6087 |
| 2471.5 | 3788 | 2994 | 1072 | 4204.5 | 1758 | 645 | 10390 | 4364 | 5985 |
| 2455 | 3726 | 3068 | 1077 | 4249 | 1761 | 655 | 11261 | 4272 | 5916 |
| 2401 | 3672 | 3056 | 1099 | 4302 | 1798 | 681 | 10017 | 4451 | 6145 |
| 2285 | 3581 | 3081 | 1113 | 4226 | 1795 | 679 | 9550 | 4415 | 6019 |
| 2243.5 | 3473.5 | 2994 | 1081 | 4152.5 | 1789 | 664 | 9245.5 | 4243 | 5811 |
| 2231.5 | 3619 | 3033 | 1118 | 4305 | 1830 | 674 | 9320 | 4411 | 6127 |
| 2239 | 4071 | 3437 | 1191 | 4742 | 1925 | 720 | 10793 | 4428 | 6854 |
| 49324.5 | 26827.5 | 8328.5 | 1373 | 4430 | 1823.5 | 654.5 | 12869 | 5093 | 6856.5 |
| 2788 | 7971 | 4370 | 1406 | 5754 | 2193 | 790 | 14711 | 5015 | 12559 |
| 2324 | 4386 | 3425 | 1203 | 4812 | 1937 | 690 | 10141 | 4404 | 7292 |
| 2268 | 3945 | 3329 | 1184 | 4653 | 1912 | 706 | 10284 | 4421 | 6674 |
| 4175 | 9717 | 4359 | 1308 | 5612 | 2026 | 661 | 14079 | 5832 | 12432 |
| 8718.5 | 19785 | 6878.5 | 1711 | 8252 | 2432 | 684 | 15421 | 10028.5 | 23153.5 |
| 8831 | 19679 | 6640 | 1643 | 7740 | 2314 | 635 | 14617 | 9646 | 22375 |

| | | | | | | | | | |
|---|---|---|---|---|---|---|---|---|---|
| 8626 | 19495 | 6402 | 1578 | 7490 | 2258 | 643 | 15182 | 9493 | 22459 |
| 3173 | 4070 | 2264 | 759 | 2993 | 1339 | 297 | 9142 | 4261 | 4993 |
| 2749 | 3772 | 2432 | 843 | 3257 | 1454 | 441 | 8576 | 3996 | 5065 |
| 2301 | 3465 | 2567 | 920 | 3450 | 1557 | 532 | 8089 | 3701 | 5145 |
| 2509 | 3605.5 | 5903.5 | 1684 | 894847 | 75539 | 615 | 9421 | 4327 | 5773 |
| 2305 | 3671 | 3500 | 1141 | 6056 | 2150 | 642 | 9182 | 4243 | 5822 |
| 2280 | 3758 | 3317 | 1160 | 5218 | 1999 | 673 | 8996 | 4354 | 6139 |
| 2286 | 3803 | 3261 | 1132.5 | 4881 | 1967 | 694 | 9109.5 | 4437 | 6297.5 |
| 2265 | 3663 | 3027 | 1089 | 4475 | 1807 | 654 | 9224 | 4190 | 5870 |
| 2397 | 3648 | 2923 | 1020 | 4097.5 | 1710.5 | 612 | 9953 | 4193 | 5706.5 |
| 2457 | 3649 | 2861.5 | 1036.5 | 4049 | 1716 | 614 | 10113 | 4207 | 5708.5 |
| 2476 | 3987 | 3196 | 1112 | 4429 | 1799 | 677 | 10890 | 4434 | 6247 |
| 2421 | 3685 | 2968 | 1076 | 4204 | 1786 | 654 | 9825.5 | 4372.5 | 6076 |
| 2279 | 3699.5 | 3043 | 1091 | 4305 | 1809 | 676 | 10045.5 | 4311 | 6056 |
| 2136 | 3503.5 | 2932 | 1041.5 | 4058 | 1741 | 646 | 8760.5 | 3986 | 5749.5 |
| 2239 | 3574 | 3072 | 1103 | 4300 | 1814 | 675 | 9713 | 4356 | 6024 |
| 2262 | 3854 | 3218 | 1123 | 4568.5 | 1885 | 683 | 9428 | 4462 | 6458.5 |
| 44213.5 | 38918 | 8509 | 1503 | 5108.5 | 2016 | 708 | 16392 | 5092.5 | 11687.5 |
| 2539.5 | 6122 | 3860 | 1269.5 | 4984.5 | 1974.5 | 713 | 11556.5 | 4577.5 | 9677.5 |
| 2417 | 5831 | 3797 | 1312 | 5222 | 2061 | 756 | 12719 | 4623 | 9659 |
| 2408.5 | 4428.5 | 3521.5 | 1206 | 4841 | 1962 | 721.5 | 10086.5 | 4650.5 | 7321.5 |
| 4985 | 12694.5 | 4987 | 1372.5 | 6021.5 | 2068.5 | 657 | 15350.5 | 6416.5 | 15732 |
| 7536 | 20447 | 7183.5 | 1780 | 8917 | 2500 | 705 | 13861 | 8756 | 23860 |
| 9099.5 | 22490 | 7258.5 | 1758 | 8403 | 2452.5 | 686 | 14372.5 | 9864 | 25243.5 |
| 8226.5 | 20212 | 6759.5 | 1643.5 | 8103.5 | 2382 | 675 | 14586 | 9179.5 | 23528.5 |
| 3160 | 4058 | 2329 | 770 | 3067 | 1336 | 308 | 9457 | 4231 | 5037 |
| 2837 | 3779 | 2481.5 | 846 | 3354 | 1445 | 440 | 8844.5 | 4103.5 | 5188.5 |
| 2638 | 3706 | 2719 | 959 | 3667 | 1616 | 549 | 8845 | 4226 | 5492 |
| 2246 | 3525.5 | 5894.5 | 1652 | 906508.5 | 74197.5 | 594 | 8643 | 3873.5 | 5522.5 |
| 2449 | 4010 | 3769 | 1240 | 6476 | 2260 | 689 | 10433 | 4520 | 6517 |
| 2398 | 3899 | 3408 | 1188 | 5490 | 2013 | 688 | 9488 | 4406 | 6419 |
| 2171 | 3510 | 3024 | 1064 | 4641 | 1819 | 640 | 8717 | 4056 | 5813 |
| 2408 | 3916 | 3253 | 1116 | 4763 | 1884 | 681 | 9823 | 4445 | 6330 |
| 2509 | 3834 | 3044 | 1055 | 4315 | 1746 | 636 | 10376 | 4292 | 5911 |
| 3358 | 4257 | 3386.5 | 1209 | 70243.5 | 7328.5 | 633 | 11267 | 4473 | 6109 |
| 2489 | 3814 | 3031 | 1062 | 4252 | 1772 | 633 | 10185 | 4325 | 5934 |
| 2594 | 3974 | 3162 | 1123 | 36476.5 | 4370.5 | 676 | 10803.5 | 4341 | 6348 |
| 2613.5 | 3759 | 3045.5 | 1090.5 | 36145 | 4429 | 670 | 9852 | 4418.5 | 6106 |
| 28319.5 | 17623.5 | 6008 | 1206.5 | 4887 | 2023.5 | 694 | 10782 | 10682.5 | 9360.5 |
| 14058.5 | 8631 | 4131 | 1155 | 4429.5 | 1883 | 736.5 | 11261 | 6117.5 | 6916 |
| 12561 | 9013 | 4404 | 1250 | 4873 | 1970 | 772 | 13147 | 5807 | 8122 |
| 50612.5 | 41644.5 | 9470.5 | 1522 | 5043.5 | 1965 | 706 | 15465 | 5260 | 10764 |
| 2734 | 7850 | 4344 | 1355 | 5503 | 2122 | 751 | 14344 | 4735 | 11989 |
| 2594 | 5668 | 3719 | 1275 | 5240 | 2030 | 730 | 10918 | 4832 | 9383 |
| 2343 | 4097 | 3344 | 1166 | 4658 | 1898 | 696 | 10629 | 4406 | 6800 |
| 5969 | 13303 | 5234 | 1480.5 | 6482 | 2178 | 709 | 15492.5 | 7831.5 | 16755 |
| 8462.5 | 19449.5 | 6544.5 | 1691 | 7806.5 | 2402.5 | 686 | 15910.5 | 9810.5 | 22731.5 |
| 8030 | 20408 | 6721 | 1660 | 8002 | 2389 | 658 | 15344 | 8920 | 23438 |

| | | | | | | | | | |
|---|---|---|---|---|---|---|---|---|---|
| 8829 | 22328 | 6946 | 1703 | 8274 | 2411 | 681 | 13874 | 9537 | 25193 |
| 3247 | 4246 | 2394 | 772 | 3179 | 1377 | 302 | 9369 | 4322 | 5112 |
| 2948 | 4051 | 2669 | 890 | 3538 | 1512 | 472 | 9279 | 4320 | 5415 |
| 2648.5 | 3858.5 | 2792 | 961.5 | 3735 | 1633 | 555 | 8902.5 | 4303 | 5636 |
| 2645 | 3932 | 6125 | 1762 | 908889 | 74324 | 632 | 9786 | 4561 | 6058 |
| 2434 | 3872.5 | 3529.5 | 1163 | 5966.5 | 2145 | 653 | 8939.5 | 4315 | 6179 |
| 2383 | 3978 | 3370 | 1161 | 5425 | 2014 | 681 | 9483 | 4333 | 6405 |
| 2114.5 | 3458 | 2918 | 1054.5 | 4500.5 | 1809 | 627 | 8460 | 3922 | 5605.5 |
| 2292 | 3662 | 3040 | 1058 | 4453 | 1788 | 636 | 9418 | 4148 | 5851 |
| 2514 | 3772 | 2998 | 1051 | 4264 | 1736 | 620 | 10137 | 4283 | 5850 |
| 2610 | 3857 | 3065 | 1077 | 4245 | 1780.5 | 628 | 10576.5 | 4343.5 | 5962 |
| 2527 | 3843 | 3044 | 1068 | 4294 | 1775 | 630 | 10570 | 4359 | 6077 |
| 2420 | 4171 | 3260 | 1153 | 4617 | 1894 | 692 | 11836 | 4325 | 6572 |
| 2258 | 3495 | 2893 | 1028 | 4072 | 1710 | 632 | 9185 | 4169 | 5773 |
| 2378 | 3611 | 2960 | 1064 | 4152 | 1744 | 644 | 8920 | 4302 | 5868.5 |
| 2316 | 3635 | 3020 | 1077 | 4198 | 1784 | 651 | 9076 | 4329 | 6014 |
| 2284 | 3883.5 | 3193.5 | 1104 | 4410.5 | 1796 | 662 | 9872 | 4281.5 | 6316 |
| 46009.5 | 35816.5 | 8920 | 1459 | 4994 | 1933 | 688 | 14775.5 | 4811.5 | 9280 |
| 2523.5 | 5318.5 | 3667 | 1158 | 4539 | 1835.5 | 635 | 10858.5 | 4251.5 | 7917 |
| 2407 | 6518 | 3718 | 1239 | 5111 | 1979 | 713 | 12003 | 4512 | 10459 |
| 2376 | 3847 | 3190 | 1121 | 4382 | 1842 | 648 | 9457 | 4326 | 6281 |
| 6191 | 13831 | 5359 | 1482 | 6589 | 2215 | 716 | 15281 | 8167 | 17423 |
| 7837 | 17647 | 6290 | 1657 | 7610 | 2358 | 667 | 13512 | 9220 | 20775 |
| 8162.5 | 18971.5 | 8395.5 | 1865 | 10074 | 2605 | 697 | 14537 | 9446.5 | 21843.5 |
| 8344 | 20734 | 6730 | 1662 | 8030 | 2357 | 652 | 12017 | 9092 | 23440 |
| 3202 | 3993 | 2279 | 762 | 3043 | 1348 | 298 | 8921 | 4172 | 4921 |
| 2883 | 3902 | 2575 | 867 | 3332 | 1466 | 486 | 8825 | 4222 | 5188 |
| 2730 | 3879 | 2855 | 992 | 3686 | 1616 | 580 | 9127 | 4414 | 5625 |
| 2664 | 3892 | 6391 | 1729 | 719743 | 68621 | 665 | 9610 | 4591 | 5997 |
| 2589 | 3935 | 3639 | 1174 | 5645 | 2113 | 694 | 9659 | 4622 | 6302 |
| 2562 | 4466.5 | 3760.5 | 1264 | 5539.5 | 2083 | 764 | 10671.5 | 4721 | 7226 |
| 2526 | 4480 | 3638 | 1244 | 5273 | 2028 | 772 | 10605 | 4684 | 7274 |
| 2566 | 4235 | 3519 | 1196 | 4940 | 1941 | 730 | 11069 | 4584.5 | 6839.5 |
| 2671 | 4125 | 3335 | 1138 | 4562 | 1841 | 703 | 11209 | 4564 | 6504 |
| 2631.5 | 4370.5 | 3448 | 1181.5 | 4603 | 1905 | 714 | 11939.5 | 4513 | 6703.5 |
| 2545 | 4198 | 3301 | 1155 | 4493 | 1837 | 705 | 12565 | 4422 | 6435 |
| 2574 | 4050 | 3315 | 1134 | 4350 | 1817 | 707 | 10799.5 | 4607.5 | 6482.5 |
| 2483 | 4389 | 3584 | 1237 | 4826 | 1967 | 761 | 12255 | 4534 | 7132 |
| 2429 | 3750 | 3141 | 1114 | 4233 | 1773 | 696 | 9867 | 4474 | 6167 |
| 2379 | 3837 | 3239 | 1159 | 4364 | 1850 | 719 | 9812 | 4462 | 6474 |
| 2402 | 6230 | 3790 | 1303 | 5171 | 2063 | 781 | 12480 | 4469 | 10441 |
| 51132 | 39798 | 9230 | 1468 | 4899 | 1922 | 664 | 15077 | 5099 | 9688 |
| 2697 | 7559 | 4157 | 1332 | 5454 | 2093 | 723 | 13281 | 4590 | 11618 |
| 2498 | 6089 | 3802 | 1282.5 | 5179.5 | 2027 | 706 | 11777 | 4544.5 | 9607.5 |
| 2507.5 | 5428 | 3648 | 1221 | 5053 | 1982 | 712 | 12012 | 4516 | 8637.5 |
| 5408 | 12432.5 | 4978 | 1381 | 6225 | 2116 | 661 | 14770 | 7183.5 | 15639.5 |
| 8123 | 20835 | 7851 | 1873 | 9542 | 2598 | 713 | 14217 | 9604 | 24085 |
| 9365.5 | 21769 | 6980.5 | 1730 | 8403 | 2419.5 | 660 | 14582.5 | 9838 | 24819 |

| | | | | | | | | | |
|---|---|---|---|---|---|---|---|---|---|
| 8039 | 21701 | 6904 | 1679 | 8169 | 2364 | 670 | 14938 | 8986 | 24786 |
| 3160 | 4181 | 2392 | 803 | 3218 | 1384 | 305 | 9939 | 4139 | 5071 |
| 3291 | 4357 | 2763 | 899 | 3784 | 1591 | 454 | 11379 | 4234 | 5571 |
| 2964 | 4255 | 3097 | 1010 | 4284 | 1726 | 565 | 10340 | 4433 | 6059 |
| 2839.5 | 4262.5 | 7935.5 | 2065.5 | 1367967 | 140501 | 635 | 9955.5 | 4630 | 6311.5 |
| 2761 | 4408 | 3961 | 1236 | 8431 | 2471 | 714 | 11458 | 4757 | 6934 |
| 2681 | 4493 | 3504 | 1182 | 7162 | 2206 | 681 | 10413 | 4442 | 6888 |
| 2689 | 4312 | 3401 | 1136 | 6414 | 2059 | 657 | 9818 | 4472 | 6454 |
| 2879.5 | 4200.5 | 3224.5 | 1081 | 6131.5 | 1923.5 | 621 | 9823 | 4486 | 6170.5 |
| 2980.5 | 4864 | 3547.5 | 1133.5 | 6264 | 2009.5 | 641 | 11932.5 | 4444 | 6811 |
| 3003 | 5918 | 3950 | 1264 | 6505 | 2125 | 653 | 15127 | 4388 | 8249 |
| 3286 | 8426 | 4075 | 1235 | 6775 | 2126 | 676 | 16261 | 4778 | 11617 |
| 3547 | 9225 | 4324.5 | 1312 | 6911 | 2245 | 714 | 15105.5 | 5080 | 12525 |
| 3691.5 | 9466 | 4647 | 1393 | 7273.5 | 2291 | 747 | 16096.5 | 4944 | 12577 |
| 7327 | 11749 | 5150 | 1323 | 6596 | 2192 | 736 | 14393 | 6071 | 12572 |
| 4069 | 9349 | 4586 | 1268 | 6672.5 | 2177 | 711 | 12612 | 5321.5 | 12248 |
| 3637 | 9768.5 | 4615 | 1360 | 6736.5 | 2231 | 740 | 14724 | 5256 | 13519 |
| 45144 | 51051 | 10026 | 1589.5 | 5949.5 | 2096.5 | 689.5 | 18974 | 5289.5 | 14375 |
| 3285 | 10240 | 4660 | 1359 | 5696 | 2063 | 693 | 14859 | 4592 | 13315 |
| 3392 | 9085 | 4346 | 1366 | 6316 | 2201 | 715 | 15012 | 5089 | 13139 |
| 3013 | 7661 | 4064 | 1291 | 5928 | 2078 | 717 | 13984 | 4721 | 11040 |
| 7438 | 16135 | 6129 | 1580 | 7793 | 2406 | 715 | 16345 | 8748 | 18882 |
| 9062 | 21876 | 7459 | 1796 | 9283 | 2568 | 701 | 14469 | 9640 | 24933 |
| 8758 | 21510.5 | 9117.5 | 1976.5 | 11087 | 2778 | 728 | 13186.5 | 9202 | 23788 |
| 8572 | 21729 | 8487 | 1893 | 10157 | 2632 | 716 | 13266 | 9282 | 24131 |
| 3602 | 4246 | 2454 | 803 | 3234 | 1379 | 297 | 10305 | 4317 | 5059 |
| 3156 | 4042 | 2621 | 860 | 3452 | 1498 | 428 | 9291 | 4276 | 5229 |
| 2975 | 3942 | 2815 | 933 | 3687 | 1584 | 522 | 9078 | 4330 | 5511 |
| 2777 | 4221 | 6156 | 1711 | 859716 | 73368 | 624 | 10336 | 4425 | 6263 |
| 2534 | 4464 | 3853 | 1214 | 6369 | 2226 | 677 | 10996 | 4302 | 6761 |
| 2517 | 4364 | 3569 | 1182 | 5459 | 2021 | 675 | 10779 | 4380 | 6712 |
| 2532 | 5387 | 3696 | 1209 | 5460 | 2027 | 710 | 11176 | 4549 | 8305 |
| 2575 | 4899 | 3572 | 1223 | 5297 | 1996 | 711 | 10693 | 4563.5 | 7711 |
| 2571.5 | 3958.5 | 3195.5 | 1069.5 | 4512.5 | 1800 | 632 | 10090 | 4460 | 6103 |
| 2626 | 3971 | 3229 | 1151 | 4350 | 1796 | 621 | 10035 | 4281 | 5982 |
| 2410 | 4108 | 3245 | 1095 | 4499 | 1827 | 653 | 10697 | 4141 | 6292 |
| 2397 | 3967 | 3165 | 1096 | 4429 | 1808 | 662 | 10703 | 4270 | 6257 |
| 2444 | 4081.5 | 3257 | 1124 | 4603 | 1853 | 689 | 9926.5 | 4430.5 | 6491 |
| 2508 | 3642 | 2965 | 1038.5 | 4111 | 1746.5 | 627 | 8872.5 | 4282.5 | 5744.5 |
| 2433 | 4005 | 3253 | 1134 | 4549.5 | 1834 | 682.5 | 10048.5 | 4444.5 | 6397 |
| 2377 | 4306 | 3300 | 1143 | 4556 | 1849 | 658 | 10287 | 4270 | 6849 |
| 53390.5 | 46672.5 | 9580.5 | 1524 | 5268 | 2007 | 698 | 16453.5 | 5472.5 | 12094.5 |
| 2554 | 6658 | 3745 | 1222 | 4909.5 | 1923 | 668 | 10975 | 4261 | 10000 |
| 2580 | 7089 | 3867 | 1279 | 5329 | 2039 | 717 | 12597 | 4624 | 11163 |
| 2291 | 4233 | 3380 | 1167 | 4674 | 1877 | 667 | 10192 | 4156 | 6712 |
| 6143 | 13768 | 5297 | 1498 | 6710 | 2236 | 708 | 15174 | 8124 | 17370 |
| 7985 | 19874 | 7981 | 1900 | 9650 | 2664 | 738 | 14258 | 9784 | 23220 |
| 8473 | 23164.5 | 8577 | 1914 | 10076 | 2626.5 | 712 | 14428 | 9604.5 | 26643 |

| | | | | | | | | | |
|---|---|---|---|---|---|---|---|---|---|
| 7957 | 22395 | 7360.5 | 1727 | 8732.5 | 2439 | 683 | 11978 | 8648 | 25697.5 |
| 3202 | 4481 | 2574 | 809 | 3457 | 1420 | 329 | 10359 | 4213 | 5579 |
| 2905 | 3973 | 2547 | 865 | 3494 | 1488 | 443 | 8796 | 4170 | 5232 |
| 2737 | 3855 | 2767 | 949 | 3740 | 1598 | 528 | 8658 | 4304 | 5529 |
| 2711 | 4047 | 6195 | 1701 | 892730 | 71293 | 617 | 9859 | 4562 | 6139 |
| 2614 | 4163 | 3724 | 1203.5 | 6185 | 2183.5 | 665 | 9833.5 | 4509.5 | 6537.5 |
| 2455 | 3669 | 3077 | 1047 | 4874.5 | 1864 | 623 | 8585 | 4282.5 | 5776 |
| 2336 | 3713 | 3140 | 1058 | 4685 | 1848 | 640 | 8780 | 4234 | 5913 |
| 2573 | 4185 | 3341 | 1131.5 | 4886.5 | 1907 | 668 | 10209 | 4474.5 | 6609.5 |
| 2570 | 4181.5 | 3200.5 | 1090.5 | 4601.5 | 1834 | 636 | 10816 | 4292.5 | 6297.5 |
| 2787.5 | 5281.5 | 3572.5 | 1232 | 4949.5 | 1981.5 | 697 | 13902.5 | 4575 | 8018.5 |
| 2552 | 4365 | 3362.5 | 1202 | 4764 | 2008.5 | 674 | 12773.5 | 4279 | 6716 |
| 2433.5 | 3816 | 3052 | 1121.5 | 4209.5 | 1929.5 | 622 | 10132.5 | 4299.5 | 5960.5 |
| 2399 | 3778 | 3015.5 | 1123.5 | 4263.5 | 1890 | 635 | 9384 | 4305 | 5949.5 |
| 2356 | 3599 | 2935 | 1050 | 4068 | 1729 | 634 | 8694 | 4044 | 5885 |
| 2499.5 | 3980 | 3253 | 1140.5 | 4429.5 | 1873 | 678 | 9090 | 4537.5 | 6352 |
| 2387 | 4041 | 3216 | 1136 | 4438 | 1832 | 658 | 9706 | 4371 | 6462 |
| 52332 | 30324 | 9051 | 1389.5 | 4550 | 1845 | 624 | 13632.5 | 4927 | 7104.5 |
| 2666 | 4465 | 3556 | 1129 | 4442 | 1782 | 614 | 10071 | 4324 | 6541 |
| 2422 | 4388 | 3352 | 1141 | 4612 | 1842 | 649 | 9888 | 4320 | 6891 |
| 2464.5 | 3966.5 | 3270 | 1097 | 4435.5 | 1801.5 | 648 | 9226 | 4457.5 | 6424 |
| 4875 | 12329 | 5069 | 1444 | 6440 | 2158 | 695 | 14752 | 6687 | 15406 |
| 8365 | 19092 | 6641.5 | 1688 | 8013.5 | 2403 | 673 | 15150 | 9975 | 22346 |
| 8751 | 20513 | 6905 | 1659 | 8023 | 2363 | 649 | 13261 | 9478 | 23428 |
| 8585.5 | 17558.5 | 5940 | 1483 | 7187.5 | 2177.5 | 608 | 13728 | 9157.5 | 20107 |
| 816 | 1068 | 845 | 575 | 1420 | 1120 | 227 | 2357 | 1455 | 1911 |
| 1194 | 1922 | 1667 | 745 | 2421 | 1334 | 423 | 4252 | 2298 | 3396 |
| 1455 | 2545 | 2217 | 880 | 3181 | 1523 | 552 | 6502 | 3004 | 4464 |
| 1603 | 2853 | 5842 | 1651 | 899707 | 81700 | 626 | 7657 | 3418 | 5150 |
| 1678 | 3035 | 3109 | 1107 | 4954 | 2137 | 652 | 7856 | 3670 | 5510 |
| 1581.5 | 2799.5 | 2617 | 977.5 | 3900 | 1729 | 612 | 7216 | 3335 | 4971.5 |
| 1578 | 2813.5 | 2602 | 995 | 3788.5 | 1698 | 623 | 7348.5 | 3402 | 5150 |
| 1748.5 | 2969 | 2618 | 982.5 | 3707 | 1664.5 | 622.5 | 8020.5 | 3506 | 5119.5 |
| 1910 | 3071 | 2541 | 946 | 3553 | 1613 | 595 | 8818 | 3488 | 5111 |
| 2361 | 3586 | 2995 | 1087 | 4171.5 | 1796 | 663 | 10602 | 4335 | 5943 |
| 2297 | 3702 | 3146 | 1148 | 4398 | 1859 | 700 | 11070 | 4421 | 6236 |
| 1963 | 3244 | 2797.5 | 1047 | 3885 | 1719 | 655 | 9100.5 | 3888.5 | 5537.5 |
| 1909 | 3302.5 | 2933 | 1092.5 | 4080 | 1780 | 685 | 8655 | 3938.5 | 5820.5 |
| 1903 | 3235.5 | 2936 | 1059 | 4071.5 | 1779 | 677 | 8561.5 | 3877 | 5758 |
| 1955 | 3344 | 2937 | 1100 | 4191 | 1835 | 707 | 9120 | 4203 | 5919 |
| 2099 | 3672 | 3273 | 1166 | 4604 | 1911.5 | 713 | 9278 | 4468 | 6549 |
| 34663 | 25068 | 7180 | 1461 | 4988 | 2024 | 739 | 14761 | 5109 | 9545 |
| 2380 | 4118.5 | 3526.5 | 1199 | 4704 | 1894 | 700.5 | 9679.5 | 4666 | 6852 |
| 2175 | 4245 | 3442 | 1228 | 4872 | 1968 | 730 | 10903 | 4560 | 7344 |
| 2131 | 3668 | 3211 | 1151 | 4516 | 1892 | 714 | 9174 | 4521 | 6467 |
| 5046.5 | 12721.5 | 4997 | 1381 | 6087 | 2063 | 683 | 15131.5 | 6627 | 16094 |
| 7300 | 19108 | 6561 | 1704 | 7963 | 2419 | 687 | 14389 | 8636 | 22593 |
| 8729 | 21376 | 7011 | 1713 | 8211 | 2426 | 675 | 14440 | 9762 | 24400 |

| | | | | | | | | | |
|---|---|---|---|---|---|---|---|---|---|
| 8502.5 | 19240.5 | 6345.5 | 1598 | 7660 | 2298 | 656 | 13134 | 9271.5 | 21953.5 |
| 3129 | 3898 | 2148 | 735 | 2805 | 1288 | 308 | 8818 | 4174 | 4779 |
| 2390 | 3569 | 2368 | 819 | 3102 | 1407 | 455 | 8231 | 3629 | 4811 |
| 2148 | 3412 | 2606 | 930 | 3537 | 1595 | 557 | 8439 | 3749 | 5235 |
| 1915 | 3148 | 5636 | 1598 | 863874 | 78759 | 603 | 8185 | 3625 | 5243 |
| 2103 | 3450.5 | 3383 | 1168 | 5453.5 | 2103 | 671 | 8736.5 | 4203 | 5908.5 |
| 1884 | 3226.5 | 2952 | 1049 | 4480 | 1824 | 656 | 8191 | 3814 | 5589.5 |
| 1811 | 3140 | 2786 | 1010.5 | 4080 | 1756 | 645 | 7957 | 3676 | 5421.5 |
| 1881.5 | 3151 | 2707 | 980 | 3909.5 | 1697 | 630 | 8485.5 | 3636 | 5297 |
| 2106 | 3318 | 2694.5 | 991.5 | 3746 | 1646 | 608 | 9224 | 3768.5 | 5396 |
| 2448 | 3632 | 2918 | 1048 | 4166 | 1786 | 647 | 10966 | 4300 | 5870 |
| 2396 | 3820.5 | 3158 | 1125 | 4412 | 1826 | 692 | 10626.5 | 4503.5 | 6284.5 |
| 2249 | 3664.5 | 3089 | 1114 | 4279 | 1822 | 699 | 10603 | 4332.5 | 6125 |
| 2149 | 3606 | 3195 | 1167 | 4522 | 1904 | 714 | 9725 | 4360 | 6214 |
| 1902 | 3176 | 2761 | 1026 | 3885 | 1694 | 647 | 8316 | 3725 | 5515 |
| 2191 | 3817 | 3388 | 1209 | 4765 | 1967 | 763 | 10323 | 4656 | 6710 |
| 2194 | 3667 | 3218 | 1144 | 4461 | 1878.5 | 699 | 9525.5 | 4458 | 6406.5 |
| 42955.5 | 29881 | 8142 | 1481.5 | 4938.5 | 1966.5 | 713 | 14783.5 | 5193 | 8933 |
| 2353 | 5151.5 | 3811 | 1254.5 | 4993.5 | 1988.5 | 724 | 11227.5 | 4535.5 | 8333 |
| 2332 | 4049 | 3404 | 1212 | 4784 | 1937 | 716 | 9832 | 4713 | 6932 |
| 2182 | 4146.5 | 3337.5 | 1213 | 4757 | 1944 | 726 | 10843 | 4516.5 | 7106 |
| 5163.5 | 11107.5 | 4577.5 | 1321 | 5709 | 2024.5 | 661 | 14230 | 6945 | 14001 |
| 7903 | 20056 | 8057 | 1929 | 9536 | 2627 | 744 | 13531 | 9304 | 23577 |
| 8587 | 22076 | 7225 | 1748 | 8456 | 2476 | 683 | 11296 | 8771 | 24869 |
| 8115 | 20178 | 6612 | 1623 | 7934 | 2341 | 667 | 14851 | 9096 | 23384 |
| 2834 | 3821.5 | 2137.5 | 738.5 | 2780 | 1304 | 296 | 8725 | 3807 | 4663 |
| 2530 | 3568 | 2301 | 821 | 3064 | 1433 | 440 | 7980 | 3684 | 4844 |
| 2276 | 3490.5 | 2585 | 921 | 3508 | 1551 | 543 | 8103.5 | 3729.5 | 5248 |
| 1927 | 3000 | 5675.5 | 1666 | 917864.5 | 80664.5 | 555.5 | 7425.5 | 3373 | 4786.5 |
| 1887 | 3121.5 | 3177 | 1070 | 6146.5 | 2190 | 596 | 7940 | 3589.5 | 5180 |
| 2208.5 | 3420 | 3256 | 1103 | 5464.5 | 2048.5 | 658 | 8875.5 | 4195 | 5655 |
| 2117 | 3353 | 2961 | 1067 | 4557 | 1830 | 653 | 8887 | 4102.5 | 5622.5 |
| 2343 | 3846 | 3284.5 | 1136 | 4873 | 1928 | 717 | 11184 | 4395 | 6447.5 |
| 2483 | 3928 | 3193 | 1134.5 | 4625 | 1875 | 693 | 11831 | 4406.5 | 6341.5 |
| 2442 | 3596 | 2939 | 1040 | 4065.5 | 1730 | 633 | 9687.5 | 4263 | 5769 |
| 2447.5 | 3762.5 | 3045 | 1099.5 | 4365 | 1810 | 674 | 11097 | 4415.5 | 6165 |
| 2283 | 3488 | 2936 | 1040 | 4113 | 1745 | 665 | 10190.5 | 4326.5 | 5882 |
| 2260 | 3580 | 3071 | 1108 | 4364 | 1840 | 692 | 9542 | 4447 | 6157 |
| 2012 | 3263 | 2801 | 1010 | 3916 | 1712 | 642 | 8553 | 3818 | 5486 |
| 1938 | 3257 | 2814 | 1029 | 4002 | 1743 | 655 | 8335 | 3822 | 5514 |
| 2202 | 3671.5 | 3143 | 1117 | 4430 | 1827 | 690 | 9619 | 4375.5 | 6270.5 |
| 51439 | 28950.5 | 8639.5 | 1423.5 | 4657.5 | 1898 | 690 | 14681.5 | 5247 | 7353.5 |
| 2531 | 5464 | 3852.5 | 1246 | 4894 | 1950 | 727 | 11910.5 | 4674 | 8676.5 |
| 2225 | 4451 | 3513 | 1229 | 4878 | 1968 | 711 | 10490 | 4381 | 7350 |
| 2276 | 3912 | 3303 | 1164 | 4602 | 1909 | 713 | 10714 | 4506 | 6644 |
| 5760 | 13009 | 5014 | 1436 | 6411 | 2151 | 716 | 15369 | 7548 | 16384 |
| 7835 | 19720 | 6826 | 1722 | 8228 | 2473 | 719 | 15683 | 9125 | 22989 |
| 9280.5 | 20527.5 | 6826 | 1687 | 8027.5 | 2395 | 680 | 15686.5 | 10031.5 | 23393.5 |

| | | | | | | | | | |
|---|---|---|---|---|---|---|---|---|---|
| 8868 | 19611.5 | 6576 | 1615 | 7814 | 2333 | 682 | 14345.5 | 9575 | 21974 |
| 3177 | 4016 | 2275 | 739 | 2947 | 1346 | 305 | 9250 | 4250 | 4895 |
| 2545 | 3560 | 2435 | 865 | 3193 | 1453 | 467 | 7927 | 3681 | 4927 |
| 2354 | 3492.5 | 2650 | 942 | 3503 | 1602 | 582.5 | 8579.5 | 3858 | 5372.5 |
| 2516 | 3714 | 6554.5 | 1760 | 1076397 | 93628.5 | 690 | 9985.5 | 4561 | 6040 |
| 2426 | 3653 | 3525 | 1180 | 5760 | 2111 | 698 | 9476 | 4519 | 6012 |
| 2519 | 4216.5 | 3705 | 1245.5 | 5676.5 | 2099.5 | 792 | 11148.5 | 4821 | 7147 |
| 2402 | 3860 | 3395 | 1194 | 5018 | 1985 | 761 | 10143 | 4717 | 6560 |
| 2429 | 3688.5 | 3112 | 1096.5 | 4492 | 1802.5 | 696 | 10096 | 4412 | 6023 |
| 2695 | 3826 | 3103 | 1076 | 4290 | 1770 | 693 | 10550 | 4598 | 6088 |
| 2728 | 3765 | 3032 | 1055 | 4138 | 1744 | 676 | 10940 | 4644 | 5952 |
| 2648 | 4035 | 3231 | 1154 | 4518 | 1845 | 736 | 12878 | 4651 | 6511 |
| 2526 | 3775 | 3151.5 | 1098 | 4237.5 | 1799.5 | 717 | 10785.5 | 4586.5 | 6159.5 |
| 2479.5 | 3956.5 | 3316 | 1173 | 4490.5 | 1886 | 761 | 10976 | 4666.5 | 6496 |
| 2418 | 3562 | 2946 | 1075 | 4045 | 1773 | 698 | 9429 | 4390 | 5874 |
| 2441 | 3649 | 3208 | 1127 | 4360 | 1838 | 740 | 9760 | 4659 | 6253 |
| 2344 | 3974 | 3341 | 1178 | 4543 | 1912 | 733 | 11093 | 4526 | 6669 |
| 57972 | 35390 | 10055 | 1514 | 4666 | 1888 | 717 | 16098 | 5482 | 8145 |
| 2632.5 | 6631 | 3719 | 1267 | 4764 | 1936 | 734 | 11925.5 | 4694.5 | 10567.5 |
| 2472.5 | 4151 | 3437.5 | 1217 | 4726 | 1926 | 736 | 11218 | 4698 | 6930 |
| 2450 | 3820 | 3164 | 1129.5 | 4250 | 1800 | 714 | 9969.5 | 4571 | 6277 |
| 5726 | 13512 | 5259 | 1424 | 6117 | 2107 | 721 | 14561 | 6791 | 16652 |
| 9797 | 20731 | 6946 | 1751 | 7835 | 2401 | 736 | 14481 | 9908 | 23462 |
| 11193.5 | 22275 | 7149 | 1709.5 | 7869.5 | 2366 | 702.5 | 15181 | 10269 | 23848 |
| 10948.5 | 20921 | 6703 | 1621.5 | 7520 | 2271 | 705 | 12619.5 | 9515.5 | 22561 |
| 3120 | 4129 | 2311 | 759 | 3056 | 1355 | 313 | 9276.5 | 4119.5 | 5038.5 |
| 2747 | 3705 | 2435.5 | 843 | 3255.5 | 1440 | 445 | 8143 | 3871 | 5068.5 |
| 2530 | 3711 | 2752 | 968 | 3630 | 1614 | 568 | 8663 | 4046 | 5530 |
| 2105 | 3250 | 5839 | 1599 | 932296 | 87275 | 578 | 8141 | 3650 | 5148 |
| 2438 | 3535 | 3366 | 1121 | 5586.5 | 2064 | 650 | 8820 | 4302.5 | 5641.5 |
| 2383 | 3597 | 3156 | 1098 | 4851 | 1869 | 670 | 8944 | 4335 | 5892 |
| 2433 | 3749 | 3148 | 1124 | 4656 | 1847 | 700 | 9887.5 | 4478.5 | 6117 |
| 2517 | 3842 | 3246 | 1102.5 | 4564 | 1881.5 | 691 | 10253 | 4454 | 6226.5 |
| 2664 | 5861 | 3737 | 1252 | 5184 | 2045 | 768 | 14886 | 4617 | 9045 |
| 2696 | 3996 | 3207 | 1122 | 4326 | 1845 | 675 | 11200 | 4511 | 6227 |
| 2638 | 4250 | 3439 | 1173 | 4685 | 1893 | 727 | 11333 | 4578 | 6741 |
| 2562.5 | 4134.5 | 3392.5 | 1195 | 4614 | 1917 | 732 | 10969.5 | 4633.5 | 6688.5 |
| 2301 | 3561 | 2909 | 1049 | 14705 | 2593 | 671 | 9500 | 4129 | 5745 |
| 2734.5 | 4230 | 3508 | 1219.5 | 5104 | 2014 | 763 | 10681.5 | 4897 | 7013 |
| 2040 | 3293 | 2744 | 997 | 3742 | 1676 | 642 | 8463 | 3730 | 5379 |
| 2403 | 3613 | 3082 | 1094 | 4180 | 1762 | 673 | 9361 | 4468 | 6047 |
| 41144 | 22501 | 7357 | 1352.5 | 4518 | 1845 | 678 | 12964 | 5206.5 | 6966 |
| 2715 | 4094.5 | 3365 | 1099 | 4228.5 | 1773 | 650 | 9879 | 4712.5 | 6381 |
| 2511 | 3830 | 3168 | 1099.5 | 4347 | 1754 | 655 | 9670.5 | 4564.5 | 6274.5 |
| 2417.5 | 4291 | 3305.5 | 1154 | 4515 | 1883 | 708 | 11135.5 | 4414 | 7126.5 |
| 4066 | 8134 | 4136 | 1216 | 5018 | 1876 | 654 | 14865 | 5234 | 10259 |
| 9117 | 19237 | 8577 | 1941 | 10049 | 2623 | 749 | 15468 | 9912 | 21902 |
| 10032 | 20337 | 7134 | 1730 | 8104 | 2407 | 689 | 14752 | 9882 | 22370 |

| | | | | | | | | | |
|---|---|---|---|---|---|---|---|---|---|
| 9510 | 20249 | 7023 | 1674 | 8027 | 2370 | 704 | 15791 | 9570 | 22764 |
| 3117 | 4007 | 2264 | 757 | 2968 | 1324 | 293 | 9250 | 4058 | 4974 |
| 2941 | 3950 | 2479 | 843 | 3295 | 1444 | 431 | 8919 | 4191 | 5158 |
| 2606 | 3880 | 2773 | 943 | 3821 | 1602 | 548 | 8855 | 4281 | 5713 |
| 2233 | 3444.5 | 5646.5 | 1546 | 883961 | 73092 | 582 | 8547.5 | 3792 | 5421 |
| 2053.5 | 3333 | 3066.5 | 1017 | 5372 | 1969 | 583 | 7982 | 3636 | 5238 |
| 2391 | 3561 | 3087 | 1057 | 4823 | 1854 | 635 | 8644 | 4290 | 5776 |
| 2344 | 3672 | 3108 | 1096 | 4662.5 | 1843.5 | 656 | 9241.5 | 4350 | 5922 |
| 2375 | 3643 | 2970 | 1035 | 4329 | 1755 | 635 | 9837 | 4250 | 5773 |
| 2707 | 6966 | 3791 | 1293 | 5330 | 2082 | 753 | 14534 | 4782 | 10817 |
| 2577 | 4147 | 3267 | 1120 | 4563 | 1837 | 668 | 11383 | 4359 | 6406 |
| 2923 | 6652.5 | 3784.5 | 1282.5 | 5384.5 | 2077.5 | 777 | 15333.5 | 5181 | 10451 |
| 2424 | 3787 | 2977 | 1048 | 4210 | 1770 | 645 | 9966 | 4346 | 5988 |
| 2365 | 4049 | 3348 | 1172 | 4752 | 1915 | 711 | 11724 | 4391 | 6688 |
| 2213 | 3516 | 2879 | 1013 | 4038 | 1728 | 630 | 8627 | 3984 | 5690 |
| 2343.5 | 3745 | 3097.5 | 1097 | 4366.5 | 1802 | 674 | 9820.5 | 4409.5 | 6221 |
| 2241.5 | 3630.5 | 2999 | 1051 | 4168.5 | 1747.5 | 633.5 | 8720.5 | 4114.5 | 5883.5 |
| 46722 | 26507 | 8264 | 1399 | 4649 | 1850 | 657 | 13689 | 5065 | 7071 |
| 2776 | 4272 | 3540 | 1109 | 4340 | 1762 | 620 | 9685 | 4647 | 6321 |
| 2364 | 4325 | 3405 | 1136 | 4693 | 1896 | 666 | 10581 | 4309 | 6940 |
| 2329 | 3510 | 2843 | 1017 | 3944 | 1694 | 612 | 8470 | 4294 | 5713 |
| 4540 | 11535 | 4523 | 1271 | 5590 | 1944 | 633 | 13943 | 5888 | 14268 |
| 8054 | 18111 | 6367 | 1624 | 7569.5 | 2334 | 667 | 13626.5 | 9241.5 | 20931.5 |
| 8708 | 20137 | 6579 | 1621 | 7815 | 2316 | 633 | 14771 | 9576 | 22896 |
| 8834 | 18885 | 6410 | 1583 | 7512 | 2279 | 644 | 15254 | 9661 | 21595 |
| 3221 | 4126.5 | 2352 | 761 | 3088.5 | 1353 | 305 | 9394.5 | 4246 | 5120 |
| 2927 | 4106.5 | 2671 | 872.5 | 3543 | 1513 | 462 | 9284.5 | 4284.5 | 5502 |
| 2690 | 3896 | 2801 | 948 | 3727 | 1591 | 550 | 9023 | 4308 | 5620 |
| 2178 | 3387 | 5186 | 1490 | 719347 | 72035 | 552 | 8168 | 3662 | 5135 |
| 2428 | 3629 | 3408 | 1103 | 5662 | 2048 | 616 | 9129 | 4281 | 5818 |
| 2361 | 3716 | 3217 | 1087 | 5017 | 1890 | 649 | 9193 | 4352 | 5915 |
| 2356 | 3663 | 3079 | 1070 | 4628 | 1833 | 654 | 9344 | 4265 | 5927 |
| 2485 | 4231.5 | 3405 | 1158 | 4928 | 1936 | 700 | 10780.5 | 4421.5 | 6647.5 |
| 2543 | 4716 | 3477 | 1194 | 5017 | 1930 | 710 | 14000.5 | 4378 | 7367 |
| 2567 | 4124 | 3245 | 1139 | 4511 | 1842 | 663 | 11807 | 4390 | 6426 |
| 2626 | 4128 | 3242 | 1126 | 4494 | 1843 | 671 | 11036 | 4545 | 6316 |
| 2482 | 4116 | 3318 | 1142 | 4576 | 1863 | 696 | 11642 | 4389 | 6545 |
| 2318 | 3692 | 2937.5 | 1047 | 4138 | 1736 | 651 | 9248 | 4197.5 | 5823 |
| 2425 | 3720.5 | 3053.5 | 1067.5 | 4233.5 | 1784 | 665 | 9498.5 | 4366 | 5993 |
| 1957 | 3215 | 2637.5 | 977 | 3674.5 | 1615 | 603 | 7811 | 3538 | 5176.5 |
| 2352 | 3675 | 3037 | 1065 | 4211 | 1772 | 642 | 9341 | 4409 | 6020 |
| 41907 | 22600 | 7302.5 | 1318 | 4495 | 1819 | 635 | 12368.5 | 4924 | 6633 |
| 2654 | 4083.5 | 3365 | 1082.5 | 4231.5 | 1759 | 615 | 9577.5 | 4523.5 | 6198.5 |
| 2436 | 4230 | 3489 | 1156 | 4748 | 1885 | 681 | 10811 | 4392 | 6857 |
| 2399 | 3602.5 | 2974.5 | 1062 | 4050.5 | 1745.5 | 624 | 8908.5 | 4330.5 | 5825.5 |
| 4497 | 11527 | 4573 | 1271 | 5617 | 1956 | 640 | 13969 | 5821 | 14430 |
| 8772.5 | 18212.5 | 6443.5 | 1657 | 7542 | 2350 | 683 | 13796.5 | 9231 | 20860 |
| 9289 | 19501 | 6588 | 1627 | 7586 | 2301 | 640 | 15706 | 9738 | 22199 |

| | | | | | | | | | |
|---|---|---|---|---|---|---|---|---|---|
| 9134.5 | 19411 | 6487 | 1598 | 7590 | 2290 | 658 | 15652 | 9592 | 21579.5 |
| 2965 | 3925 | 2205 | 736 | 2944 | 1318 | 277 | 8636.5 | 3945.5 | 4771 |
| 3030 | 4030.5 | 2628 | 858 | 3415 | 1502.5 | 453 | 8962 | 4210 | 5365.5 |
| 3005 | 4332 | 3103 | 1033 | 4179 | 1695 | 591 | 9726 | 4634 | 6164 |
| 2353 | 3528 | 6097 | 1634 | 950590 | 83688 | 571 | 8552 | 3815 | 5455 |
| 2596 | 3664 | 3385 | 1109 | 5722 | 2039 | 628 | 9132 | 4392 | 5755 |
| 2435 | 3610 | 3188 | 1070 | 5022 | 1864 | 642 | 8824 | 4281 | 5735 |
| 2493 | 3915 | 3365 | 1117 | 4899 | 1894 | 693 | 10194 | 4403 | 6276 |
| 2552 | 3855 | 3141 | 1081 | 4561 | 1806 | 656 | 10323 | 4431 | 6015 |
| 2752 | 6653 | 3798 | 1256 | 5306 | 2045 | 749 | 14659 | 4616 | 10361 |
| 2711 | 5054 | 3750.5 | 1279.5 | 5112.5 | 2033 | 735 | 14762 | 4505.5 | 7706.5 |
| 2710 | 5792 | 3716 | 1251 | 5172 | 1985 | 748 | 14940 | 4701 | 9037 |
| 2605 | 4234 | 3399 | 1205 | 4771 | 1882 | 719 | 11186.5 | 4541.5 | 6630.5 |
| 2540 | 4276.5 | 3541 | 1201.5 | 4820.5 | 1961 | 754 | 12153 | 4626.5 | 6967.5 |
| 2604 | 4180 | 3449 | 1194 | 4767 | 1942 | 741 | 11090 | 4671 | 6865 |
| 2276.5 | 3553.5 | 2963 | 1060.5 | 4037 | 1736 | 657 | 9129 | 4185 | 5725 |
| 2491 | 3930.5 | 3261 | 1139 | 4481 | 1831 | 691.5 | 10024.5 | 4661 | 6460 |
| 48257 | 27315 | 8573 | 1409 | 4681.5 | 1905 | 685 | 14130 | 5168.5 | 7204 |
| 2721 | 4125 | 3363 | 1135 | 4346 | 1787 | 642 | 9786 | 4704 | 6396 |
| 2511 | 4101 | 3387 | 1127 | 4544 | 1850 | 681 | 10402.5 | 4491 | 6642.5 |
| 2409 | 3882 | 3197 | 1109 | 4341 | 1822 | 669 | 9675 | 4337 | 6366 |
| 4567 | 11024 | 4383 | 1245 | 5342 | 1896 | 646 | 13764 | 5540 | 13750 |
| 9094 | 19768 | 6975.5 | 1736.5 | 8046.5 | 2470 | 726 | 15414.5 | 9662.5 | 22414.5 |
| 10371 | 20552.5 | 7050 | 1688 | 8052.5 | 2352.5 | 678 | 15056 | 9658.5 | 22410 |
| 10409.5 | 20444 | 7071 | 1694 | 7907.5 | 2350 | 703 | 15729 | 10000 | 22522 |
| 3360 | 4202 | 2407 | 758 | 3211.5 | 1368 | 302 | 9905 | 4338 | 5140 |
| 2926 | 3998 | 2537 | 841 | 3426 | 1468 | 437 | 8729 | 4032 | 5216 |
| 2704.5 | 3853 | 2798.5 | 934 | 3714.5 | 1593 | 532.5 | 8680.5 | 4137 | 5471.5 |
| 2432.5 | 3594.5 | 6720.5 | 1629.5 | 1253039 | 97070.5 | 565 | 8702 | 3984 | 5408 |
| 2670 | 3766 | 3386 | 1090 | 7135 | 2137 | 637 | 9062 | 4464 | 5865 |
| 2662 | 3900 | 3239 | 1113 | 6292 | 2037 | 668 | 9555 | 4485 | 6058 |
| 2569 | 4043 | 3387 | 1130 | 5858 | 1999 | 706 | 10395.5 | 4523 | 6420 |
| 2523.5 | 4205 | 3413 | 1137 | 5659.5 | 1997 | 705 | 11171.5 | 4341.5 | 6605 |
| 2770.5 | 5857 | 3614 | 1216 | 5797 | 2014 | 729 | 14504.5 | 4540 | 8891 |
| 2786 | 4455 | 3427 | 1190 | 5068 | 1951.5 | 696 | 11619.5 | 4521 | 6685 |
| 2824 | 5341 | 3688 | 1245 | 5461 | 2035 | 753 | 14617 | 4627 | 8237 |
| 2717 | 4175 | 3325 | 1137 | 4839 | 1876 | 696 | 10849 | 4604 | 6440 |
| 2639 | 4255.5 | 3416 | 1167.5 | 5003.5 | 1924 | 733 | 10676.5 | 4654 | 6791.5 |
| 2660.5 | 4142.5 | 3320 | 1151 | 4827 | 1884 | 706.5 | 10357.5 | 4560.5 | 6587 |
| 2377 | 3690 | 2969 | 1047 | 4257 | 1767 | 656 | 9097 | 4201 | 5828 |
| 2655.5 | 3860 | 3144 | 1083 | 4548 | 1815 | 666 | 9575.5 | 4564 | 6238.5 |
| 62531.5 | 34679 | 10391.5 | 1462 | 4773.5 | 1838.5 | 649 | 15116.5 | 5514.5 | 7181.5 |
| 2802.5 | 4614 | 3739 | 1176.5 | 4912.5 | 1880 | 675 | 11107.5 | 4578 | 6975 |
| 2434 | 4178 | 3249 | 1098 | 4633 | 1823 | 647 | 9748 | 4206 | 6516 |
| 2597 | 4104 | 3327 | 1129 | 4663 | 1848 | 684 | 10619 | 4524 | 6523 |
| 5415 | 12879 | 5058.5 | 1376 | 6228.5 | 2084 | 690 | 14828.5 | 6474.5 | 15943.5 |
| 9142 | 19702 | 6778 | 1677 | 8008 | 2399 | 703 | 15575 | 9655 | 22542 |
| 9919 | 21272 | 7044 | 1710 | 8182 | 2418 | 681 | 13778 | 9892 | 23501 |

| | | | | | | | | | |
|---|---|---|---|---|---|---|---|---|---|
| 9520.5 | 20611.5 | 6822 | 1635.5 | 8016.5 | 2305 | 676 | 12972 | 9516 | 22942.5 |
| 3209 | 4120 | 2333 | 770 | 3146 | 1344 | 288 | 9490 | 4195 | 4942 |
| 2771 | 3795 | 2439 | 816 | 3283 | 1421 | 404 | 8271 | 3911 | 5045 |
| 2775.5 | 3807 | 2731 | 921 | 3703 | 1585 | 531 | 8811.5 | 4279 | 5443.5 |
| 2319 | 3504 | 5713.5 | 1583 | 873842.5 | 75644 | 559 | 8447 | 3782.5 | 5351 |
| 2431 | 3964 | 3620 | 1184 | 6046 | 2203 | 635 | 9573 | 4179 | 6194 |
| 2484.5 | 4166 | 3537 | 1167 | 5497 | 2023.5 | 686 | 10527 | 4326 | 6548 |
| 2403 | 3929 | 3229 | 1106 | 4910 | 1874 | 658 | 10075 | 4264 | 6289 |
| 2453 | 5233 | 3594 | 1198 | 5271 | 2004 | 712 | 11698 | 4354 | 8206 |
| 2687 | 4902 | 3455 | 1151 | 4952 | 1915 | 683 | 12150 | 4521 | 7487 |
| 2727 | 4176 | 3267 | 1098 | 4508 | 1815 | 638 | 10986 | 4442 | 6357 |
| 2553 | 4362 | 3354 | 1136 | 4635 | 1862 | 672 | 12212 | 4346 | 6698 |
| 2587 | 4173.5 | 3353 | 1128 | 4551.5 | 1851 | 665 | 11148.5 | 4470.5 | 6538.5 |
| 2475.5 | 3762.5 | 2987.5 | 1044 | 4208 | 1754 | 631 | 9206 | 4399 | 5936.5 |
| 2546 | 3818 | 3063 | 1058 | 4170 | 1750 | 634 | 9298 | 4446 | 6014 |
| 2414.5 | 3560 | 2923 | 1024.5 | 4035.5 | 1712 | 620 | 8811.5 | 4305.5 | 5651.5 |
| 2379 | 4151.5 | 3297.5 | 1127 | 4619 | 1832 | 663 | 10300.5 | 4232.5 | 6560 |
| 46960.5 | 25757.5 | 7823.5 | 1309 | 4381.5 | 1777 | 615.5 | 12319 | 5047.5 | 6759.5 |
| 2828 | 7515 | 4002 | 1250 | 5046 | 1971 | 690 | 12458 | 4596 | 11256 |
| 2341.5 | 3965 | 3156 | 1057.5 | 4220 | 1742.5 | 609 | 9466.5 | 4029 | 6293 |
| 2411.5 | 6233 | 3527 | 1192 | 4809 | 1920 | 693.5 | 11474.5 | 4356 | 9846 |
| 4588 | 12049.5 | 4740 | 1307 | 5800 | 2039.5 | 649 | 13155.5 | 5981 | 15245.5 |
| 8235.5 | 18763.5 | 6560.5 | 1660 | 7790 | 2362 | 656.5 | 15510.5 | 9559 | 21909 |
| 8555 | 19671 | 7019 | 1614 | 8205 | 2318 | 642 | 14559 | 9339 | 22377 |
| 7748 | 19868 | 8080 | 1750 | 9669 | 2457 | 685 | 10983 | 8400 | 22899 |
| 3246 | 4168 | 2410 | 770 | 3180 | 1376 | 294 | 9623 | 4270 | 5112 |
| 2657 | 3704.5 | 2322 | 781 | 3162 | 1410 | 381 | 7986 | 3676.5 | 4799 |
| 2651 | 3634.5 | 2585.5 | 887 | 3484 | 1534 | 499 | 8334 | 4058 | 5147 |
| 2746.5 | 3870 | 5576 | 1579 | 814009.5 | 71955.5 | 590 | 9263.5 | 4416.5 | 5755 |
| 2616 | 4037 | 3561 | 1147 | 6053 | 2118 | 632 | 9941 | 4402 | 6090 |
| 2469.5 | 3673.5 | 3092 | 1040 | 4857 | 1841.5 | 612 | 8894.5 | 4293.5 | 5787 |
| 2493.5 | 4121.5 | 3325 | 1121 | 4951.5 | 1903 | 671 | 9456 | 4395.5 | 6373 |
| 2615.5 | 5583.5 | 3633 | 1214 | 5323.5 | 2031.5 | 714 | 12195.5 | 4573.5 | 8523 |
| 2478 | 4288.5 | 3246.5 | 1082 | 4639 | 1823 | 637 | 12403.5 | 4109 | 6410.5 |
| 2798 | 4500 | 3398 | 1137 | 4745 | 1845 | 692 | 13686 | 4434 | 6762 |
| 2628 | 4239.5 | 3229 | 1075 | 4544 | 1809.5 | 646 | 11756.5 | 4383 | 6374 |
| 2526 | 4094 | 3188 | 1071 | 4464.5 | 1794 | 638 | 10356 | 4290 | 6230.5 |
| 2591 | 4733.5 | 3488 | 1188 | 4851 | 1938 | 720 | 11019 | 4672 | 7327.5 |
| 2485 | 4036 | 3251 | 1107 | 4537 | 1814 | 657 | 10160 | 4337 | 6326 |
| 2555 | 3966 | 3219 | 1090 | 4392 | 1790 | 687 | 10160 | 4512 | 6273 |
| 2566 | 3832 | 3026 | 1058 | 4228 | 1747 | 628 | 9140 | 4551 | 6174 |
| 51973 | 32143 | 9267 | 1438 | 4773 | 1896 | 661 | 13573 | 5224 | 7850 |
| 3940 | 7831 | 4424 | 1322.5 | 5273.5 | 2015.5 | 909 | 16067 | 5647.5 | 10497 |
| 3647 | 7648.5 | 4107 | 1281 | 5287.5 | 2023 | 1026 | 16907 | 6245.5 | 11619 |
| 3345 | 7391.5 | 3886.5 | 1240 | 5025.5 | 1995 | 984.5 | 17731 | 5886 | 11498 |
| 9168 | 15799 | 5361 | 1428 | 6237 | 2089 | 885 | 17495 | 7861 | 16987 |
| 14981 | 22872 | 6858 | 1646 | 7561 | 2328 | 843 | 16409 | 9663 | 21993 |
| 17798.5 | 25705 | 7140.5 | 1622 | 7718.5 | 2289 | 801 | 17786.5 | 10546.5 | 22991 |

| | | | | | | | | | |
|---|---|---|---|---|---|---|---|---|---|
| 16804 | 27172 | 7691 | 1719 | 8365 | 2404 | 879 | 16565 | 10268 | 24276 |
| 839 | 1120 | 917 | 611 | 1481 | 1142 | 233 | 2450 | 1530 | 1996 |
| 1121 | 1733 | 1480 | 718 | 2196 | 1278 | 381 | 3870 | 2124 | 3090 |
| 1345 | 2239 | 1986 | 823 | 2862 | 1446.5 | 504 | 5864 | 2699 | 3990 |
| 1481.5 | 2505 | 4798 | 1433.5 | 657160 | 74122.5 | 567.5 | 6754.5 | 3066 | 4526.5 |
| 1667 | 3001 | 3073 | 1102 | 5027 | 2153 | 648 | 7992 | 3589 | 5392 |
| 1678 | 2975 | 2850 | 1064 | 4180 | 1860 | 651 | 7902 | 3553 | 5381 |
| 1865 | 3275 | 3037 | 1131 | 4430 | 1911 | 715 | 8716 | 4077 | 5984 |
| 2006 | 3334 | 2871 | 1070.5 | 4052 | 1789 | 673 | 8792.5 | 3956.5 | 5668 |
| 2261 | 3345.5 | 2694 | 970 | 3698.5 | 1643 | 606 | 9970 | 3962 | 5408.5 |
| 2530 | 3540 | 2804 | 1027 | 3840 | 1681 | 620 | 10373 | 4383 | 5614 |
| 2446 | 3796 | 3006 | 1098 | 4222 | 1802 | 673 | 12197 | 4419 | 6127 |
| 2275 | 3487 | 2981 | 1083 | 4102 | 1774 | 674 | 10149 | 4387.5 | 5873 |
| 1891 | 3126 | 2749 | 1031 | 3852 | 1714 | 658 | 8858 | 3745 | 5422 |
| 2050 | 3220 | 2871 | 1047 | 4006 | 1743 | 670 | 8786 | 3989 | 5670 |
| 1632 | 2781.5 | 2469 | 947 | 3446 | 1626.5 | 605 | 7336.5 | 3325 | 4910.5 |
| 2125 | 3755 | 3307 | 1179 | 4558 | 1925 | 733 | 10167 | 4382 | 6641 |
| 36840 | 19108.5 | 6481.5 | 1264 | 4335 | 1826 | 647 | 11183 | 4994 | 6456.5 |
| 2387 | 5792 | 3753.5 | 1248 | 4874.5 | 1963.5 | 707 | 11288.5 | 4560 | 9452.5 |
| 2219 | 4032.5 | 3408 | 1190.5 | 4755 | 1933 | 717 | 10094 | 4562.5 | 6902 |
| 2122 | 3669 | 3221 | 1157 | 4475 | 1884 | 710 | 9837 | 4442 | 6446 |
| 5509 | 12996 | 5007.5 | 1398.5 | 6176 | 2109 | 692 | 15202.5 | 7274 | 16311.5 |
| 8200 | 19199 | 6687 | 1717 | 7897 | 2392 | 704 | 15606 | 9631 | 22166 |
| 8528 | 21136 | 7573 | 1812 | 9099 | 2547 | 706 | 13804 | 9334 | 24246 |
| 8245 | 21424 | 6931 | 1713 | 8205 | 2415 | 696 | 12126 | 9259 | 24498 |
| 2682 | 3596 | 2039 | 723 | 2652 | 1262 | 291 | 8027 | 3548 | 4442 |
| 2441 | 3399 | 2281 | 823 | 3034 | 1423 | 456 | 7923 | 3536 | 4691 |
| 2117 | 3264 | 2499 | 929 | 3379 | 1564 | 563 | 8192 | 3583 | 5088 |
| 2063 | 3280 | 5447 | 1585 | 738051 | 80378 | 633 | 8388 | 3745 | 5415 |
| 2283 | 3493.5 | 3474 | 1167 | 5599 | 2167 | 708 | 9424 | 4439.5 | 6061 |
| 2016 | 3334 | 3045 | 1108 | 4476 | 1860 | 685 | 9032 | 4071 | 5799 |
| 2285 | 3715 | 3332 | 1179 | 4869.5 | 1959 | 752 | 9865.5 | 4574.5 | 6436 |
| 2252.5 | 3463 | 2872 | 1058 | 4069.5 | 1758.5 | 663 | 9494 | 4138 | 5741.5 |
| 2601 | 3727 | 2994 | 1064 | 4070 | 1731 | 659 | 10935 | 4493 | 6006 |
| 2627.5 | 3898 | 3074.5 | 1119 | 4091.5 | 1758 | 671.5 | 11724.5 | 4436.5 | 6085.5 |
| 2556 | 3535 | 2874 | 1009 | 3863 | 1694 | 654 | 10945 | 4380 | 5721 |
| 2439 | 3641 | 3100 | 1095 | 4154 | 1793 | 689 | 10843 | 4515 | 6078 |
| 2005 | 3299 | 2792 | 1047 | 3848 | 1729 | 659 | 9192 | 3891 | 5583 |
| 2016 | 3208 | 2809 | 1035 | 3826 | 1720 | 672 | 8668 | 3840 | 5556 |
| 1902 | 3096 | 2711 | 1028 | 3736 | 1674 | 654 | 8523 | 3816 | 5380 |
| 2107 | 3361 | 2941 | 1061 | 3985 | 1740 | 660 | 8913 | 4195 | 5787 |
| 37793 | 25381 | 7220 | 1401.5 | 4647 | 1890 | 699 | 13785 | 5036 | 8591 |
| 2366 | 3531.5 | 3016.5 | 1043 | 3868 | 1691 | 624 | 8993 | 4404.5 | 5805.5 |
| 2337.5 | 5243 | 3535 | 1212 | 4894 | 1980.5 | 754 | 11478.5 | 4634.5 | 9025 |
| 2165 | 3284 | 2827 | 1040 | 3818 | 1691 | 650 | 8605 | 4202.5 | 5548 |
| 6224 | 13840 | 5322 | 1435 | 6377 | 2166 | 713 | 15338 | 7717 | 17310 |
| 8159.5 | 17178 | 6173 | 1590.5 | 7134 | 2254 | 680 | 14567 | 8993 | 19676 |
| 9350 | 20247 | 7243 | 1687 | 8271 | 2377 | 661 | 15309 | 9878 | 22752 |

| | | | | | | | | | |
|---|---|---|---|---|---|---|---|---|---|
| 9406 | 18798.5 | 6184 | 1527 | 7232.5 | 2199 | 630.5 | 13532.5 | 9431 | 21647 |
| 2991 | 3943.5 | 2228 | 752 | 2889.5 | 1309.5 | 316 | 8633.5 | 3905 | 4828 |
| 2420 | 3477 | 2312 | 811 | 3098 | 1410 | 436 | 7739 | 3611 | 4829 |
| 2395 | 3537 | 2672 | 944 | 3638 | 1616 | 573 | 8532 | 4042 | 5411 |
| 1916.5 | 3025.5 | 5588.5 | 1664.5 | 853941 | 78426 | 573 | 7503 | 3415 | 4847 |
| 1917.5 | 3127.5 | 3135 | 1041 | 5644.5 | 2092.5 | 602.5 | 7736 | 3518 | 5143.5 |
| 2259 | 3462 | 3166.5 | 1105 | 4953.5 | 1935 | 671 | 9052 | 4291 | 5757.5 |
| 2164 | 3438 | 3025 | 1069 | 4559 | 1854 | 664 | 8794 | 4102 | 5692 |
| 2431 | 3782 | 3188.5 | 1119.5 | 4632.5 | 1841 | 687 | 10631 | 4553.5 | 6254 |
| 2550 | 4122.5 | 3314.5 | 1162.5 | 4630.5 | 1861 | 706 | 12987 | 4445 | 6546.5 |
| 2667 | 4154 | 3248 | 1134 | 4414 | 1840 | 683 | 13333 | 4510 | 6343 |
| 2588 | 3953 | 3177 | 1081 | 4299 | 1799 | 677 | 12272 | 4479 | 6242 |
| 2511 | 3736.5 | 3051.5 | 1108 | 4223.5 | 1823.5 | 673 | 10118 | 4558.5 | 6050 |
| 2166.5 | 3357 | 2804 | 1015 | 3905.5 | 1701.5 | 651 | 9253.5 | 4087 | 5539 |
| 2099.5 | 3364.5 | 2800 | 1018 | 3883 | 1691.5 | 645.5 | 8352.5 | 3879.5 | 5555.5 |
| 1982.5 | 3271 | 2781.5 | 1025 | 3821 | 1706 | 651 | 8507 | 3797 | 5508.5 |
| 2023 | 3323 | 2845 | 1037 | 3907 | 1707 | 633 | 8327 | 3851 | 5682 |
| 38934.5 | 21617 | 7054.5 | 1349 | 4592 | 1851.5 | 680 | 13372 | 5006.5 | 7055 |
| 2413 | 3976 | 3360 | 1110 | 4334 | 1801 | 655 | 9612 | 4482 | 6376 |
| 2411 | 4022 | 3371 | 1160 | 4726 | 1915 | 700 | 10374 | 4547 | 6696 |
| 2356 | 4164 | 3429.5 | 1190 | 4677.5 | 1964 | 727 | 10811 | 4582.5 | 6880.5 |
| 4077 | 9079 | 4431 | 1282 | 5580 | 2014 | 666 | 14581 | 5648 | 11963 |
| 8649.5 | 19050 | 7324 | 1744 | 8835 | 2481 | 708 | 15566.5 | 10009 | 21997.5 |
| 9009 | 21021 | 7208 | 1769 | 8357 | 2473 | 701 | 13991 | 9772 | 23267 |
| 8857 | 22550 | 7126 | 1743 | 8467 | 2480 | 722 | 14432 | 9546 | 25492 |
| 2967.5 | 3624.5 | 2094.5 | 734 | 2759.5 | 1315 | 304 | 8217 | 3949 | 4627 |
| 2383 | 3317 | 2269 | 817 | 2978.5 | 1425 | 446 | 7550.5 | 3444.5 | 4622 |
| 2285.5 | 3500 | 2690 | 941.5 | 3591.5 | 1585 | 579 | 8258.5 | 3789 | 5250 |
| 2214 | 3405 | 5720.5 | 1651 | 830425 | 87234.5 | 636 | 8871 | 3830.5 | 5519 |
| 2489 | 3862 | 3751 | 1245 | 6231 | 2295 | 719 | 10038 | 4510 | 6356 |
| 2411 | 3928 | 3466 | 1192 | 5216 | 2039 | 754 | 10198 | 4616 | 6487 |
| 2291 | 3541 | 3103 | 1117 | 4524 | 1821 | 703 | 9736 | 4401 | 6083 |
| 2623 | 4001 | 3368 | 1150 | 4712 | 1885 | 730 | 11352 | 4738.5 | 6538 |
| 2814.5 | 4030 | 3122 | 1078.5 | 4292.5 | 1798 | 667 | 11742.5 | 4619 | 6188.5 |
| 2795.5 | 4060.5 | 3194 | 1085 | 4190 | 1766.5 | 669 | 11118 | 4578.5 | 6126.5 |
| 2766.5 | 4191 | 3397.5 | 1153 | 4415 | 1851 | 720 | 13378 | 4615.5 | 6502 |
| 2659 | 4022.5 | 3226.5 | 1144 | 4471 | 1857.5 | 724 | 11736 | 4656 | 6469 |
| 2565 | 3739 | 3075 | 1101 | 4230 | 1787 | 699 | 10433 | 4646 | 6266 |
| 2256 | 3371 | 2849 | 1031 | 3776 | 1670 | 655 | 9078 | 4039 | 5538 |
| 2394 | 3767.5 | 3205.5 | 1123.5 | 4317.5 | 1831 | 739.5 | 10322.5 | 4416 | 6343.5 |
| 2450 | 4133.5 | 3424 | 1196.5 | 4634.5 | 1923.5 | 736 | 10771.5 | 4587 | 6789 |
| 45094 | 34394 | 8421 | 1493 | 4982 | 1972 | 735 | 15215 | 5346 | 10175 |
| 2431.5 | 7461.5 | 3915.5 | 1271 | 5050.5 | 2004 | 747.5 | 12869 | 4330 | 11758.5 |
| 2598 | 5567 | 3723 | 1274 | 5002 | 2008 | 759 | 11421 | 4831 | 9219 |
| 2530 | 5382 | 3713.5 | 1277.5 | 5021.5 | 2030.5 | 776 | 12844 | 4707.5 | 8870 |
| 5035.5 | 10375.5 | 4848.5 | 1397 | 5933.5 | 2122.5 | 721 | 14314 | 6722 | 13004.5 |
| 9410.5 | 20579.5 | 8354 | 1927.5 | 9605 | 2587.5 | 740 | 15512 | 10051 | 23162 |
| 8596 | 22611 | 7607.5 | 1821.5 | 9088.5 | 2537 | 687 | 12763 | 9359.5 | 26400.5 |

| | | | | | | | | | |
|---|---|---|---|---|---|---|---|---|---|
| 8812 | 21392.5 | 6627 | 1648 | 7898.5 | 2331.5 | 670 | 12996 | 9596.5 | 24066.5 |
| 3240 | 3971.5 | 2297.5 | 746 | 2989 | 1351.5 | 306 | 9513 | 4238.5 | 4901 |
| 2478 | 3555 | 2376 | 835 | 3089 | 1432 | 435 | 8218 | 3642 | 4791 |
| 2416 | 3521 | 2600 | 911 | 3508 | 1552 | 526 | 8403 | 3912 | 5223 |
| 2100 | 3215 | 5161 | 1498 | 713895 | 76417 | 562 | 7953 | 3608 | 5039 |
| 2374 | 3862 | 3530 | 1178 | 5615 | 2107 | 675 | 10030 | 4250 | 6294 |
| 2433 | 3852 | 3337 | 1126 | 4953 | 1930 | 692 | 9828 | 4499 | 6248 |
| 2396 | 4128 | 3446 | 1191 | 5041 | 1973 | 729 | 11006 | 4541 | 6750 |
| 2667 | 4195 | 3491 | 1178 | 4878 | 1927 | 714 | 10781 | 4779 | 6808 |
| 2672 | 4005 | 3068 | 1036 | 4203 | 1729 | 645 | 11564 | 4358 | 6070 |
| 2683 | 4080 | 3137.5 | 1088 | 4242.5 | 1780 | 652 | 11103 | 4405 | 6181 |
| 2440 | 4156 | 3264 | 1090 | 4345 | 1821 | 670 | 13133 | 4060 | 6379 |
| 2502.5 | 3924.5 | 3158 | 1130 | 4351.5 | 1819.5 | 672 | 10751.5 | 4383.5 | 6318 |
| 2368 | 3745 | 3089 | 1096 | 4221 | 1787 | 673 | 10116 | 4376.5 | 6056 |
| 2463 | 3673 | 3075 | 1061 | 4193 | 1773 | 671 | 9585 | 4438 | 5988 |
| 2437 | 3665 | 3081.5 | 1101 | 4154.5 | 1790.5 | 688.5 | 9824 | 4426.5 | 5981 |
| 2489 | 6486 | 3852 | 1303 | 5140 | 2072 | 777 | 12644 | 4623 | 10710 |
| 33557 | 19434 | 6743 | 1323 | 4512 | 1844 | 659 | 12421 | 4756 | 6877 |
| 3130 | 7994.5 | 4395 | 1432 | 5765 | 2152.5 | 779.5 | 14377 | 5481 | 12137.5 |
| 2487 | 4669 | 3619 | 1220 | 4837 | 1965 | 695 | 11156 | 4536 | 7547 |
| 2575 | 5707.5 | 3724.5 | 1290 | 5097 | 1981.5 | 757 | 11531 | 4719 | 9404.5 |
| 5804 | 13937 | 5354 | 1489 | 6493 | 2178 | 711 | 14592.5 | 7364 | 17186 |
| 8447 | 19947 | 7892 | 1873 | 9414 | 2592 | 747 | 15365 | 9904 | 23331 |
| 9290 | 22033.5 | 7775.5 | 1851 | 8940.5 | 2539 | 717 | 13615 | 9570.5 | 24895 |
| 9169 | 23039.5 | 7500 | 1765.5 | 8718 | 2463 | 718 | 14500.5 | 9872 | 25750 |
| 3130 | 4138 | 2348 | 753.5 | 3064 | 1324 | 298 | 9661.5 | 4171.5 | 5052 |
| 2712 | 3623.5 | 2385 | 830 | 3184.5 | 1429 | 433 | 8332.5 | 3880.5 | 4851 |
| 2539 | 3623 | 2660 | 907 | 3539.5 | 1565 | 535 | 8318 | 4077 | 5288.5 |
| 2596 | 3658 | 5489 | 1534 | 710709 | 73075 | 600 | 8963 | 4420 | 5655 |
| 2452 | 3816 | 3453 | 1120 | 5602 | 2056 | 642 | 9108 | 4407 | 6008 |
| 2404.5 | 4016 | 3488.5 | 1179 | 5247 | 2000 | 688 | 10206 | 4431 | 6442 |
| 2439 | 3930.5 | 3300 | 1134 | 4815 | 1892 | 680 | 10078 | 4422.5 | 6315 |
| 2758 | 5290 | 3566 | 1222.5 | 5083.5 | 1962.5 | 729 | 11034.5 | 4939 | 8383.5 |
| 2783 | 4425 | 3350 | 1127 | 4609 | 1831 | 671 | 11539 | 4568 | 6733 |
| 2672 | 4179 | 3226 | 1072 | 4381 | 1766 | 641 | 13083 | 4325 | 6328 |
| 2480 | 4037 | 3117 | 1076 | 4350 | 1773 | 649 | 11599 | 4128 | 6254 |
| 2470 | 3836 | 3009 | 1063 | 4207 | 1736 | 646 | 10836 | 4291 | 6018 |
| 2407 | 3635 | 2952.5 | 1051 | 4083 | 1745 | 640 | 9298.5 | 4403 | 5807 |
| 2416 | 3525 | 2847 | 1045 | 3944 | 1703 | 624 | 8885 | 4184 | 5544 |
| 2374 | 3688 | 3046 | 1087 | 4212 | 1763 | 659 | 9696 | 4282 | 5975 |
| 2396.5 | 3645.5 | 2997 | 1032 | 4130 | 1726 | 630 | 8788 | 4433 | 5904 |
| 42580 | 38636.5 | 8401.5 | 1531.5 | 5389 | 2066 | 746 | 17498 | 5504.5 | 12986.5 |
| 2360 | 4394 | 3535 | 1136 | 4647 | 1837 | 659 | 10135 | 4146 | 6773 |
| 2674.5 | 7074 | 3900.5 | 1298 | 5329.5 | 2051 | 754.5 | 13182.5 | 4882.5 | 11162 |
| 2317 | 4020 | 3225 | 1107 | 4385 | 1805.5 | 669 | 9641.5 | 4264 | 6394 |
| 5264 | 12561 | 5103 | 1468 | 6381 | 2152 | 722 | 14185 | 7012 | 16184 |
| 8349 | 19913 | 7586 | 1780 | 9030 | 2547 | 707 | 14425 | 9648 | 22921 |
| 8873 | 20609 | 8177 | 1792 | 9562 | 2501 | 687 | 14778 | 9709 | 23466 |

| | | | | | | | | | |
|---|---|---|---|---|---|---|---|---|---|
| 8287 | 20801 | 7575 | 1742 | 8866 | 2424 | 692 | 11905 | 8870 | 23340 |
| 3277 | 4172 | 2379 | 783 | 3112 | 1346 | 303 | 9343 | 4264 | 5116 |
| 2659.5 | 3626 | 2396 | 814 | 3181.5 | 1447 | 425 | 8082 | 3750.5 | 4903.5 |
| 2483 | 3496.5 | 2500 | 883 | 3396.5 | 1514.5 | 505 | 8086 | 3910.5 | 5112.5 |
| 2571 | 3699 | 5523.5 | 1544 | 712777.5 | 71545.5 | 596 | 9059 | 4370.5 | 5655 |
| 2505 | 3977 | 3560 | 1142 | 5584 | 2112 | 665 | 9910 | 4405 | 6261 |
| 2517 | 4258 | 3484 | 1202.5 | 5212.5 | 1979 | 709 | 10312.5 | 4545 | 6793.5 |
| 2431 | 4063 | 3387 | 1154 | 4975 | 1936 | 701 | 9451 | 4625 | 6536 |
| 2513 | 4090 | 3312 | 1118 | 4711 | 1859.5 | 661 | 11008 | 4324 | 6393 |
| 2663 | 4256 | 3232 | 1104 | 4458 | 1811 | 648 | 11166 | 4423 | 6295 |
| 2626 | 4435 | 3279 | 1123 | 4568 | 1840 | 652 | 14070 | 4213 | 6543 |
| 2707 | 3948 | 3080 | 1027 | 4126 | 1713.5 | 618 | 11278 | 4392 | 6050 |
| 2488 | 4242.5 | 3324 | 1119 | 4551.5 | 1847 | 680 | 12594 | 4246 | 6617.5 |
| 2349 | 3812 | 3043 | 1071 | 4180 | 1736 | 644 | 10008 | 4117 | 6014 |
| 2483.5 | 3792.5 | 3041 | 1074 | 4246.5 | 1766 | 647 | 9836 | 4329 | 6020 |
| 2300.5 | 3953 | 3210 | 1124 | 4393 | 1826.5 | 677 | 9899 | 4113 | 6314 |
| 2449 | 5173.5 | 3335 | 1145.5 | 4644.5 | 1859 | 674.5 | 10528.5 | 4324 | 8229 |
| 33856.5 | 28794 | 7160.5 | 1518 | 5555 | 2117 | 764 | 16373.5 | 5673.5 | 12204 |
| 2571 | 6360 | 3751 | 1195 | 4929 | 1903.5 | 677.5 | 11096.5 | 4282.5 | 9595.5 |
| 2597.5 | 6791.5 | 3704 | 1234 | 5124 | 2002 | 705 | 12025.5 | 4611.5 | 10740 |
| 2457 | 5339 | 3568 | 1209 | 4858 | 1904 | 708 | 11331 | 4453 | 8407 |
| 4954 | 12423 | 5071 | 1397 | 6126 | 2109 | 682 | 14289 | 6525 | 15665 |
| 8135 | 19655.5 | 7333.5 | 1810 | 8819.5 | 2537 | 703 | 14445.5 | 9490 | 22978 |
| 8337 | 20890 | 7154 | 1725 | 8447 | 2448 | 662 | 12861 | 9252 | 24041 |
| 9135.5 | 20615.5 | 6774.5 | 1667 | 7928.5 | 2350 | 658 | 14019 | 9552 | 23294 |
| 3270 | 3957 | 2252 | 765 | 2984 | 1351 | 298 | 9055 | 4212 | 4848 |
| 2722.5 | 3684 | 2398.5 | 820 | 3169 | 1429 | 424 | 8190 | 3775 | 4878.5 |
| 2750 | 3779 | 2719.5 | 942.5 | 3617 | 1591 | 538 | 8646 | 4301.5 | 5383 |
| 2506 | 3508 | 5312 | 1475 | 720735 | 75431 | 565 | 8536 | 4070 | 5386 |
| 2554 | 4353 | 3923 | 1246 | 6325 | 2251 | 707 | 11128 | 4439 | 6762 |
| 2532.5 | 4053 | 3432.5 | 1153 | 5125.5 | 1941 | 681 | 10132.5 | 4449 | 6331.5 |
| 2482.5 | 4232 | 3466.5 | 1167.5 | 4968.5 | 1923.5 | 701 | 10472.5 | 4353.5 | 6633 |
| 2520 | 4793 | 3570 | 1207 | 5045 | 1981 | 725 | 11978 | 4343 | 7526 |
| 2780 | 4605 | 3492.5 | 1128 | 4716.5 | 1861 | 687 | 14308.5 | 4433 | 6943.5 |
| 2724 | 4501 | 3414 | 1152 | 4614 | 1864 | 659 | 12891.5 | 4377.5 | 6708 |
| 2568.5 | 4162 | 3167.5 | 1087.5 | 4248.5 | 1750.5 | 646 | 11929 | 4137 | 6194 |
| 2675 | 4274 | 3394.5 | 1146 | 4554.5 | 1855 | 695.5 | 11519 | 4481 | 6644 |
| 2544 | 3912 | 3110.5 | 1094 | 4205.5 | 1768 | 663 | 10287 | 4437 | 6079.5 |
| 2613 | 3890 | 3078 | 1072 | 4187 | 1746 | 662 | 9565 | 4489 | 6068 |
| 2492 | 4140 | 3384 | 1161.5 | 4505 | 1883 | 695 | 10580.5 | 4438.5 | 6650.5 |
| 2515 | 6420 | 3624 | 1242 | 4977 | 1965.5 | 724 | 12131 | 4528.5 | 10173 |
| 38424 | 25289 | 7804.5 | 1403 | 4812 | 1880 | 668 | 13117 | 4802 | 7918.5 |
| 3089 | 8454 | 4384.5 | 1339 | 5475 | 2108.5 | 747 | 13722 | 4938 | 12418 |
| 2386.5 | 6372 | 3645.5 | 1185.5 | 4866.5 | 1929 | 681.5 | 10393 | 4088.5 | 9925 |
| 2564 | 5660.5 | 3627.5 | 1223 | 4986 | 1946 | 723 | 11996 | 4469 | 9052 |
| 5506 | 13655 | 5281 | 1451 | 6488 | 2201 | 710 | 14591 | 7372 | 16999 |
| 8233 | 20992 | 6993 | 1756 | 8323 | 2449 | 686 | 14696 | 9485 | 24706 |
| 8259 | 22053 | 7631 | 1824 | 8942 | 2535 | 683 | 12817 | 9131 | 24696 |

| | | | | | | | | | |
|---|---|---|---|---|---|---|---|---|---|
| 8416.5 | 21590.5 | 8182.5 | 1856 | 9388 | 2565.5 | 720 | 13217 | 8939.5 | 24820.5 |
| 3263 | 4431 | 2545 | 791 | 3347 | 1387 | 323 | 10677 | 4256 | 5364 |
| 2841 | 3833 | 2533 | 837 | 3301 | 1441 | 427 | 8376 | 4025 | 5078 |
| 2621 | 3651 | 2643 | 918 | 3548 | 1538 | 513 | 8585 | 4159 | 5294 |
| 2766 | 3993 | 5754.5 | 1614 | 745798.5 | 70523 | 616 | 9756.5 | 4538 | 6054.5 |
| 2508.5 | 4049 | 3497.5 | 1125 | 5454 | 2069 | 646 | 9827 | 4350 | 6275.5 |
| 2481.5 | 4215.5 | 3500 | 1189 | 5258 | 2019 | 705 | 10276.5 | 4399 | 6760 |
| 2518 | 3962 | 3232 | 1114 | 4786 | 1861 | 664 | 9564 | 4475 | 6265 |
| 2766 | 5287 | 3617 | 1219 | 5212 | 1983 | 717 | 11030 | 4865 | 8271 |
| 2701 | 4136 | 3103 | 1033 | 4423 | 1744 | 626 | 11299 | 4356 | 6140 |
| 2670 | 4343 | 3300 | 1127 | 4537 | 1801 | 636 | 11679 | 4249 | 6447 |
| 2653 | 4464 | 3395 | 1148 | 4728 | 1859 | 678 | 13760 | 4322 | 6651 |
| 2535 | 4064 | 3197 | 1079 | 4347 | 1781 | 648 | 11205 | 4298 | 6195 |
| 2574 | 4026 | 3215 | 1116 | 4411 | 1831 | 685 | 10843 | 4483 | 6366 |
| 2507 | 3667 | 2963 | 1026 | 4085 | 1731 | 633 | 9104 | 4299 | 5813 |
| 2519.5 | 3849 | 3134.5 | 1095 | 4386.5 | 1798 | 673 | 9555 | 4429 | 6168 |
| 2394 | 3820 | 3025 | 1064 | 4222 | 1762 | 628 | 9304 | 4285 | 6052 |
| 45474 | 41130 | 8866 | 1549 | 5420 | 2072 | 746 | 17539 | 5518 | 13063 |
| 2746 | 6788 | 3702 | 1188 | 4651 | 1879 | 660 | 11374 | 4496 | 10122 |
| 2728 | 6749 | 3867 | 1276 | 5295 | 2022 | 724 | 12134 | 4817 | 10448 |
| 2531 | 4285 | 3382 | 1147 | 4630 | 1899 | 684 | 10711 | 4409 | 6744 |
| 5748 | 13055 | 5098 | 1414 | 6282 | 2102 | 692 | 14867 | 7205 | 16130 |
| 8627 | 19773 | 8712 | 1967 | 10183 | 2678 | 765 | 14456 | 9597 | 22709 |
| 9615.5 | 20988.5 | 7992.5 | 1823.5 | 9189.5 | 2488 | 708 | 15461.5 | 9832 | 23510 |
| 9063 | 20979 | 7163 | 1736 | 8329 | 2397 | 695 | 12527 | 9099 | 23679 |
| 3294 | 4089 | 2333.5 | 769 | 3081 | 1352.5 | 289 | 9355 | 4308 | 5010 |
| 3024.5 | 4050 | 2577 | 855 | 3528 | 1468 | 445 | 9154.5 | 4308 | 5299.5 |
| 2815 | 3742.5 | 2716.5 | 943 | 3660 | 1588 | 538 | 8831 | 4313 | 5521 |
| 2691 | 3811 | 5439 | 1533 | 701550 | 70973 | 602 | 9192 | 4443 | 5892 |
| 2531.5 | 4133 | 3674 | 1173 | 5877.5 | 2113 | 665 | 10171.5 | 4362 | 6325 |
| 2500 | 4005 | 3353.5 | 1100 | 5071 | 1932.5 | 661.5 | 10130 | 4419.5 | 6344.5 |
| 2355 | 4259.5 | 3460 | 1172 | 5101 | 1948.5 | 703 | 10412.5 | 4239 | 6779 |
| 2588.5 | 4221.5 | 3386 | 1165 | 4794.5 | 1881 | 673 | 10525.5 | 4479.5 | 6654 |
| 2615 | 4482 | 3362 | 1122 | 4654 | 1870 | 661 | 13533 | 4375 | 6637 |
| 2631 | 4349 | 3293 | 1107 | 4522 | 1798 | 646 | 12619 | 4163 | 6519 |
| 2645.5 | 4026 | 3071 | 1034 | 4196 | 1740 | 623 | 11547 | 4291 | 6017 |
| 2586 | 4203.5 | 3206.5 | 1099 | 4433 | 1808.5 | 660 | 10845 | 4441 | 6349 |
| 2560 | 4045 | 3272 | 1126 | 4465 | 1803 | 673 | 10520 | 4479 | 6384 |
| 2466 | 3606.5 | 2860 | 999.5 | 3855.5 | 1696 | 612.5 | 8867.5 | 4103.5 | 5603.5 |
| 2380 | 3693 | 2970 | 1036 | 4078 | 1735 | 639 | 9117 | 4191 | 5868 |
| 2417 | 3820 | 3087.5 | 1060 | 4188.5 | 1749 | 623.5 | 9001 | 4282.5 | 6036 |
| 36943 | 22462 | 7285.5 | 1344 | 4572 | 1818.5 | 637 | 12684 | 4657 | 7103 |
| 2715 | 6577 | 3689 | 1171 | 4649 | 1860 | 649 | 10829 | 4348 | 9606 |
| 2548 | 6532 | 3561 | 1167 | 4778 | 1892 | 663 | 11156 | 4331 | 10078 |
| 2412 | 4214.5 | 3226.5 | 1090.5 | 4423.5 | 1802.5 | 643 | 10446 | 4214 | 6636 |
| 4860 | 11831 | 4804.5 | 1333 | 5897 | 2044 | 652 | 14220 | 6351.5 | 14992.5 |
| 8797 | 19632 | 7309 | 1724 | 8721 | 2453 | 685 | 14854 | 9567 | 22250 |
| 9055 | 20732 | 6711 | 1673 | 7983 | 2349 | 644 | 15348 | 9822 | 23430 |

| | | | | | | | | | |
|---|---|---|---|---|---|---|---|---|---|
| 8799 | 19345 | 6823 | 1652 | 8011.5 | 2345 | 662.5 | 14943 | 9371.5 | 22315 |
| 3149 | 3859 | 2183 | 733 | 2954 | 1310 | 275 | 8766 | 4109 | 4596 |
| 2881 | 3897 | 2471 | 830 | 3312 | 1454 | 412 | 8600.5 | 4103.5 | 5069 |
| 2538 | 3704 | 2611 | 898 | 3464 | 1528 | 499 | 8452 | 3839 | 5172 |
| 2571 | 3593 | 5206 | 1419 | 710134 | 70194 | 554 | 8680 | 4106 | 5304 |
| 2538 | 4062 | 3548 | 1137 | 5777.5 | 2115 | 638 | 9523 | 4291.5 | 6207 |
| 2632 | 3893 | 3207 | 1056 | 4751 | 1855 | 626 | 9367 | 4488 | 5964 |
| 2566 | 4173.5 | 3419 | 1162 | 4945 | 1941 | 692 | 10296 | 4434.5 | 6511.5 |
| 2531 | 6102 | 3655 | 1210 | 5147 | 1992 | 714 | 13221 | 4239 | 9210 |
| 2723 | 6213 | 3485.5 | 1132 | 4820 | 1839 | 664.5 | 14279 | 4347 | 9202.5 |
| 2825 | 5336 | 3457 | 1151 | 4711 | 1866 | 660 | 14779 | 4412 | 7767 |
| 2565 | 4546.5 | 3358.5 | 1114 | 4590 | 1817 | 660 | 14012 | 4169 | 6703.5 |
| 2644 | 4173 | 3229 | 1080 | 4469.5 | 1778 | 648 | 10767 | 4380 | 6385.5 |
| 2575 | 4366 | 3373 | 1139 | 4679 | 1864 | 693 | 10860 | 4477 | 6699 |
| 2583 | 4348 | 3384 | 1149 | 4687 | 1877 | 691 | 11478 | 4399 | 6778 |
| 2477 | 3872 | 3101 | 1062 | 4169 | 1758 | 643 | 9904 | 4306 | 6038 |
| 2513 | 4037 | 3216 | 1087 | 4352 | 1788 | 645 | 9612 | 4449 | 6360 |
| 50211.5 | 30229.5 | 9253 | 1458 | 4882.5 | 1895.5 | 720.5 | 15958 | 5419.5 | 7740.5 |
| 4059 | 6972 | 4326.5 | 1283 | 5229.5 | 2000 | 931.5 | 17219 | 6093.5 | 9722 |
| 3692 | 7603 | 4064 | 1266 | 5234 | 1982 | 1047 | 19324 | 6034 | 11617 |
| 3313 | 5316 | 3720.5 | 1206 | 4741.5 | 1884.5 | 934.5 | 15767 | 5442.5 | 7958 |
| 6147 | 11415 | 4651 | 1275 | 5482 | 1975 | 852 | 18051 | 6141 | 14003 |
| 14958 | 24303 | 7102 | 1764 | 8001 | 2423 | 803 | 15230 | 10106 | 22992 |
| 19433 | 29750 | 8158 | 1810 | 8740 | 2476 | 862 | 15917 | 10782 | 25292 |
| 16921.5 | 26646 | 7765 | 1763 | 8526.5 | 2441 | 856 | 16962 | 10853 | 24591 |
| 838 | 1133 | 958 | 595 | 1535 | 1135.5 | 239 | 2569 | 1567 | 2078 |
| 1176 | 1845 | 1636.5 | 721 | 2356 | 1317.5 | 412 | 4175 | 2261 | 3290 |
| 1461 | 2499 | 2210 | 881 | 3205 | 1530 | 543 | 6346 | 2970.5 | 4430 |
| 1557 | 2725 | 5098 | 1518 | 756396 | 77481 | 602 | 7233 | 3290 | 4930 |
| 1625 | 2862 | 2911 | 1063 | 4595 | 2019 | 632 | 7509 | 3472 | 5225 |
| 1577 | 2732 | 2579 | 981 | 3856 | 1733 | 606.5 | 7228 | 3271 | 4887 |
| 1562 | 2699.5 | 2517 | 969 | 3610 | 1650 | 607 | 7150.5 | 3275 | 4861 |
| 1733 | 2921 | 2569 | 967 | 3570 | 1642 | 610 | 7974 | 3411.5 | 5034 |
| 1959 | 3086 | 2542 | 940 | 3572 | 1621 | 602 | 8950 | 3585 | 5109 |
| 2317 | 3399 | 2799 | 1043 | 3874 | 1716 | 630.5 | 9747.5 | 4215 | 5607 |
| 2336 | 3518.5 | 2958 | 1077 | 4110.5 | 1786 | 674 | 10546.5 | 4410 | 5930 |
| 2225.5 | 3566.5 | 3109 | 1138 | 4337 | 1866.5 | 712 | 10662 | 4435 | 6195 |
| 1861 | 3159 | 2828 | 1029 | 3966 | 1742.5 | 664 | 8614 | 3720 | 5578.5 |
| 1903 | 3209 | 2835 | 1042 | 3995 | 1741 | 667 | 8596 | 3841 | 5545 |
| 1752 | 3021 | 2712.5 | 1038.5 | 3865 | 1721.5 | 654 | 8098 | 3615 | 5399 |
| 2098 | 3577.5 | 3206 | 1150 | 4480.5 | 1866 | 716 | 9298 | 4428 | 6316 |
| 39383 | 21569.5 | 7325.5 | 1389 | 4723.5 | 1888 | 690 | 12723 | 5156.5 | 6989 |
| 2356 | 4106.5 | 3584 | 1189 | 4617 | 1890.5 | 690 | 10238.5 | 4609 | 6721.5 |
| 2173 | 3600 | 3225 | 1126 | 4473 | 1844 | 681 | 9470 | 4443 | 6271 |
| 2064 | 3455 | 3064 | 1096 | 4289 | 1792 | 686 | 9122 | 4281 | 6056 |
| 3928 | 9561 | 4464 | 1320 | 5700 | 2023 | 682 | 14938 | 5381 | 12744 |
| 7598 | 17593 | 6123 | 1582 | 7385 | 2276 | 655 | 15892 | 8782 | 20393 |
| 8913 | 19610 | 6638 | 1619 | 7716 | 2331 | 649 | 15584 | 9721 | 22377 |

| | | | | | | | | | |
|---|---|---|---|---|---|---|---|---|---|
| 8485 | 18680 | 6395 | 1589 | 7505 | 2266 | 655 | 14191 | 9292 | 21676 |
| 2989 | 3763 | 2128 | 734 | 2759.5 | 1299 | 309 | 8676 | 4017 | 4643.5 |
| 2331 | 3326 | 2295 | 814 | 3009 | 1402 | 461 | 7622 | 3470 | 4661 |
| 2043 | 3153 | 2452 | 897 | 3329 | 1539 | 554 | 7600 | 3471 | 4903 |
| 2011 | 3219 | 6144 | 1657 | 959184 | 87112 | 641 | 8319 | 3788 | 5473 |
| 2091 | 3359 | 3465 | 1162 | 5646 | 2160 | 689 | 8762 | 4066 | 5789 |
| 1890 | 3170 | 2866 | 1059 | 4322 | 1812 | 662 | 8269 | 3795 | 5441 |
| 1942 | 3209 | 2924 | 1067 | 4192 | 1797 | 673 | 8359 | 3894 | 5602 |
| 2134 | 3329 | 2877 | 1030 | 4070 | 1742 | 663 | 8864 | 4040 | 5575 |
| 2228.5 | 3336 | 2762 | 990 | 3766 | 1663.5 | 633 | 9898 | 3976.5 | 5463 |
| 2480 | 3559 | 2907 | 1036 | 3993 | 1717 | 648 | 10051 | 4299 | 5665 |
| 2491 | 3862 | 3219 | 1132 | 4437 | 1832 | 722 | 12002 | 4575 | 6369 |
| 2454 | 3830 | 3219 | 1182 | 4460 | 1906 | 721 | 10384 | 4595 | 6303 |
| 2115.5 | 3417 | 2979 | 1074 | 4075.5 | 1781 | 695 | 9538.5 | 4194 | 5910 |
| 2247 | 3455 | 3010 | 1095 | 4184 | 1811 | 712 | 9625 | 4398 | 6001 |
| 2152 | 3437 | 3000 | 1122.5 | 4198.5 | 1831 | 710 | 9115.5 | 4290 | 5965 |
| 2249 | 3794.5 | 3341.5 | 1180 | 4703 | 1935.5 | 733 | 10707.5 | 4649 | 6611.5 |
| 47082 | 24851 | 7974 | 1398 | 4521 | 1830 | 688 | 12730 | 5276 | 6906 |
| 2449 | 5413 | 3911 | 1264 | 4927 | 1995 | 740 | 11979 | 4670 | 8869 |
| 2322 | 3674 | 3213 | 1120 | 4341 | 1817 | 676 | 9259 | 4644 | 6215 |
| 2210 | 3748 | 3295 | 1150 | 4546 | 1904 | 709 | 10412 | 4435 | 6476 |
| 4238 | 10431.5 | 4447.5 | 1279 | 5563 | 1976 | 651 | 14396 | 5756.5 | 13058.5 |
| 8297 | 18703 | 7583 | 1770 | 8961 | 2497.5 | 710 | 14993.5 | 9719 | 21898 |
| 8706 | 21474 | 7322 | 1758 | 8772 | 2464 | 682 | 14554 | 9876 | 24278 |
| 7949 | 19502 | 6729 | 1651 | 7949 | 2383 | 673 | 11767 | 8626 | 22263 |
| 2688 | 3574 | 1992 | 725 | 2741 | 1283 | 291 | 8114 | 3558 | 4436 |
| 2349 | 3323 | 2213 | 789 | 2952 | 1388 | 423 | 7354 | 3450 | 4598 |
| 2135 | 3314 | 2488 | 886 | 3316 | 1504 | 532 | 7764 | 3514 | 4957 |
| 1848 | 2854 | 5633 | 1615 | 935708 | 81885 | 544 | 7271 | 3305 | 4639 |
| 1904 | 3123 | 3179 | 1083 | 5995 | 2151 | 613 | 7747 | 3579 | 5191 |
| 2232 | 3374 | 3148 | 1076 | 5090 | 1957 | 673 | 9155 | 4288 | 5755 |
| 2041 | 3268 | 2879 | 1036 | 4337 | 1780 | 649 | 8682 | 3989 | 5567 |
| 2392.5 | 3797 | 3347.5 | 1173 | 5010.5 | 1954 | 716.5 | 10769 | 4552 | 6354 |
| 2495 | 3836 | 3185 | 1108 | 4585 | 1836 | 685 | 10805 | 4509 | 6212 |
| 2409.5 | 3501 | 2839 | 1039 | 3990 | 1718 | 626 | 10081.5 | 4202.5 | 5580 |
| 2479 | 3567 | 2905 | 1039 | 4042 | 1721 | 649 | 9844 | 4503 | 5760 |
| 2095 | 3369 | 2813 | 1021 | 3936.5 | 1723 | 643 | 9138.5 | 3875 | 5590.5 |
| 2056 | 3285 | 2799 | 1038 | 3972 | 1712 | 655 | 8808 | 3975 | 5580 |
| 1989 | 3220 | 2746 | 1004 | 3868 | 1713 | 644 | 8570 | 3747 | 5500 |
| 1976 | 3230.5 | 2808 | 1033 | 3960 | 1715 | 647 | 8639 | 3911 | 5484 |
| 2151 | 3396 | 2940 | 1050 | 4038 | 1724 | 656 | 8933 | 4280 | 5750 |
| 50747.5 | 26474.5 | 8202 | 1321 | 4426.5 | 1841 | 663 | 13656 | 5263 | 6740 |
| 2460 | 4131.5 | 3487 | 1184 | 4530 | 1867 | 673 | 9927.5 | 4564 | 6644 |
| 2276.5 | 3844 | 3294.5 | 1151 | 4545.5 | 1861 | 688 | 10398.5 | 4462 | 6498.5 |
| 2255 | 3882 | 3306 | 1156 | 4599 | 1917 | 705 | 10320 | 4505 | 6565 |
| 4945.5 | 10742 | 4516 | 1278.5 | 5691.5 | 2003.5 | 665 | 14217 | 6529.5 | 13882 |
| 8607 | 19453 | 6566 | 1717 | 7891 | 2416 | 719 | 15794 | 9899 | 22536 |
| 8979 | 22240 | 7545 | 1827 | 8921 | 2578 | 723 | 13538 | 9589 | 24460 |

| | | | | | | | | | |
|---|---|---|---|---|---|---|---|---|---|
| 8342 | 21149 | 6731 | 1642 | 7991 | 2394 | 679 | 15022 | 9102 | 24030 |
| 3002 | 3661 | 2115 | 754 | 2810 | 1318 | 301 | 8379 | 3951 | 4542 |
| 2382 | 3320 | 2256 | 818 | 3009 | 1423 | 453 | 7809 | 3473.5 | 4615.5 |
| 2217 | 3332 | 2547 | 923 | 3437.5 | 1557.5 | 563 | 8261 | 3647 | 5164.5 |
| 2138 | 3290 | 5846 | 1615 | 943163 | 91670 | 618 | 8646 | 3800 | 5313 |
| 2228 | 3583 | 3402 | 1128 | 5391 | 2069 | 688 | 9008 | 4133 | 5875 |
| 2360 | 3545.5 | 3153 | 1121.5 | 4748 | 1876 | 691 | 9197 | 4514 | 5843.5 |
| 2345 | 3643 | 3208.5 | 1121 | 4768 | 1925 | 722 | 9486 | 4552 | 6155.5 |
| 2520 | 3967 | 3417 | 1184 | 4833 | 1928 | 747 | 11017 | 4674 | 6634 |
| 2688 | 3946 | 3279 | 1144 | 4552 | 1863 | 712 | 11247 | 4737 | 6355 |
| 2554 | 3635 | 2895 | 1033 | 3906 | 1716 | 654 | 10294 | 4287 | 5756 |
| 2670 | 3780 | 3053 | 1062 | 4162 | 1774 | 693 | 10897 | 4614 | 5944 |
| 2529.5 | 3654 | 3066 | 1117 | 4163 | 1779 | 694 | 10468 | 4624.5 | 5933.5 |
| 2250 | 3510 | 2992 | 1069 | 4063 | 1755 | 699 | 9311 | 4277 | 5852 |
| 2158 | 3349 | 2846 | 1050 | 3943 | 1725 | 665 | 9037 | 3959 | 5528 |
| 1987 | 3242 | 2773 | 1019 | 3848 | 1675 | 652 | 8785 | 3832 | 5461 |
| 2161 | 3481 | 2929 | 1060 | 4076 | 1731 | 653 | 8698 | 4183 | 5777 |
| 49942 | 26078 | 8154 | 1318 | 4198 | 1767 | 642 | 13297 | 5211 | 6559 |
| 2481 | 3955 | 3331.5 | 1120 | 4310.5 | 1767 | 637 | 9484.5 | 4547.5 | 6251.5 |
| 2300 | 3793 | 3195 | 1105 | 4358 | 1828 | 656 | 9613 | 4357 | 6258 |
| 2178 | 3313 | 2807 | 1012 | 3837 | 1657 | 614 | 8320 | 4144 | 5562 |
| 4815.5 | 12326 | 4858 | 1362 | 5970 | 2073 | 679 | 13523 | 6431 | 15731 |
| 8049.5 | 17705 | 6028.5 | 1561 | 7260 | 2250 | 649 | 14800.5 | 9068.5 | 20768 |
| 8726 | 20496 | 6928 | 1677 | 8169 | 2366 | 656 | 14783 | 9654 | 23112 |
| 8846 | 20038 | 6486 | 1555 | 7540 | 2265 | 648 | 13148.5 | 9604.5 | 22798.5 |
| 3136 | 4027.5 | 2248.5 | 760 | 2969.5 | 1341 | 294.5 | 9301.5 | 4160 | 4849.5 |
| 2442 | 3478 | 2322 | 813 | 3096 | 1410 | 425 | 7965.5 | 3573.5 | 4779 |
| 2344 | 3468.5 | 2651 | 920 | 3726.5 | 1604 | 534 | 8268.5 | 3849 | 5209 |
| 2102 | 3243.5 | 5238.5 | 1529 | 803228 | 77817.5 | 568 | 8074 | 3668 | 5166.5 |
| 2010 | 3204 | 2983 | 1004 | 4817 | 1855 | 590 | 8035 | 3652 | 5163 |
| 2316 | 3562 | 3124 | 1081 | 4702 | 1867 | 650 | 8963 | 4380 | 5847 |
| 2276 | 3470 | 2995 | 1053 | 4451 | 1798 | 647 | 8863 | 4225 | 5687 |
| 2325.5 | 4260 | 3393 | 1154.5 | 4941 | 1919 | 717 | 12108.5 | 4269 | 6850.5 |
| 2533 | 3720 | 2992 | 1049 | 4212 | 1766 | 646 | 10129 | 4458 | 6005 |
| 2524 | 4018 | 3217 | 1118 | 4499 | 1827 | 675 | 12370 | 4358 | 6378 |
| 2442 | 3751 | 2991 | 1047 | 4189 | 1789 | 651 | 10810 | 4334 | 5912 |
| 2255 | 3467.5 | 2838.5 | 1027 | 3904.5 | 1697 | 628 | 8979.5 | 4058 | 5572 |
| 2282 | 3446 | 2903.5 | 1043 | 4040 | 1748 | 646 | 9487 | 4244 | 5712 |
| 2136 | 3259 | 2714.5 | 995.5 | 3784.5 | 1673 | 614 | 8347 | 3889 | 5347 |
| 1965 | 3159 | 2687 | 988 | 3722 | 1646 | 616 | 8224 | 3689 | 5330 |
| 2282 | 3643 | 3032 | 1098 | 4259 | 1782 | 661 | 9655 | 4420 | 6094 |
| 45001 | 24662 | 7900.5 | 1371 | 4535 | 1878 | 677 | 13173 | 5120.5 | 6924 |
| 2617 | 5017 | 3709 | 1212 | 4698 | 1911 | 688 | 10961 | 4623 | 7737 |
| 2370 | 3918 | 3260.5 | 1132 | 4454.5 | 1832 | 668 | 10220 | 4435.5 | 6481.5 |
| 2368 | 6210 | 3655 | 1258 | 5041 | 2000 | 757 | 12830 | 4509 | 10349 |
| 4472.5 | 8855 | 4703 | 1371 | 5911 | 2093 | 703 | 13787 | 6095.5 | 11713 |
| 8666.5 | 19431 | 7541.5 | 1814 | 8957 | 2519.5 | 715 | 15399.5 | 9993.5 | 22529.5 |
| 8688 | 21492 | 7769 | 1869 | 9154 | 2564 | 707 | 14042 | 9659 | 24320 |

| | | | | | | | | | |
|---|---|---|---|---|---|---|---|---|---|
| 8237 | 21738 | 7267 | 1792 | 8691 | 2480 | 712 | 11774 | 9186 | 24642 |
| 3165.5 | 4176 | 2399 | 782 | 3135.5 | 1365 | 314 | 10005 | 4182 | 5106.5 |
| 2777 | 3637 | 2323 | 807 | 3129 | 1406 | 422 | 8529 | 4060 | 4867 |
| 2618 | 3645 | 2616 | 926 | 3550 | 1571 | 534 | 8839 | 4242 | 5340 |
| 2479 | 3538 | 5899 | 1576 | 909821 | 77466 | 594 | 8855 | 4169 | 5534 |
| 2387 | 3799 | 3521 | 1176 | 5706 | 2126 | 659 | 9599 | 4374 | 6129 |
| 2381 | 3942 | 3490 | 1157.5 | 5387 | 2010 | 688 | 9709 | 4486.5 | 6389.5 |
| 2194 | 3518 | 3034.5 | 1075 | 4563 | 1813 | 647.5 | 8729.5 | 4138 | 5809.5 |
| 2440 | 3832 | 3137 | 1094 | 4553 | 1847 | 675 | 9699 | 4375 | 6160 |
| 2526 | 3727.5 | 2996 | 1039 | 4286.5 | 1752 | 635 | 10055 | 4327 | 5797.5 |
| 2491 | 3768 | 3004 | 1047 | 4204 | 1732 | 630 | 11193 | 4225 | 5891 |
| 2501 | 4087 | 3215 | 1109 | 4475 | 1855 | 679 | 12026 | 4375 | 6358 |
| 2351 | 3860 | 3175 | 1104 | 4452.5 | 1821 | 686.5 | 11320.5 | 4267 | 6283 |
| 2336 | 3614.5 | 2982 | 1075 | 4197.5 | 1759 | 662 | 9309.5 | 4375.5 | 5953 |
| 2405.5 | 3687.5 | 3126 | 1083 | 4319.5 | 1822 | 677 | 9760 | 4367.5 | 6072 |
| 2283 | 3577 | 3030.5 | 1096.5 | 4287 | 1814.5 | 666 | 8952.5 | 4333 | 6015 |
| 2275 | 3914 | 3204 | 1124 | 4523 | 1850 | 678 | 10142 | 4273 | 6488 |
| 44831 | 36024 | 8809 | 1485 | 5026 | 1977 | 698 | 14787 | 5056 | 9977 |
| 2461 | 4562 | 3782 | 1217 | 4738 | 1912 | 674 | 10498 | 4290 | 6945 |
| 2372.5 | 5144 | 3704 | 1241 | 5108 | 2008.5 | 704 | 12048 | 4536.5 | 8349.5 |
| 2365 | 3680 | 3052 | 1093 | 4230 | 1764 | 648 | 8957 | 4359 | 6088 |
| 5138.5 | 12500.5 | 5025 | 1391 | 6179.5 | 2115 | 681 | 15235 | 6802 | 15768 |
| 8302.5 | 17883 | 6358.5 | 1609 | 7632 | 2315 | 670.5 | 15391.5 | 9525.5 | 21095 |
| 8393 | 19845 | 6897 | 1689 | 8137.5 | 2348 | 648 | 14969 | 9254.5 | 22770 |
| 8395 | 19926 | 6431 | 1604 | 7601 | 2323 | 644 | 12494 | 8907 | 22421 |
| 3196 | 4215.5 | 2428 | 783 | 3203.5 | 1370 | 324 | 10326.5 | 4200 | 5226 |
| 2973 | 3878 | 2517 | 844 | 3358 | 1474 | 444 | 9110 | 4277 | 5200 |
| 2561 | 3652 | 2691.5 | 905 | 3616 | 1556 | 533 | 8790.5 | 4190.5 | 5421 |
| 2494 | 3626 | 5895 | 1593 | 897363 | 75178 | 605 | 8772 | 4306 | 5653 |
| 2447 | 3784 | 3452 | 1153 | 5732 | 2070 | 655 | 9015 | 4420.5 | 6108.5 |
| 2281 | 3565 | 3130 | 1082 | 4795 | 1884 | 639 | 8734 | 4222 | 5872 |
| 2159 | 3482 | 2967 | 1044 | 4412 | 1763 | 631 | 8610.5 | 3989.5 | 5696 |
| 2247 | 3540 | 2913 | 1026 | 4264 | 1740 | 621 | 9139 | 4096 | 5631 |
| 2527.5 | 3665 | 2918 | 1008.5 | 4146 | 1710 | 616 | 9778.5 | 4341 | 5762 |
| 2612 | 3670 | 2930 | 1034 | 4023 | 1730 | 604 | 10188 | 4330 | 5699 |
| 2459.5 | 4003 | 3132 | 1078 | 4420 | 1802.5 | 659 | 11256.5 | 4299 | 6332.5 |
| 2368 | 3438 | 2836 | 994 | 3928 | 1665 | 612 | 9479 | 4197 | 5511 |
| 2354 | 3594.5 | 2966 | 1062 | 4183 | 1758 | 651 | 9183 | 4407 | 5870.5 |
| 2242.5 | 3370 | 2775.5 | 1011 | 3884 | 1698 | 619 | 8483 | 4049.5 | 5512.5 |
| 2273 | 3698.5 | 3095 | 1089.5 | 4271 | 1814 | 672 | 9568.5 | 4268.5 | 6158.5 |
| 2292.5 | 3741 | 3130 | 1109 | 4388 | 1832 | 661 | 9583 | 4362 | 6173 |
| 41288 | 32030 | 8063 | 1408 | 4758 | 1900 | 668 | 13634 | 4772 | 9161 |
| 2571 | 6800 | 3814 | 1206.5 | 4852.5 | 1946 | 690 | 11770 | 4410 | 10388.5 |
| 2287 | 4056 | 3313 | 1111.5 | 4498 | 1835 | 654 | 9485 | 4233.5 | 6500.5 |
| 2322 | 4782 | 3501 | 1208 | 4845 | 1931 | 722 | 11116 | 4311 | 7900 |
| 3984 | 9203 | 4507 | 1301 | 5634 | 1991 | 643 | 14639 | 5481 | 11765 |
| 7813 | 18890 | 8293 | 1876 | 9832 | 2594 | 730 | 12516 | 9020 | 22246 |
| 8181 | 20883 | 8415 | 1909.5 | 9744.5 | 2623 | 709 | 11760.5 | 9137.5 | 24051 |

| | | | | | | | | | |
|---|---|---|---|---|---|---|---|---|---|
| 7437.5 | 20553 | 7324 | 1741 | 8735 | 2463 | 697 | 11604 | 8195.5 | 23708.5 |
| 3157 | 4346 | 2494 | 784 | 3306 | 1385 | 321 | 10331 | 4223 | 5325 |
| 2982 | 3995 | 2566 | 838 | 3421 | 1455 | 444 | 9071 | 4276 | 5284 |
| 2788 | 3952 | 2865 | 960 | 3875 | 1633 | 561 | 9379 | 4396 | 5727 |
| 2642 | 3848 | 6379.5 | 1607 | 921946.5 | 74111 | 638 | 9948 | 4506 | 6126.5 |
| 2483 | 3875 | 3548 | 1157 | 5751 | 2098 | 664 | 9626 | 4419 | 6140 |
| 2416 | 3854.5 | 3311 | 1141 | 5011 | 1941 | 664 | 9102 | 4359 | 6145.5 |
| 2282 | 3736 | 3144 | 1071 | 4699 | 1848 | 659 | 8819 | 4251 | 6005 |
| 2339.5 | 3644 | 2939 | 1017 | 4323.5 | 1756 | 631 | 9312.5 | 4133.5 | 5811.5 |
| 2592.5 | 3719 | 2911 | 1013 | 4169 | 1713 | 612.5 | 9740.5 | 4361 | 5771 |
| 2569 | 3881 | 3084.5 | 1074 | 4222.5 | 1778.5 | 633 | 11279 | 4288.5 | 5986 |
| 2391.5 | 3949.5 | 3121.5 | 1102 | 4319.5 | 1813 | 642 | 11650.5 | 4054.5 | 6164.5 |
| 2418 | 3864 | 3086.5 | 1084 | 4341 | 1787 | 665 | 10852.5 | 4303.5 | 6147 |
| 2458 | 3996 | 3315 | 1167 | 4637 | 1870 | 700 | 10204 | 4536 | 6490 |
| 2348 | 3423 | 2864 | 1014 | 3963 | 1667 | 624 | 8789 | 4225 | 5571 |
| 2392 | 3829 | 3158 | 1124 | 4415 | 1832 | 680 | 9658 | 4396 | 6229 |
| 2256 | 3926 | 3214.5 | 1132 | 4492 | 1834 | 668 | 10224.5 | 4283.5 | 6468.5 |
| 48692.5 | 39834.5 | 8945 | 1483.5 | 4973 | 1943 | 700 | 14960.5 | 5243 | 10525.5 |
| 2545.5 | 6286 | 3694 | 1204 | 4745 | 1907 | 665 | 10760.5 | 4358 | 9606 |
| 2420 | 5795 | 3592 | 1206 | 4884 | 1948 | 691 | 11549 | 4405 | 9469 |
| 2352 | 4342 | 3481 | 1183 | 4799 | 1921 | 706 | 11128 | 4326 | 7149 |
| 4914 | 11767 | 4826 | 1368 | 5908 | 2060 | 662 | 13739 | 6397 | 14724 |
| 8024 | 19348 | 8531 | 1915 | 10221 | 2672 | 764 | 14174 | 9272 | 22441 |
| 8016 | 20823 | 7198 | 1761 | 8645 | 2494 | 679 | 11487 | 9030 | 23553 |
| 7840 | 20271 | 6981 | 1653 | 8171 | 2395 | 668 | 12044 | 8596 | 23023 |
| 3194.5 | 4011.5 | 2221 | 755 | 2998 | 1345.5 | 294 | 9184 | 4196 | 4874.5 |
| 2908 | 3755 | 2438 | 813 | 3254 | 1440 | 425 | 8595 | 4152 | 4989 |
| 2715.5 | 3708 | 2676.5 | 920 | 3667 | 1559 | 530 | 8621 | 4230.5 | 5370 |
| 2590 | 3701 | 6045.5 | 1573 | 916970 | 73550 | 593 | 9023.5 | 4341 | 5647 |
| 2501 | 4009 | 3714 | 1203 | 6153 | 2218 | 658 | 9633 | 4430 | 6221 |
| 2368 | 3705 | 3180 | 1073 | 4847 | 1868 | 643 | 8841 | 4287 | 5896 |
| 2412 | 3907 | 3318 | 1146 | 4874 | 1915 | 679 | 9276 | 4483 | 6327 |
| 2411 | 3773 | 3059 | 1041 | 4430 | 1787 | 637 | 9443 | 4359 | 5977 |
| 2610 | 3880 | 3077 | 1045 | 4347 | 1759 | 634 | 10206 | 4428 | 6029 |
| 2635.5 | 3800 | 2998 | 1055 | 4096.5 | 1744 | 611 | 10163 | 4410 | 5781 |
| 2558 | 3900 | 3086 | 1057 | 4290 | 1758 | 639 | 11085 | 4298 | 6055 |
| 2456 | 3730 | 2980 | 1068 | 4210 | 1752 | 641 | 9976 | 4304 | 5883 |
| 2336 | 3537 | 2911.5 | 1030.5 | 4095 | 1740 | 638 | 9256.5 | 4310.5 | 5693.5 |
| 2402 | 3591 | 2896 | 1059 | 4076 | 1746 | 642 | 8856 | 4282 | 5735 |
| 2284.5 | 3415.5 | 2812 | 1002 | 3932 | 1697 | 627.5 | 8498.5 | 4147 | 5536.5 |
| 2395 | 4998 | 3413.5 | 1178.5 | 4751.5 | 1918 | 698 | 10684.5 | 4493 | 8267 |
| 45367 | 27606.5 | 8602.5 | 1434.5 | 4811 | 1896 | 665 | 13794 | 4859 | 7322.5 |
| 2591.5 | 6379.5 | 3788.5 | 1195 | 4848 | 1904 | 681 | 11103.5 | 4373 | 9474 |
| 2319 | 4327.5 | 3499 | 1170.5 | 4791 | 1919.5 | 674 | 10820.5 | 4306 | 7034 |
| 2298 | 4121 | 3338 | 1149 | 4619 | 1906 | 686 | 9713 | 4292 | 6655 |
| 4530 | 12359 | 5139 | 1453 | 6518 | 2212 | 718 | 14348 | 6227 | 15933 |
| 7607.5 | 19841.5 | 6906 | 1736 | 8370.5 | 2457.5 | 691 | 12412.5 | 8600 | 22890 |
| 8349 | 22165 | 7793.5 | 1781 | 9147.5 | 2483 | 682.5 | 13245 | 9483 | 25356 |

| | | | | | | | | | |
|---|---|---|---|---|---|---|---|---|---|
| 8538.5 | 21066 | 7241 | 1720 | 8493 | 2440 | 688 | 13993 | 9455 | 23948.5 |
| 3196 | 3896 | 2200.5 | 760 | 3019.5 | 1356 | 289 | 9038 | 4156.5 | 4808.5 |
| 2687 | 3673 | 2416 | 825 | 3200 | 1431 | 416 | 8309 | 3861 | 4918 |
| 2446.5 | 3583 | 2581.5 | 904 | 3506 | 1553 | 510.5 | 8216.5 | 3790.5 | 5169.5 |
| 2346 | 3469 | 5361 | 1558.5 | 785589.5 | 75261.5 | 576 | 8503 | 3836 | 5399 |
| 2275 | 3489 | 3041 | 1028 | 4941.5 | 1853 | 596 | 8269.5 | 3976 | 5500.5 |
| 2366 | 3521 | 3033 | 1040 | 4642 | 1798 | 623 | 8639 | 4189 | 5659 |
| 2408 | 3784 | 3158 | 1099 | 4731 | 1846 | 658 | 9430 | 4415 | 6063 |
| 2463 | 3936 | 3257 | 1099 | 4717 | 1862 | 667 | 10603 | 4350 | 6290.5 |
| 2554 | 5143 | 3369 | 1131 | 4851 | 1885 | 685 | 12601 | 4328 | 7909 |
| 2625 | 6138.5 | 3581.5 | 1187 | 4907 | 1934 | 697 | 14267.5 | 4453 | 9442 |
| 2656 | 5843 | 3548 | 1199 | 5045 | 1954 | 723 | 14190 | 4564 | 9079 |
| 2566 | 4985 | 3544 | 1213 | 5010 | 1966 | 733 | 11973 | 4652 | 7816 |
| 2433 | 3973.5 | 3244.5 | 1129 | 4539.5 | 1848 | 690 | 11092 | 4412 | 6401.5 |
| 2501 | 4084 | 3343.5 | 1152 | 4677.5 | 1898 | 706 | 10715.5 | 4524 | 6615 |
| 2344 | 3675 | 3049 | 1069 | 4257 | 1767 | 661 | 8832 | 4343 | 6026 |
| 2349 | 4095 | 3363 | 1154 | 4700 | 1859 | 678 | 10109 | 4473 | 6652 |
| 47367 | 25760 | 8038 | 1372 | 4623.5 | 1850 | 651 | 12417.5 | 5211.5 | 7074.5 |
| 2662.5 | 4392 | 3555.5 | 1154 | 4564 | 1827 | 646 | 10155 | 4506 | 6664 |
| 2507.5 | 3870 | 3211.5 | 1085 | 4419.5 | 1774 | 628 | 9291 | 4475.5 | 6235 |
| 2490 | 3973 | 3209 | 1111 | 4384 | 1819 | 660 | 9653 | 4582 | 6382 |
| 4364 | 11465.5 | 4598 | 1304.5 | 5746 | 1987 | 653 | 13077.5 | 5829 | 14655.5 |
| 7272 | 16506 | 5933 | 1536 | 7050 | 2211 | 639 | 13850 | 8195 | 19360 |
| 8378 | 18881 | 6373 | 1606 | 7567.5 | 2286 | 620 | 14229 | 8875.5 | 21236 |
| 8815 | 19065 | 6397 | 1560 | 7596 | 2270 | 640 | 14877 | 9539 | 21762 |
| 3310.5 | 4006 | 2318 | 761 | 3083.5 | 1345 | 288 | 9624 | 4230 | 4876 |
| 2512 | 3500 | 2272 | 783 | 3073 | 1391 | 381 | 7745 | 3564 | 4608 |
| 2298 | 3392 | 2440 | 868 | 3288 | 1491 | 484 | 7576 | 3577 | 4838 |
| 2628 | 3659 | 5482 | 1534 | 797396 | 72075 | 580 | 8890 | 4310 | 5483 |
| 2481 | 3850.5 | 3471 | 1119 | 5585 | 2053.5 | 636 | 9524.5 | 4267 | 5971.5 |
| 2445 | 3766 | 3162 | 1081 | 4814 | 1861 | 639 | 8922 | 4389 | 5885 |
| 2426 | 4008 | 3327 | 1122 | 4913 | 1914 | 681 | 9415 | 4406 | 6376 |
| 2477.5 | 3781 | 3085.5 | 1064 | 4480.5 | 1793.5 | 639 | 9884.5 | 4293 | 6013 |
| 2544 | 4330.5 | 3345 | 1099 | 4710.5 | 1832.5 | 667 | 12216 | 4290 | 6589.5 |
| 2550 | 4185 | 3290 | 1138 | 4529 | 1844 | 649 | 12592 | 4203.5 | 6360 |
| 2608 | 3999 | 3184 | 1100 | 4405 | 1786.5 | 648 | 10797.5 | 4376 | 6181 |
| 2556 | 3824.5 | 3046 | 1036 | 4251.5 | 1747 | 634 | 10104.5 | 4416.5 | 5911.5 |
| 2515.5 | 4119 | 3279.5 | 1146 | 4642.5 | 1877 | 698 | 10182.5 | 4617 | 6558 |
| 2540 | 3840 | 3087 | 1103 | 4311 | 1816 | 658 | 9217 | 4503 | 6045 |
| 2391 | 3784.5 | 3100 | 1089 | 4320 | 1774 | 653 | 9612.5 | 4348.5 | 6004.5 |
| 2444 | 3918 | 3218 | 1097 | 4463 | 1806 | 677 | 10267 | 4499 | 6447 |
| 56451 | 31165 | 9279 | 1410 | 4546 | 1791 | 712 | 15804 | 5707 | 7402 |
| 3448 | 4728 | 3690 | 1142 | 4476 | 1806 | 758 | 13132 | 5587 | 7069 |
| 3172 | 4623.5 | 3628 | 1157 | 4685.5 | 1878 | 898 | 14308.5 | 5725 | 7421 |
| 3058 | 6027 | 3950 | 1290 | 5162 | 2007 | 970 | 16875 | 5632 | 9692 |
| 7936 | 14569.5 | 5248 | 1360 | 6060.5 | 2046 | 849 | 16749.5 | 7022.5 | 16247.5 |
| 16386 | 24731 | 7179 | 1756 | 7860.5 | 2421.5 | 880 | 18390.5 | 11018.5 | 23285.5 |
| 21739 | 29667.5 | 7653 | 1729 | 7935 | 2362.5 | 904.5 | 17229.5 | 10947 | 23357 |

| | | | | | | | | | |
|---|---|---|---|---|---|---|---|---|---|
| 14781 | 25079 | 7254 | 1674 | 7822 | 2319 | 829 | 17700 | 9225 | 23413 |
| 828 | 1088 | 915 | 607 | 1482 | 1132 | 231 | 2423 | 1517 | 1967 |
| 1170 | 1814 | 1583 | 705 | 2343 | 1303 | 405 | 4098 | 2219 | 3244 |
| 1456 | 2564.5 | 2282.5 | 896 | 3220.5 | 1535 | 547 | 6546 | 3036 | 4499 |
| 1585 | 2780 | 5285 | 1578 | 800680 | 81691 | 617 | 7341 | 3369 | 5043 |
| 1522 | 2594.5 | 2808 | 992 | 4717 | 2075.5 | 583 | 6858 | 3165 | 4623 |
| 1557 | 2689 | 2548 | 951 | 3764 | 1700 | 603 | 7194 | 3299 | 4829 |
| 1551 | 2681 | 2457.5 | 938 | 3556 | 1649 | 608 | 7128 | 3289 | 4820 |
| 1762 | 3025 | 2767 | 1021 | 3835 | 1732 | 648 | 8409 | 3625 | 5360 |
| 2137 | 3176 | 2728 | 990 | 3772 | 1669 | 620 | 9466 | 4005 | 5376 |
| 2348 | 3477 | 2904 | 1081 | 4046 | 1765 | 666 | 10099 | 4383 | 5791 |
| 2348 | 3665 | 3139 | 1114 | 4379 | 1859 | 713 | 10684 | 4510 | 6186 |
| 1919 | 3178 | 2735.5 | 1002 | 3816 | 1695 | 645 | 8740.5 | 3801 | 5415.5 |
| 2006 | 3329 | 2952 | 1097 | 4139.5 | 1780 | 693 | 8739.5 | 4116.5 | 5804 |
| 1754.5 | 2910 | 2599 | 988 | 3596 | 1664.5 | 624 | 7745.5 | 3479 | 5082 |
| 1849 | 3221 | 2893 | 1073 | 4063 | 1749 | 684 | 8608 | 3896 | 5671 |
| 2024 | 3442 | 3089 | 1114 | 4259 | 1854 | 699 | 8769 | 4259 | 6105 |
| 35847 | 18902 | 6521 | 1276 | 4196 | 1772 | 652 | 11271 | 4851 | 6382 |
| 2277 | 3834 | 3310 | 1108 | 4393 | 1813 | 665 | 9138 | 4538 | 6300 |
| 1949 | 3326 | 2971 | 1059 | 4106 | 1750 | 645 | 8546 | 3981 | 5833 |
| 1879.5 | 3260 | 2889.5 | 1055.5 | 4057 | 1748 | 656 | 8462.5 | 3968.5 | 5804.5 |
| 4216 | 11268 | 4435 | 1231 | 5447 | 1934 | 632 | 12915.5 | 5459.5 | 14058.5 |
| 8073.5 | 17307 | 6050.5 | 1553 | 7193.5 | 2244.5 | 657 | 16370 | 9291 | 20148.5 |
| 8954 | 19224.5 | 6515.5 | 1648 | 7627 | 2305 | 648.5 | 14507 | 9586.5 | 21740.5 |
| 8121 | 19164.5 | 6283 | 1552.5 | 7429.5 | 2242 | 651 | 15259 | 8974.5 | 21910.5 |
| 2904 | 3764 | 2169.5 | 742.5 | 2783 | 1288 | 319 | 8660 | 3858 | 4653 |
| 2551 | 3556 | 2401 | 833 | 3114 | 1442 | 463 | 8164 | 3631 | 4914 |
| 2037 | 3140 | 2466 | 886 | 3355 | 1524 | 550 | 7690 | 3508 | 4932 |
| 2080.5 | 3306 | 6104.5 | 1678 | 974918.5 | 87159.5 | 651 | 8835 | 3986 | 5559 |
| 1841 | 3044 | 3000.5 | 1049 | 4813.5 | 1936 | 625 | 7947.5 | 3560.5 | 5122 |
| 1724.5 | 2851.5 | 2607 | 954 | 3867.5 | 1667.5 | 610 | 7546 | 3409.5 | 4932 |
| 1833 | 3089 | 2787 | 1018.5 | 4068.5 | 1755 | 656 | 8136.5 | 3659.5 | 5366.5 |
| 1856 | 3010 | 2596 | 962 | 3641 | 1623 | 615 | 8300 | 3573 | 5081 |
| 2417 | 3497.5 | 2911 | 1042 | 4018 | 1724.5 | 655 | 9928.5 | 4394 | 5716.5 |
| 2517 | 3701 | 3087 | 1120 | 4216 | 1822 | 685 | 11043 | 4508 | 6007 |
| 2398 | 3602 | 3051 | 1083 | 4180 | 1781 | 676 | 10463 | 4424 | 6068 |
| 2273 | 3651 | 3111 | 1127 | 4306 | 1846 | 704 | 10160 | 4493 | 6211 |
| 1942 | 3204 | 2764 | 1023 | 3913 | 1698 | 651 | 8583 | 3802 | 5448.5 |
| 2009 | 3231.5 | 2899.5 | 1054 | 3947 | 1722 | 660 | 8406.5 | 3941 | 5638 |
| 2011 | 3380 | 2959 | 1077 | 4198 | 1789 | 699 | 8909 | 4155 | 6022 |
| 2199 | 3722.5 | 3229 | 1159 | 4535 | 1873 | 719 | 9894 | 4515.5 | 6464 |
| 40137 | 21267 | 7126 | 1329 | 4414 | 1840 | 656 | 11761 | 5046 | 6637 |
| 2471 | 4094.5 | 3482 | 1135 | 4388 | 1806 | 656 | 9743 | 4592 | 6342.5 |
| 2218 | 3665 | 3162 | 1126 | 4451 | 1830 | 668.5 | 9229.5 | 4485 | 6309.5 |
| 1913 | 3307.5 | 2858 | 1049 | 3945.5 | 1720 | 641 | 8471.5 | 3872 | 5650.5 |
| 4689 | 11222 | 4612 | 1315 | 5632.5 | 1985.5 | 658 | 14639 | 6291.5 | 14095.5 |
| 8215 | 17512 | 6206 | 1568 | 7325 | 2252 | 667 | 15699 | 9474 | 19972 |
| 8729 | 19734 | 6797 | 1636 | 7816 | 2327 | 668 | 14959 | 9480 | 22640 |

| | | | | | | | | | |
|---|---|---|---|---|---|---|---|---|---|
| 9001 | 20674 | 6691 | 1645 | 7946.5 | 2346.5 | 685 | 14475 | 9656.5 | 23182 |
| 3207.5 | 3880 | 2221 | 734 | 2845 | 1298 | 296 | 8880 | 4249 | 4816 |
| 2657 | 3597 | 2320 | 815 | 3043 | 1400 | 432 | 8195 | 3830 | 4817 |
| 2171 | 3242 | 2405 | 851.5 | 3205 | 1505 | 512 | 7444 | 3480.5 | 4780 |
| 2031 | 3193.5 | 5858 | 1773 | 939341.5 | 81308.5 | 588 | 8170 | 3635 | 5223.5 |
| 2052 | 3256 | 3366 | 1123 | 6185 | 2232 | 629 | 8346 | 3769 | 5417 |
| 2343 | 3707 | 3484 | 1187 | 5624 | 2092 | 719 | 9891 | 4568 | 6346 |
| 2183 | 3557 | 3188 | 1110 | 4797 | 1938 | 696 | 8918 | 4414 | 6123 |
| 2114.5 | 3320 | 2856 | 1015 | 4144.5 | 1728 | 639 | 8896.5 | 3960 | 5572.5 |
| 2315 | 3453 | 2869 | 1017 | 4168 | 1707 | 632 | 9286 | 4184 | 5618 |
| 2468 | 3507 | 2864.5 | 1081 | 3988 | 1756.5 | 634 | 9843 | 4444 | 5584 |
| 2473 | 3735 | 3103 | 1103.5 | 4298 | 1820 | 685.5 | 10629 | 4436 | 6124 |
| 2388 | 3742 | 3133 | 1127 | 4382.5 | 1847 | 702 | 10568.5 | 4617 | 6259.5 |
| 2270 | 3642 | 3152 | 1114 | 4464 | 1838 | 726 | 10724.5 | 4528 | 6165.5 |
| 2234 | 3471 | 3058 | 1087 | 4239 | 1828 | 696 | 9041 | 4277 | 5901 |
| 2105 | 3366 | 2953 | 1065 | 4180 | 1763 | 675 | 9112 | 4209 | 5828 |
| 2325 | 3836 | 3376 | 1178 | 4776 | 1933 | 728 | 9797 | 4774 | 6692 |
| 41748 | 25064 | 7760 | 1454 | 5063 | 1984 | 744 | 14615 | 5296 | 8198 |
| 2450 | 4138 | 3439 | 1172 | 4617 | 1881 | 684 | 9571 | 4677 | 6715 |
| 2281.5 | 3866 | 3333.5 | 1148 | 4620.5 | 1866 | 699 | 9731.5 | 4502 | 6581 |
| 2176 | 3526 | 3064 | 1095 | 4236 | 1785 | 675 | 9072 | 4321 | 6036 |
| 4754 | 11728 | 4755 | 1361 | 6000 | 2051 | 685 | 15004 | 6246 | 15102 |
| 8801 | 18764 | 6397 | 1654 | 7526 | 2342 | 695 | 15552 | 9806 | 21560 |
| 8378 | 18798 | 6342 | 1595 | 7446 | 2257 | 645 | 13621 | 8773 | 21055 |
| 9071 | 19815 | 6473 | 1614 | 7618 | 2299 | 668 | 15313 | 9994 | 22428 |
| 3237 | 4180 | 2400 | 769 | 3114 | 1368 | 322 | 9396 | 4396 | 5172 |
| 2828 | 3770 | 2509 | 866 | 3313 | 1494 | 475 | 8615 | 4108 | 5118 |
| 2407 | 3481.5 | 2638 | 927 | 3505.5 | 1548.5 | 554 | 8333.5 | 3990.5 | 5281 |
| 2551 | 3869 | 6581.5 | 1944 | 994244.5 | 86339.5 | 694 | 10328 | 4678 | 6331 |
| 2198 | 3434.5 | 3440.5 | 1146 | 5663.5 | 2090 | 670 | 8668 | 4113.5 | 5682 |
| 2004 | 3237 | 2990 | 1048 | 4624 | 1825 | 644 | 8569 | 3848 | 5526.5 |
| 1869 | 3057 | 2725 | 961 | 4003 | 1697 | 626 | 7950 | 3620 | 5206 |
| 2230 | 3375 | 2899 | 1055 | 4284 | 1759 | 655 | 9533 | 4268 | 5620 |
| 2446 | 3523 | 2879 | 1018 | 4052 | 1720 | 636 | 10251 | 4390 | 5661 |
| 2528 | 3905 | 3214 | 1134 | 4399 | 1868 | 683 | 10955 | 4532.5 | 6230.5 |
| 2472 | 3953 | 3261 | 1142 | 4489 | 1862 | 702 | 11207 | 4553 | 6347 |
| 2344.5 | 3670 | 3044.5 | 1086 | 4250.5 | 1777 | 688 | 9829.5 | 4472.5 | 6047 |
| 2264 | 3633 | 3109 | 1104.5 | 4412 | 1847 | 707 | 9454.5 | 4511 | 6201 |
| 2095 | 3390 | 2913 | 1062 | 4061 | 1762 | 658 | 8590 | 4044 | 5657 |
| 2223 | 3567 | 3093 | 1103 | 4343 | 1800 | 694 | 9432.5 | 4423.5 | 6104.5 |
| 2317 | 3846 | 3328 | 1175 | 4625 | 1914 | 706 | 9274 | 4653 | 6604 |
| 44747 | 23064 | 7437 | 1306 | 4313 | 1769 | 635 | 11989 | 5011 | 6566 |
| 2538 | 4237 | 3561 | 1161 | 4680 | 1901 | 688 | 9919 | 4826 | 6761 |
| 2331 | 3744 | 3180.5 | 1113 | 4451.5 | 1831 | 666 | 9308 | 4554 | 6345 |
| 2073 | 3452.5 | 2930 | 1049 | 4043 | 1746 | 646 | 8553.5 | 4072 | 5837.5 |
| 4561 | 11521 | 4670 | 1292 | 5701 | 1966 | 645 | 14320 | 5863 | 14482 |
| 8149 | 17307 | 6189.5 | 1572 | 7168.5 | 2264 | 659 | 15562 | 9242 | 19986 |
| 8989 | 19552 | 6512.5 | 1618.5 | 7631.5 | 2296 | 655 | 14573 | 9494.5 | 21763.5 |

| | | | | | | | | | |
|---|---|---|---|---|---|---|---|---|---|
| 8960 | 19080 | 6412.5 | 1565 | 7517.5 | 2262.5 | 655 | 14484.5 | 9663 | 22088 |
| 3260 | 4233.5 | 2442 | 785 | 3176.5 | 1371 | 322 | 9557.5 | 4358 | 5255.5 |
| 2923 | 3949.5 | 2543 | 868.5 | 3420 | 1489 | 457 | 9077.5 | 4285.5 | 5360 |
| 2392 | 3562.5 | 2651 | 915.5 | 3595.5 | 1573.5 | 545 | 8516 | 3975 | 5321 |
| 2553.5 | 3799.5 | 6250.5 | 1862.5 | 920865 | 79572 | 649 | 9800.5 | 4592 | 6073 |
| 2112 | 3409 | 3211 | 1104 | 5328 | 2002 | 624 | 8162 | 3906 | 5524 |
| 1981 | 3209.5 | 2933 | 1018 | 4526 | 1783 | 618 | 8012 | 3750 | 5383.5 |
| 1916.5 | 3200 | 2767.5 | 1001.5 | 4206 | 1732.5 | 618 | 7931 | 3686.5 | 5250.5 |
| 2065 | 3302.5 | 2789 | 1005 | 4078.5 | 1693 | 614 | 8754 | 3724 | 5413.5 |
| 2340 | 3522 | 2851 | 1015 | 4069 | 1688 | 624 | 9384.5 | 4190 | 5631.5 |
| 2510 | 3709 | 3022 | 1099 | 4112 | 1744 | 640 | 10561 | 4418 | 5804 |
| 2438 | 3982 | 3195 | 1108 | 4524 | 1851 | 681 | 11333 | 4435 | 6370 |
| 2375 | 3883 | 3228 | 1126 | 4587 | 1864 | 700 | 11079 | 4464 | 6334 |
| 2286.5 | 3648 | 3139.5 | 1120 | 4366.5 | 1832 | 693 | 10195.5 | 4431.5 | 6119.5 |
| 2320 | 3643.5 | 3122 | 1122 | 4274 | 1816 | 690 | 9532 | 4409 | 6011.5 |
| 2260 | 3792.5 | 3320 | 1158 | 4588 | 1889 | 730 | 10499.5 | 4489 | 6469.5 |
| 2266 | 3785 | 3207 | 1129 | 4538 | 1857 | 691 | 9165 | 4577 | 6373 |
| 39594 | 29797 | 7833.5 | 1444 | 5113.5 | 1985 | 721 | 15144.5 | 5082 | 9737 |
| 2468.5 | 4244 | 3522.5 | 1155.5 | 4576 | 1866.5 | 675 | 9689.5 | 4640 | 6687 |
| 2300 | 4149 | 3435.5 | 1166.5 | 4787.5 | 1906.5 | 695.5 | 10446 | 4520 | 6927.5 |
| 2210 | 3516 | 3025 | 1065 | 4118 | 1763 | 648 | 8851 | 4190 | 5916 |
| 4697 | 12020 | 4734 | 1321 | 5715 | 2001 | 654 | 14387 | 6014 | 15079 |
| 8177 | 16983 | 6215.5 | 1560.5 | 7330.5 | 2262 | 657 | 16112.5 | 9637.5 | 19969.5 |
| 8836 | 19707 | 6716 | 1653 | 7902 | 2336 | 660 | 15504 | 9879 | 22841 |
| 8453 | 18769 | 6235.5 | 1567 | 7544 | 2282.5 | 656 | 12655.5 | 9116 | 21062 |
| 3076 | 4163 | 2384 | 749 | 3129 | 1349 | 301 | 9305 | 4133 | 5143 |
| 2974 | 3902 | 2587 | 858 | 3404 | 1473 | 459 | 8694 | 4271 | 5276 |
| 2585 | 3711.5 | 2720 | 943 | 3729 | 1616 | 556.5 | 8693.5 | 4270 | 5442 |
| 2543 | 3867.5 | 6526.5 | 1980 | 944553 | 79464.5 | 663.5 | 10312.5 | 4534 | 6277 |
| 2205 | 3492 | 3450 | 1142 | 5947 | 2149 | 625 | 8468 | 4119 | 5665 |
| 2162 | 3471 | 3117 | 1067 | 4907 | 1897 | 644 | 8612 | 4029 | 5719 |
| 1939 | 3154 | 2774 | 993 | 4178 | 1705 | 610 | 7881 | 3620 | 5241 |
| 2196.5 | 3330.5 | 2836 | 1003 | 4122.5 | 1676 | 611 | 8906.5 | 3964 | 5430 |
| 2499.5 | 3751.5 | 3112 | 1060.5 | 4320 | 1762 | 644 | 10473 | 4326.5 | 5936 |
| 2445 | 3920.5 | 3154 | 1125 | 4502 | 1869 | 673.5 | 11822.5 | 4302 | 6206 |
| 2559.5 | 4168 | 3387 | 1158.5 | 4724 | 1880.5 | 717 | 11069.5 | 4562 | 6672 |
| 2416 | 3842 | 3216.5 | 1119 | 4445.5 | 1824 | 687 | 10726 | 4424 | 6248 |
| 2342 | 3738 | 3194 | 1120 | 4438 | 1848 | 692 | 9625 | 4507.5 | 6181 |
| 2275.5 | 3541 | 2952 | 1053 | 4149 | 1765 | 661 | 8954.5 | 4237.5 | 5849.5 |
| 2262 | 3652 | 3153.5 | 1110 | 4350.5 | 1821 | 696 | 9455 | 4436 | 6123.5 |
| 2354 | 4058 | 3424 | 1194 | 4838 | 1934.5 | 727 | 9930.5 | 4675 | 6996 |
| 43907 | 29288 | 8384 | 1482 | 5083 | 2002 | 708 | 13968 | 5145 | 8568 |
| 2633.5 | 4446 | 3660 | 1185 | 4607 | 1837 | 666 | 10060 | 4639 | 6718.5 |
| 2342 | 4443 | 3627 | 1206 | 4925 | 1951 | 707 | 10675 | 4528 | 7457 |
| 2305 | 3653 | 3113 | 1074 | 4201 | 1774 | 648 | 9176 | 4352 | 6042 |
| 5856.5 | 13498 | 5159.5 | 1448 | 6370 | 2124 | 693.5 | 15427 | 7630 | 16909.5 |
| 8250 | 18286 | 6406 | 1645 | 7663 | 2359 | 668 | 15360 | 9641 | 21399 |
| 8202 | 20787 | 6743 | 1682 | 7906 | 2341 | 661 | 15459 | 9174 | 23429 |

| | | | | | | | | | |
|---|---|---|---|---|---|---|---|---|---|
| 8813 | 22175 | 7251.5 | 1716 | 8448 | 2425.5 | 705 | 14307 | 9741 | 25320.5 |
| 3237 | 4519 | 2622 | 796 | 3443 | 1412 | 326 | 10603 | 4327 | 5574 |
| 2870 | 3875 | 2548 | 836 | 3392 | 1464 | 449 | 8809 | 4271.5 | 5231 |
| 2724 | 3986 | 2906 | 988 | 3973 | 1660 | 569 | 9605 | 4439 | 5892 |
| 2404 | 3564 | 6000 | 1783 | 939513 | 80200 | 610 | 8572 | 4113 | 5643 |
| 2360.5 | 3698 | 3620 | 1180 | 6261 | 2235.5 | 647 | 8746.5 | 4300 | 5963 |
| 2153 | 3472.5 | 3092 | 1063.5 | 4947 | 1875 | 623.5 | 8195 | 3929 | 5562 |
| 1998 | 3279 | 2872 | 1013 | 4334 | 1756 | 611 | 8137 | 3689 | 5358 |
| 2368 | 3535.5 | 2975.5 | 1023.5 | 4323 | 1783 | 635 | 9043.5 | 4274 | 5740 |
| 2508 | 3739 | 3014 | 1057 | 4314 | 1759 | 632 | 10440 | 4364 | 5834 |
| 2597 | 4033 | 3265 | 1189 | 4551 | 1897 | 668 | 10757 | 4429 | 6312 |
| 2522 | 4084 | 3248 | 1157 | 4629 | 1856 | 681 | 11749 | 4463 | 6395 |
| 2438.5 | 4040 | 3265 | 1148 | 4639 | 1867 | 694.5 | 10275.5 | 4530.5 | 6518 |
| 2387 | 3793 | 3178 | 1127 | 4437 | 1847 | 689 | 9881 | 4519 | 6218 |
| 2336 | 3573 | 3013 | 1053 | 4204 | 1786 | 664 | 9384 | 4337 | 5939 |
| 2369 | 3961 | 3371 | 1177 | 4651 | 1916 | 726 | 10398 | 4520 | 6618 |
| 2389 | 3995 | 3380 | 1156 | 4669 | 1896 | 697 | 9958 | 4593 | 6630 |
| 38986.5 | 24268.5 | 7458.5 | 1386.5 | 4883.5 | 1949.5 | 691.5 | 12354 | 5213.5 | 7927 |
| 2725.5 | 5063 | 3928.5 | 1275 | 4974.5 | 2003 | 702 | 10740 | 4700.5 | 7497 |
| 2434 | 3985 | 3311 | 1126 | 4582.5 | 1865 | 664 | 9490.5 | 4612 | 6461 |
| 2417.5 | 3887 | 3235 | 1139 | 4520 | 1853 | 684 | 9764.5 | 4540 | 6482 |
| 4820 | 11201 | 4612 | 1266 | 5670 | 1956 | 643 | 14490 | 6421 | 14111 |
| 8488 | 19114 | 6603.5 | 1659 | 7748.5 | 2359 | 665 | 15888.5 | 10047.5 | 22248 |
| 8613 | 20550 | 7022 | 1709 | 8417 | 2430 | 668 | 12481 | 9231 | 23342 |
| 8538.5 | 19771.5 | 6571.5 | 1632 | 7813.5 | 2324.5 | 667 | 14239 | 9446.5 | 22703.5 |
| 3232 | 4352 | 2435 | 764 | 3225 | 1387 | 304 | 9589.5 | 4296 | 5286 |
| 2996 | 4294 | 2808 | 893 | 3688 | 1560 | 474 | 10056.5 | 4359 | 5744.5 |
| 2782 | 3972 | 2872 | 959 | 3907 | 1640 | 545 | 9245.5 | 4425 | 5774.5 |
| 2570 | 3878 | 6149.5 | 1876 | 918059.5 | 77460 | 628 | 9717 | 4462 | 6038 |
| 2490 | 3971.5 | 3814 | 1234.5 | 6398 | 2277 | 674.5 | 9340 | 4470 | 6241 |
| 2180.5 | 3432 | 3117 | 1062 | 4880 | 1863 | 615 | 8419 | 3934.5 | 5496 |
| 2090 | 3432 | 2912.5 | 1032 | 4385 | 1767 | 625 | 8302.5 | 3812 | 5538 |
| 2158 | 3322 | 2810 | 997 | 4080 | 1696 | 604 | 8578 | 3751 | 5320 |
| 2530 | 3787 | 3039 | 1027 | 4344 | 1766 | 633 | 10078 | 4326 | 5958 |
| 2536.5 | 3784.5 | 3077.5 | 1104.5 | 4172 | 1801 | 629 | 10527.5 | 4325 | 5814 |
| 2536 | 4505.5 | 3468 | 1187.5 | 4838.5 | 1942 | 720 | 13147.5 | 4473.5 | 7079 |
| 2460.5 | 3882 | 3170 | 1109 | 4453 | 1826 | 673.5 | 10507.5 | 4462 | 6226.5 |
| 2378.5 | 4209.5 | 3372.5 | 1167 | 4692.5 | 1900 | 718 | 10728 | 4472 | 6851 |
| 2343 | 3543 | 2984 | 1056 | 4110.5 | 1751 | 645 | 8941 | 4250 | 5764.5 |
| 2366.5 | 4066 | 3350 | 1176 | 4791.5 | 1923 | 725 | 10573.5 | 4480 | 6659.5 |
| 2330 | 4168 | 3439 | 1199 | 4795 | 1927 | 710 | 10443 | 4459 | 6884 |
| 48193 | 31817 | 8824 | 1464 | 4896 | 1951 | 698 | 14404 | 5217 | 8444 |
| 2607 | 6127.5 | 3997 | 1274 | 5201 | 2001.5 | 714 | 12044 | 4600 | 9354 |
| 2480.5 | 4295 | 3514 | 1169 | 4799.5 | 1905 | 693 | 9842.5 | 4569.5 | 6951 |
| 2356.5 | 3903 | 3237 | 1147 | 4453.5 | 1832.5 | 666 | 9668 | 4428 | 6429.5 |
| 3869 | 8545 | 4357 | 1288 | 5587 | 2006 | 655 | 13741 | 5353 | 11313 |
| 8512.5 | 19620 | 6872 | 1639 | 8108.5 | 2357.5 | 671 | 15722.5 | 9727.5 | 22705.5 |
| 9036 | 20398 | 7017 | 1690 | 8324 | 2398 | 658 | 15554 | 10073 | 23284 |

| | | | | | | | | | |
|---|---|---|---|---|---|---|---|---|---|
| 8102 | 20631.5 | 6698 | 1672.5 | 7997.5 | 2344.5 | 665 | 11730.5 | 8907.5 | 23554.5 |
| 3307 | 4149 | 2408 | 763 | 3198 | 1377 | 295 | 8872 | 4329 | 5158 |
| 3023 | 4227 | 2736 | 879 | 3667 | 1526 | 461 | 9523 | 4364 | 5700.5 |
| 2679 | 3785.5 | 2794.5 | 911 | 3710 | 1600 | 523 | 8558.5 | 4181 | 5417.5 |
| 2717 | 3964 | 6475 | 1812 | 926493 | 75791 | 634 | 9852 | 4589 | 6089 |
| 2429.5 | 3780.5 | 3654 | 1178.5 | 5995 | 2190.5 | 642.5 | 8858 | 4334 | 6012.5 |
| 2206 | 3434 | 3006 | 1038.5 | 4688 | 1816 | 608.5 | 8285.5 | 3922 | 5484 |
| 2232 | 3541 | 3082 | 1039 | 4528.5 | 1798 | 639 | 8533.5 | 4046 | 5711 |
| 2295 | 3486 | 2867 | 999 | 4139 | 1715 | 609 | 8887 | 3988 | 5498 |
| 2540 | 3786 | 3039 | 1044 | 4286 | 1737 | 632 | 10458 | 4311 | 5892 |
| 2469.5 | 4002 | 3102 | 1080.5 | 4313.5 | 1805 | 645 | 10872.5 | 4086 | 6051 |
| 2477 | 3997 | 3244 | 1081 | 4496 | 1799 | 665 | 11738 | 4271 | 6241 |
| 2335 | 4054 | 3228 | 1126 | 4581 | 1878 | 688 | 10458 | 4272.5 | 6424 |
| 2388.5 | 3740 | 3113.5 | 1089 | 4368.5 | 1803 | 665 | 9777.5 | 4352 | 6109.5 |
| 2499 | 3788 | 3142 | 1127 | 4377 | 1833 | 671 | 9224 | 4564 | 6188 |
| 2362 | 3837 | 3257 | 1108 | 4515 | 1850 | 694 | 10022 | 4483 | 6334 |
| 2317 | 4531 | 3456.5 | 1198 | 4885 | 1936 | 714 | 10059.5 | 4426 | 7318 |
| 53338 | 32079 | 9446 | 1459 | 4894 | 1927 | 677 | 14008 | 5076 | 7485 |
| 2691 | 5234 | 3988 | 1245 | 4980 | 1957 | 688 | 10547 | 4502 | 7718 |
| 2456 | 4528 | 3563 | 1207 | 4906 | 1941 | 705 | 10593 | 4501 | 7363 |
| 2441 | 4135 | 3382 | 1154 | 4713 | 1920 | 689 | 9567 | 4549 | 6642 |
| 5182.5 | 13498.5 | 5195.5 | 1436 | 6439.5 | 2131 | 684 | 14654.5 | 6654 | 16761 |
| 8012 | 18212 | 6603 | 1672 | 7898 | 2400 | 693 | 14688 | 9262 | 21351 |
| 8553 | 20902 | 7334 | 1748 | 8701 | 2453 | 673 | 15169 | 9581 | 24079 |
| 8744 | 20709 | 6740 | 1649 | 7958 | 2361 | 660 | 12989 | 9512 | 23648 |
| 3262 | 4712 | 2701 | 827 | 3560 | 1459 | 332 | 10945 | 4287 | 5728 |
| 2998.5 | 3996.5 | 2597.5 | 858.5 | 3453 | 1489 | 429 | 8926 | 4297 | 5240 |
| 2792 | 4231 | 3007 | 999 | 4091.5 | 1659 | 566 | 10138.5 | 4486.5 | 6140 |
| 2632 | 3796 | 6087.5 | 1765 | 870989.5 | 76493 | 607 | 9161 | 4517.5 | 5869 |
| 2486.5 | 3852 | 3558.5 | 1163.5 | 5997.5 | 2188 | 643 | 9117 | 4310.5 | 6015.5 |
| 2347.5 | 3740 | 3217.5 | 1083 | 4976.5 | 1910 | 641.5 | 8912.5 | 4183 | 5941 |
| 2054.5 | 3316.5 | 2775.5 | 970 | 4089.5 | 1692 | 584 | 8062.5 | 3677.5 | 5233 |
| 2481 | 3710 | 3044 | 1039 | 4346 | 1757 | 636 | 9335 | 4431 | 5864 |
| 2583.5 | 3814 | 2979 | 1033 | 4247 | 1721 | 627.5 | 10430 | 4370 | 5860.5 |
| 2541 | 4004.5 | 3208.5 | 1136 | 4370 | 1854 | 650 | 11423 | 4333 | 6191 |
| 2395 | 4273 | 3376 | 1135 | 4707 | 1876 | 687 | 12447 | 4185 | 6611 |
| 2453 | 3848 | 3101 | 1083 | 4349 | 1794 | 655 | 10366 | 4400 | 6113 |
| 2450 | 4233 | 3453 | 1204 | 4841 | 1962 | 725 | 11827 | 4509.5 | 6906.5 |
| 2391 | 3621 | 2944 | 1048 | 4091 | 1730 | 641 | 9042 | 4270 | 5796 |
| 2462 | 4300 | 3417 | 1205 | 4822 | 1926 | 732 | 10725 | 4626 | 7072 |
| 2339 | 4100.5 | 3362 | 1161 | 4717 | 1873 | 688 | 10760 | 4324 | 6785 |
| 37423 | 29097.5 | 7277.5 | 1430 | 5117 | 1981.5 | 704 | 13439 | 5128 | 10121 |
| 2767 | 6858 | 4121 | 1315 | 5366 | 2063 | 732 | 12816 | 4730 | 10216 |
| 2416 | 4216 | 3440.5 | 1142 | 4729.5 | 1882 | 667.5 | 10078 | 4440 | 6828 |
| 2406 | 4357.5 | 3506.5 | 1199 | 4830 | 1932 | 704 | 11114 | 4516.5 | 7127 |
| 4057.5 | 9312 | 4495 | 1316 | 5671 | 1992 | 667 | 14351 | 5556.5 | 12066.5 |
| 8419.5 | 20546 | 6939 | 1751.5 | 8230 | 2444 | 686 | 15607.5 | 9661.5 | 24019 |
| 8575 | 22542 | 7553.5 | 1775 | 8962.5 | 2476 | 688 | 13399.5 | 9340.5 | 25613 |

| | | | | | | | | | |
|---|---|---|---|---|---|---|---|---|---|
| 8212 | 19886 | 6596 | 1633 | 7814.5 | 2344.5 | 668.5 | 12491.5 | 8978 | 22134 |
| 3128 | 4242 | 2449 | 761 | 3238 | 1373 | 300 | 9674 | 4178 | 5188 |
| 2996 | 4143 | 2656 | 868 | 3551 | 1509 | 430 | 9401 | 4279 | 5436 |
| 2743 | 3952 | 2858 | 960 | 3844 | 1603.5 | 533.5 | 8839 | 4352.5 | 5610.5 |
| 2764 | 3998 | 6207 | 1844 | 889972 | 72149 | 618 | 9598 | 4631 | 6060 |
| 2494 | 3949 | 3677.5 | 1188 | 5939 | 2181 | 644 | 9199.5 | 4413 | 6193 |
| 2450 | 3713 | 3137 | 1062 | 4838 | 1846 | 633 | 8769 | 4395 | 5884 |
| 2269 | 3570.5 | 3013 | 1058 | 4441 | 1809.5 | 627 | 8419.5 | 4153 | 5692 |
| 2529.5 | 3645.5 | 2975.5 | 1009 | 4242 | 1742 | 613 | 9256 | 4314 | 5694 |
| 2594 | 4312 | 3319 | 1098 | 4704.5 | 1858 | 673 | 10638 | 4381 | 6555.5 |
| 2650 | 4296 | 3397 | 1146 | 4646 | 1865 | 672 | 11870 | 4418 | 6583 |
| 2535 | 4221 | 3329 | 1116 | 4605 | 1849 | 683 | 12517.5 | 4307.5 | 6513.5 |
| 2358 | 3842 | 3067 | 1059 | 4273 | 1786 | 652 | 10104 | 4160 | 5957 |
| 2433 | 3699 | 3014 | 1048 | 4235 | 1742 | 653 | 9664 | 4378 | 5918 |
| 2101 | 3354 | 2810 | 972 | 3821 | 1647 | 602 | 8149 | 3731 | 5415 |
| 2373.5 | 3739.5 | 3123 | 1093 | 4383.5 | 1817 | 668 | 9220.5 | 4431 | 6045.5 |
| 2400 | 4319 | 3547 | 1207 | 4940 | 1972 | 715 | 10667 | 4531 | 7063 |
| 47954.5 | 28672.5 | 8488.5 | 1387 | 4736.5 | 1881 | 702 | 14844.5 | 5295 | 7716.5 |
| 3320 | 6831 | 3973 | 1232 | 4828 | 1913 | 851 | 15686 | 5348 | 9772 |
| 3133 | 4556 | 3500 | 1158 | 4595 | 1825 | 925 | 14938 | 5733 | 7343 |
| 3073 | 4127 | 3218.5 | 1078 | 4203.5 | 1748 | 839 | 13259.5 | 5488 | 6503 |
| 6694 | 12239 | 4670.5 | 1253 | 5430.5 | 1904.5 | 841 | 17474.5 | 6239 | 13831 |
| 17873 | 26928 | 7562.5 | 1797 | 8239.5 | 2484 | 978 | 17521.5 | 11284 | 24604.5 |
| 21549.5 | 30792 | 8231 | 1850.5 | 8552.5 | 2501 | 973 | 16855.5 | 10965.5 | 25128.5 |
| 20719.5 | 26027 | 7095.5 | 1593 | 7505.5 | 2255 | 890 | 18923.5 | 11403.5 | 22126 |
| 960 | 1565 | 1363 | 713 | 2107 | 1318 | 394 | 3465 | 1949 | 2822 |
| 1365 | 2362 | 2089 | 897 | 3074 | 1565 | 563 | 6253 | 2838 | 4317 |
| 1535 | 2905.5 | 2648 | 1033.5 | 3853 | 1735 | 683 | 7739.5 | 3454 | 5251.5 |
| 1396 | 2579 | 4994 | 1521 | 666946 | 55392 | 639 | 6933 | 3176 | 4826 |
| 1486 | 2703 | 2935 | 1054 | 5525 | 2049 | 672 | 7413 | 3402 | 5139 |
| 1571 | 2826 | 2822 | 1055 | 4674 | 1937 | 701 | 7672 | 3399 | 5247 |
| 1573 | 2853.5 | 2708 | 1054 | 4192 | 1815 | 697 | 7656 | 3611 | 5482.5 |
| 1598 | 2628 | 2307 | 886 | 3407 | 1590 | 600 | 7576 | 3240 | 4704 |
| 2415 | 4340.5 | 3070.5 | 1134 | 4457 | 1915.5 | 746 | 10971 | 4604 | 7406 |
| 2153 | 4031 | 3057 | 1137 | 4344 | 1847 | 754 | 11246 | 4241 | 7061 |
| 2233 | 4570 | 3287 | 1286 | 4851 | 2093 | 814 | 10953 | 4754 | 8038 |
| 2083 | 4099 | 3310 | 1271 | 4777 | 2061 | 844 | 10241 | 4723 | 7522 |
| 2144 | 5505 | 3745 | 1372 | 5337 | 2198 | 917 | 12403 | 4908 | 10382 |
| 13304 | 11019 | 4641 | 1229 | 4411 | 1948 | 778 | 9518 | 7065 | 8906 |
| 2531 | 3756 | 2954 | 1064 | 4082 | 1778 | 716 | 8314 | 4318 | 6153 |
| 2214 | 3464 | 2790 | 995 | 3732 | 1691 | 627 | 7747 | 3958 | 5781 |
| 68077.5 | 34155.5 | 10067.5 | 1460 | 4262.5 | 1807 | 663.5 | 12693 | 5862 | 6924.5 |
| 2577.5 | 3838 | 3262.5 | 1076 | 4178.5 | 1773.5 | 654 | 9091 | 4625.5 | 6333 |
| 2267 | 3647 | 3122 | 1146 | 4401 | 1858.5 | 723 | 9062.5 | 4411 | 6278.5 |
| 2395 | 3707.5 | 3092.5 | 1110 | 4190.5 | 1820 | 727 | 9063 | 4419.5 | 6173.5 |
| 6381 | 14824 | 5234 | 1446 | 6454 | 2159 | 687 | 14588 | 7740 | 17138 |
| 8072 | 22013 | 8173 | 1808 | 9766 | 2555 | 722 | 12419 | 8768 | 24594 |
| 7500 | 21449 | 9035 | 1968 | 10464 | 2693 | 775 | 12440 | 8717 | 23902 |

| | | | | | | | | | |
|---|---|---|---|---|---|---|---|---|---|
| 7186.5 | 17536.5 | 8368 | 1932 | 10190 | 2761.5 | 835 | 11166 | 8559 | 19992.5 |
| 2589 | 3574 | 2225 | 768 | 2995 | 1378 | 411 | 8045 | 3800 | 4829 |
| 2453 | 3563 | 2605 | 940 | 3635 | 1622 | 582 | 8531 | 4032 | 5439 |
| 2422 | 3693 | 2856 | 1055 | 3937 | 1739 | 648 | 9547 | 4291 | 5709 |
| 2348 | 4012 | 6072 | 1621 | 684840 | 53321 | 744 | 10900 | 4314 | 6491 |
| 1965 | 3250 | 2838.5 | 995 | 6405 | 1978 | 651 | 8287.5 | 3678.5 | 5378.5 |
| 2191.5 | 3413.5 | 2894 | 1090 | 5323 | 1922 | 683 | 8971.5 | 4238.5 | 5754.5 |
| 1994 | 3308 | 2758 | 1066 | 4669 | 1847 | 686 | 8764 | 3898 | 5722 |
| 2178 | 3241 | 2699 | 986 | 4200 | 1733 | 642 | 9188 | 3970 | 5516 |
| 2475 | 3572.5 | 2788.5 | 1013 | 4202.5 | 1742.5 | 661 | 10806.5 | 4372.5 | 5841 |
| 2811.5 | 6992 | 3728.5 | 1320 | 5444 | 2152 | 852.5 | 13110.5 | 5370.5 | 11784.5 |
| 2279 | 3750.5 | 2967 | 1099 | 4259 | 1878 | 706 | 9691.5 | 4395 | 6280.5 |
| 2242.5 | 5220.5 | 3575 | 1295.5 | 5098 | 2095.5 | 826 | 11911 | 4450 | 9099.5 |
| 2388 | 5579 | 3723 | 1362 | 5343 | 2230 | 889 | 12760 | 5041 | 9815 |
| 12759 | 10035 | 4154 | 1118 | 3864 | 1770 | 704 | 8579 | 6542 | 7559 |
| 2716 | 3854 | 2906 | 1032 | 3926 | 1739 | 667 | 8156 | 4271 | 5732 |
| 2481 | 3662 | 2910 | 1003 | 3894 | 1690 | 641 | 8247 | 4316 | 5854 |
| 63138.5 | 55805.5 | 10360.5 | 1500.5 | 4603 | 1838.5 | 677 | 15684.5 | 5442 | 11889.5 |
| 3211 | 10179 | 5076 | 1640 | 6858 | 2499 | 944 | 14981 | 5514 | 16255 |
| 2630 | 5537.5 | 3670 | 1307 | 5320.5 | 2059 | 810 | 10877.5 | 4832.5 | 9255.5 |
| 2519 | 4085 | 3325 | 1177 | 4471 | 1891 | 735 | 10279 | 4374 | 6466 |
| 6552 | 17625.5 | 7147 | 1789 | 8647.5 | 2562.5 | 807 | 13994 | 8093 | 21672 |
| 8025.5 | 23304.5 | 8031.5 | 1805 | 9625.5 | 2547 | 705 | 12743.5 | 8957.5 | 26140 |
| 8482 | 19088.5 | 9179 | 1984 | 10510.5 | 2807.5 | 795 | 12563.5 | 8562.5 | 21508 |
| 7225 | 18126 | 9462 | 2128 | 11373 | 2971 | 876 | 10850 | 8173 | 21416 |
| 2800 | 3786.5 | 2314 | 793 | 2960.5 | 1429 | 400 | 8174.5 | 3868 | 4928.5 |
| 2575.5 | 3597 | 2552 | 890.5 | 3449.5 | 1581 | 545 | 8531.5 | 4096.5 | 5363 |
| 2498 | 3659 | 2840 | 1016 | 3870 | 1735 | 649 | 9575 | 4249 | 5782 |
| 2380 | 5222 | 5899.5 | 1632.5 | 646655 | 51749.5 | 736 | 10860 | 4427 | 8577.5 |
| 2195 | 3517 | 3044 | 1069 | 6994 | 2072 | 676 | 8810 | 3967 | 5672 |
| 2160 | 3442 | 2898.5 | 1089 | 5855.5 | 1997 | 701 | 8830.5 | 3962 | 5685.5 |
| 2130.5 | 3460 | 2873.5 | 1058.5 | 5103 | 1876 | 703.5 | 9162.5 | 4079 | 5873 |
| 2356 | 3586 | 2860 | 1049 | 4524 | 1819 | 684 | 9738 | 4178 | 5842 |
| 2482 | 3674 | 2848 | 1012 | 4306 | 1771 | 670 | 10463 | 4305 | 5830 |
| 2386 | 4783.5 | 3030.5 | 1162 | 4314 | 1918 | 718 | 10966.5 | 4295.5 | 7634.5 |
| 2502 | 6441 | 4019 | 1443.5 | 5818.5 | 2305.5 | 891.5 | 13287.5 | 4873 | 11079 |
| 2636.5 | 6428.5 | 3930.5 | 1429.5 | 5692.5 | 2288.5 | 896.5 | 11524.5 | 5311.5 | 11254.5 |
| 2041 | 3771.5 | 2756.5 | 1085 | 4036 | 1810 | 719 | 9355.5 | 4089.5 | 6310 |
| 12573 | 8893 | 4090 | 1077.5 | 4028.5 | 1775.5 | 698.5 | 8736 | 6533.5 | 7261.5 |
| 2530 | 3739 | 2961 | 1015 | 3758 | 1668 | 665 | 8045 | 4085 | 5679 |
| 2584 | 4222 | 3062.5 | 1058 | 4152.5 | 1816.5 | 663 | 8754 | 4400 | 6670.5 |
| 54330 | 46663 | 9896 | 1676 | 5493 | 2126 | 786 | 16754 | 6037 | 12820 |
| 2412 | 5896 | 3357 | 1175 | 4478 | 1855 | 691 | 9962 | 4134 | 9245 |
| 2422.5 | 6579.5 | 3834.5 | 1368.5 | 5495 | 2134 | 817 | 12437.5 | 4511 | 10546 |
| 2702 | 4916 | 3535 | 1239 | 4811 | 1963 | 762 | 10550 | 4756 | 7617 |
| 6479.5 | 14700.5 | 5887.5 | 1541 | 7399.5 | 2309 | 741 | 15200.5 | 7922 | 17384.5 |
| 8485 | 22658 | 8499 | 1924 | 9982 | 2697 | 765 | 13686 | 9179 | 25328 |
| 8315 | 23219 | 8533 | 1963 | 10277.5 | 2676.5 | 772 | 12464 | 8655.5 | 26124.5 |

| | | | | | | | | | |
|---|---|---|---|---|---|---|---|---|---|
| 6447 | 17252 | 9261 | 2099 | 11286 | 2978 | 892 | 11240 | 7806 | 21127 |
| 2579 | 3584 | 2215 | 779.5 | 2990 | 1397 | 394 | 7818.5 | 3630.5 | 4738 |
| 2849 | 4197 | 2983 | 994 | 4002 | 1699 | 600 | 10137 | 4605 | 6138 |
| 2828 | 4767 | 3487 | 1179 | 4844 | 1939 | 728 | 12090 | 4766 | 7552 |
| 2439.5 | 4118.5 | 5819.5 | 1611 | 643893 | 52905.5 | 725.5 | 10677 | 4434.5 | 6429 |
| 2373 | 4209.5 | 3547.5 | 1203.5 | 7656 | 2283 | 744 | 10793 | 4408 | 6873.5 |
| 2494 | 6038.5 | 3868 | 1351.5 | 6854 | 2298.5 | 842.5 | 13743.5 | 4650.5 | 10011 |
| 2332 | 3902 | 3248.5 | 1131.5 | 5377 | 2000 | 739.5 | 10682 | 4439 | 6349 |
| 2278 | 4211 | 3076 | 1135 | 4938 | 1910 | 712 | 11994 | 4058 | 6599 |
| 2823 | 7128 | 3905 | 1337 | 5865 | 2175 | 825 | 14678 | 4926 | 11510 |
| 2704 | 5871 | 3803 | 1361.5 | 5322.5 | 2143.5 | 822.5 | 13011 | 4908.5 | 9471 |
| 2375.5 | 3985.5 | 3055 | 1141 | 4427 | 1832.5 | 718 | 10290.5 | 4427 | 6655.5 |
| 2084 | 3293 | 2714 | 1024 | 3794 | 1740 | 668 | 8651 | 3916 | 5512 |
| 1940 | 3095 | 2607 | 1010 | 3692 | 1680 | 651 | 8331 | 3817 | 5355 |
| 12845 | 9116 | 4056 | 1072 | 3740 | 1689 | 643 | 8434 | 6665 | 6819 |
| 2867.5 | 4818 | 3361.5 | 1123 | 4466 | 1855.5 | 723.5 | 9775 | 4471 | 7304.5 |
| 2779 | 7445 | 3939.5 | 1291.5 | 5259 | 2047 | 777 | 12041.5 | 4891 | 11755 |
| 57684 | 56546 | 11007 | 1652 | 5346 | 2077 | 756 | 17128 | 5677 | 13663 |
| 2448 | 7715 | 3946 | 1312 | 5207 | 2034 | 759 | 11650 | 4196 | 11842 |
| 2440 | 5314.5 | 3724 | 1262.5 | 5134.5 | 2058.5 | 782.5 | 11309 | 4360 | 8386.5 |
| 2815.5 | 6724 | 3854 | 1323.5 | 5166 | 2082 | 822 | 11305 | 4841 | 10641 |
| 6788 | 16120 | 5712 | 1582 | 6983 | 2241 | 724 | 15725 | 8310 | 18446 |
| 7677.5 | 21058.5 | 10385.5 | 2274 | 11896.5 | 3026 | 838.5 | 11470.5 | 8954.5 | 23998 |
| 8286 | 21978 | 9189 | 2040 | 10454 | 2749 | 778 | 13811 | 9132 | 24490 |
| 7022 | 20918 | 8901 | 1940 | 10432 | 2682 | 788 | 9459 | 8174 | 23869 |
| 2772 | 3791 | 2312 | 781 | 3222 | 1417 | 402 | 8570 | 3931 | 4845 |
| 2798.5 | 4073 | 2913.5 | 995.5 | 3931 | 1677.5 | 584 | 9630.5 | 4364.5 | 5817.5 |
| 2708 | 4249 | 3131 | 1126 | 4297 | 1856 | 690 | 11107 | 4486 | 6412 |
| 2437.5 | 5018 | 6250.5 | 1700.5 | 704441 | 53997 | 731 | 11530.5 | 4351.5 | 8050 |
| 2397 | 3732 | 3128 | 1067 | 6413 | 2002 | 681 | 10118 | 4293 | 6053 |
| 2392 | 3553 | 2940 | 1024 | 5322 | 1867 | 679 | 9652 | 4372 | 5917 |
| 2419.5 | 3578.5 | 2829 | 1048 | 4798 | 1874 | 696 | 9601.5 | 4433 | 5864 |
| 2571 | 4034 | 3068 | 1116 | 4716 | 1885 | 717 | 11382 | 4437 | 6385 |
| 2671 | 6148 | 3403 | 1153 | 5021 | 2007 | 777 | 14018 | 4481 | 9644 |
| 2441 | 5190 | 3482 | 1239 | 4886 | 2019 | 773 | 13999 | 4317 | 8327 |
| 3201 | 7560 | 4151 | 1481 | 5992 | 2318 | 944 | 14144 | 5923 | 12219 |
| 2444.5 | 4078.5 | 3315 | 1203 | 4568 | 1928.5 | 777 | 10277.5 | 4583.5 | 6778 |
| 1979 | 3131 | 2522 | 962 | 3590 | 1682 | 658 | 8477 | 3688 | 5345 |
| 19134.5 | 13559 | 5020 | 1081.5 | 3714 | 1652 | 671.5 | 8687 | 8260 | 8005 |
| 2862.5 | 5055 | 3538.5 | 1232 | 4627 | 1929.5 | 779 | 10051 | 4548 | 7840 |
| 2619 | 5232 | 3330 | 1130 | 4445 | 1850 | 711 | 9672 | 4490 | 8217 |
| 47092 | 44228 | 9868 | 1761 | 6448 | 2376 | 878 | 16372 | 6404 | 15455 |
| 3497 | 10464 | 5060 | 1637 | 6606 | 2456 | 930 | 14280 | 6067 | 16753 |
| 2852 | 6867 | 3751.5 | 1306.5 | 5238.5 | 2098.5 | 794.5 | 12016.5 | 5056.5 | 11010.5 |
| 3228 | 8112 | 4477 | 1534 | 6184 | 2427 | 941 | 15230 | 5580 | 12828 |
| 6229 | 14243 | 5333 | 1469.5 | 6529.5 | 2185 | 704 | 15788.5 | 7557.5 | 16408 |
| 8831 | 20124 | 9796 | 2278 | 11439.5 | 3030.5 | 837.5 | 12817 | 9326.5 | 22696.5 |
| 8378 | 23390 | 8772 | 1938 | 10242 | 2670 | 788 | 13970 | 9368 | 25761 |

| | | | | | | | | | |
|---|---|---|---|---|---|---|---|---|---|
| 7804 | 20875 | 9137 | 2048 | 10874 | 2822 | 878 | 11324 | 8620 | 22997 |
| 2910 | 3824 | 2283 | 832 | 3115 | 1399.5 | 388 | 8140 | 3902.5 | 4828.5 |
| 4884 | 4813 | 3158 | 974 | 14602 | 2438 | 600 | 10328 | 4753 | 5853 |
| 4012 | 4687 | 3234 | 1096 | 11297 | 2297 | 694 | 11403 | 4786 | 6386 |
| 5164.5 | 4746.5 | 5999 | 1631 | 287837 | 27276.5 | 688 | 10617 | 4467.5 | 6053.5 |
| 3662 | 4473 | 4961 | 1360 | 441731 | 49952 | 788 | 11661 | 4785 | 6569 |
| 3641 | 4222 | 4334 | 1238 | 320052 | 36781 | 769 | 10480 | 4954 | 6330 |
| 3419 | 4512 | 3979 | 1221 | 230168 | 26466 | 788 | 11934 | 4761 | 6807 |
| 3846 | 4173 | 3650 | 1116 | 211141 | 23699 | 738 | 11692 | 4910 | 6141 |
| 3507 | 7226 | 3866 | 1250 | 154776 | 17093 | 816 | 15673 | 4732 | 10600 |
| 3282 | 4833 | 3682 | 1260 | 28654 | 3815 | 808 | 14516 | 4742 | 7281 |
| 3618 | 6633.5 | 3778.5 | 1352 | 48139.5 | 5469.5 | 885.5 | 14349.5 | 5304.5 | 10239 |
| 3566 | 7503.5 | 4456 | 1601 | 18641.5 | 3499.5 | 1002.5 | 14777 | 5472 | 12051 |
| 3715 | 6365 | 4058 | 1502 | 29473 | 4030 | 969 | 13678 | 5585 | 9995 |
| 70673 | 53063.5 | 18548.5 | 3768 | 22870.5 | 3295 | 781 | 10564 | 23221.5 | 18666.5 |
| 9322 | 9229 | 4809 | 1389 | 22391 | 3316 | 881 | 11583 | 7525 | 9855 |
| 6375 | 8909 | 4212 | 1308 | 17722 | 3028 | 839 | 12702 | 5970 | 11699 |
| 643093 | 596449 | 117601 | 8082 | 17633 | 2824 | 688 | 72154 | 22377 | 24252 |
| 8857 | 15206 | 6189 | 1913 | 20151 | 3658 | 1002 | 17758 | 6794 | 18678 |
| 3960 | 6273 | 3944 | 1319 | 27472 | 5029 | 788 | 11512 | 5106 | 8684 |
| 4737 | 8607 | 4194 | 1386 | 17157 | 3069 | 842 | 14090 | 5049 | 11173 |
| 9581 | 18889 | 9343 | 2191 | 19336 | 3636 | 947 | 14929 | 9467 | 21390 |
| 8828 | 23444.5 | 10722 | 2351 | 18254 | 3544 | 900 | 10634.5 | 8217.5 | 26199.5 |
| 10985 | 26865.5 | 9417.5 | 2052.5 | 16571 | 3203 | 839 | 14180 | 9515.5 | 28097.5 |
| 8906 | 18986 | 10046 | 2334 | 16230 | 3490 | 962 | 11560 | 8730 | 21192 |
| 3297 | 4425.5 | 2727 | 866.5 | 3678 | 1524 | 450 | 9878 | 4556.5 | 5682 |
| 2836 | 4548 | 3135 | 1051 | 4259 | 1747 | 624 | 11206 | 4380 | 6418 |
| 2767 | 4356 | 3279 | 1098 | 4463.5 | 1841 | 676 | 11305.5 | 4441 | 6542.5 |
| 2534 | 5902 | 6580 | 1724 | 626126 | 52570 | 744 | 12438 | 4331 | 9165 |
| 2586 | 6071 | 3666 | 1273 | 7662 | 2289 | 782 | 12623 | 4518 | 9578 |
| 2418 | 4197.5 | 3380 | 1188 | 5992 | 2061.5 | 731 | 11013.5 | 4353.5 | 6773 |
| 2360 | 5605 | 3492 | 1187 | 5581 | 2062 | 761 | 11670 | 4405 | 8997 |
| 2498 | 4060 | 3165 | 1095.5 | 4814.5 | 1862 | 702 | 11378 | 4335.5 | 6356 |
| 2664 | 6737 | 3669 | 1244 | 5326 | 2091 | 776 | 14558 | 4458 | 10537 |
| 2707.5 | 6865 | 3796 | 1310 | 5395.5 | 2163.5 | 801 | 15029.5 | 4699.5 | 10686.5 |
| 2772 | 6054 | 3885 | 1371 | 5489 | 2204 | 850 | 13145 | 5192 | 9649 |
| 2455 | 5238 | 3547 | 1258 | 5047 | 2035 | 797 | 11573 | 4688 | 8561 |
| 2348 | 5397 | 3340 | 1185 | 4761 | 1934 | 777 | 11130 | 4393 | 9131 |
| 14831 | 10872 | 4325 | 1058 | 3770 | 1657 | 659 | 8599 | 6742 | 7322 |
| 2964 | 5113 | 3135 | 1072 | 4136 | 1777 | 682 | 9344 | 4730 | 7588 |
| 2724 | 4276 | 3302 | 1077 | 4247 | 1760 | 665 | 9519 | 4526 | 6717 |
| 50036 | 48238 | 9913.5 | 1695 | 5801 | 2160 | 790.5 | 17504 | 5908.5 | 14024.5 |
| 2589 | 7935 | 3920 | 1263.5 | 5058.5 | 2039.5 | 748 | 12354 | 4214 | 12070.5 |
| 2878 | 8719 | 4346 | 1513 | 6277 | 2335 | 883 | 15054.5 | 5086 | 13870 |
| 2852.5 | 6156 | 3866.5 | 1235 | 5180 | 2017 | 777.5 | 11369.5 | 4719.5 | 9435.5 |
| 7736 | 18454 | 7914 | 1939 | 9642 | 2668 | 852 | 14673 | 9537 | 21518 |
| 7714.5 | 19726 | 8844 | 1984.5 | 10260 | 2739.5 | 766.5 | 12021 | 8607 | 22036 |
| 8559 | 23228 | 8501 | 1917.5 | 9797 | 2624 | 765 | 13462 | 9047 | 25701 |

| | | | | | | | | | |
|---|---|---|---|---|---|---|---|---|---|
| 7859 | 21991 | 8460.5 | 1881 | 10197 | 2584 | 796.5 | 10214 | 7940.5 | 25293.5 |
| 3151 | 4369 | 2688 | 859 | 3484 | 1523 | 439 | 10079 | 4345 | 5566 |
| 2862.5 | 4572 | 3189.5 | 1078 | 4314 | 1800 | 585 | 10301 | 4441.5 | 6786 |
| 2620 | 6244 | 3527.5 | 1253.5 | 5089 | 2094.5 | 690 | 11657 | 4716 | 10327 |
| 2456 | 6851.5 | 7239 | 1805.5 | 563771 | 47060 | 747 | 13389 | 4707 | 12192 |
| 2282 | 6256 | 3816 | 1398 | 7864 | 2578 | 726 | 11877 | 4454 | 11656 |
| 2405 | 6972.5 | 3956.5 | 1487 | 7206.5 | 2619.5 | 748 | 13192 | 4643.5 | 13184.5 |
| 2166 | 6734 | 3851 | 1458 | 6552 | 2466 | 731 | 11978 | 4216 | 13266 |
| 2241.5 | 5790.5 | 3371 | 1283 | 5596.5 | 2168 | 666 | 11850 | 4341.5 | 10739.5 |
| 2462 | 7740 | 4003 | 1469 | 6261 | 2439 | 753 | 14780 | 4638 | 13942 |
| 2742 | 7684 | 4542 | 1627.5 | 6717 | 2608.5 | 813 | 14134.5 | 5171.5 | 13749.5 |
| 2430 | 5716 | 3850 | 1420 | 5702 | 2344 | 758 | 11882 | 4766 | 10492 |
| 2446.5 | 4201.5 | 3353.5 | 1257.5 | 4926 | 2125 | 702 | 10121 | 4864 | 7762.5 |
| 2174 | 4900 | 3318 | 1240 | 4867 | 2089 | 694 | 9692 | 4388 | 9343 |
| 5898.5 | 5441.5 | 3649 | 1226 | 4591 | 1988 | 641 | 8976 | 5451.5 | 7230.5 |
| 2917.5 | 7631.5 | 4043.5 | 1445 | 5887 | 2378.5 | 768 | 12040.5 | 5008.5 | 13415.5 |
| 2497.5 | 6693 | 3722 | 1336 | 5428 | 2184 | 672 | 10345.5 | 4616 | 11734.5 |
| 53975 | 50779.5 | 10927 | 2111.5 | 8053.5 | 2950.5 | 876 | 17811.5 | 7628.5 | 19101.5 |
| 2791 | 8977 | 4599 | 1569 | 6348 | 2496 | 772 | 14432 | 4878 | 15228 |
| 3300 | 9636 | 5122 | 1821 | 7811 | 2931 | 896 | 15649 | 6235 | 17387 |
| 2908 | 7588 | 4201 | 1495 | 6199 | 2439 | 779 | 11867 | 5345 | 13296 |
| 7460 | 19783 | 7965 | 2141 | 10228.5 | 3037.5 | 801 | 13994.5 | 9121.5 | 24755.5 |
| 8104 | 19465 | 8759 | 2134 | 10400 | 3028 | 742 | 14070 | 9110 | 22820 |
| 8428 | 19738 | 8890 | 2132 | 10583 | 2957 | 757 | 12432 | 8995 | 23095 |
| 7728 | 20557 | 8970 | 2186 | 10854 | 3093 | 805 | 12512 | 9123 | 25028 |
| 3058 | 4056 | 2506 | 849 | 3373 | 1460 | 402 | 8836 | 4121 | 5103 |
| 3402.5 | 4826 | 3265.5 | 1075.5 | 6653 | 2010.5 | 581.5 | 12373 | 4389.5 | 6362.5 |
| 3147 | 5084 | 3445 | 1182 | 9334 | 2325 | 681 | 11898 | 4495 | 7142 |
| 3375 | 8969 | 7782 | 2274 | 410801 | 58584 | 768 | 14294 | 4750 | 12337 |
| 3448.5 | 9270.5 | 5279.5 | 1521 | 169778.5 | 25579.5 | 807 | 15674.5 | 4991 | 12982 |
| 3183.5 | 4747 | 3958.5 | 1243.5 | 116252.5 | 16560.5 | 692.5 | 11266 | 4666.5 | 6699.5 |
| 3547 | 8218 | 4294 | 1385 | 63471 | 8528 | 801 | 17101 | 6564 | 14589 |
| 2995 | 4921 | 3641 | 1212 | 54586 | 7208 | 716 | 13122 | 4337 | 7201 |
| 3151 | 6710 | 3905 | 1253 | 38170 | 5184 | 771 | 14146 | 4440 | 9372 |
| 3743 | 12265.5 | 7846 | 2425 | 25017.5 | 4470 | 882 | 15082.5 | 5248.5 | 15754 |
| 3410 | 9758 | 5056 | 1618 | 20426 | 3815 | 940 | 14912 | 5429 | 14178 |
| 2849.5 | 6329 | 4190.5 | 1367 | 10750.5 | 2667 | 819.5 | 12659 | 4765.5 | 9589.5 |
| 3480 | 5064.5 | 4184.5 | 1273.5 | 12139 | 2569.5 | 723 | 10129.5 | 4854.5 | 6704 |
| 30881 | 28300.5 | 13006.5 | 1834 | 11117 | 2494.5 | 789 | 9876.5 | 13509.5 | 14662 |
| 4834 | 6048.5 | 4162 | 1221 | 10201 | 2356.5 | 718 | 9808.5 | 5510 | 7785.5 |
| 4352 | 7386.5 | 3799.5 | 1181.5 | 9414.5 | 2291 | 707 | 11073 | 5356.5 | 9937.5 |
| 371607.5 | 349812.5 | 81301 | 6727 | 11421.5 | 2953 | 850.5 | 42380 | 14264.5 | 21202.5 |
| 8545.5 | 14671 | 5625.5 | 1589 | 9590.5 | 2557 | 820.5 | 15438 | 5969.5 | 14840.5 |
| 5811 | 12375 | 5572 | 1569 | 44707 | 8009 | 887 | 14977 | 6681 | 15070 |
| 7505 | 15802 | 6899 | 1937 | 10734 | 2897 | 828 | 16380 | 7102 | 15775 |
| 10037 | 23459 | 8971 | 2091 | 13527 | 3159 | 866 | 14747 | 9643 | 24549 |
| 9835 | 26362 | 9607 | 2157.5 | 14082 | 3162.5 | 771 | 11688.5 | 8731 | 27451.5 |
| 9711 | 24988 | 8747 | 1933 | 12592 | 2867 | 752 | 14339 | 9684 | 26926 |

| | | | | | | | | | |
|---|---|---|---|---|---|---|---|---|---|
| 9092 | 21264.5 | 9104 | 1948 | 13055.5 | 2870.5 | 793.5 | 10559 | 8348 | 23064 |
| 2967.5 | 3868.5 | 2446 | 826.5 | 3324.5 | 1484 | 398.5 | 9344 | 4146 | 4997 |
| 2780 | 4125 | 2984 | 1027 | 4057 | 1686 | 579 | 10389 | 4363 | 5928 |
| 2677.5 | 4565 | 3370 | 1155.5 | 4599.5 | 1891 | 707 | 12188.5 | 4476.5 | 6890.5 |
| 2508 | 3924 | 5205 | 1471 | 554852 | 49380 | 662 | 10007 | 4304 | 5998 |
| 2520 | 6051 | 3514 | 1206 | 7066 | 2218 | 737 | 11357 | 4388 | 9465 |
| 2398 | 4152 | 3484.5 | 1179 | 5875 | 2058 | 751 | 10912 | 4396 | 6729 |
| 2440 | 3761 | 3160 | 1144 | 5128 | 1930 | 711 | 9978 | 4476 | 6164 |
| 2501 | 3566 | 2807 | 994 | 4240 | 1702 | 640 | 9742 | 4486 | 5608 |
| 2641.5 | 7216.5 | 3398.5 | 1161.5 | 5092.5 | 1944.5 | 734 | 13745.5 | 4401 | 11176 |
| 2608 | 7411.5 | 3831.5 | 1371 | 5459 | 2163.5 | 797 | 14703.5 | 4579 | 11535 |
| 2448 | 4625 | 3319 | 1188 | 4709 | 1897 | 747 | 11382 | 4328 | 7521 |
| 2806 | 5963 | 3815 | 1375 | 5337 | 2210 | 850 | 12162 | 5352 | 9521 |
| 2757 | 4904 | 3530 | 1240 | 5020 | 2020 | 799 | 10903 | 5234 | 8042 |
| 13518 | 9874.5 | 4305.5 | 1115 | 4053 | 1773 | 658.5 | 8661 | 6577 | 7374.5 |
| 2867 | 3923 | 2961 | 1000 | 3918 | 1691 | 641 | 8196 | 4394 | 5847 |
| 2706 | 5651 | 3548 | 1178 | 4763 | 1864 | 696 | 10312 | 4468 | 8805 |
| 76918 | 62482.5 | 12265.5 | 1549 | 4601.5 | 1882 | 672 | 16261.5 | 5962.5 | 11686.5 |
| 3001 | 8313 | 4036 | 1279 | 5077 | 2024 | 744 | 13266 | 4772 | 12252 |
| 3042 | 8948 | 4587 | 1514 | 6287 | 2371 | 881 | 14991 | 5268 | 14567 |
| 2783 | 4270 | 3232 | 1124.5 | 4375.5 | 1797 | 684.5 | 10034.5 | 4482.5 | 6512 |
| 6988 | 18341 | 8774 | 2092 | 10600 | 2934 | 872 | 12511 | 8420 | 21667 |
| 8337 | 22868 | 8323 | 1911 | 9778 | 2607 | 727 | 11978 | 8856 | 25931 |
| 8121.5 | 19746 | 9491.5 | 2150 | 11282.5 | 2834 | 817 | 11843.5 | 8831.5 | 22405.5 |
| 7802 | 20399 | 9052 | 2096 | 10708 | 2870 | 837 | 12230 | 9464 | 23629 |
| 3220.5 | 4241 | 2555 | 843 | 3399.5 | 1468.5 | 392.5 | 9003 | 4235 | 5298 |
| 2921 | 4597.5 | 3183 | 1038.5 | 4314.5 | 1720.5 | 592 | 11565.5 | 4276.5 | 6375 |
| 2872.5 | 6655.5 | 3801 | 1247 | 5109.5 | 1991 | 741 | 14119 | 4656.5 | 9796 |
| 2531 | 4151 | 5458 | 1517 | 557639 | 50781 | 661 | 10634 | 4192 | 6201 |
| 2512 | 6571 | 3649 | 1200.5 | 6774 | 2194 | 741 | 11660 | 4124.5 | 10039 |
| 2624 | 6190 | 3726 | 1258 | 6189 | 2158 | 771 | 12856 | 4653 | 9766 |
| 2499.5 | 4218.5 | 3346 | 1195 | 5272.5 | 1986 | 739 | 11003.5 | 4344.5 | 6610.5 |
| 2612.5 | 6661.5 | 3403.5 | 1148 | 4940.5 | 1922 | 734 | 12824.5 | 4344.5 | 10353 |
| 3325.5 | 8273 | 4142 | 1373 | 5973.5 | 2168.5 | 807.5 | 14705.5 | 5549.5 | 12441.5 |
| 2595.5 | 6685.5 | 3658.5 | 1273.5 | 5051 | 2028 | 763.5 | 14857 | 4373.5 | 10246.5 |
| 2440.5 | 6144 | 3456.5 | 1188 | 4751 | 1944.5 | 727 | 13066 | 4345.5 | 9666.5 |
| 2416.5 | 3769.5 | 2979.5 | 1077.5 | 4182.5 | 1760 | 679 | 9751 | 4297 | 6044.5 |
| 2373 | 3756 | 2981 | 1085.5 | 4186 | 1770 | 688 | 9459.5 | 4314 | 6052 |
| 10861 | 8911 | 4419 | 1181 | 4496 | 1861 | 730 | 9528 | 6451 | 7925 |
| 3184.5 | 6569.5 | 3881 | 1290 | 5206.5 | 2065 | 787.5 | 12269.5 | 5005.5 | 9697 |
| 2704 | 5649 | 3487 | 1160 | 4590 | 1868 | 692 | 10020 | 4366 | 8782 |
| 66791.5 | 55755 | 11668.5 | 1838 | 6188.5 | 2307 | 823 | 18592.5 | 6828.5 | 15045.5 |
| 2705 | 7536 | 3793 | 1200 | 4787 | 1887 | 700 | 11832 | 4309 | 11429 |
| 3073 | 5175 | 3609 | 1201 | 4951 | 1968 | 728 | 10999 | 5192 | 8077 |
| 2884 | 5247 | 3673 | 1218 | 4910 | 1994 | 739 | 10822 | 4600 | 7774 |
| 6826 | 18364 | 8711 | 2087 | 10849 | 2913 | 890 | 13957.5 | 8486.5 | 22118 |
| 9432 | 21027 | 7116 | 1743 | 8307 | 2408 | 684 | 14736 | 9670 | 22328 |
| 7941 | 19448 | 8711 | 1940 | 10336 | 2676 | 769 | 12811 | 8488 | 21258 |

| | | | | | | | | | |
|---|---|---|---|---|---|---|---|---|---|
| 8661 | 18906 | 6343 | 1555 | 7234 | 2216 | 693 | 12446 | 9037 | 21475 |
| 935.5 | 1473.5 | 1322 | 646.5 | 2040 | 1230 | 352 | 3406.5 | 1939.5 | 2746.5 |
| 1326 | 2326 | 2094 | 854 | 3004 | 1487 | 528 | 6149 | 2820 | 4200 |
| 1586 | 2956 | 2737 | 1034 | 3956 | 1739 | 684 | 8145 | 3597 | 5427 |
| 1704 | 3033 | 5408 | 1626 | 665531 | 54649 | 718 | 8250 | 3887 | 5690 |
| 1678 | 2952 | 3025 | 1125 | 4721 | 1972 | 717 | 8170 | 3846 | 5587 |
| 1609 | 2835 | 2728 | 1032 | 4089 | 1834 | 695 | 7980 | 3648 | 5277 |
| 1620 | 2810 | 2669 | 1027 | 3941.5 | 1786.5 | 681 | 8050 | 3671.5 | 5227 |
| 1867 | 3012.5 | 2664 | 1028 | 3756.5 | 1733 | 657 | 8752 | 3810 | 5233.5 |
| 2244.5 | 3614 | 2996.5 | 1138 | 4178.5 | 1870.5 | 721 | 11459.5 | 4215.5 | 6087.5 |
| 2188 | 3663 | 3161 | 1237 | 4487 | 2003 | 750 | 11714 | 4370 | 6268 |
| 2074 | 3672 | 3071 | 1198 | 4452 | 1935 | 764 | 11191 | 4237 | 6439 |
| 1969 | 3412 | 3048 | 1135 | 4347 | 1914 | 753 | 10384 | 4348 | 6290 |
| 1874 | 3301 | 3013 | 1142 | 4362 | 1923 | 779 | 9325 | 4230 | 6219 |
| 15633 | 11217 | 4408 | 1070 | 3633 | 1709 | 656 | 7833 | 6790 | 6991 |
| 2323 | 3328 | 2740 | 1070 | 3789 | 1792 | 679 | 8330 | 3970 | 5547 |
| 1980 | 3066 | 2536 | 985 | 3452 | 1664 | 608 | 7321 | 3718 | 5197 |
| 66541.5 | 41141 | 10468 | 1383 | 3976.5 | 1721.5 | 631 | 13675 | 5267 | 7400 |
| 3249 | 9423 | 5067 | 1628 | 6193 | 2416 | 895 | 12935 | 5407 | 14126 |
| 2444 | 6578 | 4012 | 1397 | 5924 | 2320 | 864 | 13671 | 5060 | 11443 |
| 2311.5 | 3692.5 | 3197 | 1164 | 4363 | 1900.5 | 749.5 | 10115.5 | 4522.5 | 6417.5 |
| 7034.5 | 16659 | 6511 | 1744 | 7957 | 2465 | 819 | 14335.5 | 9269.5 | 20812.5 |
| 7482.5 | 23251 | 9042.5 | 2186 | 11142 | 2913.5 | 818 | 12176 | 8742.5 | 26789 |
| 8460 | 22705.5 | 7409 | 1719.5 | 8494 | 2430 | 696 | 13793.5 | 9070 | 25766.5 |
| 7371 | 20995.5 | 6826 | 1560 | 7961.5 | 2275 | 717.5 | 9966 | 8201 | 23707.5 |
| 2498 | 3627.5 | 2279 | 770.5 | 3032 | 1418 | 411 | 7636 | 3816.5 | 4807.5 |
| 2398 | 3411.5 | 2637.5 | 928 | 3609.5 | 1611 | 567 | 8169 | 4014 | 5240.5 |
| 2254.5 | 3437 | 2777.5 | 1016.5 | 3870 | 1691.5 | 637 | 8646.5 | 4195 | 5582.5 |
| 2222 | 3394 | 5400 | 1540 | 629033 | 53743 | 679 | 8927 | 4228 | 5717 |
| 2249 | 3575 | 3379 | 1183 | 6002 | 2172 | 745 | 10154 | 4275 | 6327 |
| 2084 | 3153 | 2785 | 1047 | 4695 | 1844 | 669 | 8841 | 4095 | 5466 |
| 1829 | 3022 | 2607 | 970 | 4118 | 1704 | 654 | 8011 | 3642 | 5196 |
| 2178 | 3160 | 2581 | 963 | 3834 | 1679 | 632 | 9120 | 3971 | 5140 |
| 2381 | 4458 | 3200 | 1141 | 4592 | 1881 | 731 | 12413 | 4239 | 7359 |
| 2414.5 | 4069 | 3320 | 1194 | 4583 | 1924 | 737 | 12290.5 | 4345 | 6623.5 |
| 2472.5 | 6025 | 3696 | 1347.5 | 5319 | 2149.5 | 835.5 | 14300 | 4725.5 | 10294.5 |
| 2366 | 3956 | 3147.5 | 1224 | 4638.5 | 1942.5 | 792 | 11984 | 4857.5 | 6938.5 |
| 2375 | 4595 | 3509 | 1301 | 5085 | 2093 | 845 | 11147 | 4880 | 7958 |
| 15744.5 | 11264 | 4657 | 1194 | 4055.5 | 1796 | 699 | 9824 | 7345.5 | 7781 |
| 2740 | 4197 | 3119 | 1129 | 4202 | 1892 | 728 | 10160 | 4623 | 6641 |
| 2259 | 3459 | 2656 | 987 | 3735.5 | 1627 | 621 | 8445.5 | 4039.5 | 5546 |
| 57304.5 | 57035.5 | 10722 | 1722 | 5767.5 | 2182.5 | 792 | 18823.5 | 5931 | 14951 |
| 3737 | 9998 | 4992.5 | 1753.5 | 6800.5 | 2601.5 | 971 | 14682 | 6612.5 | 15975.5 |
| 2912 | 7143 | 4360 | 1459.5 | 6332.5 | 2345 | 895.5 | 13617 | 5464.5 | 12004.5 |
| 2334 | 6421 | 3440 | 1237 | 4960 | 2004 | 777 | 12369 | 4207 | 10346 |
| 7199 | 16813 | 8374 | 1982 | 10340 | 2773 | 858 | 13423 | 9331 | 20032 |
| 8021 | 23897 | 7863 | 1959 | 9405 | 2717 | 782 | 11597 | 8852 | 26779 |
| 8410.5 | 23873.5 | 8217 | 1803 | 9432.5 | 2450 | 708.5 | 11931.5 | 8631.5 | 26567 |

| | | | | | | | | | |
|---|---|---|---|---|---|---|---|---|---|
| 7352 | 22350 | 9194 | 1952 | 10790 | 2744 | 818 | 9267 | 7151 | 24345 |
| 2684.5 | 3616.5 | 2250.5 | 808.5 | 2983.5 | 1388 | 393 | 8004 | 3848.5 | 4825 |
| 2433 | 3464 | 2514 | 886 | 3463 | 1551 | 548 | 8314 | 3978 | 5088 |
| 2260 | 3316 | 2599 | 985 | 3677 | 1665 | 630 | 8350 | 4040 | 5265 |
| 2333 | 4633 | 6029.5 | 1644 | 653664 | 52116.5 | 708 | 10941.5 | 4350.5 | 7673.5 |
| 2355 | 3910.5 | 3333 | 1185 | 6264.5 | 2182 | 742 | 10376.5 | 4374 | 6441 |
| 2293 | 3632 | 3100 | 1106 | 5128 | 1951 | 702 | 9651 | 4361 | 6072 |
| 2254 | 3470 | 2874 | 1070 | 4630 | 1839 | 683 | 9645 | 4267 | 5848 |
| 2381 | 3463 | 2790 | 1023 | 4208 | 1767 | 660 | 10304 | 4241 | 5760 |
| 2487 | 4441 | 3244 | 1168 | 4876 | 1936 | 745 | 13333 | 4431 | 7046 |
| 2280 | 4674 | 3355 | 1233 | 4768 | 1956 | 754 | 12907 | 4258 | 7522 |
| 2374 | 4095 | 3222 | 1182 | 4727 | 1968 | 757 | 11060 | 4493 | 6859 |
| 2284.5 | 3506.5 | 2869.5 | 1067.5 | 4080.5 | 1770 | 710 | 9802 | 4345.5 | 5980.5 |
| 2009 | 3212 | 2742 | 1007 | 3813 | 1734 | 668 | 8679 | 4003 | 5535 |
| 14214 | 10382 | 4159 | 994 | 3388 | 1559 | 624 | 8053 | 6212 | 6684 |
| 2771 | 3786 | 3010 | 1060 | 4069 | 1739 | 685 | 9419 | 4597 | 6014 |
| 2393 | 4602 | 2915 | 1076 | 3912 | 1696 | 660 | 9620 | 4113 | 7519 |
| 58136 | 50551 | 10444 | 1622 | 4904 | 1940 | 718 | 17110 | 5849 | 12142 |
| 3812 | 9744 | 5178 | 1677 | 7011 | 2539 | 936 | 14793 | 6834 | 14616 |
| 2904 | 7641 | 4109.5 | 1414 | 5974 | 2238 | 858 | 13853.5 | 5215.5 | 12484 |
| 2554.5 | 6536.5 | 3802 | 1338 | 5215.5 | 2102.5 | 817 | 13362 | 4578 | 10727 |
| 6916 | 18656 | 7202 | 1803 | 8927 | 2580 | 789 | 14356 | 8791 | 21879 |
| 8122 | 24295 | 8661 | 1990 | 10403 | 2735 | 783 | 13343 | 9080 | 27447 |
| 7729 | 22479 | 7720 | 1760 | 8993 | 2435 | 704 | 14111 | 8237 | 24852 |
| 8479 | 21966 | 7041 | 1766 | 8469 | 2515 | 758 | 10485 | 8457 | 25084 |
| 2634.5 | 3671.5 | 2325 | 784 | 3009 | 1397 | 389 | 7821 | 3562.5 | 4778.5 |
| 2700 | 3833 | 2879 | 983 | 3859 | 1677 | 586 | 9242 | 4390 | 5782 |
| 2576.5 | 3863.5 | 3011 | 1072 | 4112 | 1776.5 | 666.5 | 9982.5 | 4479 | 6005 |
| 2368 | 3627 | 5572 | 1542 | 607315 | 52975 | 655 | 9571 | 4197 | 5668 |
| 2344 | 3742 | 3154 | 1143 | 6064 | 2074 | 700 | 10134 | 4274 | 6024 |
| 2305.5 | 3435 | 2866.5 | 1039.5 | 4951 | 1839 | 667 | 9325.5 | 4177.5 | 5689 |
| 2279 | 3524.5 | 2868 | 1055.5 | 4660.5 | 1845 | 688 | 9246 | 4217 | 5892 |
| 2336 | 3435 | 2786 | 993 | 4114 | 1699 | 643 | 9820 | 4106 | 5501 |
| 2754 | 6022 | 3258 | 1131 | 4948 | 1905 | 734 | 12882 | 4696 | 9307 |
| 2521 | 6047 | 3648.5 | 1304.5 | 4981.5 | 2092.5 | 791.5 | 14268.5 | 4462 | 9373.5 |
| 2313.5 | 5772 | 3265 | 1218 | 4813.5 | 1938 | 758 | 12862 | 4187 | 9365.5 |
| 2488 | 4982.5 | 3414 | 1234.5 | 4876.5 | 2043 | 801.5 | 10923.5 | 4732 | 8395 |
| 2496.5 | 4630.5 | 3645.5 | 1329.5 | 5148 | 2105.5 | 832 | 11315 | 4794 | 8065 |
| 14044 | 10263.5 | 4245.5 | 1033 | 3517.5 | 1629 | 628 | 8172.5 | 6501.5 | 6984.5 |
| 2573 | 3679.5 | 2887.5 | 1035 | 3911 | 1708 | 658 | 8308.5 | 4084.5 | 5774.5 |
| 2617.5 | 3676.5 | 2903 | 1025 | 3947.5 | 1678.5 | 635 | 8907 | 4483 | 5888 |
| 68955 | 55474 | 11314 | 1553 | 4691 | 1922 | 697 | 16897 | 5955 | 11627 |
| 2978 | 8852 | 4145 | 1384 | 5552 | 2185 | 814 | 12984 | 5117 | 14177 |
| 3851 | 9917 | 5028 | 1734 | 7342 | 2672 | 1016 | 14230 | 6956 | 16570 |
| 2829 | 7017 | 4074 | 1419 | 5631 | 2224 | 863 | 14484 | 5029 | 11465 |
| 7412 | 17428 | 8081.5 | 1962.5 | 10038.5 | 2764.5 | 860 | 14196.5 | 9504.5 | 20581 |
| 8061.5 | 22951.5 | 7989.5 | 1857 | 9284 | 2594.5 | 726 | 14215 | 8770 | 25738.5 |
| 8462 | 22604 | 7826.5 | 1809.5 | 9258.5 | 2526 | 714 | 13536.5 | 9259.5 | 26069 |

| | | | | | | | | | |
|---|---|---|---|---|---|---|---|---|---|
| 8656 | 19894 | 6584 | 1555 | 7799 | 2330 | 713 | 11349 | 9032 | 22487 |
| 2764.5 | 3769.5 | 2348 | 795.5 | 3192 | 1430 | 386 | 7906 | 3785.5 | 4893 |
| 2872 | 4132 | 2948.5 | 1009.5 | 4014.5 | 1717 | 593 | 9883 | 4521.5 | 5915.5 |
| 2593 | 4149 | 3227 | 1119 | 4479 | 1853.5 | 698 | 10945 | 4394 | 6598.5 |
| 2483 | 4057.5 | 5591 | 1671 | 610585 | 53090.5 | 703.5 | 10467 | 4435 | 6368.5 |
| 2352 | 4516.5 | 3415 | 1189 | 6092 | 2124 | 722.5 | 11162.5 | 4228 | 7263.5 |
| 2414 | 5180 | 3464 | 1197 | 5647 | 2083 | 767 | 11447 | 4422 | 8296 |
| 2427.5 | 3679.5 | 3103.5 | 1114 | 4870 | 1911 | 731.5 | 9788 | 4514 | 6009.5 |
| 2481.5 | 3821 | 2999.5 | 1064 | 4487.5 | 1823.5 | 694 | 10582.5 | 4297 | 6099.5 |
| 2705 | 6052.5 | 3474 | 1176.5 | 4988 | 1971 | 768 | 14957 | 4518.5 | 9419 |
| 2539 | 4251 | 3395.5 | 1222 | 4659 | 1928 | 737 | 11621 | 4313.5 | 6829 |
| 2671 | 4134 | 3230 | 1141 | 4561.5 | 1895 | 766 | 10814.5 | 4770.5 | 6729 |
| 2429 | 3853 | 3136 | 1152 | 4368 | 1832 | 741 | 10273 | 4618 | 6573 |
| 1958 | 3088 | 2522 | 953.5 | 3532 | 1580 | 640 | 8475 | 3728 | 5142.5 |
| 21892 | 15079 | 5449 | 1117 | 3974 | 1729 | 689 | 9073 | 9073 | 8455 |
| 2755 | 3901 | 3055 | 1055 | 4082 | 1764 | 715 | 9395 | 4648 | 6079 |
| 2480 | 3864 | 2914 | 1031 | 3915 | 1676 | 649 | 9301 | 4496 | 6227 |
| 50517 | 48078 | 9903 | 1759 | 5823 | 2150 | 802 | 16528 | 6356 | 15153 |
| 3851 | 10056 | 5129 | 1643 | 6789 | 2544 | 952 | 14146 | 6736 | 15545 |
| 2803 | 7601 | 4411 | 1470 | 5977 | 2295 | 888 | 13885 | 5013 | 12096 |
| 2854.5 | 7446.5 | 4117.5 | 1423 | 5754.5 | 2239.5 | 871 | 14181.5 | 4895 | 12432 |
| 7143 | 15962 | 6738 | 1687 | 8573 | 2488 | 787 | 14536 | 8478 | 18857 |
| 8292 | 23278 | 8064 | 1888 | 9335 | 2595 | 733 | 14435 | 8874 | 25530 |
| 7736 | 23836 | 7949 | 1870 | 9400 | 2596 | 747 | 11134 | 8153 | 27149 |
| 8053 | 22423 | 6860 | 1722 | 8292 | 2404 | 762 | 12453 | 8703 | 25321 |
| 3158 | 4062 | 2573 | 867 | 3412 | 1489 | 411 | 8963 | 4360 | 5239 |
| 4882 | 4852 | 3285 | 1026 | 15310 | 2545 | 623 | 10232 | 4756 | 6063 |
| 4376 | 4891 | 3434 | 1170 | 16132 | 2737 | 740 | 11862 | 4890 | 6675 |
| 5487.5 | 5004.5 | 6356.5 | 1833 | 296668 | 30397.5 | 711 | 10734 | 4479 | 6210.5 |
| 3614.5 | 5929.5 | 5468 | 1484.5 | 445205 | 51385.5 | 853 | 13029 | 4700 | 8638.5 |
| 3577 | 4549 | 4414 | 1333 | 297949 | 34167 | 817 | 12039 | 4867 | 6712 |
| 3537 | 4333 | 3966.5 | 1236.5 | 227896.5 | 25977 | 798.5 | 11243 | 4842 | 6645 |
| 3730 | 6991 | 4127 | 1277 | 193356.5 | 21738 | 853 | 14939.5 | 5106 | 10312 |
| 3687 | 6546 | 4164 | 1295 | 171480 | 19034 | 853 | 16309 | 5037 | 9917 |
| 3325 | 5344 | 3801 | 1295 | 26377 | 3601 | 829 | 12849 | 4940 | 8128 |
| 3444.5 | 4589.5 | 3529.5 | 1200.5 | 56224 | 6172.5 | 826 | 13174.5 | 5024.5 | 7089.5 |
| 3476 | 4430 | 3372 | 1206 | 22397 | 3133 | 833 | 11551 | 5137 | 6875 |
| 3715 | 4212 | 3188 | 1110 | 38162 | 4312 | 788 | 10759 | 4708 | 6318 |
| 72604 | 52216 | 20541 | 4628 | 27890 | 3636 | 723 | 9970 | 23524 | 17561 |
| 11102 | 8398 | 4233 | 1099 | 33592 | 3910 | 768 | 9710 | 6547 | 7003 |
| 7236.5 | 7562.5 | 3840 | 1140.5 | 28996 | 3582 | 764 | 11393.5 | 5683 | 8401.5 |
| 747808 | 785098.5 | 177779 | 14471.5 | 23070.5 | 3364.5 | 770 | 94686 | 28662 | 32522.5 |
| 13250.5 | 14713.5 | 6399.5 | 1878 | 30634 | 4321.5 | 1042 | 16270.5 | 8229 | 16964.5 |
| 4815 | 10582 | 5513 | 1812 | 37964 | 6865 | 1053 | 16046 | 6982 | 16572 |
| 6218.5 | 8896.5 | 4622 | 1437 | 26287.5 | 3764 | 927.5 | 15592.5 | 6091 | 12196 |
| 10788.5 | 19831 | 7236 | 1881 | 24600.5 | 3778.5 | 894.5 | 16574 | 9689 | 22172 |
| 8376 | 28662.5 | 11135.5 | 2376.5 | 23034 | 3931 | 930 | 9761 | 7680 | 30301.5 |
| 11766 | 25722 | 8355 | 1875 | 16430 | 3159 | 801 | 14022 | 9728 | 25443 |

| | | | | | | | | | |
|---|---|---|---|---|---|---|---|---|---|
| 10272.5 | 22705 | 8158 | 1835 | 14501.5 | 2913.5 | 797.5 | 11503 | 8797.5 | 24034.5 |
| 3092.5 | 4165 | 2608 | 876 | 3554 | 1525 | 416.5 | 8710 | 4228.5 | 5341.5 |
| 2843.5 | 4152 | 2974 | 1017 | 4072 | 1697.5 | 589 | 9743 | 4493 | 6010.5 |
| 2650 | 4371 | 3316 | 1183 | 4534 | 1893 | 718 | 11618 | 4337 | 6688 |
| 2408 | 3912.5 | 5647.5 | 1653.5 | 587368 | 52347 | 675 | 10098.5 | 4268.5 | 6183.5 |
| 2456 | 4049.5 | 3447 | 1186.5 | 6708.5 | 2154.5 | 727 | 10792 | 4311.5 | 6477 |
| 2311 | 4000 | 3302.5 | 1139 | 5605 | 2011.5 | 714 | 10394 | 4215.5 | 6361 |
| 2172 | 5419 | 3274 | 1147 | 5202 | 1986 | 752 | 10856 | 4051 | 8956 |
| 2518.5 | 3672 | 2882 | 1016.5 | 4356.5 | 1747.5 | 657 | 10077.5 | 4312 | 5616 |
| 2619.5 | 4474 | 3218.5 | 1097.5 | 4748 | 1905.5 | 719 | 12333 | 4347.5 | 6976.5 |
| 2743.5 | 6951.5 | 3741.5 | 1333 | 5225.5 | 2098.5 | 797 | 14656 | 4826 | 11013.5 |
| 2567 | 6175 | 3638 | 1306 | 5205 | 2092 | 812 | 13915 | 4538 | 9981 |
| 2399 | 4917 | 3446 | 1198 | 4827 | 1974 | 767 | 11189 | 4460 | 8196 |
| 2672 | 4694.5 | 3501 | 1292 | 5113.5 | 2084 | 829.5 | 10709.5 | 5181.5 | 7948.5 |
| 14985 | 11403 | 4993 | 1287 | 4761 | 2001 | 765 | 10460 | 7764 | 8754 |
| 2943 | 4692 | 3311 | 1189 | 4509 | 1930 | 742 | 10377 | 4593 | 7223 |
| 2586.5 | 5002 | 3182 | 1098 | 4195.5 | 1791 | 674 | 9391 | 4453 | 7906.5 |
| 72376 | 51411 | 11484 | 1441 | 4037 | 1719 | 617 | 15130 | 5560 | 8622 |
| 3469 | 9311 | 4549 | 1536 | 6450 | 2388 | 858 | 12972 | 6114 | 14750 |
| 3461.5 | 9463 | 4890.5 | 1631.5 | 6793.5 | 2541 | 946 | 15603 | 6871 | 15387 |
| 2438 | 5229 | 3492 | 1220 | 4775 | 1963 | 743 | 10023 | 4452 | 8342 |
| 6177 | 16576 | 6738 | 1743 | 8408 | 2490 | 787 | 12427 | 7837 | 19775 |
| 7619 | 24495 | 9809 | 2214 | 11778 | 3050 | 828 | 13256 | 8943 | 28159 |
| 9009 | 22694 | 7996 | 1796 | 9271 | 2572 | 732 | 13743 | 9289 | 25582 |
| 8858.5 | 22887.5 | 7345.5 | 1744.5 | 8611.5 | 2490 | 762 | 13979.5 | 9415.5 | 26409 |
| 2909 | 3898.5 | 2550.5 | 847 | 3374 | 1489 | 418 | 8059.5 | 4133.5 | 5122 |
| 2755.5 | 4320.5 | 3285.5 | 1159 | 4540 | 1883.5 | 608 | 10573.5 | 4412 | 6600 |
| 2371 | 4139 | 3226 | 1191.5 | 4633 | 1995.5 | 654.5 | 9826 | 4348.5 | 6909.5 |
| 2239 | 5829 | 6045 | 1676 | 542482 | 49100 | 689 | 11296 | 4284 | 10469 |
| 2380 | 6165 | 3694 | 1345 | 6678 | 2425 | 713 | 11368 | 4570 | 11253 |
| 2305.5 | 6265.5 | 3555.5 | 1334 | 6150.5 | 2326 | 704 | 10651.5 | 4437 | 11805 |
| 2203 | 6277 | 3558 | 1351 | 5898 | 2320 | 717 | 11134 | 4419 | 11916 |
| 2469.5 | 6997 | 3890 | 1504 | 6085.5 | 2482 | 742 | 13160 | 4740 | 13047.5 |
| 2654 | 7147 | 4060 | 1487 | 6272 | 2464 | 749.5 | 15255 | 4992.5 | 13178 |
| 2314 | 5167 | 3614 | 1334 | 5276 | 2269 | 693 | 11575 | 4567 | 9656 |
| 2195 | 4005.5 | 3318.5 | 1260 | 5003 | 2144 | 666 | 10405.5 | 4470 | 7827.5 |
| 2115 | 3514 | 3100 | 1190.5 | 4807 | 2060 | 645 | 9744 | 4396.5 | 6969 |
| 2042 | 3117 | 2833 | 1094 | 4249 | 1867 | 605 | 8374 | 4198 | 6028 |
| 10539 | 10947.5 | 4198.5 | 1220 | 4474.5 | 1959 | 643 | 9096.5 | 6074.5 | 10200 |
| 2293 | 6513 | 3618 | 1304 | 5162 | 2146 | 701 | 10243 | 4226 | 11930 |
| 2381 | 6721 | 3781 | 1370 | 5441 | 2207 | 691 | 10367 | 4474 | 12341 |
| 53839 | 55775 | 10622 | 1868 | 6223 | 2425 | 745 | 18437 | 5898 | 15948 |
| 3285 | 10375 | 5330 | 1809 | 7664 | 2862 | 882 | 14026 | 6153 | 18526 |
| 2702 | 9388 | 4902 | 1736 | 7411 | 2883 | 876 | 14475 | 5340 | 17249 |
| 2812.5 | 7824 | 4441 | 1595 | 6411 | 2524 | 819 | 14004 | 5439 | 13633 |
| 7616 | 17376 | 8341 | 2082 | 10196 | 2965 | 811 | 14592 | 9498 | 21913 |
| 7529 | 21940.5 | 9530 | 2351 | 11760.5 | 3317 | 799 | 12803 | 9154.5 | 25889 |
| 8388.5 | 21577 | 8210.5 | 2006 | 10074 | 2816 | 696.5 | 12281 | 9003.5 | 25085 |

| | | | | | | | | | |
|---|---|---|---|---|---|---|---|---|---|
| 7414 | 18986 | 8241 | 1943 | 9875.5 | 2827 | 739 | 11751 | 8545 | 23331.5 |
| 2891 | 3838 | 2680.5 | 943 | 3721.5 | 1638.5 | 388 | 8399 | 4297.5 | 5525 |
| 3491 | 4594 | 3402 | 1098.5 | 6906 | 2152.5 | 596 | 10373.5 | 4769 | 6782 |
| 3045 | 6750 | 3752 | 1294.5 | 9694 | 2584.5 | 709 | 12741 | 4819 | 10661 |
| 3027 | 7172 | 7303 | 2297 | 357299 | 51453 | 732 | 12692 | 4507 | 10669 |
| 3355 | 8720 | 4980 | 1508 | 154255 | 23119 | 771 | 14457 | 4966 | 12574 |
| 3293 | 9181.5 | 4692.5 | 1430.5 | 111165 | 15815.5 | 787 | 14477.5 | 4943 | 13283.5 |
| 3396 | 8301 | 4355 | 1332 | 72822 | 9979 | 752 | 16197 | 5941 | 14138 |
| 3352.5 | 8394 | 4099 | 1304.5 | 66920 | 9079 | 742.5 | 14658.5 | 4793.5 | 11926.5 |
| 3653 | 9720 | 4468 | 1454 | 51963 | 7047 | 810 | 15752 | 5355 | 14086 |
| 4597 | 12158.5 | 8110.5 | 2604.5 | 37785.5 | 6012.5 | 888 | 17761.5 | 6188.5 | 15819.5 |
| 3085 | 5049 | 3739 | 1222.5 | 34884 | 4860.5 | 709.5 | 11422 | 4702 | 7365.5 |
| 2832 | 4782 | 3858 | 1295 | 16013 | 2943.5 | 738 | 10846 | 4615 | 7126 |
| 3050.5 | 4064.5 | 3242.5 | 1113.5 | 22714.5 | 3487.5 | 676 | 9686.5 | 4569 | 6073.5 |
| 38447.5 | 30982 | 26378.5 | 3391 | 18279 | 3094 | 691 | 9977.5 | 15025.5 | 13686 |
| 5506 | 11060 | 5181 | 1389 | 15588 | 3087 | 779 | 12430 | 6830 | 14436 |
| 3492 | 9159 | 4329 | 1354 | 15054 | 2950 | 749.5 | 12461 | 4843 | 13237.5 |
| 360086 | 425951 | 105645 | 9333 | 14937 | 3189 | 813 | 52600 | 13274 | 22542 |
| 5325 | 12163 | 5679.5 | 1796.5 | 14490.5 | 3224.5 | 890 | 16622.5 | 6882 | 16399.5 |
| 5178 | 11546 | 5549 | 1706 | 61889 | 11430 | 894 | 16492 | 6816 | 16378 |
| 4746 | 10362 | 5090 | 1602 | 14249 | 3153 | 845 | 14523 | 5948 | 13588 |
| 8756.5 | 18822 | 8162.5 | 1910.5 | 14336.5 | 3140 | 789 | 15400.5 | 8707.5 | 20681 |
| 8275 | 22244 | 9761 | 2314 | 14884 | 3368 | 798 | 14159 | 8567 | 24712 |
| 9498 | 24672 | 9115 | 2039 | 14141 | 3167 | 768 | 13910 | 9656 | 26585 |
| 9334 | 23990 | 8016 | 1932 | 12554 | 2946 | 808 | 13967.5 | 9609 | 26324 |
| 2785.5 | 3705.5 | 2437 | 857 | 3494.5 | 1519.5 | 403.5 | 8941 | 3909 | 5101.5 |
| 2784 | 3919 | 2819.5 | 998 | 3945.5 | 1686 | 535 | 9312 | 4347.5 | 5614.5 |
| 2542 | 3949 | 3124 | 1083.5 | 4302.5 | 1787 | 650 | 10211.5 | 4259 | 6186.5 |
| 2410 | 4031 | 5435 | 1491 | 570185 | 48800 | 663 | 9655 | 4187 | 6486 |
| 2443 | 4614 | 3374 | 1156 | 6814 | 2162 | 700 | 10742 | 4376 | 7486 |
| 2330 | 4148 | 3281 | 1157 | 5674 | 2043 | 718 | 10412 | 4255 | 6765 |
| 2269 | 3776 | 3102 | 1117 | 5190 | 1926 | 711 | 9955 | 4297 | 6359 |
| 2430 | 3782.5 | 3026.5 | 1066 | 4679 | 1845 | 671 | 10627 | 4269.5 | 6126.5 |
| 2515 | 4438 | 3293 | 1145 | 4835 | 1915 | 711 | 12529 | 4276 | 6853 |
| 2402.5 | 6097.5 | 3601 | 1396.5 | 5044 | 2090 | 725 | 13575.5 | 4256 | 9569 |
| 2394 | 7496 | 4020 | 1370 | 5785 | 2223 | 847 | 14220 | 4496 | 12628 |
| 2263 | 4355 | 3395.5 | 1223.5 | 4913 | 1947.5 | 763 | 10895 | 4452 | 7438 |
| 2288 | 3519.5 | 2940 | 1072 | 4162 | 1801 | 680 | 9261.5 | 4357.5 | 5975.5 |
| 13533 | 9942 | 4215.5 | 1018.5 | 3705 | 1636.5 | 629.5 | 8207.5 | 6135.5 | 6759.5 |
| 2768 | 3813 | 2973 | 1049 | 3979 | 1743 | 668 | 8941 | 4352 | 5966 |
| 2578.5 | 4099.5 | 3212.5 | 1091 | 4330 | 1806.5 | 657 | 9126 | 4421 | 6510 |
| 91630 | 82288 | 14759 | 1786 | 5101 | 2014 | 709 | 19324 | 6373 | 13443 |
| 3207 | 7279 | 3811 | 1294 | 5078 | 2059 | 745 | 11419 | 5112 | 10822 |
| 2873 | 8443 | 4173 | 1450 | 6045 | 2336 | 831 | 13873 | 4973 | 13811 |
| 2574 | 7098 | 3764 | 1332 | 5253 | 2074 | 787 | 12535 | 4668 | 11516 |
| 7115 | 17334 | 6726 | 1724 | 8353 | 2468 | 797 | 15384 | 8907 | 21318 |
| 8294 | 20876 | 9422 | 2152 | 10885 | 2910 | 784 | 13406 | 9564 | 23880 |
| 8453.5 | 22559 | 9268.5 | 2073 | 11049.5 | 2834 | 771.5 | 10137 | 8534.5 | 25895 |

| | | | | | | | | | |
|---|---|---|---|---|---|---|---|---|---|
| 7593 | 21285 | 6895 | 1672 | 8246 | 2407 | 720 | 10971 | 8478 | 24458 |
| 2391 | 3280 | 2214 | 807 | 3109 | 1437 | 338 | 7120 | 3422 | 4591 |
| 2617 | 3626 | 2616 | 930 | 3672.5 | 1630.5 | 514 | 8363 | 4128 | 5383.5 |
| 2599 | 3830 | 2928 | 995 | 4075 | 1726 | 615 | 9237 | 4400 | 5897 |
| 2438 | 3884 | 5835 | 1641.5 | 606089.5 | 51289.5 | 647.5 | 9554 | 4263.5 | 6223.5 |
| 2369 | 3620 | 2982 | 1038 | 5714 | 1915.5 | 639 | 8903 | 4197 | 5773 |
| 2446.5 | 5595 | 3435 | 1188.5 | 5589 | 1988.5 | 723 | 10378 | 4223 | 8711 |
| 2420 | 3753 | 3033 | 1076 | 4802 | 1886 | 674 | 9559 | 4435 | 5982 |
| 2538 | 3879 | 3036.5 | 1089 | 4646.5 | 1817.5 | 690 | 10399 | 4445.5 | 6211.5 |
| 2617 | 6193 | 3437 | 1145 | 4975 | 1934 | 712 | 13843 | 4382 | 9601 |
| 2426 | 3947 | 3094 | 1109 | 4206 | 1807 | 659 | 11309 | 4016 | 6161 |
| 2607.5 | 3999 | 3051.5 | 1087.5 | 4396 | 1875.5 | 699 | 10657 | 4542.5 | 6423 |
| 2351 | 3806 | 3013.5 | 1117 | 4287.5 | 1837.5 | 690 | 9638.5 | 4491 | 6099.5 |
| 2398 | 3996 | 3350 | 1187 | 4716 | 1949 | 744 | 9951 | 4601 | 6617 |
| 11411 | 10321 | 4928 | 1291 | 4947 | 1999 | 775 | 11344 | 6376 | 9070 |
| 2918 | 6421 | 3445 | 1163 | 4733 | 1937 | 742 | 10755 | 4594 | 9916 |
| 2712 | 4257 | 3014 | 1066 | 4026.5 | 1716 | 636 | 9393.5 | 4423.5 | 6567.5 |
| 75279.5 | 72178 | 13223.5 | 1750.5 | 5500.5 | 2156.5 | 750 | 19502 | 6074 | 14280 |
| 2513 | 7149.5 | 3548 | 1143.5 | 4680 | 1886 | 694.5 | 10737.5 | 4287 | 11029.5 |
| 2705 | 4564.5 | 3338.5 | 1165 | 4686 | 1905 | 711.5 | 9705 | 4755.5 | 7324.5 |
| 2745.5 | 5664 | 3652.5 | 1233.5 | 4874 | 1933 | 742 | 10581 | 4610.5 | 9203 |
| 7573 | 17135.5 | 8174.5 | 1962 | 9825 | 2759.5 | 835 | 14677.5 | 9399 | 20852.5 |
| 8747.5 | 22795 | 8342.5 | 1889.5 | 9683.5 | 2639 | 731.5 | 14144 | 9383 | 25269 |
| 8875.5 | 20082 | 7258.5 | 1631 | 8269.5 | 2308.5 | 664 | 13856 | 9082.5 | 22266 |
| 7778 | 19415 | 7116 | 1609 | 8424 | 2362 | 723 | 12894 | 8414 | 21540 |
| 867 | 1255 | 1120 | 634 | 1798 | 1200 | 299 | 2948 | 1751 | 2485 |
| 1302 | 2184.5 | 1979 | 829 | 2905 | 1459 | 519 | 5797.5 | 2689 | 4026 |
| 1526.5 | 2700 | 2479 | 970 | 3599 | 1690 | 641 | 7371.5 | 3301 | 5019 |
| 1615 | 2893.5 | 5139.5 | 1544 | 619516 | 56279.5 | 707 | 8013 | 3590.5 | 5401 |
| 1532.5 | 2764 | 2709 | 1026 | 4370.5 | 1881 | 674 | 7838 | 3574.5 | 5236.5 |
| 1564.5 | 2786.5 | 2749 | 1054 | 4118 | 1796 | 678 | 8029 | 3592.5 | 5177.5 |
| 1642 | 2827.5 | 2676 | 1059.5 | 4018 | 1807.5 | 702 | 8160 | 3690 | 5316.5 |
| 1758 | 2795 | 2425 | 963 | 3526 | 1656 | 653 | 8424 | 3529 | 4954 |
| 2250.5 | 3488.5 | 2866 | 1107 | 4147.5 | 1849.5 | 722 | 10709.5 | 4335.5 | 5914.5 |
| 2223 | 3582.5 | 2989.5 | 1176 | 4191.5 | 1892.5 | 727 | 10862 | 4457 | 6138 |
| 2118 | 3653 | 3288 | 1222 | 4550 | 2054 | 791 | 11077 | 4460 | 6641 |
| 2290.5 | 4518 | 3670 | 1390 | 5208 | 2227 | 887 | 10851 | 5112 | 8378.5 |
| 2101 | 4007 | 3372 | 1290 | 4803 | 2093 | 850 | 11265 | 4890 | 7373 |
| 16727 | 11899 | 4548 | 1111 | 3611 | 1718 | 670 | 7463 | 7210 | 7327 |
| 2113 | 3240 | 2766.5 | 1026.5 | 3707 | 1683 | 691 | 7971 | 3892 | 5584.5 |
| 2093 | 3372 | 2799.5 | 1011 | 3866 | 1725 | 666 | 8330 | 3973.5 | 5753.5 |
| 74717 | 39973 | 11111 | 1407 | 3822 | 1700 | 620 | 13996 | 5808 | 6758 |
| 2481 | 6562 | 3726 | 1284 | 5085 | 2075.5 | 792 | 12108 | 4841.5 | 11163 |
| 1933 | 5830.5 | 3241 | 1223 | 5040 | 2059 | 755.5 | 11241 | 3975 | 10390 |
| 2285.5 | 5387 | 3500.5 | 1276.5 | 5031.5 | 2061 | 833 | 11736 | 4625.5 | 9245 |
| 5678 | 13618 | 5701 | 1665 | 7468 | 2523 | 828 | 13093 | 7651 | 17071 |
| 8550.5 | 24057 | 8949 | 1974 | 10258.5 | 2733 | 787 | 14124.5 | 9530 | 27159.5 |
| 8177 | 21985 | 8360 | 1866 | 10023 | 2564 | 737 | 11456 | 8674 | 24475 |

| | | | | | | | | | |
|---|---|---|---|---|---|---|---|---|---|
| 7896 | 21012 | 6744 | 1639 | 7981 | 2356 | 736 | 11652 | 8625 | 24633 |
| 2105 | 2895 | 1852 | 727 | 2522 | 1303 | 320 | 6122 | 2987 | 3961 |
| 1981 | 2944 | 2251.5 | 849 | 3022 | 1498 | 497 | 7287 | 3266.5 | 4559 |
| 1975 | 3031 | 2417 | 929 | 3328 | 1574 | 589 | 7826 | 3507 | 4904 |
| 1907.5 | 3101.5 | 4952 | 1434 | 601393 | 55574 | 641.5 | 7757 | 3551 | 5151 |
| 2024 | 3273 | 2907 | 1082 | 5096 | 1934 | 684 | 8451 | 3861 | 5496 |
| 1896 | 3131 | 2789 | 1026 | 4249 | 1812 | 681 | 8138 | 3987 | 5543 |
| 1928 | 3219 | 2778 | 1032 | 4108 | 1807 | 710 | 8427 | 3927 | 5628 |
| 1869.5 | 2885 | 2413 | 913 | 3487 | 1611.5 | 625 | 8278.5 | 3521.5 | 4870.5 |
| 2483 | 3455 | 2726 | 976 | 3910 | 1720 | 659 | 10551 | 4375 | 5725 |
| 2700 | 4017 | 3141.5 | 1242.5 | 4478.5 | 2014.5 | 741 | 11072.5 | 4974.5 | 6554 |
| 2261 | 3649.5 | 2979 | 1115.5 | 4340.5 | 1848.5 | 738 | 10224.5 | 4435.5 | 6183.5 |
| 2298.5 | 3789.5 | 3257 | 1197 | 4626.5 | 1971 | 785 | 10355 | 4679.5 | 6686 |
| 2495 | 4665 | 3739 | 1369 | 5189 | 2175 | 878 | 11004 | 5409 | 8084 |
| 16209 | 12321 | 4620 | 1088 | 3609.5 | 1748 | 668 | 8133 | 7411.5 | 7842.5 |
| 2217.5 | 3200 | 2533 | 961 | 3511 | 1625 | 636.5 | 7650.5 | 3722.5 | 5260 |
| 2202.5 | 3342.5 | 2713 | 986.5 | 3744 | 1646 | 640.5 | 8480 | 3978.5 | 5624.5 |
| 71321.5 | 47930 | 11374 | 1485 | 4269 | 1845 | 674 | 14697.5 | 5665 | 8931.5 |
| 2590.5 | 7713 | 3834.5 | 1320.5 | 5299 | 2088.5 | 779.5 | 12947.5 | 4659 | 12468.5 |
| 2833 | 7734.5 | 4288.5 | 1470.5 | 6447 | 2415.5 | 912 | 14083.5 | 5630.5 | 13278.5 |
| 2368.5 | 5294 | 3300.5 | 1236 | 4737.5 | 2040.5 | 739.5 | 10261.5 | 4449.5 | 9357.5 |
| 6941 | 16149 | 7049 | 1737 | 8641 | 2517 | 810 | 14312 | 9011 | 19528 |
| 7766 | 22988.5 | 8306.5 | 1867.5 | 9441.5 | 2599 | 740 | 11828 | 8538 | 26455 |
| 8215 | 23223 | 8459 | 1843 | 10023 | 2553 | 750 | 12573 | 8707 | 25347 |
| 8235 | 21912.5 | 8230 | 1912 | 9680.5 | 2613 | 807 | 11830 | 8806.5 | 25213 |
| 2666 | 3535 | 2330 | 841 | 3210.5 | 1472 | 380 | 7898 | 3863 | 4949.5 |
| 2145 | 3257 | 2440 | 883 | 3311 | 1505 | 517 | 7785 | 3541 | 4973 |
| 2264 | 3324 | 2685 | 979 | 3667 | 1679 | 608 | 8560 | 3905 | 5202 |
| 2336 | 3434 | 5474 | 1534 | 620254 | 53302 | 661 | 9328 | 4264 | 5673 |
| 2289 | 3640.5 | 3101.5 | 1089 | 5375 | 2024.5 | 711 | 8998 | 4114.5 | 5865 |
| 2141.5 | 3324 | 2803.5 | 1025.5 | 4540.5 | 1837 | 689.5 | 8654 | 4045 | 5605 |
| 2218.5 | 3374.5 | 2948 | 1049.5 | 4425.5 | 1796 | 695 | 9172 | 4285.5 | 5589 |
| 2160 | 3320 | 2643.5 | 992.5 | 3951 | 1657 | 647.5 | 8902 | 3813 | 5367.5 |
| 2675 | 3661 | 2931 | 1058 | 4187 | 1789 | 697 | 10872 | 4617 | 5945 |
| 2465 | 4186 | 3100 | 1179 | 4352 | 1887 | 722 | 12105 | 4456 | 6702 |
| 2430 | 6125 | 3524 | 1253 | 4944 | 2031 | 821 | 13606 | 4488 | 9992 |
| 2205 | 3732 | 3109 | 1128 | 4476 | 1859 | 757 | 10242 | 4460 | 6384 |
| 2337 | 4049 | 3268 | 1155 | 4560 | 1961 | 788 | 9977 | 4755 | 6928 |
| 16211 | 11984 | 5055 | 1227 | 4431 | 1928 | 753 | 9853.5 | 7779.5 | 8487 |
| 2600.5 | 3613 | 2872 | 1044 | 3828 | 1766 | 694 | 8826 | 4228 | 5776 |
| 2414 | 3671 | 2865 | 1019 | 3889 | 1722 | 657 | 8888 | 4256 | 5931 |
| 59596 | 45184 | 9874.5 | 1324 | 3609.5 | 1621.5 | 588 | 13364.5 | 4834 | 7978 |
| 2767.5 | 7778 | 3926.5 | 1371 | 5388.5 | 2142.5 | 802 | 13493.5 | 4876 | 12218.5 |
| 2801 | 7803 | 4164 | 1467 | 6106.5 | 2359 | 868.5 | 14106 | 5319 | 12987 |
| 2955.5 | 7104.5 | 4130 | 1438.5 | 5778.5 | 2326 | 900 | 14111 | 5493.5 | 11437.5 |
| 6004 | 14355.5 | 6085 | 1721.5 | 7658 | 2471 | 839 | 14261 | 7753 | 16953.5 |
| 6658.5 | 23339.5 | 9466.5 | 2172 | 11119.5 | 2975 | 796 | 10191.5 | 7274.5 | 26774 |
| 8803.5 | 23209 | 7422.5 | 1767 | 8669.5 | 2438 | 708 | 14280.5 | 9499.5 | 25541.5 |

| | | | | | | | | | |
|---|---|---|---|---|---|---|---|---|---|
| 8754 | 20984 | 6851 | 1690 | 8056 | 2414 | 743 | 11242 | 8741 | 23271 |
| 2894 | 3737 | 2396 | 804 | 3109.5 | 1405.5 | 393 | 8312 | 4021 | 4829 |
| 2716 | 3871 | 2823 | 969 | 3696 | 1658 | 564 | 9051 | 4289 | 5569 |
| 2590 | 3737.5 | 2918.5 | 1022 | 4017.5 | 1726 | 652 | 9456.5 | 4471 | 5879 |
| 2424.5 | 3947.5 | 5724.5 | 1684 | 610582.5 | 53865 | 708.5 | 10510 | 4392.5 | 6328.5 |
| 2374 | 3677 | 3194 | 1095 | 5574 | 1992 | 688 | 9787 | 4351 | 5967 |
| 2373 | 3585 | 3072 | 1099.5 | 4876 | 1895 | 708 | 9507.5 | 4459.5 | 6029 |
| 2362 | 3607 | 2984 | 1064 | 4556 | 1836 | 711 | 9324 | 4385 | 5813 |
| 2546 | 3855 | 3022 | 1096 | 4579 | 1841 | 716 | 11063 | 4480 | 6135 |
| 2504 | 3752 | 2930 | 1001 | 4234 | 1777 | 678 | 10902 | 4331 | 5920 |
| 2588.5 | 5922.5 | 3785.5 | 1368 | 5203 | 2138.5 | 807 | 14468.5 | 4552 | 9408 |
| 2550 | 5162 | 3641 | 1279 | 5148 | 2111 | 818 | 11622 | 4656 | 8537 |
| 2407 | 4724 | 3627 | 1322 | 5125 | 2106 | 843 | 11205 | 4690 | 8169 |
| 2455 | 4299 | 3363 | 1195 | 4806 | 1968 | 800 | 10158 | 4904 | 7370 |
| 16107 | 12599.5 | 4893.5 | 1203.5 | 4143 | 1827.5 | 729 | 8983.5 | 7652.5 | 8643 |
| 2690 | 3763.5 | 2911.5 | 1038 | 3853.5 | 1745 | 686 | 8805 | 4360 | 5852.5 |
| 2538 | 3829 | 2900 | 1078 | 3896 | 1748 | 657 | 8920 | 4396 | 5998 |
| 74249 | 47580 | 11696 | 1462 | 4004 | 1714 | 632 | 15216 | 5547 | 7913 |
| 2636 | 7099 | 3596 | 1210 | 4692 | 1935 | 732 | 11496 | 4587 | 11315 |
| 2570.5 | 6135 | 3474.5 | 1198 | 4985.5 | 1947.5 | 743 | 11601 | 4665.5 | 10246 |
| 2502 | 5581 | 3412 | 1174 | 4660 | 1932 | 751 | 11469 | 4508 | 9326 |
| 6962 | 16531 | 5957 | 1644 | 7401 | 2410 | 775 | 15353 | 8898 | 20043 |
| 7881 | 23146.5 | 9050 | 2072.5 | 10591.5 | 2867 | 802 | 12207 | 8841 | 26654 |
| 8835 | 20661.5 | 8380.5 | 1765 | 9669 | 2508 | 727 | 12066 | 8927.5 | 23294.5 |
| 6951 | 20529 | 9048.5 | 1986 | 10596.5 | 2670.5 | 804 | 9459 | 7753.5 | 23450 |
| 2932 | 3763 | 2431 | 820 | 3139 | 1417 | 397 | 8544 | 4031 | 4909 |
| 2475.5 | 3494 | 2521.5 | 895.5 | 3341.5 | 1560.5 | 542 | 8055 | 3764 | 4975.5 |
| 2594.5 | 3699.5 | 2818.5 | 1007 | 3830 | 1715 | 637 | 9153 | 4391.5 | 5599 |
| 2514 | 3589.5 | 5339.5 | 1560 | 611333 | 53407.5 | 688 | 9334 | 4400.5 | 5854 |
| 2523.5 | 3722 | 3177.5 | 1121.5 | 5529 | 2017.5 | 722 | 9920 | 4580 | 5994 |
| 2400 | 3781 | 3137 | 1089 | 5039 | 1955 | 742 | 9629 | 4491 | 6255 |
| 2291 | 3527 | 2959 | 1094 | 4445 | 1795 | 709 | 9086 | 4204 | 5817 |
| 2357 | 3484 | 2747 | 968 | 4094 | 1715 | 669 | 9557 | 4082 | 5631 |
| 2538 | 4389 | 3439 | 1188 | 4895 | 1931 | 772 | 13258 | 4339 | 7006 |
| 2657.5 | 4622.5 | 3614 | 1331.5 | 4965 | 2023.5 | 793 | 13559.5 | 4594 | 7280 |
| 2569 | 3995 | 3311 | 1158 | 4586 | 1880 | 768 | 10906 | 4644 | 6631 |
| 2414 | 3924 | 3125 | 1123 | 4344 | 1865 | 755 | 9981 | 4595 | 6519 |
| 2173 | 3575 | 2888 | 1084 | 4079 | 1822 | 729 | 9060 | 4256 | 6152 |
| 21986.5 | 15810 | 5346 | 1048 | 3435 | 1575 | 661.5 | 8082 | 8914 | 8254 |
| 2811 | 3778.5 | 3060.5 | 1053 | 3938 | 1768.5 | 705 | 9410 | 4688 | 6105.5 |
| 2709 | 3781 | 2999 | 1030 | 3966 | 1704 | 684 | 9082 | 4783 | 6035 |
| 49879 | 44605.5 | 9487.5 | 1514.5 | 4866 | 1958 | 732 | 16230.5 | 5384 | 11736.5 |
| 2877 | 8327 | 4326 | 1427 | 5860 | 2264 | 865 | 14735 | 5168 | 13250 |
| 2501 | 7626 | 4038 | 1418 | 5888 | 2245 | 877 | 13741 | 4573 | 12659 |
| 2626 | 5991.5 | 4011.5 | 1325 | 5454.5 | 2161 | 859.5 | 12666.5 | 4853.5 | 9979 |
| 8305 | 18993.5 | 8058.5 | 1974.5 | 9617.5 | 2735 | 882 | 15121 | 9795.5 | 22247 |
| 7911 | 24588 | 8810 | 2110 | 10597 | 2838 | 835 | 11760 | 8778 | 27625 |
| 8823 | 22553 | 7954.5 | 1813.5 | 9049.5 | 2509 | 737 | 14698 | 9168.5 | 24659.5 |

| | | | | | | | | | |
|---|---|---|---|---|---|---|---|---|---|
| 9293.5 | 22903.5 | 7188.5 | 1775.5 | 8286 | 2501 | 794 | 13308.5 | 9408 | 24687 |
| 2785 | 3730.5 | 2328 | 802.5 | 3077 | 1431.5 | 377 | 7968 | 3800.5 | 4832.5 |
| 5630 | 4926.5 | 3193 | 999 | 16126.5 | 2589 | 621 | 9997.5 | 4640 | 5779.5 |
| 4716 | 4803 | 3334 | 1137 | 15584 | 2737 | 766 | 10865 | 5168 | 6404 |
| 6856 | 5324 | 6562 | 1899 | 300279.5 | 30427.5 | 786 | 11481.5 | 4902.5 | 6318.5 |
| 4313.5 | 4750.5 | 5211 | 1425.5 | 413182.5 | 46553.5 | 861.5 | 12106.5 | 5386.5 | 6751 |
| 3953 | 4397 | 4450 | 1240 | 304056 | 34565 | 835 | 11202 | 4890 | 6513 |
| 4039 | 4636 | 4296 | 1291 | 219109 | 24840.5 | 892 | 12114 | 5319 | 6975.5 |
| 4250 | 4489 | 3881 | 1180 | 191364 | 21328 | 851 | 12119 | 5319 | 6623 |
| 4027.5 | 5281.5 | 4080.5 | 1241 | 161184.5 | 17846.5 | 908 | 15805 | 5110.5 | 7615 |
| 3729.5 | 7467 | 4213.5 | 1454 | 34574 | 4280.5 | 951.5 | 17283 | 5531 | 11220 |
| 3785.5 | 6635 | 4149.5 | 1383 | 69181 | 7572 | 974 | 16807 | 5412 | 10376.5 |
| 3640.5 | 4968.5 | 3774.5 | 1283 | 22189 | 3384.5 | 920.5 | 12663 | 5284 | 7664.5 |
| 3985 | 5633 | 4036 | 1389 | 40483 | 4800 | 987 | 14414 | 5464 | 8626 |
| 70867 | 56577 | 23318 | 8306.5 | 28508.5 | 4022.5 | 936 | 12035 | 22961.5 | 20056 |
| 10964.5 | 9076 | 4884 | 1296 | 35462.5 | 4265.5 | 879 | 11574.5 | 7251.5 | 8718.5 |
| 8676.5 | 7570.5 | 4105.5 | 1216 | 30971.5 | 3761 | 852 | 11889.5 | 6383 | 7953 |
| 848129.5 | 597245 | 190376 | 16133.5 | 22639.5 | 3534 | 819.5 | 100853 | 32959 | 24799.5 |
| 13661 | 13211 | 4842 | 1290 | 31303 | 4079 | 883 | 13645 | 5977 | 11177 |
| 3709 | 7102 | 3831 | 1269 | 38650 | 6484 | 820 | 12755 | 4902 | 10503 |
| 6702.5 | 9392.5 | 4622 | 1428 | 27539 | 3854.5 | 1006 | 15876.5 | 5769.5 | 12235 |
| 11548 | 21730 | 7996 | 2041 | 26635 | 4093 | 997 | 16740 | 9800 | 23699 |
| 11541 | 28765.5 | 11062.5 | 2377 | 24192.5 | 4032.5 | 983.5 | 10980 | 8877 | 29357 |
| 11508 | 24113.5 | 7714 | 1786.5 | 10970 | 2648.5 | 760 | 15313 | 10003 | 24963 |
| 8994 | 23285 | 9290 | 2046 | 14518 | 3161 | 895 | 13087 | 8708 | 24701 |
| 2959.5 | 3838 | 2465 | 815.5 | 3206.5 | 1454 | 390 | 8378.5 | 4012 | 4939 |
| 2423.5 | 3459 | 2444 | 858 | 3333.5 | 1521 | 509 | 7990 | 3752 | 4899 |
| 2556 | 3601 | 2795 | 1005 | 3767 | 1642 | 615 | 9179 | 4180 | 5511 |
| 2361.5 | 3977.5 | 6258.5 | 1743 | 675352 | 53739 | 698 | 10128 | 4180 | 6162 |
| 2508 | 3795 | 3323 | 1144 | 6354 | 2097 | 692 | 9374 | 4522 | 6027 |
| 2453 | 3988 | 3276 | 1153 | 5623 | 1997 | 725 | 9875 | 4453 | 6351 |
| 2339 | 3975 | 3289 | 1151 | 5291 | 1990 | 751 | 10254 | 4401 | 6551 |
| 2609 | 3981 | 3206 | 1107 | 4862 | 1925 | 725 | 10671 | 4515 | 6387 |
| 2560 | 4330 | 3392 | 1149.5 | 4927 | 1903 | 752.5 | 12657 | 4372 | 6867 |
| 2577 | 4833.5 | 3564 | 1258 | 4953.5 | 1987.5 | 766 | 14147 | 4487 | 7724 |
| 2505 | 4727 | 3494 | 1215 | 4941 | 1982 | 774 | 10990 | 4655 | 7634 |
| 2238 | 3561 | 2936 | 1054 | 4063 | 1758 | 677 | 9224 | 4208 | 5989 |
| 2091 | 3347 | 2842 | 1004 | 3955 | 1710 | 672 | 8742 | 3919 | 5613 |
| 14977 | 10832 | 4597 | 1131 | 4087 | 1769 | 696 | 9110 | 6984 | 7569 |
| 2903 | 4427 | 3447 | 1210 | 4671 | 1939 | 764 | 10045 | 4727 | 6877 |
| 2681 | 6841.5 | 3733 | 1259.5 | 4990.5 | 1975 | 765.5 | 11825.5 | 4599 | 10926 |
| 65960 | 63870 | 12947 | 1700 | 5158 | 1968 | 732 | 16752 | 5287 | 11892 |
| 3070 | 9008 | 4590 | 1457.5 | 6084 | 2326.5 | 862 | 15168 | 5173.5 | 13964 |
| 2745 | 7864 | 4087.5 | 1390.5 | 5843.5 | 2220 | 839 | 13987.5 | 4937 | 12583.5 |
| 2625 | 6861 | 4093 | 1420 | 5826 | 2270 | 882 | 13807 | 4773 | 11211 |
| 7813 | 17923 | 6782 | 1756 | 8184 | 2457 | 804 | 14906 | 9287 | 20766 |
| 8065 | 23695 | 9196 | 2146 | 11115 | 2918 | 820 | 11650 | 9121 | 26921 |
| 8962 | 23411.5 | 7680.5 | 1827.5 | 8974.5 | 2594 | 729 | 12785.5 | 9322.5 | 26287 |

| | | | | | | | | | |
|---|---|---|---|---|---|---|---|---|---|
| 8269 | 20700 | 6942 | 1689 | 8267 | 2395 | 741 | 11982 | 9098 | 23226 |
| 2578 | 3463 | 2176 | 789 | 2914 | 1385 | 354 | 7351 | 3467 | 4523 |
| 2765 | 3740 | 2764 | 996 | 3911 | 1666.5 | 525.5 | 8533 | 4227.5 | 5575 |
| 2489 | 3813 | 3108 | 1140 | 4510.5 | 1932 | 623 | 9497 | 4430 | 6319 |
| 2389.5 | 4413.5 | 6108.5 | 1726.5 | 544365 | 50220.5 | 667.5 | 10068.5 | 4434 | 7959.5 |
| 2223 | 5929.5 | 3578.5 | 1294 | 6605.5 | 2343 | 681 | 10192 | 4422 | 11003 |
| 2145.5 | 5779.5 | 3504 | 1316 | 6002.5 | 2282 | 691 | 10472 | 4341 | 11171.5 |
| 2154.5 | 5614 | 3588.5 | 1342 | 5771.5 | 2325.5 | 691 | 9934.5 | 4395 | 11016.5 |
| 2272 | 6350 | 3649 | 1395 | 5908 | 2323 | 707 | 12050 | 4500 | 12198 |
| 2595.5 | 7102 | 3938.5 | 1463.5 | 6233.5 | 2439 | 746 | 15636 | 4976.5 | 13371.5 |
| 2367 | 6317 | 3922 | 1483 | 5975 | 2422 | 737 | 13940 | 4777 | 11935 |
| 2302 | 5928 | 3845 | 1462 | 6029 | 2412 | 745 | 13723 | 4633 | 11686 |
| 2340 | 5148 | 3769 | 1446 | 5871 | 2427 | 752 | 10845 | 5028 | 10316 |
| 2149 | 3545.5 | 3142 | 1233.5 | 4913 | 2136 | 650 | 9151.5 | 4638 | 7303.5 |
| 5824.5 | 4996.5 | 3555 | 1215 | 4574 | 1967 | 621.5 | 9018.5 | 5689 | 7092 |
| 2595 | 3783 | 3177 | 1222 | 4690 | 2080 | 642 | 8640 | 4824 | 7054 |
| 2466 | 6128 | 3451 | 1304 | 5123.5 | 2151 | 665 | 9785.5 | 4568 | 10998 |
| 38435 | 37736.5 | 7846 | 1585 | 5421.5 | 2177 | 649 | 14442.5 | 5142 | 13438 |
| 2739.5 | 8276 | 4456 | 1601 | 6578 | 2582 | 777 | 14185.5 | 5241 | 15025.5 |
| 2735 | 8097.5 | 4452 | 1606.5 | 7085 | 2669 | 830 | 14798.5 | 5326.5 | 15395 |
| 2471 | 6952 | 4121 | 1487 | 6028 | 2461 | 776 | 11848 | 4878 | 12896 |
| 6581 | 14851 | 6207.5 | 1746.5 | 7872 | 2677.5 | 744 | 14873 | 8599.5 | 20038 |
| 8204.5 | 20240.5 | 9092.5 | 2223.5 | 10897.5 | 3102 | 771 | 12586.5 | 9274 | 24371.5 |
| 8995 | 21170.5 | 8415 | 1902.5 | 9877.5 | 2676 | 693 | 12789.5 | 8957.5 | 24518 |
| 7735 | 18972 | 8253 | 1958 | 10150 | 2865 | 749 | 10520 | 7881 | 22213 |
| 2760 | 3679 | 2352.5 | 808 | 3248 | 1430 | 372 | 8016.5 | 3814 | 4847 |
| 3777 | 4507 | 3156 | 1040 | 6034 | 1911 | 584 | 10579 | 4559 | 5883 |
| 3208 | 4448 | 3496 | 1136 | 8362 | 2243 | 692 | 11260 | 4534 | 6470 |
| 2906 | 6601 | 6806.5 | 2219.5 | 323757 | 48851.5 | 702.5 | 11778.5 | 4190.5 | 9583.5 |
| 3085 | 7931 | 4800 | 1444 | 120013 | 17182 | 772 | 13178 | 4582 | 11064 |
| 3063.5 | 7810.5 | 4223 | 1280.5 | 94257 | 12884.5 | 736.5 | 12178 | 4420.5 | 11080.5 |
| 3520 | 8184 | 4267 | 1293.5 | 67020.5 | 9119 | 738.5 | 15235 | 5923 | 13812 |
| 3495 | 8977 | 4482 | 1368 | 60159 | 8064 | 781 | 15887 | 4836 | 12556 |
| 3570 | 9160.5 | 4461 | 1411 | 49757 | 6826 | 795.5 | 16666.5 | 5130 | 13089 |
| 4730 | 14761.5 | 9109 | 2951.5 | 36663.5 | 5862.5 | 931 | 17214.5 | 6169.5 | 18191.5 |
| 3249 | 6568 | 4173 | 1311 | 36633 | 5137 | 802 | 13195 | 4728 | 9097 |
| 3012 | 4908 | 3959 | 1290 | 18558 | 3168 | 725 | 11614 | 4706 | 7150 |
| 3409.5 | 4373 | 3497.5 | 1140 | 25457.5 | 3814.5 | 709 | 10644 | 4812.5 | 6433.5 |
| 43611.5 | 32930 | 28643.5 | 3438.5 | 20558.5 | 3204 | 687 | 9793.5 | 17409.5 | 14375 |
| 4639.5 | 6894.5 | 4205.5 | 1234.5 | 20073 | 3357 | 735 | 10494 | 5194 | 9101 |
| 3639 | 8464 | 3987 | 1223 | 17129 | 3074 | 716 | 11117 | 4710 | 11824 |
| 394220.5 | 418523 | 105664 | 9343.5 | 16051 | 2997 | 699.5 | 50329.5 | 13728 | 19450 |
| 4940 | 11795 | 5297 | 1520 | 15255 | 3163 | 862 | 14966 | 6124 | 15686 |
| 3813 | 9538.5 | 4838.5 | 1522.5 | 74415.5 | 13868 | 817 | 15557.5 | 5653 | 13884.5 |
| 5222 | 12149 | 5759 | 1759 | 16457 | 3531 | 965 | 15546 | 6889 | 16491 |
| 8787.5 | 21879 | 7718.5 | 1962 | 15387.5 | 3336 | 819 | 16026.5 | 9531 | 25183 |
| 8639 | 22605 | 11038 | 2526.5 | 17507 | 3695.5 | 886 | 12694 | 9071.5 | 25408.5 |
| 9875.5 | 23482 | 8663 | 1819.5 | 13956 | 2936.5 | 729 | 13834.5 | 9557.5 | 25062 |

| | | | | | | | | | |
|---|---|---|---|---|---|---|---|---|---|
| 8587 | 20495 | 9381 | 2105 | 14577.5 | 3255 | 876.5 | 11938.5 | 8681 | 23073.5 |
| 2729 | 3488 | 2179 | 777 | 3088 | 1392 | 351 | 8241 | 3353 | 4352 |
| 2818.5 | 3789.5 | 2780.5 | 920.5 | 3624.5 | 1584.5 | 527 | 8911 | 4238 | 5380.5 |
| 2753 | 4046 | 3033 | 1058 | 4186 | 1724 | 649 | 9801 | 4500 | 6051 |
| 2536 | 4186 | 5758 | 1624 | 565490 | 49048 | 697 | 10635 | 4344 | 6526 |
| 2486.5 | 3879.5 | 3274.5 | 1131.5 | 6374.5 | 2058 | 698 | 10061 | 4442.5 | 6056 |
| 2571 | 3858 | 3102 | 1116 | 5412 | 1976 | 693 | 9543 | 4605 | 6034 |
| 2424 | 3859 | 3095 | 1114 | 4929 | 1882 | 709 | 9172 | 4477 | 6152 |
| 2631.5 | 3696.5 | 3009 | 1019 | 4476 | 1793 | 666 | 9983.5 | 4516.5 | 5873 |
| 2615 | 6422 | 3371 | 1167 | 5061 | 1936 | 747 | 13745 | 4422 | 10035 |
| 2591 | 6643.5 | 4026 | 1535 | 5397 | 2314 | 807.5 | 15388.5 | 4592 | 10217.5 |
| 2480 | 4927 | 3504 | 1240 | 5038 | 2019 | 780 | 13073 | 4499 | 8135 |
| 2370 | 4497 | 3350 | 1183 | 4780 | 1961 | 755 | 10235 | 4541 | 7452 |
| 2326 | 3915.5 | 3132 | 1124 | 4415 | 1852 | 725.5 | 9646.5 | 4511 | 6544 |
| 17050.5 | 12625 | 5162 | 1231 | 4589.5 | 1900.5 | 757 | 9847.5 | 7815 | 8783.5 |
| 2789.5 | 4157 | 3145.5 | 1092.5 | 4192 | 1807 | 710 | 9078 | 4668 | 6332 |
| 2715 | 4122 | 3074 | 1044 | 4155 | 1773 | 663 | 9064 | 4536 | 6395 |
| 101408 | 84037.5 | 16054.5 | 1691 | 4458.5 | 1877.5 | 676 | 19211.5 | 6347 | 11296.5 |
| 2766 | 5556 | 3250.5 | 1106 | 4206 | 1785 | 653.5 | 9970.5 | 4371.5 | 8383 |
| 2948.5 | 7927 | 4175.5 | 1370 | 5946 | 2252 | 853 | 13885 | 5335.5 | 12317.5 |
| 2578 | 6930 | 3879 | 1384 | 5422 | 2138 | 816 | 12908 | 4636 | 10808 |
| 7396 | 18601.5 | 7673 | 1911.5 | 9522 | 2737 | 870 | 14909.5 | 9347.5 | 22229.5 |
| 8198 | 20932 | 9097 | 2078 | 10744 | 2851 | 813 | 14012 | 9301 | 23190 |
| 8074 | 23315 | 9139 | 2041 | 10770 | 2751 | 780 | 10687 | 8320 | 25905 |
| 8034.5 | 23016 | 7705 | 1826 | 9090 | 2540.5 | 763 | 12187.5 | 8719.5 | 25514.5 |
| 3182 | 4186 | 2665 | 875 | 3519 | 1499 | 411 | 8925 | 4394 | 5381 |
| 2789 | 4044 | 2897 | 964 | 3842 | 1616 | 545 | 9081 | 4293 | 5603 |
| 2637 | 3858 | 2976 | 1011 | 4062 | 1700 | 628 | 9578 | 4320 | 5878 |
| 2514 | 4280 | 5892 | 1693 | 572130 | 50762 | 722 | 10539.5 | 4223 | 6460 |
| 2532 | 4150 | 3444 | 1176 | 6159 | 2090 | 733 | 10377 | 4433 | 6380 |
| 2574.5 | 3709.5 | 3040.5 | 1088 | 4959 | 1905 | 685 | 9306 | 4585.5 | 5839 |
| 2508.5 | 3686 | 3004 | 1064.5 | 4672 | 1827 | 690 | 9368 | 4492 | 5797 |
| 2690.5 | 3906.5 | 3071.5 | 1065 | 4640 | 1816 | 680 | 10503.5 | 4507 | 6137 |
| 2669.5 | 6614 | 3619 | 1251 | 5240 | 2025.5 | 778 | 14838 | 4486.5 | 10106.5 |
| 2525 | 4231 | 3308 | 1151 | 4478 | 1863 | 714 | 12270 | 4328.5 | 6454 |
| 2672 | 4015 | 3137 | 1073 | 4392 | 1830 | 714 | 10904 | 4668 | 6354 |
| 2598 | 3750.5 | 3039 | 1090 | 4157.5 | 1801 | 705 | 9807.5 | 4639.5 | 6073 |
| 2362 | 3972 | 3238 | 1131 | 4525 | 1908 | 750 | 10061 | 4483 | 6660 |
| 12544 | 10003 | 4829 | 1251 | 4764 | 1928 | 768.5 | 10055 | 6773 | 8266.5 |
| 2838 | 4601 | 3555 | 1190 | 4737 | 1916 | 758 | 10639 | 4524 | 7125 |
| 2682.5 | 4374.5 | 3415.5 | 1145 | 4640 | 1878.5 | 709 | 9947 | 4609.5 | 7048.5 |
| 75906 | 73267 | 13491 | 1576 | 4665 | 1880 | 678 | 16315 | 5564 | 11668 |
| 2700 | 4515 | 3464 | 1122 | 4621.5 | 1835.5 | 677 | 9882 | 4496.5 | 6777 |
| 2568 | 4432.5 | 3477.5 | 1107.5 | 4801.5 | 1889 | 715 | 10024.5 | 4497.5 | 7035.5 |
| 2778 | 5719 | 3718.5 | 1280 | 5185 | 2039.5 | 803 | 10906 | 4783 | 9220 |
| 8475 | 19118 | 6468 | 1685 | 7816 | 2469 | 771 | 14731 | 9562 | 21955 |
| 8153 | 22241 | 8929 | 2023 | 10368 | 2711 | 784 | 11657 | 8829 | 24507 |
| 8401 | 21310 | 8668 | 1800 | 9602 | 2558 | 738 | 12082 | 8720 | 23289 |

| | | | | | | | | | |
|---|---|---|---|---|---|---|---|---|---|
| 8874.5 | 18955 | 6142.5 | 1517.5 | 6987 | 2157 | 687 | 14428.5 | 9344.5 | 21413.5 |
| 894 | 1344 | 1165 | 637 | 1828 | 1201 | 319 | 3143 | 1810 | 2482 |
| 1330 | 2355 | 2059 | 884 | 3070 | 1528 | 545 | 6195 | 2810 | 4272 |
| 1522 | 2723.5 | 2483 | 981.5 | 3641 | 1669.5 | 647 | 7287 | 3318.5 | 5041.5 |
| 1637 | 3009 | 5085 | 1561 | 569293 | 51998 | 710 | 8377 | 3689 | 5699 |
| 1615 | 2950 | 2926 | 1095 | 4540 | 1943 | 713 | 8093 | 3738 | 5579 |
| 1784 | 3170 | 3023 | 1123 | 4578.5 | 1988 | 761 | 8744.5 | 4099.5 | 5862 |
| 1732 | 3080 | 2918 | 1121 | 4274 | 1864 | 733 | 8562 | 3905 | 5762 |
| 1911.5 | 3074.5 | 2712 | 997 | 4004.5 | 1732.5 | 685.5 | 8968.5 | 3786.5 | 5479.5 |
| 2260 | 3723 | 3220 | 1136 | 4427.5 | 1878 | 761 | 11276.5 | 4434 | 6399.5 |
| 2344 | 3801 | 3350 | 1313 | 4628 | 2060 | 784 | 11858 | 4504 | 6673 |
| 2256 | 4400.5 | 3539.5 | 1321 | 4995 | 2116 | 856 | 12254 | 4676.5 | 7627.5 |
| 1961.5 | 3490 | 3020.5 | 1154 | 4350 | 1897 | 771.5 | 9637.5 | 4354 | 6293 |
| 1677 | 2961 | 2616 | 1027 | 3708 | 1754 | 681 | 7941 | 3690 | 5302 |
| 14921 | 10672 | 4176 | 999 | 3328 | 1632 | 623 | 7297 | 6409 | 6552 |
| 2075 | 3132 | 2812 | 1026 | 3798.5 | 1704.5 | 686 | 8036.5 | 3828.5 | 5396.5 |
| 1955 | 3141 | 2710.5 | 1009.5 | 3671.5 | 1712 | 653 | 7948.5 | 3778 | 5443 |
| 61453.5 | 45964.5 | 10143 | 1479 | 4297.5 | 1869 | 682.5 | 14942.5 | 5686 | 9860 |
| 2405 | 7344 | 4019 | 1385 | 5521 | 2188 | 841 | 13111 | 4813 | 12485 |
| 2190 | 5879 | 3612 | 1287 | 5416 | 2117 | 814 | 11287 | 4439 | 10211 |
| 2252.5 | 5176.5 | 3614 | 1330 | 5223 | 2070 | 824 | 12369 | 4665 | 9265.5 |
| 7110.5 | 17095.5 | 7340 | 1892 | 9346 | 2655.5 | 855 | 14562 | 9284 | 20608.5 |
| 7467.5 | 23259.5 | 9210 | 2157 | 10944 | 2893.5 | 827.5 | 11430.5 | 8827 | 27627.5 |
| 7922 | 22638 | 7447 | 1742 | 8990 | 2441 | 713 | 11861 | 8716 | 25876 |
| 8409 | 22263.5 | 7313 | 1734 | 8603 | 2511 | 762 | 13162 | 9275.5 | 25563 |
| 2240 | 3117 | 1946.5 | 737 | 2625 | 1290.5 | 350.5 | 6765 | 3154.5 | 4136 |
| 2084 | 3265 | 2375 | 901 | 3253.5 | 1512 | 533 | 7586.5 | 3427.5 | 4831 |
| 2202 | 3432 | 2732 | 1005 | 3752 | 1686 | 638 | 8446 | 3790 | 5368 |
| 1977 | 3179 | 5044.5 | 1468.5 | 580194.5 | 53485.5 | 669 | 8000 | 3759 | 5507 |
| 1933 | 3153 | 2915 | 1041 | 4923 | 1926 | 676 | 8381 | 3726 | 5477 |
| 2016 | 3244.5 | 2867 | 1051 | 4485 | 1876.5 | 710 | 8413 | 4020.5 | 5706.5 |
| 2054 | 3388 | 2995 | 1117 | 4517 | 1883 | 727 | 8921 | 4169 | 6052 |
| 1927 | 3069 | 2564 | 980 | 3714 | 1660 | 628 | 8406 | 3639 | 5096 |
| 2422 | 3907 | 3252 | 1157 | 4558 | 1854.5 | 731 | 11819.5 | 4396.5 | 6362.5 |
| 2498 | 3948 | 3270 | 1241 | 4536 | 1972 | 747 | 11351 | 4667 | 6484 |
| 2317 | 3477.5 | 3005 | 1135 | 4192.5 | 1829.5 | 718.5 | 10738 | 4469 | 6037 |
| 2296 | 4240 | 3411 | 1250 | 4871.5 | 2035 | 804 | 11194.5 | 4674.5 | 7511 |
| 2691 | 6872 | 4066 | 1498 | 6087 | 2388 | 965 | 14198 | 5503 | 12375 |
| 15764 | 11900.5 | 5093 | 1257 | 4579 | 1987 | 794 | 9392.5 | 7783.5 | 8864 |
| 2183 | 3347 | 2748.5 | 988 | 3649 | 1669 | 665.5 | 8262.5 | 3878 | 5461.5 |
| 2360 | 3649 | 3017 | 1091 | 4146 | 1778 | 681 | 9029 | 4442 | 6092 |
| 62487 | 33153 | 9711 | 1414 | 4193 | 1796 | 663 | 13783 | 5716 | 7133 |
| 2604 | 7274 | 3979 | 1325 | 5246 | 2110 | 798 | 12260 | 4726 | 11774 |
| 2705 | 7092 | 4096 | 1416 | 6019 | 2313 | 868 | 13956 | 5242 | 11878 |
| 2510.5 | 6141.5 | 3810 | 1333 | 5263 | 2108 | 834.5 | 12171.5 | 4868 | 10284 |
| 6500 | 14660.5 | 5669.5 | 1543 | 7058 | 2273 | 737 | 14208.5 | 8174.5 | 17207.5 |
| 8141 | 23311 | 9519 | 2154 | 10989 | 2916 | 800 | 13342 | 9327 | 26488 |
| 8020 | 22635 | 8472 | 1956 | 10096 | 2658 | 761 | 11988 | 8837 | 25655 |

| | | | | | | | | | |
|---|---|---|---|---|---|---|---|---|---|
| 7663.5 | 21987 | 6651.5 | 1651 | 7938.5 | 2383 | 731 | 10200.5 | 8086.5 | 24836.5 |
| 2438 | 3386 | 2073 | 766.5 | 2765 | 1353 | 365 | 7361.5 | 3386.5 | 4491.5 |
| 2139.5 | 3278.5 | 2405.5 | 863 | 3156.5 | 1474.5 | 500.5 | 7624 | 3408 | 4784.5 |
| 2226 | 3326 | 2627 | 973 | 3547 | 1625 | 602 | 8419 | 3765 | 5071 |
| 2139 | 3255 | 4991 | 1468 | 583617 | 53076 | 621 | 8423 | 3874 | 5419 |
| 2220.5 | 3582 | 3097 | 1098.5 | 5188.5 | 1938 | 688.5 | 9019.5 | 4150 | 5891 |
| 2197.5 | 3491.5 | 2928 | 1059.5 | 4610 | 1839.5 | 700 | 8924 | 4133.5 | 5741.5 |
| 2007 | 3188 | 2773 | 1004 | 4167.5 | 1727 | 671 | 8661.5 | 3822 | 5496 |
| 2361 | 3526 | 2875 | 1007 | 4143 | 1787 | 689 | 9635 | 4290 | 5768 |
| 2423 | 4060 | 3294 | 1175 | 4565 | 1875 | 725 | 12685 | 4285 | 6426 |
| 2443 | 5722 | 3459 | 1297 | 4861 | 2081 | 781 | 13996 | 4476 | 9264 |
| 2468.5 | 4643.5 | 3435.5 | 1222 | 4873.5 | 2021.5 | 785 | 12508 | 4558 | 7944.5 |
| 2430.5 | 4078.5 | 3270 | 1210 | 4717.5 | 1903 | 770 | 10251.5 | 4785.5 | 6874.5 |
| 2391 | 4400 | 3232.5 | 1198 | 4760 | 2025.5 | 791.5 | 9790.5 | 4795.5 | 7486.5 |
| 14708.5 | 10718.5 | 4336 | 1052.5 | 3562.5 | 1669 | 650 | 8058.5 | 6764 | 7145 |
| 2498 | 3625.5 | 2935.5 | 1047 | 3936 | 1741.5 | 693.5 | 8687 | 4152.5 | 5875 |
| 2406 | 3526 | 2808.5 | 1048 | 3911 | 1708 | 657 | 8695 | 4162 | 5956 |
| 53138 | 35480 | 9037 | 1355 | 4191 | 1752 | 644 | 13501 | 5403 | 8559 |
| 2808 | 7991 | 4171 | 1374 | 5443 | 2097 | 794 | 13477 | 4918 | 12698 |
| 2333 | 4320 | 3361 | 1171 | 4984 | 1961 | 745 | 10693 | 4246 | 7075 |
| 2595 | 6531.5 | 3819 | 1344.5 | 5347.5 | 2143.5 | 850 | 12968.5 | 4778 | 10718.5 |
| 8139 | 18003 | 7534.5 | 1883.5 | 9126.5 | 2647 | 844.5 | 14992.5 | 10162 | 22117.5 |
| 6795 | 22809 | 8505 | 2045 | 10321 | 2837 | 805 | 10061 | 7951 | 26203 |
| 8353 | 22366.5 | 7833.5 | 1737 | 8948 | 2436.5 | 702.5 | 11286 | 8746.5 | 24835.5 |
| 7351 | 22271 | 8081 | 1788 | 9605.5 | 2551 | 775 | 10295.5 | 8519.5 | 25561.5 |
| 2402 | 3317 | 2073 | 750 | 2779 | 1350 | 349 | 7268 | 3316 | 4348 |
| 2176 | 3220 | 2329 | 862 | 3202 | 1499 | 510 | 7634 | 3462 | 4741 |
| 2228 | 3345.5 | 2630 | 956 | 3568 | 1621.5 | 596 | 8469.5 | 3825 | 5219 |
| 2055 | 3275 | 4934 | 1459 | 575881 | 53161 | 615 | 8335 | 3730 | 5267 |
| 2433 | 3585 | 3156 | 1090 | 5201 | 1969 | 668 | 9093 | 4365 | 5695 |
| 2375.5 | 3552.5 | 3036 | 1096.5 | 4780.5 | 1846 | 680 | 9391.5 | 4456 | 5790 |
| 2346 | 3707 | 3173.5 | 1101 | 4669.5 | 1868.5 | 729 | 9222 | 4366.5 | 6252.5 |
| 2407 | 3496 | 2838 | 1019 | 4119 | 1691 | 659 | 9387 | 4106 | 5547 |
| 2588 | 3885 | 3001 | 1062 | 4302 | 1795 | 699 | 11783 | 4387 | 6197 |
| 2595 | 3814 | 3007 | 1162.5 | 4261.5 | 1791.5 | 686 | 11175.5 | 4463.5 | 6016.5 |
| 2423 | 6299 | 3605 | 1299 | 5257 | 2095 | 803 | 14148 | 4435 | 10292 |
| 2463 | 5762 | 3708 | 1310 | 5282.5 | 2126 | 846 | 13646 | 4685 | 9750 |
| 2412 | 4086 | 3320 | 1177.5 | 4592.5 | 1982 | 776 | 10241 | 4771 | 7026.5 |
| 14785 | 11436 | 4739 | 1222 | 4514 | 1859 | 746 | 9734 | 7386 | 8589 |
| 2689 | 3895 | 3082 | 1071 | 4247 | 1817 | 710 | 8948 | 4464 | 6109 |
| 2402.5 | 3676 | 2882.5 | 1019 | 3810 | 1708.5 | 635 | 8558.5 | 4101.5 | 5844 |
| 70797 | 37280 | 10899 | 1474 | 4448.5 | 1848 | 680 | 14641 | 6031 | 7379 |
| 2429 | 6484.5 | 3262.5 | 1110 | 4415 | 1828.5 | 682 | 10500.5 | 4244.5 | 10356.5 |
| 2352 | 5699 | 3225 | 1131 | 4576 | 1867 | 694 | 10309 | 4135 | 9569 |
| 2585 | 6119 | 3622 | 1252 | 4985 | 2007 | 776 | 11925 | 4605 | 9861 |
| 6127 | 16394 | 6157 | 1657 | 7853 | 2478 | 801 | 13034 | 7675 | 19805 |
| 6396 | 22283 | 9743 | 2209 | 11857 | 3039 | 832 | 9372 | 7248 | 25527 |
| 8350.5 | 22795.5 | 7629.5 | 1698.5 | 8798 | 2421.5 | 686 | 12815.5 | 8945 | 25270.5 |

| | | | | | | | | | |
|---|---|---|---|---|---|---|---|---|---|
| 8042.5 | 22539 | 7888.5 | 1844 | 9416 | 2557 | 769 | 12695.5 | 9045.5 | 26107.5 |
| 2564 | 3728 | 2243.5 | 822 | 3007.5 | 1394 | 376 | 7797 | 3703.5 | 4660 |
| 2388 | 3602.5 | 2599 | 892 | 3443 | 1564.5 | 543 | 8372 | 3801 | 5083 |
| 2574.5 | 3562 | 2731.5 | 975 | 3737.5 | 1604 | 618 | 9022.5 | 4310 | 5549.5 |
| 2432 | 3552 | 5876 | 1685 | 706862 | 53748 | 654 | 9504 | 4389 | 5784 |
| 2498.5 | 3746 | 3323 | 1142 | 5622.5 | 2091.5 | 737 | 9703.5 | 4513 | 6075 |
| 2588 | 3844 | 3209 | 1112 | 5025 | 1940 | 733 | 9704 | 4699 | 6139 |
| 2419 | 3569 | 3050 | 1071 | 4539 | 1828 | 725 | 8862 | 4393 | 5893 |
| 2674 | 3753 | 3119 | 1079 | 4548 | 1858 | 727 | 10361 | 4584 | 6103 |
| 2571.5 | 4663.5 | 3544.5 | 1262 | 5043 | 1959 | 792 | 14111.5 | 4403 | 7293.5 |
| 2677 | 6159.5 | 3774.5 | 1309 | 5135 | 2078.5 | 807 | 14553.5 | 4702 | 9651 |
| 2516 | 4839 | 3581 | 1257 | 4921 | 2011 | 809 | 13759 | 4595 | 7819 |
| 2636 | 4780.5 | 3587 | 1269.5 | 5048 | 2076 | 822 | 11393 | 4967 | 8041 |
| 2500 | 4041.5 | 3256 | 1185.5 | 4574.5 | 1908.5 | 782 | 10205.5 | 4665 | 6949 |
| 20514 | 14932 | 5204 | 1047 | 3418 | 1605 | 630 | 8287 | 8489 | 7722 |
| 2834 | 4053 | 3204 | 1108 | 4274 | 1839 | 756 | 10423 | 4767 | 6424 |
| 2714 | 3870 | 3139 | 1062 | 4081 | 1705 | 689 | 9778 | 4826 | 6311 |
| 44091 | 41814 | 8772 | 1520 | 4860 | 1948 | 740 | 15667 | 5332 | 12191 |
| 2844 | 8411 | 4178 | 1413 | 5685 | 2224 | 861 | 14861 | 5004 | 13496 |
| 2841.5 | 7501.5 | 4125.5 | 1431.5 | 5980.5 | 2241.5 | 872 | 14676.5 | 5158 | 12324.5 |
| 2695 | 6402.5 | 3849 | 1330 | 5362 | 2080 | 829 | 14222.5 | 4752 | 10347.5 |
| 8379 | 18498 | 7842 | 1934 | 9419 | 2688 | 854 | 15015 | 9881 | 21860 |
| 7053 | 24737 | 9328 | 2166 | 11288 | 2964 | 844 | 10178 | 8017 | 28469 |
| 9189 | 23994 | 7973 | 1869 | 9256 | 2551 | 762 | 13697 | 9078 | 25143 |
| 8541 | 21707 | 7180 | 1760 | 8203 | 2424 | 760 | 11509 | 8835 | 24503 |
| 2544 | 3551 | 2235.5 | 817 | 2930.5 | 1413 | 366 | 7812 | 3523 | 4489 |
| 5280 | 4836 | 3002.5 | 972 | 11354 | 2229 | 569 | 9638.5 | 4155 | 5339 |
| 4888 | 4709 | 3260.5 | 1104 | 13488.5 | 2528.5 | 738 | 11275.5 | 5065.5 | 6259.5 |
| 6061 | 5197.5 | 7033 | 1955 | 318724.5 | 29377 | 727 | 11365 | 4735.5 | 6320 |
| 4171.5 | 4815 | 5077.5 | 1396 | 351516 | 40177.5 | 859 | 12003.5 | 5272.5 | 7120 |
| 4236 | 4678 | 4441 | 1298 | 265797 | 30293 | 875 | 12023 | 5365 | 6727 |
| 3778 | 4548 | 3985 | 1239 | 163961 | 18700 | 851 | 11999 | 4918 | 6814 |
| 3977 | 4598 | 3892 | 1229 | 157752 | 17676.5 | 849.5 | 12763.5 | 4986 | 6629.5 |
| 3908 | 6538.5 | 4104.5 | 1324 | 123919.5 | 13641.5 | 946 | 17644.5 | 5156 | 9842.5 |
| 3634 | 6086.5 | 4153 | 1462.5 | 30014 | 4014.5 | 925 | 16899.5 | 5153.5 | 9080 |
| 3954 | 5084 | 3790 | 1259 | 61319 | 6659 | 923 | 13324 | 5683 | 7587 |
| 3717 | 4478 | 3341 | 1179 | 23967 | 3340 | 854 | 11943 | 5170 | 6716 |
| 3702 | 4005 | 3131 | 1095 | 37275 | 4247 | 836 | 11346 | 4782 | 6101 |
| 67771.5 | 50628.5 | 21597.5 | 7150 | 26332.5 | 3567.5 | 789 | 10472 | 22297.5 | 17060 |
| 10950 | 8306 | 4543 | 1209 | 29714 | 3752 | 886 | 11667 | 6907 | 7922 |
| 7658 | 6652.5 | 3923 | 1203.5 | 25528 | 3364 | 831.5 | 12367 | 6248.5 | 7658 |
| 658732 | 769460 | 166478 | 13368 | 20480 | 3153 | 773 | 97485 | 26278 | 30937 |
| 11162 | 13940 | 5419 | 1578 | 25788 | 3731 | 981 | 18355 | 6363 | 14920 |
| 3672 | 8692.5 | 4566.5 | 1468 | 37703.5 | 6526.5 | 906.5 | 15389 | 5188.5 | 13413.5 |
| 6283 | 9051 | 4526 | 1479 | 22914 | 3527 | 982 | 15593 | 5581 | 12098 |
| 12454 | 23201 | 8689 | 2090 | 22760 | 3920 | 991 | 13698 | 9916 | 24879 |
| 11022 | 25697 | 10949 | 2430.5 | 20065.5 | 3828 | 965 | 14358.5 | 10002.5 | 27782.5 |
| 11617 | 25327 | 9307 | 1957.5 | 15775 | 3162.5 | 853 | 12374.5 | 9134 | 25958.5 |

| | | | | | | | | | |
|---|---|---|---|---|---|---|---|---|---|
| 10426 | 25065 | 8838 | 1934 | 12667 | 2926 | 864 | 14150 | 9698 | 27590 |
| 2628.5 | 3684 | 2225 | 793.5 | 3041 | 1367 | 366 | 7776 | 3569 | 4665 |
| 2701 | 3684.5 | 2627 | 924 | 3557.5 | 1562.5 | 541 | 8464 | 4189 | 5202 |
| 2660.5 | 3911 | 2938 | 1062.5 | 4016.5 | 1730 | 636 | 9493.5 | 4447.5 | 5818.5 |
| 2477 | 3919 | 5659 | 1680 | 597267 | 52111 | 675 | 9099 | 4423 | 6093 |
| 2530 | 3919.5 | 3539 | 1174.5 | 6262 | 2137.5 | 712.5 | 10075 | 4482.5 | 6127.5 |
| 2417 | 3955.5 | 3309 | 1133 | 5406 | 1959 | 705 | 10246 | 4366.5 | 6327 |
| 2432 | 3781 | 3258 | 1109 | 4879 | 1902 | 716 | 9880 | 4425 | 6162 |
| 2524 | 4070 | 3254 | 1108 | 4797 | 1902 | 715 | 11090 | 4334 | 6397 |
| 2701 | 6264 | 3718 | 1275 | 5347 | 2085 | 795 | 14913 | 4639 | 9725 |
| 2692 | 5411.5 | 3869 | 1377 | 5282 | 2107.5 | 795 | 14037 | 4579.5 | 8496 |
| 2466 | 4539 | 3326 | 1182 | 4791.5 | 1943 | 748 | 10730 | 4449 | 7316 |
| 2458 | 3955 | 3066 | 1117 | 4432 | 1830 | 738 | 9915 | 4638 | 6534 |
| 2168 | 3400 | 2817 | 1030 | 4034 | 1736 | 685 | 8578 | 4039 | 5708 |
| 13613 | 10123 | 4281 | 1065 | 3777.5 | 1677 | 660 | 8443 | 6432 | 7043 |
| 2882 | 3769 | 2939 | 1033 | 3954 | 1705 | 665 | 8841 | 4542 | 5746 |
| 2643.5 | 4205.5 | 3315.5 | 1123.5 | 4449.5 | 1845.5 | 704 | 9582.5 | 4493.5 | 6575.5 |
| 71394 | 41814 | 11219 | 1421 | 4168 | 1771 | 659 | 15210 | 5744 | 7712 |
| 2763 | 7966 | 3830 | 1237 | 5117 | 2012 | 754 | 12551 | 4678 | 12245 |
| 2901 | 7287 | 4064 | 1337 | 5827 | 2182 | 817 | 13642 | 5120 | 11731 |
| 2619 | 7184.5 | 4061 | 1377.5 | 5697.5 | 2189 | 857 | 13044 | 4775 | 11418 |
| 6907 | 18065 | 6502 | 1734 | 7776 | 2446 | 785 | 13511 | 8554 | 22007 |
| 7958 | 22773.5 | 9980 | 2273 | 11856 | 3130.5 | 832 | 12988 | 9234.5 | 25724.5 |
| 8176 | 22327 | 8299 | 1858 | 9639 | 2587 | 724 | 12274 | 8757 | 25528 |
| 8072 | 22412 | 7260 | 1720 | 8588 | 2438 | 752 | 12772.5 | 9208.5 | 25724 |
| 3097.5 | 3940.5 | 2484 | 821 | 3209 | 1447 | 389 | 8557 | 4099 | 4933.5 |
| 2841 | 3867 | 2828 | 999 | 3857 | 1693 | 557 | 9114 | 4439 | 5539 |
| 2583 | 4028 | 3258 | 1116 | 4428 | 1876 | 643 | 9822 | 4502 | 6179 |
| 2309 | 4113 | 6166 | 1779 | 539205 | 48655 | 668 | 10038 | 4148 | 7385 |
| 2216 | 6179.5 | 3779 | 1400.5 | 6734.5 | 2460 | 718 | 10797.5 | 4271 | 11470 |
| 2131 | 6236 | 3716 | 1420 | 6239 | 2356 | 725 | 11016 | 4422 | 11901 |
| 2199 | 5736.5 | 3639 | 1360.5 | 5823 | 2288.5 | 714 | 10365 | 4307.5 | 10928 |
| 2295 | 6718.5 | 3899 | 1466 | 6048.5 | 2383 | 734 | 12794.5 | 4413 | 12541.5 |
| 2603 | 7307.5 | 4231.5 | 1562 | 6453.5 | 2565.5 | 770 | 15861.5 | 4954.5 | 13459 |
| 2459 | 6065 | 3962.5 | 1493 | 5892 | 2432 | 737 | 13474.5 | 4790 | 11616 |
| 2266 | 5210 | 3652.5 | 1373 | 5473 | 2258 | 693 | 10935.5 | 4595 | 10200.5 |
| 2269.5 | 4118.5 | 3472 | 1318 | 5081 | 2207 | 704 | 10377.5 | 4602 | 8051 |
| 2205 | 3502 | 2978 | 1185 | 4593 | 1972 | 653 | 8781 | 4419 | 6940 |
| 5497.5 | 5169.5 | 3673 | 1213 | 4680.5 | 2003.5 | 671.5 | 9260.5 | 5218.5 | 7005.5 |
| 2441 | 5881 | 3560 | 1286 | 5305 | 2152 | 710 | 10362 | 4572 | 10852 |
| 2373 | 6646 | 3789 | 1348 | 5435 | 2192 | 709.5 | 10818 | 4452 | 12151 |
| 38327 | 40722 | 8508 | 1656 | 5774 | 2303 | 711 | 15703 | 5064 | 14224 |
| 2986.5 | 8439 | 4704 | 1601.5 | 6615 | 2596.5 | 824 | 14545.5 | 5661.5 | 15137.5 |
| 2870 | 9115 | 4762 | 1672 | 7081 | 2694 | 862 | 14540 | 5440 | 16115 |
| 2752.5 | 7814 | 4461 | 1634 | 6647.5 | 2578.5 | 828.5 | 14646 | 5100 | 14259 |
| 6584.5 | 19482.5 | 7213.5 | 1998.5 | 9248 | 2917.5 | 789 | 14436.5 | 8821 | 24266 |
| 8117.5 | 21112 | 9395 | 2277 | 11414.5 | 3161.5 | 789 | 12020.5 | 9301 | 25002.5 |
| 8100 | 20661.5 | 9135 | 2172.5 | 11110 | 3073 | 781 | 12911 | 9199.5 | 24577.5 |

| | | | | | | | | | |
|---|---|---|---|---|---|---|---|---|---|
| 7608 | 22060 | 8192 | 1950 | 10089 | 2774 | 763 | 11294 | 8732 | 26311 |
| 2544.5 | 3552 | 2401 | 846 | 3228 | 1449 | 363 | 7812.5 | 3524 | 4768 |
| 3791 | 4244 | 3081 | 1038 | 5391 | 1853 | 554 | 10372 | 4544 | 5605 |
| 3167.5 | 4260 | 3263.5 | 1143.5 | 7567 | 2166 | 668 | 10755.5 | 4657.5 | 6223 |
| 2989 | 4358 | 6702 | 2184 | 328851 | 47605.5 | 678 | 10167 | 4247.5 | 6450.5 |
| 2905 | 6123 | 4430 | 1399 | 105610 | 14965 | 762.5 | 12329.5 | 4486.5 | 9050.5 |
| 2968 | 7418 | 4147 | 1276 | 75909 | 10106 | 752 | 12717 | 4427 | 10872 |
| 3659 | 8146 | 4382.5 | 1280 | 52373 | 7111.5 | 753 | 15730.5 | 5716.5 | 13641 |
| 3428 | 9287 | 4326 | 1324 | 51660 | 7076 | 749 | 16241 | 4648 | 12829 |
| 3525 | 10711 | 4782 | 1454 | 42504 | 5970 | 847 | 15673 | 5254 | 15012 |
| 5006 | 16348 | 12766 | 3743 | 31455.5 | 5598 | 882.5 | 17257 | 6062 | 19129 |
| 3201 | 6215 | 4086 | 1291 | 33093.5 | 4779 | 762 | 12705.5 | 4858 | 8776.5 |
| 3082.5 | 6092.5 | 4344 | 1396 | 16652 | 3194 | 764.5 | 12215 | 5010 | 8893.5 |
| 3393 | 4425 | 3536 | 1136 | 24210 | 3571 | 705 | 10618 | 4848 | 6541 |
| 39161 | 30711 | 25014 | 3016 | 19105 | 3162 | 682 | 10217 | 15591 | 13724 |
| 4393 | 9758 | 4619 | 1394 | 18695 | 3397 | 794 | 11992 | 5172 | 13281 |
| 3472.5 | 9151 | 4436.5 | 1318.5 | 16455 | 3117.5 | 761 | 11540.5 | 4666.5 | 13001.5 |
| 344833 | 390382 | 94328 | 8235 | 15196 | 2915 | 705 | 47970.5 | 12388 | 19281 |
| 5286 | 11794 | 5506 | 1721 | 14941 | 3234 | 895 | 15658 | 6342 | 15895 |
| 3547.5 | 10274.5 | 4979.5 | 1625 | 75576 | 13636.5 | 856 | 15279 | 5380.5 | 14883.5 |
| 4058 | 10916.5 | 4999.5 | 1502 | 15159 | 3169.5 | 863.5 | 16100 | 5329 | 14816 |
| 8310 | 20337 | 7755 | 1968 | 15653 | 3329 | 857 | 14636 | 9265 | 23618 |
| 8087 | 24208 | 11064 | 2372 | 16915 | 3611 | 863 | 11144 | 8475 | 27402 |
| 9128 | 24351 | 9588 | 2176 | 15192 | 3238 | 812 | 12724 | 9237 | 26513 |
| 9178 | 23744.5 | 8463 | 1930 | 13071.5 | 3040.5 | 797.5 | 14528 | 9517 | 26749 |
| 2984.5 | 3776.5 | 2439 | 842 | 3459 | 1481 | 358 | 8987.5 | 3842 | 4853.5 |
| 3038 | 3969 | 2880 | 975 | 3899 | 1651 | 536 | 9860 | 4348 | 5652 |
| 2763.5 | 4148.5 | 3195.5 | 1062 | 4356 | 1787 | 637 | 10222.5 | 4352 | 6050 |
| 2508.5 | 4167 | 5989.5 | 1625.5 | 591888 | 49008 | 689 | 10407 | 4333.5 | 6513.5 |
| 2578 | 4093 | 3595 | 1215 | 7284 | 2235 | 710 | 10285 | 4422 | 6482 |
| 2487 | 3822 | 3166 | 1099.5 | 5468 | 1929 | 697 | 9633 | 4457 | 6107 |
| 2503 | 4133 | 3408.5 | 1198.5 | 5403 | 1960 | 742 | 10535 | 4423.5 | 6613.5 |
| 2577 | 3872 | 3031 | 1048 | 4690 | 1801 | 680 | 10025 | 4467 | 5998 |
| 2580 | 6287 | 3535.5 | 1207 | 5083.5 | 2012.5 | 752 | 13996 | 4375 | 9684.5 |
| 2606 | 5653 | 3609 | 1414 | 5044 | 2112 | 723 | 13331 | 4456 | 8941 |
| 2546.5 | 6901.5 | 4089.5 | 1426.5 | 5708 | 2300.5 | 855.5 | 14950 | 4547.5 | 11272 |
| 2567 | 4920 | 3611 | 1251 | 5135 | 2034 | 804 | 11276 | 4815 | 8087 |
| 2262.5 | 3731 | 2984 | 1066 | 4233.5 | 1808.5 | 696 | 9233 | 4229 | 6177.5 |
| 14813 | 10697.5 | 4371 | 1079 | 3809 | 1678 | 652 | 8419.5 | 6656.5 | 6975 |
| 2897 | 3882 | 3075 | 1066 | 4156 | 1746 | 682 | 8691 | 4649 | 5996 |
| 2677 | 4021 | 3153 | 1064 | 4309 | 1777 | 659 | 9149 | 4520 | 6303 |
| 73831 | 75652 | 13316 | 1552 | 4595 | 1858 | 671 | 17336 | 5422 | 12424 |
| 2693 | 8562.5 | 4211.5 | 1372 | 5646.5 | 2181.5 | 795.5 | 13774 | 4655.5 | 13576.5 |
| 2744 | 7689.5 | 4130.5 | 1387 | 5863 | 2232.5 | 831.5 | 13793 | 4791 | 12437 |
| 2581.5 | 7146 | 3884.5 | 1366.5 | 5492.5 | 2123.5 | 813.5 | 13243 | 4554.5 | 11352.5 |
| 8249 | 19224.5 | 7087.5 | 1800.5 | 8565.5 | 2551 | 781 | 15512 | 9887.5 | 22374 |
| 7734.5 | 20553.5 | 9259 | 2110.5 | 10832 | 2878.5 | 798 | 11043 | 8643 | 23156.5 |
| 8822.5 | 23209.5 | 8439.5 | 1849 | 9971.5 | 2641.5 | 731 | 13616 | 9452 | 26300.5 |

| | | | | | | | | | |
|---|---|---|---|---|---|---|---|---|---|
| 8444 | 21854 | 7539 | 1796 | 9081 | 2533 | 772 | 13221 | 9213 | 25381 |
| 2622 | 3654 | 2289 | 798 | 3090 | 1390 | 334 | 7711 | 3557 | 4695 |
| 2659 | 3656 | 2692 | 928 | 3611 | 1621 | 516 | 8291 | 4083 | 5261 |
| 2674 | 4106.5 | 3071 | 1074.5 | 4248 | 1774 | 636.5 | 9934 | 4497 | 6269 |
| 2404 | 3766 | 5287 | 1612 | 547871 | 49854 | 659 | 9095 | 4283 | 6026 |
| 2510 | 3809 | 3531 | 1165 | 6205 | 2110 | 696 | 9462 | 4460 | 6256 |
| 2528 | 3942 | 3309 | 1151 | 5429 | 2018 | 695 | 9665 | 4393 | 6296 |
| 2473 | 3690 | 3111 | 1100 | 4814.5 | 1872.5 | 689 | 9459 | 4468.5 | 6055 |
| 2590 | 3941 | 3091 | 1093 | 4679 | 1840 | 680 | 10400 | 4330 | 6220 |
| 2734 | 6937 | 3551 | 1186.5 | 5142.5 | 1997 | 744 | 14542.5 | 4499 | 10725.5 |
| 2479 | 4005 | 3142 | 1163 | 4354 | 1858 | 670 | 10900 | 4254 | 6200 |
| 2555.5 | 4101 | 3236.5 | 1175.5 | 4571 | 1882.5 | 721.5 | 10956.5 | 4507.5 | 6736.5 |
| 2517 | 4559 | 3468 | 1227 | 4850 | 1972 | 773 | 10724 | 4666 | 7458 |
| 2541.5 | 5039.5 | 3687.5 | 1246 | 5091 | 2040.5 | 800 | 10502.5 | 4826.5 | 8239 |
| 13001 | 9242 | 4464 | 1173 | 4343 | 1840 | 706 | 9054 | 6981 | 7543 |
| 3023 | 4027 | 3167 | 1093.5 | 4220 | 1812 | 698 | 9044.5 | 4837 | 6230 |
| 2838 | 4020 | 3212 | 1083 | 4245 | 1800 | 672 | 9116 | 4677 | 6407 |
| 68853 | 43360 | 12106 | 1590 | 4893 | 1947 | 696 | 15492 | 5761 | 8267 |
| 2909 | 5736 | 3986 | 1276 | 5038 | 1993 | 747 | 11042 | 4780 | 8455 |
| 2779.5 | 7260 | 4016.5 | 1365 | 5746 | 2183 | 800.5 | 11846.5 | 5024 | 11305 |
| 2522 | 6927 | 3850 | 1287 | 5332 | 2107 | 796 | 12038 | 4532 | 11259 |
| 7589.5 | 19174 | 7113 | 1818.5 | 9008.5 | 2594.5 | 812 | 14756.5 | 9183.5 | 23139.5 |
| 8449.5 | 22825 | 8155 | 1902 | 9625.5 | 2583 | 727.5 | 13154.5 | 8990 | 25577 |
| 8105 | 20980.5 | 8363 | 1795.5 | 9650 | 2502 | 709 | 13003.5 | 8626.5 | 23247 |
| 8482 | 19925 | 6522 | 1582 | 7721 | 2297 | 696 | 12951 | 9032 | 22736 |
| 900 | 1301 | 1161 | 661.5 | 1881.5 | 1227 | 300 | 2917 | 1795.5 | 2565 |
| 1291 | 2156.5 | 1989 | 813 | 2906.5 | 1458 | 516 | 5896 | 2708.5 | 3945 |
| 1424 | 2519 | 2279 | 920 | 3295 | 1609 | 597 | 6789 | 3078 | 4545 |
| 1496 | 2731 | 4672 | 1475 | 571210 | 52470 | 674 | 7467 | 3381 | 5118 |
| 1541 | 2653 | 2714 | 1025 | 4544 | 1950 | 670 | 7319 | 3414 | 5060 |
| 1582 | 2915 | 2813 | 1100 | 4309 | 1857 | 708 | 7986 | 3672 | 5522 |
| 1715 | 2958 | 2737.5 | 1081 | 4157 | 1873 | 745 | 8256 | 3880 | 5731 |
| 1738 | 2877 | 2515 | 964 | 3627 | 1682 | 661 | 8274.5 | 3572 | 5111 |
| 2200.5 | 3308 | 2773 | 1039 | 3900.5 | 1720 | 684 | 9679 | 4241 | 5621 |
| 2296 | 3358 | 2975 | 1196 | 4089 | 1904 | 720 | 10101 | 4466 | 5941 |
| 1845 | 3050 | 2674 | 1038 | 3776 | 1713 | 680 | 8878 | 3739 | 5347 |
| 2054 | 3209 | 2924 | 1123 | 4241 | 1875 | 751 | 9668 | 4392 | 5964 |
| 2007 | 3539 | 3345 | 1247.5 | 4728.5 | 1992 | 818 | 10118 | 4558.5 | 6703 |
| 15630 | 11681 | 4944 | 1316 | 4839 | 2047 | 831 | 9548 | 7487 | 8939 |
| 2428.5 | 4222.5 | 3507 | 1246 | 4887.5 | 2054 | 839 | 10836.5 | 4958.5 | 7579 |
| 2626 | 4844 | 3739 | 1337 | 5342 | 2128 | 863 | 11576 | 5301 | 8443 |
| 69134 | 45472 | 11553 | 1672 | 5070 | 2055 | 780 | 17217 | 6182 | 9626 |
| 2410 | 3878 | 3233 | 1112 | 4205 | 1794 | 684 | 9644 | 4733 | 6683 |
| 2250 | 3726 | 3153 | 1126 | 4792 | 1956 | 724 | 9301 | 4523 | 6512 |
| 2238.5 | 3480 | 3011.5 | 1091 | 4253 | 1803 | 726 | 9499 | 4562 | 5980 |
| 6150 | 13340.5 | 5100.5 | 1364.5 | 6097.5 | 2100 | 699.5 | 15379 | 7379 | 15728.5 |
| 8423 | 24197 | 8902 | 2045 | 10233 | 2741 | 810 | 12161 | 9411 | 27296 |
| 9257 | 19208 | 6222 | 1526 | 7149 | 2154 | 658 | 15295 | 9313 | 21458 |

| | | | | | | | | | |
|---|---|---|---|---|---|---|---|---|---|
| 7805 | 20254 | 6965 | 1629 | 8077 | 2329 | 733 | 11551 | 8446 | 22991 |
| 2371 | 3150 | 1986 | 739 | 2699 | 1343 | 349 | 7000 | 3223 | 4208 |
| 2155.5 | 3173.5 | 2408 | 894 | 3325 | 1528.5 | 553 | 7532 | 3579 | 4903.5 |
| 2160 | 3195 | 2644 | 1003 | 3708 | 1621 | 657 | 8418 | 3948 | 5373 |
| 2162 | 3448 | 5868 | 1646 | 734922 | 64360 | 711 | 8872 | 4259.5 | 6011.5 |
| 2272 | 3727 | 3416 | 1189 | 5999 | 2221 | 800 | 9493 | 4479 | 6505 |
| 2279 | 3817 | 3305 | 1208 | 5214 | 2054 | 806 | 9908 | 4616 | 6711 |
| 2157.5 | 3440 | 3112 | 1158 | 4548.5 | 1881 | 769 | 9537 | 4419.5 | 6131 |
| 2209 | 3315 | 2826 | 1023 | 4092 | 1743 | 710 | 9512 | 4223 | 5738 |
| 2473.5 | 3598.5 | 2984 | 1065.5 | 4123 | 1765 | 737 | 10645.5 | 4523.5 | 5965 |
| 2501 | 3664.5 | 3118 | 1195.5 | 4215 | 1868 | 724 | 11383.5 | 4614 | 6055 |
| 2545.5 | 3620 | 3195.5 | 1133.5 | 4341.5 | 1849.5 | 771 | 10772 | 4717 | 6260.5 |
| 2342 | 3660 | 3230 | 1145 | 4391 | 1868 | 805 | 10013 | 4624 | 6443 |
| 2256 | 3820.5 | 3346.5 | 1232 | 4749.5 | 1990 | 814.5 | 10172 | 4803 | 6907 |
| 18089.5 | 13269 | 5258.5 | 1316 | 4671 | 1982.5 | 844.5 | 10287 | 8382.5 | 9403.5 |
| 2794 | 4275 | 3414 | 1226 | 4496 | 1973 | 819 | 9851 | 4911 | 7167 |
| 2436.5 | 3565.5 | 3053.5 | 1065 | 4032 | 1770 | 703 | 9189.5 | 4442.5 | 6207.5 |
| 84796 | 44270.5 | 12601 | 1551 | 4384.5 | 1840 | 702 | 15539 | 6295 | 7525 |
| 2729 | 4275 | 3487 | 1191 | 4540 | 1845 | 750 | 10187 | 5025 | 7094 |
| 2536 | 3879 | 3208 | 1143 | 4940 | 1943 | 748 | 9832 | 4852 | 6516 |
| 2663 | 4361 | 3532 | 1241 | 4818.5 | 2030 | 833.5 | 10884.5 | 5169 | 7436.5 |
| 7042 | 15134 | 5621 | 1530 | 6663 | 2194 | 762 | 15454 | 8402 | 17833 |
| 8138 | 23704 | 8143 | 1891 | 9528 | 2557 | 777 | 13754 | 8363 | 25387 |
| 8841 | 21512 | 7216 | 1745 | 8074 | 2367 | 739 | 13969 | 8439 | 23436 |
| 8475.5 | 18826 | 6339 | 1469.5 | 7174.5 | 2262 | 721 | 12276 | 8470.5 | 20407.5 |
| 2625 | 3598 | 2332 | 811 | 3088.5 | 1400.5 | 384 | 8057.5 | 3821 | 4937.5 |
| 2968 | 4259 | 3108 | 1075 | 4284 | 1753 | 627 | 10307 | 4822 | 6460 |
| 2649 | 4053 | 3212.5 | 1144 | 4416 | 1863.5 | 715 | 10692 | 4652.5 | 6436 |
| 2423 | 5140 | 7069 | 2192 | 718342 | 57621 | 781 | 10952 | 4601 | 8568 |
| 2470 | 4041 | 4162 | 1351 | 7987 | 2588 | 790 | 11215 | 4652 | 6828 |
| 2612.5 | 6257 | 4201.5 | 1458 | 7071 | 2435.5 | 866 | 13569 | 4835.5 | 10341 |
| 2447 | 3978.5 | 3552 | 1218 | 5516.5 | 2065.5 | 810.5 | 11246.5 | 4696 | 6757 |
| 2809 | 5174 | 3608 | 1263 | 5435 | 2084 | 828 | 11295 | 5124 | 8744 |
| 2720 | 3901 | 3129 | 1095 | 4421 | 1813 | 730 | 10666 | 4678 | 6289 |
| 2619 | 3999 | 3289 | 1419 | 4754 | 2156 | 729 | 11437 | 4564 | 6426 |
| 2476.5 | 3579.5 | 3121.5 | 1179 | 4378.5 | 1907 | 722.5 | 10194.5 | 4634 | 6107.5 |
| 2351 | 4111 | 3445 | 1254 | 4976 | 2017 | 803 | 11635 | 4650 | 6980 |
| 2377 | 5157 | 3686 | 1344 | 5346 | 2114 | 877 | 12895 | 4648 | 8998 |
| 16719 | 11642 | 5032 | 1256 | 4387 | 1839 | 753 | 9373 | 7754 | 7803 |
| 3006 | 5921 | 3926 | 1349 | 5268 | 2113 | 852 | 11951 | 5131 | 9068 |
| 2712 | 4330 | 3371 | 1181.5 | 4617.5 | 1904 | 735 | 9931 | 4847.5 | 7071.5 |
| 91824 | 54830 | 14893 | 1750 | 4906 | 1947 | 732 | 16882 | 6361 | 8615 |
| 2686.5 | 4665 | 3412.5 | 1144 | 4479.5 | 1859 | 709.5 | 10571.5 | 4665 | 7527.5 |
| 2544 | 4050 | 3365 | 1169 | 5163 | 1997 | 745 | 10061 | 4556 | 6758 |
| 2647 | 3933 | 3274 | 1147 | 4508 | 1898 | 751 | 10331 | 4671 | 6568 |
| 7956 | 16211 | 5944 | 1623 | 7279 | 2309 | 759 | 15958 | 9123 | 19394 |
| 9207 | 22105 | 7622 | 1797 | 8736 | 2489 | 747 | 14209 | 9377 | 24760 |
| 9382 | 22027 | 7320 | 1720 | 8531 | 2465 | 738 | 14078 | 9369 | 24914 |

| | | | | | | | | | |
|---|---|---|---|---|---|---|---|---|---|
| 8821 | 19997 | 6517 | 1594 | 7532 | 2310 | 744 | 14215 | 9492 | 22986 |
| 2201.5 | 3041.5 | 1841 | 740 | 2585 | 1315 | 325.5 | 6631 | 3054 | 3978 |
| 2275 | 3249 | 2367.5 | 888 | 3142.5 | 1501 | 522 | 7481.5 | 3495 | 4770 |
| 2381 | 3609 | 2792 | 998 | 3923 | 1709 | 684 | 8891 | 4266 | 5682 |
| 2437 | 3930 | 5977 | 1782 | 721705 | 61170 | 730 | 9458 | 4518 | 6501 |
| 2573.5 | 4271.5 | 4017.5 | 1325.5 | 7332.5 | 2491.5 | 803.5 | 10356.5 | 4902 | 7229.5 |
| 2549.5 | 4437 | 3712.5 | 1251.5 | 5910.5 | 2160 | 826 | 10551.5 | 4826.5 | 7233 |
| 2679 | 4835 | 3818 | 1342 | 5842 | 2177 | 859 | 12896 | 5149 | 8207 |
| 2514 | 3901 | 3083.5 | 1133 | 4640.5 | 1915 | 746 | 10524 | 4728 | 6383 |
| 2520 | 3522 | 2732.5 | 989 | 4022 | 1683 | 673 | 9998 | 4278.5 | 5557 |
| 2471 | 3509 | 2948 | 1119 | 3844 | 1775 | 693 | 9915 | 4181 | 5626 |
| 2602.5 | 3590.5 | 3020.5 | 1100 | 4150 | 1818 | 735 | 10645.5 | 4733.5 | 6008 |
| 2346 | 3847 | 3098 | 1133 | 4416 | 1902 | 754 | 10589 | 4388 | 6539 |
| 2694.5 | 4916 | 3973.5 | 1413.5 | 5465 | 2237 | 912 | 12014.5 | 5498.5 | 8670.5 |
| 17581.5 | 12610.5 | 5196 | 1254.5 | 4489 | 1921 | 798 | 9979.5 | 7998 | 8753.5 |
| 3032 | 4387 | 3477 | 1187 | 4430 | 1894 | 778 | 9745 | 5057 | 6814 |
| 2691.5 | 4055 | 3146 | 1118.5 | 4309.5 | 1852.5 | 723.5 | 9641.5 | 4758.5 | 6667.5 |
| 91862.5 | 45696 | 13582.5 | 1573.5 | 4182.5 | 1756.5 | 685 | 16003.5 | 6512 | 7274 |
| 2626 | 4679 | 3478 | 1191 | 4601 | 1867 | 745 | 10631 | 4572 | 7636 |
| 2643 | 4088 | 3344 | 1171.5 | 5037.5 | 1971 | 772.5 | 10083.5 | 4812 | 6887 |
| 2639 | 3747 | 3100 | 1068 | 4140 | 1816 | 742 | 9283 | 4778 | 6223 |
| 8995 | 18246 | 6548 | 1700 | 7802 | 2422 | 819 | 16205 | 10093 | 21214 |
| 9516.5 | 21740.5 | 8058 | 1873 | 9164.5 | 2546 | 768 | 14660.5 | 9860 | 24022 |
| 9286.5 | 23783.5 | 8800.5 | 1819 | 9911 | 2556 | 770 | 12804 | 9366.5 | 25494.5 |
| 9540.5 | 19606 | 6464 | 1547.5 | 7374 | 2216.5 | 729 | 14414 | 9539.5 | 21926.5 |
| 2454 | 3389 | 2170.5 | 786.5 | 2887 | 1396.5 | 368 | 7131.5 | 3380.5 | 4454 |
| 2642 | 3638 | 2738 | 965 | 3568 | 1635 | 573 | 8531 | 3969 | 5402 |
| 2505 | 3582.5 | 2840.5 | 1024.5 | 3804 | 1705.5 | 652 | 9080 | 4237 | 5768 |
| 2509 | 3879 | 5963 | 1850 | 674216 | 61614 | 718 | 9776 | 4522 | 6404 |
| 2673 | 4084 | 3714 | 1278 | 6529 | 2269 | 792 | 10229 | 4873 | 6761 |
| 2591 | 4048 | 3508 | 1227 | 5439 | 2062 | 803 | 10317 | 4916 | 6782 |
| 2399 | 3601 | 3160 | 1076 | 4795 | 1836 | 742 | 9724 | 4579 | 6113 |
| 2625.5 | 3691 | 3094 | 1086 | 4422 | 1806 | 738.5 | 10127 | 4534 | 6032.5 |
| 2903 | 4154 | 3320 | 1141.5 | 4703 | 1887.5 | 771 | 12004 | 4834.5 | 6513 |
| 2701.5 | 4162.5 | 3382 | 1291 | 4597.5 | 1984 | 786 | 12026.5 | 4900 | 6626 |
| 2535 | 4138.5 | 3421 | 1188 | 4719 | 1982 | 818 | 12392 | 4689 | 6831.5 |
| 2499 | 4273 | 3555.5 | 1298 | 4928 | 2058.5 | 844 | 12312 | 4755 | 7311.5 |
| 2569 | 4164 | 3332 | 1252 | 4828 | 2011 | 841 | 10701 | 5055 | 7247 |
| 17922.5 | 12533.5 | 4989.5 | 1218.5 | 4096 | 1766 | 759 | 9614.5 | 7944.5 | 8007.5 |
| 3126 | 4060.5 | 3375.5 | 1185.5 | 4365 | 1849.5 | 806.5 | 9893 | 5037 | 6608.5 |
| 2914 | 4084 | 3222 | 1119 | 4328 | 1770 | 744 | 10114 | 5119 | 6615 |
| 70636 | 37282 | 11248 | 1506 | 4389 | 1824 | 728 | 15079 | 6435 | 7554 |
| 2787 | 5314 | 3290 | 1148 | 4211 | 1770 | 721 | 10591 | 4722 | 8502 |
| 2686.5 | 3957 | 3241 | 1164 | 4663 | 1889.5 | 751 | 10369 | 4815.5 | 6612 |
| 2586 | 6303 | 3682 | 1279 | 4938 | 2026 | 848 | 11384 | 4478 | 10453 |
| 6567 | 17725 | 6674 | 1796 | 8123 | 2597 | 857 | 14530 | 7816 | 20628 |
| 10795 | 26458 | 8551 | 1925.5 | 9479 | 2625.5 | 803 | 12985 | 9652.5 | 26735.5 |
| 9591.5 | 23481.5 | 8862 | 1847.5 | 9807.5 | 2550.5 | 778.5 | 12346 | 9032 | 24888.5 |

| | | | | | | | | | |
|---|---|---|---|---|---|---|---|---|---|
| 10529 | 21344 | 6873 | 1636 | 7505 | 2381 | 779 | 14628 | 9771 | 22993 |
| 2382.5 | 3266.5 | 1998 | 755 | 2728 | 1308 | 334.5 | 6809.5 | 3198 | 4157.5 |
| 2346 | 3360 | 2413 | 864 | 3317 | 1513 | 515 | 7668 | 3576 | 4892 |
| 2430.5 | 3693.5 | 2946 | 1001.5 | 3912.5 | 1688.5 | 622 | 8988.5 | 4170.5 | 5639.5 |
| 2364 | 3670 | 5589 | 1704.5 | 603197 | 53036.5 | 679 | 8949 | 4302 | 5921.5 |
| 2496 | 3783.5 | 3408.5 | 1153 | 6188 | 2141 | 711 | 9382 | 4462 | 6070 |
| 2469.5 | 3840 | 3358.5 | 1138 | 5271.5 | 1941 | 724.5 | 9563.5 | 4566.5 | 6279.5 |
| 2491 | 3898 | 3297 | 1177 | 5095 | 1937 | 757 | 9636 | 4563 | 6409 |
| 2530 | 3686 | 3054.5 | 1047.5 | 4474 | 1791 | 701.5 | 9954 | 4444 | 6112 |
| 2593 | 3965 | 3140 | 1089 | 4495 | 1848 | 719 | 11626 | 4402 | 6267 |
| 2412 | 4215 | 3449.5 | 1297 | 4783.5 | 2012 | 754.5 | 12618.5 | 4197.5 | 6785.5 |
| 2438 | 4074.5 | 3348.5 | 1194.5 | 4676 | 1897.5 | 766 | 11606.5 | 4404.5 | 6754 |
| 2487 | 4159 | 3348 | 1209 | 4763 | 1960 | 794 | 10532 | 4734 | 6866 |
| 2469.5 | 4305.5 | 3361.5 | 1234 | 4794 | 1993.5 | 800.5 | 10058 | 4909.5 | 7183.5 |
| 14759 | 11022 | 4908 | 1228 | 4590 | 1901 | 774 | 9791 | 7487 | 8499 |
| 2899 | 4145.5 | 3210.5 | 1144 | 4387.5 | 1847.5 | 745 | 9308.5 | 4718.5 | 6559.5 |
| 2807 | 4414 | 3369 | 1136.5 | 4586.5 | 1895.5 | 731 | 9922 | 4897 | 7118.5 |
| 75340 | 42382 | 11696 | 1579 | 4683 | 1875 | 709 | 15371 | 6532 | 8144 |
| 2660 | 6008 | 3502 | 1149 | 4676 | 1850 | 712 | 10867 | 4677 | 9531 |
| 2499 | 4149.5 | 3411 | 1109.5 | 4933.5 | 1930.5 | 749 | 10088.5 | 4538.5 | 6849.5 |
| 2381 | 3993 | 3330 | 1151 | 4473 | 1873 | 748 | 9937 | 4331 | 6593 |
| 7130 | 15733.5 | 5764.5 | 1579.5 | 7162 | 2285 | 764 | 14734.5 | 8437 | 18855 |
| 8077 | 22188 | 9066 | 2048 | 10391 | 2732 | 776 | 11425 | 8739 | 24557 |
| 8835 | 22779.5 | 8301 | 1837 | 9715 | 2553.5 | 735 | 12684.5 | 8928.5 | 25029 |
| 8455 | 22647 | 7545 | 1749 | 8913 | 2479 | 763 | 13593 | 9223 | 25036 |
| 2645 | 3479 | 2183 | 793 | 2984 | 1417 | 368 | 7442 | 3474 | 4452 |
| 2382.5 | 3430.5 | 2443 | 866.5 | 3295 | 1539 | 517.5 | 7781 | 3668.5 | 4911 |
| 2431 | 3465.5 | 2708 | 981 | 3694 | 1633.5 | 631 | 8452 | 4008 | 5414 |
| 2414 | 3767 | 5655.5 | 1785.5 | 602774.5 | 54544.5 | 682 | 9071.5 | 4364.5 | 6111 |
| 2706 | 4169 | 3767.5 | 1230 | 6677.5 | 2280.5 | 762 | 10055 | 4840 | 6754 |
| 2639.5 | 4047 | 3402 | 1181 | 5488 | 2042.5 | 757 | 9892.5 | 4715 | 6572 |
| 2612 | 3879 | 3388 | 1180 | 5167 | 1990 | 750 | 9978 | 4733 | 6515 |
| 2556 | 3752 | 3062 | 1064 | 4570 | 1823 | 707 | 9955 | 4390 | 6068 |
| 2709.5 | 4226.5 | 3339.5 | 1154 | 4783 | 1877 | 744.5 | 12613 | 4533 | 6657.5 |
| 2642.5 | 5498 | 3624.5 | 1353 | 4881.5 | 2076 | 787 | 13978.5 | 4578.5 | 8780 |
| 2519 | 4345 | 3423.5 | 1210 | 4889 | 1999 | 788 | 12549 | 4488.5 | 6934.5 |
| 2556 | 4466 | 3530 | 1200 | 4868 | 1978 | 814 | 11003 | 4830 | 7522 |
| 2479 | 4071.5 | 3177 | 1192 | 4544 | 1893.5 | 794 | 10008.5 | 4807 | 6772.5 |
| 14316 | 10565 | 4749 | 1219 | 4408 | 1877 | 756 | 9526 | 7297 | 8092 |
| 3169 | 4685 | 3511.5 | 1230 | 4666.5 | 1964 | 779 | 9672.5 | 5079.5 | 7453.5 |
| 2813 | 4508 | 3325.5 | 1127.5 | 4432 | 1860.5 | 716 | 9715 | 4874 | 7254.5 |
| 67049 | 39627 | 11184 | 1579 | 4447 | 1839 | 699 | 15634 | 5925 | 7720 |
| 2518 | 4552 | 3385 | 1109 | 4435 | 1810 | 690 | 10234 | 4375 | 7130 |
| 2485 | 4485 | 3347 | 1156 | 4813.5 | 1967 | 736 | 10162.5 | 4336.5 | 7417 |
| 2568 | 4247 | 3484 | 1212 | 4796 | 1936 | 775 | 10539 | 4545 | 6983 |
| 8212 | 17007 | 6272 | 1593 | 7294 | 2343 | 778 | 16497 | 9570 | 19844 |
| 9742 | 24843 | 8278 | 1925 | 9888 | 2663 | 801 | 14487 | 9666 | 27123 |
| 9485 | 22419 | 7312 | 1718 | 8140 | 2436 | 729 | 13200 | 9184 | 24278 |

| | | | | | | | | | |
|---|---|---|---|---|---|---|---|---|---|
| 9595.5 | 20011.5 | 6292 | 1588 | 7190 | 2199.5 | 719 | 13715.5 | 9201.5 | 21833 |
| 2416 | 3342 | 2088.5 | 756 | 2863 | 1372 | 338 | 7141.5 | 3265.5 | 4364.5 |
| 2519 | 3594 | 2537 | 917 | 3358 | 1562 | 525 | 8075 | 3692 | 5068 |
| 2759 | 3841.5 | 3043.5 | 1082 | 3994.5 | 1702.5 | 654 | 9359.5 | 4464.5 | 5858 |
| 2691.5 | 4037.5 | 6207 | 1968.5 | 691875.5 | 60922 | 727 | 9861.5 | 4742.5 | 6366 |
| 2816 | 4027.5 | 3788.5 | 1256.5 | 6992.5 | 2388.5 | 758.5 | 10225 | 4912 | 6479 |
| 2774 | 4034 | 3468 | 1159.5 | 5497 | 2085 | 797.5 | 10278 | 4972 | 6513.5 |
| 2739 | 4140 | 3427 | 1189 | 5286 | 2005 | 788 | 10230 | 4967 | 6686 |
| 2807 | 3945 | 3209 | 1098 | 4714 | 1816 | 725 | 10491 | 4789 | 6241 |
| 2834 | 4321.5 | 3251 | 1122.5 | 4688 | 1857.5 | 755.5 | 12454.5 | 4628 | 6624 |
| 2666 | 6220 | 3748.5 | 1381 | 5047.5 | 2086 | 806 | 15379 | 4459 | 9850.5 |
| 2647 | 4751.5 | 3669 | 1278.5 | 5085.5 | 2064 | 822.5 | 13656 | 4706 | 7678.5 |
| 2688.5 | 4065.5 | 3484.5 | 1207 | 4760.5 | 1920 | 788 | 11509.5 | 4765.5 | 6842.5 |
| 2504 | 4659.5 | 3495 | 1268 | 5010 | 2081.5 | 855 | 12010 | 4773 | 8003 |
| 16212 | 12028 | 5095 | 1256 | 4609 | 1916 | 793 | 10148 | 7747 | 8717 |
| 3258 | 4768 | 3564 | 1207 | 4682 | 1943 | 805 | 10208 | 5289 | 7548 |
| 3120 | 4531.5 | 3395 | 1183.5 | 4546 | 1867 | 757 | 10294 | 5129 | 7083.5 |
| 87367 | 60240 | 13771 | 1825 | 5332 | 2104 | 801 | 17701 | 7392 | 11148 |
| 3006 | 4616.5 | 3674.5 | 1187 | 4661 | 1904 | 746 | 11173.5 | 4857 | 7299.5 |
| 2899 | 4451 | 3411 | 1122 | 4828 | 1947 | 755 | 10457 | 5083 | 7284 |
| 2866 | 4254 | 3394 | 1178 | 4569 | 1897 | 772 | 10369 | 4888 | 6729 |
| 8215 | 18101.5 | 6274.5 | 1646.5 | 7694 | 2340 | 778 | 16163.5 | 9071.5 | 20773.5 |
| 9776.5 | 24783.5 | 9140 | 1958 | 10194.5 | 2706.5 | 804.5 | 14548.5 | 9567 | 27068 |
| 10905 | 21424 | 6977 | 1682 | 7931 | 2339 | 733 | 14799 | 9903 | 22513 |
| 9951 | 21371 | 7681 | 1678 | 8828 | 2361 | 791 | 12784 | 8948 | 22703 |
| 2562.5 | 3522 | 2168 | 808 | 2917.5 | 1404.5 | 344 | 7389.5 | 3370.5 | 4432 |
| 2555 | 3589 | 2470 | 875 | 3761 | 1559 | 492 | 8001 | 3657 | 4956 |
| 2609 | 3734 | 2748 | 986 | 4319 | 1715 | 605 | 8630 | 4133 | 5545 |
| 3045.5 | 4471.5 | 5885.5 | 1921 | 339561.5 | 33314.5 | 676 | 9337.5 | 4615 | 6579.5 |
| 2872 | 4884 | 3780 | 1271 | 31366 | 4162 | 758 | 10178 | 4944 | 7602 |
| 2601 | 3944 | 3266.5 | 1141 | 21001 | 3121 | 722 | 9727 | 4506.5 | 6294 |
| 2658 | 4348 | 3460 | 1195 | 15659.5 | 2758 | 757 | 10036.5 | 4703 | 6904 |
| 2734 | 3906 | 3130 | 1054 | 12835 | 2417 | 687 | 10422 | 4576 | 6088 |
| 2697 | 4175.5 | 3198.5 | 1081.5 | 11194 | 2331.5 | 706 | 11314.5 | 4370 | 6282.5 |
| 2584 | 6338 | 3765.5 | 1357 | 6183 | 2213 | 773 | 14801 | 4240.5 | 9928.5 |
| 2432 | 5263 | 3693 | 1282 | 7918 | 2334 | 808 | 14310 | 4422 | 8339 |
| 2514.5 | 4264.5 | 3356 | 1194.5 | 6498.5 | 2112 | 760 | 10513.5 | 4597 | 7075.5 |
| 2478 | 3820.5 | 3107 | 1111 | 6211 | 2000 | 722 | 9845 | 4552.5 | 6266.5 |
| 11078 | 8361 | 4140 | 1134 | 6056 | 1953 | 714 | 9059 | 6281 | 7285 |
| 3037.5 | 4185.5 | 3297.5 | 1133 | 5616 | 1973.5 | 725 | 9591.5 | 4792.5 | 6512 |
| 2946 | 4264 | 3278 | 1092 | 5529 | 1926 | 688 | 9452 | 4929 | 6603 |
| 147091.5 | 135979 | 25308.5 | 2071.5 | 6189 | 2026 | 693 | 24659 | 7150 | 12992.5 |
| 2693 | 7467.5 | 3696 | 1237 | 5903 | 2073 | 747 | 11970.5 | 4388 | 11805 |
| 2427 | 6190 | 3368 | 1202 | 9849 | 2636 | 721 | 10557 | 4289 | 10446 |
| 2700.5 | 6323 | 3691 | 1208 | 5994.5 | 2103.5 | 788 | 11415.5 | 4478 | 9933 |
| 8197.5 | 19162 | 6609.5 | 1731.5 | 8879.5 | 2535.5 | 781 | 16116 | 9581 | 22732 |
| 8846 | 21883 | 9094 | 1989 | 11209 | 2851 | 789 | 13690 | 9298 | 24176 |
| 8952 | 21482 | 8744 | 1794 | 10757 | 2622 | 755 | 12326 | 8834 | 23691 |

| | | | | | | | | | |
|---|---|---|---|---|---|---|---|---|---|
| 8920 | 22864.5 | 7034.5 | 1703 | 8817 | 2445.5 | 758 | 13372 | 9310.5 | 25090.5 |
| 2543.5 | 3590.5 | 2198 | 787.5 | 3020.5 | 1412 | 350 | 7234 | 3448 | 4576 |
| 2553 | 3661.5 | 2683.5 | 916 | 3521.5 | 1538.5 | 529 | 8298.5 | 3846 | 5127 |
| 2711.5 | 3826 | 2888.5 | 1003 | 3805 | 1677.5 | 665 | 9144.5 | 4394 | 5876.5 |
| 2746.5 | 4074.5 | 5953 | 1832 | 639080 | 55289 | 719 | 10093 | 4897.5 | 6419.5 |
| 2815.5 | 4144.5 | 3707.5 | 1243 | 6558 | 2334.5 | 740.5 | 10177.5 | 4832.5 | 6559.5 |
| 2753 | 4322 | 3477 | 1231 | 5654.5 | 2070 | 784 | 10726 | 4883 | 6926 |
| 2801 | 4604 | 3708 | 1253 | 5500 | 2114 | 813 | 11323 | 4986 | 7333 |
| 2906.5 | 4167.5 | 3273 | 1109 | 4827.5 | 1879 | 751 | 11205.5 | 5079 | 6583.5 |
| 2845 | 4310 | 3260 | 1132 | 4583 | 1859 | 732 | 12751 | 4607 | 6652 |
| 2723 | 4474 | 3566 | 1297 | 4780 | 1995 | 776 | 13692 | 4598 | 7048 |
| 2533.5 | 4385.5 | 3488.5 | 1217 | 4734.5 | 1981.5 | 795.5 | 12878.5 | 4536.5 | 7012.5 |
| 2452.5 | 4816.5 | 3576 | 1264 | 4952.5 | 2028.5 | 830 | 12905 | 4543.5 | 7925 |
| 2734.5 | 5952 | 4139 | 1466 | 5842 | 2256 | 924 | 14067 | 5296.5 | 10029.5 |
| 32165 | 24010 | 7633 | 1362 | 4437 | 1837 | 759 | 10380 | 13168 | 12282 |
| 3098 | 4537 | 3419 | 1191 | 4498 | 1843 | 779 | 10354 | 5243 | 7196 |
| 2956 | 4631 | 3525 | 1165 | 4633 | 1849 | 760 | 10519 | 5085 | 7485 |
| 69518 | 43027 | 11927 | 1580 | 4745 | 1880 | 754 | 15796 | 6317 | 8706 |
| 2648 | 7710 | 3921 | 1233 | 5041 | 1973 | 780 | 12187 | 4544 | 11742 |
| 2823 | 4446 | 3455 | 1149 | 4844 | 1910 | 774 | 10427 | 4933 | 7006 |
| 2735 | 4599 | 3478 | 1151 | 4753 | 1896 | 791 | 11366 | 4703 | 7325 |
| 9067.5 | 18219.5 | 6427 | 1632 | 7538.5 | 2413 | 792 | 16762.5 | 9933 | 20574.5 |
| 9657.5 | 26730 | 9495 | 2209.5 | 10919.5 | 2938 | 855 | 14433.5 | 9770.5 | 28786.5 |
| 10793 | 22546 | 7084 | 1671 | 7906 | 2296 | 729 | 15409 | 9958 | 23974 |
| 10392 | 21079 | 6934 | 1633 | 7901 | 2324 | 768 | 14092 | 9528 | 22537 |
| 2677 | 3551 | 2298 | 808 | 3059 | 1442 | 345 | 7336 | 3500 | 4563 |
| 2743.5 | 3830.5 | 2723 | 915.5 | 3644.5 | 1573.5 | 530 | 8326 | 4070 | 5309 |
| 2750 | 3946 | 2981 | 1041 | 3967 | 1713 | 641 | 9107 | 4266 | 5897 |
| 2728 | 4239 | 6067 | 1859 | 605486 | 52373 | 693 | 9716 | 4649 | 6513 |
| 2726 | 4251 | 3696 | 1206 | 6408 | 2173 | 722 | 9848 | 4816 | 6675 |
| 2725 | 4202 | 3592 | 1233 | 5574 | 2066 | 736 | 10218 | 4754 | 6671 |
| 2752 | 4448 | 3523 | 1222 | 5449 | 1993 | 751 | 10164 | 4937 | 7034 |
| 2838 | 3825 | 3093 | 1039 | 4590 | 1799 | 702 | 10354 | 4642 | 6130 |
| 2860 | 4545 | 3393 | 1145 | 4972 | 1945 | 740 | 12415 | 4707 | 6686 |
| 2952 | 5159 | 3811 | 1347 | 5072 | 2082 | 786 | 13215 | 4798 | 8023 |
| 2763.5 | 5391.5 | 3744.5 | 1290 | 5198.5 | 2067 | 801 | 12299.5 | 4733 | 8608.5 |
| 2661 | 5059 | 3642 | 1237 | 5067 | 2043 | 807 | 11526 | 4818 | 7991 |
| 2583.5 | 4214.5 | 3545 | 1246 | 4994 | 1984 | 800 | 11102.5 | 4723 | 7132.5 |
| 13141 | 9424 | 4550 | 1160 | 4284 | 1889 | 731 | 9854 | 6972 | 7661 |
| 3079 | 4143 | 3125 | 1048 | 4135 | 1731 | 689 | 9225 | 4740 | 6165 |
| 3063 | 4775 | 3481 | 1172 | 4568 | 1859 | 722 | 10037 | 5123 | 7318 |
| 75096.5 | 61451.5 | 13743 | 1978 | 6253.5 | 2307 | 832.5 | 19330 | 6948 | 13454.5 |
| 3039 | 7079 | 4250 | 1360 | 5726 | 2141 | 828 | 12321 | 5057 | 10752 |
| 2601.5 | 7154.5 | 3758.5 | 1238 | 5441 | 2044.5 | 791.5 | 10951 | 4481.5 | 11525 |
| 2647 | 4256 | 3313 | 1164 | 4560 | 1850 | 747 | 10524 | 4416 | 6785 |
| 7370.5 | 15913.5 | 5751 | 1527 | 6762.5 | 2178 | 714 | 16191.5 | 8372.5 | 18303.5 |
| 10210.5 | 23102.5 | 7637 | 1837.5 | 8524.5 | 2547.5 | 761 | 15307 | 10159.5 | 25282.5 |
| 9577 | 23557 | 7942 | 1841 | 9042 | 2527 | 771 | 13319 | 9321 | 25635 |

| | | | | | | | | | |
|---|---|---|---|---|---|---|---|---|---|
| 8684.5 | 23348.5 | 7622.5 | 1762 | 8894 | 2554.5 | 790 | 11015.5 | 8802 | 25245.5 |
| 799 | 1158 | 1020 | 622 | 1729.5 | 1196 | 270 | 2742 | 1691 | 2309.5 |
| 1188 | 1949 | 1757 | 764 | 2611 | 1397 | 475 | 4463 | 2428 | 3603 |
| 1394 | 2468.5 | 2276 | 892 | 3166 | 1575 | 585 | 6641.5 | 3070 | 4551 |
| 1499 | 2660 | 4730 | 1410 | 580289.5 | 53768.5 | 644 | 7077.5 | 3347.5 | 4962 |
| 1564 | 2867 | 2826 | 1052 | 4475 | 1951 | 688 | 7673 | 3589 | 5282 |
| 1666 | 2948 | 2799 | 1084 | 4270 | 1846 | 715 | 8050 | 3798 | 5542 |
| 1762 | 2985 | 2846.5 | 1096 | 4272 | 1852 | 741 | 8314 | 3931 | 5776.5 |
| 1731 | 2833 | 2482 | 971 | 3600 | 1635 | 648 | 8232 | 3511 | 5059 |
| 1957 | 3075 | 2624 | 978 | 3664 | 1699 | 672 | 9224 | 3772 | 5348 |
| 2067.5 | 3287 | 2822 | 1110.5 | 3919.5 | 1840 | 697 | 9574.5 | 4010 | 5551 |
| 1971.5 | 3107.5 | 2750.5 | 1071 | 3909.5 | 1783 | 705.5 | 9317 | 4050.5 | 5569.5 |
| 1952 | 2957 | 2683.5 | 1034.5 | 3796 | 1735 | 702 | 8728.5 | 4096.5 | 5366.5 |
| 1931 | 3230 | 3005.5 | 1169.5 | 4339 | 1897.5 | 760 | 9719.5 | 4424 | 6112 |
| 16556 | 15261 | 5709 | 1517 | 5610 | 2275 | 929 | 12770 | 7883 | 12016 |
| 2426.5 | 3790 | 3208.5 | 1152.5 | 4482 | 1961 | 779 | 9037 | 4614 | 6542.5 |
| 2313 | 3849 | 3281.5 | 1185 | 4633.5 | 1927.5 | 753 | 9155 | 4650 | 6765.5 |
| 70525 | 37122 | 10981 | 1487 | 4305 | 1841 | 691 | 13998 | 5637 | 6962 |
| 2351 | 3840.5 | 3356.5 | 1118 | 4386 | 1834 | 708 | 9457.5 | 4653 | 6449 |
| 2281.5 | 3897 | 3260 | 1169 | 4957 | 2018 | 754 | 9665.5 | 4706.5 | 6874.5 |
| 2168 | 3475 | 3018.5 | 1107 | 4137 | 1818.5 | 715 | 8959.5 | 4256 | 6043 |
| 7269 | 16734 | 6647 | 1780 | 8497 | 2578 | 855 | 14532 | 9094 | 20266 |
| 7313 | 21594 | 7546 | 1802 | 8641.5 | 2554.5 | 757 | 14014 | 8109 | 24268 |
| 8258 | 21553 | 6948 | 1676 | 7787 | 2382 | 694 | 13075 | 8814 | 23912 |
| 8227 | 19015 | 6165 | 1525 | 7177 | 2221 | 693 | 11918 | 8745 | 20611 |
| 2516 | 3428.5 | 2097.5 | 767 | 2832.5 | 1359.5 | 390 | 7877.5 | 3572 | 4474.5 |
| 2489.5 | 3565 | 2586 | 945 | 3398 | 1613.5 | 572 | 8714.5 | 3856.5 | 5228 |
| 2262 | 3396 | 2750 | 1020 | 3691 | 1657 | 661 | 8395 | 3970 | 5608 |
| 2434 | 3911 | 6471 | 1793 | 839097 | 69122 | 770 | 9763 | 4725 | 6572 |
| 2395 | 3742 | 3495 | 1221 | 5831 | 2194 | 805 | 9823 | 4717 | 6525 |
| 2221 | 3613 | 3245 | 1173 | 4979 | 1995 | 780 | 9791 | 4582 | 6502 |
| 2332 | 3831.5 | 3419.5 | 1246 | 5037 | 2040.5 | 815 | 9943 | 4829.5 | 6601.5 |
| 2293.5 | 3398.5 | 2962.5 | 1077 | 4046.5 | 1793.5 | 735 | 9720 | 4289 | 5840 |
| 2619.5 | 3745.5 | 3135.5 | 1108.5 | 4201 | 1827 | 748 | 11152 | 4724 | 6189 |
| 2573 | 3763 | 3174 | 1196 | 4378 | 1888 | 751 | 11201 | 4770 | 6285 |
| 2580 | 3738 | 3222 | 1187 | 4423 | 1895 | 772 | 11036.5 | 4833 | 6336 |
| 2350 | 3472 | 2952 | 1123 | 4124 | 1820 | 766 | 9870 | 4577 | 5958 |
| 2229.5 | 3583.5 | 3191.5 | 1191 | 4418 | 1888.5 | 808.5 | 9778 | 4734 | 6457 |
| 19920 | 15727 | 5940 | 1473 | 5195 | 2148 | 885 | 11716 | 8951 | 10592 |
| 2821.5 | 4158 | 3425 | 1280.5 | 4790 | 1967.5 | 817 | 10068.5 | 5047 | 7141.5 |
| 2921 | 4702 | 3736 | 1374 | 5174 | 2106 | 845 | 10646 | 5384 | 8480 |
| 89298 | 49842 | 13760 | 1646 | 4489 | 1856 | 730 | 16333 | 6481 | 7976 |
| 2657 | 4616 | 3603 | 1218 | 4912 | 1946 | 775 | 10684 | 5014 | 7640 |
| 2490.5 | 3889 | 3285 | 1164.5 | 5042.5 | 1971 | 762 | 10007.5 | 4813 | 6719.5 |
| 2688 | 3879 | 3306 | 1187 | 4469 | 1884 | 774 | 10011 | 4879 | 6674 |
| 8023.5 | 17510 | 6542.5 | 1749.5 | 7584.5 | 2478 | 824 | 15164.5 | 9014.5 | 20701 |
| 7167 | 23103.5 | 8778.5 | 2041 | 9940 | 2788.5 | 824 | 12200 | 7539.5 | 27229 |
| 8428.5 | 21984 | 7756 | 1749.5 | 8612.5 | 2489.5 | 771 | 14657 | 7958.5 | 24024.5 |

| | | | | | | | | | |
|---|---|---|---|---|---|---|---|---|---|
| 8294 | 19291 | 6385 | 1533 | 7328 | 2233 | 742 | 15146 | 7904 | 21804 |
| 2549 | 3478 | 2271 | 782.5 | 2991.5 | 1389 | 382 | 7928.5 | 3710 | 4687.5 |
| 2777 | 3874 | 2850 | 976 | 3869 | 1610 | 592 | 9298 | 4422 | 5611 |
| 2583 | 3816 | 2926 | 1068 | 4090 | 1758 | 695 | 9472 | 4635 | 5992 |
| 2486 | 3802 | 6539 | 1993 | 783787 | 62995 | 718 | 10038 | 4379 | 6131 |
| 2427 | 5567 | 4292 | 1384 | 8016 | 2637 | 817 | 12312 | 4381 | 9192 |
| 2425 | 4243 | 3572 | 1215 | 5838 | 2162 | 785 | 11819 | 4424 | 7224 |
| 2443 | 3621 | 3104 | 1100 | 4709 | 1875 | 738 | 9884 | 4474 | 6108 |
| 2708 | 5173 | 3674 | 1343 | 5615 | 2175 | 842 | 14453 | 4728 | 8424 |
| 2787.5 | 4572.5 | 3563.5 | 1232 | 5159.5 | 2015 | 798 | 11608.5 | 4847.5 | 7356 |
| 2641 | 3812 | 3217 | 1295 | 4459 | 2020 | 720 | 10857 | 4611 | 6211 |
| 2550 | 3612 | 3069 | 1140 | 4371 | 1909 | 723 | 10315 | 4541 | 6122 |
| 2466 | 3792 | 3260 | 1170 | 4467 | 1899 | 762 | 10954 | 4616 | 6312 |
| 2477 | 4426 | 3633.5 | 1311 | 5090 | 2073 | 859 | 12788.5 | 4697.5 | 7538.5 |
| 18627.5 | 13104 | 5435 | 1305 | 4675.5 | 1986.5 | 799 | 11031.5 | 8303.5 | 8749.5 |
| 3040 | 4497 | 3508 | 1249 | 4924 | 2020 | 798 | 10317 | 5080 | 7341 |
| 2693 | 3906 | 3188 | 1116 | 4295 | 1792 | 710 | 9560 | 4746 | 6519 |
| 105660 | 56092 | 15433 | 1666 | 4540 | 1885 | 708 | 17243 | 6826 | 7907 |
| 2705 | 4244.5 | 3427.5 | 1113 | 4394.5 | 1817 | 705 | 10140 | 4667.5 | 6746.5 |
| 2537 | 4290 | 3475 | 1187 | 5234 | 2063 | 755 | 10962 | 4655 | 7124 |
| 2708.5 | 3946.5 | 3258 | 1135 | 4396.5 | 1870.5 | 741 | 10040.5 | 4685.5 | 6268.5 |
| 7346 | 15663 | 5841 | 1580 | 7138 | 2295 | 762 | 15648 | 8747 | 18699 |
| 8882 | 19913.5 | 6773 | 1730.5 | 7819.5 | 2406.5 | 723.5 | 13889 | 9440 | 22208 |
| 9431 | 22651.5 | 7186 | 1736 | 8481.5 | 2437.5 | 729 | 13348.5 | 9456.5 | 25115 |
| 8664 | 18703 | 6120.5 | 1577 | 7197.5 | 2203 | 704 | 12413 | 8761.5 | 20701.5 |
| 2478 | 3452 | 2100 | 778 | 2793 | 1330 | 355 | 7313 | 3432 | 4425 |
| 2548 | 3564 | 2563 | 930 | 3456 | 1601 | 553 | 8586 | 3884 | 5134 |
| 2610.5 | 3836.5 | 3031.5 | 1105.5 | 4104 | 1782.5 | 690 | 9725 | 4575 | 6055.5 |
| 2638.5 | 4669.5 | 6609 | 1853 | 806264 | 67341 | 783 | 10686.5 | 4840 | 7571.5 |
| 2525.5 | 4094 | 3716.5 | 1262 | 6473 | 2258 | 784 | 10231 | 4657.5 | 6669 |
| 2425.5 | 4052 | 3347 | 1178.5 | 5453.5 | 2013.5 | 779.5 | 10030.5 | 4589 | 6718 |
| 2599 | 4646 | 3694.5 | 1284.5 | 5601 | 2142.5 | 863 | 10722 | 5034.5 | 7803.5 |
| 2741 | 4211 | 3283.5 | 1142 | 4723.5 | 1935 | 788 | 10638 | 4771.5 | 6792.5 |
| 2817 | 3937 | 3249 | 1126 | 4401 | 1842 | 741 | 10928 | 4867 | 6362 |
| 2769 | 3974.5 | 3139 | 1195 | 4321.5 | 1868 | 746.5 | 10737 | 4678.5 | 6183 |
| 2647 | 3934.5 | 3159.5 | 1162 | 4466 | 1876.5 | 773 | 10692 | 4702 | 6229.5 |
| 2369 | 3569 | 3145 | 1096 | 4179 | 1802 | 750 | 9736 | 4526 | 6091 |
| 2470 | 3936 | 3349 | 1260 | 4551 | 1990 | 828 | 10880 | 4801 | 6823 |
| 17809 | 16155 | 5877 | 1451 | 5487 | 2151 | 879 | 11356 | 8234 | 11437 |
| 3140.5 | 5544 | 4102 | 1427 | 5439.5 | 2190.5 | 891 | 12694.5 | 5302 | 9025.5 |
| 2934 | 6206 | 4202 | 1448 | 5648 | 2244 | 861 | 12022 | 5297 | 10423 |
| 100266 | 61392.5 | 15195.5 | 1810 | 5113.5 | 2037.5 | 773 | 18110.5 | 7273 | 10049 |
| 2851 | 4296.5 | 3644.5 | 1150.5 | 4536 | 1880 | 730 | 10564.5 | 4965.5 | 6951 |
| 2709 | 4198.5 | 3456.5 | 1192 | 5167 | 1977.5 | 767 | 10505 | 5057 | 7027 |
| 2745.5 | 4060 | 3264 | 1157 | 4572 | 1886 | 776 | 10162.5 | 4852.5 | 6588.5 |
| 5556 | 13338 | 5399 | 1542 | 6570 | 2226 | 790 | 13236 | 6798 | 15489 |
| 8898.5 | 24564.5 | 8578.5 | 1988 | 9917.5 | 2715 | 816 | 12369 | 9115.5 | 27113.5 |
| 9893 | 21260 | 6987 | 1647 | 7940 | 2407 | 725 | 13591 | 9245 | 22630 |

| | | | | | | | | | |
|---|---|---|---|---|---|---|---|---|---|
| 9464 | 20101 | 6722 | 1632 | 7887 | 2299.5 | 754 | 13691.5 | 9257 | 22805 |
| 2715 | 3653 | 2295 | 805 | 3036 | 1426 | 366 | 7945 | 3758 | 4685 |
| 2562.5 | 3697 | 2580.5 | 929.5 | 3435 | 1562 | 550 | 8089 | 3783.5 | 5045.5 |
| 2698.5 | 3800 | 3019.5 | 1042 | 3979.5 | 1739 | 684 | 9562.5 | 4473.5 | 6111.5 |
| 2609 | 3918 | 6172 | 1869 | 729453 | 68467 | 738 | 9841 | 4561 | 6275 |
| 2777 | 4209 | 3705.5 | 1314.5 | 6179.5 | 2193.5 | 806 | 10492 | 4991 | 6763.5 |
| 2685 | 4028 | 3499 | 1223 | 5374 | 2013 | 803 | 10302 | 4855 | 6678 |
| 2752 | 4513 | 3613 | 1260 | 5259 | 2093 | 836 | 10696 | 5103 | 7401 |
| 2718 | 3841 | 3106 | 1106 | 4464 | 1813 | 748 | 10684 | 4694 | 6188 |
| 2776 | 4013 | 3190 | 1115 | 4507 | 1838 | 762 | 11827 | 4674 | 6335 |
| 2833 | 4265 | 3487 | 1343 | 4748 | 2044.5 | 778 | 12753.5 | 4771 | 6709 |
| 2698 | 4410 | 3626 | 1265 | 4981 | 2007 | 845 | 13020 | 4758 | 7057 |
| 2532 | 4925.5 | 3613.5 | 1323 | 5093 | 2072.5 | 859 | 13485.5 | 4776 | 8208.5 |
| 2601.5 | 4681.5 | 3615 | 1311 | 5053.5 | 2075 | 890 | 11162 | 5061.5 | 7962 |
| 20051.5 | 14393 | 5643 | 1295 | 4530.5 | 1922 | 789 | 10170 | 8555.5 | 8969 |
| 3310.5 | 4975.5 | 3741.5 | 1259 | 4882.5 | 2022 | 857 | 10714.5 | 5323.5 | 7972.5 |
| 3222 | 5478 | 3731 | 1291 | 5023 | 2041 | 855 | 11187 | 5583 | 8806 |
| 75952.5 | 42926 | 11752.5 | 1612 | 4761 | 1949 | 773.5 | 15973.5 | 6658.5 | 8574.5 |
| 3239 | 4715 | 3737 | 1223 | 4678 | 1919 | 791 | 11183 | 5490 | 7537 |
| 2868 | 3988 | 3298.5 | 1122 | 4545.5 | 1916.5 | 739 | 10177.5 | 4976.5 | 6425 |
| 3068 | 4386 | 3402 | 1189 | 4496 | 1909 | 802 | 10807 | 5322 | 6978 |
| 5751 | 14524 | 5961 | 1577 | 6828 | 2253 | 822 | 16457 | 6755 | 16995 |
| 6682 | 26484 | 12048 | 2633 | 14950 | 3471 | 983 | 11005 | 7461 | 30307 |
| 10864 | 20759 | 6916 | 1640 | 7519 | 2276 | 741 | 15007 | 9550 | 22031 |
| 10320.5 | 19948 | 6308 | 1535 | 6959 | 2210 | 749 | 15819 | 9946.5 | 21620 |
| 2688.5 | 3635 | 2295 | 791 | 3025 | 1395.5 | 373 | 7849 | 3684.5 | 4719 |
| 2530 | 3631.5 | 2580 | 897.5 | 3467 | 1529.5 | 521 | 8003.5 | 3809.5 | 5155 |
| 2556 | 3815.5 | 3015.5 | 1044.5 | 4087 | 1749 | 647 | 9235.5 | 4307 | 5918.5 |
| 2464 | 4162.5 | 6091.5 | 1800.5 | 655893.5 | 55198 | 719 | 9649 | 4531.5 | 6587 |
| 2505 | 4205 | 3608 | 1228 | 6178 | 2197 | 761.5 | 9969 | 4639 | 6877.5 |
| 2472.5 | 4115 | 3441 | 1174.5 | 5421 | 2096 | 748.5 | 9899 | 4546.5 | 6616.5 |
| 2604 | 4325 | 3580 | 1222 | 5280 | 2052 | 800 | 10061 | 4735 | 7251 |
| 2541.5 | 3728.5 | 3041 | 1051 | 4400 | 1774 | 697 | 9772.5 | 4474.5 | 5870 |
| 2584 | 3975.5 | 3188.5 | 1096 | 4508 | 1847.5 | 712 | 11092 | 4432.5 | 6131 |
| 2558.5 | 4261.5 | 3545 | 1286.5 | 4842 | 1987 | 758 | 12222.5 | 4406 | 6724 |
| 2366.5 | 4070.5 | 3407 | 1193 | 4634.5 | 1898 | 750 | 11591 | 4277.5 | 6565 |
| 2381.5 | 4112.5 | 3375 | 1189 | 4754.5 | 1949 | 775 | 10658 | 4566.5 | 6927 |
| 2372 | 4040.5 | 3312.5 | 1183.5 | 4721.5 | 1951 | 785 | 10264 | 4647.5 | 6844.5 |
| 15503.5 | 11068 | 4824.5 | 1184 | 4311.5 | 1819 | 735 | 9334.5 | 7363.5 | 7962 |
| 3063 | 4666 | 3610.5 | 1254.5 | 4815 | 1975.5 | 789 | 9973 | 5143.5 | 7452.5 |
| 2636.5 | 3978 | 3200.5 | 1087.5 | 4314 | 1822 | 705 | 9299.5 | 4567 | 6492 |
| 76387.5 | 40912.5 | 11928 | 1508.5 | 4437.5 | 1785 | 677 | 14959.5 | 5990.5 | 7295 |
| 2682 | 4675 | 3590.5 | 1186.5 | 4736.5 | 1927 | 717 | 10632 | 4544.5 | 7306 |
| 2501 | 3824 | 3231 | 1105 | 4672 | 1861 | 690 | 9795 | 4613 | 6198 |
| 2589 | 4086 | 3417 | 1193 | 4590 | 1897 | 751 | 10309 | 4627 | 6569 |
| 5711.5 | 14329.5 | 5503.5 | 1519 | 6736 | 2274 | 740 | 15854 | 6863 | 17721 |
| 8281 | 21950 | 7479 | 1791 | 8862 | 2539 | 737 | 12294 | 8894 | 24693 |
| 8671 | 20974 | 6930 | 1670 | 8011 | 2362 | 693 | 13586 | 9019 | 23393 |

| | | | | | | | | | |
|---|---|---|---|---|---|---|---|---|---|
| 7900 | 17523.5 | 5838.5 | 1495.5 | 6885 | 2129.5 | 665 | 12964.5 | 8130 | 19184 |
| 2606.5 | 3498.5 | 2210 | 787.5 | 2909 | 1395.5 | 353 | 7505.5 | 3469.5 | 4469.5 |
| 2384 | 3411 | 2468 | 893 | 3276 | 1465 | 510 | 7658 | 3614 | 4931 |
| 2666.5 | 3890.5 | 3051.5 | 1034 | 3996.5 | 1731 | 668 | 9320.5 | 4345 | 5941 |
| 2591 | 4144 | 6354 | 1932 | 686957 | 61190 | 726 | 9890 | 4662 | 6579 |
| 2706 | 4197 | 3714 | 1244 | 6426 | 2207 | 753 | 10097 | 4768 | 6687 |
| 2653.5 | 4426 | 3664 | 1256 | 5882.5 | 2109 | 791.5 | 10340.5 | 4739.5 | 7124.5 |
| 2669.5 | 4634 | 3676 | 1273 | 5798 | 2088 | 813 | 10558.5 | 4885 | 7503 |
| 2676 | 3802 | 3050 | 1059 | 4514 | 1783 | 726 | 10128 | 4581 | 6027 |
| 2756 | 4367 | 3487 | 1178 | 4924 | 1929 | 770 | 13149 | 4590 | 6877 |
| 2759.5 | 4783 | 3670 | 1365 | 4871.5 | 2048 | 795 | 12140.5 | 4691 | 7307 |
| 2697 | 4178 | 3396 | 1210.5 | 4638.5 | 1982 | 770 | 11929 | 4707.5 | 6707.5 |
| 2444 | 4751 | 3558.5 | 1278 | 4990.5 | 2014 | 824 | 12499 | 4574.5 | 7899 |
| 2649 | 5069.5 | 3670 | 1312 | 5326.5 | 2057 | 855 | 11529 | 4960 | 8440 |
| 17083.5 | 12514.5 | 5167.5 | 1268 | 4597 | 1928 | 793 | 10074.5 | 7915.5 | 8807.5 |
| 3159 | 4496 | 3491.5 | 1206 | 4595 | 1949 | 778 | 9849 | 4972.5 | 6908.5 |
| 3232 | 5542 | 3779 | 1295 | 5074 | 1998 | 801 | 10856 | 5581 | 8782 |
| 77877 | 47117 | 12083 | 1615 | 4668 | 1961 | 720 | 15424 | 6522 | 8886 |
| 2901 | 4400.5 | 3493 | 1170 | 4540 | 1837.5 | 726.5 | 10560.5 | 4922.5 | 7035.5 |
| 2699 | 3957 | 3178 | 1108 | 4537 | 1822 | 708 | 9911 | 4808 | 6520 |
| 2822 | 4245 | 3340 | 1136 | 4547 | 1885 | 778 | 10489 | 4871 | 6632 |
| 7699 | 16624 | 6391 | 1662 | 7484 | 2355 | 803 | 16304 | 8954 | 19575 |
| 9282 | 24759.5 | 8980.5 | 1986.5 | 9995.5 | 2758 | 808 | 14461.5 | 9774 | 27355.5 |
| 9944.5 | 21826 | 7164 | 1684 | 8104.5 | 2392 | 747 | 13434.5 | 9530 | 23924 |
| 9304 | 20402 | 6412 | 1590 | 7359.5 | 2267.5 | 740 | 14146 | 9430.5 | 22259 |
| 2574 | 3513.5 | 2199.5 | 771 | 2907 | 1387.5 | 344 | 7511 | 3503.5 | 4450 |
| 2617 | 3630.5 | 2726.5 | 941 | 3539 | 1550 | 534 | 8304.5 | 3905 | 5108 |
| 2863.5 | 4106.5 | 3150 | 1086.5 | 4138.5 | 1761 | 680 | 9482.5 | 4668 | 6311 |
| 2607.5 | 4154 | 6129 | 1819.5 | 689428.5 | 63494 | 725 | 9913 | 4577 | 6672 |
| 2780 | 4436 | 3862 | 1270 | 6539 | 2247 | 794 | 10628 | 4853 | 6947 |
| 2715 | 4707 | 3617 | 1249 | 5776 | 2133 | 799 | 10549 | 4735 | 7618 |
| 2772 | 4748.5 | 3550 | 1301 | 5527.5 | 2114.5 | 818.5 | 10843.5 | 5010.5 | 7715.5 |
| 2773 | 3844 | 3071 | 1046 | 4421 | 1807 | 717 | 10246 | 4584 | 6134 |
| 2909.5 | 4283 | 3347.5 | 1119.5 | 4609.5 | 1853 | 761.5 | 12206.5 | 4578 | 6581.5 |
| 2758 | 4441.5 | 3654.5 | 1280 | 4769 | 2015 | 768 | 13232.5 | 4683.5 | 6847 |
| 2592.5 | 4604 | 3799 | 1302 | 5063 | 2062.5 | 835 | 13902 | 4555.5 | 7309.5 |
| 2603 | 4852 | 3680.5 | 1301.5 | 5084 | 2058 | 829 | 12479 | 4704 | 8050.5 |
| 2652.5 | 4670.5 | 3569.5 | 1317 | 5091 | 2027 | 834 | 11106 | 4969.5 | 7700 |
| 18814 | 13098.5 | 5087.5 | 1199 | 4223.5 | 1801.5 | 743 | 9663 | 8029.5 | 8294.5 |
| 3213 | 4654 | 3598 | 1253 | 4667 | 1956 | 795 | 10571 | 5168 | 7464 |
| 3061 | 4859 | 3672 | 1219 | 4827 | 1954 | 776 | 10771 | 5113 | 7544 |
| 88983 | 49036 | 13956 | 1596 | 4472 | 1822 | 707 | 15992.5 | 6416 | 7835 |
| 3051.5 | 4960.5 | 3874 | 1280 | 4913.5 | 1937 | 782 | 11184.5 | 5033.5 | 7746.5 |
| 2728 | 4563 | 3556 | 1163 | 5064.5 | 1975.5 | 771 | 10857.5 | 4727 | 7243 |
| 2769 | 4361.5 | 3552 | 1167 | 4693.5 | 1855.5 | 770 | 10304.5 | 4660 | 6750 |
| 6182.5 | 15497 | 5733.5 | 1547.5 | 6891.5 | 2243.5 | 777.5 | 14979.5 | 7110 | 18537 |
| 9105.5 | 24552.5 | 8789.5 | 1970 | 9950 | 2689 | 783.5 | 13721 | 9407.5 | 27200 |
| 9344 | 21954 | 7177 | 1686 | 8317.5 | 2384.5 | 706 | 13277.5 | 9021.5 | 23651 |

| | | | | | | | | | |
|---|---|---|---|---|---|---|---|---|---|
| 8836 | 17822 | 5740 | 1423 | 6739 | 2147 | 661 | 15083 | 9122 | 19993 |
| 2888 | 3748 | 2355 | 808 | 3140 | 1437 | 355 | 7974 | 3710 | 4729 |
| 2892 | 3897 | 2763.5 | 946 | 4102 | 1641 | 534.5 | 8544 | 4097.5 | 5496.5 |
| 2698 | 4199 | 3083 | 1055 | 4754 | 1805 | 657 | 9278 | 4496 | 6254 |
| 3078 | 4318 | 5829 | 1818 | 351894 | 34652 | 675 | 9366 | 4449 | 6436 |
| 2659 | 4486 | 3689.5 | 1283.5 | 26072.5 | 3673.5 | 745.5 | 10193 | 4580 | 6933 |
| 2683 | 4745.5 | 3686.5 | 1240.5 | 19018 | 3106.5 | 786 | 10456 | 4681.5 | 7384 |
| 2701 | 4629 | 3611 | 1246 | 13981 | 2720 | 780 | 10527 | 4865 | 7273 |
| 2673 | 3909 | 3125 | 1087 | 11429 | 2321 | 688 | 10264 | 4465 | 6213 |
| 2695 | 4381 | 3276 | 1112 | 9790 | 2244 | 712 | 11442 | 4473 | 6502 |
| 2616 | 4470 | 3591 | 1276 | 5933 | 2097 | 737 | 13455 | 4384 | 6890 |
| 2528.5 | 6183.5 | 3614 | 1244 | 7380.5 | 2226 | 786 | 14170 | 4302 | 9595.5 |
| 2584 | 4888 | 3569 | 1240 | 6711 | 2193 | 818 | 11331 | 4755 | 7998 |
| 2505 | 4020 | 3297 | 1156 | 5901 | 2012 | 746 | 9879 | 4677 | 6584 |
| 11723 | 8448 | 4140.5 | 1130 | 5809 | 1896 | 694.5 | 9399.5 | 6441.5 | 7145.5 |
| 2919 | 4260 | 3263 | 1133 | 5746 | 1987 | 717 | 9502 | 4727 | 6615 |
| 2763 | 4070.5 | 3255 | 1094 | 5201.5 | 1874.5 | 664 | 9170 | 4545 | 6413.5 |
| 157467 | 119468 | 25799 | 2084 | 6100 | 1997 | 688 | 24674 | 7346 | 11028 |
| 2756 | 7257 | 3771 | 1248 | 5954 | 2014 | 723 | 11304 | 4418 | 11188 |
| 2547 | 5284 | 3600 | 1203 | 11251 | 2784 | 731 | 10833 | 4347 | 8525 |
| 2589 | 6350 | 3537 | 1195 | 5818 | 2071 | 759 | 11571 | 4362 | 9756 |
| 7433 | 16370.5 | 6101.5 | 1625 | 8091 | 2348 | 755 | 15785.5 | 8685 | 19585 |
| 8542.5 | 22058.5 | 8175 | 1893 | 10105.5 | 2725.5 | 765 | 11652 | 8933.5 | 25183.5 |
| 8822 | 20739 | 6862 | 1666 | 8502 | 2391 | 679 | 13673 | 9218 | 22778 |
| 8756.5 | 18609 | 6175 | 1495.5 | 7800 | 2243.5 | 691 | 13164.5 | 8960 | 20700.5 |
| 2744 | 3650 | 2304 | 819 | 3136 | 1438 | 365 | 7848 | 3717 | 4731 |
| 2869 | 4004 | 2725.5 | 959.5 | 3756 | 1649.5 | 567.5 | 8791.5 | 4268 | 5562.5 |
| 2934.5 | 4272.5 | 3075 | 1094 | 4275 | 1756.5 | 693 | 9910 | 4744 | 6364.5 |
| 2871 | 4806 | 6392 | 1907 | 723644 | 58662 | 762 | 10683 | 4999 | 7524 |
| 2972 | 4624 | 4002 | 1318 | 6964 | 2382 | 794 | 10964 | 5024 | 7311 |
| 2834 | 5025.5 | 3877 | 1231.5 | 5966.5 | 2127 | 818 | 11170 | 4952.5 | 8022.5 |
| 2977.5 | 4925.5 | 3762 | 1298.5 | 5733 | 2134.5 | 847 | 11158.5 | 5263.5 | 8114 |
| 2907 | 4146 | 3181 | 1133 | 4671 | 1843 | 758 | 11042 | 4916 | 6428 |
| 2767 | 4353 | 3337 | 1161 | 4674 | 1894 | 774 | 12811 | 4718 | 6771 |
| 2864 | 4961 | 3708 | 1295.5 | 5070.5 | 2058 | 802 | 13303.5 | 4819.5 | 7603.5 |
| 2463 | 4548 | 3531 | 1242 | 4940 | 1941 | 811 | 13902 | 4287 | 7141 |
| 2557 | 5817.5 | 3765.5 | 1306.5 | 5288 | 2099 | 860 | 13732.5 | 4890.5 | 9655.5 |
| 2668 | 4215 | 3470 | 1238 | 4823 | 1954 | 815 | 10948 | 5061 | 6950 |
| 37321 | 26575 | 8275 | 1306 | 4407 | 1894 | 764 | 9935 | 14621 | 12370 |
| 3093 | 4111 | 3287 | 1136 | 4289 | 1804 | 754 | 9615 | 5214 | 6598 |
| 2988 | 4477 | 3482 | 1153 | 4665 | 1859 | 759 | 10912 | 5059 | 6999 |
| 64059 | 37626 | 11294 | 1554 | 4686 | 1872 | 725.5 | 16035 | 5732.5 | 7687.5 |
| 2853.5 | 6670 | 3844.5 | 1265 | 4882.5 | 1937 | 777 | 12088 | 4662.5 | 10400.5 |
| 2495 | 7441 | 3702 | 1235 | 5287 | 2039 | 804 | 11857 | 4412 | 11928 |
| 2796 | 4688 | 3684 | 1235 | 4841.5 | 1975 | 808.5 | 11562.5 | 4827 | 7398.5 |
| 8990.5 | 18692.5 | 6671.5 | 1730.5 | 7885.5 | 2499.5 | 834 | 14963.5 | 9881 | 21778 |
| 9564 | 23638 | 8804 | 2026 | 10280 | 2734 | 825 | 14415 | 9517 | 25887 |
| 9780.5 | 23700 | 8778 | 1854 | 10101 | 2616.5 | 785.5 | 13347.5 | 9278.5 | 24429.5 |

| | | | | | | | | | |
|---|---|---|---|---|---|---|---|---|---|
| 10049.5 | 22110.5 | 7176 | 1695.5 | 8093 | 2429.5 | 793 | 13550.5 | 9132.5 | 23886.5 |
| 2734 | 3710 | 2290 | 792 | 3025 | 1397 | 341 | 7592 | 3607 | 4546 |
| 2545.5 | 3668 | 2599.5 | 900 | 3497.5 | 1535 | 496 | 7923.5 | 3785 | 5045.5 |
| 2724.5 | 4031.5 | 3060 | 1031 | 4045 | 1709 | 618 | 8885 | 4379 | 5887 |
| 2671.5 | 4247.5 | 6041 | 1814.5 | 606240.5 | 52914.5 | 694 | 9809.5 | 4500.5 | 6534 |
| 2716 | 4304 | 3682 | 1240 | 6422 | 2189 | 723 | 10029 | 4675 | 6716 |
| 2748 | 4407 | 3519 | 1223 | 5732 | 2063 | 759 | 10349 | 4743 | 6857 |
| 2791 | 4307.5 | 3500 | 1186.5 | 5430 | 2000 | 753 | 10191 | 4941.5 | 6733 |
| 2844.5 | 4019 | 3206 | 1080 | 4813.5 | 1854 | 714 | 10314.5 | 4663.5 | 6196 |
| 2906 | 4463 | 3460 | 1167 | 4878 | 1942 | 745 | 12340 | 4645 | 6835 |
| 2811.5 | 4810 | 3718.5 | 1282.5 | 4901.5 | 2013.5 | 756 | 12879 | 4602.5 | 7407 |
| 2815 | 5278 | 3672 | 1253 | 5136 | 2023 | 789 | 12291 | 4894 | 8181 |
| 2643 | 4881 | 3653 | 1260 | 5034 | 2008 | 808 | 11616 | 4779 | 7860 |
| 2484 | 4157 | 3475.5 | 1172 | 4683 | 1902 | 765 | 10911.5 | 4525 | 6746 |
| 13569 | 10709.5 | 5053.5 | 1253 | 4779.5 | 1979.5 | 777 | 10942 | 6874 | 8351 |
| 3211 | 4586 | 3578 | 1204 | 4736 | 1915 | 775 | 10279 | 4957 | 6872 |
| 3047 | 5044 | 3690 | 1244 | 4847 | 1945 | 752 | 10470 | 5027 | 7941 |
| 83348 | 67041 | 14063 | 1910 | 5750 | 2221 | 797 | 17675 | 6729 | 12794 |
| 2953 | 7096 | 4292 | 1377 | 5412 | 2105 | 808 | 12832 | 4897 | 10832 |
| 2666 | 6702 | 3603 | 1193 | 5181 | 1996 | 748 | 11447 | 4523 | 10369 |
| 2768 | 7161 | 3752 | 1271 | 5009.5 | 2058 | 787 | 12260 | 4357.5 | 11199.5 |
| 7755 | 16617 | 5899 | 1567 | 6953 | 2240 | 728 | 16628 | 8610 | 18927 |
| 10101 | 22677.5 | 7901 | 1893.5 | 9024.5 | 2632 | 762 | 14846.5 | 10116 | 24906.5 |
| 9522.5 | 22665 | 7513 | 1689.5 | 8295.5 | 2395 | 726 | 14292.5 | 9167 | 24703 |
| 8829 | 22353 | 7503 | 1759 | 8692 | 2488 | 762 | 12908 | 9388 | 25320 |
| 872 | 1296 | 1093 | 627 | 1738 | 1190 | 310 | 2889 | 1748 | 2417 |
| 1200 | 1958 | 1768 | 769 | 2618 | 1351 | 483 | 4500 | 2475 | 3645 |
| 1424 | 2359 | 2156 | 875 | 3231 | 1539 | 587 | 6565 | 3001 | 4483 |
| 1564.5 | 2804 | 4882 | 1477 | 576452.5 | 53050.5 | 684 | 7417 | 3509.5 | 5251.5 |
| 1664 | 2940.5 | 2887 | 1065.5 | 4732.5 | 1993 | 715 | 7835.5 | 3642 | 5396 |
| 1702 | 3002 | 2901 | 1110 | 4422 | 1905 | 721 | 8068 | 3861 | 5533 |
| 1758 | 3118 | 2937 | 1104 | 4356 | 1939 | 752 | 8147 | 4002 | 5879 |
| 1811 | 2976 | 2645 | 989 | 3760 | 1719 | 681 | 8585 | 3721 | 5267 |
| 2112 | 3316 | 2732 | 1051 | 4005 | 1756 | 698 | 9839 | 4056 | 5658 |
| 2293 | 3528 | 3090 | 1158 | 4272 | 1878 | 739 | 10677 | 4489 | 6068 |
| 2243 | 3526 | 3113 | 1115 | 4339 | 1892 | 755 | 10207 | 4495 | 6102 |
| 2089 | 3609 | 3274 | 1221 | 4673 | 1987 | 820 | 11020 | 4573 | 6678 |
| 2005.5 | 3453 | 3218 | 1208 | 4483 | 1951 | 813 | 9223 | 4576.5 | 6426.5 |
| 16454 | 11597 | 4906.5 | 1233.5 | 4443 | 1945 | 778 | 8975.5 | 7349 | 8260 |
| 2050 | 3181 | 2800 | 1035 | 3818 | 1764 | 711.5 | 7967 | 3978 | 5653.5 |
| 2084.5 | 3306.5 | 2879.5 | 1058 | 4028 | 1740 | 680 | 8326 | 4056.5 | 5680 |
| 69795 | 37391 | 11234 | 1502 | 4196 | 1804 | 678 | 13768 | 5652 | 6920 |
| 2417 | 4108 | 3366 | 1194 | 4524 | 1915 | 745 | 9721 | 4866 | 6890 |
| 2110 | 3384 | 2943 | 1067 | 4600 | 1855 | 694 | 8704 | 4161 | 5824 |
| 2123.5 | 3360 | 2905.5 | 1055 | 4101.5 | 1771 | 711.5 | 8751 | 4141 | 5849 |
| 7142 | 15133 | 5678 | 1494 | 6786 | 2192 | 721 | 15587 | 8921 | 18468 |
| 8264 | 20766 | 7324 | 1810 | 8432 | 2509 | 742 | 14942 | 9365 | 24588 |
| 8999 | 21878 | 6933 | 1683 | 8059 | 2377 | 701 | 13887 | 9502 | 24755 |

| | | | | | | | | | |
|---|---|---|---|---|---|---|---|---|---|
| 8578 | 17028 | 5844 | 1409 | 6841 | 2142 | 667 | 13877 | 9123 | 19761 |
| 2297 | 3217 | 1949 | 716 | 2607 | 1312 | 346 | 7200 | 3291 | 4173 |
| 2287 | 3320 | 2439 | 880 | 3328 | 1520 | 542 | 7842 | 3625 | 4780 |
| 2405 | 3620.5 | 2985 | 1071.5 | 4086 | 1764 | 689 | 9141.5 | 4417.5 | 5956.5 |
| 2214 | 3575 | 5574 | 1557 | 708194 | 62624 | 732 | 9082 | 4358 | 6095 |
| 2312 | 3672 | 3423 | 1206 | 5841 | 2169 | 777 | 9649 | 4687 | 6524 |
| 2318 | 4023 | 3416 | 1267 | 5410 | 2098 | 824 | 11151 | 4768 | 6943 |
| 2397 | 4309 | 3527 | 1297 | 5256 | 2100 | 851 | 10432 | 4767 | 7596 |
| 2310 | 3523.5 | 2968.5 | 1075 | 4307.5 | 1829 | 741 | 9866 | 4322 | 5943.5 |
| 2583.5 | 3730 | 3118.5 | 1099 | 4350.5 | 1816 | 759 | 10814 | 4693.5 | 6160 |
| 2466 | 3586 | 3016 | 1125 | 4075 | 1799 | 741 | 10543 | 4513 | 5928 |
| 2380 | 3625 | 3077 | 1140 | 4269 | 1845 | 766 | 10010 | 4539 | 6254 |
| 2285 | 3610 | 3149 | 1150 | 4480 | 1911 | 768 | 10349 | 4672 | 6306 |
| 2234 | 3669 | 3236 | 1198 | 4586 | 1958 | 798.5 | 10379 | 4712 | 6635 |
| 19059 | 13880 | 5383 | 1332 | 4783 | 2014 | 835 | 10690 | 8286 | 9240 |
| 2904.5 | 4922.5 | 3737.5 | 1365.5 | 5188 | 2154 | 872 | 10842.5 | 5264 | 8278 |
| 2718.5 | 4868.5 | 3625 | 1274.5 | 5045.5 | 2011.5 | 811 | 10565.5 | 5189 | 8028 |
| 84027 | 53440 | 13619 | 1799 | 5139 | 2097 | 798 | 16933 | 6667 | 9738 |
| 2602.5 | 4510 | 3720 | 1276.5 | 4928 | 1970 | 760 | 10352 | 5032.5 | 7388 |
| 2435 | 4341.5 | 3505.5 | 1227.5 | 5447 | 2131.5 | 801.5 | 10881 | 4756 | 7654 |
| 2673 | 4088 | 3388.5 | 1229.5 | 4775.5 | 1941 | 794.5 | 10237 | 5049.5 | 6916.5 |
| 6964 | 15706 | 7368.5 | 1874 | 9358.5 | 2681.5 | 836.5 | 13366.5 | 8908.5 | 19144.5 |
| 8233 | 24969 | 9383 | 2210 | 11221 | 2904 | 851 | 12146 | 8689 | 28184 |
| 7925.5 | 21501 | 7080.5 | 1698.5 | 8028.5 | 2405 | 710 | 14164.5 | 8505.5 | 23933.5 |
| 7089 | 17201 | 5786 | 1413 | 6655 | 2097 | 671 | 14284 | 7541 | 19440 |
| 2746.5 | 3635 | 2285.5 | 784 | 2975.5 | 1378 | 397.5 | 8418.5 | 3814.5 | 4817.5 |
| 2497 | 3469 | 2591 | 887.5 | 3491.5 | 1531 | 538 | 8231.5 | 3971 | 5146.5 |
| 2575 | 3740.5 | 2971.5 | 1046.5 | 4068.5 | 1761 | 660 | 9561 | 4412.5 | 5861 |
| 2323 | 3638 | 6038 | 1792 | 714858 | 58547 | 700 | 10275 | 4319 | 6099 |
| 2446.5 | 4270 | 3750 | 1277.5 | 6672.5 | 2323 | 793 | 11359.5 | 4549 | 6966 |
| 2425 | 4015.5 | 3531.5 | 1210 | 5603 | 2103.5 | 779 | 11369 | 4548.5 | 6778.5 |
| 2441 | 3880 | 3395 | 1139.5 | 5020.5 | 1968.5 | 767 | 10686.5 | 4641 | 6522 |
| 2618 | 4662.5 | 3571 | 1222.5 | 5278.5 | 2044.5 | 817.5 | 13127.5 | 4716 | 7710 |
| 2714 | 3976 | 3239 | 1124 | 4628 | 1865 | 736 | 10888 | 4713 | 6458 |
| 2537 | 3664 | 2973.5 | 1051 | 4178 | 1759.5 | 691 | 10075 | 4555 | 5843.5 |
| 2592 | 3756.5 | 3128.5 | 1098 | 4343 | 1853 | 734 | 10521 | 4617.5 | 6098 |
| 2441 | 3840.5 | 3244 | 1177 | 4549.5 | 1919 | 768 | 11301 | 4651 | 6353.5 |
| 2386.5 | 3837 | 3213.5 | 1197 | 4676 | 1928 | 774 | 11214.5 | 4595.5 | 6532.5 |
| 17320 | 13220 | 5490 | 1355 | 4884.5 | 2046.5 | 827 | 10915.5 | 8002.5 | 9482 |
| 2959 | 4257 | 3499 | 1213 | 4784 | 1972 | 801 | 10376 | 5019 | 6926 |
| 3115 | 5223 | 3737 | 1296 | 5224 | 2069 | 805 | 11139 | 5468 | 8522 |
| 98003 | 55725.5 | 15058 | 1674.5 | 4550.5 | 1841.5 | 706 | 16484 | 6503 | 8333 |
| 2823 | 4125 | 3330.5 | 1123.5 | 4448 | 1845.5 | 701.5 | 9899 | 4809.5 | 6596.5 |
| 2638 | 4307 | 3466 | 1212 | 5449 | 2094 | 769 | 11302 | 4627 | 7079 |
| 2697 | 3990 | 3280 | 1122 | 4493 | 1883 | 725 | 10397 | 4800 | 6441 |
| 6564 | 15010 | 5704 | 1575 | 6938 | 2239 | 764 | 15599 | 7965 | 18383 |
| 8744.5 | 21304.5 | 7346 | 1805.5 | 8348 | 2515.5 | 723 | 13841.5 | 9462.5 | 23895.5 |
| 9310 | 19819 | 6727 | 1653 | 7719 | 2315 | 697 | 15007 | 9646 | 22104 |

| | | | | | | | | | |
|---|---|---|---|---|---|---|---|---|---|
| 8834 | 18551 | 6374 | 1566 | 7384 | 2269 | 704 | 12876 | 9297 | 20680 |
| 2630 | 3562 | 2139 | 754 | 2869 | 1344 | 366 | 7656 | 3602 | 4544 |
| 2485 | 3415.5 | 2435.5 | 880.5 | 3302.5 | 1517.5 | 536.5 | 8084.5 | 3704.5 | 4988 |
| 2658 | 3866 | 2987 | 1055 | 4171 | 1794 | 676 | 9509 | 4623 | 6175 |
| 2405 | 3801 | 5829 | 1594 | 724673 | 63114 | 708 | 9531 | 4446 | 6154 |
| 2556 | 4107 | 3563 | 1191 | 6045 | 2128 | 758 | 9974 | 4674 | 6816 |
| 2545 | 4487 | 3683 | 1227 | 5622.5 | 2104 | 790 | 10639.5 | 4847.5 | 7364.5 |
| 2816.5 | 4979.5 | 3748.5 | 1280.5 | 5497.5 | 2091 | 840 | 10748.5 | 5351.5 | 8393.5 |
| 2608 | 3718 | 3061 | 1075 | 4440 | 1832 | 720 | 10305 | 4548 | 5937 |
| 2779 | 3944 | 3097 | 1087 | 4417 | 1813 | 720 | 10963 | 4652 | 6228 |
| 2620 | 3739 | 3082 | 1143 | 4118 | 1866 | 725 | 10734 | 4513 | 5972 |
| 2653 | 3749 | 3016 | 1099 | 4185 | 1810 | 717 | 10573 | 4681 | 5959 |
| 2485.5 | 3900.5 | 3247.5 | 1171 | 4457.5 | 1846 | 789 | 11018.5 | 4727 | 6494.5 |
| 2515.5 | 4363 | 3644 | 1308 | 4992.5 | 2047 | 839 | 12510.5 | 4686 | 7357.5 |
| 16469 | 15497 | 5701 | 1445 | 5375 | 2219 | 918 | 13725 | 8020 | 11758 |
| 3015.5 | 5414 | 3730 | 1325.5 | 5089.5 | 2117 | 849 | 11027 | 5091.5 | 9032.5 |
| 3268 | 6894 | 4267.5 | 1449.5 | 5854 | 2262 | 875 | 11984.5 | 5889 | 11399.5 |
| 84231 | 59329.5 | 14497 | 1784 | 5162.5 | 2001.5 | 768 | 16963 | 6327.5 | 9776.5 |
| 3052 | 6061 | 4368.5 | 1376.5 | 5394.5 | 2146.5 | 791 | 11784.5 | 5127 | 9115.5 |
| 2959 | 4925 | 3615 | 1297 | 5570 | 2168 | 791 | 11418 | 5308 | 7891 |
| 2883.5 | 4370 | 3478.5 | 1230.5 | 4794 | 1917.5 | 781 | 10678.5 | 5183 | 7211.5 |
| 6174 | 15509.5 | 6572 | 1717 | 8089 | 2517 | 851 | 13999 | 7646 | 18568 |
| 7170 | 23818.5 | 9709.5 | 2224 | 11410 | 3046.5 | 853 | 12325 | 8178.5 | 27538.5 |
| 7456 | 19817 | 6449 | 1589 | 7419 | 2188 | 666 | 13831 | 7701 | 21680 |
| 8343 | 18161 | 5932 | 1514 | 7090 | 2217 | 699 | 13624 | 8709 | 20993 |
| 2830 | 3672 | 2113 | 766 | 2816 | 1355 | 344 | 8168 | 3763 | 4597 |
| 2803.5 | 3811 | 2720 | 946 | 3541 | 1559 | 564 | 8912.5 | 4033 | 5292 |
| 2884 | 4160 | 3155 | 1099 | 4212 | 1768 | 712 | 9879 | 4793 | 6353 |
| 2631.5 | 4492 | 6231 | 1772 | 687805.5 | 62791 | 769 | 10459 | 4748 | 7135.5 |
| 2750 | 4726 | 4002 | 1338 | 6595 | 2299 | 835 | 11214 | 4916 | 7569 |
| 2728 | 5377 | 4000.5 | 1356.5 | 6056.5 | 2265 | 875.5 | 12844 | 5060.5 | 8751.5 |
| 2691 | 4573 | 3809 | 1276 | 5532 | 2092 | 833 | 11464 | 5037 | 7820 |
| 2846 | 4127 | 3284 | 1141 | 4635 | 1868 | 770 | 10764 | 4928 | 6452 |
| 2869 | 4136 | 3204 | 1116 | 4442 | 1827 | 746 | 11742 | 4639 | 6410 |
| 2604 | 4256.5 | 3527.5 | 1246 | 4599.5 | 1947.5 | 775 | 12515 | 4457 | 6741 |
| 2609 | 4145 | 3374 | 1178 | 4681 | 1898 | 794 | 12294 | 4622 | 6678 |
| 2541 | 4243 | 3534 | 1214 | 4908 | 1967 | 812 | 12505 | 4737 | 7101 |
| 2520 | 4744 | 3655 | 1315 | 5009 | 2045 | 862 | 12038 | 4856 | 8021 |
| 19430.5 | 14467 | 5639.5 | 1298.5 | 4696.5 | 1969.5 | 820 | 10533 | 8646 | 9683.5 |
| 3469.5 | 5107.5 | 3748 | 1293.5 | 4973 | 2087.5 | 841 | 10821.5 | 5400 | 7786 |
| 2968 | 4906.5 | 3623 | 1201 | 4865 | 1998 | 783 | 10460.5 | 5060.5 | 7803 |
| 70669 | 41779 | 11423 | 1568 | 4737 | 1847 | 749 | 15648 | 6232 | 8190 |
| 2796 | 4534 | 3580 | 1225 | 4683 | 1913 | 743 | 11087 | 4695 | 7084 |
| 2695.5 | 4150 | 3444.5 | 1151.5 | 4791 | 1914.5 | 754 | 10464 | 4782.5 | 6832 |
| 2710 | 4350 | 3638 | 1235 | 4702 | 1936 | 799 | 10990 | 4675 | 6969 |
| 7134 | 15351 | 5866 | 1541 | 6887 | 2239 | 790 | 15910 | 8344 | 17573 |
| 9859.5 | 23706.5 | 8405.5 | 1990 | 9575 | 2655.5 | 792 | 14356 | 9798 | 26535.5 |
| 10542 | 21147 | 7030.5 | 1719.5 | 7910.5 | 2391.5 | 745 | 14090.5 | 9682 | 22931.5 |

| | | | | | | | | | |
|---|---|---|---|---|---|---|---|---|---|
| 9628 | 19598 | 6831 | 1608 | 7899 | 2282 | 770 | 12356 | 8801 | 21290 |
| 2638 | 3604 | 2206 | 783 | 2987 | 1360 | 352 | 7713 | 3614 | 4599 |
| 2495 | 3504 | 2476 | 874 | 3338 | 1529 | 514 | 7975 | 3737 | 4951 |
| 2559 | 3792 | 2882 | 1033 | 4032 | 1734 | 637 | 8830 | 4211 | 5878 |
| 2507 | 4012 | 5761 | 1697 | 641216 | 54412.5 | 694.5 | 9287.5 | 4508.5 | 6401.5 |
| 2559.5 | 4382 | 3681.5 | 1257 | 6039 | 2174 | 748 | 10066.5 | 4596.5 | 7090.5 |
| 2524 | 4015 | 3492 | 1171 | 5332 | 2005 | 742 | 9805 | 4595 | 6581 |
| 2541 | 4097 | 3326 | 1178 | 5058 | 1942 | 755 | 9665.5 | 4642.5 | 6755 |
| 2568.5 | 3843.5 | 3094.5 | 1093 | 4516 | 1826.5 | 686 | 10031.5 | 4459.5 | 6061.5 |
| 2581 | 4189 | 3337 | 1152 | 4721 | 1873 | 728 | 12210 | 4419 | 6506 |
| 2409 | 4097 | 3377 | 1227 | 4528 | 1906 | 725 | 11846 | 4195 | 6336 |
| 2524 | 4242 | 3450 | 1182.5 | 4814 | 1960 | 765 | 12018 | 4510 | 6723 |
| 2459 | 4265 | 3391.5 | 1208.5 | 4763 | 1916 | 788 | 11498 | 4512 | 7120 |
| 2490 | 4284 | 3417 | 1193 | 4817 | 1953 | 781 | 10166 | 4821 | 7329 |
| 14138 | 9868 | 4564 | 1142 | 4218 | 1816 | 729 | 9190 | 7017 | 7707 |
| 2832 | 4162 | 3200 | 1109 | 4465 | 1847 | 730 | 9496 | 4647 | 6608 |
| 2848 | 4182 | 3271 | 1153 | 4462 | 1901 | 700 | 9595 | 4768 | 6562 |
| 78523 | 44022 | 11939 | 1536 | 4463 | 1849 | 676 | 14892 | 6359 | 7702 |
| 2646.5 | 4325 | 3494.5 | 1163 | 4683 | 1845 | 702.5 | 10294 | 4492 | 6835 |
| 2473.5 | 4305.5 | 3553 | 1203.5 | 5162 | 1992.5 | 740 | 10454.5 | 4472 | 6787 |
| 2413 | 4199.5 | 3460 | 1147 | 4605 | 1873.5 | 748 | 10246.5 | 4272.5 | 6675.5 |
| 6838 | 15917 | 5761 | 1538 | 6978 | 2221 | 742 | 14943 | 8397 | 18417 |
| 9119 | 23168 | 7786 | 1830.5 | 9002.5 | 2557 | 742 | 14480.5 | 9586 | 25587.5 |
| 9468 | 19822.5 | 6774.5 | 1639.5 | 7687.5 | 2329 | 674 | 14048 | 9282 | 22204.5 |
| 8445 | 18486 | 6131 | 1455 | 7061 | 2153 | 697 | 13124 | 8784 | 20266 |
| 2838.5 | 3643 | 2255.5 | 764 | 2931.5 | 1359.5 | 376 | 7983 | 3754 | 4650.5 |
| 2592.5 | 3671.5 | 2634 | 859.5 | 3474 | 1522 | 523 | 8043.5 | 3878.5 | 5247 |
| 2548 | 3788 | 2888 | 999 | 3942.5 | 1672 | 618 | 8849 | 4264 | 5746 |
| 2501.5 | 4219.5 | 6005 | 1802 | 634357.5 | 53593 | 730 | 9784.5 | 4519.5 | 6596.5 |
| 2602.5 | 4771 | 3974 | 1279.5 | 6750.5 | 2320.5 | 804 | 10787 | 4775 | 7684.5 |
| 2552.5 | 4885.5 | 3914 | 1301.5 | 6099 | 2148.5 | 816 | 11251 | 4624 | 7867 |
| 2529 | 4735 | 3703 | 1275 | 5639 | 2123 | 802 | 10570 | 4636 | 7729 |
| 2694 | 3885.5 | 3130.5 | 1090 | 4645 | 1821.5 | 711 | 10259.5 | 4654.5 | 6165 |
| 2689 | 4118 | 3348 | 1108 | 4643 | 1875 | 731 | 11957 | 4534 | 6431 |
| 2668 | 4328.5 | 3590 | 1266 | 4729.5 | 1975 | 749 | 12417 | 4458.5 | 6687 |
| 2644.5 | 4280 | 3428.5 | 1226 | 4791.5 | 1944.5 | 770 | 12238.5 | 4635.5 | 6738.5 |
| 2541.5 | 4684.5 | 3607.5 | 1262 | 5068 | 2019 | 809.5 | 12853.5 | 4673.5 | 7699.5 |
| 2520 | 5009 | 3702 | 1287 | 5243 | 2120 | 847 | 12119 | 4837 | 8551 |
| 15045 | 11212 | 4882 | 1264 | 4393 | 1903 | 761 | 9857 | 7424 | 8505 |
| 2971 | 4193 | 3297 | 1105 | 4347 | 1860 | 738 | 9628 | 4730 | 6577 |
| 2900.5 | 4504 | 3368 | 1176 | 4465 | 1918.5 | 705.5 | 9762.5 | 4965 | 7163.5 |
| 73129.5 | 40772.5 | 11770 | 1575 | 4544 | 1914.5 | 719 | 15259.5 | 6104 | 7734.5 |
| 2687 | 4433 | 3459 | 1157 | 4595 | 1886 | 715 | 10548.5 | 4503 | 7035.5 |
| 2650 | 4183.5 | 3454.5 | 1183.5 | 4911.5 | 1960 | 743.5 | 10223.5 | 4676 | 6857 |
| 2670.5 | 4076.5 | 3331.5 | 1134 | 4489.5 | 1902.5 | 746 | 10208 | 4650.5 | 6656 |
| 7496 | 16889 | 6249 | 1631 | 7554 | 2356 | 772 | 14944 | 8962 | 20171 |
| 8530 | 21433 | 7414 | 1811 | 8685 | 2535 | 761 | 12712 | 9033 | 23840 |
| 8211.5 | 21418.5 | 6930.5 | 1656.5 | 8124 | 2403.5 | 738 | 14039.5 | 8376.5 | 23512 |

| | | | | | | | | | |
|---|---|---|---|---|---|---|---|---|---|
| 9193 | 18270 | 5992 | 1453.5 | 6891 | 2127 | 696.5 | 14097 | 9097 | 20422.5 |
| 2919 | 3737 | 2244.5 | 769 | 3019.5 | 1411.5 | 363.5 | 7925 | 3727.5 | 4792 |
| 2861 | 3988 | 2758 | 958.5 | 3699 | 1614 | 541 | 8963 | 4242 | 5623 |
| 2913 | 4321 | 3259 | 1124 | 4470 | 1801 | 703 | 9806 | 4814 | 6602 |
| 2623 | 4514 | 6376 | 1868 | 674554 | 59374 | 736 | 10091 | 4653 | 6978 |
| 2686 | 4566.5 | 4042.5 | 1314 | 6946.5 | 2330.5 | 773 | 10526.5 | 4690.5 | 7244.5 |
| 2801.5 | 5098.5 | 3935 | 1345 | 6301 | 2252.5 | 831.5 | 11040 | 4977.5 | 8139.5 |
| 2717.5 | 4773.5 | 3849 | 1301 | 5686 | 2151 | 830 | 10939 | 4956 | 7946 |
| 2671 | 4000 | 3294 | 1109 | 4808 | 1865 | 735 | 10663 | 4573 | 6169 |
| 2837 | 4072 | 3191.5 | 1110 | 4463 | 1827.5 | 722 | 11507.5 | 4628.5 | 6298 |
| 2676 | 4297 | 3421 | 1256 | 4585 | 1958 | 751 | 12250 | 4519 | 6703 |
| 2564 | 4349.5 | 3456 | 1238.5 | 4791 | 1970.5 | 793 | 13176.5 | 4408 | 6832 |
| 2515 | 5197 | 3614 | 1264 | 5087 | 2023 | 829 | 13493 | 4647 | 8515 |
| 2479 | 4457.5 | 3578 | 1283 | 4994.5 | 2050.5 | 804 | 10809.5 | 4812.5 | 7571.5 |
| 14358 | 10817 | 4756 | 1225 | 4445 | 1911.5 | 751.5 | 9910.5 | 7338.5 | 8365 |
| 3035 | 4264.5 | 3315 | 1173 | 4508 | 1890.5 | 741.5 | 9427.5 | 4748.5 | 6596 |
| 2953 | 4780 | 3567 | 1176 | 4822.5 | 1959 | 765 | 10085.5 | 5116 | 7333 |
| 84929 | 49359.5 | 13358.5 | 1668.5 | 4956 | 1942 | 736.5 | 15675 | 6570 | 8741.5 |
| 2906.5 | 4795.5 | 3773 | 1224 | 4856.5 | 1938.5 | 735 | 10853.5 | 4795.5 | 7439 |
| 2572 | 4544 | 3520.5 | 1167 | 5055.5 | 1955.5 | 734 | 10288.5 | 4424 | 7347.5 |
| 2691 | 4264.5 | 3370 | 1172 | 4573.5 | 1896 | 736 | 10061.5 | 4725 | 6850 |
| 7680 | 16803 | 6227 | 1614 | 7223 | 2360 | 774 | 14804 | 8901 | 19920 |
| 8091 | 22067 | 7866.5 | 1885.5 | 9081.5 | 2547 | 763 | 14809.5 | 9120 | 24817 |
| 9578 | 21206 | 6978 | 1720 | 8149 | 2387 | 726 | 12013 | 8659 | 23155 |
| 8154 | 18858 | 6202 | 1491 | 7173 | 2154 | 698 | 14177 | 8701 | 20847 |
| 2955.5 | 3798.5 | 2299 | 786 | 3028 | 1394 | 373.5 | 7917 | 3788 | 4701.5 |
| 2759 | 3760 | 2621 | 889 | 3731.5 | 1591.5 | 517.5 | 8491 | 3975 | 5316.5 |
| 2688.5 | 4042 | 3020.5 | 1030.5 | 4370.5 | 1767 | 632.5 | 8960.5 | 4351.5 | 5882.5 |
| 2815 | 4267 | 5717 | 1712 | 371983 | 36936 | 686 | 9348 | 4324 | 6443 |
| 2868.5 | 5425.5 | 3864 | 1288 | 19117 | 3178 | 783 | 10471.5 | 4972 | 8328.5 |
| 2719 | 4650 | 3648 | 1263 | 13690 | 2714 | 777 | 10130 | 4703 | 7378 |
| 2611 | 4595 | 3418 | 1202 | 10649 | 2454 | 756 | 10421 | 4589 | 7282 |
| 2611 | 3875 | 3123 | 1075 | 9158 | 2172 | 701 | 10146 | 4422 | 6056 |
| 2764.5 | 3958 | 3078 | 1025 | 8055.5 | 2058 | 662 | 10750.5 | 4400.5 | 6006 |
| 2678 | 4321.5 | 3518.5 | 1227 | 5638 | 2040.5 | 728 | 12728.5 | 4343 | 6783 |
| 2542 | 4326.5 | 3424 | 1176.5 | 6558.5 | 2091 | 752.5 | 12480.5 | 4440 | 6790.5 |
| 2514 | 4472.5 | 3506 | 1177 | 6261.5 | 2096 | 778 | 12511 | 4471 | 7258 |
| 2531.5 | 4745.5 | 3524 | 1244.5 | 5973.5 | 2119 | 796 | 11449 | 4615 | 7728 |
| 12251 | 9516 | 4703 | 1240 | 5902 | 2068 | 766 | 10076 | 6890 | 8420 |
| 3254 | 5625 | 3882 | 1296 | 6009 | 2158 | 827 | 10422 | 5335 | 8823 |
| 2927 | 4949.5 | 3532 | 1193 | 5621 | 1989 | 741 | 10177.5 | 4836 | 7703 |
| 159903 | 102671 | 24518 | 2171 | 6007 | 2064 | 726 | 22420.5 | 7909 | 10235.5 |
| 2927 | 4686 | 3710 | 1197 | 5599 | 1984 | 716 | 11177 | 4866 | 7365 |
| 2660 | 4085 | 3274 | 1128 | 9651 | 2536 | 703 | 9776 | 4615 | 6399 |
| 2686 | 4246 | 3414 | 1130 | 5222 | 1947 | 734 | 10251 | 4462 | 6624 |
| 7871.5 | 17778 | 6489.5 | 1669 | 8111 | 2473 | 775.5 | 14909.5 | 9181 | 21022.5 |
| 9126.5 | 23270.5 | 7783 | 1907 | 9527 | 2650 | 756 | 14268 | 9867 | 25718 |
| 8065 | 19795 | 6601.5 | 1628 | 8092.5 | 2334 | 668.5 | 13915.5 | 8429.5 | 22179.5 |

| | | | | | | | | | |
|---|---|---|---|---|---|---|---|---|---|
| 8470.5 | 18288 | 6068 | 1518.5 | 7596 | 2255 | 680 | 12472.5 | 8874.5 | 20837 |
| 2925 | 4106 | 2567 | 825 | 3377 | 1460 | 377 | 8421 | 4126 | 5193 |
| 2936 | 4194 | 2930 | 1004 | 3917 | 1647 | 577 | 9176 | 4573 | 5899 |
| 3025 | 4781 | 3258 | 1164 | 4613 | 1853 | 704 | 10033 | 5106 | 7219 |
| 2684 | 5083 | 6549 | 1847 | 701246 | 56278 | 774 | 11107 | 4711 | 8131 |
| 2741 | 5057.5 | 4023 | 1309 | 6474.5 | 2352 | 790.5 | 11257.5 | 4725 | 8028.5 |
| 2783 | 5809 | 4135.5 | 1380 | 6175.5 | 2285.5 | 846 | 13293 | 5068.5 | 9529 |
| 2788 | 5493 | 4012.5 | 1334.5 | 5849.5 | 2193.5 | 867 | 12690 | 4974.5 | 8914.5 |
| 2801 | 4480 | 3420 | 1178 | 4991 | 1976 | 765 | 11373 | 4836 | 6895 |
| 2850 | 4144 | 3269.5 | 1112 | 4532.5 | 1825 | 729.5 | 11739 | 4755.5 | 6596 |
| 2668.5 | 4311.5 | 3458 | 1245.5 | 4631 | 1925 | 745 | 12805.5 | 4504.5 | 6723.5 |
| 2658.5 | 4340 | 3492.5 | 1192 | 4853 | 1944.5 | 785 | 12376 | 4694.5 | 6906.5 |
| 2510 | 5449 | 3710.5 | 1298 | 5188 | 2027.5 | 819 | 13910 | 4668.5 | 8829 |
| 2517 | 5221 | 3797 | 1314 | 5329 | 2139 | 866 | 12789 | 4814 | 8767 |
| 31440 | 25648 | 7824 | 1389 | 5019 | 2036 | 808 | 10916 | 12971 | 13044 |
| 2998 | 5114 | 3779 | 1235 | 4942 | 2034 | 830 | 10977 | 5090 | 8132 |
| 2840 | 5031.5 | 3740 | 1221.5 | 5031.5 | 1987 | 780.5 | 10747.5 | 5076.5 | 7946.5 |
| 64730 | 43960 | 11864 | 1652 | 5167 | 2006 | 765 | 16162 | 6000 | 9637 |
| 2805 | 4955 | 3829 | 1248 | 4954 | 1977 | 766 | 11585 | 4846 | 7651 |
| 2604.5 | 4379.5 | 3418 | 1124 | 4792.5 | 1881 | 739 | 9970.5 | 4537 | 6679.5 |
| 2768.5 | 4395 | 3367.5 | 1180 | 4626 | 1902.5 | 774.5 | 10744 | 4778 | 6869 |
| 8116 | 16876 | 6236 | 1650 | 7347 | 2336 | 804 | 15309 | 9254 | 20196 |
| 8966 | 24893 | 8881.5 | 2051 | 10152.5 | 2763 | 846 | 13033 | 9457 | 27870.5 |
| 9472 | 21842 | 7163 | 1717 | 8129 | 2336 | 733 | 13387 | 9371 | 23804 |
| 9585 | 21023.5 | 6724 | 1649 | 7590.5 | 2245 | 759.5 | 14386 | 9734.5 | 23316 |
| 2868 | 3854 | 2395 | 811 | 3114 | 1389 | 363 | 7917 | 3829 | 4795 |
| 2779 | 3864 | 2766 | 955 | 3626 | 1569 | 532 | 8442 | 4143 | 5382 |
| 2858 | 4772 | 3398 | 1145 | 4649 | 1863 | 704 | 10446 | 4786 | 7167 |
| 2563 | 4603 | 6356 | 1858 | 614533 | 51595 | 714 | 10074 | 4418 | 7082 |
| 2681 | 4821 | 3848.5 | 1272 | 6189 | 2212.5 | 751 | 10561 | 4596.5 | 7625 |
| 2673 | 5098.5 | 3906 | 1344 | 6055.5 | 2186 | 783 | 10892 | 4789.5 | 8043 |
| 2651 | 5549 | 3831 | 1331 | 5809 | 2128 | 809 | 10842 | 4828 | 8716 |
| 2683 | 3856.5 | 2990 | 1046 | 4406 | 1740 | 675 | 9893 | 4502.5 | 6030 |
| 2807.5 | 4677.5 | 3502.5 | 1146 | 4942 | 1857.5 | 728 | 12363 | 4617.5 | 7070.5 |
| 2985.5 | 6311.5 | 4003.5 | 1395 | 5405.5 | 2182.5 | 809 | 15460.5 | 5165.5 | 9567 |
| 2683 | 5446 | 3671 | 1280 | 5085 | 1996 | 794 | 13910 | 4657 | 8342 |
| 2470 | 4685 | 3664 | 1250 | 4932 | 1983 | 771 | 12812 | 4340 | 7537 |
| 2492 | 4186 | 3507 | 1219 | 4840 | 2016 | 778 | 11034 | 4676 | 6969 |
| 12902 | 9463.5 | 4584 | 1153 | 4301.5 | 1823 | 720 | 9876.5 | 6846 | 7630 |
| 3011.5 | 4531.5 | 3388 | 1182 | 4584.5 | 1883.5 | 760.5 | 9798 | 4981 | 7066 |
| 3080.5 | 4986 | 3767.5 | 1203 | 4927 | 1939.5 | 747 | 10314.5 | 5030 | 7955 |
| 73097 | 51950 | 12952 | 1707 | 5088 | 1993 | 743 | 16556 | 5992 | 9599 |
| 3029 | 7054.5 | 4190.5 | 1360.5 | 5473 | 2115.5 | 795 | 11957 | 5003.5 | 10808.5 |
| 2405.5 | 6702 | 3389 | 1181 | 5062.5 | 1922 | 746 | 10018 | 4117.5 | 10485 |
| 2767 | 4250.5 | 3383.5 | 1134.5 | 4411 | 1826.5 | 731 | 10049.5 | 4499 | 6563.5 |
| 6288 | 15312 | 5636 | 1492 | 6687 | 2190 | 729 | 14755 | 6844 | 18043 |
| 8768 | 23307 | 8566 | 1993 | 10075 | 2762 | 802 | 14029 | 9601 | 26052 |
| 8485.5 | 23506 | 8210.5 | 1922 | 9842 | 2667.5 | 764 | 11496 | 8814 | 24993.5 |

| | | | | | | | | | |
|---|---|---|---|---|---|---|---|---|---|
| 7604 | 21002 | 7736 | 1805 | 9492 | 2554 | 769 | 9852 | 8039 | 23337 |
| 831 | 1210 | 1047 | 590 | 1685 | 1150 | 293 | 2900 | 1682 | 2324 |
| 1179 | 1987.5 | 1790 | 776.5 | 2671.5 | 1387 | 482 | 4931 | 2502.5 | 3600.5 |
| 1414 | 2466 | 2247.5 | 882 | 3255 | 1553 | 603 | 6597.5 | 3086 | 4490.5 |
| 1542 | 2739 | 4735 | 1420 | 577394 | 52775 | 654 | 7303 | 3453 | 5109 |
| 1619 | 2774 | 2861 | 1057 | 4495 | 1943 | 686 | 7728 | 3551 | 5242 |
| 1641 | 2909 | 2762 | 1062 | 4142 | 1818 | 715 | 8137 | 3810 | 5453 |
| 1735 | 3012 | 2832 | 1094 | 4183 | 1829.5 | 747.5 | 8345.5 | 4003.5 | 5660 |
| 1774.5 | 2917.5 | 2593.5 | 1020.5 | 3715 | 1709 | 663 | 8310.5 | 3652.5 | 5228 |
| 2233.5 | 3343.5 | 2871 | 1055.5 | 4036 | 1769 | 708 | 9976 | 4301.5 | 5765.5 |
| 2072 | 3220 | 2717 | 1009 | 3828.5 | 1697 | 669 | 9570.5 | 3995.5 | 5455 |
| 1950 | 3153 | 2785 | 1013 | 3975 | 1758 | 704 | 9076 | 4056 | 5541 |
| 1907 | 3202 | 2966 | 1093 | 4123 | 1839 | 738 | 9214 | 4136 | 5805 |
| 1916 | 3232 | 3054 | 1123 | 4326 | 1852 | 766 | 9437 | 4365 | 6077 |
| 15303 | 10667 | 4567 | 1155 | 4059 | 1755 | 732 | 8353 | 6887 | 7287 |
| 2191.5 | 3393 | 3065.5 | 1123.5 | 4274.5 | 1879 | 758.5 | 8770.5 | 4252.5 | 6019 |
| 2320.5 | 3949.5 | 3282 | 1209 | 4590 | 1947.5 | 756 | 9228 | 4614.5 | 6960.5 |
| 77356.5 | 40577 | 11881.5 | 1515.5 | 4385.5 | 1843 | 701 | 13857 | 5722.5 | 7136.5 |
| 2276 | 3556 | 3126 | 1088 | 4194 | 1764 | 696 | 9166 | 4508 | 6173 |
| 2052 | 3331.5 | 2851.5 | 1048 | 4498.5 | 1856 | 690 | 8634.5 | 4139.5 | 5767 |
| 2088 | 3355 | 2935 | 1044 | 4096 | 1779 | 703 | 8929 | 4131 | 5796 |
| 4851 | 13509 | 5151 | 1424 | 6345 | 2117 | 700 | 15596 | 6096 | 16266 |
| 8416.5 | 21347.5 | 7566 | 1804 | 8968 | 2514 | 724 | 13439.5 | 9400.5 | 23755 |
| 8981 | 20591.5 | 6794 | 1669 | 7873 | 2360.5 | 699 | 12961 | 8756 | 22602.5 |
| 8454.5 | 18216 | 5961 | 1505 | 7067 | 2146.5 | 687 | 14405 | 9128.5 | 20897 |
| 2480 | 3429 | 2073 | 733 | 2737 | 1313 | 374 | 7796 | 3495 | 4490 |
| 2335 | 3369.5 | 2425 | 881 | 3382 | 1542 | 546 | 7924 | 3730 | 5052 |
| 2197 | 3373 | 2781 | 969 | 3836 | 1664 | 663 | 8585 | 4075 | 5531 |
| 2217.5 | 3515.5 | 5697 | 1602.5 | 689254 | 60514 | 722 | 9147.5 | 4295.5 | 5902.5 |
| 2355 | 3772 | 3606 | 1239 | 5698 | 2183 | 794 | 10043 | 4609 | 6666 |
| 2244 | 3702 | 3333 | 1191 | 5064 | 2025 | 804 | 9979 | 4656 | 6483 |
| 2271 | 3719 | 3279 | 1206 | 4809 | 1985 | 787 | 9938 | 4605 | 6498 |
| 2355.5 | 3508.5 | 3051 | 1116.5 | 4347 | 1852 | 730 | 9794.5 | 4390 | 6040.5 |
| 2493 | 3546.5 | 2913 | 1046 | 4076.5 | 1736.5 | 710 | 10360.5 | 4388.5 | 5848.5 |
| 2397 | 3499 | 2943.5 | 1124 | 3993.5 | 1819 | 726.5 | 10479 | 4354.5 | 5840 |
| 2416 | 3452 | 2996 | 1057 | 4181 | 1775 | 732 | 10153 | 4623 | 6020 |
| 2336 | 3524 | 3106 | 1152 | 4367 | 1874 | 780 | 10172 | 4592 | 6254 |
| 2283 | 3795 | 3309 | 1208 | 4828 | 1990 | 823 | 11087 | 4669 | 6702 |
| 17530 | 12238.5 | 4994 | 1246 | 4407.5 | 1854 | 783.5 | 9603.5 | 7813.5 | 8263 |
| 2651.5 | 3976.5 | 3302 | 1223 | 4561.5 | 1903 | 818 | 9746.5 | 4761 | 6808.5 |
| 2577 | 4502 | 3517 | 1233 | 4765 | 2001.5 | 796.5 | 10192 | 5023 | 7785.5 |
| 92208 | 51840 | 14221 | 1689 | 4743 | 1943 | 748 | 16158 | 6562 | 8380 |
| 2520 | 3972 | 3349 | 1174 | 4365 | 1873 | 719 | 9883 | 4658 | 6606 |
| 2408 | 3893 | 3307 | 1112 | 5041 | 1999 | 747 | 9572 | 4701 | 6590 |
| 2192 | 3494 | 3002 | 1064 | 4177.5 | 1809.5 | 718 | 9004.5 | 4332 | 6094 |
| 5123.5 | 13811.5 | 5260 | 1476.5 | 6474.5 | 2124 | 705 | 15247.5 | 6359 | 16355 |
| 8683 | 22561 | 7701 | 1808 | 8943.5 | 2557 | 745 | 14540.5 | 9538.5 | 25679 |
| 9355 | 20396 | 6725 | 1636 | 7802 | 2350 | 681 | 13661 | 9283 | 22480 |

| | | | | | | | | | |
|---|---|---|---|---|---|---|---|---|---|
| 7895.5 | 17272 | 5630 | 1418.5 | 6635 | 2101.5 | 664 | 14461 | 8313.5 | 19525.5 |
| 2777 | 3796 | 2290 | 799 | 3080 | 1404 | 410 | 8701 | 3885 | 4797 |
| 2345.5 | 3409.5 | 2458.5 | 896 | 3421.5 | 1499 | 534.5 | 7746 | 3841.5 | 5018.5 |
| 2493.5 | 3649.5 | 2949.5 | 1048 | 4119 | 1748 | 672 | 8870 | 4351.5 | 5956 |
| 2237 | 3401 | 5177 | 1499 | 567726 | 51256 | 690 | 8852 | 4318 | 5798 |
| 2331 | 3760 | 3537.5 | 1189 | 5861.5 | 2147.5 | 751 | 9840.5 | 4454.5 | 6246.5 |
| 2243 | 3494 | 3120 | 1091 | 4849 | 1873 | 718 | 9671 | 4391 | 5960 |
| 2328.5 | 3815.5 | 3408 | 1164 | 4976.5 | 1992 | 776 | 10683.5 | 4609 | 6679.5 |
| 2605 | 4588 | 3493 | 1237 | 5237 | 2054 | 802 | 12469 | 4798 | 7789 |
| 2568 | 4124 | 3247 | 1133 | 4714 | 1864 | 750 | 12391 | 4673 | 6709 |
| 2572 | 3767 | 3164.5 | 1210 | 4360.5 | 1933.5 | 713 | 10837.5 | 4578.5 | 6141.5 |
| 2500 | 3588 | 3057 | 1106 | 4301 | 1838 | 720 | 10681 | 4552 | 5965 |
| 2359 | 3471.5 | 2949 | 1061 | 4145.5 | 1778.5 | 733 | 9598 | 4513 | 5888.5 |
| 2371 | 4076 | 3443 | 1214 | 4869 | 1971 | 823 | 12080 | 4505 | 6914 |
| 16755 | 11902.5 | 5098 | 1303.5 | 4713.5 | 1950 | 799.5 | 10646 | 7809 | 8449 |
| 2826 | 4048.5 | 3404.5 | 1184.5 | 4686.5 | 1907 | 777 | 9799 | 4904.5 | 6640.5 |
| 2724 | 4582.5 | 3508.5 | 1233.5 | 4871 | 1965.5 | 788 | 10327 | 5008.5 | 7675.5 |
| 100807.5 | 60372 | 15703.5 | 1693.5 | 4448 | 1862 | 717.5 | 16816.5 | 6335.5 | 8286 |
| 2723 | 4053 | 3277 | 1117 | 4369 | 1802 | 696 | 10087 | 4849 | 6346 |
| 2531 | 4153 | 3487 | 1183 | 5312.5 | 2035.5 | 749 | 10733 | 4612.5 | 6853 |
| 2482 | 3648 | 3057 | 1079 | 4157 | 1760 | 712 | 9281 | 4479 | 5974 |
| 6682.5 | 14372 | 5422 | 1519 | 6739.5 | 2229.5 | 738 | 15521 | 8155 | 17537 |
| 8856.5 | 21581.5 | 7158.5 | 1750 | 8225.5 | 2451.5 | 724.5 | 15628 | 9408 | 23786.5 |
| 9453 | 21426.5 | 6921.5 | 1693 | 8093 | 2407.5 | 718 | 13945.5 | 9313 | 23042 |
| 8792 | 19245.5 | 6414 | 1563.5 | 7442.5 | 2249.5 | 714 | 12739 | 9147 | 21334.5 |
| 2602 | 3500 | 2165 | 756 | 2857 | 1344 | 361 | 7707 | 3470 | 4439 |
| 2524 | 3611 | 2639 | 922 | 3463 | 1560 | 550 | 8493 | 3986 | 5185 |
| 2546 | 3859 | 3009 | 1069 | 4163 | 1785 | 671 | 9702 | 4427 | 5953 |
| 2496 | 4083 | 6166.5 | 1674 | 704283 | 60455 | 756 | 10258 | 4721.5 | 6910 |
| 2462 | 4365 | 3698 | 1265 | 6147 | 2225 | 802 | 11023 | 4596 | 7440 |
| 2393 | 4222 | 3592 | 1228 | 5502 | 2046 | 803 | 11017 | 4655 | 6979 |
| 2426 | 4603.5 | 3566 | 1257 | 5241.5 | 2033 | 818.5 | 11231 | 4738 | 7970 |
| 2613.5 | 3839 | 3074.5 | 1119 | 4544 | 1833.5 | 739 | 10115.5 | 4613 | 6267 |
| 2680.5 | 4026.5 | 3174.5 | 1106.5 | 4450 | 1846.5 | 738 | 11264 | 4674.5 | 6313 |
| 2604 | 3759 | 3129 | 1161 | 4147 | 1831 | 714 | 11033 | 4707 | 6087 |
| 2493 | 3525 | 2929 | 1057 | 4094 | 1764 | 702 | 9782 | 4536 | 5876 |
| 2442.5 | 3536.5 | 3038 | 1090 | 4146 | 1826.5 | 738 | 10036.5 | 4567.5 | 5952.5 |
| 2314.5 | 4268 | 3511 | 1269 | 4991.5 | 2047.5 | 822 | 12068 | 4490.5 | 7272 |
| 14612 | 12639 | 5137 | 1361 | 5220 | 2116 | 830 | 11657 | 7456 | 10407 |
| 2676 | 4196 | 3480 | 1217 | 4620 | 1930 | 776 | 9955 | 4712 | 6945 |
| 3020 | 5937 | 3814 | 1336 | 5485 | 2155 | 823 | 11887 | 5599 | 10093 |
| 79980 | 53213 | 12981 | 1753 | 5353 | 2095 | 777 | 16037 | 6616 | 10071 |
| 2825 | 5400 | 3749 | 1280 | 5063 | 1976 | 773 | 10498 | 5048 | 8664 |
| 2564.5 | 4149.5 | 3314.5 | 1135.5 | 5001 | 1954 | 739.5 | 9622.5 | 4820 | 6814 |
| 2710.5 | 4430 | 3521 | 1237.5 | 4866 | 1978.5 | 806.5 | 10209 | 5007.5 | 7412 |
| 7727 | 16464.5 | 5961 | 1575.5 | 7105.5 | 2298 | 731 | 15766.5 | 9267.5 | 19843 |
| 6761 | 24652 | 8865 | 2071 | 10652 | 2907 | 810 | 11263 | 7565 | 28023 |
| 8749 | 21443 | 6684 | 1578 | 7662 | 2307 | 671 | 12540 | 8898 | 23491 |

| | | | | | | | | | |
|---|---|---|---|---|---|---|---|---|---|
| 8728 | 18207 | 5892 | 1452 | 6798 | 2145 | 673 | 15508 | 8829 | 19968 |
| 2790.5 | 3735 | 2374.5 | 796 | 3146.5 | 1400 | 396.5 | 7993.5 | 4060.5 | 4898.5 |
| 2818 | 3921.5 | 2873.5 | 965 | 3751.5 | 1650.5 | 578 | 9075.5 | 4297.5 | 5653.5 |
| 2828 | 4110 | 3189 | 1086 | 4329 | 1794 | 706 | 9953 | 4801 | 6538 |
| 2602 | 4414 | 6197 | 1775 | 637433 | 60128 | 789 | 10395 | 4746 | 7076 |
| 2560.5 | 4482 | 3719 | 1277 | 6154.5 | 2237 | 806.5 | 10949 | 4735 | 7252.5 |
| 2700.5 | 4628 | 3600 | 1276.5 | 5615.5 | 2109.5 | 838.5 | 10624.5 | 5018.5 | 7834 |
| 2558 | 4346.5 | 3641 | 1227 | 5385 | 2013.5 | 816 | 10741 | 4819 | 7277 |
| 2690 | 3987.5 | 3319.5 | 1121 | 4623.5 | 1841.5 | 759 | 10458.5 | 4819 | 6487.5 |
| 2870 | 4040 | 3217 | 1092 | 4317 | 1838 | 743 | 11550 | 4761 | 6411 |
| 2743.5 | 4225 | 3545 | 1309.5 | 4726 | 2021 | 770 | 12432 | 4703.5 | 6679 |
| 2406 | 4114 | 3336 | 1200 | 4710 | 1905 | 805 | 12165 | 4376 | 6725 |
| 2489 | 4119.5 | 3506 | 1267 | 4915.5 | 1960 | 833 | 11214 | 4720 | 7002.5 |
| 2368 | 3922 | 3378 | 1190 | 4628 | 1949 | 819 | 10590 | 4767 | 6712 |
| 17163 | 12082 | 5046 | 1226 | 4247 | 1837 | 762 | 9420 | 7764 | 8071 |
| 3001 | 4223.5 | 3342.5 | 1189 | 4349 | 1910 | 793.5 | 9853 | 4861 | 6689 |
| 3049.5 | 4453 | 3565 | 1199.5 | 4659 | 1967.5 | 788.5 | 10441 | 5181.5 | 7272.5 |
| 75145 | 40296 | 11752 | 1579 | 4693 | 1878 | 744 | 15534 | 6347 | 7749 |
| 2805 | 4322.5 | 3543 | 1161 | 4690.5 | 1881 | 756.5 | 10431.5 | 4950.5 | 6873.5 |
| 2656 | 4086.5 | 3354 | 1140.5 | 4789.5 | 1888.5 | 750 | 10152 | 4697 | 6518.5 |
| 2678.5 | 4077 | 3303.5 | 1144.5 | 4459.5 | 1836 | 767 | 10308.5 | 4806 | 6570 |
| 8420 | 17450.5 | 6288 | 1611.5 | 7411.5 | 2381 | 798 | 14790 | 9239 | 20641.5 |
| 9572 | 23518 | 8127 | 1866 | 9374 | 2609 | 778 | 15286 | 9921 | 25746 |
| 10481.5 | 20699.5 | 6676 | 1610.5 | 7606 | 2245 | 690 | 14795.5 | 9731 | 21906.5 |
| 9829 | 21269 | 6603 | 1622 | 7638 | 2345 | 763 | 12243 | 8984 | 22814 |
| 2802.5 | 3742.5 | 2278.5 | 782 | 2990.5 | 1374 | 370 | 8034 | 3765.5 | 4737.5 |
| 2600 | 3746.5 | 2646 | 944.5 | 3632 | 1588 | 546 | 8393 | 4050.5 | 5325.5 |
| 2633.5 | 3929 | 3061 | 1030 | 4141.5 | 1770.5 | 664.5 | 9442 | 4552 | 6160 |
| 2437.5 | 4229 | 5942.5 | 1725 | 620594.5 | 52721.5 | 723 | 9665 | 4560 | 6791.5 |
| 2646 | 4766.5 | 3871 | 1327 | 6222 | 2232 | 780 | 10172 | 4919.5 | 7812.5 |
| 2480 | 4524 | 3612 | 1268.5 | 5496.5 | 2095 | 792 | 10275 | 4643 | 7445 |
| 2487 | 4659 | 3572 | 1242 | 5331 | 2058 | 797 | 10507 | 4710 | 7731 |
| 2542 | 3816.5 | 3036.5 | 1055 | 4437 | 1780 | 701 | 9677 | 4437 | 6048.5 |
| 2550 | 4007 | 3134 | 1083 | 4427 | 1809 | 701 | 11708 | 4359 | 6287 |
| 2607.5 | 4056.5 | 3231 | 1193 | 4447 | 1904.5 | 695.5 | 11430.5 | 4461 | 6385 |
| 2501 | 4324.5 | 3515 | 1228.5 | 4886.5 | 2004 | 775 | 13406 | 4474 | 7038.5 |
| 2466.5 | 4605 | 3528 | 1262.5 | 4902 | 2014 | 810 | 12336.5 | 4680 | 7681 |
| 2332 | 4467 | 3353 | 1232 | 4894 | 1994 | 797 | 10359 | 4606 | 7368 |
| 13894 | 9853 | 4579 | 1155 | 4143 | 1797 | 722 | 9114 | 6907 | 7473 |
| 2812 | 4150.5 | 3341 | 1155 | 4525 | 1853.5 | 755 | 9414 | 4664.5 | 6477.5 |
| 2653 | 3957 | 3176 | 1122 | 4375 | 1836 | 715 | 9472 | 4680 | 6424 |
| 76659 | 42770 | 12724 | 1567 | 4545 | 1848 | 699 | 15559 | 5899 | 7525 |
| 2620 | 4341 | 3487.5 | 1185 | 4650 | 1901.5 | 727 | 10254.5 | 4615 | 6776 |
| 2538 | 3875 | 3122 | 1078 | 4669 | 1849 | 703.5 | 9317.5 | 4602 | 6250 |
| 2463.5 | 3932 | 3295 | 1100 | 4369 | 1820 | 723 | 9650 | 4533 | 6341 |
| 7084 | 15725 | 5797 | 1500 | 6891 | 2271 | 739 | 15275 | 8525 | 18431 |
| 8362 | 23949 | 8348 | 1885 | 9935 | 2658 | 764 | 13469 | 9317 | 26931 |
| 9704.5 | 21076.5 | 6730 | 1620.5 | 7899.5 | 2312.5 | 688 | 13796 | 9317 | 22201.5 |

| | | | | | | | | | |
|---|---|---|---|---|---|---|---|---|---|
| 9099 | 19467.5 | 6362 | 1606 | 7405 | 2269.5 | 699 | 13899 | 9022.5 | 21786.5 |
| 2933 | 3808 | 2385 | 790 | 3182 | 1403 | 386 | 8157 | 3856 | 4900 |
| 2623 | 3868 | 2698 | 940 | 3643.5 | 1574.5 | 525 | 8546.5 | 4066 | 5305.5 |
| 2653 | 3974 | 3017 | 1061 | 4144 | 1745 | 655 | 9394 | 4410 | 6066 |
| 2516 | 4204.5 | 5939 | 1708 | 600675.5 | 52190.5 | 698 | 9647 | 4512.5 | 6772.5 |
| 2527 | 4475.5 | 3727 | 1255.5 | 6259.5 | 2233 | 758 | 10393 | 4537.5 | 7199.5 |
| 2515 | 4717 | 3662 | 1282 | 5699 | 2125 | 798 | 10651 | 4635 | 7769 |
| 2569 | 4317 | 3588 | 1252 | 5397 | 2071 | 791 | 10511 | 4788 | 7168 |
| 2602 | 3875 | 3118 | 1074 | 4503 | 1817 | 700 | 10145 | 4442 | 5993 |
| 2777 | 4060 | 3225.5 | 1101 | 4465 | 1814 | 702 | 11029 | 4525.5 | 6325 |
| 2649 | 4169 | 3417 | 1257 | 4608 | 1982 | 724 | 11850 | 4514 | 6581 |
| 2561.5 | 4052 | 3290.5 | 1173 | 4674 | 1926 | 756 | 11586.5 | 4570.5 | 6503.5 |
| 2468 | 4355 | 3590 | 1257 | 5065 | 2021 | 811 | 12647 | 4612 | 7064 |
| 2463.5 | 4392.5 | 3430 | 1222.5 | 4876 | 1989.5 | 810 | 11325.5 | 4635.5 | 7248 |
| 14785.5 | 10514 | 4853 | 1222 | 4483.5 | 1878 | 752.5 | 9813 | 7372 | 8110 |
| 3152 | 5432 | 3818 | 1326 | 5221 | 2074 | 832 | 11790 | 5149 | 8680 |
| 2906 | 4901 | 3621 | 1220 | 4837 | 1964 | 774 | 10095 | 5080 | 7821 |
| 78099 | 48257 | 12323 | 1594 | 4691 | 1885 | 727 | 15287 | 6280 | 8537 |
| 2739 | 4539.5 | 3688.5 | 1217.5 | 4897 | 1934 | 751 | 11003 | 4556 | 7187 |
| 2664 | 3921 | 3178 | 1101.5 | 4670.5 | 1821 | 709 | 9628.5 | 4764.5 | 6445 |
| 2603 | 4206 | 3451 | 1159 | 4670 | 1913 | 756 | 10600 | 4563 | 6856 |
| 7821 | 15663 | 5803 | 1508 | 6803 | 2227 | 727 | 16279 | 9253 | 18804.5 |
| 9313 | 22220.5 | 8140.5 | 1891.5 | 9334.5 | 2656.5 | 787 | 13940.5 | 9625 | 24135 |
| 9565 | 21361 | 6897 | 1702 | 8122 | 2354 | 723 | 14205 | 9228 | 22740 |
| 9083 | 18894.5 | 6252.5 | 1528 | 7218 | 2207.5 | 706 | 12552.5 | 8975 | 20696.5 |
| 2821 | 3705 | 2292 | 786 | 3078 | 1379 | 358 | 7841 | 3803 | 4705 |
| 2869 | 3942 | 2839 | 973 | 3763 | 1641 | 556 | 9021 | 4260 | 5553 |
| 2795 | 4296 | 3227 | 1116 | 4386 | 1834 | 700 | 10354 | 4693 | 6448 |
| 2702 | 5285 | 6692 | 1939 | 665794 | 56756 | 789 | 10679 | 4803 | 8481 |
| 2700.5 | 4997 | 4061.5 | 1348 | 6721 | 2331 | 814.5 | 10961.5 | 4793 | 7843 |
| 2602 | 4942 | 3821 | 1281 | 5793 | 2165 | 809 | 10708 | 4802 | 7991 |
| 2810 | 5105 | 3732 | 1316 | 5743 | 2146 | 843 | 10816 | 5199 | 8366 |
| 2628 | 4139 | 3279 | 1133 | 4825 | 1886 | 729 | 10893 | 4590 | 6471 |
| 2795 | 3918 | 3078.5 | 1043 | 4350 | 1754.5 | 684 | 10828 | 4530.5 | 6053.5 |
| 2659 | 4232 | 3458 | 1246 | 4680 | 1995 | 738 | 12628 | 4474 | 6575 |
| 2595.5 | 4327.5 | 3490 | 1212 | 4809.5 | 1944.5 | 772 | 12521 | 4558.5 | 6751.5 |
| 2511.5 | 4239 | 3451.5 | 1199.5 | 4827 | 1956.5 | 795 | 12214.5 | 4523.5 | 6927.5 |
| 2449 | 4957 | 3584 | 1278.5 | 5210 | 2037.5 | 825 | 11500.5 | 4757 | 8303 |
| 14603 | 10784 | 4927 | 1226 | 4554 | 1901 | 770 | 9916 | 7488 | 8483 |
| 2950 | 4144 | 3243 | 1124 | 4346 | 1783 | 749 | 9505 | 4739 | 6433 |
| 2787 | 4195 | 3280 | 1139 | 4455 | 1820 | 715 | 9736 | 4800 | 6743 |
| 86195 | 49635 | 13423 | 1669 | 4846 | 1885 | 735 | 16199 | 6500 | 8282 |
| 2728 | 4699.5 | 3632 | 1213.5 | 4839 | 1901 | 732 | 10873.5 | 4627.5 | 7270 |
| 2538 | 4155 | 3306 | 1101 | 4889 | 1938 | 722 | 10258 | 4604 | 6689 |
| 2584 | 4364.5 | 3484 | 1217 | 4807 | 1960.5 | 765 | 10824 | 4527.5 | 7085 |
| 8076.5 | 16712 | 5905.5 | 1561 | 7119.5 | 2269 | 739 | 15852 | 9300 | 19590.5 |
| 9357.5 | 22292 | 7638 | 1843.5 | 8747 | 2568 | 746 | 13893 | 9369.5 | 24782.5 |
| 9375 | 21026.5 | 6899.5 | 1645 | 8103 | 2394 | 700 | 13987 | 9411 | 22560 |

| | | | | | | | | | |
|---|---|---|---|---|---|---|---|---|---|
| 8896.5 | 19087 | 6101.5 | 1536 | 7264.5 | 2238 | 695 | 13241.5 | 9133 | 20519.5 |
| 2876.5 | 3794.5 | 2370.5 | 786 | 3116 | 1386.5 | 364 | 8283 | 3935 | 4853.5 |
| 2947.5 | 4075 | 2886 | 973.5 | 3896.5 | 1645 | 545 | 8794.5 | 4319.5 | 5715.5 |
| 3007.5 | 4781 | 3383 | 1160.5 | 4857.5 | 1883.5 | 688.5 | 10025.5 | 5010.5 | 7349.5 |
| 2798.5 | 4647 | 5923 | 1758 | 368440 | 36070.5 | 711 | 9939.5 | 4444.5 | 7013.5 |
| 2646 | 5133 | 3924 | 1273 | 16025 | 2981 | 773 | 11043 | 4670 | 8104 |
| 2577 | 5137 | 3875 | 1313 | 13067 | 2733 | 786 | 11245 | 4673 | 7985 |
| 2649 | 5879 | 3988 | 1330 | 10262 | 2540 | 831 | 13225 | 4834 | 9150 |
| 2683 | 4022 | 3221 | 1104 | 8399 | 2148 | 713 | 10590 | 4505 | 6345 |
| 2751 | 4233 | 3267 | 1125 | 7508 | 2082 | 713 | 11587 | 4517 | 6541 |
| 2665 | 4178 | 3362 | 1199 | 5149 | 1967 | 711 | 11714 | 4448 | 6447 |
| 2558 | 4090 | 3301 | 1103 | 5935 | 1985 | 721 | 11608 | 4388 | 6475 |
| 2279.5 | 4060 | 3348.5 | 1177 | 5669 | 1998 | 742 | 11414 | 4140 | 6561.5 |
| 2454.5 | 4681 | 3511 | 1243 | 5746.5 | 2096.5 | 788.5 | 12245 | 4541 | 7641 |
| 11050 | 11196 | 4897 | 1315 | 6174 | 2187 | 838 | 11473 | 6642 | 10447 |
| 3025.5 | 4616.5 | 3551 | 1228 | 5459 | 2042.5 | 779.5 | 9944.5 | 5043 | 7321.5 |
| 2952 | 5382 | 3761.5 | 1256 | 5580.5 | 2036 | 767.5 | 10293.5 | 5096 | 8741.5 |
| 160813.5 | 97914.5 | 26242.5 | 2125 | 5645 | 1966.5 | 699 | 23311.5 | 7443 | 9042 |
| 2688.5 | 4578 | 3560.5 | 1169.5 | 5167.5 | 1927 | 697 | 9991.5 | 4442.5 | 6885.5 |
| 2598 | 4228 | 3316 | 1169 | 9703 | 2561 | 728 | 10136 | 4528 | 6711 |
| 2738 | 4062 | 3269 | 1122 | 4976 | 1842 | 717 | 9700 | 4500 | 6335 |
| 7984 | 16961 | 6065.5 | 1588.5 | 7743.5 | 2383 | 768 | 14739.5 | 9078 | 20037.5 |
| 8787.5 | 22832.5 | 8253 | 1886 | 9906.5 | 2640 | 755 | 13824 | 9475 | 26606 |
| 9474.5 | 20364 | 6861 | 1633.5 | 7988 | 2293 | 683 | 14704 | 9551.5 | 22395.5 |
| 9124 | 18801 | 6043 | 1510 | 7345 | 2212 | 676 | 14178 | 9228 | 20651 |
| 2939.5 | 3880 | 2472.5 | 823 | 3242 | 1434.5 | 368 | 8363.5 | 4050 | 4983 |
| 2866 | 4076.5 | 2975.5 | 965 | 3818.5 | 1643.5 | 552.5 | 8947 | 4423 | 5784 |
| 2854 | 4511 | 3266 | 1107 | 4581 | 1848 | 685 | 9792 | 4814 | 6792 |
| 2638 | 4890.5 | 6389 | 1881.5 | 678727 | 55456.5 | 747 | 10659.5 | 4581 | 7540 |
| 2744 | 5210 | 4094 | 1394 | 6681 | 2411 | 830 | 11493 | 5006 | 8356 |
| 2679.5 | 4997 | 3843 | 1275.5 | 5932 | 2136.5 | 808.5 | 11508.5 | 4826 | 8054.5 |
| 2740 | 5902 | 4092 | 1349 | 5921 | 2243 | 864 | 13793 | 5003 | 9588 |
| 2860 | 4439 | 3439 | 1210 | 4921 | 1920 | 776 | 11749 | 4901 | 7012 |
| 2801 | 4410 | 3316 | 1116 | 4694 | 1903 | 750 | 12793 | 4629 | 6826 |
| 2755 | 4523 | 3520 | 1265 | 4880 | 2001 | 762 | 13602 | 4609 | 6909 |
| 2706 | 4584 | 3630 | 1274 | 4973 | 1990 | 802 | 14103 | 4624 | 7081 |
| 2483.5 | 5417 | 3639 | 1264.5 | 5026 | 2013 | 844.5 | 13008 | 4631 | 8660 |
| 2578.5 | 5029.5 | 3687.5 | 1308.5 | 5076 | 2054 | 839 | 11485.5 | 4935 | 8276.5 |
| 32783 | 23651 | 7629 | 1330 | 4345 | 1839 | 758 | 9925 | 13257 | 11574 |
| 2888.5 | 4002 | 3257.5 | 1091.5 | 4231 | 1814.5 | 733 | 9809 | 4957 | 6434.5 |
| 2839 | 4275 | 3440 | 1160 | 4548 | 1837 | 743 | 10362 | 4924 | 6874 |
| 67612.5 | 39711.5 | 11855 | 1597 | 4776 | 1905.5 | 755 | 15973 | 5795 | 8090.5 |
| 2787 | 4557 | 3680.5 | 1179 | 4764.5 | 1875 | 754 | 11325.5 | 4665.5 | 7269.5 |
| 2709 | 4317.5 | 3359.5 | 1151 | 4888 | 1897.5 | 746 | 10697.5 | 4709.5 | 6778 |
| 2716 | 4502 | 3578 | 1194 | 4767 | 1935 | 768 | 11287 | 4588 | 6825 |
| 7835 | 17080.5 | 6042.5 | 1614.5 | 7162.5 | 2240.5 | 766.5 | 15952 | 8879.5 | 19699.5 |
| 9253 | 23658 | 8556 | 1905 | 9762 | 2614 | 806 | 12554 | 9159 | 25541 |
| 10359 | 22361 | 6980 | 1718 | 8124 | 2359 | 736 | 14150 | 9478 | 23008 |

| | | | | | | | | | |
|---|---|---|---|---|---|---|---|---|---|
| 9730 | 20257 | 6408.5 | 1570 | 7431.5 | 2227 | 739.5 | 14012 | 9460.5 | 22168 |
| 2823 | 3732 | 2348 | 808 | 3007 | 1411.5 | 357.5 | 7835 | 3791.5 | 4668.5 |
| 2776.5 | 3774.5 | 2719 | 937 | 3630.5 | 1547 | 523.5 | 8497 | 4157 | 5264.5 |
| 2756 | 4153 | 3072 | 1067 | 4171 | 1764 | 652.5 | 9283 | 4492 | 6055.5 |
| 2652 | 4681 | 6115 | 1790 | 589662 | 50557 | 706 | 9900 | 4623 | 7189 |
| 2776 | 5352 | 4095 | 1363 | 6748 | 2397 | 782 | 11307 | 4888 | 8359 |
| 2611 | 5290 | 3755 | 1245 | 5740 | 2111 | 763 | 10263 | 4699 | 8404 |
| 2619 | 5272 | 3791 | 1290 | 5676 | 2127 | 802 | 11611 | 4653 | 8373 |
| 2700 | 4308 | 3330 | 1141 | 4908 | 1922 | 743 | 10865 | 4539 | 6560 |
| 2920 | 6140 | 3815 | 1276 | 5310 | 2037 | 772 | 12776 | 4835 | 9257 |
| 2886 | 5711 | 3934 | 1366 | 5360 | 2138 | 804 | 15442 | 4778 | 8584 |
| 2679 | 6454 | 3788 | 1268 | 5271.5 | 2059 | 811 | 14756.5 | 4680 | 10061.5 |
| 2539.5 | 4489 | 3486 | 1223 | 4847 | 1954 | 772 | 12896.5 | 4480 | 7233 |
| 2466 | 4117 | 3392 | 1204 | 4656 | 1949 | 769 | 10911 | 4495 | 6713 |
| 13026 | 9408 | 4511 | 1108 | 4227 | 1785 | 702 | 9332 | 6748 | 7459 |
| 3051 | 4499 | 3352 | 1163 | 4609 | 1934 | 749 | 9529 | 4885 | 6985 |
| 2906.5 | 5060 | 3563.5 | 1163.5 | 4757 | 1941.5 | 740 | 9957.5 | 4960.5 | 7960.5 |
| 75799 | 60384 | 12864 | 1787 | 5616 | 2077 | 788 | 16421 | 6328 | 12285 |
| 2914.5 | 7874.5 | 4246.5 | 1392 | 5570.5 | 2152 | 812 | 13859.5 | 4853.5 | 12165 |
| 2507 | 6557 | 3532 | 1180 | 5189 | 2000 | 751 | 10999 | 4394 | 10277 |
| 2637 | 3909.5 | 3126 | 1089.5 | 4251 | 1759.5 | 688 | 8989 | 4371 | 6132.5 |
| 6696 | 14818 | 5431 | 1443 | 6410 | 2095 | 704 | 15487 | 7995 | 17091 |
| 8927 | 23330 | 7980 | 1885 | 9092 | 2572 | 757 | 13790 | 9472 | 26470 |
| 9063.5 | 23042 | 7801 | 1768 | 8947.5 | 2526 | 735.5 | 13685.5 | 9601.5 | 24734 |
| 8359 | 21590.5 | 7095.5 | 1705 | 8593 | 2414 | 744 | 14007.5 | 9361.5 | 24807.5 |

| VL5 | VL6 |
|---|---|
| 1361 | 1103 |
| 2096 | 1284 |
| 2777 | 1447 |
| 11533 | 3945 |
| 4162 | 1792 |
| 3287 | 1585 |
| 3144 | 1553 |
| 3300 | 1609.5 |
| 3124 | 1568 |
| 3114 | 1575 |
| 3676.5 | 1685 |
| 3308 | 1632 |
| 3683 | 1703 |
| 3315 | 1617 |
| 3112 | 1569 |
| 2698 | 1425 |
| 4336 | 1818 |
| 3889 | 1716 |
| 4230 | 1789 |
| 4789.5 | 1983 |
| 6238 | 2075 |
| 8854.5 | 2483.5 |
| 7048 | 2152.5 |
| 7636 | 2225 |
| 2582 | 1284 |
| 2527 | 1335 |
| 3042 | 1482 |
| 8001 | 3359 |
| 4683 | 1913 |
| 3521 | 1639 |
| 3414 | 1603 |
| 3122 | 1539 |
| 3082 | 1537 |
| 3501 | 1642 |
| 2974 | 1528 |
| 3234 | 1572 |
| 3527 | 1694 |
| 3579 | 1686 |
| 3553 | 1654 |
| 3633.5 | 1673 |
| 4936.5 | 1908 |
| 4730 | 1869 |
| 4776 | 1858 |
| 5076 | 1947 |
| 4941 | 1901 |
| 7498 | 2253 |
| 7027 | 2172 |

| | |
|---|---|
| 7256 | 2229 |
| 2365 | 1251 |
| 2974.5 | 1462 |
| 3572.5 | 1641 |
| 9485 | 3128 |
| 4173 | 1787 |
| 3920.5 | 1747.5 |
| 4146.5 | 1838 |
| 3994 | 1789 |
| 4132 | 1840.5 |
| 4755 | 2038 |
| 4422 | 1932 |
| 4429 | 1963 |
| 4552 | 1979 |
| 4039.5 | 1825 |
| 4105 | 1876 |
| 4484 | 1945 |
| 5052 | 2058 |
| 4639 | 1950 |
| 4952 | 2049 |
| 4516 | 1980 |
| 6289 | 2235 |
| 7522.5 | 2402 |
| 7339 | 2271 |
| 8065 | 2419 |
| 2657 | 1288 |
| 2991 | 1443 |
| 3499 | 1606 |
| 7646 | 3063 |
| 4267 | 1835 |
| 3694 | 1685 |
| 3490 | 1645 |
| 4121 | 1802 |
| 3916 | 1768 |
| 4073.5 | 1799.5 |
| 4240 | 1868 |
| 4402 | 1865 |
| 4323 | 1908 |
| 4018 | 1794 |
| 3725 | 1694 |
| 4504 | 1919 |
| 4608 | 1914 |
| 4716 | 1961 |
| 4256 | 1837 |
| 4855 | 2023 |
| 5733 | 2103 |
| 7810 | 2343 |
| 6909 | 2166 |

| | |
|---|---|
| 6760 | 2091 |
| 2437 | 1253 |
| 2883 | 1449 |
| 3448 | 1638 |
| 8220.5 | 2879 |
| 4264 | 1810 |
| 4009 | 1767 |
| 3975 | 1782 |
| 4067 | 1786 |
| 4415 | 1905 |
| 4598.5 | 1973 |
| 4715 | 2011 |
| 4259 | 1875 |
| 4031 | 1814 |
| 4636.5 | 1996 |
| 4692 | 1996 |
| 4039 | 1792 |
| 4559 | 1902 |
| 4234 | 1825 |
| 4118 | 1802 |
| 3921.5 | 1756 |
| 6289 | 2148 |
| 6772 | 2191 |
| 7862 | 2358 |
| 7370 | 2253 |
| 2377 | 1254 |
| 3039.5 | 1465 |
| 3531.5 | 1645.5 |
| 8679 | 2840 |
| 4297 | 1816 |
| 4063 | 1780 |
| 4365 | 1865.5 |
| 4087.5 | 1787 |
| 5075 | 2089 |
| 6701 | 3636 |
| 5055 | 2148 |
| 4450 | 1937 |
| 4476 | 1922 |
| 3980 | 1749 |
| 3866.5 | 1689 |
| 4293.5 | 1843.5 |
| 9555 | 2440 |
| 4918 | 1920 |
| 4452 | 1839 |
| 4496 | 1813 |
| 5591.5 | 1962 |
| 7961 | 2354 |
| 7816 | 2352 |

| | |
|---|---|
| 7608 | 2248 |
| 2477 | 1258 |
| 2771 | 1379 |
| 3082 | 1478 |
| 8428 | 2860 |
| 4338 | 1805 |
| 4044 | 1750 |
| 4465 | 1860 |
| 3803 | 1695 |
| 4290.5 | 1805 |
| 3585 | 1639 |
| 3907 | 1720 |
| 4198 | 1814 |
| 4654 | 1888 |
| 3915 | 1734 |
| 3792.5 | 1690 |
| 4576 | 1865 |
| 4514 | 1830 |
| 4717 | 1943 |
| 3969 | 1734 |
| 4899 | 1994 |
| 5224.5 | 1914 |
| 7044 | 2224 |
| 8057 | 2331 |
| 7683 | 2290 |
| 2385 | 1246 |
| 3096 | 1479 |
| 3446 | 1601 |
| 9143 | 3146 |
| 4679 | 1921 |
| 4808 | 1993 |
| 4538 | 1917 |
| 4740.5 | 2001 |
| 4298 | 1891 |
| 4117.5 | 1826 |
| 4272 | 1855 |
| 3826 | 1752 |
| 4222.5 | 1872 |
| 3984 | 1789 |
| 3720 | 1735.5 |
| 4316 | 1892 |
| 4443 | 1857 |
| 4287 | 1817 |
| 4258 | 1839 |
| 3957 | 1753 |
| 5730.5 | 2045 |
| 7285 | 2306 |
| 9481 | 2646 |

| | |
|---|---|
| 7596 | 2366 |
| 2514.5 | 1272 |
| 2928 | 1439 |
| 3287 | 1538 |
| 14074 | 4113 |
| 5951 | 2146 |
| 5468 | 2036 |
| 4736.5 | 1902 |
| 4184 | 1758 |
| 4420 | 1820.5 |
| 5168 | 2181 |
| 4623 | 1900 |
| 4580 | 1892 |
| 4521 | 1893 |
| 4453 | 1844 |
| 4839 | 1960 |
| 5236 | 2052 |
| 5906 | 2134 |
| 5238 | 2046 |
| 4876 | 1946 |
| 5142 | 2010.5 |
| 7052 | 2330 |
| 9902 | 2688 |
| 9481 | 2606.5 |
| 9520 | 2584 |
| 2639 | 1300.5 |
| 2870 | 1397 |
| 3301 | 1548 |
| 9693.5 | 3095.5 |
| 4607 | 1880 |
| 4347 | 1819 |
| 3893.5 | 1699 |
| 3759 | 1666 |
| 3652 | 1663 |
| 3687 | 1739 |
| 3391 | 1597 |
| 3613 | 1670 |
| 3696 | 1684 |
| 3265 | 1563 |
| 3623 | 1659 |
| 3592 | 1636 |
| 4505.5 | 1829 |
| 4596 | 1831 |
| 4322 | 1819 |
| 3855 | 1749.5 |
| 4993.5 | 1854 |
| 7439 | 2285 |
| 8100 | 2330 |

| | |
|---|---|
| 7783 | 2279 |
| 2700 | 1306 |
| 3119 | 1467 |
| 3520 | 1595 |
| 10058 | 3322 |
| 4357 | 1858 |
| 3868 | 1735 |
| 3496 | 1628 |
| 3962 | 1773 |
| 3936 | 1750 |
| 3516.5 | 1663 |
| 4271.5 | 1859 |
| 3582 | 1686 |
| 3777 | 1717 |
| 3466 | 1664 |
| 4339 | 1872 |
| 3953 | 1723 |
| 4234 | 1773 |
| 4861 | 1947 |
| 3999 | 1742 |
| 3657 | 1668 |
| 5470 | 1937 |
| 9401 | 2613 |
| 7807 | 2348 |
| 9381.5 | 2457 |
| 1360 | 1093 |
| 1986 | 1235 |
| 2599 | 1404 |
| 9745 | 3539 |
| 3588 | 1699 |
| 3619 | 1695 |
| 3651 | 1706 |
| 3562 | 1676 |
| 3364.5 | 1622 |
| 3220 | 1611 |
| 3255 | 1577 |
| 3465 | 1647 |
| 3547 | 1673 |
| 3322 | 1602 |
| 3577 | 1667 |
| 3672 | 1703 |
| 3708.5 | 1659 |
| 4052.5 | 1759.5 |
| 3789 | 1700 |
| 3841.5 | 1739 |
| 5440 | 1949 |
| 8106.5 | 2327 |
| 8829 | 2442 |

| | |
|---|---|
| 7010 | 2153 |
| 2234 | 1207 |
| 2562 | 1324 |
| 3108 | 1512 |
| 9863 | 3232 |
| 3584 | 1637 |
| 3427.5 | 1604 |
| 3491 | 1621 |
| 3502 | 1638 |
| 3604 | 1655 |
| 3851 | 1749 |
| 3468 | 1637 |
| 3502 | 1629 |
| 3497 | 1662 |
| 3535 | 1658 |
| 3676.5 | 1672 |
| 3630 | 1645 |
| 4046 | 1732 |
| 3625 | 1641 |
| 3840 | 1702 |
| 3499.5 | 1633 |
| 4602 | 1778 |
| 7530 | 2268 |
| 7092.5 | 2185 |
| 7596.5 | 2256 |
| 2443 | 1241 |
| 2801 | 1413 |
| 3361 | 1599 |
| 10667 | 3478 |
| 3945 | 1744 |
| 3734 | 1708 |
| 3810.5 | 1757 |
| 4044 | 1816 |
| 3982.5 | 1784 |
| 4044 | 1833 |
| 3845 | 1759 |
| 3829 | 1744 |
| 3773 | 1752 |
| 3782.5 | 1779.5 |
| 4001 | 1816 |
| 4155 | 1838 |
| 4208.5 | 1805 |
| 4157 | 1814 |
| 4234 | 1845 |
| 4009 | 1797 |
| 5000.5 | 1876 |
| 7747 | 2399.5 |
| 7924 | 2447.5 |

| | |
|---|---|
| 7683.5 | 2344 |
| 2507 | 1263 |
| 2888 | 1424 |
| 3207 | 1548 |
| 9521 | 3136 |
| 4641 | 1919 |
| 3580 | 1658 |
| 3323 | 1608 |
| 3728 | 1693 |
| 3851.5 | 1744 |
| 3861 | 1742 |
| 4026 | 1793 |
| 3766 | 1734 |
| 3947 | 1796 |
| 3659 | 1711 |
| 3984 | 1818 |
| 5007 | 2056 |
| 4084 | 1779 |
| 4005 | 1754 |
| 4138.5 | 1798 |
| 3356 | 1609 |
| 5329 | 1945 |
| 7227 | 2290 |
| 9848 | 2540 |
| 7304 | 2206 |
| 2350 | 1229.5 |
| 2868 | 1413 |
| 3190 | 1539 |
| 8166 | 2855 |
| 4041 | 1756 |
| 4391 | 1849 |
| 4236 | 1841 |
| 4090 | 1794 |
| 3933 | 1778 |
| 3702 | 1716 |
| 3912 | 1771 |
| 3811 | 1738 |
| 4134 | 1841 |
| 3564 | 1681 |
| 3684 | 1726 |
| 4350 | 1916 |
| 4506.5 | 1889 |
| 4307 | 1829.5 |
| 4020.5 | 1757 |
| 3911 | 1768 |
| 5535 | 2029.5 |
| 7878.5 | 2445 |
| 8262 | 2410 |

| | |
|---|---|
| 7605 | 2355 |
| 2465 | 1260 |
| 2891 | 1421 |
| 3436 | 1580 |
| 7874.5 | 2676 |
| 4493 | 1837 |
| 4060 | 1791 |
| 4649 | 1933 |
| 4302 | 1887 |
| 4026.5 | 1790.5 |
| 5492 | 3265 |
| 4048.5 | 1781.5 |
| 4133 | 1790 |
| 3869 | 1734 |
| 3855 | 1686 |
| 3826 | 1720 |
| 3951 | 1733 |
| 9045 | 2240 |
| 4073 | 1690 |
| 4384 | 1822 |
| 4162 | 1777.5 |
| 5484 | 1928 |
| 7771 | 2417 |
| 7433.5 | 2308 |
| 7999 | 2416 |
| 2576 | 1287 |
| 2903.5 | 1398 |
| 3396 | 1552 |
| 9303 | 3081 |
| 4633 | 1872 |
| 4116 | 1754 |
| 3558 | 1611 |
| 3913 | 1705 |
| 3297 | 1557 |
| 3626 | 1647 |
| 3951 | 1729 |
| 3971 | 1742 |
| 4276 | 1805 |
| 3588 | 1652 |
| 4044 | 1747 |
| 3599 | 1631 |
| 4240 | 1761 |
| 4078 | 1743 |
| 3764 | 1683.5 |
| 4001 | 1752 |
| 5428 | 1935 |
| 7749 | 2326 |
| 7319 | 2225 |

| | |
|---|---|
| 8299 | 2352 |
| 2437 | 1264 |
| 3433 | 1584 |
| 3615.5 | 1663 |
| 9853 | 3348.5 |
| 4582 | 1894 |
| 4008 | 1761 |
| 4581 | 1951 |
| 4211 | 1839 |
| 4536 | 1942 |
| 4317 | 1895 |
| 4088 | 1796 |
| 4509.5 | 1940 |
| 4140 | 1854 |
| 4218 | 1873 |
| 3930 | 1782 |
| 4161 | 1826 |
| 4663 | 1941 |
| 4674 | 1959 |
| 4468.5 | 1906 |
| 4283 | 1882 |
| 5841.5 | 2081 |
| 7200.5 | 2287.5 |
| 8660.5 | 2468 |
| 7940.5 | 2424 |
| 2609 | 1305 |
| 2875 | 1402 |
| 3642.5 | 1642.5 |
| 11616 | 3605 |
| 4620 | 1875 |
| 4683 | 1859 |
| 4565 | 1865 |
| 4955 | 1963 |
| 4511 | 1875.5 |
| 5022 | 2137 |
| 4162.5 | 1783 |
| 4061 | 1773 |
| 4149 | 1773 |
| 3934 | 1698 |
| 3918.5 | 1726.5 |
| 4553 | 1887 |
| 4795 | 1833 |
| 4490 | 1851 |
| 4436.5 | 1836 |
| 4140 | 1792.5 |
| 5908 | 2039 |
| 7415 | 2331 |
| 7476 | 2302 |

| | |
|---|---|
| 8404 | 2460 |
| 2644 | 1304 |
| 2944 | 1419 |
| 3278 | 1530 |
| 8194 | 2855 |
| 3862 | 1696 |
| 4089 | 1760 |
| 3591 | 1629 |
| 4127 | 1780 |
| 4285 | 1801 |
| 4232 | 1891 |
| 4400 | 1843 |
| 3679 | 1654 |
| 4253 | 1771 |
| 3370 | 1579 |
| 3413.5 | 1603 |
| 3841 | 1707.5 |
| 4196 | 1748 |
| 3784 | 1664 |
| 3708 | 1644 |
| 3983 | 1717.5 |
| 4953 | 1824 |
| 7193 | 2229 |
| 6672 | 2078 |
| 6343 | 2034 |
| 2550 | 1273 |
| 3038 | 1417 |
| 3293 | 1543 |
| 8324 | 2887 |
| 4421 | 1835 |
| 4121 | 1778 |
| 4402 | 1851 |
| 4074.5 | 1763.5 |
| 3995 | 1759 |
| 3733 | 1690 |
| 4086 | 1798 |
| 3948 | 1753 |
| 3928.5 | 1759.5 |
| 3444 | 1619.5 |
| 3798 | 1721 |
| 4386.5 | 1867 |
| 3958.5 | 1729 |
| 4011 | 1740 |
| 3832 | 1704 |
| 3757 | 1706.5 |
| 5187 | 1902 |
| 6904 | 2223 |
| 6958 | 2192 |

| | |
|---|---|
| 7124 | 2196 |
| 1222 | 1058 |
| 2117 | 1257 |
| 2622 | 1409 |
| 9432 | 3527 |
| 3582 | 1724 |
| 3574 | 1674 |
| 3743.5 | 1732 |
| 3476 | 1660 |
| 3416 | 1633 |
| 3298 | 1594 |
| 3329 | 1603 |
| 3537 | 1667 |
| 3518 | 1670 |
| 3683 | 1707 |
| 3760 | 1719 |
| 3701 | 1700 |
| 3914 | 1728 |
| 4220 | 1785 |
| 3992 | 1752 |
| 4190 | 1826 |
| 5427 | 1975 |
| 7383 | 2289 |
| 8707.5 | 2458 |
| 8134.5 | 2348 |
| 2119 | 1177 |
| 2452 | 1304 |
| 2875 | 1441 |
| 9689 | 3282 |
| 3869 | 1716 |
| 3700 | 1682 |
| 3894 | 1752 |
| 3671 | 1664 |
| 3455 | 1607 |
| 3356 | 1602 |
| 3379 | 1607 |
| 3627 | 1672 |
| 3770 | 1713 |
| 3730 | 1719 |
| 4119.5 | 1801 |
| 3774 | 1713 |
| 4247 | 1791 |
| 3997 | 1737 |
| 4384 | 1843 |
| 4282.5 | 1835 |
| 5840 | 2055 |
| 8244 | 2439 |
| 7848 | 2335 |

| | |
|---|---|
| 8552 | 2371 |
| 2370 | 1224 |
| 2849.5 | 1411 |
| 3139.5 | 1523 |
| 9262 | 3219 |
| 3979 | 1772 |
| 4029 | 1822.5 |
| 4066 | 1813 |
| 4169 | 1847 |
| 3809 | 1731 |
| 3606 | 1695 |
| 3523 | 1663 |
| 3873.5 | 1778 |
| 4120 | 1853 |
| 4055 | 1851 |
| 4253 | 1915 |
| 4091 | 1807 |
| 4350 | 1855 |
| 4672 | 1969 |
| 4420 | 1887 |
| 4481 | 1959 |
| 6052 | 2142 |
| 7904 | 2478 |
| 8652 | 2548.5 |
| 8523 | 2468 |
| 2348 | 1227.5 |
| 2808 | 1396 |
| 3186 | 1519 |
| 8334 | 3009 |
| 4008 | 1737 |
| 3932 | 1754 |
| 4013 | 1781 |
| 3798 | 1726 |
| 3977.5 | 1774 |
| 3877 | 1744 |
| 3365.5 | 1608 |
| 3719 | 1727 |
| 3574 | 1692 |
| 3800 | 1733 |
| 4037 | 1786 |
| 4003 | 1776 |
| 3954 | 1733 |
| 4221.5 | 1812 |
| 4362 | 1883 |
| 4299 | 1872 |
| 5597 | 2045 |
| 7369 | 2327 |
| 7939 | 2301 |

| | |
|---|---|
| 8728 | 2401 |
| 2364 | 1225 |
| 2742 | 1386 |
| 3075 | 1521 |
| 7898 | 2863 |
| 3780 | 1696 |
| 3824 | 1714 |
| 3944 | 1768 |
| 3813.5 | 1750 |
| 3966 | 1779 |
| 3902.5 | 1783.5 |
| 3455.5 | 1647 |
| 3533 | 1672 |
| 3435 | 1641 |
| 3613 | 1695 |
| 3658 | 1697 |
| 3799 | 1725 |
| 3913 | 1749 |
| 3912 | 1715.5 |
| 3886 | 1762 |
| 3819 | 1729 |
| 5294 | 1933 |
| 6679 | 2193 |
| 7323 | 2291 |
| 8505 | 2523 |
| 2487 | 1249 |
| 2771 | 1368 |
| 3078 | 1492 |
| 7608 | 2587 |
| 4659 | 1893.5 |
| 4345 | 1846 |
| 4156 | 1784 |
| 4109.5 | 1780 |
| 4052 | 1763 |
| 5026 | 2618 |
| 3934 | 1761 |
| 4035.5 | 1797 |
| 3945 | 1771 |
| 3943 | 1713 |
| 3646 | 1664 |
| 4478 | 1865 |
| 8527.5 | 2301 |
| 4577.5 | 1888.5 |
| 4339 | 1835 |
| 4152 | 1795 |
| 5226.5 | 1921 |
| 7784.5 | 2350 |
| 8111 | 2324 |

| | |
|---|---|
| 7866 | 2381 |
| 2486 | 1247 |
| 2753 | 1354.5 |
| 3006 | 1458 |
| 9453 | 3164 |
| 4274 | 1800 |
| 4064.5 | 1738 |
| 4064 | 1739 |
| 3908 | 1701 |
| 3633 | 1661 |
| 3397 | 1590 |
| 3777 | 1694 |
| 3605 | 1647 |
| 3714 | 1699 |
| 3460 | 1608.5 |
| 3622 | 1663 |
| 4198 | 1793.5 |
| 4426 | 1826 |
| 4221 | 1788 |
| 4159 | 1764 |
| 3858 | 1702 |
| 5240 | 1883 |
| 7210 | 2255 |
| 6912 | 2119 |
| 7945 | 2319 |
| 2537 | 1283 |
| 2956 | 1426 |
| 3200 | 1547 |
| 8207 | 2881 |
| 4774 | 1998.5 |
| 4488 | 1879 |
| 4687 | 1957 |
| 4462.5 | 1914 |
| 4209 | 1838 |
| 3718 | 1711 |
| 4214 | 1856 |
| 4402 | 1921 |
| 4417 | 1922 |
| 4013 | 1784 |
| 4066 | 1841 |
| 4931 | 2080 |
| 5063 | 2060 |
| 5017.5 | 2030 |
| 4449 | 1881 |
| 4599 | 1973 |
| 6049.5 | 2127 |
| 8434 | 2613 |
| 7923 | 2446 |

| | |
|---|---|
| 9917 | 2678 |
| 2558 | 1272 |
| 2948 | 1412 |
| 3464 | 1588.5 |
| 11898 | 3725 |
| 4748 | 1936 |
| 4448 | 1901 |
| 4323.5 | 1833 |
| 4197 | 1791 |
| 3980 | 1736 |
| 4523 | 1986 |
| 4486.5 | 1904.5 |
| 4406 | 1861 |
| 4080 | 1787 |
| 4125.5 | 1764 |
| 4231 | 1794 |
| 4591 | 1907 |
| 5231 | 2020 |
| 5095 | 1999 |
| 5087 | 1985 |
| 4842.5 | 1972 |
| 6525.5 | 2236 |
| 8331.5 | 2495 |
| 8279 | 2424 |
| 8604.5 | 2505 |
| 2509 | 1274 |
| 2743 | 1369 |
| 3128 | 1481 |
| 8576.5 | 2906 |
| 4140 | 1741 |
| 4392 | 1826 |
| 4346 | 1805 |
| 4247 | 1778 |
| 3632 | 1632 |
| 3790 | 1805 |
| 3874 | 1740 |
| 3917 | 1726 |
| 4154 | 1793 |
| 3572 | 1646 |
| 4138 | 1812 |
| 4462 | 1867.5 |
| 4769 | 1910 |
| 4571 | 1865 |
| 4420.5 | 1842 |
| 4260 | 1808 |
| 5784 | 2009 |
| 7421.5 | 2309.5 |
| 7989 | 2377 |

| | |
|---|---|
| 7504 | 2231 |
| 2538 | 1263 |
| 2709 | 1340 |
| 3082 | 1484 |
| 8174 | 2880 |
| 4044 | 1713 |
| 4162 | 1769 |
| 3923 | 1713 |
| 3971 | 1743 |
| 3773 | 1675 |
| 3749 | 1689 |
| 3867 | 1700 |
| 4063.5 | 1780 |
| 4018 | 1763 |
| 3489.5 | 1614 |
| 3720 | 1688 |
| 4228.5 | 1811 |
| 4526 | 1850 |
| 4270 | 1805 |
| 3701 | 1663.5 |
| 4157 | 1795 |
| 5357 | 1943 |
| 6659 | 2159 |
| 6687 | 2093.5 |
| 7362 | 2253.5 |
| 1287 | 1070 |
| 2151 | 1243 |
| 2729 | 1384 |
| 8502.5 | 3565 |
| 4363 | 1919 |
| 4100 | 1753 |
| 4173 | 1766 |
| 4040 | 1724 |
| 3606 | 1624 |
| 3589 | 1637 |
| 3688 | 1651 |
| 4099 | 1747 |
| 4145 | 1749 |
| 3990 | 1730 |
| 4259.5 | 1787 |
| 4203.5 | 1760 |
| 4229.5 | 1755.5 |
| 4507 | 1808 |
| 4245 | 1771 |
| 4575 | 1823 |
| 5916 | 2004 |
| 8099 | 2269 |
| 9098 | 2384 |

| | |
|---|---|
| 7311 | 2171 |
| 2361 | 1213 |
| 2649 | 1326 |
| 3155 | 1490 |
| 10131.5 | 3436.5 |
| 4523.5 | 1850 |
| 4271 | 1801 |
| 4073 | 1728 |
| 4113.5 | 1759 |
| 3640 | 1642 |
| 3622 | 1658 |
| 3670 | 1645 |
| 3943 | 1732 |
| 3875 | 1697 |
| 3835 | 1707 |
| 4169 | 1768 |
| 4017 | 1725 |
| 4250 | 1779 |
| 4352 | 1797 |
| 4200 | 1772 |
| 4466 | 1844 |
| 5592 | 1963 |
| 7409.5 | 2263 |
| 7999 | 2364 |
| 7298 | 2210 |
| 2494 | 1236 |
| 2860 | 1366.5 |
| 3506 | 1544 |
| 11578.5 | 3879 |
| 4705 | 1870 |
| 4277 | 1753 |
| 4304 | 1762 |
| 4258 | 1735 |
| 3732 | 1636 |
| 3834 | 1756 |
| 3703.5 | 1667 |
| 4033 | 1764 |
| 4218 | 1775.5 |
| 4256 | 1779 |
| 4443 | 1798 |
| 4469 | 1820 |
| 4113 | 1723 |
| 4816 | 1833 |
| 4288 | 1760.5 |
| 4267 | 1767 |
| 5341 | 1900 |
| 6475 | 2075 |
| 7478 | 2223 |

| | |
|---|---|
| 6910 | 2104 |
| 2563 | 1253 |
| 2843.5 | 1373 |
| 3301 | 1536 |
| 10475 | 3465.5 |
| 4388 | 1807 |
| 4366 | 1817 |
| 4306.5 | 1809 |
| 3872 | 1705 |
| 3968 | 1743 |
| 3795 | 1709 |
| 4167 | 1777 |
| 3816 | 1681 |
| 3800 | 1680 |
| 3764 | 1687 |
| 3751.5 | 1661 |
| 4175 | 1785 |
| 4298.5 | 1769 |
| 4048 | 1706 |
| 4516 | 1883 |
| 4308 | 1808 |
| 5317 | 1908 |
| 7350 | 2314 |
| 7583 | 2274 |
| 8741.5 | 2382 |
| 2639 | 1277 |
| 3438.5 | 1533 |
| 3713 | 1664 |
| 12002.5 | 3820 |
| 5742 | 2194 |
| 5100 | 2039 |
| 4250.5 | 1819 |
| 4585.5 | 1914 |
| 4035 | 1750 |
| 4152 | 1805 |
| 4523 | 1925 |
| 3601 | 1671 |
| 4105.5 | 1786 |
| 5046 | 2049 |
| 3737 | 1700 |
| 4536 | 1883 |
| 4675 | 1889 |
| 4588.5 | 1873 |
| 4468 | 1851 |
| 4475 | 1873 |
| 5608 | 2014 |
| 6931.5 | 2187 |
| 8700 | 2538 |

| | |
|---|---|
| 8149 | 2436 |
| 2468 | 1245 |
| 2797 | 1355 |
| 3118 | 1461 |
| 10656 | 3449.5 |
| 4276 | 1755 |
| 4091 | 1705 |
| 4203 | 1727 |
| 4292 | 1744 |
| 3744 | 1631 |
| 3655 | 1612 |
| 3771 | 1657 |
| 4081 | 1715 |
| 4077 | 1729 |
| 4074 | 1726 |
| 4512 | 1780 |
| 4421 | 1783 |
| 4631 | 1811 |
| 4463 | 1785 |
| 4539 | 1790 |
| 4413 | 1790 |
| 6302 | 2047 |
| 7911.5 | 2286 |
| 8566 | 2345 |
| 8178.5 | 2262.5 |
| 2474 | 1251 |
| 2821 | 1375 |
| 3358 | 1525 |
| 10158.5 | 3435 |
| 4748 | 1893 |
| 4509 | 1870 |
| 4278.5 | 1783.5 |
| 4334 | 1808 |
| 3904.5 | 1712 |
| 3724 | 1664 |
| 3765 | 1661 |
| 4188 | 1783 |
| 4085 | 1752 |
| 4173 | 1774 |
| 4495 | 1903 |
| 4307 | 1800 |
| 4704 | 1888 |
| 4528 | 1844 |
| 4424 | 1823 |
| 4463 | 1817 |
| 6298.5 | 2120 |
| 8023.5 | 2421 |
| 8417 | 2412 |

| | |
|---|---|
| 8523 | 2399 |
| 2473 | 1241 |
| 3095 | 1457 |
| 3490 | 1587 |
| 11890 | 3860 |
| 4871 | 1942 |
| 4641 | 1893 |
| 4730 | 1919 |
| 4620 | 1882 |
| 4242 | 1796 |
| 4125.5 | 1784.5 |
| 4064 | 1761.5 |
| 4357 | 1836 |
| 4444 | 1861.5 |
| 4217 | 1812 |
| 4762 | 1978 |
| 4632 | 1919 |
| 4667 | 1873 |
| 4989 | 1983 |
| 4616 | 1900 |
| 5021 | 2012 |
| 6165 | 2118 |
| 8632 | 2564 |
| 10318.5 | 2723 |
| 8900 | 2541 |
| 2487 | 1244 |
| 3077.5 | 1423 |
| 3300 | 1513 |
| 12181 | 3731 |
| 4510 | 1807 |
| 4343 | 1778 |
| 4267 | 1784 |
| 4493 | 1850 |
| 4156 | 1745 |
| 3880 | 1704 |
| 3960 | 1719 |
| 4198 | 1766 |
| 4106 | 1746 |
| 4214 | 1781 |
| 4557 | 1862 |
| 4645 | 1902 |
| 4798 | 1905 |
| 4722 | 1881 |
| 4607.5 | 1854 |
| 4709 | 1895 |
| 6539 | 2186 |
| 8039 | 2466 |
| 8322 | 2448 |

| | |
|---|---|
| 8695 | 2417.5 |
| 2388 | 1232 |
| 2828 | 1356 |
| 3235 | 1482 |
| 10155.5 | 3377.5 |
| 4337 | 1765 |
| 4265 | 1790 |
| 4491 | 1799 |
| 4308.5 | 1751.5 |
| 4053.5 | 1697 |
| 3729 | 1644 |
| 3760 | 1647 |
| 4017 | 1721 |
| 4139 | 1745 |
| 4073 | 1734 |
| 4287 | 1776 |
| 4276.5 | 1776 |
| 4537 | 1813 |
| 4496 | 1801 |
| 4583 | 1827 |
| 4559 | 1820 |
| 6123 | 2043 |
| 8286.5 | 2371 |
| 8736.5 | 2351.5 |
| 8474 | 2283 |
| 2519 | 1245 |
| 2765 | 1346 |
| 3142 | 1456 |
| 10645 | 3495 |
| 4470 | 1772 |
| 4171 | 1729 |
| 3917 | 1647 |
| 4238.5 | 1749 |
| 4702 | 1835 |
| 4281 | 1764 |
| 3845 | 1659 |
| 3834 | 1664 |
| 4552 | 1782 |
| 4654.5 | 1810 |
| 4720 | 1835 |
| 4484 | 1785 |
| 5027 | 1868 |
| 4956 | 1863 |
| 4718 | 1812 |
| 4660 | 1778 |
| 6088 | 2034 |
| 8031.5 | 2406 |
| 7706 | 2233 |

| | |
|---|---|
| 7218 | 2162.5 |
| 1248 | 1050 |
| 2132 | 1244 |
| 2883.5 | 1445 |
| 9776 | 3736 |
| 4213.5 | 1885 |
| 4022 | 1782 |
| 3815 | 1701 |
| 3760 | 1693 |
| 3593.5 | 1624 |
| 3433 | 1630 |
| 3481 | 1620 |
| 3803 | 1722 |
| 3792 | 1710 |
| 3897 | 1733 |
| 3718 | 1674.5 |
| 4102 | 1774 |
| 4296 | 1791 |
| 4388 | 1822 |
| 4594.5 | 1885 |
| 4720 | 1952.5 |
| 6013 | 2070 |
| 7367.5 | 2309 |
| 8182 | 2405.5 |
| 9136 | 2487 |
| 2460 | 1226.5 |
| 2698 | 1346 |
| 3042 | 1472 |
| 9630 | 3161 |
| 4623 | 1902 |
| 4006.5 | 1755 |
| 3984 | 1751 |
| 3878 | 1737 |
| 3884 | 1695 |
| 3705 | 1678 |
| 3678 | 1662 |
| 3878 | 1742 |
| 4006 | 1769 |
| 4009 | 1762 |
| 4248.5 | 1834 |
| 4178 | 1812 |
| 4350 | 1800 |
| 4534 | 1867 |
| 4188 | 1789 |
| 4503 | 1910 |
| 6345 | 2168.5 |
| 7820 | 2372 |
| 8500 | 2375 |

| | |
|---|---|
| 8436 | 2432 |
| 2493 | 1249 |
| 2840 | 1363 |
| 3149 | 1488 |
| 12591 | 4024 |
| 4291 | 1780 |
| 4196.5 | 1764 |
| 4105 | 1773 |
| 3894 | 1707 |
| 3861 | 1703 |
| 3792 | 1776 |
| 3529 | 1628 |
| 3828.5 | 1703 |
| 3848 | 1710 |
| 3799 | 1698 |
| 3944 | 1750 |
| 4167 | 1777 |
| 4094 | 1712 |
| 4769 | 1934 |
| 4778 | 1936 |
| 4747 | 1921 |
| 5549 | 1994 |
| 7955 | 2357 |
| 7418 | 2279 |
| 7814 | 2289 |
| 2721 | 1284 |
| 3087 | 1464 |
| 3532 | 1609 |
| 10809.5 | 3506 |
| 4409 | 1862 |
| 4527 | 1882 |
| 4348 | 1845 |
| 4186 | 1804 |
| 3826.5 | 1715 |
| 4001 | 1769 |
| 3917 | 1751 |
| 4431 | 1862 |
| 4318 | 1839 |
| 4506 | 1922 |
| 4350 | 1866 |
| 4595 | 1896 |
| 4752.5 | 1914 |
| 4692 | 1933 |
| 4913.5 | 1989.5 |
| 4747 | 1944 |
| 6271 | 2159.5 |
| 8244 | 2510 |
| 10309 | 2690 |

| | |
|---|---|
| 8641 | 2487 |
| 2398 | 1242 |
| 2956 | 1417 |
| 3371 | 1572 |
| 11186 | 3659 |
| 4350.5 | 1818.5 |
| 4363 | 1826 |
| 4581 | 1908 |
| 4408 | 1868 |
| 4389 | 1859 |
| 4162 | 1814 |
| 4076 | 1773.5 |
| 4559 | 1960.5 |
| 4530 | 1930 |
| 4585 | 1939 |
| 4903.5 | 2008 |
| 4691 | 1953 |
| 5031 | 2003 |
| 5229.5 | 2058 |
| 5031 | 2044 |
| 4785 | 1995 |
| 6612 | 2299 |
| 8434.5 | 2614.5 |
| 9926 | 2791 |
| 9693 | 2674 |
| 2534 | 1258 |
| 2771 | 1364.5 |
| 3133 | 1470 |
| 9570 | 3241 |
| 4448 | 1834.5 |
| 3999 | 1712 |
| 4128 | 1754 |
| 4377 | 1818 |
| 3904 | 1695.5 |
| 3493.5 | 1622 |
| 3682.5 | 1661 |
| 3737.5 | 1666 |
| 4095 | 1764 |
| 3798 | 1701 |
| 4158 | 1781 |
| 4411 | 1855 |
| 4293 | 1791 |
| 5083 | 1991 |
| 4388 | 1817 |
| 4613 | 1869 |
| 6041 | 2092 |
| 7838 | 2376 |
| 9242 | 2512 |

| | |
|---|---|
| 8516.5 | 2405 |
| 2572 | 1266 |
| 2899 | 1395 |
| 3288 | 1527 |
| 9883 | 3180 |
| 4219 | 1773 |
| 4051.5 | 1754 |
| 4144.5 | 1799 |
| 4365 | 1822.5 |
| 3970 | 1740 |
| 3966 | 1741.5 |
| 4072 | 1755 |
| 4111 | 1790 |
| 4182 | 1816 |
| 4320 | 1849 |
| 4470.5 | 1917.5 |
| 4377 | 1849 |
| 4656 | 1920 |
| 4715 | 1916 |
| 4580 | 1869 |
| 4853 | 2001.5 |
| 6299 | 2160 |
| 7640 | 2389 |
| 9245 | 2685 |
| 8817 | 2517 |
| 2434.5 | 1244.5 |
| 3130 | 1488 |
| 3675 | 1662 |
| 12046 | 3878.5 |
| 4783 | 1930 |
| 4473 | 1847.5 |
| 4610 | 1897 |
| 4618 | 1910 |
| 4805 | 2008 |
| 4273 | 1831 |
| 4381.5 | 1854.5 |
| 4389 | 1867 |
| 4636 | 1916 |
| 4592.5 | 1916.5 |
| 4584.5 | 1922 |
| 4638 | 1923 |
| 4993.5 | 1963 |
| 5354 | 2104 |
| 4929 | 2016 |
| 5193.5 | 2079.5 |
| 6560 | 2264 |
| 8929 | 2674 |
| 9221 | 2679.5 |

| | |
|---|---|
| 10075 | 2748 |
| 2546 | 1263 |
| 2945 | 1424 |
| 3287.5 | 1535.5 |
| 10984 | 3590 |
| 4570 | 1859 |
| 4397 | 1806 |
| 4367.5 | 1841 |
| 4541.5 | 1860 |
| 4318 | 1796 |
| 3957 | 1733 |
| 3975 | 1722 |
| 4105 | 1778 |
| 4250 | 1807 |
| 4034 | 1767.5 |
| 4592 | 1899 |
| 4435 | 1836 |
| 4670 | 1859 |
| 4779 | 1908 |
| 4582 | 1865.5 |
| 4942 | 1979 |
| 6327 | 2164 |
| 9552 | 2709 |
| 8876.5 | 2555.5 |
| 8905.5 | 2549 |
| 2443 | 1228 |
| 2949 | 1407 |
| 3301 | 1508 |
| 8412.5 | 3023 |
| 4292 | 1781 |
| 4018.5 | 1721 |
| 4106 | 1768 |
| 4473 | 1821 |
| 3944 | 1701 |
| 3999 | 1732 |
| 3814 | 1652 |
| 4079 | 1756 |
| 3693 | 1629 |
| 4055 | 1754 |
| 4010.5 | 1753 |
| 4153 | 1770 |
| 4416 | 1826 |
| 4454 | 1821 |
| 4565 | 1867 |
| 4439 | 1854 |
| 6274 | 2146 |
| 8077 | 2436 |
| 8595 | 2480 |

| | |
|---|---|
| 8822 | 2452.5 |
| 2450 | 1238 |
| 2847 | 1383 |
| 3150 | 1486 |
| 9360 | 3029 |
| 4051 | 1714 |
| 4219 | 1765 |
| 3882 | 1665 |
| 4325 | 1796 |
| 4375.5 | 1812 |
| 4676 | 1875 |
| 4258 | 1761 |
| 4062.5 | 1728.5 |
| 4135.5 | 1753 |
| 4676 | 1859 |
| 4338 | 1794 |
| 4244 | 1768 |
| 4902.5 | 1869 |
| 4928 | 1910 |
| 4752 | 1848 |
| 4408 | 1784 |
| 5751 | 1993 |
| 7724 | 2327 |
| 7515 | 2247 |
| 7412 | 2253 |
| 1267 | 1056 |
| 2223 | 1264 |
| 2901 | 1440 |
| 9702 | 3655 |
| 3964 | 1803 |
| 3903.5 | 1724 |
| 3488 | 1621 |
| 3202 | 1538 |
| 4024 | 1736.5 |
| 3807.5 | 1724 |
| 3626 | 1653 |
| 3389 | 1581 |
| 3652 | 1664.5 |
| 3724 | 1687 |
| 3789 | 1705 |
| 3651 | 1655 |
| 3915 | 1682 |
| 4329 | 1816 |
| 4212 | 1777 |
| 3852 | 1695 |
| 5028 | 1838.5 |
| 7521 | 2326 |
| 7891.5 | 2348 |

| | |
|---|---|
| 7503 | 2274 |
| 2587 | 1250 |
| 2705 | 1336 |
| 3403.5 | 1537 |
| 9788 | 3258 |
| 4564 | 1889 |
| 4141 | 1778 |
| 3921 | 1716 |
| 3414 | 1585 |
| 3828.5 | 1702 |
| 3531 | 1628 |
| 3529 | 1609 |
| 3551 | 1624 |
| 3352 | 1571 |
| 3816 | 1699 |
| 3789 | 1685 |
| 3863 | 1698 |
| 3947 | 1675 |
| 4091.5 | 1744 |
| 3973 | 1710 |
| 3826 | 1688 |
| 5285 | 1913 |
| 6316 | 2050 |
| 6329.5 | 2007 |
| 6621 | 2056 |
| 2509.5 | 1250 |
| 2816 | 1365 |
| 3257.5 | 1504 |
| 12291 | 3884 |
| 4812 | 1900 |
| 4406 | 1827 |
| 4148 | 1789 |
| 4230 | 1787 |
| 3931 | 1701 |
| 3786 | 1754 |
| 3805.5 | 1697 |
| 3920 | 1721 |
| 4206 | 1816 |
| 4242 | 1801 |
| 4312 | 1825 |
| 4350 | 1834.5 |
| 4139 | 1741 |
| 4263 | 1794 |
| 4258 | 1806 |
| 4219 | 1792 |
| 5475 | 1959 |
| 6739 | 2176 |
| 7097 | 2210 |

| | |
|---|---|
| 7538 | 2264 |
| 2703 | 1282 |
| 2970 | 1406 |
| 3418 | 1569 |
| 10840 | 3528 |
| 4503 | 1863 |
| 4294 | 1781 |
| 4021 | 1755 |
| 3736 | 1677 |
| 3812 | 1685 |
| 4038 | 1788 |
| 4000 | 1738 |
| 4203 | 1808 |
| 4372 | 1865 |
| 4043 | 1764 |
| 4041 | 1770 |
| 4216 | 1779 |
| 4470.5 | 1817 |
| 4428.5 | 1833 |
| 4460 | 1847 |
| 4540 | 1891.5 |
| 6035 | 2110 |
| 6927.5 | 2217 |
| 7511.5 | 2264.5 |
| 7504 | 2254 |
| 2464 | 1247 |
| 2968 | 1414 |
| 3330 | 1537 |
| 10917.5 | 3530.5 |
| 4503 | 1866.5 |
| 4612 | 1926 |
| 4277 | 1826 |
| 4504 | 1906 |
| 3934.5 | 1719 |
| 3887 | 1730 |
| 4326 | 1842 |
| 4332 | 1840 |
| 4591 | 1948 |
| 4395 | 1874 |
| 4533 | 1922 |
| 4422 | 1868 |
| 4692.5 | 1874 |
| 4660 | 1888 |
| 4498.5 | 1868.5 |
| 4676 | 1920 |
| 6452.5 | 2231 |
| 7974 | 2445.5 |
| 8424.5 | 2506 |

| | |
|---|---|
| 7846.5 | 2298 |
| 2606 | 1270 |
| 2847 | 1366 |
| 3312.5 | 1526.5 |
| 9654 | 3176 |
| 4506 | 1846 |
| 4312 | 1801 |
| 4227 | 1786 |
| 4019 | 1732 |
| 3705 | 1639 |
| 3847 | 1695 |
| 3725 | 1640 |
| 4021 | 1737 |
| 4055 | 1753 |
| 4197 | 1764 |
| 4388 | 1826 |
| 4321 | 1808 |
| 4513 | 1828 |
| 4339 | 1788 |
| 4451.5 | 1819 |
| 4447 | 1843 |
| 5897 | 2018 |
| 7288.5 | 2258 |
| 7506 | 2255 |
| 7591 | 2272 |
| 2529 | 1262 |
| 2922 | 1418 |
| 3315 | 1529 |
| 10112.5 | 3331.5 |
| 4438 | 1827 |
| 4514 | 1868 |
| 4781.5 | 1930 |
| 4051 | 1762 |
| 3885 | 1677 |
| 4088 | 1779.5 |
| 4000 | 1738 |
| 4431 | 1869 |
| 4404 | 1884 |
| 4423 | 1863 |
| 4629 | 1914 |
| 4411 | 1868 |
| 4627 | 1847 |
| 4898 | 1949 |
| 4581 | 1869 |
| 4699 | 1925 |
| 6280 | 2175 |
| 7847.5 | 2454 |
| 8285 | 2484 |

| | |
|---|---|
| 8628 | 2495 |
| 2447 | 1253 |
| 3128 | 1479 |
| 3616 | 1635.5 |
| 11509 | 3709 |
| 4997 | 1975 |
| 4708 | 1908 |
| 4789 | 1925 |
| 4734 | 1960 |
| 4078 | 1772 |
| 3963 | 1745 |
| 4174 | 1801 |
| 4164 | 1787 |
| 4405 | 1845 |
| 4481 | 1882 |
| 4746.5 | 1975 |
| 4730 | 1951 |
| 4548 | 1836 |
| 4700 | 1925 |
| 4840 | 1933 |
| 4925 | 1980 |
| 6593.5 | 2261 |
| 7830.5 | 2416 |
| 8465 | 2451 |
| 8072 | 2370 |
| 2509 | 1258 |
| 3029 | 1434 |
| 3365 | 1542 |
| 10859 | 3521 |
| 4586 | 1854 |
| 4523 | 1848 |
| 4802 | 1941 |
| 4161 | 1764 |
| 4242 | 1758 |
| 4039 | 1737 |
| 3891 | 1692.5 |
| 4300 | 1828 |
| 4175 | 1778 |
| 4422.5 | 1851 |
| 4754.5 | 1946 |
| 4611 | 1907 |
| 4885 | 1899 |
| 4785 | 1918.5 |
| 4854.5 | 1945.5 |
| 4583 | 1878 |
| 6509 | 2210 |
| 7538 | 2371 |
| 8089 | 2398 |

| | |
|---|---|
| 8224.5 | 2392 |
| 2629 | 1274 |
| 2994 | 1402 |
| 3301 | 1509 |
| 9708 | 3244 |
| 4437 | 1823 |
| 4322 | 1789.5 |
| 4376 | 1814 |
| 4042 | 1729 |
| 3774 | 1647 |
| 3984.5 | 1723 |
| 3946.5 | 1713 |
| 4323 | 1822 |
| 4279 | 1787.5 |
| 4349 | 1830 |
| 4413.5 | 1836 |
| 4401 | 1834 |
| 4518 | 1817 |
| 4540 | 1843 |
| 4472 | 1822 |
| 4582.5 | 1876.5 |
| 6048 | 2098 |
| 7625 | 2353 |
| 7932 | 2327 |
| 8145 | 2367 |
| 2531 | 1266 |
| 2883 | 1373 |
| 3138 | 1468 |
| 9047.5 | 3065 |
| 4200.5 | 1767 |
| 4469 | 1817 |
| 3875 | 1675 |
| 3868 | 1680 |
| 4034 | 1714 |
| 4118 | 1741 |
| 4155.5 | 1736 |
| 3758 | 1647 |
| 3879 | 1683 |
| 3907 | 1688 |
| 4265 | 1773 |
| 4059.5 | 1713 |
| 4231 | 1745 |
| 4361 | 1783 |
| 4396 | 1799 |
| 4392.5 | 1775 |
| 5619 | 1951 |
| 7192 | 2219 |
| 7545 | 2271.5 |

| | |
|---|---|
| 7453.5 | 2247.5 |
| 1556 | 1183 |
| 2234.5 | 1339 |
| 2913 | 1487 |
| 13721 | 4083 |
| 5057 | 2013.5 |
| 4291 | 1894.5 |
| 4112.5 | 1860 |
| 4629.5 | 1945 |
| 4422.5 | 1937 |
| 4314.5 | 1962.5 |
| 4564 | 1966 |
| 5336 | 2241 |
| 4502 | 1951 |
| 4829 | 1997 |
| 4681 | 2054.5 |
| 3830 | 1758 |
| 5917 | 2239.5 |
| 5033 | 2074 |
| 5151 | 2141.5 |
| 3615 | 1667 |
| 5297 | 1937 |
| 7854 | 2436 |
| 8122 | 2423 |
| 7862 | 2325 |
| 2595 | 1334 |
| 2815 | 1381 |
| 3199.5 | 1526.5 |
| 8817 | 2881 |
| 4240 | 1795 |
| 4022.5 | 1701.5 |
| 4155 | 1847 |
| 4260 | 1877 |
| 4297 | 1903 |
| 5614 | 2281 |
| 4990.5 | 2139 |
| 5361.5 | 2207 |
| 4827 | 2101 |
| 4543 | 1918 |
| 4465.5 | 1981 |
| 4212 | 1912 |
| 5479 | 2213 |
| 4065 | 1701 |
| 4103 | 1782 |
| 3811.5 | 1735 |
| 5238 | 1897 |
| 8447.5 | 2531.5 |
| 8532 | 2502 |

| | |
|---|---|
| 9231 | 2551 |
| 2652.5 | 1342.5 |
| 2929.5 | 1448.5 |
| 3343.5 | 1621.5 |
| 10152 | 3143.5 |
| 4549 | 1851 |
| 4531 | 1903 |
| 4363 | 1885.5 |
| 5069 | 2046 |
| 4884.5 | 1991 |
| 4636.5 | 2014 |
| 4301.5 | 1887.5 |
| 4771.5 | 2002.5 |
| 4320.5 | 1827.5 |
| 5207 | 2126 |
| 4013 | 1805 |
| 4118 | 1811 |
| 6128.5 | 2329 |
| 4793 | 1934 |
| 5300 | 2168 |
| 3964 | 1713 |
| 5875 | 2056.5 |
| 7821 | 2473.5 |
| 8416 | 2496 |
| 8062 | 2425 |
| 2456 | 1320 |
| 2743.5 | 1378 |
| 3132 | 1550 |
| 12003 | 3440 |
| 5274.5 | 2002.5 |
| 4904 | 1951 |
| 4564 | 1935 |
| 4585 | 1935 |
| 3538 | 1655 |
| 4681 | 2067 |
| 5764.5 | 2387 |
| 4643 | 1885 |
| 4966 | 2068 |
| 4168 | 1777 |
| 3553 | 1628.5 |
| 4037.5 | 1754 |
| 4438 | 1818.5 |
| 4439.5 | 1871 |
| 4186 | 1806 |
| 4426 | 1911 |
| 6028 | 2116 |
| 10730 | 3013 |
| 10954 | 2789 |

| | |
|---|---|
| 8825.5 | 2575 |
| 2678 | 1345 |
| 2889 | 1464 |
| 3961 | 1763 |
| 13980 | 3926 |
| 5839.5 | 2190 |
| 4359 | 1803 |
| 3921 | 1714 |
| 4014 | 1744 |
| 4003.5 | 1812.5 |
| 4087 | 1884 |
| 4228.5 | 1826 |
| 5298 | 2100 |
| 5662.5 | 2279 |
| 5502 | 2047 |
| 6974 | 2611 |
| 3972 | 1740 |
| 4337 | 1786 |
| 4399 | 1845.5 |
| 4503 | 1877 |
| 3840.5 | 1701.5 |
| 5976 | 2090 |
| 8411 | 2496 |
| 10587 | 2832 |
| 9459.5 | 2733 |
| 2759.5 | 1353 |
| 3400 | 1497 |
| 3870.5 | 1673 |
| 11399.5 | 3530.5 |
| 6874 | 2379 |
| 5830 | 2135 |
| 5804 | 2238 |
| 6126 | 2314.5 |
| 5559 | 2206 |
| 5179.5 | 2207.5 |
| 4972 | 2015 |
| 4660 | 1929 |
| 4847 | 2051 |
| 6367 | 2413 |
| 4707 | 1821 |
| 4610 | 1831 |
| 8415 | 2325 |
| 5237.5 | 2001.5 |
| 4550 | 1926 |
| 4669 | 1816.5 |
| 6402 | 2131 |
| 8344.5 | 2629.5 |
| 8805 | 2557 |

| | |
|---|---|
| 8378 | 2397 |
| 2850 | 1346 |
| 3072 | 1474 |
| 3550 | 1627 |
| 12533 | 3647 |
| 5068 | 1936.5 |
| 5052 | 2026 |
| 4813.5 | 1984 |
| 4701 | 1904 |
| 4361 | 1876 |
| 4001 | 1747 |
| 4095 | 1773 |
| 4114 | 1850 |
| 4006 | 1791 |
| 4601 | 1759 |
| 3854 | 1716 |
| 4603 | 1921 |
| 4549 | 1845 |
| 4282 | 1742 |
| 4838 | 1950 |
| 4440 | 1877 |
| 6320 | 2094.5 |
| 11867 | 2960 |
| 8730 | 2659 |
| 12125.5 | 3164.5 |
| 2935 | 1418 |
| 3349.5 | 1557.5 |
| 3822 | 1730 |
| 10849 | 3426 |
| 5557 | 2288 |
| 5738 | 2329.5 |
| 5429.5 | 2267.5 |
| 5516 | 2340.5 |
| 5488 | 2333.5 |
| 4857 | 2137 |
| 5068 | 2171 |
| 4716 | 2054.5 |
| 5311 | 2248.5 |
| 4803 | 2075 |
| 4253 | 1892 |
| 5449.5 | 2300 |
| 4978 | 2118 |
| 4928 | 2081 |
| 5335 | 2253 |
| 5007.5 | 2117 |
| 5948.5 | 2490.5 |
| 7531 | 2984 |
| 6144 | 2332 |

| | |
|---|---|
| 6800 | 2457 |
| 3055 | 1401 |
| 3869 | 1662 |
| 4083 | 1695.5 |
| 14302 | 4232 |
| 9254 | 2970 |
| 7014 | 2384 |
| 10805.5 | 3115.5 |
| 5978 | 2160 |
| 5550 | 2091 |
| 6875 | 2710 |
| 5963.5 | 2161.5 |
| 5385.5 | 2108 |
| 5413 | 1994 |
| 15289 | 3156 |
| 5990 | 2197 |
| 5778.5 | 2157.5 |
| 7465.5 | 2367 |
| 6096 | 2151 |
| 9797 | 3181 |
| 6018 | 2147 |
| 6112 | 2212.5 |
| 6419.5 | 2240 |
| 6328 | 2279 |
| 6773 | 2282 |
| 3066 | 1413 |
| 3358.5 | 1540.5 |
| 3855.5 | 1671.5 |
| 11123 | 3342 |
| 5559 | 2098 |
| 5184 | 2039 |
| 4340 | 1853 |
| 4422.5 | 1885 |
| 3974 | 1750 |
| 4391 | 2052 |
| 4247 | 1861 |
| 3864.5 | 1723 |
| 4207 | 1835 |
| 4272.5 | 1750 |
| 3791 | 1708 |
| 4209 | 1800 |
| 5322 | 2133 |
| 4742 | 1942 |
| 4568 | 1924 |
| 4356.5 | 1850 |
| 4482 | 1873 |
| 4776.5 | 1937.5 |
| 5389.5 | 2003.5 |

| | |
|---|---|
| 5935 | 2069 |
| 3075 | 1440 |
| 3420 | 1531 |
| 3591 | 1635 |
| 13232 | 3718 |
| 5349 | 2003 |
| 4720 | 1926 |
| 4234 | 1777 |
| 4297 | 1843 |
| 4526.5 | 1855.5 |
| 4195 | 1836 |
| 4239 | 1872 |
| 4386.5 | 1889.5 |
| 4288 | 1841.5 |
| 4107 | 1727 |
| 4238 | 1845 |
| 4309 | 1827 |
| 5024 | 1961 |
| 4347 | 1801.5 |
| 4543 | 1862 |
| 4617 | 1912 |
| 4188 | 1797 |
| 4558.5 | 1824 |
| 4716 | 1880.5 |
| 5363 | 1901.5 |
| 1385 | 1125 |
| 2092 | 1299 |
| 2767 | 1425 |
| 8907 | 3222 |
| 4320 | 1873 |
| 4208 | 1918 |
| 3980 | 1789 |
| 4062 | 1863 |
| 4133 | 1827 |
| 3881.5 | 1785.5 |
| 3555 | 1693 |
| 3496 | 1686 |
| 3538.5 | 1747.5 |
| 4021.5 | 1747 |
| 4080 | 1826 |
| 4195 | 1875 |
| 4476 | 1835 |
| 3963 | 1737 |
| 3721 | 1695 |
| 3774 | 1751 |
| 5134 | 1985 |
| 7914.5 | 2466 |
| 7473 | 2324 |

| | |
|---|---|
| 8274 | 2452.5 |
| 2717 | 1300.5 |
| 2767 | 1418 |
| 3261 | 1525 |
| 8897.5 | 2941 |
| 3989 | 1754 |
| 3753 | 1662 |
| 3856.5 | 1770.5 |
| 4223 | 1836 |
| 4027 | 1741 |
| 4264 | 1879 |
| 3681 | 1679 |
| 3927 | 1783 |
| 3589 | 1697.5 |
| 4199.5 | 1776.5 |
| 4111.5 | 1802 |
| 4168 | 1787 |
| 4384 | 1856 |
| 4340 | 1855 |
| 4402.5 | 1876.5 |
| 4410 | 1857 |
| 6159 | 2150 |
| 8007.5 | 2518 |
| 8318 | 2474.5 |
| 8369 | 2477 |
| 2774.5 | 1326.5 |
| 3066.5 | 1435.5 |
| 3515 | 1618 |
| 9261 | 2994 |
| 4034 | 1753 |
| 3998.5 | 1745.5 |
| 3773 | 1684 |
| 4123.5 | 1848.5 |
| 4214 | 1764 |
| 4422 | 1953.5 |
| 3864.5 | 1705 |
| 3883 | 1782 |
| 3967 | 1797.5 |
| 3989 | 1756.5 |
| 3967.5 | 1768.5 |
| 4224 | 1827 |
| 4278.5 | 1791 |
| 4621 | 1988 |
| 4370 | 1838 |
| 4306 | 1897 |
| 5975 | 2149 |
| 7995.5 | 2443 |
| 10840 | 2848 |

| | |
|---|---|
| 8392.5 | 2415 |
| 2848 | 1373 |
| 2867 | 1403 |
| 3302 | 1573 |
| 9100 | 2911 |
| 4220 | 1750 |
| 3886 | 1757 |
| 4194 | 1848 |
| 4066.5 | 1798.5 |
| 4208.5 | 1797 |
| 4003 | 1824 |
| 3620 | 1641 |
| 3788 | 1756 |
| 3836 | 1756 |
| 4111 | 1751.5 |
| 4078 | 1770 |
| 4081 | 1771 |
| 4470 | 1796 |
| 4372 | 1854 |
| 4366 | 1844 |
| 4331 | 1859 |
| 6235 | 2149 |
| 7888 | 2476 |
| 9855 | 2689 |
| 8719 | 2548 |
| 2903 | 1345 |
| 3127 | 1489 |
| 3578.5 | 1597 |
| 8835 | 2798 |
| 4337 | 1824.5 |
| 3869 | 1716 |
| 4269 | 1826.5 |
| 4413 | 1831 |
| 4141.5 | 1790 |
| 4256.5 | 1799 |
| 3728 | 1777 |
| 3755 | 1718 |
| 3921 | 1767 |
| 4520 | 1773 |
| 4045 | 1782.5 |
| 4268 | 1795 |
| 4668 | 1921 |
| 4216.5 | 1799.5 |
| 4295 | 1871 |
| 4407.5 | 1893.5 |
| 6245.5 | 2167 |
| 7905 | 2471 |
| 9668 | 2675 |

| | |
|---|---|
| 8607.5 | 2583 |
| 3118 | 1436.5 |
| 3341 | 1540 |
| 3877 | 1663 |
| 8671 | 2955 |
| 4947 | 1949.5 |
| 4485 | 1822 |
| 5042 | 1966 |
| 4893.5 | 2010 |
| 5147.5 | 2068 |
| 5623 | 2320 |
| 4151.5 | 1746 |
| 4237 | 1799 |
| 4110.5 | 1821 |
| 5225 | 1906 |
| 4755.5 | 1930 |
| 4753 | 1891 |
| 7289 | 2158 |
| 5096 | 2009 |
| 4615 | 1973 |
| 4977 | 2073 |
| 6882 | 2305.5 |
| 8985.5 | 2640 |
| 9535 | 2666 |
| 8933.5 | 2505 |
| 2927 | 1375.5 |
| 3118 | 1491 |
| 3599.5 | 1621 |
| 9583 | 3146 |
| 4175 | 1737 |
| 4157 | 1769 |
| 4067 | 1761 |
| 4229 | 1855 |
| 4545 | 1881 |
| 4459 | 1883 |
| 3774 | 1635 |
| 3918 | 1707 |
| 3749 | 1704 |
| 4247.5 | 1775.5 |
| 4204 | 1793 |
| 4255 | 1818 |
| 4354 | 1766 |
| 4531 | 1836 |
| 4366 | 1786 |
| 4074 | 1752 |
| 5865 | 2022 |
| 7684.5 | 2344.5 |
| 8694.5 | 2557.5 |

| | |
|---|---|
| 8616 | 2481 |
| 2815 | 1327 |
| 3305 | 1450 |
| 3891.5 | 1731.5 |
| 9207.5 | 3110 |
| 5058 | 2009 |
| 4452 | 1958 |
| 4835 | 2058 |
| 5265 | 2254 |
| 5177.5 | 2196.5 |
| 5239.5 | 2249.5 |
| 4509 | 1967 |
| 4806.5 | 2083.5 |
| 4816 | 2090 |
| 4772.5 | 2034.5 |
| 5147 | 2197 |
| 5090 | 2227 |
| 5254 | 2127 |
| 5293 | 2183 |
| 5465 | 2253.5 |
| 5525 | 2348 |
| 6970 | 2526 |
| 9106.5 | 2821 |
| 10765 | 2877 |
| 11136 | 3065 |
| 2840 | 1406 |
| 3389 | 1566 |
| 4065 | 1721.5 |
| 10511 | 3658 |
| 5951.5 | 2152 |
| 5251 | 2034 |
| 7882 | 2562 |
| 5531 | 2078 |
| 5818 | 2072.5 |
| 6872.5 | 2609 |
| 5060 | 1913 |
| 5526 | 2089 |
| 5083.5 | 1978 |
| 13340 | 2632 |
| 5290 | 1976 |
| 5046 | 1957 |
| 6978.5 | 2137.5 |
| 5222.5 | 2003.5 |
| 10575 | 3198 |
| 5274 | 2035 |
| 7160 | 2408 |
| 9266 | 2645 |
| 9118 | 2613 |

| | |
|---|---|
| 9698.5 | 2779 |
| 2731 | 1338 |
| 3268 | 1477 |
| 3793.5 | 1638 |
| 9380 | 3010 |
| 4458 | 1779 |
| 4306 | 1856 |
| 4379 | 1836.5 |
| 4433 | 1917 |
| 4571 | 1896 |
| 4445 | 2012 |
| 4013 | 1743 |
| 3972.5 | 1766.5 |
| 4046 | 1809 |
| 4388.5 | 1829.5 |
| 4202 | 1811 |
| 4182 | 1774 |
| 4617 | 1808 |
| 4315 | 1862.5 |
| 4573 | 1876 |
| 4223 | 1758 |
| 6204 | 2087 |
| 7941 | 2434 |
| 8292 | 2454 |
| 7961 | 2357 |
| 2931.5 | 1391 |
| 3121 | 1440 |
| 3495.5 | 1559.5 |
| 9593 | 3149 |
| 4169.5 | 1786.5 |
| 4438 | 1828 |
| 4499 | 1838.5 |
| 4522 | 1872 |
| 4088 | 1803 |
| 4030.5 | 1757.5 |
| 4382 | 1873.5 |
| 4571 | 1930 |
| 4040 | 1797 |
| 4060 | 1695 |
| 4345 | 1847.5 |
| 4608.5 | 1914 |
| 4799 | 1892 |
| 4391 | 1774.5 |
| 4648 | 1917 |
| 4791 | 2021 |
| 5739 | 2065 |
| 7295 | 2268 |
| 7270 | 2172 |

| | |
|---|---|
| 8074.5 | 2378.5 |
| 1368 | 1129 |
| 2382 | 1313 |
| 2947 | 1483 |
| 7657.5 | 2956.5 |
| 4633 | 2051 |
| 3921.5 | 1802 |
| 3507 | 1682 |
| 3565 | 1668 |
| 3938 | 1733 |
| 4047 | 1820 |
| 4122 | 1875 |
| 3664.5 | 1738 |
| 3862 | 1775 |
| 4477 | 1915 |
| 4284 | 1866 |
| 4064 | 1823.5 |
| 4576 | 1895 |
| 4287.5 | 1879.5 |
| 4670 | 2018 |
| 4663 | 1999.5 |
| 5980 | 2156 |
| 7871.5 | 2472 |
| 8583 | 2537 |
| 9328.5 | 2555 |
| 2626.5 | 1306 |
| 2794 | 1396 |
| 3393 | 1568.5 |
| 8562 | 2753 |
| 4168 | 1755 |
| 4276 | 1832 |
| 3938 | 1767 |
| 3703 | 1722 |
| 3688 | 1687 |
| 4190.5 | 1894 |
| 3988.5 | 1837.5 |
| 4146.5 | 1875 |
| 4057.5 | 1834 |
| 4427 | 1758 |
| 4345 | 1894 |
| 4339.5 | 1847.5 |
| 4572 | 1834 |
| 4483 | 1895 |
| 4543 | 1899 |
| 4394.5 | 1921 |
| 5758 | 2082 |
| 7343.5 | 2257 |
| 8187 | 2464 |

| | |
|---|---|
| 8576 | 2509 |
| 2758.5 | 1296.5 |
| 2973 | 1449 |
| 3462.5 | 1601.5 |
| 8777 | 2918 |
| 4833 | 1975.5 |
| 4486.5 | 1933 |
| 4094 | 1852 |
| 3889 | 1784 |
| 3852.5 | 1689.5 |
| 4086 | 1844 |
| 4334 | 1838 |
| 4221 | 1852 |
| 3941 | 1775.5 |
| 4493 | 1851.5 |
| 4041 | 1780 |
| 4353.5 | 1879 |
| 4721 | 1967 |
| 4774 | 1954 |
| 4676 | 1896 |
| 4708 | 1935 |
| 6574.5 | 2271 |
| 8095 | 2421 |
| 9525 | 2735 |
| 9407 | 2600 |
| 2678.5 | 1293.5 |
| 2826.5 | 1404 |
| 3143.5 | 1505 |
| 8425 | 2751.5 |
| 4922.5 | 2004 |
| 4286 | 1872 |
| 4121 | 1823 |
| 3944 | 1787 |
| 3876.5 | 1778.5 |
| 4100 | 1869 |
| 3989 | 1785 |
| 4379 | 1935 |
| 4188 | 1886 |
| 4500 | 1868 |
| 4580.5 | 1989.5 |
| 4533 | 1962 |
| 4434 | 1836 |
| 4651 | 1925 |
| 4557 | 1981 |
| 4706 | 1987 |
| 6371.5 | 2199.5 |
| 8160.5 | 2502 |
| 8687 | 2546.5 |

| | |
|---|---|
| 8408 | 2435.5 |
| 2977.5 | 1365.5 |
| 3158 | 1468 |
| 3472.5 | 1602 |
| 8596 | 2832 |
| 4799 | 1977 |
| 4817.5 | 1927 |
| 4318.5 | 1869 |
| 3941 | 1684.5 |
| 4057 | 1789 |
| 4021.5 | 1795.5 |
| 4196.5 | 1884.5 |
| 4253 | 1896.5 |
| 4337 | 1871 |
| 5179 | 1945 |
| 4423 | 1887 |
| 4537 | 1878 |
| 4762 | 1927 |
| 4864 | 2013 |
| 4808 | 1942 |
| 4660 | 2000 |
| 6644 | 2225 |
| 8460 | 2605 |
| 8923 | 2569 |
| 9646 | 2600 |
| 3054 | 1429 |
| 3275 | 1492.5 |
| 3796 | 1731 |
| 7680 | 2645 |
| 5662 | 2190 |
| 5519.5 | 2155 |
| 5065 | 2030 |
| 4584 | 1876 |
| 4417 | 1921 |
| 5207 | 2243 |
| 4472 | 1811 |
| 4888 | 1958 |
| 4513 | 1902 |
| 5913.5 | 2246 |
| 4937 | 2088 |
| 4888.5 | 2039 |
| 7134.5 | 2223.5 |
| 5179 | 2062 |
| 4882.5 | 2070.5 |
| 5136 | 2077 |
| 7104 | 2332.5 |
| 8559.5 | 2580.5 |
| 8957 | 2599 |

| | |
|---|---|
| 8826 | 2591 |
| 2932.5 | 1352.5 |
| 3075 | 1474 |
| 3311 | 1520 |
| 9162 | 3074 |
| 4805 | 1878 |
| 4545 | 1904 |
| 4277.5 | 1852 |
| 4124 | 1756 |
| 3891 | 1756 |
| 4084 | 1840 |
| 4100 | 1793.5 |
| 4393 | 1900 |
| 4435 | 1864 |
| 4546 | 1870 |
| 4736.5 | 1962.5 |
| 4653.5 | 1953 |
| 4882.5 | 1986.5 |
| 4888 | 1877 |
| 5147 | 2153 |
| 4831 | 2072 |
| 6356 | 2197.5 |
| 8383 | 2517 |
| 9474 | 2676 |
| 10214 | 2778 |
| 2918 | 1330 |
| 3275 | 1514 |
| 4026 | 1793 |
| 8143 | 2909 |
| 5202 | 2123 |
| 5468 | 2148 |
| 5480.5 | 2291 |
| 5024 | 2119 |
| 4678.5 | 1983.5 |
| 4838 | 2148 |
| 5197 | 2218 |
| 5375 | 2266 |
| 5436 | 2298 |
| 5515 | 2334 |
| 5737 | 2377 |
| 5498.5 | 2320.5 |
| 5761.5 | 2366.5 |
| 5923 | 2465 |
| 5689.5 | 2283 |
| 5711.5 | 2455 |
| 7195.5 | 2549.5 |
| 9275 | 3004 |
| 9781.5 | 2924 |

| | |
|---|---|
| 9795 | 2824 |
| 3181.5 | 1438 |
| 3543 | 1527 |
| 4059 | 1757 |
| 10100 | 3507 |
| 5396.5 | 2036.5 |
| 5157 | 1982 |
| 8206 | 2514 |
| 4909.5 | 1985 |
| 4721 | 1901.5 |
| 6229 | 2565 |
| 5414 | 2016 |
| 5595 | 2131 |
| 5721 | 2048 |
| 11939 | 2814 |
| 5783 | 2154.5 |
| 5526.5 | 2127.5 |
| 7739 | 2391 |
| 5833 | 2172 |
| 9904 | 2990 |
| 5898 | 2221 |
| 7369 | 2358 |
| 9671 | 2787 |
| 10229 | 2867 |
| 9970.5 | 2771.5 |
| 3082 | 1406 |
| 3311 | 1478 |
| 3644 | 1675 |
| 8448.5 | 2760.5 |
| 4846.5 | 2004.5 |
| 4563 | 1871 |
| 4535 | 1855 |
| 4263.5 | 1802 |
| 4133 | 1783 |
| 4453 | 2036 |
| 4400.5 | 1825.5 |
| 4603 | 1936 |
| 4440 | 1882.5 |
| 4647 | 1896 |
| 4597 | 1876 |
| 4631 | 1912 |
| 5164 | 2006 |
| 4779 | 1892 |
| 5195 | 2070 |
| 4914.5 | 2039 |
| 6411 | 2147 |
| 9548.5 | 2742 |
| 10601 | 2795 |

| | |
|---|---|
| 10386 | 2747 |
| 3161.5 | 1402.5 |
| 3106 | 1473 |
| 3642.5 | 1652 |
| 7718.5 | 2651.5 |
| 4588.5 | 1932 |
| 4680 | 1928.5 |
| 4455 | 1860 |
| 4161 | 1827.5 |
| 4304 | 1827 |
| 4399 | 1901 |
| 4308.5 | 1812 |
| 4080 | 1809 |
| 3966 | 1787 |
| 4339 | 1830 |
| 4313 | 1847 |
| 4301.5 | 1836 |
| 4364 | 1812 |
| 4557 | 1943.5 |
| 4686.5 | 1978.5 |
| 4494.5 | 1930.5 |
| 5658.5 | 2014 |
| 7113.5 | 2215.5 |
| 8730.5 | 2510 |
| 8966 | 2556 |
| 1362 | 1089.5 |
| 2401.5 | 1339.5 |
| 2991 | 1484 |
| 7593 | 2976 |
| 4042 | 1852 |
| 4171 | 1868 |
| 3738 | 1710 |
| 3909 | 1773 |
| 3791 | 1737 |
| 4061 | 1814 |
| 4399.5 | 1935.5 |
| 4278 | 1879 |
| 4229 | 1886 |
| 4637 | 1922 |
| 4203 | 1899 |
| 4205 | 1850 |
| 4588 | 1916 |
| 4403 | 1881 |
| 4323.5 | 1842.5 |
| 4511 | 1897 |
| 5457 | 2055.5 |
| 8050 | 2463 |
| 8677.5 | 2549.5 |

| | |
|---|---|
| 8801 | 2577 |
| 2839.5 | 1339.5 |
| 2939.5 | 1428.5 |
| 3550.5 | 1644 |
| 8968 | 2956 |
| 4242 | 1917 |
| 4858 | 2050 |
| 4530 | 1892 |
| 4071 | 1793 |
| 3955.5 | 1761 |
| 4068.5 | 1863 |
| 4180.5 | 1822.5 |
| 4696.5 | 2014.5 |
| 4650 | 1992 |
| 4975 | 1993 |
| 4841 | 2055.5 |
| 4498 | 1918 |
| 5168 | 2077 |
| 4686 | 1963 |
| 4889 | 2062 |
| 4809 | 2117 |
| 6743.5 | 2327 |
| 8540.5 | 2537.5 |
| 10469 | 2895 |
| 9264 | 2694 |
| 2853 | 1304 |
| 3070 | 1502 |
| 3438 | 1614.5 |
| 12491 | 3752.5 |
| 4900 | 1971 |
| 4856.5 | 2000.5 |
| 4513 | 1916 |
| 4717 | 1916 |
| 4200 | 1823 |
| 4040 | 1900 |
| 3844 | 1840 |
| 4273 | 1914.5 |
| 4287 | 1798 |
| 4952.5 | 1988 |
| 4762 | 2002 |
| 4815.5 | 1968 |
| 5264 | 2011 |
| 4953 | 2027 |
| 4855 | 1970 |
| 4651 | 2003 |
| 6271 | 2210 |
| 8367 | 2549 |
| 9473.5 | 2743 |

| | |
|---|---|
| 8780 | 2475 |
| 2930 | 1328 |
| 3039 | 1446 |
| 3541 | 1648 |
| 9071 | 3112 |
| 4423.5 | 1876.5 |
| 4653 | 1984 |
| 4233 | 1831 |
| 4433 | 1861 |
| 4249 | 1859 |
| 4222 | 1876 |
| 4425.5 | 1851.5 |
| 4518 | 1997 |
| 4594 | 1963 |
| 5193.5 | 2000.5 |
| 4759.5 | 2056 |
| 4774.5 | 1950.5 |
| 5037 | 2042 |
| 5299 | 2119.5 |
| 4867 | 2070 |
| 4874 | 2049 |
| 6289 | 2173 |
| 9377 | 2735 |
| 11019 | 2874 |
| 9848 | 2684.5 |
| 3026 | 1373 |
| 3181 | 1513 |
| 3423 | 1643.5 |
| 9590 | 3125 |
| 4402 | 1817 |
| 4496 | 1916 |
| 4902.5 | 2039.5 |
| 4547.5 | 1908 |
| 4227 | 1820 |
| 3993 | 1845 |
| 4183 | 1871 |
| 4344.5 | 1935.5 |
| 4649.5 | 1983.5 |
| 4644 | 1842 |
| 4817 | 2039 |
| 4821 | 2051 |
| 4895.5 | 1991 |
| 5383.5 | 2209.5 |
| 5223.5 | 2091.5 |
| 4973 | 2054 |
| 6680 | 2204.5 |
| 8723 | 2778.5 |
| 10365.5 | 2932.5 |

| | |
|---|---|
| 10153 | 2798 |
| 2978.5 | 1380 |
| 3100 | 1459 |
| 3711 | 1628 |
| 8344 | 2820 |
| 4647 | 1883 |
| 4491.5 | 1827.5 |
| 4592.5 | 1966 |
| 4488 | 1941.5 |
| 4315 | 1790 |
| 3967.5 | 1830.5 |
| 4150 | 1791 |
| 4455 | 1924 |
| 4450.5 | 1971.5 |
| 4684 | 1925 |
| 4390 | 1913 |
| 4701.5 | 1945.5 |
| 4964 | 1985 |
| 4766 | 2014.5 |
| 4913 | 1968 |
| 4872 | 2033 |
| 6578 | 2286 |
| 8967 | 2669 |
| 10829.5 | 2735 |
| 9963 | 2691 |
| 3017 | 1378 |
| 3232.5 | 1472 |
| 3556 | 1629 |
| 8864.5 | 3083.5 |
| 4551 | 1905 |
| 4965.5 | 1979 |
| 4718 | 1981 |
| 3897 | 1710 |
| 4575 | 1860.5 |
| 4813.5 | 2053.5 |
| 4336 | 1831 |
| 4363.5 | 1865.5 |
| 4391 | 1845 |
| 4932.5 | 1976.5 |
| 4546 | 1893 |
| 4667.5 | 1935 |
| 4849 | 1989 |
| 4881 | 1895 |
| 4660 | 1965 |
| 4692 | 1914 |
| 5799.5 | 2083 |
| 8146 | 2467 |
| 9390 | 2695 |

| | |
|---|---|
| 7323 | 2263 |
| 2955 | 1394 |
| 3256 | 1491 |
| 4044 | 1830 |
| 9772.5 | 3244.5 |
| 4689 | 1920 |
| 5010 | 2057 |
| 5536 | 2214.5 |
| 4694 | 1976 |
| 4098.5 | 1825.5 |
| 4773 | 2137.5 |
| 4913 | 2064 |
| 5099.5 | 2162.5 |
| 4503 | 1969.5 |
| 5384 | 2066 |
| 3850.5 | 1748 |
| 4581.5 | 1983.5 |
| 5190 | 2100 |
| 5002 | 2046 |
| 4651 | 1998.5 |
| 4611 | 1990 |
| 5666 | 1977.5 |
| 7339.5 | 2306 |
| 10957 | 2836 |
| 11231 | 2864.5 |
| 2762 | 1347 |
| 3244.5 | 1507.5 |
| 3405.5 | 1573 |
| 7686 | 2645 |
| 4702 | 1895 |
| 4355.5 | 1824 |
| 4475 | 1842 |
| 4808 | 1942 |
| 4457 | 1816 |
| 3983.5 | 1742.5 |
| 4038.5 | 1781 |
| 4041.5 | 1720 |
| 4484 | 1878 |
| 4302 | 1830 |
| 4493 | 1902 |
| 4565.5 | 1919.5 |
| 4911 | 1921 |
| 4573 | 1859 |
| 5206 | 1989.5 |
| 4697 | 1952 |
| 6317 | 2187.5 |
| 7750 | 2435 |
| 10110 | 2641.5 |

| | |
|---|---|
| 9084 | 2588 |
| 2899 | 1357 |
| 3181 | 1480.5 |
| 3432 | 1603 |
| 8971 | 3052 |
| 4142.5 | 1754.5 |
| 4668 | 1877 |
| 4455.5 | 1846 |
| 4875.5 | 2070 |
| 4603.5 | 1859.5 |
| 4241 | 1823 |
| 4263.5 | 1815 |
| 4478.5 | 1919.5 |
| 4407 | 1849 |
| 6225 | 2044 |
| 4700 | 1922 |
| 4592 | 1928 |
| 5103 | 1964 |
| 4650 | 1958 |
| 4662 | 1957 |
| 4490 | 1934.5 |
| 5608 | 1956 |
| 8222 | 2512.5 |
| 10317 | 2769.5 |
| 8136 | 2298.5 |
| 2892 | 1365.5 |
| 3084 | 1460.5 |
| 3359 | 1501 |
| 8564 | 2879 |
| 4300.5 | 1816 |
| 4549 | 1883 |
| 4326 | 1826 |
| 4596 | 1910 |
| 4810 | 2053 |
| 4490 | 1876 |
| 4039 | 1734 |
| 4396 | 1872 |
| 4437.5 | 1880 |
| 4560 | 1883 |
| 4386 | 1861.5 |
| 4727 | 1910 |
| 4970 | 1979 |
| 5025 | 2032 |
| 4298.5 | 1837.5 |
| 4453 | 1894 |
| 5719 | 2043 |
| 7955.5 | 2398.5 |
| 8287 | 2559.5 |

| | |
|---|---|
| 7890 | 2406 |
| 1445 | 1101.5 |
| 2210 | 1303 |
| 3114 | 1575 |
| 7488 | 3034 |
| 4331.5 | 1939 |
| 4071 | 1844 |
| 3633 | 1694 |
| 3638 | 1697 |
| 3686 | 1721 |
| 3908 | 1772 |
| 4026 | 1819 |
| 4064 | 1856 |
| 4024.5 | 1824 |
| 4826 | 1904.5 |
| 4312 | 1887 |
| 4291 | 1859 |
| 4318 | 1862.5 |
| 4442 | 1863 |
| 4379 | 1886 |
| 4237 | 1830.5 |
| 5967 | 2076 |
| 7264 | 2315 |
| 8252 | 2480 |
| 8064 | 2436.5 |
| 2814 | 1340 |
| 2896.5 | 1404.5 |
| 3560.5 | 1618 |
| 9213 | 3035 |
| 4882 | 1957 |
| 4564 | 1943.5 |
| 4167 | 1870 |
| 3999 | 1787 |
| 3975.5 | 1780.5 |
| 4222.5 | 1922 |
| 4338 | 1935.5 |
| 4512 | 1996 |
| 4478.5 | 1932.5 |
| 4958 | 1945.5 |
| 4457.5 | 1925.5 |
| 4696 | 1968 |
| 4900 | 1935 |
| 4905 | 1980 |
| 5158 | 2123.5 |
| 4986 | 2102 |
| 6893.5 | 2344.5 |
| 8365 | 2584 |
| 9654.5 | 2656 |

| | |
|---|---|
| 9147 | 2631 |
| 2780 | 1306.5 |
| 2792 | 1417 |
| 3562 | 1645 |
| 13794.5 | 4003.5 |
| 4890 | 1938 |
| 4566 | 1898 |
| 4775 | 2017 |
| 4239 | 1888 |
| 3837.5 | 1707.5 |
| 4059 | 1865 |
| 4036 | 1851 |
| 4466.5 | 1955 |
| 4394 | 1927 |
| 4755.5 | 1958 |
| 4622 | 2027 |
| 4610 | 1982 |
| 4793 | 1923 |
| 4618 | 1904 |
| 4617 | 1888 |
| 4549 | 1950 |
| 6143 | 2213.5 |
| 7524 | 2430 |
| 8354 | 2553 |
| 8857 | 2618 |
| 2938 | 1359 |
| 3093 | 1449 |
| 3470 | 1689 |
| 9309.5 | 3113.5 |
| 4938 | 1985 |
| 4682 | 1997 |
| 4541 | 1883 |
| 4082 | 1794.5 |
| 3927 | 1751.5 |
| 4220 | 1936 |
| 4237 | 1827 |
| 4358.5 | 1904 |
| 4569 | 1960 |
| 4812 | 1903.5 |
| 4700.5 | 1947.5 |
| 4861 | 2062 |
| 4970 | 1947 |
| 5015 | 2037 |
| 4857.5 | 2074 |
| 4739.5 | 1927 |
| 6698 | 2313 |
| 8568 | 2654 |
| 8564 | 2645 |

| | |
|---|---|
| 9756 | 2703 |
| 2670 | 1326 |
| 3118 | 1506 |
| 3522 | 1608 |
| 9027 | 3065 |
| 4878 | 2037.5 |
| 5016 | 2075 |
| 4735 | 1884.5 |
| 4364 | 1857 |
| 4344 | 1832 |
| 4290 | 1889 |
| 4206 | 1897 |
| 4648 | 2028 |
| 4503 | 1972.5 |
| 5118.5 | 1959 |
| 4962 | 2089 |
| 4637 | 1939.5 |
| 5023.5 | 2081 |
| 4995 | 2099 |
| 5118.5 | 2054.5 |
| 5225 | 2102 |
| 6628 | 2261.5 |
| 8655 | 2711 |
| 9894 | 2826 |
| 9439.5 | 2697.5 |
| 2793.5 | 1347.5 |
| 2975 | 1416 |
| 3458 | 1584 |
| 8383.5 | 2804.5 |
| 4459.5 | 1884.5 |
| 4781.5 | 1931.5 |
| 4424 | 1839 |
| 4265 | 1843 |
| 3911.5 | 1784 |
| 4114 | 1904 |
| 4416 | 1909 |
| 4415.5 | 1925 |
| 4526 | 1930 |
| 4693 | 1910 |
| 4424 | 1905 |
| 4575 | 1977 |
| 4787.5 | 1918 |
| 4883.5 | 2032.5 |
| 4949 | 2053 |
| 5059.5 | 2037.5 |
| 6276 | 2205 |
| 8306.5 | 2590 |
| 9197 | 2607 |

| | |
|---|---|
| 8772 | 2507 |
| 2976 | 1384 |
| 3074 | 1465.5 |
| 3308 | 1525 |
| 8805 | 2909 |
| 4850 | 1956.5 |
| 4822 | 1987 |
| 4632 | 1927 |
| 4391 | 1856 |
| 4109 | 1799 |
| 4177.5 | 1950.5 |
| 3938 | 1794 |
| 4528 | 1962 |
| 4364 | 1903 |
| 4895 | 1942 |
| 4575 | 2004 |
| 4687 | 1965 |
| 5270 | 2054 |
| 4805 | 1962 |
| 5057 | 2082 |
| 4562 | 1829 |
| 6551 | 2243 |
| 8747 | 2586 |
| 9492 | 2634 |
| 9959 | 2684 |
| 2810.5 | 1333 |
| 3178 | 1464 |
| 3592 | 1625 |
| 9411 | 3212 |
| 4839 | 1973 |
| 4965.5 | 2044.5 |
| 4802.5 | 2025.5 |
| 4427 | 1842.5 |
| 4357 | 1851 |
| 4202 | 1865 |
| 4194 | 1804.5 |
| 4458 | 1949 |
| 4526 | 1959 |
| 4778 | 1985 |
| 4361.5 | 1898.5 |
| 4808 | 1990 |
| 4923 | 2061 |
| 4791 | 1976 |
| 4494.5 | 1864 |
| 4944.5 | 2084 |
| 6392 | 2245 |
| 8854 | 2698 |
| 9464 | 2720.5 |

| | |
|---|---|
| 8867 | 2600 |
| 2856 | 1335 |
| 3088 | 1468 |
| 3437.5 | 1553.5 |
| 7482 | 2609 |
| 4606.5 | 1846 |
| 4766 | 1944 |
| 4570.5 | 1913 |
| 4612 | 1893 |
| 4226.5 | 1802 |
| 3982 | 1792 |
| 4234 | 1800 |
| 4321 | 1844.5 |
| 4273 | 1844 |
| 4428 | 1852 |
| 4455 | 1907 |
| 4626.5 | 1941 |
| 5206 | 1959 |
| 4663 | 1898 |
| 4857 | 1952.5 |
| 4582 | 1933 |
| 6245 | 2150.5 |
| 8366.5 | 2584.5 |
| 8763 | 2644 |
| 8610 | 2486 |
| 3009 | 1364 |
| 3197 | 1425 |
| 3513.5 | 1575.5 |
| 9022 | 2926 |
| 4608 | 1891 |
| 4724 | 1964 |
| 4663.5 | 1969.5 |
| 4640 | 1964 |
| 4598 | 1893.5 |
| 4176 | 1809 |
| 4094.5 | 1785.5 |
| 4367 | 1878 |
| 4091 | 1798 |
| 5850.5 | 1910.5 |
| 4902 | 1898.5 |
| 4649 | 1891.5 |
| 5013.5 | 2031.5 |
| 5044 | 2065 |
| 4789 | 2012 |
| 4950 | 2020 |
| 6610.5 | 2240.5 |
| 8461 | 2654.5 |
| 10225 | 2809 |

| | |
|---|---|
| 9268 | 2624.5 |
| 3024 | 1397 |
| 2982 | 1429 |
| 3343 | 1546 |
| 8460 | 2875.5 |
| 4500 | 1837 |
| 4204 | 1785.5 |
| 4307 | 1855 |
| 4274.5 | 1826 |
| 4465 | 1845 |
| 4646 | 1954 |
| 4078 | 1766 |
| 4149 | 1820.5 |
| 4251 | 1834 |
| 4624.5 | 1904.5 |
| 4173 | 1868 |
| 4248 | 1821 |
| 4879.5 | 1932 |
| 4804 | 1980 |
| 4809 | 2002.5 |
| 4318 | 1879.5 |
| 5606 | 2057 |
| 7677 | 2442 |
| 8449 | 2611 |
| 8104 | 2453 |
| 1425 | 1102 |
| 2284.5 | 1320.5 |
| 3197.5 | 1571 |
| 7745.5 | 2940.5 |
| 4088.5 | 1863.5 |
| 4155.5 | 1895 |
| 3896 | 1831 |
| 3645.5 | 1665 |
| 3813.5 | 1776.5 |
| 4088.5 | 1831.5 |
| 4260 | 1891 |
| 4195 | 1927 |
| 4218 | 1937 |
| 4573 | 1879 |
| 4215.5 | 1865 |
| 4179 | 1816 |
| 4375.5 | 1769.5 |
| 4538.5 | 1865.5 |
| 4480 | 1848 |
| 4392 | 1909 |
| 5716.5 | 2072 |
| 6806 | 2234 |
| 8121.5 | 2533.5 |

| | |
|---|---|
| 8046.5 | 2557 |
| 2840.5 | 1370 |
| 3187 | 1498.5 |
| 3797 | 1687.5 |
| 10155 | 3261.5 |
| 4651 | 1934 |
| 4871 | 2040 |
| 4234 | 1794 |
| 4045.5 | 1800 |
| 4074 | 1817 |
| 4612.5 | 2022 |
| 4693.5 | 2017.5 |
| 4386 | 1946 |
| 4590.5 | 1856.5 |
| 4846.5 | 1985 |
| 4382 | 1917 |
| 4390 | 1922 |
| 4826 | 1917 |
| 4730.5 | 1893 |
| 4586 | 1926 |
| 4666.5 | 1964 |
| 6268 | 2172 |
| 7749 | 2454 |
| 8804.5 | 2482.5 |
| 8664 | 2511 |
| 2624 | 1252.5 |
| 2891 | 1456 |
| 3335 | 1534 |
| 8826 | 2819 |
| 4242 | 1767 |
| 4013 | 1811 |
| 4490 | 1872 |
| 4336 | 1841 |
| 3910 | 1768.5 |
| 4051 | 1854 |
| 4011.5 | 1794.5 |
| 4729 | 1977.5 |
| 4225 | 1900 |
| 4938 | 2062.5 |
| 4512 | 1974 |
| 4720 | 1991 |
| 4994 | 2018.5 |
| 4767 | 1952.5 |
| 4915 | 1991 |
| 4667 | 1920 |
| 5842 | 2075.5 |
| 7039 | 2257 |
| 7950 | 2414 |

| | |
|---|---|
| 8045.5 | 2407 |
| 3080.5 | 1405 |
| 3343 | 1505.5 |
| 3575 | 1606 |
| 9956 | 3259 |
| 5158 | 2014 |
| 4889.5 | 2041 |
| 4137 | 1868 |
| 4486 | 1864 |
| 4671 | 1978 |
| 4015 | 1814.5 |
| 4444.5 | 1924.5 |
| 4204 | 1907 |
| 4037 | 1797 |
| 4631.5 | 1887.5 |
| 4193 | 1788 |
| 4171 | 1854 |
| 5134 | 2109 |
| 4890 | 2087 |
| 4811.5 | 1970.5 |
| 4550 | 1891 |
| 5758 | 2079.5 |
| 7004 | 2334 |
| 8573 | 2517 |
| 8007.5 | 2416.5 |
| 3007.5 | 1386.5 |
| 3142.5 | 1463.5 |
| 3490.5 | 1596.5 |
| 11071 | 3584.5 |
| 5642 | 2199 |
| 4435 | 1836 |
| 4710 | 1968 |
| 4874 | 2082 |
| 4678 | 1933 |
| 4261.5 | 1910 |
| 4194 | 1898 |
| 4355 | 1924 |
| 4797 | 2145 |
| 4329 | 1823 |
| 3973 | 1769 |
| 5472 | 2180 |
| 5141.5 | 2075.5 |
| 5101 | 1998 |
| 4959 | 2023.5 |
| 4674 | 2092 |
| 6182 | 2109 |
| 7381 | 2433.5 |
| 9443 | 2816 |

| | |
|---|---|
| 8860.5 | 2626 |
| 3038 | 1395 |
| 3199 | 1489 |
| 3583 | 1643 |
| 8622 | 2879.5 |
| 5082.5 | 2015.5 |
| 4441 | 1878 |
| 4281 | 1818 |
| 4019 | 1711 |
| 4320 | 1857.5 |
| 4574 | 1976 |
| 4530 | 1979 |
| 4734 | 1956.5 |
| 4516.5 | 1919 |
| 4873 | 1966 |
| 4719 | 1982 |
| 4709 | 1978 |
| 4712 | 1878 |
| 4648 | 1847 |
| 4481 | 1924.5 |
| 4604 | 1892 |
| 5766.5 | 2090 |
| 7082 | 2331 |
| 8928.5 | 2582 |
| 8638 | 2522 |
| 3053 | 1372.5 |
| 3240 | 1521 |
| 3482.5 | 1584 |
| 8834 | 2967 |
| 5185 | 2020 |
| 4610.5 | 1919.5 |
| 4751.5 | 1997.5 |
| 4273 | 1831 |
| 4123 | 1746.5 |
| 4017.5 | 1816.5 |
| 4215 | 1822 |
| 4736.5 | 1979 |
| 4408 | 1916.5 |
| 4859 | 1889 |
| 4824 | 1985 |
| 4845 | 2038 |
| 4802.5 | 1927 |
| 5310 | 2129 |
| 5047 | 2079 |
| 4983 | 2120 |
| 5766 | 2131.5 |
| 8038 | 2452 |
| 9821.5 | 2732 |

| | |
|---|---|
| 9645 | 2757 |
| 3034 | 1412 |
| 3366 | 1548 |
| 3618 | 1582 |
| 9266.5 | 3103 |
| 4830 | 1943 |
| 5430 | 2148 |
| 4837.5 | 1988.5 |
| 4337 | 1891 |
| 4168.5 | 1802 |
| 4312 | 1913 |
| 4510 | 1963 |
| 4441 | 1884 |
| 4595 | 2048 |
| 5003 | 2011 |
| 4888 | 2083 |
| 4961 | 2026 |
| 5051 | 1978 |
| 4871.5 | 2019 |
| 5060 | 2065.5 |
| 4797 | 1963 |
| 6595 | 2273.5 |
| 8182.5 | 2504 |
| 9186 | 2652 |
| 8940.5 | 2523.5 |
| 2944 | 1368 |
| 3389.5 | 1534.5 |
| 3625 | 1649.5 |
| 8388.5 | 2883.5 |
| 4887 | 1922 |
| 5020 | 2055 |
| 4563 | 1851 |
| 4287.5 | 1805 |
| 4338 | 1801 |
| 4529 | 1897 |
| 4571 | 1950 |
| 4363 | 1896 |
| 4731 | 1976 |
| 4845 | 1972 |
| 4743 | 2020 |
| 4749 | 1964 |
| 4765 | 1844 |
| 5100 | 2009 |
| 5512 | 2116.5 |
| 4908 | 1970 |
| 5799 | 2078 |
| 8381 | 2516 |
| 9214 | 2626 |

| | |
|---|---|
| 9726.5 | 2545.5 |
| 3010 | 1330 |
| 3360.5 | 1514.5 |
| 3492 | 1593 |
| 8948 | 2964 |
| 4703.5 | 1909.5 |
| 4864.5 | 1988 |
| 4477 | 1886 |
| 4463 | 1845.5 |
| 4227 | 1801 |
| 4175.5 | 1827 |
| 4444 | 1912.5 |
| 4921 | 2073.5 |
| 4587 | 1955 |
| 6643.5 | 2177.5 |
| 4934.5 | 2066 |
| 4897 | 1945 |
| 5084 | 2012 |
| 5102 | 2073 |
| 5496 | 2186.5 |
| 4895 | 2068 |
| 6157.5 | 2192.5 |
| 8412.5 | 2528.5 |
| 9814.5 | 2653.5 |
| 9626 | 2761 |
| 3072 | 1403.5 |
| 3430 | 1513 |
| 3775 | 1708 |
| 8363.5 | 2829 |
| 5296.5 | 2093 |
| 5181 | 2098 |
| 4911 | 1967 |
| 4423 | 1886 |
| 4416.5 | 1833.5 |
| 4886 | 1983 |
| 4489 | 1945 |
| 4419.5 | 1875 |
| 4304.5 | 1855 |
| 4811 | 1933 |
| 4764 | 1945.5 |
| 4412.5 | 1890.5 |
| 4857.5 | 1965.5 |
| 4998 | 2002 |
| 4906.5 | 1961 |
| 4802 | 1980 |
| 5583 | 1996 |
| 6934.5 | 2240 |
| 8451 | 2476 |

| | |
|---|---|
| 8510 | 2493.5 |
| 1173 | 1081 |
| 4180 | 1712 |
| 3127 | 1473 |
| 13118 | 4997 |
| 5742 | 2362 |
| 4346.5 | 1881.5 |
| 4337.5 | 1896 |
| 3909 | 1767 |
| 3845 | 1763 |
| 3799 | 1730 |
| 3819 | 1722.5 |
| 3541 | 1682 |
| 3905 | 1775 |
| 3616 | 1713 |
| 4083 | 1810 |
| 3559 | 1671 |
| 3733 | 1690 |
| 4070 | 1814 |
| 4127 | 1807 |
| 4156.5 | 1804.5 |
| 4972 | 1978 |
| 6958 | 2344 |
| 7959 | 2360 |
| 9860 | 2632 |
| 10519.5 | 2775.5 |
| 7827 | 2424 |
| 6554 | 2169.5 |
| 8340 | 2829 |
| 6704 | 2403 |
| 5646 | 2197 |
| 5124 | 2067 |
| 4545.5 | 1925 |
| 4441.5 | 1910 |
| 4149 | 1843 |
| 4500.5 | 1932 |
| 4278 | 1866 |
| 5087 | 2103.5 |
| 4270 | 1873 |
| 4248 | 1838 |
| 5127 | 2101 |
| 4484 | 1893 |
| 5171 | 2087 |
| 4915 | 2012.5 |
| 5442.5 | 2123 |
| 5126 | 2048 |
| 6872 | 2315 |
| 8757 | 2477 |

| | |
|---|---|
| 9695.5 | 2657 |
| 10837 | 2758.5 |
| 8508.5 | 2521 |
| 6934 | 2229 |
| 6752 | 2286 |
| 5615 | 2103 |
| 4642 | 1879 |
| 4635 | 1987 |
| 4704 | 2006 |
| 3908 | 1746 |
| 4631.5 | 1972 |
| 3840 | 1790 |
| 4745 | 2016 |
| 4723.5 | 1983 |
| 4045.5 | 1768 |
| 4112 | 1838.5 |
| 4185 | 1798 |
| 3997.5 | 1770.5 |
| 4555 | 1916 |
| 4943 | 2025 |
| 4539 | 1931 |
| 6468 | 2344 |
| 8667 | 2790 |
| 9346 | 2554 |
| 10809 | 2757 |
| 8779.5 | 2503.5 |
| 7607.5 | 2484 |
| 6820 | 2252 |
| 6877 | 2362 |
| 6071 | 2218 |
| 4607.5 | 1887 |
| 4645 | 1936 |
| 4637 | 1971 |
| 3916 | 1747 |
| 4646 | 2021 |
| 3602 | 1693 |
| 4581 | 1916 |
| 4050 | 1810.5 |
| 4601 | 1947 |
| 4189 | 1827 |
| 4350 | 1911 |
| 4151 | 1830.5 |
| 4748.5 | 1968 |
| 5119.5 | 2081 |
| 4383 | 1848 |
| 6238 | 2285 |
| 6070 | 2088 |
| 9370 | 2546 |

| | |
|---|---|
| 8210 | 2369 |
| 7062 | 2128 |
| 7794 | 2436.5 |
| 7099 | 2309 |
| 7270.5 | 2406 |
| 5410 | 2011 |
| 4623 | 1881 |
| 4135 | 1793 |
| 3585 | 1649 |
| 4060.5 | 1778.5 |
| 3347.5 | 1603 |
| 4349 | 1843.5 |
| 3788 | 1681 |
| 3679.5 | 1674 |
| 3646.5 | 1670 |
| 4064 | 1780 |
| 4431.5 | 1921 |
| 3903.5 | 1726.5 |
| 4760.5 | 1972.5 |
| 4360 | 1841 |
| 4046.5 | 1758 |
| 5281 | 2022 |
| 6336 | 2150.5 |
| 8864 | 2547.5 |
| 7892 | 2318 |
| 9523 | 2595 |
| 13356 | 4085 |
| 7388 | 2354.5 |
| 7957 | 2593 |
| 6266 | 2226 |
| 5469 | 2102 |
| 4519.5 | 1879 |
| 4463.5 | 1915.5 |
| 4165.5 | 1825 |
| 4428 | 1909.5 |
| 4724.5 | 2028 |
| 3864 | 1780 |
| 3998.5 | 1732 |
| 3850.5 | 1703.5 |
| 4183 | 1762 |
| 4175 | 1779 |
| 6486 | 2169 |
| 5243.5 | 2032.5 |
| 4950 | 1987 |
| 5313 | 2082 |
| 6096 | 2193 |
| 7746 | 2494 |
| 7901 | 2394 |

| | |
|---|---|
| 8914 | 2533.5 |
| 6665 | 2069 |
| 8400 | 2524 |
| 6247 | 2037.5 |
| 8335 | 2578 |
| 5566 | 2034 |
| 4397 | 1837 |
| 4777.5 | 1933.5 |
| 3738 | 1668 |
| 3393 | 1612 |
| 3453 | 1617 |
| 4148 | 1788 |
| 3848 | 1702 |
| 4341 | 1847.5 |
| 4339 | 1869 |
| 4180 | 1833 |
| 3763 | 1704.5 |
| 3797 | 1676 |
| 4843 | 2002 |
| 4063 | 1769.5 |
| 4369 | 1835 |
| 4091 | 1792 |
| 6690 | 2244 |
| 7917 | 2362 |
| 9466.5 | 2562 |
| 9393 | 2545 |
| 8572.5 | 2500.5 |
| 6569 | 2166 |
| 9339.5 | 2868 |
| 8963 | 2824 |
| 6035 | 2219 |
| 3767 | 1668 |
| 3924 | 1770 |
| 4219.5 | 1811.5 |
| 3572.5 | 1662.5 |
| 4211 | 1858 |
| 3656 | 1693.5 |
| 4388.5 | 1908 |
| 4305 | 1939 |
| 4435 | 1908 |
| 3821 | 1721 |
| 3474 | 1598.5 |
| 4033 | 1803 |
| 4414.5 | 1864 |
| 4550 | 1925 |
| 4846 | 1982 |
| 6704 | 2232.5 |
| 7222.5 | 2233.5 |

| | |
|---|---|
| 9316 | 2665 |
| 8897 | 2479 |
| 13245 | 4959.5 |
| 7268 | 2862.5 |
| 9887 | 3286 |
| 7122 | 2856 |
| 4926 | 2035 |
| 5626 | 2262 |
| 4454 | 1933 |
| 4250 | 1816 |
| 4712 | 2007 |
| 3960 | 1759 |
| 3979 | 1739.5 |
| 4497 | 1936 |
| 3537 | 1590 |
| 4729.5 | 1948.5 |
| 4304 | 1892 |
| 5080.5 | 2127.5 |
| 4726 | 1983 |
| 5106 | 2079.5 |
| 4821 | 2014 |
| 5060 | 1971 |
| 10442 | 2933 |
| 9264 | 2902.5 |
| 12877 | 3141 |
| 7693 | 2271 |
| 12186 | 4806.5 |
| 7948 | 3325 |
| 12726 | 4242 |
| 8131 | 3326 |
| 6677 | 2820 |
| 4941 | 2182 |
| 6235 | 2535.5 |
| 4526 | 1970 |
| 5223 | 2177 |
| 4662.5 | 2019.5 |
| 4583 | 1953 |
| 4433 | 1924 |
| 4964 | 2079 |
| 4391.5 | 1931.5 |
| 3916 | 1739 |
| 4764 | 2023 |
| 4744 | 2012 |
| 5229.5 | 2094 |
| 5041.5 | 2042.5 |
| 6293.5 | 2322 |
| 7609.5 | 2431 |
| 7840 | 2393 |

| | |
|---|---|
| 9076 | 2366 |
| 7902 | 2348 |
| 14877 | 5074 |
| 6715 | 2608 |
| 10846 | 3967.5 |
| 6807 | 2785 |
| 5273 | 2232 |
| 4689 | 2010 |
| 4814 | 2050 |
| 4237 | 1897 |
| 4618 | 1990 |
| 4069 | 1781 |
| 5036 | 2016.5 |
| 4558 | 1895 |
| 5025 | 2050 |
| 3735 | 1732 |
| 5058 | 2052 |
| 4932.5 | 2036.5 |
| 4376.5 | 1845 |
| 5048 | 2069 |
| 4270 | 1854 |
| 5360 | 2025 |
| 7220 | 2288 |
| 7912 | 2342 |
| 0 | 0 |
| 3567 | 1462 |
| 6669 | 2700 |
| 5465 | 2047 |
| 33859 | 13683.5 |
| 13087 | 4645 |
| 9305 | 3351 |
| 7383.5 | 2848.5 |
| 6792 | 2618 |
| 5437 | 2207 |
| 5293 | 2161.5 |
| 5109.5 | 2129 |
| 4464 | 1912 |
| 4716.5 | 1975 |
| 4210.5 | 1843 |
| 4534 | 1938.5 |
| 4219 | 1872 |
| 4460 | 1877 |
| 4792 | 1983 |
| 4748 | 1965 |
| 4549 | 1885 |
| 5467 | 2073 |
| 7179 | 2343 |
| 8112 | 2467 |

| | |
|---|---|
| 10036 | 2560 |
| 8850 | 2518 |
| 8522 | 2823 |
| 7201 | 2315 |
| 11430.5 | 3872 |
| 7014 | 2623 |
| 5603 | 2190.5 |
| 4902 | 2005 |
| 4358.5 | 1875.5 |
| 4203 | 1833 |
| 4097 | 1852 |
| 4135.5 | 1827 |
| 4297 | 1859 |
| 4013.5 | 1773.5 |
| 4066 | 1775 |
| 4272 | 1889.5 |
| 4006 | 1782 |
| 4269.5 | 1876 |
| 4414 | 1875 |
| 4557.5 | 1895 |
| 4463.5 | 1905 |
| 5039 | 1912 |
| 6675.5 | 2205 |
| 7794 | 2395 |
| 8751.5 | 2483.5 |
| 9054 | 2546 |
| 7421 | 2320 |
| 7173 | 2306 |
| 8311.5 | 2732 |
| 5216.5 | 1979 |
| 5402 | 2082.5 |
| 4337.5 | 1881 |
| 4094 | 1800 |
| 4092 | 1851 |
| 4121 | 1813 |
| 4441 | 1924 |
| 4089 | 1826 |
| 4439 | 1931 |
| 4249.5 | 1834 |
| 3926 | 1767 |
| 4226.5 | 1831.5 |
| 4289 | 1862 |
| 4773 | 2014 |
| 4431.5 | 1908 |
| 4589 | 1977 |
| 4749 | 1967 |
| 6799 | 2221 |
| 8695 | 2514 |

| | |
|---|---|
| 10497 | 2744 |
| 10445 | 2663 |
| 9473.5 | 2970 |
| 6871 | 2268 |
| 7831 | 2648 |
| 5695 | 2114.5 |
| 5413 | 2089 |
| 4630 | 1939 |
| 4936 | 2032 |
| 4334 | 1860 |
| 4849 | 2017 |
| 4566 | 1951 |
| 4573 | 1942 |
| 4707.5 | 1948 |
| 4481 | 1912 |
| 4442 | 1900 |
| 4608 | 1915 |
| 4656.5 | 1953 |
| 4686.5 | 1967 |
| 4934 | 1997 |
| 4610.5 | 1905.5 |
| 6351.5 | 2266 |
| 6818 | 2220 |
| 8908 | 2565 |
| 12488 | 3068 |
| 9373 | 2597 |
| 9013 | 2646 |
| 7437.5 | 2339.5 |
| 8196.5 | 2701 |
| 5692 | 2131.5 |
| 5049 | 2000 |
| 4676 | 1934 |
| 4564.5 | 1903 |
| 4425 | 1882.5 |
| 4406 | 1883 |
| 4886 | 2000.5 |
| 4649 | 1994 |
| 4855 | 1989 |
| 4402.5 | 1909 |
| 4143.5 | 1818 |
| 4559 | 1947 |
| 4340 | 1856 |
| 5559 | 2210 |
| 4428 | 1862 |
| 4702 | 1940 |
| 5022 | 1950 |
| 9398.5 | 2651.5 |
| 8017 | 2411 |

| | |
|---|---|
| 10339 | 2641 |
| 9637 | 2650 |
| 14456 | 4425 |
| 6639.5 | 2174 |
| 10981 | 3325 |
| 6550.5 | 2329 |
| 6027 | 2171 |
| 4908 | 1980 |
| 5040 | 2030 |
| 4811.5 | 1944.5 |
| 5024 | 2120 |
| 4989.5 | 2094.5 |
| 4344 | 1871.5 |
| 4783 | 1991 |
| 5061 | 1953 |
| 4968.5 | 1954.5 |
| 5035 | 2059 |
| 6535 | 2194.5 |
| 5100 | 2083 |
| 5621.5 | 2151 |
| 5110 | 2036.5 |
| 6215 | 2340 |
| 6595 | 2358 |
| 10060 | 2626 |
| 10786 | 2843 |
| 9944 | 2670 |
| 8821 | 2527 |
| 8563 | 2518 |
| 12149.5 | 3409.5 |
| 6583 | 2309.5 |
| 5766 | 2119.5 |
| 5361.5 | 2090 |
| 4608.5 | 1928 |
| 4649 | 1921 |
| 4172 | 1814 |
| 4176 | 1809 |
| 4339 | 1829 |
| 4517 | 1883 |
| 4399 | 1854 |
| 4461 | 1931 |
| 4078 | 1793 |
| 4371 | 1866 |
| 4453 | 1919 |
| 4605 | 1883 |
| 4899 | 1996 |
| 5027 | 1937 |
| 7315.5 | 2292 |
| 9213 | 2547 |

| | |
|---|---|
| 9751 | 2645 |
| 7939 | 2354 |
| 8717 | 2707 |
| 5779 | 1959 |
| 11327 | 3188 |
| 6858 | 2381 |
| 6277 | 2315 |
| 5644 | 2146 |
| 4833 | 2000.5 |
| 4583 | 1947 |
| 4453 | 1917 |
| 4886.5 | 2017 |
| 4699.5 | 1941 |
| 4351.5 | 1896 |
| 4968 | 2077 |
| 4466 | 1914 |
| 4769 | 2085 |
| 5319 | 2126 |
| 6158 | 2352 |
| 5343 | 2144 |
| 4945 | 2006 |
| 5529 | 2183 |
| 7461 | 2468.5 |
| 7870 | 2425 |
| 10070 | 2812 |
| 9370 | 2506 |
| 22049 | 7623 |
| 7834 | 2726 |
| 13859 | 3881 |
| 7335 | 2631 |
| 7036 | 2646.5 |
| 5837 | 2339 |
| 4896 | 2044 |
| 5262 | 2159 |
| 5038 | 2193 |
| 4080 | 1850 |
| 4597 | 1948 |
| 4387 | 1893 |
| 4400 | 1893 |
| 4454 | 1973 |
| 4039 | 1810 |
| 4350.5 | 1862.5 |
| 4243 | 1848 |
| 4257 | 1809 |
| 4275 | 1836 |
| 4787 | 1897 |
| 7793 | 2460 |
| 7362.5 | 2251 |

| | |
|---|---|
| 12163 | 2959 |
| 7991 | 2336 |
| 10963.5 | 4299.5 |
| 8986.5 | 3489.5 |
| 9165 | 3363 |
| 7599 | 3057 |
| 5885 | 2476 |
| 5079 | 2174 |
| 4128 | 1855 |
| 3873 | 1743 |
| 4724 | 2061.5 |
| 3968 | 1805 |
| 4048.5 | 1804 |
| 4087.5 | 1809 |
| 3489 | 1616 |
| 4250.5 | 1836.5 |
| 3940 | 1720.5 |
| 4309 | 1833 |
| 4118 | 1813 |
| 4537.5 | 1916 |
| 4040.5 | 1785 |
| 4837.5 | 1865 |
| 7544 | 2376 |
| 10123 | 2814 |
| 11082 | 2896 |
| 8148 | 2344 |
| 8502 | 2886 |
| 6955 | 2413 |
| 10442 | 3044 |
| 6402 | 2323 |
| 6236 | 2331 |
| 4819 | 1903 |
| 4188 | 1768 |
| 4256 | 1844 |
| 4296 | 1910 |
| 4220 | 1862 |
| 3991 | 1743 |
| 4802 | 2025 |
| 3694 | 1674 |
| 4382.5 | 1836 |
| 4174 | 1822 |
| 4033 | 1737 |
| 4206 | 1800 |
| 4373 | 1863 |
| 5455 | 2137 |
| 4361 | 1812 |
| 6378.5 | 2192 |
| 8108 | 2399 |

| | |
|---|---|
| 0 | 0 |
| 4021 | 1559 |
| 7185 | 2826 |
| 4711 | 1855 |
| 29425.5 | 10976.5 |
| 10185 | 3573 |
| 7181 | 2738 |
| 6527 | 2555 |
| 5359.5 | 2160 |
| 5505.5 | 2201 |
| 4671 | 1983.5 |
| 5030 | 2074 |
| 4643 | 1982 |
| 4329 | 1848 |
| 4123 | 1862 |
| 4108.5 | 1782 |
| 4245.5 | 1844 |
| 4279 | 1816 |
| 4319 | 1857 |
| 4330.5 | 1814.5 |
| 4331 | 1875 |
| 4977 | 1967 |
| 6592.5 | 2187 |
| 8755 | 2543 |
| 7955 | 2339 |
| 7888.5 | 2386.5 |
| 7072 | 2410 |
| 7223 | 2284.5 |
| 8790 | 3066 |
| 5501 | 2094 |
| 4703 | 1951 |
| 4403 | 1858.5 |
| 3991 | 1764 |
| 4059.5 | 1756 |
| 3738 | 1732.5 |
| 4039 | 1818 |
| 4019 | 1768 |
| 3981 | 1788 |
| 4098 | 1834 |
| 3882 | 1751 |
| 4076.5 | 1840 |
| 4136.5 | 1792 |
| 4320 | 1828 |
| 4038 | 1754 |
| 4538 | 1940 |
| 4680 | 1871 |
| 6421 | 2197 |
| 8013 | 2391 |

| | |
|---|---|
| 8956.5 | 2446 |
| 8317 | 2418 |
| 7803 | 2376 |
| 6580 | 2207 |
| 7572 | 2555 |
| 5351 | 2079 |
| 4562.5 | 1919.5 |
| 4526.5 | 1939 |
| 4035 | 1785 |
| 4537 | 1981.5 |
| 4063 | 1834 |
| 4263 | 1838.5 |
| 4157 | 1834 |
| 4399 | 1904 |
| 4343 | 1912 |
| 4160 | 1818 |
| 4345 | 1855 |
| 4158 | 1812 |
| 4478 | 1887 |
| 4304 | 1831 |
| 4291 | 1816 |
| 5152 | 1968 |
| 7244 | 2344 |
| 7779 | 2395 |
| 10221 | 2575 |
| 10797 | 2863 |
| 6968 | 2490 |
| 5975.5 | 1965.5 |
| 7956 | 2636 |
| 5319 | 2012 |
| 4655 | 1901 |
| 4053.5 | 1776 |
| 4195 | 1817 |
| 3847 | 1750 |
| 4032 | 1792 |
| 4060 | 1758 |
| 3875 | 1770 |
| 4406 | 1881 |
| 4281 | 1837 |
| 4073 | 1804 |
| 4314 | 1870 |
| 4605 | 1940 |
| 4388 | 1921.5 |
| 4404.5 | 1852 |
| 4767 | 1956.5 |
| 5381 | 2088 |
| 6600.5 | 2220.5 |
| 7999 | 2398 |

| | |
|---|---|
| 9058.5 | 2488 |
| 8876.5 | 2560 |
| 7610 | 2352 |
| 6544.5 | 2155 |
| 7372 | 2493 |
| 5549 | 2058 |
| 5160.5 | 2033 |
| 4019 | 1746 |
| 4264 | 1818 |
| 4009 | 1779 |
| 4069 | 1789 |
| 4044.5 | 1803.5 |
| 4029 | 1774 |
| 4001 | 1744 |
| 4299 | 1838 |
| 3876 | 1744 |
| 3899 | 1719 |
| 4033 | 1768 |
| 4418 | 1850 |
| 4022 | 1754 |
| 4773.5 | 1984 |
| 5022 | 2043 |
| 8565 | 2586 |
| 9250 | 2654 |
| 9571 | 2680 |
| 10228 | 2683 |
| 8919.5 | 2886 |
| 7633 | 2389 |
| 8066 | 2799 |
| 6277 | 2237 |
| 5618 | 2120.5 |
| 5163 | 2054.5 |
| 4785.5 | 1985 |
| 4940 | 1999.5 |
| 5187 | 2245 |
| 4559 | 1919 |
| 4218 | 1923 |
| 4914.5 | 2006 |
| 4589 | 1879 |
| 4791 | 1938.5 |
| 4271.5 | 1832.5 |
| 6317 | 2124 |
| 4844 | 1913 |
| 4873 | 1961 |
| 4955 | 1987.5 |
| 5340.5 | 2105.5 |
| 9802 | 2881 |
| 8446 | 2507 |

| | |
|---|---|
| 12106 | 2989 |
| 8439 | 2443 |
| 8371 | 2553 |
| 6594 | 2117 |
| 8561 | 2827.5 |
| 5827.5 | 2138.5 |
| 4928 | 2013 |
| 4611 | 1905 |
| 4502 | 1909 |
| 4044 | 1759 |
| 4453 | 1883.5 |
| 4368 | 1890.5 |
| 4385 | 1883 |
| 4317 | 1879 |
| 4002 | 1791 |
| 4389 | 1905 |
| 4154.5 | 1822.5 |
| 4329.5 | 1839 |
| 4721 | 1946 |
| 4584 | 1896 |
| 4955 | 2001 |
| 5065 | 2017.5 |
| 9579 | 2628 |
| 8190 | 2421 |
| 10959 | 2764 |
| 8170.5 | 2418 |
| 7813 | 2488 |
| 7128.5 | 2286 |
| 7495.5 | 2472 |
| 5908.5 | 2198 |
| 5248 | 2070.5 |
| 4732 | 1945 |
| 4074.5 | 1790.5 |
| 4814 | 1965 |
| 4877 | 2042 |
| 4140 | 1811.5 |
| 4299 | 1880 |
| 4663 | 1998 |
| 4183.5 | 1818 |
| 4614 | 1992 |
| 4453.5 | 1876.5 |
| 4229 | 1817 |
| 4472 | 1954 |
| 4591 | 1915 |
| 4937 | 2065 |
| 4656 | 1898 |
| 7497 | 2394 |
| 7672 | 2330.5 |

| | |
|---|---|
| 10233 | 2700 |
| 11108 | 2786 |
| 14483 | 5203 |
| 7508.5 | 2671.5 |
| 8252 | 2893 |
| 6266.5 | 2347 |
| 5460.5 | 2214 |
| 5141 | 2155 |
| 4672 | 2002 |
| 5128.5 | 2151 |
| 5132 | 2153 |
| 4547 | 1919 |
| 5365 | 2201 |
| 4454 | 1921 |
| 4668 | 1935.5 |
| 4540 | 1947 |
| 4440 | 1871 |
| 4749 | 1985.5 |
| 4750 | 1950.5 |
| 4534 | 1893.5 |
| 4692 | 2012.5 |
| 5029 | 2037 |
| 6813 | 2287 |
| 7906 | 2339 |
| 11201 | 2870 |
| 10706 | 2809 |
| 10741 | 4025 |
| 8942 | 3445 |
| 9177 | 4020 |
| 6838.5 | 2877 |
| 5166.5 | 2240 |
| 4558 | 2052 |
| 4227 | 1889 |
| 3841 | 1764 |
| 4108 | 1826 |
| 3952 | 1749 |
| 4155 | 1781 |
| 4238 | 1872 |
| 4185 | 1855 |
| 3993 | 1804 |
| 4573 | 1939 |
| 4389.5 | 1864 |
| 4453 | 1903 |
| 4686 | 1937 |
| 4732 | 1958 |
| 5116 | 1987 |
| 7604 | 2439 |
| 7854 | 2349 |

| | |
|---|---|
| 10250 | 2699.5 |
| 9176.5 | 2513 |
| 8349 | 2768 |
| 7412 | 2471 |
| 7428 | 2697 |
| 5932 | 2241 |
| 4680.5 | 1993 |
| 4202 | 1840 |
| 4516 | 1948 |
| 4930 | 2045 |
| 4213 | 1803 |
| 4384 | 1884 |
| 4355 | 1844 |
| 4531 | 1912 |
| 4524.5 | 1900 |
| 4006 | 1775 |
| 3871 | 1758 |
| 4523 | 1920 |
| 4544 | 1886.5 |
| 4543 | 1902 |
| 4660.5 | 1923 |
| 4906.5 | 2002.5 |
| 5478 | 2008 |
| 6919 | 2145 |
| 9643 | 2646.5 |
| 1158 | 1078 |
| 2699 | 1418 |
| 2793 | 1343 |
| 9785 | 3962 |
| 5346 | 2187 |
| 4202 | 1808 |
| 3956 | 1753 |
| 4084 | 1764 |
| 3826 | 1684 |
| 4088.5 | 1786.5 |
| 4065 | 1775 |
| 3806 | 1679 |
| 4030 | 1747 |
| 3943 | 1730 |
| 3995.5 | 1742 |
| 3887 | 1742.5 |
| 4280 | 1794 |
| 4458.5 | 1858 |
| 4408 | 1814 |
| 4374 | 1829 |
| 5399 | 1969 |
| 6570.5 | 2175.5 |
| 8444 | 2445 |

| | |
|---|---|
| 9096 | 2550 |
| 10279 | 2667 |
| 8707 | 2643 |
| 6535 | 2121 |
| 8210.5 | 2682 |
| 6270.5 | 2260 |
| 4938 | 1948 |
| 4622 | 1888 |
| 4430 | 1865 |
| 4441 | 1887 |
| 4471 | 1903 |
| 4200 | 1778 |
| 4431 | 1876.5 |
| 4209 | 1790 |
| 4312 | 1820 |
| 4229 | 1789 |
| 4455 | 1845 |
| 4355.5 | 1819 |
| 4486.5 | 1829.5 |
| 4638 | 1916 |
| 4288 | 1816 |
| 5458.5 | 1980.5 |
| 6511 | 2135 |
| 8681.5 | 2489 |
| 9854 | 2600 |
| 10912.5 | 2784 |
| 9804.5 | 2678 |
| 7228 | 2280 |
| 8001 | 2626.5 |
| 5994.5 | 2113 |
| 4951.5 | 1921 |
| 4516 | 1853 |
| 4556 | 1871 |
| 4250.5 | 1842.5 |
| 4516 | 1916.5 |
| 4899.5 | 2016 |
| 4474 | 1883 |
| 4538 | 1916 |
| 4434 | 1927 |
| 4247 | 1822 |
| 4480 | 1942.5 |
| 4467.5 | 1880 |
| 4873 | 1966 |
| 4888 | 1968.5 |
| 4655 | 1961 |
| 5041.5 | 1963 |
| 7448 | 2345 |
| 9208.5 | 2584 |

| | |
|---|---|
| 10942 | 2818 |
| 10288 | 2647 |
| 8837 | 2610 |
| 7413 | 2341 |
| 8245 | 2731 |
| 6166 | 2181 |
| 5054 | 1959 |
| 4826 | 1933 |
| 4498 | 1886 |
| 4345 | 1838 |
| 4334 | 1856 |
| 4728 | 1954 |
| 4320 | 1793 |
| 4420 | 1843 |
| 4338.5 | 1826 |
| 4293 | 1817.5 |
| 4398 | 1867.5 |
| 4467 | 1843 |
| 4590 | 1880 |
| 4813.5 | 1944 |
| 4822.5 | 1950 |
| 5518 | 2006 |
| 6761 | 2257 |
| 9103 | 2607 |
| 10963 | 2805 |
| 10100 | 2556 |
| 8838 | 2761 |
| 7284.5 | 2315 |
| 7963.5 | 2614.5 |
| 6465 | 2273 |
| 5067.5 | 1949 |
| 4794 | 1940 |
| 4520 | 1851 |
| 4491.5 | 1857 |
| 4363 | 1843 |
| 4513 | 1882 |
| 4512 | 1857 |
| 4356 | 1809 |
| 4381 | 1873 |
| 4487.5 | 1880.5 |
| 4277 | 1824 |
| 4506 | 1859 |
| 4638 | 1929 |
| 4621.5 | 1917.5 |
| 4591 | 1877.5 |
| 5096.5 | 1963 |
| 7302 | 2335 |
| 10368 | 2634 |

| | |
|---|---|
| 10655 | 2765 |
| 9931 | 2594 |
| 14603 | 4618 |
| 8076 | 2663 |
| 8565 | 2682 |
| 5877 | 2105.5 |
| 5308.5 | 2058 |
| 4757.5 | 1938 |
| 4640 | 1896 |
| 4861 | 1982 |
| 4989 | 2121 |
| 4804.5 | 1980 |
| 4593 | 1816 |
| 4442 | 1838 |
| 4748 | 1847 |
| 4588.5 | 1835 |
| 4408 | 1840 |
| 7169.5 | 2084.5 |
| 4766 | 1887 |
| 5453 | 2076 |
| 5037 | 1941 |
| 5742.5 | 2069.5 |
| 7583.5 | 2378.5 |
| 9622.5 | 2535.5 |
| 11090.5 | 2762.5 |
| 10660.5 | 2673 |
| 9107 | 2600.5 |
| 7937.5 | 2441.5 |
| 8531 | 2696 |
| 6147 | 2173.5 |
| 5211.5 | 1988.5 |
| 4766 | 1948 |
| 4487.5 | 1874 |
| 4328 | 1812 |
| 4335 | 1818 |
| 4496 | 1842 |
| 4165 | 1772 |
| 4298 | 1810 |
| 4168 | 1810 |
| 4206 | 1787.5 |
| 4350 | 1828 |
| 4542 | 1861 |
| 4556 | 1896 |
| 4608 | 1875 |
| 4601.5 | 1882.5 |
| 4946 | 1912 |
| 7326 | 2309 |
| 8993 | 2540 |

| | |
|---|---|
| 10136 | 2643 |
| 9867 | 2534 |
| 9276.5 | 2674 |
| 7912 | 2406 |
| 11823 | 3350 |
| 7078 | 2340 |
| 5782 | 2110.5 |
| 5083 | 1980 |
| 4875 | 1993 |
| 4769 | 1950 |
| 4809 | 1959 |
| 4668 | 1928 |
| 4681.5 | 1954.5 |
| 4603 | 1895 |
| 4567 | 1866 |
| 4514 | 1887 |
| 4304 | 1871 |
| 4656.5 | 1927 |
| 4822 | 1947 |
| 4967 | 2013 |
| 4914 | 1957 |
| 5111 | 2038 |
| 6469 | 2186 |
| 9398 | 2683 |
| 10393 | 2715 |
| 10953 | 2720.5 |
| 13536 | 4656 |
| 7951 | 2610 |
| 11582 | 3641 |
| 7965.5 | 2779.5 |
| 6379.5 | 2335.5 |
| 5163 | 1999 |
| 5162.5 | 2033.5 |
| 4832 | 1964 |
| 5157 | 2124 |
| 4611 | 1876 |
| 4641 | 1910.5 |
| 4631 | 1895 |
| 4774.5 | 1870.5 |
| 4725 | 1883 |
| 4717 | 1924 |
| 4936.5 | 1954.5 |
| 4876 | 1963 |
| 4875.5 | 1975.5 |
| 4623 | 1909.5 |
| 5553 | 2034 |
| 7236 | 2365 |
| 9865 | 2618 |

| | |
|---|---|
| 10928 | 2774 |
| 9876 | 2586 |
| 13856.5 | 5152.5 |
| 8146 | 3028 |
| 10598 | 3569.5 |
| 6985 | 2676.5 |
| 6024.5 | 2360 |
| 5226 | 2062 |
| 4968.5 | 2001 |
| 4748 | 1955 |
| 4599 | 1994 |
| 4527 | 1908 |
| 4491 | 1864 |
| 4449 | 1860 |
| 4268 | 1841 |
| 4387 | 1852 |
| 4185 | 1813 |
| 4521.5 | 1846 |
| 4592 | 1876 |
| 4735 | 1915 |
| 4536 | 1865 |
| 5292 | 1957.5 |
| 6176.5 | 2086 |
| 8474 | 2475 |
| 9317.5 | 2553 |
| 10368 | 2606 |
| 12340.5 | 4620.5 |
| 7154 | 2637 |
| 11177 | 3494 |
| 6404 | 2493.5 |
| 5232 | 2105 |
| 5239 | 2098 |
| 4753 | 1958 |
| 4372 | 1846 |
| 4571.5 | 1915 |
| 4420.5 | 1817 |
| 4262 | 1798 |
| 4190 | 1792 |
| 4490.5 | 1854 |
| 4181 | 1817 |
| 4272.5 | 1813 |
| 4389 | 1840 |
| 4728 | 1881 |
| 4473 | 1846 |
| 4190.5 | 1743 |
| 5430 | 1982 |
| 6749 | 2196 |
| 8155.5 | 2420 |

| | |
|---|---|
| 9747 | 2598 |
| 1020 | 1048.5 |
| 3079 | 1377 |
| 2756 | 1350 |
| 9238 | 3832 |
| 4624 | 2051 |
| 4146 | 1846 |
| 3870 | 1766 |
| 3977 | 1807 |
| 3836 | 1750 |
| 3822 | 1753 |
| 3514 | 1685 |
| 3886 | 1776 |
| 3624 | 1704.5 |
| 3599 | 1697 |
| 3566 | 1688 |
| 3604 | 1697 |
| 3704 | 1709 |
| 3534 | 1652 |
| 4083 | 1832 |
| 3932 | 1784 |
| 4145 | 1779 |
| 6599 | 2161.5 |
| 7612 | 2310 |
| 8669.5 | 2525 |
| 8226 | 2437 |
| 7755 | 2390 |
| 6753.5 | 2196 |
| 7217 | 2528 |
| 5314 | 2075 |
| 4499 | 1892.5 |
| 3866 | 1760 |
| 3846 | 1768 |
| 3609 | 1673 |
| 3858 | 1762 |
| 3625 | 1701.5 |
| 3665 | 1697 |
| 3529.5 | 1688.5 |
| 3648 | 1706 |
| 3690 | 1714 |
| 3744 | 1741 |
| 3869.5 | 1745 |
| 3965 | 1749 |
| 4094 | 1813 |
| 4131 | 1808 |
| 4783 | 1870 |
| 6149 | 2111 |
| 8068 | 2421 |

| | |
|---|---|
| 9494 | 2585 |
| 8389 | 2461 |
| 7414.5 | 2328.5 |
| 6237 | 2082 |
| 7743 | 2618 |
| 5554 | 2121 |
| 4405 | 1822 |
| 4085 | 1777 |
| 3991 | 1781 |
| 3829 | 1722 |
| 3846 | 1731 |
| 3734 | 1707 |
| 3754 | 1711 |
| 3801 | 1717 |
| 3487 | 1638 |
| 3597 | 1674 |
| 3763 | 1721 |
| 3879.5 | 1729 |
| 3723 | 1723 |
| 3997 | 1767 |
| 3911 | 1742 |
| 4590 | 1841 |
| 5843.5 | 1995 |
| 7084.5 | 2195 |
| 7797 | 2325 |
| 8843 | 2470 |
| 10197.5 | 3183.5 |
| 7394 | 2392 |
| 10231 | 3154 |
| 5647 | 2144 |
| 4893 | 1976 |
| 4555 | 1914 |
| 4265 | 1855 |
| 4017 | 1813 |
| 4073 | 1805.5 |
| 4100 | 1815 |
| 4342 | 1871 |
| 4048.5 | 1812 |
| 3982 | 1793 |
| 4080 | 1836.5 |
| 4182.5 | 1864.5 |
| 4266 | 1851 |
| 4432.5 | 1892 |
| 4350 | 1864 |
| 4285 | 1863 |
| 4580 | 1906 |
| 7044 | 2288 |
| 8551.5 | 2498.5 |

| | |
|---|---|
| 10862 | 2753 |
| 9127.5 | 2552.5 |
| 8145 | 2535 |
| 6759 | 2234.5 |
| 8031.5 | 2780.5 |
| 5082 | 2015 |
| 4562 | 1887 |
| 4622 | 1961 |
| 4021.5 | 1798 |
| 4115 | 1829 |
| 4272 | 1932 |
| 4233 | 1834 |
| 3989 | 1810 |
| 4225 | 1836 |
| 3853 | 1763 |
| 4200 | 1851 |
| 4150.5 | 1852 |
| 4352 | 1890 |
| 4451 | 1890 |
| 4580 | 1965 |
| 4263 | 1874 |
| 4458.5 | 1912 |
| 7071 | 2314 |
| 8915 | 2591 |
| 10056.5 | 2741 |
| 7621 | 2283 |
| 8774.5 | 2733 |
| 6509 | 2139 |
| 8282 | 2720 |
| 5619 | 2160.5 |
| 4953 | 1988.5 |
| 4198 | 1818 |
| 4062 | 1819 |
| 4052 | 1801.5 |
| 4049 | 1786 |
| 4125 | 1820 |
| 4091.5 | 1808.5 |
| 3984.5 | 1811 |
| 3955.5 | 1796.5 |
| 3757 | 1719 |
| 3956 | 1773.5 |
| 4155 | 1794 |
| 4320 | 1845 |
| 4483 | 1910 |
| 4321 | 1851 |
| 5171 | 1960 |
| 6257 | 2126 |
| 9439 | 2564 |

| | |
|---|---|
| 9191 | 2474 |
| 10122 | 2614 |
| 8446.5 | 2664 |
| 7299 | 2305 |
| 8090 | 2755 |
| 5636 | 2193 |
| 4675.5 | 1967 |
| 4549.5 | 1937 |
| 4500 | 1947 |
| 4345 | 1907 |
| 4128 | 1823 |
| 4221 | 1829 |
| 4229.5 | 1839 |
| 4167 | 1816 |
| 4118 | 1848 |
| 4019 | 1832 |
| 4049 | 1798 |
| 4113 | 1813 |
| 4486 | 1891 |
| 4505 | 1906.5 |
| 4752.5 | 2009 |
| 4868.5 | 1954 |
| 6364 | 2154 |
| 8270.5 | 2445 |
| 10695 | 2756.5 |
| 9187 | 2475 |
| 9834 | 2897 |
| 6569 | 2211 |
| 8728 | 2874 |
| 5670.5 | 2195.5 |
| 5017 | 2010 |
| 4432 | 1869 |
| 4317 | 1903 |
| 4515 | 1913 |
| 4241 | 1880 |
| 4105 | 1838 |
| 4450 | 1906 |
| 4297 | 1884 |
| 3815 | 1789 |
| 4195 | 1852 |
| 4142 | 1874 |
| 4359.5 | 1857.5 |
| 4380 | 1892 |
| 4553 | 1954 |
| 4726 | 1992 |
| 5130.5 | 2002 |
| 6309 | 2251 |
| 8572.5 | 2533.5 |

| | |
|---|---|
| 10586.5 | 2825.5 |
| 9877 | 2527 |
| 9361 | 2727 |
| 6764 | 2218 |
| 9083 | 2924 |
| 5317 | 2014 |
| 4429 | 1835.5 |
| 4684 | 1943.5 |
| 4178 | 1816 |
| 4207 | 1829 |
| 4063 | 1832 |
| 3911 | 1759 |
| 3815 | 1719 |
| 4021 | 1831 |
| 3878 | 1790 |
| 3824 | 1724 |
| 4050 | 1837.5 |
| 4175 | 1771 |
| 4278.5 | 1843.5 |
| 4516 | 1874.5 |
| 4431.5 | 1863 |
| 4276 | 1855.5 |
| 6406 | 2187 |
| 7932.5 | 2377.5 |
| 10058 | 2627 |
| 10371 | 2559 |
| 9458 | 2793 |
| 6969 | 2216 |
| 8187 | 2707 |
| 5271 | 2060 |
| 4770 | 1942 |
| 4274.5 | 1833 |
| 3966 | 1794 |
| 3719.5 | 1687 |
| 4069.5 | 1834 |
| 4186 | 1830 |
| 3943 | 1767 |
| 3829 | 1729 |
| 3979 | 1792 |
| 3866 | 1766 |
| 3712 | 1716 |
| 4406 | 1881 |
| 4433 | 1868.5 |
| 4450 | 1872 |
| 4240 | 1820 |
| 4943 | 1960 |
| 6239 | 2146 |
| 8576.5 | 2482 |

| | |
|---|---|
| 9013 | 2568 |
| 8715 | 2447 |
| 8155 | 2462 |
| 6742 | 2145 |
| 8852 | 2799 |
| 5476 | 2039 |
| 4653 | 1848 |
| 4571 | 1882 |
| 4311 | 1837 |
| 4166 | 1809 |
| 4230 | 1839 |
| 4207 | 1805 |
| 4046.5 | 1783.5 |
| 4209.5 | 1878 |
| 4474 | 1890 |
| 4071.5 | 1777 |
| 3734 | 1670 |
| 4125 | 1810 |
| 4532 | 1849 |
| 4302 | 1807 |
| 4091 | 1746 |
| 4855.5 | 1927 |
| 5866 | 2033 |
| 7391.5 | 2275 |
| 9871 | 2629 |
| 1056 | 1044 |
| 2362 | 1299 |
| 2580 | 1333 |
| 7278 | 3232 |
| 3851 | 1797 |
| 3406 | 1624 |
| 3549.5 | 1699 |
| 3670 | 1703 |
| 3718 | 1731 |
| 3753 | 1759 |
| 3540 | 1679 |
| 3710.5 | 1739 |
| 3721 | 1721 |
| 3726.5 | 1755 |
| 3573 | 1695 |
| 3793 | 1745 |
| 3909 | 1767 |
| 3799 | 1731 |
| 4055 | 1793 |
| 4004 | 1810 |
| 4983 | 1900 |
| 6717 | 2213 |
| 8207 | 2466 |

| | |
|---|---|
| 9880 | 2663 |
| 8645.5 | 2492 |
| 8128 | 2575.5 |
| 6308.5 | 2111 |
| 7755 | 2636 |
| 5639 | 2145 |
| 4785.5 | 1925 |
| 4298.5 | 1840 |
| 3819 | 1756 |
| 4043 | 1791 |
| 4063 | 1794 |
| 3988 | 1769.5 |
| 4041 | 1819 |
| 3905 | 1761 |
| 3961 | 1797 |
| 3866.5 | 1757 |
| 3887 | 1773 |
| 4097.5 | 1800 |
| 4429.5 | 1882 |
| 4290.5 | 1820 |
| 4328 | 1880 |
| 5221 | 1965 |
| 6749 | 2255 |
| 8455 | 2460 |
| 9921.5 | 2640 |
| 8751 | 2515 |
| 7830.5 | 2461 |
| 6418 | 2134 |
| 8972 | 2880 |
| 5502 | 2086 |
| 4885 | 1959 |
| 4247 | 1831 |
| 4067 | 1819 |
| 4081 | 1811 |
| 4151 | 1815 |
| 4037 | 1807 |
| 3955 | 1760 |
| 4085.5 | 1820 |
| 4004 | 1816 |
| 3745 | 1722 |
| 3906 | 1748 |
| 4412 | 1867 |
| 4050 | 1751 |
| 4150 | 1796 |
| 4241 | 1854 |
| 4907 | 1924 |
| 6293 | 2107 |
| 7807 | 2376 |

| | |
|---|---|
| 9308 | 2532 |
| 7889 | 2343 |
| 10182 | 3183.5 |
| 6624 | 2177 |
| 8735.5 | 2837 |
| 5209.5 | 2020 |
| 4601 | 1925 |
| 4296 | 1859 |
| 4221 | 1839 |
| 4128 | 1827 |
| 4198 | 1865 |
| 3920 | 1770 |
| 3970.5 | 1785 |
| 3666 | 1675 |
| 4058 | 1822 |
| 3851 | 1751 |
| 3853 | 1747 |
| 4038 | 1779 |
| 4025.5 | 1756.5 |
| 4247 | 1826 |
| 4166 | 1831 |
| 4669.5 | 1882 |
| 6238 | 2082 |
| 8213 | 2462 |
| 9750.5 | 2651 |
| 7947 | 2398 |
| 7517 | 2345 |
| 6404.5 | 2104.5 |
| 7324 | 2548 |
| 5394.5 | 2070 |
| 4616.5 | 1901 |
| 4257 | 1872 |
| 3953 | 1769 |
| 3850 | 1739 |
| 4098.5 | 1825 |
| 3725 | 1671 |
| 3630 | 1702 |
| 3911 | 1748 |
| 3910 | 1828 |
| 4056 | 1846.5 |
| 3751 | 1752 |
| 4094 | 1800 |
| 3913 | 1771.5 |
| 4020 | 1777 |
| 4101 | 1816 |
| 5109.5 | 1939.5 |
| 6153 | 2126 |
| 8180 | 2462 |

| | |
|---|---|
| 9046 | 2555.5 |
| 8306.5 | 2433 |
| 9147.5 | 2917 |
| 6211.5 | 2104.5 |
| 8382 | 2722 |
| 5214 | 1990.5 |
| 4568 | 1877 |
| 4162 | 1807 |
| 3975 | 1741 |
| 3894 | 1765 |
| 3852.5 | 1741 |
| 3796 | 1737.5 |
| 3774 | 1738 |
| 3651 | 1686 |
| 3535 | 1646 |
| 3759 | 1725 |
| 3752 | 1707 |
| 3973 | 1752.5 |
| 3877 | 1715 |
| 4148 | 1805 |
| 4079 | 1779.5 |
| 4272 | 1800 |
| 6132 | 2058 |
| 7547.5 | 2310 |
| 8867 | 2526 |
| 8144 | 2381 |
| 8145.5 | 2493 |
| 6325 | 2099.5 |
| 7832 | 2661 |
| 5271 | 2027 |
| 4632 | 1912 |
| 4269 | 1822 |
| 4075 | 1813 |
| 4200 | 1840 |
| 4093 | 1840 |
| 3850.5 | 1744.5 |
| 3856.5 | 1753 |
| 3825 | 1732 |
| 3861 | 1739 |
| 3919 | 1780.5 |
| 3734 | 1714 |
| 4066 | 1799 |
| 4094.5 | 1783 |
| 4063 | 1787 |
| 4116 | 1794 |
| 4357 | 1867 |
| 6257 | 2125 |
| 7675 | 2324 |

| | |
|---|---|
| 9579 | 2624 |
| 7860 | 2344 |
| 8968 | 2794 |
| 6099 | 2022 |
| 9150 | 2849 |
| 5622 | 2081 |
| 4700 | 1896 |
| 4341 | 1850 |
| 4250 | 1861 |
| 4076 | 1787 |
| 4088 | 1797 |
| 3853.5 | 1749.5 |
| 3941 | 1778 |
| 3893 | 1768 |
| 3965 | 1770 |
| 3831 | 1735 |
| 4123 | 1846.5 |
| 4244 | 1818 |
| 4469.5 | 1924.5 |
| 4314 | 1856 |
| 4310 | 1826 |
| 4790 | 1922 |
| 6362 | 2175 |
| 7922 | 2436 |
| 9043 | 2573 |
| 8414 | 2436.5 |
| 7563 | 2357 |
| 6491 | 2140 |
| 7980 | 2725 |
| 5642.5 | 2115.5 |
| 4784 | 1946 |
| 4207.5 | 1844 |
| 3988 | 1761 |
| 3812 | 1740 |
| 3990 | 1785 |
| 3783 | 1732 |
| 3860 | 1759 |
| 3718 | 1707 |
| 3820 | 1760.5 |
| 3877 | 1757 |
| 3902 | 1744 |
| 3814 | 1736 |
| 4032 | 1787 |
| 4068 | 1783 |
| 4025 | 1766 |
| 4399 | 1840 |
| 6638 | 2135 |
| 7546 | 2315 |

| | |
|---|---|
| 8226 | 2438 |
| 8161 | 2382 |
| 8756.5 | 2822.5 |
| 6524.5 | 2143 |
| 8104 | 2689 |
| 5481 | 2028 |
| 4723 | 1882 |
| 4008.5 | 1755 |
| 4162 | 1799 |
| 3902 | 1743 |
| 3858 | 1740 |
| 3752 | 1719 |
| 3867 | 1752 |
| 3597 | 1668 |
| 3701.5 | 1687 |
| 3727 | 1701 |
| 3656 | 1682 |
| 3950 | 1767.5 |
| 3883 | 1718 |
| 4089 | 1773 |
| 4063 | 1790 |
| 4524 | 1825 |
| 6056 | 2060 |
| 7352 | 2251 |
| 8091 | 2377 |
| 7469 | 2254 |
| 7587 | 2363.5 |
| 7150 | 2247 |
| 9005 | 2885 |
| 6129 | 2201 |
| 5313 | 2042 |
| 4819 | 1940 |
| 4425 | 1857 |
| 4267 | 1796 |
| 4159 | 1786 |
| 3990 | 1751 |
| 3985 | 1756.5 |
| 4342 | 1855 |
| 4267.5 | 1816 |
| 4007 | 1788 |
| 3920.5 | 1735.5 |
| 4240 | 1824 |
| 4336.5 | 1843.5 |
| 4333 | 1809 |
| 4304 | 1827 |
| 4489 | 1844 |
| 5918 | 2022.5 |
| 8266 | 2468 |

| | |
|---|---|
| 8385 | 2447 |
| 1016 | 1035 |
| 2052 | 1226 |
| 2397 | 1279 |
| 8080 | 3499 |
| 3807 | 1798 |
| 3640 | 1698 |
| 3694 | 1739 |
| 3746 | 1749 |
| 3695 | 1727 |
| 3870 | 1773 |
| 3803 | 1743 |
| 3746 | 1725 |
| 3840 | 1771 |
| 3825 | 1772 |
| 3733 | 1742 |
| 3725 | 1732 |
| 3909 | 1737 |
| 4083 | 1806 |
| 3880.5 | 1755 |
| 4067 | 1797 |
| 4302 | 1791 |
| 6255.5 | 2081 |
| 7530 | 2316 |
| 8431 | 2447 |
| 7597 | 2288 |
| 7509 | 2405.5 |
| 6104 | 2040 |
| 8428 | 2811.5 |
| 5157.5 | 2038 |
| 4249 | 1819 |
| 4042 | 1806 |
| 3775 | 1710.5 |
| 3701 | 1723 |
| 3759 | 1722 |
| 3729 | 1701 |
| 3577 | 1664 |
| 3779.5 | 1744.5 |
| 3541 | 1680 |
| 3582 | 1685 |
| 3712 | 1751 |
| 4066 | 1763 |
| 4076 | 1762 |
| 4188 | 1826 |
| 4188 | 1788.5 |
| 4621 | 1889 |
| 6259 | 2116 |
| 7992 | 2392 |

| | |
|---|---|
| 8531.5 | 2478 |
| 7561 | 2265 |
| 7486 | 2355 |
| 6299 | 2095 |
| 8551.5 | 2758 |
| 5541 | 2106 |
| 4492 | 1866 |
| 4173 | 1813.5 |
| 4074 | 1782 |
| 3939 | 1771 |
| 3930.5 | 1781 |
| 3808 | 1743 |
| 3701 | 1708 |
| 3733 | 1714 |
| 3702 | 1712 |
| 3760 | 1721 |
| 3653 | 1665 |
| 3981 | 1721 |
| 3770 | 1697 |
| 4001 | 1756 |
| 3736.5 | 1676 |
| 4634 | 1844 |
| 6045 | 2013 |
| 7309 | 2268 |
| 8261 | 2423 |
| 7511 | 2254 |
| 8126.5 | 2721 |
| 6415 | 2122 |
| 8517.5 | 2857.5 |
| 5323 | 2050.5 |
| 4545 | 1897.5 |
| 4276 | 1821 |
| 3891.5 | 1749 |
| 3965 | 1745 |
| 3853 | 1767 |
| 3624 | 1658 |
| 3831 | 1740 |
| 3791 | 1749 |
| 3618 | 1679 |
| 3621.5 | 1672 |
| 3664 | 1700 |
| 3994 | 1734 |
| 4152 | 1813.5 |
| 4161 | 1806 |
| 4052 | 1764 |
| 4389.5 | 1820 |
| 5847 | 2049 |
| 7620 | 2297 |

| | |
|---|---|
| 8505 | 2459 |
| 7452 | 2239.5 |
| 7261 | 2303 |
| 5971 | 1975 |
| 8518 | 2794 |
| 5312 | 2031 |
| 4617 | 1903 |
| 3947 | 1738 |
| 3977 | 1776 |
| 3871 | 1756 |
| 3993 | 1773.5 |
| 3827 | 1766 |
| 3899 | 1769 |
| 3942 | 1778 |
| 3858 | 1753.5 |
| 3765 | 1746 |
| 4125 | 1846 |
| 4196 | 1835 |
| 4110 | 1784 |
| 4254 | 1834 |
| 4355.5 | 1840 |
| 4924 | 1939 |
| 5701 | 2002 |
| 7938.5 | 2390 |
| 8924 | 2568 |
| 7807.5 | 2350 |
| 8126.5 | 2546 |
| 6333.5 | 2116 |
| 7612.5 | 2622 |
| 5419 | 2056 |
| 4524 | 1861 |
| 4167 | 1789 |
| 3865 | 1718 |
| 3792 | 1722 |
| 3929 | 1759.5 |
| 3776 | 1734 |
| 3616 | 1702.5 |
| 3844 | 1747.5 |
| 3604 | 1675 |
| 3720 | 1713 |
| 3697 | 1703 |
| 3864 | 1713 |
| 3901 | 1739 |
| 4107 | 1764 |
| 4026 | 1759 |
| 5012 | 1921 |
| 5742.5 | 1995 |
| 7765 | 2339.5 |

| | |
|---|---|
| 8606 | 2406 |
| 8924 | 2502 |
| 7906.5 | 2436.5 |
| 6980 | 2222 |
| 8503 | 2778 |
| 5659 | 2120 |
| 4901 | 1972 |
| 4168 | 1795 |
| 4392 | 1868 |
| 4133 | 1805 |
| 4079.5 | 1816.5 |
| 4220.5 | 1868 |
| 3968 | 1796 |
| 4073 | 1813 |
| 4026.5 | 1805 |
| 3962.5 | 1775.5 |
| 3969 | 1830 |
| 4219.5 | 1839.5 |
| 4200 | 1820 |
| 4160 | 1812 |
| 4145 | 1827 |
| 4800 | 1948 |
| 5819 | 2049 |
| 7660 | 2318 |
| 8936 | 2507 |
| 7719 | 2289 |
| 8219 | 2596 |
| 6599 | 2137 |
| 7877 | 2668.5 |
| 5826.5 | 2190 |
| 4566.5 | 1881 |
| 4308.5 | 1834 |
| 4100.5 | 1819 |
| 3982.5 | 1797 |
| 4030.5 | 1829 |
| 3803 | 1720 |
| 4005 | 1802 |
| 3981.5 | 1803.5 |
| 3940.5 | 1794 |
| 3807 | 1728 |
| 3881 | 1763 |
| 4083 | 1778 |
| 4406 | 1876 |
| 4227 | 1822 |
| 4242 | 1841 |
| 4522 | 1871 |
| 6356 | 2131 |
| 7937 | 2395 |

| | |
|---|---|
| 8909 | 2616 |
| 8039 | 2366 |
| 7524 | 2384.5 |
| 6688 | 2173 |
| 8876 | 2852 |
| 5304 | 2075.5 |
| 4364 | 1807.5 |
| 3921 | 1736 |
| 3924 | 1770 |
| 3669 | 1687 |
| 4018 | 1783 |
| 3656 | 1668 |
| 3720 | 1697 |
| 3843 | 1727 |
| 3718 | 1708 |
| 3704 | 1688 |
| 3554 | 1652 |
| 4008 | 1756 |
| 3875 | 1702 |
| 3922 | 1739 |
| 4101.5 | 1798 |
| 4782 | 1881 |
| 6051.5 | 2040 |
| 7547 | 2339 |
| 8754 | 2453 |
| 8115 | 2385 |
| 7917 | 2535.5 |
| 6537 | 2174 |
| 7750 | 2643 |
| 5745.5 | 2164 |
| 4651 | 1914 |
| 4127 | 1778 |
| 4063.5 | 1814 |
| 4052 | 1803 |
| 4091.5 | 1811 |
| 3773 | 1729 |
| 3936.5 | 1756 |
| 3681 | 1663.5 |
| 3766 | 1703 |
| 3853 | 1739 |
| 3830 | 1727 |
| 4162 | 1797 |
| 4057.5 | 1783 |
| 3956 | 1766 |
| 4084 | 1781 |
| 4518 | 1838 |
| 5774 | 2015.5 |
| 7497 | 2315 |

| | |
|---|---|
| 7679 | 2284 |
| 7617 | 2267 |
| 8187 | 2486.5 |
| 6918 | 2159 |
| 8875 | 2874 |
| 5894 | 2184 |
| 4744 | 1892 |
| 4516 | 1842 |
| 4261 | 1813 |
| 4140 | 1756 |
| 4016.5 | 1750 |
| 3794.5 | 1695 |
| 3931 | 1742 |
| 4277 | 1828 |
| 4278.5 | 1817 |
| 3937 | 1752 |
| 4065 | 1790 |
| 4457 | 1868 |
| 4220 | 1807 |
| 4138.5 | 1784 |
| 4276 | 1821 |
| 4830 | 1846 |
| 5738.5 | 1988 |
| 7741 | 2348 |
| 8828 | 2467 |
| 1019 | 1080 |
| 4656.5 | 1796 |
| 3422.5 | 1503 |
| 16914 | 6030 |
| 6801.5 | 2593.5 |
| 4566 | 1872 |
| 4233.5 | 1743 |
| 3700 | 1640 |
| 3951 | 1694 |
| 3768 | 1728 |
| 3613 | 1701 |
| 3755 | 1674 |
| 3551.5 | 1682 |
| 4449 | 1839 |
| 3689.5 | 1781.5 |
| 3880.5 | 1725 |
| 4145 | 1777 |
| 4405 | 1715 |
| 4318.5 | 1853 |
| 3829 | 1817 |
| 4521 | 1865 |
| 6702 | 2256 |
| 10451.5 | 2799.5 |

| | |
|---|---|
| 10809.5 | 2591.5 |
| 11535 | 2806 |
| 75034 | 35622 |
| 8420 | 2513 |
| 8000.5 | 2587 |
| 6307 | 2142 |
| 4684 | 1955 |
| 4146 | 1781 |
| 4182 | 1749 |
| 3979 | 1806 |
| 4270 | 1898 |
| 4191 | 1692 |
| 3890 | 1822 |
| 4170 | 1910 |
| 5053 | 1919 |
| 4159 | 1863 |
| 4148 | 1716 |
| 4347 | 1735 |
| 4121 | 1807.5 |
| 4818 | 1873 |
| 4241.5 | 1912.5 |
| 4417.5 | 1933 |
| 5919 | 2119 |
| 10510 | 2711 |
| 10847 | 2761.5 |
| 11959 | 2998 |
| 12320.5 | 3258.5 |
| 8072 | 2545 |
| 8032.5 | 2582 |
| 6376 | 2039.5 |
| 4913 | 1801 |
| 4117 | 1729 |
| 4333 | 1912 |
| 4155 | 1842 |
| 4335.5 | 1815 |
| 4007 | 1666 |
| 3917 | 1783.5 |
| 4147 | 1759 |
| 4171.5 | 1601 |
| 4183 | 1816 |
| 3909 | 1775 |
| 3962.5 | 1824 |
| 4524 | 1884 |
| 3975 | 1845 |
| 4128 | 1820 |
| 4691 | 1798.5 |
| 5349.5 | 1981 |
| 8719 | 2420 |

| | |
|---|---|
| 11960.5 | 2903 |
| 10612 | 2742 |
| 71136 | 31933 |
| 8747 | 2700 |
| 8155 | 2500 |
| 6616 | 2137.5 |
| 5786.5 | 1947.5 |
| 4478 | 1874.5 |
| 4316.5 | 1782 |
| 4610.5 | 1982.5 |
| 4158 | 1763 |
| 4327 | 1854 |
| 3862.5 | 1692 |
| 4296 | 1868 |
| 4519.5 | 1978 |
| 3821 | 1732 |
| 4144.5 | 1829 |
| 4367 | 1921 |
| 4288.5 | 1727.5 |
| 4638 | 1779 |
| 4315 | 2063.5 |
| 4656 | 1890 |
| 6186.5 | 2094.5 |
| 8929 | 2429 |
| 10327 | 2514.5 |
| 12508 | 3090 |
| 17457 | 4803 |
| 9311 | 2770 |
| 8066 | 2486.5 |
| 6657.5 | 2211.5 |
| 5179.5 | 1988 |
| 4226 | 1733 |
| 4159 | 1924.5 |
| 3955.5 | 1749 |
| 4129.5 | 1867 |
| 4423 | 1927 |
| 4009 | 1823 |
| 4121 | 1830 |
| 4367 | 1631.5 |
| 4253.5 | 1710.5 |
| 4464 | 1892 |
| 4202.5 | 1759 |
| 3743.5 | 1850.5 |
| 4347 | 1890 |
| 4673 | 1983 |
| 4669 | 1815 |
| 6172 | 2171 |
| 8566 | 2512.5 |

| | |
|---|---|
| 11885 | 2760 |
| 10368 | 2722 |
| 83800 | 39452 |
| 10404.5 | 3421 |
| 9029 | 3165 |
| 7969 | 2597 |
| 5200 | 1959 |
| 4490 | 1828 |
| 4611 | 1795 |
| 4815 | 1869 |
| 4323 | 2016 |
| 4465.5 | 1819.5 |
| 4273 | 1858 |
| 4219 | 1815 |
| 4381 | 1901 |
| 4761 | 1874 |
| 4616.5 | 2024 |
| 5161 | 1818.5 |
| 5240 | 2075.5 |
| 4775 | 1945.5 |
| 4598.5 | 1833.5 |
| 4526 | 1807 |
| 6711 | 2266 |
| 7270.5 | 2095.5 |
| 8503 | 2460.5 |
| 12994 | 2993 |
| 17687.5 | 4466.5 |
| 8973 | 2560 |
| 8688 | 2480 |
| 5974 | 2149 |
| 5072 | 1886 |
| 4462.5 | 1823 |
| 4617 | 2045 |
| 4158 | 1855.5 |
| 3719.5 | 1838 |
| 3830 | 1758 |
| 4158.5 | 1857.5 |
| 4025 | 1729 |
| 4744 | 1902 |
| 4345 | 1848 |
| 3584.5 | 1697 |
| 3971 | 1820 |
| 4328 | 1760.5 |
| 4641 | 1742 |
| 4272.5 | 1699.5 |
| 4845.5 | 1898 |
| 6250.5 | 2218.5 |
| 8519 | 2472.5 |

| | |
|---|---|
| 11150 | 2939 |
| 11920 | 2846.5 |
| 86495 | 37303 |
| 10024 | 3045 |
| 8208.5 | 2434 |
| 7103 | 2312 |
| 5718 | 2415 |
| 6402.5 | 2539 |
| 5440.5 | 2244 |
| 5283 | 2166 |
| 5087 | 2284 |
| 5016.5 | 2084.5 |
| 5401 | 2208 |
| 5481.5 | 2213.5 |
| 5238.5 | 2285.5 |
| 5421 | 2233 |
| 4757.5 | 2162.5 |
| 5452 | 2338 |
| 5539 | 2407 |
| 5768.5 | 2402 |
| 5323 | 2235 |
| 5424.5 | 2347 |
| 6843 | 2485.5 |
| 8883 | 2730 |
| 10180 | 2947.5 |
| 13533 | 3404 |
| 48725 | 14967.5 |
| 19397.5 | 5780.5 |
| 9688 | 3477 |
| 9695 | 3440 |
| 6909.5 | 2501 |
| 5088 | 2167 |
| 5238 | 2005 |
| 5016.5 | 1916 |
| 4824 | 2203 |
| 4343.5 | 1829.5 |
| 4165.5 | 1819.5 |
| 4428 | 1857 |
| 4767 | 1906 |
| 5114 | 1897.5 |
| 4647 | 1769 |
| 4967.5 | 1874.5 |
| 5207 | 1951 |
| 5031.5 | 2073 |
| 4401 | 1963 |
| 5011 | 1953.5 |
| 6716 | 2075 |
| 11094 | 2909 |

| | |
|---|---|
| 9939 | 2651 |
| 11055 | 2841 |
| 134745.5 | 54972 |
| 78290 | 32447 |
| 14012.5 | 5616.5 |
| 8301 | 3080 |
| 5806 | 2269 |
| 5754 | 2198 |
| 5023 | 1879 |
| 4854.5 | 1977 |
| 4723.5 | 1991 |
| 4695 | 2153.5 |
| 4583 | 2008 |
| 3848 | 1857 |
| 4592 | 1955 |
| 4343 | 1964 |
| 4280 | 1801 |
| 4057 | 1935.5 |
| 4199.5 | 1785.5 |
| 4455.5 | 1912 |
| 4027.5 | 1668.5 |
| 4896.5 | 1814 |
| 6645 | 2201 |
| 6566.5 | 2043 |
| 10006 | 2640 |
| 11349 | 2750 |
| 135951 | 56455 |
| 16137 | 4935 |
| 10188.5 | 3416.5 |
| 6352 | 2503 |
| 4989 | 2030 |
| 4598 | 1962 |
| 4640 | 1801.5 |
| 4339 | 1917 |
| 4010.5 | 1822 |
| 4229 | 1765 |
| 4166 | 1841 |
| 4006.5 | 1694 |
| 4405 | 1732 |
| 3858.5 | 1866.5 |
| 4080.5 | 1852.5 |
| 3713.5 | 1800.5 |
| 4295.5 | 1702.5 |
| 4523 | 1832 |
| 4326 | 1803 |
| 4020 | 1740.5 |
| 6150 | 2043 |
| 7489 | 2336 |

| | |
|---|---|
| 10370.5 | 2650.5 |
| 1974 | 1247 |
| 5665 | 2441 |
| 3922 | 1767.5 |
| 9909.5 | 3827 |
| 5121 | 2036 |
| 4511 | 1902 |
| 4413 | 1830.5 |
| 4538 | 1884 |
| 4749.5 | 1889 |
| 4582 | 2111.5 |
| 4747 | 1965 |
| 4600 | 1967 |
| 4491 | 1919 |
| 5027 | 1945 |
| 4400.5 | 1803.5 |
| 4750 | 1841 |
| 5082.5 | 1968 |
| 4642 | 1951 |
| 4671 | 1964.5 |
| 4519 | 1990 |
| 11178 | 2907.5 |
| 14080.5 | 3465.5 |
| 14262 | 3526 |
| 11554 | 3037 |
| 9777 | 2591.5 |
| 73504 | 35717 |
| 6127 | 2245 |
| 8478.5 | 2932.5 |
| 5652 | 2124 |
| 5103 | 2004 |
| 4875 | 1977 |
| 4618 | 1886 |
| 4870.5 | 1960.5 |
| 5011.5 | 2117 |
| 4842.5 | 2081.5 |
| 4592.5 | 1932.5 |
| 4588 | 1971 |
| 5493 | 2017 |
| 4417 | 1925 |
| 4976 | 2003 |
| 5253.5 | 2125 |
| 5264 | 2047 |
| 4946 | 2020 |
| 4660 | 1940.5 |
| 12219.5 | 3116 |
| 14599 | 3553 |
| 14002 | 3485 |

| | |
|---|---|
| 11952.5 | 3140.5 |
| 9919 | 2524.5 |
| 69671 | 32994 |
| 6072 | 2191 |
| 8613 | 2882 |
| 5660.5 | 2210.5 |
| 4957.5 | 2009.5 |
| 4660.5 | 1969 |
| 4956.5 | 2014.5 |
| 4663 | 1937 |
| 4805 | 1992 |
| 4630 | 1955 |
| 4566 | 1956 |
| 4708 | 1961 |
| 5217 | 1978 |
| 4622 | 2058 |
| 4895.5 | 1940 |
| 5147 | 2025.5 |
| 4976.5 | 2003 |
| 4935 | 1963 |
| 4325 | 1904 |
| 12133 | 3111 |
| 14929 | 3665 |
| 14594 | 3661 |
| 12174 | 2989.5 |
| 10063 | 2656 |
| 96167.5 | 45444.5 |
| 6497.5 | 2277.5 |
| 8799 | 2915 |
| 5764.5 | 2081.5 |
| 5012 | 2006 |
| 5028 | 2027 |
| 4800 | 1968 |
| 5070 | 1982 |
| 5081 | 1991.5 |
| 4655 | 2028 |
| 4518.5 | 1979.5 |
| 4643 | 1915 |
| 4842 | 2049 |
| 4549 | 1930 |
| 4767 | 2028 |
| 5272 | 2100 |
| 5322.5 | 2160.5 |
| 5091 | 2100 |
| 4695 | 1980 |
| 11769.5 | 2938 |
| 14451 | 3687 |
| 14735 | 3431.5 |

| | |
|---|---|
| 11968 | 3074 |
| 9532 | 2646 |
| 87177 | 40897 |
| 6116 | 2118 |
| 8961 | 2851 |
| 5917 | 2177 |
| 5186 | 2005 |
| 4721 | 1933 |
| 4681 | 1945 |
| 5089 | 2003 |
| 5228.5 | 2098.5 |
| 4778 | 1921.5 |
| 4682 | 1925 |
| 4638 | 1932 |
| 5355 | 2052 |
| 4703.5 | 1889 |
| 4937 | 1969 |
| 5186 | 2028 |
| 5042 | 2124 |
| 4948 | 2073 |
| 4862 | 1915.5 |
| 12024.5 | 3038 |
| 14712.5 | 3693 |
| 13656 | 3559 |
| 12122 | 3098 |
| 9821.5 | 2650 |
| 101333 | 47162.5 |
| 6932.5 | 2577 |
| 8584 | 3138.5 |
| 7508 | 2553 |
| 5507 | 2196 |
| 4995.5 | 1995 |
| 5043 | 2006 |
| 5287 | 2025.5 |
| 5171.5 | 2057 |
| 4938 | 2109 |
| 4738 | 2021.5 |
| 5022 | 2065 |
| 5123 | 2218 |
| 4836 | 1981 |
| 5072.5 | 2116 |
| 6196.5 | 2171.5 |
| 5493 | 2103 |
| 5570 | 2194.5 |
| 5181 | 1968.5 |
| 12449 | 3136 |
| 14485.5 | 3850 |
| 14708 | 3625 |

| | |
|---|---|
| 12575 | 3195.5 |
| 9925 | 2613 |
| 91740 | 42027.5 |
| 5997 | 2338 |
| 8937 | 2978 |
| 5879 | 2286 |
| 5411.5 | 2093 |
| 4892 | 2008 |
| 4926 | 1977 |
| 4982 | 2028 |
| 5032 | 2143 |
| 4840 | 1979 |
| 4864 | 1994 |
| 4724 | 1925 |
| 5283 | 2075 |
| 4661.5 | 1899 |
| 4924 | 2029 |
| 5139 | 2024 |
| 4855 | 1954 |
| 5142 | 2017 |
| 4838 | 2000 |
| 11848.5 | 2959 |
| 14647.5 | 3611 |
| 14721 | 3523 |
| 12025 | 2945 |
| 8606 | 2493 |
| 94024 | 44007 |
| 6492 | 2518 |
| 8976 | 3202.5 |
| 8355 | 2968 |
| 6620 | 2738 |
| 6441 | 2562 |
| 6624 | 2686 |
| 6922.5 | 2775.5 |
| 6646 | 2687 |
| 6332 | 2717 |
| 6603.5 | 2696.5 |
| 6664 | 2814 |
| 7021.5 | 2834.5 |
| 6528 | 2772 |
| 6641 | 2709 |
| 6973.5 | 2855 |
| 7264 | 2944 |
| 6809 | 2669 |
| 6535 | 2738 |
| 12187 | 3594 |
| 15547 | 3992.5 |
| 13756 | 3886 |

| | |
|---|---|
| 13088 | 3659 |
| 11080.5 | 3291 |
| 56627.5 | 17873 |
| 27749.5 | 9157.5 |
| 10262 | 3848 |
| 33701 | 6980.5 |
| 6435 | 2396 |
| 6154 | 2339 |
| 5614 | 2200 |
| 5517 | 2150 |
| 8856 | 3445 |
| 5244 | 2143 |
| 5074 | 2133.5 |
| 5126 | 1989 |
| 6370.5 | 2267.5 |
| 4925 | 2055 |
| 5259.5 | 2055 |
| 5885 | 2295 |
| 5661 | 2097 |
| 6274.5 | 2304.5 |
| 5386 | 2102 |
| 12166 | 3114 |
| 14670 | 3627 |
| 14246.5 | 3612 |
| 12298 | 3144.5 |
| 9394 | 2631.5 |
| 122336 | 52668 |
| 8009 | 3366 |
| 10663 | 3637 |
| 6188 | 2296 |
| 5727 | 2185 |
| 5130 | 2076 |
| 5301 | 2062 |
| 5268.5 | 2037.5 |
| 5509 | 2268 |
| 5057.5 | 2078 |
| 4997 | 2062.5 |
| 4993 | 2050.5 |
| 5523.5 | 2060.5 |
| 4841 | 2038 |
| 5238 | 2130.5 |
| 5747 | 2157 |
| 5758 | 2151.5 |
| 5442.5 | 2068.5 |
| 4961.5 | 2054.5 |
| 12043.5 | 3265 |
| 14954.5 | 3821.5 |
| 14273 | 3485 |

| | |
|---|---|
| 12080 | 3185.5 |
| 10312 | 2823 |
| 127726.5 | 54033 |
| 7882 | 3077 |
| 9505 | 3680 |
| 6325.5 | 2657 |
| 6255 | 2448 |
| 5700 | 2240.5 |
| 5677 | 2262 |
| 5345 | 2181 |
| 5678 | 2342 |
| 5443 | 2178 |
| 4931 | 2044 |
| 4624.5 | 1935 |
| 5271 | 2078 |
| 4984.5 | 2092 |
| 5165 | 2033 |
| 5412 | 2086 |
| 4988 | 2098 |
| 4864 | 2023 |
| 4864 | 1985 |
| 12109 | 3099 |
| 13629 | 3444.5 |
| 13072 | 3157 |
| 11076 | 2966.5 |
| 1988 | 1200 |
| 4215.5 | 1805.5 |
| 3779 | 1679.5 |
| 7429 | 2715 |
| 4656.5 | 1910.5 |
| 4499 | 1800 |
| 4176 | 1852 |
| 4601 | 1928 |
| 4361.5 | 1786 |
| 4418 | 1991 |
| 4561.5 | 1931.5 |
| 4376 | 1841 |
| 4203 | 1872 |
| 4816 | 1928 |
| 4349.5 | 1830 |
| 4702 | 1930 |
| 4868 | 1982 |
| 4565 | 1957 |
| 4551 | 1874 |
| 4410 | 1819 |
| 11173 | 2812 |
| 13964.5 | 3489.5 |
| 13256 | 3349 |

| | |
|---|---|
| 11408 | 2766 |
| 9204 | 2540 |
| 71691 | 35060 |
| 5671 | 2114 |
| 8952 | 2933 |
| 5659 | 2109 |
| 5037 | 2091 |
| 4787 | 1954.5 |
| 4626.5 | 1982 |
| 4899 | 1996 |
| 4833.5 | 2004 |
| 4796 | 1902 |
| 4797.5 | 1840.5 |
| 4490 | 1892 |
| 5065 | 1953.5 |
| 4440 | 1945 |
| 4796 | 1881 |
| 5313.5 | 2036 |
| 5025 | 2019 |
| 4645 | 1917 |
| 4473 | 1940.5 |
| 10847 | 2764 |
| 14064.5 | 3417.5 |
| 13572 | 3374 |
| 11931 | 2948 |
| 9917.5 | 2588 |
| 9606 | 3152 |
| 5711 | 2043 |
| 8613 | 2785 |
| 5822 | 2125 |
| 5054 | 2033.5 |
| 4662 | 1933.5 |
| 4812 | 1926 |
| 4516.5 | 1873.5 |
| 4394 | 1932 |
| 4417 | 1873 |
| 4433.5 | 1881.5 |
| 4473.5 | 1851.5 |
| 4699.5 | 1811.5 |
| 4384.5 | 1846.5 |
| 4677 | 1939 |
| 4845.5 | 1931 |
| 4849 | 1986.5 |
| 4637 | 1961 |
| 4396.5 | 1898.5 |
| 11070 | 2965.5 |
| 14199.5 | 3584.5 |
| 13664 | 3434 |

| | |
|---|---|
| 11441 | 2875 |
| 9410 | 2696 |
| 86548.5 | 40862 |
| 6333 | 2315 |
| 8683 | 2770 |
| 5885 | 2163 |
| 4964.5 | 1971 |
| 4767 | 1968 |
| 4688.5 | 1985.5 |
| 4591 | 1830 |
| 4720.5 | 1981.5 |
| 4422 | 1919 |
| 4465.5 | 1872 |
| 4402.5 | 1836.5 |
| 4886.5 | 1966.5 |
| 4139 | 1884.5 |
| 4498 | 1898 |
| 5332 | 2027 |
| 5034 | 2009 |
| 4880 | 1947 |
| 4457 | 1836 |
| 10508 | 2853 |
| 13864 | 3394 |
| 13454.5 | 3225.5 |
| 12242 | 2967 |
| 9465 | 2588.5 |
| 10545 | 3685 |
| 5671 | 2098 |
| 8573 | 2897.5 |
| 5940 | 2201 |
| 5147.5 | 2066.5 |
| 4781 | 2000 |
| 5057.5 | 1973 |
| 4707 | 1900 |
| 4441 | 2024 |
| 4436 | 1947.5 |
| 4436 | 1877 |
| 4360 | 1971.5 |
| 5075 | 1972 |
| 4421.5 | 1965 |
| 4833 | 1994 |
| 5006 | 1934 |
| 4982 | 1856 |
| 4839 | 1971 |
| 4495.5 | 1848 |
| 11647.5 | 3039.5 |
| 14359 | 3505 |
| 13632.5 | 3344 |

| | |
|---|---|
| 11724 | 2953 |
| 9908.5 | 2582.5 |
| 90196 | 42839 |
| 6594.5 | 2524.5 |
| 8556 | 3109 |
| 6961 | 2401.5 |
| 6086.5 | 2297.5 |
| 5444 | 2076 |
| 5326.5 | 2124 |
| 5375 | 2011 |
| 5212 | 2112 |
| 4705 | 1962 |
| 4830 | 1984 |
| 4693 | 1973 |
| 4938.5 | 2080 |
| 4903.5 | 2045.5 |
| 5149 | 1994 |
| 5854 | 2124 |
| 5513 | 2078 |
| 5248.5 | 2022 |
| 5090.5 | 1967.5 |
| 10632.5 | 2927 |
| 14341 | 3745 |
| 14408 | 3548 |
| 12496.5 | 3191 |
| 9426 | 2607 |
| 21676 | 7874 |
| 5839 | 2093.5 |
| 8768 | 2925 |
| 6072 | 2229.5 |
| 5080.5 | 2026 |
| 4771 | 1949 |
| 4808 | 1962 |
| 4631 | 1881 |
| 4528 | 1931 |
| 4338.5 | 1938 |
| 4298.5 | 1877.5 |
| 4208.5 | 1902.5 |
| 4908 | 1878.5 |
| 4380 | 1870 |
| 4375 | 1879 |
| 5054.5 | 2030 |
| 4850 | 2027.5 |
| 4679 | 1926 |
| 4493 | 1886 |
| 10547 | 2792 |
| 14387 | 3592 |
| 14163 | 3475 |

| | |
|---|---|
| 12059 | 2989 |
| 9001 | 2573 |
| 89456 | 41924 |
| 7329 | 2707 |
| 9144 | 3177 |
| 8164.5 | 2962 |
| 6386 | 2610 |
| 6429 | 2605 |
| 6242 | 2460 |
| 6030 | 2411.5 |
| 6056 | 2429 |
| 6121 | 2509 |
| 5593 | 2432 |
| 5724 | 2368 |
| 5502 | 2299.5 |
| 5135 | 2046.5 |
| 5328 | 2259 |
| 5617 | 2245 |
| 5862 | 2196 |
| 5432 | 2170 |
| 5047 | 2211 |
| 10963 | 2973 |
| 13789 | 3441 |
| 14357.5 | 3638 |
| 11777.5 | 3155 |
| 9263 | 2805 |
| 115334 | 36539 |
| 31679 | 11661.5 |
| 9064.5 | 3667 |
| 19491 | 4658 |
| 5990 | 2272 |
| 5788 | 2136 |
| 5357 | 2096 |
| 5247 | 2074.5 |
| 6831 | 2683.5 |
| 5044 | 2050 |
| 5060 | 2019.5 |
| 4805 | 2005 |
| 6172.5 | 2113 |
| 4999 | 1955 |
| 4874 | 1889 |
| 6385 | 2199 |
| 5397.5 | 2095.5 |
| 6403 | 2325 |
| 5149 | 1975 |
| 10754.5 | 2833 |
| 14565.5 | 3644 |
| 14435.5 | 3592.5 |

| | |
|---|---|
| 12006 | 2992 |
| 8160 | 2505 |
| 116281 | 49837 |
| 8095 | 3473 |
| 10183 | 3733 |
| 6284.5 | 2554.5 |
| 5112 | 2084 |
| 5069 | 2060 |
| 4916 | 2006 |
| 4827 | 2034 |
| 5261.5 | 2252 |
| 4792.5 | 1954.5 |
| 4796 | 1970 |
| 4543 | 1919.5 |
| 5069 | 1943 |
| 4601 | 1836 |
| 4835 | 2053 |
| 5332 | 2100 |
| 5142 | 2125 |
| 5032.5 | 1970.5 |
| 4848 | 1940 |
| 11156 | 2783.5 |
| 14026 | 3385 |
| 12802 | 3267 |
| 11567 | 2974.5 |
| 9115 | 2635 |
| 81318 | 35123 |
| 6841.5 | 2777.5 |
| 8659.5 | 3186 |
| 6550.5 | 2518.5 |
| 5347 | 2180 |
| 4891 | 1986 |
| 4762.5 | 1959.5 |
| 4554.5 | 1858.5 |
| 4912.5 | 2046 |
| 4932 | 2049.5 |
| 4953 | 2040.5 |
| 4822 | 1972 |
| 4959.5 | 1909 |
| 4521 | 1953.5 |
| 4862.5 | 1922.5 |
| 5590.5 | 2052 |
| 4897 | 2021.5 |
| 4785.5 | 1978 |
| 4491 | 1927 |
| 11404 | 2921 |
| 14709 | 3609.5 |
| 13934 | 3481 |

| | |
|---|---|
| 11833.5 | 2930 |
| 1906 | 1194 |
| 4082.5 | 1789 |
| 3578 | 1668 |
| 6976 | 2624 |
| 4394 | 1839 |
| 4347.5 | 1880.5 |
| 4300.5 | 1833.5 |
| 4355 | 1855 |
| 4154.5 | 1707.5 |
| 4310 | 1953 |
| 4154 | 1780 |
| 3948 | 1796 |
| 4008 | 1798 |
| 4552 | 1847 |
| 4163 | 1826 |
| 4379.5 | 1805 |
| 4731 | 1928 |
| 4367 | 1808 |
| 4476 | 1836 |
| 4215 | 1818 |
| 11028 | 2756 |
| 13471 | 3265 |
| 12987 | 3126 |
| 11132 | 2762 |
| 8245 | 2429 |
| 64547 | 31212.5 |
| 5468 | 1998 |
| 8681.5 | 2844.5 |
| 5253 | 2062 |
| 4572 | 1885.5 |
| 4398 | 1828.5 |
| 4243 | 1857 |
| 4369 | 1791 |
| 4407 | 1906.5 |
| 4239 | 1787 |
| 4108 | 1831 |
| 4117 | 1825 |
| 4639 | 1837 |
| 4043 | 1729 |
| 4616 | 1906 |
| 4793 | 1906 |
| 4623 | 1842 |
| 4421 | 1898.5 |
| 4337 | 1818 |
| 10696 | 2791 |
| 13074.5 | 3313.5 |
| 13169 | 3248 |

| | |
|---|---|
| 11337 | 2923 |
| 8288 | 2425 |
| 54155.5 | 25919 |
| 5221 | 2000.5 |
| 8488 | 2738.5 |
| 5309 | 1982 |
| 4583 | 1972.5 |
| 4555 | 1899.5 |
| 4457 | 1824 |
| 4309 | 1821 |
| 4308 | 1901 |
| 4303 | 1804 |
| 4207 | 1837 |
| 3985 | 1770 |
| 4517 | 1792 |
| 4040 | 1758 |
| 4281 | 1848 |
| 4591.5 | 1901 |
| 4542.5 | 1840 |
| 4321.5 | 1868 |
| 4220 | 1832 |
| 10929 | 2804 |
| 12746 | 3226 |
| 12542.5 | 3243.5 |
| 11316 | 2774 |
| 7612 | 2387 |
| 82427 | 38752 |
| 5312 | 2102 |
| 8329 | 2727 |
| 5504.5 | 2093.5 |
| 4761 | 1902 |
| 4604 | 1891 |
| 4283 | 1770 |
| 4364.5 | 1827.5 |
| 4632 | 1944 |
| 4291 | 1817.5 |
| 4133 | 1744.5 |
| 4232.5 | 1826.5 |
| 4686 | 1847 |
| 4197 | 1827.5 |
| 4328 | 1850 |
| 4814 | 1914 |
| 4613 | 1853.5 |
| 4484 | 1868 |
| 4146.5 | 1790.5 |
| 10589 | 2785 |
| 13586 | 3400 |
| 13349.5 | 3261 |

| | |
|---|---|
| 11331 | 2840 |
| 8387 | 2468 |
| 61654 | 28223 |
| 5477 | 2101 |
| 8222 | 2718 |
| 5625 | 2160 |
| 5005.5 | 1981 |
| 4686 | 1884.5 |
| 4680 | 1873.5 |
| 4548 | 1840 |
| 4382 | 1864.5 |
| 4443 | 1838 |
| 4297 | 1790 |
| 4140 | 1852 |
| 5009.5 | 1844 |
| 4343 | 1838 |
| 4438 | 1849 |
| 4788 | 1932 |
| 4506 | 1842 |
| 4614 | 1862 |
| 4256 | 1765 |
| 10720 | 2780 |
| 13030 | 3351.5 |
| 12864.5 | 3240.5 |
| 11141 | 2806 |
| 8879.5 | 2490 |
| 89727.5 | 41589.5 |
| 5799.5 | 2203.5 |
| 8163 | 2874 |
| 6813 | 2350 |
| 5098 | 2054 |
| 4884 | 1923.5 |
| 4690 | 1974 |
| 4598 | 1811.5 |
| 4933 | 2124 |
| 4449.5 | 1835.5 |
| 4346 | 1919 |
| 4429 | 1853 |
| 4810 | 1904 |
| 4404 | 1870 |
| 4593 | 1793 |
| 5645 | 2042 |
| 4838.5 | 1916.5 |
| 4793 | 1949 |
| 4770.5 | 1906.5 |
| 10990 | 2811 |
| 14439 | 3404 |
| 13420.5 | 3167.5 |

| | |
|---|---|
| 11674 | 2867 |
| 8716 | 2636 |
| 64119.5 | 29602 |
| 5585 | 2117.5 |
| 8566 | 2856 |
| 5537 | 2118 |
| 4728.5 | 1940.5 |
| 4526 | 1870 |
| 4562 | 1868 |
| 4469 | 1775 |
| 4429 | 1874 |
| 4412 | 1840 |
| 4213 | 1778 |
| 4201 | 1821.5 |
| 4944 | 1919 |
| 4269 | 1766 |
| 4514 | 1893 |
| 4861 | 1833 |
| 4551.5 | 1916 |
| 4697 | 1860 |
| 4310.5 | 1821.5 |
| 10617 | 2791 |
| 13578 | 3247 |
| 13266 | 3320 |
| 10657 | 2833 |
| 8137.5 | 2469 |
| 71548.5 | 33279.5 |
| 6209.5 | 2307.5 |
| 8901 | 3007 |
| 5961.5 | 2346.5 |
| 5253 | 2147 |
| 5172 | 2102 |
| 5187 | 1997 |
| 4896 | 2018 |
| 4649 | 1989 |
| 4803 | 1963 |
| 4818 | 1993.5 |
| 4792 | 1964 |
| 5084 | 2097.5 |
| 4737 | 2031 |
| 5048 | 1972.5 |
| 5582 | 2166 |
| 5108 | 2248 |
| 5001 | 2018 |
| 4765 | 1954 |
| 10849 | 2970 |
| 15442 | 3696 |
| 13265 | 3526.5 |

| | |
|---|---|
| 11998.5 | 3067.5 |
| 8539 | 2706 |
| 95175 | 33789 |
| 10105 | 4092 |
| 8024 | 3266.5 |
| 23943 | 4878 |
| 5735 | 2247 |
| 6157 | 2111 |
| 5097 | 1965 |
| 5296 | 2032 |
| 6545.5 | 2643.5 |
| 5126 | 2120 |
| 4776.5 | 2017.5 |
| 4718 | 2082 |
| 7405.5 | 2371 |
| 5379 | 2137 |
| 5104 | 2027 |
| 6036.5 | 2194.5 |
| 5301 | 2074 |
| 6788.5 | 2549.5 |
| 4883 | 2021 |
| 11976 | 3055 |
| 14458 | 3538.5 |
| 14214 | 3500 |
| 11463 | 2883 |
| 8107 | 2480 |
| 112701.5 | 48511.5 |
| 7070 | 3131 |
| 10259 | 3628.5 |
| 6015.5 | 2316.5 |
| 5110.5 | 2148 |
| 4619 | 1933 |
| 4661 | 2031 |
| 4782 | 1911 |
| 5329.5 | 2183 |
| 5076 | 2069 |
| 4513.5 | 1874 |
| 4576.5 | 1966.5 |
| 5381.5 | 2050.5 |
| 4852 | 2001 |
| 4954 | 1930.5 |
| 5218 | 2056 |
| 5309 | 2141 |
| 4878 | 1990 |
| 4695.5 | 1980 |
| 12510 | 3117.5 |
| 14620 | 3572 |
| 13212.5 | 3303 |

| | |
|---|---|
| 11530 | 2994 |
| 7828.5 | 2401.5 |
| 94252 | 42966 |
| 6077 | 2331 |
| 9009 | 3193 |
| 5869 | 2219 |
| 4814 | 2006 |
| 4963 | 2043 |
| 4725.5 | 1936 |
| 4637 | 1890 |
| 5358 | 2192 |
| 4789 | 1972.5 |
| 4846 | 2057 |
| 4982 | 2009 |
| 5205 | 2152 |
| 4984 | 2008 |
| 4590 | 1929 |
| 5451 | 1947.5 |
| 5402 | 2194.5 |
| 5425 | 2142 |
| 4753 | 2006 |
| 11471 | 2893 |
| 15846 | 3734.5 |
| 14933 | 3602 |
| 11653 | 2898 |
| 1841.5 | 1225 |
| 4015.5 | 1746.5 |
| 3447 | 1686 |
| 7201 | 2713 |
| 4423.5 | 1871.5 |
| 4085 | 1806.5 |
| 3956.5 | 1763.5 |
| 4156 | 1815 |
| 4215 | 1786.5 |
| 4222 | 1900 |
| 4146 | 1942 |
| 4113.5 | 1799 |
| 4238 | 1788.5 |
| 4668.5 | 1858.5 |
| 4150 | 1779 |
| 4273.5 | 1764.5 |
| 4678.5 | 1957 |
| 4419 | 1881.5 |
| 4321 | 1764 |
| 4130 | 1785 |
| 10721 | 2777.5 |
| 14301.5 | 3387.5 |
| 13352.5 | 3298.5 |

| | |
|---|---|
| 10823 | 2762.5 |
| 7524 | 2336 |
| 67157.5 | 32800.5 |
| 5529 | 2148 |
| 9294.5 | 3079.5 |
| 5588 | 2101 |
| 4557.5 | 1867 |
| 4468 | 1793 |
| 4490.5 | 1829 |
| 4317 | 1787 |
| 4380 | 2017 |
| 4097 | 1842.5 |
| 4339 | 1853.5 |
| 4160 | 1869.5 |
| 5021 | 1851 |
| 4347.5 | 1778 |
| 4737 | 2002 |
| 4881 | 1946 |
| 4853.5 | 1970.5 |
| 4829 | 1925 |
| 4326 | 1913 |
| 11044.5 | 2935 |
| 14178 | 3386 |
| 13230.5 | 3299 |
| 11834.5 | 2917 |
| 9023 | 2567 |
| 12894 | 5149 |
| 5571 | 1986 |
| 9655.5 | 3155 |
| 6086.5 | 2305.5 |
| 5234.5 | 2064 |
| 4752 | 1994 |
| 4713.5 | 1910.5 |
| 4470 | 1818.5 |
| 4715 | 2049.5 |
| 4568 | 1860 |
| 4469 | 1845 |
| 4275 | 1838 |
| 4573.5 | 1904.5 |
| 4342 | 1837 |
| 4623 | 1886 |
| 4892.5 | 1889 |
| 4708 | 1826 |
| 4620 | 1874.5 |
| 4195 | 1772 |
| 11166 | 2829 |
| 13244 | 3306.5 |
| 13572.5 | 3312.5 |

| | |
|---|---|
| 11237.5 | 2934 |
| 8198 | 2379 |
| 80199 | 36950.5 |
| 5882.5 | 2258.5 |
| 9016.5 | 3240.5 |
| 6415 | 2483 |
| 5190 | 2092 |
| 4752.5 | 1929 |
| 4593.5 | 1875.5 |
| 4412.5 | 1838 |
| 4550 | 1917 |
| 4386 | 1931 |
| 4295.5 | 1826.5 |
| 4278 | 1804.5 |
| 4663 | 1872 |
| 4210 | 1814.5 |
| 4732 | 1984 |
| 5048 | 1972.5 |
| 4789 | 1970 |
| 4657 | 1947 |
| 4243 | 1819 |
| 9236 | 2725 |
| 14288 | 3347 |
| 13752.5 | 3401 |
| 11303 | 2924 |
| 8772 | 2494 |
| 11001.5 | 4343 |
| 5451.5 | 2118.5 |
| 8835 | 3058 |
| 5891 | 2278 |
| 5043.5 | 1994 |
| 4725.5 | 1889.5 |
| 4647 | 1897 |
| 4577 | 1891 |
| 4513.5 | 1919.5 |
| 4585 | 1902 |
| 4168 | 1837 |
| 4620.5 | 1927.5 |
| 4942.5 | 1869 |
| 4302 | 1805 |
| 4621 | 1860.5 |
| 5108 | 1975 |
| 4677.5 | 1889.5 |
| 4778 | 1852 |
| 4704.5 | 1887 |
| 10428 | 2722 |
| 14783.5 | 3640.5 |
| 14029 | 3531 |

| | |
|---|---|
| 12001 | 3082.5 |
| 9259 | 2536 |
| 10647.5 | 4202 |
| 5766 | 2097 |
| 8424 | 2920 |
| 6006.5 | 2245 |
| 5271.5 | 2030 |
| 4833 | 1928.5 |
| 4673 | 1931 |
| 4530 | 1902.5 |
| 4666 | 2037 |
| 4452 | 1914 |
| 4444.5 | 1822 |
| 4424.5 | 1840.5 |
| 4734.5 | 1945 |
| 4508 | 1803.5 |
| 4336.5 | 1913 |
| 5088.5 | 1971.5 |
| 4795 | 1999.5 |
| 4504 | 1898 |
| 4356 | 1851 |
| 9606 | 2634 |
| 15207 | 3555 |
| 14464 | 3485 |
| 11384 | 2960 |
| 9382 | 2554.5 |
| 20410 | 7887 |
| 5769 | 2206.5 |
| 8658 | 2888 |
| 5884.5 | 2279 |
| 5231.5 | 2003 |
| 4838.5 | 2044.5 |
| 4837.5 | 1960.5 |
| 4532 | 1887 |
| 4830 | 1988.5 |
| 4609 | 1909 |
| 4410 | 1932 |
| 4599.5 | 1954.5 |
| 4944 | 1900 |
| 4566.5 | 1898.5 |
| 4727 | 2007 |
| 5122 | 1932.5 |
| 4937 | 1933 |
| 4956 | 1987 |
| 4683 | 1848 |
| 10408.5 | 2856 |
| 14827 | 3565.5 |
| 14658 | 3596 |

| | |
|---|---|
| 11667 | 2955 |
| 8256 | 2479 |
| 66422 | 30740 |
| 5808 | 2277 |
| 8904.5 | 3077 |
| 6010 | 2291 |
| 5087 | 2002.5 |
| 4928 | 2003 |
| 4634 | 1944.5 |
| 4591 | 1887 |
| 4712 | 1962 |
| 4381 | 1896 |
| 4311 | 1845 |
| 4435 | 1856 |
| 5064.5 | 1961.5 |
| 4349 | 1878 |
| 4566 | 1905.5 |
| 5013.5 | 2023 |
| 4817.5 | 1927.5 |
| 4853 | 1946 |
| 4482 | 1862 |
| 10028 | 2721 |
| 14357.5 | 3500.5 |
| 14748 | 3451 |
| 12312 | 3098 |
| 7960 | 2440 |
| 28947 | 11364 |
| 5827 | 2129 |
| 7321 | 2550 |
| 5555.5 | 2023.5 |
| 5117.5 | 2033.5 |
| 4478 | 1919.5 |
| 4535 | 1836 |
| 4323 | 1810.5 |
| 4561 | 1944 |
| 4227 | 1827 |
| 4151 | 1773.5 |
| 4264 | 1871 |
| 4371.5 | 1836.5 |
| 4195 | 1768.5 |
| 4471 | 1829 |
| 5074.5 | 1942.5 |
| 4645 | 1900 |
| 4871 | 1932 |
| 4236 | 1798 |
| 11047 | 2806 |
| 13878 | 3407 |
| 13070 | 3298.5 |

| | |
|---|---|
| 11129 | 2728 |
| 7997 | 2483 |
| 65259.5 | 30002 |
| 5687 | 2164 |
| 8456 | 2909 |
| 5824 | 2182 |
| 5089 | 2068.5 |
| 4734 | 1910 |
| 4437.5 | 1826 |
| 4466 | 1808 |
| 4607 | 1928 |
| 4387.5 | 1859.5 |
| 4367.5 | 1884 |
| 4191.5 | 1864.5 |
| 5865 | 2045.5 |
| 4350 | 1833 |
| 4645 | 1948 |
| 5209 | 1950 |
| 4853.5 | 1957 |
| 4904 | 1822 |
| 4487 | 1895 |
| 10956.5 | 2778 |
| 14290 | 3604 |
| 13959 | 3376 |
| 11696 | 2887 |
| 8762 | 2487 |
| 7764 | 2719 |
| 5579 | 2020.5 |
| 8136 | 2712 |
| 5657 | 2159 |
| 4937 | 2026.5 |
| 4709.5 | 1961.5 |
| 4550 | 1862 |
| 4212 | 1773 |
| 4685.5 | 1948.5 |
| 4747 | 1927 |
| 4766 | 2039 |
| 4554.5 | 1955.5 |
| 4826 | 1861 |
| 4371 | 1895 |
| 4828 | 1944 |
| 5253 | 2097 |
| 4944 | 2049 |
| 4701 | 1910 |
| 4431.5 | 1925 |
| 11643 | 3004 |
| 15973.5 | 3830.5 |
| 14659 | 3592 |

| | |
|---|---|
| 11210.5 | 2786.5 |
| 1724.5 | 1195.5 |
| 3783 | 1753 |
| 3559 | 1674 |
| 6820.5 | 2553.5 |
| 4288.5 | 1820 |
| 4080.5 | 1759.5 |
| 3941 | 1743 |
| 4194 | 1769 |
| 4118 | 1730 |
| 4571 | 2048.5 |
| 4387.5 | 1908.5 |
| 4041 | 1813 |
| 4379 | 1868 |
| 4380 | 1809 |
| 4273 | 1925 |
| 4447 | 1769.5 |
| 4861 | 1947 |
| 4602.5 | 1891 |
| 4611.5 | 1897.5 |
| 4209 | 1787 |
| 10256 | 2767 |
| 14919 | 3647 |
| 13927.5 | 3317 |
| 11670.5 | 3003.5 |
| 7886.5 | 2391 |
| 72928.5 | 35873 |
| 5630 | 2035 |
| 8812 | 2977 |
| 5221.5 | 2061 |
| 4633 | 1894 |
| 4120 | 1777 |
| 4144 | 1833 |
| 4281 | 1781 |
| 4373.5 | 1988.5 |
| 4555.5 | 1916 |
| 4175.5 | 1840 |
| 4091 | 1755.5 |
| 4967 | 1913 |
| 4279 | 1904 |
| 4296 | 1851 |
| 4868 | 1923.5 |
| 4559 | 1864.5 |
| 4550 | 1929 |
| 4247 | 1911 |
| 11481.5 | 2902.5 |
| 13692 | 3379 |
| 13231 | 3289 |

| | |
|---|---|
| 11183.5 | 2831 |
| 8398 | 2535.5 |
| 9085 | 3122 |
| 5545 | 2002.5 |
| 9292 | 3096.5 |
| 5885 | 2209 |
| 5075 | 1979 |
| 4666 | 1896.5 |
| 4525 | 1966 |
| 4300.5 | 1812 |
| 4373 | 1945 |
| 4304 | 1906.5 |
| 4265 | 1860 |
| 4118.5 | 1784.5 |
| 4611 | 1810 |
| 4175 | 1712 |
| 4451 | 1843.5 |
| 4692 | 1865 |
| 4480 | 1834.5 |
| 4655 | 1848 |
| 4197.5 | 1771 |
| 11159 | 2732 |
| 13457 | 3241 |
| 13556.5 | 3330 |
| 11076 | 2940 |
| 8239 | 2396 |
| 92141 | 43009.5 |
| 6006 | 2371.5 |
| 8670 | 3253 |
| 5946 | 2356 |
| 5122 | 2091 |
| 4454 | 1899 |
| 4490.5 | 1885 |
| 4292 | 1835 |
| 4451 | 1978 |
| 4320 | 1916 |
| 4329.5 | 1837 |
| 4145 | 1866 |
| 4858.5 | 1891.5 |
| 4170 | 1807.5 |
| 4461 | 1902 |
| 4908 | 1954 |
| 4668 | 1940 |
| 4664 | 1898.5 |
| 4118 | 1752 |
| 9820.5 | 2555.5 |
| 14707 | 3359 |
| 13442 | 3333.5 |

| | |
|---|---|
| 11591 | 2833 |
| 7884 | 2336 |
| 63546 | 29479.5 |
| 5180 | 2051.5 |
| 8322 | 2860 |
| 5561.5 | 2162 |
| 4899 | 1922 |
| 4547.5 | 1901.5 |
| 4584 | 1879 |
| 4219.5 | 1817 |
| 4495 | 1958 |
| 4150 | 1879 |
| 4314 | 1838 |
| 4078 | 1770 |
| 4712 | 1869 |
| 4126.5 | 1803.5 |
| 4354.5 | 1825.5 |
| 4927 | 1863 |
| 4661 | 1825 |
| 4513 | 1862 |
| 4210 | 1830 |
| 10548 | 2753 |
| 14009 | 3418 |
| 13266 | 3367 |
| 11849 | 2956.5 |
| 8489 | 2364 |
| 83607.5 | 38773.5 |
| 5496.5 | 2120 |
| 8467 | 2812 |
| 5798 | 2258 |
| 4686 | 1914 |
| 4643.5 | 1956 |
| 4349 | 1815 |
| 4406 | 1823.5 |
| 4507 | 1968 |
| 4269.5 | 1829 |
| 4147 | 1725 |
| 4078 | 1819.5 |
| 4548 | 1761 |
| 4304 | 1775 |
| 4335.5 | 1774.5 |
| 4686 | 1833 |
| 4629.5 | 1877 |
| 4492 | 1853.5 |
| 4148.5 | 1762.5 |
| 10376 | 2802 |
| 13896 | 3354 |
| 13112 | 3249 |

| | |
|---|---|
| 11205 | 2849.5 |
| 7831.5 | 2414 |
| 64853 | 29627 |
| 4969 | 2038 |
| 7704.5 | 2659.5 |
| 5402 | 2049 |
| 4983 | 1932 |
| 4548.5 | 1790 |
| 4384 | 1859 |
| 4231.5 | 1720 |
| 4824 | 1934 |
| 4291 | 1785 |
| 4102 | 1799.5 |
| 4066 | 1731 |
| 4433.5 | 1808 |
| 4052 | 1795.5 |
| 4633 | 1822 |
| 4811 | 1836.5 |
| 4490 | 1834 |
| 4620 | 1865 |
| 4336 | 1771 |
| 11391.5 | 2834 |
| 14430 | 3398 |
| 13639 | 3349.5 |
| 11237 | 2757 |
| 8301 | 2434 |
| 80419 | 37434 |
| 5805.5 | 2189.5 |
| 8309.5 | 2805 |
| 5395 | 2136 |
| 4783.5 | 1934.5 |
| 4762 | 1904 |
| 4226 | 1835 |
| 4452 | 1830 |
| 4804.5 | 2039 |
| 4565 | 1799 |
| 4536 | 1981 |
| 4256 | 1860 |
| 5223 | 1911 |
| 4353.5 | 1849.5 |
| 4757 | 1912 |
| 5128 | 2031.5 |
| 4974 | 1955 |
| 4938 | 1944 |
| 4538.5 | 1839 |
| 12026 | 3035 |
| 14687 | 3496 |
| 13460 | 3258 |

| | |
|---|---|
| 11291.5 | 2939 |
| 8315 | 2488.5 |
| 67065 | 30746.5 |
| 5578 | 2036 |
| 7645 | 2635 |
| 5116 | 2022.5 |
| 4810 | 1907 |
| 4376.5 | 1904 |
| 4512 | 1804 |
| 4253 | 1806 |
| 4526 | 1897 |
| 4412 | 1882 |
| 4198 | 1776 |
| 3937 | 1754 |
| 4576 | 1916 |
| 4191.5 | 1787 |
| 4526 | 1852 |
| 4805 | 1877 |
| 4475 | 1894 |
| 4683.5 | 1884 |
| 4283 | 1817 |
| 11729 | 2807 |
| 13796.5 | 3314 |
| 13730 | 3514 |
| 10878 | 2816 |
| 7819 | 2312 |
| 81034.5 | 37551 |
| 5601 | 2135 |
| 7955 | 2777 |
| 5347 | 2058 |
| 4830 | 1936.5 |
| 4531 | 1893.5 |
| 4481 | 1850 |
| 4470 | 1935 |
| 4640 | 1980.5 |
| 4527.5 | 2025 |
| 4552 | 1891 |
| 4410.5 | 1893.5 |
| 5565 | 1865.5 |
| 4508 | 1895 |
| 4590 | 1835 |
| 5175 | 2027 |
| 4797 | 1937 |
| 4510 | 1871 |
| 4367 | 1851 |
| 11410 | 2940.5 |
| 13644 | 3404 |
| 13770 | 3271 |

| | |
|---|---|
| 10886 | 2769 |
| 8244 | 2371 |
| 8567.5 | 2933.5 |
| 5475 | 2017 |
| 7782 | 2711 |
| 5324 | 2028 |
| 4655.5 | 1885.5 |
| 4342.5 | 1811 |
| 4275 | 1889 |
| 4435 | 1836 |
| 4828 | 2044 |
| 4598 | 1854 |
| 4498 | 1974 |
| 4590.5 | 1942 |
| 4919 | 2004.5 |
| 4486 | 1929.5 |
| 4636 | 1876.5 |
| 5011 | 2006 |
| 5126 | 2002.5 |
| 4892 | 2079 |
| 4519.5 | 1862.5 |
| 11125 | 2894 |
| 15481 | 3658 |
| 13789 | 3471 |
| 11236.5 | 2884.5 |
| 1940 | 1241 |
| 4663 | 2035.5 |
| 3598 | 1615 |
| 6823 | 2652 |
| 4053 | 1845 |
| 4111 | 1762 |
| 4097 | 1792.5 |
| 4205 | 1821 |
| 3965 | 1766 |
| 4195 | 1857 |
| 4152 | 1838.5 |
| 4098 | 1756 |
| 4060 | 1860 |
| 4339.5 | 1804 |
| 4032.5 | 1735 |
| 4303 | 1846.5 |
| 4778 | 1944 |
| 4407.5 | 1777.5 |
| 4288 | 1788.5 |
| 4332 | 1731.5 |
| 10710 | 2898 |
| 14582 | 3568 |
| 13857 | 3295 |

| | |
|---|---|
| 11160 | 2688 |
| 8015 | 2373 |
| 87323 | 42882 |
| 5504 | 2111 |
| 9201.5 | 3015.5 |
| 5240.5 | 2058.5 |
| 4743 | 1921.5 |
| 4640.5 | 1875.5 |
| 4500 | 1853 |
| 4243 | 1716 |
| 4610 | 1968 |
| 4178 | 1757 |
| 4204 | 1840 |
| 4393 | 1808 |
| 5068.5 | 1830 |
| 4211 | 1850 |
| 4536.5 | 1809.5 |
| 5072 | 1926.5 |
| 4705 | 1905 |
| 4800 | 1892 |
| 4407 | 1771 |
| 11338 | 2894 |
| 14603 | 3388 |
| 13692 | 3231 |
| 11891 | 2928 |
| 8432.5 | 2481.5 |
| 65410 | 32756 |
| 5785.5 | 2131.5 |
| 9532.5 | 3201 |
| 6374.5 | 2349 |
| 5616 | 2225 |
| 4807.5 | 1995 |
| 4844.5 | 1975.5 |
| 4389 | 1896.5 |
| 4564 | 1975 |
| 4388 | 1844 |
| 4212.5 | 1827 |
| 4442.5 | 1889.5 |
| 4742 | 1869 |
| 4274.5 | 1796 |
| 4662 | 1918 |
| 4909 | 1893 |
| 4687 | 1964 |
| 4653 | 1900 |
| 4265.5 | 1844 |
| 11476 | 2896 |
| 14409 | 3408.5 |
| 12960 | 3277 |

| | |
|---|---|
| 11875.5 | 2965.5 |
| 8221 | 2413.5 |
| 102715 | 46339 |
| 6249 | 2331 |
| 9182 | 3286.5 |
| 6193 | 2528 |
| 5343 | 2128 |
| 4834 | 2017 |
| 4625 | 1909 |
| 4212.5 | 1715 |
| 4522 | 2060 |
| 4241 | 1819 |
| 4201 | 1768 |
| 4200.5 | 1768 |
| 5151.5 | 1889 |
| 4345 | 1846 |
| 4495 | 1894 |
| 5013 | 1967 |
| 4814.5 | 1931.5 |
| 4363 | 1851 |
| 4161 | 1783 |
| 11779 | 2946 |
| 14325 | 3506.5 |
| 13465.5 | 3371 |
| 11007 | 2747.5 |
| 8308 | 2383 |
| 74705 | 34582 |
| 5518 | 2085.5 |
| 8795 | 2859.5 |
| 5772 | 2196 |
| 5053 | 1993.5 |
| 4594 | 1842 |
| 4717 | 1930 |
| 4306 | 1840 |
| 4340.5 | 1910 |
| 4511.5 | 1798.5 |
| 4240.5 | 1801 |
| 4186.5 | 1790 |
| 4522 | 1907 |
| 4163 | 1868 |
| 4449 | 1906 |
| 4959 | 1902 |
| 4698 | 1935 |
| 4695 | 1873 |
| 4364.5 | 1785.5 |
| 11468 | 2952 |
| 14250 | 3383 |
| 13889 | 3390 |

| | |
|---|---|
| 12045 | 2839.5 |
| 8323 | 2467 |
| 79444.5 | 37087 |
| 5915 | 2157 |
| 8477 | 2768 |
| 5731.5 | 2191 |
| 4864 | 1923 |
| 4556 | 1872 |
| 4559 | 1819 |
| 4177 | 1821 |
| 4496 | 1944 |
| 4312 | 1899 |
| 4066 | 1734 |
| 4070.5 | 1719 |
| 4572 | 1798 |
| 4303 | 1811 |
| 4343.5 | 1809 |
| 5025 | 1896 |
| 4538.5 | 1947.5 |
| 4370 | 1865.5 |
| 4257.5 | 1851.5 |
| 11769.5 | 2927 |
| 13920 | 3381 |
| 13749 | 3305.5 |
| 11330 | 2878 |
| 8522 | 2490.5 |
| 75457 | 34415 |
| 5567 | 2104 |
| 8470.5 | 2735.5 |
| 5711 | 2208 |
| 4899 | 1953 |
| 4672 | 1936 |
| 4604.5 | 1903 |
| 4354 | 1855 |
| 4416 | 1891 |
| 4352 | 1862 |
| 4064.5 | 1814.5 |
| 4203.5 | 1787 |
| 4559 | 1881.5 |
| 4312 | 1822 |
| 4373 | 1831 |
| 5054.5 | 1949.5 |
| 4748 | 1929 |
| 4473 | 1851 |
| 4429 | 1866 |
| 11266.5 | 3020 |
| 14056 | 3416 |
| 13435 | 3324 |

| | |
|---|---|
| 11252 | 2667 |
| 8645 | 2408 |
| 88196 | 41054 |
| 5926 | 2263.5 |
| 8963 | 2945 |
| 5698.5 | 2263 |
| 4837.5 | 2011 |
| 4707 | 1938 |
| 4621.5 | 1919.5 |
| 4314.5 | 1823.5 |
| 4594 | 1936 |
| 4410 | 1966 |
| 4175 | 1803 |
| 4465 | 1884 |
| 4824.5 | 1853.5 |
| 4319 | 1853 |
| 4647 | 1936 |
| 5155 | 2002 |
| 4857 | 1897 |
| 4837 | 1966 |
| 4583 | 1972.5 |
| 11554 | 2810.5 |
| 14509 | 3523.5 |
| 14061.5 | 3421.5 |
| 11719 | 2933 |
| 8479 | 2430 |
| 72354 | 33419 |
| 5906.5 | 2184.5 |
| 7620 | 2637.5 |
| 5209 | 2185 |
| 4886 | 1981 |
| 4379 | 1955 |
| 4447 | 1831 |
| 4253 | 1714 |
| 4562 | 1977 |
| 4243.5 | 1816.5 |
| 4261 | 1831 |
| 4088.5 | 1793 |
| 4313.5 | 1726 |
| 4254 | 1852 |
| 4579.5 | 1857 |
| 4898 | 1875 |
| 4634 | 1838.5 |
| 4847 | 1943 |
| 4266 | 1735 |
| 11251 | 2781 |
| 14627.5 | 3363 |
| 13441 | 3227 |

| | |
|---|---|
| 11139 | 2823 |
| 7914 | 2360 |
| 94289.5 | 43329 |
| 6182.5 | 2443.5 |
| 8451 | 2815.5 |
| 5798 | 2198.5 |
| 5055 | 2059.5 |
| 4763 | 1912 |
| 4560.5 | 1905 |
| 4477 | 1890.5 |
| 4605 | 1885 |
| 4375.5 | 1832.5 |
| 4154.5 | 1907 |
| 4332 | 1860 |
| 6008 | 1954 |
| 4661 | 1950 |
| 4632 | 1813 |
| 4927.5 | 1917 |
| 5051 | 1853 |
| 4782.5 | 1901.5 |
| 4228 | 1848 |
| 11151.5 | 2847.5 |
| 13269.5 | 3257 |
| 13717 | 3356 |
| 12075 | 2914 |
| 8579.5 | 2471 |
| 9432 | 3315 |
| 5340.5 | 1979 |
| 8259.5 | 2624.5 |
| 5256 | 2070 |
| 4754 | 1965 |
| 4456 | 1875.5 |
| 4411 | 1903 |
| 4308.5 | 1837 |
| 4664 | 1986 |
| 4453.5 | 1909.5 |
| 4473.5 | 1901 |
| 4768 | 1919 |
| 4795 | 1910.5 |
| 4355 | 1814 |
| 4682 | 1957 |
| 5198.5 | 1990 |
| 5341 | 2120 |
| 4985 | 1953 |
| 4609 | 1926.5 |
| 11796 | 2903 |
| 15500 | 3839 |
| 14084 | 3426 |

| | |
|---|---|
| 11088.5 | 2783.5 |
| 1902.5 | 1183 |
| 3803 | 1735.5 |
| 3690 | 1655 |
| 7569 | 2840 |
| 4556 | 1889 |
| 4151 | 1755.5 |
| 4156 | 1789 |
| 4397 | 1848 |
| 4118 | 1827.5 |
| 4328 | 1946 |
| 4207 | 1796 |
| 4062.5 | 1744 |
| 4198 | 1826.5 |
| 4435 | 1799 |
| 4112 | 1828 |
| 4326 | 1777 |
| 4709 | 1925 |
| 4518 | 1809 |
| 4433 | 1850 |
| 4248 | 1776 |
| 11028 | 2822 |
| 14326 | 3412 |
| 13473.5 | 3297 |
| 11108 | 2820 |
| 8482 | 2561.5 |
| 77601.5 | 38599.5 |
| 5877 | 2211 |
| 9639 | 3185 |
| 5435 | 2147 |
| 4560.5 | 1818.5 |
| 4522.5 | 1871 |
| 4645.5 | 1923.5 |
| 4243.5 | 1771.5 |
| 4438 | 1966.5 |
| 4505 | 1855.5 |
| 4207 | 1842 |
| 4202.5 | 1806 |
| 5070.5 | 1912 |
| 4283.5 | 1829.5 |
| 4816 | 1912 |
| 5198 | 2049 |
| 4778 | 1944 |
| 4621.5 | 1881.5 |
| 4412.5 | 1892.5 |
| 11899 | 3042 |
| 14842.5 | 3644 |
| 14498 | 3494 |

| | |
|---|---|
| 11667.5 | 2940.5 |
| 9405.5 | 2622 |
| 11302.5 | 3708 |
| 5867.5 | 2132.5 |
| 10405 | 3330.5 |
| 6695 | 2552 |
| 5357 | 2121 |
| 5132 | 1934.5 |
| 4994 | 1936 |
| 4492 | 1872 |
| 4682 | 2035.5 |
| 4668.5 | 1914.5 |
| 4356 | 1835 |
| 4398 | 1891 |
| 5091 | 1903 |
| 4406.5 | 1853 |
| 4598 | 1888 |
| 4943 | 1871 |
| 4600 | 1890.5 |
| 4508 | 1894 |
| 4423 | 1843 |
| 11589.5 | 2808 |
| 14022.5 | 3432.5 |
| 13459.5 | 3209.5 |
| 11174 | 2855.5 |
| 8955 | 2500.5 |
| 89293 | 41168 |
| 6302 | 2372 |
| 8866.5 | 3343 |
| 6203 | 2477 |
| 5185 | 2086 |
| 4700 | 2011 |
| 4722 | 1901.5 |
| 4343 | 1856 |
| 4616 | 2034 |
| 4385 | 1904 |
| 4328 | 1773 |
| 4121 | 1842 |
| 4673.5 | 1895 |
| 4175 | 1880 |
| 4441 | 1893 |
| 5264 | 1954 |
| 4655 | 1883 |
| 4492 | 1818 |
| 4233 | 1800.5 |
| 11615.5 | 2954.5 |
| 13962 | 3411.5 |
| 13524.5 | 3373 |

| | |
|---|---|
| 11715 | 2834 |
| 8816 | 2578 |
| 54394 | 25056 |
| 5860 | 2050 |
| 8885 | 2953 |
| 5793 | 2162 |
| 4863 | 1954.5 |
| 4736.5 | 1876 |
| 4616 | 1859.5 |
| 4201 | 1806.5 |
| 4755 | 1943 |
| 4580 | 1934 |
| 4255 | 1843 |
| 4254 | 1830 |
| 4735 | 1880 |
| 4575 | 1879.5 |
| 4407 | 1875 |
| 4875 | 1987.5 |
| 4616.5 | 1947 |
| 4574 | 1868 |
| 4400 | 1780 |
| 11772.5 | 3019 |
| 14766 | 3554.5 |
| 13847 | 3448 |
| 11648 | 2903 |
| 8823.5 | 2501.5 |
| 71866 | 33786 |
| 5705 | 2100 |
| 8823 | 2848 |
| 5665 | 2240 |
| 4730 | 1979 |
| 4521.5 | 1890 |
| 4418.5 | 1829 |
| 4327.5 | 1760 |
| 4404.5 | 1876 |
| 4443 | 1898.5 |
| 4190 | 1833 |
| 4283 | 1835 |
| 4830 | 1870 |
| 4457 | 1867 |
| 4498 | 1833 |
| 4973 | 1958 |
| 4653 | 1924 |
| 4594.5 | 1857 |
| 4357 | 1847.5 |
| 11898.5 | 3007.5 |
| 14360 | 3488 |
| 13884 | 3417 |

| | |
|---|---|
| 11330 | 2900 |
| 8546 | 2448 |
| 45812.5 | 20648.5 |
| 5679 | 2061.5 |
| 8563 | 2780 |
| 5504 | 2183 |
| 5168 | 2035 |
| 4796 | 1982 |
| 4677 | 1852.5 |
| 4383 | 1794 |
| 4714 | 1936.5 |
| 4404 | 1929.5 |
| 4185 | 1784 |
| 4384 | 1819.5 |
| 4515 | 1866.5 |
| 4266 | 1824 |
| 4429 | 1862 |
| 5033 | 1918 |
| 4835 | 1990.5 |
| 4428 | 1868 |
| 4347 | 1900 |
| 11101 | 2856 |
| 13898.5 | 3419.5 |
| 13749 | 3401 |
| 11735.5 | 2953.5 |
| 9046.5 | 2476.5 |
| 82495 | 38080 |
| 5961 | 2243 |
| 9279 | 3100 |
| 6292.5 | 2370 |
| 4981.5 | 2044.5 |
| 4663.5 | 1948.5 |
| 4661 | 1922 |
| 4576 | 1794 |
| 4551 | 1860 |
| 4692 | 1898 |
| 4201 | 1815 |
| 4337 | 1753 |
| 5065.5 | 1896.5 |
| 4418 | 1839 |
| 4437 | 1949 |
| 5258 | 2008 |
| 4684 | 1920 |
| 4784 | 1906 |
| 4512 | 1833 |
| 12085 | 2926 |
| 14576 | 3465.5 |
| 13844 | 3452 |

| | |
|---|---|
| 11118 | 2719 |
| 8035 | 2403 |
| 12790 | 4270 |
| 5677 | 2052 |
| 7957.5 | 2795 |
| 5392 | 2082 |
| 4646 | 1889 |
| 4471 | 1857 |
| 4502.5 | 1831 |
| 4400.5 | 1824.5 |
| 4548 | 1858 |
| 4291 | 1748 |
| 4341 | 1843 |
| 4124.5 | 1838.5 |
| 4510 | 1817.5 |
| 4197 | 1856.5 |
| 4453.5 | 1867.5 |
| 4962 | 1907 |
| 4870 | 1868 |
| 4840 | 1872 |
| 4323 | 1812 |
| 11678 | 3058 |
| 13982 | 3463.5 |
| 14207 | 3326 |
| 11347 | 2824 |
| 8617 | 2555 |
| 82029 | 38781.5 |
| 5617.5 | 2109 |
| 8684.5 | 2996 |
| 5898.5 | 2173.5 |
| 5033 | 1925 |
| 4728 | 1862.5 |
| 4764 | 1826 |
| 4534.5 | 1799 |
| 4605 | 1940 |
| 4501 | 1891 |
| 4399.5 | 1866 |
| 4164 | 1780 |
| 5208 | 1873 |
| 4586.5 | 1889 |
| 4512 | 1930 |
| 5170.5 | 1993.5 |
| 4998 | 1904.5 |
| 4619 | 1818 |
| 4466 | 1800 |
| 12042 | 2952.5 |
| 14447.5 | 3469 |
| 13906 | 3521 |

| | |
|---|---|
| 11021 | 2828 |
| 8439 | 2434 |
| 7928.5 | 2717 |
| 5194 | 1935 |
| 7979 | 2809 |
| 5374 | 2050 |
| 4735 | 1849 |
| 4512 | 1869 |
| 4451 | 1849 |
| 4424 | 1833 |
| 4778 | 1957 |
| 4507 | 1955.5 |
| 4636 | 1864 |
| 4648 | 1852 |
| 4934 | 1933 |
| 4409 | 1964 |
| 4710.5 | 1951 |
| 5344.5 | 1970.5 |
| 5125 | 2003 |
| 4838 | 1921 |
| 4571 | 1894 |
| 12329 | 3099 |
| 15839 | 3682 |
| 14446 | 3523 |
| 11621 | 2891 |
| 1457 | 1154 |
| 2385.5 | 1351 |
| 2466 | 1376 |
| 12679 | 3810 |
| 4319 | 1818.5 |
| 3855 | 1781 |
| 4664 | 2005 |
| 4220 | 1858 |
| 4836 | 2030 |
| 4533 | 1954 |
| 5333 | 2196 |
| 4649.5 | 2015 |
| 5052 | 2122.5 |
| 4227 | 1880 |
| 4792.5 | 2106 |
| 6122 | 2389 |
| 3470 | 1611.5 |
| 5144.5 | 2072 |
| 5446 | 2188.5 |
| 5974 | 2103 |
| 12996 | 3341 |
| 6804 | 2164 |
| 9304 | 2742 |

| | |
|---|---|
| 7865 | 2313 |
| 3099.5 | 1389 |
| 3358.5 | 1512.5 |
| 3058.5 | 1492 |
| 11251 | 3428 |
| 5257 | 1981 |
| 5085 | 2019 |
| 5182 | 2076.5 |
| 5513.5 | 2188 |
| 5683 | 2205 |
| 5972 | 2216 |
| 6991 | 2641 |
| 6147 | 2410 |
| 5795.5 | 2324.5 |
| 5876.5 | 2354.5 |
| 5163 | 2160 |
| 5298 | 2176 |
| 5616 | 2155 |
| 5734 | 2226 |
| 5211 | 2091 |
| 5164.5 | 2113 |
| 6685 | 2318 |
| 10341 | 2839 |
| 7476 | 2206 |
| 7656 | 2295 |
| 3355 | 1400 |
| 3367 | 1459 |
| 3817 | 1663.5 |
| 10279 | 3074 |
| 5577 | 2125 |
| 4186.5 | 1781 |
| 4873.5 | 1983 |
| 4229 | 1880 |
| 4622 | 1946 |
| 4660 | 1986 |
| 5270.5 | 2135 |
| 5274 | 2106 |
| 4764.5 | 2038.5 |
| 4803.5 | 2003 |
| 8590 | 3063 |
| 9004 | 3043.5 |
| 4593 | 1999 |
| 5832.5 | 2282.5 |
| 7031 | 2677 |
| 5354 | 2240 |
| 6069.5 | 2164.5 |
| 9011 | 2586 |
| 7309.5 | 2288.5 |

| | |
|---|---|
| 7747 | 2303 |
| 2936 | 1348 |
| 3201 | 1485 |
| 3120.5 | 1527.5 |
| 9527 | 2927 |
| 4769 | 1890.5 |
| 4456 | 1887.5 |
| 5151 | 2034 |
| 4038 | 1724 |
| 4383 | 1872 |
| 4092.5 | 1805 |
| 4798 | 2034 |
| 4140 | 1814 |
| 4248.5 | 1863.5 |
| 4131.5 | 1816 |
| 4448 | 1910 |
| 3879 | 1696 |
| 5319 | 2099 |
| 4377 | 1815 |
| 6602 | 2504 |
| 4964 | 2015 |
| 9372 | 2826 |
| 9776.5 | 2690 |
| 7606 | 2299 |
| 8008 | 2371 |
| 3218 | 1410 |
| 3629 | 1561 |
| 3247.5 | 1510 |
| 10828 | 3277 |
| 5980 | 2136 |
| 4162.5 | 1772 |
| 4516 | 1910 |
| 5802 | 2276.5 |
| 3956 | 1732 |
| 6128.5 | 2302.5 |
| 4033.5 | 1813 |
| 5344 | 2186 |
| 4653 | 1907 |
| 4506.5 | 1881.5 |
| 4775 | 2001 |
| 5835 | 2261 |
| 5097 | 2045 |
| 5484 | 2160 |
| 4960 | 1943 |
| 5883.5 | 2261 |
| 5915 | 2107 |
| 11757 | 3199 |
| 9176 | 2643 |

| | |
|---|---|
| 7974.5 | 2407 |
| 2954.5 | 1384 |
| 3503 | 1549.5 |
| 4181.5 | 1719.5 |
| 11444 | 3158 |
| 5993 | 2114.5 |
| 6405 | 2256 |
| 5882.5 | 2212.5 |
| 5702.5 | 2140.5 |
| 6798.5 | 2516.5 |
| 4716 | 1896 |
| 5777 | 2307 |
| 4561 | 1913 |
| 4679 | 1855 |
| 5319 | 2003 |
| 9096.5 | 3135.5 |
| 4360 | 1748 |
| 13729 | 3434.5 |
| 6322 | 2294 |
| 5301 | 2002.5 |
| 7714 | 2657 |
| 6310 | 2100.5 |
| 13137 | 3303 |
| 8088 | 2428 |
| 7709 | 2290 |
| 3418 | 1441 |
| 3429.5 | 1515 |
| 3771.5 | 1628 |
| 10616.5 | 3216.5 |
| 5079.5 | 1923 |
| 4570 | 1818 |
| 4443 | 1888 |
| 4640.5 | 1983 |
| 4255 | 1803 |
| 5159 | 2081 |
| 4046 | 1779 |
| 6337.5 | 2376 |
| 4048 | 1775 |
| 5879 | 2278 |
| 5152 | 2128 |
| 6030.5 | 2317.5 |
| 4924 | 1962.5 |
| 9725 | 3110 |
| 4208 | 1791.5 |
| 6895 | 2614.5 |
| 6031 | 2114 |
| 11040 | 2967 |
| 9416.5 | 2713.5 |

| | |
|---|---|
| 7546.5 | 2280.5 |
| 3172 | 1417 |
| 3754 | 1566 |
| 3380 | 1528 |
| 9623 | 2993 |
| 4590.5 | 1854 |
| 4313 | 1748 |
| 4337 | 1819 |
| 4147.5 | 1738 |
| 4274 | 1793 |
| 4017.5 | 1710.5 |
| 4326 | 1845 |
| 3561 | 1634 |
| 4345 | 1821 |
| 4952 | 2010 |
| 4577 | 1938 |
| 4338 | 1794 |
| 4519 | 1807 |
| 5893 | 2241.5 |
| 6280 | 2372 |
| 6269 | 2284 |
| 8171 | 2614 |
| 9675 | 2669.5 |
| 9260 | 2714 |
| 10072 | 2578.5 |
| 3244 | 1424 |
| 4009 | 1595 |
| 3417.5 | 1522 |
| 15073 | 4185 |
| 5886 | 2045.5 |
| 5823 | 2069 |
| 4898 | 1894 |
| 5078.5 | 1946 |
| 4746 | 1883 |
| 5663 | 2122 |
| 5143 | 1972.5 |
| 5240 | 2039 |
| 4538 | 1886 |
| 4646 | 1755 |
| 5013 | 1962.5 |
| 5167.5 | 2001 |
| 5649 | 2039 |
| 5357.5 | 2033 |
| 6085 | 2128 |
| 5549 | 2094 |
| 7896 | 2599 |
| 10435 | 2822 |
| 10049 | 2761 |

| | |
|---|---|
| 10021 | 2592 |
| 3125 | 1424.5 |
| 3510.5 | 1507 |
| 4603.5 | 1817 |
| 12519 | 3603 |
| 5363 | 1977 |
| 4804 | 1906 |
| 5395 | 2094 |
| 5080 | 2019 |
| 5076 | 2001.5 |
| 5160 | 2053 |
| 4561 | 1864 |
| 9788.5 | 3256 |
| 4992 | 2080 |
| 8114 | 2898 |
| 7007 | 2698.5 |
| 5101 | 1993 |
| 8018 | 2767 |
| 5984 | 2311 |
| 4763 | 1871 |
| 6125.5 | 2453.5 |
| 5338 | 1876 |
| 12177 | 3114.5 |
| 9213 | 2532 |
| 8726 | 2476.5 |
| 3877 | 1501 |
| 3575 | 1506 |
| 4113 | 1675 |
| 11167 | 3148 |
| 5225 | 1935 |
| 5160 | 1968 |
| 4659 | 1882 |
| 5261.5 | 2075 |
| 5562.5 | 2176 |
| 4445.5 | 1798 |
| 4844 | 1919 |
| 4651 | 1911 |
| 5375.5 | 2126 |
| 7260 | 2650 |
| 5091.5 | 2102.5 |
| 7210.5 | 2582.5 |
| 7349 | 2663 |
| 5570.5 | 2098 |
| 6780 | 2463 |
| 4926.5 | 1970 |
| 5567 | 2056.5 |
| 10525 | 2743 |
| 8136 | 2407 |

| | |
|---|---|
| 7717.5 | 2346.5 |
| 1640 | 1147 |
| 2405 | 1305 |
| 3441 | 1579 |
| 9735 | 3433 |
| 4392.5 | 1903 |
| 3661 | 1657 |
| 4068 | 1775 |
| 3975 | 1742.5 |
| 4313 | 1811 |
| 4759 | 2007 |
| 4445 | 1880 |
| 4512 | 1894 |
| 4255 | 1824 |
| 4710 | 1951 |
| 3817.5 | 1691 |
| 4413 | 1841 |
| 4984 | 1973 |
| 4285 | 1795 |
| 4814 | 1926 |
| 4170 | 1767 |
| 5734 | 1975 |
| 7547.5 | 2327 |
| 6928 | 2125 |
| 7015.5 | 2153 |
| 3236.5 | 1348.5 |
| 3404.5 | 1470 |
| 3735 | 1613 |
| 11802 | 3842 |
| 5011 | 1944 |
| 4114 | 1727 |
| 3921 | 1707 |
| 4210 | 1774 |
| 4465 | 1836 |
| 4738 | 1959 |
| 4401 | 1847 |
| 4647 | 1920 |
| 4534.5 | 1889 |
| 4232 | 1800 |
| 4168 | 1802 |
| 4183 | 1781 |
| 4308 | 1754 |
| 5230 | 2033 |
| 4497 | 1853.5 |
| 4289 | 1798 |
| 6090 | 2060 |
| 7043 | 2204 |
| 7000.5 | 2157 |

| | |
|---|---|
| 7553 | 2274 |
| 2954 | 1309 |
| 3418 | 1463 |
| 4428.5 | 1793 |
| 11929 | 3679 |
| 5038 | 1928 |
| 4317 | 1792 |
| 4292 | 1827 |
| 4178 | 1789 |
| 4600 | 1874 |
| 4568.5 | 1890.5 |
| 4383 | 1839 |
| 4563 | 1897 |
| 4471 | 1879 |
| 4388 | 1813.5 |
| 3875 | 1696 |
| 4494 | 1901 |
| 4297 | 1777 |
| 4999 | 1959 |
| 4197 | 1743 |
| 4939 | 1993 |
| 5195.5 | 1844 |
| 7959 | 2373.5 |
| 7317 | 2216 |
| 7435 | 2244.5 |
| 3265 | 1358 |
| 3480 | 1489 |
| 4125 | 1721 |
| 12304 | 3793.5 |
| 5117 | 1949 |
| 4490.5 | 1798 |
| 3930.5 | 1671.5 |
| 4054.5 | 1744 |
| 4091 | 1759 |
| 4128 | 1775 |
| 4327 | 1837.5 |
| 4240 | 1778 |
| 4628 | 1916 |
| 4830 | 1980 |
| 4611 | 1903 |
| 4692 | 1895 |
| 4885 | 1935 |
| 5275.5 | 2038 |
| 4534 | 1847 |
| 5052 | 2026 |
| 5926 | 2055 |
| 8988 | 2524 |
| 7945.5 | 2359.5 |

| | |
|---|---|
| 7526 | 2272 |
| 3274 | 1376 |
| 3187.5 | 1419 |
| 3985 | 1671 |
| 12701 | 3974 |
| 5867 | 2206 |
| 4842.5 | 1906 |
| 4388 | 1805 |
| 4035 | 1695 |
| 4072 | 1709 |
| 4565 | 1898 |
| 4388 | 1805 |
| 4642 | 1925 |
| 4334 | 1797 |
| 5397 | 2115 |
| 4270 | 1794 |
| 5065 | 1988 |
| 5028 | 1960 |
| 4579 | 1825 |
| 5376 | 2092 |
| 4653 | 1895 |
| 6481 | 2151 |
| 7733.5 | 2340 |
| 7701.5 | 2296 |
| 7480 | 2234 |
| 3481.5 | 1411 |
| 3697 | 1517 |
| 3805 | 1618 |
| 15535.5 | 4483.5 |
| 6570.5 | 2302 |
| 5524.5 | 2069.5 |
| 5465 | 2056 |
| 4906 | 1914 |
| 4538 | 1809 |
| 5814 | 2572.5 |
| 4672 | 1853 |
| 4664 | 1871 |
| 5063.5 | 1977 |
| 6016.5 | 2062.5 |
| 5697 | 2157 |
| 5554 | 2070 |
| 8961 | 2356 |
| 5844.5 | 2128 |
| 5758 | 2114 |
| 5465 | 2048 |
| 7302 | 2330.5 |
| 9434 | 2644 |
| 7922 | 2335 |

| | |
|---|---|
| 8035.5 | 2307.5 |
| 3465 | 1401 |
| 3756.5 | 1532.5 |
| 4176 | 1681 |
| 14550 | 4544 |
| 6141.5 | 2204 |
| 4932 | 1907 |
| 4170 | 1738.5 |
| 4178.5 | 1772 |
| 4150 | 1725 |
| 4905 | 1969 |
| 4354 | 1777 |
| 4680 | 1906 |
| 4578 | 1868 |
| 5088 | 2005 |
| 4671 | 1920 |
| 4697.5 | 1885 |
| 4992 | 1916 |
| 5177 | 2002 |
| 4911 | 1937 |
| 4885 | 1929 |
| 6704 | 2230 |
| 8984.5 | 2533.5 |
| 7524 | 2259 |
| 8269 | 2387 |
| 3639 | 1430 |
| 3785 | 1531 |
| 4219 | 1691 |
| 13954 | 4222 |
| 5425 | 2062 |
| 4785 | 1882 |
| 4073 | 1708 |
| 4510 | 1854 |
| 4236 | 1749 |
| 5184.5 | 2053 |
| 4792 | 1920 |
| 4107 | 1736 |
| 4523 | 1877 |
| 5034.5 | 2005 |
| 4625 | 1888 |
| 5435 | 2133 |
| 5018 | 1939 |
| 5370 | 2068 |
| 5415 | 2072 |
| 5350 | 2093 |
| 7038 | 2290 |
| 9024 | 2589 |
| 8920 | 2463.5 |

| | |
|---|---|
| 8148 | 2378 |
| 3563.5 | 1428 |
| 3790 | 1553 |
| 3840.5 | 1594 |
| 16183 | 4821.5 |
| 6560 | 2325 |
| 4933.5 | 1900 |
| 5187 | 1957.5 |
| 5110 | 1906 |
| 5202 | 1886 |
| 5492 | 2149 |
| 5853 | 2129.5 |
| 5301 | 1992 |
| 5033 | 1934 |
| 6038.5 | 2142 |
| 5979 | 2083 |
| 5454 | 1983.5 |
| 7047 | 2367 |
| 6473 | 2304 |
| 6003 | 2142 |
| 6748 | 2306 |
| 7090 | 2280 |
| 10085 | 2653 |
| 9829 | 2657 |
| 9148 | 2561 |
| 3583.5 | 1426 |
| 3698.5 | 1523 |
| 4022 | 1654 |
| 12885 | 3836 |
| 5791 | 2124 |
| 4981.5 | 1917.5 |
| 4609 | 1830 |
| 4719 | 1881 |
| 4143 | 1720 |
| 4521 | 1883 |
| 3900 | 1668 |
| 4423 | 1805 |
| 3957 | 1699 |
| 4562 | 1875.5 |
| 4386.5 | 1812 |
| 4907 | 1924 |
| 5017 | 1932 |
| 5205 | 1998 |
| 5125 | 1991 |
| 5365 | 2081 |
| 6517 | 2162 |
| 9561.5 | 2586.5 |
| 8505 | 2429 |

| | |
|---|---|
| 8867 | 2439 |
| 3247 | 1373 |
| 3467 | 1483.5 |
| 3696 | 1557 |
| 13150 | 3899.5 |
| 5139 | 1953 |
| 4923 | 1933 |
| 4238 | 1758 |
| 5097 | 1994 |
| 4208 | 1727 |
| 4888 | 1938 |
| 4237.5 | 1758 |
| 4347 | 1783 |
| 3917.5 | 1708 |
| 4699.5 | 1879.5 |
| 4395 | 1827 |
| 4760.5 | 1906 |
| 4620 | 1830 |
| 4687 | 1847 |
| 4740.5 | 1884 |
| 4420 | 1782 |
| 6238 | 2124 |
| 8612 | 2451 |
| 8804 | 2486 |
| 7190.5 | 2208 |
| 1493 | 1131 |
| 2325 | 1296 |
| 2978 | 1462 |
| 8044 | 3116 |
| 4667 | 2033.5 |
| 4173.5 | 1837.5 |
| 4224 | 1816.5 |
| 4299.5 | 1852.5 |
| 4612 | 1919 |
| 3950 | 1756 |
| 4062.5 | 1764 |
| 4394 | 1853 |
| 3811 | 1702 |
| 4298.5 | 1841.5 |
| 3783.5 | 1707 |
| 3988 | 1734 |
| 4427 | 1813 |
| 4515 | 1859 |
| 4139 | 1726.5 |
| 4099 | 1738.5 |
| 5900 | 2046 |
| 7154 | 2236 |
| 7298.5 | 2261 |

| | |
|---|---|
| 7540 | 2293 |
| 2949.5 | 1312 |
| 3337 | 1456 |
| 3646.5 | 1602 |
| 10953 | 3328 |
| 4555 | 1840.5 |
| 4142 | 1737 |
| 4360 | 1820 |
| 4717 | 1912 |
| 4347 | 1842.5 |
| 4652 | 1897 |
| 4347 | 1829 |
| 4557 | 1883.5 |
| 4207 | 1824 |
| 3782 | 1684 |
| 3965 | 1747 |
| 4112 | 1751 |
| 4700 | 1895 |
| 4036 | 1739 |
| 4516.5 | 1860 |
| 4166.5 | 1793 |
| 5685 | 1983.5 |
| 8042 | 2420 |
| 7639 | 2306 |
| 7991 | 2333 |
| 2960 | 1317 |
| 3010 | 1392 |
| 3799 | 1629 |
| 10905 | 3207.5 |
| 4630 | 1852 |
| 4490 | 1836.5 |
| 4767 | 1957 |
| 4731 | 1927 |
| 4991 | 1970 |
| 4244 | 1813 |
| 4061 | 1724 |
| 4337 | 1858.5 |
| 4051 | 1785.5 |
| 3927.5 | 1724 |
| 4052 | 1752 |
| 4110 | 1742 |
| 4591.5 | 1832.5 |
| 4420.5 | 1807.5 |
| 4884 | 1969 |
| 4068 | 1769 |
| 5949 | 2059 |
| 7876 | 2401 |
| 7735.5 | 2346 |

| | |
|---|---|
| 7974 | 2367 |
| 3107.5 | 1335 |
| 3485 | 1509 |
| 3664 | 1595 |
| 11136 | 3201 |
| 4533 | 1818 |
| 4302 | 1774 |
| 4657 | 1921 |
| 4497 | 1854 |
| 4957 | 1945 |
| 4540 | 1851 |
| 4573 | 1833 |
| 4031 | 1754 |
| 4468 | 1837 |
| 4068 | 1755 |
| 3641 | 1653 |
| 4785 | 1950 |
| 4231 | 1721 |
| 4937 | 1926 |
| 4266 | 1776 |
| 4431 | 1849 |
| 5562 | 1944 |
| 7646 | 2332 |
| 7536.5 | 2284.5 |
| 7542 | 2267 |
| 2987.5 | 1338 |
| 3389 | 1456.5 |
| 3993 | 1696 |
| 11359 | 3359 |
| 4414 | 1789 |
| 4681 | 1873 |
| 4334.5 | 1823 |
| 4911 | 1998 |
| 4617 | 1886 |
| 4633 | 1909 |
| 4244.5 | 1810.5 |
| 4456 | 1867 |
| 3990 | 1747 |
| 4031 | 1720 |
| 3757 | 1688 |
| 4648 | 1926 |
| 4191 | 1723 |
| 4855 | 1915 |
| 4254 | 1774 |
| 4612 | 1897 |
| 5754.5 | 1976.5 |
| 8897.5 | 2506 |
| 8003 | 2402 |

| | |
|---|---|
| 8468 | 2368 |
| 3142 | 1366 |
| 3227 | 1450 |
| 3979.5 | 1666 |
| 11506 | 3376 |
| 4691 | 1850 |
| 4721 | 1864 |
| 4962 | 1959 |
| 4635.5 | 1903 |
| 4805 | 1955 |
| 4478 | 1847 |
| 4375 | 1793.5 |
| 4379 | 1838.5 |
| 4085 | 1736 |
| 4497 | 1858 |
| 3744 | 1647 |
| 4354 | 1822 |
| 4605 | 1849 |
| 4361 | 1794 |
| 4458.5 | 1845 |
| 4554 | 1885 |
| 5512 | 1950 |
| 7935 | 2376 |
| 7775 | 2320 |
| 7302 | 2226 |
| 3269 | 1377.5 |
| 3378 | 1471 |
| 3988 | 1671 |
| 11295.5 | 3325 |
| 5038 | 1952.5 |
| 4030 | 1689 |
| 4387.5 | 1828.5 |
| 4480 | 1835 |
| 4737 | 1898 |
| 4761 | 1934 |
| 5110 | 2027 |
| 4619 | 1872.5 |
| 4808 | 1919 |
| 4822 | 1955 |
| 4013 | 1733 |
| 4622 | 1884 |
| 4668 | 1891 |
| 4591 | 1826 |
| 4736 | 1873 |
| 4382.5 | 1800 |
| 6403 | 2141 |
| 7153.5 | 2221.5 |
| 7585 | 2262 |

| | |
|---|---|
| 7616 | 2273 |
| 3233.5 | 1369 |
| 3630 | 1523 |
| 4624 | 1842 |
| 11265 | 3325 |
| 4786 | 1859 |
| 4433.5 | 1822 |
| 4403 | 1843 |
| 4524 | 1877 |
| 5009.5 | 1972.5 |
| 5820 | 2234 |
| 4608 | 1867 |
| 5572 | 2156.5 |
| 4971 | 2039 |
| 5242 | 2084 |
| 4855 | 2000 |
| 5014 | 1961 |
| 4914 | 1923 |
| 5882 | 2223 |
| 4676.5 | 1831.5 |
| 4773.5 | 1909 |
| 6903.5 | 2333 |
| 7667 | 2320 |
| 8581 | 2470 |
| 8266 | 2448.5 |
| 3845 | 1490 |
| 4034.5 | 1595 |
| 4496 | 1781 |
| 12906.5 | 3866 |
| 5018.5 | 1894 |
| 4205 | 1726 |
| 4928 | 1926 |
| 4479 | 1757 |
| 4768.5 | 1880 |
| 5777 | 2125 |
| 5617.5 | 2065 |
| 5181 | 1985 |
| 5927 | 2132 |
| 6489 | 2186 |
| 6157 | 2127 |
| 6077 | 2101 |
| 6100 | 2121 |
| 5497 | 1991 |
| 5336 | 2020 |
| 5343.5 | 2033 |
| 6142 | 2109 |
| 8444.5 | 2473.5 |
| 8505.5 | 2448 |

| | |
|---|---|
| 7890 | 2320 |
| 3919 | 1485 |
| 3945 | 1598 |
| 4074 | 1678 |
| 14059.5 | 4012 |
| 5314 | 1957 |
| 5185 | 1984 |
| 4435 | 1792 |
| 4340 | 1774 |
| 4176 | 1718 |
| 4596 | 1959 |
| 4479 | 1857 |
| 4293 | 1795.5 |
| 4400 | 1824 |
| 5482.5 | 2165.5 |
| 4476 | 1839 |
| 5447 | 2106.5 |
| 4887 | 1888 |
| 5348.5 | 2048 |
| 5305 | 2061 |
| 4706 | 1896.5 |
| 6221.5 | 2140.5 |
| 7954.5 | 2420 |
| 8133 | 2431 |
| 7468 | 2250 |
| 3869 | 1461 |
| 4123 | 1620 |
| 4394.5 | 1723.5 |
| 12284 | 3588 |
| 5455 | 2047 |
| 4620 | 1876 |
| 4775.5 | 1879.5 |
| 4136.5 | 1716 |
| 4624 | 1857 |
| 4788 | 1931 |
| 5091 | 1976 |
| 4401 | 1819 |
| 3898 | 1678 |
| 4329 | 1788 |
| 4118 | 1734 |
| 4909.5 | 1945 |
| 4838 | 1905.5 |
| 5091.5 | 1959 |
| 4619 | 1855.5 |
| 4447 | 1835 |
| 5798.5 | 1991 |
| 7543 | 2299 |
| 8502 | 2554.5 |

| | |
|---|---|
| 8289.5 | 2446 |
| 1601 | 1125 |
| 2440 | 1318 |
| 3165.5 | 1528 |
| 9741 | 3551 |
| 4548 | 1991 |
| 3912 | 1727 |
| 3662 | 1650 |
| 3658 | 1637 |
| 3710 | 1664.5 |
| 4193.5 | 1818 |
| 4393.5 | 1847 |
| 4177 | 1794 |
| 4208 | 1803 |
| 4023 | 1773 |
| 4349.5 | 1866 |
| 4579 | 1911.5 |
| 4986 | 1968 |
| 4480 | 1831.5 |
| 5134 | 2042 |
| 4466 | 1867 |
| 6049 | 2145 |
| 7889.5 | 2410 |
| 7759 | 2310 |
| 8117 | 2401 |
| 2749 | 1279 |
| 3155 | 1423 |
| 3467 | 1564 |
| 10027 | 3268 |
| 5147 | 2033 |
| 4337 | 1810 |
| 4078 | 1757 |
| 3739 | 1692 |
| 4053 | 1739 |
| 3725 | 1656 |
| 4584 | 1907 |
| 4008.5 | 1755 |
| 4221 | 1817 |
| 4030 | 1751 |
| 4406 | 1859 |
| 4891 | 2018 |
| 4845 | 1914 |
| 5038.5 | 2002 |
| 4741 | 1915 |
| 4978 | 2009 |
| 5739.5 | 2059.5 |
| 9240.5 | 2531.5 |
| 8168 | 2371 |

| | |
|---|---|
| 8111.5 | 2360 |
| 3017 | 1322 |
| 3128 | 1440 |
| 3610 | 1597 |
| 11772 | 3491 |
| 5151 | 1983 |
| 4380 | 1794 |
| 4127 | 1747 |
| 3873 | 1685 |
| 3901.5 | 1692 |
| 4198 | 1765 |
| 4630 | 1899 |
| 4298 | 1774 |
| 4448 | 1870.5 |
| 4021 | 1747 |
| 4226 | 1806 |
| 5132 | 2050 |
| 4790.5 | 1922.5 |
| 5392 | 2104 |
| 4944 | 1961 |
| 4819 | 1952 |
| 5881.5 | 2071.5 |
| 8495 | 2477.5 |
| 7909 | 2354 |
| 7965 | 2363 |
| 3019 | 1319 |
| 3084.5 | 1393 |
| 3726 | 1592.5 |
| 11783 | 3482 |
| 5376 | 2030 |
| 4819 | 1916 |
| 4291 | 1812 |
| 4185 | 1765 |
| 4369 | 1813 |
| 4202.5 | 1756 |
| 4278 | 1781 |
| 4392 | 1811 |
| 4288 | 1846 |
| 4227 | 1790.5 |
| 4373.5 | 1864 |
| 4748 | 1943 |
| 4623 | 1838 |
| 5790 | 2226 |
| 4774 | 1942 |
| 4738 | 1913 |
| 5686 | 2034 |
| 8282.5 | 2471.5 |
| 7831 | 2320 |

| | |
|---|---|
| 7597 | 2256 |
| 3010 | 1305 |
| 3269 | 1443 |
| 3507 | 1570 |
| 11945 | 3509 |
| 5353 | 1999 |
| 4895 | 1920 |
| 4723.5 | 1896 |
| 4337 | 1763 |
| 4006.5 | 1703 |
| 4073.5 | 1732.5 |
| 4451 | 1823 |
| 4270.5 | 1792 |
| 4310 | 1807 |
| 4080 | 1746 |
| 4297 | 1831 |
| 4613.5 | 1876 |
| 5317.5 | 2061 |
| 5082 | 2011 |
| 5320 | 2074 |
| 4921 | 1971.5 |
| 6148.5 | 2082 |
| 8843 | 2527 |
| 8566 | 2475 |
| 8162 | 2404 |
| 3025 | 1324 |
| 3331 | 1458 |
| 3745 | 1597 |
| 11610 | 3374 |
| 5705 | 2119 |
| 5065 | 1970 |
| 4408 | 1800 |
| 4649 | 1870 |
| 4230 | 1743 |
| 4662 | 1916 |
| 4176 | 1734 |
| 4846 | 1895.5 |
| 4682.5 | 1865.5 |
| 4941.5 | 1819.5 |
| 4556 | 1853 |
| 4964 | 1946 |
| 5167 | 1987 |
| 5576 | 2115 |
| 5272 | 2060 |
| 4748 | 1920 |
| 6604 | 2224.5 |
| 7977 | 2395 |
| 8045 | 2357 |

| | |
|---|---|
| 8408 | 2445 |
| 3135 | 1334 |
| 3542 | 1504 |
| 3794 | 1631 |
| 12288 | 3686 |
| 5279 | 2028 |
| 4980 | 1954 |
| 4284 | 1774 |
| 4291 | 1763 |
| 4179 | 1728 |
| 4245.5 | 1785 |
| 4264 | 1793 |
| 4591 | 1890 |
| 4043 | 1756 |
| 4154 | 1754 |
| 4273 | 1793 |
| 4486 | 1860 |
| 5057 | 1981.5 |
| 4626.5 | 1861 |
| 5092 | 2008 |
| 4415 | 1843 |
| 6753 | 2226 |
| 7816 | 2390 |
| 10071 | 2581.5 |
| 8066 | 2348 |
| 3026 | 1333 |
| 3368 | 1489 |
| 3807 | 1639 |
| 11061 | 3255 |
| 5261 | 2005 |
| 5466 | 2063 |
| 5130 | 2034 |
| 4912.5 | 1965 |
| 4605 | 1851 |
| 4768 | 1914.5 |
| 4559 | 1849 |
| 4557 | 1841 |
| 4973 | 2023 |
| 4354 | 1833 |
| 4517 | 1885 |
| 5198 | 2090 |
| 4991 | 1929 |
| 5431 | 2088 |
| 5204.5 | 2040 |
| 5098 | 2004 |
| 6262 | 2111 |
| 9575 | 2649 |
| 8451 | 2430.5 |

| | |
|---|---|
| 8197 | 2377 |
| 3159 | 1351 |
| 3613 | 1532 |
| 4131 | 1675 |
| 13921.5 | 4150 |
| 5766 | 2145 |
| 5063 | 1962 |
| 4697 | 1891 |
| 4453 | 1763 |
| 4772 | 1850.5 |
| 5189 | 1964 |
| 5329 | 2008 |
| 5619 | 2084 |
| 5692 | 2057 |
| 5687 | 1971 |
| 5559 | 1991 |
| 5879 | 2137 |
| 5809.5 | 2082.5 |
| 5652 | 2095 |
| 5825 | 2123 |
| 5479 | 2079 |
| 7216 | 2298 |
| 8707 | 2534 |
| 10478 | 2730 |
| 10045 | 2592 |
| 3185.5 | 1362 |
| 3444 | 1473 |
| 3764 | 1596 |
| 12279 | 3577 |
| 5717 | 2115 |
| 5159 | 1977 |
| 5210 | 2000 |
| 5106.5 | 1985 |
| 4345 | 1798 |
| 4312 | 1818 |
| 4457 | 1818 |
| 4398 | 1795 |
| 4616 | 1879 |
| 4135.5 | 1722 |
| 4511.5 | 1858 |
| 4532 | 1866 |
| 5362 | 2052.5 |
| 4924 | 1933 |
| 5307 | 2067 |
| 4746 | 1875 |
| 6713 | 2236 |
| 9692 | 2672 |
| 10220 | 2639 |

| | |
|---|---|
| 8662.5 | 2491.5 |
| 3420 | 1390 |
| 3492 | 1460 |
| 3767 | 1575 |
| 11622 | 3446 |
| 5425 | 2054 |
| 4495 | 1807 |
| 4509 | 1803 |
| 4733 | 1858 |
| 4496.5 | 1802 |
| 4985.5 | 1967 |
| 4706 | 1983.5 |
| 4193 | 1896.5 |
| 4202.5 | 1926 |
| 4090 | 1735 |
| 4422 | 1864 |
| 4478 | 1848 |
| 4666 | 1872 |
| 4521 | 1806 |
| 4627 | 1843 |
| 4502.5 | 1814 |
| 6476 | 2172 |
| 8097.5 | 2421.5 |
| 8103 | 2378 |
| 7223 | 2159 |
| 1454 | 1097 |
| 2511 | 1321.5 |
| 3247 | 1535 |
| 10428 | 3627 |
| 4576 | 1978 |
| 3850 | 1716 |
| 3841.5 | 1702 |
| 3718 | 1661 |
| 3618 | 1618 |
| 4239 | 1795.5 |
| 4516 | 1874 |
| 4002 | 1755 |
| 4241.5 | 1793 |
| 4226.5 | 1793 |
| 4338 | 1860 |
| 4697 | 1930.5 |
| 5210 | 2045 |
| 4803 | 1929.5 |
| 4964 | 2006 |
| 4621 | 1928 |
| 6227 | 2102 |
| 8139 | 2427 |
| 8474 | 2445 |

| | |
|---|---|
| 7610.5 | 2292 |
| 2855 | 1268 |
| 3151 | 1425 |
| 3593 | 1596 |
| 11256 | 3354 |
| 5060.5 | 1964.5 |
| 4407.5 | 1817 |
| 4026 | 1748 |
| 3851.5 | 1689 |
| 3841 | 1656 |
| 4152 | 1788 |
| 4476.5 | 1837 |
| 4399 | 1850 |
| 4546 | 1918 |
| 3951 | 1731 |
| 4858 | 1961 |
| 4597 | 1885 |
| 5040 | 2010.5 |
| 5090 | 2031 |
| 4879 | 1926 |
| 4818.5 | 1978.5 |
| 5866.5 | 2055.5 |
| 9627 | 2629 |
| 8545 | 2514 |
| 7982 | 2317 |
| 2820 | 1287 |
| 3142 | 1413 |
| 3567 | 1569 |
| 13079.5 | 3953 |
| 5133 | 1946.5 |
| 4938.5 | 1935 |
| 4349 | 1795 |
| 4789.5 | 1947 |
| 4541 | 1863 |
| 4095 | 1738 |
| 4376 | 1816.5 |
| 4154 | 1766 |
| 4380 | 1845 |
| 3937 | 1720 |
| 4048 | 1751 |
| 4527 | 1849 |
| 4798.5 | 1904.5 |
| 4988.5 | 1982 |
| 4829 | 1978 |
| 4762 | 1877 |
| 6458 | 2156 |
| 8356 | 2468 |
| 8067.5 | 2392 |

| | |
|---|---|
| 7893.5 | 2383 |
| 2984 | 1316 |
| 3288 | 1446 |
| 3637 | 1590 |
| 12010.5 | 3516 |
| 5194 | 2023 |
| 5357.5 | 2080 |
| 4907 | 1955 |
| 4367 | 1847 |
| 4290 | 1777 |
| 4126 | 1748 |
| 4605 | 1870 |
| 4336 | 1803.5 |
| 4556 | 1898.5 |
| 4143 | 1782 |
| 4416 | 1849 |
| 4684 | 1908 |
| 4923 | 1916 |
| 4954 | 1947 |
| 4820 | 1943 |
| 4349.5 | 1823 |
| 6394 | 2128 |
| 7976 | 2433 |
| 8072.5 | 2411 |
| 7740.5 | 2319.5 |
| 3050.5 | 1335 |
| 3263 | 1446 |
| 3700 | 1609 |
| 10935 | 3220 |
| 4903 | 1910 |
| 4518 | 1826 |
| 4530.5 | 1845 |
| 4450 | 1822 |
| 5210 | 2055 |
| 4379 | 1821 |
| 4691 | 1916 |
| 4718 | 1918 |
| 4267 | 1742 |
| 4899.5 | 1956.5 |
| 3811 | 1678 |
| 4219 | 1801 |
| 4626 | 1853 |
| 4408 | 1792 |
| 4393.5 | 1807 |
| 4683 | 1913.5 |
| 5079 | 1888 |
| 10154 | 2652 |
| 8330 | 2426 |

| | |
|---|---|
| 8093 | 2352 |
| 3009 | 1299 |
| 3352 | 1436 |
| 3845 | 1623 |
| 10894 | 3154 |
| 4649 | 1817 |
| 4552 | 1832 |
| 4503.5 | 1836.5 |
| 4222 | 1746 |
| 5318 | 2068 |
| 4489 | 1831 |
| 5431 | 2099 |
| 4217 | 1770 |
| 4730 | 1915 |
| 4080 | 1732 |
| 4403.5 | 1800 |
| 4208.5 | 1765 |
| 4785 | 1876 |
| 4459 | 1769 |
| 4703 | 1872 |
| 4047 | 1696 |
| 5690 | 1972 |
| 7621 | 2351 |
| 7931 | 2329 |
| 7646 | 2300 |
| 3090.5 | 1333.5 |
| 3531.5 | 1494 |
| 3829 | 1612 |
| 9408 | 2999 |
| 5018 | 1934 |
| 4699 | 1867 |
| 4438 | 1804 |
| 4861.5 | 1894 |
| 4897 | 1955 |
| 4437 | 1846 |
| 4493 | 1836 |
| 4670 | 1909 |
| 4190.5 | 1751 |
| 4261.5 | 1806.5 |
| 3709 | 1627 |
| 4253.5 | 1787 |
| 4550 | 1837 |
| 4395.5 | 1768.5 |
| 4746 | 1906 |
| 4092 | 1725 |
| 5741 | 1981 |
| 7740.5 | 2340 |
| 7678 | 2318 |

| | |
|---|---|
| 7738.5 | 2308 |
| 2913 | 1290 |
| 3470 | 1479 |
| 4214 | 1708 |
| 11378 | 3318 |
| 4966 | 1927 |
| 4621 | 1831 |
| 4739 | 1855 |
| 4365 | 1807 |
| 5256 | 2045 |
| 5129 | 2028.5 |
| 5103 | 2001 |
| 4770 | 1908 |
| 4825 | 1935 |
| 4816 | 1972 |
| 4121.5 | 1735 |
| 4593 | 1848 |
| 4929 | 1919 |
| 4499 | 1791 |
| 4670.5 | 1865.5 |
| 4399 | 1821 |
| 5425 | 1904 |
| 8251 | 2458 |
| 8158 | 2372.5 |
| 8070.5 | 2416 |
| 3178 | 1337 |
| 3398 | 1447 |
| 3731 | 1575 |
| 11237 | 3169.5 |
| 4914 | 1891 |
| 4674 | 1872 |
| 4807.5 | 1902 |
| 4813 | 1883 |
| 5105 | 1955 |
| 4752 | 1916 |
| 5012 | 2001 |
| 4598 | 1859 |
| 4795 | 1905 |
| 4585 | 1850 |
| 4128 | 1746 |
| 4323 | 1772 |
| 4713 | 1850 |
| 4841 | 1880 |
| 4480 | 1794 |
| 4574 | 1843 |
| 6212.5 | 2093 |
| 8013 | 2421 |
| 8226 | 2399 |

| | |
|---|---|
| 8087 | 2344 |
| 3077 | 1337 |
| 3275 | 1431 |
| 3691 | 1567 |
| 10603 | 3156 |
| 5362 | 2038 |
| 5143 | 1976 |
| 4680 | 1860 |
| 5037 | 1982 |
| 4848 | 1923 |
| 4491 | 1828 |
| 4651 | 1860 |
| 4534.5 | 1831 |
| 4188.5 | 1734 |
| 4221 | 1763 |
| 4043.5 | 1714 |
| 4622.5 | 1858 |
| 4606.5 | 1802 |
| 5165 | 1986 |
| 4329 | 1742 |
| 4899 | 1945.5 |
| 5905 | 2022 |
| 7881 | 2384.5 |
| 8288 | 2355 |
| 9609 | 2493 |
| 3188 | 1351 |
| 3210 | 1384 |
| 3520.5 | 1531 |
| 9545 | 3071 |
| 5206 | 1960 |
| 4505 | 1778 |
| 4773 | 1879 |
| 5160.5 | 2036 |
| 4486.5 | 1805 |
| 4660 | 1843 |
| 4467 | 1773.5 |
| 4387 | 1783 |
| 4833 | 1903.5 |
| 4497 | 1809 |
| 4408 | 1807 |
| 4230 | 1754 |
| 4988 | 1894 |
| 5496.5 | 2033 |
| 5449 | 2011 |
| 5290.5 | 1998 |
| 6412 | 2084 |
| 7606 | 2344 |
| 7898.5 | 2290 |

| | |
|---|---|
| 8633 | 2449 |
| 1535 | 1123.5 |
| 2261 | 1288 |
| 2932.5 | 1449 |
| 7192 | 2910 |
| 4532 | 1944 |
| 4173 | 1801 |
| 4467 | 1892 |
| 4166.5 | 1797 |
| 3785 | 1660 |
| 3975 | 1706 |
| 4314 | 1798 |
| 4249.5 | 1818 |
| 3970 | 1735 |
| 4084 | 1746 |
| 3567.5 | 1649 |
| 4716 | 1928 |
| 4459 | 1836.5 |
| 4997 | 1973.5 |
| 4803.5 | 1942 |
| 4686 | 1909 |
| 6371 | 2119.5 |
| 8082 | 2447 |
| 9238 | 2578 |
| 8315 | 2411 |
| 2686 | 1241 |
| 3118 | 1436 |
| 3539 | 1555 |
| 9088 | 3063 |
| 5210.5 | 2022 |
| 4473 | 1848 |
| 4812 | 1949 |
| 4150 | 1769 |
| 4242 | 1765 |
| 4211 | 1775 |
| 3982 | 1718 |
| 4312 | 1826 |
| 3954 | 1729 |
| 3970 | 1741 |
| 3828 | 1699 |
| 4123 | 1763 |
| 4846.5 | 1933 |
| 4070.5 | 1707 |
| 4944.5 | 1999 |
| 3973 | 1726.5 |
| 6538 | 2162 |
| 7287.5 | 2282 |
| 8408 | 2382 |

| | |
|---|---|
| 7392 | 2206.5 |
| 2939 | 1309 |
| 3139 | 1415 |
| 3737 | 1619 |
| 12329 | 3648 |
| 4923.5 | 1899 |
| 4699.5 | 1874 |
| 4391 | 1820.5 |
| 4545 | 1859 |
| 4705 | 1880 |
| 4499 | 1849 |
| 4420 | 1819 |
| 4294 | 1804 |
| 3952.5 | 1733.5 |
| 3992 | 1717 |
| 3957 | 1704 |
| 4028 | 1725 |
| 4773.5 | 1897.5 |
| 4460 | 1826 |
| 4725 | 1923 |
| 4845 | 1948 |
| 5677 | 2008 |
| 8914 | 2523.5 |
| 8588 | 2508 |
| 8624 | 2488 |
| 2857.5 | 1276 |
| 3068.5 | 1415.5 |
| 3646 | 1597 |
| 9215.5 | 3053 |
| 5537 | 2115 |
| 5023 | 1970 |
| 4499 | 1856 |
| 4726 | 1893 |
| 4309 | 1779 |
| 4248.5 | 1786 |
| 4545.5 | 1865 |
| 4507 | 1862 |
| 4337 | 1836 |
| 3887 | 1710 |
| 4496 | 1881 |
| 4789 | 1925.5 |
| 5164 | 1979 |
| 5202 | 2051 |
| 5203 | 2024 |
| 5194 | 2066 |
| 5996 | 2112.5 |
| 9701 | 2624 |
| 9256 | 2524 |

| | |
|---|---|
| 7984.5 | 2326 |
| 2996.5 | 1331 |
| 3176 | 1423 |
| 3580 | 1563 |
| 8594 | 2914 |
| 5238 | 1991 |
| 4823 | 1923 |
| 4979 | 1973 |
| 4814 | 1928 |
| 4278 | 1732 |
| 4322 | 1799 |
| 4450 | 1791 |
| 4435.5 | 1816 |
| 4316.5 | 1833.5 |
| 4242 | 1792 |
| 4288.5 | 1820 |
| 5248 | 2070 |
| 4645 | 1832 |
| 5841 | 2203 |
| 4933 | 1988 |
| 5216 | 2038 |
| 6593 | 2189 |
| 9460 | 2586 |
| 9199 | 2577 |
| 8982 | 2488 |
| 3071 | 1333.5 |
| 3218.5 | 1421.5 |
| 3597.5 | 1579 |
| 8930 | 2957 |
| 5120 | 1941 |
| 5075 | 1945.5 |
| 4682 | 1895 |
| 5014.5 | 1972 |
| 4612 | 1846 |
| 4367 | 1782 |
| 4377 | 1776 |
| 4241 | 1755 |
| 4141 | 1756 |
| 4011 | 1729 |
| 4303 | 1776 |
| 4207 | 1736.5 |
| 5632.5 | 2111.5 |
| 4680 | 1837 |
| 5399.5 | 2086 |
| 4526.5 | 1822 |
| 6518 | 2192 |
| 9014 | 2522 |
| 9637 | 2510 |

| | |
|---|---|
| 9007 | 2487 |
| 3109 | 1343 |
| 3242 | 1430 |
| 3465 | 1517 |
| 8843 | 2910 |
| 5163 | 1963 |
| 5067 | 1977 |
| 4862 | 1899 |
| 4592 | 1845 |
| 4515 | 1824 |
| 4586 | 1833 |
| 4116.5 | 1733 |
| 4599 | 1856 |
| 4238 | 1800 |
| 4314 | 1789 |
| 4455 | 1815 |
| 4751 | 1893.5 |
| 5672.5 | 2142 |
| 4918.5 | 1914 |
| 5170 | 1987 |
| 4921 | 1948 |
| 6317 | 2144 |
| 8931 | 2533 |
| 8538 | 2441 |
| 8001 | 2345 |
| 3007 | 1326 |
| 3185.5 | 1428 |
| 3693 | 1575 |
| 8215 | 2733 |
| 5696 | 2105 |
| 4856 | 1901 |
| 4792.5 | 1912.5 |
| 4915 | 1972 |
| 4756 | 1856 |
| 4648.5 | 1858 |
| 4246 | 1748.5 |
| 4559 | 1848.5 |
| 4254.5 | 1788 |
| 4274 | 1771 |
| 4573.5 | 1872 |
| 5037 | 2005 |
| 4944.5 | 1929 |
| 5572.5 | 2131 |
| 4891 | 1907.5 |
| 5039.5 | 1961 |
| 6587 | 2184 |
| 8375 | 2446 |
| 9123 | 2510 |

| | |
|---|---|
| 9635 | 2577 |
| 3307 | 1379 |
| 3316 | 1441 |
| 3670 | 1539 |
| 9901 | 3072 |
| 4950.5 | 1950.5 |
| 5079 | 1975 |
| 4588 | 1879 |
| 5125 | 1987 |
| 4345 | 1750 |
| 4495 | 1814 |
| 4651 | 1861 |
| 4353 | 1800 |
| 4440 | 1820 |
| 4117 | 1744 |
| 4357.5 | 1813 |
| 4218 | 1792 |
| 5559 | 2119 |
| 4798 | 1900 |
| 5225 | 2022 |
| 4702 | 1891 |
| 6326 | 2104 |
| 10257 | 2721 |
| 9436.5 | 2547 |
| 8216 | 2402 |
| 3023.5 | 1328 |
| 3484.5 | 1476.5 |
| 3717 | 1590 |
| 8804 | 2887 |
| 5367 | 2010 |
| 4842.5 | 1898 |
| 4992.5 | 1944 |
| 4760 | 1873 |
| 4688 | 1825 |
| 4451 | 1804 |
| 4222 | 1736.5 |
| 4450 | 1819.5 |
| 4486 | 1842 |
| 3992.5 | 1708 |
| 4102 | 1733 |
| 4182.5 | 1764 |
| 4808 | 1835 |
| 4794 | 1890 |
| 4913 | 1925 |
| 4517 | 1804 |
| 6042.5 | 2040.5 |
| 8791 | 2462 |
| 8054 | 2375 |

| | |
|---|---|
| 8158.5 | 2369 |
| 2917 | 1278 |
| 3316 | 1434 |
| 3525 | 1528 |
| 7821 | 2611 |
| 5124 | 1954 |
| 4481 | 1788 |
| 4851 | 1911 |
| 5037 | 1984 |
| 4742 | 1851 |
| 4722 | 1859 |
| 4649 | 1812 |
| 4443 | 1787 |
| 4715 | 1886 |
| 4664 | 1877 |
| 4248 | 1756 |
| 4440 | 1807 |
| 5051 | 1890 |
| 5452 | 2009 |
| 5481 | 1992 |
| 4983 | 1892 |
| 5699 | 1998 |
| 8166 | 2445 |
| 8829 | 2475 |
| 8822.5 | 2452 |
| 1567 | 1116 |
| 2400 | 1291 |
| 3301 | 1529 |
| 7795 | 3157 |
| 4339 | 1892 |
| 3795 | 1683 |
| 3638 | 1649 |
| 3669 | 1635 |
| 3672 | 1636 |
| 3959 | 1727 |
| 4236 | 1788 |
| 4475 | 1866 |
| 4052.5 | 1738 |
| 4104 | 1781 |
| 3972 | 1745 |
| 4574.5 | 1916.5 |
| 4844.5 | 1932.5 |
| 4744.5 | 1910 |
| 4558 | 1864 |
| 4396 | 1821 |
| 5726 | 2022 |
| 7491 | 2288 |
| 7917 | 2315 |

| | |
|---|---|
| 7622 | 2275 |
| 2812 | 1276 |
| 3093 | 1400 |
| 3400 | 1560 |
| 11047 | 3289 |
| 5065 | 2011 |
| 4257 | 1791 |
| 4267 | 1785 |
| 4063 | 1738 |
| 3839 | 1673 |
| 4048 | 1738 |
| 4520 | 1868 |
| 4546 | 1901 |
| 4188 | 1798 |
| 4299 | 1792 |
| 4314 | 1832.5 |
| 4792.5 | 1918 |
| 4774 | 1862 |
| 5116 | 2035 |
| 4433 | 1819 |
| 4674 | 1904 |
| 5692 | 1979.5 |
| 9115.5 | 2531.5 |
| 8781 | 2482 |
| 8205 | 2369 |
| 2759 | 1285 |
| 3030 | 1392 |
| 3446 | 1531 |
| 12053 | 3579 |
| 5054 | 1918 |
| 4639 | 1879 |
| 4257 | 1772 |
| 4850 | 1955 |
| 4486 | 1849 |
| 4030 | 1721 |
| 4056 | 1738 |
| 3992.5 | 1729 |
| 3987 | 1731 |
| 3914 | 1698 |
| 3945 | 1724.5 |
| 4145 | 1754 |
| 4684.5 | 1858 |
| 4672 | 1906.5 |
| 4658 | 1882.5 |
| 4665 | 1942 |
| 5718.5 | 2029.5 |
| 8107 | 2427 |
| 9055 | 2588 |

| | |
|---|---|
| 8017 | 2383 |
| 2836 | 1294 |
| 3100 | 1419 |
| 3540.5 | 1573 |
| 9447 | 3116 |
| 4963 | 1924 |
| 4576.5 | 1882.5 |
| 4617.5 | 1876 |
| 4807 | 1923 |
| 4526 | 1844 |
| 4016 | 1715 |
| 4300 | 1775 |
| 4245 | 1810 |
| 4167 | 1772 |
| 4027 | 1717 |
| 3941 | 1703 |
| 4094 | 1756 |
| 4431 | 1803 |
| 4430 | 1797 |
| 4445 | 1813 |
| 3942 | 1690 |
| 6082 | 2067 |
| 7481.5 | 2288 |
| 8248 | 2377 |
| 7636 | 2264 |
| 3010.5 | 1341 |
| 3131.5 | 1404 |
| 3699 | 1571 |
| 9276 | 3032 |
| 4409 | 1769 |
| 4532 | 1853 |
| 4355 | 1784 |
| 4852 | 1936 |
| 4226 | 1737 |
| 4538 | 1844 |
| 4220 | 1760 |
| 3912 | 1714 |
| 4127.5 | 1745 |
| 3804 | 1685 |
| 3825 | 1666 |
| 4335 | 1799 |
| 4803 | 1898 |
| 4898 | 1918 |
| 4565 | 1838 |
| 5182 | 2032 |
| 5941 | 2095.5 |
| 9016.5 | 2533 |
| 9300 | 2569 |

| | |
|---|---|
| 8719 | 2519 |
| 3163 | 1342 |
| 3169 | 1402 |
| 3623 | 1578 |
| 10427 | 3128 |
| 5181 | 2008 |
| 5065 | 1955 |
| 4410 | 1800 |
| 4513 | 1818 |
| 4225.5 | 1730.5 |
| 4189 | 1737 |
| 4556 | 1866 |
| 4440 | 1843 |
| 4249 | 1795 |
| 4356 | 1835 |
| 4270 | 1808 |
| 4575 | 1861 |
| 5155 | 2001 |
| 4829 | 1915 |
| 5159 | 2004 |
| 4311 | 1785 |
| 6253 | 2075 |
| 7671.5 | 2376 |
| 8273 | 2383 |
| 7746 | 2316 |
| 3200 | 1353 |
| 3365 | 1456 |
| 3683 | 1578.5 |
| 10884 | 3190 |
| 5194.5 | 1963 |
| 4583 | 1825 |
| 4284 | 1759 |
| 4216 | 1752 |
| 4083.5 | 1693 |
| 4025 | 1709 |
| 4435 | 1807 |
| 3952 | 1666 |
| 4194.5 | 1778.5 |
| 3934 | 1721 |
| 4335 | 1803 |
| 4365 | 1844 |
| 4834 | 1913 |
| 4979.5 | 1952 |
| 4555 | 1850 |
| 4865 | 1959 |
| 5794 | 2020 |
| 9937 | 2588 |
| 9900.5 | 2603 |

| | |
|---|---|
| 8702 | 2496 |
| 3316 | 1355 |
| 3421 | 1468 |
| 3913 | 1644 |
| 10859 | 3193 |
| 5254 | 1949 |
| 4789.5 | 1899 |
| 4534 | 1818 |
| 4220.5 | 1743.5 |
| 4106 | 1703.5 |
| 4220.5 | 1791 |
| 4309 | 1796 |
| 4364.5 | 1826 |
| 4706 | 1882 |
| 4002 | 1697 |
| 4498 | 1838 |
| 4515 | 1849 |
| 5128.5 | 1977 |
| 4811.5 | 1905.5 |
| 4933 | 1936 |
| 4857 | 1937 |
| 6030 | 2056 |
| 10456 | 2711 |
| 8843 | 2536 |
| 8224 | 2354 |
| 3029 | 1307 |
| 3242 | 1447 |
| 3678 | 1568 |
| 10990.5 | 3139 |
| 5426 | 2052 |
| 4593 | 1838 |
| 4766 | 1882 |
| 4404 | 1777 |
| 4321 | 1759 |
| 4091 | 1749.5 |
| 4303 | 1753 |
| 4181 | 1771 |
| 4084 | 1741 |
| 4119 | 1745 |
| 4044.5 | 1710 |
| 4771 | 1934.5 |
| 4945 | 1907 |
| 4931 | 1938 |
| 4804 | 1919 |
| 4691 | 1897 |
| 6587 | 2233 |
| 8386 | 2457 |
| 9236 | 2536.5 |

| | |
|---|---|
| 8671.5 | 2455 |
| 2956.5 | 1317 |
| 3221 | 1435 |
| 3495.5 | 1523 |
| 9139 | 3041.5 |
| 4454.5 | 1784 |
| 4358 | 1758 |
| 4577 | 1831 |
| 4612 | 1840 |
| 4740 | 1883 |
| 4948.5 | 1963.5 |
| 4996 | 1937 |
| 4977 | 1969 |
| 4562.5 | 1857 |
| 4709.5 | 1922 |
| 4340 | 1796 |
| 4675 | 1891 |
| 4727.5 | 1864 |
| 4710 | 1867 |
| 4398 | 1809 |
| 4484 | 1844 |
| 5839.5 | 2005.5 |
| 7209 | 2234 |
| 7583 | 2288.5 |
| 7659 | 2272 |
| 3036.5 | 1310.5 |
| 3051 | 1387 |
| 3333 | 1480 |
| 9193 | 2971 |
| 5036.5 | 1919 |
| 4590 | 1824 |
| 4785 | 1893 |
| 4358 | 1782 |
| 4598.5 | 1855.5 |
| 4521 | 1866.5 |
| 4433 | 1778 |
| 4238.5 | 1753 |
| 4657.5 | 1885 |
| 4381 | 1804 |
| 4347 | 1789.5 |
| 4534 | 1817 |
| 4739 | 1837 |
| 4685 | 1813 |
| 4960.5 | 1908 |
| 5407 | 2056 |
| 6160.5 | 2064 |
| 8182 | 2422 |
| 8126.5 | 2384 |

| | |
|---|---|
| 8066 | 2347 |
| 1516 | 1105 |
| 2398 | 1293 |
| 3309 | 1546.5 |
| 8464 | 3334 |
| 4161 | 1819 |
| 3710 | 1673 |
| 3624.5 | 1639 |
| 3896 | 1732 |
| 3891 | 1666 |
| 4147 | 1789 |
| 4476 | 1880 |
| 3902 | 1713.5 |
| 4232 | 1828 |
| 3751 | 1661.5 |
| 4174 | 1784 |
| 4437 | 1839 |
| 4398 | 1807 |
| 4515 | 1843 |
| 4192 | 1756 |
| 4178.5 | 1778.5 |
| 5529.5 | 1948 |
| 7325 | 2264.5 |
| 7793.5 | 2339 |
| 7479.5 | 2249 |
| 2831 | 1295 |
| 3196 | 1438 |
| 3451 | 1539 |
| 11677.5 | 3554.5 |
| 4487 | 1821 |
| 3805 | 1659.5 |
| 4042.5 | 1731 |
| 3672 | 1627 |
| 4096 | 1752 |
| 4293 | 1839 |
| 4281 | 1819 |
| 4432 | 1864 |
| 3975 | 1748.5 |
| 4074.5 | 1761.5 |
| 4327 | 1838 |
| 4643.5 | 1929.5 |
| 4549 | 1845 |
| 4525 | 1841.5 |
| 4508 | 1813.5 |
| 4122 | 1747 |
| 5856.5 | 2022 |
| 7384 | 2285 |
| 8017 | 2387 |

| | |
|---|---|
| 8060.5 | 2363 |
| 2918 | 1300.5 |
| 3104 | 1410 |
| 3296 | 1482 |
| 13536 | 4069 |
| 5247 | 2008 |
| 5237 | 2034 |
| 4661 | 1884 |
| 4097 | 1732 |
| 4104 | 1714 |
| 4036 | 1762 |
| 4379 | 1838.5 |
| 4508.5 | 1853 |
| 4573 | 1869 |
| 4271 | 1823 |
| 4232 | 1802 |
| 4815 | 1925 |
| 5264 | 2028 |
| 4714 | 1894 |
| 4693.5 | 1893.5 |
| 4391 | 1807 |
| 5988 | 2052 |
| 7679 | 2342 |
| 7524 | 2286 |
| 7699 | 2333 |
| 3181 | 1352 |
| 3356 | 1476 |
| 3640 | 1570 |
| 13270 | 4032.5 |
| 5107 | 1978 |
| 4392 | 1778 |
| 3957 | 1681 |
| 4180 | 1778 |
| 4035 | 1728 |
| 4485 | 1873 |
| 4544 | 1885 |
| 4338 | 1817 |
| 4447 | 1856 |
| 4151 | 1783 |
| 4436 | 1841 |
| 4759 | 1905 |
| 4504 | 1833 |
| 4765 | 1889 |
| 4545 | 1860 |
| 4236.5 | 1775 |
| 5765 | 1967 |
| 7388 | 2257 |
| 7754 | 2333 |

| | |
|---|---|
| 7673.5 | 2291 |
| 3167 | 1349.5 |
| 3477 | 1475.5 |
| 3664 | 1597.5 |
| 12439 | 3839 |
| 4784 | 1882 |
| 4266 | 1760.5 |
| 4078.5 | 1738 |
| 3976 | 1700 |
| 4051.5 | 1711 |
| 4194 | 1799 |
| 4531 | 1866 |
| 4599 | 1893 |
| 4495.5 | 1835 |
| 4396 | 1842 |
| 4728 | 1930 |
| 4587 | 1847 |
| 5246.5 | 2002 |
| 4663.5 | 1900 |
| 4824 | 1912 |
| 4198 | 1804 |
| 5882 | 2021 |
| 7322 | 2266 |
| 7919 | 2355 |
| 7633.5 | 2292.5 |
| 3193 | 1334 |
| 3392 | 1483 |
| 3788 | 1619 |
| 14091.5 | 4242 |
| 5272 | 1994 |
| 4652 | 1842 |
| 3992 | 1694 |
| 4004.5 | 1694 |
| 4301.5 | 1771 |
| 4516 | 1838.5 |
| 4717 | 1899.5 |
| 4456.5 | 1830.5 |
| 4474.5 | 1844.5 |
| 4203.5 | 1777 |
| 4440 | 1833 |
| 4926.5 | 1950 |
| 5183 | 2019 |
| 4707 | 1882 |
| 4972 | 1975 |
| 4313 | 1783 |
| 6630 | 2177 |
| 7694 | 2366 |
| 8027 | 2377 |

| | |
|---|---|
| 8631 | 2434.5 |
| 3406 | 1383 |
| 3437 | 1449 |
| 4038 | 1659 |
| 12116 | 3755 |
| 5518 | 2070.5 |
| 4596.5 | 1823.5 |
| 4176 | 1718 |
| 4209 | 1743 |
| 4239 | 1745 |
| 4568 | 1923 |
| 4607 | 1889 |
| 4645 | 1888 |
| 4472 | 1860 |
| 4220 | 1779 |
| 4844 | 1935 |
| 4774 | 1912 |
| 5010 | 1969 |
| 5118 | 1987.5 |
| 4618 | 1876 |
| 4642 | 1858 |
| 5812 | 2008 |
| 7990.5 | 2360 |
| 8363 | 2456 |
| 7923 | 2347 |
| 3215 | 1352.5 |
| 3773 | 1553 |
| 3938 | 1618 |
| 12838.5 | 3927 |
| 5772 | 2112 |
| 4562.5 | 1809 |
| 4254.5 | 1726 |
| 3958 | 1661 |
| 4257 | 1759 |
| 4238 | 1801 |
| 4892 | 1941 |
| 4424 | 1836 |
| 4780.5 | 1939 |
| 4145 | 1775 |
| 4757 | 1935 |
| 4845 | 1949 |
| 5087 | 1968 |
| 5262 | 2029 |
| 4860.5 | 1930 |
| 4527 | 1824 |
| 5644 | 2014 |
| 8207 | 2410 |
| 8327 | 2408 |

| | |
|---|---|
| 8064.5 | 2357 |
| 3149.5 | 1356 |
| 3667 | 1517.5 |
| 3704 | 1586 |
| 11582 | 3598 |
| 5346 | 2016 |
| 4349.5 | 1770.5 |
| 4404.5 | 1769 |
| 4050 | 1713 |
| 4230 | 1740 |
| 4365 | 1801 |
| 4487 | 1826 |
| 4565 | 1869.5 |
| 4347 | 1806 |
| 4401 | 1829 |
| 4593 | 1864 |
| 4918 | 1959.5 |
| 5039 | 1947 |
| 5043 | 1975 |
| 4968 | 1940 |
| 4731 | 1887 |
| 6482 | 2149 |
| 8035 | 2401 |
| 8916 | 2476 |
| 8151 | 2398 |
| 3587 | 1416 |
| 3486 | 1466 |
| 4173 | 1663.5 |
| 11199 | 3596.5 |
| 5358.5 | 2012 |
| 4708 | 1840 |
| 3985.5 | 1671 |
| 4246 | 1758 |
| 4143.5 | 1739 |
| 4436 | 1858 |
| 4688 | 1885 |
| 4404 | 1795 |
| 4873.5 | 1944.5 |
| 4130 | 1748 |
| 4870 | 1930 |
| 4770 | 1862 |
| 5255 | 1982 |
| 5426 | 2071 |
| 4782 | 1916 |
| 4908 | 1928 |
| 5709.5 | 2029.5 |
| 8304.5 | 2474 |
| 9089 | 2527.5 |

| | |
|---|---|
| 8003 | 2370.5 |
| 3218 | 1351 |
| 3577 | 1487 |
| 3842.5 | 1608 |
| 11486 | 3638 |
| 5409 | 2027 |
| 4580 | 1824 |
| 4300.5 | 1770 |
| 4132 | 1737 |
| 4607 | 1844 |
| 4681 | 1891 |
| 4666 | 1832.5 |
| 4352 | 1772 |
| 4235 | 1749 |
| 3837 | 1660 |
| 4399 | 1801 |
| 4980 | 1962 |
| 4955.5 | 1893 |
| 5004 | 1947 |
| 4767 | 1846 |
| 4419 | 1773 |
| 5691 | 1943 |
| 8501 | 2520 |
| 8809.5 | 2531.5 |
| 7702.5 | 2273.5 |
| 2198 | 1310 |
| 3221 | 1580 |
| 3973.5 | 1780 |
| 12804 | 3636 |
| 4652 | 1941 |
| 4295 | 1892 |
| 4048.5 | 1798.5 |
| 3404 | 1574 |
| 4514.5 | 1929.5 |
| 4445 | 1879 |
| 4926 | 2122 |
| 4956 | 2102 |
| 5519 | 2273 |
| 4853 | 2003 |
| 4173 | 1842 |
| 4046 | 1748 |
| 4482.5 | 1850.5 |
| 4383 | 1769 |
| 4503.5 | 1864.5 |
| 4399 | 1888 |
| 6514 | 2167 |
| 9954 | 2586 |
| 10705 | 2742 |

| | |
|---|---|
| 10491.5 | 2807 |
| 3075 | 1391 |
| 3698 | 1604 |
| 4068 | 1784 |
| 12240 | 3312 |
| 4457 | 1809.5 |
| 4312.5 | 1845 |
| 4183 | 1806 |
| 3967 | 1684 |
| 3977 | 1745.5 |
| 5435 | 2130.5 |
| 4270.5 | 1879.5 |
| 5160 | 2100 |
| 5371 | 2266 |
| 4209 | 1808 |
| 4028 | 1699 |
| 4083 | 1735 |
| 4762 | 1931 |
| 6797 | 2572 |
| 5253 | 2099.5 |
| 4610 | 1918 |
| 8652.5 | 2584 |
| 9601.5 | 2576 |
| 10791 | 2738 |
| 11571 | 3013 |
| 3098.5 | 1401 |
| 3562 | 1567 |
| 3983 | 1775 |
| 12133.5 | 3276.5 |
| 4618 | 1846 |
| 4342 | 1858.5 |
| 4291.5 | 1859.5 |
| 4164 | 1773 |
| 4074 | 1782 |
| 4471.5 | 1909.5 |
| 5831.5 | 2365 |
| 5848 | 2333 |
| 4089 | 1814.5 |
| 4466 | 1843.5 |
| 3965 | 1709 |
| 4262.5 | 1825.5 |
| 5768 | 2203 |
| 4628 | 1844 |
| 5469.5 | 2149.5 |
| 5023 | 2046 |
| 7517 | 2311.5 |
| 10031 | 2680 |
| 10316 | 2740 |

| | |
|---|---|
| 11431 | 2985 |
| 3041 | 1350 |
| 4176 | 1738 |
| 4828 | 1961 |
| 11499.5 | 3269 |
| 5395 | 2082 |
| 5600.5 | 2202.5 |
| 4646.5 | 1898 |
| 4591 | 1846 |
| 5520 | 2199 |
| 5369.5 | 2176 |
| 4473.5 | 1887 |
| 3851 | 1754 |
| 3803 | 1705 |
| 4126 | 1731 |
| 4597.5 | 1901 |
| 5324.5 | 2095.5 |
| 5504 | 2159 |
| 5248 | 2063 |
| 5216.5 | 2067.5 |
| 5332 | 2121 |
| 7035 | 2321 |
| 11903.5 | 3165 |
| 10584 | 2751 |
| 10638 | 2711 |
| 3140 | 1409 |
| 3981 | 1687 |
| 4449 | 1847 |
| 12044 | 3265.5 |
| 4579 | 1873 |
| 4344 | 1790 |
| 4297 | 1792 |
| 4509 | 1840 |
| 4874 | 1986 |
| 4913 | 2020 |
| 6106 | 2385 |
| 4698.5 | 1931.5 |
| 3759 | 1687 |
| 4374.5 | 1744 |
| 4755.5 | 1979.5 |
| 4571 | 1868 |
| 6484 | 2437 |
| 6685 | 2527 |
| 5333 | 2104.5 |
| 6309 | 2450 |
| 6764 | 2232.5 |
| 11782 | 3116.5 |
| 10474 | 2772 |

| | |
|---|---|
| 11133 | 2897 |
| 3086 | 1405.5 |
| 4033 | 1650 |
| 4414 | 1806 |
| 10459 | 2966 |
| 6452 | 2259 |
| 5665 | 2120 |
| 5610 | 2133 |
| 5055 | 1937 |
| 5540 | 2102 |
| 5093 | 2076 |
| 5740 | 2162 |
| 6486.5 | 2478.5 |
| 5966 | 2331 |
| 7087 | 2573.5 |
| 5734 | 2170 |
| 5647 | 2134 |
| 6447 | 2018 |
| 7493 | 2742 |
| 6099 | 2289 |
| 5712 | 2183 |
| 10878 | 2959 |
| 12175.5 | 3081 |
| 10571.5 | 2788 |
| 12160 | 3173 |
| 3652 | 1524 |
| 4376 | 1752 |
| 4521.5 | 1823 |
| 11422 | 3181 |
| 5513 | 2111 |
| 4942 | 2005 |
| 5020 | 2027 |
| 4479 | 1851 |
| 5129 | 2058 |
| 5436 | 2170 |
| 5602 | 2217 |
| 5100 | 2087 |
| 4836 | 1984 |
| 4214 | 1705 |
| 4334 | 1829 |
| 4509 | 1797 |
| 5846.5 | 2166 |
| 5257.5 | 2065 |
| 6278 | 2327 |
| 5269 | 2090 |
| 9816 | 2800 |
| 10391 | 2796.5 |
| 10083.5 | 2624.5 |

| | |
|---|---|
| 10083.5 | 2640.5 |
| 3532 | 1517 |
| 4442 | 1818 |
| 5134.5 | 2132.5 |
| 11838 | 3473.5 |
| 6040 | 2419 |
| 6294 | 2503.5 |
| 5910 | 2460 |
| 5330 | 2189.5 |
| 6135 | 2480 |
| 6794.5 | 2685.5 |
| 5824 | 2388 |
| 5014.5 | 2159.5 |
| 5028 | 2136 |
| 4933 | 2029.5 |
| 6095.5 | 2488.5 |
| 5536 | 2253.5 |
| 8187.5 | 2980.5 |
| 6524 | 2536 |
| 7495 | 2971 |
| 6277 | 2465 |
| 10332 | 3111.5 |
| 10633 | 3048 |
| 11109 | 3035 |
| 11006 | 3083 |
| 3361 | 1427 |
| 4368.5 | 1787.5 |
| 4748 | 1901 |
| 16775 | 4563 |
| 8252 | 2753.5 |
| 5986.5 | 2134 |
| 8965 | 2754 |
| 5532 | 2006 |
| 5608 | 2106 |
| 8232 | 3140 |
| 7057 | 2544 |
| 5749 | 2133 |
| 4924 | 1905.5 |
| 7153.5 | 2108.5 |
| 5136 | 1914.5 |
| 4994.5 | 1880 |
| 8112.5 | 2620.5 |
| 6453 | 2270.5 |
| 8610 | 2748 |
| 7105 | 2502 |
| 10108 | 2830 |
| 10751 | 2872.5 |
| 9937 | 2697 |

| | |
|---|---|
| 10628.5 | 2627.5 |
| 3275.5 | 1438 |
| 4017 | 1732 |
| 4686 | 1887 |
| 8883 | 2900 |
| 5283 | 2028 |
| 4953.5 | 1997.5 |
| 4628 | 1881 |
| 4043 | 1702 |
| 4813.5 | 1939.5 |
| 5358 | 2170 |
| 4785 | 1971 |
| 5466 | 2220 |
| 5091 | 2087 |
| 4382.5 | 1774 |
| 4013 | 1705 |
| 4895 | 1942 |
| 4922 | 1940.5 |
| 5296 | 2013 |
| 6265 | 2381 |
| 4466.5 | 1833 |
| 11027 | 2930 |
| 9631 | 2619 |
| 11495.5 | 2938.5 |
| 10901 | 2853 |
| 3423.5 | 1463 |
| 4248.5 | 1736 |
| 5022.5 | 2034 |
| 8665 | 2839 |
| 5242.5 | 2029 |
| 5372 | 2072 |
| 4797.5 | 1945 |
| 4795.5 | 1909.5 |
| 5674 | 2195.5 |
| 5121 | 2063 |
| 4810 | 1955.5 |
| 4284 | 1788 |
| 4239.5 | 1796 |
| 4819 | 1899 |
| 5352 | 2082.5 |
| 4796 | 1897 |
| 6399.5 | 2363 |
| 5020 | 1920 |
| 4953 | 1969 |
| 4927 | 1968 |
| 10861.5 | 2971 |
| 8472 | 2436 |
| 10374 | 2724 |

| | |
|---|---|
| 7549 | 2233 |
| 2104 | 1248 |
| 3119 | 1495 |
| 4035 | 1816 |
| 11381 | 3573 |
| 4525 | 1936 |
| 4027 | 1809 |
| 4002 | 1782.5 |
| 3857 | 1721.5 |
| 4338 | 1874 |
| 4522 | 2007 |
| 4525 | 1994 |
| 4525 | 1929 |
| 4516 | 1917 |
| 4138 | 1751 |
| 3929 | 1831 |
| 3655 | 1681 |
| 4285.5 | 1783 |
| 6377 | 2495 |
| 5795 | 2324 |
| 4516 | 1865.5 |
| 8077.5 | 2513 |
| 11112 | 2963.5 |
| 8698 | 2491 |
| 7920 | 2349 |
| 3124 | 1386 |
| 3635.5 | 1603 |
| 3959.5 | 1725.5 |
| 10039 | 2956 |
| 5106 | 2013 |
| 4255 | 1765 |
| 3801 | 1691 |
| 3707 | 1670 |
| 4516 | 1923 |
| 4708.5 | 1977.5 |
| 5370 | 2150 |
| 4695 | 2015 |
| 5241 | 2103 |
| 4502 | 1844 |
| 4458 | 1898 |
| 3816.5 | 1641 |
| 5805.5 | 2257 |
| 7127 | 2644 |
| 6145 | 2388.5 |
| 4905 | 2098 |
| 10257 | 2797 |
| 9640 | 2694 |
| 9623.5 | 2514 |

| | |
|---|---|
| 10415 | 2761 |
| 3067 | 1388 |
| 3514 | 1569 |
| 3722 | 1674 |
| 11349.5 | 3168.5 |
| 5099.5 | 2009.5 |
| 4508 | 1896 |
| 4283 | 1831 |
| 4082 | 1745 |
| 4712 | 1900 |
| 4835 | 1995 |
| 4808 | 1948 |
| 4287.5 | 1858 |
| 3906 | 1746 |
| 3870 | 1641 |
| 4192 | 1777 |
| 4180 | 1765 |
| 5196 | 2090 |
| 7080 | 2517 |
| 5754 | 2259 |
| 5236 | 2158 |
| 9006 | 2575 |
| 10480 | 2824 |
| 9125 | 2450 |
| 8672 | 2543 |
| 3076 | 1391 |
| 3881 | 1686 |
| 4229.5 | 1797 |
| 9360 | 2905 |
| 4704 | 1934 |
| 4147 | 1778 |
| 4179 | 1801.5 |
| 3908 | 1689 |
| 4676 | 1929 |
| 5209.5 | 2089 |
| 4814 | 1977.5 |
| 5038.5 | 2069 |
| 5275.5 | 2133.5 |
| 4060 | 1708.5 |
| 4073.5 | 1713 |
| 4103 | 1716 |
| 4943 | 1988 |
| 5845 | 2254 |
| 7263 | 2646 |
| 5651 | 2242 |
| 10299 | 2866.5 |
| 9343 | 2653.5 |
| 9483 | 2510.5 |

| | |
|---|---|
| 8063 | 2365 |
| 3185 | 1386.5 |
| 4081.5 | 1734 |
| 4491.5 | 1843 |
| 9995.5 | 3125.5 |
| 4952 | 2015.5 |
| 5039 | 2036 |
| 4408.5 | 1869.5 |
| 4273.5 | 1765 |
| 4932 | 1987.5 |
| 4706 | 1941 |
| 4621.5 | 1909 |
| 4515 | 1856 |
| 3655.5 | 1658 |
| 4589 | 1782 |
| 4255 | 1799 |
| 3980 | 1755 |
| 5984 | 2234 |
| 6890 | 2556 |
| 6067 | 2322 |
| 5720.5 | 2307 |
| 8585 | 2533 |
| 9551 | 2620 |
| 9357 | 2627 |
| 8249 | 2423 |
| 3413 | 1487 |
| 4186 | 1711 |
| 4754 | 1877 |
| 9516 | 3026 |
| 7040.5 | 2444 |
| 6181 | 2258 |
| 5543 | 2177 |
| 5862.5 | 2199.5 |
| 5970 | 2211 |
| 5280 | 2100 |
| 5266 | 2045 |
| 4953 | 1939 |
| 4619 | 1901 |
| 6970 | 2469 |
| 4626 | 1837 |
| 4737.5 | 1901 |
| 8499.5 | 2389 |
| 7922 | 2831 |
| 8509 | 2993 |
| 6256 | 2296 |
| 8393 | 2627.5 |
| 12701 | 3244 |
| 9134 | 2569 |

| | |
|---|---|
| 9479.5 | 2593.5 |
| 3479.5 | 1498 |
| 4160 | 1732.5 |
| 4738 | 1913 |
| 9371.5 | 3076 |
| 5040.5 | 1989.5 |
| 4823.5 | 1948.5 |
| 4712 | 1948 |
| 4052 | 1745.5 |
| 4557 | 1899.5 |
| 5356 | 2112 |
| 5299 | 2078 |
| 4894 | 1988 |
| 5180.5 | 2084.5 |
| 5181 | 2013 |
| 4735 | 1979 |
| 4415 | 1862 |
| 4289 | 1778 |
| 5986 | 2315 |
| 6659 | 2536.5 |
| 4939 | 1983 |
| 8696 | 2564 |
| 11832 | 3108 |
| 9396 | 2592 |
| 9012.5 | 2521.5 |
| 3366.5 | 1482.5 |
| 4621.5 | 1915 |
| 4827.5 | 2027.5 |
| 9467 | 3127 |
| 5688 | 2280 |
| 5491 | 2271.5 |
| 5520 | 2270 |
| 5967 | 2450 |
| 6177 | 2484 |
| 5528 | 2299 |
| 5219.5 | 2186.5 |
| 4973 | 2115.5 |
| 4333 | 1897 |
| 4823.5 | 2055 |
| 5341 | 2241 |
| 5544 | 2233 |
| 6535 | 2475 |
| 7720 | 2889 |
| 7340 | 2904 |
| 6711 | 2585 |
| 10500 | 3016 |
| 11945.5 | 3291 |
| 10228.5 | 2853.5 |

| | |
|---|---|
| 10386 | 2867 |
| 3782 | 1691 |
| 4714 | 1914 |
| 5440.5 | 2087.5 |
| 13615 | 4079 |
| 8100 | 2712 |
| 7270.5 | 2499.5 |
| 9022 | 2810 |
| 6332.5 | 2275.5 |
| 6754 | 2344 |
| 9276.5 | 3454.5 |
| 5325.5 | 2057.5 |
| 5317.5 | 2057 |
| 4719 | 1882.5 |
| 9533.5 | 2452.5 |
| 6245 | 2161 |
| 5827.5 | 2114.5 |
| 8029 | 2573 |
| 7324.5 | 2612.5 |
| 9376 | 3041 |
| 6920 | 2423 |
| 9699.5 | 2600 |
| 11006 | 3018 |
| 10809 | 2806 |
| 9661.5 | 2754.5 |
| 3436 | 1523.5 |
| 3939.5 | 1708.5 |
| 4312.5 | 1838.5 |
| 9367 | 2818 |
| 5002 | 1944 |
| 4797 | 1980 |
| 4592 | 1897 |
| 4339 | 1806.5 |
| 4625 | 1885 |
| 5046 | 2162.5 |
| 5746 | 2256 |
| 4898.5 | 2015 |
| 4286.5 | 1812.5 |
| 4035 | 1681 |
| 4102 | 1784 |
| 4398 | 1810 |
| 5205 | 2023 |
| 5129 | 2094 |
| 5874 | 2301 |
| 5267 | 2106 |
| 8463 | 2505 |
| 11081 | 2973 |
| 11026.5 | 2824.5 |

| | |
|---|---|
| 8284 | 2400 |
| 3073 | 1440 |
| 3776.5 | 1648 |
| 4171 | 1747 |
| 10189.5 | 3086 |
| 4392.5 | 1803.5 |
| 4872 | 1991.5 |
| 4351 | 1828 |
| 4356 | 1812.5 |
| 4848 | 1936 |
| 4280 | 1794 |
| 4347.5 | 1884.5 |
| 4315 | 1847.5 |
| 4733 | 1985 |
| 5311 | 2058 |
| 4824 | 1958 |
| 4221 | 1744 |
| 5706 | 2157.5 |
| 4780.5 | 1942 |
| 4758.5 | 1925 |
| 4968 | 1974 |
| 10130 | 2791.5 |
| 9730.5 | 2625 |
| 8488 | 2347.5 |
| 8504 | 2342 |
| 1885 | 1206 |
| 3031 | 1505 |
| 3735 | 1711.5 |
| 9240 | 3294 |
| 4264.5 | 1865 |
| 4093 | 1803.5 |
| 4106.5 | 1847 |
| 3681 | 1690 |
| 4261 | 1864.5 |
| 4350 | 1921.5 |
| 4715 | 2012 |
| 5376 | 2251 |
| 5156 | 2096 |
| 4139 | 1775 |
| 3919 | 1753.5 |
| 4102 | 1741.5 |
| 4193 | 1744 |
| 5374.5 | 2115.5 |
| 4718.5 | 1995.5 |
| 5077.5 | 2096 |
| 7695 | 2514 |
| 10532 | 2732 |
| 10024 | 2620 |

| | |
|---|---|
| 8232 | 2393 |
| 2634 | 1324 |
| 3125 | 1504.5 |
| 3466 | 1610 |
| 8002.5 | 2640.5 |
| 4411 | 1855 |
| 4117 | 1760 |
| 4148 | 1819 |
| 3524 | 1587 |
| 4004 | 1716 |
| 4696.5 | 2024 |
| 4336 | 1878 |
| 4750 | 2014 |
| 5383 | 2188 |
| 4179 | 1792.5 |
| 3641.5 | 1651 |
| 3909 | 1678 |
| 4466 | 1913 |
| 5414 | 2094.5 |
| 6168.5 | 2397 |
| 4584.5 | 2016 |
| 8735 | 2547 |
| 9595.5 | 2639.5 |
| 10028 | 2624 |
| 9813 | 2646 |
| 3229 | 1493 |
| 3450 | 1573 |
| 3778 | 1715 |
| 9275 | 2895 |
| 4661 | 1932 |
| 4158.5 | 1798 |
| 4251.5 | 1789.5 |
| 3839 | 1671 |
| 4195 | 1788 |
| 4467 | 1913 |
| 5038 | 2065 |
| 4566 | 1895 |
| 4754 | 1953 |
| 4902 | 1924.5 |
| 3977 | 1760.5 |
| 4076 | 1709 |
| 3923.5 | 1665.5 |
| 5567.5 | 2129.5 |
| 5985.5 | 2313 |
| 5957 | 2278 |
| 7800 | 2603 |
| 11195.5 | 2956 |
| 8728 | 2501 |

| | |
|---|---|
| 8203 | 2379 |
| 3124.5 | 1377 |
| 3866 | 1674 |
| 4112.5 | 1738.5 |
| 9800 | 3159.5 |
| 4668 | 1894 |
| 4412 | 1844.5 |
| 4304 | 1806 |
| 4468 | 1837 |
| 4168 | 1766 |
| 5285 | 2157.5 |
| 5140 | 2108 |
| 5269 | 2139 |
| 4955 | 2045 |
| 4626 | 1900.5 |
| 4064 | 1751.5 |
| 4020 | 1809 |
| 4258 | 1772 |
| 4931 | 1967 |
| 4828 | 1985 |
| 4861 | 1961 |
| 7546 | 2419 |
| 10972 | 2909.5 |
| 9782.5 | 2525.5 |
| 10781.5 | 2805.5 |
| 3206 | 1412 |
| 3446 | 1543 |
| 3997 | 1722 |
| 9567 | 3055.5 |
| 4776.5 | 1939.5 |
| 4577 | 1940 |
| 4214 | 1811 |
| 3961 | 1733 |
| 4876 | 1905 |
| 5032 | 2127 |
| 4731 | 1924 |
| 4530 | 1880 |
| 4216 | 1816 |
| 4238 | 1653 |
| 4198.5 | 1799 |
| 4200 | 1752 |
| 5128.5 | 1986.5 |
| 6040 | 2296 |
| 5861 | 2236 |
| 5559.5 | 2196 |
| 9784.5 | 2776 |
| 10777 | 2834 |
| 9570.5 | 2596 |

| | |
|---|---|
| 8629 | 2544.5 |
| 3091 | 1413 |
| 4028.5 | 1716 |
| 4653 | 1867 |
| 9517 | 3195.5 |
| 6476 | 2339.5 |
| 5923 | 2139 |
| 5983 | 2192 |
| 5448 | 2024 |
| 5746.5 | 2131.5 |
| 5953 | 2243 |
| 6008.5 | 2320 |
| 5431.5 | 2138 |
| 5865 | 2281 |
| 8097 | 3248.5 |
| 5403.5 | 2053 |
| 5092.5 | 1973.5 |
| 9080.5 | 2427.5 |
| 5626 | 2122 |
| 6410 | 2267 |
| 6214 | 2323.5 |
| 9269 | 2848 |
| 12642.5 | 3149 |
| 8621 | 2463.5 |
| 10939 | 2885 |
| 3255 | 1446 |
| 3412 | 1503 |
| 3926 | 1694 |
| 11286 | 3303.5 |
| 4946 | 1974 |
| 4759 | 1936 |
| 4859 | 1970 |
| 4621 | 1877 |
| 4674 | 1906 |
| 5096.5 | 2027 |
| 4943 | 2008 |
| 4194 | 1794 |
| 3956 | 1768 |
| 4484 | 1794 |
| 4833 | 1974 |
| 5137 | 2020 |
| 5352 | 2015 |
| 6184.5 | 2331 |
| 5660 | 2193 |
| 5738 | 2261 |
| 8403 | 2490 |
| 11081 | 2944 |
| 9074.5 | 2543.5 |

| | |
|---|---|
| 8403 | 2422 |
| 2918 | 1392 |
| 3993.5 | 1728.5 |
| 4606.5 | 1945.5 |
| 9561 | 3217 |
| 5662 | 2292.5 |
| 5540 | 2281 |
| 5544 | 2300 |
| 5748 | 2318 |
| 6074.5 | 2470 |
| 6161 | 2480 |
| 6082 | 2502 |
| 5972 | 2453 |
| 5123 | 2181.5 |
| 4907.5 | 2064.5 |
| 4930 | 2130 |
| 5265.5 | 2205.5 |
| 5646 | 2242 |
| 6649.5 | 2656 |
| 7038.5 | 2776.5 |
| 6211 | 2533 |
| 8072 | 2715 |
| 11206.5 | 3150 |
| 10212.5 | 2811 |
| 10350 | 2939 |
| 3176 | 1418 |
| 4162 | 1718 |
| 4656 | 1837 |
| 13481 | 4083.5 |
| 7604 | 2535 |
| 6554 | 2287 |
| 8815.5 | 2672 |
| 6432 | 2218 |
| 6476.5 | 2263 |
| 9864 | 3620 |
| 5997 | 2152 |
| 5336 | 2006 |
| 4868.5 | 1913 |
| 10177.5 | 2473 |
| 5392.5 | 2058.5 |
| 5339 | 2010 |
| 6608 | 2160.5 |
| 6655 | 2356 |
| 9009 | 2939.5 |
| 7898 | 2703 |
| 9544.5 | 2730 |
| 13167 | 3323 |
| 10366.5 | 2550.5 |

| | |
|---|---|
| 11308 | 2928.5 |
| 2929 | 1382 |
| 3680 | 1591 |
| 4235 | 1742 |
| 9874 | 2988 |
| 4916 | 1904 |
| 4604 | 1878 |
| 4516 | 1851 |
| 4215 | 1775.5 |
| 4798 | 1957 |
| 5598 | 2352.5 |
| 5057 | 2018 |
| 4758 | 1966 |
| 4488 | 1865 |
| 5021.5 | 1959.5 |
| 4338 | 1803 |
| 4321 | 1785 |
| 4819 | 1896.5 |
| 4357 | 1836.5 |
| 5775 | 2256.5 |
| 5451 | 2150 |
| 9718.5 | 2707.5 |
| 10891 | 2878 |
| 10770 | 2772 |
| 9181.5 | 2555.5 |
| 3503 | 1499 |
| 3896 | 1692 |
| 4127 | 1730 |
| 9636.5 | 3141.5 |
| 5074 | 2003 |
| 4407.5 | 1857.5 |
| 4323 | 1780 |
| 4342 | 1783.5 |
| 5142 | 2017 |
| 4670 | 1927.5 |
| 4398 | 1862 |
| 4243.5 | 1799.5 |
| 4645 | 1891 |
| 5103 | 1991 |
| 4843 | 1957 |
| 4626 | 1931 |
| 4889 | 1917 |
| 4690 | 1859.5 |
| 4712 | 1897 |
| 5278.5 | 2032.5 |
| 7960 | 2488 |
| 10578 | 2787 |
| 10236 | 2545 |

| | |
|---|---|
| 7218.5 | 2191 |
| 1897 | 1181 |
| 3200 | 1535 |
| 3735 | 1692 |
| 9513 | 3414 |
| 4440 | 1902 |
| 4402.5 | 1976 |
| 4392 | 1895 |
| 4024 | 1765 |
| 4547.5 | 1914 |
| 4813 | 2079 |
| 5193.5 | 2132.5 |
| 4447.5 | 1932 |
| 3900 | 1755 |
| 3891 | 1657 |
| 3915.5 | 1740 |
| 3982.5 | 1745 |
| 4735.5 | 1893 |
| 5810 | 2240 |
| 5235 | 2103 |
| 5260.5 | 2156 |
| 9410.5 | 2714 |
| 10891.5 | 2887.5 |
| 9251 | 2476 |
| 8647.5 | 2523 |
| 2711.5 | 1319.5 |
| 3410.5 | 1529.5 |
| 3879 | 1734 |
| 8868 | 2856 |
| 4498 | 1816 |
| 4300 | 1843 |
| 4395 | 1836 |
| 3750 | 1623 |
| 4637.5 | 1913.5 |
| 4664 | 2055 |
| 4281 | 1836 |
| 4979.5 | 2054.5 |
| 6144 | 2447 |
| 5067 | 2056.5 |
| 3888 | 1708.5 |
| 4376 | 1811 |
| 4487 | 1838 |
| 5436 | 2130 |
| 5906 | 2277 |
| 5300.5 | 2144.5 |
| 7158 | 2312 |
| 11324 | 2935 |
| 10117 | 2670 |

| | |
|---|---|
| 8077.5 | 2393.5 |
| 2899.5 | 1358 |
| 3301.5 | 1508.5 |
| 3688 | 1657 |
| 9080 | 2882 |
| 4619 | 1883 |
| 4356.5 | 1820 |
| 4083.5 | 1712 |
| 4122 | 1741 |
| 4547 | 1897 |
| 5038 | 2089 |
| 4914 | 2038 |
| 4700.5 | 1955.5 |
| 4783.5 | 1992 |
| 4045.5 | 1701.5 |
| 4152 | 1808 |
| 4041 | 1767.5 |
| 4428 | 1815 |
| 5640 | 2136 |
| 4765 | 1935 |
| 5475 | 2185.5 |
| 9253.5 | 2617 |
| 10457 | 2917 |
| 9032 | 2453.5 |
| 9611 | 2600.5 |
| 2862 | 1327 |
| 3282 | 1491 |
| 3697.5 | 1631.5 |
| 8827 | 2785 |
| 4647 | 1852 |
| 4437.5 | 1831.5 |
| 4545.5 | 1880 |
| 4081 | 1712 |
| 4408 | 1808 |
| 4309.5 | 1872 |
| 5289 | 2090 |
| 5441.5 | 2189 |
| 4846 | 2005.5 |
| 4802 | 1970 |
| 4354 | 1863 |
| 4002 | 1725.5 |
| 4664 | 1887 |
| 4484 | 1859.5 |
| 4448 | 1855 |
| 5097 | 2028 |
| 7832 | 2414 |
| 11916 | 3081 |
| 9001 | 2418.5 |

| | |
|---|---|
| 9454 | 2528 |
| 3056 | 1391 |
| 3542 | 1560.5 |
| 3821 | 1643 |
| 11718 | 3262 |
| 5002 | 1994.5 |
| 4708 | 1912 |
| 4388 | 1828 |
| 4454 | 1815 |
| 5016.5 | 2013.5 |
| 5292.5 | 2129.5 |
| 5116 | 2062 |
| 5158.5 | 2128 |
| 4625.5 | 1958 |
| 4103 | 1624 |
| 4541 | 1841 |
| 4284 | 1805 |
| 5238 | 2005 |
| 5884 | 2246 |
| 5881 | 2306.5 |
| 5521.5 | 2161 |
| 9733 | 2675 |
| 11414 | 2988 |
| 9756 | 2609 |
| 8353 | 2468 |
| 3033 | 1402.5 |
| 3720.5 | 1636 |
| 4376.5 | 1813.5 |
| 11350.5 | 3367 |
| 6755 | 2358 |
| 6105 | 2213 |
| 5732 | 2105 |
| 5579 | 2035 |
| 6049 | 2251.5 |
| 5740 | 2321.5 |
| 5553 | 2149 |
| 4744 | 1910 |
| 4469 | 1825 |
| 7063 | 2688.5 |
| 5248 | 1922 |
| 5123 | 1948 |
| 8254 | 2311 |
| 6759 | 2386 |
| 7307 | 2526 |
| 6139 | 2311 |
| 9922 | 2880 |
| 12386 | 3158.5 |
| 10597 | 2692 |

| | |
|---|---|
| 10302 | 2706 |
| 3069 | 1406.5 |
| 3621 | 1588.5 |
| 4178 | 1756 |
| 10256 | 3265 |
| 5223.5 | 1997 |
| 4835.5 | 1903 |
| 4623 | 1845 |
| 4649 | 1907 |
| 5315 | 2083 |
| 5299 | 2172 |
| 4782 | 1960.5 |
| 4567 | 1920 |
| 4099 | 1785 |
| 4190 | 1705 |
| 4096 | 1730 |
| 4657 | 1870.5 |
| 4458 | 1794 |
| 5233 | 2039 |
| 5637 | 2174 |
| 5750.5 | 2252 |
| 7960 | 2461 |
| 12000 | 3168 |
| 9676 | 2529 |
| 8716 | 2429 |
| 3193.5 | 1448 |
| 3991 | 1677 |
| 4600 | 1909 |
| 10224 | 3328 |
| 5830.5 | 2348 |
| 5915 | 2367 |
| 5630 | 2321.5 |
| 5981 | 2484.5 |
| 6548 | 2597.5 |
| 6071.5 | 2482.5 |
| 5588.5 | 2328 |
| 5366 | 2233.5 |
| 4703 | 2019 |
| 4848 | 2065 |
| 5275 | 2182 |
| 5751 | 2275.5 |
| 6102 | 2360 |
| 6773.5 | 2699.5 |
| 7006 | 2738 |
| 6616.5 | 2588 |
| 9687 | 2923.5 |
| 11624 | 3229.5 |
| 11376.5 | 3111.5 |

| | |
|---|---|
| 10520 | 2774 |
| 3128.5 | 1462 |
| 3986 | 1727 |
| 4546 | 1833 |
| 12468 | 3877 |
| 7052 | 2403.5 |
| 6552 | 2257 |
| 8355.5 | 2560 |
| 6348 | 2213 |
| 6972 | 2379 |
| 11022.5 | 4185.5 |
| 5844 | 2146.5 |
| 5826.5 | 2168 |
| 4959 | 1935 |
| 9436 | 2380 |
| 6098 | 2144 |
| 5902 | 2132 |
| 6714.5 | 2190 |
| 7210 | 2509 |
| 9147.5 | 3013 |
| 6692.5 | 2413 |
| 9560 | 2823 |
| 13158 | 3231 |
| 11196 | 2900 |
| 9862.5 | 2688 |
| 3240.5 | 1437 |
| 3843 | 1658 |
| 4385 | 1805.5 |
| 10948 | 3043.5 |
| 5248 | 2064 |
| 4706 | 1878.5 |
| 4991.5 | 1981.5 |
| 4295 | 1786 |
| 5032.5 | 1995 |
| 5075 | 2185 |
| 5803 | 2260.5 |
| 5138 | 2093 |
| 4276 | 1845.5 |
| 4191 | 1745 |
| 4306 | 1786 |
| 4421 | 1807 |
| 4893 | 1899 |
| 5744.5 | 2239.5 |
| 5819.5 | 2217 |
| 5590.5 | 2157 |
| 8688 | 2580.5 |
| 11051.5 | 2928.5 |
| 10075 | 2629 |

| | |
|---|---|
| 9050 | 2522 |
| 3116 | 1416 |
| 3642 | 1584 |
| 4280 | 1806.5 |
| 9496 | 3067 |
| 5219 | 2017 |
| 4755 | 1964 |
| 4447 | 1820.5 |
| 4349 | 1835 |
| 4994 | 1988.5 |
| 4356 | 1881 |
| 4627.5 | 1900.5 |
| 4914 | 1999 |
| 5100.5 | 2083 |
| 4713 | 1872 |
| 4443.5 | 1830 |
| 4423 | 1752 |
| 5120 | 1952 |
| 5231 | 1999 |
| 5713 | 2187.5 |
| 5265 | 2115 |
| 8902 | 2625 |
| 9864.5 | 2634.5 |
| 9993 | 2537 |
| 7748 | 2322 |
| 1962 | 1267 |
| 2996 | 1512 |
| 3464 | 1625 |
| 8716 | 3200 |
| 4140 | 1821 |
| 4306 | 1884 |
| 4251.5 | 1872.5 |
| 3759.5 | 1745 |
| 4147 | 1706 |
| 4294 | 1952 |
| 3917 | 1762 |
| 4360 | 1895 |
| 4908.5 | 2074.5 |
| 5287 | 2104 |
| 5047.5 | 2127 |
| 5429 | 2188 |
| 5357 | 2045 |
| 4363 | 1898 |
| 4626 | 1930 |
| 4366.5 | 1886 |
| 6276.5 | 2108.5 |
| 10479 | 2768 |
| 7317 | 2216 |

| | |
|---|---|
| 8298 | 2335 |
| 2771 | 1358 |
| 3393 | 1600.5 |
| 3797 | 1680 |
| 10670 | 3209.5 |
| 5280 | 2071 |
| 5011 | 2052 |
| 4526.5 | 1918.5 |
| 4145 | 1780 |
| 4281 | 1790 |
| 4471.5 | 1890.5 |
| 4411 | 1928.5 |
| 4575 | 1943 |
| 4873.5 | 2027 |
| 5341.5 | 2060.5 |
| 4911 | 2043 |
| 4273.5 | 1779 |
| 4783.5 | 1892.5 |
| 4854 | 1968 |
| 4648 | 1914 |
| 5032.5 | 2107.5 |
| 7042 | 2282 |
| 9591 | 2678 |
| 8380 | 2421 |
| 7494 | 2262.5 |
| 3241 | 1469 |
| 4375 | 1817 |
| 4456.5 | 1892 |
| 18353 | 5138 |
| 6470 | 2324 |
| 6329.5 | 2363 |
| 5152.5 | 2033 |
| 5312 | 2073 |
| 4348 | 1814 |
| 4669 | 2156 |
| 4447.5 | 1915 |
| 4955 | 2092 |
| 5424 | 2179 |
| 4764 | 1941 |
| 5306 | 2154 |
| 4801 | 1903.5 |
| 5118 | 2036 |
| 4642 | 1869 |
| 4760 | 1977 |
| 4656 | 1926 |
| 7302 | 2327 |
| 8873 | 2583 |
| 8649 | 2451 |

| | |
|---|---|
| 7620 | 2366 |
| 2665 | 1307.5 |
| 3306 | 1534 |
| 4032 | 1713 |
| 11545 | 3462 |
| 6088.5 | 2281.5 |
| 5442 | 2107.5 |
| 5500 | 2202 |
| 4601 | 1878.5 |
| 4044.5 | 1676.5 |
| 4053 | 1849 |
| 4323 | 1826 |
| 4427 | 1851 |
| 5854 | 2306 |
| 5013.5 | 1974.5 |
| 4624 | 1920 |
| 4522 | 1889.5 |
| 4595 | 1817 |
| 4720 | 1925 |
| 4900.5 | 1948.5 |
| 4361 | 1828 |
| 8003 | 2463 |
| 9168 | 2592.5 |
| 10253 | 2561 |
| 7583 | 2236 |
| 3004 | 1396 |
| 3666 | 1639 |
| 4002 | 1719 |
| 11371 | 3548 |
| 5568 | 2123 |
| 5044 | 2015 |
| 4571 | 1861 |
| 4367 | 1796 |
| 4631.5 | 1873 |
| 4800.5 | 1992.5 |
| 4822.5 | 2001 |
| 5155 | 2070 |
| 4821 | 2030 |
| 4620 | 1863 |
| 4699 | 1901.5 |
| 4542 | 1852 |
| 4750 | 1882 |
| 4373 | 1807 |
| 4603.5 | 1868 |
| 5125 | 2051 |
| 8528 | 2560 |
| 9564.5 | 2716.5 |
| 10041.5 | 2568 |

| | |
|---|---|
| 7756 | 2337 |
| 2742.5 | 1322 |
| 3387 | 1527 |
| 4016.5 | 1744.5 |
| 10259 | 3306.5 |
| 5226 | 2043 |
| 4788 | 1923.5 |
| 4808 | 1974 |
| 4336 | 1809 |
| 4492 | 1816 |
| 4943 | 2041.5 |
| 4752 | 1911.5 |
| 4845 | 1977 |
| 4924.5 | 2001.5 |
| 4998 | 1973 |
| 4554.5 | 1895.5 |
| 4620.5 | 1897 |
| 4998 | 1909 |
| 4839 | 1902 |
| 4752 | 1922 |
| 4667 | 1874 |
| 7139.5 | 2337 |
| 10669 | 2769 |
| 9854.5 | 2600 |
| 8965 | 2462 |
| 2951 | 1413 |
| 3362.5 | 1522.5 |
| 3756.5 | 1655 |
| 10278 | 3384.5 |
| 5593.5 | 2133.5 |
| 5013 | 1958 |
| 4808 | 1943 |
| 4380 | 1799 |
| 4672 | 1888 |
| 5080 | 2097 |
| 4974.5 | 2018.5 |
| 4966 | 2015 |
| 4719.5 | 1980 |
| 4939 | 1926 |
| 4770 | 1963 |
| 4570 | 1906.5 |
| 4842 | 1872 |
| 4642 | 1854 |
| 4728.5 | 1875.5 |
| 4915 | 1962 |
| 7644 | 2385 |
| 9737 | 2684 |
| 8522 | 2476 |

| | |
|---|---|
| 7583.5 | 2274 |
| 2899.5 | 1366 |
| 3477 | 1572 |
| 4112 | 1762 |
| 11653 | 3642 |
| 5713.5 | 2132.5 |
| 5049 | 1960 |
| 4874 | 1960 |
| 4488 | 1847 |
| 4607 | 1884.5 |
| 5260 | 2168 |
| 5215 | 2074.5 |
| 4858 | 1959 |
| 5235.5 | 2025.5 |
| 5093 | 1998 |
| 4857 | 1965 |
| 4751.5 | 1889.5 |
| 5732 | 2127 |
| 4853.5 | 1934 |
| 4728 | 1898 |
| 4684 | 1929 |
| 7687.5 | 2322 |
| 10200.5 | 2683 |
| 7878 | 2339 |
| 9027 | 2434 |
| 2908 | 1379 |
| 3392 | 1530 |
| 3974 | 1672 |
| 10914 | 3486 |
| 6132 | 2219 |
| 4938 | 1911.5 |
| 5187 | 2035 |
| 4500 | 1796 |
| 4624.5 | 1863.5 |
| 5230.5 | 2165 |
| 5340 | 2139 |
| 4871 | 1961.5 |
| 4578 | 1926 |
| 4535 | 1795 |
| 4569.5 | 1888.5 |
| 4602 | 1823 |
| 5131.5 | 1924.5 |
| 5161 | 1963 |
| 5058 | 2005 |
| 5212 | 2056 |
| 8048.5 | 2499 |
| 11012 | 2758 |
| 10170 | 2565 |

| | |
|---|---|
| 8442 | 2465 |
| 3044 | 1408 |
| 3584.5 | 1591 |
| 3999 | 1702 |
| 10616 | 3465.5 |
| 5571.5 | 2118 |
| 5193 | 2053.5 |
| 5304 | 2078 |
| 4712 | 1877.5 |
| 4661 | 1882 |
| 5000 | 2015 |
| 4911.5 | 1994 |
| 5108 | 2066.5 |
| 5957.5 | 2343 |
| 5455 | 1938 |
| 4708 | 1929 |
| 4905 | 1931 |
| 5194 | 1930 |
| 5193 | 2015 |
| 4910 | 1944 |
| 4864 | 1963 |
| 7716.5 | 2412.5 |
| 11488.5 | 2939 |
| 8115 | 2430 |
| 8154 | 2338 |
| 3059 | 1409 |
| 3716.5 | 1572.5 |
| 4081 | 1741 |
| 10789 | 3517 |
| 5376 | 2039 |
| 5123 | 2004 |
| 5024 | 1974 |
| 4308 | 1797 |
| 4832 | 1902 |
| 5200 | 2099 |
| 5253 | 2066 |
| 5205 | 2008 |
| 5068.5 | 2039.5 |
| 4688 | 1862 |
| 4254 | 1760 |
| 4796 | 1896 |
| 6650 | 2365.5 |
| 5762 | 2190 |
| 5371.5 | 2035.5 |
| 4634 | 1834 |
| 6983 | 2211.5 |
| 9071.5 | 2587.5 |
| 9231 | 2581 |

| | |
|---|---|
| 9111.5 | 2531.5 |
| 1833 | 1180 |
| 2732 | 1393 |
| 3383.5 | 1588 |
| 7652 | 2893 |
| 4347 | 1821 |
| 4282 | 1842 |
| 4304.5 | 1846 |
| 3666 | 1685 |
| 3923 | 1725 |
| 4106 | 1925 |
| 4086.5 | 1799.5 |
| 3940 | 1790.5 |
| 4487 | 1914 |
| 6068 | 2320 |
| 4696.5 | 1991.5 |
| 4691 | 1968 |
| 4629 | 1895 |
| 4582.5 | 1903.5 |
| 4765 | 1967.5 |
| 4321.5 | 1831 |
| 8486 | 2648 |
| 9000.5 | 2556 |
| 7973 | 2406 |
| 7428 | 2204 |
| 2952.5 | 1384 |
| 3636.5 | 1568.5 |
| 3932 | 1733 |
| 12122 | 3544 |
| 5219 | 2052 |
| 4847 | 1951 |
| 4930 | 2025.5 |
| 4137 | 1792.5 |
| 4382 | 1823 |
| 4470 | 1900 |
| 4623.5 | 1936 |
| 4304 | 1881 |
| 4685.5 | 1991 |
| 5770 | 2234 |
| 4876 | 2060 |
| 5401 | 2183 |
| 4887 | 1898 |
| 5042 | 1998 |
| 4823 | 1888.5 |
| 4724 | 1919 |
| 8016.5 | 2490.5 |
| 10115.5 | 2847 |
| 8933 | 2462.5 |

| | |
|---|---|
| 7465 | 2257 |
| 3058.5 | 1400.5 |
| 3908 | 1685 |
| 4196 | 1798 |
| 15123 | 4336 |
| 6416 | 2377 |
| 5269 | 2082 |
| 4528 | 1852 |
| 5428 | 2147 |
| 5077 | 2013.5 |
| 4558 | 2033 |
| 4319 | 1873 |
| 4576 | 1941 |
| 5193.5 | 2128.5 |
| 5198.5 | 2026 |
| 4934 | 2006 |
| 4406 | 1849 |
| 4829 | 1921 |
| 4642 | 1863.5 |
| 4838 | 1964 |
| 4597 | 1890 |
| 7228 | 2309 |
| 8069.5 | 2442.5 |
| 8474 | 2435 |
| 7254 | 2286.5 |
| 2916 | 1349 |
| 3601 | 1592 |
| 4350.5 | 1789.5 |
| 12635.5 | 3746 |
| 5488.5 | 2143.5 |
| 4995 | 2005 |
| 5394.5 | 2095 |
| 4709 | 1952 |
| 4504 | 1834 |
| 4476.5 | 1933 |
| 4407 | 1874 |
| 4325 | 1845 |
| 4822 | 2043 |
| 5838 | 2204 |
| 5521.5 | 2212 |
| 5831 | 2247 |
| 5484 | 2075 |
| 4772.5 | 1915 |
| 4838.5 | 1960 |
| 4671.5 | 1953.5 |
| 6568 | 2285 |
| 9929 | 2835.5 |
| 8294 | 2433 |

| | |
|---|---|
| 7947.5 | 2312 |
| 3060 | 1410 |
| 3469 | 1599 |
| 4154 | 1745.5 |
| 11468 | 3554 |
| 5473 | 2100.5 |
| 4984 | 1992 |
| 5236 | 2079 |
| 4457 | 1836 |
| 4531 | 1822 |
| 4882.5 | 2026.5 |
| 4921 | 2057 |
| 5289.5 | 2064.5 |
| 5196.5 | 2148.5 |
| 5057 | 1935.5 |
| 5108 | 2083 |
| 5286 | 2069 |
| 5218.5 | 2008.5 |
| 5009 | 1937 |
| 4625.5 | 1903.5 |
| 4729 | 1978 |
| 7137 | 2343 |
| 14958 | 3491 |
| 7871 | 2314 |
| 7251.5 | 2213.5 |
| 3065.5 | 1401.5 |
| 3533.5 | 1575 |
| 4231.5 | 1772 |
| 11854.5 | 3566 |
| 5425 | 2120.5 |
| 5142 | 1989 |
| 5111 | 2046 |
| 4389.5 | 1771.5 |
| 4417 | 1834 |
| 5001.5 | 2080 |
| 4700.5 | 1921.5 |
| 4890 | 1984.5 |
| 4853.5 | 2007 |
| 4813.5 | 1886 |
| 4936 | 2032 |
| 4516 | 1816.5 |
| 4671.5 | 1861 |
| 4913 | 1938 |
| 4544 | 1839 |
| 4801 | 1941 |
| 6822.5 | 2216 |
| 8931 | 2553 |
| 8161 | 2359 |

| | |
|---|---|
| 7059.5 | 2147.5 |
| 2962 | 1379 |
| 3372 | 1508 |
| 4146.5 | 1763.5 |
| 11931 | 3643 |
| 5440 | 2054 |
| 5266 | 2094 |
| 5349 | 2056 |
| 4368 | 1808 |
| 4762 | 1931 |
| 5101 | 2097.5 |
| 4892 | 1955 |
| 5100.5 | 2044.5 |
| 5277 | 2162.5 |
| 5051 | 1986.5 |
| 4881.5 | 1946 |
| 5288 | 2101 |
| 5019 | 1953 |
| 4825.5 | 1903.5 |
| 4370 | 1839 |
| 4729 | 1937 |
| 7657 | 2430 |
| 10324 | 2728.5 |
| 8129 | 2360 |
| 7545 | 2315.5 |
| 2954 | 1380.5 |
| 3529.5 | 1561 |
| 4311.5 | 1764.5 |
| 10346.5 | 3338 |
| 5577 | 2142 |
| 5360 | 2151 |
| 5278.5 | 2107 |
| 4302 | 1798 |
| 4666.5 | 1874.5 |
| 4903 | 2075 |
| 5165.5 | 2088.5 |
| 5175 | 2097.5 |
| 5008 | 2053.5 |
| 4710 | 1857 |
| 4947 | 2003 |
| 5097 | 2004 |
| 4876 | 1834.5 |
| 5160 | 1991.5 |
| 4966 | 1943.5 |
| 4742 | 1919 |
| 7074 | 2315 |
| 9937 | 2730 |
| 8535 | 2425 |

| | |
|---|---|
| 6911 | 2160 |
| 3084 | 1440 |
| 3756 | 1612 |
| 4332 | 1790 |
| 10201 | 3373 |
| 5775 | 2174.5 |
| 5521 | 2123 |
| 5397 | 2074 |
| 4572 | 1853 |
| 4777 | 1886 |
| 4951 | 2056 |
| 5171.5 | 2080.5 |
| 5210 | 2061 |
| 4688 | 1939 |
| 4585 | 1817.5 |
| 4583 | 1856 |
| 4492.5 | 1824.5 |
| 5162 | 1934 |
| 5105 | 1969 |
| 5277 | 2041 |
| 5016 | 2005 |
| 7441 | 2384 |
| 9529 | 2649.5 |
| 8109 | 2333 |
| 7447.5 | 2219 |
| 3128 | 1435 |
| 3793.5 | 1681.5 |
| 4351 | 1818.5 |
| 12243 | 3750 |
| 5930 | 2235 |
| 5655 | 2133 |
| 5447.5 | 2108.5 |
| 4650 | 1853 |
| 4773 | 1907 |
| 5151 | 2124.5 |
| 5106 | 1991 |
| 5394 | 2115.5 |
| 5004 | 2047 |
| 5476 | 1920 |
| 4460 | 1788 |
| 4793 | 1911 |
| 5001 | 1937.5 |
| 5117 | 2071.5 |
| 5103 | 2038 |
| 4997 | 2002.5 |
| 8159 | 2505 |
| 10396 | 2713 |
| 10548 | 2683 |

| | |
|---|---|
| 8402 | 2486 |
| 3001 | 1412 |
| 3464.5 | 1566 |
| 4144.5 | 1753 |
| 10748.5 | 3440 |
| 5421 | 2091 |
| 5124 | 2018 |
| 4910 | 1992 |
| 4421.5 | 1818 |
| 4772 | 1884 |
| 5018 | 2055 |
| 5218 | 2033 |
| 5133 | 2002 |
| 4766 | 1973 |
| 5081 | 2025.5 |
| 4803 | 1935 |
| 4974 | 2011 |
| 6006 | 2283 |
| 5649 | 2123 |
| 4991 | 1970 |
| 5199.5 | 2034.5 |
| 7129 | 2280 |
| 9245 | 2624 |
| 8654 | 2496 |
| 8763 | 2514 |
| 1835 | 1190 |
| 2723 | 1394 |
| 3341 | 1545 |
| 8429 | 3173 |
| 4442.5 | 1928 |
| 4348 | 1877 |
| 4386 | 1898 |
| 3902 | 1711 |
| 4133 | 1768 |
| 4491 | 1903 |
| 4579 | 1926 |
| 4816 | 1989 |
| 4778.5 | 1974.5 |
| 4937 | 1973 |
| 4079.5 | 1781 |
| 4170 | 1753 |
| 4552 | 1867 |
| 4777 | 1941 |
| 4353 | 1797 |
| 4325.5 | 1841 |
| 6921 | 2240 |
| 8882 | 2521 |
| 8073 | 2426 |

| | |
|---|---|
| 7007 | 2139 |
| 2662 | 1298 |
| 3372 | 1538 |
| 4249 | 1782 |
| 9935 | 3078 |
| 5233 | 2047 |
| 5141 | 2071 |
| 5257 | 2112 |
| 4320.5 | 1843 |
| 4451.5 | 1861 |
| 4257 | 1851 |
| 4532 | 1886 |
| 4582 | 1934 |
| 4856 | 1972 |
| 5291 | 2038 |
| 5375 | 2149.5 |
| 5220 | 2114 |
| 5524 | 2131 |
| 5130 | 2002 |
| 5021 | 2032 |
| 4949.5 | 1983.5 |
| 9217.5 | 2671 |
| 11402 | 2952 |
| 8223 | 2398.5 |
| 6950 | 2099 |
| 3102 | 1382.5 |
| 3520.5 | 1526.5 |
| 4127 | 1768 |
| 15158 | 4031 |
| 5662 | 2174 |
| 5219.5 | 2083.5 |
| 4902.5 | 1964 |
| 5184 | 2027.5 |
| 4635 | 1884 |
| 4192.5 | 1754.5 |
| 4332.5 | 1828.5 |
| 4707.5 | 1916.5 |
| 4714.5 | 1916.5 |
| 5516 | 2084.5 |
| 5015 | 2005 |
| 5393 | 2119 |
| 4898.5 | 1878 |
| 4604.5 | 1842 |
| 5059 | 1994 |
| 4554 | 1880 |
| 7050 | 2282 |
| 8523 | 2569 |
| 7933 | 2346 |

| | |
|---|---|
| 7471 | 2280 |
| 2858 | 1342 |
| 3442.5 | 1528.5 |
| 4320 | 1759 |
| 10429 | 3184 |
| 5302 | 2031 |
| 5252.5 | 2087 |
| 5408.5 | 2091 |
| 4298 | 1831 |
| 4485 | 1806 |
| 4268 | 1886 |
| 4198 | 1860 |
| 4531 | 1924.5 |
| 5050.5 | 2101 |
| 5965 | 2207 |
| 5276.5 | 2143.5 |
| 5958.5 | 2241 |
| 5486.5 | 2092 |
| 5441.5 | 2159 |
| 5159 | 2110 |
| 4913 | 1979.5 |
| 8471.5 | 2516 |
| 11569.5 | 3026.5 |
| 7496 | 2319 |
| 7321 | 2192 |
| 2932 | 1336 |
| 3722 | 1580.5 |
| 4388 | 1819 |
| 11483.5 | 3323 |
| 6022 | 2217 |
| 5932.5 | 2231 |
| 5362 | 2053 |
| 4618 | 1857 |
| 4487 | 1858 |
| 4734.5 | 2007.5 |
| 4762 | 1929 |
| 4962 | 2018 |
| 5245 | 2097 |
| 5200.5 | 1999.5 |
| 5071.5 | 2098 |
| 5014 | 1984.5 |
| 4973 | 1979 |
| 4903 | 1931 |
| 4670 | 1908 |
| 4849 | 1992 |
| 6985 | 2287 |
| 9724 | 2659.5 |
| 8244.5 | 2388 |

| | |
|---|---|
| 8036 | 2357 |
| 3033 | 1356 |
| 3450 | 1520 |
| 4104 | 1748 |
| 11236.5 | 3371 |
| 5408.5 | 2086 |
| 4927 | 1939 |
| 4773 | 1970 |
| 4480.5 | 1831 |
| 4729 | 1888 |
| 4628 | 1930 |
| 4782 | 1952.5 |
| 4936.5 | 1987 |
| 4895 | 1980 |
| 4627 | 1830 |
| 4587 | 1913 |
| 4639 | 1865 |
| 4767 | 1858 |
| 4663 | 1895 |
| 4906 | 1947 |
| 4784.5 | 1944 |
| 7246 | 2299 |
| 9383 | 2554 |
| 7873 | 2390.5 |
| 7175 | 2203 |
| 3022 | 1382.5 |
| 3541.5 | 1572 |
| 4035.5 | 1708 |
| 11565.5 | 3447.5 |
| 5985.5 | 2210.5 |
| 5593.5 | 2202 |
| 5270 | 2126 |
| 4487 | 1842.5 |
| 4532 | 1855 |
| 4878.5 | 2056.5 |
| 4869 | 1971 |
| 5196.5 | 2104 |
| 5407 | 2138 |
| 4994 | 1933 |
| 4505 | 1854 |
| 4777.5 | 1910.5 |
| 4910.5 | 1886.5 |
| 4814.5 | 1945 |
| 4775 | 1935 |
| 4732.5 | 1897 |
| 7658 | 2336 |
| 8718 | 2599 |
| 8406 | 2420.5 |

| | |
|---|---|
| 7183 | 2181 |
| 3104 | 1404 |
| 3778 | 1616 |
| 4601 | 1857 |
| 12310 | 3683 |
| 5915.5 | 2179 |
| 5760.5 | 2189.5 |
| 5399.5 | 2145 |
| 4578 | 1857 |
| 4458 | 1809 |
| 4750 | 1983 |
| 4927.5 | 1956.5 |
| 5199 | 2085 |
| 5099.5 | 2035 |
| 4931 | 1922.5 |
| 4550 | 1921.5 |
| 4983.5 | 1963 |
| 5172 | 1961 |
| 5039 | 2034 |
| 4824 | 1923 |
| 4656 | 1881.5 |
| 7352 | 2375 |
| 9348.5 | 2578.5 |
| 8278 | 2448 |
| 7427 | 2224 |
| 3022 | 1396 |
| 3660.5 | 1552 |
| 4199 | 1783 |
| 9624 | 3155 |
| 5816.5 | 2154 |
| 5413 | 2068 |
| 5098 | 2011 |
| 4547 | 1815 |
| 4342 | 1764 |
| 4935 | 2011 |
| 4801 | 1979 |
| 5035 | 2025 |
| 5080.5 | 2020.5 |
| 5168 | 2029 |
| 5374 | 2120 |
| 5091.5 | 2020.5 |
| 5489 | 2021 |
| 5086 | 1968 |
| 4989 | 1946 |
| 4713 | 1909 |
| 7877 | 2457 |
| 9236 | 2607 |
| 7673.5 | 2266.5 |

| | |
|---|---|
| 7293 | 2237.5 |
| 3455 | 1446 |
| 4032 | 1706 |
| 4783 | 1870 |
| 11707 | 3513 |
| 5963.5 | 2204.5 |
| 5899 | 2295.5 |
| 5729 | 2219 |
| 4902 | 1929 |
| 4598 | 1845.5 |
| 4745 | 1969.5 |
| 4838.5 | 1988 |
| 5264 | 2079.5 |
| 5496 | 2189 |
| 5942 | 2086 |
| 5246 | 2018 |
| 5212.5 | 2090 |
| 5523 | 2098 |
| 5235 | 2029 |
| 4675 | 1906.5 |
| 4848 | 1947.5 |
| 7633 | 2424 |
| 10055.5 | 2788.5 |
| 8454 | 2418 |
| 7714 | 2379.5 |
| 3112 | 1422 |
| 3748 | 1582 |
| 4745 | 1874 |
| 11174 | 3470 |
| 5568 | 2127 |
| 5534.5 | 2148 |
| 5553 | 2145 |
| 4323 | 1782.5 |
| 4772 | 1901 |
| 5549 | 2162.5 |
| 5197 | 2021 |
| 5182 | 2044 |
| 4961 | 1990 |
| 4706.5 | 1897 |
| 4789 | 1900.5 |
| 5167.5 | 1970 |
| 5447 | 2017 |
| 5616 | 2105 |
| 4745.5 | 1935 |
| 4622 | 1869.5 |
| 6765 | 2253 |
| 10139 | 2798 |
| 9967 | 2670.5 |

| | |
|---|---|
| 9576 | 2588 |
| 1743 | 1143 |
| 2742 | 1374.5 |
| 3327 | 1577 |
| 7779 | 2928 |
| 4278 | 1869 |
| 4191 | 1822 |
| 4313 | 1867 |
| 3907 | 1737 |
| 4231.5 | 1778 |
| 3988.5 | 1719 |
| 4052 | 1773 |
| 4308 | 1872 |
| 4503 | 1899 |
| 4539 | 1876 |
| 4474.5 | 1874 |
| 4867.5 | 2012 |
| 4697 | 1914 |
| 4425 | 1782 |
| 4230 | 1770 |
| 4192 | 1819 |
| 6444 | 2177 |
| 9003 | 2520 |
| 8003.5 | 2399 |
| 7362 | 2223 |
| 2815 | 1326 |
| 3408 | 1549 |
| 3898 | 1707 |
| 10389.5 | 3115 |
| 5341 | 2067 |
| 4990 | 2002 |
| 4826 | 1970 |
| 4380.5 | 1809.5 |
| 4236.5 | 1755 |
| 4235 | 1832 |
| 4336 | 1864 |
| 4460 | 1928 |
| 4922 | 2059 |
| 4945.5 | 1974 |
| 4761 | 2008.5 |
| 5122 | 1998.5 |
| 5163 | 2006 |
| 4664 | 1915 |
| 4825 | 1911 |
| 4359 | 1843.5 |
| 6600.5 | 2145.5 |
| 8957 | 2583.5 |
| 8005 | 2345 |

| | |
|---|---|
| 6921 | 2117 |
| 3147 | 1418 |
| 3451 | 1536 |
| 4264.5 | 1779 |
| 9388 | 2954 |
| 5292 | 2071 |
| 4635 | 1873 |
| 5034.5 | 1977 |
| 5223 | 2059 |
| 4729 | 1881 |
| 4505.5 | 1977.5 |
| 4280 | 1866 |
| 4217 | 1811.5 |
| 5016 | 2035 |
| 5194.5 | 2047.5 |
| 4722.5 | 1940 |
| 4958 | 1978.5 |
| 4770 | 1877.5 |
| 4577 | 1831 |
| 4922.5 | 1972 |
| 4272 | 1800 |
| 6848.5 | 2279.5 |
| 8498.5 | 2458 |
| 8204 | 2400 |
| 7552.5 | 2310 |
| 2936 | 1354 |
| 3588 | 1561 |
| 4219 | 1779 |
| 11125.5 | 3310.5 |
| 5541 | 2109 |
| 5257 | 2048 |
| 5257 | 2076.5 |
| 4480 | 1856 |
| 4495.5 | 1858.5 |
| 4388 | 1889 |
| 4192 | 1811 |
| 4325.5 | 1839.5 |
| 5144 | 2079 |
| 5629 | 2154 |
| 4803 | 1973 |
| 5474 | 2175 |
| 5689 | 2129 |
| 5238 | 2019 |
| 4761.5 | 1915.5 |
| 4930 | 1971.5 |
| 7285 | 2286.5 |
| 10881 | 2882 |
| 7894 | 2325 |

| | |
|---|---|
| 7044 | 2152 |
| 3237 | 1429 |
| 3916 | 1677 |
| 4509 | 1870 |
| 10581 | 3298 |
| 5604.5 | 2172 |
| 5518 | 2170 |
| 5178.5 | 2040.5 |
| 4603.5 | 1879 |
| 4520 | 1821 |
| 4795 | 2029.5 |
| 4739 | 1954 |
| 5062.5 | 2028 |
| 4774 | 1981 |
| 4763 | 1874 |
| 4685 | 1900.5 |
| 4926.5 | 1975 |
| 4910 | 1950 |
| 4797 | 1948 |
| 4662 | 1918.5 |
| 4643 | 1873.5 |
| 7561.5 | 2377.5 |
| 9659 | 2601 |
| 7623 | 2296 |
| 7773 | 2379 |
| 3069.5 | 1379 |
| 3793 | 1584 |
| 4302.5 | 1780.5 |
| 11103 | 3386.5 |
| 5761.5 | 2184.5 |
| 5306.5 | 2072 |
| 5141 | 2080 |
| 4383 | 1781 |
| 4477 | 1823 |
| 4596.5 | 1941 |
| 4981.5 | 2026 |
| 5081 | 2018 |
| 4993 | 2008 |
| 4532 | 1837 |
| 4578.5 | 1904.5 |
| 4599 | 1869 |
| 4983 | 1898 |
| 4860.5 | 1944 |
| 4443 | 1828 |
| 4458 | 1839 |
| 7097 | 2281 |
| 9961 | 2704 |
| 7878 | 2324.5 |

| | |
|---|---|
| 7672.5 | 2301 |
| 3211 | 1419 |
| 3678.5 | 1569.5 |
| 4305 | 1787 |
| 10511.5 | 3299 |
| 5722 | 2141.5 |
| 5452 | 2145 |
| 5213 | 2069 |
| 4487 | 1841 |
| 4450.5 | 1838.5 |
| 4695 | 2012 |
| 4652.5 | 1907 |
| 5056 | 2060 |
| 4912.5 | 1984 |
| 4891 | 1933 |
| 5355 | 2148 |
| 4984 | 2041 |
| 5178 | 1976 |
| 5040.5 | 1950 |
| 4538 | 1797 |
| 4808 | 1927 |
| 7132 | 2260.5 |
| 9602.5 | 2619.5 |
| 8164 | 2402 |
| 7492.5 | 2254 |
| 3050 | 1366 |
| 3765 | 1575 |
| 4516 | 1841 |
| 12120 | 3682 |
| 6030 | 2249.5 |
| 5509 | 2134 |
| 5553 | 2160 |
| 4695 | 1884 |
| 4336 | 1764 |
| 4770 | 2009 |
| 4844 | 1962 |
| 4894 | 2028.5 |
| 5192 | 2097 |
| 5004 | 1949 |
| 4554 | 1867 |
| 4604 | 1880 |
| 5191 | 1962 |
| 4974.5 | 1964.5 |
| 4737 | 1854 |
| 4962 | 1962 |
| 7303 | 2304 |
| 8934 | 2585.5 |
| 8270.5 | 2360.5 |

| | |
|---|---|
| 7311.5 | 2239 |
| 3168 | 1394 |
| 3952.5 | 1673.5 |
| 4690 | 1919 |
| 9538 | 3098.5 |
| 5939 | 2172 |
| 5629 | 2153 |
| 5706 | 2242 |
| 4685 | 1870 |
| 4654 | 1854 |
| 4639 | 1943 |
| 4568 | 1881 |
| 4691 | 1927.5 |
| 5033 | 2015 |
| 5579 | 2122 |
| 4933 | 1971 |
| 5165.5 | 2058 |
| 5268 | 1960.5 |
| 4845.5 | 1911.5 |
| 5041 | 1991 |
| 4520 | 1857 |
| 7635 | 2387 |
| 9809.5 | 2654 |
| 7933 | 2346 |
| 7287 | 2232 |
| 3324 | 1428 |
| 3899 | 1657.5 |
| 4503 | 1862 |
| 11604 | 3585 |
| 6251 | 2296 |
| 5691 | 2153 |
| 5749 | 2225 |
| 4904 | 1983 |
| 4731 | 1939 |
| 5003 | 2028 |
| 5131 | 2057 |
| 5194.5 | 2092.5 |
| 5203 | 2093 |
| 5421 | 1929 |
| 4536.5 | 1831.5 |
| 4812 | 1930 |
| 5144 | 1960 |
| 5087 | 1961 |
| 4781 | 1923.5 |
| 4924 | 1990 |
| 7471 | 2355.5 |
| 10082 | 2633 |
| 8227 | 2401 |

| | |
|---|---|
| 7633 | 2276 |
| 3033.5 | 1408 |
| 3720 | 1607.5 |
| 4319 | 1787.5 |
| 10556 | 3318 |
| 5965 | 2272 |
| 5428 | 2095 |
| 5511 | 2115 |
| 4816 | 1873 |
| 5356 | 2049 |
| 5513 | 2140 |
| 5283.5 | 2057 |
| 4998.5 | 2018.5 |
| 4789 | 1942 |
| 4564 | 1829 |
| 4725 | 1895 |
| 4974 | 1966 |
| 5826 | 2148 |
| 5853.5 | 2170 |
| 5031 | 1994 |
| 4357.5 | 1790 |
| 6553 | 2133 |
| 9247 | 2592 |
| 9154.5 | 2551 |
| 8659 | 2446.5 |
