## Supplementary table 2 for "Rapid accumulation of fluorophores and fast kill identify drugs with bactericidal effects against Gram-negative bacteria"

| Species | Antibiotic | Fluor | n | Signal.mec | Iq.signal | Events.me | Iq.events |
|---|---|---|---|---|---|---|---|
| A. baylyi | Amoxacilli | 5-Carboxyf | 4 | 2135 | [1039 - 303 | 942 | [756 - 974] |
| A. baylyi | Amoxacilli | Acridine or | 4 | 1 | [1 - 1] | 873 | [678 - 942] |
| A. baylyi | Amoxacilli | Alizarin | 4 | 1 | [1 - 1] | 900 | [726 - 923] |
| A. baylyi | Amoxacilli | Allura Red | 4 | 1 | [1 - 1] | 938 | [788 - 954] |
| A. baylyi | Amoxacilli | Amaranth | 4 | 4571 | [2552 - 206 | 878 | [605 - 1008] |
| A. baylyi | Amoxacilli | Apigenin | 4 | 303 | [1 - 8612] | 988 | [746 - 1060] |
| A. baylyi | Amoxacilli | Calcein | 4 | 3049 | [2033 - 367 | 910 | [748 - 973] |
| A. baylyi | Amoxacilli | Chlorophe | 4 | 1 | [1 - 1] | 978 | [764 - 1024] |
| A. baylyi | Amoxacilli | Congo Red | 4 | 1 | [1 - 1] | 955 | [772 - 1000] |
| A. baylyi | Amoxacilli | DiSC3(5) | 4 | 654208 | [629431 - 1 | 911 | [714 - 941] |
| A. baylyi | Amoxacilli | Eosin Y | 4 | 1794 | [1219 - 270 | 926 | [732 - 990] |
| A. baylyi | Amoxacilli | H2FDA | 4 | 1 | [1 - 1] | 950 | [747 - 978] |
| A. baylyi | Amoxacilli | Malachite ( | 4 | 1 | [1 - 1] | 926 | [753 - 969] |
| A. baylyi | Amoxacilli | Neutral Re | 4 | 1 | [1 - 1] | 971 | [806 - 982] |
| A. baylyi | Amoxacilli | Oxonol | 4 | 1 | [1 - 1] | 2500 | [1959 - 2903] |
| A. baylyi | Amoxacilli | Pyronin Y | 4 | 1 | [1 - 1] | 910 | [734 - 920] |
| A. baylyi | Amoxacilli | Quinacrine | 4 | 2295 | [903 - 1297 | 914 | [718 - 947] |
| A. baylyi | Amoxacilli | Rhodamine | 4 | 1 | [1 - 1] | 928 | [753 - 968] |
| A. baylyi | Amoxacilli | Rhodamine | 4 | 1 | [1 - 1] | 912 | [754 - 970] |
| A. baylyi | Amoxacilli | Riboflavin | 4 | 1 | [1 - 1] | 899 | [714 - 904] |
| A. baylyi | Amoxacilli | Sunset Yell | 4 | 1 | [1 - 1] | 1008 | [776 - 1072] |
| A. baylyi | Amoxacilli | SYBR Gree | 4 | 293838 | [282349 - 3 | 932 | [783 - 946] |
| A. baylyi | Amoxacilli | Thioflavin T | 4 | 7090 | [4531 - 885 | 922 | [785 - 930] |
| A. baylyi | Ampicillin | 5-Carboxyf | 4 | 11422 | [9986 - 120 | 826 | [782 - 875] |
| A. baylyi | Ampicillin | Acridine or | 4 | 1 | [1 - 1] | 812 | [802 - 819] |
| A. baylyi | Ampicillin | Alizarin | 4 | 1 | [1 - 1] | 838 | [814 - 864] |
| A. baylyi | Ampicillin | Allura Red | 4 | 1 | [1 - 1] | 931 | [894 - 982] |
| A. baylyi | Ampicillin | Amaranth | 4 | 12663 | [11832 - 13 | 782 | [736 - 810] |
| A. baylyi | Ampicillin | Apigenin | 4 | 1 | [1 - 1] | 888 | [866 - 892] |
| A. baylyi | Ampicillin | Calcein | 4 | 9541 | [7886 - 112 | 826 | [815 - 835] |
| A. baylyi | Ampicillin | Chlorophe | 4 | 1 | [1 - 1] | 860 | [814 - 892] |
| A. baylyi | Ampicillin | Congo Red | 4 | 1 | [1 - 1] | 878 | [846 - 903] |
| A. baylyi | Ampicillin | DiSC3(5) | 4 | 1024866 | [981479 - 1 | 806 | [794 - 825] |
| A. baylyi | Ampicillin | Eosin Y | 4 | 11693 | [9541 - 129 | 794 | [768 - 820] |
| A. baylyi | Ampicillin | H2FDA | 4 | 1 | [1 - 1] | 788 | [780 - 801] |
| A. baylyi | Ampicillin | Malachite ( | 4 | 1 | [1 - 1] | 847 | [828 - 879] |
| A. baylyi | Ampicillin | Neutral Re | 4 | 1 | [1 - 1] | 906 | [870 - 927] |
| A. baylyi | Ampicillin | Oxonol | 4 | 1 | [1 - 1] | 3239 | [2903 - 3442] |
| A. baylyi | Ampicillin | Pyronin Y | 4 | 1 | [1 - 1] | 758 | [734 - 784] |
| A. baylyi | Ampicillin | Quinacrine | 4 | 580 | [212 - 1378 | 783 | [757 - 822] |
| A. baylyi | Ampicillin | Rhodamine | 4 | 1 | [1 - 1] | 812 | [759 - 851] |
| A. baylyi | Ampicillin | Rhodamine | 4 | 1 | [1 - 1] | 902 | [866 - 938] |
| A. baylyi | Ampicillin | Riboflavin | 4 | 1 | [1 - 1] | 798 | [779 - 805] |
| A. baylyi | Ampicillin | Sunset Yell | 4 | 1 | [1 - 1] | 836 | [833 - 874] |
| A. baylyi | Ampicillin | SYBR Gree | 4 | 604427 | [536823 - 6 | 844 | [837 - 870] |
| A. baylyi | Ampicillin | Thioflavin T | 4 | 3113 | [1582 - 444 | 849 | [796 - 896] |
| A. baylyi | Azithromyc | 5-Carboxyf | 4 | 1794 | [568 - 3128 | 948 | [806 - 967] |

| | | | | | | | |
|---|---|---|---|---|---|---|---|
| A. baylyi | Azithromyc | Acridine or | 4 | 1 | [1 - 1] | 982 | [788 - 1018] |
| A. baylyi | Azithromyc | Alizarin | 4 | 1 | [1 - 1] | 982 | [806 - 1023] |
| A. baylyi | Azithromyc | Allura Red | 4 | 1 | [1 - 1] | 945 | [786 - 972] |
| A. baylyi | Azithromyc | Amaranth | 4 | 3091 | [1386 - 558 | 946 | [737 - 996] |
| A. baylyi | Azithromyc | Apigenin | 4 | 1 | [1 - 1917] | 1047 | [861 - 1064] |
| A. baylyi | Azithromyc | Calcein | 4 | 1598 | [1079 - 222 | 944 | [779 - 993] |
| A. baylyi | Azithromyc | Chlorophe | 4 | 1 | [1 - 1] | 1010 | [809 - 1036] |
| A. baylyi | Azithromyc | Congo Red | 4 | 1 | [1 - 1] | 964 | [811 - 986] |
| A. baylyi | Azithromyc | DiSC3(5) | 4 | 677525 | [630313 - 8 | 858 | [680 - 931] |
| A. baylyi | Azithromyc | Eosin Y | 4 | 3106 | [2000 - 772 | 928 | [722 - 978] |
| A. baylyi | Azithromyc | H2FDA | 4 | 1 | [1 - 1] | 972 | [826 - 983] |
| A. baylyi | Azithromyc | Malachite ( | 4 | 1 | [1 - 1] | 956 | [736 - 1038] |
| A. baylyi | Azithromyc | Neutral Re | 4 | 1 | [1 - 1] | 923 | [758 - 967] |
| A. baylyi | Azithromyc | Oxonol | 4 | 1 | [1 - 1] | 1624 | [1129 - 1990] |
| A. baylyi | Azithromyc | Pyronin Y | 4 | 1 | [1 - 1] | 906 | [679 - 984] |
| A. baylyi | Azithromyc | Quinacrine | 4 | 1 | [1 - 13241] | 995 | [766 - 1043] |
| A. baylyi | Azithromyc | Rhodamine | 4 | 1 | [1 - 1] | 962 | [782 - 1013] |
| A. baylyi | Azithromyc | Rhodamine | 4 | 1 | [1 - 1] | 898 | [736 - 948] |
| A. baylyi | Azithromyc | Riboflavin | 4 | 1 | [1 - 1] | 936 | [757 - 977] |
| A. baylyi | Azithromyc | Sunset Yell | 4 | 1 | [1 - 1] | 937 | [749 - 1008] |
| A. baylyi | Azithromyc | SYBR Gree | 4 | 239267 | [215456 - 2 | 926 | [723 - 979] |
| A. baylyi | Azithromyc | Thioflavin T | 4 | 4483 | [2253 - 129 | 928 | [697 - 1008] |
| A. baylyi | Bacitracin | 5-Carboxyf | 4 | 3974 | [2379 - 547 | 1095 | [1042 - 1106] |
| A. baylyi | Bacitracin | Acridine or | 4 | 1 | [1 - 1] | 1034 | [979 - 1069] |
| A. baylyi | Bacitracin | Alizarin | 4 | 1 | [1 - 1] | 1100 | [1029 - 1175] |
| A. baylyi | Bacitracin | Allura Red | 4 | 1 | [1 - 1] | 1036 | [960 - 1108] |
| A. baylyi | Bacitracin | Amaranth | 4 | 4847 | [3262 - 675 | 1026 | [945 - 1078] |
| A. baylyi | Bacitracin | Apigenin | 4 | 1 | [1 - 1] | 1167 | [1102 - 1184] |
| A. baylyi | Bacitracin | Calcein | 4 | 4110 | [2619 - 565 | 996 | [983 - 1033] |
| A. baylyi | Bacitracin | Chlorophe | 4 | 1 | [1 - 1] | 1113 | [1090 - 1116] |
| A. baylyi | Bacitracin | Congo Red | 4 | 1 | [1 - 1] | 1014 | [995 - 1036] |
| A. baylyi | Bacitracin | DiSC3(5) | 4 | 666953 | [609741 - 7 | 967 | [896 - 1017] |
| A. baylyi | Bacitracin | Eosin Y | 4 | 2190 | [1566 - 294 | 1091 | [1037 - 1147] |
| A. baylyi | Bacitracin | H2FDA | 4 | 1 | [1 - 1] | 1060 | [1024 - 1086] |
| A. baylyi | Bacitracin | Malachite ( | 4 | 1 | [1 - 1] | 1086 | [1040 - 1108] |
| A. baylyi | Bacitracin | Neutral Re | 4 | 1 | [1 - 1] | 1126 | [1061 - 1155] |
| A. baylyi | Bacitracin | Oxonol | 4 | 1 | [1 - 1] | 1573 | [1479 - 1831] |
| A. baylyi | Bacitracin | Pyronin Y | 4 | 1 | [1 - 1] | 992 | [943 - 1039] |
| A. baylyi | Bacitracin | Quinacrine | 4 | 1 | [1 - 1] | 1120 | [1078 - 1168] |
| A. baylyi | Bacitracin | Rhodamine | 4 | 1 | [1 - 1] | 1061 | [1031 - 1093] |
| A. baylyi | Bacitracin | Rhodamine | 4 | 1 | [1 - 1] | 1115 | [1055 - 1171] |
| A. baylyi | Bacitracin | Riboflavin | 4 | 1 | [1 - 1] | 1088 | [1038 - 1126] |
| A. baylyi | Bacitracin | Sunset Yell | 4 | 1 | [1 - 1] | 1105 | [1062 - 1147] |
| A. baylyi | Bacitracin | SYBR Gree | 4 | 594375 | [537100 - 6 | 978 | [952 - 1014] |
| A. baylyi | Bacitracin | Thioflavin T | 4 | 133 | [1 - 339] | 1080 | [1049 - 1099] |
| A. baylyi | Ceftriaxon | 5-Carboxyf | 4 | 6470 | [2781 - 991 | 912 | [879 - 968] |
| A. baylyi | Ceftriaxon | Acridine or | 4 | 1 | [1 - 1] | 886 | [878 - 903] |
| A. baylyi | Ceftriaxon | Alizarin | 4 | 1 | [1 - 1] | 900 | [880 - 924] |

| | | | | | | | |
|---|---|---|---|---|---|---|---|
| A. baylyi | Ceftriaxon | Allura Red | 4 | 1 | [1 - 1] | 1144 | [1087 - 1227] |
| A. baylyi | Ceftriaxon | Amaranth | 4 | 7037 | [5724 - 82 | 874 | [849 - 916] |
| A. baylyi | Ceftriaxon | Apigenin | 4 | 1 | [1 - 1] | 998 | [961 - 1041] |
| A. baylyi | Ceftriaxon | Calcein | 4 | 3937 | [3255 - 48 | 940 | [924 - 971] |
| A. baylyi | Ceftriaxon | Chlorophe | 4 | 1 | [1 - 1] | 1018 | [959 - 1066] |
| A. baylyi | Ceftriaxon | Congo Red | 4 | 1 | [1 - 1] | 919 | [864 - 965] |
| A. baylyi | Ceftriaxon | DiSC3(5) | 4 | 850441 | [791222 - | 858 | [820 - 900] |
| A. baylyi | Ceftriaxon | Eosin Y | 4 | 5685 | [4810 - 67 | 902 | [868 - 944] |
| A. baylyi | Ceftriaxon | H2FDA | 4 | 1 | [1 - 1] | 952 | [891 - 991] |
| A. baylyi | Ceftriaxon | Malachite | 4 | 1 | [1 - 1] | 978 | [949 - 996] |
| A. baylyi | Ceftriaxon | Neutral Re | 4 | 1 | [1 - 1] | 1040 | [1020 - 1053] |
| A. baylyi | Ceftriaxon | Oxonol | 4 | 1 | [1 - 1] | 4314 | [3481 - 5010] |
| A. baylyi | Ceftriaxon | Pyronin Y | 4 | 1 | [1 - 1] | 804 | [782 - 837] |
| A. baylyi | Ceftriaxon | Quinacrine | 4 | 1 | [1 - 1] | 938 | [922 - 954] |
| A. baylyi | Ceftriaxon | Rhodamine | 4 | 1 | [1 - 1] | 911 | [879 - 951] |
| A. baylyi | Ceftriaxon | Rhodamine | 4 | 1 | [1 - 1] | 940 | [884 - 1015] |
| A. baylyi | Ceftriaxon | Riboflavin | 4 | 1 | [1 - 1] | 968 | [948 - 986] |
| A. baylyi | Ceftriaxon | Sunset Yel | 4 | 1 | [1 - 1] | 983 | [934 - 1037] |
| A. baylyi | Ceftriaxon | SYBR Gree | 4 | 493565 | [454952 - | 950 | [927 - 977] |
| A. baylyi | Ceftriaxon | Thioflavin | 4 | 333 | [132 - 764] | 980 | [942 - 1000] |
| A. baylyi | Chloramph | 5-Carboxy | 4 | 3847 | [2600 - 55 | 808 | [687 - 830] |
| A. baylyi | Chloramph | Acridine or | 4 | 1 | [1 - 1] | 809 | [658 - 840] |
| A. baylyi | Chloramph | Alizarin | 4 | 1 | [1 - 1] | 785 | [648 - 815] |
| A. baylyi | Chloramph | Allura Red | 4 | 1 | [1 - 4653] | 824 | [660 - 857] |
| A. baylyi | Chloramph | Amaranth | 4 | 8161 | [5949 - 10 | 799 | [660 - 840] |
| A. baylyi | Chloramph | Apigenin | 4 | 289 | [1 - 13046] | 825 | [611 - 905] |
| A. baylyi | Chloramph | Calcein | 4 | 5758 | [2725 - 81 | 828 | [680 - 857] |
| A. baylyi | Chloramph | Chlorophe | 4 | 1 | [1 - 1] | 840 | [702 - 852] |
| A. baylyi | Chloramph | Congo Red | 4 | 1 | [1 - 1] | 758 | [598 - 816] |
| A. baylyi | Chloramph | DiSC3(5) | 4 | 731169 | [660625 - | 792 | [671 - 834] |
| A. baylyi | Chloramph | Eosin Y | 4 | 7866 | [4533 - 11 | 787 | [663 - 794] |
| A. baylyi | Chloramph | H2FDA | 4 | 1 | [1 - 1] | 860 | [726 - 894] |
| A. baylyi | Chloramph | Malachite | 4 | 1 | [1 - 1] | 839 | [684 - 850] |
| A. baylyi | Chloramph | Neutral Re | 4 | 1 | [1 - 1] | 779 | [648 - 803] |
| A. baylyi | Chloramph | Oxonol | 4 | 1 | [1 - 1] | 1556 | [1089 - 1993] |
| A. baylyi | Chloramph | Pyronin Y | 4 | 1 | [1 - 1] | 792 | [611 - 870] |
| A. baylyi | Chloramph | Quinacrine | 4 | 1654 | [1058 - 87 | 858 | [718 - 872] |
| A. baylyi | Chloramph | Rhodamine | 4 | 1 | [1 - 1] | 808 | [656 - 867] |
| A. baylyi | Chloramph | Rhodamine | 4 | 1 | [1 - 1] | 719 | [602 - 770] |
| A. baylyi | Chloramph | Riboflavin | 4 | 1 | [1 - 1] | 836 | [678 - 897] |
| A. baylyi | Chloramph | Sunset Yel | 4 | 1 | [1 - 1] | 838 | [672 - 885] |
| A. baylyi | Chloramph | SYBR Gree | 4 | 235813 | [179238 - | 868 | [727 - 896] |
| A. baylyi | Chloramph | Thioflavin | 4 | 5287 | [3433 - 11 | 870 | [729 - 877] |
| A. baylyi | Ciprofloxa | 5-Carboxy | 4 | 10578 | [7890 - 12 | 848 | [825 - 870] |
| A. baylyi | Ciprofloxa | Acridine or | 4 | 1 | [1 - 1] | 830 | [809 - 850] |
| A. baylyi | Ciprofloxa | Alizarin | 4 | 1 | [1 - 1] | 844 | [822 - 885] |
| A. baylyi | Ciprofloxa | Allura Red | 4 | 1 | [1 - 1] | 884 | [863 - 912] |
| A. baylyi | Ciprofloxa | Amaranth | 4 | 9434 | [8301 - 10 | 879 | [860 - 886] |

| | | | | | | | |
|---|---|---|---|---|---|---|---|
| A. baylyi | Ciprofloxa | Apigenin | 4 | 1 | [1 - 1] | 906 | [893 - 928] |
| A. baylyi | Ciprofloxa | Calcein | 4 | 9333 | [6962 - 11 | 859 | [836 - 887] |
| A. baylyi | Ciprofloxa | Chlorophe | 4 | 1 | [1 - 1] | 870 | [842 - 897] |
| A. baylyi | Ciprofloxa | Congo Red | 4 | 1 | [1 - 1] | 847 | [832 - 855] |
| A. baylyi | Ciprofloxa | DiSC3(5) | 4 | 802184 | [765251 - | 812 | [802 - 836] |
| A. baylyi | Ciprofloxa | Eosin Y | 4 | 11667 | [9632 - 12 | 808 | [775 - 855] |
| A. baylyi | Ciprofloxa | H2FDA | 4 | 1 | [1 - 1] | 899 | [874 - 925] |
| A. baylyi | Ciprofloxa | Malachite | 4 | 1 | [1 - 1] | 850 | [808 - 905] |
| A. baylyi | Ciprofloxa | Neutral Re | 4 | 1 | [1 - 1] | 841 | [829 - 869] |
| A. baylyi | Ciprofloxa | Oxonol | 4 | 1 | [1 - 1] | 1796 | [1354 - 2265] |
| A. baylyi | Ciprofloxa | Pyronin Y | 4 | 1 | [1 - 1] | 870 | [838 - 898] |
| A. baylyi | Ciprofloxa | Quinacrine | 4 | 1529 | [1071 - 17 | 834 | [818 - 862] |
| A. baylyi | Ciprofloxa | Rhodamine | 4 | 1 | [1 - 1] | 851 | [843 - 858] |
| A. baylyi | Ciprofloxa | Rhodamine | 4 | 1 | [1 - 1] | 891 | [876 - 912] |
| A. baylyi | Ciprofloxa | Riboflavin | 4 | 1 | [1 - 1] | 836 | [794 - 880] |
| A. baylyi | Ciprofloxa | Sunset Yel | 4 | 1 | [1 - 1] | 874 | [853 - 894] |
| A. baylyi | Ciprofloxa | SYBR Gree | 4 | 414881 | [385895 - | 841 | [829 - 854] |
| A. baylyi | Ciprofloxa | Thioflavin T | 4 | 4807 | [3757 - 51 | 830 | [820 - 860] |
| A. baylyi | Colistin | 5-Carboxyf | 4 | 50856 | [38065 - 8 | 590 | [501 - 631] |
| A. baylyi | Colistin | Acridine or | 4 | 112713 | [90861 - 1 | 636 | [520 - 658] |
| A. baylyi | Colistin | Alizarin | 4 | 53810 | [48598 - 6 | 624 | [510 - 642] |
| A. baylyi | Colistin | Allura Red | 4 | 1050 | [558 - 978 | 674 | [591 - 695] |
| A. baylyi | Colistin | Amaranth | 4 | 44492 | [38481 - 5 | 612 | [502 - 628] |
| A. baylyi | Colistin | Apigenin | 4 | 6617 | [4380 - 15 | 660 | [543 - 709] |
| A. baylyi | Colistin | Calcein | 4 | 45275 | [36544 - 8 | 613 | [503 - 623] |
| A. baylyi | Colistin | Chlorophe | 4 | 1 | [1 - 1] | 590 | [453 - 664] |
| A. baylyi | Colistin | Congo Red | 4 | 1 | [1 - 1] | 641 | [511 - 656] |
| A. baylyi | Colistin | DiSC3(5) | 4 | 284948 | [276081 - | 507 | [426 - 516] |
| A. baylyi | Colistin | Eosin Y | 4 | 41950 | [38270 - 7 | 607 | [484 - 625] |
| A. baylyi | Colistin | H2FDA | 4 | 96047 | [82801 - 1 | 624 | [508 - 646] |
| A. baylyi | Colistin | Malachite | 4 | 138810 | [124853 - | 631 | [484 - 685] |
| A. baylyi | Colistin | Neutral Re | 4 | 55597 | [51524 - 6 | 657 | [544 - 680] |
| A. baylyi | Colistin | Oxonol | 4 | 1 | [1 - 1] | 2948 | [2206 - 3326] |
| A. baylyi | Colistin | Pyronin Y | 4 | 82919 | [73773 - 9 | 597 | [464 - 641] |
| A. baylyi | Colistin | Quinacrine | 4 | 11019 | [7600 - 33 | 624 | [500 - 646] |
| A. baylyi | Colistin | Rhodamine | 4 | 235983 | [223304 - | 575 | [466 - 590] |
| A. baylyi | Colistin | Rhodamine | 4 | 66438 | [58231 - 7 | 596 | [498 - 634] |
| A. baylyi | Colistin | Riboflavin | 4 | 56587 | [45146 - 6 | 586 | [466 - 623] |
| A. baylyi | Colistin | Sunset Yel | 4 | 1 | [1 - 629] | 674 | [550 - 692] |
| A. baylyi | Colistin | SYBR Gree | 4 | 2348518 | [2084069 - | 632 | [498 - 663] |
| A. baylyi | Colistin | Thioflavin T | 4 | 14601 | [12426 - 2 | 662 | [524 - 719] |
| A. baylyi | Fosfomyci | 5-Carboxyf | 4 | 109 | [1 - 784] | 1168 | [1113 - 1183] |
| A. baylyi | Fosfomyci | Acridine or | 4 | 1 | [1 - 1] | 1132 | [1097 - 1163] |
| A. baylyi | Fosfomyci | Alizarin | 4 | 1 | [1 - 1] | 1080 | [1033 - 1129] |
| A. baylyi | Fosfomyci | Allura Red | 4 | 1 | [1 - 1] | 1160 | [1124 - 1191] |
| A. baylyi | Fosfomyci | Amaranth | 4 | 1201 | [861 - 137 | 1094 | [1067 - 1108] |
| A. baylyi | Fosfomyci | Apigenin | 4 | 1 | [1 - 1] | 1121 | [1112 - 1141] |
| A. baylyi | Fosfomyci | Calcein | 4 | 1 | [1 - 156] | 1158 | [1118 - 1197] |

| | | | | | | | |
|---|---|---|---|---|---|---|---|
| A. baylyi | Fosfomycii | Chlorophe | 4 | 1 | [1 - 1] | 1138 | [1080 - 1194] |
| A. baylyi | Fosfomycii | Congo Red | 4 | 1 | [1 - 1] | 1142 | [1103 - 1160] |
| A. baylyi | Fosfomycii | DiSC3(5) | 4 | 645680 | [617749 - 6 | 950 | [930 - 967] |
| A. baylyi | Fosfomycii | Eosin Y | 4 | 300 | [1 - 1516] | 1070 | [1021 - 1097] |
| A. baylyi | Fosfomycii | H2FDA | 4 | 1 | [1 - 1] | 1060 | [1054 - 1092] |
| A. baylyi | Fosfomycii | Malachite ( | 4 | 1 | [1 - 1] | 1099 | [1060 - 1110] |
| A. baylyi | Fosfomycii | Neutral Re | 4 | 1 | [1 - 1] | 1131 | [1113 - 1144] |
| A. baylyi | Fosfomycii | Oxonol | 4 | 1 | [1 - 1] | 2884 | [2326 - 3362] |
| A. baylyi | Fosfomycii | Pyronin Y | 4 | 1 | [1 - 1] | 998 | [970 - 1026] |
| A. baylyi | Fosfomycii | Quinacrine | 4 | 1 | [1 - 1] | 1074 | [1032 - 1110] |
| A. baylyi | Fosfomycii | Rhodamine | 4 | 1 | [1 - 1] | 1087 | [1070 - 1096] |
| A. baylyi | Fosfomycii | Rhodamine | 4 | 1 | [1 - 1] | 1068 | [1051 - 1118] |
| A. baylyi | Fosfomycii | Riboflavin | 4 | 1 | [1 - 1] | 1105 | [1044 - 1142] |
| A. baylyi | Fosfomycii | Sunset Yell | 4 | 1 | [1 - 1] | 1104 | [1047 - 1162] |
| A. baylyi | Fosfomycii | SYBR Gree | 4 | 351176 | [333895 - 3 | 1064 | [1015 - 1110] |
| A. baylyi | Fosfomycii | Thioflavin T | 4 | 1 | [1 - 1] | 1076 | [1066 - 1112] |
| A. baylyi | Gentamicii | 5-Carboxyf | 4 | 701 | [508 - 1452 | 918 | [764 - 947] |
| A. baylyi | Gentamicii | Acridine or | 4 | 1 | [1 - 1] | 875 | [716 - 924] |
| A. baylyi | Gentamicii | Alizarin | 4 | 1 | [1 - 1] | 924 | [751 - 942] |
| A. baylyi | Gentamicii | Allura Red | 4 | 1 | [1 - 1076] | 938 | [782 - 962] |
| A. baylyi | Gentamicii | Amaranth | 4 | 962 | [377 - 2308 | 899 | [744 - 943] |
| A. baylyi | Gentamicii | Apigenin | 4 | 1 | [1 - 2206] | 1004 | [828 - 1034] |
| A. baylyi | Gentamicii | Calcein | 4 | 1170 | [455 - 7330 | 906 | [731 - 912] |
| A. baylyi | Gentamicii | Chlorophe | 4 | 1 | [1 - 1] | 928 | [726 - 996] |
| A. baylyi | Gentamicii | Congo Red | 4 | 1 | [1 - 1] | 956 | [785 - 998] |
| A. baylyi | Gentamicii | DiSC3(5) | 4 | 625986 | [608121 - 7 | 926 | [773 - 944] |
| A. baylyi | Gentamicii | Eosin Y | 4 | 1 | [1 - 243] | 948 | [780 - 979] |
| A. baylyi | Gentamicii | H2FDA | 4 | 1 | [1 - 1] | 918 | [771 - 950] |
| A. baylyi | Gentamicii | Malachite ( | 4 | 1 | [1 - 1] | 974 | [801 - 994] |
| A. baylyi | Gentamicii | Neutral Re | 4 | 1 | [1 - 1] | 937 | [760 - 996] |
| A. baylyi | Gentamicii | Oxonol | 4 | 1 | [1 - 1] | 1884 | [1367 - 2428] |
| A. baylyi | Gentamicii | Pyronin Y | 4 | 1 | [1 - 1] | 874 | [660 - 988] |
| A. baylyi | Gentamicii | Quinacrine | 4 | 462 | [1 - 4590] | 967 | [778 - 1028] |
| A. baylyi | Gentamicii | Rhodamine | 4 | 1 | [1 - 1] | 852 | [714 - 922] |
| A. baylyi | Gentamicii | Rhodamine | 4 | 1 | [1 - 1] | 890 | [711 - 929] |
| A. baylyi | Gentamicii | Riboflavin | 4 | 1 | [1 - 1] | 916 | [736 - 953] |
| A. baylyi | Gentamicii | Sunset Yell | 4 | 1 | [1 - 1] | 876 | [710 - 930] |
| A. baylyi | Gentamicii | SYBR Gree | 4 | 321440 | [210555 - 3 | 944 | [700 - 1031] |
| A. baylyi | Gentamicii | Thioflavin T | 4 | 6753 | [5139 - 103 | 937 | [755 - 965] |
| A. baylyi | Nalidixic ac | 5-Carboxyf | 4 | 6031 | [4846 - 709 | 980 | [931 - 1044] |
| A. baylyi | Nalidixic ac | Acridine or | 4 | 1 | [1 - 1] | 1017 | [985 - 1050] |
| A. baylyi | Nalidixic ac | Alizarin | 4 | 1 | [1 - 1] | 958 | [930 - 1015] |
| A. baylyi | Nalidixic ac | Allura Red | 4 | 1 | [1 - 1] | 1035 | [1025 - 1053] |
| A. baylyi | Nalidixic ac | Amaranth | 4 | 7221 | [6542 - 790 | 956 | [906 - 1003] |
| A. baylyi | Nalidixic ac | Apigenin | 4 | 1 | [1 - 9] | 978 | [944 - 1042] |
| A. baylyi | Nalidixic ac | Calcein | 4 | 4340 | [3860 - 508 | 984 | [963 - 1027] |
| A. baylyi | Nalidixic ac | Chlorophe | 4 | 1 | [1 - 1] | 1022 | [1000 - 1052] |
| A. baylyi | Nalidixic ac | Congo Red | 4 | 1 | [1 - 1] | 936 | [908 - 1005] |

| | | | | | | | |
|---|---|---|---|---|---|---|---|
| A. baylyi | Nalidixic a | DiSC3(5) | 4 | 699599 | [647353 - 7 | 910 | [876 - 949] |
| A. baylyi | Nalidixic a | Eosin Y | 4 | 6976 | [6266 - 72 | 970 | [922 - 1006] |
| A. baylyi | Nalidixic a | H2FDA | 4 | 1 | [1 - 1] | 1010 | [968 - 1043] |
| A. baylyi | Nalidixic a | Malachite | 4 | 1 | [1 - 1] | 928 | [914 - 961] |
| A. baylyi | Nalidixic a | Neutral Re | 4 | 1 | [1 - 1] | 1003 | [972 - 1042] |
| A. baylyi | Nalidixic a | Oxonol | 4 | 1 | [1 - 1] | 1971 | [1668 - 2395] |
| A. baylyi | Nalidixic a | Pyronin Y | 4 | 1 | [1 - 1] | 930 | [904 - 944] |
| A. baylyi | Nalidixic a | Quinacrine | 4 | 1 | [1 - 203] | 1030 | [959 - 1098] |
| A. baylyi | Nalidixic a | Rhodamine | 4 | 1 | [1 - 1] | 997 | [955 - 1044] |
| A. baylyi | Nalidixic a | Rhodamine | 4 | 1 | [1 - 1] | 1046 | [1008 - 1081] |
| A. baylyi | Nalidixic a | Riboflavin | 4 | 1 | [1 - 1] | 926 | [906 - 978] |
| A. baylyi | Nalidixic a | Sunset Yel | 4 | 1 | [1 - 1] | 996 | [931 - 1057] |
| A. baylyi | Nalidixic a | SYBR Gree | 4 | 373370 | [350110 - 3 | 970 | [941 - 995] |
| A. baylyi | Nalidixic a | Thioflavin T | 4 | 2160 | [1438 - 273 | 1007 | [976 - 1018] |
| A. baylyi | Nitrofurant | 5-Carboxyf | 4 | 1718 | [555 - 374 | 883 | [700 - 951] |
| A. baylyi | Nitrofurant | Acridine or | 4 | 1 | [1 - 1] | 807 | [652 - 837] |
| A. baylyi | Nitrofurant | Alizarin | 4 | 1 | [1 - 1] | 744 | [600 - 802] |
| A. baylyi | Nitrofurant | Allura Red | 4 | 1 | [1 - 1] | 958 | [804 - 1023] |
| A. baylyi | Nitrofurant | Amaranth | 4 | 630 | [1 - 12262] | 854 | [637 - 960] |
| A. baylyi | Nitrofurant | Apigenin | 4 | 1250 | [239 - 113 | 808 | [607 - 942] |
| A. baylyi | Nitrofurant | Calcein | 4 | 2478 | [123 - 872 | 890 | [718 - 942] |
| A. baylyi | Nitrofurant | Chlorophe | 4 | 1 | [1 - 1] | 830 | [682 - 914] |
| A. baylyi | Nitrofurant | Congo Red | 4 | 1 | [1 - 1] | 763 | [648 - 830] |
| A. baylyi | Nitrofurant | DiSC3(5) | 4 | 469084 | [370735 - 4 | 742 | [566 - 843] |
| A. baylyi | Nitrofurant | Eosin Y | 4 | 2586 | [1583 - 60 | 794 | [667 - 814] |
| A. baylyi | Nitrofurant | H2FDA | 4 | 1 | [1 - 1] | 865 | [670 - 958] |
| A. baylyi | Nitrofurant | Malachite | 4 | 1 | [1 - 1] | 904 | [696 - 1008] |
| A. baylyi | Nitrofurant | Neutral Re | 4 | 1 | [1 - 1] | 942 | [751 - 994] |
| A. baylyi | Nitrofurant | Oxonol | 4 | 1 | [1 - 1] | 2788 | [1903 - 3418] |
| A. baylyi | Nitrofurant | Pyronin Y | 4 | 1 | [1 - 1] | 816 | [606 - 916] |
| A. baylyi | Nitrofurant | Quinacrine | 4 | 3985 | [2917 - 12 | 820 | [675 - 861] |
| A. baylyi | Nitrofurant | Rhodamine | 4 | 1 | [1 - 841] | 743 | [566 - 867] |
| A. baylyi | Nitrofurant | Rhodamine | 4 | 1 | [1 - 1] | 856 | [679 - 941] |
| A. baylyi | Nitrofurant | Riboflavin | 4 | 1 | [1 - 1] | 828 | [623 - 909] |
| A. baylyi | Nitrofurant | Sunset Yel | 4 | 1 | [1 - 1] | 896 | [660 - 1030] |
| A. baylyi | Nitrofurant | SYBR Gree | 4 | 207286 | [167533 - 2 | 905 | [736 - 983] |
| A. baylyi | Nitrofurant | Thioflavin T | 4 | 18489 | [9785 - 30 | 912 | [726 - 988] |
| A. baylyi | None | 5-Carboxyf | 8 | 17296 | [16047 - 1 | 1043 | [1014 - 1075] |
| A. baylyi | None | Acridine or | 8 | 3933 | [3561 - 43 | 1046 | [992 - 1112] |
| A. baylyi | None | Alizarin | 8 | 3267 | [3094 - 40 | 1112 | [1041 - 1154] |
| A. baylyi | None | Allura Red | 8 | 2839 | [2536 - 32 | 1596 | [1455 - 1712] |
| A. baylyi | None | Amaranth | 8 | 16687 | [15409 - 1 | 1085 | [967 - 1103] |
| A. baylyi | None | Apigenin | 8 | 6105 | [5793 - 62 | 1174 | [1135 - 1213] |
| A. baylyi | None | Calcein | 8 | 16352 | [15633 - 1 | 990 | [942 - 1122] |
| A. baylyi | None | Chlorophe | 8 | 2572 | [2350 - 29 | 1169 | [1070 - 1233] |
| A. baylyi | None | Congo Red | 8 | 2628 | [2551 - 30 | 1125 | [1039 - 1171] |
| A. baylyi | None | DiSC3(5) | 8 | 318790 | [258712 - 3 | 1395 | [1350 - 1485] |
| A. baylyi | None | Eosin Y | 8 | 17369 | [16527 - 1 | 1020 | [970 - 1107] |

| | | | | | | | |
|---|---|---|---|---|---|---|---|
| A. baylyi | None | H2FDA | 8 | 4218 | [3788 - 505 | 1196 | [1091 - 1260] |
| A. baylyi | None | Malachite ( | 8 | 3294 | [3192 - 354 | 1051 | [1008 - 1091] |
| A. baylyi | None | Neutral Re | 8 | 2811 | [2677 - 318 | 1108 | [1031 - 1183] |
| A. baylyi | None | Oxonol | 8 | 799 | [582 - 875] | 2402 | [2150 - 2686] |
| A. baylyi | None | Pyronin Y | 8 | 3301 | [3065 - 364 | 1024 | [949 - 1114] |
| A. baylyi | None | Quinacrine | 8 | 5933 | [5057 - 744 | 1282 | [1149 - 1379] |
| A. baylyi | None | Rhodamine | 8 | 4194 | [3794 - 452 | 1064 | [1016 - 1130] |
| A. baylyi | None | Rhodamine | 8 | 2559 | [2382 - 289 | 1084 | [1012 - 1189] |
| A. baylyi | None | Riboflavin | 8 | 2870 | [2667 - 324 | 1104 | [1024 - 1143] |
| A. baylyi | None | Sunset Yell | 8 | 1587 | [1418 - 197 | 1056 | [989 - 1198] |
| A. baylyi | None | SYBR Gree | 8 | 291969 | [256206 - 3 | 1078 | [998 - 1128] |
| A. baylyi | None | Thioflavin T | 8 | 5809 | [5037 - 669 | 1386 | [1270 - 1441] |
| A. baylyi | Ofloxacin | 5-Carboxyf | 4 | 11353 | [8459 - 136 | 866 | [812 - 931] |
| A. baylyi | Ofloxacin | Acridine or | 4 | 1 | [1 - 1] | 818 | [795 - 844] |
| A. baylyi | Ofloxacin | Alizarin | 4 | 1 | [1 - 1] | 855 | [842 - 893] |
| A. baylyi | Ofloxacin | Allura Red | 4 | 1 | [1 - 1] | 1003 | [993 - 1014] |
| A. baylyi | Ofloxacin | Amaranth | 4 | 9154 | [8166 - 103 | 861 | [814 - 900] |
| A. baylyi | Ofloxacin | Apigenin | 4 | 1 | [1 - 339] | 920 | [874 - 971] |
| A. baylyi | Ofloxacin | Calcein | 4 | 8937 | [8309 - 106 | 884 | [830 - 940] |
| A. baylyi | Ofloxacin | Chlorophe | 4 | 1 | [1 - 1] | 923 | [889 - 972] |
| A. baylyi | Ofloxacin | Congo Red | 4 | 1 | [1 - 1] | 881 | [870 - 908] |
| A. baylyi | Ofloxacin | DiSC3(5) | 4 | 855503 | [809857 - 8 | 808 | [784 - 830] |
| A. baylyi | Ofloxacin | Eosin Y | 4 | 10306 | [9141 - 118 | 856 | [808 - 887] |
| A. baylyi | Ofloxacin | H2FDA | 4 | 1 | [1 - 1] | 869 | [850 - 893] |
| A. baylyi | Ofloxacin | Malachite ( | 4 | 1 | [1 - 1] | 912 | [852 - 972] |
| A. baylyi | Ofloxacin | Neutral Re | 4 | 1 | [1 - 1] | 879 | [871 - 888] |
| A. baylyi | Ofloxacin | Oxonol | 4 | 1 | [1 - 1] | 3083 | [2642 - 3278] |
| A. baylyi | Ofloxacin | Pyronin Y | 4 | 1 | [1 - 1] | 806 | [775 - 845] |
| A. baylyi | Ofloxacin | Quinacrine | 4 | 1065 | [584 - 1663 | 885 | [851 - 910] |
| A. baylyi | Ofloxacin | Rhodamine | 4 | 1 | [1 - 1] | 912 | [850 - 952] |
| A. baylyi | Ofloxacin | Rhodamine | 4 | 1 | [1 - 1] | 921 | [910 - 945] |
| A. baylyi | Ofloxacin | Riboflavin | 4 | 1 | [1 - 1] | 800 | [778 - 860] |
| A. baylyi | Ofloxacin | Sunset Yell | 4 | 1 | [1 - 1] | 946 | [937 - 958] |
| A. baylyi | Ofloxacin | SYBR Gree | 4 | 480981 | [465391 - 5 | 885 | [814 - 934] |
| A. baylyi | Ofloxacin | Thioflavin T | 4 | 1631 | [1292 - 239 | 927 | [921 - 950] |
| A. baylyi | Pentamidi | 5-Carboxyf | 4 | 7723 | [5569 - 803 | 706 | [588 - 760] |
| A. baylyi | Pentamidi | Acridine or | 4 | 1 | [1 - 5059] | 696 | [555 - 750] |
| A. baylyi | Pentamidi | Alizarin | 4 | 1 | [1 - 1] | 766 | [621 - 803] |
| A. baylyi | Pentamidi | Allura Red | 4 | 9295 | [6971 - 157 | 1273 | [1032 - 1391] |
| A. baylyi | Pentamidi | Amaranth | 4 | 3639 | [1622 - 970 | 752 | [629 - 772] |
| A. baylyi | Pentamidi | Apigenin | 4 | 3327 | [1538 - 751 | 726 | [605 - 759] |
| A. baylyi | Pentamidi | Calcein | 4 | 2016 | [1087 - 295 | 734 | [672 - 810] |
| A. baylyi | Pentamidi | Chlorophe | 4 | 1 | [1 - 1] | 730 | [595 - 750] |
| A. baylyi | Pentamidi | Congo Red | 4 | 1 | [1 - 1] | 876 | [695 - 964] |
| A. baylyi | Pentamidi | DiSC3(5) | 4 | 173942 | [138567 - 1 | 776 | [590 - 858] |
| A. baylyi | Pentamidi | Eosin Y | 4 | 4384 | [973 - 1987 | 738 | [634 - 779] |
| A. baylyi | Pentamidi | H2FDA | 4 | 1 | [1 - 1] | 782 | [651 - 805] |
| A. baylyi | Pentamidi | Malachite ( | 4 | 321 | [149 - 3107 | 793 | [658 - 805] |

| | | | | | | | |
|---|---|---|---|---|---|---|---|
| A. baylyi | Pentamidi | Neutral Re | 4 | 1 | [1 - 694] | 760 | [625 - 800] |
| A. baylyi | Pentamidi | Oxonol | 4 | 1 | [1 - 1] | 10882 | [7698 - 13194] |
| A. baylyi | Pentamidi | Pyronin Y | 4 | 1 | [1 - 1] | 712 | [566 - 760] |
| A. baylyi | Pentamidi | Quinacrine | 4 | 3306 | [2100 - 88 | 741 | [625 - 809] |
| A. baylyi | Pentamidi | Rhodamine | 4 | 24893 | [20220 - 4 | 743 | [595 - 776] |
| A. baylyi | Pentamidi | Rhodamine | 4 | 1 | [1 - 1] | 752 | [625 - 804] |
| A. baylyi | Pentamidi | Riboflavin | 4 | 1 | [1 - 1] | 744 | [646 - 788] |
| A. baylyi | Pentamidi | Sunset Yel | 4 | 1 | [1 - 1] | 721 | [570 - 768] |
| A. baylyi | Pentamidi | SYBR Gree | 4 | 1128879 | [657579 - | 754 | [584 - 901] |
| A. baylyi | Pentamidi | Thioflavin | 4 | 57192 | [38086 - 9 | 726 | [614 - 750] |
| A. baylyi | Polymyxin | 5-Carboxy | 4 | 651 | [406 - 982] | 1132 | [1092 - 1195] |
| A. baylyi | Polymyxin | Acridine or | 4 | 1 | [1 - 1] | 1106 | [1089 - 1120] |
| A. baylyi | Polymyxin | Alizarin | 4 | 1 | [1 - 1] | 1138 | [1093 - 1193] |
| A. baylyi | Polymyxin | Allura Red | 4 | 1 | [1 - 1] | 1294 | [1228 - 1321] |
| A. baylyi | Polymyxin | Amaranth | 4 | 433 | [1 - 1416] | 1126 | [1086 - 1157] |
| A. baylyi | Polymyxin | Apigenin | 4 | 1 | [1 - 1] | 1258 | [1220 - 1290] |
| A. baylyi | Polymyxin | Calcein | 4 | 55 | [39 - 234] | 1184 | [1123 - 1219] |
| A. baylyi | Polymyxin | Chlorophe | 4 | 1 | [1 - 1] | 1248 | [1208 - 1265] |
| A. baylyi | Polymyxin | Congo Red | 4 | 1 | [1 - 1] | 1174 | [1136 - 1198] |
| A. baylyi | Polymyxin | DiSC3(5) | 4 | 281897 | [250576 - | 1060 | [1028 - 1098] |
| A. baylyi | Polymyxin | Eosin Y | 4 | 1236 | [867 - 133 | 1146 | [1053 - 1228] |
| A. baylyi | Polymyxin | H2FDA | 4 | 1 | [1 - 1] | 1198 | [1153 - 1238] |
| A. baylyi | Polymyxin | Malachite | 4 | 1 | [1 - 1] | 1144 | [1102 - 1158] |
| A. baylyi | Polymyxin | Neutral Re | 4 | 1 | [1 - 1] | 1165 | [1097 - 1214] |
| A. baylyi | Polymyxin | Oxonol | 4 | 1 | [1 - 1] | 2200 | [1697 - 2743] |
| A. baylyi | Polymyxin | Pyronin Y | 4 | 1 | [1 - 1] | 1109 | [1082 - 1141] |
| A. baylyi | Polymyxin | Quinacrine | 4 | 1 | [1 - 1] | 1198 | [1156 - 1201] |
| A. baylyi | Polymyxin | Rhodamine | 4 | 1 | [1 - 1] | 1182 | [1130 - 1235] |
| A. baylyi | Polymyxin | Rhodamine | 4 | 1 | [1 - 1] | 1131 | [1066 - 1199] |
| A. baylyi | Polymyxin | Riboflavin | 4 | 1 | [1 - 1] | 1153 | [1067 - 1215] |
| A. baylyi | Polymyxin | Sunset Yel | 4 | 1 | [1 - 1] | 1174 | [1154 - 1220] |
| A. baylyi | Polymyxin | SYBR Gree | 4 | 586182 | [543021 - | 1038 | [1019 - 1069] |
| A. baylyi | Polymyxin | Thioflavin | 4 | 1 | [1 - 1] | 1168 | [1137 - 1182] |
| A. baylyi | Rifampicin | 5-Carboxy | 4 | 1 | [1 - 819] | 1048 | [867 - 1118] |
| A. baylyi | Rifampicin | Acridine or | 4 | 1 | [1 - 1] | 926 | [700 - 1015] |
| A. baylyi | Rifampicin | Alizarin | 4 | 1 | [1 - 1] | 930 | [738 - 963] |
| A. baylyi | Rifampicin | Allura Red | 4 | 1 | [1 - 18540] | 1300 | [1128 - 1376] |
| A. baylyi | Rifampicin | Amaranth | 4 | 1847 | [452 - 343 | 978 | [747 - 1055] |
| A. baylyi | Rifampicin | Apigenin | 4 | 1 | [1 - 6848] | 1049 | [868 - 1062] |
| A. baylyi | Rifampicin | Calcein | 4 | 1601 | [1 - 10839] | 1022 | [824 - 1042] |
| A. baylyi | Rifampicin | Chlorophe | 4 | 1 | [1 - 1] | 1018 | [820 - 1057] |
| A. baylyi | Rifampicin | Congo Red | 4 | 1 | [1 - 1] | 972 | [768 - 992] |
| A. baylyi | Rifampicin | DiSC3(5) | 4 | 437072 | [345767 - | 1030 | [801 - 1125] |
| A. baylyi | Rifampicin | Eosin Y | 4 | 1193 | [1 - 4378] | 996 | [744 - 1091] |
| A. baylyi | Rifampicin | H2FDA | 4 | 1 | [1 - 1] | 1048 | [785 - 1118] |
| A. baylyi | Rifampicin | Malachite | 4 | 1 | [1 - 1] | 1034 | [816 - 1121] |
| A. baylyi | Rifampicin | Neutral Re | 4 | 1 | [1 - 1] | 1001 | [801 - 1079] |
| A. baylyi | Rifampicin | Oxonol | 4 | 1 | [1 - 1] | 5178 | [4336 - 5429] |

| | | | | | | | |
|---|---|---|---|---|---|---|---|
| A. baylyi | Rifampicin | Pyronin Y | 4 | 1 | [1 - 1] | 904 | [735 - 962] |
| A. baylyi | Rifampicin | Quinacrine | 4 | 1 | [1 - 5319] | 1072 | [876 - 1104] |
| A. baylyi | Rifampicin | Rhodamine | 4 | 1 | [1 - 17400] | 986 | [776 - 1026] |
| A. baylyi | Rifampicin | Rhodamine | 4 | 1 | [1 - 1] | 1029 | [857 - 1052] |
| A. baylyi | Rifampicin | Riboflavin | 4 | 1 | [1 - 1] | 949 | [767 - 1015] |
| A. baylyi | Rifampicin | Sunset Yel | 4 | 1 | [1 - 1] | 1020 | [848 - 1066] |
| A. baylyi | Rifampicin | SYBR Gree | 4 | 320103 | [229947 - 3 | 998 | [798 - 1017] |
| A. baylyi | Rifampicin | Thioflavin T | 4 | 3446 | [1448 - 627 | 985 | [808 - 1045] |
| A. baylyi | Tetracyclin | 5-Carboxyf | 4 | 6407 | [5309 - 784 | 879 | [852 - 906] |
| A. baylyi | Tetracyclin | Acridine or | 4 | 1 | [1 - 1] | 785 | [773 - 796] |
| A. baylyi | Tetracyclin | Alizarin | 4 | 1 | [1 - 1] | 850 | [800 - 884] |
| A. baylyi | Tetracyclin | Allura Red | 4 | 1 | [1 - 1] | 930 | [905 - 981] |
| A. baylyi | Tetracyclin | Amaranth | 4 | 8199 | [6640 - 10( | 740 | [730 - 792] |
| A. baylyi | Tetracyclin | Apigenin | 4 | 176 | [123 - 380] | 876 | [835 - 915] |
| A. baylyi | Tetracyclin | Calcein | 4 | 6176 | [4516 - 837 | 806 | [760 - 875] |
| A. baylyi | Tetracyclin | Chlorophe | 4 | 1 | [1 - 1] | 862 | [846 - 894] |
| A. baylyi | Tetracyclin | Congo Red | 4 | 1 | [1 - 1] | 822 | [787 - 838] |
| A. baylyi | Tetracyclin | DiSC3(5) | 4 | 752251 | [749092 - 7 | 744 | [721 - 760] |
| A. baylyi | Tetracyclin | Eosin Y | 4 | 9117 | [7769 - 987 | 784 | [769 - 798] |
| A. baylyi | Tetracyclin | H2FDA | 4 | 1 | [1 - 1] | 788 | [772 - 834] |
| A. baylyi | Tetracyclin | Malachite ( | 4 | 1 | [1 - 1] | 829 | [776 - 865] |
| A. baylyi | Tetracyclin | Neutral Re | 4 | 1 | [1 - 1] | 862 | [779 - 943] |
| A. baylyi | Tetracyclin | Oxonol | 4 | 1 | [1 - 1] | 2557 | [2377 - 2819] |
| A. baylyi | Tetracyclin | Pyronin Y | 4 | 1 | [1 - 1] | 752 | [714 - 799] |
| A. baylyi | Tetracyclin | Quinacrine | 4 | 1996 | [962 - 2712 | 849 | [806 - 880] |
| A. baylyi | Tetracyclin | Rhodamine | 4 | 1 | [1 - 1] | 820 | [798 - 837] |
| A. baylyi | Tetracyclin | Rhodamine | 4 | 1 | [1 - 1] | 862 | [801 - 923] |
| A. baylyi | Tetracyclin | Riboflavin | 4 | 1 | [1 - 1] | 764 | [714 - 813] |
| A. baylyi | Tetracyclin | Sunset Yel | 4 | 1 | [1 - 1] | 884 | [840 - 901] |
| A. baylyi | Tetracyclin | SYBR Gree | 4 | 480848 | [440387 - 5 | 870 | [852 - 876] |
| A. baylyi | Tetracyclin | Thioflavin T | 4 | 7095 | [5901 - 815 | 856 | [812 - 900] |
| A. baylyi | Trimethopr | 5-Carboxyf | 8 | 918 | [1 - 1974] | 920 | [734 - 1057] |
| A. baylyi | Trimethopr | Acridine or | 8 | 1 | [1 - 1] | 884 | [693 - 1006] |
| A. baylyi | Trimethopr | Alizarin | 8 | 1 | [1 - 1] | 913 | [744 - 950] |
| A. baylyi | Trimethopr | Allura Red | 8 | 1 | [1 - 1] | 1008 | [851 - 1055] |
| A. baylyi | Trimethopr | Amaranth | 8 | 1460 | [272 - 3307 | 895 | [722 - 956] |
| A. baylyi | Trimethopr | Apigenin | 8 | 1 | [1 - 601] | 996 | [824 - 1128] |
| A. baylyi | Trimethopr | Calcein | 8 | 132 | [1 - 1044] | 961 | [776 - 1035] |
| A. baylyi | Trimethopr | Chlorophe | 8 | 1 | [1 - 1] | 1015 | [768 - 1039] |
| A. baylyi | Trimethopr | Congo Red | 8 | 1 | [1 - 1] | 938 | [711 - 1101] |
| A. baylyi | Trimethopr | DiSC3(5) | 8 | 546636 | [489489 - 5 | 873 | [728 - 904] |
| A. baylyi | Trimethopr | Eosin Y | 8 | 731 | [1 - 3261] | 926 | [734 - 976] |
| A. baylyi | Trimethopr | H2FDA | 8 | 1 | [1 - 1] | 929 | [706 - 1021] |
| A. baylyi | Trimethopr | Malachite ( | 8 | 1 | [1 - 1] | 1012 | [828 - 1083] |
| A. baylyi | Trimethopr | Neutral Re | 8 | 1 | [1 - 1] | 953 | [801 - 1020] |
| A. baylyi | Trimethopr | Oxonol | 8 | 1 | [1 - 1] | 3086 | [2198 - 3679] |
| A. baylyi | Trimethopr | Pyronin Y | 8 | 1 | [1 - 1] | 895 | [698 - 985] |
| A. baylyi | Trimethopr | Quinacrine | 8 | 1 | [1 - 7178] | 924 | [727 - 993] |

| | | | | | | | |
|---|---|---|---|---|---|---|---|
| A. baylyi | Trimethopr | Rhodamin | 8 | 1 | [1 - 1] | 886 | [711 - 995] |
| A. baylyi | Trimethopr | Rhodamin | 8 | 1 | [1 - 1] | 912 | [745 - 1103] |
| A. baylyi | Trimethopr | Riboflavin | 8 | 1 | [1 - 1] | 870 | [708 - 972] |
| A. baylyi | Trimethopr | Sunset Yel | 8 | 1 | [1 - 1] | 950 | [800 - 986] |
| A. baylyi | Trimethopr | SYBR Gree | 8 | 359202 | [300842 - | 954 | [686 - 997] |
| A. baylyi | Trimethopr | Thioflavin | 8 | 4899 | [1814 - 51 | 913 | [760 - 948] |
| A. baylyi | Vancomyc | 5-Carboxyf | 4 | 3237 | [1380 - 51 | 1008 | [908 - 1086] |
| A. baylyi | Vancomyc | Acridine or | 4 | 1 | [1 - 1] | 1100 | [1061 - 1128] |
| A. baylyi | Vancomyc | Alizarin | 4 | 1 | [1 - 1] | 1106 | [1063 - 1146] |
| A. baylyi | Vancomyc | Allura Red | 4 | 1 | [1 - 1] | 1063 | [991 - 1134] |
| A. baylyi | Vancomyc | Amaranth | 4 | 3665 | [1628 - 60 | 1044 | [978 - 1086] |
| A. baylyi | Vancomyc | Apigenin | 4 | 1 | [1 - 1] | 1099 | [1065 - 1126] |
| A. baylyi | Vancomyc | Calcein | 4 | 1602 | [1055 - 20 | 1071 | [1061 - 1080] |
| A. baylyi | Vancomyc | Chlorophe | 4 | 1 | [1 - 1] | 1132 | [1093 - 1146] |
| A. baylyi | Vancomyc | Congo Red | 4 | 1 | [1 - 1] | 1058 | [1014 - 1120] |
| A. baylyi | Vancomyc | DiSC3(5) | 4 | 687450 | [623261 - | 928 | [884 - 982] |
| A. baylyi | Vancomyc | Eosin Y | 4 | 1 | [1 - 179] | 1031 | [982 - 1085] |
| A. baylyi | Vancomyc | H2FDA | 4 | 1 | [1 - 1] | 1092 | [1040 - 1134] |
| A. baylyi | Vancomyc | Malachite | 4 | 1 | [1 - 1] | 1042 | [992 - 1086] |
| A. baylyi | Vancomyc | Neutral Re | 4 | 1 | [1 - 1] | 1068 | [1021 - 1102] |
| A. baylyi | Vancomyc | Oxonol | 4 | 1 | [1 - 1] | 1398 | [1306 - 1436] |
| A. baylyi | Vancomyc | Pyronin Y | 4 | 1 | [1 - 1] | 972 | [962 - 993] |
| A. baylyi | Vancomyc | Quinacrin | 4 | 1 | [1 - 1] | 1124 | [1069 - 1141] |
| A. baylyi | Vancomyc | Rhodamin | 4 | 1 | [1 - 1] | 1040 | [1027 - 1061] |
| A. baylyi | Vancomyc | Rhodamin | 4 | 1 | [1 - 1] | 1122 | [1104 - 1138] |
| A. baylyi | Vancomyc | Riboflavin | 4 | 1 | [1 - 1] | 1074 | [1017 - 1123] |
| A. baylyi | Vancomyc | Sunset Yel | 4 | 1 | [1 - 1] | 1055 | [1042 - 1070] |
| A. baylyi | Vancomyc | SYBR Gree | 4 | 424419 | [374560 - | 1016 | [976 - 1049] |
| A. baylyi | Vancomyc | Thioflavin | 4 | 1 | [1 - 1] | 1086 | [1029 - 1148] |
| E. coli | Amoxacilli | 5-Carboxyf | 2 | 1 | [1 - 1] | 8913 | [8152 - 9674] |
| E. coli | Amoxacilli | Acridine or | 2 | 1 | [1 - 1] | 8624 | [8421 - 8826] |
| E. coli | Amoxacilli | Alizarin | 2 | 1 | [1 - 1] | 7915 | [7814 - 8016] |
| E. coli | Amoxacilli | Allura Red | 2 | 1 | [1 - 1] | 8674 | [8495 - 8854] |
| E. coli | Amoxacilli | Amaranth | 2 | 274 | [215 - 333] | 8944 | [8577 - 9312] |
| E. coli | Amoxacilli | Apigenin | 2 | 433 | [395 - 471] | 8578 | [8370 - 8786] |
| E. coli | Amoxacilli | Calcein | 2 | 59 | [44 - 74] | 8768 | [8405 - 9132] |
| E. coli | Amoxacilli | Chlorophe | 2 | 1 | [1 - 1] | 8930 | [8591 - 9270] |
| E. coli | Amoxacilli | Congo Red | 2 | 1 | [1 - 1] | 7948 | [7846 - 8051] |
| E. coli | Amoxacilli | DiSC3(5) | 2 | 52734 | [29588 - 75 | 6734 | [6531 - 6936] |
| E. coli | Amoxacilli | Eosin Y | 2 | 223 | [112 - 334] | 7522 | [7082 - 7963] |
| E. coli | Amoxacilli | H2FDA | 2 | 1 | [1 - 1] | 8802 | [8460 - 9145] |
| E. coli | Amoxacilli | Malachite | 2 | 1 | [1 - 1] | 8323 | [7944 - 8702] |
| E. coli | Amoxacilli | Neutral Re | 2 | 1 | [1 - 1] | 8214 | [7636 - 8792] |
| E. coli | Amoxacilli | Oxonol | 2 | 1 | [1 - 1] | 9312 | [9106 - 9517] |
| E. coli | Amoxacilli | Pyronin Y | 2 | 1 | [1 - 1] | 7904 | [7706 - 8103] |
| E. coli | Amoxacilli | Quinacrin | 2 | 394 | [373 - 414] | 9542 | [9140 - 9944] |
| E. coli | Amoxacilli | Rhodamin | 2 | 1 | [1 - 1] | 7602 | [7010 - 8193] |
| E. coli | Amoxacilli | Rhodamin | 2 | 1 | [1 - 1] | 9068 | [8780 - 9357] |

| | | | | | | | |
|---|---|---|---|---|---|---|---|
| E. coli | Amoxacillii | Riboflavin | 2 | 1 | [1 - 1] | 8700 | [8506 - 8893] |
| E. coli | Amoxacillii | Sunset Yel | 2 | 1 | [1 - 1] | 9402 | [9026 - 9779] |
| E. coli | Amoxacillii | SYBR Gree | 2 | 12593 | [11659 - 1: | 8616 | [8472 - 8761] |
| E. coli | Amoxacillii | Thioflavin T | 2 | 212 | [107 - 318] | 9076 | [8839 - 9312] |
| E. coli | Ampicillin | 5-Carboxyf | 3 | 1 | [1 - 1] | 13629 | [13058 - 14030] |
| E. coli | Ampicillin | Acridine or | 3 | 1 | [1 - 1] | 13768 | [12815 - 14560] |
| E. coli | Ampicillin | Alizarin | 3 | 1 | [1 - 1] | 14297 | [12770 - 14486] |
| E. coli | Ampicillin | Allura Red | 3 | 1 | [1 - 1] | 11456 | [11214 - 13354] |
| E. coli | Ampicillin | Amaranth | 3 | 60 | [30 - 163] | 13547 | [13124 - 13614] |
| E. coli | Ampicillin | Apigenin | 3 | 1 | [1 - 1] | 15147 | [13664 - 15196] |
| E. coli | Ampicillin | Calcein | 3 | 1 | [1 - 105] | 14749 | [13824 - 15027] |
| E. coli | Ampicillin | Chlorophe | 3 | 1 | [1 - 1] | 16171 | [15459 - 16778] |
| E. coli | Ampicillin | Congo Red | 3 | 1 | [1 - 1] | 15122 | [13612 - 15758] |
| E. coli | Ampicillin | DiSC3(5) | 3 | 48579 | [28169 - 5( | 11015 | [10890 - 12298] |
| E. coli | Ampicillin | Eosin Y | 3 | 86 | [43 - 120] | 10173 | [10010 - 12640] |
| E. coli | Ampicillin | H2FDA | 3 | 1 | [1 - 1] | 12507 | [11216 - 13870] |
| E. coli | Ampicillin | Malachite ( | 3 | 1 | [1 - 1] | 14148 | [12822 - 14279] |
| E. coli | Ampicillin | Neutral Re | 3 | 1 | [1 - 1] | 13143 | [12552 - 13758] |
| E. coli | Ampicillin | Oxonol | 3 | 1 | [1 - 19] | 13296 | [12802 - 13784] |
| E. coli | Ampicillin | Pyronin Y | 3 | 1 | [1 - 1] | 15419 | [13800 - 15964] |
| E. coli | Ampicillin | Quinacrine | 3 | 180 | [125 - 233] | 14076 | [14021 - 15065] |
| E. coli | Ampicillin | Rhodamine | 3 | 1 | [1 - 1] | 15502 | [14179 - 15588] |
| E. coli | Ampicillin | Rhodamine | 3 | 1 | [1 - 1] | 14013 | [13264 - 14348] |
| E. coli | Ampicillin | Riboflavin | 3 | 1 | [1 - 1] | 10436 | [10300 - 12352] |
| E. coli | Ampicillin | Sunset Yel | 3 | 1 | [1 - 1] | 13256 | [13005 - 13488] |
| E. coli | Ampicillin | SYBR Gree | 3 | 14021 | [10373 - 1! | 10806 | [10687 - 14196] |
| E. coli | Ampicillin | Thioflavin T | 3 | 1 | [1 - 3] | 17328 | [16758 - 17435] |
| E. coli | Azithromyc | 5-Carboxyf | 2 | 821 | [545 - 109{ | 4874 | [4769 - 4980] |
| E. coli | Azithromyc | Acridine or | 2 | 1 | [1 - 1] | 4956 | [4775 - 5138] |
| E. coli | Azithromyc | Alizarin | 2 | 1 | [1 - 1] | 4890 | [4858 - 4923] |
| E. coli | Azithromyc | Allura Red | 2 | 374 | [347 - 401] | 5846 | [5768 - 5924] |
| E. coli | Azithromyc | Amaranth | 2 | 2377 | [2152 - 26( | 4990 | [4968 - 5012] |
| E. coli | Azithromyc | Apigenin | 2 | 1031 | [962 - 110( | 6356 | [6056 - 6656] |
| E. coli | Azithromyc | Calcein | 2 | 1788 | [1518 - 20! | 5042 | [4749 - 5334] |
| E. coli | Azithromyc | Chlorophe | 2 | 1 | [1 - 1] | 5242 | [5152 - 5333] |
| E. coli | Azithromyc | Congo Red | 2 | 1 | [1 - 1] | 6078 | [5889 - 6267] |
| E. coli | Azithromyc | DiSC3(5) | 2 | 69975 | [61958 - 7; | 5318 | [4826 - 5809] |
| E. coli | Azithromyc | Eosin Y | 2 | 1124 | [901 - 134; | 5025 | [4893 - 5157] |
| E. coli | Azithromyc | H2FDA | 2 | 1 | [1 - 1] | 5768 | [5754 - 5782] |
| E. coli | Azithromyc | Malachite ( | 2 | 1 | [1 - 1] | 5722 | [5325 - 6119] |
| E. coli | Azithromyc | Neutral Re | 2 | 1 | [1 - 1] | 5412 | [5083 - 5742] |
| E. coli | Azithromyc | Oxonol | 2 | 413 | [292 - 533] | 6011 | [5606 - 6416] |
| E. coli | Azithromyc | Pyronin Y | 2 | 1 | [1 - 1] | 5562 | [5345 - 5780] |
| E. coli | Azithromyc | Quinacrine | 2 | 1554 | [1446 - 16( | 5962 | [5586 - 6339] |
| E. coli | Azithromyc | Rhodamine | 2 | 1 | [1 - 1] | 5720 | [5398 - 6042] |
| E. coli | Azithromyc | Rhodamine | 2 | 1 | [1 - 1] | 5164 | [4988 - 5341] |
| E. coli | Azithromyc | Riboflavin | 2 | 1 | [1 - 1] | 4769 | [4520 - 5018] |
| E. coli | Azithromyc | Sunset Yel | 2 | 1 | [1 - 1] | 4884 | [4448 - 5321] |

| | | | | | | | |
|---|---|---|---|---|---|---|---|
| E. coli | Azithromyc | SYBR Gree | 2 | 4059 | [2653 - 546 | 4999 | [4718 - 5280] |
| E. coli | Azithromyc | Thioflavin T | 2 | 1444 | [1399 - 149 | 5804 | [5678 - 5929] |
| E. coli | Bacitracin | 5-Carboxyf | 3 | 1 | [1 - 1] | 14451 | [13742 - 14693] |
| E. coli | Bacitracin | Acridine or | 3 | 1 | [1 - 1] | 14410 | [13154 - 15353] |
| E. coli | Bacitracin | Alizarin | 3 | 1 | [1 - 1] | 13170 | [12981 - 14110] |
| E. coli | Bacitracin | Allura Red | 3 | 1 | [1 - 1] | 10099 | [9767 - 11841] |
| E. coli | Bacitracin | Amaranth | 3 | 128 | [64 - 680] | 13153 | [12368 - 13618] |
| E. coli | Bacitracin | Apigenin | 3 | 1 | [1 - 90] | 12389 | [11844 - 13498] |
| E. coli | Bacitracin | Calcein | 3 | 1 | [1 - 20] | 14170 | [13594 - 14826] |
| E. coli | Bacitracin | Chlorophe | 3 | 1 | [1 - 1] | 14653 | [14388 - 15990] |
| E. coli | Bacitracin | Congo Red | 3 | 1 | [1 - 1] | 13430 | [13371 - 14201] |
| E. coli | Bacitracin | DiSC3(5) | 3 | 49519 | [43740 - 55 | 13284 | [12172 - 13885] |
| E. coli | Bacitracin | Eosin Y | 3 | 1 | [1 - 67] | 16710 | [13671 - 16869] |
| E. coli | Bacitracin | H2FDA | 3 | 1 | [1 - 1] | 14119 | [13682 - 14553] |
| E. coli | Bacitracin | Malachite ( | 3 | 1 | [1 - 1] | 13843 | [13820 - 14381] |
| E. coli | Bacitracin | Neutral Re | 3 | 1 | [1 - 1] | 15451 | [14010 - 15491] |
| E. coli | Bacitracin | Oxonol | 3 | 1 | [1 - 31] | 13757 | [12554 - 14326] |
| E. coli | Bacitracin | Pyronin Y | 3 | 1 | [1 - 1] | 13116 | [12894 - 14298] |
| E. coli | Bacitracin | Quinacrine | 3 | 77 | [39 - 88] | 15707 | [14592 - 16320] |
| E. coli | Bacitracin | Rhodamine | 3 | 1 | [1 - 1] | 14327 | [13645 - 15582] |
| E. coli | Bacitracin | Rhodamine | 3 | 1 | [1 - 1] | 14272 | [14035 - 15234] |
| E. coli | Bacitracin | Riboflavin | 3 | 1 | [1 - 1] | 13996 | [13326 - 14506] |
| E. coli | Bacitracin | Sunset Yell | 3 | 1 | [1 - 1] | 14232 | [13474 - 14659] |
| E. coli | Bacitracin | SYBR Gree | 3 | 11935 | [10822 - 14 | 11632 | [11330 - 13127] |
| E. coli | Bacitracin | Thioflavin T | 3 | 1 | [1 - 1] | 14615 | [14471 - 15304] |
| E. coli | Ceftriaxon | 5-Carboxyf | 3 | 1 | [1 - 1] | 10648 | [10404 - 11115] |
| E. coli | Ceftriaxon | Acridine or | 3 | 1 | [1 - 1] | 10473 | [10254 - 12072] |
| E. coli | Ceftriaxon | Alizarin | 3 | 1 | [1 - 1] | 10526 | [10213 - 10906] |
| E. coli | Ceftriaxon | Allura Red | 3 | 1 | [1 - 164] | 13411 | [12025 - 13754] |
| E. coli | Ceftriaxon | Amaranth | 3 | 864 | [623 - 1009 | 11289 | [10576 - 11849] |
| E. coli | Ceftriaxon | Apigenin | 3 | 1 | [1 - 101] | 13637 | [13584 - 15718] |
| E. coli | Ceftriaxon | Calcein | 3 | 274 | [266 - 308] | 11759 | [11382 - 12933] |
| E. coli | Ceftriaxon | Chlorophe | 3 | 1 | [1 - 1] | 11003 | [10749 - 14568] |
| E. coli | Ceftriaxon | Congo Red | 3 | 1 | [1 - 1] | 12841 | [11307 - 12964] |
| E. coli | Ceftriaxon | DiSC3(5) | 3 | 70166 | [62922 - 75 | 11443 | [11012 - 12104] |
| E. coli | Ceftriaxon | Eosin Y | 3 | 63 | [33 - 203] | 11889 | [10753 - 11928] |
| E. coli | Ceftriaxon | H2FDA | 3 | 1 | [1 - 1] | 12029 | [11270 - 12054] |
| E. coli | Ceftriaxon | Malachite ( | 3 | 1 | [1 - 1] | 13007 | [11906 - 14586] |
| E. coli | Ceftriaxon | Neutral Re | 3 | 1 | [1 - 1] | 11869 | [11062 - 12594] |
| E. coli | Ceftriaxon | Oxonol | 3 | 73 | [67 - 118] | 13842 | [12141 - 13847] |
| E. coli | Ceftriaxon | Pyronin Y | 3 | 1 | [1 - 1] | 10173 | [10124 - 12273] |
| E. coli | Ceftriaxon | Quinacrine | 3 | 232 | [194 - 279] | 13050 | [12469 - 13240] |
| E. coli | Ceftriaxon | Rhodamine | 3 | 1 | [1 - 1] | 9560 | [9409 - 10812] |
| E. coli | Ceftriaxon | Rhodamine | 3 | 1 | [1 - 1] | 11300 | [11070 - 12085] |
| E. coli | Ceftriaxon | Riboflavin | 3 | 1 | [1 - 1] | 11342 | [11256 - 11910] |
| E. coli | Ceftriaxon | Sunset Yell | 3 | 1 | [1 - 1] | 12660 | [12570 - 15276] |
| E. coli | Ceftriaxon | SYBR Gree | 3 | 11581 | [11116 - 15 | 10894 | [10394 - 12228] |
| E. coli | Ceftriaxon | Thioflavin T | 3 | 1 | [1 - 79] | 13287 | [13222 - 15478] |

| | | | | | | | |
|---|---|---|---|---|---|---|---|
| E. coli | Chloramph | 5-Carboxyf | 2 | 1 | [1 - 1] | 7726 | [7166 - 8287] |
| E. coli | Chloramph | Acridine or | 2 | 1 | [1 - 1] | 7583 | [7052 - 8114] |
| E. coli | Chloramph | Alizarin | 2 | 1 | [1 - 1] | 7721 | [7667 - 7775] |
| E. coli | Chloramph | Allura Red | 2 | 143 | [72 - 214] | 6952 | [6757 - 7147] |
| E. coli | Chloramph | Amaranth | 2 | 167 | [84 - 250] | 8040 | [7668 - 8412] |
| E. coli | Chloramph | Apigenin | 2 | 184 | [93 - 276] | 8528 | [8290 - 8766] |
| E. coli | Chloramph | Calcein | 2 | 63 | [32 - 94] | 7440 | [7156 - 7725] |
| E. coli | Chloramph | Chlorophe | 2 | 1 | [1 - 1] | 8110 | [7409 - 8811] |
| E. coli | Chloramph | Congo Red | 2 | 1 | [1 - 1] | 6604 | [6450 - 6759] |
| E. coli | Chloramph | DiSC3(5) | 2 | 67834 | [67072 - 68 | 7205 | [6890 - 7520] |
| E. coli | Chloramph | Eosin Y | 2 | 1 | [1 - 1] | 7466 | [7052 - 7881] |
| E. coli | Chloramph | H2FDA | 2 | 1 | [1 - 1] | 7345 | [6788 - 7902] |
| E. coli | Chloramph | Malachite ( | 2 | 1 | [1 - 1] | 7526 | [7135 - 7916] |
| E. coli | Chloramph | Neutral Re | 2 | 1 | [1 - 1] | 8294 | [8191 - 8398] |
| E. coli | Chloramph | Oxonol | 2 | 99 | [50 - 149] | 7230 | [6805 - 7654] |
| E. coli | Chloramph | Pyronin Y | 2 | 1 | [1 - 1] | 7586 | [6850 - 8321] |
| E. coli | Chloramph | Quinacrine | 2 | 378 | [364 - 393] | 7750 | [7607 - 7894] |
| E. coli | Chloramph | Rhodamine | 2 | 1 | [1 - 1] | 7286 | [6547 - 8024] |
| E. coli | Chloramph | Rhodamine | 2 | 1 | [1 - 1] | 7674 | [7490 - 7857] |
| E. coli | Chloramph | Riboflavin | 2 | 1 | [1 - 1] | 6727 | [6582 - 6872] |
| E. coli | Chloramph | Sunset Yell | 2 | 1 | [1 - 1] | 7149 | [6470 - 7828] |
| E. coli | Chloramph | SYBR Gree | 2 | 5360 | [4106 - 661 | 7886 | [7616 - 8155] |
| E. coli | Chloramph | Thioflavin T | 2 | 446 | [332 - 559] | 7616 | [7054 - 8178] |
| E. coli | Ciprofloxa | 5-Carboxyf | 3 | 1 | [1 - 1] | 12524 | [11079 - 12568] |
| E. coli | Ciprofloxa | Acridine or | 3 | 1 | [1 - 1] | 10431 | [10205 - 11114] |
| E. coli | Ciprofloxa | Alizarin | 3 | 1 | [1 - 1] | 10041 | [8794 - 10266] |
| E. coli | Ciprofloxa | Allura Red | 3 | 1 | [1 - 1] | 10750 | [10749 - 11870] |
| E. coli | Ciprofloxa | Amaranth | 3 | 48 | [24 - 72] | 11128 | [10866 - 11152] |
| E. coli | Ciprofloxa | Apigenin | 3 | 1 | [1 - 183] | 11117 | [10680 - 12046] |
| E. coli | Ciprofloxa | Calcein | 3 | 219 | [110 - 449] | 10320 | [10070 - 11330] |
| E. coli | Ciprofloxa | Chlorophe | 3 | 1 | [1 - 1] | 10667 | [10504 - 11131] |
| E. coli | Ciprofloxa | Congo Red | 3 | 1 | [1 - 1] | 10672 | [9254 - 12037] |
| E. coli | Ciprofloxa | DiSC3(5) | 3 | 48677 | [48232 - 55 | 12877 | [11978 - 13964] |
| E. coli | Ciprofloxa | Eosin Y | 3 | 1 | [1 - 117] | 9849 | [9459 - 10985] |
| E. coli | Ciprofloxa | H2FDA | 3 | 1 | [1 - 1] | 10264 | [10010 - 11328] |
| E. coli | Ciprofloxa | Malachite ( | 3 | 1 | [1 - 1] | 9987 | [9875 - 10876] |
| E. coli | Ciprofloxa | Neutral Re | 3 | 1 | [1 - 1] | 11741 | [11627 - 12271] |
| E. coli | Ciprofloxa | Oxonol | 3 | 1 | [1 - 1] | 12663 | [12358 - 14414] |
| E. coli | Ciprofloxa | Pyronin Y | 3 | 1 | [1 - 1] | 8835 | [8589 - 9700] |
| E. coli | Ciprofloxa | Quinacrine | 3 | 144 | [109 - 214] | 11748 | [11512 - 12808] |
| E. coli | Ciprofloxa | Rhodamine | 3 | 1 | [1 - 1] | 11114 | [9740 - 11851] |
| E. coli | Ciprofloxa | Rhodamine | 3 | 1 | [1 - 1] | 12633 | [10796 - 12648] |
| E. coli | Ciprofloxa | Riboflavin | 3 | 1 | [1 - 1] | 10615 | [10065 - 11982] |
| E. coli | Ciprofloxa | Sunset Yell | 3 | 1 | [1 - 1] | 12152 | [10730 - 12586] |
| E. coli | Ciprofloxa | SYBR Gree | 3 | 15449 | [13331 - 21 | 11959 | [10112 - 12201] |
| E. coli | Ciprofloxa | Thioflavin T | 3 | 23 | [12 - 118] | 11534 | [11430 - 12698] |
| E. coli | Colistin | 5-Carboxyf | 2 | 23817 | [20306 - 27 | 2596 | [2368 - 2823] |
| E. coli | Colistin | Acridine or | 2 | 60055 | [45961 - 74 | 2390 | [1961 - 2820] |

| | | | | | | | |
|---|---|---|---|---|---|---|---|
| E. coli | Colistin | Alizarin | 2 | 5222 | [3398 - 704 | 2772 | [2442 - 3102] |
| E. coli | Colistin | Allura Red | 2 | 5219 | [3773 - 666 | 2923 | [2600 - 3246] |
| E. coli | Colistin | Amaranth | 2 | 10444 | [9878 - 110 | 1989 | [1780 - 2198] |
| E. coli | Colistin | Apigenin | 2 | 2806 | [2697 - 291 | 3371 | [3136 - 3606] |
| E. coli | Colistin | Calcein | 2 | 10904 | [10438 - 11 | 2120 | [1957 - 2284] |
| E. coli | Colistin | Chlorophe | 2 | 1 | [1 - 1] | 3462 | [3358 - 3565] |
| E. coli | Colistin | Congo Red | 2 | 1 | [1 - 1] | 2608 | [2603 - 2613] |
| E. coli | Colistin | DiSC3(5) | 2 | 38961 | [38343 - 39 | 2337 | [2103 - 2571] |
| E. coli | Colistin | Eosin Y | 2 | 15193 | [14221 - 16 | 2186 | [2078 - 2295] |
| E. coli | Colistin | H2FDA | 2 | 7185 | [5178 - 919 | 2730 | [2629 - 2832] |
| E. coli | Colistin | Malachite ( | 2 | 43808 | [34054 - 53 | 2952 | [2616 - 3288] |
| E. coli | Colistin | Neutral Re | 2 | 3104 | [1938 - 427 | 3628 | [3423 - 3832] |
| E. coli | Colistin | Oxonol | 2 | 754 | [565 - 944] | 4888 | [4399 - 5378] |
| E. coli | Colistin | Pyronin Y | 2 | 7927 | [5229 - 106 | 2238 | [1878 - 2597] |
| E. coli | Colistin | Quinacrine | 2 | 5047 | [4584 - 551 | 2734 | [2656 - 2811] |
| E. coli | Colistin | Rhodamine | 2 | 120164 | [109765 - 1 | 2118 | [1922 - 2314] |
| E. coli | Colistin | Rhodamine | 2 | 5587 | [3398 - 777 | 2720 | [2394 - 3045] |
| E. coli | Colistin | Riboflavin | 2 | 8642 | [5896 - 113 | 2126 | [1896 - 2356] |
| E. coli | Colistin | Sunset Yell | 2 | 796 | [785 - 808] | 2532 | [2384 - 2680] |
| E. coli | Colistin | SYBR Gree | 2 | 981583 | [827617 - 1 | 2754 | [2463 - 3046] |
| E. coli | Colistin | Thioflavin T | 2 | 4966 | [4904 - 502 | 2378 | [2334 - 2421] |
| E. coli | Fosfomycin | 5-Carboxyf | 3 | 1 | [1 - 1] | 18385 | [15146 - 18781] |
| E. coli | Fosfomycin | Acridine or | 3 | 1 | [1 - 1] | 14888 | [14191 - 14940] |
| E. coli | Fosfomycin | Alizarin | 3 | 1 | [1 - 1] | 12308 | [11436 - 12780] |
| E. coli | Fosfomycin | Allura Red | 3 | 1 | [1 - 1] | 13416 | [12813 - 14320] |
| E. coli | Fosfomycin | Amaranth | 3 | 80 | [40 - 127] | 13163 | [12290 - 14982] |
| E. coli | Fosfomycin | Apigenin | 3 | 1 | [1 - 1] | 16951 | [15756 - 18255] |
| E. coli | Fosfomycin | Calcein | 3 | 187 | [94 - 305] | 16714 | [13851 - 17164] |
| E. coli | Fosfomycin | Chlorophe | 3 | 1 | [1 - 1] | 13428 | [12317 - 14320] |
| E. coli | Fosfomycin | Congo Red | 3 | 1 | [1 - 1] | 11972 | [11902 - 13746] |
| E. coli | Fosfomycin | DiSC3(5) | 3 | 53831 | [48793 - 54 | 13035 | [12348 - 13628] |
| E. coli | Fosfomycin | Eosin Y | 3 | 67 | [34 - 160] | 11886 | [10750 - 13035] |
| E. coli | Fosfomycin | H2FDA | 3 | 1 | [1 - 1] | 11288 | [11250 - 13276] |
| E. coli | Fosfomycin | Malachite ( | 3 | 1 | [1 - 1] | 12051 | [11886 - 15510] |
| E. coli | Fosfomycin | Neutral Re | 3 | 1 | [1 - 1] | 14662 | [12844 - 14750] |
| E. coli | Fosfomycin | Oxonol | 3 | 24 | [12 - 48] | 11134 | [10512 - 11991] |
| E. coli | Fosfomycin | Pyronin Y | 3 | 1 | [1 - 1] | 12650 | [12112 - 13182] |
| E. coli | Fosfomycin | Quinacrine | 3 | 350 | [271 - 500] | 12997 | [11570 - 13144] |
| E. coli | Fosfomycin | Rhodamine | 3 | 1 | [1 - 1] | 13443 | [12236 - 14378] |
| E. coli | Fosfomycin | Rhodamine | 3 | 1 | [1 - 1] | 15181 | [15156 - 15893] |
| E. coli | Fosfomycin | Riboflavin | 3 | 1 | [1 - 1] | 13802 | [13352 - 14123] |
| E. coli | Fosfomycin | Sunset Yell | 3 | 1 | [1 - 1] | 15640 | [12972 - 16180] |
| E. coli | Fosfomycin | SYBR Gree | 3 | 10975 | [8856 - 134 | 13141 | [12358 - 14578] |
| E. coli | Fosfomycin | Thioflavin T | 3 | 1 | [1 - 1] | 14534 | [14487 - 16702] |
| E. coli | Gentamicin | 5-Carboxyf | 2 | 1 | [1 - 1] | 8430 | [8022 - 8839] |
| E. coli | Gentamicin | Acridine or | 2 | 1 | [1 - 1] | 8386 | [7983 - 8790] |
| E. coli | Gentamicin | Alizarin | 2 | 1 | [1 - 1] | 8052 | [7542 - 8562] |
| E. coli | Gentamicin | Allura Red | 2 | 1 | [1 - 1] | 6826 | [6423 - 7229] |

| | | | | | | | |
|---|---|---|---|---|---|---|---|
| E. coli | Gentamici | Amaranth | 2 | 1 | [1 - 1] | 8416 | [8196 - 8637] |
| E. coli | Gentamici | Apigenin | 2 | 1 | [1 - 1] | 8776 | [8178 - 9374] |
| E. coli | Gentamici | Calcein | 2 | 1 | [1 - 1] | 8698 | [8122 - 9273] |
| E. coli | Gentamici | Chlorophe | 2 | 1 | [1 - 1] | 7974 | [7530 - 8418] |
| E. coli | Gentamici | Congo Red | 2 | 1 | [1 - 1] | 9486 | [9264 - 9709] |
| E. coli | Gentamici | DiSC3(5) | 2 | 36137 | [32527 - 3 | 8234 | [7450 - 9018] |
| E. coli | Gentamici | Eosin Y | 2 | 1 | [1 - 1] | 8230 | [7732 - 8728] |
| E. coli | Gentamici | H2FDA | 2 | 1 | [1 - 1] | 8651 | [8594 - 8708] |
| E. coli | Gentamici | Malachite | 2 | 1 | [1 - 1] | 8438 | [8332 - 8544] |
| E. coli | Gentamici | Neutral Re | 2 | 1 | [1 - 1] | 9560 | [9406 - 9715] |
| E. coli | Gentamici | Oxonol | 2 | 24 | [12 - 35] | 6530 | [5995 - 7066] |
| E. coli | Gentamici | Pyronin Y | 2 | 1 | [1 - 1] | 7746 | [7360 - 8131] |
| E. coli | Gentamici | Quinacrine | 2 | 1 | [1 - 1] | 8502 | [8169 - 8836] |
| E. coli | Gentamici | Rhodamin | 2 | 1 | [1 - 1] | 8432 | [8422 - 8443] |
| E. coli | Gentamici | Rhodamin | 2 | 1 | [1 - 1] | 8818 | [8629 - 9008] |
| E. coli | Gentamici | Riboflavin | 2 | 1 | [1 - 1] | 9226 | [8669 - 9783] |
| E. coli | Gentamici | Sunset Yel | 2 | 1 | [1 - 1] | 8782 | [8675 - 8888] |
| E. coli | Gentamici | SYBR Gree | 2 | 10781 | [6525 - 15 | 8230 | [7289 - 9172] |
| E. coli | Gentamici | Thioflavin | 2 | 1 | [1 - 1] | 10149 | [10008 - 10290] |
| E. coli | Nalidixic a | 5-Carboxy | 3 | 1 | [1 - 1] | 12024 | [11373 - 12274] |
| E. coli | Nalidixic a | Acridine or | 3 | 1 | [1 - 1] | 8894 | [8795 - 10112] |
| E. coli | Nalidixic a | Alizarin | 3 | 1 | [1 - 1] | 9786 | [9444 - 10514] |
| E. coli | Nalidixic a | Allura Red | 3 | 1 | [1 - 1] | 13508 | [13097 - 14104] |
| E. coli | Nalidixic a | Amaranth | 3 | 1 | [1 - 365] | 11248 | [10760 - 12445] |
| E. coli | Nalidixic a | Apigenin | 3 | 100 | [50 - 191] | 14071 | [13852 - 14936] |
| E. coli | Nalidixic a | Calcein | 3 | 1 | [1 - 412] | 11320 | [10728 - 11970] |
| E. coli | Nalidixic a | Chlorophe | 3 | 1 | [1 - 1] | 12174 | [11675 - 13404] |
| E. coli | Nalidixic a | Congo Red | 3 | 1 | [1 - 1] | 12884 | [11241 - 13098] |
| E. coli | Nalidixic a | DiSC3(5) | 3 | 56892 | [54719 - 5 | 11100 | [10570 - 11825] |
| E. coli | Nalidixic a | Eosin Y | 3 | 1 | [1 - 233] | 10457 | [10327 - 10809] |
| E. coli | Nalidixic a | H2FDA | 3 | 1 | [1 - 1] | 13598 | [11740 - 14886] |
| E. coli | Nalidixic a | Malachite | 3 | 1 | [1 - 1] | 11313 | [11186 - 11462] |
| E. coli | Nalidixic a | Neutral Re | 3 | 1 | [1 - 1] | 10630 | [10360 - 11534] |
| E. coli | Nalidixic a | Oxonol | 3 | 105 | [53 - 138] | 11034 | [10897 - 13526] |
| E. coli | Nalidixic a | Pyronin Y | 3 | 1 | [1 - 1] | 10580 | [10214 - 11344] |
| E. coli | Nalidixic a | Quinacrine | 3 | 368 | [335 - 495] | 12069 | [11444 - 13931] |
| E. coli | Nalidixic a | Rhodamin | 3 | 1 | [1 - 1] | 13421 | [12002 - 13729] |
| E. coli | Nalidixic a | Rhodamin | 3 | 1 | [1 - 1] | 8700 | [8572 - 10051] |
| E. coli | Nalidixic a | Riboflavin | 3 | 1 | [1 - 1] | 10281 | [10255 - 11756] |
| E. coli | Nalidixic a | Sunset Yel | 3 | 1 | [1 - 1] | 10426 | [10070 - 11334] |
| E. coli | Nalidixic a | SYBR Gree | 3 | 11243 | [8287 - 11 | 10020 | [9642 - 11126] |
| E. coli | Nalidixic a | Thioflavin | 3 | 1 | [1 - 161] | 16421 | [14772 - 17478] |
| E. coli | Nitrofurant | 5-Carboxy | 2 | 1 | [1 - 1] | 5897 | [5434 - 6360] |
| E. coli | Nitrofurant | Acridine or | 2 | 1 | [1 - 1] | 5294 | [5186 - 5402] |
| E. coli | Nitrofurant | Alizarin | 2 | 1 | [1 - 1] | 5507 | [5138 - 5876] |
| E. coli | Nitrofurant | Allura Red | 2 | 1 | [1 - 1] | 8034 | [7543 - 8526] |
| E. coli | Nitrofurant | Amaranth | 2 | 1539 | [1328 - 17 | 5244 | [5212 - 5277] |
| E. coli | Nitrofurant | Apigenin | 2 | 367 | [184 - 551] | 8559 | [8083 - 9035] |

| | | | | | | | |
|---|---|---|---|---|---|---|---|
| E. coli | Nitrofurant | Calcein | 2 | 783 | [781 - 785] | 6860 | [6371 - 7350] |
| E. coli | Nitrofurant | Chlorophe | 2 | 1 | [1 - 1] | 6924 | [6387 - 7462] |
| E. coli | Nitrofurant | Congo Red | 2 | 1 | [1 - 1] | 6608 | [6432 - 6785] |
| E. coli | Nitrofurant | DiSC3(5) | 2 | 64771 | [33080 - 9 | 5860 | [5755 - 5966] |
| E. coli | Nitrofurant | Eosin Y | 2 | 738 | [569 - 906] | 6160 | [5578 - 6742] |
| E. coli | Nitrofurant | H2FDA | 2 | 1 | [1 - 1] | 6188 | [5410 - 6966] |
| E. coli | Nitrofurant | Malachite | 2 | 1 | [1 - 1] | 5862 | [5442 - 6283] |
| E. coli | Nitrofurant | Neutral Re | 2 | 1 | [1 - 1] | 6834 | [6786 - 6883] |
| E. coli | Nitrofurant | Oxonol | 2 | 1 | [1 - 1] | 7878 | [7545 - 8211] |
| E. coli | Nitrofurant | Pyronin Y | 2 | 1 | [1 - 1] | 5540 | [4743 - 6338] |
| E. coli | Nitrofurant | Quinacrine | 2 | 1195 | [1060 - 13 | 6092 | [6056 - 6127] |
| E. coli | Nitrofurant | Rhodamine | 2 | 1 | [1 - 1] | 5983 | [5893 - 6073] |
| E. coli | Nitrofurant | Rhodamine | 2 | 1 | [1 - 1] | 7674 | [7463 - 7886] |
| E. coli | Nitrofurant | Riboflavin | 2 | 1 | [1 - 1] | 6126 | [5222 - 7030] |
| E. coli | Nitrofurant | Sunset Yel | 2 | 1 | [1 - 1] | 7081 | [6843 - 7319] |
| E. coli | Nitrofurant | SYBR Gree | 2 | 4670 | [3324 - 60 | 6750 | [6669 - 6831] |
| E. coli | Nitrofurant | Thioflavin T | 2 | 1134 | [996 - 127 | 6658 | [5890 - 7426] |
| E. coli | None | 5-Carboxyf | 5 | 824 | [608 - 118 | 13840 | [9212 - 13841] |
| E. coli | None | Acridine or | 5 | 167 | [148 - 296] | 12393 | [10432 - 13095] |
| E. coli | None | Alizarin | 5 | 284 | [202 - 360] | 11592 | [9578 - 14283] |
| E. coli | None | Allura Red | 5 | 142 | [134 - 250] | 15149 | [10527 - 15993] |
| E. coli | None | Amaranth | 5 | 1914 | [1370 - 19 | 12066 | [9609 - 13439] |
| E. coli | None | Apigenin | 5 | 477 | [447 - 667] | 13004 | [10417 - 14242] |
| E. coli | None | Calcein | 5 | 1383 | [1209 - 16 | 13798 | [10827 - 14631] |
| E. coli | None | Chlorophe | 5 | 121 | [96 - 226] | 13479 | [8875 - 15238] |
| E. coli | None | Congo Red | 5 | 176 | [114 - 218] | 10841 | [9440 - 14998] |
| E. coli | None | DiSC3(5) | 5 | 62080 | [54549 - 6 | 12883 | [10007 - 13530] |
| E. coli | None | Eosin Y | 5 | 1124 | [896 - 198 | 14001 | [8426 - 17756] |
| E. coli | None | H2FDA | 5 | 172 | [143 - 397] | 14949 | [9871 - 15993] |
| E. coli | None | Malachite | 5 | 166 | [153 - 324] | 13570 | [9873 - 14678] |
| E. coli | None | Neutral Re | 5 | 237 | [159 - 496] | 12676 | [8189 - 14392] |
| E. coli | None | Oxonol | 5 | 141 | [71 - 269] | 9633 | [9577 - 13643] |
| E. coli | None | Pyronin Y | 5 | 205 | [132 - 415] | 12264 | [8877 - 16314] |
| E. coli | None | Quinacrine | 5 | 496 | [364 - 109 | 15281 | [10352 - 15973] |
| E. coli | None | Rhodamine | 5 | 172 | [159 - 492] | 13743 | [9225 - 16445] |
| E. coli | None | Rhodamine | 5 | 167 | [100 - 273] | 13000 | [10633 - 14815] |
| E. coli | None | Riboflavin | 5 | 125 | [88 - 352] | 14191 | [7955 - 16280] |
| E. coli | None | Sunset Yel | 5 | 32 | [23 - 102] | 15480 | [9652 - 15619] |
| E. coli | None | SYBR Gree | 5 | 11705 | [8214 - 13 | 11256 | [8751 - 14939] |
| E. coli | None | Thioflavin T | 5 | 506 | [503 - 867] | 12984 | [10679 - 12987] |
| E. coli | Ofloxacin | 5-Carboxyf | 3 | 1 | [1 - 1] | 10944 | [9804 - 11238] |
| E. coli | Ofloxacin | Acridine or | 3 | 1 | [1 - 1] | 10057 | [9780 - 10366] |
| E. coli | Ofloxacin | Alizarin | 3 | 1 | [1 - 1] | 9407 | [9103 - 9624] |
| E. coli | Ofloxacin | Allura Red | 3 | 1 | [1 - 18] | 11855 | [11649 - 11862] |
| E. coli | Ofloxacin | Amaranth | 3 | 1387 | [1257 - 18 | 9096 | [8108 - 9112] |
| E. coli | Ofloxacin | Apigenin | 3 | 1 | [1 - 273] | 14952 | [12606 - 15022] |
| E. coli | Ofloxacin | Calcein | 3 | 621 | [576 - 724] | 10466 | [10218 - 10798] |
| E. coli | Ofloxacin | Chlorophe | 3 | 1 | [1 - 1] | 12728 | [12197 - 13270] |

| | | | | | | | |
|---|---|---|---|---|---|---|---|
| E. coli | Ofloxacin | Congo Red | 3 | 1 | [1 - 1] | 10032 | [9034 - 10394] |
| E. coli | Ofloxacin | DiSC3(5) | 3 | 70649 | [58116 - 81 | 10846 | [9552 - 11916] |
| E. coli | Ofloxacin | Eosin Y | 3 | 571 | [393 - 878] | 10037 | [9346 - 11116] |
| E. coli | Ofloxacin | H2FDA | 3 | 1 | [1 - 1] | 9000 | [8865 - 10292] |
| E. coli | Ofloxacin | Malachite ( | 3 | 1 | [1 - 1] | 9876 | [9588 - 11720] |
| E. coli | Ofloxacin | Neutral Re | 3 | 1 | [1 - 1] | 9506 | [9384 - 9598] |
| E. coli | Ofloxacin | Oxonol | 3 | 1 | [1 - 156] | 12485 | [11111 - 14374] |
| E. coli | Ofloxacin | Pyronin Y | 3 | 1 | [1 - 1] | 10517 | [9641 - 10566] |
| E. coli | Ofloxacin | Quinacrine | 3 | 145 | [73 - 445] | 12608 | [11680 - 15012] |
| E. coli | Ofloxacin | Rhodamine | 3 | 1 | [1 - 1] | 8880 | [8734 - 9295] |
| E. coli | Ofloxacin | Rhodamine | 3 | 1 | [1 - 1] | 12228 | [11140 - 12553] |
| E. coli | Ofloxacin | Riboflavin | 3 | 1 | [1 - 1] | 6848 | [6809 - 7824] |
| E. coli | Ofloxacin | Sunset Yel | 3 | 1 | [1 - 1] | 10809 | [10298 - 11803] |
| E. coli | Ofloxacin | SYBR Gree | 3 | 15136 | [12829 - 15 | 12796 | [12346 - 13192] |
| E. coli | Ofloxacin | Thioflavin T | 3 | 1 | [1 - 86] | 16656 | [15130 - 16882] |
| E. coli | Pentamidi | 5-Carboxyf | 2 | 1 | [1 - 1] | 6940 | [6526 - 7354] |
| E. coli | Pentamidi | Acridine or | 2 | 1 | [1 - 1] | 6724 | [6362 - 7087] |
| E. coli | Pentamidi | Alizarin | 2 | 1 | [1 - 1] | 5954 | [5907 - 6001] |
| E. coli | Pentamidi | Allura Red | 2 | 1 | [1 - 1] | 7784 | [7248 - 8319] |
| E. coli | Pentamidi | Amaranth | 2 | 176 | [89 - 264] | 7044 | [6994 - 7094] |
| E. coli | Pentamidi | Apigenin | 2 | 681 | [341 - 102 | 6898 | [6254 - 7542] |
| E. coli | Pentamidi | Calcein | 2 | 1579 | [886 - 227 | 6252 | [5269 - 7235] |
| E. coli | Pentamidi | Chlorophe | 2 | 1 | [1 - 1] | 6278 | [5894 - 6662] |
| E. coli | Pentamidi | Congo Red | 2 | 1 | [1 - 1] | 7084 | [6748 - 7420] |
| E. coli | Pentamidi | DiSC3(5) | 2 | 161636 | [127997 - 1 | 5610 | [4969 - 6251] |
| E. coli | Pentamidi | Eosin Y | 2 | 172 | [86 - 257] | 6442 | [6285 - 6598] |
| E. coli | Pentamidi | H2FDA | 2 | 1 | [1 - 1] | 6370 | [6314 - 6426] |
| E. coli | Pentamidi | Malachite ( | 2 | 1 | [1 - 1] | 6554 | [6234 - 6874] |
| E. coli | Pentamidi | Neutral Re | 2 | 1 | [1 - 1] | 5914 | [5614 - 6214] |
| E. coli | Pentamidi | Oxonol | 2 | 1 | [1 - 1] | 8460 | [8024 - 8897] |
| E. coli | Pentamidi | Pyronin Y | 2 | 1 | [1 - 1] | 6846 | [6604 - 7089] |
| E. coli | Pentamidi | Quinacrine | 2 | 840 | [421 - 126( | 6971 | [6052 - 7890] |
| E. coli | Pentamidi | Rhodamine | 2 | 1 | [1 - 1] | 6862 | [6154 - 7570] |
| E. coli | Pentamidi | Rhodamine | 2 | 1 | [1 - 1] | 5962 | [5787 - 6138] |
| E. coli | Pentamidi | Riboflavin | 2 | 1 | [1 - 1] | 5034 | [5028 - 5039] |
| E. coli | Pentamidi | Sunset Yel | 2 | 1 | [1 - 1] | 6478 | [6058 - 6897] |
| E. coli | Pentamidi | SYBR Gree | 2 | 84616 | [68920 - 10 | 5133 | [5000 - 5266] |
| E. coli | Pentamidi | Thioflavin T | 2 | 755 | [483 - 102 | 6008 | [5993 - 6024] |
| E. coli | Polymyxin | 5-Carboxyf | 3 | 1 | [1 - 1] | 11333 | [11102 - 12833] |
| E. coli | Polymyxin | Acridine or | 3 | 1 | [1 - 1] | 13432 | [13024 - 14123] |
| E. coli | Polymyxin | Alizarin | 3 | 1 | [1 - 1] | 15239 | [13989 - 15490] |
| E. coli | Polymyxin | Allura Red | 3 | 1 | [1 - 1] | 10273 | [10224 - 12130] |
| E. coli | Polymyxin | Amaranth | 3 | 166 | [83 - 1906] | 13436 | [13150 - 15460] |
| E. coli | Polymyxin | Apigenin | 3 | 1 | [1 - 262] | 13152 | [12227 - 13362] |
| E. coli | Polymyxin | Calcein | 3 | 1 | [1 - 702] | 12376 | [11895 - 13280] |
| E. coli | Polymyxin | Chlorophe | 3 | 1 | [1 - 1] | 15374 | [14796 - 15730] |
| E. coli | Polymyxin | Congo Red | 3 | 1 | [1 - 1] | 11081 | [10907 - 11481] |
| E. coli | Polymyxin | DiSC3(5) | 3 | 52844 | [49110 - 57 | 14805 | [12819 - 15980] |

| | | | | | | | |
|---|---|---|---|---|---|---|---|
| E. coli | Polymyxin | Eosin Y | 3 | 1 | [1 - 742] | 13773 | [13449 - 14244] |
| E. coli | Polymyxin | H2FDA | 3 | 1 | [1 - 1] | 12596 | [12545 - 13042] |
| E. coli | Polymyxin | Malachite ( | 3 | 1 | [1 - 1] | 14474 | [13484 - 15180] |
| E. coli | Polymyxin | Neutral Re | 3 | 1 | [1 - 1] | 14303 | [12632 - 15460] |
| E. coli | Polymyxin | Oxonol | 3 | 1 | [1 - 1] | 14803 | [13866 - 15204] |
| E. coli | Polymyxin | Pyronin Y | 3 | 1 | [1 - 1] | 12546 | [11780 - 13500] |
| E. coli | Polymyxin | Quinacrine | 3 | 152 | [77 - 205] | 16508 | [13864 - 17076] |
| E. coli | Polymyxin | Rhodamine | 3 | 1 | [1 - 1] | 14822 | [13468 - 15392] |
| E. coli | Polymyxin | Rhodamine | 3 | 1 | [1 - 1] | 13944 | [13568 - 14990] |
| E. coli | Polymyxin | Riboflavin | 3 | 1 | [1 - 1] | 11562 | [11080 - 11676] |
| E. coli | Polymyxin | Sunset Yel | 3 | 1 | [1 - 1] | 13057 | [12058 - 13677] |
| E. coli | Polymyxin | SYBR Gree | 3 | 10586 | [7909 - 10 | 13451 | [11878 - 15008] |
| E. coli | Polymyxin | Thioflavin T | 3 | 27 | [14 - 106] | 14316 | [14157 - 14450] |
| E. coli | Rifampicin | 5-Carboxyf | 2 | 123 | [62 - 185] | 6789 | [6260 - 7318] |
| E. coli | Rifampicin | Acridine or | 2 | 1 | [1 - 1] | 8326 | [8153 - 8500] |
| E. coli | Rifampicin | Alizarin | 2 | 1 | [1 - 1] | 8506 | [7916 - 9095] |
| E. coli | Rifampicin | Allura Red | 2 | 1 | [1 - 1] | 8322 | [7584 - 9060] |
| E. coli | Rifampicin | Amaranth | 2 | 521 | [261 - 781] | 8296 | [7531 - 9062] |
| E. coli | Rifampicin | Apigenin | 2 | 205 | [103 - 307] | 8335 | [7879 - 8791] |
| E. coli | Rifampicin | Calcein | 2 | 73 | [37 - 109] | 9060 | [8909 - 9211] |
| E. coli | Rifampicin | Chlorophe | 2 | 1 | [1 - 1] | 8901 | [8685 - 9117] |
| E. coli | Rifampicin | Congo Red | 2 | 1 | [1 - 1] | 7424 | [6892 - 7955] |
| E. coli | Rifampicin | DiSC3(5) | 2 | 89593 | [88797 - 9( | 6544 | [6526 - 6562] |
| E. coli | Rifampicin | Eosin Y | 2 | 1 | [1 - 1] | 8054 | [7899 - 8209] |
| E. coli | Rifampicin | H2FDA | 2 | 1 | [1 - 1] | 9089 | [8312 - 9866] |
| E. coli | Rifampicin | Malachite ( | 2 | 1 | [1 - 1] | 8516 | [7812 - 9221] |
| E. coli | Rifampicin | Neutral Re | 2 | 1 | [1 - 1] | 7692 | [7432 - 7953] |
| E. coli | Rifampicin | Oxonol | 2 | 649 | [646 - 652] | 6502 | [6296 - 6708] |
| E. coli | Rifampicin | Pyronin Y | 2 | 1 | [1 - 1] | 7789 | [7618 - 7960] |
| E. coli | Rifampicin | Quinacrine | 2 | 147 | [74 - 221] | 8768 | [8399 - 9138] |
| E. coli | Rifampicin | Rhodamine | 2 | 1 | [1 - 1] | 7800 | [7724 - 7875] |
| E. coli | Rifampicin | Rhodamine | 2 | 1 | [1 - 1] | 8140 | [7638 - 8643] |
| E. coli | Rifampicin | Riboflavin | 2 | 1 | [1 - 1] | 7797 | [7663 - 7931] |
| E. coli | Rifampicin | Sunset Yel | 2 | 1 | [1 - 1] | 8508 | [8265 - 8751] |
| E. coli | Rifampicin | SYBR Gree | 2 | 3386 | [1693 - 507 | 8503 | [7746 - 9260] |
| E. coli | Rifampicin | Thioflavin T | 2 | 72 | [37 - 108] | 10172 | [9985 - 10358] |
| E. coli | Tetracyclin | 5-Carboxyf | 3 | 1 | [1 - 1] | 14607 | [14213 - 16183] |
| E. coli | Tetracyclin | Acridine or | 3 | 1 | [1 - 1] | 13388 | [11784 - 13960] |
| E. coli | Tetracyclin | Alizarin | 3 | 1 | [1 - 1] | 11279 | [10617 - 12814] |
| E. coli | Tetracyclin | Allura Red | 3 | 1 | [1 - 1] | 13706 | [12958 - 13830] |
| E. coli | Tetracyclin | Amaranth | 3 | 36 | [19 - 287] | 10724 | [10620 - 11898] |
| E. coli | Tetracyclin | Apigenin | 3 | 1 | [1 - 1] | 17181 | [15574 - 18104] |
| E. coli | Tetracyclin | Calcein | 3 | 1 | [1 - 60] | 14288 | [13986 - 14387] |
| E. coli | Tetracyclin | Chlorophe | 3 | 1 | [1 - 1] | 13962 | [13588 - 14559] |
| E. coli | Tetracyclin | Congo Red | 3 | 1 | [1 - 1] | 10604 | [10507 - 11574] |
| E. coli | Tetracyclin | DiSC3(5) | 3 | 50894 | [49668 - 5: | 12128 | [11966 - 12914] |
| E. coli | Tetracyclin | Eosin Y | 3 | 1 | [1 - 1] | 12504 | [11616 - 13360] |
| E. coli | Tetracyclin | H2FDA | 3 | 1 | [1 - 1] | 11860 | [11848 - 12124] |

| | | | | | | | |
|---|---|---|---|---|---|---|---|
| E. coli | Tetracyclin | Malachite ( | 3 | 1 | [1 - 1] | 16125 | [15433 - 16257] |
| E. coli | Tetracyclin | Neutral Re | 3 | 1 | [1 - 1] | 15128 | [13312 - 15339] |
| E. coli | Tetracyclin | Oxonol | 3 | 16 | [8 - 37] | 11747 | [11506 - 12052] |
| E. coli | Tetracyclin | Pyronin Y | 3 | 1 | [1 - 1] | 11672 | [11268 - 11682] |
| E. coli | Tetracyclin | Quinacrine | 3 | 444 | [389 - 507] | 11392 | [10918 - 11738] |
| E. coli | Tetracyclin | Rhodamine | 3 | 1 | [1 - 1] | 11957 | [11814 - 12358] |
| E. coli | Tetracyclin | Rhodamine | 3 | 1 | [1 - 1] | 13211 | [12478 - 14940] |
| E. coli | Tetracyclin | Riboflavin | 3 | 1 | [1 - 1] | 14014 | [13144 - 14338] |
| E. coli | Tetracyclin | Sunset Yel | 3 | 1 | [1 - 1] | 13540 | [13527 - 14550] |
| E. coli | Tetracyclin | SYBR Gree | 3 | 9806 | [9487 - 10 | 12934 | [12730 - 13176] |
| E. coli | Tetracyclin | Thioflavin T | 3 | 1 | [1 - 1] | 16759 | [16013 - 17990] |
| E. coli | Trimethopr | 5-Carboxyf | 4 | 1 | [1 - 1] | 7875 | [7778 - 7969] |
| E. coli | Trimethopr | Acridine or | 4 | 1 | [1 - 1] | 8271 | [8199 - 8427] |
| E. coli | Trimethopr | Alizarin | 4 | 1 | [1 - 1] | 7308 | [7105 - 7531] |
| E. coli | Trimethopr | Allura Red | 4 | 1 | [1 - 1] | 7473 | [7280 - 7798] |
| E. coli | Trimethopr | Amaranth | 4 | NA | [117 - 397] | 7139 | [5134 - 7614] |
| E. coli | Trimethopr | Apigenin | 4 | 119 | [46 - 292] | 7986 | [7602 - 8256] |
| E. coli | Trimethopr | Calcein | 4 | 134 | [1 - 317] | 8358 | [7922 - 8616] |
| E. coli | Trimethopr | Chlorophe | 4 | 1 | [1 - 1] | 8040 | [7331 - 8251] |
| E. coli | Trimethopr | Congo Red | 4 | 1 | [1 - 1] | 6926 | [6486 - 7645] |
| E. coli | Trimethopr | DiSC3(5) | 4 | 85120 | [80756 - 9( | 7198 | [7024 - 7303] |
| E. coli | Trimethopr | Eosin Y | 4 | 96 | [1 - 206] | 6876 | [6540 - 7106] |
| E. coli | Trimethopr | H2FDA | 4 | 1 | [1 - 1] | 7780 | [7153 - 8086] |
| E. coli | Trimethopr | Malachite ( | 4 | 1 | [1 - 1] | 6735 | [6595 - 7317] |
| E. coli | Trimethopr | Neutral Re | 4 | 1 | [1 - 1] | 7832 | [7340 - 8227] |
| E. coli | Trimethopr | Oxonol | 4 | 36 | [1 - 265] | 7114 | [6262 - 7924] |
| E. coli | Trimethopr | Pyronin Y | 4 | 1 | [1 - 1] | 6778 | [6438 - 7233] |
| E. coli | Trimethopr | Quinacrine | 4 | 674 | [482 - 838] | 6845 | [6505 - 7090] |
| E. coli | Trimethopr | Rhodamine | 4 | 1 | [1 - 1] | 6649 | [5564 - 8070] |
| E. coli | Trimethopr | Rhodamine | 4 | 1 | [1 - 1] | 7891 | [7478 - 8070] |
| E. coli | Trimethopr | Riboflavin | 4 | 1 | [1 - 1] | 7053 | [6431 - 7782] |
| E. coli | Trimethopr | Sunset Yel | 4 | 1 | [1 - 1] | 7672 | [7102 - 8390] |
| E. coli | Trimethopr | SYBR Gree | 4 | 10781 | [9382 - 12( | 7368 | [7115 - 7852] |
| E. coli | Trimethopr | Thioflavin T | 4 | 332 | [218 - 411] | 7934 | [7746 - 8234] |
| E. coli | Vancomyc | 5-Carboxyf | 3 | 1 | [1 - 233] | 12580 | [11002 - 13158] |
| E. coli | Vancomyc | Acridine or | 3 | 1 | [1 - 1] | 14732 | [11932 - 15166] |
| E. coli | Vancomyc | Alizarin | 3 | 1 | [1 - 1] | 11204 | [10443 - 13194] |
| E. coli | Vancomyc | Allura Red | 3 | 531 | [481 - 627] | 10889 | [10601 - 11988] |
| E. coli | Vancomyc | Amaranth | 3 | 3117 | [2930 - 41 | 14278 | [12594 - 14448] |
| E. coli | Vancomyc | Apigenin | 3 | 700 | [520 - 718] | 13502 | [13262 - 13996] |
| E. coli | Vancomyc | Calcein | 3 | 3702 | [3543 - 38 | 11212 | [10764 - 11678] |
| E. coli | Vancomyc | Chlorophe | 3 | 1 | [1 - 1] | 12214 | [11288 - 13143] |
| E. coli | Vancomyc | Congo Red | 3 | 1 | [1 - 1] | 11309 | [11098 - 11418] |
| E. coli | Vancomyc | DiSC3(5) | 3 | 56241 | [55960 - 6 | 11672 | [11032 - 12871] |
| E. coli | Vancomyc | Eosin Y | 3 | 1614 | [1538 - 18 | 12334 | [11802 - 13723] |
| E. coli | Vancomyc | H2FDA | 3 | 1 | [1 - 1] | 11549 | [11158 - 12582] |
| E. coli | Vancomyc | Malachite ( | 3 | 1 | [1 - 1] | 12712 | [11794 - 12783] |
| E. coli | Vancomyc | Neutral Re | 3 | 1 | [1 - 1] | 10139 | [10102 - 11249] |

| | | | | | | | |
|---|---|---|---|---|---|---|---|
| E. coli | Vancomyc | Oxonol | 3 | 117 | [59 - 189] | 11518 | [11367 - 13542] |
| E. coli | Vancomyc | Pyronin Y | 3 | 1 | [1 - 1] | 12856 | [12550 - 13013] |
| E. coli | Vancomyc | Quinacrine | 3 | 889 | [707 - 949] | 13779 | [12362 - 14788] |
| E. coli | Vancomyc | Rhodamine | 3 | 1 | [1 - 1] | 11639 | [10728 - 13209] |
| E. coli | Vancomyc | Rhodamine | 3 | 1 | [1 - 1] | 10924 | [10668 - 12170] |
| E. coli | Vancomyc | Riboflavin | 3 | 1 | [1 - 1] | 13807 | [13282 - 14299] |
| E. coli | Vancomyc | Sunset Yell | 3 | 1 | [1 - 1] | 13673 | [13495 - 14438] |
| E. coli | Vancomyc | SYBR Gree | 3 | 18505 | [15886 - 18 | 11236 | [10190 - 11460] |
| E. coli | Vancomyc | Thioflavin T | 3 | 719 | [626 - 733] | 11481 | [11022 - 12586] |
