## Supplementary table 3 for "Rapid accumulation of fluorophores and fast kill identify drugs with bactericidal effects against Gram-negative bacteria"

| PID | Name |
|---|---|
| Prestw-1 | Azaguanine-8 |
| Prestw-10 | Sulfaguanidine |
| Prestw-100 | Nocodazole |
| Prestw-1000 | Lymecycline |
| Prestw-1001 | Alfadolone acetate |
| Prestw-1002 | Alfaxalone |
| Prestw-1003 | Azapropazone |
| Prestw-1004 | Meptazinol hydrochloride |
| Prestw-1005 | Apramycin |
| Prestw-1007 | Fursultiamine Hydrochloride |
| Prestw-1008 | Gabexate mesilate |
| Prestw-1009 | Pivampicillin |
| Prestw-101 | R(-) Apomorphine hydrochloride hemihydrate |
| Prestw-1011 | Flucloxacillin sodium |
| Prestw-1012 | Trapidil |
| Prestw-1013 | Deptropine citrate |
| Prestw-1014 | Sertraline |
| Prestw-1015 | Ethamsylate |
| Prestw-1016 | Moxonidine |
| Prestw-1017 | Etilefrine hydrochloride |
| Prestw-1018 | Alprostadil |
| Prestw-1019 | Tribenoside |
| Prestw-102 | Amoxapine |
| Prestw-1020 | Rimexolone |
| Prestw-1021 | Isradipine |
| Prestw-1023 | Isometheptene mucate |
| Prestw-1024 | Nifurtimox |
| Prestw-1025 | Letrozole |
| Prestw-1027 | Tocainide hydrochloride |
| Prestw-1028 | Benzathine benzylpenicillin |
| Prestw-1029 | Risperidone |
| Prestw-103 | Cyproheptadine hydrochloride |
| Prestw-1030 | Torsemide |
| Prestw-1031 | Halofantrine hydrochloride |
| Prestw-1032 | Articaine hydrochloride |
| Prestw-1033 | Nomegestrol acetate |
| Prestw-1034 | Pancuronium bromide |
| Prestw-1035 | Molindone hydrochloride |
| Prestw-1036 | Alcuronium chloride |
| Prestw-1037 | Zalcitabine |
| Prestw-1038 | Methyldopate hydrochloride |
| Prestw-1039 | Levocabastine hydrochloride |
| Prestw-104 | Famotidine |
| Prestw-1040 | Pyrvinium pamoate |

| | |
|---|---|
| Prestw-1041 | Etomidate |
| Prestw-1042 | Tridihexethyl chloride |
| Prestw-1043 | Penbutolol sulfate |
| Prestw-1044 | Prednicarbate |
| Prestw-1045 | Sertaconazole nitrate |
| Prestw-1046 | Repaglinide |
| Prestw-1047 | Piretanide |
| Prestw-1048 | Piperacetazine |
| Prestw-1049 | Oxyphenbutazone |
| Prestw-105 | Danazol |
| Prestw-1050 | Quinethazone |
| Prestw-1051 | Moricizine hydrochloride |
| Prestw-1052 | Iopanoic acid |
| Prestw-1053 | Pivmecillinam hydrochloride |
| Prestw-1054 | Levopropoxyphene napsylate |
| Prestw-1055 | Piperidolate hydrochloride |
| Prestw-1056 | Trifluridine |
| Prestw-1057 | Oxprenolol hydrochloride |
| Prestw-1058 | Ondansetron Hydrochloride |
| Prestw-1059 | Propoxycaine hydrochloride |
| Prestw-106 | Nicorandil |
| Prestw-1060 | Oxaprozin |
| Prestw-1061 | Phensuximide |
| Prestw-1062 | Ioxaglic acid |
| Prestw-1063 | Naftifine hydrochloride |
| Prestw-1064 | Meprylcaine hydrochloride |
| Prestw-1065 | Milrinone |
| Prestw-1066 | Methantheline bromide |
| Prestw-1067 | Ticarcillin sodium |
| Prestw-1068 | Thiethylperazine dimalate |
| Prestw-1069 | Mesalamine |
| Prestw-1071 | Imidurea |
| Prestw-1072 | Lansoprazole |
| Prestw-1073 | Bethanechol chloride |
| Prestw-1074 | Cyproterone acetate |
| Prestw-1075 | (R)-Propranolol hydrochloride |
| Prestw-1076 | Ciprofibrate |
| Prestw-1078 | Benzylpenicillin sodium |
| Prestw-108 | Nomifensine maleate |
| Prestw-1080 | Methiazole |
| Prestw-1081 | (S)-propranolol hydrochloride |
| Prestw-1082 | (-)-Eseroline fumarate salt |
| Prestw-1086 | D-cycloserine |
| Prestw-1088 | (+,-)-Synephrine |
| Prestw-1089 | (S)-(-)-Cycloserine |

| | |
|---|---|
| Prestw-1090 | Homosalate |
| Prestw-1091 | Spaglumic acid |
| Prestw-1092 | Ranolazine |
| Prestw-1094 | Sulfadoxine |
| Prestw-1095 | Cyclopentolate hydrochloride |
| Prestw-1096 | Estriol |
| Prestw-1097 | (-)-Isoproterenol hydrochloride |
| Prestw-1099 | Nialamide |
| Prestw-11 | Meticrane |
| Prestw-110 | Acenocoumarol |
| Prestw-1101 | Perindopril |
| Prestw-1102 | Fexofenadine hydrochloride |
| Prestw-1104 | Clonixin Lysinate |
| Prestw-1105 | Verteporfin |
| Prestw-1106 | Meropenem |
| Prestw-1107 | Ramipril |
| Prestw-1108 | Mephenytoin |
| Prestw-1109 | Rifabutin |
| Prestw-111 | Naloxone hydrochloride |
| Prestw-1110 | Parbendazole |
| Prestw-1111 | Mecamylamine hydrochloride |
| Prestw-1112 | Procarbazine hydrochloride |
| Prestw-1113 | Viomycin sulfate |
| Prestw-1114 | Saquinavir mesylate |
| Prestw-1115 | Ronidazole |
| Prestw-1116 | Dorzolamide hydrochloride |
| Prestw-1117 | Azaperone |
| Prestw-1118 | Cefepime hydrochloride |
| Prestw-1119 | Clocortolone pivalate |
| Prestw-112 | Metolazone |
| Prestw-1120 | Nadifloxacin |
| Prestw-1121 | Carbadox |
| Prestw-1125 | Oxiconazole Nitrate |
| Prestw-1127 | Acipimox |
| Prestw-1129 | Benazepril hydrochloride |
| Prestw-113 | Ciprofloxacin hydrochloride monohydrate |
| Prestw-1130 | Azelastine hydrochloride |
| Prestw-1132 | Celiprolol hydrochloride |
| Prestw-1134 | Cytarabine |
| Prestw-1136 | Doxofylline |
| Prestw-1137 | Esmolol hydrochloride |
| Prestw-1139 | Itraconazole |
| Prestw-114 | Ampicillin trihydrate |
| Prestw-1140 | Liranaftate |
| Prestw-1144 | Mirtazapine |

| | |
|---|---|
| Prestw-1147 | Modafinil |
| Prestw-115 | Haloperidol |
| Prestw-1152 | Nefazodone hydrochloride |
| Prestw-1154 | Nilvadipine |
| Prestw-1156 | Oxcarbazepine |
| Prestw-1157 | Rifapentine |
| Prestw-1158 | Ropinirole hydrochloride |
| Prestw-1159 | Sibutramine hydrochloride |
| Prestw-116 | Naltrexone hydrochloride dihydrate |
| Prestw-1161 | Stanozolol |
| Prestw-1162 | Zonisamide |
| Prestw-1165 | Acitretin |
| Prestw-1166 | Rebamipide |
| Prestw-1167 | Diacerein |
| Prestw-1169 | Miglitol |
| Prestw-117 | Chlorpheniramine maleate |
| Prestw-1170 | Venlafaxine |
| Prestw-1173 | Irsogladine maleate |
| Prestw-1174 | Acarbose |
| Prestw-1177 | Carbidopa |
| Prestw-1178 | Aniracetam |
| Prestw-1179 | Busulfan |
| Prestw-118 | Nalbuphine hydrochloride |
| Prestw-1180 | Docetaxel |
| Prestw-1181 | Tibolone |
| Prestw-1182 | Tizanidine hydrochloride |
| Prestw-1183 | Temozolomide |
| Prestw-1184 | Tioconazole |
| Prestw-1187 | Granisetron |
| Prestw-1188 | Ziprasidone Hydrochloride |
| Prestw-1189 | Montelukast |
| Prestw-119 | Picotamide monohydrate |
| Prestw-1190 | Olmesartan |
| Prestw-1192 | Oxandrolone |
| Prestw-1195 | Toltrazuril |
| Prestw-1196 | Topotecan |
| Prestw-1197 | Toremifene |
| Prestw-1198 | Tranilast |
| Prestw-1199 | Tripelennamine hydrochloride |
| Prestw-12 | Benzonatate |
| Prestw-120 | Triamcinolone |
| Prestw-1202 | 4-aminosalicylic acid |
| Prestw-1203 | 5-fluorouracil |
| Prestw-1206 | Acetylcysteine |
| Prestw-1207 | Acetylsalicylic acid |

| | |
|---|---|
| Prestw-121 | Bromocryptine mesylate |
| Prestw-1210 | Alendronate sodium |
| Prestw-1211 | Alfacalcidol |
| Prestw-1213 | Allopurinol |
| Prestw-1217 | Amisulpride |
| Prestw-1219 | Amlodipine |
| Prestw-1222 | Anastrozole |
| Prestw-1223 | Anethole-trithione |
| Prestw-1224 | Anthralin |
| Prestw-1228 | Argatroban |
| Prestw-1229 | Aripiprazole |
| Prestw-123 | Dehydrocholic acid |
| Prestw-1231 | Asenapine maleate |
| Prestw-1232 | Atorvastatin |
| Prestw-1233 | Auranofin |
| Prestw-1234 | Azithromycin |
| Prestw-1236 | Benztropine mesylate |
| Prestw-1239 | Bicalutamide |
| Prestw-1241 | Bifonazole |
| Prestw-1242 | Erlotinib |
| Prestw-1244 | Bosentan |
| Prestw-1246 | Bromhexine hydrochloride |
| Prestw-1249 | Famciclovir |
| Prestw-125 | Perphenazine |
| Prestw-1251 | Butalbital |
| Prestw-1252 | Butenafine Hydrochloride |
| Prestw-1253 | Butylscopolammonium (n-) bromide |
| Prestw-1254 | Fentiazac |
| Prestw-1256 | Caffeine |
| Prestw-1257 | Calcipotriene |
| Prestw-1258 | Candesartan |
| Prestw-1259 | Canrenone |
| Prestw-126 | Mefloquine hydrochloride |
| Prestw-1261 | Carprofen |
| Prestw-1262 | Carvedilol |
| Prestw-1263 | Cefdinir |
| Prestw-1265 | Gatifloxacin |
| Prestw-1266 | Gemcitabine |
| Prestw-1267 | Gestrinone |
| Prestw-1268 | Guaiacol |
| Prestw-1269 | Haloprogin |
| Prestw-127 | Isoconazole |
| Prestw-1270 | Gefitinib |
| Prestw-1271 | Escitalopram oxalate |
| Prestw-1274 | Emedastine |

| | |
|---|---|
| Prestw-1279 | Stavudine |
| Prestw-128 | Spironolactone |
| Prestw-1280 | Mepivacaine hydrochloride |
| Prestw-1282 | Methenamine |
| Prestw-1283 | Buspirone hydrochloride |
| Prestw-1284 | Hydroxychloroquine sulfate |
| Prestw-1285 | Ibandronate sodium |
| Prestw-1286 | Ibudilast |
| Prestw-1288 | Idebenone |
| Prestw-129 | Pirenzepine dihydrochloride |
| Prestw-1290 | Imatinib |
| Prestw-1291 | Imiquimod |
| Prestw-1292 | Ipsapirone |
| Prestw-1294 | Isosorbide mononitrate |
| Prestw-1295 | Itopride |
| Prestw-1297 | Lacidipine |
| Prestw-1298 | Lamivudine |
| Prestw-13 | Hydroflumethiazide |
| Prestw-130 | Dexamethasone acetate |
| Prestw-1303 | Pefloxacine |
| Prestw-1307 | Olopatadine hydrochloride |
| Prestw-1308 | Phentermine hydrochloride |
| Prestw-131 | Glipizide |
| Prestw-1310 | Phenylbutazone |
| Prestw-1314 | Pioglitazone |
| Prestw-1315 | Clavulanate potassium salt |
| Prestw-1316 | Pramipexole dihydrochloride |
| Prestw-1317 | Pranlukast |
| Prestw-1318 | Pranoprofen |
| Prestw-1319 | Pravastatin |
| Prestw-132 | Loxapine succinate |
| Prestw-1321 | Prothionamide |
| Prestw-1322 | Pyridostigmine iodide |
| Prestw-1323 | Quetiapine hemifumarate |
| Prestw-1325 | Raclopride |
| Prestw-1328 | Reboxetine mesylate |
| Prestw-133 | Hydroxyzine dihydrochloride |
| Prestw-1331 | Rimantadine Hydrochloride |
| Prestw-1334 | Rivastigmine |
| Prestw-1336 | Rofecoxib |
| Prestw-1337 | Rosiglitazone Hydrochloride |
| Prestw-1338 | Rufloxacin |
| Prestw-1339 | Sarafloxacin |
| Prestw-134 | Diltiazem hydrochloride |
| Prestw-1340 | Secnidazole |

| | |
|---|---|
| Prestw-1341 | Sertindole |
| Prestw-1342 | Sildenafil |
| Prestw-1343 | Sparfloxacin |
| Prestw-1345 | Sulbactam |
| Prestw-1346 | Sumatriptan succinate |
| Prestw-1349 | Tazobactam |
| Prestw-1350 | Telmisartan |
| Prestw-1351 | Tenatoprazole |
| Prestw-1352 | Tulobuterol |
| Prestw-1353 | Tylosin |
| Prestw-1356 | Vardenafil |
| Prestw-1358 | Vatalanib |
| Prestw-1359 | Vecuronium bromide |
| Prestw-136 | Astemizole |
| Prestw-1361 | Viloxazine hydrochloride |
| Prestw-1362 | Vorinostat |
| Prestw-1363 | Warfarin |
| Prestw-1364 | Zafirlukast |
| Prestw-1365 | Zileuton |
| Prestw-1367 | Zopiclone |
| Prestw-1368 | Zotepine |
| Prestw-1369 | Zaleplon |
| Prestw-137 | Clindamycin hydrochloride |
| Prestw-1371 | Celecoxib |
| Prestw-1374 | Chlormadinone acetate |
| Prestw-1376 | Cilnidipine |
| Prestw-1378 | Clarithromycin |
| Prestw-1379 | Clinafloxacin |
| Prestw-138 | Terfenadine |
| Prestw-1380 | Clobutinol hydrochloride |
| Prestw-1383 | Clofibrate |
| Prestw-1385 | Closantel |
| Prestw-1387 | Tetraethylenepentamine pentahydrochloride |
| Prestw-139 | Cefotaxime sodium salt |
| Prestw-1390 | Desloratadine |
| Prestw-1392 | Dexfenfluramine hydrochloride |
| Prestw-1393 | Dibenzepine hydrochloride |
| Prestw-1394 | Diclazuril |
| Prestw-1398 | Dopamine hydrochloride |
| Prestw-1399 | Doxycycline hydrochloride |
| Prestw-14 | Sulfacetamide sodic hydrate |
| Prestw-140 | Tetracycline hydrochloride |
| Prestw-1400 | Efavirenz |
| Prestw-1401 | Enoxacin |
| Prestw-1403 | Entacapone |

| | |
|---|---|
| Prestw-1405 | Ethinylestradiol |
| Prestw-1407 | Etofenamate |
| Prestw-1408 | Etoricoxib |
| Prestw-1409 | Etretinate |
| Prestw-141 | Verapamil hydrochloride |
| Prestw-1410 | Exemestane |
| Prestw-1414 | Fleroxacin |
| Prestw-1415 | Floxuridine |
| Prestw-1416 | Flubendazol |
| Prestw-1417 | Fluconazole |
| Prestw-1419 | Fluocinolone acetonide |
| Prestw-142 | Dipyridamole |
| Prestw-1420 | Formestane |
| Prestw-1421 | Formoterol fumarate |
| Prestw-1423 | Fosinopril |
| Prestw-1424 | Fulvestrant |
| Prestw-1427 | Levetiracetam |
| Prestw-1429 | Linezolid |
| Prestw-143 | Chlorhexidine |
| Prestw-1430 | Lofexidine |
| Prestw-1431 | Loracarbef |
| Prestw-1432 | Loratadine |
| Prestw-1433 | Losartan |
| Prestw-1435 | Melengestrol acetate |
| Prestw-144 | Loperamide hydrochloride |
| Prestw-1441 | Mevastatin |
| Prestw-1443 | Misoprostol |
| Prestw-1444 | Mitotane |
| Prestw-1446 | Moxifloxacin |
| Prestw-1447 | Nalidixic acid sodium salt |
| Prestw-1449 | Nicotinamide |
| Prestw-145 | Chlortetracycline hydrochloride |
| Prestw-1452 | Norgestimate |
| Prestw-1454 | Nylidrin |
| Prestw-1455 | Olanzapine |
| Prestw-1456 | Opipramol dihydrochloride |
| Prestw-1459 | Oxfendazol |
| Prestw-146 | Tamoxifen citrate |
| Prestw-1460 | Oxibendazol |
| Prestw-1463 | Tomoxetine hydrochloride |
| Prestw-1464 | Tosufloxacin hydrochloride |
| Prestw-1465 | Tramadol hydrochloride |
| Prestw-1467 | Troglitazone |
| Prestw-1469 | Mercaptopurine |
| Prestw-147 | Nicergoline |

| | |
|---|---|
| Prestw-1471 | Amfepramone hydrochloride |
| Prestw-1472 | Hexachlorophene |
| Prestw-1473 | Estradiol Valerate |
| Prestw-1474 | Chloroxine |
| Prestw-1475 | Oxacillin sodium |
| Prestw-1476 | Amcinonide |
| Prestw-1477 | D,L-Penicillamine |
| Prestw-1478 | Rifaximin |
| Prestw-1479 | Triclosan |
| Prestw-148 | Canrenoic acid potassium salt |
| Prestw-1480 | Racepinephrine hydrochloride |
| Prestw-1482 | Valproic acid |
| Prestw-1483 | Fludarabine |
| Prestw-1484 | Cladribine |
| Prestw-1486 | Cortisol acetate |
| Prestw-1487 | Mesna |
| Prestw-1488 | Penciclovir |
| Prestw-1489 | Amifostine |
| Prestw-149 | Thioproperazine dimesylate |
| Prestw-1490 | Nalmefene hydrochloride |
| Prestw-1491 | Pentobarbital |
| Prestw-1492 | Lamotrigine |
| Prestw-1493 | Topiramate |
| Prestw-1494 | Irinotecan hydrochloride trihydrate |
| Prestw-1495 | Rabeprazole Sodium salt |
| Prestw-1497 | Ambrisentan |
| Prestw-1498 | Camylofine chlorhydrate |
| Prestw-1499 | Fomepizole |
| Prestw-15 | Heptaminol hydrochloride |
| Prestw-150 | Dihydroergotamine tartrate |
| Prestw-1500 | Voriconazole |
| Prestw-1501 | Fenipentol |
| Prestw-1502 | Acamprosate calcium |
| Prestw-1503 | Diosmin |
| Prestw-1505 | Valacyclovir hydrochloride |
| Prestw-1506 | Mizolastine |
| Prestw-1507 | Acefylline |
| Prestw-1508 | Ibutilide fumarate |
| Prestw-1509 | Deflazacort |
| Prestw-151 | Erythromycin |
| Prestw-1510 | Dolasetron mesilate |
| Prestw-1511 | Aceclidine Hydrochloride |
| Prestw-1512 | Abacavir Sulfate |
| Prestw-1514 | Darifenacin hydrobromide |
| Prestw-1516 | Levalbuterol hydrochloride |

| | |
|---|---|
| Prestw-153 | Didanosine |
| Prestw-154 | Josamycin |
| Prestw-155 | Paclitaxel |
| Prestw-156 | Ivermectin |
| Prestw-157 | Gallamine triethiodide |
| Prestw-158 | Neomycin sulfate |
| Prestw-159 | Dihydrostreptomycin sulfate |
| Prestw-16 | Sulfathiazole |
| Prestw-160 | Gentamicine sulfate |
| Prestw-1600 | Aprepitant |
| Prestw-1601 | Pinaverium bromide |
| Prestw-1602 | Indatraline hydrochloride |
| Prestw-1603 | Pemirolast potassium |
| Prestw-1604 | (-)-Emtricitabine |
| Prestw-161 | Isoniazid |
| Prestw-162 | Pentylenetetrazole |
| Prestw-163 | Chlorzoxazone |
| Prestw-164 | Ornidazole |
| Prestw-165 | Ethosuximide |
| Prestw-166 | Mafenide hydrochloride |
| Prestw-167 | Riluzole hydrochloride |
| Prestw-168 | Nitrofurantoin |
| Prestw-169 | Hydralazine hydrochloride |
| Prestw-17 | Levodopa |
| Prestw-170 | Phenelzine sulfate |
| Prestw-1700 | Histamine dihydrochloride |
| Prestw-1701 | Clopidogrel |
| Prestw-1702 | Trimetozine |
| Prestw-1703 | 1,8-Dihydroxyanthraquinone |
| Prestw-1704 | Desonide |
| Prestw-1705 | Diatrizoic acid dihydrate |
| Prestw-1706 | Donepezil hydrochloride |
| Prestw-1707 | Doxapram hydrochloride |
| Prestw-1708 | (R)-Duloxetine hydrochloride |
| Prestw-1709 | Estramustine |
| Prestw-171 | Tranexamic acid |
| Prestw-1710 | Ethoxzolamide |
| Prestw-1711 | Felbamate |
| Prestw-1712 | Flumethasone pivalate |
| Prestw-1713 | Fenoldopam |
| Prestw-1714 | Adapalene |
| Prestw-1715 | Algestone acetophenide |
| Prestw-1716 | Altrenogest |
| Prestw-1717 | Aminacrine |
| Prestw-1718 | Amlexanox |

| | |
|---|---|
| Prestw-1719 | Amorolfine hydrochloride |
| Prestw-172 | Etofylline |
| Prestw-1720 | Anagrelide |
| Prestw-1721 | Avobenzone |
| Prestw-1722 | Azatadine maleate |
| Prestw-1723 | Benzoxiquine |
| Prestw-1726 | Cefpiramide |
| Prestw-1727 | Cefpodoxime proxetil |
| Prestw-1728 | Cefprozil |
| Prestw-1729 | Ceftibuten |
| Prestw-173 | Tranylcypromine hydrochloride |
| Prestw-1730 | Cefuroxime axetil |
| Prestw-1731 | Zoledronic acid hydrate |
| Prestw-1732 | Tolcapone |
| Prestw-1733 | Nitazoxanide |
| Prestw-1734 | Rasagiline |
| Prestw-1735 | Valsartan |
| Prestw-1736 | Irbesartan |
| Prestw-1737 | Indinavir sulfate |
| Prestw-1738 | Risedronic acid monohydrate |
| Prestw-1739 | Nevirapine |
| Prestw-174 | Alverine citrate salt |
| Prestw-1740 | Besifloxacin hydrochloride |
| Prestw-1741 | Loteprednol etabonate |
| Prestw-1743 | Dofetilide |
| Prestw-1744 | Tolterodine tartrate |
| Prestw-1745 | Cisatracurium besylate |
| Prestw-1746 | Lodoxamide |
| Prestw-1748 | Nateglinide |
| Prestw-1749 | Silodosin |
| Prestw-175 | Aceclofenac |
| Prestw-1750 | Terbinafine |
| Prestw-1751 | Brimonidine L-Tartrate |
| Prestw-1752 | Epirubicin hydrochloride |
| Prestw-1753 | Tegafur |
| Prestw-1756 | Lofepramine |
| Prestw-1758 | Pantoprazole sodium |
| Prestw-1759 | Valdecoxib |
| Prestw-176 | Iproniazide phosphate |
| Prestw-1760 | Tegaserod maleate |
| Prestw-1761 | Rizatriptan benzoate |
| Prestw-1762 | Ezetimibe |
| Prestw-1763 | Actarit |
| Prestw-1764 | Nelfinavir mesylate |
| Prestw-1765 | Latanoprost |

| | |
|---|---|
| Prestw-1766 | Benidipine hydrochloride |
| Prestw-1768 | Triclabendazole |
| Prestw-1769 | Enalaprilat dihydrate |
| Prestw-177 | Sulfamethoxazole |
| Prestw-1770 | Perospirone |
| Prestw-1771 | Carmofur |
| Prestw-1772 | Ampiroxicam |
| Prestw-1773 | Ipriflavone |
| Prestw-1774 | Nifekalant |
| Prestw-1775 | Lomerizine hydrochloride |
| Prestw-1776 | Phenprobamate |
| Prestw-1777 | Trimebutine |
| Prestw-1778 | Troxipide |
| Prestw-178 | Mephenesin |
| Prestw-1780 | Oxymetholone |
| Prestw-1781 | Alosetron hydrochloride |
| Prestw-1782 | Ritonavir |
| Prestw-1783 | Palonosetron hydrochloride |
| Prestw-1784 | Ubenimex |
| Prestw-1785 | Phenothiazine |
| Prestw-1786 | Enrofloxacin |
| Prestw-1787 | Homoveratrylamine |
| Prestw-1788 | Milnacipran hydrochloride |
| Prestw-179 | Phenformin hydrochloride |
| Prestw-1791 | Pregabalin |
| Prestw-1792 | Pidotimod |
| Prestw-1793 | Raltitrexed |
| Prestw-1794 | Pemetrexed disodium |
| Prestw-1795 | Mupirocin |
| Prestw-1796 | Delavirdine |
| Prestw-1797 | Grepafloxacin |
| Prestw-1798 | Mirabegron |
| Prestw-1799 | Tigecycline |
| Prestw-18 | Idoxuridine |
| Prestw-180 | Flutamide |
| Prestw-1800 | Zolmitriptan |
| Prestw-1801 | Ciclesonide |
| Prestw-1802 | Darunavir |
| Prestw-1803 | Methicillin sodium |
| Prestw-1804 | Nelarabine |
| Prestw-181 | Ampyrone |
| Prestw-1810 | Aliskiren hemifumarate |
| Prestw-1811 | Azilsartan kamedoxomil |
| Prestw-1812 | Lacosamide |
| Prestw-1813 | Lopinavir |

| | |
|---|---|
| Prestw-1814 | Nebivolol hydrochloride |
| Prestw-1815 | Pitavastatin calcium |
| Prestw-1816 | Ropivacaine hydrochloride |
| Prestw-1817 | Tazarotene |
| Prestw-1818 | Tirofiban hydrochloride |
| Prestw-1819 | Tolvaptan |
| Prestw-182 | Levamisole hydrochloride |
| Prestw-1820 | Amprenavir |
| Prestw-1821 | Bimatoprost |
| Prestw-1822 | Dronedarone hydrochloride |
| Prestw-1823 | Eprosartan mesylate |
| Prestw-1824 | Eszopiclone |
| Prestw-1825 | Gemifloxacin mesylate |
| Prestw-1826 | Posaconazole |
| Prestw-1827 | Paliperidone |
| Prestw-1828 | Capecitabine |
| Prestw-1829 | Levofloxacin |
| Prestw-183 | Pargyline hydrochloride |
| Prestw-1830 | Dexrazoxane hydrochloride |
| Prestw-184 | Methocarbamol |
| Prestw-185 | Aztreonam |
| Prestw-186 | Cloxacillin sodium salt |
| Prestw-188 | Pentolinium bitartrate |
| Prestw-189 | Aminopurine, 6-benzyl |
| Prestw-19 | Captopril |
| Prestw-190 | Tolbutamide |
| Prestw-191 | Midodrine hydrochloride |
| Prestw-192 | Thalidomide |
| Prestw-193 | Oxolinic acid |
| Prestw-194 | Nimesulide |
| Prestw-196 | Pentoxifylline |
| Prestw-197 | Metaraminol bitartrate |
| Prestw-198 | Salbutamol |
| Prestw-199 | Prilocaine hydrochloride |
| Prestw-2 | Allantoin |
| Prestw-20 | Minoxidil |
| Prestw-200 | Camptothecine (S,+) |
| Prestw-201 | Ranitidine hydrochloride |
| Prestw-202 | Tiratricol, 3,3',5-triiodothyroacetic acid |
| Prestw-203 | Flufenamic acid |
| Prestw-204 | Flumequine |
| Prestw-205 | Tolfenamic acid |
| Prestw-206 | Meclofenamic acid sodium salt monohydrate |
| Prestw-208 | Trimethoprim |
| Prestw-209 | Metoclopramide monohydrochloride |

| | |
|---|---|
| Prestw-21 | Sulfaphenazole |
| Prestw-210 | Fenbendazole |
| Prestw-211 | Piroxicam |
| Prestw-212 | Pyrantel tartrate |
| Prestw-213 | Fenspiride hydrochloride |
| Prestw-214 | Gemfibrozil |
| Prestw-215 | Mefexamide hydrochloride |
| Prestw-216 | Tiapride hydrochloride |
| Prestw-217 | Mebendazole |
| Prestw-218 | Fenbufen |
| Prestw-219 | Ketoprofen |
| Prestw-22 | Panthenol (D) |
| Prestw-220 | Indapamide |
| Prestw-221 | Norfloxacin |
| Prestw-222 | Antimycin A |
| Prestw-223 | Xylometazoline hydrochloride |
| Prestw-224 | Oxymetazoline hydrochloride |
| Prestw-225 | Nifenazone |
| Prestw-226 | Griseofulvin |
| Prestw-227 | Clemizole hydrochloride |
| Prestw-228 | Tropicamide |
| Prestw-229 | Nefopam hydrochloride |
| Prestw-23 | Sulfadiazine |
| Prestw-230 | Phentolamine hydrochloride |
| Prestw-231 | Etodolac |
| Prestw-232 | Scopolamin-N-oxide hydrobromide |
| Prestw-233 | Hyoscyamine (L) |
| Prestw-234 | Chlorphensin carbamate |
| Prestw-236 | Dilazep dihydrochloride |
| Prestw-237 | Ofloxacin |
| Prestw-238 | Lomefloxacin hydrochloride |
| Prestw-239 | Orphenadrine hydrochloride |
| Prestw-24 | Norethynodrel |
| Prestw-240 | Proglumide |
| Prestw-241 | Mexiletine hydrochloride |
| Prestw-242 | Flavoxate hydrochloride |
| Prestw-243 | Bufexamac |
| Prestw-244 | Glutethimide, para-amino |
| Prestw-245 | Dropropizine (R,S) |
| Prestw-246 | Pinacidil |
| Prestw-247 | Albendazole |
| Prestw-248 | Clonidine hydrochloride |
| Prestw-249 | Bupropion hydrochloride |
| Prestw-25 | Thiamphenicol |
| Prestw-250 | Alprenolol hydrochloride |

| | |
|---|---|
| Prestw-251 | Chlorothiazide |
| Prestw-252 | Diphenidol hydrochloride |
| Prestw-253 | Norethindrone |
| Prestw-254 | Nortriptyline hydrochloride |
| Prestw-255 | Niflumic acid |
| Prestw-256 | Isotretinoin |
| Prestw-257 | Retinoic acid |
| Prestw-258 | Antazoline hydrochloride |
| Prestw-259 | Ethacrynic acid |
| Prestw-26 | Cimetidine |
| Prestw-260 | Praziquantel |
| Prestw-261 | Ethisterone |
| Prestw-262 | Triprolidine hydrochloride |
| Prestw-263 | Doxepin hydrochloride |
| Prestw-264 | Dyclonine hydrochloride |
| Prestw-265 | Dimenhydrinate |
| Prestw-266 | Disopyramide |
| Prestw-267 | Clotrimazole |
| Prestw-268 | Vinpocetine |
| Prestw-269 | Clomipramine hydrochloride |
| Prestw-27 | Doxylamine succinate |
| Prestw-270 | Fendiline hydrochloride |
| Prestw-271 | Vincamine |
| Prestw-272 | Indomethacin |
| Prestw-273 | Cortisone |
| Prestw-274 | Prednisolone |
| Prestw-275 | Fenofibrate |
| Prestw-276 | Bumetanide |
| Prestw-277 | Labetalol hydrochloride |
| Prestw-278 | Cinnarizine |
| Prestw-279 | Methylprednisolone, 6-alpha |
| Prestw-28 | Ethambutol dihydrochloride |
| Prestw-280 | Quinidine hydrochloride monohydrate |
| Prestw-281 | Fludrocortisone acetate |
| Prestw-282 | Fenoterol hydrobromide |
| Prestw-283 | Homochlorcyclizine dihydrochloride |
| Prestw-284 | Diethylcarbamazine citrate |
| Prestw-285 | Chenodiol |
| Prestw-286 | Perhexiline maleate |
| Prestw-287 | Oxybutynin chloride |
| Prestw-288 | Spiperone |
| Prestw-289 | Pyrilamine maleate |
| Prestw-29 | Antipyrine |
| Prestw-290 | Sulfinpyrazone |
| Prestw-291 | Dantrolene sodium salt |

| | |
|---|---|
| Prestw-292 | Trazodone hydrochloride |
| Prestw-293 | Glafenine hydrochloride |
| Prestw-294 | Pimethixene maleate |
| Prestw-295 | Pergolide mesylate |
| Prestw-296 | Acemetacin |
| Prestw-297 | Benzydamine hydrochloride |
| Prestw-298 | Fipexide hydrochloride |
| Prestw-299 | Mifepristone |
| Prestw-3 | Acetazolamide |
| Prestw-30 | Antipyrine, 4-hydroxy |
| Prestw-300 | Diperodon hydrochloride |
| Prestw-301 | Lisinopril |
| Prestw-302 | Lincomycin hydrochloride |
| Prestw-303 | Telenzepine dihydrochloride |
| Prestw-304 | Econazole nitrate |
| Prestw-305 | Bupivacaine hydrochloride |
| Prestw-306 | Clemastine fumarate |
| Prestw-307 | Oxytetracycline dihydrate |
| Prestw-308 | Pimozide |
| Prestw-309 | Amodiaquin dihydrochloride dihydrate |
| Prestw-31 | Chloramphenicol |
| Prestw-310 | Mebeverine hydrochloride |
| Prestw-311 | Ifenprodil tartrate |
| Prestw-312 | Flunarizine dihydrochloride |
| Prestw-313 | Trifluoperazine dihydrochloride |
| Prestw-314 | Enalapril maleate |
| Prestw-315 | Minocycline hydrochloride |
| Prestw-316 | Glibenclamide |
| Prestw-317 | Guanethidine sulfate |
| Prestw-318 | Quinacrine dihydrochloride hydrate |
| Prestw-319 | Clofilium tosylate |
| Prestw-32 | Epirizole |
| Prestw-320 | Fluphenazine dihydrochloride |
| Prestw-321 | Streptomycin sulfate |
| Prestw-322 | Alfuzosin hydrochloride |
| Prestw-323 | Chlorpropamide |
| Prestw-324 | Phenylpropanolamine hydrochloride |
| Prestw-325 | Ascorbic acid |
| Prestw-326 | Methyldopa (L,-) |
| Prestw-327 | Cefoperazone dihydrate |
| Prestw-328 | Zoxazolamine |
| Prestw-329 | Tacrine hydrochloride |
| Prestw-33 | Diprophylline |
| Prestw-330 | Bisoprolol fumarate |
| Prestw-332 | Practolol |

| | |
|---|---|
| Prestw-333 | Zidovudine, AZT |
| Prestw-334 | Sulfisoxazole |
| Prestw-335 | Zaprinast |
| Prestw-336 | Chlormezanone |
| Prestw-337 | Procainamide hydrochloride |
| Prestw-338 | N6-methyladenosine |
| Prestw-339 | Guanfacine hydrochloride |
| Prestw-34 | Triamterene |
| Prestw-340 | Domperidone |
| Prestw-341 | Furosemide |
| Prestw-342 | Methapyrilene hydrochloride |
| Prestw-343 | Desipramine hydrochloride |
| Prestw-344 | Clorgyline hydrochloride |
| Prestw-345 | Clenbuterol hydrochloride |
| Prestw-346 | Maprotiline hydrochloride |
| Prestw-347 | Thioguanosine |
| Prestw-348 | Chlorprothixene hydrochloride |
| Prestw-349 | Ritodrine hydrochloride |
| Prestw-35 | Dapsone |
| Prestw-350 | Clozapine |
| Prestw-351 | Chlorthalidone |
| Prestw-352 | Dobutamine hydrochloride |
| Prestw-353 | Moclobemide |
| Prestw-354 | Clopamide |
| Prestw-355 | Hycanthone |
| Prestw-356 | Adenosine 5'-monophosphate monohydrate |
| Prestw-357 | Amoxicillin |
| Prestw-359 | Dextromethorphan hydrobromide monohydrate |
| Prestw-36 | Troleandomycin |
| Prestw-360 | Droperidol |
| Prestw-361 | Bambuterol hydrochloride |
| Prestw-362 | Betamethasone |
| Prestw-363 | Colchicine |
| Prestw-364 | Metergoline |
| Prestw-365 | Brinzolamide |
| Prestw-366 | Ambroxol hydrochloride |
| Prestw-367 | Benfluorex |
| Prestw-368 | Bepridil hydrochloride |
| Prestw-369 | Meloxicam |
| Prestw-37 | Pyrimethamine |
| Prestw-370 | Benzbromarone |
| Prestw-371 | Ketotifen fumarate |
| Prestw-372 | Debrisoquin sulfate |
| Prestw-373 | Amethopterin (R,S) |
| Prestw-374 | Methylergometrine maleate |

| | |
|---|---|
| Prestw-375 | Methiothepin maleate |
| Prestw-376 | Clofazimine |
| Prestw-377 | Nafronyl oxalate |
| Prestw-378 | Bezafibrate |
| Prestw-38 | Hexamethonium dibromide dihydrate |
| Prestw-380 | Clebopride maleate |
| Prestw-381 | Lidoflazine |
| Prestw-382 | Betaxolol hydrochloride |
| Prestw-383 | Nicardipine hydrochloride |
| Prestw-384 | Probucol |
| Prestw-385 | Mitoxantrone dihydrochloride |
| Prestw-386 | GBR 12909 dihydrochloride |
| Prestw-387 | Carbetapentane citrate |
| Prestw-388 | Dequalinium dichloride |
| Prestw-389 | Ketoconazole |
| Prestw-39 | Diflunisal |
| Prestw-390 | Fusidic acid sodium salt |
| Prestw-391 | Terbutaline hemisulfate |
| Prestw-392 | Ketanserin tartrate hydrate |
| Prestw-393 | Hemicholinium bromide |
| Prestw-394 | Kanamycin A sulfate |
| Prestw-395 | Amikacin hydrate |
| Prestw-396 | Etoposide |
| Prestw-397 | Clomiphene citrate (Z,E) |
| Prestw-398 | Oxantel pamoate |
| Prestw-399 | Prochlorperazine dimaleate |
| Prestw-4 | Metformin hydrochloride |
| Prestw-40 | Niclosamide |
| Prestw-400 | Hesperidin |
| Prestw-401 | Testosterone propionate |
| Prestw-403 | Thyroxine (L) |
| Prestw-405 | Pepstatin A |
| Prestw-407 | Adamantamine fumarate |
| Prestw-408 | Butoconazole nitrate |
| Prestw-409 | Amiodarone hydrochloride |
| Prestw-41 | Procaine hydrochloride |
| Prestw-410 | Amphotericin B |
| Prestw-411 | Androsterone |
| Prestw-413 | Carbarsone |
| Prestw-416 | Bacampicillin hydrochloride |
| Prestw-418 | Biotin |
| Prestw-419 | Bisacodyl |
| Prestw-42 | Moxisylyte hydrochoride |
| Prestw-421 | Suloctidil |
| Prestw-423 | Carisoprodol |

| | |
|---|---|
| Prestw-424 | Cephalosporanic acid, 7-amino |
| Prestw-425 | Chicago sky blue 6B |
| Prestw-426 | Buflomedil hydrochloride |
| Prestw-428 | Roxatidine Acetate hydrochloride |
| Prestw-43 | Betazole hydrochloride |
| Prestw-430 | Cisapride |
| Prestw-432 | Corticosterone |
| Prestw-433 | Cyanocobalamin |
| Prestw-434 | Cefadroxil |
| Prestw-435 | Cyclosporin A |
| Prestw-436 | Digitoxigenin |
| Prestw-437 | Digoxin |
| Prestw-438 | Doxorubicin hydrochloride |
| Prestw-439 | Carbimazole |
| Prestw-44 | Isoxicam |
| Prestw-440 | Epiandrosterone |
| Prestw-441 | Estradiol-17 beta |
| Prestw-445 | Cyclobenzaprine hydrochloride |
| Prestw-446 | Carteolol hydrochloride |
| Prestw-447 | Hydrocortisone base |
| Prestw-449 | Pilocarpine nitrate |
| Prestw-45 | Naproxen |
| Prestw-450 | Dicloxacillin sodium salt hydrate |
| Prestw-451 | Alizapride hydrochloride |
| Prestw-455 | Mebhydroline 1,5-naphtalenedisulfonate |
| Prestw-456 | Meclocycline sulfosalicylate |
| Prestw-457 | Meclozine dihydrochloride |
| Prestw-458 | Melatonin |
| Prestw-46 | Naphazoline hydrochloride |
| Prestw-460 | Dinoprost trometamol |
| Prestw-461 | Tropisetron hydrochloride |
| Prestw-462 | Cefixime |
| Prestw-463 | Metrizamide |
| Prestw-468 | Neostigmine bromide |
| Prestw-469 | Niridazole |
| Prestw-47 | Ticlopidine hydrochloride |
| Prestw-470 | Ceforanide |
| Prestw-473 | Cefotetan |
| Prestw-475 | Brompheniramine maleate |
| Prestw-476 | Primaquine diphosphate |
| Prestw-477 | Progesterone |
| Prestw-478 | Felodipine |
| Prestw-48 | Dicyclomine hydrochloride |
| Prestw-481 | Serotonin hydrochloride |
| Prestw-482 | Cefotiam hydrochloride |

| | |
|---|---|
| Prestw-484 | Benperidol |
| Prestw-485 | Cefaclor hydrate |
| Prestw-486 | Colistin sulfate |
| Prestw-487 | Daunorubicin hydrochloride |
| Prestw-488 | Dosulepin hydrochloride |
| Prestw-489 | Ceftazidime pentahydrate |
| Prestw-49 | Amyleine hydrochloride |
| Prestw-490 | Iobenguane sulfate |
| Prestw-491 | Metixene hydrochloride |
| Prestw-492 | Nitrofural |
| Prestw-493 | Omeprazole |
| Prestw-494 | Propylthiouracil |
| Prestw-495 | Terconazole |
| Prestw-496 | Tiaprofenic acid |
| Prestw-497 | Vancomycin hydrochloride |
| Prestw-498 | Artemisinin |
| Prestw-499 | Propafenone hydrochloride |
| Prestw-5 | Atracurium besylate |
| Prestw-50 | Lidocaine hydrochloride |
| Prestw-500 | Ethamivan |
| Prestw-501 | Vigabatrin hydrochloride |
| Prestw-502 | Biperiden hydrochloride |
| Prestw-503 | Cetirizine dihydrochloride |
| Prestw-504 | Etifenin |
| Prestw-505 | Metaproterenol sulfate, orciprenaline sulfate |
| Prestw-506 | Sisomicin sulfate |
| Prestw-509 | Bromperidol |
| Prestw-510 | Cyclizine hydrochloride |
| Prestw-511 | Fluoxetine hydrochloride |
| Prestw-512 | Iohexol |
| Prestw-513 | Norcyclobenzaprine |
| Prestw-514 | Pyrazinamide |
| Prestw-515 | Trimethadione |
| Prestw-516 | Lovastatin |
| Prestw-517 | Nystatine |
| Prestw-518 | Budesonide |
| Prestw-519 | Imipenem |
| Prestw-52 | Carbamazepine |
| Prestw-520 | Sulfasalazine |
| Prestw-522 | Thiostrepton |
| Prestw-524 | Tiabendazole |
| Prestw-525 | Rifampicin |
| Prestw-526 | Ethionamide |
| Prestw-527 | Tenoxicam |
| Prestw-528 | Triflusal |

| | |
|---|---|
| Prestw-529 | Mesoridazine besylate |
| Prestw-53 | Triflupromazine hydrochloride |
| Prestw-530 | Trolox |
| Prestw-531 | Pirenperone |
| Prestw-533 | Phenacetin |
| Prestw-534 | Atovaquone |
| Prestw-535 | Methoxamine hydrochloride |
| Prestw-536 | (R)-(+)-Atenolol |
| Prestw-537 | Piracetam |
| Prestw-538 | Phenindione |
| Prestw-539 | Thiocolchicoside |
| Prestw-54 | Mefenamic acid |
| Prestw-540 | Clorsulon |
| Prestw-541 | Ciclopirox ethanolamine |
| Prestw-542 | Probenecid |
| Prestw-543 | Betahistine mesylate |
| Prestw-544 | Tobramycin |
| Prestw-545 | Tetramisole hydrochloride |
| Prestw-546 | Pregnenolone |
| Prestw-547 | Molsidomine |
| Prestw-548 | Chloroquine diphosphate |
| Prestw-549 | Trimetazidine dihydrochloride |
| Prestw-55 | Acetohexamide |
| Prestw-551 | Hexetidine |
| Prestw-552 | Selegiline hydrochloride |
| Prestw-553 | Pentamidine isethionate |
| Prestw-554 | Tolazamide |
| Prestw-555 | Nifuroxazide |
| Prestw-557 | Dirithromycin |
| Prestw-558 | Gliclazide |
| Prestw-56 | Sulpiride |
| Prestw-560 | Prenylamine lactate |
| Prestw-565 | Atropine sulfate monohydrate |
| Prestw-566 | Eserine hemisulfate salt |
| Prestw-57 | Benoxinate hydrochloride |
| Prestw-571 | Tetracaïne hydrochloride |
| Prestw-572 | Mometasone furoate |
| Prestw-574 | Dacarbazine |
| Prestw-576 | Acetopromazine maleate salt |
| Prestw-58 | Oxethazaine |
| Prestw-583 | Papaverine hydrochloride |
| Prestw-584 | Yohimbine hydrochloride |
| Prestw-587 | Cilostazol |
| Prestw-588 | Galanthamine hydrobromide |
| Prestw-59 | Pheniramine maleate |

| | |
|---|---|
| Prestw-594 | Diclofenac sodium |
| Prestw-598 | Xylazine |
| Prestw-6 | Isoflupredone acetate |
| Prestw-60 | Tolazoline hydrochloride |
| Prestw-61 | Morantel tartrate |
| Prestw-616 | Demecarium bromide |
| Prestw-619 | Diflorasone Diacetate |
| Prestw-62 | Homatropine hydrobromide (R,S) |
| Prestw-623 | Pyridoxine hydrochloride |
| Prestw-626 | Racecadotril |
| Prestw-627 | Folic acid |
| Prestw-63 | Nifedipine |
| Prestw-630 | Dimethisoquin hydrochloride |
| Prestw-631 | Thiamine hydrochloride |
| Prestw-632 | Dipivefrin hydrochloride |
| Prestw-633 | Thiorphan |
| Prestw-64 | Chlorpromazine hydrochloride |
| Prestw-641 | Sulmazole |
| Prestw-643 | Flunisolide |
| Prestw-644 | N-Acetyl-DL-homocysteine Thiolactone |
| Prestw-645 | Flurandrenolide |
| Prestw-649 | Etanidazole |
| Prestw-65 | Diphenhydramine hydrochloride |
| Prestw-651 | Glimepiride |
| Prestw-652 | Picrotoxinin |
| Prestw-653 | Mepenzolate bromide |
| Prestw-654 | Benfotiamine |
| Prestw-655 | Halcinonide |
| Prestw-656 | Lanatoside C |
| Prestw-657 | Benzamil hydrochloride |
| Prestw-658 | Suxibuzone |
| Prestw-659 | 6-Furfurylaminopurine |
| Prestw-66 | Minaprine dihydrochloride |
| Prestw-660 | Avermectin B1 |
| Prestw-666 | Nisoldipine |
| Prestw-67 | Miconazole |
| Prestw-671 | Dydrogesterone |
| Prestw-676 | Beta-Escin |
| Prestw-68 | Isoxsuprine hydrochloride |
| Prestw-683 | Pempidine |
| Prestw-688 | Estropipate |
| Prestw-69 | Acebutolol hydrochloride |
| Prestw-692 | Citalopram Hydrobromide |
| Prestw-693 | Promazine hydrochloride |
| Prestw-694 | Sulfamerazine |

| | |
|---|---|
| Prestw-696 | Ethotoin |
| Prestw-697 | 3-alpha-Hydroxy-5-beta-androstan-17-one |
| Prestw-698 | Tetrahydrozoline hydrochloride |
| Prestw-699 | Hexestrol |
| Prestw-7 | Amiloride hydrochloride dihydrate |
| Prestw-70 | Tolnaftate |
| Prestw-700 | Cefmetazole sodium salt |
| Prestw-701 | Trihexyphenidyl-D,L Hydrochloride |
| Prestw-702 | Succinylsulfathiazole |
| Prestw-703 | Famprofazone |
| Prestw-704 | Bromopride |
| Prestw-705 | Methyl benzethonium chloride |
| Prestw-706 | Chlorcyclizine hydrochloride |
| Prestw-707 | Diphenylpyraline hydrochloride |
| Prestw-708 | Benzethonium chloride |
| Prestw-709 | Trioxsalen |
| Prestw-71 | Todralazine hydrochloride |
| Prestw-711 | Sulfabenzamide |
| Prestw-712 | Benzocaine |
| Prestw-713 | Dipyrone |
| Prestw-714 | Isosorbide dinitrate |
| Prestw-715 | Sulfachloropyridazine |
| Prestw-716 | Pramoxine hydrochloride |
| Prestw-717 | Finasteride |
| Prestw-718 | Fluorometholone |
| Prestw-719 | Cephalothin sodium salt |
| Prestw-72 | Imipramine hydrochloride |
| Prestw-720 | Cefuroxime sodium salt |
| Prestw-721 | Althiazide |
| Prestw-722 | Isopyrin hydrochloride |
| Prestw-723 | Phenethicillin potassium salt |
| Prestw-724 | Sulfamethoxypyridazine |
| Prestw-725 | Deferoxamine mesylate |
| Prestw-726 | Mephentermine hemisulfate |
| Prestw-728 | Sulfadimethoxine |
| Prestw-729 | Sulfanilamide |
| Prestw-73 | Sulindac |
| Prestw-730 | Balsalazide Sodium |
| Prestw-731 | Sulfaquinoxaline sodium salt |
| Prestw-732 | Streptozotocin |
| Prestw-733 | Metoprolol-(+,-) (+)-tartrate salt |
| Prestw-734 | Flumethasone |
| Prestw-735 | Flecainide acetate |
| Prestw-736 | Cefazolin sodium salt |
| Prestw-738 | Folinic acid calcium salt |

| | |
|---|---|
| Prestw-739 | Levonordefrin |
| Prestw-74 | Amitryptiline hydrochloride |
| Prestw-742 | Sulfamethizole |
| Prestw-743 | Medrysone |
| Prestw-744 | Flunixin meglumine |
| Prestw-745 | Spiramycin |
| Prestw-746 | Glycopyrrolate |
| Prestw-748 | Monensin sodium salt |
| Prestw-749 | Isoetharine mesylate salt |
| Prestw-75 | Adiphenine hydrochloride |
| Prestw-751 | Terazosin hydrochloride |
| Prestw-752 | Phenazopyridine hydrochloride |
| Prestw-753 | Demeclocycline hydrochloride |
| Prestw-754 | Fenoprofen calcium salt dihydrate |
| Prestw-755 | Piperacillin sodium salt |
| Prestw-756 | Diethylstilbestrol |
| Prestw-757 | Chlorotrianisene |
| Prestw-758 | Ribostamycin sulfate salt |
| Prestw-759 | Methacholine chloride |
| Prestw-76 | Dibucaine |
| Prestw-760 | Pipenzolate bromide |
| Prestw-761 | Butamben |
| Prestw-762 | Sulfapyridine |
| Prestw-763 | Meclofenoxate hydrochloride |
| Prestw-764 | Furaltadone hydrochloride |
| Prestw-765 | Ethoxyquin |
| Prestw-766 | Tinidazole |
| Prestw-767 | Guanadrel sulfate |
| Prestw-768 | Vidarabine |
| Prestw-769 | Sulfameter |
| Prestw-77 | Prednisone |
| Prestw-770 | Isopropamide iodide |
| Prestw-771 | Alclometasone dipropionate |
| Prestw-772 | Leflunomide |
| Prestw-773 | Norgestrel-(-)-D |
| Prestw-774 | Fluocinonide |
| Prestw-775 | Sulfamethazine sodium salt |
| Prestw-776 | Guaifenesin |
| Prestw-777 | Alexidine dihydrochloride |
| Prestw-778 | Proadifen hydrochloride |
| Prestw-779 | Zomepirac sodium salt |
| Prestw-78 | Thioridazine hydrochloride |
| Prestw-780 | Cinoxacin |
| Prestw-781 | Clobetasol propionate |
| Prestw-782 | Podophyllotoxin |

| | |
|---|---|
| Prestw-783 | Clofibric acid |
| Prestw-784 | Bendroflumethiazide |
| Prestw-785 | Dicumarol |
| Prestw-786 | Methimazole |
| Prestw-787 | Merbromin |
| Prestw-788 | Hexylcaine hydrochloride |
| Prestw-789 | Drofenine hydrochloride |
| Prestw-79 | Diphemanil methylsulfate |
| Prestw-790 | Cycloheximide |
| Prestw-791 | (R) -Naproxen sodium salt |
| Prestw-792 | Propidium iodide |
| Prestw-793 | Cloperastine hydrochloride |
| Prestw-795 | Isocarboxazid |
| Prestw-796 | Lithocholic acid |
| Prestw-797 | Methotrimeprazine maleat salt |
| Prestw-798 | Dienestrol |
| Prestw-799 | Pridinol methanesulfonate salt |
| Prestw-8 | Amprolium hydrochloride |
| Prestw-80 | Trimethobenzamide hydrochloride |
| Prestw-800 | Amrinone |
| Prestw-801 | Carbinoxamine maleate salt |
| Prestw-802 | Methazolamide |
| Prestw-803 | Pyrithyldione |
| Prestw-804 | Spectinomycin dihydrochloride |
| Prestw-805 | Piromidic acid |
| Prestw-806 | Trimipramine maleate salt |
| Prestw-807 | Chloropyramine hydrochloride |
| Prestw-808 | Furazolidone |
| Prestw-809 | Dichlorphenamide |
| Prestw-81 | Metronidazole |
| Prestw-810 | Sulconazole nitrate |
| Prestw-812 | Cromolyn disodium salt |
| Prestw-813 | Bucladesine sodium salt |
| Prestw-814 | Cefsulodin sodium salt |
| Prestw-815 | Fosfosal |
| Prestw-816 | Suprofen |
| Prestw-818 | Nadolol |
| Prestw-819 | Moxalactam disodium salt |
| Prestw-820 | Aminophylline |
| Prestw-821 | Azlocillin sodium salt |
| Prestw-822 | Clidinium bromide |
| Prestw-823 | Sulfamonomethoxine |
| Prestw-824 | Benzthiazide |
| Prestw-825 | Trichlormethiazide |
| Prestw-826 | Oxalamine citrate salt |

| | |
|---|---|
| Prestw-827 | Propantheline bromide |
| Prestw-829 | Dimethadione |
| Prestw-83 | Edrophonium chloride |
| Prestw-830 | Ethaverine hydrochloride |
| Prestw-831 | Butacaine |
| Prestw-832 | Cefoxitin sodium salt |
| Prestw-834 | Novobiocin sodium salt |
| Prestw-836 | Indoprofen |
| Prestw-837 | Carbenoxolone disodium salt |
| Prestw-838 | Iocetamic acid |
| Prestw-839 | Ganciclovir |
| Prestw-84 | Moroxidine hydrochloride |
| Prestw-840 | Ethopropazine hydrochloride |
| Prestw-842 | Trimeprazine tartrate |
| Prestw-843 | Nafcillin sodium salt monohydrate |
| Prestw-844 | Procyclidine hydrochloride |
| Prestw-845 | Amiprilose hydrochloride |
| Prestw-846 | Ethynylestradiol 3-methyl ether |
| Prestw-847 | (-) -Levobunolol hydrochloride |
| Prestw-848 | Iodixanol |
| Prestw-85 | Baclofen (R,S) |
| Prestw-850 | Equilin |
| Prestw-851 | Paroxetine Hydrochloride |
| Prestw-853 | Liothyronine |
| Prestw-854 | Roxithromycin |
| Prestw-855 | Beclomethasone dipropionate |
| Prestw-856 | Tolmetin sodium salt dihydrate |
| Prestw-857 | (+) -Levobunolol hydrochloride |
| Prestw-858 | Doxazosin mesylate |
| Prestw-859 | Fluvastatin sodium salt |
| Prestw-86 | Acyclovir |
| Prestw-860 | Methylhydantoin-5-(L) |
| Prestw-861 | Gabapentin |
| Prestw-862 | Raloxifene hydrochloride |
| Prestw-864 | Methylhydantoin-5-(D) |
| Prestw-865 | Simvastatin |
| Prestw-866 | Azacytidine-5 |
| Prestw-867 | Paromomycin sulfate |
| Prestw-868 | Acetaminophen |
| Prestw-869 | Phthalylsulfathiazole |
| Prestw-87 | Diazoxide |
| Prestw-870 | Luteolin |
| Prestw-871 | Iopamidol |
| Prestw-872 | Iopromide |
| Prestw-873 | Theophylline monohydrate |

| | |
|---|---|
| Prestw-874 | Theobromine |
| Prestw-875 | Reserpine |
| Prestw-877 | Scopolamine hydrochloride |
| Prestw-878 | Ioversol |
| Prestw-88 | Amidopyrine |
| Prestw-880 | Carbachol |
| Prestw-881 | Niacin |
| Prestw-882 | Bemegride |
| Prestw-883 | Digoxigenin |
| Prestw-884 | Meglumine |
| Prestw-886 | Clioquinol |
| Prestw-887 | Oxybenzone |
| Prestw-888 | Promethazine hydrochloride |
| Prestw-893 | Felbinac |
| Prestw-894 | Butylparaben |
| Prestw-895 | Aminohippuric acid |
| Prestw-896 | N-Acetyl-L-leucine |
| Prestw-897 | Pipemidic acid |
| Prestw-898 | Dioxybenzone |
| Prestw-899 | Adrenosterone |
| Prestw-9 | Hydrochlorothiazide |
| Prestw-90 | Pindolol |
| Prestw-900 | Methylatropine nitrate |
| Prestw-901 | Hymecromone |
| Prestw-903 | Diloxanide furoate |
| Prestw-904 | Metyrapone |
| Prestw-905 | Urapidil hydrochloride |
| Prestw-906 | Fluspirilen |
| Prestw-907 | S-(+)-ibuprofen |
| Prestw-908 | Ethynodiol diacetate |
| Prestw-909 | Nabumetone |
| Prestw-91 | Khellin |
| Prestw-910 | Nisoxetine hydrochloride |
| Prestw-911 | (+)-Isoproterenol (+)-bitartrate salt |
| Prestw-912 | Monobenzone |
| Prestw-913 | 2-Aminobenzenesulfonamide |
| Prestw-914 | Estrone |
| Prestw-915 | Lorglumide sodium salt |
| Prestw-916 | Nitrendipine |
| Prestw-917 | Flurbiprofen |
| Prestw-918 | Nimodipine |
| Prestw-919 | Bacitracin |
| Prestw-92 | Zimelidine dihydrochloride monohydrate |
| Prestw-921 | Nizatidine |
| Prestw-922 | Thioperamide maleate |

| | |
|---|---|
| Prestw-923 | Xamoterol hemifumarate |
| Prestw-925 | Thonzonium bromide |
| Prestw-926 | Idazoxan hydrochloride |
| Prestw-927 | Quinapril hydrochloride |
| Prestw-928 | Nilutamide |
| Prestw-929 | Ketorolac tromethamine |
| Prestw-93 | Azacyclonol |
| Prestw-930 | Protriptyline hydrochloride |
| Prestw-931 | Propofol |
| Prestw-932 | S(-)Eticlopride hydrochloride |
| Prestw-933 | Primidone |
| Prestw-934 | Flucytosine |
| Prestw-936 | Bephenium hydroxynaphthoate |
| Prestw-937 | Dehydroisoandosterone 3-acetate |
| Prestw-938 | Benserazide hydrochloride |
| Prestw-939 | Iodipamide |
| Prestw-94 | Azathioprine |
| Prestw-941 | Pentetic acid |
| Prestw-942 | Bretylium tosylate |
| Prestw-943 | Pralidoxime chloride |
| Prestw-944 | Phenoxybenzamine hydrochloride |
| Prestw-945 | Salmeterol |
| Prestw-946 | Altretamine |
| Prestw-947 | Prazosin hydrochloride |
| Prestw-948 | Timolol maleate salt |
| Prestw-949 | (+,-)-Octopamine hydrochloride |
| Prestw-95 | Lynestrenol |
| Prestw-951 | Crotamiton |
| Prestw-953 | (S)-(-)-Atenolol |
| Prestw-954 | Tyloxapol |
| Prestw-955 | Florfenicol |
| Prestw-956 | Megestrol acetate |
| Prestw-957 | Deoxycorticosterone |
| Prestw-958 | Urosiol |
| Prestw-959 | Proparacaine hydrochloride |
| Prestw-96 | Guanabenz acetate |
| Prestw-960 | Aminocaproic acid |
| Prestw-961 | Denatonium benzoate |
| Prestw-963 | Enilconazole |
| Prestw-964 | Methacycline hydrochloride |
| Prestw-966 | Sotalol hydrochloride |
| Prestw-968 | Decamethonium bromide |
| Prestw-97 | Disulfiram |
| Prestw-971 | Remoxipride Hydrochloride |
| Prestw-972 | THIP Hydrochloride |

| | |
|---|---|
| Prestw-973 | Pirlindole mesylate |
| Prestw-974 | Pronethalol hydrochloride |
| Prestw-975 | Naftopidil dihydrochloride |
| Prestw-976 | Tracazolate hydrochloride |
| Prestw-977 | Zardaverine |
| Prestw-978 | Memantine Hydrochloride |
| Prestw-979 | Ozagrel hydrochloride |
| Prestw-98 | Acetylsalicylsalicylic acid |
| Prestw-980 | Piribedil hydrochloride |
| Prestw-981 | Nitrocaramiphen hydrochloride |
| Prestw-982 | Nandrolone |
| Prestw-983 | Dimaprit dihydrochloride |
| Prestw-99 | Mianserine hydrochloride |
| Prestw-991 | Gliquidone |
| Prestw-992 | Pizotifen malate |
| Prestw-993 | Ribavirin |
| Prestw-994 | Cyclopenthiazide |
| Prestw-995 | Fluvoxamine maleate |
| Prestw-997 | Fluticasone propionate |
| Prestw-998 | Zuclopenthixol dihydrochloride |
| Prestw-999 | Proguanil hydrochloride |
