## Supplementary table 4 for "Rapid accumulation of fluorophores and fast kill identify drugs with bactericidal effects against Gram-negative bacteria"

| Repeat | PID | Drug | Events | Signal.minus.drug.bgk | Signal.minus.autof |
|---|---|---|---|---|---|
| 2 | Prestw-847 | (-) -Levobunolol hydrochloride | 2687 | 2779.571134 | 2750.999269 |
| 1 | Prestw-847 | (-) -Levobunolol hydrochloride | 4057 | 944.1648981 | 892.1821945 |
| 2 | Prestw-1604 | (-)-Emtricitabine | 4221 | 47.1048713 | 30.4916873 |
| 1 | Prestw-1604 | (-)-Emtricitabine | 10148 | 63.01012151 | 1 |
| 1 | Prestw-1082 | (-)-Eseroline fumarate salt | 7980.5 | 1 | 1 |
| 2 | Prestw-1082 | (-)-Eseroline fumarate salt | 2919.5 | 1 | 1 |
| 1 | Prestw-1097 | (-)-Isoproterenol hydrochloride | 8575 | 15.70498507 | 14.70498507 |
| 2 | Prestw-1097 | (-)-Isoproterenol hydrochloride | 2762.5 | 21.5940879 | 1 |
| 1 | Prestw-949 | (+,-)-Octopamine hydrochloride | 16553 | 61.77085156 | 10.36352405 |
| 2 | Prestw-949 | (+,-)-Octopamine hydrochloride | 4881 | 1 | 1 |
| 2 | Prestw-1088 | (+,-)-Synephrine | 2525 | 73.4348004 | 72.4348004 |
| 1 | Prestw-1088 | (+,-)-Synephrine | 8019.5 | 1 | 1 |
| 1 | Prestw-857 | (+) -Levobunolol hydrochloride | 16883.5 | 61.44766954 | 10.04034203 |
| 2 | Prestw-857 | (+) -Levobunolol hydrochloride | 5957.5 | 24.245372 | 4.6595403 |
| 1 | Prestw-911 | (+)-Isoproterenol (+)-bitartrate salt | 16011.5 | 2.90874876 | 1.90874876 |
| 2 | Prestw-911 | (+)-Isoproterenol (+)-bitartrate salt | 5499.5 | 1 | 1 |
| 2 | Prestw-791 | (R) -Naproxen sodium salt | 5972 | 55.511875 | 42.7551879 |
| 1 | Prestw-791 | (R) -Naproxen sodium salt | 17684 | 47.74920599 | 13.72093772 |
| 2 | Prestw-536 | (R)-(+)-Atenolol | 4359.5 | 135.3504304 | 31.8138533 |
| 1 | Prestw-536 | (R)-(+)-Atenolol | 12696 | 8.64364977 | 1 |
| 2 | Prestw-1708 | (R)-Duloxetine hydrochloride | 4333.5 | 88.9195342 | 1 |
| 1 | Prestw-1708 | (R)-Duloxetine hydrochloride | 12027 | 122.065997 | 1 |
| 1 | Prestw-1075 | (R)-Propranolol hydrochloride | 9777 | 56.81545271 | 21.56503427 |
| 2 | Prestw-1075 | (R)-Propranolol hydrochloride | 3169.5 | 1 | 1 |
| 1 | Prestw-953 | (S)-(-)-Atenolol | 20541.5 | 83.7594013 | 47.20538297 |
| 2 | Prestw-953 | (S)-(-)-Atenolol | 6812.5 | 73.5885588 | 70.3800184 |
| 1 | Prestw-1089 | (S)-(-)-Cycloserine | 7792.5 | 1 | 1 |
| 2 | Prestw-1089 | (S)-(-)-Cycloserine | 3205 | 1 | 1 |
| 2 | Prestw-1081 | (S)-propranolol hydrochloride | 2990 | 9.0939114 | 8.0939114 |
| 1 | Prestw-1081 | (S)-propranolol hydrochloride | 8031.5 | 1 | 1 |
| 2 | Prestw-1703 | 1,8-Dihydroxyanthraquinone | 5392.5 | 93.9040773 | 1 |
| 1 | Prestw-1703 | 1,8-Dihydroxyanthraquinone | 14068 | 110.1487953 | 1 |
| 1 | Prestw-913 | 2-Aminobenzenesulfonamide | 14432 | 15.98435231 | 14.98435231 |
| 2 | Prestw-913 | 2-Aminobenzenesulfonamide | 6058 | 1 | 1 |
| 1 | Prestw-697 | 3-alpha-Hydroxy-5-beta-androstan-17-one | 17281 | 41.63243291 | 1 |
| 2 | Prestw-697 | 3-alpha-Hydroxy-5-beta-androstan-17-one | 5888.5 | 19.5053391 | 1 |
| 1 | Prestw-1202 | 4-aminosalicylic acid | 8264 | 17.990575 | 12.4418223 |
| 2 | Prestw-1202 | 4-aminosalicylic acid | 3013 | 31.676074 | 1 |
| 1 | Prestw-1203 | 5-fluorouracil | 8746.5 | 28.6483055 | 23.0995528 |
| 2 | Prestw-1203 | 5-fluorouracil | 2972 | 83.0824598 | 6.7883217 |
| 1 | Prestw-659 | 6-Furfurylaminopurine | 14774.5 | 26.76606528 | 1 |
| 2 | Prestw-659 | 6-Furfurylaminopurine | 5462 | 1 | 1 |
| 1 | Prestw-1512 | Abacavir Sulfate | 9995 | 1 | 1 |
| 2 | Prestw-1512 | Abacavir Sulfate | 4207 | 1 | 1 |
| 2 | Prestw-1502 | Acamprosate calcium | 4293 | 16.1015216 | 12.6072603 |
| 1 | Prestw-1502 | Acamprosate calcium | 10870.5 | 111.0036934 | 25.47565947 |
| 1 | Prestw-1174 | Acarbose | 10127.5 | 40.46382454 | 5.2134061 |
| 2 | Prestw-1174 | Acarbose | 3374 | 1 | 1 |
| 2 | Prestw-69 | Acebutolol hydrochloride | 5438 | 112.8878426 | 81.9997662 |
| 1 | Prestw-69 | Acebutolol hydrochloride | 15976 | 47.15239168 | 15.2637636 |
| 1 | Prestw-1511 | Aceclidine Hydrochloride | 10062 | 1 | 1 |
| 2 | Prestw-1511 | Aceclidine Hydrochloride | 4625.5 | 1 | 1 |
| 2 | Prestw-175 | Aceclofenac | 3986.5 | 95.7233289 | 92.2290676 |
| 1 | Prestw-175 | Aceclofenac | 10715.5 | 134.4513105 | 48.92327663 |
| 2 | Prestw-1507 | Acefylline | 4609 | 45.6050826 | 42.1108213 |
| 1 | Prestw-1507 | Acefylline | 11831.5 | 106.67284 | 21.1448061 |
| 1 | Prestw-296 | Acemetacin | 13367 | 22.43782145 | 17.79275215 |
| 2 | Prestw-296 | Acemetacin | 4414.5 | 123.2316441 | 16.2228771 |
| 1 | Prestw-110 | Acenocoumarol | 8498.5 | 2.7464808 | 1.7464808 |
| 2 | Prestw-110 | Acenocoumarol | 3000 | 5.5254833 | 1 |
| 1 | Prestw-868 | Acetaminophen | 16489.5 | 52.57448315 | 16.02046482 |
| 2 | Prestw-868 | Acetaminophen | 5654 | 18.634196 | 15.4256556 |

| | | | | | |
|---|---|---|---|---|---|
| 1 | Prestw-3 | Acetazolamide | 11485.5 | 50.63846531 | 42.61280727 |
| 2 | Prestw-3 | Acetazolamide | 4456.5 | 136.3164392 | 1 |
| 1 | Prestw-55 | Acetohexamide | 15711 | 25.16118579 | 1 |
| 2 | Prestw-55 | Acetohexamide | 5778 | 93.81149 | 1 |
| 1 | Prestw-576 | Acetopromazine maleate salt | 12852 | 50.4775395 | 23.80046456 |
| 2 | Prestw-576 | Acetopromazine maleate salt | 4361.5 | 167.0036217 | 1 |
| 1 | Prestw-1206 | Acetylcysteine | 8389 | 19.1867332 | 13.6379805 |
| 2 | Prestw-1206 | Acetylcysteine | 3203 | 1 | 1 |
| 1 | Prestw-1207 | Acetylsalicylic acid | 8345 | 12.1048228 | 6.5560701 |
| 2 | Prestw-1207 | Acetylsalicylic acid | 2704.5 | 1 | 1 |
| 1 | Prestw-98 | Acetylsalicylsalicylic acid | 19508 | 35.4512257 | 6.20164518 |
| 2 | Prestw-98 | Acetylsalicylsalicylic acid | 5599.5 | 1 | 1 |
| 1 | Prestw-1127 | Acipimox | 11331 | 39.22788073 | 1 |
| 2 | Prestw-1127 | Acipimox | 3829 | 1 | 1 |
| 2 | Prestw-1165 | Acitretin | 3531 | 19.3389531 | 1 |
| 1 | Prestw-1165 | Acitretin | 10173 | 87.98600572 | 1 |
| 2 | Prestw-1763 | Actarit | 4279.5 | 47.7043881 | 1 |
| 1 | Prestw-1763 | Actarit | 9567.5 | 78.43165383 | 1 |
| 2 | Prestw-86 | Acyclovir | 5533 | 102.1499731 | 82.5641414 |
| 1 | Prestw-86 | Acyclovir | 14312 | 84.11517154 | 32.70784403 |
| 2 | Prestw-407 | Adamantamine fumarate | 4131.5 | 91.3139773 | 27.9638017 |
| 1 | Prestw-407 | Adamantamine fumarate | 12383 | 28.81680902 | 1 |
| 1 | Prestw-1714 | Adapalene | 14047.5 | 140.0307971 | 22.50430823 |
| 2 | Prestw-1714 | Adapalene | 5463 | 59.4690337 | 1 |
| 1 | Prestw-356 | Adenosine 5'-monophosphate monohydrate | 11178 | 23.06345083 | 9.59700128 |
| 2 | Prestw-356 | Adenosine 5'-monophosphate monohydrate | 4260 | 141.140212 | 35.7348268 |
| 1 | Prestw-75 | Adiphenine hydrochloride | 14649.5 | 1 | 1 |
| 2 | Prestw-75 | Adiphenine hydrochloride | 5300 | 1 | 1 |
| 2 | Prestw-899 | Adrenosterone | 5721.5 | 41.7074743 | 40.7074743 |
| 1 | Prestw-899 | Adrenosterone | 19485 | 41.90802253 | 12.65844201 |
| 2 | Prestw-247 | Albendazole | 4915 | 59.64726 | 1 |
| 1 | Prestw-247 | Albendazole | 14289 | 101.734437 | 1 |
| 2 | Prestw-771 | Alclometasone dipropionate | 5257.5 | 62.6380006 | 31.7499242 |
| 1 | Prestw-771 | Alclometasone dipropionate | 16752.5 | 45.32053881 | 13.43191073 |
| 1 | Prestw-1036 | Alcuronium chloride | 9153.5 | 39.476349 | 16.3322547 |
| 2 | Prestw-1036 | Alcuronium chloride | 3093.5 | 14.3950336 | 1 |
| 2 | Prestw-1210 | Alendronate sodium | 3507 | 42.2308056 | 39.1568604 |
| 1 | Prestw-1210 | Alendronate sodium | 9814.5 | 13.89027462 | 1 |
| 1 | Prestw-777 | Alexidine dihydrochloride | 19405 | 68.17522628 | 22.7958431 |
| 2 | Prestw-777 | Alexidine dihydrochloride | 5332.5 | 1 | 1 |
| 1 | Prestw-1211 | Alfacalcidol | 8901 | 9.2446554 | 1 |
| 2 | Prestw-1211 | Alfacalcidol | 3164.5 | 1 | 1 |
| 1 | Prestw-1001 | Alfadolone acetate | 7294.5 | 1 | 1 |
| 2 | Prestw-1001 | Alfadolone acetate | 3054.5 | 1 | 1 |
| 1 | Prestw-1002 | Alfaxalone | 7841.5 | 1 | 1 |
| 2 | Prestw-1002 | Alfaxalone | 3139 | 1 | 1 |
| 1 | Prestw-322 | Alfuzosin hydrochloride | 12805.5 | 54.40705835 | 41.44900385 |
| 2 | Prestw-322 | Alfuzosin hydrochloride | 4591.5 | 167.8568456 | 6.9931893 |
| 2 | Prestw-1715 | Algestone acetophenide | 5166.5 | 89.1269656 | 2.4248496 |
| 1 | Prestw-1715 | Algestone acetophenide | 13179 | 113.4350384 | 1 |
| 1 | Prestw-1810 | Aliskiren hemifumarate | 11725 | 233.5015779 | 44.22538765 |
| 2 | Prestw-1810 | Aliskiren hemifumarate | 4851.5 | 37.5966094 | 1 |
| 2 | Prestw-451 | Alizapride hydrochloride | 4168 | 123.6579227 | 31.477131 |
| 1 | Prestw-451 | Alizapride hydrochloride | 11892.5 | 10.7975102 | 1 |
| 1 | Prestw-2 | Allantoin | 10375.5 | 134.1398045 | 51.39278698 |
| 2 | Prestw-2 | Allantoin | 4533 | 56.822684 | 1 |
| 2 | Prestw-1213 | Allopurinol | 2987.5 | 97.5375604 | 94.4636152 |
| 1 | Prestw-1213 | Allopurinol | 8772.5 | 16.1436612 | 1 |
| 1 | Prestw-1781 | Alosetron hydrochloride | 11735.5 | 132.9048408 | 21.75596973 |
| 2 | Prestw-1781 | Alosetron hydrochloride | 4957.5 | 101.3511877 | 7.892725 |
| 1 | Prestw-250 | Alprenolol hydrochloride | 13231.5 | 130.8858963 | 10.97313301 |
| 2 | Prestw-250 | Alprenolol hydrochloride | 4945.5 | 63.0498034 | 1 |
| 2 | Prestw-1018 | Alprostadil | 3046.5 | 21.2103815 | 18.1046633 |

| | | | | | |
|---|---|---|---|---|---|
| 1 | Prestw-1018 | Alprostadil | 7918.5 | 6.360034 | 1 |
| 2 | Prestw-721 | Althiazide | 5977 | 14.6195704 | 13.6195704 |
| 1 | Prestw-721 | Althiazide | 18364.5 | 42.60037941 | 1 |
| 1 | Prestw-1716 | Altrenogest | 13527.5 | 149.336259 | 31.80977018 |
| 2 | Prestw-1716 | Altrenogest | 4842 | 89.3436092 | 2.6414932 |
| 1 | Prestw-946 | Altretamine | 18591 | 57.76245178 | 6.35512427 |
| 2 | Prestw-946 | Altretamine | 6130 | 33.8311801 | 14.2453484 |
| 2 | Prestw-174 | Alverine citrate salt | 4386 | 42.0845928 | 38.5903315 |
| 1 | Prestw-174 | Alverine citrate salt | 11463 | 110.2347487 | 24.70671476 |
| 2 | Prestw-1497 | Ambrisentan | 4361 | 95.3652594 | 23.3731189 |
| 1 | Prestw-1497 | Ambrisentan | 11781.5 | 187.0775509 | 43.21687956 |
| 1 | Prestw-366 | Ambroxol hydrochloride | 7286 | 62.904173 | 58.2591037 |
| 2 | Prestw-366 | Ambroxol hydrochloride | 3831 | 612.1466477 | 505.1378807 |
| 2 | Prestw-1476 | Amcinonide | 4527.5 | 153.5713992 | 66.8692832 |
| 1 | Prestw-1476 | Amcinonide | 10676.5 | 134.8248371 | 17.29834829 |
| 1 | Prestw-373 | Amethopterin (R,S) | 15248.5 | 10.73514722 | 2.70948918 |
| 2 | Prestw-373 | Amethopterin (R,S) | 5902 | 127.2583468 | 1 |
| 2 | Prestw-1471 | Amfepramone hydrochloride | 5343.5 | 83.8562291 | 1 |
| 1 | Prestw-1471 | Amfepramone hydrochloride | 13563.5 | 122.2356263 | 1 |
| 2 | Prestw-88 | Amidopyrine | 5809 | 9.9592606 | 8.9592606 |
| 1 | Prestw-88 | Amidopyrine | 17199 | 42.06920009 | 1 |
| 2 | Prestw-1489 | Amifostine | 4231.5 | 182.8277978 | 3.8528798 |
| 1 | Prestw-1489 | Amifostine | 11794 | 166.0838028 | 2.44800682 |
| 1 | Prestw-395 | Amikacin hydrate | 13418.5 | 30.2320742 | 17.2740197 |
| 2 | Prestw-395 | Amikacin hydrate | 4196.5 | 89.7934761 | 1 |
| 2 | Prestw-7 | Amiloride hydrochloride dihydrate | 4816.5 | 98.6278568 | 29.2224606 |
| 1 | Prestw-7 | Amiloride hydrochloride dihydrate | 16365.5 | 57.00531745 | 11.62593427 |
| 1 | Prestw-1717 | Aminacrine | 11861 | 153.6393982 | 36.11290931 |
| 2 | Prestw-1717 | Aminacrine | 4968 | 87.2854812 | 0.5833652 |
| 2 | Prestw-960 | Aminocaproic acid | 6587.5 | 1 | 1 |
| 1 | Prestw-960 | Aminocaproic acid | 18074.5 | 46.12843381 | 1 |
| 2 | Prestw-895 | Aminohippuric acid | 5086 | 19.6596973 | 18.6596973 |
| 1 | Prestw-895 | Aminohippuric acid | 17564.5 | 35.70256036 | 6.45297984 |
| 2 | Prestw-820 | Aminophylline | 2701 | 2510.883333 | 2509.883333 |
| 1 | Prestw-820 | Aminophylline | 2783.5 | 1139.981872 | 1125.336863 |
| 1 | Prestw-189 | Aminopurine, 6-benzyl | 10638.5 | 37.46918885 | 17.76332223 |
| 2 | Prestw-189 | Aminopurine, 6-benzyl | 4396 | 1 | 1 |
| 2 | Prestw-409 | Amiodarone hydrochloride | 4504.5 | 75.9750833 | 12.6249077 |
| 1 | Prestw-409 | Amiodarone hydrochloride | 12383.5 | 28.62274846 | 1 |
| 1 | Prestw-845 | Amiprilose hydrochloride | 17389.5 | 34.89271151 | 1 |
| 2 | Prestw-845 | Amiprilose hydrochloride | 5852 | 1 | 1 |
| 2 | Prestw-1217 | Amisulpride | 3333 | 44.4684993 | 41.3945541 |
| 1 | Prestw-1217 | Amisulpride | 8831.5 | 23.5024015 | 5.9401993 |
| 1 | Prestw-74 | Amitryptiline hydrochloride | 15711.5 | 31.23131189 | 16.58630295 |
| 2 | Prestw-74 | Amitryptiline hydrochloride | 5509.5 | 1 | 1 |
| 2 | Prestw-1718 | Amlexanox | 4987.5 | 94.1267338 | 7.4246178 |
| 1 | Prestw-1718 | Amlexanox | 13385 | 117.0457232 | 1 |
| 1 | Prestw-1219 | Amlodipine | 3122.5 | 636.7278969 | 619.1656947 |
| 2 | Prestw-1219 | Amlodipine | 1189 | 891.21271 | 888.1387648 |
| 1 | Prestw-309 | Amodiaquin dihydrochloride dihydrate | 11373.5 | 0.72672 | 1 |
| 2 | Prestw-309 | Amodiaquin dihydrochloride dihydrate | 4129 | 115.0614944 | 1 |
| 2 | Prestw-1719 | Amorolfine hydrochloride | 5063.5 | 73.6577082 | 1 |
| 1 | Prestw-1719 | Amorolfine hydrochloride | 13660 | 116.5196517 | 1 |
| 2 | Prestw-102 | Amoxapine | 2810 | 63.9609611 | 60.8552429 |
| 1 | Prestw-102 | Amoxapine | 8419.5 | 9.1526331 | 1 |
| 1 | Prestw-357 | Amoxicillin | 12907 | 13.39393086 | 1 |
| 2 | Prestw-357 | Amoxicillin | 4380.5 | 118.8668105 | 13.4614253 |
| 1 | Prestw-410 | Amphotericin B | 10780.5 | 1 | 1 |
| 2 | Prestw-410 | Amphotericin B | 3821.5 | 1 | 1 |
| 2 | Prestw-114 | Ampicillin trihydrate | 3409 | 59.6661668 | 1 |
| 1 | Prestw-114 | Ampicillin trihydrate | 10477.5 | 72.19989926 | 1 |
| 2 | Prestw-1772 | Ampiroxicam | 4994 | 46.8993243 | 1 |
| 1 | Prestw-1772 | Ampiroxicam | 13054 | 71.68101689 | 1 |

| | | | | | |
|---|---|---|---|---|---|
| 1 | Prestw-1820 | Amprenavir | 12031 | 152.5959418 | 32.68317853 |
| 2 | Prestw-1820 | Amprenavir | 4426 | 35.4228022 | 1 |
| 1 | Prestw-8 | Amprolium hydrochloride | 19600 | 42.89436387 | 1 |
| 2 | Prestw-8 | Amprolium hydrochloride | 5902 | 1 | 1 |
| 1 | Prestw-181 | Ampyrone | 10847 | 207.6520127 | 18.37582246 |
| 2 | Prestw-181 | Ampyrone | 4626 | 82.714865 | 1 |
| 2 | Prestw-800 | Amrinone | 6686.5 | 7.4216016 | 6.4216016 |
| 1 | Prestw-800 | Amrinone | 21687.5 | 31.30198353 | 1 |
| 1 | Prestw-49 | Amyleine hydrochloride | 11763 | 17.4097588 | 1 |
| 2 | Prestw-49 | Amyleine hydrochloride | 2926 | 1 | 1 |
| 2 | Prestw-1720 | Anagrelide | 5527.5 | 101.0565101 | 1 |
| 1 | Prestw-1720 | Anagrelide | 14533 | 127.6053329 | 1 |
| 2 | Prestw-1222 | Anastrozole | 2877.5 | 94.6155402 | 91.541595 |
| 1 | Prestw-1222 | Anastrozole | 9099 | 23.8287607 | 6.2665585 |
| 1 | Prestw-411 | Androsterone | 10203.5 | 1 | 1 |
| 2 | Prestw-411 | Androsterone | 3781.5 | 1 | 1 |
| 2 | Prestw-1223 | Anethole-trithione | 3336.5 | 81.5510111 | 78.4770659 |
| 1 | Prestw-1223 | Anethole-trithione | 8926 | 23.9653894 | 6.4031872 |
| 1 | Prestw-1178 | Aniracetam | 9850 | 45.83524861 | 10.58483017 |
| 2 | Prestw-1178 | Aniracetam | 3255.5 | 1 | 1 |
| 1 | Prestw-258 | Antazoline hydrochloride | 12323.5 | 88.18472269 | 25.83478167 |
| 2 | Prestw-258 | Antazoline hydrochloride | 4719.5 | 6.9376688 | 5.9376688 |
| 1 | Prestw-1224 | Anthralin | 9218.5 | 39.58991236 | 22.02771016 |
| 2 | Prestw-1224 | Anthralin | 3178.5 | 1 | 1 |
| 2 | Prestw-222 | Antimycin A | 4601.5 | 60.8994741 | 1 |
| 1 | Prestw-222 | Antimycin A | 11934.5 | 119.3921486 | 1 |
| 1 | Prestw-29 | Antipyrine | 11067.5 | 28.0740398 | 23.4289705 |
| 2 | Prestw-29 | Antipyrine | 4198.5 | 109.7379349 | 2.7291679 |
| 1 | Prestw-30 | Antipyrine, 4-hydroxy | 11947 | 36.92599391 | 28.90033587 |
| 2 | Prestw-30 | Antipyrine, 4-hydroxy | 4269 | 123.4610092 | 1 |
| 1 | Prestw-1005 | Apramycin | 7393 | 1 | 1 |
| 2 | Prestw-1005 | Apramycin | 3029.5 | 1 | 1 |
| 2 | Prestw-1600 | Aprepitant | 4206 | 80.4351595 | 63.8219755 |
| 1 | Prestw-1600 | Aprepitant | 11036.5 | 65.17435651 | 1 |
| 1 | Prestw-1228 | Argatroban | 8911 | 52.35263905 | 34.79043685 |
| 2 | Prestw-1228 | Argatroban | 3212.5 | 55.1840476 | 52.1101024 |
| 2 | Prestw-1229 | Aripiprazole | 3274 | 45.0716304 | 41.9976852 |
| 1 | Prestw-1229 | Aripiprazole | 9214.5 | 18.8099235 | 1.2477213 |
| 1 | Prestw-498 | Artemisinin | 12043 | 34.2643685 | 8.76585113 |
| 2 | Prestw-498 | Artemisinin | 4124 | 112.4610344 | 1 |
| 2 | Prestw-1032 | Articaine hydrochloride | 2988.5 | 98.0245578 | 49.8705926 |
| 1 | Prestw-1032 | Articaine hydrochloride | 8349.5 | 35.4805795 | 12.3364852 |
| 2 | Prestw-325 | Ascorbic acid | 4058.5 | 119.3179921 | 55.9678165 |
| 1 | Prestw-325 | Ascorbic acid | 11165.5 | 34.242113 | 1 |
| 2 | Prestw-1231 | Asenapine maleate | 3400 | 129.8109694 | 43.258176 |
| 1 | Prestw-1231 | Asenapine maleate | 9407.5 | 34.71475597 | 30.12683487 |
| 2 | Prestw-136 | Astemizole | 3769 | 28.8716856 | 6.715508 |
| 1 | Prestw-136 | Astemizole | 11601.5 | 29.97910348 | 1 |
| 2 | Prestw-1232 | Atorvastatin | 2954.5 | 144.3874878 | 57.8346944 |
| 1 | Prestw-1232 | Atorvastatin | 8396 | 48.4626844 | 43.8747633 |
| 1 | Prestw-534 | Atovaquone | 13164.5 | 5.58692657 | 1 |
| 2 | Prestw-534 | Atovaquone | 4677.5 | 105.176892 | 1.6403149 |
| 1 | Prestw-5 | Atracurium besylate | 10632.5 | 37.38894123 | 34.20204221 |
| 2 | Prestw-5 | Atracurium besylate | 2749 | 1 | 1 |
| 1 | Prestw-565 | Atropine sulfate monohydrate | 13285 | 48.68756351 | 22.01048857 |
| 2 | Prestw-565 | Atropine sulfate monohydrate | 4461 | 163.9736695 | 1 |
| 1 | Prestw-1233 | Auranofin | 8965.5 | 31.6040899 | 27.0161688 |
| 2 | Prestw-1233 | Auranofin | 3033.5 | 76.7559458 | 1 |
| 1 | Prestw-660 | Avermectin B1 | 13582.5 | 21.12917691 | 1 |
| 2 | Prestw-660 | Avermectin B1 | 4881.5 | 1 | 1 |
| 2 | Prestw-1721 | Avobenzone | 5627.5 | 88.044256 | 1 |
| 1 | Prestw-1721 | Avobenzone | 13842.5 | 162.5057949 | 1 |
| 2 | Prestw-93 | Azacyclonol | 5741.5 | 58.2088054 | 29.6369413 |

| | | | | | |
|---|---|---|---|---|---|
| 1 | Prestw-93 | Azacyclonol | 18627.5 | 59.84070996 | 7.85800638 |
| 2 | Prestw-866 | Azacytidine-5 | 5778.5 | 59.1950425 | 55.9865021 |
| 1 | Prestw-866 | Azacytidine-5 | 16154.5 | 38.64383893 | 2.0898206 |
| 1 | Prestw-1 | Azaguanine-8 | 7443.5 | 1 | 1 |
| 2 | Prestw-1 | Azaguanine-8 | 2912 | 1 | 1 |
| 1 | Prestw-1117 | Azaperone | 9039.5 | 31.960512 | 8.8164177 |
| 2 | Prestw-1117 | Azaperone | 3372.5 | 16.9889144 | 1 |
| 1 | Prestw-1003 | Azapropazone | 7527.5 | 6.6038288 | 5.6038288 |
| 2 | Prestw-1003 | Azapropazone | 3225.5 | 1 | 1 |
| 1 | Prestw-1722 | Azatadine maleate | 13650.5 | 168.461297 | 4.82550106 |
| 2 | Prestw-1722 | Azatadine maleate | 4844.5 | 88.1454454 | 1 |
| 2 | Prestw-94 | Azathioprine | 6148 | 103.5957384 | 84.0099067 |
| 1 | Prestw-94 | Azathioprine | 20611 | 60.28382798 | 8.87650047 |
| 1 | Prestw-1130 | Azelastine hydrochloride | 10048.5 | 33.57626035 | 1 |
| 2 | Prestw-1130 | Azelastine hydrochloride | 3524.5 | 10.7502742 | 1 |
| 2 | Prestw-1811 | Azilsartan kamedoxomil | 4869 | 49.2267768 | 1 |
| 1 | Prestw-1811 | Azilsartan kamedoxomil | 12614 | 174.1244019 | 1 |
| 1 | Prestw-1234 | Azithromycin | 8967.5 | 30.2689262 | 25.6810051 |
| 2 | Prestw-1234 | Azithromycin | 3275 | 62.1952905 | 1 |
| 1 | Prestw-821 | Azlocillin sodium salt | 17050 | 11.26873188 | 1 |
| 2 | Prestw-821 | Azlocillin sodium salt | 5533 | 1 | 1 |
| 1 | Prestw-185 | Aztreonam | 12389.5 | 74.41985622 | 12.0699152 |
| 2 | Prestw-185 | Aztreonam | 4876 | 1 | 1 |
| 1 | Prestw-416 | Bacampicillin hydrochloride | 10449 | 1 | 1 |
| 2 | Prestw-416 | Bacampicillin hydrochloride | 3796 | 0.568573 | 1 |
| 2 | Prestw-919 | Bacitracin | 4940.5 | 51.0983899 | 7.017267 |
| 1 | Prestw-919 | Bacitracin | 13763 | 30.01662037 | 1 |
| 2 | Prestw-85 | Baclofen (R,S) | 5197 | 49.2298324 | 20.6579683 |
| 1 | Prestw-85 | Baclofen (R,S) | 15804.5 | 46.97432554 | 1 |
| 2 | Prestw-730 | Balsalazide Sodium | 5364 | 1 | 1 |
| 1 | Prestw-730 | Balsalazide Sodium | 16873 | 43.26990815 | 1 |
| 1 | Prestw-361 | Bambuterol hydrochloride | 11877 | 7.59401409 | 1 |
| 2 | Prestw-361 | Bambuterol hydrochloride | 3940.5 | 87.8795538 | 1 |
| 2 | Prestw-855 | Beclomethasone dipropionate | 6020 | 73.2708073 | 44.6989432 |
| 1 | Prestw-855 | Beclomethasone dipropionate | 16320.5 | 64.43045833 | 12.44775475 |
| 2 | Prestw-882 | Bemegride | 6042 | 5.7525664 | 4.7525664 |
| 1 | Prestw-882 | Bemegride | 18753 | 32.92563987 | 1 |
| 1 | Prestw-1129 | Benazepril hydrochloride | 10944.5 | 26.88156089 | 1 |
| 2 | Prestw-1129 | Benazepril hydrochloride | 3582 | 8.8594815 | 1 |
| 1 | Prestw-784 | Bendroflumethiazide | 19557.5 | 89.9891219 | 55.96085363 |
| 2 | Prestw-784 | Bendroflumethiazide | 6298.5 | 40.039805 | 27.2831179 |
| 1 | Prestw-367 | Benfluorex | 13637.5 | 7.53447381 | 2.88940451 |
| 2 | Prestw-367 | Benfluorex | 4631.5 | 117.959529 | 10.950762 |
| 1 | Prestw-654 | Benfotiamine | 15251.5 | 1 | 1 |
| 2 | Prestw-654 | Benfotiamine | 5911.5 | 1 | 1 |
| 1 | Prestw-1766 | Benidipine hydrochloride | 9451 | 137.7401856 | 54.99316809 |
| 2 | Prestw-1766 | Benidipine hydrochloride | 4084.5 | 68.6754414 | 1 |
| 1 | Prestw-57 | Benoxinate hydrochloride | 14096.5 | 49.52005803 | 22.84298309 |
| 2 | Prestw-57 | Benoxinate hydrochloride | 4183 | 106.146552 | 1 |
| 1 | Prestw-484 | Benperidol | 11549.5 | 40.6736424 | 13.99656746 |
| 2 | Prestw-484 | Benperidol | 3931.5 | 209.1103633 | 31.5856368 |
| 2 | Prestw-938 | Benserazide hydrochloride | 6104 | 53.5713056 | 24.9994415 |
| 1 | Prestw-938 | Benserazide hydrochloride | 16821.5 | 49.59063515 | 1 |
| 1 | Prestw-657 | Benzamil hydrochloride | 14356 | 3.96617167 | 1 |
| 2 | Prestw-657 | Benzamil hydrochloride | 5049.5 | 1 | 1 |
| 1 | Prestw-1028 | Benzathine benzylpenicillin | 8728 | 19.970414 | 5.8155603 |
| 2 | Prestw-1028 | Benzathine benzylpenicillin | 3134 | 1 | 1 |
| 2 | Prestw-370 | Benzbromarone | 4172.5 | 156.417326 | 49.408559 |
| 1 | Prestw-370 | Benzbromarone | 12959 | 6.63469379 | 1.98962449 |
| 1 | Prestw-708 | Benzethonium chloride | 17006 | 69.55513691 | 35.52686864 |
| 2 | Prestw-708 | Benzethonium chloride | 5660.5 | 41.6236404 | 28.8669533 |
| 1 | Prestw-712 | Benzocaine | 17979 | 70.87002956 | 36.84176129 |
| 2 | Prestw-712 | Benzocaine | 5762.5 | 63.7348424 | 50.9781553 |

| | | | | | |
|---|---|---|---|---|---|
| 1 | Prestw-12 | Benzonatate | 8087 | 10.0285478 | 4.4797951 |
| 2 | Prestw-12 | Benzonatate | 2861 | 30.4525073 | 1 |
| 2 | Prestw-1723 | Benzoxiquine | 4959 | 87.3756974 | 1 |
| 1 | Prestw-1723 | Benzoxiquine | 14021 | 158.2146484 | 1 |
| 1 | Prestw-824 | Benzthiazide | 15525.5 | 1 | 1 |
| 2 | Prestw-824 | Benzthiazide | 5525.5 | 1 | 1 |
| 1 | Prestw-1236 | Benztropine mesylate | 9344 | 30.5880754 | 26.0001543 |
| 2 | Prestw-1236 | Benztropine mesylate | 3333.5 | 92.9396529 | 6.3868595 |
| 1 | Prestw-297 | Benzydamine hydrochloride | 14245 | 22.08606604 | 17.44099674 |
| 2 | Prestw-297 | Benzydamine hydrochloride | 4891 | 144.0619378 | 37.0531708 |
| 1 | Prestw-1078 | Benzylpenicillin sodium | 8235.5 | 1 | 1 |
| 2 | Prestw-1078 | Benzylpenicillin sodium | 3517 | 1 | 1 |
| 2 | Prestw-936 | Bephenium hydroxynaphthoate | 6528 | 46.4658607 | 17.8939966 |
| 1 | Prestw-936 | Bephenium hydroxynaphthoate | 17915 | 52.08922482 | 0.10652124 |
| 2 | Prestw-368 | Bepridil hydrochloride | 4557.5 | 122.7127392 | 15.7039722 |
| 1 | Prestw-368 | Bepridil hydrochloride | 13331 | 1.99744952 | 1 |
| 2 | Prestw-1740 | Besifloxacin hydrochloride | 5062.5 | 26.0391998 | 22.5449385 |
| 1 | Prestw-1740 | Besifloxacin hydrochloride | 12203 | 109.5326729 | 24.00463895 |
| 1 | Prestw-676 | Beta-Escin | 15350 | 40.72613872 | 8.83751064 |
| 2 | Prestw-676 | Beta-Escin | 5547.5 | 45.1049392 | 14.2168628 |
| 1 | Prestw-543 | Betahistine mesylate | 14624.5 | 7.26713 | 1 |
| 2 | Prestw-543 | Betahistine mesylate | 3770 | 1 | 1 |
| 2 | Prestw-362 | Betamethasone | 5620.5 | 139.1830199 | 32.1742529 |
| 1 | Prestw-362 | Betamethasone | 15316.5 | 4.12047631 | 1 |
| 1 | Prestw-382 | Betaxolol hydrochloride | 13999 | 39.7012858 | 2.2735216 |
| 2 | Prestw-382 | Betaxolol hydrochloride | 4926 | 124.0587657 | 1 |
| 1 | Prestw-43 | Betazole hydrochloride | 11694.5 | 3.2557928 | 1 |
| 2 | Prestw-43 | Betazole hydrochloride | 3825 | 31.7141497 | 1 |
| 1 | Prestw-1073 | Bethanechol chloride | 10035 | 48.31925842 | 13.06883998 |
| 2 | Prestw-1073 | Bethanechol chloride | 3338.5 | 1 | 1 |
| 1 | Prestw-378 | Bezafibrate | 12769.5 | 4.41294445 | 1 |
| 2 | Prestw-378 | Bezafibrate | 4546 | 145.1650897 | 1 |
| 1 | Prestw-1239 | Bicalutamide | 8930 | 53.95267972 | 49.36475862 |
| 2 | Prestw-1239 | Bicalutamide | 3469 | 69.1879514 | 1 |
| 1 | Prestw-1241 | Bifonazole | 9442 | 27.5763282 | 22.9884071 |
| 2 | Prestw-1241 | Bifonazole | 3259 | 51.589905 | 1 |
| 2 | Prestw-1821 | Bimatoprost | 4932 | 49.5711256 | 1 |
| 1 | Prestw-1821 | Bimatoprost | 13399.5 | 117.689389 | 1 |
| 1 | Prestw-418 | Biotin | 10446.5 | 1 | 1 |
| 2 | Prestw-418 | Biotin | 3184 | 1 | 1 |
| 2 | Prestw-502 | Biperiden hydrochloride | 4622 | 90.7185048 | 57.8731646 |
| 1 | Prestw-502 | Biperiden hydrochloride | 11631 | 10.91832798 | 7.73142896 |
| 1 | Prestw-419 | Bisacodyl | 10537.5 | 1 | 1 |
| 2 | Prestw-419 | Bisacodyl | 3936 | 1 | 1 |
| 2 | Prestw-330 | Bisoprolol fumarate | 4551.5 | 125.3752333 | 62.0250577 |
| 1 | Prestw-330 | Bisoprolol fumarate | 11914 | 30.81204295 | 1 |
| 1 | Prestw-1244 | Bosentan | 10579 | 32.0890315 | 27.5011104 |
| 2 | Prestw-1244 | Bosentan | 3491.5 | 1 | 1 |
| 2 | Prestw-942 | Bretylium tosylate | 6184.5 | 90.0356413 | 70.4498096 |
| 1 | Prestw-942 | Bretylium tosylate | 21157.5 | 66.39893749 | 14.99160998 |
| 1 | Prestw-1751 | Brimonidine L-Tartrate | 2748 | 256.3574968 | 236.6516302 |
| 2 | Prestw-1751 | Brimonidine L-Tartrate | 1510.5 | 33.7747988 | 1 |
| 2 | Prestw-365 | Brinzolamide | 4937 | 123.63847 | 16.629703 |
| 1 | Prestw-365 | Brinzolamide | 13395.5 | 1 | 1 |
| 2 | Prestw-1246 | Bromhexine hydrochloride | 3371.5 | 70.1335962 | 47.9774186 |
| 1 | Prestw-1246 | Bromhexine hydrochloride | 9864.5 | 54.84077543 | 1 |
| 1 | Prestw-121 | Bromocryptine mesylate | 8571.5 | 21.3522526 | 15.8034999 |
| 2 | Prestw-121 | Bromocryptine mesylate | 3218 | 4.4805683 | 1 |
| 1 | Prestw-704 | Bromopride | 17818.5 | 57.30189825 | 11.92251507 |
| 2 | Prestw-704 | Bromopride | 5833.5 | 72.4351276 | 3.0297314 |
| 2 | Prestw-509 | Bromperidol | 4177.5 | 85.9594382 | 52.368998 |
| 1 | Prestw-509 | Bromperidol | 10754 | 1 | 1 |
| 2 | Prestw-475 | Brompheniramine maleate | 4711.5 | 156.2195094 | 39.0015553 |

| | | | | | |
|---|---|---|---|---|---|
| 1 | Prestw-475 | Brompheniramine maleate | 14064.5 | 18.23117724 | 1 |
| 1 | Prestw-813 | Bucladesine sodium salt | 17437.5 | 32.66502494 | 18.020016 |
| 2 | Prestw-813 | Bucladesine sodium salt | 5975.5 | 1 | 1 |
| 1 | Prestw-518 | Budesonide | 10583.5 | 1 | 1 |
| 2 | Prestw-518 | Budesonide | 3771 | 1 | 1 |
| 1 | Prestw-243 | Bufexamac | 12868.5 | 127.6821044 | 7.76934115 |
| 2 | Prestw-243 | Bufexamac | 4487.5 | 57.7174219 | 1 |
| 1 | Prestw-426 | Buflomedil hydrochloride | 10472 | 0.2315966 | 1 |
| 2 | Prestw-426 | Buflomedil hydrochloride | 4052.5 | 41.8434098 | 1 |
| 1 | Prestw-276 | Bumetanide | 10055 | 1 | 1 |
| 2 | Prestw-276 | Bumetanide | 3704.5 | 29.3935102 | 0.8452971 |
| 1 | Prestw-305 | Bupivacaine hydrochloride | 14297.5 | 27.56469354 | 19.5390355 |
| 2 | Prestw-305 | Bupivacaine hydrochloride | 5450 | 121.9804382 | 1 |
| 2 | Prestw-249 | Bupropion hydrochloride | 5059.5 | 28.5010646 | 1 |
| 1 | Prestw-249 | Bupropion hydrochloride | 15072 | 109.6669302 | 1 |
| 2 | Prestw-1283 | Buspirone hydrochloride | 3612 | 1 | 1 |
| 1 | Prestw-1283 | Buspirone hydrochloride | 11080 | 63.48949211 | 1 |
| 1 | Prestw-1179 | Busulfan | 9305.5 | 47.66544322 | 12.41502478 |
| 2 | Prestw-1179 | Busulfan | 3387.5 | 1 | 1 |
| 1 | Prestw-831 | Butacaine | 15489.5 | 1 | 1 |
| 2 | Prestw-831 | Butacaine | 5487.5 | 1 | 1 |
| 2 | Prestw-1251 | Butalbital | 2902.5 | 133.9393589 | 111.7831813 |
| 1 | Prestw-1251 | Butalbital | 8296.5 | 65.9470274 | 1 |
| 1 | Prestw-761 | Butamben | 15175.5 | 10.42121116 | 1 |
| 2 | Prestw-761 | Butamben | 5103 | 1 | 1 |
| 2 | Prestw-1252 | Butenafine Hydrochloride | 3102.5 | 35.1781338 | 13.0219562 |
| 1 | Prestw-1252 | Butenafine Hydrochloride | 9578 | 42.19312074 | 1 |
| 1 | Prestw-408 | Butoconazole nitrate | 12300.5 | 25.01553254 | 1 |
| 2 | Prestw-408 | Butoconazole nitrate | 2766.5 | 1 | 1 |
| 1 | Prestw-894 | Butylparaben | 16962.5 | 38.07312959 | 8.82354907 |
| 2 | Prestw-894 | Butylparaben | 5241.5 | 1 | 1 |
| 2 | Prestw-1253 | Butylscopolammonium (n-) bromide | 2925 | 66.1272076 | 43.97103 |
| 1 | Prestw-1253 | Butylscopolammonium (n-) bromide | 10336.5 | 39.09563651 | 1 |
| 2 | Prestw-1256 | Caffeine | 2423.5 | 152.9738614 | 130.8176838 |
| 1 | Prestw-1256 | Caffeine | 7378 | 78.69517 | 2.4460422 |
| 2 | Prestw-1257 | Calcipotriene | 3603 | 24.0731778 | 1.9170002 |
| 1 | Prestw-1257 | Calcipotriene | 9935 | 43.57861751 | 1 |
| 1 | Prestw-200 | Camptothecine (S,+) | 10807.5 | 104.0324657 | 21.28544817 |
| 2 | Prestw-200 | Camptothecine (S,+) | 4551 | 46.948611 | 1 |
| 2 | Prestw-1498 | Camylofine chlorhydrate | 4978 | 105.6032354 | 33.6110949 |
| 1 | Prestw-1498 | Camylofine chlorhydrate | 11938.5 | 173.7696171 | 29.90894581 |
| 1 | Prestw-1258 | Candesartan | 10232 | 34.65660454 | 1 |
| 2 | Prestw-1258 | Candesartan | 3539 | 1 | 1 |
| 1 | Prestw-148 | Canrenoic acid potassium salt | 14413 | 127.1080246 | 9.58153577 |
| 2 | Prestw-148 | Canrenoic acid potassium salt | 5677.5 | 67.1714864 | 1 |
| 2 | Prestw-1259 | Canrenone | 3199 | 102.8875337 | 47.7960121 |
| 1 | Prestw-1259 | Canrenone | 9910 | 79.33006944 | 1 |
| 2 | Prestw-1828 | Capecitabine | 4514.5 | 16.3010804 | 15.3010804 |
| 1 | Prestw-1828 | Capecitabine | 11581 | 82.56485804 | 20.21491702 |
| 1 | Prestw-19 | Captopril | 10818.5 | 29.05546228 | 9.34959566 |
| 2 | Prestw-19 | Captopril | 4228.5 | 36.6837617 | 1 |
| 2 | Prestw-880 | Carbachol | 5409.5 | 43.8299225 | 42.8299225 |
| 1 | Prestw-880 | Carbachol | 16444.5 | 44.2488854 | 1 |
| 1 | Prestw-1121 | Carbadox | 9847 | 27.5006821 | 1 |
| 2 | Prestw-1121 | Carbadox | 3788 | 1 | 1 |
| 1 | Prestw-52 | Carbamazepine | 11899.5 | 2.8337991 | 1 |
| 2 | Prestw-52 | Carbamazepine | 4330.5 | 72.5324961 | 1 |
| 1 | Prestw-413 | Carbarsone | 10229 | 1 | 1 |
| 2 | Prestw-413 | Carbarsone | 3631.5 | 1 | 1 |
| 1 | Prestw-837 | Carbenoxolone disodium salt | 14304 | 30.91399395 | 1 |
| 2 | Prestw-837 | Carbenoxolone disodium salt | 4944.5 | 7.3634357 | 1 |
| 2 | Prestw-387 | Carbetapentane citrate | 4201 | 289.7718543 | 114.1312871 |
| 1 | Prestw-387 | Carbetapentane citrate | 10892.5 | 58.3583539 | 20.9305897 |

| | | | | | |
|---|---|---|---|---|---|
| 1 | Prestw-1177 | Carbidopa | 9576 | 40.43225313 | 5.18183469 |
| 2 | Prestw-1177 | Carbidopa | 3672.5 | 1 | 1 |
| 1 | Prestw-439 | Carbimazole | 10807 | 30.5050762 | 17.6529497 |
| 2 | Prestw-439 | Carbimazole | 2519.5 | 1 | 1 |
| 1 | Prestw-801 | Carbinoxamine maleate salt | 18317.5 | 38.40442336 | 1 |
| 2 | Prestw-801 | Carbinoxamine maleate salt | 6091.5 | 1 | 1 |
| 1 | Prestw-423 | Carisoprodol | 9445 | 1 | 1 |
| 2 | Prestw-423 | Carisoprodol | 3642.5 | 33.6768422 | 0.086402 |
| 1 | Prestw-1771 | Carmofur | 10988.5 | 114.3380208 | 31.59100326 |
| 2 | Prestw-1771 | Carmofur | 4596 | 81.9723667 | 1 |
| 2 | Prestw-1261 | Carprofen | 3220.5 | 28.8191949 | 1 |
| 1 | Prestw-1261 | Carprofen | 10183.5 | 75.01428296 | 1 |
| 1 | Prestw-446 | Carteolol hydrochloride | 11601.5 | 20.28776079 | 7.43563429 |
| 2 | Prestw-446 | Carteolol hydrochloride | 4449.5 | 155.6726632 | 52.1360861 |
| 2 | Prestw-1262 | Carvedilol | 3143 | 117.9663643 | 62.8748427 |
| 1 | Prestw-1262 | Carvedilol | 9600.5 | 85.52131757 | 3.58348281 |
| 1 | Prestw-485 | Cefaclor hydrate | 12545 | 49.18976336 | 22.51268842 |
| 2 | Prestw-485 | Cefaclor hydrate | 4067.5 | 125.9052033 | 1 |
| 1 | Prestw-434 | Cefadroxil | 11617 | 0.5375625 | 1 |
| 2 | Prestw-434 | Cefadroxil | 4141.5 | 67.9023578 | 1 |
| 2 | Prestw-736 | Cefazolin sodium salt | 5442 | 1 | 1 |
| 1 | Prestw-736 | Cefazolin sodium salt | 15796.5 | 14.71709805 | 0.07208911 |
| 1 | Prestw-1263 | Cefdinir | 9301 | 89.63509883 | 7.69726407 |
| 2 | Prestw-1263 | Cefdinir | 3295 | 25.4172512 | 1 |
| 1 | Prestw-1118 | Cefepime hydrochloride | 9284 | 34.11009385 | 10.96599955 |
| 2 | Prestw-1118 | Cefepime hydrochloride | 3192.5 | 31.8419249 | 1 |
| 1 | Prestw-462 | Cefixime | 12151 | 30.85933452 | 1 |
| 2 | Prestw-462 | Cefixime | 4834.5 | 118.2671468 | 1.0491927 |
| 2 | Prestw-700 | Cefmetazole sodium salt | 5509 | 86.3384688 | 16.9330726 |
| 1 | Prestw-700 | Cefmetazole sodium salt | 17432 | 46.41729408 | 1.0379109 |
| 2 | Prestw-327 | Cefoperazone dihydrate | 3625.5 | 164.8392065 | 101.4890309 |
| 1 | Prestw-327 | Cefoperazone dihydrate | 10915.5 | 39.71012078 | 1 |
| 1 | Prestw-470 | Ceforanide | 12275.5 | 37.30156511 | 3.38486987 |
| 2 | Prestw-470 | Ceforanide | 3501 | 1 | 1 |
| 2 | Prestw-139 | Cefotaxime sodium salt | 3600.5 | 1 | 1 |
| 1 | Prestw-139 | Cefotaxime sodium salt | 11206 | 55.42870105 | 1 |
| 1 | Prestw-473 | Cefotetan | 13139 | 41.07197775 | 7.15528251 |
| 2 | Prestw-473 | Cefotetan | 4795 | 141.1282606 | 23.9103065 |
| 1 | Prestw-482 | Cefotiam hydrochloride | 11652 | 44.8181385 | 18.14106356 |
| 2 | Prestw-482 | Cefotiam hydrochloride | 3881.5 | 143.4237016 | 1 |
| 1 | Prestw-832 | Cefoxitin sodium salt | 15232.5 | 1 | 1 |
| 2 | Prestw-832 | Cefoxitin sodium salt | 5466 | 1 | 1 |
| 2 | Prestw-1726 | Cefpiramide | 3337.5 | 266.0300303 | 87.0551123 |
| 1 | Prestw-1726 | Cefpiramide | 8182.5 | 239.6534959 | 76.0177 |
| 2 | Prestw-1727 | Cefpodoxime proxetil | 4541 | 171.6542385 | 1 |
| 1 | Prestw-1727 | Cefpodoxime proxetil | 12937.5 | 144.034355 | 1 |
| 1 | Prestw-1728 | Cefprozil | 12231 | 184.030418 | 20.39462202 |
| 2 | Prestw-1728 | Cefprozil | 4785.5 | 116.5467894 | 1 |
| 1 | Prestw-814 | Cefsulodin sodium salt | 17916.5 | 24.01108849 | 9.36607955 |
| 2 | Prestw-814 | Cefsulodin sodium salt | 5630.5 | 1 | 1 |
| 1 | Prestw-489 | Ceftazidime pentahydrate | 14200.5 | 37.43239513 | 10.75532019 |
| 2 | Prestw-489 | Ceftazidime pentahydrate | 4802 | 163.8823221 | 1 |
| 2 | Prestw-1729 | Ceftibuten | 5105 | 129.1282379 | 1 |
| 1 | Prestw-1729 | Ceftibuten | 14613.5 | 135.8285564 | 1 |
| 2 | Prestw-1730 | Cefuroxime axetil | 5167 | 92.0132088 | 20.0210683 |
| 1 | Prestw-1730 | Cefuroxime axetil | 13028.5 | 170.0383254 | 26.17765406 |
| 2 | Prestw-720 | Cefuroxime sodium salt | 5630 | 48.7591707 | 47.7591707 |
| 1 | Prestw-720 | Cefuroxime sodium salt | 16899 | 38.57866207 | 1 |
| 2 | Prestw-1371 | Celecoxib | 3402 | 31.8875086 | 1 |
| 1 | Prestw-1371 | Celecoxib | 10822.5 | 59.08908192 | 1 |
| 1 | Prestw-1132 | Celiprolol hydrochloride | 9847 | 29.3458641 | 1 |
| 2 | Prestw-1132 | Celiprolol hydrochloride | 3583.5 | 27.516529 | 1 |
| 1 | Cells | Cells | 14520.5 | 1 | 1 |

| | | | | | |
|---|---|---|---|---|---|
| 1 | Cells | Cells | 15157 | 1 | 1 |
| 1 | Cells | Cells | 11410.5 | 13.46644955 | 1 |
| 1 | Cells | Cells | 10411.5 | 10.0923656 | 1 |
| 1 | Cells | Cells | 10148.5 | 1 | 1 |
| 1 | Cells | Cells | 9431 | 1 | 1 |
| 1 | Cells | Cells | 10191 | 1 | 1 |
| 1 | Cells | Cells | 7868.5 | 1 | 1 |
| 1 | Cells | Cells | 11777 | 4.6450693 | 1 |
| 2 | Cells | Cells | 5317.5 | 1 | 1 |
| 1 | Cells | Cells | 10523.5 | 12.8521265 | 1 |
| 1 | Cells | Cells | 8040 | 1 | 1 |
| 2 | Cells | Cells | 3293 | 15.5031276 | 1 |
| 1 | Cells | Cells | 12562.5 | 33.91669524 | 1 |
| 1 | Cells | Cells | 12485.5 | 8.02565804 | 1 |
| 1 | Cells | Cells | 12012 | 3.18689902 | 1 |
| 2 | Cells | Cells | 3754.5 | 1 | 1 |
| 1 | Cells | Cells | 10405.5 | 1 | 1 |
| 2 | Cells | Cells | 4951 | 1 | 1 |
| 1 | Cells | Cells | 11655 | 12.9580545 | 1 |
| 1 | Cells | Cells | 10881.5 | 1.95550112 | 1 |
| 1 | Cells | Cells | 16281.5 | 10.93488149 | 1 |
| 1 | Cells | Cells | 11535 | 25.49851737 | 1 |
| 1 | Cells | Cells | 11998 | 25.85254565 | 1 |
| 1 | Cells | Cells | 8036 | 1 | 1 |
| 1 | Cells | Cells | 11240 | 40.12874414 | 1 |
| 1 | Cells | Cells | 11622.5 | 37.4277642 | 1 |
| 2 | Cells | Cells | 3689 | 33.5904402 | 1 |
| 1 | Cells | Cells | 14176 | 38.03047291 | 1 |
| 1 | Cells | Cells | 8266.5 | 5.5487527 | 1 |
| 2 | Cells | Cells | 3145.5 | 59.5933451 | 1 |
| 1 | Cells | Cells | 9294.5 | 35.25041844 | 1 |
| 2 | Cells | Cells | 3909.5 | 1 | 1 |
| 1 | Cells | Cells | 8778.5 | 17.5622022 | 1 |
| 1 | Cells | Cells | 12181.5 | 26.67707494 | 1 |
| 2 | Cells | Cells | 2957 | 55.0915216 | 1 |
| 2 | Cells | Cells | 5127 | 20.5794015 | 1 |
| 1 | Cells | Cells | 9847 | 19.70586662 | 1 |
| 2 | Cells | Cells | 2762 | 1 | 1 |
| 2 | Cells | Cells | 2786 | 1 | 1 |
| 2 | Cells | Cells | 3132.5 | 52.5252085 | 1 |
| 2 | Cells | Cells | 6109 | 1 | 1 |
| 2 | Cells | Cells | 2710.5 | 76.2941381 | 1 |
| 1 | Cells | Cells | 8945.5 | 23.1440943 | 1 |
| 1 | Cells | Cells | 15892 | 51.40732751 | 1 |
| 1 | Cells | Cells | 8639 | 14.1548537 | 1 |
| 2 | Cells | Cells | 3814 | 65.3728526 | 1 |
| 2 | Cells | Cells | 4982.5 | 1 | 1 |
| 2 | Cells | Cells | 3058.5 | 1 | 1 |
| 2 | Cells | Cells | 4207.5 | 32.8453402 | 1 |
| 2 | Cells | Cells | 4379.5 | 67.4940618 | 1 |
| 1 | Cells | Cells | 17067.5 | 29.24958052 | 1 |
| 2 | Cells | Cells | 4125 | 28.5482131 | 1 |
| 1 | Cells | Cells | 9174.5 | 52.135354 | 1 |
| 2 | Cells | Cells | 4356.5 | 3.4942613 | 1 |
| 2 | Cells | Cells | 3005.5 | 60.8740933 | 1 |
| 1 | Cells | Cells | 15113 | 40.24036095 | 1 |
| 2 | Cells | Cells | 4607.5 | 1 | 1 |
| 2 | Cells | Cells | 4338.5 | 16.613184 | 1 |
| 1 | Cells | Cells | 9379.5 | 4.5879211 | 1 |
| 2 | Cells | Cells | 4021 | 1 | 1 |
| 1 | Cells | Cells | 16659.5 | 14.64500894 | 1 |
| 2 | Cells | Cells | 4715.5 | 93.4584627 | 1 |
| 2 | Cells | Cells | 6060.5 | 1 | 1 |

| | | | | | |
|---|---|---|---|---|---|
| 2 | Cells | Cells | 3021 | 3.0739452 | 1 |
| 2 | Cells | Cells | 3003 | 48.1539652 | 1 |
| 2 | Cells | Cells | 3912.5 | 139.1288273 | 1 |
| 1 | Cells | Cells | 16771.5 | 31.88862808 | 1 |
| 2 | Cells | Cells | 3064.5 | 152.9449772 | 1 |
| 1 | Cells | Cells | 9276.5 | 83.91545542 | 1 |
| 1 | Cells | Cells | 17506.5 | 48.53293508 | 1 |
| 2 | Cells | Cells | 3052 | 3.1057182 | 1 |
| 1 | Cells | Cells | 15564.5 | 51.98270358 | 1 |
| 1 | Cells | Cells | 8950 | 76.2491278 | 1 |
| 1 | Cells | Cells | 11609 | 67.81179385 | 1 |
| 1 | Cells | Cells | 15586.5 | 34.02826827 | 1 |
| 2 | Cells | Cells | 4481.5 | 63.5924085 | 1 |
| 2 | Cells | Cells | 3067 | 86.5527934 | 1 |
| 1 | Cells | Cells | 12381 | 62.34994102 | 1 |
| 2 | Cells | Cells | 6065.5 | 1 | 1 |
| 1 | Cells | Cells | 16674.5 | 36.55401833 | 1 |
| 1 | Cells | Cells | 9428.5 | 81.93783476 | 1 |
| 1 | Cells | Cells | 9274 | 102.5064791 | 1 |
| 2 | Cells | Cells | 4084 | 93.6353447 | 1 |
| 2 | Cells | Cells | 5095.5 | 1 | 1 |
| 2 | Cells | Cells | 3361.5 | 22.1561776 | 1 |
| 1 | Cells | Cells | 16709 | 45.37938318 | 1 |
| 2 | Cells | Cells | 3904.5 | 98.9008145 | 1 |
| 2 | Cells | Cells | 4397 | 112.4538381 | 1 |
| 1 | Cells | Cells | 11429.5 | 82.74701755 | 1 |
| 2 | Cells | Cells | 6169.5 | 28.5718641 | 1 |
| 1 | Cells | Cells | 9117 | 106.0689324 | 1 |
| 1 | Cells | Cells | 16867.5 | 47.75596792 | 1 |
| 2 | Cells | Cells | 4948 | 103.4815422 | 1 |
| 1 | Cells | Cells | 17654.5 | 44.12780876 | 1 |
| 2 | Cells | Cells | 4341 | 157.3951968 | 1 |
| 2 | Cells | Cells | 4533 | 136.5984955 | 1 |
| 2 | Cells | Cells | 4136 | 63.3501756 | 1 |
| 2 | Cells | Cells | 5456 | 44.0811229 | 1 |
| 2 | Cells | Cells | 4387.5 | 71.9921405 | 1 |
| 2 | Cells | Cells | 5588 | 1 | 1 |
| 2 | Cells | Cells | 4885.5 | 117.2179541 | 1 |
| 2 | Cells | Cells | 3656 | 105.4053852 | 1 |
| 2 | Cells | Cells | 5748 | 19.5858317 | 1 |
| 2 | Cells | Cells | 3926 | 177.5247265 | 1 |
| 2 | Cells | Cells | 4150.5 | 154.1661667 | 1 |
| 1 | Cells | Cells | 11478 | 104.3823129 | 1 |
| 1 | Cells | Cells | 12040.5 | 131.6744348 | 1 |
| 2 | Cells | Cells | 4609.5 | 86.702116 | 1 |
| 1 | Cells | Cells | 11829 | 111.1488711 | 1 |
| 1 | Cells | Cells | 11217 | 85.5280339 | 1 |
| 2 | Cells | Cells | 3980.5 | 103.5365771 | 1 |
| 2 | Cells | Cells | 3406.5 | 175.6405672 | 1 |
| 2 | Cells | Cells | 4301 | 107.008767 | 1 |
| 1 | Cells | Cells | 11513 | 160.0402442 | 1 |
| 2 | Cells | Cells | 5693 | 3.2085404 | 1 |
| 2 | Cells | Cells | 4325.5 | 178.974918 | 1 |
| 2 | Cells | Cells | 5267 | 30.8880764 | 1 |
| 1 | Cells | Cells | 12420.5 | 117.5264888 | 1 |
| 1 | Cells | Cells | 12573 | 119.9127633 | 1 |
| 2 | Cells | Cells | 5303 | 69.4053962 | 1 |
| 2 | Cells | Cells | 5491.5 | 12.7566871 | 1 |
| 2 | Cells | Cells | 3787 | 160.8636563 | 1 |
| 2 | Cells | Cells | 4278 | 92.1807917 | 1 |
| 1 | Cells | Cells | 12927.5 | 173.2425215 | 1 |
| 1 | Cells | Cells | 11879.5 | 189.2761902 | 1 |
| 1 | Cells | Cells | 12249.5 | 143.8606713 | 1 |

| | | | | | |
|---|---|---|---|---|---|
| 1 | Cells | Cells | 12319 | 163.6357959 | 1 |
| 1 | Prestw-424 | Cephalosporanic acid, 7-amino | 10729 | 1 | 1 |
| 2 | Prestw-424 | Cephalosporanic acid, 7-amino | 3738.5 | 20.2129957 | 1 |
| 2 | Prestw-719 | Cephalothin sodium salt | 5171 | 40.8710791 | 39.8710791 |
| 1 | Prestw-719 | Cephalothin sodium salt | 17128.5 | 36.7607114 | 1 |
| 2 | Prestw-503 | Cetirizine dihydrochloride | 4032 | 79.9976052 | 47.152265 |
| 1 | Prestw-503 | Cetirizine dihydrochloride | 11459 | 8.39428012 | 5.2073811 |
| 1 | Prestw-285 | Chenodiol | 10881 | 20.280591 | 6.81414145 |
| 2 | Prestw-285 | Chenodiol | 3820 | 102.601199 | 1 |
| 1 | Prestw-425 | Chicago sky blue 6B | 11476.5 | 1 | 1 |
| 2 | Prestw-425 | Chicago sky blue 6B | 3814 | 31.032301 | 1 |
| 1 | Prestw-31 | Chloramphenicol | 11558 | 44.19513 | 6.7673658 |
| 2 | Prestw-31 | Chloramphenicol | 4115 | 185.0079511 | 9.3673839 |
| 1 | Prestw-706 | Chlorcyclizine hydrochloride | 17210.5 | 59.53517594 | 25.50690767 |
| 2 | Prestw-706 | Chlorcyclizine hydrochloride | 5929.5 | 51.7322574 | 38.9755703 |
| 2 | Prestw-143 | Chlorhexidine | 1912 | 527.4408078 | 510.8276238 |
| 1 | Prestw-143 | Chlorhexidine | 3534.5 | 417.9255606 | 350.1137668 |
| 2 | Prestw-1374 | Chlormadinone acetate | 3490 | 1 | 1 |
| 1 | Prestw-1374 | Chlormadinone acetate | 10211 | 79.20238974 | 1 |
| 1 | Prestw-336 | Chlormezanone | 10835.5 | 1 | 1 |
| 2 | Prestw-336 | Chlormezanone | 3815.5 | 1 | 1 |
| 2 | Prestw-807 | Chloropyramine hydrochloride | 4052 | 126.6231129 | 125.6231129 |
| 1 | Prestw-807 | Chloropyramine hydrochloride | 12621 | 58.65423039 | 10.12129531 |
| 2 | Prestw-548 | Chloroquine diphosphate | 3348.5 | 998.5158057 | 906.335014 |
| 1 | Prestw-548 | Chloroquine diphosphate | 4789.5 | 6.5715868 | 1 |
| 1 | Prestw-251 | Chlorothiazide | 12465 | 128.3118852 | 8.39912194 |
| 2 | Prestw-251 | Chlorothiazide | 3924 | 22.165454 | 1 |
| 1 | Prestw-757 | Chlorotrianisene | 14998 | 17.37397661 | 1 |
| 2 | Prestw-757 | Chlorotrianisene | 5053 | 1 | 1 |
| 1 | Prestw-1474 | Chloroxine | 11037.5 | 137.1193569 | 19.59286806 |
| 2 | Prestw-1474 | Chloroxine | 4651 | 49.3272281 | 1 |
| 2 | Prestw-117 | Chlorpheniramine maleate | 3369.5 | 23.2687207 | 1 |
| 1 | Prestw-117 | Chlorpheniramine maleate | 9612 | 69.6062403 | 1 |
| 2 | Prestw-234 | Chlorphensin carbamate | 5051 | 48.576416 | 1 |
| 1 | Prestw-234 | Chlorphensin carbamate | 13319 | 153.3963218 | 1 |
| 1 | Prestw-64 | Chlorpromazine hydrochloride | 13467.5 | 1 | 1 |
| 2 | Prestw-64 | Chlorpromazine hydrochloride | 5241.5 | 1 | 1 |
| 1 | Prestw-323 | Chlorpropamide | 13818.5 | 40.11519005 | 27.15713555 |
| 2 | Prestw-323 | Chlorpropamide | 4287 | 143.350235 | 1 |
| 1 | Prestw-348 | Chlorprothixene hydrochloride | 9614.5 | 1 | 1 |
| 2 | Prestw-348 | Chlorprothixene hydrochloride | 3546.5 | 1 | 1 |
| 1 | Prestw-145 | Chlortetracycline hydrochloride | 11143.5 | 107.0001124 | 2.61779945 |
| 2 | Prestw-145 | Chlortetracycline hydrochloride | 4495 | 93.2017942 | 1 |
| 1 | Prestw-351 | Chlorthalidone | 10716 | 1 | 1 |
| 2 | Prestw-351 | Chlorthalidone | 3975 | 22.592419 | 1 |
| 1 | Prestw-163 | Chlorzoxazone | 11066.5 | 120.255816 | 15.87350311 |
| 2 | Prestw-163 | Chlorzoxazone | 4807 | 103.9314456 | 10.2961009 |
| 2 | Prestw-1801 | Ciclesonide | 4329.5 | 79.0888034 | 1 |
| 1 | Prestw-1801 | Ciclesonide | 12536.5 | 158.6177719 | 1 |
| 2 | Prestw-541 | Ciclopirox ethanolamine | 5914.5 | 109.42437 | 17.2435783 |
| 1 | Prestw-541 | Ciclopirox ethanolamine | 15239 | 15.30883393 | 1 |
| 2 | Prestw-1376 | Cilnidipine | 3663.5 | 3.5003938 | 1 |
| 1 | Prestw-1376 | Cilnidipine | 11310.5 | 52.20166321 | 1 |
| 1 | Prestw-587 | Cilostazol | 13548 | 32.08413403 | 6.58561666 |
| 2 | Prestw-587 | Cilostazol | 4718.5 | 110.8011627 | 1 |
| 1 | Prestw-26 | Cimetidine | 11816 | 104.884775 | 42.534834 |
| 2 | Prestw-26 | Cimetidine | 4827 | 1 | 1 |
| 1 | Prestw-278 | Cinnarizine | 10970 | 1 | 1 |
| 2 | Prestw-278 | Cinnarizine | 3672.5 | 1 | 1 |
| 2 | Prestw-780 | Cinoxacin | 5811.5 | 78.2787205 | 8.8733243 |
| 1 | Prestw-780 | Cinoxacin | 18354 | 48.10508843 | 2.72570525 |
| 1 | Prestw-1076 | Ciprofibrate | 7960.5 | 1 | 1 |
| 2 | Prestw-1076 | Ciprofibrate | 3397.5 | 1 | 1 |

| | | | | | |
|---|---|---|---|---|---|
| 1 | Prestw-113 | Ciprofloxacin hydrochloride monohydrate | 10597 | 40.25291977 | 1 |
| 2 | Prestw-113 | Ciprofloxacin hydrochloride monohydrate | 3547.5 | 2.6589595 | 1 |
| 1 | Prestw-430 | Cisapride | 10760 | 7.1695204 | 1 |
| 2 | Prestw-430 | Cisapride | 4053.5 | 26.8132621 | 1 |
| 2 | Prestw-1745 | Cisatracurium besylate | 4332.5 | 61.4552585 | 57.9609972 |
| 1 | Prestw-1745 | Cisatracurium besylate | 12950.5 | 106.3411489 | 20.813115 |
| 2 | Prestw-692 | Citalopram Hydrobromide | 5361 | 67.04164 | 36.1535636 |
| 1 | Prestw-692 | Citalopram Hydrobromide | 15055.5 | 45.47865284 | 13.59002476 |
| 2 | Prestw-1484 | Cladribine | 4494 | 76.7260896 | 1 |
| 1 | Prestw-1484 | Cladribine | 11206.5 | 158.8782526 | 1 |
| 2 | Prestw-1378 | Clarithromycin | 3597 | 1 | 1 |
| 1 | Prestw-1378 | Clarithromycin | 10638 | 57.7344474 | 1 |
| 1 | Prestw-1315 | Clavulanate potassium salt | 8523.5 | 1 | 1 |
| 2 | Prestw-1315 | Clavulanate potassium salt | 2701.5 | 1 | 1 |
| 1 | Prestw-380 | Clebopride maleate | 13068 | 16.65102512 | 8.62536708 |
| 2 | Prestw-380 | Clebopride maleate | 4696 | 99.6890347 | 1 |
| 1 | Prestw-306 | Clemastine fumarate | 12525 | 21.44677534 | 1 |
| 2 | Prestw-306 | Clemastine fumarate | 4164.5 | 165.3580391 | 1 |
| 2 | Prestw-227 | Clemizole hydrochloride | 4936.5 | 135.2685383 | 31.7869961 |
| 1 | Prestw-227 | Clemizole hydrochloride | 14914.5 | 113.0251261 | 1 |
| 1 | Prestw-345 | Clenbuterol hydrochloride | 10511.5 | 8.1576063 | 7.1576063 |
| 2 | Prestw-345 | Clenbuterol hydrochloride | 3485 | 1 | 1 |
| 2 | Prestw-822 | Clidinium bromide | 4878.5 | 85.2974815 | 84.2974815 |
| 1 | Prestw-822 | Clidinium bromide | 11551 | 56.7902422 | 42.14523326 |
| 2 | Prestw-1379 | Clinafloxacin | 3463 | 1 | 1 |
| 1 | Prestw-1379 | Clinafloxacin | 10332 | 57.43191734 | 1 |
| 2 | Prestw-137 | Clindamycin hydrochloride | 3542 | 51.2197612 | 1 |
| 1 | Prestw-137 | Clindamycin hydrochloride | 10294 | 74.21349097 | 1 |
| 2 | Prestw-886 | Clioquinol | 6563 | 26.2742199 | 25.2742199 |
| 1 | Prestw-886 | Clioquinol | 20068.5 | 41.33570519 | 1 |
| 1 | Prestw-781 | Clobetasol propionate | 17559 | 51.58528566 | 6.20590248 |
| 2 | Prestw-781 | Clobetasol propionate | 5477.5 | 73.8328749 | 4.4274787 |
| 2 | Prestw-1380 | Clobutinol hydrochloride | 3584.5 | 1 | 1 |
| 1 | Prestw-1380 | Clobutinol hydrochloride | 10582 | 75.41018189 | 1 |
| 2 | Prestw-1119 | Clocortolone pivalate | 3229.5 | 56.6929752 | 8.53901 |
| 1 | Prestw-1119 | Clocortolone pivalate | 9410.5 | 28.4229215 | 5.2788272 |
| 1 | Prestw-376 | Clofazimine | 13440 | 19.71686323 | 11.69120519 |
| 2 | Prestw-376 | Clofazimine | 4992 | 139.7822882 | 1 |
| 1 | Prestw-1383 | Clofibrate | 9537.5 | 86.44027298 | 4.50243822 |
| 2 | Prestw-1383 | Clofibrate | 3312 | 1 | 1 |
| 1 | Prestw-783 | Clofibric acid | 17639 | 49.8949134 | 4.51553022 |
| 2 | Prestw-783 | Clofibric acid | 5647 | 55.7955021 | 1 |
| 1 | Prestw-319 | Clofilium tosylate | 11652 | 31.8150288 | 18.8569743 |
| 2 | Prestw-319 | Clofilium tosylate | 4236.5 | 169.3122432 | 8.4485869 |
| 2 | Prestw-397 | Clomiphene citrate (Z,E) | 4622 | 588.6540271 | 427.7903708 |
| 1 | Prestw-397 | Clomiphene citrate (Z,E) | 14421 | 1 | 1 |
| 1 | Prestw-269 | Clomipramine hydrochloride | 9543 | 1 | 1 |
| 2 | Prestw-269 | Clomipramine hydrochloride | 3558.5 | 1 | 1 |
| 2 | Prestw-248 | Clonidine hydrochloride | 4836 | 52.2485202 | 1 |
| 1 | Prestw-248 | Clonidine hydrochloride | 13943 | 113.5425413 | 1 |
| 1 | Prestw-1104 | Clonixin Lysinate | 9491.5 | 13.773089 | 1 |
| 2 | Prestw-1104 | Clonixin Lysinate | 3151 | 1 | 1 |
| 1 | Prestw-354 | Clopamide | 12975.5 | 15.0082032 | 1.54175365 |
| 2 | Prestw-354 | Clopamide | 4117 | 130.2560998 | 24.8507146 |
| 1 | Prestw-793 | Cloperastine hydrochloride | 20379 | 34.31391982 | 1 |
| 2 | Prestw-793 | Cloperastine hydrochloride | 6250 | 1 | 1 |
| 2 | Prestw-1701 | Clopidogrel | 2297.5 | 681.8921014 | 545.2936059 |
| 1 | Prestw-1701 | Clopidogrel | 5023.5 | 627.3541066 | 495.6796718 |
| 2 | Prestw-344 | Clorgyline hydrochloride | 3371 | 46.9938449 | 18.4456318 |
| 1 | Prestw-344 | Clorgyline hydrochloride | 9733 | 1 | 1 |
| 2 | Prestw-540 | Clorsulon | 5704.5 | 150.8461746 | 58.6653829 |
| 1 | Prestw-540 | Clorsulon | 14214 | 24.16906678 | 1 |
| 2 | Prestw-1385 | Closantel | 3516 | 1 | 1 |

| | | | | | |
|---|---|---|---|---|---|
| 1 | Prestw-1385 | Closantel | 9818.5 | 81.4049894 | 1 |
| 2 | Prestw-267 | Clotrimazole | 4080 | 28.6001592 | 27.6001592 |
| 1 | Prestw-267 | Clotrimazole | 10249.5 | 1 | 1 |
| 1 | Prestw-186 | Cloxacillin sodium salt | 10317.5 | 59.33839399 | 39.63252737 |
| 2 | Prestw-186 | Cloxacillin sodium salt | 4134.5 | 1 | 1 |
| 2 | Prestw-350 | Clozapine | 3919 | 38.2924593 | 9.7442462 |
| 1 | Prestw-350 | Clozapine | 10446.5 | 1 | 1 |
| 1 | Prestw-363 | Colchicine | 14987 | 11.33189516 | 6.68682586 |
| 2 | Prestw-363 | Colchicine | 5282 | 124.5555739 | 17.5468069 |
| 1 | Colistin | Colistin | 2774.5 | 68.9330331 | 67.9330331 |
| 1 | Colistin | Colistin | 2838 | 136.6661221 | 135.6661221 |
| 1 | Colistin | Colistin | 2922 | 270.4444589 | 269.4444589 |
| 1 | Colistin | Colistin | 3013.5 | 333.6650211 | 328.1162684 |
| 1 | Colistin | Colistin | 2844 | 428.0442057 | 413.889352 |
| 1 | Colistin | Colistin | 2973 | 361.3226105 | 343.7604083 |
| 1 | Colistin | Colistin | 2889.5 | 406.9403708 | 383.7962765 |
| 1 | Colistin | Colistin | 3028.5 | 482.0669054 | 477.4789843 |
| 1 | Colistin | Colistin | 2869 | 500.5739298 | 448.4385758 |
| 1 | Colistin | Colistin | 3060 | 662.800831 | 586.5517032 |
| 1 | Colistin | Colistin | 2689 | 867.6561022 | 761.5871698 |
| 1 | Colistin | Colistin | 2816.5 | 754.5264876 | 672.5886528 |
| 1 | Colistin | Colistin | 2781 | 779.6342764 | 677.1277973 |
| 1 | Colistin | Colistin | 2757.5 | 648.1976039 | 564.2821485 |
| 1 | Colistin | Colistin | 2719 | 366.0766769 | 330.8262585 |
| 1 | Colistin | Colistin | 2703 | 237.8955714 | 235.9400703 |
| 1 | Colistin | Colistin | 2300 | 188.1414531 | 187.1414531 |
| 1 | Colistin | Colistin | 2332.5 | 378.0663583 | 358.3604917 |
| 1 | Colistin | Colistin | 2176 | 663.0471765 | 595.2353827 |
| 1 | Colistin | Colistin | 2352.5 | 833.8169335 | 751.069916 |
| 1 | Colistin | Colistin | 2018 | 1105.648824 | 1001.266511 |
| 1 | Colistin | Colistin | 2263.5 | 1145.998127 | 1034.849256 |
| 1 | Colistin | Colistin | 2295 | 1149.072597 | 1017.398162 |
| 1 | Colistin | Colistin | 2271 | 1275.142589 | 1115.102345 |
| 1 | Colistin | Colistin | 2259 | 1337.438553 | 1219.912065 |
| 1 | Colistin | Colistin | 2313.5 | 1630.88506 | 1457.642538 |
| 1 | Colistin | Colistin | 2097 | 1943.958379 | 1780.322583 |
| 1 | Colistin | Colistin | 2187.5 | 1681.170746 | 1491.894556 |
| 1 | Colistin | Colistin | 1886.5 | 1894.53024 | 1750.669569 |
| 1 | Colistin | Colistin | 2149.5 | 1363.624151 | 1243.711388 |
| 1 | Colistin | Colistin | 2048.5 | 1123.241438 | 1037.713404 |
| 1 | Colistin | Colistin | 2150.5 | 701.0793374 | 638.7293964 |
| 1 | Colistin | Colistin | 2245 | 59.4636179 | 58.4636179 |
| 1 | Colistin | Colistin | 2002.5 | 59.4636179 | 58.4636179 |
| 1 | Colistin | Colistin | 2078 | 59.4636179 | 58.4636179 |
| 1 | Colistin | Colistin | 2043 | 59.4636179 | 49.3712523 |
| 1 | Colistin | Colistin | 1951.5 | 59.4636179 | 45.99716835 |
| 1 | Colistin | Colistin | 2283.5 | 140.1151627 | 127.2630362 |
| 1 | Colistin | Colistin | 2005.5 | 129.4247386 | 124.7796693 |
| 1 | Colistin | Colistin | 1844 | 294.4661585 | 268.6136129 |
| 1 | Colistin | Colistin | 2062.5 | 165.7801388 | 157.7544808 |
| 1 | Colistin | Colistin | 2135 | 191.1577431 | 157.2410479 |
| 1 | Colistin | Colistin | 1885 | 337.3635047 | 299.9357405 |
| 1 | Colistin | Colistin | 1898 | 375.0804644 | 348.4033895 |
| 1 | Colistin | Colistin | 1935 | 241.2224908 | 228.2644363 |
| 1 | Colistin | Colistin | 1939 | 289.2853876 | 263.7868702 |
| 1 | Colistin | Colistin | 1928 | 314.9597968 | 274.8310527 |
| 1 | Colistin | Colistin | 1764.5 | 166.4949436 | 163.3080446 |
| 1 | Colistin | Colistin | 2804 | 27.0677937 | 26.0677937 |
| 1 | Colistin | Colistin | 2867.5 | 27.0677937 | 26.0677937 |
| 1 | Colistin | Colistin | 2782.5 | 350.6702153 | 312.6397424 |
| 1 | Colistin | Colistin | 2961.5 | 508.7010339 | 468.460673 |
| 1 | Colistin | Colistin | 2941 | 567.9618502 | 536.0732221 |
| 1 | Colistin | Colistin | 2891 | 628.8149273 | 576.8322237 |

| | | | | | |
|---|---|---|---|---|---|
| 1 | Colistin | Colistin | 2693 | 753.7605006 | 708.3811174 |
| 1 | Colistin | Colistin | 3071.5 | 443.1944074 | 391.7870799 |
| 1 | Colistin | Colistin | 2618.5 | 638.112473 | 604.0842047 |
| 1 | Colistin | Colistin | 2990.5 | 678.9244935 | 642.3704752 |
| 1 | Colistin | Colistin | 2638 | 829.0941439 | 784.9663351 |
| 1 | Colistin | Colistin | 2588.5 | 812.622413 | 764.8664451 |
| 1 | Colistin | Colistin | 2727 | 613.9225126 | 565.3895775 |
| 1 | Colistin | Colistin | 2694 | 469.9403054 | 440.6907249 |
| 1 | Colistin | Colistin | 2267 | 501.8096917 | 487.1646828 |
| 1 | Colistin | Colistin | 2587.5 | 253.4503345 | 242.515453 |
| 2 | Colistin | Colistin | 1237.5 | 361.8713762 | 360.8713762 |
| 2 | Colistin | Colistin | 1172 | 361.8713762 | 360.8713762 |
| 2 | Colistin | Colistin | 1196.5 | 388.0176746 | 328.4243295 |
| 2 | Colistin | Colistin | 1178.5 | 456.0321002 | 379.7379621 |
| 2 | Colistin | Colistin | 1138 | 569.0835427 | 565.9778245 |
| 2 | Colistin | Colistin | 1182.5 | 537.4980142 | 534.424069 |
| 2 | Colistin | Colistin | 1228 | 583.1745404 | 535.0205752 |
| 2 | Colistin | Colistin | 1235 | 712.3180328 | 625.7652394 |
| 2 | Colistin | Colistin | 1214 | 700.8628906 | 547.9179134 |
| 2 | Colistin | Colistin | 1154 | 726.1104149 | 703.9542373 |
| 2 | Colistin | Colistin | 1073.5 | 527.2983118 | 466.4242185 |
| 2 | Colistin | Colistin | 1109.5 | 409.9470994 | 354.8555778 |
| 2 | Colistin | Colistin | 1109.5 | 423.6522968 | 371.1270883 |
| 2 | Colistin | Colistin | 1059.5 | 423.6522968 | 422.6522968 |
| 2 | Colistin | Colistin | 1034 | 167.985966 | 152.4828384 |
| 2 | Colistin | Colistin | 1290.5 | 167.985966 | 166.985966 |
| 2 | Colistin | Colistin | 1435.5 | 483.5488228 | 482.5488228 |
| 2 | Colistin | Colistin | 1200.5 | 483.5488228 | 418.1759702 |
| 2 | Colistin | Colistin | 1219.5 | 485.8900857 | 469.2769017 |
| 2 | Colistin | Colistin | 1283.5 | 675.0541123 | 535.925285 |
| 2 | Colistin | Colistin | 1182 | 779.3017559 | 685.6664112 |
| 2 | Colistin | Colistin | 1218 | 615.5775853 | 522.1191226 |
| 2 | Colistin | Colistin | 1169 | 976.0929961 | 839.4945006 |
| 2 | Colistin | Colistin | 1225 | 681.2177448 | 617.6253363 |
| 2 | Colistin | Colistin | 1097 | 1118.853856 | 1032.15174 |
| 2 | Colistin | Colistin | 1158.5 | 873.1502575 | 769.6687153 |
| 2 | Colistin | Colistin | 1069 | 1333.73169 | 1154.756772 |
| 2 | Colistin | Colistin | 1098 | 863.2858039 | 750.8319658 |
| 2 | Colistin | Colistin | 992 | 942.9947378 | 871.0025973 |
| 2 | Colistin | Colistin | 1200.5 | 506.8532322 | 439.3591704 |
| 2 | Colistin | Colistin | 1026.5 | 464.7366468 | 461.2423855 |
| 2 | Colistin | Colistin | 1108.5 | 469.8903217 | 468.8903217 |
| 2 | Colistin | Colistin | 1167.5 | 341.7876238 | 340.7876238 |
| 2 | Colistin | Colistin | 1062.5 | 341.7876238 | 308.1971836 |
| 2 | Colistin | Colistin | 1045.5 | 476.9816231 | 448.43341 |
| 2 | Colistin | Colistin | 1134.5 | 820.9955112 | 722.0946967 |
| 2 | Colistin | Colistin | 1024.5 | 1046.248283 | 940.8428976 |
| 2 | Colistin | Colistin | 1053.5 | 1159.175284 | 1055.638707 |
| 2 | Colistin | Colistin | 955.5 | 1207.775041 | 1100.766274 |
| 2 | Colistin | Colistin | 930.5 | 1524.634022 | 1432.45323 |
| 2 | Colistin | Colistin | 1009.5 | 988.7903893 | 831.3951925 |
| 2 | Colistin | Colistin | 921.5 | 1040.75658 | 923.5386258 |
| 2 | Colistin | Colistin | 1006.5 | 1254.199938 | 1078.559371 |
| 2 | Colistin | Colistin | 980.5 | 1146.074269 | 968.5495425 |
| 2 | Colistin | Colistin | 914 | 1551.631355 | 1390.767699 |
| 2 | Colistin | Colistin | 862 | 1126.787692 | 972.6215253 |
| 2 | Colistin | Colistin | 885 | 920.7518929 | 857.4017173 |
| 2 | Colistin | Colistin | 861 | 466.512768 | 433.6674278 |
| 2 | Colistin | Colistin | 1352.5 | 126.0570253 | 125.0570253 |
| 2 | Colistin | Colistin | 1270 | 218.9199689 | 217.9199689 |
| 2 | Colistin | Colistin | 1454 | 377.0031012 | 356.4236997 |
| 2 | Colistin | Colistin | 1193.5 | 909.6789468 | 865.5978239 |
| 2 | Colistin | Colistin | 1188 | 1210.561522 | 1179.673445 |

| | | | | | |
|---|---|---|---|---|---|
| 2 | Colistin | Colistin | 1339.5 | 789.4907801 | 760.918916 |
| 2 | Colistin | Colistin | 1257 | 1348.603357 | 1279.197961 |
| 2 | Colistin | Colistin | 1148.5 | 968.6660651 | 949.0802334 |
| 2 | Colistin | Colistin | 1186.5 | 1318.002163 | 1305.245476 |
| 2 | Colistin | Colistin | 1212.5 | 1148.971052 | 1145.762512 |
| 2 | Colistin | Colistin | 1223.5 | 888.4582239 | 887.4582239 |
| 2 | Colistin | Colistin | 1170 | 695.921053 | 694.921053 |
| 2 | Colistin | Colistin | 1161.5 | 419.6782008 | 418.6782008 |
| 2 | Colistin | Colistin | 1098.5 | 642.5018509 | 641.5018509 |
| 2 | Colistin | Colistin | 1120 | 526.5182885 | 525.5182885 |
| 2 | Colistin | Colistin | 1088.5 | 372.8926646 | 371.8926646 |
| 1 | Prestw-486 | Colistin sulfate | 11680 | 71.81633958 | 45.13926464 |
| 2 | Prestw-486 | Colistin sulfate | 3793 | 189.3499816 | 11.8252551 |
| 1 | Prestw-432 | Corticosterone | 11113 | 0.9216787 | 1 |
| 2 | Prestw-432 | Corticosterone | 3983.5 | 59.3085826 | 1 |
| 1 | Prestw-1486 | Cortisol acetate | 10952.5 | 166.3610508 | 2.72525487 |
| 2 | Prestw-1486 | Cortisol acetate | 4705 | 91.1224298 | 1 |
| 1 | Prestw-273 | Cortisone | 9521 | 1 | 1 |
| 2 | Prestw-273 | Cortisone | 3661 | 1 | 1 |
| 1 | Prestw-812 | Cromolyn disodium salt | 16707.5 | 29.63774223 | 14.99273329 |
| 2 | Prestw-812 | Cromolyn disodium salt | 5327.5 | 1 | 1 |
| 2 | Prestw-951 | Crotamiton | 6580 | 66.528579 | 63.3200386 |
| 1 | Prestw-951 | Crotamiton | 20690 | 45.58100239 | 9.02698406 |
| 1 | Prestw-433 | Cyanocobalamin | 11821 | 0.9669709 | 1 |
| 2 | Prestw-433 | Cyanocobalamin | 4059.5 | 89.9343483 | 1 |
| 2 | Prestw-510 | Cyclizine hydrochloride | 3838 | 58.1716465 | 24.5812063 |
| 1 | Prestw-510 | Cyclizine hydrochloride | 10698.5 | 1 | 1 |
| 1 | Prestw-445 | Cyclobenzaprine hydrochloride | 12173.5 | 32.58138483 | 19.72925833 |
| 2 | Prestw-445 | Cyclobenzaprine hydrochloride | 4376.5 | 149.8279069 | 46.2913298 |
| 2 | Prestw-790 | Cycloheximide | 5987.5 | 45.314429 | 32.5577419 |
| 1 | Prestw-790 | Cycloheximide | 18877 | 46.54653323 | 12.51826496 |
| 1 | Prestw-994 | Cyclopenthiazide | 16826 | 8.18716596 | 1 |
| 2 | Prestw-994 | Cyclopenthiazide | 6276 | 1 | 1 |
| 1 | Prestw-1095 | Cyclopentolate hydrochloride | 8048 | 16.2749921 | 15.2749921 |
| 2 | Prestw-1095 | Cyclopentolate hydrochloride | 3056.5 | 1 | 1 |
| 2 | Prestw-435 | Cyclosporin A | 2809 | 1037.845501 | 938.9446863 |
| 1 | Prestw-435 | Cyclosporin A | 4697.5 | 1 | 1 |
| 2 | Prestw-103 | Cyproheptadine hydrochloride | 3293.5 | 53.7073836 | 5.5534184 |
| 1 | Prestw-103 | Cyproheptadine hydrochloride | 8856 | 25.0694581 | 1.9253638 |
| 1 | Prestw-1074 | Cyproterone acetate | 9668.5 | 58.55685893 | 23.30644049 |
| 2 | Prestw-1074 | Cyproterone acetate | 3473.5 | 1 | 1 |
| 1 | Prestw-1134 | Cytarabine | 9538 | 55.13370679 | 2.99835279 |
| 2 | Prestw-1134 | Cytarabine | 3443 | 57.0570141 | 1 |
| 1 | Prestw-1086 | D-cycloserine | 7880 | 1 | 1 |
| 2 | Prestw-1086 | D-cycloserine | 3110.5 | 1 | 1 |
| 2 | Prestw-1477 | D,L-Penicillamine | 4640.5 | 144.4128444 | 57.7107284 |
| 1 | Prestw-1477 | D,L-Penicillamine | 12829 | 146.9787943 | 29.4523055 |
| 1 | Prestw-574 | Dacarbazine | 14609 | 31.14365704 | 4.4665821 |
| 2 | Prestw-574 | Dacarbazine | 4857 | 135.5912703 | 1 |
| 2 | Prestw-105 | Danazol | 2911.5 | 59.6197108 | 1 |
| 1 | Prestw-105 | Danazol | 8916.5 | 100.3350532 | 1 |
| 1 | Prestw-291 | Dantrolene sodium salt | 11217 | 30.9084474 | 26.2633781 |
| 2 | Prestw-291 | Dantrolene sodium salt | 3787 | 110.2200299 | 3.2112629 |
| 1 | Prestw-35 | Dapsone | 10272.5 | 1 | 1 |
| 2 | Prestw-35 | Dapsone | 3546.5 | 17.9912876 | 1 |
| 1 | Prestw-1514 | Darifenacin hydrobromide | 10246 | 1 | 1 |
| 2 | Prestw-1514 | Darifenacin hydrobromide | 4255 | 1 | 1 |
| 2 | Prestw-1802 | Darunavir | 2031 | 793.9772944 | 681.5234563 |
| 1 | Prestw-1802 | Darunavir | 4346 | 1121.577344 | 932.3011537 |
| 1 | Prestw-487 | Daunorubicin hydrochloride | 12830.5 | 39.96929971 | 13.29222477 |
| 2 | Prestw-487 | Daunorubicin hydrochloride | 4245.5 | 199.6543834 | 22.1296569 |
| 1 | Prestw-372 | Debrisoquin sulfate | 15085 | 18.53362284 | 10.5079648 |
| 2 | Prestw-372 | Debrisoquin sulfate | 5829 | 114.6798002 | 1 |

| | | | | | |
|---|---|---|---|---|---|
| 2 | Prestw-968 | Decamethonium bromide | 6018 | 31.2923845 | 30.2923845 |
| 1 | Prestw-968 | Decamethonium bromide | 19363.5 | 58.51450419 | 10.75853627 |
| 2 | Prestw-725 | Deferoxamine mesylate | 4745 | 1 | 1 |
| 1 | Prestw-725 | Deferoxamine mesylate | 16734.5 | 37.08670927 | 1 |
| 2 | Prestw-1509 | Deflazacort | 4630 | 0.6343321 | 1 |
| 1 | Prestw-1509 | Deflazacort | 13036.5 | 86.63414086 | 1.10610696 |
| 1 | Prestw-123 | Dehydrocholic acid | 9448 | 20.0667336 | 15.4788125 |
| 2 | Prestw-123 | Dehydrocholic acid | 3115 | 102.2966701 | 15.7438767 |
| 2 | Prestw-937 | Dehydroisoandosterone 3-acetate | 5514 | 62.1360941 | 33.56423 |
| 1 | Prestw-937 | Dehydroisoandosterone 3-acetate | 17038.5 | 41.69581291 | 1 |
| 2 | Prestw-1796 | Delavirdine | 4457 | 45.0200204 | 1 |
| 1 | Prestw-1796 | Delavirdine | 11196.5 | 168.5879956 | 1 |
| 1 | Prestw-616 | Demecarium bromide | 11908.5 | 13.73975309 | 1 |
| 2 | Prestw-616 | Demecarium bromide | 4082 | 118.7484016 | 1 |
| 1 | Prestw-753 | Demeclocycline hydrochloride | 15556.5 | 1 | 1 |
| 2 | Prestw-753 | Demeclocycline hydrochloride | 5172 | 1 | 1 |
| 2 | Prestw-961 | Denatonium benzoate | 6126.5 | 30.5525446 | 29.5525446 |
| 1 | Prestw-961 | Denatonium benzoate | 18995.5 | 49.42617797 | 1.67021005 |
| 2 | Prestw-957 | Deoxycorticosterone | 5848 | 67.9444988 | 64.7359584 |
| 1 | Prestw-957 | Deoxycorticosterone | 18750 | 42.46504393 | 5.9110256 |
| 1 | Prestw-1013 | Deptropine citrate | 7934.5 | 1 | 1 |
| 2 | Prestw-1013 | Deptropine citrate | 2964 | 43.1544598 | 1 |
| 1 | Prestw-388 | Dequalinium dichloride | 11224.5 | 60.77615281 | 23.34838861 |
| 2 | Prestw-388 | Dequalinium dichloride | 3159 | 1 | 1 |
| 1 | Prestw-343 | Desipramine hydrochloride | 11200 | 1 | 1 |
| 2 | Prestw-343 | Desipramine hydrochloride | 3723.5 | 18.0491847 | 1 |
| 2 | Prestw-1390 | Desloratadine | 3483 | 1 | 1 |
| 1 | Prestw-1390 | Desloratadine | 10473.5 | 51.18711255 | 1 |
| 2 | Prestw-1704 | Desonide | 5155.5 | 96.635172 | 1 |
| 1 | Prestw-1704 | Desonide | 12650 | 128.0677104 | 1 |
| 1 | Prestw-130 | Dexamethasone acetate | 7935 | 1 | 1 |
| 2 | Prestw-130 | Dexamethasone acetate | 3116.5 | 1 | 1 |
| 2 | Prestw-1392 | Dexfenfluramine hydrochloride | 3561 | 1 | 1 |
| 1 | Prestw-1392 | Dexfenfluramine hydrochloride | 10769 | 77.11691325 | 1 |
| 2 | Prestw-1830 | Dexrazoxane hydrochloride | 4870 | 16.496789 | 15.496789 |
| 1 | Prestw-1830 | Dexrazoxane hydrochloride | 13039.5 | 77.60022139 | 15.25028037 |
| 1 | Prestw-359 | Dextromethorphan hydrobromide monohydrat | 11721.5 | 9.63752024 | 1 |
| 2 | Prestw-359 | Dextromethorphan hydrobromide monohydrat | 3857 | 101.5464881 | 1 |
| 2 | Prestw-1167 | Diacerein | 3291 | 1 | 1 |
| 1 | Prestw-1167 | Diacerein | 10293.5 | 93.29689131 | 1 |
| 2 | Prestw-1705 | Diatrizoic acid dihydrate | 4938 | 82.7125129 | 1 |
| 1 | Prestw-1705 | Diatrizoic acid dihydrate | 13210 | 119.2993211 | 1 |
| 2 | Prestw-87 | Diazoxide | 5119.5 | 108.6008397 | 105.3922993 |
| 1 | Prestw-87 | Diazoxide | 16638 | 67.38858795 | 30.83456962 |
| 2 | Prestw-1393 | Dibenzepine hydrochloride | 3379.5 | 1 | 1 |
| 1 | Prestw-1393 | Dibenzepine hydrochloride | 10395 | 59.95314848 | 1 |
| 1 | Prestw-76 | Dibucaine | 15063 | 12.73496093 | 1 |
| 2 | Prestw-76 | Dibucaine | 5150.5 | 1 | 1 |
| 2 | Prestw-809 | Dichlorphenamide | 5553 | 1 | 1 |
| 1 | Prestw-809 | Dichlorphenamide | 20489.5 | 28.29669786 | 1 |
| 2 | Prestw-1394 | Diclazuril | 3008 | 1 | 1 |
| 1 | Prestw-1394 | Diclazuril | 9919.5 | 84.14836247 | 0.23290705 |
| 1 | Prestw-594 | Diclofenac sodium | 13959 | 31.31320116 | 5.81468379 |
| 2 | Prestw-594 | Diclofenac sodium | 4900.5 | 102.6705533 | 1 |
| 1 | Prestw-450 | Dicloxacillin sodium salt hydrate | 12681.5 | 20.09894979 | 1 |
| 2 | Prestw-450 | Dicloxacillin sodium salt hydrate | 4379 | 82.4136846 | 1 |
| 2 | Prestw-785 | Dicumarol | 6546.5 | 87.7007417 | 74.9440546 |
| 1 | Prestw-785 | Dicumarol | 20821.5 | 66.90849594 | 32.88022767 |
| 1 | Prestw-48 | Dicyclomine hydrochloride | 13136 | 31.3122804 | 4.63520546 |
| 2 | Prestw-48 | Dicyclomine hydrochloride | 4434 | 165.5204766 | 1 |
| 1 | Prestw-153 | Didanosine | 10452.5 | 0.05847109 | 1 |
| 2 | Prestw-153 | Didanosine | 4123 | 1 | 1 |
| 2 | Prestw-798 | Dienestrol | 5300 | 67.1729191 | 66.1729191 |

| | | | | | |
|---|---|---|---|---|---|
| 1 | Prestw-798 | Dienestrol | 19283 | 56.52686764 | 12.39905888 |
| 1 | Prestw-284 | Diethylcarbamazine citrate | 11430.5 | 19.43445012 | 5.96800057 |
| 2 | Prestw-284 | Diethylcarbamazine citrate | 4340.5 | 90.0260192 | 1 |
| 1 | Prestw-756 | Diethylstilbestrol | 15881.5 | 1 | 1 |
| 2 | Prestw-756 | Diethylstilbestrol | 5135 | 1 | 1 |
| 1 | Prestw-619 | Diflorasone Diacetate | 11651.5 | 26.34116373 | 23.15426471 |
| 2 | Prestw-619 | Diflorasone Diacetate | 3969.5 | 53.4994019 | 20.6540617 |
| 2 | Prestw-39 | Diflunisal | 4738 | 184.2302254 | 8.5896582 |
| 1 | Prestw-39 | Diflunisal | 12545.5 | 24.49459604 | 1 |
| 1 | Prestw-436 | Digitoxigenin | 11088 | 8.6465727 | 1 |
| 2 | Prestw-436 | Digitoxigenin | 4267 | 99.8532459 | 0.9524314 |
| 1 | Prestw-883 | Digoxigenin | 17693.5 | 49.81155851 | 2.05559059 |
| 2 | Prestw-883 | Digoxigenin | 5334 | 1 | 1 |
| 1 | Prestw-437 | Digoxin | 11406.5 | 14.08925336 | 1.23712686 |
| 2 | Prestw-437 | Digoxin | 4316.5 | 106.9050852 | 3.3685081 |
| 2 | Prestw-150 | Dihydroergotamine tartrate | 4547.5 | 104.2209863 | 32.2288458 |
| 1 | Prestw-150 | Dihydroergotamine tartrate | 12929.5 | 173.2516248 | 29.39095351 |
| 2 | Prestw-159 | Dihydrostreptomycin sulfate | 4172.5 | 87.3320675 | 70.7188835 |
| 1 | Prestw-159 | Dihydrostreptomycin sulfate | 10023 | 88.78793333 | 20.97613948 |
| 2 | Prestw-236 | Dilazep dihydrochloride | 5425 | 32.5829218 | 1 |
| 1 | Prestw-236 | Dilazep dihydrochloride | 14972.5 | 145.0330654 | 1 |
| 1 | Prestw-903 | Diloxanide furoate | 11630.5 | 38.69463021 | 27.75974872 |
| 2 | Prestw-903 | Diloxanide furoate | 4825.5 | 35.4696867 | 34.4696867 |
| 2 | Prestw-134 | Diltiazem hydrochloride | 3081.5 | 40.5991998 | 37.5252546 |
| 1 | Prestw-134 | Diltiazem hydrochloride | 9080 | 21.4833106 | 3.9211084 |
| 1 | Prestw-983 | Dimaprit dihydrochloride | 17596.5 | 9.2714659 | 1 |
| 2 | Prestw-983 | Dimaprit dihydrochloride | 6850 | 1 | 1 |
| 1 | Prestw-265 | Dimenhydrinate | 9951.5 | 1 | 1 |
| 2 | Prestw-265 | Dimenhydrinate | 3692 | 1 | 1 |
| 1 | Prestw-829 | Dimethadione | 14764 | 1 | 1 |
| 2 | Prestw-829 | Dimethadione | 4942.5 | 1 | 1 |
| 2 | Prestw-630 | Dimethisoquin hydrochloride | 4023 | 93.1214958 | 60.2761556 |
| 1 | Prestw-630 | Dimethisoquin hydrochloride | 12382 | 9.07622262 | 5.8893236 |
| 2 | Prestw-460 | Dinoprost trometamol | 4980 | 141.50532 | 49.3245283 |
| 1 | Prestw-460 | Dinoprost trometamol | 13880.5 | 25.43402245 | 1 |
| 2 | Prestw-1503 | Diosmin | 4509 | 47.2221153 | 43.727854 |
| 1 | Prestw-1503 | Diosmin | 10841.5 | 126.6601653 | 41.13213139 |
| 1 | Prestw-898 | Dioxybenzone | 17919.5 | 34.95473976 | 5.70515924 |
| 2 | Prestw-898 | Dioxybenzone | 6398 | 6.7387506 | 5.7387506 |
| 1 | Prestw-300 | Diperodon hydrochloride | 11006.5 | 41.12494204 | 33.099284 |
| 2 | Prestw-300 | Diperodon hydrochloride | 4093.5 | 127.2884561 | 1 |
| 2 | Prestw-79 | Diphemanil methylsulfate | 6223.5 | 44.6018171 | 31.84513 |
| 1 | Prestw-79 | Diphemanil methylsulfate | 18391 | 48.10489474 | 14.07662647 |
| 1 | Prestw-65 | Diphenhydramine hydrochloride | 15243.5 | 1 | 1 |
| 2 | Prestw-65 | Diphenhydramine hydrochloride | 5672 | 1 | 1 |
| 2 | Prestw-252 | Diphenidol hydrochloride | 4867.5 | 10.267559 | 9.267559 |
| 1 | Prestw-252 | Diphenidol hydrochloride | 12441 | 65.67424438 | 3.32430336 |
| 1 | Prestw-707 | Diphenylpyraline hydrochloride | 16289 | 62.62924549 | 28.60097722 |
| 2 | Prestw-707 | Diphenylpyraline hydrochloride | 5439 | 54.2788285 | 41.5221414 |
| 2 | Prestw-632 | Dipivefrin hydrochloride | 4182.5 | 77.248062 | 44.4027218 |
| 1 | Prestw-632 | Dipivefrin hydrochloride | 11752.5 | 1.51641068 | 1 |
| 2 | Prestw-33 | Diprophylline | 4101 | 112.3038722 | 48.9536966 |
| 1 | Prestw-33 | Diprophylline | 11212.5 | 42.50738859 | 2.37864445 |
| 1 | Prestw-142 | Dipyridamole | 10255.5 | 28.89963648 | 27.89963648 |
| 2 | Prestw-142 | Dipyridamole | 4592 | 1 | 1 |
| 2 | Prestw-713 | Dipyrone | 5996 | 111.7713006 | 99.0146135 |
| 1 | Prestw-713 | Dipyrone | 19216 | 76.7802251 | 42.75195683 |
| 2 | Prestw-557 | Dirithromycin | 4372.5 | 191.0400593 | 73.8221052 |
| 1 | Prestw-557 | Dirithromycin | 12382.5 | 29.67349463 | 1 |
| 1 | Prestw-266 | Disopyramide | 10301 | 1 | 1 |
| 2 | Prestw-266 | Disopyramide | 4104 | 4.1459782 | 3.1459782 |
| 1 | Prestw-97 | Disulfiram | 18975.5 | 73.92395016 | 26.16798224 |
| 2 | Prestw-97 | Disulfiram | 6120 | 12.7599478 | 11.7599478 |

| | | | | | |
|---|---|---|---|---|---|
| 2 | Prestw-352 | Dobutamine hydrochloride | 4041 | 154.5811961 | 49.1758109 |
| 1 | Prestw-352 | Dobutamine hydrochloride | 12022 | 5.89661685 | 1 |
| 1 | Prestw-1180 | Docetaxel | 9239.5 | 39.39520898 | 4.14479054 |
| 2 | Prestw-1180 | Docetaxel | 2991.5 | 1 | 1 |
| 2 | Prestw-1743 | Dofetilide | 4707 | 38.6606589 | 35.1663976 |
| 1 | Prestw-1743 | Dofetilide | 13326 | 91.47523539 | 5.94720149 |
| 2 | Prestw-1510 | Dolasetron mesilate | 4458 | 11.6586138 | 8.1643525 |
| 1 | Prestw-1510 | Dolasetron mesilate | 13349 | 81.04707621 | 1 |
| 1 | Prestw-340 | Domperidone | 9709.5 | 1 | 1 |
| 2 | Prestw-340 | Domperidone | 3873.5 | 1 | 1 |
| 2 | Prestw-1706 | Donepezil hydrochloride | 5142.5 | 63.4565198 | 1 |
| 1 | Prestw-1706 | Donepezil hydrochloride | 13413 | 110.0825705 | 1 |
| 2 | Prestw-1398 | Dopamine hydrochloride | 3313.5 | 6.2844276 | 5.2844276 |
| 1 | Prestw-1398 | Dopamine hydrochloride | 9381.5 | 82.23989062 | 1 |
| 2 | Prestw-1116 | Dorzolamide hydrochloride | 3274.5 | 55.0037004 | 6.8497352 |
| 1 | Prestw-1116 | Dorzolamide hydrochloride | 9612.5 | 22.7771139 | 1 |
| 1 | Prestw-488 | Dosulepin hydrochloride | 12932 | 52.21860903 | 25.54153409 |
| 2 | Prestw-488 | Dosulepin hydrochloride | 3758 | 149.230778 | 1 |
| 2 | Prestw-1707 | Doxapram hydrochloride | 5109.5 | 88.9366302 | 1 |
| 1 | Prestw-1707 | Doxapram hydrochloride | 12915.5 | 120.8031251 | 1 |
| 2 | Prestw-858 | Doxazosin mesylate | 5712 | 67.7401827 | 48.154351 |
| 1 | Prestw-858 | Doxazosin mesylate | 17518 | 54.02305302 | 2.61572551 |
| 1 | Prestw-263 | Doxepin hydrochloride | 10447 | 1 | 1 |
| 2 | Prestw-263 | Doxepin hydrochloride | 4059.5 | 1 | 1 |
| 1 | Prestw-1136 | Doxofylline | 9393 | 53.08776859 | 0.95241459 |
| 2 | Prestw-1136 | Doxofylline | 3296 | 17.100151 | 1 |
| 1 | Prestw-438 | Doxorubicin hydrochloride | 11518.5 | 14.8459413 | 1.9938148 |
| 2 | Prestw-438 | Doxorubicin hydrochloride | 3970 | 105.8883815 | 2.3518044 |
| 1 | Prestw-1399 | Doxycycline hydrochloride | 9313.5 | 90.42753724 | 6.51208182 |
| 2 | Prestw-1399 | Doxycycline hydrochloride | 3279.5 | 1 | 1 |
| 2 | Prestw-27 | Doxylamine succinate | 4022 | 48.7446936 | 47.7446936 |
| 1 | Prestw-27 | Doxylamine succinate | 11032 | 1 | 1 |
| 2 | Prestw-789 | Drofenine hydrochloride | 6250.5 | 41.7785686 | 29.0218815 |
| 1 | Prestw-789 | Drofenine hydrochloride | 20698 | 42.44765476 | 8.41938649 |
| 1 | Prestw-1822 | Dronedarone hydrochloride | 12580.5 | 150.8286498 | 30.91588656 |
| 2 | Prestw-1822 | Dronedarone hydrochloride | 4776 | 69.7598101 | 2.2657483 |
| 2 | Prestw-360 | Droperidol | 4189.5 | 133.9177881 | 28.5124029 |
| 1 | Prestw-360 | Droperidol | 12545 | 1 | 1 |
| 2 | Prestw-245 | Dropropizine (R,S) | 5170 | 22.2005594 | 1 |
| 1 | Prestw-245 | Dropropizine (R,S) | 14263 | 112.3101 | 1 |
| 1 | Prestw-264 | Dyclonine hydrochloride | 10314.5 | 1 | 1 |
| 2 | Prestw-264 | Dyclonine hydrochloride | 3916 | 1 | 1 |
| 1 | Prestw-671 | Dydrogesterone | 15920 | 17.70280889 | 1 |
| 2 | Prestw-671 | Dydrogesterone | 5114 | 1 | 1 |
| 1 | Prestw-304 | Econazole nitrate | 14691 | 23.62031317 | 15.59465513 |
| 2 | Prestw-304 | Econazole nitrate | 5200 | 149.93959 | 1 |
| 1 | Prestw-83 | Edrophonium chloride | 15205 | 1 | 1 |
| 2 | Prestw-83 | Edrophonium chloride | 5396 | 1 | 1 |
| 1 | Prestw-1400 | Efavirenz | 10129 | 14.43259323 | 12.47709211 |
| 2 | Prestw-1400 | Efavirenz | 3497.5 | 1 | 1 |
| 2 | Prestw-1274 | Emedastine | 3253 | 28.6343057 | 27.6343057 |
| 1 | Prestw-1274 | Emedastine | 9544 | 75.49700393 | 1 |
| 1 | Prestw-314 | Enalapril maleate | 11438.5 | 78.35348115 | 40.92571695 |
| 2 | Prestw-314 | Enalapril maleate | 3760.5 | 183.2614561 | 7.6208889 |
| 1 | Prestw-1769 | Enalaprilat dihydrate | 10181.5 | 115.8882204 | 33.14120286 |
| 2 | Prestw-1769 | Enalaprilat dihydrate | 4292 | 96.3272066 | 1 |
| 2 | Prestw-963 | Enilconazole | 6350 | 26.9054023 | 25.9054023 |
| 1 | Prestw-963 | Enilconazole | 20107.5 | 55.00793224 | 7.25196432 |
| 1 | Prestw-1401 | Enoxacin | 9998.5 | 9.38974257 | 7.43424145 |
| 2 | Prestw-1401 | Enoxacin | 3592 | 1 | 1 |
| 2 | Prestw-1786 | Enrofloxacin | 4641 | 122.9928943 | 59.4004858 |
| 1 | Prestw-1786 | Enrofloxacin | 11548.5 | 174.5699914 | 14.52974723 |
| 2 | Prestw-1403 | Entacapone | 3740.5 | 5.665006 | 4.665006 |

| | | | | | |
|---|---|---|---|---|---|
| 1 | Prestw-1403 | Entacapone | 10925 | 7.35816454 | 5.40266342 |
| 1 | Prestw-440 | Epiandrosterone | 11495 | 29.20744998 | 16.35532348 |
| 2 | Prestw-440 | Epiandrosterone | 4057 | 77.0572066 | 1 |
| 1 | Prestw-32 | Epirizole | 12920 | 60.76052818 | 47.80247368 |
| 2 | Prestw-32 | Epirizole | 2864.5 | 1 | 1 |
| 1 | Prestw-1752 | Epirubicin hydrochloride | 10009 | 22.12071578 | 2.41484916 |
| 2 | Prestw-1752 | Epirubicin hydrochloride | 4324 | 1 | 1 |
| 1 | Prestw-1823 | Eprosartan mesylate | 13588.5 | 127.5604978 | 7.64773448 |
| 2 | Prestw-1823 | Eprosartan mesylate | 5315 | 59.3312797 | 1 |
| 2 | Prestw-850 | Equilin | 5060 | 73.1694983 | 44.5976342 |
| 1 | Prestw-850 | Equilin | 17798.5 | 47.30621415 | 1 |
| 1 | Prestw-1242 | Erlotinib | 10202.5 | 28.11632997 | 23.52840887 |
| 2 | Prestw-1242 | Erlotinib | 3285 | 66.2077201 | 1 |
| 2 | Prestw-151 | Erythromycin | 4340 | 64.5089059 | 61.0146446 |
| 1 | Prestw-151 | Erythromycin | 11963 | 124.9858153 | 39.45778135 |
| 2 | Prestw-1271 | Escitalopram oxalate | 3308 | 22.2991609 | 21.2991609 |
| 1 | Prestw-1271 | Escitalopram oxalate | 9221 | 88.99652758 | 5.08107216 |
| 1 | Prestw-566 | Eserine hemisulfate salt | 14335.5 | 33.04164227 | 6.36456733 |
| 2 | Prestw-566 | Eserine hemisulfate salt | 5094.5 | 77.7189932 | 1 |
| 1 | Prestw-1137 | Esmolol hydrochloride | 9494.5 | 42.0198194 | 1 |
| 2 | Prestw-1137 | Esmolol hydrochloride | 3218 | 132.6990062 | 1 |
| 1 | Prestw-1473 | Estradiol Valerate | 11263 | 154.693469 | 37.16698014 |
| 2 | Prestw-1473 | Estradiol Valerate | 4711 | 94.5368231 | 7.8347071 |
| 1 | Prestw-441 | Estradiol-17 beta | 12486.5 | 26.98609213 | 14.13396563 |
| 2 | Prestw-441 | Estradiol-17 beta | 4335.5 | 111.4110307 | 7.8744536 |
| 2 | Prestw-1709 | Estramustine | 4822 | 81.4083934 | 1 |
| 1 | Prestw-1709 | Estramustine | 13093 | 106.5072766 | 1 |
| 1 | Prestw-1096 | Estriol | 8081 | 3.0305223 | 2.0305223 |
| 2 | Prestw-1096 | Estriol | 3105.5 | 1 | 1 |
| 1 | Prestw-914 | Estrone | 15255 | 0.71965451 | 1 |
| 2 | Prestw-914 | Estrone | 5181.5 | 1 | 1 |
| 1 | Prestw-688 | Estropipate | 15656.5 | 44.47729271 | 12.58866463 |
| 2 | Prestw-688 | Estropipate | 4968 | 17.5472352 | 1 |
| 1 | Prestw-1824 | Eszopiclone | 11876.5 | 99.74247172 | 37.3925307 |
| 2 | Prestw-1824 | Eszopiclone | 5096.5 | 10.6997554 | 9.6997554 |
| 1 | Prestw-649 | Etanidazole | 14530.5 | 0.06174593 | 1 |
| 2 | Prestw-649 | Etanidazole | 5380 | 1 | 1 |
| 1 | Prestw-259 | Ethacrynic acid | 12206.5 | 65.79618536 | 3.44624434 |
| 2 | Prestw-259 | Ethacrynic acid | 4903 | 1 | 1 |
| 1 | Prestw-28 | Ethambutol dihydrochloride | 11456.5 | 1 | 1 |
| 2 | Prestw-28 | Ethambutol dihydrochloride | 3951 | 56.2006786 | 1 |
| 1 | Prestw-500 | Ethamivan | 10933.5 | 19.89082744 | 16.70392842 |
| 2 | Prestw-500 | Ethamivan | 3469 | 51.1236118 | 18.2782716 |
| 1 | Prestw-1015 | Ethamsylate | 8389.5 | 1 | 1 |
| 2 | Prestw-1015 | Ethamsylate | 3068 | 1 | 1 |
| 1 | Prestw-830 | Ethaverine hydrochloride | 15110.5 | 1 | 1 |
| 2 | Prestw-830 | Ethaverine hydrochloride | 4651.5 | 1 | 1 |
| 1 | Prestw-1405 | Ethinylestradiol | 9561 | 18.75237057 | 16.79686945 |
| 2 | Prestw-1405 | Ethinylestradiol | 3534 | 1 | 1 |
| 1 | Prestw-526 | Ethionamide | 11350 | 1 | 1 |
| 2 | Prestw-526 | Ethionamide | 3459.5 | 55.1510338 | 1 |
| 1 | Prestw-261 | Ethisterone | 9629.5 | 1 | 1 |
| 2 | Prestw-261 | Ethisterone | 3847 | 4.5674806 | 3.5674806 |
| 2 | Prestw-840 | Ethopropazine hydrochloride | 5054 | 48.8782868 | 4.7971639 |
| 1 | Prestw-840 | Ethopropazine hydrochloride | 16086.5 | 28.35433844 | 1 |
| 2 | Prestw-165 | Ethosuximide | 4439 | 140.1845708 | 46.5492261 |
| 1 | Prestw-165 | Ethosuximide | 11235.5 | 132.9523136 | 28.57000062 |
| 1 | Prestw-696 | Ethotoin | 16808 | 49.38019082 | 17.49156274 |
| 2 | Prestw-696 | Ethotoin | 5451.5 | 37.5933134 | 6.705237 |
| 1 | Prestw-765 | Ethoxyquin | 13865 | 19.05743972 | 1 |
| 2 | Prestw-765 | Ethoxyquin | 4797.5 | 0.6241247 | 1 |
| 2 | Prestw-1710 | Ethoxzolamide | 5517 | 88.7732669 | 2.0711509 |
| 1 | Prestw-1710 | Ethoxzolamide | 14259.5 | 116.5205192 | 1 |

| | | | | | |
|---|---|---|---|---|---|
| 1 | Prestw-908 | Ethynodiol diacetate | 15477.5 | 16.53198261 | 5.59710112 |
| 2 | Prestw-908 | Ethynodiol diacetate | 5977 | 1 | 1 |
| 1 | Prestw-846 | Ethynylestradiol 3-methyl ether | 16295 | 35.94825859 | 1 |
| 2 | Prestw-846 | Ethynylestradiol 3-methyl ether | 5684 | 29.5204483 | 0.9485842 |
| 1 | Prestw-504 | Etifenin | 10427.5 | 33.9709079 | 30.78400888 |
| 2 | Prestw-504 | Etifenin | 4324 | 111.1040877 | 78.2587475 |
| 1 | Prestw-1017 | Etilefrine hydrochloride | 8317 | 2.3554897 | 1.3554897 |
| 2 | Prestw-1017 | Etilefrine hydrochloride | 3319.5 | 15.8153905 | 1 |
| 2 | Prestw-231 | Etodolac | 4597.5 | 149.6466917 | 46.1651495 |
| 1 | Prestw-231 | Etodolac | 13113.5 | 130.1172894 | 1 |
| 1 | Prestw-1407 | Etofenamate | 9792 | 22.06335234 | 20.10785122 |
| 2 | Prestw-1407 | Etofenamate | 3892.5 | 1 | 1 |
| 1 | Prestw-172 | Etofylline | 12794.5 | 178.7930574 | 15.15726149 |
| 2 | Prestw-172 | Etofylline | 4853 | 163.0508663 | 1 |
| 1 | Prestw-1041 | Etomidate | 9262.5 | 39.7946728 | 1 |
| 2 | Prestw-1041 | Etomidate | 3202.5 | 130.5522474 | 1 |
| 2 | Prestw-396 | Etoposide | 4335.5 | 378.3599149 | 217.4962586 |
| 1 | Prestw-396 | Etoposide | 13245 | 1 | 1 |
| 1 | Prestw-1408 | Etoricoxib | 9461.5 | 18.24706993 | 16.29156881 |
| 2 | Prestw-1408 | Etoricoxib | 3309.5 | 1 | 1 |
| 1 | Prestw-1409 | Etretinate | 10041 | 5.8148437 | 3.85934258 |
| 2 | Prestw-1409 | Etretinate | 3239.5 | 1 | 1 |
| 1 | Prestw-1410 | Exemestane | 8974.5 | 22.04223984 | 20.08673872 |
| 2 | Prestw-1410 | Exemestane | 3889.5 | 1 | 1 |
| 2 | Prestw-1762 | Ezetimibe | 4616.5 | 44.3921429 | 1 |
| 1 | Prestw-1762 | Ezetimibe | 11068.5 | 59.73225769 | 1 |
| 2 | Prestw-1249 | Famciclovir | 3530.5 | 105.5471673 | 83.3909897 |
| 1 | Prestw-1249 | Famciclovir | 9385 | 48.78495788 | 1 |
| 1 | Prestw-104 | Famotidine | 9157 | 30.6710705 | 1 |
| 2 | Prestw-104 | Famotidine | 3274 | 58.9684901 | 1 |
| 1 | Prestw-703 | Famprofazone | 17724.5 | 74.21006417 | 28.83068099 |
| 2 | Prestw-703 | Famprofazone | 5636.5 | 98.2986854 | 28.8932892 |
| 2 | Prestw-1711 | Felbamate | 5688 | 86.4901737 | 1 |
| 1 | Prestw-1711 | Felbamate | 15193 | 112.2729461 | 1 |
| 2 | Prestw-893 | Felbinac | 5483.5 | 1 | 1 |
| 1 | Prestw-893 | Felbinac | 14982.5 | 29.3387822 | 0.08920168 |
| 1 | Prestw-478 | Felodipine | 11748.5 | 31.8370817 | 5.16000676 |
| 2 | Prestw-478 | Felodipine | 4111.5 | 104.478424 | 1 |
| 2 | Prestw-210 | Fenbendazole | 4811.5 | 101.7872194 | 8.3287567 |
| 1 | Prestw-210 | Fenbendazole | 11972.5 | 116.1860289 | 5.03715783 |
| 2 | Prestw-218 | Fenbufen | 5741 | 74.248531 | 10.6561225 |
| 1 | Prestw-218 | Fenbufen | 13397.5 | 135.9837949 | 1 |
| 1 | Prestw-270 | Fendiline hydrochloride | 10975 | 1 | 1 |
| 2 | Prestw-270 | Fendiline hydrochloride | 3840.5 | 1 | 1 |
| 2 | Prestw-1501 | Fenipentol | 2797.5 | 36.0282046 | 32.5339433 |
| 1 | Prestw-1501 | Fenipentol | 10889 | 99.73038432 | 14.20235042 |
| 1 | Prestw-275 | Fenofibrate | 10032.5 | 1 | 1 |
| 2 | Prestw-275 | Fenofibrate | 3948.5 | 24.7997094 | 1 |
| 1 | Prestw-1713 | Fenoldopam | 14305.5 | 135.8413064 | 18.31481753 |
| 2 | Prestw-1713 | Fenoldopam | 5717.5 | 59.0596628 | 1 |
| 1 | Prestw-754 | Fenoprofen calcium salt dihydrate | 14716.5 | 1 | 1 |
| 2 | Prestw-754 | Fenoprofen calcium salt dihydrate | 5243.5 | 1 | 1 |
| 1 | Prestw-282 | Fenoterol hydrobromide | 10536 | 37.43666591 | 23.97021636 |
| 2 | Prestw-282 | Fenoterol hydrobromide | 3787.5 | 78.3496602 | 1 |
| 1 | Prestw-213 | Fenspiride hydrochloride | 11817.5 | 129.5023652 | 18.35349414 |
| 2 | Prestw-213 | Fenspiride hydrochloride | 4688.5 | 82.455189 | 1 |
| 1 | Prestw-1254 | Fentiazac | 9342.5 | 63.03253433 | 1 |
| 2 | Prestw-1254 | Fentiazac | 3388 | 1 | 1 |
| 1 | Prestw-1102 | Fexofenadine hydrochloride | 7916.5 | 50.4439644 | 36.2891107 |
| 2 | Prestw-1102 | Fexofenadine hydrochloride | 3053 | 11.8559535 | 8.7502353 |
| 2 | Prestw-717 | Finasteride | 5735.5 | 17.0939196 | 16.0939196 |
| 1 | Prestw-717 | Finasteride | 17568 | 22.47300981 | 1 |
| 1 | Prestw-298 | Fipexide hydrochloride | 11985 | 31.56533352 | 23.53967548 |

| | | | | | |
|---|---|---|---|---|---|
| 2 | Prestw-298 | Fipexide hydrochloride | 4392.5 | 93.1724836 | 1 |
| 2 | Prestw-242 | Flavoxate hydrochloride | 4686.5 | 118.7560634 | 6.3022253 |
| 1 | Prestw-242 | Flavoxate hydrochloride | 13841.5 | 155.7367186 | 1 |
| 1 | Prestw-735 | Flecainide acetate | 13703 | 23.72348335 | 9.07847441 |
| 2 | Prestw-735 | Flecainide acetate | 5183 | 6.8503773 | 5.8503773 |
| 1 | Prestw-1414 | Fleroxacin | 9607.5 | 1 | 1 |
| 2 | Prestw-1414 | Fleroxacin | 3643 | 1 | 1 |
| 2 | Prestw-955 | Florfenicol | 6314 | 32.1988552 | 28.9903148 |
| 1 | Prestw-955 | Florfenicol | 20490 | 43.4216477 | 6.86762937 |
| 1 | Prestw-1415 | Floxuridine | 9943 | 8.42422557 | 7.42422557 |
| 2 | Prestw-1415 | Floxuridine | 4189.5 | 1 | 1 |
| 1 | Prestw-1416 | Flubendazol | 10558.5 | 18.66318245 | 17.66318245 |
| 2 | Prestw-1416 | Flubendazol | 4430 | 1 | 1 |
| 1 | Prestw-1011 | Flucloxacillin sodium | 7798 | 1 | 1 |
| 2 | Prestw-1011 | Flucloxacillin sodium | 3088.5 | 59.4775068 | 1 |
| 1 | Prestw-1417 | Fluconazole | 9967 | 29.95803956 | 28.95803956 |
| 2 | Prestw-1417 | Fluconazole | 4415 | 1 | 1 |
| 2 | Prestw-934 | Flucytosine | 6188 | 46.0944723 | 17.5226082 |
| 1 | Prestw-934 | Flucytosine | 18403.5 | 54.67408287 | 2.69137929 |
| 2 | Prestw-1483 | Fludarabine | 4943.5 | 87.7661271 | 1 |
| 1 | Prestw-1483 | Fludarabine | 12640 | 144.0921149 | 1 |
| 1 | Prestw-281 | Fludrocortisone acetate | 10111 | 10.4340248 | 1 |
| 2 | Prestw-281 | Fludrocortisone acetate | 3835 | 62.4132789 | 1 |
| 1 | Prestw-203 | Flufenamic acid | 10873.5 | 87.80144328 | 5.05442573 |
| 2 | Prestw-203 | Flufenamic acid | 4293 | 68.5529411 | 1 |
| 1 | Prestw-204 | Flumequine | 10637.5 | 84.15527223 | 1.40825468 |
| 2 | Prestw-204 | Flumequine | 4048.5 | 93.9997095 | 1 |
| 1 | Prestw-734 | Flumethasone | 15558 | 26.43281586 | 11.78780692 |
| 2 | Prestw-734 | Flumethasone | 5043.5 | 1 | 1 |
| 1 | Prestw-1712 | Flumethasone pivalate | 14651.5 | 138.8839384 | 21.35744952 |
| 2 | Prestw-1712 | Flumethasone pivalate | 6309 | 86.4906649 | 1 |
| 2 | Prestw-312 | Flunarizine dihydrochloride | 4560 | 189.9305556 | 14.2899884 |
| 1 | Prestw-312 | Flunarizine dihydrochloride | 12146 | 38.93345316 | 1.50568896 |
| 1 | Prestw-643 | Flunisolide | 16284 | 1 | 1 |
| 2 | Prestw-643 | Flunisolide | 5442 | 1 | 1 |
| 1 | Prestw-744 | Flunixin meglumine | 16486 | 29.09799324 | 14.4529843 |
| 2 | Prestw-744 | Flunixin meglumine | 5454.5 | 1 | 1 |
| 1 | Prestw-1419 | Fluocinolone acetonide | 10132.5 | 28.48607771 | 27.48607771 |
| 2 | Prestw-1419 | Fluocinolone acetonide | 4280.5 | 1 | 1 |
| 1 | Prestw-774 | Fluocinonide | 16389.5 | 45.12130992 | 13.23268184 |
| 2 | Prestw-774 | Fluocinonide | 5211.5 | 19.6027103 | 1 |
| 2 | Prestw-718 | Fluorometholone | 5653.5 | 18.9955927 | 17.9955927 |
| 1 | Prestw-718 | Fluorometholone | 16228.5 | 37.07379982 | 1 |
| 1 | Prestw-511 | Fluoxetine hydrochloride | 10489.5 | 1 | 1 |
| 2 | Prestw-511 | Fluoxetine hydrochloride | 3670 | 35.0813781 | 1.4909379 |
| 1 | Prestw-320 | Fluphenazine dihydrochloride | 12474 | 46.13782249 | 33.17976799 |
| 2 | Prestw-320 | Fluphenazine dihydrochloride | 4734.5 | 166.0603178 | 5.1966615 |
| 1 | Prestw-645 | Flurandrenolide | 9336.5 | 21.4452561 | 20.4452561 |
| 2 | Prestw-645 | Flurandrenolide | 3883.5 | 57.6067855 | 56.6067855 |
| 1 | Prestw-917 | Flurbiprofen | 15093 | 1 | 1 |
| 2 | Prestw-917 | Flurbiprofen | 5485 | 1 | 1 |
| 1 | Prestw-906 | Fluspirilen | 16273.5 | 10.34120344 | 1 |
| 2 | Prestw-906 | Fluspirilen | 6582 | 1 | 1 |
| 2 | Prestw-180 | Flutamide | 5334 | 64.0910075 | 1 |
| 1 | Prestw-180 | Flutamide | 13999 | 124.4911444 | 1 |
| 1 | Prestw-997 | Fluticasone propionate | 16709 | 1 | 1 |
| 2 | Prestw-997 | Fluticasone propionate | 6013 | 1 | 1 |
| 2 | Prestw-859 | Fluvastatin sodium salt | 5608 | 76.5317699 | 56.9459382 |
| 1 | Prestw-859 | Fluvastatin sodium salt | 16473.5 | 63.18949705 | 11.78216954 |
| 1 | Prestw-995 | Fluvoxamine maleate | 17091.5 | 1 | 1 |
| 2 | Prestw-995 | Fluvoxamine maleate | 6230 | 1 | 1 |
| 2 | Prestw-627 | Folic acid | 4274.5 | 94.9394406 | 62.0941004 |
| 1 | Prestw-627 | Folic acid | 11669 | 0.21550624 | 1 |

| | | | | | |
|---|---|---|---|---|---|
| 1 | Prestw-738 | Folinic acid calcium salt | 14867.5 | 34.28030218 | 19.63529324 |
| 2 | Prestw-738 | Folinic acid calcium salt | 5452.5 | 1 | 1 |
| 2 | Prestw-1499 | Fomepizole | 5109.5 | 101.1675773 | 29.1754368 |
| 1 | Prestw-1499 | Fomepizole | 12481.5 | 179.8948426 | 36.03417135 |
| 1 | Prestw-1420 | Formestane | 2780 | 478.4608786 | 477.4608786 |
| 2 | Prestw-1420 | Formestane | 1898 | 284.0515288 | 283.0515288 |
| 2 | Prestw-1421 | Formoterol fumarate | 3917.5 | 105.432725 | 104.432725 |
| 1 | Prestw-1421 | Formoterol fumarate | 10003.5 | 22.42500485 | 21.42500485 |
| 1 | Prestw-815 | Fosfosal | 17722.5 | 26.42118692 | 11.77617798 |
| 2 | Prestw-815 | Fosfosal | 5895 | 1 | 1 |
| 1 | Prestw-1423 | Fosinopril | 10140.5 | 20.75194371 | 19.75194371 |
| 2 | Prestw-1423 | Fosinopril | 4422.5 | 1 | 1 |
| 1 | Prestw-1424 | Fulvestrant | 10073 | 22.76252889 | 21.76252889 |
| 2 | Prestw-1424 | Fulvestrant | 4113.5 | 1 | 1 |
| 1 | Prestw-764 | Furaltadone hydrochloride | 14291 | 9.11675572 | 1 |
| 2 | Prestw-764 | Furaltadone hydrochloride | 4797 | 1 | 1 |
| 2 | Prestw-808 | Furazolidone | 5334 | 1 | 1 |
| 1 | Prestw-808 | Furazolidone | 19190 | 38.39334956 | 1 |
| 1 | Prestw-341 | Furosemide | 10540 | 1 | 1 |
| 2 | Prestw-341 | Furosemide | 3988.5 | 2.431825 | 1.431825 |
| 1 | Prestw-1007 | Fursultiamine Hydrochloride | 7108.5 | 1 | 1 |
| 2 | Prestw-1007 | Fursultiamine Hydrochloride | 3010 | 1 | 1 |
| 1 | Prestw-390 | Fusidic acid sodium salt | 13313.5 | 44.32847149 | 31.37041699 |
| 2 | Prestw-390 | Fusidic acid sodium salt | 4337.5 | 153.9218579 | 1 |
| 2 | Prestw-861 | Gabapentin | 5957 | 95.197836 | 75.6120043 |
| 1 | Prestw-861 | Gabapentin | 17585 | 74.62840877 | 23.22108126 |
| 1 | Prestw-1008 | Gabexate mesilate | 8446.5 | 1 | 1 |
| 2 | Prestw-1008 | Gabexate mesilate | 3335.5 | 30.7816149 | 1 |
| 1 | Prestw-588 | Galanthamine hydrobromide | 13321 | 26.11779376 | 0.61927639 |
| 2 | Prestw-588 | Galanthamine hydrobromide | 4685.5 | 125.8951571 | 1 |
| 1 | Prestw-157 | Gallamine triethiodide | 10166 | 1 | 1 |
| 2 | Prestw-157 | Gallamine triethiodide | 4082.5 | 1 | 1 |
| 1 | Prestw-839 | Ganciclovir | 15849.5 | 27.05845407 | 1 |
| 2 | Prestw-839 | Ganciclovir | 4948.5 | 18.1682704 | 1 |
| 1 | Prestw-1265 | Gatifloxacin | 9494.5 | 88.92413554 | 6.98630078 |
| 2 | Prestw-1265 | Gatifloxacin | 3575.5 | 25.9575396 | 1 |
| 1 | Prestw-386 | GBR 12909 dihydrochloride | 13723.5 | 44.51000489 | 7.08224069 |
| 2 | Prestw-386 | GBR 12909 dihydrochloride | 4664 | 114.1146951 | 1 |
| 1 | Prestw-1270 | Gefitinib | 9277 | 87.69215013 | 3.77669471 |
| 2 | Prestw-1270 | Gefitinib | 3267.5 | 1 | 1 |
| 2 | Prestw-1266 | Gemcitabine | 3318 | 61.6842571 | 6.5927355 |
| 1 | Prestw-1266 | Gemcitabine | 9586 | 88.05171327 | 6.11387851 |
| 2 | Prestw-214 | Gemfibrozil | 4758 | 84.8610918 | 1 |
| 1 | Prestw-214 | Gemfibrozil | 11717 | 103.8691126 | 1 |
| 2 | Prestw-1825 | Gemifloxacin mesylate | 4635.5 | 42.5593371 | 41.5593371 |
| 1 | Prestw-1825 | Gemifloxacin mesylate | 11672.5 | 93.3668516 | 31.01691058 |
| 1 | Prestw-160 | Gentamicine sulfate | 10410 | 91.04353254 | 23.23173869 |
| 2 | Prestw-160 | Gentamicine sulfate | 4180 | 30.9992378 | 14.3860538 |
| 2 | Prestw-1267 | Gestrinone | 3534 | 37.1550109 | 1 |
| 1 | Prestw-1267 | Gestrinone | 10116 | 80.79897805 | 1 |
| 1 | Prestw-293 | Glafenine hydrochloride | 12221.5 | 31.17728704 | 26.53221774 |
| 2 | Prestw-293 | Glafenine hydrochloride | 4076.5 | 64.2327815 | 1 |
| 1 | Prestw-316 | Glibenclamide | 11369 | 23.9871876 | 11.0291331 |
| 2 | Prestw-316 | Glibenclamide | 4169 | 143.3576138 | 1 |
| 2 | Prestw-558 | Gliclazide | 4389 | 211.6493677 | 94.4314136 |
| 1 | Prestw-558 | Gliclazide | 14473 | 28.29934683 | 1 |
| 1 | Prestw-651 | Glimepiride | 15652.5 | 5.81095923 | 4.81095923 |
| 2 | Prestw-651 | Glimepiride | 5484.5 | 1 | 1 |
| 2 | Prestw-131 | Glipizide | 3001 | 11.4967401 | 10.4967401 |
| 1 | Prestw-131 | Glipizide | 7828 | 1 | 1 |
| 1 | Prestw-991 | Gliquidone | 17979.5 | 4.27287545 | 1 |
| 2 | Prestw-991 | Gliquidone | 6297.5 | 1 | 1 |
| 2 | Prestw-244 | Glutethimide, para-amino | 4688.5 | 29.2093871 | 1 |

| | | | | | |
|---|---|---|---|---|---|
| 1 | Prestw-244 | Glutethimide, para-amino | 13668 | 117.3045936 | 1 |
| 1 | Prestw-746 | Glycopyrrolate | 15050 | 3.1632915 | 2.1632915 |
| 2 | Prestw-746 | Glycopyrrolate | 5062 | 1 | 1 |
| 2 | Prestw-1187 | Granisetron | 3276.5 | 5.3150857 | 4.3150857 |
| 1 | Prestw-1187 | Granisetron | 7287 | 1 | 1 |
| 2 | Prestw-1797 | Grepafloxacin | 4482 | 114.1091219 | 10.6275797 |
| 1 | Prestw-1797 | Grepafloxacin | 10524.5 | 182.2009413 | 8.95841973 |
| 2 | Prestw-226 | Griseofulvin | 6093.5 | 58.0402675 | 1 |
| 1 | Prestw-226 | Griseofulvin | 15761.5 | 99.01653302 | 1 |
| 2 | Prestw-1268 | Guaiacol | 3237 | 51.630864 | 1 |
| 1 | Prestw-1268 | Guaiacol | 10188 | 67.15803079 | 1 |
| 2 | Prestw-776 | Guaifenesin | 6178 | 82.9068613 | 13.5014651 |
| 1 | Prestw-776 | Guaifenesin | 18630 | 49.4375076 | 4.05812442 |
| 2 | Prestw-96 | Guanabenz acetate | 6259.5 | 45.9222485 | 42.7137081 |
| 1 | Prestw-96 | Guanabenz acetate | 19976 | 35.74385813 | 1 |
| 1 | Prestw-767 | Guanadrel sulfate | 16525.5 | 54.50732317 | 22.61869509 |
| 2 | Prestw-767 | Guanadrel sulfate | 5700 | 72.7618563 | 41.8737799 |
| 1 | Prestw-317 | Guanethidine sulfate | 11092.5 | 32.2284914 | 19.2704369 |
| 2 | Prestw-317 | Guanethidine sulfate | 4138.5 | 146.8838742 | 1 |
| 1 | Prestw-339 | Guanfacine hydrochloride | 9726.5 | 1 | 1 |
| 2 | Prestw-339 | Guanfacine hydrochloride | 4152 | 1 | 1 |
| 1 | Prestw-655 | Halcinonide | 15376.5 | 1 | 1 |
| 2 | Prestw-655 | Halcinonide | 5206 | 1 | 1 |
| 1 | Prestw-1031 | Halofantrine hydrochloride | 8940.5 | 15.2562277 | 1 |
| 2 | Prestw-1031 | Halofantrine hydrochloride | 2665.5 | 1 | 1 |
| 2 | Prestw-115 | Haloperidol | 3346 | 1 | 1 |
| 1 | Prestw-115 | Haloperidol | 10629 | 73.01415397 | 1 |
| 2 | Prestw-1269 | Haloprogin | 3210 | 57.870747 | 2.7792254 |
| 1 | Prestw-1269 | Haloprogin | 10240.5 | 61.93680798 | 1 |
| 1 | Prestw-393 | Hemicholinium bromide | 14577.5 | 33.46363785 | 20.50558335 |
| 2 | Prestw-393 | Hemicholinium bromide | 4392.5 | 193.9840793 | 33.120423 |
| 2 | Prestw-15 | Heptaminol hydrochloride | 4777.5 | 101.631341 | 29.6392005 |
| 1 | Prestw-15 | Heptaminol hydrochloride | 12097 | 192.3837178 | 48.52304652 |
| 2 | Prestw-400 | Hesperidin | 4763.5 | 92.1533552 | 28.8031796 |
| 1 | Prestw-400 | Hesperidin | 12607 | 25.41093084 | 1 |
| 1 | Prestw-1472 | Hexachlorophene | 11402 | 127.124244 | 9.59775516 |
| 2 | Prestw-1472 | Hexachlorophene | 4695 | 84.3753813 | 1 |
| 1 | Prestw-38 | Hexamethonium dibromide dihydrate | 12199 | 22.60614092 | 14.58048288 |
| 2 | Prestw-38 | Hexamethonium dibromide dihydrate | 4601.5 | 115.1290255 | 1 |
| 1 | Prestw-699 | Hexestrol | 16115.5 | 66.66702893 | 21.28764575 |
| 2 | Prestw-699 | Hexestrol | 5031 | 55.5828433 | 1 |
| 1 | Prestw-551 | Hexetidine | 15178 | 32.03986749 | 1 |
| 2 | Prestw-551 | Hexetidine | 5410 | 98.987073 | 1 |
| 2 | Prestw-788 | Hexylcaine hydrochloride | 6043.5 | 119.6243828 | 106.8676957 |
| 1 | Prestw-788 | Hexylcaine hydrochloride | 19255 | 80.47621216 | 46.44794389 |
| 2 | Prestw-1700 | Histamine dihydrochloride | 4468 | 109.4245374 | 15.7891927 |
| 1 | Prestw-1700 | Histamine dihydrochloride | 11335.5 | 102.4774768 | 1 |
| 1 | Prestw-62 | Homatropine hydrobromide (R,S) | 12149.5 | 10.19565583 | 7.00875681 |
| 2 | Prestw-62 | Homatropine hydrobromide (R,S) | 4382 | 48.3428966 | 15.4975564 |
| 1 | Prestw-283 | Homochlorcyclizine dihydrochloride | 11223 | 16.8064983 | 3.34004875 |
| 2 | Prestw-283 | Homochlorcyclizine dihydrochloride | 4020.5 | 119.1764214 | 13.7710362 |
| 1 | Prestw-1090 | Homosalate | 7433.5 | 1 | 1 |
| 2 | Prestw-1090 | Homosalate | 3231.5 | 1 | 1 |
| 2 | Prestw-1787 | Homoveratrylamine | 5141.5 | 113.5057544 | 49.9133459 |
| 1 | Prestw-1787 | Homoveratrylamine | 12146.5 | 179.4980804 | 19.45783621 |
| 1 | Prestw-355 | Hycanthone | 12060 | 8.6662534 | 1 |
| 2 | Prestw-355 | Hycanthone | 4157 | 113.635789 | 8.2304038 |
| 1 | Prestw-169 | Hydralazine hydrochloride | 11057 | 121.3598511 | 16.97753816 |
| 2 | Prestw-169 | Hydralazine hydrochloride | 4800 | 70.8879463 | 1 |
| 1 | Prestw-9 | Hydrochlorothiazide | 17411 | 39.54437527 | 10.29479475 |
| 2 | Prestw-9 | Hydrochlorothiazide | 5733.5 | 1 | 1 |
| 2 | Prestw-447 | Hydrocortisone base | 4381.5 | 139.8843255 | 36.3477484 |
| 1 | Prestw-447 | Hydrocortisone base | 13081 | 7.69025925 | 1 |

| | | | | | |
|---|---|---|---|---|---|
| 1 | Prestw-13 | Hydroflumethiazide | 8201.5 | 1 | 1 |
| 2 | Prestw-13 | Hydroflumethiazide | 2954.5 | 1 | 1 |
| 2 | Prestw-1284 | Hydroxychloroquine sulfate | 3525.5 | 1 | 1 |
| 1 | Prestw-1284 | Hydroxychloroquine sulfate | 9873.5 | 67.95647797 | 1 |
| 1 | Prestw-133 | Hydroxyzine dihydrochloride | 9214 | 49.47740114 | 31.91519894 |
| 2 | Prestw-133 | Hydroxyzine dihydrochloride | 3515.5 | 1 | 1 |
| 1 | Prestw-901 | Hymecromone | 15203.5 | 7.97921395 | 1 |
| 2 | Prestw-901 | Hymecromone | 5422 | 1 | 1 |
| 2 | Prestw-233 | Hyoscyamine (L) | 4898 | 148.8219486 | 36.3681105 |
| 1 | Prestw-233 | Hyoscyamine (L) | 12521 | 176.8244282 | 1 |
| 1 | Prestw-1285 | Ibandronate sodium | 9587 | 30.06514354 | 28.10964242 |
| 2 | Prestw-1285 | Ibandronate sodium | 3507.5 | 12.1226393 | 11.1226393 |
| 1 | Prestw-1286 | Ibudilast | 9843 | 25.22735038 | 23.27184926 |
| 2 | Prestw-1286 | Ibudilast | 3606.5 | 1 | 1 |
| 2 | Prestw-1508 | Ibutilide fumarate | 4498.5 | 29.0491293 | 25.554868 |
| 1 | Prestw-1508 | Ibutilide fumarate | 11684 | 94.70975665 | 9.18172275 |
| 1 | Prestw-926 | Idazoxan hydrochloride | 16095.5 | 38.44142164 | 1 |
| 2 | Prestw-926 | Idazoxan hydrochloride | 5466 | 11.3104597 | 1 |
| 1 | Prestw-1288 | Idebenone | 9719 | 31.51614242 | 29.5606413 |
| 2 | Prestw-1288 | Idebenone | 3401.5 | 1 | 1 |
| 2 | Prestw-18 | Idoxuridine | 5250.5 | 77.1543165 | 1 |
| 1 | Prestw-18 | Idoxuridine | 12950 | 120.4638069 | 1 |
| 2 | Prestw-311 | Ifenprodil tartrate | 4240.5 | 233.4322734 | 57.7917062 |
| 1 | Prestw-311 | Ifenprodil tartrate | 11979 | 33.6889459 | 1 |
| 1 | Prestw-1290 | Imatinib | 9381 | 27.25736268 | 25.30186156 |
| 2 | Prestw-1290 | Imatinib | 3529 | 1 | 1 |
| 1 | Prestw-1071 | Imidurea | 9152.5 | 54.68212681 | 19.43170837 |
| 2 | Prestw-1071 | Imidurea | 3616 | 5.7475232 | 1 |
| 1 | Prestw-519 | Imipenem | 12217.5 | 1 | 1 |
| 2 | Prestw-519 | Imipenem | 4360 | 90.2590349 | 1 |
| 2 | Prestw-72 | Imipramine hydrochloride | 6042.5 | 18.6535946 | 17.6535946 |
| 1 | Prestw-72 | Imipramine hydrochloride | 18534.5 | 32.71889585 | 1 |
| 1 | Prestw-1291 | Imiquimod | 10352 | 18.65949232 | 16.7039912 |
| 2 | Prestw-1291 | Imiquimod | 3585 | 1 | 1 |
| 2 | Prestw-220 | Indapamide | 5103 | 114.8678789 | 51.2754704 |
| 1 | Prestw-220 | Indapamide | 14026.5 | 167.7121875 | 7.67194328 |
| 2 | Prestw-1602 | Indatraline hydrochloride | 4204 | 72.7967997 | 56.1836157 |
| 1 | Prestw-1602 | Indatraline hydrochloride | 9916.5 | 82.24061549 | 14.42882164 |
| 2 | Prestw-1737 | Indinavir sulfate | 5169.5 | 90.8933147 | 18.9011742 |
| 1 | Prestw-1737 | Indinavir sulfate | 13875 | 169.5794085 | 25.71873717 |
| 1 | Prestw-272 | Indomethacin | 10934.5 | 1 | 1 |
| 2 | Prestw-272 | Indomethacin | 3893.5 | 1 | 1 |
| 1 | Prestw-836 | Indoprofen | 15902 | 25.08563256 | 1 |
| 2 | Prestw-836 | Indoprofen | 5606 | 1 | 1 |
| 1 | Prestw-490 | Iobenguane sulfate | 11623.5 | 29.5272685 | 4.02875113 |
| 2 | Prestw-490 | Iobenguane sulfate | 4108.5 | 142.12559 | 1 |
| 1 | Prestw-838 | Iocetamic acid | 15484 | 33.74272872 | 1 |
| 2 | Prestw-838 | Iocetamic acid | 5310 | 0.2578954 | 1 |
| 1 | Prestw-939 | Iodipamide | 16477 | 33.70381686 | 1 |
| 2 | Prestw-939 | Iodipamide | 5849.5 | 29.2483939 | 0.6765298 |
| 2 | Prestw-848 | Iodixanol | 5395 | 33.7154017 | 5.1435376 |
| 1 | Prestw-848 | Iodixanol | 15232.5 | 52.91015866 | 0.92745508 |
| 1 | Prestw-512 | Iohexol | 9672 | 1 | 1 |
| 2 | Prestw-512 | Iohexol | 3940.5 | 8.0219187 | 1 |
| 2 | Prestw-871 | Iopamidol | 5188 | 70.0485135 | 66.8399731 |
| 1 | Prestw-871 | Iopamidol | 17925 | 50.4557524 | 13.90173407 |
| 2 | Prestw-1052 | Iopanoic acid | 2880.5 | 52.4250105 | 1 |
| 1 | Prestw-1052 | Iopanoic acid | 9623 | 71.80409158 | 1 |
| 2 | Prestw-872 | Iopromide | 5927 | 56.0437759 | 52.8352355 |
| 1 | Prestw-872 | Iopromide | 17987.5 | 48.04173349 | 11.48771516 |
| 2 | Prestw-878 | Ioversol | 5641.5 | 1 | 1 |
| 1 | Prestw-878 | Ioversol | 17112 | 33.11212849 | 1 |
| 2 | Prestw-1062 | Ioxaglic acid | 3416 | 29.7832776 | 1 |

| | | | | | |
|---|---|---|---|---|---|
| 1 | Prestw-1062 | Ioxaglic acid | 10049.5 | 65.10849532 | 1 |
| 2 | Prestw-1773 | Ipriflavone | 4769 | 71.556148 | 1 |
| 1 | Prestw-1773 | Ipriflavone | 10280 | 104.765665 | 1 |
| 1 | Prestw-176 | Iproniazide phosphate | 10253 | 31.41288955 | 11.70702293 |
| 2 | Prestw-176 | Iproniazide phosphate | 4330.5 | 22.3486429 | 1 |
| 1 | Prestw-1292 | Ipsapirone | 4587.5 | 266.0129775 | 264.0574764 |
| 2 | Prestw-1292 | Ipsapirone | 1627.5 | 150.3720431 | 149.3720431 |
| 1 | Prestw-1736 | Irbesartan | 13627 | 167.206519 | 23.34584773 |
| 2 | Prestw-1736 | Irbesartan | 4786.5 | 49.1360874 | 1 |
| 1 | Prestw-1494 | Irinotecan hydrochloride trihydrate | 11909 | 159.099068 | 15.23839674 |
| 2 | Prestw-1494 | Irinotecan hydrochloride trihydrate | 4373 | 73.85656 | 1.8644195 |
| 1 | Prestw-1173 | Irsogladine maleate | 10286 | 43.40735864 | 8.1569402 |
| 2 | Prestw-1173 | Irsogladine maleate | 3297.5 | 1 | 1 |
| 2 | Prestw-795 | Isocarboxazid | 6279 | 32.6298675 | 31.6298675 |
| 1 | Prestw-795 | Isocarboxazid | 19292 | 44.27927192 | 0.15146316 |
| 2 | Prestw-127 | Isoconazole | 1564.5 | 487.8493279 | 486.8493279 |
| 1 | Prestw-127 | Isoconazole | 4665.5 | 471.376849 | 387.4613936 |
| 1 | Prestw-749 | Isoetharine mesylate salt | 15535 | 1 | 1 |
| 2 | Prestw-749 | Isoetharine mesylate salt | 5688.5 | 1 | 1 |
| 1 | Prestw-6 | Isoflupredone acetate | 12931.5 | 40.88837494 | 15.38985757 |
| 2 | Prestw-6 | Isoflupredone acetate | 4963 | 126.4263679 | 1 |
| 2 | Prestw-1023 | Isometheptene mucate | 3113 | 86.4337321 | 83.3280139 |
| 1 | Prestw-1023 | Isometheptene mucate | 8381.5 | 45.5593213 | 31.4044676 |
| 2 | Prestw-161 | Isoniazid | 4156.5 | 47.8214291 | 31.2082451 |
| 1 | Prestw-161 | Isoniazid | 10912 | 60.37997329 | 1 |
| 1 | Prestw-770 | Isopropamide iodide | 18159 | 41.08114851 | 9.19252043 |
| 2 | Prestw-770 | Isopropamide iodide | 5670.5 | 25.2196453 | 1 |
| 2 | Prestw-722 | Isopyrin hydrochloride | 5227.5 | 65.5429608 | 64.5429608 |
| 1 | Prestw-722 | Isopyrin hydrochloride | 18097.5 | 42.09417263 | 1 |
| 1 | Prestw-714 | Isosorbide dinitrate | 19473.5 | 63.0774192 | 29.04915093 |
| 2 | Prestw-714 | Isosorbide dinitrate | 6357 | 66.3720004 | 53.6153133 |
| 2 | Prestw-1294 | Isosorbide mononitrate | 1539.5 | 226.3047364 | 225.3047364 |
| 1 | Prestw-1294 | Isosorbide mononitrate | 4227 | 259.796522 | 257.8410209 |
| 1 | Prestw-256 | Isotretinoin | 13963 | 65.3428216 | 2.99288058 |
| 2 | Prestw-256 | Isotretinoin | 4666.5 | 1 | 1 |
| 1 | Prestw-44 | Isoxicam | 11461.5 | 22.58672329 | 9.73459679 |
| 2 | Prestw-44 | Isoxicam | 4010 | 68.4546913 | 1 |
| 1 | Prestw-68 | Isoxsuprine hydrochloride | 14933.5 | 45.30484128 | 13.4162132 |
| 2 | Prestw-68 | Isoxsuprine hydrochloride | 4986 | 52.576673 | 21.6885966 |
| 1 | Prestw-1021 | Isradipine | 8563 | 30.787007 | 16.6321533 |
| 2 | Prestw-1021 | Isradipine | 3187.5 | 1 | 1 |
| 1 | Prestw-1295 | Itopride | 9560.5 | 22.04129448 | 20.08579336 |
| 2 | Prestw-1295 | Itopride | 3314 | 1 | 1 |
| 2 | Prestw-1139 | Itraconazole | 3430.5 | 57.6232785 | 1 |
| 1 | Prestw-1139 | Itraconazole | 9475 | 80.5672148 | 1 |
| 1 | Prestw-156 | Ivermectin | 9629.5 | 1 | 1 |
| 2 | Prestw-156 | Ivermectin | 3871 | 1 | 1 |
| 1 | Prestw-154 | Josamycin | 10189 | 1 | 1 |
| 2 | Prestw-154 | Josamycin | 3778.5 | 1 | 1 |
| 1 | Prestw-394 | Kanamycin A sulfate | 14143.5 | 35.02629941 | 22.06824491 |
| 2 | Prestw-394 | Kanamycin A sulfate | 4553 | 165.5993535 | 4.7356972 |
| 1 | Prestw-392 | Ketanserin tartrate hydrate | 13600.5 | 39.90616845 | 26.94811395 |
| 2 | Prestw-392 | Ketanserin tartrate hydrate | 4794.5 | 113.8728545 | 1 |
| 2 | Prestw-389 | Ketoconazole | 5268.5 | 99.8713976 | 1 |
| 1 | Prestw-389 | Ketoconazole | 13671.5 | 37.46592813 | 0.03816393 |
| 2 | Prestw-219 | Ketoprofen | 4812.5 | 116.1615565 | 52.569148 |
| 1 | Prestw-219 | Ketoprofen | 14367 | 129.6731409 | 1 |
| 2 | Prestw-929 | Ketorolac tromethamine | 5004 | 66.5739339 | 22.492811 |
| 1 | Prestw-929 | Ketorolac tromethamine | 15245.5 | 43.77047185 | 3.5301109 |
| 1 | Prestw-371 | Ketotifen fumarate | 14554.5 | 17.68983336 | 9.66417532 |
| 2 | Prestw-371 | Ketotifen fumarate | 5094.5 | 124.2373308 | 1 |
| 1 | Prestw-91 | Khellin | 14896 | 1 | 1 |
| 2 | Prestw-91 | Khellin | 5311.5 | 1 | 1 |

| | | | | | |
|---|---|---|---|---|---|
| 2 | Prestw-277 | Labetalol hydrochloride | 3765 | 77.7437962 | 49.1955831 |
| 1 | Prestw-277 | Labetalol hydrochloride | 10112 | 6.0502993 | 5.0502993 |
| 1 | Prestw-1297 | Lacidipine | 10031.5 | 23.02558559 | 21.07008447 |
| 2 | Prestw-1297 | Lacidipine | 3347 | 1 | 1 |
| 2 | Prestw-1812 | Lacosamide | 4725.5 | 98.3716834 | 1 |
| 1 | Prestw-1812 | Lacosamide | 12564.5 | 180.6490463 | 1 |
| 1 | Prestw-1298 | Lamivudine | 8388.5 | 1 | 1 |
| 2 | Prestw-1298 | Lamivudine | 2900 | 1 | 1 |
| 2 | Prestw-1492 | Lamotrigine | 4661.5 | 102.3058802 | 30.3137397 |
| 1 | Prestw-1492 | Lamotrigine | 11310 | 193.4337382 | 49.5730669 |
| 1 | Prestw-656 | Lanatoside C | 14666.5 | 4.57398586 | 1 |
| 2 | Prestw-656 | Lanatoside C | 5031 | 1 | 1 |
| 1 | Prestw-1072 | Lansoprazole | 9071.5 | 53.92743125 | 18.67701281 |
| 2 | Prestw-1072 | Lansoprazole | 3304 | 1 | 1 |
| 1 | Prestw-1765 | Latanoprost | 10891 | 87.08253805 | 4.3355205 |
| 2 | Prestw-1765 | Latanoprost | 4773.5 | 44.1239976 | 1 |
| 1 | Prestw-772 | Leflunomide | 16631.5 | 53.02043017 | 21.13180209 |
| 2 | Prestw-772 | Leflunomide | 5097.5 | 45.0830086 | 14.1949322 |
| 1 | Prestw-1025 | Letrozole | 8664 | 23.136023 | 8.9811693 |
| 2 | Prestw-1025 | Letrozole | 3086.5 | 10.6728312 | 7.567113 |
| 1 | Prestw-1516 | Levalbuterol hydrochloride | 9925.5 | 1 | 1 |
| 2 | Prestw-1516 | Levalbuterol hydrochloride | 4154 | 1 | 1 |
| 1 | Prestw-182 | Levamisole hydrochloride | 11316.5 | 197.9421856 | 78.02942234 |
| 2 | Prestw-182 | Levamisole hydrochloride | 4943 | 39.8901295 | 1 |
| 2 | Prestw-1427 | Levetiracetam | 4969.5 | 1 | 1 |
| 1 | Prestw-1427 | Levetiracetam | 10719 | 28.33212063 | 1 |
| 1 | Prestw-1039 | Levocabastine hydrochloride | 9127 | 32.5808482 | 1 |
| 2 | Prestw-1039 | Levocabastine hydrochloride | 3387.5 | 80.2385059 | 1 |
| 1 | Prestw-17 | Levodopa | 11273 | 114.4155231 | 10.03321012 |
| 2 | Prestw-17 | Levodopa | 4392 | 91.177228 | 1 |
| 1 | Prestw-1829 | Levofloxacin | 12616.5 | 78.51177039 | 16.16182937 |
| 2 | Prestw-1829 | Levofloxacin | 4639.5 | 1 | 1 |
| 1 | Prestw-739 | Levonordefrin | 14845.5 | 32.91961839 | 18.27460945 |
| 2 | Prestw-739 | Levonordefrin | 5284.5 | 1 | 1 |
| 2 | Prestw-1054 | Levopropoxyphene napsylate | 3465.5 | 33.0156879 | 1 |
| 1 | Prestw-1054 | Levopropoxyphene napsylate | 9815.5 | 84.41197077 | 1 |
| 1 | Prestw-50 | Lidocaine hydrochloride | 10509.5 | 44.60423893 | 41.41733991 |
| 2 | Prestw-50 | Lidocaine hydrochloride | 3971.5 | 60.3703453 | 27.5250051 |
| 1 | Prestw-381 | Lidoflazine | 13872 | 41.89464971 | 4.46688551 |
| 2 | Prestw-381 | Lidoflazine | 4745 | 153.1386432 | 1 |
| 1 | Prestw-302 | Lincomycin hydrochloride | 11983 | 36.73485921 | 28.70920117 |
| 2 | Prestw-302 | Lincomycin hydrochloride | 4755 | 159.0203724 | 1.6251756 |
| 2 | Prestw-1429 | Linezolid | 4460.5 | 1 | 1 |
| 1 | Prestw-1429 | Linezolid | 10239.5 | 39.03720035 | 1 |
| 2 | Prestw-853 | Liothyronine | 5474.5 | 56.8745913 | 28.3027272 |
| 1 | Prestw-853 | Liothyronine | 17510.5 | 43.04151775 | 1 |
| 2 | Prestw-1140 | Liranaftate | 3589 | 1 | 1 |
| 1 | Prestw-1140 | Liranaftate | 10503 | 63.23305791 | 1 |
| 1 | Prestw-301 | Lisinopril | 13394.5 | 39.75370812 | 31.72805008 |
| 2 | Prestw-301 | Lisinopril | 4433.5 | 141.6162298 | 1 |
| 1 | Prestw-796 | Lithocholic acid | 20331.5 | 46.70131652 | 2.57350776 |
| 2 | Prestw-796 | Lithocholic acid | 6284.5 | 1 | 1 |
| 2 | Prestw-1746 | Lodoxamide | 4054.5 | 33.5450143 | 30.050753 |
| 1 | Prestw-1746 | Lodoxamide | 12774.5 | 97.93654369 | 12.40850979 |
| 1 | Prestw-1756 | Lofepramine | 10199 | 22.71906063 | 3.01319401 |
| 2 | Prestw-1756 | Lofepramine | 4253.5 | 1 | 1 |
| 2 | Prestw-1430 | Lofexidine | 4089.5 | 27.0122573 | 10.3990733 |
| 1 | Prestw-1430 | Lofexidine | 8782.5 | 64.40335457 | 1 |
| 2 | Prestw-238 | Lomefloxacin hydrochloride | 5371 | 27.4261783 | 1 |
| 1 | Prestw-238 | Lomefloxacin hydrochloride | 14917.5 | 139.1754144 | 1 |
| 1 | Prestw-1775 | Lomerizine hydrochloride | 11085 | 126.9673075 | 15.81843644 |
| 2 | Prestw-1775 | Lomerizine hydrochloride | 4594 | 77.9776166 | 1 |
| 2 | Prestw-144 | Loperamide hydrochloride | 4047 | 55.0451424 | 38.4319584 |

| | | | | | |
|---|---|---|---|---|---|
| 1 | Prestw-144 | Loperamide hydrochloride | 9440 | 98.71869217 | 30.90689832 |
| 1 | Prestw-1813 | Lopinavir | 12351 | 201.6687409 | 12.39255063 |
| 2 | Prestw-1813 | Lopinavir | 4613.5 | 56.8635544 | 1 |
| 2 | Prestw-1431 | Loracarbef | 4145.5 | 43.3999694 | 26.7867854 |
| 1 | Prestw-1431 | Loracarbef | 10126 | 73.62156692 | 5.80977307 |
| 1 | Prestw-1432 | Loratadine | 10046.5 | 77.3216175 | 9.50982365 |
| 2 | Prestw-1432 | Loratadine | 4234.5 | 20.0466562 | 3.4334722 |
| 1 | Prestw-915 | Lorglumide sodium salt | 15177 | 8.64594386 | 7.64594386 |
| 2 | Prestw-915 | Lorglumide sodium salt | 5511.5 | 1 | 1 |
| 1 | Prestw-1433 | Losartan | 9558 | 107.740133 | 39.92833919 |
| 2 | Prestw-1433 | Losartan | 4184.5 | 14.4181915 | 1 |
| 2 | Prestw-1741 | Loteprednol etabonate | 1276 | 1280.589623 | 1277.095361 |
| 1 | Prestw-1741 | Loteprednol etabonate | 2956.5 | 2021.593413 | 1936.065379 |
| 1 | Prestw-516 | Lovastatin | 9433 | 1 | 1 |
| 2 | Prestw-516 | Lovastatin | 3164 | 1 | 1 |
| 1 | Prestw-132 | Loxapine succinate | 8432.5 | 12.5282473 | 6.9794946 |
| 2 | Prestw-132 | Loxapine succinate | 3147 | 1 | 1 |
| 2 | Prestw-870 | Luteolin | 3228 | 1163.380558 | 1160.172018 |
| 1 | Prestw-870 | Luteolin | 6662.5 | 371.2293277 | 334.6753094 |
| 1 | Prestw-1000 | Lymecycline | 8524 | 1 | 1 |
| 2 | Prestw-1000 | Lymecycline | 2875 | 1 | 1 |
| 1 | Prestw-95 | Lynestrenol | 19536 | 76.06398496 | 39.50996663 |
| 2 | Prestw-95 | Lynestrenol | 6580.5 | 36.1030456 | 32.8945052 |
| 1 | Prestw-166 | Mafenide hydrochloride | 11636 | 120.0129466 | 15.63063368 |
| 2 | Prestw-166 | Mafenide hydrochloride | 4405 | 61.6820619 | 1 |
| 1 | Prestw-346 | Maprotiline hydrochloride | 10626.5 | 1 | 1 |
| 2 | Prestw-346 | Maprotiline hydrochloride | 3964 | 28.2268017 | 1 |
| 2 | Prestw-217 | Mebendazole | 5586.5 | 66.6295066 | 3.0370981 |
| 1 | Prestw-217 | Mebendazole | 13636.5 | 135.9839538 | 1 |
| 1 | Prestw-310 | Mebeverine hydrochloride | 11988.5 | 47.53885609 | 10.11109189 |
| 2 | Prestw-310 | Mebeverine hydrochloride | 4285.5 | 178.3710882 | 2.730521 |
| 1 | Prestw-455 | Mebhydroline 1,5-naphtalenedisulfonate | 11642 | 33.84275571 | 7.99021006 |
| 2 | Prestw-455 | Mebhydroline 1,5-naphtalenedisulfonate | 4280 | 104.4118109 | 12.2310192 |
| 1 | Prestw-1111 | Mecamylamine hydrochloride | 8741.5 | 19.1288094 | 4.9739557 |
| 2 | Prestw-1111 | Mecamylamine hydrochloride | 3195.5 | 7.2390604 | 4.1333422 |
| 2 | Prestw-456 | Meclocycline sulfosalicylate | 3817 | 112.3368809 | 20.1560892 |
| 1 | Prestw-456 | Meclocycline sulfosalicylate | 12067.5 | 18.761513 | 1 |
| 2 | Prestw-206 | Meclofenamic acid sodium salt monohydrate | 5516 | 67.0216182 | 1 |
| 1 | Prestw-206 | Meclofenamic acid sodium salt monohydrate | 13378.5 | 101.8104482 | 1 |
| 1 | Prestw-763 | Meclofenoxate hydrochloride | 13627.5 | 1 | 1 |
| 2 | Prestw-763 | Meclofenoxate hydrochloride | 4872 | 1 | 1 |
| 1 | Prestw-457 | Meclozine dihydrochloride | 11471 | 27.2025668 | 1.35002115 |
| 2 | Prestw-457 | Meclozine dihydrochloride | 3148 | 1 | 1 |
| 1 | Prestw-743 | Medrysone | 16841 | 31.1184373 | 16.47342836 |
| 2 | Prestw-743 | Medrysone | 6091 | 1 | 1 |
| 2 | Prestw-54 | Mefenamic acid | 5463.5 | 155.4650843 | 63.2842926 |
| 1 | Prestw-54 | Mefenamic acid | 15204 | 11.12616434 | 1 |
| 2 | Prestw-215 | Mefexamide hydrochloride | 5995.5 | 65.2495504 | 1.6571419 |
| 1 | Prestw-215 | Mefexamide hydrochloride | 14927 | 128.614996 | 1 |
| 1 | Prestw-126 | Mefloquine hydrochloride | 10375.5 | 87.14574505 | 5.20791029 |
| 2 | Prestw-126 | Mefloquine hydrochloride | 3556.5 | 37.6199374 | 1 |
| 1 | Prestw-956 | Megestrol acetate | 18159.5 | 68.61653632 | 32.06251799 |
| 2 | Prestw-956 | Megestrol acetate | 6554 | 51.32913 | 48.1205896 |
| 1 | Prestw-884 | Meglumine | 18793 | 58.09741085 | 10.34144293 |
| 2 | Prestw-884 | Meglumine | 6417.5 | 1 | 1 |
| 1 | Prestw-458 | Melatonin | 13138 | 34.81630747 | 8.96376182 |
| 2 | Prestw-458 | Melatonin | 4512.5 | 125.9645935 | 33.7838018 |
| 2 | Prestw-1435 | Melengestrol acetate | 4287.5 | 60.9749172 | 44.3617332 |
| 1 | Prestw-1435 | Melengestrol acetate | 9948.5 | 93.52636211 | 25.71456826 |
| 1 | Prestw-369 | Meloxicam | 11632.5 | 22.06465798 | 17.41958868 |
| 2 | Prestw-369 | Meloxicam | 4478 | 139.0849017 | 32.0761347 |
| 1 | Prestw-978 | Memantine Hydrochloride | 20024 | 37.70885979 | 8.45927927 |
| 2 | Prestw-978 | Memantine Hydrochloride | 6347 | 1 | 1 |

| | | | | | |
|---|---|---|---|---|---|
| 1 | Prestw-653 | Mepenzolate bromide | 15578.5 | 3.93281116 | 2.93281116 |
| 2 | Prestw-653 | Mepenzolate bromide | 5170.5 | 1 | 1 |
| 2 | Prestw-178 | Mephenesin | 4842.5 | 109.4108518 | 15.9523891 |
| 1 | Prestw-178 | Mephenesin | 11401 | 114.8018917 | 3.65302062 |
| 2 | Prestw-726 | Mephentermine hemisulfate | 5536.5 | 6.5029351 | 5.5029351 |
| 1 | Prestw-726 | Mephentermine hemisulfate | 16657.5 | 26.39974197 | 1 |
| 2 | Prestw-1108 | Mephenytoin | 2960.5 | 148.9942512 | 145.888533 |
| 1 | Prestw-1108 | Mephenytoin | 8292 | 41.1426523 | 26.9877986 |
| 1 | Prestw-1280 | Mepivacaine hydrochloride | 9823 | 87.52506095 | 3.60960553 |
| 2 | Prestw-1280 | Mepivacaine hydrochloride | 3562.5 | 1 | 1 |
| 2 | Prestw-1064 | Meprylcaine hydrochloride | 3550 | 20.6357968 | 1 |
| 1 | Prestw-1064 | Meprylcaine hydrochloride | 10724 | 75.09458856 | 1 |
| 2 | Prestw-1004 | Meptazinol hydrochloride | 2801 | 11.1196962 | 10.1196962 |
| 1 | Prestw-1004 | Meptazinol hydrochloride | 7852 | 1 | 1 |
| 2 | Prestw-787 | Merbromin | 6387.5 | 92.8007159 | 80.0440288 |
| 1 | Prestw-787 | Merbromin | 20459.5 | 46.82574743 | 12.79747916 |
| 2 | Prestw-1469 | Mercaptopurine | 4807.5 | 89.5214702 | 1 |
| 1 | Prestw-1469 | Mercaptopurine | 11831 | 131.9253213 | 0.25088646 |
| 2 | Prestw-1106 | Meropenem | 3060.5 | 75.3661598 | 72.2604416 |
| 1 | Prestw-1106 | Meropenem | 8242 | 37.6569548 | 23.5021011 |
| 1 | Prestw-1069 | Mesalamine | 8744.5 | 74.38208972 | 39.13167128 |
| 2 | Prestw-1069 | Mesalamine | 3500 | 1 | 1 |
| 1 | Prestw-1487 | Mesna | 11708.5 | 186.1747467 | 22.53895072 |
| 2 | Prestw-1487 | Mesna | 4383 | 114.5822029 | 1 |
| 2 | Prestw-529 | Mesoridazine besylate | 3837.5 | 129.1808723 | 30.2800578 |
| 1 | Prestw-529 | Mesoridazine besylate | 11028.5 | 0.0444154 | 1 |
| 2 | Prestw-505 | Metaproterenol sulfate, orciprenaline sulfate | 4030 | 97.7421223 | 64.8967821 |
| 1 | Prestw-505 | Metaproterenol sulfate, orciprenaline sulfate | 11792 | 17.99367485 | 14.80677583 |
| 1 | Prestw-197 | Metaraminol bitartrate | 10945 | 105.4901101 | 22.74309256 |
| 2 | Prestw-197 | Metaraminol bitartrate | 4717.5 | 44.0102284 | 1 |
| 1 | Prestw-364 | Metergoline | 7107.5 | 31.9406498 | 27.2955805 |
| 2 | Prestw-364 | Metergoline | 4086 | 347.3486728 | 240.3399058 |
| 2 | Prestw-4 | Metformin hydrochloride | 3800.5 | 161.5291848 | 98.1790092 |
| 1 | Prestw-4 | Metformin hydrochloride | 11339.5 | 36.80933121 | 1 |
| 2 | Prestw-759 | Methacholine chloride | 4296 | 46.5100814 | 25.9306799 |
| 1 | Prestw-759 | Methacholine chloride | 11615.5 | 29.14537247 | 1 |
| 2 | Prestw-964 | Methacycline hydrochloride | 6222 | 1 | 1 |
| 1 | Prestw-964 | Methacycline hydrochloride | 20973.5 | 48.23574117 | 0.47977325 |
| 1 | Prestw-1066 | Methantheline bromide | 9066 | 88.66592373 | 53.41550529 |
| 2 | Prestw-1066 | Methantheline bromide | 2876.5 | 36.6231901 | 21.1200625 |
| 1 | Prestw-342 | Methapyrilene hydrochloride | 10986 | 1 | 1 |
| 2 | Prestw-342 | Methapyrilene hydrochloride | 4307 | 1 | 1 |
| 2 | Prestw-802 | Methazolamide | 5989 | 1 | 1 |
| 1 | Prestw-802 | Methazolamide | 17893 | 47.20810794 | 1 |
| 2 | Prestw-1282 | Methenamine | 3679.5 | 1 | 1 |
| 1 | Prestw-1282 | Methenamine | 10559.5 | 67.10866086 | 1 |
| 1 | Prestw-1080 | Methiazole | 8385 | 1 | 1 |
| 2 | Prestw-1080 | Methiazole | 3244.5 | 1 | 1 |
| 2 | Prestw-1803 | Methicillin sodium | 4748.5 | 85.8185841 | 1 |
| 1 | Prestw-1803 | Methicillin sodium | 11559.5 | 162.5900464 | 1 |
| 1 | Prestw-786 | Methimazole | 20832 | 86.63265793 | 52.60438966 |
| 2 | Prestw-786 | Methimazole | 6808.5 | 72.0660512 | 59.3093641 |
| 1 | Prestw-375 | Methiothepin maleate | 14652.5 | 4.58315908 | 1 |
| 2 | Prestw-375 | Methiothepin maleate | 4997 | 117.7220194 | 1 |
| 1 | Prestw-184 | Methocarbamol | 12070 | 73.05106871 | 10.70112769 |
| 2 | Prestw-184 | Methocarbamol | 4453 | 1 | 1 |
| 1 | Prestw-797 | Methotrimeprazine maleat salt | 21030 | 58.11835478 | 13.99054602 |
| 2 | Prestw-797 | Methotrimeprazine maleat salt | 5838.5 | 1 | 1 |
| 1 | Prestw-535 | Methoxamine hydrochloride | 12350.5 | 16.30711845 | 3.45499195 |
| 2 | Prestw-535 | Methoxamine hydrochloride | 4742 | 120.8500284 | 17.3134513 |
| 1 | Prestw-705 | Methyl benzethonium chloride | 16687.5 | 58.91418405 | 24.88591578 |
| 2 | Prestw-705 | Methyl benzethonium chloride | 5701.5 | 46.4983686 | 33.7416815 |
| 2 | Prestw-900 | Methylatropine nitrate | 3011 | 917.0736953 | 916.0736953 |

| | | | | | |
|---|---|---|---|---|---|
| 1 | Prestw-900 | Methylatropine nitrate | 4183 | 305.5878917 | 294.6530102 |
| 2 | Prestw-326 | Methyldopa (L,-) | 3703 | 164.0744563 | 100.7242807 |
| 1 | Prestw-326 | Methyldopa (L,-) | 10616.5 | 26.8556778 | 1 |
| 1 | Prestw-1038 | Methyldopate hydrochloride | 9532 | 34.7662529 | 1 |
| 2 | Prestw-1038 | Methyldopate hydrochloride | 3286.5 | 44.8666389 | 1 |
| 1 | Prestw-374 | Methylergometrine maleate | 13743.5 | 9.25269762 | 1.22703958 |
| 2 | Prestw-374 | Methylergometrine maleate | 5434.5 | 137.1103236 | 1 |
| 1 | Prestw-864 | Methylhydantoin-5-(D) | 18672 | 95.47315457 | 44.06582706 |
| 2 | Prestw-864 | Methylhydantoin-5-(D) | 6247.5 | 66.2562144 | 46.6703827 |
| 2 | Prestw-860 | Methylhydantoin-5-(L) | 3563.5 | 826.8191264 | 807.2332947 |
| 1 | Prestw-860 | Methylhydantoin-5-(L) | 6742.5 | 383.6642012 | 332.2568737 |
| 2 | Prestw-279 | Methylprednisolone, 6-alpha | 4154 | 59.9865046 | 31.4382915 |
| 1 | Prestw-279 | Methylprednisolone, 6-alpha | 11035 | 1 | 1 |
| 1 | Prestw-11 | Meticrane | 8193.5 | 1 | 1 |
| 2 | Prestw-11 | Meticrane | 3074.5 | 1 | 1 |
| 1 | Prestw-491 | Metixene hydrochloride | 11038 | 33.617831 | 8.11931363 |
| 2 | Prestw-491 | Metixene hydrochloride | 4340 | 116.3879884 | 1 |
| 2 | Prestw-209 | Metoclopramide monohydrochloride | 5200.5 | 75.2751012 | 1 |
| 1 | Prestw-209 | Metoclopramide monohydrochloride | 13289 | 107.3090101 | 1 |
| 2 | Prestw-112 | Metolazone | 3195 | 60.7434766 | 12.5895114 |
| 1 | Prestw-112 | Metolazone | 9454.5 | 22.8257937 | 1 |
| 2 | Prestw-733 | Metoprolol-(+,-) (+)-tartrate salt | 6071.5 | 1 | 1 |
| 1 | Prestw-733 | Metoprolol-(+,-) (+)-tartrate salt | 19985 | 41.4043336 | 1 |
| 2 | Prestw-463 | Metrizamide | 4296.5 | 130.6316123 | 13.4136582 |
| 1 | Prestw-463 | Metrizamide | 12861 | 27.39503962 | 1 |
| 2 | Prestw-81 | Metronidazole | 5380.5 | 1 | 1 |
| 1 | Prestw-81 | Metronidazole | 18905 | 43.64057496 | 1 |
| 1 | Prestw-904 | Metyrapone | 16221.5 | 18.22318389 | 7.2883024 |
| 2 | Prestw-904 | Metyrapone | 5776.5 | 1 | 1 |
| 1 | Prestw-1441 | Mevastatin | 9177.5 | 141.2703782 | 73.45858439 |
| 2 | Prestw-1441 | Mevastatin | 3882 | 64.9200139 | 48.3068299 |
| 2 | Prestw-241 | Mexiletine hydrochloride | 5784.5 | 51.0299997 | 1 |
| 1 | Prestw-241 | Mexiletine hydrochloride | 13615 | 152.5860985 | 1 |
| 2 | Prestw-99 | Mianserine hydrochloride | 6049.5 | 4.546714 | 3.546714 |
| 1 | Prestw-99 | Mianserine hydrochloride | 17974.5 | 9.68675194 | 1 |
| 1 | Prestw-67 | Miconazole | 15917 | 12.86313238 | 1 |
| 2 | Prestw-67 | Miconazole | 4758.5 | 1 | 1 |
| 1 | Prestw-191 | Midodrine hydrochloride | 10269 | 26.39409655 | 6.68822993 |
| 2 | Prestw-191 | Midodrine hydrochloride | 4120.5 | 1 | 1 |
| 1 | Prestw-299 | Mifepristone | 12151.5 | 23.93068343 | 15.90502539 |
| 2 | Prestw-299 | Mifepristone | 4506 | 129.2773086 | 1 |
| 2 | Prestw-1169 | Miglitol | 3474 | 1 | 1 |
| 1 | Prestw-1169 | Miglitol | 10305.5 | 92.84732669 | 1 |
| 2 | Prestw-1788 | Milnacipran hydrochloride | 4866.5 | 148.1952792 | 84.6028707 |
| 1 | Prestw-1788 | Milnacipran hydrochloride | 13452 | 123.5931957 | 1 |
| 2 | Prestw-1065 | Milrinone | 2421 | 74.3048571 | 58.8017295 |
| 1 | Prestw-1065 | Milrinone | 9132.5 | 91.05916895 | 55.80875051 |
| 1 | Prestw-66 | Minaprine dihydrochloride | 15027.5 | 23.91741689 | 1 |
| 2 | Prestw-66 | Minaprine dihydrochloride | 4573.5 | 1 | 1 |
| 1 | Prestw-315 | Minocycline hydrochloride | 11339.5 | 31.5785214 | 18.6204669 |
| 2 | Prestw-315 | Minocycline hydrochloride | 4135.5 | 94.2839836 | 1 |
| 1 | Prestw-20 | Minoxidil | 11196 | 87.50465091 | 4.75763336 |
| 2 | Prestw-20 | Minoxidil | 4194.5 | 105.4916445 | 1 |
| 2 | Prestw-1798 | Mirabegron | 5194 | 110.2943579 | 6.8128157 |
| 1 | Prestw-1798 | Mirabegron | 12357 | 120.9992382 | 1 |
| 2 | Prestw-1144 | Mirtazapine | 3515 | 36.7684419 | 1 |
| 1 | Prestw-1144 | Mirtazapine | 10656 | 63.54768534 | 1 |
| 2 | Prestw-1443 | Misoprostol | 4625.5 | 33.3629533 | 1 |
| 1 | Prestw-1443 | Misoprostol | 11717 | 69.16944628 | 1 |
| 2 | Prestw-1444 | Mitotane | 4004.5 | 123.0985914 | 29.4632467 |
| 1 | Prestw-1444 | Mitotane | 10541.5 | 94.2909963 | 1 |
| 2 | Prestw-385 | Mitoxantrone dihydrochloride | 4443.5 | 232.3800717 | 56.7395045 |
| 1 | Prestw-385 | Mitoxantrone dihydrochloride | 13854.5 | 42.52189252 | 5.09412832 |

| | | | | | |
|---|---|---|---|---|---|
| 1 | Prestw-1506 | Mizolastine | 10696.5 | 124.5733712 | 39.04533725 |
| 2 | Prestw-1506 | Mizolastine | 4481 | 19.513299 | 16.0190377 |
| 1 | Prestw-353 | Moclobemide | 12446.5 | 22.29044501 | 8.82399546 |
| 2 | Prestw-353 | Moclobemide | 4522.5 | 103.0211715 | 1 |
| 2 | Prestw-1147 | Modafinil | 3601.5 | 47.5887223 | 1 |
| 1 | Prestw-1147 | Modafinil | 10284 | 63.98783231 | 1 |
| 1 | Prestw-1035 | Molindone hydrochloride | 8639 | 41.6819421 | 18.5378478 |
| 2 | Prestw-1035 | Molindone hydrochloride | 3368.5 | 41.710658 | 1 |
| 2 | Prestw-547 | Molsidomine | 4563.5 | 160.2412964 | 68.0605047 |
| 1 | Prestw-547 | Molsidomine | 13952 | 24.22032289 | 1 |
| 1 | Prestw-572 | Mometasone furoate | 13775.5 | 43.51139271 | 16.83431777 |
| 2 | Prestw-572 | Mometasone furoate | 4904 | 132.1940796 | 1 |
| 1 | Prestw-748 | Monensin sodium salt | 13956.5 | 2.824897 | 1.824897 |
| 2 | Prestw-748 | Monensin sodium salt | 5306.5 | 1 | 1 |
| 1 | Prestw-912 | Monobenzone | 15134.5 | 1.35930876 | 0.35930876 |
| 2 | Prestw-912 | Monobenzone | 5152 | 1 | 1 |
| 2 | Prestw-1189 | Montelukast | 3153.5 | 14.6789804 | 13.6789804 |
| 1 | Prestw-1189 | Montelukast | 7895.5 | 1 | 1 |
| 1 | Prestw-61 | Morantel tartrate | 12876 | 14.36547234 | 1 |
| 2 | Prestw-61 | Morantel tartrate | 4717 | 122.450297 | 1 |
| 2 | Prestw-1051 | Moricizine hydrochloride | 3205.5 | 99.7188279 | 38.8447346 |
| 1 | Prestw-1051 | Moricizine hydrochloride | 9607 | 96.77523335 | 1 |
| 1 | Prestw-84 | Moroxidine hydrochloride | 15334 | 48.74379043 | 8.50342948 |
| 2 | Prestw-84 | Moroxidine hydrochloride | 5435 | 22.3563032 | 1 |
| 1 | Prestw-819 | Moxalactam disodium salt | 16922.5 | 25.24961524 | 10.6046063 |
| 2 | Prestw-819 | Moxalactam disodium salt | 6436 | 1 | 1 |
| 2 | Prestw-1446 | Moxifloxacin | 4161.5 | 106.4269251 | 12.7915804 |
| 1 | Prestw-1446 | Moxifloxacin | 10239 | 103.5208842 | 1 |
| 1 | Prestw-42 | Moxisylyte hydrochoride | 10021.5 | 1 | 1 |
| 2 | Prestw-42 | Moxisylyte hydrochoride | 3796.5 | 2.5186265 | 1 |
| 2 | Prestw-1016 | Moxonidine | 2850 | 65.4339736 | 5.8406285 |
| 1 | Prestw-1016 | Moxonidine | 7610.5 | 1 | 1 |
| 2 | Prestw-1795 | Mupirocin | 5089 | 63.6071983 | 1 |
| 1 | Prestw-1795 | Mupirocin | 11754 | 144.7121858 | 1 |
| 1 | Prestw-644 | N-Acetyl-DL-homocysteine Thiolactone | 10707.5 | 11.52200929 | 10.52200929 |
| 2 | Prestw-644 | N-Acetyl-DL-homocysteine Thiolactone | 4604.5 | 1 | 1 |
| 2 | Prestw-896 | N-Acetyl-L-leucine | 5455 | 50.510489 | 49.510489 |
| 1 | Prestw-896 | N-Acetyl-L-leucine | 15638.5 | 17.08428515 | 1 |
| 2 | Prestw-338 | N6-methyladenosine | 3940.5 | 48.5911983 | 47.5911983 |
| 1 | Prestw-338 | N6-methyladenosine | 10766 | 1 | 1 |
| 1 | Prestw-909 | Nabumetone | 14873.5 | 17.15544873 | 6.22056724 |
| 2 | Prestw-909 | Nabumetone | 5956.5 | 1 | 1 |
| 1 | Prestw-1120 | Nadifloxacin | 9207.5 | 29.0637988 | 5.9197045 |
| 2 | Prestw-1120 | Nadifloxacin | 3407.5 | 41.4697791 | 1 |
| 1 | Prestw-818 | Nadolol | 17635 | 25.58474063 | 10.93973169 |
| 2 | Prestw-818 | Nadolol | 5812 | 1 | 1 |
| 1 | Prestw-843 | Nafcillin sodium salt monohydrate | 15831 | 37.38752878 | 1 |
| 2 | Prestw-843 | Nafcillin sodium salt monohydrate | 4121.5 | 1 | 1 |
| 1 | Prestw-377 | Nafronyl oxalate | 13186.5 | 19.73260678 | 11.70694874 |
| 2 | Prestw-377 | Nafronyl oxalate | 4757.5 | 119.975862 | 1 |
| 2 | Prestw-1063 | Naftifine hydrochloride | 3145.5 | 10.5673745 | 1 |
| 1 | Prestw-1063 | Naftifine hydrochloride | 9355 | 93.65391744 | 1 |
| 2 | Prestw-975 | Naftopidil dihydrochloride | 6269 | 4.8692442 | 3.8692442 |
| 1 | Prestw-975 | Naftopidil dihydrochloride | 18787 | 27.78550624 | 1 |
| 1 | Prestw-118 | Nalbuphine hydrochloride | 9733 | 42.88500919 | 7.63459075 |
| 2 | Prestw-118 | Nalbuphine hydrochloride | 3507.5 | 1 | 1 |
| 2 | Prestw-1447 | Nalidixic acid sodium salt | 4183.5 | 97.3524435 | 3.7170988 |
| 1 | Prestw-1447 | Nalidixic acid sodium salt | 10719 | 107.4316325 | 3.0493196 |
| 1 | Prestw-1490 | Nalmefene hydrochloride | 13670.5 | 169.2576009 | 5.62180497 |
| 2 | Prestw-1490 | Nalmefene hydrochloride | 4695 | 106.2530853 | 1 |
| 1 | Prestw-111 | Naloxone hydrochloride | 8365.5 | 30.150511 | 15.9956573 |
| 2 | Prestw-111 | Naloxone hydrochloride | 3185.5 | 20.112654 | 17.0069358 |
| 2 | Prestw-116 | Naltrexone hydrochloride dihydrate | 3473.5 | 11.6982727 | 1 |

| | | | | | |
|---|---|---|---|---|---|
| 1 | Prestw-116 | Naltrexone hydrochloride dihydrate | 9690 | 88.23240064 | 1 |
| 2 | Prestw-982 | Nandrolone | 5910.5 | 8.8603823 | 7.8603823 |
| 1 | Prestw-982 | Nandrolone | 16428 | 20.98818829 | 1 |
| 1 | Prestw-46 | Naphazoline hydrochloride | 12983.5 | 39.65674636 | 13.80420071 |
| 2 | Prestw-46 | Naphazoline hydrochloride | 4892.5 | 143.1890391 | 51.0082474 |
| 2 | Prestw-45 | Naproxen | 4139 | 108.2689226 | 16.0881309 |
| 1 | Prestw-45 | Naproxen | 12614.5 | 8.15554536 | 1 |
| 1 | Prestw-1748 | Nateglinide | 12210 | 114.4451959 | 28.91716202 |
| 2 | Prestw-1748 | Nateglinide | 4537.5 | 13.2681018 | 9.7738405 |
| 2 | Prestw-1814 | Nebivolol hydrochloride | 4869.5 | 169.8002052 | 57.3463671 |
| 1 | Prestw-1814 | Nebivolol hydrochloride | 12803 | 168.0250864 | 1 |
| 2 | Prestw-1152 | Nefazodone hydrochloride | 3338.5 | 31.4696275 | 1 |
| 1 | Prestw-1152 | Nefazodone hydrochloride | 9944 | 76.17376401 | 1 |
| 2 | Prestw-229 | Nefopam hydrochloride | 5161.5 | 121.437625 | 17.9560828 |
| 1 | Prestw-229 | Nefopam hydrochloride | 13387.5 | 124.3092496 | 1 |
| 2 | Prestw-1804 | Nelarabine | 4487 | 136.2635884 | 23.8097503 |
| 1 | Prestw-1804 | Nelarabine | 11914 | 157.6346644 | 1 |
| 1 | Prestw-1764 | Nelfinavir mesylate | 10309 | 93.80952493 | 11.06250738 |
| 2 | Prestw-1764 | Nelfinavir mesylate | 4303.5 | 40.8442568 | 1 |
| 1 | Prestw-158 | Neomycin sulfate | 10485 | 1 | 1 |
| 2 | Prestw-158 | Neomycin sulfate | 4069 | 1 | 1 |
| 1 | Prestw-468 | Neostigmine bromide | 12115.5 | 22.4387659 | 1 |
| 2 | Prestw-468 | Neostigmine bromide | 4103.5 | 76.8576553 | 1 |
| 2 | Prestw-1739 | Nevirapine | 2870 | 319.4198279 | 247.4276874 |
| 1 | Prestw-1739 | Nevirapine | 8693 | 426.5408895 | 282.6802182 |
| 1 | Prestw-881 | Niacin | 18029.5 | 52.4160812 | 4.66011328 |
| 2 | Prestw-881 | Niacin | 5889.5 | 1 | 1 |
| 1 | Prestw-1099 | Nialamide | 8102.5 | 8.8100515 | 7.8100515 |
| 2 | Prestw-1099 | Nialamide | 3202 | 19.7540705 | 1 |
| 1 | Prestw-383 | Nicardipine hydrochloride | 12902.5 | 59.83782811 | 22.41006391 |
| 2 | Prestw-383 | Nicardipine hydrochloride | 4738.5 | 139.1856719 | 1 |
| 1 | Prestw-147 | Nicergoline | 12507 | 143.8868623 | 12.21242749 |
| 2 | Prestw-147 | Nicergoline | 4900.5 | 112.4965942 | 1 |
| 2 | Prestw-40 | Niclosamide | 4224 | 124.185982 | 60.8358064 |
| 1 | Prestw-40 | Niclosamide | 12726.5 | 31.91913848 | 1 |
| 2 | Prestw-106 | Nicorandil | 3344 | 37.671128 | 1 |
| 1 | Prestw-106 | Nicorandil | 8855.5 | 61.3633215 | 1 |
| 1 | Prestw-1449 | Nicotinamide | 10362 | 118.7848365 | 14.40252356 |
| 2 | Prestw-1449 | Nicotinamide | 4405.5 | 75.3638254 | 1 |
| 2 | Prestw-63 | Nifedipine | 4359.5 | 75.5535053 | 42.7081651 |
| 1 | Prestw-63 | Nifedipine | 12506.5 | 10.82787024 | 7.64097122 |
| 2 | Prestw-1774 | Nifekalant | 4657 | 94.8894386 | 1.4309759 |
| 1 | Prestw-1774 | Nifekalant | 11353 | 105.146477 | 1 |
| 2 | Prestw-225 | Nifenazone | 5940 | 98.9858238 | 1 |
| 1 | Prestw-225 | Nifenazone | 15148.5 | 115.8148937 | 1 |
| 2 | Prestw-255 | Niflumic acid | 4466 | 18.8647959 | 17.8647959 |
| 1 | Prestw-255 | Niflumic acid | 12478 | 85.06500819 | 22.71506717 |
| 2 | Prestw-555 | Nifuroxazide | 4798 | 144.17256 | 26.9546059 |
| 1 | Prestw-555 | Nifuroxazide | 13333 | 25.6699801 | 1 |
| 2 | Prestw-1024 | Nifurtimox | 2976 | 69.6283202 | 66.522602 |
| 1 | Prestw-1024 | Nifurtimox | 8100 | 27.0345861 | 12.8797324 |
| 1 | Prestw-928 | Nilutamide | 14791 | 36.52842709 | 1 |
| 2 | Prestw-928 | Nilutamide | 5482.5 | 1 | 1 |
| 2 | Prestw-1154 | Nilvadipine | 3356 | 30.3708137 | 1 |
| 1 | Prestw-1154 | Nilvadipine | 10100 | 73.7350427 | 1 |
| 1 | Prestw-194 | Nimesulide | 10532.5 | 23.79737937 | 4.09151275 |
| 2 | Prestw-194 | Nimesulide | 4236 | 4.8484205 | 1 |
| 1 | Prestw-918 | Nimodipine | 14355 | 1 | 1 |
| 2 | Prestw-918 | Nimodipine | 5411 | 1 | 1 |
| 1 | Prestw-469 | Niridazole | 12415.5 | 35.28008189 | 1.36338665 |
| 2 | Prestw-469 | Niridazole | 4342.5 | 111.6441546 | 1 |
| 1 | Prestw-666 | Nisoldipine | 15171 | 24.87577346 | 1 |
| 2 | Prestw-666 | Nisoldipine | 5035 | 1 | 1 |

| | | | | | |
|---|---|---|---|---|---|
| 1 | Prestw-910 | Nisoxetine hydrochloride | 14550 | 1 | 1 |
| 2 | Prestw-910 | Nisoxetine hydrochloride | 5298 | 1 | 1 |
| 2 | Prestw-1733 | Nitazoxanide | 4781.5 | 95.6934389 | 23.7012984 |
| 1 | Prestw-1733 | Nitazoxanide | 13906.5 | 156.2556564 | 12.39498515 |
| 2 | Prestw-916 | Nitrendipine | 3148.5 | 442.3872421 | 441.3872421 |
| 1 | Prestw-916 | Nitrendipine | 5344 | 1 | 1 |
| 2 | Prestw-981 | Nitrocaramiphen hydrochloride | 5962 | 1 | 1 |
| 1 | Prestw-981 | Nitrocaramiphen hydrochloride | 18647 | 29.27308238 | 0.02350186 |
| 1 | Prestw-492 | Nitrofural | 11074 | 41.4294508 | 15.93093343 |
| 2 | Prestw-492 | Nitrofural | 3782.5 | 149.8768111 | 1 |
| 1 | Prestw-168 | Nitrofurantoin | 11819.5 | 112.8371472 | 8.45483429 |
| 2 | Prestw-168 | Nitrofurantoin | 4837.5 | 71.7414262 | 1 |
| 1 | Prestw-921 | Nizatidine | 16999 | 36.57733604 | 1 |
| 2 | Prestw-921 | Nizatidine | 5690.5 | 1 | 1 |
| 1 | Prestw-100 | Nocodazole | 7401.5 | 1 | 1 |
| 2 | Prestw-100 | Nocodazole | 3059 | 1 | 1 |
| 1 | Prestw-1033 | Nomegestrol acetate | 8788.5 | 42.5727669 | 19.4286726 |
| 2 | Prestw-1033 | Nomegestrol acetate | 3082 | 69.4980808 | 21.3441156 |
| 1 | Prestw-108 | Nomifensine maleate | 7719.5 | 1 | 1 |
| 2 | Prestw-108 | Nomifensine maleate | 3298 | 1 | 1 |
| 1 | Prestw-513 | Norcyclobenzaprine | 10851.5 | 1 | 1 |
| 2 | Prestw-513 | Norcyclobenzaprine | 3971 | 34.2891445 | 0.6987043 |
| 1 | Prestw-253 | Norethindrone | 11913.5 | 83.65386726 | 21.30392624 |
| 2 | Prestw-253 | Norethindrone | 5369 | 1 | 1 |
| 2 | Prestw-24 | Norethynodrel | 5482.5 | 66.6597795 | 1 |
| 1 | Prestw-24 | Norethynodrel | 13772 | 176.9760972 | 1 |
| 2 | Prestw-221 | Norfloxacin | 4978.5 | 42.0166094 | 1 |
| 1 | Prestw-221 | Norfloxacin | 12258.5 | 148.9855962 | 1 |
| 2 | Prestw-1452 | Norgestimate | 4111.5 | 174.195771 | 80.5604263 |
| 1 | Prestw-1452 | Norgestimate | 10345 | 139.703248 | 35.32093503 |
| 1 | Prestw-773 | Norgestrel-(-)-D | 16677.5 | 47.42990647 | 15.54127839 |
| 2 | Prestw-773 | Norgestrel-(-)-D | 5563.5 | 28.7696207 | 1 |
| 2 | Prestw-254 | Nortriptyline hydrochloride | 4534 | 15.5631777 | 14.5631777 |
| 1 | Prestw-254 | Nortriptyline hydrochloride | 13113.5 | 70.13103759 | 7.78109657 |
| 1 | Prestw-834 | Novobiocin sodium salt | 14819.5 | 20.54610283 | 1 |
| 2 | Prestw-834 | Novobiocin sodium salt | 5572.5 | 1 | 1 |
| 2 | Prestw-1454 | Nylidrin | 4284 | 98.9762003 | 5.3408556 |
| 1 | Prestw-1454 | Nylidrin | 11593.5 | 105.1996302 | 0.81731726 |
| 1 | Prestw-517 | Nystatine | 10644.5 | 1 | 1 |
| 2 | Prestw-517 | Nystatine | 3930.5 | 1 | 1 |
| 2 | Prestw-237 | Ofloxacin | 5431.5 | 35.5041039 | 1 |
| 1 | Prestw-237 | Ofloxacin | 13045 | 178.874755 | 1 |
| 2 | Prestw-1455 | Olanzapine | 4462 | 130.2662937 | 36.630949 |
| 1 | Prestw-1455 | Olanzapine | 11126 | 113.074218 | 8.69190504 |
| 2 | Prestw-1190 | Olmesartan | 3308 | 25.3683976 | 24.3683976 |
| 1 | Prestw-1190 | Olmesartan | 7638 | 1 | 1 |
| 1 | Prestw-1307 | Olopatadine hydrochloride | 7805.5 | 1 | 1 |
| 2 | Prestw-1307 | Olopatadine hydrochloride | 2712 | 1 | 1 |
| 1 | Prestw-493 | Omeprazole | 11692 | 35.2138505 | 9.71533313 |
| 2 | Prestw-493 | Omeprazole | 4172 | 163.8235657 | 9.657399 |
| 2 | Prestw-1058 | Ondansetron Hydrochloride | 3009 | 1 | 1 |
| 1 | Prestw-1058 | Ondansetron Hydrochloride | 9604 | 60.14151025 | 1 |
| 1 | Prestw-1456 | Opipramol dihydrochloride | 10832 | 136.7893338 | 32.40702083 |
| 2 | Prestw-1456 | Opipramol dihydrochloride | 4441.5 | 86.4811949 | 1 |
| 2 | Prestw-164 | Ornidazole | 4729.5 | 100.9555378 | 7.3201931 |
| 1 | Prestw-164 | Ornidazole | 12729 | 93.78348197 | 1 |
| 2 | Prestw-239 | Orphenadrine hydrochloride | 5202 | 59.1180517 | 1 |
| 1 | Prestw-239 | Orphenadrine hydrochloride | 14110 | 123.9416165 | 1 |
| 1 | Prestw-1475 | Oxacillin sodium | 10837.5 | 184.8653616 | 67.3388728 |
| 2 | Prestw-1475 | Oxacillin sodium | 4856 | 100.841886 | 14.13977 |
| 1 | Prestw-826 | Oxalamine citrate salt | 15476 | 0.80456373 | 1 |
| 2 | Prestw-826 | Oxalamine citrate salt | 5494 | 1 | 1 |
| 2 | Prestw-1192 | Oxandrolone | 2969 | 76.5242806 | 75.5242806 |

| | | | | | |
|---|---|---|---|---|---|
| 1 | Prestw-1192 | Oxandrolone | 8044 | 1 | 1 |
| 2 | Prestw-398 | Oxantel pamoate | 4349 | 398.0144091 | 237.1507528 |
| 1 | Prestw-398 | Oxantel pamoate | 15032 | 1 | 1 |
| 2 | Prestw-1060 | Oxaprozin | 3332.5 | 1 | 1 |
| 1 | Prestw-1060 | Oxaprozin | 9434 | 77.01618939 | 1 |
| 1 | Prestw-1156 | Oxcarbazepine | 3772.5 | 1029.179317 | 923.1103846 |
| 2 | Prestw-1156 | Oxcarbazepine | 1341 | 389.2011669 | 328.3270736 |
| 1 | Prestw-58 | Oxethazaine | 13763 | 32.20348168 | 5.52640674 |
| 2 | Prestw-58 | Oxethazaine | 3436.5 | 6.6824098 | 1 |
| 2 | Prestw-1459 | Oxfendazol | 4716.5 | 95.7710683 | 1 |
| 1 | Prestw-1459 | Oxfendazol | 11510.5 | 115.6670357 | 1 |
| 1 | Prestw-1460 | Oxibendazol | 11489 | 134.5830747 | 2.90863992 |
| 2 | Prestw-1460 | Oxibendazol | 4592 | 118.5258964 | 1 |
| 1 | Prestw-1125 | Oxiconazole Nitrate | 11617.5 | 17.15652708 | 1 |
| 2 | Prestw-1125 | Oxiconazole Nitrate | 3990.5 | 1 | 1 |
| 1 | Prestw-193 | Oxolinic acid | 9871 | 34.45747695 | 14.75161033 |
| 2 | Prestw-193 | Oxolinic acid | 4105 | 6.3366349 | 1 |
| 2 | Prestw-1057 | Oxprenolol hydrochloride | 3342 | 27.5739458 | 1 |
| 1 | Prestw-1057 | Oxprenolol hydrochloride | 9870.5 | 57.13931295 | 1 |
| 2 | Prestw-887 | Oxybenzone | 2894.5 | 1087.176344 | 1086.176344 |
| 1 | Prestw-887 | Oxybenzone | 4589.5 | 422.9958203 | 393.7462398 |
| 1 | Prestw-287 | Oxybutynin chloride | 11834 | 18.87518062 | 5.40873107 |
| 2 | Prestw-287 | Oxybutynin chloride | 3982.5 | 101.3492752 | 1 |
| 2 | Prestw-224 | Oxymetazoline hydrochloride | 5724 | 96.8505342 | 1 |
| 1 | Prestw-224 | Oxymetazoline hydrochloride | 13884.5 | 118.0383894 | 1 |
| 2 | Prestw-1780 | Oxymetholone | 4859.5 | 123.2481397 | 29.789677 |
| 1 | Prestw-1780 | Oxymetholone | 12432.5 | 122.0911616 | 10.94229051 |
| 2 | Prestw-1049 | Oxyphenbutazone | 3117.5 | 98.0372671 | 37.1631738 |
| 1 | Prestw-1049 | Oxyphenbutazone | 9710 | 78.30743656 | 1 |
| 1 | Prestw-307 | Oxytetracycline dihydrate | 12295 | 13.57696385 | 1 |
| 2 | Prestw-307 | Oxytetracycline dihydrate | 4209.5 | 122.8298431 | 1 |
| 1 | Prestw-979 | Ozagrel hydrochloride | 19886 | 20.93524921 | 1 |
| 2 | Prestw-979 | Ozagrel hydrochloride | 6028 | 1 | 1 |
| 1 | Prestw-155 | Paclitaxel | 10165 | 1 | 1 |
| 2 | Prestw-155 | Paclitaxel | 4335 | 1 | 1 |
| 1 | Prestw-1827 | Paliperidone | 10950.5 | 94.10933034 | 31.75938932 |
| 2 | Prestw-1827 | Paliperidone | 4467.5 | 6.1545271 | 5.1545271 |
| 2 | Prestw-1783 | Palonosetron hydrochloride | 4704.5 | 93.2843971 | 29.6919886 |
| 1 | Prestw-1783 | Palonosetron hydrochloride | 12123 | 138.2142782 | 1 |
| 2 | Prestw-1034 | Pancuronium bromide | 3058 | 72.7363121 | 24.5823469 |
| 1 | Prestw-1034 | Pancuronium bromide | 8816.5 | 32.2489073 | 9.104813 |
| 2 | Prestw-22 | Panthenol (D) | 5000.5 | 86.3557715 | 22.763363 |
| 1 | Prestw-22 | Panthenol (D) | 13408 | 149.398633 | 1 |
| 1 | Prestw-1758 | Pantoprazole sodium | 2173 | 338.774401 | 319.0685344 |
| 2 | Prestw-1758 | Pantoprazole sodium | 1274 | 298.7187268 | 233.3458742 |
| 1 | Prestw-583 | Papaverine hydrochloride | 12785 | 38.31061803 | 11.63354309 |
| 2 | Prestw-583 | Papaverine hydrochloride | 4799.5 | 152.7080998 | 1 |
| 2 | Prestw-1110 | Parbendazole | 3180 | 122.9895271 | 119.8838089 |
| 1 | Prestw-1110 | Parbendazole | 8615.5 | 23.6055213 | 9.4506676 |
| 1 | Prestw-183 | Pargyline hydrochloride | 12128 | 72.17447969 | 9.82453867 |
| 2 | Prestw-183 | Pargyline hydrochloride | 4490.5 | 1 | 1 |
| 1 | Prestw-867 | Paromomycin sulfate | 17839 | 44.75313263 | 8.1991143 |
| 2 | Prestw-867 | Paromomycin sulfate | 5944.5 | 3.4528644 | 0.244324 |
| 2 | Prestw-851 | Paroxetine Hydrochloride | 5747.5 | 44.6212073 | 16.0493432 |
| 1 | Prestw-851 | Paroxetine Hydrochloride | 17701 | 46.25896392 | 1 |
| 1 | Prestw-1303 | Pefloxacine | 8105 | 1 | 1 |
| 2 | Prestw-1303 | Pefloxacine | 3044 | 1 | 1 |
| 2 | Prestw-1794 | Pemetrexed disodium | 5068 | 126.9752904 | 23.4937482 |
| 1 | Prestw-1794 | Pemetrexed disodium | 11919.5 | 183.7373539 | 10.49483233 |
| 1 | Prestw-1603 | Pemirolast potassium | 9930 | 86.96554246 | 19.15374861 |
| 2 | Prestw-1603 | Pemirolast potassium | 4110.5 | 1 | 1 |
| 2 | Prestw-683 | Pempidine | 4941 | 66.2759955 | 35.3879191 |
| 1 | Prestw-683 | Pempidine | 16095 | 41.24247682 | 9.35384874 |

| | | | | | |
|---|---|---|---|---|---|
| 1 | Prestw-1043 | Penbutolol sulfate | 9272.5 | 34.2754486 | 1 |
| 2 | Prestw-1043 | Penbutolol sulfate | 3473 | 20.7540003 | 1 |
| 1 | Prestw-1488 | Penciclovir | 11585 | 200.7641082 | 37.12831224 |
| 2 | Prestw-1488 | Penciclovir | 5018.5 | 83.2844936 | 1 |
| 1 | Prestw-553 | Pentamidine isethionate | 14670 | 30.54832668 | 1 |
| 2 | Prestw-553 | Pentamidine isethionate | 5281.5 | 101.1197324 | 1 |
| 2 | Prestw-941 | Pentetic acid | 6751.5 | 75.6336946 | 56.0478629 |
| 1 | Prestw-941 | Pentetic acid | 20566.5 | 59.92083035 | 8.51350284 |
| 2 | Prestw-1491 | Pentobarbital | 5110 | 144.8106737 | 1 |
| 1 | Prestw-1491 | Pentobarbital | 13546 | 147.682872 | 1 |
| 1 | Prestw-188 | Pentolinium bitartrate | 10214 | 48.09166002 | 28.3857934 |
| 2 | Prestw-188 | Pentolinium bitartrate | 4023 | 1 | 1 |
| 1 | Prestw-196 | Pentoxifylline | 9691 | 23.90518178 | 4.19931516 |
| 2 | Prestw-196 | Pentoxifylline | 4175 | 1 | 1 |
| 2 | Prestw-162 | Pentylenetetrazole | 4079 | 54.2423999 | 37.6292159 |
| 1 | Prestw-162 | Pentylenetetrazole | 10063.5 | 68.32979545 | 0.5180016 |
| 2 | Prestw-405 | Pepstatin A | 4481 | 91.1772617 | 27.8270861 |
| 1 | Prestw-405 | Pepstatin A | 12315.5 | 38.32259037 | 1 |
| 1 | Prestw-295 | Pergolide mesylate | 12776.5 | 35.29303274 | 30.64796344 |
| 2 | Prestw-295 | Pergolide mesylate | 4345 | 133.8929608 | 26.8841938 |
| 1 | Prestw-286 | Perhexiline maleate | 11260 | 25.70728604 | 12.24083649 |
| 2 | Prestw-286 | Perhexiline maleate | 3880.5 | 47.8970015 | 1 |
| 2 | Prestw-1101 | Perindopril | 2891 | 76.4523216 | 16.8589765 |
| 1 | Prestw-1101 | Perindopril | 8368.5 | 1 | 1 |
| 1 | Prestw-1770 | Perospirone | 11197.5 | 94.31566164 | 11.56864409 |
| 2 | Prestw-1770 | Perospirone | 4180.5 | 80.7612695 | 1 |
| 2 | Prestw-125 | Perphenazine | 3446 | 107.1906404 | 85.0344628 |
| 1 | Prestw-125 | Perphenazine | 9772.5 | 45.55373128 | 1 |
| 1 | Prestw-533 | Phenacetin | 13015.5 | 16.51038897 | 3.65826247 |
| 2 | Prestw-533 | Phenacetin | 4582.5 | 105.3040205 | 1.7674434 |
| 1 | Prestw-752 | Phenazopyridine hydrochloride | 14249.5 | 1.25253925 | 0.25253925 |
| 2 | Prestw-752 | Phenazopyridine hydrochloride | 5202.5 | 1 | 1 |
| 2 | Prestw-170 | Phenelzine sulfate | 4637 | 93.0130631 | 1 |
| 1 | Prestw-170 | Phenelzine sulfate | 11839 | 96.98821136 | 1 |
| 1 | Prestw-723 | Phenethicillin potassium salt | 18959.5 | 59.35151099 | 15.22370223 |
| 2 | Prestw-723 | Phenethicillin potassium salt | 5643 | 1 | 1 |
| 2 | Prestw-179 | Phenformin hydrochloride | 5142 | 82.195004 | 18.6025955 |
| 1 | Prestw-179 | Phenformin hydrochloride | 13092.5 | 137.9132545 | 1 |
| 1 | Prestw-538 | Phenindione | 11867 | 20.18867026 | 7.33654376 |
| 2 | Prestw-538 | Phenindione | 4249 | 127.1453044 | 23.6087273 |
| 1 | Prestw-59 | Pheniramine maleate | 13852.5 | 39.2094782 | 13.71096083 |
| 2 | Prestw-59 | Pheniramine maleate | 4880.5 | 135.6698366 | 1 |
| 2 | Prestw-1785 | Phenothiazine | 4707.5 | 133.6825224 | 70.0901139 |
| 1 | Prestw-1785 | Phenothiazine | 11121.5 | 164.9906208 | 4.9503766 |
| 2 | Prestw-944 | Phenoxybenzamine hydrochloride | 3728 | 777.6317477 | 758.045916 |
| 1 | Prestw-944 | Phenoxybenzamine hydrochloride | 9566 | 229.3511885 | 177.943861 |
| 1 | Prestw-1776 | Phenprobamate | 11140 | 123.2175148 | 12.06864376 |
| 2 | Prestw-1776 | Phenprobamate | 4358.5 | 90.0378987 | 1 |
| 2 | Prestw-1061 | Phensuximide | 3294.5 | 25.0377735 | 1 |
| 1 | Prestw-1061 | Phensuximide | 9882.5 | 61.5692054 | 1 |
| 2 | Prestw-1308 | Phentermine hydrochloride | 2980.5 | 5.9092714 | 4.9092714 |
| 1 | Prestw-1308 | Phentermine hydrochloride | 7723.5 | 1 | 1 |
| 2 | Prestw-230 | Phentolamine hydrochloride | 5054.5 | 90.8069017 | 1 |
| 1 | Prestw-230 | Phentolamine hydrochloride | 13244.5 | 119.5967616 | 1 |
| 2 | Prestw-1310 | Phenylbutazone | 2811.5 | 50.3932712 | 49.3932712 |
| 1 | Prestw-1310 | Phenylbutazone | 7586 | 1 | 1 |
| 2 | Prestw-324 | Phenylpropanolamine hydrochloride | 2115.5 | 259.3387471 | 195.9885715 |
| 1 | Prestw-324 | Phenylpropanolamine hydrochloride | 10338.5 | 20.876832 | 1 |
| 2 | Prestw-869 | Phthalylsulfathiazole | 5359.5 | 52.0865287 | 48.8779883 |
| 1 | Prestw-869 | Phthalylsulfathiazole | 17723 | 56.60220385 | 20.04818552 |
| 1 | Prestw-119 | Picotamide monohydrate | 7114 | 1 | 1 |
| 2 | Prestw-119 | Picotamide monohydrate | 3057 | 1 | 1 |
| 1 | Prestw-652 | Picrotoxinin | 15413.5 | 6.60326861 | 5.60326861 |

| | | | | | |
|---|---|---|---|---|---|
| 2 | Prestw-652 | Picrotoxinin | 5711.5 | 1 | 1 |
| 2 | Prestw-1792 | Pidotimod | 5384.5 | 115.2558667 | 51.6634582 |
| 1 | Prestw-1792 | Pidotimod | 13982 | 131.237014 | 1 |
| 1 | Prestw-449 | Pilocarpine nitrate | 12463.5 | 22.94027631 | 10.08814981 |
| 2 | Prestw-449 | Pilocarpine nitrate | 4430.5 | 122.9540611 | 19.417484 |
| 1 | Prestw-294 | Pimethixene maleate | 11451.5 | 31.94347922 | 27.29840992 |
| 2 | Prestw-294 | Pimethixene maleate | 4260 | 179.0711026 | 72.0623356 |
| 1 | Prestw-308 | Pimozide | 11198 | 15.2119149 | 1 |
| 2 | Prestw-308 | Pimozide | 4321 | 130.2823157 | 1 |
| 2 | Prestw-246 | Pinacidil | 5170.5 | 56.4276076 | 1 |
| 1 | Prestw-246 | Pinacidil | 14218.5 | 98.7217987 | 1 |
| 2 | Prestw-1601 | Pinaverium bromide | 4023.5 | 44.4589894 | 27.8458054 |
| 1 | Prestw-1601 | Pinaverium bromide | 9376 | 102.2054898 | 34.39369592 |
| 2 | Prestw-90 | Pindolol | 5961 | 1 | 1 |
| 1 | Prestw-90 | Pindolol | 16003.5 | 11.41053886 | 0.47565737 |
| 1 | Prestw-1314 | Pioglitazone | 8181 | 1 | 1 |
| 2 | Prestw-1314 | Pioglitazone | 2966.5 | 3.2703481 | 2.2703481 |
| 1 | Prestw-897 | Pipemidic acid | 16977 | 31.03652507 | 1.78694455 |
| 2 | Prestw-897 | Pipemidic acid | 5949 | 0.5561614 | 1 |
| 1 | Prestw-760 | Pipenzolate bromide | 16006.5 | 5.20071126 | 1 |
| 2 | Prestw-760 | Pipenzolate bromide | 5495.5 | 1 | 1 |
| 2 | Prestw-1048 | Piperacetazine | 3262.5 | 85.8064161 | 24.9323228 |
| 1 | Prestw-1048 | Piperacetazine | 8888.5 | 73.1447335 | 1 |
| 1 | Prestw-755 | Piperacillin sodium salt | 15854.5 | 1 | 1 |
| 2 | Prestw-755 | Piperacillin sodium salt | 5330 | 1 | 1 |
| 2 | Prestw-1055 | Piperidolate hydrochloride | 3309.5 | 52.3301412 | 1 |
| 1 | Prestw-1055 | Piperidolate hydrochloride | 9880 | 69.64601813 | 1 |
| 2 | Prestw-537 | Piracetam | 4298 | 144.4647202 | 40.9281431 |
| 1 | Prestw-537 | Piracetam | 11811 | 11.2932041 | 1 |
| 2 | Prestw-531 | Pirenperone | 4485.5 | 141.8930123 | 38.3564352 |
| 1 | Prestw-531 | Pirenperone | 13060.5 | 7.29269382 | 1 |
| 1 | Prestw-129 | Pirenzepine dihydrochloride | 9233 | 34.00411521 | 32.04861409 |
| 2 | Prestw-129 | Pirenzepine dihydrochloride | 3730.5 | 1 | 1 |
| 2 | Prestw-1047 | Piretanide | 3262.5 | 1 | 1 |
| 1 | Prestw-1047 | Piretanide | 9334.5 | 76.14684712 | 1 |
| 1 | Prestw-980 | Piribedil hydrochloride | 17841.5 | 39.31097906 | 10.06139854 |
| 2 | Prestw-980 | Piribedil hydrochloride | 5665.5 | 1 | 1 |
| 2 | Prestw-973 | Pirlindole mesylate | 5442.5 | 17.7471867 | 16.7471867 |
| 1 | Prestw-973 | Pirlindole mesylate | 17814.5 | 51.58808768 | 3.83211976 |
| 2 | Prestw-805 | Piromidic acid | 6037.5 | 1 | 1 |
| 1 | Prestw-805 | Piromidic acid | 19919.5 | 41.90055897 | 1 |
| 2 | Prestw-211 | Piroxicam | 5165 | 90.4084455 | 1 |
| 1 | Prestw-211 | Piroxicam | 12373.5 | 103.6328449 | 1 |
| 1 | Prestw-1815 | Pitavastatin calcium | 11486 | 175.5868315 | 55.67406818 |
| 2 | Prestw-1815 | Pitavastatin calcium | 4358 | 80.8196964 | 13.3256346 |
| 1 | Prestw-1009 | Pivampicillin | 7718.5 | 1 | 1 |
| 2 | Prestw-1009 | Pivampicillin | 2888 | 32.1004412 | 1 |
| 2 | Prestw-1053 | Pivmecillinam hydrochloride | 3442 | 5.1137709 | 1 |
| 1 | Prestw-1053 | Pivmecillinam hydrochloride | 9237 | 78.0488557 | 1 |
| 2 | Prestw-992 | Pizotifen malate | 3011 | 1216.181495 | 1215.181495 |
| 1 | Prestw-992 | Pizotifen malate | 5750.5 | 212.2104626 | 201.2755811 |
| 1 | Prestw-782 | Podophyllotoxin | 16595 | 63.47113306 | 18.09174988 |
| 2 | Prestw-782 | Podophyllotoxin | 5800.5 | 31.9007199 | 1 |
| 1 | Prestw-1826 | Posaconazole | 11372 | 83.37641708 | 21.02647606 |
| 2 | Prestw-1826 | Posaconazole | 4169 | 1 | 1 |
| 2 | Prestw-332 | Practolol | 4073.5 | 199.8256863 | 136.4755107 |
| 1 | Prestw-332 | Practolol | 12946 | 29.93597262 | 1 |
| 1 | Prestw-943 | Pralidoxime chloride | 19828.5 | 72.46203682 | 21.05470931 |
| 2 | Prestw-943 | Pralidoxime chloride | 6283 | 42.0629414 | 22.4771097 |
| 1 | Prestw-1316 | Pramipexole dihydrochloride | 8880 | 1 | 1 |
| 2 | Prestw-1316 | Pramipexole dihydrochloride | 2970.5 | 1 | 1 |
| 2 | Prestw-716 | Pramoxine hydrochloride | 5422 | 41.1206507 | 40.1206507 |
| 1 | Prestw-716 | Pramoxine hydrochloride | 16217.5 | 22.2690139 | 1 |

| | | | | | |
|---|---|---|---|---|---|
| 1 | Prestw-1317 | Pranlukast | 8669 | 28.1236051 | 22.5748524 |
| 2 | Prestw-1317 | Pranlukast | 3148 | 1 | 1 |
| 1 | Prestw-1318 | Pranoprofen | 8869 | 8.6014816 | 3.0527289 |
| 2 | Prestw-1318 | Pranoprofen | 3243 | 1 | 1 |
| 1 | Prestw-1319 | Pravastatin | 8515 | 10.8107001 | 5.2619474 |
| 2 | Prestw-1319 | Pravastatin | 2841.5 | 1 | 1 |
| 2 | Prestw-260 | Praziquantel | 4556.5 | 29.6438378 | 28.6438378 |
| 1 | Prestw-260 | Praziquantel | 11895.5 | 64.87582418 | 2.52588316 |
| 2 | Prestw-947 | Prazosin hydrochloride | 5450.5 | 94.2606002 | 74.6747685 |
| 1 | Prestw-947 | Prazosin hydrochloride | 17914.5 | 42.56958448 | 1 |
| 1 | Prestw-1044 | Prednicarbate | 9800 | 35.31728171 | 1 |
| 2 | Prestw-1044 | Prednicarbate | 3354 | 52.5493781 | 1 |
| 1 | Prestw-274 | Prednisolone | 10485 | 1 | 1 |
| 2 | Prestw-274 | Prednisolone | 3840 | 17.5431742 | 1 |
| 1 | Prestw-77 | Prednisone | 16690.5 | 51.76211353 | 19.87348545 |
| 2 | Prestw-77 | Prednisone | 5685 | 56.4710863 | 25.5830099 |
| 2 | Prestw-1791 | Pregabalin | 5637 | 68.6273282 | 5.0349197 |
| 1 | Prestw-1791 | Pregabalin | 13620 | 127.370701 | 1 |
| 2 | Prestw-546 | Pregnenolone | 4417 | 128.4225675 | 36.2417758 |
| 1 | Prestw-546 | Pregnenolone | 13269 | 20.53241743 | 1 |
| 1 | Prestw-560 | Prenylamine lactate | 13325.5 | 50.98449165 | 24.30741671 |
| 2 | Prestw-560 | Prenylamine lactate | 4187 | 173.6283199 | 1 |
| 2 | Prestw-799 | Pridinol methanesulfonate salt | 6155.5 | 7.1325659 | 6.1325659 |
| 1 | Prestw-799 | Pridinol methanesulfonate salt | 19236.5 | 48.9464774 | 4.81866864 |
| 1 | Prestw-199 | Prilocaine hydrochloride | 11030.5 | 99.65165136 | 16.90463381 |
| 2 | Prestw-199 | Prilocaine hydrochloride | 4228.5 | 88.5941966 | 1 |
| 2 | Prestw-476 | Primaquine diphosphate | 3454.5 | 1147.396273 | 1030.178319 |
| 1 | Prestw-476 | Primaquine diphosphate | 5212 | 160.7100183 | 126.7933231 |
| 2 | Prestw-933 | Primidone | 5866 | 32.9619671 | 4.390103 |
| 1 | Prestw-933 | Primidone | 17499 | 52.85048223 | 0.86777865 |
| 1 | Prestw-778 | Proadifen hydrochloride | 18688 | 88.08566048 | 42.7062773 |
| 2 | Prestw-778 | Proadifen hydrochloride | 5893 | 52.0055724 | 1 |
| 1 | Prestw-542 | Probenecid | 15044.5 | 17.00339137 | 1 |
| 2 | Prestw-542 | Probenecid | 5182.5 | 78.2147133 | 1 |
| 1 | Prestw-384 | Probucol | 14177.5 | 35.49248348 | 1 |
| 2 | Prestw-384 | Probucol | 5136 | 120.7641987 | 1 |
| 2 | Prestw-337 | Procainamide hydrochloride | 3851 | 7.0139842 | 6.0139842 |
| 1 | Prestw-337 | Procainamide hydrochloride | 10711.5 | 1 | 1 |
| 1 | Prestw-41 | Procaine hydrochloride | 11041 | 42.65520901 | 2.52646487 |
| 2 | Prestw-41 | Procaine hydrochloride | 3523.5 | 1 | 1 |
| 1 | Prestw-1112 | Procarbazine hydrochloride | 9574 | 18.3857459 | 1 |
| 2 | Prestw-1112 | Procarbazine hydrochloride | 3171 | 3.5871831 | 1 |
| 2 | Prestw-399 | Prochlorperazine dimaleate | 4235 | 254.9953877 | 94.1317314 |
| 1 | Prestw-399 | Prochlorperazine dimaleate | 13008 | 7.1065213 | 1 |
| 1 | Prestw-844 | Procyclidine hydrochloride | 15972 | 59.8398694 | 19.59950845 |
| 2 | Prestw-844 | Procyclidine hydrochloride | 5275.5 | 38.8966676 | 1 |
| 1 | Prestw-477 | Progesterone | 14866.5 | 27.35778059 | 1 |
| 2 | Prestw-477 | Progesterone | 5284.5 | 107.5011822 | 1 |
| 2 | Prestw-240 | Proglumide | 4928.5 | 65.7435736 | 1 |
| 1 | Prestw-240 | Proglumide | 13384.5 | 173.434081 | 1 |
| 1 | Prestw-999 | Proguanil hydrochloride | 15291.5 | 1 | 1 |
| 2 | Prestw-999 | Proguanil hydrochloride | 5928 | 1 | 1 |
| 1 | Prestw-693 | Promazine hydrochloride | 16469.5 | 44.8186076 | 12.92997952 |
| 2 | Prestw-693 | Promazine hydrochloride | 5494 | 55.2890838 | 24.4010074 |
| 1 | Prestw-888 | Promethazine hydrochloride | 16012 | 41.6252662 | 12.37568568 |
| 2 | Prestw-888 | Promethazine hydrochloride | 5661 | 1 | 1 |
| 1 | Prestw-974 | Pronethalol hydrochloride | 17730.5 | 35.3573135 | 6.10773298 |
| 2 | Prestw-974 | Pronethalol hydrochloride | 5514 | 3.2203517 | 2.2203517 |
| 1 | Prestw-499 | Propafenone hydrochloride | 11478 | 50.74680149 | 47.55990247 |
| 2 | Prestw-499 | Propafenone hydrochloride | 4002 | 112.8907586 | 80.0454184 |
| 1 | Prestw-827 | Propantheline bromide | 6469 | 59.3935718 | 58.3935718 |
| 2 | Prestw-827 | Propantheline bromide | 3120 | 360.0709618 | 359.0709618 |
| 1 | Prestw-959 | Proparacaine hydrochloride | 18316.5 | 57.22523578 | 20.67121745 |

| | | | | | |
|---|---|---|---|---|---|
| 2 | Prestw-959 | Proparacaine hydrochloride | 5857 | 29.0601462 | 25.8516058 |
| 2 | Prestw-792 | Propidium iodide | 6111 | 51.5341489 | 38.7774618 |
| 1 | Prestw-792 | Propidium iodide | 18049.5 | 35.63624721 | 1.60797894 |
| 2 | Prestw-931 | Propofol | 5955 | 47.7869727 | 19.2151086 |
| 1 | Prestw-931 | Propofol | 16450 | 66.23875138 | 14.2560478 |
| 2 | Prestw-1059 | Propoxycaine hydrochloride | 3251.5 | 9.4216255 | 1 |
| 1 | Prestw-1059 | Propoxycaine hydrochloride | 8960 | 66.8296871 | 1 |
| 1 | Prestw-494 | Propylthiouracil | 11617.5 | 41.12595281 | 15.62743544 |
| 2 | Prestw-494 | Propylthiouracil | 4186 | 142.0287468 | 1 |
| 1 | Prestw-1321 | Prothionamide | 8272 | 4.8463095 | 1 |
| 2 | Prestw-1321 | Prothionamide | 3222.5 | 8.8914817 | 1 |
| 2 | Prestw-930 | Protriptyline hydrochloride | 6399 | 76.2577817 | 47.6859176 |
| 1 | Prestw-930 | Protriptyline hydrochloride | 18870 | 57.72120136 | 5.73849778 |
| 2 | Prestw-212 | Pyrantel tartrate | 4521 | 76.766888 | 1 |
| 1 | Prestw-212 | Pyrantel tartrate | 12182.5 | 108.1981772 | 1 |
| 1 | Prestw-514 | Pyrazinamide | 10932 | 1 | 1 |
| 2 | Prestw-514 | Pyrazinamide | 3567.5 | 1 | 1 |
| 1 | Prestw-1322 | Pyridostigmine iodide | 8585.5 | 12.0295799 | 6.4808272 |
| 2 | Prestw-1322 | Pyridostigmine iodide | 3040.5 | 38.6825626 | 1 |
| 2 | Prestw-623 | Pyridoxine hydrochloride | 4521 | 60.9172173 | 28.0718771 |
| 1 | Prestw-623 | Pyridoxine hydrochloride | 12196 | 8.88726106 | 5.70036204 |
| 1 | Prestw-289 | Pyrilamine maleate | 11279 | 21.15960615 | 16.51453685 |
| 2 | Prestw-289 | Pyrilamine maleate | 4346 | 84.0392905 | 1 |
| 2 | Prestw-37 | Pyrimethamine | 4404.5 | 157.2809219 | 50.2721549 |
| 1 | Prestw-37 | Pyrimethamine | 11931 | 5.5218181 | 0.8767488 |
| 2 | Prestw-803 | Pyrithyldione | 6004.5 | 1 | 1 |
| 1 | Prestw-803 | Pyrithyldione | 16508.5 | 43.6945395 | 1 |
| 1 | Prestw-1040 | Pyrvinium pamoate | 8648.5 | 47.504934 | 1 |
| 2 | Prestw-1040 | Pyrvinium pamoate | 3389 | 71.5658317 | 1 |
| 1 | Prestw-1323 | Quetiapine hemifumarate | 7965.5 | 11.1071491 | 5.5583964 |
| 2 | Prestw-1323 | Quetiapine hemifumarate | 3043.5 | 12.6523999 | 1 |
| 1 | Prestw-318 | Quinacrine dihydrochloride hydrate | 11317.5 | 38.0325078 | 25.0744533 |
| 2 | Prestw-318 | Quinacrine dihydrochloride hydrate | 4054 | 153.1588518 | 1 |
| 1 | Prestw-927 | Quinapril hydrochloride | 15369.5 | 44.10001307 | 3.85965212 |
| 2 | Prestw-927 | Quinapril hydrochloride | 5041 | 1 | 1 |
| 2 | Prestw-1050 | Quinethazone | 3328.5 | 40.1508763 | 1 |
| 1 | Prestw-1050 | Quinethazone | 9102.5 | 80.56575545 | 1 |
| 1 | Prestw-280 | Quinidine hydrochloride monohydrate | 11691.5 | 7.542586 | 1 |
| 2 | Prestw-280 | Quinidine hydrochloride monohydrate | 4181 | 63.5990561 | 1 |
| 1 | Prestw-101 | R(-) Apomorphine hydrochloride hemihydrate | 7761.5 | 1 | 1 |
| 2 | Prestw-101 | R(-) Apomorphine hydrochloride hemihydrate | 2909.5 | 1 | 1 |
| 1 | Prestw-1495 | Rabeprazole Sodium salt | 11195.5 | 174.9263739 | 31.06570257 |
| 2 | Prestw-1495 | Rabeprazole Sodium salt | 4681.5 | 65.7144082 | 1 |
| 1 | Prestw-626 | Racecadotril | 12609 | 11.03549457 | 7.84859555 |
| 2 | Prestw-626 | Racecadotril | 3030.5 | 1 | 1 |
| 2 | Prestw-1480 | Racepinephrine hydrochloride | 5625 | 130.5737566 | 43.8716406 |
| 1 | Prestw-1480 | Racepinephrine hydrochloride | 14445 | 125.5592039 | 8.03271507 |
| 1 | Prestw-1325 | Raclopride | 8693.5 | 26.4614428 | 20.9126901 |
| 2 | Prestw-1325 | Raclopride | 3008 | 31.9389697 | 1 |
| 2 | Prestw-862 | Raloxifene hydrochloride | 5744 | 77.8980212 | 58.3121895 |
| 1 | Prestw-862 | Raloxifene hydrochloride | 19130 | 73.94396335 | 22.53663584 |
| 2 | Prestw-1793 | Raltitrexed | 5323 | 77.2586706 | 1 |
| 1 | Prestw-1793 | Raltitrexed | 12053 | 123.3057966 | 1 |
| 2 | Prestw-1107 | Ramipril | 3017.5 | 29.8068752 | 26.701157 |
| 1 | Prestw-1107 | Ramipril | 9161 | 33.30231698 | 19.14746328 |
| 2 | Prestw-201 | Ranitidine hydrochloride | 3979 | 182.4323709 | 43.3035436 |
| 1 | Prestw-201 | Ranitidine hydrochloride | 10580.5 | 115.8223485 | 33.07533091 |
| 1 | Prestw-1092 | Ranolazine | 8031 | 8.3250673 | 7.3250673 |
| 2 | Prestw-1092 | Ranolazine | 2946 | 60.8115825 | 1.2182374 |
| 2 | Prestw-1734 | Rasagiline | 2297.5 | 473.9889767 | 401.9968362 |
| 1 | Prestw-1734 | Rasagiline | 5141 | 791.8096208 | 647.9489495 |
| 2 | Prestw-1166 | Rebamipide | 3190.5 | 1 | 1 |
| 1 | Prestw-1166 | Rebamipide | 10709 | 68.35851729 | 1 |

| | | | | | |
|---|---|---|---|---|---|
| 1 | Prestw-1328 | Reboxetine mesylate | 8433 | 11.3424217 | 5.793669 |
| 2 | Prestw-1328 | Reboxetine mesylate | 2970 | 1 | 1 |
| 2 | Prestw-971 | Remoxipride Hydrochloride | 6268.5 | 1 | 1 |
| 1 | Prestw-971 | Remoxipride Hydrochloride | 19821.5 | 46.62855475 | 1 |
| 1 | Prestw-1046 | Repaglinide | 10974 | 11.64062171 | 1 |
| 2 | Prestw-1046 | Repaglinide | 3672 | 1 | 1 |
| 2 | Prestw-875 | Reserpine | 5752 | 1 | 1 |
| 1 | Prestw-875 | Reserpine | 17781.5 | 43.01595825 | 1 |
| 2 | Prestw-257 | Retinoic acid | 5014 | 14.0975286 | 13.0975286 |
| 1 | Prestw-257 | Retinoic acid | 13136 | 68.15534918 | 5.80540816 |
| 1 | Prestw-993 | Ribavirin | 18630 | 4.91671491 | 1 |
| 2 | Prestw-993 | Ribavirin | 6226 | 1 | 1 |
| 1 | Prestw-758 | Ribostamycin sulfate salt | 15039 | 10.10356907 | 1 |
| 2 | Prestw-758 | Ribostamycin sulfate salt | 4686.5 | 1 | 1 |
| 1 | Prestw-1109 | Rifabutin | 8423 | 21.1152055 | 6.9603518 |
| 2 | Prestw-1109 | Rifabutin | 3109 | 9.3136676 | 6.2079494 |
| 1 | Prestw-525 | Rifampicin | 11451 | 2.87491198 | 1 |
| 2 | Prestw-525 | Rifampicin | 4032 | 93.9304506 | 1 |
| 2 | Prestw-1157 | Rifapentine | 3018 | 74.6234154 | 13.7493221 |
| 1 | Prestw-1157 | Rifapentine | 8782 | 99.1143378 | 1 |
| 2 | Prestw-1478 | Rifaximin | 4877.5 | 143.8974099 | 57.1952939 |
| 1 | Prestw-1478 | Rifaximin | 13225 | 132.1007054 | 14.57421655 |
| 1 | Prestw-167 | Riluzole hydrochloride | 10985.5 | 126.6205249 | 22.23821199 |
| 2 | Prestw-167 | Riluzole hydrochloride | 4538 | 73.691792 | 1 |
| 1 | Prestw-1331 | Rimantadine Hydrochloride | 10047.5 | 24.80425524 | 7.24205304 |
| 2 | Prestw-1331 | Rimantadine Hydrochloride | 3537.5 | 3.2576397 | 0.1836945 |
| 2 | Prestw-1020 | Rimexolone | 3100 | 58.1419521 | 55.0362339 |
| 1 | Prestw-1020 | Rimexolone | 8701.5 | 1 | 1 |
| 1 | Prestw-1738 | Risedronic acid monohydrate | 13404.5 | 147.0737856 | 3.21311426 |
| 2 | Prestw-1738 | Risedronic acid monohydrate | 4896.5 | 29.944743 | 1 |
| 1 | Prestw-1029 | Risperidone | 8571 | 35.4042666 | 12.2601723 |
| 2 | Prestw-1029 | Risperidone | 2990.5 | 43.606984 | 1 |
| 1 | Prestw-349 | Ritodrine hydrochloride | 9861.5 | 13.0918987 | 12.0918987 |
| 2 | Prestw-349 | Ritodrine hydrochloride | 3539 | 37.3562692 | 8.8080561 |
| 2 | Prestw-1782 | Ritonavir | 4900 | 100.4549997 | 36.8625912 |
| 1 | Prestw-1782 | Ritonavir | 12031.5 | 102.0642195 | 1 |
| 1 | Prestw-1334 | Rivastigmine | 8876.5 | 33.4064988 | 15.8442966 |
| 2 | Prestw-1334 | Rivastigmine | 3174.5 | 1 | 1 |
| 1 | Prestw-1761 | Rizatriptan benzoate | 9514 | 46.66806813 | 26.96220151 |
| 2 | Prestw-1761 | Rizatriptan benzoate | 4006.5 | 20.9022313 | 1 |
| 2 | Prestw-1336 | Rofecoxib | 3310 | 24.6918201 | 21.6178749 |
| 1 | Prestw-1336 | Rofecoxib | 9208.5 | 25.7114811 | 8.1492789 |
| 1 | Prestw-1115 | Ronidazole | 9502.5 | 26.3117214 | 3.1676271 |
| 2 | Prestw-1115 | Ronidazole | 3404 | 1 | 1 |
| 2 | Prestw-1158 | Ropinirole hydrochloride | 3373 | 1 | 1 |
| 1 | Prestw-1158 | Ropinirole hydrochloride | 10074 | 81.87644972 | 1 |
| 1 | Prestw-1816 | Ropivacaine hydrochloride | 12037.5 | 138.7115662 | 18.7988029 |
| 2 | Prestw-1816 | Ropivacaine hydrochloride | 4487 | 71.0471463 | 3.5530845 |
| 2 | Prestw-1337 | Rosiglitazone Hydrochloride | 3052 | 34.9733614 | 31.8994162 |
| 1 | Prestw-1337 | Rosiglitazone Hydrochloride | 9431 | 36.74353938 | 19.18133718 |
| 1 | Prestw-428 | Roxatidine Acetate hydrochloride | 10565.5 | 17.8358363 | 7.7434707 |
| 2 | Prestw-428 | Roxatidine Acetate hydrochloride | 3812 | 16.3654898 | 1 |
| 2 | Prestw-854 | Roxithromycin | 5887.5 | 89.4918638 | 60.9199997 |
| 1 | Prestw-854 | Roxithromycin | 17124 | 46.78166158 | 1 |
| 2 | Prestw-1338 | Rufloxacin | 3393.5 | 30.3176639 | 27.2437187 |
| 1 | Prestw-1338 | Rufloxacin | 9516 | 17.46261223 | 1 |
| 1 | Prestw-907 | S-(+)-ibuprofen | 16633.5 | 12.67725778 | 1.74237629 |
| 2 | Prestw-907 | S-(+)-ibuprofen | 5898.5 | 1 | 1 |
| 2 | Prestw-932 | S(-)Eticlopride hydrochloride | 6073.5 | 67.7238389 | 39.1519748 |
| 1 | Prestw-932 | S(-)Eticlopride hydrochloride | 18174 | 54.53665888 | 2.5539553 |
| 1 | Prestw-198 | Salbutamol | 12302.5 | 88.40436322 | 5.65734567 |
| 2 | Prestw-198 | Salbutamol | 4578 | 63.3040281 | 1 |
| 2 | Prestw-945 | Salmeterol | 5643.5 | 65.3673933 | 45.7815616 |

| | | | | | |
|---|---|---|---|---|---|
| 1 | Prestw-945 | Salmeterol | 19375 | 42.73294771 | 1 |
| 1 | Prestw-1114 | Saquinavir mesylate | 7933 | 64.7526394 | 41.6085451 |
| 2 | Prestw-1114 | Saquinavir mesylate | 2588 | 37.4351723 | 1 |
| 2 | Prestw-1339 | Sarafloxacin | 3399.5 | 20.1659183 | 17.0919731 |
| 1 | Prestw-1339 | Sarafloxacin | 9254 | 21.3137955 | 3.7515933 |
| 2 | Prestw-232 | Scopolamin-N-oxide hydrobromide | 5064 | 68.7449457 | 1 |
| 1 | Prestw-232 | Scopolamin-N-oxide hydrobromide | 13835.5 | 145.8151371 | 1 |
| 2 | Prestw-877 | Scopolamine hydrochloride | 6588 | 5.248098 | 4.248098 |
| 1 | Prestw-877 | Scopolamine hydrochloride | 17852.5 | 36.75553427 | 1 |
| 2 | Prestw-1340 | Secnidazole | 3308.5 | 31.3649234 | 28.2909782 |
| 1 | Prestw-1340 | Secnidazole | 9218.5 | 17.2138219 | 1 |
| 2 | Prestw-552 | Selegiline hydrochloride | 5073.5 | 127.1726508 | 9.9546967 |
| 1 | Prestw-552 | Selegiline hydrochloride | 16117 | 23.44042179 | 1 |
| 1 | Prestw-481 | Serotonin hydrochloride | 11411 | 38.371223 | 11.69414806 |
| 2 | Prestw-481 | Serotonin hydrochloride | 4112.5 | 107.5168831 | 1 |
| 1 | Prestw-1045 | Sertaconazole nitrate | 10619 | 23.09809691 | 1 |
| 2 | Prestw-1045 | Sertaconazole nitrate | 3658 | 6.419858 | 1 |
| 2 | Prestw-1341 | Sertindole | 3080 | 31.4533243 | 28.3793791 |
| 1 | Prestw-1341 | Sertindole | 9076.5 | 31.84217293 | 14.27997073 |
| 2 | Prestw-1014 | Sertraline | 3117 | 82.7959878 | 23.2026427 |
| 1 | Prestw-1014 | Sertraline | 7926.5 | 1 | 1 |
| 2 | Prestw-1159 | Sibutramine hydrochloride | 3764.5 | 1 | 1 |
| 1 | Prestw-1159 | Sibutramine hydrochloride | 10231.5 | 76.66546102 | 1 |
| 1 | Prestw-1342 | Sildenafil | 10169.5 | 26.11177093 | 21.52384983 |
| 2 | Prestw-1342 | Sildenafil | 3472 | 1 | 1 |
| 2 | Prestw-1749 | Silodosin | 4765.5 | 19.7866443 | 16.292383 |
| 1 | Prestw-1749 | Silodosin | 12474.5 | 117.2142846 | 31.68625067 |
| 2 | Prestw-865 | Simvastatin | 6037.5 | 93.0173666 | 73.4315349 |
| 1 | Prestw-865 | Simvastatin | 19402 | 70.19939621 | 18.7920687 |
| 2 | Prestw-506 | Sisomicin sulfate | 4417.5 | 109.4988098 | 76.6534696 |
| 1 | Prestw-506 | Sisomicin sulfate | 11351.5 | 6.8255279 | 3.63862888 |
| 2 | Prestw-966 | Sotalol hydrochloride | 6383 | 1 | 1 |
| 1 | Prestw-966 | Sotalol hydrochloride | 19181 | 44.02965157 | 1 |
| 1 | Prestw-1091 | Spaglumic acid | 8004 | 30.1768588 | 29.1768588 |
| 2 | Prestw-1091 | Spaglumic acid | 3188.5 | 1 | 1 |
| 1 | Prestw-1343 | Sparfloxacin | 10495.5 | 22.48843779 | 17.90051669 |
| 2 | Prestw-1343 | Sparfloxacin | 3608.5 | 1 | 1 |
| 2 | Prestw-804 | Spectinomycin dihydrochloride | 6371.5 | 5.5187961 | 4.5187961 |
| 1 | Prestw-804 | Spectinomycin dihydrochloride | 21388 | 33.70017688 | 1 |
| 1 | Prestw-288 | Spiperone | 11201 | 33.69414835 | 20.2276988 |
| 2 | Prestw-288 | Spiperone | 4180 | 138.6987942 | 33.293409 |
| 1 | Prestw-745 | Spiramycin | 17893.5 | 30.66700816 | 16.02199922 |
| 2 | Prestw-745 | Spiramycin | 5247.5 | 1 | 1 |
| 1 | Prestw-128 | Spironolactone | 9158.5 | 98.09557664 | 14.18012122 |
| 2 | Prestw-128 | Spironolactone | 3454.5 | 1 | 1 |
| 2 | Prestw-1161 | Stanozolol | 3408.5 | 1 | 1 |
| 1 | Prestw-1161 | Stanozolol | 10858.5 | 95.98931537 | 1 |
| 1 | Prestw-1279 | Stavudine | 9797.5 | 87.03706257 | 3.12160715 |
| 2 | Prestw-1279 | Stavudine | 3209 | 2.2734799 | 1.2734799 |
| 1 | Prestw-321 | Streptomycin sulfate | 13195.5 | 38.26331377 | 25.30525927 |
| 2 | Prestw-321 | Streptomycin sulfate | 4368.5 | 150.0684534 | 1 |
| 2 | Prestw-732 | Streptozotocin | 5470.5 | 8.8859176 | 7.8859176 |
| 1 | Prestw-732 | Streptozotocin | 17059.5 | 31.35219517 | 1 |
| 1 | Prestw-702 | Succinylsulfathiazole | 18370.5 | 65.45618156 | 20.07679838 |
| 2 | Prestw-702 | Succinylsulfathiazole | 5912.5 | 97.6948807 | 28.2894845 |
| 1 | Prestw-1345 | Sulbactam | 10934 | 29.01885213 | 24.43093103 |
| 2 | Prestw-1345 | Sulbactam | 3944.5 | 4.8831907 | 1 |
| 2 | Prestw-810 | Sulconazole nitrate | 5572.5 | 1 | 1 |
| 1 | Prestw-810 | Sulconazole nitrate | 20387.5 | 30.710399 | 1 |
| 1 | Prestw-711 | Sulfabenzamide | 17789 | 99.66674617 | 65.6384779 |
| 2 | Prestw-711 | Sulfabenzamide | 5732.5 | 107.8859806 | 95.1292935 |
| 2 | Prestw-14 | Sulfacetamide sodic hydrate | 3144 | 1 | 1 |
| 1 | Prestw-14 | Sulfacetamide sodic hydrate | 9568 | 83.22787199 | 1 |

| | | | | | |
|---|---|---|---|---|---|
| 1 | Prestw-715 | Sulfachloropyridazine | 16746 | 53.84869233 | 9.72088357 |
| 2 | Prestw-715 | Sulfachloropyridazine | 6205 | 1 | 1 |
| 2 | Prestw-23 | Sulfadiazine | 4527 | 118.8122354 | 15.3306932 |
| 1 | Prestw-23 | Sulfadiazine | 13729.5 | 110.7271419 | 1 |
| 2 | Prestw-728 | Sulfadimethoxine | 5216 | 1 | 1 |
| 1 | Prestw-728 | Sulfadimethoxine | 15834 | 49.79706639 | 1.26413131 |
| 1 | Prestw-1094 | Sulfadoxine | 8122 | 1 | 1 |
| 2 | Prestw-1094 | Sulfadoxine | 2966 | 22.5925275 | 1 |
| 1 | Prestw-10 | Sulfaguanidine | 7688.5 | 1 | 1 |
| 2 | Prestw-10 | Sulfaguanidine | 2843.5 | 1 | 1 |
| 2 | Prestw-694 | Sulfamerazine | 5150 | 69.60093 | 38.7128536 |
| 1 | Prestw-694 | Sulfamerazine | 16851.5 | 37.93229076 | 6.04366268 |
| 1 | Prestw-769 | Sulfameter | 16528 | 54.36465451 | 22.47602643 |
| 2 | Prestw-769 | Sulfameter | 5019 | 61.1341077 | 30.2460313 |
| 1 | Prestw-775 | Sulfamethazine sodium salt | 18951.5 | 59.13098281 | 13.75159963 |
| 2 | Prestw-775 | Sulfamethazine sodium salt | 6501.5 | 90.3242355 | 20.9188393 |
| 1 | Prestw-742 | Sulfamethizole | 16450 | 29.18289412 | 14.53788518 |
| 2 | Prestw-742 | Sulfamethizole | 5586.5 | 1 | 1 |
| 1 | Prestw-177 | Sulfamethoxazole | 10667.5 | 117.6401973 | 34.89317979 |
| 2 | Prestw-177 | Sulfamethoxazole | 4081.5 | 103.8530551 | 1 |
| 2 | Prestw-724 | Sulfamethoxypyridazine | 5140 | 1 | 1 |
| 1 | Prestw-724 | Sulfamethoxypyridazine | 16738 | 41.64331193 | 1 |
| 1 | Prestw-823 | Sulfamonomethoxine | 16077.5 | 1 | 1 |
| 2 | Prestw-823 | Sulfamonomethoxine | 5858.5 | 1 | 1 |
| 2 | Prestw-729 | Sulfanilamide | 5161.5 | 3.7207379 | 2.7207379 |
| 1 | Prestw-729 | Sulfanilamide | 16122.5 | 37.92554274 | 1 |
| 1 | Prestw-21 | Sulfaphenazole | 11578 | 129.1477009 | 17.99882981 |
| 2 | Prestw-21 | Sulfaphenazole | 4997 | 72.8997618 | 1 |
| 1 | Prestw-762 | Sulfapyridine | 14930 | 13.78143177 | 1 |
| 2 | Prestw-762 | Sulfapyridine | 5118.5 | 1 | 1 |
| 2 | Prestw-731 | Sulfaquinoxaline sodium salt | 5850.5 | 8.9264977 | 7.9264977 |
| 1 | Prestw-731 | Sulfaquinoxaline sodium salt | 17934.5 | 54.63819293 | 6.10525785 |
| 1 | Prestw-520 | Sulfasalazine | 11690.5 | 1.28521663 | 1 |
| 2 | Prestw-520 | Sulfasalazine | 4115.5 | 94.5138471 | 1 |
| 1 | Prestw-16 | Sulfathiazole | 10416.5 | 89.21392578 | 21.40213193 |
| 2 | Prestw-16 | Sulfathiazole | 4349.5 | 14.8104323 | 1 |
| 1 | Prestw-290 | Sulfinpyrazone | 11548.5 | 14.7760121 | 10.1309428 |
| 2 | Prestw-290 | Sulfinpyrazone | 4134 | 118.6176727 | 11.6089057 |
| 1 | Prestw-334 | Sulfisoxazole | 10509 | 1 | 1 |
| 2 | Prestw-334 | Sulfisoxazole | 4080 | 1 | 1 |
| 2 | Prestw-73 | Sulindac | 2919.5 | 1118.933649 | 1117.933649 |
| 1 | Prestw-73 | Sulindac | 4914.5 | 434.3235191 | 385.790584 |
| 1 | Prestw-641 | Sulmazole | 15072.5 | 1 | 1 |
| 2 | Prestw-641 | Sulmazole | 6039 | 1 | 1 |
| 2 | Prestw-421 | Suloctidil | 3352.5 | 141.8040365 | 108.2135963 |
| 1 | Prestw-421 | Suloctidil | 7598.5 | 1 | 1 |
| 2 | Prestw-56 | Sulpiride | 4521 | 138.8938117 | 21.6758576 |
| 1 | Prestw-56 | Sulpiride | 13714 | 11.07478956 | 1 |
| 1 | Prestw-1346 | Sumatriptan succinate | 10273.5 | 29.01167063 | 24.42374953 |
| 2 | Prestw-1346 | Sumatriptan succinate | 3537 | 16.9229226 | 1 |
| 1 | Prestw-816 | Suprofen | 19070 | 18.00434164 | 3.3593327 |
| 2 | Prestw-816 | Suprofen | 5351.5 | 1 | 1 |
| 1 | Prestw-658 | Suxibuzone | 15655 | 11.27424516 | 1 |
| 2 | Prestw-658 | Suxibuzone | 4834.5 | 1 | 1 |
| 2 | Prestw-329 | Tacrine hydrochloride | 4344 | 106.8994676 | 43.549292 |
| 1 | Prestw-329 | Tacrine hydrochloride | 10889.5 | 26.8256824 | 1 |
| 2 | Prestw-146 | Tamoxifen citrate | 4687 | 72.6027024 | 1 |
| 1 | Prestw-146 | Tamoxifen citrate | 11265.5 | 128.9100056 | 1 |
| 1 | Prestw-1817 | Tazarotene | 12236 | 132.3388355 | 12.42607219 |
| 2 | Prestw-1817 | Tazarotene | 4724.5 | 54.6481443 | 1 |
| 1 | Prestw-1349 | Tazobactam | 10155.5 | 21.96641096 | 17.37848986 |
| 2 | Prestw-1349 | Tazobactam | 3394.5 | 22.3458153 | 1 |
| 1 | Prestw-1753 | Tegafur | 9531 | 51.58612853 | 31.88026191 |

| | | | | | |
|---|---|---|---|---|---|
| 2 | Prestw-1753 | Tegafur | 4303 | 6.9213385 | 1 |
| 1 | Prestw-1760 | Tegaserod maleate | 3821.5 | 381.8252904 | 362.1194238 |
| 2 | Prestw-1760 | Tegaserod maleate | 1942.5 | 276.1897527 | 210.8169001 |
| 1 | Prestw-303 | Telenzepine dihydrochloride | 13519.5 | 41.85066168 | 33.82500364 |
| 2 | Prestw-303 | Telenzepine dihydrochloride | 4896.5 | 180.2195307 | 22.8243339 |
| 1 | Prestw-1350 | Telmisartan | 10333 | 20.9009728 | 16.3130517 |
| 2 | Prestw-1350 | Telmisartan | 3296 | 30.7607754 | 1 |
| 2 | Prestw-1183 | Temozolomide | 2932 | 25.9404823 | 24.9404823 |
| 1 | Prestw-1183 | Temozolomide | 8458 | 1 | 1 |
| 1 | Prestw-1351 | Tenatoprazole | 10097 | 16.829528 | 12.2416069 |
| 2 | Prestw-1351 | Tenatoprazole | 3524.5 | 74.6270408 | 1 |
| 1 | Prestw-527 | Tenoxicam | 10381 | 25.63571436 | 15.54334876 |
| 2 | Prestw-527 | Tenoxicam | 4042.5 | 110.5081688 | 11.6073543 |
| 1 | Prestw-751 | Terazosin hydrochloride | 13748.5 | 5.35760045 | 4.35760045 |
| 2 | Prestw-751 | Terazosin hydrochloride | 4662.5 | 1 | 1 |
| 1 | Prestw-1750 | Terbinafine | 9830 | 12.8133638 | 1 |
| 2 | Prestw-1750 | Terbinafine | 4462.5 | 1 | 1 |
| 1 | Prestw-391 | Terbutaline hemisulfate | 13025 | 32.32906725 | 19.37101275 |
| 2 | Prestw-391 | Terbutaline hemisulfate | 4567 | 177.6261326 | 16.7624763 |
| 1 | Prestw-495 | Terconazole | 12478.5 | 40.56892088 | 15.07040351 |
| 2 | Prestw-495 | Terconazole | 4204 | 140.7953491 | 1 |
| 2 | Prestw-138 | Terfenadine | 3088 | 10.2466288 | 1 |
| 1 | Prestw-138 | Terfenadine | 10115.5 | 71.06984951 | 1 |
| 2 | Prestw-401 | Testosterone propionate | 4810.5 | 100.221022 | 36.8708464 |
| 1 | Prestw-401 | Testosterone propionate | 13773.5 | 15.3388232 | 1 |
| 1 | Prestw-571 | Tetracaïne hydrochloride | 13522.5 | 48.42807623 | 21.75100129 |
| 2 | Prestw-571 | Tetracaïne hydrochloride | 4689.5 | 165.7367295 | 1 |
| 1 | Prestw-140 | Tetracycline hydrochloride | 10293.5 | 24.78045388 | 22.82495276 |
| 2 | Prestw-140 | Tetracycline hydrochloride | 3625 | 1 | 1 |
| 2 | Prestw-1387 | Tetraethylenepentamine pentahydrochloride | 3537 | 1 | 1 |
| 1 | Prestw-1387 | Tetraethylenepentamine pentahydrochloride | 10300.5 | 75.31220809 | 1 |
| 1 | Prestw-698 | Tetrahydrozoline hydrochloride | 15461.5 | 56.85757299 | 11.47818981 |
| 2 | Prestw-698 | Tetrahydrozoline hydrochloride | 5115.5 | 33.2048926 | 1 |
| 2 | Prestw-545 | Tetramisole hydrochloride | 4614.5 | 129.7205015 | 37.5397098 |
| 1 | Prestw-545 | Tetramisole hydrochloride | 13126 | 12.95153043 | 1 |
| 1 | Prestw-192 | Thalidomide | 9868.5 | 29.46938762 | 9.763521 |
| 2 | Prestw-192 | Thalidomide | 4042 | 1 | 1 |
| 2 | Prestw-874 | Theobromine | 6552.5 | 61.3305021 | 58.1219617 |
| 1 | Prestw-874 | Theobromine | 20630.5 | 49.06237679 | 12.50835846 |
| 1 | Prestw-873 | Theophylline monohydrate | 19401.5 | 73.83746614 | 37.28344781 |
| 2 | Prestw-873 | Theophylline monohydrate | 5066.5 | 1 | 1 |
| 2 | Prestw-631 | Thiamine hydrochloride | 4372 | 69.5067171 | 36.6613769 |
| 1 | Prestw-631 | Thiamine hydrochloride | 12924.5 | 4.17410824 | 0.98720922 |
| 2 | Prestw-25 | Thiamphenicol | 4784.5 | 35.1058578 | 1 |
| 1 | Prestw-25 | Thiamphenicol | 13887.5 | 111.223359 | 1 |
| 1 | Prestw-1068 | Thiethylperazine dimalate | 9088 | 80.35428655 | 45.10386811 |
| 2 | Prestw-1068 | Thiethylperazine dimalate | 3227.5 | 1 | 1 |
| 1 | Prestw-539 | Thiocolchicoside | 12221 | 14.87948668 | 2.02736018 |
| 2 | Prestw-539 | Thiocolchicoside | 4307 | 107.1438444 | 3.6072673 |
| 1 | Prestw-347 | Thioguanosine | 9737.5 | 7.4293309 | 6.4293309 |
| 2 | Prestw-347 | Thioguanosine | 3887.5 | 30.9691484 | 2.4209353 |
| 1 | Prestw-922 | Thioperamide maleate | 15110 | 48.36277562 | 8.12241467 |
| 2 | Prestw-922 | Thioperamide maleate | 4898.5 | 32.1728808 | 1 |
| 1 | Prestw-149 | Thioproperazine dimesylate | 11778.5 | 199.6331425 | 35.99734656 |
| 2 | Prestw-149 | Thioproperazine dimesylate | 4124.5 | 147.5862444 | 1 |
| 1 | Prestw-78 | Thioridazine hydrochloride | 17960.5 | 48.16898019 | 2.78959701 |
| 2 | Prestw-78 | Thioridazine hydrochloride | 5952.5 | 55.4889987 | 1 |
| 2 | Prestw-633 | Thiorphan | 4108 | 62.0495512 | 29.204211 |
| 1 | Prestw-633 | Thiorphan | 10942 | 6.33755951 | 3.15066049 |
| 1 | Prestw-522 | Thiostrepton | 10632 | 6.9302512 | 1 |
| 2 | Prestw-522 | Thiostrepton | 4099 | 39.2844404 | 1 |
| 2 | Prestw-972 | THIP Hydrochloride | 4257.5 | 345.055853 | 344.055853 |
| 1 | Prestw-972 | THIP Hydrochloride | 6881 | 410.2579636 | 362.5019957 |

| | | | | | |
|---|---|---|---|---|---|
| 1 | Prestw-925 | Thonzonium bromide | 17440.5 | 29.64640036 | 1 |
| 2 | Prestw-925 | Thonzonium bromide | 4978 | 1 | 1 |
| 2 | Prestw-403 | Thyroxine (L) | 4309.5 | 106.0552568 | 42.7050812 |
| 1 | Prestw-403 | Thyroxine (L) | 12667 | 33.23287631 | 1 |
| 1 | Prestw-524 | Tiabendazole | 11548.5 | 1 | 1 |
| 2 | Prestw-524 | Tiabendazole | 4555 | 106.9038961 | 8.0030816 |
| 2 | Prestw-216 | Tiapride hydrochloride | 6138.5 | 49.752785 | 1 |
| 1 | Prestw-216 | Tiapride hydrochloride | 15204.5 | 130.6555414 | 1 |
| 1 | Prestw-496 | Tiaprofenic acid | 12809.5 | 45.51244078 | 20.01392341 |
| 2 | Prestw-496 | Tiaprofenic acid | 4256 | 118.4775922 | 1 |
| 2 | Prestw-1181 | Tibolone | 3246.5 | 1 | 1 |
| 1 | Prestw-1181 | Tibolone | 9537 | 31.0111282 | 1 |
| 1 | Prestw-1067 | Ticarcillin sodium | 8837 | 70.41665859 | 35.16624015 |
| 2 | Prestw-1067 | Ticarcillin sodium | 3469.5 | 19.2063768 | 3.7032492 |
| 2 | Prestw-47 | Ticlopidine hydrochloride | 4039 | 125.3096364 | 8.0916823 |
| 1 | Prestw-47 | Ticlopidine hydrochloride | 12871.5 | 17.45436648 | 1 |
| 2 | Prestw-1799 | Tigecycline | 5021.5 | 109.6982437 | 6.2167015 |
| 1 | Prestw-1799 | Tigecycline | 13168 | 139.6413834 | 1 |
| 1 | Prestw-948 | Timolol maleate salt | 16433 | 58.95220833 | 7.54488082 |
| 2 | Prestw-948 | Timolol maleate salt | 5970 | 35.0265066 | 15.4406749 |
| 1 | Prestw-766 | Tinidazole | 15400 | 60.49507322 | 28.60644514 |
| 2 | Prestw-766 | Tinidazole | 5528 | 65.3851129 | 34.4970365 |
| 1 | Prestw-1184 | Tioconazole | 8432.5 | 1 | 1 |
| 2 | Prestw-1184 | Tioconazole | 3159.5 | 1 | 1 |
| 1 | Prestw-202 | Tiratricol, 3,3',5-triiodothyroacetic acid | 11012.5 | 98.05599509 | 15.30897754 |
| 2 | Prestw-202 | Tiratricol, 3,3',5-triiodothyroacetic acid | 4055.5 | 72.3584584 | 1 |
| 1 | Prestw-1818 | Tirofiban hydrochloride | 11311 | 134.2708975 | 14.35813418 |
| 2 | Prestw-1818 | Tirofiban hydrochloride | 4009 | 64.2416071 | 1 |
| 1 | Prestw-1182 | Tizanidine hydrochloride | 9292 | 47.34827147 | 12.09785303 |
| 2 | Prestw-1182 | Tizanidine hydrochloride | 3123.5 | 1 | 1 |
| 2 | Prestw-544 | Tobramycin | 4777.5 | 112.0236856 | 19.8428939 |
| 1 | Prestw-544 | Tobramycin | 13705.5 | 5.31100022 | 1 |
| 1 | Prestw-1027 | Tocainide hydrochloride | 8815.5 | 26.9936442 | 12.8387905 |
| 2 | Prestw-1027 | Tocainide hydrochloride | 3264 | 1 | 1 |
| 2 | Prestw-71 | Todralazine hydrochloride | 6093 | 64.1682011 | 51.411514 |
| 1 | Prestw-71 | Todralazine hydrochloride | 17366 | 54.15257816 | 20.12430989 |
| 1 | Prestw-554 | Tolazamide | 14375.5 | 34.51608579 | 0.59939055 |
| 2 | Prestw-554 | Tolazamide | 4822 | 117.9602814 | 0.7423273 |
| 1 | Prestw-60 | Tolazoline hydrochloride | 13501 | 33.22218892 | 7.72367155 |
| 2 | Prestw-60 | Tolazoline hydrochloride | 4417.5 | 116.1614578 | 1 |
| 1 | Prestw-190 | Tolbutamide | 9481.5 | 54.4331551 | 34.72728848 |
| 2 | Prestw-190 | Tolbutamide | 4024.5 | 1 | 1 |
| 2 | Prestw-1732 | Tolcapone | 5549 | 105.771762 | 33.7796215 |
| 1 | Prestw-1732 | Tolcapone | 14123.5 | 161.1955504 | 17.33487907 |
| 2 | Prestw-205 | Tolfenamic acid | 5186.5 | 64.7483415 | 1 |
| 1 | Prestw-205 | Tolfenamic acid | 12912 | 99.47717513 | 1 |
| 2 | Prestw-856 | Tolmetin sodium salt dihydrate | 5688 | 76.604662 | 57.0188303 |
| 1 | Prestw-856 | Tolmetin sodium salt dihydrate | 16422 | 53.12424611 | 1.7169186 |
| 1 | Prestw-70 | Tolnaftate | 15629 | 72.64023652 | 27.26085334 |
| 2 | Prestw-70 | Tolnaftate | 5241 | 53.809618 | 1 |
| 1 | Prestw-1744 | Tolterodine tartrate | 12816 | 100.1540832 | 14.62604934 |
| 2 | Prestw-1744 | Tolterodine tartrate | 4368 | 1 | 1 |
| 2 | Prestw-1195 | Toltrazuril | 3157 | 19.652714 | 18.652714 |
| 1 | Prestw-1195 | Toltrazuril | 7748.5 | 1 | 1 |
| 1 | Prestw-1819 | Tolvaptan | 12636 | 136.3110042 | 16.39824092 |
| 2 | Prestw-1819 | Tolvaptan | 4911.5 | 52.5277552 | 1 |
| 1 | Prestw-1463 | Tomoxetine hydrochloride | 10792 | 146.0413531 | 14.36691834 |
| 2 | Prestw-1463 | Tomoxetine hydrochloride | 4685.5 | 90.3165317 | 1 |
| 2 | Prestw-1493 | Topiramate | 4581.5 | 106.5084589 | 34.5163184 |
| 1 | Prestw-1493 | Topiramate | 11739 | 181.6762415 | 37.81557019 |
| 2 | Prestw-1196 | Topotecan | 2703.5 | 76.0758622 | 75.0758622 |
| 1 | Prestw-1196 | Topotecan | 8133.5 | 1 | 1 |
| 1 | Prestw-1197 | Toremifene | 8788 | 1 | 1 |

| | | | | | |
|---|---|---|---|---|---|
| 2 | Prestw-1197 | Toremifene | 3155 | 33.7418372 | 1 |
| 1 | Prestw-1030 | Torsemide | 8336 | 15.3726471 | 1 |
| 2 | Prestw-1030 | Torsemide | 2987 | 38.662463 | 1 |
| 1 | Prestw-1464 | Tosufloxacin hydrochloride | 11141 | 157.5656639 | 25.89122906 |
| 2 | Prestw-1464 | Tosufloxacin hydrochloride | 4360 | 89.8304047 | 1 |
| 1 | Prestw-976 | Tracazolate hydrochloride | 19863 | 43.6021481 | 14.35256758 |
| 2 | Prestw-976 | Tracazolate hydrochloride | 5934 | 1 | 1 |
| 1 | Prestw-1465 | Tramadol hydrochloride | 11040.5 | 143.1847893 | 11.51035449 |
| 2 | Prestw-1465 | Tramadol hydrochloride | 4523 | 114.5499868 | 1 |
| 2 | Prestw-171 | Tranexamic acid | 4718.5 | 79.5436608 | 1 |
| 1 | Prestw-171 | Tranexamic acid | 12062.5 | 118.6575965 | 1 |
| 1 | Prestw-1198 | Tranilast | 8245.5 | 1 | 1 |
| 2 | Prestw-1198 | Tranilast | 2928 | 61.5648901 | 1 |
| 2 | Prestw-173 | Tranylcypromine hydrochloride | 4686 | 72.1031344 | 1 |
| 1 | Prestw-173 | Tranylcypromine hydrochloride | 13851 | 150.1357391 | 1 |
| 2 | Prestw-1012 | Trapidil | 2898 | 79.898301 | 20.3049559 |
| 1 | Prestw-1012 | Trapidil | 7866 | 3.6346731 | 2.6346731 |
| 1 | Prestw-292 | Trazodone hydrochloride | 11029.5 | 42.05953021 | 37.41446091 |
| 2 | Prestw-292 | Trazodone hydrochloride | 3852.5 | 137.3517236 | 30.3429566 |
| 1 | Prestw-120 | Triamcinolone | 8241 | 11.43101 | 5.8822573 |
| 2 | Prestw-120 | Triamcinolone | 3012.5 | 81.705406 | 5.4112679 |
| 1 | Prestw-34 | Triamterene | 10921.5 | 1 | 1 |
| 2 | Prestw-34 | Triamterene | 4041.5 | 1 | 1 |
| 2 | Prestw-1019 | Tribenoside | 3277 | 22.2819933 | 19.1762751 |
| 1 | Prestw-1019 | Tribenoside | 9349.5 | 0.8706675 | 1 |
| 1 | Prestw-825 | Trichlormethiazide | 14715 | 1 | 1 |
| 2 | Prestw-825 | Trichlormethiazide | 4899.5 | 1 | 1 |
| 1 | Prestw-1768 | Triclabendazole | 10859 | 99.59898778 | 16.85197023 |
| 2 | Prestw-1768 | Triclabendazole | 4449.5 | 45.4026163 | 1 |
| 2 | Prestw-1479 | Triclosan | 4896 | 128.9526731 | 42.2505571 |
| 1 | Prestw-1479 | Triclosan | 13251 | 147.4818565 | 29.95536765 |
| 1 | Prestw-1042 | Tridihexethyl chloride | 9338 | 25.4721899 | 1 |
| 2 | Prestw-1042 | Tridihexethyl chloride | 3089 | 43.6988722 | 1 |
| 1 | Prestw-313 | Trifluoperazine dihydrochloride | 12540 | 57.52666001 | 20.09889581 |
| 2 | Prestw-313 | Trifluoperazine dihydrochloride | 4604 | 132.2840345 | 1 |
| 2 | Prestw-53 | Triflupromazine hydrochloride | 4425.5 | 182.3200948 | 78.7835177 |
| 1 | Prestw-53 | Triflupromazine hydrochloride | 14372 | 14.05360318 | 1.20147668 |
| 2 | Prestw-1056 | Trifluridine | 3373.5 | 67.1655025 | 14.640294 |
| 1 | Prestw-1056 | Trifluridine | 9706.5 | 46.90150589 | 1 |
| 1 | Prestw-528 | Triflusal | 11846 | 2.73650878 | 1 |
| 2 | Prestw-528 | Triflusal | 3812.5 | 100.3457407 | 1.4449262 |
| 1 | Prestw-701 | Trihexyphenidyl-D,L Hydrochloride | 15911.5 | 66.49008212 | 21.11069894 |
| 2 | Prestw-701 | Trihexyphenidyl-D,L Hydrochloride | 5506.5 | 93.1196839 | 23.7142877 |
| 1 | Prestw-1777 | Trimebutine | 11082.5 | 118.2733507 | 7.12447963 |
| 2 | Prestw-1777 | Trimebutine | 4874.5 | 83.5124152 | 1 |
| 1 | Prestw-842 | Trimeprazine tartrate | 15858.5 | 32.31866119 | 1 |
| 2 | Prestw-842 | Trimeprazine tartrate | 4931.5 | 10.9784333 | 1 |
| 1 | Prestw-549 | Trimetazidine dihydrochloride | 14713.5 | 22.94691194 | 1 |
| 2 | Prestw-549 | Trimetazidine dihydrochloride | 5436 | 108.615479 | 1 |
| 1 | Prestw-515 | Trimethadione | 10223 | 1 | 1 |
| 2 | Prestw-515 | Trimethadione | 3630.5 | 16.6041028 | 1 |
| 2 | Prestw-80 | Trimethobenzamide hydrochloride | 3282 | 995.8213066 | 994.8213066 |
| 1 | Prestw-80 | Trimethobenzamide hydrochloride | 5911 | 389.5541901 | 345.4263813 |
| 2 | Prestw-208 | Trimethoprim | 4797.5 | 74.8976 | 1 |
| 1 | Prestw-208 | Trimethoprim | 12530 | 109.5496894 | 1 |
| 2 | Prestw-1702 | Trimetozine | 5694.5 | 91.0712828 | 1 |
| 1 | Prestw-1702 | Trimetozine | 13691 | 114.3774057 | 1 |
| 2 | Prestw-806 | Trimipramine maleate salt | 5490 | 38.4757363 | 37.4757363 |
| 1 | Prestw-806 | Trimipramine maleate salt | 19645.5 | 44.6565903 | 1 |
| 2 | Prestw-709 | Trioxsalen | 5196.5 | 89.4680872 | 76.7114001 |
| 1 | Prestw-709 | Trioxsalen | 16468.5 | 57.98161673 | 23.95334846 |
| 1 | Prestw-1199 | Tripelennamine hydrochloride | 8322 | 1 | 1 |
| 2 | Prestw-1199 | Tripelennamine hydrochloride | 2973.5 | 50.5317748 | 1 |

| | | | | | |
|---|---|---|---|---|---|
| 1 | Prestw-262 | Triprolidine hydrochloride | 10122 | 1 | 1 |
| 2 | Prestw-262 | Triprolidine hydrochloride | 3819.5 | 1 | 1 |
| 1 | Prestw-1467 | Troglitazone | 11083.5 | 143.7233523 | 12.04891747 |
| 2 | Prestw-1467 | Troglitazone | 4898 | 102.7990742 | 1 |
| 1 | Prestw-36 | Troleandomycin | 11478 | 1 | 1 |
| 2 | Prestw-36 | Troleandomycin | 4186.5 | 96.6346704 | 1 |
| 1 | Prestw-530 | Trolox | 13116 | 17.19870704 | 4.34658054 |
| 2 | Prestw-530 | Trolox | 4883.5 | 131.6447313 | 28.1081542 |
| 2 | Prestw-228 | Tropicamide | 5952.5 | 79.7321649 | 1 |
| 1 | Prestw-228 | Tropicamide | 14721.5 | 116.0806268 | 1 |
| 2 | Prestw-461 | Tropisetron hydrochloride | 5516 | 127.7982545 | 35.6174628 |
| 1 | Prestw-461 | Tropisetron hydrochloride | 13609.5 | 20.00321084 | 1 |
| 1 | Prestw-1778 | Troxipide | 11346.5 | 119.1525718 | 8.0037007 |
| 2 | Prestw-1778 | Troxipide | 4408 | 74.0398946 | 1 |
| 2 | Prestw-1352 | Tulobuterol | 2756 | 165.1061517 | 78.5533583 |
| 1 | Prestw-1352 | Tulobuterol | 10147 | 24.23305029 | 19.64512919 |
| 1 | Prestw-1353 | Tylosin | 9838 | 40.19080369 | 35.60288259 |
| 2 | Prestw-1353 | Tylosin | 3394 | 96.826527 | 10.2737336 |
| 2 | Prestw-954 | Tyloxapol | 6421.5 | 101.2756026 | 98.0670622 |
| 1 | Prestw-954 | Tyloxapol | 20185 | 61.99771162 | 25.44369329 |
| 2 | Prestw-1784 | Ubenimex | 4628 | 105.3733008 | 41.7808923 |
| 1 | Prestw-1784 | Ubenimex | 11019 | 144.6927429 | 1 |
| 2 | Prestw-905 | Urapidil hydrochloride | 5861.5 | 1 | 1 |
| 1 | Prestw-905 | Urapidil hydrochloride | 16951 | 11.57980322 | 0.64492173 |
| 1 | Prestw-958 | Urosiol | 17703.5 | 55.64416702 | 19.09014869 |
| 2 | Prestw-958 | Urosiol | 6147 | 33.363489 | 30.1549486 |
| 2 | Prestw-1505 | Valacyclovir hydrochloride | 4144 | 52.6988182 | 49.2045569 |
| 1 | Prestw-1505 | Valacyclovir hydrochloride | 11465.5 | 128.3883658 | 42.86033193 |
| 1 | Prestw-1759 | Valdecoxib | 10013.5 | 18.77949161 | 1 |
| 2 | Prestw-1759 | Valdecoxib | 4256.5 | 1 | 1 |
| 1 | Prestw-1482 | Valproic acid | 12128 | 176.1524279 | 12.51663199 |
| 2 | Prestw-1482 | Valproic acid | 4585 | 87.7791636 | 1 |
| 2 | Prestw-1735 | Valsartan | 4348 | 96.7271692 | 24.7350287 |
| 1 | Prestw-1735 | Valsartan | 13665 | 159.7769746 | 15.91630329 |
| 1 | Prestw-497 | Vancomycin hydrochloride | 13785 | 28.9389447 | 3.44042733 |
| 2 | Prestw-497 | Vancomycin hydrochloride | 4538.5 | 149.7226401 | 1 |
| 1 | Prestw-1356 | Vardenafil | 9494.5 | 47.99939486 | 43.41147376 |
| 2 | Prestw-1356 | Vardenafil | 3250 | 52.4114729 | 1 |
| 1 | Prestw-1358 | Vatalanib | 11270 | 25.31220271 | 1 |
| 2 | Prestw-1358 | Vatalanib | 3736 | 1 | 1 |
| 1 | Prestw-1359 | Vecuronium bromide | 10885 | 31.75447139 | 1 |
| 2 | Prestw-1359 | Vecuronium bromide | 3984 | 1 | 1 |
| 1 | Prestw-1170 | Venlafaxine | 9340 | 56.09154048 | 20.84112204 |
| 2 | Prestw-1170 | Venlafaxine | 3074.5 | 1 | 1 |
| 1 | Prestw-141 | Verapamil hydrochloride | 9824 | 30.79058827 | 28.83508715 |
| 2 | Prestw-141 | Verapamil hydrochloride | 3420.5 | 1 | 1 |
| 1 | Prestw-1105 | Verteporfin | 9352 | 26.5342605 | 12.3794068 |
| 2 | Prestw-1105 | Verteporfin | 3048.5 | 19.7842752 | 16.678557 |
| 2 | Prestw-768 | Vidarabine | 4947 | 80.7648321 | 49.8767557 |
| 1 | Prestw-768 | Vidarabine | 17929.5 | 52.17699357 | 20.28836549 |
| 1 | Prestw-501 | Vigabatrin hydrochloride | 11617 | 24.82338545 | 21.63648643 |
| 2 | Prestw-501 | Vigabatrin hydrochloride | 4475.5 | 75.5536711 | 42.7083309 |
| 1 | Prestw-1361 | Viloxazine hydrochloride | 11599.5 | 21.86422791 | 1 |
| 2 | Prestw-1361 | Viloxazine hydrochloride | 3620.5 | 1 | 1 |
| 1 | Prestw-271 | Vincamine | 10353.5 | 1 | 1 |
| 2 | Prestw-271 | Vincamine | 3762.5 | 1 | 1 |
| 2 | Prestw-268 | Vinpocetine | 4039 | 24.7123189 | 23.7123189 |
| 1 | Prestw-268 | Vinpocetine | 9500 | 1 | 1 |
| 1 | Prestw-1113 | Viomycin sulfate | 10437.5 | 18.92845673 | 1 |
| 2 | Prestw-1113 | Viomycin sulfate | 3320.5 | 1 | 1 |
| 1 | Prestw-1500 | Voriconazole | 12765.5 | 181.9778618 | 38.11719051 |
| 2 | Prestw-1500 | Voriconazole | 4884 | 86.2656889 | 14.2735484 |
| 1 | Prestw-1362 | Vorinostat | 9775.5 | 57.29918794 | 1 |

| | | | | | |
|---|---|---|---|---|---|
| 2 | Prestw-1362 | Vorinostat | 3746.5 | 1 | 1 |
| 2 | Prestw-1363 | Warfarin | 3545 | 25.3520286 | 3.195851 |
| 1 | Prestw-1363 | Warfarin | 10720 | 35.22771899 | 1 |
| 1 | Prestw-923 | Xamoterol hemifumarate | 16982 | 32.23400938 | 1 |
| 2 | Prestw-923 | Xamoterol hemifumarate | 5460 | 12.3199107 | 1 |
| 2 | Prestw-598 | Xylazine | 4044.5 | 202.8107785 | 48.6446118 |
| 1 | Prestw-598 | Xylazine | 13477.5 | 32.92641004 | 7.42789267 |
| 2 | Prestw-223 | Xylometazoline hydrochloride | 4568 | 88.7539087 | 25.1615002 |
| 1 | Prestw-223 | Xylometazoline hydrochloride | 12980 | 142.3248151 | 1 |
| 1 | Prestw-584 | Yohimbine hydrochloride | 12766 | 42.93570585 | 17.43718848 |
| 2 | Prestw-584 | Yohimbine hydrochloride | 4038.5 | 108.6088016 | 1 |
| 2 | Prestw-1364 | Zafirlukast | 3763 | 47.1728397 | 25.0166621 |
| 1 | Prestw-1364 | Zafirlukast | 10221 | 52.10393725 | 1 |
| 1 | Prestw-1037 | Zalcitabine | 9284 | 26.856685 | 3.7125907 |
| 2 | Prestw-1037 | Zalcitabine | 3603.5 | 52.0091755 | 3.8552103 |
| 2 | Prestw-1369 | Zaleplon | 3523.5 | 6.7764655 | 1 |
| 1 | Prestw-1369 | Zaleplon | 10018.5 | 67.57602725 | 1 |
| 1 | Prestw-335 | Zaprinast | 9319.5 | 1 | 1 |
| 2 | Prestw-335 | Zaprinast | 3925 | 1 | 1 |
| 1 | Prestw-977 | Zardaverine | 21520.5 | 30.88608877 | 1.63650825 |
| 2 | Prestw-977 | Zardaverine | 5889 | 1 | 1 |
| 1 | Prestw-333 | Zidovudine, AZT | 9906.5 | 60.4622768 | 20.33353266 |
| 2 | Prestw-333 | Zidovudine, AZT | 4023.5 | 112.7667364 | 49.4165608 |
| 2 | Prestw-1365 | Zileuton | 3524.5 | 31.8986141 | 9.7424365 |
| 1 | Prestw-1365 | Zileuton | 10041.5 | 76.37580314 | 0.12667534 |
| 1 | Prestw-92 | Zimelidine dihydrochloride monohydrate | 16173 | 51.50921883 | 11.26885788 |
| 2 | Prestw-92 | Zimelidine dihydrochloride monohydrate | 5944 | 39.0582167 | 1 |
| 1 | Prestw-1188 | Ziprasidone Hydrochloride | 7880.5 | 1 | 1 |
| 2 | Prestw-1188 | Ziprasidone Hydrochloride | 2931 | 1 | 1 |
| 2 | Prestw-1731 | Zoledronic acid hydrate | 1407 | 2038.156366 | 1966.164225 |
| 1 | Prestw-1731 | Zoledronic acid hydrate | 3174.5 | 2958.669089 | 2814.808418 |
| 2 | Prestw-1800 | Zolmitriptan | 5785 | 94.2948787 | 1 |
| 1 | Prestw-1800 | Zolmitriptan | 14878.5 | 123.9056445 | 1 |
| 1 | Prestw-779 | Zomepirac sodium salt | 18502.5 | 42.01642022 | 1 |
| 2 | Prestw-779 | Zomepirac sodium salt | 5424.5 | 1 | 1 |
| 2 | Prestw-1162 | Zonisamide | 2598 | 83.6012036 | 31.0759951 |
| 1 | Prestw-1162 | Zonisamide | 8415.5 | 116.4739411 | 13.96746197 |
| 2 | Prestw-1367 | Zopiclone | 3468.5 | 55.0864611 | 32.9302835 |
| 1 | Prestw-1367 | Zopiclone | 9544 | 59.0982427 | 1 |
| 1 | Prestw-1368 | Zotepine | 9993.5 | 62.22560055 | 1 |
| 2 | Prestw-1368 | Zotepine | 3533 | 18.5573096 | 1 |
| 2 | Prestw-328 | Zoxazolamine | 4053 | 104.9625326 | 41.612357 |
| 1 | Prestw-328 | Zoxazolamine | 9948 | 19.2202112 | 1 |
| 2 | Prestw-998 | Zuclopenthixol dihydrochloride | 5090 | 3.9536515 | 2.9536515 |
| 1 | Prestw-998 | Zuclopenthixol dihydrochloride | 15458.5 | 1 | 1 |

| Signal.ratio.over.Colistin | Events.ratio.no.drug |
|---|---|
| 3.615364544 | 0.435529622 |
| 1.546692709 | 0.260657265 |
| 0.064975896 | 0.972916907 |
| 0.001680008 | 0.874149367 |
| 0.014720379 | 1.014233971 |
| 0.00277107 | 1.057023896 |
| 0.054575199 | 1.067073171 |
| 0.003044841 | 0.878238754 |
| 0.026451929 | 1.041593254 |
| 0.001053652 | 0.849164927 |
| 0.200721934 | 0.914192614 |
| 0.014720379 | 1.019190443 |
| 0.025627037 | 1.062389882 |
| 0.004909533 | 1.03644746 |
| 0.07322249 | 1.056376592 |
| 0.004588841 | 1.1107857 |
| 0.032756434 | 1.087498862 |
| 0.022713618 | 1.134571584 |
| 0.030137066 | 1.095214169 |
| 0.007857741 | 1.206442723 |
| 0.001191193 | 0.955989411 |
| 0.000982899 | 0.998878784 |
| 0.065185377 | 1.051912421 |
| 0.006558115 | 0.962496204 |
| 0.073486228 | 1.231911002 |
| 0.061426358 | 1.196645003 |
| 0.014720379 | 0.990341234 |
| 0.00277107 | 1.160391021 |
| 0.022428799 | 1.082548878 |
| 0.014720379 | 1.020715511 |
| 0.001191193 | 1.18960953 |
| 0.000982899 | 1.168390017 |
| 0.574822422 | 0.952167315 |
| 0.004588841 | 1.223591194 |
| 0.001411669 | 1.034233048 |
| 0.00078174 | 1.110409202 |
| 0.037918944 | 0.999697575 |
| 0.002633395 | 1.111603025 |
| 0.070400511 | 1.058065687 |
| 0.017876332 | 1.096476665 |
| 0.00319857 | 1.042219244 |
| 0.00280565 | 1.065340355 |
| 0.005343552 | 0.960549709 |
| 0.002072329 | 1.04625715 |
| 0.027333265 | 0.985424079 |
| 0.024549803 | 0.969109388 |
| 0.015758743 | 1.089622895 |
| 0.006558115 | 1.024597631 |
| 0.069510564 | 1.0324663 |
| 0.028473281 | 0.952568345 |
| 0.005343552 | 0.966988612 |
| 0.002072329 | 1.150335737 |
| 0.199957919 | 0.915069436 |
| 0.047145268 | 0.955291076 |
| 0.091298681 | 1.057959371 |
| 0.020376345 | 1.054782919 |
| 0.142593359 | 1.135008916 |
| 0.014737804 | 1.026389212 |
| 0.006481784 | 1.057553509 |
| 0.003044841 | 0.953743443 |
| 0.024939603 | 0.988905215 |
| 0.013463222 | 0.993149482 |

| | |
|---|---|
| 0.270121058 | 0.919907092 |
| 0.001202797 | 1.026606773 |
| 0.006359663 | 1.250626866 |
| 0.001082792 | 1.182683451 |
| 0.068312954 | 1.055042482 |
| 0.001032472 | 1.110927152 |
| 0.041564475 | 1.014818847 |
| 0.002633395 | 1.181700793 |
| 0.019980936 | 1.009496159 |
| 0.002633395 | 0.997786386 |
| 0.014072557 | 1.142991065 |
| 0.001558842 | 0.923172039 |
| 0.00222996 | 1.235053681 |
| 0.001825091 | 1.249469734 |
| 0.002694495 | 1.127214685 |
| 0.001476826 | 1.096937675 |
| 0.001865932 | 1.093801917 |
| 0.001331434 | 0.837088237 |
| 0.086993848 | 0.962595685 |
| 0.083483723 | 0.900578908 |
| 0.032614586 | 0.998911992 |
| 0.003638599 | 1.101690391 |
| 0.018447484 | 1.130993116 |
| 0.00096885 | 1.18516108 |
| 0.208643306 | 0.97962403 |
| 0.037981715 | 1.165207877 |
| 0.038361513 | 1.008883992 |
| 0.007996352 | 0.99670898 |
| 0.063456519 | 0.943285797 |
| 0.028724094 | 1.141643474 |
| 0.002276042 | 1.122274232 |
| 0.000804045 | 1.13648294 |
| 0.026914164 | 0.998196317 |
| 0.025056112 | 0.998867126 |
| 0.04255449 | 1.023251914 |
| 0.001869087 | 1.03013653 |
| 0.073269268 | 1.160873883 |
| 0.002909003 | 1.118015606 |
| 0.032180196 | 1.161350171 |
| 0.00078174 | 1.005562889 |
| 0.002909003 | 1.013954548 |
| 0.001871173 | 1.047500828 |
| 0.014720379 | 0.927050899 |
| 0.00277107 | 1.105901521 |
| 0.014720379 | 0.996568596 |
| 0.00277107 | 1.136495293 |
| 0.181583275 | 1.098712999 |
| 0.005028294 | 1.212437285 |
| 0.002349315 | 1.120837401 |
| 0.000819731 | 1.061068395 |
| 0.029643776 | 0.986994402 |
| 0.001331856 | 1.103365931 |
| 0.021974282 | 0.97428705 |
| 0.00372282 | 0.991206868 |
| 0.068426102 | 0.907782493 |
| 0.001865932 | 1.158594249 |
| 0.176757786 | 0.988910957 |
| 0.002909003 | 0.999316512 |
| 0.021023323 | 0.992095697 |
| 0.015116713 | 1.051320115 |
| 0.008822893 | 1.052374135 |
| 0.002276042 | 1.129238498 |
| 0.031988291 | 0.998197903 |

| | |
|---|---|
| 0.002416105 | 0.916599143 |
| 0.015346717 | 1.069613457 |
| 0.00127394 | 1.040216375 |
| 0.026075462 | 1.089126847 |
| 0.00255921 | 1.05043931 |
| 0.016220862 | 1.169833879 |
| 0.015009636 | 1.066457898 |
| 0.083666057 | 1.006771491 |
| 0.023808804 | 1.021930998 |
| 0.026834729 | 0.993960114 |
| 0.024685915 | 0.961794359 |
| 0.466895801 | 0.618663497 |
| 0.458896582 | 0.890723088 |
| 0.064786291 | 0.982210652 |
| 0.014179996 | 0.859586973 |
| 0.017175355 | 1.221296704 |
| 0.001202797 | 1.359594563 |
| 0.001191193 | 1.178799912 |
| 0.000982899 | 1.126489764 |
| 0.012892487 | 1.140025513 |
| 0.001307418 | 1.019653179 |
| 0.003336529 | 0.978268408 |
| 0.001375036 | 0.957382904 |
| 0.075675475 | 1.151308451 |
| 0.000719027 | 1.108133087 |
| 0.022844361 | 0.908259476 |
| 0.016411977 | 0.979442217 |
| 0.029602879 | 0.954953504 |
| 0.000565193 | 1.077774162 |
| 0.001439012 | 1.292807379 |
| 0.001307418 | 1.071557729 |
| 0.029087519 | 0.838512901 |
| 0.014642876 | 1.029119672 |
| 4.776015199 | 0.445672799 |
| 2.309972177 | 0.167081845 |
| 0.049568305 | 1.080379811 |
| 0.002391338 | 1.1525957 |
| 0.014724612 | 1.089095745 |
| 0.003638599 | 1.101734875 |
| 0.001733606 | 1.117254008 |
| 0.0013142 | 0.948537159 |
| 0.077456381 | 1.103277061 |
| 0.017280057 | 1.006037478 |
| 0.034046604 | 0.943095531 |
| 0.001902883 | 0.909083409 |
| 0.007193339 | 1.082004556 |
| 0.000819731 | 1.077653879 |
| 1.801154757 | 0.355698582 |
| 1.661861462 | 0.393578285 |
| 0.003334047 | 0.978576038 |
| 0.000927163 | 1.212094525 |
| 0.00096885 | 1.098492244 |
| 0.000819731 | 1.099794694 |
| 0.10752231 | 0.920707733 |
| 0.002416105 | 0.974591967 |
| 0.021740469 | 1.131151133 |
| 0.014307835 | 1.198167396 |
| 0.017104655 | 1.143091931 |
| 0.003244676 | 1.035917593 |
| 0.002143971 | 1.134253868 |
| 0.001313047 | 1.149226719 |
| 0.001915272 | 1.059060545 |
| 0.000966324 | 1.10355905 |

| | |
|---|---|
| 0.026278748 | 0.956891752 |
| 0.002276042 | 1.01061765 |
| 0.00127394 | 1.110198533 |
| 0.001126814 | 1.05619184 |
| 0.012317105 | 0.913085568 |
| 0.001331856 | 1.052080964 |
| 0.007235948 | 1.196581961 |
| 0.00127394 | 1.228440341 |
| 0.003790939 | 1.01976593 |
| 0.001028149 | 0.704975304 |
| 0.000865983 | 1.277886949 |
| 0.000561696 | 1.17972238 |
| 0.17129018 | 0.952499172 |
| 0.018229436 | 1.036509654 |
| 0.017104655 | 1.08191072 |
| 0.003244676 | 1.025074546 |
| 0.146844183 | 1.104435617 |
| 0.01862689 | 1.016802415 |
| 0.031995133 | 1.059766529 |
| 0.006558115 | 0.988612208 |
| 0.040447147 | 0.995355787 |
| 0.012663236 | 1.024308193 |
| 0.064078671 | 1.050122458 |
| 0.001871173 | 1.052135055 |
| 0.001619105 | 1.026776749 |
| 0.000896779 | 1.036610788 |
| 0.187762723 | 0.939755456 |
| 0.002479335 | 0.976168333 |
| 0.183198193 | 0.956869969 |
| 0.001202797 | 0.98341396 |
| 0.014720379 | 0.939569168 |
| 0.00277107 | 1.096850109 |
| 0.136000675 | 0.969459491 |
| 0.001680008 | 0.950684814 |
| 0.101205479 | 1.015093695 |
| 0.097507028 | 1.063389606 |
| 0.078584943 | 1.083747104 |
| 0.003629625 | 1.0496668 |
| 0.033230809 | 1.044039879 |
| 0.001028149 | 0.993615227 |
| 0.093212476 | 0.995171495 |
| 0.032143317 | 0.933374322 |
| 0.065276072 | 0.981262089 |
| 0.003638599 | 0.993371886 |
| 0.069128442 | 1.108575155 |
| 0.063095625 | 1.002985234 |
| 0.009539694 | 1.121225643 |
| 0.00170488 | 1.296256983 |
| 0.092422351 | 0.963319204 |
| 0.091888365 | 0.895143664 |
| 0.007857741 | 1.250962132 |
| 0.00155386 | 1.17510363 |
| 0.209432685 | 0.88515651 |
| 0.002305914 | 0.6533571 |
| 0.0631753 | 1.090588187 |
| 0.001032472 | 1.136271014 |
| 0.056580854 | 0.955861187 |
| 0.001598043 | 0.989077274 |
| 0.00319857 | 0.958133465 |
| 0.00280565 | 0.952116247 |
| 0.000865983 | 1.301005664 |
| 0.000561696 | 1.123670752 |
| 0.038948882 | 0.930626469 |

| | |
|---|---|
| 0.01362269 | 1.196793986 |
| 0.048863967 | 1.015018444 |
| 0.003253295 | 0.968814657 |
| 0.014720379 | 0.945987164 |
| 0.00277107 | 1.054308472 |
| 0.022971608 | 1.010508077 |
| 0.001869087 | 1.123043623 |
| 0.082490484 | 0.956662642 |
| 0.00277107 | 1.167813179 |
| 0.002710464 | 1.108085072 |
| 0.000865983 | 1.119986129 |
| 0.088517181 | 1.069589422 |
| 0.02265644 | 1.296941858 |
| 0.00222996 | 1.095264047 |
| 0.001825091 | 1.150106053 |
| 0.001331856 | 1.107345918 |
| 0.000670289 | 1.061829202 |
| 0.053784577 | 0.956074418 |
| 0.001598043 | 1.067818715 |
| 0.002052694 | 1.023440079 |
| 0.001902883 | 0.912960977 |
| 0.018896759 | 1.000686536 |
| 0.002132695 | 1.058274552 |
| 0.017104655 | 1.107941894 |
| 0.003244676 | 1.02900515 |
| 0.008106845 | 0.905516862 |
| 0.002134651 | 0.910672931 |
| 0.027148712 | 0.842369722 |
| 0.001733606 | 1.015419705 |
| 0.002388469 | 1.076567988 |
| 0.001768692 | 0.963813441 |
| 0.021740469 | 1.040883397 |
| 0.001062877 | 1.077817287 |
| 0.058743372 | 0.97576789 |
| 0.021579507 | 1.048572071 |
| 0.006839002 | 1.185752134 |
| 0.001307418 | 1.111783015 |
| 0.00222996 | 1.192926045 |
| 0.001825091 | 1.16886931 |
| 0.092637505 | 1.254771758 |
| 0.020902672 | 1.146954384 |
| 0.023156052 | 1.157977414 |
| 0.009948308 | 1.076842595 |
| 0.00319857 | 1.075867664 |
| 0.00280565 | 1.153013458 |
| 0.073219772 | 0.826895315 |
| 0.001865932 | 1.043961661 |
| 0.065564756 | 1.157205599 |
| 0.001032472 | 1.065461029 |
| 0.040173454 | 0.948118048 |
| 0.032611276 | 1.001400917 |
| 0.032854278 | 0.989383256 |
| 0.001733606 | 1.080760705 |
| 0.00319857 | 1.012697517 |
| 0.00280565 | 0.984883948 |
| 0.014051002 | 1.010302118 |
| 0.001766854 | 1.026867628 |
| 0.044885604 | 0.970123227 |
| 0.015945101 | 1.100365118 |
| 0.05881112 | 1.091072402 |
| 0.022116111 | 1.030774834 |
| 0.060987791 | 1.15349822 |
| 0.039056374 | 1.049348994 |

| | |
|---|---|
| 0.013653072 | 0.978285853 |
| 0.002633395 | 1.055524811 |
| 0.000865983 | 1.146457057 |
| 0.000561696 | 1.138160565 |
| 0.038361513 | 1.024312199 |
| 0.004588841 | 1.116037164 |
| 0.054452982 | 0.99621515 |
| 0.010206479 | 1.086892729 |
| 0.139774347 | 1.209561009 |
| 0.033661252 | 1.137177401 |
| 0.014720379 | 1.046641672 |
| 0.00277107 | 1.273352643 |
| 0.023516299 | 1.058108437 |
| 0.000184666 | 1.151016737 |
| 0.0142664 | 1.059637294 |
| 0.008014126 | 1.13195211 |
| 0.048878722 | 1.162056697 |
| 0.023132243 | 1.087902291 |
| 0.016485641 | 0.915243121 |
| 0.012051524 | 1.053256123 |
| 0.00372282 | 1.218911485 |
| 0.000698103 | 0.881252922 |
| 0.02922896 | 1.306789119 |
| 0.008014126 | 1.300543432 |
| 0.007580029 | 1.20447408 |
| 0.000927163 | 1.446059005 |
| 0.020254702 | 1.123229122 |
| 0.00138486 | 0.979638878 |
| 0.039503636 | 1.079670773 |
| 0.006558115 | 1.013817188 |
| 0.006338964 | 1.022746386 |
| 0.001202797 | 1.047224142 |
| 0.103386244 | 0.952076337 |
| 0.001598043 | 1.131072709 |
| 0.04814538 | 1.006663468 |
| 0.001598043 | 1.062601891 |
| 0.002276042 | 1.126155954 |
| 0.000804045 | 1.065736101 |
| 0.017104655 | 1.107676811 |
| 0.003244676 | 0.863106533 |
| 0.133450568 | 1.098514557 |
| 0.047342609 | 0.968281718 |
| 0.017104655 | 1.11732584 |
| 0.003244676 | 1.066955815 |
| 0.07234072 | 1.100459381 |
| 0.003638599 | 1.059964413 |
| 0.057596483 | 1.127885282 |
| 0.001598043 | 1.138408869 |
| 0.074229562 | 1.075939457 |
| 0.038264687 | 1.331330229 |
| 0.660373104 | 0.279069767 |
| 0.002391338 | 0.396040902 |
| 0.015107388 | 1.147872588 |
| 0.008014126 | 1.137428887 |
| 0.068154173 | 1.002974862 |
| 0.00170488 | 1.102178771 |
| 0.048164329 | 1.036895905 |
| 0.002633395 | 1.187234828 |
| 0.016830651 | 1.066401341 |
| 0.002368462 | 1.100037715 |
| 0.16992043 | 1.13242071 |
| 0.017104655 | 1.140282049 |
| 0.042230562 | 0.964384403 |

| | |
|---|---|
| 0.006359663 | 1.119562189 |
| 0.036989578 | 1.046700081 |
| 0.001902883 | 0.985974755 |
| 0.017104655 | 1.122203372 |
| 0.003244676 | 1.022228246 |
| 0.0062469 | 1.023502744 |
| 0.002276042 | 1.024660349 |
| 0.020254702 | 1.005810882 |
| 0.00138486 | 1.037904981 |
| 0.017104655 | 0.986654892 |
| 0.001885 | 0.898060606 |
| 0.123857246 | 1.145128349 |
| 0.001202797 | 1.25547109 |
| 0.002276042 | 1.155268866 |
| 0.000804045 | 1.198759246 |
| 0.002366011 | 1.180971064 |
| 0.001772163 | 1.194415997 |
| 0.037527326 | 1.001183496 |
| 0.006558115 | 1.028697237 |
| 0.038361513 | 1.021937059 |
| 0.004588841 | 1.108361947 |
| 0.15879325 | 0.863453815 |
| 0.00170488 | 0.92698324 |
| 0.00319857 | 1.07050649 |
| 0.00280565 | 0.9953189 |
| 0.018498299 | 0.922951064 |
| 0.00170488 | 1.070167598 |
| 0.003638599 | 1.094350534 |
| 0.001166314 | 0.668882979 |
| 0.020022089 | 0.993847957 |
| 0.001558842 | 0.864149699 |
| 0.06246291 | 0.870147256 |
| 0.00170488 | 1.154916201 |
| 0.185832653 | 0.720957906 |
| 0.004170207 | 0.824357542 |
| 0.002723189 | 1.071842927 |
| 0.00170488 | 1.110055866 |
| 0.028340169 | 0.945579422 |
| 0.001865932 | 1.163194888 |
| 0.038588972 | 1.134586895 |
| 0.01708429 | 0.974611209 |
| 0.00170488 | 1.143240223 |
| 0.001420547 | 1.052803808 |
| 0.007854284 | 1.160420273 |
| 0.00096885 | 1.231695412 |
| 0.134691449 | 1.081839702 |
| 0.001486793 | 1.051068569 |
| 0.032632536 | 0.979815518 |
| 0.03164864 | 0.935384864 |
| 0.026089917 | 1.09865949 |
| 0.002391338 | 1.108678553 |
| 0.061632789 | 1.061623001 |
| 0.001307418 | 0.974922188 |
| 0.00222996 | 1.073300997 |
| 0.001825091 | 1.236090716 |
| 0.020254702 | 1.142918888 |
| 0.00138486 | 1.109104879 |
| 0.017104655 | 1.084614569 |
| 0.003244676 | 0.98441312 |
| 0.002134651 | 0.946469927 |
| 0.001155271 | 0.90625 |
| 0.10581827 | 1.233230589 |
| 0.06978358 | 0.937190794 |

| | |
|---|---|
| 0.015663311 | 1.030286729 |
| 0.006558115 | 1.115244458 |
| 0.13871231 | 1.026939706 |
| 0.000947294 | 0.632960683 |
| 0.00127394 | 1.037554165 |
| 0.001126814 | 1.090103794 |
| 0.017104655 | 1.001484466 |
| 0.000280346 | 0.987394958 |
| 0.042061335 | 0.961415635 |
| 0.001865932 | 1.174696486 |
| 0.002818048 | 1.089110585 |
| 0.001486793 | 1.080076364 |
| 0.05842729 | 1.102437402 |
| 0.049388191 | 1.117824394 |
| 0.177184316 | 1.062901589 |
| 0.005327897 | 1.018242562 |
| 0.064616732 | 1.029840332 |
| 0.001032472 | 1.036041773 |
| 0.020254702 | 1.11578543 |
| 0.00138486 | 1.060699193 |
| 0.001902883 | 0.897945714 |
| 0.000147977 | 0.948197725 |
| 0.011444237 | 0.98647717 |
| 0.002818048 | 1.114305039 |
| 0.028572449 | 1.037840255 |
| 0.001869087 | 1.063103563 |
| 0.006359663 | 0.967243781 |
| 0.001136057 | 0.989560946 |
| 0.013237257 | 1.038845936 |
| 0.001465187 | 1.043270094 |
| 0.118368122 | 0.876571567 |
| 0.003638599 | 0.971129893 |
| 0.02152663 | 0.977154229 |
| 0.001082792 | 0.716610378 |
| 0.002366011 | 1.177211051 |
| 0.001772163 | 1.207998706 |
| 0.045505182 | 1.045890547 |
| 0.025889882 | 0.981475796 |
| 0.052069136 | 0.956532447 |
| 0.001032472 | 0.988665308 |
| 0.038361513 | 1.004981197 |
| 0.004588841 | 1.10401939 |
| 0.075388267 | 0.7715871 |
| 0.042698835 | 0.664217875 |
| 0.000865983 | 1.04982083 |
| 0.000561696 | 1.050206997 |
| 0.011455577 | 0.992856563 |
| 0.000865983 | 1.106346087 |
| 0.019225695 | 1.075452445 |
| 0.001902883 | 0.929048758 |
| 0.030870309 | 1.165743135 |
| 0.001032472 | 1.223127866 |
| 0.000865983 | 1.18021038 |
| 0.000561696 | 1.186257001 |
| 0.022986233 | 1.177663818 |
| 0.014952938 | 1.063594432 |
| 0.053815683 | 1.007516106 |
| 0.00127394 | 0.957206378 |
| 0.002818048 | 1.150490362 |
| 0.001486793 | 1.147849605 |
| 0.00222996 | 1.073300997 |
| 0.001825091 | 1.169358786 |
| 0.038361513 | 1 |

| | |
|---|---|
| 0.038361513 | 1 |
| 0.021740469 | 1 |
| 0.020254702 | 1 |
| 0.017104655 | 1 |
| 0.017104655 | 1 |
| 0.017104655 | 1 |
| 0.014720379 | 1 |
| 0.008014126 | 1 |
| 0.007996352 | 1 |
| 0.007857741 | 1 |
| 0.007371037 | 1 |
| 0.006558115 | 1 |
| 0.006359663 | 1 |
| 0.006338964 | 1 |
| 0.006123397 | 1 |
| 0.005988527 | 1 |
| 0.005343552 | 1 |
| 0.004588841 | 1 |
| 0.004380884 | 1 |
| 0.004238364 | 1 |
| 0.004123449 | 1 |
| 0.003790939 | 1 |
| 0.00372282 | 1 |
| 0.00371134 | 1 |
| 0.003638599 | 1 |
| 0.003334047 | 1 |
| 0.003244676 | 1 |
| 0.00319857 | 1 |
| 0.0030477 | 1 |
| 0.003044841 | 1 |
| 0.003022735 | 1 |
| 0.002934379 | 1 |
| 0.002909003 | 1 |
| 0.002870236 | 1 |
| 0.002818048 | 1 |
| 0.00280565 | 1 |
| 0.002790486 | 1 |
| 0.00277107 | 1 |
| 0.00277107 | 1 |
| 0.002694495 | 1 |
| 0.002688948 | 1 |
| 0.002633395 | 1 |
| 0.002605549 | 1 |
| 0.002552407 | 1 |
| 0.002416105 | 1 |
| 0.002391338 | 1 |
| 0.002388469 | 1 |
| 0.002366011 | 1 |
| 0.002305914 | 1 |
| 0.002276042 | 1 |
| 0.002269165 | 1 |
| 0.002229985 | 1 |
| 0.00222996 | 1 |
| 0.002168057 | 1 |
| 0.002143971 | 1 |
| 0.002134651 | 1 |
| 0.002132695 | 1 |
| 0.002130938 | 1 |
| 0.002094333 | 1 |
| 0.002072329 | 1 |
| 0.002052694 | 1 |
| 0.001915272 | 1 |
| 0.001902883 | 1 |

| | |
|---|---|
| 0.001871173 | 1 |
| 0.001869087 | 1 |
| 0.001865932 | 1 |
| 0.001865417 | 1 |
| 0.001825091 | 1 |
| 0.001772163 | 1 |
| 0.001768692 | 1 |
| 0.001766854 | 1 |
| 0.001733606 | 1 |
| 0.00170488 | 1 |
| 0.001680008 | 1 |
| 0.001655398 | 1 |
| 0.001619105 | 1 |
| 0.001598043 | 1 |
| 0.001565608 | 1 |
| 0.001558842 | 1 |
| 0.001556734 | 1 |
| 0.001486793 | 1 |
| 0.001476826 | 1 |
| 0.001458435 | 1 |
| 0.001439012 | 1 |
| 0.001420547 | 1 |
| 0.001411669 | 1 |
| 0.00138486 | 1 |
| 0.001331856 | 1 |
| 0.001331434 | 1 |
| 0.0013142 | 1 |
| 0.001313047 | 1 |
| 0.001307418 | 1 |
| 0.00129926 | 1 |
| 0.00127394 | 1 |
| 0.001202797 | 1 |
| 0.001191193 | 1 |
| 0.001166314 | 1 |
| 0.001155271 | 1 |
| 0.001148102 | 1 |
| 0.001126814 | 1 |
| 0.001082792 | 1 |
| 0.001062877 | 1 |
| 0.001053652 | 1 |
| 0.001032472 | 1 |
| 0.001028149 | 1 |
| 0.000998735 | 1 |
| 0.000982899 | 1 |
| 0.00096885 | 1 |
| 0.000966324 | 1 |
| 0.000963657 | 1 |
| 0.000947294 | 1 |
| 0.000927163 | 1 |
| 0.000908458 | 1 |
| 0.000896779 | 1 |
| 0.000872781 | 1 |
| 0.000865983 | 1 |
| 0.000847692 | 1 |
| 0.000819731 | 1 |
| 0.000804045 | 1 |
| 0.00078174 | 1 |
| 0.000766139 | 1 |
| 0.000719027 | 1 |
| 0.000698103 | 1 |
| 0.000686039 | 1 |
| 0.000670289 | 1 |
| 0.00057121 | 1 |

| | |
|---|---|
| 0.000561696 | 1 |
| 0.017104655 | 1.137631216 |
| 0.003244676 | 1.013418271 |
| 0.044927274 | 0.925375805 |
| 0.00127394 | 0.970205897 |
| 0.10872909 | 0.95828877 |
| 0.031886862 | 0.953962704 |
| 0.148142629 | 0.953595373 |
| 0.001062877 | 1.044857768 |
| 0.020254702 | 1.102290736 |
| 0.00138486 | 0.976821616 |
| 0.022562719 | 0.994450419 |
| 0.008685089 | 1.207984735 |
| 0.042224093 | 1.104192731 |
| 0.02986072 | 1.079759629 |
| 1.088542014 | 0.440705313 |
| 0.588193809 | 0.304462055 |
| 0.002818048 | 1.180250254 |
| 0.001486793 | 1.082993053 |
| 0.017104655 | 1.067694733 |
| 0.002934379 | 0.975956005 |
| 0.30004694 | 0.813246362 |
| 0.017901454 | 0.720932225 |
| 0.632715257 | 0.782725573 |
| 0.00372282 | 0.399191532 |
| 0.006753273 | 0.991410165 |
| 0.002276042 | 0.895992693 |
| 0.00319857 | 1.057985327 |
| 0.00280565 | 0.985566608 |
| 0.016060886 | 0.888651826 |
| 0.00096885 | 1.009003146 |
| 0.002694495 | 1.07565842 |
| 0.001476826 | 1.036445978 |
| 0.001331856 | 1.148737776 |
| 0.000670289 | 1.121175134 |
| 0.038361513 | 0.927481836 |
| 0.007996352 | 0.985707569 |
| 0.118972259 | 1.185628486 |
| 0.000719027 | 1.132030631 |
| 0.017104655 | 0.943430478 |
| 0.002229985 | 0.859757576 |
| 0.002614488 | 0.970857292 |
| 0.001458435 | 1.100636631 |
| 0.017104655 | 1.051516044 |
| 0.002229985 | 0.963636364 |
| 0.015853425 | 0.964148806 |
| 0.015016196 | 1.177032321 |
| 0.001331856 | 0.984648624 |
| 0.000670289 | 1.055305358 |
| 0.012037795 | 1.382538569 |
| 0.00372282 | 1.270128355 |
| 0.002818048 | 1.238924586 |
| 0.001486793 | 1.199607573 |
| 0.024965673 | 1.174512354 |
| 0.001028149 | 1.136850982 |
| 0.066592886 | 0.95436556 |
| 0.002132695 | 1.047639718 |
| 0.017104655 | 1.076439996 |
| 0.002229985 | 0.89030303 |
| 0.006936631 | 1.095889119 |
| 0.003847795 | 1.098449937 |
| 0.014720379 | 1.01169219 |
| 0.00277107 | 1.230086894 |

| | |
|---|---|
| 0.00222996 | 1.155049321 |
| 0.001825091 | 1.157611356 |
| 0.020254702 | 1.033472602 |
| 0.00138486 | 1.038161096 |
| 0.125662773 | 0.99449099 |
| 0.020056708 | 1.154542213 |
| 0.030647095 | 1.017846972 |
| 0.025351061 | 0.89768357 |
| 0.000865983 | 1.038955034 |
| 0.000561696 | 0.909692345 |
| 0.002818048 | 1.216435577 |
| 0.001486793 | 1.128281275 |
| 0.007371037 | 1.060136816 |
| 0.00277107 | 0.969669777 |
| 0.054675893 | 1.046654119 |
| 0.001202797 | 1.081778392 |
| 0.003334047 | 1.077651108 |
| 0.000927163 | 1.222515779 |
| 0.041299582 | 0.997675829 |
| 0.000686039 | 1.153703346 |
| 0.122428385 | 1.031449318 |
| 0.002229985 | 0.844848485 |
| 0.160408274 | 0.804966587 |
| 0.086511266 | 0.693358144 |
| 0.002818048 | 1.171119378 |
| 0.001486793 | 1.095826484 |
| 0.002818048 | 1.197835644 |
| 0.001486793 | 1.09179615 |
| 0.036369915 | 1.287999215 |
| 0.001307418 | 1.189773233 |
| 0.008760683 | 1.050870788 |
| 0.003461136 | 1.032905902 |
| 0.002818048 | 1.212208319 |
| 0.001486793 | 1.122341836 |
| 0.015960153 | 1.075424575 |
| 0.013754243 | 1.051981443 |
| 0.074110131 | 1.07644868 |
| 0.001202797 | 1.149965446 |
| 0.006694193 | 1.011560694 |
| 0.002818048 | 1.120054109 |
| 0.006374436 | 1.055658627 |
| 0.00078174 | 1.064868942 |
| 0.082610216 | 0.9997426 |
| 0.006074765 | 1.118695537 |
| 0.307592973 | 1.220491154 |
| 0.004380884 | 1.237323037 |
| 0.017104655 | 0.94033601 |
| 0.002934379 | 0.910218698 |
| 0.002276042 | 1.104235643 |
| 0.000804045 | 1.108963652 |
| 0.002416105 | 1.098680403 |
| 0.001766854 | 1.032437746 |
| 0.033518447 | 1.137154375 |
| 0.026413246 | 1.126094092 |
| 0.00127394 | 1.15432326 |
| 0.001126814 | 1.118468146 |
| 0.649549944 | 0.506838738 |
| 0.487203231 | 0.417216893 |
| 0.041133491 | 0.817212121 |
| 0.017104655 | 0.955058385 |
| 0.040954484 | 1.33345021 |
| 0.00372282 | 1.18469745 |
| 0.002366011 | 1.149583129 |

| | |
|---|---|
| 0.001772163 | 1.058427209 |
| 0.080989324 | 1.043611715 |
| 0.017104655 | 1.00995221 |
| 0.11059402 | 1.04778105 |
| 0.002391338 | 1.084032512 |
| 0.021729528 | 0.950060606 |
| 0.017104655 | 1.025071141 |
| 0.053589065 | 1.272565169 |
| 0.015940538 | 1.228086492 |
| 1 | 0.352608502 |
| 1 | 0.352985075 |
| 1 | 0.363613738 |
| 1 | 0.36454364 |
| 1 | 0.329204769 |
| 1 | 0.338668337 |
| 1 | 0.32301157 |
| 1 | 0.322885015 |
| 1 | 0.312714589 |
| 1 | 0.341899441 |
| 1 | 0.294943512 |
| 1 | 0.29872196 |
| 1 | 0.299870606 |
| 1 | 0.297256508 |
| 1 | 0.292538598 |
| 1 | 0.248403253 |
| 1 | 0.221036952 |
| 1 | 0.236874175 |
| 1 | 0.187440779 |
| 1 | 0.205827027 |
| 1 | 0.175814602 |
| 1 | 0.191351763 |
| 1 | 0.190606702 |
| 1 | 0.197255277 |
| 1 | 0.181876736 |
| 1 | 0.178959582 |
| 1 | 0.170224856 |
| 1 | 0.184140747 |
| 1 | 0.154006286 |
| 1 | 0.170961584 |
| 1 | 0.182624588 |
| 1 | 0.173693563 |
| 1 | 0.221214958 |
| 1 | 0.212331672 |
| 1 | 0.203905407 |
| 1 | 0.196225328 |
| 1 | 0.171026686 |
| 1 | 0.216990545 |
| 1 | 0.170289547 |
| 1 | 0.153692282 |
| 1 | 0.165191622 |
| 1 | 0.169950249 |
| 1 | 0.162185416 |
| 1 | 0.15581004 |
| 1 | 0.166023166 |
| 1 | 0.168097096 |
| 1 | 0.171530249 |
| 1 | 0.146894772 |
| 1 | 0.193106298 |
| 1 | 0.189186514 |
| 1 | 0.196282449 |
| 1 | 0.195957123 |
| 1 | 0.175357004 |
| 1 | 0.185743198 |

| | |
|---|---|
| 1 | 0.161170627 |
| 1 | 0.193273345 |
| 1 | 0.167997947 |
| 1 | 0.179345708 |
| 1 | 0.14942366 |
| 1 | 0.153460797 |
| 1 | 0.155770714 |
| 1 | 0.157843855 |
| 1 | 0.136078514 |
| 1 | 0.158922704 |
| 1 | 0.448044895 |
| 1 | 0.420674803 |
| 1 | 0.380384677 |
| 1 | 0.434790629 |
| 1 | 0.372870249 |
| 1 | 0.39142668 |
| 1 | 0.408924409 |
| 1 | 0.402673622 |
| 1 | 0.396149453 |
| 1 | 0.343299122 |
| 1 | 0.357178506 |
| 1 | 0.375211363 |
| 1 | 0.354189944 |
| 1 | 0.34641164 |
| 1 | 0.313999393 |
| 1 | 0.343720868 |
| 1 | 0.357000746 |
| 1 | 0.314761405 |
| 1 | 0.281087934 |
| 1 | 0.328051118 |
| 1 | 0.289422135 |
| 1 | 0.258297105 |
| 1 | 0.257886609 |
| 1 | 0.273345978 |
| 1 | 0.237986766 |
| 1 | 0.234135004 |
| 1 | 0.247139059 |
| 1 | 0.249715715 |
| 1 | 0.226096866 |
| 1 | 0.27411805 |
| 1 | 0.235624928 |
| 1 | 0.240586001 |
| 1 | 0.298631539 |
| 1 | 0.288018433 |
| 1 | 0.253454545 |
| 1 | 0.290562172 |
| 1 | 0.280224289 |
| 1 | 0.264665243 |
| 1 | 0.222157638 |
| 1 | 0.217508181 |
| 1 | 0.232550104 |
| 1 | 0.188619384 |
| 1 | 0.295464553 |
| 1 | 0.249745288 |
| 1 | 0.241351994 |
| 1 | 0.207685821 |
| 1 | 0.213974855 |
| 1 | 0.204634581 |
| 1 | 0.254348848 |
| 1 | 0.256513836 |
| 1 | 0.283596645 |
| 1 | 0.21875 |
| 1 | 0.225555345 |

| | |
|---|---|
| 1 | 0.21711646 |
| 1 | 0.23703564 |
| 1 | 0.199808629 |
| 1 | 0.216061185 |
| 1 | 0.212980854 |
| 1 | 0.218951324 |
| 1 | 0.229614366 |
| 1 | 0.233115906 |
| 1 | 0.181106257 |
| 1 | 0.184803234 |
| 1 | 0.178179735 |
| 0.129560349 | 0.958831014 |
| 0.012209241 | 0.966123281 |
| 0.020254702 | 1.067377419 |
| 0.00138486 | 1.020233064 |
| 0.001530765 | 0.889073788 |
| 0.000865983 | 1.087735522 |
| 0.017104655 | 0.934255716 |
| 0.002229985 | 0.887515152 |
| 0.030775493 | 1.002881239 |
| 0.001902883 | 0.879052883 |
| 0.05526454 | 1.155805375 |
| 0.014052614 | 1.240816816 |
| 0.020254702 | 1.135379148 |
| 0.00138486 | 1.039697785 |
| 0.07975805 | 1.04039035 |
| 0.017104655 | 1.134397201 |
| 0.155027406 | 1.156791942 |
| 0.04385149 | 1.099484989 |
| 0.024943769 | 1.090321406 |
| 0.020722715 | 1.21111218 |
| 0.004123449 | 1.033442865 |
| 0.002688948 | 1.027336716 |
| 0.056690689 | 1.00149328 |
| 0.003044841 | 0.971705611 |
| 1.300306858 | 0.719426303 |
| 0.020254702 | 0.451183787 |
| 0.010379822 | 1.096736597 |
| 0.00501663 | 0.98999497 |
| 0.070449186 | 1.040238851 |
| 0.006558115 | 1.05481324 |
| 0.006686206 | 1.039620688 |
| 0.001825091 | 1.123511176 |
| 0.014720379 | 1.001461524 |
| 0.00277107 | 1.126176684 |
| 0.055913027 | 1.006725241 |
| 0.024142974 | 1.032889175 |
| 0.012820145 | 1.199277593 |
| 0.001032472 | 1.237137035 |
| 0.002143971 | 0.968724006 |
| 0.001313047 | 0.978008117 |
| 0.210478023 | 0.95244969 |
| 0.002917298 | 0.880492909 |
| 0.017104655 | 1.007997252 |
| 0.002229985 | 0.859757576 |
| 0.005343552 | 0.984671568 |
| 0.002072329 | 1.058194479 |
| 0.907691051 | 0.461905845 |
| 0.624910889 | 0.365840313 |
| 0.038151824 | 1.053277511 |
| 0.022848245 | 1.08138054 |
| 0.066609612 | 1.208201514 |
| 0.001202797 | 1.342778162 |

| | |
|---|---|
| 0.043591116 | 1.181042096 |
| 0.014065902 | 1.147976879 |
| 0.002388469 | 0.952333166 |
| 0.001768692 | 0.955902094 |
| 0.002168057 | 1.062779754 |
| 0.001065908 | 1.162209147 |
| 0.032417788 | 1.007303161 |
| 0.025159398 | 1.015650473 |
| 0.044110127 | 0.89375152 |
| 0.001733606 | 1.094702689 |
| 0.00129926 | 0.900767987 |
| 0.000686039 | 0.866099401 |
| 0.003790939 | 1.032379714 |
| 0.001028149 | 0.983495964 |
| 0.038361513 | 1.071347405 |
| 0.007996352 | 0.972637518 |
| 0.042526478 | 1.202335394 |
| 0.002183662 | 1.126159775 |
| 0.056500329 | 1.027226418 |
| 0.009201895 | 1.124471498 |
| 0.00371134 | 0.987369338 |
| 0.003044841 | 0.942298522 |
| 0.077844636 | 0.965756077 |
| 0.000927163 | 0.927344782 |
| 0.017104655 | 1.099008929 |
| 0.002229985 | 0.902666667 |
| 0.002366011 | 1.138793526 |
| 0.001772163 | 1.129035735 |
| 0.001191193 | 1.137326274 |
| 0.000982899 | 1.050620821 |
| 0.007371037 | 0.986940299 |
| 0.00277107 | 1.118628859 |
| 0.002366011 | 1.164296224 |
| 0.001772163 | 1.160890422 |
| 0.033049923 | 1.056972328 |
| 0.023875964 | 1.053186334 |
| 0.021740469 | 1.027255598 |
| 0.001062877 | 1.054978118 |
| 0.002694495 | 1.050598563 |
| 0.001476826 | 1.10993099 |
| 0.001191193 | 1.089344805 |
| 0.000982899 | 1.097130518 |
| 0.091984419 | 0.899262252 |
| 0.048001225 | 0.997811029 |
| 0.002366011 | 1.104953409 |
| 0.001772163 | 1.120573492 |
| 0.00319857 | 1.062570542 |
| 0.00280565 | 1.004583577 |
| 0.002388469 | 1.114500753 |
| 0.001768692 | 1.170393854 |
| 0.002366011 | 0.983488638 |
| 0.000412749 | 1.069314936 |
| 0.022043113 | 1.210143043 |
| 0.001028149 | 1.18070112 |
| 0.00372282 | 1.056967828 |
| 0.000698103 | 1.023609163 |
| 0.057417594 | 1.192115087 |
| 0.054429875 | 1.335867578 |
| 0.013304134 | 1.078356524 |
| 0.001032472 | 1.129393785 |
| 0.005343552 | 1.004516842 |
| 0.002072329 | 1.025366824 |
| 0.074564545 | 0.948460988 |

| | |
|---|---|
| 0.015795657 | 1.092242771 |
| 0.12974713 | 1.001752772 |
| 0.001062877 | 1.187226477 |
| 0.038361513 | 1.093729555 |
| 0.007996352 | 0.965679361 |
| 0.141782756 | 0.969988345 |
| 0.0476265 | 0.943434343 |
| 0.007964011 | 1.390870395 |
| 0.003334047 | 1.079414928 |
| 0.020254702 | 1.064976228 |
| 0.001318984 | 1.092841593 |
| 0.002687516 | 1.048969913 |
| 0.001439012 | 1.046806005 |
| 0.009721023 | 1.083907445 |
| 0.003190967 | 1.084411506 |
| 0.037002009 | 1.036467236 |
| 0.016788407 | 1.05551247 |
| 0.150697559 | 0.961737928 |
| 0.035240075 | 0.863381859 |
| 0.001331856 | 1.23379577 |
| 0.000670289 | 1.260364493 |
| 0.114465896 | 0.714338359 |
| 0.092687192 | 0.789900147 |
| 0.070216251 | 1.020026481 |
| 0.011406515 | 1.034345275 |
| 0.004123449 | 1.080766514 |
| 0.002688948 | 1.121296448 |
| 0.017104655 | 0.980588264 |
| 0.002934379 | 0.944366287 |
| 0.038361513 | 0.974071386 |
| 0.004588841 | 0.998283175 |
| 0.13899166 | 0.956149733 |
| 0.036062667 | 1.030802531 |
| 0.034433605 | 1.164095372 |
| 0.00372282 | 1.15690115 |
| 0.094804501 | 1.035005165 |
| 0.039637275 | 0.966524026 |
| 0.012945948 | 1.049919438 |
| 0.008945805 | 1.054818234 |
| 0.209815175 | 0.881542589 |
| 0.001202797 | 0.942985487 |
| 0.024397809 | 1.133296913 |
| 0.023302424 | 1.179931351 |
| 0.038361513 | 1.049791674 |
| 0.007996352 | 1.066666667 |
| 0.019764876 | 1.056429734 |
| 0.005204557 | 1.004846135 |
| 0.047346011 | 1.045071055 |
| 0.031811749 | 0.990439771 |
| 0.102388879 | 0.994058229 |
| 0.006123397 | 0.978396603 |
| 0.057095403 | 0.991537718 |
| 0.008654933 | 0.997553381 |
| 0.149083145 | 0.985584547 |
| 0.002072329 | 1.142004476 |
| 0.075858998 | 1.091869252 |
| 0.070771519 | 1.232861771 |
| 0.079933966 | 0.894995395 |
| 0.006359663 | 0.985671642 |
| 0.017104655 | 1.015026851 |
| 0.009231492 | 1.049750607 |
| 0.034212486 | 1.124974063 |
| 0.01692271 | 1.201059759 |

| | |
|---|---|
| 0.052267824 | 1.105306346 |
| 0.021740469 | 1.053590991 |
| 0.012528602 | 0.994082522 |
| 0.006558115 | 0.90844215 |
| 0.076242771 | 1.080454493 |
| 0.005731064 | 1.188018187 |
| 0.017700785 | 1.023298519 |
| 0.000963657 | 1.190068646 |
| 0.017104655 | 0.956742376 |
| 0.002934379 | 0.990791661 |
| 0.001191193 | 1.134458416 |
| 0.000982899 | 1.113990283 |
| 0.012503014 | 1.083374203 |
| 0.001772163 | 1.011318924 |
| 0.012802751 | 1.09040959 |
| 0.002605549 | 1.074562629 |
| 0.073310234 | 1.061609818 |
| 0.001032472 | 0.957208355 |
| 0.001191193 | 1.127178469 |
| 0.000982899 | 1.072671401 |
| 0.050737914 | 0.993736952 |
| 0.006676396 | 1.102315631 |
| 0.017104655 | 1.029413214 |
| 0.002934379 | 1.038368078 |
| 0.002123846 | 1.023816012 |
| 0.001825091 | 1.075542503 |
| 0.015666881 | 1.094550292 |
| 0.00222785 | 0.99736214 |
| 0.011540471 | 1.003988573 |
| 0.002366011 | 1.072257643 |
| 0.14010102 | 1.028776058 |
| 0.017104655 | 1.087057201 |
| 0.022234807 | 1.138213603 |
| 0.013937439 | 1.327944054 |
| 0.024857766 | 1.000596516 |
| 0.005156939 | 1.090535449 |
| 0.030305169 | 1.145924508 |
| 0.021740469 | 1.099425967 |
| 0.002276042 | 1.180500057 |
| 0.000804045 | 1.134415016 |
| 0.017104655 | 1.016357097 |
| 0.002934379 | 1.001662617 |
| 0.001865417 | 0.949229347 |
| 0.000847692 | 0.970951206 |
| 0.09885396 | 1.176644908 |
| 0.001202797 | 1.197880673 |
| 0.038361513 | 1.003166854 |
| 0.004588841 | 1.089880832 |
| 0.052882463 | 0.930845931 |
| 0.005988527 | 0.931548808 |
| 0.065383072 | 1.063593265 |
| 0.001772163 | 1.028836307 |
| 0.136448283 | 0.984168638 |
| 0.007065804 | 1.103918978 |
| 0.044125323 | 0.890808872 |
| 0.001865932 | 1.096996805 |
| 0.037278195 | 1.246197625 |
| 0.009481347 | 1.192085371 |
| 0.031509024 | 0.918853099 |
| 0.005988527 | 0.956718604 |
| 0.096175598 | 1.035590762 |
| 0.013029967 | 1.003083471 |
| 0.027936515 | 0.996271141 |

| | |
|---|---|
| 0.022898456 | 1.003997611 |
| 0.128515899 | 1.092317195 |
| 0.000947294 | 1.019218691 |
| 0.209417089 | 1.108537109 |
| 0.000719027 | 0.756403486 |
| 0.006738603 | 1.016451711 |
| 0.002391338 | 1.133717881 |
| 0.006149123 | 1.080768313 |
| 0.002276042 | 1.213608859 |
| 0.058610232 | 0.820163709 |
| 0.001733606 | 1.143531755 |
| 0.049276323 | 1.087744549 |
| 0.001598043 | 1.071079231 |
| 0.132283256 | 0.996212556 |
| 0.038023775 | 1.066506196 |
| 0.05039405 | 1.081575936 |
| 0.009004489 | 0.99401714 |
| 0.018267811 | 1.176825514 |
| 0.001032472 | 1.297631177 |
| 0.00222996 | 1.034879285 |
| 0.001825091 | 1.050089737 |
| 0.030466934 | 0.906807294 |
| 0.007590654 | 1.022019742 |
| 0.111061044 | 1.186534898 |
| 0.007459421 | 1.089184776 |
| 0.001191193 | 1.063754688 |
| 0.000982899 | 1.087413313 |
| 0.007535959 | 1.005599801 |
| 0.003044841 | 0.987283421 |
| 0.038361513 | 1.006465659 |
| 0.004588841 | 1.046556251 |
| 0.023483107 | 0.933518171 |
| 0.000847692 | 0.943231441 |
| 0.058542054 | 0.95925208 |
| 0.020686619 | 1.106131308 |
| 0.038361513 | 1.000688682 |
| 0.007996352 | 1.011753644 |
| 0.005395469 | 0.985905823 |
| 0.002132695 | 1.064134563 |
| 0.021740469 | 1.004031375 |
| 0.001062877 | 1.080689278 |
| 0.102284786 | 0.910214785 |
| 0.042148131 | 0.824480095 |
| 0.00371134 | 1.043989547 |
| 0.003044841 | 0.975361628 |
| 0.038361513 | 0.996932111 |
| 0.004588841 | 0.93950717 |
| 0.071191254 | 0.878647245 |
| 0.005988527 | 0.941270475 |
| 0.020254702 | 1.09014071 |
| 0.00138486 | 0.886028941 |
| 0.017104655 | 0.948859437 |
| 0.01046834 | 0.984013301 |
| 0.005542024 | 0.926319648 |
| 0.002134651 | 1.064414742 |
| 0.067889028 | 1.086924584 |
| 0.028533862 | 0.978872626 |
| 0.032629055 | 1.002176311 |
| 0.005683977 | 1.035029429 |
| 0.00319857 | 0.978061512 |
| 0.00280565 | 0.935732397 |
| 0.002006634 | 1.196876017 |
| 0.000819731 | 1.148061672 |

| | |
|---|---|
| 0.023079359 | 0.9506188 |
| 0.002688948 | 0.978392536 |
| 0.001733606 | 1.046933727 |
| 0.00124663 | 0.921306427 |
| 0.188502709 | 0.868090243 |
| 0.180457979 | 1.027688651 |
| 0.005030683 | 1.034967646 |
| 0.003044841 | 1.05531712 |
| 0.059980546 | 0.929163298 |
| 0.000686039 | 1.014387933 |
| 0.085224401 | 0.899875936 |
| 0.005988527 | 1.036755893 |
| 0.008513773 | 1.038598912 |
| 0.000865983 | 1.12195122 |
| 0.00222996 | 1.009591803 |
| 0.001825091 | 1.045031816 |
| 0.156385756 | 1.144837602 |
| 0.004380884 | 1.136422136 |
| 0.069049606 | 0.869503285 |
| 0.005988527 | 0.881475563 |
| 0.0163573 | 0.922758811 |
| 0.005988527 | 0.862831269 |
| 0.085134919 | 0.824748426 |
| 0.005988527 | 1.035956852 |
| 0.001865932 | 1.179936102 |
| 0.001331434 | 0.968415066 |
| 0.118460811 | 1.050275175 |
| 0.00170488 | 1.048603352 |
| 0.00222996 | 0.998092539 |
| 0.001825091 | 1.068363518 |
| 0.040699392 | 1.06077563 |
| 0.022587035 | 1.062888931 |
| 0.00096885 | 1.233973316 |
| 0.000819731 | 1.223219677 |
| 0.001558842 | 0.904047482 |
| 0.000202413 | 0.877837996 |
| 0.014810438 | 0.964454295 |
| 0.001032472 | 1.047249109 |
| 0.015951832 | 1.020358393 |
| 0.004867528 | 1.012131203 |
| 0.017253377 | 1.281044293 |
| 0.000896779 | 1.163684531 |
| 0.017104655 | 1.076930625 |
| 0.002229985 | 0.931030303 |
| 0.070535459 | 0.642143923 |
| 0.013686197 | 0.97075867 |
| 0.017104655 | 0.984447061 |
| 0.002229985 | 0.957212121 |
| 0.015013228 | 1.151765227 |
| 0.00096885 | 1.240373142 |
| 0.038361513 | 1.013498158 |
| 0.007996352 | 0.986083686 |
| 0.521123739 | 0.923360063 |
| 0.001062877 | 1.035968271 |
| 0.017735428 | 0.999027813 |
| 0.001915272 | 0.994274202 |
| 0.00170488 | 1.043854749 |
| 0.001420547 | 1.007883385 |
| 0.08767829 | 0.916367635 |
| 0.015460385 | 1.000327654 |
| 0.018134848 | 1.026395848 |
| 0.00127394 | 0.995100399 |
| 0.149217159 | 0.9599135 |

| | |
|---|---|
| 0.001202797 | 1.011863626 |
| 0.008393656 | 1.065840346 |
| 0.000670289 | 1.165158466 |
| 0.01863533 | 0.822533689 |
| 0.011132586 | 0.855209966 |
| 0.005343552 | 0.923309788 |
| 0.002072329 | 0.905993534 |
| 0.025302202 | 1.109081328 |
| 0.010691073 | 1.228822453 |
| 0.039671732 | 0.955552352 |
| 0.002072329 | 1.041904999 |
| 0.094384126 | 1.014703762 |
| 0.002072329 | 1.101715991 |
| 0.00371134 | 0.970383275 |
| 0.003044841 | 0.981878875 |
| 0.154738777 | 0.957858825 |
| 0.002072329 | 1.097985576 |
| 0.02302822 | 1.002998622 |
| 0.004665792 | 1.182402262 |
| 0.000865983 | 1.142873656 |
| 0.000561696 | 1.02605731 |
| 0.021740469 | 0.886113667 |
| 0.001062877 | 1.048960613 |
| 0.006729634 | 0.951353952 |
| 0.001865932 | 1.097252396 |
| 0.001874998 | 0.93070563 |
| 0.001865932 | 1.034760383 |
| 0.02419676 | 0.933881569 |
| 0.001902883 | 0.832192063 |
| 0.017507368 | 1.179622398 |
| 0.00096885 | 1.368695086 |
| 0.013249144 | 1.338617349 |
| 0.005020038 | 1.045041945 |
| 0.038361513 | 1.121448986 |
| 0.007996352 | 1.023413258 |
| 0.029667554 | 0.989585522 |
| 0.001902883 | 0.90000825 |
| 0.146873273 | 0.973763875 |
| 0.002072329 | 1.064536185 |
| 0.024684467 | 0.977223266 |
| 0.000847692 | 0.989462692 |
| 0.020277679 | 1.011721546 |
| 0.00127394 | 0.919227392 |
| 0.017104655 | 1.112236242 |
| 0.00483761 | 0.994849553 |
| 0.145356712 | 1.07027027 |
| 0.003736542 | 1.250198046 |
| 0.784310952 | 0.6429875 |
| 0.452647785 | 0.730324401 |
| 0.038361513 | 0.995777529 |
| 0.004588841 | 1.107856999 |
| 0.004123449 | 0.999508645 |
| 0.002688948 | 1.077426747 |
| 0.00129926 | 1.078011318 |
| 0.000686039 | 1.082885322 |
| 0.004123449 | 1.026256795 |
| 0.002688948 | 0.98428548 |
| 0.060001185 | 0.975643702 |
| 0.030072889 | 1.036590737 |
| 0.004123449 | 1.049749716 |
| 0.002688948 | 1.019806842 |
| 0.143183685 | 1.015923945 |
| 0.006123397 | 0.971445221 |

| | |
|---|---|
| 0.040305248 | 0.892433747 |
| 0.001902883 | 0.899678244 |
| 0.033496383 | 1.164558405 |
| 0.02058308 | 1.018939549 |
| 2.551336813 | 0.267166402 |
| 0.586575939 | 0.472021885 |
| 0.216418982 | 0.974260134 |
| 0.114485618 | 0.961366585 |
| 0.024172889 | 1.063807437 |
| 0.001902883 | 0.972692022 |
| 0.105545529 | 0.974532699 |
| 0.002072329 | 1.099850783 |
| 0.116289195 | 0.968045745 |
| 0.002072329 | 1.023004228 |
| 0.00319857 | 1.008112302 |
| 0.00280565 | 0.935634874 |
| 0.002388469 | 1.070546914 |
| 0.001768692 | 1.096164282 |
| 0.017104655 | 1.03857713 |
| 0.004201517 | 1.020207188 |
| 0.014720379 | 0.90341234 |
| 0.00277107 | 1.089790007 |
| 0.137430156 | 1.142299442 |
| 0.000719027 | 1.145365725 |
| 0.079668717 | 1.036360473 |
| 0.059269645 | 1.106531588 |
| 0.00371134 | 1.051082628 |
| 0.003044841 | 1.060403751 |
| 0.002347639 | 1.154833117 |
| 0.001028149 | 1.128900133 |
| 0.005343552 | 0.976983326 |
| 0.002072329 | 1.015294703 |
| 0.002134651 | 1.048732879 |
| 0.001155271 | 0.906983138 |
| 0.010387182 | 1.007000053 |
| 0.002818048 | 1.209164694 |
| 0.023612527 | 1.180770058 |
| 0.000927163 | 1.369147219 |
| 0.006692919 | 1.0000539 |
| 0.002366011 | 1.068334151 |
| 0.018578644 | 1.122083192 |
| 0.009090071 | 1.016704672 |
| 0.001915272 | 1.00901283 |
| 0.000966324 | 0.990531744 |
| 0.088633386 | 1.006077048 |
| 0.04856033 | 0.94277522 |
| 0.039029499 | 0.896718064 |
| 0.030655789 | 0.963466636 |
| 0.002818048 | 1.1951302 |
| 0.001486793 | 1.072917219 |
| 0.212632538 | 1.037743059 |
| 0.000908458 | 0.947802837 |
| 0.048317352 | 0.975461175 |
| 0.000719027 | 1.100871402 |
| 0.102249555 | 0.898372736 |
| 0.006359663 | 1.152079602 |
| 0.184555674 | 1.077958748 |
| 0.007996352 | 1.031405736 |
| 0.029087206 | 1.077171572 |
| 0.007371037 | 0.973631841 |
| 0.004123449 | 1.104290145 |
| 0.002688948 | 1.030856114 |
| 0.002276042 | 1.070556 |

| | |
|---|---|
| 0.000804045 | 1.087091386 |
| 0.082987134 | 1.036465686 |
| 0.007996352 | 0.951951105 |
| 0.011957406 | 1.176058866 |
| 0.007371037 | 0.906343284 |
| 0.013807992 | 0.905820534 |
| 0.006145828 | 0.814117192 |
| 0.00129926 | 1.23150768 |
| 0.000686039 | 1.219222588 |
| 0.002818048 | 1.094690565 |
| 0.001486793 | 1.080553641 |
| 0.010554633 | 1.165000943 |
| 0.00572873 | 1.114967981 |
| 0.037279722 | 1.099508168 |
| 0.001556734 | 1.197996941 |
| 0.042193294 | 0.98533226 |
| 0.035496077 | 1.082209987 |
| 0.084421547 | 0.951737452 |
| 0.000719027 | 1.092817534 |
| 0.017104655 | 0.9584175 |
| 0.002934379 | 1.062028392 |
| 0.00319857 | 1.084685384 |
| 0.00280565 | 1.015408621 |
| 0.002605549 | 0.99944106 |
| 0.001869087 | 0.887612388 |
| 0.002143971 | 1.113292297 |
| 0.001313047 | 1.165844028 |
| 0.00783199 | 1.085559689 |
| 0.001486793 | 1.086121865 |
| 0.089832581 | 1.250750751 |
| 0.02381449 | 1.159889094 |
| 0.034028831 | 1.088888889 |
| 0.02771685 | 0.987550512 |
| 0.033593564 | 1.151716634 |
| 0.003638599 | 1.121619217 |
| 0.00786758 | 0.91799847 |
| 0.00096885 | 1.01854865 |
| 0.092425158 | 0.977053382 |
| 0.001202797 | 1.060009214 |
| 0.03005112 | 0.96448022 |
| 0.00078174 | 0.948708278 |
| 0.006359663 | 1.208199005 |
| 0.001082792 | 1.10735851 |
| 0.081875553 | 1.100518984 |
| 0.07688985 | 1.235363937 |
| 0.023027514 | 1.094025465 |
| 0.000998735 | 0.987584945 |
| 0.042917401 | 1.011446886 |
| 0.03573604 | 1.041473559 |
| 0.072614225 | 0.983567767 |
| 0.014636914 | 1.099699125 |
| 0.014720379 | 0.944716274 |
| 0.00277107 | 1.169985518 |
| 0.080814926 | 1.147272119 |
| 0.017449373 | 1.055024755 |
| 0.021740469 | 1.056921257 |
| 0.008747904 | 1.137035011 |
| 0.016956063 | 0.963321136 |
| 0.001458435 | 1.175318315 |
| 0.023360589 | 1.02012597 |
| 0.001558842 | 0.945264199 |
| 0.034431997 | 1.100741113 |
| 0.007857741 | 1.24302751 |

| | |
|---|---|
| 0.007371037 | 1.020087065 |
| 0.00277107 | 1.060480976 |
| 0.002366011 | 1.152689227 |
| 0.001772163 | 1.064356169 |
| 0.092841404 | 1.049609842 |
| 0.001871173 | 1.163687521 |
| 0.004123449 | 0.933789884 |
| 0.002688948 | 0.887542969 |
| 0.048437083 | 1.113941324 |
| 0.000670289 | 1.054000589 |
| 0.119138908 | 0.881036622 |
| 0.066608228 | 0.934212279 |
| 0.098634578 | 0.90456279 |
| 0.005988527 | 0.960580637 |
| 0.055404422 | 1.032594973 |
| 0.008848033 | 1.041633235 |
| 0.002134651 | 1.065010256 |
| 0.001155271 | 1.001832845 |
| 0.12528877 | 0.893167302 |
| 0.005988527 | 0.905979491 |
| 0.00129926 | 1.061135812 |
| 0.000686039 | 1.001740476 |
| 0.053582313 | 1.244826068 |
| 0.003334047 | 1.030673263 |
| 0.10723851 | 0.862105408 |
| 0.005988527 | 0.93993874 |
| 0.058736899 | 0.984722148 |
| 0.006558115 | 1.098086851 |
| 0.020254702 | 1.173462037 |
| 0.00138486 | 1.116660264 |
| 0.019892311 | 1.081335004 |
| 0.00127394 | 1.049845648 |
| 0.070797602 | 0.951339429 |
| 0.005988527 | 0.954854175 |
| 0.083020348 | 1.138681245 |
| 0.006880035 | 1.218318423 |
| 0.119723804 | 0.968998502 |
| 0.024240531 | 0.854207942 |
| 0.02170048 | 1.178233618 |
| 0.0146908 | 1.132699294 |
| 0.017104655 | 1.07295653 |
| 0.002229985 | 0.943878788 |
| 0.002134651 | 1.052206709 |
| 0.001155271 | 1.027492669 |
| 0.015272751 | 1.007672302 |
| 0.001028149 | 0.989880737 |
| 0.002134651 | 1.024548402 |
| 0.001155271 | 0.973240469 |
| 0.001733606 | 1.058627004 |
| 0.000889096 | 0.948131939 |
| 0.00675964 | 0.874463085 |
| 0.001607842 | 0.978669408 |
| 0.017104655 | 1.025554024 |
| 0.003244676 | 1.068175657 |
| 0.058336673 | 0.911294572 |
| 0.021641303 | 1.074994752 |
| 0.002143971 | 0.958409582 |
| 0.001313047 | 1.055500713 |
| 0.046113601 | 1.041103109 |
| 0.017883317 | 1.078742991 |
| 0.001439012 | 1.107153371 |
| 0.001307418 | 1.014495331 |
| 0.002694495 | 1.090502793 |

| | |
|---|---|
| 0.001476826 | 1.083620876 |
| 0.001915272 | 1.011345563 |
| 0.000966324 | 0.869050638 |
| 0.032668286 | 1.041230832 |
| 0.002391338 | 1.135422129 |
| 1.119171814 | 0.421587097 |
| 0.894518544 | 0.433479824 |
| 0.013335382 | 1.11245357 |
| 0.001148102 | 1.090940171 |
| 0.008704325 | 0.972202947 |
| 0.002140544 | 0.996695157 |
| 0.024656266 | 1.106675991 |
| 0.006558115 | 1.001366535 |
| 0.035640965 | 1.123657838 |
| 0.000192955 | 1.092752556 |
| 1.151890884 | 0.511525257 |
| 0.686644783 | 0.50293753 |
| 0.038361513 | 1.06986674 |
| 0.007996352 | 1.069769629 |
| 0.058342015 | 1.12106632 |
| 0.001028149 | 1.195759547 |
| 0.147228408 | 1.019986894 |
| 0.075876481 | 0.970193309 |
| 0.066502837 | 0.958050017 |
| 0.001680008 | 0.939960376 |
| 0.017147882 | 1.082729631 |
| 0.000847692 | 1.076609075 |
| 0.072727886 | 0.935486757 |
| 0.00127394 | 1.025092753 |
| 0.048087917 | 1.249382478 |
| 0.041076805 | 1.157607211 |
| 1.349243543 | 0.410041284 |
| 1.09282421 | 0.388457474 |
| 0.004685678 | 1.127776432 |
| 0.002132695 | 1.012805209 |
| 0.076491942 | 1.089133843 |
| 0.000947294 | 1.007411129 |
| 0.02502683 | 0.890409325 |
| 0.018385255 | 0.946648946 |
| 0.040185023 | 0.991202685 |
| 0.001766854 | 1.044397117 |
| 0.085130912 | 0.878601296 |
| 0.005988527 | 0.882674124 |
| 0.002143971 | 1.14140742 |
| 0.001313047 | 1.039267303 |
| 0.005343552 | 0.925424055 |
| 0.002072329 | 0.962695847 |
| 0.005343552 | 0.979193696 |
| 0.002072329 | 0.939691619 |
| 0.096678419 | 1.213513514 |
| 0.003405096 | 1.202270927 |
| 0.118056559 | 1.166924067 |
| 0.000719027 | 1.266041722 |
| 0.000927163 | 1.546602084 |
| 0.00012724 | 1.176295978 |
| 0.085114947 | 1.073859199 |
| 0.000896779 | 1.247893685 |
| 0.025985291 | 0.917155425 |
| 0.007535554 | 1.008767286 |
| 0.061260861 | 1.165712226 |
| 0.001202797 | 1.173577517 |
| 0.038361513 | 0.982780234 |
| 0.004588841 | 1.072813573 |

| | |
|---|---|
| 0.109705437 | 0.912727273 |
| 0.086383626 | 0.992248062 |
| 0.089302696 | 0.921885769 |
| 0.005988527 | 0.891463577 |
| 0.001331856 | 1.07471003 |
| 0.000670289 | 1.05766236 |
| 0.007371037 | 1.043345771 |
| 0.00277107 | 1.04091888 |
| 0.034803271 | 1.062450142 |
| 0.028316633 | 0.923302992 |
| 0.00319857 | 1.034600734 |
| 0.00280565 | 0.9812756 |
| 0.056455654 | 0.976007316 |
| 0.006558115 | 1.003340419 |
| 0.005772459 | 0.952885078 |
| 0.001865932 | 1.220063898 |
| 0.039419619 | 0.991652506 |
| 0.012032934 | 0.967818493 |
| 0.021699445 | 1.002893853 |
| 0.013369981 | 1.011304063 |
| 0.005343552 | 0.953870549 |
| 0.002072329 | 1.033076349 |
| 0.062739172 | 0.900063628 |
| 0.002276042 | 1.128667656 |
| 0.002130938 | 1.145441973 |
| 0.001680008 | 0.923335343 |
| 0.00222996 | 0.994822606 |
| 0.001825091 | 1.105400555 |
| 0.010020519 | 0.982139746 |
| 0.001458435 | 1.075416259 |
| 0.025303093 | 1.019021081 |
| 0.002132695 | 1.006945198 |
| 0.03751218 | 0.891113179 |
| 0.001902883 | 0.871957759 |
| 0.002143971 | 1.153052737 |
| 0.001313047 | 1.076615115 |
| 0.253614817 | 0.87491675 |
| 0.063470308 | 0.943909685 |
| 0.014892808 | 1.193546999 |
| 0.000927163 | 1.39292529 |
| 0.181986597 | 0.959753314 |
| 0.001954757 | 1.09536973 |
| 0.002130938 | 1.028120318 |
| 0.001680008 | 0.882031183 |
| 0.037195457 | 0.887349056 |
| 0.001733606 | 1.125028109 |
| 0.002143971 | 1.194144069 |
| 0.001313047 | 1.152023692 |
| 0.201122972 | 1.072804453 |
| 0.001202797 | 1.021308454 |
| 0.003278494 | 1.151632728 |
| 0.001126814 | 1.12464209 |
| 0.06515176 | 0.930678297 |
| 0.01195755 | 1.138851743 |
| 0.008408276 | 1.035746928 |
| 0.002391338 | 1.115233351 |
| 0.022159781 | 0.942606892 |
| 0.001680008 | 0.75652511 |
| 0.001331856 | 1.221514669 |
| 0.000670289 | 1.255734669 |
| 0.01528574 | 0.937103728 |
| 0.001915272 | 0.974233909 |
| 0.081896122 | 0.932810879 |

| | |
|---|---|
| 0.051923826 | 0.813162202 |
| 0.008306586 | 1.039690223 |
| 0.001331856 | 1.049238117 |
| 0.05708098 | 0.955514579 |
| 0.009760463 | 0.872254285 |
| 0.015976577 | 0.86540615 |
| 0.007316517 | 0.976028581 |
| 0.293309973 | 1.001319522 |
| 0.004588841 | 1.113209453 |
| 0.067079916 | 0.823326729 |
| 0.002130938 | 0.964503861 |
| 2.76881614 | 0.292895673 |
| 1.865703354 | 0.263573148 |
| 0.017104655 | 1.000212067 |
| 0.003244676 | 0.857685009 |
| 0.021271407 | 1.02008105 |
| 0.002633395 | 1.161040398 |
| 1.012576346 | 0.56701212 |
| 0.521000454 | 0.399562206 |
| 0.014720379 | 1.083306856 |
| 0.00277107 | 1.040912382 |
| 0.061506511 | 1.171609344 |
| 0.028709706 | 1.155893202 |
| 0.015610862 | 1.013765464 |
| 0.001458435 | 1.078599412 |
| 0.017104655 | 1.042733785 |
| 0.002229985 | 0.960969697 |
| 0.004917379 | 1.246569229 |
| 0.000896779 | 1.184443672 |
| 0.03371086 | 1.031490643 |
| 0.002531637 | 1.258036107 |
| 0.02974611 | 0.970328388 |
| 0.008538512 | 1.000467508 |
| 0.012017598 | 1.011864799 |
| 0.007303011 | 1.047018349 |
| 0.014071028 | 0.892239364 |
| 0.00372282 | 1.005792632 |
| 0.001915272 | 1.169759304 |
| 0.000966324 | 1.130991631 |
| 0.00319857 | 0.961307844 |
| 0.00280565 | 0.950263312 |
| 0.005025885 | 0.956076013 |
| 0.000698103 | 0.735857878 |
| 0.033814907 | 1.010894685 |
| 0.001902883 | 1.005032588 |
| 0.044178959 | 1.277115475 |
| 0.00372282 | 1.267211202 |
| 0.002683086 | 1.337833315 |
| 0.000896779 | 1.296534352 |
| 0.007743084 | 1.100440155 |
| 0.002818048 | 1.202739263 |
| 0.049912814 | 1.089058143 |
| 0.041998747 | 1.151238363 |
| 0.013520586 | 1.114154439 |
| 0.001439012 | 1.259444608 |
| 0.033370467 | 1.095015836 |
| 0.023584576 | 1.054815334 |
| 0.094532105 | 0.988244785 |
| 0.043200672 | 0.856964424 |
| 0.13960278 | 0.987730322 |
| 0.029139823 | 1.04115322 |
| 0.019195501 | 1.173223964 |
| 0.001558842 | 1.046410024 |

| | |
|---|---|
| 0.112507073 | 1.072862505 |
| 0.007996352 | 0.97235543 |
| 0.03055316 | 1.026932457 |
| 0.003530003 | 0.963817736 |
| 0.013143591 | 1.111189162 |
| 0.001768692 | 0.951503727 |
| 0.257763691 | 0.970019659 |
| 0.065205346 | 0.959833314 |
| 0.00639681 | 1.058912305 |
| 0.002366011 | 1.16478666 |
| 0.002694495 | 1.133280128 |
| 0.001476826 | 1.156351089 |
| 0.028042391 | 1.014120203 |
| 0.014720379 | 0.997903031 |
| 0.061324885 | 1.163161249 |
| 0.021184926 | 1.312642351 |
| 0.001191193 | 1.060555923 |
| 0.000246596 | 0.98260039 |
| 0.127673627 | 1.002785059 |
| 0.056783536 | 0.954045607 |
| 0.118284659 | 0.940825219 |
| 0.006558115 | 1.062860613 |
| 0.012660038 | 0.950442406 |
| 0.000865983 | 1.013293261 |
| 0.041933638 | 0.982840312 |
| 0.020254702 | 1.059261394 |
| 0.149646429 | 0.957813428 |
| 0.090667768 | 0.981684982 |
| 0.030280926 | 0.957609694 |
| 0.001865932 | 1.205750799 |
| 0.218750223 | 0.603506835 |
| 0.218338726 | 0.950011625 |
| 0.114507596 | 0.918882979 |
| 0.003638599 | 1.008852313 |
| 0.072752401 | 0.83791691 |
| 0.00319857 | 0.819377822 |
| 0.001439012 | 1.221077421 |
| 0.000627264 | 1.243426708 |
| 0.1614609 | 0.975415568 |
| 0.1385078 | 0.873519587 |
| 0.017104655 | 1.082524511 |
| 0.002934379 | 1.101675406 |
| 0.002388469 | 1.202007025 |
| 0.001768692 | 1.022077514 |
| 0.002366011 | 1.203040706 |
| 0.001772163 | 1.138306473 |
| 0.014720379 | 1.065641482 |
| 0.00277107 | 1.174692252 |
| 0.001331856 | 1.079940869 |
| 0.000670289 | 0.973062839 |
| 0.08708122 | 1.336541238 |
| 0.045439241 | 1.239825184 |
| 0.006338964 | 1.173561331 |
| 0.001202797 | 1.151117254 |
| 0.016753774 | 0.974880866 |
| 0.002132695 | 0.966467716 |
| 0.017823116 | 1.191197712 |
| 0.001126814 | 1.044828203 |
| 0.027148433 | 1.173611441 |
| 0.016400925 | 1.191307625 |
| 0.041196104 | 1.070638052 |
| 0.025850832 | 1.038240918 |
| 2.46327444 | 0.492879358 |

| | |
|---|---|
| 1.214986536 | 0.25691736 |
| 0.117476183 | 0.895309478 |
| 0.003638599 | 0.94452847 |
| 0.00222996 | 1.038966701 |
| 0.001825091 | 1.072442487 |
| 0.00777816 | 1.100756878 |
| 0.001202797 | 1.251900484 |
| 0.112473916 | 1.174930783 |
| 0.049174328 | 1.086899791 |
| 0.850542732 | 0.619954767 |
| 0.848054698 | 0.424270073 |
| 0.070106934 | 1.007030303 |
| 0.017104655 | 1.082818173 |
| 0.00371134 | 1.019599303 |
| 0.003044841 | 0.977428072 |
| 0.030779825 | 0.956913741 |
| 0.001028149 | 1.04565715 |
| 0.001915272 | 1.102852296 |
| 0.000966324 | 1.12342548 |
| 0.023530892 | 1.063936064 |
| 0.002605549 | 1.056900117 |
| 0.002388469 | 1.218564977 |
| 0.001768692 | 1.141575986 |
| 0.014524198 | 0.879439157 |
| 0.006359663 | 1.023761194 |
| 0.002388469 | 1.079879579 |
| 0.001768692 | 1.079884614 |
| 0.03005294 | 0.996314836 |
| 0.002688948 | 0.945572107 |
| 0.123410984 | 0.790550435 |
| 0.102938861 | 0.894779302 |
| 0.001331856 | 1.315556061 |
| 0.000670289 | 1.146092007 |
| 0.009536929 | 0.990260272 |
| 0.004123449 | 1.103983048 |
| 0.00319857 | 1.122813205 |
| 0.00280565 | 0.92812561 |
| 0.018663413 | 1.042855692 |
| 0.002391338 | 1.080361825 |
| 0.100821386 | 0.973248969 |
| 0.001202797 | 1.038009675 |
| 0.002694495 | 1.109018356 |
| 0.001476826 | 1.11122493 |
| 0.136980894 | 1.085908736 |
| 0.000896779 | 1.16841831 |
| 0.385628508 | 0.73519587 |
| 0.168695045 | 0.982570337 |
| 0.00319857 | 1.060066309 |
| 0.00280565 | 0.89204213 |
| 0.081574104 | 0.972930073 |
| 0.000719027 | 1.09202535 |
| 0.006334475 | 0.97957041 |
| 0.001865932 | 1.072076677 |
| 0.008851621 | 1.049717057 |
| 0.000686039 | 0.955869271 |
| 0.002143971 | 1.169522542 |
| 0.001313047 | 1.168805528 |
| 0.001458435 | 1.132590597 |
| 0.000998735 | 1.020822443 |
| 0.042970235 | 0.98053379 |
| 0.000998735 | 0.918409131 |
| 0.052606751 | 1.304418024 |
| 0.016984066 | 1.192041299 |

| | |
|---|---|
| 0.037626321 | 0.953597219 |
| 0.034730194 | 1.028577987 |
| 0.191837797 | 1.090793567 |
| 0.001062877 | 1.237007659 |
| 0.002143971 | 1.198303111 |
| 0.001313047 | 1.128002632 |
| 0.048301271 | 0.965736963 |
| 0.001869087 | 1.121711622 |
| 0.047513247 | 1.066736793 |
| 0.00372282 | 1.162860477 |
| 0.048318467 | 1.130854164 |
| 0.001032472 | 1.249108507 |
| 0.07000581 | 0.961158362 |
| 0.007996352 | 0.997931359 |
| 0.013783628 | 0.998515537 |
| 0.004588841 | 1.040597859 |
| 0.037905418 | 1.131909548 |
| 0.007371037 | 0.982027363 |
| 0.003790939 | 1.116254876 |
| 0.001028149 | 1.13648958 |
| 0.083281985 | 1.066544668 |
| 0.001313047 | 1.05374575 |
| 0.018151853 | 1.014623172 |
| 0.001155271 | 0.996151026 |
| 0.021768011 | 1.015786788 |
| 0.001902883 | 1.061958584 |
| 0.018655691 | 1.018976494 |
| 0.000998735 | 0.892054365 |
| 0.017104655 | 1.06261266 |
| 0.003244676 | 1.029140689 |
| 0.017783788 | 0.906056271 |
| 0.00371134 | 0.947050772 |
| 0.00129926 | 1.028496362 |
| 0.000686039 | 0.909224521 |
| 0.403640193 | 0.737405737 |
| 0.007996352 | 0.865914433 |
| 0.077179028 | 0.899348776 |
| 0.002269165 | 0.916273619 |
| 0.139650606 | 1.007929403 |
| 0.017104655 | 1.060846431 |
| 0.025650189 | 0.913521481 |
| 0.002688948 | 0.975036831 |
| 0.01542408 | 1.029288469 |
| 0.001869087 | 1.134698635 |
| 0.022455921 | 1.058555179 |
| 0.001902883 | 0.958996782 |
| 0.002134651 | 1.047508767 |
| 0.001155271 | 0.755406891 |
| 0.074209929 | 1.056145128 |
| 0.001202797 | 1.095945635 |
| 0.002694495 | 1.00415004 |
| 0.001476826 | 1.008734095 |
| 0.00603154 | 1.033550408 |
| 0.002269165 | 1.100747034 |
| 0.023077342 | 1.047178439 |
| 0.006558115 | 1.065138172 |
| 0.005421148 | 1.024363369 |
| 0.003045462 | 0.933873497 |
| 0.003157745 | 1.10970858 |
| 0.000865983 | 1.08542365 |
| 0.038647182 | 0.968341243 |
| 0.030048767 | 1.043741809 |
| 0.002694495 | 1.108858739 |

| | |
|---|---|
| 0.001476826 | 1.044856588 |
| 0.012253094 | 0.974445635 |
| 0.002269165 | 0.962531126 |
| 0.051390548 | 1.08213869 |
| 0.035609014 | 1.143641889 |
| 0.011231174 | 0.967508181 |
| 0.00372282 | 1.051383564 |
| 0.027866232 | 1.088526344 |
| 0.021190248 | 1.041547114 |
| 0.076377099 | 1.107459632 |
| 0.000670289 | 1.077738962 |
| 0.002143971 | 1.110796872 |
| 0.001313047 | 1.090709663 |
| 0.023329625 | 1.043148747 |
| 0.000686039 | 1.035583059 |
| 0.031711157 | 1.020468501 |
| 0.000670289 | 1.002904163 |
| 0.014728998 | 0.901964215 |
| 0.001865932 | 1.099936102 |
| 0.005343552 | 1.00764019 |
| 0.002072329 | 1.011937329 |
| 0.006359663 | 0.96441791 |
| 0.001082792 | 0.8399345 |
| 0.284072273 | 0.654131054 |
| 0.161469773 | 0.709661619 |
| 0.006092715 | 1.068889877 |
| 0.001439012 | 1.155823766 |
| 0.028985757 | 1.008275261 |
| 0.003044841 | 1.017962168 |
| 0.074716217 | 1.110131211 |
| 0.000927163 | 1.391017173 |
| 0.012003587 | 1.038744238 |
| 0.001191193 | 1.081072138 |
| 0.070953679 | 1.021276596 |
| 0.003638599 | 1.13225089 |
| 0.002694495 | 1.067517957 |
| 0.001476826 | 0.954873841 |
| 0.014384306 | 0.902770518 |
| 0.001458435 | 1.078721841 |
| 0.098481376 | 1.036125966 |
| 0.046788701 | 1.041167166 |
| 0.002740708 | 0.987594105 |
| 0.000966324 | 0.959759912 |
| 0.00129926 | 1.200485044 |
| 0.000686039 | 1.171804293 |
| 0.038100159 | 0.969289202 |
| 0.035562896 | 1.007834585 |
| 0.029186225 | 0.982089858 |
| 0.006359663 | 1.061333333 |
| 0.117535704 | 0.975098296 |
| 0.031118782 | 0.93760852 |
| 0.002134651 | 0.97869384 |
| 0.001155271 | 1.004857038 |
| 0.002143971 | 1.116619531 |
| 0.001313047 | 1.107820555 |
| 0.01141731 | 1.069615111 |
| 0.002391338 | 1.110644992 |
| 0.038361513 | 0.947087154 |
| 0.004588841 | 1.092910523 |
| 0.008670679 | 0.988298507 |
| 0.001082792 | 0.888854774 |
| 0.00319857 | 1.070189052 |
| 0.00280565 | 0.982055783 |

| | |
|---|---|
| 0.038361513 | 0.959952497 |
| 0.004588841 | 1.070086851 |
| 0.027211513 | 1.08980057 |
| 0.00708014 | 1.135270827 |
| 2.025455695 | 0.635932135 |
| 0.038361513 | 0.352576367 |
| 0.001558842 | 0.982936279 |
| 5.33E-05 | 1.092544309 |
| 0.060393201 | 0.960034677 |
| 0.001028149 | 0.911335984 |
| 0.00844414 | 1.02975257 |
| 0.001458435 | 1.18450049 |
| 0.002134651 | 1.124793224 |
| 0.001155271 | 1.042980205 |
| 0.014720379 | 0.940649425 |
| 0.00277107 | 1.107530775 |
| 0.050622358 | 0.982449276 |
| 0.039894009 | 1.026307026 |
| 0.014720379 | 0.981063735 |
| 0.00277107 | 1.194062274 |
| 0.017104655 | 1.150620295 |
| 0.002267069 | 1.076443481 |
| 0.033353602 | 0.96224053 |
| 0.002132695 | 1.16527401 |
| 0.001331856 | 1.246872868 |
| 0.000670289 | 1.159308052 |
| 0.001619105 | 1.110900368 |
| 0.000896779 | 1.064752888 |
| 0.117492158 | 1.006733595 |
| 0.035276257 | 0.901289423 |
| 0.028990962 | 0.994395254 |
| 0.000847692 | 1.056293905 |
| 0.031058815 | 0.984047748 |
| 0.012182149 | 1.059163234 |
| 0.002134651 | 0.980579633 |
| 0.001155271 | 1.021352639 |
| 0.007789292 | 1.048971596 |
| 0.000816283 | 1.010062729 |
| 0.017104655 | 1.128671403 |
| 0.003244676 | 1.065464896 |
| 0.001331856 | 1.23527405 |
| 0.000670289 | 1.09811019 |
| 0.053423864 | 1.092556317 |
| 0.008680911 | 0.969332636 |
| 0.067526546 | 1.187365398 |
| 0.007371037 | 0.95 |
| 0.007371037 | 0.970833333 |
| 0.00277107 | 0.973438622 |
| 0.036830238 | 1.01361075 |
| 0.009929247 | 1.005180099 |
| 0.002694495 | 0.960574621 |
| 0.001476826 | 1.035583351 |
| 0.032366029 | 0.943718418 |
| 0.001458435 | 1.087536729 |
| 0.010676027 | 1.158055828 |
| 0.000998735 | 1.108991113 |
| 0.001331856 | 1.183079372 |
| 0.000670289 | 1.187760428 |
| 0.055199776 | 0.872549414 |
| 0.013699313 | 1.053476516 |
| 0.038361513 | 1.021046381 |
| 0.004588841 | 1.109674813 |
| 0.209283101 | 1.065685571 |

| | |
|---|---|
| 0.007371037 | 1.000497512 |
| 0.170517875 | 1.148402429 |
| 0.004380884 | 1.28974689 |
| 0.002694495 | 1.063846768 |
| 0.001476826 | 1.017252534 |
| 1.212087626 | 0.41378743 |
| 0.703923726 | 0.446182 |
| 0.015862092 | 1.129828018 |
| 0.001032472 | 0.87531839 |
| 0.001191193 | 1.040480918 |
| 0.000982899 | 0.955981894 |
| 0.0028589 | 0.954196254 |
| 0.001191193 | 1.013015663 |
| 0.00222996 | 1.266281541 |
| 0.001825091 | 1.302170011 |
| 0.041164165 | 1.002437291 |
| 0.002391338 | 1.07629785 |
| 0.002694495 | 1.066879489 |
| 0.001476826 | 1.064319603 |
| 1.693177256 | 0.477207155 |
| 0.893475214 | 0.268902886 |
| 0.117588349 | 1.037114938 |
| 0.001062877 | 1.089305252 |
| 0.00129926 | 1.156831043 |
| 0.000686039 | 1.074028234 |
| 0.057055326 | 1.030537589 |
| 0.010573801 | 1.051018683 |
| 0.079676767 | 1.037265014 |
| 0.001313047 | 1.065043326 |
| 0.003334047 | 1.057861906 |
| 0.000927163 | 1.235725818 |
| 0.002269165 | 1.165138421 |
| 0.001558842 | 0.993817492 |
| 0.005343552 | 0.976887223 |
| 0.002072329 | 1.078090027 |
| 0.049722761 | 0.88446006 |
| 0.010993034 | 0.969614759 |
| 0.048074434 | 1.049760125 |
| 0.000896779 | 1.052983584 |
| 0.045946545 | 1.018315018 |
| 0.023723036 | 0.985579342 |
| 0.036856265 | 1.115809439 |
| 0.000896779 | 1.164596543 |
| 0.890356336 | 0.220676348 |
| 0.558008807 | 0.334032512 |
| 0.033391016 | 1.049542339 |
| 0.001032472 | 1.222491085 |
| 0.211817149 | 1.041939712 |
| 0.022833802 | 0.997279778 |
| 0.015381379 | 0.979565463 |
| 0.002132695 | 0.97460662 |
| 0.012763841 | 1.069837177 |
| 0.000213241 | 1.04417706 |
| 0.021092054 | 0.931598995 |
| 0.001733606 | 1.1372675 |
| 0.007371037 | 1.008084577 |
| 0.00277107 | 1.092605887 |
| 0.030524494 | 1.024252223 |
| 0.007199867 | 0.922026687 |
| 0.032178444 | 0.855370833 |
| 0.002130938 | 0.947447274 |
| 0.029998064 | 0.938105183 |
| 0.017448827 | 0.959663715 |

| | |
|---|---|
| 0.00222996 | 1.010681781 |
| 0.001825091 | 1.133300702 |
| 0.020854823 | 0.940417242 |
| 0.000865983 | 1.160212692 |
| 0.006359663 | 1.167761194 |
| 0.001082792 | 1.081056187 |
| 0.059054926 | 1.174582463 |
| 0.021729922 | 1.294141707 |
| 0.000865983 | 1.181366316 |
| 0.000561696 | 1.09960224 |
| 0.079210164 | 1.037270235 |
| 0.002391338 | 1.054798112 |
| 0.011718131 | 0.984157611 |
| 0.002391338 | 1.094651285 |
| 0.080185527 | 0.9401867 |
| 0.000870247 | 0.866870531 |
| 0.032455132 | 1.083413926 |
| 0.003638599 | 1.095685053 |
| 0.245616643 | 1.084868812 |
| 0.024423163 | 1.010230179 |
| 0.266121523 | 0.986810394 |
| 0.001062877 | 1.061405908 |
| 0.05133291 | 0.919090765 |
| 0.00371134 | 1.041376307 |
| 0.015402886 | 0.979701649 |
| 0.001865932 | 1.068498403 |
| 0.120795441 | 1.025137587 |
| 0.00170488 | 1.091899441 |
| 0.02874568 | 1.236803345 |
| 0.001674288 | 1.151237282 |
| 0.009687788 | 0.981336731 |
| 0.007996352 | 0.978373296 |
| 0.001458435 | 1.135406464 |
| 0.000998735 | 1.031451472 |
| 0.019394083 | 1.073918831 |
| 0.001126814 | 1.00984252 |
| 0.030119547 | 1.147383688 |
| 0.000896779 | 1.137192739 |
| 0.057648662 | 1.127666651 |
| 0.022364401 | 1.067453837 |
| 0.05197742 | 1.200910273 |
| 0.001028149 | 1.175882424 |
| 0.113483223 | 1.050429544 |
| 0.004439392 | 0.965994962 |
| 0.798716367 | 0.648573417 |
| 0.454185118 | 0.601938082 |
| 0.011662224 | 0.941753318 |
| 0.001915272 | 0.924292228 |
| 0.002694495 | 1.051715882 |
| 0.001476826 | 1.065613543 |
| 0.013603937 | 1.069813352 |
| 0.007371037 | 0.960634328 |
| 0.00129926 | 1.021523848 |
| 0.000686039 | 1.024521369 |
| 0.136872233 | 1.009152907 |
| 0.007371037 | 0.943532338 |
| 0.2285843 | 0.511484526 |
| 0.003638599 | 0.919795374 |
| 0.04265979 | 0.941419287 |
| 0.031209693 | 1.062880446 |
| 0.007371037 | 0.884825871 |
| 0.00277107 | 1.097272075 |
| 0.21494986 | 1.06149926 |

| | |
|---|---|
| 0.007996352 | 1.074094969 |
| 0.083648541 | 1.201495035 |
| 0.000896779 | 1.214453227 |
| 0.07927007 | 1.184349313 |
| 0.018394062 | 1.113051124 |
| 0.218772898 | 0.972361382 |
| 0.06546561 | 0.990467333 |
| 0.003334047 | 0.963476016 |
| 0.000927163 | 1.268457361 |
| 0.002276042 | 1.180614225 |
| 0.000804045 | 1.130875686 |
| 0.059337686 | 0.927394261 |
| 0.057781672 | 0.807649238 |
| 0.002688948 | 0.975773449 |
| 0.001961349 | 0.982925406 |
| 0.007371037 | 1.017537313 |
| 0.006291294 | 1.064788227 |
| 0.004054872 | 0.994697525 |
| 0.001558842 | 0.98079301 |
| 0.00319857 | 1.129126693 |
| 0.00280565 | 1.07187439 |
| 0.053454177 | 1.085509899 |
| 0.001313047 | 0.974936931 |
| 0.038361513 | 1.091870115 |
| 0.007996352 | 1.002350729 |
| 0.002143971 | 1.101147896 |
| 0.001313047 | 1.08368981 |
| 0.038770976 | 1.079763849 |
| 0.007857741 | 1.122345227 |
| 0.036334813 | 1.126868484 |
| 0.007857741 | 1.241079489 |
| 0.135833706 | 0.848504342 |
| 0.005988527 | 0.993607671 |
| 0.002143971 | 1.085509899 |
| 0.001313047 | 1.023856532 |
| 0.022830974 | 1.045349348 |
| 0.001558842 | 0.934053252 |
| 0.024099409 | 1.068099303 |
| 0.005010182 | 1.056143471 |
| 0.002388469 | 1.211741094 |
| 0.001768692 | 1.137834519 |
| 0.001915272 | 1.095323932 |
| 0.000966324 | 1.046030941 |
| 0.04476446 | 0.913544898 |
| 0.030329706 | 0.995090764 |
| 0.00371134 | 0.960490294 |
| 0.003044841 | 0.918137021 |
| 0.002143971 | 1.145233738 |
| 0.001313047 | 1.013162224 |
| 3.267559731 | 0.492879358 |
| 0.829949509 | 0.353192273 |
| 0.025539571 | 0.99317733 |
| 0.00078174 | 1.093814822 |
| 0.032919224 | 0.91850416 |
| 0.002132695 | 0.904829083 |
| 0.159173358 | 0.984888781 |
| 0.003638599 | 1.151779359 |
| 0.053740183 | 1.247703247 |
| 0.023683045 | 1.093075852 |
| 0.0030477 | 1.074215206 |
| 0.002633395 | 1.095923261 |
| 0.045208495 | 0.970293486 |
| 0.00127394 | 0.918604322 |

| | |
|---|---|
| 0.068801381 | 1.048690498 |
| 0.002633395 | 1.161409334 |
| 0.009303802 | 1.072884534 |
| 0.002633395 | 1.196458218 |
| 0.016036838 | 1.03006109 |
| 0.002633395 | 1.048330566 |
| 0.061088567 | 0.988931091 |
| 0.003954543 | 0.960786689 |
| 0.078681197 | 0.948242867 |
| 0.002552407 | 1.127265291 |
| 0.00222996 | 1.068178102 |
| 0.001825091 | 1.094468918 |
| 0.017104655 | 1.028848984 |
| 0.002229985 | 0.930909091 |
| 0.037072334 | 0.995170378 |
| 0.021686518 | 1.079362066 |
| 0.008152061 | 1.257837778 |
| 0.000896779 | 1.18301051 |
| 0.025300495 | 1.032491819 |
| 0.00372282 | 1.105934322 |
| 0.069768026 | 1.093912901 |
| 0.001032472 | 1.066479878 |
| 0.006910259 | 1.101556908 |
| 0.006138695 | 1.089608882 |
| 0.022507404 | 0.965090336 |
| 0.001865932 | 1.080766773 |
| 1.115468579 | 0.707092416 |
| 0.806362746 | 0.414885572 |
| 0.005769475 | 0.950806386 |
| 0.001504387 | 1.124289248 |
| 0.060287148 | 1.118439165 |
| 0.00078174 | 1.111257779 |
| 0.00372282 | 1.25391732 |
| 0.000698103 | 1.211430575 |
| 0.003334047 | 1.219832222 |
| 0.000927163 | 1.507705856 |
| 0.017647308 | 0.98503645 |
| 0.017104655 | 1.055476179 |
| 0.009192793 | 0.982295374 |
| 0.001166314 | 0.851910058 |
| 0.002605549 | 1.070258789 |
| 0.001869087 | 1.055944056 |
| 0.067683288 | 1.118299445 |
| 0.004380884 | 1.116087516 |
| 0.041838108 | 1.056838483 |
| 0.001155271 | 0.966917155 |
| 0.006359663 | 1.183402985 |
| 0.001082792 | 1.081670249 |
| 0.001331856 | 1.120877871 |
| 0.000670289 | 1.126688834 |
| 0.004123449 | 0.939194792 |
| 0.002688948 | 0.970371583 |
| 0.024119801 | 0.981993262 |
| 0.020684544 | 1.043098538 |
| 0.028082474 | 0.938157317 |
| 0.001558842 | 0.933311351 |
| 0.013859454 | 1.038845759 |
| 0.003461177 | 0.909075921 |
| 0.291228167 | 0.955544456 |
| 0.184577889 | 0.951158645 |
| 2.240065748 | 0.426799499 |
| 1.647719406 | 0.630175722 |
| 0.032179588 | 1.098473717 |

| | |
|---|---|
| 0.022562796 | 1.028807307 |
| 0.029708942 | 1.112810707 |
| 0.002661846 | 1.158021365 |
| 0.025252505 | 0.965232191 |
| 0.024714375 | 1.056892287 |
| 0.002694495 | 1.037988827 |
| 0.001476826 | 0.966141902 |
| 0.059242658 | 1.007152146 |
| 0.001028149 | 1.008553186 |
| 0.0030477 | 1.000665336 |
| 0.002633395 | 1.188895038 |
| 0.062668856 | 1.037199125 |
| 0.009948296 | 1.212374313 |
| 0.001915272 | 0.958753048 |
| 0.000966324 | 1.029884183 |
| 0.017104655 | 1.159155975 |
| 0.003244676 | 0.967064245 |
| 0.019751618 | 1.038589488 |
| 0.002633395 | 1.121748755 |
| 0.064731348 | 1.074509804 |
| 0.03490558 | 1.015318015 |
| 0.13234958 | 0.957714189 |
| 0.000908458 | 1.010462683 |
| 0.045670145 | 1.024064171 |
| 0.007026375 | 1.013076335 |
| 0.002388469 | 1.205117913 |
| 0.001768692 | 0.942992603 |
| 0.00222996 | 0.942667175 |
| 0.001825091 | 1.105890031 |
| 0.016940326 | 0.963587976 |
| 0.002633395 | 1.122855562 |
| 0.109848269 | 0.971042471 |
| 0.000719027 | 1.070504357 |
| 0.00823901 | 1.016972143 |
| 0.001155271 | 0.92393695 |
| 0.002143971 | 1.107469639 |
| 0.001313047 | 0.998409565 |
| 0.021740469 | 1.024626441 |
| 0.001062877 | 1.143599562 |
| 0.00371134 | 0.965841215 |
| 0.003044841 | 0.924972182 |
| 0.017745041 | 0.913955672 |
| 0.001148102 | 1.067008547 |
| 0.048060067 | 1.0497003 |
| 0.002305914 | 0.720261438 |
| 0.04250503 | 1.22030589 |
| 0.006584667 | 1.162996659 |
| 0.063735609 | 1.051654267 |
| 0.002633395 | 1.109758347 |
| 0.061440738 | 0.999304106 |
| 0.057522662 | 1.203750315 |
| 0.00129926 | 1.075788197 |
| 0.000686039 | 0.93235351 |
| 0.047177037 | 0.988695937 |
| 0.046262276 | 1.06042366 |
| 0.080801456 | 1.016996805 |
| 0.044037619 | 0.925718535 |
| 0.027185815 | 0.9993778 |
| 0.00370934 | 0.936576061 |
| 0.461533453 | 0.523646724 |
| 0.370114933 | 0.4196906 |
| 0.002694495 | 1.018515563 |
| 0.001476826 | 1.154733664 |

| | |
|---|---|
| 0.017657366 | 1.020141535 |
| 0.002633395 | 1.095738794 |
| 0.001439012 | 1.23020312 |
| 0.001307418 | 1.175129687 |
| 0.00222996 | 1.196141479 |
| 0.001825091 | 1.198237885 |
| 0.001439012 | 1.128839172 |
| 0.001307418 | 1.054187046 |
| 0.027933033 | 1.088225719 |
| 0.009088995 | 1.060980535 |
| 0.004123449 | 1.144243467 |
| 0.002688948 | 1.019152071 |
| 0.00319857 | 1.06087754 |
| 0.00280565 | 0.914082309 |
| 0.016816939 | 0.974997106 |
| 0.010968538 | 1.018676278 |
| 0.020254702 | 1.099841521 |
| 0.00138486 | 1.032654629 |
| 0.029478148 | 1.004159042 |
| 0.001313047 | 0.963255457 |
| 0.055413649 | 1.058140796 |
| 0.01194694 | 1.06477195 |
| 0.022210083 | 0.957091828 |
| 0.001458435 | 1.111165524 |
| 0.021067153 | 1.144557726 |
| 0.000343724 | 1.170969878 |
| 0.097240972 | 1.015727392 |
| 0.002416105 | 1.007234634 |
| 0.001835363 | 1.094289563 |
| 0.001148102 | 1.116011396 |
| 0.03194448 | 0.958135375 |
| 0.001869087 | 0.995837496 |
| 0.206827753 | 0.96766755 |
| 0.019641837 | 0.857939394 |
| 0.05968439 | 1.093383912 |
| 0.000896779 | 1.045036046 |
| 0.046091104 | 1.011163638 |
| 0.001871173 | 1.05081099 |
| 0.075237651 | 0.966182594 |
| 0.002391338 | 1.050471945 |
| 0.040450788 | 1.095663688 |
| 0.023706275 | 1.048983311 |
| 0.008253408 | 1.062265944 |
| 0.001869087 | 1.133533134 |
| 0.002694495 | 1.076775738 |
| 0.001476826 | 1.08626267 |
| 0.015115085 | 0.957408733 |
| 0.00808697 | 1.024546181 |
| 0.059689333 | 1.010261503 |
| 0.055798564 | 1.074329327 |
| 0.156841691 | 1.014791337 |
| 0.00138486 | 0.976309387 |
| 0.080061092 | 0.954291272 |
| 0.001733606 | 1.100195959 |
| 0.050977717 | 1.123303542 |
| 0.002909003 | 1.084012075 |
| 0.007184599 | 1.02161963 |
| 0.002688948 | 0.965542642 |
| 0.051453544 | 0.984439582 |
| 0.004427553 | 1.167657169 |
| 0.007532382 | 1.076381294 |
| 0.001865932 | 1.170095847 |
| 0.04823782 | 0.981819763 |

| | |
|---|---|
| 0.002552407 | 1.219166876 |
| 0.108413103 | 0.8868146 |
| 0.001869087 | 0.861804862 |
| 0.031982042 | 1.125289639 |
| 0.010913396 | 1.054166429 |
| 0.00129926 | 1.023443816 |
| 0.000686039 | 1.070237865 |
| 0.006113066 | 1.292905505 |
| 0.001307418 | 1.058396324 |
| 0.05293732 | 1.095167163 |
| 0.002909003 | 1.050122458 |
| 0.010778863 | 1.03848122 |
| 0.006359663 | 1.282945274 |
| 0.033564966 | 0.936748348 |
| 0.001032472 | 1.047503821 |
| 0.00222996 | 1.157447272 |
| 0.001825091 | 1.19366944 |
| 0.053102734 | 1.019529957 |
| 0.041540475 | 1.033946574 |
| 0.070648367 | 0.990939437 |
| 0.00371134 | 0.986373818 |
| 0.002694495 | 1.201755786 |
| 0.001476826 | 1.103245633 |
| 0.045078109 | 1.084226238 |
| 0.001598043 | 1.132050864 |
| 0.035322823 | 1.093882704 |
| 0.030534684 | 1.112106624 |
| 0.077371261 | 1.050365344 |
| 0.047965004 | 1.220865844 |
| 0.176756345 | 1.049910873 |
| 0.022280769 | 0.94501332 |
| 0.001439012 | 1.252673928 |
| 0.001307418 | 1.137157255 |
| 0.108285243 | 0.996017919 |
| 0.003044841 | 1.013670323 |
| 0.037489643 | 1.118982888 |
| 0.001598043 | 1.176556896 |
| 0.010793005 | 1.278775715 |
| 0.001768692 | 1.221717648 |
| 0.439759653 | 0.981639718 |
| 0.035386789 | 1.143326039 |
| 0.032888261 | 1.074071851 |
| 0.001902883 | 0.865852652 |
| 0.025129488 | 0.987279685 |
| 0.002366011 | 1.129475233 |
| 0.002694495 | 1.08810854 |
| 0.001476826 | 1.170854 |
| 0.005531997 | 1.056163424 |
| 0.003013068 | 1.049207128 |
| 0.110859404 | 1.132175032 |
| 0.000719027 | 1.153551624 |
| 0.018835272 | 1.0979428 |
| 0.001768692 | 0.974466627 |
| 0.028341803 | 1.099437429 |
| 0.022115017 | 1.114934942 |
| 0.051166505 | 1.165733781 |
| 0.001598043 | 1.286110205 |
| 0.002388469 | 1.118414451 |
| 0.001768692 | 1.164567446 |
| 0.108657828 | 1.141308183 |
| 0.072882301 | 1.043886006 |
| 0.002366011 | 1.02795488 |
| 0.001772163 | 1.031423489 |

| | |
|---|---|
| 0.012383822 | 0.948540032 |
| 0.001126814 | 1.110415175 |
| 0.019918561 | 0.914915117 |
| 0.000686039 | 1.06203829 |
| 0.002388469 | 1.046864024 |
| 0.002235859 | 0.904464056 |
| 0.00371134 | 1.010701842 |
| 0.003044841 | 0.942934351 |
| 0.014720379 | 0.977123975 |
| 0.00277107 | 1.029507603 |
| 0.032816585 | 0.977786216 |
| 0.01127395 | 1.004769997 |
| 0.041927158 | 0.985481322 |
| 0.025639325 | 0.952914373 |
| 0.019412713 | 1.134209109 |
| 0.01635309 | 1.226004149 |
| 0.029841829 | 0.987424593 |
| 0.001902883 | 0.921788631 |
| 0.046457965 | 0.933330417 |
| 0.001865932 | 1.043194888 |
| 0.002388469 | 1.031610637 |
| 0.001768692 | 0.956102019 |
| 0.038361513 | 1.060731015 |
| 0.004588841 | 1.183296304 |
| 0.006498399 | 1.03592574 |
| 0.001768692 | 0.92094365 |
| 0.017392707 | 0.978780962 |
| 0.001915272 | 1.059696745 |
| 0.00319857 | 1.053188488 |
| 0.00280565 | 0.99834211 |
| 0.018932196 | 1.174209734 |
| 0.01079832 | 1.024448062 |
| 0.020254702 | 1.122844931 |
| 0.00138486 | 1.05404021 |
| 0.035955745 | 0.897277974 |
| 0.002130938 | 1.002535439 |
| 0.081190653 | 0.980597775 |
| 0.010546204 | 0.961171821 |
| 0.017104655 | 1.035522491 |
| 0.002934379 | 1.043611715 |
| 2.670150122 | 0.585950828 |
| 0.682344704 | 0.280724302 |
| 0.038361513 | 1.03801522 |
| 0.007996352 | 1.135684062 |
| 0.351118057 | 0.908782868 |
| 0.017104655 | 0.805693988 |
| 0.02347044 | 0.925391465 |
| 0.006359663 | 1.091661692 |
| 0.051151465 | 1.095314249 |
| 0.001598043 | 1.153244213 |
| 0.006895682 | 1.144692218 |
| 0.001902883 | 0.883012953 |
| 0.00319857 | 1.104331264 |
| 0.00280565 | 0.942949093 |
| 0.050792168 | 1.050290135 |
| 0.003638599 | 0.968816726 |
| 0.001191193 | 1.033973086 |
| 0.000982899 | 0.935633902 |
| 0.009991122 | 0.973196532 |
| 0.002276042 | 1.078776116 |
| 0.036396345 | 1.082733621 |
| 0.001598043 | 1.106781872 |
| 0.08896143 | 0.967909008 |

| | |
|---|---|
| 0.002391338 | 1.128211851 |
| 1.010489248 | 0.388087742 |
| 0.50413442 | 0.509307813 |
| 0.214415486 | 1.082816067 |
| 0.02745305 | 1.127965906 |
| 0.034164963 | 1.101657871 |
| 0.001598043 | 1.074665797 |
| 0.069111833 | 1.052404882 |
| 0.007371037 | 1.05199005 |
| 0.025638001 | 1.076496615 |
| 0.001598043 | 1.149168569 |
| 0.314825896 | 0.997070547 |
| 0.01607456 | 1.035343834 |
| 0.167164145 | 0.946833787 |
| 0.007996352 | 0.876821815 |
| 0.002790486 | 0.998273586 |
| 0.002391338 | 1.170031463 |
| 0.084862158 | 1.117546118 |
| 0.012052679 | 1.205967785 |
| 0.057130984 | 1.081794538 |
| 0.001028149 | 1.012890013 |
| 0.002818048 | 1.044301657 |
| 0.001486793 | 1.072864188 |
| 0.043003 | 1.163080271 |
| 0.003638599 | 1.225400356 |
| 0.06243051 | 1.110084965 |
| 0.001032472 | 1.194472746 |
| 0.096740468 | 0.945963332 |
| 0.005988527 | 0.965508057 |
| 0.002366011 | 1.15644924 |
| 0.001772163 | 1.11038646 |
| 0.01620341 | 0.925339637 |
| 0.00078174 | 0.964642655 |
| 0.026206587 | 1.078658252 |
| 0.00372282 | 1.094015669 |
| 0.02724497 | 1.002183406 |
| 0.002391338 | 1.059779759 |
| 0.050727757 | 1.150974881 |
| 0.019472188 | 1.237248493 |
| 0.058040413 | 1.163543135 |
| 0.000872781 | 0.889952573 |
| 0.084538 | 1.039096851 |
| 0.006045074 | 1.075965701 |
| 0.002276042 | 1.09247631 |
| 0.000804045 | 1.104549431 |
| 0.136337026 | 0.97778256 |
| 0.006558115 | 0.980109323 |
| 0.015930472 | 1.161305649 |
| 0.003417142 | 1.082024871 |
| 0.109971485 | 0.955499951 |
| 0.005398651 | 0.942424242 |
| 0.01733852 | 0.999801495 |
| 0.001155271 | 0.897818915 |
| 0.020219564 | 0.956124685 |
| 0.000865983 | 0.953531384 |
| 0.003937989 | 1.074899755 |
| 0.00078174 | 1.122477843 |
| 0.067342413 | 0.976351753 |
| 0.019292745 | 0.910922411 |
| 0.020254702 | 1.021178505 |
| 0.00138486 | 1.049814317 |
| 0.495100633 | 0.835541164 |
| 0.473941559 | 0.407944272 |

| | |
|---|---|
| 0.002134651 | 1.154006484 |
| 0.001155271 | 0.912390029 |
| 0.049807553 | 1.041948743 |
| 0.003638599 | 1.126957295 |
| 0.020254702 | 1.109206166 |
| 0.011083147 | 1.166602638 |
| 0.001619105 | 1.369742274 |
| 0.000896779 | 1.32063754 |
| 0.075871568 | 1.110489814 |
| 0.001028149 | 1.025418624 |
| 0.006558115 | 0.985879138 |
| 0.003022735 | 1.026090699 |
| 0.106298213 | 0.950777341 |
| 0.024286334 | 1.053598542 |
| 0.008761607 | 0.826732167 |
| 0.006359663 | 1.024597015 |
| 0.008077113 | 1.014854487 |
| 0.000686039 | 1.018603752 |
| 0.019257605 | 1.034042285 |
| 0.016269093 | 1.038622129 |
| 0.053362944 | 0.918224369 |
| 0.029242869 | 1.049553826 |
| 0.007371037 | 1.048818408 |
| 0.00277107 | 1.134063173 |
| 0.020382893 | 0.963515464 |
| 0.001865932 | 1.036549521 |
| 0.011544587 | 0.899626183 |
| 0.002276042 | 0.915401302 |
| 0.0365686 | 0.999731024 |
| 0.006558115 | 0.948527179 |
| 0.013852385 | 1.116760168 |
| 0.00372282 | 1.142315386 |
| 0.031019862 | 1.020430605 |
| 0.001766854 | 1.069462647 |
| 0.039388387 | 1.109532915 |
| 0.03331375 | 1.114169313 |
| 0.003811922 | 1.144318408 |
| 0.000803786 | 0.987002354 |
| 0.02927997 | 1.170437798 |
| 0.001028149 | 1.064329599 |
| 0.096906019 | 0.962882096 |
| 0.002391338 | 1.0551914 |
| 0.038782458 | 1.264729345 |
| 0.009901857 | 1.152985836 |
| 0.001915272 | 1.099883363 |
| 0.000966324 | 1.091554654 |
| 0.060077987 | 0.989561587 |
| 0.004382275 | 1.033350113 |
| 0.038483315 | 0.935364175 |
| 0.00078174 | 0.988308505 |
| 0.014094498 | 1.142551484 |
| 0.002168057 | 1.002639734 |
| 0.051687984 | 1.133165829 |
| 0.007371037 | 0.963743781 |
| 0.013184925 | 1.005010737 |
| 0.002276042 | 1.121475054 |
| 0.014121235 | 0.896308293 |
| 0.001191193 | 1.03364218 |
| 0.039628261 | 1.044216524 |
| 0.021600633 | 0.95832483 |
| 0.208040502 | 0.970387653 |
| 0.007371037 | 1.011629353 |
| 0.0030477 | 1.063085949 |

| | |
|---|---|
| 0.002633395 | 1.163991883 |
| 0.002605549 | 0.931865184 |
| 0.001869087 | 0.994671995 |
| 0.025448472 | 0.9252938 |
| 0.001191193 | 0.961835429 |
| 0.032568345 | 1.163790831 |
| 0.001558842 | 0.978320007 |
| 0.01131352 | 0.916946971 |
| 0.001191193 | 0.997793955 |
| 0.001191193 | 1.040922127 |
| 0.000982899 | 1.001827167 |
| 0.0030477 | 0.997459626 |
| 0.002633395 | 1.080243498 |
| 0.000865983 | 1.083342966 |
| 0.000561696 | 1.124360744 |
| 0.061825371 | 0.921316166 |
| 0.009778168 | 0.978845197 |
| 0.299844206 | 0.936528827 |
| 0.027565304 | 0.895721925 |
| 0.017927356 | 0.99691526 |
| 0.014250005 | 1.111418557 |
| 0.017104655 | 1.076168892 |
| 0.002934379 | 1.033763908 |
| 0.033881672 | 1.073722149 |
| 0.002416105 | 1.082243315 |
| 0.038361513 | 0.970838556 |
| 0.004588841 | 0.989598061 |
| 0.022437286 | 0.950085306 |
| 0.001865932 | 1.137252396 |
| 0.040934444 | 1.062154247 |
| 0.02455535 | 1.066865263 |
| 0.00222996 | 1.017821135 |
| 0.001825091 | 1.007994779 |
| 0.067010673 | 1.078941708 |
| 0.000927163 | 1.351533832 |
| 0.074631138 | 1.111795001 |
| 0.009440893 | 1.365705326 |
| 0.039448196 | 1.076935355 |
| 0.001476826 | 1.046635756 |
| 0.020254702 | 1.137780339 |
| 0.00200102 | 0.976437444 |
| 0.029801329 | 0.952271231 |
| 0.018538403 | 1.038374505 |
| 0.006884558 | 0.936892383 |
| 0.001915272 | 1.033718588 |
| 0.002134651 | 1.049328393 |
| 0.001155271 | 0.903867302 |
| 0.006359663 | 1.171223881 |
| 0.001082792 | 1.112680381 |
| 0.017104655 | 1.083978369 |
| 0.003244676 | 0.984142044 |
| 1.120978182 | 0.587329993 |
| 0.440052479 | 0.334815486 |
| 0.001915272 | 1.01738946 |
| 0.000966324 | 1.059261138 |
| 0.001191193 | 1.256232076 |
| 0.000982899 | 1.137079025 |
| 0.089509643 | 1.101856498 |
| 0.001768692 | 1.122183189 |
| 0.058771627 | 0.946280615 |
| 0.039652334 | 1.056587431 |
| 0.0030477 | 1.006713845 |
| 0.002633395 | 1.097030068 |

| | |
|---|---|
| 0.017104655 | 0.997388777 |
| 0.002934379 | 0.976979153 |
| 0.011842873 | 0.920518251 |
| 0.001191193 | 1.080520627 |
| 0.021740469 | 1.005915604 |
| 0.001062877 | 1.145103939 |
| 0.034154305 | 1.2463534 |
| 0.02662668 | 1.226855923 |
| 0.00129926 | 1.203011318 |
| 0.000686039 | 1.138773932 |
| 0.02486466 | 1.289387564 |
| 0.00372282 | 1.134314052 |
| 0.007734171 | 0.959210415 |
| 0.001915272 | 0.934789524 |
| 0.125531674 | 0.898597978 |
| 0.041143443 | 1.08182739 |
| 0.074564292 | 1.048883203 |
| 0.016417872 | 1.106618846 |
| 0.08559109 | 1.127964167 |
| 0.039609064 | 1.21053105 |
| 0.067647633 | 1.032689948 |
| 0.000896779 | 0.957091983 |
| 0.002688948 | 0.959486004 |
| 0.002659302 | 1.04112029 |
| 0.029718285 | 1.061710996 |
| 0.026318673 | 1.079747058 |
| 0.106678307 | 0.951222311 |
| 0.041302668 | 1.022153874 |
| 0.002790486 | 1.016908703 |
| 0.002391338 | 1.116019927 |
| 0.007030542 | 0.984495495 |
| 0.000865983 | 1.059993064 |
| 0.028398341 | 0.990997151 |
| 0.009091552 | 1.115555737 |
| 0.013042451 | 1.195058518 |
| 0.001028149 | 1.093482713 |
| 0.090918083 | 1.012260781 |
| 0.001598043 | 1.059667427 |
| 0.00170488 | 1.259217877 |
| 0.001420547 | 1.111408597 |
| 0.00170488 | 1.216201117 |
| 0.001420547 | 1.185185185 |
| 0.062997182 | 1.004895368 |
| 0.006558115 | 0.93364713 |
| 0.122213608 | 0.902816707 |
| 0.005988527 | 0.911040085 |
| 0.029909943 | 1.082532701 |
| 0.02946857 | 0.998853211 |
| 0.042280138 | 0.939244352 |
| 0.037846258 | 1.069045703 |
| 0.132488797 | 0.967116217 |
| 0.098481759 | 1.063695781 |
| 0.00170488 | 1.29603352 |
| 0.001420547 | 1.077048936 |
| 0.017104655 | 1.015945442 |
| 0.002229985 | 0.912121212 |
| 0.069580927 | 1.03312444 |
| 0.017104655 | 0.936098931 |
| 0.002605549 | 1.16678777 |
| 0.001869087 | 1.105727606 |
| 0.021772921 | 1.042124168 |
| 0.016387492 | 1.113162393 |
| 0.00170488 | 1.092234637 |

| | |
|---|---|
| 0.001420547 | 1.114532203 |
| 0.004539856 | 1.054588725 |
| 0.00170488 | 1.197765363 |
| 0.002134651 | 1.123668365 |
| 0.001155271 | 1.000733138 |
| 0.050013917 | 0.974460908 |
| 0.02815869 | 1.16840052 |
| 0.0407391 | 1.019301573 |
| 0.000896779 | 1.127421176 |
| 0.066103322 | 1.106718682 |
| 0.001028149 | 0.973015299 |
| 0.035537341 | 1.119440726 |
| 0.00170488 | 1.142011173 |
| 0.009673337 | 1.037840255 |
| 0.007205723 | 1.1999667 |
| 0.002818048 | 1.191579303 |
| 0.001486793 | 1.062576232 |
| 0.017104655 | 0.918313051 |
| 0.002934379 | 1.003964701 |
| 0.003713507 | 1.260905229 |
| 0.001558842 | 0.970900997 |
| 0.073985572 | 0.88136121 |
| 0.057635248 | 0.972799807 |
| 0.013839588 | 1.048490257 |
| 0.000215966 | 1.121955307 |
| 0.024055078 | 1.070138292 |
| 0.001155271 | 1.089442815 |
| 0.007371037 | 0.980161692 |
| 0.00277107 | 1.052045944 |
| 2.257357477 | 0.320683761 |
| 1.607846774 | 0.259153435 |
| 0.00129926 | 1.169159256 |
| 0.000686039 | 1.150918584 |
| 0.001411669 | 1.107337363 |
| 0.00078174 | 1.022911559 |
| 0.083734106 | 0.829369513 |
| 0.020627512 | 0.907429372 |
| 0.046779012 | 1.031831028 |
| 0.00170488 | 1.066368715 |
| 0.00170488 | 1.116592179 |
| 0.001420547 | 1.05101889 |
| 0.048533093 | 0.979932302 |
| 0.003638599 | 0.885053381 |
| 0.007942215 | 0.833196923 |
| 0.004123449 | 0.949451832 |
