## Supplementary figures and images for "Rapid accumulation of fluorophores and fast kill identify drugs with bactericidal effects against Gram-negative bacteria"

### Supplementary Figure 1

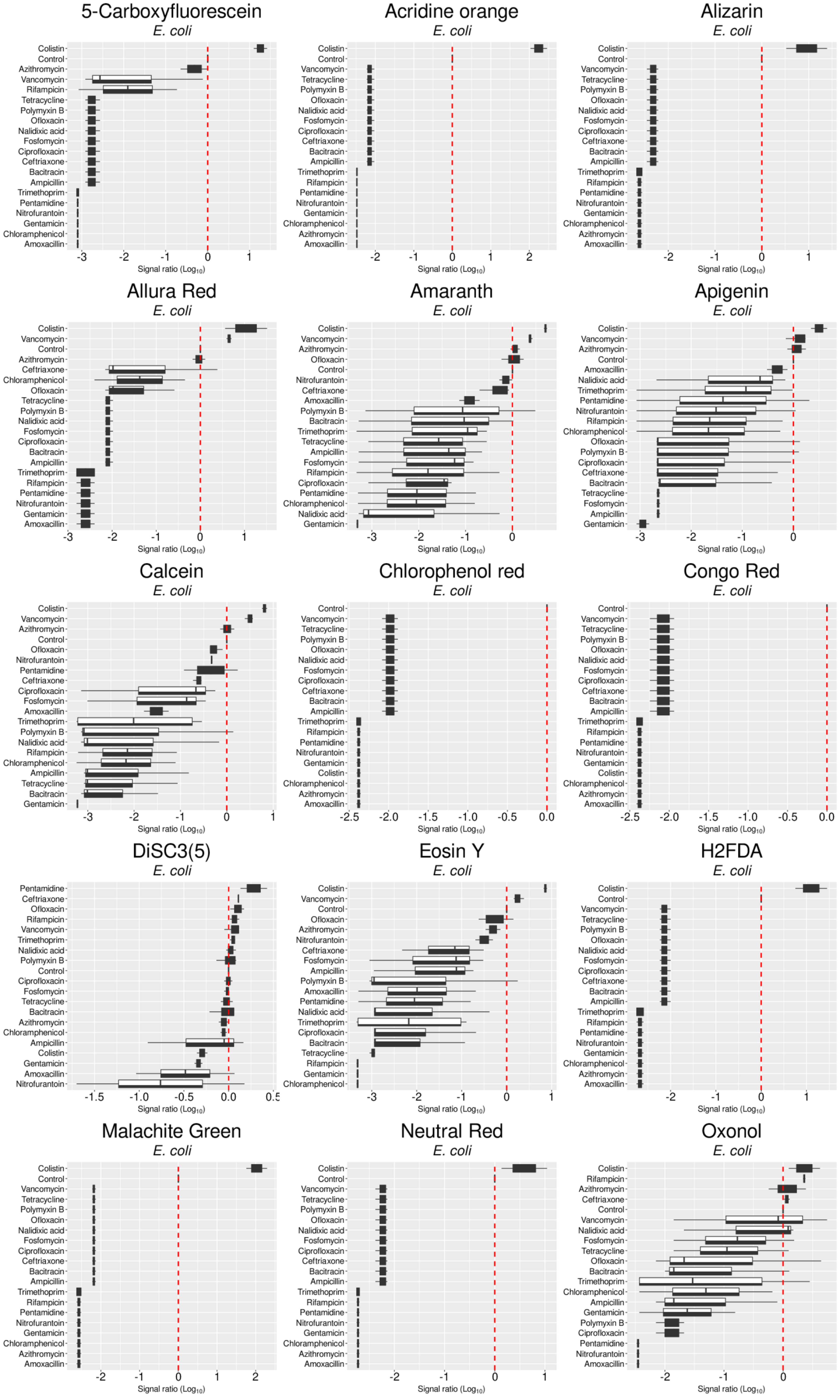

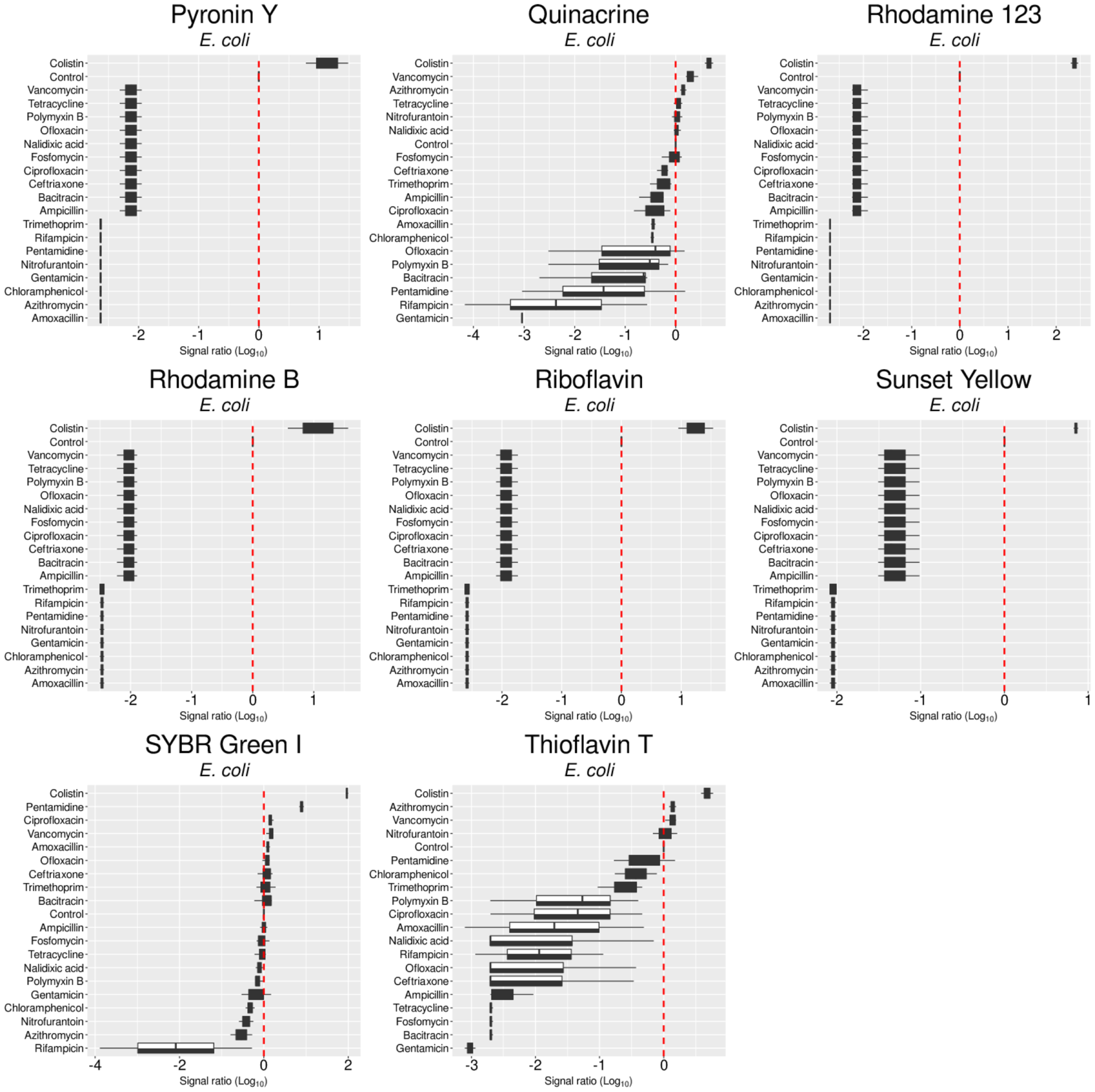

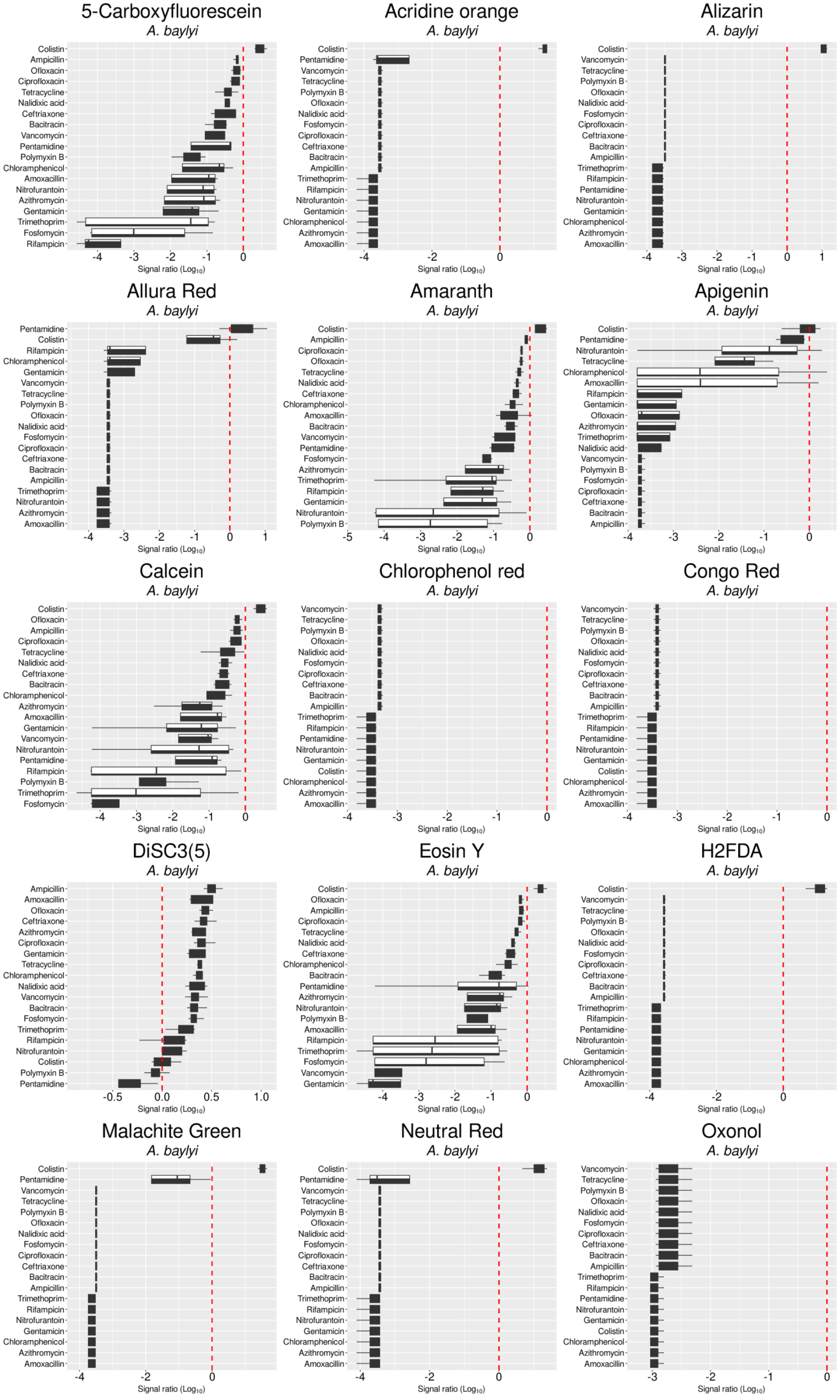

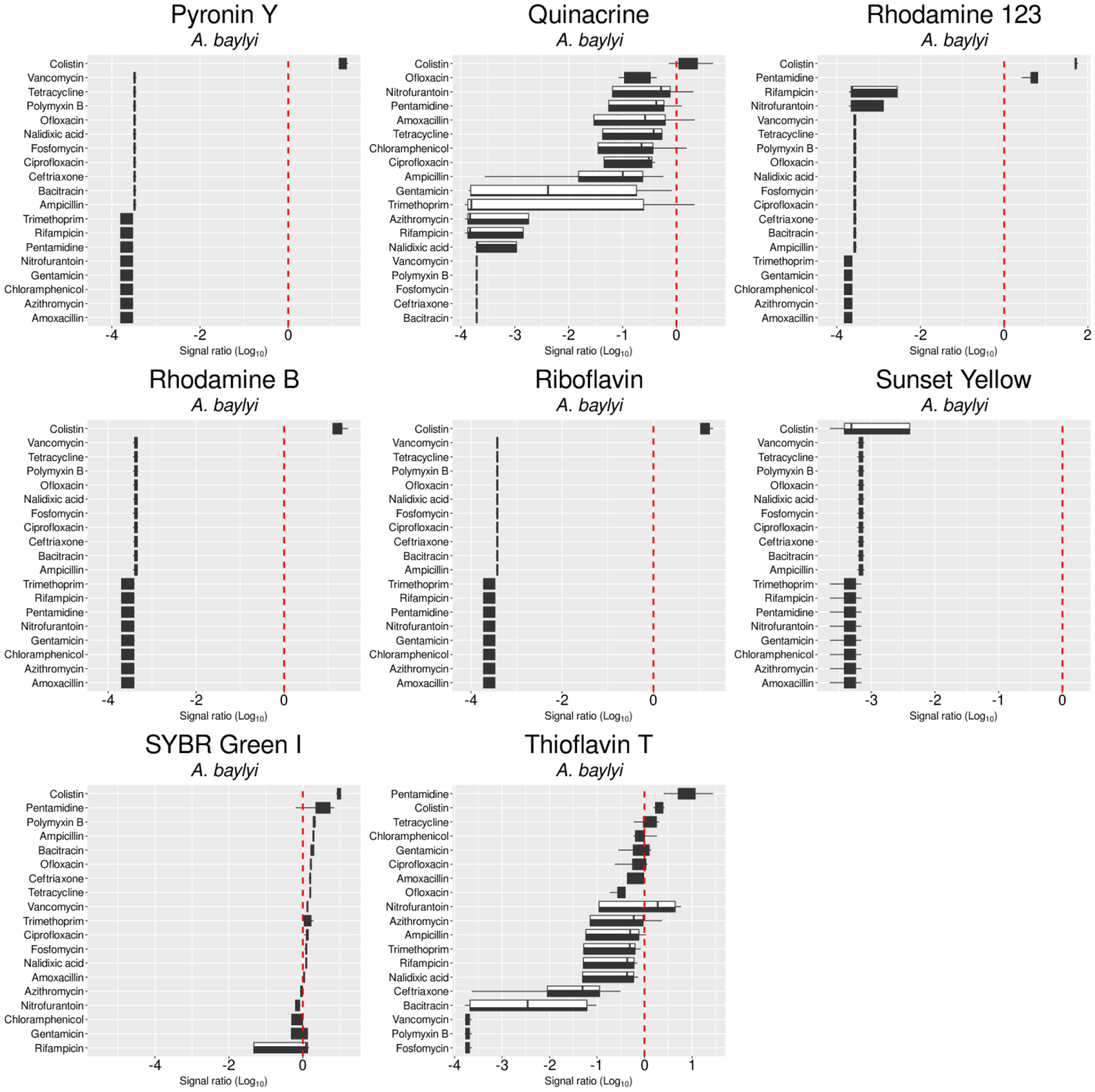
